# Ubiquitous, B_12_-dependent virioplankton utilizing ribonucleotide triphosphate reductase demonstrate interseasonal dynamics and associate with a diverse range of bacterial hosts in the pelagic ocean

**DOI:** 10.1101/2023.03.13.532061

**Authors:** Ling-Yi Wu, Gonçalo J. Piedade, Ryan M. Moore, Amelia O. Harrison, Ana M. Martins, Kay D. Bidle, Shawn W. Polson, Eric Sakowski, Jozef I. Nissimov, Jacob T. Dums, Barbra D. Ferrell, K. Eric Wommack

## Abstract

Through infection and lysis of their coexisting bacterial hosts, viruses impact the biogeochemical cycles sustaining globally significant pelagic oceanic ecosystems. Currently, little is known of the ecological interactions between lytic viruses and their bacterial hosts underlying these biogeochemical impacts at ecosystem scales. This study focused on populations of lytic viruses carrying the B_12_- dependent Class II monomeric ribonucleotide reductase (RNR) gene, ribonucleotide triphosphate reductase (RTPR), documenting seasonal changes in pelagic virioplankton and bacterioplankton using amplicon sequences of RTPR and the 16S rRNA gene, respectively. Amplicon sequence libraries were analyzed using compositional data analysis tools that account for the compositional nature of these data. Both virio- and bacterioplankton communities responded to environmental changes typically seen across seasonal cycles as well as shorter term upwelling–downwelling events. Defining RTPR-carrying viral populations according to major phylogenetic clades proved a more robust means of exploring virioplankton ecology than operational taxonomic units defined by percent sequence homology. Virioplankton RTPR populations showed positive associations with a broad phylogenetic diversity of bacterioplankton including dominant taxa within pelagic oceanic ecosystems such as *Prochlorococcus* and SAR11. Temporal changes in RTPR-virioplankton, occurring as both free viruses and within infected cells, indicated possible viral–host pairs undergoing sustained infection and lysis cycles throughout the seasonal study. Phylogenetic relationships inferred from RTPR sequences mirrored ecological patterns in virio- and bacterioplankton populations demonstrating possible genome to phenome associations for an essential viral replication gene.

## INTRODUCTION

The subtropical pelagic ocean gyres are extensive, coherent regions that collectively represent the largest ecosystems on Earth, accounting for 40% of the earth’s surface ^1^. Circulation within the upper kilometer of subtropical gyres is primarily wind driven ^2^. Due to their size, the horizontal and vertical motion of water within this upper layer of the gyres plays an outsized role in regulating global nutrient cycles and the size of the atmospheric carbon pool. Both of these issues are critical concerns for understanding anthropogenic impacts on fisheries, oceanic general circulation, and global climate. Ultimately, the activities of and the interactions between oceanic microbial communities—composed of protists, phytoplankton, bacterioplankton, and virioplankton—regulate the flow of carbon and nutrient elements within the pelagic ocean gyres ^3,4^. Truly appreciating how oceans will respond to environmental changes brought on by anthropogenic activities requires detailed, mechanistic understanding of oceanic microbial communities ^5–8^. This study focused on specific interactions between important and ubiquitous populations of lytic viruses ^9,10^ and bacterioplankton, the most abundant microbial host group within the low nutrient ecosystems that dominate much of the pelagic ocean.

Viral cell lysis is among the microbial community interactions most directly responsible for the conversion of cellular biomass into dissolved and particulate organic carbon ^11^. This process is one of the major top-down controls on bacterioplankton productivity, responsible for between 10 and 40% of daily bacterioplankton mortality in the ocean ^12^. Upon lysis, cell contents not consumed by viral replication are released, contributing to dissolved nutrient pools within marine ecosystems. Bacterioplankton cell lysis comprises the largest known carbon flux process between biomass and the dissolved organic matter (DOM) pool amounting to ∼150 gigatons C/yr ^6,13^. The conversion of biomass to DOM from viral lysis is five times greater than that performed by other biological mechanisms such as grazing and programmed cell death ^6,13^. This flux from biomass to DOM sustains nutrient cycles, providing essential growth substrates for bacterioplankton and phytoplankton productivity. The availability and concentrations of different nutrients within the DOM pool comprise the bottom-up controls on bacterioplankton productivity ^6,13,14^. Besides influencing the composition, diversity, and productivity of marine bacterial communities through top-down and bottom-up controls, viruses can influence bacterial physiology, improving fitness in the face of environmental changes ^15,16^ by altering metabolic pathways ^17,18^ through horizontal gene transfer; and through the expression of metabolic genes carried within viral genomes during viral infection ^19,20^. However, the behavior of individual virioplankton populations within ecosystems and their detailed interactions with bacterioplankton host populations remain largely unknown. This study examined both virioplankton and bacterioplankton dynamics within the North Atlantic Subtropical Gyre across time with the objective of observing interactions at the community and population levels.

For some time, microbial ecologists have investigated bacterial populations and communities utilizing 16S rRNA gene amplicon sequences, a universal gene marker for bacterial diversity ^21^. Due to their polyphyletic origins, viruses do not have a universally conserved marker gene that could be similarly used for tracing viral evolutionary history and community ecology investigations. Nevertheless, there are a few widespread genes exhibiting robust evolutionary histories that can be used as markers for viral community ecology studies within natural environments. Genes such as the DNA polymerase A gene (encoding DNA polymerase I) ^22,23^, terminase gp20 gene ^24–27^, major capsid protein gp23 gene of T4-like phages ^28–30^, and ribonucleotide reductase (RNR) genes ^9^ have all informed studies assessing the diversity of abundant viral populations. This study used amplicons of the ribonucleotide triphosphate reductase (RTPR) gene for following the dynamics of virioplankton populations.

RNRs are the only known enzymes capable of reducing ribonucleotides to deoxyribonucleotides, essential substrates for DNA synthesis ^31,32^. RNRs are ancient enzymes that were essential to the emergence of DNA ^33^. As a consequence, RNR genes are under rigorous evolutionary selection pressure, and among the most abundantly identified genes in marine virome libraries ^34^. Importantly, RNR genes are present in tailed phages within the class Caudoviricetes and have been identified within the genomes of viruses infecting hosts within all three domains of life. Generally, lytic viruses carry RNR genes, which ensures a steady supply of deoxyribonucleotides for DNA synthesis ^35^. Therefore, RNRs are most commonly observed within the genomes of lytic phages.

RNRs are biologically informative as this group of enzymes consists of three physiological classes according to reactivity with oxygen. Class I RNRs are oxygen-dependent. Class II RNRs are oxygen-independent and rely upon an adenosylcobalamin (vitamin B_12_) cofactor for ribonucleotide reduction. Class II RNRs are further divided according to structure and substrate utilization. Monomeric Class II RTPR reduces ribonucleotide triphosphate to deoxyribonucleotide triphosphate (dNTPs), the substrate for DNA synthesis. Dimeric Class II reduces ribonucleotide diphosphate to deoxyribonucleotide diphosphate thus requiring a dinucleotide kinase for the final step in forming dNTPs. Class III RNRs are sensitive to oxygen. As such, Class I and Class II RNRs, not Class III RNRs, are commonly present within virioplankton communities found in oxygenated ocean water ^10,32^. This study leveraged the predominance of RTPR genes within viruses for examining the ecological dynamics of virioplankton populations. RTPR and 16S rRNA amplicon sequence data were analyzed against a backdrop of microbial and oceanographic observations using the latest compositional data analysis (CoDA) tools ^36,37 38,39 40,41^ for assessing how lytic, B_12_-dependent viral populations responded to seasonal dynamic changes within the euphotic zone of a subtropical oceanic gyre, a region critically important within the global carbon cycle.

## MATERIALS AND METHODS

### Study Site and Sample Collection

Seawater samples were collected from a fixed station at South of Channel Faial-Pico islands (SCFP; 38°23’11.67”N, 28°35’6.14”W), located in the northeastern Atlantic, 6.4 nautical miles off São Mateus (Pico Island) in the Azores archipelago (ICES subdivision Xa2). The station is representative of a larger open ocean ecosystem, the Mid-Atlantic Ridge ecoregion, corresponding to the International Council for the Exploration of the Seas (ICES) areas X and XII, which includes a high proportion of high seas ^42^. The Azores are located along the northern boundary of the North Atlantic Subtropical Gyre, which constitutes a barrier between superficial cold waters of Nordic provenance and warmer waters of the gyre. Due to the influence of large scale currents such as the Gulf Stream, the North Atlantic Current, and the Azores Current, the islands are mostly exposed to the transport of warm waters of equatorial and tropical origin. These currents result in high salinity, sharp horizontal temperature gradients, and high sea surface temperatures (on average 15–16 °C in winter and 22–24 °C in summer). These physical conditions result in low nutrient regimes, and correspond with low ecosystem productivity ^43^. Nevertheless, frontal regions and underwater features (seamounts, eddies, upwelling areas) can enhance local productivity ^44,45^.

Water samples were collected in acid-rinsed (0.1 M HCl) and seawater-flushed carboys between 9:30 and 11:00 local time from 0, 5, 25, 50, and 75 m depths during seven sampling events on May 12th, June 8th, June 22nd, July 11th, July 27th, August 5th, and September 8th of 2016. Surface seawater (0 m depth) was collected by directly submerging a carboy. Seawater at other depths was collected using a 2.5 L Niskin bottle and subsequently transferred to a 5 L carboy. Seawater samples were transported inside a light-tight cooler and laboratory-processed within 2 h of collection. Subsamples for flow cytometric enumeration of phytoplankton were fixed immediately using glutaraldehyde (0.5% final concentration). Flow cytometry samples were transported on ice (1–2 h) and stored at -80 °C until further processing.

Water column conductivity, temperature, and depth (CTD) data was collected down to 150 m using a CTD instrument (MIDAS Valeport, Totnes, United Kingdom) (**Supplementary Table S1)**. Salinity (unitless, on the Practical Salinity Scale PSS-78) was expressed as a function of conductivity. Density (kg/m³) was expressed as a function of pressure. CTD profiles were visualized using the cmocean colour palettes ^46^ on Ocean Data View 5.3.0 ^47^ (https://odv.awi.de). Temperature and salinity values between sampling days were interpolated using the weighted average grid method ^47^. Temperature, salinity, and density were determined by averaging all values 0.5 m above and below the targeted sampling depths.

Equipment malfunction prevented CTD data collection on September 8th, thus, temperature, salinity, and density metadata were estimated from the nearest time, pixel, and depth from the Marine Copernicus Global Ocean Physics model ^48^ (GLOBAL_ANALYSISFORECAST_PHY_CPL_001_015, http://marine.copernicus.eu/). Average values for chlorophyll a (mg/m^3^), sea surface temperature (SST, °C), particulate organic carbon (POC, mg/m^3^), and photosynthetically active radiation (PAR, Einstein/m^2^d) data in 2016 were obtained from the NASA Ocean Color website (http://oceancolor.gsfc.nasa.gov/) at 1 km resolution. Satellite data was mapped using SeaDAS 7 ^49^ for the 9 km^2^ area surrounding sampling station SCFP.

### Phytoplankton Enumeration using Flow Cytometry

Phytoplankton cells were enumerated using a BD Influx Mariner 209s Flow Cytometer (BD Biosciences, San Jose, CA) equipped with a 488 nm argon laser and 525 ± 15, 542 ± 27, 585 ± 40, and 692 ± 40 nm emission filters with a nozzle size of 70 μm. After thawing and gentle vortexing, samples were run for 2 min ^50^. A blank consisting of 0.22 µm filtered Milli-Q water was run before and after counting. The flow rate was determined by running Milli-Q water for 2 min and dividing the mass difference (determined with a scale) by time; mass was converted to volume using the density of 1 kg/L. Phytoplankton cells were identified based on chlorophyll fluorescence (692 ± 40 nm) from 488 nm excitation (**Supplementary Table S1**).

### Virio- and Bacterioplankton Biomass Collection

The cellular (>0.22 µm) and subcellular (0.02–0.22 µm) fractions, biologically defined in this study as bacterioplankton and virioplankton, respectively, were separated through two consecutive syringe filtrations using 0.22 and 0.02 µm filters. All tubing and containers were acid-washed and deionized water-rinsed prior to sample filtration. The bacterioplankton fraction was collected by filtering 5 L of seawater through a 0.22 µm pore-size Sterivex filter (Merck Millipore, Darmstadt, Germany) under a vacuum pressure system linked to a filtration ramp (Merck Millipore, Burlington, MA). The sample collection flask was rinsed twice using the first 200 mL of 0.22 µm filtrate. Subsequently, the virioplankton fraction was collected by filtering 1 L of 0.22 µm filtrate through a 0.02 µm pore size Anotop filter (Whatman, Maidstone, United Kingdom) ^51,52^ connected to a 60 mL syringe, using a caulk gun which generated a steady pressure on the syringe. Both 0.22 µm and 0.02 µm filters with bacterio- or virioplankton biomass were parafilm sealed and stored at -80 °C until DNA extraction.

### DNA Extraction from the Virioplankton Fraction and Tag-Encoded RTPR Amplicon Sequencing

In brief, DNA was extracted from the 0.02 µm Anotop filters (virioplankton fraction) and quantified using previously reported protocols ^52^. RTPR PCR amplicons of ∼750 bp were obtained using virioplankton template DNA and degenerate primers designed using sequence alignments derived from uncultivated virus populations observed within marine viromes. After magnetic bead purification, amplicon molecules were tagged with a unique barcode sequence using ligation. Purified, barcoded amplicons were then subjected to a limited cycle PCR with barcode sequences as template targets. Enriched barcoded amplicons from each sample were then purified, pooled in equal proportions, and sequenced using a PacBio RSII sequencer (Pacific Biosciences, Menlo Park, CA). Detailed methods describing DNA extraction, amplification of RTPR target genes, and amplicon sequencing are provided in **Supplementary Methods**.

### DNA Extraction from the Bacterioplankton Fraction, and 16S rRNA Amplicon and Tag-Encoded RTPR Sequencing

DNA was extracted from the 0.22 µm Sterivex filters (bacterioplankton fraction) using the phenol/chloroform method ^53^. The V3–V4 hypervariable region of the 16S rRNA gene was PCR-amplified and sequenced from the bacterioplankton fraction on the Illumina MiSeq (Illumina, San Diego, CA) using a Nextera XT DNA Library Preparation Kit (Illumina), which exploits a dual-indexing strategy for multiplexed sequencing ^54^. RTPR genes within the bacterioplankton fraction were PCR-amplified, barcoded, and enriched using the same protocol as described for virioplankton. Detailed methods describing DNA extraction, amplification of 16S rRNA and RTPR target genes, and amplicon sequencing are provided in **Supplementary Methods**.

### RTPR amplicon quality control

General and detailed bioinformatics analysis pipelines are summarized in **Fig. 1** and **Supplementary Fig. S1**, respectively. Briefly, RTPR gene amplicon sequences from the virioplankton fraction (PacBio RSII sequencer) and the bacterioplankton fraction (PacBio Sequel sequencer) were initially screened for low quality bases and read length. Circular consensus sequence (CCS) reads were generated. Reads with less than three full passes, less than 98% minimum predicted accuracy, and with a length shorter than 250 bp or longer than 5,000 bp were excluded. Subsequently, sequence reads were demultiplexed, and primer and barcode sequences were detected and removed. Reads were translated into predicted amino acid sequences using a custom frameshift polishing pipeline (https://github.com/dnasko/frameshift_polisher). Key catalytic residues within translated amplicons were used for validating sequences as true RTPRs using the Protein Active Site Validation (PASV) (https://github.com/mooreryan/pasv) pipeline ^55^. Detailed methods describing bioinformatic processing of RTPR amplicon sequences are provided in **Supplementary Methods**.

**Figure 1.**
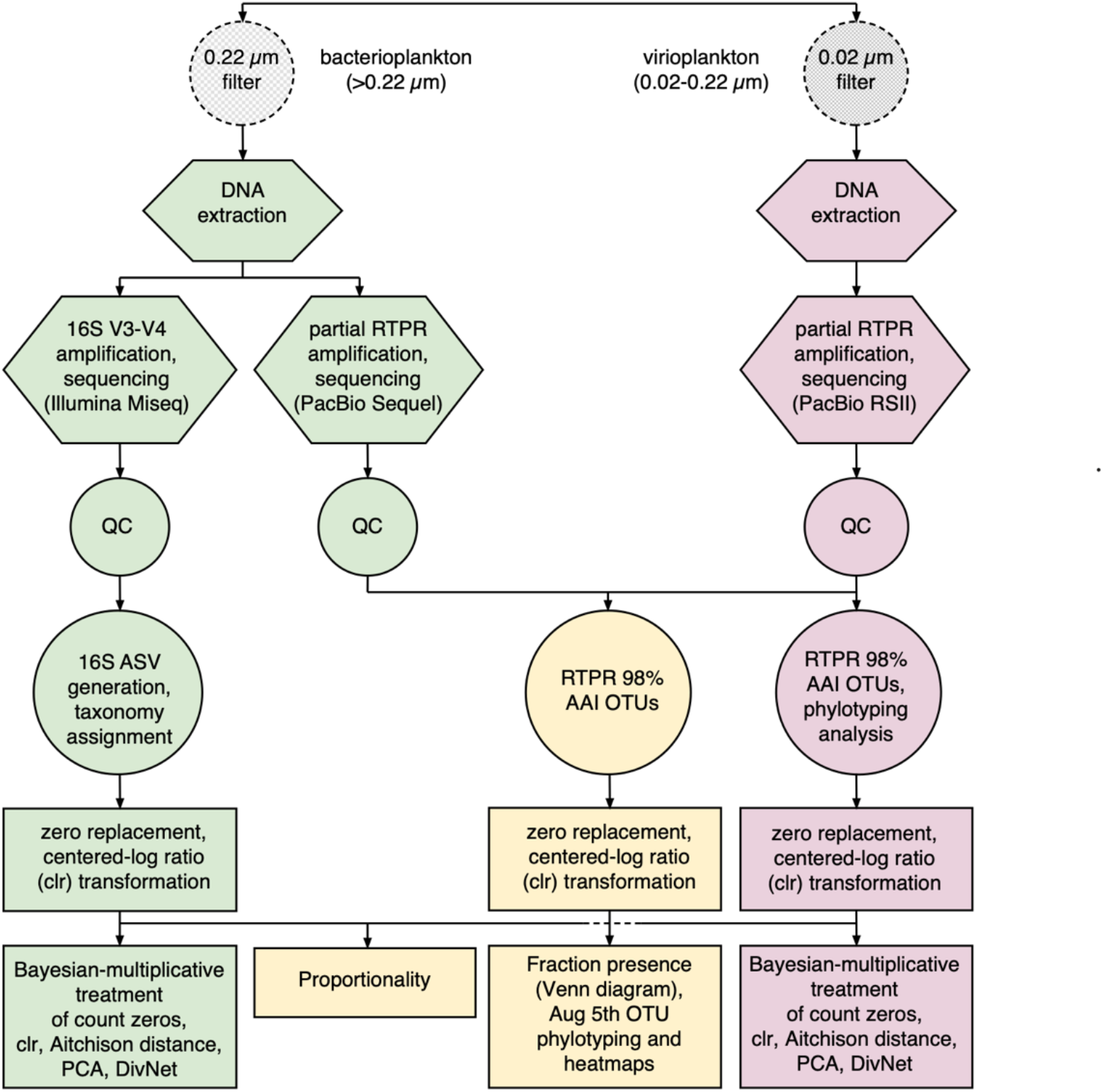
Overview of experimental methods and analysis pipeline. The 0.22 µm filter captured bacterioplankton cells. The 0.02 µm filter captured virioplankton particles. Hexagons represent molecular genetic experimental methods. QC stands for a collection of quality control steps specific to the type of next-generation sequencing technology used in sequencing amplicon libraries. Circles represent conventional data analysis methods. Squares represent compositional data analysis (CoDA) methods.

### Virioplankton RTPR 98% OTUs and phylogenetic clades

Quality-controlled virioplankton RTPR amino acid sequences were run through 11 *de novo* operational taxonomic unit (OTUs) clustering assessments varying the percent sequence identity (90– 100% amino acid sequence identity) using the *cluster_fast* command with default settings in USEARCH (version 11.0) ^56^. This heuristic process identified 98% OTU clusters as a balance between optimal lumping and splitting of OTUs (**Supplementary Fig. S2**). Virioplankton RTPR 98% OTU amino acid sequences were taxonomically divided into clades using a phylotyping method. OTUs were aligned using MAFFT (version 7.450) with the ginsi method. A phylogenetic tree of virioplankton RTPR 98% OTUs was built using FastTree (version 2.1.11) ^57^ and visualized using Iroki ^58^. Virioplankton RTPR phylogenetic clades were defined manually.

### 16S quality control and classification

General and detailed analysis pipelines are summarized in **Fig. 1** and **Supplementary Fig. S1**. Demultiplexed, paired-end 16S rRNA reads from the bacterioplankton fraction were imported into QIIME2 (QIIME2 2019.1) ^59^ for quality filtering and generation of amplicon sequence variants (ASVs) using DADA2 ^60^. Forward and reverse reads were truncated at positions 35 and 300 and at positions 35 and 295, respectively. 16S ASVs were taxonomically identified (**Supplementary Table S2**) using the Silva 132 QIIME-compatible release (full length, seven-level taxonomy) ^61^ as a reference database in QIIME2. Sequences taxonomically assigned to Archaea (22 ASVs) and chloroplasts (230 ASVs) were removed prior to downstream analysis.

For proportionality tests with bacterioplankton 16S ASVs and virioplankton RTPR 98% OTUs, ASVs were restricted to only those that occurred in the 29 samples with virioplankton RTPR amplicons. From that subset of 29 samples, only ASVs with a total count greater than 100 and with at least 10 counts in at least 10 samples were retained for downstream analysis. This avoided the risk of spurious associations based on rare populations observed in few samples.

### RTPR and 16S community and population analyses

Virioplankton RTPR 98% OTUs, virioplankton RTPR phylogenetic clades, and bacterioplankton 16S ASVs were used in community and population analyses such as alpha and beta diversity, and population proportionality. The influence of environmental conditions on virio- and bacterioplankton communities were determined based on complete-linkage hierarchical clustering of RTPR 98% OTUs, RTPR phylogenetic clades, or 16S ASV community profiles using the hclust method in R. Tested environmental features include sampling day, depth, temperature, density, current velocity, salinity, nanoeukaryote abundance, picoeukaryote abundance, *Prochlorococcus* abundance, *Synechococcus* abundance, total autotroph abundance, viral abundance, and bacterial abundance. Clustering of virio- and bacterioplankton communities was based on the Aitchison distance between the May 12th 5 m sample and every other sample. Resulting dendrograms were visualized with Iroki. Correlations between virio- or bacterioplankton communities and associated environmental features were tested using the *qiime diversity mantel* ^*65*^ command in QIIME2 (permutation times: 999). Significant correlations were defined as *p* <0.05. Associated environmental features that showed significant correlations with sample community composition were plotted next to the sample community composition dendrogram.

Alpha diversity of the virio- or bacterioplankton communities in each sample was estimated based on the Shannon index using the R package DivNet (version 0.3.2) ^66^ and displayed using ggplot2 (version 3.3.0) ^67^. Significance of alpha diversity differences was tested using betta function in the R package DivNet breakaway (version 4.6.14) ^68^. Estimated Shannon indices were converted to the effective number of species (ENS, Hill numbers) ^69^. Correlations of ENS estimates based on 98% OTUs or RTPR phylogenetic clades were tested using the cor.test method in R. Beta diversity was visualized through principal component analysis (PCA) plots of sample community compositions using the R package compositions (version 1.40.3) ^70^. Significance of beta diversity differences between sampling day or depth groups was tested using *beta-group-significance* command (permanova) method ^71^ in QIIME2.

Associations within or between virioplankton RTPR phylogenetic clades and bacterioplankton 16S ASV populations were explored using proportionality (*rho*, ρ) tests, with -1 < ρ < 1 ^72^. Only positive ρ values reflecting a positive association between two populations were considered, as negative ρ values cannot be conclusively explained ^73^. Only those associations with false discovery rates (FDRs) lower than 0.05 were retained for further analysis. Proportionality and FDR tests were performed using the R package propr (version 4.2.8). Associations with FDR <0.05 were extracted and plotted in a heatmap using ComplexHeatmap ^74^. Columns and rows of the heatmap were clustered based on phylogenetic relations of virio- and bacterioplankton, respectively. Virioplankton RTPR clades were arranged by their topological order on the 98% OTU phylogenetic tree. Bacterioplankton 16S ASVs were arranged by their 16S phylogenetic placement or association patterns.

### Virio- and bacterioplankton population analysis using common RTPR 98% OTUs

In 27 of 35 samples, RTPR amplicons were obtained from both virio- and bacterioplankton fractions yielding 47,997 quality-controlled RTPR amino acid sequences. Quality-controlled virio- and bacterioplankton RTPR amino acid sequences were run through 11 *de novo* operational taxonomic unit (OTU) clustering assessments varying the percent sequence identity (90–100% amino acid sequence identity) using the *cluster_fast* command with default settings in USEARCH. This heuristic process identified 5,411 98% OTU clusters as a balance between optimal lumping and splitting of OTUs (**Supplementary Fig. S3**). OTUs with less than ten counts were removed prior to downstream analysis. The COUNTIF function in Excel was used for binning the remaining 254 RTPR 98% OTUs into those containing sequences from both fractions, only the virioplankton fraction, or only the bacterioplankton fraction. RTPR 98% OTU counts were calculated using the SUM function in Excel prior to zero replacement and clr transformation. An arbitrary clr value of 5 or -5 was assigned to those common 98% OTUs found exclusively in the virio- or bacterioplankton fraction, respectively. The difference in clr abundance between fractions of each RTPR 98% OTU was calculated using the subtraction formula in Excel. Venn diagrams were plotted to show the total clr abundance of each OTU and the distribution of each OTU (difference in clr abundance) in the virio- and/or bacterioplankton fraction.

Only the August 5th sampling date yielded RTPR amplicons from virio- and bacterioplankton fractions at all depths. Thus, this date was chosen for assessing depth-resolved distribution patterns of RTPR virioplankton populations within the virio- and bacterioplankton fractions. Calculation of clr abundance values for the August 5th OTUs was performed by removing any OTU not present in at least one August 5th sample, leaving 1,477 of the 5,411 OTUs observed in the combined virio- and bacterioplankton datasets. Twenty-nine August 5th OTUs contained more than 50 sequences and had two or fewer zero replacements, and were retained for further analysis. Similarity between the phylogenetic distance of these 29 RTPR 98% OTUs and their depth-resolved distribution patterns within the virioplankton fraction, the bacterioplankton fraction, and the combined virio- and bacterioplankton fraction were tested by plotting the cophenetic distance versus Aithison distance for all combinations of 29 OTUs. Combinations rather than permutations of OTUs were used, as the distances used are symmetric (e.g., d(A, B) = d(B, A), where d is the distance function, and A and B are OTUs). Self-distances (e.g., d(A, A), d(B, B)) were not included. Linear models and significance tests were performed with the lm function in R.

### Data transformation

Compositional data analysis (CoDA) methods were applied to all analyses of community diversity and population interactions. Zero values in the feature tables [virioplankton RTPR 98% OTUs (**Supplementary Table S3**); RTPR phylogenetic clade (**Supplementary Table S4**); bacterioplankton 16S ASVs (**Supplementary Table S5**); bacterioplankton 16S ASVs retained in the 29 samples with RTPR amplicons (**Supplementary Table S6**); and common RTPR 98% OTUs found in both the bacterio- and virioplankton fractions (**Supplementary Tables S7 & S8**)] were replaced by imputed values using the count zero multiplicative method from the R package zCompositions (version 1.3.3.1) ^62^ in R (version 3.5.3) ^63^. The centered log-ratio (clr) transform was applied to the zero-replaced data set to produce clr abundance using the R package CoDaSeq (version 0.99.4) ^64^. Within the August 5th subset (sampling date with RTPR amplicons in both fractions at all depths) of common RTPR 98% OTUs (**Supplementary Table S9**), zero count values were replaced using *cmultRepl* with the method count zero multiplicative (CZM) in the zCompositions package (version 1.3.4) of R (version 4.0.2). Subsequently, the clr transformation was performed using base 2 in the R package CoDaSeq.

### RTPR 98% OTUs within global RNR context

RTPR 98% OTUs from both virio- and bacterioplankton fractions, RTPRs identified through stringent homology search of the *Tara* Ocean dataset ^75^, and RNRdb ^76^ databases were aligned using Geneious 10.0.9 MAFFT alignment FFT-NS-2 and trimmed to the region of interest (H346 to S643 in *Lactobacillus leichmannii* monomeric class 2 adenosylcobalamin-dependent ribonucleotide-triphosphate reductase). A phylogenetic tree of trimmed sequences was built using FastTree and visualized with Iroki.

### Data Deposition

RTPR and 16S rRNA gene sequences were deposited in NCBI BioProject under the accession number PRJNA842570 and Zenodo under DOI 10.5281/zenodo.7313881 (https://doi.org/10.5281/zenodo.7313881).

## RESULTS

### Water Column Environment

Temperature of surface waters at station SCFP increased from May 12th (16.4 ºC) to August 5th (22.0 ºC) (**Fig. 2A**), agreeing with broader sea surface temperature (SST) data derived from satellite imagery (**Supplementary Fig. S4A**). Surface salinity (using the Practical Salinity Scale) increased from May 12th (35.9) to July 11th (36.1), decreased by 0.1 on July 27th, and increased again to 36.1 on August 5th (**Fig. 2B**). Surface waters were well mixed down to 50 m as indicated by temperature and salinity contours prior to June (**Fig. 2**). Water column stratification intensified throughout the sampling period, accompanying the mixed layer depth rising from 43 m (May 12th) to 5 m (August 5th). Salinity showed distinct sub-surface increases during the study period, indicating substantial water movement in the sampling region. Salinity changes in the sub-surface were centered at the 75 m depth, from June 8th to July 27th, especially on July 11th. The observations suggested the presence of an anticyclonic (downwelling) vortex at the sampling location around July 11th. Uplifts of isotherms and isohalines found in temperature and salinity profiles between sampling dates suggested the presence of cyclonic (upwelling) vortices, on June 22nd (more pronounced) and on July 27th (less pronounced) extending to August 5th. Satellite-derived data in 2016 (**Supplementary Fig. S4A**) showed one phytoplankton bloom, based on chlorophyll a and particulate organic carbon (POC), that occurred between February and April, with its maximum in March.

**Figure 2.**
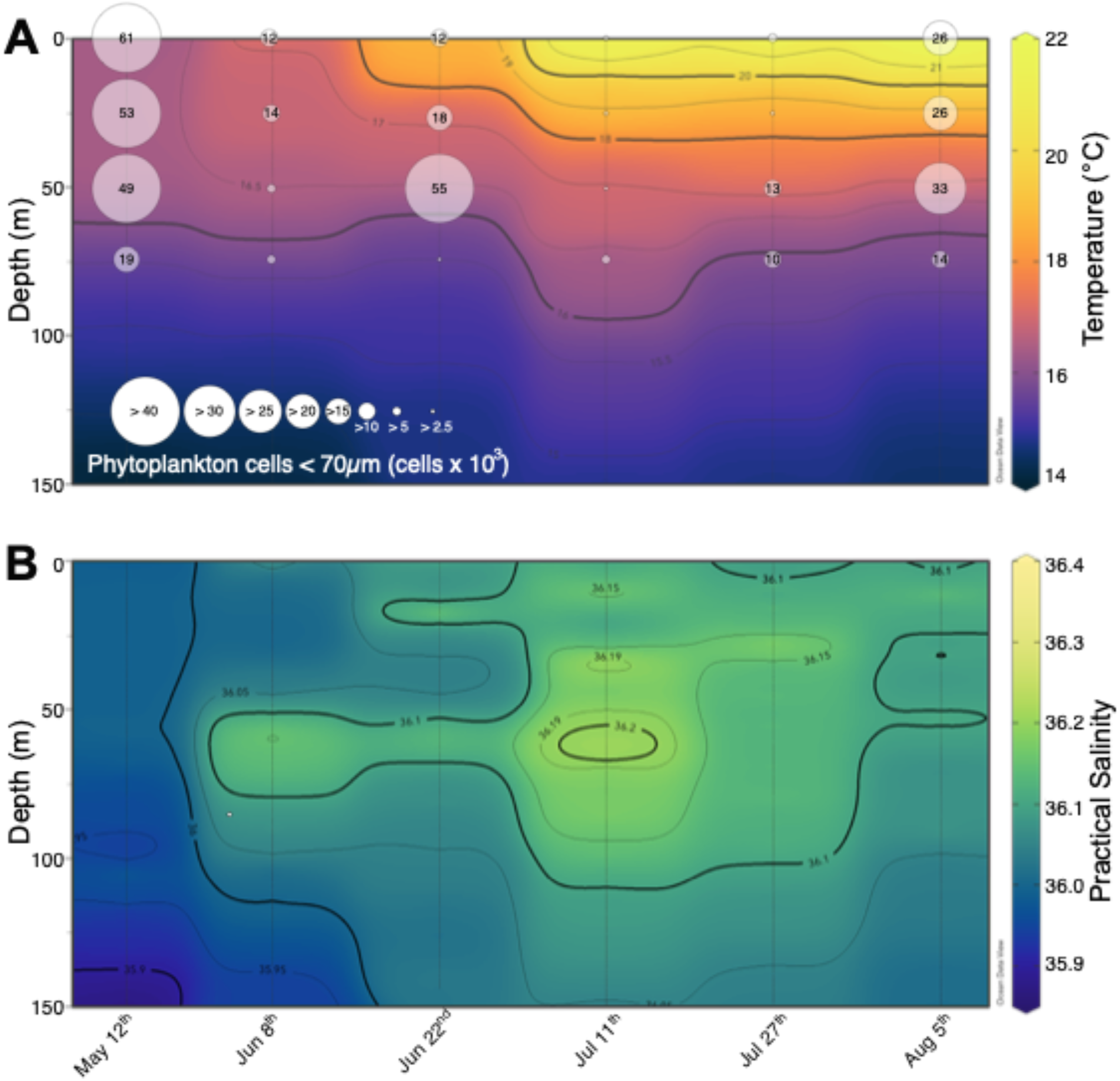
Vertical profile contours of **(A)** temperature and phytoplankton cell numbers, and **(B)** salinity at station South of Channel Faial-Pico islands (SCFP) from May 12th to August 5th, 2016. Data was not collected on September 8th due to equipment malfunction. Vertical gray lines indicate the sampling date and the maximum depth of data collection. Temperature and salinity data between sampling dates was interpolated using the weighted average grid method. Circle sizes indicate phytoplankton abundances at each sampled depth (surface, 25, 50 and 75 m). Surface values were the average abundance at the 0 and 5 m depths.

The sampling period started after the seasonal spring phytoplankton bloom and spanned the environmental and biological conditions observed from late spring to late summer. Observed changes in phytoplankton abundance corresponded with seasonal environmental changes during the sampling period (**Fig. 2A** and **Supplementary Fig. S4B**). High phytoplankton abundance (19–61 × 10^3^ cells/mL) was observed throughout the water column from 0 to 75 m depth on May 12th, indicating the end of the annual spring bloom. Subsequently, phytoplankton abundance fell below 2 × 10^3^ cells/mL with the exception of a sub-surface peak of ∼55 × 10^3^ cells/mL at 50 m on June 22nd, coinciding with the observed cyclonic upwelling event (**Supplementary Table S1**). The lowest phytoplankton abundance was observed on July 11th, coinciding with the anticyclonic downwelling event. Another increase in phytoplankton abundance coincided with the second, albeit less pronounced, cyclonic upwelling event during the subsequent two sampling dates, July 27th and August 5th.

### Changing environmental conditions and virioplankton and bacterioplankton community dynamics

For unknown reasons, only 29 of 35 virioplankton samples yielded RTPR amplicons (**Supplementary Tables S3 & S10**). From a total of 30,473 virioplankton RTPR amino acid sequences, 3,697 RTPR 98% OTUs were identified (**Supplementary Fig. S1 and S2**). The RTPR 98% OTUs clustered into 17 phylogenetic clades as manually determined by phylogenetic tree topology (**Fig. 3**). The three clades topologically located between clades 12 and 13 (gray color branches in **Fig. 3**) were excluded from downstream analyses due to their low abundance (7 OTUs, 12 amino acid sequences) and distant phylogenetic relations to other clades, leaving 3,690 98% OTUs (30,461 amino acid sequences, **Supplementary Table S3**) in 17 clades (**Supplementary Table S4**). Phylogenetic clade 5 was the most abundant based on both amino acid sequence counts (3,860) and clr abundance (1.07). The 17 RTPR phylogenetic clades were used in subsequent compositional data analyses, including community diversity and population interactions analyses, as clades of related OTUs provided clearer ecological signals than the larger collection of more highly resolved RTPR 98% OTUs. However, as shown in supplemental analyses, RTPR 98% OTU community diversity within and between samples was also determined for comparison with the phylogenetic approach.

**Figure 3.**
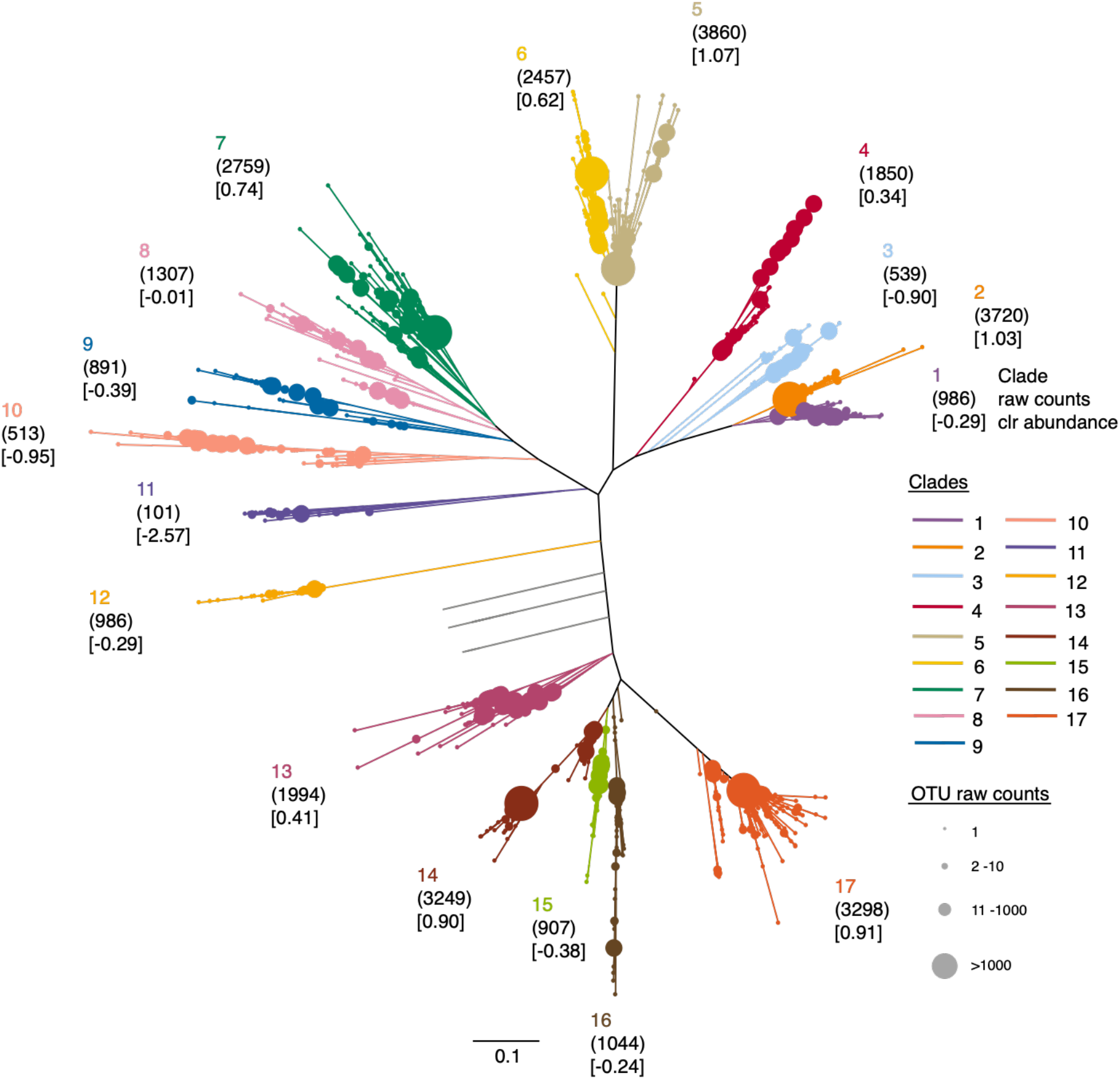
Approximate maximum likelihood tree of virioplankton RTPR 98% OTU representative sequences. Shared branch, node, and clade number colors indicate RTPR phylogenetic clade manual assignment. Seven OTUs (between clades 12 and 13) with low abundance and distant relationships with other phylogenetic clades are represented by gray branches and were excluded from downstream analysis. Node sizes indicate 98% OTU counts. Each phylogenetic clade’s total amino acid sequence counts and clr abundance are indicated in rounded and squared parentheses, respectively. Scale bar indicates the average number of amino acid substitutions per site.

The V3–V4 region of the 16S rRNA gene was PCR-amplified from 35 bacterioplankton fraction samples and sequenced (**Supplementary Tables S2 & S10**). After sequence quality filtering, 1,779 amplicon sequence variants (ASVs) were identified from a total of 918,792 sequences across all samples (**Supplementary Table S5**). Complete-linkage hierarchical clustering based on the Aitchison distance of clr-transformed sequence abundance between the first (May 12th 5 m) water sample and every other sample was performed for each of three datasets (virioplankton RTPR 98% OTUs, RTPR phylogenetic clades, or bacterioplankton 16S ASVs). Mantel tests assessed each Aitchison distance matrix with distance matrices of all collected environmental metadata (**Supplementary Table S11**) to identify environmental factors correlated with and potentially driving the clustering of virio- or bacterioplankton communities. Sample depth, collection day (displayed as Julian day), temperature, and water density were significantly (Spearman and Pearson *p*-value <0.02) correlated with virioplankton community composition (RTPR phylogenetic clades and RTPR 98% OTUs) and bacterioplankton community composition. For the significant environmental variables, correlation coefficients with virioplankton populations were generally higher when using the RTPR phylogenetic clades.

Hierarchical clustering of virioplankton communities based on RTPR phylogenetic clades formed four large groups (vA, vB, vC, and vD) (**Fig. 4A**) based on their distance from the May 12th 5 m sample. Community groups vA/vB and groups vC/vD were organized into equally distant supergroups. Within each group, communities had similar profiles for depth/density and for temperature/day. Groups vA/vB contained communities generally occurring at deeper depths with lower water temperatures, whereas groups vC/vD contained communities generally collected at shallower depths with higher water temperatures. However, for all groups, there were outliers likely reflecting some of the low rank correlation coefficients (0.19–0.51) between community structure and environmental metadata (**Supplementary Table S11**). Bacterioplankton communities formed three groups (bA, bB, and bC) (**Fig. 4B**) based on their distance to the May 12th 5 m sample. Communities in groups bA and bB were observed at lower water temperatures, whereas group bC communities were found in higher water temperatures at shallow depths (<25 m). The division between groups bA and bB was more subtle and may have been forced by depth as group bB contained communities in only 50 and 75 m samples.

**Figure 4.**
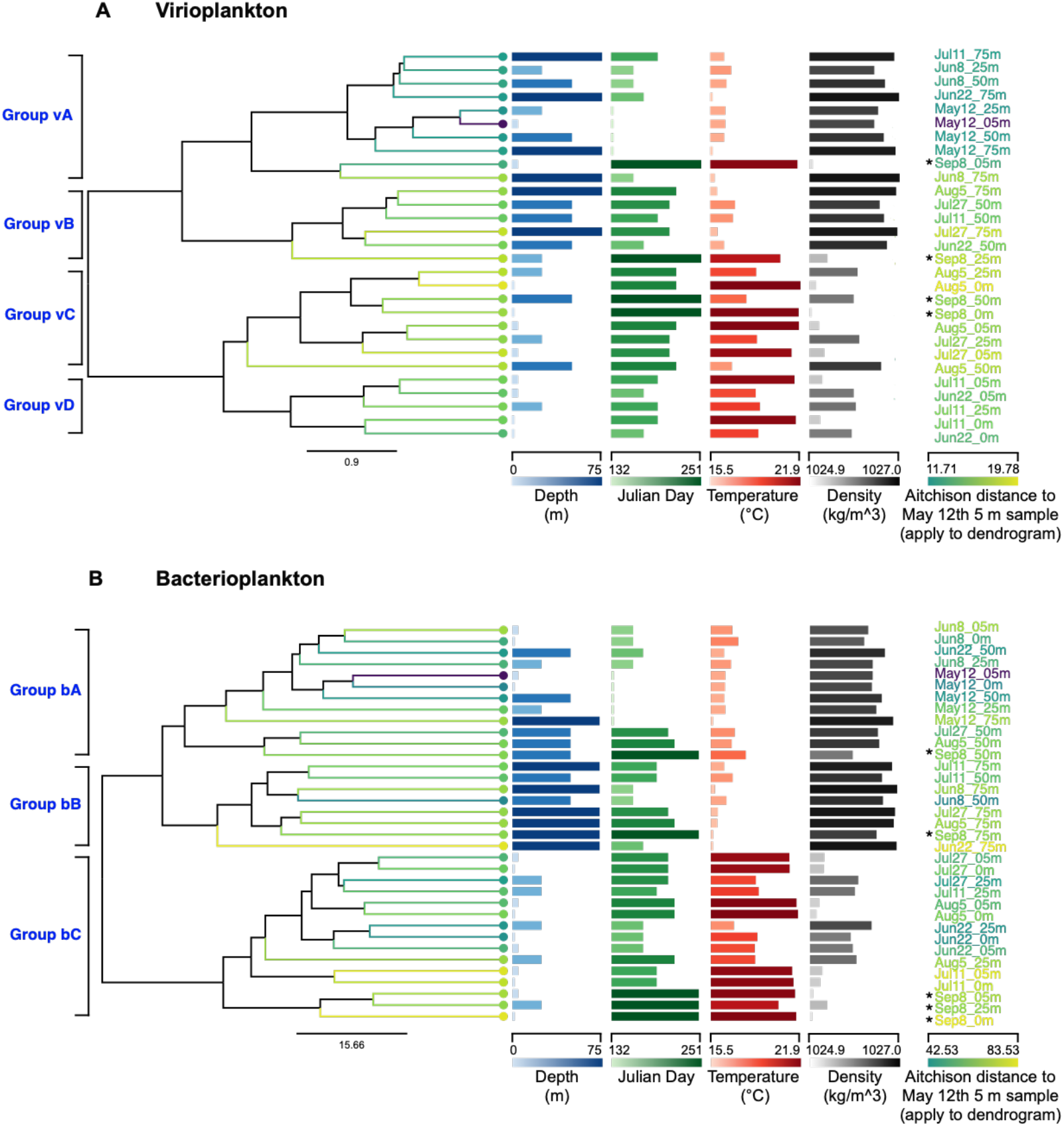
Phylogenetic clustering of virio- and bacterioplankton communities (dendrograms) based on **(A)** virioplankton RTPR phylogenetic clades and **(B)** bacterioplankton 16S ASVs, alongside environmental variables (columns of horizontal bars) demonstrating significant correlations (Spearman and Pearson tests, *p* <.05). Dendrograms are based on complete-linkage hierarchical clustering of each sample’s Aitchison distance to the sample collected May 12th at 5 m (May12_05m) based on centered-log ratio (clr) abundance. Scale bar indicates Aitchison distance between samples. Major groups of virio- or bacterioplankton communities are labeled to the left of each dendrogram. Environmental variables are represented in columns, with horizontal bars indicating the measurement for each sample both by color gradient and by bar length compared to the numerical scale along each column’s horizontal axis [Depth (blue), Day (green), Temperature (red), and Density (gray)]. Sample labels with asterisks (*) indicate samples where temperature and density were estimated using the Marine Copernicus Global Ocean Physics model.

The effects of sampling date (**Fig. 5A**) and depth (**Supplementary Fig. S5A**) on viral and bacterial ENS were significant (betta <0.05; data not shown); therefore, these environmental parameters were selected for analysis of their interactions with alpha and beta diversity of the virio- and bacterioplankton communities. Over the interseasonal sampling period, virioplankton alpha diversity according to RTPR phylogenetic clades (**Fig. 5A)** fluctuated slightly reaching a maximum around the July 11th anticyclonic downwelling event (**Fig. 2)**. This peak was shouldered by minimum diversity estimates on June 8th and July 27th corresponding with cyclonic upwelling events. Virioplankton alpha diversity fluctuated slightly with depth (**Supplementary Fig. S5A)**. Maximal and minimal diversity estimates occurred at 25 and 75 m, respectively. ENS estimates of alpha diversity based on 98% OTUs (**Supplementary Fig. S6A and B**) were three to six-fold higher than ENS estimates for virioplankton based on RTPR phylogenetic clades (**Fig. 5A and Supplementary Fig. S5A**) as there were thousands of 98% OTUs versus only 17 RTPR phylogenetic clades. The general trends of ENS estimates based on 98% OTUs and RTPR phylogenetic clades were consistent (Spearman’s rank correlation: ρ = 0.82, *p* value = 0.034; Pearson’s product-moment correlation: *r* =0.78, *p* value = 0.037). Increases in RTPR 98% OTU ENS (**Supplementary Fig. S6A**) were observed on June 22nd and July 11th, corresponding with the observed cyclonic upwelling event. The maximum ENS was observed on September 8th; unfortunately its association with an upwelling event is unknown as equipment failure prevented temperature measurements on this date. The maximum and minimum virioplankton RTPR 98% OTU ENS in samples by depth were 5 and 50 m, respectively (**Supplementary Fig. S6B**). Bacterioplankton alpha diversity steadily declined from its May 12th peak to a minimum value on July 11th followed by a slight increase over the remaining sampling dates (**Fig. 5A**). In contrast to virioplankton, bacterioplankton alpha diversity did change dramatically with depth demonstrating steady increases from minimal diversity estimates in surface waters to the maximal diversity estimates observed at 75 m (**Supplementary Fig. S5A**).

**Figure 5.**
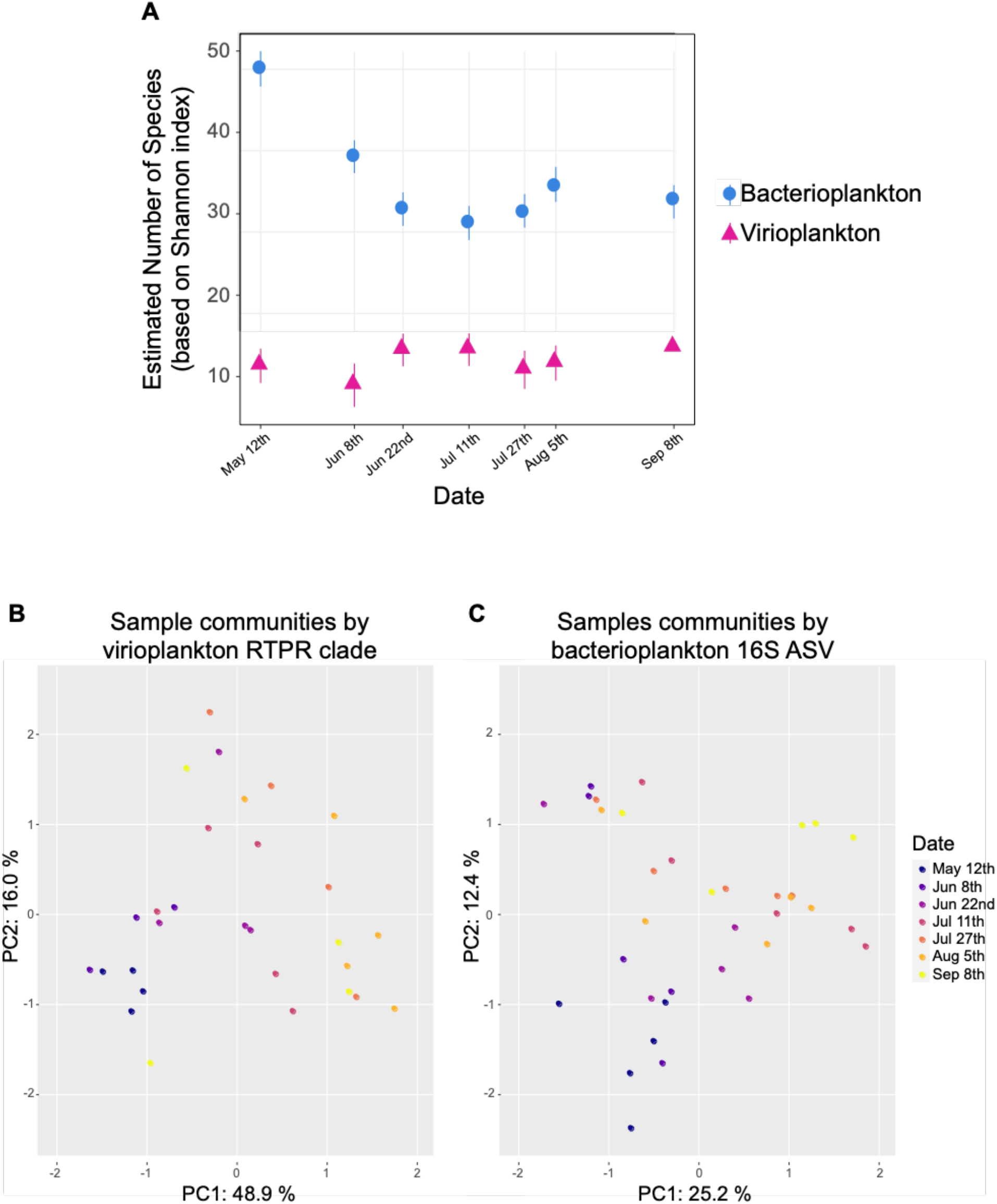
Sampling date influenced virioplankton (RTPR phylogenetic clade) and bacterioplankton (16S ASV) community alpha and beta diversity. **(A)** Effective number of species (ENS, based on Shannon indices) of virioplankton RTPR phylogenetic clades and bacterioplankton 16S communities by sampling date. Error bars represent two standard deviations of the estimates. Principal component analysis (PCA) plots of beta diversity based on centered-log ratio (clr) transformed abundance of **(B)** virioplankton RPTR phylogenetic clade communities or **(C)** bacterioplankton 16S ASV communities. Samples (circles) are colored according to sampling date.

Relationships between virioplankton RTPR phylogenetic clade communities were examined using principal components analysis (**Fig. 5B)**. Sixty-five percent of the variability between communities was explained by the first two principal components (PC1 48.9%, PC2 16.0%), and sampling date (**Fig. 5B)** seemed to explain this variability more than sample depth (**Supplementary Fig. S5B**). Using the 98% OTU populations, ∼35% of the variability between communities was explained by the first two principal components (PC1 22.8%, PC2 12.1%), and was associated with both sampling date and depth (**Supplementary Fig. S6C and D**). The variability in virioplankton communities explained by the RTPR 98% OTUs was approximately half of that explained by phylogenetic clades. This trend was attributed to the higher number and granularity of RTPR 98% OTU populations in describing virioplankton community structure. For either means of defining virioplankton communities, PCA analyses showed that the May 12th virioplankton communities grouped closer to the origins of both the PC1 and PC2 axes, apart from most other sampling dates, and the August 5th samples tended to place toward the ends of PC1 and PC2 axes. Virioplankton communities as defined by RTPR phylogenetic clades demonstrated significant differences based on date and depth (PERMANOVA *p* value <0.05; **Supplementary Table S12A & D)**, largely agreeing with observations of 98% OTUs **(Supplementary Table S12B & E**). May 12th virioplankton communities differed from most other timepoints. June 8th communities differed from July 27th, and August 5th communities differed from those on June 8th, June 22nd, and July 11th. Depth-based virioplankton community structure (**Supplementary Fig. S5B)** did not group as strongly on the PCA plots as date-based communities (**Fig. 5B**). Nevertheless, surface communities (0 m) were different from those at 50 and 75 m, and 5 m communities differed from 75 m communities.

Relationships between bacterioplankton 16S ASV communities were examined using principal components analysis as well (**Fig. 5C)**. Approximately 38% of the variability between communities was explained by the first two principal components (PC1 25.2%, PC2 12.4%), and was associated with both sampling date (**Fig. 5C**) and depth (**Supplementary Fig. S5C**). As seen with viral communities defined by either RTPR phylogenetic clades or 98% OTUs, the May 12th bacterioplankton samples grouped toward the origin of both PC1 and PC2, apart from most other sampling dates. The August 5th bacterioplankton communities showed a greater spread across PC1 than the virioplankton communities for this sampling date. These observations were statistically confirmed. Bacterioplankton communities on May 12th and June 8th were different from those observed on all other sampling dates, except the pair of samples from June 8th and June 22nd (PERMANOVA *p* value and/or *q* value <0.05; **Supplementary Table S12C**). Bacterial 16S ASV communities were also separated by depth, with 50 m and 75 m communities occurring along the left side of PC1 (**Supplementary Fig. S5C**). Surface bacterioplankton communities at 0 and 5 m significantly diverged from the deep communities at 50 and 75 m; 25 m communities diverged from the 75 m communities; and 50 m communities diverged from the 75 m communities (PERMANOVA *p* value and/or *q* value <0.05; **Supplementary Table S12F**).

### Associations between virioplankton RTPR phylogenetic clades and bacterioplankton 16S ASVs

The proportionality test was used for comparing changes in the clr abundance of 17 virioplankton RTPR populations (as defined by phylogenetic clades on **Fig. 3**) and the subset of 221 bacterioplankton 16S ASV populations in 27 samples with both virioplankton RTPR and bacterioplankton 16S amplicons (**Supplementary Table S13**). Virio- and bacterioplankton population pairs with ρ values ≥0.48 had positive associations as the false discovery rates were ≤0.05 (**Supplementary Fig. S7**). Nine of the 17 virioplankton RTPR phylogenetic clades had positive associations (ρ ≥.48) with 39 of 221 bacterioplankton ASV populations (**Fig. 6**). Associating virioplankton clades were phylogenetically diverse and were observed at both high (clade 2 and 5; clr 1.03 and 1.07, respectively) and low abundance (clade 3 and 11; clr -0.90 and -2.57, respectively) (**Fig. 3**). Virio- and bacterioplankton abundance did not appear to drive these associations as low-abundance RTPR virioplankton associated with high-abundance bacterioplankton and vice versa (**Supplementary Fig. S8)**. The phylogenetic diversity of positively-associating bacterioplankton populations was broad, encompassing seven bacterial phyla observed among the included 16S ASV populations. The bacterioplankton 16S ASV populations ranged in clr abundance from the most abundant taxon 3.65 (Prochlorococcus_MIT9313) to one of the least abundant taxon -1.99 (SAR116_clade).

**Figure 6.**
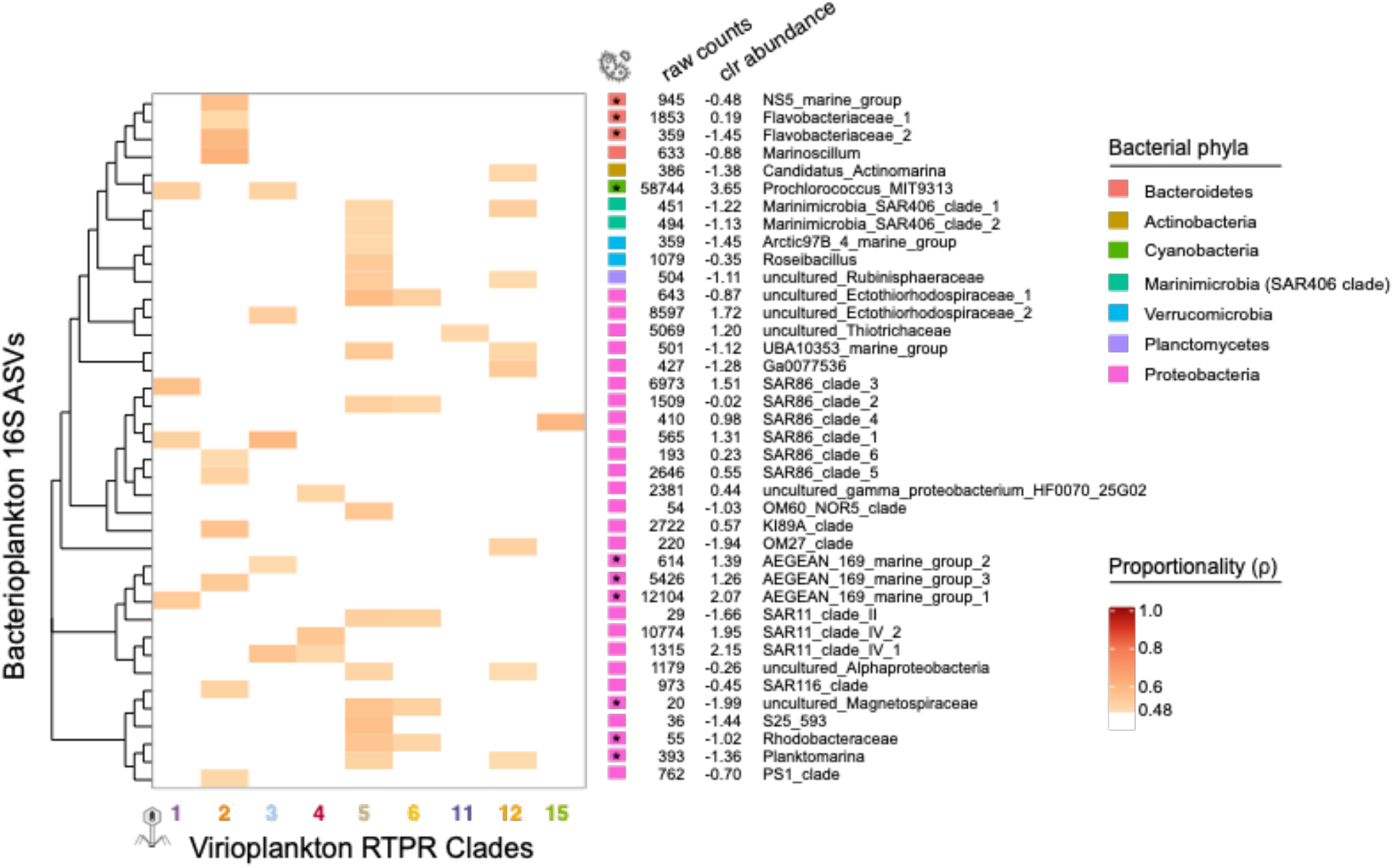
Matrix (ρ proportionality) of significantly associated (ρ ≥.48) virioplankton RTPR phylogenetic clades (x-axis) and bacterioplankton 16S ASVs (y-axis) (**Supplementary Table S13**). Virioplankton RTPR clades are colored and arranged by their topological order on the 98% OTU phylogenetic tree (**Fig. 3**). Bacterioplankton 16S ASVs are arranged according to phylogeny. Bacterioplankton 16S ASV phylum (colored block), raw sequence count, clr abundance, and taxonomic assignment are shown on the right. Asterisk indicates those ASVs within orders, families, or genera containing species capable of B_12_ synthesis according to a pangenomic survey by Heal *et al*. 2017 ^90^.

Virioplankton RTPR phylogenetic clades 1 and 3 were found to be associated with the most abundant bacterioplankton 16S ASV, Prochlorococcus_MIT9313, and several 16S ASVs from Proteobacteria, such as SAR86, SAR11, and the Aegan169 marine group, well known to be abundant within oceanic ecosystems. Virioplankton clade 2 showed associations with multiple bacterioplankton ASVs from Proteobacteria and Bacteroidetes. Interestingly, RTPR phylogenetic clade 2 was not associated with any of the same 16S ASVs as clades 1 and 3, despite its close phylogenetic relationship with these clades (**Fig. 3**). Phylogenetic clades 3 and 4 were both associated with SAR11_clade_IV_1, and clade 4 was also associated with SAR11_clade_IV_2. Virioplankton clades 11 and 15 were each only associated with a single bacterioplankton 16S ASV, uncultured_Thiotrichaceae and SAR86_clade_4, respectively. Virioplankton clade 5, the most abundant RTPR phylogenetic clade (clr 1.07, **Fig. 3**), showed the most positive associations across a wide phylogenetic breadth of bacterioplankton populations, encompassing four of the nine bacterial phyla and 15 of the 39 total positively-associated 16S ASVs, including SAR11_clade_II.

Fourteen out of 136 non-self comparisons between virioplankton RTPR populations showed a positive association (**Supplementary Fig. S9, Supplementary Table S13**). These associations between virioplankton clades did not follow a pattern according to phylogenetic distance as only three of the 14 associations were between neighboring RTPR clades (clades 2 and 3; clades 5 and 6; and clades 8 and 9). Of the 221 bacterioplankton 16S ASV populations, 36 showed positive, non-self associations with other ASV populations (**Supplementary Fig. S10, Supplementary Table S13**). Overall, there were 390 positive non-self associations among the 36 ASVs, ∼5% of all the non-self comparisons (1260).

Positively associating bacterioplankton 16S ASV populations were clearly divided by their association with either SAR11_clade_II or Prochlorococcus_MIT9313 (red boxes in **Supplementary Fig. S10**). No 16S ASVs associated with both SAR11_clade_II and Prochlorococcus_MIT9313. The SAR_clade_II population was associated with only other low abundance (clr <0) ASVs, whereas Prochlorococcus_MIT9313 was associated with mostly high abundance (clr >0) ASVs. Beyond the delineation of SAR11_clade_II from Prochlorococcus_MIT9313, positively associating bacterioplankton ASV populations could be divided into three larger groups (**Supplementary Fig. S10** x-axis). Nearly all of the ASV populations within group 1 all showed positive associations with SAR11_clade_II and nine other taxa (clr values ranging from -1.38 to -0.35) across five phyla (Marinimicrobia: Marinimicrobia_SAR406_clades 1 (clr -1.22) & 2 (clr -1.13); Actinobacteria: Candidatus_Actinomarina (clr -1.38); Planctomycetes: uncultured_Rubinisphaeraceae (clr -1.11); Verrucomicrobia: Roseibacillus (clr - 0.35); Proteobacteria: UBA10353_marine_group (clr -1.12), Ga0077536 (clr -1.28), OM60_NOR5_clade (clr -1.03), and Rhodobacteraceae (clr -1.02)). ASV populations within group 2 were defined by their consistent association with the most dominant ASV taxa Cyanobacteria: Prochlorococcus_MIT9313 (clr 3.65). In addition to this singular representative of the Cyanobacteria, group 2 consisted of 13 *Proteobacteria* (clr values ranging from -0.45 to 2.15) and three *Bacteriodetes* (clr values ranging from - 1.45 to 0.19). Several of the ASV taxa in group 2, such as Proteobacteria: uncultured_Ectothiorhodospiraceae_2 (clr 1.72), Proteobacteria: SAR11_clade_IV_1 (clr 2.15) & 2 (clr 1.95), and Proteobacteria: SAR86_clade 3 (clr 1.51) were the most abundant bacterioplankton populations observed in the study. ASV populations within group 3, which consisted of only three Proteobacteria taxa, showed no consistent associations within the group and few associations overall.

### Depth- and time-resolved viral–host associations

Spatiotemporal patterns in bacterioplankton associations with clade 5, the most abundant RTPR virioplankton population, were explored by plotting changes in clr abundance at each sampling depth over time (**Supplementary Fig. S11**). Clade 5 RTPR viral populations showed their highest abundances at sampling depths of 25 m and deeper, with consistently high abundances occurring at 75 m. Deeper samples also contained a greater number and diversity of bacterioplankton ASV taxa showing associations with clade 5 RTPR virioplankton. Among the sampling dates, July 11th was noteworthy as several of the ASV taxa associated with clade 5 RTPR virioplankton showed dramatic reductions in clr abundance corresponding with the anticyclonic downwelling event. Correspondingly, the clr abundance of clade 5 virioplankton held steady or declined on the two subsequent sampling dates (July 27th and August 5th). A similar spatiotemporal analysis of associations between virio- and bacterioplankton populations was conducted for abundant bacterioplankton populations, Prochlorococcus_MIT9313 and three SAR11 clades, and four associating RTPR virioplankton populations (**Supplementary Fig. S12**). Clades 1 and 3 RTPR virioplankton populations showed associations with Prochlorococcus_MIT9313 (**Fig. 6**). Within a sampled depth across time, the clr abundance of clade 1 and 3 viruses was relatively steady, however, clr abundance of these two populations decreased in the 75 m samples as did the abundance of their associated Prochlorococcus_MIT9313 population. RTPR clade 3 was also associated with SAR11_clade_IV_1 which, like Prochlorococcus_MIT9313, was most abundant in surface waters. RTPR clade 4 showed positive associations with SAR11_clade_IV_1 and _2 **(Fig. 6)**, which were the second and third most abundant bacterioplankton taxa showing associations with RTPR virioplankton. While these SAR11 populations were most abundant at the surface (clr values of around 3.75), they were also abundant throughout the water column and changed only modestly with time. The behavior of RTPR clade 4 showed a similar pattern as it was most abundant in surface waters (clr values ≥ 1) and then lower abundance at 50 and 75 m (clr values ∼ 0). RTPR clade 5 associated with SAR11_clade_II which was the rarest of the SAR11 taxa showing associations with an RTPR virioplankton population. SAR11_clade_II populations increased with depth, showing their highest abundances (clr values ∼ 0) at 75 m. Similarly, RTPR clade 5 was less abundant at shallow depths (clr values < 1) and then increased to its highest abundances at 75 m (clr values > 1).

### Free and actively replicating viral populations: RTPR 98% OTUs occurring in virio- and bacterioplankton fractions

Twenty-nine virioplankton and 31 bacterioplankton samples provided RTPR amplicons with an overlap of 27 samples providing amplicons from both. These 27 paired samples provided 47,997 RTPR quality-filtered amino acid sequences, which were clustered *de novo* into 5,411 RTPR common 98% OTUs (**Supplementary Table S7**). After removing 98% RTPR OTUs with less than ten sequences, 21 OTUs were observed only in the virioplankton fraction, 40 only in the bacterioplankton fraction, and 193 in both fractions (**Fig. 7** and **Supplementary Table S8**). An encompassing phylogenetic analysis of these RTPR 98% OTUs, sequences from the RNR database, and viromes from the *Tara* Oceans expedition showed that the RTPR 98% OTUs from the virio- and bacterioplankton fractions occurred within a section of the tree containing few sequences from known bacteria or viruses (RNRdb) but several sequences from *Tara* Oceans viromes (**Supplementary Fig. S13**). The most abundant OTUs were primarily found in the shared fraction (**Fig. 7A**). Less abundant OTUs were largely found in either virio- or bacterioplankton fractions only. Examining the relative contribution of virio- or bacterioplankton amplicon sequences to RTPR 98% OTUs required assignment of an arbitrary clr of 5 and -5 to those OTUs found only within the virio- or bacterioplankton fractions, respectively. The difference in clr abundance within each RTPR 98% OTU (clr(virioplankton)-clr(bacterioplankton), **Fig. 7B**) found that within most of the shared OTUs, virioplankton contributed a greater number of amplicons (66% of all shared OTUs (128 of 193), red colored OTUs above the dashed line in **Fig. 7B**).

**Figure 7.**
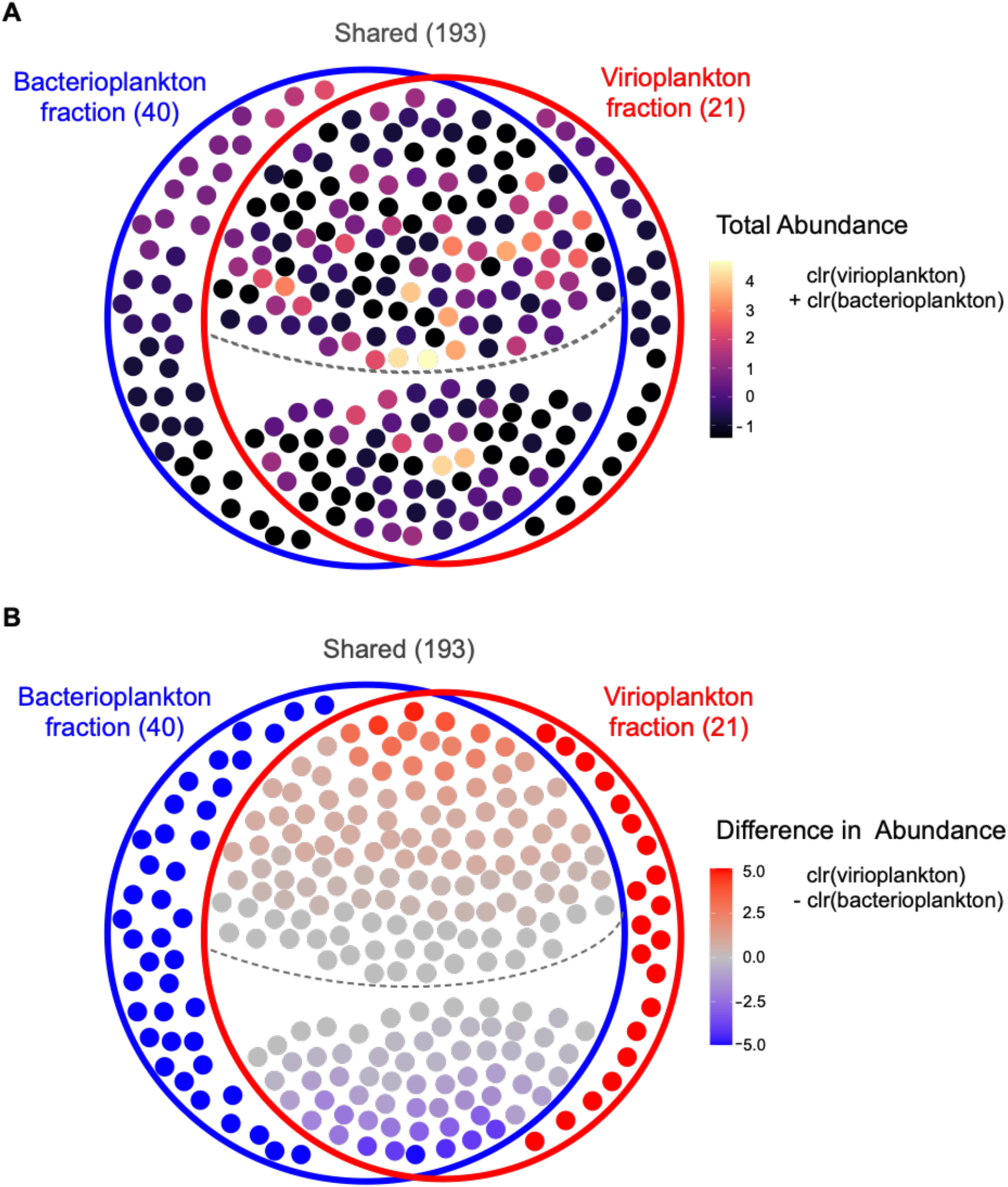
RTPR 98% OTUs (n=254) identified from amino acid sequences present in 27 paired samples with amplicons from the virio- and bacterioplankton fractions. RTPR 98% OTUs (filled circles) were observed in either only the bacterioplankton fraction (blue circle, n=40), only the virioplankton fraction (red circle, n=21), or both (Shared, n=193). The dashed line in the overlap region divides the shared OTUs more abundant in the virioplankton fraction (above the line, n=128) from those more abundant in the bacterioplankton fraction (below the line, n=65). **(A)** RTPR 98% OTUs are colored by their total centered log ratio (clr) abundance in both fractions (clr[virioplankton]+clr[bacterioplankton]). **(B)** RTPR 98% OTUs are colored by the difference between clr abundance in the virioplankton and bacterioplankton fractions (clr[virioplankton] - clr[bacterioplankton]). Red colors indicate greater abundance of that OTU in the virioplankton fraction (values > 0, where 5 is an arbitrary maximum representing OTU presence in the virioplankton only). Blue colors indicate greater abundance of that OTU in the bacterioplankton fraction (values < 0, where -5 is an arbitrary minimum representing OTU presence in the bacterioplankton only).

All samples (fractions and depths) from the August 5th sampling date provided RTPR amplicon sequences, enabling coordinate examination of virioplankton populations observed both as free viruses (i.e., within the virioplankton fraction) and viruses replicating within cells (i.e., within the bacterioplankton fraction). Twenty-nine RTPR 98% OTU populations were selected, each containing greater than 50 total sequences with two or fewer zero-replaced samples (**Supplementary Table S9**). These heuristics prevented the inclusion of rare and inconsistently observed virioplankton populations. Phylogenetic analysis of these 29 RTPR 98% OTUs alongside observations of clr abundance of each OTU within the virio- and bacterioplankton fractions addressed the hypothesis that the evolutionary distance between virioplankton populations reflects similarity in their ecological behavior (**Fig. 8**). Statistical testing comparing the cophenetic distance of the OTUs with the Aitchison distance of the OTUs according to their clr abundances across the August 5th samples demonstrated a significant (*p* = 1.6 e^-5^), weak (R^*2*^ = 0.045), positive correlation (correlation r = 0.212) between evolutionary distance and clr abundance in the bacterioplankton fraction (**Supplementary Fig. S14**). There was no significant relationship between cophenetic distance and clr abundance in the virioplankton fraction or when combining the virio- and bacterioplankton fractions (data not shown). Thus, more closely related RTPR virioplankton populations demonstrated similar ecological behavior in terms of abundance as viruses replicating within cells (i.e., the bacterioplankton fraction), but not as free viruses.

**Figure 8.**
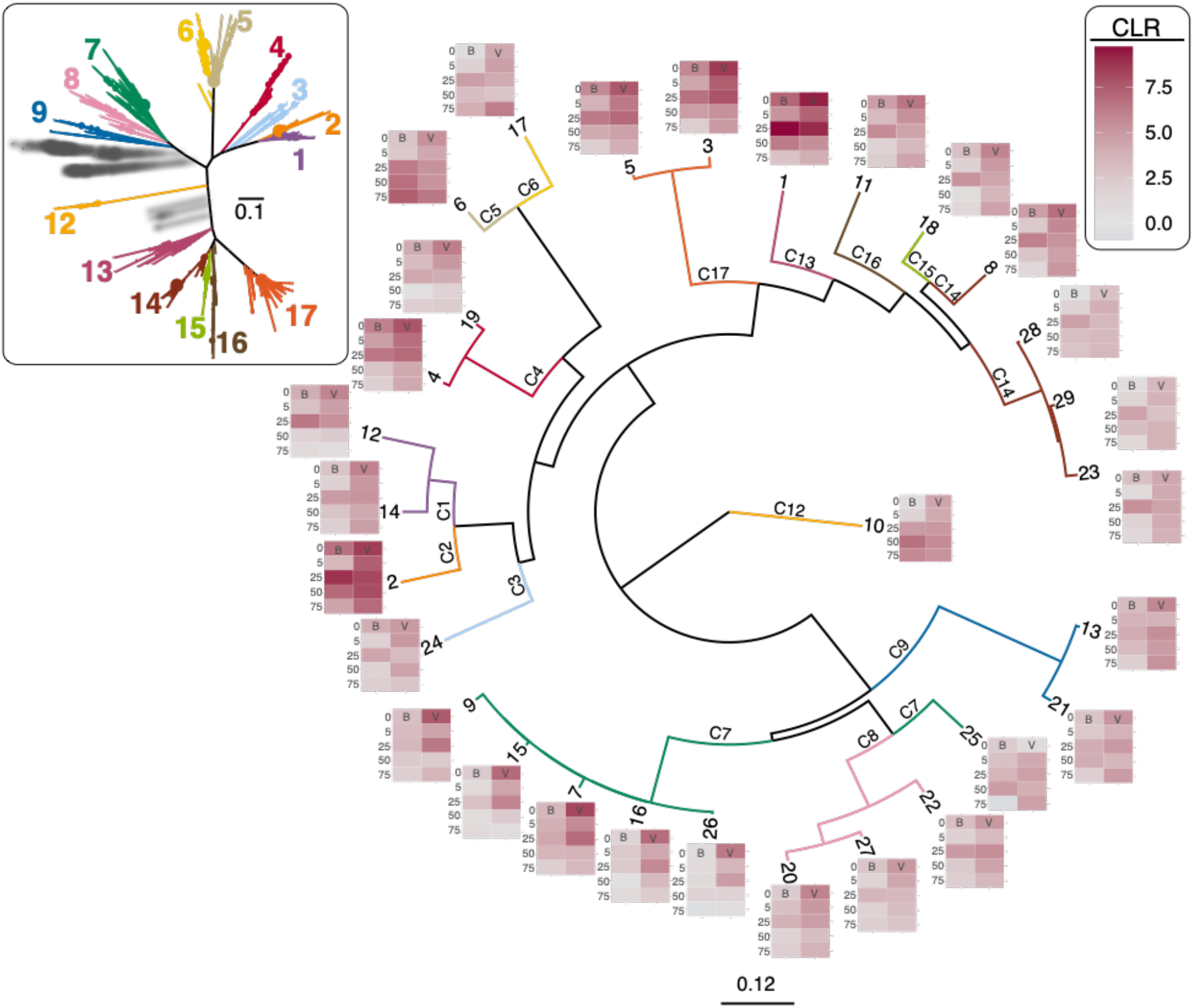
Phylogram of 29 RTPR 98% OTUs occurring within both the bacterioplankton (B) and virioplankton (V) fractions on the August 5th sampling date, with heat maps at nodes representing OTU centered-log ratio (clr) abundance in each sample (fraction and depth). Node labels indicate OTU number. Branches are colored and labeled according to membership of the OTUs in the RTPR clades (C1–C9; C12–C17) identified in the inset tree (**Fig. 3**). Scale bar indicates the average number of amino acid substitutions per site.

The 29 August 5th RTPR 98% OTUs occurred across nearly all of the major RTPR clades identified in **Fig. 3**, with the exception of clades 10 and 11 which were among the rarest clades observed in the study (clr values -0.95 and -2.57, respectively). Clade 7 was the most represented among the August 5th OTUs. These OTUs demonstrated a distinctive evolutionary split with a group of closely related OTUs (9, 15, 7, 16 and 26) clustering away from OTU 25 (**Fig. 8**). The clr abundance heat maps of the closely related group were similar with the majority of sequences occurring in the virioplankton fraction above 25 meters. In contrast, OTU 25 sequences were more abundant below 25 m demonstrating high abundance in the bacterioplankton fraction at 50 m. By and large, closely related August 5th OTUs showed similar clr abundance patterns supporting the finding of a significant relationship between evolutionary distance of RTPR virioplankton populations and Aithison distance.

## DISCUSSION

### Ribonucleotide triphosphate reductase: a hallmark gene within viruses

As the only enzyme capable of reducing ribonucleotides to deoxyribonucleotides, RNR effectively controls the rate of DNA synthesis ^33^. Thus, it makes intuitive sense that RNR genes would be common within the genomes of lytic viruses as speed and efficiency in genome replication is under positive selection pressure for viruses exhibiting a virulent lifecycle. Indeed, RNR genes are among the most commonly observed genes within dsDNA viral genomes and viromes, lending support to this hypothesis ^34,77–79^. RNRs are ancient proteins as enzymatic ribonucleotide reduction was a critical step from the RNA world to the emergent dominance of DNA as the principle information-carrying molecule for life on Earth ^80–82^. Thus, it is possible that RNRs existed within viral genomes not long after the emergence of DNA. Phylogenetic analysis of known and environmental RTPRs (**Supplementary Fig. S13**) indicated that viral RTPR proteins diverged from cellular RTPRs and are as deeply branching, reflecting both the unique selective pressure on viral RTPRs and their long evolutionary history. However, the tree also shows the emergence of cellular RTPRs from clades dominated by viral sequences and vice versa. These instances may represent viral to cellular (and cellular to viral) gene transfer events. Throughout the long evolutionary history of RNRs there is evidence of gene transfer between cellular kingdoms, most notably the transfer of Class I RNRs from bacteria to archaea and ultimately eukaryotes ^31^ where Class I RNRs predominate (**Supplementary Fig. S15A**). In the case of nucleotide metabolism proteins, such as RNRs, it is possible that greater levels of biochemical innovation occur within lytic viruses than within cells, as viruses typically carry one allele of a particular nucleotide metabolism gene which is under intense selective pressure for fast and efficient DNA replication. Certainly, at the level of RNR class, it is clear that selective processes may influence the distribution of RNR classes across the tree of life, where viruses demonstrate a distinctly high frequency of Class II RTPRs in comparison with cellular kingdoms. Among known viruses, Class II RTPRs demonstrate a distinctive skew in taxonomic distribution favoring tailed phages having a myoviridae or siphoviridae morphology (**Supplementary Figs. S15A & B**).

Interestingly, a breadth of microbial diversity was associated with several of the RTPR viral phylogenetic clades (**Fig. 6**). There are a few possible explanations for this. One possibility is that physiochemical factors were stronger drivers of microbial dynamics than viral-induced mortality, resulting in positive associations between RTPR viral clades and a sub-community of similarly adapted organisms, both host and non-host alike. However, it is also possible that the diverse taxa positively associated with several of the RTPR viral clades are a reflection of the hosts for viruses within these clades. Abundant marine phages, like 37-F6, have been predicted to infect hosts spanning different phyla ^83^, while direct linkages between viral RNR genes and 16S rRNA genes from diverse bacterial taxa have been observed ^84^. In the latter case, viruses associated with diverse taxa were less efficient at infecting these hosts than viruses with narrow host ranges ^84^, but even infrequent cross-taxa infections could provide ample opportunities for horizontal gene transfer (HGT) of RNR genes between viral populations, resulting in similar RNR sequences across a diverse group of viruses infecting different hosts. Furthermore, horizontal gene transfer of RNRs across all three domains of life ^31^, as well as between phage populations ^34^, is well-established, suggesting that HGT has been critical to shaping viral ecology and cobalamin cycling in marine environments. Answering questions surrounding cellular–viral gene exchanges and their contribution towards the emergence of biochemical innovation will improve understanding of the evolutionary mechanisms behind adaptation in microbial communities.

### Biochemistry constrains RTPR virioplankton population ecology

RNR is an essential gene for cellular life as all cells contain dsDNA genomes. However, the frequency of RNR types varies substantially among known cellular organisms and viruses (**Supplementary Fig. S15A**). Meta-analysis of RNR protein sequences within the RNRdb ^76^ along with sequences in the UniProt knowledgebase ^85^ showed the high frequency of RTPR (i.e., Class II monomeric RNR) within known viruses (∼22%) as compared to cellular kingdoms (2–3%). Within the Caudovirales most RTPR genes occurred within siphophages (**Supplementary Fig. S15B**). In contrast, Archaea and Bacteria favor Class II dimeric or Class III RNRs for anaerobic ribonucleotide reduction, a biochemical strategy strikingly different from viruses. In the case of RTPR, this monomeric Class II RNR requires a adensyolcobalamin (B_12_) co-factor (like all Class II RNRs), can perform ribonucleotide reduction under aerobic or anaerobic conditions, and, along with Class III RNRs, utilizes a ribonucleotide triphosphate (NTP) substrate ^86^. The B_12_ requirement and use of NTP substrates have intriguing implications for the ecology of viruses carrying RTPR. Since cobamide nutrient (including B_12_) synthesis is complex, specialized, and energy intensive, few marine bacterial and archaeal phyla possess the entire synthesis pathway ^87,88^. Most bacteria and phytoplankton are auxotrophic for B_12_ and are thus dependent on the few taxa capable of B_12_ production. RTPR virioplankton populations must infect a host that can either synthesize B_12_ or assimilate and subsequently modify a B_12_ precursor containing the corrin ring.

Moreover, B_12_ has several chemical analogs depending on the combination of ligands at the *ɑ* and *β* sites of the corrin ring ^89,90^.

There is a growing recognition that selectivity in B_12_ analogs may drive interaction networks within microbial communities as cobamide specificity is commonly observed in microorganisms ^91^. For example, pseudocobalamin, produced by cyanobacteria such as *Prochlorococcus* (the dominant bacterioplankton taxa observed in this study), is poorly utilized by B_12_ auxotrophic microalgae ^90,92^. Specificity in B_12_ requirements among bacterioplankton taxa may explain the separation observed between bacterioplankton populations associating with either SAR11_clade_II populations (**Supplementary Fig. S10**, Group 1) or Prochlorococcus_MIT9313 (**Supplementary Fig. S10**, Group 2). Positive association patterns between virioplankton RTPR phylotype populations and bacterioplankton 16S ASV populations reflected the split seen between SAR11 and *Prochlorococcus* associations. RTPR virioplankton clades 5 and 6 associated with SAR11_clade_II and 14 other bacterioplankton populations (**Fig. 6**), three of which (Magnetospiraceae, Rhodobacteraceae, Planktomarina) may be capable of B_12_ production ^90^. Eleven of these bacterioplankton populations are associated with SAR11_clade_II (**Supplementary Fig. S10**, Group 1). Similarly, RTPR virioplankton clades 1 and 3 associated with Prochlorococcus_MIT9313, and six other bacterioplankton populations that also demonstrated associations with *Prochlorococcus* (**Supplementary Fig. S9**, Group 2). Three of these seven (AEGEAN_169_marine_group_1, AEGEAN_169_marine_group_2, and Prochlorococcus_MIT9313) occur in an order, family, or genera containing species capable of B_12_ production (**Fig. 6**).

Time- and depth-resolved observations of RTPR virioplankton clades 1, 3, and 5 (**Supplementary Figs. S11** and **S12**) indicate that changes in the abundance of these virioplankton populations followed those of their associated bacterioplankton populations throughout the euphotic zone. It seems unlikely that the parallels in association patterns observed between bacterioplankton populations (**Supplementary Fig. S10**) and those observed between RTPR virioplankton and bacterioplankton populations (**Fig. 6**) are happenstance. It is likely that these positive virio- to bacterioplankton association patterns reflect the B_12_-driven association patterns among their bacterioplankton hosts ^91^. It is likely that a few heterotrophic bacterial taxa and *Prochlorococcus* were the principal B_12_-producing populations in the euphotic surface waters of the Azores. Although Thaumarchaeota are also important oceanic B_12_ producers, metagenomic biogeographic surveys indicate that these archaeal populations predominate at high latitudes or in mesopelagic waters ^88^.

It has been hypothesized that virioplankton influence oceanic primary and secondary productivity by controlling the flux of limiting trace nutrients, such as iron ^93^, through cell lysis. The common presence and dynamic nature of RTPR virioplankton populations now places viral–host interactions as another important component within the vitamin traffic pathways of this trace nutrient in the oligotrophic ocean ^94^. While technically B_12_ is not a nutrient as it is not incorporated into cellular biomass, it displays oceanic distribution patterns reminiscent of nutrients such as nitrogen. Within surface waters of the oligotrophic oceanic gyres, B_12_ occurs in vanishingly low concentrations of less than 1.75 picomolar ^90,95^. However, deeper waters typically demonstrate higher B_12_ concentrations ^96^. Depth-resolved dissolved B_12_ concentrations occurring at our study site were likely highest below 25 m, similar to those recorded for in April and August at the western-most station of a recent study of eastern north Atlantic central waters off the Iberian peninsula ^95^. While RTPR virioplankton populations did show differences in their depth and time distribution (**Fig. 8, Supplementary Figs. S11** and **S12**), the abundance of these populations as a fraction of overall virioplankton abundance could not be discerned using an amplicon approach. Nevertheless, as RTPR viral populations depend on B_12_ for a steady supply of deoxyribonucleotides, it is possible that these virioplankton regulate or enhance B_12_ uptake or production during infection. Subsequently, lysis by RTPR-utilizing virioplankton could increase the flux of B_12_ pools disproportionately over virioplankton populations that do not utilize B_12_-dependent proteins. New quantitative approaches for observations of viral–host interactions based on detecting single viral genes using polonies ^97–99^ could provide data on the *in situ* frequency and production rate of RTPR virioplankton populations, as has been recently demonstrated for pelagic cyanophage populations ^100^.

Unlike Class I and dimeric Class II RNRs that use ribonucleotide diphosphate (NDP) substrates, RTPR uses ribonucleotide triphosphate (NTP) substrates. This difference in substrates may correlate with significant differences in viral population ecology. Virioplankton populations utilizing a Class I or dimeric Class II RNR would be dependent on NDP production from RNA digestion ^101^ as well as an NDP kinase for producing dNTPs ^102^. In contrast, RTPR virioplankton populations have direct and immediate access to the cellular pool of NTPs (which is substantially larger than the NDP pool ^103^) for DNA synthesis. These biochemical features of RTPR would favor a rapid lytic cycle. It is possible that RTPR-carrying virioplankton are highly lytic viruses specifically infecting actively growing bacterioplankton populations containing large intercellular pools of rNTP substrates.

However, there are possible costs to an RTPR strategy in terms of DNA synthesis regulation. Improved control of dNTP substrate levels for DNA synthesis and allosteric regulation of RNR enzyme activity have been proposed as possible advantages of NDP versus NTP reduction ^82^. Thus, RTPR-carrying viruses may pay a cost for faster DNA synthesis with lower fidelity as imbalances or excesses in dNTP levels from runaway RTPR activity can increase mutagenesis rates ^102,104^. The fact that RTPRs are common within the virioplankton ^34,77,78^ and are proportionally more represented within viruses than within cellular genomes (**Supplementary Fig. S15**) indicates that the possible fitness costs of RTPR do not outweigh the benefits of utilizing this gene for dNTP synthesis. Nevertheless, it is possible that the frequency of RTPR-carrying viruses may depend on environmental context as the majority of known viruses carry a Class I or Class II dimeric RNR (**Supplementary Fig. S15**).

### RTPR virioplankton phylotypes demonstrated dynamic responses to ecosystem change

At the community and population scale, RTPR-carrying virioplankton demonstrated dynamic behavior throughout the four month study, responding to seasonal and oceanographic changes influencing the activity of co-occurring phyto- and bacterioplankton communities. In this sense, RTPR virioplankton populations demonstrated community dynamic behavior reminiscent of that seen from studies utilizing amplicons of structural genes such as T4 major capsid protein ^28–30^, portal protein ^24,105^, and terminase ^25– 27^. A unique aspect of this study was that primer sets targeting viral RTPR genes were used for amplifying RTPR from both the virioplankton (0.02–0.22 µm) and bacterioplankton (>0.22 µm) fractions. While viral genes have been amplified and sequenced from bacterioplankton fractions in past studies ^29^, to our knowledge none have examined coordinate occurrence of OTUs across extracellular (i.e., virioplankton fraction) and intracellular (i.e., bacterioplankton fraction) virioplankton populations. The majority of RTPR OTUs observed in both fractions showed a greater abundance in the virioplankton fraction (**Fig. 7**) indicating host populations undergoing active lysis. These OTUs had the highest clr abundance of all RTPR virioplankton populations observed in the study. This observation makes sense as each viral particle released contains an RTPR gene copy and dozens of viral particles are released for each lysed cell. RTPR OTUs observed solely within either the virio- or bacterioplankton fraction were less frequent (8% virioplankton only; 16% bacterioplankton only) and typically showed low clr abundance. Virioplankton-only RTPR OTUs likely represented viral populations either no longer actively infecting hosts or somehow displaced from their host populations. Bacterioplankton-only RTPR OTUs may have represented lysogenic viruses existing as prophages, free viral genomic DNA within the >0.22 µm size fraction, some lytic viral populations yet to lyse infected cells, or inadvertent amplification of a bacterial RTPR gene. Distinguishing between these four possibilities would be difficult; nevertheless, it is clear that the majority of observed RTPR OTUs (84%) occurred within the virioplankton.

Individual RTPR virioplankton populations were actively infecting, replicating, and subsequently disappearing from the pelagic euphotic zone. Using an amplicon approach provided the advantage of deeply sampling RTPR virioplankton; it was less clear how these data should be partitioned into ecologically meaningful populations. Community and population ecology analyses conducted using either a heuristic approach (binning amplicons into 98% amino acid identity OTUs) or an evolutionary approach (defining major phylogenetic clades of RTPR amplicons) demonstrated the superiority of the evolutionary approach. Community beta diversity analyses based on clr-transformed OTU or clade abundance showed that the evolutionary approach (clades, **Fig. 5B** and **Supplementary Fig. S5B**) captured a greater proportion of variability in virioplankton communities than the heuristic approach (98% OTUs **Supplementary Fig. S6C and D**). The evolutionary approach lumped RTPR virioplankton populations into 17 clades as opposed to thousands of 98% OTUs, likely providing a greater signal to noise ratio for observing virioplankton ecological behavior. Indeed, virioplankton evolutionary relationships gleaned from RTPR phylogeny demonstrated a significant connection with the ecological behavior of virioplankton populations on the August 5th sampling date (**Fig. 8**). The RTPR genes carried within each clade population likely occurred within a diversity of genomic backgrounds. Thus, it was surprising that this single gene showed a statistically significant positive relationship between evolutionary distance (RTPR phylogenetic tree) and patterns of clr abundance within the bacterioplankton fraction (Aitchison distance) **(Supplementary Fig. S14)**. The significant correlation of cophenetic distance (RTPR phylogeny) and ecological behavior (Aitchison distance) of RTPR OTUs identified within the bacterioplankton and not within free viruses is interesting as these populations represented those RTPR viral populations undergoing active replication.

Examinations of connections between gene-based viral phylogeny and viral ecology have been limited, in large part due to the widely acknowledged mosaicism within viral genomes ^106^. Nevertheless, when considering the influence of enzyme biochemistry or the presence/absence of particular auxiliary metabolic genes ^107^, it is clear that single genes have a dramatic influence on a virus’ ecology. Single-gene amplicon approaches provide not only an evolutionary framework for understanding viral ecology, but also have practical advantages over metagenomic approaches in terms of cost, depth and breadth of sampling, and computational complexity ^108^. Prior work examining genetic connections to phage host range phenotypes found that while whole genome networks provided better predictive capability, single gene groups could also show strong connections ^109^. Interestingly, DNA replication and phage replication were among those Rapid Annotation Subsystem gene groups ^110^ showing a stronger predictive ability for host range. This might not be surprising when considering that phage genome and replication processes may require intimate interaction with host anabolic systems. Examining possible connections between single genes and whole genomes and other important phage phenotypes, especially those describing infection dynamics, should ultimately improve the predictive ability of metagenome data for ecosystem modeling of phage impacts on ecosystem processes.

## AUTHOR CONTRIBUTIONS

GP designed the study, performed processing of water samples and DNA extraction following water samples collection and part of the RTPR library preparation. LW conducted the majority of the RTPR library preparation and all of the 16S rDNA library preparation, performed all of the bioinformatics and biostatistical analyses following sequence processing, and wrote and edited the manuscript. RM, CL, AH, JD, BF, AM, KB, SP, JN and EW contributed to the study design, protocol development, data analysis and interpretation, and manuscript preparation. All authors read, edited, and approved the final manuscript.

## ACKNOWLEDGEMENTS

We would like to thank, by order of involvement, Clara M. Loureiro, Catharina Pieper, Ricardo Fernandes, Christien Laber, Prasanna Joglekar, Joana Botelho, Ana Pavon, and Marilia Olio for the help collecting and processing the samples. We would like to thank Dr. Daniel Nasko for input on library preparation and statistical analysis. We would like to acknowledge the skipper Renato Bettencourt, and all laboratory and facility management personnel from Departamento de Oceanografia e P, OKEANOS Institute of Marine Research, and IMAR-Instituto do Mar from the University of the Azores, DMCS-RU and DBI-UD.

