## supplementary tables for "Ubiquitous, B_12_-dependent virioplankton utilizing ribonucleotide triphosphate reductase demonstrate interseasonal dynamics and associate with a diverse range of bacterial hosts in the pelagic ocean"

**Supplementary Table S1**. Environmental and phytoplankton abundance metadata

| SampleID | Date | Day | Depth | Density(kg*m | Temperature | Salinity(PSU) | Current_head | Current vol (r | Sat_Chl(mg* | Sat_POC(mg | Sat_PIC(mg* | Sat_PAR( Ein |
|---|---|---|---|---|---|---|---|---|---|---|---|---|
| May12_0m | May 12th, 20 | 132 | 0 | 1026.4152 | 16.3931936 | 35.983086 | 287.421505 | 0.30090738 | 0.22685016 | 66.6776831 | 1.2109E-05 | 59.9592967 |
| May12_05m | May 12th, 20 | 132 | 5 | 1026.43446 | 16.4162 | 35.9969333 | 113.2 | 0.06062379 | 0.22685016 | 66.6776831 | 1.2109E-05 | 59.9592967 |
| May12_25m | May 12th, 20 | 132 | 25 | 1026.5237 | 16.417625 | 35.9944167 | 189.175 | 0.04826189 | 0.22685016 | 66.6776831 | 1.2109E-05 | 59.9592967 |
| May12_50m | May 12th, 20 | 132 | 50 | 1026.6575 | 16.33725 | 35.994625 | 159.375 | 0.06194113 | 0.22685016 | 66.6776831 | 1.2109E-05 | 59.9592967 |
| May12_75m | May 12th, 20 | 132 | 75 | 1026.93417 | 15.4928333 | 35.9576667 | 199.575 | 0.03842525 | 0.22685016 | 66.6776831 | 1.2109E-05 | 59.9592967 |
| Jun8_0m | June 8th, 201 | 159 | 0 | 1026.22833 | 17.4017333 | 36.0534667 | 164.76 | 0.17237898 | 0.23117758 | 68.1554612 | 1.2887E-05 | 63.4369999 |
| Jun8_05m | June 8th, 201 | 159 | 5 | 1026.32763 | 16.9490625 | 36.0135 | 62.40625 | 0.25205537 | 0.23117758 | 68.1554612 | 1.2887E-05 | 63.4369999 |
| Jun8_25m | June 8th, 201 | 159 | 25 | 1026.43517 | 16.84885 | 36.0052167 | 150.143333 | 0.07261246 | 0.23117758 | 68.1554612 | 1.2887E-05 | 63.4369999 |
| Jun8_50m | June 8th, 201 | 159 | 50 | 1026.68606 | 16.5063889 | 36.0846111 | 239.077778 | 0.10410502 | 0.23117758 | 68.1554612 | 1.2887E-05 | 63.4369999 |
| Jun8_75m | June 8th, 201 | 159 | 75 | 1027.03215 | 15.6404615 | 36.1294615 | 90.3692308 | 0.15200257 | 0.23117758 | 68.1554612 | 1.2887E-05 | 63.4369999 |
| Jun22_0m | June 22nd, 20 | 173 | 0 | 1025.90309 | 18.8472727 | 36.0994546 | 124.345455 | 0.05122903 | 0.21196049 | 64.1110166 | 1.7109E-05 | 63.5183609 |
| Jun22_05m | June 22nd, 20 | 173 | 5 | 1025.95092 | 18.65125 | 36.0686667 | 252.9 | 0.13518858 | 0.21196049 | 64.1110166 | 1.7109E-05 | 63.5183609 |
| Jun22_25m | June 22nd, 20 | 173 | 25 | 1026.4088 | 17.07192 | 36.0416 | 233.824 | 0.26230657 | 0.21196049 | 64.1110166 | 1.7109E-05 | 63.5183609 |
| Jun22_50m | June 22nd, 20 | 173 | 50 | 1026.73255 | 16.3304 | 36.0901 | 168.57 | 0.5715793 | 0.21196049 | 64.1110166 | 1.7109E-05 | 63.5183609 |
| Jun22_75m | June 22nd, 20 | 173 | 75 | 1027.01775 | 15.4870625 | 36.065625 | 163.95625 | 0.38000683 | 0.21196049 | 64.1110166 | 1.7109E-05 | 63.5183609 |
| Jul11_0m | July 11st, 201 | 192 | 0 | 1025.16006 | 21.5934375 | 36.0871875 | 295.8375 | 0.23252226 | 0.11430367 | 43.7110163 | 0.00015111 | 62.6303995 |
| Jul11_05m | July 11st, 201 | 192 | 5 | 1025.21167 | 21.4927778 | 36.09 | 298.811111 | 0.09218614 | 0.11430367 | 43.7110163 | 0.00015111 | 62.6303995 |
| Jul11_25m | July 11st, 201 | 192 | 25 | 1026.00083 | 18.9545833 | 36.1210417 | 284.1625 | 0.03366411 | 0.11430367 | 43.7110163 | 0.00015111 | 62.6303995 |
| Jul11_50m | July 11st, 201 | 192 | 50 | 1026.66276 | 16.977 | 36.2 | 219.682353 | 0.01638465 | 0.11430367 | 43.7110163 | 0.00015111 | 62.6303995 |
| Jul11_75m | July 11st, 201 | 192 | 75 | 1026.90562 | 16.3356154 | 36.1741539 | 118.984615 | 0.04290101 | 0.11430367 | 43.7110163 | 0.00015111 | 62.6303995 |
| Jul27_0m | July 27th, 201 | 208 | 0 | 1025.24355 | 21.2976364 | 36.0894546 | 61.0636364 | 0.07375283 | 0.13663873 | 48.4499049 | 0.000177 | 60.3527671 |
| Jul27_05m | July 27th, 201 | 208 | 5 | 1025.25945 | 21.2777273 | 36.0740909 | 44.7909091 | 0.20369214 | 0.13663873 | 48.4499049 | 0.000177 | 60.3527671 |
| Jul27_25m | July 27th, 201 | 208 | 25 | 1026.08281 | 18.737875 | 36.1536875 | 184.2625 | 0.24727257 | 0.13663873 | 48.4499049 | 0.000177 | 60.3527671 |
| Jul27_50m | July 27th, 201 | 208 | 50 | 1026.56178 | 17.1333333 | 36.1173333 | 132.055556 | 0.18117153 | 0.13663873 | 48.4499049 | 0.000177 | 60.3527671 |
| Jul27_75m | July 27th, 201 | 208 | 75 | 1026.979 | 15.85175 | 36.12275 | 307.925 | 0.08543276 | 0.13663873 | 48.4499049 | 0.000177 | 60.3527671 |
| Aug5_0m | August 5th, 2 | 217 | 0 | 1025.06785 | 21.9334615 | 36.0909231 | 313.584615 | 0.22378113 | 0.14524373 | 51.1999054 | 0.000182 | 58.6197134 |
| Aug5_05m | August 5th, 2 | 217 | 5 | 1025.13681 | 21.8215 | 36.112125 | 71.49375 | 0.20292673 | 0.14524373 | 51.1999054 | 0.000182 | 58.6197134 |
| Aug5_25m | August 5th, 2 | 217 | 25 | 1026.04113 | 18.6854667 | 36.0836 | 54.82 | 0.03887101 | 0.14524373 | 51.1999054 | 0.000182 | 58.6197134 |
| Aug5_50m | August 5th, 2 | 217 | 50 | 1026.59569 | 16.901625 | 36.089 | 235.675 | 0.05768753 | 0.14524373 | 51.1999054 | 0.000182 | 58.6197134 |
| Aug5_75m | August 5th, 2 | 217 | 75 | 1026.94953 | 15.8157059 | 36.0729412 | 173.464706 | 0.23643618 | 0.14524373 | 51.1999054 | 0.000182 | 58.6197134 |
| Sep8_0m* | September 8 | 251 | 0 | 1024.965 | 21.80276 | 35.872982 | NA | 0.08259867 | 0.11669481 | 42.8443498 | 1.27E-05 | 48.5509349 |
| Sep8_05m* | September 8 | 251 | 5 | 1024.99 | 21.709738 | 35.871456 | NA | 0.10939939 | 0.11669481 | 42.8443498 | 1.27E-05 | 48.5509349 |
| Sep8_25m* | September 8 | 251 | 25 | 1025.333 | 20.454329 | 35.86993 | NA | 0.14939488 | 0.11669481 | 42.8443498 | 1.27E-05 | 48.5509349 |
| Sep8_50m* | September 8 | 251 | 50 | 1025.95 | 17.96768 | 35.837887 | NA | 0.12136702 | 0.11669481 | 42.8443498 | 1.27E-05 | 48.5509349 |
| Sep8_75m* | September 8 | 251 | 75 | 1026.531 | 15.499344 | 35.83331 | NA | 0.10140599 | 0.11669481 | 42.8443498 | 1.27E-05 | 48.5509349 |

*Equipment malfunction prevented CTD data collection on September 8, thus, temperature, salinity, and density metadata was estimated from the nearest time, pixel, and depth

**Supplemen**

| SampleID | Sat_SST (°C) | Total Autotro | Syn (Cells/ml | Proc (Cells/m | PikoEuk (Cells | NanoEuk (Cel | Virus x10^6 ( | Bacteria x10^5 (Cells/mL) |
|---|---|---|---|---|---|---|---|---|
| May12_0m | 17.4449997 | 64107.74 | 36756.86 | 4260.03 | 18303.03 | 3091.35 | 6900372.67 | 807972.39 |
| May12_05m | 17.4449997 | 58490.54 | 33260.25 | 3996.13 | 17275.73 | 2818.03 | 6714223.6 | 797454.27 |
| May12_25m | 17.4449997 | 52873.33 | 29763.63 | 3732.24 | 16248.42 | 2544.71 | 6736024.84 | 790102.9 |
| May12_50m | 17.4449997 | 49367.29 | 28123.71 | 2601.25 | 15739.48 | 2393.91 | 7259254.66 | 731518.12 |
| May12_75m | 17.4449997 | 19000.47 | 11309.8 | 1319.48 | 5032.86 | 527.79 | 4427888.2 | 450695.69 |
| Jun8_0m | 17.1299992 | 10942.35 | 1302.89 | 8042.68 | 871.86 | 440.83 | 1149145.78 | 136263.13 |
| Jun8_05m | 17.1299992 | 13185.68 | 1508.61 | 9717.83 | 1009.01 | 626.96 | 1337482.83 | 163750.44 |
| Jun8_25m | 17.1299992 | 13695.08 | 4124.2 | 6230.38 | 1988.63 | 773.9 | 1166820.48 | 156827.28 |
| Jun8_50m | 17.1299992 | 6769.17 | 1430.24 | 3134.78 | 2018.02 | 58.78 | 622961.02 | 57761.49 |
| Jun8_75m | 17.1299992 | 6230.38 | 1459.63 | 2458.85 | 2096.39 | 107.76 | 597173.33 | 56382.72 |
| Jun22_0m | 19.2950001 | 11718.11 | 41.15 | 8004.12 | 2551.44 | 751.03 | 4642514.47 | 349484.6 |
| Jun22_05m | 19.2950001 | 11481.48 | 0 | 6965.02 | 2890.95 | 915.64 | 4405624.48 | 294567.7 |
| Jun22_25m | 19.2950001 | 17860.08 | 72.02 | 12500 | 3312.76 | 1306.58 | 4919768.4 | 331475 |
| Jun22_50m | 19.2950001 | 55102.88 | 39670.78 | 3497.94 | 9104.94 | 2150.21 | 4938957.82 | 551882.6 |
| Jun22_75m | 19.2950001 | 4434.16 | 1481.48 | 1255.14 | 1121.4 | 92.59 | 3084201.82 | 191789.7 |
| Jul11_0m | 20.6700001 | 2938.75 | 996.36 | 905.78 | 523.34 | 281.8 | 1468000 | 763873.53 |
| Jul11_05m | 20.6700001 | 3753.95 | 774.94 | 986.29 | 734.69 | 664.24 | 1759111.11 | 856021.44 |
| Jul11_25m | 20.6700001 | 3190.35 | 291.86 | 946.04 | 1378.8 | 352.25 | 1953333.33 | 942795.06 |
| Jul11_50m | 20.6700001 | 7558.22 | 1046.68 | 1710.92 | 2254.38 | 1127.19 | 2322666.67 | 1096022.65 |
| Jul11_75m | 20.6700001 | 8212.4 | 956.1 | 1922.26 | 4418.19 | 241.54 | 1739111.11 | 874861.64 |
| Jul27_0m | 23.3850002 | 3849.15 | 119.22 | 2622.87 | 187.35 | | 2551086.96 | 280689.78 |
| Jul27_05m | 23.3850002 | 6378.35 | 25.55 | 2861.31 | 85.16 | 459.85 | 2169347.83 | 261478.1 |
| Jul27_25m | 23.3850002 | 4164.23 | 17.03 | 2537.71 | 851.58 | 391.73 | 2868043.48 | 310759.12 |
| Jul27_50m | 23.3850002 | 13046.23 | 2946.47 | 3593.67 | 5279.81 | 630.17 | 2538695.65 | 297372.26 |
| Jul27_75m | 23.3850002 | 10278.59 | 1388.08 | 5194.65 | 3287.1 | 187.35 | 1636521.74 | 120941.61 |
| Aug5_0m | 23.0949993 | 17666.89 | 6832.56 | 1568.87 | 7617 | 989.07 | 1047399.96 | 412682.12 |
| Aug5_05m | 23.0949993 | 24738.19 | 9583.77 | 1898.57 | 11232.23 | 1261.92 | 1245522.98 | 1049406.29 |
| Aug5_25m | 23.0949993 | 26398.01 | 12016.67 | 1978.15 | 10231.79 | 1625.72 | 1414941.56 | 601867.88 |
| Aug5_50m | 23.0949993 | 33196.47 | 16723.29 | 6468.76 | 8594.7 | 886.75 | 1565639.86 | 673695.03 |
| Aug5_75m | 23.0949993 | 14369.98 | 3717.55 | 5491.06 | 3421.96 | 454.75 | 807780.31 | 160741.39 |
| Sep8_0m* | 22.8850002 | 9730.79 | 6288.42 | 704.74 | 1825.09 | 722.81 | 3557722.93 | 252401.5 |
| Sep8_05m* | 22.8850002 | 9685.61 | 6180 | 614.39 | 1797.98 | 731.84 | 3603403.66 | 285722 |
| Sep8_25m* | 22.8850002 | 9007.98 | 4987.37 | 876.4 | 1969.65 | 650.53 | 3384136.15 | 313326 |
| Sep8_50m* | 22.8850002 | 15892.72 | 4589.82 | 2132.28 | 5899.91 | 1165.53 | 5558807.72 | 509695.5 |
| Sep8_75m* | 22.8850002 | 9450.7 | 1852.19 | 3532.72 | 3523.68 | 252.98 | 3650965.37 | 383778 |

*Equipment n from the Marine Copernicus Global Ocean Physics model

**Supplementary Table S2**. 16S ASV taxonomic identification

| Feature_ID | Kingdom | Phylum | Class | Order | Family | Genus | Species | Confidence |
|---|---|---|---|---|---|---|---|---|
| e5cad0f83c5da3a8e4f138aac9423460 | Bacteria | Cyanobacteria | Oxyphotobacteria | Synechococcales | Cyanobiaceae | Prochlorococcus MIT9313 | | 1.000 |
| 36b651dfb1ecf5381fe35ea3c4a1061f | Bacteria | Proteobacteria | Alphaproteobacteria | SAR11 clade | Clade I | Clade Ia | | 0.982 |
| 1d72f3f992bcd714b2a0c1b37d614dd2 | Bacteria | Cyanobacteria | Oxyphotobacteria | Synechococcales | Cyanobiaceae | Synechococcus CC9902 | | 0.999 |
| 936be975fb97fd3b6b5829e1c83be6bd | Bacteria | Proteobacteria | Alphaproteobacteria | Puniceispirillales | SAR116 clade | alpha proteobacterium SCGC AAA015-N04 | alpha proteobacterium SCGC AAA015-N04 | 0.815 |
| a6a79b8a486f3cdaf899162d34b8035a | Bacteria | Proteobacteria | Alphaproteobacteria | SAR11 clade | Clade I | Clade Ia | | 0.819 |
| a3d4f90910b762e33dd4d344f63357f8 | Bacteria | Proteobacteria | Alphaproteobacteria | SAR11 clade | Clade I | Clade Ia | | 0.906 |
| 1122e6b7ea8ece945016e945302ee1ad | Bacteria | Proteobacteria | Alphaproteobacteria | SAR11 clade | Clade II | | | 0.996 |
| a756a146af3f113055fcd3b5ebebe18f | Bacteria | Proteobacteria | Alphaproteobacteria | SAR11 clade | Clade I | Clade Ia | | 0.876 |
| b77bf87ad1c17314baa9d7c9acc10a3b | Bacteria | Proteobacteria | Alphaproteobacteria | SAR11 clade | Clade IV | Ambiguous_taxa | Ambiguous_taxa | 0.785 |
| 2d4bbf1e16d0492f979b8c5510decc4c | Bacteria | Proteobacteria | Alphaproteobacteria | SAR11 clade | Clade II | | | 0.997 |
| 5f2b8d3eb00c873859709a32681cae6a | Bacteria | Cyanobacteria | Oxyphotobacteria | Chloroplast | marine metagenome | marine metagenome | marine metagenome | 1.000 |
| 0194fcb71fb1f171d26db7ddaeb20811 | Bacteria | Proteobacteria | Alphaproteobacteria | Rhodospirillales | AEGEAN-169 marine group | Ambiguous_taxa | Ambiguous_taxa | 0.999 |
| 2b7c6eeca2efe4e826759346836efc05 | Bacteria | Cyanobacteria | Oxyphotobacteria | Synechococcales | Cyanobiaceae | Prochlorococcus MIT9313 | | 0.999 |
| c1b1851dde5aaca57cc59dc350c37be7 | Bacteria | Proteobacteria | Alphaproteobacteria | SAR11 clade | Clade IV | | | 1.000 |
| c7d8e6d1fd5ae641fff2b96fa1112d7b | Bacteria | Marinimicrobia (SAR406 clade) | Alteromonas macleodii | Alteromonas macleodii | Alteromonas macleodii | Alteromonas macleodii | Alteromonas macleodii | 0.858 |
| 81092899f1c0c2db02ec2ab6e65d86f2 | Bacteria | Proteobacteria | Alphaproteobacteria | SAR11 clade | Clade I | Clade Ia | | 0.849 |
| 8b0192de06c3b10f85bc482236c92366 | Bacteria | Cyanobacteria | Oxyphotobacteria | Synechococcales | Cyanobiaceae | Synechococcus CC9902 | | 0.998 |
| b4aaa30f8f64f72ba6cfd139f2023a8d | Bacteria | Proteobacteria | Alphaproteobacteria | Rhodospirillales | AEGEAN-169 marine group | uncultured marine bacterium | uncultured marine bacterium | 0.803 |
| 1c54ed87c1705610cad2896671526efe | Bacteria | Cyanobacteria | Oxyphotobacteria | Synechococcales | Cyanobiaceae | Prochlorococcus MIT9313 | | 1.000 |
| a5e6de786770e07687674cc32d097319 | Bacteria | Proteobacteria | Gammaproteobacteria | Ectothiorhodospirales | Ectothiorhodospiraceae | uncultured | uncultured marine bacterium | 0.970 |
| a35da92fe92a502bc206941a91a633bc | Bacteria | Proteobacteria | Alphaproteobacteria | Rhodobacterales | Rhodobacteraceae | uncultured | Rhodobacteraceae bacterium REDSEA-S03_B4 | 0.744 |
| e43703ac625ad9fda149e97e827b0553 | Bacteria | Marinimicrobia (SAR406 clade) | Ambiguous_taxa | Ambiguous_taxa | Ambiguous_taxa | Ambiguous_taxa | Ambiguous_taxa | 0.909 |
| 3ab38fc647166c931ad18a473daa0560 | Bacteria | Proteobacteria | Alphaproteobacteria | Rhodobacterales | Rhodobacteraceae | Ascidiaceihabitans | | 0.780 |
| 91f687fdb62026d040baa83be4ad4556 | Bacteria | Cyanobacteria | Oxyphotobacteria | Synechococcales | Cyanobiaceae | Prochlorococcus MIT9313 | | 1.000 |
| 5234f65d442af2c647f49e161bb35ffa | Bacteria | Verrucomicrobia | Verrucomicrobiae | Verrucomicrobiales | Rubritaleaceae | Roseibacillus | uncultured marine bacterium | 0.948 |
| 8941b912fff03d55d958a89a8510b9c2 | Bacteria | Proteobacteria | Gammaproteobacteria | SAR86 clade | | | | 1.000 |
| 1b980624ac540cbc5036300a8919d6da | Bacteria | Proteobacteria | Alphaproteobacteria | Rhodospirillales | AEGEAN-169 marine group | Ambiguous_taxa | Ambiguous_taxa | 1.000 |
| 46eb9145a13d564c00f01e905866be62 | Bacteria | Actinobacteria | Acidimicrobiia | Actinomarinales | Actinomarinaceae | Candidatus Actinomarina | uncultured marine bacterium | 0.737 |
| 05932a0c91ddcdbca8bcd2c6ad0849b7 | Bacteria | Proteobacteria | Alphaproteobacteria | Rhodospirillales | AEGEAN-169 marine group | marine metagenome | marine metagenome | 0.893 |
| de6ce53e1a0ea2d20425f04a7b4e733f | Bacteria | Proteobacteria | Gammaproteobacteria | SAR86 clade | uncultured gamma proteobacterium KTc1121 | uncultured gamma proteobacterium KTc1121 | uncultured gamma proteobacterium KTc1121 | 0.767 |
| 539970532548c674f88ff4566f27590b | Bacteria | Proteobacteria | Alphaproteobacteria | Puniceispirillales | SAR116 clade | | | 1.000 |
| bb26c60c17351a7aef92a8c89d3b4dd9 | Bacteria | Proteobacteria | Alphaproteobacteria | Rhodobacterales | Rhodobacteraceae | | | 1.000 |
| 2c6e619dbaab11dccebfaa0c0bc5d1d3 | Bacteria | Proteobacteria | Alphaproteobacteria | SAR11 clade | Clade IV | | | 1.000 |
| 08223140d04d57bdcd77795646bbb117 | Bacteria | Proteobacteria | Alphaproteobacteria | Rhodospirillales | AEGEAN-169 marine group | Ambiguous_taxa | Ambiguous_taxa | 0.999 |
| e057f66ccacf6c694b1b69b11f19533f | Bacteria | Proteobacteria | Gammaproteobacteria | SAR86 clade | | | | 1.000 |
| 7290b3cdf5690aa2bd532a5dc67d77ad | Bacteria | Proteobacteria | Gammaproteobacteria | SAR86 clade | uncultured SAR86 cluster bacterium | uncultured SAR86 cluster bacterium | uncultured SAR86 cluster bacterium | 0.886 |
| a6a7d35754f59891dba1d67c6c7144bf | Bacteria | Proteobacteria | Gammaproteobacteria | Cellvibrionales | Porticoccaceae | SAR92 clade | | 1.000 |
| b85ed7a908f42de3eea96eea6301f7f4 | Bacteria | Marinimicrobia (SAR406 clade) | unidentified marine bacterioplankton | unidentified marine bacterioplankton | unidentified marine bacterioplankton | unidentified marine bacterioplankton | unidentified marine bacterioplankton | 0.985 |
| 9eb3f4b8a7fb033b51fb84fdc95e4775 | Bacteria | Proteobacteria | Gammaproteobacteria | Thiotrichales | Thiotrichaceae | uncultured | uncultured marine bacterium | 1.000 |
| 7db24e7c57b22d168f4d3b7bba68a3ea | Bacteria | Bacteroidetes | Bacteroidia | Flavobacteriales | Flavobacteriaceae | NS2b marine group | uncultured marine bacterium | 0.786 |
| b1c4d3c70082054ddc4a26eb17563fbc | Bacteria | Proteobacteria | Alphaproteobacteria | Parvibaculales | OCS116 clade | | | 1.000 |
| c95be457c741775ab35521676639b910 | Bacteria | Actinobacteria | Acidimicrobiia | Actinomarinales | Actinomarinaceae | Candidatus Actinomarina | Ambiguous_taxa | 0.828 |
| 191d7e7da61a2dc4cbe8c2f898d070c4 | Bacteria | Proteobacteria | Gammaproteobacteria | SAR86 clade | | | | 1.000 |
| 584ade95e9d1b271b83a5f45e85a4152 | Bacteria | Proteobacteria | Alphaproteobacteria | SAR11 clade | Clade II | | | 0.993 |
| 5472553891fee5a529081d99d11dea18 | Bacteria | Actinobacteria | Acidimicrobiia | Actinomarinales | Actinomarinaceae | Candidatus Actinomarina | | 1.000 |
| 477382dd44cb51c8fe5d476693824054 | Bacteria | Proteobacteria | Gammaproteobacteria | Cellvibrionales | Porticoccaceae | SAR92 clade | uncultured marine bacterium | 0.979 |
| 0b3451bfacccfe3f45e24b486da99a9c | Bacteria | Bacteroidetes | Bacteroidia | Flavobacteriales | Flavobacteriaceae | NS4 marine group | | 1.000 |
| 5d51ef8a47229f789328b35ea0b5b39a | Bacteria | Proteobacteria | Gammaproteobacteria | SAR86 clade | | | | 1.000 |
| dbd6a30965f3370841c1850f121f6c4e | Bacteria | Proteobacteria | Gammaproteobacteria | Betaproteobacteriales | Methylophilaceae | OM43 clade | Ambiguous_taxa | 0.805 |
| b3ede366cfbb90e778be76dcbd29aa11 | Bacteria | Proteobacteria | Alphaproteobacteria | Rhodobacterales | Rhodobacteraceae | Ascidiaceihabitans | | 0.903 |
| bbaa4707c6e1d69a212df3828254e70f | Bacteria | Proteobacteria | Alphaproteobacteria | Rickettsiales | S25-593 | uncultured marine bacterium | uncultured marine bacterium | 0.748 |
| 802c4a4234381e27c4ea1730da37a783 | Bacteria | Proteobacteria | Gammaproteobacteria | SAR86 clade | uncultured proteobacterium OCS44 | uncultured proteobacterium OCS44 | uncultured proteobacterium OCS44 | 0.769 |
| b97409614c1176bec58b078d86169ffb | Bacteria | Proteobacteria | Gammaproteobacteria | Thiomicrospirales | Thioglobaceae | SUP05 cluster | | 1.000 |
| 3582e1beb44986ccf8fed703ecd0a314 | Bacteria | Proteobacteria | Gammaproteobacteria | SAR86 clade | | | | 1.000 |
| f67951137456a41df3608df7675ae032 | Bacteria | Proteobacteria | Alphaproteobacteria | SAR11 clade | Clade II | | | 0.994 |
| 9f4e97a31dd378385a259d3f043220af | Bacteria | Proteobacteria | Alphaproteobacteria | SAR11 clade | Clade I | | | 0.998 |
| 35b220edde7a107dfd6e8c88f1e36ec5 | Bacteria | Cyanobacteria | Oxyphotobacteria | Synechococcales | Cyanobiaceae | Prochlorococcus MIT9313 | | 1.000 |
| 44c88775a796caf1fc5f29bf7bc73033 | Bacteria | Proteobacteria | Alphaproteobacteria | SAR11 clade | Clade I | Clade Ib | | 0.754 |
| fe8f740f4d7f9ae55b4eda61d15dc311 | Bacteria | Proteobacteria | Alphaproteobacteria | Puniceispirillales | SAR116 clade | Candidatus Puniceispirillum | | 0.848 |
| a8f894d55f7034b905cb45a15df31bc8 | Bacteria | Proteobacteria | Alphaproteobacteria | SAR11 clade | Clade I | | | 1.000 |
| 10884bc8cb1cbbbd39749627ef5e323f | Bacteria | Proteobacteria | Alphaproteobacteria | Rhodobacterales | Rhodobacteraceae | | | 1.000 |
| 9e7dfa8ee4c2408adfd9f088ed34ecd5 | Bacteria | Bacteroidetes | Bacteroidia | Flavobacteriales | Flavobacteriaceae | NS4 marine group | | 1.000 |
| f4c50f1ab46aaf3860b962135d8ac239 | Bacteria | Bacteroidetes | Bacteroidia | Cytophagales | Cyclobacteriaceae | Marinoscillum | | 1.000 |
| 47ff44a42565e4d0db3a806b626dd11f | Bacteria | Proteobacteria | Gammaproteobacteria | SAR86 clade | | | | 1.000 |
| 6d99895665fb90d94a10f0204333500d | Bacteria | Proteobacteria | Gammaproteobacteria | SAR86 clade | | | | 1.000 |
| d077a35cd9abcb1074362008c48f0cd3 | Bacteria | Cyanobacteria | Oxyphotobacteria | Synechococcales | Cyanobiaceae | Prochlorococcus MIT9313 | | 1.000 |
| daac7046e78b2c355f0518e9d7cc5692 | Bacteria | Proteobacteria | Deltaproteobacteria | SAR324 clade(Marine group B) | | | | 1.000 |
| 26746171a93079b0b9d8fad0b5468038 | Bacteria | Cyanobacteria | Oxyphotobacteria | Synechococcales | Cyanobiaceae | Synechococcus CC9902 | | 0.999 |
| c582f5e706adebf9fa8aa9dba825e821 | Bacteria | Proteobacteria | Gammaproteobacteria | SAR86 clade | | | | 1.000 |
| 992fe20d2957d584c944da77375e4496 | Bacteria | Bacteroidetes | Bacteroidia | Flavobacteriales | Flavobacteriaceae | Formosa | uncultured marine bacterium | 0.822 |
| a10d9d5d48bef4fab665520e8981884d | Bacteria | Verrucomicrobia | Verrucomicrobiae | Opitutales | Puniceicoccaceae | Lentimonas | Ambiguous_taxa | 0.808 |
| b516660d69b8590d4917b26e8a493dcf | Bacteria | Proteobacteria | Gammaproteobacteria | KI89A clade | uncultured marine bacterium | uncultured marine bacterium | uncultured marine bacterium | 0.877 |
| 04a389e30cb0917b65770980426f47e9 | Bacteria | Bacteroidetes | Bacteroidia | Flavobacteriales | Flavobacteriaceae | NS5 marine group | Ambiguous_taxa | 0.745 |
| 6cf2f2a5b633ef318ec491e8379f02fe | Bacteria | Proteobacteria | Gammaproteobacteria | SAR86 clade | uncultured gamma proteobacterium HF0070_25G02 | uncultured gamma proteobacterium HF0070_25G02 | uncultured gamma proteobacterium HF0070_25G02 | 0.951 |
| d054169d4bf17f483a2af0da2aa2d686 | Bacteria | Bacteroidetes | Bacteroidia | Flavobacteriales | Flavobacteriaceae | NS4 marine group | | 1.000 |
| 3c59436521bef1ebffc31d2c98c50dd8 | Bacteria | Proteobacteria | Gammaproteobacteria | SAR86 clade | | | | 1.000 |

| Feature_ID | Kingdom | Phylum | Class | Order | Family | Genus | Species | Confidence |
|---|---|---|---|---|---|---|---|---|
| 252554d217904f1bd746191daf86ff98 | Bacteria | Proteobacteria | Gammaproteobacteria | SAR86 clade | uncultured gamma proteobacterium HF0070_25G02 | uncultured gamma proteobacterium HF0070_25G02 | uncultured gamma proteobacterium HF0070_25G02 | 1.000 |
| 9ed82d76e0e6c7b93614e35870c76787 | Bacteria | Proteobacteria | Alphaproteobacteria | Rhodospirillales | AEGEAN-169 marine group | uncultured marine bacterium | uncultured marine bacterium | 0.744 |
| 6b31940a8beebbbf26dc9f013108f1d5 | Bacteria | Proteobacteria | Alphaproteobacteria | Parvibaculales | OCS116 clade | | | 1.000 |
| 4a1f5b9c4d1ae591ffaa0b264dd24d60 | Bacteria | Proteobacteria | Alphaproteobacteria | Rhodospirillales | AEGEAN-169 marine group | Ambiguous_taxa | Ambiguous_taxa | 0.999 |
| b17f25f6a3ce44c2802197794cc61846 | Bacteria | Proteobacteria | Gammaproteobacteria | SAR86 clade | | | | 1.000 |
| 3cc0470e5ae9eecee24011e0f9708b45 | Bacteria | Proteobacteria | Gammaproteobacteria | SAR86 clade | uncultured gamma proteobacterium KTc1121 | uncultured gamma proteobacterium KTc1121 | uncultured gamma proteobacterium KTc1121 | 0.759 |
| 77643ca079d7ae633ff198b7b17d75cf | Bacteria | Proteobacteria | Alphaproteobacteria | Thalassobaculales | Nisaeaceae | OM75 clade | | 1.000 |
| db13c5b283352a7f288dd617c0adb847 | Bacteria | Bacteroidetes | Bacteroidia | Flavobacteriales | Cryomorphaceae | uncultured | uncultured Owenweeksia sp. | 0.907 |
| b2a544438dab82c2a06dbeb43ef7137f | Bacteria | Cyanobacteria | Oxyphotobacteria | Chloroplast | Phalacroma mitra | Phalacroma mitra | Phalacroma mitra | 0.972 |
| 931b5b8dcc30783ebb945f1bf7008850 | Bacteria | Proteobacteria | Alphaproteobacteria | uncultured | uncultured marine bacterium | uncultured marine bacterium | uncultured marine bacterium | 0.960 |
| 88c221ea1bba372ca003d01321272911 | Bacteria | Proteobacteria | Gammaproteobacteria | HOC36 | Ambiguous_taxa | Ambiguous_taxa | Ambiguous_taxa | 0.927 |
| 289dd5fdf6016e756a0ace95d394acd8 | Bacteria | Proteobacteria | Alphaproteobacteria | Rhodospirillales | AEGEAN-169 marine group | | | 1.000 |
| 5ca72a029c2aff7c4ee9e6d7b0787e24 | Bacteria | Proteobacteria | Alphaproteobacteria | Rickettsiales | S25-593 | | | 1.000 |
| 781bf210724ebca9606340c2ce9fa190 | Bacteria | Proteobacteria | Alphaproteobacteria | Puniceispirillales | SAR116 clade | Ambiguous_taxa | Ambiguous_taxa | 0.761 |
| ed85ffc35c0865b3bc572cdf5fda97b8 | Bacteria | Bacteroidetes | Bacteroidia | Flavobacteriales | Flavobacteriaceae | NS2b marine group | | 1.000 |
| bde0f7cede6a0f51d7c7e78513f6cca2 | Bacteria | Proteobacteria | Alphaproteobacteria | SAR11 clade | Clade II | | | 0.990 |
| 7dcb42dd3b53586e69d0d52a42ad9519 | Bacteria | Proteobacteria | Alphaproteobacteria | SAR11 clade | Clade I | Clade Ia | | 0.860 |
| ce5f5fefa9a0156716cc33690a335dcb | Bacteria | Proteobacteria | Alphaproteobacteria | Rhodospirillales | AEGEAN-169 marine group | | | 1.000 |
| e0a62ac838662cb1b9fa1caba62aad66 | Bacteria | Bacteroidetes | Bacteroidia | Flavobacteriales | NS9 marine group | uncultured Flavobacteriia bacterium | uncultured Flavobacteriia bacterium | 0.970 |
| f8b4478984e7ecd3a28c667cc9d29100 | Bacteria | Proteobacteria | Gammaproteobacteria | SAR86 clade | | | | 1.000 |
| 95aac9dadcf8629bed8e50619d7bf87c | Bacteria | Proteobacteria | Gammaproteobacteria | SAR86 clade | | | | 1.000 |
| ec847826ecd243a4384951e5fa668c5e | Bacteria | Cyanobacteria | Oxyphotobacteria | Chloroplast | uncultured marine eukaryote | uncultured marine eukaryote | uncultured marine eukaryote | 0.902 |
| d4025c348719cedb2ea9c148e1d8837f | Bacteria | Bacteroidetes | Bacteroidia | Flavobacteriales | Flavobacteriaceae | | | 1.000 |
| 4e1167fbbbc5e772fad1530d2b3d2b3d | Bacteria | Bacteroidetes | Bacteroidia | Flavobacteriales | NS7 marine group | Ambiguous_taxa | Ambiguous_taxa | 0.740 |
| 63b47a5c24b84d4bcce61a8e4d500bd7 | Bacteria | Bacteroidetes | Bacteroidia | Flavobacteriales | Crocinitomicaceae | Fluviicola | uncultured marine bacterium | 0.845 |
| 291deb67818a7f99faadc6ddaacb2842 | Bacteria | Proteobacteria | Alphaproteobacteria | Parvibaculales | OCS116 clade | uncultured marine bacterium | uncultured marine bacterium | 0.840 |
| 076922ad9957104dbe93a1dc92877fa1 | Bacteria | Cyanobacteria | Oxyphotobacteria | Chloroplast | Phalacroma mitra | Phalacroma mitra | Phalacroma mitra | 0.979 |
| 05794f49d61313bcf77f6880ae90cfa0 | Bacteria | Verrucomicrobia | Verrucomicrobiae | Pedosphaerales | Pedosphaeraceae | SCGC AAA164-E04 | | 1.000 |
| c45b037c35d97b0afa741dbad837674f | Bacteria | Proteobacteria | Gammaproteobacteria | SAR86 clade | | | | 1.000 |
| e46b4d859dc6df8f7c07cd3cf053284f | Bacteria | Bacteroidetes | Bacteroidia | Flavobacteriales | Flavobacteriaceae | Formosa | uncultured marine bacterium | 0.889 |
| f9730793fd8ff31c44cb686796bb6188 | Bacteria | Proteobacteria | Gammaproteobacteria | SAR86 clade | | | | 1.000 |
| 85631eb6b9bc924e5ed6053447181c3e | Bacteria | Bacteroidetes | Bacteroidia | Flavobacteriales | NS9 marine group | Ambiguous_taxa | Ambiguous_taxa | 0.860 |
| 71f6bf6f8f5090b414c3ea8dd9eb713b | Bacteria | Proteobacteria | Gammaproteobacteria | Steroidobacterales | Woeseiaceae | Woeseia | uncultured marine bacterium | 1.000 |
| f4dfaada866a3f67086adc50567f5065 | Bacteria | Proteobacteria | Alphaproteobacteria | Puniceispirillales | SAR116 clade | Candidatus Puniceispirillum | | 0.952 |
| a441d96a218fb7aa31a3d7b1404a698b | Bacteria | Proteobacteria | Alphaproteobacteria | Rhodobacterales | Rhodobacteraceae | Amylibacter | | 0.949 |
| f4a6e50e16e7c6f7a8d4ebb1229e1326 | Bacteria | Bacteroidetes | Bacteroidia | Flavobacteriales | Flavobacteriaceae | NS4 marine group | | 1.000 |
| 0d04c71dcf004a6321b5a41cb75f12fa | Bacteria | Proteobacteria | Alphaproteobacteria | Puniceispirillales | SAR116 clade | | | 1.000 |
| b9ec28fefa62f2e28a7189169b9c5734 | Bacteria | Planctomycetes | Planctomycetacia | Pirellulales | Pirellulaceae | Rhodopirellula | marine metagenome | 0.998 |
| b7f557adfdc461b578dc4f47ea4bbcd0 | Bacteria | Proteobacteria | Alphaproteobacteria | Puniceispirillales | SAR116 clade | Candidatus Puniceispirillum | | 0.942 |
| 1b7e6b1555ef9b7c5aba76214e21bfff | Bacteria | Bacteroidetes | Bacteroidia | Flavobacteriales | NS9 marine group | uncultured Flavobacteriia bacterium | uncultured Flavobacteriia bacterium | 0.975 |
| 843859b79828f1386a1f9ae0f630774b | Bacteria | Proteobacteria | Gammaproteobacteria | Ectothiorhodospirales | Ectothiorhodospiraceae | uncultured | Ambiguous_taxa | 0.904 |
| db06db4029f8cb3b6f899ffbe76c03aa | Bacteria | Marinimicrobia (SAR406 clade) | Ambiguous_taxa | Ambiguous_taxa | Ambiguous_taxa | Ambiguous_taxa | Ambiguous_taxa | 0.831 |
| 9d240954c03c9e53333e3a4d2f10d6e6 | Bacteria | Verrucomicrobia | Verrucomicrobiae | Verrucomicrobiales | DEV007 | | | 1.000 |
| 1a6128b13c51f1f35cd10c0456a6b8e2 | Bacteria | Bacteroidetes | Bacteroidia | Flavobacteriales | Flavobacteriaceae | NS4 marine group | uncultured marine bacterium | 0.895 |
| 5afd2e8cb777b78f3999b91d97dd305a | Bacteria | Proteobacteria | Alphaproteobacteria | uncultured | | | | 1.000 |
| 5cbccd30f9dd319ee8f0f6e2b8dfec37 | Bacteria | Bacteroidetes | Bacteroidia | Flavobacteriales | Flavobacteriaceae | NS4 marine group | | 1.000 |
| 6efc1f46a7713ccbf6592587f88c5d9e | Bacteria | Cyanobacteria | Oxyphotobacteria | Synechococcales | Cyanobiaceae | | | 1.000 |
| 3c5c4c699d122f12e342f531f0a57159 | Bacteria | Proteobacteria | Alphaproteobacteria | SAR11 clade | Clade II | | | 0.801 |
| 9e893d8a87f8fb7eade73e5095b731c9 | Bacteria | Cyanobacteria | Oxyphotobacteria | Synechococcales | Cyanobiaceae | Synechococcus CC9902 | | 0.993 |
| 07f203cb593fa3e98a355dc445235d40 | Bacteria | Bacteroidetes | Bacteroidia | Flavobacteriales | Flavobacteriaceae | NS5 marine group | | 1.000 |
| dfc67b35046703839f39ea644d857bbd | Bacteria | Proteobacteria | Alphaproteobacteria | SAR11 clade | Clade II | unidentified marine bacterioplankton | unidentified marine bacterioplankton | 0.739 |
| 15c5581ecf3c9229c2c206379d488ce6 | Bacteria | Bacteroidetes | Bacteroidia | Flavobacteriales | Flavobacteriaceae | NS2b marine group | uncultured marine bacterium | 0.913 |
| 21767e83f19921d3cb152934ab5b10bb | Bacteria | Cyanobacteria | Oxyphotobacteria | Chloroplast | uncultured prymnesiophyte C8704 | uncultured prymnesiophyte C8704 | uncultured prymnesiophyte C8704 | 0.996 |
| efe10567db1cbd5febc8ffb808575e98 | Bacteria | Proteobacteria | Gammaproteobacteria | SAR86 clade | uncultured gamma proteobacterium HF0070_25G02 | uncultured gamma proteobacterium HF0070_25G02 | uncultured gamma proteobacterium HF0070_25G02 | 1.000 |
| 4bf887a6d1b93c8b0b13ca9a677800a1 | Bacteria | Proteobacteria | Alphaproteobacteria | Puniceispirillales | SAR116 clade | Candidatus Puniceispirillum | uncultured bacterium | 0.748 |
| 08f5c4a8d1100d6ed3b234abbc6ef0b1 | Bacteria | Actinobacteria | Acidimicrobiia | Actinomarinales | Actinomarinaceae | Candidatus Actinomarina | uncultured marine bacterium | 0.789 |
| 82636c13aeb820d1aebf7dedaecf8cd2 | Bacteria | Verrucomicrobia | Verrucomicrobiae | Verrucomicrobiales | Rubritaleaceae | Roseibacillus | uncultured marine bacterium | 0.999 |
| 85809ab6d10133261b503338a1f1cd60 | Bacteria | Proteobacteria | Alphaproteobacteria | Parvibaculales | Parvibaculaceae | uncultured | Ambiguous_taxa | 0.998 |
| b6e8e35011fbccc0d4c70b5870f9961b | Bacteria | Cyanobacteria | Oxyphotobacteria | Synechococcales | Cyanobiaceae | Synechococcus CC9902 | | 0.995 |
| 664af6a633547b3dd5b07372f47ee95b | Bacteria | Proteobacteria | Gammaproteobacteria | Oceanospirillales | Pseudohongiellaceae | Pseudohongiella | | 1.000 |
| 3f669b23c56f9b7cc0bae37b9458c22a | Bacteria | Cyanobacteria | Oxyphotobacteria | Chloroplast | uncultured Cryptomonadaceae | uncultured Cryptomonadaceae | uncultured Cryptomonadaceae | 0.975 |
| f4fe3ea99e3849bdad1e4e7bf559eb2c | Bacteria | Cyanobacteria | Oxyphotobacteria | Chloroplast | | | | 1.000 |
| e71e2cdbc8f65de93e08721c92b89c90 | Bacteria | Verrucomicrobia | Verrucomicrobiae | Verrucomicrobiales | Rubritaleaceae | Roseibacillus | marine metagenome | 0.947 |
| 82d8d63aec8f3d3860586defded5be32 | Bacteria | Marinimicrobia (SAR406 clade) | | | | | | 1.000 |
| 55fba4512510e83ea3af0871e3cbd291 | Bacteria | Bacteroidetes | Bacteroidia | Flavobacteriales | Cryomorphaceae | uncultured | Ambiguous_taxa | 0.867 |
| e787656029734838905049 54a9bdc9b0 | Bacteria | Bacteroidetes | Bacteroidia | Flavobacteriales | Flavobacteriaceae | NS4 marine group | | 1.000 |
| aa4e24fcdfd89ec545f1d14d1d3e9300 | Bacteria | Chloroflexi | Dehalococcoidia | SAR202 clade | Ambiguous_taxa | Ambiguous_taxa | Ambiguous_taxa | 0.997 |
| 7c5a0fd8a87be2cb862f42d207bfbcb0 | Bacteria | Proteobacteria | Alphaproteobacteria | Rhodobacterales | Rhodobacteraceae | | | 1.000 |
| ef58a82c6abb005e535d37100f1ca82a | Bacteria | Proteobacteria | Deltaproteobacteria | SAR324 clade(Marine group B) | Ambiguous_taxa | Ambiguous_taxa | Ambiguous_taxa | 0.965 |
| 282999fee21149dccedbab19c2449394 | Bacteria | Cyanobacteria | Oxyphotobacteria | Chloroplast | marine metagenome | marine metagenome | marine metagenome | 0.978 |
| e30d19fe5b0ebf9554c052840c4d956a | Bacteria | Proteobacteria | Alphaproteobacteria | SAR11 clade | Clade I | Clade Ia | | 0.950 |
| 29baa5ec8562d9147686ed0145ed2b61 | Bacteria | Cyanobacteria | Oxyphotobacteria | Chloroplast | | | | 1.000 |
| 753197e4f87bd09a6ffc97956daa03a8 | Bacteria | Bacteroidetes | Bacteroidia | Flavobacteriales | Flavobacteriaceae | NS5 marine group | | 1.000 |
| 605559dba01eee0ccfdfb68e4dd35cb8 | Bacteria | Proteobacteria | Alphaproteobacteria | SAR11 clade | Clade I | Clade Ib | | 0.706 |
| bcbbb314a8fa5891dae2d2ac6cfe667e | Bacteria | Bacteroidetes | Bacteroidia | Flavobacteriales | Flavobacteriaceae | NS4 marine group | Ambiguous_taxa | 0.712 |

| Feature_ID | Kingdom | Phylum | Class | Order | Family | Genus | Species | Confidence |
|---|---|---|---|---|---|---|---|---|
| 508de5bb11bb82fa44239af13f00ad82 | Bacteria | Proteobacteria | Alphaproteobacteria | Rhodospirillales | AEGEAN-169 marine group | | | 1.000 |
| de392404ac3ef9a07e3629ed62484d85 | Bacteria | Proteobacteria | Gammaproteobacteria | SAR86 clade | | | | 1.000 |
| cc092684f2a7f76179a466e844d29f23 | Bacteria | Marinimicrobia (SAR406 clade) | Marinimicrobia bacterium JGI 0000113-D11 | Marinimicrobia bacterium JGI 0000113-D11 | Marinimicrobia bacterium JGI 0000113-D11 | Marinimicrobia bacterium JGI 0000113-D11 | Marinimicrobia bacterium JGI 0000113-D11 | 0.785 |
| 3e8607ee4c179ad7f18201b67f76863a | Bacteria | Proteobacteria | Alphaproteobacteria | Puniceispirillales | SAR116 clade | Ambiguous_taxa | Ambiguous_taxa | 0.721 |
| 8a13fa187cbfd57eeba9e8067576048a | Bacteria | Proteobacteria | Alphaproteobacteria | SAR11 clade | Clade I | Clade Ib | | 0.719 |
| c5034d5ed01af74802cdae21bda00b7a | Bacteria | Proteobacteria | Gammaproteobacteria | SAR86 clade | uncultured gamma proteobacterium HF0070_25G02 | uncultured gamma proteobacterium HF0070_25G02 | uncultured gamma proteobacterium HF0070_25G02 | 1.000 |
| d2f397f5b7eaa4e05f0f54e9e9b19611 | Bacteria | Verrucomicrobia | Verrucomicrobiae | Opitutales | Puniceicoccaceae | Coraliomargarita | uncultured marine bacterium | 0.763 |
| 98c93109ee2ee98c7b1fabbf0075f0e5 | Bacteria | Proteobacteria | Gammaproteobacteria | SAR86 clade | | | | 1.000 |
| 5ef547d6704a6fb5bc89f620b8b8cef8 | Bacteria | Cyanobacteria | Oxyphotobacteria | Chloroplast | uncultured diatom | uncultured diatom | uncultured diatom | 0.740 |
| c23de50eb911e9febd9dfc8c6515faa5 | Bacteria | Bacteroidetes | Bacteroidia | Cytophagales | Cyclobacteriaceae | Marinoscillum | marine metagenome | 0.827 |
| 73bb08decc3b8de17a42365a3e530dad | Bacteria | Proteobacteria | Alphaproteobacteria | SAR11 clade | Clade III | | | 1.000 |
| 118632c3c8e0d6461ce3af52e785dd3e | Bacteria | Cyanobacteria | Oxyphotobacteria | Chloroplast | Phalacroma mitra | Phalacroma mitra | Phalacroma mitra | 0.996 |
| b6b738a3d99d9e95203548f7dfba8e32 | Bacteria | Verrucomicrobia | Verrucomicrobiae | Opitutales | Puniceicoccaceae | | | 1.000 |
| 599689144411ce9c897d565b1439ed0f | Bacteria | Proteobacteria | Gammaproteobacteria | Ectothiorhodospirales | Ectothiorhodospiraceae | uncultured | uncultured marine bacterium | 0.950 |
| 19f01309d32a8c51acb7fcc6ac03ab0a | Bacteria | Cyanobacteria | Oxyphotobacteria | Synechococcales | Cyanobiaceae | Synechococcus CC9902 | | 0.999 |
| c01b1b54d3ec820b9861c26638955c30 | Bacteria | Proteobacteria | Gammaproteobacteria | Oceanospirillales | Pseudohongiellaceae | Pseudohongiella | | 1.000 |
| ff20761d24c2c2160f37ca8d4914e5c5 | Bacteria | Nitrospinae | Nitrospinia | Nitrospinales | Nitrospinaceae | LS-NOB | | 1.000 |
| ef5d8bc94e7e90cc298c54b439ce0bc0 | Bacteria | Cyanobacteria | Oxyphotobacteria | Chloroplast | Phalacroma mitra | Phalacroma mitra | Phalacroma mitra | 0.914 |
| c5bf382008838fe10ad58441a530a80a | Archaea | Euryarchaeota | Thermoplasmata | Marine Group II | uncultured marine archaeon | uncultured marine archaeon | uncultured marine archaeon | 0.828 |
| 6887783221d5d3c0067d9d9582ad443b | Bacteria | Proteobacteria | Alphaproteobacteria | Puniceispirillales | SAR116 clade | Candidatus Puniceispirillum | Ambiguous_taxa | 0.734 |
| 03a52768936e3d07ef55f97972de1bc7 | Bacteria | Bacteroidetes | Bacteroidia | Flavobacteriales | NS9 marine group | | | 1.000 |
| ff02653b16980833c506a2b99a8820c8 | Bacteria | Proteobacteria | Gammaproteobacteria | SAR86 clade | | | | 1.000 |
| f21c48ae04d886c87524f1afeee09f97 | Bacteria | Proteobacteria | Alphaproteobacteria | Puniceispirillales | SAR116 clade | | | 1.000 |
| 4487839f1123763907bc1ff10e85833e | Bacteria | Cyanobacteria | Oxyphotobacteria | Chloroplast | | | | 1.000 |
| b772e3ea4020555e3cd29722b277d5d1 | Bacteria | Bacteroidetes | Bacteroidia | Flavobacteriales | Flavobacteriaceae | NS2b marine group | | 1.000 |
| 4d072cfea4d0d51d223a1de459017a90 | Bacteria | Proteobacteria | Alphaproteobacteria | Parvibaculales | PS1 clade | | | 1.000 |
| 71362904ece78f66227f6d6a23402ecf | Bacteria | Bacteroidetes | Bacteroidia | Flavobacteriales | Flavobacteriaceae | NS4 marine group | | 1.000 |
| c5dbf2ea44cadec702ee97926cfe8184 | Bacteria | Proteobacteria | Alphaproteobacteria | uncultured | uncultured marine bacterium | uncultured marine bacterium | uncultured marine bacterium | 0.839 |
| fe37c03cee5f1b60dc84416bbfec8213 | Bacteria | Proteobacteria | Gammaproteobacteria | SAR86 clade | | | | 1.000 |
| bf18173ef06b0e8cd0a55ce86358c75f | Bacteria | Chloroflexi | Dehalococcoidia | SAR202 clade | metagenome | metagenome | metagenome | 1.000 |
| 44b6d0d63263b2f4004cbc43da70130b | Bacteria | Bacteroidetes | Bacteroidia | Flavobacteriales | Cryomorphaceae | uncultured | | 1.000 |
| de986b628b6ef085bf3b6e57f5f505bf | Bacteria | Proteobacteria | Alphaproteobacteria | Rickettsiales | S25-593 | | | 1.000 |
| dbff2a1f6252b3e862810065f0a47c96 | Bacteria | Nitrospinae | Nitrospinia | Nitrospinales | Nitrospinaceae | LS-NOB | | 1.000 |
| 2f6a5b789dc79c84a6f981c1166ae302 | Bacteria | Proteobacteria | Gammaproteobacteria | KI89A clade | uncultured marine bacterium | uncultured marine bacterium | uncultured marine bacterium | 0.909 |
| c53dad5262ba5e488c5284f6a083e9b3 | Bacteria | Bacteroidetes | Bacteroidia | Flavobacteriales | NS7 marine group | uncultured marine bacterium | uncultured marine bacterium | 0.789 |
| d3cdda9df90471774e94fad3fa66968a | Bacteria | Verrucomicrobia | Verrucomicrobiae | Opitutales | Puniceicoccaceae | Pelagicoccus | uncultured marine bacterium | 0.985 |
| 6b3d10862ba35a1b4736215214f4e0ae | Bacteria | Actinobacteria | Acidimicrobiia | Microtrichales | Microtrichaceae | Sva0996 marine group | uncultured marine bacterium | 0.896 |
| b6c0d89b177b3dd41142025330da6cba | Bacteria | Proteobacteria | Gammaproteobacteria | Ectothiorhodospirales | Ectothiorhodospiraceae | uncultured | | 1.000 |
| c895c3255c9cb80d01b06f4d4ea099bb | Bacteria | Bacteroidetes | Bacteroidia | Flavobacteriales | NS9 marine group | uncultured Flavobacteriia bacterium | uncultured Flavobacteriia bacterium | 0.948 |
| b5cf59752a56d6d4fcfa57ab815e9f6b | Bacteria | Proteobacteria | Gammaproteobacteria | Cellvibrionales | Halieaceae | OM60(NOR5) clade | Ambiguous_taxa | 0.888 |
| 1c42fd5161b8a27b461f68058f035fab | Bacteria | Bacteroidetes | Bacteroidia | Flavobacteriales | NS9 marine group | uncultured Flavobacteriia bacterium | uncultured Flavobacteriia bacterium | 0.842 |
| ed1f814cf7dececf59c956fb9e7091b0 | Bacteria | Proteobacteria | Alphaproteobacteria | SAR11 clade | Clade IV | | | 1.000 |
| a8d6e8aaa8fdfc9e88461c8f4c5a694f | Bacteria | Bacteroidetes | Bacteroidia | Flavobacteriales | NS9 marine group | uncultured marine bacterium | uncultured marine bacterium | 0.965 |
| 31686a4eece3e2575abefc8cdbefac08 | Bacteria | Proteobacteria | Alphaproteobacteria | SAR11 clade | Clade I | Clade Ib | | 0.700 |
| e582bb515b125c554a9fd3589d39aa8f | Bacteria | Proteobacteria | Alphaproteobacteria | Rhodospirillales | AEGEAN-169 marine group | | | 1.000 |
| a67d31cadaa0705b60077cea1b01eb3c | Bacteria | Bacteroidetes | Bacteroidia | Cytophagales | Cyclobacteriaceae | Marinoscillum | uncultured Bacteroidetes bacterium | 0.754 |
| 09e533a38f7cea3dbdc6ddc568385026 | Bacteria | Proteobacteria | Alphaproteobacteria | Rhodovibrionales | Kiloniellaceae | Kiloniella | | 0.923 |
| 0c74eae4dbf73f8e229295b0106c102a | Bacteria | Proteobacteria | Alphaproteobacteria | Rhodobacterales | Rhodobacteraceae | uncultured | | 0.981 |
| 68f8a4ce3475ff0ca3babe811577e2c6 | Bacteria | Bacteroidetes | Bacteroidia | Flavobacteriales | NS9 marine group | uncultured Flavobacteriales bacterium | uncultured Flavobacteriales bacterium | 0.843 |
| a5555d4fb3c468646e2ccbc60debf2eb | Bacteria | Proteobacteria | Alphaproteobacteria | Parvibaculales | Parvibaculaceae | uncultured | Ambiguous_taxa | 0.996 |
| df3a0e0d9593bdee1fb0bfe9a3551fd7 | Bacteria | Bacteroidetes | Bacteroidia | Flavobacteriales | NS7 marine group | | | 1.000 |
| 215cb8c53fd190b0093e97f0d2ba0f05 | Bacteria | Proteobacteria | Gammaproteobacteria | Ectothiorhodospirales | Ectothiorhodospiraceae | uncultured | uncultured marine bacterium | 0.983 |
| 269c56d6f6d510fca9e695dd4ef3aded | Bacteria | Verrucomicrobia | Verrucomicrobiae | Verrucomicrobiales | DEV007 | uncultured Verrucomicrobia bacterium | uncultured Verrucomicrobia bacterium | 0.839 |
| a47d1a3ccb217e8a847cf3ffe67d583c | Bacteria | Cyanobacteria | Oxyphotobacteria | Chloroplast | Phalacroma mitra | Phalacroma mitra | Phalacroma mitra | 0.999 |
| 635b862162c7602ce467943c39341ec6 | Bacteria | Proteobacteria | Gammaproteobacteria | Ectothiorhodospirales | Ectothiorhodospiraceae | uncultured | uncultured marine bacterium | 0.924 |
| 07915a1137f699de14f72c85f5bbd2a6 | Bacteria | Proteobacteria | Gammaproteobacteria | OM182 clade | Ambiguous_taxa | Ambiguous_taxa | Ambiguous_taxa | 0.702 |
| 62131078606f33fa2aee4fb1e24d1325 | Bacteria | Bacteroidetes | Bacteroidia | Flavobacteriales | Flavobacteriaceae | NS5 marine group | | 1.000 |
| 30f098ac7a82cddb029ee14ef9099c3f | Bacteria | Cyanobacteria | Oxyphotobacteria | Chloroplast | marine metagenome | marine metagenome | marine metagenome | 0.733 |
| 4756f9f656811df6fd09118ba685b062 | Bacteria | Proteobacteria | Alphaproteobacteria | Rhodobacterales | Rhodobacteraceae | | | 1.000 |
| 868e7facae9a1b8f62b269fa7bfaa730 | Bacteria | Verrucomicrobia | Verrucomicrobiae | Opitutales | Puniceicoccaceae | MB11C04 marine group | Ambiguous_taxa | 0.925 |
| cd803d1bccf1637b1b70fd30b8e8fd1c | Bacteria | Proteobacteria | Gammaproteobacteria | Cellvibrionales | Halieaceae | OM60(NOR5) clade | | 1.000 |
| acf6b2c4a4df518a1bd69c54d7521a66 | Bacteria | Proteobacteria | Alphaproteobacteria | Puniceispirillales | SAR116 clade | | | 1.000 |
| bdc0a8e3e90a5e0985147bf5144416be | Bacteria | Dadabacteria | Dadabacteriia | Dadabacteriales | metagenome | metagenome | metagenome | 0.999 |
| addcc2812e993a95129c169e0873386c | Bacteria | Proteobacteria | Deltaproteobacteria | Bdellovibrionales | Bdellovibrionaceae | OM27 clade | uncultured marine bacterium | 0.972 |
| 2eb2a18ea8b945b538211ad4c0be4b3f | Bacteria | Cyanobacteria | Oxyphotobacteria | Chloroplast | marine metagenome | marine metagenome | marine metagenome | 0.981 |
| f078b26f395a12e1eba3c71c485b4250 | Bacteria | Proteobacteria | Alphaproteobacteria | Parvibaculales | PS1 clade | | | 1.000 |
| b2969b12e4a530fde932492f44706d40 | Bacteria | Bacteroidetes | Bacteroidia | Flavobacteriales | Flavobacteriaceae | NS4 marine group | | 1.000 |
| cc020cdf107dc095a23b764f390a2293 | Bacteria | Cyanobacteria | Oxyphotobacteria | Chloroplast | Phalacroma mitra | Phalacroma mitra | Phalacroma mitra | 0.977 |
| 35871cbc31c90c10072c12c2f33d5238 | Bacteria | Cyanobacteria | Oxyphotobacteria | Chloroplast | unidentified marine bacterioplankton | unidentified marine bacterioplankton | unidentified marine bacterioplankton | 1.000 |
| 2219cd2896936f3ef4554239084b8846 | Bacteria | Proteobacteria | Deltaproteobacteria | SAR324 clade(Marine group B) | uncultured marine bacterium | uncultured marine bacterium | uncultured marine bacterium | 0.912 |
| ce330d07615fe7d3ec50c3016a5633f4 | Bacteria | Bacteroidetes | Bacteroidia | Flavobacteriales | NS9 marine group | Ambiguous_taxa | Ambiguous_taxa | 0.930 |
| 46dcd9314a0ff27108e1825c812d2eb0 | Bacteria | Marinimicrobia (SAR406 clade) | Ambiguous_taxa | Ambiguous_taxa | Ambiguous_taxa | Ambiguous_taxa | Ambiguous_taxa | 0.793 |
| 50c80c9baee4fa5da977674a08447452 | Bacteria | Proteobacteria | Gammaproteobacteria | Cellvibrionales | Spongiibacteraceae | BD1-7 clade | uncultured gamma proteobacterium | 0.803 |
| a1c2281577778d0172e74978fa84ed67 | Bacteria | Bacteroidetes | Bacteroidia | Flavobacteriales | NS9 marine group | Ambiguous_taxa | Ambiguous_taxa | 0.753 |
| 8fffad6a6296ad2edc7ca2c798df3792 | Bacteria | Bacteroidetes | Bacteroidia | Flavobacteriales | Cryomorphaceae | uncultured | | 1.000 |
| 38e67f409674a5d8f3df37ddbbb771a0 | Bacteria | Marinimicrobia (SAR406 clade) | uncultured marine bacterium | uncultured marine bacterium | uncultured marine bacterium | uncultured marine bacterium | uncultured marine bacterium | 0.943 |

| Feature_ID | Kingdom | Phylum | Class | Order | Family | Genus | Species | Confidence |
|---|---|---|---|---|---|---|---|---|
| 314b54066917a95d63f9354217b50cb5 | Bacteria | Chloroflexi | Dehalococcoidia | SAR202 clade | marine metagenome | marine metagenome | marine metagenome | 1.000 |
| c2ffe73bcebceeeb92743ada0e34d4fb | Bacteria | Bacteroidetes | Bacteroidia | Flavobacteriales | NS9 marine group | uncultured Flavobacteriia bacterium | uncultured Flavobacteriia bacterium | 0.913 |
| e08fbb4b67723c99b3d2d43e56045915 | Bacteria | Planctomycetes | Planctomycetacia | Pirellulales | Pirellulaceae | Blastopirellula | | 1.000 |
| 5a8040f87477bb12b890b5887c52c2fb | Bacteria | Dadabacteria | Dadabacteriia | Dadabacteriales | uncultured marine bacterium | uncultured marine bacterium | uncultured marine bacterium | 0.993 |
| 3c618a9049bc88d4ac4eff7acb13eda4 | Bacteria | Proteobacteria | Gammaproteobacteria | UBA10353 marine group | Ambiguous_taxa | Ambiguous_taxa | Ambiguous_taxa | 0.943 |
| dcb5ea7fb45ddcc0b93680167a8b7c44 | Bacteria | Verrucomicrobia | Verrucomicrobiae | Opitutales | Puniceicoccaceae | MB11C04 marine group | Ambiguous_taxa | 0.913 |
| b121e808a19922f89690f50bdf4ec19d | Bacteria | Bacteroidetes | Rhodothermia | Balneolales | Balneolaceae | Balneola | uncultured bacterium | 0.931 |
| c6079ff7dbdc0cd1fada9d054207a754 | Bacteria | Bacteroidetes | Bacteroidia | Flavobacteriales | Flavobacteriaceae | NS4 marine group | | 1.000 |
| e57d8fdef0d6b6588e27684430cfa57c | Bacteria | Planctomycetes | Planctomycetacia | Planctomycetales | Rubinisphaeraceae | uncultured | uncultured marine bacterium | 0.749 |
| 576e7ca7e6605d3309c5bf7adb453b8c | Bacteria | Proteobacteria | Gammaproteobacteria | Ectothiorhodospirales | Ectothiorhodospiraceae | uncultured | Ambiguous_taxa | 0.830 |
| c2de020fd6003f4489484e3e0d69db71 | Bacteria | Proteobacteria | Gammaproteobacteria | Ga0077536 | Ambiguous_taxa | Ambiguous_taxa | Ambiguous_taxa | 0.984 |
| 0b2c428b3e51956fff53742bc5eaa987 | Bacteria | Cyanobacteria | Oxyphotobacteria | Chloroplast | | | | 1.000 |
| b014d868ab6ea2b0ef47d3a7c8c8ae90 | Bacteria | Proteobacteria | Gammaproteobacteria | Oceanospirillales | Litoricolaceae | Litoricola | uncultured bacterium | 0.992 |
| e7ba9027d00b4757a5c590c5a3854b5e | Bacteria | Proteobacteria | Gammaproteobacteria | SAR86 clade | Ambiguous_taxa | Ambiguous_taxa | Ambiguous_taxa | 0.928 |
| 661f6d8a8debecb2adb641ddf8492956 | Bacteria | Verrucomicrobia | Verrucomicrobiae | Opitutales | Puniceicoccaceae | MB11C04 marine group | | 1.000 |
| bfea4a71b894ef01085b645ec34c4aa8 | Bacteria | Bacteroidetes | Bacteroidia | Flavobacteriales | NS9 marine group | uncultured Flavobacteriia bacterium | uncultured Flavobacteriia bacterium | 0.997 |
| 24c0daead3c6eb74983e67a111a6d5b7 | Bacteria | Proteobacteria | Alphaproteobacteria | Rickettsiales | S25-593 | uncultured marine bacterium | uncultured marine bacterium | 0.968 |
| 7574ca5823843687003b85b37ded3ce1 | Bacteria | Proteobacteria | Gammaproteobacteria | Thiotrichales | Thiotrichaceae | uncultured | uncultured marine bacterium | 1.000 |
| 75bd04ce5abd401ad49f6d6a41635e1b | Bacteria | Proteobacteria | Gammaproteobacteria | Betaproteobacteriales | Burkholderiaceae | MWH-UniP1 aquatic group | uncultured marine bacterium | 0.999 |
| f67f8c6729bafe9f2f6ce460dab0cc16 | Bacteria | Cyanobacteria | Oxyphotobacteria | Chloroplast | uncultured Cryptomonadaceae | uncultured Cryptomonadaceae | uncultured Cryptomonadaceae | 0.998 |
| 1bed6912133387cfebeb22b9970014ec | Bacteria | Proteobacteria | Alphaproteobacteria | SAR11 clade | Clade IV | | | 1.000 |
| 743d8bbb6cc5be08e50f1ab343f7b7a1 | Bacteria | Cyanobacteria | Oxyphotobacteria | Chloroplast | | | | 1.000 |
| 7ae0274f9c137205c5bfe756d1f8afb1 | Bacteria | Bacteroidetes | Bacteroidia | Flavobacteriales | Flavobacteriaceae | NS4 marine group | | 1.000 |
| e1a3d5864576c71c8d39697ad2342caf | Bacteria | Cyanobacteria | Oxyphotobacteria | Chloroplast | marine metagenome | marine metagenome | marine metagenome | 0.998 |
| 79b19ccb9d64ce35b4e4528c44ea7b0d | Bacteria | Proteobacteria | Gammaproteobacteria | Cellvibrionales | Porticoccaceae | SAR92 clade | | 1.000 |
| c7763753d3f0e889277fcdae148eb371 | Bacteria | Cyanobacteria | Oxyphotobacteria | Nostocales | Microcystaceae | Atelocyanobacterium (UCYN-A) | Ambiguous_taxa | 1.000 |
| f4a251958883c58397ad4ba8175c930d | Bacteria | Actinobacteria | Acidimicrobiia | Actinomarinales | Actinomarinaceae | Candidatus Actinomarina | | 1.000 |
| a5b834320abae36afa162bf9ea064f9a | Bacteria | Proteobacteria | Alphaproteobacteria | Rhodobacterales | Rhodobacteraceae | Planktomarina | Ambiguous_taxa | 0.745 |
| 0774d9e02be877dc80ca29d200c55c4b | Bacteria | Marinimicrobia (SAR406 clade) | metagenome | metagenome | metagenome | metagenome | metagenome | 0.867 |
| 7ca9503d5f097e8043782f5c59be4a06 | Bacteria | Bacteroidetes | Bacteroidia | Flavobacteriales | NS7 marine group | Ambiguous_taxa | Ambiguous_taxa | 0.746 |
| d04ca5488d9bc098653e3e767ad669a8 | Bacteria | Bacteroidetes | Bacteroidia | Flavobacteriales | NS9 marine group | | | 1.000 |
| a4f1edf6207afee5cede60c8a78ab396 | Bacteria | Cyanobacteria | Oxyphotobacteria | Synechococcales | Cyanobiaceae | Synechococcus CC9902 | | 0.991 |
| 52774904c524c524c46822631aec7615 | Bacteria | Cyanobacteria | Oxyphotobacteria | Chloroplast | Phalacroma mitra | Phalacroma mitra | Phalacroma mitra | 0.999 |
| d7716b0e4b28cf56e18343f29efe53b3 | Bacteria | Proteobacteria | Gammaproteobacteria | Cellvibrionales | Halieaceae | OM60(NOR5) clade | | 0.999 |
| a80ce521498d598368b58e246b4202ae | Bacteria | Proteobacteria | Alphaproteobacteria | Sphingomonadales | Sphingomonadaceae | | | 1.000 |
| 9fa3553e1f858086b01c3e1277682a7b | Bacteria | Actinobacteria | Acidimicrobiia | Actinomarinales | Actinomarinaceae | Candidatus Actinomarina | uncultured marine eukaryote | 1.000 |
| e6b625b62f2d1399d4704ababeee004f | Bacteria | Cyanobacteria | Oxyphotobacteria | Synechococcales | Cyanobiaceae | Synechococcus CC9902 | | 0.999 |
| 2e3f2c75bd357a5066741f97ea465f18 | Bacteria | Bacteroidetes | Bacteroidia | Flavobacteriales | Flavobacteriaceae | NS5 marine group | | 1.000 |
| 5b10a9bd60ddd45023e745da09053f52 | Bacteria | Verrucomicrobia | Verrucomicrobiae | Opitutales | Puniceicoccaceae | MB11C04 marine group | | 1.000 |
| 959dbedb4ef1b825ed569c48e49f7f79 | Bacteria | Proteobacteria | Deltaproteobacteria | SAR324 clade(Marine group B) | | | | 1.000 |
| b393a6979af18f43c36502f2cbdb96f1 | Bacteria | Verrucomicrobia | Verrucomicrobiae | Arctic97B-4 marine group | uncultured marine bacterium | uncultured marine bacterium | uncultured marine bacterium | 0.976 |
| 5da7c5389fac9c8aa4f254d7864439e5 | Bacteria | Proteobacteria | Gammaproteobacteria | SAR86 clade | Ambiguous_taxa | Ambiguous_taxa | Ambiguous_taxa | 0.922 |
| f06900dc9acb0a6894ff5db2bb7ec67e | Bacteria | Bacteroidetes | Bacteroidia | Flavobacteriales | Flavobacteriaceae | | | 1.000 |
| 94b8e4ad007f32353a2ace0106a57f80 | Bacteria | Proteobacteria | Gammaproteobacteria | SAR86 clade | uncultured gamma proteobacterium | uncultured gamma proteobacterium | uncultured gamma proteobacterium | 0.928 |
| 77ccdc9de7f2053dadebbb04c863c012 | Bacteria | Bacteroidetes | Bacteroidia | Flavobacteriales | Flavobacteriaceae | NS5 marine group | | 1.000 |
| 4a1c9b400c293e6639ee864ef90732df | Bacteria | Bacteroidetes | Bacteroidia | Flavobacteriales | Flavobacteriaceae | NS5 marine group | | 1.000 |
| 622c0dc6e0db0561ea646406466d6478 | Bacteria | Proteobacteria | Alphaproteobacteria | SAR11 clade | Clade I | Clade Ib | | 0.898 |
| 2b167da5985a57acbc0f508b7839170e | Bacteria | Proteobacteria | Alphaproteobacteria | SAR11 clade | Clade I | Clade Ib | | 0.975 |
| 911a0b11f962659a5f292a13f46eff9e | Bacteria | Dadabacteria | Dadabacteriia | Dadabacteriales | uncultured marine bacterium | uncultured marine bacterium | uncultured marine bacterium | 0.838 |
| 3c7c9c6e7201cfc573bc0641aedee109 | Bacteria | Marinimicrobia (SAR406 clade) | hydrothermal vent metagenome | hydrothermal vent metagenome | hydrothermal vent metagenome | hydrothermal vent metagenome | hydrothermal vent metagenome | 0.999 |
| 9ec12434c1f8c66b0e4b010fb2759252 | Bacteria | Planctomycetes | OM190 | Ambiguous_taxa | Ambiguous_taxa | Ambiguous_taxa | Ambiguous_taxa | 0.914 |
| c074ffe7bde46fb34a5fb3d12b1d16f1 | Bacteria | Proteobacteria | Gammaproteobacteria | SAR86 clade | metagenome | metagenome | metagenome | 0.972 |
| 16cf0460d6cd711c823a97642f23c7c8 | Bacteria | Verrucomicrobia | Verrucomicrobiae | Arctic97B-4 marine group | metagenome | metagenome | metagenome | 0.878 |
| 4ff9a81b1cd1216940c572029e4b0ff3 | Bacteria | Proteobacteria | Alphaproteobacteria | Rhodospirillales | AEGEAN-169 marine group | unidentified marine bacterioplankton | unidentified marine bacterioplankton | 0.983 |
| e5fa824335930cce17c586a78a3ffdf0 | Bacteria | Chloroflexi | Dehalococcoidia | SAR202 clade | metagenome | metagenome | metagenome | 0.876 |
| 00e9343d8fc364bc3c93a300efef122a | Bacteria | Proteobacteria | Alphaproteobacteria | Rickettsiales | S25-593 | uncultured marine bacterium | uncultured marine bacterium | 0.795 |
| f07714adb6189b375fa21a85940c5592 | Bacteria | Proteobacteria | Gammaproteobacteria | Ectothiorhodospirales | Ectothiorhodospiraceae | uncultured | uncultured marine bacterium | 0.991 |
| 30f7b76d702e7a619d7d84a8c5b8d21e | Bacteria | Proteobacteria | Gammaproteobacteria | Alteromonadales | Idiomarinaceae | Idiomarina | | 1.000 |
| d772fc4bf76190ab1f52ab252da575a2 | Bacteria | Proteobacteria | Alphaproteobacteria | Rhodobacterales | Rhodobacteraceae | Ascidiaceihabitans | | 0.992 |
| ad06170a0751836a5d6af3c0fcb4ac48 | Bacteria | Proteobacteria | Alphaproteobacteria | Puniceispirillales | SAR116 clade | Ambiguous_taxa | Ambiguous_taxa | 0.820 |
| 3ba46e5569172c5a1f6f0b5051d7c280 | Bacteria | Bacteroidetes | Bacteroidia | Flavobacteriales | NS9 marine group | Ambiguous_taxa | Ambiguous_taxa | 0.987 |
| 204e689938a2d8b443b249866bd369b6 | Bacteria | Bacteroidetes | Bacteroidia | Flavobacteriales | Cryomorphaceae | uncultured | uncultured Sphingobacterium sp. EB080_L08E11 | 0.879 |
| fe80e81f7f407fd78769dabebeb49cbd | Bacteria | Bacteroidetes | Bacteroidia | Flavobacteriales | Flavobacteriaceae | NS5 marine group | | 1.000 |
| 960cacb58c173c89bea7572d45d37361 | Bacteria | Cyanobacteria | Oxyphotobacteria | Synechococcales | Cyanobiaceae | Prochlorococcus MIT9313 | | 1.000 |
| 4cf34a75a4156783701a3a2c722ef874 | Bacteria | Verrucomicrobia | Verrucomicrobiae | Opitutales | Puniceicoccaceae | MB11C04 marine group | | 1.000 |
| 80942f5b2b50a0a0066512d64f78f246 | Bacteria | Bacteroidetes | Bacteroidia | Flavobacteriales | NS9 marine group | uncultured Flavobacteriia bacterium | uncultured Flavobacteriia bacterium | 0.972 |
| 2548fb397f3e7864863220b3c057c21e | Bacteria | Proteobacteria | Alphaproteobacteria | SAR11 clade | Clade I | Clade Ib | | 0.970 |
| 7b7819fef86dfa73b226800b43cd63f9 | Bacteria | Bacteroidetes | Bacteroidia | Flavobacteriales | Flavobacteriaceae | NS2b marine group | uncultured marine bacterium | 0.704 |
| 54312722d8893ce472adee8afb643f98 | Archaea | Euryarchaeota | Thermoplasmata | Marine Group II | uncultured marine group II euryarchaeote HF70_89B11 | uncultured marine group II euryarchaeote HF70_89B11 | uncultured marine group II euryarchaeote HF70_89B11 | 0.755 |
| 8cebec5a31a2bad840a6f58d37a6dc16 | Bacteria | Proteobacteria | Gammaproteobacteria | SAR86 clade | | | | 1.000 |
| d1af6b6bc0d37351821bc3652777aaa4 | Bacteria | Proteobacteria | Gammaproteobacteria | SAR86 clade | metagenome | metagenome | metagenome | 0.943 |
| bca465403b6d93fff384bccfb567f5c5 | Bacteria | Proteobacteria | Gammaproteobacteria | Alteromonadales | Alteromonadaceae | Alteromonas | | 0.943 |
| 11a6b07ab2d79d3dbbbad77d1c992448 | Bacteria | Bacteroidetes | Bacteroidia | Flavobacteriales | NS9 marine group | uncultured Flavobacteriia bacterium | uncultured Flavobacteriia bacterium | 0.880 |
| 63d8db74a2f9118f15242f0f04cace91 | Bacteria | Verrucomicrobia | Verrucomicrobiae | Opitutales | Puniceicoccaceae | uncultured | marine metagenome | 1.000 |

| Feature_ID | Kingdom | Phylum | Class | Order | Family | Genus | Species | Confidence |
|---|---|---|---|---|---|---|---|---|
| 143db17801b690d35b3b46f1c3ff348f | Bacteria | Proteobacteria | Alphaproteobacteria | uncultured | uncultured marine bacterium | uncultured marine bacterium | uncultured marine bacterium | 0.866 |
| e0714bc55e746c0a4e9e0e19648d367c | Bacteria | Bacteroidetes | Bacteroidia | Chitinophagales | uncultured | Ambiguous_taxa | Ambiguous_taxa | 1.000 |
| 0a233c064695292c966162c237d28200 | Bacteria | Cyanobacteria | Oxyphotobacteria | Chloroplast | Phalacroma mitra | Phalacroma mitra | Phalacroma mitra | 0.963 |
| 4088787ee607d0020d0e932c8903442e | Bacteria | Proteobacteria | Deltaproteobacteria | SAR324 clade(Marine group B) | uncultured SAR324 cluster bacterium | uncultured SAR324 cluster bacterium | uncultured SAR324 cluster bacterium | 0.744 |
| 115951f203f4294ad187121b08976bfa | Bacteria | Bacteroidetes | Bacteroidia | Flavobacteriales | NS9 marine group | | | 1.000 |
| a0fa10c9d5441215c28904385d35c43b | Bacteria | Proteobacteria | Gammaproteobacteria | Oceanospirillales | Pseudohongiellaceae | Pseudohongiella | | 1.000 |
| c0333556d049a08bad538681720e010e | Bacteria | Cyanobacteria | Oxyphotobacteria | Chloroplast | | | | 1.000 |
| 813fca6e40120aa92ead0e7596a9ce17 | Bacteria | Proteobacteria | Alphaproteobacteria | SAR11 clade | Clade I | Clade Ib | | 0.712 |
| 3b521bfe4081d4e6a5f952bc58deee87 | Bacteria | Bacteroidetes | Bacteroidia | Flavobacteriales | Flavobacteriaceae | NS5 marine group | uncultured marine bacterium | 0.728 |
| 32fe8473a9a604eb82e09f1096e13a86 | Bacteria | Cyanobacteria | Oxyphotobacteria | Chloroplast | | | | 1.000 |
| 907e0d9e928232d66afbdaeefb1f0213 | Bacteria | Marinimicrobia (SAR406 clade) | unidentified marine bacterioplankton | unidentified marine bacterioplankton | unidentified marine bacterioplankton | unidentified marine bacterioplankton | unidentified marine bacterioplankton | 0.971 |
| 1213d0907ccaacf7e640c612dbb5da1d | Bacteria | Bacteroidetes | Bacteroidia | Flavobacteriales | Cryomorphaceae | NS10 marine group | uncultured marine bacterium | 0.987 |
| 63448f9947173da993aecd18b86f6812 | Bacteria | Proteobacteria | Gammaproteobacteria | Oceanospirillales | Alcanivoracaceae | Alcanivorax | | 1.000 |
| 68c91a31fb541719277ec47133244630 | Bacteria | Bacteroidetes | Bacteroidia | Flavobacteriales | NS9 marine group | uncultured Flavobacteriia bacterium | uncultured Flavobacteriia bacterium | 0.932 |
| 3203cbd461967fea767f3fa3176c97cd | Bacteria | Bacteroidetes | Bacteroidia | Cytophagales | Cyclobacteriaceae | Marinoscillum | uncultured marine bacterium | 0.890 |
| ef08dbff58adabf9fb8e78614c9c9d83 | Bacteria | Marinimicrobia (SAR406 clade) | uncultured marine bacterium | uncultured marine bacterium | uncultured marine bacterium | uncultured marine bacterium | uncultured marine bacterium | 0.989 |
| c786dc70f37af8d7eacf501d0c0d0319 | Bacteria | Proteobacteria | Alphaproteobacteria | SAR11 clade | Clade I | Clade Ia | | 0.983 |
| adb2b9855cf752b255960c6c19323266 | Bacteria | Bacteroidetes | Bacteroidia | Flavobacteriales | Flavobacteriaceae | Aurantivirga | Ambiguous_taxa | 0.990 |
| b70002d086310254f3de3e75ab244179 | Bacteria | Proteobacteria | Gammaproteobacteria | Vibrionales | Vibrionaceae | Photobacterium | | 1.000 |
| dba6da18688461248a5e43144412a1be | Bacteria | Actinobacteria | Acidimicrobiia | Microtrichales | Microtrichaceae | Sva0996 marine group | Ambiguous_taxa | 0.949 |
| 53918459d75e5cb5a3153ad250b468a6 | Bacteria | Cyanobacteria | Oxyphotobacteria | Chloroplast | | | | 1.000 |
| 34c14db13812a643d994433f4b34ab3d | Bacteria | Proteobacteria | Alphaproteobacteria | Thalassobaculales | uncultured | Ambiguous_taxa | Ambiguous_taxa | 0.913 |
| a50962f849c1ba4479cdf46a8210487c | Bacteria | Bacteroidetes | Bacteroidia | Flavobacteriales | Flavobacteriaceae | NS4 marine group | uncultured marine bacterium | 0.957 |
| e5288906456af9668fe428eb598fe498 | Bacteria | Proteobacteria | Alphaproteobacteria | Parvibaculales | PS1 clade | | | 1.000 |
| 9c66d37b8847f69ceb8cfd2f8f198689 | Bacteria | Cyanobacteria | Oxyphotobacteria | Chloroplast | | | | 1.000 |
| b6e1ed0f35f11f020c01991572b1b82c | Bacteria | Proteobacteria | Gammaproteobacteria | SAR86 clade | | | | 1.000 |
| 671b031173df10dbc6b4a2ae58d35709 | Bacteria | Chloroflexi | Dehalococcoidia | SAR202 clade | metagenome | metagenome | metagenome | 1.000 |
| 0399040dcb298884a3c4ad72bf922fad | Bacteria | Proteobacteria | Gammaproteobacteria | Betaproteobacteriales | Burkholderiaceae | MWH-UniP1 aquatic group | uncultured bacterium | 0.848 |
| f8c92c975389e2b8865c4c9491f10ea5 | Bacteria | Proteobacteria | Alphaproteobacteria | SAR11 clade | Clade II | | | 0.921 |
| 0c4013f3f1e92ea31f4f88c95ab4cf5c | Bacteria | Bacteroidetes | Bacteroidia | Flavobacteriales | Flavobacteriaceae | NS5 marine group | uncultured marine bacterium | 0.886 |
| 76a0cbc88131e0e43e3cabf757e37d96 | Archaea | Euryarchaeota | Thermoplasmata | Marine Group II | uncultured crenarchaeote | uncultured crenarchaeote | uncultured crenarchaeote | 0.994 |
| 9e153487f61544ea8fa29e0f620a4a88 | Bacteria | Bacteroidetes | Bacteroidia | Flavobacteriales | NS9 marine group | Ambiguous_taxa | Ambiguous_taxa | 0.918 |
| 09508e6050ac255206b48a33c69138bd | Bacteria | Proteobacteria | Gammaproteobacteria | Cellvibrionales | Halieaceae | OM60(NOR5) clade | Ambiguous_taxa | 0.875 |
| 7fbf48227c4b9352edcb6ccff23ce4ac | Bacteria | Bacteroidetes | Bacteroidia | Flavobacteriales | Flavobacteriaceae | NS5 marine group | | 1.000 |
| c47a1935d92cf41f81f4a5bfc760646c | Bacteria | Verrucomicrobia | Verrucomicrobiae | Opitutales | Puniceicoccaceae | Coraliomargarita | Ambiguous_taxa | 0.901 |
| e2d47798ea14e8cfd9aca01c23f03f2b | Bacteria | Kiritimatiellaeota | Kiritimatiellae | Kiritimatiellales | Kiritimatiellaceae | R76-B128 | marine metagenome | 0.994 |
| 0183fa601627e66c81dbbfb04bc34dd5 | Bacteria | Proteobacteria | Deltaproteobacteria | Bdellovibrionales | Bdellovibrionaceae | OM27 clade | uncultured marine bacterium | 0.970 |
| 8f8971517c3b9ba638f1c16893f7c5e5 | Bacteria | Proteobacteria | Gammaproteobacteria | HOC36 | uncultured marine bacterium | uncultured marine bacterium | uncultured marine bacterium | 0.999 |
| 109ab8dc170b0e876ebfb7b035d732a8 | Bacteria | Proteobacteria | Gammaproteobacteria | Cellvibrionales | Halieaceae | OM60(NOR5) clade | Ambiguous_taxa | 0.774 |
| c39a297dd4bb432d36bb58efa4459136 | Bacteria | Bacteroidetes | Bacteroidia | Flavobacteriales | Flavobacteriaceae | NS4 marine group | | 1.000 |
| f81a41c7e17ffea136fa5918a45487ad | Bacteria | Cyanobacteria | Oxyphotobacteria | Chloroplast | Phalacroma mitra | Phalacroma mitra | Phalacroma mitra | 0.965 |
| 18582b991c92644a21af1c370302c751 | Bacteria | Bacteroidetes | Bacteroidia | Flavobacteriales | Flavobacteriaceae | NS5 marine group | | 1.000 |
| f6aad746ffbe240a1a557d63d93213d8 | Bacteria | Planctomycetes | Phycisphaerae | Phycisphaerales | Phycisphaeraceae | Urania-1B-19 marine sediment group | Ambiguous_taxa | 1.000 |
| 60b8d3b8a6b413f71c162b62ce4834af | Bacteria | Proteobacteria | Alphaproteobacteria | uncultured | uncultured marine bacterium | uncultured marine bacterium | uncultured marine bacterium | 0.860 |
| d4c50a4f1df4d200b425cfe6f53e1e11 | Bacteria | Proteobacteria | Gammaproteobacteria | Cellvibrionales | Halieaceae | OM60(NOR5) clade | Ambiguous_taxa | 0.925 |
| e64da0dab06bc13809e495c0bbc848c5 | Archaea | Euryarchaeota | Thermoplasmata | Marine Group II | unidentified archaeon | unidentified archaeon | unidentified archaeon | 0.883 |
| 0b3737e9a96def03842568bf3a9a7812 | Bacteria | Proteobacteria | Alphaproteobacteria | SAR11 clade | Clade I | Clade Ia | | 0.876 |
| 6361b5166b308289706c741177f925da | Bacteria | Cyanobacteria | Oxyphotobacteria | Chloroplast | uncultured marine microorganism | uncultured marine microorganism | uncultured marine microorganism | 0.759 |
| c2a36810118e967801413ff3e8a66740 | Bacteria | Bacteroidetes | Bacteroidia | Flavobacteriales | NS7 marine group | uncultured Bacteroidetes bacterium | uncultured Bacteroidetes bacterium | 0.783 |
| ff40d8496e342401901cc4330e3118bc | Bacteria | Proteobacteria | Alphaproteobacteria | uncultured | uncultured marine bacterium | uncultured marine bacterium | uncultured marine bacterium | 0.996 |
| 69ecd66736e63361a229650c557fe5c4 | Bacteria | Marinimicrobia (SAR406 clade) | marine metagenome | marine metagenome | marine metagenome | marine metagenome | marine metagenome | 0.997 |
| e1d62bb8c36eb390ca125cabe55de8f9 | Bacteria | Bacteroidetes | Bacteroidia | Flavobacteriales | Flavobacteriaceae | NS5 marine group | | 1.000 |
| d8dc217567f18be6d10da68c7001a5c1 | Bacteria | Proteobacteria | Gammaproteobacteria | Alteromonadales | Pseudoalteromonadaceae | Pseudoalteromonas | | 0.994 |
| 7d83cdc24ba825ba451cf37b3a951c4b | Bacteria | Cyanobacteria | Oxyphotobacteria | Synechococcales | Cyanobiaceae | Prochlorococcus MIT9313 | | 0.999 |
| f1cf5556ba5ae779fb4a4ce255648b2f | Bacteria | Nitrospinae | Nitrospinia | Nitrospinales | Nitrospinaceae | LS-NOB | | 1.000 |
| 6ef9372cea995ed5a8eabf66e2f39185 | Bacteria | Bacteroidetes | Bacteroidia | Flavobacteriales | NS9 marine group | | | 1.000 |
| eeec7f5124ace2a069883082159a62c3 | Bacteria | Proteobacteria | Alphaproteobacteria | SAR11 clade | Clade I | Clade Ia | | 0.818 |
| c728fde5a5f19c94efce2372645cc585 | Bacteria | Proteobacteria | Gammaproteobacteria | Cellvibrionales | Porticoccaceae | SAR92 clade | uncultured marine bacterium | 0.978 |
| 22f859ea1ee356e0a6f34f7377384f21 | Bacteria | Proteobacteria | Alphaproteobacteria | Rickettsiales | S25-593 | uncultured marine bacterium | uncultured marine bacterium | 0.996 |
| 519bbbb8d914fde6e7b88bca185fc74f | Bacteria | Proteobacteria | Alphaproteobacteria | SAR11 clade | Clade I | Clade Ia | | 0.879 |
| 84593f6b964b1877c9bb0fb44c5b0c98 | Bacteria | Proteobacteria | Gammaproteobacteria | Cellvibrionales | Porticoccaceae | SAR92 clade | uncultured marine bacterium | 0.982 |
| 2f5bd880f7bd639de3221f12f6aefc19 | Bacteria | Proteobacteria | Alphaproteobacteria | Rhodobacterales | Rhodobacteraceae | Ascidiaceihabitans | | 0.990 |
| 279a085f1a6de8c0b976ce0dece5a72a | Bacteria | Planctomycetes | BD7-11 | metagenome | metagenome | metagenome | metagenome | 1.000 |
| 5d6bd15b0b80e19bfe563fc39acaac9c | Bacteria | Verrucomicrobia | Verrucomicrobiae | Verrucomicrobiales | DEV007 | Ambiguous_taxa | Ambiguous_taxa | 0.940 |
| d38047a1749bc3322bad6e66197cf3a9 | Bacteria | Proteobacteria | Alphaproteobacteria | Rhodospirillales | AEGEAN-169 marine group | uncultured marine bacterium | uncultured marine bacterium | 0.754 |
| 91df06809dde0e0fb06cda1f8300d16d | Bacteria | Proteobacteria | Gammaproteobacteria | Thiomicrospirales | Thioglobaceae | SUP05 cluster | | 1.000 |
| e3cda4357a7d5bff4372ebece2458911 | Bacteria | Proteobacteria | Gammaproteobacteria | Oceanospirillales | Halomonadaceae | Halomonas | | 1.000 |
| ecfc9cf5774ed0fe7f28a4a722c3a500 | Bacteria | Verrucomicrobia | Verrucomicrobiae | Opitutales | Puniceicoccaceae | MB11C04 marine group | | 1.000 |
| 793c0bb9242212f0a754cd687ac7cc00 | Bacteria | Proteobacteria | Gammaproteobacteria | Steroidobacterales | Woeseiaceae | Woeseia | uncultured marine bacterium | 1.000 |
| 5b467f21a97309d3b35f9cd90691a62e | Bacteria | Proteobacteria | Alphaproteobacteria | Rhodobacterales | Rhodobacteraceae | uncultured | | 0.982 |
| dd1936c4d663b6564aa4d99707c77565 | Bacteria | Proteobacteria | Alphaproteobacteria | Rhodospirillales | AEGEAN-169 marine group | | | 1.000 |
| ae08999425247cf343c495c5da35b8a7 | Bacteria | Actinobacteria | Acidimicrobiia | Microtrichales | Microtrichaceae | Sva0996 marine group | Ambiguous_taxa | 0.882 |
| b718cd41d94f25bb1e717f061bc8e015 | Bacteria | Proteobacteria | Gammaproteobacteria | SAR86 clade | uncultured bacterium | uncultured bacterium | uncultured bacterium | 0.749 |
| 7d6b1e56ef74b824d9f8385d2a4347bf | Bacteria | Bacteroidetes | Bacteroidia | Flavobacteriales | NS9 marine group | hydrothermal vent metagenome | hydrothermal vent metagenome | 1.000 |
| 59f2f4cc5c43362afd6a37dac554e6cd | Bacteria | Cyanobacteria | Oxyphotobacteria | Chloroplast | | | | 1.000 |
| b3a4e7f7a8e676839126c78d23367ce9 | Bacteria | Proteobacteria | Deltaproteobacteria | Bdellovibrionales | Bdellovibrionaceae | OM27 clade | Ambiguous_taxa | 0.994 |

| Feature_ID | Kingdom | Phylum | Class | Order | Family | Genus | Species | Confidence |
|---|---|---|---|---|---|---|---|---|
| 3965e4dde16b4284dc71c970c6ab6b29 | Bacteria | Cyanobacteria | Oxyphotobacteria | Chloroplast | | | | 1.000 |
| 8cba59be471be99c970f382a80c0e73b | Bacteria | Proteobacteria | Alphaproteobacteria | SAR11 clade | Clade III | uncultured marine bacterium | uncultured marine bacterium | 0.949 |
| efd3b8f4a8fb27585152cd819d2c7a1f | Bacteria | Proteobacteria | Alphaproteobacteria | Rickettsiales | S25-593 | | | 1.000 |
| db6a43a54997935c94a0ab38c03cf77c | Bacteria | Verrucomicrobia | Verrucomicrobiae | Verrucomicrobiales | DEV007 | hydrothermal vent metagenome | hydrothermal vent metagenome | 0.956 |
| 9880e34d3fc7ebb5a8ec39440e6f58e1 | Bacteria | Bacteroidetes | Bacteroidia | Flavobacteriales | Cryomorphaceae | NS10 marine group | marine metagenome | 0.812 |
| 181ecd927f4a6a0d11dd2f8a21679701 | Bacteria | Cyanobacteria | Oxyphotobacteria | Chloroplast | marine metagenome | marine metagenome | marine metagenome | 0.980 |
| d19c5daff6e47acadaff7a609bcf900f | Bacteria | Actinobacteria | Acidimicrobiia | Microtrichales | Microtrichaceae | Sva0996 marine group | uncultured bacterium | 0.942 |
| 8381bfa1787dd0d5fbfa1a5108465af9 | Bacteria | Proteobacteria | Alphaproteobacteria | Rhodospirillales | Magnetospiraceae | uncultured | hydrothermal vent metagenome | 0.997 |
| 476a9e90f22d21911ed55e1bf39cb63c | Bacteria | Marinimicrobia (SAR406 clade) | | | | | | 1.000 |
| af0620d9127ad460ea001a57c69c45ee | Bacteria | Bacteroidetes | Bacteroidia | Flavobacteriales | Flavobacteriaceae | NS2b marine group | | 1.000 |
| dd3de45077fc66df0414ae85313bb861 | Bacteria | Proteobacteria | Alphaproteobacteria | Puniceispirillales | SAR116 clade | alpha proteobacterium SCGC AAA015-N04 | alpha proteobacterium SCGC AAA015-N04 | 0.864 |
| 7ba1b164eece4ac4ec3cd37dceb033cc | Bacteria | Proteobacteria | Gammaproteobacteria | OM182 clade | | | | 1.000 |
| 7db426ecf20394ecff2e04290207d90a | Bacteria | Cyanobacteria | Oxyphotobacteria | Synechococcales | Cyanobiaceae | Prochlorococcus MIT9313 | | 1.000 |
| a4e4bd6a602c189491cd53e6d9005e4b | Bacteria | Proteobacteria | Gammaproteobacteria | Betaproteobacteriales | Methylophilaceae | OM43 clade | Ambiguous_taxa | 0.900 |
| 761d583ba5e18e2ca7cbbde95e794211 | Bacteria | Cyanobacteria | Oxyphotobacteria | Chloroplast | Phalacroma mitra | Phalacroma mitra | Phalacroma mitra | 0.826 |
| cb049fad99506ef1618856b1379dd33c | Bacteria | Cyanobacteria | Oxyphotobacteria | Synechococcales | Cyanobiaceae | Prochlorococcus MIT9313 | | 0.974 |
| 4831ee76ffe1eddaddccd612333d560d | Bacteria | Cyanobacteria | Oxyphotobacteria | Chloroplast | Bathycoccus prasinos | Bathycoccus prasinos | Bathycoccus prasinos | 1.000 |
| 1090f885623e3b30f61b79ef7d460499 | Bacteria | Cyanobacteria | Oxyphotobacteria | Synechococcales | Cyanobiaceae | Prochlorococcus MIT9313 | | 0.999 |
| 2a9ae138a4b2a2444170b139bba24a1e | Bacteria | Bacteroidetes | Bacteroidia | Chitinophagales | uncultured | Ambiguous_taxa | Ambiguous_taxa | 1.000 |
| 687cbd7f96e71438cbbe66e5815c0d0f | Bacteria | Proteobacteria | Gammaproteobacteria | Alteromonadales | Pseudoalteromonadaceae | Pseudoalteromonas | | 1.000 |
| e1664283d0598e4bb8cb4ae585c64682 | Bacteria | Marinimicrobia (SAR406 clade) | uncultured Fibrobacteres/Acidobacteria group bacterium | uncultured Fibrobacteres/Acidobacteria group bacterium | uncultured Fibrobacteres/Acidobacteria group bacterium | uncultured Fibrobacteres/Acidobacteria group bacterium | uncultured Fibrobacteres/Acidobacteria group bacterium | 0.926 |
| 5ed4b360e428d248c2768acd9a1d24c0 | Archaea | Euryarchaeota | Thermoplasmata | Marine Group II | | | | 1.000 |
| 52d0f2934d086aa298ed0029c5406289 | Bacteria | Bacteroidetes | Bacteroidia | Flavobacteriales | NS7 marine group | | | 1.000 |
| 26e2f2042251004de245a74911c75056 | Bacteria | Proteobacteria | Alphaproteobacteria | SAR11 clade | Clade I | | | 1.000 |
| ef82db7a784194691c8b8fecc41b156a | Bacteria | Bacteroidetes | Bacteroidia | Flavobacteriales | Cryomorphaceae | uncultured | | 1.000 |
| 95d0c087a9ba84341ed21e80b3277f4c | Bacteria | Proteobacteria | Gammaproteobacteria | SAR86 clade | uncultured gamma proteobacterium KTc1121 | uncultured gamma proteobacterium KTc1121 | uncultured gamma proteobacterium KTc1121 | 0.728 |
| 17378062be25460666bac995a3d7422a | Bacteria | Proteobacteria | Deltaproteobacteria | Bdellovibrionales | Bdellovibrionaceae | OM27 clade | uncultured marine bacterium | 0.989 |
| 05e651042e4aed4750673b6bc095e742 | Bacteria | Proteobacteria | Gammaproteobacteria | SAR86 clade | uncultured gamma proteobacterium KTc1121 | uncultured gamma proteobacterium KTc1121 | uncultured gamma proteobacterium KTc1121 | 0.795 |
| f837b692fe04e44a18ff1f6beca3a304 | Bacteria | Proteobacteria | Gammaproteobacteria | OM182 clade | uncultured gamma proteobacterium | uncultured gamma proteobacterium | uncultured gamma proteobacterium | 1.000 |
| e773509c44bfc8d4dc539473d6d207ef | Bacteria | Proteobacteria | Gammaproteobacteria | SAR86 clade | uncultured SAR86 cluster bacterium | uncultured SAR86 cluster bacterium | uncultured SAR86 cluster bacterium | 0.812 |
| 454a402f1fe882e9331f60eb969990ff | Bacteria | Proteobacteria | Gammaproteobacteria | Ectothiorhodospirales | Ectothiorhodospiraceae | uncultured | uncultured marine bacterium | 0.983 |
| 3e9b7bc1b40e8d02d12d76820c7dbd27 | Bacteria | Proteobacteria | Alphaproteobacteria | Puniceispirillales | SAR116 clade | marine metagenome | marine metagenome | 0.994 |
| eb7d813dc7982fd1efb8d1b52b420ac5 | Bacteria | Proteobacteria | Gammaproteobacteria | Alteromonadales | Pseudoalteromonadaceae | Pseudoalteromonas | | 0.998 |
| d126165e4c921e1da5f8580ef4b88c60 | Bacteria | Bacteroidetes | Bacteroidia | Flavobacteriales | NS9 marine group | uncultured marine bacterium | uncultured marine bacterium | 0.994 |
| 5fc0c1dab4b1a256250b7e460dbcb14e | Bacteria | Proteobacteria | Deltaproteobacteria | PB19 | uncultured marine microorganism | uncultured marine microorganism | uncultured marine microorganism | 0.974 |
| 7b741115d3b2b52b5f3428f399658f2b | Bacteria | Proteobacteria | Gammaproteobacteria | Betaproteobacteriales | Burkholderiaceae | MWH-UniP1 aquatic group | uncultured bacterium | 0.980 |
| bcd22c817f400c5a6bba8fdf1a77ca09 | Bacteria | Proteobacteria | Alphaproteobacteria | Rickettsiales | S25-593 | uncultured marine bacterium | uncultured marine bacterium | 0.889 |
| 7549e57eaad565c09cfab6cd0a99bd72 | Bacteria | Proteobacteria | Deltaproteobacteria | Bdellovibrionales | Bdellovibrionaceae | OM27 clade | uncultured marine microorganism | 1.000 |
| 4247391984c4f10ae4e4f01d1dac9da7 | Bacteria | Bacteroidetes | Bacteroidia | Flavobacteriales | NS9 marine group | uncultured marine bacterium | uncultured marine bacterium | 0.995 |
| 5a7e96a9e3ad4156b4df47bf6b3ecf99 | Bacteria | Proteobacteria | Alphaproteobacteria | Rhodobacterales | Rhodobacteraceae | | | 1.000 |
| 12e271fae4693221effe71b5607f6727 | Bacteria | Proteobacteria | Gammaproteobacteria | OM182 clade | uncultured gamma proteobacterium | uncultured gamma proteobacterium | uncultured gamma proteobacterium | 0.999 |
| 1fb9f17bde5c2c5c910a3ca6f8396270 | Bacteria | Cyanobacteria | Oxyphotobacteria | Chloroplast | Ostreococcus tauri | Ostreococcus tauri | Ostreococcus tauri | 0.929 |
| 3a91733367e0cd9c6ec2d6da668886ac | Bacteria | Proteobacteria | Gammaproteobacteria | Cellvibrionales | Halieaceae | OM60(NOR5) clade | Ambiguous_taxa | 0.724 |
| 6ce9a49ca2fbb47045727f39fcf2133c | Bacteria | Proteobacteria | Gammaproteobacteria | Nitrosococcales | Methylophagaceae | Methylophaga | | 1.000 |
| b2c68127cdc77dd97b46ebfd4e431f23 | Bacteria | Cyanobacteria | Oxyphotobacteria | Chloroplast | | | | 1.000 |
| 1557d232fa3f9a321dc2f9e6a09d44e3 | Bacteria | Bacteroidetes | Bacteroidia | Flavobacteriales | NS9 marine group | uncultured marine bacterium | uncultured marine bacterium | 0.991 |
| 3a1ebbcfec4cef5125e161a21dc6c2e5 | Bacteria | Bacteroidetes | Bacteroidia | Flavobacteriales | Crocinitomicaceae | Fluviicola | | 1.000 |
| 27593cb90ba89d4d5ce558f5480d532b | Bacteria | Cyanobacteria | Oxyphotobacteria | Synechococcales | Cyanobiaceae | Synechococcus CC9902 | | 0.853 |
| a1a8af62eb4a2bfd344fddf895e99ebc | Bacteria | Proteobacteria | Gammaproteobacteria | Oceanospirillales | Pseudohongiellaceae | Pseudohongiella | | 1.000 |
| 5119344990960b88d08ffb94476a773d | Bacteria | Bacteroidetes | Bacteroidia | Flavobacteriales | Flavobacteriaceae | NS4 marine group | | 1.000 |
| 00959df1876e5f1060cd36231d6dba28 | Bacteria | Proteobacteria | Alphaproteobacteria | SAR11 clade | Clade I | uncultured | | 0.799 |
| 9b9c1b1fb9e71df47db6b098db3c3a8f | Bacteria | Proteobacteria | Gammaproteobacteria | Cellvibrionales | Halieaceae | OM60(NOR5) clade | Ambiguous_taxa | 0.787 |
| 2f6655c5d2d73ff441db8af0caa31a1c | Bacteria | Cyanobacteria | Oxyphotobacteria | Chloroplast | marine metagenome | marine metagenome | marine metagenome | 0.986 |
| dc18af6b42a55969fbd79055ab19413f | Bacteria | Cyanobacteria | Oxyphotobacteria | Chloroplast | | | | 1.000 |
| 94b18f0c88b7f338ca85b8229d9331b2 | Bacteria | Proteobacteria | Gammaproteobacteria | HOC36 | Candidatus Thioglobus sp. REDSEA-S12_B1 | Candidatus Thioglobus sp. REDSEA-S12_B1 | Candidatus Thioglobus sp. REDSEA-S12_B1 | 1.000 |
| 788e4ec70767db30d51973b83124fdc4 | Bacteria | Proteobacteria | Gammaproteobacteria | Ectothiorhodospirales | Ectothiorhodospiraceae | uncultured | uncultured marine bacterium | 0.930 |
| d225904882df61c5e7f13c370df2b59a | Bacteria | Bacteroidetes | Bacteroidia | Flavobacteriales | NS9 marine group | | | 1.000 |
| 98c815507ac78088ded3997fad6945c2 | Bacteria | Bacteroidetes | Bacteroidia | Chitinophagales | Saprospiraceae | uncultured | unidentified marine eubacterium | 1.000 |
| be9d6040f58aa6bf2e34f435083e68ce | Bacteria | Proteobacteria | Gammaproteobacteria | Cellvibrionales | Halieaceae | OM60(NOR5) clade | Ambiguous_taxa | 0.952 |
| 8f7715fdc5d711ffc6b1d0835995ed06 | Bacteria | Proteobacteria | Gammaproteobacteria | OM182 clade | | | | 1.000 |
| 6d78d68afe425ef7969b917bc6fff9da | Bacteria | Cyanobacteria | Oxyphotobacteria | Chloroplast | Phalacroma mitra | Phalacroma mitra | Phalacroma mitra | 0.744 |
| e98a2eeb39347ef4470121821d56762c | Bacteria | Proteobacteria | Gammaproteobacteria | Betaproteobacteriales | Methylophilaceae | OM43 clade | Ambiguous_taxa | 0.962 |
| dc13bae2b5ebe13bb53f537d385abd34 | Bacteria | Proteobacteria | Deltaproteobacteria | Bdellovibrionales | Bdellovibrionaceae | OM27 clade | | 1.000 |
| 1fff2c958e5d34bad7dd270d6a95a2c1 | Bacteria | Cyanobacteria | Oxyphotobacteria | Chloroplast | | | | 1.000 |
| 703e9daecc082479f6778bd1086d2567 | Bacteria | Cyanobacteria | Oxyphotobacteria | Chloroplast | unidentified marine bacterioplankton | unidentified marine bacterioplankton | unidentified marine bacterioplankton | 0.978 |
| 2100177b2369bd065753dcf742fefb7c | Bacteria | Bacteroidetes | Bacteroidia | Flavobacteriales | NS9 marine group | uncultured Flavobacteriia bacterium | uncultured Flavobacteriia bacterium | 0.989 |
| b947f336bfc0d1577fbdab76c84dd5e6 | Bacteria | Bacteroidetes | Bacteroidia | Flavobacteriales | NS9 marine group | uncultured marine bacterium | uncultured marine bacterium | 0.936 |
| 47cdd08d5cb118b8c5e02018326917ad | Bacteria | Bacteroidetes | Bacteroidia | Chitinophagales | Saprospiraceae | uncultured | Ambiguous_taxa | 0.994 |
| b9bcda3d6e2486b03ac6d8738eda35a7 | Bacteria | Proteobacteria | Gammaproteobacteria | UBA10353 marine group | Ambiguous_taxa | Ambiguous_taxa | Ambiguous_taxa | 0.932 |
| b4881312c25a93bd996f6e4f5c83af72 | Bacteria | Bacteroidetes | Bacteroidia | Flavobacteriales | Cryomorphaceae | NS10 marine group | uncultured marine bacterium | 0.983 |
| 1d447bcacab9dccafb0042146672bc29 | Bacteria | Cyanobacteria | Oxyphotobacteria | Chloroplast | | | | 1.000 |
| 30c84d5d82cdae04d8aa3d5635cd1668 | Bacteria | Proteobacteria | Alphaproteobacteria | Rhodospirillales | Magnetospiraceae | uncultured | | 1.000 |
| 1d4406231f90416ad02f4a90255fe1b7 | Bacteria | Proteobacteria | Gammaproteobacteria | Thiohalorhabdales | Thiohalorhabdaceae | uncultured | Ambiguous_taxa | 0.985 |

| Feature_ID | Kingdom | Phylum | Class | Order | Family | Genus | Species | Confidence |
|---|---|---|---|---|---|---|---|---|
| 11aa54de56bbdb5f712df193d8c1f387 | Bacteria | Bacteroidetes | Bacteroidia | Flavobacteriales | NS9 marine group | uncultured bacterium | uncultured bacterium | 0.977 |
| 872fca46f3da5bc5a13812cbe8c90916 | Bacteria | Marinimicrobia (SAR406 clade) | uncultured bacterium | uncultured bacterium | uncultured bacterium | uncultured bacterium | uncultured bacterium | 0.966 |
| 2ac7e83bad1a6532bdffb1ee46bb972c | Bacteria | Bacteroidetes | Bacteroidia | Flavobacteriales | Flavobacteriaceae | NS5 marine group | | 1.000 |
| 43c9cfaeaa171d99ef4c1b7903ed7b4e | Bacteria | Cyanobacteria | Oxyphotobacteria | Chloroplast | | | | 1.000 |
| f1d1e5a2e3ee2e178d1b0505eefe2bb8 | Bacteria | Proteobacteria | Gammaproteobacteria | SAR86 clade | | | | 1.000 |
| 21584f179042f46294a407e9fa2a5c33 | Bacteria | Proteobacteria | Gammaproteobacteria | SAR86 clade | | | | 1.000 |
| 0dde1473306732b68bf2f5850473db22 | Bacteria | Cyanobacteria | Oxyphotobacteria | Chloroplast | uncultured Cryptomonadaceae | uncultured Cryptomonadaceae | uncultured Cryptomonadaceae | 0.999 |
| 735be70943879eb8c58ae2dfe9cdc2be | Bacteria | Cyanobacteria | Oxyphotobacteria | Chloroplast | | | | 1.000 |
| a6028017c7a5ac7620288c31299c1d3f | Bacteria | Proteobacteria | Gammaproteobacteria | SAR86 clade | uncultured proteobacterium OCS44 | uncultured proteobacterium OCS44 | uncultured proteobacterium OCS44 | 0.758 |
| 386e761d37a480eec0fefff5ad237238 | Bacteria | Bacteroidetes | Bacteroidia | Flavobacteriales | Flavobacteriaceae | NS2b marine group | | 1.000 |
| 55f9ddd0a55d21d36855eb0ac350eb1e | Bacteria | Proteobacteria | Gammaproteobacteria | KI89A clade | uncultured marine bacterium | uncultured marine bacterium | uncultured marine bacterium | 0.918 |
| afeb7677ee9a9c58a829c478d976c684 | Bacteria | Proteobacteria | Deltaproteobacteria | PB19 | uncultured marine bacterium | uncultured marine bacterium | uncultured marine bacterium | 0.922 |
| cf79c866027f0504bafd7b17f368c680 | Bacteria | Proteobacteria | Deltaproteobacteria | Bdellovibrionales | Bdellovibrionaceae | OM27 clade | uncultured marine microorganism | 0.991 |
| e8a2a6b71377024d827e801d0a96f59f | Bacteria | Cyanobacteria | Oxyphotobacteria | Chloroplast | Phalacroma mitra | Phalacroma mitra | Phalacroma mitra | 0.895 |
| adbc904a96d3f7374951e78838a78e73 | Bacteria | Bacteroidetes | Bacteroidia | Flavobacteriales | Flavobacteriaceae | NS5 marine group | | 1.000 |
| 3be56f97905eb61efaed31bb61b2b27b | Bacteria | Cyanobacteria | Oxyphotobacteria | Chloroplast | | | | 1.000 |
| 8cc0eafcb297ffc4102c34ed82dda261 | Bacteria | Proteobacteria | Gammaproteobacteria | Coxiellales | Coxiellaceae | Coxiella | | 1.000 |
| 997910b2ca10daca8e44fc35c048cf16 | Bacteria | WPS-2 | Ambiguous_taxa | Ambiguous_taxa | Ambiguous_taxa | Ambiguous_taxa | Ambiguous_taxa | 1.000 |
| c10d11b0f648e102467303bf277aee80 | Bacteria | Cyanobacteria | Oxyphotobacteria | Chloroplast | marine metagenome | marine metagenome | marine metagenome | 0.985 |
| 653732bd7b346fa2cd5e890185be2df4 | Bacteria | Proteobacteria | Gammaproteobacteria | Ectothiorhodospirales | Ectothiorhodospiraceae | uncultured | uncultured marine bacterium | 0.954 |
| af935abb3901414fa7d5f0b7f94524d9 | Bacteria | Marinimicrobia (SAR406 clade) | | | | | | 1.000 |
| 1fea099b7995754082857e9f09a18bcc | Bacteria | Bacteroidetes | Bacteroidia | Flavobacteriales | Flavobacteriaceae | NS4 marine group | | 1.000 |
| 2fc3478a790ed02f9b51bd62e7d085a8 | Bacteria | Proteobacteria | Gammaproteobacteria | UBA10353 marine group | | | | 1.000 |
| d16445c268193aeea8e232f32a0958a1 | Bacteria | Proteobacteria | Alphaproteobacteria | SAR11 clade | Clade II | | | 0.910 |
| d0465cdb3fd1e2f7053e8403b58f7080 | Bacteria | Marinimicrobia (SAR406 clade) | uncultured Fibrobacteres/Acidobacteria group bacterium | uncultured Fibrobacteres/Acidobacteria group bacterium | uncultured Fibrobacteres/Acidobacteria group bacterium | uncultured Fibrobacteres/Acidobacteria group bacterium | uncultured Fibrobacteres/Acidobacteria group bacterium | 1.000 |
| ebb62d3089cc18ec91cdcc27f31946ee | Bacteria | Bacteroidetes | Bacteroidia | Flavobacteriales | Crocinitomicaceae | Fluviicola | | 1.000 |
| 7bb1a4b17607ac5694f3810866ed6171 | Bacteria | Cyanobacteria | Oxyphotobacteria | Chloroplast | uncultured diatom | uncultured diatom | uncultured diatom | 0.878 |
| aaa9c7cf85d077f62c94e73bf478fb70 | Bacteria | Bacteroidetes | Bacteroidia | Flavobacteriales | NS9 marine group | | | 1.000 |
| 8cb7619b459585e05071471fa0c7cc08 | Bacteria | Bacteroidetes | Bacteroidia | Flavobacteriales | NS9 marine group | hydrothermal vent metagenome | hydrothermal vent metagenome | 1.000 |
| 3d038b0c2c0fb1082e65589eedffaf89 | Bacteria | Bacteroidetes | Bacteroidia | Flavobacteriales | Flavobacteriaceae | NS5 marine group | | 1.000 |
| b4789b7fd53aff4db2da7e2bfdcab2f3 | Bacteria | Proteobacteria | Deltaproteobacteria | SAR324 clade(Marine group B) | | | | 1.000 |
| acf57007ceb65cafb1878d270ade8a07 | Bacteria | Proteobacteria | Deltaproteobacteria | Bdellovibrionales | Bdellovibrionaceae | OM27 clade | unidentified marine bacterioplankton | 0.999 |
| 734013662141e128c2e80d600f2c0c34 | Bacteria | Proteobacteria | Alphaproteobacteria | Rhodobacterales | Rhodobacteraceae | | | 1.000 |
| 8617f557db300c492dbc1f7c63ab890c | Bacteria | Proteobacteria | Alphaproteobacteria | SAR11 clade | Clade II | | | 0.923 |
| 59095dbe42d8898c1ba3c36332f92413 | Bacteria | Proteobacteria | Alphaproteobacteria | Rickettsiales | S25-593 | | | 1.000 |
| 1744e6c70f0956cd14bb9d51502723b7 | Bacteria | Bacteroidetes | Bacteroidia | Flavobacteriales | NS9 marine group | Ambiguous_taxa | Ambiguous_taxa | 0.763 |
| baf96a8b9cdd13c5e845ef9301953ee0 | Bacteria | Bacteroidetes | Bacteroidia | Flavobacteriales | NS9 marine group | uncultured marine bacterium | uncultured marine bacterium | 0.997 |
| bebbae5872888d156411954d22113fe7 | Bacteria | Cyanobacteria | Oxyphotobacteria | Synechococcales | Cyanobiaceae | Prochlorococcus MIT9313 | Ambiguous_taxa | 0.744 |
| 3f3eda52b690d79809a9a3dc82c3b4f4 | Bacteria | Bacteroidetes | Bacteroidia | Flavobacteriales | Flavobacteriaceae | NS4 marine group | | 1.000 |
| 618875f933901ddf7bb4861ef80152fd | Bacteria | Proteobacteria | Alphaproteobacteria | SAR11 clade | Clade I | Clade Ib | | 0.761 |
| e8706f8a8396b8e188f0df8ddb93f578 | Bacteria | Proteobacteria | Deltaproteobacteria | Oligoflexales | Oligoflexaceae | uncultured | uncultured bacterium | 1.000 |
| 6aa1fe4a28c722eb339e995c65cc830e | Bacteria | Marinimicrobia (SAR406 clade) | marine metagenome | marine metagenome | marine metagenome | marine metagenome | marine metagenome | 0.890 |
| 6a05b15945eec7af4ca9f50e502fd09d | Bacteria | Proteobacteria | Gammaproteobacteria | Vibrionales | Vibrionaceae | Photobacterium | | 0.999 |
| 42e5e5b596055786676327f0f3d44c92 | Bacteria | Chloroflexi | Dehalococcoidia | SAR202 clade | uncultured bacterium | uncultured bacterium | uncultured bacterium | 0.726 |
| 578c0f91a1b2aa15042b6debfd5c9622 | Bacteria | Cyanobacteria | Oxyphotobacteria | Chloroplast | Epipyxis sp. PR26KG | Epipyxis sp. PR26KG | Epipyxis sp. PR26KG | 0.901 |
| 92d82058dd082469df0891638930c0c9 | Bacteria | Bacteroidetes | Bacteroidia | Flavobacteriales | NS7 marine group | Ambiguous_taxa | Ambiguous_taxa | 0.939 |
| 4f6a759f559a9368ed02ffba9b3f015a | Bacteria | Bacteroidetes | Bacteroidia | Flavobacteriales | Flavobacteriaceae | Formosa | uncultured marine bacterium | 0.925 |
| 435a4b0cafdb283687f497e8e8b81535 | Bacteria | Actinobacteria | Acidimicrobiia | Actinomarinales | Actinomarinaceae | Candidatus Actinomarina | uncultured marine bacterium | 0.833 |
| 27ff93acae11247e39f1723121c50d1d | Bacteria | Bacteroidetes | Bacteroidia | Flavobacteriales | Cryomorphaceae | uncultured | uncultured Owenweeksia sp. | 0.955 |
| 066eac4266e76faaaa82fa09ce7634e3 | Bacteria | Bacteroidetes | Bacteroidia | Flavobacteriales | Flavobacteriaceae | NS2b marine group | uncultured marine bacterium | 0.786 |
| 51a13138db3fcff88c2826b4bd8069aa | Bacteria | Bacteroidetes | Bacteroidia | Flavobacteriales | NS9 marine group | uncultured Flavobacteriia bacterium | uncultured Flavobacteriia bacterium | 0.973 |
| ee70504f852ac061e0a62b77f3140f2a | Bacteria | Dadabacteria | Dadabacteriia | Dadabacteriales | uncultured marine bacterium | uncultured marine bacterium | uncultured marine bacterium | 0.995 |
| fa74446f4295dac546c352f6b8bb005c | Bacteria | Proteobacteria | Alphaproteobacteria | Puniceispirillales | SAR116 clade | Candidatus Puniceispirillum | | 0.807 |
| 09bbdda11f384bb7adc2d5abc17ac7be | Bacteria | Bacteroidetes | Bacteroidia | Flavobacteriales | Flavobacteriaceae | NS4 marine group | | 1.000 |
| 4ee7d437fd17c1e71a45402a7703ce4e | Bacteria | Bacteroidetes | Bacteroidia | Flavobacteriales | NS9 marine group | uncultured marine bacterium | uncultured marine bacterium | 0.976 |
| 6e367f752ccff4c9725ffc30b268c417 | Bacteria | Chloroflexi | Dehalococcoidia | SAR202 clade | marine metagenome | marine metagenome | marine metagenome | 1.000 |
| 3616465ab65721d774e4dfeb74ceea09 | Bacteria | Proteobacteria | Alphaproteobacteria | Puniceispirillales | SAR116 clade | Candidatus Puniceispirillum | uncultured bacterium | 0.987 |
| 05ffc4cd84c947a6d4aea5535afaf70c | Archaea | Euryarchaeota | Thermoplasmata | Marine Group II | marine metagenome | marine metagenome | marine metagenome | 0.849 |
| b263c550540161d6282b050e663c7e91 | Bacteria | Bacteroidetes | Bacteroidia | Flavobacteriales | Flavobacteriaceae | | | 1.000 |
| 166b894a06a4408204d1737ec7057877 | Bacteria | Cyanobacteria | Oxyphotobacteria | Chloroplast | | | | 1.000 |
| f708204d5c243dfd81951f43b0f85af2 | Bacteria | Bacteroidetes | Bacteroidia | Flavobacteriales | NS9 marine group | uncultured marine bacterium | uncultured marine bacterium | 0.996 |
| afdfbebea593f028408a4171fbb64347 | Bacteria | Cyanobacteria | Oxyphotobacteria | Chloroplast | uncultured bacterium ARCTIC13_E_12 | uncultured bacterium ARCTIC13_E_12 | uncultured bacterium ARCTIC13_E_12 | 0.795 |
| 0a2463f6dd1229442ec2aea45a54b5b9 | Bacteria | Bacteroidetes | Bacteroidia | Flavobacteriales | Flavobacteriaceae | NS2b marine group | uncultured marine bacterium | 0.821 |
| cf915c661e4a061e2793fc8ed1430399 | Bacteria | Proteobacteria | Alphaproteobacteria | SAR11 clade | Clade II | | | 0.714 |
| a06a4e8db2076fff72fcbca835066262 | Bacteria | Proteobacteria | Gammaproteobacteria | KI89A clade | uncultured marine bacterium | uncultured marine bacterium | uncultured marine bacterium | 0.873 |
| 0c84384e6722111c020aac8b7fbda142 | Bacteria | Bacteroidetes | Bacteroidia | Flavobacteriales | NS9 marine group | uncultured Flavobacteriia bacterium | uncultured Flavobacteriia bacterium | 0.987 |
| d3893ebcfc1920ccd1bc3d9d98b83e2c | Bacteria | Bacteroidetes | Bacteroidia | Flavobacteriales | Crocinitomicaceae | Fluviicola | | 1.000 |
| c33858cc31aaa71b939e48b2890b39fa | Bacteria | Proteobacteria | Gammaproteobacteria | Cellvibrionales | Halieaceae | OM60(NOR5) clade | Ambiguous_taxa | 0.925 |
| acac251921d52037b96e1597edc68c88 | Bacteria | Bacteroidetes | Bacteroidia | Flavobacteriales | NS9 marine group | uncultured Flavobacteriia bacterium | uncultured Flavobacteriia bacterium | 0.896 |
| d444fec272ce5980c0adcca448ee5d86 | Bacteria | Marinimicrobia (SAR406 clade) | Alteromonas macleodii | Alteromonas macleodii | Alteromonas macleodii | Alteromonas macleodii | Alteromonas macleodii | 0.976 |
| a69a75d5ee08d78bd8c71bcf504e0595 | Bacteria | Proteobacteria | Gammaproteobacteria | SAR86 clade | uncultured bacterium | uncultured bacterium | uncultured bacterium | 0.741 |

| Feature_ID | Kingdom | Phylum | Class | Order | Family | Genus | Species | Confidence |
|---|---|---|---|---|---|---|---|---|
| df64d0b5d48450d6fb060b22e12a6d9e | Bacteria | Proteobacteria | Alphaproteobacteria | Puniceispirillales | SAR116 clade | Candidatus Puniceispirillum | uncultured bacterium | 0.971 |
| 109ca30c2be2933970f662b78cb8bd5f | Bacteria | Proteobacteria | Alphaproteobacteria | Puniceispirillales | SAR116 clade | | | 1.000 |
| 60d6dd425dfa0a6d45761b918a36b771 | Bacteria | Cyanobacteria | Oxyphotobacteria | Chloroplast | marine metagenome | marine metagenome | marine metagenome | 0.911 |
| 83c24c6a57efff56a44f89c4dc028029 | Bacteria | Marinimicrobia (SAR406 clade) | Marinimicrobia bacterium SCGC AAA003-L08 | Marinimicrobia bacterium SCGC AAA003-L08 | Marinimicrobia bacterium SCGC AAA003-L08 | Marinimicrobia bacterium SCGC AAA003-L08 | Marinimicrobia bacterium SCGC AAA003-L08 | 1.000 |
| e9d803cb50b88b4bf18ccbe7d2b3d28b | Bacteria | Proteobacteria | Alphaproteobacteria | SAR11 clade | Clade I | Clade Ia | | 0.962 |
| edea52bf051cf317ecea1df86a0a361f | Bacteria | Proteobacteria | Deltaproteobacteria | Bdellovibrionales | Bdellovibrionaceae | OM27 clade | Ambiguous_taxa | 0.993 |
| 490b60bbce7028dfa0dc5fcc5c861c0b | Bacteria | Proteobacteria | Gammaproteobacteria | SAR86 clade | | | | 1.000 |
| bfa4ab900ba91b96587d09efcb9e690e | Bacteria | Proteobacteria | Deltaproteobacteria | Bdellovibrionales | Bacteriovoracaceae | uncultured | marine metagenome | 0.946 |
| 51dc000b86ea1bf3ed2c68e314aceef1 | Bacteria | Cyanobacteria | Oxyphotobacteria | Chloroplast | | | | 1.000 |
| 764f5a270afa9cec1e520c872eb53b46 | Bacteria | Cyanobacteria | Oxyphotobacteria | Synechococcales | Cyanobiaceae | Synechococcus CC9902 | | 0.995 |
| e8e59dc192aa41449f1733e846c9cedb | Bacteria | Proteobacteria | Alphaproteobacteria | Rhodospirillales | AEGEAN-169 marine group | uncultured alpha proteobacterium | uncultured alpha proteobacterium | 0.744 |
| 438927eaf1aa85db349f08730025030d | Bacteria | Proteobacteria | Gammaproteobacteria | KI89A clade | Ambiguous_taxa | Ambiguous_taxa | Ambiguous_taxa | 0.701 |
| 4779e8ffc760750e779f185f7dc018c7 | Bacteria | Proteobacteria | Gammaproteobacteria | Cellvibrionales | Spongiibacteraceae | BD1-7 clade | Ambiguous_taxa | 0.936 |
| 67b03cc55c2d03d5238253fc74ae69ae | Bacteria | Bacteroidetes | Bacteroidia | Flavobacteriales | Crocinitomicaceae | Fluviicola | | 1.000 |
| 707cea640b10f610ee5e226ee5cde8ae | Bacteria | Bacteroidetes | Bacteroidia | Flavobacteriales | Flavobacteriaceae | Formosa | | 0.995 |
| b4c3ec664912855ce7f90bec277919cc | Bacteria | Planctomycetes | Planctomycetacia | Pirellulales | Pirellulaceae | Rubripirellula | | 1.000 |
| 6c892d490d7a40ffa2a60bc99b72dc20 | Bacteria | Bacteroidetes | Bacteroidia | Flavobacteriales | NS9 marine group | uncultured Flavobacteriia bacterium | uncultured Flavobacteriia bacterium | 0.912 |
| 9e54883092ada57a33865076a45b2525 | Bacteria | Bacteroidetes | Bacteroidia | Flavobacteriales | NS9 marine group | uncultured marine bacterium | uncultured marine bacterium | 0.952 |
| 1a98ab03c1fd435eb11d6937d35d9f48 | Bacteria | Cyanobacteria | Oxyphotobacteria | Chloroplast | | | | 1.000 |
| e2a8fe5c36b4b352191f25b265be144b | Bacteria | Proteobacteria | Gammaproteobacteria | KI89A clade | | | | 1.000 |
| 25ace10ce6d4d75a0fe11ebcf33f91c3 | Bacteria | Proteobacteria | Gammaproteobacteria | HOC36 | Candidatus Thioglobus sp. REDSEA-S12_B1 | Candidatus Thioglobus sp. REDSEA-S12_B1 | Candidatus Thioglobus sp. REDSEA-S12_B1 | 0.999 |
| 6dd86130168bcc3876f2aabb906a5934 | Bacteria | Cyanobacteria | Oxyphotobacteria | Chloroplast | Prasinoderma coloniale | Prasinoderma coloniale | Prasinoderma coloniale | 1.000 |
| d295eae8eeebd9125ca1d1787cfb2066 | Bacteria | Proteobacteria | Gammaproteobacteria | Vibrionales | Vibrionaceae | Vibrio | | 0.999 |
| 6a9308a3d4b6be920b44f03b3f80f832 | Bacteria | Proteobacteria | Gammaproteobacteria | Thiomicrospirales | Thioglobaceae | SUP05 cluster | | 0.998 |
| f0fec588b2b57fa6da085e782bfb80e3 | Bacteria | Proteobacteria | Deltaproteobacteria | SAR324 clade(Marine group B) | | | | 1.000 |
| e08b5550e729154028140570e43a147c | Bacteria | Proteobacteria | Alphaproteobacteria | SAR11 clade | Clade II | | | 0.920 |
| e8980bb8dc27afa9264051725c463489 | Bacteria | Verrucomicrobia | Verrucomicrobiae | Opitutales | Puniceicoccaceae | Coraliomargarita | Ambiguous_taxa | 0.922 |
| 1bc1d3897b046fc022d03617c4935f4c | Bacteria | Verrucomicrobia | Verrucomicrobiae | Opitutales | Puniceicoccaceae | MB11C04 marine group | | 1.000 |
| b0e3b77912ab2763fd39818553d851fa | Bacteria | WPS-2 | Ambiguous_taxa | Ambiguous_taxa | Ambiguous_taxa | Ambiguous_taxa | Ambiguous_taxa | 1.000 |
| 1950553995cfd481756d5466a2286744 | Bacteria | Proteobacteria | Gammaproteobacteria | Cellvibrionales | Halieaceae | OM60(NOR5) clade | Ambiguous_taxa | 0.751 |
| 741fcf923ccd4e5d89e9ca2670eb4ea7 | Bacteria | Proteobacteria | Alphaproteobacteria | SAR11 clade | Clade II | | | 0.986 |
| 80cd4f2b322ab6bfee0b1b37928c0906 | Bacteria | Bacteroidetes | Bacteroidia | Flavobacteriales | NS9 marine group | Ambiguous_taxa | Ambiguous_taxa | 0.959 |
| 2610fd16edf6db0e82333aec225a71dc | Bacteria | Proteobacteria | Gammaproteobacteria | Vibrionales | Vibrionaceae | uncultured | Ambiguous_taxa | 0.899 |
| 95ee4bda57ccbfef6aeb1e85401a91d7 | Bacteria | Marinimicrobia (SAR406 clade) | Ambiguous_taxa | Ambiguous_taxa | Ambiguous_taxa | Ambiguous_taxa | Ambiguous_taxa | 0.733 |
| 583af40a1f36b9a80c32636005c1272e | Bacteria | Proteobacteria | Deltaproteobacteria | Bdellovibrionales | Bdellovibrionaceae | OM27 clade | uncultured marine microorganism | 1.000 |
| 953bf575ad5f467cc5d188c920f7edc1 | Bacteria | Proteobacteria | Gammaproteobacteria | Ectothiorhodospirales | Ectothiorhodospiraceae | uncultured | uncultured marine bacterium | 0.966 |
| b5d4643929e9855e7aed51249c3f3ff5 | Bacteria | Bacteroidetes | Bacteroidia | Flavobacteriales | Flavobacteriaceae | | | 1.000 |
| 1160896bd5e218ed85652f62084c1cae | Bacteria | Proteobacteria | Gammaproteobacteria | Alteromonadales | Marinobacteraceae | Marinobacter | | 1.000 |
| 7333eff94312512fc6598937e48adad4 | Bacteria | Proteobacteria | Gammaproteobacteria | SAR86 clade | uncultured gamma proteobacterium HF0070_25G02 | uncultured gamma proteobacterium HF0070_25G02 | uncultured gamma proteobacterium HF0070_25G02 | 1.000 |
| acc650be2c9c7ef74ee782f4cec342ea | Bacteria | Proteobacteria | Gammaproteobacteria | Cellvibrionales | Halieaceae | Parahaliea | uncultured marine bacterium | 0.999 |
| 4ff740aace27a1d0c81ecbec439acab6 | Bacteria | Marinimicrobia (SAR406 clade) | | | | | | 1.000 |
| 696635c4819a7f013309c252229954a7 | Bacteria | Cyanobacteria | Oxyphotobacteria | Chloroplast | Chaetoceros affinis | Chaetoceros affinis | Chaetoceros affinis | 0.751 |
| 0545daa49207838ca7a4b88fe149359f | Bacteria | Proteobacteria | Alphaproteobacteria | Rhodospirillales | AEGEAN-169 marine group | Ambiguous_taxa | Ambiguous_taxa | 0.998 |
| dd26b396d97a6ea2a87c8f094a0bbdb2 | Bacteria | Verrucomicrobia | Verrucomicrobiae | Opitutales | Puniceicoccaceae | MB11C04 marine group | | 1.000 |
| 84b495a19bc220e89f24a1050bdb2811 | Bacteria | Proteobacteria | Gammaproteobacteria | | | | | 1.000 |
| f8f42cf8dc3d3201a1f2941391b1e5c5 | Bacteria | Proteobacteria | Alphaproteobacteria | Rhodovibrionales | Kiloniellaceae | uncultured | uncultured beta proteobacterium | 0.995 |
| f42d180a8f895bee1b150c680ec39441 | Archaea | Euryarchaeota | Thermoplasmata | Marine Group II | unidentified archaeon | unidentified archaeon | unidentified archaeon | 0.989 |
| 1c6b4a698fddfd978f7a170d5b7fc44a | Bacteria | Marinimicrobia (SAR406 clade) | | | | | | 1.000 |
| ab6d977125509c30e9517cdf91d20fec | Bacteria | Bacteroidetes | Bacteroidia | Flavobacteriales | NS9 marine group | uncultured Flavobacteriales bacterium | uncultured Flavobacteriales bacterium | 0.809 |
| b193c299f6f39e5a45cf0c400d591aa4 | Bacteria | Bacteroidetes | Bacteroidia | Flavobacteriales | NS9 marine group | uncultured marine bacterium | uncultured marine bacterium | 0.981 |
| 8b96d2c683aacd1638d5e162d286dc22 | Bacteria | Planctomycetes | OM190 | Ambiguous_taxa | Ambiguous_taxa | Ambiguous_taxa | Ambiguous_taxa | 0.920 |
| c007cbfea90d1d3ce82a604bd899b8f8 | Bacteria | Bacteroidetes | Bacteroidia | Flavobacteriales | Cryomorphaceae | NS10 marine group | uncultured Bacteroidetes/Chlorobi group bacterium | 0.898 |
| b2c416efeade8d6cf941e29d51e96a29 | Bacteria | Proteobacteria | Deltaproteobacteria | SAR324 clade(Marine group B) | | | | 1.000 |
| dbe59cce2795c14f3f932124e96fbc52 | Bacteria | Planctomycetes | Planctomycetacia | Pirellulales | Pirellulaceae | uncultured | | 0.971 |
| c38fa7fc466f7365132face254f24025 | Bacteria | Proteobacteria | Deltaproteobacteria | Bdellovibrionales | Bdellovibrionaceae | OM27 clade | uncultured bacterium | 0.751 |
| c56d0651acaf1f91a5a23914d6e44395 | Bacteria | Bacteroidetes | Bacteroidia | Flavobacteriales | NS7 marine group | | | 1.000 |
| 4b0db1d7135e7871e025da6a7674e300 | Bacteria | Proteobacteria | Gammaproteobacteria | KI89A clade | uncultured bacterium | uncultured bacterium | uncultured bacterium | 0.912 |
| 3b660f8a4f16bbc1f794576f53f86c50 | Bacteria | Bacteroidetes | Bacteroidia | Flavobacteriales | NS9 marine group | uncultured marine bacterium | uncultured marine bacterium | 0.972 |
| 1eb35efbb8bf646d238c3abbad909e28 | Bacteria | Bacteroidetes | Bacteroidia | Flavobacteriales | Flavobacteriaceae | NS5 marine group | | 1.000 |
| 8a3a7875c5271e6034bd82cd2dafec75 | Bacteria | Proteobacteria | Gammaproteobacteria | Ectothiorhodospirales | Ectothiorhodospiraceae | uncultured | uncultured marine bacterium | 0.980 |
| 3eac82e0aaee53b75ce74b2c63bbdd9c | Bacteria | Verrucomicrobia | Verrucomicrobiae | Opitutales | Puniceicoccaceae | MB11C04 marine group | Ambiguous_taxa | 0.962 |
| 4eb74ff4aa61d2b5fe256006c831a349 | Bacteria | Cyanobacteria | Oxyphotobacteria | Synechococcales | Cyanobiaceae | Prochlorococcus MIT9313 | Ambiguous_taxa | 0.794 |
| b967e649393e9f2dc6efb3eb6bcec102 | Bacteria | Proteobacteria | Alphaproteobacteria | SAR11 clade | Clade I | uncultured | | 0.727 |
| f11bc49d4ac43fc3267f1f5d0de157d6 | Bacteria | Proteobacteria | Alphaproteobacteria | Puniceispirillales | SAR116 clade | uncultured Oceanibaculum sp. | uncultured Oceanibaculum sp. | 0.961 |
| 14f30649d0c2397658289522bdecef38 | Bacteria | Bacteroidetes | Bacteroidia | Cytophagales | Cyclobacteriaceae | Marinoscillum | | 1.000 |
| b5baf3d173d1d6459bd4f91c6c5999e2 | Bacteria | Proteobacteria | Alphaproteobacteria | Rhodospirillales | Magnetospiraceae | uncultured | uncultured Rhodospirillaceae bacterium | 0.831 |
| 83cedb510097a430a123bd2a35663c7a | Bacteria | Verrucomicrobia | Verrucomicrobiae | Verrucomicrobiales | Rubritaleaceae | Rubritalea | uncultured bacterium | 1.000 |
| 6551125ac9d66702b81b14df675c0c2f | Bacteria | Bacteroidetes | Bacteroidia | Flavobacteriales | NS9 marine group | | | 1.000 |
| 65b9a59ab09b8c11d448b5d6a60f4146 | Bacteria | Proteobacteria | Gammaproteobacteria | SAR86 clade | | | | 1.000 |
| fc8ab582b19fdb8013da173f308af0ca | Bacteria | Bacteroidetes | Bacteroidia | Flavobacteriales | Flavobacteriaceae | NS5 marine group | uncultured Bacteroidetes bacterium | 0.703 |
| 34855835fa93bfacba38577b657fd6f4 | Bacteria | Cyanobacteria | Oxyphotobacteria | Chloroplast | | | | 1.000 |
| f089527968cb352321cecc0e8f488a94 | Bacteria | Bacteroidetes | Bacteroidia | Flavobacteriales | Flavobacteriaceae | NS4 marine group | | 1.000 |
| 04c9d96f9a15fcc80a82c38e826ef601 | Bacteria | Planctomycetes | Phycisphaerae | Phycisphaerales | Phycisphaeraceae | CL500-3 | Ambiguous_taxa | 1.000 |
| fe8d5a20d41835d4f91d6cc6045949ee | Bacteria | Cyanobacteria | Oxyphotobacteria | Chloroplast | | | | 1.000 |

| Feature_ID | Kingdom | Phylum | Class | Order | Family | Genus | Species | Confidence |
|---|---|---|---|---|---|---|---|---|
| e6b2e4b665d631ef395efe3e7af8a857 | Bacteria | Proteobacteria | Gammaproteobacteria | KI89A clade | uncultured marine bacterium | uncultured marine bacterium | uncultured marine bacterium | 0.910 |
| c1be7dc01420dc4e57d33742e32f2a64 | Bacteria | Proteobacteria | Deltaproteobacteria | Bdellovibrionales | Bdellovibrionaceae | OM27 clade | Ambiguous_taxa | 0.987 |
| 0618434dab7767fd79535af457c280a8 | Bacteria | Cyanobacteria | Oxyphotobacteria | Chloroplast | Phalacroma mitra | Phalacroma mitra | Phalacroma mitra | 0.928 |
| 6c1b110dda53f48f54613f3356675c5a | Bacteria | Cyanobacteria | Oxyphotobacteria | Chloroplast | Epipyxis sp. PR26KG | Epipyxis sp. PR26KG | Epipyxis sp. PR26KG | 0.757 |
| d71e24176473157cf83bc8427aa93c01 | Bacteria | Proteobacteria | Alphaproteobacteria | Sphingomonadales | Sphingomonadaceae | Sphingobium | | 0.998 |
| b9e2ecb08a556c40daa75c0952e4c339 | Bacteria | Proteobacteria | Alphaproteobacteria | SAR11 clade | Clade I | Clade Ia | | 0.834 |
| a1a058b4498dfd8d9bd5a7b96ddfaf4e | Bacteria | Bacteroidetes | Bacteroidia | Flavobacteriales | NS9 marine group | uncultured Flavobacteriia bacterium | uncultured Flavobacteriia bacterium | 0.995 |
| 90e62fbd8d59d93b9bcdcbd7c0c995cd | Bacteria | Bacteroidetes | Bacteroidia | Flavobacteriales | Flavobacteriaceae | Formosa | | 0.892 |
| 3b91ffa8b3acb07761d92436e1ded491 | Bacteria | Proteobacteria | Alphaproteobacteria | SAR11 clade | Clade II | | | 0.856 |
| d0d0f02008f5cf3316b9fc3b938a7ff2 | Bacteria | Bacteroidetes | Bacteroidia | Cytophagales | Cyclobacteriaceae | uncultured | uncultured marine bacterium | 1.000 |
| ed6ece5c674d250cf879fbc7b614ef36 | Bacteria | Cyanobacteria | Oxyphotobacteria | Chloroplast | | | | 1.000 |
| 05a88fdc0f88be8b42e82f21b33849ee | Bacteria | Bacteroidetes | Bacteroidia | Flavobacteriales | NS9 marine group | uncultured Flavobacteriia bacterium | uncultured Flavobacteriia bacterium | 0.993 |
| c0a2c1a45a6a4b91c6b83fa4ea0dd44c | Bacteria | Planctomycetes | Planctomycetacia | Pirellulales | Pirellulaceae | uncultured | | 0.999 |
| 1bd6beae6c81232c64257bd36195f29a | Bacteria | Proteobacteria | Alphaproteobacteria | Rhodobacterales | Rhodobacteraceae | Ascidiaceihabitans | | 0.955 |
| 0e563142cb8e40de8ece72fb81df70cc | Bacteria | Cyanobacteria | Oxyphotobacteria | Chloroplast | Epipyxis sp. PR26KG | Epipyxis sp. PR26KG | Epipyxis sp. PR26KG | 0.807 |
| 4d6a47f6ee2ad4d1cccdd6f33544d360 | Bacteria | Proteobacteria | Gammaproteobacteria | Ectothiorhodospirales | Ectothiorhodospiraceae | uncultured | uncultured marine bacterium | 0.968 |
| 7f959926ff2ad151c3e5e6358c2fd443 | Bacteria | Proteobacteria | Gammaproteobacteria | Cellvibrionales | Halieaceae | OM60(NOR5) clade | Ambiguous_taxa | 0.926 |
| a95a7d56ac96f044ee872a0bd83a2364 | Bacteria | Cyanobacteria | Oxyphotobacteria | Chloroplast | Virgulinella fragilis | Virgulinella fragilis | Virgulinella fragilis | 0.723 |
| 2a08a47582ef129964bbd5158d29ccc3 | Bacteria | Marinimicrobia (SAR406 clade) | marine metagenome | marine metagenome | marine metagenome | marine metagenome | marine metagenome | 0.878 |
| 0197714ed0a1106b893866905a492004 | Bacteria | Proteobacteria | Alphaproteobacteria | Thalassobaculales | Nisaeaceae | OM75 clade | | 1.000 |
| 52dbb7439eeba96a1c2829f5b505bf5b | Bacteria | Bacteroidetes | Bacteroidia | Flavobacteriales | NS9 marine group | uncultured Flavobacteriia bacterium | uncultured Flavobacteriia bacterium | 0.993 |
| 66f775d06f38671095ccab4be4ce728c | Bacteria | Verrucomicrobia | Verrucomicrobiae | Arctic97B-4 marine group | uncultured marine bacterium | uncultured marine bacterium | uncultured marine bacterium | 0.981 |
| 55edac7ccbfaa44ee4000e412c41dadf | Bacteria | Cyanobacteria | Oxyphotobacteria | Chloroplast | marine metagenome | marine metagenome | marine metagenome | 0.702 |
| fb5509311f749334f29f271df933c651 | Bacteria | Actinobacteria | Acidimicrobiia | Microtrichales | Microtrichaceae | Sva0996 marine group | Ambiguous_taxa | 0.982 |
| c557eb0f4fad212f420561d38db2ebf1 | Bacteria | Bacteroidetes | Bacteroidia | Flavobacteriales | Crocinitomicaceae | Fluviicola | uncultured marine bacterium | 0.806 |
| 47354d572da1facadb38c900097a5a20 | Bacteria | Proteobacteria | Deltaproteobacteria | PB19 | Ambiguous_taxa | Ambiguous_taxa | Ambiguous_taxa | 0.869 |
| 5ae95cdb613d1123e3deda2c9a00c879 | Bacteria | Verrucomicrobia | Verrucomicrobiae | Arctic97B-4 marine group | | | | 1.000 |
| ce91478373bf9c8bd0531cfea26928cb | Bacteria | Cyanobacteria | Oxyphotobacteria | Chloroplast | Phalacroma mitra | Phalacroma mitra | Phalacroma mitra | 0.884 |
| 4e64935fcd01c436fc6f7c7d0a48d636 | Bacteria | Proteobacteria | Alphaproteobacteria | SAR11 clade | | | | 1.000 |
| f0e88f855c1c5849ad439b1a1b942f94 | Bacteria | Proteobacteria | Gammaproteobacteria | KI89A clade | Ambiguous_taxa | Ambiguous_taxa | Ambiguous_taxa | 0.937 |
| 1711ad8be5919385e302b3fe53e49bd9 | Bacteria | Proteobacteria | Alphaproteobacteria | Puniceispirillales | SAR116 clade | Candidatus Puniceispirillum | | 0.724 |
| dc4dae920a565377ec418a79da23c6d1 | Bacteria | Cyanobacteria | Oxyphotobacteria | Chloroplast | Phalacroma mitra | Phalacroma mitra | Phalacroma mitra | 0.983 |
| 2f49cfd4834a95c8b3a246b698750661 | Bacteria | Bacteroidetes | Bacteroidia | Flavobacteriales | Cryomorphaceae | uncultured | uncultured Owenweeksia sp. | 0.779 |
| ab5305c1e15fb1df58ce822e78d29b82 | Bacteria | Proteobacteria | Gammaproteobacteria | Cellvibrionales | Porticoccaceae | SAR92 clade | uncultured marine bacterium | 0.893 |
| 9c609ecc137025ff8e78deb827a4549e | Bacteria | Proteobacteria | Alphaproteobacteria | Thalassobaculales | Nisaeaceae | OM75 clade | | 1.000 |
| c8e0c1a307726d4f9dc3726837eaaaaa | Bacteria | Cyanobacteria | Oxyphotobacteria | Synechococcales | Cyanobiaceae | Prochlorococcus MIT9313 | | 0.998 |
| d06c0c3c746d75a5e0ff799ec160cae2 | Bacteria | Proteobacteria | Alphaproteobacteria | Rhodospirillales | Magnetospiraceae | uncultured | uncultured nuHF2 cluster bacterium HF0130_29D04 | 0.999 |
| dafcab2d646767ee5c3fcc019648ca46 | Bacteria | Bacteroidetes | Bacteroidia | Flavobacteriales | Flavobacteriaceae | NS5 marine group | uncultured marine bacterium | 0.920 |
| 7c911fee228a7ddc4f8c394027613329 | Bacteria | Bacteroidetes | Bacteroidia | Flavobacteriales | NS9 marine group | uncultured Flavobacteriia bacterium | uncultured Flavobacteriia bacterium | 0.989 |
| dd322adb9a196d775eb20363f1220892 | Bacteria | Cyanobacteria | Oxyphotobacteria | Chloroplast | | | | 1.000 |
| 0b536f8854e11d193387ae39b7d08579 | Archaea | Euryarchaeota | Thermoplasmata | Marine Group II | | | | 1.000 |
| f19c29319a758454c02a87cc8a493ce9 | Bacteria | Proteobacteria | Gammaproteobacteria | Alteromonadales | Idiomarinaceae | Idiomarina | | 1.000 |
| 549f2f4cecba95049d30ce6f0886c6b4 | Bacteria | Verrucomicrobia | Verrucomicrobiae | Verrucomicrobiales | DEV007 | uncultured Verrucomicrobiaceae bacterium | uncultured Verrucomicrobiaceae bacterium | 1.000 |
| c779e047db8920984b7495a22a773243 | Bacteria | Cyanobacteria | Oxyphotobacteria | Chloroplast | marine metagenome | marine metagenome | marine metagenome | 0.867 |
| ac178abd52ee0f4f3ec9133bd18df952 | Bacteria | Marinimicrobia (SAR406 clade) | unidentified marine bacterioplankton | unidentified marine bacterioplankton | unidentified marine bacterioplankton | unidentified marine bacterioplankton | unidentified marine bacterioplankton | 0.872 |
| 8ee5980ab9d6562886e98207f38f77ed | Bacteria | Proteobacteria | Gammaproteobacteria | OM182 clade | uncultured gamma proteobacterium | uncultured gamma proteobacterium | uncultured gamma proteobacterium | 0.756 |
| 2ed972fdbc5f2c2f8b31d768b4bec3bb | Bacteria | Proteobacteria | Alphaproteobacteria | Rickettsiales | Mitochondria | uncultured bacterium | uncultured bacterium | 0.998 |
| a291edca74ee6a661f7337ff14d6cdac | Bacteria | Proteobacteria | Deltaproteobacteria | PB19 | Ambiguous_taxa | Ambiguous_taxa | Ambiguous_taxa | 0.868 |
| 092805d73c5687ed3575a5e60e0a197e | Bacteria | Cyanobacteria | Oxyphotobacteria | Chloroplast | Epipyxis sp. PR26KG | Epipyxis sp. PR26KG | Epipyxis sp. PR26KG | 0.868 |
| 183c25e0ca904c246dc4c9c2cccc8497 | Bacteria | Proteobacteria | Alphaproteobacteria | Rhodospirillales | Magnetospiraceae | uncultured | uncultured marine bacterium | 0.952 |
| 37bc07d963ba97d0017a3fe2818ee9af | Bacteria | Proteobacteria | Alphaproteobacteria | Rhodospirillales | AEGEAN-169 marine group | | | 1.000 |
| 51e8dc03f8773502475a842214173542 | Bacteria | Bacteroidetes | Bacteroidia | Flavobacteriales | Flavobacteriaceae | uncultured | uncultured marine bacterium | 0.924 |
| d4a3dd6950897f44dc38e4cc61d2ba08 | Bacteria | Bacteroidetes | Bacteroidia | Flavobacteriales | Flavobacteriaceae | NS5 marine group | | 1.000 |
| 7ca6b80c0a19aff33c547bafefcb7d57 | Bacteria | Cyanobacteria | Oxyphotobacteria | Chloroplast | uncultured soil bacterium | uncultured soil bacterium | uncultured soil bacterium | 0.972 |
| 0e703419c9e0c4c447cbb29ab3443324 | Bacteria | Cyanobacteria | Oxyphotobacteria | Chloroplast | Phalacroma mitra | Phalacroma mitra | Phalacroma mitra | 0.995 |
| 545a8c7991b2cdca012b52ff315b2a87 | Bacteria | Proteobacteria | Gammaproteobacteria | Ectothiorhodospirales | Ectothiorhodospiraceae | uncultured | uncultured marine bacterium | 0.953 |
| 20a7d89e3104a1b8562ffc0b216bbf87 | Bacteria | Proteobacteria | Deltaproteobacteria | PB19 | uncultured delta proteobacterium | uncultured delta proteobacterium | uncultured delta proteobacterium | 0.980 |
| fb122356537261dc96ed534a428dc9cc | Bacteria | Proteobacteria | Gammaproteobacteria | KI89A clade | uncultured bacterium | uncultured bacterium | uncultured bacterium | 0.751 |
| 1c3a2a0910f2b514330be5e8ef493cb1 | Bacteria | Bacteroidetes | Bacteroidia | Flavobacteriales | Flavobacteriaceae | NS5 marine group | | 1.000 |
| 88acfab92ed933b887cfb2721ca078ea | Bacteria | Bacteroidetes | Bacteroidia | Flavobacteriales | Cryomorphaceae | uncultured | uncultured Flavobacterium sp. | 0.828 |
| 017b83e3cb9536619714f41ed6d6d594 | Bacteria | Proteobacteria | Gammaproteobacteria | SAR86 clade | | | | 1.000 |
| a68a11708036b30d87ab564bf1650e22 | Bacteria | Proteobacteria | Alphaproteobacteria | SAR11 clade | Clade I | Clade Ib | | 0.809 |
| ba1f0129b1a627ee8516ce97ee3177d4 | Bacteria | Proteobacteria | Alphaproteobacteria | Rickettsiales | S25-593 | | | 1.000 |
| 84dbc43ff05fc49d790f888b908de64a | Bacteria | Bacteroidetes | Bacteroidia | Chitinophagales | Saprospiraceae | Aureispira | uncultured marine bacterium | 0.816 |
| ec731edd3b79a4b2fb69d3292e414d5a | Bacteria | Cyanobacteria | Oxyphotobacteria | Chloroplast | marine metagenome | marine metagenome | marine metagenome | 0.966 |
| 8bc0014fd986d2426e3f1eedd8c82e27 | Bacteria | Planctomycetes | Planctomycetacia | Pirellulales | Pirellulaceae | uncultured | | 1.000 |
| 052ee499e583ef9e8fa91f3005006edf | Bacteria | Marinimicrobia (SAR406 clade) | unidentified marine bacterioplankton | unidentified marine bacterioplankton | unidentified marine bacterioplankton | unidentified marine bacterioplankton | unidentified marine bacterioplankton | 0.958 |
| 5aab5c0a105bf1377b1ded69ed6cb985 | Archaea | Euryarchaeota | Thermoplasmata | Marine Group II | metagenome | metagenome | metagenome | 1.000 |
| fd74bfe395348d8f527fd3030664b0bb | Bacteria | Bacteroidetes | Bacteroidia | Flavobacteriales | Flavobacteriaceae | | | 1.000 |
| ea8b0afe862b4b6d2287895db88647f6 | Bacteria | Margulisbacteria | uncultured marine cyanobacterium | uncultured marine cyanobacterium | uncultured marine cyanobacterium | uncultured marine cyanobacterium | uncultured marine cyanobacterium | 1.000 |
| 409bb8c4521da5952f1d9aab3de8d8d8 | Bacteria | Proteobacteria | Alphaproteobacteria | SAR11 clade | Clade II | | | 0.881 |
| c6738714274a3eba8008a61098e604c8 | Bacteria | Proteobacteria | Gammaproteobacteria | SAR86 clade | | | | 1.000 |
| c8cf89d98fc51716912700553e661eda | Bacteria | Proteobacteria | Deltaproteobacteria | Bdellovibrionales | Bdellovibrionaceae | OM27 clade | uncultured marine bacterium | 0.991 |

| Feature_ID | Kingdom | Phylum | Class | Order | Family | Genus | Species | Confidence |
|---|---|---|---|---|---|---|---|---|
| 4c2378bd8d7bb48fc8b19a62b77ed00c | Bacteria | Cyanobacteria | Oxyphotobacteria | Synechococcales | Cyanobiaceae | Prochlorococcus MIT9313 | | 0.980 |
| 742483a385e6d8bcfc0bc44298783d96 | Bacteria | Proteobacteria | Gammaproteobacteria | Cellvibrionales | Spongiibacteraceae | BD1-7 clade | uncultured gamma proteobacterium | 0.957 |
| e2c9ba2b677a88bda505b8ddef62b784 | Bacteria | Proteobacteria | Gammaproteobacteria | HgCo23 | uncultured marine bacterium | uncultured marine bacterium | uncultured marine bacterium | 0.990 |
| 7b1d62f61e7e3de5b0f330a603980a79 | Bacteria | Verrucomicrobia | Verrucomicrobiae | Opitutales | Puniceicoccaceae | | | 1.000 |
| c87c523c6fefef01dfffc92da489a0ad | Bacteria | Proteobacteria | Gammaproteobacteria | Pseudomonadales | Moraxellaceae | Psychrobacter | | 1.000 |
| 874d9e136f552dbd708db51efe352b6b | Bacteria | Proteobacteria | Gammaproteobacteria | Alteromonadales | Pseudoalteromonadaceae | Pseudoalteromonas | | 0.987 |
| 9ac20859a7c74ee65993bdfe260f6824 | Bacteria | Marinimicrobia (SAR406 clade) | Alteromonas macleodii | Alteromonas macleodii | Alteromonas macleodii | Alteromonas macleodii | Alteromonas macleodii | 1.000 |
| 245260e4d0fdef9d3446d42053e858b9 | Bacteria | Proteobacteria | Gammaproteobacteria | SAR86 clade | uncultured gamma proteobacterium HF0070_25G02 | uncultured gamma proteobacterium HF0070_25G02 | uncultured gamma proteobacterium HF0070_25G02 | 0.999 |
| e2c21fcedcd026dfd695839df7ced4b9 | Archaea | Euryarchaeota | Thermoplasmata | Marine Group II | | | | 1.000 |
| 0d8977fdc2a364a6522825b2bbf5209c | Bacteria | Marinimicrobia (SAR406 clade) | hydrothermal vent metagenome | hydrothermal vent metagenome | hydrothermal vent metagenome | hydrothermal vent metagenome | hydrothermal vent metagenome | 0.996 |
| 9c44eec10b6576a9b66c450a18cb14cc | Bacteria | Marinimicrobia (SAR406 clade) | marine metagenome | marine metagenome | marine metagenome | marine metagenome | marine metagenome | 0.704 |
| 919cb4708a7a2cf89b1198e5bec4e12c | Bacteria | Proteobacteria | Alphaproteobacteria | Rhodospirillales | AEGEAN-169 marine group | metagenome | metagenome | 0.998 |
| 1bb16fe2b96655065c488559217ebe59 | Bacteria | Proteobacteria | Gammaproteobacteria | UBA10353 marine group | Ambiguous_taxa | Ambiguous_taxa | Ambiguous_taxa | 0.924 |
| 761a173d79e70868cca44afb8ebcb1c6 | Bacteria | Chloroflexi | JG30-KF-CM66 | marine metagenome | marine metagenome | marine metagenome | marine metagenome | 1.000 |
| fdf2a5cac1fbf537da59850e9c8bc88a | Bacteria | Proteobacteria | Gammaproteobacteria | OM182 clade | | | | 1.000 |
| 0c65eea8e9074423132d9d946d19cf93 | Bacteria | Proteobacteria | Deltaproteobacteria | Myxococcales | P3OB-42 | uncultured marine bacterium | uncultured marine bacterium | 1.000 |
| cd306f4f534d90f752e17556f3e04011 | Bacteria | Proteobacteria | Deltaproteobacteria | Bdellovibrionales | Bdellovibrionaceae | OM27 clade | uncultured delta proteobacterium | 0.999 |
| ff59c48ddbe153708c0e45e41da60ad3 | Bacteria | Proteobacteria | Alphaproteobacteria | Sphingomonadales | Sphingomonadaceae | Sphingobium | | 1.000 |
| 0d5094dfea127a2c2ce1caa0f387b3a2 | Bacteria | Bacteroidetes | Bacteroidia | Flavobacteriales | Cryomorphaceae | uncultured | | 1.000 |
| adcb0e67dfd880b1e6227cb506d294ee | Bacteria | Bacteroidetes | Bacteroidia | Flavobacteriales | Flavobacteriaceae | NS2b marine group | Ambiguous_taxa | 0.786 |
| 155c12b0a3a87120411d46627f7f4b7a | Bacteria | Proteobacteria | Gammaproteobacteria | Alteromonadales | Pseudoalteromonadaceae | Pseudoalteromonas | | 0.991 |
| 7ac5166f9380a4c18232004fd7a2da4c | Bacteria | Proteobacteria | Deltaproteobacteria | Bdellovibrionales | Bdellovibrionaceae | OM27 clade | | 1.000 |
| c5a0edf440b9cb64d8657b950937120c | Bacteria | Verrucomicrobia | Verrucomicrobiae | Opitutales | Puniceicoccaceae | Lentimonas | Ambiguous_taxa | 0.794 |
| 6eda1a6288cf233b3aba1835e3620506 | Bacteria | Bacteroidetes | Bacteroidia | Flavobacteriales | NS9 marine group | uncultured Flavobacteriia bacterium | uncultured Flavobacteriia bacterium | 0.933 |
| 4729d63a3816597ce8eb96512ca0b1c8 | Bacteria | Cyanobacteria | Oxyphotobacteria | Chloroplast | Phalacroma mitra | Phalacroma mitra | Phalacroma mitra | 0.937 |
| d61cf02c7e220952dab4b9828a0d0a41 | Bacteria | Bacteroidetes | Bacteroidia | Flavobacteriales | Flavobacteriaceae | NS5 marine group | | 1.000 |
| 252e439ccbdca22c735fa29d4271e9e9 | Bacteria | Proteobacteria | Gammaproteobacteria | Cellvibrionales | Spongiibacteraceae | BD1-7 clade | uncultured gamma proteobacterium | 0.874 |
| 6b1e35bd3080f862d4150e5df8b474ac | Bacteria | Proteobacteria | Gammaproteobacteria | Betaproteobacteriales | Methylophilaceae | OM43 clade | | 1.000 |
| 859ab5a6e6e6f2df91544acc0950a4c4 | Bacteria | Bacteroidetes | Bacteroidia | Flavobacteriales | NS9 marine group | Ambiguous_taxa | Ambiguous_taxa | 0.998 |
| f7b2a213bf9a691b2b61df129eeafcb9 | Bacteria | Proteobacteria | Gammaproteobacteria | Ectothiorhodospirales | Ectothiorhodospiraceae | uncultured | uncultured marine bacterium | 0.996 |
| 32bc678776f916ca3001805fac6f6482 | Bacteria | Proteobacteria | Gammaproteobacteria | Cellvibrionales | Halieaceae | OM60(NOR5) clade | Ambiguous_taxa | 0.880 |
| 79fa7199885be179a3b093762d928125 | Bacteria | Cyanobacteria | Oxyphotobacteria | Chloroplast | Cymbomonas tetramitiformis | Cymbomonas tetramitiformis | Cymbomonas tetramitiformis | 0.976 |
| a774191f4bd981dd110fd5abc0ec667d | Bacteria | Proteobacteria | Deltaproteobacteria | PB19 | Ambiguous_taxa | Ambiguous_taxa | Ambiguous_taxa | 0.995 |
| 8fbe91a176e7dd7935cbd873a926dd80 | Bacteria | Bacteroidetes | Bacteroidia | Flavobacteriales | Flavobacteriaceae | NS5 marine group | uncultured marine bacterium | 0.813 |
| 692890348a075c0845fdb30e8707edee | Bacteria | Proteobacteria | Gammaproteobacteria | SAR86 clade | | | | 1.000 |
| da9d4a63e9bff3156e80c95e6ebf3085 | Bacteria | Proteobacteria | Gammaproteobacteria | Coxiellales | Coxiellaceae | Coxiella | | 1.000 |
| 6452c5dbadde2b8693ac7abe232592d0 | Bacteria | Proteobacteria | Gammaproteobacteria | SAR86 clade | | | | 1.000 |
| 32adea307c0da7150d5abe0208c6e16b | Bacteria | Proteobacteria | Gammaproteobacteria | EPR3968-O8a-Bc78 | uncultured gamma proteobacterium | uncultured gamma proteobacterium | uncultured gamma proteobacterium | 0.996 |
| d1baec9ea84c26fc15be9229a3496488 | Bacteria | Bacteroidetes | Bacteroidia | Flavobacteriales | Flavobacteriaceae | NS5 marine group | Ambiguous_taxa | 0.855 |
| 54f5e3b3abe03e4920fd5e9372bf3490 | Bacteria | Marinimicrobia (SAR406 clade) | uncultured bacterium | uncultured bacterium | uncultured bacterium | uncultured bacterium | uncultured bacterium | 0.992 |
| b761abc16f93da22268f2fbb19412ef2 | Bacteria | Margulisbacteria | metagenome | metagenome | metagenome | metagenome | metagenome | 0.830 |
| 3c9e6449d55561e2924625e44e4a5691 | Bacteria | Bacteroidetes | Bacteroidia | Flavobacteriales | NS7 marine group | Ambiguous_taxa | Ambiguous_taxa | 0.980 |
| 1725f2fa7f32b27d507ff91a3fd851f9 | Bacteria | Proteobacteria | Alphaproteobacteria | SAR11 clade | Clade II | | | 0.989 |
| 84b557603c26e79eb618048375bf27dd | Bacteria | Cyanobacteria | Oxyphotobacteria | Chloroplast | marine metagenome | marine metagenome | marine metagenome | 0.983 |
| 6d570e6a2bda42f16fa5f6bc138a9252 | Bacteria | Proteobacteria | Deltaproteobacteria | Bdellovibrionales | Bdellovibrionaceae | OM27 clade | | 1.000 |
| c66920b0f261138f92f8d7c5abf47832 | Bacteria | Proteobacteria | Gammaproteobacteria | Cellvibrionales | Porticoccaceae | SAR92 clade | | 1.000 |
| 500ab9e56663fed86d81ea30e00fbfec | Bacteria | Proteobacteria | Deltaproteobacteria | Bdellovibrionales | Bdellovibrionaceae | OM27 clade | unidentified marine bacterioplankton | 1.000 |
| f6502f81e1a9ec0794c30b424047c432 | Bacteria | Bacteroidetes | Bacteroidia | Flavobacteriales | Cryomorphaceae | uncultured | uncultured Flavobacterium sp. | 0.727 |
| 4122b4f14ca5c928b97d9b4beda24aff | Bacteria | Bacteroidetes | Bacteroidia | Flavobacteriales | NS9 marine group | hydrothermal vent metagenome | hydrothermal vent metagenome | 1.000 |
| 098cca0005074921c2cd828564aaa85b | Bacteria | Bacteroidetes | Bacteroidia | Flavobacteriales | Crocinitomicaceae | Fluviicola | uncultured marine bacterium | 0.771 |
| 36c205347e8a79f3f7e94c8b7db5325f | Bacteria | Planctomycetes | Planctomycetacia | Pirellulales | Pirellulaceae | Blastopirellula | uncultured bacterium | 0.876 |
| ab9cafa035f9b76e30ccd95d342d7020 | Bacteria | Cyanobacteria | Oxyphotobacteria | Chloroplast | Prasinophyceae sp. CCMP1205 | Prasinophyceae sp. CCMP1205 | Prasinophyceae sp. CCMP1205 | 1.000 |
| 2e23cbb77c33137b704a828c747c2d2d | Bacteria | Proteobacteria | Alphaproteobacteria | Rhodospirillales | AEGEAN-169 marine group | | | 1.000 |
| 4cb87ce1035934817464e64fe9ff8e76 | Bacteria | Proteobacteria | Alphaproteobacteria | Rhodospirillales | Magnetospiraceae | uncultured | | 0.918 |
| 36aa2d54e7141490ab0bf50e64e896dd | Bacteria | Proteobacteria | Gammaproteobacteria | JTB23 | uncultured marine microorganism | uncultured marine microorganism | uncultured marine microorganism | 0.783 |
| 5a83fb0df7c3c67a97f224353badb1bd | Bacteria | Bacteroidetes | Bacteroidia | Chitinophagales | Saprospiraceae | uncultured | uncultured marine bacterium | 0.992 |
| fd84c514cc4d48dc63f65c8e2911c337 | Bacteria | Bacteroidetes | Bacteroidia | Flavobacteriales | Flavobacteriaceae | NS5 marine group | | 1.000 |
| c3f089bed9742584bc47b98c4fb2271d | Bacteria | Bacteroidetes | Bacteroidia | Flavobacteriales | NS7 marine group | Ambiguous_taxa | Ambiguous_taxa | 0.752 |
| c7524bb695bc9ff58468eb2b41db7c23 | Bacteria | Proteobacteria | Alphaproteobacteria | uncultured | uncultured marine bacterium | uncultured marine bacterium | uncultured marine bacterium | 0.832 |
| e6f6f772608bdf40d30b88cdab922c57 | Bacteria | Marinimicrobia (SAR406 clade) | unidentified marine bacterioplankton | unidentified marine bacterioplankton | unidentified marine bacterioplankton | unidentified marine bacterioplankton | unidentified marine bacterioplankton | 0.905 |
| c418a7de7b19e944befbd9812a46dbbc | Bacteria | Proteobacteria | Gammaproteobacteria | Cellvibrionales | Halieaceae | Parahaliea | | 0.998 |
| 96cb90db2b8c045b07e5ae06e8fb2be7 | Bacteria | Proteobacteria | Deltaproteobacteria | Bdellovibrionales | Bdellovibrionaceae | OM27 clade | uncultured delta proteobacterium | 0.874 |
| f6d026b76000d326e3c106b1ed1aeb58 | Bacteria | Proteobacteria | Gammaproteobacteria | UBA10353 marine group | Ambiguous_taxa | Ambiguous_taxa | Ambiguous_taxa | 0.994 |
| a2d386eb6e2c6e769f94800164c14b37 | Bacteria | Bacteroidetes | Bacteroidia | Flavobacteriales | Flavobacteriaceae | NS4 marine group | | 1.000 |
| 36a62faf56eaa19ea2dcad578e032bfb | Bacteria | Verrucomicrobia | Verrucomicrobiae | Opitutales | Puniceicoccaceae | A714019 | uncultured bacterium | 0.996 |
| aa24c87bfa55411297865dadd4ca78f6 | Bacteria | Proteobacteria | Alphaproteobacteria | Rhodospirillales | Rhodospirillaceae | uncultured | uncultured alpha proteobacterium | 0.996 |
| 4e7fb19f71d3adb01fe91ff4115dfd0f | Bacteria | Proteobacteria | Alphaproteobacteria | SAR11 clade | Clade IV | Ambiguous_taxa | Ambiguous_taxa | 0.881 |
| 1ad1f2a089b88988c7a9aa615b54cae8 | Bacteria | Proteobacteria | Gammaproteobacteria | HOC36 | uncultured marine bacterium | uncultured marine bacterium | uncultured marine bacterium | 0.996 |
| c4380c96be50a1e72af3a42ef3fe5807 | Bacteria | Bacteroidetes | Bacteroidia | Chitinophagales | Saprospiraceae | uncultured | marine metagenome | 1.000 |
| 938f5c6eda24b0e792eb6240c0a114ef | Bacteria | Proteobacteria | Alphaproteobacteria | Rhodospirillales | Magnetospiraceae | Magnetospira | uncultured alpha proteobacterium | 0.951 |

| Feature_ID | Kingdom | Phylum | Class | Order | Family | Genus | Species | Confidence |
|---|---|---|---|---|---|---|---|---|
| 6d819599793bfc0a051dcddc44a24895 | Bacteria | Proteobacteria | Alphaproteobacteria | Puniceispirillales | SAR116 clade | alpha proteobacterium SCGC AAA015-N04 | alpha proteobacterium SCGC AAA015-N04 | 0.904 |
| a19d7650f6581c8bd5ca189b3e0d0d3f | Bacteria | Proteobacteria | Gammaproteobacteria | Oceanospirillales | Halomonadaceae | Salinicola | Ambiguous_taxa | 0.725 |
| 0576236553062a4957d64a26a94409e3 | Bacteria | Marinimicrobia (SAR406 clade) | | | | | | 1.000 |
| b582eb48adb2998e16f87695a0d9663b | Bacteria | Bacteroidetes | Bacteroidia | Flavobacteriales | Flavobacteriaceae | Aurantivirga | Ambiguous_taxa | 0.990 |
| 8a02577ea757407d9484f9f14d7c8dde | Bacteria | Cyanobacteria | Oxyphotobacteria | Chloroplast | | | | 0.988 |
| 9970e1e5f6c4a3b14fbb44ca24846e83 | Bacteria | Proteobacteria | Gammaproteobacteria | Xanthomonadales | Xanthomonadaceae | Stenotrophomonas | | 1.000 |
| 7956ff617d71e0a98c575dee33bbc251 | Bacteria | Proteobacteria | Gammaproteobacteria | Salinisphaerales | Salinisphaeraceae | Salinisphaera | Ambiguous_taxa | 0.958 |
| 7f7481008cf6fe648fbc909e72cbda85 | Bacteria | Proteobacteria | Gammaproteobacteria | Alteromonadales | Pseudoalteromonadaceae | Pseudoalteromonas | | 0.996 |
| 1dfb182a50f16ec1a5a1c37053e04467 | Bacteria | Proteobacteria | Alphaproteobacteria | SAR11 clade | Clade I | Clade Ib | | 0.759 |
| d6c65e37a93b8f6bbd8759dc083bdab8 | Bacteria | Cyanobacteria | Oxyphotobacteria | Chloroplast | Phalacroma mitra | Phalacroma mitra | Phalacroma mitra | 0.723 |
| 20ec9ae8c9a5ccda6ac7e522e146f64b | Archaea | Euryarchaeota | Thermoplasmata | Marine Group II | | | | 1.000 |
| 1346daf56a7e3633222e9bbb0abe19f7 | Bacteria | Proteobacteria | Alphaproteobacteria | Rhodospirillales | Magnetospiraceae | uncultured | marine metagenome | 0.997 |
| fe06c9d61f1a99567c659af731b04bfe | Bacteria | Cyanobacteria | Oxyphotobacteria | Chloroplast | uncultured bacterium | uncultured bacterium | uncultured bacterium | 0.928 |
| 6720d6d947ab8fdf2bd70b0337c15928 | Bacteria | Verrucomicrobia | Verrucomicrobiae | Arctic97B-4 marine group | | | | 1.000 |
| bd25eb9fcb0661f9b637c1e1850633c9 | Bacteria | Proteobacteria | Deltaproteobacteria | Bdellovibrionales | Bdellovibrionaceae | OM27 clade | Ambiguous_taxa | 0.987 |
| e8a35fee05dc471741f3434fdf99a98c | Bacteria | Proteobacteria | Deltaproteobacteria | SAR324 clade(Marine group B) | uncultured bacterium | uncultured bacterium | uncultured bacterium | 0.853 |
| bdf6912cb0477e9b6bf8b88a1e226840 | Bacteria | Marinimicrobia (SAR406 clade) | uncultured bacterium | uncultured bacterium | uncultured bacterium | uncultured bacterium | uncultured bacterium | 0.829 |
| a077ecf7545c82b3523b4437ac273406 | Bacteria | Proteobacteria | Gammaproteobacteria | SAR86 clade | uncultured SAR86 cluster bacterium | uncultured SAR86 cluster bacterium | uncultured SAR86 cluster bacterium | 0.789 |
| cb134f230b9469ae4fd0fc73e699557a | Bacteria | Proteobacteria | Gammaproteobacteria | SAR86 clade | | | | 1.000 |
| 4e9abbf3b2477e4547e4115ffa4d9bda | Bacteria | Proteobacteria | Deltaproteobacteria | Bdellovibrionales | Bacteriovoracaceae | uncultured | marine metagenome | 1.000 |
| 3db31e37eb2def3876dedbffc09ad6ed | Bacteria | Bacteroidetes | Bacteroidia | Flavobacteriales | NS9 marine group | uncultured Flavobacteriales bacterium | uncultured Flavobacteriales bacterium | 0.811 |
| f85f3e981b1b820047fa00c384414905 | Bacteria | Bacteroidetes | Bacteroidia | Flavobacteriales | NS7 marine group | uncultured bacterium | uncultured bacterium | 0.997 |
| dd9012533ca8fd1c74a0ca0414d4a9ad | Bacteria | Bacteroidetes | Bacteroidia | Flavobacteriales | Crocinitomicaceae | Salinirepens | Ambiguous_taxa | 0.997 |
| b33b631d66b50c1afc97615c54337f41 | Bacteria | Bacteroidetes | Bacteroidia | Flavobacteriales | Flavobacteriaceae | | | 1.000 |
| ba7c83ec470314f4213a58e428fca582 | Bacteria | Cyanobacteria | Oxyphotobacteria | Chloroplast | | | | 1.000 |
| 6766ad8f12879b15c6e8fc6cdedab85f | Bacteria | Cyanobacteria | Oxyphotobacteria | Chloroplast | Phalacroma mitra | Phalacroma mitra | Phalacroma mitra | 0.998 |
| 7bd01c648a6bdd62b802b3b24b8fa58d | Bacteria | Cyanobacteria | Oxyphotobacteria | Synechococcales | Cyanobiaceae | Synechococcus CC9902 | | 0.996 |
| 3e89520d137d665367d6f049ce4c72b6 | Bacteria | Proteobacteria | Deltaproteobacteria | Bdellovibrionales | Bdellovibrionaceae | OM27 clade | | 1.000 |
| 41e4cb63bebc0f7b08da26bd8eb82fb6 | Bacteria | Chloroflexi | Dehalococcoidia | SAR202 clade | Ambiguous_taxa | Ambiguous_taxa | Ambiguous_taxa | 0.999 |
| 10eab941eaa795e010b0f2eb14af6e11 | Bacteria | Bacteroidetes | Bacteroidia | Flavobacteriales | Flavobacteriaceae | Croceibacter | Ambiguous_taxa | 0.999 |
| 69087073e43695e81e7b165d2ba1be49 | Bacteria | Proteobacteria | Alphaproteobacteria | uncultured | unidentified marine bacterioplankton | unidentified marine bacterioplankton | unidentified marine bacterioplankton | 0.919 |
| 379a21bedd7b13ec636e765b5f412eeb | Bacteria | Cyanobacteria | Oxyphotobacteria | Chloroplast | | | | 1.000 |
| 6938d6ebc15f951e64a0ab34b36b4f50 | Bacteria | Bacteroidetes | Bacteroidia | Flavobacteriales | NS9 marine group | uncultured marine bacterium | uncultured marine bacterium | 0.975 |
| b2c68312e48a93811a34f0c7f1df2e17 | Bacteria | Cyanobacteria | Oxyphotobacteria | Chloroplast | uncultured marine eukaryote | uncultured marine eukaryote | uncultured marine eukaryote | 0.869 |
| 507ed460400113404367b7e077f69bb2 | Bacteria | Bacteroidetes | Bacteroidia | Flavobacteriales | NS9 marine group | uncultured Flavobacteriia bacterium | uncultured Flavobacteriia bacterium | 0.897 |
| 284c18945802255b2b5113daee125657 | Bacteria | Cyanobacteria | Oxyphotobacteria | Chloroplast | unidentified marine bacterioplankton | unidentified marine bacterioplankton | unidentified marine bacterioplankton | 0.921 |
| 1aa53435b5a8e93d74144a31aa938830 | Bacteria | Marinimicrobia (SAR406 clade) | uncultured Fibrobacteres/Acidobacteria group bacterium | uncultured Fibrobacteres/Acidobacteria group bacterium | uncultured Fibrobacteres/Acidobacteria group bacterium | uncultured Fibrobacteres/Acidobacteria group bacterium | uncultured Fibrobacteres/Acidobacteria group bacterium | 0.976 |
| d24db56ca957b8e88d4e05201186da9d | Bacteria | Marinimicrobia (SAR406 clade) | marine metagenome | marine metagenome | marine metagenome | marine metagenome | marine metagenome | 0.995 |
| 5603ed8c3e539916f650ec8743f6745f | Bacteria | Cyanobacteria | Oxyphotobacteria | Synechococcales | Cyanobiaceae | | | 1.000 |
| f5cf85ac34af6fbbe8467c6b8595bdc4 | Bacteria | Proteobacteria | Alphaproteobacteria | Sphingomonadales | Sphingomonadaceae | Sphingomonas | | 0.923 |
| ce20c85816fa8785a16f02420bc2b6fd | Bacteria | Proteobacteria | Gammaproteobacteria | Betaproteobacteriales | Methylophilaceae | OM43 clade | | 1.000 |
| 7284bc721fac6ea21182cce1ee717004 | Bacteria | Proteobacteria | Gammaproteobacteria | Pseudomonadales | Moraxellaceae | Psychrobacter | | 1.000 |
| 3dd10745ff94ea3ca85b8ba24d456c48 | Bacteria | Proteobacteria | Alphaproteobacteria | Rhodospirillales | Rhodospirillaceae | uncultured | uncultured alpha proteobacterium | 0.998 |
| 23acebf2fe6e76f3433071e97a82d4fb | Bacteria | Proteobacteria | Deltaproteobacteria | PB19 | uncultured delta proteobacterium | uncultured delta proteobacterium | uncultured delta proteobacterium | 0.800 |
| 4d5354dd57633c0f74fa0c02a9cd9cf4 | Bacteria | Marinimicrobia (SAR406 clade) | Ambiguous_taxa | Ambiguous_taxa | Ambiguous_taxa | Ambiguous_taxa | Ambiguous_taxa | 0.714 |
| 0b048f047591ca2c4eb56091e74f9a72 | Bacteria | Gemmatimonadetes | BD2-11 terrestrial group | Ambiguous_taxa | Ambiguous_taxa | Ambiguous_taxa | Ambiguous_taxa | 0.992 |
| 2401ffde2674954c10184462993f67b3 | Bacteria | Actinobacteria | Acidimicrobiia | Microtrichales | Microtrichaceae | Sva0996 marine group | | 1.000 |
| e3347c171f10ca0bc57df2a5526e0c54 | Bacteria | Bacteroidetes | Bacteroidia | Flavobacteriales | NS9 marine group | uncultured marine bacterium | uncultured marine bacterium | 0.997 |
| bfd6ebbd1a2857bc0838f160b41a120c | Bacteria | Proteobacteria | Alphaproteobacteria | Puniceispirillales | SAR116 clade | Candidatus Puniceispirillum | Ambiguous_taxa | 0.841 |
| ffbd0c01dd69853f36c624f8101f504a | Bacteria | Fibrobacteres | Fibrobacteria | Fibrobacterales | Fibrobacteraceae | uncultured | uncultured bacterium | 0.972 |
| 559015c4570044b7a109be7b6498f739 | Bacteria | Proteobacteria | Deltaproteobacteria | PB19 | | | | 1.000 |
| ffb6efb8c674fe69a5028ee06cc867bc | Bacteria | Proteobacteria | Gammaproteobacteria | SAR86 clade | uncultured bacterium | uncultured bacterium | uncultured bacterium | 0.943 |
| eef2ba0e710187e4a2f88341b32ad66c | Bacteria | Bacteroidetes | Bacteroidia | Flavobacteriales | Flavobacteriaceae | NS4 marine group | | 1.000 |
| b72d76feb701353e5d58c174b8453c7b | Bacteria | | | | | | | 1.000 |
| 62e3ce37bb87fbd042c3f0ce46aa97c5 | Bacteria | Bacteroidetes | Bacteroidia | Flavobacteriales | NS9 marine group | uncultured bacterium | uncultured bacterium | 0.954 |
| cccc004d306e6aa2a7e369d73e6d92c7 | Bacteria | Proteobacteria | Gammaproteobacteria | Cellvibrionales | Porticoccaceae | SAR92 clade | | 1.000 |
| 4f0c41bbf3cf4584c8bced52e0ebb1ef | Bacteria | Proteobacteria | Alphaproteobacteria | Rhodospirillales | Magnetospiraceae | uncultured | uncultured bacterium | 0.773 |
| 2477d6a3ed9c86ef58538e576e572046 | Bacteria | Bacteroidetes | Bacteroidia | Cytophagales | Cyclobacteriaceae | Marinoscillum | | 1.000 |
| 7d3a370630c3d31bca8aadb36874de47 | Bacteria | Cyanobacteria | Oxyphotobacteria | Chloroplast | | | | 1.000 |
| 136e1e07b41914727e04d24f8b7080ae | Bacteria | Proteobacteria | Deltaproteobacteria | Oligoflexales | Oligoflexaceae | uncultured | uncultured bacterium | 1.000 |
| 1e10d369d4bd3144de4d87380ecafd22 | Bacteria | Proteobacteria | Gammaproteobacteria | Ga0077536 | uncultured bacterium | uncultured bacterium | uncultured bacterium | 0.913 |
| 3610432799d779ca62115b4cad254bb8 | Bacteria | Proteobacteria | Gammaproteobacteria | SAR86 clade | | | | 1.000 |
| ee8cbabfee85de6de441dbbaddd215e3 | Bacteria | Cyanobacteria | Oxyphotobacteria | Chloroplast | | | | 1.000 |
| 399bbfd7c02e91837eee973f8dbd915f | Bacteria | Bacteroidetes | Bacteroidia | Flavobacteriales | NS9 marine group | | | 1.000 |
| 795d21bba0b44d09cc1777593f7b6a7e | Archaea | Euryarchaeota | Thermoplasmata | Marine Group II | marine metagenome | marine metagenome | marine metagenome | 0.858 |
| ca544d3c90a84336cae1297722dc3a50 | Bacteria | Proteobacteria | Gammaproteobacteria | Alteromonadales | Marinobacteraceae | Marinobacter | | 1.000 |
| f333d706d83753fd34566aa843200754 | Bacteria | Cyanobacteria | Oxyphotobacteria | Chloroplast | Cymbomonas tetramitiformis | Cymbomonas tetramitiformis | Cymbomonas tetramitiformis | 0.973 |
| a55361bfacb8322774a22320a37a11cd | Bacteria | Proteobacteria | Deltaproteobacteria | PB19 | uncultured marine bacterium | uncultured marine bacterium | uncultured marine bacterium | 0.994 |
| f54e22d74b4bf7bdbcac607ea8093c93 | Bacteria | Cyanobacteria | Oxyphotobacteria | Chloroplast | | | | 1.000 |
| 7479fa23f086238f9a098e756d821767 | Bacteria | Proteobacteria | Alphaproteobacteria | SAR11 clade | Clade I | Clade Ia | | 0.973 |
| 52bace6ee6ba168ec3b3cd77c5b278b6 | Bacteria | Proteobacteria | Gammaproteobacteria | Vibrionales | Vibrionaceae | Vibrio | | 0.996 |
| 402ee8edfe9534a85cc4bee939b8b4d2 | Bacteria | Proteobacteria | Alphaproteobacteria | Rickettsiales | S25-593 | marine metagenome | marine metagenome | 0.996 |
| 9b494b22b5ea13244eba4ace4d6cc61a | Bacteria | Bacteroidetes | Bacteroidia | Cytophagales | Cyclobacteriaceae | Marinoscillum | | 1.000 |
| 1ddffd158d4ef764a4876dcee271813b | Bacteria | Bacteroidetes | Bacteroidia | Flavobacteriales | Flavobacteriaceae | NS5 marine group | | 1.000 |
| d20d8dc000987d40328f788af4a2be51 | Bacteria | Bacteroidetes | Bacteroidia | Flavobacteriales | Flavobacteriaceae | Salegentibacter | Ambiguous_taxa | 0.801 |
| ac5cc2d0174be98688c242dbaf421230 | Bacteria | Proteobacteria | Deltaproteobacteria | PB19 | uncultured bacterium | uncultured bacterium | uncultured bacterium | 0.889 |
| 06dfcef1d90509eb81c8e6c7ab4da60a | Bacteria | Proteobacteria | Gammaproteobacteria | Alteromonadales | Alteromonadaceae | Alteromonas | | 0.951 |
| 60a976eedf1cc46cba152c5ccee4fcf7 | Bacteria | Cyanobacteria | Oxyphotobacteria | Chloroplast | uncultured marine eukaryote | uncultured marine eukaryote | uncultured marine eukaryote | 0.944 |
| 059ab9b4a35762f3d177703561869fbc | Bacteria | Proteobacteria | Deltaproteobacteria | PB19 | Ambiguous_taxa | Ambiguous_taxa | Ambiguous_taxa | 0.998 |

| Feature_ID | Kingdom | Phylum | Class | Order | Family | Genus | Species | Confidence |
|---|---|---|---|---|---|---|---|---|
| 403f3512c4d6b1ec996efa04d6d9e61f | Bacteria | Cyanobacteria | Oxyphotobacteria | Chloroplast | bacterium WHC4-2 | bacterium WHC4-2 | bacterium WHC4-2 | 0.960 |
| 6b9feab1259cc3059a2a4c4152e9f2ad | Bacteria | Proteobacteria | Alphaproteobacteria | SAR11 clade | Clade II | | | 0.834 |
| 6a8885ddd92b2b5609daf6f5da3d7e7a | Bacteria | Proteobacteria | Alphaproteobacteria | Rhodobacterales | Rhodobacteraceae | | | 1.000 |
| 3a579139b071b67c7672735bbf99f042 | Bacteria | Proteobacteria | Gammaproteobacteria | Cellvibrionales | Porticoccaceae | SAR92 clade | | 1.000 |
| d9358f6e983aa612273437d249dd8fd2 | Bacteria | Proteobacteria | Alphaproteobacteria | SAR11 clade | Clade II | | | 0.874 |
| 36dd501e70e91938af5b0708fb47fbfc | Bacteria | Proteobacteria | Deltaproteobacteria | Bdellovibrionales | Bdellovibrionaceae | OM27 clade | | 1.000 |
| fea477a0ba2762112dec1d651368ec77 | Bacteria | Verrucomicrobia | Verrucomicrobiae | Verrucomicrobiales | Rubritaleaceae | Roseibacillus | uncultured marine bacterium | 0.910 |
| 28d669ac294ae5fbae9d54154fb276e5 | Bacteria | Cyanobacteria | Oxyphotobacteria | Chloroplast | Prasinophyceae sp. MBIC10622 | Prasinophyceae sp. MBIC10622 | Prasinophyceae sp. MBIC10622 | 1.000 |
| 29cabdfd807d2812109a1cac9a8b8ae2 | Bacteria | Bacteroidetes | Bacteroidia | Flavobacteriales | Flavobacteriaceae | NS5 marine group | Ambiguous_taxa | 0.747 |
| 3086c4fc3fd78479c80f4bc32e391009 | Bacteria | Cyanobacteria | Oxyphotobacteria | Chloroplast | Phalacroma mitra | Phalacroma mitra | Phalacroma mitra | 0.910 |
| ec4a28fe927717ec73bef0714356c901 | Bacteria | Proteobacteria | Gammaproteobacteria | Coxiellales | Coxiellaceae | Coxiella | uncultured bacterium | 0.976 |
| 788f21dfa7cadfe2cb92bcf13a402b47 | Bacteria | Bacteroidetes | Bacteroidia | Flavobacteriales | NS9 marine group | uncultured marine bacterium | uncultured marine bacterium | 0.955 |
| e17c3bcdf295ecdb250c4dc46240c27f | Bacteria | Proteobacteria | Alphaproteobacteria | Rhodospirillales | AEGEAN-169 marine group | | | 1.000 |
| a7a99613af34480141dec4cad7e8e388 | Bacteria | Cyanobacteria | Oxyphotobacteria | Chloroplast | | | | 0.983 |
| bfe56699d0d7697de49c1e6e612a0078 | Bacteria | Proteobacteria | Gammaproteobacteria | Vibrionales | Vibrionaceae | Vibrio | | 0.877 |
| 1d7e23ca83ad12d1b985f3361de47cd7 | Bacteria | Proteobacteria | Alphaproteobacteria | Rickettsiales | S25-593 | | | 1.000 |
| ab48a8acbe9cc25dce8a8779642b5f5d | Bacteria | Proteobacteria | Deltaproteobacteria | Bdellovibrionales | Bacteriovoracaceae | uncultured | marine metagenome | 1.000 |
| c420d283c4b6a94130cf985f63cac23b | Bacteria | Poribacteria | metagenome | metagenome | metagenome | metagenome | metagenome | 1.000 |
| 6810507a21d7aab9e87d9fc5cdf1aff5 | Bacteria | Cyanobacteria | Oxyphotobacteria | Chloroplast | | | | 1.000 |
| 89ff4b670b08a73f5fdec27395ae3d0f | Bacteria | Proteobacteria | Deltaproteobacteria | PB19 | Ambiguous_taxa | Ambiguous_taxa | Ambiguous_taxa | 0.994 |
| 6270e792a6ac0051e785be9eb522935d | Bacteria | Chloroflexi | Dehalococcoidia | SAR202 clade | uncultured bacterium | uncultured bacterium | uncultured bacterium | 0.768 |
| 0dddbe3104d4ed3502f3f16c4adeeb73 | Bacteria | Proteobacteria | Alphaproteobacteria | Puniceispirillales | SAR116 clade | marine metagenome | marine metagenome | 0.994 |
| 3d94f17abb64e5f1183ecaad951124a9 | Bacteria | Proteobacteria | Gammaproteobacteria | Betaproteobacteriales | Burkholderiaceae | MWH-UniP1 aquatic group | uncultured marine bacterium | 0.999 |
| 09f42b53f97dcd197a49d362b8cca2ce | Bacteria | Proteobacteria | Alphaproteobacteria | SAR11 clade | Clade I | Clade Ia | | 0.819 |
| eda4911fd79a17c624c06c73624fca09 | Bacteria | Proteobacteria | Gammaproteobacteria | Oceanospirillales | Halomonadaceae | Halomonas | | 0.952 |
| 943f12db9377b7417175f1fad0b68943 | Bacteria | Kiritimatiellaeota | Kiritimatiellae | Kiritimatiellales | Kiritimatiellaceae | R76-B128 | marine metagenome | 0.985 |
| add63ee675e77105f8e4b6d0bc33c5b5 | Bacteria | Bacteroidetes | Bacteroidia | Flavobacteriales | Cryomorphaceae | uncultured | | 1.000 |
| 0423b95d027f8ec68dcb16792a478a03 | Bacteria | Cyanobacteria | Oxyphotobacteria | Chloroplast | uncultured Cryptomonadaceae | uncultured Cryptomonadaceae | uncultured Cryptomonadaceae | 0.955 |
| edfb0ff2d79eac08d15aff6cf7f8244c | Bacteria | Bacteroidetes | Bacteroidia | Flavobacteriales | NS9 marine group | Ambiguous_taxa | Ambiguous_taxa | 0.996 |
| 37d0f8c647939d5187161d5073014fb1 | Bacteria | Cyanobacteria | Oxyphotobacteria | Chloroplast | uncultured bacterium | uncultured bacterium | uncultured bacterium | 0.708 |
| c617caf3d30c56c60290bdc804b4aa75 | Bacteria | Proteobacteria | Alphaproteobacteria | Rhodospirillales | AEGEAN-169 marine group | | | 1.000 |
| a7338f29f3c13fd3394eff71d3e6397f | Bacteria | Proteobacteria | Deltaproteobacteria | Bdellovibrionales | Bacteriovoracaceae | uncultured | uncultured Bacteriovoracaceae bacterium | 0.993 |
| f09630dfd032360bdefbebb01acd093b | Bacteria | Proteobacteria | Alphaproteobacteria | Puniceispirillales | SAR116 clade | Candidatus Puniceispirillum | | 0.796 |
| 6316342e2f80d6d37f9affaa69f2e5b8 | Bacteria | Bacteroidetes | Bacteroidia | Flavobacteriales | NS7 marine group | | | 1.000 |
| ef9e3487ed35499459c1d33d36b958e1 | Bacteria | Proteobacteria | Gammaproteobacteria | SAR86 clade | | | | 1.000 |
| 5a2d247249bac5b6f5992101fd10885b | Bacteria | Cyanobacteria | Oxyphotobacteria | Chloroplast | | | | 1.000 |
| a4ec7c4f7b8a811e2e9db71452bff424 | Bacteria | Bacteroidetes | Bacteroidia | Flavobacteriales | Flavobacteriaceae | uncultured | uncultured marine bacterium | 0.939 |
| 1cf8eb2d8b552f7fbf69f7a76bc82f5b | Archaea | Euryarchaeota | Thermoplasmata | Marine Group III | Ambiguous_taxa | Ambiguous_taxa | Ambiguous_taxa | 0.928 |
| 4d5510a36e9e663a18e3760fd733bb89 | Bacteria | Proteobacteria | Alphaproteobacteria | SAR11 clade | Clade II | | | 0.905 |
| 3b15897490d8c2ebc860907371423f57 | Bacteria | Bacteroidetes | Bacteroidia | Flavobacteriales | Flavobacteriaceae | NS2b marine group | uncultured marine bacterium | 0.912 |
| 6f9d2f842a1e9583d1c0d77af83fe972 | Bacteria | Proteobacteria | Alphaproteobacteria | Thalassobaculales | uncultured | Ambiguous_taxa | Ambiguous_taxa | 0.989 |
| 6bb2bbb3f39683f256cdc16fedf4abcc | Archaea | Euryarchaeota | Thermoplasmata | Marine Group II | | | | 1.000 |
| 8a3f418340868a85d4c96e39bbe62f18 | Bacteria | Proteobacteria | Alphaproteobacteria | | | | | 1.000 |
| 0f269ae1a3247c55a95b147fa51070b5 | Bacteria | Proteobacteria | Deltaproteobacteria | PB19 | Ambiguous_taxa | Ambiguous_taxa | Ambiguous_taxa | 0.776 |
| 75484a0c70aac472f6e8e0a92af3fa5f | Bacteria | Proteobacteria | Alphaproteobacteria | Puniceispirillales | SAR116 clade | Candidatus Puniceispirillum | | 0.820 |
| 0998d947a5580debbdd89084c90c0f96 | Bacteria | Proteobacteria | Deltaproteobacteria | Bdellovibrionales | Bdellovibrionaceae | OM27 clade | uncultured marine bacterium | 0.997 |
| ef522f083e80a50f25ff65d44287b58a | Bacteria | Cyanobacteria | Oxyphotobacteria | Chloroplast | Ochromonas sp. CCMP1393 | Ochromonas sp. CCMP1393 | Ochromonas sp. CCMP1393 | 0.724 |
| 10adfc865553280ab106b2ea4f00f53c | Bacteria | Marinimicrobia (SAR406 clade) | uncultured bacterium | uncultured bacterium | uncultured bacterium | uncultured bacterium | uncultured bacterium | 0.999 |
| d40abcedfd40a26ffc29cad86982c0e5 | Bacteria | Verrucomicrobia | Verrucomicrobiae | Verrucomicrobiales | DEV007 | uncultured Verrucomicrobiaceae bacterium | uncultured Verrucomicrobiaceae bacterium | 0.991 |
| 2c360f7bc9ccc4b9b81cab119f4721bf | Bacteria | Proteobacteria | Alphaproteobacteria | Rhodospirillales | AEGEAN-169 marine group | | | 1.000 |
| ceff91f9dd377b5c81f7dd1f56885d31 | Bacteria | Proteobacteria | Deltaproteobacteria | Bdellovibrionales | Bdellovibrionaceae | OM27 clade | unidentified marine bacterioplankton | 0.997 |
| 645c030e89d66e484c63c92cd76fcac4 | Bacteria | Cyanobacteria | Oxyphotobacteria | Synechococcales | Cyanobiaceae | Prochlorococcus MIT9313 | | 0.856 |
| 82e97640a9f189f986e929badadf6b33 | Bacteria | Proteobacteria | Gammaproteobacteria | Vibrionales | Vibrionaceae | Vibrio | | 0.927 |
| 2d1c8b8679937f346a0e25171516a53f | Bacteria | Bacteroidetes | Bacteroidia | Flavobacteriales | NS9 marine group | | | 1.000 |
| ac0afe30f7b1e6adc020738560823c2f | Bacteria | Cyanobacteria | Oxyphotobacteria | Chloroplast | Phalacroma mitra | Phalacroma mitra | Phalacroma mitra | 0.964 |
| 6e6ff17e226695fb54905aeff02b0a5c | Bacteria | Cyanobacteria | Oxyphotobacteria | Chloroplast | uncultured Cryptomonadaceae | uncultured Cryptomonadaceae | uncultured Cryptomonadaceae | 0.991 |
| 9ec07b621e5c39707fca59c775c01ba8 | Bacteria | Proteobacteria | Alphaproteobacteria | Caulobacterales | Caulobacteraceae | Caulobacter | | 1.000 |
| 9b7b685ce0aa1b8ea665d1e4f65dc26d | Bacteria | Cyanobacteria | Oxyphotobacteria | Chloroplast | marine metagenome | marine metagenome | marine metagenome | 0.917 |
| b82e2014805dd0172f2752c6b42117ab | Bacteria | Proteobacteria | Gammaproteobacteria | Cellvibrionales | Halieaceae | OM60(NOR5) clade | Ambiguous_taxa | 0.879 |
| c5a6b7f671a4ba6fb3da11ebc93db10c | Bacteria | Bacteroidetes | Bacteroidia | Flavobacteriales | NS9 marine group | uncultured marine bacterium | uncultured marine bacterium | 0.993 |
| 007d3c256f5dd3f66157e11fc1ef2b84 | Bacteria | Cyanobacteria | Oxyphotobacteria | Chloroplast | Phalacroma mitra | Phalacroma mitra | Phalacroma mitra | 0.996 |
| 4559531c6e44bba3a292663065c0169d | Bacteria | Proteobacteria | Gammaproteobacteria | OM182 clade | uncultured gamma proteobacterium | uncultured gamma proteobacterium | uncultured gamma proteobacterium | 0.999 |
| 3a370d2e2de7eb3c8fc777925e15d5d2 | Bacteria | Proteobacteria | Alphaproteobacteria | uncultured | unidentified marine bacterioplankton | unidentified marine bacterioplankton | unidentified marine bacterioplankton | 0.958 |
| eb47536874b17614e23359145b2ce17d | Bacteria | Proteobacteria | Gammaproteobacteria | Alteromonadales | Colwelliaceae | Colwellia | uncultured marine bacterium | 0.764 |
| f94195e38396e6fc026795ce9c0a909a | Bacteria | Cyanobacteria | Oxyphotobacteria | Chloroplast | | | | 1.000 |
| f85cac35de5da1926cc3a6bca4505cd3 | Bacteria | Proteobacteria | Gammaproteobacteria | Vibrionales | Vibrionaceae | Grimontia | Ambiguous_taxa | 0.884 |
| 884ac7da7fafadbcc83a2d1fbcf04dbf | Archaea | Euryarchaeota | Thermoplasmata | Marine Group II | | | | 1.000 |
| 3ce64164af3dced54c914846e658a539 | Bacteria | Bacteroidetes | Bacteroidia | Flavobacteriales | Flavobacteriaceae | NS5 marine group | | 1.000 |
| cf8a13d68056ef677bb2b3b8c767ca00 | Bacteria | Proteobacteria | Gammaproteobacteria | Oceanospirillales | Litoricolaceae | Litoricola | uncultured bacterium | 0.992 |
| 9d10f9a491680e42f44577a6f61cc98a | Bacteria | Cyanobacteria | Oxyphotobacteria | Chloroplast | Phalacroma mitra | Phalacroma mitra | Phalacroma mitra | 0.968 |
| dc4edb34d73b342615d766eb3f7b1f9f | Bacteria | Verrucomicrobia | Verrucomicrobiae | Arctic97B-4 marine group | uncultured marine bacterium | uncultured marine bacterium | uncultured marine bacterium | 0.979 |
| 35e4263003039c8a71765b6b349a7f3b | Bacteria | Cyanobacteria | Oxyphotobacteria | Chloroplast | uncultured bacterium | uncultured bacterium | uncultured bacterium | 0.715 |
| ceecbd04b484e7c09d0cbdbed9f2d619 | Bacteria | Gemmatimonadetes | BD2-11 terrestrial group | Ambiguous_taxa | Ambiguous_taxa | Ambiguous_taxa | Ambiguous_taxa | 0.986 |
| 06b7076c4e52263cb56c2825bdd670f9 | Bacteria | Proteobacteria | Gammaproteobacteria | Ectothiorhodospirales | Ectothiorhodospiraceae | uncultured | uncultured marine bacterium | 0.939 |
| 2c291c318a22d880979ca3c31b5e9020 | Bacteria | Bacteroidetes | Bacteroidia | Flavobacteriales | Flavobacteriaceae | NS5 marine group | | 1.000 |
| 99bd4b0129ee50825546135981da6b2d | Bacteria | Bacteroidetes | Bacteroidia | Flavobacteriales | Flavobacteriaceae | NS5 marine group | uncultured Flavobacteriaceae bacterium | 0.882 |
| a100291f0a1641433508441cde92c313 | Bacteria | Bacteroidetes | Bacteroidia | Flavobacteriales | Flavobacteriaceae | NS4 marine group | | 1.000 |

| Feature_ID | Kingdom | Phylum | Class | Order | Family | Genus | Species | Confidence |
|---|---|---|---|---|---|---|---|---|
| b263ca3a884c152249379b71f77f9ce9 | Bacteria | Cyanobacteria | Oxyphotobacteria | Chloroplast | | | | 1.000 |
| 33b037990c3a01e072da843c85f72835 | Bacteria | Bacteroidetes | Bacteroidia | Flavobacteriales | Flavobacteriaceae | NS2b marine group | | 1.000 |
| 8a117a94fe2a1518a70e7f367ebb0d2b | Bacteria | Bacteroidetes | Bacteroidia | Flavobacteriales | Flavobacteriaceae | Mesonia | Ambiguous_taxa | 0.999 |
| 1842462a3eabf3aa148026cbcb2dea24 | Bacteria | Proteobacteria | Alphaproteobacteria | Rickettsiales | S25-593 | uncultured marine bacterium | uncultured marine bacterium | 0.996 |
| efd641491199a46b60bc7177660bb019 | Bacteria | Proteobacteria | Alphaproteobacteria | Rickettsiales | Mitochondria | Proteobacteria bacterium JGI 0000113-D06 | Proteobacteria bacterium JGI 0000113-D06 | 0.712 |
| 27513b6877183e6762674232300b5fab | Bacteria | Proteobacteria | Gammaproteobacteria | Ectothiorhodospirales | Ectothiorhodospiraceae | uncultured | uncultured marine bacterium | 0.963 |
| 906a43b16efad5007c2f6c205fc83b36 | Bacteria | Bacteroidetes | Bacteroidia | Flavobacteriales | Flavobacteriaceae | | | 1.000 |
| 8aac3aa49c80f843f0d856f0cbfa21b7 | Bacteria | Verrucomicrobia | Verrucomicrobiae | Opitutales | Puniceicoccaceae | MB11C04 marine group | | 1.000 |
| ef5bc45d1cd1bde46eb1213c12f09dc7 | Bacteria | Proteobacteria | Alphaproteobacteria | Rickettsiales | Mitochondria | uncultured bacterium | uncultured bacterium | 0.960 |
| ad7795885ac78faa2cceab8f506599d4 | Bacteria | Bacteroidetes | Bacteroidia | Flavobacteriales | NS9 marine group | | | 1.000 |
| d9d2f8b40aac80acb2359078d021f802 | Bacteria | Proteobacteria | Gammaproteobacteria | KI89A clade | uncultured marine bacterium | uncultured marine bacterium | uncultured marine bacterium | 0.797 |
| 28941a14797f40a565c11789339091f7 | Bacteria | Proteobacteria | Deltaproteobacteria | Bdellovibrionales | Bdellovibrionaceae | OM27 clade | uncultured marine bacterium | 0.774 |
| a75cb8cb9915ece52d9deffedd2fa017 | Bacteria | Cyanobacteria | Oxyphotobacteria | Chloroplast | uncultured Cryptomonadaceae | uncultured Cryptomonadaceae | uncultured Cryptomonadaceae | 0.999 |
| fb410157f67230f2fba616c8760a1b3f | Bacteria | Proteobacteria | Alphaproteobacteria | Rhodospirillales | AEGEAN-169 marine group | uncultured bacterium | uncultured bacterium | 0.996 |
| 36e28bc583126a618d65ce5df0210f1c | Bacteria | Marinimicrobia (SAR406 clade) | Ambiguous_taxa | Ambiguous_taxa | Ambiguous_taxa | Ambiguous_taxa | Ambiguous_taxa | 0.792 |
| 600d3a9b14608d34fc67c76a3751d1a8 | Bacteria | Proteobacteria | Alphaproteobacteria | SAR11 clade | Clade II | | | 0.968 |
| d3c4242dd33e85ee902b833b78b9624a | Bacteria | Cyanobacteria | Oxyphotobacteria | Chloroplast | uncultured marine eukaryote | uncultured marine eukaryote | uncultured marine eukaryote | 0.943 |
| 1794f8c96ca30e7466df78e09e696d81 | Bacteria | Proteobacteria | Gammaproteobacteria | Alteromonadales | Idiomarinaceae | Idiomarina | | 1.000 |
| 674ddf111dca6ef8a5220d9af6af2ce5 | Bacteria | Proteobacteria | Alphaproteobacteria | uncultured | uncultured marine bacterium | uncultured marine bacterium | uncultured marine bacterium | 0.998 |
| e090cf29c31ef128b0d654c754fe7cbc | Bacteria | Proteobacteria | Gammaproteobacteria | SAR86 clade | Ambiguous_taxa | Ambiguous_taxa | Ambiguous_taxa | 0.960 |
| b9de90e19d3c2d167720964a93f2e98e | Bacteria | Cyanobacteria | Oxyphotobacteria | Chloroplast | Phalacroma mitra | Phalacroma mitra | Phalacroma mitra | 0.837 |
| aa5493b0951aaf80fc6d84f57e261e23 | Bacteria | Bacteroidetes | Bacteroidia | Flavobacteriales | NS9 marine group | | | 1.000 |
| b13b606cb96fed3f237b1fc0bd10bd2a | Bacteria | Cyanobacteria | Oxyphotobacteria | Synechococcales | Cyanobiaceae | Prochlorococcus MIT9313 | | 0.957 |
| 86172db1d4c4d3285d770a69c05a2a0d | Bacteria | Proteobacteria | Alphaproteobacteria | SAR11 clade | Clade I | Clade Ib | | 0.985 |
| 58969e6e509b2655b712e7ecceb9711a | Bacteria | Verrucomicrobia | Verrucomicrobiae | Arctic97B-4 marine group | uncultured marine bacterium | uncultured marine bacterium | uncultured marine bacterium | 0.990 |
| df3adaa210f476e1b738eb8fdaf97f38 | Bacteria | Proteobacteria | Gammaproteobacteria | EPR3968-O8a-Bc78 | Ambiguous_taxa | Ambiguous_taxa | Ambiguous_taxa | 1.000 |
| 4f40256d33ce0b8f6921d557161fb1a8 | Bacteria | Kiritimatiellaeota | Kiritimatiellae | Kiritimatiellales | Kiritimatiellaceae | R76-B128 | uncultured bacterium | 0.869 |
| d8f872ff12f209b874fc79f045a8797e | Bacteria | Marinimicrobia (SAR406 clade) | uncultured marine microorganism | uncultured marine microorganism | uncultured marine microorganism | uncultured marine microorganism | uncultured marine microorganism | 0.999 |
| ee1fbe16af52df6ee86c1a5fd67f3089 | Bacteria | Proteobacteria | Gammaproteobacteria | Vibrionales | Vibrionaceae | Grimontia | Ambiguous_taxa | 0.921 |
| c6d7ed89bca593e0ebe90592597ad7de | Bacteria | Proteobacteria | Gammaproteobacteria | Vibrionales | Vibrionaceae | Vibrio | | 0.990 |
| 183722946bc3c942bde9beab5de2daab | Bacteria | Planctomycetes | Planctomycetacia | Planctomycetales | Gimesiaceae | uncultured | Ambiguous_taxa | 1.000 |
| 58daeddf7b2aadf8106ce6e555cb76d8 | Bacteria | Proteobacteria | Alphaproteobacteria | SAR11 clade | | | | 1.000 |
| 633a1a4fc546a667c8087dd78e382c7e | Bacteria | Bacteroidetes | Bacteroidia | Flavobacteriales | Crocinitomicaceae | Fluviicola | uncultured marine bacterium | 0.738 |
| f2a5c7c217e2349ca24fe0fe9cd6a9e9 | Bacteria | Proteobacteria | Deltaproteobacteria | Myxococcales | P3OB-42 | uncultured marine bacterium | uncultured marine bacterium | 1.000 |
| 31133f9969f3484e3df7be17a59599ef | Bacteria | Cyanobacteria | Oxyphotobacteria | Chloroplast | | | | 1.000 |
| 29685b6d9e5657869f319ccf1d44268b | Bacteria | Proteobacteria | Gammaproteobacteria | SAR86 clade | Ambiguous_taxa | Ambiguous_taxa | Ambiguous_taxa | 0.977 |
| 7200261ab7c9ea64651d6f4372966531 | Bacteria | Proteobacteria | Alphaproteobacteria | Puniceispirillales | SAR116 clade | Candidatus Puniceispirillum | Ambiguous_taxa | 0.849 |
| 5f07fc450df756f36b4bb001ce3b6a02 | Bacteria | Bacteroidetes | Bacteroidia | Flavobacteriales | NS9 marine group | Ambiguous_taxa | Ambiguous_taxa | 0.937 |
| 678cbd6ed8d41f279ba07b1772e92c71 | Bacteria | Planctomycetes | Phycisphaerae | Phycisphaerales | Phycisphaeraceae | CL500-3 | uncultured marine bacterium | 1.000 |
| 9a6775272ac7e083d960795661bcb390 | Bacteria | Bacteroidetes | Bacteroidia | Flavobacteriales | NS9 marine group | Ambiguous_taxa | Ambiguous_taxa | 0.743 |
| b37c475b60c82495ddc22b2d69ee15ba | Bacteria | Proteobacteria | Alphaproteobacteria | Rhodospirillales | AEGEAN-169 marine group | | | 1.000 |
| 7520ecf4798e5ea244df6d92ad7c81cd | Bacteria | Cyanobacteria | Oxyphotobacteria | Chloroplast | | | | 1.000 |
| cd8fe65bdebd14e3d6998be03b400e1e | Bacteria | Bacteroidetes | Bacteroidia | Flavobacteriales | Cryomorphaceae | uncultured | uncultured Owenweeksia sp. | 0.905 |
| 2893165dcb93179cdce6bd7208275ce1 | Bacteria | Proteobacteria | Gammaproteobacteria | Vibrionales | Vibrionaceae | uncultured | Ambiguous_taxa | 0.788 |
| 390ee06cb481079746410ffa563e8147 | Bacteria | Proteobacteria | Alphaproteobacteria | Rhodospirillales | Magnetospiraceae | Magnetospira | uncultured alpha proteobacterium | 0.848 |
| 76fa3adee60554b089b6bba2e98f2cad | Bacteria | Bacteroidetes | Bacteroidia | Flavobacteriales | Flavobacteriaceae | Tenacibaculum | Ambiguous_taxa | 0.846 |
| f6380181180883a3384a7e984f432a24 | Bacteria | Bacteroidetes | Bacteroidia | Flavobacteriales | Flavobacteriaceae | NS5 marine group | | 1.000 |
| 264fb7729d11d482103be8340d10f43f | Bacteria | Proteobacteria | Alphaproteobacteria | Rhodospirillales | AEGEAN-169 marine group | unidentified marine bacterioplankton | unidentified marine bacterioplankton | 0.989 |
| ffeb4e1ae0736450f932a80f39b058fb | Bacteria | Bacteroidetes | Bacteroidia | Cytophagales | Cyclobacteriaceae | Marinoscillum | | 1.000 |
| 05da9fd5392d5fe48c14595919e3dfce | Bacteria | Bacteroidetes | Bacteroidia | Flavobacteriales | NS9 marine group | uncultured Flavobacteriia bacterium | uncultured Flavobacteriia bacterium | 0.901 |
| cd7a1bb3f0324a8ff5b8678375ef6372 | Bacteria | Proteobacteria | Gammaproteobacteria | SAR86 clade | uncultured bacterium | uncultured bacterium | uncultured bacterium | 0.818 |
| 6d2cf7f8b39cb8594fcc3a7ff768f8ee | Bacteria | Proteobacteria | Alphaproteobacteria | Sphingomonadales | Sphingomonadaceae | Stakelama | Ambiguous_taxa | 0.994 |
| 2a5c1f50e248635d823a780156b73362 | Bacteria | Bacteroidetes | Bacteroidia | Flavobacteriales | NS9 marine group | uncultured Flavobacteriia bacterium | uncultured Flavobacteriia bacterium | 0.992 |
| 0e78e0242ec250f63fe4fa0280725a83 | Bacteria | Proteobacteria | Deltaproteobacteria | PB19 | uncultured marine microorganism | uncultured marine microorganism | uncultured marine microorganism | 0.851 |
| 7a54d3f3d9ad99f1396ba5cfc666b724 | Bacteria | Cyanobacteria | Oxyphotobacteria | Chloroplast | uncultured bacterium | uncultured bacterium | uncultured bacterium | 0.916 |
| c8aa00ef3ea9ac280f5d569682304e58 | Bacteria | Proteobacteria | Deltaproteobacteria | Bdellovibrionales | Bdellovibrionaceae | OM27 clade | | 1.000 |
| 2c2472fd81ab76ae8db0877ed805ca6a | Bacteria | Cyanobacteria | Oxyphotobacteria | Chloroplast | | | | 1.000 |
| af7bd3e90e6157da864e01aaab657990 | Bacteria | Proteobacteria | Alphaproteobacteria | Rhizobiales | Rhizobiaceae | | | 1.000 |
| 8610ed9c52d4b3c43cc394e3b98d4a54 | Bacteria | Proteobacteria | Alphaproteobacteria | SAR11 clade | Clade I | Clade Ia | | 0.861 |
| fd161277b49fb8718a9a66788e9cfabe | Bacteria | Bacteroidetes | Bacteroidia | Flavobacteriales | NS9 marine group | uncultured Flavobacteriia bacterium | uncultured Flavobacteriia bacterium | 0.984 |
| 2ac36fa4b4bb697fae28e66ae4806d8f | Bacteria | Cyanobacteria | Oxyphotobacteria | Chloroplast | | | | 1.000 |
| 688001dd2fb8cee5bf2fbf6bc908af1d | Bacteria | Proteobacteria | Deltaproteobacteria | SAR324 clade(Marine group B) | | | | 1.000 |
| 0fa7065c65455e5d0a4d9861fd04c953 | Bacteria | Marinimicrobia (SAR406 clade) | uncultured marine microorganism | uncultured marine microorganism | uncultured marine microorganism | uncultured marine microorganism | uncultured marine microorganism | 0.992 |
| 1eb1da33226fbd6dfff646373ee1079b | Bacteria | Proteobacteria | Gammaproteobacteria | Thiomicrospirales | Thioglobaceae | SUP05 cluster | | 0.999 |
| e3d76972ac790bb92ffff3afdf05abdd | Bacteria | Bacteroidetes | Bacteroidia | Flavobacteriales | NS9 marine group | uncultured marine bacterium | uncultured marine bacterium | 0.993 |
| 6c2be4e95eb066d84141a5e3e59ecca1 | Bacteria | Proteobacteria | Alphaproteobacteria | Rickettsiales | Midichloriaceae | MD3-55 | uncultured alpha proteobacterium | 0.835 |
| 10006b208ccb6988dd4b8f02ac0f9040 | Bacteria | Proteobacteria | Gammaproteobacteria | Thiomicrospirales | Thioglobaceae | SUP05 cluster | | 1.000 |
| f05b1386fdb55c65a7622c8aeba21ef3 | Bacteria | Bacteroidetes | Bacteroidia | Flavobacteriales | Flavobacteriaceae | NS4 marine group | | 1.000 |
| e31241d8eb3c94aac7fcf4f94980e7c4 | Bacteria | Proteobacteria | Deltaproteobacteria | Bdellovibrionales | Bdellovibrionaceae | OM27 clade | unidentified marine bacterioplankton | 1.000 |
| 0b9bb4a0e000955919cacd0cb535c459 | Bacteria | Cyanobacteria | Oxyphotobacteria | Chloroplast | | | | 1.000 |
| 0481fc952cc4e413f80a75e3195f7a44 | Bacteria | Cyanobacteria | Oxyphotobacteria | Chloroplast | Epipyxis sp. PR26KG | Epipyxis sp. PR26KG | Epipyxis sp. PR26KG | 0.826 |
| 50b4087cc1b6c678359ef79a31560fb4 | Bacteria | Proteobacteria | Deltaproteobacteria | Myxococcales | Blfdi19 | Ambiguous_taxa | Ambiguous_taxa | 1.000 |
| 0ea6807cfbd229bdb90a2cc69096c96e | Bacteria | Proteobacteria | Alphaproteobacteria | uncultured | | | | 1.000 |
| 9ac94f2186f68c25f52ded40a6cb1a78 | Bacteria | Proteobacteria | Alphaproteobacteria | Puniceispirillales | SAR116 clade | Ambiguous_taxa | Ambiguous_taxa | 0.903 |

| Feature_ID | Kingdom | Phylum | Class | Order | Family | Genus | Species | Confidence |
|---|---|---|---|---|---|---|---|---|
| 3eaa11d4571a5423fe23f0fb25c7e07c | Bacteria | Cyanobacteria | Oxyphotobacteria | Chloroplast | | | | 1.000 |
| 63cba9f02bdbbb9d63a7ed8f40e76d4d | Bacteria | Marinimicrobia (SAR406 clade) | | | | | | 1.000 |
| 7b1c1b2c651a16bffa3eab3367ef8e0d | Bacteria | Proteobacteria | Gammaproteobacteria | KI89A clade | | | | 1.000 |
| 64534c5c9aa3a7575a34bcc33007ab84 | Bacteria | Cyanobacteria | Oxyphotobacteria | Chloroplast | Phalacroma mitra | Phalacroma mitra | Phalacroma mitra | 0.830 |
| c32dca4586fdbb79e6245dd8a6a97a22 | Bacteria | Proteobacteria | Deltaproteobacteria | SAR324 clade(Marine group B) | | | | 1.000 |
| e08c6823fd9536f248fd6fdb99e9207c | Bacteria | Proteobacteria | Deltaproteobacteria | PB19 | Ambiguous_taxa | Ambiguous_taxa | Ambiguous_taxa | 0.708 |
| dfc250acebc32e454acaf152ab233172 | Bacteria | Proteobacteria | Deltaproteobacteria | NB1-j | uncultured prokaryote | uncultured prokaryote | uncultured prokaryote | 0.837 |
| 0d7caaeae72516a70dcb3b57dbbfcc5a | Bacteria | Proteobacteria | Alphaproteobacteria | Rickettsiales | S25-593 | uncultured marine bacterium | uncultured marine bacterium | 0.703 |
| ff9333e98d509d20f0f183ffe965a410 | Bacteria | Proteobacteria | Alphaproteobacteria | Puniceispirillales | SAR116 clade | | | 1.000 |
| 72b5518687c102c26151b9034e15d264 | Bacteria | Cyanobacteria | Oxyphotobacteria | Chloroplast | Phalacroma mitra | Phalacroma mitra | Phalacroma mitra | 0.763 |
| 1b156a8899662c0e4e1df7946ca7a072 | Bacteria | Chloroflexi | Dehalococcoidia | SAR202 clade | uncultured Chloroflexi bacterium | uncultured Chloroflexi bacterium | uncultured Chloroflexi bacterium | 0.947 |
| ad50aadf132d31c1db49a5554f9fe7e1 | Bacteria | Verrucomicrobia | Verrucomicrobiae | Opitutales | Puniceicoccaceae | Coraliomargarita | | 0.995 |
| c8e79e64c10b78a2dcc2e0e9f46871b5 | Bacteria | Bacteroidetes | Bacteroidia | Flavobacteriales | Flavobacteriaceae | NS2b marine group | | 1.000 |
| 220a0f7aaf106aa7fa6e2d77315a5a67 | Bacteria | Proteobacteria | Gammaproteobacteria | Vibrionales | Vibrionaceae | Vibrio | | 0.999 |
| a6f8819eb9e3765cc985d6389d41d7b9 | Bacteria | Planctomycetes | Phycisphaerae | Phycisphaerales | Phycisphaeraceae | Urania-1B-19 marine sediment group | Ambiguous_taxa | 1.000 |
| b6970bf611c0d72299d098b23d00c601 | Bacteria | Proteobacteria | Gammaproteobacteria | Thiotrichales | Thiotrichaceae | uncultured | uncultured marine bacterium | 1.000 |
| 388045997bbe90966cae3f044db6ec40 | Bacteria | Proteobacteria | Alphaproteobacteria | uncultured | unidentified marine bacterioplankton | unidentified marine bacterioplankton | unidentified marine bacterioplankton | 0.965 |
| fb05c383e039e97cbe277b5ed000729a | Bacteria | Proteobacteria | Gammaproteobacteria | Betaproteobacteriales | Burkholderiaceae | MWH-UniP1 aquatic group | uncultured marine bacterium | 0.987 |
| 88cd7aa61edb6556a55cc48b2ae9fed7 | Bacteria | Proteobacteria | Gammaproteobacteria | Oceanospirillales | Halomonadaceae | Salinicola | | 1.000 |
| 9f66e6c672a6dec59a2f9ac846525448 | Bacteria | Verrucomicrobia | Verrucomicrobiae | Verrucomicrobiales | Rubritaleaceae | Roseibacillus | uncultured marine bacterium | 0.962 |
| c95a8f8b0c5899b80bff68debb92a298 | Bacteria | Proteobacteria | Alphaproteobacteria | uncultured | unidentified marine bacterioplankton | unidentified marine bacterioplankton | unidentified marine bacterioplankton | 0.940 |
| a7e605d7248513cba8f445bf1ddf389c | Bacteria | Epsilonbacteraeota | Campylobacteria | Campylobacterales | Arcobacteraceae | Arcobacter | uncultured bacterium | 0.971 |
| 239ba189230f81170e8f127ec44ddade | Bacteria | Proteobacteria | Deltaproteobacteria | Desulfarculales | Desulfarculaceae | uncultured | uncultured delta proteobacterium | 1.000 |
| 6687d307cbf76d544c2ce5f730214e78 | Bacteria | Bacteroidetes | Bacteroidia | Flavobacteriales | Flavobacteriaceae | NS5 marine group | Ambiguous_taxa | 0.734 |
| 9414f21cd74d30c55dfd4e1722835236 | Bacteria | Cyanobacteria | Oxyphotobacteria | Chloroplast | Phalacroma mitra | Phalacroma mitra | Phalacroma mitra | 0.871 |
| dee3cd038831e45b444ddce5206a7d18 | Bacteria | Proteobacteria | Alphaproteobacteria | Rhodospirillales | Magnetospiraceae | uncultured | uncultured Rhodospirillaceae bacterium | 0.718 |
| 8e8dde2d3b5b6b20f62754849d02f66d | Bacteria | Proteobacteria | Gammaproteobacteria | Thiomicrospirales | Thioglobaceae | SUP05 cluster | uncultured marine bacterium | 0.780 |
| 0bcc04aceaef8b5f35e42adae728149e | Bacteria | Proteobacteria | Gammaproteobacteria | UBA10353 marine group | Ambiguous_taxa | Ambiguous_taxa | Ambiguous_taxa | 0.987 |
| d9b0bb6a47ec18c7e5e0435ed3b15dbe | Bacteria | Cyanobacteria | Oxyphotobacteria | Chloroplast | Epipyxis sp. PR26KG | Epipyxis sp. PR26KG | Epipyxis sp. PR26KG | 0.712 |
| d5f19299f9d0049719548317dd53bcc8 | Bacteria | Proteobacteria | Gammaproteobacteria | KI89A clade | Ambiguous_taxa | Ambiguous_taxa | Ambiguous_taxa | 0.934 |
| 34606f52693436d00f15fed60f92f990 | Bacteria | Proteobacteria | Alphaproteobacteria | Rhodobacterales | Rhodobacteraceae | uncultured | | 0.960 |
| a35fa0e36211b8c532c11e3912663742 | Bacteria | Proteobacteria | Alphaproteobacteria | Rhodospirillales | AEGEAN-169 marine group | | | 1.000 |
| d59742a48c41856d1bb241940c9824ed | Bacteria | Verrucomicrobia | Verrucomicrobiae | Opitutales | Puniceicoccaceae | MB11C04 marine group | | 1.000 |
| 0aa35bb423ff0b679e9bbc3118e38238 | Bacteria | Bacteroidetes | Bacteroidia | Flavobacteriales | Flavobacteriaceae | NS5 marine group | | 0.833 |
| 1b7981b73440119740be00aa57a15ffb | Bacteria | Proteobacteria | Deltaproteobacteria | NB1-j | uncultured marine bacterium | uncultured marine bacterium | uncultured marine bacterium | 0.999 |
| b95b5f64e7d69286efcb393af1459743 | Bacteria | Bacteroidetes | Bacteroidia | Flavobacteriales | Flavobacteriaceae | Flavicella | uncultured bacterium | 0.992 |
| b9abee5f6a84c79a844d2559cf468992 | Bacteria | Proteobacteria | Alphaproteobacteria | Rickettsiales | S25-593 | uncultured marine bacterium | uncultured marine bacterium | 0.854 |
| 43dac0ed1e740bffb10d3155cb781a2c | Bacteria | Bacteroidetes | Bacteroidia | Flavobacteriales | NS9 marine group | uncultured marine bacterium | uncultured marine bacterium | 0.987 |
| 39666b64f1359f6e9cf2d89c00fc3eb2 | Bacteria | Proteobacteria | Gammaproteobacteria | Alteromonadales | Pseudoalteromonadaceae | Pseudoalteromonas | | 1.000 |
| 8957bb3386795e1251684fa5904255b3 | Bacteria | Bacteroidetes | Bacteroidia | Flavobacteriales | Flavobacteriaceae | Aquibacter | Ambiguous_taxa | 0.988 |
| 3ed40e38620c9d25aa427651d0366110 | Bacteria | Cyanobacteria | Oxyphotobacteria | Chloroplast | uncultured diatom | uncultured diatom | uncultured diatom | 0.865 |
| 8a2f347867e5521ca27c6e063ae87414 | Bacteria | Bacteroidetes | Bacteroidia | Flavobacteriales | Flavobacteriaceae | Flavicella | uncultured Flavobacteriaceae bacterium | 0.841 |
| 59f65c6f8a89ea793ed529d79170f19f | Bacteria | Proteobacteria | Deltaproteobacteria | Bdellovibrionales | Bdellovibrionaceae | OM27 clade | uncultured marine bacterium | 0.775 |
| f5c5cb218d1e635bd299562ec6ef20fe | Bacteria | Cyanobacteria | Oxyphotobacteria | Chloroplast | uncultured prymnesiophyte C5574 | uncultured prymnesiophyte C5574 | uncultured prymnesiophyte C5574 | 0.792 |
| a02578e4deb6b1fdfb3482731ff6bfaa | Bacteria | Planctomycetes | Planctomycetacia | Pirellulales | Pirellulaceae | Blastopirellula | uncultured marine bacterium | 0.939 |
| e58f94718a486e75a8882088a4e933b2 | Bacteria | Cyanobacteria | Oxyphotobacteria | Chloroplast | uncultured bacterium | uncultured bacterium | uncultured bacterium | 0.788 |
| ca9e427c68a4edb76f10f2d1e946ed93 | Bacteria | Proteobacteria | Alphaproteobacteria | Rhodospirillales | Thalassospiraceae | Thalassospira | | 0.997 |
| 30a762fd54aeea1651783f9be433f90a | Archaea | Euryarchaeota | Thermoplasmata | Marine Group II | marine metagenome | marine metagenome | marine metagenome | 0.951 |
| 8602c264af08d04c8f2ae249be302f28 | Bacteria | Proteobacteria | Deltaproteobacteria | Myxococcales | Blfdi19 | Ambiguous_taxa | Ambiguous_taxa | 1.000 |
| c57b019ae5ef2a6ed2500f52334612fa | Bacteria | Proteobacteria | Gammaproteobacteria | Coxiellales | Coxiellaceae | Coxiella | uncultured bacterium | 0.989 |
| d4c70d3cdb480b123d8852f87399dcbd | Bacteria | Cyanobacteria | Oxyphotobacteria | Chloroplast | | | | 1.000 |
| 7cf008337c9da74f49339bc1f0e0f638 | Bacteria | Proteobacteria | Deltaproteobacteria | SAR324 clade(Marine group B) | | | | 1.000 |
| 0025d4e026e55b3aab0e7cb7c3f58a9c | Bacteria | Proteobacteria | Alphaproteobacteria | Caulobacterales | Caulobacteraceae | Brevundimonas | | 1.000 |
| df2c834919c479cfdc315d2f6c66176e | Bacteria | Cyanobacteria | Oxyphotobacteria | Chloroplast | uncultured marine eukaryote | uncultured marine eukaryote | uncultured marine eukaryote | 0.829 |
| 7157ed43df2ff14eeadc91537f662d79 | Bacteria | Proteobacteria | Deltaproteobacteria | Bdellovibrionales | Bdellovibrionaceae | OM27 clade | | 1.000 |
| aaf9cfef609895e3ee54126a19296d91 | Bacteria | Marinimicrobia (SAR406 clade) | uncultured marine bacterium | uncultured marine bacterium | uncultured marine bacterium | uncultured marine bacterium | uncultured marine bacterium | 0.946 |
| 6bc7512f71287465a5050495ab7b079e | Bacteria | Proteobacteria | Gammaproteobacteria | Alteromonadales | | | | 0.981 |
| e3e519c01594b121dee8dd03cbd2b201 | Bacteria | Bacteroidetes | Bacteroidia | Flavobacteriales | Flavobacteriaceae | NS5 marine group | | 1.000 |
| dc034c38f5f3c7fad8bdd3932ad6450b | Bacteria | Proteobacteria | Gammaproteobacteria | Alteromonadales | Idiomarinaceae | Idiomarina | | 1.000 |
| f08120a4f490c4d29ea6952a941b7d27 | Bacteria | Proteobacteria | Gammaproteobacteria | Ectothiorhodospirales | Ectothiorhodospiraceae | uncultured | uncultured marine bacterium | 0.935 |
| a0bfae3e3ba805032859313f80519a12 | Bacteria | Nitrospinae | P9X2b3D02 | Candidatus Tectomicrobia bacterium RIFCSPLOWO2_12_FULL_69_37 | Candidatus Tectomicrobia bacterium RIFCSPLOWO2_12_FULL_69_37 | Candidatus Tectomicrobia bacterium RIFCSPLOWO2_12_FULL_69_37 | Candidatus Tectomicrobia bacterium RIFCSPLOWO2_12_FULL_69_37 | 0.999 |
| e0fa2fd0a5bad4ad7b812eddb7eaa78e | Bacteria | Proteobacteria | Alphaproteobacteria | SAR11 clade | Clade II | | | 0.840 |
| fb6ec7a317dd602dff316484c2be9abf | Bacteria | Marinimicrobia (SAR406 clade) | uncultured bacterium | uncultured bacterium | uncultured bacterium | uncultured bacterium | uncultured bacterium | 0.938 |
| 7cdd7a6f6a2118fdf61c22899f31b5d7 | Bacteria | Cyanobacteria | Oxyphotobacteria | Chloroplast | Phalacroma mitra | Phalacroma mitra | Phalacroma mitra | 0.939 |
| 580db4d4bd0cf21c127ec2b011e0a0bd | Bacteria | Proteobacteria | Deltaproteobacteria | Bdellovibrionales | Bdellovibrionaceae | OM27 clade | | 1.000 |
| a4fecb717e471eaac52e143d2379573d | Bacteria | Marinimicrobia (SAR406 clade) | uncultured bacterium | uncultured bacterium | uncultured bacterium | uncultured bacterium | uncultured bacterium | 0.748 |
| 9f513d43bdf034d4adb1c2a8324d2d3f | Bacteria | Proteobacteria | Gammaproteobacteria | SAR86 clade | | | | 1.000 |
| ca46261eee68bd963df824c3a3d92139 | Bacteria | Patescibacteria | Gracilibacteria | Candidatus Peregrinibacteria | uncultured bacterium | uncultured bacterium | uncultured bacterium | 0.999 |

| Feature_ID | Kingdom | Phylum | Class | Order | Family | Genus | Species | Confidence |
|---|---|---|---|---|---|---|---|---|
| f162459a1558d8b818b4744e9eca5186 | Bacteria | Cyanobacteria | Oxyphotobacteria | Chloroplast | | | | 1.000 |
| b851e7a5c19adc9e1d13ce7541f92a22 | Bacteria | Proteobacteria | Alphaproteobacteria | Rickettsiales | Mitochondria | uncultured bacterium | uncultured bacterium | 0.886 |
| f80ed8ce09d5c92170bbfca86fdc60dc | Bacteria | Proteobacteria | Deltaproteobacteria | Oligoflexales | Oligoflexaceae | uncultured | uncultured marine bacterium | 0.925 |
| af74e689e59b7dca53e3124153475820 | Bacteria | Bacteroidetes | Bacteroidia | Flavobacteriales | Flavobacteriaceae | NS5 marine group | | 1.000 |
| 6fedacc082b8ca9d6a3bb420dc75e639 | Bacteria | Cyanobacteria | Oxyphotobacteria | Chloroplast | Phalacroma mitra | Phalacroma mitra | Phalacroma mitra | 0.937 |
| 637d25313b426e43102f213ef262bc3e | Bacteria | Proteobacteria | Alphaproteobacteria | uncultured | uncultured marine bacterium | uncultured marine bacterium | uncultured marine bacterium | 0.956 |
| d6eff63ee5dda4a2a45c0965480d9f24 | Bacteria | Proteobacteria | Deltaproteobacteria | Oligoflexales | Oligoflexaceae | uncultured | uncultured bacterium | 1.000 |
| 771805b755687efc965b188e88440a84 | Bacteria | Marinimicrobia (SAR406 clade) | | | | | | 1.000 |
| cc1c4244ac5cd5835a0164d94b5e3373 | Bacteria | Chloroflexi | Dehalococcoidia | SAR202 clade | uncultured Chloroflexi bacterium HF4000_28F02 | uncultured Chloroflexi bacterium HF4000_28F02 | uncultured Chloroflexi bacterium HF4000_28F02 | 1.000 |
| 88e8503f537f9e169ffdc83985042ee2 | Bacteria | Bacteroidetes | Bacteroidia | Flavobacteriales | NS9 marine group | uncultured Flavobacteriia bacterium | uncultured Flavobacteriia bacterium | 0.991 |
| 6b4faa79fdf4a7e509a421ddeb16d37e | Bacteria | Proteobacteria | Gammaproteobacteria | OM182 clade | uncultured bacterium | uncultured bacterium | uncultured bacterium | 0.895 |
| ee763c29f15b2a475081c27a81f372e2 | Bacteria | Marinimicrobia (SAR406 clade) | unidentified marine bacterioplankton | unidentified marine bacterioplankton | unidentified marine bacterioplankton | unidentified marine bacterioplankton | unidentified marine bacterioplankton | 0.977 |
| e290d868f19d48e068affd0982444191 | Bacteria | Verrucomicrobia | Verrucomicrobiae | Opitutales | Puniceicoccaceae | MB11C04 marine group | | 1.000 |
| edca081e502f2fb05c1515d43fb0520f | Bacteria | Cyanobacteria | Oxyphotobacteria | Chloroplast | uncultured marine eukaryote | uncultured marine eukaryote | uncultured marine eukaryote | 0.878 |
| cb4e56f878a02942c3ebc105a19fb27f | Bacteria | Verrucomicrobia | Verrucomicrobiae | Opitutales | Puniceicoccaceae | MB11C04 marine group | | 1.000 |
| 9b4e34ffda88c9cf44066d920afd299c | Bacteria | Bacteroidetes | Bacteroidia | Flavobacteriales | Cryomorphaceae | uncultured | metagenome | 1.000 |
| 31ed31c06a369469e1bc5be9f171ef9a | Bacteria | Marinimicrobia (SAR406 clade) | | | | | | 1.000 |
| 027a873fcb6b062795b762d5530bdd8d | Bacteria | Proteobacteria | Gammaproteobacteria | Coxiellales | Coxiellaceae | Coxiella | uncultured Legionellales bacterium | 0.856 |
| 6228d1ad2221fba614c514cd6c491283 | Bacteria | Proteobacteria | Alphaproteobacteria | Rickettsiales | S25-593 | uncultured marine bacterium | uncultured marine bacterium | 0.975 |
| ee1e4f823448199e4530f6da89491365 | Bacteria | Cyanobacteria | Oxyphotobacteria | Synechococcales | Cyanobiaceae | Cyanobium PCC-6307 | | 1.000 |
| 00c181aba51117925b1ef5d41bdb2894 | Bacteria | Proteobacteria | Gammaproteobacteria | HOC36 | Candidatus Thioglobus sp. REDSEA-S12_B1 | Candidatus Thioglobus sp. REDSEA-S12_B1 | Candidatus Thioglobus sp. REDSEA-S12_B1 | 1.000 |
| 4b145ed5300b3671fe4c469872f6e7e2 | Bacteria | Proteobacteria | Alphaproteobacteria | SAR11 clade | Clade II | | | 0.876 |
| 88a94fba34bacac07d76902215ea8373 | Bacteria | Proteobacteria | Alphaproteobacteria | Rhodospirillales | AEGEAN-169 marine group | uncultured bacterium | uncultured bacterium | 1.000 |
| 38cb37725dfacb60970083a288351883 | Bacteria | Proteobacteria | Gammaproteobacteria | Cellvibrionales | Halieaceae | OM60(NOR5) clade | | 0.882 |
| 0f0b97057a67130d2e47ba1dd24c4d5f | Bacteria | Bacteroidetes | Bacteroidia | Flavobacteriales | NS9 marine group | | | 1.000 |
| 25d7d6b9575def80901aa1527dca456c | Bacteria | Proteobacteria | Gammaproteobacteria | Alteromonadales | Pseudoalteromonadaceae | Pseudoalteromonas | | 1.000 |
| 6517d7c89575ed21d516a34b791d61ae | Bacteria | Bacteroidetes | Bacteroidia | Flavobacteriales | Flavobacteriaceae | Pseudofulvibacter | | 0.975 |
| 620d7368c912e4ff0e74270d877f0c99 | Bacteria | Proteobacteria | Deltaproteobacteria | Bdellovibrionales | Bdellovibrionaceae | OM27 clade | uncultured marine bacterium | 0.985 |
| 07d4cd7ab52f9d37be844ad18261173b | Bacteria | Cyanobacteria | Oxyphotobacteria | Chloroplast | Phalacroma mitra | Phalacroma mitra | Phalacroma mitra | 0.968 |
| 88e5ba79e63b5563a1bf8261ead3612e | Bacteria | Bacteroidetes | Bacteroidia | Flavobacteriales | NS9 marine group | uncultured marine bacterium | uncultured marine bacterium | 0.991 |
| 451eda6352291763ca4a5ffe434611d7 | Bacteria | Bacteroidetes | Bacteroidia | Flavobacteriales | Flavobacteriaceae | | | 1.000 |
| c2164bd99dc5c74f0034724a044f5111 | Bacteria | Verrucomicrobia | Verrucomicrobiae | Arctic97B-4 marine group | metagenome | metagenome | metagenome | 0.982 |
| c29a30bed46a3f896ba42d082b5c3276 | Bacteria | Verrucomicrobia | Verrucomicrobiae | Arctic97B-4 marine group | | | | 1.000 |
| 6e0f61eeffdbec98093160499d2d6617 | Bacteria | Proteobacteria | Alphaproteobacteria | uncultured | uncultured marine bacterium | uncultured marine bacterium | uncultured marine bacterium | 0.923 |
| a5845fe981a4dec16fa256ba66166f0d | Bacteria | Bacteroidetes | Bacteroidia | Flavobacteriales | Flavobacteriaceae | NS5 marine group | uncultured bacterium | 0.779 |
| f1878dc4551507d4eb7ce32ab19ed4b1 | Bacteria | Marinimicrobia (SAR406 clade) | uncultured bacterium | uncultured bacterium | uncultured bacterium | uncultured bacterium | uncultured bacterium | 0.883 |
| 8fd69e37e3d162a32d0060c44876aa65 | Bacteria | Proteobacteria | Deltaproteobacteria | Bdellovibrionales | Bdellovibrionaceae | OM27 clade | uncultured marine bacterium | 0.975 |
| e4167abfa3a7e3f4cc679131bde17c07 | Bacteria | Cyanobacteria | Oxyphotobacteria | Chloroplast | Aureoumbra lagunensis | Aureoumbra lagunensis | Aureoumbra lagunensis | 0.739 |
| d374259f696caa3d2d186213f65bf358 | Bacteria | Chloroflexi | Dehalococcoidia | SAR202 clade | hydrothermal vent metagenome | hydrothermal vent metagenome | hydrothermal vent metagenome | 0.998 |
| 32dd08a35d3d54d6ac3ec41e00d2463f | Bacteria | Proteobacteria | Alphaproteobacteria | SAR11 clade | Clade II | unidentified marine bacterioplankton | unidentified marine bacterioplankton | 0.740 |
| 9086635cdc238f32f1c29767b6af575c | Bacteria | Proteobacteria | Alphaproteobacteria | SAR11 clade | Clade II | | | 0.910 |
| ae052211ae38f61e97050cd585d50c8e | Bacteria | Bacteroidetes | Bacteroidia | Chitinophagales | uncultured | Ambiguous_taxa | Ambiguous_taxa | 1.000 |
| 42d4a272cbf36bbf4c61ee5d048eca0a | Bacteria | Bacteroidetes | Bacteroidia | Flavobacteriales | NS7 marine group | | | 1.000 |
| e4bf9639d8597708045bc47b479fd3ec | Bacteria | Bacteroidetes | Bacteroidia | Flavobacteriales | Flavobacteriaceae | NS4 marine group | | 1.000 |
| 9127d993e81bdac48f41aa11e10ab8da | Bacteria | Bacteroidetes | Bacteroidia | Flavobacteriales | Cryomorphaceae | uncultured | uncultured Owenweeksia sp. | 0.960 |
| d0fb0be18a4aa56b9fae7fdd3f14f2d8 | Bacteria | Proteobacteria | Deltaproteobacteria | Myxococcales | P3OB-42 | uncultured marine bacterium | uncultured marine bacterium | 1.000 |
| 0dcac9c9a7c4da86de187e4154d96bbf | Bacteria | Bacteroidetes | Bacteroidia | Flavobacteriales | NS9 marine group | | | 0.867 |
| cc5a556f71fc75e1ef452eaf7f299de8 | Bacteria | Proteobacteria | Gammaproteobacteria | Gammaproteobacteria Incertae Sedis | Unknown Family | uncultured marine bacterium | uncultured marine bacterium | 1.000 |
| 6ed2144335fd13a44ea88ad1647c14e1 | Bacteria | Margulisbacteria | marine metagenome | marine metagenome | marine metagenome | marine metagenome | marine metagenome | 1.000 |
| 1c3b99535e970758789ca0d3917d3407 | Bacteria | Proteobacteria | Gammaproteobacteria | OM182 clade | uncultured gamma proteobacterium | uncultured gamma proteobacterium | uncultured gamma proteobacterium | 0.999 |
| a1abf6d2e203cc96084ceae2ab70ed07 | Bacteria | Cyanobacteria | Oxyphotobacteria | Chloroplast | uncultured bacterium | uncultured bacterium | uncultured bacterium | 0.778 |
| b76b8b9404d2545eb85227a25a40c01c | Bacteria | Proteobacteria | Gammaproteobacteria | Cellvibrionales | Halieaceae | Parahaliea | uncultured marine bacterium | 0.993 |
| 81df44c26a3077c53d93af81be9c72b2 | Bacteria | Chloroflexi | Dehalococcoidia | SAR202 clade | Ambiguous_taxa | Ambiguous_taxa | Ambiguous_taxa | 0.999 |
| 3fcb84cdb86b6d60a2067e3a3975a5e2 | Bacteria | Proteobacteria | Alphaproteobacteria | Rhodospirillales | Magnetospiraceae | uncultured | | 1.000 |
| 63d4c67c563ddcc06ce4250b3dd5dd7c | Bacteria | Bacteroidetes | Bacteroidia | Flavobacteriales | NS9 marine group | uncultured marine bacterium | uncultured marine bacterium | 0.772 |
| 0ed5ec5fcda63e9ce8541d59cb7bab97 | Bacteria | Cyanobacteria | Oxyphotobacteria | Chloroplast | Phalacroma mitra | Phalacroma mitra | Phalacroma mitra | 0.737 |
| 5da424385c3728994cf9ef85b841c43a | Bacteria | Chloroflexi | Dehalococcoidia | SAR202 clade | Ambiguous_taxa | Ambiguous_taxa | Ambiguous_taxa | 1.000 |
| b527099585068309fd2fe03ca738a52e | Bacteria | Marinimicrobia (SAR406 clade) | uncultured marine bacterium | uncultured marine bacterium | uncultured marine bacterium | uncultured marine bacterium | uncultured marine bacterium | 0.870 |
| 2266168a6e92d138ec6617f4fc43c4c4 | Bacteria | Firmicutes | Clostridia | Clostridiales | Ruminococcaceae | Ruminococcus 2 | | 1.000 |
| 2917e02786dbb9fa5529c90489d369b4 | Bacteria | Proteobacteria | Alphaproteobacteria | Puniceispirillales | SAR116 clade | | | 1.000 |
| d13f6e32063aa6e9ef787e7f45e0c381 | Bacteria | Proteobacteria | Deltaproteobacteria | Oligoflexales | Oligoflexaceae | uncultured | uncultured marine bacterium | 0.996 |
| 9b8029d59106a1eebe1ae917a928d813 | Bacteria | Proteobacteria | Gammaproteobacteria | | | | | 1.000 |
| b5179407010d40364159775445436411 | Bacteria | Marinimicrobia (SAR406 clade) | marine metagenome | marine metagenome | marine metagenome | marine metagenome | marine metagenome | 0.880 |
| 40e30ba7bf07eaca5ecde4a9537f1793 | Bacteria | Bacteroidetes | Bacteroidia | Flavobacteriales | Flavobacteriaceae | NS4 marine group | | 1.000 |
| ebba845db68505a5c2ca03c99d913afc | Bacteria | Proteobacteria | Gammaproteobacteria | Cellvibrionales | Halieaceae | Parahaliea | uncultured gamma proteobacterium | 0.914 |
| 726b8e13e3fa04830245dfee0c8e0e43 | Bacteria | Cyanobacteria | Oxyphotobacteria | Chloroplast | uncultured marine eukaryote | uncultured marine eukaryote | uncultured marine eukaryote | 0.785 |
| 455d144a95f5df6e89797c0ba9d1a8f8 | Bacteria | Bacteroidetes | Bacteroidia | Flavobacteriales | Flavobacteriaceae | Pseudofulvibacter | uncultured bacterium | 0.770 |
| 1826547cb4f9a711ac6ce0c4813cd62a | Bacteria | Bacteroidetes | Bacteroidia | Flavobacteriales | NS9 marine group | uncultured marine bacterium | uncultured marine bacterium | 0.950 |
| 22eb0b7ab3d074ab16074d9319572543 | Bacteria | Bacteroidetes | Bacteroidia | Flavobacteriales | NS9 marine group | uncultured marine bacterium | uncultured marine bacterium | 0.990 |

| Feature_ID | Kingdom | Phylum | Class | Order | Family | Genus | Species | Confidence |
|---|---|---|---|---|---|---|---|---|
| 4894ce1485e4097610534ff85c58107f | Bacteria | Marinimicrobia (SAR406 clade) | uncultured eubacterium OCS307 | uncultured eubacterium OCS307 | uncultured eubacterium OCS307 | uncultured eubacterium OCS307 | uncultured eubacterium OCS307 | 1.000 |
| a2934032a92509a20ee4a6a828e680ba | Bacteria | Bacteroidetes | Bacteroidia | Flavobacteriales | Cryomorphaceae | NS10 marine group | | 1.000 |
| 80c777f382d668114b2a202528eede79 | Bacteria | Marinimicrobia (SAR406 clade) | uncultured marine bacterium | uncultured marine bacterium | uncultured marine bacterium | uncultured marine bacterium | uncultured marine bacterium | 0.989 |
| f1e4d87808760a9f1c3623bde9791889 | Bacteria | Proteobacteria | Gammaproteobacteria | UBA10353 marine group | Ambiguous_taxa | Ambiguous_taxa | Ambiguous_taxa | 0.998 |
| 85aaba025c8d933e4669bf383e8c48cb | Bacteria | Bacteroidetes | Bacteroidia | Flavobacteriales | Flavobacteriaceae | NS5 marine group | | 1.000 |
| c56cee2a8d5aa9e9a1f8e5e570e0fda6 | Bacteria | Proteobacteria | Gammaproteobacteria | Vibrionales | Vibrionaceae | Photobacterium | | 0.999 |
| bedff13cff8cf4bc1edc1ea136268541 | Bacteria | Proteobacteria | Deltaproteobacteria | SAR324 clade(Marine group B) | uncultured bacterium | uncultured bacterium | uncultured bacterium | 0.819 |
| e7e36d3ba9a67f9338ae78bf1694a6e5 | Bacteria | Proteobacteria | Gammaproteobacteria | | | | | 1.000 |
| 506b89e9c4e106a4e9f44ff399b131f4 | Bacteria | Proteobacteria | Alphaproteobacteria | Rickettsiales | S25-593 | | | 1.000 |
| 614bfd70d138c55999f1a85ef1de7822 | Bacteria | Bacteroidetes | Bacteroidia | Flavobacteriales | Crocinitomicaceae | Salinirepens | Ambiguous_taxa | 0.986 |
| 069529b71f2d7f4aa2c873f425e6a9de | Bacteria | Proteobacteria | Gammaproteobacteria | Cellvibrionales | Spongiibacteraceae | BD1-7 clade | uncultured gamma proteobacterium | 0.999 |
| c16038743097bb323a839e8b6f1bdb7e | Bacteria | Proteobacteria | Gammaproteobacteria | Coxiellales | Coxiellaceae | Coxiella | | 1.000 |
| 9ec187f02620af7343e43e2e4a4a2634 | Bacteria | Bacteroidetes | Bacteroidia | Flavobacteriales | Flavobacteriaceae | uncultured | uncultured Flavobacteriales bacterium | 0.774 |
| 960a44e0d9d81685fc52c43cf25ca58f | Bacteria | Proteobacteria | Gammaproteobacteria | EPR3968-O8a-Bc78 | uncultured marine bacterium | uncultured marine bacterium | uncultured marine bacterium | 1.000 |
| 1c1f30a08a71956b5f5b8c5ee204cd08 | Bacteria | Bacteroidetes | Bacteroidia | Flavobacteriales | Flavobacteriaceae | NS5 marine group | | 1.000 |
| 88968e43eec79a677d51c7863bb839bb | Bacteria | Bacteroidetes | Bacteroidia | Flavobacteriales | NS9 marine group | uncultured marine bacterium | uncultured marine bacterium | 0.972 |
| 3b0a8656f57e74f096a2788c033fc5be | Bacteria | Proteobacteria | Deltaproteobacteria | Bdellovibrionales | Bdellovibrionaceae | OM27 clade | uncultured marine bacterium | 0.962 |
| 113cf225927980de5a96ba247fbaee96 | Bacteria | Planctomycetes | Planctomycetacia | Pirellulales | Pirellulaceae | Rhodopirellula | hydrothermal vent metagenome | 0.995 |
| 258ef60630a4b8f8b4b6e564a4f99102 | Bacteria | Verrucomicrobia | Verrucomicrobiae | Arctic97B-4 marine group | | | | 1.000 |
| 9ac2bce222bd253c77af2751142f4f47 | Bacteria | Verrucomicrobia | Verrucomicrobiae | Arctic97B-4 marine group | Ambiguous_taxa | Ambiguous_taxa | Ambiguous_taxa | 0.998 |
| 13bebf1e665953a4be970862ebffd837 | Bacteria | Verrucomicrobia | Verrucomicrobiae | Opitutales | Puniceicoccaceae | Pelagicoccus | uncultured marine bacterium | 0.999 |
| 00fc329ff5060ebf8b997f77edc0093c | Bacteria | Bacteroidetes | Bacteroidia | Flavobacteriales | NS9 marine group | | | 1.000 |
| 71ac71a7ad69a78af36bfe17775ad65c | Bacteria | Bacteroidetes | Bacteroidia | Flavobacteriales | NS9 marine group | uncultured Cytophagaceae bacterium | uncultured Cytophagaceae bacterium | 0.872 |
| 5b41fc61b067b7d335dc2beb8465b375 | Bacteria | Bacteroidetes | Bacteroidia | Flavobacteriales | NS9 marine group | uncultured marine bacterium | uncultured marine bacterium | 0.974 |
| a636832e0bc8d549b0d604b9291133c3 | Bacteria | Proteobacteria | Deltaproteobacteria | Bdellovibrionales | Bdellovibrionaceae | OM27 clade | | 1.000 |
| 2a397771272ab7b3ef0774e6be2cabd9 | Bacteria | Bacteroidetes | Bacteroidia | Flavobacteriales | NS9 marine group | Ambiguous_taxa | Ambiguous_taxa | 0.826 |
| f7f4d2293a1c32b3662687bc2c01ee90 | Bacteria | Cyanobacteria | Oxyphotobacteria | Chloroplast | | | | 1.000 |
| 7dd3449b4a59f1b4935e78e10c5562a4 | Bacteria | Proteobacteria | Deltaproteobacteria | Myxococcales | Sandaracinaceae | uncultured | uncultured marine bacterium | 1.000 |
| 71ed703262a4b122e89dd7b50c4cf8fe | Bacteria | Proteobacteria | Gammaproteobacteria | Gammaproteobacteria Incertae Sedis | Unknown Family | uncultured marine bacterium | uncultured marine bacterium | 1.000 |
| 90cad10504185809da7fa842dcc5724d | Bacteria | Proteobacteria | Alphaproteobacteria | SAR11 clade | | | | 1.000 |
| 9724d2faa939dad49f6a6452304feed1 | Bacteria | Bacteroidetes | Bacteroidia | Flavobacteriales | Flavobacteriaceae | NS5 marine group | uncultured Bacteroidetes bacterium | 0.796 |
| 95920d5edeaca8b3d15edb278f9d92be | Bacteria | Cyanobacteria | Oxyphotobacteria | Chloroplast | Prasinophyceae sp. CCMP1205 | Prasinophyceae sp. CCMP1205 | Prasinophyceae sp. CCMP1205 | 1.000 |
| 4a920fd9646420ec7af245c937a8adc8 | Bacteria | Proteobacteria | Gammaproteobacteria | Oceanospirillales | Halomonadaceae | Halomonas | | 1.000 |
| 76c7ab14ae61d10cc1fcc75171be5d06 | Bacteria | Verrucomicrobia | Verrucomicrobiae | Arctic97B-4 marine group | uncultured Verrucomicrobia bacterium | uncultured Verrucomicrobia bacterium | uncultured Verrucomicrobia bacterium | 0.889 |
| 883f963b133759bc47cb69db69273e00 | Bacteria | Cyanobacteria | Oxyphotobacteria | Chloroplast | Phalacroma mitra | Phalacroma mitra | Phalacroma mitra | 0.989 |
| d5c306e14c90cded95c265c57913f1a8 | Bacteria | Proteobacteria | Alphaproteobacteria | SAR11 clade | Clade II | | | 0.888 |
| 64cf84b7200c858fe591463c6a568494 | Bacteria | Proteobacteria | Alphaproteobacteria | Rickettsiales | S25-593 | marine metagenome | marine metagenome | 0.994 |
| eaefb3e09d049747b784dfbb6681df5c | Bacteria | Proteobacteria | Gammaproteobacteria | JTB23 | uncultured marine microorganism | uncultured marine microorganism | uncultured marine microorganism | 0.967 |
| 4128e2dd0f81f9e1e275dab32c676271 | Bacteria | Chloroflexi | Dehalococcoidia | SAR202 clade | hydrothermal vent metagenome | hydrothermal vent metagenome | hydrothermal vent metagenome | 0.979 |
| 2ee85a51dff0dda58b8698ddd73139ca | Bacteria | Bacteroidetes | Bacteroidia | Cytophagales | Cyclobacteriaceae | uncultured | uncultured marine bacterium | 1.000 |
| d15181c78bfd695e82830217bad091ca | Bacteria | Planctomycetes | Phycisphaerae | Phycisphaerales | Phycisphaeraceae | CL500-3 | Ambiguous_taxa | 0.982 |
| a048e53366d02ba31fd7db52a2b11630 | Bacteria | Bacteroidetes | Bacteroidia | Flavobacteriales | Flavobacteriaceae | NS5 marine group | | 1.000 |
| 6a9c0583ee93123f5e4b019922320328 | Bacteria | Proteobacteria | Alphaproteobacteria | Rhodobacterales | Rhodobacteraceae | Paracoccus | | 0.999 |
| f9a29a30d9f8b57f0adac25494538000 | Archaea | Euryarchaeota | Thermoplasmata | Marine Group II | marine metagenome | marine metagenome | marine metagenome | 0.806 |
| 81d321d2a6079631aa7cea94105e07ba | Bacteria | Proteobacteria | Alphaproteobacteria | Puniceispirillales | SAR116 clade | Candidatus Puniceispirillum | | 0.937 |
| e705e1eac9a072b1d92093fc55c3dc21 | Bacteria | Proteobacteria | Gammaproteobacteria | Cellvibrionales | Halieaceae | OM60(NOR5) clade | | 0.996 |
| 6abff36b05cbb0ffd8918c9895e1e23c | Bacteria | Bacteroidetes | Bacteroidia | Flavobacteriales | NS7 marine group | | | 1.000 |
| 0b6e5f9a4e9f4b950184f9a734adf1eb | Bacteria | Kiritimatiellaeota | Kiritimatiellae | Kiritimatiellales | Kiritimatiellaceae | R76-B128 | | 1.000 |
| 8fcb01aba87231a77d087e66fc7726cd | Bacteria | Proteobacteria | Gammaproteobacteria | Coxiellales | Coxiellaceae | Coxiella | uncultured bacterium | 0.908 |
| 8dd8b95a5447a89020880b9bf8575f0a | Bacteria | Proteobacteria | Gammaproteobacteria | Betaproteobacteriales | Burkholderiaceae | MWH-UniP1 aquatic group | uncultured marine bacterium | 0.982 |
| 15ff669914e8a6237f663872ecc52031 | Bacteria | Proteobacteria | Gammaproteobacteria | | | | | 1.000 |
| 9bfa918711ab999d86291def4d390d98 | Bacteria | Marinimicrobia (SAR406 clade) | | | | | | 1.000 |
| 466deaca70e770235ca8a5c91ab4dd88 | Bacteria | Proteobacteria | Alphaproteobacteria | Rhodospirillales | AEGEAN-169 marine group | | | 1.000 |
| 8d424edbd58ce46385af53a88ed1d2b7 | Bacteria | Marinimicrobia (SAR406 clade) | uncultured marine bacterium | uncultured marine bacterium | uncultured marine bacterium | uncultured marine bacterium | uncultured marine bacterium | 0.934 |
| 0d510288bd374edfc141b3bf0deb79fb | Bacteria | Marinimicrobia (SAR406 clade) | | | | | | 1.000 |
| 18ba657a31199cb69d98f53ed906f4cc | Bacteria | Chloroflexi | Dehalococcoidia | SAR202 clade | | | | 1.000 |
| 93de0b378643d09fc936358553d6d859 | Bacteria | Verrucomicrobia | Verrucomicrobiae | Opitutales | Puniceicoccaceae | Coraliomargarita | uncultured marine bacterium | 1.000 |
| c149a6e394387bfd3deebb246110e628 | Bacteria | Marinimicrobia (SAR406 clade) | uncultured Fibrobacteres/Acidobacteria group bacterium | uncultured Fibrobacteres/Acidobacteria group bacterium | uncultured Fibrobacteres/Acidobacteria group bacterium | uncultured Fibrobacteres/Acidobacteria group bacterium | uncultured Fibrobacteres/Acidobacteria group bacterium | 0.968 |
| f19bc9d4642c6f5a7c46b6bd0e4c7738 | Bacteria | Bacteroidetes | Bacteroidia | Cytophagales | Cyclobacteriaceae | Marinoscillum | uncultured Bacteroidetes bacterium | 0.745 |
| 113bef0f47c237002039949ec9656a1a | Bacteria | Bacteroidetes | Bacteroidia | Flavobacteriales | Cryomorphaceae | uncultured | uncultured Owenweeksia sp. | 0.936 |
| 6c524597d23595a47ec9228fd1bd5147 | Bacteria | Proteobacteria | Deltaproteobacteria | Oligoflexales | Oligoflexaceae | uncultured | | 1.000 |
| e19ffd6d48beceb057d0c01696dc2f2c | Bacteria | Marinimicrobia (SAR406 clade) | | | | | | 1.000 |
| 1f9784a5b95cfc2a60308846c58f01fd | Bacteria | Chloroflexi | Dehalococcoidia | SAR202 clade | hydrothermal vent metagenome | hydrothermal vent metagenome | hydrothermal vent metagenome | 0.994 |
| a41ea6398c29bf4aca87ee2f86b0e952 | Bacteria | Bacteroidetes | Bacteroidia | Chitinophagales | Saprospiraceae | uncultured | unidentified marine eubacterium | 1.000 |
| 1649df8989783bf5fc53666fbe1fffed | Bacteria | Proteobacteria | Gammaproteobacteria | Vibrionales | Vibrionaceae | Vibrio | | 1.000 |
| 19522f299389440fc748c41354f14d33 | Bacteria | Proteobacteria | Gammaproteobacteria | Cellvibrionales | Halieaceae | Parahaliea | uncultured marine bacterium | 0.844 |
| 9799f032f7139a687a2b1af8d0fdb0f9 | Bacteria | Proteobacteria | Gammaproteobacteria | Oceanospirillales | Halomonadaceae | Salinicola | Ambiguous_taxa | 0.842 |

| Feature_ID | Kingdom | Phylum | Class | Order | Family | Genus | Species | Confidence |
|---|---|---|---|---|---|---|---|---|
| 8f91d9967191d5cc3543b11145156336 | Bacteria | Cyanobacteria | Oxyphotobacteria | Chloroplast | | | | 1.000 |
| f00e595d02d58e2f5a1cf9b37d5998bf | Bacteria | Bacteroidetes | Bacteroidia | Flavobacteriales | NS9 marine group | | | 1.000 |
| 8b3ee47d66e1d3cc5d0ab6b822e5beaf | Bacteria | Bacteroidetes | Bacteroidia | Flavobacteriales | Flavobacteriaceae | Gramella | Ambiguous_taxa | 0.963 |
| 242e410bd5e857abfba5f5e62dfd75c7 | Bacteria | Proteobacteria | Gammaproteobacteria | | | | | 1.000 |
| 1b4b18c328209dc0afe6e9d0c6785b6b | Bacteria | Proteobacteria | Gammaproteobacteria | Vibrionales | Vibrionaceae | Vibrio | | 0.987 |
| 53b1d6533607400a44e56018f8f7c229 | Bacteria | Proteobacteria | Alphaproteobacteria | uncultured | uncultured marine bacterium | uncultured marine bacterium | uncultured marine bacterium | 0.746 |
| 10c2e08bb7c87897d8aaf55e7243d67a | Bacteria | Verrucomicrobia | Verrucomicrobiae | Arctic97B-4 marine group | Ambiguous_taxa | Ambiguous_taxa | Ambiguous_taxa | 0.890 |
| 934d5a60529e32e60eaf011708bda184 | Bacteria | Proteobacteria | Alphaproteobacteria | Rhodospirillales | AEGEAN-169 marine group | uncultured bacterium | uncultured bacterium | 0.992 |
| 3cf3e1861113d8faad912034e0077dd6 | Bacteria | Proteobacteria | Gammaproteobacteria | OM182 clade | | | | 1.000 |
| 109ec420feca589e50e9412da4289d66 | Bacteria | Cyanobacteria | Oxyphotobacteria | Chloroplast | | | | 1.000 |
| acd070ed8e99fa6456a2c6c5cd652acf | Bacteria | Cyanobacteria | Oxyphotobacteria | Chloroplast | | | | 1.000 |
| 64cbd96da71203e36b714636a7e54bf0 | Bacteria | Margulisbacteria | uncultured bacterium | uncultured bacterium | uncultured bacterium | uncultured bacterium | uncultured bacterium | 0.966 |
| 3b8edc60c17c2e71d27fd8151e57b553 | Bacteria | Proteobacteria | Deltaproteobacteria | Oligoflexales | Oligoflexaceae | uncultured | marine metagenome | 1.000 |
| f93e5daac2d2202f8dff217a7b06363f | Bacteria | Proteobacteria | Alphaproteobacteria | Rickettsiales | | | | 1.000 |
| 5878eee334440bf972ee8b0a0f9d9e3f | Bacteria | Bacteroidetes | Bacteroidia | Flavobacteriales | Flavobacteriaceae | Pseudofulvibacter | uncultured Bacteroidetes bacterium | 0.808 |
| ef44e064fa6629bb99b6ccabfba9ab2f | Bacteria | Bacteroidetes | Bacteroidia | Flavobacteriales | Flavobacteriaceae | NS5 marine group | | 1.000 |
| 6b291baa9e0c11a04b22c5e62f3a4874 | Bacteria | Proteobacteria | Gammaproteobacteria | Vibrionales | Vibrionaceae | Vibrio | | 0.927 |
| ec5527dcd577c4dc849f4b17323496d3 | Bacteria | Proteobacteria | Alphaproteobacteria | uncultured | | | | 1.000 |
| 1aebeafe2e1830cf907be36a6618199e | Bacteria | Marinimicrobia (SAR406 clade) | | | | | | 1.000 |
| 2debcf34b919dc7062a36c54469e5f7f | Bacteria | Bacteroidetes | Bacteroidia | Flavobacteriales | NS9 marine group | Ambiguous_taxa | Ambiguous_taxa | 0.774 |
| 9a03cc45e25cb5a8407aa84059e484fa | Bacteria | Bacteroidetes | Bacteroidia | Flavobacteriales | NS9 marine group | uncultured Flavobacteriia bacterium | uncultured Flavobacteriia bacterium | 0.927 |
| 5dc579902e511bf63c41608f333db905 | Bacteria | Proteobacteria | Alphaproteobacteria | | | | | 1.000 |
| 0b4c0501d3fa6c035b8f513b1c0afa9e | Bacteria | Proteobacteria | Gammaproteobacteria | SAR86 clade | | | | 1.000 |
| 674c3042c46e748ca9da22479ca173b0 | Bacteria | Bacteroidetes | Bacteroidia | Flavobacteriales | NS9 marine group | uncultured Flavobacteriia bacterium | uncultured Flavobacteriia bacterium | 0.886 |
| c1c1f1fe56e143bb89cea4c69851a5e2 | Bacteria | Marinimicrobia (SAR406 clade) | | | | | | 1.000 |
| 1411131900f309c76068f482e219d76f | Bacteria | Cyanobacteria | Oxyphotobacteria | Chloroplast | | | | 1.000 |
| 98ef073af44a1f987c04c5d0087fcb1c | Bacteria | Cyanobacteria | Oxyphotobacteria | Chloroplast | | | | 1.000 |
| 6a148edba636e8d6d3af9f5249f95d38 | Bacteria | Margulisbacteria | metagenome | metagenome | metagenome | metagenome | metagenome | 0.997 |
| b2f015f98dc5d1cf9e63fe266a3fe5ff | Bacteria | Proteobacteria | Gammaproteobacteria | Vibrionales | Vibrionaceae | Vibrio | | 0.998 |
| 6d95c423bbd15da2b1c50fdeed503c4f | Bacteria | Bacteroidetes | Bacteroidia | Flavobacteriales | NS9 marine group | | | 0.999 |
| 7a9313e2d8263fc1f2ae1f85c1e529e3 | Bacteria | Proteobacteria | Gammaproteobacteria | Pseudomonadales | Pseudomonadaceae | Pseudomonas | | 1.000 |
| 420b912eeb1748b6395cbb8a66ed2829 | Bacteria | Proteobacteria | Gammaproteobacteria | Oceanospirillales | Pseudohongiellaceae | Pseudohongiella | | 1.000 |
| 7fb0f37f4c87925de92b0802475ce081 | Bacteria | Proteobacteria | Alphaproteobacteria | Sneathiellales | Sneathiellaceae | AT-s3-44 | uncultured bacterium | 0.869 |
| 01780edb0abafb4055ba2127de54267b | Bacteria | Chloroflexi | Dehalococcoidia | SAR202 clade | | | | 1.000 |
| 35e02a58869d9d396e07002fc92be69e | Bacteria | Cyanobacteria | Oxyphotobacteria | Chloroplast | uncultured diatom | uncultured diatom | uncultured diatom | 0.762 |
| edec9507817b6e548d2fdd21b6d55be2 | Bacteria | Cyanobacteria | Oxyphotobacteria | Chloroplast | marine metagenome | marine metagenome | marine metagenome | 0.817 |
| 3413244e718c5b28f53fdae23b625fec | Bacteria | Proteobacteria | Alphaproteobacteria | SAR11 clade | Clade II | | | 0.962 |
| c83a07183f934ecd41b3de1fa76fbcc8 | Bacteria | Proteobacteria | Alphaproteobacteria | uncultured | hydrothermal vent metagenome | hydrothermal vent metagenome | hydrothermal vent metagenome | 0.738 |
| 7c2b94b80c08548a8da35f492d7298fa | Bacteria | Bacteroidetes | Bacteroidia | Flavobacteriales | Flavobacteriaceae | | | 1.000 |
| 2b1673ad9db2e798c0b4cafd34ba339c | Bacteria | Proteobacteria | Gammaproteobacteria | Coxiellales | Coxiellaceae | Coxiella | | 1.000 |
| 91795ad59366c22f2678695b123f625a | Bacteria | Proteobacteria | Gammaproteobacteria | Cellvibrionales | Porticoccaceae | SAR92 clade | | 1.000 |
| 3fdc3646c1d51844cba8f9e362a0ef3b | Bacteria | Proteobacteria | Gammaproteobacteria | SAR86 clade | | | | 1.000 |
| eb2d8cd665871a846060efa7d2f5b1f5 | Bacteria | Proteobacteria | Alphaproteobacteria | Rickettsiales | Rickettsiaceae | uncultured | uncultured bacterium | 0.989 |
| 3fb9aab89b211b8970d125e19361309d | Bacteria | Proteobacteria | Alphaproteobacteria | Caulobacterales | Caulobacteraceae | Phenylobacterium | | 0.866 |
| 039d12c3a75924ebdca5370aa403164e | Bacteria | Proteobacteria | Alphaproteobacteria | uncultured | unidentified marine bacterioplankton | unidentified marine bacterioplankton | unidentified marine bacterioplankton | 0.993 |
| 32cc406e27d9dab097cb599f3b80d708 | Bacteria | Proteobacteria | Deltaproteobacteria | Bdellovibrionales | Bdellovibrionaceae | OM27 clade | | 1.000 |
| 1c88d362891cef109ebadb147709de6d | Bacteria | Proteobacteria | Alphaproteobacteria | Rickettsiales | S25-593 | uncultured marine bacterium | uncultured marine bacterium | 0.820 |
| 779040b594019ca993388916964dbd2f | Bacteria | Bacteroidetes | Bacteroidia | Flavobacteriales | NS9 marine group | uncultured Flavobacteriales bacterium | uncultured Flavobacteriales bacterium | 0.742 |
| be8e3cb9a0a0cefb775f0a214d1dc33d | Bacteria | Bacteroidetes | Bacteroidia | Flavobacteriales | NS9 marine group | | | 1.000 |
| b6b2f5043f133c8528a403bb434a58fe | Bacteria | Proteobacteria | Gammaproteobacteria | Gammaproteobacteria Incertae Sedis | Unknown Family | uncultured marine bacterium | uncultured marine bacterium | 1.000 |
| e6411ee5941ee104ec9116bd3cc3686e | Bacteria | Verrucomicrobia | Verrucomicrobiae | Opitutales | Puniceicoccaceae | MB11C04 marine group | | 1.000 |
| aaa9bf0307bc79f5bf9589c36d998f20 | Bacteria | Proteobacteria | Gammaproteobacteria | Thiohalorhabdales | Thiohalorhabdaceae | uncultured | | 0.915 |
| e4c38ff9819dea40b8877e8d1c7bbd5b | Bacteria | Proteobacteria | Gammaproteobacteria | Oceanospirillales | Saccharospirillaceae | uncultured deep-sea bacterium | uncultured deep-sea bacterium | 1.000 |
| 6ec45751ec571ab29fc808c33b5044a5 | Bacteria | Bacteroidetes | Bacteroidia | Chitinophagales | Saprospiraceae | uncultured | | 0.812 |
| 6fc4c89b73a3dcb015836601296a7152 | Bacteria | Proteobacteria | Alphaproteobacteria | Rhodobacterales | Rhodobacteraceae | Ascidiaceihabitans | | 0.775 |
| 5b5578251cd19f073302171fe4c6611d | Bacteria | Verrucomicrobia | Verrucomicrobiae | Opitutales | Puniceicoccaceae | MB11C04 marine group | uncultured marine microorganism | 0.938 |
| ebce346d2886742517e4cf309c8b1360 | Bacteria | Chloroflexi | Dehalococcoidia | SAR202 clade | | | | 1.000 |
| 27c3e42d1738c926b9a14de3da9ad292 | Bacteria | Proteobacteria | Alphaproteobacteria | Rhodospirillales | AEGEAN-169 marine group | uncultured marine bacterium | uncultured marine bacterium | 0.763 |
| 8c823447f4042c1e3f81d260edfc76e9 | Bacteria | Marinimicrobia (SAR406 clade) | | | | | | 1.000 |
| 7a71b71d2637db332653a9488671ecd5 | Bacteria | Marinimicrobia (SAR406 clade) | uncultured marine microorganism | uncultured marine microorganism | uncultured marine microorganism | uncultured marine microorganism | uncultured marine microorganism | 1.000 |
| 7ccb862a0ebd9c55823caa45deff3bff | Bacteria | Chloroflexi | Dehalococcoidia | SAR202 clade | uncultured marine microorganism | uncultured marine microorganism | uncultured marine microorganism | 0.999 |
| 7d026221a04a6922abc6f0b7621094a4 | Bacteria | Bacteroidetes | Bacteroidia | Flavobacteriales | Flavobacteriaceae | NS5 marine group | uncultured marine bacterium | 0.790 |
| 5526ae2ea4b0e35126c2d72c0a6f9e47 | Bacteria | Proteobacteria | Gammaproteobacteria | SAR86 clade | uncultured bacterium | uncultured bacterium | uncultured bacterium | 0.820 |
| 00a0d0f543eab5fd5b33cc504d254d17 | Bacteria | Proteobacteria | Gammaproteobacteria | Oceanospirillales | Pseudohongiellaceae | Pseudohongiella | Ambiguous_taxa | 0.861 |
| bfce84f0234d7dd647f0e7a34bc02283 | Bacteria | Proteobacteria | Deltaproteobacteria | Bdellovibrionales | Bdellovibrionaceae | OM27 clade | Ambiguous_taxa | 0.987 |
| 56935a78569bcca4af9742d98e995448 | Bacteria | Verrucomicrobia | Verrucomicrobiae | Arctic97B-4 marine group | uncultured marine bacterium | uncultured marine bacterium | uncultured marine bacterium | 0.986 |
| aa6ebc72e62352638109e6dc0dde8d0e | Bacteria | Proteobacteria | Gammaproteobacteria | Coxiellales | Coxiellaceae | Coxiella | uncultured gamma proteobacterium | 0.794 |
| 127a0c1ce8288c17ed00ff92f06fd2bf | Bacteria | Proteobacteria | Gammaproteobacteria | Oceanospirillales | Halomonadaceae | Halomonas | | 1.000 |
| 69b79e3aab5e84f0f5ea6dae2bc9b191 | Bacteria | Margulisbacteria | uncultured marine cyanobacterium | uncultured marine cyanobacterium | uncultured marine cyanobacterium | uncultured marine cyanobacterium | uncultured marine cyanobacterium | 0.994 |
| 4cd9af307d4f6b6233ac4854010a50b2 | Bacteria | Proteobacteria | Alphaproteobacteria | Rickettsiales | S25-593 | marine metagenome | marine metagenome | 0.993 |
| dcaa50c4ae0791ec92b0fcfa485db2d8 | Bacteria | Bacteroidetes | Bacteroidia | Flavobacteriales | NS9 marine group | | | 1.000 |
| 8fd2e72fb268b46c811756c7ad0ec8d3 | Bacteria | Bacteroidetes | Bacteroidia | Flavobacteriales | Flavobacteriaceae | NS2b marine group | | 1.000 |
| 6ab430a4044fc25e2e4baf3ec65b8e4a | Bacteria | Proteobacteria | Alphaproteobacteria | Rhodospirillales | uncultured | uncultured bacterium | uncultured bacterium | 1.000 |
| ff66de6490e449485656ce330bdd96a6 | Bacteria | Bacteroidetes | Bacteroidia | Flavobacteriales | Flavobacteriaceae | NS4 marine group | | 1.000 |
| 74db8edd11200f8dfa6895aaf97f0807 | Bacteria | Proteobacteria | Alphaproteobacteria | Rickettsiales | Midichloriaceae | MD3-55 | uncultured proteobacterium | 0.987 |
| 27f0cdf902357cd1b90b30b3962621e0 | Bacteria | Bacteroidetes | Bacteroidia | Flavobacteriales | Crocinitomicaceae | Salinirepens | uncultured marine bacterium | 0.955 |
| b4a0a8ea69e5ad8e5970d02746c351d6 | Bacteria | Proteobacteria | Deltaproteobacteria | Bdellovibrionales | Bdellovibrionaceae | OM27 clade | | 1.000 |
| c99f9c438c35cb7b518843cda91196b3 | Bacteria | Proteobacteria | Gammaproteobacteria | Alteromonadales | Alteromonadaceae | Paraglaciecola | Ambiguous_taxa | 0.757 |

| Feature_ID | Kingdom | Phylum | Class | Order | Family | Genus | Species | Confidence |
|---|---|---|---|---|---|---|---|---|
| 0f2405a72c541eeba8cbcda9f5b444e0 | Bacteria | Marinimicrobia (SAR406 clade) | Marinimicrobia bacterium JGI 0000113-D11 | Marinimicrobia bacterium JGI 0000113-D11 | Marinimicrobia bacterium JGI 0000113-D11 | Marinimicrobia bacterium JGI 0000113-D11 | Marinimicrobia bacterium JGI 0000113-D11 | 0.803 |
| da6cd99773764cd9272f81ebc9472ec5 | Bacteria | Proteobacteria | Alphaproteobacteria | Puniceispirillales | SAR116 clade | Candidatus Puniceispirillum | Ambiguous_taxa | 0.790 |
| 8dabe0d786933279d9318c6aa9c72276 | Bacteria | Proteobacteria | Gammaproteobacteria | Alteromonadales | Alteromonadaceae | | | 1.000 |
| 65dc92e561a3bbdf7f098d20b356d29e | Bacteria | Proteobacteria | Deltaproteobacteria | Bdellovibrionales | Bacteriovoracaceae | uncultured | uncultured marine bacterium | 0.998 |
| 07ae0e04f58fce205e946fbfa3118d00 | Bacteria | Cyanobacteria | Oxyphotobacteria | Chloroplast | uncultured marine eukaryote | uncultured marine eukaryote | uncultured marine eukaryote | 0.934 |
| eb9c74886ea7d9808bac1f860d89664d | Bacteria | Proteobacteria | Alphaproteobacteria | Rhodospirillales | Magnetospiraceae | uncultured | uncultured bacterium | 0.812 |
| f34326e80f8ce3c25ec5ea3e1fc5534a | Bacteria | Bacteroidetes | Bacteroidia | Flavobacteriales | Cryomorphaceae | uncultured | uncultured Owenweeksia sp. | 0.927 |
| 22606a3a126cb8506775d1a0665ed3e6 | Bacteria | Proteobacteria | Gammaproteobacteria | | | | | 0.996 |
| b61a70e09618762451f6204a4236d742 | Bacteria | Proteobacteria | Deltaproteobacteria | SAR324 clade(Marine group B) | uncultured bacterium | uncultured bacterium | uncultured bacterium | 0.909 |
| 51f3434f8d941d6d4762c6f67fbe6675 | Bacteria | Proteobacteria | Gammaproteobacteria | Alteromonadales | Alteromonadaceae | Aestuariibacter | uncultured bacterium | 0.751 |
| 9063ece8e9fed2580066030d040e3a95 | Bacteria | Bacteroidetes | Bacteroidia | Flavobacteriales | Cryomorphaceae | uncultured | Ambiguous_taxa | 0.865 |
| e2f108d9908f3004813330d999e24d64 | Bacteria | Cyanobacteria | Oxyphotobacteria | Chloroplast | | | | 1.000 |
| 819c6125778e7d4f8e2c2afc1b09673a | Bacteria | Bacteroidetes | Bacteroidia | Chitinophagales | Saprospiraceae | Aureispira | uncultured Saprospiraceae bacterium | 0.896 |
| a4f63be19364a7d2a6de6a42ecc3c400 | Bacteria | Proteobacteria | Gammaproteobacteria | Betaproteobacteriales | Burkholderiaceae | MWH-UniP1 aquatic group | uncultured bacterium | 0.997 |
| 0996ed3ed6bd9152724d9e1cbb0e3fca | Bacteria | Proteobacteria | Alphaproteobacteria | Rhodospirillales | AEGEAN-169 marine group | uncultured bacterium | uncultured bacterium | 0.960 |
| 3f338bff81ad6542370e256a45c97ff5 | Bacteria | Proteobacteria | Gammaproteobacteria | OM182 clade | uncultured bacterium | uncultured bacterium | uncultured bacterium | 0.741 |
| b7b614066e23569ff8e9d5e3caa40f7b | Bacteria | Marinimicrobia (SAR406 clade) | uncultured bacterium | uncultured bacterium | uncultured bacterium | uncultured bacterium | uncultured bacterium | 0.770 |
| 7225bc4d50a6f57ec520f4ed8d12e2d4 | Bacteria | Chloroflexi | Dehalococcoidia | SAR202 clade | uncultured Chloroflexi bacterium | uncultured Chloroflexi bacterium | uncultured Chloroflexi bacterium | 0.910 |
| 00e747f4d30b7924005c3b34c4c68f15 | Bacteria | Cyanobacteria | Oxyphotobacteria | Chloroplast | marine metagenome | marine metagenome | marine metagenome | 0.962 |
| d87828eba5eb9851ccab4532200a8c37 | Bacteria | Proteobacteria | Gammaproteobacteria | Piscirickettsiales | Piscirickettsiaceae | Candidatus Endoecteinascidia | Ambiguous_taxa | 0.787 |
| 55c0cbce3271fe391f870b520b2d6c82 | Bacteria | Proteobacteria | Gammaproteobacteria | Oceanospirillales | Pseudohongiellaceae | Pseudohongiella | | 1.000 |
| e3bacd0bceb66498b385ea52fdc907f5 | Bacteria | Bacteroidetes | Bacteroidia | Flavobacteriales | Flavobacteriaceae | Flavicella | uncultured bacterium | 0.992 |
| ebfd1f70b6173e816e1556dc80407da9 | Bacteria | Bacteroidetes | Bacteroidia | Flavobacteriales | Flavobacteriaceae | NS2b marine group | uncultured marine bacterium | 0.930 |
| e17de8149c20f0f7a843f0c760864dcf | Bacteria | Proteobacteria | Gammaproteobacteria | Cellvibrionales | Halieaceae | OM60(NOR5) clade | Ambiguous_taxa | 0.891 |
| 9592044e2d42cccf0c9a114bfa9d1b2e | Bacteria | Bacteroidetes | Bacteroidia | Flavobacteriales | NS9 marine group | uncultured marine bacterium | uncultured marine bacterium | 0.999 |
| 8efd03f85e12d953815a8ded41775c31 | Bacteria | Bacteroidetes | Bacteroidia | Cytophagales | Cyclobacteriaceae | Marinoscillum | | 1.000 |
| c0a09bdf990420b37a732a9c7b29af09 | Bacteria | Proteobacteria | Alphaproteobacteria | Rickettsiales | Mitochondria | | | 0.995 |
| 2942c9b812d98529cf017cfdaab3a4f1 | Bacteria | Proteobacteria | Deltaproteobacteria | PB19 | | | | 1.000 |
| 58d621afa51805e4b1d0ba74da49b39d | Bacteria | Proteobacteria | Alphaproteobacteria | Rickettsiales | S25-593 | marine metagenome | marine metagenome | 0.968 |
| ea23e76aeb2e53e7a9ecdd13c5afebce | Bacteria | Proteobacteria | Deltaproteobacteria | Myxococcales | Blfdi19 | | | 1.000 |
| d5b9f90404e41e20a29e8f5ec35dbca6 | Bacteria | Proteobacteria | Alphaproteobacteria | SAR11 clade | Clade II | | | 0.850 |
| 7368aabac8e6a3566c9e9f1ada315055 | Archaea | Euryarchaeota | Thermoplasmata | Marine Group II | uncultured marine microorganism HF200_10-06-02 | uncultured marine microorganism HF200_10-06-02 | uncultured marine microorganism HF200_10-06-02 | 0.901 |
| ccee959e431c6aba9891913c12e3e4c3 | Bacteria | Proteobacteria | Gammaproteobacteria | SAR86 clade | uncultured bacterium | uncultured bacterium | uncultured bacterium | 0.962 |
| 10f0b07aad1421776d9a9e9f654bf50e | Bacteria | Bacteroidetes | Bacteroidia | Flavobacteriales | NS9 marine group | | | 1.000 |
| 3bfee15642e1097c6a183c8948244720 | Bacteria | Cyanobacteria | Oxyphotobacteria | Chloroplast | Virgulinella fragilis | Virgulinella fragilis | Virgulinella fragilis | 0.982 |
| f67c2eeced00990a00ec20cfa7441682 | Bacteria | Cyanobacteria | Oxyphotobacteria | Chloroplast | | | | 1.000 |
| 52766904bbb796864e53328a0e66ecdf | Bacteria | Bacteroidetes | Bacteroidia | Flavobacteriales | NS9 marine group | Ambiguous_taxa | Ambiguous_taxa | 0.756 |
| 39ad46665d820262f9c9b74d2377e8ca | Bacteria | Proteobacteria | Alphaproteobacteria | uncultured | uncultured marine bacterium | uncultured marine bacterium | uncultured marine bacterium | 0.952 |
| 2a45dc931ebb5357ce53558f4d16afd8 | Bacteria | Proteobacteria | Deltaproteobacteria | Oligoflexales | Oligoflexaceae | uncultured | uncultured bacterium | 1.000 |
| dfc288635a89458900abda2eed28ed61 | Bacteria | Proteobacteria | Gammaproteobacteria | Betaproteobacteriales | Methylophilaceae | OM43 clade | Ambiguous_taxa | 0.975 |
| 5c337f48cfaab2233d22592f22fb3cb9 | Bacteria | Marinimicrobia (SAR406 clade) | | | | | | 1.000 |
| 1a265652ef04f2fddd66fdeeb1449d73 | Bacteria | Proteobacteria | Alphaproteobacteria | SAR11 clade | | | | 1.000 |
| a74512558fdb2939dd64a4ce30f3c544 | Bacteria | Kiritimatiellaeota | Kiritimatiellae | Kiritimatiellales | Kiritimatiellaceae | R76-B128 | Ambiguous_taxa | 1.000 |
| 034f7a43009080b35358f715ff3e6e25 | Bacteria | Proteobacteria | Alphaproteobacteria | Puniceispirillales | SAR116 clade | alpha proteobacterium SCGC AAA015-N04 | alpha proteobacterium SCGC AAA015-N04 | 0.985 |
| 0056be93807dd5c15df627f478f05c4d | Bacteria | Cyanobacteria | Oxyphotobacteria | Nostocales | Nostocaceae | Richelia HH01 | | 0.908 |
| a968ca643083ccf6f939d03dff997b87 | Bacteria | Proteobacteria | Deltaproteobacteria | PB19 | Ambiguous_taxa | Ambiguous_taxa | Ambiguous_taxa | 0.862 |
| c8a9e0a31e4031863a5b13a5cde0a02d | Bacteria | Proteobacteria | Alphaproteobacteria | Rickettsiales | S25-593 | marine metagenome | marine metagenome | 1.000 |
| 9507052d9a42124864d919d3efbc461a | Bacteria | Cyanobacteria | Oxyphotobacteria | Chloroplast | Pyramimonas parkeae | Pyramimonas parkeae | Pyramimonas parkeae | 0.999 |
| 91f1c19d43f14a32283c20b7adc6b3ed | Bacteria | Proteobacteria | Deltaproteobacteria | PB19 | Ambiguous_taxa | Ambiguous_taxa | Ambiguous_taxa | 0.997 |
| 7c191ba791c2265f96d93048dbf6586e | Bacteria | Proteobacteria | Deltaproteobacteria | Myxococcales | P3OB-42 | uncultured bacterium | uncultured bacterium | 0.729 |
| 8de8dbfc853fab0e246e7374b2a1936d | Bacteria | Gemmatimonadetes | BD2-11 terrestrial group | Ambiguous_taxa | Ambiguous_taxa | Ambiguous_taxa | Ambiguous_taxa | 0.989 |
| 61317041500d17631ab0f6cbb3e817a5 | Bacteria | Firmicutes | Clostridia | Clostridiales | Ruminococcaceae | Ruminococcus 2 | | 1.000 |
| 6f4eb9151ebf33e973cf2d54487269f2 | Bacteria | Proteobacteria | Gammaproteobacteria | Nitrosococcales | Methylophagaceae | Methylophaga | | 1.000 |
| 928e2cdbd60a5c68e73815a93669cf6b | Bacteria | Bacteroidetes | Bacteroidia | Chitinophagales | Saprospiraceae | uncultured | unidentified marine eubacterium | 1.000 |
| d4b598d8ff697ac14b1e75f944eaf5c1 | Bacteria | Proteobacteria | Alphaproteobacteria | uncultured | unidentified marine bacterioplankton | unidentified marine bacterioplankton | unidentified marine bacterioplankton | 0.873 |
| 5770edaa5784b93ee9164638583db0e7 | Bacteria | Proteobacteria | Gammaproteobacteria | Oceanospirillales | Nitrincolaceae | uncultured | uncultured bacterium | 0.824 |
| 2924406489859b3c0477c2d402a50b31 | Bacteria | Marinimicrobia (SAR406 clade) | Ambiguous_taxa | Ambiguous_taxa | Ambiguous_taxa | Ambiguous_taxa | Ambiguous_taxa | 0.756 |
| 4ffeafa7fe10dab30d93074012305c1d | Bacteria | Proteobacteria | Alphaproteobacteria | Rhodospirillales | Magnetospiraceae | uncultured | uncultured bacterium | 0.797 |
| c9c77276627b37ce078ec9526aaf01d1 | Bacteria | Epsilonbacteraeota | Campylobacteria | Campylobacterales | Arcobacteraceae | Arcobacter | Arcobacter sp. EP1 | 0.953 |
| df8facef76b20cf0c9c92e7389555c0e | Bacteria | Chloroflexi | Dehalococcoidia | SAR202 clade | marine metagenome | marine metagenome | marine metagenome | 1.000 |
| bdc7f0de3785a216e23f635afe9b4990 | Bacteria | Bacteroidetes | Bacteroidia | Flavobacteriales | NS9 marine group | uncultured marine bacterium | uncultured marine bacterium | 0.995 |
| 837c2680dc200a2fe6775d373957b4ba | Bacteria | Marinimicrobia (SAR406 clade) | marine metagenome | marine metagenome | marine metagenome | marine metagenome | marine metagenome | 0.859 |
| c5589bf3a56bb27ceb8bd2d6fa2beb09 | Bacteria | Proteobacteria | Alphaproteobacteria | uncultured | unidentified marine bacterioplankton | unidentified marine bacterioplankton | unidentified marine bacterioplankton | 0.986 |
| 91b7b5980e6c71840511f893834ac1fd | Bacteria | Proteobacteria | Alphaproteobacteria | Rhodospirillales | Magnetospiraceae | Magnetospira | uncultured nuHF1 cluster bacterium HF0770_35I22 | 0.974 |
| da9d3dcbb6499fc0749249c872d334f6 | Bacteria | Bacteroidetes | Bacteroidia | Flavobacteriales | Flavobacteriaceae | | | 1.000 |
| c1338c53aa4ca64c42cc4180ac1ad3e4 | Bacteria | Planctomycetes | Planctomycetacia | Pirellulales | Pirellulaceae | Rhodopirellula | uncultured marine microorganism | 1.000 |
| b28e037a0e8cc75b7129f37ce619fcc2 | Bacteria | Marinimicrobia (SAR406 clade) | uncultured bacterium | uncultured bacterium | uncultured bacterium | uncultured bacterium | uncultured bacterium | 0.711 |
| 95104cd4b3bf8838a6c82a1f08b10a4b | Bacteria | Cyanobacteria | Oxyphotobacteria | Chloroplast | | | | 1.000 |
| 28033eccf64be4f2a0195ea681a8dc74 | Bacteria | Margulisbacteria | uncultured bacterium | uncultured bacterium | uncultured bacterium | uncultured bacterium | uncultured bacterium | 0.847 |
| 7cdff66257319b627d474b60dc0a6e9b | Bacteria | Proteobacteria | Gammaproteobacteria | KI89A clade | | | | 1.000 |
| f78737c77d691d4d980e30f3961fcc61 | Bacteria | Proteobacteria | Gammaproteobacteria | Gammaproteobacteria Incertae Sedis | Unknown Family | uncultured marine bacterium | uncultured marine bacterium | 1.000 |
| f19bebb06f1c88140d0551e9b92df422 | Bacteria | Proteobacteria | Gammaproteobacteria | Alteromonadales | Pseudoalteromonadaceae | Pseudoalteromonas | | 1.000 |
| 87d48901d273c86525561c5e7de8427a | Bacteria | Bacteroidetes | Bacteroidia | Flavobacteriales | Flavobacteriaceae | NS4 marine group | | 1.000 |
| c48362c4d1a72c2b616b38260c0eae53 | Bacteria | Proteobacteria | Deltaproteobacteria | PB19 | Ambiguous_taxa | Ambiguous_taxa | Ambiguous_taxa | 0.998 |

| Feature_ID | Kingdom | Phylum | Class | Order | Family | Genus | Species | Confidence |
|---|---|---|---|---|---|---|---|---|
| db07348ff5f05e43aaa43cbd702f3da9 | Bacteria | Bacteroidetes | Bacteroidia | Flavobacteriales | Cryomorphaceae | uncultured | uncultured Owenweeksia sp. | 0.820 |
| d9a230acb012bf5bf00adf14650bf064 | Bacteria | Proteobacteria | Alphaproteobacteria | Rhodospirillales | Magnetospiraceae | uncultured | Ambiguous_taxa | 0.701 |
| 1ed9ac0dbe585389b452bcb15f2d226c | Bacteria | Proteobacteria | Deltaproteobacteria | Bdellovibrionales | Bdellovibrionaceae | OM27 clade | unidentified marine bacterioplankton | 1.000 |
| 0242e24810311ddcade2fd7a2d8dfe3c | Bacteria | Proteobacteria | Gammaproteobacteria | | | | | 1.000 |
| 3fda4837da1f326f15b2824599fa48a8 | Bacteria | Proteobacteria | Deltaproteobacteria | PB19 | | | | 1.000 |
| 505890a7b867642bc38eaee26bd28fe5 | Bacteria | Proteobacteria | Deltaproteobacteria | Bdellovibrionales | Bdellovibrionaceae | OM27 clade | | 1.000 |
| 78bf063117de2d7f8643ba18a0d87325 | Bacteria | Bacteroidetes | Bacteroidia | Flavobacteriales | NS9 marine group | Ambiguous_taxa | Ambiguous_taxa | 0.742 |
| e495b390f80436ee4e5aab08f622e1dc | Bacteria | Proteobacteria | Gammaproteobacteria | Nitrosococcales | Nitrosococcaceae | Cm1-21 | uncultured bacterium | 0.720 |
| 58e025e8b700699d091844183890a496 | Bacteria | Cyanobacteria | Oxyphotobacteria | Chloroplast | | | | 1.000 |
| 40dd3c7b96b092bf07570bc2be2f0894 | Bacteria | Bacteroidetes | Bacteroidia | Flavobacteriales | NS9 marine group | uncultured Flavobacteriales bacterium | uncultured Flavobacteriales bacterium | 0.756 |
| 164a0cba1d3f1f16e25ecce79272b9f7 | Bacteria | Proteobacteria | Gammaproteobacteria | Alteromonadales | Marinobacteraceae | Marinobacter | | 1.000 |
| 82d8e38d3cf3c5a210b7415f99f735a4 | Bacteria | Bacteroidetes | Bacteroidia | Chitinophagales | Saprospiraceae | uncultured | uncultured bacterium | 0.878 |
| 20b193755613010e88acd22d4862bc04 | Bacteria | Cyanobacteria | Oxyphotobacteria | Chloroplast | Proboscia indica | Proboscia indica | Proboscia indica | 0.930 |
| 174c97734e591a4bf141667dc6f3e648 | Bacteria | Margulisbacteria | metagenome | metagenome | metagenome | metagenome | metagenome | 0.997 |
| c490573917992ad569826e2915eb7722 | Bacteria | Proteobacteria | Alphaproteobacteria | uncultured | unidentified marine bacterioplankton | unidentified marine bacterioplankton | unidentified marine bacterioplankton | 0.825 |
| 0a1a78504742154754324da0445a8458 | Bacteria | Cyanobacteria | Oxyphotobacteria | Chloroplast | uncultured Cryptomonadaceae | uncultured Cryptomonadaceae | uncultured Cryptomonadaceae | 0.973 |
| fc2406b82069df9c4a63188ceec675be | Bacteria | Proteobacteria | Deltaproteobacteria | Bdellovibrionales | Bdellovibrionaceae | OM27 clade | unidentified marine bacterioplankton | 1.000 |
| bd3b15486f3c1c00a1d3a70350040558 | Bacteria | Proteobacteria | Gammaproteobacteria | Coxiellales | Coxiellaceae | Coxiella | uncultured marine bacterium | 0.880 |
| 243f934f46e5a4330c430249c34b9da2 | Bacteria | Proteobacteria | Alphaproteobacteria | Rhodobacterales | Rhodobacteraceae | Ascidiaceihabitans | uncultured bacterium | 0.776 |
| 4539c99762e5ee56961a72c88976029c | Bacteria | Proteobacteria | Gammaproteobacteria | SAR86 clade | uncultured gamma proteobacterium HF4000_19M20 | uncultured gamma proteobacterium HF4000_19M20 | uncultured gamma proteobacterium HF4000_19M20 | 1.000 |
| 477889c36e471f777e7a2bdbe0031d85 | Bacteria | Chloroflexi | Dehalococcoidia | SAR202 clade | | | | 1.000 |
| b348bb98c3d2a0fad1489024240a6bfd | Bacteria | Chloroflexi | Dehalococcoidia | SAR202 clade | Ambiguous_taxa | Ambiguous_taxa | Ambiguous_taxa | 0.996 |
| 48e2a9d6f9c9beb8a4fac9428cf00cdc | Bacteria | Cyanobacteria | Oxyphotobacteria | Chloroplast | | | | 1.000 |
| 389066fa02c4216d54c9d73644e713a7 | Bacteria | Proteobacteria | Deltaproteobacteria | Bdellovibrionales | Bdellovibrionaceae | OM27 clade | uncultured marine bacterium | 0.999 |
| 15885bb18320facec0de5cb894309af3 | Bacteria | Bacteroidetes | Bacteroidia | Flavobacteriales | Flavobacteriaceae | NS5 marine group | | 1.000 |
| 088a9758f9e5da4adaf4e29ea86e6a5b | Bacteria | Proteobacteria | Alphaproteobacteria | SAR11 clade | | | | 1.000 |
| a44dea011b7cd27d7b35b0aa92123cc8 | Bacteria | Bacteroidetes | Bacteroidia | Cytophagales | Cyclobacteriaceae | uncultured | uncultured Bacteroidetes bacterium | 0.858 |
| 95986efe6abb250f94b5db516751e56d | Bacteria | Proteobacteria | Deltaproteobacteria | Bdellovibrionales | Bdellovibrionaceae | OM27 clade | | 1.000 |
| c5e03ed3783b06e0676f452a2b89bf64 | Bacteria | Proteobacteria | Deltaproteobacteria | NB1-j | uncultured marine bacterium | uncultured marine bacterium | uncultured marine bacterium | 0.992 |
| 704f4a03f6937995a459747c1111a1eb | Bacteria | Proteobacteria | Deltaproteobacteria | Bradymonadales | uncultured bacterium | uncultured bacterium | uncultured bacterium | 0.998 |
| 8cb126c23f73c0fe1c9bbdd421f51f39 | Bacteria | Bacteroidetes | Bacteroidia | Flavobacteriales | NS9 marine group | | | 1.000 |
| 08e50b782be6edd7ffdcf52b2c01769e | Bacteria | Proteobacteria | Gammaproteobacteria | Alteromonadales | Alteromonadaceae | | | 1.000 |
| 1d01169d8c7eeee52d3edd346978a8aa | Bacteria | Bacteroidetes | Bacteroidia | Flavobacteriales | NS9 marine group | hydrothermal vent metagenome | hydrothermal vent metagenome | 1.000 |
| 6fd148441ffe78592ca4c3dc0d301ac8 | Bacteria | Bacteroidetes | Bacteroidia | Flavobacteriales | NS9 marine group | uncultured Flavobacteriales bacterium | uncultured Flavobacteriales bacterium | 0.746 |
| 3bf79ebc3c7e8ee32a8fcc25f18c429e | Bacteria | Bacteroidetes | Bacteroidia | Flavobacteriales | NS9 marine group | uncultured Flavobacteriia bacterium | uncultured Flavobacteriia bacterium | 0.748 |
| a08034cfb7b79bf9fb7545edceac7e71 | Bacteria | Proteobacteria | Alphaproteobacteria | Micavibrionales | Micavibrionaceae | uncultured | uncultured bacterium | 0.981 |
| c9a9f40efb94587e51d9d62d09844698 | Bacteria | Proteobacteria | Gammaproteobacteria | Betaproteobacteriales | Burkholderiaceae | MWH-UniP1 aquatic group | uncultured marine bacterium | 0.987 |
| 8e6e50c6ce00858a391ebf57fd7d9e5b | Bacteria | Proteobacteria | Gammaproteobacteria | Coxiellales | Coxiellaceae | Coxiella | uncultured bacterium | 0.984 |
| 564bf8d5970f39dd46de6693fe02f911 | Bacteria | Proteobacteria | Alphaproteobacteria | Rickettsiales | AB1 | uncultured alpha proteobacterium | uncultured alpha proteobacterium | 0.827 |
| 2afb47ba5ef359f27c10d898b5b243b8 | Bacteria | Proteobacteria | Deltaproteobacteria | SAR324 clade(Marine group B) | | | | 1.000 |
| 06f09aed5e3b7a83d07352415c1c04be | Bacteria | Bacteroidetes | Bacteroidia | Flavobacteriales | Flavobacteriaceae | NS5 marine group | | 1.000 |
| eb6c7926524cf5345b6b6c043192849e | Bacteria | Proteobacteria | Gammaproteobacteria | Vibrionales | Vibrionaceae | Vibrio | | 1.000 |
| e026ba5f41ce4323c4efb70065e18af4 | Bacteria | Proteobacteria | Alphaproteobacteria | SAR11 clade | Clade I | Clade Ib | | 0.997 |
| eb44ac888b7cc18167cd2fff31c1455e | Bacteria | Proteobacteria | Alphaproteobacteria | SAR11 clade | Clade II | | | 0.702 |
| db495bd28b4bf328ee567c3db452fdcb | Bacteria | Proteobacteria | Gammaproteobacteria | | | | | 1.000 |
| 2248d33b5b487ce0db56407efb7665c9 | Bacteria | Verrucomicrobia | Verrucomicrobiae | Arctic97B-4 marine group | metagenome | metagenome | metagenome | 0.746 |
| 8da95e55c8d353206e7a837f0555fdc1 | Bacteria | Cyanobacteria | Oxyphotobacteria | Chloroplast | | | | 1.000 |
| 82dd528f90d5062d4a46deff7f2e2c40 | Bacteria | Proteobacteria | Gammaproteobacteria | Ectothiorhodospirales | Ectothiorhodospiraceae | uncultured | | 1.000 |
| 3377f176fcf21ab023a17802b091cb01 | Bacteria | Proteobacteria | Gammaproteobacteria | SAR86 clade | Ambiguous_taxa | Ambiguous_taxa | Ambiguous_taxa | 0.937 |
| fc2c768a1b8bb117063781e28b7e5d14 | Bacteria | Proteobacteria | Alphaproteobacteria | Rhodospirillales | Magnetospiraceae | uncultured | uncultured alpha proteobacterium | 0.786 |
| d65f0c644337272bf24bec18a180b30e | Bacteria | Proteobacteria | Gammaproteobacteria | Alteromonadales | Alteromonadaceae | uncultured | | 0.738 |
| 27ba8f5586f04f3414b3bfd925553231 | Bacteria | Marinimicrobia (SAR406 clade) | uncultured bacterium | uncultured bacterium | uncultured bacterium | uncultured bacterium | uncultured bacterium | 0.966 |
| 84a6628e19d94e146b0438ee91040754 | Bacteria | Planctomycetes | Phycisphaerae | Phycisphaerales | Phycisphaeraceae | FS140-16B-02 marine group | uncultured bacterium | 1.000 |
| c34ac2be3bde10d39ef6104e29a4cae9 | Bacteria | Chloroflexi | Dehalococcoidia | SAR202 clade | uncultured marine microorganism | uncultured marine microorganism | uncultured marine microorganism | 0.709 |
| 4609b3fdee7a2db9e8cac99e6d0fe992 | Bacteria | Marinimicrobia (SAR406 clade) | marine metagenome | marine metagenome | marine metagenome | marine metagenome | marine metagenome | 0.875 |
| c27727ce5053c2f3475b3ce2921579ed | Bacteria | Proteobacteria | Gammaproteobacteria | Thiotrichales | Thiotrichaceae | uncultured | uncultured marine bacterium | 0.851 |
| 91e74e2fc1dc499905a8d59c5f741048 | Bacteria | Proteobacteria | Gammaproteobacteria | Coxiellales | Coxiellaceae | Coxiella | uncultured bacterium | 0.982 |
| 3a101da250415d86317fad6f8b03be2f | Bacteria | Bacteroidetes | Bacteroidia | Flavobacteriales | Cryomorphaceae | uncultured | uncultured Owenweeksia sp. | 0.920 |
| 22caf10c09d69dc90994eec1dd48f872 | Bacteria | Proteobacteria | Gammaproteobacteria | Vibrionales | Vibrionaceae | Vibrio | | 0.998 |
| 8707c0dd0667f9a226a700d5a835bb65 | Bacteria | Bacteroidetes | Bacteroidia | Flavobacteriales | Flavobacteriaceae | NS5 marine group | | 1.000 |
| f820c9a395454c8b4327d34beaf12c2b | Bacteria | Margulisbacteria | uncultured marine cyanobacterium | uncultured marine cyanobacterium | uncultured marine cyanobacterium | uncultured marine cyanobacterium | uncultured marine cyanobacterium | 1.000 |
| 0aeda5d66cc192abf9f47d6d6dd11f8d | Bacteria | Proteobacteria | Gammaproteobacteria | Vibrionales | Vibrionaceae | Vibrio | uncultured marine bacterium | 0.705 |
| 1ad3a013ca39802c694a38bb03eeea1d | Bacteria | Verrucomicrobia | Verrucomicrobiae | Opitutales | Puniceicoccaceae | MB11C04 marine group | | 1.000 |
| fb4c659afc2f990337827c997413acdc | Bacteria | Proteobacteria | Alphaproteobacteria | SAR11 clade | | | | 1.000 |
| 819d506138c655db3983972f18d3a509 | Bacteria | Marinimicrobia (SAR406 clade) | uncultured marine bacterium | uncultured marine bacterium | uncultured marine bacterium | uncultured marine bacterium | uncultured marine bacterium | 0.892 |
| c45482a5b650d5a2a8ca4158a647e844 | Bacteria | Proteobacteria | Deltaproteobacteria | Bdellovibrionales | Bdellovibrionaceae | OM27 clade | | 1.000 |
| 1f883e40a2e31708d6cc982de8030403 | Bacteria | Proteobacteria | Deltaproteobacteria | Bdellovibrionales | Bdellovibrionaceae | OM27 clade | uncultured bacterium | 0.952 |
| 5309c5dfc97d6c5c7a5d0bb86585c999 | Bacteria | Cyanobacteria | Oxyphotobacteria | Chloroplast | uncultured diatom | uncultured diatom | uncultured diatom | 0.801 |
| 4746d060dcfd5cd475d942dff026c51a | Bacteria | Bacteroidetes | Bacteroidia | Flavobacteriales | Flavobacteriaceae | NS5 marine group | uncultured Flavobacteriaceae bacterium | 0.870 |
| 695676ad9e9bf8fc440dfbe8b21c3be5 | Bacteria | Proteobacteria | Alphaproteobacteria | uncultured | uncultured marine bacterium | uncultured marine bacterium | uncultured marine bacterium | 0.953 |

| Feature_ID | Kingdom | Phylum | Class | Order | Family | Genus | Species | Confidence |
|---|---|---|---|---|---|---|---|---|
| 08f7d9154e1d2cfcd090a9d3404a8240 | Bacteria | Actinobacteria | Acidimicrobiia | Microtrichales | Microtrichaceae | Sva0996 marine group | uncultured marine bacterium | 0.810 |
| 26841b4cd9e2d82e0e0713d78ac67cc5 | Bacteria | Proteobacteria | Alphaproteobacteria | Rickettsiales | Midichloriaceae | MD3-55 | | 1.000 |
| 8fdc5ad59a290e03a95ba210fa220f34 | Bacteria | Proteobacteria | Deltaproteobacteria | Bdellovibrionales | Bacteriovoracaceae | uncultured | marine metagenome | 1.000 |
| 9381196598889b85411a88db5c4ecda1 | Bacteria | Bacteroidetes | Bacteroidia | Sphingobacteriales | Lentimicrobiaceae | | | 0.822 |
| 674ad4fd0093e4e9af52e4ac7c9861a7 | Bacteria | Planctomycetes | Phycisphaerae | Phycisphaerales | Phycisphaeraceae | JL-ETNP-F27 | | 1.000 |
| 1d2312249d67dd4cb080857859d359cc | Bacteria | Proteobacteria | Alphaproteobacteria | Rhodobacterales | Rhodobacteraceae | | | 1.000 |
| a8c0a1705820ee05019345f7bd12a48a | Bacteria | Proteobacteria | Deltaproteobacteria | Oligoflexales | Oligoflexaceae | uncultured | uncultured bacterium | 1.000 |
| 06731b15d250abacac01f893c9eefee1 | Bacteria | Bacteroidetes | Bacteroidia | Flavobacteriales | Flavobacteriaceae | Joostella | Ambiguous_taxa | 0.981 |
| 830d8ebb1edcf6fcd0147221d4dab143 | Bacteria | Proteobacteria | Gammaproteobacteria | Alteromonadales | Alteromonadaceae | | | 1.000 |
| 42f32014a4d814b49a564fa5c9c6f9cb | Bacteria | Proteobacteria | Deltaproteobacteria | PB19 | uncultured bacterium | uncultured bacterium | uncultured bacterium | 0.738 |
| e2fab2a594d422476d9139cd7198c5e4 | Bacteria | Proteobacteria | Gammaproteobacteria | Nitrosococcales | Methylophagaceae | Methylophaga | | 1.000 |
| face6e33b73eba34df912fc3c1b28a97 | Bacteria | Cyanobacteria | Oxyphotobacteria | Chloroplast | uncultured bacterium | uncultured bacterium | uncultured bacterium | 0.886 |
| 17bc9a76366330dfb98fd0e357b18ba9 | Bacteria | Proteobacteria | Gammaproteobacteria | EPR3968-O8a-Bc78 | uncultured gamma proteobacterium | uncultured gamma proteobacterium | uncultured gamma proteobacterium | 0.981 |
| 3f4a04a3231869ed49cef439cbbe7397 | Bacteria | Cyanobacteria | Oxyphotobacteria | Chloroplast | | | | 1.000 |
| a610a6301d44bf7a84daf879e9a92a4b | Bacteria | Cyanobacteria | Oxyphotobacteria | Chloroplast | | | | 1.000 |
| 60d93e044dacf3de599b65a9d1cbbe59 | Bacteria | Proteobacteria | Deltaproteobacteria | Bdellovibrionales | Bdellovibrionaceae | OM27 clade | uncultured marine bacterium | 0.935 |
| 3de83fe4c9c3709ec7af5204fc41f74d | Bacteria | Cyanobacteria | Oxyphotobacteria | Nostocales | Phormidiaceae | Trichodesmium IMS101 | Ambiguous_taxa | 0.999 |
| b8de4d844df0e19ba8eddf1e4b44bd62 | Bacteria | Cyanobacteria | Oxyphotobacteria | Chloroplast | | | | 1.000 |
| 95b53fb7d0fc6023a1a921cf00e13cc5 | Bacteria | Proteobacteria | Gammaproteobacteria | Cellvibrionales | Cellvibrionaceae | Gilvimarinus | uncultured bacterium | 0.777 |
| 0cfd5a997e489407f7436413f3604085 | Bacteria | Proteobacteria | Gammaproteobacteria | Gammaproteobacteria Incertae Sedis | Unknown Family | uncultured marine bacterium | uncultured marine bacterium | 1.000 |
| 95e44c0cfc0081858400b5e8e9cf729d | Bacteria | Proteobacteria | Deltaproteobacteria | Myxococcales | P3OB-42 | uncultured marine bacterium | uncultured marine bacterium | 1.000 |
| 3258f6ac7487ad5051a4672a89a011ff | Bacteria | Margulisbacteria | | | | | | 1.000 |
| c0cce68eb3c19a2d2c312d0595cef7ed | Bacteria | Proteobacteria | Alphaproteobacteria | Rickettsiales | S25-593 | uncultured marine bacterium | uncultured marine bacterium | 0.736 |
| 3404f9299340c28afecaf392ecfdd1c4 | Bacteria | Bacteroidetes | Bacteroidia | Flavobacteriales | Flavobacteriaceae | NS4 marine group | | 1.000 |
| cdfbc000e9585911547e376adf8c6d40 | Bacteria | Proteobacteria | Deltaproteobacteria | Oligoflexales | Oligoflexaceae | uncultured | uncultured bacterium | 1.000 |
| 177aaf11037d8ea3f4ec8bb0b37e9a93 | Archaea | Euryarchaeota | Thermoplasmata | Marine Group III | uncultured marine archaeon | uncultured marine archaeon | uncultured marine archaeon | 0.886 |
| 8a9f2ed191788d8008bc416cedc34c2a | Bacteria | Marinimicrobia (SAR406 clade) | uncultured Fibrobacteres/Acidobacteria group bacterium | uncultured Fibrobacteres/Acidobacteria group bacterium | uncultured Fibrobacteres/Acidobacteria group bacterium | uncultured Fibrobacteres/Acidobacteria group bacterium | uncultured Fibrobacteres/Acidobacteria group bacterium | 0.997 |
| 4d5c8a8090fb3872444b7fec20642a31 | Bacteria | Proteobacteria | Gammaproteobacteria | SAR86 clade | | | | 1.000 |
| 4d56acb3808d1d3f000db3318c40f83e | Bacteria | Cyanobacteria | Oxyphotobacteria | Nostocales | Microcystaceae | Atelocyanobacterium (UCYN-A) | Ambiguous_taxa | 1.000 |
| 4c4f92bf3807cd8c84b6a31bf9dde0d9 | Bacteria | Cyanobacteria | Oxyphotobacteria | Chloroplast | uncultured marine eukaryote | uncultured marine eukaryote | uncultured marine eukaryote | 0.930 |
| 95af87e0bd799ae45d49df55f73e6954 | Bacteria | Proteobacteria | Gammaproteobacteria | Cellvibrionales | Cellvibrionaceae | uncultured | uncultured gamma proteobacterium | 0.865 |
| 124813d7efd84d0fb2053eea0391c5dc | Bacteria | Cyanobacteria | Oxyphotobacteria | Chloroplast | Cylindrotheca closterium | Cylindrotheca closterium | Cylindrotheca closterium | 0.988 |
| 04aef29ff0a4e1fc0a82b01ddb1a4e40 | Bacteria | Verrucomicrobia | Verrucomicrobiae | Arctic97B-4 marine group | uncultured marine bacterium | uncultured marine bacterium | uncultured marine bacterium | 0.977 |
| b15c1b61bf901ec81957f583a04224f3 | Bacteria | Proteobacteria | Deltaproteobacteria | Bdellovibrionales | Bdellovibrionaceae | OM27 clade | unidentified marine bacterioplankton | 0.999 |
| 49aeb496d206146c69ede52850843148 | Bacteria | Cyanobacteria | Oxyphotobacteria | Chloroplast | | | | 1.000 |
| ea6bc720f965aa3d0e196aca79787c12 | Bacteria | Proteobacteria | Gammaproteobacteria | Alteromonadales | Shewanellaceae | Shewanella | | 1.000 |
| 3e9e854ae4fc8c0e2b869f9e73bbe7c4 | Bacteria | Proteobacteria | Gammaproteobacteria | KI89A clade | | | | 1.000 |
| 38f7c51a29ebb6421963c79aa84a321e | Bacteria | Proteobacteria | Alphaproteobacteria | Rhodospirillales | AEGEAN-169 marine group | | | 1.000 |
| 5cbbb3191e720136b04253284fa52c69 | Bacteria | Proteobacteria | Deltaproteobacteria | Bdellovibrionales | Bdellovibrionaceae | OM27 clade | | 1.000 |
| a631f22bbf6ad3330eb6e5508ba4e37e | Bacteria | Cyanobacteria | Oxyphotobacteria | Chloroplast | | | | 1.000 |
| 15ad8638c39176ce52b676bedc5dada1 | Bacteria | Cyanobacteria | Oxyphotobacteria | Chloroplast | | | | 1.000 |
| 3835d0369abd8f8d64d96db200164d77 | Bacteria | Proteobacteria | Alphaproteobacteria | Sneathiellales | Sneathiellaceae | AT-s3-44 | uncultured marine bacterium | 0.986 |
| 062c5c26200a970015d23571f4f4e97c | Bacteria | Marinimicrobia (SAR406 clade) | uncultured bacterium | uncultured bacterium | uncultured bacterium | uncultured bacterium | uncultured bacterium | 0.783 |
| dcb98b9f9b6edee73f0409e7c2b91f8f | Bacteria | Proteobacteria | Gammaproteobacteria | Pseudomonadales | Moraxellaceae | Acinetobacter | | 1.000 |
| 564c3c6610a1d7b46b32972d5c363fda | Bacteria | Proteobacteria | Alphaproteobacteria | Rhodospirillales | Magnetospiraceae | uncultured | uncultured marine bacterium | 0.966 |
| 8675c76a37afa539c4cbdf680bf9e776 | Bacteria | Cyanobacteria | Oxyphotobacteria | Chloroplast | marine metagenome | marine metagenome | marine metagenome | 0.999 |
| a5e1d9afda67ebb3082a5ee93c5e8faa | Bacteria | Proteobacteria | Alphaproteobacteria | Rickettsiales | S25-593 | | | 1.000 |
| f9c91207f762e3f6b5c678a48a4e4790 | Bacteria | Proteobacteria | Alphaproteobacteria | Rhodospirillales | Magnetospiraceae | uncultured | marine metagenome | 0.998 |
| ebe708b42df1fb5343799dd02e143c10 | Bacteria | Marinimicrobia (SAR406 clade) | metagenome | metagenome | metagenome | metagenome | metagenome | 0.868 |
| 6598be55fe50a2139cfba55d3ee622fd | Bacteria | Planctomycetes | OM190 | uncultured Planctomycetaceae bacterium | uncultured Planctomycetaceae bacterium | uncultured Planctomycetaceae bacterium | uncultured Planctomycetaceae bacterium | 0.915 |
| 1921cf951a8b48e97d556fa59188b1e1 | Bacteria | Proteobacteria | Deltaproteobacteria | Oligoflexales | Oligoflexaceae | uncultured | uncultured bacterium | 0.977 |
| 72478d44ccfe95f06b2db611a268391b | Bacteria | Proteobacteria | Deltaproteobacteria | Bdellovibrionales | Bdellovibrionaceae | OM27 clade | Ambiguous_taxa | 0.983 |
| c77a164cd2429a25febf7648c932a4f3 | Bacteria | Proteobacteria | Alphaproteobacteria | | | | | 1.000 |
| e34ef8095dd1e5b5affde91cb3f07327 | Bacteria | Proteobacteria | Deltaproteobacteria | Bdellovibrionales | Bdellovibrionaceae | OM27 clade | | 0.966 |
| bb157bfccbd11ae30c7449b2ac0279a8 | Bacteria | Bacteroidetes | Bacteroidia | Flavobacteriales | Flavobacteriaceae | NS5 marine group | | 1.000 |
| ce6531e8c56ad6900d6c31e9339913e6 | Bacteria | Bacteroidetes | Bacteroidia | Chitinophagales | Saprospiraceae | uncultured | unidentified marine eubacterium | 1.000 |
| e7987a6fcb37c925f38b08b88214c9e6 | Bacteria | Bacteroidetes | Bacteroidia | Flavobacteriales | NS9 marine group | | | 1.000 |
| 1c54604c687153f81612b4d10e481e77 | Bacteria | Cyanobacteria | Oxyphotobacteria | Nostocales | Microcystaceae | Atelocyanobacterium (UCYN-A) | Ambiguous_taxa | 1.000 |
| 580190655efb478a9783c9f56245e774 | Bacteria | Bacteroidetes | Bacteroidia | Flavobacteriales | Cryomorphaceae | uncultured | | 0.759 |
| 2ac69cc152adf9c108cdc2adca94c2c4 | Bacteria | Proteobacteria | Gammaproteobacteria | Oceanospirillales | Alcanivoracaceae | Alcanivorax | uncultured Alcanivorax sp. | 0.849 |
| 108581dd41a110366795f9cd45e9ef9e | Bacteria | Proteobacteria | Gammaproteobacteria | Oceanospirillales | Halomonadaceae | Halomonas | | 1.000 |
| 6e4f053a5105f5c1c2a47a5e63057595 | Bacteria | Bacteroidetes | Bacteroidia | Cytophagales | Cyclobacteriaceae | uncultured | uncultured bacterium | 0.775 |
| 2d56d9d44ff80b6419a42f7cd240b01e | Bacteria | Proteobacteria | Gammaproteobacteria | Alteromonadales | Alteromonadaceae | Alteromonas | | 0.930 |
| ac3694e776b7854f0eab79d3a3cea64d | Bacteria | Proteobacteria | Deltaproteobacteria | PB19 | uncultured delta proteobacterium | uncultured delta proteobacterium | uncultured delta proteobacterium | 0.802 |
| 3554e46b91e25e22492d6ff951ff9132 | Bacteria | Proteobacteria | Alphaproteobacteria | | | | | 1.000 |
| d26cd7f6bdb174b5f292ede6e5f66b83 | Bacteria | Proteobacteria | Deltaproteobacteria | Oligoflexales | Oligoflexaceae | uncultured | | 1.000 |
| 726cf59156ce5cc81212a5781b7b51ee | Bacteria | Verrucomicrobia | Verrucomicrobiae | Opitutales | Puniceicoccaceae | MB11C04 marine group | uncultured Verrucomicrobia bacterium | 0.898 |
| f7cba05fcaeeb04512f63d9dc07df9d7 | Bacteria | Verrucomicrobia | Verrucomicrobiae | Opitutales | Puniceicoccaceae | MB11C04 marine group | uncultured marine microorganism | 0.972 |
| 72aa622a6806dbd010efc5162ddd45a0 | Bacteria | Proteobacteria | Deltaproteobacteria | Bdellovibrionales | Bacteriovoracaceae | uncultured | uncultured marine bacterium | 0.794 |
| fec41ffee3bf29a2dbb32158baaba256 | Bacteria | Proteobacteria | Deltaproteobacteria | PB19 | Ambiguous_taxa | Ambiguous_taxa | Ambiguous_taxa | 0.880 |
| 263c7a8bb12e13ebca4accfee8ab75c5 | Bacteria | Bacteroidetes | Bacteroidia | Flavobacteriales | Flavobacteriaceae | | | 1.000 |
| 8a78b2a83c2261a0af9cdf8dc5166e6b | Bacteria | Proteobacteria | Gammaproteobacteria | Oceanospirillales | Alcanivoracaceae | Alcanivorax | | 1.000 |
| 943b298ff2de3289f1a030f28cae8cd7 | Bacteria | Bacteroidetes | Bacteroidia | Flavobacteriales | Flavobacteriaceae | Leeuwenhoekiella | Ambiguous_taxa | 0.991 |
| baa60b1c0933dd8ea7dd4fcb169ea545 | Bacteria | Proteobacteria | Deltaproteobacteria | Bdellovibrionales | Bdellovibrionaceae | OM27 clade | | 1.000 |

| Feature_ID | Kingdom | Phylum | Class | Order | Family | Genus | Species | Confidence |
|---|---|---|---|---|---|---|---|---|
| 3ab2f61acc234dbf7e15decdd063bda2 | Bacteria | Proteobacteria | Deltaproteobacteria | Bdellovibrionales | Bacteriovoracaceae | uncultured | uncultured marine bacterium | 0.788 |
| ecbe8a8ccb118a09b551ede83d9bd156 | Bacteria | Proteobacteria | Alphaproteobacteria | Rickettsiales | S25-593 | uncultured marine bacterium | uncultured marine bacterium | 0.985 |
| f333eb1f71a39bbfb57fc69e28889302 | Bacteria | Proteobacteria | Gammaproteobacteria | Vibrionales | Vibrionaceae | Vibrio | | 0.999 |
| 59cdf625fe552ea257c8448a1be68a71 | Bacteria | Proteobacteria | Gammaproteobacteria | Coxiellales | Coxiellaceae | Coxiella | uncultured bacterium | 0.769 |
| 15f5ea541d3f680e1c5261a8f976ec36 | Bacteria | Bacteroidetes | Bacteroidia | Flavobacteriales | Crocinitomicaceae | Fluviicola | unidentified marine bacterioplankton | 0.922 |
| 6a6bbedb9479860b6ad9e339c3afc02d | Bacteria | Cyanobacteria | Oxyphotobacteria | Chloroplast | | | | 1.000 |
| 9fdc6db7f88f1e100435b617bb90abb4 | Bacteria | Proteobacteria | Deltaproteobacteria | Bdellovibrionales | Bdellovibrionaceae | OM27 clade | uncultured marine bacterium | 0.920 |
| 6e9fb661da416eaad867b6bec08c384c | Bacteria | Bacteroidetes | Bacteroidia | Flavobacteriales | NS9 marine group | uncultured marine bacterium | uncultured marine bacterium | 0.998 |
| 98f11dae43bd94330ad8bcc7c81f64b7 | Bacteria | Proteobacteria | Gammaproteobacteria | Thiotrichales | Thiotrichaceae | uncultured | | 1.000 |
| 8c9a461cb0181764183ce27a326f7312 | Bacteria | Cyanobacteria | Oxyphotobacteria | Chloroplast | | | | 1.000 |
| 246b7de821c45dcb1c6aa7354d9234be | Bacteria | Proteobacteria | Deltaproteobacteria | Bdellovibrionales | Bdellovibrionaceae | OM27 clade | uncultured marine bacterium | 0.952 |
| b7b973f5199a63612e67c50f4d3f6956 | Bacteria | Proteobacteria | Deltaproteobacteria | Bdellovibrionales | Bdellovibrionaceae | OM27 clade | uncultured marine bacterium | 0.994 |
| 6e54cec1baba407238a8a50b2182e6ce | Bacteria | Proteobacteria | Alphaproteobacteria | Rickettsiales | Midichloriaceae | MD3-55 | | 1.000 |
| b7c31f68ffdb36fded29daff92f61ac6 | Bacteria | Proteobacteria | Gammaproteobacteria | Coxiellales | Coxiellaceae | Coxiella | uncultured marine bacterium | 0.920 |
| 9a40dc9e0f4026f7a7cdf205b122dcf4 | Bacteria | Proteobacteria | Deltaproteobacteria | Bdellovibrionales | Bdellovibrionaceae | OM27 clade | uncultured marine bacterium | 0.977 |
| 0fae4a961394e190e538ed045712f96b | Bacteria | Acidobacteria | Subgroup 6 | uncultured bacterium AD248-D7-1A | uncultured bacterium AD248-D7-1A | uncultured bacterium AD248-D7-1A | uncultured bacterium AD248-D7-1A | 1.000 |
| 509d18c6dbf5f602454babb84ccb7a08 | Bacteria | Proteobacteria | Gammaproteobacteria | Alteromonadales | | | | 0.895 |
| 3a5cd96f54d5d5e2438f65901797579d | Bacteria | Marinimicrobia (SAR406 clade) | uncultured bacterium | uncultured bacterium | uncultured bacterium | uncultured bacterium | uncultured bacterium | 0.976 |
| f5568c692638dad4055d068e7ac42f4b | Bacteria | Marinimicrobia (SAR406 clade) | uncultured marine bacterium | uncultured marine bacterium | uncultured marine bacterium | uncultured marine bacterium | uncultured marine bacterium | 0.979 |
| a3b2da88ac893b68ca4118fa22a9be4d | Bacteria | Proteobacteria | Gammaproteobacteria | Alteromonadales | Colwelliaceae | Thalassotalea | | 1.000 |
| c70043f9bb0a913d8511927cf3ad75df | Bacteria | Proteobacteria | Deltaproteobacteria | SAR324 clade(Marine group B) | | | | 1.000 |
| 489760bf3d8630c338b9555185c85393 | Bacteria | Proteobacteria | Alphaproteobacteria | SAR11 clade | Clade I | Clade Ia | unidentified marine eubacterium | 0.852 |
| 66f5120c0c6b6c37ecf4c68b9ccbd84c | Bacteria | Bacteroidetes | Bacteroidia | Chitinophagales | Saprospiraceae | Aureispira | uncultured Saprospiraceae bacterium | 0.932 |
| 3fc41c8a7634c1db166ac76b215ae08f | Bacteria | Cyanobacteria | Oxyphotobacteria | Chloroplast | | | | 1.000 |
| 8e8214b1c5b5a374b0fc694198b26892 | Bacteria | Proteobacteria | Deltaproteobacteria | PB19 | uncultured delta proteobacterium | uncultured delta proteobacterium | uncultured delta proteobacterium | 0.998 |
| 33e705ea1eba802a738eb2d83c971545 | Bacteria | Proteobacteria | Gammaproteobacteria | Cellvibrionales | Cellvibrionaceae | Umboniibacter | uncultured bacterium | 0.993 |
| a651f5af1c2290b9f98427ad86dad014 | Bacteria | Bacteroidetes | Bacteroidia | Cytophagales | Cyclobacteriaceae | uncultured | uncultured marine bacterium | 1.000 |
| 0e9b8b829dbc8224db268ccb40ac244e | Bacteria | Proteobacteria | Gammaproteobacteria | SAR86 clade | uncultured marine bacterium | uncultured marine bacterium | uncultured marine bacterium | 0.725 |
| d3c66e77494f1ca7509c81285eb0bbeb | Bacteria | Cyanobacteria | Oxyphotobacteria | Chloroplast | uncultured prymnesiophyte C5574 | uncultured prymnesiophyte C5574 | uncultured prymnesiophyte C5574 | 0.850 |
| 97e80f21afbfa600efc7d5e1a1ec841a | Bacteria | Bacteroidetes | Bacteroidia | Flavobacteriales | Flavobacteriaceae | Leeuwenhoekiella | Ambiguous_taxa | 0.996 |
| d070a05a7ab0838ceb97040c56f8c826 | Bacteria | | | | | | | 0.998 |
| 43d1a2865083679a9e704e21e15c1e97 | Bacteria | Cyanobacteria | Oxyphotobacteria | Chloroplast | | | | 1.000 |
| a5413ebe87428ad9c522b477b96a9814 | Bacteria | Cyanobacteria | Oxyphotobacteria | Chloroplast | | | | 1.000 |
| 9edef7d55154ea788972b95ad557e425 | Bacteria | Bacteroidetes | Bacteroidia | Flavobacteriales | Cryomorphaceae | uncultured | uncultured organism | 0.906 |
| 2e4be662f07eb90fc93d4227815035f4 | Bacteria | Proteobacteria | Gammaproteobacteria | Betaproteobacteriales | Methylophilaceae | OM43 clade | Ambiguous_taxa | 0.916 |
| a808d3512350b552d836b87b7847f05d | Bacteria | Proteobacteria | Deltaproteobacteria | Bdellovibrionales | Bdellovibrionaceae | OM27 clade | | 1.000 |
| 2ba659a32f325695bb8a15542c316bc2 | Bacteria | Firmicutes | Bacilli | Lactobacillales | Streptococcaceae | Streptococcus | Streptococcus salivarius subsp. thermophilus | 0.913 |
| 87e65411867e8f3ff99f7a3b84c7bd85 | Bacteria | Proteobacteria | Deltaproteobacteria | Myxococcales | Sandaracinaceae | uncultured | uncultured delta proteobacterium | 0.838 |
| 94a0ddc0e963a3710f1ddb1e92752acf | Bacteria | Proteobacteria | Alphaproteobacteria | Parvibaculales | OCS116 clade | uncultured marine bacterium | uncultured marine bacterium | 0.973 |
| 53d4e655ba7bd47ab0584caf454c6701 | Bacteria | Proteobacteria | Gammaproteobacteria | Coxiellales | Coxiellaceae | Coxiella | uncultured bacterium | 0.709 |
| 62d822227d0dd754c5a2a9f910ba9ac9 | Bacteria | Proteobacteria | Alphaproteobacteria | Rhodospirillales | Magnetospiraceae | Varunaivibrio | uncultured marine bacterium | 1.000 |
| ac0f3d11539f32d50d4c766e70ddda39 | Bacteria | Proteobacteria | Gammaproteobacteria | OM182 clade | uncultured marine microorganism | uncultured marine microorganism | uncultured marine microorganism | 1.000 |
| ab20d637985d91aa1d1a1ae4618a81e6 | Bacteria | Bacteroidetes | Bacteroidia | Flavobacteriales | NS9 marine group | uncultured marine bacterium | uncultured marine bacterium | 0.943 |
| dba9f72c944f23bf1064240c9c0addab | Bacteria | Bacteroidetes | Bacteroidia | Flavobacteriales | Cryomorphaceae | uncultured | uncultured Owenweeksia sp. | 0.829 |
| 1393f76bb109e2cb04f0fa618d93acba | Bacteria | Proteobacteria | Deltaproteobacteria | Bdellovibrionales | Bacteriovoracaceae | uncultured | uncultured marine bacterium | 1.000 |
| f682a9827a51759f9e6521e43738ae03 | Bacteria | Planctomycetes | Phycisphaerae | Phycisphaerales | Phycisphaeraceae | FS140-16B-02 marine group | Ambiguous_taxa | 0.828 |
| 922f28646c7468d0ba06ae438a3f4c38 | Bacteria | Bacteroidetes | Bacteroidia | Cytophagales | Cyclobacteriaceae | uncultured | uncultured bacterium | 0.949 |
| 23cd2b04141187e066e74330d91295a0 | Bacteria | Marinimicrobia (SAR406 clade) | unidentified marine bacterioplankton | unidentified marine bacterioplankton | unidentified marine bacterioplankton | unidentified marine bacterioplankton | unidentified marine bacterioplankton | 0.984 |
| 7f0420aa9f2555976c432cf19fedcb0e | Bacteria | Proteobacteria | Gammaproteobacteria | Ga0077536 | uncultured bacterium | uncultured bacterium | uncultured bacterium | 0.955 |
| d949487c027f8702ae7f2934bb047524 | Bacteria | Proteobacteria | Gammaproteobacteria | | | | | 1.000 |
| 793d5df649983fe2c6e6b805e8a82599 | Bacteria | Margulisbacteria | metagenome | metagenome | metagenome | metagenome | metagenome | 0.984 |
| 05ba0090032caa922f6586e0d6110d03 | Bacteria | Cyanobacteria | Oxyphotobacteria | Chloroplast | uncultured marine eukaryote | uncultured marine eukaryote | uncultured marine eukaryote | 0.981 |
| 3d651a342f804d73db76e69ca097a5c3 | Bacteria | Proteobacteria | Gammaproteobacteria | Betaproteobacteriales | Burkholderiaceae | MWH-UniP1 aquatic group | uncultured bacterium | 0.940 |
| 9e7e8df738a103cc431e0d989c3e1cc6 | Bacteria | Bacteroidetes | Bacteroidia | Flavobacteriales | Cryomorphaceae | NS10 marine group | uncultured marine bacterium | 0.972 |
| 001de51c1b99e92f09510a30248fa0fc | Bacteria | Proteobacteria | Gammaproteobacteria | KI89A clade | | | | 1.000 |
| 8cd7b7a8c140912813bb62d2b4695487 | Bacteria | Proteobacteria | Deltaproteobacteria | SAR324 clade(Marine group B) | uncultured bacterium | uncultured bacterium | uncultured bacterium | 0.894 |
| a44e518aa92d1f0007fece9fe55a7588 | Bacteria | Proteobacteria | Gammaproteobacteria | SAR86 clade | metagenome | metagenome | metagenome | 0.998 |
| 66e1ef493960e6e492820ea2f6381282 | Bacteria | Proteobacteria | Deltaproteobacteria | Oligoflexales | Oligoflexaceae | uncultured | uncultured bacterium | 1.000 |
| 7d3e1f71754a5d2a13420ff415486f30 | Bacteria | Proteobacteria | Gammaproteobacteria | Oceanospirillales | Marinomonadaceae | Marinomonas | Achromobacter cholinophagum | 0.955 |
| ca0727f74f0cc400c4aa687b5664573f | Bacteria | Marinimicrobia (SAR406 clade) | uncultured bacterium MB13C05 | uncultured bacterium MB13C05 | uncultured bacterium MB13C05 | uncultured bacterium MB13C05 | uncultured bacterium MB13C05 | 0.953 |
| 4d94478a3714e0dc7df0a17bbbcf3095 | Bacteria | Proteobacteria | Deltaproteobacteria | Myxococcales | P3OB-42 | uncultured bacterium | uncultured bacterium | 0.928 |
| 200462cdae28b75673fa86076de5eb9f | Bacteria | Bacteroidetes | Bacteroidia | Cytophagales | Cyclobacteriaceae | uncultured | uncultured Bacteroidetes bacterium | 0.730 |
| e4298f09f6a24096c64f84c04a4e0e1d | Bacteria | Proteobacteria | Gammaproteobacteria | Gammaproteobacteria Incertae Sedis | Unknown Family | uncultured marine bacterium | uncultured marine bacterium | 1.000 |
| 2276e0df2a2758aa75c2fef7278264dd | Bacteria | Proteobacteria | Gammaproteobacteria | Oceanospirillales | Oleiphilaceae | Oleiphilus | uncultured bacterium | 0.858 |
| b0dc5f42017e58175f9c63f4a2d7f738 | Bacteria | Bacteroidetes | Bacteroidia | Chitinophagales | Saprospiraceae | uncultured | uncultured bacterium | 0.919 |

| Feature_ID | Kingdom | Phylum | Class | Order | Family | Genus | Species | Confidence |
|---|---|---|---|---|---|---|---|---|
| 5caa316ccb110285eb4ef5e8a848fc0e | Bacteria | Proteobacteria | Gammaproteobacteria | Coxiellales | Coxiellaceae | Coxiella | Ambiguous_taxa | 0.955 |
| 0eb26f6155f1348e4f4a4159d9d5e183 | Bacteria | Proteobacteria | Gammaproteobacteria | Alteromonadales | Alteromonadaceae | Aestuariibacter | | 0.875 |
| f9ca9b4717452789fc9277d5fb6805f9 | Bacteria | Proteobacteria | Deltaproteobacteria | SAR324 clade(Marine group B) | uncultured marine bacterium | uncultured marine bacterium | uncultured marine bacterium | 0.753 |
| fe2e9c776a53a048453e2af7905842b3 | Bacteria | Bacteroidetes | Bacteroidia | Chitinophagales | Saprospiraceae | uncultured | | 0.999 |
| 9621167ddea9bf14cfaf69e9aa64adc5 | Bacteria | Bacteroidetes | Bacteroidia | Cytophagales | Cyclobacteriaceae | uncultured | uncultured marine bacterium | 1.000 |
| 49fd7322a61a4e1f0e563e194ca91bef | Bacteria | Proteobacteria | Alphaproteobacteria | Rhodospirillales | Magnetospiraceae | uncultured | | 1.000 |
| 9e8488ea1b111a5d7f51b67c73187fda | Bacteria | Cyanobacteria | Oxyphotobacteria | Chloroplast | | | | 1.000 |
| c3f312932b7c3ed5454ead5114e708c9 | Bacteria | Cyanobacteria | Oxyphotobacteria | Chloroplast | | | | 1.000 |
| a88539f77639d0d912e4f05c896aa553 | Bacteria | Verrucomicrobia | Verrucomicrobiae | Arctic97B-4 marine group | uncultured organism | uncultured organism | uncultured organism | 0.896 |
| 0e4dc486dd016bc0ed9559944379f437 | Bacteria | Proteobacteria | Alphaproteobacteria | Rickettsiales | S25-593 | uncultured bacterium | uncultured bacterium | 0.928 |
| c9f02f1cb795a8a03b4318e3ec4afffe | Bacteria | Bacteroidetes | Bacteroidia | Flavobacteriales | Flavobacteriaceae | Gramella | uncultured bacterium | 0.784 |
| 3a2337d28291dc6d1d90d8dcc6fcc11a | Bacteria | Bacteroidetes | Bacteroidia | Flavobacteriales | NS7 marine group | Ambiguous_taxa | Ambiguous_taxa | 0.942 |
| 42f24da52fcf1f82c0452d6aec8f43c2 | Bacteria | Marinimicrobia (SAR406 clade) | Marinimicrobia bacterium JGI 0000113-D11 | Marinimicrobia bacterium JGI 0000113-D11 | Marinimicrobia bacterium JGI 0000113-D11 | Marinimicrobia bacterium JGI 0000113-D11 | Marinimicrobia bacterium JGI 0000113-D11 | 0.965 |
| 3036c113bed51ae4ce64c2101df060a1 | Bacteria | Chloroflexi | Dehalococcoidia | SAR202 clade | | | | 1.000 |
| 74edab1156c1e9d8dc1a93b4e581340c | Bacteria | Proteobacteria | Gammaproteobacteria | OM182 clade | uncultured marine microorganism | uncultured marine microorganism | uncultured marine microorganism | 1.000 |
| bdcf1f0f9d60f20de4e2e13519239b7e | Bacteria | Firmicutes | Bacilli | Bacillales | Staphylococcaceae | Staphylococcus | | 1.000 |
| af42e6e4fd9ce265eac5fbf5be1e53fe | Bacteria | Margulisbacteria | uncultured bacterium | uncultured bacterium | uncultured bacterium | uncultured bacterium | uncultured bacterium | 0.957 |
| 49a3ac4f56939e8cb917c275f2f658c3 | Bacteria | Verrucomicrobia | Verrucomicrobiae | Opitutales | Puniceicoccaceae | MB11C04 marine group | uncultured Verrucomicrobia bacterium | 0.745 |
| 05ed747393e56a4ac09caa3d55dcd5e0 | Archaea | Euryarchaeota | Thermoplasmata | Marine Group III | Ambiguous_taxa | Ambiguous_taxa | Ambiguous_taxa | 0.896 |
| d4929e92dd032b69e53d422a6b7d67c4 | Bacteria | Proteobacteria | Alphaproteobacteria | Rhodospirillales | AEGEAN-169 marine group | uncultured bacterium | uncultured bacterium | 0.982 |
| 6cc38a14c3d89a96bc4e7eace391ee46 | Bacteria | Chloroflexi | Dehalococcoidia | SAR202 clade | uncultured Chloroflexi bacterium HF4000_28F02 | uncultured Chloroflexi bacterium HF4000_28F02 | uncultured Chloroflexi bacterium HF4000_28F02 | 0.999 |
| 71ef1116b46ad80e44b4addf828fb6e5 | Bacteria | Chloroflexi | Dehalococcoidia | SAR202 clade | uncultured SAR202 cluster bacterium | uncultured SAR202 cluster bacterium | uncultured SAR202 cluster bacterium | 0.796 |
| d770a3bc8c48882c5a09855445637762 | Bacteria | Proteobacteria | Deltaproteobacteria | Bdellovibrionales | Bacteriovoracaceae | uncultured | marine metagenome | 0.848 |
| 66eec867ffd555c10a1eb61e4822f1a3 | Bacteria | Cyanobacteria | Oxyphotobacteria | Chloroplast | uncultured marine eukaryote | uncultured marine eukaryote | uncultured marine eukaryote | 0.872 |
| 8d59216f6a8027463789bb6a3be53c00 | Bacteria | Margulisbacteria | uncultured marine bacterium | uncultured marine bacterium | uncultured marine bacterium | uncultured marine bacterium | uncultured marine bacterium | 0.999 |
| 93843b3fbf03c8a0d144d88bfb670d38 | Bacteria | Proteobacteria | Deltaproteobacteria | Oligoflexales | 053A03-B-DI-P58 | | | 0.951 |
| 23e5352a7b4a5f887afc8f4976521f56 | Bacteria | Proteobacteria | Gammaproteobacteria | Ectothiorhodospirales | Ectothiorhodospiraceae | uncultured | | 1.000 |
| 2785a702537affce697fc936c2adb1b7 | Bacteria | Proteobacteria | Deltaproteobacteria | Myxococcales | Eel-36e1D6 | uncultured delta proteobacterium | uncultured delta proteobacterium | 1.000 |
| c20d4ad50c059e48ecf84e0ce5f6b22c | Bacteria | Proteobacteria | Alphaproteobacteria | Rhodospirillales | AEGEAN-169 marine group | | | 1.000 |
| afe5bed304b45d993f5bb92da51e4c04 | Bacteria | Proteobacteria | Deltaproteobacteria | Bradymonadales | uncultured delta proteobacterium | uncultured delta proteobacterium | uncultured delta proteobacterium | 1.000 |
| 47d479b609afd0946aecbd9abed79a09 | Archaea | Thaumarchaeota | Nitrososphaeria | Nitrosopumilales | Nitrosopumilaceae | Candidatus Nitrosopelagicus | uncultured marine archaeon | 0.927 |
| d4e873350e95ce20129bc6636b3f7ecd | Bacteria | Marinimicrobia (SAR406 clade) | | | | | | 1.000 |
| 0bcab4ff927bf216f2c3de77dda28434 | Bacteria | Proteobacteria | Gammaproteobacteria | Vibrionales | Vibrionaceae | Vibrio | | 0.948 |
| b13a6e99d54f6c1105b0182debd65121 | Bacteria | Cyanobacteria | Oxyphotobacteria | Chloroplast | uncultured bacterium | uncultured bacterium | uncultured bacterium | 0.792 |
| 398b7a80dc26d77f1028fb5d631d7355 | Bacteria | Proteobacteria | Gammaproteobacteria | SAR86 clade | | | | 1.000 |
| 3f08bb4aa92817a4684bcccf54b9ce08 | Bacteria | Proteobacteria | Gammaproteobacteria | Alteromonadales | Alteromonadaceae | | | 1.000 |
| 6f3554b8e49cc2a513d9cd3c94bb16a4 | Bacteria | Proteobacteria | Alphaproteobacteria | Rickettsiales | Midichloriaceae | MD3-55 | Ambiguous_taxa | 0.990 |
| 8fc4f917919c0e03531c463916c49cc2 | Bacteria | Proteobacteria | Gammaproteobacteria | Steroidobacterales | Woeseiaceae | Woeseia | uncultured marine bacterium | 1.000 |
| 5c1723f82a8ac43ccba713ad7b08fb1c | Bacteria | Proteobacteria | Deltaproteobacteria | PB19 | | | | 1.000 |
| 6150f78b7fa8370afc64e9a76d91b807 | Bacteria | Margulisbacteria | uncultured marine cyanobacterium | uncultured marine cyanobacterium | uncultured marine cyanobacterium | uncultured marine cyanobacterium | uncultured marine cyanobacterium | 0.999 |
| c8eed146433dcc71fa4dfa89d2e8fa05 | Bacteria | Proteobacteria | Gammaproteobacteria | Piscirickettsiales | Piscirickettsiaceae | Candidatus Endoecteinascidia | uncultured marine bacterium | 1.000 |
| 48b1d0d4a44acc2206f4b152abeedf0a | Bacteria | Bacteroidetes | Bacteroidia | Flavobacteriales | Flavobacteriaceae | NS5 marine group | uncultured Flavobacteriaceae bacterium | 0.916 |
| 2193f1a58afd44d536db4fbac52e78e7 | Bacteria | Marinimicrobia (SAR406 clade) | uncultured marine bacterium | uncultured marine bacterium | uncultured marine bacterium | uncultured marine bacterium | uncultured marine bacterium | 0.943 |
| 1176e7dc97e8723fc8461b18b78dc510 | Bacteria | Bacteroidetes | Bacteroidia | Flavobacteriales | Flavobacteriaceae | Flavobacterium | Flavobacterium sp. SM1502 | 0.915 |
| 89a258b55e5f7d8621f92089614e02bd | Bacteria | Proteobacteria | Deltaproteobacteria | Oligoflexales | Oligoflexaceae | uncultured | uncultured marine bacterium | 1.000 |
| 3fc88169d2f320339ac71dcb12e4e00c | Bacteria | Cyanobacteria | Oxyphotobacteria | Chloroplast | | | | 1.000 |
| aa215cef5ee303af70e3d53aa11102bc | Bacteria | Verrucomicrobia | Verrucomicrobiae | Opitutales | Puniceicoccaceae | Coraliomargarita | uncultured Verrucomicrobia bacterium | 0.952 |
| 001f650db10d041f58889d89c7b568d1 | Bacteria | Cyanobacteria | Oxyphotobacteria | Chloroplast | uncultured bacterium | uncultured bacterium | uncultured bacterium | 0.869 |
| 171bbc6a554d9c341a669bded9b11c0a | Bacteria | Margulisbacteria | uncultured marine cyanobacterium | uncultured marine cyanobacterium | uncultured marine cyanobacterium | uncultured marine cyanobacterium | uncultured marine cyanobacterium | 0.995 |
| cd8c09391fa8e30a173e06e3972d5dae | Bacteria | Bacteroidetes | Bacteroidia | Flavobacteriales | Flavobacteriaceae | Ulvibacter | uncultured marine bacterium | 0.974 |
| 0e6ac77416b8cfb9d8c10727dfd3a526 | Bacteria | Proteobacteria | Alphaproteobacteria | SAR11 clade | Clade II | uncultured Candidatus Pelagibacter sp. | uncultured Candidatus Pelagibacter sp. | 0.730 |
| e2c9a0e5e622f6a0154451da7d7921c6 | Bacteria | Bacteroidetes | Bacteroidia | Flavobacteriales | NS9 marine group | uncultured Flavobacteriia bacterium | uncultured Flavobacteriia bacterium | 0.943 |
| 1b9bd4a7185432d3f42a79b6bc76a82a | Bacteria | Marinimicrobia (SAR406 clade) | uncultured marine microorganism | uncultured marine microorganism | uncultured marine microorganism | uncultured marine microorganism | uncultured marine microorganism | 1.000 |
| 6313cb7de0be333c38fab205a680ba5b | Bacteria | Proteobacteria | Deltaproteobacteria | Bdellovibrionales | Bdellovibrionaceae | OM27 clade | uncultured alpha proteobacterium | 0.972 |
| 9a0cc027d73c3315610d1fd1b24d2bc1 | Bacteria | Proteobacteria | Deltaproteobacteria | Bdellovibrionales | Bdellovibrionaceae | OM27 clade | | 1.000 |
| aa225ae35df636d92ed55b79d4141eaf | Bacteria | Bacteroidetes | Bacteroidia | Flavobacteriales | Flavobacteriaceae | NS5 marine group | uncultured marine bacterium | 0.710 |
| d9b94b24f2feab12b6e65903e1756929 | Bacteria | Bacteroidetes | Bacteroidia | Flavobacteriales | Cryomorphaceae | uncultured | | 1.000 |
| 24dcc3462a9bed82d0362bb1b1cef52a | Bacteria | Proteobacteria | Alphaproteobacteria | | | | | 1.000 |
| c6b4b1189dc35ab21ddc6c47d921b02b | Bacteria | Proteobacteria | Alphaproteobacteria | Rhodobacterales | Rhodobacteraceae | uncultured | | 0.865 |
| e226f1251b916d179d59037f823b84a2 | Bacteria | Cyanobacteria | Oxyphotobacteria | Chloroplast | Virgulinella fragilis | Virgulinella fragilis | Virgulinella fragilis | 0.717 |
| 15928f46e09fc8b13d28534468b2a6bb | Bacteria | Proteobacteria | Gammaproteobacteria | Enterobacteriales | Enterobacteriaceae | | | 1.000 |
| 1eec9b16acc18632cb9aa20c11784484 | Bacteria | Bacteroidetes | Bacteroidia | Flavobacteriales | Flavobacteriaceae | Tenacibaculum | Ambiguous_taxa | 0.966 |
| 8e45207056ee62d480c45c63602507a1 | Bacteria | Dadabacteria | Dadabacteriia | Dadabacteriales | metagenome | metagenome | metagenome | 0.999 |
| 24f27c51e96e3ce699dbf1ae8d0b3b35 | Bacteria | Proteobacteria | Alphaproteobacteria | Puniceispirillales | SAR116 clade | | | 1.000 |
| 520391183ab71056132188d1a34bbe8d | Bacteria | Verrucomicrobia | Verrucomicrobiae | Arctic97B-4 marine group | uncultured marine bacterium | uncultured marine bacterium | uncultured marine bacterium | 0.986 |
| fb6d8e6ae6afd378d3edca0c7d68df08 | Bacteria | Proteobacteria | Alphaproteobacteria | Rickettsiales | Midichloriaceae | MD3-55 | uncultured proteobacterium | 1.000 |
| 65b974e2d7fdbb174be55f45f2bc46d9 | Bacteria | Proteobacteria | Deltaproteobacteria | SAR324 clade(Marine group B) | uncultured bacterium | uncultured bacterium | uncultured bacterium | 0.807 |
| e9df451896a5489bcab7a68d6cf4cf76 | Bacteria | Proteobacteria | Gammaproteobacteria | Coxiellales | Coxiellaceae | Coxiella | uncultured Legionellales bacterium | 0.870 |

| Feature_ID | Kingdom | Phylum | Class | Order | Family | Genus | Species | Confidence |
|---|---|---|---|---|---|---|---|---|
| 62e1e6a5e256c486f543f5e2b5cf7332 | Bacteria | Bacteroidetes | Bacteroidia | Flavobacteriales | NS9 marine group | hydrothermal vent metagenome | hydrothermal vent metagenome | 1.000 |
| 4103ac66d2abddfe9ff7397b6781d837 | Bacteria | Proteobacteria | Gammaproteobacteria | Coxiellales | Coxiellaceae | Coxiella | | 0.995 |
| 278156bf3df2e32adeffbe915c862035 | Bacteria | Proteobacteria | Gammaproteobacteria | Francisellales | Francisellaceae | Francisella | | 1.000 |
| 8fda521c8234eb40f32d22b4388eb752 | Bacteria | Planctomycetes | Planctomycetacia | Pirellulales | Pirellulaceae | Blastopirellula | uncultured marine bacterium | 0.900 |
| 75c45a45ff494c1a828941cf043aa9d8 | Bacteria | Proteobacteria | Deltaproteobacteria | PB19 | uncultured bacterium | uncultured bacterium | uncultured bacterium | 0.998 |
| 6e30705e019f71f615444a952120a3c8 | Bacteria | Proteobacteria | Deltaproteobacteria | Bdellovibrionales | Bacteriovoracaceae | Peredibacter | uncultured bacterium | 0.923 |
| 8dfd63f862faf069b981a643b6aa914b | Bacteria | Proteobacteria | Deltaproteobacteria | SAR324 clade(Marine group B) | uncultured marine bacterium | uncultured marine bacterium | uncultured marine bacterium | 0.797 |
| 3bd052bf05a5402143aa0a1a965f6ced | Bacteria | Cyanobacteria | Oxyphotobacteria | Chloroplast | | | | 1.000 |
| b8a0949c76ba19e6aa43d3708fd619bf | Bacteria | Proteobacteria | Gammaproteobacteria | Oceanospirillales | Oleiphilaceae | Oleiphilus | Ambiguous_taxa | 0.989 |
| 43b5d7c5feabdfa57219b2640a689204 | Bacteria | Marinimicrobia (SAR406 clade) | uncultured bacterium | uncultured bacterium | uncultured bacterium | uncultured bacterium | uncultured bacterium | 0.971 |
| 0db1f51505067f53eff3118d1945a5ac | Bacteria | Marinimicrobia (SAR406 clade) | | | | | | 1.000 |
| 8d6345af37291471adbb7bdbaae7ea60 | Bacteria | Cyanobacteria | Oxyphotobacteria | Chloroplast | | | | 1.000 |
| 325ac32109fb7d68db9c5ae4ff158209 | Bacteria | Proteobacteria | Alphaproteobacteria | Rhodospirillales | Magnetospiraceae | uncultured | uncultured alpha proteobacterium | 0.802 |
| 7ebe7391dbd3988c5fe920b02cc8853b | Bacteria | Proteobacteria | Alphaproteobacteria | Rickettsiales | uncultured bacterium | uncultured bacterium | uncultured bacterium | 1.000 |
| 8e9e1c3f4720eadf67ce35cfa7c2903c | Bacteria | Chloroflexi | N9D0 | marine metagenome | marine metagenome | marine metagenome | marine metagenome | 1.000 |
| 5397d90c9cd07104c90c2e8d6ff3f338 | Bacteria | Proteobacteria | Gammaproteobacteria | Coxiellales | Coxiellaceae | Coxiella | | 1.000 |
| c252e2ba6c3f8c9d27b07cd922baed92 | Bacteria | Cyanobacteria | Oxyphotobacteria | | | | | 0.972 |
| 96a513a3e8ba4bf5e13d98bfc658f536 | Bacteria | Proteobacteria | Alphaproteobacteria | Rickettsiales | Midichloriaceae | MD3-55 | uncultured proteobacterium | 1.000 |
| bd39bf6933afee365dd00081c8ea2400 | Bacteria | Patescibacteria | Gracilibacteria | JGI 0000069-P22 | uncultured bacterium | uncultured bacterium | uncultured bacterium | 0.840 |
| 4eefa79728b7a964df56c16710f0871e | Bacteria | Chloroflexi | Dehalococcoidia | SAR202 clade | Ambiguous_taxa | Ambiguous_taxa | Ambiguous_taxa | 1.000 |
| 7b67152335dd4ee11eb7b575d6571347 | Bacteria | Bacteroidetes | Bacteroidia | Flavobacteriales | NS9 marine group | uncultured marine bacterium | uncultured marine bacterium | 0.807 |
| 105102e9216534ff50d72aef49f9a067 | Bacteria | Fibrobacteres | Fibrobacteria | Fibrobacterales | Fibrobacteraceae | uncultured | uncultured bacterium | 0.964 |
| cff1003e1d0ae7c71be54ff0d2a4ada4 | Bacteria | Cyanobacteria | Oxyphotobacteria | Chloroplast | | | | 1.000 |
| 6f52ad7b849a5420973ed3de06b6e770 | Bacteria | Planctomycetes | Phycisphaerae | Phycisphaerales | Phycisphaeraceae | FS140-16B-02 marine group | | 1.000 |
| 76ddac6b0bb8ef390412ba76c602db79 | Bacteria | Chloroflexi | Dehalococcoidia | SAR202 clade | hydrothermal vent metagenome | hydrothermal vent metagenome | hydrothermal vent metagenome | 0.885 |
| 528e6043f6cb9d717e085884df09003e | Bacteria | Proteobacteria | Gammaproteobacteria | SAR86 clade | | | | 1.000 |
| 10390c2c16e6dd7567f0d93c0b18fb14 | Bacteria | Bacteroidetes | Bacteroidia | Flavobacteriales | Flavobacteriaceae | Pseudofulvibacter | uncultured Bacteroidetes bacterium | 0.846 |
| 3d0e8c71d8de69a154557dabdbaee4f1 | Bacteria | Proteobacteria | Deltaproteobacteria | Bdellovibrionales | Bacteriovoracaceae | uncultured | | 1.000 |
| 9a08cb6c81b0704d6ceb0928e229187b | Bacteria | Bacteroidetes | Bacteroidia | Chitinophagales | uncultured | uncultured Bacteroidetes/Chlorobi group bacterium | uncultured Bacteroidetes/Chlorobi group bacterium | 0.998 |
| 5e34faab5973bcb278757f4541637c70 | Bacteria | Actinobacteria | Actinobacteria | PeM15 | uncultured Actinomycetales bacterium | uncultured Actinomycetales bacterium | uncultured Actinomycetales bacterium | 0.848 |
| 243337808edfd7522422e3d267343a70 | Bacteria | Proteobacteria | Gammaproteobacteria | Cellvibrionales | Spongiibacteraceae | BD1-7 clade | uncultured bacterium | 0.962 |
| 34d3b687c4d943195214d71151965aee | Bacteria | Proteobacteria | Gammaproteobacteria | Coxiellales | Coxiellaceae | Coxiella | uncultured gamma proteobacterium | 0.958 |
| 90669f55d647b98ea6a6c785aa7c1a70 | Bacteria | Proteobacteria | Gammaproteobacteria | Alteromonadales | Pseudoalteromonadaceae | Pseudoalteromonas | | 1.000 |
| 04e1a13c3759654c528852df6177138c | Bacteria | Planctomycetes | Phycisphaerae | Phycisphaerales | Phycisphaeraceae | CL500-3 | uncultured Phycisphaeraceae bacterium | 0.974 |
| 3354179ec2f50428225698d082ccd4e2 | Bacteria | Proteobacteria | Deltaproteobacteria | Bdellovibrionales | Bdellovibrionaceae | OM27 clade | uncultured marine bacterium | 0.936 |
| 526515e306335cc684711beaa2fac9c8 | Bacteria | Marinimicrobia (SAR406 clade) | uncultured Fibrobacteres/Acidobacteria group bacterium | uncultured Fibrobacteres/Acidobacteria group bacterium | uncultured Fibrobacteres/Acidobacteria group bacterium | uncultured Fibrobacteres/Acidobacteria group bacterium | uncultured Fibrobacteres/Acidobacteria group bacterium | 0.997 |
| 35b3ee3c85c4c22109f2eef76692605f | Bacteria | Proteobacteria | Deltaproteobacteria | Oligoflexales | Oligoflexaceae | uncultured | marine metagenome | 0.741 |
| d03872c3a7f58a4fb9ba9a58d9bcfc21 | Bacteria | Verrucomicrobia | Verrucomicrobiae | Opitutales | Puniceicoccaceae | MB11C04 marine group | uncultured Verrucomicrobia bacterium | 0.987 |
| de1c7559fd02330f766a8d54cbe4cff0 | Bacteria | Bacteroidetes | Bacteroidia | Flavobacteriales | Flavobacteriaceae | Muricauda | Muricauda aquimarina | 0.781 |
| c79df9f8a1051056e47349f81c455ecb | Bacteria | Chloroflexi | Dehalococcoidia | SAR202 clade | uncultured Chloroflexi bacterium HF4000_28F02 | uncultured Chloroflexi bacterium HF4000_28F02 | uncultured Chloroflexi bacterium HF4000_28F02 | 1.000 |
| b30ec1714cfcbb5654a140c222f89d95 | Bacteria | Proteobacteria | Gammaproteobacteria | Ectothiorhodospirales | Ectothiorhodospiraceae | uncultured | uncultured marine bacterium | 0.936 |
| 689ec83d9bd2521b17fa36d976c3bc8f | Bacteria | Proteobacteria | Gammaproteobacteria | Coxiellales | Coxiellaceae | Coxiella | uncultured marine bacterium | 0.727 |
| 09efea11e0dc746a2891cd8fbbc9c0bf | Bacteria | Proteobacteria | Deltaproteobacteria | Bdellovibrionales | Bdellovibrionaceae | OM27 clade | uncultured marine bacterium | 0.994 |
| cece916144b6694706d4b68147b93029 | Bacteria | Bacteroidetes | Bacteroidia | Cytophagales | Cyclobacteriaceae | uncultured | | 0.974 |
| 901f7d86a89f0017bebe819b465d2708 | Bacteria | Proteobacteria | Alphaproteobacteria | Rickettsiales | Rickettsiaceae | uncultured | uncultured bacterium | 0.945 |
| f2b2bbb61cf3a80f0ef687eb9e234557 | Bacteria | Cyanobacteria | Oxyphotobacteria | Chloroplast | | | | 1.000 |
| de3f928b57fcbd8c1add376a86238f12 | Bacteria | Proteobacteria | Gammaproteobacteria | Coxiellales | Coxiellaceae | Coxiella | uncultured bacterium | 0.914 |
| 79226c5333a6cfde17c4eead2a6a8d8a | Bacteria | Cyanobacteria | Oxyphotobacteria | Chloroplast | Epipyxis sp. PR26KG | Epipyxis sp. PR26KG | Epipyxis sp. PR26KG | 0.832 |
| 2d8cdefe87d8592b11811dde1c9bdafc | Bacteria | Proteobacteria | Alphaproteobacteria | uncultured | uncultured marine bacterium | uncultured marine bacterium | uncultured marine bacterium | 0.926 |
| 5db00182d76885e5275ae0959d6ac59e | Bacteria | Chlamydiae | Chlamydiae | Chlamydiales | Simkaniaceae | Candidatus Fritschea | Candidatus Syngnamydia salmonis | 0.877 |
| 8bcbb796a33f4d24c1500262fb166027 | Bacteria | Proteobacteria | Gammaproteobacteria | SAR86 clade | Ambiguous_taxa | Ambiguous_taxa | Ambiguous_taxa | 0.981 |
| 1445692ac0f1e0ccbb37b90c729b00fc | Bacteria | Proteobacteria | Alphaproteobacteria | Rickettsiales | S25-593 | | | 1.000 |
| 84f4194acd73a885343a94e9079ca0b0 | Bacteria | Proteobacteria | Deltaproteobacteria | PB19 | uncultured marine bacterium | uncultured marine bacterium | uncultured marine bacterium | 0.999 |
| f231f9253aa760706fa7cf76625a569f | Bacteria | Proteobacteria | Gammaproteobacteria | Vibrionales | Vibrionaceae | Enterovibrio | uncultured bacterium | 0.974 |
| 7cb6ffbadfa93cc06922e1318fd47097 | Bacteria | Cyanobacteria | Oxyphotobacteria | Chloroplast | Epipyxis sp. PR26KG | Epipyxis sp. PR26KG | Epipyxis sp. PR26KG | 0.868 |
| a5968078fe6fd98cffce5d691f89c3f5 | Bacteria | Proteobacteria | Gammaproteobacteria | Cellvibrionales | Spongiibacteraceae | BD1-7 clade | | 0.964 |
| 7be9301973d5ca479ce74dc307e4b3eb | Bacteria | Proteobacteria | Alphaproteobacteria | Rickettsiales | Mitochondria | | | 1.000 |
| 814123085f30789c3846da5db2408947 | Bacteria | Cyanobacteria | Oxyphotobacteria | Chloroplast | | | | 0.887 |
| b4f8d588bab5ed2fa2600166254435f5 | Bacteria | Proteobacteria | Deltaproteobacteria | Bdellovibrionales | Bdellovibrionaceae | OM27 clade | uncultured marine bacterium | 0.754 |
| 53f247034169a941e8aac06e48c573f5 | Bacteria | Cyanobacteria | Oxyphotobacteria | | | | | 0.994 |
| 9eee8f818ee25fe170c5a2183832817d | Bacteria | Proteobacteria | Alphaproteobacteria | Rickettsiales | Mitochondria | uncultured bacterium | uncultured bacterium | 0.945 |
| 0bc9da7ebfaaa1f5f053b4fbd076e306 | Bacteria | Proteobacteria | Deltaproteobacteria | Myxococcales | P3OB-42 | uncultured marine bacterium | uncultured marine bacterium | 1.000 |
| 14408a7f00373c8ec34d1182fb81d027 | Bacteria | Margulisbacteria | uncultured bacterium | uncultured bacterium | uncultured bacterium | uncultured bacterium | uncultured bacterium | 0.882 |
| f05c96b76bb6b8856de5632315d1918e | Bacteria | Proteobacteria | Gammaproteobacteria | Cellvibrionales | Cellvibrionaceae | Gilvimarinus | uncultured bacterium | 0.809 |
| 17e3fe1edd7843f5333c5f633d4ffa71 | Bacteria | Bacteroidetes | Bacteroidia | Cytophagales | Cyclobacteriaceae | Algoriphagus | | 1.000 |
| 8b2414e92a8eb825c102a338b7c4f3f9 | Bacteria | Proteobacteria | Gammaproteobacteria | Oceanospirillales | Halomonadaceae | Halomonas | | 1.000 |
| 178639bcfd43ea0e76ea5664b5feff78 | Bacteria | Proteobacteria | Alphaproteobacteria | Rhodobacterales | Rhodobacteraceae | Sulfitobacter | | 0.983 |
| 672c650be25f41fd17ab570894b22a48 | Bacteria | Bacteroidetes | Bacteroidia | Flavobacteriales | Flavobacteriaceae | Tenacibaculum | Ambiguous_taxa | 0.954 |
| cee750c9da03be487136468043fdfcbb | Bacteria | Proteobacteria | Gammaproteobacteria | Gammaproteobacteria Incertae Sedis | Unknown Family | Candidatus Berkiella | | 1.000 |
| 9be3e6827037097f35e3f4039d9a4b75 | Bacteria | Proteobacteria | Deltaproteobacteria | Bdellovibrionales | Bdellovibrionaceae | OM27 clade | uncultured marine bacterium | 0.999 |

| Feature_ID | Kingdom | Phylum | Class | Order | Family | Genus | Species | Confidence |
|---|---|---|---|---|---|---|---|---|
| fc4f9cc518e1e47a64273583e9447023 | Bacteria | Bacteroidetes | Bacteroidia | Chitinophagales | Saprospiraceae | uncultured | unidentified marine eubacterium | 1.000 |
| 44fa2b32fe41b860055fac1781f0ba42 | Bacteria | Proteobacteria | Gammaproteobacteria | Nitrosococcales | Methylophagaceae | uncultured | uncultured marine bacterium | 0.995 |
| ab3630d2ff5936f4a0b2e58e37c03c4c | Bacteria | Proteobacteria | Gammaproteobacteria | Coxiellales | Coxiellaceae | Coxiella | uncultured marine bacterium | 0.724 |
| fbd60c0f8826ab44a3407ebe139ea0ec | Bacteria | Bacteroidetes | Bacteroidia | Flavobacteriales | Cryomorphaceae | uncultured | uncultured Owenweeksia sp. | 0.890 |
| 3615f1a7df592eb3b6f4a2f5a8640ffc | Bacteria | Margulisbacteria | uncultured bacterium | uncultured bacterium | uncultured bacterium | uncultured bacterium | uncultured bacterium | 0.988 |
| 587bb6802824b1790d5442957b3c651e | Bacteria | Proteobacteria | Gammaproteobacteria | Coxiellales | Coxiellaceae | Coxiella | uncultured bacterium | 0.931 |
| 49a86b3a708040af7c222095eebe24e9 | Bacteria | Chloroflexi | Dehalococcoidia | SAR202 clade | | | | 0.972 |
| 8334ac344c0f15794d5fdc1cad11dac9 | Bacteria | Marinimicrobia (SAR406 clade) | | | | | | 1.000 |
| b457abff0c04f689d87b9a4572859d66 | Bacteria | Chloroflexi | Dehalococcoidia | SAR202 clade | Ambiguous_taxa | Ambiguous_taxa | Ambiguous_taxa | 0.990 |
| 6953c7da1a5a3ac279e35569a4af471e | Bacteria | Proteobacteria | Alphaproteobacteria | Rhodobacterales | Rhodobacteraceae | uncultured | | 0.754 |
| dd2a8a748f25215259c7664e3e53ff44 | Bacteria | Cyanobacteria | Oxyphotobacteria | Chloroplast | | | | 1.000 |
| 2f1416e0cb2c00d579fd5531e873ea64 | Bacteria | Proteobacteria | Gammaproteobacteria | SAR86 clade | | | | 1.000 |
| 440e78141b19e5c4d9737b14c0c7ab4b | Bacteria | Bacteroidetes | Bacteroidia | Chitinophagales | Saprospiraceae | Saprospira | uncultured bacterium | 1.000 |
| 6da8c172e82883d5ea687fae198a5f76 | Bacteria | Cyanobacteria | Oxyphotobacteria | Chloroplast | uncultured marine rappemonad | uncultured marine rappemonad | uncultured marine rappemonad | 1.000 |
| 0fabb1c309a6341e07badd27e991a9d9 | Bacteria | Bacteroidetes | Bacteroidia | Cytophagales | Cyclobacteriaceae | Marinoscillum | uncultured marine bacterium | 0.990 |
| d60c0fa2a9bcc75181bd8272786ed7d7 | Bacteria | Proteobacteria | Alphaproteobacteria | Rhodospirillales | Magnetospiraceae | uncultured | | 0.852 |
| 6fa2d5478f3ed88f1d7ebc18a0eb12d7 | Bacteria | Marinimicrobia (SAR406 clade) | | | | | | 1.000 |
| df431af4616e0cc266b68e9ada7b28cd | Bacteria | Chloroflexi | Dehalococcoidia | SAR202 clade | | | | 1.000 |
| 80da5f0b030114ad7e02c2099c79dd3a | Bacteria | Bacteroidetes | Bacteroidia | Flavobacteriales | Flavobacteriaceae | Winogradskyella | uncultured bacterium | 0.897 |
| da1fcf4c87e0e46080df28df1ddd7b74 | Bacteria | Bacteroidetes | Bacteroidia | Cytophagales | Cyclobacteriaceae | uncultured | uncultured Bacteroidetes bacterium | 0.818 |
| da78a9d089f55db0f56bc378d1cd17a1 | Bacteria | Cyanobacteria | Oxyphotobacteria | | | | | 0.980 |
| eddf0245aedcd394a5d05866700222e3 | Bacteria | Proteobacteria | Gammaproteobacteria | Vibrionales | Vibrionaceae | Vibrio | | 0.995 |
| 9f6d839784a47fc7a1e0c955bad35fea | Bacteria | Cyanobacteria | Oxyphotobacteria | Chloroplast | | | | 0.992 |
| 007609a10926db5d44eb19ab8b909813 | Bacteria | Marinimicrobia (SAR406 clade) | uncultured bacterium | uncultured bacterium | uncultured bacterium | uncultured bacterium | uncultured bacterium | 0.991 |
| c37ce5ba81967cd15c472df5a6349288 | Bacteria | Proteobacteria | Gammaproteobacteria | Oceanospirillales | Pseudohongiellaceae | Pseudohongiella | | 1.000 |
| 20e4f3f6724aa9cd8855be803d59ff71 | Bacteria | Proteobacteria | Gammaproteobacteria | Ectothiorhodospirales | Ectothiorhodospiraceae | uncultured | Ambiguous_taxa | 0.905 |
| 2c96e1f2b667aeadfe5af8e5b905e08c | Bacteria | Bacteroidetes | Bacteroidia | Flavobacteriales | NS9 marine group | uncultured Flavobacteriia bacterium | uncultured Flavobacteriia bacterium | 0.871 |
| 199e7e3b09d52fe2256fb8794f9a10e4 | Bacteria | Proteobacteria | Alphaproteobacteria | SAR11 clade | Clade II | uncultured Candidatus Pelagibacter sp. | uncultured Candidatus Pelagibacter sp. | 0.702 |
| 8dc8e2bcd7c9c1a891e72450af5945df | Bacteria | Cyanobacteria | Oxyphotobacteria | | | | | 0.999 |
| 40a33eb515cc704462eb1bcd35e3c989 | Bacteria | Cyanobacteria | Oxyphotobacteria | Chloroplast | uncultured marine eukaryote | uncultured marine eukaryote | uncultured marine eukaryote | 0.974 |
| 818898f8bffcbc265b6d083b120fe51e | Bacteria | Proteobacteria | Alphaproteobacteria | Rickettsiales | Rickettsiaceae | Candidatus Megaira | | 1.000 |
| a1d78803716a9028d4b2e48fe6f7b4c5 | Bacteria | Bacteroidetes | Bacteroidia | Chitinophagales | uncultured | Ambiguous_taxa | Ambiguous_taxa | 0.997 |
| a54c2dd068c3c7f2d7bef432709383bf | Bacteria | Bacteroidetes | Bacteroidia | Flavobacteriales | NS9 marine group | marine metagenome | marine metagenome | 0.801 |
| ab04c03d9ef9c5f51dc78ca2ea4df2a2 | Bacteria | Proteobacteria | Deltaproteobacteria | Bdellovibrionales | Bdellovibrionaceae | OM27 clade | uncultured marine microorganism | 0.987 |
| 6a2c29d16a603987adea935189ab51b1 | Bacteria | Proteobacteria | Alphaproteobacteria | | | | | 1.000 |
| 12350db43e3d0752c3b3c85edc00f950 | Bacteria | Proteobacteria | Deltaproteobacteria | NB1-j | uncultured delta proteobacterium | uncultured delta proteobacterium | uncultured delta proteobacterium | 0.812 |
| 5b8092bd1e71c93238c7841cf5126b3d | Bacteria | Proteobacteria | Gammaproteobacteria | Pseudomonadales | Pseudomonadaceae | Pseudomonas | Pseudomonas oryzihabitans | 0.924 |
| e1274bc2a7bea010a4fba797a261a014 | Bacteria | Proteobacteria | Deltaproteobacteria | Oligoflexales | Oligoflexaceae | uncultured | | 1.000 |
| 9b2582585c83cb97c3faaff6b4a23f3d | Bacteria | Proteobacteria | Gammaproteobacteria | Nitrosococcales | Methylophagaceae | Methylophaga | | 1.000 |
| a9f80c705a56ba457bc11079b066ac06 | Bacteria | Proteobacteria | Alphaproteobacteria | uncultured | uncultured marine bacterium | uncultured marine bacterium | uncultured marine bacterium | 0.810 |
| 4914d60eb347837dac7331e1616e365e | Bacteria | Proteobacteria | Alphaproteobacteria | Puniceispirillales | SAR116 clade | marine metagenome | marine metagenome | 0.993 |
| 5a58f0c92ff2a486e0906f347b97316c | Bacteria | Cyanobacteria | Oxyphotobacteria | Chloroplast | | | | 1.000 |
| 81c6ed55e029d9af6a02901c9e0ce44b | Bacteria | Proteobacteria | Alphaproteobacteria | Rhodospirillales | AEGEAN-169 marine group | unidentified marine bacterioplankton | unidentified marine bacterioplankton | 0.990 |
| b9565621e581887520ec11b371e78f44 | Bacteria | Marinimicrobia (SAR406 clade) | | | | | | 1.000 |
| b5102527585dcc887fce894ccce3433f | Bacteria | Proteobacteria | Gammaproteobacteria | Oceanospirillales | Halomonadaceae | Salinicola | Ambiguous_taxa | 0.937 |
| 7128721f0a3f5f2732138ebd96b7cdc0 | Bacteria | Planctomycetes | BD7-11 | metagenome | metagenome | metagenome | metagenome | 1.000 |
| 1f83d9e9221043716663bb5f72d5c423 | Bacteria | Proteobacteria | Deltaproteobacteria | Bdellovibrionales | Bdellovibrionaceae | OM27 clade | unidentified marine bacterioplankton | 1.000 |
| 73f9c4cf86ef43304f311bed3b6100b9 | Bacteria | Bacteroidetes | Bacteroidia | Cytophagales | Cyclobacteriaceae | Roseivirga | Ambiguous_taxa | 0.927 |
| 0207e404beade20d260c178b7a0b48d9 | Bacteria | Proteobacteria | Gammaproteobacteria | Alteromonadales | Shewanellaceae | Shewanella | | 1.000 |
| eda8a677029353f504dd7764d9d68be3 | Bacteria | Bacteroidetes | Bacteroidia | Flavobacteriales | Flavobacteriaceae | Muricauda | Ambiguous_taxa | 0.812 |
| 9a442e6daf46f3b21b9d7a5c0e6af5ba | Bacteria | Proteobacteria | Alphaproteobacteria | Rhodospirillales | Magnetospiraceae | uncultured | | 0.847 |
| 173014114c720db39cd0b67d5fd5dcc4 | Bacteria | Margulisbacteria | uncultured bacterium | uncultured bacterium | uncultured bacterium | uncultured bacterium | uncultured bacterium | 0.812 |
| f328237b63388075cf60d22e6ec2d968 | Bacteria | Chloroflexi | Dehalococcoidia | SAR202 clade | metagenome | metagenome | metagenome | 1.000 |
| 06b802ac5ad16494488f7f89012fe143 | Bacteria | Bacteroidetes | Bacteroidia | Flavobacteriales | Flavobacteriaceae | NS4 marine group | | 1.000 |
| 3646a09ac8f1317ce3453b7ca6a9b7d4 | Bacteria | Proteobacteria | Gammaproteobacteria | Oceanospirillales | Nitrincolaceae | uncultured | | 0.994 |
| 243344b04b8a75ae61331223e24fef5d | Bacteria | Bacteroidetes | Bacteroidia | Flavobacteriales | Flavobacteriaceae | Pseudofulvibacter | uncultured bacterium | 0.997 |
| 3ca6b8c53d10682a06100f438d99e06a | Bacteria | Proteobacteria | Deltaproteobacteria | Bdellovibrionales | Bdellovibrionaceae | OM27 clade | uncultured delta proteobacterium | 0.868 |
| 865d95bfe65904fa4acc82fa6dcf7e8b | Bacteria | Proteobacteria | Gammaproteobacteria | Betaproteobacteriales | Burkholderiaceae | MWH-UniP1 aquatic group | uncultured marine bacterium | 0.998 |
| 4334a619479aad06fad95a086019e491 | Bacteria | Cyanobacteria | Oxyphotobacteria | Chloroplast | uncultured marine eukaryote | uncultured marine eukaryote | uncultured marine eukaryote | 0.856 |
| 90316d25d47f8c8e5be067170fc14fb4 | Bacteria | Proteobacteria | Gammaproteobacteria | Coxiellales | Coxiellaceae | Coxiella | | 1.000 |
| 973207b51ade60a64c0812280739269e | Bacteria | Proteobacteria | Gammaproteobacteria | Coxiellales | Coxiellaceae | Coxiella | uncultured marine bacterium | 0.900 |
| adbd1f75c2669f7436c25aeab65a92f1 | Bacteria | Cyanobacteria | Oxyphotobacteria | Chloroplast | | | | 1.000 |
| 8fe6daf570f851673d4dd68a78deb5ea | Bacteria | Proteobacteria | Deltaproteobacteria | Oligoflexales | Oligoflexaceae | Pseudobacteriovorax | uncultured delta proteobacterium | 0.991 |
| c7b34b91b512a0b57d9cbca40a193824 | Bacteria | Cyanobacteria | Oxyphotobacteria | Chloroplast | uncultured bacterium | uncultured bacterium | uncultured bacterium | 0.948 |
| cf50ac719721ceb3aa67b51b7ce51cde | Bacteria | Marinimicrobia (SAR406 clade) | | | | | | 1.000 |
| c82305cfe51886f190537c68a5ade9f3 | Bacteria | Bacteroidetes | Bacteroidia | Flavobacteriales | Flavobacteriaceae | Salegentibacter | Ambiguous_taxa | 0.859 |
| f1cf3687e81473580eb4582e2402a55b | Bacteria | Proteobacteria | Alphaproteobacteria | Rhodospirillales | AEGEAN-169 marine group | marine metagenome | marine metagenome | 0.935 |
| 8a63c5e82d07611dd5bfca55740d94ef | Archaea | Euryarchaeota | Thermoplasmata | Marine Group II | Ambiguous_taxa | Ambiguous_taxa | Ambiguous_taxa | 0.989 |
| 8166a4ff88b68f6ab56a677404f1630d | Bacteria | Cyanobacteria | Oxyphotobacteria | Chloroplast | uncultured marine rappemonad | uncultured marine rappemonad | uncultured marine rappemonad | 1.000 |
| 20e2bc59d7911ca70fa88de37cafda6f | Bacteria | Proteobacteria | Gammaproteobacteria | Coxiellales | Coxiellaceae | Coxiella | | 1.000 |
| 069be6b7ab40cac991b68c2b6461a0af | Bacteria | Cyanobacteria | Oxyphotobacteria | Chloroplast | | | | 1.000 |
| 3e617551815055fa8d1e3f60cd72b835 | Bacteria | Bacteroidetes | Bacteroidia | Flavobacteriales | Flavobacteriaceae | NS4 marine group | marine metagenome | 0.965 |
| 3659fa8e677aeebdb502090adad0561c | Bacteria | Proteobacteria | Alphaproteobacteria | uncultured | uncultured marine bacterium | uncultured marine bacterium | uncultured marine bacterium | 0.973 |

| Feature_ID | Kingdom | Phylum | Class | Order | Family | Genus | Species | Confidence |
|---|---|---|---|---|---|---|---|---|
| d7ea75c97d8856f2e491e55adf787535 | Bacteria | Proteobacteria | Gammaproteobacteria | SAR86 clade | Ambiguous_taxa | Ambiguous_taxa | Ambiguous_taxa | 0.898 |
| 91bb9516bef340053f23b2ee24371039 | Bacteria | Cyanobacteria | Oxyphotobacteria | Chloroplast | | | | 1.000 |
| a28515c4aba81ff4f6a07457a7d9e075 | Bacteria | Cyanobacteria | Oxyphotobacteria | Chloroplast | Virgulinella fragilis | Virgulinella fragilis | Virgulinella fragilis | 0.782 |
| 428b28ac9837a14c8f000adfa09d797a | Bacteria | Cyanobacteria | Oxyphotobacteria | Chloroplast | | | | 1.000 |
| e610873f7fee94ef26f5ba9cdeed24e6 | Bacteria | Cyanobacteria | Oxyphotobacteria | Chloroplast | | | | 1.000 |
| 6bf2daf6e156869751cb3894b7256d72 | Bacteria | Proteobacteria | Gammaproteobacteria | Steroidobacterales | Woeseiaceae | Woeseia | uncultured marine microorganism | 0.752 |
| 2bea4e2bd82d9a9468f816db244c9164 | Bacteria | Bacteroidetes | Bacteroidia | Flavobacteriales | NS9 marine group | | | 1.000 |
| b0cc16ef0aa7efb5d23e1acb8daa4ce3 | Bacteria | Bacteroidetes | Bacteroidia | Flavobacteriales | NS7 marine group | uncultured bacterium | uncultured bacterium | 0.971 |
| 61cdf20ff2d92b30e582779f2fc02110 | Bacteria | Cyanobacteria | Oxyphotobacteria | Chloroplast | | | | 1.000 |
| 99a2b7b9b67164e14b29b9abe1b70df1 | Bacteria | Chloroflexi | Dehalococcoidia | SAR202 clade | uncultured marine microorganism | uncultured marine microorganism | uncultured marine microorganism | 1.000 |
| 616129302f3bc96117136df0df370784 | Bacteria | Cyanobacteria | Oxyphotobacteria | Chloroplast | | | | 1.000 |
| 885b34a83261aba755e006ff5b44f158 | Bacteria | Bacteroidetes | Bacteroidia | Flavobacteriales | Flavobacteriaceae | | | 1.000 |
| 3939bbb1803f9995031619b8340125db | Bacteria | Proteobacteria | Deltaproteobacteria | Oligoflexales | Oligoflexaceae | uncultured | uncultured bacterium | 1.000 |
| 372b0e58df16ef710b2a6dc41c976b13 | Bacteria | Bacteroidetes | Bacteroidia | Flavobacteriales | NS7 marine group | seawater metagenome | seawater metagenome | 0.969 |
| ab2163ba3594b8f12987b23193acb882 | Bacteria | Margulisbacteria | marine metagenome | marine metagenome | marine metagenome | marine metagenome | marine metagenome | 1.000 |
| 2a860a97764198a71697cb8e5f9f89be | Bacteria | Proteobacteria | Deltaproteobacteria | Oligoflexales | Oligoflexaceae | uncultured | uncultured marine bacterium | 0.991 |
| 889187307fa3b4625e3163c98aae20bf | Bacteria | Bacteroidetes | Bacteroidia | Sphingobacteriales | Lentimicrobiaceae | | | 0.876 |
| a0d3971a66c84efc7547eb8a6cc8ebd9 | Bacteria | Bacteroidetes | Bacteroidia | Flavobacteriales | Weeksellaceae | Chryseobacterium | | 1.000 |
| c0535dfac7304e11bec310ece5f8c1af | Bacteria | Proteobacteria | Alphaproteobacteria | Rickettsiales | Mitochondria | uncultured bacterium | uncultured bacterium | 0.725 |
| bf8c7e778f8046ec32c585970b9271a1 | Bacteria | Proteobacteria | Gammaproteobacteria | Alteromonadales | Psychromonadaceae | Psychromonas | | 0.774 |
| 9e84a367e991b2f5ecc5baff597f51a0 | Bacteria | Chloroflexi | Dehalococcoidia | SAR202 clade | uncultured marine microorganism | uncultured marine microorganism | uncultured marine microorganism | 0.775 |
| 55e87a00a72fe0e18497e20e29c59bff | Bacteria | Marinimicrobia (SAR406 clade) | | | | | | 1.000 |
| 34bdb4d45e8dbcff28436627eaf64c58 | Bacteria | Proteobacteria | Alphaproteobacteria | SAR11 clade | Clade III | | | 1.000 |
| df1e0a4f564f1c6f4a98c75729d7baf6 | Bacteria | Chloroflexi | Dehalococcoidia | SAR202 clade | | | | 1.000 |
| 5242c543602e38181e8d985e21a3ecab | Bacteria | Bacteroidetes | Bacteroidia | Cytophagales | Cyclobacteriaceae | uncultured | | 0.999 |
| a2070fda7c93d647f044fcfbca708abf | Bacteria | Cyanobacteria | Oxyphotobacteria | Chloroplast | unidentified rhodophyte PRD01a010B | unidentified rhodophyte PRD01a010B | unidentified rhodophyte PRD01a010B | 0.807 |
| 8caefd97aca4503b3c4d55ddd364e948 | Bacteria | Cyanobacteria | Oxyphotobacteria | Chloroplast | | | | 0.793 |
| 2454fdcb79a9ce3e7e584aca5f6ebb28 | Bacteria | Bacteroidetes | Bacteroidia | Flavobacteriales | Flavobacteriaceae | NS5 marine group | uncultured Flavobacteriia bacterium | 0.921 |
| e8483dc5ea4e2fff54b1b33989c8817f | Bacteria | Margulisbacteria | uncultured bacterium | uncultured bacterium | uncultured bacterium | uncultured bacterium | uncultured bacterium | 0.980 |
| e79caaf336d49f27c3aa85ea660b7d4b | Bacteria | Actinobacteria | Actinobacteria | PeM15 | uncultured Actinomycetales bacterium | uncultured Actinomycetales bacterium | uncultured Actinomycetales bacterium | 0.807 |
| 131101715377ae010fce3565ee486f40 | Bacteria | Proteobacteria | Alphaproteobacteria | Rickettsiales | Rickettsiaceae | uncultured | uncultured bacterium | 0.967 |
| 71d30286b6f5cdf26718ba721c1ded8f | Bacteria | Proteobacteria | Alphaproteobacteria | Rhodospirillales | Magnetospiraceae | uncultured | uncultured bacterium | 0.784 |
| 4d0ece734c54be2729898074bfda295e | Bacteria | Proteobacteria | Alphaproteobacteria | SAR11 clade | Clade I | | | 0.894 |
| d6698271c81f5713ffe7bc2c744b175d | Bacteria | Proteobacteria | Alphaproteobacteria | Rhodospirillales | AEGEAN-169 marine group | Ambiguous_taxa | Ambiguous_taxa | 0.967 |
| d4e3121ddb0ac740110f1e6a667bac10 | Bacteria | Proteobacteria | Alphaproteobacteria | Rickettsiales | Mitochondria | | | 0.999 |
| c5ee1fcd89e7cdf736a91203b48447e8 | Bacteria | Bacteroidetes | Bacteroidia | Flavobacteriales | Flavobacteriaceae | Mesonia | | 1.000 |
| 441919abe886b2ca53488e9d6378804a | Bacteria | Proteobacteria | Alphaproteobacteria | Rhizobiales | Xanthobacteraceae | Bradyrhizobium | | 0.738 |
| d4c9e63d6b4c939623938a1b68ab4539 | Bacteria | Verrucomicrobia | Verrucomicrobiae | Arctic97B-4 marine group | uncultured bacterium | uncultured bacterium | uncultured bacterium | 1.000 |
| 4a79d27b813e39d1e7a820c2ebfda80d | Bacteria | Proteobacteria | Deltaproteobacteria | Bdellovibrionales | Bacteriovoracaceae | Peredibacter | uncultured organism | 0.739 |
| 39604659583f9b6c8ce705dfc42be7ae | Bacteria | Proteobacteria | Gammaproteobacteria | Cellvibrionales | Porticoccaceae | C1-B045 | | 0.853 |
| 0ff69bf5a7eff24d27efe4fd84899fa0 | Bacteria | Proteobacteria | Gammaproteobacteria | KI89A clade | | | | 1.000 |
| df52e9427f7e132d85cc9938a9d778db | Bacteria | Proteobacteria | Alphaproteobacteria | Rickettsiales | S25-593 | | | 1.000 |
| 4209435f8077944eecebebb74013fc40 | Bacteria | Proteobacteria | Alphaproteobacteria | Rickettsiales | Mitochondria | uncultured bacterium | uncultured bacterium | 0.963 |
| c98092e9ee7ac56f8f27968ae18324eb | Bacteria | Kiritimatiellaeota | Kiritimatiellae | Kiritimatiellales | Kiritimatiellaceae | R76-B128 | uncultured bacterium | 0.834 |
| e657989a25a0c6bee5049305ea7e7230 | Bacteria | Planctomycetes | Planctomycetacia | Pirellulales | Pirellulaceae | Rubripirellula | | 1.000 |
| b76d5b554464d7c729677ece2d8f512e | Bacteria | Planctomycetes | Phycisphaerae | Phycisphaerales | Phycisphaeraceae | CL500-3 | uncultured Phycisphaeraceae bacterium | 0.996 |
| 92a5cf6d0f6abbba30b0fec62671cdc8 | Bacteria | Proteobacteria | Deltaproteobacteria | PB19 | uncultured delta proteobacterium DeepAnt-32C6 | uncultured delta proteobacterium DeepAnt-32C6 | uncultured delta proteobacterium DeepAnt-32C6 | 1.000 |
| 18b5a613386c1f6e4c59a1c3127ccc44 | Bacteria | Proteobacteria | Deltaproteobacteria | Bdellovibrionales | Bdellovibrionaceae | OM27 clade | | 1.000 |
| f1d297fd71ae0b707cf47b14acb6849b | Bacteria | Proteobacteria | Deltaproteobacteria | PB19 | Ambiguous_taxa | Ambiguous_taxa | Ambiguous_taxa | 0.999 |
| 9c62f33cc4fe3178364d5acf7cb2bc98 | Bacteria | Cyanobacteria | Oxyphotobacteria | Chloroplast | uncultured bacterium | uncultured bacterium | uncultured bacterium | 0.780 |
| 7b9ecdcf8513468c17795c60245694f2 | Bacteria | Bacteroidetes | Bacteroidia | Flavobacteriales | NS9 marine group | uncultured marine bacterium | uncultured marine bacterium | 0.963 |
| d8a292b392a5dcd5f4d20e8d0135bb9a | Bacteria | Proteobacteria | Deltaproteobacteria | Myxococcales | P3OB-42 | uncultured bacterium | uncultured bacterium | 0.999 |
| ffba9d1c736b8beb1383181d7a7ab53c | Bacteria | Proteobacteria | Gammaproteobacteria | | | | | 1.000 |
| ab9a06144b6b5b97a55e8c2721e58df4 | Bacteria | Proteobacteria | Gammaproteobacteria | Pseudomonadales | Pseudomonadaceae | Pseudomonas | | 0.999 |
| ea9ae875c76b5d3c66068789c4f7c43e | Bacteria | Proteobacteria | Deltaproteobacteria | Oligoflexales | 053A03-B-DI-P58 | | | 0.754 |
| 8b749642527ab9f2a65c9407bb635278 | Bacteria | Proteobacteria | Deltaproteobacteria | Oligoflexales | Oligoflexaceae | uncultured | uncultured bacterium | 0.769 |
| c48f41767b13bea73c71f81ab410d6e5 | Bacteria | Proteobacteria | Deltaproteobacteria | SAR324 clade(Marine group B) | uncultured bacterium | uncultured bacterium | uncultured bacterium | 0.852 |
| f6f24174dbc30d50ee043ce9e52af60f | Bacteria | Actinobacteria | Actinobacteria | Micrococcales | Microbacteriaceae | Microbacterium | | 0.993 |
| 700766015f60e7166d87b7ce56a54804 | Bacteria | Proteobacteria | Gammaproteobacteria | | | | | 1.000 |
| 0dccdbb119b701769060b6fa3f31bc34 | Bacteria | Proteobacteria | Gammaproteobacteria | Gammaproteobacteria Incertae Sedis | Unknown Family | uncultured marine bacterium | uncultured marine bacterium | 1.000 |
| 1df81a0fc27ad29683ab62f2e5c41f79 | Bacteria | Proteobacteria | Gammaproteobacteria | Vibrionales | Vibrionaceae | Vibrio | Vibrio halioticoli | 0.976 |
| 4691037dc9393528b778152a04f83dfc | Bacteria | Verrucomicrobia | Verrucomicrobiae | Arctic97B-4 marine group | uncultured marine bacterium | uncultured marine bacterium | uncultured marine bacterium | 0.989 |
| ba7bb52bc701ee50f6d62a94e604e5d3 | Bacteria | Proteobacteria | Alphaproteobacteria | Rickettsiales | Midichloriaceae | MD3-55 | | 0.997 |
| f0ecd50f752ae26722aa239485ff723c | Bacteria | Proteobacteria | | | | | | 0.885 |
| b64938b7e6251d23aa9af43d59f0f4a3 | Bacteria | Proteobacteria | Alphaproteobacteria | Rhodospirillales | Thalassospiraceae | Thalassospira | | 0.998 |
| 4d4566311554bccc9cb8dd95de1af5a1 | Bacteria | Proteobacteria | Alphaproteobacteria | Rhodospirillales | Magnetospiraceae | Magnetospira | uncultured alpha proteobacterium | 0.993 |
| 44b08a0a9378e175946bbed1ddf9232a | Bacteria | Bacteroidetes | Bacteroidia | Cytophagales | Cyclobacteriaceae | uncultured | | 0.988 |
| 862114615c3f87dcefaf1e9e1738a61c | Bacteria | Proteobacteria | Deltaproteobacteria | PB19 | uncultured bacterium | uncultured bacterium | uncultured bacterium | 0.975 |
| e45f610b2ea63b298276b3002f05be42 | Bacteria | Proteobacteria | Gammaproteobacteria | Piscirickettsiales | Piscirickettsiaceae | Candidatus Endoecteinascidia | Ambiguous_taxa | 1.000 |
| 55ddb1d5456c91d1c38e1dfd684c1a25 | Bacteria | Proteobacteria | Gammaproteobacteria | | | | | 0.982 |
| a328ebe3a24b340af7bfa5147c4a1fe3 | Bacteria | Proteobacteria | Alphaproteobacteria | Rickettsiales | Mitochondria | uncultured bacterium | uncultured bacterium | 0.999 |
| 1f1b47cc9d41a8b313be0c5d305934bc | Bacteria | Proteobacteria | Alphaproteobacteria | Puniceispirillales | SAR116 clade | alpha proteobacterium SCGC AAA015-N04 | alpha proteobacterium SCGC AAA015-N04 | 1.000 |
| b994cbc19e95fcfd2b4cd417b91cd4e2 | Bacteria | Verrucomicrobia | Verrucomicrobiae | Opitutales | Puniceicoccaceae | Cerasicoccus | uncultured Verrucomicrobia bacterium | 0.960 |
| 3bee117d5dadf5f578502e37e2de03ef | Bacteria | Proteobacteria | Alphaproteobacteria | Rhodospirillales | Thalassospiraceae | Thalassospira | | 0.997 |
| dc630db72ca98919c1f4d6e71951c057 | Bacteria | Proteobacteria | Gammaproteobacteria | Coxiellales | Coxiellaceae | Coxiella | uncultured bacterium | 0.892 |
| 3797c84fb3aca015fc5efd67a52ea6e4 | Bacteria | Proteobacteria | Alphaproteobacteria | Rhodobacterales | Rhodobacteraceae | | | 1.000 |
| 71d2cac4c42dcb3f7ccb3e1c5d8658d0 | Bacteria | Kiritimatiellaeota | Kiritimatiellae | Kiritimatiellales | Kiritimatiellaceae | R76-B128 | marine metagenome | 0.996 |
| 2cfca4b42781188e1d2f440f110d5cab | Bacteria | Proteobacteria | Gammaproteobacteria | Vibrionales | Vibrionaceae | Vibrio | | 0.963 |
| 720a4420b88125a4e38f19ad466d95c9 | Bacteria | Proteobacteria | Alphaproteobacteria | Rickettsiales | Mitochondria | Proteobacteria bacterium JGI 0000113-D06 | Proteobacteria bacterium JGI 0000113-D06 | 1.000 |

| Feature_ID | Kingdom | Phylum | Class | Order | Family | Genus | Species | Confidence |
|---|---|---|---|---|---|---|---|---|
| 271ae02863ae0666db7f57ac2434ac69 | Bacteria | Proteobacteria | Gammaproteobacteria | UBA10353 marine group | Ambiguous_taxa | Ambiguous_taxa | Ambiguous_taxa | 0.854 |
| f11bd7b1f30f110f80b81a8bd0cb7b9a | Bacteria | Proteobacteria | Alphaproteobacteria | Rhodospirillales | AEGEAN-169 marine group | Ambiguous_taxa | Ambiguous_taxa | 0.928 |
| 7b6cccf3c22cf7c0b1e9d02aa4444d7d | Bacteria | Proteobacteria | Gammaproteobacteria | Betaproteobacteriales | Nitrosomonadaceae | oc32 | uncultured Rhodocyclaceae bacterium | 0.848 |
| 20c2a3a2c1c41e4834f4abcbb2a8718b | Bacteria | Proteobacteria | Alphaproteobacteria | SAR11 clade | Clade II | | | 0.971 |
| de4b641533d829c51c21fc0f554e302d | Bacteria | Proteobacteria | Deltaproteobacteria | Bdellovibrionales | Bacteriovoracaceae | Peredibacter | uncultured bacterium | 0.817 |
| 74a382c0aa3d7b37174e599a1482a8b2 | Bacteria | Proteobacteria | Alphaproteobacteria | Puniceispirillales | SAR116 clade | uncultured bacterium | uncultured bacterium | 0.889 |
| 7b631e625a5c8ee9c5a2f5ea02750b63 | Bacteria | Proteobacteria | Alphaproteobacteria | Rickettsiales | Mitochondria | Proteobacteria bacterium JGI 0000113-D06 | Proteobacteria bacterium JGI 0000113-D06 | 0.930 |
| 22d1c875fe677df7ac94e0d5cb9319ff | Bacteria | Proteobacteria | Alphaproteobacteria | SAR11 clade | Clade I | Clade Ia | unidentified marine eubacterium | 0.944 |
| 2d7f0eab395ddc395dff65f7400f7934 | Bacteria | Marinimicrobia (SAR406 clade) | uncultured bacterium | uncultured bacterium | uncultured bacterium | uncultured bacterium | uncultured bacterium | 0.999 |
| ac6ae6c0a3b285517bb1eb20a643ec7d | Bacteria | Cyanobacteria | Oxyphotobacteria | Chloroplast | | | | 1.000 |
| 795565ed0227ef2808fdf4fb0b6f7151 | Bacteria | Margulisbacteria | uncultured bacterium | uncultured bacterium | uncultured bacterium | uncultured bacterium | uncultured bacterium | 1.000 |
| 7bca88288746a3b7cb70aa7aecef2623 | Bacteria | Cyanobacteria | Oxyphotobacteria | Chloroplast | | | | 1.000 |
| 9b45962bf31c586fecdbc1d62b00e673 | Bacteria | Proteobacteria | Alphaproteobacteria | SAR11 clade | Clade IV | | | 0.946 |
| d30a36e9fcfe8a553cd7f37a2b19151e | Bacteria | Proteobacteria | Gammaproteobacteria | Oceanospirillales | Nitrincolaceae | uncultured | uncultured bacterium | 0.871 |
| 48cbc0483fe4a48c15179cad32be7fba | Bacteria | Proteobacteria | Alphaproteobacteria | Rickettsiales | Rickettsiaceae | | | 0.772 |
| ad5d7acebf4ce29f604d85d069ebfa48 | Bacteria | Proteobacteria | Gammaproteobacteria | Nitrosococcales | Methylophagaceae | uncultured | uncultured marine bacterium | 0.873 |
| db58c6514d11269384e70c960020acd2 | Bacteria | Cyanobacteria | Oxyphotobacteria | Chloroplast | unidentified rhodophyte PRD01a010B | unidentified rhodophyte PRD01a010B | unidentified rhodophyte PRD01a010B | 0.945 |
| ddc478f8f4e75ca2e0a9c408eca8d5f2 | Bacteria | Cyanobacteria | Oxyphotobacteria | Chloroplast | | | | 1.000 |
| c94e14eead29c37508244ae20a80d233 | Bacteria | Bacteroidetes | Bacteroidia | Chitinophagales | Saprospiraceae | uncultured | uncultured marine bacterium | 0.999 |
| b65d8b1e41bbc6f4fcfa5028f9732118 | Bacteria | Bacteroidetes | Bacteroidia | Flavobacteriales | NS9 marine group | uncultured marine bacterium | uncultured marine bacterium | 0.969 |
| db8a20878bd1dfcdb6247c9250f9f253 | Bacteria | Firmicutes | Bacilli | Bacillales | Bacillaceae | Marinococcus | Marinococcus halophilus | 0.743 |
| 0e8a2ce7a56cbe8d650edf191a121d0f | Bacteria | Proteobacteria | Alphaproteobacteria | uncultured | | | | 0.765 |
| cbb82d9f44ced97837434e8ae2d70bdc | Bacteria | Planctomycetes | Phycisphaerae | Phycisphaerales | Phycisphaeraceae | CL500-3 | uncultured planctomycete | 0.835 |
| 1ca133cbfe1474dd8da269e711ba1103 | Bacteria | Proteobacteria | Deltaproteobacteria | Bdellovibrionales | Bdellovibrionaceae | OM27 clade | uncultured delta proteobacterium | 0.730 |
| 640ebce32e65e763c60a0bd707083eab | Bacteria | Proteobacteria | Alphaproteobacteria | Rickettsiales | Mitochondria | uncultured Rickettsiales bacterium | uncultured Rickettsiales bacterium | 0.741 |
| 041e39e9319e3b43da1ebb3127b12d3a | Bacteria | Bacteroidetes | Bacteroidia | Flavobacteriales | NS9 marine group | | | 0.999 |
| 57fe72114711d713f73124abbc2e4e16 | Bacteria | Bacteroidetes | Bacteroidia | Flavobacteriales | Cryomorphaceae | uncultured | marine metagenome | 0.869 |
| 0d1771f0238c75198678d929eebd43db | Bacteria | Proteobacteria | Deltaproteobacteria | Oligoflexales | Oligoflexaceae | uncultured | uncultured marine bacterium | 1.000 |
| 1e58ed425f1fe6931428f38066be69b7 | Bacteria | Proteobacteria | Gammaproteobacteria | Gammaproteobacteria Incertae Sedis | Unknown Family | uncultured bacterium | uncultured bacterium | 0.992 |
| 41c5a6971518cd156fb72350eaed3d5f | Bacteria | Proteobacteria | Gammaproteobacteria | Chromatiales | Sedimenticolaceae | Sedimenticola | Anodontia fragilis gill symbiont | 0.723 |
| 467f23910e6c1566e461fdece5e2d5f9 | Bacteria | Proteobacteria | Deltaproteobacteria | PB19 | Ambiguous_taxa | Ambiguous_taxa | Ambiguous_taxa | 0.944 |
| eb1f76bfcd72504629e62902629a229a | Bacteria | Bacteroidetes | Bacteroidia | Cytophagales | Cyclobacteriaceae | uncultured | uncultured bacterium | 0.821 |
| ceff7e2f6423173ad221c93d8beb1082 | Bacteria | Proteobacteria | Gammaproteobacteria | UBA10353 marine group | uncultured gamma proteobacterium HF0200_24F15 | uncultured gamma proteobacterium HF0200_24F15 | uncultured gamma proteobacterium HF0200_24F15 | 0.969 |
| 060be68394ed806ebd4ac9fa8ab72eb2 | Bacteria | Proteobacteria | Gammaproteobacteria | Alteromonadales | Moritellaceae | Moritella | uncultured bacterium | 0.969 |
| 3953c46d66987793e229eeffdef816ad | Bacteria | Proteobacteria | Alphaproteobacteria | Rhodospirillales | AEGEAN-169 marine group | marine metagenome | marine metagenome | 0.990 |
| 8bbaec5dba9743b14ac46b38b93dce12 | Bacteria | Actinobacteria | Acidimicrobiia | Microtrichales | Ilumatobacteraceae | uncultured | Ambiguous_taxa | 0.974 |
| 1660941eec313e30e4c253bc65d61096 | Bacteria | Proteobacteria | Alphaproteobacteria | Rickettsiales | Anaplasmataceae | Wolbachia | | 1.000 |
| ee64c77c94771b2ed6c0593b2af042fe | Bacteria | Verrucomicrobia | Verrucomicrobiae | Opitutales | Puniceicoccaceae | uncultured | marine metagenome | 1.000 |
| 6f1e4320d1587f40c5be49a181b11f80 | Bacteria | Proteobacteria | Alphaproteobacteria | SAR11 clade | | | | 1.000 |
| 8efd07f86b0a83ad48ba8fd9b4c35a0b | Bacteria | Proteobacteria | Alphaproteobacteria | SAR11 clade | Clade I | | | 0.998 |
| ac2146746003b211e8fbe8a53171c277 | Bacteria | Proteobacteria | Deltaproteobacteria | NB1-j | uncultured marine bacterium | uncultured marine bacterium | uncultured marine bacterium | 0.995 |
| 5eec1b46e052532bad718c343d3d3a49 | Bacteria | Proteobacteria | Alphaproteobacteria | Rickettsiales | Mitochondria | uncultured bacterium | uncultured bacterium | 0.991 |
| 211a4522cb189758bc1c2fd1afc37149 | Bacteria | Proteobacteria | Alphaproteobacteria | Caulobacterales | Hyphomonadaceae | Henriciella | Ambiguous_taxa | 0.994 |
| 4fb3beb75df8a08a9f878e850da82c6d | Bacteria | Proteobacteria | Alphaproteobacteria | Rickettsiales | Mitochondria | Proteobacteria bacterium JGI 0000113-D06 | Proteobacteria bacterium JGI 0000113-D06 | 0.930 |
| 79b0cf973f2f15bec9686a19d7097019 | Bacteria | Bacteroidetes | Bacteroidia | Flavobacteriales | Crocinitomicaceae | Crocinitomix | | 1.000 |
| f555a90112bd1cdc8aa46c73b4ebd765 | Bacteria | Proteobacteria | Alphaproteobacteria | Rickettsiales | Mitochondria | | | 1.000 |
| 95b582d787137d763bab0706a918d826 | Bacteria | Lentisphaerae | Lentisphaeria | Lentisphaerales | Lentisphaeraceae | Lentisphaera | Ambiguous_taxa | 0.999 |
| b3ff176f9e9d4f43557b1032f180fb6d | Bacteria | Bacteroidetes | Bacteroidia | Flavobacteriales | Cryomorphaceae | uncultured | | 1.000 |
| bbcb32b8ed4df6ec5828d01c2676cfa0 | Bacteria | Proteobacteria | Deltaproteobacteria | | | | | 0.977 |
| 7aaf58488391c09316c457bf5a0b60fa | Bacteria | Cyanobacteria | Oxyphotobacteria | Chloroplast | | | | 1.000 |
| ab89265ceb6a9f19668565df4684157f | Bacteria | Proteobacteria | Deltaproteobacteria | Myxococcales | Nannocystaceae | uncultured | uncultured marine bacterium | 1.000 |
| cabbe95cc39c2340a9724141e9ffdd8a | Bacteria | Chloroflexi | Dehalococcoidia | SAR202 clade | | | | 1.000 |
| 2c6075d66e634f35b381cab617592c4c | Bacteria | Marinimicrobia (SAR406 clade) | | | | | | 1.000 |
| 46739cf0e795f8bdc0152f49dac85124 | Bacteria | PAUC34f | uncultured marine bacterium | uncultured marine bacterium | uncultured marine bacterium | uncultured marine bacterium | uncultured marine bacterium | 0.855 |
| f8817219b8e3c232c105151768a5f881 | Bacteria | Proteobacteria | Deltaproteobacteria | PB19 | uncultured bacterium | uncultured bacterium | uncultured bacterium | 0.976 |
| 06e4826650d46a7c52a86c399a047ddf | Bacteria | Bacteroidetes | Bacteroidia | Cytophagales | Flammeovirgaceae | uncultured bacterium | uncultured bacterium | 0.996 |
| 194c9098513248666b674801f1c31f32 | Bacteria | Patescibacteria | Gracilibacteria | JGI 0000069-P22 | uncultured bacterium | uncultured bacterium | uncultured bacterium | 0.921 |
| 83af8db305f08b8249b4309df6dad3cc | Bacteria | Cyanobacteria | Oxyphotobacteria | Chloroplast | uncultured marine eukaryote | uncultured marine eukaryote | uncultured marine eukaryote | 0.836 |
| 72d6d1a9083c147ef35cda526cce5ef9 | Bacteria | Proteobacteria | Deltaproteobacteria | Bdellovibrionales | Bacteriovoracaceae | Peredibacter | | 1.000 |
| b567ad23661637ee76224fcca8899595 | Bacteria | Bacteroidetes | Bacteroidia | Cytophagales | Cyclobacteriaceae | uncultured | | 0.905 |
| 52cf327326d098fe00a5f3d67a6610b5 | Bacteria | Proteobacteria | Deltaproteobacteria | Bdellovibrionales | Bacteriovoracaceae | Halobacteriovorax | Ambiguous_taxa | 0.962 |
| eaba607e55478219a4dffb8212a59219 | Bacteria | Proteobacteria | Gammaproteobacteria | Oceanospirillales | Pseudohongiellaceae | Pseudohongiella | | 0.998 |
| 83144d5ea68560b6a34b62dd77e90a93 | Bacteria | Proteobacteria | Gammaproteobacteria | Vibrionales | Vibrionaceae | uncultured | Ambiguous_taxa | 0.904 |
| 502ff238dd72c4282405c5d4c606b9c5 | Bacteria | Proteobacteria | Gammaproteobacteria | Francisellales | Francisellaceae | [Caedibacter] taeniospiralis group | uncultured bacterium | 0.964 |
| 987c9175dc6cb29f1c6d0c0c7b36dafb | Bacteria | Proteobacteria | Gammaproteobacteria | Alteromonadales | Alteromonadaceae | | | 1.000 |
| 1b2ea70c0f68c51f73069bdcbd44023e | Bacteria | Proteobacteria | Gammaproteobacteria | OM182 clade | | | | 1.000 |
| 4e88736f7eb48a39270742b786e54db3 | Bacteria | Proteobacteria | Gammaproteobacteria | Salinisphaerales | Salinisphaeraceae | Salinisphaera | Ambiguous_taxa | 0.996 |
| 75a25d3f98c339cfe6fe46684475f796 | Bacteria | Cyanobacteria | Oxyphotobacteria | Chloroplast | uncultured marine eukaryote | uncultured marine eukaryote | uncultured marine eukaryote | 0.938 |
| 9f7e6f2877fa6cdffb1b048880e2953a | Bacteria | Proteobacteria | Alphaproteobacteria | Rhodospirillales | Magnetospiraceae | Magnetospira | uncultured alpha proteobacterium | 0.816 |
| 527dec942f359745bf26782d154d73b8 | Bacteria | Proteobacteria | Gammaproteobacteria | | | | | 0.994 |
| 063f45b16093c0c7995b2c3f07e45c25 | Bacteria | Margulisbacteria | uncultured bacterium | uncultured bacterium | uncultured bacterium | uncultured bacterium | uncultured bacterium | 1.000 |
| 8d26e8a6c10844cfdd479f1bc2d9ebe2 | Bacteria | Proteobacteria | Gammaproteobacteria | Pseudomonadales | Pseudomonadaceae | Pseudomonas | | 0.956 |
| ea1b1a98376b80f47c50ee80e01f1543 | Bacteria | Bacteroidetes | Bacteroidia | Flavobacteriales | Flavobacteriaceae | Salegentibacter | | 1.000 |
| 69da0c771e90024641c3f209ba18b4ae | Bacteria | Proteobacteria | Gammaproteobacteria | Francisellales | Francisellaceae | | | 0.998 |
| 50a31f6b1968a2196b966812a975671b | Bacteria | Cyanobacteria | Oxyphotobacteria | Synechococcales | Cyanobiaceae | | | 0.976 |
| ce22e0a48c35e050bff1cee23e9dad16 | Bacteria | Cyanobacteria | Oxyphotobacteria | Chloroplast | | | | 0.898 |

| Feature_ID | Kingdom | Phylum | Class | Order | Family | Genus | Species | Confidence |
|---|---|---|---|---|---|---|---|---|
| 95034f4bcb05ea3ba5b85315a6dc6676 | Bacteria | Cyanobacteria | Oxyphotobacteria | Chloroplast | | | | 1.000 |
| 193e863ef3d7d270fe6811cb5b8287f6 | Bacteria | Bacteroidetes | Bacteroidia | Flavobacteriales | Flavobacteriaceae | Tenacibaculum | | 0.964 |
| 06e730d3fa6726ce14aae5a7c50e4a66 | Bacteria | Proteobacteria | Gammaproteobacteria | Thiohalorhabdales | Thiohalorhabdaceae | uncultured | uncultured Sinobacteraceae bacterium | 0.784 |
| 461e5f49467b6566962e5976cf346fa7 | Bacteria | Proteobacteria | Alphaproteobacteria | SAR11 clade | Clade II | uncultured Candidatus Pelagibacter sp. | uncultured Candidatus Pelagibacter sp. | 0.886 |
| 2630ab298a78fed6183abe894fbd7605 | Bacteria | Proteobacteria | Gammaproteobacteria | Oceanospirillales | Halomonadaceae | Halomonas | | 1.000 |
| f38ddf60368f14a50dc16fa67b22da00 | Bacteria | Proteobacteria | Gammaproteobacteria | SAR86 clade | uncultured bacterium | uncultured bacterium | uncultured bacterium | 0.810 |
| cbfd06baa4e95cb45bb65ff6da5d4883 | Bacteria | Bacteroidetes | Bacteroidia | Flavobacteriales | Flavobacteriaceae | | | 1.000 |
| 2b2dad8ddae34a19c64db383241f915f | Bacteria | Proteobacteria | Deltaproteobacteria | Bdellovibrionales | Bdellovibrionaceae | OM27 clade | uncultured delta proteobacterium | 0.795 |
| 55793d1d78c620668a41aebc3ee388ea | Bacteria | Bacteroidetes | Bacteroidia | Flavobacteriales | NS7 marine group | uncultured Bacteroidetes bacterium | uncultured Bacteroidetes bacterium | 0.864 |
| dc1e524db5ba5268c9ce9e24342864be | Bacteria | Epsilonbacteraeota | Campylobacteria | Campylobacterales | Arcobacteraceae | Arcobacter | | 1.000 |
| daf3e8d6a1d6eed3ec1cc79e8a5768f1 | Bacteria | | | | | | | 0.951 |
| 63ec8b0c3dd3d45c074f2dee8317d215 | Bacteria | Proteobacteria | Gammaproteobacteria | | | | | 1.000 |
| 4d07490bc411474fb17925d4c5e088b7 | Bacteria | Bacteroidetes | Bacteroidia | Flavobacteriales | Cryomorphaceae | Luteibaculum | uncultured bacterium | 1.000 |
| 00836362a8f932c0fee8019b2a26b7d1 | Bacteria | Proteobacteria | Alphaproteobacteria | Caulobacterales | Hyphomonadaceae | uncultured | uncultured bacterium | 0.958 |
| fe5dfe6da0d9d5ee04b99c04dc74f165 | Bacteria | | | | | | | 1.000 |
| f5e881da9a0592f169a3caccdb4d20dc | Bacteria | Proteobacteria | Alphaproteobacteria | Rhodobacterales | Rhodobacteraceae | uncultured | uncultured bacterium GRIST14 | 0.853 |
| 0a5c89a348813885ad37925bb467454d | Bacteria | Cyanobacteria | Oxyphotobacteria | Chloroplast | | | | 0.998 |
| 3286a679a2c01bc04566ff6f6777a913 | Bacteria | Proteobacteria | Gammaproteobacteria | | | | | 0.966 |
| d95bdb9087239a5e00dd7c5c2110fb29 | Bacteria | Bacteroidetes | Bacteroidia | Flavobacteriales | NS7 marine group | | | 0.998 |
| be557549dcc5eb659cf76a470e695c74 | Bacteria | Proteobacteria | Deltaproteobacteria | Bdellovibrionales | Bdellovibrionaceae | OM27 clade | | 1.000 |
| af2dc8ee24b834ce6732072cbe824d48 | Bacteria | Proteobacteria | Alphaproteobacteria | SAR11 clade | Clade I | | | 0.999 |
| 3ed5e39ce71075c88bd4850cb9833239 | Bacteria | Cyanobacteria | Oxyphotobacteria | Chloroplast | | | | 0.963 |
| 2075c3dcd08d54458db7adc568078514 | Bacteria | Proteobacteria | Alphaproteobacteria | Rickettsiales | AB1 | uncultured bacterium | uncultured bacterium | 0.834 |
| 377ba6e5a430b22a2d48cda47dd1363f | Bacteria | Proteobacteria | Gammaproteobacteria | KI89A clade | obligately oligotrophic bacterium KI89C | obligately oligotrophic bacterium KI89C | obligately oligotrophic bacterium KI89C | 0.961 |
| afdf3a5808ade10a577ef7580156f874 | Bacteria | Proteobacteria | Alphaproteobacteria | SAR11 clade | Clade I | | | 1.000 |
| 7dbe029ba4e2ddd7ce18659368a47252 | Bacteria | Proteobacteria | Alphaproteobacteria | SAR11 clade | Clade IV | | | 1.000 |
| ac0250c03da863239501e5aacefd33a9 | Bacteria | Bacteroidetes | Bacteroidia | Flavobacteriales | NS9 marine group | | | 0.931 |
| ce294127f9aeaedfcc717689b8328443 | Bacteria | Cyanobacteria | Oxyphotobacteria | Synechococcales | Cyanobiaceae | Prochlorococcus MIT9313 | | 0.710 |
| a0eb637a7067a8d5b6341db30381bb78 | Bacteria | Cyanobacteria | Oxyphotobacteria | Chloroplast | uncultured marine rappemonad | uncultured marine rappemonad | uncultured marine rappemonad | 1.000 |
| 0890ca596fba39fb10dc44146f67dc6b | Bacteria | Proteobacteria | Gammaproteobacteria | SAR86 clade | uncultured gamma proteobacterium HF0070_25G02 | uncultured gamma proteobacterium HF0070_25G02 | uncultured gamma proteobacterium HF0070_25G02 | 0.999 |
| a35eff4dfbc0828f6f791fc20ea1f69d | Bacteria | Verrucomicrobia | Verrucomicrobiae | Opitutales | Puniceicoccaceae | uncultured | marine metagenome | 1.000 |
| fc026e2c89a6a12a861abe0fce7e0b97 | Bacteria | Proteobacteria | Alphaproteobacteria | Thalassobaculales | Nisaeaceae | OM75 clade | uncultured marine bacterium | 0.781 |
| 0b6c7cb65337ee606309242eb842a1bc | Bacteria | Proteobacteria | Alphaproteobacteria | Rickettsiales | S25-593 | metagenome | metagenome | 0.858 |
| b2b8a8c9475a2e53d861790f592629de | Bacteria | Proteobacteria | Alphaproteobacteria | SAR11 clade | | | | 0.809 |
| 573115e964a1e74c071e71526fb01660 | Bacteria | Cyanobacteria | Oxyphotobacteria | | | | | 0.999 |
| db122da0233bcbd3acbe017ff2a81cf7 | Bacteria | Proteobacteria | Gammaproteobacteria | SAR86 clade | uncultured gamma proteobacterium | uncultured gamma proteobacterium | uncultured gamma proteobacterium | 0.725 |
| a7fa870a72f4567135e186c656bd9dcc | Bacteria | Proteobacteria | Gammaproteobacteria | SAR86 clade | uncultured gamma proteobacterium | uncultured gamma proteobacterium | uncultured gamma proteobacterium | 0.735 |
| 02553af81ed0a26dec2b8cd96b26973c | Bacteria | Cyanobacteria | Oxyphotobacteria | Synechococcales | Cyanobiaceae | | | 0.848 |
| 936d27e23f844df5a2380087b165d7a4 | Bacteria | Proteobacteria | Deltaproteobacteria | Bdellovibrionales | Bdellovibrionaceae | OM27 clade | uncultured bacterium | 0.977 |
| 5a576b5f62a7631d660bc7b64da791d0 | Bacteria | Proteobacteria | Alphaproteobacteria | uncultured | | | | 0.946 |
| ed5bf1ff620ea094b9ab3692449eb320 | Bacteria | Bacteroidetes | Bacteroidia | Flavobacteriales | NS9 marine group | uncultured marine bacterium | uncultured marine bacterium | 0.956 |
| d7d214b1152e6e151478c261411f2a2f | Bacteria | Bacteroidetes | Bacteroidia | Flavobacteriales | Flavobacteriaceae | | | 1.000 |
| d500d56d74f8b9aeeac20872a0e89f6f | Bacteria | Proteobacteria | Alphaproteobacteria | Rickettsiales | Mitochondria | uncultured bacterium | uncultured bacterium | 0.896 |
| 4e85fdab57a66def75a6e7607da3cb63 | Bacteria | Dadabacteria | Dadabacteriia | Dadabacteriales | uncultured marine bacterium | uncultured marine bacterium | uncultured marine bacterium | 0.992 |
| 502d45c3e4fd7f4aa73841d6a5a776b0 | Bacteria | Cyanobacteria | Oxyphotobacteria | Chloroplast | | | | 1.000 |
| d16a1386b99c01202eb4b1aa078b6b86 | Bacteria | Bacteroidetes | Bacteroidia | Chitinophagales | Saprospiraceae | uncultured | Ambiguous_taxa | 0.776 |
| 656ea0e46f324ee5f662c10447bc0466 | Bacteria | Proteobacteria | Alphaproteobacteria | SAR11 clade | Clade II | | | 0.959 |
| 6c59936d3fa685a313ff794d7792c16f | Bacteria | Proteobacteria | Gammaproteobacteria | Vibrionales | Vibrionaceae | Vibrio | uncultured marine bacterium | 0.904 |
| 7d5294ecf2ccb39c3580fabd3fcd6f44 | Bacteria | Proteobacteria | Gammaproteobacteria | Gammaproteobacteria Incertae Sedis | Unknown Family | uncultured bacterium | uncultured bacterium | 0.999 |
| 7725306f309fab615cec5176135dcbf8 | Bacteria | Cyanobacteria | Oxyphotobacteria | Chloroplast | | | | 0.990 |
| 2c26ffbff0cc9f658c38cc7def2eddce | Bacteria | Cyanobacteria | Oxyphotobacteria | Chloroplast | | | | 0.978 |
| 0a3b2ecf7f780fcf7dfc61ed0f080219 | Bacteria | Proteobacteria | Deltaproteobacteria | Oligoflexales | Oligoflexaceae | uncultured | marine metagenome | 1.000 |
| b41f4d87061ad27192997fb39779bcd3 | Bacteria | Actinobacteria | Actinobacteria | Micrococcales | Microbacteriaceae | | | 1.000 |
| 9e47775fe27ba6de15af5057a46e4d57 | Bacteria | Bacteroidetes | Bacteroidia | Flavobacteriales | Cryomorphaceae | Luteibaculum | uncultured bacterium | 1.000 |
| 41ca5ce8d28e1df4ac044265fd4f3757 | Bacteria | Proteobacteria | Alphaproteobacteria | Rhodobacterales | Rhodobacteraceae | Ascidiaceihabitans | uncultured Roseobacter NAC1-2 | 0.718 |
| 8463537d406629ed67edb1ead1f1b46a | Bacteria | Nitrospinae | Nitrospinia | Nitrospinales | Nitrospinaceae | LS-NOB | | 1.000 |
| 500e1a71fab496d1fb78affa15c442cc | Bacteria | Marinimicrobia (SAR406 clade) | Alteromonas macleodii | Alteromonas macleodii | Alteromonas macleodii | Alteromonas macleodii | Alteromonas macleodii | 0.849 |
| f95ef8dc3b93df3335d6711cd2e725de | Bacteria | Marinimicrobia (SAR406 clade) | Alteromonas macleodii | Alteromonas macleodii | Alteromonas macleodii | Alteromonas macleodii | Alteromonas macleodii | 0.925 |
| b8348d8601c0a34964a77e270b2da3d1 | Bacteria | Proteobacteria | Alphaproteobacteria | Parvibaculales | OCS116 clade | uncultured alpha proteobacterium | uncultured alpha proteobacterium | 0.709 |
| 5c6c50dd5bb2f2c5546fc905128be7da | Bacteria | Proteobacteria | Alphaproteobacteria | Rickettsiales | Mitochondria | | | 1.000 |
| 9ca2cf18be3ce6d78a7b1dd6e6556595 | Bacteria | Cyanobacteria | Oxyphotobacteria | Chloroplast | | | | 1.000 |
| 1a555dba75553e90ce1f60e3f2a54f37 | Bacteria | Proteobacteria | Gammaproteobacteria | Enterobacteriales | Enterobacteriaceae | | | 0.999 |
| f97dbc45358a8aa620622d3d482304d8 | Bacteria | Proteobacteria | Alphaproteobacteria | Rickettsiales | Midichloriaceae | MD3-55 | uncultured proteobacterium | 1.000 |
| bc18e10bb9fdae5aafeb8cb091aa0840 | Bacteria | Cyanobacteria | Oxyphotobacteria | Chloroplast | | | | 1.000 |
| 2d8b0db2bd0fc0ce33e252327cece26c | Bacteria | Proteobacteria | Gammaproteobacteria | Coxiellales | Coxiellaceae | Coxiella | uncultured bacterium | 0.923 |
| e4dc9940e2d8877a81e83d35a583500b | Bacteria | Proteobacteria | Gammaproteobacteria | Alteromonadales | Psychromonadaceae | Psychromonas | | 0.965 |
| 28d11587b7355b07322c2d81eaa92db5 | Bacteria | Proteobacteria | Alphaproteobacteria | uncultured | | | | 0.999 |
| de721b107711e949b16e928427d202ba | Bacteria | Proteobacteria | Gammaproteobacteria | Enterobacteriales | Enterobacteriaceae | Erwinia | | 0.734 |
| eab6d4a131c05e95abf744a0b10610ff | Bacteria | Fibrobacteres | Fibrobacteria | Fibrobacterales | Fibrobacteraceae | uncultured | uncultured bacterium | 1.000 |
| 0293224acd74642d8896df9e4cbcb4f0 | Bacteria | Proteobacteria | Gammaproteobacteria | Betaproteobacteriales | Burkholderiaceae | Delftia | | 1.000 |
| e01085ec4f0ac927f12d9a01691e0857 | Bacteria | Bacteroidetes | Bacteroidia | Flavobacteriales | NS9 marine group | uncultured marine bacterium | uncultured marine bacterium | 0.930 |
| aad97b97b6dbea3f201adb6bcae25be2 | Bacteria | Proteobacteria | Gammaproteobacteria | Betaproteobacteriales | Nitrosomonadaceae | Nitrosomonas | | 1.000 |
| fbacaf25ec023ba1db5335303011cead | Bacteria | Bacteroidetes | Bacteroidia | Flavobacteriales | NS9 marine group | marine metagenome | marine metagenome | 0.701 |
| 316cc31f48635bcfd5c653a0fef7e6cb | Bacteria | Bacteroidetes | Bacteroidia | Flavobacteriales | Flavobacteriaceae | Flavicella | | 0.724 |
| 985b3ce30953ce34f3ea958e420754b6 | Bacteria | Bacteroidetes | Bacteroidia | Flavobacteriales | Flavobacteriaceae | NS2b marine group | uncultured Bacteroidetes bacterium | 0.739 |
| ef6514c3180c1632d301e91d88170b2e | Bacteria | Bacteroidetes | Bacteroidia | Flavobacteriales | Crocinitomicaceae | Fluviicola | marine metagenome | 0.986 |
| a62a9e8d5925a2ae970fb58006491773 | Bacteria | Proteobacteria | Alphaproteobacteria | Rickettsiales | Mitochondria | uncultured bacterium | uncultured bacterium | 0.731 |
| 376f506f23241f47ad2ed3e71fc2bccf | Bacteria | Proteobacteria | Alphaproteobacteria | Rickettsiales | Rickettsiaceae | Candidatus Megaira | | 0.991 |

| Feature_ID | Kingdom | Phylum | Class | Order | Family | Genus | Species | Confidence |
|---|---|---|---|---|---|---|---|---|
| a094d9f986844f32634753310f95464e | Bacteria | Proteobacteria | Deltaproteobacteria | Bdellovibrionales | Bdellovibrionaceae | OM27 clade | uncultured bacterium | 0.921 |
| e1e5809eb353b2c6a463c213e5fae102 | Bacteria | Proteobacteria | Gammaproteobacteria | Alteromonadales | | | | 0.936 |
| 54eccb5abc9eabc5af9c0eaac29c2572 | Bacteria | Cyanobacteria | Oxyphotobacteria | Chloroplast | uncultured marine bacterium | uncultured marine bacterium | uncultured marine bacterium | 0.929 |
| 2bcdb0f74ff8a99c9dd1c9f136b2bec3 | Bacteria | Proteobacteria | Alphaproteobacteria | Sphingomonadales | Sphingomonadaceae | Erythrobacter | | 0.888 |
| 155d7c348f373b4885df3e8c975ceb56 | Bacteria | Proteobacteria | Deltaproteobacteria | SAR324 clade(Marine group B) | metagenome | metagenome | metagenome | 1.000 |
| 4a30345adaf66cd5f45eebac19f97b72 | Bacteria | Proteobacteria | Gammaproteobacteria | Cellvibrionales | | | | 0.724 |
| 8a33676217d1c488a34c7b553f6ae0dd | Bacteria | Planctomycetes | vadinHA49 | metagenome | metagenome | metagenome | metagenome | 1.000 |
| ca25f1b88c33a55f450b79c4304ace7c | Bacteria | Proteobacteria | Alphaproteobacteria | SAR11 clade | Clade IV | | | 1.000 |
| ebec26a136b56d3e674c189888625c25 | Bacteria | Cyanobacteria | Oxyphotobacteria | Chloroplast | Crustomastix stigmatica | Crustomastix stigmatica | Crustomastix stigmatica | 1.000 |
| 6aa70ce7d904bb80c5b1c6dd2ed3f891 | Bacteria | Planctomycetes | BD7-11 | metagenome | metagenome | metagenome | metagenome | 0.996 |
| 89c861363bf481a3a9b68c696bd6e430 | Bacteria | Proteobacteria | Alphaproteobacteria | Rhodobacterales | Rhodobacteraceae | uncultured | | 0.903 |
| 02d9429323e69ee3a21df9dbaea92148 | Bacteria | Proteobacteria | Alphaproteobacteria | Rickettsiales | AB1 | uncultured alpha proteobacterium | uncultured alpha proteobacterium | 0.852 |
| c0c27820c72e6986fdca1da7e24ad2ba | Bacteria | Bacteroidetes | Bacteroidia | Flavobacteriales | Crocinitomicaceae | Crocinitomix | uncultured Flavobacteriales bacterium | 0.822 |
| 563fce8b9367a23051bf9938ee8c3b01 | Bacteria | Proteobacteria | Gammaproteobacteria | Coxiellales | Coxiellaceae | Coxiella | Ambiguous_taxa | 0.932 |
| bc70c55e3fae378a0e233dcfe3c625e2 | Bacteria | Proteobacteria | Alphaproteobacteria | SAR11 clade | Clade I | | | 1.000 |
| c067a20175575b96769242d0e93af7a8 | Bacteria | Proteobacteria | Gammaproteobacteria | Steroidobacterales | Woeseiaceae | Woeseia | uncultured marine bacterium | 0.966 |
| 1caab32ff200b6926c99925c6348c41a | Bacteria | Proteobacteria | Alphaproteobacteria | Rhodospirillales | Magnetospiraceae | uncultured | | 0.783 |
| c7752b2f352f50b3fd911867bb47e2f3 | Bacteria | Proteobacteria | Gammaproteobacteria | Salinisphaerales | Salinisphaeraceae | Salinisphaera | Ambiguous_taxa | 0.904 |
| 351ff60066fc06ca0648211621431b9f | Bacteria | Proteobacteria | Alphaproteobacteria | Rhodospirillales | Magnetospiraceae | uncultured | uncultured marine bacterium | 0.999 |
| 85c7bc111a31308dc6069d427c48a907 | Bacteria | Marinimicrobia (SAR406 clade) | marine metagenome | marine metagenome | marine metagenome | marine metagenome | marine metagenome | 0.966 |
| ea2ffaf32d4e63268411bfb102e2b23e | Bacteria | Planctomycetes | OM190 | uncultured bacterium | uncultured bacterium | uncultured bacterium | uncultured bacterium | 0.995 |
| b4d2191952ef6eeed45b037a1dfa6af5 | Bacteria | Bacteroidetes | Bacteroidia | Flavobacteriales | NS9 marine group | uncultured marine bacterium | uncultured marine bacterium | 0.992 |
| 818f93a7f90c508a4aa706babeb80870 | Bacteria | Proteobacteria | Gammaproteobacteria | KI89A clade | | | | 0.998 |
| 1ecf1353a5dc5b3ff7e5be4e46069f71 | Bacteria | Proteobacteria | Gammaproteobacteria | Enterobacteriales | Enterobacteriaceae | Pantoea | uncultured marine bacterium | 1.000 |
| d9caf2767956ffb4af2095396ae9357f | Bacteria | Proteobacteria | Gammaproteobacteria | Oceanospirillales | Nitrincolaceae | uncultured | uncultured bacterium | 0.954 |
| 757ff74929537f5d6e7b8c0f091f66da | Bacteria | Proteobacteria | Gammaproteobacteria | Vibrionales | Vibrionaceae | Vibrio | | 0.870 |
| 319e508ce3111e0228cc65142e84c9a5 | Bacteria | Bacteroidetes | Bacteroidia | Flavobacteriales | Flavobacteriaceae | | | 1.000 |
| 88a322662a85dc4431c7c9b3c5ce5918 | Bacteria | Proteobacteria | Gammaproteobacteria | SAR86 clade | | | | 1.000 |
| 8445de4328ede24e6b09707c4653e198 | Bacteria | Bacteroidetes | Bacteroidia | Flavobacteriales | Cryomorphaceae | uncultured | | 0.996 |
| 47b8a40cbc51ba17902745210e46b142 | Bacteria | Marinimicrobia (SAR406 clade) | Ambiguous_taxa | Ambiguous_taxa | Ambiguous_taxa | Ambiguous_taxa | Ambiguous_taxa | 0.727 |
| b3f1eebb5a61f444d67917f43383ecd3 | Bacteria | Marinimicrobia (SAR406 clade) | Ambiguous_taxa | Ambiguous_taxa | Ambiguous_taxa | Ambiguous_taxa | Ambiguous_taxa | 0.706 |
| be55a4adbb1205e74f77b8d78687412a | Bacteria | Proteobacteria | Gammaproteobacteria | Betaproteobacteriales | Neisseriaceae | uncultured | uncultured beta proteobacterium | 0.997 |
| e5dc7b37b1611a955f4207f00b49db61 | Bacteria | Bacteroidetes | Bacteroidia | Flavobacteriales | Flavobacteriaceae | NS4 marine group | | 0.999 |
| a71f6d2e3330c83b7412500ab2801c11 | Bacteria | Proteobacteria | Deltaproteobacteria | SAR324 clade(Marine group B) | uncultured bacterium | uncultured bacterium | uncultured bacterium | 0.790 |
| 18dfde1fcfd0c4a7f80533f0c4f13b5c | Bacteria | Verrucomicrobia | Verrucomicrobiae | Opitutales | Puniceicoccaceae | uncultured | uncultured deep-sea bacterium | 0.777 |
| fa787dac5fb8025f368b04268cda9334 | Bacteria | Bacteroidetes | Bacteroidia | Flavobacteriales | NS9 marine group | | | 0.937 |
| 92ad9e749720d468fa16b755cc5fe389 | Bacteria | Verrucomicrobia | Verrucomicrobiae | Arctic97B-4 marine group | | | | 1.000 |
| 3827820c615ab8eee488774e38b8b94e | Bacteria | Proteobacteria | Gammaproteobacteria | Ectothiorhodospirales | Ectothiorhodospiraceae | uncultured | metagenome | 0.914 |
| 99178e8d3012181eaf83f7389bfd3fea | Bacteria | Proteobacteria | Gammaproteobacteria | Thiotrichales | Thiotrichaceae | uncultured | uncultured marine bacterium | 0.985 |
| feb5892ebbe208a61011ac04a07a286f | Bacteria | Cyanobacteria | Oxyphotobacteria | Chloroplast | uncultured bacterium | uncultured bacterium | uncultured bacterium | 0.778 |
| aa5a62173f28e00961e9a5e58fee74a1 | Bacteria | Proteobacteria | Alphaproteobacteria | Rhodospirillales | AEGEAN-169 marine group | unidentified marine bacterioplankton | unidentified marine bacterioplankton | 0.991 |
| 2037353a4493f18ea25ca6ccaaa4e3c5 | Bacteria | Bacteroidetes | Bacteroidia | Flavobacteriales | Cryomorphaceae | uncultured | uncultured bacterium | 0.956 |
| 5af725f575405bbae470dc97247499df | Bacteria | Proteobacteria | Gammaproteobacteria | Oceanospirillales | Nitrincolaceae | uncultured | metagenome | 0.843 |
| d80b58c5235f0608ac448d1577963289 | Bacteria | Proteobacteria | Deltaproteobacteria | Bdellovibrionales | Bdellovibrionaceae | OM27 clade | | 1.000 |
| de9a2905a6425b78b2cbe80a7aa8adae | Bacteria | Proteobacteria | Alphaproteobacteria | Rickettsiales | Rickettsiaceae | Candidatus Cryptoprodotis | uncultured bacterium | 1.000 |
| e67f3df98f4b2d74b20a33754008fd19 | Bacteria | Proteobacteria | Gammaproteobacteria | Pseudomonadales | Moraxellaceae | Acinetobacter | Acinetobacter lwoffii | 0.879 |
| cea09f9f364d59684c86fbdc348b15c6 | Bacteria | Proteobacteria | Alphaproteobacteria | Rhodospirillales | AEGEAN-169 marine group | marine metagenome | marine metagenome | 0.997 |
| a03f77553daec241407186b671f0573b | Bacteria | Bacteroidetes | Bacteroidia | Cytophagales | Cyclobacteriaceae | Marinoscillum | | 1.000 |
| 27a816f6ed96d6abcaf41291462f3d7e | Bacteria | Proteobacteria | Deltaproteobacteria | PB19 | uncultured bacterium | uncultured bacterium | uncultured bacterium | 0.999 |
| 06f31d9b89ddd034fe1b4c127afc2434 | Bacteria | Margulisbacteria | uncultured bacterium | uncultured bacterium | uncultured bacterium | uncultured bacterium | uncultured bacterium | 0.888 |
| 077ec792baa7dcf315493973b1657218 | Bacteria | Proteobacteria | Alphaproteobacteria | SAR11 clade | Clade I | Clade Ia | | 0.965 |
| 7556c38714d7c73cc85d7562d37209cb | Bacteria | Bacteroidetes | Bacteroidia | Flavobacteriales | NS9 marine group | uncultured Flavobacteriia bacterium | uncultured Flavobacteriia bacterium | 0.903 |
| b1c785e8fd5dca5540fbf9e3498da62d | Bacteria | Bacteroidetes | Bacteroidia | Flavobacteriales | NS9 marine group | uncultured marine bacterium | uncultured marine bacterium | 0.981 |
| be50a2ea6349b983d7a6d764ea1bd627 | Bacteria | Bacteroidetes | Bacteroidia | Flavobacteriales | NS7 marine group | Ambiguous_taxa | Ambiguous_taxa | 0.726 |

**Supplementary Table S3**. Virioplankton RTPR 98% OTU feature table

|Feature_ID|654|May12_05m|May12_25m|May12_50m|May12_75m|Jun8_25m|Jun8_50m|Jun8_75m|Jun22_0m|Jun22_05m|Jun22_50m|Jun22_75m|Jul11_0m|Jul11_05m|Jul11_25m|Jul11_50m|Jul11_75m|Jul27_05m|Jul27_25m|Jul27_50m|Jul27_75m|Aug5_0m|Aug5_05m|Aug5_25m|Aug5_50m|Aug5_75m|Sep8_0m|Sep8_05m|Sep8_25m|Sep8_50m|
|---|---|---|---|---|---|---|---|---|---|---|---|---|---|---|---|---|---|---|---|---|---|---|---|---|---|---|---|---|---|---|
|Jul11.5v_1468|156|5|1|0|5|1|3|0|7|1|2|1|2|9|1|4|1|12|7|5|0|31|11|21|0|0|8|2|1|15|
|Aug5.75v_39888|1|0|0|0|0|0|0|0|0|0|0|0|0|0|0|0|0|0|0|0|0|0|0|0|0|1|0|0|0|0|
|Aug5.25v_22287|1|0|0|0|0|0|0|0|0|0|0|0|0|0|0|0|0|0|0|0|0|0|0|1|0|0|0|0|0|0|
|Jul27.25v_13759|1|0|0|0|0|0|0|0|0|0|0|0|0|0|0|0|0|0|1|0|0|0|0|0|0|0|0|0|0|0|
|Jun8.50v_686|1|0|0|0|0|0|1|0|0|0|0|0|0|0|0|0|0|0|0|0|0|0|0|0|0|0|0|0|0|0|
|Sep8.0v_1101|2|0|0|0|0|0|0|0|0|0|0|0|0|0|0|0|0|0|0|0|0|1|0|0|0|0|1|0|0|0|
|Aug5.5v_28183|277|7|10|7|3|3|9|0|6|2|10|7|6|4|5|5|4|12|9|12|1|34|17|19|8|15|4|3|3|52|
|Sep8.50v_25890|2|0|0|0|0|0|0|0|0|0|0|0|0|0|0|0|0|0|0|0|0|0|0|0|1|0|0|0|0|1|
|Jul11.0v_11656|1|0|0|0|0|0|0|0|0|0|0|0|1|0|0|0|0|0|0|0|0|0|0|0|0|0|0|0|0|0|
|Aug5.0v_1978|1|0|0|0|0|0|0|0|0|0|0|0|0|0|0|0|0|0|0|0|0|1|0|0|0|0|0|0|0|0|
|May12.50v_24230|1|0|0|1|0|0|0|0|0|0|0|0|0|0|0|0|0|0|0|0|0|0|0|0|0|0|0|0|0|0|
|Sep8.25v_20738|1|0|0|0|0|0|0|0|0|0|0|0|0|0|0|0|0|0|0|0|0|0|0|0|0|0|0|0|1|0|
|Sep8.50v_21432|18|0|0|1|1|0|0|0|1|0|0|0|0|1|0|2|0|1|0|0|0|3|1|2|0|0|1|0|0|4|
|Sep8.50v_29250|1|0|0|0|0|0|0|0|0|0|0|0|0|0|0|0|0|0|0|0|0|0|0|0|0|0|0|0|0|1|
|Jul11.25v_13450|5|0|0|0|0|0|0|0|1|0|0|1|0|0|2|0|0|0|0|0|0|0|1|0|0|0|0|0|0|0|
|Sep8.0v_8165|1|0|0|0|0|0|0|0|0|0|0|0|0|0|0|0|0|0|0|0|0|0|0|0|0|0|1|0|0|0|
|Jun22.0v_13723|6|0|0|0|0|0|0|0|1|0|0|0|0|1|1|0|0|0|0|0|0|0|0|1|0|1|0|0|0|1|
|Sep8.50v_36517|1|0|0|0|0|0|0|0|0|0|0|0|0|0|0|0|0|0|0|0|0|0|0|0|0|0|0|0|0|1|
|Aug5.5v_13719|1|0|0|0|0|0|0|0|0|0|0|0|0|0|0|0|0|0|0|0|0|0|1|0|0|0|0|0|0|0|
|Sep8.50v_34751|1|0|0|0|0|0|0|0|0|0|0|0|0|0|0|0|0|0|0|0|0|0|0|0|0|0|0|0|0|1|
|Sep8.0v_8944|1|0|0|0|0|0|0|0|0|0|0|0|0|0|0|0|0|0|0|0|0|0|0|0|0|0|1|0|0|0|
|May12.50v_16901|1|0|0|1|0|0|0|0|0|0|0|0|0|0|0|0|0|0|0|0|0|0|0|0|0|0|0|0|0|0|
|Sep8.50v_29894|1|0|0|0|0|0|0|0|0|0|0|0|0|0|0|0|0|0|0|0|0|0|0|0|0|0|0|0|0|1|
|Sep8.50v_37785|1|0|0|0|0|0|0|0|0|0|0|0|0|0|0|0|0|0|0|0|0|0|0|0|0|0|0|0|0|1|
|Sep8.25v_17295|1|0|0|0|0|0|0|0|0|0|0|0|0|0|0|0|0|0|0|0|0|0|0|0|0|0|0|0|1|0|
|Aug5.50v_35605|1|0|0|0|0|0|0|0|0|0|0|0|0|0|0|0|0|0|0|0|0|0|0|0|1|0|0|0|0|0|
|Jul27.5v_6261|1|0|0|0|0|0|0|0|0|0|0|0|0|0|0|0|0|1|0|0|0|0|0|0|0|0|0|0|0|0|
|May12.5v_4693|1|1|0|0|0|0|0|0|0|0|0|0|0|0|0|0|0|0|0|0|0|0|0|0|0|0|0|0|0|0|
|May12.50v_5666|1|0|0|1|0|0|0|0|0|0|0|0|0|0|0|0|0|0|0|0|0|0|0|0|0|0|0|0|0|0|
|Sep8.50v_9662|1|0|0|0|0|0|0|0|0|0|0|0|0|0|0|0|0|0|0|0|0|0|0|0|0|0|0|0|0|1|
|Jun22.50v_34570|1|0|0|0|0|0|0|0|0|0|1|0|0|0|0|0|0|0|0|0|0|0|0|0|0|0|0|0|0|0|
|May12.50v_10549|1|0|0|1|0|0|0|0|0|0|0|0|0|0|0|0|0|0|0|0|0|0|0|0|0|0|0|0|0|0|
|Jul11.0v_8727|2|0|0|0|0|0|0|0|0|0|0|0|1|0|0|0|0|0|0|0|0|0|0|0|0|0|0|0|0|1|
|May12.25v_1530|5|0|1|0|0|0|0|0|0|0|0|0|0|0|0|1|1|1|0|0|0|0|0|1|0|0|0|0|0|0|
|Aug5.0v_1320|28|1|0|1|1|1|4|0|2|2|2|2|3|0|0|0|1|0|0|0|0|2|3|0|1|0|1|0|0|1|
|Jul27.5v_8356|3|0|0|0|0|0|0|0|0|0|0|0|0|0|0|0|0|2|0|1|0|0|0|0|0|0|0|0|0|0|
|Jun8.50v_10797|184|1|4|5|8|0|4|2|1|4|2|2|3|10|4|1|0|20|10|1|0|34|17|23|3|1|14|4|0|6|
|Sep8.0v_16261|2|0|0|0|0|0|0|0|0|0|0|0|0|0|0|0|0|0|0|0|0|0|0|0|0|0|2|0|0|0|
|Jun22.50v_32786|35|2|2|0|0|5|1|0|1|2|2|1|1|1|0|0|0|3|0|0|0|3|1|1|0|0|1|4|0|4|
|Jul27.50v_14259|10|0|0|0|0|0|0|0|0|1|0|0|1|0|3|0|0|0|0|2|0|2|0|0|0|1|0|0|0|0|
|May12.75v_113|1|0|0|0|1|0|0|0|0|0|0|0|0|0|0|0|0|0|0|0|0|0|0|0|0|0|0|0|0|0|
|Aug5.0v_2075|2|0|0|0|0|0|0|0|0|0|0|0|0|0|0|0|0|0|0|0|0|1|0|0|0|0|1|0|0|0|
|Jun22.75v_5791|1|0|0|0|0|0|0|0|0|0|0|1|0|0|0|0|0|0|0|0|0|0|0|0|0|0|0|0|0|0|
|Jul27.5v_13640|1|0|0|0|0|0|0|0|0|0|0|0|0|0|0|0|0|1|0|0|0|0|0|0|0|0|0|0|0|0|
|May12.75v_37898|1|0|0|0|1|0|0|0|0|0|0|0|0|0|0|0|0|0|0|0|0|0|0|0|0|0|0|0|0|0|
|Aug5.5v_14779|1|0|0|0|0|0|0|0|0|0|0|0|0|0|0|0|0|0|0|0|0|0|1|0|0|0|0|0|0|0|
|Aug5.5v_24741|29|0|6|3|0|1|0|0|1|1|1|1|1|0|0|0|0|2|0|0|0|2|2|3|0|2|1|0|1|1|
|Sep8.0v_8244|1|0|0|0|0|0|0|0|0|0|0|0|0|0|0|0|0|0|0|0|0|0|0|0|0|0|1|0|0|0|
|Jun8.50v_19908|2|0|0|0|0|0|1|0|0|1|0|0|0|0|0|0|0|0|0|0|0|0|0|0|0|0|0|0|0|0|
|Aug5.0v_1182|25|1|1|0|0|0|0|0|0|1|1|0|1|1|0|0|0|2|1|0|0|7|1|3|0|0|5|0|0|0|
|Jul27.5v_17596|21|2|0|0|1|1|0|0|1|1|1|2|1|1|1|0|0|4|0|0|0|2|1|0|0|0|0|0|0|2|
|Aug5.25v_23298|4|0|0|0|0|0|0|0|0|1|0|0|0|0|0|0|0|0|0|0|0|0|0|2|0|0|0|0|0|1|
|Aug5.25v_25628|1|0|0|0|0|0|0|0|0|0|0|0|0|0|0|0|0|0|0|0|0|0|0|1|0|0|0|0|0|0|
|Aug5.25v_30997|1|0|0|0|0|0|0|0|0|0|0|0|0|0|0|0|0|0|0|0|0|0|0|1|0|0|0|0|0|0|
|Aug5.25v_38436|1|0|0|0|0|0|0|0|0|0|0|0|0|0|0|0|0|0|0|0|0|0|0|1|0|0|0|0|0|0|
|Aug5.0v_4761|1|0|0|0|0|0|0|0|0|0|0|0|0|0|0|0|0|0|0|0|0|1|0|0|0|0|0|0|0|0|
|Aug5.5v_19590|1|0|0|0|0|0|0|0|0|0|0|0|0|0|0|0|0|0|0|0|0|0|1|0|0|0|0|0|0|0|
|Sep8.0v_6624|2|0|0|0|0|0|0|0|0|0|1|0|0|0|0|0|0|0|0|0|0|0|0|0|0|0|1|0|0|0|
|Jul27.5v_12484|2|0|0|0|0|0|0|0|0|0|0|0|0|0|0|0|0|1|1|0|0|0|0|0|0|0|0|0|0|0|
|Jun22.5v_7264|1|0|0|0|0|0|0|0|0|1|0|0|0|0|0|0|0|0|0|0|0|0|0|0|0|0|0|0|0|0|
|Sep8.25v_6070|1|0|0|0|0|0|0|0|0|0|0|0|0|0|0|0|0|0|0|0|0|0|0|0|0|0|0|0|1|0|
|Sep8.5v_616|1|0|0|0|0|0|0|0|0|0|0|0|0|0|0|0|0|0|0|0|0|0|0|0|0|0|0|1|0|0|
|Jul27.5v_179|1|0|0|0|0|0|0|0|0|0|0|0|0|0|0|0|0|1|0|0|0|0|0|0|0|0|0|0|0|0|
|Jul27.5v_3646|1|0|0|0|0|0|0|0|0|0|0|0|0|0|0|0|0|1|0|0|0|0|0|0|0|0|0|0|0|0|
|Aug5.0v_4228|1|0|0|0|0|0|0|0|0|0|0|0|0|0|0|0|0|0|0|0|0|1|0|0|0|0|0|0|0|0|
|Jun22.0v_11126|1|0|0|0|0|0|0|0|1|0|0|0|0|0|0|0|0|0|0|0|0|0|0|0|0|0|0|0|0|0|
|May12.25v_18241|1|0|1|0|0|0|0|0|0|0|0|0|0|0|0|0|0|0|0|0|0|0|0|0|0|0|0|0|0|0|
|Jun22.75v_20936|1|0|0|0|0|0|0|0|0|0|0|1|0|0|0|0|0|0|0|0|0|0|0|0|0|0|0|0|0|0|
|Jul27.5v_2985|1|0|0|0|0|0|0|0|0|0|0|0|0|0|0|0|0|1|0|0|0|0|0|0|0|0|0|0|0|0|
|May12.5v_30272|1|1|0|0|0|0|0|0|0|0|0|0|0|0|0|0|0|0|0|0|0|0|0|0|0|0|0|0|0|0|
|Aug5.25v_8358|1|0|0|0|0|0|0|0|0|0|0|0|0|0|0|0|0|0|0|0|0|0|0|1|0|0|0|0|0|0|
|Jun8.50v_12566|1|0|0|0|0|0|1|0|0|0|0|0|0|0|0|0|0|0|0|0|0|0|0|0|0|0|0|0|0|0|
|Jul27.25v_31892|1|0|0|0|0|0|0|0|0|0|0|0|0|0|0|0|0|0|1|0|0|0|0|0|0|0|0|0|0|0|
|Aug5.0v_2155|1|0|0|0|0|0|0|0|0|0|0|0|0|0|0|0|0|0|0|0|0|1|0|0|0|0|0|0|0|0|
|Aug5.75v_6381|1|0|0|0|0|0|0|0|0|0|0|0|0|0|0|0|0|0|0|0|0|0|0|0|0|1|0|0|0|0|
|Jul27.5v_20083|1|0|0|0|0|0|0|0|0|0|0|0|0|0|0|0|0|1|0|0|0|0|0|0|0|0|0|0|0|0|
|Jul27.5v_7708|1|0|0|0|0|0|0|0|0|0|0|0|0|0|0|0|0|1|0|0|0|0|0|0|0|0|0|0|0|0|
|Aug5.25v_27584|1|0|0|0|0|0|0|0|0|0|0|0|0|0|0|0|0|0|0|0|0|0|0|1|0|0|0|0|0|0|
|Jul11.75v_537|1|0|0|0|0|0|0|0|0|0|0|0|0|0|0|0|1|0|0|0|0|0|0|0|0|0|0|0|0|0|
|Jun22.5v_17560|2|0|0|0|0|0|0|0|0|1|0|0|0|0|0|0|0|0|0|0|0|0|0|1|0|0|0|0|0|0|
|Jun22.5v_90|3|0|0|0|0|0|0|0|0|1|0|0|0|0|0|0|0|0|0|0|0|1|0|0|0|0|0|0|0|1|
|Jul27.5v_5739|1|0|0|0|0|0|0|0|0|0|0|0|0|0|0|0|0|1|0|0|0|0|0|0|0|0|0|0|0|0|
|Jul27.5v_15916|1|0|0|0|0|0|0|0|0|0|0|0|0|0|0|0|0|1|0|0|0|0|0|0|0|0|0|0|0|0|
|Aug5.50v_720|1|0|0|0|0|0|0|0|0|0|0|0|0|0|0|0|0|0|0|0|0|0|0|0|1|0|0|0|0|0|
|Jul11.5v_3506|4|1|0|0|0|0|0|0|0|0|0|0|0|3|0|0|0|0|0|0|0|0|0|0|0|0|0|0|0|0|
|Jun22.0v_20743|1|0|0|0|0|0|0|0|1|0|0|0|0|0|0|0|0|0|0|0|0|0|0|0|0|0|0|0|0|0|
|Jun8.50v_4303|2|0|0|0|0|1|1|0|0|0|0|0|0|0|0|0|0|0|0|0|0|0|0|0|0|0|0|0|0|0|
|Jun22.0v_10421|1|0|0|0|0|0|0|0|1|0|0|0|0|0|0|0|0|0|0|0|0|0|0|0|0|0|0|0|0|0|
|Aug5.0v_1855|1|0|0|0|0|0|0|0|0|0|0|0|0|0|0|0|0|0|0|0|0|1|0|0|0|0|0|0|0|0|
|Jun22.75v_25000|1|0|0|0|0|0|0|0|0|0|0|1|0|0|0|0|0|0|0|0|0|0|0|0|0|0|0|0|0|0|
|Sep8.5v_13874|1|0|0|0|0|0|0|0|0|0|0|0|0|0|0|0|0|0|0|0|0|0|0|0|0|0|0|1|0|0|
|Sep8.50v_3217|1|0|0|0|0|0|0|0|0|0|0|0|0|0|0|0|0|0|0|0|0|0|0|0|0|0|0|0|0|1|
|Jul27.25v_26842|1|0|0|0|0|0|0|0|0|0|0|0|0|0|0|0|0|0|1|0|0|0|0|0|0|0|0|0|0|0|
|Jul11.25v_13332|1|0|0|0|0|0|0|0|0|0|0|0|0|0|1|0|0|0|0|0|0|0|0|0|0|0|0|0|0|0|
|Jul11.5v_17515|1|0|0|0|0|0|0|0|0|0|0|0|0|1|0|0|0|0|0|0|0|0|0|0|0|0|0|0|0|0|
|May12.5v_20244|1|1|0|0|0|0|0|0|0|0|0|0|0|0|0|0|0|0|0|0|0|0|0|0|0|0|0|0|0|0|
|Aug5.0v_4926|1|0|0|0|0|0|0|0|0|0|0|0|0|0|0|0|0|0|0|0|0|1|0|0|0|0|0|0|0|0|
|Aug5.0v_5390|1|0|0|0|0|0|0|0|0|0|0|0|0|0|0|0|0|0|0|0|0|1|0|0|0|0|0|0|0|0|
|Aug5.5v_357|1|0|0|0|0|0|0|0|0|0|0|0|0|0|0|0|0|0|0|0|0|0|1|0|0|0|0|0|0|0|
|Aug5.0v_809|1|0|0|0|0|0|0|0|0|0|0|0|0|0|0|0|0|0|0|0|0|1|0|0|0|0|0|0|0|0|
|Jul11.0v_1613|1|0|0|0|0|0|0|0|0|0|0|0|1|0|0|0|0|0|0|0|0|0|0|0|0|0|0|0|0|0|
|Aug5.75v_24062|1|0|0|0|0|0|0|0|0|0|0|0|0|0|0|0|0|0|0|0|0|0|0|0|0|1|0|0|0|0|
|Jul11.50v_14199|1|0|0|0|0|0|0|0|0|0|0|0|0|0|0|1|0|0|0|0|0|0|0|0|0|0|0|0|0|0|
|Aug5.75v_3902|1|0|0|0|0|0|0|0|0|0|0|0|0|0|0|0|0|0|0|0|0|0|0|0|0|1|0|0|0|0|
|Jul27.25v_11877|1|0|0|0|0|0|0|0|0|0|0|0|0|0|0|0|0|0|1|0|0|0|0|0|0|0|0|0|0|0|
|Jul27.5v_11560|1|0|0|0|0|0|0|0|0|0|0|0|0|0|0|0|0|1|0|0|0|0|0|0|0|0|0|0|0|0|
|Sep8.50v_11308|1|0|0|0|0|0|0|0|0|0|0|0|0|0|0|0|0|0|0|0|0|0|0|0|0|0|0|0|0|1|
|Jun22.50v_34629|1|0|0|0|0|0|0|0|0|0|1|0|0|0|0|0|0|0|0|0|0|0|0|0|0|0|0|0|0|0|
|Sep8.50v_15084|1|0|0|0|0|0|0|0|0|0|0|0|0|0|0|0|0|0|0|0|0|0|0|0|0|0|0|0|0|1|
|Aug5.25v_24273|1|0|0|0|0|0|0|0|0|0|0|0|0|0|0|0|0|0|0|0|0|0|0|1|0|0|0|0|0|0|
|Aug5.5v_40425|1|0|0|0|0|0|0|0|0|0|0|0|0|0|0|0|0|0|0|0|0|0|1|0|0|0|0|0|0|0|
|Sep8.0v_6231|2|0|0|0|0|0|1|0|0|0|0|0|0|0|0|0|0|0|0|0|0|0|0|0|0|0|1|0|0|0|
|Aug5.0v_2957|1|0|0|0|0|0|0|0|0|0|0|0|0|0|0|0|0|0|0|0|0|1|0|0|0|0|0|0|0|0|
|Jul27.5v_1048|1|0|0|0|0|0|0|0|0|0|0|0|0|0|0|0|0|1|0|0|0|0|0|0|0|0|0|0|0|0|
|Sep8.50v_15779|1|0|0|0|0|0|0|0|0|0|0|0|0|0|0|0|0|0|0|0|0|0|0|0|0|0|0|0|0|1|
|Jul11.5v_4595|1|0|0|0|0|0|0|0|0|0|0|0|0|1|0|0|0|0|0|0|0|0|0|0|0|0|0|0|0|0|
|Aug5.0v_305|2|0|0|0|1|0|0|0|0|0|0|0|0|0|0|0|0|0|0|0|0|1|0|0|0|0|0|0|0|0|
|May12.25v_3553|1|0|1|0|0|0|0|0|0|0|0|0|0|0|0|0|0|0|0|0|0|0|0|0|0|0|0|0|0|0|
|Jul27.5v_12271|1|0|0|0|0|0|0|0|0|0|0|0|0|0|0|0|0|1|0|0|0|0|0|0|0|0|0|0|0|0|
|May12.5v_8029|1|1|0|0|0|0|0|0|0|0|0|0|0|0|0|0|0|0|0|0|0|0|0|0|0|0|0|0|0|0|
|Jul27.5v_2154|1|0|0|0|0|0|0|0|0|0|0|0|0|0|0|0|0|1|0|0|0|0|0|0|0|0|0|0|0|0|
|Jun22.0v_6729|1|0|0|0|0|0|0|0|1|0|0|0|0|0|0|0|0|0|0|0|0|0|0|0|0|0|0|0|0|0|
|Aug5.5v_11015|1|0|0|0|0|0|0|0|0|0|0|0|0|0|0|0|0|0|0|0|0|0|1|0|0|0|0|0|0|0|
|Aug5.5v_23987|1|0|0|0|0|0|0|0|0|0|0|0|0|0|0|0|0|0|0|0|0|0|1|0|0|0|0|0|0|0|
|Jul27.5v_2009|1|0|0|0|0|0|0|0|0|0|0|0|0|0|0|0|0|1|0|0|0|0|0|0|0|0|0|0|0|0|
|Aug5.5v_23135|2|0|0|0|0|0|0|0|0|0|0|0|0|0|0|1|0|0|0|0|0|0|1|0|0|0|0|0|0|0|
|Jun8.25v_26408|1|0|0|0|0|1|0|0|0|0|0|0|0|0|0|0|0|0|0|0|0|0|0|0|0|0|0|0|0|0|
|Jun8.50v_16697|1|0|0|0|0|0|1|0|0|0|0|0|0|0|0|0|0|0|0|0|0|0|0|0|0|0|0|0|0|0|
|Jul11.75v_7569|1|0|0|0|0|0|0|0|0|0|0|0|0|0|0|0|1|0|0|0|0|0|0|0|0|0|0|0|0|0|
|Aug5.75v_26509|1|0|0|0|0|0|0|0|0|0|0|0|0|0|0|0|0|0|0|0|0|0|0|0|0|1|0|0|0|0|
|Aug5.50v_709|1|0|0|0|0|0|0|0|0|0|0|0|0|0|0|0|0|0|0|0|0|0|0|0|1|0|0|0|0|0|
|Sep8.5v_20924|1|0|0|0|0|0|0|0|0|0|0|0|0|0|0|0|0|0|0|0|0|0|0|0|0|0|0|1|0|0|
|Aug5.25v_1084|1|0|0|0|0|0|0|0|0|0|0|0|0|0|0|0|0|0|0|0|0|0|0|1|0|0|0|0|0|0|
|Jul27.25v_34099|1|0|0|0|0|0|0|0|0|0|0|0|0|0|0|0|0|0|1|0|0|0|0|0|0|0|0|0|0|0|
|May12.25v_13980|1|0|1|0|0|0|0|0|0|0|0|0|0|0|0|0|0|0|0|0|0|0|0|0|0|0|0|0|0|0|
|Aug5.25v_21957|1|0|0|0|0|0|0|0|0|0|0|0|0|0|0|0|0|0|0|0|0|0|0|1|0|0|0|0|0|0|
|Aug5.0v_3013|2|0|0|0|0|0|0|0|0|0|0|0|0|0|0|0|0|0|0|0|0|2|0|0|0|0|0|0|0|0|
|Jun22.5v_1792|1|0|0|0|0|0|0|0|0|1|0|0|0|0|0|0|0|0|0|0|0|0|0|0|0|0|0|0|0|0|
|Jun8.75v_2661|1|0|0|0|0|0|0|1|0|0|0|0|0|0|0|0|0|0|0|0|0|0|0|0|0|0|0|0|0|0|
|Aug5.5v_8128|6|0|0|0|0|0|0|0|0|0|0|0|0|1|0|0|0|0|0|0|0|0|4|0|0|0|0|0|0|1|
|Jul27.5v_6482|1|0|0|0|0|0|0|0|0|0|0|0|0|0|0|0|0|1|0|0|0|0|0|0|0|0|0|0|0|0|
|May12.5v_29257|2|1|0|0|0|0|0|0|0|0|0|0|0|0|0|0|0|0|0|0|0|0|0|0|0|0|0|0|0|1|
|Sep8.5v_4513|1|0|0|0|0|0|0|0|0|0|0|0|0|0|0|0|0|0|0|0|0|0|0|0|0|0|0|1|0|0|
|Sep8.0v_9488|1|0|0|0|0|0|0|0|0|0|0|0|0|0|0|0|0|0|0|0|0|0|0|0|0|0|1|0|0|0|
|Jul27.5v_8800|1|0|0|0|0|0|0|0|0|0|0|0|0|0|0|0|0|1|0|0|0|0|0|0|0|0|0|0|0|0|
|Aug5.0v_2246|1|0|0|0|0|0|0|0|0|0|0|0|0|0|0|0|0|0|0|0|0|1|0|0|0|0|0|0|0|0|
|Jul27.5v_13488|1|0|0|0|0|0|0|0|0|0|0|0|0|0|0|0|0|1|0|0|0|0|0|0|0|0|0|0|0|0|
|Sep8.50v_5872|1|0|0|0|0|0|0|0|0|0|0|0|0|0|0|0|0|0|0|0|0|0|0|0|0|0|0|0|0|1|
|Aug5.0v_1688|1|0|0|0|0|0|0|0|0|0|0|0|0|0|0|0|0|0|0|0|0|1|0|0|0|0|0|0|0|0|
|Jun8.75v_10966|1|0|0|0|0|0|0|1|0|0|0|0|0|0|0|0|0|0|0|0|0|0|0|0|0|0|0|0|0|0|
|May12.5v_13906|1|1|0|0|0|0|0|0|0|0|0|0|0|0|0|0|0|0|0|0|0|0|0|0|0|0|0|0|0|0|
|May12.50v_7190|1|0|0|1|0|0|0|0|0|0|0|0|0|0|0|0|0|0|0|0|0|0|0|0|0|0|0|0|0|0|
|Sep8.0v_17533|1|0|0|0|0|0|0|0|0|0|0|0|0|0|0|0|0|0|0|0|0|0|0|0|0|0|1|0|0|0|
|Jul27.5v_9780|1|0|0|0|0|0|0|0|0|0|0|0|0|0|0|0|0|1|0|0|0|0|0|0|0|0|0|0|0|0|
|Jul11.25v_3735|1|0|0|0|0|0|0|0|0|0|0|0|0|0|1|0|0|0|0|0|0|0|0|0|0|0|0|0|0|0|
|Jun22.0v_9832|1|0|0|0|0|0|0|0|1|0|0|0|0|0|0|0|0|0|0|0|0|0|0|0|0|0|0|0|0|0|
|Aug5.25v_36318|1|0|0|0|0|0|0|0|0|0|0|0|0|0|0|0|0|0|0|0|0|0|0|1|0|0|0|0|0|0|
|Jul11.0v_2038|1|0|0|0|0|0|0|0|0|0|0|0|1|0|0|0|0|0|0|0|0|0|0|0|0|0|0|0|0|0|
|Jul27.25v_40180|1|0|0|0|0|0|0|0|0|0|0|0|0|0|0|0|0|0|1|0|0|0|0|0|0|0|0|0|0|0|
|Aug5.25v_86|1|0|0|0|0|0|0|0|0|0|0|0|0|0|0|0|0|0|0|0|0|0|0|1|0|0|0|0|0|0|
|Aug5.25v_40792|2|0|0|0|0|0|0|0|0|0|0|0|0|0|0|0|0|0|0|0|0|0|0|1|0|0|0|0|0|1|
|Sep8.5v_4344|1|0|0|0|0|0|0|0|0|0|0|0|0|0|0|0|0|0|0|0|0|0|0|0|0|0|0|1|0|0|
|Jul27.5v_10681|1|0|0|0|0|0|0|0|0|0|0|0|0|0|0|0|0|1|0|0|0|0|0|0|0|0|0|0|0|0|
|Sep8.0v_2419|1|0|0|0|0|0|0|0|0|0|0|0|0|0|0|0|0|0|0|0|0|0|0|0|0|0|1|0|0|0|
|Jul27.5v_14056|1|0|0|0|0|0|0|0|0|0|0|0|0|0|0|0|0|1|0|0|0|0|0|0|0|0|0|0|0|0|
|Jun8.50v_8853|1|0|0|0|0|0|1|0|0|0|0|0|0|0|0|0|0|0|0|0|0|0|0|0|0|0|0|0|0|0|
|Jul27.5v_13619|1|0|0|0|0|0|0|0|0|0|0|0|0|0|0|0|0|1|0|0|0|0|0|0|0|0|0|0|0|0|
|Aug5.0v_1242|1|0|0|0|0|0|0|0|0|0|0|0|0|0|0|0|0|0|0|0|0|1|0|0|0|0|0|0|0|0|
|Aug5.0v_2151|1|0|0|0|0|0|0|0|0|0|0|0|0|0|0|0|0|0|0|0|0|1|0|0|0|0|0|0|0|0|
|Aug5.0v_2401|3435|5|8|10|4|12|18|1|37|91|353|27|24|131|157|60|14|335|290|359|44|285|161|249|224|117|66|5|38|310|
|Aug5.0v_1166|1|0|0|0|0|0|0|0|0|0|0|0|0|0|0|0|0|0|0|0|0|1|0|0|0|0|0|0|0|0|
|Aug5.25v_30841|6|0|0|0|0|0|0|0|0|1|0|0|1|1|0|0|0|0|0|0|0|0|0|1|0|0|1|1|0|0|
|Jul27.50v_30921|22|0|0|0|0|0|1|0|1|2|2|0|1|3|1|0|0|1|2|3|1|2|0|1|0|0|0|0|0|1|
|Sep8.50v_32597|7|0|0|0|0|0|0|0|0|1|0|1|0|0|2|0|0|0|0|0|1|0|0|0|1|0|0|0|0|1|
|Jun22.50v_12093|1|0|0|0|0|0|0|0|0|0|1|0|0|0|0|0|0|0|0|0|0|0|0|0|0|0|0|0|0|0|
|Jul27.5v_5032|1|0|0|0|0|0|0|0|0|0|0|0|0|0|0|0|0|1|0|0|0|0|0|0|0|0|0|0|0|0|
|Sep8.50v_8612|4|0|0|0|0|0|0|0|0|0|0|0|0|0|1|0|0|0|0|0|0|0|0|0|0|0|1|0|0|2|
|Aug5.0v_4045|1|0|0|0|0|0|0|0|0|0|0|0|0|0|0|0|0|0|0|0|0|1|0|0|0|0|0|0|0|0|
|Aug5.50v_13702|61|0|0|0|0|0|0|0|1|5|4|0|2|2|4|3|1|14|4|4|1|0|3|3|3|1|1|1|0|4|
|Jul27.50v_11414|10|0|0|0|0|0|0|0|0|1|2|0|0|0|1|1|0|0|0|2|0|1|0|1|1|0|0|0|0|0|
|Sep8.50v_16464|4|0|0|0|0|0|0|0|1|0|0|0|0|0|0|0|0|1|0|1|0|0|0|0|0|0|0|0|0|1|
|Jul27.5v_10641|1|0|0|0|0|0|0|0|0|0|0|0|0|0|0|0|0|1|0|0|0|0|0|0|0|0|0|0|0|0|
|Jul11.5v_6885|1|0|0|0|0|0|0|0|0|0|0|0|0|1|0|0|0|0|0|0|0|0|0|0|0|0|0|0|0|0|
|Jun22.5v_19613|2|0|0|0|0|0|0|0|0|1|0|0|0|0|1|0|0|0|0|0|0|0|0|0|0|0|0|0|0|0|
|Aug5.75v_14295|1|0|0|0|0|0|0|0|0|0|0|0|0|0|0|0|0|0|0|0|0|0|0|0|0|1|0|0|0|0|
|Sep8.50v_23834|1|0|0|0|0|0|0|0|0|0|0|0|0|0|0|0|0|0|0|0|0|0|0|0|0|0|0|0|0|1|
|Jul11.25v_20568|3|0|0|0|0|0|0|0|0|0|0|0|0|0|1|0|0|0|0|0|0|0|0|1|0|0|0|0|0|1|
|Jun22.50v_16304|3|0|0|0|0|0|0|0|0|0|1|0|0|0|0|0|0|1|1|0|0|0|0|0|0|0|0|0|0|0|
|Jul11.5v_13808|1|0|0|0|0|0|0|0|0|0|0|0|0|1|0|0|0|0|0|0|0|0|0|0|0|0|0|0|0|0|
|Jul11.25v_3211|1|0|0|0|0|0|0|0|0|0|0|0|0|0|1|0|0|0|0|0|0|0|0|0|0|0|0|0|0|0|
|Jul27.25v_35958|1|0|0|0|0|0|0|0|0|0|0|0|0|0|0|0|0|0|1|0|0|0|0|0|0|0|0|0|0|0|
|Aug5.0v_5214|1|0|0|0|0|0|0|0|0|0|0|0|0|0|0|0|0|0|0|0|0|1|0|0|0|0|0|0|0|0|
|Aug5.25v_13924|1|0|0|0|0|0|0|0|0|0|0|0|0|0|0|0|0|0|0|0|0|0|0|1|0|0|0|0|0|0|
|Jul11.25v_16201|2|0|0|0|0|0|0|0|0|1|0|0|0|0|1|0|0|0|0|0|0|0|0|0|0|0|0|0|0|0|
|Jun22.50v_23024|1|0|0|0|0|0|0|0|0|0|1|0|0|0|0|0|0|0|0|0|0|0|0|0|0|0|0|0|0|0|
|Aug5.50v_32970|1|0|0|0|0|0|0|0|0|0|0|0|0|0|0|0|0|0|0|0|0|0|0|0|1|0|0|0|0|0|
|Jul27.5v_12604|8|0|0|0|0|0|0|0|1|0|0|0|0|2|0|0|0|4|0|1|0|0|0|0|0|0|0|0|0|0|
|Jun22.5v_7140|1|0|0|0|0|0|0|0|0|1|0|0|0|0|0|0|0|0|0|0|0|0|0|0|0|0|0|0|0|0|
|Jul27.25v_9384|1|0|0|0|0|0|0|0|0|0|0|0|0|0|0|0|0|0|1|0|0|0|0|0|0|0|0|0|0|0|
|Jun22.5v_15315|1|0|0|0|0|0|0|0|0|1|0|0|0|0|0|0|0|0|0|0|0|0|0|0|0|0|0|0|0|0|
|Jul27.25v_3573|1|0|0|0|0|0|0|0|0|0|0|0|0|0|0|0|0|0|1|0|0|0|0|0|0|0|0|0|0|0|
|Aug5.75v_19722|1|0|0|0|0|0|0|0|0|0|0|0|0|0|0|0|0|0|0|0|0|0|0|0|0|1|0|0|0|0|
|Aug5.50v_37660|1|0|0|0|0|0|0|0|0|0|0|0|0|0|0|0|0|0|0|0|0|0|0|0|1|0|0|0|0|0|
|Aug5.0v_4777|1|0|0|0|0|0|0|0|0|0|0|0|0|0|0|0|0|0|0|0|0|1|0|0|0|0|0|0|0|0|
|Sep8.50v_18336|1|0|0|0|0|0|0|0|0|0|0|0|0|0|0|0|0|0|0|0|0|0|0|0|0|0|0|0|0|1|
|Jul27.5v_18355|1|0|0|0|0|0|0|0|0|0|0|0|0|0|0|0|0|1|0|0|0|0|0|0|0|0|0|0|0|0|
|Aug5.0v_5311|1|0|0|0|0|0|0|0|0|0|0|0|0|0|0|0|0|0|0|0|0|1|0|0|0|0|0|0|0|0|
|Jun22.5v_11053|1|0|0|0|0|0|0|0|0|1|0|0|0|0|0|0|0|0|0|0|0|0|0|0|0|0|0|0|0|0|
|Jun22.50v_15780|1|0|0|0|0|0|0|0|0|0|1|0|0|0|0|0|0|0|0|0|0|0|0|0|0|0|0|0|0|0|
|Jul11.5v_12061|2|0|0|0|0|0|0|0|0|0|0|0|0|1|0|0|0|0|0|0|0|0|0|1|0|0|0|0|0|0|

| Feature_ID | 654 | May12_05m | May12_25m | May12_50m | May12_75m | Jun8_25m | Jun8_50m | Jun8_75m | Jun22_0m | Jun22_05m | Jun22_50m | Jun22_75m | Jul11_0m | Jul11_05m | Jul11_25m | Jul11_50m | Jul11_75m | Jul27_05m | Jul27_25m | Jul27_50m | Jul27_75m | Aug5_0m | Aug5_05m | Aug5_25m | Aug5_50m | Aug5_75m | Sep8_0m | Sep8_05m | Sep8_25m | Sep8_50m |
|---|---|---|---|---|---|---|---|---|---|---|---|---|---|---|---|---|---|---|---|---|---|---|---|---|---|---|---|---|---|---|
| Sep8.0v_7899 | 1 | 0 | 0 | 0 | 0 | 0 | 0 | 0 | 0 | 0 | 0 | 0 | 0 | 0 | 0 | 0 | 0 | 0 | 0 | 0 | 0 | 0 | 0 | 0 | 0 | 0 | 1 | 0 | 0 | 0 |
| Jun22.5v_9520 | 1 | 0 | 0 | 0 | 0 | 0 | 0 | 0 | 0 | 1 | 0 | 0 | 0 | 0 | 0 | 0 | 0 | 0 | 0 | 0 | 0 | 0 | 0 | 0 | 0 | 0 | 0 | 0 | 0 | 0 |
| Sep8.50v_6763 | 2 | 0 | 0 | 0 | 0 | 0 | 0 | 0 | 0 | 0 | 0 | 0 | 0 | 0 | 0 | 0 | 0 | 0 | 0 | 0 | 0 | 1 | 0 | 0 | 0 | 0 | 0 | 0 | 0 | 1 |
| Aug5.50v_15857 | 1 | 0 | 0 | 0 | 0 | 0 | 0 | 0 | 0 | 0 | 0 | 0 | 0 | 0 | 0 | 0 | 0 | 0 | 0 | 0 | 0 | 0 | 0 | 0 | 1 | 0 | 0 | 0 | 0 | 0 |
| Jun22.0v_5371 | 1 | 0 | 0 | 0 | 0 | 0 | 0 | 0 | 1 | 0 | 0 | 0 | 0 | 0 | 0 | 0 | 0 | 0 | 0 | 0 | 0 | 0 | 0 | 0 | 0 | 0 | 0 | 0 | 0 | 0 |
| Jul11.50v_14475 | 1 | 0 | 0 | 0 | 0 | 0 | 0 | 0 | 0 | 0 | 0 | 0 | 0 | 0 | 0 | 1 | 0 | 0 | 0 | 0 | 0 | 0 | 0 | 0 | 0 | 0 | 0 | 0 | 0 | 0 |
| Aug5.50v_10921 | 1 | 0 | 0 | 0 | 0 | 0 | 0 | 0 | 0 | 0 | 0 | 0 | 0 | 0 | 0 | 0 | 0 | 0 | 0 | 0 | 0 | 0 | 0 | 0 | 1 | 0 | 0 | 0 | 0 | 0 |
| Jul27.5v_12791 | 1 | 0 | 0 | 0 | 0 | 0 | 0 | 0 | 0 | 0 | 0 | 0 | 0 | 0 | 0 | 0 | 0 | 1 | 0 | 0 | 0 | 0 | 0 | 0 | 0 | 0 | 0 | 0 | 0 | 0 |
| Jul27.5v_417 | 2 | 0 | 0 | 0 | 0 | 0 | 0 | 0 | 0 | 0 | 0 | 0 | 0 | 0 | 0 | 0 | 0 | 1 | 0 | 0 | 0 | 0 | 0 | 0 | 0 | 0 | 0 | 0 | 0 | 1 |
| Sep8.50v_37938 | 1 | 0 | 0 | 0 | 0 | 0 | 0 | 0 | 0 | 0 | 0 | 0 | 0 | 0 | 0 | 0 | 0 | 0 | 0 | 0 | 0 | 0 | 0 | 0 | 0 | 0 | 0 | 0 | 0 | 1 |
| Jul27.50v_34186 | 1 | 0 | 0 | 0 | 0 | 0 | 0 | 0 | 0 | 0 | 0 | 0 | 0 | 0 | 0 | 0 | 0 | 0 | 0 | 1 | 0 | 0 | 0 | 0 | 0 | 0 | 0 | 0 | 0 | 0 |
| Jul27.25v_32416 | 1 | 0 | 0 | 0 | 0 | 0 | 0 | 0 | 0 | 0 | 0 | 0 | 0 | 0 | 0 | 0 | 0 | 0 | 1 | 0 | 0 | 0 | 0 | 0 | 0 | 0 | 0 | 0 | 0 | 0 |
| Jun22.50v_1073 | 1 | 0 | 0 | 0 | 0 | 0 | 0 | 0 | 0 | 0 | 1 | 0 | 0 | 0 | 0 | 0 | 0 | 0 | 0 | 0 | 0 | 0 | 0 | 0 | 0 | 0 | 0 | 0 | 0 | 0 |
| Jun22.75v_22534 | 1 | 0 | 0 | 0 | 0 | 0 | 0 | 0 | 0 | 0 | 0 | 1 | 0 | 0 | 0 | 0 | 0 | 0 | 0 | 0 | 0 | 0 | 0 | 0 | 0 | 0 | 0 | 0 | 0 | 0 |
| Jul27.5v_3869 | 1 | 0 | 0 | 0 | 0 | 0 | 0 | 0 | 0 | 0 | 0 | 0 | 0 | 0 | 0 | 0 | 0 | 1 | 0 | 0 | 0 | 0 | 0 | 0 | 0 | 0 | 0 | 0 | 0 | 0 |
| Aug5.75v_17202 | 1 | 0 | 0 | 0 | 0 | 0 | 0 | 0 | 0 | 0 | 0 | 0 | 0 | 0 | 0 | 0 | 0 | 0 | 0 | 0 | 0 | 0 | 0 | 0 | 0 | 1 | 0 | 0 | 0 | 0 |
| Jul11.25v_8845 | 1 | 0 | 0 | 0 | 0 | 0 | 0 | 0 | 0 | 0 | 0 | 0 | 0 | 0 | 1 | 0 | 0 | 0 | 0 | 0 | 0 | 0 | 0 | 0 | 0 | 0 | 0 | 0 | 0 | 0 |
| Jul11.0v_12290 | 1 | 0 | 0 | 0 | 0 | 0 | 0 | 0 | 0 | 0 | 0 | 0 | 1 | 0 | 0 | 0 | 0 | 0 | 0 | 0 | 0 | 0 | 0 | 0 | 0 | 0 | 0 | 0 | 0 | 0 |
| Jul27.5v_6577 | 1 | 0 | 0 | 0 | 0 | 0 | 0 | 0 | 0 | 0 | 0 | 0 | 0 | 0 | 0 | 0 | 0 | 1 | 0 | 0 | 0 | 0 | 0 | 0 | 0 | 0 | 0 | 0 | 0 | 0 |
| Sep8.0v_18372 | 1 | 0 | 0 | 0 | 0 | 0 | 0 | 0 | 0 | 0 | 0 | 0 | 0 | 0 | 0 | 0 | 0 | 0 | 0 | 0 | 0 | 0 | 0 | 0 | 0 | 0 | 1 | 0 | 0 | 0 |
| Sep8.0v_17221 | 1 | 0 | 0 | 0 | 0 | 0 | 0 | 0 | 0 | 0 | 0 | 0 | 0 | 0 | 0 | 0 | 0 | 0 | 0 | 0 | 0 | 0 | 0 | 0 | 0 | 0 | 1 | 0 | 0 | 0 |
| Jun22.5v_14889 | 1 | 0 | 0 | 0 | 0 | 0 | 0 | 0 | 0 | 1 | 0 | 0 | 0 | 0 | 0 | 0 | 0 | 0 | 0 | 0 | 0 | 0 | 0 | 0 | 0 | 0 | 0 | 0 | 0 | 0 |
| Aug5.0v_3883 | 1 | 0 | 0 | 0 | 0 | 0 | 0 | 0 | 0 | 0 | 0 | 0 | 0 | 0 | 0 | 0 | 0 | 0 | 0 | 0 | 0 | 1 | 0 | 0 | 0 | 0 | 0 | 0 | 0 | 0 |
| Jul27.25v_39770 | 1 | 0 | 0 | 0 | 0 | 0 | 0 | 0 | 0 | 0 | 0 | 0 | 0 | 0 | 0 | 0 | 0 | 0 | 1 | 0 | 0 | 0 | 0 | 0 | 0 | 0 | 0 | 0 | 0 | 0 |
| Aug5.50v_40182 | 1 | 0 | 0 | 0 | 0 | 0 | 0 | 0 | 0 | 0 | 0 | 0 | 0 | 0 | 0 | 0 | 0 | 0 | 0 | 0 | 0 | 0 | 0 | 0 | 1 | 0 | 0 | 0 | 0 | 0 |
| Jul27.5v_20206 | 1 | 0 | 0 | 0 | 0 | 0 | 0 | 0 | 0 | 0 | 0 | 0 | 0 | 0 | 0 | 0 | 0 | 1 | 0 | 0 | 0 | 0 | 0 | 0 | 0 | 0 | 0 | 0 | 0 | 0 |
| Jun22.5v_2907 | 1 | 0 | 0 | 0 | 0 | 0 | 0 | 0 | 0 | 1 | 0 | 0 | 0 | 0 | 0 | 0 | 0 | 0 | 0 | 0 | 0 | 0 | 0 | 0 | 0 | 0 | 0 | 0 | 0 | 0 |
| Jul27.5v_3024 | 1 | 0 | 0 | 0 | 0 | 0 | 0 | 0 | 0 | 0 | 0 | 0 | 0 | 0 | 0 | 0 | 0 | 1 | 0 | 0 | 0 | 0 | 0 | 0 | 0 | 0 | 0 | 0 | 0 | 0 |
| Sep8.50v_37960 | 2 | 0 | 0 | 0 | 0 | 0 | 0 | 0 | 0 | 0 | 0 | 0 | 0 | 0 | 0 | 0 | 0 | 0 | 0 | 0 | 0 | 0 | 0 | 0 | 0 | 0 | 0 | 0 | 0 | 2 |
| Aug5.25v_31270 | 4 | 0 | 0 | 0 | 0 | 0 | 0 | 0 | 0 | 0 | 0 | 0 | 0 | 0 | 0 | 0 | 0 | 0 | 1 | 0 | 0 | 0 | 0 | 2 | 0 | 0 | 0 | 0 | 0 | 1 |
| Jul27.25v_39795 | 1 | 0 | 0 | 0 | 0 | 0 | 0 | 0 | 0 | 0 | 0 | 0 | 0 | 0 | 0 | 0 | 0 | 0 | 1 | 0 | 0 | 0 | 0 | 0 | 0 | 0 | 0 | 0 | 0 | 0 |
| Aug5.5v_14858 | 6 | 0 | 0 | 0 | 0 | 0 | 0 | 0 | 1 | 0 | 0 | 0 | 0 | 0 | 1 | 0 | 0 | 0 | 1 | 0 | 0 | 1 | 1 | 0 | 0 | 0 | 0 | 0 | 0 | 1 |
| Aug5.0v_519 | 1 | 0 | 0 | 0 | 0 | 0 | 0 | 0 | 0 | 0 | 0 | 0 | 0 | 0 | 0 | 0 | 0 | 0 | 0 | 0 | 0 | 1 | 0 | 0 | 0 | 0 | 0 | 0 | 0 | 0 |
| Jul27.25v_6981 | 1 | 0 | 0 | 0 | 0 | 0 | 0 | 0 | 0 | 0 | 0 | 0 | 0 | 0 | 0 | 0 | 0 | 0 | 1 | 0 | 0 | 0 | 0 | 0 | 0 | 0 | 0 | 0 | 0 | 0 |
| Aug5.50v_7705 | 2 | 0 | 0 | 0 | 0 | 0 | 0 | 0 | 0 | 0 | 0 | 0 | 0 | 0 | 1 | 0 | 0 | 0 | 0 | 0 | 0 | 0 | 0 | 0 | 1 | 0 | 0 | 0 | 0 | 0 |
| Jul11.25v_20780 | 1 | 0 | 0 | 0 | 0 | 0 | 0 | 0 | 0 | 0 | 0 | 0 | 0 | 0 | 1 | 0 | 0 | 0 | 0 | 0 | 0 | 0 | 0 | 0 | 0 | 0 | 0 | 0 | 0 | 0 |
| Jul27.75v_898 | 1 | 0 | 0 | 0 | 0 | 0 | 0 | 0 | 0 | 0 | 0 | 0 | 0 | 0 | 0 | 0 | 0 | 0 | 0 | 0 | 1 | 0 | 0 | 0 | 0 | 0 | 0 | 0 | 0 | 0 |
| Aug5.0v_1237 | 1 | 0 | 0 | 0 | 0 | 0 | 0 | 0 | 0 | 0 | 0 | 0 | 0 | 0 | 0 | 0 | 0 | 0 | 0 | 0 | 0 | 1 | 0 | 0 | 0 | 0 | 0 | 0 | 0 | 0 |
| Jun22.5v_4882 | 1 | 0 | 0 | 0 | 0 | 0 | 0 | 0 | 0 | 1 | 0 | 0 | 0 | 0 | 0 | 0 | 0 | 0 | 0 | 0 | 0 | 0 | 0 | 0 | 0 | 0 | 0 | 0 | 0 | 0 |
| Sep8.50v_28128 | 1 | 0 | 0 | 0 | 0 | 0 | 0 | 0 | 0 | 0 | 0 | 0 | 0 | 0 | 0 | 0 | 0 | 0 | 0 | 0 | 0 | 0 | 0 | 0 | 0 | 0 | 0 | 0 | 0 | 1 |
| Jul27.5v_18574 | 1 | 0 | 0 | 0 | 0 | 0 | 0 | 0 | 0 | 0 | 0 | 0 | 0 | 0 | 0 | 0 | 0 | 1 | 0 | 0 | 0 | 0 | 0 | 0 | 0 | 0 | 0 | 0 | 0 | 0 |
| Jul27.25v_26285 | 1 | 0 | 0 | 0 | 0 | 0 | 0 | 0 | 0 | 0 | 0 | 0 | 0 | 0 | 0 | 0 | 0 | 0 | 1 | 0 | 0 | 0 | 0 | 0 | 0 | 0 | 0 | 0 | 0 | 0 |
| Jul11.5v_13541 | 1 | 0 | 0 | 0 | 0 | 0 | 0 | 0 | 0 | 0 | 0 | 0 | 0 | 1 | 0 | 0 | 0 | 0 | 0 | 0 | 0 | 0 | 0 | 0 | 0 | 0 | 0 | 0 | 0 | 0 |
| Jun22.0v_1189 | 1 | 0 | 0 | 0 | 0 | 0 | 0 | 0 | 1 | 0 | 0 | 0 | 0 | 0 | 0 | 0 | 0 | 0 | 0 | 0 | 0 | 0 | 0 | 0 | 0 | 0 | 0 | 0 | 0 | 0 |
| Jul27.5v_6098 | 1 | 0 | 0 | 0 | 0 | 0 | 0 | 0 | 0 | 0 | 0 | 0 | 0 | 0 | 0 | 0 | 0 | 1 | 0 | 0 | 0 | 0 | 0 | 0 | 0 | 0 | 0 | 0 | 0 | 0 |
| Jul11.25v_1628 | 1 | 0 | 0 | 0 | 0 | 0 | 0 | 0 | 0 | 0 | 0 | 0 | 0 | 0 | 1 | 0 | 0 | 0 | 0 | 0 | 0 | 0 | 0 | 0 | 0 | 0 | 0 | 0 | 0 | 0 |
| Jul27.5v_14174 | 1 | 0 | 0 | 0 | 0 | 0 | 0 | 0 | 0 | 0 | 0 | 0 | 0 | 0 | 0 | 0 | 0 | 1 | 0 | 0 | 0 | 0 | 0 | 0 | 0 | 0 | 0 | 0 | 0 | 0 |
| Jul27.5v_13580 | 1 | 0 | 0 | 0 | 0 | 0 | 0 | 0 | 0 | 0 | 0 | 0 | 0 | 0 | 0 | 0 | 0 | 1 | 0 | 0 | 0 | 0 | 0 | 0 | 0 | 0 | 0 | 0 | 0 | 0 |
| Jul11.5v_505 | 1 | 0 | 0 | 0 | 0 | 0 | 0 | 0 | 0 | 0 | 0 | 0 | 0 | 1 | 0 | 0 | 0 | 0 | 0 | 0 | 0 | 0 | 0 | 0 | 0 | 0 | 0 | 0 | 0 | 0 |
| Jul27.25v_21996 | 1 | 0 | 0 | 0 | 0 | 0 | 0 | 0 | 0 | 0 | 0 | 0 | 0 | 0 | 0 | 0 | 0 | 0 | 1 | 0 | 0 | 0 | 0 | 0 | 0 | 0 | 0 | 0 | 0 | 0 |
| Jul27.5v_21101 | 1 | 0 | 0 | 0 | 0 | 0 | 0 | 0 | 0 | 0 | 0 | 0 | 0 | 0 | 0 | 0 | 0 | 1 | 0 | 0 | 0 | 0 | 0 | 0 | 0 | 0 | 0 | 0 | 0 | 0 |
| Aug5.50v_21180 | 1 | 0 | 0 | 0 | 0 | 0 | 0 | 0 | 0 | 0 | 0 | 0 | 0 | 0 | 0 | 0 | 0 | 0 | 0 | 0 | 0 | 0 | 0 | 0 | 1 | 0 | 0 | 0 | 0 | 0 |
| Jun22.50v_41054 | 1 | 0 | 0 | 0 | 0 | 0 | 0 | 0 | 0 | 0 | 1 | 0 | 0 | 0 | 0 | 0 | 0 | 0 | 0 | 0 | 0 | 0 | 0 | 0 | 0 | 0 | 0 | 0 | 0 | 0 |
| Jul11.25v_12389 | 1 | 0 | 0 | 0 | 0 | 0 | 0 | 0 | 0 | 0 | 0 | 0 | 0 | 0 | 1 | 0 | 0 | 0 | 0 | 0 | 0 | 0 | 0 | 0 | 0 | 0 | 0 | 0 | 0 | 0 |
| Jul27.25v_22225 | 1 | 0 | 0 | 0 | 0 | 0 | 0 | 0 | 0 | 0 | 0 | 0 | 0 | 0 | 0 | 0 | 0 | 0 | 1 | 0 | 0 | 0 | 0 | 0 | 0 | 0 | 0 | 0 | 0 | 0 |
| Jul11.50v_8236 | 1 | 0 | 0 | 0 | 0 | 0 | 0 | 0 | 0 | 0 | 0 | 0 | 0 | 0 | 0 | 1 | 0 | 0 | 0 | 0 | 0 | 0 | 0 | 0 | 0 | 0 | 0 | 0 | 0 | 0 |
| Aug5.25v_19259 | 1 | 0 | 0 | 0 | 0 | 0 | 0 | 0 | 0 | 0 | 0 | 0 | 0 | 0 | 0 | 0 | 0 | 0 | 0 | 0 | 0 | 0 | 0 | 1 | 0 | 0 | 0 | 0 | 0 | 0 |
| Aug5.25v_8038 | 1 | 0 | 0 | 0 | 0 | 0 | 0 | 0 | 0 | 0 | 0 | 0 | 0 | 0 | 0 | 0 | 0 | 0 | 0 | 0 | 0 | 0 | 0 | 1 | 0 | 0 | 0 | 0 | 0 | 0 |
| Jul11.50v_9303 | 1 | 0 | 0 | 0 | 0 | 0 | 0 | 0 | 0 | 0 | 0 | 0 | 0 | 0 | 0 | 1 | 0 | 0 | 0 | 0 | 0 | 0 | 0 | 0 | 0 | 0 | 0 | 0 | 0 | 0 |
| Aug5.25v_29883 | 1 | 0 | 0 | 0 | 0 | 0 | 0 | 0 | 0 | 0 | 0 | 0 | 0 | 0 | 0 | 0 | 0 | 0 | 0 | 0 | 0 | 0 | 0 | 1 | 0 | 0 | 0 | 0 | 0 | 0 |
| Aug5.0v_4956 | 1 | 0 | 0 | 0 | 0 | 0 | 0 | 0 | 0 | 0 | 0 | 0 | 0 | 0 | 0 | 0 | 0 | 0 | 0 | 0 | 0 | 1 | 0 | 0 | 0 | 0 | 0 | 0 | 0 | 0 |
| Jul27.5v_8906 | 1 | 0 | 0 | 0 | 0 | 0 | 0 | 0 | 0 | 0 | 0 | 0 | 0 | 0 | 0 | 0 | 0 | 1 | 0 | 0 | 0 | 0 | 0 | 0 | 0 | 0 | 0 | 0 | 0 | 0 |
| Jul27.75v_10776 | 1 | 0 | 0 | 0 | 0 | 0 | 0 | 0 | 0 | 0 | 0 | 0 | 0 | 0 | 0 | 0 | 0 | 0 | 0 | 0 | 1 | 0 | 0 | 0 | 0 | 0 | 0 | 0 | 0 | 0 |
| Jul11.50v_9468 | 1 | 0 | 0 | 0 | 0 | 0 | 0 | 0 | 0 | 0 | 0 | 0 | 0 | 0 | 0 | 1 | 0 | 0 | 0 | 0 | 0 | 0 | 0 | 0 | 0 | 0 | 0 | 0 | 0 | 0 |
| Jul11.25v_5616 | 1 | 0 | 0 | 0 | 0 | 0 | 0 | 0 | 0 | 0 | 0 | 0 | 0 | 0 | 1 | 0 | 0 | 0 | 0 | 0 | 0 | 0 | 0 | 0 | 0 | 0 | 0 | 0 | 0 | 0 |
| Sep8.50v_40086 | 1 | 0 | 0 | 0 | 0 | 0 | 0 | 0 | 0 | 0 | 0 | 0 | 0 | 0 | 0 | 0 | 0 | 0 | 0 | 0 | 0 | 0 | 0 | 0 | 0 | 0 | 0 | 0 | 0 | 1 |
| Jul27.25v_32554 | 1 | 0 | 0 | 0 | 0 | 0 | 0 | 0 | 0 | 0 | 0 | 0 | 0 | 0 | 0 | 0 | 0 | 0 | 1 | 0 | 0 | 0 | 0 | 0 | 0 | 0 | 0 | 0 | 0 | 0 |
| Jul11.25v_17097 | 1 | 0 | 0 | 0 | 0 | 0 | 0 | 0 | 0 | 0 | 0 | 0 | 0 | 0 | 1 | 0 | 0 | 0 | 0 | 0 | 0 | 0 | 0 | 0 | 0 | 0 | 0 | 0 | 0 | 0 |
| Aug5.25v_18561 | 1 | 0 | 0 | 0 | 0 | 0 | 0 | 0 | 0 | 0 | 0 | 0 | 0 | 0 | 0 | 0 | 0 | 0 | 0 | 0 | 0 | 0 | 0 | 1 | 0 | 0 | 0 | 0 | 0 | 0 |
| Jul27.50v_28599 | 1 | 0 | 0 | 0 | 0 | 0 | 0 | 0 | 0 | 0 | 0 | 0 | 0 | 0 | 0 | 0 | 0 | 0 | 0 | 1 | 0 | 0 | 0 | 0 | 0 | 0 | 0 | 0 | 0 | 0 |
| May12.50v_32397 | 1 | 0 | 0 | 1 | 0 | 0 | 0 | 0 | 0 | 0 | 0 | 0 | 0 | 0 | 0 | 0 | 0 | 0 | 0 | 0 | 0 | 0 | 0 | 0 | 0 | 0 | 0 | 0 | 0 | 0 |
| Jul11.0v_14239 | 1 | 0 | 0 | 0 | 0 | 0 | 0 | 0 | 0 | 0 | 0 | 0 | 1 | 0 | 0 | 0 | 0 | 0 | 0 | 0 | 0 | 0 | 0 | 0 | 0 | 0 | 0 | 0 | 0 | 0 |
| Sep8.25v_4207 | 1 | 0 | 0 | 0 | 0 | 0 | 0 | 0 | 0 | 0 | 0 | 0 | 0 | 0 | 0 | 0 | 0 | 0 | 0 | 0 | 0 | 0 | 0 | 0 | 0 | 0 | 0 | 0 | 1 | 0 |
| Sep8.0v_6998 | 1 | 0 | 0 | 0 | 0 | 0 | 0 | 0 | 0 | 0 | 0 | 0 | 0 | 0 | 0 | 0 | 0 | 0 | 0 | 0 | 0 | 0 | 0 | 0 | 0 | 0 | 1 | 0 | 0 | 0 |
| Jul11.25v_12958 | 1 | 0 | 0 | 0 | 0 | 0 | 0 | 0 | 0 | 0 | 0 | 0 | 0 | 0 | 1 | 0 | 0 | 0 | 0 | 0 | 0 | 0 | 0 | 0 | 0 | 0 | 0 | 0 | 0 | 0 |
| Jul27.5v_12864 | 1 | 0 | 0 | 0 | 0 | 0 | 0 | 0 | 0 | 0 | 0 | 0 | 0 | 0 | 0 | 0 | 0 | 1 | 0 | 0 | 0 | 0 | 0 | 0 | 0 | 0 | 0 | 0 | 0 | 0 |
| Jul27.50v_33834 | 1 | 0 | 0 | 0 | 0 | 0 | 0 | 0 | 0 | 0 | 0 | 0 | 0 | 0 | 0 | 0 | 0 | 0 | 0 | 1 | 0 | 0 | 0 | 0 | 0 | 0 | 0 | 0 | 0 | 0 |
| Aug5.50v_11588 | 2 | 0 | 0 | 0 | 0 | 0 | 0 | 0 | 0 | 1 | 0 | 0 | 0 | 0 | 0 | 0 | 0 | 0 | 0 | 0 | 0 | 0 | 0 | 0 | 1 | 0 | 0 | 0 | 0 | 0 |
| Jul27.50v_35593 | 1 | 0 | 0 | 0 | 0 | 0 | 0 | 0 | 0 | 0 | 0 | 0 | 0 | 0 | 0 | 0 | 0 | 0 | 0 | 1 | 0 | 0 | 0 | 0 | 0 | 0 | 0 | 0 | 0 | 0 |
| Jun22.0v_20897 | 1 | 0 | 0 | 0 | 0 | 0 | 0 | 0 | 1 | 0 | 0 | 0 | 0 | 0 | 0 | 0 | 0 | 0 | 0 | 0 | 0 | 0 | 0 | 0 | 0 | 0 | 0 | 0 | 0 | 0 |
| Jul11.25v_17868 | 1 | 0 | 0 | 0 | 0 | 0 | 0 | 0 | 0 | 0 | 0 | 0 | 0 | 0 | 1 | 0 | 0 | 0 | 0 | 0 | 0 | 0 | 0 | 0 | 0 | 0 | 0 | 0 | 0 | 0 |
| Jul27.5v_8443 | 1 | 0 | 0 | 0 | 0 | 0 | 0 | 0 | 0 | 0 | 0 | 0 | 0 | 0 | 0 | 0 | 0 | 1 | 0 | 0 | 0 | 0 | 0 | 0 | 0 | 0 | 0 | 0 | 0 | 0 |
| Jul11.25v_14388 | 1 | 0 | 0 | 0 | 0 | 0 | 0 | 0 | 0 | 0 | 0 | 0 | 0 | 0 | 1 | 0 | 0 | 0 | 0 | 0 | 0 | 0 | 0 | 0 | 0 | 0 | 0 | 0 | 0 | 0 |
| Jul27.5v_14695 | 1 | 0 | 0 | 0 | 0 | 0 | 0 | 0 | 0 | 0 | 0 | 0 | 0 | 0 | 0 | 0 | 0 | 1 | 0 | 0 | 0 | 0 | 0 | 0 | 0 | 0 | 0 | 0 | 0 | 0 |
| Aug5.0v_2912 | 1 | 0 | 0 | 0 | 0 | 0 | 0 | 0 | 0 | 0 | 0 | 0 | 0 | 0 | 0 | 0 | 0 | 0 | 0 | 0 | 0 | 1 | 0 | 0 | 0 | 0 | 0 | 0 | 0 | 0 |
| Jun22.0v_16908 | 1 | 0 | 0 | 0 | 0 | 0 | 0 | 0 | 1 | 0 | 0 | 0 | 0 | 0 | 0 | 0 | 0 | 0 | 0 | 0 | 0 | 0 | 0 | 0 | 0 | 0 | 0 | 0 | 0 | 0 |
| Aug5.0v_2789 | 1 | 0 | 0 | 0 | 0 | 0 | 0 | 0 | 0 | 0 | 0 | 0 | 0 | 0 | 0 | 0 | 0 | 0 | 0 | 0 | 0 | 1 | 0 | 0 | 0 | 0 | 0 | 0 | 0 | 0 |
| Jul27.25v_21411 | 1 | 0 | 0 | 0 | 0 | 0 | 0 | 0 | 0 | 0 | 0 | 0 | 0 | 0 | 0 | 0 | 0 | 0 | 1 | 0 | 0 | 0 | 0 | 0 | 0 | 0 | 0 | 0 | 0 | 0 |
| Aug5.5v_3223 | 1 | 0 | 0 | 0 | 0 | 0 | 0 | 0 | 0 | 0 | 0 | 0 | 0 | 0 | 0 | 0 | 0 | 0 | 0 | 0 | 0 | 0 | 1 | 0 | 0 | 0 | 0 | 0 | 0 | 0 |
| Aug5.0v_845 | 1 | 0 | 0 | 0 | 0 | 0 | 0 | 0 | 0 | 0 | 0 | 0 | 0 | 0 | 0 | 0 | 0 | 0 | 0 | 0 | 0 | 1 | 0 | 0 | 0 | 0 | 0 | 0 | 0 | 0 |
| Jul27.5v_20227 | 1 | 0 | 0 | 0 | 0 | 0 | 0 | 0 | 0 | 0 | 0 | 0 | 0 | 0 | 0 | 0 | 0 | 1 | 0 | 0 | 0 | 0 | 0 | 0 | 0 | 0 | 0 | 0 | 0 | 0 |
| Aug5.5v_21429 | 1 | 0 | 0 | 0 | 0 | 0 | 0 | 0 | 0 | 0 | 0 | 0 | 0 | 0 | 0 | 0 | 0 | 0 | 0 | 0 | 0 | 0 | 1 | 0 | 0 | 0 | 0 | 0 | 0 | 0 |
| Jul27.50v_33776 | 1 | 0 | 0 | 0 | 0 | 0 | 0 | 0 | 0 | 0 | 0 | 0 | 0 | 0 | 0 | 0 | 0 | 0 | 0 | 1 | 0 | 0 | 0 | 0 | 0 | 0 | 0 | 0 | 0 | 0 |
| Jul11.25v_7137 | 3 | 0 | 0 | 0 | 0 | 0 | 0 | 0 | 0 | 0 | 1 | 0 | 0 | 0 | 1 | 0 | 0 | 1 | 0 | 0 | 0 | 0 | 0 | 0 | 0 | 0 | 0 | 0 | 0 | 0 |
| Jul27.5v_4197 | 1 | 0 | 0 | 0 | 0 | 0 | 0 | 0 | 0 | 0 | 0 | 0 | 0 | 0 | 0 | 0 | 0 | 1 | 0 | 0 | 0 | 0 | 0 | 0 | 0 | 0 | 0 | 0 | 0 | 0 |
| Jul27.25v_22199 | 1 | 0 | 0 | 0 | 0 | 0 | 0 | 0 | 0 | 0 | 0 | 0 | 0 | 0 | 0 | 0 | 0 | 0 | 1 | 0 | 0 | 0 | 0 | 0 | 0 | 0 | 0 | 0 | 0 | 0 |
| Jul11.25v_17506 | 1 | 0 | 0 | 0 | 0 | 0 | 0 | 0 | 0 | 0 | 0 | 0 | 0 | 0 | 1 | 0 | 0 | 0 | 0 | 0 | 0 | 0 | 0 | 0 | 0 | 0 | 0 | 0 | 0 | 0 |
| Jul27.5v_8536 | 1 | 0 | 0 | 0 | 0 | 0 | 0 | 0 | 0 | 0 | 0 | 0 | 0 | 0 | 0 | 0 | 0 | 1 | 0 | 0 | 0 | 0 | 0 | 0 | 0 | 0 | 0 | 0 | 0 | 0 |
| Sep8.0v_5444 | 1 | 0 | 0 | 0 | 0 | 0 | 0 | 0 | 0 | 0 | 0 | 0 | 0 | 0 | 0 | 0 | 0 | 0 | 0 | 0 | 0 | 0 | 0 | 0 | 0 | 0 | 1 | 0 | 0 | 0 |
| Jul27.50v_31914 | 1 | 0 | 0 | 0 | 0 | 0 | 0 | 0 | 0 | 0 | 0 | 0 | 0 | 0 | 0 | 0 | 0 | 0 | 0 | 1 | 0 | 0 | 0 | 0 | 0 | 0 | 0 | 0 | 0 | 0 |
| Jul11.25v_11011 | 1 | 0 | 0 | 0 | 0 | 0 | 0 | 0 | 0 | 0 | 0 | 0 | 0 | 0 | 1 | 0 | 0 | 0 | 0 | 0 | 0 | 0 | 0 | 0 | 0 | 0 | 0 | 0 | 0 | 0 |
| Jun22.5v_20249 | 1 | 0 | 0 | 0 | 0 | 0 | 0 | 0 | 0 | 1 | 0 | 0 | 0 | 0 | 0 | 0 | 0 | 0 | 0 | 0 | 0 | 0 | 0 | 0 | 0 | 0 | 0 | 0 | 0 | 0 |
| Jul27.50v_15416 | 1 | 0 | 0 | 0 | 0 | 0 | 0 | 0 | 0 | 0 | 0 | 0 | 0 | 0 | 0 | 0 | 0 | 0 | 0 | 1 | 0 | 0 | 0 | 0 | 0 | 0 | 0 | 0 | 0 | 0 |
| Jul27.75v_4388 | 1 | 0 | 0 | 0 | 0 | 0 | 0 | 0 | 0 | 0 | 0 | 0 | 0 | 0 | 0 | 0 | 0 | 0 | 0 | 0 | 1 | 0 | 0 | 0 | 0 | 0 | 0 | 0 | 0 | 0 |
| Aug5.0v_348 | 1 | 0 | 0 | 0 | 0 | 0 | 0 | 0 | 0 | 0 | 0 | 0 | 0 | 0 | 0 | 0 | 0 | 0 | 0 | 0 | 0 | 1 | 0 | 0 | 0 | 0 | 0 | 0 | 0 | 0 |
| Jun22.5v_2753 | 1 | 0 | 0 | 0 | 0 | 0 | 0 | 0 | 0 | 1 | 0 | 0 | 0 | 0 | 0 | 0 | 0 | 0 | 0 | 0 | 0 | 0 | 0 | 0 | 0 | 0 | 0 | 0 | 0 | 0 |
| Jun22.75v_36860 | 1 | 0 | 0 | 0 | 0 | 0 | 0 | 0 | 0 | 0 | 0 | 1 | 0 | 0 | 0 | 0 | 0 | 0 | 0 | 0 | 0 | 0 | 0 | 0 | 0 | 0 | 0 | 0 | 0 | 0 |
| Sep8.25v_18733 | 1 | 0 | 0 | 0 | 0 | 0 | 0 | 0 | 0 | 0 | 0 | 0 | 0 | 0 | 0 | 0 | 0 | 0 | 0 | 0 | 0 | 0 | 0 | 0 | 0 | 0 | 0 | 0 | 1 | 0 |
| May12.50v_6736 | 1 | 0 | 0 | 1 | 0 | 0 | 0 | 0 | 0 | 0 | 0 | 0 | 0 | 0 | 0 | 0 | 0 | 0 | 0 | 0 | 0 | 0 | 0 | 0 | 0 | 0 | 0 | 0 | 0 | 0 |
| Jun22.0v_18752 | 1 | 0 | 0 | 0 | 0 | 0 | 0 | 0 | 1 | 0 | 0 | 0 | 0 | 0 | 0 | 0 | 0 | 0 | 0 | 0 | 0 | 0 | 0 | 0 | 0 | 0 | 0 | 0 | 0 | 0 |
| Sep8.0v_3893 | 1 | 0 | 0 | 0 | 0 | 0 | 0 | 0 | 0 | 0 | 0 | 0 | 0 | 0 | 0 | 0 | 0 | 0 | 0 | 0 | 0 | 0 | 0 | 0 | 0 | 0 | 1 | 0 | 0 | 0 |
| Aug5.25v_29513 | 1 | 0 | 0 | 0 | 0 | 0 | 0 | 0 | 0 | 0 | 0 | 0 | 0 | 0 | 0 | 0 | 0 | 0 | 0 | 0 | 0 | 0 | 0 | 1 | 0 | 0 | 0 | 0 | 0 | 0 |
| Jul11.5v_9333 | 242 | 0 | 0 | 0 | 0 | 0 | 0 | 0 | 0 | 10 | 13 | 0 | 3 | 31 | 29 | 1 | 0 | 34 | 18 | 19 | 2 | 20 | 26 | 5 | 13 | 4 | 4 | 0 | 1 | 9 |
| Jul27.25v_39146 | 5 | 0 | 0 | 0 | 0 | 0 | 0 | 0 | 0 | 1 | 0 | 0 | 0 | 0 | 0 | 1 | 0 | 1 | 1 | 0 | 0 | 1 | 0 | 0 | 0 | 0 | 0 | 0 | 0 | 0 |
| Aug5.50v_7694 | 1 | 0 | 0 | 0 | 0 | 0 | 0 | 0 | 0 | 0 | 0 | 0 | 0 | 0 | 0 | 0 | 0 | 0 | 0 | 0 | 0 | 0 | 0 | 0 | 1 | 0 | 0 | 0 | 0 | 0 |
| Jul27.5v_9148 | 14 | 0 | 0 | 0 | 0 | 0 | 0 | 0 | 1 | 0 | 1 | 0 | 0 | 1 | 4 | 0 | 0 | 4 | 0 | 1 | 1 | 0 | 0 | 0 | 0 | 0 | 0 | 0 | 0 | 1 |
| Jul27.5v_13178 | 2 | 0 | 0 | 0 | 0 | 0 | 0 | 0 | 0 | 0 | 0 | 0 | 0 | 0 | 0 | 0 | 0 | 1 | 0 | 0 | 0 | 0 | 0 | 1 | 0 | 0 | 0 | 0 | 0 | 0 |
| Sep8.0v_16990 | 2 | 0 | 0 | 0 | 0 | 0 | 0 | 0 | 0 | 0 | 0 | 0 | 0 | 0 | 0 | 0 | 0 | 0 | 1 | 0 | 0 | 0 | 0 | 0 | 0 | 0 | 1 | 0 | 0 | 0 |
| Jul11.5v_15366 | 1 | 0 | 0 | 0 | 0 | 0 | 0 | 0 | 0 | 0 | 0 | 0 | 0 | 1 | 0 | 0 | 0 | 0 | 0 | 0 | 0 | 0 | 0 | 0 | 0 | 0 | 0 | 0 | 0 | 0 |
| Aug5.25v_9357 | 1 | 0 | 0 | 0 | 0 | 0 | 0 | 0 | 0 | 0 | 0 | 0 | 0 | 0 | 0 | 0 | 0 | 0 | 0 | 0 | 0 | 0 | 0 | 1 | 0 | 0 | 0 | 0 | 0 | 0 |
| Jul11.25v_20446 | 1 | 0 | 0 | 0 | 0 | 0 | 0 | 0 | 0 | 0 | 0 | 0 | 0 | 0 | 1 | 0 | 0 | 0 | 0 | 0 | 0 | 0 | 0 | 0 | 0 | 0 | 0 | 0 | 0 | 0 |
| Jul27.5v_8478 | 1 | 0 | 0 | 0 | 0 | 0 | 0 | 0 | 0 | 0 | 0 | 0 | 0 | 0 | 0 | 0 | 0 | 1 | 0 | 0 | 0 | 0 | 0 | 0 | 0 | 0 | 0 | 0 | 0 | 0 |
| Aug5.75v_33481 | 3 | 0 | 0 | 0 | 0 | 0 | 0 | 0 | 0 | 0 | 0 | 0 | 0 | 0 | 0 | 0 | 0 | 0 | 0 | 0 | 0 | 0 | 0 | 1 | 1 | 1 | 0 | 0 | 0 | 0 |
| Jul11.25v_4514 | 1 | 0 | 0 | 0 | 0 | 0 | 0 | 0 | 0 | 0 | 0 | 0 | 0 | 0 | 1 | 0 | 0 | 0 | 0 | 0 | 0 | 0 | 0 | 0 | 0 | 0 | 0 | 0 | 0 | 0 |
| Jul27.5v_16802 | 3 | 0 | 0 | 0 | 0 | 0 | 0 | 0 | 0 | 0 | 0 | 0 | 0 | 0 | 0 | 0 | 0 | 1 | 0 | 0 | 1 | 0 | 0 | 0 | 1 | 0 | 0 | 0 | 0 | 0 |
| Aug5.5v_11011 | 1 | 0 | 0 | 0 | 0 | 0 | 0 | 0 | 0 | 0 | 0 | 0 | 0 | 0 | 0 | 0 | 0 | 0 | 0 | 0 | 0 | 0 | 1 | 0 | 0 | 0 | 0 | 0 | 0 | 0 |
| Jul27.5v_8937 | 1 | 0 | 0 | 0 | 0 | 0 | 0 | 0 | 0 | 0 | 0 | 0 | 0 | 0 | 0 | 0 | 0 | 1 | 0 | 0 | 0 | 0 | 0 | 0 | 0 | 0 | 0 | 0 | 0 | 0 |
| Jun22.50v_20864 | 1 | 0 | 0 | 0 | 0 | 0 | 0 | 0 | 0 | 0 | 1 | 0 | 0 | 0 | 0 | 0 | 0 | 0 | 0 | 0 | 0 | 0 | 0 | 0 | 0 | 0 | 0 | 0 | 0 | 0 |
| Aug5.5v_20692 | 1 | 0 | 0 | 0 | 0 | 0 | 0 | 0 | 0 | 0 | 0 | 0 | 0 | 0 | 0 | 0 | 0 | 0 | 0 | 0 | 0 | 0 | 1 | 0 | 0 | 0 | 0 | 0 | 0 | 0 |
| Jul11.25v_20696 | 39 | 0 | 0 | 0 | 0 | 0 | 0 | 0 | 0 | 0 | 2 | 0 | 2 | 1 | 7 | 0 | 0 | 2 | 5 | 3 | 2 | 3 | 5 | 1 | 1 | 2 | 1 | 0 | 0 | 2 |
| Jun22.5v_6252 | 1 | 0 | 0 | 0 | 0 | 0 | 0 | 0 | 0 | 1 | 0 | 0 | 0 | 0 | 0 | 0 | 0 | 0 | 0 | 0 | 0 | 0 | 0 | 0 | 0 | 0 | 0 | 0 | 0 | 0 |
| Jul27.5v_364 | 1 | 0 | 0 | 0 | 0 | 0 | 0 | 0 | 0 | 0 | 0 | 0 | 0 | 0 | 0 | 0 | 0 | 1 | 0 | 0 | 0 | 0 | 0 | 0 | 0 | 0 | 0 | 0 | 0 | 0 |
| Jul11.5v_12724 | 1 | 0 | 0 | 0 | 0 | 0 | 0 | 0 | 0 | 0 | 0 | 0 | 0 | 1 | 0 | 0 | 0 | 0 | 0 | 0 | 0 | 0 | 0 | 0 | 0 | 0 | 0 | 0 | 0 | 0 |
| Jul11.5v_3453 | 20 | 1 | 1 | 0 | 2 | 0 | 1 | 0 | 0 | 0 | 1 | 0 | 0 | 2 | 1 | 3 | 2 | 0 | 1 | 0 | 1 | 0 | 0 | 1 | 0 | 1 | 0 | 2 | 0 | 0 |
| Jul11.0v_1381 | 27 | 1 | 3 | 0 | 2 | 0 | 1 | 3 | 0 | 4 | 2 | 0 | 1 | 1 | 2 | 0 | 0 | 1 | 0 | 0 | 0 | 0 | 0 | 0 | 0 | 1 | 1 | 3 | 0 | 1 |
| Jun22.50v_11549 | 1 | 0 | 0 | 0 | 0 | 0 | 0 | 0 | 0 | 0 | 1 | 0 | 0 | 0 | 0 | 0 | 0 | 0 | 0 | 0 | 0 | 0 | 0 | 0 | 0 | 0 | 0 | 0 | 0 | 0 |
| Sep8.5v_15643 | 1 | 0 | 0 | 0 | 0 | 0 | 0 | 0 | 0 | 0 | 0 | 0 | 0 | 0 | 0 | 0 | 0 | 0 | 0 | 0 | 0 | 0 | 0 | 0 | 0 | 0 | 0 | 1 | 0 | 0 |
| Jul11.5v_20935 | 1 | 0 | 0 | 0 | 0 | 0 | 0 | 0 | 0 | 0 | 0 | 0 | 0 | 1 | 0 | 0 | 0 | 0 | 0 | 0 | 0 | 0 | 0 | 0 | 0 | 0 | 0 | 0 | 0 | 0 |
| Jun22.50v_38736 | 1 | 0 | 0 | 0 | 0 | 0 | 0 | 0 | 0 | 0 | 1 | 0 | 0 | 0 | 0 | 0 | 0 | 0 | 0 | 0 | 0 | 0 | 0 | 0 | 0 | 0 | 0 | 0 | 0 | 0 |
| Jun22.0v_10946 | 9 | 0 | 1 | 0 | 0 | 2 | 0 | 0 | 2 | 0 | 0 | 0 | 0 | 0 | 1 | 1 | 0 | 0 | 0 | 0 | 0 | 0 | 0 | 0 | 1 | 1 | 0 | 0 | 0 | 0 |
| Jul11.25v_14000 | 12 | 0 | 0 | 0 | 0 | 1 | 1 | 0 | 0 | 0 | 0 | 0 | 0 | 1 | 2 | 0 | 3 | 0 | 2 | 0 | 0 | 0 | 0 | 1 | 0 | 0 | 0 | 0 | 1 | 0 |
| Aug5.75v_13845 | 1 | 0 | 0 | 0 | 0 | 0 | 0 | 0 | 0 | 0 | 0 | 0 | 0 | 0 | 0 | 0 | 0 | 0 | 0 | 0 | 0 | 0 | 0 | 0 | 0 | 1 | 0 | 0 | 0 | 0 |
| Jun22.0v_5034 | 12 | 1 | 1 | 0 | 0 | 0 | 0 | 0 | 1 | 0 | 1 | 0 | 0 | 0 | 0 | 0 | 1 | 1 | 0 | 0 | 1 | 1 | 0 | 1 | 0 | 0 | 0 | 1 | 1 | 1 |
| Aug5.5v_36607 | 1 | 0 | 0 | 0 | 0 | 0 | 0 | 0 | 0 | 0 | 0 | 0 | 0 | 0 | 0 | 0 | 0 | 0 | 0 | 0 | 0 | 0 | 1 | 0 | 0 | 0 | 0 | 0 | 0 | 0 |
| Aug5.5v_9607 | 1 | 0 | 0 | 0 | 0 | 0 | 0 | 0 | 0 | 0 | 0 | 0 | 0 | 0 | 0 | 0 | 0 | 0 | 0 | 0 | 0 | 0 | 1 | 0 | 0 | 0 | 0 | 0 | 0 | 0 |
| Jun22.0v_8658 | 1 | 0 | 0 | 0 | 0 | 0 | 0 | 0 | 1 | 0 | 0 | 0 | 0 | 0 | 0 | 0 | 0 | 0 | 0 | 0 | 0 | 0 | 0 | 0 | 0 | 0 | 0 | 0 | 0 | 0 |
| Aug5.75v_17768 | 1 | 0 | 0 | 0 | 0 | 0 | 0 | 0 | 0 | 0 | 0 | 0 | 0 | 0 | 0 | 0 | 0 | 0 | 0 | 0 | 0 | 0 | 0 | 0 | 0 | 1 | 0 | 0 | 0 | 0 |
| May12.5v_33502 | 39 | 7 | 2 | 5 | 5 | 1 | 1 | 1 | 0 | 1 | 3 | 2 | 0 | 2 | 0 | 1 | 2 | 0 | 0 | 2 | 0 | 0 | 0 | 2 | 0 | 0 | 1 | 0 | 0 | 1 |
| Aug5.5v_34126 | 1 | 0 | 0 | 0 | 0 | 0 | 0 | 0 | 0 | 0 | 0 | 0 | 0 | 0 | 0 | 0 | 0 | 0 | 0 | 0 | 0 | 0 | 1 | 0 | 0 | 0 | 0 | 0 | 0 | 0 |
| Jun22.5v_6872 | 3 | 0 | 0 | 0 | 1 | 0 | 0 | 0 | 0 | 2 | 0 | 0 | 0 | 0 | 0 | 0 | 0 | 0 | 0 | 0 | 0 | 0 | 0 | 0 | 0 | 0 | 0 | 0 | 0 | 0 |
| Jul11.50v_12164 | 7 | 0 | 0 | 0 | 0 | 0 | 0 | 0 | 0 | 0 | 0 | 0 | 1 | 0 | 0 | 1 | 0 | 0 | 1 | 0 | 0 | 1 | 0 | 1 | 0 | 2 | 0 | 0 | 0 | 0 |
| May12.25v_6827 | 24 | 1 | 2 | 3 | 1 | 0 | 0 | 0 | 0 | 0 | 1 | 1 | 0 | 0 | 2 | 3 | 1 | 0 | 0 | 0 | 1 | 1 | 2 | 1 | 0 | 1 | 2 | 0 | 0 | 1 |
| Jul11.75v_5099 | 1 | 0 | 0 | 0 | 0 | 0 | 0 | 0 | 0 | 0 | 0 | 0 | 0 | 0 | 0 | 0 | 1 | 0 | 0 | 0 | 0 | 0 | 0 | 0 | 0 | 0 | 0 | 0 | 0 | 0 |
| Jun8.50v_30605 | 8 | 0 | 0 | 0 | 0 | 1 | 1 | 0 | 1 | 0 | 0 | 0 | 1 | 0 | 1 | 0 | 0 | 0 | 0 | 1 | 0 | 0 | 0 | 1 | 0 | 0 | 0 | 1 | 0 | 0 |
| Jul27.50v_28876 | 1 | 0 | 0 | 0 | 0 | 0 | 0 | 0 | 0 | 0 | 0 | 0 | 0 | 0 | 0 | 0 | 0 | 0 | 0 | 1 | 0 | 0 | 0 | 0 | 0 | 0 | 0 | 0 | 0 | 0 |
| Sep8.50v_15929 | 1 | 0 | 0 | 0 | 0 | 0 | 0 | 0 | 0 | 0 | 0 | 0 | 0 | 0 | 0 | 0 | 0 | 0 | 0 | 0 | 0 | 0 | 0 | 0 | 0 | 0 | 0 | 0 | 0 | 1 |
| Jul11.5v_6807 | 2 | 0 | 0 | 0 | 0 | 0 | 0 | 0 | 0 | 0 | 0 | 0 | 0 | 1 | 0 | 0 | 0 | 0 | 0 | 0 | 0 | 0 | 0 | 1 | 0 | 0 | 0 | 0 | 0 | 0 |
| May12.50v_4477 | 1 | 0 | 0 | 1 | 0 | 0 | 0 | 0 | 0 | 0 | 0 | 0 | 0 | 0 | 0 | 0 | 0 | 0 | 0 | 0 | 0 | 0 | 0 | 0 | 0 | 0 | 0 | 0 | 0 | 0 |
| Jul11.0v_3574 | 1 | 0 | 0 | 0 | 0 | 0 | 0 | 0 | 0 | 0 | 0 | 0 | 1 | 0 | 0 | 0 | 0 | 0 | 0 | 0 | 0 | 0 | 0 | 0 | 0 | 0 | 0 | 0 | 0 | 0 |
| Jul27.25v_34443 | 1 | 0 | 0 | 0 | 0 | 0 | 0 | 0 | 0 | 0 | 0 | 0 | 0 | 0 | 0 | 0 | 0 | 0 | 1 | 0 | 0 | 0 | 0 | 0 | 0 | 0 | 0 | 0 | 0 | 0 |
| Sep8.50v_13262 | 1 | 0 | 0 | 0 | 0 | 0 | 0 | 0 | 0 | 0 | 0 | 0 | 0 | 0 | 0 | 0 | 0 | 0 | 0 | 0 | 0 | 0 | 0 | 0 | 0 | 0 | 0 | 0 | 0 | 1 |
| Aug5.0v_2268 | 1 | 0 | 0 | 0 | 0 | 0 | 0 | 0 | 0 | 0 | 0 | 0 | 0 | 0 | 0 | 0 | 0 | 0 | 0 | 0 | 0 | 1 | 0 | 0 | 0 | 0 | 0 | 0 | 0 | 0 |
| Jul27.5v_4921 | 1 | 0 | 0 | 0 | 0 | 0 | 0 | 0 | 0 | 0 | 0 | 0 | 0 | 0 | 0 | 0 | 0 | 1 | 0 | 0 | 0 | 0 | 0 | 0 | 0 | 0 | 0 | 0 | 0 | 0 |
| Aug5.5v_36555 | 1 | 0 | 0 | 0 | 0 | 0 | 0 | 0 | 0 | 0 | 0 | 0 | 0 | 0 | 0 | 0 | 0 | 0 | 0 | 0 | 0 | 0 | 1 | 0 | 0 | 0 | 0 | 0 | 0 | 0 |
| Sep8.50v_24010 | 1 | 0 | 0 | 0 | 0 | 0 | 0 | 0 | 0 | 0 | 0 | 0 | 0 | 0 | 0 | 0 | 0 | 0 | 0 | 0 | 0 | 0 | 0 | 0 | 0 | 0 | 0 | 0 | 0 | 1 |
| Jul11.5v_15655 | 1 | 0 | 0 | 0 | 0 | 0 | 0 | 0 | 0 | 0 | 0 | 0 | 0 | 1 | 0 | 0 | 0 | 0 | 0 | 0 | 0 | 0 | 0 | 0 | 0 | 0 | 0 | 0 | 0 | 0 |
| Aug5.25v_38987 | 1 | 0 | 0 | 0 | 0 | 0 | 0 | 0 | 0 | 0 | 0 | 0 | 0 | 0 | 0 | 0 | 0 | 0 | 0 | 0 | 0 | 0 | 0 | 1 | 0 | 0 | 0 | 0 | 0 | 0 |
| Aug5.25v_24775 | 1 | 0 | 0 | 0 | 0 | 0 | 0 | 0 | 0 | 0 | 0 | 0 | 0 | 0 | 0 | 0 | 0 | 0 | 0 | 0 | 0 | 0 | 0 | 1 | 0 | 0 | 0 | 0 | 0 | 0 |
| Jul27.50v_27383 | 1 | 0 | 0 | 0 | 0 | 0 | 0 | 0 | 0 | 0 | 0 | 0 | 0 | 0 | 0 | 0 | 0 | 0 | 0 | 1 | 0 | 0 | 0 | 0 | 0 | 0 | 0 | 0 | 0 | 0 |
| Aug5.50v_32846 | 1 | 0 | 0 | 0 | 0 | 0 | 0 | 0 | 0 | 0 | 0 | 0 | 0 | 0 | 0 | 0 | 0 | 0 | 0 | 0 | 0 | 0 | 0 | 0 | 1 | 0 | 0 | 0 | 0 | 0 |
| Jul27.5v_10453 | 1 | 0 | 0 | 0 | 0 | 0 | 0 | 0 | 0 | 0 | 0 | 0 | 0 | 0 | 0 | 0 | 0 | 1 | 0 | 0 | 0 | 0 | 0 | 0 | 0 | 0 | 0 | 0 | 0 | 0 |
| Jul27.5v_3540 | 1 | 0 | 0 | 0 | 0 | 0 | 0 | 0 | 0 | 0 | 0 | 0 | 0 | 0 | 0 | 0 | 0 | 1 | 0 | 0 | 0 | 0 | 0 | 0 | 0 | 0 | 0 | 0 | 0 | 0 |
| Aug5.5v_22887 | 1 | 0 | 0 | 0 | 0 | 0 | 0 | 0 | 0 | 0 | 0 | 0 | 0 | 0 | 0 | 0 | 0 | 0 | 0 | 0 | 0 | 0 | 1 | 0 | 0 | 0 | 0 | 0 | 0 | 0 |
| Aug5.0v_2052 | 1 | 0 | 0 | 0 | 0 | 0 | 0 | 0 | 0 | 0 | 0 | 0 | 0 | 0 | 0 | 0 | 0 | 0 | 0 | 0 | 0 | 1 | 0 | 0 | 0 | 0 | 0 | 0 | 0 | 0 |
| Jul27.5v_1182 | 1 | 0 | 0 | 0 | 0 | 0 | 0 | 0 | 0 | 0 | 0 | 0 | 0 | 0 | 0 | 0 | 0 | 1 | 0 | 0 | 0 | 0 | 0 | 0 | 0 | 0 | 0 | 0 | 0 | 0 |
| Jul11.5v_3073 | 1 | 0 | 0 | 0 | 0 | 0 | 0 | 0 | 0 | 0 | 0 | 0 | 0 | 1 | 0 | 0 | 0 | 0 | 0 | 0 | 0 | 0 | 0 | 0 | 0 | 0 | 0 | 0 | 0 | 0 |
| Jul11.25v_7357 | 1 | 0 | 0 | 0 | 0 | 0 | 0 | 0 | 0 | 0 | 0 | 0 | 0 | 0 | 1 | 0 | 0 | 0 | 0 | 0 | 0 | 0 | 0 | 0 | 0 | 0 | 0 | 0 | 0 | 0 |
| Jul11.5v_4412 | 1 | 0 | 0 | 0 | 0 | 0 | 0 | 0 | 0 | 0 | 0 | 0 | 0 | 1 | 0 | 0 | 0 | 0 | 0 | 0 | 0 | 0 | 0 | 0 | 0 | 0 | 0 | 0 | 0 | 0 |
| Jul27.5v_17045 | 1 | 0 | 0 | 0 | 0 | 0 | 0 | 0 | 0 | 0 | 0 | 0 | 0 | 0 | 0 | 0 | 0 | 1 | 0 | 0 | 0 | 0 | 0 | 0 | 0 | 0 | 0 | 0 | 0 | 0 |
| Jul27.75v_1714 | 1 | 0 | 0 | 0 | 0 | 0 | 0 | 0 | 0 | 0 | 0 | 0 | 0 | 0 | 0 | 0 | 0 | 0 | 0 | 0 | 1 | 0 | 0 | 0 | 0 | 0 | 0 | 0 | 0 | 0 |
| Jun8.50v_21260 | 1 | 0 | 0 | 0 | 0 | 0 | 1 | 0 | 0 | 0 | 0 | 0 | 0 | 0 | 0 | 0 | 0 | 0 | 0 | 0 | 0 | 0 | 0 | 0 | 0 | 0 | 0 | 0 | 0 | 0 |
| Jul11.5v_412 | 1 | 0 | 0 | 0 | 0 | 0 | 0 | 0 | 0 | 0 | 0 | 0 | 0 | 1 | 0 | 0 | 0 | 0 | 0 | 0 | 0 | 0 | 0 | 0 | 0 | 0 | 0 | 0 | 0 | 0 |
| Jul11.0v_20942 | 1 | 0 | 0 | 0 | 0 | 0 | 0 | 0 | 0 | 0 | 0 | 0 | 1 | 0 | 0 | 0 | 0 | 0 | 0 | 0 | 0 | 0 | 0 | 0 | 0 | 0 | 0 | 0 | 0 | 0 |
| May12.5v_7184 | 1 | 1 | 0 | 0 | 0 | 0 | 0 | 0 | 0 | 0 | 0 | 0 | 0 | 0 | 0 | 0 | 0 | 0 | 0 | 0 | 0 | 0 | 0 | 0 | 0 | 0 | 0 | 0 | 0 | 0 |
| Aug5.0v_422 | 1 | 0 | 0 | 0 | 0 | 0 | 0 | 0 | 0 | 0 | 0 | 0 | 0 | 0 | 0 | 0 | 0 | 0 | 0 | 0 | 0 | 1 | 0 | 0 | 0 | 0 | 0 | 0 | 0 | 0 |
| Jun22.0v_9242 | 1 | 0 | 0 | 0 | 0 | 0 | 0 | 0 | 1 | 0 | 0 | 0 | 0 | 0 | 0 | 0 | 0 | 0 | 0 | 0 | 0 | 0 | 0 | 0 | 0 | 0 | 0 | 0 | 0 | 0 |
| Jul11.25v_6769 | 1 | 0 | 0 | 0 | 0 | 0 | 0 | 0 | 0 | 0 | 0 | 0 | 0 | 0 | 1 | 0 | 0 | 0 | 0 | 0 | 0 | 0 | 0 | 0 | 0 | 0 | 0 | 0 | 0 | 0 |
| Jun22.75v_25850 | 1 | 0 | 0 | 0 | 0 | 0 | 0 | 0 | 0 | 0 | 0 | 1 | 0 | 0 | 0 | 0 | 0 | 0 | 0 | 0 | 0 | 0 | 0 | 0 | 0 | 0 | 0 | 0 | 0 | 0 |
| Jun22.5v_5002 | 1 | 0 | 0 | 0 | 0 | 0 | 0 | 0 | 0 | 1 | 0 | 0 | 0 | 0 | 0 | 0 | 0 | 0 | 0 | 0 | 0 | 0 | 0 | 0 | 0 | 0 | 0 | 0 | 0 | 0 |
| Sep8.50v_33778 | 2 | 0 | 0 | 0 | 0 | 0 | 0 | 0 | 0 | 0 | 0 | 1 | 0 | 0 | 0 | 0 | 0 | 0 | 0 | 0 | 0 | 0 | 0 | 0 | 0 | 0 | 0 | 0 | 0 | 1 |
| Jul11.25v_2817 | 1 | 0 | 0 | 0 | 0 | 0 | 0 | 0 | 0 | 0 | 0 | 0 | 0 | 0 | 1 | 0 | 0 | 0 | 0 | 0 | 0 | 0 | 0 | 0 | 0 | 0 | 0 | 0 | 0 | 0 |
| Jun22.5v_706 | 1 | 0 | 0 | 0 | 0 | 0 | 0 | 0 | 0 | 1 | 0 | 0 | 0 | 0 | 0 | 0 | 0 | 0 | 0 | 0 | 0 | 0 | 0 | 0 | 0 | 0 | 0 | 0 | 0 | 0 |
| Sep8.0v_11196 | 1 | 0 | 0 | 0 | 0 | 0 | 0 | 0 | 0 | 0 | 0 | 0 | 0 | 0 | 0 | 0 | 0 | 0 | 0 | 0 | 0 | 0 | 0 | 0 | 0 | 0 | 1 | 0 | 0 | 0 |
| Jul11.0v_11308 | 1 | 0 | 0 | 0 | 0 | 0 | 0 | 0 | 0 | 0 | 0 | 0 | 1 | 0 | 0 | 0 | 0 | 0 | 0 | 0 | 0 | 0 | 0 | 0 | 0 | 0 | 0 | 0 | 0 | 0 |
| Aug5.25v_18348 | 1 | 0 | 0 | 0 | 0 | 0 | 0 | 0 | 0 | 0 | 0 | 0 | 0 | 0 | 0 | 0 | 0 | 0 | 0 | 0 | 0 | 0 | 0 | 1 | 0 | 0 | 0 | 0 | 0 | 0 |
| May12.5v_34960 | 31 | 2 | 0 | 1 | 1 | 0 | 0 | 0 | 1 | 1 | 1 | 1 | 1 | 1 | 2 | 3 | 1 | 0 | 1 | 0 | 2 | 5 | 2 | 2 | 0 | 0 | 2 | 0 | 0 | 1 |
| Aug5.0v_1911 | 971 | 27 | 32 | 32 | 26 | 14 | 29 | 8 | 24 | 22 | 29 | 19 | 38 | 60 | 61 | 17 | 18 | 67 | 29 | 24 | 5 | 115 | 61 | 64 | 11 | 12 | 57 | 8 | 4 | 58 |
| Jun22.75v_27590 | 5 | 0 | 0 | 0 | 0 | 0 | 0 | 0 | 0 | 0 | 0 | 1 | 0 | 0 | 1 | 0 | 0 | 0 | 0 | 0 | 0 | 0 | 0 | 0 | 1 | 0 | 0 | 0 | 0 | 2 |
| Jun22.75v_28044 | 1 | 0 | 0 | 0 | 0 | 0 | 0 | 0 | 0 | 0 | 0 | 1 | 0 | 0 | 0 | 0 | 0 | 0 | 0 | 0 | 0 | 0 | 0 | 0 | 0 | 0 | 0 | 0 | 0 | 0 |
| Jun8.50v_8834 | 1 | 0 | 0 | 0 | 0 | 0 | 1 | 0 | 0 | 0 | 0 | 0 | 0 | 0 | 0 | 0 | 0 | 0 | 0 | 0 | 0 | 0 | 0 | 0 | 0 | 0 | 0 | 0 | 0 | 0 |
| Sep8.5v_3819 | 1 | 0 | 0 | 0 | 0 | 0 | 0 | 0 | 0 | 0 | 0 | 0 | 0 | 0 | 0 | 0 | 0 | 0 | 0 | 0 | 0 | 0 | 0 | 0 | 0 | 0 | 0 | 1 | 0 | 0 |
| Aug5.5v_31730 | 1 | 0 | 0 | 0 | 0 | 0 | 0 | 0 | 0 | 0 | 0 | 0 | 0 | 0 | 0 | 0 | 0 | 0 | 0 | 0 | 0 | 0 | 1 | 0 | 0 | 0 | 0 | 0 | 0 | 0 |
| Sep8.50v_29542 | 3 | 0 | 0 | 0 | 0 | 0 | 0 | 0 | 1 | 0 | 0 | 0 | 1 | 0 | 0 | 0 | 0 | 0 | 0 | 0 | 0 | 0 | 0 | 0 | 0 | 0 | 0 | 0 | 0 | 1 |
| May12.25v_9204 | 1 | 0 | 1 | 0 | 0 | 0 | 0 | 0 | 0 | 0 | 0 | 0 | 0 | 0 | 0 | 0 | 0 | 0 | 0 | 0 | 0 | 0 | 0 | 0 | 0 | 0 | 0 | 0 | 0 | 0 |
| Jul27.50v_26221 | 1 | 0 | 0 | 0 | 0 | 0 | 0 | 0 | 0 | 0 | 0 | 0 | 0 | 0 | 0 | 0 | 0 | 0 | 0 | 1 | 0 | 0 | 0 | 0 | 0 | 0 | 0 | 0 | 0 | 0 |
| Aug5.0v_3190 | 1 | 0 | 0 | 0 | 0 | 0 | 0 | 0 | 0 | 0 | 0 | 0 | 0 | 0 | 0 | 0 | 0 | 0 | 0 | 0 | 0 | 1 | 0 | 0 | 0 | 0 | 0 | 0 | 0 | 0 |
| May12.5v_19292 | 5 | 1 | 0 | 2 | 0 | 0 | 0 | 0 | 0 | 0 | 0 | 0 | 0 | 0 | 0 | 0 | 1 | 0 | 0 | 0 | 0 | 0 | 0 | 0 | 0 | 0 | 0 | 1 | 0 | 0 |
| Aug5.0v_2709 | 300 | 9 | 3 | 12 | 12 | 4 | 5 | 0 | 10 | 5 | 8 | 7 | 12 | 21 | 10 | 6 | 3 | 24 | 14 | 8 | 0 | 53 | 27 | 11 | 2 | 3 | 9 | 1 | 0 | 21 |
| Sep8.5v_8500 | 2 | 0 | 0 | 0 | 0 | 0 | 0 | 0 | 0 | 0 | 0 | 0 | 0 | 0 | 0 | 0 | 0 | 1 | 0 | 0 | 0 | 0 | 0 | 0 | 0 | 0 | 0 | 1 | 0 | 0 |
| Aug5.25v_24445 | 1 | 0 | 0 | 0 | 0 | 0 | 0 | 0 | 0 | 0 | 0 | 0 | 0 | 0 | 0 | 0 | 0 | 0 | 0 | 0 | 0 | 0 | 0 | 1 | 0 | 0 | 0 | 0 | 0 | 0 |
| Aug5.0v_3222 | 1 | 0 | 0 | 0 | 0 | 0 | 0 | 0 | 0 | 0 | 0 | 0 | 0 | 0 | 0 | 0 | 0 | 0 | 0 | 0 | 0 | 1 | 0 | 0 | 0 | 0 | 0 | 0 | 0 | 0 |

| Feature_ID | 654 | May12_05m | May12_25m | May12_50m | May12_75m | Jun8_25m | Jun8_50m | Jun8_75m | Jun22_0m | Jun22_05m | Jun22_50m | Jun22_75m | Jul11_0m | Jul11_05m | Jul11_25m | Jul11_50m | Jul11_75m | Jul27_05m | Jul27_25m | Jul27_50m | Jul27_75m | Aug5_0m | Aug5_05m | Aug5_25m | Aug5_50m | Aug5_75m | Sep8_0m | Sep8_05m | Sep8_25m | Sep8_50m |
|---|---|---|---|---|---|---|---|---|---|---|---|---|---|---|---|---|---|---|---|---|---|---|---|---|---|---|---|---|---|---|
| Jul27.50v_15266 | 1 | 0 | 0 | 0 | 0 | 0 | 0 | 0 | 0 | 0 | 0 | 0 | 0 | 0 | 0 | 0 | 0 | 0 | 0 | 1 | 0 | 0 | 0 | 0 | 0 | 0 | 0 | 0 | 0 | 0 |
| May12.50v_7012 | 1 | 0 | 0 | 1 | 0 | 0 | 0 | 0 | 0 | 0 | 0 | 0 | 0 | 0 | 0 | 0 | 0 | 0 | 0 | 0 | 0 | 0 | 0 | 0 | 0 | 0 | 0 | 0 | 0 | 0 |
| Jul27.5v_3767 | 1 | 0 | 0 | 0 | 0 | 0 | 0 | 0 | 0 | 0 | 0 | 0 | 0 | 0 | 0 | 0 | 0 | 1 | 0 | 0 | 0 | 0 | 0 | 0 | 0 | 0 | 0 | 0 | 0 | 0 |
| Sep8.0v_13128 | 1 | 0 | 0 | 0 | 0 | 0 | 0 | 0 | 0 | 0 | 0 | 0 | 0 | 0 | 0 | 0 | 0 | 0 | 0 | 0 | 0 | 0 | 0 | 0 | 0 | 0 | 1 | 0 | 0 | 0 |
| Jul11.5v_2104 | 1 | 0 | 0 | 0 | 0 | 0 | 0 | 0 | 0 | 0 | 0 | 0 | 0 | 1 | 0 | 0 | 0 | 0 | 0 | 0 | 0 | 0 | 0 | 0 | 0 | 0 | 0 | 0 | 0 | 0 |
| Jul11.0v_20214 | 1 | 0 | 0 | 0 | 0 | 0 | 0 | 0 | 0 | 0 | 0 | 0 | 1 | 0 | 0 | 0 | 0 | 0 | 0 | 0 | 0 | 0 | 0 | 0 | 0 | 0 | 0 | 0 | 0 | 0 |
| May12.25v_40130 | 1 | 0 | 1 | 0 | 0 | 0 | 0 | 0 | 0 | 0 | 0 | 0 | 0 | 0 | 0 | 0 | 0 | 0 | 0 | 0 | 0 | 0 | 0 | 0 | 0 | 0 | 0 | 0 | 0 | 0 |
| Jun22.75v_37026 | 1 | 0 | 0 | 0 | 0 | 0 | 0 | 0 | 0 | 0 | 0 | 1 | 0 | 0 | 0 | 0 | 0 | 0 | 0 | 0 | 0 | 0 | 0 | 0 | 0 | 0 | 0 | 0 | 0 | 0 |
| May12.50v_18378 | 1 | 0 | 0 | 1 | 0 | 0 | 0 | 0 | 0 | 0 | 0 | 0 | 0 | 0 | 0 | 0 | 0 | 0 | 0 | 0 | 0 | 0 | 0 | 0 | 0 | 0 | 0 | 0 | 0 | 0 |
| Jul27.5v_17680 | 3 | 0 | 0 | 0 | 0 | 0 | 0 | 0 | 0 | 0 | 0 | 1 | 0 | 1 | 0 | 0 | 0 | 1 | 0 | 0 | 0 | 0 | 0 | 0 | 0 | 0 | 0 | 0 | 0 | 0 |
| Jul11.25v_14091 | 1 | 0 | 0 | 0 | 0 | 0 | 0 | 0 | 0 | 0 | 0 | 0 | 0 | 0 | 1 | 0 | 0 | 0 | 0 | 0 | 0 | 0 | 0 | 0 | 0 | 0 | 0 | 0 | 0 | 0 |
| Jul27.5v_13084 | 1 | 0 | 0 | 0 | 0 | 0 | 0 | 0 | 0 | 0 | 0 | 0 | 0 | 0 | 0 | 0 | 0 | 1 | 0 | 0 | 0 | 0 | 0 | 0 | 0 | 0 | 0 | 0 | 0 | 0 |
| Jul11.25v_12490 | 1 | 0 | 0 | 0 | 0 | 0 | 0 | 0 | 0 | 0 | 0 | 0 | 0 | 0 | 1 | 0 | 0 | 0 | 0 | 0 | 0 | 0 | 0 | 0 | 0 | 0 | 0 | 0 | 0 | 0 |
| Jul27.75v_3872 | 1 | 0 | 0 | 0 | 0 | 0 | 0 | 0 | 0 | 0 | 0 | 0 | 0 | 0 | 0 | 0 | 0 | 0 | 0 | 0 | 1 | 0 | 0 | 0 | 0 | 0 | 0 | 0 | 0 | 0 |
| Jul27.5v_19079 | 1 | 0 | 0 | 0 | 0 | 0 | 0 | 0 | 0 | 0 | 0 | 0 | 0 | 0 | 0 | 0 | 0 | 1 | 0 | 0 | 0 | 0 | 0 | 0 | 0 | 0 | 0 | 0 | 0 | 0 |
| Jul27.5v_11463 | 1 | 0 | 0 | 0 | 0 | 0 | 0 | 0 | 0 | 0 | 0 | 0 | 0 | 0 | 0 | 0 | 0 | 1 | 0 | 0 | 0 | 0 | 0 | 0 | 0 | 0 | 0 | 0 | 0 | 0 |
| Jun22.50v_33242 | 1 | 0 | 0 | 0 | 0 | 0 | 0 | 0 | 0 | 0 | 1 | 0 | 0 | 0 | 0 | 0 | 0 | 0 | 0 | 0 | 0 | 0 | 0 | 0 | 0 | 0 | 0 | 0 | 0 | 0 |
| Jun8.25v_35482 | 1 | 0 | 0 | 0 | 0 | 1 | 0 | 0 | 0 | 0 | 0 | 0 | 0 | 0 | 0 | 0 | 0 | 0 | 0 | 0 | 0 | 0 | 0 | 0 | 0 | 0 | 0 | 0 | 0 | 0 |
| Sep8.50v_7969 | 1 | 0 | 0 | 0 | 0 | 0 | 0 | 0 | 0 | 0 | 0 | 0 | 0 | 0 | 0 | 0 | 0 | 0 | 0 | 0 | 0 | 0 | 0 | 0 | 0 | 0 | 0 | 0 | 0 | 1 |
| Jun22.5v_3552 | 1 | 0 | 0 | 0 | 0 | 0 | 0 | 0 | 0 | 1 | 0 | 0 | 0 | 0 | 0 | 0 | 0 | 0 | 0 | 0 | 0 | 0 | 0 | 0 | 0 | 0 | 0 | 0 | 0 | 0 |
| Aug5.50v_3350 | 1 | 0 | 0 | 0 | 0 | 0 | 0 | 0 | 0 | 0 | 0 | 0 | 0 | 0 | 0 | 0 | 0 | 0 | 0 | 0 | 0 | 0 | 0 | 0 | 1 | 0 | 0 | 0 | 0 | 0 |
| Sep8.0v_9852 | 1 | 0 | 0 | 0 | 0 | 0 | 0 | 0 | 0 | 0 | 0 | 0 | 0 | 0 | 0 | 0 | 0 | 0 | 0 | 0 | 0 | 0 | 0 | 0 | 0 | 0 | 1 | 0 | 0 | 0 |
| Jul11.0v_14761 | 2 | 1 | 0 | 0 | 0 | 0 | 0 | 0 | 0 | 0 | 0 | 0 | 1 | 0 | 0 | 0 | 0 | 0 | 0 | 0 | 0 | 0 | 0 | 0 | 0 | 0 | 0 | 0 | 0 | 0 |
| Sep8.50v_20345 | 1 | 0 | 0 | 0 | 0 | 0 | 0 | 0 | 0 | 0 | 0 | 0 | 0 | 0 | 0 | 0 | 0 | 0 | 0 | 0 | 0 | 0 | 0 | 0 | 0 | 0 | 0 | 0 | 0 | 1 |
| Jul11.25v_6485 | 1 | 0 | 0 | 0 | 0 | 0 | 0 | 0 | 0 | 0 | 0 | 0 | 0 | 0 | 1 | 0 | 0 | 0 | 0 | 0 | 0 | 0 | 0 | 0 | 0 | 0 | 0 | 0 | 0 | 0 |
| Jul27.25v_16951 | 2 | 0 | 0 | 0 | 0 | 0 | 0 | 0 | 0 | 0 | 0 | 0 | 0 | 0 | 0 | 0 | 0 | 0 | 1 | 0 | 0 | 0 | 0 | 0 | 0 | 0 | 1 | 0 | 0 | 0 |
| Jul11.0v_19513 | 1 | 0 | 0 | 0 | 0 | 0 | 0 | 0 | 0 | 0 | 0 | 0 | 1 | 0 | 0 | 0 | 0 | 0 | 0 | 0 | 0 | 0 | 0 | 0 | 0 | 0 | 0 | 0 | 0 | 0 |
| Aug5.0v_2290 | 1 | 0 | 0 | 0 | 0 | 0 | 0 | 0 | 0 | 0 | 0 | 0 | 0 | 0 | 0 | 0 | 0 | 0 | 0 | 0 | 0 | 1 | 0 | 0 | 0 | 0 | 0 | 0 | 0 | 0 |
| Aug5.25v_11480 | 1 | 0 | 0 | 0 | 0 | 0 | 0 | 0 | 0 | 0 | 0 | 0 | 0 | 0 | 0 | 0 | 0 | 0 | 0 | 0 | 0 | 0 | 0 | 1 | 0 | 0 | 0 | 0 | 0 | 0 |
| May12.25v_26490 | 1 | 0 | 1 | 0 | 0 | 0 | 0 | 0 | 0 | 0 | 0 | 0 | 0 | 0 | 0 | 0 | 0 | 0 | 0 | 0 | 0 | 0 | 0 | 0 | 0 | 0 | 0 | 0 | 0 | 0 |
| Jun8.25v_16559 | 1 | 0 | 0 | 0 | 0 | 1 | 0 | 0 | 0 | 0 | 0 | 0 | 0 | 0 | 0 | 0 | 0 | 0 | 0 | 0 | 0 | 0 | 0 | 0 | 0 | 0 | 0 | 0 | 0 | 0 |
| Aug5.5v_41029 | 1 | 0 | 0 | 0 | 0 | 0 | 0 | 0 | 0 | 0 | 0 | 0 | 0 | 0 | 0 | 0 | 0 | 0 | 0 | 0 | 0 | 0 | 1 | 0 | 0 | 0 | 0 | 0 | 0 | 0 |
| May12.5v_19934 | 1 | 1 | 0 | 0 | 0 | 0 | 0 | 0 | 0 | 0 | 0 | 0 | 0 | 0 | 0 | 0 | 0 | 0 | 0 | 0 | 0 | 0 | 0 | 0 | 0 | 0 | 0 | 0 | 0 | 0 |
| Aug5.0v_789 | 1 | 0 | 0 | 0 | 0 | 0 | 0 | 0 | 0 | 0 | 0 | 0 | 0 | 0 | 0 | 0 | 0 | 0 | 0 | 0 | 0 | 1 | 0 | 0 | 0 | 0 | 0 | 0 | 0 | 0 |
| Jul27.25v_14799 | 3 | 0 | 0 | 0 | 0 | 0 | 0 | 0 | 0 | 0 | 0 | 0 | 0 | 0 | 0 | 0 | 0 | 0 | 1 | 1 | 0 | 0 | 1 | 0 | 0 | 0 | 0 | 0 | 0 | 0 |
| Jun8.25v_468 | 1 | 0 | 0 | 0 | 0 | 1 | 0 | 0 | 0 | 0 | 0 | 0 | 0 | 0 | 0 | 0 | 0 | 0 | 0 | 0 | 0 | 0 | 0 | 0 | 0 | 0 | 0 | 0 | 0 | 0 |
| Jul27.5v_519 | 1 | 0 | 0 | 0 | 0 | 0 | 0 | 0 | 0 | 0 | 0 | 0 | 0 | 0 | 0 | 0 | 0 | 1 | 0 | 0 | 0 | 0 | 0 | 0 | 0 | 0 | 0 | 0 | 0 | 0 |
| Jul11.0v_542 | 1 | 0 | 0 | 0 | 0 | 0 | 0 | 0 | 0 | 0 | 0 | 0 | 1 | 0 | 0 | 0 | 0 | 0 | 0 | 0 | 0 | 0 | 0 | 0 | 0 | 0 | 0 | 0 | 0 | 0 |
| Jun22.50v_39821 | 1 | 0 | 0 | 0 | 0 | 0 | 0 | 0 | 0 | 0 | 1 | 0 | 0 | 0 | 0 | 0 | 0 | 0 | 0 | 0 | 0 | 0 | 0 | 0 | 0 | 0 | 0 | 0 | 0 | 0 |
| Aug5.0v_708 | 1 | 0 | 0 | 0 | 0 | 0 | 0 | 0 | 0 | 0 | 0 | 0 | 0 | 0 | 0 | 0 | 0 | 0 | 0 | 0 | 0 | 1 | 0 | 0 | 0 | 0 | 0 | 0 | 0 | 0 |
| Jul27.25v_33044 | 1 | 0 | 0 | 0 | 0 | 0 | 0 | 0 | 0 | 0 | 0 | 0 | 0 | 0 | 0 | 0 | 0 | 0 | 1 | 0 | 0 | 0 | 0 | 0 | 0 | 0 | 0 | 0 | 0 | 0 |
| Sep8.0v_12732 | 1 | 0 | 0 | 0 | 0 | 0 | 0 | 0 | 0 | 0 | 0 | 0 | 0 | 0 | 0 | 0 | 0 | 0 | 0 | 0 | 0 | 0 | 0 | 0 | 0 | 0 | 1 | 0 | 0 | 0 |
| May12.25v_13063 | 1 | 0 | 1 | 0 | 0 | 0 | 0 | 0 | 0 | 0 | 0 | 0 | 0 | 0 | 0 | 0 | 0 | 0 | 0 | 0 | 0 | 0 | 0 | 0 | 0 | 0 | 0 | 0 | 0 | 0 |
| Aug5.25v_18767 | 1 | 0 | 0 | 0 | 0 | 0 | 0 | 0 | 0 | 0 | 0 | 0 | 0 | 0 | 0 | 0 | 0 | 0 | 0 | 0 | 0 | 0 | 0 | 1 | 0 | 0 | 0 | 0 | 0 | 0 |
| May12.25v_21905 | 1 | 0 | 1 | 0 | 0 | 0 | 0 | 0 | 0 | 0 | 0 | 0 | 0 | 0 | 0 | 0 | 0 | 0 | 0 | 0 | 0 | 0 | 0 | 0 | 0 | 0 | 0 | 0 | 0 | 0 |
| Jul27.50v_27527 | 1 | 0 | 0 | 0 | 0 | 0 | 0 | 0 | 0 | 0 | 0 | 0 | 0 | 0 | 0 | 0 | 0 | 0 | 0 | 1 | 0 | 0 | 0 | 0 | 0 | 0 | 0 | 0 | 0 | 0 |
| May12.25v_34745 | 1 | 0 | 1 | 0 | 0 | 0 | 0 | 0 | 0 | 0 | 0 | 0 | 0 | 0 | 0 | 0 | 0 | 0 | 0 | 0 | 0 | 0 | 0 | 0 | 0 | 0 | 0 | 0 | 0 | 0 |
| Aug5.0v_4754 | 1 | 0 | 0 | 0 | 0 | 0 | 0 | 0 | 0 | 0 | 0 | 0 | 0 | 0 | 0 | 0 | 0 | 0 | 0 | 0 | 0 | 1 | 0 | 0 | 0 | 0 | 0 | 0 | 0 | 0 |
| Sep8.0v_4419 | 1 | 0 | 0 | 0 | 0 | 0 | 0 | 0 | 0 | 0 | 0 | 0 | 0 | 0 | 0 | 0 | 0 | 0 | 0 | 0 | 0 | 0 | 0 | 0 | 0 | 0 | 1 | 0 | 0 | 0 |
| Sep8.50v_1458 | 1 | 0 | 0 | 0 | 0 | 0 | 0 | 0 | 0 | 0 | 0 | 0 | 0 | 0 | 0 | 0 | 0 | 0 | 0 | 0 | 0 | 0 | 0 | 0 | 0 | 0 | 0 | 0 | 0 | 1 |
| Sep8.0v_19482 | 1 | 0 | 0 | 0 | 0 | 0 | 0 | 0 | 0 | 0 | 0 | 0 | 0 | 0 | 0 | 0 | 0 | 0 | 0 | 0 | 0 | 0 | 0 | 0 | 0 | 0 | 1 | 0 | 0 | 0 |
| Aug5.75v_11363 | 3 | 0 | 0 | 0 | 0 | 0 | 0 | 0 | 0 | 1 | 0 | 0 | 0 | 0 | 0 | 0 | 0 | 1 | 0 | 0 | 0 | 0 | 0 | 0 | 0 | 1 | 0 | 0 | 0 | 0 |
| Jun22.50v_11217 | 1 | 0 | 0 | 0 | 0 | 0 | 0 | 0 | 0 | 0 | 1 | 0 | 0 | 0 | 0 | 0 | 0 | 0 | 0 | 0 | 0 | 0 | 0 | 0 | 0 | 0 | 0 | 0 | 0 | 0 |
| May12.25v_32088 | 1 | 0 | 1 | 0 | 0 | 0 | 0 | 0 | 0 | 0 | 0 | 0 | 0 | 0 | 0 | 0 | 0 | 0 | 0 | 0 | 0 | 0 | 0 | 0 | 0 | 0 | 0 | 0 | 0 | 0 |
| Jul27.5v_5430 | 1 | 0 | 0 | 0 | 0 | 0 | 0 | 0 | 0 | 0 | 0 | 0 | 0 | 0 | 0 | 0 | 0 | 1 | 0 | 0 | 0 | 0 | 0 | 0 | 0 | 0 | 0 | 0 | 0 | 0 |
| Jul11.25v_19568 | 3 | 0 | 0 | 0 | 0 | 0 | 0 | 0 | 0 | 0 | 0 | 0 | 0 | 0 | 1 | 0 | 0 | 0 | 1 | 0 | 0 | 0 | 1 | 0 | 0 | 0 | 0 | 0 | 0 | 0 |
| Jun8.50v_37130 | 1 | 0 | 0 | 0 | 0 | 0 | 1 | 0 | 0 | 0 | 0 | 0 | 0 | 0 | 0 | 0 | 0 | 0 | 0 | 0 | 0 | 0 | 0 | 0 | 0 | 0 | 0 | 0 | 0 | 0 |
| Jul27.5v_15103 | 1 | 0 | 0 | 0 | 0 | 0 | 0 | 0 | 0 | 0 | 0 | 0 | 0 | 0 | 0 | 0 | 0 | 1 | 0 | 0 | 0 | 0 | 0 | 0 | 0 | 0 | 0 | 0 | 0 | 0 |
| Jun8.50v_13861 | 1 | 0 | 0 | 0 | 0 | 0 | 1 | 0 | 0 | 0 | 0 | 0 | 0 | 0 | 0 | 0 | 0 | 0 | 0 | 0 | 0 | 0 | 0 | 0 | 0 | 0 | 0 | 0 | 0 | 0 |
| Jul27.50v_12301 | 1 | 0 | 0 | 0 | 0 | 0 | 0 | 0 | 0 | 0 | 0 | 0 | 0 | 0 | 0 | 0 | 0 | 0 | 0 | 1 | 0 | 0 | 0 | 0 | 0 | 0 | 0 | 0 | 0 | 0 |
| May12.75v_33047 | 1 | 0 | 0 | 0 | 1 | 0 | 0 | 0 | 0 | 0 | 0 | 0 | 0 | 0 | 0 | 0 | 0 | 0 | 0 | 0 | 0 | 0 | 0 | 0 | 0 | 0 | 0 | 0 | 0 | 0 |
| Jun22.0v_16126 | 1 | 0 | 0 | 0 | 0 | 0 | 0 | 0 | 1 | 0 | 0 | 0 | 0 | 0 | 0 | 0 | 0 | 0 | 0 | 0 | 0 | 0 | 0 | 0 | 0 | 0 | 0 | 0 | 0 | 0 |
| Aug5.0v_2318 | 3 | 0 | 0 | 0 | 0 | 0 | 0 | 0 | 1 | 0 | 0 | 0 | 1 | 0 | 0 | 0 | 0 | 0 | 0 | 0 | 0 | 1 | 0 | 0 | 0 | 0 | 0 | 0 | 0 | 0 |
| Aug5.5v_22369 | 1 | 0 | 0 | 0 | 0 | 0 | 0 | 0 | 0 | 0 | 0 | 0 | 0 | 0 | 0 | 0 | 0 | 0 | 0 | 0 | 0 | 0 | 1 | 0 | 0 | 0 | 0 | 0 | 0 | 0 |
| May12.25v_31397 | 1 | 0 | 1 | 0 | 0 | 0 | 0 | 0 | 0 | 0 | 0 | 0 | 0 | 0 | 0 | 0 | 0 | 0 | 0 | 0 | 0 | 0 | 0 | 0 | 0 | 0 | 0 | 0 | 0 | 0 |
| Jun22.5v_13263 | 26 | 3 | 1 | 2 | 2 | 0 | 1 | 0 | 1 | 1 | 1 | 1 | 2 | 1 | 0 | 0 | 0 | 2 | 2 | 0 | 0 | 1 | 2 | 0 | 0 | 0 | 2 | 0 | 0 | 1 |
| Jun22.75v_9026 | 1 | 0 | 0 | 0 | 0 | 0 | 0 | 0 | 0 | 0 | 0 | 1 | 0 | 0 | 0 | 0 | 0 | 0 | 0 | 0 | 0 | 0 | 0 | 0 | 0 | 0 | 0 | 0 | 0 | 0 |
| Aug5.0v_3863 | 2 | 0 | 0 | 0 | 0 | 0 | 0 | 0 | 0 | 0 | 0 | 0 | 0 | 1 | 0 | 0 | 0 | 0 | 0 | 0 | 0 | 1 | 0 | 0 | 0 | 0 | 0 | 0 | 0 | 0 |
| May12.75v_26086 | 4 | 0 | 0 | 2 | 1 | 0 | 0 | 0 | 0 | 1 | 0 | 0 | 0 | 0 | 0 | 0 | 0 | 0 | 0 | 0 | 0 | 0 | 0 | 0 | 0 | 0 | 0 | 0 | 0 | 0 |
| Jul11.0v_7048 | 7 | 0 | 0 | 0 | 0 | 0 | 0 | 0 | 0 | 0 | 0 | 0 | 2 | 0 | 0 | 0 | 0 | 0 | 0 | 1 | 0 | 2 | 1 | 0 | 0 | 0 | 1 | 0 | 0 | 0 |
| Jul27.5v_17032 | 2 | 1 | 0 | 0 | 0 | 0 | 0 | 0 | 0 | 0 | 0 | 0 | 0 | 0 | 0 | 0 | 0 | 1 | 0 | 0 | 0 | 0 | 0 | 0 | 0 | 0 | 0 | 0 | 0 | 0 |
| Jun22.0v_1940 | 16 | 0 | 0 | 1 | 1 | 0 | 0 | 0 | 2 | 0 | 0 | 1 | 1 | 0 | 2 | 2 | 0 | 0 | 0 | 0 | 1 | 1 | 2 | 1 | 0 | 1 | 0 | 0 | 0 | 0 |
| Aug5.0v_2459 | 3 | 0 | 0 | 0 | 0 | 0 | 0 | 0 | 0 | 0 | 0 | 0 | 1 | 0 | 0 | 0 | 1 | 0 | 0 | 0 | 0 | 1 | 0 | 0 | 0 | 0 | 0 | 0 | 0 | 0 |
| Jun8.50v_40458 | 1 | 0 | 0 | 0 | 0 | 0 | 1 | 0 | 0 | 0 | 0 | 0 | 0 | 0 | 0 | 0 | 0 | 0 | 0 | 0 | 0 | 0 | 0 | 0 | 0 | 0 | 0 | 0 | 0 | 0 |
| Sep8.0v_17751 | 1 | 0 | 0 | 0 | 0 | 0 | 0 | 0 | 0 | 0 | 0 | 0 | 0 | 0 | 0 | 0 | 0 | 0 | 0 | 0 | 0 | 0 | 0 | 0 | 0 | 0 | 1 | 0 | 0 | 0 |
| Jun22.75v_15046 | 13 | 1 | 0 | 2 | 1 | 1 | 0 | 0 | 0 | 0 | 1 | 1 | 1 | 0 | 1 | 1 | 0 | 0 | 0 | 0 | 0 | 0 | 1 | 0 | 0 | 1 | 0 | 0 | 0 | 1 |
| Jul11.25v_13598 | 5 | 0 | 0 | 0 | 1 | 0 | 0 | 0 | 0 | 0 | 0 | 0 | 0 | 0 | 1 | 1 | 0 | 0 | 0 | 1 | 0 | 0 | 0 | 0 | 1 | 0 | 0 | 0 | 0 | 0 |
| Jun22.50v_6477 | 5 | 0 | 0 | 0 | 1 | 0 | 0 | 0 | 0 | 0 | 1 | 2 | 0 | 1 | 0 | 0 | 0 | 0 | 0 | 0 | 0 | 0 | 0 | 0 | 0 | 0 | 0 | 0 | 0 | 0 |
| Jul11.5v_1191 | 1 | 0 | 0 | 0 | 0 | 0 | 0 | 0 | 0 | 0 | 0 | 0 | 0 | 1 | 0 | 0 | 0 | 0 | 0 | 0 | 0 | 0 | 0 | 0 | 0 | 0 | 0 | 0 | 0 | 0 |
| Aug5.5v_10878 | 9 | 0 | 0 | 1 | 1 | 0 | 0 | 0 | 0 | 0 | 1 | 0 | 0 | 2 | 2 | 0 | 0 | 0 | 0 | 1 | 0 | 0 | 1 | 0 | 0 | 0 | 0 | 0 | 0 | 0 |
| Sep8.50v_38833 | 9 | 1 | 0 | 0 | 0 | 0 | 0 | 0 | 0 | 1 | 0 | 0 | 0 | 0 | 0 | 0 | 0 | 2 | 0 | 2 | 0 | 1 | 1 | 0 | 0 | 0 | 0 | 0 | 0 | 1 |
| Jul11.75v_7480 | 51 | 3 | 7 | 2 | 2 | 1 | 0 | 0 | 1 | 3 | 3 | 4 | 2 | 4 | 2 | 0 | 1 | 4 | 2 | 1 | 0 | 0 | 2 | 1 | 1 | 0 | 2 | 2 | 0 | 1 |
| Jul11.5v_7560 | 1 | 0 | 0 | 0 | 0 | 0 | 0 | 0 | 0 | 0 | 0 | 0 | 0 | 1 | 0 | 0 | 0 | 0 | 0 | 0 | 0 | 0 | 0 | 0 | 0 | 0 | 0 | 0 | 0 | 0 |
| Sep8.0v_14462 | 26 | 2 | 1 | 0 | 3 | 3 | 3 | 0 | 0 | 1 | 0 | 3 | 1 | 0 | 0 | 2 | 0 | 2 | 0 | 3 | 0 | 0 | 1 | 0 | 0 | 0 | 1 | 0 | 0 | 0 |
| Jun8.50v_5126 | 1 | 0 | 0 | 0 | 0 | 0 | 1 | 0 | 0 | 0 | 0 | 0 | 0 | 0 | 0 | 0 | 0 | 0 | 0 | 0 | 0 | 0 | 0 | 0 | 0 | 0 | 0 | 0 | 0 | 0 |
| Jul27.5v_18255 | 1 | 0 | 0 | 0 | 0 | 0 | 0 | 0 | 0 | 0 | 0 | 0 | 0 | 0 | 0 | 0 | 0 | 1 | 0 | 0 | 0 | 0 | 0 | 0 | 0 | 0 | 0 | 0 | 0 | 0 |
| Aug5.0v_4066 | 1 | 0 | 0 | 0 | 0 | 0 | 0 | 0 | 0 | 0 | 0 | 0 | 0 | 0 | 0 | 0 | 0 | 0 | 0 | 0 | 0 | 1 | 0 | 0 | 0 | 0 | 0 | 0 | 0 | 0 |
| Sep8.0v_13637 | 4 | 0 | 0 | 0 | 0 | 0 | 0 | 0 | 0 | 0 | 1 | 0 | 0 | 0 | 1 | 0 | 0 | 0 | 1 | 0 | 0 | 0 | 0 | 0 | 0 | 0 | 1 | 0 | 0 | 0 |
| Jul11.5v_12100 | 1 | 0 | 0 | 0 | 0 | 0 | 0 | 0 | 0 | 0 | 0 | 0 | 0 | 1 | 0 | 0 | 0 | 0 | 0 | 0 | 0 | 0 | 0 | 0 | 0 | 0 | 0 | 0 | 0 | 0 |
| Aug5.0v_433 | 19 | 3 | 0 | 2 | 2 | 0 | 1 | 0 | 0 | 0 | 0 | 1 | 1 | 0 | 0 | 1 | 0 | 0 | 1 | 0 | 0 | 3 | 0 | 3 | 0 | 0 | 1 | 0 | 0 | 0 |
| Jul27.5v_18031 | 1 | 0 | 0 | 0 | 0 | 0 | 0 | 0 | 0 | 0 | 0 | 0 | 0 | 0 | 0 | 0 | 0 | 1 | 0 | 0 | 0 | 0 | 0 | 0 | 0 | 0 | 0 | 0 | 0 | 0 |
| May12.50v_15696 | 4 | 0 | 0 | 1 | 1 | 0 | 0 | 0 | 0 | 0 | 0 | 0 | 0 | 0 | 0 | 0 | 0 | 0 | 0 | 0 | 0 | 0 | 0 | 0 | 0 | 2 | 0 | 0 | 0 | 0 |
| Jul27.50v_31991 | 1 | 0 | 0 | 0 | 0 | 0 | 0 | 0 | 0 | 0 | 0 | 0 | 0 | 0 | 0 | 0 | 0 | 0 | 0 | 1 | 0 | 0 | 0 | 0 | 0 | 0 | 0 | 0 | 0 | 0 |
| Jul11.0v_6292 | 35 | 1 | 1 | 0 | 3 | 1 | 2 | 0 | 4 | 1 | 0 | 0 | 3 | 3 | 0 | 2 | 1 | 3 | 1 | 0 | 0 | 3 | 1 | 1 | 0 | 1 | 1 | 0 | 0 | 2 |
| Jun22.0v_12525 | 1 | 0 | 0 | 0 | 0 | 0 | 0 | 0 | 1 | 0 | 0 | 0 | 0 | 0 | 0 | 0 | 0 | 0 | 0 | 0 | 0 | 0 | 0 | 0 | 0 | 0 | 0 | 0 | 0 | 0 |
| Jul27.5v_15809 | 1 | 0 | 0 | 0 | 0 | 0 | 0 | 0 | 0 | 0 | 0 | 0 | 0 | 0 | 0 | 0 | 0 | 1 | 0 | 0 | 0 | 0 | 0 | 0 | 0 | 0 | 0 | 0 | 0 | 0 |
| Jul27.5v_14805 | 1 | 0 | 0 | 0 | 0 | 0 | 0 | 0 | 0 | 0 | 0 | 0 | 0 | 0 | 0 | 0 | 0 | 1 | 0 | 0 | 0 | 0 | 0 | 0 | 0 | 0 | 0 | 0 | 0 | 0 |
| Jul11.5v_14896 | 1 | 0 | 0 | 0 | 0 | 0 | 0 | 0 | 0 | 0 | 0 | 0 | 0 | 1 | 0 | 0 | 0 | 0 | 0 | 0 | 0 | 0 | 0 | 0 | 0 | 0 | 0 | 0 | 0 | 0 |
| May12.25v_26384 | 1 | 0 | 1 | 0 | 0 | 0 | 0 | 0 | 0 | 0 | 0 | 0 | 0 | 0 | 0 | 0 | 0 | 0 | 0 | 0 | 0 | 0 | 0 | 0 | 0 | 0 | 0 | 0 | 0 | 0 |
| Aug5.5v_23640 | 15 | 0 | 2 | 1 | 0 | 2 | 0 | 0 | 0 | 0 | 2 | 0 | 2 | 0 | 0 | 0 | 0 | 2 | 1 | 0 | 0 | 0 | 3 | 0 | 0 | 0 | 0 | 0 | 0 | 0 |
| Aug5.5v_15451 | 1 | 0 | 0 | 0 | 0 | 0 | 0 | 0 | 0 | 0 | 0 | 0 | 0 | 0 | 0 | 0 | 0 | 0 | 0 | 0 | 0 | 0 | 1 | 0 | 0 | 0 | 0 | 0 | 0 | 0 |
| Aug5.0v_335 | 4 | 0 | 0 | 0 | 0 | 0 | 0 | 0 | 0 | 0 | 0 | 0 | 0 | 3 | 0 | 0 | 0 | 0 | 0 | 0 | 0 | 1 | 0 | 0 | 0 | 0 | 0 | 0 | 0 | 0 |
| May12.25v_1813 | 1 | 0 | 1 | 0 | 0 | 0 | 0 | 0 | 0 | 0 | 0 | 0 | 0 | 0 | 0 | 0 | 0 | 0 | 0 | 0 | 0 | 0 | 0 | 0 | 0 | 0 | 0 | 0 | 0 | 0 |
| Aug5.25v_7670 | 18 | 1 | 2 | 0 | 0 | 0 | 1 | 1 | 0 | 1 | 0 | 0 | 1 | 1 | 1 | 0 | 1 | 0 | 2 | 0 | 1 | 1 | 2 | 1 | 0 | 0 | 0 | 0 | 0 | 1 |
| Aug5.0v_659 | 2 | 0 | 0 | 0 | 0 | 0 | 0 | 0 | 0 | 0 | 0 | 0 | 0 | 0 | 0 | 0 | 0 | 1 | 0 | 0 | 0 | 1 | 0 | 0 | 0 | 0 | 0 | 0 | 0 | 0 |
| May12.50v_31645 | 42 | 2 | 4 | 1 | 2 | 1 | 1 | 0 | 4 | 2 | 2 | 2 | 0 | 3 | 5 | 0 | 0 | 4 | 2 | 1 | 0 | 0 | 3 | 2 | 0 | 0 | 0 | 1 | 0 | 0 |
| Jul27.5v_6125 | 1 | 0 | 0 | 0 | 0 | 0 | 0 | 0 | 0 | 0 | 0 | 0 | 0 | 0 | 0 | 0 | 0 | 1 | 0 | 0 | 0 | 0 | 0 | 0 | 0 | 0 | 0 | 0 | 0 | 0 |
| May12.5v_5978 | 1 | 1 | 0 | 0 | 0 | 0 | 0 | 0 | 0 | 0 | 0 | 0 | 0 | 0 | 0 | 0 | 0 | 0 | 0 | 0 | 0 | 0 | 0 | 0 | 0 | 0 | 0 | 0 | 0 | 0 |
| Jul27.5v_1171 | 1 | 0 | 0 | 0 | 0 | 0 | 0 | 0 | 0 | 0 | 0 | 0 | 0 | 0 | 0 | 0 | 0 | 1 | 0 | 0 | 0 | 0 | 0 | 0 | 0 | 0 | 0 | 0 | 0 | 0 |
| Aug5.25v_10634 | 1 | 0 | 0 | 0 | 0 | 0 | 0 | 0 | 0 | 0 | 0 | 0 | 0 | 0 | 0 | 0 | 0 | 0 | 0 | 0 | 0 | 0 | 0 | 1 | 0 | 0 | 0 | 0 | 0 | 0 |
| Jul11.25v_13785 | 1 | 0 | 0 | 0 | 0 | 0 | 0 | 0 | 0 | 0 | 0 | 0 | 0 | 0 | 1 | 0 | 0 | 0 | 0 | 0 | 0 | 0 | 0 | 0 | 0 | 0 | 0 | 0 | 0 | 0 |
| Jul11.25v_914 | 1 | 0 | 0 | 0 | 0 | 0 | 0 | 0 | 0 | 0 | 0 | 0 | 0 | 0 | 1 | 0 | 0 | 0 | 0 | 0 | 0 | 0 | 0 | 0 | 0 | 0 | 0 | 0 | 0 | 0 |
| Jul11.25v_12713 | 1 | 0 | 0 | 0 | 0 | 0 | 0 | 0 | 0 | 0 | 0 | 0 | 0 | 0 | 1 | 0 | 0 | 0 | 0 | 0 | 0 | 0 | 0 | 0 | 0 | 0 | 0 | 0 | 0 | 0 |
| Jul27.50v_25724 | 1 | 0 | 0 | 0 | 0 | 0 | 0 | 0 | 0 | 0 | 0 | 0 | 0 | 0 | 0 | 0 | 0 | 0 | 0 | 1 | 0 | 0 | 0 | 0 | 0 | 0 | 0 | 0 | 0 | 0 |
| May12.25v_32267 | 1 | 0 | 1 | 0 | 0 | 0 | 0 | 0 | 0 | 0 | 0 | 0 | 0 | 0 | 0 | 0 | 0 | 0 | 0 | 0 | 0 | 0 | 0 | 0 | 0 | 0 | 0 | 0 | 0 | 0 |
| Aug5.50v_38181 | 3 | 0 | 0 | 1 | 0 | 0 | 0 | 0 | 0 | 0 | 0 | 0 | 0 | 0 | 0 | 0 | 0 | 0 | 0 | 0 | 0 | 0 | 0 | 0 | 1 | 0 | 0 | 0 | 0 | 1 |
| Jul27.75v_3174 | 1 | 0 | 0 | 0 | 0 | 0 | 0 | 0 | 0 | 0 | 0 | 0 | 0 | 0 | 0 | 0 | 0 | 0 | 0 | 0 | 1 | 0 | 0 | 0 | 0 | 0 | 0 | 0 | 0 | 0 |
| Jun22.0v_15566 | 1 | 0 | 0 | 0 | 0 | 0 | 0 | 0 | 1 | 0 | 0 | 0 | 0 | 0 | 0 | 0 | 0 | 0 | 0 | 0 | 0 | 0 | 0 | 0 | 0 | 0 | 0 | 0 | 0 | 0 |
| Sep8.50v_22677 | 1 | 0 | 0 | 0 | 0 | 0 | 0 | 0 | 0 | 0 | 0 | 0 | 0 | 0 | 0 | 0 | 0 | 0 | 0 | 0 | 0 | 0 | 0 | 0 | 0 | 0 | 0 | 0 | 0 | 1 |
| Aug5.0v_4322 | 1 | 0 | 0 | 0 | 0 | 0 | 0 | 0 | 0 | 0 | 0 | 0 | 0 | 0 | 0 | 0 | 0 | 0 | 0 | 0 | 0 | 1 | 0 | 0 | 0 | 0 | 0 | 0 | 0 | 0 |
| Jul11.25v_9931 | 1 | 0 | 0 | 0 | 0 | 0 | 0 | 0 | 0 | 0 | 0 | 0 | 0 | 0 | 1 | 0 | 0 | 0 | 0 | 0 | 0 | 0 | 0 | 0 | 0 | 0 | 0 | 0 | 0 | 0 |
| Jul11.50v_8038 | 1 | 0 | 0 | 0 | 0 | 0 | 0 | 0 | 0 | 0 | 0 | 0 | 0 | 0 | 0 | 1 | 0 | 0 | 0 | 0 | 0 | 0 | 0 | 0 | 0 | 0 | 0 | 0 | 0 | 0 |
| Aug5.5v_4540 | 1 | 0 | 0 | 0 | 0 | 0 | 0 | 0 | 0 | 0 | 0 | 0 | 0 | 0 | 0 | 0 | 0 | 0 | 0 | 0 | 0 | 0 | 1 | 0 | 0 | 0 | 0 | 0 | 0 | 0 |
| Aug5.25v_28677 | 1 | 0 | 0 | 0 | 0 | 0 | 0 | 0 | 0 | 0 | 0 | 0 | 0 | 0 | 0 | 0 | 0 | 0 | 0 | 0 | 0 | 0 | 0 | 1 | 0 | 0 | 0 | 0 | 0 | 0 |
| Jul11.0v_18769 | 2 | 0 | 0 | 0 | 0 | 0 | 0 | 0 | 0 | 0 | 0 | 0 | 1 | 0 | 0 | 0 | 0 | 0 | 0 | 0 | 0 | 0 | 1 | 0 | 0 | 0 | 0 | 0 | 0 | 0 |
| Sep8.0v_19664 | 1 | 0 | 0 | 0 | 0 | 0 | 0 | 0 | 0 | 0 | 0 | 0 | 0 | 0 | 0 | 0 | 0 | 0 | 0 | 0 | 0 | 0 | 0 | 0 | 0 | 0 | 1 | 0 | 0 | 0 |
| Jul11.25v_400 | 1 | 0 | 0 | 0 | 0 | 0 | 0 | 0 | 0 | 0 | 0 | 0 | 0 | 0 | 1 | 0 | 0 | 0 | 0 | 0 | 0 | 0 | 0 | 0 | 0 | 0 | 0 | 0 | 0 | 0 |
| Aug5.75v_27553 | 2 | 0 | 0 | 0 | 0 | 0 | 0 | 0 | 0 | 0 | 0 | 0 | 0 | 0 | 0 | 0 | 0 | 0 | 0 | 0 | 1 | 0 | 0 | 0 | 0 | 1 | 0 | 0 | 0 | 0 |
| Jul11.0v_4124 | 1 | 0 | 0 | 0 | 0 | 0 | 0 | 0 | 0 | 0 | 0 | 0 | 1 | 0 | 0 | 0 | 0 | 0 | 0 | 0 | 0 | 0 | 0 | 0 | 0 | 0 | 0 | 0 | 0 | 0 |
| May12.5v_35782 | 1 | 1 | 0 | 0 | 0 | 0 | 0 | 0 | 0 | 0 | 0 | 0 | 0 | 0 | 0 | 0 | 0 | 0 | 0 | 0 | 0 | 0 | 0 | 0 | 0 | 0 | 0 | 0 | 0 | 0 |
| Sep8.50v_17827 | 1 | 0 | 0 | 0 | 0 | 0 | 0 | 0 | 0 | 0 | 0 | 0 | 0 | 0 | 0 | 0 | 0 | 0 | 0 | 0 | 0 | 0 | 0 | 0 | 0 | 0 | 0 | 0 | 0 | 1 |
| Sep8.0v_3288 | 1 | 0 | 0 | 0 | 0 | 0 | 0 | 0 | 0 | 0 | 0 | 0 | 0 | 0 | 0 | 0 | 0 | 0 | 0 | 0 | 0 | 0 | 0 | 0 | 0 | 0 | 1 | 0 | 0 | 0 |
| Sep8.5v_7952 | 1 | 0 | 0 | 0 | 0 | 0 | 0 | 0 | 0 | 0 | 0 | 0 | 0 | 0 | 0 | 0 | 0 | 0 | 0 | 0 | 0 | 0 | 0 | 0 | 0 | 0 | 0 | 1 | 0 | 0 |
| Jul11.25v_4155 | 1 | 0 | 0 | 0 | 0 | 0 | 0 | 0 | 0 | 0 | 0 | 0 | 0 | 0 | 1 | 0 | 0 | 0 | 0 | 0 | 0 | 0 | 0 | 0 | 0 | 0 | 0 | 0 | 0 | 0 |
| Sep8.0v_14734 | 1 | 0 | 0 | 0 | 0 | 0 | 0 | 0 | 0 | 0 | 0 | 0 | 0 | 0 | 0 | 0 | 0 | 0 | 0 | 0 | 0 | 0 | 0 | 0 | 0 | 0 | 1 | 0 | 0 | 0 |
| May12.50v_2088 | 1 | 0 | 0 | 1 | 0 | 0 | 0 | 0 | 0 | 0 | 0 | 0 | 0 | 0 | 0 | 0 | 0 | 0 | 0 | 0 | 0 | 0 | 0 | 0 | 0 | 0 | 0 | 0 | 0 | 0 |
| May12.75v_21511 | 1 | 0 | 0 | 0 | 1 | 0 | 0 | 0 | 0 | 0 | 0 | 0 | 0 | 0 | 0 | 0 | 0 | 0 | 0 | 0 | 0 | 0 | 0 | 0 | 0 | 0 | 0 | 0 | 0 | 0 |
| Aug5.0v_1537 | 1 | 0 | 0 | 0 | 0 | 0 | 0 | 0 | 0 | 0 | 0 | 0 | 0 | 0 | 0 | 0 | 0 | 0 | 0 | 0 | 0 | 1 | 0 | 0 | 0 | 0 | 0 | 0 | 0 | 0 |
| Jul27.5v_19626 | 1 | 0 | 0 | 0 | 0 | 0 | 0 | 0 | 0 | 0 | 0 | 0 | 0 | 0 | 0 | 0 | 0 | 1 | 0 | 0 | 0 | 0 | 0 | 0 | 0 | 0 | 0 | 0 | 0 | 0 |
| Jul27.5v_18499 | 1 | 0 | 0 | 0 | 0 | 0 | 0 | 0 | 0 | 0 | 0 | 0 | 0 | 0 | 0 | 0 | 0 | 1 | 0 | 0 | 0 | 0 | 0 | 0 | 0 | 0 | 0 | 0 | 0 | 0 |
| Aug5.5v_36992 | 1 | 0 | 0 | 0 | 0 | 0 | 0 | 0 | 0 | 0 | 0 | 0 | 0 | 0 | 0 | 0 | 0 | 0 | 0 | 0 | 0 | 0 | 1 | 0 | 0 | 0 | 0 | 0 | 0 | 0 |
| Jul27.5v_12825 | 1 | 0 | 0 | 0 | 0 | 0 | 0 | 0 | 0 | 0 | 0 | 0 | 0 | 0 | 0 | 0 | 0 | 1 | 0 | 0 | 0 | 0 | 0 | 0 | 0 | 0 | 0 | 0 | 0 | 0 |
| Jul11.50v_6014 | 1 | 0 | 0 | 0 | 0 | 0 | 0 | 0 | 0 | 0 | 0 | 0 | 0 | 0 | 0 | 1 | 0 | 0 | 0 | 0 | 0 | 0 | 0 | 0 | 0 | 0 | 0 | 0 | 0 | 0 |
| May12.5v_41032 | 1 | 1 | 0 | 0 | 0 | 0 | 0 | 0 | 0 | 0 | 0 | 0 | 0 | 0 | 0 | 0 | 0 | 0 | 0 | 0 | 0 | 0 | 0 | 0 | 0 | 0 | 0 | 0 | 0 | 0 |
| Aug5.25v_17243 | 1 | 0 | 0 | 0 | 0 | 0 | 0 | 0 | 0 | 0 | 0 | 0 | 0 | 0 | 0 | 0 | 0 | 0 | 0 | 0 | 0 | 0 | 0 | 1 | 0 | 0 | 0 | 0 | 0 | 0 |
| Jul27.5v_1734 | 1 | 0 | 0 | 0 | 0 | 0 | 0 | 0 | 0 | 0 | 0 | 0 | 0 | 0 | 0 | 0 | 0 | 1 | 0 | 0 | 0 | 0 | 0 | 0 | 0 | 0 | 0 | 0 | 0 | 0 |
| Jul27.25v_27739 | 1 | 0 | 0 | 0 | 0 | 0 | 0 | 0 | 0 | 0 | 0 | 0 | 0 | 0 | 0 | 0 | 0 | 0 | 1 | 0 | 0 | 0 | 0 | 0 | 0 | 0 | 0 | 0 | 0 | 0 |
| Jul27.75v_18485 | 1 | 0 | 0 | 0 | 0 | 0 | 0 | 0 | 0 | 0 | 0 | 0 | 0 | 0 | 0 | 0 | 0 | 0 | 0 | 0 | 1 | 0 | 0 | 0 | 0 | 0 | 0 | 0 | 0 | 0 |
| May12.50v_41150 | 1 | 0 | 0 | 1 | 0 | 0 | 0 | 0 | 0 | 0 | 0 | 0 | 0 | 0 | 0 | 0 | 0 | 0 | 0 | 0 | 0 | 0 | 0 | 0 | 0 | 0 | 0 | 0 | 0 | 0 |
| Aug5.0v_2941 | 1 | 0 | 0 | 0 | 0 | 0 | 0 | 0 | 0 | 0 | 0 | 0 | 0 | 0 | 0 | 0 | 0 | 0 | 0 | 0 | 0 | 1 | 0 | 0 | 0 | 0 | 0 | 0 | 0 | 0 |
| Jul27.5v_4775 | 1 | 0 | 0 | 0 | 0 | 0 | 0 | 0 | 0 | 0 | 0 | 0 | 0 | 0 | 0 | 0 | 0 | 1 | 0 | 0 | 0 | 0 | 0 | 0 | 0 | 0 | 0 | 0 | 0 | 0 |
| Jul27.75v_6963 | 1 | 0 | 0 | 0 | 0 | 0 | 0 | 0 | 0 | 0 | 0 | 0 | 0 | 0 | 0 | 0 | 0 | 0 | 0 | 0 | 1 | 0 | 0 | 0 | 0 | 0 | 0 | 0 | 0 | 0 |
| Jul11.0v_7008 | 1 | 0 | 0 | 0 | 0 | 0 | 0 | 0 | 0 | 0 | 0 | 0 | 1 | 0 | 0 | 0 | 0 | 0 | 0 | 0 | 0 | 0 | 0 | 0 | 0 | 0 | 0 | 0 | 0 | 0 |
| Sep8.0v_3366 | 1 | 0 | 0 | 0 | 0 | 0 | 0 | 0 | 0 | 0 | 0 | 0 | 0 | 0 | 0 | 0 | 0 | 0 | 0 | 0 | 0 | 0 | 0 | 0 | 0 | 0 | 1 | 0 | 0 | 0 |
| Jun8.50v_20991 | 1 | 0 | 0 | 0 | 0 | 0 | 1 | 0 | 0 | 0 | 0 | 0 | 0 | 0 | 0 | 0 | 0 | 0 | 0 | 0 | 0 | 0 | 0 | 0 | 0 | 0 | 0 | 0 | 0 | 0 |
| Jul27.5v_213 | 1 | 0 | 0 | 0 | 0 | 0 | 0 | 0 | 0 | 0 | 0 | 0 | 0 | 0 | 0 | 0 | 0 | 1 | 0 | 0 | 0 | 0 | 0 | 0 | 0 | 0 | 0 | 0 | 0 | 0 |
| Jul11.0v_17580 | 1 | 0 | 0 | 0 | 0 | 0 | 0 | 0 | 0 | 0 | 0 | 0 | 1 | 0 | 0 | 0 | 0 | 0 | 0 | 0 | 0 | 0 | 0 | 0 | 0 | 0 | 0 | 0 | 0 | 0 |
| Jul11.25v_6271 | 1 | 0 | 0 | 0 | 0 | 0 | 0 | 0 | 0 | 0 | 0 | 0 | 0 | 0 | 1 | 0 | 0 | 0 | 0 | 0 | 0 | 0 | 0 | 0 | 0 | 0 | 0 | 0 | 0 | 0 |
| Jul27.5v_15851 | 1 | 0 | 0 | 0 | 0 | 0 | 0 | 0 | 0 | 0 | 0 | 0 | 0 | 0 | 0 | 0 | 0 | 1 | 0 | 0 | 0 | 0 | 0 | 0 | 0 | 0 | 0 | 0 | 0 | 0 |
| Jun8.50v_5068 | 1 | 0 | 0 | 0 | 0 | 0 | 1 | 0 | 0 | 0 | 0 | 0 | 0 | 0 | 0 | 0 | 0 | 0 | 0 | 0 | 0 | 0 | 0 | 0 | 0 | 0 | 0 | 0 | 0 | 0 |
| Jul11.50v_9390 | 1 | 0 | 0 | 0 | 0 | 0 | 0 | 0 | 0 | 0 | 0 | 0 | 0 | 0 | 0 | 1 | 0 | 0 | 0 | 0 | 0 | 0 | 0 | 0 | 0 | 0 | 0 | 0 | 0 | 0 |
| Aug5.0v_4206 | 1 | 0 | 0 | 0 | 0 | 0 | 0 | 0 | 0 | 0 | 0 | 0 | 0 | 0 | 0 | 0 | 0 | 0 | 0 | 0 | 0 | 1 | 0 | 0 | 0 | 0 | 0 | 0 | 0 | 0 |
| Aug5.0v_5228 | 1 | 0 | 0 | 0 | 0 | 0 | 0 | 0 | 0 | 0 | 0 | 0 | 0 | 0 | 0 | 0 | 0 | 0 | 0 | 0 | 0 | 1 | 0 | 0 | 0 | 0 | 0 | 0 | 0 | 0 |
| Jun8.50v_13809 | 2 | 1 | 0 | 0 | 0 | 0 | 1 | 0 | 0 | 0 | 0 | 0 | 0 | 0 | 0 | 0 | 0 | 0 | 0 | 0 | 0 | 0 | 0 | 0 | 0 | 0 | 0 | 0 | 0 | 0 |
| Jun22.50v_9466 | 1 | 0 | 0 | 0 | 0 | 0 | 0 | 0 | 0 | 0 | 1 | 0 | 0 | 0 | 0 | 0 | 0 | 0 | 0 | 0 | 0 | 0 | 0 | 0 | 0 | 0 | 0 | 0 | 0 | 0 |
| Jul27.50v_6950 | 1 | 0 | 0 | 0 | 0 | 0 | 0 | 0 | 0 | 0 | 0 | 0 | 0 | 0 | 0 | 0 | 0 | 0 | 0 | 1 | 0 | 0 | 0 | 0 | 0 | 0 | 0 | 0 | 0 | 0 |
| Jun22.50v_1537 | 1 | 0 | 0 | 0 | 0 | 0 | 0 | 0 | 0 | 0 | 1 | 0 | 0 | 0 | 0 | 0 | 0 | 0 | 0 | 0 | 0 | 0 | 0 | 0 | 0 | 0 | 0 | 0 | 0 | 0 |
| Jul11.50v_1478 | 1 | 0 | 0 | 0 | 0 | 0 | 0 | 0 | 0 | 0 | 0 | 0 | 0 | 0 | 0 | 1 | 0 | 0 | 0 | 0 | 0 | 0 | 0 | 0 | 0 | 0 | 0 | 0 | 0 | 0 |
| Jul27.25v_33018 | 1 | 0 | 0 | 0 | 0 | 0 | 0 | 0 | 0 | 0 | 0 | 0 | 0 | 0 | 0 | 0 | 0 | 0 | 1 | 0 | 0 | 0 | 0 | 0 | 0 | 0 | 0 | 0 | 0 | 0 |
| May12.5v_33832 | 1 | 1 | 0 | 0 | 0 | 0 | 0 | 0 | 0 | 0 | 0 | 0 | 0 | 0 | 0 | 0 | 0 | 0 | 0 | 0 | 0 | 0 | 0 | 0 | 0 | 0 | 0 | 0 | 0 | 0 |
| Aug5.0v_1609 | 1 | 0 | 0 | 0 | 0 | 0 | 0 | 0 | 0 | 0 | 0 | 0 | 0 | 0 | 0 | 0 | 0 | 0 | 0 | 0 | 0 | 1 | 0 | 0 | 0 | 0 | 0 | 0 | 0 | 0 |
| Jul27.50v_1924 | 2 | 0 | 0 | 0 | 0 | 0 | 0 | 0 | 1 | 0 | 0 | 0 | 0 | 0 | 0 | 0 | 0 | 0 | 0 | 1 | 0 | 0 | 0 | 0 | 0 | 0 | 0 | 0 | 0 | 0 |
| Jun22.75v_3936 | 1 | 0 | 0 | 0 | 0 | 0 | 0 | 0 | 0 | 0 | 0 | 1 | 0 | 0 | 0 | 0 | 0 | 0 | 0 | 0 | 0 | 0 | 0 | 0 | 0 | 0 | 0 | 0 | 0 | 0 |
| Aug5.0v_4514 | 1 | 0 | 0 | 0 | 0 | 0 | 0 | 0 | 0 | 0 | 0 | 0 | 0 | 0 | 0 | 0 | 0 | 0 | 0 | 0 | 0 | 1 | 0 | 0 | 0 | 0 | 0 | 0 | 0 | 0 |
| Aug5.25v_4754 | 1 | 0 | 0 | 0 | 0 | 0 | 0 | 0 | 0 | 0 | 0 | 0 | 0 | 0 | 0 | 0 | 0 | 0 | 0 | 0 | 0 | 0 | 0 | 1 | 0 | 0 | 0 | 0 | 0 | 0 |
| Jun22.50v_28817 | 3 | 0 | 0 | 0 | 0 | 0 | 0 | 0 | 0 | 0 | 1 | 1 | 0 | 0 | 0 | 0 | 0 | 0 | 0 | 0 | 0 | 0 | 0 | 0 | 0 | 0 | 0 | 0 | 0 | 1 |
| Sep8.50v_36214 | 1 | 0 | 0 | 0 | 0 | 0 | 0 | 0 | 0 | 0 | 0 | 0 | 0 | 0 | 0 | 0 | 0 | 0 | 0 | 0 | 0 | 0 | 0 | 0 | 0 | 0 | 0 | 0 | 0 | 1 |
| Sep8.50v_11419 | 1 | 0 | 0 | 0 | 0 | 0 | 0 | 0 | 0 | 0 | 0 | 0 | 0 | 0 | 0 | 0 | 0 | 0 | 0 | 0 | 0 | 0 | 0 | 0 | 0 | 0 | 0 | 0 | 0 | 1 |
| Jun22.75v_37444 | 1 | 0 | 0 | 0 | 0 | 0 | 0 | 0 | 0 | 0 | 0 | 1 | 0 | 0 | 0 | 0 | 0 | 0 | 0 | 0 | 0 | 0 | 0 | 0 | 0 | 0 | 0 | 0 | 0 | 0 |
| Aug5.5v_10746 | 1 | 0 | 0 | 0 | 0 | 0 | 0 | 0 | 0 | 0 | 0 | 0 | 0 | 0 | 0 | 0 | 0 | 0 | 0 | 0 | 0 | 0 | 1 | 0 | 0 | 0 | 0 | 0 | 0 | 0 |
| May12.75v_26962 | 1 | 0 | 0 | 0 | 1 | 0 | 0 | 0 | 0 | 0 | 0 | 0 | 0 | 0 | 0 | 0 | 0 | 0 | 0 | 0 | 0 | 0 | 0 | 0 | 0 | 0 | 0 | 0 | 0 | 0 |
| May12.75v_37974 | 1 | 0 | 0 | 0 | 1 | 0 | 0 | 0 | 0 | 0 | 0 | 0 | 0 | 0 | 0 | 0 | 0 | 0 | 0 | 0 | 0 | 0 | 0 | 0 | 0 | 0 | 0 | 0 | 0 | 0 |
| Jul11.0v_583 | 1 | 0 | 0 | 0 | 0 | 0 | 0 | 0 | 0 | 0 | 0 | 0 | 1 | 0 | 0 | 0 | 0 | 0 | 0 | 0 | 0 | 0 | 0 | 0 | 0 | 0 | 0 | 0 | 0 | 0 |
| May12.75v_26613 | 1 | 0 | 0 | 0 | 1 | 0 | 0 | 0 | 0 | 0 | 0 | 0 | 0 | 0 | 0 | 0 | 0 | 0 | 0 | 0 | 0 | 0 | 0 | 0 | 0 | 0 | 0 | 0 | 0 | 0 |
| Jul11.5v_3195 | 1 | 0 | 0 | 0 | 0 | 0 | 0 | 0 | 0 | 0 | 0 | 0 | 0 | 1 | 0 | 0 | 0 | 0 | 0 | 0 | 0 | 0 | 0 | 0 | 0 | 0 | 0 | 0 | 0 | 0 |
| Aug5.25v_33983 | 1 | 0 | 0 | 0 | 0 | 0 | 0 | 0 | 0 | 0 | 0 | 0 | 0 | 0 | 0 | 0 | 0 | 0 | 0 | 0 | 0 | 0 | 0 | 1 | 0 | 0 | 0 | 0 | 0 | 0 |
| Jul11.0v_7815 | 1 | 0 | 0 | 0 | 0 | 0 | 0 | 0 | 0 | 0 | 0 | 0 | 1 | 0 | 0 | 0 | 0 | 0 | 0 | 0 | 0 | 0 | 0 | 0 | 0 | 0 | 0 | 0 | 0 | 0 |
| Jul27.5v_354 | 1 | 0 | 0 | 0 | 0 | 0 | 0 | 0 | 0 | 0 | 0 | 0 | 0 | 0 | 0 | 0 | 0 | 1 | 0 | 0 | 0 | 0 | 0 | 0 | 0 | 0 | 0 | 0 | 0 | 0 |
| May12.5v_13033 | 1 | 1 | 0 | 0 | 0 | 0 | 0 | 0 | 0 | 0 | 0 | 0 | 0 | 0 | 0 | 0 | 0 | 0 | 0 | 0 | 0 | 0 | 0 | 0 | 0 | 0 | 0 | 0 | 0 | 0 |
| Sep8.0v_14400 | 1 | 0 | 0 | 0 | 0 | 0 | 0 | 0 | 0 | 0 | 0 | 0 | 0 | 0 | 0 | 0 | 0 | 0 | 0 | 0 | 0 | 0 | 0 | 0 | 0 | 0 | 1 | 0 | 0 | 0 |
| Jul27.5v_8588 | 1 | 0 | 0 | 0 | 0 | 0 | 0 | 0 | 0 | 0 | 0 | 0 | 0 | 0 | 0 | 0 | 0 | 1 | 0 | 0 | 0 | 0 | 0 | 0 | 0 | 0 | 0 | 0 | 0 | 0 |
| Jun22.0v_19636 | 1 | 0 | 0 | 0 | 0 | 0 | 0 | 0 | 1 | 0 | 0 | 0 | 0 | 0 | 0 | 0 | 0 | 0 | 0 | 0 | 0 | 0 | 0 | 0 | 0 | 0 | 0 | 0 | 0 | 0 |
| Sep8.0v_4929 | 1 | 0 | 0 | 0 | 0 | 0 | 0 | 0 | 0 | 0 | 0 | 0 | 0 | 0 | 0 | 0 | 0 | 0 | 0 | 0 | 0 | 0 | 0 | 0 | 0 | 0 | 1 | 0 | 0 | 0 |
| Sep8.50v_28342 | 1 | 0 | 0 | 0 | 0 | 0 | 0 | 0 | 0 | 0 | 0 | 0 | 0 | 0 | 0 | 0 | 0 | 0 | 0 | 0 | 0 | 0 | 0 | 0 | 0 | 0 | 0 | 0 | 0 | 1 |
| Jun8.50v_17015 | 1 | 0 | 0 | 0 | 0 | 0 | 1 | 0 | 0 | 0 | 0 | 0 | 0 | 0 | 0 | 0 | 0 | 0 | 0 | 0 | 0 | 0 | 0 | 0 | 0 | 0 | 0 | 0 | 0 | 0 |
| Sep8.50v_9900 | 1 | 0 | 0 | 0 | 0 | 0 | 0 | 0 | 0 | 0 | 0 | 0 | 0 | 0 | 0 | 0 | 0 | 0 | 0 | 0 | 0 | 0 | 0 | 0 | 0 | 0 | 0 | 0 | 0 | 1 |
| Jun22.5v_10791 | 1 | 0 | 0 | 0 | 0 | 0 | 0 | 0 | 0 | 1 | 0 | 0 | 0 | 0 | 0 | 0 | 0 | 0 | 0 | 0 | 0 | 0 | 0 | 0 | 0 | 0 | 0 | 0 | 0 | 0 |
| Aug5.0v_5312 | 1 | 0 | 0 | 0 | 0 | 0 | 0 | 0 | 0 | 0 | 0 | 0 | 0 | 0 | 0 | 0 | 0 | 0 | 0 | 0 | 0 | 1 | 0 | 0 | 0 | 0 | 0 | 0 | 0 | 0 |
| Sep8.50v_25933 | 1 | 0 | 0 | 0 | 0 | 0 | 0 | 0 | 0 | 0 | 0 | 0 | 0 | 0 | 0 | 0 | 0 | 0 | 0 | 0 | 0 | 0 | 0 | 0 | 0 | 0 | 0 | 0 | 0 | 1 |
| Jul11.5v_11755 | 1 | 0 | 0 | 0 | 0 | 0 | 0 | 0 | 0 | 0 | 0 | 0 | 0 | 1 | 0 | 0 | 0 | 0 | 0 | 0 | 0 | 0 | 0 | 0 | 0 | 0 | 0 | 0 | 0 | 0 |
| May12.5v_13330 | 1 | 1 | 0 | 0 | 0 | 0 | 0 | 0 | 0 | 0 | 0 | 0 | 0 | 0 | 0 | 0 | 0 | 0 | 0 | 0 | 0 | 0 | 0 | 0 | 0 | 0 | 0 | 0 | 0 | 0 |
| Jul11.50v_4788 | 1 | 0 | 0 | 0 | 0 | 0 | 0 | 0 | 0 | 0 | 0 | 0 | 0 | 0 | 0 | 1 | 0 | 0 | 0 | 0 | 0 | 0 | 0 | 0 | 0 | 0 | 0 | 0 | 0 | 0 |
| May12.5v_31792 | 1 | 1 | 0 | 0 | 0 | 0 | 0 | 0 | 0 | 0 | 0 | 0 | 0 | 0 | 0 | 0 | 0 | 0 | 0 | 0 | 0 | 0 | 0 | 0 | 0 | 0 | 0 | 0 | 0 | 0 |
| Jun8.25v_21043 | 1 | 0 | 0 | 0 | 0 | 1 | 0 | 0 | 0 | 0 | 0 | 0 | 0 | 0 | 0 | 0 | 0 | 0 | 0 | 0 | 0 | 0 | 0 | 0 | 0 | 0 | 0 | 0 | 0 | 0 |
| May12.50v_24086 | 1 | 0 | 0 | 1 | 0 | 0 | 0 | 0 | 0 | 0 | 0 | 0 | 0 | 0 | 0 | 0 | 0 | 0 | 0 | 0 | 0 | 0 | 0 | 0 | 0 | 0 | 0 | 0 | 0 | 0 |
| May12.75v_17992 | 1 | 0 | 0 | 0 | 1 | 0 | 0 | 0 | 0 | 0 | 0 | 0 | 0 | 0 | 0 | 0 | 0 | 0 | 0 | 0 | 0 | 0 | 0 | 0 | 0 | 0 | 0 | 0 | 0 | 0 |
| Jul27.5v_6385 | 139 | 6 | 3 | 15 | 10 | 2 | 17 | 6 | 1 | 4 | 8 | 6 | 0 | 4 | 3 | 8 | 8 | 1 | 1 | 2 | 2 | 0 | 0 | 0 | 0 | 4 | 1 | 2 | 23 | 2 |
| May12.50v_29396 | 4 | 0 | 0 | 2 | 0 | 0 | 0 | 0 | 0 | 0 | 0 | 0 | 0 | 0 | 0 | 0 | 1 | 0 | 0 | 0 | 0 | 0 | 0 | 0 | 0 | 0 | 0 | 0 | 1 | 0 |
| May12.50v_18440 | 13 | 0 | 0 | 2 | 1 | 0 | 2 | 1 | 0 | 1 | 0 | 0 | 0 | 0 | 0 | 0 | 0 | 0 | 0 | 0 | 0 | 0 | 0 | 1 | 0 | 2 | 0 | 0 | 3 | 0 |
| Jun22.50v_25542 | 2671 | 96 | 117 | 231 | 381 | 75 | 308 | 115 | 15 | 37 | 232 | 249 | 9 | 22 | 49 | 122 | 101 | 14 | 28 | 83 | 21 | 21 | 10 | 20 | 22 | 46 | 5 | 45 | 172 | 25 |
| Jun22.75v_6874 | 136 | 3 | 6 | 7 | 13 | 2 | 11 | 11 | 3 | 3 | 7 | 14 | 0 | 5 | 4 | 10 | 3 | 1 | 1 | 0 | 3 | 0 | 0 | 2 | 0 | 2 | 1 | 4 | 18 | 2 |

| Feature_ID | 654 | May12_05m | May12_25m | May12_50m | May12_75m | Jun8_25m | Jun8_50m | Jun8_75m | Jun22_0m | Jun22_05m | Jun22_50m | Jun22_75m | Jul11_0m | Jul11_05m | Jul11_25m | Jul11_50m | Jul11_75m | Jul27_05m | Jul27_25m | Jul27_50m | Jul27_75m | Aug5_0m | Aug5_05m | Aug5_25m | Aug5_50m | Aug5_75m | Sep8_0m | Sep8_05m | Sep8_25m | Sep8_50m |
|---|---|---|---|---|---|---|---|---|---|---|---|---|---|---|---|---|---|---|---|---|---|---|---|---|---|---|---|---|---|---|
| Jun8.50v_16840 | 2 | 0 | 0 | 0 | 0 | 0 | 1 | 0 | 0 | 0 | 0 | 1 | 0 | 0 | 0 | 0 | 0 | 0 | 0 | 0 | 0 | 0 | 0 | 0 | 0 | 0 | 0 | 0 | 0 | 0 |
| Sep8.25v_13893 | 1 | 0 | 0 | 0 | 0 | 0 | 0 | 0 | 0 | 0 | 0 | 0 | 0 | 0 | 0 | 0 | 0 | 0 | 0 | 0 | 0 | 0 | 0 | 0 | 0 | 0 | 0 | 0 | 1 | 0 |
| Sep8.25v_18005 | 7 | 0 | 0 | 0 | 0 | 0 | 1 | 0 | 0 | 0 | 1 | 1 | 0 | 0 | 0 | 0 | 1 | 0 | 0 | 0 | 0 | 0 | 0 | 0 | 0 | 0 | 0 | 0 | 3 | 0 |
| Jul11.50v_14872 | 1 | 0 | 0 | 0 | 0 | 0 | 0 | 0 | 0 | 0 | 0 | 0 | 0 | 0 | 0 | 1 | 0 | 0 | 0 | 0 | 0 | 0 | 0 | 0 | 0 | 0 | 0 | 0 | 0 | 0 |
| Jun22.75v_30304 | 1 | 0 | 0 | 0 | 0 | 0 | 0 | 0 | 0 | 0 | 0 | 1 | 0 | 0 | 0 | 0 | 0 | 0 | 0 | 0 | 0 | 0 | 0 | 0 | 0 | 0 | 0 | 0 | 0 | 0 |
| Jun8.50v_34262 | 1 | 0 | 0 | 0 | 0 | 0 | 1 | 0 | 0 | 0 | 0 | 0 | 0 | 0 | 0 | 0 | 0 | 0 | 0 | 0 | 0 | 0 | 0 | 0 | 0 | 0 | 0 | 0 | 0 | 0 |
| May12.75v_6261 | 1 | 0 | 0 | 0 | 1 | 0 | 0 | 0 | 0 | 0 | 0 | 0 | 0 | 0 | 0 | 0 | 0 | 0 | 0 | 0 | 0 | 0 | 0 | 0 | 0 | 0 | 0 | 0 | 0 | 0 |
| Jun22.75v_7618 | 4 | 0 | 0 | 0 | 0 | 0 | 1 | 0 | 0 | 0 | 0 | 1 | 0 | 0 | 1 | 0 | 0 | 0 | 0 | 0 | 0 | 0 | 0 | 0 | 0 | 0 | 0 | 0 | 1 | 0 |
| May12.5v_33427 | 2 | 1 | 0 | 0 | 0 | 0 | 0 | 0 | 0 | 0 | 0 | 1 | 0 | 0 | 0 | 0 | 0 | 0 | 0 | 0 | 0 | 0 | 0 | 0 | 0 | 0 | 0 | 0 | 0 | 0 |
| May12.50v_517 | 14 | 0 | 1 | 5 | 2 | 0 | 2 | 0 | 0 | 0 | 3 | 0 | 0 | 0 | 0 | 0 | 1 | 0 | 0 | 0 | 0 | 0 | 0 | 0 | 0 | 0 | 0 | 0 | 0 | 0 |
| Jun8.25v_20851 | 65 | 3 | 2 | 3 | 11 | 3 | 4 | 1 | 0 | 0 | 5 | 12 | 0 | 3 | 0 | 2 | 1 | 0 | 2 | 3 | 0 | 0 | 1 | 0 | 0 | 2 | 1 | 1 | 1 | 4 |
| Jun22.50v_541 | 70 | 0 | 0 | 2 | 11 | 1 | 7 | 4 | 0 | 0 | 10 | 7 | 0 | 0 | 3 | 5 | 6 | 0 | 0 | 2 | 0 | 0 | 1 | 1 | 0 | 1 | 0 | 0 | 8 | 1 |
| Jul27.75v_16246 | 5 | 0 | 0 | 0 | 0 | 0 | 1 | 0 | 0 | 1 | 0 | 0 | 0 | 0 | 0 | 1 | 0 | 0 | 0 | 0 | 1 | 1 | 0 | 0 | 0 | 0 | 0 | 0 | 0 | 0 |
| Jun8.25v_22848 | 1 | 0 | 0 | 0 | 0 | 1 | 0 | 0 | 0 | 0 | 0 | 0 | 0 | 0 | 0 | 0 | 0 | 0 | 0 | 0 | 0 | 0 | 0 | 0 | 0 | 0 | 0 | 0 | 0 | 0 |
| Jul27.75v_9932 | 2 | 0 | 0 | 0 | 0 | 0 | 0 | 0 | 0 | 0 | 0 | 0 | 0 | 0 | 0 | 0 | 0 | 0 | 0 | 0 | 1 | 0 | 0 | 0 | 0 | 0 | 0 | 0 | 1 | 0 |
| Jun22.50v_9359 | 1 | 0 | 0 | 0 | 0 | 0 | 0 | 0 | 0 | 0 | 1 | 0 | 0 | 0 | 0 | 0 | 0 | 0 | 0 | 0 | 0 | 0 | 0 | 0 | 0 | 0 | 0 | 0 | 0 | 0 |
| Jul11.50v_17524 | 1 | 0 | 0 | 0 | 0 | 0 | 0 | 0 | 0 | 0 | 0 | 0 | 0 | 0 | 0 | 1 | 0 | 0 | 0 | 0 | 0 | 0 | 0 | 0 | 0 | 0 | 0 | 0 | 0 | 0 |
| Jun22.50v_4319 | 1 | 0 | 0 | 0 | 0 | 0 | 0 | 0 | 0 | 0 | 1 | 0 | 0 | 0 | 0 | 0 | 0 | 0 | 0 | 0 | 0 | 0 | 0 | 0 | 0 | 0 | 0 | 0 | 0 | 0 |
| Jul11.75v_10703 | 1 | 0 | 0 | 0 | 0 | 0 | 0 | 0 | 0 | 0 | 0 | 0 | 0 | 0 | 0 | 0 | 1 | 0 | 0 | 0 | 0 | 0 | 0 | 0 | 0 | 0 | 0 | 0 | 0 | 0 |
| May12.75v_33699 | 1 | 0 | 0 | 0 | 1 | 0 | 0 | 0 | 0 | 0 | 0 | 0 | 0 | 0 | 0 | 0 | 0 | 0 | 0 | 0 | 0 | 0 | 0 | 0 | 0 | 0 | 0 | 0 | 0 | 0 |
| Sep8.25v_20096 | 1 | 0 | 0 | 0 | 0 | 0 | 0 | 0 | 0 | 0 | 0 | 0 | 0 | 0 | 0 | 0 | 0 | 0 | 0 | 0 | 0 | 0 | 0 | 0 | 0 | 0 | 0 | 0 | 1 | 0 |
| Jun8.50v_25735 | 1 | 0 | 0 | 0 | 0 | 0 | 1 | 0 | 0 | 0 | 0 | 0 | 0 | 0 | 0 | 0 | 0 | 0 | 0 | 0 | 0 | 0 | 0 | 0 | 0 | 0 | 0 | 0 | 0 | 0 |
| Jun8.50v_18944 | 1 | 0 | 0 | 0 | 0 | 0 | 1 | 0 | 0 | 0 | 0 | 0 | 0 | 0 | 0 | 0 | 0 | 0 | 0 | 0 | 0 | 0 | 0 | 0 | 0 | 0 | 0 | 0 | 0 | 0 |
| Sep8.5v_8909 | 1 | 0 | 0 | 0 | 0 | 0 | 0 | 0 | 0 | 0 | 0 | 0 | 0 | 0 | 0 | 0 | 0 | 0 | 0 | 0 | 0 | 0 | 0 | 0 | 0 | 0 | 0 | 1 | 0 | 0 |
| Jul11.50v_5203 | 1 | 0 | 0 | 0 | 0 | 0 | 0 | 0 | 0 | 0 | 0 | 0 | 0 | 0 | 0 | 1 | 0 | 0 | 0 | 0 | 0 | 0 | 0 | 0 | 0 | 0 | 0 | 0 | 0 | 0 |
| May12.5v_11768 | 1 | 1 | 0 | 0 | 0 | 0 | 0 | 0 | 0 | 0 | 0 | 0 | 0 | 0 | 0 | 0 | 0 | 0 | 0 | 0 | 0 | 0 | 0 | 0 | 0 | 0 | 0 | 0 | 0 | 0 |
| Jun22.75v_19953 | 1 | 0 | 0 | 0 | 0 | 0 | 0 | 0 | 0 | 0 | 0 | 1 | 0 | 0 | 0 | 0 | 0 | 0 | 0 | 0 | 0 | 0 | 0 | 0 | 0 | 0 | 0 | 0 | 0 | 0 |
| Sep8.25v_5364 | 1 | 0 | 0 | 0 | 0 | 0 | 0 | 0 | 0 | 0 | 0 | 0 | 0 | 0 | 0 | 0 | 0 | 0 | 0 | 0 | 0 | 0 | 0 | 0 | 0 | 0 | 0 | 0 | 1 | 0 |
| Jun8.25v_27165 | 1 | 0 | 0 | 0 | 0 | 1 | 0 | 0 | 0 | 0 | 0 | 0 | 0 | 0 | 0 | 0 | 0 | 0 | 0 | 0 | 0 | 0 | 0 | 0 | 0 | 0 | 0 | 0 | 0 | 0 |
| Jun22.75v_20573 | 8 | 0 | 1 | 1 | 0 | 0 | 0 | 1 | 0 | 0 | 0 | 1 | 0 | 1 | 0 | 2 | 0 | 0 | 1 | 0 | 0 | 0 | 0 | 0 | 0 | 0 | 0 | 0 | 0 | 0 |
| Jun8.50v_17840 | 1 | 0 | 0 | 0 | 0 | 0 | 1 | 0 | 0 | 0 | 0 | 0 | 0 | 0 | 0 | 0 | 0 | 0 | 0 | 0 | 0 | 0 | 0 | 0 | 0 | 0 | 0 | 0 | 0 | 0 |
| Sep8.5v_5247 | 1 | 0 | 0 | 0 | 0 | 0 | 0 | 0 | 0 | 0 | 0 | 0 | 0 | 0 | 0 | 0 | 0 | 0 | 0 | 0 | 0 | 0 | 0 | 0 | 0 | 0 | 0 | 1 | 0 | 0 |
| Sep8.25v_9814 | 2 | 0 | 0 | 0 | 0 | 0 | 0 | 0 | 0 | 0 | 0 | 0 | 0 | 0 | 0 | 0 | 0 | 0 | 0 | 0 | 0 | 0 | 0 | 0 | 0 | 0 | 0 | 0 | 2 | 0 |
| Jun22.5v_10198 | 1 | 0 | 0 | 0 | 0 | 0 | 0 | 0 | 0 | 1 | 0 | 0 | 0 | 0 | 0 | 0 | 0 | 0 | 0 | 0 | 0 | 0 | 0 | 0 | 0 | 0 | 0 | 0 | 0 | 0 |
| Sep8.25v_9026 | 1 | 0 | 0 | 0 | 0 | 0 | 0 | 0 | 0 | 0 | 0 | 0 | 0 | 0 | 0 | 0 | 0 | 0 | 0 | 0 | 0 | 0 | 0 | 0 | 0 | 0 | 0 | 0 | 1 | 0 |
| Aug5.50v_26854 | 1 | 0 | 0 | 0 | 0 | 0 | 0 | 0 | 0 | 0 | 0 | 0 | 0 | 0 | 0 | 0 | 0 | 0 | 0 | 0 | 0 | 0 | 0 | 0 | 1 | 0 | 0 | 0 | 0 | 0 |
| Jul11.50v_5185 | 2 | 0 | 0 | 0 | 0 | 0 | 0 | 0 | 0 | 0 | 0 | 1 | 0 | 0 | 0 | 1 | 0 | 0 | 0 | 0 | 0 | 0 | 0 | 0 | 0 | 0 | 0 | 0 | 0 | 0 |
| Sep8.0v_1177 | 1 | 0 | 0 | 0 | 0 | 0 | 0 | 0 | 0 | 0 | 0 | 0 | 0 | 0 | 0 | 0 | 0 | 0 | 0 | 0 | 0 | 0 | 0 | 0 | 0 | 0 | 1 | 0 | 0 | 0 |
| May12.50v_8251 | 1 | 0 | 0 | 1 | 0 | 0 | 0 | 0 | 0 | 0 | 0 | 0 | 0 | 0 | 0 | 0 | 0 | 0 | 0 | 0 | 0 | 0 | 0 | 0 | 0 | 0 | 0 | 0 | 0 | 0 |
| May12.50v_19565 | 1 | 0 | 0 | 1 | 0 | 0 | 0 | 0 | 0 | 0 | 0 | 0 | 0 | 0 | 0 | 0 | 0 | 0 | 0 | 0 | 0 | 0 | 0 | 0 | 0 | 0 | 0 | 0 | 0 | 0 |
| May12.50v_13050 | 3 | 0 | 0 | 1 | 0 | 0 | 0 | 0 | 0 | 0 | 0 | 1 | 0 | 0 | 0 | 0 | 0 | 0 | 0 | 0 | 0 | 0 | 0 | 0 | 0 | 0 | 0 | 0 | 1 | 0 |
| Jun22.75v_9284 | 1 | 0 | 0 | 0 | 0 | 0 | 0 | 0 | 0 | 0 | 0 | 1 | 0 | 0 | 0 | 0 | 0 | 0 | 0 | 0 | 0 | 0 | 0 | 0 | 0 | 0 | 0 | 0 | 0 | 0 |
| Jun22.75v_35587 | 1 | 0 | 0 | 0 | 0 | 0 | 0 | 0 | 0 | 0 | 0 | 1 | 0 | 0 | 0 | 0 | 0 | 0 | 0 | 0 | 0 | 0 | 0 | 0 | 0 | 0 | 0 | 0 | 0 | 0 |
| Sep8.25v_5813 | 3 | 0 | 1 | 0 | 0 | 0 | 0 | 0 | 0 | 0 | 0 | 0 | 0 | 0 | 0 | 0 | 1 | 0 | 0 | 0 | 0 | 0 | 0 | 0 | 0 | 0 | 0 | 0 | 1 | 0 |
| May12.25v_6517 | 1 | 0 | 1 | 0 | 0 | 0 | 0 | 0 | 0 | 0 | 0 | 0 | 0 | 0 | 0 | 0 | 0 | 0 | 0 | 0 | 0 | 0 | 0 | 0 | 0 | 0 | 0 | 0 | 0 | 0 |
| Jul11.50v_11498 | 1 | 0 | 0 | 0 | 0 | 0 | 0 | 0 | 0 | 0 | 0 | 0 | 0 | 0 | 0 | 1 | 0 | 0 | 0 | 0 | 0 | 0 | 0 | 0 | 0 | 0 | 0 | 0 | 0 | 0 |
| Jun8.75v_18911 | 1 | 0 | 0 | 0 | 0 | 0 | 0 | 1 | 0 | 0 | 0 | 0 | 0 | 0 | 0 | 0 | 0 | 0 | 0 | 0 | 0 | 0 | 0 | 0 | 0 | 0 | 0 | 0 | 0 | 0 |
| Jul27.5v_5955 | 1 | 0 | 0 | 0 | 0 | 0 | 0 | 0 | 0 | 0 | 0 | 0 | 0 | 0 | 0 | 0 | 0 | 1 | 0 | 0 | 0 | 0 | 0 | 0 | 0 | 0 | 0 | 0 | 0 | 0 |
| Jul11.75v_23511 | 1 | 0 | 0 | 0 | 0 | 0 | 0 | 0 | 0 | 0 | 0 | 0 | 0 | 0 | 0 | 0 | 1 | 0 | 0 | 0 | 0 | 0 | 0 | 0 | 0 | 0 | 0 | 0 | 0 | 0 |
| Jul11.25v_8380 | 1 | 0 | 0 | 0 | 0 | 0 | 0 | 0 | 0 | 0 | 0 | 0 | 0 | 0 | 1 | 0 | 0 | 0 | 0 | 0 | 0 | 0 | 0 | 0 | 0 | 0 | 0 | 0 | 0 | 0 |
| May12.75v_20042 | 6 | 1 | 0 | 0 | 2 | 0 | 0 | 0 | 0 | 0 | 2 | 0 | 0 | 0 | 0 | 0 | 0 | 0 | 0 | 1 | 0 | 0 | 0 | 0 | 0 | 0 | 0 | 0 | 0 | 0 |
| May12.50v_16267 | 1 | 0 | 0 | 1 | 0 | 0 | 0 | 0 | 0 | 0 | 0 | 0 | 0 | 0 | 0 | 0 | 0 | 0 | 0 | 0 | 0 | 0 | 0 | 0 | 0 | 0 | 0 | 0 | 0 | 0 |
| May12.50v_40642 | 1 | 0 | 0 | 1 | 0 | 0 | 0 | 0 | 0 | 0 | 0 | 0 | 0 | 0 | 0 | 0 | 0 | 0 | 0 | 0 | 0 | 0 | 0 | 0 | 0 | 0 | 0 | 0 | 0 | 0 |
| Jun8.25v_22631 | 1 | 0 | 0 | 0 | 0 | 1 | 0 | 0 | 0 | 0 | 0 | 0 | 0 | 0 | 0 | 0 | 0 | 0 | 0 | 0 | 0 | 0 | 0 | 0 | 0 | 0 | 0 | 0 | 0 | 0 |
| Jun8.50v_22877 | 1 | 0 | 0 | 0 | 0 | 0 | 1 | 0 | 0 | 0 | 0 | 0 | 0 | 0 | 0 | 0 | 0 | 0 | 0 | 0 | 0 | 0 | 0 | 0 | 0 | 0 | 0 | 0 | 0 | 0 |
| Jun22.50v_31296 | 1 | 0 | 0 | 0 | 0 | 0 | 0 | 0 | 0 | 0 | 1 | 0 | 0 | 0 | 0 | 0 | 0 | 0 | 0 | 0 | 0 | 0 | 0 | 0 | 0 | 0 | 0 | 0 | 0 | 0 |
| Jun22.75v_35672 | 1 | 0 | 0 | 0 | 0 | 0 | 0 | 0 | 0 | 0 | 0 | 1 | 0 | 0 | 0 | 0 | 0 | 0 | 0 | 0 | 0 | 0 | 0 | 0 | 0 | 0 | 0 | 0 | 0 | 0 |
| Jul11.5v_18748 | 1 | 0 | 0 | 0 | 0 | 0 | 0 | 0 | 0 | 0 | 0 | 0 | 0 | 1 | 0 | 0 | 0 | 0 | 0 | 0 | 0 | 0 | 0 | 0 | 0 | 0 | 0 | 0 | 0 | 0 |
| Sep8.25v_9252 | 1 | 0 | 0 | 0 | 0 | 0 | 0 | 0 | 0 | 0 | 0 | 0 | 0 | 0 | 0 | 0 | 0 | 0 | 0 | 0 | 0 | 0 | 0 | 0 | 0 | 0 | 0 | 0 | 1 | 0 |
| May12.50v_254 | 1 | 0 | 0 | 1 | 0 | 0 | 0 | 0 | 0 | 0 | 0 | 0 | 0 | 0 | 0 | 0 | 0 | 0 | 0 | 0 | 0 | 0 | 0 | 0 | 0 | 0 | 0 | 0 | 0 | 0 |
| Jul11.75v_20708 | 1 | 0 | 0 | 0 | 0 | 0 | 0 | 0 | 0 | 0 | 0 | 0 | 0 | 0 | 0 | 0 | 1 | 0 | 0 | 0 | 0 | 0 | 0 | 0 | 0 | 0 | 0 | 0 | 0 | 0 |
| Jul11.50v_20336 | 1 | 0 | 0 | 0 | 0 | 0 | 0 | 0 | 0 | 0 | 0 | 0 | 0 | 0 | 0 | 1 | 0 | 0 | 0 | 0 | 0 | 0 | 0 | 0 | 0 | 0 | 0 | 0 | 0 | 0 |
| Jun8.25v_2122 | 1 | 0 | 0 | 0 | 0 | 1 | 0 | 0 | 0 | 0 | 0 | 0 | 0 | 0 | 0 | 0 | 0 | 0 | 0 | 0 | 0 | 0 | 0 | 0 | 0 | 0 | 0 | 0 | 0 | 0 |
| Jul11.5v_11935 | 6 | 0 | 0 | 1 | 0 | 0 | 0 | 0 | 0 | 0 | 0 | 0 | 0 | 1 | 0 | 0 | 1 | 0 | 1 | 0 | 0 | 0 | 0 | 0 | 0 | 0 | 0 | 0 | 2 | 0 |
| Jun22.75v_11180 | 1 | 0 | 0 | 0 | 0 | 0 | 0 | 0 | 0 | 0 | 0 | 1 | 0 | 0 | 0 | 0 | 0 | 0 | 0 | 0 | 0 | 0 | 0 | 0 | 0 | 0 | 0 | 0 | 0 | 0 |
| Jul11.75v_32550 | 1 | 0 | 0 | 0 | 0 | 0 | 0 | 0 | 0 | 0 | 0 | 0 | 0 | 0 | 0 | 0 | 1 | 0 | 0 | 0 | 0 | 0 | 0 | 0 | 0 | 0 | 0 | 0 | 0 | 0 |
| Jul11.5v_14058 | 2 | 0 | 0 | 0 | 0 | 0 | 1 | 0 | 0 | 0 | 0 | 0 | 0 | 1 | 0 | 0 | 0 | 0 | 0 | 0 | 0 | 0 | 0 | 0 | 0 | 0 | 0 | 0 | 0 | 0 |
| Jul27.50v_10128 | 3 | 0 | 0 | 1 | 0 | 0 | 0 | 0 | 0 | 0 | 0 | 0 | 0 | 0 | 0 | 0 | 0 | 0 | 0 | 1 | 0 | 0 | 0 | 0 | 0 | 0 | 0 | 0 | 1 | 0 |
| May12.25v_30513 | 1 | 0 | 1 | 0 | 0 | 0 | 0 | 0 | 0 | 0 | 0 | 0 | 0 | 0 | 0 | 0 | 0 | 0 | 0 | 0 | 0 | 0 | 0 | 0 | 0 | 0 | 0 | 0 | 0 | 0 |
| Jun22.75v_12312 | 1 | 0 | 0 | 0 | 0 | 0 | 0 | 0 | 0 | 0 | 0 | 1 | 0 | 0 | 0 | 0 | 0 | 0 | 0 | 0 | 0 | 0 | 0 | 0 | 0 | 0 | 0 | 0 | 0 | 0 |
| Jun22.75v_33405 | 1 | 0 | 0 | 0 | 0 | 0 | 0 | 0 | 0 | 0 | 0 | 1 | 0 | 0 | 0 | 0 | 0 | 0 | 0 | 0 | 0 | 0 | 0 | 0 | 0 | 0 | 0 | 0 | 0 | 0 |
| May12.75v_22671 | 1 | 0 | 0 | 0 | 1 | 0 | 0 | 0 | 0 | 0 | 0 | 0 | 0 | 0 | 0 | 0 | 0 | 0 | 0 | 0 | 0 | 0 | 0 | 0 | 0 | 0 | 0 | 0 | 0 | 0 |
| May12.50v_2606 | 1 | 0 | 0 | 1 | 0 | 0 | 0 | 0 | 0 | 0 | 0 | 0 | 0 | 0 | 0 | 0 | 0 | 0 | 0 | 0 | 0 | 0 | 0 | 0 | 0 | 0 | 0 | 0 | 0 | 0 |
| Jun8.50v_29977 | 1 | 0 | 0 | 0 | 0 | 0 | 1 | 0 | 0 | 0 | 0 | 0 | 0 | 0 | 0 | 0 | 0 | 0 | 0 | 0 | 0 | 0 | 0 | 0 | 0 | 0 | 0 | 0 | 0 | 0 |
| Jun22.50v_35476 | 1 | 0 | 0 | 0 | 0 | 0 | 0 | 0 | 0 | 0 | 1 | 0 | 0 | 0 | 0 | 0 | 0 | 0 | 0 | 0 | 0 | 0 | 0 | 0 | 0 | 0 | 0 | 0 | 0 | 0 |
| Jul27.50v_27881 | 5 | 0 | 2 | 0 | 0 | 0 | 0 | 0 | 0 | 0 | 0 | 1 | 0 | 0 | 0 | 0 | 0 | 0 | 0 | 1 | 0 | 0 | 0 | 0 | 0 | 0 | 0 | 0 | 1 | 0 |
| Jul11.5v_14940 | 1 | 0 | 0 | 0 | 0 | 0 | 0 | 0 | 0 | 0 | 0 | 0 | 0 | 1 | 0 | 0 | 0 | 0 | 0 | 0 | 0 | 0 | 0 | 0 | 0 | 0 | 0 | 0 | 0 | 0 |
| Jun22.75v_14341 | 2 | 0 | 0 | 0 | 0 | 0 | 0 | 0 | 0 | 0 | 0 | 1 | 0 | 0 | 0 | 0 | 0 | 0 | 0 | 0 | 0 | 0 | 0 | 0 | 0 | 0 | 0 | 0 | 1 | 0 |
| May12.50v_8118 | 1 | 0 | 0 | 1 | 0 | 0 | 0 | 0 | 0 | 0 | 0 | 0 | 0 | 0 | 0 | 0 | 0 | 0 | 0 | 0 | 0 | 0 | 0 | 0 | 0 | 0 | 0 | 0 | 0 | 0 |
| Sep8.25v_3777 | 1 | 0 | 0 | 0 | 0 | 0 | 0 | 0 | 0 | 0 | 0 | 0 | 0 | 0 | 0 | 0 | 0 | 0 | 0 | 0 | 0 | 0 | 0 | 0 | 0 | 0 | 0 | 0 | 1 | 0 |
| Aug5.75v_3732 | 1 | 0 | 0 | 0 | 0 | 0 | 0 | 0 | 0 | 0 | 0 | 0 | 0 | 0 | 0 | 0 | 0 | 0 | 0 | 0 | 0 | 0 | 0 | 0 | 0 | 1 | 0 | 0 | 0 | 0 |
| May12.50v_39093 | 1 | 0 | 0 | 1 | 0 | 0 | 0 | 0 | 0 | 0 | 0 | 0 | 0 | 0 | 0 | 0 | 0 | 0 | 0 | 0 | 0 | 0 | 0 | 0 | 0 | 0 | 0 | 0 | 0 | 0 |
| Sep8.50v_36039 | 1 | 0 | 0 | 0 | 0 | 0 | 0 | 0 | 0 | 0 | 0 | 0 | 0 | 0 | 0 | 0 | 0 | 0 | 0 | 0 | 0 | 0 | 0 | 0 | 0 | 0 | 0 | 0 | 0 | 1 |
| May12.75v_31874 | 1 | 0 | 0 | 0 | 1 | 0 | 0 | 0 | 0 | 0 | 0 | 0 | 0 | 0 | 0 | 0 | 0 | 0 | 0 | 0 | 0 | 0 | 0 | 0 | 0 | 0 | 0 | 0 | 0 | 0 |
| Jun8.50v_24242 | 1 | 0 | 0 | 0 | 0 | 0 | 1 | 0 | 0 | 0 | 0 | 0 | 0 | 0 | 0 | 0 | 0 | 0 | 0 | 0 | 0 | 0 | 0 | 0 | 0 | 0 | 0 | 0 | 0 | 0 |
| Jun8.50v_14260 | 1 | 0 | 0 | 0 | 0 | 0 | 1 | 0 | 0 | 0 | 0 | 0 | 0 | 0 | 0 | 0 | 0 | 0 | 0 | 0 | 0 | 0 | 0 | 0 | 0 | 0 | 0 | 0 | 0 | 0 |
| Jun8.25v_5246 | 1 | 0 | 0 | 0 | 0 | 1 | 0 | 0 | 0 | 0 | 0 | 0 | 0 | 0 | 0 | 0 | 0 | 0 | 0 | 0 | 0 | 0 | 0 | 0 | 0 | 0 | 0 | 0 | 0 | 0 |
| Jun8.25v_20772 | 1 | 0 | 0 | 0 | 0 | 1 | 0 | 0 | 0 | 0 | 0 | 0 | 0 | 0 | 0 | 0 | 0 | 0 | 0 | 0 | 0 | 0 | 0 | 0 | 0 | 0 | 0 | 0 | 0 | 0 |
| Jun22.75v_36577 | 1 | 0 | 0 | 0 | 0 | 0 | 0 | 0 | 0 | 0 | 0 | 1 | 0 | 0 | 0 | 0 | 0 | 0 | 0 | 0 | 0 | 0 | 0 | 0 | 0 | 0 | 0 | 0 | 0 | 0 |
| May12.75v_16591 | 1 | 0 | 0 | 0 | 1 | 0 | 0 | 0 | 0 | 0 | 0 | 0 | 0 | 0 | 0 | 0 | 0 | 0 | 0 | 0 | 0 | 0 | 0 | 0 | 0 | 0 | 0 | 0 | 0 | 0 |
| Sep8.25v_14972 | 8 | 1 | 0 | 1 | 2 | 0 | 0 | 0 | 0 | 0 | 0 | 1 | 0 | 0 | 1 | 0 | 0 | 0 | 0 | 0 | 0 | 0 | 0 | 0 | 0 | 0 | 0 | 0 | 2 | 0 |
| Jun8.50v_3611 | 1 | 0 | 0 | 0 | 0 | 0 | 1 | 0 | 0 | 0 | 0 | 0 | 0 | 0 | 0 | 0 | 0 | 0 | 0 | 0 | 0 | 0 | 0 | 0 | 0 | 0 | 0 | 0 | 0 | 0 |
| May12.75v_24812 | 1 | 0 | 0 | 0 | 1 | 0 | 0 | 0 | 0 | 0 | 0 | 0 | 0 | 0 | 0 | 0 | 0 | 0 | 0 | 0 | 0 | 0 | 0 | 0 | 0 | 0 | 0 | 0 | 0 | 0 |
| Sep8.25v_2068 | 1 | 0 | 0 | 0 | 0 | 0 | 0 | 0 | 0 | 0 | 0 | 0 | 0 | 0 | 0 | 0 | 0 | 0 | 0 | 0 | 0 | 0 | 0 | 0 | 0 | 0 | 0 | 0 | 1 | 0 |
| May12.50v_35328 | 1 | 0 | 0 | 1 | 0 | 0 | 0 | 0 | 0 | 0 | 0 | 0 | 0 | 0 | 0 | 0 | 0 | 0 | 0 | 0 | 0 | 0 | 0 | 0 | 0 | 0 | 0 | 0 | 0 | 0 |
| May12.75v_33261 | 1 | 0 | 0 | 0 | 1 | 0 | 0 | 0 | 0 | 0 | 0 | 0 | 0 | 0 | 0 | 0 | 0 | 0 | 0 | 0 | 0 | 0 | 0 | 0 | 0 | 0 | 0 | 0 | 0 | 0 |
| May12.75v_23095 | 1 | 0 | 0 | 0 | 1 | 0 | 0 | 0 | 0 | 0 | 0 | 0 | 0 | 0 | 0 | 0 | 0 | 0 | 0 | 0 | 0 | 0 | 0 | 0 | 0 | 0 | 0 | 0 | 0 | 0 |
| May12.50v_11431 | 1 | 0 | 0 | 1 | 0 | 0 | 0 | 0 | 0 | 0 | 0 | 0 | 0 | 0 | 0 | 0 | 0 | 0 | 0 | 0 | 0 | 0 | 0 | 0 | 0 | 0 | 0 | 0 | 0 | 0 |
| May12.50v_17053 | 1 | 0 | 0 | 1 | 0 | 0 | 0 | 0 | 0 | 0 | 0 | 0 | 0 | 0 | 0 | 0 | 0 | 0 | 0 | 0 | 0 | 0 | 0 | 0 | 0 | 0 | 0 | 0 | 0 | 0 |
| Jul11.50v_15915 | 6 | 0 | 0 | 2 | 1 | 0 | 1 | 0 | 0 | 1 | 0 | 0 | 0 | 0 | 0 | 1 | 0 | 0 | 0 | 0 | 0 | 0 | 0 | 0 | 0 | 0 | 0 | 0 | 0 | 0 |
| Jun8.50v_27214 | 2 | 0 | 0 | 1 | 0 | 0 | 1 | 0 | 0 | 0 | 0 | 0 | 0 | 0 | 0 | 0 | 0 | 0 | 0 | 0 | 0 | 0 | 0 | 0 | 0 | 0 | 0 | 0 | 0 | 0 |
| Jun8.50v_22104 | 1 | 0 | 0 | 0 | 0 | 0 | 1 | 0 | 0 | 0 | 0 | 0 | 0 | 0 | 0 | 0 | 0 | 0 | 0 | 0 | 0 | 0 | 0 | 0 | 0 | 0 | 0 | 0 | 0 | 0 |
| Jun8.50v_17308 | 1 | 0 | 0 | 0 | 0 | 0 | 1 | 0 | 0 | 0 | 0 | 0 | 0 | 0 | 0 | 0 | 0 | 0 | 0 | 0 | 0 | 0 | 0 | 0 | 0 | 0 | 0 | 0 | 0 | 0 |
| May12.50v_27103 | 1 | 0 | 0 | 1 | 0 | 0 | 0 | 0 | 0 | 0 | 0 | 0 | 0 | 0 | 0 | 0 | 0 | 0 | 0 | 0 | 0 | 0 | 0 | 0 | 0 | 0 | 0 | 0 | 0 | 0 |
| Jun22.75v_307 | 5 | 0 | 0 | 1 | 1 | 0 | 0 | 0 | 0 | 0 | 1 | 2 | 0 | 0 | 0 | 0 | 0 | 0 | 0 | 0 | 0 | 0 | 0 | 0 | 0 | 0 | 0 | 0 | 0 | 0 |
| May12.75v_16633 | 4 | 0 | 0 | 0 | 1 | 0 | 0 | 0 | 0 | 0 | 2 | 0 | 0 | 0 | 0 | 0 | 0 | 0 | 0 | 0 | 0 | 0 | 0 | 0 | 0 | 1 | 0 | 0 | 0 | 0 |
| May12.5v_26224 | 1 | 1 | 0 | 0 | 0 | 0 | 0 | 0 | 0 | 0 | 0 | 0 | 0 | 0 | 0 | 0 | 0 | 0 | 0 | 0 | 0 | 0 | 0 | 0 | 0 | 0 | 0 | 0 | 0 | 0 |
| Jun22.75v_40035 | 2 | 0 | 0 | 0 | 0 | 0 | 0 | 0 | 0 | 0 | 0 | 1 | 0 | 0 | 0 | 0 | 0 | 0 | 0 | 0 | 0 | 0 | 0 | 0 | 0 | 0 | 0 | 0 | 0 | 1 |
| Jun22.75v_8553 | 10 | 1 | 1 | 1 | 2 | 1 | 0 | 0 | 0 | 0 | 0 | 2 | 0 | 0 | 0 | 0 | 0 | 0 | 0 | 0 | 0 | 0 | 0 | 0 | 0 | 0 | 0 | 0 | 2 | 0 |
| Jun22.50v_16165 | 1 | 0 | 0 | 0 | 0 | 0 | 0 | 0 | 0 | 0 | 1 | 0 | 0 | 0 | 0 | 0 | 0 | 0 | 0 | 0 | 0 | 0 | 0 | 0 | 0 | 0 | 0 | 0 | 0 | 0 |
| May12.75v_38123 | 1 | 0 | 0 | 0 | 1 | 0 | 0 | 0 | 0 | 0 | 0 | 0 | 0 | 0 | 0 | 0 | 0 | 0 | 0 | 0 | 0 | 0 | 0 | 0 | 0 | 0 | 0 | 0 | 0 | 0 |
| May12.75v_18089 | 1 | 0 | 0 | 0 | 1 | 0 | 0 | 0 | 0 | 0 | 0 | 0 | 0 | 0 | 0 | 0 | 0 | 0 | 0 | 0 | 0 | 0 | 0 | 0 | 0 | 0 | 0 | 0 | 0 | 0 |
| Jul11.75v_38552 | 1 | 0 | 0 | 0 | 0 | 0 | 0 | 0 | 0 | 0 | 0 | 0 | 0 | 0 | 0 | 0 | 1 | 0 | 0 | 0 | 0 | 0 | 0 | 0 | 0 | 0 | 0 | 0 | 0 | 0 |
| Jun22.50v_15849 | 1 | 0 | 0 | 0 | 0 | 0 | 0 | 0 | 0 | 0 | 1 | 0 | 0 | 0 | 0 | 0 | 0 | 0 | 0 | 0 | 0 | 0 | 0 | 0 | 0 | 0 | 0 | 0 | 0 | 0 |
| Jul11.50v_12800 | 2 | 0 | 0 | 0 | 0 | 0 | 0 | 0 | 0 | 0 | 0 | 0 | 0 | 0 | 0 | 1 | 0 | 0 | 0 | 1 | 0 | 0 | 0 | 0 | 0 | 0 | 0 | 0 | 0 | 0 |
| Jun22.75v_11022 | 1 | 0 | 0 | 0 | 0 | 0 | 0 | 0 | 0 | 0 | 0 | 1 | 0 | 0 | 0 | 0 | 0 | 0 | 0 | 0 | 0 | 0 | 0 | 0 | 0 | 0 | 0 | 0 | 0 | 0 |
| Jul11.50v_10932 | 1 | 0 | 0 | 0 | 0 | 0 | 0 | 0 | 0 | 0 | 0 | 0 | 0 | 0 | 0 | 1 | 0 | 0 | 0 | 0 | 0 | 0 | 0 | 0 | 0 | 0 | 0 | 0 | 0 | 0 |
| May12.75v_32405 | 1 | 0 | 0 | 0 | 1 | 0 | 0 | 0 | 0 | 0 | 0 | 0 | 0 | 0 | 0 | 0 | 0 | 0 | 0 | 0 | 0 | 0 | 0 | 0 | 0 | 0 | 0 | 0 | 0 | 0 |
| May12.50v_30157 | 1 | 0 | 0 | 1 | 0 | 0 | 0 | 0 | 0 | 0 | 0 | 0 | 0 | 0 | 0 | 0 | 0 | 0 | 0 | 0 | 0 | 0 | 0 | 0 | 0 | 0 | 0 | 0 | 0 | 0 |
| May12.75v_10377 | 1 | 0 | 0 | 0 | 1 | 0 | 0 | 0 | 0 | 0 | 0 | 0 | 0 | 0 | 0 | 0 | 0 | 0 | 0 | 0 | 0 | 0 | 0 | 0 | 0 | 0 | 0 | 0 | 0 | 0 |
| May12.75v_23551 | 1 | 0 | 0 | 0 | 1 | 0 | 0 | 0 | 0 | 0 | 0 | 0 | 0 | 0 | 0 | 0 | 0 | 0 | 0 | 0 | 0 | 0 | 0 | 0 | 0 | 0 | 0 | 0 | 0 | 0 |
| May12.75v_15769 | 1 | 0 | 0 | 0 | 1 | 0 | 0 | 0 | 0 | 0 | 0 | 0 | 0 | 0 | 0 | 0 | 0 | 0 | 0 | 0 | 0 | 0 | 0 | 0 | 0 | 0 | 0 | 0 | 0 | 0 |
| Jun8.75v_3585 | 1 | 0 | 0 | 0 | 0 | 0 | 0 | 1 | 0 | 0 | 0 | 0 | 0 | 0 | 0 | 0 | 0 | 0 | 0 | 0 | 0 | 0 | 0 | 0 | 0 | 0 | 0 | 0 | 0 | 0 |
| Jun22.75v_34153 | 1 | 0 | 0 | 0 | 0 | 0 | 0 | 0 | 0 | 0 | 0 | 1 | 0 | 0 | 0 | 0 | 0 | 0 | 0 | 0 | 0 | 0 | 0 | 0 | 0 | 0 | 0 | 0 | 0 | 0 |
| May12.50v_23212 | 3 | 0 | 0 | 1 | 0 | 0 | 0 | 0 | 0 | 0 | 0 | 1 | 0 | 0 | 0 | 0 | 0 | 0 | 0 | 0 | 0 | 0 | 0 | 0 | 0 | 1 | 0 | 0 | 0 | 0 |
| May12.75v_29693 | 1 | 0 | 0 | 0 | 1 | 0 | 0 | 0 | 0 | 0 | 0 | 0 | 0 | 0 | 0 | 0 | 0 | 0 | 0 | 0 | 0 | 0 | 0 | 0 | 0 | 0 | 0 | 0 | 0 | 0 |
| Jun22.50v_18819 | 1 | 0 | 0 | 0 | 0 | 0 | 0 | 0 | 0 | 0 | 1 | 0 | 0 | 0 | 0 | 0 | 0 | 0 | 0 | 0 | 0 | 0 | 0 | 0 | 0 | 0 | 0 | 0 | 0 | 0 |
| Sep8.25v_13890 | 1 | 0 | 0 | 0 | 0 | 0 | 0 | 0 | 0 | 0 | 0 | 0 | 0 | 0 | 0 | 0 | 0 | 0 | 0 | 0 | 0 | 0 | 0 | 0 | 0 | 0 | 0 | 0 | 1 | 0 |
| Jun22.0v_4056 | 1 | 0 | 0 | 0 | 0 | 0 | 0 | 0 | 1 | 0 | 0 | 0 | 0 | 0 | 0 | 0 | 0 | 0 | 0 | 0 | 0 | 0 | 0 | 0 | 0 | 0 | 0 | 0 | 0 | 0 |
| Jul11.75v_7232 | 1 | 0 | 0 | 0 | 0 | 0 | 0 | 0 | 0 | 0 | 0 | 0 | 0 | 0 | 0 | 0 | 1 | 0 | 0 | 0 | 0 | 0 | 0 | 0 | 0 | 0 | 0 | 0 | 0 | 0 |
| May12.75v_40381 | 4 | 0 | 1 | 0 | 1 | 0 | 0 | 0 | 0 | 0 | 0 | 0 | 0 | 0 | 0 | 1 | 0 | 1 | 0 | 0 | 0 | 0 | 0 | 0 | 0 | 0 | 0 | 0 | 0 | 0 |
| May12.5v_8472 | 1 | 1 | 0 | 0 | 0 | 0 | 0 | 0 | 0 | 0 | 0 | 0 | 0 | 0 | 0 | 0 | 0 | 0 | 0 | 0 | 0 | 0 | 0 | 0 | 0 | 0 | 0 | 0 | 0 | 0 |
| Jul27.50v_6705 | 1 | 0 | 0 | 0 | 0 | 0 | 0 | 0 | 0 | 0 | 0 | 0 | 0 | 0 | 0 | 0 | 0 | 0 | 0 | 1 | 0 | 0 | 0 | 0 | 0 | 0 | 0 | 0 | 0 | 0 |
| Jun22.75v_37712 | 1 | 0 | 0 | 0 | 0 | 0 | 0 | 0 | 0 | 0 | 0 | 1 | 0 | 0 | 0 | 0 | 0 | 0 | 0 | 0 | 0 | 0 | 0 | 0 | 0 | 0 | 0 | 0 | 0 | 0 |
| May12.50v_36826 | 2 | 0 | 0 | 1 | 0 | 0 | 0 | 0 | 0 | 0 | 0 | 0 | 0 | 0 | 0 | 0 | 0 | 0 | 0 | 0 | 1 | 0 | 0 | 0 | 0 | 0 | 0 | 0 | 0 | 0 |
| Aug5.75v_26312 | 1 | 0 | 0 | 0 | 0 | 0 | 0 | 0 | 0 | 0 | 0 | 0 | 0 | 0 | 0 | 0 | 0 | 0 | 0 | 0 | 0 | 0 | 0 | 0 | 0 | 1 | 0 | 0 | 0 | 0 |
| Jun22.50v_40415 | 1 | 0 | 0 | 0 | 0 | 0 | 0 | 0 | 0 | 0 | 1 | 0 | 0 | 0 | 0 | 0 | 0 | 0 | 0 | 0 | 0 | 0 | 0 | 0 | 0 | 0 | 0 | 0 | 0 | 0 |
| Jun22.75v_35110 | 1 | 0 | 0 | 0 | 0 | 0 | 0 | 0 | 0 | 0 | 0 | 1 | 0 | 0 | 0 | 0 | 0 | 0 | 0 | 0 | 0 | 0 | 0 | 0 | 0 | 0 | 0 | 0 | 0 | 0 |
| Jun8.25v_36017 | 1 | 0 | 0 | 0 | 0 | 1 | 0 | 0 | 0 | 0 | 0 | 0 | 0 | 0 | 0 | 0 | 0 | 0 | 0 | 0 | 0 | 0 | 0 | 0 | 0 | 0 | 0 | 0 | 0 | 0 |
| Jun22.50v_9394 | 1 | 0 | 0 | 0 | 0 | 0 | 0 | 0 | 0 | 0 | 1 | 0 | 0 | 0 | 0 | 0 | 0 | 0 | 0 | 0 | 0 | 0 | 0 | 0 | 0 | 0 | 0 | 0 | 0 | 0 |
| May12.75v_32293 | 1 | 0 | 0 | 0 | 1 | 0 | 0 | 0 | 0 | 0 | 0 | 0 | 0 | 0 | 0 | 0 | 0 | 0 | 0 | 0 | 0 | 0 | 0 | 0 | 0 | 0 | 0 | 0 | 0 | 0 |
| Jun8.25v_23058 | 1 | 0 | 0 | 0 | 0 | 1 | 0 | 0 | 0 | 0 | 0 | 0 | 0 | 0 | 0 | 0 | 0 | 0 | 0 | 0 | 0 | 0 | 0 | 0 | 0 | 0 | 0 | 0 | 0 | 0 |
| Jul11.50v_15682 | 1 | 0 | 0 | 0 | 0 | 0 | 0 | 0 | 0 | 0 | 0 | 0 | 0 | 0 | 0 | 1 | 0 | 0 | 0 | 0 | 0 | 0 | 0 | 0 | 0 | 0 | 0 | 0 | 0 | 0 |
| Sep8.25v_11089 | 1 | 0 | 0 | 0 | 0 | 0 | 0 | 0 | 0 | 0 | 0 | 0 | 0 | 0 | 0 | 0 | 0 | 0 | 0 | 0 | 0 | 0 | 0 | 0 | 0 | 0 | 0 | 0 | 1 | 0 |
| Jul11.75v_743 | 1 | 0 | 0 | 0 | 0 | 0 | 0 | 0 | 0 | 0 | 0 | 0 | 0 | 0 | 0 | 0 | 1 | 0 | 0 | 0 | 0 | 0 | 0 | 0 | 0 | 0 | 0 | 0 | 0 | 0 |
| Jul27.50v_13551 | 1 | 0 | 0 | 0 | 0 | 0 | 0 | 0 | 0 | 0 | 0 | 0 | 0 | 0 | 0 | 0 | 0 | 0 | 0 | 1 | 0 | 0 | 0 | 0 | 0 | 0 | 0 | 0 | 0 | 0 |
| May12.75v_17754 | 1 | 0 | 0 | 0 | 1 | 0 | 0 | 0 | 0 | 0 | 0 | 0 | 0 | 0 | 0 | 0 | 0 | 0 | 0 | 0 | 0 | 0 | 0 | 0 | 0 | 0 | 0 | 0 | 0 | 0 |
| Jul11.50v_5842 | 1 | 0 | 0 | 0 | 0 | 0 | 0 | 0 | 0 | 0 | 0 | 0 | 0 | 0 | 0 | 1 | 0 | 0 | 0 | 0 | 0 | 0 | 0 | 0 | 0 | 0 | 0 | 0 | 0 | 0 |
| Jun8.50v_13238 | 1 | 0 | 0 | 0 | 0 | 0 | 1 | 0 | 0 | 0 | 0 | 0 | 0 | 0 | 0 | 0 | 0 | 0 | 0 | 0 | 0 | 0 | 0 | 0 | 0 | 0 | 0 | 0 | 0 | 0 |
| Sep8.25v_5311 | 1 | 0 | 0 | 0 | 0 | 0 | 0 | 0 | 0 | 0 | 0 | 0 | 0 | 0 | 0 | 0 | 0 | 0 | 0 | 0 | 0 | 0 | 0 | 0 | 0 | 0 | 0 | 0 | 1 | 0 |
| May12.75v_14086 | 1 | 0 | 0 | 0 | 1 | 0 | 0 | 0 | 0 | 0 | 0 | 0 | 0 | 0 | 0 | 0 | 0 | 0 | 0 | 0 | 0 | 0 | 0 | 0 | 0 | 0 | 0 | 0 | 0 | 0 |
| Jul11.50v_14411 | 1 | 0 | 0 | 0 | 0 | 0 | 0 | 0 | 0 | 0 | 0 | 0 | 0 | 0 | 0 | 1 | 0 | 0 | 0 | 0 | 0 | 0 | 0 | 0 | 0 | 0 | 0 | 0 | 0 | 0 |
| May12.75v_4068 | 1 | 0 | 0 | 0 | 1 | 0 | 0 | 0 | 0 | 0 | 0 | 0 | 0 | 0 | 0 | 0 | 0 | 0 | 0 | 0 | 0 | 0 | 0 | 0 | 0 | 0 | 0 | 0 | 0 | 0 |
| Sep8.25v_8224 | 1 | 0 | 0 | 0 | 0 | 0 | 0 | 0 | 0 | 0 | 0 | 0 | 0 | 0 | 0 | 0 | 0 | 0 | 0 | 0 | 0 | 0 | 0 | 0 | 0 | 0 | 0 | 0 | 1 | 0 |
| Jul11.50v_13741 | 1 | 0 | 0 | 0 | 0 | 0 | 0 | 0 | 0 | 0 | 0 | 0 | 0 | 0 | 0 | 1 | 0 | 0 | 0 | 0 | 0 | 0 | 0 | 0 | 0 | 0 | 0 | 0 | 0 | 0 |
| Jun8.75v_20610 | 1 | 0 | 0 | 0 | 0 | 0 | 0 | 1 | 0 | 0 | 0 | 0 | 0 | 0 | 0 | 0 | 0 | 0 | 0 | 0 | 0 | 0 | 0 | 0 | 0 | 0 | 0 | 0 | 0 | 0 |
| Jun22.50v_32202 | 1 | 0 | 0 | 0 | 0 | 0 | 0 | 0 | 0 | 0 | 1 | 0 | 0 | 0 | 0 | 0 | 0 | 0 | 0 | 0 | 0 | 0 | 0 | 0 | 0 | 0 | 0 | 0 | 0 | 0 |
| May12.50v_35624 | 1 | 0 | 0 | 1 | 0 | 0 | 0 | 0 | 0 | 0 | 0 | 0 | 0 | 0 | 0 | 0 | 0 | 0 | 0 | 0 | 0 | 0 | 0 | 0 | 0 | 0 | 0 | 0 | 0 | 0 |
| Jul11.50v_16447 | 1 | 0 | 0 | 0 | 0 | 0 | 0 | 0 | 0 | 0 | 0 | 0 | 0 | 0 | 0 | 1 | 0 | 0 | 0 | 0 | 0 | 0 | 0 | 0 | 0 | 0 | 0 | 0 | 0 | 0 |
| Jul11.50v_14281 | 1 | 0 | 0 | 0 | 0 | 0 | 0 | 0 | 0 | 0 | 0 | 0 | 0 | 0 | 0 | 1 | 0 | 0 | 0 | 0 | 0 | 0 | 0 | 0 | 0 | 0 | 0 | 0 | 0 | 0 |
| Jun8.75v_17411 | 1 | 0 | 0 | 0 | 0 | 0 | 0 | 1 | 0 | 0 | 0 | 0 | 0 | 0 | 0 | 0 | 0 | 0 | 0 | 0 | 0 | 0 | 0 | 0 | 0 | 0 | 0 | 0 | 0 | 0 |
| May12.50v_2422 | 6 | 0 | 0 | 1 | 0 | 0 | 2 | 0 | 0 | 0 | 0 | 0 | 0 | 0 | 0 | 1 | 1 | 0 | 0 | 0 | 1 | 0 | 0 | 0 | 0 | 0 | 0 | 0 | 0 | 0 |
| Jun22.75v_3974 | 231 | 7 | 20 | 3 | 25 | 11 | 24 | 3 | 4 | 4 | 12 | 31 | 4 | 4 | 2 | 3 | 16 | 4 | 9 | 4 | 1 | 8 | 10 | 4 | 2 | 7 | 1 | 3 | 0 | 5 |
| Jun8.50v_27870 | 1 | 0 | 0 | 0 | 0 | 0 | 1 | 0 | 0 | 0 | 0 | 0 | 0 | 0 | 0 | 0 | 0 | 0 | 0 | 0 | 0 | 0 | 0 | 0 | 0 | 0 | 0 | 0 | 0 | 0 |
| Jul11.25v_18199 | 1 | 0 | 0 | 0 | 0 | 0 | 0 | 0 | 0 | 0 | 0 | 0 | 0 | 0 | 1 | 0 | 0 | 0 | 0 | 0 | 0 | 0 | 0 | 0 | 0 | 0 | 0 | 0 | 0 | 0 |
| May12.25v_34244 | 1 | 0 | 1 | 0 | 0 | 0 | 0 | 0 | 0 | 0 | 0 | 0 | 0 | 0 | 0 | 0 | 0 | 0 | 0 | 0 | 0 | 0 | 0 | 0 | 0 | 0 | 0 | 0 | 0 | 0 |
| Aug5.75v_5078 | 1 | 0 | 0 | 0 | 0 | 0 | 0 | 0 | 0 | 0 | 0 | 0 | 0 | 0 | 0 | 0 | 0 | 0 | 0 | 0 | 0 | 0 | 0 | 0 | 0 | 1 | 0 | 0 | 0 | 0 |
| Jul11.25v_16748 | 1 | 0 | 0 | 0 | 0 | 0 | 0 | 0 | 0 | 0 | 0 | 0 | 0 | 0 | 1 | 0 | 0 | 0 | 0 | 0 | 0 | 0 | 0 | 0 | 0 | 0 | 0 | 0 | 0 | 0 |
| Jun8.50v_24602 | 1 | 0 | 0 | 0 | 0 | 0 | 1 | 0 | 0 | 0 | 0 | 0 | 0 | 0 | 0 | 0 | 0 | 0 | 0 | 0 | 0 | 0 | 0 | 0 | 0 | 0 | 0 | 0 | 0 | 0 |
| Jun8.25v_35951 | 1 | 0 | 0 | 0 | 0 | 1 | 0 | 0 | 0 | 0 | 0 | 0 | 0 | 0 | 0 | 0 | 0 | 0 | 0 | 0 | 0 | 0 | 0 | 0 | 0 | 0 | 0 | 0 | 0 | 0 |
| Jun8.50v_22775 | 1 | 0 | 0 | 0 | 0 | 0 | 1 | 0 | 0 | 0 | 0 | 0 | 0 | 0 | 0 | 0 | 0 | 0 | 0 | 0 | 0 | 0 | 0 | 0 | 0 | 0 | 0 | 0 | 0 | 0 |
| Jun22.0v_13049 | 1 | 0 | 0 | 0 | 0 | 0 | 0 | 0 | 1 | 0 | 0 | 0 | 0 | 0 | 0 | 0 | 0 | 0 | 0 | 0 | 0 | 0 | 0 | 0 | 0 | 0 | 0 | 0 | 0 | 0 |
| May12.75v_5955 | 1 | 0 | 0 | 0 | 1 | 0 | 0 | 0 | 0 | 0 | 0 | 0 | 0 | 0 | 0 | 0 | 0 | 0 | 0 | 0 | 0 | 0 | 0 | 0 | 0 | 0 | 0 | 0 | 0 | 0 |
| Jul11.5v_16457 | 32 | 0 | 1 | 0 | 3 | 0 | 5 | 5 | 0 | 1 | 3 | 0 | 0 | 3 | 1 | 2 | 2 | 0 | 1 | 0 | 1 | 0 | 0 | 0 | 0 | 0 | 0 | 4 | 0 | 0 |
| Jul11.0v_7805 | 12 | 0 | 0 | 0 | 0 | 1 | 4 | 4 | 0 | 0 | 0 | 0 | 1 | 1 | 0 | 0 | 0 | 0 | 0 | 0 | 0 | 0 | 0 | 0 | 0 | 0 | 0 | 1 | 0 | 0 |
| Jul27.5v_5862 | 1 | 0 | 0 | 0 | 0 | 0 | 0 | 0 | 0 | 0 | 0 | 0 | 0 | 0 | 0 | 0 | 0 | 1 | 0 | 0 | 0 | 0 | 0 | 0 | 0 | 0 | 0 | 0 | 0 | 0 |
| Jun8.75v_20644 | 1 | 0 | 0 | 0 | 0 | 0 | 0 | 1 | 0 | 0 | 0 | 0 | 0 | 0 | 0 | 0 | 0 | 0 | 0 | 0 | 0 | 0 | 0 | 0 | 0 | 0 | 0 | 0 | 0 | 0 |
| Sep8.50v_14560 | 1 | 0 | 0 | 0 | 0 | 0 | 0 | 0 | 0 | 0 | 0 | 0 | 0 | 0 | 0 | 0 | 0 | 0 | 0 | 0 | 0 | 0 | 0 | 0 | 0 | 0 | 0 | 0 | 0 | 1 |
| May12.75v_14700 | 1 | 0 | 0 | 0 | 1 | 0 | 0 | 0 | 0 | 0 | 0 | 0 | 0 | 0 | 0 | 0 | 0 | 0 | 0 | 0 | 0 | 0 | 0 | 0 | 0 | 0 | 0 | 0 | 0 | 0 |
| Sep8.25v_19340 | 1 | 0 | 0 | 0 | 0 | 0 | 0 | 0 | 0 | 0 | 0 | 0 | 0 | 0 | 0 | 0 | 0 | 0 | 0 | 0 | 0 | 0 | 0 | 0 | 0 | 0 | 0 | 0 | 1 | 0 |
| Jun22.50v_36324 | 1 | 0 | 0 | 0 | 0 | 0 | 0 | 0 | 0 | 0 | 1 | 0 | 0 | 0 | 0 | 0 | 0 | 0 | 0 | 0 | 0 | 0 | 0 | 0 | 0 | 0 | 0 | 0 | 0 | 0 |
| Aug5.25v_19585 | 1 | 0 | 0 | 0 | 0 | 0 | 0 | 0 | 0 | 0 | 0 | 0 | 0 | 0 | 0 | 0 | 0 | 0 | 0 | 0 | 0 | 0 | 0 | 1 | 0 | 0 | 0 | 0 | 0 | 0 |
| May12.75v_17207 | 1 | 0 | 0 | 0 | 1 | 0 | 0 | 0 | 0 | 0 | 0 | 0 | 0 | 0 | 0 | 0 | 0 | 0 | 0 | 0 | 0 | 0 | 0 | 0 | 0 | 0 | 0 | 0 | 0 | 0 |
| Jun22.50v_37442 | 1 | 0 | 0 | 0 | 0 | 0 | 0 | 0 | 0 | 0 | 1 | 0 | 0 | 0 | 0 | 0 | 0 | 0 | 0 | 0 | 0 | 0 | 0 | 0 | 0 | 0 | 0 | 0 | 0 | 0 |
| Jul27.50v_9795 | 2 | 0 | 0 | 1 | 0 | 0 | 0 | 0 | 0 | 0 | 0 | 0 | 0 | 0 | 0 | 0 | 0 | 0 | 0 | 1 | 0 | 0 | 0 | 0 | 0 | 0 | 0 | 0 | 0 | 0 |
| Sep8.25v_18163 | 1 | 0 | 0 | 0 | 0 | 0 | 0 | 0 | 0 | 0 | 0 | 0 | 0 | 0 | 0 | 0 | 0 | 0 | 0 | 0 | 0 | 0 | 0 | 0 | 0 | 0 | 0 | 0 | 1 | 0 |
| Sep8.50v_28215 | 1 | 0 | 0 | 0 | 0 | 0 | 0 | 0 | 0 | 0 | 0 | 0 | 0 | 0 | 0 | 0 | 0 | 0 | 0 | 0 | 0 | 0 | 0 | 0 | 0 | 0 | 0 | 0 | 0 | 1 |
| May12.50v_5150 | 1 | 0 | 0 | 1 | 0 | 0 | 0 | 0 | 0 | 0 | 0 | 0 | 0 | 0 | 0 | 0 | 0 | 0 | 0 | 0 | 0 | 0 | 0 | 0 | 0 | 0 | 0 | 0 | 0 | 0 |
| Jul11.75v_33942 | 1 | 0 | 0 | 0 | 0 | 0 | 0 | 0 | 0 | 0 | 0 | 0 | 0 | 0 | 0 | 0 | 1 | 0 | 0 | 0 | 0 | 0 | 0 | 0 | 0 | 0 | 0 | 0 | 0 | 0 |
| Jun22.75v_35948 | 1 | 0 | 0 | 0 | 0 | 0 | 0 | 0 | 0 | 0 | 0 | 1 | 0 | 0 | 0 | 0 | 0 | 0 | 0 | 0 | 0 | 0 | 0 | 0 | 0 | 0 | 0 | 0 | 0 | 0 |
| Jun22.75v_2986 | 1 | 0 | 0 | 0 | 0 | 0 | 0 | 0 | 0 | 0 | 0 | 1 | 0 | 0 | 0 | 0 | 0 | 0 | 0 | 0 | 0 | 0 | 0 | 0 | 0 | 0 | 0 | 0 | 0 | 0 |
| Jul11.25v_7261 | 1 | 0 | 0 | 0 | 0 | 0 | 0 | 0 | 0 | 0 | 0 | 0 | 0 | 0 | 1 | 0 | 0 | 0 | 0 | 0 | 0 | 0 | 0 | 0 | 0 | 0 | 0 | 0 | 0 | 0 |
| Jun8.75v_15662 | 1 | 0 | 0 | 0 | 0 | 0 | 0 | 1 | 0 | 0 | 0 | 0 | 0 | 0 | 0 | 0 | 0 | 0 | 0 | 0 | 0 | 0 | 0 | 0 | 0 | 0 | 0 | 0 | 0 | 0 |
| May12.25v_33105 | 1 | 0 | 1 | 0 | 0 | 0 | 0 | 0 | 0 | 0 | 0 | 0 | 0 | 0 | 0 | 0 | 0 | 0 | 0 | 0 | 0 | 0 | 0 | 0 | 0 | 0 | 0 | 0 | 0 | 0 |
| Jun8.50v_5512 | 1 | 0 | 0 | 0 | 0 | 0 | 1 | 0 | 0 | 0 | 0 | 0 | 0 | 0 | 0 | 0 | 0 | 0 | 0 | 0 | 0 | 0 | 0 | 0 | 0 | 0 | 0 | 0 | 0 | 0 |
| Aug5.75v_31614 | 1 | 0 | 0 | 0 | 0 | 0 | 0 | 0 | 0 | 0 | 0 | 0 | 0 | 0 | 0 | 0 | 0 | 0 | 0 | 0 | 0 | 0 | 0 | 0 | 0 | 1 | 0 | 0 | 0 | 0 |
| Jun8.50v_28076 | 4 | 0 | 0 | 0 | 1 | 0 | 1 | 0 | 0 | 0 | 0 | 1 | 0 | 0 | 1 | 0 | 0 | 0 | 0 | 0 | 0 | 0 | 0 | 0 | 0 | 0 | 0 | 0 | 0 | 0 |
| Jun22.50v_29480 | 1 | 0 | 0 | 0 | 0 | 0 | 0 | 0 | 0 | 0 | 1 | 0 | 0 | 0 | 0 | 0 | 0 | 0 | 0 | 0 | 0 | 0 | 0 | 0 | 0 | 0 | 0 | 0 | 0 | 0 |
| May12.50v_16362 | 3 | 0 | 1 | 1 | 0 | 0 | 0 | 0 | 0 | 0 | 0 | 0 | 0 | 0 | 0 | 0 | 0 | 0 | 0 | 0 | 0 | 0 | 0 | 0 | 0 | 1 | 0 | 0 | 0 | 0 |
| May12.50v_5278 | 1 | 0 | 0 | 1 | 0 | 0 | 0 | 0 | 0 | 0 | 0 | 0 | 0 | 0 | 0 | 0 | 0 | 0 | 0 | 0 | 0 | 0 | 0 | 0 | 0 | 0 | 0 | 0 | 0 | 0 |
| Jun8.50v_7115 | 1 | 0 | 0 | 0 | 0 | 0 | 1 | 0 | 0 | 0 | 0 | 0 | 0 | 0 | 0 | 0 | 0 | 0 | 0 | 0 | 0 | 0 | 0 | 0 | 0 | 0 | 0 | 0 | 0 | 0 |
| May12.25v_9922 | 1 | 0 | 1 | 0 | 0 | 0 | 0 | 0 | 0 | 0 | 0 | 0 | 0 | 0 | 0 | 0 | 0 | 0 | 0 | 0 | 0 | 0 | 0 | 0 | 0 | 0 | 0 | 0 | 0 | 0 |
| May12.75v_7263 | 1 | 0 | 0 | 0 | 1 | 0 | 0 | 0 | 0 | 0 | 0 | 0 | 0 | 0 | 0 | 0 | 0 | 0 | 0 | 0 | 0 | 0 | 0 | 0 | 0 | 0 | 0 | 0 | 0 | 0 |
| Jul11.25v_19468 | 1 | 0 | 0 | 0 | 0 | 0 | 0 | 0 | 0 | 0 | 0 | 0 | 0 | 0 | 1 | 0 | 0 | 0 | 0 | 0 | 0 | 0 | 0 | 0 | 0 | 0 | 0 | 0 | 0 | 0 |
| May12.75v_31989 | 1 | 0 | 0 | 0 | 1 | 0 | 0 | 0 | 0 | 0 | 0 | 0 | 0 | 0 | 0 | 0 | 0 | 0 | 0 | 0 | 0 | 0 | 0 | 0 | 0 | 0 | 0 | 0 | 0 | 0 |
| Aug5.25v_30934 | 1 | 0 | 0 | 0 | 0 | 0 | 0 | 0 | 0 | 0 | 0 | 0 | 0 | 0 | 0 | 0 | 0 | 0 | 0 | 0 | 0 | 0 | 0 | 1 | 0 | 0 | 0 | 0 | 0 | 0 |
| Jul27.75v_15687 | 1 | 0 | 0 | 0 | 0 | 0 | 0 | 0 | 0 | 0 | 0 | 0 | 0 | 0 | 0 | 0 | 0 | 0 | 0 | 0 | 1 | 0 | 0 | 0 | 0 | 0 | 0 | 0 | 0 | 0 |
| Sep8.50v_35096 | 1 | 0 | 0 | 0 | 0 | 0 | 0 | 0 | 0 | 0 | 0 | 0 | 0 | 0 | 0 | 0 | 0 | 0 | 0 | 0 | 0 | 0 | 0 | 0 | 0 | 0 | 0 | 0 | 0 | 1 |
| May12.25v_8993 | 1 | 0 | 1 | 0 | 0 | 0 | 0 | 0 | 0 | 0 | 0 | 0 | 0 | 0 | 0 | 0 | 0 | 0 | 0 | 0 | 0 | 0 | 0 | 0 | 0 | 0 | 0 | 0 | 0 | 0 |
| May12.50v_26155 | 3 | 0 | 0 | 1 | 2 | 0 | 0 | 0 | 0 | 0 | 0 | 0 | 0 | 0 | 0 | 0 | 0 | 0 | 0 | 0 | 0 | 0 | 0 | 0 | 0 | 0 | 0 | 0 | 0 | 0 |
| May12.50v_16743 | 1 | 0 | 0 | 1 | 0 | 0 | 0 | 0 | 0 | 0 | 0 | 0 | 0 | 0 | 0 | 0 | 0 | 0 | 0 | 0 | 0 | 0 | 0 | 0 | 0 | 0 | 0 | 0 | 0 | 0 |

| Feature_ID | 654 | May12_05m | May12_25m | May12_50m | May12_75m | Jun8_25m | Jun8_50m | Jun8_75m | Jun22_0m | Jun22_05m | Jun22_50m | Jun22_75m | Jul11_0m | Jul11_05m | Jul11_25m | Jul11_50m | Jul11_75m | Jul27_05m | Jul27_25m | Jul27_50m | Jul27_75m | Aug5_0m | Aug5_05m | Aug5_25m | Aug5_50m | Aug5_75m | Sep8_0m | Sep8_05m | Sep8_25m | Sep8_50m |
|---|---|---|---|---|---|---|---|---|---|---|---|---|---|---|---|---|---|---|---|---|---|---|---|---|---|---|---|---|---|---|
| Jul11.75v_31337 | 1 | 0 | 0 | 0 | 0 | 0 | 0 | 0 | 0 | 0 | 0 | 0 | 0 | 0 | 0 | 0 | 1 | 0 | 0 | 0 | 0 | 0 | 0 | 0 | 0 | 0 | 0 | 0 | 0 | 0 |
| May12.5v_16869 | 1 | 1 | 0 | 0 | 0 | 0 | 0 | 0 | 0 | 0 | 0 | 0 | 0 | 0 | 0 | 0 | 0 | 0 | 0 | 0 | 0 | 0 | 0 | 0 | 0 | 0 | 0 | 0 | 0 | 0 |
| Jun8.50v_28682 | 2 | 0 | 0 | 0 | 0 | 0 | 1 | 0 | 0 | 0 | 0 | 0 | 0 | 0 | 0 | 0 | 1 | 0 | 0 | 0 | 0 | 0 | 0 | 0 | 0 | 0 | 0 | 0 | 0 | 0 |
| Jun22.75v_35460 | 1 | 0 | 0 | 0 | 0 | 0 | 0 | 0 | 0 | 0 | 0 | 1 | 0 | 0 | 0 | 0 | 0 | 0 | 0 | 0 | 0 | 0 | 0 | 0 | 0 | 0 | 0 | 0 | 0 | 0 |
| Jun8.50v_14843 | 1 | 0 | 0 | 0 | 0 | 0 | 1 | 0 | 0 | 0 | 0 | 0 | 0 | 0 | 0 | 0 | 0 | 0 | 0 | 0 | 0 | 0 | 0 | 0 | 0 | 0 | 0 | 0 | 0 | 0 |
| Jun8.50v_8716 | 1 | 0 | 0 | 0 | 0 | 0 | 1 | 0 | 0 | 0 | 0 | 0 | 0 | 0 | 0 | 0 | 0 | 0 | 0 | 0 | 0 | 0 | 0 | 0 | 0 | 0 | 0 | 0 | 0 | 0 |
| Jun22.50v_40701 | 1 | 0 | 0 | 0 | 0 | 0 | 0 | 0 | 0 | 0 | 1 | 0 | 0 | 0 | 0 | 0 | 0 | 0 | 0 | 0 | 0 | 0 | 0 | 0 | 0 | 0 | 0 | 0 | 0 | 0 |
| May12.75v_13877 | 1 | 0 | 0 | 0 | 1 | 0 | 0 | 0 | 0 | 0 | 0 | 0 | 0 | 0 | 0 | 0 | 0 | 0 | 0 | 0 | 0 | 0 | 0 | 0 | 0 | 0 | 0 | 0 | 0 | 0 |
| Aug5.0v_570 | 1 | 0 | 0 | 0 | 0 | 0 | 0 | 0 | 0 | 0 | 0 | 0 | 0 | 0 | 0 | 0 | 0 | 0 | 0 | 0 | 0 | 1 | 0 | 0 | 0 | 0 | 0 | 0 | 0 | 0 |
| Jun8.50v_1347 | 1 | 0 | 0 | 0 | 0 | 0 | 1 | 0 | 0 | 0 | 0 | 0 | 0 | 0 | 0 | 0 | 0 | 0 | 0 | 0 | 0 | 0 | 0 | 0 | 0 | 0 | 0 | 0 | 0 | 0 |
| May12.75v_8282 | 5 | 0 | 1 | 0 | 2 | 0 | 1 | 0 | 0 | 0 | 0 | 0 | 0 | 0 | 0 | 0 | 0 | 0 | 0 | 0 | 0 | 0 | 0 | 0 | 0 | 0 | 0 | 0 | 0 | 1 |
| May12.75v_29124 | 1 | 0 | 0 | 0 | 1 | 0 | 0 | 0 | 0 | 0 | 0 | 0 | 0 | 0 | 0 | 0 | 0 | 0 | 0 | 0 | 0 | 0 | 0 | 0 | 0 | 0 | 0 | 0 | 0 | 0 |
| May12.75v_29360 | 1 | 0 | 0 | 0 | 1 | 0 | 0 | 0 | 0 | 0 | 0 | 0 | 0 | 0 | 0 | 0 | 0 | 0 | 0 | 0 | 0 | 0 | 0 | 0 | 0 | 0 | 0 | 0 | 0 | 0 |
| Jun8.50v_28675 | 1 | 0 | 0 | 0 | 0 | 0 | 1 | 0 | 0 | 0 | 0 | 0 | 0 | 0 | 0 | 0 | 0 | 0 | 0 | 0 | 0 | 0 | 0 | 0 | 0 | 0 | 0 | 0 | 0 | 0 |
| Jun22.75v_12019 | 1 | 0 | 0 | 0 | 0 | 0 | 0 | 0 | 0 | 0 | 0 | 1 | 0 | 0 | 0 | 0 | 0 | 0 | 0 | 0 | 0 | 0 | 0 | 0 | 0 | 0 | 0 | 0 | 0 | 0 |
| Jul11.50v_10110 | 1 | 0 | 0 | 0 | 0 | 0 | 0 | 0 | 0 | 0 | 0 | 0 | 0 | 0 | 0 | 1 | 0 | 0 | 0 | 0 | 0 | 0 | 0 | 0 | 0 | 0 | 0 | 0 | 0 | 0 |
| Jun22.75v_7798 | 1 | 0 | 0 | 0 | 0 | 0 | 0 | 0 | 0 | 0 | 0 | 1 | 0 | 0 | 0 | 0 | 0 | 0 | 0 | 0 | 0 | 0 | 0 | 0 | 0 | 0 | 0 | 0 | 0 | 0 |
| Jun8.75v_17613 | 1 | 0 | 0 | 0 | 0 | 0 | 0 | 1 | 0 | 0 | 0 | 0 | 0 | 0 | 0 | 0 | 0 | 0 | 0 | 0 | 0 | 0 | 0 | 0 | 0 | 0 | 0 | 0 | 0 | 0 |
| Jun22.5v_16084 | 1 | 0 | 0 | 0 | 0 | 0 | 0 | 0 | 0 | 1 | 0 | 0 | 0 | 0 | 0 | 0 | 0 | 0 | 0 | 0 | 0 | 0 | 0 | 0 | 0 | 0 | 0 | 0 | 0 | 0 |
| May12.75v_39250 | 1 | 0 | 0 | 0 | 1 | 0 | 0 | 0 | 0 | 0 | 0 | 0 | 0 | 0 | 0 | 0 | 0 | 0 | 0 | 0 | 0 | 0 | 0 | 0 | 0 | 0 | 0 | 0 | 0 | 0 |
| Sep8.0v_13501 | 2 | 0 | 0 | 0 | 0 | 0 | 0 | 0 | 0 | 0 | 0 | 0 | 0 | 0 | 1 | 0 | 0 | 0 | 0 | 0 | 0 | 0 | 0 | 0 | 0 | 0 | 1 | 0 | 0 | 0 |
| Sep8.25v_7191 | 1 | 0 | 0 | 0 | 0 | 0 | 0 | 0 | 0 | 0 | 0 | 0 | 0 | 0 | 0 | 0 | 0 | 0 | 0 | 0 | 0 | 0 | 0 | 0 | 0 | 0 | 0 | 0 | 1 | 0 |
| Jul11.75v_11591 | 1 | 0 | 0 | 0 | 0 | 0 | 0 | 0 | 0 | 0 | 0 | 0 | 0 | 0 | 0 | 0 | 1 | 0 | 0 | 0 | 0 | 0 | 0 | 0 | 0 | 0 | 0 | 0 | 0 | 0 |
| May12.50v_7135 | 1 | 0 | 0 | 1 | 0 | 0 | 0 | 0 | 0 | 0 | 0 | 0 | 0 | 0 | 0 | 0 | 0 | 0 | 0 | 0 | 0 | 0 | 0 | 0 | 0 | 0 | 0 | 0 | 0 | 0 |
| Jul27.50v_27422 | 1 | 0 | 0 | 0 | 0 | 0 | 0 | 0 | 0 | 0 | 0 | 0 | 0 | 0 | 0 | 0 | 0 | 0 | 0 | 1 | 0 | 0 | 0 | 0 | 0 | 0 | 0 | 0 | 0 | 0 |
| Sep8.5v_11860 | 1 | 0 | 0 | 0 | 0 | 0 | 0 | 0 | 0 | 0 | 0 | 0 | 0 | 0 | 0 | 0 | 0 | 0 | 0 | 0 | 0 | 0 | 0 | 0 | 0 | 0 | 0 | 1 | 0 | 0 |
| Jun8.50v_35792 | 2 | 0 | 0 | 0 | 0 | 0 | 1 | 0 | 0 | 0 | 1 | 0 | 0 | 0 | 0 | 0 | 0 | 0 | 0 | 0 | 0 | 0 | 0 | 0 | 0 | 0 | 0 | 0 | 0 | 0 |
| May12.25v_35238 | 1 | 0 | 1 | 0 | 0 | 0 | 0 | 0 | 0 | 0 | 0 | 0 | 0 | 0 | 0 | 0 | 0 | 0 | 0 | 0 | 0 | 0 | 0 | 0 | 0 | 0 | 0 | 0 | 0 | 0 |
| Jul27.50v_36180 | 1 | 0 | 0 | 0 | 0 | 0 | 0 | 0 | 0 | 0 | 0 | 0 | 0 | 0 | 0 | 0 | 0 | 0 | 0 | 1 | 0 | 0 | 0 | 0 | 0 | 0 | 0 | 0 | 0 | 0 |
| Jun8.75v_20642 | 1 | 0 | 0 | 0 | 0 | 0 | 0 | 1 | 0 | 0 | 0 | 0 | 0 | 0 | 0 | 0 | 0 | 0 | 0 | 0 | 0 | 0 | 0 | 0 | 0 | 0 | 0 | 0 | 0 | 0 |
| May12.75v_31817 | 1 | 0 | 0 | 0 | 1 | 0 | 0 | 0 | 0 | 0 | 0 | 0 | 0 | 0 | 0 | 0 | 0 | 0 | 0 | 0 | 0 | 0 | 0 | 0 | 0 | 0 | 0 | 0 | 0 | 0 |
| Sep8.25v_3583 | 1 | 0 | 0 | 0 | 0 | 0 | 0 | 0 | 0 | 0 | 0 | 0 | 0 | 0 | 0 | 0 | 0 | 0 | 0 | 0 | 0 | 0 | 0 | 0 | 0 | 0 | 0 | 0 | 1 | 0 |
| May12.50v_32600 | 1 | 0 | 0 | 1 | 0 | 0 | 0 | 0 | 0 | 0 | 0 | 0 | 0 | 0 | 0 | 0 | 0 | 0 | 0 | 0 | 0 | 0 | 0 | 0 | 0 | 0 | 0 | 0 | 0 | 0 |
| Jun22.0v_8746 | 1 | 0 | 0 | 0 | 0 | 0 | 0 | 0 | 1 | 0 | 0 | 0 | 0 | 0 | 0 | 0 | 0 | 0 | 0 | 0 | 0 | 0 | 0 | 0 | 0 | 0 | 0 | 0 | 0 | 0 |
| May12.75v_18171 | 1 | 0 | 0 | 0 | 1 | 0 | 0 | 0 | 0 | 0 | 0 | 0 | 0 | 0 | 0 | 0 | 0 | 0 | 0 | 0 | 0 | 0 | 0 | 0 | 0 | 0 | 0 | 0 | 0 | 0 |
| May12.50v_6912 | 1 | 0 | 0 | 1 | 0 | 0 | 0 | 0 | 0 | 0 | 0 | 0 | 0 | 0 | 0 | 0 | 0 | 0 | 0 | 0 | 0 | 0 | 0 | 0 | 0 | 0 | 0 | 0 | 0 | 0 |
| Jun8.25v_23465 | 1 | 0 | 0 | 0 | 0 | 1 | 0 | 0 | 0 | 0 | 0 | 0 | 0 | 0 | 0 | 0 | 0 | 0 | 0 | 0 | 0 | 0 | 0 | 0 | 0 | 0 | 0 | 0 | 0 | 0 |
| May12.75v_20799 | 1 | 0 | 0 | 0 | 1 | 0 | 0 | 0 | 0 | 0 | 0 | 0 | 0 | 0 | 0 | 0 | 0 | 0 | 0 | 0 | 0 | 0 | 0 | 0 | 0 | 0 | 0 | 0 | 0 | 0 |
| Jul27.50v_39912 | 1 | 0 | 0 | 0 | 0 | 0 | 0 | 0 | 0 | 0 | 0 | 0 | 0 | 0 | 0 | 0 | 0 | 0 | 0 | 1 | 0 | 0 | 0 | 0 | 0 | 0 | 0 | 0 | 0 | 0 |
| Jul27.50v_23584 | 2 | 0 | 0 | 0 | 0 | 0 | 0 | 0 | 0 | 0 | 1 | 0 | 0 | 0 | 0 | 0 | 0 | 0 | 0 | 1 | 0 | 0 | 0 | 0 | 0 | 0 | 0 | 0 | 0 | 0 |
| May12.75v_38256 | 1 | 0 | 0 | 0 | 1 | 0 | 0 | 0 | 0 | 0 | 0 | 0 | 0 | 0 | 0 | 0 | 0 | 0 | 0 | 0 | 0 | 0 | 0 | 0 | 0 | 0 | 0 | 0 | 0 | 0 |
| May12.50v_26028 | 1 | 0 | 0 | 1 | 0 | 0 | 0 | 0 | 0 | 0 | 0 | 0 | 0 | 0 | 0 | 0 | 0 | 0 | 0 | 0 | 0 | 0 | 0 | 0 | 0 | 0 | 0 | 0 | 0 | 0 |
| Jul27.50v_31625 | 2 | 0 | 0 | 0 | 1 | 0 | 0 | 0 | 0 | 0 | 0 | 0 | 0 | 0 | 0 | 0 | 0 | 0 | 0 | 1 | 0 | 0 | 0 | 0 | 0 | 0 | 0 | 0 | 0 | 0 |
| Jun22.75v_8632 | 1 | 0 | 0 | 0 | 0 | 0 | 0 | 0 | 0 | 0 | 0 | 1 | 0 | 0 | 0 | 0 | 0 | 0 | 0 | 0 | 0 | 0 | 0 | 0 | 0 | 0 | 0 | 0 | 0 | 0 |
| Sep8.25v_5114 | 1 | 0 | 0 | 0 | 0 | 0 | 0 | 0 | 0 | 0 | 0 | 0 | 0 | 0 | 0 | 0 | 0 | 0 | 0 | 0 | 0 | 0 | 0 | 0 | 0 | 0 | 0 | 0 | 1 | 0 |
| May12.75v_8932 | 1 | 0 | 0 | 0 | 1 | 0 | 0 | 0 | 0 | 0 | 0 | 0 | 0 | 0 | 0 | 0 | 0 | 0 | 0 | 0 | 0 | 0 | 0 | 0 | 0 | 0 | 0 | 0 | 0 | 0 |
| May12.75v_10506 | 1 | 0 | 0 | 0 | 1 | 0 | 0 | 0 | 0 | 0 | 0 | 0 | 0 | 0 | 0 | 0 | 0 | 0 | 0 | 0 | 0 | 0 | 0 | 0 | 0 | 0 | 0 | 0 | 0 | 0 |
| Jun22.0v_18741 | 1 | 0 | 0 | 0 | 0 | 0 | 0 | 0 | 1 | 0 | 0 | 0 | 0 | 0 | 0 | 0 | 0 | 0 | 0 | 0 | 0 | 0 | 0 | 0 | 0 | 0 | 0 | 0 | 0 | 0 |
| Jun22.75v_23240 | 1 | 0 | 0 | 0 | 0 | 0 | 0 | 0 | 0 | 0 | 0 | 1 | 0 | 0 | 0 | 0 | 0 | 0 | 0 | 0 | 0 | 0 | 0 | 0 | 0 | 0 | 0 | 0 | 0 | 0 |
| May12.75v_10859 | 1 | 0 | 0 | 0 | 1 | 0 | 0 | 0 | 0 | 0 | 0 | 0 | 0 | 0 | 0 | 0 | 0 | 0 | 0 | 0 | 0 | 0 | 0 | 0 | 0 | 0 | 0 | 0 | 0 | 0 |
| Aug5.5v_33657 | 1 | 0 | 0 | 0 | 0 | 0 | 0 | 0 | 0 | 0 | 0 | 0 | 0 | 0 | 0 | 0 | 0 | 0 | 0 | 0 | 0 | 0 | 1 | 0 | 0 | 0 | 0 | 0 | 0 | 0 |
| Jun22.75v_23329 | 1 | 0 | 0 | 0 | 0 | 0 | 0 | 0 | 0 | 0 | 0 | 1 | 0 | 0 | 0 | 0 | 0 | 0 | 0 | 0 | 0 | 0 | 0 | 0 | 0 | 0 | 0 | 0 | 0 | 0 |
| Jul11.25v_3495 | 1 | 0 | 0 | 0 | 0 | 0 | 0 | 0 | 0 | 0 | 0 | 0 | 0 | 0 | 1 | 0 | 0 | 0 | 0 | 0 | 0 | 0 | 0 | 0 | 0 | 0 | 0 | 0 | 0 | 0 |
| Jun8.50v_40922 | 1 | 0 | 0 | 0 | 0 | 0 | 1 | 0 | 0 | 0 | 0 | 0 | 0 | 0 | 0 | 0 | 0 | 0 | 0 | 0 | 0 | 0 | 0 | 0 | 0 | 0 | 0 | 0 | 0 | 0 |
| Sep8.25v_8497 | 1 | 0 | 0 | 0 | 0 | 0 | 0 | 0 | 0 | 0 | 0 | 0 | 0 | 0 | 0 | 0 | 0 | 0 | 0 | 0 | 0 | 0 | 0 | 0 | 0 | 0 | 0 | 0 | 1 | 0 |
| Jun8.75v_14242 | 1 | 0 | 0 | 0 | 0 | 0 | 0 | 1 | 0 | 0 | 0 | 0 | 0 | 0 | 0 | 0 | 0 | 0 | 0 | 0 | 0 | 0 | 0 | 0 | 0 | 0 | 0 | 0 | 0 | 0 |
| Sep8.50v_27680 | 1 | 0 | 0 | 0 | 0 | 0 | 0 | 0 | 0 | 0 | 0 | 0 | 0 | 0 | 0 | 0 | 0 | 0 | 0 | 0 | 0 | 0 | 0 | 0 | 0 | 0 | 0 | 0 | 0 | 1 |
| Jul27.50v_9836 | 1 | 0 | 0 | 0 | 0 | 0 | 0 | 0 | 0 | 0 | 0 | 0 | 0 | 0 | 0 | 0 | 0 | 0 | 0 | 1 | 0 | 0 | 0 | 0 | 0 | 0 | 0 | 0 | 0 | 0 |
| May12.75v_26799 | 1 | 0 | 0 | 0 | 1 | 0 | 0 | 0 | 0 | 0 | 0 | 0 | 0 | 0 | 0 | 0 | 0 | 0 | 0 | 0 | 0 | 0 | 0 | 0 | 0 | 0 | 0 | 0 | 0 | 0 |
| Jun8.50v_27147 | 1 | 0 | 0 | 0 | 0 | 0 | 1 | 0 | 0 | 0 | 0 | 0 | 0 | 0 | 0 | 0 | 0 | 0 | 0 | 0 | 0 | 0 | 0 | 0 | 0 | 0 | 0 | 0 | 0 | 0 |
| May12.75v_13301 | 1 | 0 | 0 | 0 | 1 | 0 | 0 | 0 | 0 | 0 | 0 | 0 | 0 | 0 | 0 | 0 | 0 | 0 | 0 | 0 | 0 | 0 | 0 | 0 | 0 | 0 | 0 | 0 | 0 | 0 |
| May12.50v_31084 | 1 | 0 | 0 | 1 | 0 | 0 | 0 | 0 | 0 | 0 | 0 | 0 | 0 | 0 | 0 | 0 | 0 | 0 | 0 | 0 | 0 | 0 | 0 | 0 | 0 | 0 | 0 | 0 | 0 | 0 |
| Sep8.25v_20688 | 1 | 0 | 0 | 0 | 0 | 0 | 0 | 0 | 0 | 0 | 0 | 0 | 0 | 0 | 0 | 0 | 0 | 0 | 0 | 0 | 0 | 0 | 0 | 0 | 0 | 0 | 0 | 0 | 1 | 0 |
| May12.50v_20272 | 1 | 0 | 0 | 1 | 0 | 0 | 0 | 0 | 0 | 0 | 0 | 0 | 0 | 0 | 0 | 0 | 0 | 0 | 0 | 0 | 0 | 0 | 0 | 0 | 0 | 0 | 0 | 0 | 0 | 0 |
| Jul11.75v_36647 | 1 | 0 | 0 | 0 | 0 | 0 | 0 | 0 | 0 | 0 | 0 | 0 | 0 | 0 | 0 | 0 | 1 | 0 | 0 | 0 | 0 | 0 | 0 | 0 | 0 | 0 | 0 | 0 | 0 | 0 |
| Jul11.50v_13333 | 1 | 0 | 0 | 0 | 0 | 0 | 0 | 0 | 0 | 0 | 0 | 0 | 0 | 0 | 0 | 1 | 0 | 0 | 0 | 0 | 0 | 0 | 0 | 0 | 0 | 0 | 0 | 0 | 0 | 0 |
| Jun8.25v_8782 | 1 | 0 | 0 | 0 | 0 | 1 | 0 | 0 | 0 | 0 | 0 | 0 | 0 | 0 | 0 | 0 | 0 | 0 | 0 | 0 | 0 | 0 | 0 | 0 | 0 | 0 | 0 | 0 | 0 | 0 |
| May12.50v_41099 | 1 | 0 | 0 | 1 | 0 | 0 | 0 | 0 | 0 | 0 | 0 | 0 | 0 | 0 | 0 | 0 | 0 | 0 | 0 | 0 | 0 | 0 | 0 | 0 | 0 | 0 | 0 | 0 | 0 | 0 |
| Sep8.25v_20861 | 1 | 0 | 0 | 0 | 0 | 0 | 0 | 0 | 0 | 0 | 0 | 0 | 0 | 0 | 0 | 0 | 0 | 0 | 0 | 0 | 0 | 0 | 0 | 0 | 0 | 0 | 0 | 0 | 1 | 0 |
| Jul11.50v_3570 | 1 | 0 | 0 | 0 | 0 | 0 | 0 | 0 | 0 | 0 | 0 | 0 | 0 | 0 | 0 | 1 | 0 | 0 | 0 | 0 | 0 | 0 | 0 | 0 | 0 | 0 | 0 | 0 | 0 | 0 |
| May12.75v_31407 | 1 | 0 | 0 | 0 | 1 | 0 | 0 | 0 | 0 | 0 | 0 | 0 | 0 | 0 | 0 | 0 | 0 | 0 | 0 | 0 | 0 | 0 | 0 | 0 | 0 | 0 | 0 | 0 | 0 | 0 |
| Jul11.5v_15561 | 1 | 0 | 0 | 0 | 0 | 0 | 0 | 0 | 0 | 0 | 0 | 0 | 0 | 1 | 0 | 0 | 0 | 0 | 0 | 0 | 0 | 0 | 0 | 0 | 0 | 0 | 0 | 0 | 0 | 0 |
| Jun22.50v_28951 | 1 | 0 | 0 | 0 | 0 | 0 | 0 | 0 | 0 | 0 | 1 | 0 | 0 | 0 | 0 | 0 | 0 | 0 | 0 | 0 | 0 | 0 | 0 | 0 | 0 | 0 | 0 | 0 | 0 | 0 |
| May12.75v_24328 | 1 | 0 | 0 | 0 | 1 | 0 | 0 | 0 | 0 | 0 | 0 | 0 | 0 | 0 | 0 | 0 | 0 | 0 | 0 | 0 | 0 | 0 | 0 | 0 | 0 | 0 | 0 | 0 | 0 | 0 |
| May12.75v_4038 | 1 | 0 | 0 | 0 | 1 | 0 | 0 | 0 | 0 | 0 | 0 | 0 | 0 | 0 | 0 | 0 | 0 | 0 | 0 | 0 | 0 | 0 | 0 | 0 | 0 | 0 | 0 | 0 | 0 | 0 |
| May12.50v_40416 | 1 | 0 | 0 | 1 | 0 | 0 | 0 | 0 | 0 | 0 | 0 | 0 | 0 | 0 | 0 | 0 | 0 | 0 | 0 | 0 | 0 | 0 | 0 | 0 | 0 | 0 | 0 | 0 | 0 | 0 |
| Jun8.50v_21295 | 1 | 0 | 0 | 0 | 0 | 0 | 1 | 0 | 0 | 0 | 0 | 0 | 0 | 0 | 0 | 0 | 0 | 0 | 0 | 0 | 0 | 0 | 0 | 0 | 0 | 0 | 0 | 0 | 0 | 0 |
| May12.25v_25260 | 1 | 0 | 1 | 0 | 0 | 0 | 0 | 0 | 0 | 0 | 0 | 0 | 0 | 0 | 0 | 0 | 0 | 0 | 0 | 0 | 0 | 0 | 0 | 0 | 0 | 0 | 0 | 0 | 0 | 0 |
| May12.5v_39071 | 1 | 1 | 0 | 0 | 0 | 0 | 0 | 0 | 0 | 0 | 0 | 0 | 0 | 0 | 0 | 0 | 0 | 0 | 0 | 0 | 0 | 0 | 0 | 0 | 0 | 0 | 0 | 0 | 0 | 0 |
| May12.50v_14520 | 1 | 0 | 0 | 1 | 0 | 0 | 0 | 0 | 0 | 0 | 0 | 0 | 0 | 0 | 0 | 0 | 0 | 0 | 0 | 0 | 0 | 0 | 0 | 0 | 0 | 0 | 0 | 0 | 0 | 0 |
| May12.75v_2819 | 1 | 0 | 0 | 0 | 1 | 0 | 0 | 0 | 0 | 0 | 0 | 0 | 0 | 0 | 0 | 0 | 0 | 0 | 0 | 0 | 0 | 0 | 0 | 0 | 0 | 0 | 0 | 0 | 0 | 0 |
| Sep8.25v_12326 | 1 | 0 | 0 | 0 | 0 | 0 | 0 | 0 | 0 | 0 | 0 | 0 | 0 | 0 | 0 | 0 | 0 | 0 | 0 | 0 | 0 | 0 | 0 | 0 | 0 | 0 | 0 | 0 | 1 | 0 |
| Jun8.75v_19828 | 1 | 0 | 0 | 0 | 0 | 0 | 0 | 1 | 0 | 0 | 0 | 0 | 0 | 0 | 0 | 0 | 0 | 0 | 0 | 0 | 0 | 0 | 0 | 0 | 0 | 0 | 0 | 0 | 0 | 0 |
| May12.50v_20309 | 1 | 0 | 0 | 1 | 0 | 0 | 0 | 0 | 0 | 0 | 0 | 0 | 0 | 0 | 0 | 0 | 0 | 0 | 0 | 0 | 0 | 0 | 0 | 0 | 0 | 0 | 0 | 0 | 0 | 0 |
| May12.50v_30530 | 1 | 0 | 0 | 1 | 0 | 0 | 0 | 0 | 0 | 0 | 0 | 0 | 0 | 0 | 0 | 0 | 0 | 0 | 0 | 0 | 0 | 0 | 0 | 0 | 0 | 0 | 0 | 0 | 0 | 0 |
| Jun22.75v_4550 | 1 | 0 | 0 | 0 | 0 | 0 | 0 | 0 | 0 | 0 | 0 | 1 | 0 | 0 | 0 | 0 | 0 | 0 | 0 | 0 | 0 | 0 | 0 | 0 | 0 | 0 | 0 | 0 | 0 | 0 |
| May12.75v_23165 | 1 | 0 | 0 | 0 | 1 | 0 | 0 | 0 | 0 | 0 | 0 | 0 | 0 | 0 | 0 | 0 | 0 | 0 | 0 | 0 | 0 | 0 | 0 | 0 | 0 | 0 | 0 | 0 | 0 | 0 |
| Jun8.50v_2855 | 1 | 0 | 0 | 0 | 0 | 0 | 1 | 0 | 0 | 0 | 0 | 0 | 0 | 0 | 0 | 0 | 0 | 0 | 0 | 0 | 0 | 0 | 0 | 0 | 0 | 0 | 0 | 0 | 0 | 0 |
| Jul27.50v_39869 | 1 | 0 | 0 | 0 | 0 | 0 | 0 | 0 | 0 | 0 | 0 | 0 | 0 | 0 | 0 | 0 | 0 | 0 | 0 | 1 | 0 | 0 | 0 | 0 | 0 | 0 | 0 | 0 | 0 | 0 |
| Jul11.50v_10799 | 1 | 0 | 0 | 0 | 0 | 0 | 0 | 0 | 0 | 0 | 0 | 0 | 0 | 0 | 0 | 1 | 0 | 0 | 0 | 0 | 0 | 0 | 0 | 0 | 0 | 0 | 0 | 0 | 0 | 0 |
| Jun22.50v_3360 | 1 | 0 | 0 | 0 | 0 | 0 | 0 | 0 | 0 | 0 | 1 | 0 | 0 | 0 | 0 | 0 | 0 | 0 | 0 | 0 | 0 | 0 | 0 | 0 | 0 | 0 | 0 | 0 | 0 | 0 |
| May12.25v_35885 | 1 | 0 | 1 | 0 | 0 | 0 | 0 | 0 | 0 | 0 | 0 | 0 | 0 | 0 | 0 | 0 | 0 | 0 | 0 | 0 | 0 | 0 | 0 | 0 | 0 | 0 | 0 | 0 | 0 | 0 |
| May12.50v_29146 | 1 | 0 | 0 | 1 | 0 | 0 | 0 | 0 | 0 | 0 | 0 | 0 | 0 | 0 | 0 | 0 | 0 | 0 | 0 | 0 | 0 | 0 | 0 | 0 | 0 | 0 | 0 | 0 | 0 | 0 |
| May12.50v_27974 | 1 | 0 | 0 | 1 | 0 | 0 | 0 | 0 | 0 | 0 | 0 | 0 | 0 | 0 | 0 | 0 | 0 | 0 | 0 | 0 | 0 | 0 | 0 | 0 | 0 | 0 | 0 | 0 | 0 | 0 |
| Jun8.50v_17880 | 1 | 0 | 0 | 0 | 0 | 0 | 1 | 0 | 0 | 0 | 0 | 0 | 0 | 0 | 0 | 0 | 0 | 0 | 0 | 0 | 0 | 0 | 0 | 0 | 0 | 0 | 0 | 0 | 0 | 0 |
| May12.50v_8873 | 1 | 0 | 0 | 1 | 0 | 0 | 0 | 0 | 0 | 0 | 0 | 0 | 0 | 0 | 0 | 0 | 0 | 0 | 0 | 0 | 0 | 0 | 0 | 0 | 0 | 0 | 0 | 0 | 0 | 0 |
| Jun8.50v_36178 | 1 | 0 | 0 | 0 | 0 | 0 | 1 | 0 | 0 | 0 | 0 | 0 | 0 | 0 | 0 | 0 | 0 | 0 | 0 | 0 | 0 | 0 | 0 | 0 | 0 | 0 | 0 | 0 | 0 | 0 |
| May12.75v_21233 | 1 | 0 | 0 | 0 | 1 | 0 | 0 | 0 | 0 | 0 | 0 | 0 | 0 | 0 | 0 | 0 | 0 | 0 | 0 | 0 | 0 | 0 | 0 | 0 | 0 | 0 | 0 | 0 | 0 | 0 |
| Sep8.25v_20896 | 2 | 0 | 0 | 0 | 0 | 0 | 0 | 0 | 0 | 0 | 0 | 0 | 0 | 0 | 0 | 0 | 0 | 0 | 0 | 0 | 0 | 0 | 0 | 0 | 0 | 0 | 0 | 0 | 2 | 0 |
| Sep8.25v_3018 | 1 | 0 | 0 | 0 | 0 | 0 | 0 | 0 | 0 | 0 | 0 | 0 | 0 | 0 | 0 | 0 | 0 | 0 | 0 | 0 | 0 | 0 | 0 | 0 | 0 | 0 | 0 | 0 | 1 | 0 |
| Jun22.75v_35385 | 1 | 0 | 0 | 0 | 0 | 0 | 0 | 0 | 0 | 0 | 0 | 1 | 0 | 0 | 0 | 0 | 0 | 0 | 0 | 0 | 0 | 0 | 0 | 0 | 0 | 0 | 0 | 0 | 0 | 0 |
| Jun8.50v_35836 | 1 | 0 | 0 | 0 | 0 | 0 | 1 | 0 | 0 | 0 | 0 | 0 | 0 | 0 | 0 | 0 | 0 | 0 | 0 | 0 | 0 | 0 | 0 | 0 | 0 | 0 | 0 | 0 | 0 | 0 |
| May12.25v_6494 | 1 | 0 | 1 | 0 | 0 | 0 | 0 | 0 | 0 | 0 | 0 | 0 | 0 | 0 | 0 | 0 | 0 | 0 | 0 | 0 | 0 | 0 | 0 | 0 | 0 | 0 | 0 | 0 | 0 | 0 |
| May12.75v_6936 | 1 | 0 | 0 | 0 | 1 | 0 | 0 | 0 | 0 | 0 | 0 | 0 | 0 | 0 | 0 | 0 | 0 | 0 | 0 | 0 | 0 | 0 | 0 | 0 | 0 | 0 | 0 | 0 | 0 | 0 |
| Jun22.50v_33089 | 1 | 0 | 0 | 0 | 0 | 0 | 0 | 0 | 0 | 0 | 1 | 0 | 0 | 0 | 0 | 0 | 0 | 0 | 0 | 0 | 0 | 0 | 0 | 0 | 0 | 0 | 0 | 0 | 0 | 0 |
| Jun8.75v_16443 | 1 | 0 | 0 | 0 | 0 | 0 | 0 | 1 | 0 | 0 | 0 | 0 | 0 | 0 | 0 | 0 | 0 | 0 | 0 | 0 | 0 | 0 | 0 | 0 | 0 | 0 | 0 | 0 | 0 | 0 |
| May12.5v_35100 | 1 | 1 | 0 | 0 | 0 | 0 | 0 | 0 | 0 | 0 | 0 | 0 | 0 | 0 | 0 | 0 | 0 | 0 | 0 | 0 | 0 | 0 | 0 | 0 | 0 | 0 | 0 | 0 | 0 | 0 |
| Jun22.75v_5416 | 1 | 0 | 0 | 0 | 0 | 0 | 0 | 0 | 0 | 0 | 0 | 1 | 0 | 0 | 0 | 0 | 0 | 0 | 0 | 0 | 0 | 0 | 0 | 0 | 0 | 0 | 0 | 0 | 0 | 0 |
| May12.50v_10911 | 1 | 0 | 0 | 1 | 0 | 0 | 0 | 0 | 0 | 0 | 0 | 0 | 0 | 0 | 0 | 0 | 0 | 0 | 0 | 0 | 0 | 0 | 0 | 0 | 0 | 0 | 0 | 0 | 0 | 0 |
| May12.75v_39261 | 1 | 0 | 0 | 0 | 1 | 0 | 0 | 0 | 0 | 0 | 0 | 0 | 0 | 0 | 0 | 0 | 0 | 0 | 0 | 0 | 0 | 0 | 0 | 0 | 0 | 0 | 0 | 0 | 0 | 0 |
| Jun8.50v_663 | 1 | 0 | 0 | 0 | 0 | 0 | 1 | 0 | 0 | 0 | 0 | 0 | 0 | 0 | 0 | 0 | 0 | 0 | 0 | 0 | 0 | 0 | 0 | 0 | 0 | 0 | 0 | 0 | 0 | 0 |
| Jun8.50v_14959 | 1 | 0 | 0 | 0 | 0 | 0 | 1 | 0 | 0 | 0 | 0 | 0 | 0 | 0 | 0 | 0 | 0 | 0 | 0 | 0 | 0 | 0 | 0 | 0 | 0 | 0 | 0 | 0 | 0 | 0 |
| Sep8.25v_10270 | 1 | 0 | 0 | 0 | 0 | 0 | 0 | 0 | 0 | 0 | 0 | 0 | 0 | 0 | 0 | 0 | 0 | 0 | 0 | 0 | 0 | 0 | 0 | 0 | 0 | 0 | 0 | 0 | 1 | 0 |
| Sep8.25v_21004 | 1 | 0 | 0 | 0 | 0 | 0 | 0 | 0 | 0 | 0 | 0 | 0 | 0 | 0 | 0 | 0 | 0 | 0 | 0 | 0 | 0 | 0 | 0 | 0 | 0 | 0 | 0 | 0 | 1 | 0 |
| May12.75v_30830 | 1 | 0 | 0 | 0 | 1 | 0 | 0 | 0 | 0 | 0 | 0 | 0 | 0 | 0 | 0 | 0 | 0 | 0 | 0 | 0 | 0 | 0 | 0 | 0 | 0 | 0 | 0 | 0 | 0 | 0 |
| Jun22.50v_19633 | 1 | 0 | 0 | 0 | 0 | 0 | 0 | 0 | 0 | 0 | 1 | 0 | 0 | 0 | 0 | 0 | 0 | 0 | 0 | 0 | 0 | 0 | 0 | 0 | 0 | 0 | 0 | 0 | 0 | 0 |
| Sep8.25v_8066 | 1 | 0 | 0 | 0 | 0 | 0 | 0 | 0 | 0 | 0 | 0 | 0 | 0 | 0 | 0 | 0 | 0 | 0 | 0 | 0 | 0 | 0 | 0 | 0 | 0 | 0 | 0 | 0 | 1 | 0 |
| Jun8.50v_30966 | 1 | 0 | 0 | 0 | 0 | 0 | 1 | 0 | 0 | 0 | 0 | 0 | 0 | 0 | 0 | 0 | 0 | 0 | 0 | 0 | 0 | 0 | 0 | 0 | 0 | 0 | 0 | 0 | 0 | 0 |
| Jun22.50v_23468 | 1 | 0 | 0 | 0 | 0 | 0 | 0 | 0 | 0 | 0 | 1 | 0 | 0 | 0 | 0 | 0 | 0 | 0 | 0 | 0 | 0 | 0 | 0 | 0 | 0 | 0 | 0 | 0 | 0 | 0 |
| Jul11.50v_16973 | 1 | 0 | 0 | 0 | 0 | 0 | 0 | 0 | 0 | 0 | 0 | 0 | 0 | 0 | 0 | 1 | 0 | 0 | 0 | 0 | 0 | 0 | 0 | 0 | 0 | 0 | 0 | 0 | 0 | 0 |
| May12.75v_39997 | 1 | 0 | 0 | 0 | 1 | 0 | 0 | 0 | 0 | 0 | 0 | 0 | 0 | 0 | 0 | 0 | 0 | 0 | 0 | 0 | 0 | 0 | 0 | 0 | 0 | 0 | 0 | 0 | 0 | 0 |
| Jun22.5v_15148 | 2 | 0 | 0 | 0 | 1 | 0 | 0 | 0 | 0 | 1 | 0 | 0 | 0 | 0 | 0 | 0 | 0 | 0 | 0 | 0 | 0 | 0 | 0 | 0 | 0 | 0 | 0 | 0 | 0 | 0 |
| Jun8.50v_37894 | 1 | 0 | 0 | 0 | 0 | 0 | 1 | 0 | 0 | 0 | 0 | 0 | 0 | 0 | 0 | 0 | 0 | 0 | 0 | 0 | 0 | 0 | 0 | 0 | 0 | 0 | 0 | 0 | 0 | 0 |
| May12.25v_25031 | 1 | 0 | 1 | 0 | 0 | 0 | 0 | 0 | 0 | 0 | 0 | 0 | 0 | 0 | 0 | 0 | 0 | 0 | 0 | 0 | 0 | 0 | 0 | 0 | 0 | 0 | 0 | 0 | 0 | 0 |
| May12.5v_30341 | 1 | 1 | 0 | 0 | 0 | 0 | 0 | 0 | 0 | 0 | 0 | 0 | 0 | 0 | 0 | 0 | 0 | 0 | 0 | 0 | 0 | 0 | 0 | 0 | 0 | 0 | 0 | 0 | 0 | 0 |
| Jul27.75v_821 | 1 | 0 | 0 | 0 | 0 | 0 | 0 | 0 | 0 | 0 | 0 | 0 | 0 | 0 | 0 | 0 | 0 | 0 | 0 | 0 | 1 | 0 | 0 | 0 | 0 | 0 | 0 | 0 | 0 | 0 |
| May12.5v_9996 | 1 | 1 | 0 | 0 | 0 | 0 | 0 | 0 | 0 | 0 | 0 | 0 | 0 | 0 | 0 | 0 | 0 | 0 | 0 | 0 | 0 | 0 | 0 | 0 | 0 | 0 | 0 | 0 | 0 | 0 |
| Jun22.75v_29778 | 1 | 0 | 0 | 0 | 0 | 0 | 0 | 0 | 0 | 0 | 0 | 1 | 0 | 0 | 0 | 0 | 0 | 0 | 0 | 0 | 0 | 0 | 0 | 0 | 0 | 0 | 0 | 0 | 0 | 0 |
| Aug5.50v_7119 | 1 | 0 | 0 | 0 | 0 | 0 | 0 | 0 | 0 | 0 | 0 | 0 | 0 | 0 | 0 | 0 | 0 | 0 | 0 | 0 | 0 | 0 | 0 | 0 | 1 | 0 | 0 | 0 | 0 | 0 |
| May12.5v_37184 | 1 | 1 | 0 | 0 | 0 | 0 | 0 | 0 | 0 | 0 | 0 | 0 | 0 | 0 | 0 | 0 | 0 | 0 | 0 | 0 | 0 | 0 | 0 | 0 | 0 | 0 | 0 | 0 | 0 | 0 |
| Jun22.0v_7074 | 1 | 0 | 0 | 0 | 0 | 0 | 0 | 0 | 1 | 0 | 0 | 0 | 0 | 0 | 0 | 0 | 0 | 0 | 0 | 0 | 0 | 0 | 0 | 0 | 0 | 0 | 0 | 0 | 0 | 0 |
| Jul11.75v_11827 | 1 | 0 | 0 | 0 | 0 | 0 | 0 | 0 | 0 | 0 | 0 | 0 | 0 | 0 | 0 | 0 | 1 | 0 | 0 | 0 | 0 | 0 | 0 | 0 | 0 | 0 | 0 | 0 | 0 | 0 |
| May12.50v_34468 | 1 | 0 | 0 | 1 | 0 | 0 | 0 | 0 | 0 | 0 | 0 | 0 | 0 | 0 | 0 | 0 | 0 | 0 | 0 | 0 | 0 | 0 | 0 | 0 | 0 | 0 | 0 | 0 | 0 | 0 |
| Jun8.50v_32615 | 1 | 0 | 0 | 0 | 0 | 0 | 1 | 0 | 0 | 0 | 0 | 0 | 0 | 0 | 0 | 0 | 0 | 0 | 0 | 0 | 0 | 0 | 0 | 0 | 0 | 0 | 0 | 0 | 0 | 0 |
| May12.75v_13829 | 1 | 0 | 0 | 0 | 1 | 0 | 0 | 0 | 0 | 0 | 0 | 0 | 0 | 0 | 0 | 0 | 0 | 0 | 0 | 0 | 0 | 0 | 0 | 0 | 0 | 0 | 0 | 0 | 0 | 0 |
| May12.5v_9250 | 1 | 1 | 0 | 0 | 0 | 0 | 0 | 0 | 0 | 0 | 0 | 0 | 0 | 0 | 0 | 0 | 0 | 0 | 0 | 0 | 0 | 0 | 0 | 0 | 0 | 0 | 0 | 0 | 0 | 0 |
| May12.5v_10726 | 1 | 1 | 0 | 0 | 0 | 0 | 0 | 0 | 0 | 0 | 0 | 0 | 0 | 0 | 0 | 0 | 0 | 0 | 0 | 0 | 0 | 0 | 0 | 0 | 0 | 0 | 0 | 0 | 0 | 0 |
| Jun8.50v_13174 | 2 | 0 | 0 | 0 | 1 | 0 | 1 | 0 | 0 | 0 | 0 | 0 | 0 | 0 | 0 | 0 | 0 | 0 | 0 | 0 | 0 | 0 | 0 | 0 | 0 | 0 | 0 | 0 | 0 | 0 |
| Sep8.0v_3627 | 1 | 0 | 0 | 0 | 0 | 0 | 0 | 0 | 0 | 0 | 0 | 0 | 0 | 0 | 0 | 0 | 0 | 0 | 0 | 0 | 0 | 0 | 0 | 0 | 0 | 0 | 1 | 0 | 0 | 0 |
| May12.75v_8633 | 1 | 0 | 0 | 0 | 1 | 0 | 0 | 0 | 0 | 0 | 0 | 0 | 0 | 0 | 0 | 0 | 0 | 0 | 0 | 0 | 0 | 0 | 0 | 0 | 0 | 0 | 0 | 0 | 0 | 0 |
| Aug5.25v_32847 | 1 | 0 | 0 | 0 | 0 | 0 | 0 | 0 | 0 | 0 | 0 | 0 | 0 | 0 | 0 | 0 | 0 | 0 | 0 | 0 | 0 | 0 | 0 | 1 | 0 | 0 | 0 | 0 | 0 | 0 |
| May12.75v_8217 | 1 | 0 | 0 | 0 | 1 | 0 | 0 | 0 | 0 | 0 | 0 | 0 | 0 | 0 | 0 | 0 | 0 | 0 | 0 | 0 | 0 | 0 | 0 | 0 | 0 | 0 | 0 | 0 | 0 | 0 |
| Sep8.25v_13433 | 1 | 0 | 0 | 0 | 0 | 0 | 0 | 0 | 0 | 0 | 0 | 0 | 0 | 0 | 0 | 0 | 0 | 0 | 0 | 0 | 0 | 0 | 0 | 0 | 0 | 0 | 0 | 0 | 1 | 0 |
| Jun8.50v_5606 | 1 | 0 | 0 | 0 | 0 | 0 | 1 | 0 | 0 | 0 | 0 | 0 | 0 | 0 | 0 | 0 | 0 | 0 | 0 | 0 | 0 | 0 | 0 | 0 | 0 | 0 | 0 | 0 | 0 | 0 |
| Aug5.50v_39142 | 1 | 0 | 0 | 0 | 0 | 0 | 0 | 0 | 0 | 0 | 0 | 0 | 0 | 0 | 0 | 0 | 0 | 0 | 0 | 0 | 0 | 0 | 0 | 0 | 1 | 0 | 0 | 0 | 0 | 0 |
| Jun22.75v_12338 | 1 | 0 | 0 | 0 | 0 | 0 | 0 | 0 | 0 | 0 | 0 | 1 | 0 | 0 | 0 | 0 | 0 | 0 | 0 | 0 | 0 | 0 | 0 | 0 | 0 | 0 | 0 | 0 | 0 | 0 |
| Jun22.50v_39505 | 1 | 0 | 0 | 0 | 0 | 0 | 0 | 0 | 0 | 0 | 1 | 0 | 0 | 0 | 0 | 0 | 0 | 0 | 0 | 0 | 0 | 0 | 0 | 0 | 0 | 0 | 0 | 0 | 0 | 0 |
| May12.5v_33563 | 1 | 1 | 0 | 0 | 0 | 0 | 0 | 0 | 0 | 0 | 0 | 0 | 0 | 0 | 0 | 0 | 0 | 0 | 0 | 0 | 0 | 0 | 0 | 0 | 0 | 0 | 0 | 0 | 0 | 0 |
| Jul11.75v_18370 | 1 | 0 | 0 | 0 | 0 | 0 | 0 | 0 | 0 | 0 | 0 | 0 | 0 | 0 | 0 | 0 | 1 | 0 | 0 | 0 | 0 | 0 | 0 | 0 | 0 | 0 | 0 | 0 | 0 | 0 |
| Jun22.50v_19128 | 1 | 0 | 0 | 0 | 0 | 0 | 0 | 0 | 0 | 0 | 1 | 0 | 0 | 0 | 0 | 0 | 0 | 0 | 0 | 0 | 0 | 0 | 0 | 0 | 0 | 0 | 0 | 0 | 0 | 0 |
| Jun8.50v_13689 | 1 | 0 | 0 | 0 | 0 | 0 | 1 | 0 | 0 | 0 | 0 | 0 | 0 | 0 | 0 | 0 | 0 | 0 | 0 | 0 | 0 | 0 | 0 | 0 | 0 | 0 | 0 | 0 | 0 | 0 |
| Jul11.75v_18794 | 1 | 0 | 0 | 0 | 0 | 0 | 0 | 0 | 0 | 0 | 0 | 0 | 0 | 0 | 0 | 0 | 1 | 0 | 0 | 0 | 0 | 0 | 0 | 0 | 0 | 0 | 0 | 0 | 0 | 0 |
| Jul11.75v_3084 | 1 | 0 | 0 | 0 | 0 | 0 | 0 | 0 | 0 | 0 | 0 | 0 | 0 | 0 | 0 | 0 | 1 | 0 | 0 | 0 | 0 | 0 | 0 | 0 | 0 | 0 | 0 | 0 | 0 | 0 |
| Jun8.75v_451 | 1 | 0 | 0 | 0 | 0 | 0 | 0 | 1 | 0 | 0 | 0 | 0 | 0 | 0 | 0 | 0 | 0 | 0 | 0 | 0 | 0 | 0 | 0 | 0 | 0 | 0 | 0 | 0 | 0 | 0 |
| Jul11.5v_16593 | 1 | 0 | 0 | 0 | 0 | 0 | 0 | 0 | 0 | 0 | 0 | 0 | 0 | 1 | 0 | 0 | 0 | 0 | 0 | 0 | 0 | 0 | 0 | 0 | 0 | 0 | 0 | 0 | 0 | 0 |
| Jul11.50v_4166 | 1 | 0 | 0 | 0 | 0 | 0 | 0 | 0 | 0 | 0 | 0 | 0 | 0 | 0 | 0 | 1 | 0 | 0 | 0 | 0 | 0 | 0 | 0 | 0 | 0 | 0 | 0 | 0 | 0 | 0 |
| Jun8.25v_24590 | 1 | 0 | 0 | 0 | 0 | 1 | 0 | 0 | 0 | 0 | 0 | 0 | 0 | 0 | 0 | 0 | 0 | 0 | 0 | 0 | 0 | 0 | 0 | 0 | 0 | 0 | 0 | 0 | 0 | 0 |
| May12.50v_36443 | 2 | 0 | 0 | 1 | 0 | 0 | 0 | 0 | 0 | 0 | 1 | 0 | 0 | 0 | 0 | 0 | 0 | 0 | 0 | 0 | 0 | 0 | 0 | 0 | 0 | 0 | 0 | 0 | 0 | 0 |
| May12.50v_14452 | 1 | 0 | 0 | 1 | 0 | 0 | 0 | 0 | 0 | 0 | 0 | 0 | 0 | 0 | 0 | 0 | 0 | 0 | 0 | 0 | 0 | 0 | 0 | 0 | 0 | 0 | 0 | 0 | 0 | 0 |
| Jun22.75v_24388 | 1 | 0 | 0 | 0 | 0 | 0 | 0 | 0 | 0 | 0 | 0 | 1 | 0 | 0 | 0 | 0 | 0 | 0 | 0 | 0 | 0 | 0 | 0 | 0 | 0 | 0 | 0 | 0 | 0 | 0 |
| Aug5.75v_39058 | 2 | 0 | 0 | 1 | 0 | 0 | 0 | 0 | 0 | 0 | 0 | 0 | 0 | 0 | 0 | 0 | 0 | 0 | 0 | 0 | 0 | 0 | 0 | 0 | 0 | 1 | 0 | 0 | 0 | 0 |
| May12.25v_5937 | 4 | 0 | 1 | 0 | 1 | 0 | 0 | 0 | 0 | 0 | 0 | 1 | 0 | 0 | 0 | 0 | 0 | 0 | 1 | 0 | 0 | 0 | 0 | 0 | 0 | 0 | 0 | 0 | 0 | 0 |
| May12.75v_22117 | 1205 | 97 | 98 | 101 | 174 | 21 | 150 | 5 | 21 | 16 | 33 | 121 | 14 | 20 | 11 | 29 | 70 | 17 | 22 | 8 | 31 | 11 | 16 | 11 | 4 | 46 | 4 | 30 | 12 | 12 |
| Jun22.75v_2628 | 1 | 0 | 0 | 0 | 0 | 0 | 0 | 0 | 0 | 0 | 0 | 1 | 0 | 0 | 0 | 0 | 0 | 0 | 0 | 0 | 0 | 0 | 0 | 0 | 0 | 0 | 0 | 0 | 0 | 0 |
| May12.50v_36914 | 1 | 0 | 0 | 1 | 0 | 0 | 0 | 0 | 0 | 0 | 0 | 0 | 0 | 0 | 0 | 0 | 0 | 0 | 0 | 0 | 0 | 0 | 0 | 0 | 0 | 0 | 0 | 0 | 0 | 0 |
| Sep8.5v_15115 | 455 | 29 | 19 | 8 | 57 | 19 | 53 | 26 | 3 | 16 | 20 | 39 | 1 | 13 | 17 | 10 | 12 | 3 | 2 | 22 | 1 | 1 | 1 | 0 | 2 | 3 | 0 | 72 | 3 | 3 |
| Jun22.50v_13394 | 1 | 0 | 0 | 0 | 0 | 0 | 0 | 0 | 0 | 0 | 1 | 0 | 0 | 0 | 0 | 0 | 0 | 0 | 0 | 0 | 0 | 0 | 0 | 0 | 0 | 0 | 0 | 0 | 0 | 0 |
| Jun8.75v_19582 | 116 | 8 | 7 | 3 | 16 | 0 | 11 | 11 | 0 | 6 | 3 | 16 | 1 | 2 | 3 | 4 | 3 | 1 | 0 | 4 | 1 | 1 | 0 | 0 | 0 | 1 | 1 | 13 | 0 | 0 |
| Jun8.75v_18380 | 33 | 1 | 1 | 0 | 2 | 1 | 3 | 6 | 0 | 3 | 1 | 1 | 0 | 2 | 1 | 2 | 0 | 0 | 0 | 1 | 1 | 0 | 0 | 0 | 0 | 0 | 0 | 7 | 0 | 0 |
| May12.5v_40693 | 1 | 1 | 0 | 0 | 0 | 0 | 0 | 0 | 0 | 0 | 0 | 0 | 0 | 0 | 0 | 0 | 0 | 0 | 0 | 0 | 0 | 0 | 0 | 0 | 0 | 0 | 0 | 0 | 0 | 0 |
| Jul11.25v_1821 | 1 | 0 | 0 | 0 | 0 | 0 | 0 | 0 | 0 | 0 | 0 | 0 | 0 | 0 | 1 | 0 | 0 | 0 | 0 | 0 | 0 | 0 | 0 | 0 | 0 | 0 | 0 | 0 | 0 | 0 |
| May12.50v_26320 | 1 | 0 | 0 | 1 | 0 | 0 | 0 | 0 | 0 | 0 | 0 | 0 | 0 | 0 | 0 | 0 | 0 | 0 | 0 | 0 | 0 | 0 | 0 | 0 | 0 | 0 | 0 | 0 | 0 | 0 |
| Aug5.25v_10994 | 1 | 0 | 0 | 0 | 0 | 0 | 0 | 0 | 0 | 0 | 0 | 0 | 0 | 0 | 0 | 0 | 0 | 0 | 0 | 0 | 0 | 0 | 0 | 1 | 0 | 0 | 0 | 0 | 0 | 0 |
| Jun8.25v_20883 | 1 | 0 | 0 | 0 | 0 | 1 | 0 | 0 | 0 | 0 | 0 | 0 | 0 | 0 | 0 | 0 | 0 | 0 | 0 | 0 | 0 | 0 | 0 | 0 | 0 | 0 | 0 | 0 | 0 | 0 |
| Jun8.75v_18908 | 1 | 0 | 0 | 0 | 0 | 0 | 0 | 1 | 0 | 0 | 0 | 0 | 0 | 0 | 0 | 0 | 0 | 0 | 0 | 0 | 0 | 0 | 0 | 0 | 0 | 0 | 0 | 0 | 0 | 0 |
| May12.5v_7454 | 1 | 1 | 0 | 0 | 0 | 0 | 0 | 0 | 0 | 0 | 0 | 0 | 0 | 0 | 0 | 0 | 0 | 0 | 0 | 0 | 0 | 0 | 0 | 0 | 0 | 0 | 0 | 0 | 0 | 0 |
| Jun22.75v_28013 | 1 | 0 | 0 | 0 | 0 | 0 | 0 | 0 | 0 | 0 | 0 | 1 | 0 | 0 | 0 | 0 | 0 | 0 | 0 | 0 | 0 | 0 | 0 | 0 | 0 | 0 | 0 | 0 | 0 | 0 |
| Jul11.50v_6203 | 16 | 0 | 0 | 0 | 0 | 0 | 0 | 1 | 0 | 1 | 0 | 2 | 0 | 0 | 2 | 2 | 0 | 1 | 0 | 0 | 0 | 0 | 0 | 0 | 0 | 0 | 0 | 6 | 1 | 0 |
| Jul27.50v_15882 | 3 | 0 | 0 | 0 | 1 | 0 | 0 | 0 | 0 | 0 | 0 | 0 | 0 | 0 | 0 | 0 | 0 | 0 | 0 | 1 | 0 | 0 | 0 | 0 | 1 | 0 | 0 | 0 | 0 | 0 |
| May12.5v_1939 | 2 | 1 | 0 | 0 | 0 | 0 | 1 | 0 | 0 | 0 | 0 | 0 | 0 | 0 | 0 | 0 | 0 | 0 | 0 | 0 | 0 | 0 | 0 | 0 | 0 | 0 | 0 | 0 | 0 | 0 |
| Jun22.50v_4774 | 1 | 0 | 0 | 0 | 0 | 0 | 0 | 0 | 0 | 0 | 1 | 0 | 0 | 0 | 0 | 0 | 0 | 0 | 0 | 0 | 0 | 0 | 0 | 0 | 0 | 0 | 0 | 0 | 0 | 0 |
| Jul11.50v_3752 | 2 | 0 | 0 | 0 | 0 | 1 | 0 | 0 | 0 | 0 | 0 | 0 | 0 | 0 | 0 | 1 | 0 | 0 | 0 | 0 | 0 | 0 | 0 | 0 | 0 | 0 | 0 | 0 | 0 | 0 |
| May12.25v_24101 | 1 | 0 | 1 | 0 | 0 | 0 | 0 | 0 | 0 | 0 | 0 | 0 | 0 | 0 | 0 | 0 | 0 | 0 | 0 | 0 | 0 | 0 | 0 | 0 | 0 | 0 | 0 | 0 | 0 | 0 |
| Jul11.25v_19914 | 1 | 0 | 0 | 0 | 0 | 0 | 0 | 0 | 0 | 0 | 0 | 0 | 0 | 0 | 1 | 0 | 0 | 0 | 0 | 0 | 0 | 0 | 0 | 0 | 0 | 0 | 0 | 0 | 0 | 0 |
| Jul27.5v_1426 | 2 | 0 | 0 | 0 | 0 | 0 | 0 | 0 | 0 | 1 | 0 | 0 | 0 | 0 | 0 | 0 | 0 | 1 | 0 | 0 | 0 | 0 | 0 | 0 | 0 | 0 | 0 | 0 | 0 | 0 |
| Jun8.50v_27280 | 2 | 0 | 0 | 0 | 0 | 0 | 1 | 0 | 0 | 0 | 0 | 0 | 0 | 0 | 1 | 0 | 0 | 0 | 0 | 0 | 0 | 0 | 0 | 0 | 0 | 0 | 0 | 0 | 0 | 0 |
| Sep8.5v_6515 | 2 | 0 | 0 | 0 | 0 | 0 | 0 | 0 | 0 | 0 | 0 | 0 | 0 | 0 | 1 | 0 | 0 | 0 | 0 | 0 | 0 | 0 | 0 | 0 | 0 | 0 | 0 | 1 | 0 | 0 |
| May12.50v_27028 | 1 | 0 | 0 | 1 | 0 | 0 | 0 | 0 | 0 | 0 | 0 | 0 | 0 | 0 | 0 | 0 | 0 | 0 | 0 | 0 | 0 | 0 | 0 | 0 | 0 | 0 | 0 | 0 | 0 | 0 |
| Sep8.5v_19629 | 1 | 0 | 0 | 0 | 0 | 0 | 0 | 0 | 0 | 0 | 0 | 0 | 0 | 0 | 0 | 0 | 0 | 0 | 0 | 0 | 0 | 0 | 0 | 0 | 0 | 0 | 0 | 1 | 0 | 0 |
| Sep8.25v_16285 | 1 | 0 | 0 | 0 | 0 | 0 | 0 | 0 | 0 | 0 | 0 | 0 | 0 | 0 | 0 | 0 | 0 | 0 | 0 | 0 | 0 | 0 | 0 | 0 | 0 | 0 | 0 | 0 | 1 | 0 |
| Jul11.25v_8611 | 1 | 0 | 0 | 0 | 0 | 0 | 0 | 0 | 0 | 0 | 0 | 0 | 0 | 0 | 1 | 0 | 0 | 0 | 0 | 0 | 0 | 0 | 0 | 0 | 0 | 0 | 0 | 0 | 0 | 0 |
| May12.50v_9895 | 1 | 0 | 0 | 1 | 0 | 0 | 0 | 0 | 0 | 0 | 0 | 0 | 0 | 0 | 0 | 0 | 0 | 0 | 0 | 0 | 0 | 0 | 0 | 0 | 0 | 0 | 0 | 0 | 0 | 0 |
| May12.25v_741 | 1 | 0 | 1 | 0 | 0 | 0 | 0 | 0 | 0 | 0 | 0 | 0 | 0 | 0 | 0 | 0 | 0 | 0 | 0 | 0 | 0 | 0 | 0 | 0 | 0 | 0 | 0 | 0 | 0 | 0 |
| Jun8.50v_4633 | 1 | 0 | 0 | 0 | 0 | 0 | 1 | 0 | 0 | 0 | 0 | 0 | 0 | 0 | 0 | 0 | 0 | 0 | 0 | 0 | 0 | 0 | 0 | 0 | 0 | 0 | 0 | 0 | 0 | 0 |
| Jun8.75v_1576 | 1 | 0 | 0 | 0 | 0 | 0 | 0 | 1 | 0 | 0 | 0 | 0 | 0 | 0 | 0 | 0 | 0 | 0 | 0 | 0 | 0 | 0 | 0 | 0 | 0 | 0 | 0 | 0 | 0 | 0 |
| Jun22.5v_10829 | 1 | 0 | 0 | 0 | 0 | 0 | 0 | 0 | 0 | 1 | 0 | 0 | 0 | 0 | 0 | 0 | 0 | 0 | 0 | 0 | 0 | 0 | 0 | 0 | 0 | 0 | 0 | 0 | 0 | 0 |
| Jul11.5v_2027 | 1 | 0 | 0 | 0 | 0 | 0 | 0 | 0 | 0 | 0 | 0 | 0 | 0 | 1 | 0 | 0 | 0 | 0 | 0 | 0 | 0 | 0 | 0 | 0 | 0 | 0 | 0 | 0 | 0 | 0 |
| Jul11.50v_14820 | 1 | 0 | 0 | 0 | 0 | 0 | 0 | 0 | 0 | 0 | 0 | 0 | 0 | 0 | 0 | 1 | 0 | 0 | 0 | 0 | 0 | 0 | 0 | 0 | 0 | 0 | 0 | 0 | 0 | 0 |
| May12.75v_28437 | 2 | 0 | 0 | 0 | 1 | 0 | 0 | 0 | 0 | 0 | 0 | 1 | 0 | 0 | 0 | 0 | 0 | 0 | 0 | 0 | 0 | 0 | 0 | 0 | 0 | 0 | 0 | 0 | 0 | 0 |
| Jun22.75v_28096 | 1 | 0 | 0 | 0 | 0 | 0 | 0 | 0 | 0 | 0 | 0 | 1 | 0 | 0 | 0 | 0 | 0 | 0 | 0 | 0 | 0 | 0 | 0 | 0 | 0 | 0 | 0 | 0 | 0 | 0 |
| May12.75v_29897 | 1 | 0 | 0 | 0 | 1 | 0 | 0 | 0 | 0 | 0 | 0 | 0 | 0 | 0 | 0 | 0 | 0 | 0 | 0 | 0 | 0 | 0 | 0 | 0 | 0 | 0 | 0 | 0 | 0 | 0 |
| Sep8.25v_16511 | 1 | 0 | 0 | 0 | 0 | 0 | 0 | 0 | 0 | 0 | 0 | 0 | 0 | 0 | 0 | 0 | 0 | 0 | 0 | 0 | 0 | 0 | 0 | 0 | 0 | 0 | 0 | 0 | 1 | 0 |
| Jul27.25v_16175 | 8 | 1 | 1 | 0 | 2 | 0 | 2 | 0 | 0 | 0 | 0 | 0 | 0 | 0 | 0 | 0 | 0 | 0 | 2 | 0 | 0 | 0 | 0 | 0 | 0 | 0 | 0 | 0 | 0 | 0 |
| May12.25v_18938 | 1 | 0 | 1 | 0 | 0 | 0 | 0 | 0 | 0 | 0 | 0 | 0 | 0 | 0 | 0 | 0 | 0 | 0 | 0 | 0 | 0 | 0 | 0 | 0 | 0 | 0 | 0 | 0 | 0 | 0 |
| May12.25v_22625 | 1 | 0 | 1 | 0 | 0 | 0 | 0 | 0 | 0 | 0 | 0 | 0 | 0 | 0 | 0 | 0 | 0 | 0 | 0 | 0 | 0 | 0 | 0 | 0 | 0 | 0 | 0 | 0 | 0 | 0 |
| Jul27.75v_7636 | 248 | 6 | 8 | 2 | 11 | 15 | 14 | 15 | 0 | 10 | 11 | 27 | 2 | 31 | 26 | 18 | 11 | 6 | 5 | 6 | 1 | 0 | 4 | 2 | 0 | 1 | 0 | 9 | 3 | 4 |
| Jun8.50v_3688 | 18 | 1 | 1 | 0 | 0 | 1 | 2 | 0 | 0 | 1 | 0 | 0 | 0 | 2 | 5 | 1 | 0 | 2 | 0 | 0 | 0 | 0 | 0 | 1 | 0 | 0 | 0 | 1 | 0 | 0 |
| Jul27.75v_283 | 3 | 0 | 0 | 0 | 0 | 0 | 0 | 0 | 0 | 1 | 0 | 0 | 0 | 1 | 0 | 0 | 0 | 0 | 0 | 0 | 1 | 0 | 0 | 0 | 0 | 0 | 0 | 0 | 0 | 0 |
| Jul11.5v_7794 | 87 | 2 | 4 | 2 | 4 | 3 | 8 | 5 | 2 | 6 | 1 | 7 | 1 | 6 | 4 | 14 | 3 | 5 | 0 | 3 | 1 | 0 | 1 | 0 | 1 | 1 | 0 | 2 | 0 | 1 |
| Jul11.25v_7446 | 1 | 0 | 0 | 0 | 0 | 0 | 0 | 0 | 0 | 0 | 0 | 0 | 0 | 0 | 1 | 0 | 0 | 0 | 0 | 0 | 0 | 0 | 0 | 0 | 0 | 0 | 0 | 0 | 0 | 0 |
| Jul11.25v_1406 | 1 | 0 | 0 | 0 | 0 | 0 | 0 | 0 | 0 | 0 | 0 | 0 | 0 | 0 | 1 | 0 | 0 | 0 | 0 | 0 | 0 | 0 | 0 | 0 | 0 | 0 | 0 | 0 | 0 | 0 |

| Feature_ID | 654 | May12_05m | May12_25m | May12_50m | May12_75m | Jun8_25m | Jun8_50m | Jun8_75m | Jun22_0m | Jun22_05m | Jun22_50m | Jun22_75m | Jul11_0m | Jul11_05m | Jul11_25m | Jul11_50m | Jul11_75m | Jul27_05m | Jul27_25m | Jul27_50m | Jul27_75m | Aug5_0m | Aug5_05m | Aug5_25m | Aug5_50m | Aug5_75m | Sep8_0m | Sep8_05m | Sep8_25m | Sep8_50m |
|---|---|---|---|---|---|---|---|---|---|---|---|---|---|---|---|---|---|---|---|---|---|---|---|---|---|---|---|---|---|---|
| Jul11.5v_811 | 4 | 0 | 0 | 0 | 0 | 0 | 1 | 0 | 0 | 0 | 0 | 0 | 0 | 2 | 1 | 0 | 0 | 0 | 0 | 0 | 0 | 0 | 0 | 0 | 0 | 0 | 0 | 0 | 0 | 0 |
| Jul27.25v_41111 | 1 | 0 | 0 | 0 | 0 | 0 | 0 | 0 | 0 | 0 | 0 | 0 | 0 | 0 | 0 | 0 | 0 | 0 | 1 | 0 | 0 | 0 | 0 | 0 | 0 | 0 | 0 | 0 | 0 | 0 |
| May12.5v_38399 | 1 | 1 | 0 | 0 | 0 | 0 | 0 | 0 | 0 | 0 | 0 | 0 | 0 | 0 | 0 | 0 | 0 | 0 | 0 | 0 | 0 | 0 | 0 | 0 | 0 | 0 | 0 | 0 | 0 | 0 |
| Jul27.5v_17608 | 4 | 0 | 0 | 0 | 0 | 1 | 0 | 0 | 0 | 0 | 0 | 0 | 0 | 1 | 0 | 0 | 0 | 2 | 0 | 0 | 0 | 0 | 0 | 0 | 0 | 0 | 0 | 0 | 0 | 0 |
| Jul11.5v_9282 | 10 | 0 | 0 | 0 | 3 | 0 | 0 | 0 | 0 | 0 | 1 | 2 | 0 | 2 | 0 | 0 | 0 | 0 | 0 | 1 | 0 | 0 | 0 | 0 | 0 | 0 | 0 | 1 | 0 | 0 |
| May12.75v_38832 | 3 | 0 | 0 | 0 | 1 | 1 | 0 | 0 | 0 | 0 | 0 | 0 | 0 | 0 | 1 | 0 | 0 | 0 | 0 | 0 | 0 | 0 | 0 | 0 | 0 | 0 | 0 | 0 | 0 | 0 |
| Jul11.5v_18487 | 1 | 0 | 0 | 0 | 0 | 0 | 0 | 0 | 0 | 0 | 0 | 0 | 0 | 1 | 0 | 0 | 0 | 0 | 0 | 0 | 0 | 0 | 0 | 0 | 0 | 0 | 0 | 0 | 0 | 0 |
| Jul27.5v_7935 | 1 | 0 | 0 | 0 | 0 | 0 | 0 | 0 | 0 | 0 | 0 | 0 | 0 | 0 | 0 | 0 | 0 | 1 | 0 | 0 | 0 | 0 | 0 | 0 | 0 | 0 | 0 | 0 | 0 | 0 |
| May12.75v_12425 | 6 | 0 | 0 | 0 | 2 | 0 | 0 | 0 | 0 | 0 | 0 | 0 | 0 | 2 | 0 | 0 | 0 | 1 | 1 | 0 | 0 | 0 | 0 | 0 | 0 | 0 | 0 | 0 | 0 | 0 |
| Jun22.0v_7656 | 1 | 0 | 0 | 0 | 0 | 0 | 0 | 0 | 1 | 0 | 0 | 0 | 0 | 0 | 0 | 0 | 0 | 0 | 0 | 0 | 0 | 0 | 0 | 0 | 0 | 0 | 0 | 0 | 0 | 0 |
| Jun8.50v_40750 | 1 | 0 | 0 | 0 | 0 | 0 | 1 | 0 | 0 | 0 | 0 | 0 | 0 | 0 | 0 | 0 | 0 | 0 | 0 | 0 | 0 | 0 | 0 | 0 | 0 | 0 | 0 | 0 | 0 | 0 |
| Jul11.5v_17790 | 1 | 0 | 0 | 0 | 0 | 0 | 0 | 0 | 0 | 0 | 0 | 0 | 0 | 1 | 0 | 0 | 0 | 0 | 0 | 0 | 0 | 0 | 0 | 0 | 0 | 0 | 0 | 0 | 0 | 0 |
| Jul11.5v_15011 | 2 | 0 | 0 | 0 | 0 | 0 | 0 | 0 | 0 | 0 | 0 | 0 | 0 | 2 | 0 | 0 | 0 | 0 | 0 | 0 | 0 | 0 | 0 | 0 | 0 | 0 | 0 | 0 | 0 | 0 |
| Jul11.5v_12620 | 1 | 0 | 0 | 0 | 0 | 0 | 0 | 0 | 0 | 0 | 0 | 0 | 0 | 1 | 0 | 0 | 0 | 0 | 0 | 0 | 0 | 0 | 0 | 0 | 0 | 0 | 0 | 0 | 0 | 0 |
| Jul11.50v_9849 | 1 | 0 | 0 | 0 | 0 | 0 | 0 | 0 | 0 | 0 | 0 | 0 | 0 | 0 | 0 | 1 | 0 | 0 | 0 | 0 | 0 | 0 | 0 | 0 | 0 | 0 | 0 | 0 | 0 | 0 |
| May12.75v_22525 | 2 | 0 | 1 | 0 | 1 | 0 | 0 | 0 | 0 | 0 | 0 | 0 | 0 | 0 | 0 | 0 | 0 | 0 | 0 | 0 | 0 | 0 | 0 | 0 | 0 | 0 | 0 | 0 | 0 | 0 |
| Jun22.75v_35061 | 1 | 0 | 0 | 0 | 0 | 0 | 0 | 0 | 0 | 0 | 0 | 1 | 0 | 0 | 0 | 0 | 0 | 0 | 0 | 0 | 0 | 0 | 0 | 0 | 0 | 0 | 0 | 0 | 0 | 0 |
| Jun22.75v_16753 | 5 | 1 | 1 | 0 | 1 | 0 | 0 | 0 | 0 | 0 | 0 | 1 | 0 | 0 | 0 | 0 | 0 | 0 | 1 | 0 | 0 | 0 | 0 | 0 | 0 | 0 | 0 | 0 | 0 | 0 |
| May12.75v_20751 | 1 | 0 | 0 | 0 | 1 | 0 | 0 | 0 | 0 | 0 | 0 | 0 | 0 | 0 | 0 | 0 | 0 | 0 | 0 | 0 | 0 | 0 | 0 | 0 | 0 | 0 | 0 | 0 | 0 | 0 |
| Aug5.0v_4694 | 3 | 0 | 0 | 0 | 1 | 0 | 0 | 0 | 0 | 0 | 0 | 0 | 0 | 0 | 0 | 1 | 0 | 0 | 0 | 0 | 0 | 1 | 0 | 0 | 0 | 0 | 0 | 0 | 0 | 0 |
| Aug5.75v_13224 | 7 | 2 | 1 | 2 | 1 | 0 | 0 | 0 | 0 | 0 | 0 | 0 | 0 | 0 | 0 | 0 | 0 | 0 | 0 | 0 | 0 | 0 | 0 | 0 | 0 | 1 | 0 | 0 | 0 | 0 |
| May12.50v_24282 | 1 | 0 | 0 | 1 | 0 | 0 | 0 | 0 | 0 | 0 | 0 | 0 | 0 | 0 | 0 | 0 | 0 | 0 | 0 | 0 | 0 | 0 | 0 | 0 | 0 | 0 | 0 | 0 | 0 | 0 |
| Jun8.50v_36234 | 3 | 0 | 0 | 0 | 2 | 0 | 1 | 0 | 0 | 0 | 0 | 0 | 0 | 0 | 0 | 0 | 0 | 0 | 0 | 0 | 0 | 0 | 0 | 0 | 0 | 0 | 0 | 0 | 0 | 0 |
| May12.50v_5143 | 1 | 0 | 0 | 1 | 0 | 0 | 0 | 0 | 0 | 0 | 0 | 0 | 0 | 0 | 0 | 0 | 0 | 0 | 0 | 0 | 0 | 0 | 0 | 0 | 0 | 0 | 0 | 0 | 0 | 0 |
| Sep8.5v_2354 | 1 | 0 | 0 | 0 | 0 | 0 | 0 | 0 | 0 | 0 | 0 | 0 | 0 | 0 | 0 | 0 | 0 | 0 | 0 | 0 | 0 | 0 | 0 | 0 | 0 | 0 | 0 | 1 | 0 | 0 |
| May12.5v_22134 | 2 | 1 | 0 | 1 | 0 | 0 | 0 | 0 | 0 | 0 | 0 | 0 | 0 | 0 | 0 | 0 | 0 | 0 | 0 | 0 | 0 | 0 | 0 | 0 | 0 | 0 | 0 | 0 | 0 | 0 |
| Jun8.50v_24075 | 2 | 0 | 0 | 0 | 0 | 0 | 1 | 0 | 1 | 0 | 0 | 0 | 0 | 0 | 0 | 0 | 0 | 0 | 0 | 0 | 0 | 0 | 0 | 0 | 0 | 0 | 0 | 0 | 0 | 0 |
| May12.50v_25386 | 1 | 0 | 0 | 1 | 0 | 0 | 0 | 0 | 0 | 0 | 0 | 0 | 0 | 0 | 0 | 0 | 0 | 0 | 0 | 0 | 0 | 0 | 0 | 0 | 0 | 0 | 0 | 0 | 0 | 0 |
| May12.25v_2423 | 1 | 0 | 1 | 0 | 0 | 0 | 0 | 0 | 0 | 0 | 0 | 0 | 0 | 0 | 0 | 0 | 0 | 0 | 0 | 0 | 0 | 0 | 0 | 0 | 0 | 0 | 0 | 0 | 0 | 0 |
| Jun22.5v_12098 | 1 | 0 | 0 | 0 | 0 | 0 | 0 | 0 | 0 | 1 | 0 | 0 | 0 | 0 | 0 | 0 | 0 | 0 | 0 | 0 | 0 | 0 | 0 | 0 | 0 | 0 | 0 | 0 | 0 | 0 |
| Jun8.50v_5830 | 2 | 0 | 0 | 0 | 0 | 0 | 1 | 0 | 0 | 0 | 0 | 0 | 0 | 0 | 0 | 0 | 0 | 0 | 0 | 0 | 0 | 0 | 0 | 0 | 0 | 0 | 0 | 1 | 0 | 0 |
| May12.5v_1781 | 1 | 1 | 0 | 0 | 0 | 0 | 0 | 0 | 0 | 0 | 0 | 0 | 0 | 0 | 0 | 0 | 0 | 0 | 0 | 0 | 0 | 0 | 0 | 0 | 0 | 0 | 0 | 0 | 0 | 0 |
| May12.5v_29532 | 1 | 1 | 0 | 0 | 0 | 0 | 0 | 0 | 0 | 0 | 0 | 0 | 0 | 0 | 0 | 0 | 0 | 0 | 0 | 0 | 0 | 0 | 0 | 0 | 0 | 0 | 0 | 0 | 0 | 0 |
| May12.75v_13232 | 1 | 0 | 0 | 0 | 1 | 0 | 0 | 0 | 0 | 0 | 0 | 0 | 0 | 0 | 0 | 0 | 0 | 0 | 0 | 0 | 0 | 0 | 0 | 0 | 0 | 0 | 0 | 0 | 0 | 0 |
| Jul11.75v_36621 | 3 | 0 | 0 | 0 | 2 | 0 | 0 | 0 | 0 | 0 | 0 | 0 | 0 | 0 | 0 | 0 | 1 | 0 | 0 | 0 | 0 | 0 | 0 | 0 | 0 | 0 | 0 | 0 | 0 | 0 |
| Jul11.5v_16225 | 1 | 0 | 0 | 0 | 0 | 0 | 0 | 0 | 0 | 0 | 0 | 0 | 0 | 1 | 0 | 0 | 0 | 0 | 0 | 0 | 0 | 0 | 0 | 0 | 0 | 0 | 0 | 0 | 0 | 0 |
| Jun8.50v_28378 | 1 | 0 | 0 | 0 | 0 | 0 | 1 | 0 | 0 | 0 | 0 | 0 | 0 | 0 | 0 | 0 | 0 | 0 | 0 | 0 | 0 | 0 | 0 | 0 | 0 | 0 | 0 | 0 | 0 | 0 |
| Jul11.50v_9853 | 1 | 0 | 0 | 0 | 0 | 0 | 0 | 0 | 0 | 0 | 0 | 0 | 0 | 0 | 0 | 1 | 0 | 0 | 0 | 0 | 0 | 0 | 0 | 0 | 0 | 0 | 0 | 0 | 0 | 0 |
| May12.25v_35881 | 1 | 0 | 1 | 0 | 0 | 0 | 0 | 0 | 0 | 0 | 0 | 0 | 0 | 0 | 0 | 0 | 0 | 0 | 0 | 0 | 0 | 0 | 0 | 0 | 0 | 0 | 0 | 0 | 0 | 0 |
| Jun8.50v_4437 | 1 | 0 | 0 | 0 | 0 | 0 | 1 | 0 | 0 | 0 | 0 | 0 | 0 | 0 | 0 | 0 | 0 | 0 | 0 | 0 | 0 | 0 | 0 | 0 | 0 | 0 | 0 | 0 | 0 | 0 |
| Jun22.5v_1197 | 1 | 0 | 0 | 0 | 0 | 0 | 0 | 0 | 0 | 1 | 0 | 0 | 0 | 0 | 0 | 0 | 0 | 0 | 0 | 0 | 0 | 0 | 0 | 0 | 0 | 0 | 0 | 0 | 0 | 0 |
| May12.5v_21702 | 1 | 1 | 0 | 0 | 0 | 0 | 0 | 0 | 0 | 0 | 0 | 0 | 0 | 0 | 0 | 0 | 0 | 0 | 0 | 0 | 0 | 0 | 0 | 0 | 0 | 0 | 0 | 0 | 0 | 0 |
| May12.75v_35295 | 1 | 0 | 0 | 0 | 1 | 0 | 0 | 0 | 0 | 0 | 0 | 0 | 0 | 0 | 0 | 0 | 0 | 0 | 0 | 0 | 0 | 0 | 0 | 0 | 0 | 0 | 0 | 0 | 0 | 0 |
| Jun22.50v_31867 | 1 | 0 | 0 | 0 | 0 | 0 | 0 | 0 | 0 | 0 | 1 | 0 | 0 | 0 | 0 | 0 | 0 | 0 | 0 | 0 | 0 | 0 | 0 | 0 | 0 | 0 | 0 | 0 | 0 | 0 |
| May12.5v_14498 | 1 | 1 | 0 | 0 | 0 | 0 | 0 | 0 | 0 | 0 | 0 | 0 | 0 | 0 | 0 | 0 | 0 | 0 | 0 | 0 | 0 | 0 | 0 | 0 | 0 | 0 | 0 | 0 | 0 | 0 |
| Jul27.75v_8129 | 1 | 0 | 0 | 0 | 0 | 0 | 0 | 0 | 0 | 0 | 0 | 0 | 0 | 0 | 0 | 0 | 0 | 0 | 0 | 0 | 1 | 0 | 0 | 0 | 0 | 0 | 0 | 0 | 0 | 0 |
| Jul27.75v_20235 | 3 | 0 | 0 | 1 | 1 | 0 | 0 | 0 | 0 | 0 | 0 | 0 | 0 | 0 | 0 | 0 | 0 | 0 | 0 | 0 | 1 | 0 | 0 | 0 | 0 | 0 | 0 | 0 | 0 | 0 |
| Jun8.50v_32444 | 1 | 0 | 0 | 0 | 0 | 0 | 1 | 0 | 0 | 0 | 0 | 0 | 0 | 0 | 0 | 0 | 0 | 0 | 0 | 0 | 0 | 0 | 0 | 0 | 0 | 0 | 0 | 0 | 0 | 0 |
| Jun22.50v_12192 | 2 | 0 | 0 | 0 | 1 | 0 | 0 | 0 | 0 | 0 | 1 | 0 | 0 | 0 | 0 | 0 | 0 | 0 | 0 | 0 | 0 | 0 | 0 | 0 | 0 | 0 | 0 | 0 | 0 | 0 |
| May12.50v_33817 | 1 | 0 | 0 | 1 | 0 | 0 | 0 | 0 | 0 | 0 | 0 | 0 | 0 | 0 | 0 | 0 | 0 | 0 | 0 | 0 | 0 | 0 | 0 | 0 | 0 | 0 | 0 | 0 | 0 | 0 |
| Jul11.5v_2359 | 1 | 0 | 0 | 0 | 0 | 0 | 0 | 0 | 0 | 0 | 0 | 0 | 0 | 1 | 0 | 0 | 0 | 0 | 0 | 0 | 0 | 0 | 0 | 0 | 0 | 0 | 0 | 0 | 0 | 0 |
| Jul27.50v_31841 | 1 | 0 | 0 | 0 | 0 | 0 | 0 | 0 | 0 | 0 | 0 | 0 | 0 | 0 | 0 | 0 | 0 | 0 | 0 | 1 | 0 | 0 | 0 | 0 | 0 | 0 | 0 | 0 | 0 | 0 |
| Jul27.5v_9135 | 5 | 0 | 0 | 0 | 0 | 0 | 1 | 0 | 0 | 0 | 0 | 1 | 0 | 1 | 0 | 0 | 0 | 1 | 0 | 0 | 0 | 0 | 0 | 0 | 0 | 0 | 0 | 1 | 0 | 0 |
| Jun22.50v_40868 | 1 | 0 | 0 | 0 | 0 | 0 | 0 | 0 | 0 | 0 | 1 | 0 | 0 | 0 | 0 | 0 | 0 | 0 | 0 | 0 | 0 | 0 | 0 | 0 | 0 | 0 | 0 | 0 | 0 | 0 |
| Jun8.75v_6003 | 2 | 0 | 0 | 0 | 0 | 0 | 0 | 1 | 0 | 0 | 0 | 0 | 0 | 0 | 0 | 0 | 0 | 0 | 0 | 1 | 0 | 0 | 0 | 0 | 0 | 0 | 0 | 0 | 0 | 0 |
| May12.5v_31990 | 1 | 1 | 0 | 0 | 0 | 0 | 0 | 0 | 0 | 0 | 0 | 0 | 0 | 0 | 0 | 0 | 0 | 0 | 0 | 0 | 0 | 0 | 0 | 0 | 0 | 0 | 0 | 0 | 0 | 0 |
| Sep8.5v_8082 | 2 | 0 | 0 | 0 | 0 | 0 | 0 | 1 | 0 | 0 | 0 | 0 | 0 | 0 | 0 | 0 | 0 | 0 | 0 | 0 | 0 | 0 | 0 | 0 | 0 | 0 | 0 | 1 | 0 | 0 |
| Jul11.25v_14119 | 1 | 0 | 0 | 0 | 0 | 0 | 0 | 0 | 0 | 0 | 0 | 0 | 0 | 0 | 1 | 0 | 0 | 0 | 0 | 0 | 0 | 0 | 0 | 0 | 0 | 0 | 0 | 0 | 0 | 0 |
| Jul11.5v_6463 | 2 | 0 | 0 | 0 | 0 | 0 | 0 | 0 | 0 | 0 | 0 | 0 | 0 | 1 | 0 | 0 | 0 | 0 | 0 | 1 | 0 | 0 | 0 | 0 | 0 | 0 | 0 | 0 | 0 | 0 |
| Jun22.5v_21048 | 1 | 0 | 0 | 0 | 0 | 0 | 0 | 0 | 0 | 1 | 0 | 0 | 0 | 0 | 0 | 0 | 0 | 0 | 0 | 0 | 0 | 0 | 0 | 0 | 0 | 0 | 0 | 0 | 0 | 0 |
| Jul27.5v_20218 | 1 | 0 | 0 | 0 | 0 | 0 | 0 | 0 | 0 | 0 | 0 | 0 | 0 | 0 | 0 | 0 | 0 | 1 | 0 | 0 | 0 | 0 | 0 | 0 | 0 | 0 | 0 | 0 | 0 | 0 |
| Jun8.50v_30352 | 1 | 0 | 0 | 0 | 0 | 0 | 1 | 0 | 0 | 0 | 0 | 0 | 0 | 0 | 0 | 0 | 0 | 0 | 0 | 0 | 0 | 0 | 0 | 0 | 0 | 0 | 0 | 0 | 0 | 0 |
| May12.25v_13023 | 1 | 0 | 1 | 0 | 0 | 0 | 0 | 0 | 0 | 0 | 0 | 0 | 0 | 0 | 0 | 0 | 0 | 0 | 0 | 0 | 0 | 0 | 0 | 0 | 0 | 0 | 0 | 0 | 0 | 0 |
| May12.5v_29920 | 1 | 1 | 0 | 0 | 0 | 0 | 0 | 0 | 0 | 0 | 0 | 0 | 0 | 0 | 0 | 0 | 0 | 0 | 0 | 0 | 0 | 0 | 0 | 0 | 0 | 0 | 0 | 0 | 0 | 0 |
| Jun22.50v_36714 | 1 | 0 | 0 | 0 | 0 | 0 | 0 | 0 | 0 | 0 | 1 | 0 | 0 | 0 | 0 | 0 | 0 | 0 | 0 | 0 | 0 | 0 | 0 | 0 | 0 | 0 | 0 | 0 | 0 | 0 |
| May12.50v_40245 | 1 | 0 | 0 | 1 | 0 | 0 | 0 | 0 | 0 | 0 | 0 | 0 | 0 | 0 | 0 | 0 | 0 | 0 | 0 | 0 | 0 | 0 | 0 | 0 | 0 | 0 | 0 | 0 | 0 | 0 |
| Jun22.75v_22208 | 1 | 0 | 0 | 0 | 0 | 0 | 0 | 0 | 0 | 0 | 0 | 1 | 0 | 0 | 0 | 0 | 0 | 0 | 0 | 0 | 0 | 0 | 0 | 0 | 0 | 0 | 0 | 0 | 0 | 0 |
| May12.75v_7025 | 1 | 0 | 0 | 0 | 1 | 0 | 0 | 0 | 0 | 0 | 0 | 0 | 0 | 0 | 0 | 0 | 0 | 0 | 0 | 0 | 0 | 0 | 0 | 0 | 0 | 0 | 0 | 0 | 0 | 0 |
| Jun22.75v_1757 | 1 | 0 | 0 | 0 | 0 | 0 | 0 | 0 | 0 | 0 | 0 | 1 | 0 | 0 | 0 | 0 | 0 | 0 | 0 | 0 | 0 | 0 | 0 | 0 | 0 | 0 | 0 | 0 | 0 | 0 |
| Sep8.5v_17398 | 1 | 0 | 0 | 0 | 0 | 0 | 0 | 0 | 0 | 0 | 0 | 0 | 0 | 0 | 0 | 0 | 0 | 0 | 0 | 0 | 0 | 0 | 0 | 0 | 0 | 0 | 0 | 1 | 0 | 0 |
| May12.75v_5824 | 1 | 0 | 0 | 0 | 1 | 0 | 0 | 0 | 0 | 0 | 0 | 0 | 0 | 0 | 0 | 0 | 0 | 0 | 0 | 0 | 0 | 0 | 0 | 0 | 0 | 0 | 0 | 0 | 0 | 0 |
| Jun22.75v_18459 | 1 | 0 | 0 | 0 | 0 | 0 | 0 | 0 | 0 | 0 | 0 | 1 | 0 | 0 | 0 | 0 | 0 | 0 | 0 | 0 | 0 | 0 | 0 | 0 | 0 | 0 | 0 | 0 | 0 | 0 |
| Jun8.75v_15985 | 1 | 0 | 0 | 0 | 0 | 0 | 0 | 1 | 0 | 0 | 0 | 0 | 0 | 0 | 0 | 0 | 0 | 0 | 0 | 0 | 0 | 0 | 0 | 0 | 0 | 0 | 0 | 0 | 0 | 0 |
| May12.50v_11507 | 1 | 0 | 0 | 1 | 0 | 0 | 0 | 0 | 0 | 0 | 0 | 0 | 0 | 0 | 0 | 0 | 0 | 0 | 0 | 0 | 0 | 0 | 0 | 0 | 0 | 0 | 0 | 0 | 0 | 0 |
| Jul11.75v_38413 | 2 | 0 | 0 | 0 | 0 | 0 | 0 | 0 | 0 | 0 | 0 | 0 | 0 | 0 | 0 | 0 | 1 | 0 | 0 | 0 | 0 | 0 | 0 | 0 | 0 | 1 | 0 | 0 | 0 | 0 |
| Jul11.75v_36883 | 1 | 0 | 0 | 0 | 0 | 0 | 0 | 0 | 0 | 0 | 0 | 0 | 0 | 0 | 0 | 0 | 1 | 0 | 0 | 0 | 0 | 0 | 0 | 0 | 0 | 0 | 0 | 0 | 0 | 0 |
| May12.25v_34956 | 1 | 0 | 1 | 0 | 0 | 0 | 0 | 0 | 0 | 0 | 0 | 0 | 0 | 0 | 0 | 0 | 0 | 0 | 0 | 0 | 0 | 0 | 0 | 0 | 0 | 0 | 0 | 0 | 0 | 0 |
| Sep8.5v_2219 | 1 | 0 | 0 | 0 | 0 | 0 | 0 | 0 | 0 | 0 | 0 | 0 | 0 | 0 | 0 | 0 | 0 | 0 | 0 | 0 | 0 | 0 | 0 | 0 | 0 | 0 | 0 | 1 | 0 | 0 |
| May12.25v_18011 | 1 | 0 | 1 | 0 | 0 | 0 | 0 | 0 | 0 | 0 | 0 | 0 | 0 | 0 | 0 | 0 | 0 | 0 | 0 | 0 | 0 | 0 | 0 | 0 | 0 | 0 | 0 | 0 | 0 | 0 |
| Jun8.50v_6958 | 1 | 0 | 0 | 0 | 0 | 0 | 1 | 0 | 0 | 0 | 0 | 0 | 0 | 0 | 0 | 0 | 0 | 0 | 0 | 0 | 0 | 0 | 0 | 0 | 0 | 0 | 0 | 0 | 0 | 0 |
| May12.5v_20341 | 1 | 1 | 0 | 0 | 0 | 0 | 0 | 0 | 0 | 0 | 0 | 0 | 0 | 0 | 0 | 0 | 0 | 0 | 0 | 0 | 0 | 0 | 0 | 0 | 0 | 0 | 0 | 0 | 0 | 0 |
| Sep8.5v_10759 | 1 | 0 | 0 | 0 | 0 | 0 | 0 | 0 | 0 | 0 | 0 | 0 | 0 | 0 | 0 | 0 | 0 | 0 | 0 | 0 | 0 | 0 | 0 | 0 | 0 | 0 | 0 | 1 | 0 | 0 |
| Jun8.25v_24785 | 1 | 0 | 0 | 0 | 0 | 1 | 0 | 0 | 0 | 0 | 0 | 0 | 0 | 0 | 0 | 0 | 0 | 0 | 0 | 0 | 0 | 0 | 0 | 0 | 0 | 0 | 0 | 0 | 0 | 0 |
| Jun22.75v_4346 | 1 | 0 | 0 | 0 | 0 | 0 | 0 | 0 | 0 | 0 | 0 | 1 | 0 | 0 | 0 | 0 | 0 | 0 | 0 | 0 | 0 | 0 | 0 | 0 | 0 | 0 | 0 | 0 | 0 | 0 |
| Sep8.5v_553 | 1 | 0 | 0 | 0 | 0 | 0 | 0 | 0 | 0 | 0 | 0 | 0 | 0 | 0 | 0 | 0 | 0 | 0 | 0 | 0 | 0 | 0 | 0 | 0 | 0 | 0 | 0 | 1 | 0 | 0 |
| May12.25v_16354 | 1 | 0 | 1 | 0 | 0 | 0 | 0 | 0 | 0 | 0 | 0 | 0 | 0 | 0 | 0 | 0 | 0 | 0 | 0 | 0 | 0 | 0 | 0 | 0 | 0 | 0 | 0 | 0 | 0 | 0 |
| Jun22.75v_38535 | 1 | 0 | 0 | 0 | 0 | 0 | 0 | 0 | 0 | 0 | 0 | 1 | 0 | 0 | 0 | 0 | 0 | 0 | 0 | 0 | 0 | 0 | 0 | 0 | 0 | 0 | 0 | 0 | 0 | 0 |
| Aug5.5v_32033 | 1 | 0 | 0 | 0 | 0 | 0 | 0 | 0 | 0 | 0 | 0 | 0 | 0 | 0 | 0 | 0 | 0 | 0 | 0 | 0 | 0 | 0 | 1 | 0 | 0 | 0 | 0 | 0 | 0 | 0 |
| Jul11.50v_18812 | 1 | 0 | 0 | 0 | 0 | 0 | 0 | 0 | 0 | 0 | 0 | 0 | 0 | 0 | 0 | 1 | 0 | 0 | 0 | 0 | 0 | 0 | 0 | 0 | 0 | 0 | 0 | 0 | 0 | 0 |
| Jun8.50v_502 | 1 | 0 | 0 | 0 | 0 | 0 | 1 | 0 | 0 | 0 | 0 | 0 | 0 | 0 | 0 | 0 | 0 | 0 | 0 | 0 | 0 | 0 | 0 | 0 | 0 | 0 | 0 | 0 | 0 | 0 |
| Jul27.50v_22531 | 1 | 0 | 0 | 0 | 0 | 0 | 0 | 0 | 0 | 0 | 0 | 0 | 0 | 0 | 0 | 0 | 0 | 0 | 0 | 1 | 0 | 0 | 0 | 0 | 0 | 0 | 0 | 0 | 0 | 0 |
| May12.75v_30198 | 1 | 0 | 0 | 0 | 1 | 0 | 0 | 0 | 0 | 0 | 0 | 0 | 0 | 0 | 0 | 0 | 0 | 0 | 0 | 0 | 0 | 0 | 0 | 0 | 0 | 0 | 0 | 0 | 0 | 0 |
| Jun8.75v_17650 | 1 | 0 | 0 | 0 | 0 | 0 | 0 | 1 | 0 | 0 | 0 | 0 | 0 | 0 | 0 | 0 | 0 | 0 | 0 | 0 | 0 | 0 | 0 | 0 | 0 | 0 | 0 | 0 | 0 | 0 |
| Jul11.75v_28798 | 1 | 0 | 0 | 0 | 0 | 0 | 0 | 0 | 0 | 0 | 0 | 0 | 0 | 0 | 0 | 0 | 1 | 0 | 0 | 0 | 0 | 0 | 0 | 0 | 0 | 0 | 0 | 0 | 0 | 0 |
| Sep8.5v_5564 | 1 | 0 | 0 | 0 | 0 | 0 | 0 | 0 | 0 | 0 | 0 | 0 | 0 | 0 | 0 | 0 | 0 | 0 | 0 | 0 | 0 | 0 | 0 | 0 | 0 | 0 | 0 | 1 | 0 | 0 |
| Jun8.75v_3911 | 5 | 0 | 0 | 0 | 0 | 0 | 1 | 2 | 0 | 0 | 0 | 1 | 0 | 0 | 1 | 0 | 0 | 0 | 0 | 0 | 0 | 0 | 0 | 0 | 0 | 0 | 0 | 0 | 0 | 0 |
| Sep8.5v_18285 | 1 | 0 | 0 | 0 | 0 | 0 | 0 | 0 | 0 | 0 | 0 | 0 | 0 | 0 | 0 | 0 | 0 | 0 | 0 | 0 | 0 | 0 | 0 | 0 | 0 | 0 | 0 | 1 | 0 | 0 |
| Jun8.75v_2736 | 1 | 0 | 0 | 0 | 0 | 0 | 0 | 1 | 0 | 0 | 0 | 0 | 0 | 0 | 0 | 0 | 0 | 0 | 0 | 0 | 0 | 0 | 0 | 0 | 0 | 0 | 0 | 0 | 0 | 0 |
| Jun8.50v_13937 | 1 | 0 | 0 | 0 | 0 | 0 | 1 | 0 | 0 | 0 | 0 | 0 | 0 | 0 | 0 | 0 | 0 | 0 | 0 | 0 | 0 | 0 | 0 | 0 | 0 | 0 | 0 | 0 | 0 | 0 |
| Jun22.5v_18240 | 1 | 0 | 0 | 0 | 0 | 0 | 0 | 0 | 0 | 1 | 0 | 0 | 0 | 0 | 0 | 0 | 0 | 0 | 0 | 0 | 0 | 0 | 0 | 0 | 0 | 0 | 0 | 0 | 0 | 0 |
| Jun22.0v_838 | 1 | 0 | 0 | 0 | 0 | 0 | 0 | 0 | 1 | 0 | 0 | 0 | 0 | 0 | 0 | 0 | 0 | 0 | 0 | 0 | 0 | 0 | 0 | 0 | 0 | 0 | 0 | 0 | 0 | 0 |
| May12.75v_28554 | 1 | 0 | 0 | 0 | 1 | 0 | 0 | 0 | 0 | 0 | 0 | 0 | 0 | 0 | 0 | 0 | 0 | 0 | 0 | 0 | 0 | 0 | 0 | 0 | 0 | 0 | 0 | 0 | 0 | 0 |
| Jun8.50v_39939 | 2 | 0 | 0 | 0 | 0 | 0 | 2 | 0 | 0 | 0 | 0 | 0 | 0 | 0 | 0 | 0 | 0 | 0 | 0 | 0 | 0 | 0 | 0 | 0 | 0 | 0 | 0 | 0 | 0 | 0 |
| Jul11.5v_1952 | 1 | 0 | 0 | 0 | 0 | 0 | 0 | 0 | 0 | 0 | 0 | 0 | 0 | 1 | 0 | 0 | 0 | 0 | 0 | 0 | 0 | 0 | 0 | 0 | 0 | 0 | 0 | 0 | 0 | 0 |
| Jun22.75v_40825 | 1 | 0 | 0 | 0 | 0 | 0 | 0 | 0 | 0 | 0 | 0 | 1 | 0 | 0 | 0 | 0 | 0 | 0 | 0 | 0 | 0 | 0 | 0 | 0 | 0 | 0 | 0 | 0 | 0 | 0 |
| Jul11.5v_14609 | 1 | 0 | 0 | 0 | 0 | 0 | 0 | 0 | 0 | 0 | 0 | 0 | 0 | 1 | 0 | 0 | 0 | 0 | 0 | 0 | 0 | 0 | 0 | 0 | 0 | 0 | 0 | 0 | 0 | 0 |
| May12.75v_16871 | 1 | 0 | 0 | 0 | 1 | 0 | 0 | 0 | 0 | 0 | 0 | 0 | 0 | 0 | 0 | 0 | 0 | 0 | 0 | 0 | 0 | 0 | 0 | 0 | 0 | 0 | 0 | 0 | 0 | 0 |
| Jun22.75v_31125 | 1 | 0 | 0 | 0 | 0 | 0 | 0 | 0 | 0 | 0 | 0 | 1 | 0 | 0 | 0 | 0 | 0 | 0 | 0 | 0 | 0 | 0 | 0 | 0 | 0 | 0 | 0 | 0 | 0 | 0 |
| Jun8.75v_6517 | 1 | 0 | 0 | 0 | 0 | 0 | 0 | 1 | 0 | 0 | 0 | 0 | 0 | 0 | 0 | 0 | 0 | 0 | 0 | 0 | 0 | 0 | 0 | 0 | 0 | 0 | 0 | 0 | 0 | 0 |
| Sep8.5v_4658 | 1 | 0 | 0 | 0 | 0 | 0 | 0 | 0 | 0 | 0 | 0 | 0 | 0 | 0 | 0 | 0 | 0 | 0 | 0 | 0 | 0 | 0 | 0 | 0 | 0 | 0 | 0 | 1 | 0 | 0 |
| May12.25v_28165 | 1 | 0 | 1 | 0 | 0 | 0 | 0 | 0 | 0 | 0 | 0 | 0 | 0 | 0 | 0 | 0 | 0 | 0 | 0 | 0 | 0 | 0 | 0 | 0 | 0 | 0 | 0 | 0 | 0 | 0 |
| Jul27.5v_4018 | 1 | 0 | 0 | 0 | 0 | 0 | 0 | 0 | 0 | 0 | 0 | 0 | 0 | 0 | 0 | 0 | 0 | 1 | 0 | 0 | 0 | 0 | 0 | 0 | 0 | 0 | 0 | 0 | 0 | 0 |
| Jul11.0v_19251 | 1 | 0 | 0 | 0 | 0 | 0 | 0 | 0 | 0 | 0 | 0 | 0 | 1 | 0 | 0 | 0 | 0 | 0 | 0 | 0 | 0 | 0 | 0 | 0 | 0 | 0 | 0 | 0 | 0 | 0 |
| Jun8.75v_3555 | 1 | 0 | 0 | 0 | 0 | 0 | 0 | 1 | 0 | 0 | 0 | 0 | 0 | 0 | 0 | 0 | 0 | 0 | 0 | 0 | 0 | 0 | 0 | 0 | 0 | 0 | 0 | 0 | 0 | 0 |
| Jun22.75v_39617 | 1 | 0 | 0 | 0 | 0 | 0 | 0 | 0 | 0 | 0 | 0 | 1 | 0 | 0 | 0 | 0 | 0 | 0 | 0 | 0 | 0 | 0 | 0 | 0 | 0 | 0 | 0 | 0 | 0 | 0 |
| Jul11.5v_11372 | 1 | 0 | 0 | 0 | 0 | 0 | 0 | 0 | 0 | 0 | 0 | 0 | 0 | 1 | 0 | 0 | 0 | 0 | 0 | 0 | 0 | 0 | 0 | 0 | 0 | 0 | 0 | 0 | 0 | 0 |
| Jun8.75v_115 | 1 | 0 | 0 | 0 | 0 | 0 | 0 | 1 | 0 | 0 | 0 | 0 | 0 | 0 | 0 | 0 | 0 | 0 | 0 | 0 | 0 | 0 | 0 | 0 | 0 | 0 | 0 | 0 | 0 | 0 |
| Jun8.25v_13954 | 1 | 0 | 0 | 0 | 0 | 1 | 0 | 0 | 0 | 0 | 0 | 0 | 0 | 0 | 0 | 0 | 0 | 0 | 0 | 0 | 0 | 0 | 0 | 0 | 0 | 0 | 0 | 0 | 0 | 0 |
| Aug5.75v_5167 | 4 | 0 | 1 | 0 | 1 | 0 | 0 | 0 | 0 | 0 | 0 | 0 | 0 | 0 | 0 | 0 | 0 | 0 | 0 | 0 | 0 | 0 | 0 | 0 | 0 | 2 | 0 | 0 | 0 | 0 |
| Jul11.75v_6584 | 1 | 0 | 0 | 0 | 0 | 0 | 0 | 0 | 0 | 0 | 0 | 0 | 0 | 0 | 0 | 0 | 1 | 0 | 0 | 0 | 0 | 0 | 0 | 0 | 0 | 0 | 0 | 0 | 0 | 0 |
| May12.75v_35955 | 1 | 0 | 0 | 0 | 1 | 0 | 0 | 0 | 0 | 0 | 0 | 0 | 0 | 0 | 0 | 0 | 0 | 0 | 0 | 0 | 0 | 0 | 0 | 0 | 0 | 0 | 0 | 0 | 0 | 0 |
| Jun22.50v_8676 | 1 | 0 | 0 | 0 | 0 | 0 | 0 | 0 | 0 | 0 | 1 | 0 | 0 | 0 | 0 | 0 | 0 | 0 | 0 | 0 | 0 | 0 | 0 | 0 | 0 | 0 | 0 | 0 | 0 | 0 |
| Jun22.75v_33788 | 1 | 0 | 0 | 0 | 0 | 0 | 0 | 0 | 0 | 0 | 0 | 1 | 0 | 0 | 0 | 0 | 0 | 0 | 0 | 0 | 0 | 0 | 0 | 0 | 0 | 0 | 0 | 0 | 0 | 0 |
| May12.5v_37173 | 2 | 1 | 0 | 0 | 0 | 0 | 0 | 0 | 0 | 0 | 0 | 0 | 0 | 0 | 0 | 0 | 1 | 0 | 0 | 0 | 0 | 0 | 0 | 0 | 0 | 0 | 0 | 0 | 0 | 0 |
| Jul11.25v_3200 | 1 | 0 | 0 | 0 | 0 | 0 | 0 | 0 | 0 | 0 | 0 | 0 | 0 | 0 | 1 | 0 | 0 | 0 | 0 | 0 | 0 | 0 | 0 | 0 | 0 | 0 | 0 | 0 | 0 | 0 |
| May12.25v_19002 | 1 | 0 | 1 | 0 | 0 | 0 | 0 | 0 | 0 | 0 | 0 | 0 | 0 | 0 | 0 | 0 | 0 | 0 | 0 | 0 | 0 | 0 | 0 | 0 | 0 | 0 | 0 | 0 | 0 | 0 |
| May12.50v_9880 | 1 | 0 | 0 | 1 | 0 | 0 | 0 | 0 | 0 | 0 | 0 | 0 | 0 | 0 | 0 | 0 | 0 | 0 | 0 | 0 | 0 | 0 | 0 | 0 | 0 | 0 | 0 | 0 | 0 | 0 |
| Jul11.50v_11281 | 1 | 0 | 0 | 0 | 0 | 0 | 0 | 0 | 0 | 0 | 0 | 0 | 0 | 0 | 0 | 1 | 0 | 0 | 0 | 0 | 0 | 0 | 0 | 0 | 0 | 0 | 0 | 0 | 0 | 0 |
| Jul11.5v_3734 | 1 | 0 | 0 | 0 | 0 | 0 | 0 | 0 | 0 | 0 | 0 | 0 | 0 | 1 | 0 | 0 | 0 | 0 | 0 | 0 | 0 | 0 | 0 | 0 | 0 | 0 | 0 | 0 | 0 | 0 |
| Jul11.25v_11137 | 1 | 0 | 0 | 0 | 0 | 0 | 0 | 0 | 0 | 0 | 0 | 0 | 0 | 0 | 1 | 0 | 0 | 0 | 0 | 0 | 0 | 0 | 0 | 0 | 0 | 0 | 0 | 0 | 0 | 0 |
| Jun22.75v_29454 | 1 | 0 | 0 | 0 | 0 | 0 | 0 | 0 | 0 | 0 | 0 | 1 | 0 | 0 | 0 | 0 | 0 | 0 | 0 | 0 | 0 | 0 | 0 | 0 | 0 | 0 | 0 | 0 | 0 | 0 |
| Jun8.50v_2049 | 1 | 0 | 0 | 0 | 0 | 0 | 1 | 0 | 0 | 0 | 0 | 0 | 0 | 0 | 0 | 0 | 0 | 0 | 0 | 0 | 0 | 0 | 0 | 0 | 0 | 0 | 0 | 0 | 0 | 0 |
| Jul11.75v_26829 | 1 | 0 | 0 | 0 | 0 | 0 | 0 | 0 | 0 | 0 | 0 | 0 | 0 | 0 | 0 | 0 | 1 | 0 | 0 | 0 | 0 | 0 | 0 | 0 | 0 | 0 | 0 | 0 | 0 | 0 |
| May12.75v_1319 | 1 | 0 | 0 | 0 | 1 | 0 | 0 | 0 | 0 | 0 | 0 | 0 | 0 | 0 | 0 | 0 | 0 | 0 | 0 | 0 | 0 | 0 | 0 | 0 | 0 | 0 | 0 | 0 | 0 | 0 |
| Jul11.75v_37192 | 1 | 0 | 0 | 0 | 0 | 0 | 0 | 0 | 0 | 0 | 0 | 0 | 0 | 0 | 0 | 0 | 1 | 0 | 0 | 0 | 0 | 0 | 0 | 0 | 0 | 0 | 0 | 0 | 0 | 0 |
| Jul11.75v_21494 | 3 | 0 | 0 | 0 | 0 | 0 | 1 | 0 | 0 | 0 | 1 | 0 | 0 | 0 | 0 | 0 | 1 | 0 | 0 | 0 | 0 | 0 | 0 | 0 | 0 | 0 | 0 | 0 | 0 | 0 |
| Jul11.75v_4234 | 1 | 0 | 0 | 0 | 0 | 0 | 0 | 0 | 0 | 0 | 0 | 0 | 0 | 0 | 0 | 0 | 1 | 0 | 0 | 0 | 0 | 0 | 0 | 0 | 0 | 0 | 0 | 0 | 0 | 0 |
| Jun8.50v_16334 | 1 | 0 | 0 | 0 | 0 | 0 | 1 | 0 | 0 | 0 | 0 | 0 | 0 | 0 | 0 | 0 | 0 | 0 | 0 | 0 | 0 | 0 | 0 | 0 | 0 | 0 | 0 | 0 | 0 | 0 |
| Jun22.0v_17047 | 1 | 0 | 0 | 0 | 0 | 0 | 0 | 0 | 1 | 0 | 0 | 0 | 0 | 0 | 0 | 0 | 0 | 0 | 0 | 0 | 0 | 0 | 0 | 0 | 0 | 0 | 0 | 0 | 0 | 0 |
| May12.25v_32287 | 1 | 0 | 1 | 0 | 0 | 0 | 0 | 0 | 0 | 0 | 0 | 0 | 0 | 0 | 0 | 0 | 0 | 0 | 0 | 0 | 0 | 0 | 0 | 0 | 0 | 0 | 0 | 0 | 0 | 0 |
| Jun22.0v_16886 | 1 | 0 | 0 | 0 | 0 | 0 | 0 | 0 | 1 | 0 | 0 | 0 | 0 | 0 | 0 | 0 | 0 | 0 | 0 | 0 | 0 | 0 | 0 | 0 | 0 | 0 | 0 | 0 | 0 | 0 |
| Jul11.50v_19926 | 1 | 0 | 0 | 0 | 0 | 0 | 0 | 0 | 0 | 0 | 0 | 0 | 0 | 0 | 0 | 1 | 0 | 0 | 0 | 0 | 0 | 0 | 0 | 0 | 0 | 0 | 0 | 0 | 0 | 0 |
| Jul11.25v_5123 | 1 | 0 | 0 | 0 | 0 | 0 | 0 | 0 | 0 | 0 | 0 | 0 | 0 | 0 | 1 | 0 | 0 | 0 | 0 | 0 | 0 | 0 | 0 | 0 | 0 | 0 | 0 | 0 | 0 | 0 |
| Jul11.25v_1151 | 1 | 0 | 0 | 0 | 0 | 0 | 0 | 0 | 0 | 0 | 0 | 0 | 0 | 0 | 1 | 0 | 0 | 0 | 0 | 0 | 0 | 0 | 0 | 0 | 0 | 0 | 0 | 0 | 0 | 0 |
| Jul27.5v_19951 | 1 | 0 | 0 | 0 | 0 | 0 | 0 | 0 | 0 | 0 | 0 | 0 | 0 | 0 | 0 | 0 | 0 | 1 | 0 | 0 | 0 | 0 | 0 | 0 | 0 | 0 | 0 | 0 | 0 | 0 |
| Jul11.25v_15347 | 1 | 0 | 0 | 0 | 0 | 0 | 0 | 0 | 0 | 0 | 0 | 0 | 0 | 0 | 1 | 0 | 0 | 0 | 0 | 0 | 0 | 0 | 0 | 0 | 0 | 0 | 0 | 0 | 0 | 0 |
| Aug5.25v_14483 | 179 | 7 | 3 | 6 | 1 | 5 | 4 | 0 | 13 | 4 | 19 | 10 | 5 | 2 | 3 | 7 | 6 | 1 | 8 | 7 | 2 | 1 | 11 | 14 | 9 | 16 | 2 | 2 | 1 | 10 |
| May12.50v_39718 | 1 | 0 | 0 | 1 | 0 | 0 | 0 | 0 | 0 | 0 | 0 | 0 | 0 | 0 | 0 | 0 | 0 | 0 | 0 | 0 | 0 | 0 | 0 | 0 | 0 | 0 | 0 | 0 | 0 | 0 |
| Aug5.75v_9669 | 45 | 1 | 1 | 2 | 2 | 3 | 2 | 0 | 4 | 1 | 6 | 1 | 3 | 0 | 1 | 0 | 1 | 1 | 2 | 2 | 0 | 0 | 0 | 4 | 2 | 1 | 0 | 1 | 0 | 4 |
| Jul11.75v_12927 | 1 | 0 | 0 | 0 | 0 | 0 | 0 | 0 | 0 | 0 | 0 | 0 | 0 | 0 | 0 | 0 | 1 | 0 | 0 | 0 | 0 | 0 | 0 | 0 | 0 | 0 | 0 | 0 | 0 | 0 |
| May12.25v_29365 | 7 | 1 | 1 | 0 | 0 | 0 | 0 | 0 | 0 | 0 | 0 | 0 | 0 | 0 | 0 | 0 | 1 | 0 | 0 | 0 | 0 | 1 | 1 | 0 | 0 | 0 | 0 | 0 | 0 | 2 |
| Aug5.0v_1161 | 38 | 2 | 1 | 2 | 1 | 1 | 3 | 0 | 3 | 0 | 0 | 1 | 1 | 0 | 1 | 4 | 4 | 0 | 1 | 1 | 0 | 3 | 2 | 1 | 0 | 0 | 2 | 1 | 0 | 3 |
| Jun22.75v_32131 | 6 | 0 | 0 | 0 | 0 | 0 | 0 | 0 | 0 | 0 | 2 | 2 | 0 | 0 | 0 | 0 | 0 | 0 | 0 | 0 | 0 | 0 | 1 | 1 | 0 | 0 | 0 | 0 | 0 | 0 |
| Jun22.50v_33321 | 1 | 0 | 0 | 0 | 0 | 0 | 0 | 0 | 0 | 0 | 1 | 0 | 0 | 0 | 0 | 0 | 0 | 0 | 0 | 0 | 0 | 0 | 0 | 0 | 0 | 0 | 0 | 0 | 0 | 0 |
| Aug5.75v_25916 | 82 | 11 | 12 | 3 | 23 | 2 | 2 | 0 | 2 | 0 | 0 | 2 | 1 | 1 | 1 | 3 | 4 | 2 | 0 | 2 | 0 | 3 | 3 | 0 | 1 | 2 | 0 | 0 | 0 | 2 |
| Jun22.50v_20925 | 3 | 0 | 0 | 0 | 1 | 0 | 0 | 0 | 0 | 0 | 1 | 0 | 0 | 0 | 0 | 1 | 0 | 0 | 0 | 0 | 0 | 0 | 0 | 0 | 0 | 0 | 0 | 0 | 0 | 0 |
| Jun22.75v_24468 | 3 | 1 | 0 | 0 | 0 | 0 | 0 | 0 | 0 | 0 | 0 | 1 | 1 | 0 | 0 | 0 | 0 | 0 | 0 | 0 | 0 | 0 | 0 | 0 | 0 | 0 | 0 | 0 | 0 | 0 |
| May12.5v_30061 | 1 | 1 | 0 | 0 | 0 | 0 | 0 | 0 | 0 | 0 | 0 | 0 | 0 | 0 | 0 | 0 | 0 | 0 | 0 | 0 | 0 | 0 | 0 | 0 | 0 | 0 | 0 | 0 | 0 | 0 |
| Jun22.0v_13574 | 1 | 0 | 0 | 0 | 0 | 0 | 0 | 0 | 1 | 0 | 0 | 0 | 0 | 0 | 0 | 0 | 0 | 0 | 0 | 0 | 0 | 0 | 0 | 0 | 0 | 0 | 0 | 0 | 0 | 0 |
| May12.25v_8321 | 1 | 0 | 1 | 0 | 0 | 0 | 0 | 0 | 0 | 0 | 0 | 0 | 0 | 0 | 0 | 0 | 0 | 0 | 0 | 0 | 0 | 0 | 0 | 0 | 0 | 0 | 0 | 0 | 0 | 0 |
| Jun22.50v_3231 | 122 | 9 | 9 | 14 | 14 | 2 | 12 | 0 | 10 | 0 | 4 | 15 | 1 | 0 | 0 | 1 | 6 | 0 | 3 | 1 | 1 | 0 | 1 | 1 | 3 | 4 | 2 | 3 | 1 | 5 |
| May12.75v_3844 | 2 | 0 | 0 | 0 | 2 | 0 | 0 | 0 | 0 | 0 | 0 | 0 | 0 | 0 | 0 | 0 | 0 | 0 | 0 | 0 | 0 | 0 | 0 | 0 | 0 | 0 | 0 | 0 | 0 | 0 |
| Aug5.50v_25476 | 1 | 0 | 0 | 0 | 0 | 0 | 0 | 0 | 0 | 0 | 0 | 0 | 0 | 0 | 0 | 0 | 0 | 0 | 0 | 0 | 0 | 0 | 0 | 0 | 1 | 0 | 0 | 0 | 0 | 0 |
| Jul11.75v_25191 | 3 | 0 | 0 | 0 | 0 | 0 | 0 | 0 | 0 | 0 | 0 | 0 | 0 | 0 | 0 | 0 | 3 | 0 | 0 | 0 | 0 | 0 | 0 | 0 | 0 | 0 | 0 | 0 | 0 | 0 |
| May12.25v_37667 | 1 | 0 | 1 | 0 | 0 | 0 | 0 | 0 | 0 | 0 | 0 | 0 | 0 | 0 | 0 | 0 | 0 | 0 | 0 | 0 | 0 | 0 | 0 | 0 | 0 | 0 | 0 | 0 | 0 | 0 |
| Jun22.75v_21972 | 1 | 0 | 0 | 0 | 0 | 0 | 0 | 0 | 0 | 0 | 0 | 1 | 0 | 0 | 0 | 0 | 0 | 0 | 0 | 0 | 0 | 0 | 0 | 0 | 0 | 0 | 0 | 0 | 0 | 0 |
| Aug5.50v_13857 | 3 | 0 | 0 | 0 | 0 | 0 | 0 | 0 | 0 | 0 | 0 | 0 | 0 | 0 | 1 | 0 | 0 | 0 | 0 | 0 | 0 | 0 | 0 | 1 | 1 | 0 | 0 | 0 | 0 | 0 |
| Jun22.5v_8141 | 1 | 0 | 0 | 0 | 0 | 0 | 0 | 0 | 0 | 1 | 0 | 0 | 0 | 0 | 0 | 0 | 0 | 0 | 0 | 0 | 0 | 0 | 0 | 0 | 0 | 0 | 0 | 0 | 0 | 0 |
| Jul27.5v_14499 | 1 | 0 | 0 | 0 | 0 | 0 | 0 | 0 | 0 | 0 | 0 | 0 | 0 | 0 | 0 | 0 | 0 | 1 | 0 | 0 | 0 | 0 | 0 | 0 | 0 | 0 | 0 | 0 | 0 | 0 |
| Aug5.75v_14261 | 2 | 0 | 0 | 0 | 0 | 0 | 0 | 0 | 0 | 0 | 0 | 0 | 0 | 0 | 1 | 0 | 0 | 0 | 0 | 0 | 0 | 0 | 0 | 0 | 0 | 1 | 0 | 0 | 0 | 0 |
| Jun22.0v_11947 | 1 | 0 | 0 | 0 | 0 | 0 | 0 | 0 | 1 | 0 | 0 | 0 | 0 | 0 | 0 | 0 | 0 | 0 | 0 | 0 | 0 | 0 | 0 | 0 | 0 | 0 | 0 | 0 | 0 | 0 |
| Sep8.50v_2515 | 1 | 0 | 0 | 0 | 0 | 0 | 0 | 0 | 0 | 0 | 0 | 0 | 0 | 0 | 0 | 0 | 0 | 0 | 0 | 0 | 0 | 0 | 0 | 0 | 0 | 0 | 0 | 0 | 0 | 1 |
| Aug5.25v_8671 | 5 | 0 | 0 | 0 | 0 | 0 | 0 | 0 | 1 | 0 | 0 | 0 | 0 | 0 | 0 | 0 | 0 | 1 | 0 | 0 | 0 | 1 | 0 | 2 | 0 | 0 | 0 | 0 | 0 | 0 |
| Jun22.0v_3429 | 1 | 0 | 0 | 0 | 0 | 0 | 0 | 0 | 1 | 0 | 0 | 0 | 0 | 0 | 0 | 0 | 0 | 0 | 0 | 0 | 0 | 0 | 0 | 0 | 0 | 0 | 0 | 0 | 0 | 0 |
| Aug5.25v_35003 | 1 | 0 | 0 | 0 | 0 | 0 | 0 | 0 | 0 | 0 | 0 | 0 | 0 | 0 | 0 | 0 | 0 | 0 | 0 | 0 | 0 | 0 | 0 | 1 | 0 | 0 | 0 | 0 | 0 | 0 |
| Aug5.25v_20101 | 1 | 0 | 0 | 0 | 0 | 0 | 0 | 0 | 0 | 0 | 0 | 0 | 0 | 0 | 0 | 0 | 0 | 0 | 0 | 0 | 0 | 0 | 0 | 1 | 0 | 0 | 0 | 0 | 0 | 0 |
| Aug5.25v_3366 | 1 | 0 | 0 | 0 | 0 | 0 | 0 | 0 | 0 | 0 | 0 | 0 | 0 | 0 | 0 | 0 | 0 | 0 | 0 | 0 | 0 | 0 | 0 | 1 | 0 | 0 | 0 | 0 | 0 | 0 |
| Jul11.5v_18498 | 1466 | 2 | 1 | 0 | 1 | 0 | 0 | 0 | 5 | 4 | 0 | 0 | 77 | 82 | 30 | 1 | 0 | 116 | 105 | 19 | 0 | 543 | 93 | 236 | 23 | 22 | 40 | 0 | 3 | 63 |
| Aug5.0v_5277 | 3 | 0 | 0 | 0 | 0 | 0 | 0 | 0 | 0 | 0 | 0 | 0 | 0 | 0 | 0 | 0 | 0 | 0 | 0 | 0 | 0 | 1 | 0 | 0 | 0 | 1 | 0 | 0 | 0 | 1 |
| Aug5.0v_829 | 1 | 0 | 0 | 0 | 0 | 0 | 0 | 0 | 0 | 0 | 0 | 0 | 0 | 0 | 0 | 0 | 0 | 0 | 0 | 0 | 0 | 1 | 0 | 0 | 0 | 0 | 0 | 0 | 0 | 0 |
| Jul27.25v_861 | 1 | 0 | 0 | 0 | 0 | 0 | 0 | 0 | 0 | 0 | 0 | 0 | 0 | 0 | 0 | 0 | 0 | 0 | 1 | 0 | 0 | 0 | 0 | 0 | 0 | 0 | 0 | 0 | 0 | 0 |
| Sep8.0v_15334 | 1 | 0 | 0 | 0 | 0 | 0 | 0 | 0 | 0 | 0 | 0 | 0 | 0 | 0 | 0 | 0 | 0 | 0 | 0 | 0 | 0 | 0 | 0 | 0 | 0 | 0 | 1 | 0 | 0 | 0 |
| Aug5.5v_31869 | 1 | 0 | 0 | 0 | 0 | 0 | 0 | 0 | 0 | 0 | 0 | 0 | 0 | 0 | 0 | 0 | 0 | 0 | 0 | 0 | 0 | 0 | 1 | 0 | 0 | 0 | 0 | 0 | 0 | 0 |
| Aug5.0v_3711 | 1 | 0 | 0 | 0 | 0 | 0 | 0 | 0 | 0 | 0 | 0 | 0 | 0 | 0 | 0 | 0 | 0 | 0 | 0 | 0 | 0 | 1 | 0 | 0 | 0 | 0 | 0 | 0 | 0 | 0 |
| Aug5.25v_10907 | 1 | 0 | 0 | 0 | 0 | 0 | 0 | 0 | 0 | 0 | 0 | 0 | 0 | 0 | 0 | 0 | 0 | 0 | 0 | 0 | 0 | 0 | 0 | 1 | 0 | 0 | 0 | 0 | 0 | 0 |
| Aug5.25v_37357 | 1 | 0 | 0 | 0 | 0 | 0 | 0 | 0 | 0 | 0 | 0 | 0 | 0 | 0 | 0 | 0 | 0 | 0 | 0 | 0 | 0 | 0 | 0 | 1 | 0 | 0 | 0 | 0 | 0 | 0 |
| Aug5.5v_32649 | 1 | 0 | 0 | 0 | 0 | 0 | 0 | 0 | 0 | 0 | 0 | 0 | 0 | 0 | 0 | 0 | 0 | 0 | 0 | 0 | 0 | 0 | 1 | 0 | 0 | 0 | 0 | 0 | 0 | 0 |
| Aug5.0v_722 | 1 | 0 | 0 | 0 | 0 | 0 | 0 | 0 | 0 | 0 | 0 | 0 | 0 | 0 | 0 | 0 | 0 | 0 | 0 | 0 | 0 | 1 | 0 | 0 | 0 | 0 | 0 | 0 | 0 | 0 |
| Aug5.25v_34955 | 1 | 0 | 0 | 0 | 0 | 0 | 0 | 0 | 0 | 0 | 0 | 0 | 0 | 0 | 0 | 0 | 0 | 0 | 0 | 0 | 0 | 0 | 0 | 1 | 0 | 0 | 0 | 0 | 0 | 0 |
| Aug5.0v_3014 | 2 | 0 | 0 | 0 | 0 | 0 | 0 | 0 | 0 | 0 | 0 | 0 | 0 | 0 | 0 | 0 | 0 | 0 | 0 | 0 | 0 | 1 | 0 | 0 | 0 | 0 | 0 | 0 | 0 | 1 |
| Jul27.5v_19824 | 1 | 0 | 0 | 0 | 0 | 0 | 0 | 0 | 0 | 0 | 0 | 0 | 0 | 0 | 0 | 0 | 0 | 1 | 0 | 0 | 0 | 0 | 0 | 0 | 0 | 0 | 0 | 0 | 0 | 0 |
| Jul27.5v_17437 | 1 | 0 | 0 | 0 | 0 | 0 | 0 | 0 | 0 | 0 | 0 | 0 | 0 | 0 | 0 | 0 | 0 | 1 | 0 | 0 | 0 | 0 | 0 | 0 | 0 | 0 | 0 | 0 | 0 | 0 |
| Jul11.5v_17079 | 2 | 0 | 0 | 0 | 0 | 0 | 0 | 0 | 0 | 0 | 0 | 0 | 0 | 1 | 0 | 0 | 0 | 0 | 1 | 0 | 0 | 0 | 0 | 0 | 0 | 0 | 0 | 0 | 0 | 0 |
| Jul27.5v_15609 | 1 | 0 | 0 | 0 | 0 | 0 | 0 | 0 | 0 | 0 | 0 | 0 | 0 | 0 | 0 | 0 | 0 | 1 | 0 | 0 | 0 | 0 | 0 | 0 | 0 | 0 | 0 | 0 | 0 | 0 |
| Sep8.0v_19152 | 1 | 0 | 0 | 0 | 0 | 0 | 0 | 0 | 0 | 0 | 0 | 0 | 0 | 0 | 0 | 0 | 0 | 0 | 0 | 0 | 0 | 0 | 0 | 0 | 0 | 0 | 1 | 0 | 0 | 0 |
| Jul27.5v_11613 | 1 | 0 | 0 | 0 | 0 | 0 | 0 | 0 | 0 | 0 | 0 | 0 | 0 | 0 | 0 | 0 | 0 | 1 | 0 | 0 | 0 | 0 | 0 | 0 | 0 | 0 | 0 | 0 | 0 | 0 |
| Jul11.0v_3981 | 1 | 0 | 0 | 0 | 0 | 0 | 0 | 0 | 0 | 0 | 0 | 0 | 1 | 0 | 0 | 0 | 0 | 0 | 0 | 0 | 0 | 0 | 0 | 0 | 0 | 0 | 0 | 0 | 0 | 0 |
| Jul11.0v_5391 | 1 | 0 | 0 | 0 | 0 | 0 | 0 | 0 | 0 | 0 | 0 | 0 | 1 | 0 | 0 | 0 | 0 | 0 | 0 | 0 | 0 | 0 | 0 | 0 | 0 | 0 | 0 | 0 | 0 | 0 |
| Aug5.0v_4361 | 1 | 0 | 0 | 0 | 0 | 0 | 0 | 0 | 0 | 0 | 0 | 0 | 0 | 0 | 0 | 0 | 0 | 0 | 0 | 0 | 0 | 1 | 0 | 0 | 0 | 0 | 0 | 0 | 0 | 0 |
| Aug5.0v_4136 | 1 | 0 | 0 | 0 | 0 | 0 | 0 | 0 | 0 | 0 | 0 | 0 | 0 | 0 | 0 | 0 | 0 | 0 | 0 | 0 | 0 | 1 | 0 | 0 | 0 | 0 | 0 | 0 | 0 | 0 |
| Aug5.25v_1114 | 1 | 0 | 0 | 0 | 0 | 0 | 0 | 0 | 0 | 0 | 0 | 0 | 0 | 0 | 0 | 0 | 0 | 0 | 0 | 0 | 0 | 0 | 0 | 1 | 0 | 0 | 0 | 0 | 0 | 0 |
| Aug5.50v_41091 | 1 | 0 | 0 | 0 | 0 | 0 | 0 | 0 | 0 | 0 | 0 | 0 | 0 | 0 | 0 | 0 | 0 | 0 | 0 | 0 | 0 | 0 | 0 | 0 | 1 | 0 | 0 | 0 | 0 | 0 |
| Aug5.25v_638 | 1 | 0 | 0 | 0 | 0 | 0 | 0 | 0 | 0 | 0 | 0 | 0 | 0 | 0 | 0 | 0 | 0 | 0 | 0 | 0 | 0 | 0 | 0 | 1 | 0 | 0 | 0 | 0 | 0 | 0 |
| Aug5.25v_19141 | 1 | 0 | 0 | 0 | 0 | 0 | 0 | 0 | 0 | 0 | 0 | 0 | 0 | 0 | 0 | 0 | 0 | 0 | 0 | 0 | 0 | 0 | 0 | 1 | 0 | 0 | 0 | 0 | 0 | 0 |
| Aug5.0v_4464 | 1 | 0 | 0 | 0 | 0 | 0 | 0 | 0 | 0 | 0 | 0 | 0 | 0 | 0 | 0 | 0 | 0 | 0 | 0 | 0 | 0 | 1 | 0 | 0 | 0 | 0 | 0 | 0 | 0 | 0 |
| Aug5.5v_7503 | 1 | 0 | 0 | 0 | 0 | 0 | 0 | 0 | 0 | 0 | 0 | 0 | 0 | 0 | 0 | 0 | 0 | 0 | 0 | 0 | 0 | 0 | 1 | 0 | 0 | 0 | 0 | 0 | 0 | 0 |
| Aug5.0v_2804 | 1 | 0 | 0 | 0 | 0 | 0 | 0 | 0 | 0 | 0 | 0 | 0 | 0 | 0 | 0 | 0 | 0 | 0 | 0 | 0 | 0 | 1 | 0 | 0 | 0 | 0 | 0 | 0 | 0 | 0 |
| Jul11.5v_2297 | 1 | 0 | 0 | 0 | 0 | 0 | 0 | 0 | 0 | 0 | 0 | 0 | 0 | 1 | 0 | 0 | 0 | 0 | 0 | 0 | 0 | 0 | 0 | 0 | 0 | 0 | 0 | 0 | 0 | 0 |
| Aug5.25v_23454 | 1 | 0 | 0 | 0 | 0 | 0 | 0 | 0 | 0 | 0 | 0 | 0 | 0 | 0 | 0 | 0 | 0 | 0 | 0 | 0 | 0 | 0 | 0 | 1 | 0 | 0 | 0 | 0 | 0 | 0 |
| Aug5.0v_4276 | 1 | 0 | 0 | 0 | 0 | 0 | 0 | 0 | 0 | 0 | 0 | 0 | 0 | 0 | 0 | 0 | 0 | 0 | 0 | 0 | 0 | 1 | 0 | 0 | 0 | 0 | 0 | 0 | 0 | 0 |
| Jul11.5v_9652 | 1 | 0 | 0 | 0 | 0 | 0 | 0 | 0 | 0 | 0 | 0 | 0 | 0 | 1 | 0 | 0 | 0 | 0 | 0 | 0 | 0 | 0 | 0 | 0 | 0 | 0 | 0 | 0 | 0 | 0 |

| Feature_ID | 654 | May12_05m | May12_25m | May12_50m | May12_75m | Jun8_25m | Jun8_50m | Jun8_75m | Jun22_0m | Jun22_05m | Jun22_50m | Jun22_75m | Jul11_0m | Jul11_05m | Jul11_25m | Jul11_50m | Jul11_75m | Jul27_05m | Jul27_25m | Jul27_50m | Jul27_75m | Aug5_0m | Aug5_05m | Aug5_25m | Aug5_50m | Aug5_75m | Sep8_0m | Sep8_05m | Sep8_25m | Sep8_50m |
|---|---|---|---|---|---|---|---|---|---|---|---|---|---|---|---|---|---|---|---|---|---|---|---|---|---|---|---|---|---|---|
| Aug5.5v_12554 | 5 | 0 | 0 | 0 | 0 | 0 | 0 | 0 | 0 | 0 | 0 | 0 | 1 | 1 | 0 | 0 | 0 | 1 | 0 | 0 | 0 | 0 | 1 | 1 | 0 | 0 | 0 | 0 | 0 | 0 |
| Jul27.25v_12717 | 1 | 0 | 0 | 0 | 0 | 0 | 0 | 0 | 0 | 0 | 0 | 0 | 0 | 0 | 0 | 0 | 0 | 0 | 1 | 0 | 0 | 0 | 0 | 0 | 0 | 0 | 0 | 0 | 0 | 0 |
| Aug5.25v_21303 | 1 | 0 | 0 | 0 | 0 | 0 | 0 | 0 | 0 | 0 | 0 | 0 | 0 | 0 | 0 | 0 | 0 | 0 | 0 | 0 | 0 | 0 | 0 | 1 | 0 | 0 | 0 | 0 | 0 | 0 |
| Aug5.0v_2845 | 2 | 0 | 0 | 0 | 0 | 0 | 0 | 0 | 0 | 0 | 0 | 0 | 0 | 0 | 0 | 0 | 0 | 1 | 0 | 0 | 0 | 1 | 0 | 0 | 0 | 0 | 0 | 0 | 0 | 0 |
| Jul11.0v_12253 | 1 | 0 | 0 | 0 | 0 | 0 | 0 | 0 | 0 | 0 | 0 | 0 | 1 | 0 | 0 | 0 | 0 | 0 | 0 | 0 | 0 | 0 | 0 | 0 | 0 | 0 | 0 | 0 | 0 | 0 |
| Jul11.0v_17430 | 1 | 0 | 0 | 0 | 0 | 0 | 0 | 0 | 0 | 0 | 0 | 0 | 1 | 0 | 0 | 0 | 0 | 0 | 0 | 0 | 0 | 0 | 0 | 0 | 0 | 0 | 0 | 0 | 0 | 0 |
| Aug5.25v_14023 | 1 | 0 | 0 | 0 | 0 | 0 | 0 | 0 | 0 | 0 | 0 | 0 | 0 | 0 | 0 | 0 | 0 | 0 | 0 | 0 | 0 | 0 | 0 | 1 | 0 | 0 | 0 | 0 | 0 | 0 |
| Sep8.50v_12933 | 9 | 0 | 1 | 0 | 2 | 0 | 0 | 0 | 0 | 0 | 0 | 0 | 0 | 0 | 0 | 0 | 0 | 0 | 0 | 0 | 0 | 2 | 0 | 1 | 0 | 1 | 1 | 0 | 0 | 1 |
| Jul11.75v_38959 | 1 | 0 | 0 | 0 | 0 | 0 | 0 | 0 | 0 | 0 | 0 | 0 | 0 | 0 | 0 | 0 | 1 | 0 | 0 | 0 | 0 | 0 | 0 | 0 | 0 | 0 | 0 | 0 | 0 | 0 |
| Jul11.0v_9962 | 1 | 0 | 0 | 0 | 0 | 0 | 0 | 0 | 0 | 0 | 0 | 0 | 1 | 0 | 0 | 0 | 0 | 0 | 0 | 0 | 0 | 0 | 0 | 0 | 0 | 0 | 0 | 0 | 0 | 0 |
| Jun22.0v_568 | 11 | 0 | 1 | 0 | 1 | 1 | 1 | 0 | 2 | 0 | 0 | 1 | 0 | 1 | 0 | 0 | 0 | 1 | 1 | 0 | 0 | 0 | 0 | 1 | 0 | 0 | 0 | 0 | 0 | 0 |
| Jun8.25v_15884 | 1 | 0 | 0 | 0 | 0 | 1 | 0 | 0 | 0 | 0 | 0 | 0 | 0 | 0 | 0 | 0 | 0 | 0 | 0 | 0 | 0 | 0 | 0 | 0 | 0 | 0 | 0 | 0 | 0 | 0 |
| May12.75v_20279 | 135 | 0 | 5 | 2 | 6 | 1 | 1 | 0 | 1 | 1 | 9 | 6 | 7 | 7 | 2 | 5 | 2 | 4 | 8 | 3 | 2 | 21 | 9 | 11 | 6 | 6 | 4 | 2 | 1 | 3 |
| Jul11.25v_11920 | 25 | 0 | 0 | 0 | 1 | 0 | 0 | 0 | 4 | 1 | 1 | 0 | 0 | 1 | 1 | 2 | 0 | 3 | 0 | 1 | 0 | 3 | 2 | 0 | 1 | 4 | 0 | 0 | 0 | 0 |
| Jun22.0v_11186 | 1 | 0 | 0 | 0 | 0 | 0 | 0 | 0 | 1 | 0 | 0 | 0 | 0 | 0 | 0 | 0 | 0 | 0 | 0 | 0 | 0 | 0 | 0 | 0 | 0 | 0 | 0 | 0 | 0 | 0 |
| May12.75v_1135 | 1 | 0 | 0 | 0 | 1 | 0 | 0 | 0 | 0 | 0 | 0 | 0 | 0 | 0 | 0 | 0 | 0 | 0 | 0 | 0 | 0 | 0 | 0 | 0 | 0 | 0 | 0 | 0 | 0 | 0 |
| Aug5.5v_17722 | 1 | 0 | 0 | 0 | 0 | 0 | 0 | 0 | 0 | 0 | 0 | 0 | 0 | 0 | 0 | 0 | 0 | 0 | 0 | 0 | 0 | 0 | 1 | 0 | 0 | 0 | 0 | 0 | 0 | 0 |
| Aug5.0v_3862 | 1 | 0 | 0 | 0 | 0 | 0 | 0 | 0 | 0 | 0 | 0 | 0 | 0 | 0 | 0 | 0 | 0 | 0 | 0 | 0 | 0 | 1 | 0 | 0 | 0 | 0 | 0 | 0 | 0 | 0 |
| Jul11.75v_30859 | 1 | 0 | 0 | 0 | 0 | 0 | 0 | 0 | 0 | 0 | 0 | 0 | 0 | 0 | 0 | 0 | 1 | 0 | 0 | 0 | 0 | 0 | 0 | 0 | 0 | 0 | 0 | 0 | 0 | 0 |
| Jul27.25v_13167 | 2 | 0 | 0 | 0 | 0 | 0 | 0 | 0 | 0 | 0 | 0 | 0 | 0 | 0 | 0 | 0 | 0 | 0 | 1 | 0 | 0 | 0 | 0 | 0 | 0 | 0 | 1 | 0 | 0 | 0 |
| Jul11.0v_5373 | 2 | 0 | 0 | 0 | 0 | 0 | 0 | 0 | 0 | 0 | 0 | 0 | 1 | 0 | 0 | 0 | 0 | 1 | 0 | 0 | 0 | 0 | 0 | 0 | 0 | 0 | 0 | 0 | 0 | 0 |
| Jul11.50v_15972 | 5 | 0 | 0 | 0 | 0 | 0 | 0 | 0 | 1 | 0 | 0 | 0 | 1 | 0 | 0 | 1 | 0 | 0 | 0 | 0 | 0 | 0 | 1 | 0 | 0 | 1 | 0 | 0 | 0 | 0 |
| Jul27.25v_14674 | 1 | 0 | 0 | 0 | 0 | 0 | 0 | 0 | 0 | 0 | 0 | 0 | 0 | 0 | 0 | 0 | 0 | 0 | 1 | 0 | 0 | 0 | 0 | 0 | 0 | 0 | 0 | 0 | 0 | 0 |
| Aug5.5v_13145 | 2 | 0 | 0 | 0 | 0 | 0 | 0 | 0 | 0 | 0 | 0 | 0 | 0 | 0 | 0 | 0 | 0 | 0 | 0 | 0 | 0 | 0 | 1 | 0 | 1 | 0 | 0 | 0 | 0 | 0 |
| Aug5.0v_981 | 3 | 0 | 0 | 0 | 0 | 0 | 0 | 0 | 0 | 0 | 0 | 0 | 0 | 0 | 0 | 0 | 0 | 0 | 0 | 1 | 0 | 2 | 0 | 0 | 0 | 0 | 0 | 0 | 0 | 0 |
| Aug5.5v_26601 | 1 | 0 | 0 | 0 | 0 | 0 | 0 | 0 | 0 | 0 | 0 | 0 | 0 | 0 | 0 | 0 | 0 | 0 | 0 | 0 | 0 | 0 | 1 | 0 | 0 | 0 | 0 | 0 | 0 | 0 |
| Aug5.5v_22996 | 1 | 0 | 0 | 0 | 0 | 0 | 0 | 0 | 0 | 0 | 0 | 0 | 0 | 0 | 0 | 0 | 0 | 0 | 0 | 0 | 0 | 0 | 1 | 0 | 0 | 0 | 0 | 0 | 0 | 0 |
| Jul11.5v_3420 | 1 | 0 | 0 | 0 | 0 | 0 | 0 | 0 | 0 | 0 | 0 | 0 | 0 | 1 | 0 | 0 | 0 | 0 | 0 | 0 | 0 | 0 | 0 | 0 | 0 | 0 | 0 | 0 | 0 | 0 |
| Jul11.50v_53 | 1 | 0 | 0 | 0 | 0 | 0 | 0 | 0 | 0 | 0 | 0 | 0 | 0 | 0 | 0 | 1 | 0 | 0 | 0 | 0 | 0 | 0 | 0 | 0 | 0 | 0 | 0 | 0 | 0 | 0 |
| May12.25v_5181 | 6 | 0 | 3 | 1 | 0 | 0 | 0 | 1 | 0 | 0 | 0 | 0 | 0 | 0 | 0 | 0 | 0 | 0 | 0 | 0 | 0 | 0 | 1 | 0 | 0 | 0 | 0 | 0 | 0 | 0 |
| Aug5.5v_28796 | 1 | 0 | 0 | 0 | 0 | 0 | 0 | 0 | 0 | 0 | 0 | 0 | 0 | 0 | 0 | 0 | 0 | 0 | 0 | 0 | 0 | 0 | 1 | 0 | 0 | 0 | 0 | 0 | 0 | 0 |
| Jul27.50v_37796 | 1 | 0 | 0 | 0 | 0 | 0 | 0 | 0 | 0 | 0 | 0 | 0 | 0 | 0 | 0 | 0 | 0 | 0 | 0 | 1 | 0 | 0 | 0 | 0 | 0 | 0 | 0 | 0 | 0 | 0 |
| Aug5.75v_33388 | 1 | 0 | 0 | 0 | 0 | 0 | 0 | 0 | 0 | 0 | 0 | 0 | 0 | 0 | 0 | 0 | 0 | 0 | 0 | 0 | 0 | 0 | 0 | 0 | 0 | 1 | 0 | 0 | 0 | 0 |
| Jun22.0v_19176 | 1 | 0 | 0 | 0 | 0 | 0 | 0 | 0 | 1 | 0 | 0 | 0 | 0 | 0 | 0 | 0 | 0 | 0 | 0 | 0 | 0 | 0 | 0 | 0 | 0 | 0 | 0 | 0 | 0 | 0 |
| Sep8.25v_4126 | 1 | 0 | 0 | 0 | 0 | 0 | 0 | 0 | 0 | 0 | 0 | 0 | 0 | 0 | 0 | 0 | 0 | 0 | 0 | 0 | 0 | 0 | 0 | 0 | 0 | 0 | 0 | 0 | 1 | 0 |
| Jun8.50v_40442 | 10 | 0 | 2 | 0 | 3 | 0 | 1 | 0 | 1 | 0 | 0 | 0 | 0 | 0 | 0 | 0 | 0 | 0 | 1 | 1 | 0 | 0 | 1 | 0 | 0 | 0 | 0 | 0 | 0 | 0 |
| May12.75v_11957 | 1 | 0 | 0 | 0 | 1 | 0 | 0 | 0 | 0 | 0 | 0 | 0 | 0 | 0 | 0 | 0 | 0 | 0 | 0 | 0 | 0 | 0 | 0 | 0 | 0 | 0 | 0 | 0 | 0 | 0 |
| Jul11.25v_17316 | 1 | 0 | 0 | 0 | 0 | 0 | 0 | 0 | 0 | 0 | 0 | 0 | 0 | 0 | 1 | 0 | 0 | 0 | 0 | 0 | 0 | 0 | 0 | 0 | 0 | 0 | 0 | 0 | 0 | 0 |
| May12.50v_35971 | 1 | 0 | 0 | 1 | 0 | 0 | 0 | 0 | 0 | 0 | 0 | 0 | 0 | 0 | 0 | 0 | 0 | 0 | 0 | 0 | 0 | 0 | 0 | 0 | 0 | 0 | 0 | 0 | 0 | 0 |
| Jun22.5v_7980 | 1 | 0 | 0 | 0 | 0 | 0 | 0 | 0 | 0 | 1 | 0 | 0 | 0 | 0 | 0 | 0 | 0 | 0 | 0 | 0 | 0 | 0 | 0 | 0 | 0 | 0 | 0 | 0 | 0 | 0 |
| Jun22.75v_32988 | 1 | 0 | 0 | 0 | 0 | 0 | 0 | 0 | 0 | 0 | 0 | 1 | 0 | 0 | 0 | 0 | 0 | 0 | 0 | 0 | 0 | 0 | 0 | 0 | 0 | 0 | 0 | 0 | 0 | 0 |
| May12.75v_12970 | 1 | 0 | 0 | 0 | 1 | 0 | 0 | 0 | 0 | 0 | 0 | 0 | 0 | 0 | 0 | 0 | 0 | 0 | 0 | 0 | 0 | 0 | 0 | 0 | 0 | 0 | 0 | 0 | 0 | 0 |
| Aug5.25v_34856 | 3 | 0 | 0 | 0 | 0 | 0 | 0 | 0 | 0 | 0 | 0 | 0 | 0 | 0 | 0 | 0 | 0 | 0 | 0 | 0 | 0 | 0 | 1 | 1 | 1 | 0 | 0 | 0 | 0 | 0 |
| Jul27.5v_12865 | 44 | 5 | 5 | 2 | 3 | 2 | 1 | 0 | 2 | 1 | 1 | 2 | 3 | 1 | 0 | 1 | 1 | 2 | 1 | 1 | 0 | 3 | 1 | 2 | 0 | 0 | 1 | 0 | 1 | 2 |
| Jul11.25v_19237 | 1 | 0 | 0 | 0 | 0 | 0 | 0 | 0 | 0 | 0 | 0 | 0 | 0 | 0 | 1 | 0 | 0 | 0 | 0 | 0 | 0 | 0 | 0 | 0 | 0 | 0 | 0 | 0 | 0 | 0 |
| Aug5.25v_18580 | 1 | 0 | 0 | 0 | 0 | 0 | 0 | 0 | 0 | 0 | 0 | 0 | 0 | 0 | 0 | 0 | 0 | 0 | 0 | 0 | 0 | 0 | 0 | 1 | 0 | 0 | 0 | 0 | 0 | 0 |
| Sep8.50v_40365 | 2 | 0 | 0 | 0 | 0 | 0 | 0 | 0 | 0 | 0 | 0 | 0 | 0 | 0 | 0 | 0 | 0 | 0 | 1 | 0 | 0 | 0 | 0 | 0 | 0 | 0 | 0 | 0 | 0 | 1 |
| Jun8.50v_11785 | 7 | 0 | 0 | 0 | 1 | 1 | 2 | 1 | 0 | 0 | 0 | 0 | 0 | 0 | 0 | 0 | 0 | 0 | 1 | 0 | 0 | 0 | 0 | 0 | 0 | 0 | 0 | 0 | 0 | 1 |
| May12.5v_29401 | 1 | 1 | 0 | 0 | 0 | 0 | 0 | 0 | 0 | 0 | 0 | 0 | 0 | 0 | 0 | 0 | 0 | 0 | 0 | 0 | 0 | 0 | 0 | 0 | 0 | 0 | 0 | 0 | 0 | 0 |
| Aug5.75v_35530 | 2 | 0 | 0 | 1 | 0 | 0 | 0 | 0 | 0 | 0 | 0 | 0 | 0 | 0 | 0 | 0 | 0 | 0 | 0 | 0 | 0 | 0 | 0 | 0 | 0 | 1 | 0 | 0 | 0 | 0 |
| May12.75v_2831 | 1 | 0 | 0 | 0 | 1 | 0 | 0 | 0 | 0 | 0 | 0 | 0 | 0 | 0 | 0 | 0 | 0 | 0 | 0 | 0 | 0 | 0 | 0 | 0 | 0 | 0 | 0 | 0 | 0 | 0 |
| Jun22.50v_28667 | 2 | 0 | 1 | 0 | 0 | 0 | 0 | 0 | 0 | 0 | 1 | 0 | 0 | 0 | 0 | 0 | 0 | 0 | 0 | 0 | 0 | 0 | 0 | 0 | 0 | 0 | 0 | 0 | 0 | 0 |
| Jul11.0v_10133 | 1 | 0 | 0 | 0 | 0 | 0 | 0 | 0 | 0 | 0 | 0 | 0 | 1 | 0 | 0 | 0 | 0 | 0 | 0 | 0 | 0 | 0 | 0 | 0 | 0 | 0 | 0 | 0 | 0 | 0 |
| May12.5v_34438 | 1 | 1 | 0 | 0 | 0 | 0 | 0 | 0 | 0 | 0 | 0 | 0 | 0 | 0 | 0 | 0 | 0 | 0 | 0 | 0 | 0 | 0 | 0 | 0 | 0 | 0 | 0 | 0 | 0 | 0 |
| Jun8.25v_10081 | 1 | 0 | 0 | 0 | 0 | 1 | 0 | 0 | 0 | 0 | 0 | 0 | 0 | 0 | 0 | 0 | 0 | 0 | 0 | 0 | 0 | 0 | 0 | 0 | 0 | 0 | 0 | 0 | 0 | 0 |
| May12.25v_26376 | 1 | 0 | 1 | 0 | 0 | 0 | 0 | 0 | 0 | 0 | 0 | 0 | 0 | 0 | 0 | 0 | 0 | 0 | 0 | 0 | 0 | 0 | 0 | 0 | 0 | 0 | 0 | 0 | 0 | 0 |
| Jul11.5v_19872 | 1 | 0 | 0 | 0 | 0 | 0 | 0 | 0 | 0 | 0 | 0 | 0 | 0 | 1 | 0 | 0 | 0 | 0 | 0 | 0 | 0 | 0 | 0 | 0 | 0 | 0 | 0 | 0 | 0 | 0 |
| May12.25v_935 | 3 | 0 | 1 | 0 | 0 | 0 | 0 | 0 | 0 | 0 | 0 | 0 | 0 | 0 | 0 | 1 | 1 | 0 | 0 | 0 | 0 | 0 | 0 | 0 | 0 | 0 | 0 | 0 | 0 | 0 |
| Aug5.25v_38816 | 1 | 0 | 0 | 0 | 0 | 0 | 0 | 0 | 0 | 0 | 0 | 0 | 0 | 0 | 0 | 0 | 0 | 0 | 0 | 0 | 0 | 0 | 0 | 1 | 0 | 0 | 0 | 0 | 0 | 0 |
| May12.75v_6881 | 1 | 0 | 0 | 0 | 1 | 0 | 0 | 0 | 0 | 0 | 0 | 0 | 0 | 0 | 0 | 0 | 0 | 0 | 0 | 0 | 0 | 0 | 0 | 0 | 0 | 0 | 0 | 0 | 0 | 0 |
| Jul27.5v_19797 | 41 | 1 | 2 | 0 | 1 | 3 | 8 | 1 | 1 | 3 | 0 | 3 | 0 | 1 | 1 | 3 | 4 | 3 | 0 | 0 | 1 | 0 | 2 | 1 | 1 | 0 | 1 | 0 | 0 | 0 |
| Jul11.50v_9019 | 1 | 0 | 0 | 0 | 0 | 0 | 0 | 0 | 0 | 0 | 0 | 0 | 0 | 0 | 0 | 1 | 0 | 0 | 0 | 0 | 0 | 0 | 0 | 0 | 0 | 0 | 0 | 0 | 0 | 0 |
| Aug5.0v_719 | 6 | 1 | 0 | 0 | 0 | 0 | 3 | 0 | 0 | 0 | 0 | 0 | 0 | 0 | 0 | 0 | 0 | 0 | 0 | 0 | 0 | 2 | 0 | 0 | 0 | 0 | 0 | 0 | 0 | 0 |
| Jul11.5v_4199 | 1 | 0 | 0 | 0 | 0 | 0 | 0 | 0 | 0 | 0 | 0 | 0 | 0 | 1 | 0 | 0 | 0 | 0 | 0 | 0 | 0 | 0 | 0 | 0 | 0 | 0 | 0 | 0 | 0 | 0 |
| Jun22.0v_2092 | 1 | 0 | 0 | 0 | 0 | 0 | 0 | 0 | 1 | 0 | 0 | 0 | 0 | 0 | 0 | 0 | 0 | 0 | 0 | 0 | 0 | 0 | 0 | 0 | 0 | 0 | 0 | 0 | 0 | 0 |
| Jun22.75v_8296 | 2 | 0 | 0 | 0 | 0 | 0 | 0 | 0 | 0 | 0 | 0 | 1 | 0 | 1 | 0 | 0 | 0 | 0 | 0 | 0 | 0 | 0 | 0 | 0 | 0 | 0 | 0 | 0 | 0 | 0 |
| May12.5v_3677 | 1 | 1 | 0 | 0 | 0 | 0 | 0 | 0 | 0 | 0 | 0 | 0 | 0 | 0 | 0 | 0 | 0 | 0 | 0 | 0 | 0 | 0 | 0 | 0 | 0 | 0 | 0 | 0 | 0 | 0 |
| May12.25v_12260 | 1 | 0 | 1 | 0 | 0 | 0 | 0 | 0 | 0 | 0 | 0 | 0 | 0 | 0 | 0 | 0 | 0 | 0 | 0 | 0 | 0 | 0 | 0 | 0 | 0 | 0 | 0 | 0 | 0 | 0 |
| Aug5.0v_1938 | 7 | 1 | 1 | 2 | 0 | 0 | 0 | 0 | 1 | 0 | 0 | 1 | 0 | 0 | 0 | 0 | 0 | 0 | 0 | 0 | 0 | 1 | 0 | 0 | 0 | 0 | 0 | 0 | 0 | 0 |
| Jun22.50v_22265 | 114 | 3 | 4 | 2 | 8 | 1 | 21 | 1 | 4 | 2 | 3 | 2 | 4 | 2 | 2 | 10 | 19 | 0 | 5 | 1 | 0 | 0 | 2 | 4 | 2 | 2 | 1 | 2 | 0 | 7 |
| Jul11.75v_8152 | 53 | 3 | 1 | 2 | 4 | 3 | 5 | 0 | 4 | 1 | 2 | 7 | 0 | 0 | 0 | 4 | 6 | 0 | 2 | 1 | 0 | 1 | 0 | 0 | 3 | 1 | 0 | 1 | 1 | 1 |
| Jul27.25v_18805 | 4 | 0 | 0 | 1 | 0 | 0 | 2 | 0 | 0 | 0 | 0 | 0 | 0 | 0 | 0 | 0 | 0 | 0 | 1 | 0 | 0 | 0 | 0 | 0 | 0 | 0 | 0 | 0 | 0 | 0 |
| Jul11.50v_11809 | 1 | 0 | 0 | 0 | 0 | 0 | 0 | 0 | 0 | 0 | 0 | 0 | 0 | 0 | 0 | 1 | 0 | 0 | 0 | 0 | 0 | 0 | 0 | 0 | 0 | 0 | 0 | 0 | 0 | 0 |
| May12.25v_21467 | 16 | 1 | 4 | 0 | 2 | 3 | 2 | 0 | 0 | 0 | 0 | 2 | 0 | 0 | 0 | 1 | 1 | 0 | 0 | 0 | 0 | 0 | 0 | 0 | 0 | 0 | 0 | 0 | 0 | 0 |
| Jul11.75v_958 | 2 | 0 | 0 | 0 | 0 | 0 | 1 | 0 | 0 | 0 | 0 | 0 | 0 | 0 | 0 | 0 | 1 | 0 | 0 | 0 | 0 | 0 | 0 | 0 | 0 | 0 | 0 | 0 | 0 | 0 |
| Jul11.50v_14684 | 1 | 0 | 0 | 0 | 0 | 0 | 0 | 0 | 0 | 0 | 0 | 0 | 0 | 0 | 0 | 1 | 0 | 0 | 0 | 0 | 0 | 0 | 0 | 0 | 0 | 0 | 0 | 0 | 0 | 0 |
| Jul11.5v_18582 | 1 | 0 | 0 | 0 | 0 | 0 | 0 | 0 | 0 | 0 | 0 | 0 | 0 | 1 | 0 | 0 | 0 | 0 | 0 | 0 | 0 | 0 | 0 | 0 | 0 | 0 | 0 | 0 | 0 | 0 |
| May12.25v_40561 | 1 | 0 | 1 | 0 | 0 | 0 | 0 | 0 | 0 | 0 | 0 | 0 | 0 | 0 | 0 | 0 | 0 | 0 | 0 | 0 | 0 | 0 | 0 | 0 | 0 | 0 | 0 | 0 | 0 | 0 |
| Jul11.0v_13563 | 1 | 0 | 0 | 0 | 0 | 0 | 0 | 0 | 0 | 0 | 0 | 0 | 1 | 0 | 0 | 0 | 0 | 0 | 0 | 0 | 0 | 0 | 0 | 0 | 0 | 0 | 0 | 0 | 0 | 0 |
| Aug5.5v_12014 | 29 | 2 | 0 | 0 | 0 | 0 | 0 | 0 | 3 | 0 | 1 | 0 | 0 | 1 | 1 | 4 | 0 | 0 | 2 | 2 | 0 | 3 | 1 | 2 | 1 | 1 | 0 | 0 | 0 | 5 |
| Jul11.50v_1249 | 1 | 0 | 0 | 0 | 0 | 0 | 0 | 0 | 0 | 0 | 0 | 0 | 0 | 0 | 0 | 1 | 0 | 0 | 0 | 0 | 0 | 0 | 0 | 0 | 0 | 0 | 0 | 0 | 0 | 0 |
| Jul27.25v_626 | 1 | 0 | 0 | 0 | 0 | 0 | 0 | 0 | 0 | 0 | 0 | 0 | 0 | 0 | 0 | 0 | 0 | 0 | 1 | 0 | 0 | 0 | 0 | 0 | 0 | 0 | 0 | 0 | 0 | 0 |
| May12.5v_38517 | 1 | 1 | 0 | 0 | 0 | 0 | 0 | 0 | 0 | 0 | 0 | 0 | 0 | 0 | 0 | 0 | 0 | 0 | 0 | 0 | 0 | 0 | 0 | 0 | 0 | 0 | 0 | 0 | 0 | 0 |
| Jun22.50v_18262 | 1 | 0 | 0 | 0 | 0 | 0 | 0 | 0 | 0 | 0 | 1 | 0 | 0 | 0 | 0 | 0 | 0 | 0 | 0 | 0 | 0 | 0 | 0 | 0 | 0 | 0 | 0 | 0 | 0 | 0 |
| Aug5.0v_4614 | 1 | 0 | 0 | 0 | 0 | 0 | 0 | 0 | 0 | 0 | 0 | 0 | 0 | 0 | 0 | 0 | 0 | 0 | 0 | 0 | 0 | 1 | 0 | 0 | 0 | 0 | 0 | 0 | 0 | 0 |
| Jul11.50v_9013 | 1 | 0 | 0 | 0 | 0 | 0 | 0 | 0 | 0 | 0 | 0 | 0 | 0 | 0 | 0 | 1 | 0 | 0 | 0 | 0 | 0 | 0 | 0 | 0 | 0 | 0 | 0 | 0 | 0 | 0 |
| Jul11.75v_26150 | 1 | 0 | 0 | 0 | 0 | 0 | 0 | 0 | 0 | 0 | 0 | 0 | 0 | 0 | 0 | 0 | 1 | 0 | 0 | 0 | 0 | 0 | 0 | 0 | 0 | 0 | 0 | 0 | 0 | 0 |
| May12.50v_7282 | 1 | 0 | 0 | 1 | 0 | 0 | 0 | 0 | 0 | 0 | 0 | 0 | 0 | 0 | 0 | 0 | 0 | 0 | 0 | 0 | 0 | 0 | 0 | 0 | 0 | 0 | 0 | 0 | 0 | 0 |
| Jul11.75v_23737 | 1 | 0 | 0 | 0 | 0 | 0 | 0 | 0 | 0 | 0 | 0 | 0 | 0 | 0 | 0 | 0 | 1 | 0 | 0 | 0 | 0 | 0 | 0 | 0 | 0 | 0 | 0 | 0 | 0 | 0 |
| Jul27.25v_13389 | 1 | 0 | 0 | 0 | 0 | 0 | 0 | 0 | 0 | 0 | 0 | 0 | 0 | 0 | 0 | 0 | 0 | 0 | 1 | 0 | 0 | 0 | 0 | 0 | 0 | 0 | 0 | 0 | 0 | 0 |
| Aug5.25v_17057 | 1 | 0 | 0 | 0 | 0 | 0 | 0 | 0 | 0 | 0 | 0 | 0 | 0 | 0 | 0 | 0 | 0 | 0 | 0 | 0 | 0 | 0 | 0 | 1 | 0 | 0 | 0 | 0 | 0 | 0 |
| Aug5.25v_33587 | 1 | 0 | 0 | 0 | 0 | 0 | 0 | 0 | 0 | 0 | 0 | 0 | 0 | 0 | 0 | 0 | 0 | 0 | 0 | 0 | 0 | 0 | 0 | 1 | 0 | 0 | 0 | 0 | 0 | 0 |
| Jul11.50v_8132 | 1 | 0 | 0 | 0 | 0 | 0 | 0 | 0 | 0 | 0 | 0 | 0 | 0 | 0 | 0 | 1 | 0 | 0 | 0 | 0 | 0 | 0 | 0 | 0 | 0 | 0 | 0 | 0 | 0 | 0 |
| Aug5.75v_19284 | 1 | 0 | 0 | 0 | 0 | 0 | 0 | 0 | 0 | 0 | 0 | 0 | 0 | 0 | 0 | 0 | 0 | 0 | 0 | 0 | 0 | 0 | 0 | 0 | 0 | 1 | 0 | 0 | 0 | 0 |
| May12.5v_29202 | 1 | 1 | 0 | 0 | 0 | 0 | 0 | 0 | 0 | 0 | 0 | 0 | 0 | 0 | 0 | 0 | 0 | 0 | 0 | 0 | 0 | 0 | 0 | 0 | 0 | 0 | 0 | 0 | 0 | 0 |
| May12.25v_22797 | 1 | 0 | 1 | 0 | 0 | 0 | 0 | 0 | 0 | 0 | 0 | 0 | 0 | 0 | 0 | 0 | 0 | 0 | 0 | 0 | 0 | 0 | 0 | 0 | 0 | 0 | 0 | 0 | 0 | 0 |
| May12.25v_13925 | 1 | 0 | 1 | 0 | 0 | 0 | 0 | 0 | 0 | 0 | 0 | 0 | 0 | 0 | 0 | 0 | 0 | 0 | 0 | 0 | 0 | 0 | 0 | 0 | 0 | 0 | 0 | 0 | 0 | 0 |
| Jun8.50v_19258 | 1 | 0 | 0 | 0 | 0 | 0 | 1 | 0 | 0 | 0 | 0 | 0 | 0 | 0 | 0 | 0 | 0 | 0 | 0 | 0 | 0 | 0 | 0 | 0 | 0 | 0 | 0 | 0 | 0 | 0 |
| Jun8.50v_20182 | 1 | 0 | 0 | 0 | 0 | 0 | 1 | 0 | 0 | 0 | 0 | 0 | 0 | 0 | 0 | 0 | 0 | 0 | 0 | 0 | 0 | 0 | 0 | 0 | 0 | 0 | 0 | 0 | 0 | 0 |
| May12.50v_28331 | 2 | 0 | 0 | 1 | 0 | 0 | 0 | 0 | 0 | 0 | 0 | 0 | 0 | 0 | 0 | 0 | 0 | 0 | 0 | 0 | 0 | 0 | 0 | 0 | 0 | 1 | 0 | 0 | 0 | 0 |
| Jul27.5v_6689 | 1 | 0 | 0 | 0 | 0 | 0 | 0 | 0 | 0 | 0 | 0 | 0 | 0 | 0 | 0 | 0 | 0 | 1 | 0 | 0 | 0 | 0 | 0 | 0 | 0 | 0 | 0 | 0 | 0 | 0 |
| Aug5.0v_4892 | 1 | 0 | 0 | 0 | 0 | 0 | 0 | 0 | 0 | 0 | 0 | 0 | 0 | 0 | 0 | 0 | 0 | 0 | 0 | 0 | 0 | 1 | 0 | 0 | 0 | 0 | 0 | 0 | 0 | 0 |
| Jul11.50v_19297 | 1 | 0 | 0 | 0 | 0 | 0 | 0 | 0 | 0 | 0 | 0 | 0 | 0 | 0 | 0 | 1 | 0 | 0 | 0 | 0 | 0 | 0 | 0 | 0 | 0 | 0 | 0 | 0 | 0 | 0 |
| Aug5.25v_17930 | 1 | 0 | 0 | 0 | 0 | 0 | 0 | 0 | 0 | 0 | 0 | 0 | 0 | 0 | 0 | 0 | 0 | 0 | 0 | 0 | 0 | 0 | 0 | 1 | 0 | 0 | 0 | 0 | 0 | 0 |
| Jun22.50v_27084 | 1 | 0 | 0 | 0 | 0 | 0 | 0 | 0 | 0 | 0 | 1 | 0 | 0 | 0 | 0 | 0 | 0 | 0 | 0 | 0 | 0 | 0 | 0 | 0 | 0 | 0 | 0 | 0 | 0 | 0 |
| Sep8.5v_8152 | 1 | 0 | 0 | 0 | 0 | 0 | 0 | 0 | 0 | 0 | 0 | 0 | 0 | 0 | 0 | 0 | 0 | 0 | 0 | 0 | 0 | 0 | 0 | 0 | 0 | 0 | 0 | 1 | 0 | 0 |
| May12.75v_7621 | 1 | 0 | 0 | 0 | 1 | 0 | 0 | 0 | 0 | 0 | 0 | 0 | 0 | 0 | 0 | 0 | 0 | 0 | 0 | 0 | 0 | 0 | 0 | 0 | 0 | 0 | 0 | 0 | 0 | 0 |
| Jun22.75v_20711 | 1 | 0 | 0 | 0 | 0 | 0 | 0 | 0 | 0 | 0 | 0 | 1 | 0 | 0 | 0 | 0 | 0 | 0 | 0 | 0 | 0 | 0 | 0 | 0 | 0 | 0 | 0 | 0 | 0 | 0 |
| Jun22.0v_8194 | 1 | 0 | 0 | 0 | 0 | 0 | 0 | 0 | 1 | 0 | 0 | 0 | 0 | 0 | 0 | 0 | 0 | 0 | 0 | 0 | 0 | 0 | 0 | 0 | 0 | 0 | 0 | 0 | 0 | 0 |
| Sep8.0v_205 | 1 | 0 | 0 | 0 | 0 | 0 | 0 | 0 | 0 | 0 | 0 | 0 | 0 | 0 | 0 | 0 | 0 | 0 | 0 | 0 | 0 | 0 | 0 | 0 | 0 | 0 | 1 | 0 | 0 | 0 |
| Aug5.25v_40304 | 1 | 0 | 0 | 0 | 0 | 0 | 0 | 0 | 0 | 0 | 0 | 0 | 0 | 0 | 0 | 0 | 0 | 0 | 0 | 0 | 0 | 0 | 0 | 1 | 0 | 0 | 0 | 0 | 0 | 0 |
| Jul27.25v_20852 | 1 | 0 | 0 | 0 | 0 | 0 | 0 | 0 | 0 | 0 | 0 | 0 | 0 | 0 | 0 | 0 | 0 | 0 | 1 | 0 | 0 | 0 | 0 | 0 | 0 | 0 | 0 | 0 | 0 | 0 |
| Aug5.0v_960 | 2 | 0 | 0 | 0 | 0 | 0 | 0 | 0 | 0 | 0 | 0 | 0 | 0 | 0 | 0 | 0 | 0 | 0 | 0 | 0 | 0 | 2 | 0 | 0 | 0 | 0 | 0 | 0 | 0 | 0 |
| Aug5.0v_1224 | 1 | 0 | 0 | 0 | 0 | 0 | 0 | 0 | 0 | 0 | 0 | 0 | 0 | 0 | 0 | 0 | 0 | 0 | 0 | 0 | 0 | 1 | 0 | 0 | 0 | 0 | 0 | 0 | 0 | 0 |
| Aug5.0v_5270 | 1 | 0 | 0 | 0 | 0 | 0 | 0 | 0 | 0 | 0 | 0 | 0 | 0 | 0 | 0 | 0 | 0 | 0 | 0 | 0 | 0 | 1 | 0 | 0 | 0 | 0 | 0 | 0 | 0 | 0 |
| Jul27.25v_35277 | 1 | 0 | 0 | 0 | 0 | 0 | 0 | 0 | 0 | 0 | 0 | 0 | 0 | 0 | 0 | 0 | 0 | 0 | 1 | 0 | 0 | 0 | 0 | 0 | 0 | 0 | 0 | 0 | 0 | 0 |
| Sep8.50v_24060 | 1 | 0 | 0 | 0 | 0 | 0 | 0 | 0 | 0 | 0 | 0 | 0 | 0 | 0 | 0 | 0 | 0 | 0 | 0 | 0 | 0 | 0 | 0 | 0 | 0 | 0 | 0 | 0 | 0 | 1 |
| Jun22.75v_36044 | 1 | 0 | 0 | 0 | 0 | 0 | 0 | 0 | 0 | 0 | 0 | 1 | 0 | 0 | 0 | 0 | 0 | 0 | 0 | 0 | 0 | 0 | 0 | 0 | 0 | 0 | 0 | 0 | 0 | 0 |
| Jul27.5v_9686 | 1 | 0 | 0 | 0 | 0 | 0 | 0 | 0 | 0 | 0 | 0 | 0 | 0 | 0 | 0 | 0 | 0 | 1 | 0 | 0 | 0 | 0 | 0 | 0 | 0 | 0 | 0 | 0 | 0 | 0 |
| Jul27.75v_18104 | 1 | 0 | 0 | 0 | 0 | 0 | 0 | 0 | 0 | 0 | 0 | 0 | 0 | 0 | 0 | 0 | 0 | 0 | 0 | 0 | 1 | 0 | 0 | 0 | 0 | 0 | 0 | 0 | 0 | 0 |
| Aug5.0v_4642 | 1 | 0 | 0 | 0 | 0 | 0 | 0 | 0 | 0 | 0 | 0 | 0 | 0 | 0 | 0 | 0 | 0 | 0 | 0 | 0 | 0 | 1 | 0 | 0 | 0 | 0 | 0 | 0 | 0 | 0 |
| Aug5.25v_37860 | 2 | 0 | 0 | 0 | 0 | 0 | 0 | 0 | 1 | 0 | 0 | 0 | 0 | 0 | 0 | 0 | 0 | 0 | 0 | 0 | 0 | 0 | 0 | 1 | 0 | 0 | 0 | 0 | 0 | 0 |
| Aug5.5v_37490 | 1 | 0 | 0 | 0 | 0 | 0 | 0 | 0 | 0 | 0 | 0 | 0 | 0 | 0 | 0 | 0 | 0 | 0 | 0 | 0 | 0 | 0 | 1 | 0 | 0 | 0 | 0 | 0 | 0 | 0 |
| Jun22.0v_628 | 1 | 0 | 0 | 0 | 0 | 0 | 0 | 0 | 1 | 0 | 0 | 0 | 0 | 0 | 0 | 0 | 0 | 0 | 0 | 0 | 0 | 0 | 0 | 0 | 0 | 0 | 0 | 0 | 0 | 0 |
| Aug5.0v_4969 | 1 | 0 | 0 | 0 | 0 | 0 | 0 | 0 | 0 | 0 | 0 | 0 | 0 | 0 | 0 | 0 | 0 | 0 | 0 | 0 | 0 | 1 | 0 | 0 | 0 | 0 | 0 | 0 | 0 | 0 |
| Jul11.0v_5545 | 1 | 0 | 0 | 0 | 0 | 0 | 0 | 0 | 0 | 0 | 0 | 0 | 1 | 0 | 0 | 0 | 0 | 0 | 0 | 0 | 0 | 0 | 0 | 0 | 0 | 0 | 0 | 0 | 0 | 0 |
| Aug5.50v_27756 | 1 | 0 | 0 | 0 | 0 | 0 | 0 | 0 | 0 | 0 | 0 | 0 | 0 | 0 | 0 | 0 | 0 | 0 | 0 | 0 | 0 | 0 | 0 | 0 | 1 | 0 | 0 | 0 | 0 | 0 |
| Aug5.0v_3249 | 1 | 0 | 0 | 0 | 0 | 0 | 0 | 0 | 0 | 0 | 0 | 0 | 0 | 0 | 0 | 0 | 0 | 0 | 0 | 0 | 0 | 1 | 0 | 0 | 0 | 0 | 0 | 0 | 0 | 0 |
| Sep8.0v_16579 | 1 | 0 | 0 | 0 | 0 | 0 | 0 | 0 | 0 | 0 | 0 | 0 | 0 | 0 | 0 | 0 | 0 | 0 | 0 | 0 | 0 | 0 | 0 | 0 | 0 | 0 | 1 | 0 | 0 | 0 |
| Jul27.25v_37043 | 1 | 0 | 0 | 0 | 0 | 0 | 0 | 0 | 0 | 0 | 0 | 0 | 0 | 0 | 0 | 0 | 0 | 0 | 1 | 0 | 0 | 0 | 0 | 0 | 0 | 0 | 0 | 0 | 0 | 0 |
| Jul27.5v_10375 | 1 | 0 | 0 | 0 | 0 | 0 | 0 | 0 | 0 | 0 | 0 | 0 | 0 | 0 | 0 | 0 | 0 | 1 | 0 | 0 | 0 | 0 | 0 | 0 | 0 | 0 | 0 | 0 | 0 | 0 |
| Jun22.0v_15267 | 1 | 0 | 0 | 0 | 0 | 0 | 0 | 0 | 1 | 0 | 0 | 0 | 0 | 0 | 0 | 0 | 0 | 0 | 0 | 0 | 0 | 0 | 0 | 0 | 0 | 0 | 0 | 0 | 0 | 0 |
| Jul27.25v_21311 | 1 | 0 | 0 | 0 | 0 | 0 | 0 | 0 | 0 | 0 | 0 | 0 | 0 | 0 | 0 | 0 | 0 | 0 | 1 | 0 | 0 | 0 | 0 | 0 | 0 | 0 | 0 | 0 | 0 | 0 |
| Aug5.0v_5038 | 1 | 0 | 0 | 0 | 0 | 0 | 0 | 0 | 0 | 0 | 0 | 0 | 0 | 0 | 0 | 0 | 0 | 0 | 0 | 0 | 0 | 1 | 0 | 0 | 0 | 0 | 0 | 0 | 0 | 0 |
| Aug5.5v_28046 | 2 | 0 | 0 | 0 | 0 | 0 | 0 | 0 | 0 | 0 | 0 | 0 | 1 | 0 | 0 | 0 | 0 | 0 | 0 | 0 | 0 | 0 | 1 | 0 | 0 | 0 | 0 | 0 | 0 | 0 |
| Jul11.5v_11138 | 1 | 0 | 0 | 0 | 0 | 0 | 0 | 0 | 0 | 0 | 0 | 0 | 0 | 1 | 0 | 0 | 0 | 0 | 0 | 0 | 0 | 0 | 0 | 0 | 0 | 0 | 0 | 0 | 0 | 0 |
| Jun22.0v_20016 | 1 | 0 | 0 | 0 | 0 | 0 | 0 | 0 | 1 | 0 | 0 | 0 | 0 | 0 | 0 | 0 | 0 | 0 | 0 | 0 | 0 | 0 | 0 | 0 | 0 | 0 | 0 | 0 | 0 | 0 |
| Jul27.5v_19719 | 1 | 0 | 0 | 0 | 0 | 0 | 0 | 0 | 0 | 0 | 0 | 0 | 0 | 0 | 0 | 0 | 0 | 1 | 0 | 0 | 0 | 0 | 0 | 0 | 0 | 0 | 0 | 0 | 0 | 0 |
| Aug5.0v_1656 | 1 | 0 | 0 | 0 | 0 | 0 | 0 | 0 | 0 | 0 | 0 | 0 | 0 | 0 | 0 | 0 | 0 | 0 | 0 | 0 | 0 | 1 | 0 | 0 | 0 | 0 | 0 | 0 | 0 | 0 |
| Aug5.25v_9868 | 1 | 0 | 0 | 0 | 0 | 0 | 0 | 0 | 0 | 0 | 0 | 0 | 0 | 0 | 0 | 0 | 0 | 0 | 0 | 0 | 0 | 0 | 0 | 1 | 0 | 0 | 0 | 0 | 0 | 0 |
| Aug5.25v_18338 | 1 | 0 | 0 | 0 | 0 | 0 | 0 | 0 | 0 | 0 | 0 | 0 | 0 | 0 | 0 | 0 | 0 | 0 | 0 | 0 | 0 | 0 | 0 | 1 | 0 | 0 | 0 | 0 | 0 | 0 |
| Jul11.0v_17969 | 1 | 0 | 0 | 0 | 0 | 0 | 0 | 0 | 0 | 0 | 0 | 0 | 1 | 0 | 0 | 0 | 0 | 0 | 0 | 0 | 0 | 0 | 0 | 0 | 0 | 0 | 0 | 0 | 0 | 0 |
| Jul11.5v_16835 | 1 | 0 | 0 | 0 | 0 | 0 | 0 | 0 | 0 | 0 | 0 | 0 | 0 | 1 | 0 | 0 | 0 | 0 | 0 | 0 | 0 | 0 | 0 | 0 | 0 | 0 | 0 | 0 | 0 | 0 |
| Sep8.0v_12412 | 1 | 0 | 0 | 0 | 0 | 0 | 0 | 0 | 0 | 0 | 0 | 0 | 0 | 0 | 0 | 0 | 0 | 0 | 0 | 0 | 0 | 0 | 0 | 0 | 0 | 0 | 1 | 0 | 0 | 0 |
| Aug5.25v_11212 | 1 | 0 | 0 | 0 | 0 | 0 | 0 | 0 | 0 | 0 | 0 | 0 | 0 | 0 | 0 | 0 | 0 | 0 | 0 | 0 | 0 | 0 | 0 | 1 | 0 | 0 | 0 | 0 | 0 | 0 |
| Jul27.5v_10258 | 1 | 0 | 0 | 0 | 0 | 0 | 0 | 0 | 0 | 0 | 0 | 0 | 0 | 0 | 0 | 0 | 0 | 1 | 0 | 0 | 0 | 0 | 0 | 0 | 0 | 0 | 0 | 0 | 0 | 0 |
| Jul27.5v_5152 | 1 | 0 | 0 | 0 | 0 | 0 | 0 | 0 | 0 | 0 | 0 | 0 | 0 | 0 | 0 | 0 | 0 | 1 | 0 | 0 | 0 | 0 | 0 | 0 | 0 | 0 | 0 | 0 | 0 | 0 |
| Aug5.25v_33144 | 1 | 0 | 0 | 0 | 0 | 0 | 0 | 0 | 0 | 0 | 0 | 0 | 0 | 0 | 0 | 0 | 0 | 0 | 0 | 0 | 0 | 0 | 0 | 1 | 0 | 0 | 0 | 0 | 0 | 0 |
| Sep8.50v_8381 | 1 | 0 | 0 | 0 | 0 | 0 | 0 | 0 | 0 | 0 | 0 | 0 | 0 | 0 | 0 | 0 | 0 | 0 | 0 | 0 | 0 | 0 | 0 | 0 | 0 | 0 | 0 | 0 | 0 | 1 |
| Aug5.25v_17104 | 1 | 0 | 0 | 0 | 0 | 0 | 0 | 0 | 0 | 0 | 0 | 0 | 0 | 0 | 0 | 0 | 0 | 0 | 0 | 0 | 0 | 0 | 0 | 1 | 0 | 0 | 0 | 0 | 0 | 0 |
| Aug5.25v_31155 | 1 | 0 | 0 | 0 | 0 | 0 | 0 | 0 | 0 | 0 | 0 | 0 | 0 | 0 | 0 | 0 | 0 | 0 | 0 | 0 | 0 | 0 | 0 | 1 | 0 | 0 | 0 | 0 | 0 | 0 |
| Jul27.5v_10429 | 1 | 0 | 0 | 0 | 0 | 0 | 0 | 0 | 0 | 0 | 0 | 0 | 0 | 0 | 0 | 0 | 0 | 1 | 0 | 0 | 0 | 0 | 0 | 0 | 0 | 0 | 0 | 0 | 0 | 0 |
| Jul27.5v_7080 | 1 | 0 | 0 | 0 | 0 | 0 | 0 | 0 | 0 | 0 | 0 | 0 | 0 | 0 | 0 | 0 | 0 | 1 | 0 | 0 | 0 | 0 | 0 | 0 | 0 | 0 | 0 | 0 | 0 | 0 |
| Jul11.5v_13320 | 1 | 0 | 0 | 0 | 0 | 0 | 0 | 0 | 0 | 0 | 0 | 0 | 0 | 1 | 0 | 0 | 0 | 0 | 0 | 0 | 0 | 0 | 0 | 0 | 0 | 0 | 0 | 0 | 0 | 0 |
| Aug5.25v_41255 | 1 | 0 | 0 | 0 | 0 | 0 | 0 | 0 | 0 | 0 | 0 | 0 | 0 | 0 | 0 | 0 | 0 | 0 | 0 | 0 | 0 | 0 | 0 | 1 | 0 | 0 | 0 | 0 | 0 | 0 |
| Sep8.0v_14852 | 1 | 0 | 0 | 0 | 0 | 0 | 0 | 0 | 0 | 0 | 0 | 0 | 0 | 0 | 0 | 0 | 0 | 0 | 0 | 0 | 0 | 0 | 0 | 0 | 0 | 0 | 1 | 0 | 0 | 0 |
| Sep8.0v_17805 | 1 | 0 | 0 | 0 | 0 | 0 | 0 | 0 | 0 | 0 | 0 | 0 | 0 | 0 | 0 | 0 | 0 | 0 | 0 | 0 | 0 | 0 | 0 | 0 | 0 | 0 | 1 | 0 | 0 | 0 |
| Aug5.0v_1188 | 1 | 0 | 0 | 0 | 0 | 0 | 0 | 0 | 0 | 0 | 0 | 0 | 0 | 0 | 0 | 0 | 0 | 0 | 0 | 0 | 0 | 1 | 0 | 0 | 0 | 0 | 0 | 0 | 0 | 0 |
| Jul11.5v_4525 | 1 | 0 | 0 | 0 | 0 | 0 | 0 | 0 | 0 | 0 | 0 | 0 | 0 | 1 | 0 | 0 | 0 | 0 | 0 | 0 | 0 | 0 | 0 | 0 | 0 | 0 | 0 | 0 | 0 | 0 |
| Aug5.5v_4974 | 1 | 0 | 0 | 0 | 0 | 0 | 0 | 0 | 0 | 0 | 0 | 0 | 0 | 0 | 0 | 0 | 0 | 0 | 0 | 0 | 0 | 0 | 1 | 0 | 0 | 0 | 0 | 0 | 0 | 0 |
| Jul11.0v_9478 | 1 | 0 | 0 | 0 | 0 | 0 | 0 | 0 | 0 | 0 | 0 | 0 | 1 | 0 | 0 | 0 | 0 | 0 | 0 | 0 | 0 | 0 | 0 | 0 | 0 | 0 | 0 | 0 | 0 | 0 |
| Jun22.0v_15281 | 1 | 0 | 0 | 0 | 0 | 0 | 0 | 0 | 1 | 0 | 0 | 0 | 0 | 0 | 0 | 0 | 0 | 0 | 0 | 0 | 0 | 0 | 0 | 0 | 0 | 0 | 0 | 0 | 0 | 0 |
| Jul27.5v_14291 | 1 | 0 | 0 | 0 | 0 | 0 | 0 | 0 | 0 | 0 | 0 | 0 | 0 | 0 | 0 | 0 | 0 | 1 | 0 | 0 | 0 | 0 | 0 | 0 | 0 | 0 | 0 | 0 | 0 | 0 |
| Jul11.50v_1938 | 1 | 0 | 0 | 0 | 0 | 0 | 0 | 0 | 0 | 0 | 0 | 0 | 0 | 0 | 0 | 1 | 0 | 0 | 0 | 0 | 0 | 0 | 0 | 0 | 0 | 0 | 0 | 0 | 0 | 0 |
| Aug5.5v_35015 | 1 | 0 | 0 | 0 | 0 | 0 | 0 | 0 | 0 | 0 | 0 | 0 | 0 | 0 | 0 | 0 | 0 | 0 | 0 | 0 | 0 | 0 | 1 | 0 | 0 | 0 | 0 | 0 | 0 | 0 |
| May12.75v_1619 | 1 | 0 | 0 | 0 | 1 | 0 | 0 | 0 | 0 | 0 | 0 | 0 | 0 | 0 | 0 | 0 | 0 | 0 | 0 | 0 | 0 | 0 | 0 | 0 | 0 | 0 | 0 | 0 | 0 | 0 |
| Aug5.5v_9825 | 2 | 0 | 0 | 0 | 0 | 0 | 0 | 0 | 0 | 0 | 1 | 0 | 0 | 0 | 0 | 0 | 0 | 0 | 0 | 0 | 0 | 0 | 1 | 0 | 0 | 0 | 0 | 0 | 0 | 0 |
| Aug5.0v_294 | 1 | 0 | 0 | 0 | 0 | 0 | 0 | 0 | 0 | 0 | 0 | 0 | 0 | 0 | 0 | 0 | 0 | 0 | 0 | 0 | 0 | 1 | 0 | 0 | 0 | 0 | 0 | 0 | 0 | 0 |
| Aug5.0v_4814 | 1 | 0 | 0 | 0 | 0 | 0 | 0 | 0 | 0 | 0 | 0 | 0 | 0 | 0 | 0 | 0 | 0 | 0 | 0 | 0 | 0 | 1 | 0 | 0 | 0 | 0 | 0 | 0 | 0 | 0 |
| Aug5.25v_26838 | 1 | 0 | 0 | 0 | 0 | 0 | 0 | 0 | 0 | 0 | 0 | 0 | 0 | 0 | 0 | 0 | 0 | 0 | 0 | 0 | 0 | 0 | 0 | 1 | 0 | 0 | 0 | 0 | 0 | 0 |
| Jul27.5v_20216 | 1 | 0 | 0 | 0 | 0 | 0 | 0 | 0 | 0 | 0 | 0 | 0 | 0 | 0 | 0 | 0 | 0 | 1 | 0 | 0 | 0 | 0 | 0 | 0 | 0 | 0 | 0 | 0 | 0 | 0 |
| Aug5.0v_4466 | 1 | 0 | 0 | 0 | 0 | 0 | 0 | 0 | 0 | 0 | 0 | 0 | 0 | 0 | 0 | 0 | 0 | 0 | 0 | 0 | 0 | 1 | 0 | 0 | 0 | 0 | 0 | 0 | 0 | 0 |
| May12.50v_39082 | 1 | 0 | 0 | 1 | 0 | 0 | 0 | 0 | 0 | 0 | 0 | 0 | 0 | 0 | 0 | 0 | 0 | 0 | 0 | 0 | 0 | 0 | 0 | 0 | 0 | 0 | 0 | 0 | 0 | 0 |
| Aug5.0v_2599 | 1 | 0 | 0 | 0 | 0 | 0 | 0 | 0 | 0 | 0 | 0 | 0 | 0 | 0 | 0 | 0 | 0 | 0 | 0 | 0 | 0 | 1 | 0 | 0 | 0 | 0 | 0 | 0 | 0 | 0 |
| Aug5.5v_38006 | 1 | 0 | 0 | 0 | 0 | 0 | 0 | 0 | 0 | 0 | 0 | 0 | 0 | 0 | 0 | 0 | 0 | 0 | 0 | 0 | 0 | 0 | 1 | 0 | 0 | 0 | 0 | 0 | 0 | 0 |
| Jun8.50v_19106 | 1 | 0 | 0 | 0 | 0 | 0 | 1 | 0 | 0 | 0 | 0 | 0 | 0 | 0 | 0 | 0 | 0 | 0 | 0 | 0 | 0 | 0 | 0 | 0 | 0 | 0 | 0 | 0 | 0 | 0 |
| Jun22.0v_19488 | 1 | 0 | 0 | 0 | 0 | 0 | 0 | 0 | 1 | 0 | 0 | 0 | 0 | 0 | 0 | 0 | 0 | 0 | 0 | 0 | 0 | 0 | 0 | 0 | 0 | 0 | 0 | 0 | 0 | 0 |
| Jun8.75v_18738 | 1 | 0 | 0 | 0 | 0 | 0 | 0 | 1 | 0 | 0 | 0 | 0 | 0 | 0 | 0 | 0 | 0 | 0 | 0 | 0 | 0 | 0 | 0 | 0 | 0 | 0 | 0 | 0 | 0 | 0 |
| Jun22.0v_1817 | 3 | 0 | 0 | 0 | 0 | 0 | 0 | 0 | 1 | 0 | 0 | 0 | 0 | 0 | 0 | 1 | 0 | 0 | 0 | 0 | 0 | 0 | 1 | 0 | 0 | 0 | 0 | 0 | 0 | 0 |
| Jul27.25v_6615 | 1 | 0 | 0 | 0 | 0 | 0 | 0 | 0 | 0 | 0 | 0 | 0 | 0 | 0 | 0 | 0 | 0 | 0 | 1 | 0 | 0 | 0 | 0 | 0 | 0 | 0 | 0 | 0 | 0 | 0 |
| Jun8.50v_24007 | 3 | 0 | 0 | 0 | 0 | 0 | 1 | 0 | 0 | 0 | 0 | 0 | 1 | 0 | 0 | 0 | 1 | 0 | 0 | 0 | 0 | 0 | 0 | 0 | 0 | 0 | 0 | 0 | 0 | 0 |
| Sep8.5v_15067 | 1 | 0 | 0 | 0 | 0 | 0 | 0 | 0 | 0 | 0 | 0 | 0 | 0 | 0 | 0 | 0 | 0 | 0 | 0 | 0 | 0 | 0 | 0 | 0 | 0 | 0 | 0 | 1 | 0 | 0 |
| Jun8.50v_6226 | 1 | 0 | 0 | 0 | 0 | 0 | 1 | 0 | 0 | 0 | 0 | 0 | 0 | 0 | 0 | 0 | 0 | 0 | 0 | 0 | 0 | 0 | 0 | 0 | 0 | 0 | 0 | 0 | 0 | 0 |
| May12.75v_26000 | 1 | 0 | 0 | 0 | 1 | 0 | 0 | 0 | 0 | 0 | 0 | 0 | 0 | 0 | 0 | 0 | 0 | 0 | 0 | 0 | 0 | 0 | 0 | 0 | 0 | 0 | 0 | 0 | 0 | 0 |
| May12.25v_26714 | 1 | 0 | 1 | 0 | 0 | 0 | 0 | 0 | 0 | 0 | 0 | 0 | 0 | 0 | 0 | 0 | 0 | 0 | 0 | 0 | 0 | 0 | 0 | 0 | 0 | 0 | 0 | 0 | 0 | 0 |
| Jun8.25v_24974 | 1 | 0 | 0 | 0 | 0 | 1 | 0 | 0 | 0 | 0 | 0 | 0 | 0 | 0 | 0 | 0 | 0 | 0 | 0 | 0 | 0 | 0 | 0 | 0 | 0 | 0 | 0 | 0 | 0 | 0 |
| Jul11.0v_1929 | 1 | 0 | 0 | 0 | 0 | 0 | 0 | 0 | 0 | 0 | 0 | 0 | 1 | 0 | 0 | 0 | 0 | 0 | 0 | 0 | 0 | 0 | 0 | 0 | 0 | 0 | 0 | 0 | 0 | 0 |
| Sep8.50v_27169 | 1 | 0 | 0 | 0 | 0 | 0 | 0 | 0 | 0 | 0 | 0 | 0 | 0 | 0 | 0 | 0 | 0 | 0 | 0 | 0 | 0 | 0 | 0 | 0 | 0 | 0 | 0 | 0 | 0 | 1 |
| Jun22.75v_1417 | 2 | 0 | 0 | 0 | 0 | 0 | 0 | 0 | 0 | 0 | 1 | 1 | 0 | 0 | 0 | 0 | 0 | 0 | 0 | 0 | 0 | 0 | 0 | 0 | 0 | 0 | 0 | 0 | 0 | 0 |
| Aug5.5v_36926 | 1 | 0 | 0 | 0 | 0 | 0 | 0 | 0 | 0 | 0 | 0 | 0 | 0 | 0 | 0 | 0 | 0 | 0 | 0 | 0 | 0 | 0 | 1 | 0 | 0 | 0 | 0 | 0 | 0 | 0 |
| Jun8.75v_8412 | 1 | 0 | 0 | 0 | 0 | 0 | 0 | 1 | 0 | 0 | 0 | 0 | 0 | 0 | 0 | 0 | 0 | 0 | 0 | 0 | 0 | 0 | 0 | 0 | 0 | 0 | 0 | 0 | 0 | 0 |
| Jul27.75v_2284 | 1 | 0 | 0 | 0 | 0 | 0 | 0 | 0 | 0 | 0 | 0 | 0 | 0 | 0 | 0 | 0 | 0 | 0 | 0 | 0 | 1 | 0 | 0 | 0 | 0 | 0 | 0 | 0 | 0 | 0 |
| Jun22.0v_2785 | 1 | 0 | 0 | 0 | 0 | 0 | 0 | 0 | 1 | 0 | 0 | 0 | 0 | 0 | 0 | 0 | 0 | 0 | 0 | 0 | 0 | 0 | 0 | 0 | 0 | 0 | 0 | 0 | 0 | 0 |
| Jul11.50v_6448 | 1 | 0 | 0 | 0 | 0 | 0 | 0 | 0 | 0 | 0 | 0 | 0 | 0 | 0 | 0 | 1 | 0 | 0 | 0 | 0 | 0 | 0 | 0 | 0 | 0 | 0 | 0 | 0 | 0 | 0 |
| Jun22.50v_17142 | 1 | 0 | 0 | 0 | 0 | 0 | 0 | 0 | 0 | 0 | 1 | 0 | 0 | 0 | 0 | 0 | 0 | 0 | 0 | 0 | 0 | 0 | 0 | 0 | 0 | 0 | 0 | 0 | 0 | 0 |
| Jul11.75v_37858 | 3 | 0 | 0 | 0 | 0 | 0 | 0 | 0 | 0 | 0 | 1 | 0 | 0 | 0 | 0 | 0 | 1 | 0 | 0 | 0 | 1 | 0 | 0 | 0 | 0 | 0 | 0 | 0 | 0 | 0 |
| Jul11.75v_22505 | 1 | 0 | 0 | 0 | 0 | 0 | 0 | 0 | 0 | 0 | 0 | 0 | 0 | 0 | 0 | 0 | 1 | 0 | 0 | 0 | 0 | 0 | 0 | 0 | 0 | 0 | 0 | 0 | 0 | 0 |
| Jul11.0v_5945 | 1 | 0 | 0 | 0 | 0 | 0 | 0 | 0 | 0 | 0 | 0 | 0 | 1 | 0 | 0 | 0 | 0 | 0 | 0 | 0 | 0 | 0 | 0 | 0 | 0 | 0 | 0 | 0 | 0 | 0 |
| Jun8.25v_4189 | 1 | 0 | 0 | 0 | 0 | 1 | 0 | 0 | 0 | 0 | 0 | 0 | 0 | 0 | 0 | 0 | 0 | 0 | 0 | 0 | 0 | 0 | 0 | 0 | 0 | 0 | 0 | 0 | 0 | 0 |
| Jun22.0v_8397 | 1 | 0 | 0 | 0 | 0 | 0 | 0 | 0 | 1 | 0 | 0 | 0 | 0 | 0 | 0 | 0 | 0 | 0 | 0 | 0 | 0 | 0 | 0 | 0 | 0 | 0 | 0 | 0 | 0 | 0 |
| Jun8.50v_32175 | 1 | 0 | 0 | 0 | 0 | 0 | 1 | 0 | 0 | 0 | 0 | 0 | 0 | 0 | 0 | 0 | 0 | 0 | 0 | 0 | 0 | 0 | 0 | 0 | 0 | 0 | 0 | 0 | 0 | 0 |
| May12.75v_23945 | 1 | 0 | 0 | 0 | 1 | 0 | 0 | 0 | 0 | 0 | 0 | 0 | 0 | 0 | 0 | 0 | 0 | 0 | 0 | 0 | 0 | 0 | 0 | 0 | 0 | 0 | 0 | 0 | 0 | 0 |
| Aug5.0v_3442 | 1 | 0 | 0 | 0 | 0 | 0 | 0 | 0 | 0 | 0 | 0 | 0 | 0 | 0 | 0 | 0 | 0 | 0 | 0 | 0 | 0 | 1 | 0 | 0 | 0 | 0 | 0 | 0 | 0 | 0 |
| Jul11.25v_19545 | 1 | 0 | 0 | 0 | 0 | 0 | 0 | 0 | 0 | 0 | 0 | 0 | 0 | 0 | 1 | 0 | 0 | 0 | 0 | 0 | 0 | 0 | 0 | 0 | 0 | 0 | 0 | 0 | 0 | 0 |
| Aug5.0v_1512 | 1 | 0 | 0 | 0 | 0 | 0 | 0 | 0 | 0 | 0 | 0 | 0 | 0 | 0 | 0 | 0 | 0 | 0 | 0 | 0 | 0 | 1 | 0 | 0 | 0 | 0 | 0 | 0 | 0 | 0 |
| Aug5.0v_3818 | 584 | 8 | 5 | 11 | 1 | 8 | 4 | 0 | 28 | 8 | 79 | 11 | 22 | 5 | 25 | 24 | 12 | 6 | 17 | 25 | 6 | 83 | 51 | 44 | 13 | 27 | 8 | 5 | 2 | 46 |
| Aug5.5v_2335 | 2 | 0 | 0 | 0 | 0 | 0 | 0 | 0 | 0 | 0 | 1 | 0 | 0 | 0 | 0 | 0 | 0 | 0 | 0 | 0 | 0 | 0 | 1 | 0 | 0 | 0 | 0 | 0 | 0 | 0 |
| Aug5.0v_470 | 3 | 0 | 0 | 0 | 0 | 0 | 0 | 0 | 0 | 0 | 0 | 0 | 0 | 0 | 0 | 0 | 0 | 0 | 0 | 0 | 0 | 1 | 2 | 0 | 0 | 0 | 0 | 0 | 0 | 0 |
| May12.50v_34908 | 1 | 0 | 0 | 1 | 0 | 0 | 0 | 0 | 0 | 0 | 0 | 0 | 0 | 0 | 0 | 0 | 0 | 0 | 0 | 0 | 0 | 0 | 0 | 0 | 0 | 0 | 0 | 0 | 0 | 0 |
| Aug5.75v_739 | 2 | 0 | 0 | 0 | 0 | 0 | 0 | 0 | 0 | 0 | 0 | 0 | 0 | 0 | 0 | 0 | 0 | 0 | 0 | 0 | 0 | 0 | 0 | 0 | 0 | 1 | 1 | 0 | 0 | 0 |
| Aug5.5v_9076 | 1 | 0 | 0 | 0 | 0 | 0 | 0 | 0 | 0 | 0 | 0 | 0 | 0 | 0 | 0 | 0 | 0 | 0 | 0 | 0 | 0 | 0 | 1 | 0 | 0 | 0 | 0 | 0 | 0 | 0 |

| Feature_ID | 654 | May12_05m | May12_25m | May12_50m | May12_75m | Jun8_25m | Jun8_50m | Jun8_75m | Jun22_0m | Jun22_05m | Jun22_50m | Jun22_75m | Jul11_0m | Jul11_05m | Jul11_25m | Jul11_50m | Jul11_75m | Jul27_05m | Jul27_25m | Jul27_50m | Jul27_75m | Aug5_0m | Aug5_05m | Aug5_25m | Aug5_50m | Aug5_75m | Sep8_0m | Sep8_05m | Sep8_25m | Sep8_50m |
|---|---|---|---|---|---|---|---|---|---|---|---|---|---|---|---|---|---|---|---|---|---|---|---|---|---|---|---|---|---|---|
| Sep8.0v_10243 | 7 | 0 | 0 | 0 | 0 | 0 | 0 | 0 | 1 | 0 | 1 | 0 | 0 | 0 | 1 | 0 | 0 | 1 | 0 | 0 | 1 | 0 | 1 | 0 | 0 | 0 | 1 | 0 | 0 | 0 |
| Jul27.50v_26402 | 1 | 0 | 0 | 0 | 0 | 0 | 0 | 0 | 0 | 0 | 0 | 0 | 0 | 0 | 0 | 0 | 0 | 0 | 0 | 1 | 0 | 0 | 0 | 0 | 0 | 0 | 0 | 0 | 0 | 0 |
| Aug5.0v_5330 | 1 | 0 | 0 | 0 | 0 | 0 | 0 | 0 | 0 | 0 | 0 | 0 | 0 | 0 | 0 | 0 | 0 | 0 | 0 | 0 | 0 | 1 | 0 | 0 | 0 | 0 | 0 | 0 | 0 | 0 |
| Jun22.50v_8292 | 2 | 0 | 0 | 0 | 0 | 0 | 0 | 0 | 0 | 0 | 1 | 0 | 0 | 0 | 0 | 0 | 0 | 0 | 0 | 0 | 0 | 0 | 0 | 1 | 0 | 0 | 0 | 0 | 0 | 0 |
| Aug5.0v_968 | 2 | 0 | 0 | 0 | 0 | 0 | 0 | 0 | 0 | 0 | 0 | 0 | 0 | 0 | 0 | 0 | 0 | 0 | 0 | 0 | 0 | 1 | 0 | 0 | 0 | 0 | 0 | 0 | 0 | 1 |
| Aug5.25v_5611 | 1 | 0 | 0 | 0 | 0 | 0 | 0 | 0 | 0 | 0 | 0 | 0 | 0 | 0 | 0 | 0 | 0 | 0 | 0 | 0 | 0 | 0 | 0 | 1 | 0 | 0 | 0 | 0 | 0 | 0 |
| Aug5.5v_20294 | 1 | 0 | 0 | 0 | 0 | 0 | 0 | 0 | 0 | 0 | 0 | 0 | 0 | 0 | 0 | 0 | 0 | 0 | 0 | 0 | 0 | 0 | 1 | 0 | 0 | 0 | 0 | 0 | 0 | 0 |
| Jun22.5v_4664 | 2 | 0 | 0 | 0 | 0 | 0 | 0 | 0 | 0 | 1 | 0 | 0 | 0 | 0 | 1 | 0 | 0 | 0 | 0 | 0 | 0 | 0 | 0 | 0 | 0 | 0 | 0 | 0 | 0 | 0 |
| Aug5.0v_3852 | 1 | 0 | 0 | 0 | 0 | 0 | 0 | 0 | 0 | 0 | 0 | 0 | 0 | 0 | 0 | 0 | 0 | 0 | 0 | 0 | 0 | 1 | 0 | 0 | 0 | 0 | 0 | 0 | 0 | 0 |
| Sep8.0v_17222 | 1 | 0 | 0 | 0 | 0 | 0 | 0 | 0 | 0 | 0 | 0 | 0 | 0 | 0 | 0 | 0 | 0 | 0 | 0 | 0 | 0 | 0 | 0 | 0 | 0 | 0 | 1 | 0 | 0 | 0 |
| Sep8.50v_24863 | 1 | 0 | 0 | 0 | 0 | 0 | 0 | 0 | 0 | 0 | 0 | 0 | 0 | 0 | 0 | 0 | 0 | 0 | 0 | 0 | 0 | 0 | 0 | 0 | 0 | 0 | 0 | 0 | 0 | 1 |
| Jun22.75v_32931 | 1 | 0 | 0 | 0 | 0 | 0 | 0 | 0 | 0 | 0 | 0 | 1 | 0 | 0 | 0 | 0 | 0 | 0 | 0 | 0 | 0 | 0 | 0 | 0 | 0 | 0 | 0 | 0 | 0 | 0 |
| Sep8.50v_13124 | 1 | 0 | 0 | 0 | 0 | 0 | 0 | 0 | 0 | 0 | 0 | 0 | 0 | 0 | 0 | 0 | 0 | 0 | 0 | 0 | 0 | 0 | 0 | 0 | 0 | 0 | 0 | 0 | 0 | 1 |
| Jul27.50v_20682 | 16 | 0 | 0 | 0 | 1 | 0 | 0 | 0 | 0 | 1 | 1 | 0 | 0 | 1 | 0 | 0 | 0 | 0 | 0 | 2 | 0 | 4 | 1 | 1 | 0 | 3 | 0 | 0 | 0 | 1 |
| Aug5.75v_2507 | 1 | 0 | 0 | 0 | 0 | 0 | 0 | 0 | 0 | 0 | 0 | 0 | 0 | 0 | 0 | 0 | 0 | 0 | 0 | 0 | 0 | 0 | 0 | 0 | 0 | 1 | 0 | 0 | 0 | 0 |
| Jun22.0v_17581 | 1 | 0 | 0 | 0 | 0 | 0 | 0 | 0 | 1 | 0 | 0 | 0 | 0 | 0 | 0 | 0 | 0 | 0 | 0 | 0 | 0 | 0 | 0 | 0 | 0 | 0 | 0 | 0 | 0 | 0 |
| Jun22.5v_12968 | 1 | 0 | 0 | 0 | 0 | 0 | 0 | 0 | 0 | 1 | 0 | 0 | 0 | 0 | 0 | 0 | 0 | 0 | 0 | 0 | 0 | 0 | 0 | 0 | 0 | 0 | 0 | 0 | 0 | 0 |
| Jul11.50v_18630 | 1 | 0 | 0 | 0 | 0 | 0 | 0 | 0 | 0 | 0 | 0 | 0 | 0 | 0 | 0 | 1 | 0 | 0 | 0 | 0 | 0 | 0 | 0 | 0 | 0 | 0 | 0 | 0 | 0 | 0 |
| Aug5.0v_3944 | 1 | 0 | 0 | 0 | 0 | 0 | 0 | 0 | 0 | 0 | 0 | 0 | 0 | 0 | 0 | 0 | 0 | 0 | 0 | 0 | 0 | 1 | 0 | 0 | 0 | 0 | 0 | 0 | 0 | 0 |
| Aug5.0v_910 | 1 | 0 | 0 | 0 | 0 | 0 | 0 | 0 | 0 | 0 | 0 | 0 | 0 | 0 | 0 | 0 | 0 | 0 | 0 | 0 | 0 | 1 | 0 | 0 | 0 | 0 | 0 | 0 | 0 | 0 |
| Aug5.25v_27987 | 1 | 0 | 0 | 0 | 0 | 0 | 0 | 0 | 0 | 0 | 0 | 0 | 0 | 0 | 0 | 0 | 0 | 0 | 0 | 0 | 0 | 0 | 0 | 1 | 0 | 0 | 0 | 0 | 0 | 0 |
| Jul11.25v_6383 | 1 | 0 | 0 | 0 | 0 | 0 | 0 | 0 | 0 | 0 | 0 | 0 | 0 | 0 | 1 | 0 | 0 | 0 | 0 | 0 | 0 | 0 | 0 | 0 | 0 | 0 | 0 | 0 | 0 | 0 |
| Sep8.50v_39897 | 1 | 0 | 0 | 0 | 0 | 0 | 0 | 0 | 0 | 0 | 0 | 0 | 0 | 0 | 0 | 0 | 0 | 0 | 0 | 0 | 0 | 0 | 0 | 0 | 0 | 0 | 0 | 0 | 0 | 1 |
| Aug5.5v_27059 | 1 | 0 | 0 | 0 | 0 | 0 | 0 | 0 | 0 | 0 | 0 | 0 | 0 | 0 | 0 | 0 | 0 | 0 | 0 | 0 | 0 | 0 | 1 | 0 | 0 | 0 | 0 | 0 | 0 | 0 |
| Jun22.50v_7192 | 1 | 0 | 0 | 0 | 0 | 0 | 0 | 0 | 0 | 0 | 1 | 0 | 0 | 0 | 0 | 0 | 0 | 0 | 0 | 0 | 0 | 0 | 0 | 0 | 0 | 0 | 0 | 0 | 0 | 0 |
| Jun22.50v_25905 | 1 | 0 | 0 | 0 | 0 | 0 | 0 | 0 | 0 | 0 | 1 | 0 | 0 | 0 | 0 | 0 | 0 | 0 | 0 | 0 | 0 | 0 | 0 | 0 | 0 | 0 | 0 | 0 | 0 | 0 |
| Sep8.0v_16923 | 1 | 0 | 0 | 0 | 0 | 0 | 0 | 0 | 0 | 0 | 0 | 0 | 0 | 0 | 0 | 0 | 0 | 0 | 0 | 0 | 0 | 0 | 0 | 0 | 0 | 0 | 1 | 0 | 0 | 0 |
| Aug5.25v_22567 | 1 | 0 | 0 | 0 | 0 | 0 | 0 | 0 | 0 | 0 | 0 | 0 | 0 | 0 | 0 | 0 | 0 | 0 | 0 | 0 | 0 | 0 | 0 | 1 | 0 | 0 | 0 | 0 | 0 | 0 |
| Jul27.25v_58 | 1 | 0 | 0 | 0 | 0 | 0 | 0 | 0 | 0 | 0 | 0 | 0 | 0 | 0 | 0 | 0 | 0 | 0 | 1 | 0 | 0 | 0 | 0 | 0 | 0 | 0 | 0 | 0 | 0 | 0 |
| Jun22.50v_6918 | 1 | 0 | 0 | 0 | 0 | 0 | 0 | 0 | 0 | 0 | 1 | 0 | 0 | 0 | 0 | 0 | 0 | 0 | 0 | 0 | 0 | 0 | 0 | 0 | 0 | 0 | 0 | 0 | 0 | 0 |
| Aug5.25v_31117 | 5 | 0 | 0 | 0 | 0 | 0 | 0 | 0 | 0 | 0 | 0 | 0 | 0 | 0 | 0 | 0 | 1 | 0 | 1 | 0 | 0 | 0 | 0 | 2 | 0 | 1 | 0 | 0 | 0 | 0 |
| Aug5.0v_5183 | 1 | 0 | 0 | 0 | 0 | 0 | 0 | 0 | 0 | 0 | 0 | 0 | 0 | 0 | 0 | 0 | 0 | 0 | 0 | 0 | 0 | 1 | 0 | 0 | 0 | 0 | 0 | 0 | 0 | 0 |
| Aug5.25v_31091 | 2 | 0 | 0 | 0 | 0 | 0 | 0 | 0 | 0 | 0 | 0 | 1 | 0 | 0 | 0 | 0 | 0 | 0 | 0 | 0 | 0 | 0 | 0 | 1 | 0 | 0 | 0 | 0 | 0 | 0 |
| May12.50v_8235 | 1 | 0 | 0 | 1 | 0 | 0 | 0 | 0 | 0 | 0 | 0 | 0 | 0 | 0 | 0 | 0 | 0 | 0 | 0 | 0 | 0 | 0 | 0 | 0 | 0 | 0 | 0 | 0 | 0 | 0 |
| Aug5.0v_642 | 45 | 2 | 1 | 1 | 0 | 1 | 2 | 0 | 3 | 1 | 2 | 3 | 1 | 0 | 0 | 0 | 0 | 0 | 0 | 2 | 1 | 6 | 2 | 3 | 2 | 9 | 0 | 0 | 0 | 3 |
| Aug5.0v_4784 | 1 | 0 | 0 | 0 | 0 | 0 | 0 | 0 | 0 | 0 | 0 | 0 | 0 | 0 | 0 | 0 | 0 | 0 | 0 | 0 | 0 | 1 | 0 | 0 | 0 | 0 | 0 | 0 | 0 | 0 |
| Aug5.0v_131 | 1 | 0 | 0 | 0 | 0 | 0 | 0 | 0 | 0 | 0 | 0 | 0 | 0 | 0 | 0 | 0 | 0 | 0 | 0 | 0 | 0 | 1 | 0 | 0 | 0 | 0 | 0 | 0 | 0 | 0 |
| May12.50v_37932 | 1 | 0 | 0 | 1 | 0 | 0 | 0 | 0 | 0 | 0 | 0 | 0 | 0 | 0 | 0 | 0 | 0 | 0 | 0 | 0 | 0 | 0 | 0 | 0 | 0 | 0 | 0 | 0 | 0 | 0 |
| Sep8.50v_7661 | 2 | 0 | 0 | 0 | 0 | 0 | 0 | 0 | 0 | 0 | 1 | 0 | 0 | 0 | 0 | 0 | 0 | 0 | 0 | 0 | 0 | 0 | 0 | 0 | 0 | 0 | 0 | 0 | 0 | 1 |
| Jul11.75v_1147 | 1 | 0 | 0 | 0 | 0 | 0 | 0 | 0 | 0 | 0 | 0 | 0 | 0 | 0 | 0 | 0 | 1 | 0 | 0 | 0 | 0 | 0 | 0 | 0 | 0 | 0 | 0 | 0 | 0 | 0 |
| Aug5.50v_8711 | 1 | 0 | 0 | 0 | 0 | 0 | 0 | 0 | 0 | 0 | 0 | 0 | 0 | 0 | 0 | 0 | 0 | 0 | 0 | 0 | 0 | 0 | 0 | 0 | 1 | 0 | 0 | 0 | 0 | 0 |
| Jun22.75v_2967 | 1 | 0 | 0 | 0 | 0 | 0 | 0 | 0 | 0 | 0 | 0 | 1 | 0 | 0 | 0 | 0 | 0 | 0 | 0 | 0 | 0 | 0 | 0 | 0 | 0 | 0 | 0 | 0 | 0 | 0 |
| Jun22.75v_11744 | 1 | 0 | 0 | 0 | 0 | 0 | 0 | 0 | 0 | 0 | 0 | 1 | 0 | 0 | 0 | 0 | 0 | 0 | 0 | 0 | 0 | 0 | 0 | 0 | 0 | 0 | 0 | 0 | 0 | 0 |
| Jul27.5v_19879 | 1 | 0 | 0 | 0 | 0 | 0 | 0 | 0 | 0 | 0 | 0 | 0 | 0 | 0 | 0 | 0 | 0 | 1 | 0 | 0 | 0 | 0 | 0 | 0 | 0 | 0 | 0 | 0 | 0 | 0 |
| Jun22.75v_15214 | 1 | 0 | 0 | 0 | 0 | 0 | 0 | 0 | 0 | 0 | 0 | 1 | 0 | 0 | 0 | 0 | 0 | 0 | 0 | 0 | 0 | 0 | 0 | 0 | 0 | 0 | 0 | 0 | 0 | 0 |
| Jun22.50v_18064 | 1 | 0 | 0 | 0 | 0 | 0 | 0 | 0 | 0 | 0 | 1 | 0 | 0 | 0 | 0 | 0 | 0 | 0 | 0 | 0 | 0 | 0 | 0 | 0 | 0 | 0 | 0 | 0 | 0 | 0 |
| Jul27.25v_14698 | 1 | 0 | 0 | 0 | 0 | 0 | 0 | 0 | 0 | 0 | 0 | 0 | 0 | 0 | 0 | 0 | 0 | 0 | 1 | 0 | 0 | 0 | 0 | 0 | 0 | 0 | 0 | 0 | 0 | 0 |
| May12.5v_23281 | 1 | 1 | 0 | 0 | 0 | 0 | 0 | 0 | 0 | 0 | 0 | 0 | 0 | 0 | 0 | 0 | 0 | 0 | 0 | 0 | 0 | 0 | 0 | 0 | 0 | 0 | 0 | 0 | 0 | 0 |
| Jul27.50v_18643 | 1 | 0 | 0 | 0 | 0 | 0 | 0 | 0 | 0 | 0 | 0 | 0 | 0 | 0 | 0 | 0 | 0 | 0 | 0 | 1 | 0 | 0 | 0 | 0 | 0 | 0 | 0 | 0 | 0 | 0 |
| Aug5.0v_2334 | 1 | 0 | 0 | 0 | 0 | 0 | 0 | 0 | 0 | 0 | 0 | 0 | 0 | 0 | 0 | 0 | 0 | 0 | 0 | 0 | 0 | 1 | 0 | 0 | 0 | 0 | 0 | 0 | 0 | 0 |
| Jun22.50v_1497 | 2 | 0 | 0 | 0 | 0 | 0 | 0 | 0 | 0 | 0 | 1 | 1 | 0 | 0 | 0 | 0 | 0 | 0 | 0 | 0 | 0 | 0 | 0 | 0 | 0 | 0 | 0 | 0 | 0 | 0 |
| May12.5v_18492 | 1 | 1 | 0 | 0 | 0 | 0 | 0 | 0 | 0 | 0 | 0 | 0 | 0 | 0 | 0 | 0 | 0 | 0 | 0 | 0 | 0 | 0 | 0 | 0 | 0 | 0 | 0 | 0 | 0 | 0 |
| Sep8.50v_39402 | 1 | 0 | 0 | 0 | 0 | 0 | 0 | 0 | 0 | 0 | 0 | 0 | 0 | 0 | 0 | 0 | 0 | 0 | 0 | 0 | 0 | 0 | 0 | 0 | 0 | 0 | 0 | 0 | 0 | 1 |
| Jul27.50v_36250 | 1 | 0 | 0 | 0 | 0 | 0 | 0 | 0 | 0 | 0 | 0 | 0 | 0 | 0 | 0 | 0 | 0 | 0 | 0 | 1 | 0 | 0 | 0 | 0 | 0 | 0 | 0 | 0 | 0 | 0 |
| Jun22.50v_14587 | 1 | 0 | 0 | 0 | 0 | 0 | 0 | 0 | 0 | 0 | 1 | 0 | 0 | 0 | 0 | 0 | 0 | 0 | 0 | 0 | 0 | 0 | 0 | 0 | 0 | 0 | 0 | 0 | 0 | 0 |
| Jul11.50v_15797 | 1 | 0 | 0 | 0 | 0 | 0 | 0 | 0 | 0 | 0 | 0 | 0 | 0 | 0 | 0 | 1 | 0 | 0 | 0 | 0 | 0 | 0 | 0 | 0 | 0 | 0 | 0 | 0 | 0 | 0 |
| Jun8.50v_26836 | 1 | 0 | 0 | 0 | 0 | 0 | 1 | 0 | 0 | 0 | 0 | 0 | 0 | 0 | 0 | 0 | 0 | 0 | 0 | 0 | 0 | 0 | 0 | 0 | 0 | 0 | 0 | 0 | 0 | 0 |
| Jul11.25v_7999 | 1 | 0 | 0 | 0 | 0 | 0 | 0 | 0 | 0 | 0 | 0 | 0 | 0 | 0 | 1 | 0 | 0 | 0 | 0 | 0 | 0 | 0 | 0 | 0 | 0 | 0 | 0 | 0 | 0 | 0 |
| Aug5.75v_34146 | 1 | 0 | 0 | 0 | 0 | 0 | 0 | 0 | 0 | 0 | 0 | 0 | 0 | 0 | 0 | 0 | 0 | 0 | 0 | 0 | 0 | 0 | 0 | 0 | 0 | 1 | 0 | 0 | 0 | 0 |
| Aug5.5v_6412 | 1 | 0 | 0 | 0 | 0 | 0 | 0 | 0 | 0 | 0 | 0 | 0 | 0 | 0 | 0 | 0 | 0 | 0 | 0 | 0 | 0 | 0 | 1 | 0 | 0 | 0 | 0 | 0 | 0 | 0 |
| Jun22.75v_31458 | 1 | 0 | 0 | 0 | 0 | 0 | 0 | 0 | 0 | 0 | 0 | 1 | 0 | 0 | 0 | 0 | 0 | 0 | 0 | 0 | 0 | 0 | 0 | 0 | 0 | 0 | 0 | 0 | 0 | 0 |
| Jul27.5v_13698 | 2 | 0 | 0 | 0 | 0 | 0 | 0 | 0 | 1 | 0 | 0 | 0 | 0 | 0 | 0 | 0 | 0 | 1 | 0 | 0 | 0 | 0 | 0 | 0 | 0 | 0 | 0 | 0 | 0 | 0 |
| Aug5.50v_33137 | 1 | 0 | 0 | 0 | 0 | 0 | 0 | 0 | 0 | 0 | 0 | 0 | 0 | 0 | 0 | 0 | 0 | 0 | 0 | 0 | 0 | 0 | 0 | 0 | 1 | 0 | 0 | 0 | 0 | 0 |
| Jul27.25v_20090 | 1 | 0 | 0 | 0 | 0 | 0 | 0 | 0 | 0 | 0 | 0 | 0 | 0 | 0 | 0 | 0 | 0 | 0 | 1 | 0 | 0 | 0 | 0 | 0 | 0 | 0 | 0 | 0 | 0 | 0 |
| Jun8.25v_22300 | 1 | 0 | 0 | 0 | 0 | 1 | 0 | 0 | 0 | 0 | 0 | 0 | 0 | 0 | 0 | 0 | 0 | 0 | 0 | 0 | 0 | 0 | 0 | 0 | 0 | 0 | 0 | 0 | 0 | 0 |
| Aug5.75v_21779 | 1 | 0 | 0 | 0 | 0 | 0 | 0 | 0 | 0 | 0 | 0 | 0 | 0 | 0 | 0 | 0 | 0 | 0 | 0 | 0 | 0 | 0 | 0 | 0 | 0 | 1 | 0 | 0 | 0 | 0 |
| Sep8.50v_35890 | 1 | 0 | 0 | 0 | 0 | 0 | 0 | 0 | 0 | 0 | 0 | 0 | 0 | 0 | 0 | 0 | 0 | 0 | 0 | 0 | 0 | 0 | 0 | 0 | 0 | 0 | 0 | 0 | 0 | 1 |
| Jun22.0v_9662 | 1 | 0 | 0 | 0 | 0 | 0 | 0 | 0 | 1 | 0 | 0 | 0 | 0 | 0 | 0 | 0 | 0 | 0 | 0 | 0 | 0 | 0 | 0 | 0 | 0 | 0 | 0 | 0 | 0 | 0 |
| Jul11.25v_4477 | 1 | 0 | 0 | 0 | 0 | 0 | 0 | 0 | 0 | 0 | 0 | 0 | 0 | 0 | 1 | 0 | 0 | 0 | 0 | 0 | 0 | 0 | 0 | 0 | 0 | 0 | 0 | 0 | 0 | 0 |
| May12.50v_20803 | 1 | 0 | 0 | 1 | 0 | 0 | 0 | 0 | 0 | 0 | 0 | 0 | 0 | 0 | 0 | 0 | 0 | 0 | 0 | 0 | 0 | 0 | 0 | 0 | 0 | 0 | 0 | 0 | 0 | 0 |
| Jul11.50v_17530 | 1 | 0 | 0 | 0 | 0 | 0 | 0 | 0 | 0 | 0 | 0 | 0 | 0 | 0 | 0 | 1 | 0 | 0 | 0 | 0 | 0 | 0 | 0 | 0 | 0 | 0 | 0 | 0 | 0 | 0 |
| Sep8.5v_18579 | 9 | 1 | 1 | 0 | 1 | 1 | 0 | 0 | 0 | 1 | 0 | 1 | 0 | 0 | 0 | 0 | 1 | 0 | 0 | 0 | 0 | 0 | 1 | 0 | 0 | 0 | 0 | 1 | 0 | 0 |
| Aug5.0v_2801 | 1 | 0 | 0 | 0 | 0 | 0 | 0 | 0 | 0 | 0 | 0 | 0 | 0 | 0 | 0 | 0 | 0 | 0 | 0 | 0 | 0 | 1 | 0 | 0 | 0 | 0 | 0 | 0 | 0 | 0 |
| Jul11.75v_23690 | 143 | 4 | 4 | 6 | 0 | 0 | 0 | 1 | 6 | 3 | 8 | 1 | 4 | 3 | 5 | 5 | 3 | 0 | 5 | 9 | 0 | 16 | 6 | 24 | 10 | 6 | 2 | 0 | 1 | 11 |
| Aug5.50v_25289 | 1 | 0 | 0 | 0 | 0 | 0 | 0 | 0 | 0 | 0 | 0 | 0 | 0 | 0 | 0 | 0 | 0 | 0 | 0 | 0 | 0 | 0 | 0 | 0 | 1 | 0 | 0 | 0 | 0 | 0 |
| Jul27.25v_19311 | 1 | 0 | 0 | 0 | 0 | 0 | 0 | 0 | 0 | 0 | 0 | 0 | 0 | 0 | 0 | 0 | 0 | 0 | 1 | 0 | 0 | 0 | 0 | 0 | 0 | 0 | 0 | 0 | 0 | 0 |
| Aug5.25v_12048 | 1 | 0 | 0 | 0 | 0 | 0 | 0 | 0 | 0 | 0 | 0 | 0 | 0 | 0 | 0 | 0 | 0 | 0 | 0 | 0 | 0 | 0 | 0 | 1 | 0 | 0 | 0 | 0 | 0 | 0 |
| Aug5.75v_20510 | 1 | 0 | 0 | 0 | 0 | 0 | 0 | 0 | 0 | 0 | 0 | 0 | 0 | 0 | 0 | 0 | 0 | 0 | 0 | 0 | 0 | 0 | 0 | 0 | 0 | 1 | 0 | 0 | 0 | 0 |
| Aug5.0v_169 | 1 | 0 | 0 | 0 | 0 | 0 | 0 | 0 | 0 | 0 | 0 | 0 | 0 | 0 | 0 | 0 | 0 | 0 | 0 | 0 | 0 | 1 | 0 | 0 | 0 | 0 | 0 | 0 | 0 | 0 |
| Jul11.50v_12927 | 1 | 0 | 0 | 0 | 0 | 0 | 0 | 0 | 0 | 0 | 0 | 0 | 0 | 0 | 0 | 1 | 0 | 0 | 0 | 0 | 0 | 0 | 0 | 0 | 0 | 0 | 0 | 0 | 0 | 0 |
| Jul11.5v_4074 | 4 | 0 | 1 | 0 | 0 | 0 | 0 | 0 | 1 | 0 | 0 | 0 | 0 | 1 | 0 | 0 | 0 | 1 | 0 | 0 | 0 | 0 | 0 | 0 | 0 | 0 | 0 | 0 | 0 | 0 |
| Aug5.50v_17781 | 1 | 0 | 0 | 0 | 0 | 0 | 0 | 0 | 0 | 0 | 0 | 0 | 0 | 0 | 0 | 0 | 0 | 0 | 0 | 0 | 0 | 0 | 0 | 0 | 1 | 0 | 0 | 0 | 0 | 0 |
| Aug5.0v_879 | 1 | 0 | 0 | 0 | 0 | 0 | 0 | 0 | 0 | 0 | 0 | 0 | 0 | 0 | 0 | 0 | 0 | 0 | 0 | 0 | 0 | 1 | 0 | 0 | 0 | 0 | 0 | 0 | 0 | 0 |
| Sep8.50v_39605 | 1 | 0 | 0 | 0 | 0 | 0 | 0 | 0 | 0 | 0 | 0 | 0 | 0 | 0 | 0 | 0 | 0 | 0 | 0 | 0 | 0 | 0 | 0 | 0 | 0 | 0 | 0 | 0 | 0 | 1 |
| Jun8.50v_22514 | 9 | 0 | 0 | 1 | 2 | 1 | 1 | 0 | 0 | 0 | 0 | 1 | 0 | 0 | 2 | 0 | 0 | 0 | 0 | 0 | 0 | 0 | 0 | 0 | 0 | 1 | 0 | 0 | 0 | 0 |
| May12.50v_14405 | 1 | 0 | 0 | 1 | 0 | 0 | 0 | 0 | 0 | 0 | 0 | 0 | 0 | 0 | 0 | 0 | 0 | 0 | 0 | 0 | 0 | 0 | 0 | 0 | 0 | 0 | 0 | 0 | 0 | 0 |
| Jul27.50v_10744 | 1 | 0 | 0 | 0 | 0 | 0 | 0 | 0 | 0 | 0 | 0 | 0 | 0 | 0 | 0 | 0 | 0 | 0 | 0 | 1 | 0 | 0 | 0 | 0 | 0 | 0 | 0 | 0 | 0 | 0 |
| May12.25v_17147 | 1 | 0 | 1 | 0 | 0 | 0 | 0 | 0 | 0 | 0 | 0 | 0 | 0 | 0 | 0 | 0 | 0 | 0 | 0 | 0 | 0 | 0 | 0 | 0 | 0 | 0 | 0 | 0 | 0 | 0 |
| Jun22.0v_980 | 1 | 0 | 0 | 0 | 0 | 0 | 0 | 0 | 1 | 0 | 0 | 0 | 0 | 0 | 0 | 0 | 0 | 0 | 0 | 0 | 0 | 0 | 0 | 0 | 0 | 0 | 0 | 0 | 0 | 0 |
| Jun22.50v_12714 | 1 | 0 | 0 | 0 | 0 | 0 | 0 | 0 | 0 | 0 | 1 | 0 | 0 | 0 | 0 | 0 | 0 | 0 | 0 | 0 | 0 | 0 | 0 | 0 | 0 | 0 | 0 | 0 | 0 | 0 |
| Jul11.50v_15150 | 1 | 0 | 0 | 0 | 0 | 0 | 0 | 0 | 0 | 0 | 0 | 0 | 0 | 0 | 0 | 1 | 0 | 0 | 0 | 0 | 0 | 0 | 0 | 0 | 0 | 0 | 0 | 0 | 0 | 0 |
| Aug5.0v_5247 | 1 | 0 | 0 | 0 | 0 | 0 | 0 | 0 | 0 | 0 | 0 | 0 | 0 | 0 | 0 | 0 | 0 | 0 | 0 | 0 | 0 | 1 | 0 | 0 | 0 | 0 | 0 | 0 | 0 | 0 |
| Aug5.5v_12363 | 1 | 0 | 0 | 0 | 0 | 0 | 0 | 0 | 0 | 0 | 0 | 0 | 0 | 0 | 0 | 0 | 0 | 0 | 0 | 0 | 0 | 0 | 1 | 0 | 0 | 0 | 0 | 0 | 0 | 0 |
| Aug5.5v_36765 | 82 | 3 | 4 | 4 | 2 | 1 | 2 | 0 | 3 | 0 | 12 | 1 | 4 | 4 | 1 | 2 | 2 | 1 | 2 | 11 | 2 | 3 | 2 | 1 | 1 | 3 | 1 | 3 | 0 | 7 |
| Jul27.50v_7288 | 3 | 0 | 0 | 0 | 0 | 0 | 0 | 0 | 0 | 0 | 0 | 0 | 0 | 0 | 0 | 0 | 0 | 0 | 0 | 1 | 0 | 0 | 0 | 0 | 0 | 0 | 0 | 0 | 0 | 2 |
| Aug5.75v_37349 | 1 | 0 | 0 | 0 | 0 | 0 | 0 | 0 | 0 | 0 | 0 | 0 | 0 | 0 | 0 | 0 | 0 | 0 | 0 | 0 | 0 | 0 | 0 | 0 | 0 | 1 | 0 | 0 | 0 | 0 |
| Aug5.0v_3972 | 3 | 0 | 0 | 0 | 0 | 0 | 0 | 0 | 0 | 0 | 1 | 0 | 0 | 0 | 1 | 0 | 0 | 0 | 0 | 0 | 0 | 1 | 0 | 0 | 0 | 0 | 0 | 0 | 0 | 0 |
| Jul11.50v_1579 | 2 | 0 | 0 | 1 | 0 | 0 | 0 | 0 | 0 | 0 | 0 | 0 | 0 | 0 | 0 | 1 | 0 | 0 | 0 | 0 | 0 | 0 | 0 | 0 | 0 | 0 | 0 | 0 | 0 | 0 |
| Aug5.50v_18806 | 1 | 0 | 0 | 0 | 0 | 0 | 0 | 0 | 0 | 0 | 0 | 0 | 0 | 0 | 0 | 0 | 0 | 0 | 0 | 0 | 0 | 0 | 0 | 0 | 1 | 0 | 0 | 0 | 0 | 0 |
| Aug5.25v_5433 | 2 | 0 | 0 | 0 | 0 | 0 | 0 | 0 | 0 | 0 | 0 | 0 | 0 | 0 | 1 | 0 | 0 | 0 | 0 | 0 | 0 | 0 | 0 | 1 | 0 | 0 | 0 | 0 | 0 | 0 |
| Sep8.50v_39031 | 66 | 2 | 9 | 6 | 6 | 0 | 7 | 1 | 4 | 0 | 1 | 4 | 1 | 0 | 0 | 1 | 3 | 2 | 4 | 0 | 3 | 3 | 1 | 3 | 0 | 3 | 0 | 1 | 0 | 1 |
| Sep8.50v_38882 | 1 | 0 | 0 | 0 | 0 | 0 | 0 | 0 | 0 | 0 | 0 | 0 | 0 | 0 | 0 | 0 | 0 | 0 | 0 | 0 | 0 | 0 | 0 | 0 | 0 | 0 | 0 | 0 | 0 | 1 |
| Aug5.0v_3570 | 1 | 0 | 0 | 0 | 0 | 0 | 0 | 0 | 0 | 0 | 0 | 0 | 0 | 0 | 0 | 0 | 0 | 0 | 0 | 0 | 0 | 1 | 0 | 0 | 0 | 0 | 0 | 0 | 0 | 0 |
| May12.50v_35113 | 1 | 0 | 0 | 1 | 0 | 0 | 0 | 0 | 0 | 0 | 0 | 0 | 0 | 0 | 0 | 0 | 0 | 0 | 0 | 0 | 0 | 0 | 0 | 0 | 0 | 0 | 0 | 0 | 0 | 0 |
| May12.75v_19610 | 1 | 0 | 0 | 0 | 1 | 0 | 0 | 0 | 0 | 0 | 0 | 0 | 0 | 0 | 0 | 0 | 0 | 0 | 0 | 0 | 0 | 0 | 0 | 0 | 0 | 0 | 0 | 0 | 0 | 0 |
| Jul11.0v_16432 | 1 | 0 | 0 | 0 | 0 | 0 | 0 | 0 | 0 | 0 | 0 | 0 | 1 | 0 | 0 | 0 | 0 | 0 | 0 | 0 | 0 | 0 | 0 | 0 | 0 | 0 | 0 | 0 | 0 | 0 |
| Jun22.75v_37745 | 49 | 2 | 5 | 2 | 2 | 2 | 6 | 1 | 1 | 1 | 0 | 6 | 0 | 3 | 2 | 3 | 2 | 0 | 2 | 1 | 0 | 2 | 1 | 0 | 0 | 1 | 1 | 2 | 0 | 1 |
| May12.75v_38521 | 1 | 0 | 0 | 0 | 1 | 0 | 0 | 0 | 0 | 0 | 0 | 0 | 0 | 0 | 0 | 0 | 0 | 0 | 0 | 0 | 0 | 0 | 0 | 0 | 0 | 0 | 0 | 0 | 0 | 0 |
| Aug5.0v_1297 | 1 | 0 | 0 | 0 | 0 | 0 | 0 | 0 | 0 | 0 | 0 | 0 | 0 | 0 | 0 | 0 | 0 | 0 | 0 | 0 | 0 | 1 | 0 | 0 | 0 | 0 | 0 | 0 | 0 | 0 |
| Jun22.5v_10316 | 1 | 0 | 0 | 0 | 0 | 0 | 0 | 0 | 0 | 1 | 0 | 0 | 0 | 0 | 0 | 0 | 0 | 0 | 0 | 0 | 0 | 0 | 0 | 0 | 0 | 0 | 0 | 0 | 0 | 0 |
| Jul11.25v_1885 | 3 | 0 | 0 | 0 | 0 | 0 | 0 | 0 | 0 | 0 | 0 | 0 | 1 | 1 | 1 | 0 | 0 | 0 | 0 | 0 | 0 | 0 | 0 | 0 | 0 | 0 | 0 | 0 | 0 | 0 |
| May12.25v_16212 | 1 | 0 | 1 | 0 | 0 | 0 | 0 | 0 | 0 | 0 | 0 | 0 | 0 | 0 | 0 | 0 | 0 | 0 | 0 | 0 | 0 | 0 | 0 | 0 | 0 | 0 | 0 | 0 | 0 | 0 |
| Jul11.25v_20394 | 23 | 2 | 2 | 1 | 4 | 1 | 0 | 1 | 0 | 1 | 0 | 4 | 0 | 0 | 2 | 1 | 0 | 0 | 1 | 0 | 0 | 0 | 0 | 1 | 1 | 0 | 0 | 1 | 0 | 0 |
| Jun22.0v_17818 | 1 | 0 | 0 | 0 | 0 | 0 | 0 | 0 | 1 | 0 | 0 | 0 | 0 | 0 | 0 | 0 | 0 | 0 | 0 | 0 | 0 | 0 | 0 | 0 | 0 | 0 | 0 | 0 | 0 | 0 |
| Jun8.25v_8544 | 33 | 1 | 0 | 4 | 4 | 1 | 3 | 2 | 0 | 0 | 1 | 3 | 0 | 1 | 0 | 1 | 0 | 1 | 1 | 1 | 0 | 3 | 1 | 0 | 0 | 0 | 1 | 2 | 0 | 2 |
| Jul11.5v_9486 | 1 | 0 | 0 | 0 | 0 | 0 | 0 | 0 | 0 | 0 | 0 | 0 | 0 | 1 | 0 | 0 | 0 | 0 | 0 | 0 | 0 | 0 | 0 | 0 | 0 | 0 | 0 | 0 | 0 | 0 |
| Aug5.0v_282 | 1 | 0 | 0 | 0 | 0 | 0 | 0 | 0 | 0 | 0 | 0 | 0 | 0 | 0 | 0 | 0 | 0 | 0 | 0 | 0 | 0 | 1 | 0 | 0 | 0 | 0 | 0 | 0 | 0 | 0 |
| Jul11.75v_40073 | 1 | 0 | 0 | 0 | 0 | 0 | 0 | 0 | 0 | 0 | 0 | 0 | 0 | 0 | 0 | 0 | 1 | 0 | 0 | 0 | 0 | 0 | 0 | 0 | 0 | 0 | 0 | 0 | 0 | 0 |
| Jun22.50v_27917 | 1 | 0 | 0 | 0 | 0 | 0 | 0 | 0 | 0 | 0 | 1 | 0 | 0 | 0 | 0 | 0 | 0 | 0 | 0 | 0 | 0 | 0 | 0 | 0 | 0 | 0 | 0 | 0 | 0 | 0 |
| Sep8.0v_7698 | 1 | 0 | 0 | 0 | 0 | 0 | 0 | 0 | 0 | 0 | 0 | 0 | 0 | 0 | 0 | 0 | 0 | 0 | 0 | 0 | 0 | 0 | 0 | 0 | 0 | 0 | 1 | 0 | 0 | 0 |
| Jun22.50v_29457 | 1 | 0 | 0 | 0 | 0 | 0 | 0 | 0 | 0 | 0 | 1 | 0 | 0 | 0 | 0 | 0 | 0 | 0 | 0 | 0 | 0 | 0 | 0 | 0 | 0 | 0 | 0 | 0 | 0 | 0 |
| Aug5.5v_36929 | 1 | 0 | 0 | 0 | 0 | 0 | 0 | 0 | 0 | 0 | 0 | 0 | 0 | 0 | 0 | 0 | 0 | 0 | 0 | 0 | 0 | 0 | 1 | 0 | 0 | 0 | 0 | 0 | 0 | 0 |
| Jul11.50v_17405 | 1 | 0 | 0 | 0 | 0 | 0 | 0 | 0 | 0 | 0 | 0 | 0 | 0 | 0 | 0 | 1 | 0 | 0 | 0 | 0 | 0 | 0 | 0 | 0 | 0 | 0 | 0 | 0 | 0 | 0 |
| Aug5.0v_2181 | 1 | 0 | 0 | 0 | 0 | 0 | 0 | 0 | 0 | 0 | 0 | 0 | 0 | 0 | 0 | 0 | 0 | 0 | 0 | 0 | 0 | 1 | 0 | 0 | 0 | 0 | 0 | 0 | 0 | 0 |
| Jun22.50v_9933 | 1 | 0 | 0 | 0 | 0 | 0 | 0 | 0 | 0 | 0 | 1 | 0 | 0 | 0 | 0 | 0 | 0 | 0 | 0 | 0 | 0 | 0 | 0 | 0 | 0 | 0 | 0 | 0 | 0 | 0 |
| Aug5.5v_25954 | 1 | 0 | 0 | 0 | 0 | 0 | 0 | 0 | 0 | 0 | 0 | 0 | 0 | 0 | 0 | 0 | 0 | 0 | 0 | 0 | 0 | 0 | 1 | 0 | 0 | 0 | 0 | 0 | 0 | 0 |
| Aug5.0v_2672 | 1 | 0 | 0 | 0 | 0 | 0 | 0 | 0 | 0 | 0 | 0 | 0 | 0 | 0 | 0 | 0 | 0 | 0 | 0 | 0 | 0 | 1 | 0 | 0 | 0 | 0 | 0 | 0 | 0 | 0 |
| Jul27.25v_3424 | 1 | 0 | 0 | 0 | 0 | 0 | 0 | 0 | 0 | 0 | 0 | 0 | 0 | 0 | 0 | 0 | 0 | 0 | 1 | 0 | 0 | 0 | 0 | 0 | 0 | 0 | 0 | 0 | 0 | 0 |
| Jun22.0v_15322 | 1 | 0 | 0 | 0 | 0 | 0 | 0 | 0 | 1 | 0 | 0 | 0 | 0 | 0 | 0 | 0 | 0 | 0 | 0 | 0 | 0 | 0 | 0 | 0 | 0 | 0 | 0 | 0 | 0 | 0 |
| Jul11.50v_14052 | 1 | 0 | 0 | 0 | 0 | 0 | 0 | 0 | 0 | 0 | 0 | 0 | 0 | 0 | 0 | 1 | 0 | 0 | 0 | 0 | 0 | 0 | 0 | 0 | 0 | 0 | 0 | 0 | 0 | 0 |
| Jun22.0v_7652 | 1 | 0 | 0 | 0 | 0 | 0 | 0 | 0 | 1 | 0 | 0 | 0 | 0 | 0 | 0 | 0 | 0 | 0 | 0 | 0 | 0 | 0 | 0 | 0 | 0 | 0 | 0 | 0 | 0 | 0 |
| Jul11.25v_20956 | 1 | 0 | 0 | 0 | 0 | 0 | 0 | 0 | 0 | 0 | 0 | 0 | 0 | 0 | 1 | 0 | 0 | 0 | 0 | 0 | 0 | 0 | 0 | 0 | 0 | 0 | 0 | 0 | 0 | 0 |
| Jul27.75v_11058 | 1 | 0 | 0 | 0 | 0 | 0 | 0 | 0 | 0 | 0 | 0 | 0 | 0 | 0 | 0 | 0 | 0 | 0 | 0 | 0 | 1 | 0 | 0 | 0 | 0 | 0 | 0 | 0 | 0 | 0 |
| May12.50v_3419 | 1 | 0 | 0 | 1 | 0 | 0 | 0 | 0 | 0 | 0 | 0 | 0 | 0 | 0 | 0 | 0 | 0 | 0 | 0 | 0 | 0 | 0 | 0 | 0 | 0 | 0 | 0 | 0 | 0 | 0 |
| Aug5.5v_27701 | 1 | 0 | 0 | 0 | 0 | 0 | 0 | 0 | 0 | 0 | 0 | 0 | 0 | 0 | 0 | 0 | 0 | 0 | 0 | 0 | 0 | 0 | 1 | 0 | 0 | 0 | 0 | 0 | 0 | 0 |
| Aug5.75v_26667 | 1 | 0 | 0 | 0 | 0 | 0 | 0 | 0 | 0 | 0 | 0 | 0 | 0 | 0 | 0 | 0 | 0 | 0 | 0 | 0 | 0 | 0 | 0 | 0 | 0 | 1 | 0 | 0 | 0 | 0 |
| Aug5.5v_21254 | 1 | 0 | 0 | 0 | 0 | 0 | 0 | 0 | 0 | 0 | 0 | 0 | 0 | 0 | 0 | 0 | 0 | 0 | 0 | 0 | 0 | 0 | 1 | 0 | 0 | 0 | 0 | 0 | 0 | 0 |
| Jun22.50v_22954 | 1 | 0 | 0 | 0 | 0 | 0 | 0 | 0 | 0 | 0 | 1 | 0 | 0 | 0 | 0 | 0 | 0 | 0 | 0 | 0 | 0 | 0 | 0 | 0 | 0 | 0 | 0 | 0 | 0 | 0 |
| Aug5.50v_41265 | 2 | 0 | 0 | 0 | 0 | 0 | 0 | 0 | 0 | 0 | 0 | 0 | 0 | 0 | 0 | 0 | 0 | 0 | 0 | 0 | 0 | 0 | 0 | 0 | 1 | 0 | 0 | 0 | 0 | 1 |
| Sep8.50v_22977 | 1 | 0 | 0 | 0 | 0 | 0 | 0 | 0 | 0 | 0 | 0 | 0 | 0 | 0 | 0 | 0 | 0 | 0 | 0 | 0 | 0 | 0 | 0 | 0 | 0 | 0 | 0 | 0 | 0 | 1 |
| Aug5.75v_37750 | 1 | 0 | 0 | 0 | 0 | 0 | 0 | 0 | 0 | 0 | 0 | 0 | 0 | 0 | 0 | 0 | 0 | 0 | 0 | 0 | 0 | 0 | 0 | 0 | 0 | 1 | 0 | 0 | 0 | 0 |
| May12.50v_22477 | 1 | 0 | 0 | 1 | 0 | 0 | 0 | 0 | 0 | 0 | 0 | 0 | 0 | 0 | 0 | 0 | 0 | 0 | 0 | 0 | 0 | 0 | 0 | 0 | 0 | 0 | 0 | 0 | 0 | 0 |
| Aug5.25v_14374 | 1 | 0 | 0 | 0 | 0 | 0 | 0 | 0 | 0 | 0 | 0 | 0 | 0 | 0 | 0 | 0 | 0 | 0 | 0 | 0 | 0 | 0 | 0 | 1 | 0 | 0 | 0 | 0 | 0 | 0 |
| Aug5.25v_9034 | 1 | 0 | 0 | 0 | 0 | 0 | 0 | 0 | 0 | 0 | 0 | 0 | 0 | 0 | 0 | 0 | 0 | 0 | 0 | 0 | 0 | 0 | 0 | 1 | 0 | 0 | 0 | 0 | 0 | 0 |
| Jul27.50v_17843 | 1 | 0 | 0 | 0 | 0 | 0 | 0 | 0 | 0 | 0 | 0 | 0 | 0 | 0 | 0 | 0 | 0 | 0 | 0 | 1 | 0 | 0 | 0 | 0 | 0 | 0 | 0 | 0 | 0 | 0 |
| Jun22.50v_8283 | 1 | 0 | 0 | 0 | 0 | 0 | 0 | 0 | 0 | 0 | 1 | 0 | 0 | 0 | 0 | 0 | 0 | 0 | 0 | 0 | 0 | 0 | 0 | 0 | 0 | 0 | 0 | 0 | 0 | 0 |
| Aug5.25v_4284 | 1 | 0 | 0 | 0 | 0 | 0 | 0 | 0 | 0 | 0 | 0 | 0 | 0 | 0 | 0 | 0 | 0 | 0 | 0 | 0 | 0 | 0 | 0 | 1 | 0 | 0 | 0 | 0 | 0 | 0 |
| Jul27.25v_18026 | 1 | 0 | 0 | 0 | 0 | 0 | 0 | 0 | 0 | 0 | 0 | 0 | 0 | 0 | 0 | 0 | 0 | 0 | 1 | 0 | 0 | 0 | 0 | 0 | 0 | 0 | 0 | 0 | 0 | 0 |
| Jun22.50v_20221 | 1 | 0 | 0 | 0 | 0 | 0 | 0 | 0 | 0 | 0 | 1 | 0 | 0 | 0 | 0 | 0 | 0 | 0 | 0 | 0 | 0 | 0 | 0 | 0 | 0 | 0 | 0 | 0 | 0 | 0 |
| Jul27.25v_1251 | 1 | 0 | 0 | 0 | 0 | 0 | 0 | 0 | 0 | 0 | 0 | 0 | 0 | 0 | 0 | 0 | 0 | 0 | 1 | 0 | 0 | 0 | 0 | 0 | 0 | 0 | 0 | 0 | 0 | 0 |
| Jul11.0v_10117 | 1 | 0 | 0 | 0 | 0 | 0 | 0 | 0 | 0 | 0 | 0 | 0 | 1 | 0 | 0 | 0 | 0 | 0 | 0 | 0 | 0 | 0 | 0 | 0 | 0 | 0 | 0 | 0 | 0 | 0 |
| Jun22.50v_28404 | 1 | 0 | 0 | 0 | 0 | 0 | 0 | 0 | 0 | 0 | 1 | 0 | 0 | 0 | 0 | 0 | 0 | 0 | 0 | 0 | 0 | 0 | 0 | 0 | 0 | 0 | 0 | 0 | 0 | 0 |
| Jun22.0v_5863 | 1 | 0 | 0 | 0 | 0 | 0 | 0 | 0 | 1 | 0 | 0 | 0 | 0 | 0 | 0 | 0 | 0 | 0 | 0 | 0 | 0 | 0 | 0 | 0 | 0 | 0 | 0 | 0 | 0 | 0 |
| Aug5.25v_2547 | 1 | 0 | 0 | 0 | 0 | 0 | 0 | 0 | 0 | 0 | 0 | 0 | 0 | 0 | 0 | 0 | 0 | 0 | 0 | 0 | 0 | 0 | 0 | 1 | 0 | 0 | 0 | 0 | 0 | 0 |
| Jul11.50v_1941 | 1 | 0 | 0 | 0 | 0 | 0 | 0 | 0 | 0 | 0 | 0 | 0 | 0 | 0 | 0 | 1 | 0 | 0 | 0 | 0 | 0 | 0 | 0 | 0 | 0 | 0 | 0 | 0 | 0 | 0 |
| Jul11.50v_12103 | 1 | 0 | 0 | 0 | 0 | 0 | 0 | 0 | 0 | 0 | 0 | 0 | 0 | 0 | 0 | 1 | 0 | 0 | 0 | 0 | 0 | 0 | 0 | 0 | 0 | 0 | 0 | 0 | 0 | 0 |
| Aug5.75v_31744 | 1 | 0 | 0 | 0 | 0 | 0 | 0 | 0 | 0 | 0 | 0 | 0 | 0 | 0 | 0 | 0 | 0 | 0 | 0 | 0 | 0 | 0 | 0 | 0 | 0 | 1 | 0 | 0 | 0 | 0 |
| Sep8.50v_30609 | 1 | 0 | 0 | 0 | 0 | 0 | 0 | 0 | 0 | 0 | 0 | 0 | 0 | 0 | 0 | 0 | 0 | 0 | 0 | 0 | 0 | 0 | 0 | 0 | 0 | 0 | 0 | 0 | 0 | 1 |
| Jul11.50v_8700 | 1 | 0 | 0 | 0 | 0 | 0 | 0 | 0 | 0 | 0 | 0 | 0 | 0 | 0 | 0 | 1 | 0 | 0 | 0 | 0 | 0 | 0 | 0 | 0 | 0 | 0 | 0 | 0 | 0 | 0 |
| Jul11.50v_6100 | 1 | 0 | 0 | 0 | 0 | 0 | 0 | 0 | 0 | 0 | 0 | 0 | 0 | 0 | 0 | 1 | 0 | 0 | 0 | 0 | 0 | 0 | 0 | 0 | 0 | 0 | 0 | 0 | 0 | 0 |
| Jul11.0v_14910 | 1 | 0 | 0 | 0 | 0 | 0 | 0 | 0 | 0 | 0 | 0 | 0 | 1 | 0 | 0 | 0 | 0 | 0 | 0 | 0 | 0 | 0 | 0 | 0 | 0 | 0 | 0 | 0 | 0 | 0 |
| Aug5.5v_39132 | 1 | 0 | 0 | 0 | 0 | 0 | 0 | 0 | 0 | 0 | 0 | 0 | 0 | 0 | 0 | 0 | 0 | 0 | 0 | 0 | 0 | 0 | 1 | 0 | 0 | 0 | 0 | 0 | 0 | 0 |
| Aug5.50v_17273 | 1 | 0 | 0 | 0 | 0 | 0 | 0 | 0 | 0 | 0 | 0 | 0 | 0 | 0 | 0 | 0 | 0 | 0 | 0 | 0 | 0 | 0 | 0 | 0 | 1 | 0 | 0 | 0 | 0 | 0 |
| May12.25v_8483 | 1 | 0 | 1 | 0 | 0 | 0 | 0 | 0 | 0 | 0 | 0 | 0 | 0 | 0 | 0 | 0 | 0 | 0 | 0 | 0 | 0 | 0 | 0 | 0 | 0 | 0 | 0 | 0 | 0 | 0 |
| Jul11.75v_3912 | 1 | 0 | 0 | 0 | 0 | 0 | 0 | 0 | 0 | 0 | 0 | 0 | 0 | 0 | 0 | 0 | 1 | 0 | 0 | 0 | 0 | 0 | 0 | 0 | 0 | 0 | 0 | 0 | 0 | 0 |
| Jul11.50v_5261 | 1 | 0 | 0 | 0 | 0 | 0 | 0 | 0 | 0 | 0 | 0 | 0 | 0 | 0 | 0 | 1 | 0 | 0 | 0 | 0 | 0 | 0 | 0 | 0 | 0 | 0 | 0 | 0 | 0 | 0 |
| Aug5.25v_27445 | 1 | 0 | 0 | 0 | 0 | 0 | 0 | 0 | 0 | 0 | 0 | 0 | 0 | 0 | 0 | 0 | 0 | 0 | 0 | 0 | 0 | 0 | 0 | 1 | 0 | 0 | 0 | 0 | 0 | 0 |
| Jun22.0v_11106 | 1 | 0 | 0 | 0 | 0 | 0 | 0 | 0 | 1 | 0 | 0 | 0 | 0 | 0 | 0 | 0 | 0 | 0 | 0 | 0 | 0 | 0 | 0 | 0 | 0 | 0 | 0 | 0 | 0 | 0 |
| Aug5.50v_5185 | 1 | 0 | 0 | 0 | 0 | 0 | 0 | 0 | 0 | 0 | 0 | 0 | 0 | 0 | 0 | 0 | 0 | 0 | 0 | 0 | 0 | 0 | 0 | 0 | 1 | 0 | 0 | 0 | 0 | 0 |
| Jun22.0v_17840 | 1 | 0 | 0 | 0 | 0 | 0 | 0 | 0 | 1 | 0 | 0 | 0 | 0 | 0 | 0 | 0 | 0 | 0 | 0 | 0 | 0 | 0 | 0 | 0 | 0 | 0 | 0 | 0 | 0 | 0 |
| Jun22.50v_8223 | 1 | 0 | 0 | 0 | 0 | 0 | 0 | 0 | 0 | 0 | 1 | 0 | 0 | 0 | 0 | 0 | 0 | 0 | 0 | 0 | 0 | 0 | 0 | 0 | 0 | 0 | 0 | 0 | 0 | 0 |
| May12.50v_18671 | 1 | 0 | 0 | 1 | 0 | 0 | 0 | 0 | 0 | 0 | 0 | 0 | 0 | 0 | 0 | 0 | 0 | 0 | 0 | 0 | 0 | 0 | 0 | 0 | 0 | 0 | 0 | 0 | 0 | 0 |
| Sep8.5v_7188 | 1 | 0 | 0 | 0 | 0 | 0 | 0 | 0 | 0 | 0 | 0 | 0 | 0 | 0 | 0 | 0 | 0 | 0 | 0 | 0 | 0 | 0 | 0 | 0 | 0 | 0 | 0 | 1 | 0 | 0 |
| Jul27.50v_7770 | 1 | 0 | 0 | 0 | 0 | 0 | 0 | 0 | 0 | 0 | 0 | 0 | 0 | 0 | 0 | 0 | 0 | 0 | 0 | 1 | 0 | 0 | 0 | 0 | 0 | 0 | 0 | 0 | 0 | 0 |
| Jul27.50v_4112 | 1 | 0 | 0 | 0 | 0 | 0 | 0 | 0 | 0 | 0 | 0 | 0 | 0 | 0 | 0 | 0 | 0 | 0 | 0 | 1 | 0 | 0 | 0 | 0 | 0 | 0 | 0 | 0 | 0 | 0 |
| Jun22.50v_8425 | 1 | 0 | 0 | 0 | 0 | 0 | 0 | 0 | 0 | 0 | 1 | 0 | 0 | 0 | 0 | 0 | 0 | 0 | 0 | 0 | 0 | 0 | 0 | 0 | 0 | 0 | 0 | 0 | 0 | 0 |
| Jul27.50v_1242 | 2 | 0 | 0 | 0 | 0 | 0 | 1 | 0 | 0 | 0 | 0 | 0 | 0 | 0 | 0 | 0 | 0 | 0 | 0 | 1 | 0 | 0 | 0 | 0 | 0 | 0 | 0 | 0 | 0 | 0 |
| Jun8.25v_25903 | 1 | 0 | 0 | 0 | 0 | 1 | 0 | 0 | 0 | 0 | 0 | 0 | 0 | 0 | 0 | 0 | 0 | 0 | 0 | 0 | 0 | 0 | 0 | 0 | 0 | 0 | 0 | 0 | 0 | 0 |
| May12.50v_21136 | 1 | 0 | 0 | 1 | 0 | 0 | 0 | 0 | 0 | 0 | 0 | 0 | 0 | 0 | 0 | 0 | 0 | 0 | 0 | 0 | 0 | 0 | 0 | 0 | 0 | 0 | 0 | 0 | 0 | 0 |
| Jul11.50v_11059 | 1 | 0 | 0 | 0 | 0 | 0 | 0 | 0 | 0 | 0 | 0 | 0 | 0 | 0 | 0 | 1 | 0 | 0 | 0 | 0 | 0 | 0 | 0 | 0 | 0 | 0 | 0 | 0 | 0 | 0 |
| Jun22.0v_19116 | 1 | 0 | 0 | 0 | 0 | 0 | 0 | 0 | 1 | 0 | 0 | 0 | 0 | 0 | 0 | 0 | 0 | 0 | 0 | 0 | 0 | 0 | 0 | 0 | 0 | 0 | 0 | 0 | 0 | 0 |
| May12.50v_5341 | 1 | 0 | 0 | 1 | 0 | 0 | 0 | 0 | 0 | 0 | 0 | 0 | 0 | 0 | 0 | 0 | 0 | 0 | 0 | 0 | 0 | 0 | 0 | 0 | 0 | 0 | 0 | 0 | 0 | 0 |
| Aug5.0v_2536 | 1 | 0 | 0 | 0 | 0 | 0 | 0 | 0 | 0 | 0 | 0 | 0 | 0 | 0 | 0 | 0 | 0 | 0 | 0 | 0 | 0 | 1 | 0 | 0 | 0 | 0 | 0 | 0 | 0 | 0 |
| Aug5.5v_26523 | 1 | 0 | 0 | 0 | 0 | 0 | 0 | 0 | 0 | 0 | 0 | 0 | 0 | 0 | 0 | 0 | 0 | 0 | 0 | 0 | 0 | 0 | 1 | 0 | 0 | 0 | 0 | 0 | 0 | 0 |
| Jul11.0v_9971 | 1 | 0 | 0 | 0 | 0 | 0 | 0 | 0 | 0 | 0 | 0 | 0 | 1 | 0 | 0 | 0 | 0 | 0 | 0 | 0 | 0 | 0 | 0 | 0 | 0 | 0 | 0 | 0 | 0 | 0 |
| Jul11.0v_17606 | 1 | 0 | 0 | 0 | 0 | 0 | 0 | 0 | 0 | 0 | 0 | 0 | 1 | 0 | 0 | 0 | 0 | 0 | 0 | 0 | 0 | 0 | 0 | 0 | 0 | 0 | 0 | 0 | 0 | 0 |
| Jul27.50v_34735 | 1 | 0 | 0 | 0 | 0 | 0 | 0 | 0 | 0 | 0 | 0 | 0 | 0 | 0 | 0 | 0 | 0 | 0 | 0 | 1 | 0 | 0 | 0 | 0 | 0 | 0 | 0 | 0 | 0 | 0 |
| Sep8.50v_29960 | 2 | 0 | 0 | 0 | 0 | 0 | 0 | 0 | 0 | 0 | 0 | 0 | 0 | 0 | 0 | 0 | 0 | 0 | 0 | 0 | 0 | 0 | 0 | 0 | 1 | 0 | 0 | 0 | 0 | 1 |
| Jun8.25v_14743 | 1 | 0 | 0 | 0 | 0 | 1 | 0 | 0 | 0 | 0 | 0 | 0 | 0 | 0 | 0 | 0 | 0 | 0 | 0 | 0 | 0 | 0 | 0 | 0 | 0 | 0 | 0 | 0 | 0 | 0 |
| Aug5.0v_5001 | 1 | 0 | 0 | 0 | 0 | 0 | 0 | 0 | 0 | 0 | 0 | 0 | 0 | 0 | 0 | 0 | 0 | 0 | 0 | 0 | 0 | 1 | 0 | 0 | 0 | 0 | 0 | 0 | 0 | 0 |
| Aug5.25v_9442 | 1 | 0 | 0 | 0 | 0 | 0 | 0 | 0 | 0 | 0 | 0 | 0 | 0 | 0 | 0 | 0 | 0 | 0 | 0 | 0 | 0 | 0 | 0 | 1 | 0 | 0 | 0 | 0 | 0 | 0 |
| Aug5.0v_1816 | 1 | 0 | 0 | 0 | 0 | 0 | 0 | 0 | 0 | 0 | 0 | 0 | 0 | 0 | 0 | 0 | 0 | 0 | 0 | 0 | 0 | 1 | 0 | 0 | 0 | 0 | 0 | 0 | 0 | 0 |
| Aug5.5v_14907 | 1 | 0 | 0 | 0 | 0 | 0 | 0 | 0 | 0 | 0 | 0 | 0 | 0 | 0 | 0 | 0 | 0 | 0 | 0 | 0 | 0 | 0 | 1 | 0 | 0 | 0 | 0 | 0 | 0 | 0 |
| Jul11.50v_10755 | 1 | 0 | 0 | 0 | 0 | 0 | 0 | 0 | 0 | 0 | 0 | 0 | 0 | 0 | 0 | 1 | 0 | 0 | 0 | 0 | 0 | 0 | 0 | 0 | 0 | 0 | 0 | 0 | 0 | 0 |
| Aug5.0v_2235 | 7 | 0 | 0 | 0 | 0 | 0 | 0 | 0 | 0 | 0 | 3 | 0 | 0 | 0 | 0 | 1 | 0 | 0 | 0 | 0 | 0 | 1 | 0 | 0 | 0 | 1 | 0 | 0 | 0 | 1 |
| Aug5.50v_26673 | 1 | 0 | 0 | 0 | 0 | 0 | 0 | 0 | 0 | 0 | 0 | 0 | 0 | 0 | 0 | 0 | 0 | 0 | 0 | 0 | 0 | 0 | 0 | 0 | 1 | 0 | 0 | 0 | 0 | 0 |
| May12.50v_22484 | 1 | 0 | 0 | 1 | 0 | 0 | 0 | 0 | 0 | 0 | 0 | 0 | 0 | 0 | 0 | 0 | 0 | 0 | 0 | 0 | 0 | 0 | 0 | 0 | 0 | 0 | 0 | 0 | 0 | 0 |
| Jul27.5v_650 | 1 | 0 | 0 | 0 | 0 | 0 | 0 | 0 | 0 | 0 | 0 | 0 | 0 | 0 | 0 | 0 | 0 | 1 | 0 | 0 | 0 | 0 | 0 | 0 | 0 | 0 | 0 | 0 | 0 | 0 |
| Jun8.50v_35174 | 2 | 0 | 0 | 0 | 0 | 0 | 1 | 0 | 0 | 0 | 0 | 0 | 0 | 0 | 1 | 0 | 0 | 0 | 0 | 0 | 0 | 0 | 0 | 0 | 0 | 0 | 0 | 0 | 0 | 0 |
| Jul27.75v_10122 | 1 | 0 | 0 | 0 | 0 | 0 | 0 | 0 | 0 | 0 | 0 | 0 | 0 | 0 | 0 | 0 | 0 | 0 | 0 | 0 | 1 | 0 | 0 | 0 | 0 | 0 | 0 | 0 | 0 | 0 |
| Jun22.50v_22685 | 1 | 0 | 0 | 0 | 0 | 0 | 0 | 0 | 0 | 0 | 1 | 0 | 0 | 0 | 0 | 0 | 0 | 0 | 0 | 0 | 0 | 0 | 0 | 0 | 0 | 0 | 0 | 0 | 0 | 0 |
| Jun22.5v_12000 | 1 | 0 | 0 | 0 | 0 | 0 | 0 | 0 | 0 | 1 | 0 | 0 | 0 | 0 | 0 | 0 | 0 | 0 | 0 | 0 | 0 | 0 | 0 | 0 | 0 | 0 | 0 | 0 | 0 | 0 |
| Jun22.75v_36613 | 1 | 0 | 0 | 0 | 0 | 0 | 0 | 0 | 0 | 0 | 0 | 1 | 0 | 0 | 0 | 0 | 0 | 0 | 0 | 0 | 0 | 0 | 0 | 0 | 0 | 0 | 0 | 0 | 0 | 0 |
| Jul11.50v_19569 | 1 | 0 | 0 | 0 | 0 | 0 | 0 | 0 | 0 | 0 | 0 | 0 | 0 | 0 | 0 | 1 | 0 | 0 | 0 | 0 | 0 | 0 | 0 | 0 | 0 | 0 | 0 | 0 | 0 | 0 |
| Jun22.0v_17979 | 1 | 0 | 0 | 0 | 0 | 0 | 0 | 0 | 1 | 0 | 0 | 0 | 0 | 0 | 0 | 0 | 0 | 0 | 0 | 0 | 0 | 0 | 0 | 0 | 0 | 0 | 0 | 0 | 0 | 0 |
| May12.25v_18546 | 1 | 0 | 1 | 0 | 0 | 0 | 0 | 0 | 0 | 0 | 0 | 0 | 0 | 0 | 0 | 0 | 0 | 0 | 0 | 0 | 0 | 0 | 0 | 0 | 0 | 0 | 0 | 0 | 0 | 0 |
| May12.75v_28458 | 1 | 0 | 0 | 0 | 1 | 0 | 0 | 0 | 0 | 0 | 0 | 0 | 0 | 0 | 0 | 0 | 0 | 0 | 0 | 0 | 0 | 0 | 0 | 0 | 0 | 0 | 0 | 0 | 0 | 0 |
| May12.50v_11953 | 1 | 0 | 0 | 1 | 0 | 0 | 0 | 0 | 0 | 0 | 0 | 0 | 0 | 0 | 0 | 0 | 0 | 0 | 0 | 0 | 0 | 0 | 0 | 0 | 0 | 0 | 0 | 0 | 0 | 0 |
| Jul27.25v_28745 | 1 | 0 | 0 | 0 | 0 | 0 | 0 | 0 | 0 | 0 | 0 | 0 | 0 | 0 | 0 | 0 | 0 | 0 | 1 | 0 | 0 | 0 | 0 | 0 | 0 | 0 | 0 | 0 | 0 | 0 |
| Jul11.5v_10461 | 8 | 0 | 1 | 2 | 1 | 0 | 0 | 0 | 0 | 0 | 0 | 2 | 0 | 1 | 0 | 0 | 0 | 0 | 1 | 0 | 0 | 0 | 0 | 0 | 0 | 0 | 0 | 0 | 0 | 0 |
| Sep8.25v_19723 | 1 | 0 | 0 | 0 | 0 | 0 | 0 | 0 | 0 | 0 | 0 | 0 | 0 | 0 | 0 | 0 | 0 | 0 | 0 | 0 | 0 | 0 | 0 | 0 | 0 | 0 | 0 | 0 | 1 | 0 |

| Feature_ID | 654 | May12_05m | May12_25m | May12_50m | May12_75m | Jun8_25m | Jun8_50m | Jun8_75m | Jun22_0m | Jun22_05m | Jun22_50m | Jun22_75m | Jul11_0m | Jul11_05m | Jul11_25m | Jul11_50m | Jul11_75m | Jul27_05m | Jul27_25m | Jul27_50m | Jul27_75m | Aug5_0m | Aug5_05m | Aug5_25m | Aug5_50m | Aug5_75m | Sep8_0m | Sep8_05m | Sep8_25m | Sep8_50m |
|---|---|---|---|---|---|---|---|---|---|---|---|---|---|---|---|---|---|---|---|---|---|---|---|---|---|---|---|---|---|---|
| Jun8.25v_30752 | 10 | 1 | 0 | 1 | 0 | 1 | 0 | 0 | 0 | 0 | 0 | 2 | 0 | 1 | 0 | 0 | 0 | 0 | 0 | 1 | 0 | 1 | 0 | 0 | 0 | 1 | 0 | 0 | 0 | 1 |
| Jun22.75v_32291 | 1 | 0 | 0 | 0 | 0 | 0 | 0 | 0 | 0 | 0 | 0 | 1 | 0 | 0 | 0 | 0 | 0 | 0 | 0 | 0 | 0 | 0 | 0 | 0 | 0 | 0 | 0 | 0 | 0 | 0 |
| May12.25v_37913 | 1 | 0 | 1 | 0 | 0 | 0 | 0 | 0 | 0 | 0 | 0 | 0 | 0 | 0 | 0 | 0 | 0 | 0 | 0 | 0 | 0 | 0 | 0 | 0 | 0 | 0 | 0 | 0 | 0 | 0 |
| Aug5.50v_26485 | 2 | 0 | 0 | 1 | 0 | 0 | 0 | 0 | 0 | 0 | 0 | 0 | 0 | 0 | 0 | 0 | 0 | 0 | 0 | 0 | 0 | 0 | 0 | 0 | 1 | 0 | 0 | 0 | 0 | 0 |
| Jul27.75v_8888 | 1 | 0 | 0 | 0 | 0 | 0 | 0 | 0 | 0 | 0 | 0 | 0 | 0 | 0 | 0 | 0 | 0 | 0 | 0 | 0 | 1 | 0 | 0 | 0 | 0 | 0 | 0 | 0 | 0 | 0 |
| Jun8.50v_32285 | 1 | 0 | 0 | 0 | 0 | 0 | 1 | 0 | 0 | 0 | 0 | 0 | 0 | 0 | 0 | 0 | 0 | 0 | 0 | 0 | 0 | 0 | 0 | 0 | 0 | 0 | 0 | 0 | 0 | 0 |
| May12.25v_1198 | 1 | 0 | 1 | 0 | 0 | 0 | 0 | 0 | 0 | 0 | 0 | 0 | 0 | 0 | 0 | 0 | 0 | 0 | 0 | 0 | 0 | 0 | 0 | 0 | 0 | 0 | 0 | 0 | 0 | 0 |
| May12.75v_7270 | 4 | 0 | 0 | 0 | 1 | 0 | 0 | 0 | 0 | 0 | 1 | 1 | 0 | 0 | 0 | 0 | 0 | 0 | 0 | 0 | 0 | 0 | 0 | 1 | 0 | 0 | 0 | 0 | 0 | 0 |
| Sep8.50v_37486 | 1 | 0 | 0 | 0 | 0 | 0 | 0 | 0 | 0 | 0 | 0 | 0 | 0 | 0 | 0 | 0 | 0 | 0 | 0 | 0 | 0 | 0 | 0 | 0 | 0 | 0 | 0 | 0 | 0 | 1 |
| Aug5.25v_14889 | 3 | 0 | 0 | 0 | 0 | 0 | 0 | 0 | 0 | 0 | 0 | 0 | 0 | 0 | 0 | 0 | 1 | 0 | 1 | 0 | 0 | 0 | 0 | 1 | 0 | 0 | 0 | 0 | 0 | 0 |
| May12.75v_24636 | 296 | 4 | 4 | 8 | 8 | 2 | 6 | 1 | 7 | 2 | 41 | 14 | 10 | 1 | 10 | 15 | 5 | 2 | 6 | 22 | 4 | 25 | 8 | 23 | 10 | 24 | 5 | 1 | 1 | 27 |
| May12.50v_11314 | 295 | 8 | 5 | 10 | 11 | 1 | 3 | 0 | 8 | 2 | 33 | 18 | 5 | 4 | 4 | 18 | 7 | 2 | 7 | 14 | 4 | 32 | 8 | 23 | 15 | 23 | 4 | 2 | 2 | 22 |
| Jul27.50v_5285 | 2 | 0 | 0 | 0 | 0 | 0 | 0 | 0 | 0 | 0 | 0 | 0 | 0 | 0 | 0 | 0 | 0 | 0 | 0 | 1 | 0 | 1 | 0 | 0 | 0 | 0 | 0 | 0 | 0 | 0 |
| Jun22.50v_3841 | 2 | 0 | 0 | 0 | 0 | 0 | 0 | 0 | 0 | 0 | 2 | 0 | 0 | 0 | 0 | 0 | 0 | 0 | 0 | 0 | 0 | 0 | 0 | 0 | 0 | 0 | 0 | 0 | 0 | 0 |
| Jul11.75v_2531 | 1 | 0 | 0 | 0 | 0 | 0 | 0 | 0 | 0 | 0 | 0 | 0 | 0 | 0 | 0 | 0 | 1 | 0 | 0 | 0 | 0 | 0 | 0 | 0 | 0 | 0 | 0 | 0 | 0 | 0 |
| Aug5.50v_2277 | 1 | 0 | 0 | 0 | 0 | 0 | 0 | 0 | 0 | 0 | 0 | 0 | 0 | 0 | 0 | 0 | 0 | 0 | 0 | 0 | 0 | 0 | 0 | 0 | 1 | 0 | 0 | 0 | 0 | 0 |
| Aug5.75v_38140 | 12 | 1 | 0 | 0 | 1 | 1 | 2 | 1 | 1 | 0 | 0 | 0 | 0 | 0 | 0 | 1 | 1 | 0 | 0 | 0 | 1 | 0 | 0 | 1 | 0 | 1 | 0 | 0 | 0 | 0 |
| Jun22.50v_4162 | 2 | 0 | 0 | 0 | 0 | 0 | 0 | 0 | 0 | 0 | 1 | 0 | 0 | 0 | 0 | 1 | 0 | 0 | 0 | 0 | 0 | 0 | 0 | 0 | 0 | 0 | 0 | 0 | 0 | 0 |
| Jun22.75v_4010 | 4 | 0 | 0 | 0 | 0 | 0 | 0 | 0 | 0 | 0 | 0 | 1 | 0 | 0 | 0 | 2 | 0 | 0 | 0 | 0 | 0 | 0 | 0 | 1 | 0 | 0 | 0 | 0 | 0 | 0 |
| Aug5.75v_38675 | 1 | 0 | 0 | 0 | 0 | 0 | 0 | 0 | 0 | 0 | 0 | 0 | 0 | 0 | 0 | 0 | 0 | 0 | 0 | 0 | 0 | 0 | 0 | 0 | 0 | 1 | 0 | 0 | 0 | 0 |
| May12.75v_14789 | 1 | 0 | 0 | 0 | 1 | 0 | 0 | 0 | 0 | 0 | 0 | 0 | 0 | 0 | 0 | 0 | 0 | 0 | 0 | 0 | 0 | 0 | 0 | 0 | 0 | 0 | 0 | 0 | 0 | 0 |
| Jul11.5v_17536 | 1 | 0 | 0 | 0 | 0 | 0 | 0 | 0 | 0 | 0 | 0 | 0 | 0 | 1 | 0 | 0 | 0 | 0 | 0 | 0 | 0 | 0 | 0 | 0 | 0 | 0 | 0 | 0 | 0 | 0 |
| Aug5.5v_12255 | 1 | 0 | 0 | 0 | 0 | 0 | 0 | 0 | 0 | 0 | 0 | 0 | 0 | 0 | 0 | 0 | 0 | 0 | 0 | 0 | 0 | 0 | 1 | 0 | 0 | 0 | 0 | 0 | 0 | 0 |
| Sep8.25v_8232 | 1 | 0 | 0 | 0 | 0 | 0 | 0 | 0 | 0 | 0 | 0 | 0 | 0 | 0 | 0 | 0 | 0 | 0 | 0 | 0 | 0 | 0 | 0 | 0 | 0 | 0 | 0 | 0 | 1 | 0 |
| Sep8.0v_1105 | 1 | 0 | 0 | 0 | 0 | 0 | 0 | 0 | 0 | 0 | 0 | 0 | 0 | 0 | 0 | 0 | 0 | 0 | 0 | 0 | 0 | 0 | 0 | 0 | 0 | 0 | 1 | 0 | 0 | 0 |
| Aug5.0v_987 | 1 | 0 | 0 | 0 | 0 | 0 | 0 | 0 | 0 | 0 | 0 | 0 | 0 | 0 | 0 | 0 | 0 | 0 | 0 | 0 | 0 | 1 | 0 | 0 | 0 | 0 | 0 | 0 | 0 | 0 |
| Sep8.50v_22198 | 1 | 0 | 0 | 0 | 0 | 0 | 0 | 0 | 0 | 0 | 0 | 0 | 0 | 0 | 0 | 0 | 0 | 0 | 0 | 0 | 0 | 0 | 0 | 0 | 0 | 0 | 0 | 0 | 0 | 1 |
| Jun22.50v_9386 | 1 | 0 | 0 | 0 | 0 | 0 | 0 | 0 | 0 | 0 | 1 | 0 | 0 | 0 | 0 | 0 | 0 | 0 | 0 | 0 | 0 | 0 | 0 | 0 | 0 | 0 | 0 | 0 | 0 | 0 |
| Sep8.50v_40089 | 1 | 0 | 0 | 0 | 0 | 0 | 0 | 0 | 0 | 0 | 0 | 0 | 0 | 0 | 0 | 0 | 0 | 0 | 0 | 0 | 0 | 0 | 0 | 0 | 0 | 0 | 0 | 0 | 0 | 1 |
| May12.5v_19860 | 1 | 1 | 0 | 0 | 0 | 0 | 0 | 0 | 0 | 0 | 0 | 0 | 0 | 0 | 0 | 0 | 0 | 0 | 0 | 0 | 0 | 0 | 0 | 0 | 0 | 0 | 0 | 0 | 0 | 0 |
| Jul11.50v_2621 | 1 | 0 | 0 | 0 | 0 | 0 | 0 | 0 | 0 | 0 | 0 | 0 | 0 | 0 | 0 | 1 | 0 | 0 | 0 | 0 | 0 | 0 | 0 | 0 | 0 | 0 | 0 | 0 | 0 | 0 |
| Aug5.25v_24147 | 1 | 0 | 0 | 0 | 0 | 0 | 0 | 0 | 0 | 0 | 0 | 0 | 0 | 0 | 0 | 0 | 0 | 0 | 0 | 0 | 0 | 0 | 0 | 1 | 0 | 0 | 0 | 0 | 0 | 0 |
| Jul11.25v_5561 | 1 | 0 | 0 | 0 | 0 | 0 | 0 | 0 | 0 | 0 | 0 | 0 | 0 | 0 | 1 | 0 | 0 | 0 | 0 | 0 | 0 | 0 | 0 | 0 | 0 | 0 | 0 | 0 | 0 | 0 |
| Jul11.50v_14336 | 1 | 0 | 0 | 0 | 0 | 0 | 0 | 0 | 0 | 0 | 0 | 0 | 0 | 0 | 0 | 1 | 0 | 0 | 0 | 0 | 0 | 0 | 0 | 0 | 0 | 0 | 0 | 0 | 0 | 0 |
| Jul27.75v_18162 | 1 | 0 | 0 | 0 | 0 | 0 | 0 | 0 | 0 | 0 | 0 | 0 | 0 | 0 | 0 | 0 | 0 | 0 | 0 | 0 | 1 | 0 | 0 | 0 | 0 | 0 | 0 | 0 | 0 | 0 |
| Jun22.50v_36947 | 1 | 0 | 0 | 0 | 0 | 0 | 0 | 0 | 0 | 0 | 1 | 0 | 0 | 0 | 0 | 0 | 0 | 0 | 0 | 0 | 0 | 0 | 0 | 0 | 0 | 0 | 0 | 0 | 0 | 0 |
| Jun22.5v_2493 | 1 | 0 | 0 | 0 | 0 | 0 | 0 | 0 | 0 | 1 | 0 | 0 | 0 | 0 | 0 | 0 | 0 | 0 | 0 | 0 | 0 | 0 | 0 | 0 | 0 | 0 | 0 | 0 | 0 | 0 |
| Jun8.25v_994 | 1 | 0 | 0 | 0 | 0 | 1 | 0 | 0 | 0 | 0 | 0 | 0 | 0 | 0 | 0 | 0 | 0 | 0 | 0 | 0 | 0 | 0 | 0 | 0 | 0 | 0 | 0 | 0 | 0 | 0 |
| Jun22.50v_30599 | 17 | 1 | 0 | 1 | 1 | 0 | 0 | 0 | 2 | 0 | 2 | 1 | 1 | 0 | 0 | 0 | 0 | 0 | 0 | 0 | 0 | 1 | 0 | 2 | 0 | 4 | 1 | 0 | 0 | 0 |
| Aug5.0v_276 | 10 | 0 | 0 | 0 | 0 | 1 | 0 | 0 | 0 | 0 | 1 | 2 | 1 | 0 | 0 | 0 | 0 | 0 | 0 | 0 | 0 | 1 | 0 | 1 | 0 | 2 | 1 | 0 | 0 | 0 |
| Jun22.75v_39635 | 6 | 0 | 0 | 1 | 1 | 0 | 0 | 0 | 0 | 0 | 0 | 1 | 0 | 0 | 0 | 0 | 1 | 0 | 0 | 0 | 0 | 0 | 0 | 2 | 0 | 0 | 0 | 0 | 0 | 0 |
| May12.75v_13142 | 107 | 5 | 12 | 4 | 7 | 4 | 12 | 3 | 0 | 0 | 9 | 15 | 0 | 2 | 0 | 4 | 9 | 1 | 0 | 6 | 0 | 0 | 4 | 3 | 2 | 3 | 1 | 0 | 0 | 1 |
| Jun22.50v_11598 | 1 | 0 | 0 | 0 | 0 | 0 | 0 | 0 | 0 | 0 | 1 | 0 | 0 | 0 | 0 | 0 | 0 | 0 | 0 | 0 | 0 | 0 | 0 | 0 | 0 | 0 | 0 | 0 | 0 | 0 |
| May12.25v_18789 | 16 | 0 | 5 | 1 | 1 | 0 | 1 | 0 | 0 | 1 | 1 | 1 | 0 | 0 | 0 | 1 | 0 | 0 | 0 | 1 | 1 | 0 | 0 | 0 | 1 | 0 | 1 | 0 | 0 | 0 |
| Jul11.25v_10897 | 1 | 0 | 0 | 0 | 0 | 0 | 0 | 0 | 0 | 0 | 0 | 0 | 0 | 0 | 1 | 0 | 0 | 0 | 0 | 0 | 0 | 0 | 0 | 0 | 0 | 0 | 0 | 0 | 0 | 0 |
| Jul11.75v_24726 | 9 | 0 | 0 | 0 | 1 | 1 | 2 | 0 | 0 | 0 | 2 | 0 | 0 | 0 | 0 | 0 | 1 | 1 | 0 | 1 | 0 | 0 | 0 | 0 | 0 | 0 | 0 | 0 | 0 | 0 |
| Jun22.75v_39688 | 1 | 0 | 0 | 0 | 0 | 0 | 0 | 0 | 0 | 0 | 0 | 1 | 0 | 0 | 0 | 0 | 0 | 0 | 0 | 0 | 0 | 0 | 0 | 0 | 0 | 0 | 0 | 0 | 0 | 0 |
| Sep8.25v_293 | 1 | 0 | 0 | 0 | 0 | 0 | 0 | 0 | 0 | 0 | 0 | 0 | 0 | 0 | 0 | 0 | 0 | 0 | 0 | 0 | 0 | 0 | 0 | 0 | 0 | 0 | 0 | 0 | 1 | 0 |
| Jun22.5v_6570 | 1 | 0 | 0 | 0 | 0 | 0 | 0 | 0 | 0 | 1 | 0 | 0 | 0 | 0 | 0 | 0 | 0 | 0 | 0 | 0 | 0 | 0 | 0 | 0 | 0 | 0 | 0 | 0 | 0 | 0 |
| Jun22.0v_3466 | 1 | 0 | 0 | 0 | 0 | 0 | 0 | 0 | 1 | 0 | 0 | 0 | 0 | 0 | 0 | 0 | 0 | 0 | 0 | 0 | 0 | 0 | 0 | 0 | 0 | 0 | 0 | 0 | 0 | 0 |
| Jul27.75v_18645 | 4 | 0 | 1 | 0 | 1 | 0 | 1 | 0 | 0 | 0 | 0 | 0 | 0 | 0 | 0 | 0 | 0 | 0 | 0 | 0 | 1 | 0 | 0 | 0 | 0 | 0 | 0 | 0 | 0 | 0 |
| Sep8.0v_14474 | 1 | 0 | 0 | 0 | 0 | 0 | 0 | 0 | 0 | 0 | 0 | 0 | 0 | 0 | 0 | 0 | 0 | 0 | 0 | 0 | 0 | 0 | 0 | 0 | 0 | 0 | 1 | 0 | 0 | 0 |
| May12.75v_5309 | 1 | 0 | 0 | 0 | 1 | 0 | 0 | 0 | 0 | 0 | 0 | 0 | 0 | 0 | 0 | 0 | 0 | 0 | 0 | 0 | 0 | 0 | 0 | 0 | 0 | 0 | 0 | 0 | 0 | 0 |
| Jul27.25v_27525 | 1 | 0 | 0 | 0 | 0 | 0 | 0 | 0 | 0 | 0 | 0 | 0 | 0 | 0 | 0 | 0 | 0 | 0 | 1 | 0 | 0 | 0 | 0 | 0 | 0 | 0 | 0 | 0 | 0 | 0 |
| Sep8.0v_15001 | 1 | 0 | 0 | 0 | 0 | 0 | 0 | 0 | 0 | 0 | 0 | 0 | 0 | 0 | 0 | 0 | 0 | 0 | 0 | 0 | 0 | 0 | 0 | 0 | 0 | 0 | 1 | 0 | 0 | 0 |
| Aug5.50v_28901 | 1 | 0 | 0 | 0 | 0 | 0 | 0 | 0 | 0 | 0 | 0 | 0 | 0 | 0 | 0 | 0 | 0 | 0 | 0 | 0 | 0 | 0 | 0 | 0 | 1 | 0 | 0 | 0 | 0 | 0 |
| Jun22.50v_22219 | 1 | 0 | 0 | 0 | 0 | 0 | 0 | 0 | 0 | 0 | 1 | 0 | 0 | 0 | 0 | 0 | 0 | 0 | 0 | 0 | 0 | 0 | 0 | 0 | 0 | 0 | 0 | 0 | 0 | 0 |
| Jun22.50v_27535 | 1 | 0 | 0 | 0 | 0 | 0 | 0 | 0 | 0 | 0 | 1 | 0 | 0 | 0 | 0 | 0 | 0 | 0 | 0 | 0 | 0 | 0 | 0 | 0 | 0 | 0 | 0 | 0 | 0 | 0 |
| Jun22.75v_21281 | 1 | 0 | 0 | 0 | 0 | 0 | 0 | 0 | 0 | 0 | 0 | 1 | 0 | 0 | 0 | 0 | 0 | 0 | 0 | 0 | 0 | 0 | 0 | 0 | 0 | 0 | 0 | 0 | 0 | 0 |
| Aug5.75v_33715 | 1 | 0 | 0 | 0 | 0 | 0 | 0 | 0 | 0 | 0 | 0 | 0 | 0 | 0 | 0 | 0 | 0 | 0 | 0 | 0 | 0 | 0 | 0 | 0 | 0 | 1 | 0 | 0 | 0 | 0 |
| Aug5.25v_32541 | 1 | 0 | 0 | 0 | 0 | 0 | 0 | 0 | 0 | 0 | 0 | 0 | 0 | 0 | 0 | 0 | 0 | 0 | 0 | 0 | 0 | 0 | 0 | 1 | 0 | 0 | 0 | 0 | 0 | 0 |
| Jul27.50v_32602 | 1 | 0 | 0 | 0 | 0 | 0 | 0 | 0 | 0 | 0 | 0 | 0 | 0 | 0 | 0 | 0 | 0 | 0 | 0 | 1 | 0 | 0 | 0 | 0 | 0 | 0 | 0 | 0 | 0 | 0 |
| Jul11.25v_9154 | 1 | 0 | 0 | 0 | 0 | 0 | 0 | 0 | 0 | 0 | 0 | 0 | 0 | 0 | 1 | 0 | 0 | 0 | 0 | 0 | 0 | 0 | 0 | 0 | 0 | 0 | 0 | 0 | 0 | 0 |
| Jul27.5v_16702 | 1 | 0 | 0 | 0 | 0 | 0 | 0 | 0 | 0 | 0 | 0 | 0 | 0 | 0 | 0 | 0 | 0 | 1 | 0 | 0 | 0 | 0 | 0 | 0 | 0 | 0 | 0 | 0 | 0 | 0 |
| Aug5.50v_22303 | 1 | 0 | 0 | 0 | 0 | 0 | 0 | 0 | 0 | 0 | 0 | 0 | 0 | 0 | 0 | 0 | 0 | 0 | 0 | 0 | 0 | 0 | 0 | 0 | 1 | 0 | 0 | 0 | 0 | 0 |
| Sep8.50v_28066 | 1 | 0 | 0 | 0 | 0 | 0 | 0 | 0 | 0 | 0 | 0 | 0 | 0 | 0 | 0 | 0 | 0 | 0 | 0 | 0 | 0 | 0 | 0 | 0 | 0 | 0 | 0 | 0 | 0 | 1 |
| May12.5v_33029 | 1 | 1 | 0 | 0 | 0 | 0 | 0 | 0 | 0 | 0 | 0 | 0 | 0 | 0 | 0 | 0 | 0 | 0 | 0 | 0 | 0 | 0 | 0 | 0 | 0 | 0 | 0 | 0 | 0 | 0 |
| Jul11.50v_1408 | 1 | 0 | 0 | 0 | 0 | 0 | 0 | 0 | 0 | 0 | 0 | 0 | 0 | 0 | 0 | 1 | 0 | 0 | 0 | 0 | 0 | 0 | 0 | 0 | 0 | 0 | 0 | 0 | 0 | 0 |
| Aug5.75v_30233 | 1 | 0 | 0 | 0 | 0 | 0 | 0 | 0 | 0 | 0 | 0 | 0 | 0 | 0 | 0 | 0 | 0 | 0 | 0 | 0 | 0 | 0 | 0 | 0 | 0 | 1 | 0 | 0 | 0 | 0 |
| Aug5.25v_4313 | 1 | 0 | 0 | 0 | 0 | 0 | 0 | 0 | 0 | 0 | 0 | 0 | 0 | 0 | 0 | 0 | 0 | 0 | 0 | 0 | 0 | 0 | 0 | 1 | 0 | 0 | 0 | 0 | 0 | 0 |
| May12.25v_2201 | 1 | 0 | 1 | 0 | 0 | 0 | 0 | 0 | 0 | 0 | 0 | 0 | 0 | 0 | 0 | 0 | 0 | 0 | 0 | 0 | 0 | 0 | 0 | 0 | 0 | 0 | 0 | 0 | 0 | 0 |
| Sep8.50v_32411 | 1 | 0 | 0 | 0 | 0 | 0 | 0 | 0 | 0 | 0 | 0 | 0 | 0 | 0 | 0 | 0 | 0 | 0 | 0 | 0 | 0 | 0 | 0 | 0 | 0 | 0 | 0 | 0 | 0 | 1 |
| Aug5.25v_22083 | 1 | 0 | 0 | 0 | 0 | 0 | 0 | 0 | 0 | 0 | 0 | 0 | 0 | 0 | 0 | 0 | 0 | 0 | 0 | 0 | 0 | 0 | 0 | 1 | 0 | 0 | 0 | 0 | 0 | 0 |
| Jun22.5v_13338 | 1 | 0 | 0 | 0 | 0 | 0 | 0 | 0 | 0 | 1 | 0 | 0 | 0 | 0 | 0 | 0 | 0 | 0 | 0 | 0 | 0 | 0 | 0 | 0 | 0 | 0 | 0 | 0 | 0 | 0 |
| Aug5.0v_1201 | 1 | 0 | 0 | 0 | 0 | 0 | 0 | 0 | 0 | 0 | 0 | 0 | 0 | 0 | 0 | 0 | 0 | 0 | 0 | 0 | 0 | 1 | 0 | 0 | 0 | 0 | 0 | 0 | 0 | 0 |
| Jul11.25v_4961 | 1 | 0 | 0 | 0 | 0 | 0 | 0 | 0 | 0 | 0 | 0 | 0 | 0 | 0 | 1 | 0 | 0 | 0 | 0 | 0 | 0 | 0 | 0 | 0 | 0 | 0 | 0 | 0 | 0 | 0 |
| Aug5.0v_2450 | 7 | 0 | 0 | 0 | 0 | 0 | 1 | 0 | 0 | 0 | 0 | 1 | 0 | 0 | 0 | 0 | 1 | 0 | 0 | 0 | 0 | 1 | 0 | 0 | 0 | 1 | 0 | 0 | 0 | 2 |
| Jun22.75v_32241 | 1 | 0 | 0 | 0 | 0 | 0 | 0 | 0 | 0 | 0 | 0 | 1 | 0 | 0 | 0 | 0 | 0 | 0 | 0 | 0 | 0 | 0 | 0 | 0 | 0 | 0 | 0 | 0 | 0 | 0 |
| Aug5.75v_14254 | 1 | 0 | 0 | 0 | 0 | 0 | 0 | 0 | 0 | 0 | 0 | 0 | 0 | 0 | 0 | 0 | 0 | 0 | 0 | 0 | 0 | 0 | 0 | 0 | 0 | 1 | 0 | 0 | 0 | 0 |
| May12.50v_3975 | 1 | 0 | 0 | 1 | 0 | 0 | 0 | 0 | 0 | 0 | 0 | 0 | 0 | 0 | 0 | 0 | 0 | 0 | 0 | 0 | 0 | 0 | 0 | 0 | 0 | 0 | 0 | 0 | 0 | 0 |
| May12.75v_1555 | 1 | 0 | 0 | 0 | 1 | 0 | 0 | 0 | 0 | 0 | 0 | 0 | 0 | 0 | 0 | 0 | 0 | 0 | 0 | 0 | 0 | 0 | 0 | 0 | 0 | 0 | 0 | 0 | 0 | 0 |
| Jun8.25v_5811 | 1 | 0 | 0 | 0 | 0 | 1 | 0 | 0 | 0 | 0 | 0 | 0 | 0 | 0 | 0 | 0 | 0 | 0 | 0 | 0 | 0 | 0 | 0 | 0 | 0 | 0 | 0 | 0 | 0 | 0 |
| Jul11.50v_14384 | 1 | 0 | 0 | 0 | 0 | 0 | 0 | 0 | 0 | 0 | 0 | 0 | 0 | 0 | 0 | 1 | 0 | 0 | 0 | 0 | 0 | 0 | 0 | 0 | 0 | 0 | 0 | 0 | 0 | 0 |
| Jul11.75v_7915 | 1 | 0 | 0 | 0 | 0 | 0 | 0 | 0 | 0 | 0 | 0 | 0 | 0 | 0 | 0 | 0 | 1 | 0 | 0 | 0 | 0 | 0 | 0 | 0 | 0 | 0 | 0 | 0 | 0 | 0 |
| Jun22.50v_30078 | 1 | 0 | 0 | 0 | 0 | 0 | 0 | 0 | 0 | 0 | 1 | 0 | 0 | 0 | 0 | 0 | 0 | 0 | 0 | 0 | 0 | 0 | 0 | 0 | 0 | 0 | 0 | 0 | 0 | 0 |
| Aug5.75v_1000 | 1 | 0 | 0 | 0 | 0 | 0 | 0 | 0 | 0 | 0 | 0 | 0 | 0 | 0 | 0 | 0 | 0 | 0 | 0 | 0 | 0 | 0 | 0 | 0 | 0 | 1 | 0 | 0 | 0 | 0 |
| Jul11.75v_37018 | 1 | 0 | 0 | 0 | 0 | 0 | 0 | 0 | 0 | 0 | 0 | 0 | 0 | 0 | 0 | 0 | 1 | 0 | 0 | 0 | 0 | 0 | 0 | 0 | 0 | 0 | 0 | 0 | 0 | 0 |
| Jul11.50v_9145 | 1 | 0 | 0 | 0 | 0 | 0 | 0 | 0 | 0 | 0 | 0 | 0 | 0 | 0 | 0 | 1 | 0 | 0 | 0 | 0 | 0 | 0 | 0 | 0 | 0 | 0 | 0 | 0 | 0 | 0 |
| Aug5.0v_4824 | 1 | 0 | 0 | 0 | 0 | 0 | 0 | 0 | 0 | 0 | 0 | 0 | 0 | 0 | 0 | 0 | 0 | 0 | 0 | 0 | 0 | 1 | 0 | 0 | 0 | 0 | 0 | 0 | 0 | 0 |
| May12.25v_8994 | 1 | 0 | 1 | 0 | 0 | 0 | 0 | 0 | 0 | 0 | 0 | 0 | 0 | 0 | 0 | 0 | 0 | 0 | 0 | 0 | 0 | 0 | 0 | 0 | 0 | 0 | 0 | 0 | 0 | 0 |
| Jun8.50v_19047 | 1 | 0 | 0 | 0 | 0 | 0 | 1 | 0 | 0 | 0 | 0 | 0 | 0 | 0 | 0 | 0 | 0 | 0 | 0 | 0 | 0 | 0 | 0 | 0 | 0 | 0 | 0 | 0 | 0 | 0 |
| Jun8.25v_887 | 1 | 0 | 0 | 0 | 0 | 1 | 0 | 0 | 0 | 0 | 0 | 0 | 0 | 0 | 0 | 0 | 0 | 0 | 0 | 0 | 0 | 0 | 0 | 0 | 0 | 0 | 0 | 0 | 0 | 0 |
| Jun8.25v_21223 | 1 | 0 | 0 | 0 | 0 | 1 | 0 | 0 | 0 | 0 | 0 | 0 | 0 | 0 | 0 | 0 | 0 | 0 | 0 | 0 | 0 | 0 | 0 | 0 | 0 | 0 | 0 | 0 | 0 | 0 |
| Jul11.75v_20354 | 4 | 0 | 1 | 0 | 0 | 0 | 1 | 0 | 0 | 0 | 0 | 1 | 0 | 0 | 0 | 0 | 1 | 0 | 0 | 0 | 0 | 0 | 0 | 0 | 0 | 0 | 0 | 0 | 0 | 0 |
| Sep8.25v_13110 | 2 | 0 | 0 | 0 | 0 | 0 | 0 | 0 | 0 | 0 | 0 | 0 | 0 | 0 | 0 | 0 | 0 | 0 | 0 | 0 | 0 | 0 | 0 | 0 | 0 | 0 | 0 | 0 | 1 | 1 |
| May12.5v_38052 | 1 | 1 | 0 | 0 | 0 | 0 | 0 | 0 | 0 | 0 | 0 | 0 | 0 | 0 | 0 | 0 | 0 | 0 | 0 | 0 | 0 | 0 | 0 | 0 | 0 | 0 | 0 | 0 | 0 | 0 |
| May12.50v_18269 | 7 | 0 | 0 | 2 | 1 | 0 | 0 | 0 | 0 | 0 | 0 | 1 | 0 | 0 | 0 | 0 | 1 | 0 | 0 | 0 | 0 | 1 | 0 | 0 | 1 | 0 | 0 | 0 | 0 | 0 |
| May12.25v_31660 | 2 | 0 | 1 | 0 | 0 | 1 | 0 | 0 | 0 | 0 | 0 | 0 | 0 | 0 | 0 | 0 | 0 | 0 | 0 | 0 | 0 | 0 | 0 | 0 | 0 | 0 | 0 | 0 | 0 | 0 |
| Jul27.25v_19471 | 7 | 1 | 0 | 0 | 1 | 0 | 0 | 0 | 0 | 0 | 0 | 0 | 1 | 0 | 0 | 2 | 0 | 0 | 1 | 0 | 0 | 0 | 0 | 0 | 0 | 0 | 0 | 0 | 0 | 1 |
| Aug5.75v_10553 | 3 | 0 | 0 | 0 | 0 | 0 | 1 | 0 | 0 | 0 | 0 | 0 | 0 | 0 | 0 | 0 | 1 | 0 | 0 | 0 | 0 | 0 | 0 | 0 | 0 | 1 | 0 | 0 | 0 | 0 |
| Sep8.5v_930 | 44 | 2 | 4 | 1 | 2 | 1 | 7 | 1 | 0 | 1 | 0 | 5 | 3 | 2 | 2 | 2 | 1 | 1 | 0 | 1 | 0 | 0 | 0 | 0 | 1 | 0 | 1 | 6 | 0 | 0 |
| Jun22.75v_13651 | 1 | 0 | 0 | 0 | 0 | 0 | 0 | 0 | 0 | 0 | 0 | 1 | 0 | 0 | 0 | 0 | 0 | 0 | 0 | 0 | 0 | 0 | 0 | 0 | 0 | 0 | 0 | 0 | 0 | 0 |
| Aug5.0v_2324 | 17 | 1 | 0 | 2 | 2 | 0 | 0 | 0 | 0 | 0 | 1 | 0 | 0 | 0 | 0 | 0 | 0 | 0 | 1 | 1 | 0 | 2 | 3 | 2 | 0 | 0 | 0 | 1 | 0 | 1 |
| Aug5.0v_5212 | 1 | 0 | 0 | 0 | 0 | 0 | 0 | 0 | 0 | 0 | 0 | 0 | 0 | 0 | 0 | 0 | 0 | 0 | 0 | 0 | 0 | 1 | 0 | 0 | 0 | 0 | 0 | 0 | 0 | 0 |
| Jul11.75v_5240 | 1 | 0 | 0 | 0 | 0 | 0 | 0 | 0 | 0 | 0 | 0 | 0 | 0 | 0 | 0 | 0 | 1 | 0 | 0 | 0 | 0 | 0 | 0 | 0 | 0 | 0 | 0 | 0 | 0 | 0 |
| Jun22.75v_3480 | 2 | 0 | 0 | 0 | 0 | 0 | 0 | 0 | 0 | 0 | 0 | 1 | 0 | 0 | 0 | 0 | 0 | 0 | 0 | 0 | 0 | 0 | 0 | 0 | 0 | 0 | 1 | 0 | 0 | 0 |
| Jun22.75v_19113 | 1 | 0 | 0 | 0 | 0 | 0 | 0 | 0 | 0 | 0 | 0 | 1 | 0 | 0 | 0 | 0 | 0 | 0 | 0 | 0 | 0 | 0 | 0 | 0 | 0 | 0 | 0 | 0 | 0 | 0 |
| Sep8.0v_1275 | 2 | 0 | 0 | 0 | 0 | 0 | 0 | 0 | 0 | 1 | 0 | 0 | 0 | 0 | 0 | 0 | 0 | 0 | 0 | 0 | 0 | 0 | 0 | 0 | 0 | 0 | 1 | 0 | 0 | 0 |
| Jul11.5v_11629 | 1 | 0 | 0 | 0 | 0 | 0 | 0 | 0 | 0 | 0 | 0 | 0 | 0 | 1 | 0 | 0 | 0 | 0 | 0 | 0 | 0 | 0 | 0 | 0 | 0 | 0 | 0 | 0 | 0 | 0 |
| Jul11.75v_23390 | 1 | 0 | 0 | 0 | 0 | 0 | 0 | 0 | 0 | 0 | 0 | 0 | 0 | 0 | 0 | 0 | 1 | 0 | 0 | 0 | 0 | 0 | 0 | 0 | 0 | 0 | 0 | 0 | 0 | 0 |
| Jun22.0v_5496 | 45 | 2 | 1 | 3 | 3 | 3 | 3 | 2 | 1 | 0 | 2 | 6 | 0 | 1 | 1 | 2 | 4 | 0 | 0 | 4 | 0 | 0 | 0 | 0 | 0 | 2 | 0 | 2 | 0 | 3 |
| Jul11.5v_20599 | 4 | 0 | 0 | 1 | 0 | 0 | 1 | 0 | 0 | 0 | 0 | 0 | 0 | 1 | 0 | 1 | 0 | 0 | 0 | 0 | 0 | 0 | 0 | 0 | 0 | 0 | 0 | 0 | 0 | 0 |
| Jun8.50v_36988 | 1 | 0 | 0 | 0 | 0 | 0 | 1 | 0 | 0 | 0 | 0 | 0 | 0 | 0 | 0 | 0 | 0 | 0 | 0 | 0 | 0 | 0 | 0 | 0 | 0 | 0 | 0 | 0 | 0 | 0 |
| May12.5v_11069 | 7 | 1 | 1 | 1 | 0 | 0 | 0 | 0 | 0 | 0 | 1 | 0 | 0 | 0 | 0 | 0 | 2 | 0 | 0 | 1 | 0 | 0 | 0 | 0 | 0 | 0 | 0 | 0 | 0 | 0 |
| Jul11.50v_15751 | 9 | 0 | 0 | 0 | 0 | 0 | 0 | 0 | 0 | 0 | 1 | 0 | 0 | 1 | 0 | 2 | 1 | 2 | 0 | 0 | 0 | 0 | 0 | 0 | 1 | 1 | 0 | 0 | 0 | 0 |
| Sep8.50v_22833 | 1 | 0 | 0 | 0 | 0 | 0 | 0 | 0 | 0 | 0 | 0 | 0 | 0 | 0 | 0 | 0 | 0 | 0 | 0 | 0 | 0 | 0 | 0 | 0 | 0 | 0 | 0 | 0 | 0 | 1 |
| Jul27.50v_35901 | 6 | 0 | 0 | 0 | 0 | 0 | 1 | 0 | 0 | 0 | 0 | 0 | 0 | 1 | 0 | 1 | 1 | 0 | 0 | 1 | 0 | 0 | 0 | 0 | 0 | 0 | 0 | 0 | 0 | 1 |
| May12.75v_2430 | 2 | 0 | 0 | 0 | 1 | 0 | 0 | 0 | 0 | 0 | 0 | 0 | 0 | 0 | 0 | 1 | 0 | 0 | 0 | 0 | 0 | 0 | 0 | 0 | 0 | 0 | 0 | 0 | 0 | 0 |
| Jul27.25v_16514 | 1 | 0 | 0 | 0 | 0 | 0 | 0 | 0 | 0 | 0 | 0 | 0 | 0 | 0 | 0 | 0 | 0 | 0 | 1 | 0 | 0 | 0 | 0 | 0 | 0 | 0 | 0 | 0 | 0 | 0 |
| Aug5.25v_29117 | 2 | 0 | 0 | 0 | 0 | 1 | 0 | 0 | 0 | 0 | 0 | 0 | 0 | 0 | 0 | 0 | 0 | 0 | 0 | 0 | 0 | 0 | 0 | 1 | 0 | 0 | 0 | 0 | 0 | 0 |
| Jul11.75v_28208 | 1 | 0 | 0 | 0 | 0 | 0 | 0 | 0 | 0 | 0 | 0 | 0 | 0 | 0 | 0 | 0 | 1 | 0 | 0 | 0 | 0 | 0 | 0 | 0 | 0 | 0 | 0 | 0 | 0 | 0 |
| Jul11.50v_8859 | 5 | 0 | 0 | 0 | 0 | 0 | 2 | 0 | 1 | 0 | 0 | 0 | 0 | 0 | 0 | 1 | 1 | 0 | 0 | 0 | 0 | 0 | 0 | 0 | 0 | 0 | 0 | 0 | 0 | 0 |
| May12.50v_34881 | 3 | 0 | 0 | 1 | 0 | 0 | 0 | 0 | 0 | 0 | 0 | 0 | 0 | 0 | 0 | 0 | 1 | 0 | 0 | 0 | 0 | 0 | 1 | 0 | 0 | 0 | 0 | 0 | 0 | 0 |
| Jun8.25v_25702 | 31 | 1 | 0 | 3 | 1 | 2 | 0 | 0 | 0 | 0 | 0 | 3 | 2 | 2 | 1 | 0 | 1 | 0 | 2 | 0 | 0 | 5 | 1 | 1 | 0 | 0 | 1 | 1 | 0 | 4 |
| Jul11.25v_1916 | 5 | 0 | 0 | 0 | 0 | 0 | 0 | 0 | 0 | 0 | 0 | 0 | 0 | 0 | 1 | 0 | 1 | 0 | 0 | 1 | 0 | 0 | 1 | 0 | 0 | 0 | 0 | 0 | 0 | 1 |
| Sep8.50v_24750 | 3 | 0 | 1 | 0 | 0 | 0 | 1 | 0 | 0 | 0 | 0 | 0 | 0 | 0 | 0 | 0 | 0 | 0 | 0 | 0 | 0 | 0 | 0 | 0 | 0 | 0 | 0 | 0 | 0 | 1 |
| Jun22.50v_11727 | 1 | 0 | 0 | 0 | 0 | 0 | 0 | 0 | 0 | 0 | 1 | 0 | 0 | 0 | 0 | 0 | 0 | 0 | 0 | 0 | 0 | 0 | 0 | 0 | 0 | 0 | 0 | 0 | 0 | 0 |
| Jul11.75v_12470 | 1 | 0 | 0 | 0 | 0 | 0 | 0 | 0 | 0 | 0 | 0 | 0 | 0 | 0 | 0 | 0 | 1 | 0 | 0 | 0 | 0 | 0 | 0 | 0 | 0 | 0 | 0 | 0 | 0 | 0 |
| May12.50v_11557 | 1 | 0 | 0 | 1 | 0 | 0 | 0 | 0 | 0 | 0 | 0 | 0 | 0 | 0 | 0 | 0 | 0 | 0 | 0 | 0 | 0 | 0 | 0 | 0 | 0 | 0 | 0 | 0 | 0 | 0 |
| Sep8.0v_9401 | 17 | 2 | 1 | 1 | 1 | 0 | 1 | 0 | 1 | 0 | 0 | 2 | 0 | 1 | 0 | 0 | 0 | 0 | 0 | 1 | 0 | 1 | 0 | 1 | 1 | 1 | 1 | 1 | 0 | 0 |
| Jul11.50v_3450 | 3 | 0 | 0 | 0 | 0 | 0 | 0 | 0 | 0 | 0 | 0 | 0 | 0 | 0 | 0 | 1 | 0 | 0 | 0 | 0 | 0 | 0 | 0 | 1 | 0 | 0 | 0 | 1 | 0 | 0 |
| Jun22.50v_4776 | 3 | 0 | 0 | 0 | 0 | 0 | 1 | 0 | 0 | 0 | 1 | 1 | 0 | 0 | 0 | 0 | 0 | 0 | 0 | 0 | 0 | 0 | 0 | 0 | 0 | 0 | 0 | 0 | 0 | 0 |
| May12.5v_40762 | 1 | 1 | 0 | 0 | 0 | 0 | 0 | 0 | 0 | 0 | 0 | 0 | 0 | 0 | 0 | 0 | 0 | 0 | 0 | 0 | 0 | 0 | 0 | 0 | 0 | 0 | 0 | 0 | 0 | 0 |
| May12.5v_13173 | 69 | 5 | 4 | 8 | 5 | 2 | 5 | 1 | 1 | 3 | 3 | 2 | 3 | 2 | 2 | 0 | 0 | 1 | 1 | 2 | 1 | 3 | 5 | 0 | 1 | 0 | 2 | 3 | 0 | 4 |
| Jul11.25v_5085 | 2 | 0 | 0 | 0 | 0 | 0 | 0 | 0 | 0 | 0 | 0 | 0 | 0 | 0 | 1 | 0 | 0 | 0 | 1 | 0 | 0 | 0 | 0 | 0 | 0 | 0 | 0 | 0 | 0 | 0 |
| Sep8.50v_16868 | 1 | 0 | 0 | 0 | 0 | 0 | 0 | 0 | 0 | 0 | 0 | 0 | 0 | 0 | 0 | 0 | 0 | 0 | 0 | 0 | 0 | 0 | 0 | 0 | 0 | 0 | 0 | 0 | 0 | 1 |
| Jul11.50v_5024 | 1 | 0 | 0 | 0 | 0 | 0 | 0 | 0 | 0 | 0 | 0 | 0 | 0 | 0 | 0 | 1 | 0 | 0 | 0 | 0 | 0 | 0 | 0 | 0 | 0 | 0 | 0 | 0 | 0 | 0 |
| May12.25v_3798 | 1 | 0 | 1 | 0 | 0 | 0 | 0 | 0 | 0 | 0 | 0 | 0 | 0 | 0 | 0 | 0 | 0 | 0 | 0 | 0 | 0 | 0 | 0 | 0 | 0 | 0 | 0 | 0 | 0 | 0 |
| May12.25v_4729 | 2 | 0 | 2 | 0 | 0 | 0 | 0 | 0 | 0 | 0 | 0 | 0 | 0 | 0 | 0 | 0 | 0 | 0 | 0 | 0 | 0 | 0 | 0 | 0 | 0 | 0 | 0 | 0 | 0 | 0 |
| Aug5.75v_25496 | 1 | 0 | 0 | 0 | 0 | 0 | 0 | 0 | 0 | 0 | 0 | 0 | 0 | 0 | 0 | 0 | 0 | 0 | 0 | 0 | 0 | 0 | 0 | 0 | 0 | 1 | 0 | 0 | 0 | 0 |
| Jul11.0v_12151 | 98 | 10 | 5 | 8 | 1 | 2 | 2 | 0 | 3 | 3 | 2 | 4 | 2 | 1 | 3 | 2 | 5 | 0 | 4 | 8 | 0 | 4 | 4 | 5 | 0 | 1 | 4 | 1 | 0 | 14 |
| Sep8.50v_15464 | 4 | 0 | 0 | 0 | 1 | 0 | 0 | 0 | 0 | 0 | 0 | 0 | 0 | 0 | 0 | 0 | 0 | 0 | 0 | 0 | 0 | 0 | 0 | 0 | 0 | 0 | 0 | 1 | 0 | 2 |
| May12.75v_24867 | 9 | 1 | 1 | 0 | 1 | 0 | 0 | 0 | 0 | 1 | 0 | 0 | 1 | 0 | 0 | 0 | 0 | 1 | 0 | 0 | 0 | 1 | 0 | 0 | 1 | 0 | 1 | 0 | 0 | 0 |
| Jul27.25v_14231 | 3 | 0 | 0 | 0 | 0 | 0 | 0 | 0 | 0 | 0 | 0 | 0 | 0 | 0 | 0 | 0 | 0 | 0 | 2 | 1 | 0 | 0 | 0 | 0 | 0 | 0 | 0 | 0 | 0 | 0 |
| Jun22.50v_12875 | 1 | 0 | 0 | 0 | 0 | 0 | 0 | 0 | 0 | 0 | 1 | 0 | 0 | 0 | 0 | 0 | 0 | 0 | 0 | 0 | 0 | 0 | 0 | 0 | 0 | 0 | 0 | 0 | 0 | 0 |
| Sep8.50v_21263 | 1 | 0 | 0 | 0 | 0 | 0 | 0 | 0 | 0 | 0 | 0 | 0 | 0 | 0 | 0 | 0 | 0 | 0 | 0 | 0 | 0 | 0 | 0 | 0 | 0 | 0 | 0 | 0 | 0 | 1 |
| Jul11.50v_9863 | 1 | 0 | 0 | 0 | 0 | 0 | 0 | 0 | 0 | 0 | 0 | 0 | 0 | 0 | 0 | 1 | 0 | 0 | 0 | 0 | 0 | 0 | 0 | 0 | 0 | 0 | 0 | 0 | 0 | 0 |
| Jul11.25v_15692 | 1 | 0 | 0 | 0 | 0 | 0 | 0 | 0 | 0 | 0 | 0 | 0 | 0 | 0 | 1 | 0 | 0 | 0 | 0 | 0 | 0 | 0 | 0 | 0 | 0 | 0 | 0 | 0 | 0 | 0 |
| Jun22.75v_7718 | 6 | 1 | 0 | 0 | 1 | 0 | 1 | 0 | 1 | 0 | 0 | 2 | 0 | 0 | 0 | 0 | 0 | 0 | 0 | 0 | 0 | 0 | 0 | 0 | 0 | 0 | 0 | 0 | 0 | 0 |
| Jul11.0v_13768 | 11 | 0 | 0 | 1 | 1 | 0 | 0 | 0 | 2 | 0 | 0 | 1 | 2 | 0 | 0 | 1 | 0 | 0 | 0 | 1 | 0 | 0 | 0 | 0 | 2 | 0 | 0 | 0 | 0 | 0 |
| Jul11.75v_2729 | 5 | 0 | 2 | 0 | 0 | 1 | 0 | 0 | 0 | 0 | 0 | 0 | 1 | 0 | 0 | 0 | 1 | 0 | 0 | 0 | 0 | 0 | 0 | 0 | 0 | 0 | 0 | 0 | 0 | 0 |
| Sep8.50v_9279 | 3 | 0 | 0 | 0 | 0 | 0 | 0 | 0 | 0 | 0 | 0 | 0 | 0 | 0 | 0 | 0 | 1 | 0 | 0 | 1 | 0 | 0 | 0 | 0 | 0 | 0 | 0 | 0 | 0 | 1 |
| Sep8.50v_25302 | 1 | 0 | 0 | 0 | 0 | 0 | 0 | 0 | 0 | 0 | 0 | 0 | 0 | 0 | 0 | 0 | 0 | 0 | 0 | 0 | 0 | 0 | 0 | 0 | 0 | 0 | 0 | 0 | 0 | 1 |
| May12.25v_6591 | 1 | 0 | 1 | 0 | 0 | 0 | 0 | 0 | 0 | 0 | 0 | 0 | 0 | 0 | 0 | 0 | 0 | 0 | 0 | 0 | 0 | 0 | 0 | 0 | 0 | 0 | 0 | 0 | 0 | 0 |
| Jul11.50v_3146 | 1 | 0 | 0 | 0 | 0 | 0 | 0 | 0 | 0 | 0 | 0 | 0 | 0 | 0 | 0 | 1 | 0 | 0 | 0 | 0 | 0 | 0 | 0 | 0 | 0 | 0 | 0 | 0 | 0 | 0 |
| May12.75v_8355 | 1 | 0 | 0 | 0 | 1 | 0 | 0 | 0 | 0 | 0 | 0 | 0 | 0 | 0 | 0 | 0 | 0 | 0 | 0 | 0 | 0 | 0 | 0 | 0 | 0 | 0 | 0 | 0 | 0 | 0 |
| Jul11.50v_14197 | 1 | 0 | 0 | 0 | 0 | 0 | 0 | 0 | 0 | 0 | 0 | 0 | 0 | 0 | 0 | 1 | 0 | 0 | 0 | 0 | 0 | 0 | 0 | 0 | 0 | 0 | 0 | 0 | 0 | 0 |
| May12.25v_24482 | 1 | 0 | 1 | 0 | 0 | 0 | 0 | 0 | 0 | 0 | 0 | 0 | 0 | 0 | 0 | 0 | 0 | 0 | 0 | 0 | 0 | 0 | 0 | 0 | 0 | 0 | 0 | 0 | 0 | 0 |
| Aug5.25v_40327 | 1 | 0 | 0 | 0 | 0 | 0 | 0 | 0 | 0 | 0 | 0 | 0 | 0 | 0 | 0 | 0 | 0 | 0 | 0 | 0 | 0 | 0 | 0 | 1 | 0 | 0 | 0 | 0 | 0 | 0 |
| Jun22.5v_16516 | 1 | 0 | 0 | 0 | 0 | 0 | 0 | 0 | 0 | 1 | 0 | 0 | 0 | 0 | 0 | 0 | 0 | 0 | 0 | 0 | 0 | 0 | 0 | 0 | 0 | 0 | 0 | 0 | 0 | 0 |
| Sep8.5v_18197 | 1 | 0 | 0 | 0 | 0 | 0 | 0 | 0 | 0 | 0 | 0 | 0 | 0 | 0 | 0 | 0 | 0 | 0 | 0 | 0 | 0 | 0 | 0 | 0 | 0 | 0 | 0 | 1 | 0 | 0 |
| Sep8.50v_37801 | 1 | 0 | 0 | 0 | 0 | 0 | 0 | 0 | 0 | 0 | 0 | 0 | 0 | 0 | 0 | 0 | 0 | 0 | 0 | 0 | 0 | 0 | 0 | 0 | 0 | 0 | 0 | 0 | 0 | 1 |
| Jul11.50v_6169 | 1 | 0 | 0 | 0 | 0 | 0 | 0 | 0 | 0 | 0 | 0 | 0 | 0 | 0 | 0 | 1 | 0 | 0 | 0 | 0 | 0 | 0 | 0 | 0 | 0 | 0 | 0 | 0 | 0 | 0 |
| Jul11.50v_1658 | 1 | 0 | 0 | 0 | 0 | 0 | 0 | 0 | 0 | 0 | 0 | 0 | 0 | 0 | 0 | 1 | 0 | 0 | 0 | 0 | 0 | 0 | 0 | 0 | 0 | 0 | 0 | 0 | 0 | 0 |
| Jul11.5v_2529 | 1 | 0 | 0 | 0 | 0 | 0 | 0 | 0 | 0 | 0 | 0 | 0 | 0 | 1 | 0 | 0 | 0 | 0 | 0 | 0 | 0 | 0 | 0 | 0 | 0 | 0 | 0 | 0 | 0 | 0 |
| Jun8.50v_36205 | 1 | 0 | 0 | 0 | 0 | 0 | 1 | 0 | 0 | 0 | 0 | 0 | 0 | 0 | 0 | 0 | 0 | 0 | 0 | 0 | 0 | 0 | 0 | 0 | 0 | 0 | 0 | 0 | 0 | 0 |
| Jun22.50v_1920 | 1 | 0 | 0 | 0 | 0 | 0 | 0 | 0 | 0 | 0 | 1 | 0 | 0 | 0 | 0 | 0 | 0 | 0 | 0 | 0 | 0 | 0 | 0 | 0 | 0 | 0 | 0 | 0 | 0 | 0 |
| Jul11.75v_26801 | 1 | 0 | 0 | 0 | 0 | 0 | 0 | 0 | 0 | 0 | 0 | 0 | 0 | 0 | 0 | 0 | 1 | 0 | 0 | 0 | 0 | 0 | 0 | 0 | 0 | 0 | 0 | 0 | 0 | 0 |
| May12.75v_5202 | 1 | 0 | 0 | 0 | 1 | 0 | 0 | 0 | 0 | 0 | 0 | 0 | 0 | 0 | 0 | 0 | 0 | 0 | 0 | 0 | 0 | 0 | 0 | 0 | 0 | 0 | 0 | 0 | 0 | 0 |
| Sep8.50v_32747 | 1 | 0 | 0 | 0 | 0 | 0 | 0 | 0 | 0 | 0 | 0 | 0 | 0 | 0 | 0 | 0 | 0 | 0 | 0 | 0 | 0 | 0 | 0 | 0 | 0 | 0 | 0 | 0 | 0 | 1 |
| Aug5.5v_11623 | 1 | 0 | 0 | 0 | 0 | 0 | 0 | 0 | 0 | 0 | 0 | 0 | 0 | 0 | 0 | 0 | 0 | 0 | 0 | 0 | 0 | 0 | 1 | 0 | 0 | 0 | 0 | 0 | 0 | 0 |
| Jul11.50v_5900 | 1 | 0 | 0 | 0 | 0 | 0 | 0 | 0 | 0 | 0 | 0 | 0 | 0 | 0 | 0 | 1 | 0 | 0 | 0 | 0 | 0 | 0 | 0 | 0 | 0 | 0 | 0 | 0 | 0 | 0 |
| Sep8.50v_25830 | 1 | 0 | 0 | 0 | 0 | 0 | 0 | 0 | 0 | 0 | 0 | 0 | 0 | 0 | 0 | 0 | 0 | 0 | 0 | 0 | 0 | 0 | 0 | 0 | 0 | 0 | 0 | 0 | 0 | 1 |
| May12.5v_5817 | 1 | 1 | 0 | 0 | 0 | 0 | 0 | 0 | 0 | 0 | 0 | 0 | 0 | 0 | 0 | 0 | 0 | 0 | 0 | 0 | 0 | 0 | 0 | 0 | 0 | 0 | 0 | 0 | 0 | 0 |
| Jul27.5v_18023 | 1 | 0 | 0 | 0 | 0 | 0 | 0 | 0 | 0 | 0 | 0 | 0 | 0 | 0 | 0 | 0 | 0 | 1 | 0 | 0 | 0 | 0 | 0 | 0 | 0 | 0 | 0 | 0 | 0 | 0 |
| May12.75v_31618 | 1 | 0 | 0 | 0 | 1 | 0 | 0 | 0 | 0 | 0 | 0 | 0 | 0 | 0 | 0 | 0 | 0 | 0 | 0 | 0 | 0 | 0 | 0 | 0 | 0 | 0 | 0 | 0 | 0 | 0 |
| Jul11.0v_16850 | 1 | 0 | 0 | 0 | 0 | 0 | 0 | 0 | 0 | 0 | 0 | 0 | 1 | 0 | 0 | 0 | 0 | 0 | 0 | 0 | 0 | 0 | 0 | 0 | 0 | 0 | 0 | 0 | 0 | 0 |
| Jul11.50v_242 | 1 | 0 | 0 | 0 | 0 | 0 | 0 | 0 | 0 | 0 | 0 | 0 | 0 | 0 | 0 | 1 | 0 | 0 | 0 | 0 | 0 | 0 | 0 | 0 | 0 | 0 | 0 | 0 | 0 | 0 |
| Aug5.0v_4963 | 3 | 0 | 1 | 0 | 0 | 0 | 0 | 1 | 0 | 0 | 0 | 0 | 0 | 0 | 0 | 0 | 0 | 0 | 0 | 0 | 0 | 1 | 0 | 0 | 0 | 0 | 0 | 0 | 0 | 0 |
| Jul27.25v_2860 | 1 | 0 | 0 | 0 | 0 | 0 | 0 | 0 | 0 | 0 | 0 | 0 | 0 | 0 | 0 | 0 | 0 | 0 | 1 | 0 | 0 | 0 | 0 | 0 | 0 | 0 | 0 | 0 | 0 | 0 |
| Jul27.5v_8055 | 1 | 0 | 0 | 0 | 0 | 0 | 0 | 0 | 0 | 0 | 0 | 0 | 0 | 0 | 0 | 0 | 0 | 1 | 0 | 0 | 0 | 0 | 0 | 0 | 0 | 0 | 0 | 0 | 0 | 0 |
| Sep8.0v_15059 | 1 | 0 | 0 | 0 | 0 | 0 | 0 | 0 | 0 | 0 | 0 | 0 | 0 | 0 | 0 | 0 | 0 | 0 | 0 | 0 | 0 | 0 | 0 | 0 | 0 | 0 | 1 | 0 | 0 | 0 |
| Jul27.5v_15380 | 1 | 0 | 0 | 0 | 0 | 0 | 0 | 0 | 0 | 0 | 0 | 0 | 0 | 0 | 0 | 0 | 0 | 1 | 0 | 0 | 0 | 0 | 0 | 0 | 0 | 0 | 0 | 0 | 0 | 0 |
| Sep8.0v_15450 | 1 | 0 | 0 | 0 | 0 | 0 | 0 | 0 | 0 | 0 | 0 | 0 | 0 | 0 | 0 | 0 | 0 | 0 | 0 | 0 | 0 | 0 | 0 | 0 | 0 | 0 | 1 | 0 | 0 | 0 |
| Jul11.50v_15921 | 1 | 0 | 0 | 0 | 0 | 0 | 0 | 0 | 0 | 0 | 0 | 0 | 0 | 0 | 0 | 1 | 0 | 0 | 0 | 0 | 0 | 0 | 0 | 0 | 0 | 0 | 0 | 0 | 0 | 0 |
| Jul11.5v_9419 | 1 | 0 | 0 | 0 | 0 | 0 | 0 | 0 | 0 | 0 | 0 | 0 | 0 | 1 | 0 | 0 | 0 | 0 | 0 | 0 | 0 | 0 | 0 | 0 | 0 | 0 | 0 | 0 | 0 | 0 |
| May12.75v_13775 | 1 | 0 | 0 | 0 | 1 | 0 | 0 | 0 | 0 | 0 | 0 | 0 | 0 | 0 | 0 | 0 | 0 | 0 | 0 | 0 | 0 | 0 | 0 | 0 | 0 | 0 | 0 | 0 | 0 | 0 |
| Aug5.25v_13513 | 1 | 0 | 0 | 0 | 0 | 0 | 0 | 0 | 0 | 0 | 0 | 0 | 0 | 0 | 0 | 0 | 0 | 0 | 0 | 0 | 0 | 0 | 0 | 1 | 0 | 0 | 0 | 0 | 0 | 0 |
| Jun8.50v_17712 | 1 | 0 | 0 | 0 | 0 | 0 | 1 | 0 | 0 | 0 | 0 | 0 | 0 | 0 | 0 | 0 | 0 | 0 | 0 | 0 | 0 | 0 | 0 | 0 | 0 | 0 | 0 | 0 | 0 | 0 |
| Jun22.0v_13380 | 1 | 0 | 0 | 0 | 0 | 0 | 0 | 0 | 1 | 0 | 0 | 0 | 0 | 0 | 0 | 0 | 0 | 0 | 0 | 0 | 0 | 0 | 0 | 0 | 0 | 0 | 0 | 0 | 0 | 0 |
| Sep8.0v_12226 | 1 | 0 | 0 | 0 | 0 | 0 | 0 | 0 | 0 | 0 | 0 | 0 | 0 | 0 | 0 | 0 | 0 | 0 | 0 | 0 | 0 | 0 | 0 | 0 | 0 | 0 | 1 | 0 | 0 | 0 |
| Jun22.75v_31544 | 1 | 0 | 0 | 0 | 0 | 0 | 0 | 0 | 0 | 0 | 0 | 1 | 0 | 0 | 0 | 0 | 0 | 0 | 0 | 0 | 0 | 0 | 0 | 0 | 0 | 0 | 0 | 0 | 0 | 0 |
| Aug5.25v_21003 | 1 | 0 | 0 | 0 | 0 | 0 | 0 | 0 | 0 | 0 | 0 | 0 | 0 | 0 | 0 | 0 | 0 | 0 | 0 | 0 | 0 | 0 | 0 | 1 | 0 | 0 | 0 | 0 | 0 | 0 |
| Jun8.50v_3803 | 5 | 0 | 0 | 0 | 2 | 0 | 1 | 0 | 0 | 0 | 0 | 2 | 0 | 0 | 0 | 0 | 0 | 0 | 0 | 0 | 0 | 0 | 0 | 0 | 0 | 0 | 0 | 0 | 0 | 0 |
| Jul27.50v_36091 | 2 | 0 | 0 | 0 | 0 | 0 | 0 | 0 | 0 | 0 | 0 | 0 | 0 | 0 | 0 | 0 | 0 | 0 | 1 | 1 | 0 | 0 | 0 | 0 | 0 | 0 | 0 | 0 | 0 | 0 |
| May12.25v_781 | 2 | 1 | 1 | 0 | 0 | 0 | 0 | 0 | 0 | 0 | 0 | 0 | 0 | 0 | 0 | 0 | 0 | 0 | 0 | 0 | 0 | 0 | 0 | 0 | 0 | 0 | 0 | 0 | 0 | 0 |
| Jun22.50v_5990 | 1 | 0 | 0 | 0 | 0 | 0 | 0 | 0 | 0 | 0 | 1 | 0 | 0 | 0 | 0 | 0 | 0 | 0 | 0 | 0 | 0 | 0 | 0 | 0 | 0 | 0 | 0 | 0 | 0 | 0 |
| May12.25v_38886 | 5 | 0 | 3 | 0 | 0 | 1 | 0 | 0 | 0 | 0 | 0 | 1 | 0 | 0 | 0 | 0 | 0 | 0 | 0 | 0 | 0 | 0 | 0 | 0 | 0 | 0 | 0 | 0 | 0 | 0 |
| Jul11.0v_2723 | 2 | 0 | 0 | 0 | 0 | 0 | 0 | 0 | 0 | 0 | 0 | 0 | 1 | 0 | 0 | 0 | 0 | 0 | 0 | 0 | 0 | 0 | 0 | 0 | 0 | 1 | 0 | 0 | 0 | 0 |
| Jul27.5v_13960 | 1 | 0 | 0 | 0 | 0 | 0 | 0 | 0 | 0 | 0 | 0 | 0 | 0 | 0 | 0 | 0 | 0 | 1 | 0 | 0 | 0 | 0 | 0 | 0 | 0 | 0 | 0 | 0 | 0 | 0 |
| May12.25v_33564 | 1 | 0 | 1 | 0 | 0 | 0 | 0 | 0 | 0 | 0 | 0 | 0 | 0 | 0 | 0 | 0 | 0 | 0 | 0 | 0 | 0 | 0 | 0 | 0 | 0 | 0 | 0 | 0 | 0 | 0 |
| Jul11.75v_22172 | 1 | 0 | 0 | 0 | 0 | 0 | 0 | 0 | 0 | 0 | 0 | 0 | 0 | 0 | 0 | 0 | 1 | 0 | 0 | 0 | 0 | 0 | 0 | 0 | 0 | 0 | 0 | 0 | 0 | 0 |
| Jul11.25v_17732 | 1 | 0 | 0 | 0 | 0 | 0 | 0 | 0 | 0 | 0 | 0 | 0 | 0 | 0 | 1 | 0 | 0 | 0 | 0 | 0 | 0 | 0 | 0 | 0 | 0 | 0 | 0 | 0 | 0 | 0 |
| Jul27.50v_27916 | 6 | 0 | 0 | 0 | 0 | 0 | 0 | 0 | 1 | 0 | 0 | 0 | 1 | 1 | 0 | 2 | 0 | 0 | 0 | 1 | 0 | 0 | 0 | 0 | 0 | 0 | 0 | 0 | 0 | 0 |
| Aug5.5v_17365 | 9 | 0 | 1 | 0 | 0 | 0 | 0 | 0 | 1 | 0 | 0 | 0 | 0 | 0 | 0 | 1 | 0 | 1 | 0 | 1 | 0 | 0 | 1 | 0 | 0 | 1 | 2 | 0 | 0 | 0 |
| Aug5.5v_35201 | 2 | 0 | 0 | 0 | 0 | 0 | 0 | 0 | 0 | 0 | 0 | 1 | 0 | 0 | 0 | 0 | 0 | 0 | 0 | 0 | 0 | 0 | 1 | 0 | 0 | 0 | 0 | 0 | 0 | 0 |
| Aug5.75v_10659 | 3 | 0 | 0 | 0 | 0 | 0 | 0 | 0 | 0 | 0 | 0 | 0 | 0 | 0 | 0 | 0 | 0 | 0 | 0 | 0 | 0 | 1 | 0 | 0 | 0 | 1 | 0 | 0 | 0 | 1 |
| Jul11.0v_5366 | 5 | 0 | 0 | 0 | 0 | 0 | 0 | 0 | 1 | 1 | 0 | 0 | 1 | 0 | 1 | 0 | 0 | 0 | 0 | 0 | 0 | 0 | 0 | 0 | 0 | 0 | 1 | 0 | 0 | 0 |
| May12.50v_23630 | 4 | 0 | 0 | 2 | 1 | 0 | 0 | 0 | 0 | 0 | 0 | 0 | 0 | 1 | 0 | 0 | 0 | 0 | 0 | 0 | 0 | 0 | 0 | 0 | 0 | 0 | 0 | 0 | 0 | 0 |
| Jun8.50v_11725 | 23 | 1 | 0 | 2 | 2 | 2 | 9 | 0 | 0 | 0 | 1 | 0 | 0 | 0 | 0 | 0 | 1 | 0 | 1 | 0 | 0 | 0 | 0 | 1 | 0 | 2 | 0 | 0 | 0 | 1 |
| May12.75v_30345 | 2 | 0 | 0 | 0 | 1 | 0 | 0 | 0 | 0 | 0 | 1 | 0 | 0 | 0 | 0 | 0 | 0 | 0 | 0 | 0 | 0 | 0 | 0 | 0 | 0 | 0 | 0 | 0 | 0 | 0 |

| Feature_ID | 654 | May12_05m | May12_25m | May12_50m | May12_75m | Jun8_25m | Jun8_50m | Jun8_75m | Jun22_0m | Jun22_05m | Jun22_50m | Jun22_75m | Jul11_0m | Jul11_05m | Jul11_25m | Jul11_50m | Jul11_75m | Jul27_05m | Jul27_25m | Jul27_50m | Jul27_75m | Aug5_0m | Aug5_05m | Aug5_25m | Aug5_50m | Aug5_75m | Sep8_0m | Sep8_05m | Sep8_25m | Sep8_50m |
|---|---|---|---|---|---|---|---|---|---|---|---|---|---|---|---|---|---|---|---|---|---|---|---|---|---|---|---|---|---|---|
| Sep8.25v_11003 | 1 | 0 | 0 | 0 | 0 | 0 | 0 | 0 | 0 | 0 | 0 | 0 | 0 | 0 | 0 | 0 | 0 | 0 | 0 | 0 | 0 | 0 | 0 | 0 | 0 | 0 | 0 | 0 | 1 | 0 |
| Jul27.25v_38635 | 9 | 1 | 0 | 0 | 1 | 1 | 1 | 0 | 0 | 0 | 0 | 1 | 0 | 0 | 0 | 0 | 1 | 0 | 1 | 0 | 1 | 0 | 0 | 0 | 0 | 1 | 0 | 0 | 0 | 0 |
| Jun8.50v_7194 | 1 | 0 | 0 | 0 | 0 | 0 | 1 | 0 | 0 | 0 | 0 | 0 | 0 | 0 | 0 | 0 | 0 | 0 | 0 | 0 | 0 | 0 | 0 | 0 | 0 | 0 | 0 | 0 | 0 | 0 |
| Jun22.50v_1966 | 1 | 0 | 0 | 0 | 0 | 0 | 0 | 0 | 0 | 0 | 1 | 0 | 0 | 0 | 0 | 0 | 0 | 0 | 0 | 0 | 0 | 0 | 0 | 0 | 0 | 0 | 0 | 0 | 0 | 0 |
| Jun22.0v_2549 | 1 | 0 | 0 | 0 | 0 | 0 | 0 | 0 | 1 | 0 | 0 | 0 | 0 | 0 | 0 | 0 | 0 | 0 | 0 | 0 | 0 | 0 | 0 | 0 | 0 | 0 | 0 | 0 | 0 | 0 |
| Jun22.50v_16142 | 1 | 0 | 0 | 0 | 0 | 0 | 0 | 0 | 0 | 0 | 1 | 0 | 0 | 0 | 0 | 0 | 0 | 0 | 0 | 0 | 0 | 0 | 0 | 0 | 0 | 0 | 0 | 0 | 0 | 0 |
| Jul27.5v_11919 | 1 | 0 | 0 | 0 | 0 | 0 | 0 | 0 | 0 | 0 | 0 | 0 | 0 | 0 | 0 | 0 | 0 | 1 | 0 | 0 | 0 | 0 | 0 | 0 | 0 | 0 | 0 | 0 | 0 | 0 |
| Jun8.25v_37379 | 1 | 0 | 0 | 0 | 0 | 1 | 0 | 0 | 0 | 0 | 0 | 0 | 0 | 0 | 0 | 0 | 0 | 0 | 0 | 0 | 0 | 0 | 0 | 0 | 0 | 0 | 0 | 0 | 0 | 0 |
| Jul27.50v_7789 | 3 | 0 | 0 | 0 | 0 | 0 | 0 | 0 | 0 | 0 | 0 | 0 | 1 | 0 | 0 | 1 | 0 | 0 | 0 | 1 | 0 | 0 | 0 | 0 | 0 | 0 | 0 | 0 | 0 | 0 |
| May12.5v_4910 | 1 | 1 | 0 | 0 | 0 | 0 | 0 | 0 | 0 | 0 | 0 | 0 | 0 | 0 | 0 | 0 | 0 | 0 | 0 | 0 | 0 | 0 | 0 | 0 | 0 | 0 | 0 | 0 | 0 | 0 |
| May12.75v_25155 | 1 | 0 | 0 | 0 | 1 | 0 | 0 | 0 | 0 | 0 | 0 | 0 | 0 | 0 | 0 | 0 | 0 | 0 | 0 | 0 | 0 | 0 | 0 | 0 | 0 | 0 | 0 | 0 | 0 | 0 |
| Aug5.75v_24643 | 1 | 0 | 0 | 0 | 0 | 0 | 0 | 0 | 0 | 0 | 0 | 0 | 0 | 0 | 0 | 0 | 0 | 0 | 0 | 0 | 0 | 0 | 0 | 0 | 0 | 1 | 0 | 0 | 0 | 0 |
| May12.5v_7008 | 3 | 1 | 0 | 0 | 0 | 0 | 0 | 0 | 1 | 0 | 0 | 0 | 0 | 0 | 0 | 0 | 0 | 0 | 0 | 0 | 0 | 0 | 0 | 0 | 0 | 0 | 0 | 0 | 1 | 0 |
| Jun22.50v_19008 | 1 | 0 | 0 | 0 | 0 | 0 | 0 | 0 | 0 | 0 | 1 | 0 | 0 | 0 | 0 | 0 | 0 | 0 | 0 | 0 | 0 | 0 | 0 | 0 | 0 | 0 | 0 | 0 | 0 | 0 |
| May12.50v_22100 | 881 | 57 | 37 | 91 | 55 | 19 | 63 | 7 | 23 | 5 | 187 | 20 | 1 | 10 | 39 | 21 | 50 | 6 | 9 | 37 | 8 | 12 | 10 | 19 | 29 | 23 | 6 | 19 | 5 | 13 |
| May12.5v_17393 | 6 | 1 | 0 | 1 | 0 | 0 | 1 | 0 | 1 | 0 | 0 | 0 | 0 | 0 | 0 | 0 | 1 | 0 | 0 | 0 | 0 | 0 | 0 | 0 | 0 | 1 | 0 | 0 | 0 | 0 |
| Aug5.75v_16033 | 4 | 0 | 0 | 1 | 0 | 0 | 0 | 0 | 1 | 0 | 1 | 0 | 0 | 0 | 0 | 0 | 0 | 0 | 0 | 0 | 0 | 0 | 0 | 0 | 0 | 1 | 0 | 0 | 0 | 0 |
| Jun22.50v_24594 | 1 | 0 | 0 | 0 | 0 | 0 | 0 | 0 | 0 | 0 | 1 | 0 | 0 | 0 | 0 | 0 | 0 | 0 | 0 | 0 | 0 | 0 | 0 | 0 | 0 | 0 | 0 | 0 | 0 | 0 |
| Jun22.0v_9677 | 5 | 0 | 0 | 0 | 1 | 0 | 0 | 0 | 1 | 0 | 1 | 0 | 0 | 0 | 0 | 0 | 1 | 0 | 0 | 0 | 0 | 0 | 0 | 0 | 0 | 0 | 0 | 0 | 0 | 1 |
| Jul11.50v_17822 | 6 | 0 | 0 | 0 | 0 | 0 | 0 | 0 | 0 | 0 | 1 | 0 | 0 | 0 | 0 | 1 | 2 | 0 | 0 | 1 | 0 | 0 | 0 | 0 | 0 | 1 | 0 | 0 | 0 | 0 |
| May12.50v_15950 | 1 | 0 | 0 | 1 | 0 | 0 | 0 | 0 | 0 | 0 | 0 | 0 | 0 | 0 | 0 | 0 | 0 | 0 | 0 | 0 | 0 | 0 | 0 | 0 | 0 | 0 | 0 | 0 | 0 | 0 |
| Sep8.50v_19831 | 3 | 1 | 0 | 0 | 0 | 0 | 0 | 0 | 0 | 0 | 0 | 0 | 0 | 0 | 0 | 0 | 1 | 0 | 0 | 0 | 0 | 0 | 0 | 0 | 0 | 0 | 0 | 0 | 0 | 1 |
| Jul11.25v_5043 | 1 | 0 | 0 | 0 | 0 | 0 | 0 | 0 | 0 | 0 | 0 | 0 | 0 | 0 | 1 | 0 | 0 | 0 | 0 | 0 | 0 | 0 | 0 | 0 | 0 | 0 | 0 | 0 | 0 | 0 |
| Jul11.25v_9234 | 1 | 0 | 0 | 0 | 0 | 0 | 0 | 0 | 0 | 0 | 0 | 0 | 0 | 0 | 1 | 0 | 0 | 0 | 0 | 0 | 0 | 0 | 0 | 0 | 0 | 0 | 0 | 0 | 0 | 0 |
| Jul11.50v_1396 | 1 | 0 | 0 | 0 | 0 | 0 | 0 | 0 | 0 | 0 | 0 | 0 | 0 | 0 | 0 | 1 | 0 | 0 | 0 | 0 | 0 | 0 | 0 | 0 | 0 | 0 | 0 | 0 | 0 | 0 |
| Jun8.25v_23729 | 1 | 0 | 0 | 0 | 0 | 1 | 0 | 0 | 0 | 0 | 0 | 0 | 0 | 0 | 0 | 0 | 0 | 0 | 0 | 0 | 0 | 0 | 0 | 0 | 0 | 0 | 0 | 0 | 0 | 0 |
| May12.25v_17687 | 1 | 0 | 1 | 0 | 0 | 0 | 0 | 0 | 0 | 0 | 0 | 0 | 0 | 0 | 0 | 0 | 0 | 0 | 0 | 0 | 0 | 0 | 0 | 0 | 0 | 0 | 0 | 0 | 0 | 0 |
| May12.50v_21071 | 1 | 0 | 0 | 1 | 0 | 0 | 0 | 0 | 0 | 0 | 0 | 0 | 0 | 0 | 0 | 0 | 0 | 0 | 0 | 0 | 0 | 0 | 0 | 0 | 0 | 0 | 0 | 0 | 0 | 0 |
| Jul11.75v_23605 | 1 | 0 | 0 | 0 | 0 | 0 | 0 | 0 | 0 | 0 | 0 | 0 | 0 | 0 | 0 | 0 | 1 | 0 | 0 | 0 | 0 | 0 | 0 | 0 | 0 | 0 | 0 | 0 | 0 | 0 |
| Jul27.5v_17923 | 1 | 0 | 0 | 0 | 0 | 0 | 0 | 0 | 0 | 0 | 0 | 0 | 0 | 0 | 0 | 0 | 0 | 1 | 0 | 0 | 0 | 0 | 0 | 0 | 0 | 0 | 0 | 0 | 0 | 0 |
| Jun22.50v_31215 | 1 | 0 | 0 | 0 | 0 | 0 | 0 | 0 | 0 | 0 | 1 | 0 | 0 | 0 | 0 | 0 | 0 | 0 | 0 | 0 | 0 | 0 | 0 | 0 | 0 | 0 | 0 | 0 | 0 | 0 |
| May12.5v_39722 | 3 | 1 | 0 | 1 | 0 | 0 | 1 | 0 | 0 | 0 | 0 | 0 | 0 | 0 | 0 | 0 | 0 | 0 | 0 | 0 | 0 | 0 | 0 | 0 | 0 | 0 | 0 | 0 | 0 | 0 |
| Jul11.25v_18375 | 1 | 0 | 0 | 0 | 0 | 0 | 0 | 0 | 0 | 0 | 0 | 0 | 0 | 0 | 1 | 0 | 0 | 0 | 0 | 0 | 0 | 0 | 0 | 0 | 0 | 0 | 0 | 0 | 0 | 0 |
| Jul11.50v_9942 | 1 | 0 | 0 | 0 | 0 | 0 | 0 | 0 | 0 | 0 | 0 | 0 | 0 | 0 | 0 | 1 | 0 | 0 | 0 | 0 | 0 | 0 | 0 | 0 | 0 | 0 | 0 | 0 | 0 | 0 |
| Jun22.50v_394 | 2 | 0 | 0 | 0 | 0 | 0 | 0 | 0 | 0 | 0 | 2 | 0 | 0 | 0 | 0 | 0 | 0 | 0 | 0 | 0 | 0 | 0 | 0 | 0 | 0 | 0 | 0 | 0 | 0 | 0 |
| May12.75v_17877 | 3 | 0 | 0 | 0 | 1 | 0 | 0 | 0 | 0 | 0 | 0 | 0 | 0 | 0 | 0 | 0 | 0 | 0 | 0 | 0 | 0 | 0 | 0 | 0 | 0 | 1 | 0 | 0 | 1 | 0 |
| Jul27.50v_17338 | 9 | 0 | 0 | 2 | 0 | 0 | 1 | 0 | 0 | 0 | 2 | 0 | 0 | 0 | 1 | 1 | 1 | 0 | 0 | 1 | 0 | 0 | 0 | 0 | 0 | 0 | 0 | 0 | 0 | 0 |
| Jun22.75v_7134 | 1 | 0 | 0 | 0 | 0 | 0 | 0 | 0 | 0 | 0 | 0 | 1 | 0 | 0 | 0 | 0 | 0 | 0 | 0 | 0 | 0 | 0 | 0 | 0 | 0 | 0 | 0 | 0 | 0 | 0 |
| Aug5.50v_31053 | 1 | 0 | 0 | 0 | 0 | 0 | 0 | 0 | 0 | 0 | 0 | 0 | 0 | 0 | 0 | 0 | 0 | 0 | 0 | 0 | 0 | 0 | 0 | 0 | 1 | 0 | 0 | 0 | 0 | 0 |
| Aug5.75v_18641 | 1 | 0 | 0 | 0 | 0 | 0 | 0 | 0 | 0 | 0 | 0 | 0 | 0 | 0 | 0 | 0 | 0 | 0 | 0 | 0 | 0 | 0 | 0 | 0 | 0 | 1 | 0 | 0 | 0 | 0 |
| Jul27.75v_735 | 1 | 0 | 0 | 0 | 0 | 0 | 0 | 0 | 0 | 0 | 0 | 0 | 0 | 0 | 0 | 0 | 0 | 0 | 0 | 0 | 1 | 0 | 0 | 0 | 0 | 0 | 0 | 0 | 0 | 0 |
| May12.5v_10900 | 1 | 1 | 0 | 0 | 0 | 0 | 0 | 0 | 0 | 0 | 0 | 0 | 0 | 0 | 0 | 0 | 0 | 0 | 0 | 0 | 0 | 0 | 0 | 0 | 0 | 0 | 0 | 0 | 0 | 0 |
| Jul11.75v_38733 | 1 | 0 | 0 | 0 | 0 | 0 | 0 | 0 | 0 | 0 | 0 | 0 | 0 | 0 | 0 | 0 | 1 | 0 | 0 | 0 | 0 | 0 | 0 | 0 | 0 | 0 | 0 | 0 | 0 | 0 |
| Jun22.50v_30159 | 1 | 0 | 0 | 0 | 0 | 0 | 0 | 0 | 0 | 0 | 1 | 0 | 0 | 0 | 0 | 0 | 0 | 0 | 0 | 0 | 0 | 0 | 0 | 0 | 0 | 0 | 0 | 0 | 0 | 0 |
| Jun8.75v_19896 | 7 | 0 | 0 | 0 | 0 | 1 | 1 | 1 | 1 | 0 | 1 | 0 | 0 | 0 | 0 | 0 | 0 | 0 | 0 | 0 | 0 | 0 | 0 | 0 | 1 | 0 | 0 | 1 | 0 | 0 |
| Jun8.50v_35343 | 1 | 0 | 0 | 0 | 0 | 0 | 1 | 0 | 0 | 0 | 0 | 0 | 0 | 0 | 0 | 0 | 0 | 0 | 0 | 0 | 0 | 0 | 0 | 0 | 0 | 0 | 0 | 0 | 0 | 0 |
| Jul11.75v_27593 | 3 | 0 | 0 | 0 | 1 | 0 | 0 | 0 | 0 | 0 | 0 | 0 | 0 | 0 | 0 | 0 | 2 | 0 | 0 | 0 | 0 | 0 | 0 | 0 | 0 | 0 | 0 | 0 | 0 | 0 |
| Jun22.50v_35089 | 1 | 0 | 0 | 0 | 0 | 0 | 0 | 0 | 0 | 0 | 1 | 0 | 0 | 0 | 0 | 0 | 0 | 0 | 0 | 0 | 0 | 0 | 0 | 0 | 0 | 0 | 0 | 0 | 0 | 0 |
| May12.50v_1250 | 1 | 0 | 0 | 1 | 0 | 0 | 0 | 0 | 0 | 0 | 0 | 0 | 0 | 0 | 0 | 0 | 0 | 0 | 0 | 0 | 0 | 0 | 0 | 0 | 0 | 0 | 0 | 0 | 0 | 0 |
| Jun22.0v_8989 | 1 | 0 | 0 | 0 | 0 | 0 | 0 | 0 | 1 | 0 | 0 | 0 | 0 | 0 | 0 | 0 | 0 | 0 | 0 | 0 | 0 | 0 | 0 | 0 | 0 | 0 | 0 | 0 | 0 | 0 |
| May12.25v_10234 | 1 | 0 | 1 | 0 | 0 | 0 | 0 | 0 | 0 | 0 | 0 | 0 | 0 | 0 | 0 | 0 | 0 | 0 | 0 | 0 | 0 | 0 | 0 | 0 | 0 | 0 | 0 | 0 | 0 | 0 |
| Aug5.5v_6301 | 1 | 0 | 0 | 0 | 0 | 0 | 0 | 0 | 0 | 0 | 0 | 0 | 0 | 0 | 0 | 0 | 0 | 0 | 0 | 0 | 0 | 0 | 1 | 0 | 0 | 0 | 0 | 0 | 0 | 0 |
| May12.50v_8074 | 1 | 0 | 0 | 1 | 0 | 0 | 0 | 0 | 0 | 0 | 0 | 0 | 0 | 0 | 0 | 0 | 0 | 0 | 0 | 0 | 0 | 0 | 0 | 0 | 0 | 0 | 0 | 0 | 0 | 0 |
| Jul11.5v_18687 | 1 | 0 | 0 | 0 | 0 | 0 | 0 | 0 | 0 | 0 | 0 | 0 | 0 | 1 | 0 | 0 | 0 | 0 | 0 | 0 | 0 | 0 | 0 | 0 | 0 | 0 | 0 | 0 | 0 | 0 |
| Jun22.50v_205 | 2 | 0 | 0 | 0 | 0 | 0 | 0 | 0 | 0 | 0 | 1 | 0 | 0 | 0 | 0 | 0 | 0 | 0 | 1 | 0 | 0 | 0 | 0 | 0 | 0 | 0 | 0 | 0 | 0 | 0 |
| Jun22.50v_26457 | 1 | 0 | 0 | 0 | 0 | 0 | 0 | 0 | 0 | 0 | 1 | 0 | 0 | 0 | 0 | 0 | 0 | 0 | 0 | 0 | 0 | 0 | 0 | 0 | 0 | 0 | 0 | 0 | 0 | 0 |
| Jul11.75v_5792 | 1 | 0 | 0 | 0 | 0 | 0 | 0 | 0 | 0 | 0 | 0 | 0 | 0 | 0 | 0 | 0 | 1 | 0 | 0 | 0 | 0 | 0 | 0 | 0 | 0 | 0 | 0 | 0 | 0 | 0 |
| Jun22.75v_30527 | 1 | 0 | 0 | 0 | 0 | 0 | 0 | 0 | 0 | 0 | 0 | 1 | 0 | 0 | 0 | 0 | 0 | 0 | 0 | 0 | 0 | 0 | 0 | 0 | 0 | 0 | 0 | 0 | 0 | 0 |
| Sep8.50v_13150 | 1 | 0 | 0 | 0 | 0 | 0 | 0 | 0 | 0 | 0 | 0 | 0 | 0 | 0 | 0 | 0 | 0 | 0 | 0 | 0 | 0 | 0 | 0 | 0 | 0 | 0 | 0 | 0 | 0 | 1 |
| Jul11.25v_18292 | 1 | 0 | 0 | 0 | 0 | 0 | 0 | 0 | 0 | 0 | 0 | 0 | 0 | 0 | 1 | 0 | 0 | 0 | 0 | 0 | 0 | 0 | 0 | 0 | 0 | 0 | 0 | 0 | 0 | 0 |
| Aug5.50v_78 | 1 | 0 | 0 | 0 | 0 | 0 | 0 | 0 | 0 | 0 | 0 | 0 | 0 | 0 | 0 | 0 | 0 | 0 | 0 | 0 | 0 | 0 | 0 | 0 | 1 | 0 | 0 | 0 | 0 | 0 |
| Jun22.50v_9618 | 1 | 0 | 0 | 0 | 0 | 0 | 0 | 0 | 0 | 0 | 1 | 0 | 0 | 0 | 0 | 0 | 0 | 0 | 0 | 0 | 0 | 0 | 0 | 0 | 0 | 0 | 0 | 0 | 0 | 0 |
| May12.75v_37892 | 1 | 0 | 0 | 0 | 1 | 0 | 0 | 0 | 0 | 0 | 0 | 0 | 0 | 0 | 0 | 0 | 0 | 0 | 0 | 0 | 0 | 0 | 0 | 0 | 0 | 0 | 0 | 0 | 0 | 0 |
| Jul11.75v_34355 | 1 | 0 | 0 | 0 | 0 | 0 | 0 | 0 | 0 | 0 | 0 | 0 | 0 | 0 | 0 | 0 | 1 | 0 | 0 | 0 | 0 | 0 | 0 | 0 | 0 | 0 | 0 | 0 | 0 | 0 |
| Sep8.50v_17356 | 1 | 0 | 0 | 0 | 0 | 0 | 0 | 0 | 0 | 0 | 0 | 0 | 0 | 0 | 0 | 0 | 0 | 0 | 0 | 0 | 0 | 0 | 0 | 0 | 0 | 0 | 0 | 0 | 0 | 1 |
| Jun22.50v_10484 | 1 | 0 | 0 | 0 | 0 | 0 | 0 | 0 | 0 | 0 | 1 | 0 | 0 | 0 | 0 | 0 | 0 | 0 | 0 | 0 | 0 | 0 | 0 | 0 | 0 | 0 | 0 | 0 | 0 | 0 |
| Jul11.50v_19785 | 1 | 0 | 0 | 0 | 0 | 0 | 0 | 0 | 0 | 0 | 0 | 0 | 0 | 0 | 0 | 1 | 0 | 0 | 0 | 0 | 0 | 0 | 0 | 0 | 0 | 0 | 0 | 0 | 0 | 0 |
| Jul11.50v_10085 | 1 | 0 | 0 | 0 | 0 | 0 | 0 | 0 | 0 | 0 | 0 | 0 | 0 | 0 | 0 | 1 | 0 | 0 | 0 | 0 | 0 | 0 | 0 | 0 | 0 | 0 | 0 | 0 | 0 | 0 |
| Jun22.50v_3703 | 1 | 0 | 0 | 0 | 0 | 0 | 0 | 0 | 0 | 0 | 1 | 0 | 0 | 0 | 0 | 0 | 0 | 0 | 0 | 0 | 0 | 0 | 0 | 0 | 0 | 0 | 0 | 0 | 0 | 0 |
| May12.5v_21973 | 1 | 1 | 0 | 0 | 0 | 0 | 0 | 0 | 0 | 0 | 0 | 0 | 0 | 0 | 0 | 0 | 0 | 0 | 0 | 0 | 0 | 0 | 0 | 0 | 0 | 0 | 0 | 0 | 0 | 0 |
| Jun22.50v_4782 | 1 | 0 | 0 | 0 | 0 | 0 | 0 | 0 | 0 | 0 | 1 | 0 | 0 | 0 | 0 | 0 | 0 | 0 | 0 | 0 | 0 | 0 | 0 | 0 | 0 | 0 | 0 | 0 | 0 | 0 |
| Aug5.50v_23556 | 1 | 0 | 0 | 0 | 0 | 0 | 0 | 0 | 0 | 0 | 0 | 0 | 0 | 0 | 0 | 0 | 0 | 0 | 0 | 0 | 0 | 0 | 0 | 0 | 1 | 0 | 0 | 0 | 0 | 0 |
| May12.25v_3047 | 1 | 0 | 1 | 0 | 0 | 0 | 0 | 0 | 0 | 0 | 0 | 0 | 0 | 0 | 0 | 0 | 0 | 0 | 0 | 0 | 0 | 0 | 0 | 0 | 0 | 0 | 0 | 0 | 0 | 0 |
| Jul11.5v_3229 | 1 | 0 | 0 | 0 | 0 | 0 | 0 | 0 | 0 | 0 | 0 | 0 | 0 | 1 | 0 | 0 | 0 | 0 | 0 | 0 | 0 | 0 | 0 | 0 | 0 | 0 | 0 | 0 | 0 | 0 |
| Jun22.50v_28387 | 1 | 0 | 0 | 0 | 0 | 0 | 0 | 0 | 0 | 0 | 1 | 0 | 0 | 0 | 0 | 0 | 0 | 0 | 0 | 0 | 0 | 0 | 0 | 0 | 0 | 0 | 0 | 0 | 0 | 0 |
| May12.50v_30347 | 1 | 0 | 0 | 1 | 0 | 0 | 0 | 0 | 0 | 0 | 0 | 0 | 0 | 0 | 0 | 0 | 0 | 0 | 0 | 0 | 0 | 0 | 0 | 0 | 0 | 0 | 0 | 0 | 0 | 0 |
| Jul11.0v_20973 | 1 | 0 | 0 | 0 | 0 | 0 | 0 | 0 | 0 | 0 | 0 | 0 | 1 | 0 | 0 | 0 | 0 | 0 | 0 | 0 | 0 | 0 | 0 | 0 | 0 | 0 | 0 | 0 | 0 | 0 |
| May12.75v_15365 | 1 | 0 | 0 | 0 | 1 | 0 | 0 | 0 | 0 | 0 | 0 | 0 | 0 | 0 | 0 | 0 | 0 | 0 | 0 | 0 | 0 | 0 | 0 | 0 | 0 | 0 | 0 | 0 | 0 | 0 |
| Jun22.50v_40677 | 1 | 0 | 0 | 0 | 0 | 0 | 0 | 0 | 0 | 0 | 1 | 0 | 0 | 0 | 0 | 0 | 0 | 0 | 0 | 0 | 0 | 0 | 0 | 0 | 0 | 0 | 0 | 0 | 0 | 0 |
| Aug5.25v_22167 | 1 | 0 | 0 | 0 | 0 | 0 | 0 | 0 | 0 | 0 | 0 | 0 | 0 | 0 | 0 | 0 | 0 | 0 | 0 | 0 | 0 | 0 | 0 | 1 | 0 | 0 | 0 | 0 | 0 | 0 |
| Sep8.50v_30762 | 3 | 0 | 0 | 0 | 0 | 0 | 0 | 0 | 0 | 0 | 0 | 0 | 0 | 0 | 0 | 0 | 0 | 0 | 0 | 0 | 0 | 2 | 0 | 0 | 0 | 0 | 0 | 0 | 0 | 1 |
| Aug5.25v_2561 | 2 | 0 | 0 | 0 | 0 | 0 | 0 | 0 | 0 | 0 | 0 | 0 | 0 | 0 | 0 | 0 | 0 | 0 | 0 | 0 | 0 | 1 | 0 | 1 | 0 | 0 | 0 | 0 | 0 | 0 |
| May12.75v_1587 | 9 | 0 | 0 | 2 | 1 | 2 | 0 | 0 | 0 | 0 | 0 | 0 | 0 | 0 | 1 | 0 | 0 | 0 | 1 | 0 | 0 | 0 | 0 | 1 | 0 | 0 | 0 | 0 | 1 | 0 |
| Aug5.0v_3516 | 1 | 0 | 0 | 0 | 0 | 0 | 0 | 0 | 0 | 0 | 0 | 0 | 0 | 0 | 0 | 0 | 0 | 0 | 0 | 0 | 0 | 1 | 0 | 0 | 0 | 0 | 0 | 0 | 0 | 0 |
| Sep8.0v_1225 | 490 | 1 | 0 | 1 | 2 | 0 | 0 | 0 | 5 | 3 | 8 | 0 | 2 | 8 | 0 | 0 | 0 | 56 | 30 | 3 | 1 | 151 | 28 | 113 | 8 | 5 | 22 | 0 | 0 | 43 |
| Jul27.5v_16350 | 1 | 0 | 0 | 0 | 0 | 0 | 0 | 0 | 0 | 0 | 0 | 0 | 0 | 0 | 0 | 0 | 0 | 1 | 0 | 0 | 0 | 0 | 0 | 0 | 0 | 0 | 0 | 0 | 0 | 0 |
| Sep8.0v_3905 | 1 | 0 | 0 | 0 | 0 | 0 | 0 | 0 | 0 | 0 | 0 | 0 | 0 | 0 | 0 | 0 | 0 | 0 | 0 | 0 | 0 | 0 | 0 | 0 | 0 | 0 | 1 | 0 | 0 | 0 |
| Jul11.5v_5256 | 1 | 0 | 0 | 0 | 0 | 0 | 0 | 0 | 0 | 0 | 0 | 0 | 0 | 1 | 0 | 0 | 0 | 0 | 0 | 0 | 0 | 0 | 0 | 0 | 0 | 0 | 0 | 0 | 0 | 0 |
| Sep8.0v_13182 | 1 | 0 | 0 | 0 | 0 | 0 | 0 | 0 | 0 | 0 | 0 | 0 | 0 | 0 | 0 | 0 | 0 | 0 | 0 | 0 | 0 | 0 | 0 | 0 | 0 | 0 | 1 | 0 | 0 | 0 |
| Jul27.5v_4148 | 412 | 0 | 0 | 2 | 2 | 0 | 0 | 0 | 0 | 0 | 1 | 0 | 1 | 4 | 0 | 0 | 0 | 55 | 17 | 2 | 0 | 131 | 35 | 104 | 11 | 3 | 16 | 0 | 1 | 27 |
| Jul27.5v_12440 | 256 | 0 | 0 | 0 | 0 | 0 | 0 | 0 | 0 | 0 | 1 | 1 | 0 | 1 | 0 | 0 | 1 | 40 | 11 | 1 | 0 | 61 | 24 | 72 | 5 | 2 | 14 | 0 | 0 | 22 |
| Aug5.25v_16100 | 1 | 0 | 0 | 0 | 0 | 0 | 0 | 0 | 0 | 0 | 0 | 0 | 0 | 0 | 0 | 0 | 0 | 0 | 0 | 0 | 0 | 0 | 0 | 1 | 0 | 0 | 0 | 0 | 0 | 0 |
| Aug5.25v_5018 | 1 | 0 | 0 | 0 | 0 | 0 | 0 | 0 | 0 | 0 | 0 | 0 | 0 | 0 | 0 | 0 | 0 | 0 | 0 | 0 | 0 | 0 | 0 | 1 | 0 | 0 | 0 | 0 | 0 | 0 |
| Sep8.50v_1099 | 1 | 0 | 0 | 0 | 0 | 0 | 0 | 0 | 0 | 0 | 0 | 0 | 0 | 0 | 0 | 0 | 0 | 0 | 0 | 0 | 0 | 0 | 0 | 0 | 0 | 0 | 0 | 0 | 0 | 1 |
| Jul27.25v_20006 | 9 | 0 | 0 | 0 | 0 | 0 | 0 | 0 | 0 | 0 | 0 | 0 | 0 | 0 | 0 | 0 | 0 | 1 | 1 | 0 | 0 | 2 | 1 | 2 | 1 | 0 | 0 | 0 | 0 | 1 |
| Jul27.5v_8469 | 1 | 0 | 0 | 0 | 0 | 0 | 0 | 0 | 0 | 0 | 0 | 0 | 0 | 0 | 0 | 0 | 0 | 1 | 0 | 0 | 0 | 0 | 0 | 0 | 0 | 0 | 0 | 0 | 0 | 0 |
| Aug5.25v_21406 | 2 | 0 | 0 | 0 | 0 | 0 | 0 | 0 | 0 | 0 | 0 | 0 | 0 | 0 | 0 | 0 | 0 | 0 | 1 | 0 | 0 | 0 | 0 | 1 | 0 | 0 | 0 | 0 | 0 | 0 |
| Aug5.25v_34875 | 1 | 0 | 0 | 0 | 0 | 0 | 0 | 0 | 0 | 0 | 0 | 0 | 0 | 0 | 0 | 0 | 0 | 0 | 0 | 0 | 0 | 0 | 0 | 1 | 0 | 0 | 0 | 0 | 0 | 0 |
| Aug5.25v_11501 | 1 | 0 | 0 | 0 | 0 | 0 | 0 | 0 | 0 | 0 | 0 | 0 | 0 | 0 | 0 | 0 | 0 | 0 | 0 | 0 | 0 | 0 | 0 | 1 | 0 | 0 | 0 | 0 | 0 | 0 |
| Sep8.50v_10945 | 1 | 0 | 0 | 0 | 0 | 0 | 0 | 0 | 0 | 0 | 0 | 0 | 0 | 0 | 0 | 0 | 0 | 0 | 0 | 0 | 0 | 0 | 0 | 0 | 0 | 0 | 0 | 0 | 0 | 1 |
| Sep8.0v_20244 | 1 | 0 | 0 | 0 | 0 | 0 | 0 | 0 | 0 | 0 | 0 | 0 | 0 | 0 | 0 | 0 | 0 | 0 | 0 | 0 | 0 | 0 | 0 | 0 | 0 | 0 | 1 | 0 | 0 | 0 |
| Aug5.0v_1708 | 1 | 0 | 0 | 0 | 0 | 0 | 0 | 0 | 0 | 0 | 0 | 0 | 0 | 0 | 0 | 0 | 0 | 0 | 0 | 0 | 0 | 1 | 0 | 0 | 0 | 0 | 0 | 0 | 0 | 0 |
| Aug5.0v_4554 | 1 | 0 | 0 | 0 | 0 | 0 | 0 | 0 | 0 | 0 | 0 | 0 | 0 | 0 | 0 | 0 | 0 | 0 | 0 | 0 | 0 | 1 | 0 | 0 | 0 | 0 | 0 | 0 | 0 | 0 |
| Jul27.5v_18500 | 1 | 0 | 0 | 0 | 0 | 0 | 0 | 0 | 0 | 0 | 0 | 0 | 0 | 0 | 0 | 0 | 0 | 1 | 0 | 0 | 0 | 0 | 0 | 0 | 0 | 0 | 0 | 0 | 0 | 0 |
| Sep8.0v_18291 | 1 | 0 | 0 | 0 | 0 | 0 | 0 | 0 | 0 | 0 | 0 | 0 | 0 | 0 | 0 | 0 | 0 | 0 | 0 | 0 | 0 | 0 | 0 | 0 | 0 | 0 | 1 | 0 | 0 | 0 |
| Jul27.50v_16265 | 1 | 0 | 0 | 0 | 0 | 0 | 0 | 0 | 0 | 0 | 0 | 0 | 0 | 0 | 0 | 0 | 0 | 0 | 0 | 1 | 0 | 0 | 0 | 0 | 0 | 0 | 0 | 0 | 0 | 0 |
| Jul27.25v_33364 | 1 | 0 | 0 | 0 | 0 | 0 | 0 | 0 | 0 | 0 | 0 | 0 | 0 | 0 | 0 | 0 | 0 | 0 | 1 | 0 | 0 | 0 | 0 | 0 | 0 | 0 | 0 | 0 | 0 | 0 |
| Sep8.0v_9317 | 1 | 0 | 0 | 0 | 0 | 0 | 0 | 0 | 0 | 0 | 0 | 0 | 0 | 0 | 0 | 0 | 0 | 0 | 0 | 0 | 0 | 0 | 0 | 0 | 0 | 0 | 1 | 0 | 0 | 0 |
| Aug5.50v_5434 | 1 | 0 | 0 | 0 | 0 | 0 | 0 | 0 | 0 | 0 | 0 | 0 | 0 | 0 | 0 | 0 | 0 | 0 | 0 | 0 | 0 | 0 | 0 | 0 | 1 | 0 | 0 | 0 | 0 | 0 |
| Aug5.0v_3420 | 1 | 0 | 0 | 0 | 0 | 0 | 0 | 0 | 0 | 0 | 0 | 0 | 0 | 0 | 0 | 0 | 0 | 0 | 0 | 0 | 0 | 1 | 0 | 0 | 0 | 0 | 0 | 0 | 0 | 0 |
| Aug5.25v_28674 | 2 | 0 | 0 | 0 | 0 | 0 | 0 | 0 | 0 | 0 | 0 | 0 | 0 | 0 | 0 | 0 | 0 | 0 | 1 | 0 | 0 | 0 | 0 | 1 | 0 | 0 | 0 | 0 | 0 | 0 |
| Sep8.50v_18977 | 1 | 0 | 0 | 0 | 0 | 0 | 0 | 0 | 0 | 0 | 0 | 0 | 0 | 0 | 0 | 0 | 0 | 0 | 0 | 0 | 0 | 0 | 0 | 0 | 0 | 0 | 0 | 0 | 0 | 1 |
| Sep8.50v_39493 | 1 | 0 | 0 | 0 | 0 | 0 | 0 | 0 | 0 | 0 | 0 | 0 | 0 | 0 | 0 | 0 | 0 | 0 | 0 | 0 | 0 | 0 | 0 | 0 | 0 | 0 | 0 | 0 | 0 | 1 |
| Aug5.0v_5059 | 1 | 0 | 0 | 0 | 0 | 0 | 0 | 0 | 0 | 0 | 0 | 0 | 0 | 0 | 0 | 0 | 0 | 0 | 0 | 0 | 0 | 1 | 0 | 0 | 0 | 0 | 0 | 0 | 0 | 0 |
| Aug5.0v_1737 | 22 | 0 | 0 | 0 | 0 | 0 | 0 | 0 | 1 | 0 | 0 | 0 | 0 | 0 | 0 | 0 | 0 | 8 | 3 | 0 | 0 | 4 | 1 | 2 | 1 | 0 | 1 | 0 | 0 | 1 |
| Aug5.25v_21562 | 2 | 0 | 0 | 0 | 0 | 0 | 0 | 0 | 0 | 0 | 0 | 0 | 0 | 0 | 0 | 0 | 0 | 0 | 0 | 0 | 0 | 0 | 0 | 2 | 0 | 0 | 0 | 0 | 0 | 0 |
| Jul27.5v_16972 | 1 | 0 | 0 | 0 | 0 | 0 | 0 | 0 | 0 | 0 | 0 | 0 | 0 | 0 | 0 | 0 | 0 | 1 | 0 | 0 | 0 | 0 | 0 | 0 | 0 | 0 | 0 | 0 | 0 | 0 |
| Aug5.0v_586 | 3 | 0 | 0 | 0 | 0 | 0 | 0 | 0 | 0 | 0 | 0 | 0 | 0 | 0 | 0 | 0 | 0 | 0 | 0 | 0 | 0 | 3 | 0 | 0 | 0 | 0 | 0 | 0 | 0 | 0 |
| Aug5.25v_10477 | 1 | 0 | 0 | 0 | 0 | 0 | 0 | 0 | 0 | 0 | 0 | 0 | 0 | 0 | 0 | 0 | 0 | 0 | 0 | 0 | 0 | 0 | 0 | 1 | 0 | 0 | 0 | 0 | 0 | 0 |
| Jul27.5v_20148 | 1 | 0 | 0 | 0 | 0 | 0 | 0 | 0 | 0 | 0 | 0 | 0 | 0 | 0 | 0 | 0 | 0 | 1 | 0 | 0 | 0 | 0 | 0 | 0 | 0 | 0 | 0 | 0 | 0 | 0 |
| Aug5.0v_397 | 1 | 0 | 0 | 0 | 0 | 0 | 0 | 0 | 0 | 0 | 0 | 0 | 0 | 0 | 0 | 0 | 0 | 0 | 0 | 0 | 0 | 1 | 0 | 0 | 0 | 0 | 0 | 0 | 0 | 0 |
| Jul27.5v_7216 | 1 | 0 | 0 | 0 | 0 | 0 | 0 | 0 | 0 | 0 | 0 | 0 | 0 | 0 | 0 | 0 | 0 | 1 | 0 | 0 | 0 | 0 | 0 | 0 | 0 | 0 | 0 | 0 | 0 | 0 |
| Aug5.25v_27923 | 1 | 0 | 0 | 0 | 0 | 0 | 0 | 0 | 0 | 0 | 0 | 0 | 0 | 0 | 0 | 0 | 0 | 0 | 0 | 0 | 0 | 0 | 0 | 1 | 0 | 0 | 0 | 0 | 0 | 0 |
| Jun22.50v_32982 | 1 | 0 | 0 | 0 | 0 | 0 | 0 | 0 | 0 | 0 | 1 | 0 | 0 | 0 | 0 | 0 | 0 | 0 | 0 | 0 | 0 | 0 | 0 | 0 | 0 | 0 | 0 | 0 | 0 | 0 |
| Sep8.0v_9092 | 1 | 0 | 0 | 0 | 0 | 0 | 0 | 0 | 0 | 0 | 0 | 0 | 0 | 0 | 0 | 0 | 0 | 0 | 0 | 0 | 0 | 0 | 0 | 0 | 0 | 0 | 1 | 0 | 0 | 0 |
| Aug5.0v_4629 | 1 | 0 | 0 | 0 | 0 | 0 | 0 | 0 | 0 | 0 | 0 | 0 | 0 | 0 | 0 | 0 | 0 | 0 | 0 | 0 | 0 | 1 | 0 | 0 | 0 | 0 | 0 | 0 | 0 | 0 |
| May12.25v_20560 | 11 | 0 | 1 | 0 | 0 | 0 | 0 | 0 | 0 | 0 | 0 | 0 | 0 | 0 | 0 | 0 | 0 | 3 | 2 | 0 | 0 | 1 | 0 | 1 | 0 | 0 | 2 | 0 | 0 | 1 |
| Aug5.25v_11687 | 1 | 0 | 0 | 0 | 0 | 0 | 0 | 0 | 0 | 0 | 0 | 0 | 0 | 0 | 0 | 0 | 0 | 0 | 0 | 0 | 0 | 0 | 0 | 1 | 0 | 0 | 0 | 0 | 0 | 0 |
| Aug5.0v_2120 | 1 | 0 | 0 | 0 | 0 | 0 | 0 | 0 | 0 | 0 | 0 | 0 | 0 | 0 | 0 | 0 | 0 | 0 | 0 | 0 | 0 | 1 | 0 | 0 | 0 | 0 | 0 | 0 | 0 | 0 |
| Aug5.0v_2441 | 1 | 0 | 0 | 0 | 0 | 0 | 0 | 0 | 0 | 0 | 0 | 0 | 0 | 0 | 0 | 0 | 0 | 0 | 0 | 0 | 0 | 1 | 0 | 0 | 0 | 0 | 0 | 0 | 0 | 0 |
| Aug5.50v_6581 | 1 | 0 | 0 | 0 | 0 | 0 | 0 | 0 | 0 | 0 | 0 | 0 | 0 | 0 | 0 | 0 | 0 | 0 | 0 | 0 | 0 | 0 | 0 | 0 | 1 | 0 | 0 | 0 | 0 | 0 |
| Jun22.5v_15552 | 1 | 0 | 0 | 0 | 0 | 0 | 0 | 0 | 0 | 1 | 0 | 0 | 0 | 0 | 0 | 0 | 0 | 0 | 0 | 0 | 0 | 0 | 0 | 0 | 0 | 0 | 0 | 0 | 0 | 0 |
| Jul27.5v_6529 | 2 | 0 | 0 | 0 | 0 | 0 | 0 | 0 | 0 | 0 | 0 | 0 | 0 | 0 | 0 | 0 | 0 | 1 | 0 | 0 | 0 | 0 | 0 | 0 | 0 | 0 | 0 | 0 | 0 | 1 |
| Sep8.0v_4880 | 3 | 0 | 0 | 0 | 0 | 0 | 0 | 0 | 0 | 0 | 0 | 0 | 0 | 0 | 0 | 0 | 0 | 0 | 0 | 0 | 0 | 1 | 0 | 0 | 0 | 0 | 2 | 0 | 0 | 0 |
| Aug5.25v_19568 | 1 | 0 | 0 | 0 | 0 | 0 | 0 | 0 | 0 | 0 | 0 | 0 | 0 | 0 | 0 | 0 | 0 | 0 | 0 | 0 | 0 | 0 | 0 | 1 | 0 | 0 | 0 | 0 | 0 | 0 |
| Jul27.50v_24668 | 1 | 0 | 0 | 0 | 0 | 0 | 0 | 0 | 0 | 0 | 0 | 0 | 0 | 0 | 0 | 0 | 0 | 0 | 0 | 1 | 0 | 0 | 0 | 0 | 0 | 0 | 0 | 0 | 0 | 0 |
| Aug5.0v_5204 | 1 | 0 | 0 | 0 | 0 | 0 | 0 | 0 | 0 | 0 | 0 | 0 | 0 | 0 | 0 | 0 | 0 | 0 | 0 | 0 | 0 | 1 | 0 | 0 | 0 | 0 | 0 | 0 | 0 | 0 |
| Aug5.0v_3270 | 1 | 0 | 0 | 0 | 0 | 0 | 0 | 0 | 0 | 0 | 0 | 0 | 0 | 0 | 0 | 0 | 0 | 0 | 0 | 0 | 0 | 1 | 0 | 0 | 0 | 0 | 0 | 0 | 0 | 0 |
| Aug5.0v_4191 | 1 | 0 | 0 | 0 | 0 | 0 | 0 | 0 | 0 | 0 | 0 | 0 | 0 | 0 | 0 | 0 | 0 | 0 | 0 | 0 | 0 | 1 | 0 | 0 | 0 | 0 | 0 | 0 | 0 | 0 |
| Aug5.0v_1430 | 3 | 0 | 0 | 0 | 0 | 0 | 0 | 0 | 0 | 0 | 0 | 0 | 0 | 0 | 0 | 0 | 0 | 1 | 0 | 0 | 0 | 1 | 0 | 0 | 1 | 0 | 0 | 0 | 0 | 0 |
| Aug5.0v_2811 | 1 | 0 | 0 | 0 | 0 | 0 | 0 | 0 | 0 | 0 | 0 | 0 | 0 | 0 | 0 | 0 | 0 | 0 | 0 | 0 | 0 | 1 | 0 | 0 | 0 | 0 | 0 | 0 | 0 | 0 |
| Aug5.0v_1079 | 1 | 0 | 0 | 0 | 0 | 0 | 0 | 0 | 0 | 0 | 0 | 0 | 0 | 0 | 0 | 0 | 0 | 0 | 0 | 0 | 0 | 1 | 0 | 0 | 0 | 0 | 0 | 0 | 0 | 0 |
| Jul27.5v_1864 | 1 | 0 | 0 | 0 | 0 | 0 | 0 | 0 | 0 | 0 | 0 | 0 | 0 | 0 | 0 | 0 | 0 | 1 | 0 | 0 | 0 | 0 | 0 | 0 | 0 | 0 | 0 | 0 | 0 | 0 |
| Jul27.5v_19187 | 1 | 0 | 0 | 0 | 0 | 0 | 0 | 0 | 0 | 0 | 0 | 0 | 0 | 0 | 0 | 0 | 0 | 1 | 0 | 0 | 0 | 0 | 0 | 0 | 0 | 0 | 0 | 0 | 0 | 0 |
| Jul27.25v_8556 | 8 | 0 | 0 | 0 | 0 | 0 | 0 | 0 | 0 | 0 | 0 | 0 | 0 | 1 | 0 | 0 | 0 | 1 | 1 | 1 | 0 | 0 | 1 | 1 | 1 | 0 | 0 | 0 | 0 | 1 |
| Jul11.5v_4614 | 1 | 0 | 0 | 0 | 0 | 0 | 0 | 0 | 0 | 0 | 0 | 0 | 0 | 1 | 0 | 0 | 0 | 0 | 0 | 0 | 0 | 0 | 0 | 0 | 0 | 0 | 0 | 0 | 0 | 0 |
| Aug5.75v_19036 | 1 | 0 | 0 | 0 | 0 | 0 | 0 | 0 | 0 | 0 | 0 | 0 | 0 | 0 | 0 | 0 | 0 | 0 | 0 | 0 | 0 | 0 | 0 | 0 | 0 | 1 | 0 | 0 | 0 | 0 |
| Jul11.50v_15423 | 1 | 0 | 0 | 0 | 0 | 0 | 0 | 0 | 0 | 0 | 0 | 0 | 0 | 0 | 0 | 1 | 0 | 0 | 0 | 0 | 0 | 0 | 0 | 0 | 0 | 0 | 0 | 0 | 0 | 0 |
| Jun22.5v_15287 | 1 | 0 | 0 | 0 | 0 | 0 | 0 | 0 | 0 | 1 | 0 | 0 | 0 | 0 | 0 | 0 | 0 | 0 | 0 | 0 | 0 | 0 | 0 | 0 | 0 | 0 | 0 | 0 | 0 | 0 |
| Sep8.5v_8288 | 20 | 0 | 0 | 0 | 0 | 0 | 0 | 0 | 0 | 0 | 1 | 2 | 0 | 0 | 2 | 0 | 0 | 1 | 0 | 0 | 0 | 0 | 0 | 2 | 0 | 1 | 6 | 1 | 0 | 4 |
| Jun22.75v_20740 | 8 | 0 | 1 | 0 | 0 | 0 | 2 | 0 | 0 | 0 | 0 | 1 | 0 | 0 | 0 | 0 | 1 | 1 | 0 | 0 | 0 | 1 | 0 | 0 | 0 | 0 | 0 | 1 | 0 | 0 |
| Jun22.75v_21291 | 1 | 0 | 0 | 0 | 0 | 0 | 0 | 0 | 0 | 0 | 0 | 1 | 0 | 0 | 0 | 0 | 0 | 0 | 0 | 0 | 0 | 0 | 0 | 0 | 0 | 0 | 0 | 0 | 0 | 0 |
| Aug5.75v_21894 | 1 | 0 | 0 | 0 | 0 | 0 | 0 | 0 | 0 | 0 | 0 | 0 | 0 | 0 | 0 | 0 | 0 | 0 | 0 | 0 | 0 | 0 | 0 | 0 | 0 | 1 | 0 | 0 | 0 | 0 |
| Sep8.0v_3935 | 48 | 0 | 2 | 1 | 1 | 2 | 2 | 0 | 1 | 0 | 0 | 6 | 1 | 5 | 2 | 4 | 1 | 2 | 2 | 2 | 0 | 2 | 2 | 1 | 0 | 0 | 6 | 1 | 0 | 2 |
| Jul11.25v_1083 | 75 | 3 | 1 | 4 | 7 | 3 | 3 | 0 | 3 | 0 | 1 | 6 | 1 | 6 | 5 | 5 | 2 | 3 | 0 | 4 | 0 | 1 | 3 | 2 | 0 | 1 | 1 | 1 | 1 | 8 |
| Jul11.75v_40569 | 1 | 0 | 0 | 0 | 0 | 0 | 0 | 0 | 0 | 0 | 0 | 0 | 0 | 0 | 0 | 0 | 1 | 0 | 0 | 0 | 0 | 0 | 0 | 0 | 0 | 0 | 0 | 0 | 0 | 0 |
| Jun8.50v_3281 | 1 | 0 | 0 | 0 | 0 | 0 | 1 | 0 | 0 | 0 | 0 | 0 | 0 | 0 | 0 | 0 | 0 | 0 | 0 | 0 | 0 | 0 | 0 | 0 | 0 | 0 | 0 | 0 | 0 | 0 |
| Jul27.5v_16264 | 1 | 0 | 0 | 0 | 0 | 0 | 0 | 0 | 0 | 0 | 0 | 0 | 0 | 0 | 0 | 0 | 0 | 1 | 0 | 0 | 0 | 0 | 0 | 0 | 0 | 0 | 0 | 0 | 0 | 0 |
| Jul27.25v_32502 | 1 | 0 | 0 | 0 | 0 | 0 | 0 | 0 | 0 | 0 | 0 | 0 | 0 | 0 | 0 | 0 | 0 | 0 | 1 | 0 | 0 | 0 | 0 | 0 | 0 | 0 | 0 | 0 | 0 | 0 |
| Jun22.0v_13800 | 1 | 0 | 0 | 0 | 0 | 0 | 0 | 0 | 1 | 0 | 0 | 0 | 0 | 0 | 0 | 0 | 0 | 0 | 0 | 0 | 0 | 0 | 0 | 0 | 0 | 0 | 0 | 0 | 0 | 0 |
| Jun8.50v_31585 | 1 | 0 | 0 | 0 | 0 | 0 | 1 | 0 | 0 | 0 | 0 | 0 | 0 | 0 | 0 | 0 | 0 | 0 | 0 | 0 | 0 | 0 | 0 | 0 | 0 | 0 | 0 | 0 | 0 | 0 |
| Jul11.75v_21918 | 207 | 13 | 11 | 9 | 11 | 10 | 34 | 1 | 4 | 2 | 6 | 23 | 1 | 10 | 1 | 3 | 17 | 3 | 7 | 9 | 1 | 3 | 4 | 4 | 2 | 4 | 3 | 7 | 1 | 3 |
| Jun22.75v_2594 | 1 | 0 | 0 | 0 | 0 | 0 | 0 | 0 | 0 | 0 | 0 | 1 | 0 | 0 | 0 | 0 | 0 | 0 | 0 | 0 | 0 | 0 | 0 | 0 | 0 | 0 | 0 | 0 | 0 | 0 |
| Sep8.5v_5673 | 6 | 0 | 0 | 0 | 1 | 0 | 0 | 0 | 1 | 0 | 0 | 0 | 0 | 0 | 0 | 0 | 1 | 0 | 0 | 0 | 0 | 0 | 0 | 0 | 0 | 0 | 0 | 2 | 1 | 0 |
| May12.50v_11845 | 1 | 0 | 0 | 1 | 0 | 0 | 0 | 0 | 0 | 0 | 0 | 0 | 0 | 0 | 0 | 0 | 0 | 0 | 0 | 0 | 0 | 0 | 0 | 0 | 0 | 0 | 0 | 0 | 0 | 0 |
| Jul11.5v_3575 | 2 | 0 | 0 | 0 | 0 | 0 | 1 | 0 | 0 | 0 | 0 | 0 | 0 | 1 | 0 | 0 | 0 | 0 | 0 | 0 | 0 | 0 | 0 | 0 | 0 | 0 | 0 | 0 | 0 | 0 |
| Jun22.75v_19461 | 1 | 0 | 0 | 0 | 0 | 0 | 0 | 0 | 0 | 0 | 0 | 1 | 0 | 0 | 0 | 0 | 0 | 0 | 0 | 0 | 0 | 0 | 0 | 0 | 0 | 0 | 0 | 0 | 0 | 0 |
| Jul11.25v_10088 | 1 | 0 | 0 | 0 | 0 | 0 | 0 | 0 | 0 | 0 | 0 | 0 | 0 | 0 | 1 | 0 | 0 | 0 | 0 | 0 | 0 | 0 | 0 | 0 | 0 | 0 | 0 | 0 | 0 | 0 |
| Jun22.75v_3347 | 1 | 0 | 0 | 0 | 0 | 0 | 0 | 0 | 0 | 0 | 0 | 1 | 0 | 0 | 0 | 0 | 0 | 0 | 0 | 0 | 0 | 0 | 0 | 0 | 0 | 0 | 0 | 0 | 0 | 0 |
| Sep8.5v_17440 | 1 | 0 | 0 | 0 | 0 | 0 | 0 | 0 | 0 | 0 | 0 | 0 | 0 | 0 | 0 | 0 | 0 | 0 | 0 | 0 | 0 | 0 | 0 | 0 | 0 | 0 | 0 | 1 | 0 | 0 |
| Jun8.25v_23481 | 4 | 0 | 2 | 0 | 0 | 1 | 0 | 0 | 0 | 0 | 0 | 0 | 0 | 0 | 0 | 0 | 0 | 0 | 1 | 0 | 0 | 0 | 0 | 0 | 0 | 0 | 0 | 0 | 0 | 0 |
| Jul27.50v_8582 | 10 | 0 | 1 | 1 | 1 | 0 | 1 | 0 | 0 | 1 | 0 | 0 | 0 | 1 | 0 | 0 | 2 | 0 | 0 | 1 | 0 | 0 | 1 | 0 | 0 | 0 | 0 | 0 | 0 | 0 |
| Jul11.75v_9660 | 13 | 1 | 0 | 0 | 1 | 0 | 2 | 0 | 1 | 0 | 0 | 1 | 1 | 0 | 0 | 1 | 3 | 0 | 0 | 0 | 0 | 1 | 1 | 0 | 0 | 0 | 0 | 0 | 0 | 0 |
| Sep8.50v_33206 | 1 | 0 | 0 | 0 | 0 | 0 | 0 | 0 | 0 | 0 | 0 | 0 | 0 | 0 | 0 | 0 | 0 | 0 | 0 | 0 | 0 | 0 | 0 | 0 | 0 | 0 | 0 | 0 | 0 | 1 |
| Jun8.50v_31595 | 44 | 0 | 1 | 3 | 7 | 1 | 2 | 0 | 1 | 0 | 1 | 1 | 3 | 5 | 0 | 3 | 1 | 3 | 1 | 1 | 0 | 2 | 2 | 1 | 0 | 1 | 1 | 0 | 1 | 2 |
| Jun8.50v_8532 | 1 | 0 | 0 | 0 | 0 | 0 | 1 | 0 | 0 | 0 | 0 | 0 | 0 | 0 | 0 | 0 | 0 | 0 | 0 | 0 | 0 | 0 | 0 | 0 | 0 | 0 | 0 | 0 | 0 | 0 |
| Aug5.25v_11359 | 1 | 0 | 0 | 0 | 0 | 0 | 0 | 0 | 0 | 0 | 0 | 0 | 0 | 0 | 0 | 0 | 0 | 0 | 0 | 0 | 0 | 0 | 0 | 1 | 0 | 0 | 0 | 0 | 0 | 0 |
| Jul11.0v_5281 | 1 | 0 | 0 | 0 | 0 | 0 | 0 | 0 | 0 | 0 | 0 | 0 | 1 | 0 | 0 | 0 | 0 | 0 | 0 | 0 | 0 | 0 | 0 | 0 | 0 | 0 | 0 | 0 | 0 | 0 |
| Jul27.5v_16984 | 1 | 0 | 0 | 0 | 0 | 0 | 0 | 0 | 0 | 0 | 0 | 0 | 0 | 0 | 0 | 0 | 0 | 1 | 0 | 0 | 0 | 0 | 0 | 0 | 0 | 0 | 0 | 0 | 0 | 0 |
| Jun22.75v_11856 | 1 | 0 | 0 | 0 | 0 | 0 | 0 | 0 | 0 | 0 | 0 | 1 | 0 | 0 | 0 | 0 | 0 | 0 | 0 | 0 | 0 | 0 | 0 | 0 | 0 | 0 | 0 | 0 | 0 | 0 |
| Jul27.50v_18350 | 7 | 0 | 1 | 0 | 0 | 0 | 0 | 0 | 0 | 0 | 0 | 0 | 0 | 1 | 1 | 0 | 1 | 0 | 0 | 1 | 0 | 0 | 0 | 0 | 0 | 1 | 0 | 1 | 0 | 0 |
| Jul27.5v_12086 | 11 | 0 | 0 | 0 | 0 | 0 | 0 | 0 | 0 | 0 | 0 | 0 | 0 | 0 | 0 | 0 | 0 | 5 | 0 | 1 | 0 | 0 | 0 | 0 | 0 | 2 | 0 | 3 | 0 | 0 |
| Jul11.50v_17700 | 1 | 0 | 0 | 0 | 0 | 0 | 0 | 0 | 0 | 0 | 0 | 0 | 0 | 0 | 0 | 1 | 0 | 0 | 0 | 0 | 0 | 0 | 0 | 0 | 0 | 0 | 0 | 0 | 0 | 0 |
| Jun22.50v_7359 | 1 | 0 | 0 | 0 | 0 | 0 | 0 | 0 | 0 | 0 | 1 | 0 | 0 | 0 | 0 | 0 | 0 | 0 | 0 | 0 | 0 | 0 | 0 | 0 | 0 | 0 | 0 | 0 | 0 | 0 |
| Aug5.75v_19491 | 32 | 0 | 0 | 0 | 0 | 0 | 1 | 0 | 1 | 2 | 2 | 0 | 0 | 1 | 2 | 0 | 0 | 3 | 0 | 1 | 0 | 4 | 2 | 6 | 3 | 1 | 1 | 0 | 0 | 2 |
| Aug5.0v_4925 | 2 | 0 | 0 | 0 | 0 | 0 | 0 | 0 | 0 | 0 | 0 | 0 | 0 | 0 | 0 | 0 | 0 | 0 | 0 | 0 | 0 | 1 | 0 | 0 | 0 | 1 | 0 | 0 | 0 | 0 |
| Aug5.5v_15591 | 3 | 0 | 0 | 1 | 0 | 0 | 0 | 0 | 0 | 0 | 0 | 0 | 0 | 0 | 0 | 0 | 0 | 1 | 0 | 0 | 0 | 0 | 1 | 0 | 0 | 0 | 0 | 0 | 0 | 0 |
| Jul27.25v_412 | 1 | 0 | 0 | 0 | 0 | 0 | 0 | 0 | 0 | 0 | 0 | 0 | 0 | 0 | 0 | 0 | 0 | 0 | 1 | 0 | 0 | 0 | 0 | 0 | 0 | 0 | 0 | 0 | 0 | 0 |
| Jun8.50v_37590 | 1 | 0 | 0 | 0 | 0 | 0 | 1 | 0 | 0 | 0 | 0 | 0 | 0 | 0 | 0 | 0 | 0 | 0 | 0 | 0 | 0 | 0 | 0 | 0 | 0 | 0 | 0 | 0 | 0 | 0 |
| Sep8.5v_1908 | 1 | 0 | 0 | 0 | 0 | 0 | 0 | 0 | 0 | 0 | 0 | 0 | 0 | 0 | 0 | 0 | 0 | 0 | 0 | 0 | 0 | 0 | 0 | 0 | 0 | 0 | 0 | 1 | 0 | 0 |
| Aug5.0v_40 | 28 | 3 | 2 | 0 | 1 | 1 | 1 | 0 | 1 | 1 | 5 | 1 | 1 | 0 | 0 | 0 | 1 | 0 | 0 | 3 | 0 | 3 | 1 | 1 | 0 | 1 | 1 | 0 | 0 | 0 |
| Aug5.5v_15572 | 12 | 0 | 1 | 0 | 1 | 0 | 2 | 0 | 1 | 0 | 1 | 0 | 0 | 0 | 0 | 0 | 0 | 0 | 1 | 1 | 0 | 0 | 3 | 0 | 0 | 0 | 0 | 0 | 0 | 1 |
| May12.25v_38507 | 2 | 0 | 1 | 0 | 0 | 0 | 0 | 0 | 0 | 0 | 0 | 0 | 0 | 0 | 0 | 0 | 0 | 0 | 0 | 0 | 0 | 0 | 0 | 0 | 0 | 0 | 0 | 1 | 0 | 0 |
| Jun22.50v_22846 | 12 | 0 | 1 | 0 | 2 | 0 | 0 | 0 | 0 | 0 | 1 | 0 | 1 | 1 | 0 | 0 | 0 | 0 | 2 | 2 | 0 | 0 | 1 | 0 | 0 | 0 | 0 | 0 | 0 | 1 |
| Jul27.5v_16884 | 1 | 0 | 0 | 0 | 0 | 0 | 0 | 0 | 0 | 0 | 0 | 0 | 0 | 0 | 0 | 0 | 0 | 1 | 0 | 0 | 0 | 0 | 0 | 0 | 0 | 0 | 0 | 0 | 0 | 0 |
| Sep8.50v_19119 | 1 | 0 | 0 | 0 | 0 | 0 | 0 | 0 | 0 | 0 | 0 | 0 | 0 | 0 | 0 | 0 | 0 | 0 | 0 | 0 | 0 | 0 | 0 | 0 | 0 | 0 | 0 | 0 | 0 | 1 |
| Jul27.25v_31095 | 1 | 0 | 0 | 0 | 0 | 0 | 0 | 0 | 0 | 0 | 0 | 0 | 0 | 0 | 0 | 0 | 0 | 0 | 1 | 0 | 0 | 0 | 0 | 0 | 0 | 0 | 0 | 0 | 0 | 0 |
| Jul11.5v_9158 | 9 | 0 | 0 | 0 | 0 | 0 | 0 | 0 | 0 | 0 | 0 | 0 | 1 | 1 | 0 | 0 | 0 | 3 | 0 | 1 | 0 | 2 | 0 | 0 | 0 | 0 | 1 | 0 | 0 | 0 |
| Jul11.5v_15657 | 2 | 0 | 0 | 0 | 0 | 0 | 0 | 0 | 0 | 0 | 0 | 0 | 1 | 1 | 0 | 0 | 0 | 0 | 0 | 0 | 0 | 0 | 0 | 0 | 0 | 0 | 0 | 0 | 0 | 0 |
| Aug5.0v_2367 | 1 | 0 | 0 | 0 | 0 | 0 | 0 | 0 | 0 | 0 | 0 | 0 | 0 | 0 | 0 | 0 | 0 | 0 | 0 | 0 | 0 | 1 | 0 | 0 | 0 | 0 | 0 | 0 | 0 | 0 |
| Aug5.5v_18027 | 12 | 0 | 0 | 0 | 0 | 0 | 0 | 0 | 0 | 0 | 0 | 0 | 0 | 0 | 0 | 0 | 0 | 2 | 1 | 1 | 0 | 4 | 2 | 0 | 1 | 0 | 1 | 0 | 0 | 0 |
| Aug5.0v_1733 | 2 | 0 | 0 | 0 | 0 | 0 | 0 | 0 | 0 | 0 | 0 | 0 | 0 | 0 | 0 | 0 | 0 | 0 | 0 | 0 | 0 | 1 | 0 | 0 | 0 | 0 | 1 | 0 | 0 | 0 |
| Aug5.0v_2129 | 1 | 0 | 0 | 0 | 0 | 0 | 0 | 0 | 0 | 0 | 0 | 0 | 0 | 0 | 0 | 0 | 0 | 0 | 0 | 0 | 0 | 1 | 0 | 0 | 0 | 0 | 0 | 0 | 0 | 0 |
| Sep8.0v_2763 | 1 | 0 | 0 | 0 | 0 | 0 | 0 | 0 | 0 | 0 | 0 | 0 | 0 | 0 | 0 | 0 | 0 | 0 | 0 | 0 | 0 | 0 | 0 | 0 | 0 | 0 | 1 | 0 | 0 | 0 |
| Jul11.5v_18322 | 1 | 0 | 0 | 0 | 0 | 0 | 0 | 0 | 0 | 0 | 0 | 0 | 0 | 1 | 0 | 0 | 0 | 0 | 0 | 0 | 0 | 0 | 0 | 0 | 0 | 0 | 0 | 0 | 0 | 0 |
| Sep8.0v_11863 | 1 | 0 | 0 | 0 | 0 | 0 | 0 | 0 | 0 | 0 | 0 | 0 | 0 | 0 | 0 | 0 | 0 | 0 | 0 | 0 | 0 | 0 | 0 | 0 | 0 | 0 | 1 | 0 | 0 | 0 |
| Sep8.0v_4725 | 1 | 0 | 0 | 0 | 0 | 0 | 0 | 0 | 0 | 0 | 0 | 0 | 0 | 0 | 0 | 0 | 0 | 0 | 0 | 0 | 0 | 0 | 0 | 0 | 0 | 0 | 1 | 0 | 0 | 0 |
| Jul27.5v_13117 | 1 | 0 | 0 | 0 | 0 | 0 | 0 | 0 | 0 | 0 | 0 | 0 | 0 | 0 | 0 | 0 | 0 | 1 | 0 | 0 | 0 | 0 | 0 | 0 | 0 | 0 | 0 | 0 | 0 | 0 |
| Aug5.0v_2568 | 1 | 0 | 0 | 0 | 0 | 0 | 0 | 0 | 0 | 0 | 0 | 0 | 0 | 0 | 0 | 0 | 0 | 0 | 0 | 0 | 0 | 1 | 0 | 0 | 0 | 0 | 0 | 0 | 0 | 0 |
| Jul27.5v_12011 | 1 | 0 | 0 | 0 | 0 | 0 | 0 | 0 | 0 | 0 | 0 | 0 | 0 | 0 | 0 | 0 | 0 | 1 | 0 | 0 | 0 | 0 | 0 | 0 | 0 | 0 | 0 | 0 | 0 | 0 |
| Jul27.5v_12788 | 1 | 0 | 0 | 0 | 0 | 0 | 0 | 0 | 0 | 0 | 0 | 0 | 0 | 0 | 0 | 0 | 0 | 1 | 0 | 0 | 0 | 0 | 0 | 0 | 0 | 0 | 0 | 0 | 0 | 0 |

| Feature_ID | 654 | May12_05m | May12_25m | May12_50m | May12_75m | Jun8_25m | Jun8_50m | Jun8_75m | Jun22_0m | Jun22_05m | Jun22_50m | Jun22_75m | Jul11_0m | Jul11_05m | Jul11_25m | Jul11_50m | Jul11_75m | Jul27_05m | Jul27_25m | Jul27_50m | Jul27_75m | Aug5_0m | Aug5_05m | Aug5_25m | Aug5_50m | Aug5_75m | Sep8_0m | Sep8_05m | Sep8_25m | Sep8_50m |
|---|---|---|---|---|---|---|---|---|---|---|---|---|---|---|---|---|---|---|---|---|---|---|---|---|---|---|---|---|---|---|
| Aug5.0v_3538 | 1 | 0 | 0 | 0 | 0 | 0 | 0 | 0 | 0 | 0 | 0 | 0 | 0 | 0 | 0 | 0 | 0 | 0 | 0 | 0 | 0 | 1 | 0 | 0 | 0 | 0 | 0 | 0 | 0 | 0 |
| Aug5.0v_3323 | 1 | 0 | 0 | 0 | 0 | 0 | 0 | 0 | 0 | 0 | 0 | 0 | 0 | 0 | 0 | 0 | 0 | 0 | 0 | 0 | 0 | 1 | 0 | 0 | 0 | 0 | 0 | 0 | 0 | 0 |
| Aug5.25v_5891 | 1 | 0 | 0 | 0 | 0 | 0 | 0 | 0 | 0 | 0 | 0 | 0 | 0 | 0 | 0 | 0 | 0 | 0 | 0 | 0 | 0 | 0 | 0 | 1 | 0 | 0 | 0 | 0 | 0 | 0 |
| Aug5.0v_3010 | 1 | 0 | 0 | 0 | 0 | 0 | 0 | 0 | 0 | 0 | 0 | 0 | 0 | 0 | 0 | 0 | 0 | 0 | 0 | 0 | 0 | 1 | 0 | 0 | 0 | 0 | 0 | 0 | 0 | 0 |
| Aug5.0v_3099 | 3 | 0 | 0 | 0 | 0 | 0 | 0 | 0 | 0 | 0 | 0 | 0 | 0 | 0 | 0 | 0 | 0 | 0 | 0 | 0 | 0 | 2 | 0 | 1 | 0 | 0 | 0 | 0 | 0 | 0 |
| Aug5.0v_930 | 1 | 0 | 0 | 0 | 0 | 0 | 0 | 0 | 0 | 0 | 0 | 0 | 0 | 0 | 0 | 0 | 0 | 0 | 0 | 0 | 0 | 1 | 0 | 0 | 0 | 0 | 0 | 0 | 0 | 0 |
| Aug5.25v_37604 | 1 | 0 | 0 | 0 | 0 | 0 | 0 | 0 | 0 | 0 | 0 | 0 | 0 | 0 | 0 | 0 | 0 | 0 | 0 | 0 | 0 | 0 | 0 | 1 | 0 | 0 | 0 | 0 | 0 | 0 |
| Aug5.0v_888 | 1 | 0 | 0 | 0 | 0 | 0 | 0 | 0 | 0 | 0 | 0 | 0 | 0 | 0 | 0 | 0 | 0 | 0 | 0 | 0 | 0 | 1 | 0 | 0 | 0 | 0 | 0 | 0 | 0 | 0 |
| Jul11.25v_8754 | 1 | 0 | 0 | 0 | 0 | 0 | 0 | 0 | 0 | 0 | 0 | 0 | 0 | 0 | 1 | 0 | 0 | 0 | 0 | 0 | 0 | 0 | 0 | 0 | 0 | 0 | 0 | 0 | 0 | 0 |
| Sep8.0v_20712 | 1 | 0 | 0 | 0 | 0 | 0 | 0 | 0 | 0 | 0 | 0 | 0 | 0 | 0 | 0 | 0 | 0 | 0 | 0 | 0 | 0 | 0 | 0 | 0 | 0 | 0 | 1 | 0 | 0 | 0 |
| Aug5.5v_169 | 1 | 0 | 0 | 0 | 0 | 0 | 0 | 0 | 0 | 0 | 0 | 0 | 0 | 0 | 0 | 0 | 0 | 0 | 0 | 0 | 0 | 0 | 1 | 0 | 0 | 0 | 0 | 0 | 0 | 0 |
| Jul11.0v_2455 | 1 | 0 | 0 | 0 | 0 | 0 | 0 | 0 | 0 | 0 | 0 | 0 | 1 | 0 | 0 | 0 | 0 | 0 | 0 | 0 | 0 | 0 | 0 | 0 | 0 | 0 | 0 | 0 | 0 | 0 |
| Aug5.25v_12101 | 1 | 0 | 0 | 0 | 0 | 0 | 0 | 0 | 0 | 0 | 0 | 0 | 0 | 0 | 0 | 0 | 0 | 0 | 0 | 0 | 0 | 0 | 0 | 1 | 0 | 0 | 0 | 0 | 0 | 0 |
| Aug5.25v_6691 | 1 | 0 | 0 | 0 | 0 | 0 | 0 | 0 | 0 | 0 | 0 | 0 | 0 | 0 | 0 | 0 | 0 | 0 | 0 | 0 | 0 | 0 | 0 | 1 | 0 | 0 | 0 | 0 | 0 | 0 |
| Aug5.0v_4410 | 1 | 0 | 0 | 0 | 0 | 0 | 0 | 0 | 0 | 0 | 0 | 0 | 0 | 0 | 0 | 0 | 0 | 0 | 0 | 0 | 0 | 1 | 0 | 0 | 0 | 0 | 0 | 0 | 0 | 0 |
| Sep8.50v_36930 | 1 | 0 | 0 | 0 | 0 | 0 | 0 | 0 | 0 | 0 | 0 | 0 | 0 | 0 | 0 | 0 | 0 | 0 | 0 | 0 | 0 | 0 | 0 | 0 | 0 | 0 | 0 | 0 | 0 | 1 |
| Jul27.5v_13776 | 1 | 0 | 0 | 0 | 0 | 0 | 0 | 0 | 0 | 0 | 0 | 0 | 0 | 0 | 0 | 0 | 0 | 1 | 0 | 0 | 0 | 0 | 0 | 0 | 0 | 0 | 0 | 0 | 0 | 0 |
| Aug5.25v_33312 | 1 | 0 | 0 | 0 | 0 | 0 | 0 | 0 | 0 | 0 | 0 | 0 | 0 | 0 | 0 | 0 | 0 | 0 | 0 | 0 | 0 | 0 | 0 | 1 | 0 | 0 | 0 | 0 | 0 | 0 |
| Aug5.0v_4533 | 1 | 0 | 0 | 0 | 0 | 0 | 0 | 0 | 0 | 0 | 0 | 0 | 0 | 0 | 0 | 0 | 0 | 0 | 0 | 0 | 0 | 1 | 0 | 0 | 0 | 0 | 0 | 0 | 0 | 0 |
| Jul27.5v_8173 | 1 | 0 | 0 | 0 | 0 | 0 | 0 | 0 | 0 | 0 | 0 | 0 | 0 | 0 | 0 | 0 | 0 | 1 | 0 | 0 | 0 | 0 | 0 | 0 | 0 | 0 | 0 | 0 | 0 | 0 |
| Aug5.25v_38021 | 1 | 0 | 0 | 0 | 0 | 0 | 0 | 0 | 0 | 0 | 0 | 0 | 0 | 0 | 0 | 0 | 0 | 0 | 0 | 0 | 0 | 0 | 0 | 1 | 0 | 0 | 0 | 0 | 0 | 0 |
| Jul27.5v_21090 | 1 | 0 | 0 | 0 | 0 | 0 | 0 | 0 | 0 | 0 | 0 | 0 | 0 | 0 | 0 | 0 | 0 | 1 | 0 | 0 | 0 | 0 | 0 | 0 | 0 | 0 | 0 | 0 | 0 | 0 |
| Aug5.25v_19230 | 1 | 0 | 0 | 0 | 0 | 0 | 0 | 0 | 0 | 0 | 0 | 0 | 0 | 0 | 0 | 0 | 0 | 0 | 0 | 0 | 0 | 0 | 0 | 1 | 0 | 0 | 0 | 0 | 0 | 0 |
| Aug5.0v_2813 | 1 | 0 | 0 | 0 | 0 | 0 | 0 | 0 | 0 | 0 | 0 | 0 | 0 | 0 | 0 | 0 | 0 | 0 | 0 | 0 | 0 | 1 | 0 | 0 | 0 | 0 | 0 | 0 | 0 | 0 |
| Aug5.0v_390 | 1 | 0 | 0 | 0 | 0 | 0 | 0 | 0 | 0 | 0 | 0 | 0 | 0 | 0 | 0 | 0 | 0 | 0 | 0 | 0 | 0 | 1 | 0 | 0 | 0 | 0 | 0 | 0 | 0 | 0 |
| Jul27.5v_20102 | 1 | 0 | 0 | 0 | 0 | 0 | 0 | 0 | 0 | 0 | 0 | 0 | 0 | 0 | 0 | 0 | 0 | 1 | 0 | 0 | 0 | 0 | 0 | 0 | 0 | 0 | 0 | 0 | 0 | 0 |
| Aug5.25v_9058 | 1 | 0 | 0 | 0 | 0 | 0 | 0 | 0 | 0 | 0 | 0 | 0 | 0 | 0 | 0 | 0 | 0 | 0 | 0 | 0 | 0 | 0 | 0 | 1 | 0 | 0 | 0 | 0 | 0 | 0 |
| Jul27.5v_4379 | 1 | 0 | 0 | 0 | 0 | 0 | 0 | 0 | 0 | 0 | 0 | 0 | 0 | 0 | 0 | 0 | 0 | 1 | 0 | 0 | 0 | 0 | 0 | 0 | 0 | 0 | 0 | 0 | 0 | 0 |
| Aug5.25v_24047 | 1 | 0 | 0 | 0 | 0 | 0 | 0 | 0 | 0 | 0 | 0 | 0 | 0 | 0 | 0 | 0 | 0 | 0 | 0 | 0 | 0 | 0 | 0 | 1 | 0 | 0 | 0 | 0 | 0 | 0 |
| Aug5.0v_3450 | 1 | 0 | 0 | 0 | 0 | 0 | 0 | 0 | 0 | 0 | 0 | 0 | 0 | 0 | 0 | 0 | 0 | 0 | 0 | 0 | 0 | 1 | 0 | 0 | 0 | 0 | 0 | 0 | 0 | 0 |
| Aug5.25v_40869 | 1 | 0 | 0 | 0 | 0 | 0 | 0 | 0 | 0 | 0 | 0 | 0 | 0 | 0 | 0 | 0 | 0 | 0 | 0 | 0 | 0 | 0 | 0 | 1 | 0 | 0 | 0 | 0 | 0 | 0 |
| Aug5.0v_5325 | 1 | 0 | 0 | 0 | 0 | 0 | 0 | 0 | 0 | 0 | 0 | 0 | 0 | 0 | 0 | 0 | 0 | 0 | 0 | 0 | 0 | 1 | 0 | 0 | 0 | 0 | 0 | 0 | 0 | 0 |
| Jul11.0v_10567 | 1 | 0 | 0 | 0 | 0 | 0 | 0 | 0 | 0 | 0 | 0 | 0 | 1 | 0 | 0 | 0 | 0 | 0 | 0 | 0 | 0 | 0 | 0 | 0 | 0 | 0 | 0 | 0 | 0 | 0 |
| Jul27.5v_11008 | 1 | 0 | 0 | 0 | 0 | 0 | 0 | 0 | 0 | 0 | 0 | 0 | 0 | 0 | 0 | 0 | 0 | 1 | 0 | 0 | 0 | 0 | 0 | 0 | 0 | 0 | 0 | 0 | 0 | 0 |
| Aug5.0v_2748 | 1 | 0 | 0 | 0 | 0 | 0 | 0 | 0 | 0 | 0 | 0 | 0 | 0 | 0 | 0 | 0 | 0 | 0 | 0 | 0 | 0 | 1 | 0 | 0 | 0 | 0 | 0 | 0 | 0 | 0 |
| Aug5.0v_2994 | 1 | 0 | 0 | 0 | 0 | 0 | 0 | 0 | 0 | 0 | 0 | 0 | 0 | 0 | 0 | 0 | 0 | 0 | 0 | 0 | 0 | 1 | 0 | 0 | 0 | 0 | 0 | 0 | 0 | 0 |
| Jul27.5v_20949 | 1 | 0 | 0 | 0 | 0 | 0 | 0 | 0 | 0 | 0 | 0 | 0 | 0 | 0 | 0 | 0 | 0 | 1 | 0 | 0 | 0 | 0 | 0 | 0 | 0 | 0 | 0 | 0 | 0 | 0 |
| Jul27.5v_11134 | 1 | 0 | 0 | 0 | 0 | 0 | 0 | 0 | 0 | 0 | 0 | 0 | 0 | 0 | 0 | 0 | 0 | 1 | 0 | 0 | 0 | 0 | 0 | 0 | 0 | 0 | 0 | 0 | 0 | 0 |
| Aug5.0v_1769 | 1 | 0 | 0 | 0 | 0 | 0 | 0 | 0 | 0 | 0 | 0 | 0 | 0 | 0 | 0 | 0 | 0 | 0 | 0 | 0 | 0 | 1 | 0 | 0 | 0 | 0 | 0 | 0 | 0 | 0 |
| Aug5.5v_24471 | 1 | 0 | 0 | 0 | 0 | 0 | 0 | 0 | 0 | 0 | 0 | 0 | 0 | 0 | 0 | 0 | 0 | 0 | 0 | 0 | 0 | 0 | 1 | 0 | 0 | 0 | 0 | 0 | 0 | 0 |
| Jul27.25v_25399 | 1 | 0 | 0 | 0 | 0 | 0 | 0 | 0 | 0 | 0 | 0 | 0 | 0 | 0 | 0 | 0 | 0 | 0 | 1 | 0 | 0 | 0 | 0 | 0 | 0 | 0 | 0 | 0 | 0 | 0 |
| Aug5.25v_28937 | 1 | 0 | 0 | 0 | 0 | 0 | 0 | 0 | 0 | 0 | 0 | 0 | 0 | 0 | 0 | 0 | 0 | 0 | 0 | 0 | 0 | 0 | 0 | 1 | 0 | 0 | 0 | 0 | 0 | 0 |
| Jul27.5v_6090 | 1 | 0 | 0 | 0 | 0 | 0 | 0 | 0 | 0 | 0 | 0 | 0 | 0 | 0 | 0 | 0 | 0 | 1 | 0 | 0 | 0 | 0 | 0 | 0 | 0 | 0 | 0 | 0 | 0 | 0 |
| Aug5.25v_24413 | 1 | 0 | 0 | 0 | 0 | 0 | 0 | 0 | 0 | 0 | 0 | 0 | 0 | 0 | 0 | 0 | 0 | 0 | 0 | 0 | 0 | 0 | 0 | 1 | 0 | 0 | 0 | 0 | 0 | 0 |
| May12.5v_14325 | 1 | 1 | 0 | 0 | 0 | 0 | 0 | 0 | 0 | 0 | 0 | 0 | 0 | 0 | 0 | 0 | 0 | 0 | 0 | 0 | 0 | 0 | 0 | 0 | 0 | 0 | 0 | 0 | 0 | 0 |
| Aug5.25v_16298 | 1 | 0 | 0 | 0 | 0 | 0 | 0 | 0 | 0 | 0 | 0 | 0 | 0 | 0 | 0 | 0 | 0 | 0 | 0 | 0 | 0 | 0 | 0 | 1 | 0 | 0 | 0 | 0 | 0 | 0 |
| Aug5.0v_3220 | 1 | 0 | 0 | 0 | 0 | 0 | 0 | 0 | 0 | 0 | 0 | 0 | 0 | 0 | 0 | 0 | 0 | 0 | 0 | 0 | 0 | 1 | 0 | 0 | 0 | 0 | 0 | 0 | 0 | 0 |
| Jul27.5v_4002 | 1 | 0 | 0 | 0 | 0 | 0 | 0 | 0 | 0 | 0 | 0 | 0 | 0 | 0 | 0 | 0 | 0 | 1 | 0 | 0 | 0 | 0 | 0 | 0 | 0 | 0 | 0 | 0 | 0 | 0 |
| Jul11.0v_13147 | 1 | 0 | 0 | 0 | 0 | 0 | 0 | 0 | 0 | 0 | 0 | 0 | 1 | 0 | 0 | 0 | 0 | 0 | 0 | 0 | 0 | 0 | 0 | 0 | 0 | 0 | 0 | 0 | 0 | 0 |
| Jul27.50v_22655 | 1 | 0 | 0 | 0 | 0 | 0 | 0 | 0 | 0 | 0 | 0 | 0 | 0 | 0 | 0 | 0 | 0 | 0 | 0 | 1 | 0 | 0 | 0 | 0 | 0 | 0 | 0 | 0 | 0 | 0 |
| May12.75v_34836 | 1 | 0 | 0 | 0 | 1 | 0 | 0 | 0 | 0 | 0 | 0 | 0 | 0 | 0 | 0 | 0 | 0 | 0 | 0 | 0 | 0 | 0 | 0 | 0 | 0 | 0 | 0 | 0 | 0 | 0 |
| Sep8.0v_15950 | 1 | 0 | 0 | 0 | 0 | 0 | 0 | 0 | 0 | 0 | 0 | 0 | 0 | 0 | 0 | 0 | 0 | 0 | 0 | 0 | 0 | 0 | 0 | 0 | 0 | 0 | 1 | 0 | 0 | 0 |
| May12.50v_9190 | 1 | 0 | 0 | 1 | 0 | 0 | 0 | 0 | 0 | 0 | 0 | 0 | 0 | 0 | 0 | 0 | 0 | 0 | 0 | 0 | 0 | 0 | 0 | 0 | 0 | 0 | 0 | 0 | 0 | 0 |
| Sep8.0v_11708 | 1 | 0 | 0 | 0 | 0 | 0 | 0 | 0 | 0 | 0 | 0 | 0 | 0 | 0 | 0 | 0 | 0 | 0 | 0 | 0 | 0 | 0 | 0 | 0 | 0 | 0 | 1 | 0 | 0 | 0 |
| Jul27.5v_1870 | 3 | 0 | 0 | 1 | 0 | 0 | 0 | 0 | 0 | 0 | 0 | 1 | 0 | 0 | 0 | 0 | 0 | 1 | 0 | 0 | 0 | 0 | 0 | 0 | 0 | 0 | 0 | 0 | 0 | 0 |
| Sep8.50v_6231 | 1 | 0 | 0 | 0 | 0 | 0 | 0 | 0 | 0 | 0 | 0 | 0 | 0 | 0 | 0 | 0 | 0 | 0 | 0 | 0 | 0 | 0 | 0 | 0 | 0 | 0 | 0 | 0 | 0 | 1 |
| Jul11.25v_14332 | 1 | 0 | 0 | 0 | 0 | 0 | 0 | 0 | 0 | 0 | 0 | 0 | 0 | 0 | 1 | 0 | 0 | 0 | 0 | 0 | 0 | 0 | 0 | 0 | 0 | 0 | 0 | 0 | 0 | 0 |
| Sep8.5v_9010 | 1 | 0 | 0 | 0 | 0 | 0 | 0 | 0 | 0 | 0 | 0 | 0 | 0 | 0 | 0 | 0 | 0 | 0 | 0 | 0 | 0 | 0 | 0 | 0 | 0 | 0 | 0 | 1 | 0 | 0 |
| Jul27.5v_7648 | 1 | 0 | 0 | 0 | 0 | 0 | 0 | 0 | 0 | 0 | 0 | 0 | 0 | 0 | 0 | 0 | 0 | 1 | 0 | 0 | 0 | 0 | 0 | 0 | 0 | 0 | 0 | 0 | 0 | 0 |
| Jun8.75v_2130 | 1 | 0 | 0 | 0 | 0 | 0 | 0 | 1 | 0 | 0 | 0 | 0 | 0 | 0 | 0 | 0 | 0 | 0 | 0 | 0 | 0 | 0 | 0 | 0 | 0 | 0 | 0 | 0 | 0 | 0 |
| Sep8.0v_6813 | 1 | 0 | 0 | 0 | 0 | 0 | 0 | 0 | 0 | 0 | 0 | 0 | 0 | 0 | 0 | 0 | 0 | 0 | 0 | 0 | 0 | 0 | 0 | 0 | 0 | 0 | 1 | 0 | 0 | 0 |
| Jun22.75v_12846 | 1 | 0 | 0 | 0 | 0 | 0 | 0 | 0 | 0 | 0 | 0 | 1 | 0 | 0 | 0 | 0 | 0 | 0 | 0 | 0 | 0 | 0 | 0 | 0 | 0 | 0 | 0 | 0 | 0 | 0 |
| Jul11.25v_11317 | 1 | 0 | 0 | 0 | 0 | 0 | 0 | 0 | 0 | 0 | 0 | 0 | 0 | 0 | 1 | 0 | 0 | 0 | 0 | 0 | 0 | 0 | 0 | 0 | 0 | 0 | 0 | 0 | 0 | 0 |
| May12.5v_24933 | 1 | 1 | 0 | 0 | 0 | 0 | 0 | 0 | 0 | 0 | 0 | 0 | 0 | 0 | 0 | 0 | 0 | 0 | 0 | 0 | 0 | 0 | 0 | 0 | 0 | 0 | 0 | 0 | 0 | 0 |
| Jul27.50v_32021 | 1 | 0 | 0 | 0 | 0 | 0 | 0 | 0 | 0 | 0 | 0 | 0 | 0 | 0 | 0 | 0 | 0 | 0 | 0 | 1 | 0 | 0 | 0 | 0 | 0 | 0 | 0 | 0 | 0 | 0 |
| Jul11.25v_17692 | 1 | 0 | 0 | 0 | 0 | 0 | 0 | 0 | 0 | 0 | 0 | 0 | 0 | 0 | 1 | 0 | 0 | 0 | 0 | 0 | 0 | 0 | 0 | 0 | 0 | 0 | 0 | 0 | 0 | 0 |
| Jun8.50v_12412 | 1 | 0 | 0 | 0 | 0 | 0 | 1 | 0 | 0 | 0 | 0 | 0 | 0 | 0 | 0 | 0 | 0 | 0 | 0 | 0 | 0 | 0 | 0 | 0 | 0 | 0 | 0 | 0 | 0 | 0 |
| Jul11.5v_20187 | 2 | 0 | 0 | 1 | 0 | 0 | 0 | 0 | 0 | 0 | 0 | 0 | 0 | 1 | 0 | 0 | 0 | 0 | 0 | 0 | 0 | 0 | 0 | 0 | 0 | 0 | 0 | 0 | 0 | 0 |
| Sep8.0v_7000 | 1 | 0 | 0 | 0 | 0 | 0 | 0 | 0 | 0 | 0 | 0 | 0 | 0 | 0 | 0 | 0 | 0 | 0 | 0 | 0 | 0 | 0 | 0 | 0 | 0 | 0 | 1 | 0 | 0 | 0 |
| Jul11.25v_14963 | 1 | 0 | 0 | 0 | 0 | 0 | 0 | 0 | 0 | 0 | 0 | 0 | 0 | 0 | 1 | 0 | 0 | 0 | 0 | 0 | 0 | 0 | 0 | 0 | 0 | 0 | 0 | 0 | 0 | 0 |
| Aug5.0v_329 | 1 | 0 | 0 | 0 | 0 | 0 | 0 | 0 | 0 | 0 | 0 | 0 | 0 | 0 | 0 | 0 | 0 | 0 | 0 | 0 | 0 | 1 | 0 | 0 | 0 | 0 | 0 | 0 | 0 | 0 |
| Jun8.25v_8775 | 1 | 0 | 0 | 0 | 0 | 1 | 0 | 0 | 0 | 0 | 0 | 0 | 0 | 0 | 0 | 0 | 0 | 0 | 0 | 0 | 0 | 0 | 0 | 0 | 0 | 0 | 0 | 0 | 0 | 0 |
| Jul11.0v_7583 | 2 | 0 | 0 | 0 | 0 | 0 | 0 | 0 | 0 | 0 | 0 | 0 | 1 | 0 | 0 | 0 | 0 | 0 | 0 | 0 | 0 | 0 | 0 | 0 | 0 | 0 | 1 | 0 | 0 | 0 |
| Jun22.0v_10752 | 1 | 0 | 0 | 0 | 0 | 0 | 0 | 0 | 1 | 0 | 0 | 0 | 0 | 0 | 0 | 0 | 0 | 0 | 0 | 0 | 0 | 0 | 0 | 0 | 0 | 0 | 0 | 0 | 0 | 0 |
| Jul11.25v_858 | 1 | 0 | 0 | 0 | 0 | 0 | 0 | 0 | 0 | 0 | 0 | 0 | 0 | 0 | 1 | 0 | 0 | 0 | 0 | 0 | 0 | 0 | 0 | 0 | 0 | 0 | 0 | 0 | 0 | 0 |
| May12.75v_16696 | 1 | 0 | 0 | 0 | 1 | 0 | 0 | 0 | 0 | 0 | 0 | 0 | 0 | 0 | 0 | 0 | 0 | 0 | 0 | 0 | 0 | 0 | 0 | 0 | 0 | 0 | 0 | 0 | 0 | 0 |
| Jul27.5v_18332 | 1 | 0 | 0 | 0 | 0 | 0 | 0 | 0 | 0 | 0 | 0 | 0 | 0 | 0 | 0 | 0 | 0 | 1 | 0 | 0 | 0 | 0 | 0 | 0 | 0 | 0 | 0 | 0 | 0 | 0 |
| Aug5.5v_15043 | 1 | 0 | 0 | 0 | 0 | 0 | 0 | 0 | 0 | 0 | 0 | 0 | 0 | 0 | 0 | 0 | 0 | 0 | 0 | 0 | 0 | 0 | 1 | 0 | 0 | 0 | 0 | 0 | 0 | 0 |
| Jun8.75v_12670 | 1 | 0 | 0 | 0 | 0 | 0 | 0 | 1 | 0 | 0 | 0 | 0 | 0 | 0 | 0 | 0 | 0 | 0 | 0 | 0 | 0 | 0 | 0 | 0 | 0 | 0 | 0 | 0 | 0 | 0 |
| May12.50v_19340 | 1 | 0 | 0 | 1 | 0 | 0 | 0 | 0 | 0 | 0 | 0 | 0 | 0 | 0 | 0 | 0 | 0 | 0 | 0 | 0 | 0 | 0 | 0 | 0 | 0 | 0 | 0 | 0 | 0 | 0 |
| Aug5.5v_17227 | 1 | 0 | 0 | 0 | 0 | 0 | 0 | 0 | 0 | 0 | 0 | 0 | 0 | 0 | 0 | 0 | 0 | 0 | 0 | 0 | 0 | 0 | 1 | 0 | 0 | 0 | 0 | 0 | 0 | 0 |
| Jul27.5v_6357 | 1 | 0 | 0 | 0 | 0 | 0 | 0 | 0 | 0 | 0 | 0 | 0 | 0 | 0 | 0 | 0 | 0 | 1 | 0 | 0 | 0 | 0 | 0 | 0 | 0 | 0 | 0 | 0 | 0 | 0 |
| Aug5.0v_2990 | 1 | 0 | 0 | 0 | 0 | 0 | 0 | 0 | 0 | 0 | 0 | 0 | 0 | 0 | 0 | 0 | 0 | 0 | 0 | 0 | 0 | 1 | 0 | 0 | 0 | 0 | 0 | 0 | 0 | 0 |
| Aug5.25v_36100 | 1 | 0 | 0 | 0 | 0 | 0 | 0 | 0 | 0 | 0 | 0 | 0 | 0 | 0 | 0 | 0 | 0 | 0 | 0 | 0 | 0 | 0 | 0 | 1 | 0 | 0 | 0 | 0 | 0 | 0 |
| Jul27.25v_28250 | 1 | 0 | 0 | 0 | 0 | 0 | 0 | 0 | 0 | 0 | 0 | 0 | 0 | 0 | 0 | 0 | 0 | 0 | 1 | 0 | 0 | 0 | 0 | 0 | 0 | 0 | 0 | 0 | 0 | 0 |
| Jul27.50v_10491 | 1 | 0 | 0 | 0 | 0 | 0 | 0 | 0 | 0 | 0 | 0 | 0 | 0 | 0 | 0 | 0 | 0 | 0 | 0 | 1 | 0 | 0 | 0 | 0 | 0 | 0 | 0 | 0 | 0 | 0 |
| Jul11.5v_12056 | 1 | 0 | 0 | 0 | 0 | 0 | 0 | 0 | 0 | 0 | 0 | 0 | 0 | 1 | 0 | 0 | 0 | 0 | 0 | 0 | 0 | 0 | 0 | 0 | 0 | 0 | 0 | 0 | 0 | 0 |
| Jul11.0v_7341 | 57 | 0 | 1 | 2 | 1 | 2 | 6 | 2 | 1 | 4 | 2 | 4 | 2 | 6 | 2 | 1 | 2 | 2 | 1 | 3 | 0 | 0 | 1 | 1 | 1 | 0 | 0 | 7 | 0 | 3 |
| Jul11.50v_14675 | 1 | 0 | 0 | 0 | 0 | 0 | 0 | 0 | 0 | 0 | 0 | 0 | 0 | 0 | 0 | 1 | 0 | 0 | 0 | 0 | 0 | 0 | 0 | 0 | 0 | 0 | 0 | 0 | 0 | 0 |
| Jun22.75v_23848 | 114 | 6 | 6 | 5 | 2 | 2 | 15 | 4 | 3 | 2 | 6 | 9 | 0 | 4 | 1 | 6 | 7 | 1 | 3 | 3 | 1 | 0 | 3 | 3 | 2 | 3 | 1 | 6 | 3 | 7 |
| Aug5.25v_38784 | 16 | 0 | 0 | 0 | 0 | 0 | 1 | 0 | 0 | 1 | 2 | 0 | 1 | 3 | 1 | 1 | 0 | 1 | 0 | 1 | 0 | 0 | 2 | 0 | 1 | 0 | 1 | 0 | 0 | 0 |
| Jul11.50v_9436 | 20 | 0 | 1 | 2 | 1 | 0 | 2 | 0 | 1 | 1 | 3 | 0 | 1 | 0 | 1 | 1 | 0 | 1 | 0 | 1 | 0 | 0 | 1 | 0 | 0 | 0 | 1 | 1 | 0 | 1 |
| Jul11.50v_9430 | 1 | 0 | 0 | 0 | 0 | 0 | 0 | 0 | 0 | 0 | 0 | 0 | 0 | 0 | 0 | 1 | 0 | 0 | 0 | 0 | 0 | 0 | 0 | 0 | 0 | 0 | 0 | 0 | 0 | 0 |
| Aug5.50v_13703 | 1178 | 40 | 39 | 82 | 43 | 28 | 176 | 26 | 22 | 21 | 46 | 65 | 11 | 45 | 31 | 47 | 92 | 42 | 29 | 37 | 17 | 21 | 24 | 21 | 24 | 21 | 8 | 47 | 6 | 67 |
| Jun8.50v_12898 | 20 | 1 | 0 | 2 | 1 | 1 | 4 | 0 | 0 | 1 | 1 | 2 | 0 | 0 | 0 | 0 | 1 | 0 | 1 | 1 | 0 | 0 | 0 | 1 | 1 | 0 | 0 | 1 | 0 | 1 |
| Jul27.25v_32053 | 5 | 1 | 0 | 0 | 0 | 0 | 1 | 0 | 0 | 0 | 0 | 0 | 0 | 0 | 0 | 0 | 1 | 0 | 1 | 0 | 0 | 0 | 0 | 0 | 0 | 0 | 0 | 0 | 1 | 0 |
| Jun22.75v_26866 | 3 | 1 | 0 | 0 | 0 | 1 | 0 | 0 | 0 | 0 | 0 | 1 | 0 | 0 | 0 | 0 | 0 | 0 | 0 | 0 | 0 | 0 | 0 | 0 | 0 | 0 | 0 | 0 | 0 | 0 |
| Jun8.25v_10218 | 9 | 0 | 0 | 2 | 0 | 1 | 2 | 0 | 0 | 0 | 0 | 1 | 0 | 0 | 0 | 0 | 1 | 0 | 0 | 1 | 0 | 1 | 0 | 0 | 0 | 0 | 0 | 0 | 0 | 0 |
| Aug5.75v_34284 | 2 | 0 | 0 | 0 | 0 | 1 | 0 | 0 | 0 | 0 | 0 | 0 | 0 | 0 | 0 | 0 | 0 | 0 | 0 | 0 | 0 | 0 | 0 | 0 | 0 | 1 | 0 | 0 | 0 | 0 |
| Aug5.75v_22191 | 8 | 0 | 1 | 0 | 0 | 1 | 1 | 0 | 0 | 1 | 0 | 1 | 0 | 0 | 0 | 0 | 1 | 0 | 0 | 0 | 0 | 0 | 0 | 0 | 0 | 1 | 0 | 1 | 0 | 0 |
| Jun8.50v_10897 | 10 | 1 | 0 | 0 | 1 | 0 | 2 | 0 | 0 | 0 | 0 | 2 | 0 | 0 | 0 | 0 | 1 | 0 | 0 | 0 | 0 | 1 | 0 | 1 | 0 | 1 | 0 | 0 | 0 | 0 |
| Aug5.50v_1415 | 1 | 0 | 0 | 0 | 0 | 0 | 0 | 0 | 0 | 0 | 0 | 0 | 0 | 0 | 0 | 0 | 0 | 0 | 0 | 0 | 0 | 0 | 0 | 0 | 1 | 0 | 0 | 0 | 0 | 0 |
| May12.50v_31025 | 4 | 0 | 0 | 2 | 0 | 0 | 0 | 0 | 0 | 2 | 0 | 0 | 0 | 0 | 0 | 0 | 0 | 0 | 0 | 0 | 0 | 0 | 0 | 0 | 0 | 0 | 0 | 0 | 0 | 0 |
| Jul27.5v_5266 | 3 | 0 | 0 | 0 | 0 | 0 | 1 | 0 | 0 | 0 | 0 | 0 | 0 | 0 | 0 | 1 | 0 | 1 | 0 | 0 | 0 | 0 | 0 | 0 | 0 | 0 | 0 | 0 | 0 | 0 |
| Jul11.5v_2894 | 3 | 0 | 0 | 0 | 0 | 0 | 0 | 0 | 0 | 0 | 0 | 0 | 0 | 1 | 0 | 0 | 0 | 0 | 1 | 0 | 1 | 0 | 0 | 0 | 0 | 0 | 0 | 0 | 0 | 0 |
| Jun22.50v_40894 | 2 | 0 | 0 | 0 | 0 | 0 | 0 | 0 | 0 | 0 | 1 | 1 | 0 | 0 | 0 | 0 | 0 | 0 | 0 | 0 | 0 | 0 | 0 | 0 | 0 | 0 | 0 | 0 | 0 | 0 |
| Jun22.50v_8372 | 8 | 0 | 0 | 0 | 0 | 1 | 1 | 0 | 0 | 0 | 1 | 1 | 0 | 0 | 0 | 0 | 1 | 0 | 0 | 0 | 0 | 0 | 0 | 1 | 0 | 0 | 0 | 1 | 1 | 0 |
| Jul11.50v_7997 | 1 | 0 | 0 | 0 | 0 | 0 | 0 | 0 | 0 | 0 | 0 | 0 | 0 | 0 | 0 | 1 | 0 | 0 | 0 | 0 | 0 | 0 | 0 | 0 | 0 | 0 | 0 | 0 | 0 | 0 |
| Jul11.25v_8557 | 1 | 0 | 0 | 0 | 0 | 0 | 0 | 0 | 0 | 0 | 0 | 0 | 0 | 0 | 1 | 0 | 0 | 0 | 0 | 0 | 0 | 0 | 0 | 0 | 0 | 0 | 0 | 0 | 0 | 0 |
| Jul11.25v_16022 | 1 | 0 | 0 | 0 | 0 | 0 | 0 | 0 | 0 | 0 | 0 | 0 | 0 | 0 | 1 | 0 | 0 | 0 | 0 | 0 | 0 | 0 | 0 | 0 | 0 | 0 | 0 | 0 | 0 | 0 |
| Jul11.5v_4670 | 1 | 0 | 0 | 0 | 0 | 0 | 0 | 0 | 0 | 0 | 0 | 0 | 0 | 1 | 0 | 0 | 0 | 0 | 0 | 0 | 0 | 0 | 0 | 0 | 0 | 0 | 0 | 0 | 0 | 0 |
| Jul27.25v_25782 | 1 | 0 | 0 | 0 | 0 | 0 | 0 | 0 | 0 | 0 | 0 | 0 | 0 | 0 | 0 | 0 | 0 | 0 | 1 | 0 | 0 | 0 | 0 | 0 | 0 | 0 | 0 | 0 | 0 | 0 |
| Jun8.50v_5417 | 1 | 0 | 0 | 0 | 0 | 0 | 1 | 0 | 0 | 0 | 0 | 0 | 0 | 0 | 0 | 0 | 0 | 0 | 0 | 0 | 0 | 0 | 0 | 0 | 0 | 0 | 0 | 0 | 0 | 0 |
| Jul27.75v_3620 | 1 | 0 | 0 | 0 | 0 | 0 | 0 | 0 | 0 | 0 | 0 | 0 | 0 | 0 | 0 | 0 | 0 | 0 | 0 | 0 | 1 | 0 | 0 | 0 | 0 | 0 | 0 | 0 | 0 | 0 |
| Jun22.75v_37263 | 1 | 0 | 0 | 0 | 0 | 0 | 0 | 0 | 0 | 0 | 0 | 1 | 0 | 0 | 0 | 0 | 0 | 0 | 0 | 0 | 0 | 0 | 0 | 0 | 0 | 0 | 0 | 0 | 0 | 0 |
| Jul11.5v_19033 | 1 | 0 | 0 | 0 | 0 | 0 | 0 | 0 | 0 | 0 | 0 | 0 | 0 | 1 | 0 | 0 | 0 | 0 | 0 | 0 | 0 | 0 | 0 | 0 | 0 | 0 | 0 | 0 | 0 | 0 |
| Jul27.50v_14123 | 1 | 0 | 0 | 0 | 0 | 0 | 0 | 0 | 0 | 0 | 0 | 0 | 0 | 0 | 0 | 0 | 0 | 0 | 0 | 1 | 0 | 0 | 0 | 0 | 0 | 0 | 0 | 0 | 0 | 0 |
| May12.75v_19401 | 1 | 0 | 0 | 0 | 1 | 0 | 0 | 0 | 0 | 0 | 0 | 0 | 0 | 0 | 0 | 0 | 0 | 0 | 0 | 0 | 0 | 0 | 0 | 0 | 0 | 0 | 0 | 0 | 0 | 0 |
| Aug5.25v_27480 | 1 | 0 | 0 | 0 | 0 | 0 | 0 | 0 | 0 | 0 | 0 | 0 | 0 | 0 | 0 | 0 | 0 | 0 | 0 | 0 | 0 | 0 | 0 | 1 | 0 | 0 | 0 | 0 | 0 | 0 |
| Jul11.25v_19730 | 1 | 0 | 0 | 0 | 0 | 0 | 0 | 0 | 0 | 0 | 0 | 0 | 0 | 0 | 1 | 0 | 0 | 0 | 0 | 0 | 0 | 0 | 0 | 0 | 0 | 0 | 0 | 0 | 0 | 0 |
| Jul11.50v_19933 | 1 | 0 | 0 | 0 | 0 | 0 | 0 | 0 | 0 | 0 | 0 | 0 | 0 | 0 | 0 | 1 | 0 | 0 | 0 | 0 | 0 | 0 | 0 | 0 | 0 | 0 | 0 | 0 | 0 | 0 |
| Aug5.0v_3240 | 3 | 0 | 0 | 0 | 0 | 0 | 0 | 0 | 0 | 0 | 0 | 0 | 0 | 0 | 0 | 0 | 1 | 0 | 0 | 0 | 0 | 1 | 0 | 0 | 0 | 0 | 0 | 1 | 0 | 0 |
| May12.50v_3436 | 6 | 0 | 0 | 1 | 0 | 2 | 1 | 0 | 0 | 0 | 1 | 0 | 0 | 1 | 0 | 0 | 0 | 0 | 0 | 0 | 0 | 0 | 0 | 0 | 0 | 0 | 0 | 0 | 0 | 0 |
| Jun22.50v_38199 | 1 | 0 | 0 | 0 | 0 | 0 | 0 | 0 | 0 | 0 | 1 | 0 | 0 | 0 | 0 | 0 | 0 | 0 | 0 | 0 | 0 | 0 | 0 | 0 | 0 | 0 | 0 | 0 | 0 | 0 |
| Jun22.5v_17504 | 1 | 0 | 0 | 0 | 0 | 0 | 0 | 0 | 0 | 1 | 0 | 0 | 0 | 0 | 0 | 0 | 0 | 0 | 0 | 0 | 0 | 0 | 0 | 0 | 0 | 0 | 0 | 0 | 0 | 0 |
| Jul11.75v_27674 | 1 | 0 | 0 | 0 | 0 | 0 | 0 | 0 | 0 | 0 | 0 | 0 | 0 | 0 | 0 | 0 | 1 | 0 | 0 | 0 | 0 | 0 | 0 | 0 | 0 | 0 | 0 | 0 | 0 | 0 |
| Jun22.75v_16724 | 1 | 0 | 0 | 0 | 0 | 0 | 0 | 0 | 0 | 0 | 0 | 1 | 0 | 0 | 0 | 0 | 0 | 0 | 0 | 0 | 0 | 0 | 0 | 0 | 0 | 0 | 0 | 0 | 0 | 0 |
| Jun22.5v_3821 | 2 | 0 | 1 | 0 | 0 | 0 | 0 | 0 | 0 | 1 | 0 | 0 | 0 | 0 | 0 | 0 | 0 | 0 | 0 | 0 | 0 | 0 | 0 | 0 | 0 | 0 | 0 | 0 | 0 | 0 |
| Jun8.25v_16614 | 1 | 0 | 0 | 0 | 0 | 1 | 0 | 0 | 0 | 0 | 0 | 0 | 0 | 0 | 0 | 0 | 0 | 0 | 0 | 0 | 0 | 0 | 0 | 0 | 0 | 0 | 0 | 0 | 0 | 0 |
| May12.5v_28504 | 1 | 1 | 0 | 0 | 0 | 0 | 0 | 0 | 0 | 0 | 0 | 0 | 0 | 0 | 0 | 0 | 0 | 0 | 0 | 0 | 0 | 0 | 0 | 0 | 0 | 0 | 0 | 0 | 0 | 0 |
| Jun8.50v_39951 | 1 | 0 | 0 | 0 | 0 | 0 | 1 | 0 | 0 | 0 | 0 | 0 | 0 | 0 | 0 | 0 | 0 | 0 | 0 | 0 | 0 | 0 | 0 | 0 | 0 | 0 | 0 | 0 | 0 | 0 |
| Jun8.50v_14664 | 3 | 0 | 0 | 0 | 0 | 0 | 2 | 0 | 0 | 0 | 0 | 0 | 0 | 0 | 0 | 1 | 0 | 0 | 0 | 0 | 0 | 0 | 0 | 0 | 0 | 0 | 0 | 0 | 0 | 0 |
| May12.50v_14301 | 1 | 0 | 0 | 1 | 0 | 0 | 0 | 0 | 0 | 0 | 0 | 0 | 0 | 0 | 0 | 0 | 0 | 0 | 0 | 0 | 0 | 0 | 0 | 0 | 0 | 0 | 0 | 0 | 0 | 0 |
| Jul11.50v_7237 | 1 | 0 | 0 | 0 | 0 | 0 | 0 | 0 | 0 | 0 | 0 | 0 | 0 | 0 | 0 | 1 | 0 | 0 | 0 | 0 | 0 | 0 | 0 | 0 | 0 | 0 | 0 | 0 | 0 | 0 |
| Jul27.75v_3897 | 1 | 0 | 0 | 0 | 0 | 0 | 0 | 0 | 0 | 0 | 0 | 0 | 0 | 0 | 0 | 0 | 0 | 0 | 0 | 0 | 1 | 0 | 0 | 0 | 0 | 0 | 0 | 0 | 0 | 0 |
| Jun22.75v_40555 | 2 | 0 | 0 | 0 | 0 | 0 | 1 | 0 | 0 | 0 | 0 | 1 | 0 | 0 | 0 | 0 | 0 | 0 | 0 | 0 | 0 | 0 | 0 | 0 | 0 | 0 | 0 | 0 | 0 | 0 |
| Jun8.50v_16312 | 1 | 0 | 0 | 0 | 0 | 0 | 1 | 0 | 0 | 0 | 0 | 0 | 0 | 0 | 0 | 0 | 0 | 0 | 0 | 0 | 0 | 0 | 0 | 0 | 0 | 0 | 0 | 0 | 0 | 0 |
| Jun22.50v_13684 | 1 | 0 | 0 | 0 | 0 | 0 | 0 | 0 | 0 | 0 | 1 | 0 | 0 | 0 | 0 | 0 | 0 | 0 | 0 | 0 | 0 | 0 | 0 | 0 | 0 | 0 | 0 | 0 | 0 | 0 |
| May12.5v_32445 | 2 | 1 | 0 | 0 | 0 | 0 | 0 | 0 | 0 | 0 | 0 | 0 | 0 | 0 | 0 | 0 | 0 | 0 | 0 | 0 | 0 | 0 | 0 | 0 | 0 | 0 | 1 | 0 | 0 | 0 |
| Jun22.75v_17325 | 3 | 0 | 0 | 0 | 1 | 0 | 1 | 0 | 0 | 0 | 0 | 1 | 0 | 0 | 0 | 0 | 0 | 0 | 0 | 0 | 0 | 0 | 0 | 0 | 0 | 0 | 0 | 0 | 0 | 0 |
| Sep8.50v_21840 | 1 | 0 | 0 | 0 | 0 | 0 | 0 | 0 | 0 | 0 | 0 | 0 | 0 | 0 | 0 | 0 | 0 | 0 | 0 | 0 | 0 | 0 | 0 | 0 | 0 | 0 | 0 | 0 | 0 | 1 |
| Jun22.75v_27788 | 1 | 0 | 0 | 0 | 0 | 0 | 0 | 0 | 0 | 0 | 0 | 1 | 0 | 0 | 0 | 0 | 0 | 0 | 0 | 0 | 0 | 0 | 0 | 0 | 0 | 0 | 0 | 0 | 0 | 0 |
| Jul11.75v_230 | 1 | 0 | 0 | 0 | 0 | 0 | 0 | 0 | 0 | 0 | 0 | 0 | 0 | 0 | 0 | 0 | 1 | 0 | 0 | 0 | 0 | 0 | 0 | 0 | 0 | 0 | 0 | 0 | 0 | 0 |
| Jun22.50v_38892 | 1 | 0 | 0 | 0 | 0 | 0 | 0 | 0 | 0 | 0 | 1 | 0 | 0 | 0 | 0 | 0 | 0 | 0 | 0 | 0 | 0 | 0 | 0 | 0 | 0 | 0 | 0 | 0 | 0 | 0 |
| Jun22.0v_12049 | 2 | 0 | 0 | 0 | 0 | 0 | 0 | 0 | 1 | 0 | 0 | 0 | 0 | 0 | 0 | 0 | 0 | 0 | 0 | 0 | 0 | 0 | 0 | 0 | 0 | 0 | 0 | 1 | 0 | 0 |
| May12.25v_31200 | 1 | 0 | 1 | 0 | 0 | 0 | 0 | 0 | 0 | 0 | 0 | 0 | 0 | 0 | 0 | 0 | 0 | 0 | 0 | 0 | 0 | 0 | 0 | 0 | 0 | 0 | 0 | 0 | 0 | 0 |
| May12.75v_36161 | 3 | 1 | 0 | 0 | 1 | 0 | 0 | 0 | 0 | 0 | 0 | 1 | 0 | 0 | 0 | 0 | 0 | 0 | 0 | 0 | 0 | 0 | 0 | 0 | 0 | 0 | 0 | 0 | 0 | 0 |
| Jun8.50v_3759 | 1 | 0 | 0 | 0 | 0 | 0 | 1 | 0 | 0 | 0 | 0 | 0 | 0 | 0 | 0 | 0 | 0 | 0 | 0 | 0 | 0 | 0 | 0 | 0 | 0 | 0 | 0 | 0 | 0 | 0 |
| May12.5v_25221 | 1 | 1 | 0 | 0 | 0 | 0 | 0 | 0 | 0 | 0 | 0 | 0 | 0 | 0 | 0 | 0 | 0 | 0 | 0 | 0 | 0 | 0 | 0 | 0 | 0 | 0 | 0 | 0 | 0 | 0 |
| Jul11.75v_38785 | 1 | 0 | 0 | 0 | 0 | 0 | 0 | 0 | 0 | 0 | 0 | 0 | 0 | 0 | 0 | 0 | 1 | 0 | 0 | 0 | 0 | 0 | 0 | 0 | 0 | 0 | 0 | 0 | 0 | 0 |
| Jun8.75v_11582 | 1 | 0 | 0 | 0 | 0 | 0 | 0 | 1 | 0 | 0 | 0 | 0 | 0 | 0 | 0 | 0 | 0 | 0 | 0 | 0 | 0 | 0 | 0 | 0 | 0 | 0 | 0 | 0 | 0 | 0 |
| Jul11.25v_15572 | 1 | 0 | 0 | 0 | 0 | 0 | 0 | 0 | 0 | 0 | 0 | 0 | 0 | 0 | 1 | 0 | 0 | 0 | 0 | 0 | 0 | 0 | 0 | 0 | 0 | 0 | 0 | 0 | 0 | 0 |
| Jun8.50v_23587 | 1 | 0 | 0 | 0 | 0 | 0 | 1 | 0 | 0 | 0 | 0 | 0 | 0 | 0 | 0 | 0 | 0 | 0 | 0 | 0 | 0 | 0 | 0 | 0 | 0 | 0 | 0 | 0 | 0 | 0 |
| Sep8.50v_3182 | 5 | 0 | 0 | 1 | 0 | 0 | 1 | 0 | 0 | 0 | 0 | 0 | 0 | 0 | 0 | 0 | 1 | 0 | 0 | 0 | 0 | 1 | 0 | 0 | 0 | 0 | 0 | 0 | 0 | 1 |
| Jul27.50v_38659 | 1 | 0 | 0 | 0 | 0 | 0 | 0 | 0 | 0 | 0 | 0 | 0 | 0 | 0 | 0 | 0 | 0 | 0 | 0 | 1 | 0 | 0 | 0 | 0 | 0 | 0 | 0 | 0 | 0 | 0 |
| Jun22.75v_20600 | 1 | 0 | 0 | 0 | 0 | 0 | 0 | 0 | 0 | 0 | 0 | 1 | 0 | 0 | 0 | 0 | 0 | 0 | 0 | 0 | 0 | 0 | 0 | 0 | 0 | 0 | 0 | 0 | 0 | 0 |
| Jul11.75v_27949 | 1 | 0 | 0 | 0 | 0 | 0 | 0 | 0 | 0 | 0 | 0 | 0 | 0 | 0 | 0 | 0 | 1 | 0 | 0 | 0 | 0 | 0 | 0 | 0 | 0 | 0 | 0 | 0 | 0 | 0 |
| Jun22.5v_20131 | 1 | 0 | 0 | 0 | 0 | 0 | 0 | 0 | 0 | 1 | 0 | 0 | 0 | 0 | 0 | 0 | 0 | 0 | 0 | 0 | 0 | 0 | 0 | 0 | 0 | 0 | 0 | 0 | 0 | 0 |
| Jun22.5v_13612 | 3 | 0 | 0 | 0 | 0 | 0 | 1 | 0 | 0 | 1 | 0 | 1 | 0 | 0 | 0 | 0 | 0 | 0 | 0 | 0 | 0 | 0 | 0 | 0 | 0 | 0 | 0 | 0 | 0 | 0 |
| Jul11.50v_18882 | 6 | 0 | 0 | 1 | 0 | 0 | 2 | 0 | 0 | 0 | 0 | 0 | 1 | 0 | 0 | 1 | 0 | 0 | 0 | 0 | 0 | 0 | 0 | 0 | 0 | 1 | 0 | 0 | 0 | 0 |
| Sep8.5v_11546 | 1 | 0 | 0 | 0 | 0 | 0 | 0 | 0 | 0 | 0 | 0 | 0 | 0 | 0 | 0 | 0 | 0 | 0 | 0 | 0 | 0 | 0 | 0 | 0 | 0 | 0 | 0 | 1 | 0 | 0 |
| Jul27.75v_19318 | 2 | 0 | 1 | 0 | 0 | 0 | 0 | 0 | 0 | 0 | 0 | 0 | 0 | 0 | 0 | 0 | 0 | 0 | 0 | 0 | 1 | 0 | 0 | 0 | 0 | 0 | 0 | 0 | 0 | 0 |
| Jul27.5v_8922 | 1 | 0 | 0 | 0 | 0 | 0 | 0 | 0 | 0 | 0 | 0 | 0 | 0 | 0 | 0 | 0 | 0 | 1 | 0 | 0 | 0 | 0 | 0 | 0 | 0 | 0 | 0 | 0 | 0 | 0 |
| Jun22.75v_3511 | 1 | 0 | 0 | 0 | 0 | 0 | 0 | 0 | 0 | 0 | 0 | 1 | 0 | 0 | 0 | 0 | 0 | 0 | 0 | 0 | 0 | 0 | 0 | 0 | 0 | 0 | 0 | 0 | 0 | 0 |
| Jun22.5v_5977 | 1 | 0 | 0 | 0 | 0 | 0 | 0 | 0 | 0 | 1 | 0 | 0 | 0 | 0 | 0 | 0 | 0 | 0 | 0 | 0 | 0 | 0 | 0 | 0 | 0 | 0 | 0 | 0 | 0 | 0 |
| Jun8.25v_1416 | 4 | 0 | 0 | 0 | 0 | 1 | 0 | 0 | 0 | 0 | 0 | 2 | 0 | 0 | 0 | 0 | 0 | 0 | 0 | 1 | 0 | 0 | 0 | 0 | 0 | 0 | 0 | 0 | 0 | 0 |
| May12.25v_28535 | 16 | 0 | 1 | 1 | 1 | 0 | 1 | 0 | 1 | 0 | 0 | 1 | 0 | 3 | 1 | 1 | 1 | 1 | 0 | 0 | 0 | 0 | 0 | 0 | 0 | 0 | 0 | 1 | 0 | 2 |
| Jun22.50v_27550 | 1 | 0 | 0 | 0 | 0 | 0 | 0 | 0 | 0 | 0 | 1 | 0 | 0 | 0 | 0 | 0 | 0 | 0 | 0 | 0 | 0 | 0 | 0 | 0 | 0 | 0 | 0 | 0 | 0 | 0 |
| Sep8.5v_5681 | 1 | 0 | 0 | 0 | 0 | 0 | 0 | 0 | 0 | 0 | 0 | 0 | 0 | 0 | 0 | 0 | 0 | 0 | 0 | 0 | 0 | 0 | 0 | 0 | 0 | 0 | 0 | 1 | 0 | 0 |
| Sep8.25v_18280 | 1 | 0 | 0 | 0 | 0 | 0 | 0 | 0 | 0 | 0 | 0 | 0 | 0 | 0 | 0 | 0 | 0 | 0 | 0 | 0 | 0 | 0 | 0 | 0 | 0 | 0 | 0 | 0 | 1 | 0 |
| Jul11.75v_38619 | 3 | 0 | 0 | 0 | 0 | 0 | 0 | 1 | 0 | 0 | 1 | 0 | 0 | 0 | 0 | 0 | 1 | 0 | 0 | 0 | 0 | 0 | 0 | 0 | 0 | 0 | 0 | 0 | 0 | 0 |
| Jun8.50v_37882 | 1 | 0 | 0 | 0 | 0 | 0 | 1 | 0 | 0 | 0 | 0 | 0 | 0 | 0 | 0 | 0 | 0 | 0 | 0 | 0 | 0 | 0 | 0 | 0 | 0 | 0 | 0 | 0 | 0 | 0 |
| Aug5.5v_1620 | 1 | 0 | 0 | 0 | 0 | 0 | 0 | 0 | 0 | 0 | 0 | 0 | 0 | 0 | 0 | 0 | 0 | 0 | 0 | 0 | 0 | 0 | 1 | 0 | 0 | 0 | 0 | 0 | 0 | 0 |
| Jun22.50v_25362 | 1 | 0 | 0 | 0 | 0 | 0 | 0 | 0 | 0 | 0 | 1 | 0 | 0 | 0 | 0 | 0 | 0 | 0 | 0 | 0 | 0 | 0 | 0 | 0 | 0 | 0 | 0 | 0 | 0 | 0 |
| Aug5.5v_6969 | 1 | 0 | 0 | 0 | 0 | 0 | 0 | 0 | 0 | 0 | 0 | 0 | 0 | 0 | 0 | 0 | 0 | 0 | 0 | 0 | 0 | 0 | 1 | 0 | 0 | 0 | 0 | 0 | 0 | 0 |
| Jun22.75v_25721 | 1 | 0 | 0 | 0 | 0 | 0 | 0 | 0 | 0 | 0 | 0 | 1 | 0 | 0 | 0 | 0 | 0 | 0 | 0 | 0 | 0 | 0 | 0 | 0 | 0 | 0 | 0 | 0 | 0 | 0 |
| Jun22.50v_39859 | 1 | 0 | 0 | 0 | 0 | 0 | 0 | 0 | 0 | 0 | 1 | 0 | 0 | 0 | 0 | 0 | 0 | 0 | 0 | 0 | 0 | 0 | 0 | 0 | 0 | 0 | 0 | 0 | 0 | 0 |
| Aug5.25v_40325 | 1 | 0 | 0 | 0 | 0 | 0 | 0 | 0 | 0 | 0 | 0 | 0 | 0 | 0 | 0 | 0 | 0 | 0 | 0 | 0 | 0 | 0 | 0 | 1 | 0 | 0 | 0 | 0 | 0 | 0 |
| Jul11.25v_13802 | 1 | 0 | 0 | 0 | 0 | 0 | 0 | 0 | 0 | 0 | 0 | 0 | 0 | 0 | 1 | 0 | 0 | 0 | 0 | 0 | 0 | 0 | 0 | 0 | 0 | 0 | 0 | 0 | 0 | 0 |
| Aug5.0v_3017 | 34 | 1 | 0 | 0 | 1 | 1 | 2 | 1 | 0 | 2 | 1 | 1 | 1 | 5 | 2 | 1 | 6 | 0 | 2 | 0 | 0 | 2 | 0 | 0 | 0 | 2 | 0 | 0 | 0 | 3 |
| Aug5.0v_4813 | 1 | 0 | 0 | 0 | 0 | 0 | 0 | 0 | 0 | 0 | 0 | 0 | 0 | 0 | 0 | 0 | 0 | 0 | 0 | 0 | 0 | 1 | 0 | 0 | 0 | 0 | 0 | 0 | 0 | 0 |
| Jul11.5v_4917 | 1 | 0 | 0 | 0 | 0 | 0 | 0 | 0 | 0 | 0 | 0 | 0 | 0 | 1 | 0 | 0 | 0 | 0 | 0 | 0 | 0 | 0 | 0 | 0 | 0 | 0 | 0 | 0 | 0 | 0 |
| Jul11.5v_673 | 1 | 0 | 0 | 0 | 0 | 0 | 0 | 0 | 0 | 0 | 0 | 0 | 0 | 1 | 0 | 0 | 0 | 0 | 0 | 0 | 0 | 0 | 0 | 0 | 0 | 0 | 0 | 0 | 0 | 0 |
| Jul11.5v_8136 | 1 | 0 | 0 | 0 | 0 | 0 | 0 | 0 | 0 | 0 | 0 | 0 | 0 | 1 | 0 | 0 | 0 | 0 | 0 | 0 | 0 | 0 | 0 | 0 | 0 | 0 | 0 | 0 | 0 | 0 |
| Jul11.50v_18378 | 1 | 0 | 0 | 0 | 0 | 0 | 0 | 0 | 0 | 0 | 0 | 0 | 0 | 0 | 0 | 1 | 0 | 0 | 0 | 0 | 0 | 0 | 0 | 0 | 0 | 0 | 0 | 0 | 0 | 0 |
| Aug5.0v_4098 | 1 | 0 | 0 | 0 | 0 | 0 | 0 | 0 | 0 | 0 | 0 | 0 | 0 | 0 | 0 | 0 | 0 | 0 | 0 | 0 | 0 | 1 | 0 | 0 | 0 | 0 | 0 | 0 | 0 | 0 |
| Sep8.50v_38749 | 1 | 0 | 0 | 0 | 0 | 0 | 0 | 0 | 0 | 0 | 0 | 0 | 0 | 0 | 0 | 0 | 0 | 0 | 0 | 0 | 0 | 0 | 0 | 0 | 0 | 0 | 0 | 0 | 0 | 1 |
| Jul11.25v_1142 | 1 | 0 | 0 | 0 | 0 | 0 | 0 | 0 | 0 | 0 | 0 | 0 | 0 | 0 | 1 | 0 | 0 | 0 | 0 | 0 | 0 | 0 | 0 | 0 | 0 | 0 | 0 | 0 | 0 | 0 |
| Sep8.0v_13962 | 1 | 0 | 0 | 0 | 0 | 0 | 0 | 0 | 0 | 0 | 0 | 0 | 0 | 0 | 0 | 0 | 0 | 0 | 0 | 0 | 0 | 0 | 0 | 0 | 0 | 0 | 1 | 0 | 0 | 0 |
| May12.50v_1873 | 1 | 0 | 0 | 1 | 0 | 0 | 0 | 0 | 0 | 0 | 0 | 0 | 0 | 0 | 0 | 0 | 0 | 0 | 0 | 0 | 0 | 0 | 0 | 0 | 0 | 0 | 0 | 0 | 0 | 0 |
| Jul27.50v_14719 | 1 | 0 | 0 | 0 | 0 | 0 | 0 | 0 | 0 | 0 | 0 | 0 | 0 | 0 | 0 | 0 | 0 | 0 | 0 | 1 | 0 | 0 | 0 | 0 | 0 | 0 | 0 | 0 | 0 | 0 |
| Jul11.75v_19905 | 1 | 0 | 0 | 0 | 0 | 0 | 0 | 0 | 0 | 0 | 0 | 0 | 0 | 0 | 0 | 0 | 1 | 0 | 0 | 0 | 0 | 0 | 0 | 0 | 0 | 0 | 0 | 0 | 0 | 0 |
| Aug5.0v_4275 | 4 | 1 | 0 | 0 | 1 | 0 | 0 | 0 | 0 | 0 | 0 | 1 | 0 | 0 | 0 | 0 | 0 | 0 | 0 | 0 | 0 | 1 | 0 | 0 | 0 | 0 | 0 | 0 | 0 | 0 |
| Jul11.25v_15786 | 1 | 0 | 0 | 0 | 0 | 0 | 0 | 0 | 0 | 0 | 0 | 0 | 0 | 0 | 1 | 0 | 0 | 0 | 0 | 0 | 0 | 0 | 0 | 0 | 0 | 0 | 0 | 0 | 0 | 0 |
| Jul11.5v_15098 | 1 | 0 | 0 | 0 | 0 | 0 | 0 | 0 | 0 | 0 | 0 | 0 | 0 | 1 | 0 | 0 | 0 | 0 | 0 | 0 | 0 | 0 | 0 | 0 | 0 | 0 | 0 | 0 | 0 | 0 |
| Jun8.25v_17246 | 1 | 0 | 0 | 0 | 0 | 1 | 0 | 0 | 0 | 0 | 0 | 0 | 0 | 0 | 0 | 0 | 0 | 0 | 0 | 0 | 0 | 0 | 0 | 0 | 0 | 0 | 0 | 0 | 0 | 0 |
| May12.75v_22093 | 1 | 0 | 0 | 0 | 1 | 0 | 0 | 0 | 0 | 0 | 0 | 0 | 0 | 0 | 0 | 0 | 0 | 0 | 0 | 0 | 0 | 0 | 0 | 0 | 0 | 0 | 0 | 0 | 0 | 0 |
| Jul11.75v_6442 | 9 | 0 | 1 | 0 | 0 | 1 | 1 | 1 | 0 | 0 | 0 | 0 | 0 | 0 | 0 | 0 | 1 | 0 | 1 | 1 | 0 | 0 | 0 | 0 | 0 | 0 | 0 | 0 | 0 | 2 |
| Jul11.0v_19001 | 1 | 0 | 0 | 0 | 0 | 0 | 0 | 0 | 0 | 0 | 0 | 0 | 1 | 0 | 0 | 0 | 0 | 0 | 0 | 0 | 0 | 0 | 0 | 0 | 0 | 0 | 0 | 0 | 0 | 0 |
| May12.5v_34551 | 1 | 1 | 0 | 0 | 0 | 0 | 0 | 0 | 0 | 0 | 0 | 0 | 0 | 0 | 0 | 0 | 0 | 0 | 0 | 0 | 0 | 0 | 0 | 0 | 0 | 0 | 0 | 0 | 0 | 0 |
| Jul27.50v_14852 | 1 | 0 | 0 | 0 | 0 | 0 | 0 | 0 | 0 | 0 | 0 | 0 | 0 | 0 | 0 | 0 | 0 | 0 | 0 | 1 | 0 | 0 | 0 | 0 | 0 | 0 | 0 | 0 | 0 | 0 |
| May12.50v_36021 | 1 | 0 | 0 | 1 | 0 | 0 | 0 | 0 | 0 | 0 | 0 | 0 | 0 | 0 | 0 | 0 | 0 | 0 | 0 | 0 | 0 | 0 | 0 | 0 | 0 | 0 | 0 | 0 | 0 | 0 |
| Sep8.5v_10494 | 1 | 0 | 0 | 0 | 0 | 0 | 0 | 0 | 0 | 0 | 0 | 0 | 0 | 0 | 0 | 0 | 0 | 0 | 0 | 0 | 0 | 0 | 0 | 0 | 0 | 0 | 0 | 1 | 0 | 0 |
| Sep8.50v_11476 | 1 | 0 | 0 | 0 | 0 | 0 | 0 | 0 | 0 | 0 | 0 | 0 | 0 | 0 | 0 | 0 | 0 | 0 | 0 | 0 | 0 | 0 | 0 | 0 | 0 | 0 | 0 | 0 | 0 | 1 |
| Jun22.5v_12665 | 5 | 1 | 0 | 0 | 1 | 0 | 0 | 0 | 0 | 1 | 0 | 1 | 0 | 0 | 0 | 0 | 0 | 0 | 0 | 0 | 0 | 1 | 0 | 0 | 0 | 0 | 0 | 0 | 0 | 0 |
| Aug5.0v_1346 | 3 | 0 | 0 | 1 | 0 | 0 | 0 | 0 | 0 | 0 | 0 | 0 | 0 | 0 | 0 | 0 | 0 | 0 | 0 | 0 | 0 | 1 | 0 | 0 | 0 | 0 | 0 | 0 | 0 | 1 |
| Sep8.50v_20658 | 1 | 0 | 0 | 0 | 0 | 0 | 0 | 0 | 0 | 0 | 0 | 0 | 0 | 0 | 0 | 0 | 0 | 0 | 0 | 0 | 0 | 0 | 0 | 0 | 0 | 0 | 0 | 0 | 0 | 1 |
| Jul11.50v_8882 | 1 | 0 | 0 | 0 | 0 | 0 | 0 | 0 | 0 | 0 | 0 | 0 | 0 | 0 | 0 | 1 | 0 | 0 | 0 | 0 | 0 | 0 | 0 | 0 | 0 | 0 | 0 | 0 | 0 | 0 |
| Jun22.0v_7134 | 1 | 0 | 0 | 0 | 0 | 0 | 0 | 0 | 1 | 0 | 0 | 0 | 0 | 0 | 0 | 0 | 0 | 0 | 0 | 0 | 0 | 0 | 0 | 0 | 0 | 0 | 0 | 0 | 0 | 0 |
| Jul11.75v_35753 | 1 | 0 | 0 | 0 | 0 | 0 | 0 | 0 | 0 | 0 | 0 | 0 | 0 | 0 | 0 | 0 | 1 | 0 | 0 | 0 | 0 | 0 | 0 | 0 | 0 | 0 | 0 | 0 | 0 | 0 |
| May12.50v_4323 | 105 | 5 | 5 | 5 | 10 | 1 | 2 | 1 | 7 | 7 | 10 | 1 | 6 | 5 | 2 | 4 | 0 | 8 | 2 | 3 | 1 | 1 | 1 | 1 | 1 | 1 | 9 | 3 | 2 | 1 |

| Feature_ID | 654 | May12_05m | May12_25m | May12_50m | May12_75m | Jun8_25m | Jun8_50m | Jun8_75m | Jun22_0m | Jun22_05m | Jun22_50m | Jun22_75m | Jul11_0m | Jul11_05m | Jul11_25m | Jul11_50m | Jul11_75m | Jul27_05m | Jul27_25m | Jul27_50m | Jul27_75m | Aug5_0m | Aug5_05m | Aug5_25m | Aug5_50m | Aug5_75m | Sep8_0m | Sep8_05m | Sep8_25m | Sep8_50m |
|---|---|---|---|---|---|---|---|---|---|---|---|---|---|---|---|---|---|---|---|---|---|---|---|---|---|---|---|---|---|---|
| Jul11.50v_5319 | 3 | 0 | 0 | 0 | 0 | 0 | 0 | 0 | 0 | 0 | 0 | 0 | 0 | 0 | 0 | 1 | 0 | 1 | 0 | 0 | 0 | 0 | 0 | 0 | 0 | 0 | 0 | 1 | 0 | 0 |
| Jun22.0v_11636 | 1 | 0 | 0 | 0 | 0 | 0 | 0 | 0 | 1 | 0 | 0 | 0 | 0 | 0 | 0 | 0 | 0 | 0 | 0 | 0 | 0 | 0 | 0 | 0 | 0 | 0 | 0 | 1 | 0 | 0 |
| Jul27.5v_10860 | 2 | 0 | 0 | 1 | 0 | 0 | 0 | 0 | 0 | 0 | 0 | 0 | 0 | 0 | 0 | 0 | 0 | 1 | 0 | 0 | 0 | 0 | 0 | 0 | 0 | 0 | 0 | 0 | 0 | 0 |
| Jun22.5v_6858 | 750 | 45 | 46 | 98 | 36 | 14 | 12 | 4 | 49 | 20 | 72 | 22 | 48 | 32 | 17 | 11 | 5 | 21 | 24 | 19 | 1 | 40 | 20 | 11 | 9 | 12 | 23 | 23 | 3 | 13 |
| Sep8.5v_20060 | 7 | 0 | 2 | 1 | 0 | 0 | 0 | 0 | 0 | 0 | 0 | 0 | 2 | 0 | 0 | 0 | 0 | 0 | 0 | 0 | 0 | 0 | 0 | 0 | 0 | 0 | 1 | 1 | 0 | 0 |
| Jun22.5v_956 | 291 | 14 | 22 | 28 | 12 | 5 | 12 | 0 | 14 | 9 | 31 | 5 | 20 | 11 | 3 | 4 | 2 | 10 | 9 | 9 | 0 | 17 | 9 | 7 | 3 | 7 | 9 | 9 | 1 | 9 |
| May12.75v_17110 | 3 | 0 | 0 | 0 | 1 | 0 | 0 | 0 | 1 | 0 | 1 | 0 | 0 | 0 | 0 | 0 | 0 | 0 | 0 | 0 | 0 | 0 | 0 | 0 | 0 | 0 | 0 | 0 | 0 | 0 |
| May12.75v_33730 | 1 | 0 | 0 | 0 | 1 | 0 | 0 | 0 | 0 | 0 | 0 | 0 | 0 | 0 | 0 | 0 | 0 | 0 | 0 | 0 | 0 | 0 | 0 | 0 | 0 | 0 | 0 | 0 | 0 | 0 |
| Jul11.5v_5665 | 1 | 0 | 0 | 0 | 0 | 0 | 0 | 0 | 0 | 0 | 0 | 0 | 0 | 1 | 0 | 0 | 0 | 0 | 0 | 0 | 0 | 0 | 0 | 0 | 0 | 0 | 0 | 0 | 0 | 0 |
| Aug5.5v_32288 | 1 | 0 | 0 | 0 | 0 | 0 | 0 | 0 | 0 | 0 | 0 | 0 | 0 | 0 | 0 | 0 | 0 | 0 | 0 | 0 | 0 | 0 | 1 | 0 | 0 | 0 | 0 | 0 | 0 | 0 |
| Jul27.75v_17339 | 1 | 0 | 0 | 0 | 0 | 0 | 0 | 0 | 0 | 0 | 0 | 0 | 0 | 0 | 0 | 0 | 0 | 0 | 0 | 0 | 1 | 0 | 0 | 0 | 0 | 0 | 0 | 0 | 0 | 0 |
| Jun22.50v_32309 | 1 | 0 | 0 | 0 | 0 | 0 | 0 | 0 | 0 | 0 | 1 | 0 | 0 | 0 | 0 | 0 | 0 | 0 | 0 | 0 | 0 | 0 | 0 | 0 | 0 | 0 | 0 | 0 | 0 | 0 |
| May12.50v_8304 | 3 | 0 | 0 | 1 | 0 | 0 | 0 | 0 | 0 | 0 | 0 | 0 | 0 | 1 | 0 | 0 | 0 | 0 | 0 | 0 | 0 | 1 | 0 | 0 | 0 | 0 | 0 | 0 | 0 | 0 |
| Jul11.0v_9287 | 1 | 0 | 0 | 0 | 0 | 0 | 0 | 0 | 0 | 0 | 0 | 0 | 1 | 0 | 0 | 0 | 0 | 0 | 0 | 0 | 0 | 0 | 0 | 0 | 0 | 0 | 0 | 0 | 0 | 0 |
| Jul11.25v_16658 | 1 | 0 | 0 | 0 | 0 | 0 | 0 | 0 | 0 | 0 | 0 | 0 | 0 | 0 | 1 | 0 | 0 | 0 | 0 | 0 | 0 | 0 | 0 | 0 | 0 | 0 | 0 | 0 | 0 | 0 |
| Jun22.0v_12875 | 1 | 0 | 0 | 0 | 0 | 0 | 0 | 0 | 1 | 0 | 0 | 0 | 0 | 0 | 0 | 0 | 0 | 0 | 0 | 0 | 0 | 0 | 0 | 0 | 0 | 0 | 0 | 0 | 0 | 0 |
| Jun22.75v_23913 | 5 | 0 | 0 | 0 | 0 | 0 | 0 | 0 | 1 | 0 | 1 | 1 | 0 | 0 | 0 | 0 | 1 | 0 | 0 | 0 | 0 | 0 | 1 | 0 | 0 | 0 | 0 | 0 | 0 | 0 |
| May12.25v_14665 | 1 | 0 | 1 | 0 | 0 | 0 | 0 | 0 | 0 | 0 | 0 | 0 | 0 | 0 | 0 | 0 | 0 | 0 | 0 | 0 | 0 | 0 | 0 | 0 | 0 | 0 | 0 | 0 | 0 | 0 |
| Jun8.50v_4198 | 6 | 0 | 0 | 1 | 0 | 0 | 1 | 0 | 1 | 0 | 0 | 0 | 0 | 2 | 0 | 0 | 0 | 0 | 0 | 0 | 0 | 0 | 0 | 0 | 0 | 1 | 0 | 0 | 0 | 0 |
| May12.5v_39905 | 3 | 1 | 0 | 0 | 0 | 0 | 0 | 0 | 0 | 0 | 0 | 0 | 0 | 0 | 0 | 0 | 0 | 0 | 0 | 0 | 0 | 1 | 0 | 0 | 0 | 0 | 0 | 1 | 0 | 0 |
| Sep8.0v_14349 | 1 | 0 | 0 | 0 | 0 | 0 | 0 | 0 | 0 | 0 | 0 | 0 | 0 | 0 | 0 | 0 | 0 | 0 | 0 | 0 | 0 | 0 | 0 | 0 | 0 | 0 | 1 | 0 | 0 | 0 |
| Jun22.50v_13180 | 2 | 0 | 0 | 0 | 0 | 0 | 0 | 0 | 0 | 0 | 1 | 0 | 0 | 0 | 0 | 1 | 0 | 0 | 0 | 0 | 0 | 0 | 0 | 0 | 0 | 0 | 0 | 0 | 0 | 0 |
| Sep8.5v_13109 | 6 | 0 | 0 | 0 | 1 | 0 | 0 | 0 | 0 | 1 | 0 | 0 | 1 | 0 | 0 | 0 | 0 | 0 | 0 | 0 | 0 | 0 | 0 | 0 | 0 | 0 | 1 | 1 | 0 | 1 |
| Jun8.50v_40694 | 1 | 0 | 0 | 0 | 0 | 0 | 1 | 0 | 0 | 0 | 0 | 0 | 0 | 0 | 0 | 0 | 0 | 0 | 0 | 0 | 0 | 0 | 0 | 0 | 0 | 0 | 0 | 0 | 0 | 0 |
| Jul11.5v_20105 | 2 | 0 | 0 | 1 | 0 | 0 | 0 | 0 | 0 | 0 | 0 | 0 | 0 | 1 | 0 | 0 | 0 | 0 | 0 | 0 | 0 | 0 | 0 | 0 | 0 | 0 | 0 | 0 | 0 | 0 |
| Jul27.5v_11599 | 3 | 0 | 0 | 2 | 0 | 0 | 0 | 0 | 0 | 0 | 0 | 0 | 0 | 0 | 0 | 0 | 0 | 1 | 0 | 0 | 0 | 0 | 0 | 0 | 0 | 0 | 0 | 0 | 0 | 0 |
| May12.50v_27008 | 1 | 0 | 0 | 1 | 0 | 0 | 0 | 0 | 0 | 0 | 0 | 0 | 0 | 0 | 0 | 0 | 0 | 0 | 0 | 0 | 0 | 0 | 0 | 0 | 0 | 0 | 0 | 0 | 0 | 0 |
| Jul27.50v_10631 | 1 | 0 | 0 | 0 | 0 | 0 | 0 | 0 | 0 | 0 | 0 | 0 | 0 | 0 | 0 | 0 | 0 | 0 | 0 | 1 | 0 | 0 | 0 | 0 | 0 | 0 | 0 | 0 | 0 | 0 |
| May12.50v_437 | 4 | 0 | 2 | 1 | 0 | 0 | 0 | 0 | 0 | 0 | 0 | 0 | 0 | 0 | 0 | 0 | 0 | 1 | 0 | 0 | 0 | 0 | 0 | 0 | 0 | 0 | 0 | 0 | 0 | 0 |
| Jun22.5v_3114 | 1 | 0 | 0 | 0 | 0 | 0 | 0 | 0 | 0 | 1 | 0 | 0 | 0 | 0 | 0 | 0 | 0 | 0 | 0 | 0 | 0 | 0 | 0 | 0 | 0 | 0 | 0 | 0 | 0 | 0 |
| Jul27.50v_4443 | 1 | 0 | 0 | 0 | 0 | 0 | 0 | 0 | 0 | 0 | 0 | 0 | 0 | 0 | 0 | 0 | 0 | 0 | 0 | 1 | 0 | 0 | 0 | 0 | 0 | 0 | 0 | 0 | 0 | 0 |
| Jun22.50v_14931 | 1 | 0 | 0 | 0 | 0 | 0 | 0 | 0 | 0 | 0 | 1 | 0 | 0 | 0 | 0 | 0 | 0 | 0 | 0 | 0 | 0 | 0 | 0 | 0 | 0 | 0 | 0 | 0 | 0 | 0 |
| May12.50v_9415 | 3 | 1 | 0 | 1 | 0 | 0 | 0 | 0 | 0 | 0 | 0 | 0 | 0 | 0 | 0 | 0 | 1 | 0 | 0 | 0 | 0 | 0 | 0 | 0 | 0 | 0 | 0 | 0 | 0 | 0 |
| Jul11.25v_1365 | 1 | 0 | 0 | 0 | 0 | 0 | 0 | 0 | 0 | 0 | 0 | 0 | 0 | 0 | 1 | 0 | 0 | 0 | 0 | 0 | 0 | 0 | 0 | 0 | 0 | 0 | 0 | 0 | 0 | 0 |
| May12.50v_21301 | 1 | 0 | 0 | 1 | 0 | 0 | 0 | 0 | 0 | 0 | 0 | 0 | 0 | 0 | 0 | 0 | 0 | 0 | 0 | 0 | 0 | 0 | 0 | 0 | 0 | 0 | 0 | 0 | 0 | 0 |
| Jun22.0v_556 | 1 | 0 | 0 | 0 | 0 | 0 | 0 | 0 | 1 | 0 | 0 | 0 | 0 | 0 | 0 | 0 | 0 | 0 | 0 | 0 | 0 | 0 | 0 | 0 | 0 | 0 | 0 | 0 | 0 | 0 |
| May12.5v_30346 | 3 | 1 | 0 | 0 | 0 | 0 | 0 | 0 | 0 | 0 | 0 | 0 | 0 | 1 | 0 | 0 | 0 | 0 | 0 | 0 | 0 | 0 | 0 | 0 | 0 | 0 | 0 | 1 | 0 | 0 |
| May12.5v_16991 | 1 | 1 | 0 | 0 | 0 | 0 | 0 | 0 | 0 | 0 | 0 | 0 | 0 | 0 | 0 | 0 | 0 | 0 | 0 | 0 | 0 | 0 | 0 | 0 | 0 | 0 | 0 | 0 | 0 | 0 |
| Jun22.50v_4255 | 5 | 0 | 0 | 2 | 0 | 1 | 0 | 0 | 0 | 0 | 2 | 0 | 0 | 0 | 0 | 0 | 0 | 0 | 0 | 0 | 0 | 0 | 0 | 0 | 0 | 0 | 0 | 0 | 0 | 0 |
| Sep8.0v_15598 | 4 | 0 | 0 | 0 | 0 | 0 | 0 | 0 | 0 | 0 | 0 | 0 | 0 | 0 | 0 | 0 | 1 | 0 | 0 | 0 | 0 | 0 | 0 | 0 | 0 | 0 | 0 | 2 | 0 | 1 |
| Jun22.50v_34549 | 1 | 0 | 0 | 0 | 0 | 0 | 0 | 0 | 0 | 0 | 1 | 0 | 0 | 0 | 0 | 0 | 0 | 0 | 0 | 0 | 0 | 0 | 0 | 0 | 0 | 0 | 0 | 0 | 0 | 0 |
| Jun22.50v_38417 | 1 | 0 | 0 | 0 | 0 | 0 | 0 | 0 | 0 | 0 | 1 | 0 | 0 | 0 | 0 | 0 | 0 | 0 | 0 | 0 | 0 | 0 | 0 | 0 | 0 | 0 | 0 | 0 | 0 | 0 |
| Aug5.0v_1608 | 1 | 0 | 0 | 0 | 0 | 0 | 0 | 0 | 0 | 0 | 0 | 0 | 0 | 0 | 0 | 0 | 0 | 0 | 0 | 0 | 0 | 1 | 0 | 0 | 0 | 0 | 0 | 0 | 0 | 0 |
| Aug5.0v_3795 | 1 | 0 | 0 | 0 | 0 | 0 | 0 | 0 | 0 | 0 | 0 | 0 | 0 | 0 | 0 | 0 | 0 | 0 | 0 | 0 | 0 | 1 | 0 | 0 | 0 | 0 | 0 | 0 | 0 | 0 |
| Aug5.0v_1940 | 1 | 0 | 0 | 0 | 0 | 0 | 0 | 0 | 0 | 0 | 0 | 0 | 0 | 0 | 0 | 0 | 0 | 0 | 0 | 0 | 0 | 1 | 0 | 0 | 0 | 0 | 0 | 0 | 0 | 0 |
| Jul11.50v_760 | 1 | 0 | 0 | 0 | 0 | 0 | 0 | 0 | 0 | 0 | 0 | 0 | 0 | 0 | 0 | 1 | 0 | 0 | 0 | 0 | 0 | 0 | 0 | 0 | 0 | 0 | 0 | 0 | 0 | 0 |
| May12.75v_26082 | 3 | 0 | 0 | 1 | 2 | 0 | 0 | 0 | 0 | 0 | 0 | 0 | 0 | 0 | 0 | 0 | 0 | 0 | 0 | 0 | 0 | 0 | 0 | 0 | 0 | 0 | 0 | 0 | 0 | 0 |
| Aug5.25v_25218 | 1 | 0 | 0 | 0 | 0 | 0 | 0 | 0 | 0 | 0 | 0 | 0 | 0 | 0 | 0 | 0 | 0 | 0 | 0 | 0 | 0 | 0 | 0 | 1 | 0 | 0 | 0 | 0 | 0 | 0 |
| May12.75v_3667 | 1 | 0 | 0 | 0 | 1 | 0 | 0 | 0 | 0 | 0 | 0 | 0 | 0 | 0 | 0 | 0 | 0 | 0 | 0 | 0 | 0 | 0 | 0 | 0 | 0 | 0 | 0 | 0 | 0 | 0 |
| Sep8.5v_1860 | 1 | 0 | 0 | 0 | 0 | 0 | 0 | 0 | 0 | 0 | 0 | 0 | 0 | 0 | 0 | 0 | 0 | 0 | 0 | 0 | 0 | 0 | 0 | 0 | 0 | 0 | 0 | 1 | 0 | 0 |
| May12.50v_7840 | 1 | 0 | 0 | 1 | 0 | 0 | 0 | 0 | 0 | 0 | 0 | 0 | 0 | 0 | 0 | 0 | 0 | 0 | 0 | 0 | 0 | 0 | 0 | 0 | 0 | 0 | 0 | 0 | 0 | 0 |
| Jul11.75v_33809 | 1 | 0 | 0 | 0 | 0 | 0 | 0 | 0 | 0 | 0 | 0 | 0 | 0 | 0 | 0 | 0 | 1 | 0 | 0 | 0 | 0 | 0 | 0 | 0 | 0 | 0 | 0 | 0 | 0 | 0 |
| Jun8.50v_3994 | 1 | 0 | 0 | 0 | 0 | 0 | 1 | 0 | 0 | 0 | 0 | 0 | 0 | 0 | 0 | 0 | 0 | 0 | 0 | 0 | 0 | 0 | 0 | 0 | 0 | 0 | 0 | 0 | 0 | 0 |
| Jun22.5v_10311 | 1 | 0 | 0 | 0 | 0 | 0 | 0 | 0 | 0 | 1 | 0 | 0 | 0 | 0 | 0 | 0 | 0 | 0 | 0 | 0 | 0 | 0 | 0 | 0 | 0 | 0 | 0 | 0 | 0 | 0 |
| May12.5v_1395 | 1 | 1 | 0 | 0 | 0 | 0 | 0 | 0 | 0 | 0 | 0 | 0 | 0 | 0 | 0 | 0 | 0 | 0 | 0 | 0 | 0 | 0 | 0 | 0 | 0 | 0 | 0 | 0 | 0 | 0 |
| Jul27.75v_13946 | 1 | 0 | 0 | 0 | 0 | 0 | 0 | 0 | 0 | 0 | 0 | 0 | 0 | 0 | 0 | 0 | 0 | 0 | 0 | 0 | 1 | 0 | 0 | 0 | 0 | 0 | 0 | 0 | 0 | 0 |
| Jun22.50v_15153 | 1 | 0 | 0 | 0 | 0 | 0 | 0 | 0 | 0 | 0 | 1 | 0 | 0 | 0 | 0 | 0 | 0 | 0 | 0 | 0 | 0 | 0 | 0 | 0 | 0 | 0 | 0 | 0 | 0 | 0 |
| Aug5.5v_26911 | 1 | 0 | 0 | 0 | 0 | 0 | 0 | 0 | 0 | 0 | 0 | 0 | 0 | 0 | 0 | 0 | 0 | 0 | 0 | 0 | 0 | 0 | 1 | 0 | 0 | 0 | 0 | 0 | 0 | 0 |
| Jul27.5v_20671 | 1 | 0 | 0 | 0 | 0 | 0 | 0 | 0 | 0 | 0 | 0 | 0 | 0 | 0 | 0 | 0 | 0 | 1 | 0 | 0 | 0 | 0 | 0 | 0 | 0 | 0 | 0 | 0 | 0 | 0 |
| Jun22.0v_1414 | 1 | 0 | 0 | 0 | 0 | 0 | 0 | 0 | 1 | 0 | 0 | 0 | 0 | 0 | 0 | 0 | 0 | 0 | 0 | 0 | 0 | 0 | 0 | 0 | 0 | 0 | 0 | 0 | 0 | 0 |
| Aug5.25v_10962 | 1 | 0 | 0 | 0 | 0 | 0 | 0 | 0 | 0 | 0 | 0 | 0 | 0 | 0 | 0 | 0 | 0 | 0 | 0 | 0 | 0 | 0 | 0 | 1 | 0 | 0 | 0 | 0 | 0 | 0 |
| Sep8.0v_14042 | 3 | 0 | 0 | 1 | 0 | 0 | 0 | 0 | 0 | 0 | 1 | 0 | 0 | 0 | 0 | 0 | 0 | 0 | 0 | 0 | 0 | 0 | 0 | 0 | 0 | 0 | 1 | 0 | 0 | 0 |
| Sep8.50v_19712 | 1 | 0 | 0 | 0 | 0 | 0 | 0 | 0 | 0 | 0 | 0 | 0 | 0 | 0 | 0 | 0 | 0 | 0 | 0 | 0 | 0 | 0 | 0 | 0 | 0 | 0 | 0 | 0 | 0 | 1 |
| Aug5.0v_1456 | 1 | 0 | 0 | 0 | 0 | 0 | 0 | 0 | 0 | 0 | 0 | 0 | 0 | 0 | 0 | 0 | 0 | 0 | 0 | 0 | 0 | 1 | 0 | 0 | 0 | 0 | 0 | 0 | 0 | 0 |
| Jul11.0v_17021 | 1 | 0 | 0 | 0 | 0 | 0 | 0 | 0 | 0 | 0 | 0 | 0 | 1 | 0 | 0 | 0 | 0 | 0 | 0 | 0 | 0 | 0 | 0 | 0 | 0 | 0 | 0 | 0 | 0 | 0 |
| Sep8.0v_8999 | 2 | 0 | 1 | 0 | 0 | 0 | 0 | 0 | 0 | 0 | 0 | 0 | 0 | 0 | 0 | 0 | 0 | 0 | 0 | 0 | 0 | 0 | 0 | 0 | 0 | 0 | 1 | 0 | 0 | 0 |
| May12.75v_24216 | 1 | 0 | 0 | 0 | 1 | 0 | 0 | 0 | 0 | 0 | 0 | 0 | 0 | 0 | 0 | 0 | 0 | 0 | 0 | 0 | 0 | 0 | 0 | 0 | 0 | 0 | 0 | 0 | 0 | 0 |
| Jun22.0v_12932 | 1 | 0 | 0 | 0 | 0 | 0 | 0 | 0 | 1 | 0 | 0 | 0 | 0 | 0 | 0 | 0 | 0 | 0 | 0 | 0 | 0 | 0 | 0 | 0 | 0 | 0 | 0 | 0 | 0 | 0 |
| Jun22.5v_5566 | 1 | 0 | 0 | 0 | 0 | 0 | 0 | 0 | 0 | 1 | 0 | 0 | 0 | 0 | 0 | 0 | 0 | 0 | 0 | 0 | 0 | 0 | 0 | 0 | 0 | 0 | 0 | 0 | 0 | 0 |
| Jul27.5v_13610 | 1 | 0 | 0 | 0 | 0 | 0 | 0 | 0 | 0 | 0 | 0 | 0 | 0 | 0 | 0 | 0 | 0 | 1 | 0 | 0 | 0 | 0 | 0 | 0 | 0 | 0 | 0 | 0 | 0 | 0 |
| Jul27.25v_28837 | 1 | 0 | 0 | 0 | 0 | 0 | 0 | 0 | 0 | 0 | 0 | 0 | 0 | 0 | 0 | 0 | 0 | 0 | 1 | 0 | 0 | 0 | 0 | 0 | 0 | 0 | 0 | 0 | 0 | 0 |
| Jul27.25v_21526 | 1 | 0 | 0 | 0 | 0 | 0 | 0 | 0 | 0 | 0 | 0 | 0 | 0 | 0 | 0 | 0 | 0 | 1 | 0 | 0 | 0 | 0 | 0 | 0 | 0 | 0 | 0 | 0 | 0 | 0 |
| May12.25v_9039 | 1 | 0 | 1 | 0 | 0 | 0 | 0 | 0 | 0 | 0 | 0 | 0 | 0 | 0 | 0 | 0 | 0 | 0 | 0 | 0 | 0 | 0 | 0 | 0 | 0 | 0 | 0 | 0 | 0 | 0 |
| May12.50v_13126 | 1 | 0 | 0 | 1 | 0 | 0 | 0 | 0 | 0 | 0 | 0 | 0 | 0 | 0 | 0 | 0 | 0 | 0 | 0 | 0 | 0 | 0 | 0 | 0 | 0 | 0 | 0 | 0 | 0 | 0 |
| Jul11.5v_2794 | 1 | 0 | 0 | 0 | 0 | 0 | 0 | 0 | 0 | 0 | 0 | 0 | 0 | 1 | 0 | 0 | 0 | 0 | 0 | 0 | 0 | 0 | 0 | 0 | 0 | 0 | 0 | 0 | 0 | 0 |
| Aug5.50v_26932 | 8 | 0 | 1 | 1 | 0 | 0 | 0 | 0 | 2 | 1 | 1 | 0 | 0 | 0 | 0 | 0 | 0 | 0 | 0 | 0 | 0 | 0 | 0 | 0 | 0 | 1 | 1 | 0 | 0 | 0 |
| Jun22.0v_17872 | 1 | 0 | 0 | 0 | 0 | 0 | 0 | 0 | 1 | 0 | 0 | 0 | 0 | 0 | 0 | 0 | 0 | 0 | 0 | 0 | 0 | 0 | 0 | 0 | 0 | 0 | 0 | 0 | 0 | 0 |
| Jul27.5v_19245 | 1 | 0 | 0 | 0 | 0 | 0 | 0 | 0 | 0 | 0 | 0 | 0 | 0 | 0 | 0 | 0 | 0 | 1 | 0 | 0 | 0 | 0 | 0 | 0 | 0 | 0 | 0 | 0 | 0 | 0 |
| Jul11.0v_3892 | 4 | 0 | 0 | 0 | 0 | 0 | 1 | 0 | 0 | 0 | 0 | 0 | 1 | 0 | 0 | 0 | 0 | 0 | 0 | 0 | 0 | 0 | 0 | 0 | 0 | 0 | 0 | 2 | 0 | 0 |
| May12.25v_36674 | 1 | 0 | 1 | 0 | 0 | 0 | 0 | 0 | 0 | 0 | 0 | 0 | 0 | 0 | 0 | 0 | 0 | 0 | 0 | 0 | 0 | 0 | 0 | 0 | 0 | 0 | 0 | 0 | 0 | 0 |
| Jul11.5v_10009 | 1 | 0 | 0 | 0 | 0 | 0 | 0 | 0 | 0 | 0 | 0 | 0 | 0 | 1 | 0 | 0 | 0 | 0 | 0 | 0 | 0 | 0 | 0 | 0 | 0 | 0 | 0 | 0 | 0 | 0 |
| Jun22.50v_19938 | 1 | 0 | 0 | 0 | 0 | 0 | 0 | 0 | 0 | 0 | 1 | 0 | 0 | 0 | 0 | 0 | 0 | 0 | 0 | 0 | 0 | 0 | 0 | 0 | 0 | 0 | 0 | 0 | 0 | 0 |
| Jun22.0v_12982 | 1 | 0 | 0 | 0 | 0 | 0 | 0 | 0 | 1 | 0 | 0 | 0 | 0 | 0 | 0 | 0 | 0 | 0 | 0 | 0 | 0 | 0 | 0 | 0 | 0 | 0 | 0 | 0 | 0 | 0 |
| Jun22.50v_15338 | 1 | 0 | 0 | 0 | 0 | 0 | 0 | 0 | 0 | 0 | 1 | 0 | 0 | 0 | 0 | 0 | 0 | 0 | 0 | 0 | 0 | 0 | 0 | 0 | 0 | 0 | 0 | 0 | 0 | 0 |
| May12.50v_6436 | 1 | 0 | 0 | 1 | 0 | 0 | 0 | 0 | 0 | 0 | 0 | 0 | 0 | 0 | 0 | 0 | 0 | 0 | 0 | 0 | 0 | 0 | 0 | 0 | 0 | 0 | 0 | 0 | 0 | 0 |
| Jul11.50v_9111 | 1 | 0 | 0 | 0 | 0 | 0 | 0 | 0 | 0 | 0 | 0 | 0 | 0 | 0 | 0 | 1 | 0 | 0 | 0 | 0 | 0 | 0 | 0 | 0 | 0 | 0 | 0 | 0 | 0 | 0 |
| Sep8.0v_3140 | 1 | 0 | 0 | 0 | 0 | 0 | 0 | 0 | 0 | 0 | 0 | 0 | 0 | 0 | 0 | 0 | 0 | 0 | 0 | 0 | 0 | 0 | 0 | 0 | 0 | 0 | 1 | 0 | 0 | 0 |
| May12.5v_11516 | 1 | 1 | 0 | 0 | 0 | 0 | 0 | 0 | 0 | 0 | 0 | 0 | 0 | 0 | 0 | 0 | 0 | 0 | 0 | 0 | 0 | 0 | 0 | 0 | 0 | 0 | 0 | 0 | 0 | 0 |
| Jul11.50v_32 | 1 | 0 | 0 | 0 | 0 | 0 | 0 | 0 | 0 | 0 | 0 | 0 | 0 | 0 | 0 | 1 | 0 | 0 | 0 | 0 | 0 | 0 | 0 | 0 | 0 | 0 | 0 | 0 | 0 | 0 |
| Jul11.5v_8795 | 9 | 1 | 0 | 1 | 1 | 0 | 0 | 0 | 2 | 0 | 0 | 0 | 0 | 2 | 1 | 0 | 0 | 0 | 0 | 0 | 0 | 0 | 0 | 0 | 0 | 1 | 0 | 0 | 0 | 0 |
| Jul11.5v_2249 | 1 | 0 | 0 | 0 | 0 | 0 | 0 | 0 | 0 | 0 | 0 | 0 | 0 | 1 | 0 | 0 | 0 | 0 | 0 | 0 | 0 | 0 | 0 | 0 | 0 | 0 | 0 | 0 | 0 | 0 |
| Jun22.0v_1050 | 1 | 0 | 0 | 0 | 0 | 0 | 0 | 0 | 1 | 0 | 0 | 0 | 0 | 0 | 0 | 0 | 0 | 0 | 0 | 0 | 0 | 0 | 0 | 0 | 0 | 0 | 0 | 0 | 0 | 0 |
| Sep8.50v_576 | 1 | 0 | 0 | 0 | 0 | 0 | 0 | 0 | 0 | 0 | 0 | 0 | 0 | 0 | 0 | 0 | 0 | 0 | 0 | 0 | 0 | 0 | 0 | 0 | 0 | 0 | 0 | 0 | 0 | 1 |
| Jul11.5v_20584 | 1 | 0 | 0 | 0 | 0 | 0 | 0 | 0 | 0 | 0 | 0 | 0 | 0 | 1 | 0 | 0 | 0 | 0 | 0 | 0 | 0 | 0 | 0 | 0 | 0 | 0 | 0 | 0 | 0 | 0 |
| Jun22.5v_9207 | 1 | 0 | 0 | 0 | 0 | 0 | 0 | 0 | 0 | 1 | 0 | 0 | 0 | 0 | 0 | 0 | 0 | 0 | 0 | 0 | 0 | 0 | 0 | 0 | 0 | 0 | 0 | 0 | 0 | 0 |
| Aug5.0v_4047 | 1 | 0 | 0 | 0 | 0 | 0 | 0 | 0 | 0 | 0 | 0 | 0 | 0 | 0 | 0 | 0 | 0 | 0 | 0 | 0 | 0 | 1 | 0 | 0 | 0 | 0 | 0 | 0 | 0 | 0 |
| Jul11.0v_19739 | 5 | 0 | 0 | 0 | 1 | 0 | 0 | 0 | 1 | 0 | 0 | 0 | 1 | 0 | 1 | 0 | 0 | 0 | 0 | 0 | 0 | 0 | 0 | 0 | 0 | 0 | 0 | 0 | 0 | 1 |
| Aug5.0v_2530 | 4 | 0 | 1 | 0 | 1 | 0 | 0 | 0 | 0 | 0 | 1 | 0 | 0 | 0 | 0 | 0 | 0 | 0 | 0 | 0 | 0 | 1 | 0 | 0 | 0 | 0 | 0 | 0 | 0 | 0 |
| Jul11.25v_8965 | 1 | 0 | 0 | 0 | 0 | 0 | 0 | 0 | 0 | 0 | 0 | 0 | 0 | 0 | 1 | 0 | 0 | 0 | 0 | 0 | 0 | 0 | 0 | 0 | 0 | 0 | 0 | 0 | 0 | 0 |
| May12.75v_36395 | 1 | 0 | 0 | 0 | 1 | 0 | 0 | 0 | 0 | 0 | 0 | 0 | 0 | 0 | 0 | 0 | 0 | 0 | 0 | 0 | 0 | 0 | 0 | 0 | 0 | 0 | 0 | 0 | 0 | 0 |
| Jun8.50v_30353 | 1 | 0 | 0 | 0 | 0 | 0 | 1 | 0 | 0 | 0 | 0 | 0 | 0 | 0 | 0 | 0 | 0 | 0 | 0 | 0 | 0 | 0 | 0 | 0 | 0 | 0 | 0 | 0 | 0 | 0 |
| Jul11.5v_8070 | 1 | 0 | 0 | 0 | 0 | 0 | 0 | 0 | 0 | 0 | 0 | 0 | 0 | 1 | 0 | 0 | 0 | 0 | 0 | 0 | 0 | 0 | 0 | 0 | 0 | 0 | 0 | 0 | 0 | 0 |
| Aug5.0v_3417 | 1 | 0 | 0 | 0 | 0 | 0 | 0 | 0 | 0 | 0 | 0 | 0 | 0 | 0 | 0 | 0 | 0 | 0 | 0 | 0 | 0 | 1 | 0 | 0 | 0 | 0 | 0 | 0 | 0 | 0 |
| May12.75v_24178 | 1 | 0 | 0 | 0 | 1 | 0 | 0 | 0 | 0 | 0 | 0 | 0 | 0 | 0 | 0 | 0 | 0 | 0 | 0 | 0 | 0 | 0 | 0 | 0 | 0 | 0 | 0 | 0 | 0 | 0 |
| Jul27.25v_37935 | 1 | 0 | 0 | 0 | 0 | 0 | 0 | 0 | 0 | 0 | 0 | 0 | 0 | 0 | 0 | 0 | 0 | 0 | 1 | 0 | 0 | 0 | 0 | 0 | 0 | 0 | 0 | 0 | 0 | 0 |
| Jul27.5v_2526 | 1 | 0 | 0 | 0 | 0 | 0 | 0 | 0 | 0 | 0 | 0 | 0 | 0 | 0 | 0 | 0 | 0 | 1 | 0 | 0 | 0 | 0 | 0 | 0 | 0 | 0 | 0 | 0 | 0 | 0 |
| Jul27.25v_35747 | 11 | 0 | 0 | 0 | 0 | 0 | 1 | 0 | 1 | 1 | 0 | 2 | 0 | 0 | 0 | 0 | 0 | 0 | 2 | 0 | 0 | 0 | 0 | 0 | 0 | 0 | 1 | 1 | 0 | 2 |
| Jul11.0v_503 | 1 | 0 | 0 | 0 | 0 | 0 | 0 | 0 | 0 | 0 | 0 | 0 | 1 | 0 | 0 | 0 | 0 | 0 | 0 | 0 | 0 | 0 | 0 | 0 | 0 | 0 | 0 | 0 | 0 | 0 |
| Jun22.5v_19576 | 1 | 0 | 0 | 0 | 0 | 0 | 0 | 0 | 0 | 1 | 0 | 0 | 0 | 0 | 0 | 0 | 0 | 0 | 0 | 0 | 0 | 0 | 0 | 0 | 0 | 0 | 0 | 0 | 0 | 0 |
| Jul11.75v_26271 | 2 | 0 | 0 | 0 | 0 | 0 | 0 | 0 | 0 | 0 | 0 | 1 | 0 | 0 | 0 | 0 | 1 | 0 | 0 | 0 | 0 | 0 | 0 | 0 | 0 | 0 | 0 | 0 | 0 | 0 |
| Jul11.0v_19780 | 1 | 0 | 0 | 0 | 0 | 0 | 0 | 0 | 0 | 0 | 0 | 0 | 1 | 0 | 0 | 0 | 0 | 0 | 0 | 0 | 0 | 0 | 0 | 0 | 0 | 0 | 0 | 0 | 0 | 0 |
| Jun8.50v_38580 | 1 | 0 | 0 | 0 | 0 | 0 | 1 | 0 | 0 | 0 | 0 | 0 | 0 | 0 | 0 | 0 | 0 | 0 | 0 | 0 | 0 | 0 | 0 | 0 | 0 | 0 | 0 | 0 | 0 | 0 |
| Jun8.50v_2066 | 1 | 0 | 0 | 0 | 0 | 0 | 1 | 0 | 0 | 0 | 0 | 0 | 0 | 0 | 0 | 0 | 0 | 0 | 0 | 0 | 0 | 0 | 0 | 0 | 0 | 0 | 0 | 0 | 0 | 0 |
| May12.75v_22583 | 1 | 0 | 0 | 0 | 1 | 0 | 0 | 0 | 0 | 0 | 0 | 0 | 0 | 0 | 0 | 0 | 0 | 0 | 0 | 0 | 0 | 0 | 0 | 0 | 0 | 0 | 0 | 0 | 0 | 0 |
| Sep8.0v_18205 | 1 | 0 | 0 | 0 | 0 | 0 | 0 | 0 | 0 | 0 | 0 | 0 | 0 | 0 | 0 | 0 | 0 | 0 | 0 | 0 | 0 | 0 | 0 | 0 | 0 | 0 | 1 | 0 | 0 | 0 |
| Aug5.75v_31853 | 1 | 0 | 0 | 0 | 0 | 0 | 0 | 0 | 0 | 0 | 0 | 0 | 0 | 0 | 0 | 0 | 0 | 0 | 0 | 0 | 0 | 0 | 0 | 0 | 0 | 1 | 0 | 0 | 0 | 0 |
| Jul27.50v_5858 | 1 | 0 | 0 | 0 | 0 | 0 | 0 | 0 | 0 | 0 | 0 | 0 | 0 | 0 | 0 | 0 | 0 | 0 | 0 | 1 | 0 | 0 | 0 | 0 | 0 | 0 | 0 | 0 | 0 | 0 |
| Jun8.75v_596 | 1 | 0 | 0 | 0 | 0 | 0 | 0 | 1 | 0 | 0 | 0 | 0 | 0 | 0 | 0 | 0 | 0 | 0 | 0 | 0 | 0 | 0 | 0 | 0 | 0 | 0 | 0 | 0 | 0 | 0 |
| Jul11.75v_23409 | 1 | 0 | 0 | 0 | 0 | 0 | 0 | 0 | 0 | 0 | 0 | 0 | 0 | 0 | 0 | 0 | 1 | 0 | 0 | 0 | 0 | 0 | 0 | 0 | 0 | 0 | 0 | 0 | 0 | 0 |
| Aug5.25v_29627 | 1 | 0 | 0 | 0 | 0 | 0 | 0 | 0 | 0 | 0 | 0 | 0 | 0 | 0 | 0 | 0 | 0 | 0 | 0 | 0 | 0 | 0 | 0 | 1 | 0 | 0 | 0 | 0 | 0 | 0 |
| May12.25v_14815 | 1 | 0 | 1 | 0 | 0 | 0 | 0 | 0 | 0 | 0 | 0 | 0 | 0 | 0 | 0 | 0 | 0 | 0 | 0 | 0 | 0 | 0 | 0 | 0 | 0 | 0 | 0 | 0 | 0 | 0 |
| Jul11.75v_13608 | 3 | 0 | 0 | 0 | 0 | 0 | 0 | 0 | 0 | 0 | 0 | 0 | 0 | 1 | 0 | 0 | 1 | 0 | 0 | 0 | 0 | 0 | 0 | 0 | 0 | 0 | 0 | 0 | 0 | 1 |
| May12.50v_25372 | 1 | 0 | 0 | 1 | 0 | 0 | 0 | 0 | 0 | 0 | 0 | 0 | 0 | 0 | 0 | 0 | 0 | 0 | 0 | 0 | 0 | 0 | 0 | 0 | 0 | 0 | 0 | 0 | 0 | 0 |
| Jun22.50v_39965 | 1 | 0 | 0 | 0 | 0 | 0 | 0 | 0 | 0 | 0 | 1 | 0 | 0 | 0 | 0 | 0 | 0 | 0 | 0 | 0 | 0 | 0 | 0 | 0 | 0 | 0 | 0 | 0 | 0 | 0 |
| Sep8.0v_19519 | 1 | 0 | 0 | 0 | 0 | 0 | 0 | 0 | 0 | 0 | 0 | 0 | 0 | 0 | 0 | 0 | 0 | 0 | 0 | 0 | 0 | 0 | 0 | 0 | 0 | 0 | 1 | 0 | 0 | 0 |
| Sep8.5v_4102 | 1 | 0 | 0 | 0 | 0 | 0 | 0 | 0 | 0 | 0 | 0 | 0 | 0 | 0 | 0 | 0 | 0 | 0 | 0 | 0 | 0 | 0 | 0 | 0 | 0 | 0 | 0 | 1 | 0 | 0 |
| Aug5.0v_4149 | 1 | 0 | 0 | 0 | 0 | 0 | 0 | 0 | 0 | 0 | 0 | 0 | 0 | 0 | 0 | 0 | 0 | 0 | 0 | 0 | 0 | 1 | 0 | 0 | 0 | 0 | 0 | 0 | 0 | 0 |
| Jul11.5v_18843 | 1 | 0 | 0 | 0 | 0 | 0 | 0 | 0 | 0 | 0 | 0 | 0 | 0 | 1 | 0 | 0 | 0 | 0 | 0 | 0 | 0 | 0 | 0 | 0 | 0 | 0 | 0 | 0 | 0 | 0 |
| Jun8.75v_20755 | 6 | 0 | 0 | 1 | 0 | 0 | 1 | 1 | 1 | 0 | 0 | 0 | 0 | 0 | 0 | 0 | 1 | 0 | 0 | 0 | 1 | 0 | 0 | 0 | 0 | 0 | 0 | 0 | 0 | 0 |
| May12.5v_28360 | 1 | 1 | 0 | 0 | 0 | 0 | 0 | 0 | 0 | 0 | 0 | 0 | 0 | 0 | 0 | 0 | 0 | 0 | 0 | 0 | 0 | 0 | 0 | 0 | 0 | 0 | 0 | 0 | 0 | 0 |
| May12.50v_14924 | 1 | 0 | 0 | 1 | 0 | 0 | 0 | 0 | 0 | 0 | 0 | 0 | 0 | 0 | 0 | 0 | 0 | 0 | 0 | 0 | 0 | 0 | 0 | 0 | 0 | 0 | 0 | 0 | 0 | 0 |
| Jun8.50v_27475 | 1 | 0 | 0 | 0 | 0 | 0 | 1 | 0 | 0 | 0 | 0 | 0 | 0 | 0 | 0 | 0 | 0 | 0 | 0 | 0 | 0 | 0 | 0 | 0 | 0 | 0 | 0 | 0 | 0 | 0 |
| Jun22.75v_16253 | 1 | 0 | 0 | 0 | 0 | 0 | 0 | 0 | 0 | 0 | 0 | 1 | 0 | 0 | 0 | 0 | 0 | 0 | 0 | 0 | 0 | 0 | 0 | 0 | 0 | 0 | 0 | 0 | 0 | 0 |
| May12.75v_37531 | 1 | 0 | 0 | 0 | 1 | 0 | 0 | 0 | 0 | 0 | 0 | 0 | 0 | 0 | 0 | 0 | 0 | 0 | 0 | 0 | 0 | 0 | 0 | 0 | 0 | 0 | 0 | 0 | 0 | 0 |
| Jun8.50v_8635 | 1 | 0 | 0 | 0 | 0 | 0 | 1 | 0 | 0 | 0 | 0 | 0 | 0 | 0 | 0 | 0 | 0 | 0 | 0 | 0 | 0 | 0 | 0 | 0 | 0 | 0 | 0 | 0 | 0 | 0 |
| Aug5.0v_4831 | 1 | 0 | 0 | 0 | 0 | 0 | 0 | 0 | 0 | 0 | 0 | 0 | 0 | 0 | 0 | 0 | 0 | 0 | 0 | 0 | 0 | 1 | 0 | 0 | 0 | 0 | 0 | 0 | 0 | 0 |
| Jul27.75v_2820 | 1 | 0 | 0 | 0 | 0 | 0 | 0 | 0 | 0 | 0 | 0 | 0 | 0 | 0 | 0 | 0 | 0 | 0 | 0 | 0 | 1 | 0 | 0 | 0 | 0 | 0 | 0 | 0 | 0 | 0 |
| Jul11.75v_39594 | 1 | 0 | 0 | 0 | 0 | 0 | 0 | 0 | 0 | 0 | 0 | 0 | 0 | 0 | 0 | 0 | 1 | 0 | 0 | 0 | 0 | 0 | 0 | 0 | 0 | 0 | 0 | 0 | 0 | 0 |
| May12.5v_31219 | 1 | 1 | 0 | 0 | 0 | 0 | 0 | 0 | 0 | 0 | 0 | 0 | 0 | 0 | 0 | 0 | 0 | 0 | 0 | 0 | 0 | 0 | 0 | 0 | 0 | 0 | 0 | 0 | 0 | 0 |
| Jun8.50v_24700 | 1 | 0 | 0 | 0 | 0 | 0 | 1 | 0 | 0 | 0 | 0 | 0 | 0 | 0 | 0 | 0 | 0 | 0 | 0 | 0 | 0 | 0 | 0 | 0 | 0 | 0 | 0 | 0 | 0 | 0 |
| Jul11.50v_17682 | 1 | 0 | 0 | 0 | 0 | 0 | 0 | 0 | 0 | 0 | 0 | 0 | 0 | 0 | 0 | 1 | 0 | 0 | 0 | 0 | 0 | 0 | 0 | 0 | 0 | 0 | 0 | 0 | 0 | 0 |
| Jun22.5v_2873 | 1 | 0 | 0 | 0 | 0 | 0 | 0 | 0 | 0 | 1 | 0 | 0 | 0 | 0 | 0 | 0 | 0 | 0 | 0 | 0 | 0 | 0 | 0 | 0 | 0 | 0 | 0 | 0 | 0 | 0 |
| Sep8.50v_25168 | 1 | 0 | 0 | 0 | 0 | 0 | 0 | 0 | 0 | 0 | 0 | 0 | 0 | 0 | 0 | 0 | 0 | 0 | 0 | 0 | 0 | 0 | 0 | 0 | 0 | 0 | 0 | 0 | 0 | 1 |
| Aug5.0v_5323 | 1 | 0 | 0 | 0 | 0 | 0 | 0 | 0 | 0 | 0 | 0 | 0 | 0 | 0 | 0 | 0 | 0 | 0 | 0 | 0 | 0 | 1 | 0 | 0 | 0 | 0 | 0 | 0 | 0 | 0 |
| Aug5.75v_11390 | 1 | 0 | 0 | 0 | 0 | 0 | 0 | 0 | 0 | 0 | 0 | 0 | 0 | 0 | 0 | 0 | 0 | 0 | 0 | 0 | 0 | 0 | 0 | 0 | 0 | 1 | 0 | 0 | 0 | 0 |
| Jul11.5v_6464 | 1 | 0 | 0 | 0 | 0 | 0 | 0 | 0 | 0 | 0 | 0 | 0 | 0 | 1 | 0 | 0 | 0 | 0 | 0 | 0 | 0 | 0 | 0 | 0 | 0 | 0 | 0 | 0 | 0 | 0 |
| Aug5.5v_20459 | 1 | 0 | 0 | 0 | 0 | 0 | 0 | 0 | 0 | 0 | 0 | 0 | 0 | 0 | 0 | 0 | 0 | 0 | 0 | 0 | 0 | 0 | 1 | 0 | 0 | 0 | 0 | 0 | 0 | 0 |
| Jun22.75v_27749 | 1 | 0 | 0 | 0 | 0 | 0 | 0 | 0 | 0 | 0 | 0 | 1 | 0 | 0 | 0 | 0 | 0 | 0 | 0 | 0 | 0 | 0 | 0 | 0 | 0 | 0 | 0 | 0 | 0 | 0 |
| Jul11.75v_24996 | 1 | 0 | 0 | 0 | 0 | 0 | 0 | 0 | 0 | 0 | 0 | 0 | 0 | 0 | 0 | 0 | 1 | 0 | 0 | 0 | 0 | 0 | 0 | 0 | 0 | 0 | 0 | 0 | 0 | 0 |
| Jul27.75v_20954 | 1 | 0 | 0 | 0 | 0 | 0 | 0 | 0 | 0 | 0 | 0 | 0 | 0 | 0 | 0 | 0 | 0 | 0 | 0 | 0 | 1 | 0 | 0 | 0 | 0 | 0 | 0 | 0 | 0 | 0 |
| Jun8.50v_7917 | 1 | 0 | 0 | 0 | 0 | 0 | 1 | 0 | 0 | 0 | 0 | 0 | 0 | 0 | 0 | 0 | 0 | 0 | 0 | 0 | 0 | 0 | 0 | 0 | 0 | 0 | 0 | 0 | 0 | 0 |
| Jun22.0v_9080 | 1 | 0 | 0 | 0 | 0 | 0 | 0 | 0 | 1 | 0 | 0 | 0 | 0 | 0 | 0 | 0 | 0 | 0 | 0 | 0 | 0 | 0 | 0 | 0 | 0 | 0 | 0 | 0 | 0 | 0 |
| Jun8.50v_40474 | 1 | 0 | 0 | 0 | 0 | 0 | 1 | 0 | 0 | 0 | 0 | 0 | 0 | 0 | 0 | 0 | 0 | 0 | 0 | 0 | 0 | 0 | 0 | 0 | 0 | 0 | 0 | 0 | 0 | 0 |
| Aug5.0v_4945 | 1 | 0 | 0 | 0 | 0 | 0 | 0 | 0 | 0 | 0 | 0 | 0 | 0 | 0 | 0 | 0 | 0 | 0 | 0 | 0 | 0 | 1 | 0 | 0 | 0 | 0 | 0 | 0 | 0 | 0 |
| Sep8.50v_11296 | 1 | 0 | 0 | 0 | 0 | 0 | 0 | 0 | 0 | 0 | 0 | 0 | 0 | 0 | 0 | 0 | 0 | 0 | 0 | 0 | 0 | 0 | 0 | 0 | 0 | 0 | 0 | 0 | 0 | 1 |
| Jul27.25v_5603 | 1 | 0 | 0 | 0 | 0 | 0 | 0 | 0 | 0 | 0 | 0 | 0 | 0 | 0 | 0 | 0 | 0 | 0 | 1 | 0 | 0 | 0 | 0 | 0 | 0 | 0 | 0 | 0 | 0 | 0 |
| Aug5.0v_4861 | 1 | 0 | 0 | 0 | 0 | 0 | 0 | 0 | 0 | 0 | 0 | 0 | 0 | 0 | 0 | 0 | 0 | 0 | 0 | 0 | 0 | 1 | 0 | 0 | 0 | 0 | 0 | 0 | 0 | 0 |
| Aug5.0v_17 | 1 | 0 | 0 | 0 | 0 | 0 | 0 | 0 | 0 | 0 | 0 | 0 | 0 | 0 | 0 | 0 | 0 | 0 | 0 | 0 | 0 | 1 | 0 | 0 | 0 | 0 | 0 | 0 | 0 | 0 |
| Jul27.5v_8138 | 1 | 0 | 0 | 0 | 0 | 0 | 0 | 0 | 0 | 0 | 0 | 0 | 0 | 0 | 0 | 0 | 0 | 1 | 0 | 0 | 0 | 0 | 0 | 0 | 0 | 0 | 0 | 0 | 0 | 0 |
| Aug5.25v_7175 | 1 | 0 | 0 | 0 | 0 | 0 | 0 | 0 | 0 | 0 | 0 | 0 | 0 | 0 | 0 | 0 | 0 | 0 | 0 | 0 | 0 | 0 | 0 | 1 | 0 | 0 | 0 | 0 | 0 | 0 |
| Sep8.50v_12628 | 1 | 0 | 0 | 0 | 0 | 0 | 0 | 0 | 0 | 0 | 0 | 0 | 0 | 0 | 0 | 0 | 0 | 0 | 0 | 0 | 0 | 0 | 0 | 0 | 0 | 0 | 0 | 0 | 0 | 1 |
| Jun8.50v_26042 | 1 | 0 | 0 | 0 | 0 | 0 | 1 | 0 | 0 | 0 | 0 | 0 | 0 | 0 | 0 | 0 | 0 | 0 | 0 | 0 | 0 | 0 | 0 | 0 | 0 | 0 | 0 | 0 | 0 | 0 |
| Jun8.50v_1852 | 1 | 0 | 0 | 0 | 0 | 0 | 1 | 0 | 0 | 0 | 0 | 0 | 0 | 0 | 0 | 0 | 0 | 0 | 0 | 0 | 0 | 0 | 0 | 0 | 0 | 0 | 0 | 0 | 0 | 0 |
| May12.50v_23648 | 1 | 0 | 0 | 1 | 0 | 0 | 0 | 0 | 0 | 0 | 0 | 0 | 0 | 0 | 0 | 0 | 0 | 0 | 0 | 0 | 0 | 0 | 0 | 0 | 0 | 0 | 0 | 0 | 0 | 0 |
| Jul11.75v_3327 | 1 | 0 | 0 | 0 | 0 | 0 | 0 | 0 | 0 | 0 | 0 | 0 | 0 | 0 | 0 | 0 | 1 | 0 | 0 | 0 | 0 | 0 | 0 | 0 | 0 | 0 | 0 | 0 | 0 | 0 |
| Jun8.25v_4654 | 1 | 0 | 0 | 0 | 0 | 1 | 0 | 0 | 0 | 0 | 0 | 0 | 0 | 0 | 0 | 0 | 0 | 0 | 0 | 0 | 0 | 0 | 0 | 0 | 0 | 0 | 0 | 0 | 0 | 0 |
| Jun22.75v_23111 | 1 | 0 | 0 | 0 | 0 | 0 | 0 | 0 | 0 | 0 | 0 | 1 | 0 | 0 | 0 | 0 | 0 | 0 | 0 | 0 | 0 | 0 | 0 | 0 | 0 | 0 | 0 | 0 | 0 | 0 |
| Jun22.50v_15762 | 2 | 0 | 0 | 0 | 0 | 0 | 0 | 0 | 0 | 0 | 1 | 0 | 0 | 0 | 0 | 0 | 0 | 1 | 0 | 0 | 0 | 0 | 0 | 0 | 0 | 0 | 0 | 0 | 0 | 0 |
| Jul27.5v_16438 | 1 | 0 | 0 | 0 | 0 | 0 | 0 | 0 | 0 | 0 | 0 | 0 | 0 | 0 | 0 | 0 | 0 | 1 | 0 | 0 | 0 | 0 | 0 | 0 | 0 | 0 | 0 | 0 | 0 | 0 |
| Sep8.50v_22684 | 1 | 0 | 0 | 0 | 0 | 0 | 0 | 0 | 0 | 0 | 0 | 0 | 0 | 0 | 0 | 0 | 0 | 0 | 0 | 0 | 0 | 0 | 0 | 0 | 0 | 0 | 0 | 0 | 0 | 1 |
| May12.5v_8804 | 1 | 1 | 0 | 0 | 0 | 0 | 0 | 0 | 0 | 0 | 0 | 0 | 0 | 0 | 0 | 0 | 0 | 0 | 0 | 0 | 0 | 0 | 0 | 0 | 0 | 0 | 0 | 0 | 0 | 0 |
| Jul11.25v_3684 | 1 | 0 | 0 | 0 | 0 | 0 | 0 | 0 | 0 | 0 | 0 | 0 | 0 | 0 | 1 | 0 | 0 | 0 | 0 | 0 | 0 | 0 | 0 | 0 | 0 | 0 | 0 | 0 | 0 | 0 |
| Sep8.5v_11217 | 1 | 0 | 0 | 0 | 0 | 0 | 0 | 0 | 0 | 0 | 0 | 0 | 0 | 0 | 0 | 0 | 0 | 0 | 0 | 0 | 0 | 0 | 0 | 0 | 0 | 0 | 0 | 1 | 0 | 0 |
| Aug5.5v_38688 | 1 | 0 | 0 | 0 | 0 | 0 | 0 | 0 | 0 | 0 | 0 | 0 | 0 | 0 | 0 | 0 | 0 | 0 | 0 | 0 | 0 | 0 | 1 | 0 | 0 | 0 | 0 | 0 | 0 | 0 |
| Jul27.25v_33508 | 1 | 0 | 0 | 0 | 0 | 0 | 0 | 0 | 0 | 0 | 0 | 0 | 0 | 0 | 0 | 0 | 0 | 0 | 1 | 0 | 0 | 0 | 0 | 0 | 0 | 0 | 0 | 0 | 0 | 0 |
| Jun22.75v_23423 | 1 | 0 | 0 | 0 | 0 | 0 | 0 | 0 | 0 | 0 | 0 | 1 | 0 | 0 | 0 | 0 | 0 | 0 | 0 | 0 | 0 | 0 | 0 | 0 | 0 | 0 | 0 | 0 | 0 | 0 |
| Jul11.5v_15825 | 1 | 0 | 0 | 0 | 0 | 0 | 0 | 0 | 0 | 0 | 0 | 0 | 0 | 1 | 0 | 0 | 0 | 0 | 0 | 0 | 0 | 0 | 0 | 0 | 0 | 0 | 0 | 0 | 0 | 0 |
| May12.50v_33183 | 1 | 0 | 0 | 1 | 0 | 0 | 0 | 0 | 0 | 0 | 0 | 0 | 0 | 0 | 0 | 0 | 0 | 0 | 0 | 0 | 0 | 0 | 0 | 0 | 0 | 0 | 0 | 0 | 0 | 0 |
| Jul11.5v_17741 | 1 | 0 | 0 | 0 | 0 | 0 | 0 | 0 | 0 | 0 | 0 | 0 | 0 | 1 | 0 | 0 | 0 | 0 | 0 | 0 | 0 | 0 | 0 | 0 | 0 | 0 | 0 | 0 | 0 | 0 |
| Jun8.75v_18903 | 1 | 0 | 0 | 0 | 0 | 0 | 0 | 1 | 0 | 0 | 0 | 0 | 0 | 0 | 0 | 0 | 0 | 0 | 0 | 0 | 0 | 0 | 0 | 0 | 0 | 0 | 0 | 0 | 0 | 0 |
| Aug5.5v_16378 | 1 | 0 | 0 | 0 | 0 | 0 | 0 | 0 | 0 | 0 | 0 | 0 | 0 | 0 | 0 | 0 | 0 | 0 | 0 | 0 | 0 | 0 | 1 | 0 | 0 | 0 | 0 | 0 | 0 | 0 |
| Jun8.50v_20410 | 1 | 0 | 0 | 0 | 0 | 0 | 1 | 0 | 0 | 0 | 0 | 0 | 0 | 0 | 0 | 0 | 0 | 0 | 0 | 0 | 0 | 0 | 0 | 0 | 0 | 0 | 0 | 0 | 0 | 0 |
| Jul11.75v_27124 | 1 | 0 | 0 | 0 | 0 | 0 | 0 | 0 | 0 | 0 | 0 | 0 | 0 | 0 | 0 | 0 | 1 | 0 | 0 | 0 | 0 | 0 | 0 | 0 | 0 | 0 | 0 | 0 | 0 | 0 |
| Jun8.25v_39198 | 1 | 0 | 0 | 0 | 0 | 1 | 0 | 0 | 0 | 0 | 0 | 0 | 0 | 0 | 0 | 0 | 0 | 0 | 0 | 0 | 0 | 0 | 0 | 0 | 0 | 0 | 0 | 0 | 0 | 0 |
| May12.75v_39704 | 1 | 0 | 0 | 0 | 1 | 0 | 0 | 0 | 0 | 0 | 0 | 0 | 0 | 0 | 0 | 0 | 0 | 0 | 0 | 0 | 0 | 0 | 0 | 0 | 0 | 0 | 0 | 0 | 0 | 0 |
| Jun8.50v_29123 | 1 | 0 | 0 | 0 | 0 | 0 | 1 | 0 | 0 | 0 | 0 | 0 | 0 | 0 | 0 | 0 | 0 | 0 | 0 | 0 | 0 | 0 | 0 | 0 | 0 | 0 | 0 | 0 | 0 | 0 |
| Sep8.0v_10367 | 1 | 0 | 0 | 0 | 0 | 0 | 0 | 0 | 0 | 0 | 0 | 0 | 0 | 0 | 0 | 0 | 0 | 0 | 0 | 0 | 0 | 0 | 0 | 0 | 0 | 0 | 1 | 0 | 0 | 0 |
| Sep8.5v_460 | 1 | 0 | 0 | 0 | 0 | 0 | 0 | 0 | 0 | 0 | 0 | 0 | 0 | 0 | 0 | 0 | 0 | 0 | 0 | 0 | 0 | 0 | 0 | 0 | 0 | 0 | 0 | 1 | 0 | 0 |
| Jul11.0v_5433 | 1 | 0 | 0 | 0 | 0 | 0 | 0 | 0 | 0 | 0 | 0 | 0 | 1 | 0 | 0 | 0 | 0 | 0 | 0 | 0 | 0 | 0 | 0 | 0 | 0 | 0 | 0 | 0 | 0 | 0 |
| Jul11.5v_6230 | 1 | 0 | 0 | 0 | 0 | 0 | 0 | 0 | 0 | 0 | 0 | 0 | 0 | 1 | 0 | 0 | 0 | 0 | 0 | 0 | 0 | 0 | 0 | 0 | 0 | 0 | 0 | 0 | 0 | 0 |
| Jul27.25v_32275 | 1 | 0 | 0 | 0 | 0 | 0 | 0 | 0 | 0 | 0 | 0 | 0 | 0 | 0 | 0 | 0 | 0 | 0 | 1 | 0 | 0 | 0 | 0 | 0 | 0 | 0 | 0 | 0 | 0 | 0 |
| May12.75v_7417 | 1 | 0 | 0 | 0 | 1 | 0 | 0 | 0 | 0 | 0 | 0 | 0 | 0 | 0 | 0 | 0 | 0 | 0 | 0 | 0 | 0 | 0 | 0 | 0 | 0 | 0 | 0 | 0 | 0 | 0 |
| Sep8.25v_10438 | 1 | 0 | 0 | 0 | 0 | 0 | 0 | 0 | 0 | 0 | 0 | 0 | 0 | 0 | 0 | 0 | 0 | 0 | 0 | 0 | 0 | 0 | 0 | 0 | 0 | 0 | 0 | 0 | 1 | 0 |
| Jul27.25v_6801 | 1 | 0 | 0 | 0 | 0 | 0 | 0 | 0 | 0 | 0 | 0 | 0 | 0 | 0 | 0 | 0 | 0 | 0 | 1 | 0 | 0 | 0 | 0 | 0 | 0 | 0 | 0 | 0 | 0 | 0 |
| Jun8.50v_28327 | 1 | 0 | 0 | 0 | 0 | 0 | 1 | 0 | 0 | 0 | 0 | 0 | 0 | 0 | 0 | 0 | 0 | 0 | 0 | 0 | 0 | 0 | 0 | 0 | 0 | 0 | 0 | 0 | 0 | 0 |
| Jul11.5v_165 | 1 | 0 | 0 | 0 | 0 | 0 | 0 | 0 | 0 | 0 | 0 | 0 | 0 | 1 | 0 | 0 | 0 | 0 | 0 | 0 | 0 | 0 | 0 | 0 | 0 | 0 | 0 | 0 | 0 | 0 |
| Jun22.50v_9120 | 1 | 0 | 0 | 0 | 0 | 0 | 0 | 0 | 0 | 0 | 1 | 0 | 0 | 0 | 0 | 0 | 0 | 0 | 0 | 0 | 0 | 0 | 0 | 0 | 0 | 0 | 0 | 0 | 0 | 0 |
| Jul11.75v_13355 | 1 | 0 | 0 | 0 | 0 | 0 | 0 | 0 | 0 | 0 | 0 | 0 | 0 | 0 | 0 | 0 | 1 | 0 | 0 | 0 | 0 | 0 | 0 | 0 | 0 | 0 | 0 | 0 | 0 | 0 |
| May12.5v_39318 | 1 | 1 | 0 | 0 | 0 | 0 | 0 | 0 | 0 | 0 | 0 | 0 | 0 | 0 | 0 | 0 | 0 | 0 | 0 | 0 | 0 | 0 | 0 | 0 | 0 | 0 | 0 | 0 | 0 | 0 |
| Aug5.25v_21512 | 1 | 0 | 0 | 0 | 0 | 0 | 0 | 0 | 0 | 0 | 0 | 0 | 0 | 0 | 0 | 0 | 0 | 0 | 0 | 0 | 0 | 0 | 0 | 1 | 0 | 0 | 0 | 0 | 0 | 0 |
| Jun8.25v_9804 | 2 | 0 | 0 | 0 | 0 | 1 | 0 | 0 | 0 | 0 | 0 | 0 | 0 | 0 | 0 | 0 | 0 | 0 | 0 | 0 | 0 | 0 | 0 | 0 | 0 | 0 | 1 | 0 | 0 | 0 |
| Jul11.5v_2601 | 1 | 0 | 0 | 0 | 0 | 0 | 0 | 0 | 0 | 0 | 0 | 0 | 0 | 1 | 0 | 0 | 0 | 0 | 0 | 0 | 0 | 0 | 0 | 0 | 0 | 0 | 0 | 0 | 0 | 0 |
| Jun22.5v_16841 | 1 | 0 | 0 | 0 | 0 | 0 | 0 | 0 | 0 | 1 | 0 | 0 | 0 | 0 | 0 | 0 | 0 | 0 | 0 | 0 | 0 | 0 | 0 | 0 | 0 | 0 | 0 | 0 | 0 | 0 |
| Jul11.75v_35859 | 1 | 0 | 0 | 0 | 0 | 0 | 0 | 0 | 0 | 0 | 0 | 0 | 0 | 0 | 0 | 0 | 1 | 0 | 0 | 0 | 0 | 0 | 0 | 0 | 0 | 0 | 0 | 0 | 0 | 0 |
| Sep8.5v_19493 | 1 | 0 | 0 | 0 | 0 | 0 | 0 | 0 | 0 | 0 | 0 | 0 | 0 | 0 | 0 | 0 | 0 | 0 | 0 | 0 | 0 | 0 | 0 | 0 | 0 | 0 | 0 | 1 | 0 | 0 |
| Aug5.50v_25014 | 1 | 0 | 0 | 0 | 0 | 0 | 0 | 0 | 0 | 0 | 0 | 0 | 0 | 0 | 0 | 0 | 0 | 0 | 0 | 0 | 0 | 0 | 0 | 0 | 1 | 0 | 0 | 0 | 0 | 0 |
| Jun8.50v_29052 | 1 | 0 | 0 | 0 | 0 | 0 | 1 | 0 | 0 | 0 | 0 | 0 | 0 | 0 | 0 | 0 | 0 | 0 | 0 | 0 | 0 | 0 | 0 | 0 | 0 | 0 | 0 | 0 | 0 | 0 |
| Jul27.25v_10141 | 1 | 0 | 0 | 0 | 0 | 0 | 0 | 0 | 0 | 0 | 0 | 0 | 0 | 0 | 0 | 0 | 0 | 0 | 1 | 0 | 0 | 0 | 0 | 0 | 0 | 0 | 0 | 0 | 0 | 0 |
| May12.25v_13959 | 1 | 0 | 1 | 0 | 0 | 0 | 0 | 0 | 0 | 0 | 0 | 0 | 0 | 0 | 0 | 0 | 0 | 0 | 0 | 0 | 0 | 0 | 0 | 0 | 0 | 0 | 0 | 0 | 0 | 0 |
| Jul11.25v_670 | 1 | 0 | 0 | 0 | 0 | 0 | 0 | 0 | 0 | 0 | 0 | 0 | 0 | 0 | 1 | 0 | 0 | 0 | 0 | 0 | 0 | 0 | 0 | 0 | 0 | 0 | 0 | 0 | 0 | 0 |
| Jul11.50v_8009 | 1 | 0 | 0 | 0 | 0 | 0 | 0 | 0 | 0 | 0 | 0 | 0 | 0 | 0 | 0 | 1 | 0 | 0 | 0 | 0 | 0 | 0 | 0 | 0 | 0 | 0 | 0 | 0 | 0 | 0 |

| Feature_ID | 654 | May12_05m | May12_25m | May12_50m | May12_75m | Jun8_25m | Jun8_50m | Jun8_75m | Jun22_0m | Jun22_05m | Jun22_50m | Jun22_75m | Jul11_0m | Jul11_05m | Jul11_25m | Jul11_50m | Jul11_75m | Jul27_05m | Jul27_25m | Jul27_50m | Jul27_75m | Aug5_0m | Aug5_05m | Aug5_25m | Aug5_50m | Aug5_75m | Sep8_0m | Sep8_05m | Sep8_25m | Sep8_50m |
|---|---|---|---|---|---|---|---|---|---|---|---|---|---|---|---|---|---|---|---|---|---|---|---|---|---|---|---|---|---|---|
| Jun22.0v_13511 | 1 | 0 | 0 | 0 | 0 | 0 | 0 | 0 | 1 | 0 | 0 | 0 | 0 | 0 | 0 | 0 | 0 | 0 | 0 | 0 | 0 | 0 | 0 | 0 | 0 | 0 | 0 | 0 | 0 | 0 |
| Jul11.75v_20801 | 1 | 0 | 0 | 0 | 0 | 0 | 0 | 0 | 0 | 0 | 0 | 0 | 0 | 0 | 0 | 0 | 1 | 0 | 0 | 0 | 0 | 0 | 0 | 0 | 0 | 0 | 0 | 0 | 0 | 0 |
| Jul11.5v_4145 | 1 | 0 | 0 | 0 | 0 | 0 | 0 | 0 | 0 | 0 | 0 | 0 | 0 | 1 | 0 | 0 | 0 | 0 | 0 | 0 | 0 | 0 | 0 | 0 | 0 | 0 | 0 | 0 | 0 | 0 |
| May12.75v_27240 | 1 | 0 | 0 | 0 | 1 | 0 | 0 | 0 | 0 | 0 | 0 | 0 | 0 | 0 | 0 | 0 | 0 | 0 | 0 | 0 | 0 | 0 | 0 | 0 | 0 | 0 | 0 | 0 | 0 | 0 |
| Jun8.50v_40980 | 1 | 0 | 0 | 0 | 0 | 0 | 1 | 0 | 0 | 0 | 0 | 0 | 0 | 0 | 0 | 0 | 0 | 0 | 0 | 0 | 0 | 0 | 0 | 0 | 0 | 0 | 0 | 0 | 0 | 0 |
| Sep8.50v_4345 | 1 | 0 | 0 | 0 | 0 | 0 | 0 | 0 | 0 | 0 | 0 | 0 | 0 | 0 | 0 | 0 | 0 | 0 | 0 | 0 | 0 | 0 | 0 | 0 | 0 | 0 | 0 | 0 | 0 | 1 |
| Sep8.0v_8381 | 1 | 0 | 0 | 0 | 0 | 0 | 0 | 0 | 0 | 0 | 0 | 0 | 0 | 0 | 0 | 0 | 0 | 0 | 0 | 0 | 0 | 0 | 0 | 0 | 0 | 0 | 1 | 0 | 0 | 0 |
| Jul11.25v_2818 | 1 | 0 | 0 | 0 | 0 | 0 | 0 | 0 | 0 | 0 | 0 | 0 | 0 | 0 | 1 | 0 | 0 | 0 | 0 | 0 | 0 | 0 | 0 | 0 | 0 | 0 | 0 | 0 | 0 | 0 |
| Jun22.75v_36894 | 1 | 0 | 0 | 0 | 0 | 0 | 0 | 0 | 0 | 0 | 0 | 1 | 0 | 0 | 0 | 0 | 0 | 0 | 0 | 0 | 0 | 0 | 0 | 0 | 0 | 0 | 0 | 0 | 0 | 0 |
| May12.50v_23050 | 1 | 0 | 0 | 1 | 0 | 0 | 0 | 0 | 0 | 0 | 0 | 0 | 0 | 0 | 0 | 0 | 0 | 0 | 0 | 0 | 0 | 0 | 0 | 0 | 0 | 0 | 0 | 0 | 0 | 0 |
| May12.5v_12219 | 1 | 1 | 0 | 0 | 0 | 0 | 0 | 0 | 0 | 0 | 0 | 0 | 0 | 0 | 0 | 0 | 0 | 0 | 0 | 0 | 0 | 0 | 0 | 0 | 0 | 0 | 0 | 0 | 0 | 0 |
| Jul11.5v_7635 | 1 | 0 | 0 | 0 | 0 | 0 | 0 | 0 | 0 | 0 | 0 | 0 | 0 | 1 | 0 | 0 | 0 | 0 | 0 | 0 | 0 | 0 | 0 | 0 | 0 | 0 | 0 | 0 | 0 | 0 |
| Jun8.50v_1501 | 1 | 0 | 0 | 0 | 0 | 0 | 1 | 0 | 0 | 0 | 0 | 0 | 0 | 0 | 0 | 0 | 0 | 0 | 0 | 0 | 0 | 0 | 0 | 0 | 0 | 0 | 0 | 0 | 0 | 0 |
| Jun22.75v_34611 | 1 | 0 | 0 | 0 | 0 | 0 | 0 | 0 | 0 | 0 | 0 | 1 | 0 | 0 | 0 | 0 | 0 | 0 | 0 | 0 | 0 | 0 | 0 | 0 | 0 | 0 | 0 | 0 | 0 | 0 |
| Aug5.25v_32838 | 1 | 0 | 0 | 0 | 0 | 0 | 0 | 0 | 0 | 0 | 0 | 0 | 0 | 0 | 0 | 0 | 0 | 0 | 0 | 0 | 0 | 0 | 0 | 1 | 0 | 0 | 0 | 0 | 0 | 0 |
| Jun22.50v_24249 | 1 | 0 | 0 | 0 | 0 | 0 | 0 | 0 | 0 | 0 | 1 | 0 | 0 | 0 | 0 | 0 | 0 | 0 | 0 | 0 | 0 | 0 | 0 | 0 | 0 | 0 | 0 | 0 | 0 | 0 |
| Jul27.5v_1729 | 1 | 0 | 0 | 0 | 0 | 0 | 0 | 0 | 0 | 0 | 0 | 0 | 0 | 0 | 0 | 0 | 0 | 1 | 0 | 0 | 0 | 0 | 0 | 0 | 0 | 0 | 0 | 0 | 0 | 0 |
| Jul11.25v_3011 | 1 | 0 | 0 | 0 | 0 | 0 | 0 | 0 | 0 | 0 | 0 | 0 | 0 | 0 | 1 | 0 | 0 | 0 | 0 | 0 | 0 | 0 | 0 | 0 | 0 | 0 | 0 | 0 | 0 | 0 |
| Jul27.5v_20756 | 1 | 0 | 0 | 0 | 0 | 0 | 0 | 0 | 0 | 0 | 0 | 0 | 0 | 0 | 0 | 0 | 0 | 1 | 0 | 0 | 0 | 0 | 0 | 0 | 0 | 0 | 0 | 0 | 0 | 0 |
| Jul11.0v_16249 | 1 | 0 | 0 | 0 | 0 | 0 | 0 | 0 | 0 | 0 | 0 | 0 | 1 | 0 | 0 | 0 | 0 | 0 | 0 | 0 | 0 | 0 | 0 | 0 | 0 | 0 | 0 | 0 | 0 | 0 |
| Aug5.25v_32890 | 1 | 0 | 0 | 0 | 0 | 0 | 0 | 0 | 0 | 0 | 0 | 0 | 0 | 0 | 0 | 0 | 0 | 0 | 0 | 0 | 0 | 0 | 0 | 1 | 0 | 0 | 0 | 0 | 0 | 0 |
| May12.50v_22672 | 1 | 0 | 0 | 1 | 0 | 0 | 0 | 0 | 0 | 0 | 0 | 0 | 0 | 0 | 0 | 0 | 0 | 0 | 0 | 0 | 0 | 0 | 0 | 0 | 0 | 0 | 0 | 0 | 0 | 0 |
| May12.75v_41197 | 1 | 0 | 0 | 0 | 1 | 0 | 0 | 0 | 0 | 0 | 0 | 0 | 0 | 0 | 0 | 0 | 0 | 0 | 0 | 0 | 0 | 0 | 0 | 0 | 0 | 0 | 0 | 0 | 0 | 0 |
| Aug5.75v_13101 | 1 | 0 | 0 | 0 | 0 | 0 | 0 | 0 | 0 | 0 | 0 | 0 | 0 | 0 | 0 | 0 | 0 | 0 | 0 | 0 | 0 | 0 | 0 | 0 | 0 | 1 | 0 | 0 | 0 | 0 |
| Aug5.50v_819 | 1 | 0 | 0 | 0 | 0 | 0 | 0 | 0 | 0 | 0 | 0 | 0 | 0 | 0 | 0 | 0 | 0 | 0 | 0 | 0 | 0 | 0 | 0 | 0 | 1 | 0 | 0 | 0 | 0 | 0 |
| Jun22.50v_24243 | 1 | 0 | 0 | 0 | 0 | 0 | 0 | 0 | 0 | 0 | 1 | 0 | 0 | 0 | 0 | 0 | 0 | 0 | 0 | 0 | 0 | 0 | 0 | 0 | 0 | 0 | 0 | 0 | 0 | 0 |
| Jun22.0v_13167 | 1 | 0 | 0 | 0 | 0 | 0 | 0 | 0 | 1 | 0 | 0 | 0 | 0 | 0 | 0 | 0 | 0 | 0 | 0 | 0 | 0 | 0 | 0 | 0 | 0 | 0 | 0 | 0 | 0 | 0 |
| Jun22.75v_20691 | 1 | 0 | 0 | 0 | 0 | 0 | 0 | 0 | 0 | 0 | 0 | 1 | 0 | 0 | 0 | 0 | 0 | 0 | 0 | 0 | 0 | 0 | 0 | 0 | 0 | 0 | 0 | 0 | 0 | 0 |
| Aug5.25v_10776 | 1 | 0 | 0 | 0 | 0 | 0 | 0 | 0 | 0 | 0 | 0 | 0 | 0 | 0 | 0 | 0 | 0 | 0 | 0 | 0 | 0 | 0 | 0 | 1 | 0 | 0 | 0 | 0 | 0 | 0 |
| May12.50v_9068 | 1 | 0 | 0 | 1 | 0 | 0 | 0 | 0 | 0 | 0 | 0 | 0 | 0 | 0 | 0 | 0 | 0 | 0 | 0 | 0 | 0 | 0 | 0 | 0 | 0 | 0 | 0 | 0 | 0 | 0 |
| Jun22.50v_15527 | 1 | 0 | 0 | 0 | 0 | 0 | 0 | 0 | 0 | 0 | 1 | 0 | 0 | 0 | 0 | 0 | 0 | 0 | 0 | 0 | 0 | 0 | 0 | 0 | 0 | 0 | 0 | 0 | 0 | 0 |
| Aug5.5v_2741 | 1 | 0 | 0 | 0 | 0 | 0 | 0 | 0 | 0 | 0 | 0 | 0 | 0 | 0 | 0 | 0 | 0 | 0 | 0 | 0 | 0 | 0 | 1 | 0 | 0 | 0 | 0 | 0 | 0 | 0 |
| Jun22.5v_5666 | 3 | 1 | 0 | 0 | 0 | 0 | 0 | 0 | 0 | 1 | 1 | 0 | 0 | 0 | 0 | 0 | 0 | 0 | 0 | 0 | 0 | 0 | 0 | 0 | 0 | 0 | 0 | 0 | 0 | 0 |
| Jul27.5v_18654 | 1 | 0 | 0 | 0 | 0 | 0 | 0 | 0 | 0 | 0 | 0 | 0 | 0 | 0 | 0 | 0 | 0 | 1 | 0 | 0 | 0 | 0 | 0 | 0 | 0 | 0 | 0 | 0 | 0 | 0 |
| Aug5.5v_37637 | 1 | 0 | 0 | 0 | 0 | 0 | 0 | 0 | 0 | 0 | 0 | 0 | 0 | 0 | 0 | 0 | 0 | 0 | 0 | 0 | 0 | 0 | 1 | 0 | 0 | 0 | 0 | 0 | 0 | 0 |
| Jun22.50v_34065 | 1 | 0 | 0 | 0 | 0 | 0 | 0 | 0 | 0 | 0 | 1 | 0 | 0 | 0 | 0 | 0 | 0 | 0 | 0 | 0 | 0 | 0 | 0 | 0 | 0 | 0 | 0 | 0 | 0 | 0 |
| Jun22.50v_8056 | 1 | 0 | 0 | 0 | 0 | 0 | 0 | 0 | 0 | 0 | 1 | 0 | 0 | 0 | 0 | 0 | 0 | 0 | 0 | 0 | 0 | 0 | 0 | 0 | 0 | 0 | 0 | 0 | 0 | 0 |
| Aug5.0v_544 | 1 | 0 | 0 | 0 | 0 | 0 | 0 | 0 | 0 | 0 | 0 | 0 | 0 | 0 | 0 | 0 | 0 | 0 | 0 | 0 | 0 | 1 | 0 | 0 | 0 | 0 | 0 | 0 | 0 | 0 |
| May12.50v_36503 | 3 | 0 | 0 | 1 | 0 | 0 | 1 | 0 | 1 | 0 | 0 | 0 | 0 | 0 | 0 | 0 | 0 | 0 | 0 | 0 | 0 | 0 | 0 | 0 | 0 | 0 | 0 | 0 | 0 | 0 |
| Jul27.50v_20154 | 1 | 0 | 0 | 0 | 0 | 0 | 0 | 0 | 0 | 0 | 0 | 0 | 0 | 0 | 0 | 0 | 0 | 0 | 0 | 1 | 0 | 0 | 0 | 0 | 0 | 0 | 0 | 0 | 0 | 0 |
| Jul11.5v_10738 | 1 | 0 | 0 | 0 | 0 | 0 | 0 | 0 | 0 | 0 | 0 | 0 | 0 | 1 | 0 | 0 | 0 | 0 | 0 | 0 | 0 | 0 | 0 | 0 | 0 | 0 | 0 | 0 | 0 | 0 |
| Jun22.50v_18709 | 1 | 0 | 0 | 0 | 0 | 0 | 0 | 0 | 0 | 0 | 1 | 0 | 0 | 0 | 0 | 0 | 0 | 0 | 0 | 0 | 0 | 0 | 0 | 0 | 0 | 0 | 0 | 0 | 0 | 0 |
| May12.50v_33944 | 1 | 0 | 0 | 1 | 0 | 0 | 0 | 0 | 0 | 0 | 0 | 0 | 0 | 0 | 0 | 0 | 0 | 0 | 0 | 0 | 0 | 0 | 0 | 0 | 0 | 0 | 0 | 0 | 0 | 0 |
| Jun22.50v_3935 | 1 | 0 | 0 | 0 | 0 | 0 | 0 | 0 | 0 | 0 | 1 | 0 | 0 | 0 | 0 | 0 | 0 | 0 | 0 | 0 | 0 | 0 | 0 | 0 | 0 | 0 | 0 | 0 | 0 | 0 |
| May12.50v_2546 | 1 | 0 | 0 | 1 | 0 | 0 | 0 | 0 | 0 | 0 | 0 | 0 | 0 | 0 | 0 | 0 | 0 | 0 | 0 | 0 | 0 | 0 | 0 | 0 | 0 | 0 | 0 | 0 | 0 | 0 |
| May12.75v_7716 | 1 | 0 | 0 | 0 | 1 | 0 | 0 | 0 | 0 | 0 | 0 | 0 | 0 | 0 | 0 | 0 | 0 | 0 | 0 | 0 | 0 | 0 | 0 | 0 | 0 | 0 | 0 | 0 | 0 | 0 |
| May12.50v_24025 | 1 | 0 | 0 | 1 | 0 | 0 | 0 | 0 | 0 | 0 | 0 | 0 | 0 | 0 | 0 | 0 | 0 | 0 | 0 | 0 | 0 | 0 | 0 | 0 | 0 | 0 | 0 | 0 | 0 | 0 |
| May12.25v_36750 | 1 | 0 | 1 | 0 | 0 | 0 | 0 | 0 | 0 | 0 | 0 | 0 | 0 | 0 | 0 | 0 | 0 | 0 | 0 | 0 | 0 | 0 | 0 | 0 | 0 | 0 | 0 | 0 | 0 | 0 |
| Aug5.0v_2249 | 1 | 0 | 0 | 0 | 0 | 0 | 0 | 0 | 0 | 0 | 0 | 0 | 0 | 0 | 0 | 0 | 0 | 0 | 0 | 0 | 0 | 1 | 0 | 0 | 0 | 0 | 0 | 0 | 0 | 0 |
| May12.5v_34493 | 1 | 1 | 0 | 0 | 0 | 0 | 0 | 0 | 0 | 0 | 0 | 0 | 0 | 0 | 0 | 0 | 0 | 0 | 0 | 0 | 0 | 0 | 0 | 0 | 0 | 0 | 0 | 0 | 0 | 0 |
| Jul11.0v_19379 | 1 | 0 | 0 | 0 | 0 | 0 | 0 | 0 | 0 | 0 | 0 | 0 | 1 | 0 | 0 | 0 | 0 | 0 | 0 | 0 | 0 | 0 | 0 | 0 | 0 | 0 | 0 | 0 | 0 | 0 |
| May12.25v_18305 | 1 | 0 | 1 | 0 | 0 | 0 | 0 | 0 | 0 | 0 | 0 | 0 | 0 | 0 | 0 | 0 | 0 | 0 | 0 | 0 | 0 | 0 | 0 | 0 | 0 | 0 | 0 | 0 | 0 | 0 |
| Jun8.50v_18325 | 1 | 0 | 0 | 0 | 0 | 0 | 1 | 0 | 0 | 0 | 0 | 0 | 0 | 0 | 0 | 0 | 0 | 0 | 0 | 0 | 0 | 0 | 0 | 0 | 0 | 0 | 0 | 0 | 0 | 0 |
| Jun22.75v_25108 | 1 | 0 | 0 | 0 | 0 | 0 | 0 | 0 | 0 | 0 | 0 | 1 | 0 | 0 | 0 | 0 | 0 | 0 | 0 | 0 | 0 | 0 | 0 | 0 | 0 | 0 | 0 | 0 | 0 | 0 |
| May12.25v_12305 | 1 | 0 | 1 | 0 | 0 | 0 | 0 | 0 | 0 | 0 | 0 | 0 | 0 | 0 | 0 | 0 | 0 | 0 | 0 | 0 | 0 | 0 | 0 | 0 | 0 | 0 | 0 | 0 | 0 | 0 |
| Jul27.5v_6721 | 1 | 0 | 0 | 0 | 0 | 0 | 0 | 0 | 0 | 0 | 0 | 0 | 0 | 0 | 0 | 0 | 0 | 1 | 0 | 0 | 0 | 0 | 0 | 0 | 0 | 0 | 0 | 0 | 0 | 0 |
| May12.50v_26530 | 1 | 0 | 0 | 1 | 0 | 0 | 0 | 0 | 0 | 0 | 0 | 0 | 0 | 0 | 0 | 0 | 0 | 0 | 0 | 0 | 0 | 0 | 0 | 0 | 0 | 0 | 0 | 0 | 0 | 0 |
| May12.75v_131 | 1 | 0 | 0 | 0 | 1 | 0 | 0 | 0 | 0 | 0 | 0 | 0 | 0 | 0 | 0 | 0 | 0 | 0 | 0 | 0 | 0 | 0 | 0 | 0 | 0 | 0 | 0 | 0 | 0 | 0 |
| May12.50v_16193 | 1 | 0 | 0 | 1 | 0 | 0 | 0 | 0 | 0 | 0 | 0 | 0 | 0 | 0 | 0 | 0 | 0 | 0 | 0 | 0 | 0 | 0 | 0 | 0 | 0 | 0 | 0 | 0 | 0 | 0 |
| Jul11.0v_12640 | 1 | 0 | 0 | 0 | 0 | 0 | 0 | 0 | 0 | 0 | 0 | 0 | 1 | 0 | 0 | 0 | 0 | 0 | 0 | 0 | 0 | 0 | 0 | 0 | 0 | 0 | 0 | 0 | 0 | 0 |
| May12.50v_32124 | 3 | 0 | 0 | 1 | 1 | 0 | 0 | 0 | 0 | 0 | 0 | 0 | 0 | 0 | 0 | 0 | 0 | 0 | 0 | 0 | 0 | 0 | 0 | 0 | 0 | 0 | 1 | 0 | 0 | 0 |
| Jun22.75v_27325 | 1 | 0 | 0 | 0 | 0 | 0 | 0 | 0 | 0 | 0 | 0 | 1 | 0 | 0 | 0 | 0 | 0 | 0 | 0 | 0 | 0 | 0 | 0 | 0 | 0 | 0 | 0 | 0 | 0 | 0 |
| Jul27.5v_5597 | 1 | 0 | 0 | 0 | 0 | 0 | 0 | 0 | 0 | 0 | 0 | 0 | 0 | 0 | 0 | 0 | 0 | 1 | 0 | 0 | 0 | 0 | 0 | 0 | 0 | 0 | 0 | 0 | 0 | 0 |
| May12.25v_29819 | 1 | 0 | 1 | 0 | 0 | 0 | 0 | 0 | 0 | 0 | 0 | 0 | 0 | 0 | 0 | 0 | 0 | 0 | 0 | 0 | 0 | 0 | 0 | 0 | 0 | 0 | 0 | 0 | 0 | 0 |
| Jun22.0v_11123 | 1 | 0 | 0 | 0 | 0 | 0 | 0 | 0 | 1 | 0 | 0 | 0 | 0 | 0 | 0 | 0 | 0 | 0 | 0 | 0 | 0 | 0 | 0 | 0 | 0 | 0 | 0 | 0 | 0 | 0 |
| Sep8.5v_14369 | 1 | 0 | 0 | 0 | 0 | 0 | 0 | 0 | 0 | 0 | 0 | 0 | 0 | 0 | 0 | 0 | 0 | 0 | 0 | 0 | 0 | 0 | 0 | 0 | 0 | 0 | 0 | 1 | 0 | 0 |
| May12.50v_7332 | 1 | 0 | 0 | 1 | 0 | 0 | 0 | 0 | 0 | 0 | 0 | 0 | 0 | 0 | 0 | 0 | 0 | 0 | 0 | 0 | 0 | 0 | 0 | 0 | 0 | 0 | 0 | 0 | 0 | 0 |
| May12.50v_24857 | 1 | 0 | 0 | 1 | 0 | 0 | 0 | 0 | 0 | 0 | 0 | 0 | 0 | 0 | 0 | 0 | 0 | 0 | 0 | 0 | 0 | 0 | 0 | 0 | 0 | 0 | 0 | 0 | 0 | 0 |
| May12.50v_14006 | 1 | 0 | 0 | 1 | 0 | 0 | 0 | 0 | 0 | 0 | 0 | 0 | 0 | 0 | 0 | 0 | 0 | 0 | 0 | 0 | 0 | 0 | 0 | 0 | 0 | 0 | 0 | 0 | 0 | 0 |
| May12.50v_32268 | 1 | 0 | 0 | 1 | 0 | 0 | 0 | 0 | 0 | 0 | 0 | 0 | 0 | 0 | 0 | 0 | 0 | 0 | 0 | 0 | 0 | 0 | 0 | 0 | 0 | 0 | 0 | 0 | 0 | 0 |
| May12.50v_9493 | 2 | 0 | 0 | 1 | 1 | 0 | 0 | 0 | 0 | 0 | 0 | 0 | 0 | 0 | 0 | 0 | 0 | 0 | 0 | 0 | 0 | 0 | 0 | 0 | 0 | 0 | 0 | 0 | 0 | 0 |
| Jul11.0v_10255 | 1 | 0 | 0 | 0 | 0 | 0 | 0 | 0 | 0 | 0 | 0 | 0 | 1 | 0 | 0 | 0 | 0 | 0 | 0 | 0 | 0 | 0 | 0 | 0 | 0 | 0 | 0 | 0 | 0 | 0 |
| Jun8.50v_5121 | 1 | 0 | 0 | 0 | 0 | 0 | 1 | 0 | 0 | 0 | 0 | 0 | 0 | 0 | 0 | 0 | 0 | 0 | 0 | 0 | 0 | 0 | 0 | 0 | 0 | 0 | 0 | 0 | 0 | 0 |
| Jul11.0v_20941 | 1 | 0 | 0 | 0 | 0 | 0 | 0 | 0 | 0 | 0 | 0 | 0 | 1 | 0 | 0 | 0 | 0 | 0 | 0 | 0 | 0 | 0 | 0 | 0 | 0 | 0 | 0 | 0 | 0 | 0 |
| Jul27.5v_10199 | 1 | 0 | 0 | 0 | 0 | 0 | 0 | 0 | 0 | 0 | 0 | 0 | 0 | 0 | 0 | 0 | 0 | 1 | 0 | 0 | 0 | 0 | 0 | 0 | 0 | 0 | 0 | 0 | 0 | 0 |
| Jul11.50v_18621 | 1 | 0 | 0 | 0 | 0 | 0 | 0 | 0 | 0 | 0 | 0 | 0 | 0 | 0 | 0 | 1 | 0 | 0 | 0 | 0 | 0 | 0 | 0 | 0 | 0 | 0 | 0 | 0 | 0 | 0 |
| Jun22.50v_21051 | 1 | 0 | 0 | 0 | 0 | 0 | 0 | 0 | 0 | 0 | 1 | 0 | 0 | 0 | 0 | 0 | 0 | 0 | 0 | 0 | 0 | 0 | 0 | 0 | 0 | 0 | 0 | 0 | 0 | 0 |
| May12.75v_943 | 1 | 0 | 0 | 0 | 1 | 0 | 0 | 0 | 0 | 0 | 0 | 0 | 0 | 0 | 0 | 0 | 0 | 0 | 0 | 0 | 0 | 0 | 0 | 0 | 0 | 0 | 0 | 0 | 0 | 0 |
| Jun22.50v_9657 | 2 | 0 | 1 | 0 | 0 | 0 | 0 | 0 | 0 | 0 | 1 | 0 | 0 | 0 | 0 | 0 | 0 | 0 | 0 | 0 | 0 | 0 | 0 | 0 | 0 | 0 | 0 | 0 | 0 | 0 |
| Aug5.50v_4328 | 1 | 0 | 0 | 0 | 0 | 0 | 0 | 0 | 0 | 0 | 0 | 0 | 0 | 0 | 0 | 0 | 0 | 0 | 0 | 0 | 0 | 0 | 0 | 0 | 1 | 0 | 0 | 0 | 0 | 0 |
| Aug5.5v_33951 | 1 | 0 | 0 | 0 | 0 | 0 | 0 | 0 | 0 | 0 | 0 | 0 | 0 | 0 | 0 | 0 | 0 | 0 | 0 | 0 | 0 | 0 | 1 | 0 | 0 | 0 | 0 | 0 | 0 | 0 |
| Jul11.5v_13543 | 1 | 0 | 0 | 0 | 0 | 0 | 0 | 0 | 0 | 0 | 0 | 0 | 0 | 1 | 0 | 0 | 0 | 0 | 0 | 0 | 0 | 0 | 0 | 0 | 0 | 0 | 0 | 0 | 0 | 0 |
| Aug5.75v_14297 | 1 | 0 | 0 | 0 | 0 | 0 | 0 | 0 | 0 | 0 | 0 | 0 | 0 | 0 | 0 | 0 | 0 | 0 | 0 | 0 | 0 | 0 | 0 | 0 | 0 | 1 | 0 | 0 | 0 | 0 |
| Sep8.50v_36971 | 1 | 0 | 0 | 0 | 0 | 0 | 0 | 0 | 0 | 0 | 0 | 0 | 0 | 0 | 0 | 0 | 0 | 0 | 0 | 0 | 0 | 0 | 0 | 0 | 0 | 0 | 0 | 0 | 0 | 1 |
| May12.50v_19478 | 1 | 0 | 0 | 1 | 0 | 0 | 0 | 0 | 0 | 0 | 0 | 0 | 0 | 0 | 0 | 0 | 0 | 0 | 0 | 0 | 0 | 0 | 0 | 0 | 0 | 0 | 0 | 0 | 0 | 0 |
| May12.25v_25279 | 1 | 0 | 1 | 0 | 0 | 0 | 0 | 0 | 0 | 0 | 0 | 0 | 0 | 0 | 0 | 0 | 0 | 0 | 0 | 0 | 0 | 0 | 0 | 0 | 0 | 0 | 0 | 0 | 0 | 0 |
| May12.25v_23907 | 1 | 0 | 1 | 0 | 0 | 0 | 0 | 0 | 0 | 0 | 0 | 0 | 0 | 0 | 0 | 0 | 0 | 0 | 0 | 0 | 0 | 0 | 0 | 0 | 0 | 0 | 0 | 0 | 0 | 0 |
| Sep8.5v_4351 | 1 | 0 | 0 | 0 | 0 | 0 | 0 | 0 | 0 | 0 | 0 | 0 | 0 | 0 | 0 | 0 | 0 | 0 | 0 | 0 | 0 | 0 | 0 | 0 | 0 | 0 | 0 | 1 | 0 | 0 |
| May12.25v_25908 | 1 | 0 | 1 | 0 | 0 | 0 | 0 | 0 | 0 | 0 | 0 | 0 | 0 | 0 | 0 | 0 | 0 | 0 | 0 | 0 | 0 | 0 | 0 | 0 | 0 | 0 | 0 | 0 | 0 | 0 |
| May12.50v_38567 | 1 | 0 | 0 | 1 | 0 | 0 | 0 | 0 | 0 | 0 | 0 | 0 | 0 | 0 | 0 | 0 | 0 | 0 | 0 | 0 | 0 | 0 | 0 | 0 | 0 | 0 | 0 | 0 | 0 | 0 |
| Jun22.0v_17081 | 1 | 0 | 0 | 0 | 0 | 0 | 0 | 0 | 1 | 0 | 0 | 0 | 0 | 0 | 0 | 0 | 0 | 0 | 0 | 0 | 0 | 0 | 0 | 0 | 0 | 0 | 0 | 0 | 0 | 0 |
| Jun22.75v_26421 | 1 | 0 | 0 | 0 | 0 | 0 | 0 | 0 | 0 | 0 | 0 | 1 | 0 | 0 | 0 | 0 | 0 | 0 | 0 | 0 | 0 | 0 | 0 | 0 | 0 | 0 | 0 | 0 | 0 | 0 |
| Aug5.5v_38100 | 1 | 0 | 0 | 0 | 0 | 0 | 0 | 0 | 0 | 0 | 0 | 0 | 0 | 0 | 0 | 0 | 0 | 0 | 0 | 0 | 0 | 0 | 1 | 0 | 0 | 0 | 0 | 0 | 0 | 0 |
| Jun22.50v_1167 | 1 | 0 | 0 | 0 | 0 | 0 | 0 | 0 | 0 | 0 | 1 | 0 | 0 | 0 | 0 | 0 | 0 | 0 | 0 | 0 | 0 | 0 | 0 | 0 | 0 | 0 | 0 | 0 | 0 | 0 |
| May12.50v_19033 | 1 | 0 | 0 | 1 | 0 | 0 | 0 | 0 | 0 | 0 | 0 | 0 | 0 | 0 | 0 | 0 | 0 | 0 | 0 | 0 | 0 | 0 | 0 | 0 | 0 | 0 | 0 | 0 | 0 | 0 |
| Jul27.50v_24094 | 1 | 0 | 0 | 0 | 0 | 0 | 0 | 0 | 0 | 0 | 0 | 0 | 0 | 0 | 0 | 0 | 0 | 0 | 0 | 1 | 0 | 0 | 0 | 0 | 0 | 0 | 0 | 0 | 0 | 0 |
| Jul27.75v_4233 | 1 | 0 | 0 | 0 | 0 | 0 | 0 | 0 | 0 | 0 | 0 | 0 | 0 | 0 | 0 | 0 | 0 | 0 | 0 | 0 | 1 | 0 | 0 | 0 | 0 | 0 | 0 | 0 | 0 | 0 |
| Jun22.50v_5212 | 1 | 0 | 0 | 0 | 0 | 0 | 0 | 0 | 0 | 0 | 1 | 0 | 0 | 0 | 0 | 0 | 0 | 0 | 0 | 0 | 0 | 0 | 0 | 0 | 0 | 0 | 0 | 0 | 0 | 0 |
| Jun22.0v_19212 | 1 | 0 | 0 | 0 | 0 | 0 | 0 | 0 | 1 | 0 | 0 | 0 | 0 | 0 | 0 | 0 | 0 | 0 | 0 | 0 | 0 | 0 | 0 | 0 | 0 | 0 | 0 | 0 | 0 | 0 |
| May12.50v_24998 | 1 | 0 | 0 | 1 | 0 | 0 | 0 | 0 | 0 | 0 | 0 | 0 | 0 | 0 | 0 | 0 | 0 | 0 | 0 | 0 | 0 | 0 | 0 | 0 | 0 | 0 | 0 | 0 | 0 | 0 |
| Jun22.0v_14147 | 1 | 0 | 0 | 0 | 0 | 0 | 0 | 0 | 1 | 0 | 0 | 0 | 0 | 0 | 0 | 0 | 0 | 0 | 0 | 0 | 0 | 0 | 0 | 0 | 0 | 0 | 0 | 0 | 0 | 0 |
| May12.75v_7026 | 1 | 0 | 0 | 0 | 1 | 0 | 0 | 0 | 0 | 0 | 0 | 0 | 0 | 0 | 0 | 0 | 0 | 0 | 0 | 0 | 0 | 0 | 0 | 0 | 0 | 0 | 0 | 0 | 0 | 0 |
| Aug5.5v_17357 | 1 | 0 | 0 | 0 | 0 | 0 | 0 | 0 | 0 | 0 | 0 | 0 | 0 | 0 | 0 | 0 | 0 | 0 | 0 | 0 | 0 | 0 | 1 | 0 | 0 | 0 | 0 | 0 | 0 | 0 |
| Jun22.5v_9724 | 1 | 0 | 0 | 0 | 0 | 0 | 0 | 0 | 0 | 1 | 0 | 0 | 0 | 0 | 0 | 0 | 0 | 0 | 0 | 0 | 0 | 0 | 0 | 0 | 0 | 0 | 0 | 0 | 0 | 0 |
| May12.50v_22449 | 1 | 0 | 0 | 1 | 0 | 0 | 0 | 0 | 0 | 0 | 0 | 0 | 0 | 0 | 0 | 0 | 0 | 0 | 0 | 0 | 0 | 0 | 0 | 0 | 0 | 0 | 0 | 0 | 0 | 0 |
| Jul11.5v_1464 | 1 | 0 | 0 | 0 | 0 | 0 | 0 | 0 | 0 | 0 | 0 | 0 | 0 | 1 | 0 | 0 | 0 | 0 | 0 | 0 | 0 | 0 | 0 | 0 | 0 | 0 | 0 | 0 | 0 | 0 |
| Sep8.50v_11399 | 1 | 0 | 0 | 0 | 0 | 0 | 0 | 0 | 0 | 0 | 0 | 0 | 0 | 0 | 0 | 0 | 0 | 0 | 0 | 0 | 0 | 0 | 0 | 0 | 0 | 0 | 0 | 0 | 0 | 1 |
| May12.75v_38391 | 1 | 0 | 0 | 0 | 1 | 0 | 0 | 0 | 0 | 0 | 0 | 0 | 0 | 0 | 0 | 0 | 0 | 0 | 0 | 0 | 0 | 0 | 0 | 0 | 0 | 0 | 0 | 0 | 0 | 0 |
| Aug5.0v_945 | 3 | 0 | 1 | 0 | 0 | 0 | 0 | 0 | 0 | 0 | 0 | 0 | 0 | 0 | 0 | 0 | 0 | 0 | 0 | 0 | 0 | 2 | 0 | 0 | 0 | 0 | 0 | 0 | 0 | 0 |
| Aug5.0v_3301 | 1 | 0 | 0 | 0 | 0 | 0 | 0 | 0 | 0 | 0 | 0 | 0 | 0 | 0 | 0 | 0 | 0 | 0 | 0 | 0 | 0 | 1 | 0 | 0 | 0 | 0 | 0 | 0 | 0 | 0 |
| Aug5.0v_583 | 1 | 0 | 0 | 0 | 0 | 0 | 0 | 0 | 0 | 0 | 0 | 0 | 0 | 0 | 0 | 0 | 0 | 0 | 0 | 0 | 0 | 1 | 0 | 0 | 0 | 0 | 0 | 0 | 0 | 0 |
| Jun8.50v_25764 | 1 | 0 | 0 | 0 | 0 | 0 | 1 | 0 | 0 | 0 | 0 | 0 | 0 | 0 | 0 | 0 | 0 | 0 | 0 | 0 | 0 | 0 | 0 | 0 | 0 | 0 | 0 | 0 | 0 | 0 |
| May12.50v_15364 | 1 | 0 | 0 | 1 | 0 | 0 | 0 | 0 | 0 | 0 | 0 | 0 | 0 | 0 | 0 | 0 | 0 | 0 | 0 | 0 | 0 | 0 | 0 | 0 | 0 | 0 | 0 | 0 | 0 | 0 |
| Aug5.5v_31142 | 1 | 0 | 0 | 0 | 0 | 0 | 0 | 0 | 0 | 0 | 0 | 0 | 0 | 0 | 0 | 0 | 0 | 0 | 0 | 0 | 0 | 0 | 1 | 0 | 0 | 0 | 0 | 0 | 0 | 0 |
| Jul11.0v_8246 | 1 | 0 | 0 | 0 | 0 | 0 | 0 | 0 | 0 | 0 | 0 | 0 | 1 | 0 | 0 | 0 | 0 | 0 | 0 | 0 | 0 | 0 | 0 | 0 | 0 | 0 | 0 | 0 | 0 | 0 |
| Jul11.5v_11175 | 1 | 0 | 0 | 0 | 0 | 0 | 0 | 0 | 0 | 0 | 0 | 0 | 0 | 1 | 0 | 0 | 0 | 0 | 0 | 0 | 0 | 0 | 0 | 0 | 0 | 0 | 0 | 0 | 0 | 0 |
| Jun22.75v_36373 | 1 | 0 | 0 | 0 | 0 | 0 | 0 | 0 | 0 | 0 | 0 | 1 | 0 | 0 | 0 | 0 | 0 | 0 | 0 | 0 | 0 | 0 | 0 | 0 | 0 | 0 | 0 | 0 | 0 | 0 |
| Jun22.5v_13565 | 1 | 0 | 0 | 0 | 0 | 0 | 0 | 0 | 0 | 1 | 0 | 0 | 0 | 0 | 0 | 0 | 0 | 0 | 0 | 0 | 0 | 0 | 0 | 0 | 0 | 0 | 0 | 0 | 0 | 0 |
| Sep8.50v_28177 | 1 | 0 | 0 | 0 | 0 | 0 | 0 | 0 | 0 | 0 | 0 | 0 | 0 | 0 | 0 | 0 | 0 | 0 | 0 | 0 | 0 | 0 | 0 | 0 | 0 | 0 | 0 | 0 | 0 | 1 |
| May12.50v_30988 | 1 | 0 | 0 | 1 | 0 | 0 | 0 | 0 | 0 | 0 | 0 | 0 | 0 | 0 | 0 | 0 | 0 | 0 | 0 | 0 | 0 | 0 | 0 | 0 | 0 | 0 | 0 | 0 | 0 | 0 |
| May12.75v_39352 | 2 | 0 | 0 | 0 | 1 | 0 | 0 | 0 | 0 | 0 | 0 | 0 | 0 | 0 | 0 | 0 | 0 | 0 | 0 | 0 | 0 | 0 | 1 | 0 | 0 | 0 | 0 | 0 | 0 | 0 |
| Sep8.5v_3741 | 1 | 0 | 0 | 0 | 0 | 0 | 0 | 0 | 0 | 0 | 0 | 0 | 0 | 0 | 0 | 0 | 0 | 0 | 0 | 0 | 0 | 0 | 0 | 0 | 0 | 0 | 0 | 1 | 0 | 0 |
| Jul11.0v_19453 | 18 | 0 | 1 | 2 | 0 | 0 | 0 | 0 | 0 | 1 | 2 | 2 | 2 | 1 | 0 | 0 | 0 | 1 | 0 | 1 | 0 | 0 | 0 | 0 | 1 | 1 | 0 | 0 | 2 | 1 |
| Sep8.25v_3817 | 5 | 0 | 0 | 2 | 0 | 0 | 0 | 0 | 0 | 0 | 0 | 0 | 0 | 0 | 0 | 0 | 0 | 0 | 0 | 0 | 0 | 1 | 0 | 0 | 0 | 0 | 0 | 1 | 1 | 0 |
| May12.50v_13189 | 4 | 0 | 1 | 2 | 0 | 0 | 0 | 0 | 0 | 0 | 0 | 0 | 0 | 0 | 0 | 0 | 0 | 0 | 0 | 0 | 0 | 0 | 0 | 0 | 0 | 0 | 0 | 1 | 0 | 0 |
| Aug5.5v_24892 | 278 | 34 | 24 | 31 | 8 | 6 | 2 | 4 | 8 | 20 | 23 | 9 | 2 | 10 | 6 | 1 | 3 | 16 | 2 | 7 | 0 | 11 | 9 | 5 | 3 | 5 | 1 | 22 | 1 | 5 |
| Jun22.5v_13870 | 4 | 0 | 0 | 0 | 0 | 0 | 0 | 0 | 0 | 1 | 0 | 0 | 0 | 0 | 0 | 0 | 0 | 1 | 0 | 0 | 0 | 0 | 0 | 0 | 0 | 0 | 0 | 2 | 0 | 0 |
| May12.75v_20408 | 1 | 0 | 0 | 0 | 1 | 0 | 0 | 0 | 0 | 0 | 0 | 0 | 0 | 0 | 0 | 0 | 0 | 0 | 0 | 0 | 0 | 0 | 0 | 0 | 0 | 0 | 0 | 0 | 0 | 0 |
| Aug5.25v_15238 | 422 | 34 | 46 | 50 | 15 | 4 | 7 | 2 | 7 | 26 | 24 | 9 | 11 | 42 | 12 | 3 | 7 | 22 | 6 | 6 | 1 | 23 | 10 | 7 | 3 | 9 | 6 | 24 | 0 | 6 |
| Jun8.75v_6822 | 5 | 0 | 0 | 0 | 0 | 0 | 0 | 0 | 1 | 1 | 1 | 0 | 0 | 0 | 0 | 0 | 0 | 1 | 0 | 0 | 0 | 1 | 0 | 0 | 0 | 0 | 0 | 0 | 0 | 0 |
| May12.50v_12855 | 2 | 0 | 0 | 2 | 0 | 0 | 0 | 0 | 0 | 0 | 0 | 0 | 0 | 0 | 0 | 0 | 0 | 0 | 0 | 0 | 0 | 0 | 0 | 0 | 0 | 0 | 0 | 0 | 0 | 0 |
| Jul11.5v_13492 | 1 | 0 | 0 | 0 | 0 | 0 | 0 | 0 | 0 | 0 | 0 | 0 | 0 | 1 | 0 | 0 | 0 | 0 | 0 | 0 | 0 | 0 | 0 | 0 | 0 | 0 | 0 | 0 | 0 | 0 |
| Sep8.5v_19756 | 24 | 1 | 3 | 3 | 1 | 1 | 1 | 1 | 2 | 3 | 0 | 0 | 0 | 2 | 0 | 0 | 1 | 0 | 0 | 0 | 1 | 0 | 0 | 1 | 0 | 0 | 0 | 3 | 0 | 0 |
| Jun22.0v_12960 | 1 | 0 | 0 | 0 | 0 | 0 | 0 | 0 | 1 | 0 | 0 | 0 | 0 | 0 | 0 | 0 | 0 | 0 | 0 | 0 | 0 | 0 | 0 | 0 | 0 | 0 | 0 | 0 | 0 | 0 |
| May12.50v_20096 | 1 | 0 | 0 | 1 | 0 | 0 | 0 | 0 | 0 | 0 | 0 | 0 | 0 | 0 | 0 | 0 | 0 | 0 | 0 | 0 | 0 | 0 | 0 | 0 | 0 | 0 | 0 | 0 | 0 | 0 |
| May12.50v_35559 | 1 | 0 | 0 | 1 | 0 | 0 | 0 | 0 | 0 | 0 | 0 | 0 | 0 | 0 | 0 | 0 | 0 | 0 | 0 | 0 | 0 | 0 | 0 | 0 | 0 | 0 | 0 | 0 | 0 | 0 |
| Sep8.5v_1832 | 3 | 0 | 0 | 0 | 0 | 0 | 0 | 0 | 0 | 0 | 0 | 2 | 0 | 0 | 0 | 0 | 0 | 0 | 0 | 0 | 0 | 0 | 0 | 0 | 0 | 0 | 0 | 1 | 0 | 0 |
| May12.50v_40192 | 1 | 0 | 0 | 1 | 0 | 0 | 0 | 0 | 0 | 0 | 0 | 0 | 0 | 0 | 0 | 0 | 0 | 0 | 0 | 0 | 0 | 0 | 0 | 0 | 0 | 0 | 0 | 0 | 0 | 0 |
| Jul11.5v_13628 | 1 | 0 | 0 | 0 | 0 | 0 | 0 | 0 | 0 | 0 | 0 | 0 | 0 | 1 | 0 | 0 | 0 | 0 | 0 | 0 | 0 | 0 | 0 | 0 | 0 | 0 | 0 | 0 | 0 | 0 |
| Jul27.25v_21341 | 1 | 0 | 0 | 0 | 0 | 0 | 0 | 0 | 0 | 0 | 0 | 0 | 0 | 0 | 0 | 0 | 0 | 0 | 1 | 0 | 0 | 0 | 0 | 0 | 0 | 0 | 0 | 0 | 0 | 0 |
| Jul11.50v_16194 | 1 | 0 | 0 | 0 | 0 | 0 | 0 | 0 | 0 | 0 | 0 | 0 | 0 | 0 | 0 | 1 | 0 | 0 | 0 | 0 | 0 | 0 | 0 | 0 | 0 | 0 | 0 | 0 | 0 | 0 |
| Jul11.5v_1898 | 1 | 0 | 0 | 0 | 0 | 0 | 0 | 0 | 0 | 0 | 0 | 0 | 0 | 1 | 0 | 0 | 0 | 0 | 0 | 0 | 0 | 0 | 0 | 0 | 0 | 0 | 0 | 0 | 0 | 0 |
| Aug5.5v_33097 | 26 | 0 | 0 | 4 | 3 | 0 | 0 | 0 | 1 | 1 | 0 | 3 | 0 | 0 | 0 | 0 | 3 | 1 | 0 | 0 | 1 | 1 | 2 | 0 | 0 | 0 | 0 | 6 | 0 | 0 |
| Jul11.75v_38853 | 1 | 0 | 0 | 0 | 0 | 0 | 0 | 0 | 0 | 0 | 0 | 0 | 0 | 0 | 0 | 0 | 1 | 0 | 0 | 0 | 0 | 0 | 0 | 0 | 0 | 0 | 0 | 0 | 0 | 0 |
| Aug5.75v_28382 | 1 | 0 | 0 | 0 | 0 | 0 | 0 | 0 | 0 | 0 | 0 | 0 | 0 | 0 | 0 | 0 | 0 | 0 | 0 | 0 | 0 | 0 | 0 | 0 | 0 | 1 | 0 | 0 | 0 | 0 |
| Jun22.5v_2296 | 1 | 0 | 0 | 0 | 0 | 0 | 0 | 0 | 0 | 1 | 0 | 0 | 0 | 0 | 0 | 0 | 0 | 0 | 0 | 0 | 0 | 0 | 0 | 0 | 0 | 0 | 0 | 0 | 0 | 0 |
| May12.75v_11710 | 1 | 0 | 0 | 0 | 1 | 0 | 0 | 0 | 0 | 0 | 0 | 0 | 0 | 0 | 0 | 0 | 0 | 0 | 0 | 0 | 0 | 0 | 0 | 0 | 0 | 0 | 0 | 0 | 0 | 0 |
| Jul27.5v_13706 | 1 | 0 | 0 | 0 | 0 | 0 | 0 | 0 | 0 | 0 | 0 | 0 | 0 | 0 | 0 | 0 | 0 | 1 | 0 | 0 | 0 | 0 | 0 | 0 | 0 | 0 | 0 | 0 | 0 | 0 |
| Jul27.5v_6013 | 1 | 0 | 0 | 0 | 0 | 0 | 0 | 0 | 0 | 0 | 0 | 0 | 0 | 0 | 0 | 0 | 0 | 1 | 0 | 0 | 0 | 0 | 0 | 0 | 0 | 0 | 0 | 0 | 0 | 0 |
| Jun22.50v_18588 | 1 | 0 | 0 | 0 | 0 | 0 | 0 | 0 | 0 | 0 | 1 | 0 | 0 | 0 | 0 | 0 | 0 | 0 | 0 | 0 | 0 | 0 | 0 | 0 | 0 | 0 | 0 | 0 | 0 | 0 |
| Jun22.75v_4608 | 1 | 0 | 0 | 0 | 0 | 0 | 0 | 0 | 0 | 0 | 0 | 1 | 0 | 0 | 0 | 0 | 0 | 0 | 0 | 0 | 0 | 0 | 0 | 0 | 0 | 0 | 0 | 0 | 0 | 0 |
| Sep8.25v_19816 | 1 | 0 | 0 | 0 | 0 | 0 | 0 | 0 | 0 | 0 | 0 | 0 | 0 | 0 | 0 | 0 | 0 | 0 | 0 | 0 | 0 | 0 | 0 | 0 | 0 | 0 | 0 | 0 | 1 | 0 |
| May12.50v_13076 | 1 | 0 | 0 | 1 | 0 | 0 | 0 | 0 | 0 | 0 | 0 | 0 | 0 | 0 | 0 | 0 | 0 | 0 | 0 | 0 | 0 | 0 | 0 | 0 | 0 | 0 | 0 | 0 | 0 | 0 |
| May12.50v_12913 | 1 | 0 | 0 | 1 | 0 | 0 | 0 | 0 | 0 | 0 | 0 | 0 | 0 | 0 | 0 | 0 | 0 | 0 | 0 | 0 | 0 | 0 | 0 | 0 | 0 | 0 | 0 | 0 | 0 | 0 |
| Sep8.5v_8155 | 1 | 0 | 0 | 0 | 0 | 0 | 0 | 0 | 0 | 0 | 0 | 0 | 0 | 0 | 0 | 0 | 0 | 0 | 0 | 0 | 0 | 0 | 0 | 0 | 0 | 0 | 0 | 1 | 0 | 0 |
| Jul11.0v_3123 | 1 | 0 | 0 | 0 | 0 | 0 | 0 | 0 | 0 | 0 | 0 | 0 | 1 | 0 | 0 | 0 | 0 | 0 | 0 | 0 | 0 | 0 | 0 | 0 | 0 | 0 | 0 | 0 | 0 | 0 |
| Jul11.75v_40626 | 1 | 0 | 0 | 0 | 0 | 0 | 0 | 0 | 0 | 0 | 0 | 0 | 0 | 0 | 0 | 0 | 1 | 0 | 0 | 0 | 0 | 0 | 0 | 0 | 0 | 0 | 0 | 0 | 0 | 0 |
| Sep8.5v_9968 | 1 | 0 | 0 | 0 | 0 | 0 | 0 | 0 | 0 | 0 | 0 | 0 | 0 | 0 | 0 | 0 | 0 | 0 | 0 | 0 | 0 | 0 | 0 | 0 | 0 | 0 | 0 | 1 | 0 | 0 |
| Aug5.75v_16573 | 3 | 0 | 0 | 0 | 0 | 0 | 0 | 0 | 0 | 1 | 0 | 0 | 0 | 0 | 0 | 0 | 0 | 0 | 0 | 0 | 0 | 0 | 0 | 0 | 0 | 1 | 0 | 1 | 0 | 0 |
| Sep8.0v_18564 | 1 | 0 | 0 | 0 | 0 | 0 | 0 | 0 | 0 | 0 | 0 | 0 | 0 | 0 | 0 | 0 | 0 | 0 | 0 | 0 | 0 | 0 | 0 | 0 | 0 | 0 | 1 | 0 | 0 | 0 |
| Sep8.5v_6144 | 1 | 0 | 0 | 0 | 0 | 0 | 0 | 0 | 0 | 0 | 0 | 0 | 0 | 0 | 0 | 0 | 0 | 0 | 0 | 0 | 0 | 0 | 0 | 0 | 0 | 0 | 0 | 1 | 0 | 0 |
| Jun22.0v_8802 | 1 | 0 | 0 | 0 | 0 | 0 | 0 | 0 | 1 | 0 | 0 | 0 | 0 | 0 | 0 | 0 | 0 | 0 | 0 | 0 | 0 | 0 | 0 | 0 | 0 | 0 | 0 | 0 | 0 | 0 |
| May12.75v_20699 | 1 | 0 | 0 | 0 | 1 | 0 | 0 | 0 | 0 | 0 | 0 | 0 | 0 | 0 | 0 | 0 | 0 | 0 | 0 | 0 | 0 | 0 | 0 | 0 | 0 | 0 | 0 | 0 | 0 | 0 |
| May12.25v_20561 | 1 | 0 | 1 | 0 | 0 | 0 | 0 | 0 | 0 | 0 | 0 | 0 | 0 | 0 | 0 | 0 | 0 | 0 | 0 | 0 | 0 | 0 | 0 | 0 | 0 | 0 | 0 | 0 | 0 | 0 |
| May12.5v_30925 | 1 | 1 | 0 | 0 | 0 | 0 | 0 | 0 | 0 | 0 | 0 | 0 | 0 | 0 | 0 | 0 | 0 | 0 | 0 | 0 | 0 | 0 | 0 | 0 | 0 | 0 | 0 | 0 | 0 | 0 |
| May12.50v_39645 | 1 | 0 | 0 | 1 | 0 | 0 | 0 | 0 | 0 | 0 | 0 | 0 | 0 | 0 | 0 | 0 | 0 | 0 | 0 | 0 | 0 | 0 | 0 | 0 | 0 | 0 | 0 | 0 | 0 | 0 |
| Jun22.5v_12423 | 1 | 0 | 0 | 0 | 0 | 0 | 0 | 0 | 0 | 1 | 0 | 0 | 0 | 0 | 0 | 0 | 0 | 0 | 0 | 0 | 0 | 0 | 0 | 0 | 0 | 0 | 0 | 0 | 0 | 0 |
| May12.50v_36683 | 1 | 0 | 0 | 1 | 0 | 0 | 0 | 0 | 0 | 0 | 0 | 0 | 0 | 0 | 0 | 0 | 0 | 0 | 0 | 0 | 0 | 0 | 0 | 0 | 0 | 0 | 0 | 0 | 0 | 0 |
| Sep8.5v_8003 | 1 | 0 | 0 | 0 | 0 | 0 | 0 | 0 | 0 | 0 | 0 | 0 | 0 | 0 | 0 | 0 | 0 | 0 | 0 | 0 | 0 | 0 | 0 | 0 | 0 | 0 | 0 | 1 | 0 | 0 |
| Jul11.25v_19139 | 3 | 1 | 0 | 0 | 1 | 0 | 0 | 0 | 0 | 0 | 0 | 0 | 0 | 0 | 1 | 0 | 0 | 0 | 0 | 0 | 0 | 0 | 0 | 0 | 0 | 0 | 0 | 0 | 0 | 0 |
| Sep8.5v_17749 | 3 | 0 | 0 | 0 | 0 | 0 | 0 | 0 | 0 | 1 | 0 | 1 | 0 | 0 | 0 | 0 | 0 | 0 | 0 | 0 | 0 | 0 | 0 | 0 | 0 | 0 | 0 | 1 | 0 | 0 |
| Jul27.5v_8684 | 1 | 0 | 0 | 0 | 0 | 0 | 0 | 0 | 0 | 0 | 0 | 0 | 0 | 0 | 0 | 0 | 0 | 1 | 0 | 0 | 0 | 0 | 0 | 0 | 0 | 0 | 0 | 0 | 0 | 0 |
| Jul27.5v_17149 | 1 | 0 | 0 | 0 | 0 | 0 | 0 | 0 | 0 | 0 | 0 | 0 | 0 | 0 | 0 | 0 | 0 | 1 | 0 | 0 | 0 | 0 | 0 | 0 | 0 | 0 | 0 | 0 | 0 | 0 |
| May12.75v_40775 | 1 | 0 | 0 | 0 | 1 | 0 | 0 | 0 | 0 | 0 | 0 | 0 | 0 | 0 | 0 | 0 | 0 | 0 | 0 | 0 | 0 | 0 | 0 | 0 | 0 | 0 | 0 | 0 | 0 | 0 |
| Jul11.75v_30569 | 1 | 0 | 0 | 0 | 0 | 0 | 0 | 0 | 0 | 0 | 0 | 0 | 0 | 0 | 0 | 0 | 1 | 0 | 0 | 0 | 0 | 0 | 0 | 0 | 0 | 0 | 0 | 0 | 0 | 0 |
| Jun22.0v_9128 | 1 | 0 | 0 | 0 | 0 | 0 | 0 | 0 | 1 | 0 | 0 | 0 | 0 | 0 | 0 | 0 | 0 | 0 | 0 | 0 | 0 | 0 | 0 | 0 | 0 | 0 | 0 | 0 | 0 | 0 |
| Jun22.75v_39233 | 1 | 0 | 0 | 0 | 0 | 0 | 0 | 0 | 0 | 0 | 0 | 1 | 0 | 0 | 0 | 0 | 0 | 0 | 0 | 0 | 0 | 0 | 0 | 0 | 0 | 0 | 0 | 0 | 0 | 0 |
| May12.5v_40681 | 4 | 1 | 1 | 1 | 0 | 0 | 0 | 0 | 0 | 0 | 1 | 0 | 0 | 0 | 0 | 0 | 0 | 0 | 0 | 0 | 0 | 0 | 0 | 0 | 0 | 0 | 0 | 0 | 0 | 0 |
| May12.50v_37498 | 1 | 0 | 0 | 1 | 0 | 0 | 0 | 0 | 0 | 0 | 0 | 0 | 0 | 0 | 0 | 0 | 0 | 0 | 0 | 0 | 0 | 0 | 0 | 0 | 0 | 0 | 0 | 0 | 0 | 0 |
| Jun22.0v_6617 | 1 | 0 | 0 | 0 | 0 | 0 | 0 | 0 | 1 | 0 | 0 | 0 | 0 | 0 | 0 | 0 | 0 | 0 | 0 | 0 | 0 | 0 | 0 | 0 | 0 | 0 | 0 | 0 | 0 | 0 |
| May12.75v_11326 | 12 | 0 | 1 | 1 | 1 | 0 | 1 | 0 | 0 | 0 | 2 | 1 | 1 | 0 | 0 | 1 | 0 | 0 | 1 | 0 | 0 | 0 | 0 | 1 | 0 | 0 | 0 | 1 | 0 | 0 |
| Jul11.25v_12551 | 5 | 0 | 0 | 0 | 1 | 0 | 0 | 0 | 1 | 0 | 0 | 0 | 0 | 0 | 3 | 0 | 0 | 0 | 0 | 0 | 0 | 0 | 0 | 0 | 0 | 0 | 0 | 0 | 0 | 0 |
| May12.75v_30308 | 1 | 0 | 0 | 0 | 1 | 0 | 0 | 0 | 0 | 0 | 0 | 0 | 0 | 0 | 0 | 0 | 0 | 0 | 0 | 0 | 0 | 0 | 0 | 0 | 0 | 0 | 0 | 0 | 0 | 0 |
| Aug5.0v_5071 | 1 | 0 | 0 | 0 | 0 | 0 | 0 | 0 | 0 | 0 | 0 | 0 | 0 | 0 | 0 | 0 | 0 | 0 | 0 | 0 | 0 | 1 | 0 | 0 | 0 | 0 | 0 | 0 | 0 | 0 |
| Jun22.50v_29255 | 1 | 0 | 0 | 0 | 0 | 0 | 0 | 0 | 0 | 0 | 1 | 0 | 0 | 0 | 0 | 0 | 0 | 0 | 0 | 0 | 0 | 0 | 0 | 0 | 0 | 0 | 0 | 0 | 0 | 0 |
| Sep8.5v_8644 | 1 | 0 | 0 | 0 | 0 | 0 | 0 | 0 | 0 | 0 | 0 | 0 | 0 | 0 | 0 | 0 | 0 | 0 | 0 | 0 | 0 | 0 | 0 | 0 | 0 | 0 | 0 | 1 | 0 | 0 |
| May12.75v_16053 | 1 | 0 | 0 | 0 | 1 | 0 | 0 | 0 | 0 | 0 | 0 | 0 | 0 | 0 | 0 | 0 | 0 | 0 | 0 | 0 | 0 | 0 | 0 | 0 | 0 | 0 | 0 | 0 | 0 | 0 |
| Jun22.75v_12929 | 1 | 0 | 0 | 0 | 0 | 0 | 0 | 0 | 0 | 0 | 0 | 1 | 0 | 0 | 0 | 0 | 0 | 0 | 0 | 0 | 0 | 0 | 0 | 0 | 0 | 0 | 0 | 0 | 0 | 0 |
| May12.5v_20899 | 2 | 1 | 0 | 0 | 0 | 0 | 0 | 0 | 0 | 0 | 1 | 0 | 0 | 0 | 0 | 0 | 0 | 0 | 0 | 0 | 0 | 0 | 0 | 0 | 0 | 0 | 0 | 0 | 0 | 0 |
| Sep8.50v_25818 | 1 | 0 | 0 | 0 | 0 | 0 | 0 | 0 | 0 | 0 | 0 | 0 | 0 | 0 | 0 | 0 | 0 | 0 | 0 | 0 | 0 | 0 | 0 | 0 | 0 | 0 | 0 | 0 | 0 | 1 |
| Sep8.5v_20490 | 1 | 0 | 0 | 0 | 0 | 0 | 0 | 0 | 0 | 0 | 0 | 0 | 0 | 0 | 0 | 0 | 0 | 0 | 0 | 0 | 0 | 0 | 0 | 0 | 0 | 0 | 0 | 1 | 0 | 0 |
| Jul27.25v_17052 | 1 | 0 | 0 | 0 | 0 | 0 | 0 | 0 | 0 | 0 | 0 | 0 | 0 | 0 | 0 | 0 | 0 | 0 | 1 | 0 | 0 | 0 | 0 | 0 | 0 | 0 | 0 | 0 | 0 | 0 |
| Jun22.0v_6855 | 1 | 0 | 0 | 0 | 0 | 0 | 0 | 0 | 1 | 0 | 0 | 0 | 0 | 0 | 0 | 0 | 0 | 0 | 0 | 0 | 0 | 0 | 0 | 0 | 0 | 0 | 0 | 0 | 0 | 0 |
| Jun22.5v_20858 | 1 | 0 | 0 | 0 | 0 | 0 | 0 | 0 | 0 | 1 | 0 | 0 | 0 | 0 | 0 | 0 | 0 | 0 | 0 | 0 | 0 | 0 | 0 | 0 | 0 | 0 | 0 | 0 | 0 | 0 |
| Jun22.5v_17657 | 1 | 0 | 0 | 0 | 0 | 0 | 0 | 0 | 0 | 1 | 0 | 0 | 0 | 0 | 0 | 0 | 0 | 0 | 0 | 0 | 0 | 0 | 0 | 0 | 0 | 0 | 0 | 0 | 0 | 0 |
| Jul11.5v_3354 | 1 | 0 | 0 | 0 | 0 | 0 | 0 | 0 | 0 | 0 | 0 | 0 | 0 | 1 | 0 | 0 | 0 | 0 | 0 | 0 | 0 | 0 | 0 | 0 | 0 | 0 | 0 | 0 | 0 | 0 |
| Aug5.75v_865 | 1 | 0 | 0 | 0 | 0 | 0 | 0 | 0 | 0 | 0 | 0 | 0 | 0 | 0 | 0 | 0 | 0 | 0 | 0 | 0 | 0 | 0 | 0 | 0 | 0 | 1 | 0 | 0 | 0 | 0 |
| Jul11.50v_1692 | 1 | 0 | 0 | 0 | 0 | 0 | 0 | 0 | 0 | 0 | 0 | 0 | 0 | 0 | 0 | 1 | 0 | 0 | 0 | 0 | 0 | 0 | 0 | 0 | 0 | 0 | 0 | 0 | 0 | 0 |
| Sep8.5v_14931 | 1 | 0 | 0 | 0 | 0 | 0 | 0 | 0 | 0 | 0 | 0 | 0 | 0 | 0 | 0 | 0 | 0 | 0 | 0 | 0 | 0 | 0 | 0 | 0 | 0 | 0 | 0 | 1 | 0 | 0 |
| May12.50v_24689 | 1 | 0 | 0 | 1 | 0 | 0 | 0 | 0 | 0 | 0 | 0 | 0 | 0 | 0 | 0 | 0 | 0 | 0 | 0 | 0 | 0 | 0 | 0 | 0 | 0 | 0 | 0 | 0 | 0 | 0 |
| Sep8.0v_11931 | 1 | 0 | 0 | 0 | 0 | 0 | 0 | 0 | 0 | 0 | 0 | 0 | 0 | 0 | 0 | 0 | 0 | 0 | 0 | 0 | 0 | 0 | 0 | 0 | 0 | 0 | 1 | 0 | 0 | 0 |
| Aug5.0v_2756 | 1 | 0 | 0 | 0 | 0 | 0 | 0 | 0 | 0 | 0 | 0 | 0 | 0 | 0 | 0 | 0 | 0 | 0 | 0 | 0 | 0 | 1 | 0 | 0 | 0 | 0 | 0 | 0 | 0 | 0 |
| Jul11.25v_2017 | 1 | 0 | 0 | 0 | 0 | 0 | 0 | 0 | 0 | 0 | 0 | 0 | 0 | 0 | 1 | 0 | 0 | 0 | 0 | 0 | 0 | 0 | 0 | 0 | 0 | 0 | 0 | 0 | 0 | 0 |
| May12.5v_4193 | 1 | 1 | 0 | 0 | 0 | 0 | 0 | 0 | 0 | 0 | 0 | 0 | 0 | 0 | 0 | 0 | 0 | 0 | 0 | 0 | 0 | 0 | 0 | 0 | 0 | 0 | 0 | 0 | 0 | 0 |
| May12.25v_8747 | 1 | 0 | 1 | 0 | 0 | 0 | 0 | 0 | 0 | 0 | 0 | 0 | 0 | 0 | 0 | 0 | 0 | 0 | 0 | 0 | 0 | 0 | 0 | 0 | 0 | 0 | 0 | 0 | 0 | 0 |
| Jul11.5v_15535 | 1 | 0 | 0 | 0 | 0 | 0 | 0 | 0 | 0 | 0 | 0 | 0 | 0 | 1 | 0 | 0 | 0 | 0 | 0 | 0 | 0 | 0 | 0 | 0 | 0 | 0 | 0 | 0 | 0 | 0 |
| Jul11.25v_16750 | 2 | 0 | 0 | 0 | 0 | 0 | 0 | 0 | 0 | 0 | 0 | 0 | 0 | 1 | 1 | 0 | 0 | 0 | 0 | 0 | 0 | 0 | 0 | 0 | 0 | 0 | 0 | 0 | 0 | 0 |
| Jun22.0v_12683 | 1 | 0 | 0 | 0 | 0 | 0 | 0 | 0 | 1 | 0 | 0 | 0 | 0 | 0 | 0 | 0 | 0 | 0 | 0 | 0 | 0 | 0 | 0 | 0 | 0 | 0 | 0 | 0 | 0 | 0 |
| Jul11.5v_14679 | 1 | 0 | 0 | 0 | 0 | 0 | 0 | 0 | 0 | 0 | 0 | 0 | 0 | 1 | 0 | 0 | 0 | 0 | 0 | 0 | 0 | 0 | 0 | 0 | 0 | 0 | 0 | 0 | 0 | 0 |
| Jun22.5v_17483 | 1 | 0 | 0 | 0 | 0 | 0 | 0 | 0 | 0 | 1 | 0 | 0 | 0 | 0 | 0 | 0 | 0 | 0 | 0 | 0 | 0 | 0 | 0 | 0 | 0 | 0 | 0 | 0 | 0 | 0 |
| Sep8.5v_19912 | 1 | 0 | 0 | 0 | 0 | 0 | 0 | 0 | 0 | 0 | 0 | 0 | 0 | 0 | 0 | 0 | 0 | 0 | 0 | 0 | 0 | 0 | 0 | 0 | 0 | 0 | 0 | 1 | 0 | 0 |

| Feature_ID | 654 | May12_05m | May12_25m | May12_50m | May12_75m | Jun8_25m | Jun8_50m | Jun8_75m | Jun22_0m | Jun22_05m | Jun22_50m | Jun22_75m | Jul11_0m | Jul11_05m | Jul11_25m | Jul11_50m | Jul11_75m | Jul27_05m | Jul27_25m | Jul27_50m | Jul27_75m | Aug5_0m | Aug5_05m | Aug5_25m | Aug5_50m | Aug5_75m | Sep8_0m | Sep8_05m | Sep8_25m | Sep8_50m |
|---|---|---|---|---|---|---|---|---|---|---|---|---|---|---|---|---|---|---|---|---|---|---|---|---|---|---|---|---|---|---|
| Sep8.5v_11686 | 1 | 0 | 0 | 0 | 0 | 0 | 0 | 0 | 0 | 0 | 0 | 0 | 0 | 0 | 0 | 0 | 0 | 0 | 0 | 0 | 0 | 0 | 0 | 0 | 0 | 0 | 0 | 1 | 0 | 0 |
| Jun22.5v_20297 | 1 | 0 | 0 | 0 | 0 | 0 | 0 | 0 | 0 | 1 | 0 | 0 | 0 | 0 | 0 | 0 | 0 | 0 | 0 | 0 | 0 | 0 | 0 | 0 | 0 | 0 | 0 | 0 | 0 | 0 |
| May12.50v_18083 | 1 | 0 | 0 | 1 | 0 | 0 | 0 | 0 | 0 | 0 | 0 | 0 | 0 | 0 | 0 | 0 | 0 | 0 | 0 | 0 | 0 | 0 | 0 | 0 | 0 | 0 | 0 | 0 | 0 | 0 |
| Jun22.75v_40971 | 1 | 0 | 0 | 0 | 0 | 0 | 0 | 0 | 0 | 0 | 0 | 1 | 0 | 0 | 0 | 0 | 0 | 0 | 0 | 0 | 0 | 0 | 0 | 0 | 0 | 0 | 0 | 0 | 0 | 0 |
| Jun8.50v_16534 | 1 | 0 | 0 | 0 | 0 | 0 | 1 | 0 | 0 | 0 | 0 | 0 | 0 | 0 | 0 | 0 | 0 | 0 | 0 | 0 | 0 | 0 | 0 | 0 | 0 | 0 | 0 | 0 | 0 | 0 |
| May12.50v_15038 | 1 | 0 | 0 | 1 | 0 | 0 | 0 | 0 | 0 | 0 | 0 | 0 | 0 | 0 | 0 | 0 | 0 | 0 | 0 | 0 | 0 | 0 | 0 | 0 | 0 | 0 | 0 | 0 | 0 | 0 |
| Jun22.5v_19086 | 1 | 0 | 0 | 0 | 0 | 0 | 0 | 0 | 0 | 1 | 0 | 0 | 0 | 0 | 0 | 0 | 0 | 0 | 0 | 0 | 0 | 0 | 0 | 0 | 0 | 0 | 0 | 0 | 0 | 0 |
| May12.50v_7981 | 1 | 0 | 0 | 1 | 0 | 0 | 0 | 0 | 0 | 0 | 0 | 0 | 0 | 0 | 0 | 0 | 0 | 0 | 0 | 0 | 0 | 0 | 0 | 0 | 0 | 0 | 0 | 0 | 0 | 0 |
| Jul27.5v_13223 | 1 | 0 | 0 | 0 | 0 | 0 | 0 | 0 | 0 | 0 | 0 | 0 | 0 | 0 | 0 | 0 | 0 | 1 | 0 | 0 | 0 | 0 | 0 | 0 | 0 | 0 | 0 | 0 | 0 | 0 |
| Jun22.0v_12841 | 1 | 0 | 0 | 0 | 0 | 0 | 0 | 0 | 1 | 0 | 0 | 0 | 0 | 0 | 0 | 0 | 0 | 0 | 0 | 0 | 0 | 0 | 0 | 0 | 0 | 0 | 0 | 0 | 0 | 0 |
| Sep8.50v_15806 | 1 | 0 | 0 | 0 | 0 | 0 | 0 | 0 | 0 | 0 | 0 | 0 | 0 | 0 | 0 | 0 | 0 | 0 | 0 | 0 | 0 | 0 | 0 | 0 | 0 | 0 | 0 | 0 | 0 | 1 |
| Sep8.50v_11061 | 1 | 0 | 0 | 0 | 0 | 0 | 0 | 0 | 0 | 0 | 0 | 0 | 0 | 0 | 0 | 0 | 0 | 0 | 0 | 0 | 0 | 0 | 0 | 0 | 0 | 0 | 0 | 0 | 0 | 1 |
| May12.50v_5052 | 6 | 1 | 0 | 3 | 0 | 0 | 0 | 0 | 0 | 0 | 0 | 0 | 0 | 0 | 0 | 0 | 0 | 0 | 0 | 0 | 0 | 2 | 0 | 0 | 0 | 0 | 0 | 0 | 0 | 0 |
| May12.75v_7726 | 720 | 28 | 34 | 26 | 24 | 9 | 60 | 6 | 11 | 17 | 34 | 62 | 11 | 20 | 11 | 8 | 30 | 25 | 13 | 12 | 3 | 50 | 22 | 13 | 6 | 14 | 14 | 32 | 2 | 123 |
| May12.50v_12845 | 1 | 0 | 0 | 1 | 0 | 0 | 0 | 0 | 0 | 0 | 0 | 0 | 0 | 0 | 0 | 0 | 0 | 0 | 0 | 0 | 0 | 0 | 0 | 0 | 0 | 0 | 0 | 0 | 0 | 0 |
| Sep8.5v_2658 | 1 | 0 | 0 | 0 | 0 | 0 | 0 | 0 | 0 | 0 | 0 | 0 | 0 | 0 | 0 | 0 | 0 | 0 | 0 | 0 | 0 | 0 | 0 | 0 | 0 | 0 | 0 | 1 | 0 | 0 |
| Jul27.5v_17300 | 158 | 7 | 2 | 3 | 5 | 0 | 6 | 3 | 7 | 7 | 6 | 10 | 4 | 14 | 6 | 10 | 4 | 13 | 0 | 2 | 1 | 0 | 7 | 3 | 1 | 1 | 5 | 10 | 6 | 15 |
| Jul11.75v_8009 | 3 | 0 | 1 | 0 | 0 | 0 | 0 | 0 | 0 | 0 | 0 | 0 | 0 | 0 | 0 | 0 | 1 | 0 | 0 | 0 | 0 | 0 | 0 | 0 | 0 | 1 | 0 | 0 | 0 | 0 |
| Jun8.50v_3571 | 1 | 0 | 0 | 0 | 0 | 0 | 1 | 0 | 0 | 0 | 0 | 0 | 0 | 0 | 0 | 0 | 0 | 0 | 0 | 0 | 0 | 0 | 0 | 0 | 0 | 0 | 0 | 0 | 0 | 0 |
| Sep8.50v_34025 | 1 | 0 | 0 | 0 | 0 | 0 | 0 | 0 | 0 | 0 | 0 | 0 | 0 | 0 | 0 | 0 | 0 | 0 | 0 | 0 | 0 | 0 | 0 | 0 | 0 | 0 | 0 | 0 | 0 | 1 |
| Aug5.0v_482 | 1 | 0 | 0 | 0 | 0 | 0 | 0 | 0 | 0 | 0 | 0 | 0 | 0 | 0 | 0 | 0 | 0 | 0 | 0 | 0 | 0 | 1 | 0 | 0 | 0 | 0 | 0 | 0 | 0 | 0 |
| Jul11.50v_17503 | 1 | 0 | 0 | 0 | 0 | 0 | 0 | 0 | 0 | 0 | 0 | 0 | 0 | 0 | 0 | 1 | 0 | 0 | 0 | 0 | 0 | 0 | 0 | 0 | 0 | 0 | 0 | 0 | 0 | 0 |
| Aug5.5v_16045 | 2 | 0 | 0 | 0 | 0 | 0 | 0 | 0 | 1 | 0 | 0 | 0 | 0 | 0 | 0 | 0 | 0 | 0 | 0 | 0 | 0 | 0 | 1 | 0 | 0 | 0 | 0 | 0 | 0 | 0 |
| May12.75v_31281 | 1 | 0 | 0 | 0 | 1 | 0 | 0 | 0 | 0 | 0 | 0 | 0 | 0 | 0 | 0 | 0 | 0 | 0 | 0 | 0 | 0 | 0 | 0 | 0 | 0 | 0 | 0 | 0 | 0 | 0 |
| Sep8.0v_5684 | 1 | 0 | 0 | 0 | 0 | 0 | 0 | 0 | 0 | 0 | 0 | 0 | 0 | 0 | 0 | 0 | 0 | 0 | 0 | 0 | 0 | 0 | 0 | 0 | 0 | 0 | 1 | 0 | 0 | 0 |
| Jul27.5v_16976 | 5 | 0 | 0 | 0 | 0 | 0 | 1 | 0 | 0 | 0 | 0 | 1 | 0 | 2 | 0 | 0 | 0 | 1 | 0 | 0 | 0 | 0 | 0 | 0 | 0 | 0 | 0 | 0 | 0 | 0 |
| Jul27.75v_14932 | 1 | 0 | 0 | 0 | 0 | 0 | 0 | 0 | 0 | 0 | 0 | 0 | 0 | 0 | 0 | 0 | 0 | 0 | 0 | 0 | 1 | 0 | 0 | 0 | 0 | 0 | 0 | 0 | 0 | 0 |
| Jul27.25v_1425 | 1 | 0 | 0 | 0 | 0 | 0 | 0 | 0 | 0 | 0 | 0 | 0 | 0 | 0 | 0 | 0 | 0 | 0 | 1 | 0 | 0 | 0 | 0 | 0 | 0 | 0 | 0 | 0 | 0 | 0 |
| May12.50v_26875 | 2 | 0 | 0 | 1 | 0 | 0 | 0 | 0 | 0 | 0 | 0 | 0 | 0 | 0 | 0 | 0 | 0 | 0 | 0 | 0 | 0 | 0 | 0 | 0 | 0 | 0 | 0 | 0 | 0 | 1 |
| Sep8.0v_18054 | 2 | 0 | 0 | 0 | 0 | 0 | 0 | 0 | 0 | 0 | 0 | 0 | 0 | 0 | 0 | 0 | 0 | 0 | 0 | 0 | 0 | 0 | 0 | 0 | 0 | 0 | 1 | 1 | 0 | 0 |
| Jun8.75v_12607 | 1 | 0 | 0 | 0 | 0 | 0 | 0 | 1 | 0 | 0 | 0 | 0 | 0 | 0 | 0 | 0 | 0 | 0 | 0 | 0 | 0 | 0 | 0 | 0 | 0 | 0 | 0 | 0 | 0 | 0 |
| Jul11.25v_13188 | 1 | 0 | 0 | 0 | 0 | 0 | 0 | 0 | 0 | 0 | 0 | 0 | 0 | 0 | 1 | 0 | 0 | 0 | 0 | 0 | 0 | 0 | 0 | 0 | 0 | 0 | 0 | 0 | 0 | 0 |
| Jun22.5v_13056 | 1 | 0 | 0 | 0 | 0 | 0 | 0 | 0 | 0 | 1 | 0 | 0 | 0 | 0 | 0 | 0 | 0 | 0 | 0 | 0 | 0 | 0 | 0 | 0 | 0 | 0 | 0 | 0 | 0 | 0 |
| May12.5v_14099 | 2 | 2 | 0 | 0 | 0 | 0 | 0 | 0 | 0 | 0 | 0 | 0 | 0 | 0 | 0 | 0 | 0 | 0 | 0 | 0 | 0 | 0 | 0 | 0 | 0 | 0 | 0 | 0 | 0 | 0 |
| May12.25v_17629 | 1 | 0 | 1 | 0 | 0 | 0 | 0 | 0 | 0 | 0 | 0 | 0 | 0 | 0 | 0 | 0 | 0 | 0 | 0 | 0 | 0 | 0 | 0 | 0 | 0 | 0 | 0 | 0 | 0 | 0 |
| Jul11.75v_17389 | 38 | 1 | 2 | 1 | 1 | 0 | 1 | 1 | 1 | 1 | 1 | 0 | 0 | 1 | 3 | 1 | 1 | 6 | 1 | 1 | 0 | 0 | 0 | 1 | 1 | 0 | 1 | 7 | 1 | 3 |
| Jun22.5v_11255 | 1 | 0 | 0 | 0 | 0 | 0 | 0 | 0 | 0 | 1 | 0 | 0 | 0 | 0 | 0 | 0 | 0 | 0 | 0 | 0 | 0 | 0 | 0 | 0 | 0 | 0 | 0 | 0 | 0 | 0 |
| Jul11.75v_32281 | 1 | 0 | 0 | 0 | 0 | 0 | 0 | 0 | 0 | 0 | 0 | 0 | 0 | 0 | 0 | 0 | 1 | 0 | 0 | 0 | 0 | 0 | 0 | 0 | 0 | 0 | 0 | 0 | 0 | 0 |
| Jul11.5v_19652 | 1 | 0 | 0 | 0 | 0 | 0 | 0 | 0 | 0 | 0 | 0 | 0 | 0 | 1 | 0 | 0 | 0 | 0 | 0 | 0 | 0 | 0 | 0 | 0 | 0 | 0 | 0 | 0 | 0 | 0 |
| Aug5.5v_1027 | 1 | 0 | 0 | 0 | 0 | 0 | 0 | 0 | 0 | 0 | 0 | 0 | 0 | 0 | 0 | 0 | 0 | 0 | 0 | 0 | 0 | 0 | 1 | 0 | 0 | 0 | 0 | 0 | 0 | 0 |
| Jul27.5v_8304 | 2 | 0 | 0 | 0 | 0 | 0 | 0 | 0 | 0 | 0 | 0 | 0 | 0 | 0 | 0 | 1 | 0 | 1 | 0 | 0 | 0 | 0 | 0 | 0 | 0 | 0 | 0 | 0 | 0 | 0 |
| Jul11.5v_11867 | 1 | 0 | 0 | 0 | 0 | 0 | 0 | 0 | 0 | 0 | 0 | 0 | 0 | 1 | 0 | 0 | 0 | 0 | 0 | 0 | 0 | 0 | 0 | 0 | 0 | 0 | 0 | 0 | 0 | 0 |
| Jul11.50v_12998 | 1 | 0 | 0 | 0 | 0 | 0 | 0 | 0 | 0 | 0 | 0 | 0 | 0 | 0 | 0 | 1 | 0 | 0 | 0 | 0 | 0 | 0 | 0 | 0 | 0 | 0 | 0 | 0 | 0 | 0 |
| Jul27.25v_28362 | 1 | 0 | 0 | 0 | 0 | 0 | 0 | 0 | 0 | 0 | 0 | 0 | 0 | 0 | 0 | 0 | 0 | 0 | 1 | 0 | 0 | 0 | 0 | 0 | 0 | 0 | 0 | 0 | 0 | 0 |
| Jul11.5v_11265 | 1 | 0 | 0 | 0 | 0 | 0 | 0 | 0 | 0 | 0 | 0 | 0 | 0 | 1 | 0 | 0 | 0 | 0 | 0 | 0 | 0 | 0 | 0 | 0 | 0 | 0 | 0 | 0 | 0 | 0 |
| May12.75v_17774 | 2 | 0 | 0 | 0 | 1 | 0 | 0 | 0 | 0 | 0 | 0 | 0 | 0 | 0 | 0 | 0 | 0 | 0 | 0 | 0 | 0 | 0 | 0 | 0 | 0 | 0 | 0 | 0 | 0 | 1 |
| Sep8.5v_8748 | 3 | 0 | 0 | 0 | 0 | 0 | 0 | 0 | 0 | 0 | 0 | 1 | 0 | 0 | 0 | 0 | 0 | 0 | 0 | 0 | 0 | 0 | 0 | 0 | 0 | 1 | 0 | 1 | 0 | 0 |
| May12.25v_33418 | 1 | 0 | 1 | 0 | 0 | 0 | 0 | 0 | 0 | 0 | 0 | 0 | 0 | 0 | 0 | 0 | 0 | 0 | 0 | 0 | 0 | 0 | 0 | 0 | 0 | 0 | 0 | 0 | 0 | 0 |
| Jun22.0v_17853 | 1 | 0 | 0 | 0 | 0 | 0 | 0 | 0 | 1 | 0 | 0 | 0 | 0 | 0 | 0 | 0 | 0 | 0 | 0 | 0 | 0 | 0 | 0 | 0 | 0 | 0 | 0 | 0 | 0 | 0 |
| Jul11.5v_13527 | 2 | 0 | 0 | 0 | 0 | 0 | 0 | 0 | 0 | 0 | 0 | 0 | 0 | 1 | 0 | 0 | 0 | 1 | 0 | 0 | 0 | 0 | 0 | 0 | 0 | 0 | 0 | 0 | 0 | 0 |
| May12.75v_9003 | 1 | 0 | 0 | 0 | 1 | 0 | 0 | 0 | 0 | 0 | 0 | 0 | 0 | 0 | 0 | 0 | 0 | 0 | 0 | 0 | 0 | 0 | 0 | 0 | 0 | 0 | 0 | 0 | 0 | 0 |
| Jun22.5v_10036 | 22 | 1 | 1 | 0 | 0 | 0 | 1 | 0 | 1 | 2 | 0 | 1 | 0 | 2 | 0 | 1 | 0 | 1 | 1 | 1 | 0 | 0 | 1 | 0 | 0 | 1 | 0 | 2 | 0 | 5 |
| Jul11.5v_4210 | 1 | 0 | 0 | 0 | 0 | 0 | 0 | 0 | 0 | 0 | 0 | 0 | 0 | 1 | 0 | 0 | 0 | 0 | 0 | 0 | 0 | 0 | 0 | 0 | 0 | 0 | 0 | 0 | 0 | 0 |
| Jul11.0v_14923 | 1 | 0 | 0 | 0 | 0 | 0 | 0 | 0 | 0 | 0 | 0 | 0 | 1 | 0 | 0 | 0 | 0 | 0 | 0 | 0 | 0 | 0 | 0 | 0 | 0 | 0 | 0 | 0 | 0 | 0 |
| May12.50v_31451 | 1 | 0 | 0 | 1 | 0 | 0 | 0 | 0 | 0 | 0 | 0 | 0 | 0 | 0 | 0 | 0 | 0 | 0 | 0 | 0 | 0 | 0 | 0 | 0 | 0 | 0 | 0 | 0 | 0 | 0 |
| Sep8.5v_16613 | 1 | 0 | 0 | 0 | 0 | 0 | 0 | 0 | 0 | 0 | 0 | 0 | 0 | 0 | 0 | 0 | 0 | 0 | 0 | 0 | 0 | 0 | 0 | 0 | 0 | 0 | 0 | 1 | 0 | 0 |
| Aug5.0v_589 | 1 | 0 | 0 | 0 | 0 | 0 | 0 | 0 | 0 | 0 | 0 | 0 | 0 | 0 | 0 | 0 | 0 | 0 | 0 | 0 | 0 | 1 | 0 | 0 | 0 | 0 | 0 | 0 | 0 | 0 |
| May12.50v_4824 | 1 | 0 | 0 | 1 | 0 | 0 | 0 | 0 | 0 | 0 | 0 | 0 | 0 | 0 | 0 | 0 | 0 | 0 | 0 | 0 | 0 | 0 | 0 | 0 | 0 | 0 | 0 | 0 | 0 | 0 |
| Jul27.5v_10211 | 1 | 0 | 0 | 0 | 0 | 0 | 0 | 0 | 0 | 0 | 0 | 0 | 0 | 0 | 0 | 0 | 0 | 1 | 0 | 0 | 0 | 0 | 0 | 0 | 0 | 0 | 0 | 0 | 0 | 0 |
| Sep8.50v_4978 | 1 | 0 | 0 | 0 | 0 | 0 | 0 | 0 | 0 | 0 | 0 | 0 | 0 | 0 | 0 | 0 | 0 | 0 | 0 | 0 | 0 | 0 | 0 | 0 | 0 | 0 | 0 | 0 | 0 | 1 |
| Sep8.50v_3315 | 1 | 0 | 0 | 0 | 0 | 0 | 0 | 0 | 0 | 0 | 0 | 0 | 0 | 0 | 0 | 0 | 0 | 0 | 0 | 0 | 0 | 0 | 0 | 0 | 0 | 0 | 0 | 0 | 0 | 1 |
| Sep8.50v_32912 | 1 | 0 | 0 | 0 | 0 | 0 | 0 | 0 | 0 | 0 | 0 | 0 | 0 | 0 | 0 | 0 | 0 | 0 | 0 | 0 | 0 | 0 | 0 | 0 | 0 | 0 | 0 | 0 | 0 | 1 |
| Sep8.0v_3910 | 1 | 0 | 0 | 0 | 0 | 0 | 0 | 0 | 0 | 0 | 0 | 0 | 0 | 0 | 0 | 0 | 0 | 0 | 0 | 0 | 0 | 0 | 0 | 0 | 0 | 0 | 1 | 0 | 0 | 0 |
| Sep8.50v_24505 | 1 | 0 | 0 | 0 | 0 | 0 | 0 | 0 | 0 | 0 | 0 | 0 | 0 | 0 | 0 | 0 | 0 | 0 | 0 | 0 | 0 | 0 | 0 | 0 | 0 | 0 | 0 | 0 | 0 | 1 |
| Jun8.50v_17949 | 1 | 0 | 0 | 0 | 0 | 0 | 1 | 0 | 0 | 0 | 0 | 0 | 0 | 0 | 0 | 0 | 0 | 0 | 0 | 0 | 0 | 0 | 0 | 0 | 0 | 0 | 0 | 0 | 0 | 0 |
| May12.5v_9551 | 1 | 1 | 0 | 0 | 0 | 0 | 0 | 0 | 0 | 0 | 0 | 0 | 0 | 0 | 0 | 0 | 0 | 0 | 0 | 0 | 0 | 0 | 0 | 0 | 0 | 0 | 0 | 0 | 0 | 0 |
| Jul27.50v_24256 | 1 | 0 | 0 | 0 | 0 | 0 | 0 | 0 | 0 | 0 | 0 | 0 | 0 | 0 | 0 | 0 | 0 | 0 | 0 | 1 | 0 | 0 | 0 | 0 | 0 | 0 | 0 | 0 | 0 | 0 |
| Jul11.0v_5508 | 1 | 0 | 0 | 0 | 0 | 0 | 0 | 0 | 0 | 0 | 0 | 0 | 1 | 0 | 0 | 0 | 0 | 0 | 0 | 0 | 0 | 0 | 0 | 0 | 0 | 0 | 0 | 0 | 0 | 0 |
| Sep8.50v_21946 | 1 | 0 | 0 | 0 | 0 | 0 | 0 | 0 | 0 | 0 | 0 | 0 | 0 | 0 | 0 | 0 | 0 | 0 | 0 | 0 | 0 | 0 | 0 | 0 | 0 | 0 | 0 | 0 | 0 | 1 |
| Sep8.50v_3268 | 1 | 0 | 0 | 0 | 0 | 0 | 0 | 0 | 0 | 0 | 0 | 0 | 0 | 0 | 0 | 0 | 0 | 0 | 0 | 0 | 0 | 0 | 0 | 0 | 0 | 0 | 0 | 0 | 0 | 1 |
| Sep8.5v_19322 | 1 | 0 | 0 | 0 | 0 | 0 | 0 | 0 | 0 | 0 | 0 | 0 | 0 | 0 | 0 | 0 | 0 | 0 | 0 | 0 | 0 | 0 | 0 | 0 | 0 | 0 | 0 | 1 | 0 | 0 |
| Aug5.0v_3973 | 1 | 0 | 0 | 0 | 0 | 0 | 0 | 0 | 0 | 0 | 0 | 0 | 0 | 0 | 0 | 0 | 0 | 0 | 0 | 0 | 0 | 1 | 0 | 0 | 0 | 0 | 0 | 0 | 0 | 0 |
| Jul27.25v_38302 | 1 | 0 | 0 | 0 | 0 | 0 | 0 | 0 | 0 | 0 | 0 | 0 | 0 | 0 | 0 | 0 | 0 | 0 | 1 | 0 | 0 | 0 | 0 | 0 | 0 | 0 | 0 | 0 | 0 | 0 |
| Jun22.75v_28430 | 1 | 0 | 0 | 0 | 0 | 0 | 0 | 0 | 0 | 0 | 0 | 1 | 0 | 0 | 0 | 0 | 0 | 0 | 0 | 0 | 0 | 0 | 0 | 0 | 0 | 0 | 0 | 0 | 0 | 0 |
| May12.5v_16513 | 1 | 1 | 0 | 0 | 0 | 0 | 0 | 0 | 0 | 0 | 0 | 0 | 0 | 0 | 0 | 0 | 0 | 0 | 0 | 0 | 0 | 0 | 0 | 0 | 0 | 0 | 0 | 0 | 0 | 0 |
| Sep8.5v_19487 | 1 | 0 | 0 | 0 | 0 | 0 | 0 | 0 | 0 | 0 | 0 | 0 | 0 | 0 | 0 | 0 | 0 | 0 | 0 | 0 | 0 | 0 | 0 | 0 | 0 | 0 | 0 | 1 | 0 | 0 |
| Sep8.50v_12252 | 1 | 0 | 0 | 0 | 0 | 0 | 0 | 0 | 0 | 0 | 0 | 0 | 0 | 0 | 0 | 0 | 0 | 0 | 0 | 0 | 0 | 0 | 0 | 0 | 0 | 0 | 0 | 0 | 0 | 1 |
| Aug5.5v_14172 | 1 | 0 | 0 | 0 | 0 | 0 | 0 | 0 | 0 | 0 | 0 | 0 | 0 | 0 | 0 | 0 | 0 | 0 | 0 | 0 | 0 | 0 | 1 | 0 | 0 | 0 | 0 | 0 | 0 | 0 |
| May12.5v_26333 | 1 | 1 | 0 | 0 | 0 | 0 | 0 | 0 | 0 | 0 | 0 | 0 | 0 | 0 | 0 | 0 | 0 | 0 | 0 | 0 | 0 | 0 | 0 | 0 | 0 | 0 | 0 | 0 | 0 | 0 |
| Jul11.5v_9693 | 1 | 0 | 0 | 0 | 0 | 0 | 0 | 0 | 0 | 0 | 0 | 0 | 0 | 1 | 0 | 0 | 0 | 0 | 0 | 0 | 0 | 0 | 0 | 0 | 0 | 0 | 0 | 0 | 0 | 0 |
| Jul27.75v_14092 | 1 | 0 | 0 | 0 | 0 | 0 | 0 | 0 | 0 | 0 | 0 | 0 | 0 | 0 | 0 | 0 | 0 | 0 | 0 | 0 | 1 | 0 | 0 | 0 | 0 | 0 | 0 | 0 | 0 | 0 |
| Jun22.5v_2393 | 1 | 0 | 0 | 0 | 0 | 0 | 0 | 0 | 0 | 1 | 0 | 0 | 0 | 0 | 0 | 0 | 0 | 0 | 0 | 0 | 0 | 0 | 0 | 0 | 0 | 0 | 0 | 0 | 0 | 0 |
| Jul11.0v_14307 | 1 | 0 | 0 | 0 | 0 | 0 | 0 | 0 | 0 | 0 | 0 | 0 | 1 | 0 | 0 | 0 | 0 | 0 | 0 | 0 | 0 | 0 | 0 | 0 | 0 | 0 | 0 | 0 | 0 | 0 |
| Sep8.5v_13376 | 1 | 0 | 0 | 0 | 0 | 0 | 0 | 0 | 0 | 0 | 0 | 0 | 0 | 0 | 0 | 0 | 0 | 0 | 0 | 0 | 0 | 0 | 0 | 0 | 0 | 0 | 0 | 1 | 0 | 0 |
| May12.75v_31866 | 1 | 0 | 0 | 0 | 1 | 0 | 0 | 0 | 0 | 0 | 0 | 0 | 0 | 0 | 0 | 0 | 0 | 0 | 0 | 0 | 0 | 0 | 0 | 0 | 0 | 0 | 0 | 0 | 0 | 0 |
| Jul27.5v_10725 | 1 | 0 | 0 | 0 | 0 | 0 | 0 | 0 | 0 | 0 | 0 | 0 | 0 | 0 | 0 | 0 | 0 | 1 | 0 | 0 | 0 | 0 | 0 | 0 | 0 | 0 | 0 | 0 | 0 | 0 |
| May12.50v_5753 | 1 | 0 | 0 | 1 | 0 | 0 | 0 | 0 | 0 | 0 | 0 | 0 | 0 | 0 | 0 | 0 | 0 | 0 | 0 | 0 | 0 | 0 | 0 | 0 | 0 | 0 | 0 | 0 | 0 | 0 |
| Jul11.75v_17405 | 1 | 0 | 0 | 0 | 0 | 0 | 0 | 0 | 0 | 0 | 0 | 0 | 0 | 0 | 0 | 0 | 1 | 0 | 0 | 0 | 0 | 0 | 0 | 0 | 0 | 0 | 0 | 0 | 0 | 0 |
| Jun22.0v_19381 | 1 | 0 | 0 | 0 | 0 | 0 | 0 | 0 | 1 | 0 | 0 | 0 | 0 | 0 | 0 | 0 | 0 | 0 | 0 | 0 | 0 | 0 | 0 | 0 | 0 | 0 | 0 | 0 | 0 | 0 |
| Jul11.5v_18128 | 1 | 0 | 0 | 0 | 0 | 0 | 0 | 0 | 0 | 0 | 0 | 0 | 0 | 1 | 0 | 0 | 0 | 0 | 0 | 0 | 0 | 0 | 0 | 0 | 0 | 0 | 0 | 0 | 0 | 0 |
| May12.50v_30643 | 1 | 0 | 0 | 1 | 0 | 0 | 0 | 0 | 0 | 0 | 0 | 0 | 0 | 0 | 0 | 0 | 0 | 0 | 0 | 0 | 0 | 0 | 0 | 0 | 0 | 0 | 0 | 0 | 0 | 0 |
| Jun22.0v_1661 | 1 | 0 | 0 | 0 | 0 | 0 | 0 | 0 | 1 | 0 | 0 | 0 | 0 | 0 | 0 | 0 | 0 | 0 | 0 | 0 | 0 | 0 | 0 | 0 | 0 | 0 | 0 | 0 | 0 | 0 |
| Jun22.0v_13622 | 1 | 0 | 0 | 0 | 0 | 0 | 0 | 0 | 1 | 0 | 0 | 0 | 0 | 0 | 0 | 0 | 0 | 0 | 0 | 0 | 0 | 0 | 0 | 0 | 0 | 0 | 0 | 0 | 0 | 0 |
| May12.50v_3821 | 1 | 0 | 0 | 1 | 0 | 0 | 0 | 0 | 0 | 0 | 0 | 0 | 0 | 0 | 0 | 0 | 0 | 0 | 0 | 0 | 0 | 0 | 0 | 0 | 0 | 0 | 0 | 0 | 0 | 0 |
| Jun22.0v_8025 | 1 | 0 | 0 | 0 | 0 | 0 | 0 | 0 | 1 | 0 | 0 | 0 | 0 | 0 | 0 | 0 | 0 | 0 | 0 | 0 | 0 | 0 | 0 | 0 | 0 | 0 | 0 | 0 | 0 | 0 |
| Jun22.50v_21417 | 1 | 0 | 0 | 0 | 0 | 0 | 0 | 0 | 0 | 0 | 1 | 0 | 0 | 0 | 0 | 0 | 0 | 0 | 0 | 0 | 0 | 0 | 0 | 0 | 0 | 0 | 0 | 0 | 0 | 0 |
| Sep8.50v_24150 | 1 | 0 | 0 | 0 | 0 | 0 | 0 | 0 | 0 | 0 | 0 | 0 | 0 | 0 | 0 | 0 | 0 | 0 | 0 | 0 | 0 | 0 | 0 | 0 | 0 | 0 | 0 | 0 | 0 | 1 |
| May12.75v_26924 | 1 | 0 | 0 | 0 | 1 | 0 | 0 | 0 | 0 | 0 | 0 | 0 | 0 | 0 | 0 | 0 | 0 | 0 | 0 | 0 | 0 | 0 | 0 | 0 | 0 | 0 | 0 | 0 | 0 | 0 |
| Aug5.0v_468 | 1 | 0 | 0 | 0 | 0 | 0 | 0 | 0 | 0 | 0 | 0 | 0 | 0 | 0 | 0 | 0 | 0 | 0 | 0 | 0 | 0 | 1 | 0 | 0 | 0 | 0 | 0 | 0 | 0 | 0 |
| Sep8.0v_6390 | 1424 | 23 | 32 | 109 | 27 | 10 | 47 | 3 | 42 | 15 | 74 | 35 | 30 | 54 | 55 | 29 | 17 | 76 | 36 | 58 | 6 | 203 | 94 | 109 | 22 | 19 | 36 | 16 | 6 | 141 |
| Jun8.50v_34579 | 1 | 0 | 0 | 0 | 0 | 0 | 1 | 0 | 0 | 0 | 0 | 0 | 0 | 0 | 0 | 0 | 0 | 0 | 0 | 0 | 0 | 0 | 0 | 0 | 0 | 0 | 0 | 0 | 0 | 0 |
| Jun8.25v_29878 | 508 | 26 | 21 | 59 | 26 | 10 | 18 | 2 | 17 | 10 | 12 | 19 | 16 | 23 | 22 | 14 | 15 | 29 | 18 | 12 | 0 | 35 | 14 | 28 | 5 | 3 | 18 | 8 | 1 | 27 |
| Jul11.25v_16470 | 4 | 0 | 0 | 2 | 0 | 0 | 0 | 0 | 0 | 0 | 0 | 0 | 0 | 0 | 1 | 0 | 0 | 0 | 0 | 0 | 0 | 0 | 0 | 0 | 0 | 0 | 0 | 0 | 0 | 1 |
| Jul11.25v_3761 | 1 | 0 | 0 | 0 | 0 | 0 | 0 | 0 | 0 | 0 | 0 | 0 | 0 | 0 | 1 | 0 | 0 | 0 | 0 | 0 | 0 | 0 | 0 | 0 | 0 | 0 | 0 | 0 | 0 | 0 |
| Jun22.50v_37108 | 6 | 0 | 1 | 0 | 0 | 0 | 0 | 0 | 0 | 0 | 1 | 0 | 1 | 0 | 0 | 0 | 0 | 1 | 0 | 0 | 0 | 0 | 0 | 0 | 0 | 0 | 0 | 0 | 0 | 2 |
| Jul11.0v_4604 | 6 | 0 | 0 | 0 | 0 | 0 | 0 | 0 | 0 | 0 | 0 | 0 | 1 | 0 | 0 | 0 | 0 | 0 | 0 | 0 | 0 | 2 | 0 | 1 | 0 | 1 | 1 | 0 | 0 | 0 |
| May12.50v_39712 | 1 | 0 | 0 | 1 | 0 | 0 | 0 | 0 | 0 | 0 | 0 | 0 | 0 | 0 | 0 | 0 | 0 | 0 | 0 | 0 | 0 | 0 | 0 | 0 | 0 | 0 | 0 | 0 | 0 | 0 |
| Jun22.50v_4458 | 5 | 0 | 0 | 0 | 0 | 0 | 0 | 0 | 1 | 0 | 1 | 0 | 0 | 0 | 0 | 0 | 0 | 0 | 0 | 0 | 0 | 1 | 0 | 1 | 0 | 0 | 0 | 0 | 0 | 1 |
| Jun22.0v_15447 | 3 | 0 | 0 | 0 | 2 | 0 | 0 | 0 | 1 | 0 | 0 | 0 | 0 | 0 | 0 | 0 | 0 | 0 | 0 | 0 | 0 | 0 | 0 | 0 | 0 | 0 | 0 | 0 | 0 | 0 |
| Aug5.25v_12783 | 1 | 0 | 0 | 0 | 0 | 0 | 0 | 0 | 0 | 0 | 0 | 0 | 0 | 0 | 0 | 0 | 0 | 0 | 0 | 0 | 0 | 0 | 0 | 1 | 0 | 0 | 0 | 0 | 0 | 0 |
| Sep8.0v_9273 | 2 | 0 | 0 | 0 | 0 | 0 | 0 | 0 | 0 | 0 | 0 | 0 | 0 | 0 | 0 | 0 | 0 | 0 | 0 | 0 | 0 | 0 | 0 | 0 | 0 | 0 | 1 | 0 | 0 | 1 |
| Jul27.5v_11238 | 3 | 0 | 0 | 1 | 0 | 0 | 0 | 0 | 0 | 0 | 0 | 1 | 0 | 0 | 0 | 0 | 0 | 1 | 0 | 0 | 0 | 0 | 0 | 0 | 0 | 0 | 0 | 0 | 0 | 0 |
| Jun22.50v_1060 | 2 | 0 | 0 | 0 | 0 | 0 | 0 | 0 | 0 | 0 | 1 | 0 | 0 | 0 | 0 | 0 | 0 | 0 | 1 | 0 | 0 | 0 | 0 | 0 | 0 | 0 | 0 | 0 | 0 | 0 |
| Jul11.5v_15679 | 1 | 0 | 0 | 0 | 0 | 0 | 0 | 0 | 0 | 0 | 0 | 0 | 0 | 1 | 0 | 0 | 0 | 0 | 0 | 0 | 0 | 0 | 0 | 0 | 0 | 0 | 0 | 0 | 0 | 0 |
| Jul27.25v_11119 | 1 | 0 | 0 | 0 | 0 | 0 | 0 | 0 | 0 | 0 | 0 | 0 | 0 | 0 | 0 | 0 | 0 | 0 | 1 | 0 | 0 | 0 | 0 | 0 | 0 | 0 | 0 | 0 | 0 | 0 |
| Aug5.0v_4121 | 1 | 0 | 0 | 0 | 0 | 0 | 0 | 0 | 0 | 0 | 0 | 0 | 0 | 0 | 0 | 0 | 0 | 0 | 0 | 0 | 0 | 1 | 0 | 0 | 0 | 0 | 0 | 0 | 0 | 0 |
| Sep8.0v_14702 | 1 | 0 | 0 | 0 | 0 | 0 | 0 | 0 | 0 | 0 | 0 | 0 | 0 | 0 | 0 | 0 | 0 | 0 | 0 | 0 | 0 | 0 | 0 | 0 | 0 | 0 | 1 | 0 | 0 | 0 |
| Jun22.0v_20826 | 1 | 0 | 0 | 0 | 0 | 0 | 0 | 0 | 1 | 0 | 0 | 0 | 0 | 0 | 0 | 0 | 0 | 0 | 0 | 0 | 0 | 0 | 0 | 0 | 0 | 0 | 0 | 0 | 0 | 0 |
| Jul11.50v_1622 | 1 | 0 | 0 | 0 | 0 | 0 | 0 | 0 | 0 | 0 | 0 | 0 | 0 | 0 | 0 | 1 | 0 | 0 | 0 | 0 | 0 | 0 | 0 | 0 | 0 | 0 | 0 | 0 | 0 | 0 |
| May12.50v_33214 | 1 | 0 | 0 | 1 | 0 | 0 | 0 | 0 | 0 | 0 | 0 | 0 | 0 | 0 | 0 | 0 | 0 | 0 | 0 | 0 | 0 | 0 | 0 | 0 | 0 | 0 | 0 | 0 | 0 | 0 |
| Jun8.50v_8403 | 1 | 0 | 0 | 0 | 0 | 0 | 1 | 0 | 0 | 0 | 0 | 0 | 0 | 0 | 0 | 0 | 0 | 0 | 0 | 0 | 0 | 0 | 0 | 0 | 0 | 0 | 0 | 0 | 0 | 0 |
| May12.50v_21298 | 1 | 0 | 0 | 1 | 0 | 0 | 0 | 0 | 0 | 0 | 0 | 0 | 0 | 0 | 0 | 0 | 0 | 0 | 0 | 0 | 0 | 0 | 0 | 0 | 0 | 0 | 0 | 0 | 0 | 0 |
| Jun22.0v_19526 | 1 | 0 | 0 | 0 | 0 | 0 | 0 | 0 | 1 | 0 | 0 | 0 | 0 | 0 | 0 | 0 | 0 | 0 | 0 | 0 | 0 | 0 | 0 | 0 | 0 | 0 | 0 | 0 | 0 | 0 |
| Jun22.50v_1467 | 4 | 1 | 0 | 0 | 0 | 0 | 0 | 0 | 0 | 0 | 1 | 0 | 1 | 0 | 0 | 0 | 0 | 0 | 1 | 0 | 0 | 0 | 0 | 0 | 0 | 0 | 0 | 0 | 0 | 0 |
| Aug5.50v_4994 | 1 | 0 | 0 | 0 | 0 | 0 | 0 | 0 | 0 | 0 | 0 | 0 | 0 | 0 | 0 | 0 | 0 | 0 | 0 | 0 | 0 | 0 | 0 | 0 | 1 | 0 | 0 | 0 | 0 | 0 |
| Jun22.75v_731 | 1 | 0 | 0 | 0 | 0 | 0 | 0 | 0 | 0 | 0 | 0 | 1 | 0 | 0 | 0 | 0 | 0 | 0 | 0 | 0 | 0 | 0 | 0 | 0 | 0 | 0 | 0 | 0 | 0 | 0 |
| May12.50v_24948 | 1 | 0 | 0 | 1 | 0 | 0 | 0 | 0 | 0 | 0 | 0 | 0 | 0 | 0 | 0 | 0 | 0 | 0 | 0 | 0 | 0 | 0 | 0 | 0 | 0 | 0 | 0 | 0 | 0 | 0 |
| Jul11.0v_11429 | 1 | 0 | 0 | 0 | 0 | 0 | 0 | 0 | 0 | 0 | 0 | 0 | 1 | 0 | 0 | 0 | 0 | 0 | 0 | 0 | 0 | 0 | 0 | 0 | 0 | 0 | 0 | 0 | 0 | 0 |
| Aug5.5v_26014 | 1 | 0 | 0 | 0 | 0 | 0 | 0 | 0 | 0 | 0 | 0 | 0 | 0 | 0 | 0 | 0 | 0 | 0 | 0 | 0 | 0 | 0 | 1 | 0 | 0 | 0 | 0 | 0 | 0 | 0 |
| Sep8.50v_14142 | 1 | 0 | 0 | 0 | 0 | 0 | 0 | 0 | 0 | 0 | 0 | 0 | 0 | 0 | 0 | 0 | 0 | 0 | 0 | 0 | 0 | 0 | 0 | 0 | 0 | 0 | 0 | 0 | 0 | 1 |
| Jul11.0v_12835 | 1 | 0 | 0 | 0 | 0 | 0 | 0 | 0 | 0 | 0 | 0 | 0 | 1 | 0 | 0 | 0 | 0 | 0 | 0 | 0 | 0 | 0 | 0 | 0 | 0 | 0 | 0 | 0 | 0 | 0 |
| Jun22.75v_8800 | 1 | 0 | 0 | 0 | 0 | 0 | 0 | 0 | 0 | 0 | 0 | 1 | 0 | 0 | 0 | 0 | 0 | 0 | 0 | 0 | 0 | 0 | 0 | 0 | 0 | 0 | 0 | 0 | 0 | 0 |
| Aug5.0v_2333 | 6 | 0 | 0 | 0 | 0 | 0 | 0 | 0 | 0 | 0 | 0 | 0 | 0 | 0 | 1 | 0 | 0 | 1 | 0 | 0 | 0 | 3 | 0 | 1 | 0 | 0 | 0 | 0 | 0 | 0 |
| Jun22.0v_14316 | 1 | 0 | 0 | 0 | 0 | 0 | 0 | 0 | 1 | 0 | 0 | 0 | 0 | 0 | 0 | 0 | 0 | 0 | 0 | 0 | 0 | 0 | 0 | 0 | 0 | 0 | 0 | 0 | 0 | 0 |
| Jul11.25v_9244 | 4 | 0 | 0 | 0 | 0 | 0 | 0 | 0 | 0 | 0 | 0 | 0 | 0 | 0 | 2 | 0 | 0 | 1 | 0 | 0 | 0 | 0 | 0 | 1 | 0 | 0 | 0 | 0 | 0 | 0 |
| Sep8.5v_19205 | 1 | 0 | 0 | 0 | 0 | 0 | 0 | 0 | 0 | 0 | 0 | 0 | 0 | 0 | 0 | 0 | 0 | 0 | 0 | 0 | 0 | 0 | 0 | 0 | 0 | 0 | 0 | 1 | 0 | 0 |
| Sep8.0v_595 | 1 | 0 | 0 | 0 | 0 | 0 | 0 | 0 | 0 | 0 | 0 | 0 | 0 | 0 | 0 | 0 | 0 | 0 | 0 | 0 | 0 | 0 | 0 | 0 | 0 | 0 | 1 | 0 | 0 | 0 |
| Jun22.50v_4841 | 1 | 0 | 0 | 0 | 0 | 0 | 0 | 0 | 0 | 0 | 1 | 0 | 0 | 0 | 0 | 0 | 0 | 0 | 0 | 0 | 0 | 0 | 0 | 0 | 0 | 0 | 0 | 0 | 0 | 0 |
| May12.50v_10400 | 1 | 0 | 0 | 1 | 0 | 0 | 0 | 0 | 0 | 0 | 0 | 0 | 0 | 0 | 0 | 0 | 0 | 0 | 0 | 0 | 0 | 0 | 0 | 0 | 0 | 0 | 0 | 0 | 0 | 0 |
| May12.50v_32540 | 1 | 0 | 0 | 1 | 0 | 0 | 0 | 0 | 0 | 0 | 0 | 0 | 0 | 0 | 0 | 0 | 0 | 0 | 0 | 0 | 0 | 0 | 0 | 0 | 0 | 0 | 0 | 0 | 0 | 0 |
| May12.50v_4359 | 1 | 0 | 0 | 1 | 0 | 0 | 0 | 0 | 0 | 0 | 0 | 0 | 0 | 0 | 0 | 0 | 0 | 0 | 0 | 0 | 0 | 0 | 0 | 0 | 0 | 0 | 0 | 0 | 0 | 0 |
| Jul11.5v_15222 | 1 | 0 | 0 | 0 | 0 | 0 | 0 | 0 | 0 | 0 | 0 | 0 | 0 | 1 | 0 | 0 | 0 | 0 | 0 | 0 | 0 | 0 | 0 | 0 | 0 | 0 | 0 | 0 | 0 | 0 |
| Sep8.50v_17423 | 1 | 0 | 0 | 0 | 0 | 0 | 0 | 0 | 0 | 0 | 0 | 0 | 0 | 0 | 0 | 0 | 0 | 0 | 0 | 0 | 0 | 0 | 0 | 0 | 0 | 0 | 0 | 0 | 0 | 1 |
| May12.50v_19193 | 1 | 0 | 0 | 1 | 0 | 0 | 0 | 0 | 0 | 0 | 0 | 0 | 0 | 0 | 0 | 0 | 0 | 0 | 0 | 0 | 0 | 0 | 0 | 0 | 0 | 0 | 0 | 0 | 0 | 0 |
| Jul11.5v_9168 | 1 | 0 | 0 | 0 | 0 | 0 | 0 | 0 | 0 | 0 | 0 | 0 | 0 | 1 | 0 | 0 | 0 | 0 | 0 | 0 | 0 | 0 | 0 | 0 | 0 | 0 | 0 | 0 | 0 | 0 |
| May12.75v_22749 | 1 | 0 | 0 | 0 | 1 | 0 | 0 | 0 | 0 | 0 | 0 | 0 | 0 | 0 | 0 | 0 | 0 | 0 | 0 | 0 | 0 | 0 | 0 | 0 | 0 | 0 | 0 | 0 | 0 | 0 |
| Jul27.5v_19384 | 5 | 0 | 0 | 1 | 0 | 0 | 0 | 0 | 0 | 0 | 0 | 0 | 0 | 1 | 0 | 1 | 0 | 2 | 0 | 0 | 0 | 0 | 0 | 0 | 0 | 0 | 0 | 0 | 0 | 0 |
| May12.5v_28141 | 1 | 1 | 0 | 0 | 0 | 0 | 0 | 0 | 0 | 0 | 0 | 0 | 0 | 0 | 0 | 0 | 0 | 0 | 0 | 0 | 0 | 0 | 0 | 0 | 0 | 0 | 0 | 0 | 0 | 0 |
| Jun22.50v_29789 | 1 | 0 | 0 | 0 | 0 | 0 | 0 | 0 | 0 | 0 | 1 | 0 | 0 | 0 | 0 | 0 | 0 | 0 | 0 | 0 | 0 | 0 | 0 | 0 | 0 | 0 | 0 | 0 | 0 | 0 |
| Jul27.50v_12475 | 1 | 0 | 0 | 0 | 0 | 0 | 0 | 0 | 0 | 0 | 0 | 0 | 0 | 0 | 0 | 0 | 0 | 0 | 0 | 1 | 0 | 0 | 0 | 0 | 0 | 0 | 0 | 0 | 0 | 0 |
| Jun22.75v_32731 | 1 | 0 | 0 | 0 | 0 | 0 | 0 | 0 | 0 | 0 | 0 | 1 | 0 | 0 | 0 | 0 | 0 | 0 | 0 | 0 | 0 | 0 | 0 | 0 | 0 | 0 | 0 | 0 | 0 | 0 |
| Aug5.25v_32934 | 2 | 0 | 0 | 0 | 0 | 0 | 0 | 0 | 0 | 0 | 0 | 0 | 0 | 0 | 0 | 1 | 0 | 0 | 0 | 0 | 0 | 0 | 0 | 1 | 0 | 0 | 0 | 0 | 0 | 0 |
| Aug5.25v_31315 | 2 | 0 | 0 | 0 | 0 | 0 | 0 | 0 | 0 | 0 | 0 | 0 | 0 | 0 | 0 | 0 | 0 | 0 | 0 | 0 | 0 | 1 | 0 | 1 | 0 | 0 | 0 | 0 | 0 | 0 |
| Jun22.75v_17912 | 5 | 0 | 0 | 0 | 0 | 0 | 0 | 0 | 0 | 0 | 0 | 1 | 0 | 0 | 0 | 0 | 0 | 2 | 0 | 1 | 0 | 0 | 0 | 0 | 0 | 0 | 1 | 0 | 0 | 0 |
| May12.5v_11757 | 1 | 1 | 0 | 0 | 0 | 0 | 0 | 0 | 0 | 0 | 0 | 0 | 0 | 0 | 0 | 0 | 0 | 0 | 0 | 0 | 0 | 0 | 0 | 0 | 0 | 0 | 0 | 0 | 0 | 0 |
| Jun22.50v_18504 | 1 | 0 | 0 | 0 | 0 | 0 | 0 | 0 | 0 | 0 | 1 | 0 | 0 | 0 | 0 | 0 | 0 | 0 | 0 | 0 | 0 | 0 | 0 | 0 | 0 | 0 | 0 | 0 | 0 | 0 |
| May12.5v_13885 | 6 | 2 | 0 | 0 | 1 | 0 | 0 | 0 | 0 | 0 | 0 | 1 | 0 | 0 | 0 | 0 | 0 | 0 | 1 | 0 | 0 | 0 | 0 | 0 | 0 | 0 | 1 | 0 | 0 | 0 |
| May12.50v_31663 | 5 | 0 | 1 | 1 | 0 | 0 | 0 | 0 | 0 | 0 | 1 | 0 | 0 | 0 | 0 | 0 | 0 | 0 | 0 | 0 | 0 | 1 | 0 | 0 | 0 | 0 | 0 | 1 | 0 | 0 |
| Aug5.5v_1643 | 4 | 0 | 0 | 0 | 0 | 0 | 0 | 0 | 0 | 0 | 2 | 0 | 0 | 0 | 0 | 0 | 0 | 0 | 0 | 0 | 0 | 1 | 1 | 0 | 0 | 0 | 0 | 0 | 0 | 0 |
| Sep8.50v_33281 | 1 | 0 | 0 | 0 | 0 | 0 | 0 | 0 | 0 | 0 | 0 | 0 | 0 | 0 | 0 | 0 | 0 | 0 | 0 | 0 | 0 | 0 | 0 | 0 | 0 | 0 | 0 | 0 | 0 | 1 |
| Jul27.5v_7308 | 3 | 0 | 0 | 1 | 0 | 0 | 0 | 0 | 0 | 0 | 0 | 0 | 0 | 0 | 0 | 0 | 0 | 1 | 0 | 0 | 0 | 0 | 0 | 0 | 0 | 0 | 0 | 1 | 0 | 0 |
| Sep8.0v_8298 | 1 | 0 | 0 | 0 | 0 | 0 | 0 | 0 | 0 | 0 | 0 | 0 | 0 | 0 | 0 | 0 | 0 | 0 | 0 | 0 | 0 | 0 | 0 | 0 | 0 | 0 | 1 | 0 | 0 | 0 |
| Aug5.50v_40956 | 1 | 0 | 0 | 0 | 0 | 0 | 0 | 0 | 0 | 0 | 0 | 0 | 0 | 0 | 0 | 0 | 0 | 0 | 0 | 0 | 0 | 0 | 0 | 0 | 1 | 0 | 0 | 0 | 0 | 0 |
| May12.50v_34891 | 1 | 0 | 0 | 1 | 0 | 0 | 0 | 0 | 0 | 0 | 0 | 0 | 0 | 0 | 0 | 0 | 0 | 0 | 0 | 0 | 0 | 0 | 0 | 0 | 0 | 0 | 0 | 0 | 0 | 0 |
| Aug5.5v_20820 | 1 | 0 | 0 | 0 | 0 | 0 | 0 | 0 | 0 | 0 | 0 | 0 | 0 | 0 | 0 | 0 | 0 | 0 | 0 | 0 | 0 | 0 | 1 | 0 | 0 | 0 | 0 | 0 | 0 | 0 |
| Aug5.25v_32394 | 1 | 0 | 0 | 0 | 0 | 0 | 0 | 0 | 0 | 0 | 0 | 0 | 0 | 0 | 0 | 0 | 0 | 0 | 0 | 0 | 0 | 0 | 0 | 1 | 0 | 0 | 0 | 0 | 0 | 0 |
| Aug5.25v_26265 | 1 | 0 | 0 | 0 | 0 | 0 | 0 | 0 | 0 | 0 | 0 | 0 | 0 | 0 | 0 | 0 | 0 | 0 | 0 | 0 | 0 | 0 | 0 | 1 | 0 | 0 | 0 | 0 | 0 | 0 |
| Jul27.5v_1452 | 1 | 0 | 0 | 0 | 0 | 0 | 0 | 0 | 0 | 0 | 0 | 0 | 0 | 0 | 0 | 0 | 0 | 1 | 0 | 0 | 0 | 0 | 0 | 0 | 0 | 0 | 0 | 0 | 0 | 0 |
| Sep8.50v_29029 | 1 | 0 | 0 | 0 | 0 | 0 | 0 | 0 | 0 | 0 | 0 | 0 | 0 | 0 | 0 | 0 | 0 | 0 | 0 | 0 | 0 | 0 | 0 | 0 | 0 | 0 | 0 | 0 | 0 | 1 |
| May12.50v_26228 | 1 | 0 | 0 | 1 | 0 | 0 | 0 | 0 | 0 | 0 | 0 | 0 | 0 | 0 | 0 | 0 | 0 | 0 | 0 | 0 | 0 | 0 | 0 | 0 | 0 | 0 | 0 | 0 | 0 | 0 |
| May12.75v_37726 | 1 | 0 | 0 | 0 | 1 | 0 | 0 | 0 | 0 | 0 | 0 | 0 | 0 | 0 | 0 | 0 | 0 | 0 | 0 | 0 | 0 | 0 | 0 | 0 | 0 | 0 | 0 | 0 | 0 | 0 |
| Jul27.5v_9809 | 5 | 0 | 1 | 0 | 0 | 0 | 0 | 0 | 0 | 1 | 0 | 0 | 1 | 0 | 0 | 0 | 0 | 1 | 0 | 0 | 0 | 0 | 0 | 1 | 0 | 0 | 0 | 0 | 0 | 0 |
| Jul11.50v_18754 | 1 | 0 | 0 | 0 | 0 | 0 | 0 | 0 | 0 | 0 | 0 | 0 | 0 | 0 | 0 | 1 | 0 | 0 | 0 | 0 | 0 | 0 | 0 | 0 | 0 | 0 | 0 | 0 | 0 | 0 |
| Jun8.75v_4637 | 6 | 0 | 0 | 0 | 0 | 1 | 0 | 1 | 0 | 0 | 0 | 0 | 0 | 0 | 0 | 2 | 0 | 0 | 0 | 0 | 0 | 0 | 0 | 1 | 0 | 0 | 0 | 0 | 0 | 1 |
| Jun22.50v_38128 | 1 | 0 | 0 | 0 | 0 | 0 | 0 | 0 | 0 | 0 | 1 | 0 | 0 | 0 | 0 | 0 | 0 | 0 | 0 | 0 | 0 | 0 | 0 | 0 | 0 | 0 | 0 | 0 | 0 | 0 |
| Jul11.75v_37275 | 1 | 0 | 0 | 0 | 0 | 0 | 0 | 0 | 0 | 0 | 0 | 0 | 0 | 0 | 0 | 0 | 1 | 0 | 0 | 0 | 0 | 0 | 0 | 0 | 0 | 0 | 0 | 0 | 0 | 0 |
| May12.50v_4819 | 1 | 0 | 0 | 1 | 0 | 0 | 0 | 0 | 0 | 0 | 0 | 0 | 0 | 0 | 0 | 0 | 0 | 0 | 0 | 0 | 0 | 0 | 0 | 0 | 0 | 0 | 0 | 0 | 0 | 0 |
| Sep8.50v_103 | 1 | 0 | 0 | 0 | 0 | 0 | 0 | 0 | 0 | 0 | 0 | 0 | 0 | 0 | 0 | 0 | 0 | 0 | 0 | 0 | 0 | 0 | 0 | 0 | 0 | 0 | 0 | 0 | 0 | 1 |
| May12.25v_7055 | 1 | 0 | 1 | 0 | 0 | 0 | 0 | 0 | 0 | 0 | 0 | 0 | 0 | 0 | 0 | 0 | 0 | 0 | 0 | 0 | 0 | 0 | 0 | 0 | 0 | 0 | 0 | 0 | 0 | 0 |
| Aug5.25v_40635 | 4 | 0 | 0 | 0 | 0 | 0 | 0 | 0 | 0 | 0 | 1 | 1 | 0 | 0 | 0 | 0 | 1 | 0 | 0 | 0 | 0 | 0 | 0 | 1 | 0 | 0 | 0 | 0 | 0 | 0 |
| Jul11.25v_757 | 1 | 0 | 0 | 0 | 0 | 0 | 0 | 0 | 0 | 0 | 0 | 0 | 0 | 0 | 1 | 0 | 0 | 0 | 0 | 0 | 0 | 0 | 0 | 0 | 0 | 0 | 0 | 0 | 0 | 0 |
| Jul27.5v_9579 | 1 | 0 | 0 | 0 | 0 | 0 | 0 | 0 | 0 | 0 | 0 | 0 | 0 | 0 | 0 | 0 | 0 | 1 | 0 | 0 | 0 | 0 | 0 | 0 | 0 | 0 | 0 | 0 | 0 | 0 |
| May12.25v_9339 | 1 | 0 | 1 | 0 | 0 | 0 | 0 | 0 | 0 | 0 | 0 | 0 | 0 | 0 | 0 | 0 | 0 | 0 | 0 | 0 | 0 | 0 | 0 | 0 | 0 | 0 | 0 | 0 | 0 | 0 |
| May12.75v_39552 | 1 | 0 | 0 | 0 | 1 | 0 | 0 | 0 | 0 | 0 | 0 | 0 | 0 | 0 | 0 | 0 | 0 | 0 | 0 | 0 | 0 | 0 | 0 | 0 | 0 | 0 | 0 | 0 | 0 | 0 |
| May12.50v_11510 | 1 | 0 | 0 | 1 | 0 | 0 | 0 | 0 | 0 | 0 | 0 | 0 | 0 | 0 | 0 | 0 | 0 | 0 | 0 | 0 | 0 | 0 | 0 | 0 | 0 | 0 | 0 | 0 | 0 | 0 |
| May12.25v_18318 | 1 | 0 | 1 | 0 | 0 | 0 | 0 | 0 | 0 | 0 | 0 | 0 | 0 | 0 | 0 | 0 | 0 | 0 | 0 | 0 | 0 | 0 | 0 | 0 | 0 | 0 | 0 | 0 | 0 | 0 |
| Aug5.0v_753 | 5 | 0 | 0 | 0 | 0 | 0 | 0 | 0 | 0 | 0 | 0 | 0 | 1 | 0 | 0 | 0 | 0 | 0 | 0 | 0 | 0 | 1 | 1 | 1 | 0 | 0 | 0 | 0 | 0 | 1 |
| Aug5.25v_15437 | 1 | 0 | 0 | 0 | 0 | 0 | 0 | 0 | 0 | 0 | 0 | 0 | 0 | 0 | 0 | 0 | 0 | 0 | 0 | 0 | 0 | 0 | 0 | 1 | 0 | 0 | 0 | 0 | 0 | 0 |
| Aug5.50v_25883 | 1 | 0 | 0 | 0 | 0 | 0 | 0 | 0 | 0 | 0 | 0 | 0 | 0 | 0 | 0 | 0 | 0 | 0 | 0 | 0 | 0 | 0 | 0 | 0 | 1 | 0 | 0 | 0 | 0 | 0 |
| Sep8.50v_3860 | 1 | 0 | 0 | 0 | 0 | 0 | 0 | 0 | 0 | 0 | 0 | 0 | 0 | 0 | 0 | 0 | 0 | 0 | 0 | 0 | 0 | 0 | 0 | 0 | 0 | 0 | 0 | 0 | 0 | 1 |
| Aug5.50v_12302 | 1 | 0 | 0 | 0 | 0 | 0 | 0 | 0 | 0 | 0 | 0 | 0 | 0 | 0 | 0 | 0 | 0 | 0 | 0 | 0 | 0 | 0 | 0 | 0 | 1 | 0 | 0 | 0 | 0 | 0 |
| Jul11.0v_19603 | 1 | 0 | 0 | 0 | 0 | 0 | 0 | 0 | 0 | 0 | 0 | 0 | 1 | 0 | 0 | 0 | 0 | 0 | 0 | 0 | 0 | 0 | 0 | 0 | 0 | 0 | 0 | 0 | 0 | 0 |
| Sep8.0v_16220 | 1 | 0 | 0 | 0 | 0 | 0 | 0 | 0 | 0 | 0 | 0 | 0 | 0 | 0 | 0 | 0 | 0 | 0 | 0 | 0 | 0 | 0 | 0 | 0 | 0 | 0 | 1 | 0 | 0 | 0 |
| Jul11.0v_4203 | 6 | 0 | 0 | 1 | 0 | 0 | 0 | 0 | 0 | 0 | 0 | 0 | 1 | 0 | 1 | 0 | 0 | 1 | 0 | 0 | 0 | 0 | 1 | 1 | 0 | 0 | 0 | 0 | 0 | 0 |
| Sep8.0v_12989 | 1 | 0 | 0 | 0 | 0 | 0 | 0 | 0 | 0 | 0 | 0 | 0 | 0 | 0 | 0 | 0 | 0 | 0 | 0 | 0 | 0 | 0 | 0 | 0 | 0 | 0 | 1 | 0 | 0 | 0 |
| Aug5.0v_214 | 1 | 0 | 0 | 0 | 0 | 0 | 0 | 0 | 0 | 0 | 0 | 0 | 0 | 0 | 0 | 0 | 0 | 0 | 0 | 0 | 0 | 1 | 0 | 0 | 0 | 0 | 0 | 0 | 0 | 0 |
| Sep8.5v_19937 | 2 | 0 | 0 | 1 | 0 | 0 | 0 | 0 | 0 | 0 | 0 | 0 | 0 | 0 | 0 | 0 | 0 | 0 | 0 | 0 | 0 | 0 | 0 | 0 | 0 | 0 | 0 | 1 | 0 | 0 |
| May12.50v_9925 | 1 | 0 | 0 | 1 | 0 | 0 | 0 | 0 | 0 | 0 | 0 | 0 | 0 | 0 | 0 | 0 | 0 | 0 | 0 | 0 | 0 | 0 | 0 | 0 | 0 | 0 | 0 | 0 | 0 | 0 |
| May12.5v_19120 | 1 | 1 | 0 | 0 | 0 | 0 | 0 | 0 | 0 | 0 | 0 | 0 | 0 | 0 | 0 | 0 | 0 | 0 | 0 | 0 | 0 | 0 | 0 | 0 | 0 | 0 | 0 | 0 | 0 | 0 |
| Jun22.50v_32459 | 1 | 0 | 0 | 0 | 0 | 0 | 0 | 0 | 0 | 0 | 1 | 0 | 0 | 0 | 0 | 0 | 0 | 0 | 0 | 0 | 0 | 0 | 0 | 0 | 0 | 0 | 0 | 0 | 0 | 0 |
| Sep8.0v_3393 | 2 | 0 | 0 | 0 | 0 | 0 | 0 | 0 | 0 | 0 | 0 | 0 | 0 | 0 | 0 | 0 | 0 | 0 | 0 | 0 | 0 | 0 | 1 | 0 | 0 | 0 | 1 | 0 | 0 | 0 |
| Aug5.0v_2554 | 1 | 0 | 0 | 0 | 0 | 0 | 0 | 0 | 0 | 0 | 0 | 0 | 0 | 0 | 0 | 0 | 0 | 0 | 0 | 0 | 0 | 1 | 0 | 0 | 0 | 0 | 0 | 0 | 0 | 0 |
| Sep8.5v_498 | 1 | 0 | 0 | 0 | 0 | 0 | 0 | 0 | 0 | 0 | 0 | 0 | 0 | 0 | 0 | 0 | 0 | 0 | 0 | 0 | 0 | 0 | 0 | 0 | 0 | 0 | 0 | 1 | 0 | 0 |
| Jul11.5v_1837 | 1 | 0 | 0 | 0 | 0 | 0 | 0 | 0 | 0 | 0 | 0 | 0 | 0 | 1 | 0 | 0 | 0 | 0 | 0 | 0 | 0 | 0 | 0 | 0 | 0 | 0 | 0 | 0 | 0 | 0 |
| Sep8.0v_12375 | 1 | 0 | 0 | 0 | 0 | 0 | 0 | 0 | 0 | 0 | 0 | 0 | 0 | 0 | 0 | 0 | 0 | 0 | 0 | 0 | 0 | 0 | 0 | 0 | 0 | 0 | 1 | 0 | 0 | 0 |
| May12.50v_19154 | 1 | 0 | 0 | 1 | 0 | 0 | 0 | 0 | 0 | 0 | 0 | 0 | 0 | 0 | 0 | 0 | 0 | 0 | 0 | 0 | 0 | 0 | 0 | 0 | 0 | 0 | 0 | 0 | 0 | 0 |
| Sep8.0v_14253 | 1 | 0 | 0 | 0 | 0 | 0 | 0 | 0 | 0 | 0 | 0 | 0 | 0 | 0 | 0 | 0 | 0 | 0 | 0 | 0 | 0 | 0 | 0 | 0 | 0 | 0 | 1 | 0 | 0 | 0 |
| Aug5.5v_27474 | 1 | 0 | 0 | 0 | 0 | 0 | 0 | 0 | 0 | 0 | 0 | 0 | 0 | 0 | 0 | 0 | 0 | 0 | 0 | 0 | 0 | 0 | 1 | 0 | 0 | 0 | 0 | 0 | 0 | 0 |
| May12.50v_12239 | 1 | 0 | 0 | 1 | 0 | 0 | 0 | 0 | 0 | 0 | 0 | 0 | 0 | 0 | 0 | 0 | 0 | 0 | 0 | 0 | 0 | 0 | 0 | 0 | 0 | 0 | 0 | 0 | 0 | 0 |
| May12.50v_29767 | 1 | 0 | 0 | 1 | 0 | 0 | 0 | 0 | 0 | 0 | 0 | 0 | 0 | 0 | 0 | 0 | 0 | 0 | 0 | 0 | 0 | 0 | 0 | 0 | 0 | 0 | 0 | 0 | 0 | 0 |
| Jul27.50v_39825 | 4 | 0 | 0 | 0 | 0 | 0 | 0 | 0 | 0 | 0 | 1 | 0 | 0 | 0 | 0 | 0 | 0 | 0 | 0 | 1 | 0 | 1 | 0 | 0 | 0 | 0 | 0 | 0 | 0 | 1 |

| Feature_ID | 654 | May12_05m | May12_25m | May12_50m | May12_75m | Jun8_25m | Jun8_50m | Jun8_75m | Jun22_0m | Jun22_05m | Jun22_50m | Jun22_75m | Jul11_0m | Jul11_05m | Jul11_25m | Jul11_50m | Jul11_75m | Jul27_05m | Jul27_25m | Jul27_50m | Jul27_75m | Aug5_0m | Aug5_05m | Aug5_25m | Aug5_50m | Aug5_75m | Sep8_0m | Sep8_05m | Sep8_25m | Sep8_50m |
|---|---|---|---|---|---|---|---|---|---|---|---|---|---|---|---|---|---|---|---|---|---|---|---|---|---|---|---|---|---|---|
| Jun8.25v_2075 | 1 | 0 | 0 | 0 | 0 | 1 | 0 | 0 | 0 | 0 | 0 | 0 | 0 | 0 | 0 | 0 | 0 | 0 | 0 | 0 | 0 | 0 | 0 | 0 | 0 | 0 | 0 | 0 | 0 | 0 |
| Jul11.50v_10808 | 1 | 0 | 0 | 0 | 0 | 0 | 0 | 0 | 0 | 0 | 0 | 0 | 0 | 0 | 0 | 1 | 0 | 0 | 0 | 0 | 0 | 0 | 0 | 0 | 0 | 0 | 0 | 0 | 0 | 0 |
| Sep8.50v_16259 | 1 | 0 | 0 | 0 | 0 | 0 | 0 | 0 | 0 | 0 | 0 | 0 | 0 | 0 | 0 | 0 | 0 | 0 | 0 | 0 | 0 | 0 | 0 | 0 | 0 | 0 | 0 | 0 | 0 | 1 |
| May12.50v_19130 | 1 | 0 | 0 | 1 | 0 | 0 | 0 | 0 | 0 | 0 | 0 | 0 | 0 | 0 | 0 | 0 | 0 | 0 | 0 | 0 | 0 | 0 | 0 | 0 | 0 | 0 | 0 | 0 | 0 | 0 |
| Aug5.0v_334 | 2 | 0 | 0 | 0 | 0 | 0 | 0 | 0 | 0 | 0 | 0 | 0 | 0 | 0 | 0 | 0 | 0 | 0 | 0 | 0 | 0 | 1 | 0 | 1 | 0 | 0 | 0 | 0 | 0 | 0 |
| Aug5.75v_3629 | 1 | 0 | 0 | 0 | 0 | 0 | 0 | 0 | 0 | 0 | 0 | 0 | 0 | 0 | 0 | 0 | 0 | 0 | 0 | 0 | 0 | 0 | 0 | 0 | 0 | 1 | 0 | 0 | 0 | 0 |
| Jul11.5v_17803 | 1 | 0 | 0 | 0 | 0 | 0 | 0 | 0 | 0 | 0 | 0 | 0 | 0 | 1 | 0 | 0 | 0 | 0 | 0 | 0 | 0 | 0 | 0 | 0 | 0 | 0 | 0 | 0 | 0 | 0 |
| Jun22.0v_8369 | 1 | 0 | 0 | 0 | 0 | 0 | 0 | 0 | 1 | 0 | 0 | 0 | 0 | 0 | 0 | 0 | 0 | 0 | 0 | 0 | 0 | 0 | 0 | 0 | 0 | 0 | 0 | 0 | 0 | 0 |
| May12.50v_1812 | 2 | 0 | 0 | 1 | 0 | 0 | 0 | 0 | 0 | 1 | 0 | 0 | 0 | 0 | 0 | 0 | 0 | 0 | 0 | 0 | 0 | 0 | 0 | 0 | 0 | 0 | 0 | 0 | 0 | 0 |
| Jul11.25v_15141 | 1 | 0 | 0 | 0 | 0 | 0 | 0 | 0 | 0 | 0 | 0 | 0 | 0 | 0 | 1 | 0 | 0 | 0 | 0 | 0 | 0 | 0 | 0 | 0 | 0 | 0 | 0 | 0 | 0 | 0 |
| Jul27.25v_34509 | 1 | 0 | 0 | 0 | 0 | 0 | 0 | 0 | 0 | 0 | 0 | 0 | 0 | 0 | 0 | 0 | 0 | 0 | 1 | 0 | 0 | 0 | 0 | 0 | 0 | 0 | 0 | 0 | 0 | 0 |
| Sep8.50v_24605 | 2 | 0 | 0 | 0 | 0 | 0 | 0 | 0 | 0 | 0 | 0 | 0 | 0 | 0 | 1 | 0 | 0 | 0 | 0 | 0 | 0 | 0 | 0 | 0 | 0 | 0 | 0 | 0 | 0 | 1 |
| Aug5.0v_623 | 1 | 0 | 0 | 0 | 0 | 0 | 0 | 0 | 0 | 0 | 0 | 0 | 0 | 0 | 0 | 0 | 0 | 0 | 0 | 0 | 0 | 1 | 0 | 0 | 0 | 0 | 0 | 0 | 0 | 0 |
| Jul11.5v_8037 | 1 | 0 | 0 | 0 | 0 | 0 | 0 | 0 | 0 | 0 | 0 | 0 | 0 | 1 | 0 | 0 | 0 | 0 | 0 | 0 | 0 | 0 | 0 | 0 | 0 | 0 | 0 | 0 | 0 | 0 |
| Jun8.50v_4355 | 1 | 0 | 0 | 0 | 0 | 0 | 1 | 0 | 0 | 0 | 0 | 0 | 0 | 0 | 0 | 0 | 0 | 0 | 0 | 0 | 0 | 0 | 0 | 0 | 0 | 0 | 0 | 0 | 0 | 0 |
| Jul11.0v_14736 | 1 | 0 | 0 | 0 | 0 | 0 | 0 | 0 | 0 | 0 | 0 | 0 | 1 | 0 | 0 | 0 | 0 | 0 | 0 | 0 | 0 | 0 | 0 | 0 | 0 | 0 | 0 | 0 | 0 | 0 |
| Jun8.50v_36668 | 2 | 0 | 0 | 0 | 0 | 0 | 1 | 0 | 0 | 0 | 0 | 1 | 0 | 0 | 0 | 0 | 0 | 0 | 0 | 0 | 0 | 0 | 0 | 0 | 0 | 0 | 0 | 0 | 0 | 0 |
| Jul11.50v_5480 | 1 | 0 | 0 | 0 | 0 | 0 | 0 | 0 | 0 | 0 | 0 | 0 | 0 | 0 | 0 | 1 | 0 | 0 | 0 | 0 | 0 | 0 | 0 | 0 | 0 | 0 | 0 | 0 | 0 | 0 |
| May12.25v_31975 | 1 | 0 | 1 | 0 | 0 | 0 | 0 | 0 | 0 | 0 | 0 | 0 | 0 | 0 | 0 | 0 | 0 | 0 | 0 | 0 | 0 | 0 | 0 | 0 | 0 | 0 | 0 | 0 | 0 | 0 |
| May12.75v_15066 | 1 | 0 | 0 | 0 | 1 | 0 | 0 | 0 | 0 | 0 | 0 | 0 | 0 | 0 | 0 | 0 | 0 | 0 | 0 | 0 | 0 | 0 | 0 | 0 | 0 | 0 | 0 | 0 | 0 | 0 |
| Jun22.75v_30843 | 1 | 0 | 0 | 0 | 0 | 0 | 0 | 0 | 0 | 0 | 0 | 1 | 0 | 0 | 0 | 0 | 0 | 0 | 0 | 0 | 0 | 0 | 0 | 0 | 0 | 0 | 0 | 0 | 0 | 0 |
| Aug5.0v_2944 | 1 | 0 | 0 | 0 | 0 | 0 | 0 | 0 | 0 | 0 | 0 | 0 | 0 | 0 | 0 | 0 | 0 | 0 | 0 | 0 | 0 | 1 | 0 | 0 | 0 | 0 | 0 | 0 | 0 | 0 |
| Jun22.5v_1143 | 1 | 0 | 0 | 0 | 0 | 0 | 0 | 0 | 0 | 1 | 0 | 0 | 0 | 0 | 0 | 0 | 0 | 0 | 0 | 0 | 0 | 0 | 0 | 0 | 0 | 0 | 0 | 0 | 0 | 0 |
| Sep8.25v_17217 | 1 | 0 | 0 | 0 | 0 | 0 | 0 | 0 | 0 | 0 | 0 | 0 | 0 | 0 | 0 | 0 | 0 | 0 | 0 | 0 | 0 | 0 | 0 | 0 | 0 | 0 | 0 | 0 | 1 | 0 |
| Sep8.0v_11663 | 1 | 0 | 0 | 0 | 0 | 0 | 0 | 0 | 0 | 0 | 0 | 0 | 0 | 0 | 0 | 0 | 0 | 0 | 0 | 0 | 0 | 0 | 0 | 0 | 0 | 0 | 1 | 0 | 0 | 0 |
| Jul27.25v_32817 | 1 | 0 | 0 | 0 | 0 | 0 | 0 | 0 | 0 | 0 | 0 | 0 | 0 | 0 | 0 | 0 | 0 | 0 | 1 | 0 | 0 | 0 | 0 | 0 | 0 | 0 | 0 | 0 | 0 | 0 |
| Jul11.25v_11955 | 2 | 0 | 0 | 0 | 0 | 0 | 0 | 0 | 0 | 0 | 0 | 0 | 0 | 0 | 1 | 0 | 0 | 0 | 0 | 0 | 0 | 0 | 0 | 0 | 0 | 0 | 0 | 0 | 0 | 1 |
| Jul27.5v_18307 | 3 | 0 | 0 | 0 | 0 | 0 | 0 | 0 | 0 | 0 | 0 | 1 | 0 | 0 | 0 | 0 | 0 | 1 | 0 | 0 | 1 | 0 | 0 | 0 | 0 | 0 | 0 | 0 | 0 | 0 |
| Jun22.75v_20208 | 1 | 0 | 0 | 0 | 0 | 0 | 0 | 0 | 0 | 0 | 0 | 1 | 0 | 0 | 0 | 0 | 0 | 0 | 0 | 0 | 0 | 0 | 0 | 0 | 0 | 0 | 0 | 0 | 0 | 0 |
| May12.25v_4863 | 4 | 0 | 1 | 0 | 0 | 1 | 0 | 0 | 0 | 0 | 0 | 0 | 0 | 0 | 0 | 0 | 0 | 0 | 0 | 1 | 0 | 1 | 0 | 0 | 0 | 0 | 0 | 0 | 0 | 0 |
| Jul11.5v_17463 | 1 | 0 | 0 | 0 | 0 | 0 | 0 | 0 | 0 | 0 | 0 | 0 | 0 | 1 | 0 | 0 | 0 | 0 | 0 | 0 | 0 | 0 | 0 | 0 | 0 | 0 | 0 | 0 | 0 | 0 |
| Jun8.75v_3226 | 1 | 0 | 0 | 0 | 0 | 0 | 0 | 1 | 0 | 0 | 0 | 0 | 0 | 0 | 0 | 0 | 0 | 0 | 0 | 0 | 0 | 0 | 0 | 0 | 0 | 0 | 0 | 0 | 0 | 0 |
| Jun22.75v_9430 | 1 | 0 | 0 | 0 | 0 | 0 | 0 | 0 | 0 | 0 | 0 | 1 | 0 | 0 | 0 | 0 | 0 | 0 | 0 | 0 | 0 | 0 | 0 | 0 | 0 | 0 | 0 | 0 | 0 | 0 |
| May12.25v_14596 | 1 | 0 | 1 | 0 | 0 | 0 | 0 | 0 | 0 | 0 | 0 | 0 | 0 | 0 | 0 | 0 | 0 | 0 | 0 | 0 | 0 | 0 | 0 | 0 | 0 | 0 | 0 | 0 | 0 | 0 |
| Jun22.5v_6696 | 1 | 0 | 0 | 0 | 0 | 0 | 0 | 0 | 0 | 1 | 0 | 0 | 0 | 0 | 0 | 0 | 0 | 0 | 0 | 0 | 0 | 0 | 0 | 0 | 0 | 0 | 0 | 0 | 0 | 0 |
| May12.50v_34575 | 1 | 0 | 0 | 1 | 0 | 0 | 0 | 0 | 0 | 0 | 0 | 0 | 0 | 0 | 0 | 0 | 0 | 0 | 0 | 0 | 0 | 0 | 0 | 0 | 0 | 0 | 0 | 0 | 0 | 0 |
| Sep8.0v_20500 | 1 | 0 | 0 | 0 | 0 | 0 | 0 | 0 | 0 | 0 | 0 | 0 | 0 | 0 | 0 | 0 | 0 | 0 | 0 | 0 | 0 | 0 | 0 | 0 | 0 | 0 | 1 | 0 | 0 | 0 |
| Jul11.25v_16320 | 1 | 0 | 0 | 0 | 0 | 0 | 0 | 0 | 0 | 0 | 0 | 0 | 0 | 0 | 1 | 0 | 0 | 0 | 0 | 0 | 0 | 0 | 0 | 0 | 0 | 0 | 0 | 0 | 0 | 0 |
| Jul11.50v_13993 | 1 | 0 | 0 | 0 | 0 | 0 | 0 | 0 | 0 | 0 | 0 | 0 | 0 | 0 | 0 | 1 | 0 | 0 | 0 | 0 | 0 | 0 | 0 | 0 | 0 | 0 | 0 | 0 | 0 | 0 |
| May12.5v_40204 | 1 | 1 | 0 | 0 | 0 | 0 | 0 | 0 | 0 | 0 | 0 | 0 | 0 | 0 | 0 | 0 | 0 | 0 | 0 | 0 | 0 | 0 | 0 | 0 | 0 | 0 | 0 | 0 | 0 | 0 |
| Jul11.25v_1782 | 1 | 0 | 0 | 0 | 0 | 0 | 0 | 0 | 0 | 0 | 0 | 0 | 0 | 0 | 1 | 0 | 0 | 0 | 0 | 0 | 0 | 0 | 0 | 0 | 0 | 0 | 0 | 0 | 0 | 0 |
| Jul11.50v_7198 | 1 | 0 | 0 | 0 | 0 | 0 | 0 | 0 | 0 | 0 | 0 | 0 | 0 | 0 | 0 | 1 | 0 | 0 | 0 | 0 | 0 | 0 | 0 | 0 | 0 | 0 | 0 | 0 | 0 | 0 |
| Jun22.0v_20292 | 1 | 0 | 0 | 0 | 0 | 0 | 0 | 0 | 1 | 0 | 0 | 0 | 0 | 0 | 0 | 0 | 0 | 0 | 0 | 0 | 0 | 0 | 0 | 0 | 0 | 0 | 0 | 0 | 0 | 0 |
| Jun22.50v_8978 | 1 | 0 | 0 | 0 | 0 | 0 | 0 | 0 | 0 | 0 | 1 | 0 | 0 | 0 | 0 | 0 | 0 | 0 | 0 | 0 | 0 | 0 | 0 | 0 | 0 | 0 | 0 | 0 | 0 | 0 |
| Aug5.0v_1443 | 18 | 0 | 2 | 6 | 3 | 0 | 0 | 0 | 0 | 1 | 0 | 0 | 0 | 0 | 1 | 0 | 0 | 0 | 0 | 0 | 0 | 1 | 0 | 0 | 1 | 1 | 0 | 0 | 1 | 1 |
| May12.75v_38498 | 1 | 0 | 0 | 0 | 1 | 0 | 0 | 0 | 0 | 0 | 0 | 0 | 0 | 0 | 0 | 0 | 0 | 0 | 0 | 0 | 0 | 0 | 0 | 0 | 0 | 0 | 0 | 0 | 0 | 0 |
| Jul27.5v_13690 | 9 | 1 | 1 | 2 | 0 | 1 | 0 | 1 | 0 | 0 | 0 | 0 | 0 | 0 | 0 | 0 | 0 | 1 | 2 | 0 | 0 | 0 | 0 | 0 | 0 | 0 | 0 | 0 | 0 | 0 |
| Jul11.5v_6240 | 2 | 0 | 0 | 0 | 0 | 0 | 0 | 0 | 0 | 1 | 0 | 0 | 0 | 1 | 0 | 0 | 0 | 0 | 0 | 0 | 0 | 0 | 0 | 0 | 0 | 0 | 0 | 0 | 0 | 0 |
| Jun8.50v_2793 | 3 | 0 | 0 | 0 | 1 | 0 | 1 | 0 | 0 | 0 | 0 | 0 | 0 | 0 | 0 | 0 | 0 | 0 | 0 | 0 | 0 | 0 | 1 | 0 | 0 | 0 | 0 | 0 | 0 | 0 |
| May12.50v_26920 | 1 | 0 | 0 | 1 | 0 | 0 | 0 | 0 | 0 | 0 | 0 | 0 | 0 | 0 | 0 | 0 | 0 | 0 | 0 | 0 | 0 | 0 | 0 | 0 | 0 | 0 | 0 | 0 | 0 | 0 |
| Jun8.50v_24104 | 1 | 0 | 0 | 0 | 0 | 0 | 1 | 0 | 0 | 0 | 0 | 0 | 0 | 0 | 0 | 0 | 0 | 0 | 0 | 0 | 0 | 0 | 0 | 0 | 0 | 0 | 0 | 0 | 0 | 0 |
| Jul11.5v_20509 | 1 | 0 | 0 | 0 | 0 | 0 | 0 | 0 | 0 | 0 | 0 | 0 | 0 | 1 | 0 | 0 | 0 | 0 | 0 | 0 | 0 | 0 | 0 | 0 | 0 | 0 | 0 | 0 | 0 | 0 |
| Sep8.0v_1085 | 1 | 0 | 0 | 0 | 0 | 0 | 0 | 0 | 0 | 0 | 0 | 0 | 0 | 0 | 0 | 0 | 0 | 0 | 0 | 0 | 0 | 0 | 0 | 0 | 0 | 0 | 1 | 0 | 0 | 0 |
| May12.50v_22692 | 1 | 0 | 0 | 1 | 0 | 0 | 0 | 0 | 0 | 0 | 0 | 0 | 0 | 0 | 0 | 0 | 0 | 0 | 0 | 0 | 0 | 0 | 0 | 0 | 0 | 0 | 0 | 0 | 0 | 0 |
| Jul27.50v_9739 | 1 | 0 | 0 | 0 | 0 | 0 | 0 | 0 | 0 | 0 | 0 | 0 | 0 | 0 | 0 | 0 | 0 | 0 | 0 | 1 | 0 | 0 | 0 | 0 | 0 | 0 | 0 | 0 | 0 | 0 |
| May12.50v_30104 | 1 | 0 | 0 | 1 | 0 | 0 | 0 | 0 | 0 | 0 | 0 | 0 | 0 | 0 | 0 | 0 | 0 | 0 | 0 | 0 | 0 | 0 | 0 | 0 | 0 | 0 | 0 | 0 | 0 | 0 |
| May12.75v_7337 | 1 | 0 | 0 | 0 | 1 | 0 | 0 | 0 | 0 | 0 | 0 | 0 | 0 | 0 | 0 | 0 | 0 | 0 | 0 | 0 | 0 | 0 | 0 | 0 | 0 | 0 | 0 | 0 | 0 | 0 |
| May12.75v_11814 | 1 | 0 | 0 | 0 | 1 | 0 | 0 | 0 | 0 | 0 | 0 | 0 | 0 | 0 | 0 | 0 | 0 | 0 | 0 | 0 | 0 | 0 | 0 | 0 | 0 | 0 | 0 | 0 | 0 | 0 |
| Jul27.5v_2057 | 1 | 0 | 0 | 0 | 0 | 0 | 0 | 0 | 0 | 0 | 0 | 0 | 0 | 0 | 0 | 0 | 0 | 1 | 0 | 0 | 0 | 0 | 0 | 0 | 0 | 0 | 0 | 0 | 0 | 0 |
| Jun22.0v_9431 | 6 | 2 | 0 | 1 | 0 | 0 | 0 | 0 | 1 | 0 | 0 | 0 | 0 | 0 | 0 | 0 | 0 | 0 | 0 | 0 | 0 | 0 | 0 | 0 | 0 | 0 | 0 | 0 | 0 | 1 |
| Jul27.5v_13199 | 1 | 0 | 0 | 0 | 0 | 0 | 0 | 0 | 0 | 0 | 0 | 0 | 0 | 0 | 0 | 0 | 0 | 1 | 0 | 0 | 0 | 0 | 0 | 0 | 0 | 0 | 0 | 0 | 0 | 0 |
| Aug5.0v_4192 | 1 | 0 | 0 | 0 | 0 | 0 | 0 | 0 | 0 | 0 | 0 | 0 | 0 | 0 | 0 | 0 | 0 | 0 | 0 | 0 | 0 | 1 | 0 | 0 | 0 | 0 | 0 | 0 | 0 | 0 |
| May12.50v_12530 | 1 | 0 | 0 | 1 | 0 | 0 | 0 | 0 | 0 | 0 | 0 | 0 | 0 | 0 | 0 | 0 | 0 | 0 | 0 | 0 | 0 | 0 | 0 | 0 | 0 | 0 | 0 | 0 | 0 | 0 |
| Aug5.25v_23850 | 1 | 0 | 0 | 0 | 0 | 0 | 0 | 0 | 0 | 0 | 0 | 0 | 0 | 0 | 0 | 0 | 0 | 0 | 0 | 0 | 0 | 0 | 0 | 1 | 0 | 0 | 0 | 0 | 0 | 0 |
| Sep8.0v_18270 | 1 | 0 | 0 | 0 | 0 | 0 | 0 | 0 | 0 | 0 | 0 | 0 | 0 | 0 | 0 | 0 | 0 | 0 | 0 | 0 | 0 | 0 | 0 | 0 | 0 | 0 | 1 | 0 | 0 | 0 |
| May12.75v_32791 | 2 | 0 | 0 | 0 | 1 | 0 | 0 | 0 | 0 | 0 | 0 | 0 | 0 | 0 | 0 | 0 | 0 | 0 | 0 | 0 | 0 | 1 | 0 | 0 | 0 | 0 | 0 | 0 | 0 | 0 |
| Jul27.5v_384 | 1 | 0 | 0 | 0 | 0 | 0 | 0 | 0 | 0 | 0 | 0 | 0 | 0 | 0 | 0 | 0 | 0 | 1 | 0 | 0 | 0 | 0 | 0 | 0 | 0 | 0 | 0 | 0 | 0 | 0 |
| Aug5.5v_23764 | 1 | 0 | 0 | 0 | 0 | 0 | 0 | 0 | 0 | 0 | 0 | 0 | 0 | 0 | 0 | 0 | 0 | 0 | 0 | 0 | 0 | 0 | 1 | 0 | 0 | 0 | 0 | 0 | 0 | 0 |
| Jul11.5v_16566 | 1 | 0 | 0 | 0 | 0 | 0 | 0 | 0 | 0 | 0 | 0 | 0 | 0 | 1 | 0 | 0 | 0 | 0 | 0 | 0 | 0 | 0 | 0 | 0 | 0 | 0 | 0 | 0 | 0 | 0 |
| Jul27.5v_12906 | 1 | 0 | 0 | 0 | 0 | 0 | 0 | 0 | 0 | 0 | 0 | 0 | 0 | 0 | 0 | 0 | 0 | 1 | 0 | 0 | 0 | 0 | 0 | 0 | 0 | 0 | 0 | 0 | 0 | 0 |
| May12.25v_8821 | 1 | 0 | 1 | 0 | 0 | 0 | 0 | 0 | 0 | 0 | 0 | 0 | 0 | 0 | 0 | 0 | 0 | 0 | 0 | 0 | 0 | 0 | 0 | 0 | 0 | 0 | 0 | 0 | 0 | 0 |
| Aug5.0v_3552 | 1 | 0 | 0 | 0 | 0 | 0 | 0 | 0 | 0 | 0 | 0 | 0 | 0 | 0 | 0 | 0 | 0 | 0 | 0 | 0 | 0 | 1 | 0 | 0 | 0 | 0 | 0 | 0 | 0 | 0 |
| Jun22.75v_15025 | 1 | 0 | 0 | 0 | 0 | 0 | 0 | 0 | 0 | 0 | 0 | 1 | 0 | 0 | 0 | 0 | 0 | 0 | 0 | 0 | 0 | 0 | 0 | 0 | 0 | 0 | 0 | 0 | 0 | 0 |
| Jun8.50v_27724 | 1 | 0 | 0 | 0 | 0 | 0 | 1 | 0 | 0 | 0 | 0 | 0 | 0 | 0 | 0 | 0 | 0 | 0 | 0 | 0 | 0 | 0 | 0 | 0 | 0 | 0 | 0 | 0 | 0 | 0 |
| Jul11.50v_12041 | 5 | 0 | 0 | 2 | 0 | 0 | 1 | 0 | 1 | 0 | 0 | 0 | 0 | 0 | 0 | 1 | 0 | 0 | 0 | 0 | 0 | 0 | 0 | 0 | 0 | 0 | 0 | 0 | 0 | 0 |
| Jul11.75v_4957 | 1 | 0 | 0 | 0 | 0 | 0 | 0 | 0 | 0 | 0 | 0 | 0 | 0 | 0 | 0 | 0 | 1 | 0 | 0 | 0 | 0 | 0 | 0 | 0 | 0 | 0 | 0 | 0 | 0 | 0 |
| Jul27.5v_1204 | 3 | 1 | 0 | 0 | 0 | 0 | 0 | 0 | 0 | 0 | 0 | 0 | 0 | 0 | 0 | 0 | 0 | 2 | 0 | 0 | 0 | 0 | 0 | 0 | 0 | 0 | 0 | 0 | 0 | 0 |
| Jun8.50v_20158 | 5 | 0 | 1 | 1 | 0 | 0 | 1 | 0 | 1 | 0 | 0 | 0 | 0 | 0 | 0 | 0 | 0 | 0 | 0 | 0 | 0 | 0 | 1 | 0 | 0 | 0 | 0 | 0 | 0 | 0 |
| Jul11.75v_34996 | 1 | 0 | 0 | 0 | 0 | 0 | 0 | 0 | 0 | 0 | 0 | 0 | 0 | 0 | 0 | 0 | 1 | 0 | 0 | 0 | 0 | 0 | 0 | 0 | 0 | 0 | 0 | 0 | 0 | 0 |
| Jul27.5v_60 | 1 | 0 | 0 | 0 | 0 | 0 | 0 | 0 | 0 | 0 | 0 | 0 | 0 | 0 | 0 | 0 | 0 | 1 | 0 | 0 | 0 | 0 | 0 | 0 | 0 | 0 | 0 | 0 | 0 | 0 |
| May12.50v_35446 | 1 | 0 | 0 | 1 | 0 | 0 | 0 | 0 | 0 | 0 | 0 | 0 | 0 | 0 | 0 | 0 | 0 | 0 | 0 | 0 | 0 | 0 | 0 | 0 | 0 | 0 | 0 | 0 | 0 | 0 |
| Jun8.50v_31950 | 1 | 0 | 0 | 0 | 0 | 0 | 1 | 0 | 0 | 0 | 0 | 0 | 0 | 0 | 0 | 0 | 0 | 0 | 0 | 0 | 0 | 0 | 0 | 0 | 0 | 0 | 0 | 0 | 0 | 0 |
| May12.75v_1476 | 1 | 0 | 0 | 0 | 1 | 0 | 0 | 0 | 0 | 0 | 0 | 0 | 0 | 0 | 0 | 0 | 0 | 0 | 0 | 0 | 0 | 0 | 0 | 0 | 0 | 0 | 0 | 0 | 0 | 0 |
| Aug5.25v_36270 | 1 | 0 | 0 | 0 | 0 | 0 | 0 | 0 | 0 | 0 | 0 | 0 | 0 | 0 | 0 | 0 | 0 | 0 | 0 | 0 | 0 | 0 | 0 | 1 | 0 | 0 | 0 | 0 | 0 | 0 |
| Aug5.0v_5007 | 1 | 0 | 0 | 0 | 0 | 0 | 0 | 0 | 0 | 0 | 0 | 0 | 0 | 0 | 0 | 0 | 0 | 0 | 0 | 0 | 0 | 1 | 0 | 0 | 0 | 0 | 0 | 0 | 0 | 0 |
| Jul11.25v_13991 | 1 | 0 | 0 | 0 | 0 | 0 | 0 | 0 | 0 | 0 | 0 | 0 | 0 | 0 | 1 | 0 | 0 | 0 | 0 | 0 | 0 | 0 | 0 | 0 | 0 | 0 | 0 | 0 | 0 | 0 |
| Jul27.50v_16992 | 1 | 0 | 0 | 0 | 0 | 0 | 0 | 0 | 0 | 0 | 0 | 0 | 0 | 0 | 0 | 0 | 0 | 0 | 0 | 1 | 0 | 0 | 0 | 0 | 0 | 0 | 0 | 0 | 0 | 0 |
| Jul27.25v_22994 | 1 | 0 | 0 | 0 | 0 | 0 | 0 | 0 | 0 | 0 | 0 | 0 | 0 | 0 | 0 | 0 | 0 | 0 | 1 | 0 | 0 | 0 | 0 | 0 | 0 | 0 | 0 | 0 | 0 | 0 |
| May12.25v_27301 | 1 | 0 | 1 | 0 | 0 | 0 | 0 | 0 | 0 | 0 | 0 | 0 | 0 | 0 | 0 | 0 | 0 | 0 | 0 | 0 | 0 | 0 | 0 | 0 | 0 | 0 | 0 | 0 | 0 | 0 |
| Aug5.25v_21879 | 1 | 0 | 0 | 0 | 0 | 0 | 0 | 0 | 0 | 0 | 0 | 0 | 0 | 0 | 0 | 0 | 0 | 0 | 0 | 0 | 0 | 0 | 0 | 1 | 0 | 0 | 0 | 0 | 0 | 0 |
| May12.25v_22028 | 1 | 0 | 1 | 0 | 0 | 0 | 0 | 0 | 0 | 0 | 0 | 0 | 0 | 0 | 0 | 0 | 0 | 0 | 0 | 0 | 0 | 0 | 0 | 0 | 0 | 0 | 0 | 0 | 0 | 0 |
| Aug5.0v_549 | 1 | 0 | 0 | 0 | 0 | 0 | 0 | 0 | 0 | 0 | 0 | 0 | 0 | 0 | 0 | 0 | 0 | 0 | 0 | 0 | 0 | 1 | 0 | 0 | 0 | 0 | 0 | 0 | 0 | 0 |
| Aug5.0v_2093 | 1 | 0 | 0 | 0 | 0 | 0 | 0 | 0 | 0 | 0 | 0 | 0 | 0 | 0 | 0 | 0 | 0 | 0 | 0 | 0 | 0 | 1 | 0 | 0 | 0 | 0 | 0 | 0 | 0 | 0 |
| Jun22.75v_18605 | 1 | 0 | 0 | 0 | 0 | 0 | 0 | 0 | 0 | 0 | 0 | 1 | 0 | 0 | 0 | 0 | 0 | 0 | 0 | 0 | 0 | 0 | 0 | 0 | 0 | 0 | 0 | 0 | 0 | 0 |
| Jun22.0v_13007 | 1 | 0 | 0 | 0 | 0 | 0 | 0 | 0 | 1 | 0 | 0 | 0 | 0 | 0 | 0 | 0 | 0 | 0 | 0 | 0 | 0 | 0 | 0 | 0 | 0 | 0 | 0 | 0 | 0 | 0 |
| Jul11.75v_9993 | 1 | 0 | 0 | 0 | 0 | 0 | 0 | 0 | 0 | 0 | 0 | 0 | 0 | 0 | 0 | 0 | 1 | 0 | 0 | 0 | 0 | 0 | 0 | 0 | 0 | 0 | 0 | 0 | 0 | 0 |
| Jun8.50v_8433 | 1 | 0 | 0 | 0 | 0 | 0 | 1 | 0 | 0 | 0 | 0 | 0 | 0 | 0 | 0 | 0 | 0 | 0 | 0 | 0 | 0 | 0 | 0 | 0 | 0 | 0 | 0 | 0 | 0 | 0 |
| Aug5.25v_21242 | 1 | 0 | 0 | 0 | 0 | 0 | 0 | 0 | 0 | 0 | 0 | 0 | 0 | 0 | 0 | 0 | 0 | 0 | 0 | 0 | 0 | 0 | 0 | 1 | 0 | 0 | 0 | 0 | 0 | 0 |
| Aug5.0v_3688 | 1 | 0 | 0 | 0 | 0 | 0 | 0 | 0 | 0 | 0 | 0 | 0 | 0 | 0 | 0 | 0 | 0 | 0 | 0 | 0 | 0 | 1 | 0 | 0 | 0 | 0 | 0 | 0 | 0 | 0 |
| Jun8.25v_21699 | 1 | 0 | 0 | 0 | 0 | 1 | 0 | 0 | 0 | 0 | 0 | 0 | 0 | 0 | 0 | 0 | 0 | 0 | 0 | 0 | 0 | 0 | 0 | 0 | 0 | 0 | 0 | 0 | 0 | 0 |
| Jul11.50v_20995 | 1 | 0 | 0 | 0 | 0 | 0 | 0 | 0 | 0 | 0 | 0 | 0 | 0 | 0 | 0 | 1 | 0 | 0 | 0 | 0 | 0 | 0 | 0 | 0 | 0 | 0 | 0 | 0 | 0 | 0 |
| May12.50v_12016 | 1 | 0 | 0 | 1 | 0 | 0 | 0 | 0 | 0 | 0 | 0 | 0 | 0 | 0 | 0 | 0 | 0 | 0 | 0 | 0 | 0 | 0 | 0 | 0 | 0 | 0 | 0 | 0 | 0 | 0 |
| Sep8.0v_1257 | 1 | 0 | 0 | 0 | 0 | 0 | 0 | 0 | 0 | 0 | 0 | 0 | 0 | 0 | 0 | 0 | 0 | 0 | 0 | 0 | 0 | 0 | 0 | 0 | 0 | 0 | 1 | 0 | 0 | 0 |
| May12.25v_2353 | 1 | 0 | 1 | 0 | 0 | 0 | 0 | 0 | 0 | 0 | 0 | 0 | 0 | 0 | 0 | 0 | 0 | 0 | 0 | 0 | 0 | 0 | 0 | 0 | 0 | 0 | 0 | 0 | 0 | 0 |
| Jun22.0v_4843 | 1 | 0 | 0 | 0 | 0 | 0 | 0 | 0 | 1 | 0 | 0 | 0 | 0 | 0 | 0 | 0 | 0 | 0 | 0 | 0 | 0 | 0 | 0 | 0 | 0 | 0 | 0 | 0 | 0 | 0 |
| Aug5.5v_21792 | 1 | 0 | 0 | 0 | 0 | 0 | 0 | 0 | 0 | 0 | 0 | 0 | 0 | 0 | 0 | 0 | 0 | 0 | 0 | 0 | 0 | 0 | 1 | 0 | 0 | 0 | 0 | 0 | 0 | 0 |
| Jun22.75v_5319 | 1 | 0 | 0 | 0 | 0 | 0 | 0 | 0 | 0 | 0 | 0 | 1 | 0 | 0 | 0 | 0 | 0 | 0 | 0 | 0 | 0 | 0 | 0 | 0 | 0 | 0 | 0 | 0 | 0 | 0 |
| Jun8.50v_17340 | 1 | 0 | 0 | 0 | 0 | 0 | 1 | 0 | 0 | 0 | 0 | 0 | 0 | 0 | 0 | 0 | 0 | 0 | 0 | 0 | 0 | 0 | 0 | 0 | 0 | 0 | 0 | 0 | 0 | 0 |
| Sep8.0v_15142 | 2 | 0 | 0 | 0 | 0 | 0 | 0 | 0 | 0 | 0 | 0 | 0 | 0 | 0 | 0 | 0 | 0 | 0 | 0 | 0 | 0 | 0 | 0 | 1 | 0 | 0 | 1 | 0 | 0 | 0 |
| Jul27.5v_9777 | 1 | 0 | 0 | 0 | 0 | 0 | 0 | 0 | 0 | 0 | 0 | 0 | 0 | 0 | 0 | 0 | 0 | 1 | 0 | 0 | 0 | 0 | 0 | 0 | 0 | 0 | 0 | 0 | 0 | 0 |
| May12.5v_229 | 1 | 1 | 0 | 0 | 0 | 0 | 0 | 0 | 0 | 0 | 0 | 0 | 0 | 0 | 0 | 0 | 0 | 0 | 0 | 0 | 0 | 0 | 0 | 0 | 0 | 0 | 0 | 0 | 0 | 0 |
| Aug5.50v_30254 | 1 | 0 | 0 | 0 | 0 | 0 | 0 | 0 | 0 | 0 | 0 | 0 | 0 | 0 | 0 | 0 | 0 | 0 | 0 | 0 | 0 | 0 | 0 | 0 | 1 | 0 | 0 | 0 | 0 | 0 |
| May12.25v_38963 | 2 | 0 | 1 | 0 | 0 | 0 | 0 | 0 | 0 | 0 | 0 | 0 | 0 | 0 | 0 | 1 | 0 | 0 | 0 | 0 | 0 | 0 | 0 | 0 | 0 | 0 | 0 | 0 | 0 | 0 |
| Sep8.50v_32743 | 1 | 0 | 0 | 0 | 0 | 0 | 0 | 0 | 0 | 0 | 0 | 0 | 0 | 0 | 0 | 0 | 0 | 0 | 0 | 0 | 0 | 0 | 0 | 0 | 0 | 0 | 0 | 0 | 0 | 1 |
| Jul11.0v_130 | 1 | 0 | 0 | 0 | 0 | 0 | 0 | 0 | 0 | 0 | 0 | 0 | 1 | 0 | 0 | 0 | 0 | 0 | 0 | 0 | 0 | 0 | 0 | 0 | 0 | 0 | 0 | 0 | 0 | 0 |
| Jul27.5v_7634 | 1 | 0 | 0 | 0 | 0 | 0 | 0 | 0 | 0 | 0 | 0 | 0 | 0 | 0 | 0 | 0 | 0 | 1 | 0 | 0 | 0 | 0 | 0 | 0 | 0 | 0 | 0 | 0 | 0 | 0 |
| Jul11.0v_9887 | 1 | 0 | 0 | 0 | 0 | 0 | 0 | 0 | 0 | 0 | 0 | 0 | 1 | 0 | 0 | 0 | 0 | 0 | 0 | 0 | 0 | 0 | 0 | 0 | 0 | 0 | 0 | 0 | 0 | 0 |
| Sep8.0v_6185 | 1 | 0 | 0 | 0 | 0 | 0 | 0 | 0 | 0 | 0 | 0 | 0 | 0 | 0 | 0 | 0 | 0 | 0 | 0 | 0 | 0 | 0 | 0 | 0 | 0 | 0 | 1 | 0 | 0 | 0 |
| Sep8.0v_6621 | 1 | 0 | 0 | 0 | 0 | 0 | 0 | 0 | 0 | 0 | 0 | 0 | 0 | 0 | 0 | 0 | 0 | 0 | 0 | 0 | 0 | 0 | 0 | 0 | 0 | 0 | 1 | 0 | 0 | 0 |
| Jun8.50v_13375 | 1 | 0 | 0 | 0 | 0 | 0 | 1 | 0 | 0 | 0 | 0 | 0 | 0 | 0 | 0 | 0 | 0 | 0 | 0 | 0 | 0 | 0 | 0 | 0 | 0 | 0 | 0 | 0 | 0 | 0 |
| May12.50v_19765 | 1 | 0 | 0 | 1 | 0 | 0 | 0 | 0 | 0 | 0 | 0 | 0 | 0 | 0 | 0 | 0 | 0 | 0 | 0 | 0 | 0 | 0 | 0 | 0 | 0 | 0 | 0 | 0 | 0 | 0 |
| May12.50v_29669 | 1 | 0 | 0 | 1 | 0 | 0 | 0 | 0 | 0 | 0 | 0 | 0 | 0 | 0 | 0 | 0 | 0 | 0 | 0 | 0 | 0 | 0 | 0 | 0 | 0 | 0 | 0 | 0 | 0 | 0 |
| Jul27.5v_8945 | 1 | 0 | 0 | 0 | 0 | 0 | 0 | 0 | 0 | 0 | 0 | 0 | 0 | 0 | 0 | 0 | 0 | 1 | 0 | 0 | 0 | 0 | 0 | 0 | 0 | 0 | 0 | 0 | 0 | 0 |
| Jun22.0v_6931 | 1 | 0 | 0 | 0 | 0 | 0 | 0 | 0 | 1 | 0 | 0 | 0 | 0 | 0 | 0 | 0 | 0 | 0 | 0 | 0 | 0 | 0 | 0 | 0 | 0 | 0 | 0 | 0 | 0 | 0 |
| Aug5.0v_1840 | 1 | 0 | 0 | 0 | 0 | 0 | 0 | 0 | 0 | 0 | 0 | 0 | 0 | 0 | 0 | 0 | 0 | 0 | 0 | 0 | 0 | 1 | 0 | 0 | 0 | 0 | 0 | 0 | 0 | 0 |
| Jul11.50v_6507 | 1 | 0 | 0 | 0 | 0 | 0 | 0 | 0 | 0 | 0 | 0 | 0 | 0 | 0 | 0 | 1 | 0 | 0 | 0 | 0 | 0 | 0 | 0 | 0 | 0 | 0 | 0 | 0 | 0 | 0 |
| Jul27.5v_17778 | 1 | 0 | 0 | 0 | 0 | 0 | 0 | 0 | 0 | 0 | 0 | 0 | 0 | 0 | 0 | 0 | 0 | 1 | 0 | 0 | 0 | 0 | 0 | 0 | 0 | 0 | 0 | 0 | 0 | 0 |
| Aug5.0v_2189 | 1 | 0 | 0 | 0 | 0 | 0 | 0 | 0 | 0 | 0 | 0 | 0 | 0 | 0 | 0 | 0 | 0 | 0 | 0 | 0 | 0 | 1 | 0 | 0 | 0 | 0 | 0 | 0 | 0 | 0 |
| Jun8.50v_29234 | 1 | 0 | 0 | 0 | 0 | 0 | 1 | 0 | 0 | 0 | 0 | 0 | 0 | 0 | 0 | 0 | 0 | 0 | 0 | 0 | 0 | 0 | 0 | 0 | 0 | 0 | 0 | 0 | 0 | 0 |
| Aug5.5v_466 | 1 | 0 | 0 | 0 | 0 | 0 | 0 | 0 | 0 | 0 | 0 | 0 | 0 | 0 | 0 | 0 | 0 | 0 | 0 | 0 | 0 | 0 | 1 | 0 | 0 | 0 | 0 | 0 | 0 | 0 |
| Jul11.0v_1742 | 1 | 0 | 0 | 0 | 0 | 0 | 0 | 0 | 0 | 0 | 0 | 0 | 1 | 0 | 0 | 0 | 0 | 0 | 0 | 0 | 0 | 0 | 0 | 0 | 0 | 0 | 0 | 0 | 0 | 0 |
| Jun22.0v_17823 | 1 | 0 | 0 | 0 | 0 | 0 | 0 | 0 | 1 | 0 | 0 | 0 | 0 | 0 | 0 | 0 | 0 | 0 | 0 | 0 | 0 | 0 | 0 | 0 | 0 | 0 | 0 | 0 | 0 | 0 |
| Jul11.5v_8177 | 1 | 0 | 0 | 0 | 0 | 0 | 0 | 0 | 0 | 0 | 0 | 0 | 0 | 1 | 0 | 0 | 0 | 0 | 0 | 0 | 0 | 0 | 0 | 0 | 0 | 0 | 0 | 0 | 0 | 0 |
| May12.50v_24694 | 4 | 0 | 0 | 2 | 1 | 0 | 0 | 0 | 0 | 0 | 0 | 0 | 0 | 0 | 0 | 0 | 0 | 0 | 0 | 0 | 0 | 0 | 0 | 1 | 0 | 0 | 0 | 0 | 0 | 0 |
| Jun22.5v_4415 | 1 | 0 | 0 | 0 | 0 | 0 | 0 | 0 | 0 | 1 | 0 | 0 | 0 | 0 | 0 | 0 | 0 | 0 | 0 | 0 | 0 | 0 | 0 | 0 | 0 | 0 | 0 | 0 | 0 | 0 |
| Jun22.75v_2966 | 1 | 0 | 0 | 0 | 0 | 0 | 0 | 0 | 0 | 0 | 0 | 1 | 0 | 0 | 0 | 0 | 0 | 0 | 0 | 0 | 0 | 0 | 0 | 0 | 0 | 0 | 0 | 0 | 0 | 0 |
| Jul11.25v_7750 | 1 | 0 | 0 | 0 | 0 | 0 | 0 | 0 | 0 | 0 | 0 | 0 | 0 | 0 | 1 | 0 | 0 | 0 | 0 | 0 | 0 | 0 | 0 | 0 | 0 | 0 | 0 | 0 | 0 | 0 |
| Aug5.5v_35563 | 1 | 0 | 0 | 0 | 0 | 0 | 0 | 0 | 0 | 0 | 0 | 0 | 0 | 0 | 0 | 0 | 0 | 0 | 0 | 0 | 0 | 0 | 1 | 0 | 0 | 0 | 0 | 0 | 0 | 0 |
| May12.50v_35098 | 1 | 0 | 0 | 1 | 0 | 0 | 0 | 0 | 0 | 0 | 0 | 0 | 0 | 0 | 0 | 0 | 0 | 0 | 0 | 0 | 0 | 0 | 0 | 0 | 0 | 0 | 0 | 0 | 0 | 0 |
| Aug5.25v_20741 | 2 | 0 | 0 | 0 | 0 | 0 | 0 | 0 | 0 | 0 | 0 | 0 | 0 | 0 | 0 | 0 | 0 | 0 | 0 | 0 | 0 | 0 | 0 | 1 | 0 | 0 | 1 | 0 | 0 | 0 |
| Sep8.0v_20158 | 1 | 0 | 0 | 0 | 0 | 0 | 0 | 0 | 0 | 0 | 0 | 0 | 0 | 0 | 0 | 0 | 0 | 0 | 0 | 0 | 0 | 0 | 0 | 0 | 0 | 0 | 1 | 0 | 0 | 0 |
| Jul11.75v_32918 | 1 | 0 | 0 | 0 | 0 | 0 | 0 | 0 | 0 | 0 | 0 | 0 | 0 | 0 | 0 | 0 | 1 | 0 | 0 | 0 | 0 | 0 | 0 | 0 | 0 | 0 | 0 | 0 | 0 | 0 |
| May12.50v_17662 | 5 | 0 | 1 | 1 | 0 | 0 | 0 | 0 | 0 | 0 | 0 | 0 | 0 | 0 | 0 | 0 | 0 | 1 | 0 | 0 | 0 | 1 | 1 | 0 | 0 | 0 | 0 | 0 | 0 | 0 |
| Jul27.50v_969 | 1 | 0 | 0 | 0 | 0 | 0 | 0 | 0 | 0 | 0 | 0 | 0 | 0 | 0 | 0 | 0 | 0 | 0 | 0 | 1 | 0 | 0 | 0 | 0 | 0 | 0 | 0 | 0 | 0 | 0 |
| Jul27.5v_18250 | 1 | 0 | 0 | 0 | 0 | 0 | 0 | 0 | 0 | 0 | 0 | 0 | 0 | 0 | 0 | 0 | 0 | 1 | 0 | 0 | 0 | 0 | 0 | 0 | 0 | 0 | 0 | 0 | 0 | 0 |
| Aug5.25v_21486 | 1 | 0 | 0 | 0 | 0 | 0 | 0 | 0 | 0 | 0 | 0 | 0 | 0 | 0 | 0 | 0 | 0 | 0 | 0 | 0 | 0 | 0 | 0 | 1 | 0 | 0 | 0 | 0 | 0 | 0 |
| Aug5.0v_182 | 159 | 5 | 3 | 6 | 3 | 0 | 0 | 0 | 6 | 2 | 8 | 2 | 10 | 6 | 10 | 2 | 1 | 3 | 5 | 1 | 0 | 34 | 15 | 15 | 1 | 1 | 3 | 2 | 0 | 15 |
| Sep8.0v_14906 | 2 | 0 | 0 | 0 | 0 | 0 | 0 | 0 | 0 | 0 | 0 | 0 | 0 | 0 | 0 | 0 | 0 | 0 | 0 | 0 | 0 | 0 | 0 | 1 | 0 | 0 | 1 | 0 | 0 | 0 |
| Jul27.5v_16258 | 1 | 0 | 0 | 0 | 0 | 0 | 0 | 0 | 0 | 0 | 0 | 0 | 0 | 0 | 0 | 0 | 0 | 1 | 0 | 0 | 0 | 0 | 0 | 0 | 0 | 0 | 0 | 0 | 0 | 0 |
| Aug5.0v_3612 | 1 | 0 | 0 | 0 | 0 | 0 | 0 | 0 | 0 | 0 | 0 | 0 | 0 | 0 | 0 | 0 | 0 | 0 | 0 | 0 | 0 | 1 | 0 | 0 | 0 | 0 | 0 | 0 | 0 | 0 |
| Jul27.25v_1869 | 1 | 0 | 0 | 0 | 0 | 0 | 0 | 0 | 0 | 0 | 0 | 0 | 0 | 0 | 0 | 0 | 0 | 0 | 1 | 0 | 0 | 0 | 0 | 0 | 0 | 0 | 0 | 0 | 0 | 0 |
| Jul27.25v_11628 | 1 | 0 | 0 | 0 | 0 | 0 | 0 | 0 | 0 | 0 | 0 | 0 | 0 | 0 | 0 | 0 | 0 | 0 | 1 | 0 | 0 | 0 | 0 | 0 | 0 | 0 | 0 | 0 | 0 | 0 |
| Aug5.0v_847 | 1 | 0 | 0 | 0 | 0 | 0 | 0 | 0 | 0 | 0 | 0 | 0 | 0 | 0 | 0 | 0 | 0 | 0 | 0 | 0 | 0 | 1 | 0 | 0 | 0 | 0 | 0 | 0 | 0 | 0 |
| May12.5v_6661 | 2 | 1 | 1 | 0 | 0 | 0 | 0 | 0 | 0 | 0 | 0 | 0 | 0 | 0 | 0 | 0 | 0 | 0 | 0 | 0 | 0 | 0 | 0 | 0 | 0 | 0 | 0 | 0 | 0 | 0 |
| Aug5.50v_21987 | 1 | 0 | 0 | 0 | 0 | 0 | 0 | 0 | 0 | 0 | 0 | 0 | 0 | 0 | 0 | 0 | 0 | 0 | 0 | 0 | 0 | 0 | 0 | 0 | 1 | 0 | 0 | 0 | 0 | 0 |
| Aug5.5v_6543 | 1 | 0 | 0 | 0 | 0 | 0 | 0 | 0 | 0 | 0 | 0 | 0 | 0 | 0 | 0 | 0 | 0 | 0 | 0 | 0 | 0 | 0 | 1 | 0 | 0 | 0 | 0 | 0 | 0 | 0 |
| Aug5.0v_3610 | 1 | 0 | 0 | 0 | 0 | 0 | 0 | 0 | 0 | 0 | 0 | 0 | 0 | 0 | 0 | 0 | 0 | 0 | 0 | 0 | 0 | 1 | 0 | 0 | 0 | 0 | 0 | 0 | 0 | 0 |
| Jul27.5v_20577 | 1 | 0 | 0 | 0 | 0 | 0 | 0 | 0 | 0 | 0 | 0 | 0 | 0 | 0 | 0 | 0 | 0 | 1 | 0 | 0 | 0 | 0 | 0 | 0 | 0 | 0 | 0 | 0 | 0 | 0 |
| Jul27.5v_8831 | 1 | 0 | 0 | 0 | 0 | 0 | 0 | 0 | 0 | 0 | 0 | 0 | 0 | 0 | 0 | 0 | 0 | 1 | 0 | 0 | 0 | 0 | 0 | 0 | 0 | 0 | 0 | 0 | 0 | 0 |
| Sep8.50v_19042 | 1 | 0 | 0 | 0 | 0 | 0 | 0 | 0 | 0 | 0 | 0 | 0 | 0 | 0 | 0 | 0 | 0 | 0 | 0 | 0 | 0 | 0 | 0 | 0 | 0 | 0 | 0 | 0 | 0 | 1 |
| May12.50v_6472 | 1 | 0 | 0 | 1 | 0 | 0 | 0 | 0 | 0 | 0 | 0 | 0 | 0 | 0 | 0 | 0 | 0 | 0 | 0 | 0 | 0 | 0 | 0 | 0 | 0 | 0 | 0 | 0 | 0 | 0 |
| Aug5.0v_5107 | 1 | 0 | 0 | 0 | 0 | 0 | 0 | 0 | 0 | 0 | 0 | 0 | 0 | 0 | 0 | 0 | 0 | 0 | 0 | 0 | 0 | 1 | 0 | 0 | 0 | 0 | 0 | 0 | 0 | 0 |
| Aug5.5v_8109 | 1 | 0 | 0 | 0 | 0 | 0 | 0 | 0 | 0 | 0 | 0 | 0 | 0 | 0 | 0 | 0 | 0 | 0 | 0 | 0 | 0 | 0 | 1 | 0 | 0 | 0 | 0 | 0 | 0 | 0 |
| Aug5.25v_39661 | 2 | 0 | 0 | 0 | 0 | 0 | 0 | 0 | 0 | 0 | 0 | 1 | 0 | 0 | 0 | 0 | 0 | 0 | 0 | 0 | 0 | 0 | 0 | 1 | 0 | 0 | 0 | 0 | 0 | 0 |
| Jul11.0v_15554 | 1 | 0 | 0 | 0 | 0 | 0 | 0 | 0 | 0 | 0 | 0 | 0 | 1 | 0 | 0 | 0 | 0 | 0 | 0 | 0 | 0 | 0 | 0 | 0 | 0 | 0 | 0 | 0 | 0 | 0 |
| Aug5.5v_34196 | 1 | 0 | 0 | 0 | 0 | 0 | 0 | 0 | 0 | 0 | 0 | 0 | 0 | 0 | 0 | 0 | 0 | 0 | 0 | 0 | 0 | 0 | 1 | 0 | 0 | 0 | 0 | 0 | 0 | 0 |
| Jul11.50v_1319 | 2 | 0 | 0 | 0 | 0 | 0 | 0 | 0 | 0 | 0 | 1 | 0 | 0 | 0 | 0 | 1 | 0 | 0 | 0 | 0 | 0 | 0 | 0 | 0 | 0 | 0 | 0 | 0 | 0 | 0 |
| Aug5.25v_35953 | 1 | 0 | 0 | 0 | 0 | 0 | 0 | 0 | 0 | 0 | 0 | 0 | 0 | 0 | 0 | 0 | 0 | 0 | 0 | 0 | 0 | 0 | 0 | 1 | 0 | 0 | 0 | 0 | 0 | 0 |
| Jul27.25v_7166 | 1 | 0 | 0 | 0 | 0 | 0 | 0 | 0 | 0 | 0 | 0 | 0 | 0 | 0 | 0 | 0 | 0 | 0 | 1 | 0 | 0 | 0 | 0 | 0 | 0 | 0 | 0 | 0 | 0 | 0 |
| Aug5.5v_18292 | 2 | 1 | 0 | 0 | 0 | 0 | 0 | 0 | 0 | 0 | 0 | 0 | 0 | 0 | 0 | 0 | 0 | 0 | 0 | 0 | 0 | 0 | 1 | 0 | 0 | 0 | 0 | 0 | 0 | 0 |
| Jul27.5v_15209 | 1 | 0 | 0 | 0 | 0 | 0 | 0 | 0 | 0 | 0 | 0 | 0 | 0 | 0 | 0 | 0 | 0 | 1 | 0 | 0 | 0 | 0 | 0 | 0 | 0 | 0 | 0 | 0 | 0 | 0 |
| Aug5.5v_7533 | 2 | 0 | 0 | 0 | 0 | 0 | 0 | 0 | 0 | 0 | 0 | 0 | 0 | 0 | 0 | 0 | 0 | 0 | 0 | 0 | 0 | 0 | 1 | 0 | 0 | 1 | 0 | 0 | 0 | 0 |
| Jun22.0v_8431 | 1 | 0 | 0 | 0 | 0 | 0 | 0 | 0 | 1 | 0 | 0 | 0 | 0 | 0 | 0 | 0 | 0 | 0 | 0 | 0 | 0 | 0 | 0 | 0 | 0 | 0 | 0 | 0 | 0 | 0 |
| Sep8.50v_25832 | 1 | 0 | 0 | 0 | 0 | 0 | 0 | 0 | 0 | 0 | 0 | 0 | 0 | 0 | 0 | 0 | 0 | 0 | 0 | 0 | 0 | 0 | 0 | 0 | 0 | 0 | 0 | 0 | 0 | 1 |
| Jul11.50v_1558 | 1 | 0 | 0 | 0 | 0 | 0 | 0 | 0 | 0 | 0 | 0 | 0 | 0 | 0 | 0 | 1 | 0 | 0 | 0 | 0 | 0 | 0 | 0 | 0 | 0 | 0 | 0 | 0 | 0 | 0 |
| Jul27.50v_13842 | 1 | 0 | 0 | 0 | 0 | 0 | 0 | 0 | 0 | 0 | 0 | 0 | 0 | 0 | 0 | 0 | 0 | 0 | 0 | 1 | 0 | 0 | 0 | 0 | 0 | 0 | 0 | 0 | 0 | 0 |
| Jul11.5v_11436 | 1 | 0 | 0 | 0 | 0 | 0 | 0 | 0 | 0 | 0 | 0 | 0 | 0 | 1 | 0 | 0 | 0 | 0 | 0 | 0 | 0 | 0 | 0 | 0 | 0 | 0 | 0 | 0 | 0 | 0 |
| Aug5.5v_9481 | 1 | 0 | 0 | 0 | 0 | 0 | 0 | 0 | 0 | 0 | 0 | 0 | 0 | 0 | 0 | 0 | 0 | 0 | 0 | 0 | 0 | 0 | 1 | 0 | 0 | 0 | 0 | 0 | 0 | 0 |
| Aug5.0v_2478 | 2 | 0 | 0 | 0 | 0 | 0 | 0 | 0 | 0 | 0 | 0 | 0 | 0 | 0 | 0 | 0 | 0 | 0 | 0 | 0 | 0 | 1 | 1 | 0 | 0 | 0 | 0 | 0 | 0 | 0 |
| May12.25v_17390 | 1 | 0 | 1 | 0 | 0 | 0 | 0 | 0 | 0 | 0 | 0 | 0 | 0 | 0 | 0 | 0 | 0 | 0 | 0 | 0 | 0 | 0 | 0 | 0 | 0 | 0 | 0 | 0 | 0 | 0 |
| Jul11.75v_29651 | 1 | 0 | 0 | 0 | 0 | 0 | 0 | 0 | 0 | 0 | 0 | 0 | 0 | 0 | 0 | 0 | 1 | 0 | 0 | 0 | 0 | 0 | 0 | 0 | 0 | 0 | 0 | 0 | 0 | 0 |
| May12.50v_15242 | 1 | 0 | 0 | 1 | 0 | 0 | 0 | 0 | 0 | 0 | 0 | 0 | 0 | 0 | 0 | 0 | 0 | 0 | 0 | 0 | 0 | 0 | 0 | 0 | 0 | 0 | 0 | 0 | 0 | 0 |
| Aug5.0v_2301 | 1 | 0 | 0 | 0 | 0 | 0 | 0 | 0 | 0 | 0 | 0 | 0 | 0 | 0 | 0 | 0 | 0 | 0 | 0 | 0 | 0 | 1 | 0 | 0 | 0 | 0 | 0 | 0 | 0 | 0 |
| Aug5.5v_12701 | 3 | 0 | 0 | 0 | 0 | 0 | 0 | 0 | 0 | 0 | 0 | 0 | 0 | 0 | 0 | 0 | 0 | 1 | 0 | 0 | 0 | 0 | 1 | 1 | 0 | 0 | 0 | 0 | 0 | 0 |
| Jul11.5v_307 | 1 | 0 | 0 | 0 | 0 | 0 | 0 | 0 | 0 | 0 | 0 | 0 | 0 | 1 | 0 | 0 | 0 | 0 | 0 | 0 | 0 | 0 | 0 | 0 | 0 | 0 | 0 | 0 | 0 | 0 |
| Jun22.50v_19600 | 1 | 0 | 0 | 0 | 0 | 0 | 0 | 0 | 0 | 0 | 1 | 0 | 0 | 0 | 0 | 0 | 0 | 0 | 0 | 0 | 0 | 0 | 0 | 0 | 0 | 0 | 0 | 0 | 0 | 0 |
| May12.25v_32402 | 1 | 0 | 1 | 0 | 0 | 0 | 0 | 0 | 0 | 0 | 0 | 0 | 0 | 0 | 0 | 0 | 0 | 0 | 0 | 0 | 0 | 0 | 0 | 0 | 0 | 0 | 0 | 0 | 0 | 0 |
| May12.50v_33693 | 1 | 0 | 0 | 1 | 0 | 0 | 0 | 0 | 0 | 0 | 0 | 0 | 0 | 0 | 0 | 0 | 0 | 0 | 0 | 0 | 0 | 0 | 0 | 0 | 0 | 0 | 0 | 0 | 0 | 0 |
| Aug5.75v_29650 | 1 | 0 | 0 | 0 | 0 | 0 | 0 | 0 | 0 | 0 | 0 | 0 | 0 | 0 | 0 | 0 | 0 | 0 | 0 | 0 | 0 | 0 | 0 | 0 | 0 | 1 | 0 | 0 | 0 | 0 |
| Aug5.0v_3308 | 1 | 0 | 0 | 0 | 0 | 0 | 0 | 0 | 0 | 0 | 0 | 0 | 0 | 0 | 0 | 0 | 0 | 0 | 0 | 0 | 0 | 1 | 0 | 0 | 0 | 0 | 0 | 0 | 0 | 0 |
| Jul11.25v_17043 | 1 | 0 | 0 | 0 | 0 | 0 | 0 | 0 | 0 | 0 | 0 | 0 | 0 | 0 | 1 | 0 | 0 | 0 | 0 | 0 | 0 | 0 | 0 | 0 | 0 | 0 | 0 | 0 | 0 | 0 |
| Jun22.0v_219 | 1 | 0 | 0 | 0 | 0 | 0 | 0 | 0 | 1 | 0 | 0 | 0 | 0 | 0 | 0 | 0 | 0 | 0 | 0 | 0 | 0 | 0 | 0 | 0 | 0 | 0 | 0 | 0 | 0 | 0 |
| Jul11.25v_8972 | 1 | 0 | 0 | 0 | 0 | 0 | 0 | 0 | 0 | 0 | 0 | 0 | 0 | 0 | 1 | 0 | 0 | 0 | 0 | 0 | 0 | 0 | 0 | 0 | 0 | 0 | 0 | 0 | 0 | 0 |
| Aug5.0v_3647 | 1 | 0 | 0 | 0 | 0 | 0 | 0 | 0 | 0 | 0 | 0 | 0 | 0 | 0 | 0 | 0 | 0 | 0 | 0 | 0 | 0 | 1 | 0 | 0 | 0 | 0 | 0 | 0 | 0 | 0 |
| Jul11.25v_4783 | 1 | 0 | 0 | 0 | 0 | 0 | 0 | 0 | 0 | 0 | 0 | 0 | 0 | 0 | 1 | 0 | 0 | 0 | 0 | 0 | 0 | 0 | 0 | 0 | 0 | 0 | 0 | 0 | 0 | 0 |
| Jun22.50v_13859 | 1 | 0 | 0 | 0 | 0 | 0 | 0 | 0 | 0 | 0 | 1 | 0 | 0 | 0 | 0 | 0 | 0 | 0 | 0 | 0 | 0 | 0 | 0 | 0 | 0 | 0 | 0 | 0 | 0 | 0 |
| Jul11.75v_38022 | 1 | 0 | 0 | 0 | 0 | 0 | 0 | 0 | 0 | 0 | 0 | 0 | 0 | 0 | 0 | 0 | 1 | 0 | 0 | 0 | 0 | 0 | 0 | 0 | 0 | 0 | 0 | 0 | 0 | 0 |
| Sep8.50v_40266 | 1 | 0 | 0 | 0 | 0 | 0 | 0 | 0 | 0 | 0 | 0 | 0 | 0 | 0 | 0 | 0 | 0 | 0 | 0 | 0 | 0 | 0 | 0 | 0 | 0 | 0 | 0 | 0 | 0 | 1 |
| Jul27.50v_11999 | 1 | 0 | 0 | 0 | 0 | 0 | 0 | 0 | 0 | 0 | 0 | 0 | 0 | 0 | 0 | 0 | 0 | 0 | 0 | 1 | 0 | 0 | 0 | 0 | 0 | 0 | 0 | 0 | 0 | 0 |
| Jul11.5v_2538 | 2 | 0 | 0 | 0 | 0 | 0 | 0 | 0 | 0 | 0 | 0 | 0 | 0 | 1 | 0 | 0 | 0 | 0 | 0 | 0 | 0 | 0 | 0 | 1 | 0 | 0 | 0 | 0 | 0 | 0 |
| Jul11.25v_11013 | 1 | 0 | 0 | 0 | 0 | 0 | 0 | 0 | 0 | 0 | 0 | 0 | 0 | 0 | 1 | 0 | 0 | 0 | 0 | 0 | 0 | 0 | 0 | 0 | 0 | 0 | 0 | 0 | 0 | 0 |
| Aug5.0v_672 | 1 | 0 | 0 | 0 | 0 | 0 | 0 | 0 | 0 | 0 | 0 | 0 | 0 | 0 | 0 | 0 | 0 | 0 | 0 | 0 | 0 | 1 | 0 | 0 | 0 | 0 | 0 | 0 | 0 | 0 |
| Jul27.75v_11303 | 1 | 0 | 0 | 0 | 0 | 0 | 0 | 0 | 0 | 0 | 0 | 0 | 0 | 0 | 0 | 0 | 0 | 0 | 0 | 0 | 1 | 0 | 0 | 0 | 0 | 0 | 0 | 0 | 0 | 0 |
| Aug5.0v_1338 | 2 | 0 | 0 | 0 | 0 | 0 | 0 | 0 | 0 | 0 | 0 | 0 | 0 | 0 | 0 | 0 | 0 | 0 | 1 | 0 | 0 | 1 | 0 | 0 | 0 | 0 | 0 | 0 | 0 | 0 |
| May12.75v_15177 | 1 | 0 | 0 | 0 | 1 | 0 | 0 | 0 | 0 | 0 | 0 | 0 | 0 | 0 | 0 | 0 | 0 | 0 | 0 | 0 | 0 | 0 | 0 | 0 | 0 | 0 | 0 | 0 | 0 | 0 |
| Jul11.5v_19242 | 1 | 0 | 0 | 0 | 0 | 0 | 0 | 0 | 0 | 0 | 0 | 0 | 0 | 1 | 0 | 0 | 0 | 0 | 0 | 0 | 0 | 0 | 0 | 0 | 0 | 0 | 0 | 0 | 0 | 0 |
| Jul27.5v_5524 | 1 | 0 | 0 | 0 | 0 | 0 | 0 | 0 | 0 | 0 | 0 | 0 | 0 | 0 | 0 | 0 | 0 | 1 | 0 | 0 | 0 | 0 | 0 | 0 | 0 | 0 | 0 | 0 | 0 | 0 |
| Sep8.0v_634 | 1 | 0 | 0 | 0 | 0 | 0 | 0 | 0 | 0 | 0 | 0 | 0 | 0 | 0 | 0 | 0 | 0 | 0 | 0 | 0 | 0 | 0 | 0 | 0 | 0 | 0 | 1 | 0 | 0 | 0 |
| Aug5.0v_3945 | 1 | 0 | 0 | 0 | 0 | 0 | 0 | 0 | 0 | 0 | 0 | 0 | 0 | 0 | 0 | 0 | 0 | 0 | 0 | 0 | 0 | 1 | 0 | 0 | 0 | 0 | 0 | 0 | 0 | 0 |
| Sep8.0v_14899 | 1 | 0 | 0 | 0 | 0 | 0 | 0 | 0 | 0 | 0 | 0 | 0 | 0 | 0 | 0 | 0 | 0 | 0 | 0 | 0 | 0 | 0 | 0 | 0 | 0 | 0 | 1 | 0 | 0 | 0 |
| Aug5.5v_1185 | 1 | 0 | 0 | 0 | 0 | 0 | 0 | 0 | 0 | 0 | 0 | 0 | 0 | 0 | 0 | 0 | 0 | 0 | 0 | 0 | 0 | 0 | 1 | 0 | 0 | 0 | 0 | 0 | 0 | 0 |

| Feature_ID | 654 | May12_05m | May12_25m | May12_50m | May12_75m | Jun8_25m | Jun8_50m | Jun8_75m | Jun22_0m | Jun22_05m | Jun22_50m | Jun22_75m | Jul11_0m | Jul11_05m | Jul11_25m | Jul11_50m | Jul11_75m | Jul27_05m | Jul27_25m | Jul27_50m | Jul27_75m | Aug5_0m | Aug5_05m | Aug5_25m | Aug5_50m | Aug5_75m | Sep8_0m | Sep8_05m | Sep8_25m | Sep8_50m |
|---|---|---|---|---|---|---|---|---|---|---|---|---|---|---|---|---|---|---|---|---|---|---|---|---|---|---|---|---|---|---|
| Sep8.50v_26757 | 1 | 0 | 0 | 0 | 0 | 0 | 0 | 0 | 0 | 0 | 0 | 0 | 0 | 0 | 0 | 0 | 0 | 0 | 0 | 0 | 0 | 0 | 0 | 0 | 0 | 0 | 0 | 0 | 0 | 1 |
| Aug5.0v_1009 | 1 | 0 | 0 | 0 | 0 | 0 | 0 | 0 | 0 | 0 | 0 | 0 | 0 | 0 | 0 | 0 | 0 | 0 | 0 | 0 | 0 | 1 | 0 | 0 | 0 | 0 | 0 | 0 | 0 | 0 |
| Aug5.5v_32509 | 1 | 0 | 0 | 0 | 0 | 0 | 0 | 0 | 0 | 0 | 0 | 0 | 0 | 0 | 0 | 0 | 0 | 0 | 0 | 0 | 0 | 0 | 1 | 0 | 0 | 0 | 0 | 0 | 0 | 0 |
| Aug5.5v_16762 | 2 | 0 | 0 | 0 | 0 | 0 | 0 | 0 | 0 | 0 | 0 | 0 | 0 | 0 | 0 | 0 | 0 | 0 | 0 | 0 | 0 | 0 | 1 | 1 | 0 | 0 | 0 | 0 | 0 | 0 |
| Jul27.50v_20409 | 1 | 0 | 0 | 0 | 0 | 0 | 0 | 0 | 0 | 0 | 0 | 0 | 0 | 0 | 0 | 0 | 0 | 0 | 0 | 1 | 0 | 0 | 0 | 0 | 0 | 0 | 0 | 0 | 0 | 0 |
| Jul11.50v_6212 | 1 | 0 | 0 | 0 | 0 | 0 | 0 | 0 | 0 | 0 | 0 | 0 | 0 | 0 | 0 | 1 | 0 | 0 | 0 | 0 | 0 | 0 | 0 | 0 | 0 | 0 | 0 | 0 | 0 | 0 |
| Jul11.0v_20479 | 1 | 0 | 0 | 0 | 0 | 0 | 0 | 0 | 0 | 0 | 0 | 0 | 1 | 0 | 0 | 0 | 0 | 0 | 0 | 0 | 0 | 0 | 0 | 0 | 0 | 0 | 0 | 0 | 0 | 0 |
| Jul27.75v_6139 | 1 | 0 | 0 | 0 | 0 | 0 | 0 | 0 | 0 | 0 | 0 | 0 | 0 | 0 | 0 | 0 | 0 | 0 | 0 | 0 | 1 | 0 | 0 | 0 | 0 | 0 | 0 | 0 | 0 | 0 |
| Jul11.0v_12552 | 1 | 0 | 0 | 0 | 0 | 0 | 0 | 0 | 0 | 0 | 0 | 0 | 1 | 0 | 0 | 0 | 0 | 0 | 0 | 0 | 0 | 0 | 0 | 0 | 0 | 0 | 0 | 0 | 0 | 0 |
| Sep8.50v_5030 | 1 | 0 | 0 | 0 | 0 | 0 | 0 | 0 | 0 | 0 | 0 | 0 | 0 | 0 | 0 | 0 | 0 | 0 | 0 | 0 | 0 | 0 | 0 | 0 | 0 | 0 | 0 | 0 | 0 | 1 |
| Jul11.0v_1687 | 1 | 0 | 0 | 0 | 0 | 0 | 0 | 0 | 0 | 0 | 0 | 0 | 1 | 0 | 0 | 0 | 0 | 0 | 0 | 0 | 0 | 0 | 0 | 0 | 0 | 0 | 0 | 0 | 0 | 0 |
| Jul27.50v_1043 | 1 | 0 | 0 | 0 | 0 | 0 | 0 | 0 | 0 | 0 | 0 | 0 | 0 | 0 | 0 | 0 | 0 | 0 | 0 | 1 | 0 | 0 | 0 | 0 | 0 | 0 | 0 | 0 | 0 | 0 |
| Jul27.50v_15286 | 1 | 0 | 0 | 0 | 0 | 0 | 0 | 0 | 0 | 0 | 0 | 0 | 0 | 0 | 0 | 0 | 0 | 0 | 0 | 1 | 0 | 0 | 0 | 0 | 0 | 0 | 0 | 0 | 0 | 0 |
| Jun22.50v_17698 | 1 | 0 | 0 | 0 | 0 | 0 | 0 | 0 | 0 | 0 | 1 | 0 | 0 | 0 | 0 | 0 | 0 | 0 | 0 | 0 | 0 | 0 | 0 | 0 | 0 | 0 | 0 | 0 | 0 | 0 |
| Aug5.0v_393 | 1 | 0 | 0 | 0 | 0 | 0 | 0 | 0 | 0 | 0 | 0 | 0 | 0 | 0 | 0 | 0 | 0 | 0 | 0 | 0 | 0 | 1 | 0 | 0 | 0 | 0 | 0 | 0 | 0 | 0 |
| Aug5.0v_3577 | 3 | 0 | 0 | 0 | 0 | 0 | 0 | 0 | 0 | 0 | 0 | 0 | 0 | 0 | 0 | 0 | 0 | 1 | 0 | 0 | 0 | 1 | 0 | 0 | 0 | 0 | 0 | 0 | 0 | 1 |
| Sep8.0v_18039 | 1 | 0 | 0 | 0 | 0 | 0 | 0 | 0 | 0 | 0 | 0 | 0 | 0 | 0 | 0 | 0 | 0 | 0 | 0 | 0 | 0 | 0 | 0 | 0 | 0 | 0 | 1 | 0 | 0 | 0 |
| Jul27.5v_15470 | 1 | 0 | 0 | 0 | 0 | 0 | 0 | 0 | 0 | 0 | 0 | 0 | 0 | 0 | 0 | 0 | 0 | 1 | 0 | 0 | 0 | 0 | 0 | 0 | 0 | 0 | 0 | 0 | 0 | 0 |
| Aug5.25v_10464 | 1 | 0 | 0 | 0 | 0 | 0 | 0 | 0 | 0 | 0 | 0 | 0 | 0 | 0 | 0 | 0 | 0 | 0 | 0 | 0 | 0 | 0 | 0 | 1 | 0 | 0 | 0 | 0 | 0 | 0 |
| Jul11.75v_23749 | 1 | 0 | 0 | 0 | 0 | 0 | 0 | 0 | 0 | 0 | 0 | 0 | 0 | 0 | 0 | 0 | 1 | 0 | 0 | 0 | 0 | 0 | 0 | 0 | 0 | 0 | 0 | 0 | 0 | 0 |
| May12.25v_39988 | 1 | 0 | 1 | 0 | 0 | 0 | 0 | 0 | 0 | 0 | 0 | 0 | 0 | 0 | 0 | 0 | 0 | 0 | 0 | 0 | 0 | 0 | 0 | 0 | 0 | 0 | 0 | 0 | 0 | 0 |
| Jul11.25v_5451 | 1 | 0 | 0 | 0 | 0 | 0 | 0 | 0 | 0 | 0 | 0 | 0 | 0 | 0 | 1 | 0 | 0 | 0 | 0 | 0 | 0 | 0 | 0 | 0 | 0 | 0 | 0 | 0 | 0 | 0 |
| Aug5.25v_19083 | 1 | 0 | 0 | 0 | 0 | 0 | 0 | 0 | 0 | 0 | 0 | 0 | 0 | 0 | 0 | 0 | 0 | 0 | 0 | 0 | 0 | 0 | 0 | 1 | 0 | 0 | 0 | 0 | 0 | 0 |
| Jul11.75v_28567 | 1 | 0 | 0 | 0 | 0 | 0 | 0 | 0 | 0 | 0 | 0 | 0 | 0 | 0 | 0 | 0 | 1 | 0 | 0 | 0 | 0 | 0 | 0 | 0 | 0 | 0 | 0 | 0 | 0 | 0 |
| Jun8.25v_11433 | 1 | 0 | 0 | 0 | 0 | 1 | 0 | 0 | 0 | 0 | 0 | 0 | 0 | 0 | 0 | 0 | 0 | 0 | 0 | 0 | 0 | 0 | 0 | 0 | 0 | 0 | 0 | 0 | 0 | 0 |
| Jun8.25v_15820 | 1 | 0 | 0 | 0 | 0 | 1 | 0 | 0 | 0 | 0 | 0 | 0 | 0 | 0 | 0 | 0 | 0 | 0 | 0 | 0 | 0 | 0 | 0 | 0 | 0 | 0 | 0 | 0 | 0 | 0 |
| Sep8.50v_35251 | 2 | 0 | 0 | 0 | 0 | 0 | 0 | 0 | 0 | 0 | 0 | 1 | 0 | 0 | 0 | 0 | 0 | 0 | 0 | 0 | 0 | 0 | 0 | 0 | 0 | 0 | 0 | 0 | 0 | 1 |
| Aug5.0v_581 | 1 | 0 | 0 | 0 | 0 | 0 | 0 | 0 | 0 | 0 | 0 | 0 | 0 | 0 | 0 | 0 | 0 | 0 | 0 | 0 | 0 | 1 | 0 | 0 | 0 | 0 | 0 | 0 | 0 | 0 |
| Jun8.25v_2418 | 1 | 0 | 0 | 0 | 0 | 1 | 0 | 0 | 0 | 0 | 0 | 0 | 0 | 0 | 0 | 0 | 0 | 0 | 0 | 0 | 0 | 0 | 0 | 0 | 0 | 0 | 0 | 0 | 0 | 0 |
| Jul11.5v_18340 | 1 | 0 | 0 | 0 | 0 | 0 | 0 | 0 | 0 | 0 | 0 | 0 | 0 | 1 | 0 | 0 | 0 | 0 | 0 | 0 | 0 | 0 | 0 | 0 | 0 | 0 | 0 | 0 | 0 | 0 |
| Aug5.0v_1153 | 1 | 0 | 0 | 0 | 0 | 0 | 0 | 0 | 0 | 0 | 0 | 0 | 0 | 0 | 0 | 0 | 0 | 0 | 0 | 0 | 0 | 1 | 0 | 0 | 0 | 0 | 0 | 0 | 0 | 0 |
| Sep8.0v_6064 | 1 | 0 | 0 | 0 | 0 | 0 | 0 | 0 | 0 | 0 | 0 | 0 | 0 | 0 | 0 | 0 | 0 | 0 | 0 | 0 | 0 | 0 | 0 | 0 | 0 | 0 | 1 | 0 | 0 | 0 |
| Jun22.75v_2969 | 1 | 0 | 0 | 0 | 0 | 0 | 0 | 0 | 0 | 0 | 0 | 1 | 0 | 0 | 0 | 0 | 0 | 0 | 0 | 0 | 0 | 0 | 0 | 0 | 0 | 0 | 0 | 0 | 0 | 0 |
| Aug5.0v_388 | 1 | 0 | 0 | 0 | 0 | 0 | 0 | 0 | 0 | 0 | 0 | 0 | 0 | 0 | 0 | 0 | 0 | 0 | 0 | 0 | 0 | 1 | 0 | 0 | 0 | 0 | 0 | 0 | 0 | 0 |
| Aug5.5v_427 | 1 | 0 | 0 | 0 | 0 | 0 | 0 | 0 | 0 | 0 | 0 | 0 | 0 | 0 | 0 | 0 | 0 | 0 | 0 | 0 | 0 | 0 | 1 | 0 | 0 | 0 | 0 | 0 | 0 | 0 |
| Jul11.50v_924 | 4 | 0 | 0 | 0 | 0 | 0 | 0 | 0 | 0 | 0 | 0 | 0 | 0 | 0 | 0 | 1 | 0 | 0 | 0 | 0 | 0 | 1 | 1 | 0 | 0 | 0 | 0 | 0 | 0 | 1 |
| May12.5v_23953 | 1 | 1 | 0 | 0 | 0 | 0 | 0 | 0 | 0 | 0 | 0 | 0 | 0 | 0 | 0 | 0 | 0 | 0 | 0 | 0 | 0 | 0 | 0 | 0 | 0 | 0 | 0 | 0 | 0 | 0 |
| Aug5.25v_29335 | 2 | 0 | 0 | 0 | 0 | 0 | 0 | 0 | 0 | 0 | 0 | 0 | 0 | 0 | 0 | 1 | 0 | 0 | 0 | 0 | 0 | 0 | 0 | 1 | 0 | 0 | 0 | 0 | 0 | 0 |
| Jun22.75v_18829 | 4 | 0 | 0 | 0 | 0 | 0 | 0 | 0 | 0 | 0 | 0 | 1 | 0 | 0 | 0 | 0 | 0 | 1 | 0 | 0 | 0 | 0 | 1 | 0 | 0 | 0 | 0 | 0 | 0 | 1 |
| Jul11.5v_15540 | 1 | 0 | 0 | 0 | 0 | 0 | 0 | 0 | 0 | 0 | 0 | 0 | 0 | 1 | 0 | 0 | 0 | 0 | 0 | 0 | 0 | 0 | 0 | 0 | 0 | 0 | 0 | 0 | 0 | 0 |
| Aug5.0v_1583 | 1 | 0 | 0 | 0 | 0 | 0 | 0 | 0 | 0 | 0 | 0 | 0 | 0 | 0 | 0 | 0 | 0 | 0 | 0 | 0 | 0 | 1 | 0 | 0 | 0 | 0 | 0 | 0 | 0 | 0 |
| Jun8.50v_15301 | 1 | 0 | 0 | 0 | 0 | 0 | 1 | 0 | 0 | 0 | 0 | 0 | 0 | 0 | 0 | 0 | 0 | 0 | 0 | 0 | 0 | 0 | 0 | 0 | 0 | 0 | 0 | 0 | 0 | 0 |
| Aug5.0v_3105 | 1 | 0 | 0 | 0 | 0 | 0 | 0 | 0 | 0 | 0 | 0 | 0 | 0 | 0 | 0 | 0 | 0 | 0 | 0 | 0 | 0 | 1 | 0 | 0 | 0 | 0 | 0 | 0 | 0 | 0 |
| Sep8.0v_18634 | 1 | 0 | 0 | 0 | 0 | 0 | 0 | 0 | 0 | 0 | 0 | 0 | 0 | 0 | 0 | 0 | 0 | 0 | 0 | 0 | 0 | 0 | 0 | 0 | 0 | 0 | 1 | 0 | 0 | 0 |
| May12.50v_12696 | 1 | 0 | 0 | 1 | 0 | 0 | 0 | 0 | 0 | 0 | 0 | 0 | 0 | 0 | 0 | 0 | 0 | 0 | 0 | 0 | 0 | 0 | 0 | 0 | 0 | 0 | 0 | 0 | 0 | 0 |
| May12.50v_39241 | 1 | 0 | 0 | 1 | 0 | 0 | 0 | 0 | 0 | 0 | 0 | 0 | 0 | 0 | 0 | 0 | 0 | 0 | 0 | 0 | 0 | 0 | 0 | 0 | 0 | 0 | 0 | 0 | 0 | 0 |
| Sep8.50v_32558 | 1 | 0 | 0 | 0 | 0 | 0 | 0 | 0 | 0 | 0 | 0 | 0 | 0 | 0 | 0 | 0 | 0 | 0 | 0 | 0 | 0 | 0 | 0 | 0 | 0 | 0 | 0 | 0 | 0 | 1 |
| Jun22.0v_17340 | 1 | 0 | 0 | 0 | 0 | 0 | 0 | 0 | 1 | 0 | 0 | 0 | 0 | 0 | 0 | 0 | 0 | 0 | 0 | 0 | 0 | 0 | 0 | 0 | 0 | 0 | 0 | 0 | 0 | 0 |
| Aug5.25v_35301 | 1 | 0 | 0 | 0 | 0 | 0 | 0 | 0 | 0 | 0 | 0 | 0 | 0 | 0 | 0 | 0 | 0 | 0 | 0 | 0 | 0 | 0 | 0 | 1 | 0 | 0 | 0 | 0 | 0 | 0 |
| Jul27.5v_11104 | 1 | 0 | 0 | 0 | 0 | 0 | 0 | 0 | 0 | 0 | 0 | 0 | 0 | 0 | 0 | 0 | 0 | 1 | 0 | 0 | 0 | 0 | 0 | 0 | 0 | 0 | 0 | 0 | 0 | 0 |
| Jun22.0v_839 | 1 | 0 | 0 | 0 | 0 | 0 | 0 | 0 | 1 | 0 | 0 | 0 | 0 | 0 | 0 | 0 | 0 | 0 | 0 | 0 | 0 | 0 | 0 | 0 | 0 | 0 | 0 | 0 | 0 | 0 |
| Jun22.0v_15014 | 1 | 0 | 0 | 0 | 0 | 0 | 0 | 0 | 1 | 0 | 0 | 0 | 0 | 0 | 0 | 0 | 0 | 0 | 0 | 0 | 0 | 0 | 0 | 0 | 0 | 0 | 0 | 0 | 0 | 0 |
| May12.50v_39522 | 1 | 0 | 0 | 1 | 0 | 0 | 0 | 0 | 0 | 0 | 0 | 0 | 0 | 0 | 0 | 0 | 0 | 0 | 0 | 0 | 0 | 0 | 0 | 0 | 0 | 0 | 0 | 0 | 0 | 0 |
| Jul27.5v_13266 | 3 | 0 | 0 | 0 | 0 | 0 | 0 | 0 | 0 | 0 | 0 | 0 | 0 | 0 | 1 | 0 | 0 | 2 | 0 | 0 | 0 | 0 | 0 | 0 | 0 | 0 | 0 | 0 | 0 | 0 |
| Aug5.25v_21421 | 1 | 0 | 0 | 0 | 0 | 0 | 0 | 0 | 0 | 0 | 0 | 0 | 0 | 0 | 0 | 0 | 0 | 0 | 0 | 0 | 0 | 0 | 0 | 1 | 0 | 0 | 0 | 0 | 0 | 0 |
| Jun22.0v_862 | 2 | 0 | 0 | 0 | 0 | 0 | 0 | 0 | 1 | 0 | 0 | 0 | 0 | 1 | 0 | 0 | 0 | 0 | 0 | 0 | 0 | 0 | 0 | 0 | 0 | 0 | 0 | 0 | 0 | 0 |
| Aug5.25v_5053 | 1 | 0 | 0 | 0 | 0 | 0 | 0 | 0 | 0 | 0 | 0 | 0 | 0 | 0 | 0 | 0 | 0 | 0 | 0 | 0 | 0 | 0 | 0 | 1 | 0 | 0 | 0 | 0 | 0 | 0 |
| Jun22.50v_31012 | 1 | 0 | 0 | 0 | 0 | 0 | 0 | 0 | 0 | 0 | 1 | 0 | 0 | 0 | 0 | 0 | 0 | 0 | 0 | 0 | 0 | 0 | 0 | 0 | 0 | 0 | 0 | 0 | 0 | 0 |
| Aug5.0v_4075 | 2 | 0 | 0 | 0 | 0 | 0 | 0 | 0 | 0 | 0 | 0 | 0 | 0 | 0 | 0 | 0 | 0 | 1 | 0 | 0 | 0 | 1 | 0 | 0 | 0 | 0 | 0 | 0 | 0 | 0 |
| Jul11.0v_2941 | 2 | 0 | 0 | 0 | 0 | 0 | 0 | 0 | 0 | 0 | 0 | 0 | 2 | 0 | 0 | 0 | 0 | 0 | 0 | 0 | 0 | 0 | 0 | 0 | 0 | 0 | 0 | 0 | 0 | 0 |
| Aug5.25v_22964 | 1 | 0 | 0 | 0 | 0 | 0 | 0 | 0 | 0 | 0 | 0 | 0 | 0 | 0 | 0 | 0 | 0 | 0 | 0 | 0 | 0 | 0 | 0 | 1 | 0 | 0 | 0 | 0 | 0 | 0 |
| Sep8.50v_16320 | 1 | 0 | 0 | 0 | 0 | 0 | 0 | 0 | 0 | 0 | 0 | 0 | 0 | 0 | 0 | 0 | 0 | 0 | 0 | 0 | 0 | 0 | 0 | 0 | 0 | 0 | 0 | 0 | 0 | 1 |
| Jul11.0v_12503 | 1 | 0 | 0 | 0 | 0 | 0 | 0 | 0 | 0 | 0 | 0 | 0 | 1 | 0 | 0 | 0 | 0 | 0 | 0 | 0 | 0 | 0 | 0 | 0 | 0 | 0 | 0 | 0 | 0 | 0 |
| Aug5.0v_4499 | 1 | 0 | 0 | 0 | 0 | 0 | 0 | 0 | 0 | 0 | 0 | 0 | 0 | 0 | 0 | 0 | 0 | 0 | 0 | 0 | 0 | 1 | 0 | 0 | 0 | 0 | 0 | 0 | 0 | 0 |
| Aug5.25v_23422 | 8 | 1 | 0 | 1 | 0 | 2 | 0 | 0 | 0 | 0 | 0 | 0 | 0 | 0 | 0 | 0 | 0 | 0 | 1 | 0 | 0 | 0 | 1 | 2 | 0 | 0 | 0 | 0 | 0 | 0 |
| Aug5.75v_27255 | 2 | 0 | 0 | 0 | 0 | 0 | 0 | 0 | 0 | 0 | 0 | 0 | 0 | 0 | 0 | 0 | 0 | 0 | 0 | 0 | 0 | 0 | 0 | 0 | 0 | 2 | 0 | 0 | 0 | 0 |
| Jun22.0v_2007 | 3 | 0 | 0 | 0 | 0 | 0 | 0 | 0 | 1 | 0 | 0 | 0 | 0 | 0 | 0 | 1 | 0 | 0 | 0 | 0 | 0 | 0 | 0 | 0 | 0 | 0 | 0 | 0 | 0 | 1 |
| Aug5.25v_23672 | 1 | 0 | 0 | 0 | 0 | 0 | 0 | 0 | 0 | 0 | 0 | 0 | 0 | 0 | 0 | 0 | 0 | 0 | 0 | 0 | 0 | 0 | 0 | 1 | 0 | 0 | 0 | 0 | 0 | 0 |
| Jul11.50v_7178 | 2 | 0 | 0 | 1 | 0 | 0 | 0 | 0 | 0 | 0 | 0 | 0 | 0 | 0 | 0 | 1 | 0 | 0 | 0 | 0 | 0 | 0 | 0 | 0 | 0 | 0 | 0 | 0 | 0 | 0 |
| Aug5.0v_4434 | 101 | 2 | 6 | 4 | 7 | 2 | 11 | 0 | 9 | 0 | 3 | 8 | 6 | 3 | 2 | 0 | 1 | 2 | 3 | 1 | 1 | 15 | 0 | 4 | 1 | 2 | 3 | 0 | 0 | 5 |
| Jun8.50v_9271 | 1 | 0 | 0 | 0 | 0 | 0 | 1 | 0 | 0 | 0 | 0 | 0 | 0 | 0 | 0 | 0 | 0 | 0 | 0 | 0 | 0 | 0 | 0 | 0 | 0 | 0 | 0 | 0 | 0 | 0 |
| Jul11.5v_21037 | 1 | 0 | 0 | 0 | 0 | 0 | 0 | 0 | 0 | 0 | 0 | 0 | 0 | 1 | 0 | 0 | 0 | 0 | 0 | 0 | 0 | 0 | 0 | 0 | 0 | 0 | 0 | 0 | 0 | 0 |
| Jun22.75v_14616 | 1 | 0 | 0 | 0 | 0 | 0 | 0 | 0 | 0 | 0 | 0 | 1 | 0 | 0 | 0 | 0 | 0 | 0 | 0 | 0 | 0 | 0 | 0 | 0 | 0 | 0 | 0 | 0 | 0 | 0 |
| Jul11.50v_3228 | 1 | 0 | 0 | 0 | 0 | 0 | 0 | 0 | 0 | 0 | 0 | 0 | 0 | 0 | 0 | 1 | 0 | 0 | 0 | 0 | 0 | 0 | 0 | 0 | 0 | 0 | 0 | 0 | 0 | 0 |
| Jul11.50v_15968 | 1 | 0 | 0 | 0 | 0 | 0 | 0 | 0 | 0 | 0 | 0 | 0 | 0 | 0 | 0 | 1 | 0 | 0 | 0 | 0 | 0 | 0 | 0 | 0 | 0 | 0 | 0 | 0 | 0 | 0 |
| Aug5.25v_8939 | 1 | 0 | 0 | 0 | 0 | 0 | 0 | 0 | 0 | 0 | 0 | 0 | 0 | 0 | 0 | 0 | 0 | 0 | 0 | 0 | 0 | 0 | 0 | 1 | 0 | 0 | 0 | 0 | 0 | 0 |
| May12.25v_13968 | 1 | 0 | 1 | 0 | 0 | 0 | 0 | 0 | 0 | 0 | 0 | 0 | 0 | 0 | 0 | 0 | 0 | 0 | 0 | 0 | 0 | 0 | 0 | 0 | 0 | 0 | 0 | 0 | 0 | 0 |
| Jun22.0v_8769 | 17 | 0 | 1 | 1 | 0 | 0 | 0 | 0 | 3 | 0 | 0 | 2 | 0 | 0 | 0 | 2 | 0 | 2 | 1 | 0 | 0 | 3 | 0 | 0 | 1 | 0 | 1 | 0 | 0 | 0 |
| Aug5.25v_6480 | 1 | 0 | 0 | 0 | 0 | 0 | 0 | 0 | 0 | 0 | 0 | 0 | 0 | 0 | 0 | 0 | 0 | 0 | 0 | 0 | 0 | 0 | 0 | 1 | 0 | 0 | 0 | 0 | 0 | 0 |
| May12.25v_6399 | 2 | 0 | 1 | 0 | 0 | 0 | 0 | 0 | 0 | 0 | 0 | 0 | 0 | 0 | 0 | 0 | 0 | 0 | 0 | 0 | 0 | 0 | 0 | 0 | 0 | 0 | 0 | 0 | 0 | 1 |
| Jul11.50v_16904 | 1 | 0 | 0 | 0 | 0 | 0 | 0 | 0 | 0 | 0 | 0 | 0 | 0 | 0 | 0 | 1 | 0 | 0 | 0 | 0 | 0 | 0 | 0 | 0 | 0 | 0 | 0 | 0 | 0 | 0 |
| Jul11.5v_9989 | 1 | 0 | 0 | 0 | 0 | 0 | 0 | 0 | 0 | 0 | 0 | 0 | 0 | 1 | 0 | 0 | 0 | 0 | 0 | 0 | 0 | 0 | 0 | 0 | 0 | 0 | 0 | 0 | 0 | 0 |
| Aug5.75v_31515 | 3 | 0 | 0 | 0 | 0 | 0 | 0 | 0 | 0 | 0 | 0 | 0 | 0 | 0 | 0 | 0 | 0 | 0 | 0 | 0 | 0 | 0 | 0 | 0 | 0 | 2 | 0 | 1 | 0 | 0 |
| Aug5.75v_23533 | 1 | 0 | 0 | 0 | 0 | 0 | 0 | 0 | 0 | 0 | 0 | 0 | 0 | 0 | 0 | 0 | 0 | 0 | 0 | 0 | 0 | 0 | 0 | 0 | 0 | 1 | 0 | 0 | 0 | 0 |
| May12.75v_26104 | 2 | 1 | 0 | 0 | 1 | 0 | 0 | 0 | 0 | 0 | 0 | 0 | 0 | 0 | 0 | 0 | 0 | 0 | 0 | 0 | 0 | 0 | 0 | 0 | 0 | 0 | 0 | 0 | 0 | 0 |
| May12.5v_26165 | 1 | 1 | 0 | 0 | 0 | 0 | 0 | 0 | 0 | 0 | 0 | 0 | 0 | 0 | 0 | 0 | 0 | 0 | 0 | 0 | 0 | 0 | 0 | 0 | 0 | 0 | 0 | 0 | 0 | 0 |
| Jul11.50v_2116 | 1 | 0 | 0 | 0 | 0 | 0 | 0 | 0 | 0 | 0 | 0 | 0 | 0 | 0 | 0 | 1 | 0 | 0 | 0 | 0 | 0 | 0 | 0 | 0 | 0 | 0 | 0 | 0 | 0 | 0 |
| Jun22.50v_24832 | 1 | 0 | 0 | 0 | 0 | 0 | 0 | 0 | 0 | 0 | 1 | 0 | 0 | 0 | 0 | 0 | 0 | 0 | 0 | 0 | 0 | 0 | 0 | 0 | 0 | 0 | 0 | 0 | 0 | 0 |
| May12.50v_29691 | 1 | 0 | 0 | 1 | 0 | 0 | 0 | 0 | 0 | 0 | 0 | 0 | 0 | 0 | 0 | 0 | 0 | 0 | 0 | 0 | 0 | 0 | 0 | 0 | 0 | 0 | 0 | 0 | 0 | 0 |
| Jun22.50v_1308 | 1 | 0 | 0 | 0 | 0 | 0 | 0 | 0 | 0 | 0 | 1 | 0 | 0 | 0 | 0 | 0 | 0 | 0 | 0 | 0 | 0 | 0 | 0 | 0 | 0 | 0 | 0 | 0 | 0 | 0 |
| Aug5.75v_34269 | 1 | 0 | 0 | 0 | 0 | 0 | 0 | 0 | 0 | 0 | 0 | 0 | 0 | 0 | 0 | 0 | 0 | 0 | 0 | 0 | 0 | 0 | 0 | 0 | 0 | 1 | 0 | 0 | 0 | 0 |
| Aug5.25v_22388 | 1 | 0 | 0 | 0 | 0 | 0 | 0 | 0 | 0 | 0 | 0 | 0 | 0 | 0 | 0 | 0 | 0 | 0 | 0 | 0 | 0 | 0 | 0 | 1 | 0 | 0 | 0 | 0 | 0 | 0 |
| Aug5.25v_27579 | 1 | 0 | 0 | 0 | 0 | 0 | 0 | 0 | 0 | 0 | 0 | 0 | 0 | 0 | 0 | 0 | 0 | 0 | 0 | 0 | 0 | 0 | 0 | 1 | 0 | 0 | 0 | 0 | 0 | 0 |
| Jul11.25v_11270 | 1 | 0 | 0 | 0 | 0 | 0 | 0 | 0 | 0 | 0 | 0 | 0 | 0 | 0 | 1 | 0 | 0 | 0 | 0 | 0 | 0 | 0 | 0 | 0 | 0 | 0 | 0 | 0 | 0 | 0 |
| Jun22.0v_12106 | 1 | 0 | 0 | 0 | 0 | 0 | 0 | 0 | 1 | 0 | 0 | 0 | 0 | 0 | 0 | 0 | 0 | 0 | 0 | 0 | 0 | 0 | 0 | 0 | 0 | 0 | 0 | 0 | 0 | 0 |
| Aug5.5v_16686 | 1 | 0 | 0 | 0 | 0 | 0 | 0 | 0 | 0 | 0 | 0 | 0 | 0 | 0 | 0 | 0 | 0 | 0 | 0 | 0 | 0 | 0 | 1 | 0 | 0 | 0 | 0 | 0 | 0 | 0 |
| Jun22.50v_25683 | 1 | 0 | 0 | 0 | 0 | 0 | 0 | 0 | 0 | 0 | 1 | 0 | 0 | 0 | 0 | 0 | 0 | 0 | 0 | 0 | 0 | 0 | 0 | 0 | 0 | 0 | 0 | 0 | 0 | 0 |
| Sep8.50v_18146 | 1 | 0 | 0 | 0 | 0 | 0 | 0 | 0 | 0 | 0 | 0 | 0 | 0 | 0 | 0 | 0 | 0 | 0 | 0 | 0 | 0 | 0 | 0 | 0 | 0 | 0 | 0 | 0 | 0 | 1 |
| Jul11.0v_1841 | 2 | 0 | 0 | 0 | 0 | 0 | 0 | 0 | 0 | 0 | 0 | 0 | 1 | 0 | 0 | 0 | 0 | 0 | 0 | 0 | 0 | 0 | 0 | 0 | 0 | 0 | 1 | 0 | 0 | 0 |
| Jun22.5v_16706 | 1 | 0 | 0 | 0 | 0 | 0 | 0 | 0 | 0 | 1 | 0 | 0 | 0 | 0 | 0 | 0 | 0 | 0 | 0 | 0 | 0 | 0 | 0 | 0 | 0 | 0 | 0 | 0 | 0 | 0 |
| Jun22.50v_6586 | 1 | 0 | 0 | 0 | 0 | 0 | 0 | 0 | 0 | 0 | 1 | 0 | 0 | 0 | 0 | 0 | 0 | 0 | 0 | 0 | 0 | 0 | 0 | 0 | 0 | 0 | 0 | 0 | 0 | 0 |
| Jul27.5v_10419 | 2 | 0 | 0 | 0 | 0 | 0 | 0 | 0 | 0 | 0 | 0 | 0 | 0 | 0 | 0 | 0 | 0 | 2 | 0 | 0 | 0 | 0 | 0 | 0 | 0 | 0 | 0 | 0 | 0 | 0 |
| Sep8.0v_344 | 1 | 0 | 0 | 0 | 0 | 0 | 0 | 0 | 0 | 0 | 0 | 0 | 0 | 0 | 0 | 0 | 0 | 0 | 0 | 0 | 0 | 0 | 0 | 0 | 0 | 0 | 1 | 0 | 0 | 0 |
| Aug5.5v_34422 | 1 | 0 | 0 | 0 | 0 | 0 | 0 | 0 | 0 | 0 | 0 | 0 | 0 | 0 | 0 | 0 | 0 | 0 | 0 | 0 | 0 | 0 | 1 | 0 | 0 | 0 | 0 | 0 | 0 | 0 |
| Jul11.25v_4960 | 1 | 0 | 0 | 0 | 0 | 0 | 0 | 0 | 0 | 0 | 0 | 0 | 0 | 0 | 1 | 0 | 0 | 0 | 0 | 0 | 0 | 0 | 0 | 0 | 0 | 0 | 0 | 0 | 0 | 0 |
| Aug5.0v_5281 | 1 | 0 | 0 | 0 | 0 | 0 | 0 | 0 | 0 | 0 | 0 | 0 | 0 | 0 | 0 | 0 | 0 | 0 | 0 | 0 | 0 | 1 | 0 | 0 | 0 | 0 | 0 | 0 | 0 | 0 |
| Jun8.50v_17419 | 1 | 0 | 0 | 0 | 0 | 0 | 1 | 0 | 0 | 0 | 0 | 0 | 0 | 0 | 0 | 0 | 0 | 0 | 0 | 0 | 0 | 0 | 0 | 0 | 0 | 0 | 0 | 0 | 0 | 0 |
| Jul11.25v_16349 | 1 | 0 | 0 | 0 | 0 | 0 | 0 | 0 | 0 | 0 | 0 | 0 | 0 | 0 | 1 | 0 | 0 | 0 | 0 | 0 | 0 | 0 | 0 | 0 | 0 | 0 | 0 | 0 | 0 | 0 |
| Sep8.0v_6668 | 1 | 0 | 0 | 0 | 0 | 0 | 0 | 0 | 0 | 0 | 0 | 0 | 0 | 0 | 0 | 0 | 0 | 0 | 0 | 0 | 0 | 0 | 0 | 0 | 0 | 0 | 1 | 0 | 0 | 0 |
| Jul11.25v_89 | 1 | 0 | 0 | 0 | 0 | 0 | 0 | 0 | 0 | 0 | 0 | 0 | 0 | 0 | 1 | 0 | 0 | 0 | 0 | 0 | 0 | 0 | 0 | 0 | 0 | 0 | 0 | 0 | 0 | 0 |
| Aug5.25v_29501 | 1 | 0 | 0 | 0 | 0 | 0 | 0 | 0 | 0 | 0 | 0 | 0 | 0 | 0 | 0 | 0 | 0 | 0 | 0 | 0 | 0 | 0 | 0 | 1 | 0 | 0 | 0 | 0 | 0 | 0 |
| Sep8.0v_14652 | 1 | 0 | 0 | 0 | 0 | 0 | 0 | 0 | 0 | 0 | 0 | 0 | 0 | 0 | 0 | 0 | 0 | 0 | 0 | 0 | 0 | 0 | 0 | 0 | 0 | 0 | 1 | 0 | 0 | 0 |
| May12.75v_35105 | 1 | 0 | 0 | 0 | 1 | 0 | 0 | 0 | 0 | 0 | 0 | 0 | 0 | 0 | 0 | 0 | 0 | 0 | 0 | 0 | 0 | 0 | 0 | 0 | 0 | 0 | 0 | 0 | 0 | 0 |
| Jul11.75v_33366 | 1 | 0 | 0 | 0 | 0 | 0 | 0 | 0 | 0 | 0 | 0 | 0 | 0 | 0 | 0 | 0 | 1 | 0 | 0 | 0 | 0 | 0 | 0 | 0 | 0 | 0 | 0 | 0 | 0 | 0 |
| Jul27.5v_9745 | 1 | 0 | 0 | 0 | 0 | 0 | 0 | 0 | 0 | 0 | 0 | 0 | 0 | 0 | 0 | 0 | 0 | 1 | 0 | 0 | 0 | 0 | 0 | 0 | 0 | 0 | 0 | 0 | 0 | 0 |
| Jul11.50v_9688 | 1 | 0 | 0 | 0 | 0 | 0 | 0 | 0 | 0 | 0 | 0 | 0 | 0 | 0 | 0 | 1 | 0 | 0 | 0 | 0 | 0 | 0 | 0 | 0 | 0 | 0 | 0 | 0 | 0 | 0 |
| Aug5.0v_2799 | 1 | 0 | 0 | 0 | 0 | 0 | 0 | 0 | 0 | 0 | 0 | 0 | 0 | 0 | 0 | 0 | 0 | 0 | 0 | 0 | 0 | 1 | 0 | 0 | 0 | 0 | 0 | 0 | 0 | 0 |
| Aug5.0v_4127 | 1 | 0 | 0 | 0 | 0 | 0 | 0 | 0 | 0 | 0 | 0 | 0 | 0 | 0 | 0 | 0 | 0 | 0 | 0 | 0 | 0 | 1 | 0 | 0 | 0 | 0 | 0 | 0 | 0 | 0 |
| Sep8.5v_14137 | 1 | 0 | 0 | 0 | 0 | 0 | 0 | 0 | 0 | 0 | 0 | 0 | 0 | 0 | 0 | 0 | 0 | 0 | 0 | 0 | 0 | 0 | 0 | 0 | 0 | 0 | 0 | 1 | 0 | 0 |
| Aug5.0v_3413 | 1 | 0 | 0 | 0 | 0 | 0 | 0 | 0 | 0 | 0 | 0 | 0 | 0 | 0 | 0 | 0 | 0 | 0 | 0 | 0 | 0 | 1 | 0 | 0 | 0 | 0 | 0 | 0 | 0 | 0 |
| Aug5.5v_35539 | 1 | 0 | 0 | 0 | 0 | 0 | 0 | 0 | 0 | 0 | 0 | 0 | 0 | 0 | 0 | 0 | 0 | 0 | 0 | 0 | 0 | 0 | 1 | 0 | 0 | 0 | 0 | 0 | 0 | 0 |
| Aug5.0v_3749 | 1 | 0 | 0 | 0 | 0 | 0 | 0 | 0 | 0 | 0 | 0 | 0 | 0 | 0 | 0 | 0 | 0 | 0 | 0 | 0 | 0 | 1 | 0 | 0 | 0 | 0 | 0 | 0 | 0 | 0 |
| Jul11.50v_18833 | 1 | 0 | 0 | 0 | 0 | 0 | 0 | 0 | 0 | 0 | 0 | 0 | 0 | 0 | 0 | 1 | 0 | 0 | 0 | 0 | 0 | 0 | 0 | 0 | 0 | 0 | 0 | 0 | 0 | 0 |
| Jun8.50v_27437 | 2 | 0 | 0 | 0 | 0 | 0 | 1 | 0 | 0 | 0 | 0 | 0 | 0 | 0 | 0 | 0 | 0 | 0 | 0 | 0 | 0 | 0 | 0 | 0 | 0 | 0 | 0 | 0 | 0 | 1 |
| Jul11.50v_514 | 1 | 0 | 0 | 0 | 0 | 0 | 0 | 0 | 0 | 0 | 0 | 0 | 0 | 0 | 0 | 1 | 0 | 0 | 0 | 0 | 0 | 0 | 0 | 0 | 0 | 0 | 0 | 0 | 0 | 0 |
| Jun22.50v_31467 | 1 | 0 | 0 | 0 | 0 | 0 | 0 | 0 | 0 | 0 | 1 | 0 | 0 | 0 | 0 | 0 | 0 | 0 | 0 | 0 | 0 | 0 | 0 | 0 | 0 | 0 | 0 | 0 | 0 | 0 |
| Jul27.5v_7354 | 1 | 0 | 0 | 0 | 0 | 0 | 0 | 0 | 0 | 0 | 0 | 0 | 0 | 0 | 0 | 0 | 0 | 1 | 0 | 0 | 0 | 0 | 0 | 0 | 0 | 0 | 0 | 0 | 0 | 0 |
| May12.50v_3367 | 1 | 0 | 0 | 1 | 0 | 0 | 0 | 0 | 0 | 0 | 0 | 0 | 0 | 0 | 0 | 0 | 0 | 0 | 0 | 0 | 0 | 0 | 0 | 0 | 0 | 0 | 0 | 0 | 0 | 0 |
| Jul11.0v_2018 | 1 | 0 | 0 | 0 | 0 | 0 | 0 | 0 | 0 | 0 | 0 | 0 | 1 | 0 | 0 | 0 | 0 | 0 | 0 | 0 | 0 | 0 | 0 | 0 | 0 | 0 | 0 | 0 | 0 | 0 |
| May12.5v_14706 | 1 | 1 | 0 | 0 | 0 | 0 | 0 | 0 | 0 | 0 | 0 | 0 | 0 | 0 | 0 | 0 | 0 | 0 | 0 | 0 | 0 | 0 | 0 | 0 | 0 | 0 | 0 | 0 | 0 | 0 |
| Sep8.0v_304 | 1 | 0 | 0 | 0 | 0 | 0 | 0 | 0 | 0 | 0 | 0 | 0 | 0 | 0 | 0 | 0 | 0 | 0 | 0 | 0 | 0 | 0 | 0 | 0 | 0 | 0 | 1 | 0 | 0 | 0 |
| Aug5.0v_3175 | 1 | 0 | 0 | 0 | 0 | 0 | 0 | 0 | 0 | 0 | 0 | 0 | 0 | 0 | 0 | 0 | 0 | 0 | 0 | 0 | 0 | 1 | 0 | 0 | 0 | 0 | 0 | 0 | 0 | 0 |
| Jul11.75v_1633 | 1 | 0 | 0 | 0 | 0 | 0 | 0 | 0 | 0 | 0 | 0 | 0 | 0 | 0 | 0 | 0 | 1 | 0 | 0 | 0 | 0 | 0 | 0 | 0 | 0 | 0 | 0 | 0 | 0 | 0 |
| Jul11.0v_16514 | 2 | 0 | 1 | 0 | 0 | 0 | 0 | 0 | 0 | 0 | 0 | 0 | 1 | 0 | 0 | 0 | 0 | 0 | 0 | 0 | 0 | 0 | 0 | 0 | 0 | 0 | 0 | 0 | 0 | 0 |
| May12.75v_40568 | 1 | 0 | 0 | 0 | 1 | 0 | 0 | 0 | 0 | 0 | 0 | 0 | 0 | 0 | 0 | 0 | 0 | 0 | 0 | 0 | 0 | 0 | 0 | 0 | 0 | 0 | 0 | 0 | 0 | 0 |
| Aug5.25v_31446 | 1 | 0 | 0 | 0 | 0 | 0 | 0 | 0 | 0 | 0 | 0 | 0 | 0 | 0 | 0 | 0 | 0 | 0 | 0 | 0 | 0 | 0 | 0 | 1 | 0 | 0 | 0 | 0 | 0 | 0 |
| Jun8.25v_36411 | 1 | 0 | 0 | 0 | 0 | 1 | 0 | 0 | 0 | 0 | 0 | 0 | 0 | 0 | 0 | 0 | 0 | 0 | 0 | 0 | 0 | 0 | 0 | 0 | 0 | 0 | 0 | 0 | 0 | 0 |
| May12.50v_9597 | 1 | 0 | 0 | 1 | 0 | 0 | 0 | 0 | 0 | 0 | 0 | 0 | 0 | 0 | 0 | 0 | 0 | 0 | 0 | 0 | 0 | 0 | 0 | 0 | 0 | 0 | 0 | 0 | 0 | 0 |
| Aug5.25v_21388 | 2 | 0 | 0 | 0 | 0 | 0 | 0 | 0 | 0 | 0 | 0 | 0 | 0 | 0 | 0 | 1 | 0 | 0 | 0 | 0 | 0 | 0 | 0 | 1 | 0 | 0 | 0 | 0 | 0 | 0 |
| Aug5.0v_3075 | 1 | 0 | 0 | 0 | 0 | 0 | 0 | 0 | 0 | 0 | 0 | 0 | 0 | 0 | 0 | 0 | 0 | 0 | 0 | 0 | 0 | 1 | 0 | 0 | 0 | 0 | 0 | 0 | 0 | 0 |
| Jul11.5v_12428 | 2 | 0 | 0 | 0 | 0 | 0 | 0 | 0 | 0 | 0 | 0 | 0 | 0 | 1 | 0 | 0 | 1 | 0 | 0 | 0 | 0 | 0 | 0 | 0 | 0 | 0 | 0 | 0 | 0 | 0 |
| May12.75v_8791 | 1 | 0 | 0 | 0 | 1 | 0 | 0 | 0 | 0 | 0 | 0 | 0 | 0 | 0 | 0 | 0 | 0 | 0 | 0 | 0 | 0 | 0 | 0 | 0 | 0 | 0 | 0 | 0 | 0 | 0 |
| Sep8.0v_511 | 1 | 0 | 0 | 0 | 0 | 0 | 0 | 0 | 0 | 0 | 0 | 0 | 0 | 0 | 0 | 0 | 0 | 0 | 0 | 0 | 0 | 0 | 0 | 0 | 0 | 0 | 1 | 0 | 0 | 0 |
| Jun22.75v_36676 | 3 | 0 | 0 | 0 | 0 | 0 | 0 | 0 | 0 | 0 | 0 | 1 | 0 | 0 | 0 | 0 | 1 | 0 | 1 | 0 | 0 | 0 | 0 | 0 | 0 | 0 | 0 | 0 | 0 | 0 |
| Jun22.5v_2729 | 1 | 0 | 0 | 0 | 0 | 0 | 0 | 0 | 0 | 1 | 0 | 0 | 0 | 0 | 0 | 0 | 0 | 0 | 0 | 0 | 0 | 0 | 0 | 0 | 0 | 0 | 0 | 0 | 0 | 0 |
| Jul11.0v_14537 | 1 | 0 | 0 | 0 | 0 | 0 | 0 | 0 | 0 | 0 | 0 | 0 | 1 | 0 | 0 | 0 | 0 | 0 | 0 | 0 | 0 | 0 | 0 | 0 | 0 | 0 | 0 | 0 | 0 | 0 |
| Jul27.5v_9232 | 1 | 0 | 0 | 0 | 0 | 0 | 0 | 0 | 0 | 0 | 0 | 0 | 0 | 0 | 0 | 0 | 0 | 1 | 0 | 0 | 0 | 0 | 0 | 0 | 0 | 0 | 0 | 0 | 0 | 0 |
| Jul11.25v_19167 | 1 | 0 | 0 | 0 | 0 | 0 | 0 | 0 | 0 | 0 | 0 | 0 | 0 | 0 | 1 | 0 | 0 | 0 | 0 | 0 | 0 | 0 | 0 | 0 | 0 | 0 | 0 | 0 | 0 | 0 |
| May12.50v_21848 | 1 | 0 | 0 | 1 | 0 | 0 | 0 | 0 | 0 | 0 | 0 | 0 | 0 | 0 | 0 | 0 | 0 | 0 | 0 | 0 | 0 | 0 | 0 | 0 | 0 | 0 | 0 | 0 | 0 | 0 |
| Jul27.75v_19890 | 1 | 0 | 0 | 0 | 0 | 0 | 0 | 0 | 0 | 0 | 0 | 0 | 0 | 0 | 0 | 0 | 0 | 0 | 0 | 0 | 1 | 0 | 0 | 0 | 0 | 0 | 0 | 0 | 0 | 0 |
| Jul11.25v_2829 | 1 | 0 | 0 | 0 | 0 | 0 | 0 | 0 | 0 | 0 | 0 | 0 | 0 | 0 | 1 | 0 | 0 | 0 | 0 | 0 | 0 | 0 | 0 | 0 | 0 | 0 | 0 | 0 | 0 | 0 |
| Jun22.5v_20492 | 1 | 0 | 0 | 0 | 0 | 0 | 0 | 0 | 0 | 1 | 0 | 0 | 0 | 0 | 0 | 0 | 0 | 0 | 0 | 0 | 0 | 0 | 0 | 0 | 0 | 0 | 0 | 0 | 0 | 0 |
| Jun22.0v_3122 | 1 | 0 | 0 | 0 | 0 | 0 | 0 | 0 | 1 | 0 | 0 | 0 | 0 | 0 | 0 | 0 | 0 | 0 | 0 | 0 | 0 | 0 | 0 | 0 | 0 | 0 | 0 | 0 | 0 | 0 |
| Jul11.50v_10452 | 1 | 0 | 0 | 0 | 0 | 0 | 0 | 0 | 0 | 0 | 0 | 0 | 0 | 0 | 0 | 1 | 0 | 0 | 0 | 0 | 0 | 0 | 0 | 0 | 0 | 0 | 0 | 0 | 0 | 0 |
| Aug5.25v_24050 | 1 | 0 | 0 | 0 | 0 | 0 | 0 | 0 | 0 | 0 | 0 | 0 | 0 | 0 | 0 | 0 | 0 | 0 | 0 | 0 | 0 | 0 | 0 | 1 | 0 | 0 | 0 | 0 | 0 | 0 |
| Aug5.25v_21016 | 2 | 0 | 0 | 0 | 0 | 0 | 0 | 0 | 0 | 0 | 1 | 0 | 0 | 0 | 0 | 0 | 0 | 0 | 0 | 0 | 0 | 0 | 0 | 1 | 0 | 0 | 0 | 0 | 0 | 0 |
| Jul27.50v_38480 | 1 | 0 | 0 | 0 | 0 | 0 | 0 | 0 | 0 | 0 | 0 | 0 | 0 | 0 | 0 | 0 | 0 | 0 | 0 | 1 | 0 | 0 | 0 | 0 | 0 | 0 | 0 | 0 | 0 | 0 |
| May12.50v_13686 | 1 | 0 | 0 | 1 | 0 | 0 | 0 | 0 | 0 | 0 | 0 | 0 | 0 | 0 | 0 | 0 | 0 | 0 | 0 | 0 | 0 | 0 | 0 | 0 | 0 | 0 | 0 | 0 | 0 | 0 |
| May12.50v_29153 | 1 | 0 | 0 | 1 | 0 | 0 | 0 | 0 | 0 | 0 | 0 | 0 | 0 | 0 | 0 | 0 | 0 | 0 | 0 | 0 | 0 | 0 | 0 | 0 | 0 | 0 | 0 | 0 | 0 | 0 |
| Jul27.50v_33591 | 1 | 0 | 0 | 0 | 0 | 0 | 0 | 0 | 0 | 0 | 0 | 0 | 0 | 0 | 0 | 0 | 0 | 0 | 0 | 1 | 0 | 0 | 0 | 0 | 0 | 0 | 0 | 0 | 0 | 0 |
| Jul27.5v_16814 | 1 | 0 | 0 | 0 | 0 | 0 | 0 | 0 | 0 | 0 | 0 | 0 | 0 | 0 | 0 | 0 | 0 | 1 | 0 | 0 | 0 | 0 | 0 | 0 | 0 | 0 | 0 | 0 | 0 | 0 |
| May12.50v_16371 | 1 | 0 | 0 | 1 | 0 | 0 | 0 | 0 | 0 | 0 | 0 | 0 | 0 | 0 | 0 | 0 | 0 | 0 | 0 | 0 | 0 | 0 | 0 | 0 | 0 | 0 | 0 | 0 | 0 | 0 |
| May12.5v_897 | 1 | 1 | 0 | 0 | 0 | 0 | 0 | 0 | 0 | 0 | 0 | 0 | 0 | 0 | 0 | 0 | 0 | 0 | 0 | 0 | 0 | 0 | 0 | 0 | 0 | 0 | 0 | 0 | 0 | 0 |
| Aug5.0v_1806 | 1 | 0 | 0 | 0 | 0 | 0 | 0 | 0 | 0 | 0 | 0 | 0 | 0 | 0 | 0 | 0 | 0 | 0 | 0 | 0 | 0 | 1 | 0 | 0 | 0 | 0 | 0 | 0 | 0 | 0 |
| Jul11.0v_18936 | 1 | 0 | 0 | 0 | 0 | 0 | 0 | 0 | 0 | 0 | 0 | 0 | 1 | 0 | 0 | 0 | 0 | 0 | 0 | 0 | 0 | 0 | 0 | 0 | 0 | 0 | 0 | 0 | 0 | 0 |
| May12.5v_17691 | 1 | 1 | 0 | 0 | 0 | 0 | 0 | 0 | 0 | 0 | 0 | 0 | 0 | 0 | 0 | 0 | 0 | 0 | 0 | 0 | 0 | 0 | 0 | 0 | 0 | 0 | 0 | 0 | 0 | 0 |
| May12.25v_30482 | 1 | 0 | 1 | 0 | 0 | 0 | 0 | 0 | 0 | 0 | 0 | 0 | 0 | 0 | 0 | 0 | 0 | 0 | 0 | 0 | 0 | 0 | 0 | 0 | 0 | 0 | 0 | 0 | 0 | 0 |
| Jul11.75v_35862 | 1 | 0 | 0 | 0 | 0 | 0 | 0 | 0 | 0 | 0 | 0 | 0 | 0 | 0 | 0 | 0 | 1 | 0 | 0 | 0 | 0 | 0 | 0 | 0 | 0 | 0 | 0 | 0 | 0 | 0 |
| May12.5v_19922 | 1 | 1 | 0 | 0 | 0 | 0 | 0 | 0 | 0 | 0 | 0 | 0 | 0 | 0 | 0 | 0 | 0 | 0 | 0 | 0 | 0 | 0 | 0 | 0 | 0 | 0 | 0 | 0 | 0 | 0 |
| May12.75v_7338 | 1 | 0 | 0 | 0 | 1 | 0 | 0 | 0 | 0 | 0 | 0 | 0 | 0 | 0 | 0 | 0 | 0 | 0 | 0 | 0 | 0 | 0 | 0 | 0 | 0 | 0 | 0 | 0 | 0 | 0 |
| May12.75v_188 | 1 | 0 | 0 | 0 | 1 | 0 | 0 | 0 | 0 | 0 | 0 | 0 | 0 | 0 | 0 | 0 | 0 | 0 | 0 | 0 | 0 | 0 | 0 | 0 | 0 | 0 | 0 | 0 | 0 | 0 |
| May12.5v_33866 | 1 | 1 | 0 | 0 | 0 | 0 | 0 | 0 | 0 | 0 | 0 | 0 | 0 | 0 | 0 | 0 | 0 | 0 | 0 | 0 | 0 | 0 | 0 | 0 | 0 | 0 | 0 | 0 | 0 | 0 |
| Jul27.5v_18282 | 1 | 0 | 0 | 0 | 0 | 0 | 0 | 0 | 0 | 0 | 0 | 0 | 0 | 0 | 0 | 0 | 0 | 1 | 0 | 0 | 0 | 0 | 0 | 0 | 0 | 0 | 0 | 0 | 0 | 0 |
| Jul27.25v_32390 | 1 | 0 | 0 | 0 | 0 | 0 | 0 | 0 | 0 | 0 | 0 | 0 | 0 | 0 | 0 | 0 | 0 | 0 | 1 | 0 | 0 | 0 | 0 | 0 | 0 | 0 | 0 | 0 | 0 | 0 |
| Aug5.5v_14912 | 1 | 0 | 0 | 0 | 0 | 0 | 0 | 0 | 0 | 0 | 0 | 0 | 0 | 0 | 0 | 0 | 0 | 0 | 0 | 0 | 0 | 0 | 1 | 0 | 0 | 0 | 0 | 0 | 0 | 0 |
| Sep8.50v_4677 | 1 | 0 | 0 | 0 | 0 | 0 | 0 | 0 | 0 | 0 | 0 | 0 | 0 | 0 | 0 | 0 | 0 | 0 | 0 | 0 | 0 | 0 | 0 | 0 | 0 | 0 | 0 | 0 | 0 | 1 |
| Jul27.25v_13523 | 1 | 0 | 0 | 0 | 0 | 0 | 0 | 0 | 0 | 0 | 0 | 0 | 0 | 0 | 0 | 0 | 0 | 0 | 1 | 0 | 0 | 0 | 0 | 0 | 0 | 0 | 0 | 0 | 0 | 0 |
| Jul11.75v_37277 | 1 | 0 | 0 | 0 | 0 | 0 | 0 | 0 | 0 | 0 | 0 | 0 | 0 | 0 | 0 | 0 | 1 | 0 | 0 | 0 | 0 | 0 | 0 | 0 | 0 | 0 | 0 | 0 | 0 | 0 |
| May12.50v_35285 | 1 | 0 | 0 | 1 | 0 | 0 | 0 | 0 | 0 | 0 | 0 | 0 | 0 | 0 | 0 | 0 | 0 | 0 | 0 | 0 | 0 | 0 | 0 | 0 | 0 | 0 | 0 | 0 | 0 | 0 |
| Jul27.75v_7816 | 1 | 0 | 0 | 0 | 0 | 0 | 0 | 0 | 0 | 0 | 0 | 0 | 0 | 0 | 0 | 0 | 0 | 0 | 0 | 0 | 1 | 0 | 0 | 0 | 0 | 0 | 0 | 0 | 0 | 0 |
| Jul11.5v_13471 | 1 | 0 | 0 | 0 | 0 | 0 | 0 | 0 | 0 | 0 | 0 | 0 | 0 | 1 | 0 | 0 | 0 | 0 | 0 | 0 | 0 | 0 | 0 | 0 | 0 | 0 | 0 | 0 | 0 | 0 |
| May12.50v_22108 | 1 | 0 | 0 | 1 | 0 | 0 | 0 | 0 | 0 | 0 | 0 | 0 | 0 | 0 | 0 | 0 | 0 | 0 | 0 | 0 | 0 | 0 | 0 | 0 | 0 | 0 | 0 | 0 | 0 | 0 |
| Aug5.25v_8025 | 1 | 0 | 0 | 0 | 0 | 0 | 0 | 0 | 0 | 0 | 0 | 0 | 0 | 0 | 0 | 0 | 0 | 0 | 0 | 0 | 0 | 0 | 0 | 1 | 0 | 0 | 0 | 0 | 0 | 0 |
| Sep8.50v_984 | 1 | 0 | 0 | 0 | 0 | 0 | 0 | 0 | 0 | 0 | 0 | 0 | 0 | 0 | 0 | 0 | 0 | 0 | 0 | 0 | 0 | 0 | 0 | 0 | 0 | 0 | 0 | 0 | 0 | 1 |
| Jul11.5v_20375 | 1 | 0 | 0 | 0 | 0 | 0 | 0 | 0 | 0 | 0 | 0 | 0 | 0 | 1 | 0 | 0 | 0 | 0 | 0 | 0 | 0 | 0 | 0 | 0 | 0 | 0 | 0 | 0 | 0 | 0 |
| Jul11.5v_4005 | 1 | 0 | 0 | 0 | 0 | 0 | 0 | 0 | 0 | 0 | 0 | 0 | 0 | 1 | 0 | 0 | 0 | 0 | 0 | 0 | 0 | 0 | 0 | 0 | 0 | 0 | 0 | 0 | 0 | 0 |
| Sep8.50v_30300 | 1 | 0 | 0 | 0 | 0 | 0 | 0 | 0 | 0 | 0 | 0 | 0 | 0 | 0 | 0 | 0 | 0 | 0 | 0 | 0 | 0 | 0 | 0 | 0 | 0 | 0 | 0 | 0 | 0 | 1 |
| Aug5.25v_36829 | 1 | 0 | 0 | 0 | 0 | 0 | 0 | 0 | 0 | 0 | 0 | 0 | 0 | 0 | 0 | 0 | 0 | 0 | 0 | 0 | 0 | 0 | 0 | 1 | 0 | 0 | 0 | 0 | 0 | 0 |
| May12.5v_29738 | 1 | 1 | 0 | 0 | 0 | 0 | 0 | 0 | 0 | 0 | 0 | 0 | 0 | 0 | 0 | 0 | 0 | 0 | 0 | 0 | 0 | 0 | 0 | 0 | 0 | 0 | 0 | 0 | 0 | 0 |
| Jul11.25v_15024 | 1 | 0 | 0 | 0 | 0 | 0 | 0 | 0 | 0 | 0 | 0 | 0 | 0 | 0 | 1 | 0 | 0 | 0 | 0 | 0 | 0 | 0 | 0 | 0 | 0 | 0 | 0 | 0 | 0 | 0 |
| Jun22.0v_17652 | 1 | 0 | 0 | 0 | 0 | 0 | 0 | 0 | 1 | 0 | 0 | 0 | 0 | 0 | 0 | 0 | 0 | 0 | 0 | 0 | 0 | 0 | 0 | 0 | 0 | 0 | 0 | 0 | 0 | 0 |
| May12.50v_4276 | 1 | 0 | 0 | 1 | 0 | 0 | 0 | 0 | 0 | 0 | 0 | 0 | 0 | 0 | 0 | 0 | 0 | 0 | 0 | 0 | 0 | 0 | 0 | 0 | 0 | 0 | 0 | 0 | 0 | 0 |
| Aug5.0v_4551 | 1 | 0 | 0 | 0 | 0 | 0 | 0 | 0 | 0 | 0 | 0 | 0 | 0 | 0 | 0 | 0 | 0 | 0 | 0 | 0 | 0 | 1 | 0 | 0 | 0 | 0 | 0 | 0 | 0 | 0 |
| Jul11.25v_5581 | 5 | 0 | 0 | 0 | 0 | 0 | 0 | 0 | 0 | 0 | 0 | 0 | 1 | 0 | 1 | 0 | 0 | 1 | 0 | 0 | 0 | 0 | 1 | 1 | 0 | 0 | 0 | 0 | 0 | 0 |
| Sep8.50v_34557 | 1 | 0 | 0 | 0 | 0 | 0 | 0 | 0 | 0 | 0 | 0 | 0 | 0 | 0 | 0 | 0 | 0 | 0 | 0 | 0 | 0 | 0 | 0 | 0 | 0 | 0 | 0 | 0 | 0 | 1 |
| Jul11.50v_3968 | 1 | 0 | 0 | 0 | 0 | 0 | 0 | 0 | 0 | 0 | 0 | 0 | 0 | 0 | 0 | 1 | 0 | 0 | 0 | 0 | 0 | 0 | 0 | 0 | 0 | 0 | 0 | 0 | 0 | 0 |
| May12.50v_40959 | 1 | 0 | 0 | 1 | 0 | 0 | 0 | 0 | 0 | 0 | 0 | 0 | 0 | 0 | 0 | 0 | 0 | 0 | 0 | 0 | 0 | 0 | 0 | 0 | 0 | 0 | 0 | 0 | 0 | 0 |
| Aug5.0v_1519 | 1 | 0 | 0 | 0 | 0 | 0 | 0 | 0 | 0 | 0 | 0 | 0 | 0 | 0 | 0 | 0 | 0 | 0 | 0 | 0 | 0 | 1 | 0 | 0 | 0 | 0 | 0 | 0 | 0 | 0 |
| Jul27.5v_18638 | 1 | 0 | 0 | 0 | 0 | 0 | 0 | 0 | 0 | 0 | 0 | 0 | 0 | 0 | 0 | 0 | 0 | 1 | 0 | 0 | 0 | 0 | 0 | 0 | 0 | 0 | 0 | 0 | 0 | 0 |
| Jul27.5v_5273 | 1 | 0 | 0 | 0 | 0 | 0 | 0 | 0 | 0 | 0 | 0 | 0 | 0 | 0 | 0 | 0 | 0 | 1 | 0 | 0 | 0 | 0 | 0 | 0 | 0 | 0 | 0 | 0 | 0 | 0 |
| Sep8.0v_15130 | 1 | 0 | 0 | 0 | 0 | 0 | 0 | 0 | 0 | 0 | 0 | 0 | 0 | 0 | 0 | 0 | 0 | 0 | 0 | 0 | 0 | 0 | 0 | 0 | 0 | 0 | 1 | 0 | 0 | 0 |
| Jul11.0v_2254 | 1 | 0 | 0 | 0 | 0 | 0 | 0 | 0 | 0 | 0 | 0 | 0 | 1 | 0 | 0 | 0 | 0 | 0 | 0 | 0 | 0 | 0 | 0 | 0 | 0 | 0 | 0 | 0 | 0 | 0 |
| Jun22.50v_3598 | 1 | 0 | 0 | 0 | 0 | 0 | 0 | 0 | 0 | 0 | 1 | 0 | 0 | 0 | 0 | 0 | 0 | 0 | 0 | 0 | 0 | 0 | 0 | 0 | 0 | 0 | 0 | 0 | 0 | 0 |
| Aug5.25v_22660 | 1 | 0 | 0 | 0 | 0 | 0 | 0 | 0 | 0 | 0 | 0 | 0 | 0 | 0 | 0 | 0 | 0 | 0 | 0 | 0 | 0 | 0 | 0 | 1 | 0 | 0 | 0 | 0 | 0 | 0 |
| Jul11.0v_18950 | 1 | 0 | 0 | 0 | 0 | 0 | 0 | 0 | 0 | 0 | 0 | 0 | 1 | 0 | 0 | 0 | 0 | 0 | 0 | 0 | 0 | 0 | 0 | 0 | 0 | 0 | 0 | 0 | 0 | 0 |
| Jul11.0v_8773 | 1 | 0 | 0 | 0 | 0 | 0 | 0 | 0 | 0 | 0 | 0 | 0 | 1 | 0 | 0 | 0 | 0 | 0 | 0 | 0 | 0 | 0 | 0 | 0 | 0 | 0 | 0 | 0 | 0 | 0 |
| Aug5.0v_2782 | 1 | 0 | 0 | 0 | 0 | 0 | 0 | 0 | 0 | 0 | 0 | 0 | 0 | 0 | 0 | 0 | 0 | 0 | 0 | 0 | 0 | 1 | 0 | 0 | 0 | 0 | 0 | 0 | 0 | 0 |
| May12.50v_26459 | 1 | 0 | 0 | 1 | 0 | 0 | 0 | 0 | 0 | 0 | 0 | 0 | 0 | 0 | 0 | 0 | 0 | 0 | 0 | 0 | 0 | 0 | 0 | 0 | 0 | 0 | 0 | 0 | 0 | 0 |
| Jul11.0v_19050 | 1 | 0 | 0 | 0 | 0 | 0 | 0 | 0 | 0 | 0 | 0 | 0 | 1 | 0 | 0 | 0 | 0 | 0 | 0 | 0 | 0 | 0 | 0 | 0 | 0 | 0 | 0 | 0 | 0 | 0 |
| Jul11.25v_8482 | 1 | 0 | 0 | 0 | 0 | 0 | 0 | 0 | 0 | 0 | 0 | 0 | 0 | 0 | 1 | 0 | 0 | 0 | 0 | 0 | 0 | 0 | 0 | 0 | 0 | 0 | 0 | 0 | 0 | 0 |
| Aug5.25v_4744 | 1 | 0 | 0 | 0 | 0 | 0 | 0 | 0 | 0 | 0 | 0 | 0 | 0 | 0 | 0 | 0 | 0 | 0 | 0 | 0 | 0 | 0 | 0 | 1 | 0 | 0 | 0 | 0 | 0 | 0 |

| Feature_ID | 654 | May12_05m | May12_25m | May12_50m | May12_75m | Jun8_25m | Jun8_50m | Jun8_75m | Jun22_0m | Jun22_05m | Jun22_50m | Jun22_75m | Jul11_0m | Jul11_05m | Jul11_25m | Jul11_50m | Jul11_75m | Jul27_05m | Jul27_25m | Jul27_50m | Jul27_75m | Aug5_0m | Aug5_05m | Aug5_25m | Aug5_50m | Aug5_75m | Sep8_0m | Sep8_05m | Sep8_25m | Sep8_50m |
|---|---|---|---|---|---|---|---|---|---|---|---|---|---|---|---|---|---|---|---|---|---|---|---|---|---|---|---|---|---|---|
| Sep8.0v_4218 | 1 | 0 | 0 | 0 | 0 | 0 | 0 | 0 | 0 | 0 | 0 | 0 | 0 | 0 | 0 | 0 | 0 | 0 | 0 | 0 | 0 | 0 | 0 | 0 | 0 | 0 | 1 | 0 | 0 | 0 |
| Jul11.50v_5386 | 1 | 0 | 0 | 0 | 0 | 0 | 0 | 0 | 0 | 0 | 0 | 0 | 0 | 0 | 0 | 1 | 0 | 0 | 0 | 0 | 0 | 0 | 0 | 0 | 0 | 0 | 0 | 0 | 0 | 0 |
| Jun22.0v_7637 | 1 | 0 | 0 | 0 | 0 | 0 | 0 | 0 | 1 | 0 | 0 | 0 | 0 | 0 | 0 | 0 | 0 | 0 | 0 | 0 | 0 | 0 | 0 | 0 | 0 | 0 | 0 | 0 | 0 | 0 |
| Aug5.5v_3201 | 6 | 0 | 0 | 0 | 0 | 0 | 0 | 0 | 1 | 0 | 0 | 0 | 0 | 0 | 1 | 0 | 0 | 0 | 0 | 0 | 1 | 0 | 1 | 1 | 0 | 0 | 0 | 1 | 0 | 0 |
| Jul11.0v_14863 | 1 | 0 | 0 | 0 | 0 | 0 | 0 | 0 | 0 | 0 | 0 | 0 | 1 | 0 | 0 | 0 | 0 | 0 | 0 | 0 | 0 | 0 | 0 | 0 | 0 | 0 | 0 | 0 | 0 | 0 |
| Jul27.25v_35410 | 1 | 0 | 0 | 0 | 0 | 0 | 0 | 0 | 0 | 0 | 0 | 0 | 0 | 0 | 0 | 0 | 0 | 0 | 1 | 0 | 0 | 0 | 0 | 0 | 0 | 0 | 0 | 0 | 0 | 0 |
| Jul11.0v_20475 | 1 | 0 | 0 | 0 | 0 | 0 | 0 | 0 | 0 | 0 | 0 | 0 | 1 | 0 | 0 | 0 | 0 | 0 | 0 | 0 | 0 | 0 | 0 | 0 | 0 | 0 | 0 | 0 | 0 | 0 |
| Jul11.25v_11801 | 1 | 0 | 0 | 0 | 0 | 0 | 0 | 0 | 0 | 0 | 0 | 0 | 0 | 0 | 1 | 0 | 0 | 0 | 0 | 0 | 0 | 0 | 0 | 0 | 0 | 0 | 0 | 0 | 0 | 0 |
| Jul11.50v_10028 | 1 | 0 | 0 | 0 | 0 | 0 | 0 | 0 | 0 | 0 | 0 | 0 | 0 | 0 | 0 | 1 | 0 | 0 | 0 | 0 | 0 | 0 | 0 | 0 | 0 | 0 | 0 | 0 | 0 | 0 |
| Jul11.5v_2887 | 1 | 0 | 0 | 0 | 0 | 0 | 0 | 0 | 0 | 0 | 0 | 0 | 0 | 1 | 0 | 0 | 0 | 0 | 0 | 0 | 0 | 0 | 0 | 0 | 0 | 0 | 0 | 0 | 0 | 0 |
| May12.5v_25334 | 1 | 1 | 0 | 0 | 0 | 0 | 0 | 0 | 0 | 0 | 0 | 0 | 0 | 0 | 0 | 0 | 0 | 0 | 0 | 0 | 0 | 0 | 0 | 0 | 0 | 0 | 0 | 0 | 0 | 0 |
| Jul27.5v_4796 | 1 | 0 | 0 | 0 | 0 | 0 | 0 | 0 | 0 | 0 | 0 | 0 | 0 | 0 | 0 | 0 | 0 | 1 | 0 | 0 | 0 | 0 | 0 | 0 | 0 | 0 | 0 | 0 | 0 | 0 |
| Jun22.75v_19770 | 1 | 0 | 0 | 0 | 0 | 0 | 0 | 0 | 0 | 0 | 0 | 1 | 0 | 0 | 0 | 0 | 0 | 0 | 0 | 0 | 0 | 0 | 0 | 0 | 0 | 0 | 0 | 0 | 0 | 0 |
| Aug5.25v_16202 | 1 | 0 | 0 | 0 | 0 | 0 | 0 | 0 | 0 | 0 | 0 | 0 | 0 | 0 | 0 | 0 | 0 | 0 | 0 | 0 | 0 | 0 | 0 | 1 | 0 | 0 | 0 | 0 | 0 | 0 |
| Jun8.50v_38512 | 1 | 0 | 0 | 0 | 0 | 0 | 1 | 0 | 0 | 0 | 0 | 0 | 0 | 0 | 0 | 0 | 0 | 0 | 0 | 0 | 0 | 0 | 0 | 0 | 0 | 0 | 0 | 0 | 0 | 0 |
| Sep8.50v_34639 | 1 | 0 | 0 | 0 | 0 | 0 | 0 | 0 | 0 | 0 | 0 | 0 | 0 | 0 | 0 | 0 | 0 | 0 | 0 | 0 | 0 | 0 | 0 | 0 | 0 | 0 | 0 | 0 | 0 | 1 |
| Sep8.50v_37469 | 1 | 0 | 0 | 0 | 0 | 0 | 0 | 0 | 0 | 0 | 0 | 0 | 0 | 0 | 0 | 0 | 0 | 0 | 0 | 0 | 0 | 0 | 0 | 0 | 0 | 0 | 0 | 0 | 0 | 1 |
| Jul11.5v_6076 | 1 | 0 | 0 | 0 | 0 | 0 | 0 | 0 | 0 | 0 | 0 | 0 | 0 | 1 | 0 | 0 | 0 | 0 | 0 | 0 | 0 | 0 | 0 | 0 | 0 | 0 | 0 | 0 | 0 | 0 |
| May12.75v_11018 | 1 | 0 | 0 | 0 | 1 | 0 | 0 | 0 | 0 | 0 | 0 | 0 | 0 | 0 | 0 | 0 | 0 | 0 | 0 | 0 | 0 | 0 | 0 | 0 | 0 | 0 | 0 | 0 | 0 | 0 |
| Jul11.50v_7785 | 1 | 0 | 0 | 0 | 0 | 0 | 0 | 0 | 0 | 0 | 0 | 0 | 0 | 0 | 0 | 1 | 0 | 0 | 0 | 0 | 0 | 0 | 0 | 0 | 0 | 0 | 0 | 0 | 0 | 0 |
| Jun22.50v_40102 | 1 | 0 | 0 | 0 | 0 | 0 | 0 | 0 | 0 | 0 | 1 | 0 | 0 | 0 | 0 | 0 | 0 | 0 | 0 | 0 | 0 | 0 | 0 | 0 | 0 | 0 | 0 | 0 | 0 | 0 |
| Jul27.5v_21001 | 1 | 0 | 0 | 0 | 0 | 0 | 0 | 0 | 0 | 0 | 0 | 0 | 0 | 0 | 0 | 0 | 0 | 1 | 0 | 0 | 0 | 0 | 0 | 0 | 0 | 0 | 0 | 0 | 0 | 0 |
| Sep8.50v_9986 | 1 | 0 | 0 | 0 | 0 | 0 | 0 | 0 | 0 | 0 | 0 | 0 | 0 | 0 | 0 | 0 | 0 | 0 | 0 | 0 | 0 | 0 | 0 | 0 | 0 | 0 | 0 | 0 | 0 | 1 |
| Jun22.75v_24641 | 1 | 0 | 0 | 0 | 0 | 0 | 0 | 0 | 0 | 0 | 0 | 1 | 0 | 0 | 0 | 0 | 0 | 0 | 0 | 0 | 0 | 0 | 0 | 0 | 0 | 0 | 0 | 0 | 0 | 0 |
| Jun8.25v_12131 | 1 | 0 | 0 | 0 | 0 | 1 | 0 | 0 | 0 | 0 | 0 | 0 | 0 | 0 | 0 | 0 | 0 | 0 | 0 | 0 | 0 | 0 | 0 | 0 | 0 | 0 | 0 | 0 | 0 | 0 |
| May12.50v_12409 | 1 | 0 | 0 | 1 | 0 | 0 | 0 | 0 | 0 | 0 | 0 | 0 | 0 | 0 | 0 | 0 | 0 | 0 | 0 | 0 | 0 | 0 | 0 | 0 | 0 | 0 | 0 | 0 | 0 | 0 |
| Jul27.25v_14906 | 1 | 0 | 0 | 0 | 0 | 0 | 0 | 0 | 0 | 0 | 0 | 0 | 0 | 0 | 0 | 0 | 0 | 0 | 1 | 0 | 0 | 0 | 0 | 0 | 0 | 0 | 0 | 0 | 0 | 0 |
| Jun8.25v_39919 | 1 | 0 | 0 | 0 | 0 | 1 | 0 | 0 | 0 | 0 | 0 | 0 | 0 | 0 | 0 | 0 | 0 | 0 | 0 | 0 | 0 | 0 | 0 | 0 | 0 | 0 | 0 | 0 | 0 | 0 |
| May12.5v_35381 | 1 | 1 | 0 | 0 | 0 | 0 | 0 | 0 | 0 | 0 | 0 | 0 | 0 | 0 | 0 | 0 | 0 | 0 | 0 | 0 | 0 | 0 | 0 | 0 | 0 | 0 | 0 | 0 | 0 | 0 |
| Jun8.50v_34172 | 1 | 0 | 0 | 0 | 0 | 0 | 1 | 0 | 0 | 0 | 0 | 0 | 0 | 0 | 0 | 0 | 0 | 0 | 0 | 0 | 0 | 0 | 0 | 0 | 0 | 0 | 0 | 0 | 0 | 0 |
| Jun22.0v_5725 | 1 | 0 | 0 | 0 | 0 | 0 | 0 | 0 | 1 | 0 | 0 | 0 | 0 | 0 | 0 | 0 | 0 | 0 | 0 | 0 | 0 | 0 | 0 | 0 | 0 | 0 | 0 | 0 | 0 | 0 |
| Aug5.5v_37426 | 1 | 0 | 0 | 0 | 0 | 0 | 0 | 0 | 0 | 0 | 0 | 0 | 0 | 0 | 0 | 0 | 0 | 0 | 0 | 0 | 0 | 0 | 1 | 0 | 0 | 0 | 0 | 0 | 0 | 0 |
| Sep8.50v_27352 | 1 | 0 | 0 | 0 | 0 | 0 | 0 | 0 | 0 | 0 | 0 | 0 | 0 | 0 | 0 | 0 | 0 | 0 | 0 | 0 | 0 | 0 | 0 | 0 | 0 | 0 | 0 | 0 | 0 | 1 |
| Jul11.50v_3881 | 1 | 0 | 0 | 0 | 0 | 0 | 0 | 0 | 0 | 0 | 0 | 0 | 0 | 0 | 0 | 1 | 0 | 0 | 0 | 0 | 0 | 0 | 0 | 0 | 0 | 0 | 0 | 0 | 0 | 0 |
| Jun22.0v_19850 | 1 | 0 | 0 | 0 | 0 | 0 | 0 | 0 | 1 | 0 | 0 | 0 | 0 | 0 | 0 | 0 | 0 | 0 | 0 | 0 | 0 | 0 | 0 | 0 | 0 | 0 | 0 | 0 | 0 | 0 |
| Jun22.75v_4521 | 1 | 0 | 0 | 0 | 0 | 0 | 0 | 0 | 0 | 0 | 0 | 1 | 0 | 0 | 0 | 0 | 0 | 0 | 0 | 0 | 0 | 0 | 0 | 0 | 0 | 0 | 0 | 0 | 0 | 0 |
| May12.75v_5902 | 1 | 0 | 0 | 0 | 1 | 0 | 0 | 0 | 0 | 0 | 0 | 0 | 0 | 0 | 0 | 0 | 0 | 0 | 0 | 0 | 0 | 0 | 0 | 0 | 0 | 0 | 0 | 0 | 0 | 0 |
| Aug5.0v_2119 | 1 | 0 | 0 | 0 | 0 | 0 | 0 | 0 | 0 | 0 | 0 | 0 | 0 | 0 | 0 | 0 | 0 | 0 | 0 | 0 | 0 | 1 | 0 | 0 | 0 | 0 | 0 | 0 | 0 | 0 |
| May12.50v_31777 | 1 | 0 | 0 | 1 | 0 | 0 | 0 | 0 | 0 | 0 | 0 | 0 | 0 | 0 | 0 | 0 | 0 | 0 | 0 | 0 | 0 | 0 | 0 | 0 | 0 | 0 | 0 | 0 | 0 | 0 |
| Aug5.5v_92 | 1 | 0 | 0 | 0 | 0 | 0 | 0 | 0 | 0 | 0 | 0 | 0 | 0 | 0 | 0 | 0 | 0 | 0 | 0 | 0 | 0 | 0 | 1 | 0 | 0 | 0 | 0 | 0 | 0 | 0 |
| Aug5.5v_3819 | 1 | 0 | 0 | 0 | 0 | 0 | 0 | 0 | 0 | 0 | 0 | 0 | 0 | 0 | 0 | 0 | 0 | 0 | 0 | 0 | 0 | 0 | 1 | 0 | 0 | 0 | 0 | 0 | 0 | 0 |
| Aug5.0v_4562 | 1 | 0 | 0 | 0 | 0 | 0 | 0 | 0 | 0 | 0 | 0 | 0 | 0 | 0 | 0 | 0 | 0 | 0 | 0 | 0 | 0 | 1 | 0 | 0 | 0 | 0 | 0 | 0 | 0 | 0 |
| Aug5.0v_4310 | 1 | 0 | 0 | 0 | 0 | 0 | 0 | 0 | 0 | 0 | 0 | 0 | 0 | 0 | 0 | 0 | 0 | 0 | 0 | 0 | 0 | 1 | 0 | 0 | 0 | 0 | 0 | 0 | 0 | 0 |
| Jul27.5v_2477 | 1 | 0 | 0 | 0 | 0 | 0 | 0 | 0 | 0 | 0 | 0 | 0 | 0 | 0 | 0 | 0 | 0 | 1 | 0 | 0 | 0 | 0 | 0 | 0 | 0 | 0 | 0 | 0 | 0 | 0 |
| Aug5.25v_24171 | 1 | 0 | 0 | 0 | 0 | 0 | 0 | 0 | 0 | 0 | 0 | 0 | 0 | 0 | 0 | 0 | 0 | 0 | 0 | 0 | 0 | 0 | 0 | 1 | 0 | 0 | 0 | 0 | 0 | 0 |
| Aug5.5v_26095 | 1 | 0 | 0 | 0 | 0 | 0 | 0 | 0 | 0 | 0 | 0 | 0 | 0 | 0 | 0 | 0 | 0 | 0 | 0 | 0 | 0 | 0 | 1 | 0 | 0 | 0 | 0 | 0 | 0 | 0 |
| Jun22.50v_19017 | 1 | 0 | 0 | 0 | 0 | 0 | 0 | 0 | 0 | 0 | 1 | 0 | 0 | 0 | 0 | 0 | 0 | 0 | 0 | 0 | 0 | 0 | 0 | 0 | 0 | 0 | 0 | 0 | 0 | 0 |
| May12.5v_5981 | 1 | 1 | 0 | 0 | 0 | 0 | 0 | 0 | 0 | 0 | 0 | 0 | 0 | 0 | 0 | 0 | 0 | 0 | 0 | 0 | 0 | 0 | 0 | 0 | 0 | 0 | 0 | 0 | 0 | 0 |
| Aug5.25v_33505 | 1 | 0 | 0 | 0 | 0 | 0 | 0 | 0 | 0 | 0 | 0 | 0 | 0 | 0 | 0 | 0 | 0 | 0 | 0 | 0 | 0 | 0 | 0 | 1 | 0 | 0 | 0 | 0 | 0 | 0 |
| May12.50v_28989 | 1 | 0 | 0 | 1 | 0 | 0 | 0 | 0 | 0 | 0 | 0 | 0 | 0 | 0 | 0 | 0 | 0 | 0 | 0 | 0 | 0 | 0 | 0 | 0 | 0 | 0 | 0 | 0 | 0 | 0 |
| May12.75v_22155 | 1 | 0 | 0 | 0 | 1 | 0 | 0 | 0 | 0 | 0 | 0 | 0 | 0 | 0 | 0 | 0 | 0 | 0 | 0 | 0 | 0 | 0 | 0 | 0 | 0 | 0 | 0 | 0 | 0 | 0 |
| Aug5.25v_21975 | 1 | 0 | 0 | 0 | 0 | 0 | 0 | 0 | 0 | 0 | 0 | 0 | 0 | 0 | 0 | 0 | 0 | 0 | 0 | 0 | 0 | 0 | 0 | 1 | 0 | 0 | 0 | 0 | 0 | 0 |
| Jul27.5v_19539 | 1 | 0 | 0 | 0 | 0 | 0 | 0 | 0 | 0 | 0 | 0 | 0 | 0 | 0 | 0 | 0 | 0 | 1 | 0 | 0 | 0 | 0 | 0 | 0 | 0 | 0 | 0 | 0 | 0 | 0 |
| May12.5v_23524 | 1 | 1 | 0 | 0 | 0 | 0 | 0 | 0 | 0 | 0 | 0 | 0 | 0 | 0 | 0 | 0 | 0 | 0 | 0 | 0 | 0 | 0 | 0 | 0 | 0 | 0 | 0 | 0 | 0 | 0 |
| Sep8.50v_1272 | 1 | 0 | 0 | 0 | 0 | 0 | 0 | 0 | 0 | 0 | 0 | 0 | 0 | 0 | 0 | 0 | 0 | 0 | 0 | 0 | 0 | 0 | 0 | 0 | 0 | 0 | 0 | 0 | 0 | 1 |
| Jul11.0v_15398 | 1 | 0 | 0 | 0 | 0 | 0 | 0 | 0 | 0 | 0 | 0 | 0 | 1 | 0 | 0 | 0 | 0 | 0 | 0 | 0 | 0 | 0 | 0 | 0 | 0 | 0 | 0 | 0 | 0 | 0 |
| Jun22.0v_17705 | 1 | 0 | 0 | 0 | 0 | 0 | 0 | 0 | 1 | 0 | 0 | 0 | 0 | 0 | 0 | 0 | 0 | 0 | 0 | 0 | 0 | 0 | 0 | 0 | 0 | 0 | 0 | 0 | 0 | 0 |
| Jul27.25v_23752 | 1 | 0 | 0 | 0 | 0 | 0 | 0 | 0 | 0 | 0 | 0 | 0 | 0 | 0 | 0 | 0 | 0 | 0 | 1 | 0 | 0 | 0 | 0 | 0 | 0 | 0 | 0 | 0 | 0 | 0 |
| May12.50v_34294 | 1 | 0 | 0 | 1 | 0 | 0 | 0 | 0 | 0 | 0 | 0 | 0 | 0 | 0 | 0 | 0 | 0 | 0 | 0 | 0 | 0 | 0 | 0 | 0 | 0 | 0 | 0 | 0 | 0 | 0 |
| May12.50v_9460 | 1 | 0 | 0 | 1 | 0 | 0 | 0 | 0 | 0 | 0 | 0 | 0 | 0 | 0 | 0 | 0 | 0 | 0 | 0 | 0 | 0 | 0 | 0 | 0 | 0 | 0 | 0 | 0 | 0 | 0 |
| Aug5.0v_1535 | 2 | 0 | 0 | 1 | 0 | 0 | 0 | 0 | 0 | 0 | 0 | 0 | 0 | 0 | 0 | 0 | 0 | 0 | 0 | 0 | 0 | 1 | 0 | 0 | 0 | 0 | 0 | 0 | 0 | 0 |
| Sep8.0v_20558 | 1 | 0 | 0 | 0 | 0 | 0 | 0 | 0 | 0 | 0 | 0 | 0 | 0 | 0 | 0 | 0 | 0 | 0 | 0 | 0 | 0 | 0 | 0 | 0 | 0 | 0 | 1 | 0 | 0 | 0 |
| Jul27.50v_16825 | 1 | 0 | 0 | 0 | 0 | 0 | 0 | 0 | 0 | 0 | 0 | 0 | 0 | 0 | 0 | 0 | 0 | 0 | 0 | 1 | 0 | 0 | 0 | 0 | 0 | 0 | 0 | 0 | 0 | 0 |
| Jul11.50v_6262 | 1 | 0 | 0 | 0 | 0 | 0 | 0 | 0 | 0 | 0 | 0 | 0 | 0 | 0 | 0 | 1 | 0 | 0 | 0 | 0 | 0 | 0 | 0 | 0 | 0 | 0 | 0 | 0 | 0 | 0 |
| Jul11.5v_21084 | 1 | 0 | 0 | 0 | 0 | 0 | 0 | 0 | 0 | 0 | 0 | 0 | 0 | 1 | 0 | 0 | 0 | 0 | 0 | 0 | 0 | 0 | 0 | 0 | 0 | 0 | 0 | 0 | 0 | 0 |
| Aug5.0v_961 | 10 | 0 | 0 | 0 | 0 | 1 | 0 | 0 | 0 | 1 | 0 | 0 | 0 | 1 | 0 | 1 | 1 | 1 | 0 | 0 | 0 | 1 | 1 | 1 | 0 | 0 | 0 | 0 | 0 | 1 |
| May12.5v_13498 | 1 | 1 | 0 | 0 | 0 | 0 | 0 | 0 | 0 | 0 | 0 | 0 | 0 | 0 | 0 | 0 | 0 | 0 | 0 | 0 | 0 | 0 | 0 | 0 | 0 | 0 | 0 | 0 | 0 | 0 |
| Jul11.0v_2933 | 1 | 0 | 0 | 0 | 0 | 0 | 0 | 0 | 0 | 0 | 0 | 0 | 1 | 0 | 0 | 0 | 0 | 0 | 0 | 0 | 0 | 0 | 0 | 0 | 0 | 0 | 0 | 0 | 0 | 0 |
| Jul11.25v_10183 | 1 | 0 | 0 | 0 | 0 | 0 | 0 | 0 | 0 | 0 | 0 | 0 | 0 | 0 | 1 | 0 | 0 | 0 | 0 | 0 | 0 | 0 | 0 | 0 | 0 | 0 | 0 | 0 | 0 | 0 |
| May12.5v_31277 | 1 | 1 | 0 | 0 | 0 | 0 | 0 | 0 | 0 | 0 | 0 | 0 | 0 | 0 | 0 | 0 | 0 | 0 | 0 | 0 | 0 | 0 | 0 | 0 | 0 | 0 | 0 | 0 | 0 | 0 |
| Jul27.50v_6232 | 1 | 0 | 0 | 0 | 0 | 0 | 0 | 0 | 0 | 0 | 0 | 0 | 0 | 0 | 0 | 0 | 0 | 0 | 0 | 1 | 0 | 0 | 0 | 0 | 0 | 0 | 0 | 0 | 0 | 0 |
| Jun8.25v_19347 | 1 | 0 | 0 | 0 | 0 | 1 | 0 | 0 | 0 | 0 | 0 | 0 | 0 | 0 | 0 | 0 | 0 | 0 | 0 | 0 | 0 | 0 | 0 | 0 | 0 | 0 | 0 | 0 | 0 | 0 |
| Jul11.0v_14118 | 1 | 0 | 0 | 0 | 0 | 0 | 0 | 0 | 0 | 0 | 0 | 0 | 1 | 0 | 0 | 0 | 0 | 0 | 0 | 0 | 0 | 0 | 0 | 0 | 0 | 0 | 0 | 0 | 0 | 0 |
| Sep8.50v_20804 | 1 | 0 | 0 | 0 | 0 | 0 | 0 | 0 | 0 | 0 | 0 | 0 | 0 | 0 | 0 | 0 | 0 | 0 | 0 | 0 | 0 | 0 | 0 | 0 | 0 | 0 | 0 | 0 | 0 | 1 |
| May12.50v_37033 | 1 | 0 | 0 | 1 | 0 | 0 | 0 | 0 | 0 | 0 | 0 | 0 | 0 | 0 | 0 | 0 | 0 | 0 | 0 | 0 | 0 | 0 | 0 | 0 | 0 | 0 | 0 | 0 | 0 | 0 |
| Jun22.0v_7901 | 1 | 0 | 0 | 0 | 0 | 0 | 0 | 0 | 1 | 0 | 0 | 0 | 0 | 0 | 0 | 0 | 0 | 0 | 0 | 0 | 0 | 0 | 0 | 0 | 0 | 0 | 0 | 0 | 0 | 0 |
| Jun22.75v_13780 | 1 | 0 | 0 | 0 | 0 | 0 | 0 | 0 | 0 | 0 | 0 | 1 | 0 | 0 | 0 | 0 | 0 | 0 | 0 | 0 | 0 | 0 | 0 | 0 | 0 | 0 | 0 | 0 | 0 | 0 |
| Jul27.25v_13805 | 1 | 0 | 0 | 0 | 0 | 0 | 0 | 0 | 0 | 0 | 0 | 0 | 0 | 0 | 0 | 0 | 0 | 0 | 1 | 0 | 0 | 0 | 0 | 0 | 0 | 0 | 0 | 0 | 0 | 0 |
| Aug5.75v_6864 | 1 | 0 | 0 | 0 | 0 | 0 | 0 | 0 | 0 | 0 | 0 | 0 | 0 | 0 | 0 | 0 | 0 | 0 | 0 | 0 | 0 | 0 | 0 | 0 | 0 | 1 | 0 | 0 | 0 | 0 |
| Jul11.50v_2488 | 1 | 0 | 0 | 0 | 0 | 0 | 0 | 0 | 0 | 0 | 0 | 0 | 0 | 0 | 0 | 1 | 0 | 0 | 0 | 0 | 0 | 0 | 0 | 0 | 0 | 0 | 0 | 0 | 0 | 0 |
| May12.25v_23638 | 1 | 0 | 1 | 0 | 0 | 0 | 0 | 0 | 0 | 0 | 0 | 0 | 0 | 0 | 0 | 0 | 0 | 0 | 0 | 0 | 0 | 0 | 0 | 0 | 0 | 0 | 0 | 0 | 0 | 0 |
| Jul11.25v_18977 | 1 | 0 | 0 | 0 | 0 | 0 | 0 | 0 | 0 | 0 | 0 | 0 | 0 | 0 | 1 | 0 | 0 | 0 | 0 | 0 | 0 | 0 | 0 | 0 | 0 | 0 | 0 | 0 | 0 | 0 |
| May12.75v_8277 | 1 | 0 | 0 | 0 | 1 | 0 | 0 | 0 | 0 | 0 | 0 | 0 | 0 | 0 | 0 | 0 | 0 | 0 | 0 | 0 | 0 | 0 | 0 | 0 | 0 | 0 | 0 | 0 | 0 | 0 |
| Jul27.5v_13326 | 1 | 0 | 0 | 0 | 0 | 0 | 0 | 0 | 0 | 0 | 0 | 0 | 0 | 0 | 0 | 0 | 0 | 1 | 0 | 0 | 0 | 0 | 0 | 0 | 0 | 0 | 0 | 0 | 0 | 0 |
| Aug5.75v_33594 | 1 | 0 | 0 | 0 | 0 | 0 | 0 | 0 | 0 | 0 | 0 | 0 | 0 | 0 | 0 | 0 | 0 | 0 | 0 | 0 | 0 | 0 | 0 | 0 | 0 | 1 | 0 | 0 | 0 | 0 |
| Jul11.5v_14395 | 1 | 0 | 0 | 0 | 0 | 0 | 0 | 0 | 0 | 0 | 0 | 0 | 0 | 1 | 0 | 0 | 0 | 0 | 0 | 0 | 0 | 0 | 0 | 0 | 0 | 0 | 0 | 0 | 0 | 0 |
| Sep8.50v_10872 | 1 | 0 | 0 | 0 | 0 | 0 | 0 | 0 | 0 | 0 | 0 | 0 | 0 | 0 | 0 | 0 | 0 | 0 | 0 | 0 | 0 | 0 | 0 | 0 | 0 | 0 | 0 | 0 | 0 | 1 |
| Jul27.25v_3198 | 1 | 0 | 0 | 0 | 0 | 0 | 0 | 0 | 0 | 0 | 0 | 0 | 0 | 0 | 0 | 0 | 0 | 0 | 1 | 0 | 0 | 0 | 0 | 0 | 0 | 0 | 0 | 0 | 0 | 0 |
| Aug5.25v_19561 | 1 | 0 | 0 | 0 | 0 | 0 | 0 | 0 | 0 | 0 | 0 | 0 | 0 | 0 | 0 | 0 | 0 | 0 | 0 | 0 | 0 | 0 | 0 | 1 | 0 | 0 | 0 | 0 | 0 | 0 |
| Jul11.5v_7961 | 1 | 0 | 0 | 0 | 0 | 0 | 0 | 0 | 0 | 0 | 0 | 0 | 0 | 1 | 0 | 0 | 0 | 0 | 0 | 0 | 0 | 0 | 0 | 0 | 0 | 0 | 0 | 0 | 0 | 0 |
| Sep8.50v_31633 | 1 | 0 | 0 | 0 | 0 | 0 | 0 | 0 | 0 | 0 | 0 | 0 | 0 | 0 | 0 | 0 | 0 | 0 | 0 | 0 | 0 | 0 | 0 | 0 | 0 | 0 | 0 | 0 | 0 | 1 |
| Jul27.5v_1937 | 1 | 0 | 0 | 0 | 0 | 0 | 0 | 0 | 0 | 0 | 0 | 0 | 0 | 0 | 0 | 0 | 0 | 1 | 0 | 0 | 0 | 0 | 0 | 0 | 0 | 0 | 0 | 0 | 0 | 0 |
| Jul27.5v_13352 | 1 | 0 | 0 | 0 | 0 | 0 | 0 | 0 | 0 | 0 | 0 | 0 | 0 | 0 | 0 | 0 | 0 | 1 | 0 | 0 | 0 | 0 | 0 | 0 | 0 | 0 | 0 | 0 | 0 | 0 |
| Jul11.5v_2015 | 1 | 0 | 0 | 0 | 0 | 0 | 0 | 0 | 0 | 0 | 0 | 0 | 0 | 1 | 0 | 0 | 0 | 0 | 0 | 0 | 0 | 0 | 0 | 0 | 0 | 0 | 0 | 0 | 0 | 0 |
| Jun22.0v_17184 | 1 | 0 | 0 | 0 | 0 | 0 | 0 | 0 | 1 | 0 | 0 | 0 | 0 | 0 | 0 | 0 | 0 | 0 | 0 | 0 | 0 | 0 | 0 | 0 | 0 | 0 | 0 | 0 | 0 | 0 |
| Jul11.0v_4374 | 1 | 0 | 0 | 0 | 0 | 0 | 0 | 0 | 0 | 0 | 0 | 0 | 1 | 0 | 0 | 0 | 0 | 0 | 0 | 0 | 0 | 0 | 0 | 0 | 0 | 0 | 0 | 0 | 0 | 0 |
| May12.50v_26716 | 1 | 0 | 0 | 1 | 0 | 0 | 0 | 0 | 0 | 0 | 0 | 0 | 0 | 0 | 0 | 0 | 0 | 0 | 0 | 0 | 0 | 0 | 0 | 0 | 0 | 0 | 0 | 0 | 0 | 0 |
| Jul27.50v_30796 | 1 | 0 | 0 | 0 | 0 | 0 | 0 | 0 | 0 | 0 | 0 | 0 | 0 | 0 | 0 | 0 | 0 | 0 | 0 | 1 | 0 | 0 | 0 | 0 | 0 | 0 | 0 | 0 | 0 | 0 |
| Aug5.50v_38541 | 1 | 0 | 0 | 0 | 0 | 0 | 0 | 0 | 0 | 0 | 0 | 0 | 0 | 0 | 0 | 0 | 0 | 0 | 0 | 0 | 0 | 0 | 0 | 0 | 1 | 0 | 0 | 0 | 0 | 0 |
| Jun22.5v_6802 | 1 | 0 | 0 | 0 | 0 | 0 | 0 | 0 | 0 | 1 | 0 | 0 | 0 | 0 | 0 | 0 | 0 | 0 | 0 | 0 | 0 | 0 | 0 | 0 | 0 | 0 | 0 | 0 | 0 | 0 |
| May12.50v_22872 | 1 | 0 | 0 | 1 | 0 | 0 | 0 | 0 | 0 | 0 | 0 | 0 | 0 | 0 | 0 | 0 | 0 | 0 | 0 | 0 | 0 | 0 | 0 | 0 | 0 | 0 | 0 | 0 | 0 | 0 |
| Jul11.75v_4915 | 1 | 0 | 0 | 0 | 0 | 0 | 0 | 0 | 0 | 0 | 0 | 0 | 0 | 0 | 0 | 0 | 1 | 0 | 0 | 0 | 0 | 0 | 0 | 0 | 0 | 0 | 0 | 0 | 0 | 0 |
| Jun22.75v_13115 | 1 | 0 | 0 | 0 | 0 | 0 | 0 | 0 | 0 | 0 | 0 | 1 | 0 | 0 | 0 | 0 | 0 | 0 | 0 | 0 | 0 | 0 | 0 | 0 | 0 | 0 | 0 | 0 | 0 | 0 |
| Aug5.0v_3535 | 1 | 0 | 0 | 0 | 0 | 0 | 0 | 0 | 0 | 0 | 0 | 0 | 0 | 0 | 0 | 0 | 0 | 0 | 0 | 0 | 0 | 1 | 0 | 0 | 0 | 0 | 0 | 0 | 0 | 0 |
| Aug5.5v_29326 | 1 | 0 | 0 | 0 | 0 | 0 | 0 | 0 | 0 | 0 | 0 | 0 | 0 | 0 | 0 | 0 | 0 | 0 | 0 | 0 | 0 | 0 | 1 | 0 | 0 | 0 | 0 | 0 | 0 | 0 |
| May12.50v_38650 | 1 | 0 | 0 | 1 | 0 | 0 | 0 | 0 | 0 | 0 | 0 | 0 | 0 | 0 | 0 | 0 | 0 | 0 | 0 | 0 | 0 | 0 | 0 | 0 | 0 | 0 | 0 | 0 | 0 | 0 |
| Jun22.50v_16177 | 1 | 0 | 0 | 0 | 0 | 0 | 0 | 0 | 0 | 0 | 1 | 0 | 0 | 0 | 0 | 0 | 0 | 0 | 0 | 0 | 0 | 0 | 0 | 0 | 0 | 0 | 0 | 0 | 0 | 0 |
| Jul27.25v_25398 | 1 | 0 | 0 | 0 | 0 | 0 | 0 | 0 | 0 | 0 | 0 | 0 | 0 | 0 | 0 | 0 | 0 | 0 | 1 | 0 | 0 | 0 | 0 | 0 | 0 | 0 | 0 | 0 | 0 | 0 |
| Aug5.25v_552 | 1 | 0 | 0 | 0 | 0 | 0 | 0 | 0 | 0 | 0 | 0 | 0 | 0 | 0 | 0 | 0 | 0 | 0 | 0 | 0 | 0 | 0 | 0 | 1 | 0 | 0 | 0 | 0 | 0 | 0 |
| Jul11.0v_2963 | 1 | 0 | 0 | 0 | 0 | 0 | 0 | 0 | 0 | 0 | 0 | 0 | 1 | 0 | 0 | 0 | 0 | 0 | 0 | 0 | 0 | 0 | 0 | 0 | 0 | 0 | 0 | 0 | 0 | 0 |
| Jun8.25v_3003 | 1 | 0 | 0 | 0 | 0 | 1 | 0 | 0 | 0 | 0 | 0 | 0 | 0 | 0 | 0 | 0 | 0 | 0 | 0 | 0 | 0 | 0 | 0 | 0 | 0 | 0 | 0 | 0 | 0 | 0 |
| Jul11.5v_14859 | 1 | 0 | 0 | 0 | 0 | 0 | 0 | 0 | 0 | 0 | 0 | 0 | 0 | 1 | 0 | 0 | 0 | 0 | 0 | 0 | 0 | 0 | 0 | 0 | 0 | 0 | 0 | 0 | 0 | 0 |
| Aug5.5v_23129 | 1 | 0 | 0 | 0 | 0 | 0 | 0 | 0 | 0 | 0 | 0 | 0 | 0 | 0 | 0 | 0 | 0 | 0 | 0 | 0 | 0 | 0 | 1 | 0 | 0 | 0 | 0 | 0 | 0 | 0 |
| Jul27.25v_22688 | 1 | 0 | 0 | 0 | 0 | 0 | 0 | 0 | 0 | 0 | 0 | 0 | 0 | 0 | 0 | 0 | 0 | 0 | 1 | 0 | 0 | 0 | 0 | 0 | 0 | 0 | 0 | 0 | 0 | 0 |
| Jul11.75v_40291 | 1 | 0 | 0 | 0 | 0 | 0 | 0 | 0 | 0 | 0 | 0 | 0 | 0 | 0 | 0 | 0 | 1 | 0 | 0 | 0 | 0 | 0 | 0 | 0 | 0 | 0 | 0 | 0 | 0 | 0 |
| Jul11.5v_9420 | 1 | 0 | 0 | 0 | 0 | 0 | 0 | 0 | 0 | 0 | 0 | 0 | 0 | 1 | 0 | 0 | 0 | 0 | 0 | 0 | 0 | 0 | 0 | 0 | 0 | 0 | 0 | 0 | 0 | 0 |
| Jul11.50v_11036 | 1 | 0 | 0 | 0 | 0 | 0 | 0 | 0 | 0 | 0 | 0 | 0 | 0 | 0 | 0 | 1 | 0 | 0 | 0 | 0 | 0 | 0 | 0 | 0 | 0 | 0 | 0 | 0 | 0 | 0 |
| Jul11.0v_18060 | 1 | 0 | 0 | 0 | 0 | 0 | 0 | 0 | 0 | 0 | 0 | 0 | 1 | 0 | 0 | 0 | 0 | 0 | 0 | 0 | 0 | 0 | 0 | 0 | 0 | 0 | 0 | 0 | 0 | 0 |
| Jun22.75v_2016 | 1 | 0 | 0 | 0 | 0 | 0 | 0 | 0 | 0 | 0 | 0 | 1 | 0 | 0 | 0 | 0 | 0 | 0 | 0 | 0 | 0 | 0 | 0 | 0 | 0 | 0 | 0 | 0 | 0 | 0 |
| Sep8.0v_4467 | 1 | 0 | 0 | 0 | 0 | 0 | 0 | 0 | 0 | 0 | 0 | 0 | 0 | 0 | 0 | 0 | 0 | 0 | 0 | 0 | 0 | 0 | 0 | 0 | 0 | 0 | 1 | 0 | 0 | 0 |
| Jul11.5v_15683 | 1 | 0 | 0 | 0 | 0 | 0 | 0 | 0 | 0 | 0 | 0 | 0 | 0 | 1 | 0 | 0 | 0 | 0 | 0 | 0 | 0 | 0 | 0 | 0 | 0 | 0 | 0 | 0 | 0 | 0 |
| Jun22.0v_14446 | 1 | 0 | 0 | 0 | 0 | 0 | 0 | 0 | 1 | 0 | 0 | 0 | 0 | 0 | 0 | 0 | 0 | 0 | 0 | 0 | 0 | 0 | 0 | 0 | 0 | 0 | 0 | 0 | 0 | 0 |
| May12.5v_35952 | 1 | 1 | 0 | 0 | 0 | 0 | 0 | 0 | 0 | 0 | 0 | 0 | 0 | 0 | 0 | 0 | 0 | 0 | 0 | 0 | 0 | 0 | 0 | 0 | 0 | 0 | 0 | 0 | 0 | 0 |
| Jul11.25v_12881 | 1 | 0 | 0 | 0 | 0 | 0 | 0 | 0 | 0 | 0 | 0 | 0 | 0 | 0 | 1 | 0 | 0 | 0 | 0 | 0 | 0 | 0 | 0 | 0 | 0 | 0 | 0 | 0 | 0 | 0 |
| Jun8.50v_9706 | 1 | 0 | 0 | 0 | 0 | 0 | 1 | 0 | 0 | 0 | 0 | 0 | 0 | 0 | 0 | 0 | 0 | 0 | 0 | 0 | 0 | 0 | 0 | 0 | 0 | 0 | 0 | 0 | 0 | 0 |
| Jun22.50v_9863 | 1 | 0 | 0 | 0 | 0 | 0 | 0 | 0 | 0 | 0 | 1 | 0 | 0 | 0 | 0 | 0 | 0 | 0 | 0 | 0 | 0 | 0 | 0 | 0 | 0 | 0 | 0 | 0 | 0 | 0 |
| Jun22.0v_4319 | 1 | 0 | 0 | 0 | 0 | 0 | 0 | 0 | 1 | 0 | 0 | 0 | 0 | 0 | 0 | 0 | 0 | 0 | 0 | 0 | 0 | 0 | 0 | 0 | 0 | 0 | 0 | 0 | 0 | 0 |
| Jul27.5v_43 | 1 | 0 | 0 | 0 | 0 | 0 | 0 | 0 | 0 | 0 | 0 | 0 | 0 | 0 | 0 | 0 | 0 | 1 | 0 | 0 | 0 | 0 | 0 | 0 | 0 | 0 | 0 | 0 | 0 | 0 |
| Sep8.0v_20692 | 1 | 0 | 0 | 0 | 0 | 0 | 0 | 0 | 0 | 0 | 0 | 0 | 0 | 0 | 0 | 0 | 0 | 0 | 0 | 0 | 0 | 0 | 0 | 0 | 0 | 0 | 1 | 0 | 0 | 0 |
| Jun22.0v_11812 | 1 | 0 | 0 | 0 | 0 | 0 | 0 | 0 | 1 | 0 | 0 | 0 | 0 | 0 | 0 | 0 | 0 | 0 | 0 | 0 | 0 | 0 | 0 | 0 | 0 | 0 | 0 | 0 | 0 | 0 |
| May12.50v_16415 | 1 | 0 | 0 | 1 | 0 | 0 | 0 | 0 | 0 | 0 | 0 | 0 | 0 | 0 | 0 | 0 | 0 | 0 | 0 | 0 | 0 | 0 | 0 | 0 | 0 | 0 | 0 | 0 | 0 | 0 |
| Aug5.25v_20583 | 1 | 0 | 0 | 0 | 0 | 0 | 0 | 0 | 0 | 0 | 0 | 0 | 0 | 0 | 0 | 0 | 0 | 0 | 0 | 0 | 0 | 0 | 0 | 1 | 0 | 0 | 0 | 0 | 0 | 0 |
| Aug5.5v_9065 | 1 | 0 | 0 | 0 | 0 | 0 | 0 | 0 | 0 | 0 | 0 | 0 | 0 | 0 | 0 | 0 | 0 | 0 | 0 | 0 | 0 | 0 | 1 | 0 | 0 | 0 | 0 | 0 | 0 | 0 |
| Jun22.50v_1244 | 1 | 0 | 0 | 0 | 0 | 0 | 0 | 0 | 0 | 0 | 1 | 0 | 0 | 0 | 0 | 0 | 0 | 0 | 0 | 0 | 0 | 0 | 0 | 0 | 0 | 0 | 0 | 0 | 0 | 0 |
| May12.25v_32964 | 1 | 0 | 1 | 0 | 0 | 0 | 0 | 0 | 0 | 0 | 0 | 0 | 0 | 0 | 0 | 0 | 0 | 0 | 0 | 0 | 0 | 0 | 0 | 0 | 0 | 0 | 0 | 0 | 0 | 0 |
| Aug5.50v_8035 | 3 | 0 | 0 | 0 | 0 | 0 | 0 | 0 | 0 | 0 | 0 | 0 | 0 | 0 | 0 | 0 | 0 | 2 | 0 | 0 | 0 | 0 | 0 | 0 | 1 | 0 | 0 | 0 | 0 | 0 |
| Jun8.75v_11039 | 1 | 0 | 0 | 0 | 0 | 0 | 0 | 1 | 0 | 0 | 0 | 0 | 0 | 0 | 0 | 0 | 0 | 0 | 0 | 0 | 0 | 0 | 0 | 0 | 0 | 0 | 0 | 0 | 0 | 0 |
| Aug5.25v_39806 | 1 | 0 | 0 | 0 | 0 | 0 | 0 | 0 | 0 | 0 | 0 | 0 | 0 | 0 | 0 | 0 | 0 | 0 | 0 | 0 | 0 | 0 | 0 | 1 | 0 | 0 | 0 | 0 | 0 | 0 |
| Jul27.5v_8865 | 1 | 0 | 0 | 0 | 0 | 0 | 0 | 0 | 0 | 0 | 0 | 0 | 0 | 0 | 0 | 0 | 0 | 1 | 0 | 0 | 0 | 0 | 0 | 0 | 0 | 0 | 0 | 0 | 0 | 0 |
| Aug5.0v_4319 | 1 | 0 | 0 | 0 | 0 | 0 | 0 | 0 | 0 | 0 | 0 | 0 | 0 | 0 | 0 | 0 | 0 | 0 | 0 | 0 | 0 | 1 | 0 | 0 | 0 | 0 | 0 | 0 | 0 | 0 |
| Jul27.25v_27040 | 1 | 0 | 0 | 0 | 0 | 0 | 0 | 0 | 0 | 0 | 0 | 0 | 0 | 0 | 0 | 0 | 0 | 0 | 1 | 0 | 0 | 0 | 0 | 0 | 0 | 0 | 0 | 0 | 0 | 0 |
| Jul27.50v_16370 | 1 | 0 | 0 | 0 | 0 | 0 | 0 | 0 | 0 | 0 | 0 | 0 | 0 | 0 | 0 | 0 | 0 | 0 | 0 | 1 | 0 | 0 | 0 | 0 | 0 | 0 | 0 | 0 | 0 | 0 |
| May12.50v_26768 | 1 | 0 | 0 | 1 | 0 | 0 | 0 | 0 | 0 | 0 | 0 | 0 | 0 | 0 | 0 | 0 | 0 | 0 | 0 | 0 | 0 | 0 | 0 | 0 | 0 | 0 | 0 | 0 | 0 | 0 |
| Aug5.5v_7020 | 1 | 0 | 0 | 0 | 0 | 0 | 0 | 0 | 0 | 0 | 0 | 0 | 0 | 0 | 0 | 0 | 0 | 0 | 0 | 0 | 0 | 0 | 1 | 0 | 0 | 0 | 0 | 0 | 0 | 0 |
| May12.50v_25860 | 1 | 0 | 0 | 1 | 0 | 0 | 0 | 0 | 0 | 0 | 0 | 0 | 0 | 0 | 0 | 0 | 0 | 0 | 0 | 0 | 0 | 0 | 0 | 0 | 0 | 0 | 0 | 0 | 0 | 0 |
| Aug5.25v_34291 | 1 | 0 | 0 | 0 | 0 | 0 | 0 | 0 | 0 | 0 | 0 | 0 | 0 | 0 | 0 | 0 | 0 | 0 | 0 | 0 | 0 | 0 | 0 | 1 | 0 | 0 | 0 | 0 | 0 | 0 |
| Jul27.50v_17451 | 1 | 0 | 0 | 0 | 0 | 0 | 0 | 0 | 0 | 0 | 0 | 0 | 0 | 0 | 0 | 0 | 0 | 0 | 0 | 1 | 0 | 0 | 0 | 0 | 0 | 0 | 0 | 0 | 0 | 0 |
| Aug5.0v_341 | 5 | 0 | 0 | 0 | 0 | 0 | 0 | 0 | 0 | 1 | 0 | 0 | 0 | 2 | 0 | 0 | 0 | 0 | 0 | 0 | 0 | 1 | 0 | 0 | 0 | 0 | 0 | 0 | 0 | 1 |
| Aug5.25v_11221 | 1 | 0 | 0 | 0 | 0 | 0 | 0 | 0 | 0 | 0 | 0 | 0 | 0 | 0 | 0 | 0 | 0 | 0 | 0 | 0 | 0 | 0 | 0 | 1 | 0 | 0 | 0 | 0 | 0 | 0 |
| Jun22.50v_26209 | 1 | 0 | 0 | 0 | 0 | 0 | 0 | 0 | 0 | 0 | 1 | 0 | 0 | 0 | 0 | 0 | 0 | 0 | 0 | 0 | 0 | 0 | 0 | 0 | 0 | 0 | 0 | 0 | 0 | 0 |
| Aug5.5v_27837 | 1 | 0 | 0 | 0 | 0 | 0 | 0 | 0 | 0 | 0 | 0 | 0 | 0 | 0 | 0 | 0 | 0 | 0 | 0 | 0 | 0 | 0 | 1 | 0 | 0 | 0 | 0 | 0 | 0 | 0 |
| Jul27.5v_17127 | 1 | 0 | 0 | 0 | 0 | 0 | 0 | 0 | 0 | 0 | 0 | 0 | 0 | 0 | 0 | 0 | 0 | 1 | 0 | 0 | 0 | 0 | 0 | 0 | 0 | 0 | 0 | 0 | 0 | 0 |
| Aug5.0v_677 | 2 | 0 | 0 | 0 | 0 | 0 | 0 | 0 | 0 | 0 | 0 | 0 | 0 | 0 | 0 | 0 | 0 | 0 | 0 | 0 | 0 | 1 | 0 | 0 | 1 | 0 | 0 | 0 | 0 | 0 |
| May12.50v_29032 | 1 | 0 | 0 | 1 | 0 | 0 | 0 | 0 | 0 | 0 | 0 | 0 | 0 | 0 | 0 | 0 | 0 | 0 | 0 | 0 | 0 | 0 | 0 | 0 | 0 | 0 | 0 | 0 | 0 | 0 |
| Aug5.25v_9232 | 1 | 0 | 0 | 0 | 0 | 0 | 0 | 0 | 0 | 0 | 0 | 0 | 0 | 0 | 0 | 0 | 0 | 0 | 0 | 0 | 0 | 0 | 0 | 1 | 0 | 0 | 0 | 0 | 0 | 0 |
| Aug5.75v_8654 | 1 | 0 | 0 | 0 | 0 | 0 | 0 | 0 | 0 | 0 | 0 | 0 | 0 | 0 | 0 | 0 | 0 | 0 | 0 | 0 | 0 | 0 | 0 | 0 | 0 | 1 | 0 | 0 | 0 | 0 |
| Aug5.0v_1373 | 17 | 2 | 0 | 1 | 0 | 0 | 0 | 0 | 0 | 0 | 1 | 0 | 0 | 1 | 0 | 1 | 1 | 0 | 1 | 0 | 0 | 3 | 1 | 1 | 2 | 0 | 0 | 0 | 0 | 2 |
| May12.5v_20125 | 1 | 1 | 0 | 0 | 0 | 0 | 0 | 0 | 0 | 0 | 0 | 0 | 0 | 0 | 0 | 0 | 0 | 0 | 0 | 0 | 0 | 0 | 0 | 0 | 0 | 0 | 0 | 0 | 0 | 0 |
| Aug5.0v_920 | 1 | 0 | 0 | 0 | 0 | 0 | 0 | 0 | 0 | 0 | 0 | 0 | 0 | 0 | 0 | 0 | 0 | 0 | 0 | 0 | 0 | 1 | 0 | 0 | 0 | 0 | 0 | 0 | 0 | 0 |
| Aug5.50v_20206 | 2 | 0 | 0 | 0 | 0 | 0 | 0 | 0 | 0 | 0 | 0 | 0 | 0 | 1 | 0 | 0 | 0 | 0 | 0 | 0 | 0 | 0 | 0 | 0 | 1 | 0 | 0 | 0 | 0 | 0 |
| Jul11.5v_5854 | 1 | 0 | 0 | 0 | 0 | 0 | 0 | 0 | 0 | 0 | 0 | 0 | 0 | 1 | 0 | 0 | 0 | 0 | 0 | 0 | 0 | 0 | 0 | 0 | 0 | 0 | 0 | 0 | 0 | 0 |
| Sep8.50v_27094 | 1 | 0 | 0 | 0 | 0 | 0 | 0 | 0 | 0 | 0 | 0 | 0 | 0 | 0 | 0 | 0 | 0 | 0 | 0 | 0 | 0 | 0 | 0 | 0 | 0 | 0 | 0 | 0 | 0 | 1 |
| Sep8.50v_7399 | 1 | 0 | 0 | 0 | 0 | 0 | 0 | 0 | 0 | 0 | 0 | 0 | 0 | 0 | 0 | 0 | 0 | 0 | 0 | 0 | 0 | 0 | 0 | 0 | 0 | 0 | 0 | 0 | 0 | 1 |
| Jul27.5v_14706 | 1 | 0 | 0 | 0 | 0 | 0 | 0 | 0 | 0 | 0 | 0 | 0 | 0 | 0 | 0 | 0 | 0 | 1 | 0 | 0 | 0 | 0 | 0 | 0 | 0 | 0 | 0 | 0 | 0 | 0 |
| Aug5.25v_4597 | 1 | 0 | 0 | 0 | 0 | 0 | 0 | 0 | 0 | 0 | 0 | 0 | 0 | 0 | 0 | 0 | 0 | 0 | 0 | 0 | 0 | 0 | 0 | 1 | 0 | 0 | 0 | 0 | 0 | 0 |
| Jun22.75v_24684 | 1 | 0 | 0 | 0 | 0 | 0 | 0 | 0 | 0 | 0 | 0 | 1 | 0 | 0 | 0 | 0 | 0 | 0 | 0 | 0 | 0 | 0 | 0 | 0 | 0 | 0 | 0 | 0 | 0 | 0 |
| Aug5.75v_36520 | 1 | 0 | 0 | 0 | 0 | 0 | 0 | 0 | 0 | 0 | 0 | 0 | 0 | 0 | 0 | 0 | 0 | 0 | 0 | 0 | 0 | 0 | 0 | 0 | 0 | 1 | 0 | 0 | 0 | 0 |
| Sep8.0v_6359 | 1 | 0 | 0 | 0 | 0 | 0 | 0 | 0 | 0 | 0 | 0 | 0 | 0 | 0 | 0 | 0 | 0 | 0 | 0 | 0 | 0 | 0 | 0 | 0 | 0 | 0 | 1 | 0 | 0 | 0 |
| Aug5.0v_4874 | 1 | 0 | 0 | 0 | 0 | 0 | 0 | 0 | 0 | 0 | 0 | 0 | 0 | 0 | 0 | 0 | 0 | 0 | 0 | 0 | 0 | 1 | 0 | 0 | 0 | 0 | 0 | 0 | 0 | 0 |
| Aug5.0v_3911 | 1 | 0 | 0 | 0 | 0 | 0 | 0 | 0 | 0 | 0 | 0 | 0 | 0 | 0 | 0 | 0 | 0 | 0 | 0 | 0 | 0 | 1 | 0 | 0 | 0 | 0 | 0 | 0 | 0 | 0 |
| Sep8.50v_37124 | 1 | 0 | 0 | 0 | 0 | 0 | 0 | 0 | 0 | 0 | 0 | 0 | 0 | 0 | 0 | 0 | 0 | 0 | 0 | 0 | 0 | 0 | 0 | 0 | 0 | 0 | 0 | 0 | 0 | 1 |
| Aug5.75v_17539 | 1 | 0 | 0 | 0 | 0 | 0 | 0 | 0 | 0 | 0 | 0 | 0 | 0 | 0 | 0 | 0 | 0 | 0 | 0 | 0 | 0 | 0 | 0 | 0 | 0 | 1 | 0 | 0 | 0 | 0 |
| Jul27.50v_14831 | 1 | 0 | 0 | 0 | 0 | 0 | 0 | 0 | 0 | 0 | 0 | 0 | 0 | 0 | 0 | 0 | 0 | 0 | 0 | 1 | 0 | 0 | 0 | 0 | 0 | 0 | 0 | 0 | 0 | 0 |
| Aug5.0v_998 | 1 | 0 | 0 | 0 | 0 | 0 | 0 | 0 | 0 | 0 | 0 | 0 | 0 | 0 | 0 | 0 | 0 | 0 | 0 | 0 | 0 | 1 | 0 | 0 | 0 | 0 | 0 | 0 | 0 | 0 |
| Aug5.25v_15109 | 1 | 0 | 0 | 0 | 0 | 0 | 0 | 0 | 0 | 0 | 0 | 0 | 0 | 0 | 0 | 0 | 0 | 0 | 0 | 0 | 0 | 0 | 0 | 1 | 0 | 0 | 0 | 0 | 0 | 0 |
| Jun22.50v_32060 | 1 | 0 | 0 | 0 | 0 | 0 | 0 | 0 | 0 | 0 | 1 | 0 | 0 | 0 | 0 | 0 | 0 | 0 | 0 | 0 | 0 | 0 | 0 | 0 | 0 | 0 | 0 | 0 | 0 | 0 |
| Aug5.25v_7450 | 1 | 0 | 0 | 0 | 0 | 0 | 0 | 0 | 0 | 0 | 0 | 0 | 0 | 0 | 0 | 0 | 0 | 0 | 0 | 0 | 0 | 0 | 0 | 1 | 0 | 0 | 0 | 0 | 0 | 0 |
| Sep8.50v_5249 | 1 | 0 | 0 | 0 | 0 | 0 | 0 | 0 | 0 | 0 | 0 | 0 | 0 | 0 | 0 | 0 | 0 | 0 | 0 | 0 | 0 | 0 | 0 | 0 | 0 | 0 | 0 | 0 | 0 | 1 |
| Jun22.5v_994 | 1 | 0 | 0 | 0 | 0 | 0 | 0 | 0 | 0 | 1 | 0 | 0 | 0 | 0 | 0 | 0 | 0 | 0 | 0 | 0 | 0 | 0 | 0 | 0 | 0 | 0 | 0 | 0 | 0 | 0 |
| Aug5.75v_21234 | 1 | 0 | 0 | 0 | 0 | 0 | 0 | 0 | 0 | 0 | 0 | 0 | 0 | 0 | 0 | 0 | 0 | 0 | 0 | 0 | 0 | 0 | 0 | 0 | 0 | 1 | 0 | 0 | 0 | 0 |
| Aug5.0v_2382 | 1 | 0 | 0 | 0 | 0 | 0 | 0 | 0 | 0 | 0 | 0 | 0 | 0 | 0 | 0 | 0 | 0 | 0 | 0 | 0 | 0 | 1 | 0 | 0 | 0 | 0 | 0 | 0 | 0 | 0 |
| Jul27.5v_17228 | 1 | 0 | 0 | 0 | 0 | 0 | 0 | 0 | 0 | 0 | 0 | 0 | 0 | 0 | 0 | 0 | 0 | 1 | 0 | 0 | 0 | 0 | 0 | 0 | 0 | 0 | 0 | 0 | 0 | 0 |
| Jun22.0v_14164 | 1 | 0 | 0 | 0 | 0 | 0 | 0 | 0 | 1 | 0 | 0 | 0 | 0 | 0 | 0 | 0 | 0 | 0 | 0 | 0 | 0 | 0 | 0 | 0 | 0 | 0 | 0 | 0 | 0 | 0 |
| Aug5.0v_3834 | 1 | 0 | 0 | 0 | 0 | 0 | 0 | 0 | 0 | 0 | 0 | 0 | 0 | 0 | 0 | 0 | 0 | 0 | 0 | 0 | 0 | 1 | 0 | 0 | 0 | 0 | 0 | 0 | 0 | 0 |
| Aug5.0v_2296 | 1 | 0 | 0 | 0 | 0 | 0 | 0 | 0 | 0 | 0 | 0 | 0 | 0 | 0 | 0 | 0 | 0 | 0 | 0 | 0 | 0 | 1 | 0 | 0 | 0 | 0 | 0 | 0 | 0 | 0 |
| Aug5.25v_6733 | 1 | 0 | 0 | 0 | 0 | 0 | 0 | 0 | 0 | 0 | 0 | 0 | 0 | 0 | 0 | 0 | 0 | 0 | 0 | 0 | 0 | 0 | 0 | 1 | 0 | 0 | 0 | 0 | 0 | 0 |
| Jul11.50v_17433 | 1 | 0 | 0 | 0 | 0 | 0 | 0 | 0 | 0 | 0 | 0 | 0 | 0 | 0 | 0 | 1 | 0 | 0 | 0 | 0 | 0 | 0 | 0 | 0 | 0 | 0 | 0 | 0 | 0 | 0 |
| Jul27.5v_6474 | 1 | 0 | 0 | 0 | 0 | 0 | 0 | 0 | 0 | 0 | 0 | 0 | 0 | 0 | 0 | 0 | 0 | 1 | 0 | 0 | 0 | 0 | 0 | 0 | 0 | 0 | 0 | 0 | 0 | 0 |
| Aug5.25v_18150 | 2 | 0 | 0 | 0 | 0 | 0 | 0 | 0 | 0 | 0 | 0 | 0 | 0 | 0 | 0 | 0 | 0 | 0 | 0 | 0 | 0 | 0 | 0 | 1 | 0 | 0 | 1 | 0 | 0 | 0 |
| Jul27.5v_15137 | 1 | 0 | 0 | 0 | 0 | 0 | 0 | 0 | 0 | 0 | 0 | 0 | 0 | 0 | 0 | 0 | 0 | 1 | 0 | 0 | 0 | 0 | 0 | 0 | 0 | 0 | 0 | 0 | 0 | 0 |
| Aug5.25v_37408 | 1 | 0 | 0 | 0 | 0 | 0 | 0 | 0 | 0 | 0 | 0 | 0 | 0 | 0 | 0 | 0 | 0 | 0 | 0 | 0 | 0 | 0 | 0 | 1 | 0 | 0 | 0 | 0 | 0 | 0 |
| Jun8.50v_7739 | 1 | 0 | 0 | 0 | 0 | 0 | 1 | 0 | 0 | 0 | 0 | 0 | 0 | 0 | 0 | 0 | 0 | 0 | 0 | 0 | 0 | 0 | 0 | 0 | 0 | 0 | 0 | 0 | 0 | 0 |
| Aug5.0v_5413 | 1 | 0 | 0 | 0 | 0 | 0 | 0 | 0 | 0 | 0 | 0 | 0 | 0 | 0 | 0 | 0 | 0 | 0 | 0 | 0 | 0 | 1 | 0 | 0 | 0 | 0 | 0 | 0 | 0 | 0 |
| Jul27.5v_11837 | 1 | 0 | 0 | 0 | 0 | 0 | 0 | 0 | 0 | 0 | 0 | 0 | 0 | 0 | 0 | 0 | 0 | 1 | 0 | 0 | 0 | 0 | 0 | 0 | 0 | 0 | 0 | 0 | 0 | 0 |
| Aug5.5v_27177 | 1 | 0 | 0 | 0 | 0 | 0 | 0 | 0 | 0 | 0 | 0 | 0 | 0 | 0 | 0 | 0 | 0 | 0 | 0 | 0 | 0 | 0 | 1 | 0 | 0 | 0 | 0 | 0 | 0 | 0 |
| Aug5.25v_37407 | 1 | 0 | 0 | 0 | 0 | 0 | 0 | 0 | 0 | 0 | 0 | 0 | 0 | 0 | 0 | 0 | 0 | 0 | 0 | 0 | 0 | 0 | 0 | 1 | 0 | 0 | 0 | 0 | 0 | 0 |
| Sep8.50v_17601 | 1 | 0 | 0 | 0 | 0 | 0 | 0 | 0 | 0 | 0 | 0 | 0 | 0 | 0 | 0 | 0 | 0 | 0 | 0 | 0 | 0 | 0 | 0 | 0 | 0 | 0 | 0 | 0 | 0 | 1 |
| Jun22.5v_9229 | 1 | 0 | 0 | 0 | 0 | 0 | 0 | 0 | 0 | 1 | 0 | 0 | 0 | 0 | 0 | 0 | 0 | 0 | 0 | 0 | 0 | 0 | 0 | 0 | 0 | 0 | 0 | 0 | 0 | 0 |
| Aug5.0v_504 | 1 | 0 | 0 | 0 | 0 | 0 | 0 | 0 | 0 | 0 | 0 | 0 | 0 | 0 | 0 | 0 | 0 | 0 | 0 | 0 | 0 | 1 | 0 | 0 | 0 | 0 | 0 | 0 | 0 | 0 |
| Jul11.75v_33051 | 1 | 0 | 0 | 0 | 0 | 0 | 0 | 0 | 0 | 0 | 0 | 0 | 0 | 0 | 0 | 0 | 1 | 0 | 0 | 0 | 0 | 0 | 0 | 0 | 0 | 0 | 0 | 0 | 0 | 0 |
| May12.5v_29430 | 1 | 1 | 0 | 0 | 0 | 0 | 0 | 0 | 0 | 0 | 0 | 0 | 0 | 0 | 0 | 0 | 0 | 0 | 0 | 0 | 0 | 0 | 0 | 0 | 0 | 0 | 0 | 0 | 0 | 0 |
| Sep8.50v_32867 | 1 | 0 | 0 | 0 | 0 | 0 | 0 | 0 | 0 | 0 | 0 | 0 | 0 | 0 | 0 | 0 | 0 | 0 | 0 | 0 | 0 | 0 | 0 | 0 | 0 | 0 | 0 | 0 | 0 | 1 |
| Aug5.75v_30832 | 1 | 0 | 0 | 0 | 0 | 0 | 0 | 0 | 0 | 0 | 0 | 0 | 0 | 0 | 0 | 0 | 0 | 0 | 0 | 0 | 0 | 0 | 0 | 0 | 0 | 1 | 0 | 0 | 0 | 0 |
| Aug5.0v_3441 | 1 | 0 | 0 | 0 | 0 | 0 | 0 | 0 | 0 | 0 | 0 | 0 | 0 | 0 | 0 | 0 | 0 | 0 | 0 | 0 | 0 | 1 | 0 | 0 | 0 | 0 | 0 | 0 | 0 | 0 |
| Sep8.50v_20943 | 1 | 0 | 0 | 0 | 0 | 0 | 0 | 0 | 0 | 0 | 0 | 0 | 0 | 0 | 0 | 0 | 0 | 0 | 0 | 0 | 0 | 0 | 0 | 0 | 0 | 0 | 0 | 0 | 0 | 1 |
| May12.50v_1523 | 1 | 0 | 0 | 1 | 0 | 0 | 0 | 0 | 0 | 0 | 0 | 0 | 0 | 0 | 0 | 0 | 0 | 0 | 0 | 0 | 0 | 0 | 0 | 0 | 0 | 0 | 0 | 0 | 0 | 0 |
| Sep8.50v_24755 | 1 | 0 | 0 | 0 | 0 | 0 | 0 | 0 | 0 | 0 | 0 | 0 | 0 | 0 | 0 | 0 | 0 | 0 | 0 | 0 | 0 | 0 | 0 | 0 | 0 | 0 | 0 | 0 | 0 | 1 |
| Jul27.5v_251 | 1 | 0 | 0 | 0 | 0 | 0 | 0 | 0 | 0 | 0 | 0 | 0 | 0 | 0 | 0 | 0 | 0 | 1 | 0 | 0 | 0 | 0 | 0 | 0 | 0 | 0 | 0 | 0 | 0 | 0 |
| Aug5.0v_233 | 1 | 0 | 0 | 0 | 0 | 0 | 0 | 0 | 0 | 0 | 0 | 0 | 0 | 0 | 0 | 0 | 0 | 0 | 0 | 0 | 0 | 1 | 0 | 0 | 0 | 0 | 0 | 0 | 0 | 0 |
| Aug5.25v_20934 | 1 | 0 | 0 | 0 | 0 | 0 | 0 | 0 | 0 | 0 | 0 | 0 | 0 | 0 | 0 | 0 | 0 | 0 | 0 | 0 | 0 | 0 | 0 | 1 | 0 | 0 | 0 | 0 | 0 | 0 |
| Jul11.25v_4897 | 1 | 0 | 0 | 0 | 0 | 0 | 0 | 0 | 0 | 0 | 0 | 0 | 0 | 0 | 1 | 0 | 0 | 0 | 0 | 0 | 0 | 0 | 0 | 0 | 0 | 0 | 0 | 0 | 0 | 0 |

| Feature_ID | 654 | May12_05m | May12_25m | May12_50m | May12_75m | Jun8_25m | Jun8_50m | Jun8_75m | Jun22_0m | Jun22_05m | Jun22_50m | Jun22_75m | Jul11_0m | Jul11_05m | Jul11_25m | Jul11_50m | Jul11_75m | Jul27_05m | Jul27_25m | Jul27_50m | Jul27_75m | Aug5_0m | Aug5_05m | Aug5_25m | Aug5_50m | Aug5_75m | Sep8_0m | Sep8_05m | Sep8_25m | Sep8_50m |
|---|---|---|---|---|---|---|---|---|---|---|---|---|---|---|---|---|---|---|---|---|---|---|---|---|---|---|---|---|---|---|
| Jul11.50v_6384 | 1 | 0 | 0 | 0 | 0 | 0 | 0 | 0 | 0 | 0 | 0 | 0 | 0 | 0 | 0 | 1 | 0 | 0 | 0 | 0 | 0 | 0 | 0 | 0 | 0 | 0 | 0 | 0 | 0 | 0 |
| Jul27.25v_36019 | 1 | 0 | 0 | 0 | 0 | 0 | 0 | 0 | 0 | 0 | 0 | 0 | 0 | 0 | 0 | 0 | 0 | 0 | 1 | 0 | 0 | 0 | 0 | 0 | 0 | 0 | 0 | 0 | 0 | 0 |
| Jul11.25v_19966 | 1 | 0 | 0 | 0 | 0 | 0 | 0 | 0 | 0 | 0 | 0 | 0 | 0 | 0 | 1 | 0 | 0 | 0 | 0 | 0 | 0 | 0 | 0 | 0 | 0 | 0 | 0 | 0 | 0 | 0 |
| Jul11.5v_10859 | 1 | 0 | 0 | 0 | 0 | 0 | 0 | 0 | 0 | 0 | 0 | 0 | 0 | 1 | 0 | 0 | 0 | 0 | 0 | 0 | 0 | 0 | 0 | 0 | 0 | 0 | 0 | 0 | 0 | 0 |
| Aug5.0v_2502 | 1 | 0 | 0 | 0 | 0 | 0 | 0 | 0 | 0 | 0 | 0 | 0 | 0 | 0 | 0 | 0 | 0 | 0 | 0 | 0 | 0 | 1 | 0 | 0 | 0 | 0 | 0 | 0 | 0 | 0 |
| Aug5.0v_3252 | 2 | 0 | 0 | 0 | 0 | 0 | 0 | 0 | 1 | 0 | 0 | 0 | 0 | 0 | 0 | 0 | 0 | 0 | 0 | 0 | 0 | 1 | 0 | 0 | 0 | 0 | 0 | 0 | 0 | 0 |
| Aug5.0v_1977 | 1 | 0 | 0 | 0 | 0 | 0 | 0 | 0 | 0 | 0 | 0 | 0 | 0 | 0 | 0 | 0 | 0 | 0 | 0 | 0 | 0 | 1 | 0 | 0 | 0 | 0 | 0 | 0 | 0 | 0 |
| Jul11.75v_30837 | 1 | 0 | 0 | 0 | 0 | 0 | 0 | 0 | 0 | 0 | 0 | 0 | 0 | 0 | 0 | 0 | 1 | 0 | 0 | 0 | 0 | 0 | 0 | 0 | 0 | 0 | 0 | 0 | 0 | 0 |
| Aug5.0v_4177 | 1 | 0 | 0 | 0 | 0 | 0 | 0 | 0 | 0 | 0 | 0 | 0 | 0 | 0 | 0 | 0 | 0 | 0 | 0 | 0 | 0 | 1 | 0 | 0 | 0 | 0 | 0 | 0 | 0 | 0 |
| Jun22.50v_1730 | 1 | 0 | 0 | 0 | 0 | 0 | 0 | 0 | 0 | 0 | 1 | 0 | 0 | 0 | 0 | 0 | 0 | 0 | 0 | 0 | 0 | 0 | 0 | 0 | 0 | 0 | 0 | 0 | 0 | 0 |
| Jun22.50v_35217 | 1 | 0 | 0 | 0 | 0 | 0 | 0 | 0 | 0 | 0 | 1 | 0 | 0 | 0 | 0 | 0 | 0 | 0 | 0 | 0 | 0 | 0 | 0 | 0 | 0 | 0 | 0 | 0 | 0 | 0 |
| Aug5.0v_5271 | 1 | 0 | 0 | 0 | 0 | 0 | 0 | 0 | 0 | 0 | 0 | 0 | 0 | 0 | 0 | 0 | 0 | 0 | 0 | 0 | 0 | 1 | 0 | 0 | 0 | 0 | 0 | 0 | 0 | 0 |
| Jul11.0v_1557 | 1 | 0 | 0 | 0 | 0 | 0 | 0 | 0 | 0 | 0 | 0 | 0 | 1 | 0 | 0 | 0 | 0 | 0 | 0 | 0 | 0 | 0 | 0 | 0 | 0 | 0 | 0 | 0 | 0 | 0 |
| Jun22.0v_8077 | 1 | 0 | 0 | 0 | 0 | 0 | 0 | 0 | 1 | 0 | 0 | 0 | 0 | 0 | 0 | 0 | 0 | 0 | 0 | 0 | 0 | 0 | 0 | 0 | 0 | 0 | 0 | 0 | 0 | 0 |
| Sep8.50v_8700 | 1 | 0 | 0 | 0 | 0 | 0 | 0 | 0 | 0 | 0 | 0 | 0 | 0 | 0 | 0 | 0 | 0 | 0 | 0 | 0 | 0 | 0 | 0 | 0 | 0 | 0 | 0 | 0 | 0 | 1 |
| Jul11.25v_4174 | 1 | 0 | 0 | 0 | 0 | 0 | 0 | 0 | 0 | 0 | 0 | 0 | 0 | 0 | 1 | 0 | 0 | 0 | 0 | 0 | 0 | 0 | 0 | 0 | 0 | 0 | 0 | 0 | 0 | 0 |
| Sep8.50v_2826 | 1 | 0 | 0 | 0 | 0 | 0 | 0 | 0 | 0 | 0 | 0 | 0 | 0 | 0 | 0 | 0 | 0 | 0 | 0 | 0 | 0 | 0 | 0 | 0 | 0 | 0 | 0 | 0 | 0 | 1 |
| Jul11.75v_4989 | 1 | 0 | 0 | 0 | 0 | 0 | 0 | 0 | 0 | 0 | 0 | 0 | 0 | 0 | 0 | 0 | 1 | 0 | 0 | 0 | 0 | 0 | 0 | 0 | 0 | 0 | 0 | 0 | 0 | 0 |
| Aug5.0v_2215 | 1 | 0 | 0 | 0 | 0 | 0 | 0 | 0 | 0 | 0 | 0 | 0 | 0 | 0 | 0 | 0 | 0 | 0 | 0 | 0 | 0 | 1 | 0 | 0 | 0 | 0 | 0 | 0 | 0 | 0 |
| Sep8.50v_36245 | 1 | 0 | 0 | 0 | 0 | 0 | 0 | 0 | 0 | 0 | 0 | 0 | 0 | 0 | 0 | 0 | 0 | 0 | 0 | 0 | 0 | 0 | 0 | 0 | 0 | 0 | 0 | 0 | 0 | 1 |
| Jul27.5v_3834 | 1 | 0 | 0 | 0 | 0 | 0 | 0 | 0 | 0 | 0 | 0 | 0 | 0 | 0 | 0 | 0 | 0 | 1 | 0 | 0 | 0 | 0 | 0 | 0 | 0 | 0 | 0 | 0 | 0 | 0 |
| Sep8.0v_9670 | 1 | 0 | 0 | 0 | 0 | 0 | 0 | 0 | 0 | 0 | 0 | 0 | 0 | 0 | 0 | 0 | 0 | 0 | 0 | 0 | 0 | 0 | 0 | 0 | 0 | 0 | 1 | 0 | 0 | 0 |
| Jun22.50v_2620 | 1 | 0 | 0 | 0 | 0 | 0 | 0 | 0 | 0 | 0 | 1 | 0 | 0 | 0 | 0 | 0 | 0 | 0 | 0 | 0 | 0 | 0 | 0 | 0 | 0 | 0 | 0 | 0 | 0 | 0 |
| Sep8.50v_27087 | 1 | 0 | 0 | 0 | 0 | 0 | 0 | 0 | 0 | 0 | 0 | 0 | 0 | 0 | 0 | 0 | 0 | 0 | 0 | 0 | 0 | 0 | 0 | 0 | 0 | 0 | 0 | 0 | 0 | 1 |
| Aug5.5v_23712 | 2 | 0 | 0 | 0 | 0 | 0 | 0 | 0 | 0 | 0 | 0 | 0 | 0 | 0 | 0 | 0 | 0 | 1 | 0 | 0 | 0 | 0 | 1 | 0 | 0 | 0 | 0 | 0 | 0 | 0 |
| Aug5.0v_1669 | 1 | 0 | 0 | 0 | 0 | 0 | 0 | 0 | 0 | 0 | 0 | 0 | 0 | 0 | 0 | 0 | 0 | 0 | 0 | 0 | 0 | 1 | 0 | 0 | 0 | 0 | 0 | 0 | 0 | 0 |
| Jul11.25v_16800 | 1 | 0 | 0 | 0 | 0 | 0 | 0 | 0 | 0 | 0 | 0 | 0 | 0 | 0 | 1 | 0 | 0 | 0 | 0 | 0 | 0 | 0 | 0 | 0 | 0 | 0 | 0 | 0 | 0 | 0 |
| May12.25v_33619 | 1 | 0 | 1 | 0 | 0 | 0 | 0 | 0 | 0 | 0 | 0 | 0 | 0 | 0 | 0 | 0 | 0 | 0 | 0 | 0 | 0 | 0 | 0 | 0 | 0 | 0 | 0 | 0 | 0 | 0 |
| Jul27.5v_6131 | 1 | 0 | 0 | 0 | 0 | 0 | 0 | 0 | 0 | 0 | 0 | 0 | 0 | 0 | 0 | 0 | 0 | 1 | 0 | 0 | 0 | 0 | 0 | 0 | 0 | 0 | 0 | 0 | 0 | 0 |
| Jul27.25v_4130 | 1 | 0 | 0 | 0 | 0 | 0 | 0 | 0 | 0 | 0 | 0 | 0 | 0 | 0 | 0 | 0 | 0 | 0 | 1 | 0 | 0 | 0 | 0 | 0 | 0 | 0 | 0 | 0 | 0 | 0 |
| Jul27.50v_9144 | 1 | 0 | 0 | 0 | 0 | 0 | 0 | 0 | 0 | 0 | 0 | 0 | 0 | 0 | 0 | 0 | 0 | 0 | 0 | 1 | 0 | 0 | 0 | 0 | 0 | 0 | 0 | 0 | 0 | 0 |
| Jul11.50v_4694 | 1 | 0 | 0 | 0 | 0 | 0 | 0 | 0 | 0 | 0 | 0 | 0 | 0 | 0 | 0 | 1 | 0 | 0 | 0 | 0 | 0 | 0 | 0 | 0 | 0 | 0 | 0 | 0 | 0 | 0 |
| Aug5.0v_1178 | 1 | 0 | 0 | 0 | 0 | 0 | 0 | 0 | 0 | 0 | 0 | 0 | 0 | 0 | 0 | 0 | 0 | 0 | 0 | 0 | 0 | 1 | 0 | 0 | 0 | 0 | 0 | 0 | 0 | 0 |
| Jul11.25v_4567 | 1 | 0 | 0 | 0 | 0 | 0 | 0 | 0 | 0 | 0 | 0 | 0 | 0 | 0 | 1 | 0 | 0 | 0 | 0 | 0 | 0 | 0 | 0 | 0 | 0 | 0 | 0 | 0 | 0 | 0 |
| Sep8.50v_38342 | 1 | 0 | 0 | 0 | 0 | 0 | 0 | 0 | 0 | 0 | 0 | 0 | 0 | 0 | 0 | 0 | 0 | 0 | 0 | 0 | 0 | 0 | 0 | 0 | 0 | 0 | 0 | 0 | 0 | 1 |
| Aug5.0v_1451 | 1 | 0 | 0 | 0 | 0 | 0 | 0 | 0 | 0 | 0 | 0 | 0 | 0 | 0 | 0 | 0 | 0 | 0 | 0 | 0 | 0 | 1 | 0 | 0 | 0 | 0 | 0 | 0 | 0 | 0 |
| Aug5.50v_27568 | 1 | 0 | 0 | 0 | 0 | 0 | 0 | 0 | 0 | 0 | 0 | 0 | 0 | 0 | 0 | 0 | 0 | 0 | 0 | 0 | 0 | 0 | 0 | 0 | 1 | 0 | 0 | 0 | 0 | 0 |
| Jun22.50v_12097 | 1 | 0 | 0 | 0 | 0 | 0 | 0 | 0 | 0 | 0 | 1 | 0 | 0 | 0 | 0 | 0 | 0 | 0 | 0 | 0 | 0 | 0 | 0 | 0 | 0 | 0 | 0 | 0 | 0 | 0 |
| Aug5.5v_449 | 1 | 0 | 0 | 0 | 0 | 0 | 0 | 0 | 0 | 0 | 0 | 0 | 0 | 0 | 0 | 0 | 0 | 0 | 0 | 0 | 0 | 0 | 1 | 0 | 0 | 0 | 0 | 0 | 0 | 0 |
| Jul27.5v_16345 | 1 | 0 | 0 | 0 | 0 | 0 | 0 | 0 | 0 | 0 | 0 | 0 | 0 | 0 | 0 | 0 | 0 | 1 | 0 | 0 | 0 | 0 | 0 | 0 | 0 | 0 | 0 | 0 | 0 | 0 |
| Jun8.50v_17108 | 1 | 0 | 0 | 0 | 0 | 0 | 1 | 0 | 0 | 0 | 0 | 0 | 0 | 0 | 0 | 0 | 0 | 0 | 0 | 0 | 0 | 0 | 0 | 0 | 0 | 0 | 0 | 0 | 0 | 0 |
| Jun22.5v_6995 | 1 | 0 | 0 | 0 | 0 | 0 | 0 | 0 | 0 | 1 | 0 | 0 | 0 | 0 | 0 | 0 | 0 | 0 | 0 | 0 | 0 | 0 | 0 | 0 | 0 | 0 | 0 | 0 | 0 | 0 |
| Jun22.5v_594 | 1 | 0 | 0 | 0 | 0 | 0 | 0 | 0 | 0 | 1 | 0 | 0 | 0 | 0 | 0 | 0 | 0 | 0 | 0 | 0 | 0 | 0 | 0 | 0 | 0 | 0 | 0 | 0 | 0 | 0 |
| May12.50v_7837 | 1 | 0 | 0 | 1 | 0 | 0 | 0 | 0 | 0 | 0 | 0 | 0 | 0 | 0 | 0 | 0 | 0 | 0 | 0 | 0 | 0 | 0 | 0 | 0 | 0 | 0 | 0 | 0 | 0 | 0 |
| Jun22.50v_14067 | 1 | 0 | 0 | 0 | 0 | 0 | 0 | 0 | 0 | 0 | 1 | 0 | 0 | 0 | 0 | 0 | 0 | 0 | 0 | 0 | 0 | 0 | 0 | 0 | 0 | 0 | 0 | 0 | 0 | 0 |
| Jul27.50v_38632 | 1 | 0 | 0 | 0 | 0 | 0 | 0 | 0 | 0 | 0 | 0 | 0 | 0 | 0 | 0 | 0 | 0 | 0 | 0 | 1 | 0 | 0 | 0 | 0 | 0 | 0 | 0 | 0 | 0 | 0 |
| May12.5v_14281 | 1 | 1 | 0 | 0 | 0 | 0 | 0 | 0 | 0 | 0 | 0 | 0 | 0 | 0 | 0 | 0 | 0 | 0 | 0 | 0 | 0 | 0 | 0 | 0 | 0 | 0 | 0 | 0 | 0 | 0 |
| Jul11.50v_7910 | 1 | 0 | 0 | 0 | 0 | 0 | 0 | 0 | 0 | 0 | 0 | 0 | 0 | 0 | 0 | 1 | 0 | 0 | 0 | 0 | 0 | 0 | 0 | 0 | 0 | 0 | 0 | 0 | 0 | 0 |
| May12.75v_2781 | 1 | 0 | 0 | 0 | 1 | 0 | 0 | 0 | 0 | 0 | 0 | 0 | 0 | 0 | 0 | 0 | 0 | 0 | 0 | 0 | 0 | 0 | 0 | 0 | 0 | 0 | 0 | 0 | 0 | 0 |
| Jul11.0v_10988 | 1 | 0 | 0 | 0 | 0 | 0 | 0 | 0 | 0 | 0 | 0 | 0 | 1 | 0 | 0 | 0 | 0 | 0 | 0 | 0 | 0 | 0 | 0 | 0 | 0 | 0 | 0 | 0 | 0 | 0 |
| Aug5.25v_12315 | 1 | 0 | 0 | 0 | 0 | 0 | 0 | 0 | 0 | 0 | 0 | 0 | 0 | 0 | 0 | 0 | 0 | 0 | 0 | 0 | 0 | 0 | 0 | 1 | 0 | 0 | 0 | 0 | 0 | 0 |
| Jul11.25v_18940 | 1 | 0 | 0 | 0 | 0 | 0 | 0 | 0 | 0 | 0 | 0 | 0 | 0 | 0 | 1 | 0 | 0 | 0 | 0 | 0 | 0 | 0 | 0 | 0 | 0 | 0 | 0 | 0 | 0 | 0 |
| Aug5.5v_33291 | 1 | 0 | 0 | 0 | 0 | 0 | 0 | 0 | 0 | 0 | 0 | 0 | 0 | 0 | 0 | 0 | 0 | 0 | 0 | 0 | 0 | 0 | 1 | 0 | 0 | 0 | 0 | 0 | 0 | 0 |
| Jul11.5v_16378 | 1 | 0 | 0 | 0 | 0 | 0 | 0 | 0 | 0 | 0 | 0 | 0 | 0 | 1 | 0 | 0 | 0 | 0 | 0 | 0 | 0 | 0 | 0 | 0 | 0 | 0 | 0 | 0 | 0 | 0 |
| Jul11.5v_19885 | 1 | 0 | 0 | 0 | 0 | 0 | 0 | 0 | 0 | 0 | 0 | 0 | 0 | 1 | 0 | 0 | 0 | 0 | 0 | 0 | 0 | 0 | 0 | 0 | 0 | 0 | 0 | 0 | 0 | 0 |
| Aug5.0v_3520 | 1 | 0 | 0 | 0 | 0 | 0 | 0 | 0 | 0 | 0 | 0 | 0 | 0 | 0 | 0 | 0 | 0 | 0 | 0 | 0 | 0 | 1 | 0 | 0 | 0 | 0 | 0 | 0 | 0 | 0 |
| Jul27.5v_15204 | 1 | 0 | 0 | 0 | 0 | 0 | 0 | 0 | 0 | 0 | 0 | 0 | 0 | 0 | 0 | 0 | 0 | 1 | 0 | 0 | 0 | 0 | 0 | 0 | 0 | 0 | 0 | 0 | 0 | 0 |
| Sep8.50v_2637 | 1 | 0 | 0 | 0 | 0 | 0 | 0 | 0 | 0 | 0 | 0 | 0 | 0 | 0 | 0 | 0 | 0 | 0 | 0 | 0 | 0 | 0 | 0 | 0 | 0 | 0 | 0 | 0 | 0 | 1 |
| Aug5.0v_4936 | 1 | 0 | 0 | 0 | 0 | 0 | 0 | 0 | 0 | 0 | 0 | 0 | 0 | 0 | 0 | 0 | 0 | 0 | 0 | 0 | 0 | 1 | 0 | 0 | 0 | 0 | 0 | 0 | 0 | 0 |
| Aug5.25v_18399 | 1 | 0 | 0 | 0 | 0 | 0 | 0 | 0 | 0 | 0 | 0 | 0 | 0 | 0 | 0 | 0 | 0 | 0 | 0 | 0 | 0 | 0 | 0 | 1 | 0 | 0 | 0 | 0 | 0 | 0 |
| Jul11.50v_18898 | 1 | 0 | 0 | 0 | 0 | 0 | 0 | 0 | 0 | 0 | 0 | 0 | 0 | 0 | 0 | 1 | 0 | 0 | 0 | 0 | 0 | 0 | 0 | 0 | 0 | 0 | 0 | 0 | 0 | 0 |
| Sep8.0v_15947 | 1 | 0 | 0 | 0 | 0 | 0 | 0 | 0 | 0 | 0 | 0 | 0 | 0 | 0 | 0 | 0 | 0 | 0 | 0 | 0 | 0 | 0 | 0 | 0 | 0 | 0 | 1 | 0 | 0 | 0 |
| Sep8.50v_6326 | 1 | 0 | 0 | 0 | 0 | 0 | 0 | 0 | 0 | 0 | 0 | 0 | 0 | 0 | 0 | 0 | 0 | 0 | 0 | 0 | 0 | 0 | 0 | 0 | 0 | 0 | 0 | 0 | 0 | 1 |
| Aug5.0v_3985 | 1 | 0 | 0 | 0 | 0 | 0 | 0 | 0 | 0 | 0 | 0 | 0 | 0 | 0 | 0 | 0 | 0 | 0 | 0 | 0 | 0 | 1 | 0 | 0 | 0 | 0 | 0 | 0 | 0 | 0 |
| Sep8.0v_6319 | 1 | 0 | 0 | 0 | 0 | 0 | 0 | 0 | 0 | 0 | 0 | 0 | 0 | 0 | 0 | 0 | 0 | 0 | 0 | 0 | 0 | 0 | 0 | 0 | 0 | 0 | 1 | 0 | 0 | 0 |
| Aug5.0v_3625 | 1 | 0 | 0 | 0 | 0 | 0 | 0 | 0 | 0 | 0 | 0 | 0 | 0 | 0 | 0 | 0 | 0 | 0 | 0 | 0 | 0 | 1 | 0 | 0 | 0 | 0 | 0 | 0 | 0 | 0 |
| Jul11.75v_37165 | 1 | 0 | 0 | 0 | 0 | 0 | 0 | 0 | 0 | 0 | 0 | 0 | 0 | 0 | 0 | 0 | 1 | 0 | 0 | 0 | 0 | 0 | 0 | 0 | 0 | 0 | 0 | 0 | 0 | 0 |
| May12.50v_38219 | 1 | 0 | 0 | 1 | 0 | 0 | 0 | 0 | 0 | 0 | 0 | 0 | 0 | 0 | 0 | 0 | 0 | 0 | 0 | 0 | 0 | 0 | 0 | 0 | 0 | 0 | 0 | 0 | 0 | 0 |
| Aug5.25v_8581 | 1 | 0 | 0 | 0 | 0 | 0 | 0 | 0 | 0 | 0 | 0 | 0 | 0 | 0 | 0 | 0 | 0 | 0 | 0 | 0 | 0 | 0 | 0 | 1 | 0 | 0 | 0 | 0 | 0 | 0 |
| Aug5.0v_2718 | 1 | 0 | 0 | 0 | 0 | 0 | 0 | 0 | 0 | 0 | 0 | 0 | 0 | 0 | 0 | 0 | 0 | 0 | 0 | 0 | 0 | 1 | 0 | 0 | 0 | 0 | 0 | 0 | 0 | 0 |
| Jul27.5v_843 | 1 | 0 | 0 | 0 | 0 | 0 | 0 | 0 | 0 | 0 | 0 | 0 | 0 | 0 | 0 | 0 | 0 | 1 | 0 | 0 | 0 | 0 | 0 | 0 | 0 | 0 | 0 | 0 | 0 | 0 |
| Jun22.0v_19925 | 1 | 0 | 0 | 0 | 0 | 0 | 0 | 0 | 1 | 0 | 0 | 0 | 0 | 0 | 0 | 0 | 0 | 0 | 0 | 0 | 0 | 0 | 0 | 0 | 0 | 0 | 0 | 0 | 0 | 0 |
| Aug5.0v_4674 | 1 | 0 | 0 | 0 | 0 | 0 | 0 | 0 | 0 | 0 | 0 | 0 | 0 | 0 | 0 | 0 | 0 | 0 | 0 | 0 | 0 | 1 | 0 | 0 | 0 | 0 | 0 | 0 | 0 | 0 |
| May12.50v_11531 | 1 | 0 | 0 | 1 | 0 | 0 | 0 | 0 | 0 | 0 | 0 | 0 | 0 | 0 | 0 | 0 | 0 | 0 | 0 | 0 | 0 | 0 | 0 | 0 | 0 | 0 | 0 | 0 | 0 | 0 |
| Jul27.5v_380 | 1 | 0 | 0 | 0 | 0 | 0 | 0 | 0 | 0 | 0 | 0 | 0 | 0 | 0 | 0 | 0 | 0 | 1 | 0 | 0 | 0 | 0 | 0 | 0 | 0 | 0 | 0 | 0 | 0 | 0 |
| Jul11.75v_25343 | 1 | 0 | 0 | 0 | 0 | 0 | 0 | 0 | 0 | 0 | 0 | 0 | 0 | 0 | 0 | 0 | 1 | 0 | 0 | 0 | 0 | 0 | 0 | 0 | 0 | 0 | 0 | 0 | 0 | 0 |
| Jun8.50v_2489 | 1 | 0 | 0 | 0 | 0 | 0 | 1 | 0 | 0 | 0 | 0 | 0 | 0 | 0 | 0 | 0 | 0 | 0 | 0 | 0 | 0 | 0 | 0 | 0 | 0 | 0 | 0 | 0 | 0 | 0 |
| Jul11.5v_13921 | 1 | 0 | 0 | 0 | 0 | 0 | 0 | 0 | 0 | 0 | 0 | 0 | 0 | 1 | 0 | 0 | 0 | 0 | 0 | 0 | 0 | 0 | 0 | 0 | 0 | 0 | 0 | 0 | 0 | 0 |
| Sep8.0v_10103 | 1 | 0 | 0 | 0 | 0 | 0 | 0 | 0 | 0 | 0 | 0 | 0 | 0 | 0 | 0 | 0 | 0 | 0 | 0 | 0 | 0 | 0 | 0 | 0 | 0 | 0 | 1 | 0 | 0 | 0 |
| Jul11.25v_6604 | 1 | 0 | 0 | 0 | 0 | 0 | 0 | 0 | 0 | 0 | 0 | 0 | 0 | 0 | 1 | 0 | 0 | 0 | 0 | 0 | 0 | 0 | 0 | 0 | 0 | 0 | 0 | 0 | 0 | 0 |
| Jun8.50v_32582 | 1 | 0 | 0 | 0 | 0 | 0 | 1 | 0 | 0 | 0 | 0 | 0 | 0 | 0 | 0 | 0 | 0 | 0 | 0 | 0 | 0 | 0 | 0 | 0 | 0 | 0 | 0 | 0 | 0 | 0 |
| Aug5.0v_1080 | 1 | 0 | 0 | 0 | 0 | 0 | 0 | 0 | 0 | 0 | 0 | 0 | 0 | 0 | 0 | 0 | 0 | 0 | 0 | 0 | 0 | 1 | 0 | 0 | 0 | 0 | 0 | 0 | 0 | 0 |
| Aug5.5v_35840 | 1 | 0 | 0 | 0 | 0 | 0 | 0 | 0 | 0 | 0 | 0 | 0 | 0 | 0 | 0 | 0 | 0 | 0 | 0 | 0 | 0 | 0 | 1 | 0 | 0 | 0 | 0 | 0 | 0 | 0 |
| Aug5.5v_18676 | 1 | 0 | 0 | 0 | 0 | 0 | 0 | 0 | 0 | 0 | 0 | 0 | 0 | 0 | 0 | 0 | 0 | 0 | 0 | 0 | 0 | 0 | 1 | 0 | 0 | 0 | 0 | 0 | 0 | 0 |
| Jul11.50v_9034 | 1 | 0 | 0 | 0 | 0 | 0 | 0 | 0 | 0 | 0 | 0 | 0 | 0 | 0 | 0 | 1 | 0 | 0 | 0 | 0 | 0 | 0 | 0 | 0 | 0 | 0 | 0 | 0 | 0 | 0 |
| Sep8.0v_11110 | 1 | 0 | 0 | 0 | 0 | 0 | 0 | 0 | 0 | 0 | 0 | 0 | 0 | 0 | 0 | 0 | 0 | 0 | 0 | 0 | 0 | 0 | 0 | 0 | 0 | 0 | 1 | 0 | 0 | 0 |
| Aug5.0v_4348 | 1 | 0 | 0 | 0 | 0 | 0 | 0 | 0 | 0 | 0 | 0 | 0 | 0 | 0 | 0 | 0 | 0 | 0 | 0 | 0 | 0 | 1 | 0 | 0 | 0 | 0 | 0 | 0 | 0 | 0 |
| Jun8.25v_20975 | 1 | 0 | 0 | 0 | 0 | 1 | 0 | 0 | 0 | 0 | 0 | 0 | 0 | 0 | 0 | 0 | 0 | 0 | 0 | 0 | 0 | 0 | 0 | 0 | 0 | 0 | 0 | 0 | 0 | 0 |
| Jun22.75v_36530 | 1 | 0 | 0 | 0 | 0 | 0 | 0 | 0 | 0 | 0 | 0 | 1 | 0 | 0 | 0 | 0 | 0 | 0 | 0 | 0 | 0 | 0 | 0 | 0 | 0 | 0 | 0 | 0 | 0 | 0 |
| May12.50v_25185 | 1 | 0 | 0 | 1 | 0 | 0 | 0 | 0 | 0 | 0 | 0 | 0 | 0 | 0 | 0 | 0 | 0 | 0 | 0 | 0 | 0 | 0 | 0 | 0 | 0 | 0 | 0 | 0 | 0 | 0 |
| Aug5.0v_4043 | 1 | 0 | 0 | 0 | 0 | 0 | 0 | 0 | 0 | 0 | 0 | 0 | 0 | 0 | 0 | 0 | 0 | 0 | 0 | 0 | 0 | 1 | 0 | 0 | 0 | 0 | 0 | 0 | 0 | 0 |
| Sep8.0v_13681 | 1 | 0 | 0 | 0 | 0 | 0 | 0 | 0 | 0 | 0 | 0 | 0 | 0 | 0 | 0 | 0 | 0 | 0 | 0 | 0 | 0 | 0 | 0 | 0 | 0 | 0 | 1 | 0 | 0 | 0 |
| Sep8.25v_7208 | 1 | 0 | 0 | 0 | 0 | 0 | 0 | 0 | 0 | 0 | 0 | 0 | 0 | 0 | 0 | 0 | 0 | 0 | 0 | 0 | 0 | 0 | 0 | 0 | 0 | 0 | 0 | 0 | 1 | 0 |
| Jun22.0v_6413 | 1 | 0 | 0 | 0 | 0 | 0 | 0 | 0 | 1 | 0 | 0 | 0 | 0 | 0 | 0 | 0 | 0 | 0 | 0 | 0 | 0 | 0 | 0 | 0 | 0 | 0 | 0 | 0 | 0 | 0 |
| May12.25v_288 | 1 | 0 | 1 | 0 | 0 | 0 | 0 | 0 | 0 | 0 | 0 | 0 | 0 | 0 | 0 | 0 | 0 | 0 | 0 | 0 | 0 | 0 | 0 | 0 | 0 | 0 | 0 | 0 | 0 | 0 |
| Jul11.0v_18988 | 1 | 0 | 0 | 0 | 0 | 0 | 0 | 0 | 0 | 0 | 0 | 0 | 1 | 0 | 0 | 0 | 0 | 0 | 0 | 0 | 0 | 0 | 0 | 0 | 0 | 0 | 0 | 0 | 0 | 0 |
| Aug5.0v_3248 | 1 | 0 | 0 | 0 | 0 | 0 | 0 | 0 | 0 | 0 | 0 | 0 | 0 | 0 | 0 | 0 | 0 | 0 | 0 | 0 | 0 | 1 | 0 | 0 | 0 | 0 | 0 | 0 | 0 | 0 |
| Aug5.0v_3389 | 1 | 0 | 0 | 0 | 0 | 0 | 0 | 0 | 0 | 0 | 0 | 0 | 0 | 0 | 0 | 0 | 0 | 0 | 0 | 0 | 0 | 1 | 0 | 0 | 0 | 0 | 0 | 0 | 0 | 0 |
| Jun22.0v_19010 | 1 | 0 | 0 | 0 | 0 | 0 | 0 | 0 | 1 | 0 | 0 | 0 | 0 | 0 | 0 | 0 | 0 | 0 | 0 | 0 | 0 | 0 | 0 | 0 | 0 | 0 | 0 | 0 | 0 | 0 |
| Aug5.25v_8334 | 1 | 0 | 0 | 0 | 0 | 0 | 0 | 0 | 0 | 0 | 0 | 0 | 0 | 0 | 0 | 0 | 0 | 0 | 0 | 0 | 0 | 0 | 0 | 1 | 0 | 0 | 0 | 0 | 0 | 0 |
| Jun8.25v_29954 | 1 | 0 | 0 | 0 | 0 | 1 | 0 | 0 | 0 | 0 | 0 | 0 | 0 | 0 | 0 | 0 | 0 | 0 | 0 | 0 | 0 | 0 | 0 | 0 | 0 | 0 | 0 | 0 | 0 | 0 |
| Sep8.25v_2810 | 1 | 0 | 0 | 0 | 0 | 0 | 0 | 0 | 0 | 0 | 0 | 0 | 0 | 0 | 0 | 0 | 0 | 0 | 0 | 0 | 0 | 0 | 0 | 0 | 0 | 0 | 0 | 0 | 1 | 0 |
| Jul11.50v_8950 | 1 | 0 | 0 | 0 | 0 | 0 | 0 | 0 | 0 | 0 | 0 | 0 | 0 | 0 | 0 | 1 | 0 | 0 | 0 | 0 | 0 | 0 | 0 | 0 | 0 | 0 | 0 | 0 | 0 | 0 |
| Jul11.50v_1076 | 1 | 0 | 0 | 0 | 0 | 0 | 0 | 0 | 0 | 0 | 0 | 0 | 0 | 0 | 0 | 1 | 0 | 0 | 0 | 0 | 0 | 0 | 0 | 0 | 0 | 0 | 0 | 0 | 0 | 0 |
| Total 98% RTPR OTUs | 30461 | | | | | | | | | | | | | | | | | | | | | | | | | | | | | |

**Supplementary Table S4**. Virioplankton RTPR phylogenetic clade feature table

| clade | May12_05m | May12_25m | May12_50m | May12_75m | Jun8_25m | Jun8_50m | Jun8_75m | Jun22_0m | Jun22_05m | Jun22_50m | Jun22_75m | Jul11_0m | Jul11_05m | Jul11_25m | Jul11_50m | Jul11_75m |
|---|---|---|---|---|---|---|---|---|---|---|---|---|---|---|---|---|
| 1 | 26 | 28 | 22 | 22 | 14 | 28 | 4 | 27 | 21 | 24 | 20 | 23 | 34 | 19 | 15 | 9 |
| 2 | 5 | 8 | 12 | 4 | 12 | 19 | 1 | 46 | 113 | 368 | 30 | 30 | 144 | 184 | 68 | 15 |
| 3 | 12 | 10 | 9 | 11 | 5 | 6 | 4 | 8 | 21 | 27 | 5 | 10 | 47 | 54 | 11 | 10 |
| 4 | 67 | 64 | 67 | 65 | 31 | 52 | 9 | 54 | 42 | 55 | 51 | 80 | 110 | 100 | 40 | 27 |
| 5 | 131 | 169 | 320 | 526 | 108 | 433 | 163 | 29 | 54 | 309 | 362 | 14 | 49 | 72 | 178 | 159 |
| 6 | 161 | 157 | 134 | 297 | 66 | 265 | 81 | 31 | 66 | 79 | 235 | 20 | 98 | 85 | 88 | 109 |
| 7 | 57 | 67 | 45 | 86 | 32 | 77 | 6 | 78 | 20 | 57 | 62 | 120 | 112 | 48 | 59 | 68 |
| 8 | 27 | 36 | 50 | 26 | 19 | 28 | 6 | 59 | 20 | 129 | 43 | 38 | 20 | 48 | 54 | 26 |
| 9 | 22 | 32 | 30 | 37 | 15 | 30 | 5 | 19 | 8 | 99 | 63 | 17 | 10 | 18 | 47 | 29 |
| 10 | 30 | 28 | 33 | 26 | 13 | 30 | 5 | 11 | 10 | 14 | 32 | 16 | 15 | 12 | 25 | 28 |
| 11 | 5 | 6 | 4 | 8 | 5 | 12 | 0 | 5 | 1 | 6 | 5 | 4 | 2 | 2 | 4 | 3 |
| 12 | 62 | 40 | 101 | 60 | 21 | 68 | 8 | 28 | 5 | 207 | 22 | 2 | 12 | 44 | 27 | 62 |
| 13 | 23 | 26 | 29 | 40 | 21 | 56 | 3 | 22 | 11 | 30 | 49 | 19 | 52 | 21 | 18 | 32 |
| 14 | 136 | 147 | 283 | 140 | 69 | 283 | 44 | 126 | 90 | 214 | 146 | 108 | 149 | 77 | 95 | 146 |
| 15 | 74 | 79 | 110 | 37 | 11 | 12 | 8 | 26 | 62 | 55 | 29 | 19 | 62 | 24 | 7 | 17 |
| 16 | 43 | 42 | 40 | 35 | 9 | 71 | 11 | 27 | 30 | 42 | 76 | 18 | 46 | 21 | 23 | 38 |
| 17 | 92 | 94 | 270 | 92 | 40 | 101 | 9 | 117 | 44 | 137 | 96 | 103 | 130 | 131 | 95 | 59 |

**Supplementa**

| clade | Jul27_05m | Jul27_25m | Jul27_50m | Jul27_75m | Aug5_0m | Aug5_05m | Aug5_25m | Aug5_50m | Aug5_75m | Sep8_0m | Sep8_05m | Sep8_25m | Sep8_50m | Total RTPR sequences |
|---|---|---|---|---|---|---|---|---|---|---|---|---|---|---|
| 1 | 81 | 34 | 21 | 1 | 139 | 67 | 89 | 16 | 25 | 47 | 18 | 8 | 104 | 986 |
| 2 | 378 | 312 | 377 | 50 | 303 | 167 | 264 | 237 | 121 | 74 | 7 | 40 | 331 | 3720 |
| 3 | 53 | 30 | 28 | 10 | 30 | 40 | 21 | 19 | 15 | 12 | 8 | 3 | 20 | 539 |
| 4 | 137 | 62 | 51 | 14 | 205 | 120 | 95 | 18 | 22 | 90 | 16 | 4 | 102 | 1850 |
| 5 | 23 | 44 | 108 | 34 | 31 | 23 | 31 | 27 | 71 | 12 | 63 | 271 | 46 | 3860 |
| 6 | 44 | 35 | 50 | 39 | 14 | 23 | 15 | 8 | 57 | 5 | 154 | 21 | 20 | 2457 |
| 7 | 152 | 154 | 44 | 9 | 619 | 150 | 311 | 58 | 67 | 62 | 14 | 9 | 116 | 2759 |
| 8 | 16 | 40 | 62 | 15 | 145 | 82 | 92 | 37 | 64 | 18 | 16 | 3 | 88 | 1307 |
| 9 | 7 | 16 | 47 | 13 | 65 | 21 | 62 | 32 | 64 | 16 | 3 | 6 | 58 | 891 |
| 10 | 8 | 14 | 23 | 1 | 19 | 16 | 14 | 6 | 9 | 15 | 18 | 1 | 41 | 513 |
| 11 | 3 | 3 | 4 | 1 | 1 | 2 | 1 | 0 | 7 | 3 | 0 | 2 | 2 | 101 |
| 12 | 7 | 9 | 40 | 9 | 12 | 11 | 19 | 33 | 28 | 6 | 20 | 6 | 17 | 986 |
| 13 | 221 | 88 | 41 | 2 | 418 | 118 | 343 | 38 | 23 | 97 | 18 | 6 | 129 | 1994 |
| 14 | 106 | 85 | 89 | 30 | 111 | 73 | 59 | 47 | 57 | 69 | 120 | 19 | 131 | 3249 |
| 15 | 47 | 11 | 14 | 3 | 39 | 22 | 14 | 7 | 18 | 9 | 73 | 5 | 13 | 907 |
| 16 | 50 | 18 | 17 | 6 | 56 | 33 | 17 | 8 | 18 | 23 | 58 | 9 | 159 | 1044 |
| 17 | 190 | 90 | 96 | 12 | 395 | 174 | 237 | 44 | 43 | 108 | 34 | 11 | 254 | 3298 |
| | | | | | | | | | | | | total RTPR sequences | | 30461 |

**Supplementary Table S5**. Bacterioplankton 16S rRNA ASV feature table

ASVs: 1779

Seqs: 918792

| #ASV_ID | Feature_ID | total seqs | May12_0m | May12_05m | May12_25m | May12_50m | May12_75m | Jun8_0m | Jun8_05m | Jun8_25m | Jun8_50m | Jun8_75m | Jun22_0m | Jun22_05m | Jun22_25m | Jun22_50m | Jun22_75m | Jul11_0m | Jul11_05m | Jul11_25m | Jul11_50m | Jul11_75m | Jul27_0m | Jul27_05m | Jul27_25m | Jul27_50m | Jul27_75m | Aug5_0m | Aug5_05m | Aug5_25m | Aug5_50m | Aug5_75m | Sep8_0m | Sep8_05m | Sep8_25m | Sep8_50m | Sep8_75m |
|---|---|---|---|---|---|---|---|---|---|---|---|---|---|---|---|---|---|---|---|---|---|---|---|---|---|---|---|---|---|---|---|---|---|---|---|---|---|
| ASV1 | e5cad0f83c5c | 65511 | 831 | 832 | 1593 | 1024 | 671 | 1072 | 416 | 1078 | 209 | 542 | 1126 | 2924 | 1469 | 363 | 208 | 5720 | 8003 | 3002 | 951 | 74 | 2673 | 2024 | 1872 | 2203 | 423 | 3306 | 2156 | 6601 | 4155 | 394 | 3140 | 1602 | 1321 | 1227 | 306 |
| ASV2 | 36b651dfb1e | 41523 | 1283 | 646 | 2439 | 1325 | 2190 | 4643 | 2141 | 1557 | 647 | 448 | 594 | 2486 | 1235 | 1481 | 1183 | 2081 | 2012 | 1025 | 643 | 294 | 1596 | 1082 | 769 | 533 | 719 | 927 | 547 | 1325 | 978 | 908 | 375 | 315 | 190 | 526 | 380 |
| ASV3 | 1d72f3f992bc | 23646 | 1290 | 1378 | 2375 | 1395 | 1116 | 2826 | 1029 | 1559 | 248 | 410 | 417 | 943 | 1161 | 2791 | 366 | 254 | 404 | 54 | 39 | 19 | 112 | 108 | 93 | 176 | 155 | 217 | 102 | 868 | 1151 | 322 | 116 | 0 | 21 | 110 | 21 |
| ASV4 | 936be975fb9 | 19022 | 352 | 299 | 768 | 465 | 306 | 763 | 257 | 263 | 76 | 113 | 315 | 1035 | 328 | 127 | 129 | 1679 | 2895 | 474 | 213 | 63 | 714 | 440 | 174 | 179 | 117 | 1385 | 624 | 829 | 271 | 123 | 1709 | 759 | 496 | 181 | 101 |
| ASV5 | a6a79b8a486 | 18642 | 382 | 298 | 861 | 481 | 758 | 467 | 200 | 395 | 311 | 176 | 99 | 492 | 250 | 163 | 389 | 1480 | 1603 | 423 | 304 | 213 | 963 | 686 | 404 | 181 | 202 | 1357 | 804 | 987 | 328 | 224 | 1030 | 697 | 465 | 385 | 184 |
| ASV6 | a3d4f90910b | 17900 | 614 | 336 | 1063 | 609 | 997 | 848 | 471 | 674 | 699 | 419 | 121 | 511 | 245 | 286 | 767 | 618 | 691 | 440 | 511 | 395 | 504 | 389 | 356 | 391 | 717 | 647 | 395 | 601 | 419 | 439 | 420 | 271 | 231 | 467 | 338 |
| ASV7 | 1122e6b7ea8 | 17389 | 252 | 98 | 549 | 284 | 402 | 702 | 350 | 343 | 394 | 114 | 179 | 750 | 291 | 323 | 532 | 1044 | 1064 | 657 | 410 | 212 | 542 | 413 | 345 | 312 | 385 | 562 | 413 | 499 | 331 | 325 | 1680 | 1160 | 883 | 369 | 220 |
| ASV8 | a756a146af3 | 16271 | 469 | 387 | 1027 | 602 | 1100 | 706 | 317 | 494 | 454 | 306 | 115 | 507 | 238 | 236 | 515 | 664 | 827 | 413 | 423 | 245 | 544 | 402 | 328 | 288 | 363 | 713 | 484 | 661 | 408 | 337 | 491 | 360 | 247 | 364 | 236 |
| ASV9 | b77bf87ad1c | 15311 | 259 | 282 | 676 | 399 | 317 | 422 | 172 | 168 | 40 | 66 | 370 | 1077 | 442 | 220 | 98 | 1335 | 1894 | 596 | 190 | 32 | 701 | 491 | 373 | 323 | 170 | 778 | 489 | 1263 | 804 | 310 | 147 | 46 | 45 | 155 | 161 |
| ASV10 | 2d4bbf1e16d | 14853 | 182 | 114 | 398 | 191 | 377 | 452 | 248 | 328 | 244 | 185 | 172 | 604 | 285 | 227 | 424 | 759 | 898 | 523 | 292 | 135 | 433 | 338 | 251 | 247 | 421 | 458 | 294 | 430 | 304 | 231 | 1832 | 1294 | 884 | 239 | 159 |
| ASV11 | 0194fcb71fb1 | 13492 | 131 | 90 | 394 | 236 | 236 | 292 | 113 | 271 | 74 | 106 | 134 | 408 | 268 | 98 | 148 | 1133 | 1995 | 197 | 117 | 56 | 472 | 279 | 116 | 79 | 46 | 740 | 384 | 834 | 192 | 114 | 2099 | 811 | 468 | 249 | 112 |
| ASV12 | 2b7c6eeca2e | 12730 | 199 | 126 | 382 | 279 | 157 | 178 | 83 | 200 | 61 | 111 | 250 | 634 | 321 | 82 | 0 | 1000 | 1534 | 627 | 172 | 0 | 481 | 311 | 416 | 517 | 104 | 526 | 284 | 1368 | 987 | 0 | 630 | 213 | 232 | 198 | 67 |
| ASV13 | c1b1851dde5 | 12452 | 372 | 356 | 792 | 530 | 610 | 250 | 103 | 228 | 121 | 147 | 153 | 561 | 244 | 61 | 200 | 719 | 1009 | 407 | 298 | 92 | 380 | 226 | 265 | 269 | 144 | 648 | 397 | 973 | 465 | 162 | 308 | 147 | 112 | 374 | 329 |
| ASV14 | c7d8e6d1fd5 | 12316 | 447 | 325 | 1387 | 837 | 1416 | 820 | 351 | 601 | 320 | 308 | 34 | 193 | 97 | 315 | 751 | 35 | 58 | 133 | 259 | 151 | 146 | 84 | 137 | 277 | 401 | 87 | 57 | 531 | 894 | 445 | 13 | 0 | 12 | 174 | 220 |
| ASV15 | 81092899f1c | 11391 | 335 | 189 | 651 | 389 | 646 | 483 | 281 | 428 | 365 | 195 | 78 | 324 | 197 | 167 | 436 | 545 | 627 | 296 | 300 | 217 | 334 | 236 | 192 | 209 | 316 | 462 | 290 | 414 | 257 | 218 | 350 | 254 | 200 | 260 | 250 |
| ASV16 | 8b0192de06c | 11258 | 71 | 148 | 263 | 122 | 79 | 128 | 0 | 126 | 0 | 66 | 101 | 110 | 79 | 123 | 0 | 307 | 421 | 71 | 0 | 0 | 217 | 204 | 118 | 107 | 0 | 422 | 237 | 857 | 471 | 88 | 3400 | 1113 | 976 | 695 | 138 |
| ASV17 | b4aaa30f8f6 | 10550 | 228 | 182 | 581 | 427 | 630 | 643 | 291 | 367 | 264 | 224 | 136 | 369 | 198 | 345 | 693 | 207 | 381 | 147 | 169 | 129 | 306 | 200 | 170 | 245 | 320 | 239 | 139 | 602 | 683 | 384 | 78 | 53 | 54 | 155 | 311 |
| ASV18 | 1c54ed87c17 | 10124 | 0 | 0 | 52 | 38 | 51 | 75 | 0 | 620 | 945 | 1950 | 0 | 0 | 0 | 53 | 491 | 0 | 0 | 0 | 634 | 416 | 0 | 0 | 89 | 1126 | 1011 | 0 | 0 | 232 | 1034 | 479 | 0 | 0 | 0 | 243 | 585 |
| ASV19 | a5e6de78677 | 9873 | 112 | 105 | 245 | 189 | 183 | 309 | 173 | 201 | 48 | 48 | 190 | 538 | 218 | 105 | 86 | 817 | 1273 | 537 | 219 | 64 | 314 | 198 | 203 | 220 | 135 | 420 | 318 | 839 | 461 | 99 | 340 | 226 | 135 | 155 | 150 |
| ASV20 | a35da92fe92 | 9644 | 232 | 204 | 605 | 393 | 535 | 936 | 366 | 440 | 213 | 325 | 71 | 236 | 198 | 368 | 348 | 127 | 218 | 81 | 190 | 116 | 216 | 117 | 117 | 315 | 320 | 158 | 81 | 490 | 513 | 249 | 318 | 122 | 129 | 151 | 146 |
| ASV21 | e43703ac625 | 9053 | 139 | 103 | 501 | 249 | 397 | 190 | 69 | 225 | 205 | 219 | 20 | 141 | 60 | 107 | 262 | 166 | 209 | 192 | 277 | 161 | 427 | 236 | 175 | 215 | 250 | 326 | 195 | 1120 | 637 | 312 | 373 | 214 | 200 | 281 | 200 |
| ASV22 | 3ab38fc6471 | 8312 | 284 | 186 | 587 | 321 | 164 | 583 | 203 | 122 | 0 | 0 | 281 | 907 | 509 | 174 | 29 | 331 | 789 | 186 | 20 | 0 | 214 | 132 | 61 | 33 | 9 | 330 | 179 | 415 | 118 | 18 | 558 | 188 | 173 | 172 | 36 |
| ASV23 | 91f687fdb62 | 8016 | 0 | 0 | 0 | 0 | 0 | 0 | 0 | 252 | 445 | 1060 | 0 | 0 | 33 | 0 | 293 | 0 | 0 | 51 | 627 | 434 | 0 | 0 | 100 | 1175 | 1059 | 0 | 0 | 131 | 889 | 375 | 0 | 0 | 0 | 267 | 825 |
| ASV24 | 5234f65d442 | 8012 | 357 | 244 | 1484 | 637 | 459 | 311 | 79 | 103 | 15 | 34 | 26 | 51 | 36 | 64 | 111 | 59 | 257 | 18 | 15 | 0 | 104 | 54 | 13 | 28 | 21 | 205 | 165 | 684 | 395 | 104 | 857 | 366 | 260 | 333 | 63 |
| ASV25 | 8941b912fff0 | 7792 | 120 | 84 | 369 | 213 | 140 | 120 | 63 | 128 | 20 | 28 | 76 | 257 | 116 | 57 | 64 | 384 | 641 | 108 | 85 | 18 | 342 | 187 | 95 | 41 | 29 | 573 | 354 | 664 | 140 | 36 | 1028 | 535 | 388 | 231 | 58 |
| ASV26 | 1b980624ac5 | 7700 | 119 | 102 | 310 | 162 | 159 | 628 | 181 | 226 | 27 | 67 | 161 | 573 | 288 | 378 | 169 | 481 | 857 | 206 | 64 | 29 | 268 | 194 | 68 | 120 | 121 | 271 | 173 | 437 | 264 | 180 | 121 | 64 | 27 | 131 | 74 |
| ASV27 | 46eb9145a13 | 7639 | 156 | 179 | 441 | 290 | 397 | 548 | 94 | 201 | 130 | 282 | 40 | 118 | 100 | 411 | 339 | 58 | 190 | 62 | 105 | 65 | 114 | 111 | 63 | 337 | 357 | 32 | 38 | 324 | 793 | 701 | 59 | 0 | 0 | 250 | 254 |
| ASV28 | 05932a0c91d | 7561 | 185 | 121 | 377 | 286 | 416 | 516 | 216 | 191 | 97 | 101 | 108 | 354 | 181 | 225 | 379 | 261 | 451 | 143 | 75 | 57 | 256 | 185 | 125 | 137 | 176 | 243 | 120 | 555 | 438 | 207 | 64 | 29 | 27 | 111 | 148 |
| ASV29 | de6ce53e1a0 | 6917 | 195 | 158 | 504 | 357 | 659 | 639 | 260 | 205 | 186 | 246 | 0 | 177 | 135 | 393 | 439 | 0 | 0 | 86 | 156 | 92 | 72 | 40 | 103 | 261 | 241 | 0 | 0 | 290 | 511 | 319 | 0 | 0 | 0 | 51 | 142 |
| ASV30 | 53997053254 | 6894 | 329 | 264 | 765 | 427 | 279 | 629 | 185 | 118 | 0 | 0 | 183 | 593 | 222 | 80 | 51 | 417 | 762 | 211 | 68 | 0 | 244 | 119 | 70 | 110 | 47 | 180 | 92 | 249 | 136 | 0 | 0 | 0 | 0 | 48 | 16 |
| ASV31 | bb26c60c173 | 6703 | 315 | 329 | 735 | 472 | 564 | 700 | 267 | 194 | 39 | 74 | 111 | 306 | 168 | 355 | 167 | 99 | 268 | 86 | 57 | 8 | 102 | 70 | 52 | 79 | 91 | 77 | 64 | 262 | 272 | 135 | 26 | 0 | 16 | 68 | 75 |
| ASV32 | 2c6e619dbaa | 6696 | 335 | 281 | 712 | 434 | 471 | 229 | 92 | 161 | 11 | 103 | 113 | 242 | 97 | 106 | 156 | 166 | 238 | 142 | 164 | 54 | 169 | 122 | 107 | 198 | 108 | 172 | 77 | 370 | 311 | 128 | 99 | 42 | 51 | 256 | 179 |
| ASV33 | 08223140d04 | 6654 | 69 | 76 | 192 | 108 | 134 | 451 | 124 | 124 | 0 | 0 | 147 | 409 | 221 | 303 | 100 | 408 | 745 | 198 | 67 | 0 | 268 | 202 | 74 | 152 | 115 | 330 | 135 | 438 | 235 | 211 | 231 | 69 | 72 | 151 | 95 |
| ASV34 | e057f66ccacf | 6652 | 90 | 75 | 253 | 166 | 110 | 173 | 68 | 62 | 53 | 84 | 35 | 231 | 150 | 171 | 64 | 329 | 529 | 124 | 70 | 35 | 447 | 335 | 184 | 172 | 87 | 213 | 169 | 867 | 424 | 172 | 291 | 132 | 103 | 119 | 65 |
| ASV35 | 7290b3cdf56 | 6458 | 301 | 224 | 854 | 496 | 810 | 941 | 393 | 178 | 92 | 151 | 0 | 126 | 81 | 446 | 519 | 0 | 0 | 0 | 76 | 31 | 0 | 0 | 0 | 143 | 181 | 0 | 0 | 0 | 209 | 118 | 0 | 0 | 0 | 28 | 60 |
| ASV36 | a6a7d35754f | 6424 | 365 | 292 | 1272 | 636 | 356 | 796 | 307 | 187 | 40 | 44 | 98 | 354 | 229 | 280 | 128 | 134 | 209 | 140 | 58 | 0 | 21 | 14 | 24 | 65 | 31 | 19 | 16 | 86 | 113 | 24 | 10 | 12 | 0 | 47 | 17 |
| ASV37 | b85ed7a908f | 6026 | 161 | 162 | 340 | 332 | 583 | 337 | 179 | 271 | 228 | 258 | 7 | 0 | 41 | 98 | 568 | 45 | 36 | 69 | 240 | 85 | 115 | 47 | 76 | 312 | 328 | 31 | 35 | 281 | 342 | 134 | 39 | 16 | 0 | 72 | 158 |
| ASV38 | 9eb3f4b8a7f | 5943 | 138 | 128 | 425 | 225 | 404 | 325 | 121 | 155 | 71 | 114 | 71 | 295 | 123 | 184 | 166 | 350 | 595 | 210 | 336 | 77 | 61 | 36 | 74 | 166 | 150 | 85 | 48 | 154 | 172 | 141 | 41 | 0 | 51 | 145 | 106 |
| ASV39 | 7db24e7c57b | 5892 | 89 | 28 | 182 | 123 | 123 | 157 | 55 | 302 | 66 | 36 | 16 | 137 | 119 | 368 | 91 | 95 | 120 | 89 | 206 | 61 | 151 | 169 | 116 | 186 | 202 | 89 | 85 | 313 | 375 | 314 | 288 | 260 | 225 | 479 | 177 |
| ASV40 | b1c4d3c7008 | 5487 | 105 | 132 | 292 | 257 | 451 | 480 | 185 | 222 | 128 | 203 | 30 | 111 | 59 | 285 | 384 | 0 | 0 | 50 | 109 | 81 | 45 | 0 | 70 | 238 | 239 | 47 | 6 | 225 | 565 | 274 | 0 | 0 | 0 | 56 | 158 |
| ASV41 | c95be457c74 | 5374 | 51 | 29 | 189 | 107 | 208 | 114 | 53 | 204 | 103 | 278 | 8 | 37 | 37 | 96 | 245 | 0 | 126 | 42 | 124 | 103 | 70 | 92 | 54 | 316 | 313 | 42 | 32 | 352 | 632 | 492 | 166 | 60 | 17 | 300 | 282 |
| ASV42 | 191d7e7da61 | 5262 | 198 | 154 | 588 | 346 | 408 | 491 | 212 | 120 | 59 | 61 | 67 | 195 | 107 | 185 | 136 | 179 | 280 | 41 | 79 | 19 | 132 | 72 | 18 | 25 | 28 | 104 | 72 | 224 | 70 | 44 | 288 | 107 | 90 | 43 | 20 |
| ASV43 | 584ade95e9d | 5235 | 108 | 46 | 248 | 129 | 263 | 204 | 147 | 200 | 284 | 155 | 0 | 158 | 87 | 125 | 377 | 86 | 75 | 135 | 244 | 183 | 72 | 71 | 91 | 111 | 253 | 76 | 62 | 88 | 120 | 201 | 252 | 145 | 161 | 162 | 116 |
| ASV44 | 5472553891f | 5161 | 68 | 59 | 244 | 123 | 173 | 72 | 24 | 117 | 74 | 167 | 15 | 52 | 50 | 52 | 92 | 74 | 309 | 44 | 145 | 57 | 133 | 146 | 89 | 177 | 136 | 77 | 75 | 489 | 426 | 290 | 278 | 134 | 92 | 388 | 220 |
| ASV45 | 477382dd44c | 5119 | 75 | 57 | 295 | 133 | 28 | 166 | 70 | 171 | 0 | 14 | 98 | 325 | 156 | 28 | 16 | 430 | 669 | 146 | 61 | 0 | 117 | 69 | 43 | 36 | 10 | 330 | 204 | 246 | 88 | 20 | 491 | 205 | 154 | 136 | 32 |
| ASV46 | 0b3451bfacc | 5058 | 253 | 153 | 977 | 408 | 636 | 392 | 104 | 183 | 72 | 95 | 52 | 145 | 124 | 334 | 149 | 99 | 228 | 51 | 10 | 16 | 62 | 39 | 14 | 9 | 31 | 51 | 20 | 121 | 70 | 74 | 0 | 14 | 0 | 49 | 23 |
| ASV47 | 5d51ef8a472 | 4966 | 34 | 0 | 75 | 58 | 39 | 265 | 105 | 30 | 0 | 0 | 134 | 422 | 205 | 95 | 33 | 605 | 984 | 242 | 39 | 0 | 192 | 141 | 74 | 26 | 0 | 271 | 218 | 414 | 105 | 22 | 70 | 0 | 0 | 68 | 0 |
| ASV48 | dbd6a30965f | 4944 | 225 | 158 | 546 | 370 | 430 | 326 | 127 | 67 | 34 | 48 | 69 | 239 | 117 | 114 | 133 | 510 | 687 | 91 | 31 | 16 | 61 | 33 | 25 | 21 | 49 | 76 | 36 | 114 | 78 | 47 | 0 | 0 | 4 | 18 | 44 |
| ASV49 | b3ede366cfb | 4602 | 319 | 248 | 746 | 435 | 435 | 136 | 60 | 59 | 13 | 11 | 99 | 212 | 130 | 61 | 27 | 132 | 302 | 85 | 7 | 0 | 85 | 33 | 32 | 22 | 0 | 203 | 93 | 221 | 47 | 27 | 41 | 13 | 20 | 181 | 67 |
| ASV50 | bbaa4707c6e | 4454 | 44 | 36 | 115 | 48 | 68 | 117 | 129 | 142 | 101 | 86 | 41 | 113 | 116 | 89 | 90 | 179 | 306 | 67 | 115 | 70 | 68 | 14 | 110 | 208 | 146 | 132 | 71 | 168 | 205 | 133 | 459 | 260 | 213 | 105 | 90 |
| ASV51 | 802c4a42343 | 4444 | 105 | 95 | 337 | 187 | 109 | 277 | 93 | 37 | 0 | 0 | 153 | 373 | 222 | 206 | 33 | 113 | 218 | 116 | 70 | 0 | 145 | 130 | 106 | 187 | 42 | 48 | 52 | 410 | 348 | 148 | 0 | 0 | 0 | 43 | 41 |
| ASV52 | b97409614c1 | 4408 | 71 | 47 | 164 | 178 | 515 | 403 | 245 | 190 | 238 | 212 | 7 | 43 | 37 | 210 | 713 | 0 | 0 | 5 | 45 | 84 | 0 | 0 | 25 | 107 | 170 | 10 | 11 | 81 | 268 | 173 | 0 | 0 | 0 | 20 | 136 |
| ASV53 | 3582e1beb44 | 4149 | 153 | 169 | 344 | 251 | 213 | 370 | 137 | 68 | 41 | 43 | 86 | 220 | 111 | 147 | 80 | 136 | 239 | 74 | 97 | 0 | 141 | 83 | 40 | 97 | 69 | 71 | 49 | 206 | 147 | 57 | 46 | 0 | 0 | 101 | 63 |
| ASV54 | f6795113745 | 4135 | 95 | 0 | 156 | 90 | 147 | 135 | 75 | 148 | 136 | 52 | 0 | 164 | 59 | 66 | 197 | 150 | 206 | 115 | 123 | 85 | 100 | 72 | 89 | 60 | 138 | 93 | 70 | 112 | 109 | 79 | 389 | 230 | 226 | 99 | 70 |
| ASV55 | 9f4e97a31dd | 4036 | 119 | 80 | 132 | 83 | 125 | 95 | 68 | 114 | 91 | 69 | 27 | 113 | 66 | 71 | 128 | 227 | 286 | 166 | 110 | 73 | 61 | 72 | 66 | 110 | 85 | 79 | 85 | 72 | 55 | 45 | 341 | 278 | 246 | 124 | 74 |
| ASV56 | 35b220edde7 | 3974 | 0 | 0 | 0 | 0 | 0 | 0 | 0 | 218 | 321 | 816 | 0 | 0 | 0 | 0 | 311 | 0 | 0 | 0 | 184 | 246 | 0 | 0 | 0 | 452 | 415 | 0 | 0 | 119 | 390 | 136 | 0 | 0 | 0 | 122 | 244 |
| ASV57 | 44c88775a79 | 3942 | 106 | 0 | 238 | 120 | 281 | 84 | 0 | 121 | 77 | 0 | 0 | 151 | 95 | 0 | 83 | 296 | 347 | 175 | 115 | 67 | 136 | 102 | 142 | 0 | 47 | 160 | 71 | 183 | 75 | 49 | 241 | 143 | 106 | 131 | 0 |
| ASV58 | fe8f740f4d7f | 3809 | 61 | 89 | 248 | 88 | 75 | 416 | 211 | 197 | 92 | 0 | 103 | 137 | 151 | 74 | 96 | 118 | 243 | 108 | 112 | 0 | 139 | 187 | 150 | 222 | 105 | 0 | 0 | 129 | 208 | 50 | 0 | 0 | 0 | 0 | 0 |
| ASV59 | a8f894d55f7 | 3806 | 127 | 133 | 227 | 91 | 111 | 93 | 115 | 93 | 147 | 109 | 49 | 78 | 59 | 118 | 162 | 174 | 179 | 136 | 119 | 104 | 106 | 47 | 87 | 116 | 79 | 102 | 68 | 74 | 174 | 91 | 81 | 71 | 45 | 108 | 133 |
| ASV60 | 10884bc8cb1 | 3754 | 66 | 61 | 174 | 124 | 126 | 82 | 31 | 61 | 39 | 56 | 14 | 71 | 45 | 48 | 49 | 182 | 314 | 65 | 43 | 18 | 93 | 69 | 33 | 49 | 36 | 123 | 49 | 248 | 108 | 47 | 657 | 225 | 224 | 82 | 42 |
| ASV61 | 9e7dfa8ee4c | 3677 | 0 | 0 | 31 | 15 | 0 | 0 | 0 | 53 | 0 | 28 | 0 | 33 | 24 | 0 | 0 | 171 | 487 | 37 | 90 | 33 | 235 | 153 | 51 | 69 | 0 | 155 | 137 | 479 | 172 | 76 | 393 | 136 | 151 | 370 | 98 |
| ASV62 | f4c50f1ab46 | 3656 | 17 | 16 | 96 | 55 | 91 | 308 | 126 | 387 | 242 | 279 | 0 | 29 | 31 | 138 | 617 | 0 | 0 | 10 | 57 | 85 | 5 | 0 | 18 | 99 | 260 | 0 | 0 | 75 | 192 | 291 | 0 | 0 | 0 | 51 | 81 |
| ASV63 | 47ff44a4256 | 3475 | 98 | 39 | 187 | 104 | 164 | 110 | 41 | 80 | 0 | 0 | 79 | 147 | 74 | 41 | 0 | 249 | 366 | 96 | 0 | 0 | 155 | 50 | 59 | 0 | 0 | 208 | 147 | 253 | 102 | 0 | 244 | 142 | 117 | 86 | 37 |
| ASV64 | 6d99895665f | 3379 | 93 | 103 | 271 | 238 | 235 | 177 | 68 | 40 | 19 | 34 | 55 | 154 | 87 | 67 | 77 | 149 | 230 | 68 | 61 | 13 | 106 | 41 | 29 | 68 | 46 | 79 | 80 | 179 | 85 | 44 | 93 | 20 | 42 | 145 | 83 |
| ASV65 | d077a35cd9a | 3358 | 0 | 0 | 0 | 0 | 0 | 0 | 0 | 0 | 0 | 0 | 0 | 166 | 0 | 0 | 0 | 343 | 613 | 220 | 0 | 0 | 162 | 102 | 129 | 147 | 0 | 177 | 0 | 586 | 387 | 0 | 231 | 95 | 0 | 0 | 0 |
| ASV66 | daac7046e78 | 3338 | 9 | 11 | 40 | 38 | 184 | 24 | 16 | 117 | 320 | 334 | 0 | 0 | 0 | 24 | 817 | 0 | 0 | 0 | 118 | 184 | 0 | 0 | 24 | 178 | 360 | 0 | 0 | 19 | 115 | 141 | 0 | 0 | 0 | 42 | 223 |
| ASV67 | 26746171a93 | 3306 | 74 | 187 | 191 | 142 | 168 | 427 | 189 | 210 | 61 | 130 | 0 | 76 | 94 | 436 | 140 | 0 | 0 | 0 | 30 | 0 | 0 | 0 | 37 | 25 | 0 | 0 | 0 | 242 | 314 | 47 | 0 | 0 | 0 | 86 | 0 |
| ASV68 | c582f5e706a | 3280 | 47 | 0 | 112 | 41 | 79 | 64 | 43 | 31 | 0 | 0 | 0 | 79 | 42 | 30 | 35 | 199 | 284 | 37 | 67 | 45 | 101 | 62 | 39 | 0 | 28 | 190 | 110 | 266 | 60 | 23 | 654 | 196 | 223 | 60 | 33 |
| ASV69 | 992fe20d295 | 3236 | 61 | 0 | 133 | 46 | 0 | 141 | 38 | 53 | 0 | 9 | 205 | 796 | 445 | 111 | 0 | 119 | 327 | 92 | 0 | 0 | 0 | 0 | 13 | 0 | 0 | 123 | 53 | 138 | 31 | 0 | 126 | 33 | 0 | 116 | 27 |
| ASV70 | a10d9d5d48b | 3208 | 241 | 110 | 795 | 282 | 63 | 89 | 50 | 65 | 9 | 0 | 33 | 178 | 67 | 8 | 0 | 246 | 99 | 42 | 32 | 0 | 21 | 8 | 18 | 5 | 0 | 160 | 90 | 71 | 9 | 0 | 135 | 152 | 74 | 49 | 7 |
| ASV71 | b516660d69b | 2974 | 11 | 0 | 123 | 47 | 32 | 60 | 0 | 31 | 0 | 0 | 34 | 161 | 62 | 24 | 13 | 262 | 396 | 108 | 57 | 0 | 92 | 64 | 37 | 49 | 15 | 203 | 128 | 191 | 73 | 24 | 277 | 136 | 122 | 115 | 27 |
| ASV72 | 04a389e30cb | 2932 | 109 | 68 | 371 | 182 | 184 | 109 | 26 | 132 | 20 | 18 | 14 | 36 | 26 | 76 | 41 | 149 | 176 | 58 | 63 | 22 | 52 | 34 | 31 | 78 | 56 | 28 | 0 | 102 | 113 | 87 | 78 | 53 | 32 | 251 | 57 |
| ASV73 | 6cf2f2a5b63 | 2906 | 93 | 58 | 262 | 130 | 49 | 183 | 72 | 114 | 16 | 22 | 30 | 144 | 80 | 52 | 33 | 157 | 311 | 84 | 139 | 35 | 72 | 66 | 47 | 72 | 56 | 82 | 70 | 146 | 83 | 32 | 35 | 9 | 0 | 47 | 25 |
| ASV74 | d054169d4bf | 2848 | 156 | 116 | 552 | 222 | 272 | 250 | 71 | 207 | 43 | 36 | 20 | 74 | 66 | 265 | 62 | 0 | 60 | 41 | 52 | 0 | 0 | 0 | 25 | 0 | 41 | 0 | 0 | 26 | 99 | 67 | 0 | 0 | 0 | 25 | 0 |
| ASV75 | 3c59436521b | 2843 | 0 | 0 | 0 | 0 | 0 | 0 | 0 | 0 | 5 | 0 | 28 | 73 | 58 | 0 | 0 | 461 | 583 | 148 | 74 | 12 | 105 | 101 | 111 | 69 | 5 | 95 | 59 | 233 | 94 | 9 | 221 | 116 | 90 | 59 | 34 |
| ASV76 | 252554d2179 | 2823 | 49 | 45 | 180 | 81 | 57 | 258 | 130 | 121 | 0 | 0 | 42 | 154 | 97 | 156 | 80 | 64 | 152 | 77 | 207 | 28 | 112 | 54 | 58 | 123 | 110 | 7 | 0 | 70 | 139 | 95 | 53 | 0 | 0 | 0 | 24 |
| ASV77 | 9ed82d76e0e | 2784 | 54 | 28 | 91 | 78 | 341 | 64 | 49 | 80 | 238 | 230 | 0 | 0 | 0 | 58 | 330 | 0 | 0 | 0 | 61 | 126 | 0 | 0 | 111 | 188 | 0 | 0 | 0 | 64 | 151 | 197 | 0 | 0 | 0 | 61 | 184 |
| ASV78 | 6b31940a8be | 2718 | 43 | 46 | 99 | 89 | 180 | 158 | 57 | 102 | 78 | 147 | 0 | 50 | 29 | 75 | 96 | 57 | 103 | 48 | 87 | 33 | 76 | 60 | 34 | 45 | 71 | 8 | 37 | 164 | 193 | 101 | 117 | 56 | 36 | 70 | 73 |
| ASV79 | 4a1f5b9c4d1 | 2686 | 68 | 41 | 120 | 89 | 82 | 210 | 78 | 99 | 42 | 35 | 53 | 142 | 72 | 96 | 44 | 97 | 200 | 56 | 37 | 26 | 78 | 65 | 33 | 39 | 69 | 98 | 43 | 164 | 135 | 69 | 54 | 34 | 34 | 28 | 56 |
| ASV80 | b17f25f6a3c | 2644 | 109 | 107 | 332 | 229 | 286 | 254 | 98 | 125 | 105 | 119 | 13 | 57 | 51 | 62 | 222 | 0 | 22 | 24 | 95 | 24 | 0 | 0 | 18 | 33 | 44 | 0 | 0 | 47 | 52 | 36 | 14 | 0 | 0 | 37 | 29 |
| ASV81 | 3cc0470e5ae | 2473 | 73 | 68 | 231 | 116 | 93 | 159 | 74 | 37 | 26 | 0 | 44 | 151 | 67 | 60 | 42 | 152 | 282 | 101 | 28 | 0 | 93 | 67 | 37 | 0 | 0 | 89 | 63 | 185 | 46 | 0 | 34 | 0 | 0 | 36 | 19 |
| ASV82 | 77643ca079d | 2467 | 97 | 106 | 158 | 159 | 219 | 105 | 42 | 60 | 52 | 68 | 20 | 63 | 0 | 75 | 209 | 3 | 76 | 59 | 54 | 29 | 23 | 22 | 47 | 61 | 69 | 0 | 39 | 154 | 145 | 74 | 40 | 5 | 0 | 59 | 75 |
| ASV83 | db13c5b2833 | 2422 | 18 | 9 | 84 | 31 | 73 | 137 | 30 | 30 | 4 | 39 | 0 | 0 | 0 | 0 | 37 | 42 | 91 | 10 | 74 | 29 | 69 | 28 | 39 | 197 | 129 | 68 | 58 | 272 | 428 | 214 | 0 | 0 | 0 | 134 | 48 |
| ASV84 | 931b5b8dcc3 | 2377 | 85 | 104 | 182 | 121 | 93 | 132 | 63 | 76 | 29 | 31 | 41 | 82 | 44 | 65 | 63 | 102 | 156 | 80 | 39 | 19 | 60 | 53 | 30 | 87 | 46 | 47 | 24 | 129 | 111 | 41 | 24 | 17 | 0 | 57 | 44 |
| ASV85 | 88c221ea1bb | 2371 | 25 | 19 | 73 | 92 | 411 | 24 | 7 | 72 | 141 | 142 | 0 | 0 | 0 | 26 | 360 | 0 | 0 | 0 | 26 | 85 | 0 | 0 | 0 | 53 | 141 | 5 | 0 | 33 | 168 | 219 | 0 | 0 | 0 | 31 | 218 |
| ASV86 | 5ca72a029c2 | 2370 | 0 | 0 | 0 | 0 | 0 | 109 | 37 | 37 | 0 | 53 | 0 | 70 | 0 | 0 | 87 | 228 | 323 | 65 | 34 | 0 | 93 | 105 | 0 | 0 | 0 | 105 | 103 | 107 | 31 | 0 | 424 | 201 | 117 | 41 | 0 |
| ASV87 | 289dd5fdf60 | 2370 | 40 | 19 | 116 | 89 | 176 | 41 | 0 | 53 | 0 | 0 | 17 | 73 | 55 | 0 | 85 | 101 | 122 | 0 | 34 | 0 | 70 | 53 | 48 | 21 | 0 | 119 | 92 | 186 | 134 | 69 | 230 | 98 | 57 | 98 | 74 |
| ASV88 | 781bf210724 | 2352 | 31 | 37 | 76 | 45 | 70 | 24 | 22 | 43 | 13 | 17 | 15 | 51 | 18 | 7 | 46 | 93 | 171 | 68 | 73 | 17 | 90 | 35 | 52 | 98 | 62 | 188 | 97 | 233 | 138 | 34 | 135 | 66 | 43 | 67 | 77 |
| ASV89 | ed85ffc35c0 | 2312 | 58 | 35 | 252 | 100 | 163 | 249 | 75 | 163 | 32 | 28 | 24 | 59 | 60 | 499 | 91 | 8 | 10 | 10 | 32 | 0 | 0 | 8 | 8 | 54 | 58 | 3 | 0 | 16 | 97 | 78 | 0 | 0 | 0 | 20 | 22 |
| ASV90 | bde0f7cede6 | 2282 | 44 | 0 | 105 | 62 | 97 | 55 | 35 | 75 | 94 | 95 | 0 | 66 | 31 | 33 | 132 | 70 | 55 | 62 | 83 | 70 | 34 | 0 | 44 | 28 | 85 | 51 | 40 | 35 | 42 | 56 | 206 | 129 | 111 | 99 | 58 |
| ASV91 | 7dcb42dd3b5 | 2257 | 193 | 0 | 206 | 111 | 237 | 142 | 110 | 98 | 95 | 62 | 0 | 86 | 0 | 110 | 94 | 64 | 58 | 53 | 71 | 76 | 68 | 0 | 0 | 0 | 70 | 73 | 0 | 58 | 57 | 65 | 0 | 0 | 0 | 0 | 0 |
| ASV92 | ce5f5fefa9a0 | 2219 | 64 | 31 | 134 | 77 | 146 | 126 | 55 | 56 | 65 | 49 | 19 | 95 | 40 | 59 | 151 | 49 | 86 | 37 | 36 | 31 | 77 | 26 | 36 | 54 | 62 | 54 | 32 | 144 | 150 | 71 | 20 | 12 | 0 | 23 | 52 |
| ASV93 | e0a62ac8386 | 2177 | 80 | 24 | 261 | 111 | 177 | 136 | 45 | 158 | 39 | 13 | 11 | 47 | 70 | 154 | 103 | 0 | 0 | 0 | 69 | 13 | 0 | 0 | 28 | 58 | 58 | 0 | 15 | 0 | 91 | 117 | 0 | 0 | 0 | 174 | 125 |
| ASV94 | f8b4478984e | 2169 | 87 | 34 | 256 | 102 | 112 | 71 | 55 | 35 | 0 | 15 | 26 | 138 | 68 | 56 | 25 | 117 | 213 | 56 | 31 | 0 | 51 | 51 | 29 | 0 | 0 | 89 | 36 | 157 | 61 | 0 | 80 | 28 | 0 | 42 | 48 |
| ASV95 | 95aac9dadcf | 2150 | 21 | 14 | 64 | 29 | 19 | 0 | 16 | 15 | 0 | 0 | 22 | 49 | 37 | 6 | 0 | 170 | 244 | 74 | 19 | 0 | 104 | 73 | 53 | 18 | 3 | 246 | 149 | 302 | 90 | 24 | 59 | 41 | 21 | 129 | 39 |
| ASV96 | d4025c34871 | 2065 | 0 | 0 | 0 | 0 | 0 | 0 | 0 | 0 | 0 | 0 | 0 | 28 | 33 | 0 | 0 | 129 | 343 | 51 | 0 | 0 | 179 | 97 | 22 | 0 | 7 | 101 | 80 | 408 | 94 | 31 | 255 | 86 | 76 | 45 | 0 |
| ASV97 | 4e1167fbbbc | 2023 | 56 | 69 | 270 | 90 | 45 | 88 | 39 | 63 | 0 | 0 | 26 | 109 | 93 | 37 | 6 | 172 | 261 | 37 | 28 | 0 | 0 | 0 | 16 | 22 | 10 | 107 | 47 | 119 | 19 | 0 | 69 | 27 | 18 | 80 | 0 |
| ASV98 | 63b47a5c24b | 1970 | 146 | 54 | 461 | 187 | 102 | 175 | 44 | 113 | 0 | 10 | 4 | 43 | 48 | 77 | 17 | 41 | 19 | 33 | 17 | 0 | 17 | 11 | 24 | 27 | 18 | 41 | 30 | 66 | 37 | 10 | 0 | 12 | 10 | 70 | 6 |
| ASV99 | 291deb67818 | 1955 | 14 | 0 | 54 | 54 | 205 | 6 | 16 | 49 | 146 | 227 | 0 | 0 | 0 | 12 | 252 | 0 | 0 | 0 | 110 | 147 | 0 | 0 | 0 | 81 | 175 | 0 | 0 | 38 | 83 | 92 | 0 | 0 | 0 | 22 | 172 |
| ASV100 | 05794f49d61 | 1885 | 138 | 117 | 492 | 253 | 230 | 22 | 7 | 48 | 0 | 9 | 0 | 0 | 0 | 21 | 79 | 0 | 0 | 0 | 18 | 0 | 0 | 0 | 0 | 54 | 14 | 5 | 0 | 26 | 161 | 75 | 0 | 0 | 4 | 67 | 45 |
| ASV101 | c45b037c35d | 1802 | 66 | 45 | 154 | 112 | 133 | 53 | 26 | 49 | 43 | 89 | 7 | 38 | 25 | 13 | 96 | 25 | 39 | 18 | 83 | 41 | 35 | 16 | 27 | 65 | 42 | 8 | 10 | 58 | 82 | 54 | 38 | 17 | 21 | 86 | 88 |
| ASV102 | e46b4d859dc | 1798 | 46 | 32 | 294 | 119 | 83 | 51 | 0 | 0 | 11 | 15 | 71 | 379 | 183 | 130 | 25 | 55 | 124 | 0 | 44 | 0 | 0 | 0 | 0 | 0 | 0 | 35 | 0 | 44 | 0 | 0 | 0 | 0 | 0 | 51 | 6 |
| ASV103 | f9730793fd8 | 1760 | 109 | 85 | 235 | 152 | 133 | 188 | 66 | 31 | 0 | 0 | 21 | 73 | 66 | 56 | 50 | 28 | 46 | 28 | 34 | 0 | 42 | 19 | 6 | 0 | 47 | 24 | 25 | 58 | 44 | 25 | 16 | 0 | 0 | 20 | 33 |
| ASV104 | 85631eb6b9b | 1727 | 44 | 16 | 159 | 63 | 24 | 120 | 34 | 18 | 0 | 6 | 10 | 0 | 41 | 23 | 0 | 94 | 302 | 0 | 0 | 0 | 179 | 106 | 37 | 13 | 0 | 88 | 62 | 146 | 85 | 40 | 0 | 0 | 0 | 12 | 5 |
| ASV105 | 71f6bf6f8f50 | 1716 | 34 | 32 | 74 | 85 | 167 | 126 | 55 | 84 | 124 | 111 | 0 | 5 | 12 | 70 | 173 | 0 | 0 | 0 | 46 | 44 | 0 | 0 | 16 | 47 | 94 | 7 | 0 | 11 | 109 | 109 | 0 | 0 | 0 | 20 | 61 |
| ASV106 | f4dfaada866 | 1710 | 0 | 0 | 0 | 0 | 0 | 55 | 0 | 0 | 0 | 0 | 34 | 127 | 51 | 0 | 0 | 435 | 515 | 162 | 0 | 0 | 58 | 35 | 0 | 0 | 0 | 0 | 42 | 33 | 38 | 83 | 42 | 0 | 0 | 0 | 0 |
| ASV107 | a441d96a218 | 1706 | 152 | 118 | 356 | 251 | 349 | 262 | 114 | 17 | 0 | 0 | 0 | 19 | 23 | 32 | 13 | 0 | 0 | 0 | 0 | 0 | 0 | 0 | 0 | 0 | 0 | 0 | 0 | 0 | 0 | 0 | 0 | 0 | 0 | 0 | 0 |
| ASV108 | f4a6e50e16e | 1637 | 51 | 0 | 180 | 66 | 96 | 136 | 41 | 98 | 0 | 0 | 9 | 53 | 49 | 162 | 64 | 29 | 0 | 0 | 0 | 0 | 31 | 16 | 0 | 75 | 55 | 0 | 0 | 75 | 101 | 116 | 0 | 0 | 0 | 73 | 61 |
| ASV109 | 0d04c71dcf0 | 1576 | 20 | 0 | 32 | 0 | 0 | 232 | 81 | 78 | 0 | 21 | 57 | 152 | 74 | 66 | 0 | 94 | 148 | 43 | 32 | 0 | 74 | 48 | 0 | 0 | 28 | 0 | 0 | 101 | 31 | 0 | 107 | 57 | 0 | 0 | 0 |
| ASV110 | b9ec28fefa6 | 1547 | 11 | 13 | 28 | 27 | 214 | 16 | 5 | 34 | 33 | 70 | 0 | 0 | 6 | 12 | 230 | 0 | 0 | 0 | 23 | 26 | 0 | 0 | 13 | 92 | 77 | 11 | 6 | 51 | 284 | 138 | 0 | 0 | 0 | 23 | 104 |
| ASV111 | b7f557adfdc | 1541 | 0 | 0 | 0 | 0 | 0 | 0 | 0 | 0 | 0 | 0 | 0 | 93 | 0 | 0 | 0 | 291 | 546 | 0 | 0 | 0 | 0 | 0 | 0 | 0 | 0 | 268 | 80 | 76 | 0 | 0 | 131 | 56 | 0 | 0 | 0 |
| ASV112 | 1b7e6b1555e | 1538 | 56 | 0 | 147 | 94 | 160 | 43 | 0 | 112 | 119 | 79 | 0 | 0 | 0 | 161 | 165 | 0 | 0 | 0 | 59 | 56 | 0 | 0 | 0 | 127 | 78 | 0 | 0 | 0 | 0 | 82 | 0 | 0 | 0 | 0 | 0 |
| ASV113 | 843859b7982 | 1511 | 46 | 0 | 69 | 87 | 89 | 90 | 99 | 62 | 41 | 25 | 9 | 73 | 48 | 48 | 71 | 0 | 27 | 45 | 77 | 37 | 29 | 0 | 21 | 58 | 78 | 22 | 0 | 32 | 107 | 66 | 10 | 0 | 0 | 21 | 24 |
| ASV114 | db06db4029f | 1511 | 92 | 57 | 279 | 196 | 176 | 50 | 0 | 38 | 24 | 0 | 0 | 31 | 27 | 21 | 54 | 0 | 12 | 0 | 13 | 0 | 28 | 0 | 34 | 14 | 23 | 0 | 0 | 96 | 114 | 60 | 0 | 0 | 0 | 29 | 43 |
| ASV115 | 9d240954c03 | 1491 | 29 | 14 | 88 | 57 | 98 | 31 | 6 | 51 | 0 | 26 | 0 | 6 | 0 | 33 | 93 | 0 | 0 | 0 | 35 | 0 | 0 | 0 | 88 | 70 | 0 | 0 | 0 | 33 | 265 | 190 | 0 | 0 | 0 | 166 | 112 |
| ASV116 | 1a6128b13c5 | 1488 | 79 | 50 | 261 | 100 | 85 | 63 | 12 | 24 | 0 | 0 | 17 | 65 | 88 | 52 | 0 | 0 | 76 | 24 | 6 | 0 | 24 | 20 | 13 | 0 | 0 | 41 | 35 | 96 | 9 | 0 | 65 | 31 | 19 | 119 | 14 |
| ASV117 | 5afd2e8cb77 | 1486 | 38 | 45 | 92 | 80 | 211 | 103 | 60 | 73 | 47 | 38 | 0 | 10 | 16 | 97 | 160 | 0 | 0 | 7 | 21 | 0 | 0 | 0 | 0 | 28 | 58 | 0 | 0 | 19 | 64 | 111 | 0 | 0 | 0 | 18 | 90 |
| ASV118 | 5cbccd30f9d | 1473 | 0 | 14 | 32 | 22 | 0 | 15 | 0 | 0 | 12 | 0 | 0 | 0 | 0 | 0 | 0 | 33 | 45 | 13 | 0 | 0 | 107 | 49 | 29 | 0 | 0 | 22 | 0 | 128 | 42 | 0 | 429 | 203 | 164 | 114 | 0 |
| ASV119 | 6efc1f46a771 | 1424 | 0 | 55 | 84 | 24 | 35 | 129 | 64 | 67 | 64 | 65 | 0 | 0 | 36 | 297 | 150 | 25 | 39 | 0 | 0 | 0 | 0 | 0 | 0 | 40 | 25 | 0 | 0 | 30 | 87 | 27 | 39 | 0 | 0 | 42 | 0 |
| ASV120 | 3c5c4c699d1 | 1414 | 36 | 0 | 50 | 0 | 0 | 0 | 33 | 132 | 198 | 182 | 0 | 0 | 0 | 0 | 209 | 0 | 0 | 0 | 92 | 112 | 0 | 0 | 0 | 0 | 145 | 0 | 0 | 0 | 0 | 126 | 0 | 0 | 0 | 0 | 99 |
| ASV121 | 9e893d8a87f | 1411 | 137 | 90 | 238 | 130 | 146 | 168 | 60 | 150 | 0 | 0 | 0 | 55 | 56 | 102 | 25 | 0 | 0 | 0 | 0 | 0 | 0 | 0 | 0 | 0 | 24 | 0 | 0 | 0 | 30 | 0 | 0 | 0 | 0 | 0 | 0 |
| ASV122 | 07f203cb593 | 1401 | 59 | 29 | 227 | 99 | 170 | 102 | 37 | 122 | 16 | 0 | 0 | 36 | 34 | 163 | 84 | 0 | 0 | 0 | 0 | 0 | 0 | 0 | 22 | 27 | 0 | 0 | 0 | 29 | 38 | 41 | 0 | 0 | 0 | 44 | 22 |
| ASV123 | dfc67b35046 | 1400 | 0 | 0 | 0 | 0 | 0 | 49 | 0 | 0 | 0 | 0 | 0 | 0 | 0 | 0 | 0 | 166 | 143 | 89 | 0 | 0 | 44 | 49 | 0 | 0 | 0 | 67 | 39 | 50 | 0 | 0 | 332 | 210 | 162 | 0 | 0 |
| ASV124 | 15c5581ecf3 | 1380 | 0 | 0 | 16 | 0 | 0 | 0 | 0 | 16 | 0 | 0 | 4 | 27 | 25 | 0 | 0 | 110 | 94 | 27 | 39 | 0 | 47 | 34 | 26 | 0 | 0 | 90 | 65 | 119 | 16 | 6 | 157 | 166 | 134 | 149 | 13 |
| ASV125 | efe10567db1 | 1342 | 23 | 15 | 54 | 18 | 0 | 40 | 33 | 40 | 11 | 0 | 8 | 62 | 27 | 16 | 0 | 75 | 104 | 38 | 48 | 0 | 60 | 73 | 0 | 0 | 0 | 48 | 30 | 95 | 24 | 0 | 178 | 122 | 75 | 25 | 0 |
| ASV126 | 4bf887a6d1b | 1330 | 0 | 0 | 0 | 0 | 0 | 0 | 0 | 0 | 0 | 0 | 0 | 73 | 27 | 0 | 0 | 188 | 346 | 68 | 0 | 0 | 75 | 0 | 27 | 0 | 0 | 87 | 107 | 98 | 0 | 0 | 112 | 41 | 0 | 81 | 0 |
| ASV127 | 08f5c4a8d11 | 1320 | 0 | 0 | 88 | 44 | 126 | 68 | 29 | 52 | 0 | 42 | 0 | 0 | 0 | 72 | 46 | 0 | 38 | 0 | 0 | 0 | 0 | 0 | 0 | 72 | 53 | 0 | 0 | 101 | 195 | 167 | 0 | 11 | 11 | 44 | 61 |
| ASV128 | 82636c13aeb | 1272 | 31 | 27 | 118 | 73 | 78 | 179 | 41 | 85 | 16 | 18 | 8 | 18 | 17 | 64 | 52 | 0 | 0 | 0 | 31 | 0 | 7 | 0 | 8 | 21 | 11 | 18 | 8 | 83 | 115 | 41 | 15 | 11 | 7 | 57 | 14 |
| ASV129 | 85809ab6d10 | 1267 | 0 | 32 | 70 | 41 | 64 | 73 | 35 | 24 | 0 | 0 | 31 | 82 | 41 | 53 | 106 | 106 | 129 | 43 | 20 | 0 | 73 | 37 | 13 | 6 | 11 | 40 | 25 | 64 | 26 | 12 | 0 | 0 | 0 | 0 | 10 |
| ASV130 | b6e8e35011f | 1263 | 44 | 93 | 92 | 73 | 36 | 115 | 34 | 100 | 0 | 0 | 0 | 46 | 65 | 109 | 0 | 0 | 0 | 0 | 0 | 0 | 0 | 0 | 0 | 31 | 0 | 0 | 0 | 40 | 108 | 39 | 0 | 0 | 0 | 238 | 0 |
| ASV131 | 664af6a6335 | 1229 | 9 | 16 | 51 | 42 | 95 | 11 | 0 | 28 | 30 | 34 | 3 | 10 | 7 | 26 | 110 | 0 | 0 | 0 | 15 | 17 | 7 | 0 | 20 | 50 | 48 | 17 | 12 | 84 | 148 | 130 | 0 | 0 | 0 | 73 | 136 |
| ASV132 | e71e2cdbc8f | 1186 | 24 | 28 | 142 | 89 | 204 | 6 | 5 | 22 | 24 | 46 | 0 | 4 | 0 | 9 | 119 | 0 | 0 | 0 | 7 | 9 | 0 | 0 | 2 | 29 | 32 | 0 | 0 | 10 | 80 | 95 | 4 | 0 | 3 | 121 | 72 |
| ASV133 | 82d8d63aec8 | 1182 | 34 | 31 | 77 | 83 | 151 | 38 | 19 | 48 | 32 | 40 | 0 | 4 | 10 | 10 | 120 | 0 | 0 | 8 | 58 | 35 | 0 | 0 | 14 | 28 | 52 | 0 | 10 | 54 | 77 | 51 | 0 | 0 | 0 | 37 | 61 |
| ASV134 | 55fba451251 | 1181 | 24 | 19 | 133 | 69 | 119 | 83 | 32 | 22 | 6 | 0 | 11 | 57 | 37 | 118 | 15 | 47 | 76 | 49 | 24 | 0 | 23 | 15 | 17 | 8 | 21 | 7 | 12 | 64 | 33 | 17 | 0 | 0 | 0 | 13 | 10 |
| ASV135 | e7876560297 | 1162 | 29 | 15 | 70 | 39 | 41 | 44 | 23 | 50 | 0 | 0 | 0 | 12 | 30 | 105 | 0 | 14 | 28 | 0 | 0 | 0 | 52 | 31 | 0 | 48 | 37 | 23 | 0 | 88 | 146 | 103 | 0 | 0 | 0 | 92 | 42 |
| ASV136 | aa4e24fcdfd | 1150 | 7 | 6 | 13 | 11 | 113 | 0 | 5 | 31 | 85 | 54 | 0 | 0 | 0 | 0 | 290 | 0 | 0 | 0 | 11 | 36 | 0 | 0 | 20 | 30 | 94 | 4 | 0 | 35 | 92 | 89 | 0 | 0 | 0 | 18 | 106 |
| ASV137 | 7c5a0fd8a87 | 1144 | 23 | 0 | 44 | 59 | 147 | 121 | 50 | 62 | 64 | 66 | 0 | 0 | 0 | 53 | 200 | 0 | 0 | 0 | 0 | 0 | 0 | 0 | 0 | 32 | 75 | 0 | 0 | 16 | 82 | 50 | 0 | 0 | 0 | 0 | 0 |
| ASV138 | ef58a82c6ab | 1129 | 0 | 0 | 8 | 0 | 0 | 0 | 0 | 14 | 32 | 69 | 0 | 22 | 0 | 0 | 117 | 6 | 17 | 49 | 95 | 38 | 22 | 12 | 29 | 85 | 46 | 10 | 0 | 62 | 91 | 27 | 87 | 44 | 31 | 48 | 68 |
| ASV139 | e30d19fe5b0 | 1101 | 0 | 0 | 92 | 60 | 72 | 63 | 0 | 64 | 0 | 0 | 0 | 45 | 0 | 0 | 77 | 88 | 104 | 58 | 0 | 0 | 46 | 0 | 0 | 0 | 63 | 57 | 47 | 0 | 0 | 0 | 69 | 55 | 41 | 0 | 0 |
| ASV140 | 753197e4f87 | 1091 | 0 | 0 | 33 | 11 | 8 | 32 | 14 | 53 | 0 | 18 | 0 | 12 | 7 | 56 | 18 | 58 | 78 | 19 | 30 | 12 | 56 | 21 | 17 | 51 | 59 | 25 | 9 | 64 | 89 | 60 | 28 | 15 | 18 | 83 | 37 |
| ASV141 | 605559dba01 | 1084 | 0 | 0 | 82 | 55 | 86 | 0 | 0 | 30 | 100 | 93 | 0 | 24 | 0 | 0 | 159 | 0 | 0 | 0 | 60 | 66 | 14 | 0 | 38 | 0 | 75 | 14 | 0 | 17 | 0 | 87 | 18 | 8 | 8 | 0 | 50 |
| ASV142 | bcbbb314a8f | 1077 | 0 | 0 | 0 | 0 | 0 | 0 | 0 | 0 | 0 | 0 | 0 | 0 | 0 | 0 | 0 | 93 | 222 | 19 | 0 | 0 | 49 | 38 | 0 | 0 | 0 | 60 | 31 | 117 | 0 | 0 | 266 | 87 | 95 | 0 | 0 |
| ASV143 | 508de5bb11b | 1077 | 0 | 0 | 0 | 0 | 0 | 0 | 0 | 0 | 0 | 0 | 0 | 0 | 0 | 0 | 0 | 91 | 166 | 54 | 0 | 0 | 70 | 47 | 0 | 0 | 0 | 63 | 0 | 135 | 0 | 0 | 196 | 139 | 116 | 0 | 0 |
| ASV144 | de392404ac3 | 1050 | 0 | 0 | 0 | 0 | 0 | 0 | 0 | 0 | 0 | 4 | 0 | 44 | 0 | 0 | 11 | 248 | 324 | 57 | 38 | 0 | 31 | 16 | 23 | 0 | 0 | 0 | 13 | 77 | 0 | 0 | 101 | 35 | 28 | 0 | 0 |
| ASV145 | cc092684f2a | 1040 | 26 | 16 | 39 | 70 | 254 | 22 | 14 | 26 | 58 | 35 | 0 | 0 | 0 | 0 | 187 | 0 | 0 | 0 | 0 | 32 | 0 | 0 | 0 | 0 | 61 | 0 | 0 | 19 | 39 | 37 | 0 | 0 | 0 | 0 | 105 |
| ASV146 | 3e8607ee4c1 | 1036 | 0 | 0 | 0 | 0 | 0 | 0 | 0 | 0 | 0 | 0 | 0 | 0 | 0 | 0 | 0 | 94 | 210 | 46 | 0 | 0 | 53 | 43 | 21 | 8 | 0 | 134 | 121 | 149 | 15 | 0 | 54 | 0 | 32 | 46 | 10 |
| ASV147 | 8a13fa187cb | 1010 | 0 | 0 | 61 | 31 | 63 | 0 | 0 | 38 | 41 | 42 | 0 | 39 | 0 | 0 | 53 | 0 | 0 | 0 | 51 | 43 | 35 | 0 | 25 | 0 | 34 | 36 | 0 | 33 | 0 | 34 | 156 | 96 | 83 | 0 | 16 |
| ASV148 | c5034d5ed01 | 1001 | 41 | 0 | 96 | 48 | 0 | 64 | 36 | 59 | 14 | 28 | 0 | 65 | 27 | 39 | 0 | 24 | 82 | 33 | 82 | 0 | 40 | 29 | 0 | 0 | 0 | 16 | 17 | 63 | 24 | 37 | 18 | 10 | 0 | 9 | 0 |
| ASV149 | d2f397f5b7e | 989 | 0 | 0 | 0 | 0 | 0 | 21 | 4 | 0 | 0 | 0 | 63 | 102 | 60 | 0 | 0 | 151 | 286 | 22 | 18 | 0 | 0 | 0 | 0 | 0 | 0 | 53 | 38 | 53 | 7 | 0 | 60 | 21 | 16 | 14 | 0 |
| ASV150 | 98c93109ee2 | 989 | 0 | 0 | 0 | 0 | 0 | 13 | 0 | 0 | 0 | 0 | 15 | 31 | 0 | 0 | 0 | 190 | 279 | 75 | 0 | 0 | 28 | 0 | 0 | 0 | 0 | 75 | 21 | 88 | 0 | 0 | 94 | 35 | 0 | 45 | 0 |
| ASV151 | c23de50eb91 | 967 | 0 | 0 | 0 | 0 | 0 | 0 | 0 | 81 | 121 | 157 | 0 | 0 | 0 | 0 | 177 | 0 | 0 | 0 | 35 | 48 | 0 | 0 | 0 | 0 | 114 | 0 | 0 | 0 | 44 | 146 | 0 | 0 | 0 | 0 | 44 |
| ASV152 | 73bb08decc3 | 964 | 2 | 0 | 0 | 0 | 0 | 10 | 5 | 7 | 0 | 0 | 5 | 27 | 3 | 8 | 0 | 79 | 88 | 36 | 0 | 6 | 31 | 25 | 25 | 6 | 9 | 53 | 26 | 45 | 2 | 0 | 246 | 124 | 88 | 8 | 0 |
| ASV153 | b6b738a3d99 | 937 | 0 | 0 | 56 | 18 | 0 | 113 | 31 | 60 | 0 | 11 | 42 | 70 | 34 | 0 | 0 | 86 | 198 | 25 | 17 | 0 | 0 | 8 | 0 | 0 | 0 | 49 | 34 | 52 | 33 | 0 | 0 | 0 | 0 | 0 | 0 |
| ASV154 | 59968914441 | 928 | 0 | 0 | 39 | 29 | 0 | 0 | 31 | 0 | 0 | 0 | 0 | 93 | 36 | 29 | 0 | 68 | 156 | 23 | 39 | 0 | 21 | 40 | 15 | 0 | 0 | 26 | 50 | 53 | 108 | 0 | 42 | 0 | 30 | 0 | 0 |
| ASV155 | 19f01309d32 | 925 | 91 | 126 | 147 | 113 | 46 | 101 | 41 | 73 | 0 | 24 | 0 | 29 | 30 | 68 | 0 | 0 | 0 | 0 | 0 | 0 | 0 | 0 | 0 | 0 | 0 | 0 | 0 | 0 | 36 | 0 | 0 | 0 | 0 | 0 | 0 |
| ASV156 | c01b1b54d3e | 921 | 0 | 0 | 0 | 0 | 120 | 28 | 0 | 19 | 79 | 30 | 0 | 0 | 0 | 45 | 364 | 0 | 0 | 0 | 0 | 11 | 0 | 0 | 0 | 0 | 38 | 0 | 0 | 0 | 36 | 110 | 0 | 0 | 0 | 0 | 41 |
| ASV157 | ff20761d24c2 | 919 | 0 | 0 | 0 | 19 | 217 | 0 | 0 | 0 | 70 | 57 | 0 | 0 | 0 | 0 | 426 | 0 | 0 | 0 | 0 | 0 | 0 | 0 | 0 | 0 | 27 | 0 | 0 | 0 | 0 | 26 | 0 | 0 | 0 | 0 | 77 |
| ASV158 | 6887783221d | 886 | 0 | 0 | 202 | 135 | 0 | 114 | 0 | 0 | 0 | 0 | 0 | 0 | 0 | 0 | 0 | 170 | 166 | 0 | 0 | 0 | 0 | 0 | 0 | 0 | 0 | 0 | 0 | 99 | 0 | 0 | 0 | 0 | 0 | 0 | 0 |
| ASV159 | 03a52768936 | 876 | 0 | 0 | 31 | 19 | 0 | 27 | 0 | 0 | 0 | 0 | 22 | 40 | 0 | 0 | 0 | 107 | 234 | 0 | 0 | 0 | 25 | 0 | 0 | 0 | 0 | 78 | 54 | 129 | 45 | 0 | 31 | 0 | 34 | 0 | 0 |
| ASV160 | ff02653b169 | 854 | 24 | 8 | 77 | 35 | 52 | 165 | 68 | 20 | 0 | 0 | 22 | 95 | 80 | 108 | 22 | 0 | 0 | 0 | 0 | 0 | 0 | 0 | 0 | 0 | 0 | 0 | 0 | 52 | 26 | 0 | 0 | 0 | 0 | 0 | 0 |
| ASV161 | f21c48ae04d | 833 | 0 | 0 | 0 | 0 | 0 | 0 | 0 | 0 | 0 | 0 | 0 | 0 | 0 | 0 | 0 | 66 | 127 | 0 | 0 | 0 | 0 | 0 | 0 | 0 | 0 | 57 | 45 | 0 | 0 | 0 | 267 | 148 | 123 | 0 | 0 |
| ASV162 | b772e3ea402 | 830 | 0 | 0 | 0 | 0 | 0 | 0 | 0 | 0 | 0 | 0 | 0 | 0 | 0 | 0 | 0 | 0 | 0 | 0 | 0 | 0 | 0 | 0 | 0 | 0 | 0 | 0 | 0 | 0 | 0 | 0 | 349 | 284 | 176 | 21 | 0 |
| ASV163 | 4d072cfea4d | 828 | 0 | 0 | 19 | 0 | 12 | 0 | 0 | 0 | 0 | 0 | 19 | 20 | 7 | 0 | 0 | 78 | 163 | 23 | 15 | 0 | 47 | 31 | 28 | 16 | 0 | 71 | 48 | 52 | 15 | 5 | 102 | 26 | 20 | 11 | 0 |
| ASV164 | 71362904ece | 826 | 0 | 0 | 0 | 0 | 0 | 0 | 0 | 0 | 0 | 0 | 0 | 0 | 0 | 0 | 0 | 71 | 191 | 0 | 0 | 0 | 30 | 29 | 0 | 0 | 0 | 20 | 20 | 106 | 0 | 0 | 195 | 77 | 51 | 36 | 0 |
| ASV165 | c5dbf2ea44c | 825 | 0 | 0 | 50 | 0 | 31 | 0 | 0 | 0 | 0 | 0 | 0 | 28 | 13 | 0 | 0 | 69 | 138 | 23 | 0 | 0 | 32 | 30 | 24 | 0 | 0 | 50 | 29 | 65 | 0 | 15 | 64 | 76 | 42 | 46 | 0 |
| ASV166 | fe37c03cee5 | 822 | 29 | 6 | 45 | 24 | 23 | 33 | 15 | 9 | 0 | 0 | 27 | 45 | 19 | 21 | 10 | 44 | 109 | 13 | 19 | 0 | 42 | 18 | 11 | 7 | 0 | 40 | 21 | 75 | 29 | 15 | 34 | 0 | 8 | 31 | 0 |
| ASV167 | bf18173ef06 | 821 | 0 | 5 | 18 | 22 | 62 | 15 | 0 | 26 | 0 | 18 | 0 | 7 | 0 | 0 | 71 | 0 | 7 | 16 | 56 | 33 | 14 | 0 | 51 | 52 | 0 | 10 | 18 | 111 | 104 | 0 | 31 | 3 | 10 | 0 | 61 |
| ASV168 | 44b6d0d6326 | 813 | 0 | 0 | 0 | 0 | 0 | 0 | 0 | 0 | 0 | 0 | 0 | 0 | 0 | 0 | 0 | 0 | 0 | 0 | 0 | 0 | 0 | 0 | 0 | 0 | 0 | 200 | 177 | 145 | 0 | 0 | 90 | 86 | 98 | 11 | 6 |
| ASV169 | de986b628b6 | 808 | 0 | 0 | 0 | 0 | 0 | 0 | 0 | 0 | 0 | 0 | 10 | 31 | 0 | 0 | 0 | 82 | 138 | 0 | 0 | 0 | 0 | 0 | 0 | 0 | 0 | 59 | 22 | 0 | 0 | 0 | 288 | 112 | 66 | 0 | 0 |
| ASV170 | dbff2a1f6252 | 785 | 0 | 0 | 19 | 0 | 202 | 0 | 0 | 0 | 66 | 55 | 0 | 0 | 0 | 0 | 320 | 0 | 0 | 0 | 0 | 14 | 0 | 0 | 0 | 0 | 25 | 0 | 0 | 0 | 0 | 35 | 0 | 0 | 0 | 0 | 49 |
| ASV171 | 2f6a5b789dc | 779 | 82 | 51 | 241 | 133 | 86 | 110 | 50 | 0 | 8 | 14 | 0 | 0 | 0 | 4 | 0 | 0 | 0 | 0 | 0 | 0 | 0 | 0 | 0 | 0 | 0 | 0 | 0 | 0 | 0 | 0 | 0 | 0 | 0 | 0 | 0 |
| ASV172 | c53dad5262b | 767 | 0 | 0 | 42 | 0 | 17 | 0 | 0 | 0 | 0 | 0 | 7 | 32 | 28 | 0 | 0 | 86 | 195 | 36 | 24 | 0 | 0 | 0 | 0 | 11 | 0 | 35 | 27 | 34 | 0 | 0 | 44 | 38 | 34 | 60 | 17 |
| ASV173 | d3cdda9df90 | 756 | 3 | 0 | 14 | 10 | 0 | 0 | 0 | 5 | 4 | 0 | 0 | 0 | 0 | 28 | 11 | 0 | 0 | 0 | 9 | 0 | 6 | 0 | 7 | 41 | 21 | 15 | 5 | 97 | 223 | 110 | 5 | 5 | 8 | 79 | 50 |
| ASV174 | 6b3d10862ba | 743 | 0 | 0 | 0 | 0 | 22 | 11 | 0 | 71 | 36 | 60 | 0 | 0 | 0 | 18 | 69 | 0 | 0 | 0 | 0 | 19 | 0 | 0 | 0 | 72 | 78 | 0 | 0 | 26 | 62 | 112 | 0 | 0 | 0 | 54 | 33 |
| ASV175 | b6c0d89b177 | 733 | 25 | 15 | 56 | 47 | 75 | 20 | 21 | 31 | 56 | 61 | 0 | 9 | 7 | 17 | 76 | 0 | 0 | 10 | 65 | 44 | 0 | 0 | 4 | 12 | 38 | 0 | 0 | 5 | 22 | 0 | 0 | 0 | 0 | 0 | 17 |
| ASV176 | c895c3255c9 | 733 | 38 | 39 | 141 | 32 | 51 | 100 | 58 | 69 | 0 | 38 | 0 | 0 | 0 | 9 | 0 | 0 | 0 | 0 | 0 | 0 | 0 | 0 | 0 | 0 | 0 | 0 | 0 | 0 | 21 | 47 | 0 | 0 | 0 | 63 | 27 |
| ASV177 | 1c42fd5161b | 732 | 90 | 63 | 252 | 151 | 176 | 0 | 0 | 0 | 0 | 0 | 0 | 0 | 0 | 0 | 0 | 0 | 0 | 0 | 0 | 0 | 0 | 0 | 0 | 0 | 0 | 0 | 0 | 0 | 0 | 0 | 0 | 0 | 0 | 0 | 0 |
| ASV178 | b5cf59752a5 | 732 | 0 | 0 | 0 | 0 | 0 | 0 | 0 | 0 | 0 | 0 | 0 | 0 | 0 | 0 | 0 | 49 | 95 | 0 | 0 | 0 | 72 | 0 | 0 | 0 | 0 | 39 | 47 | 70 | 0 | 0 | 184 | 92 | 74 | 10 | 0 |
| ASV179 | ed1f814cf7d | 730 | 47 | 0 | 176 | 109 | 82 | 0 | 0 | 0 | 0 | 0 | 54 | 27 | 0 | 0 | 0 | 27 | 0 | 0 | 0 | 0 | 0 | 0 | 0 | 0 | 0 | 31 | 0 | 84 | 60 | 0 | 0 | 0 | 0 | 0 | 33 |
| ASV180 | a8d6e8aaa8f | 728 | 0 | 3 | 12 | 11 | 33 | 0 | 0 | 35 | 39 | 56 | 0 | 0 | 0 | 0 | 67 | 0 | 0 | 0 | 27 | 12 | 0 | 0 | 0 | 62 | 70 | 0 | 0 | 0 | 102 | 122 | 0 | 0 | 0 | 42 | 35 |
| ASV181 | 31686a4eece | 728 | 0 | 0 | 0 | 0 | 58 | 0 | 0 | 33 | 107 | 102 | 0 | 0 | 0 | 0 | 133 | 0 | 0 | 0 | 68 | 82 | 0 | 0 | 0 | 0 | 72 | 0 | 0 | 0 | 0 | 39 | 0 | 0 | 0 | 0 | 34 |
| ASV182 | e582bb515b1 | 723 | 0 | 0 | 0 | 0 | 60 | 0 | 0 | 0 | 105 | 101 | 0 | 0 | 0 | 0 | 104 | 0 | 0 | 0 | 30 | 80 | 0 | 0 | 0 | 0 | 56 | 0 | 0 | 0 | 32 | 105 | 0 | 0 | 0 | 0 | 50 |
| ASV183 | a67d31cadaa | 722 | 0 | 0 | 21 | 6 | 0 | 18 | 0 | 36 | 0 | 0 | 11 | 41 | 27 | 21 | 10 | 64 | 101 | 50 | 11 | 0 | 33 | 25 | 5 | 20 | 8 | 16 | 14 | 20 | 19 | 23 | 35 | 17 | 0 | 59 | 11 |
| ASV184 | 09e533a38f7 | 712 | 52 | 26 | 131 | 69 | 23 | 57 | 18 | 4 | 0 | 0 | 21 | 62 | 44 | 9 | 0 | 33 | 49 | 15 | 3 | 0 | 21 | 7 | 12 | 0 | 0 | 27 | 0 | 19 | 7 | 3 | 0 | 0 | 0 | 0 | 0 |
| ASV185 | 68f8a4ce347 | 712 | 0 | 0 | 0 | 0 | 0 | 0 | 0 | 0 | 0 | 0 | 0 | 0 | 0 | 0 | 0 | 37 | 56 | 0 | 0 | 0 | 36 | 8 | 0 | 0 | 0 | 0 | 0 | 20 | 0 | 0 | 313 | 122 | 120 | 0 | 0 |
| ASV186 | 0c74eae4dbf | 712 | 29 | 0 | 64 | 67 | 67 | 21 | 7 | 22 | 0 | 17 | 0 | 28 | 19 | 15 | 31 | 0 | 22 | 0 | 26 | 0 | 0 | 0 | 46 | 17 | 18 | 0 | 0 | 27 | 0 | 31 | 0 | 8 | 0 | 93 | 37 |
| ASV187 | a5555d4fb3c | 711 | 15 | 0 | 20 | 25 | 18 | 16 | 0 | 19 | 9 | 0 | 0 | 40 | 0 | 0 | 21 | 95 | 150 | 21 | 0 | 0 | 29 | 21 | 0 | 0 | 0 | 69 | 41 | 49 | 13 | 0 | 24 | 8 | 0 | 0 | 0 |
| ASV188 | df3a0e0d959 | 707 | 77 | 15 | 165 | 82 | 77 | 111 | 48 | 69 | 0 | 0 | 0 | 28 | 0 | 35 | 0 | 0 | 0 | 0 | 0 | 0 | 0 | 0 | 0 | 0 | 0 | 0 | 0 | 0 | 0 | 0 | 0 | 8 | 8 | 0 | 0 |
| ASV189 | 215cb8c53fd | 704 | 0 | 0 | 0 | 0 | 0 | 0 | 0 | 0 | 0 | 0 | 0 | 0 | 0 | 0 | 0 | 113 | 179 | 69 | 0 | 0 | 57 | 0 | 0 | 50 | 0 | 68 | 0 | 168 | 0 | 0 | 0 | 0 | 0 | 0 | 0 |
| ASV190 | 269c56d6f6d | 703 | 34 | 33 | 154 | 88 | 85 | 21 | 5 | 22 | 0 | 0 | 0 | 3 | 8 | 23 | 0 | 0 | 0 | 0 | 0 | 0 | 0 | 15 | 7 | 0 | 0 | 0 | 12 | 55 | 27 | 0 | 0 | 0 | 0 | 77 | 34 |
| ASV191 | 635b862162c | 685 | 35 | 34 | 32 | 34 | 62 | 22 | 24 | 35 | 23 | 50 | 12 | 22 | 0 | 0 | 27 | 19 | 50 | 22 | 49 | 0 | 0 | 0 | 14 | 0 | 32 | 0 | 0 | 14 | 48 | 0 | 12 | 13 | 0 | 0 | 0 |
| ASV192 | 07915a1137f | 668 | 0 | 6 | 35 | 21 | 30 | 0 | 24 | 26 | 28 | 0 | 13 | 0 | 0 | 0 | 0 | 18 | 37 | 18 | 31 | 19 | 20 | 11 | 17 | 46 | 0 | 14 | 13 | 55 | 19 | 32 | 29 | 22 | 18 | 60 | 6 |
| ASV193 | 62131078606 | 656 | 0 | 0 | 44 | 32 | 54 | 32 | 12 | 74 | 45 | 56 | 3 | 0 | 0 | 42 | 48 | 0 | 0 | 0 | 22 | 22 | 0 | 0 | 12 | 0 | 24 | 0 | 0 | 0 | 48 | 65 | 0 | 0 | 0 | 21 | 0 |
| ASV194 | 4756f9f6568 | 643 | 0 | 10 | 35 | 31 | 135 | 57 | 23 | 25 | 48 | 25 | 0 | 8 | 0 | 32 | 135 | 0 | 0 | 0 | 0 | 0 | 0 | 0 | 0 | 0 | 16 | 0 | 0 | 0 | 39 | 15 | 0 | 0 | 0 | 0 | 9 |
| ASV195 | 868e7facae9 | 639 | 0 | 0 | 53 | 0 | 0 | 80 | 21 | 27 | 0 | 0 | 23 | 54 | 56 | 0 | 0 | 31 | 73 | 63 | 0 | 0 | 0 | 0 | 17 | 4 | 0 | 0 | 23 | 0 | 0 | 7 | 76 | 12 | 19 | 0 | 0 |
| ASV196 | cd803d1bccf | 636 | 0 | 0 | 65 | 33 | 108 | 61 | 30 | 20 | 34 | 58 | 0 | 0 | 0 | 105 | 63 | 0 | 0 | 0 | 0 | 0 | 0 | 0 | 0 | 0 | 29 | 0 | 0 | 0 | 19 | 11 | 0 | 0 | 0 | 0 | 0 |
| ASV197 | acf6b2c4a4d | 630 | 0 | 0 | 0 | 0 | 0 | 0 | 0 | 0 | 0 | 0 | 0 | 0 | 0 | 0 | 0 | 0 | 0 | 0 | 0 | 0 | 0 | 0 | 0 | 0 | 0 | 49 | 0 | 0 | 0 | 0 | 316 | 88 | 177 | 0 | 0 |
| ASV198 | bdc0a8e3e90 | 628 | 0 | 0 | 18 | 6 | 19 | 25 | 10 | 30 | 0 | 0 | 0 | 10 | 8 | 43 | 22 | 7 | 0 | 16 | 12 | 0 | 16 | 12 | 27 | 11 | 0 | 9 | 8 | 54 | 66 | 30 | 40 | 26 | 21 | 55 | 27 |
| ASV199 | addcc2812e9 | 628 | 99 | 55 | 207 | 133 | 56 | 15 | 0 | 10 | 0 | 0 | 0 | 0 | 14 | 22 | 0 | 0 | 0 | 4 | 0 | 0 | 0 | 0 | 0 | 4 | 0 | 0 | 0 | 0 | 0 | 0 | 0 | 0 | 0 | 9 | 0 |
| ASV200 | b2969b12e4a | 614 | 24 | 19 | 120 | 31 | 114 | 26 | 102 | 0 | 0 | 0 | 13 | 31 | 49 | 56 | 8 | 0 | 0 | 21 | 0 | 0 | 0 | 0 | 0 | 0 | 0 | 0 | 0 | 0 | 0 | 0 | 0 | 0 | 0 | 0 | 0 |
| ASV201 | f078b26f395 | 614 | 38 | 61 | 160 | 87 | 54 | 81 | 31 | 0 | 0 | 0 | 0 | 31 | 0 | 0 | 4 | 15 | 18 | 9 | 0 | 0 | 0 | 0 | 8 | 0 | 0 | 0 | 0 | 9 | 8 | 0 | 0 | 0 | 0 | 0 | 0 |
| ASV202 | 2219cd28969 | 607 | 34 | 27 | 52 | 45 | 72 | 8 | 0 | 5 | 0 | 0 | 0 | 14 | 0 | 0 | 20 | 19 | 50 | 21 | 0 | 0 | 14 | 0 | 7 | 11 | 0 | 44 | 27 | 57 | 22 | 4 | 14 | 0 | 0 | 15 | 25 |
| ASV203 | ce330d07615 | 606 | 0 | 0 | 0 | 0 | 5 | 0 | 0 | 0 | 0 | 0 | 0 | 0 | 0 | 0 | 0 | 61 | 98 | 31 | 19 | 0 | 66 | 51 | 0 | 0 | 0 | 49 | 61 | 52 | 0 | 0 | 48 | 0 | 0 | 65 | 0 |
| ASV204 | 46dcd9314a0 | 606 | 0 | 0 | 78 | 53 | 138 | 0 | 0 | 44 | 60 | 47 | 0 | 0 | 0 | 0 | 51 | 0 | 0 | 44 | 0 | 0 | 0 | 0 | 0 | 0 | 0 | 0 | 0 | 0 | 35 | 0 | 0 | 0 | 0 | 26 | 30 |
| ASV205 | 50c80c9baee | 600 | 45 | 34 | 107 | 72 | 42 | 71 | 38 | 26 | 5 | 7 | 0 | 0 | 11 | 0 | 6 | 0 | 0 | 33 | 6 | 0 | 0 | 0 | 17 | 3 | 0 | 0 | 0 | 18 | 17 | 0 | 14 | 11 | 0 | 11 | 6 |
| ASV206 | a1c22815777 | 598 | 9 | 0 | 40 | 0 | 0 | 216 | 57 | 28 | 0 | 0 | 0 | 13 | 5 | 0 | 0 | 58 | 160 | 0 | 0 | 0 | 12 | 0 | 0 | 0 | 0 | 0 | 0 | 0 | 0 | 0 | 0 | 0 | 0 | 0 | 0 |
| ASV207 | [illegible] | [illegible] | [illegible] | [illegible] | [illegible] | [illegible] | [illegible] | [illegible] | [illegible] | [illegible] | [illegible] | [illegible] | [illegible] | [illegible] | [illegible] | [illegible] | [illegible] | [illegible] | [illegible] | [illegible] | [illegible] | [illegible] | [illegible] | [illegible] | [illegible] | [illegible] | [illegible] | [illegible] | [illegible] | [illegible] | [illegible] | [illegible] | [illegible] | [illegible] | [illegible] | [illegible] | [illegible] | [illegible] |
| ASV208 | [illegible] | [illegible] | [illegible] | [illegible] | [illegible] | [illegible] | [illegible] | [illegible] | [illegible] | [illegible] | [illegible] | [illegible] | [illegible] | [illegible] | [illegible] | [illegible] | [illegible] | [illegible] | [illegible] | [illegible] | [illegible] | [illegible] | [illegible] | [illegible] | [illegible] | [illegible] | [illegible] | [illegible] | [illegible] | [illegible] | [illegible] | [illegible] | [illegible] | [illegible] | [illegible] | [illegible] | [illegible] | [illegible] |
| ASV209 | [illegible] | [illegible] | [illegible] | [illegible] | [illegible] | [illegible] | [illegible] | [illegible] | [illegible] | [illegible] | [illegible] | [illegible] | [illegible] | [illegible] | [illegible] | [illegible] | [illegible] | [illegible] | [illegible] | [illegible] | [illegible] | [illegible] | [illegible] | [illegible] | [illegible] | [illegible] | [illegible] | [illegible] | [illegible] | [illegible] | [illegible] | [illegible] | [illegible] | [illegible] | [illegible] | [illegible] | [illegible] | [illegible] |
| ASV210 | [illegible] | [illegible] | [illegible] | [illegible] | [illegible] | [illegible] | [illegible] | [illegible] | [illegible] | [illegible] | [illegible] | [illegible] | [illegible] | [illegible] | [illegible] | [illegible] | [illegible] | [illegible] | [illegible] | [illegible] | [illegible] | [illegible] | [illegible] | [illegible] | [illegible] | [illegible] | [illegible] | [illegible] | [illegible] | [illegible] | [illegible] | [illegible] | [illegible] | [illegible] | [illegible] | [illegible] | [illegible] | [illegible] |
| ASV211 | [illegible] | [illegible] | [illegible] | [illegible] | [illegible] | [illegible] | [illegible] | [illegible] | [illegible] | [illegible] | [illegible] | [illegible] | [illegible] | [illegible] | [illegible] | [illegible] | [illegible] | [illegible] | [illegible] | [illegible] | [illegible] | [illegible] | [illegible] | [illegible] | [illegible] | [illegible] | [illegible] | [illegible] | [illegible] | [illegible] | [illegible] | [illegible] | [illegible] | [illegible] | [illegible] | [illegible] | [illegible] | [illegible] |
| ASV212 | [illegible] | [illegible] | [illegible] | [illegible] | [illegible] | [illegible] | [illegible] | [illegible] | [illegible] | [illegible] | [illegible] | [illegible] | [illegible] | [illegible] | [illegible] | [illegible] | [illegible] | [illegible] | [illegible] | [illegible] | [illegible] | [illegible] | [illegible] | [illegible] | [illegible] | [illegible] | [illegible] | [illegible] | [illegible] | [illegible] | [illegible] | [illegible] | [illegible] | [illegible] | [illegible] | [illegible] | [illegible] | [illegible] |
| ASV213 | [illegible] | [illegible] | [illegible] | [illegible] | [illegible] | [illegible] | [illegible] | [illegible] | [illegible] | [illegible] | [illegible] | [illegible] | [illegible] | [illegible] | [illegible] | [illegible] | [illegible] | [illegible] | [illegible] | [illegible] | [illegible] | [illegible] | [illegible] | [illegible] | [illegible] | [illegible] | [illegible] | [illegible] | [illegible] | [illegible] | [illegible] | [illegible] | [illegible] | [illegible] | [illegible] | [illegible] | [illegible] | [illegible] |
| ASV214 | [illegible] | [illegible] | [illegible] | [illegible] | [illegible] | [illegible] | [illegible] | [illegible] | [illegible] | [illegible] | [illegible] | [illegible] | [illegible] | [illegible] | [illegible] | [illegible] | [illegible] | [illegible] | [illegible] | [illegible] | [illegible] | [illegible] | [illegible] | [illegible] | [illegible] | [illegible] | [illegible] | [illegible] | [illegible] | [illegible] | [illegible] | [illegible] | [illegible] | [illegible] | [illegible] | [illegible] | [illegible] | [illegible] |
| ASV215 | [illegible] | [illegible] | [illegible] | [illegible] | [illegible] | [illegible] | [illegible] | [illegible] | [illegible] | [illegible] | [illegible] | [illegible] | [illegible] | [illegible] | [illegible] | [illegible] | [illegible] | [illegible] | [illegible] | [illegible] | [illegible] | [illegible] | [illegible] | [illegible] | [illegible] | [illegible] | [illegible] | [illegible] | [illegible] | [illegible] | [illegible] | [illegible] | [illegible] | [illegible] | [illegible] | [illegible] | [illegible] | [illegible] |
| ASV216 | [illegible] | [illegible] | [illegible] | [illegible] | [illegible] | [illegible] | [illegible] | [illegible] | [illegible] | [illegible] | [illegible] | [illegible] | [illegible] | [illegible] | [illegible] | [illegible] | [illegible] | [illegible] | [illegible] | [illegible] | [illegible] | [illegible] | [illegible] | [illegible] | [illegible] | [illegible] | [illegible] | [illegible] | [illegible] | [illegible] | [illegible] | [illegible] | [illegible] | [illegible] | [illegible] | [illegible] | [illegible] | [illegible] |
| ASV217 | [illegible] | [illegible] | [illegible] | [illegible] | [illegible] | [illegible] | [illegible] | [illegible] | [illegible] | [illegible] | [illegible] | [illegible] | [illegible] | [illegible] | [illegible] | [illegible] | [illegible] | [illegible] | [illegible] | [illegible] | [illegible] | [illegible] | [illegible] | [illegible] | [illegible] | [illegible] | [illegible] | [illegible] | [illegible] | [illegible] | [illegible] | [illegible] | [illegible] | [illegible] | [illegible] | [illegible] | [illegible] | [illegible] |
| ASV218 | [illegible] | [illegible] | [illegible] | [illegible] | [illegible] | [illegible] | [illegible] | [illegible] | [illegible] | [illegible] | [illegible] | [illegible] | [illegible] | [illegible] | [illegible] | [illegible] | [illegible] | [illegible] | [illegible] | [illegible] | [illegible] | [illegible] | [illegible] | [illegible] | [illegible] | [illegible] | [illegible] | [illegible] | [illegible] | [illegible] | [illegible] | [illegible] | [illegible] | [illegible] | [illegible] | [illegible] | [illegible] | [illegible] |
| ASV219 | [illegible] | [illegible] | [illegible] | [illegible] | [illegible] | [illegible] | [illegible] | [illegible] | [illegible] | [illegible] | [illegible] | [illegible] | [illegible] | [illegible] | [illegible] | [illegible] | [illegible] | [illegible] | [illegible] | [illegible] | [illegible] | [illegible] | [illegible] | [illegible] | [illegible] | [illegible] | [illegible] | [illegible] | [illegible] | [illegible] | [illegible] | [illegible] | [illegible] | [illegible] | [illegible] | [illegible] | [illegible] | [illegible] |
| ASV220 | [illegible] | [illegible] | [illegible] | [illegible] | [illegible] | [illegible] | [illegible] | [illegible] | [illegible] | [illegible] | [illegible] | [illegible] | [illegible] | [illegible] | [illegible] | [illegible] | [illegible] | [illegible] | [illegible] | [illegible] | [illegible] | [illegible] | [illegible] | [illegible] | [illegible] | [illegible] | [illegible] | [illegible] | [illegible] | [illegible] | [illegible] | [illegible] | [illegible] | [illegible] | [illegible] | [illegible] | [illegible] | [illegible] |
| ASV221 | [illegible] | [illegible] | [illegible] | [illegible] | [illegible] | [illegible] | [illegible] | [illegible] | [illegible] | [illegible] | [illegible] | [illegible] | [illegible] | [illegible] | [illegible] | [illegible] | [illegible] | [illegible] | [illegible] | [illegible] | [illegible] | [illegible] | [illegible] | [illegible] | [illegible] | [illegible] | [illegible] | [illegible] | [illegible] | [illegible] | [illegible] | [illegible] | [illegible] | [illegible] | [illegible] | [illegible] | [illegible] | [illegible] |
| ASV222 | [illegible] | [illegible] | [illegible] | [illegible] | [illegible] | [illegible] | [illegible] | [illegible] | [illegible] | [illegible] | [illegible] | [illegible] | [illegible] | [illegible] | [illegible] | [illegible] | [illegible] | [illegible] | [illegible] | [illegible] | [illegible] | [illegible] | [illegible] | [illegible] | [illegible] | [illegible] | [illegible] | [illegible] | [illegible] | [illegible] | [illegible] | [illegible] | [illegible] | [illegible] | [illegible] | [illegible] | [illegible] | [illegible] |
| ASV223 | [illegible] | [illegible] | [illegible] | [illegible] | [illegible] | [illegible] | [illegible] | [illegible] | [illegible] | [illegible] | [illegible] | [illegible] | [illegible] | [illegible] | [illegible] | [illegible] | [illegible] | [illegible] | [illegible] | [illegible] | [illegible] | [illegible] | [illegible] | [illegible] | [illegible] | [illegible] | [illegible] | [illegible] | [illegible] | [illegible] | [illegible] | [illegible] | [illegible] | [illegible] | [illegible] | [illegible] | [illegible] | [illegible] |
| ASV224 | [illegible] | [illegible] | [illegible] | [illegible] | [illegible] | [illegible] | [illegible] | [illegible] | [illegible] | [illegible] | [illegible] | [illegible] | [illegible] | [illegible] | [illegible] | [illegible] | [illegible] | [illegible] | [illegible] | [illegible] | [illegible] | [illegible] | [illegible] | [illegible] | [illegible] | [illegible] | [illegible] | [illegible] | [illegible] | [illegible] | [illegible] | [illegible] | [illegible] | [illegible] | [illegible] | [illegible] | [illegible] | [illegible] |
| ASV225 | [illegible] | [illegible] | [illegible] | [illegible] | [illegible] | [illegible] | [illegible] | [illegible] | [illegible] | [illegible] | [illegible] | [illegible] | [illegible] | [illegible] | [illegible] | [illegible] | [illegible] | [illegible] | [illegible] | [illegible] | [illegible] | [illegible] | [illegible] | [illegible] | [illegible] | [illegible] | [illegible] | [illegible] | [illegible] | [illegible] | [illegible] | [illegible] | [illegible] | [illegible] | [illegible] | [illegible] | [illegible] | [illegible] |
| ASV226 | [illegible] | [illegible] | [illegible] | [illegible] | [illegible] | [illegible] | [illegible] | [illegible] | [illegible] | [illegible] | [illegible] | [illegible] | [illegible] | [illegible] | [illegible] | [illegible] | [illegible] | [illegible] | [illegible] | [illegible] | [illegible] | [illegible] | [illegible] | [illegible] | [illegible] | [illegible] | [illegible] | [illegible] | [illegible] | [illegible] | [illegible] | [illegible] | [illegible] | [illegible] | [illegible] | [illegible] | [illegible] | [illegible] |
| ASV227 | [illegible] | [illegible] | [illegible] | [illegible] | [illegible] | [illegible] | [illegible] | [illegible] | [illegible] | [illegible] | [illegible] | [illegible] | [illegible] | [illegible] | [illegible] | [illegible] | [illegible] | [illegible] | [illegible] | [illegible] | [illegible] | [illegible] | [illegible] | [illegible] | [illegible] | [illegible] | [illegible] | [illegible] | [illegible] | [illegible] | [illegible] | [illegible] | [illegible] | [illegible] | [illegible] | [illegible] | [illegible] | [illegible] |
| ASV228 | [illegible] | [illegible] | [illegible] | [illegible] | [illegible] | [illegible] | [illegible] | [illegible] | [illegible] | [illegible] | [illegible] | [illegible] | [illegible] | [illegible] | [illegible] | [illegible] | [illegible] | [illegible] | [illegible] | [illegible] | [illegible] | [illegible] | [illegible] | [illegible] | [illegible] | [illegible] | [illegible] | [illegible] | [illegible] | [illegible] | [illegible] | [illegible] | [illegible] | [illegible] | [illegible] | [illegible] | [illegible] | [illegible] |
| ASV229 | [illegible] | [illegible] | [illegible] | [illegible] | [illegible] | [illegible] | [illegible] | [illegible] | [illegible] | [illegible] | [illegible] | [illegible] | [illegible] | [illegible] | [illegible] | [illegible] | [illegible] | [illegible] | [illegible] | [illegible] | [illegible] | [illegible] | [illegible] | [illegible] | [illegible] | [illegible] | [illegible] | [illegible] | [illegible] | [illegible] | [illegible] | [illegible] | [illegible] | [illegible] | [illegible] | [illegible] | [illegible] | [illegible] |
| ASV230 | [illegible] | [illegible] | [illegible] | [illegible] | [illegible] | [illegible] | [illegible] | [illegible] | [illegible] | [illegible] | [illegible] | [illegible] | [illegible] | [illegible] | [illegible] | [illegible] | [illegible] | [illegible] | [illegible] | [illegible] | [illegible] | [illegible] | [illegible] | [illegible] | [illegible] | [illegible] | [illegible] | [illegible] | [illegible] | [illegible] | [illegible] | [illegible] | [illegible] | [illegible] | [illegible] | [illegible] | [illegible] | [illegible] |
| ASV231 | [illegible] | [illegible] | [illegible] | [illegible] | [illegible] | [illegible] | [illegible] | [illegible] | [illegible] | [illegible] | [illegible] | [illegible] | [illegible] | [illegible] | [illegible] | [illegible] | [illegible] | [illegible] | [illegible] | [illegible] | [illegible] | [illegible] | [illegible] | [illegible] | [illegible] | [illegible] | [illegible] | [illegible] | [illegible] | [illegible] | [illegible] | [illegible] | [illegible] | [illegible] | [illegible] | [illegible] | [illegible] | [illegible] |
| ASV232 | [illegible] | [illegible] | [illegible] | [illegible] | [illegible] | [illegible] | [illegible] | [illegible] | [illegible] | [illegible] | [illegible] | [illegible] | [illegible] | [illegible] | [illegible] | [illegible] | [illegible] | [illegible] | [illegible] | [illegible] | [illegible] | [illegible] | [illegible] | [illegible] | [illegible] | [illegible] | [illegible] | [illegible] | [illegible] | [illegible] | [illegible] | [illegible] | [illegible] | [illegible] | [illegible] | [illegible] | [illegible] | [illegible] |
| ASV233 | [illegible] | [illegible] | [illegible] | [illegible] | [illegible] | [illegible] | [illegible] | [illegible] | [illegible] | [illegible] | [illegible] | [illegible] | [illegible] | [illegible] | [illegible] | [illegible] | [illegible] | [illegible] | [illegible] | [illegible] | [illegible] | [illegible] | [illegible] | [illegible] | [illegible] | [illegible] | [illegible] | [illegible] | [illegible] | [illegible] | [illegible] | [illegible] | [illegible] | [illegible] | [illegible] | [illegible] | [illegible] | [illegible] |
| ASV234 | [illegible] | [illegible] | [illegible] | [illegible] | [illegible] | [illegible] | [illegible] | [illegible] | [illegible] | [illegible] | [illegible] | [illegible] | [illegible] | [illegible] | [illegible] | [illegible] | [illegible] | [illegible] | [illegible] | [illegible] | [illegible] | [illegible] | [illegible] | [illegible] | [illegible] | [illegible] | [illegible] | [illegible] | [illegible] | [illegible] | [illegible] | [illegible] | [illegible] | [illegible] | [illegible] | [illegible] | [illegible] | [illegible] |
| ASV235 | [illegible] | [illegible] | [illegible] | [illegible] | [illegible] | [illegible] | [illegible] | [illegible] | [illegible] | [illegible] | [illegible] | [illegible] | [illegible] | [illegible] | [illegible] | [illegible] | [illegible] | [illegible] | [illegible] | [illegible] | [illegible] | [illegible] | [illegible] | [illegible] | [illegible] | [illegible] | [illegible] | [illegible] | [illegible] | [illegible] | [illegible] | [illegible] | [illegible] | [illegible] | [illegible] | [illegible] | [illegible] | [illegible] |
| ASV236 | [illegible] | [illegible] | [illegible] | [illegible] | [illegible] | [illegible] | [illegible] | [illegible] | [illegible] | [illegible] | [illegible] | [illegible] | [illegible] | [illegible] | [illegible] | [illegible] | [illegible] | [illegible] | [illegible] | [illegible] | [illegible] | [illegible] | [illegible] | [illegible] | [illegible] | [illegible] | [illegible] | [illegible] | [illegible] | [illegible] | [illegible] | [illegible] | [illegible] | [illegible] | [illegible] | [illegible] | [illegible] | [illegible] |
| ASV237 | [illegible] | [illegible] | [illegible] | [illegible] | [illegible] | [illegible] | [illegible] | [illegible] | [illegible] | [illegible] | [illegible] | [illegible] | [illegible] | [illegible] | [illegible] | [illegible] | [illegible] | [illegible] | [illegible] | [illegible] | [illegible] | [illegible] | [illegible] | [illegible] | [illegible] | [illegible] | [illegible] | [illegible] | [illegible] | [illegible] | [illegible] | [illegible] | [illegible] | [illegible] | [illegible] | [illegible] | [illegible] | [illegible] |
| ASV238 | [illegible] | [illegible] | [illegible] | [illegible] | [illegible] | [illegible] | [illegible] | [illegible] | [illegible] | [illegible] | [illegible] | [illegible] | [illegible] | [illegible] | [illegible] | [illegible] | [illegible] | [illegible] | [illegible] | [illegible] | [illegible] | [illegible] | [illegible] | [illegible] | [illegible] | [illegible] | [illegible] | [illegible] | [illegible] | [illegible] | [illegible] | [illegible] | [illegible] | [illegible] | [illegible] | [illegible] | [illegible] | [illegible] |
| ASV239 | [illegible] | [illegible] | [illegible] | [illegible] | [illegible] | [illegible] | [illegible] | [illegible] | [illegible] | [illegible] | [illegible] | [illegible] | [illegible] | [illegible] | [illegible] | [illegible] | [illegible] | [illegible] | [illegible] | [illegible] | [illegible] | [illegible] | [illegible] | [illegible] | [illegible] | [illegible] | [illegible] | [illegible] | [illegible] | [illegible] | [illegible] | [illegible] | [illegible] | [illegible] | [illegible] | [illegible] | [illegible] | [illegible] |
| ASV240 | [illegible] | [illegible] | [illegible] | [illegible] | [illegible] | [illegible] | [illegible] | [illegible] | [illegible] | [illegible] | [illegible] | [illegible] | [illegible] | [illegible] | [illegible] | [illegible] | [illegible] | [illegible] | [illegible] | [illegible] | [illegible] | [illegible] | [illegible] | [illegible] | [illegible] | [illegible] | [illegible] | [illegible] | [illegible] | [illegible] | [illegible] | [illegible] | [illegible] | [illegible] | [illegible] | [illegible] | [illegible] | [illegible] |
| ASV241 | [illegible] | [illegible] | [illegible] | [illegible] | [illegible] | [illegible] | [illegible] | [illegible] | [illegible] | [illegible] | [illegible] | [illegible] | [illegible] | [illegible] | [illegible] | [illegible] | [illegible] | [illegible] | [illegible] | [illegible] | [illegible] | [illegible] | [illegible] | [illegible] | [illegible] | [illegible] | [illegible] | [illegible] | [illegible] | [illegible] | [illegible] | [illegible] | [illegible] | [illegible] | [illegible] | [illegible] | [illegible] | [illegible] |
| ASV242 | [illegible] | [illegible] | [illegible] | [illegible] | [illegible] | [illegible] | [illegible] | [illegible] | [illegible] | [illegible] | [illegible] | [illegible] | [illegible] | [illegible] | [illegible] | [illegible] | [illegible] | [illegible] | [illegible] | [illegible] | [illegible] | [illegible] | [illegible] | [illegible] | [illegible] | [illegible] | [illegible] | [illegible] | [illegible] | [illegible] | [illegible] | [illegible] | [illegible] | [illegible] | [illegible] | [illegible] | [illegible] | [illegible] |
| ASV243 | [illegible] | [illegible] | [illegible] | [illegible] | [illegible] | [illegible] | [illegible] | [illegible] | [illegible] | [illegible] | [illegible] | [illegible] | [illegible] | [illegible] | [illegible] | [illegible] | [illegible] | [illegible] | [illegible] | [illegible] | [illegible] | [illegible] | [illegible] | [illegible] | [illegible] | [illegible] | [illegible] | [illegible] | [illegible] | [illegible] | [illegible] | [illegible] | [illegible] | [illegible] | [illegible] | [illegible] | [illegible] | [illegible] |
| ASV244 | [illegible] | [illegible] | [illegible] | [illegible] | [illegible] | [illegible] | [illegible] | [illegible] | [illegible] | [illegible] | [illegible] | [illegible] | [illegible] | [illegible] | [illegible] | [illegible] | [illegible] | [illegible] | [illegible] | [illegible] | [illegible] | [illegible] | [illegible] | [illegible] | [illegible] | [illegible] | [illegible] | [illegible] | [illegible] | [illegible] | [illegible] | [illegible] | [illegible] | [illegible] | [illegible] | [illegible] | [illegible] | [illegible] |
| ASV245 | [illegible] | [illegible] | [illegible] | [illegible] | [illegible] | [illegible] | [illegible] | [illegible] | [illegible] | [illegible] | [illegible] | [illegible] | [illegible] | [illegible] | [illegible] | [illegible] | [illegible] | [illegible] | [illegible] | [illegible] | [illegible] | [illegible] | [illegible] | [illegible] | [illegible] | [illegible] | [illegible] | [illegible] | [illegible] | [illegible] | [illegible] | [illegible] | [illegible] | [illegible] | [illegible] | [illegible] | [illegible] | [illegible] |
| ASV246 | [illegible] | [illegible] | [illegible] | [illegible] | [illegible] | [illegible] | [illegible] | [illegible] | [illegible] | [illegible] | [illegible] | [illegible] | [illegible] | [illegible] | [illegible] | [illegible] | [illegible] | [illegible] | [illegible] | [illegible] | [illegible] | [illegible] | [illegible] | [illegible] | [illegible] | [illegible] | [illegible] | [illegible] | [illegible] | [illegible] | [illegible] | [illegible] | [illegible] | [illegible] | [illegible] | [illegible] | [illegible] | [illegible] |
| ASV247 | [illegible] | [illegible] | [illegible] | [illegible] | [illegible] | [illegible] | [illegible] | [illegible] | [illegible] | [illegible] | [illegible] | [illegible] | [illegible] | [illegible] | [illegible] | [illegible] | [illegible] | [illegible] | [illegible] | [illegible] | [illegible] | [illegible] | [illegible] | [illegible] | [illegible] | [illegible] | [illegible] | [illegible] | [illegible] | [illegible] | [illegible] | [illegible] | [illegible] | [illegible] | [illegible] | [illegible] | [illegible] | [illegible] |
| ASV248 | [illegible] | [illegible] | [illegible] | [illegible] | [illegible] | [illegible] | [illegible] | [illegible] | [illegible] | [illegible] | [illegible] | [illegible] | [illegible] | [illegible] | [illegible] | [illegible] | [illegible] | [illegible] | [illegible] | [illegible] | [illegible] | [illegible] | [illegible] | [illegible] | [illegible] | [illegible] | [illegible] | [illegible] | [illegible] | [illegible] | [illegible] | [illegible] | [illegible] | [illegible] | [illegible] | [illegible] | [illegible] | [illegible] |
| ASV249 | [illegible] | [illegible] | [illegible] | [illegible] | [illegible] | [illegible] | [illegible] | [illegible] | [illegible] | [illegible] | [illegible] | [illegible] | [illegible] | [illegible] | [illegible] | [illegible] | [illegible] | [illegible] | [illegible] | [illegible] | [illegible] | [illegible] | [illegible] | [illegible] | [illegible] | [illegible] | [illegible] | [illegible] | [illegible] | [illegible] | [illegible] | [illegible] | [illegible] | [illegible] | [illegible] | [illegible] | [illegible] | [illegible] |
| ASV250 | [illegible] | [illegible] | [illegible] | [illegible] | [illegible] | [illegible] | [illegible] | [illegible] | [illegible] | [illegible] | [illegible] | [illegible] | [illegible] | [illegible] | [illegible] | [illegible] | [illegible] | [illegible] | [illegible] | [illegible] | [illegible] | [illegible] | [illegible] | [illegible] | [illegible] | [illegible] | [illegible] | [illegible] | [illegible] | [illegible] | [illegible] | [illegible] | [illegible] | [illegible] | [illegible] | [illegible] | [illegible] | [illegible] |
| ASV251 | [illegible] | [illegible] | [illegible] | [illegible] | [illegible] | [illegible] | [illegible] | [illegible] | [illegible] | [illegible] | [illegible] | [illegible] | [illegible] | [illegible] | [illegible] | [illegible] | [illegible] | [illegible] | [illegible] | [illegible] | [illegible] | [illegible] | [illegible] | [illegible] | [illegible] | [illegible] | [illegible] | [illegible] | [illegible] | [illegible] | [illegible] | [illegible] | [illegible] | [illegible] | [illegible] | [illegible] | [illegible] | [illegible] |
| ASV252 | [illegible] | [illegible] | [illegible] | [illegible] | [illegible] | [illegible] | [illegible] | [illegible] | [illegible] | [illegible] | [illegible] | [illegible] | [illegible] | [illegible] | [illegible] | [illegible] | [illegible] | [illegible] | [illegible] | [illegible] | [illegible] | [illegible] | [illegible] | [illegible] | [illegible] | [illegible] | [illegible] | [illegible] | [illegible] | [illegible] | [illegible] | [illegible] | [illegible] | [illegible] | [illegible] | [illegible] | [illegible] | [illegible] |
| ASV253 | [illegible] | [illegible] | [illegible] | [illegible] | [illegible] | [illegible] | [illegible] | [illegible] | [illegible] | [illegible] | [illegible] | [illegible] | [illegible] | [illegible] | [illegible] | [illegible] | [illegible] | [illegible] | [illegible] | [illegible] | [illegible] | [illegible] | [illegible] | [illegible] | [illegible] | [illegible] | [illegible] | [illegible] | [illegible] | [illegible] | [illegible] | [illegible] | [illegible] | [illegible] | [illegible] | [illegible] | [illegible] | [illegible] |

**Supplementary Table S5**. Bacterioplankton 16S rRNA ASV feature table
ASVs: 1779
Seqs: 918792

| #ASV_ID | Feature_ID | total seqs | May12_0m | May12_05m | May12_25m | May12_50m | May12_75m | Jun8_0m | Jun8_05m | Jun8_25m | Jun8_50m | Jun8_75m | Jun22_0m | Jun22_05m | Jun22_25m | Jun22_50m | Jun22_75m | Jul11_0m | Jul11_05m | Jul11_25m | Jul11_50m | Jul11_75m | Jul27_0m | Jul27_05m | Jul27_25m | Jul27_50m | Jul27_75m | Aug5_0m | Aug5_05m | Aug5_25m | Aug5_50m | Aug5_75m | Sep8_0m | Sep8_05m | Sep8_25m | Sep8_50m | Sep8_75m |
|---|---|---|---|---|---|---|---|---|---|---|---|---|---|---|---|---|---|---|---|---|---|---|---|---|---|---|---|---|---|---|---|---|---|---|---|---|---|
| ASV254 | 9ec12434c1f8 | 441 | 43 | 49 | 135 | 76 | 132 | 0 | 0 | 0 | 0 | 0 | 0 | 0 | 0 | 3 | 0 | 0 | 0 | 0 | 0 | 0 | 0 | 0 | 0 | 0 | 0 | 0 | 0 | 0 | 0 | 0 | 0 | 0 | 0 | 3 | 0 |
| ASV255 | c074ffe7bde4 | 433 | 0 | 0 | 34 | 8 | 8 | 24 | 0 | 0 | 0 | 0 | 0 | 21 | 3 | 37 | 0 | 32 | 35 | 16 | 0 | 0 | 47 | 31 | 14 | 7 | 0 | 38 | 13 | 46 | 12 | 0 | 0 | 0 | 0 | 7 | 0 |
| ASV256 | 16cf0460d6cc | 429 | 23 | 0 | 31 | 43 | 96 | 47 | 0 | 32 | 7 | 33 | 0 | 0 | 0 | 14 | 20 | 0 | 0 | 0 | 0 | 17 | 0 | 0 | 14 | 14 | 0 | 0 | 0 | 0 | 26 | 12 | 0 | 0 | 0 | 0 | 0 |
| ASV257 | 4ff9a81b1cd1 | 426 | 0 | 0 | 0 | 0 | 0 | 0 | 0 | 0 | 0 | 0 | 0 | 0 | 0 | 0 | 0 | 35 | 84 | 19 | 0 | 0 | 33 | 0 | 0 | 0 | 0 | 28 | 0 | 51 | 0 | 0 | 98 | 42 | 36 | 0 | 0 |
| ASV258 | 00e9343d8fc | 416 | 0 | 0 | 17 | 11 | 10 | 30 | 0 | 13 | 18 | 35 | 0 | 8 | 7 | 6 | 33 | 0 | 0 | 13 | 33 | 18 | 0 | 0 | 0 | 28 | 32 | 0 | 0 | 0 | 16 | 18 | 13 | 17 | 10 | 16 | 14 |
| ASV259 | e5fa8243359 | 416 | 8 | 0 | 0 | 0 | 57 | 0 | 0 | 0 | 30 | 31 | 0 | 0 | 2 | 0 | 146 | 0 | 0 | 0 | 7 | 20 | 0 | 0 | 0 | 8 | 41 | 0 | 0 | 0 | 17 | 20 | 0 | 0 | 0 | 0 | 29 |
| ASV260 | f07714adb61 | 413 | 0 | 6 | 22 | 0 | 43 | 36 | 19 | 16 | 9 | 15 | 0 | 32 | 0 | 21 | 23 | 0 | 7 | 17 | 20 | 0 | 0 | 19 | 11 | 21 | 22 | 0 | 0 | 15 | 24 | 0 | 0 | 0 | 0 | 0 | 15 |
| ASV261 | 30f7b76d702 | 411 | 0 | 0 | 0 | 0 | 0 | 0 | 0 | 0 | 0 | 0 | 0 | 0 | 0 | 0 | 2 | 0 | 0 | 0 | 0 | 0 | 14 | 3 | 0 | 0 | 100 | 0 | 0 | 7 | 0 | 138 | 0 | 11 | 16 | 33 | 87 |
| ASV262 | d772fc4bf761 | 408 | 27 | 17 | 95 | 46 | 82 | 36 | 0 | 21 | 0 | 14 | 0 | 10 | 0 | 9 | 0 | 0 | 15 | 0 | 0 | 0 | 0 | 0 | 0 | 0 | 0 | 15 | 21 | 0 | 0 | 0 | 0 | 0 | 0 | 0 | 0 |
| ASV263 | ad06170a075 | 405 | 0 | 0 | 0 | 0 | 0 | 0 | 0 | 0 | 0 | 0 | 4 | 18 | 0 | 0 | 0 | 71 | 91 | 0 | 0 | 0 | 22 | 0 | 0 | 0 | 0 | 57 | 0 | 0 | 9 | 0 | 93 | 34 | 6 | 0 | 0 |
| ASV264 | 3ba46e55691 | 404 | 22 | 21 | 127 | 56 | 18 | 87 | 28 | 20 | 0 | 0 | 0 | 7 | 0 | 18 | 0 | 0 | 0 | 0 | 0 | 0 | 0 | 0 | 0 | 0 | 0 | 0 | 0 | 0 | 0 | 0 | 0 | 0 | 0 | 0 | 0 |
| ASV265 | 204e689938a | 394 | 25 | 11 | 84 | 31 | 23 | 41 | 13 | 8 | 0 | 0 | 11 | 17 | 37 | 23 | 0 | 0 | 0 | 0 | 0 | 0 | 0 | 0 | 0 | 0 | 0 | 22 | 17 | 25 | 0 | 0 | 0 | 0 | 6 | 0 | 0 |
| ASV266 | fe80e81f7f40 | 394 | 0 | 0 | 0 | 0 | 0 | 0 | 0 | 0 | 0 | 0 | 0 | 0 | 0 | 0 | 0 | 17 | 32 | 0 | 0 | 0 | 8 | 8 | 0 | 0 | 0 | 6 | 0 | 16 | 0 | 0 | 128 | 72 | 43 | 64 | 0 |
| ASV267 | 960cacb58c1 | 391 | 0 | 0 | 0 | 0 | 0 | 0 | 0 | 0 | 0 | 0 | 0 | 0 | 0 | 0 | 0 | 137 | 169 | 0 | 0 | 0 | 0 | 0 | 0 | 0 | 0 | 0 | 0 | 0 | 85 | 0 | 0 | 0 | 0 | 0 | 0 |
| ASV268 | 2548fb397f3 | 387 | 28 | 0 | 37 | 24 | 77 | 0 | 0 | 0 | 48 | 59 | 0 | 0 | 0 | 0 | 71 | 0 | 0 | 0 | 0 | 0 | 0 | 0 | 0 | 0 | 0 | 0 | 0 | 0 | 0 | 0 | 0 | 0 | 0 | 26 | 17 |
| ASV269 | 4cf34a75a41 | 387 | 42 | 0 | 0 | 29 | 119 | 0 | 0 | 31 | 15 | 0 | 0 | 0 | 0 | 0 | 42 | 0 | 0 | 0 | 0 | 6 | 0 | 0 | 0 | 19 | 0 | 0 | 0 | 0 | 46 | 38 | 0 | 0 | 0 | 0 | 0 |
| ASV270 | 80942f5b2b5 | 387 | 0 | 0 | 0 | 15 | 27 | 0 | 0 | 71 | 56 | 42 | 0 | 0 | 0 | 0 | 119 | 0 | 0 | 0 | 20 | 0 | 0 | 0 | 0 | 0 | 0 | 0 | 0 | 0 | 0 | 37 | 0 | 0 | 0 | 0 | 0 |
| ASV271 | 7b7819fef86 | 384 | 0 | 0 | 74 | 0 | 0 | 0 | 0 | 0 | 0 | 0 | 0 | 0 | 0 | 89 | 0 | 0 | 0 | 0 | 0 | 0 | 0 | 0 | 0 | 0 | 0 | 0 | 0 | 66 | 70 | 0 | 0 | 0 | 0 | 85 | 0 |
| ASV272 | 8cebec5a31a | 382 | 0 | 0 | 0 | 0 | 0 | 0 | 0 | 0 | 0 | 0 | 0 | 20 | 17 | 0 | 0 | 2 | 39 | 26 | 0 | 0 | 27 | 15 | 0 | 0 | 0 | 22 | 20 | 61 | 0 | 0 | 71 | 20 | 21 | 21 | 0 |
| ASV273 | 11a6b07ab2d | 379 | 0 | 0 | 63 | 0 | 94 | 0 | 0 | 97 | 53 | 38 | 0 | 0 | 0 | 0 | 34 | 0 | 0 | 0 | 0 | 0 | 0 | 0 | 0 | 0 | 0 | 0 | 0 | 0 | 0 | 0 | 0 | 0 | 0 | 0 | 0 |
| ASV274 | bca465403b6 | 379 | 0 | 0 | 0 | 0 | 0 | 0 | 0 | 0 | 0 | 0 | 0 | 0 | 0 | 0 | 0 | 0 | 0 | 0 | 0 | 5 | 0 | 0 | 0 | 0 | 84 | 0 | 0 | 7 | 13 | 161 | 0 | 0 | 0 | 28 | 81 |
| ASV275 | d1af6b6bc0d | 379 | 14 | 4 | 0 | 0 | 0 | 20 | 32 | 18 | 0 | 0 | 17 | 0 | 0 | 0 | 0 | 18 | 0 | 0 | 17 | 0 | 43 | 23 | 0 | 0 | 13 | 31 | 14 | 12 | 0 | 10 | 45 | 25 | 23 | 0 | 0 |
| ASV276 | 63d8db74a2f | 378 | 26 | 31 | 61 | 26 | 0 | 19 | 8 | 0 | 0 | 0 | 11 | 29 | 11 | 10 | 0 | 31 | 63 | 6 | 0 | 6 | 0 | 0 | 0 | 0 | 0 | 19 | 5 | 13 | 0 | 0 | 0 | 0 | 3 | 0 | 0 |
| ASV277 | e0714bc55e7 | 377 | 51 | 27 | 175 | 84 | 40 | 0 | 0 | 0 | 0 | 0 | 0 | 0 | 0 | 0 | 0 | 0 | 0 | 0 | 0 | 0 | 0 | 0 | 0 | 0 | 0 | 0 | 0 | 0 | 0 | 0 | 0 | 0 | 0 | 0 | 0 |
| ASV278 | 143db17801b | 377 | 0 | 0 | 7 | 0 | 57 | 0 | 0 | 6 | 15 | 27 | 0 | 0 | 0 | 0 | 86 | 0 | 0 | 0 | 3 | 8 | 0 | 0 | 0 | 11 | 16 | 0 | 0 | 6 | 37 | 38 | 0 | 0 | 0 | 15 | 45 |
| ASV279 | 4088787ee60 | 375 | 0 | 0 | 0 | 0 | 0 | 0 | 0 | 0 | 129 | 0 | 0 | 0 | 0 | 0 | 246 | 0 | 0 | 0 | 0 | 0 | 0 | 0 | 0 | 0 | 0 | 0 | 0 | 0 | 0 | 0 | 0 | 0 | 0 | 0 | 0 |
| ASV280 | 115951f203f | 375 | 25 | 27 | 64 | 42 | 30 | 152 | 35 | 0 | 0 | 0 | 0 | 0 | 0 | 0 | 0 | 0 | 0 | 0 | 0 | 0 | 0 | 0 | 0 | 0 | 0 | 0 | 0 | 0 | 0 | 0 | 0 | 0 | 0 | 0 | 0 |
| ASV281 | a0fa10c9d54 | 373 | 0 | 0 | 30 | 21 | 60 | 0 | 0 | 0 | 0 | 55 | 0 | 0 | 0 | 0 | 207 | 0 | 0 | 0 | 0 | 0 | 0 | 0 | 0 | 0 | 0 | 0 | 0 | 0 | 0 | 0 | 0 | 0 | 0 | 0 | 0 |
| ASV282 | 813fca6e401 | 370 | 0 | 0 | 0 | 0 | 0 | 0 | 0 | 0 | 0 | 0 | 0 | 0 | 0 | 0 | 0 | 0 | 0 | 0 | 0 | 0 | 0 | 0 | 0 | 0 | 0 | 38 | 0 | 0 | 0 | 0 | 157 | 89 | 86 | 0 | 0 |
| ASV283 | 3b521bfe408 | 366 | 0 | 0 | 0 | 0 | 15 | 0 | 0 | 11 | 18 | 28 | 0 | 0 | 0 | 0 | 43 | 0 | 0 | 0 | 15 | 37 | 0 | 0 | 0 | 13 | 80 | 0 | 0 | 0 | 17 | 51 | 0 | 0 | 0 | 19 | 19 |
| ASV284 | 907e0d9e928 | 361 | 0 | 0 | 79 | 0 | 78 | 0 | 0 | 0 | 0 | 0 | 0 | 72 | 0 | 0 | 60 | 0 | 0 | 0 | 0 | 0 | 0 | 0 | 0 | 0 | 0 | 12 | 0 | 0 | 60 | 0 | 0 | 0 | 0 | 0 | 0 |
| ASV285 | 1213d0907cc | 358 | 21 | 0 | 55 | 23 | 47 | 23 | 12 | 27 | 36 | 12 | 0 | 0 | 0 | 0 | 0 | 0 | 0 | 0 | 8 | 24 | 0 | 0 | 7 | 0 | 0 | 0 | 0 | 0 | 0 | 0 | 0 | 0 | 0 | 34 | 29 |
| ASV286 | 63448f99471 | 357 | 0 | 0 | 3 | 7 | 93 | 0 | 0 | 0 | 0 | 0 | 0 | 0 | 0 | 0 | 0 | 81 | 0 | 0 | 0 | 0 | 47 | 0 | 0 | 0 | 0 | 17 | 0 | 0 | 0 | 4 | 0 | 8 | 0 | 3 | 94 |
| ASV287 | 68c91a31fb5 | 355 | 0 | 0 | 49 | 0 | 39 | 0 | 0 | 65 | 0 | 0 | 0 | 0 | 0 | 0 | 0 | 0 | 0 | 0 | 0 | 0 | 0 | 0 | 0 | 0 | 0 | 0 | 0 | 0 | 109 | 0 | 0 | 0 | 0 | 64 | 29 |
| ASV288 | 3203cbd4619 | 353 | 0 | 0 | 12 | 0 | 9 | 75 | 36 | 52 | 0 | 33 | 0 | 0 | 0 | 16 | 81 | 0 | 0 | 0 | 12 | 16 | 0 | 0 | 0 | 0 | 11 | 0 | 0 | 0 | 0 | 0 | 0 | 0 | 0 | 0 | 0 |
| ASV289 | ef08dbff58ad | 343 | 0 | 0 | 19 | 0 | 42 | 0 | 0 | 0 | 0 | 27 | 0 | 0 | 0 | 0 | 53 | 0 | 0 | 0 | 47 | 14 | 0 | 0 | 0 | 21 | 31 | 0 | 0 | 6 | 41 | 8 | 0 | 0 | 0 | 10 | 24 |
| ASV290 | c786dc70f37a | 342 | 0 | 0 | 0 | 0 | 0 | 224 | 118 | 0 | 0 | 0 | 0 | 0 | 0 | 0 | 0 | 0 | 0 | 0 | 0 | 0 | 0 | 0 | 0 | 0 | 0 | 0 | 0 | 0 | 0 | 0 | 0 | 0 | 0 | 0 | 0 |
| ASV291 | adb2b9855cf | 337 | 21 | 6 | 47 | 7 | 33 | 52 | 0 | 31 | 0 | 0 | 16 | 44 | 39 | 28 | 0 | 0 | 0 | 0 | 0 | 0 | 0 | 0 | 0 | 0 | 0 | 0 | 3 | 10 | 0 | 0 | 0 | 0 | 0 | 0 | 0 |
| ASV292 | b70002d0863 | 327 | 0 | 0 | 0 | 0 | 0 | 0 | 0 | 0 | 0 | 0 | 0 | 0 | 0 | 0 | 0 | 0 | 0 | 0 | 0 | 0 | 0 | 0 | 0 | 0 | 0 | 0 | 0 | 0 | 0 | 0 | 195 | 88 | 44 | 0 | 0 |
| ASV293 | dba6da18688 | 325 | 0 | 0 | 0 | 0 | 27 | 2 | 3 | 20 | 0 | 16 | 0 | 0 | 0 | 27 | 70 | 0 | 0 | 0 | 0 | 5 | 0 | 0 | 0 | 14 | 23 | 0 | 0 | 0 | 23 | 50 | 0 | 0 | 0 | 16 | 29 |
| ASV294 | 34c14db1381 | 313 | 0 | 0 | 0 | 0 | 0 | 0 | 0 | 0 | 0 | 0 | 0 | 0 | 0 | 0 | 0 | 17 | 34 | 0 | 0 | 0 | 6 | 12 | 0 | 0 | 0 | 30 | 26 | 29 | 0 | 0 | 91 | 40 | 28 | 0 | 0 |
| ASV295 | e5288906456 | 311 | 0 | 0 | 0 | 0 | 0 | 0 | 0 | 0 | 0 | 0 | 0 | 0 | 0 | 0 | 0 | 0 | 15 | 0 | 0 | 0 | 36 | 5 | 5 | 0 | 0 | 22 | 9 | 36 | 0 | 0 | 95 | 43 | 39 | 6 | 0 |
| ASV296 | a50962f849c | 311 | 0 | 0 | 60 | 0 | 0 | 0 | 0 | 0 | 0 | 0 | 0 | 0 | 0 | 43 | 0 | 76 | 122 | 0 | 0 | 0 | 0 | 0 | 0 | 0 | 0 | 10 | 0 | 0 | 0 | 0 | 0 | 0 | 0 | 0 | 0 |
| ASV297 | b6e1ed0f35f | 309 | 0 | 0 | 0 | 0 | 0 | 0 | 0 | 0 | 0 | 0 | 0 | 0 | 0 | 0 | 0 | 154 | 155 | 0 | 0 | 0 | 0 | 0 | 0 | 0 | 0 | 0 | 0 | 0 | 0 | 0 | 0 | 0 | 0 | 0 | 0 |
| ASV298 | 671b031173d | 305 | 0 | 0 | 0 | 0 | 0 | 0 | 7 | 0 | 0 | 0 | 0 | 0 | 0 | 15 | 40 | 0 | 0 | 0 | 0 | 0 | 0 | 0 | 0 | 0 | 58 | 0 | 0 | 0 | 90 | 47 | 0 | 0 | 0 | 8 | 40 |
| ASV299 | 0399040dcb2 | 303 | 0 | 7 | 9 | 26 | 0 | 0 | 0 | 0 | 6 | 0 | 0 | 0 | 13 | 0 | 9 | 26 | 56 | 0 | 25 | 0 | 14 | 0 | 9 | 11 | 0 | 22 | 0 | 21 | 13 | 0 | 0 | 0 | 0 | 36 | 0 |
| ASV300 | f8c92c97538 | 303 | 11 | 0 | 15 | 31 | 48 | 0 | 0 | 15 | 37 | 32 | 0 | 0 | 0 | 0 | 54 | 0 | 0 | 0 | 0 | 0 | 0 | 0 | 0 | 0 | 33 | 0 | 0 | 10 | 0 | 17 | 0 | 0 | 0 | 0 | 0 |
| ASV301 | 0c4013f3f1e | 303 | 0 | 0 | 13 | 9 | 37 | 0 | 0 | 45 | 21 | 7 | 0 | 0 | 0 | 0 | 64 | 0 | 0 | 0 | 0 | 22 | 0 | 0 | 0 | 14 | 26 | 0 | 0 | 0 | 0 | 27 | 0 | 0 | 0 | 0 | 18 |
| ASV302 | 9e153487f61 | 298 | 0 | 0 | 38 | 0 | 0 | 0 | 0 | 0 | 0 | 0 | 0 | 0 | 0 | 0 | 0 | 18 | 85 | 0 | 0 | 0 | 15 | 7 | 0 | 0 | 0 | 23 | 4 | 43 | 39 | 0 | 0 | 15 | 11 | 0 | 0 |
| ASV303 | 09508e6050a | 295 | 0 | 0 | 0 | 0 | 0 | 0 | 0 | 0 | 0 | 0 | 0 | 48 | 0 | 0 | 0 | 44 | 104 | 0 | 0 | 0 | 0 | 0 | 17 | 0 | 0 | 13 | 0 | 52 | 12 | 0 | 0 | 0 | 0 | 0 | 5 |
| ASV304 | 7fbf48227c4 | 294 | 35 | 0 | 96 | 28 | 17 | 43 | 0 | 51 | 12 | 0 | 0 | 0 | 0 | 0 | 12 | 0 | 0 | 0 | 0 | 0 | 0 | 0 | 0 | 0 | 0 | 0 | 0 | 0 | 0 | 0 | 0 | 0 | 0 | 0 | 0 |
| ASV305 | c47a1935d92 | 290 | 0 | 0 | 0 | 0 | 0 | 0 | 0 | 0 | 0 | 0 | 0 | 0 | 26 | 0 | 0 | 76 | 44 | 42 | 0 | 0 | 0 | 0 | 11 | 0 | 0 | 0 | 38 | 47 | 0 | 0 | 0 | 0 | 0 | 6 | 0 |
| ASV306 | e2d47798ea1 | 286 | 0 | 0 | 9 | 0 | 0 | 11 | 0 | 20 | 0 | 0 | 0 | 17 | 20 | 7 | 3 | 14 | 57 | 25 | 60 | 0 | 0 | 0 | 5 | 8 | 0 | 0 | 0 | 26 | 0 | 0 | 0 | 0 | 4 | 0 | 0 |
| ASV307 | 0183fa60162 | 285 | 68 | 0 | 109 | 68 | 0 | 0 | 28 | 0 | 0 | 0 | 0 | 0 | 0 | 0 | 12 | 0 | 0 | 0 | 0 | 0 | 0 | 0 | 0 | 0 | 0 | 0 | 0 | 0 | 0 | 0 | 0 | 0 | 0 | 0 | 0 |
| ASV308 | 8f8971517c3 | 284 | 0 | 0 | 0 | 0 | 41 | 0 | 0 | 0 | 38 | 28 | 0 | 0 | 0 | 0 | 34 | 0 | 0 | 0 | 0 | 30 | 0 | 0 | 0 | 0 | 19 | 0 | 0 | 0 | 15 | 43 | 0 | 0 | 0 | 0 | 36 |
| ASV309 | 109ab8dc170 | 282 | 0 | 0 | 35 | 0 | 52 | 0 | 0 | 0 | 0 | 0 | 0 | 44 | 0 | 0 | 0 | 0 | 0 | 0 | 0 | 0 | 0 | 0 | 0 | 0 | 0 | 0 | 0 | 0 | 52 | 0 | 27 | 0 | 0 | 72 | 0 |
| ASV310 | c39a297dd4b | 280 | 0 | 0 | 0 | 0 | 43 | 0 | 0 | 30 | 36 | 32 | 0 | 0 | 0 | 0 | 34 | 0 | 0 | 0 | 28 | 0 | 0 | 0 | 0 | 0 | 25 | 0 | 0 | 0 | 22 | 30 | 0 | 0 | 0 | 0 | 0 |
| ASV311 | 18582b991c9 | 278 | 0 | 0 | 32 | 13 | 51 | 0 | 0 | 0 | 0 | 0 | 0 | 0 | 0 | 0 | 0 | 0 | 0 | 0 | 0 | 0 | 0 | 0 | 0 | 33 | 0 | 0 | 0 | 45 | 21 | 12 | 0 | 0 | 0 | 50 | 21 |
| ASV312 | f6aad746ffbe | 277 | 3 | 6 | 0 | 0 | 0 | 0 | 0 | 0 | 0 | 0 | 0 | 0 | 0 | 0 | 0 | 0 | 0 | 0 | 0 | 0 | 0 | 0 | 0 | 0 | 0 | 9 | 0 | 0 | 0 | 5 | 174 | 39 | 14 | 27 | 0 |
| ASV313 | 60b8d3b8a6b | 275 | 0 | 0 | 36 | 0 | 0 | 0 | 0 | 0 | 0 | 0 | 0 | 41 | 0 | 0 | 0 | 45 | 51 | 0 | 0 | 0 | 62 | 23 | 0 | 0 | 0 | 0 | 0 | 0 | 0 | 17 | 0 | 0 | 0 | 0 | 0 |
| ASV314 | d4c50a4f1df4 | 274 | 0 | 0 | 0 | 0 | 0 | 11 | 0 | 0 | 0 | 0 | 0 | 48 | 17 | 0 | 0 | 76 | 101 | 21 | 0 | 0 | 0 | 0 | 0 | 0 | 0 | 0 | 0 | 0 | 0 | 0 | 0 | 0 | 0 | 0 | 0 |
| ASV315 | 0b3737e9a96 | 266 | 0 | 0 | 0 | 0 | 23 | 0 | 0 | 0 | 0 | 0 | 0 | 0 | 0 | 0 | 0 | 72 | 66 | 0 | 0 | 0 | 27 | 0 | 0 | 0 | 0 | 50 | 0 | 28 | 0 | 0 | 0 | 0 | 0 | 0 | 0 |
| ASV316 | c2a36810118 | 264 | 0 | 0 | 0 | 0 | 0 | 0 | 0 | 0 | 0 | 0 | 0 | 0 | 0 | 0 | 0 | 0 | 0 | 0 | 0 | 0 | 0 | 0 | 0 | 26 | 28 | 0 | 0 | 12 | 114 | 55 | 0 | 0 | 0 | 29 | 0 |
| ASV317 | ff40d8496e3 | 263 | 0 | 0 | 0 | 0 | 0 | 19 | 0 | 26 | 15 | 0 | 0 | 0 | 0 | 0 | 27 | 0 | 41 | 0 | 13 | 0 | 0 | 0 | 0 | 16 | 36 | 0 | 0 | 11 | 0 | 0 | 34 | 9 | 0 | 16 | 0 |
| ASV318 | 69ecd66736e | 261 | 8 | 8 | 30 | 13 | 21 | 7 | 0 | 14 | 0 | 7 | 0 | 0 | 0 | 0 | 0 | 0 | 0 | 0 | 0 | 0 | 0 | 0 | 0 | 9 | 29 | 0 | 0 | 0 | 54 | 13 | 0 | 0 | 0 | 26 | 22 |
| ASV319 | e1d62bb8c36 | 261 | 19 | 19 | 67 | 43 | 68 | 0 | 0 | 0 | 21 | 0 | 0 | 0 | 0 | 0 | 0 | 0 | 0 | 0 | 0 | 0 | 0 | 0 | 0 | 0 | 0 | 0 | 0 | 0 | 0 | 6 | 0 | 0 | 0 | 10 | 8 |
| ASV320 | d8dc217567f | 259 | 0 | 0 | 0 | 0 | 0 | 0 | 0 | 0 | 0 | 0 | 0 | 0 | 0 | 0 | 0 | 0 | 3 | 24 | 15 | 58 | 0 | 0 | 0 | 19 | 77 | 0 | 0 | 6 | 0 | 50 | 0 | 0 | 0 | 0 | 7 |
| ASV321 | 7d83cdc24ba | 258 | 0 | 0 | 0 | 0 | 0 | 0 | 0 | 0 | 0 | 88 | 0 | 0 | 0 | 0 | 0 | 0 | 0 | 0 | 0 | 0 | 0 | 0 | 0 | 0 | 83 | 0 | 0 | 0 | 87 | 0 | 0 | 0 | 0 | 0 | 0 |
| ASV322 | 6ef9372cea9 | 257 | 27 | 30 | 85 | 57 | 21 | 37 | 0 | 0 | 0 | 0 | 0 | 0 | 0 | 0 | 0 | 0 | 0 | 0 | 0 | 0 | 0 | 0 | 0 | 0 | 0 | 0 | 0 | 0 | 0 | 0 | 0 | 0 | 0 | 0 | 0 |
| ASV323 | f1cf5556ba5a | 257 | 8 | 0 | 0 | 0 | 64 | 0 | 0 | 0 | 23 | 10 | 0 | 0 | 0 | 0 | 117 | 0 | 0 | 0 | 0 | 9 | 0 | 0 | 0 | 0 | 10 | 0 | 0 | 0 | 0 | 0 | 0 | 0 | 0 | 0 | 16 |
| ASV324 | eeec7f5124a | 256 | 0 | 0 | 0 | 0 | 0 | 0 | 0 | 0 | 0 | 0 | 0 | 0 | 0 | 0 | 0 | 69 | 60 | 0 | 0 | 0 | 48 | 34 | 0 | 0 | 0 | 45 | 0 | 0 | 0 | 0 | 0 | 0 | 0 | 0 | 0 |
| ASV325 | c728fde5a5f1 | 255 | 18 | 0 | 38 | 56 | 46 | 0 | 0 | 0 | 0 | 0 | 0 | 0 | 8 | 0 | 0 | 0 | 0 | 0 | 10 | 4 | 0 | 0 | 0 | 0 | 0 | 0 | 0 | 0 | 0 | 0 | 0 | 0 | 0 | 62 | 13 |
| ASV326 | 22f859ea1ee | 251 | 0 | 0 | 0 | 0 | 0 | 0 | 0 | 0 | 0 | 0 | 0 | 0 | 0 | 0 | 0 | 43 | 75 | 0 | 0 | 0 | 0 | 0 | 0 | 0 | 0 | 38 | 0 | 13 | 0 | 0 | 53 | 19 | 10 | 0 | 0 |
| ASV327 | 519bbbb8d91 | 249 | 0 | 0 | 0 | 0 | 0 | 0 | 0 | 0 | 0 | 0 | 0 | 0 | 0 | 0 | 0 | 58 | 30 | 0 | 0 | 0 | 42 | 51 | 0 | 0 | 0 | 68 | 0 | 0 | 0 | 0 | 0 | 0 | 0 | 0 | 0 |
| ASV328 | 84593f6b964 | 249 | 0 | 0 | 0 | 0 | 0 | 0 | 0 | 0 | 0 | 0 | 0 | 42 | 0 | 0 | 0 | 64 | 98 | 0 | 0 | 0 | 0 | 0 | 0 | 0 | 0 | 45 | 0 | 0 | 0 | 0 | 0 | 0 | 0 | 0 | 0 |
| ASV329 | 2f5bd880f7b | 248 | 20 | 16 | 62 | 39 | 28 | 0 | 0 | 0 | 0 | 0 | 0 | 0 | 0 | 0 | 0 | 0 | 0 | 0 | 0 | 0 | 0 | 0 | 0 | 0 | 0 | 0 | 0 | 0 | 0 | 0 | 0 | 0 | 0 | 65 | 18 |
| ASV330 | 279a085f1a6 | 247 | 19 | 45 | 74 | 63 | 31 | 0 | 0 | 3 | 0 | 0 | 0 | 0 | 0 | 0 | 0 | 0 | 0 | 0 | 0 | 0 | 0 | 0 | 0 | 4 | 0 | 0 | 0 | 0 | 8 | 0 | 0 | 0 | 0 | 0 | 0 |
| ASV331 | d38047a1749 | 246 | 0 | 0 | 0 | 0 | 0 | 0 | 0 | 0 | 0 | 0 | 0 | 0 | 0 | 0 | 0 | 0 | 80 | 0 | 0 | 0 | 0 | 0 | 0 | 0 | 0 | 0 | 0 | 0 | 0 | 0 | 166 | 0 | 0 | 0 | 0 |
| ASV332 | 91df06809dd | 246 | 0 | 0 | 0 | 0 | 69 | 0 | 0 | 0 | 45 | 0 | 0 | 0 | 0 | 0 | 132 | 0 | 0 | 0 | 0 | 0 | 0 | 0 | 0 | 0 | 0 | 0 | 0 | 0 | 0 | 0 | 0 | 0 | 0 | 0 | 0 |
| ASV333 | 5d6bd15b0b8 | 246 | 7 | 0 | 12 | 0 | 47 | 0 | 0 | 0 | 0 | 24 | 0 | 0 | 0 | 0 | 31 | 0 | 0 | 0 | 0 | 0 | 0 | 0 | 0 | 0 | 14 | 0 | 6 | 0 | 41 | 50 | 0 | 0 | 0 | 14 | 0 |
| ASV334 | e3cda4357a7 | 244 | 0 | 0 | 0 | 0 | 0 | 0 | 0 | 0 | 0 | 0 | 0 | 0 | 0 | 0 | 0 | 0 | 0 | 0 | 0 | 0 | 0 | 0 | 0 | 0 | 50 | 0 | 0 | 0 | 0 | 77 | 0 | 0 | 17 | 20 | 80 |
| ASV335 | ecfc9cf5774e | 243 | 0 | 0 | 0 | 0 | 0 | 0 | 0 | 0 | 0 | 0 | 0 | 0 | 0 | 0 | 0 | 0 | 0 | 0 | 0 | 0 | 0 | 0 | 0 | 0 | 0 | 0 | 0 | 0 | 0 | 0 | 129 | 53 | 61 | 0 | 0 |
| ASV336 | 793c0bb9242 | 242 | 0 | 0 | 29 | 0 | 53 | 0 | 0 | 25 | 0 | 0 | 0 | 0 | 0 | 29 | 56 | 0 | 0 | 3 | 0 | 0 | 0 | 0 | 0 | 0 | 0 | 0 | 0 | 15 | 0 | 28 | 0 | 0 | 0 | 0 | 4 |
| ASV337 | 5b467f21a97 | 240 | 0 | 0 | 0 | 0 | 0 | 0 | 0 | 0 | 0 | 0 | 0 | 0 | 0 | 0 | 0 | 22 | 13 | 21 | 0 | 0 | 51 | 15 | 24 | 0 | 0 | 0 | 11 | 9 | 15 | 0 | 24 | 11 | 24 | 0 | 0 |
| ASV338 | ae089994252 | 238 | 0 | 0 | 0 | 0 | 0 | 0 | 0 | 18 | 16 | 62 | 0 | 0 | 0 | 0 | 36 | 0 | 0 | 0 | 0 | 20 | 0 | 0 | 0 | 0 | 47 | 0 | 0 | 0 | 0 | 16 | 0 | 0 | 0 | 0 | 23 |
| ASV339 | dd1936c4d66 | 238 | 0 | 0 | 0 | 0 | 63 | 34 | 0 | 0 | 34 | 0 | 0 | 0 | 0 | 0 | 49 | 0 | 0 | 0 | 0 | 0 | 0 | 0 | 0 | 0 | 0 | 0 | 0 | 0 | 0 | 28 | 0 | 0 | 0 | 0 | 30 |
| ASV340 | b718cd41d94 | 237 | 0 | 0 | 75 | 62 | 51 | 49 | 0 | 0 | 0 | 0 | 0 | 0 | 0 | 0 | 0 | 0 | 0 | 0 | 0 | 0 | 0 | 0 | 0 | 0 | 0 | 0 | 0 | 0 | 0 | 0 | 0 | 0 | 0 | 0 | 0 |
| ASV341 | 7d6b1e56ef7 | 236 | 0 | 0 | 0 | 0 | 0 | 0 | 0 | 28 | 0 | 18 | 0 | 0 | 0 | 7 | 0 | 0 | 0 | 0 | 28 | 11 | 4 | 0 | 0 | 27 | 0 | 0 | 0 | 38 | 46 | 29 | 0 | 0 | 0 | 0 | 0 |
| ASV342 | b3a4e7f7a8e | 235 | 6 | 5 | 24 | 10 | 13 | 0 | 0 | 18 | 0 | 0 | 0 | 0 | 0 | 22 | 11 | 0 | 0 | 0 | 0 | 0 | 0 | 0 | 0 | 13 | 18 | 0 | 0 | 0 | 43 | 43 | 0 | 0 | 0 | 0 | 9 |
| ASV343 | 8cba59be471 | 233 | 0 | 0 | 0 | 0 | 0 | 0 | 0 | 0 | 0 | 0 | 0 | 0 | 0 | 0 | 4 | 0 | 26 | 0 | 0 | 0 | 0 | 0 | 0 | 0 | 0 | 22 | 0 | 0 | 0 | 7 | 86 | 45 | 43 | 0 | 0 |
| ASV344 | db6a43a5499 | 232 | 0 | 0 | 0 | 0 | 0 | 0 | 0 | 0 | 19 | 40 | 0 | 0 | 0 | 0 | 107 | 0 | 0 | 0 | 0 | 9 | 0 | 0 | 0 | 0 | 0 | 0 | 0 | 0 | 0 | 57 | 0 | 0 | 0 | 0 | 0 |
| ASV345 | efd3b8f4a8fb | 232 | 0 | 0 | 16 | 13 | 0 | 41 | 0 | 0 | 0 | 0 | 0 | 15 | 0 | 0 | 30 | 0 | 32 | 15 | 23 | 0 | 0 | 0 | 0 | 0 | 0 | 23 | 0 | 0 | 0 | 0 | 24 | 0 | 0 | 0 | 0 |
| ASV346 | 9880e34d3fc | 231 | 0 | 0 | 0 | 0 | 0 | 0 | 0 | 0 | 0 | 0 | 0 | 0 | 0 | 59 | 19 | 0 | 0 | 0 | 0 | 0 | 0 | 0 | 0 | 15 | 28 | 0 | 0 | 31 | 40 | 39 | 0 | 0 | 0 | 0 | 0 |
| ASV347 | d19c5daff6e4 | 230 | 0 | 0 | 0 | 0 | 0 | 0 | 0 | 3 | 0 | 0 | 0 | 0 | 0 | 0 | 0 | 0 | 10 | 8 | 0 | 0 | 0 | 0 | 10 | 3 | 0 | 7 | 7 | 30 | 11 | 0 | 44 | 11 | 0 | 86 | 0 |
| ASV348 | dd3de45077f | 229 | 0 | 0 | 47 | 0 | 59 | 47 | 0 | 0 | 0 | 0 | 0 | 0 | 0 | 0 | 33 | 0 | 0 | 0 | 43 | 0 | 0 | 0 | 0 | 0 | 0 | 0 | 0 | 0 | 0 | 0 | 0 | 0 | 0 | 0 | 0 |
| ASV349 | 8381bfa1787 | 229 | 0 | 0 | 9 | 18 | 33 | 0 | 0 | 14 | 13 | 10 | 0 | 0 | 0 | 0 | 41 | 0 | 0 | 0 | 16 | 0 | 0 | 0 | 0 | 14 | 16 | 0 | 0 | 5 | 0 | 20 | 0 | 0 | 0 | 0 | 20 |
| ASV350 | 476a9e90f22 | 229 | 0 | 0 | 0 | 0 | 0 | 0 | 0 | 0 | 39 | 36 | 0 | 0 | 0 | 0 | 68 | 0 | 0 | 0 | 0 | 22 | 0 | 0 | 0 | 0 | 25 | 0 | 0 | 0 | 4 | 11 | 0 | 0 | 0 | 0 | 24 |
| ASV351 | af0620d9127 | 229 | 0 | 0 | 0 | 0 | 0 | 0 | 0 | 0 | 69 | 44 | 0 | 0 | 0 | 0 | 98 | 0 | 0 | 0 | 0 | 0 | 0 | 0 | 0 | 0 | 0 | 0 | 0 | 0 | 0 | 18 | 0 | 0 | 0 | 0 | 0 |
| ASV352 | 7ba1b164eec | 229 | 19 | 14 | 72 | 41 | 56 | 21 | 0 | 6 | 0 | 0 | 0 | 0 | 0 | 0 | 0 | 0 | 0 | 0 | 0 | 0 | 0 | 0 | 0 | 0 | 0 | 0 | 0 | 0 | 0 | 0 | 0 | 0 | 0 | 0 | 0 |
| ASV353 | 7db426ecf20 | 228 | 0 | 0 | 0 | 0 | 0 | 0 | 0 | 0 | 0 | 0 | 0 | 0 | 0 | 0 | 0 | 0 | 107 | 0 | 0 | 0 | 0 | 0 | 0 | 0 | 0 | 0 | 0 | 0 | 121 | 0 | 0 | 0 | 0 | 0 | 0 |
| ASV354 | a4e4bd6a602 | 226 | 0 | 0 | 0 | 0 | 89 | 0 | 0 | 0 | 27 | 21 | 0 | 0 | 0 | 0 | 44 | 0 | 0 | 0 | 0 | 0 | 0 | 0 | 0 | 0 | 13 | 0 | 0 | 0 | 0 | 11 | 0 | 0 | 0 | 0 | 21 |
| ASV355 | cb049fad995 | 226 | 0 | 0 | 0 | 0 | 0 | 0 | 0 | 0 | 43 | 0 | 0 | 0 | 0 | 0 | 0 | 0 | 0 | 0 | 0 | 40 | 0 | 0 | 0 | 14 | 99 | 0 | 0 | 0 | 0 | 0 | 0 | 0 | 0 | 0 | 30 |
| ASV356 | 1090f885623 | 225 | 0 | 0 | 0 | 14 | 0 | 0 | 0 | 0 | 0 | 0 | 0 | 22 | 0 | 0 | 0 | 38 | 41 | 0 | 0 | 0 | 0 | 17 | 0 | 14 | 0 | 15 | 0 | 25 | 39 | 0 | 0 | 0 | 0 | 0 | 0 |
| ASV357 | 2a9ae138a4b | 221 | 28 | 0 | 62 | 38 | 21 | 22 | 0 | 20 | 0 | 0 | 0 | 0 | 0 | 0 | 0 | 0 | 0 | 0 | 0 | 0 | 0 | 0 | 0 | 4 | 0 | 0 | 0 | 5 | 0 | 0 | 4 | 0 | 2 | 15 | 0 |
| ASV358 | 687cbd7f96e | 221 | 0 | 0 | 0 | 0 | 0 | 0 | 0 | 0 | 0 | 0 | 0 | 0 | 0 | 0 | 0 | 0 | 0 | 0 | 0 | 0 | 0 | 0 | 0 | 0 | 0 | 0 | 0 | 0 | 0 | 0 | 150 | 25 | 46 | 0 | 0 |
| ASV359 | e1664283d05 | 220 | 0 | 0 | 0 | 0 | 41 | 0 | 0 | 0 | 25 | 0 | 0 | 0 | 0 | 0 | 78 | 0 | 0 | 0 | 0 | 0 | 0 | 0 | 0 | 0 | 48 | 0 | 0 | 0 | 0 | 0 | 0 | 0 | 0 | 0 | 28 |
| ASV360 | 52d0f2934d0 | 217 | 0 | 0 | 0 | 0 | 0 | 0 | 0 | 0 | 0 | 0 | 0 | 33 | 0 | 0 | 0 | 53 | 61 | 21 | 0 | 0 | 17 | 32 | 0 | 0 | 0 | 0 | 0 | 0 | 0 | 0 | 0 | 0 | 0 | 0 | 0 |
| ASV361 | 26e2f204225 | 215 | 0 | 0 | 0 | 0 | 54 | 0 | 0 | 0 | 62 | 0 | 0 | 0 | 0 | 0 | 62 | 0 | 0 | 0 | 0 | 0 | 0 | 0 | 0 | 0 | 37 | 0 | 0 | 0 | 0 | 0 | 0 | 0 | 0 | 0 | 0 |
| ASV362 | ef82db7a784 | 213 | 0 | 0 | 0 | 0 | 32 | 0 | 0 | 0 | 0 | 0 | 0 | 0 | 0 | 0 | 0 | 0 | 0 | 0 | 0 | 0 | 0 | 0 | 0 | 22 | 10 | 0 | 0 | 0 | 74 | 54 | 0 | 0 | 0 | 21 | 0 |
| ASV363 | 95d0c087a9b | 211 | 1 | 26 | 17 | 0 | 2 | 7 | 0 | 1 | 18 | 0 | 33 | 7 | 0 | 1 | 27 | 0 | 36 | 0 | 31 | 0 | 0 | 0 | 0 | 0 | 0 | 0 | 0 | 1 | 3 | 0 | 0 | 0 | 0 | 0 | 0 |
| ASV364 | 17378062be2 | 210 | 0 | 0 | 0 | 0 | 0 | 0 | 0 | 0 | 0 | 0 | 0 | 0 | 0 | 0 | 0 | 49 | 114 | 15 | 0 | 0 | 5 | 0 | 0 | 0 | 0 | 7 | 7 | 13 | 0 | 0 | 0 | 0 | 0 | 0 | 0 |
| ASV365 | 05e651042e4 | 209 | 3 | 11 | 4 | 3 | 17 | 10 | 5 | 7 | 7 | 24 | 0 | 2 | 1 | 4 | 16 | 0 | 0 | 0 | 4 | 0 | 3 | 1 | 0 | 15 | 22 | 0 | 0 | 6 | 3 | 30 | 0 | 0 | 0 | 2 | 9 |
| ASV366 | f837b692fe0 | 207 | 0 | 0 | 0 | 0 | 0 | 0 | 0 | 0 | 0 | 0 | 0 | 0 | 0 | 0 | 0 | 0 | 0 | 0 | 9 | 0 | 0 | 0 | 0 | 36 | 28 | 0 | 0 | 0 | 53 | 33 | 0 | 0 | 0 | 24 | 24 |
| ASV367 | e773509c44b | 206 | 1 | 8 | 2 | 1 | 0 | 16 | 2 | 1 | 7 | 1 | 31 | 0 | 0 | 2 | 6 | 8 | 10 | 11 | 14 | 0 | 11 | 8 | 15 | 2 | 0 | 7 | 11 | 20 | 0 | 3 | 0 | 0 | 0 | 8 | 0 |
| ASV368 | eb7d813dc79 | 204 | 0 | 0 | 0 | 0 | 0 | 0 | 0 | 0 | 0 | 0 | 0 | 0 | 0 | 0 | 0 | 0 | 23 | 0 | 8 | 56 | 0 | 0 | 0 | 18 | 36 | 0 | 0 | 0 | 16 | 40 | 0 | 0 | 0 | 0 | 7 |
| ASV369 | 3e9b7bc1b40 | 204 | 7 | 0 | 10 | 7 | 14 | 8 | 0 | 0 | 12 | 21 | 0 | 0 | 0 | 0 | 0 | 0 | 12 | 5 | 23 | 8 | 4 | 0 | 0 | 0 | 12 | 0 | 0 | 13 | 7 | 7 | 18 | 13 | 0 | 3 | 0 |
| ASV370 | 454a402f1fe | 204 | 0 | 0 | 0 | 0 | 0 | 0 | 0 | 0 | 0 | 0 | 0 | 0 | 0 | 0 | 0 | 44 | 70 | 14 | 0 | 0 | 0 | 0 | 0 | 0 | 12 | 18 | 14 | 18 | 14 | 0 | 0 | 0 | 0 | 0 | 0 |
| ASV371 | d126165e4c9 | 203 | 0 | 0 | 46 | 60 | 53 | 0 | 0 | 0 | 0 | 40 | 0 | 0 | 0 | 0 | 0 | 0 | 0 | 0 | 0 | 0 | 0 | 0 | 0 | 0 | 0 | 0 | 0 | 4 | 0 | 0 | 0 | 0 | 0 | 0 | 0 |
| ASV372 | 7b741115d3b | 201 | 0 | 0 | 43 | 9 | 0 | 17 | 14 | 16 | 0 | 7 | 0 | 28 | 0 | 0 | 0 | 0 | 0 | 24 | 0 | 0 | 0 | 0 | 0 | 10 | 0 | 0 | 0 | 0 | 20 | 0 | 13 | 0 | 0 | 0 | 0 |
| ASV373 | bcd22c817f4 | 201 | 0 | 0 | 0 | 0 | 0 | 24 | 0 | 0 | 0 | 0 | 0 | 21 | 0 | 0 | 0 | 34 | 56 | 27 | 0 | 0 | 0 | 0 | 0 | 0 | 0 | 0 | 0 | 0 | 0 | 0 | 39 | 0 | 0 | 0 | 0 |
| ASV374 | 5fc0c1dab4b | 201 | 11 | 0 | 34 | 10 | 0 | 8 | 3 | 12 | 0 | 0 | 3 | 5 | 6 | 20 | 0 | 12 | 21 | 0 | 0 | 0 | 0 | 0 | 0 | 0 | 0 | 0 | 0 | 15 | 41 | 0 | 0 | 0 | 0 | 0 | 0 |
| ASV375 | 4247391984c | 200 | 42 | 0 | 35 | 26 | 97 | 0 | 0 | 0 | 0 | 0 | 0 | 0 | 0 | 0 | 0 | 0 | 0 | 0 | 0 | 0 | 0 | 0 | 0 | 0 | 0 | 0 | 0 | 0 | 0 | 0 | 0 | 0 | 0 | 0 | 0 |
| ASV376 | 7549e57eaad | 200 | 0 | 0 | 22 | 0 | 0 | 0 | 0 | 4 | 0 | 0 | 0 | 0 | 0 | 0 | 0 | 0 | 75 | 0 | 0 | 0 | 0 | 0 | 0 | 11 | 12 | 0 | 0 | 42 | 17 | 4 | 0 | 0 | 0 | 13 | 0 |
| ASV377 | 5a7e96a9e3a | 199 | 0 | 0 | 0 | 0 | 42 | 68 | 0 | 0 | 0 | 0 | 0 | 0 | 0 | 27 | 62 | 0 | 0 | 0 | 0 | 0 | 0 | 0 | 0 | 0 | 0 | 0 | 0 | 0 | 0 | 0 | 0 | 0 | 0 | 0 | 0 |
| ASV378 | 12e271fae46 | 199 | 0 | 0 | 0 | 8 | 8 | 0 | 0 | 14 | 17 | 26 | 0 | 0 | 0 | 0 | 126 | 0 | 0 | 0 | 0 | 0 | 0 | 0 | 0 | 0 | 0 | 0 | 0 | 0 | 0 | 0 | 0 | 0 | 0 | 0 | 0 |
| ASV379 | 3a91733367e | 197 | 0 | 0 | 0 | 0 | 0 | 0 | 0 | 0 | 0 | 0 | 0 | 0 | 30 | 0 | 0 | 10 | 72 | 0 | 0 | 0 | 0 | 33 | 0 | 0 | 0 | 0 | 0 | 52 | 0 | 0 | 0 | 0 | 0 | 0 | 0 |
| ASV380 | 6ce9a49ca2f | 197 | 0 | 0 | 0 | 0 | 0 | 0 | 0 | 0 | 0 | 0 | 0 | 0 | 0 | 0 | 0 | 0 | 0 | 0 | 0 | 0 | 0 | 0 | 0 | 0 | 42 | 0 | 0 | 0 | 3 | 71 | 0 | 10 | 0 | 19 | 52 |
| ASV381 | 1557d232fa3 | 194 | 16 | 5 | 24 | 33 | 70 | 0 | 0 | 14 | 0 | 32 | 0 | 0 | 0 | 0 | 0 | 0 | 0 | 0 | 0 | 0 | 0 | 0 | 0 | 0 | 0 | 0 | 0 | 0 | 0 | 0 | 0 | 0 | 0 | 0 | 0 |
| ASV382 | 3a1ebbcfec4 | 193 | 9 | 0 | 18 | 9 | 28 | 15 | 0 | 17 | 12 | 0 | 0 | 0 | 0 | 11 | 0 | 0 | 0 | 0 | 0 | 0 | 0 | 0 | 0 | 15 | 22 | 0 | 0 | 0 | 14 | 23 | 0 | 0 | 0 | 0 | 0 |
| ASV383 | 27593cb90ba | 193 | 0 | 0 | 0 | 0 | 0 | 0 | 0 | 0 | 0 | 0 | 0 | 0 | 0 | 0 | 0 | 33 | 33 | 0 | 0 | 0 | 0 | 0 | 0 | 0 | 0 | 0 | 0 | 0 | 0 | 0 | 97 | 30 | 0 | 0 | 0 |
| ASV384 | a1a8af62eb4 | 193 | 0 | 0 | 0 | 0 | 0 | 0 | 0 | 0 | 0 | 0 | 0 | 0 | 0 | 0 | 0 | 14 | 25 | 20 | 0 | 0 | 0 | 0 | 0 | 0 | 0 | 0 | 0 | 0 | 0 | 0 | 84 | 24 | 26 | 0 | 0 |
| ASV385 | 00959df1876 | 192 | 9 | 0 | 0 | 13 | 24 | 0 | 0 | 18 | 27 | 14 | 0 | 0 | 0 | 0 | 38 | 0 | 0 | 0 | 10 | 11 | 0 | 0 | 0 | 0 | 14 | 0 | 0 | 0 | 0 | 0 | 0 | 0 | 0 | 0 | 14 |
| ASV386 | 9b9c1b1fb9e | 192 | 31 | 22 | 47 | 44 | 48 | 0 | 0 | 0 | 0 | 0 | 0 | 0 | 0 | 0 | 0 | 0 | 0 | 0 | 0 | 0 | 0 | 0 | 0 | 0 | 0 | 0 | 0 | 0 | 0 | 0 | 0 | 0 | 0 | 0 | 0 |
| ASV387 | 51193449909 | 192 | 10 | 0 | 0 | 0 | 0 | 0 | 0 | 0 | 0 | 0 | 0 | 13 | 0 | 8 | 0 | 0 | 50 | 0 | 0 | 0 | 0 | 0 | 16 | 0 | 0 | 0 | 0 | 33 | 11 | 0 | 0 | 0 | 0 | 51 | 0 |
| ASV388 | 94b18f0c88b | 189 | 0 | 0 | 0 | 0 | 0 | 0 | 0 | 0 | 25 | 36 | 0 | 0 | 0 | 0 | 83 | 0 | 0 | 0 | 0 | 0 | 0 | 0 | 0 | 0 | 11 | 0 | 0 | 0 | 0 | 15 | 0 | 0 | 0 | 0 | 19 |
| ASV389 | 788e4ec7076 | 188 | 0 | 7 | 18 | 14 | 23 | 9 | 14 | 16 | 0 | 11 | 0 | 8 | 0 | 0 | 15 | 0 | 0 | 8 | 11 | 18 | 0 | 0 | 0 | 0 | 0 | 0 | 0 | 0 | 8 | 0 | 0 | 0 | 0 | 0 | 8 |
| ASV390 | d225904882d | 186 | 0 | 0 | 0 | 0 | 20 | 0 | 0 | 18 | 26 | 37 | 0 | 0 | 0 | 0 | 33 | 0 | 0 | 0 | 7 | 14 | 0 | 0 | 0 | 5 | 6 | 0 | 0 | 0 | 4 | 12 | 0 | 0 | 0 | 0 | 4 |
| ASV391 | 98c815507ac | 185 | 0 | 0 | 6 | 8 | 0 | 33 | 6 | 18 | 14 | 30 | 2 | 0 | 0 | 6 | 7 | 8 | 32 | 0 | 15 | 0 | 0 | 0 | 0 | 0 | 0 | 0 | 0 | 0 | 0 | 0 | 0 | 0 | 0 | 0 | 0 |
| ASV392 | be9d6040f58 | 184 | 0 | 0 | 0 | 0 | 0 | 30 | 10 | 10 | 0 | 0 | 19 | 65 | 27 | 0 | 0 | 0 | 15 | 0 | 0 | 0 | 0 | 0 | 0 | 0 | 0 | 0 | 0 | 8 | 0 | 0 | 0 | 0 | 0 | 0 | 0 |
| ASV393 | e98a2eeb393 | 184 | 15 | 12 | 43 | 22 | 30 | 0 | 0 | 16 | 0 | 7 | 0 | 0 | 0 | 0 | 24 | 0 | 0 | 0 | 0 | 15 | 0 | 0 | 0 | 0 | 0 | 0 | 0 | 0 | 0 | 0 | 0 | 0 | 0 | 0 | 0 |
| ASV394 | 8f7715fdc5d7 | 184 | 14 | 6 | 0 | 0 | 0 | 15 | 8 | 0 | 0 | 0 | 0 | 0 | 0 | 0 | 42 | 0 | 0 | 0 | 0 | 0 | 0 | 0 | 0 | 0 | 29 | 0 | 0 | 0 | 40 | 0 | 0 | 0 | 0 | 0 | 30 |
| ASV395 | dc13bae2b5e | 183 | 6 | 3 | 11 | 14 | 58 | 0 | 0 | 7 | 8 | 12 | 0 | 0 | 0 | 0 | 11 | 0 | 0 | 0 | 0 | 0 | 0 | 0 | 0 | 0 | 8 | 0 | 0 | 3 | 10 | 23 | 0 | 0 | 0 | 0 | 9 |
| ASV396 | 2100177b236 | 181 | 17 | 0 | 45 | 33 | 66 | 0 | 0 | 0 | 11 | 9 | 0 | 0 | 0 | 0 | 0 | 0 | 0 | 0 | 0 | 0 | 0 | 0 | 0 | 0 | 0 | 0 | 0 | 0 | 0 | 0 | 0 | 0 | 0 | 0 | 0 |
| ASV397 | b947f336bfc | 180 | 15 | 0 | 35 | 36 | 0 | 13 | 0 | 41 | 7 | 0 | 0 | 0 | 0 | 0 | 0 | 0 | 0 | 0 | 0 | 0 | 0 | 0 | 7 | 0 | 0 | 0 | 0 | 15 | 0 | 0 | 0 | 0 | 0 | 11 | 0 |
| ASV398 | 47cdd08d5cb | 180 | 6 | 8 | 23 | 11 | 10 | 6 | 6 | 10 | 0 | 0 | 0 | 0 | 0 | 25 | 0 | 0 | 0 | 0 | 0 | 0 | 0 | 0 | 0 | 0 | 0 | 0 | 0 | 8 | 51 | 0 | 0 | 0 | 0 | 11 | 5 |
| ASV399 | b9bcda3d6e2 | 180 | 13 | 0 | 0 | 0 | 23 | 0 | 0 | 0 | 6 | 15 | 0 | 0 | 0 | 0 | 62 | 0 | 0 | 0 | 0 | 0 | 0 | 0 | 0 | 0 | 0 | 0 | 0 | 0 | 0 | 61 | 0 | 0 | 0 | 0 | 0 |
| ASV400 | b4881312c25 | 179 | 0 | 0 | 0 | 0 | 0 | 0 | 0 | 17 | 17 | 8 | 0 | 0 | 0 | 0 | 20 | 0 | 0 | 0 | 0 | 0 | 0 | 0 | 0 | 8 | 15 | 0 | 0 | 26 | 22 | 18 | 0 | 0 | 0 | 15 | 13 |
| ASV401 | 30c84d5d82c | 176 | 11 | 0 | 11 | 15 | 6 | 0 | 0 | 8 | 0 | 0 | 0 | 0 | 0 | 0 | 56 | 0 | 0 | 0 | 0 | 0 | 0 | 0 | 11 | 6 | 0 | 0 | 5 | 15 | 18 | 14 | 0 | 0 | 0 | 0 | 0 |
| ASV402 | 1d4406231f9 | 175 | 0 | 0 | 7 | 0 | 10 | 0 | 0 | 0 | 0 | 0 | 0 | 0 | 0 | 0 | 0 | 0 | 4 | 0 | 11 | 4 | 6 | 9 | 10 | 25 | 9 | 0 | 0 | 11 | 40 | 13 | 0 | 0 | 0 | 7 | 9 |
| ASV403 | 11aa54de56b | 173 | 4 | 0 | 6 | 0 | 0 | 31 | 5 | 0 | 0 | 0 | 4 | 0 | 0 | 0 | 0 | 0 | 0 | 0 | 0 | 0 | 25 | 14 | 0 | 0 | 0 | 5 | 9 | 21 | 9 | 0 | 13 | 7 | 14 | 6 | 0 |
| ASV404 | 872fca46f3da | 173 | 7 | 0 | 0 | 0 | 22 | 0 | 0 | 10 | 22 | 21 | 0 | 0 | 0 | 0 | 41 | 0 | 0 | 0 | 0 | 0 | 0 | 0 | 0 | 5 | 13 | 0 | 0 | 0 | 11 | 21 | 0 | 0 | 0 | 0 | 0 |
| ASV405 | 2ac7e83bad1 | 173 | 0 | 0 | 124 | 49 | 0 | 0 | 0 | 0 | 0 | 0 | 0 | 0 | 0 | 0 | 0 | 0 | 0 | 0 | 0 | 0 | 0 | 0 | 0 | 0 | 0 | 0 | 0 | 0 | 0 | 0 | 0 | 0 | 0 | 0 | 0 |
| ASV406 | f1d1e5a2e3e | 172 | 0 | 0 | 0 | 0 | 0 | 0 | 0 | 0 | 0 | 0 | 0 | 24 | 0 | 0 | 0 | 0 | 0 | 0 | 0 | 0 | 0 | 0 | 20 | 0 | 0 | 0 | 0 | 70 | 58 | 0 | 0 | 0 | 0 | 0 | 0 |
| ASV407 | 21584f17904 | 172 | 0 | 0 | 0 | 0 | 0 | 0 | 0 | 0 | 0 | 0 | 0 | 42 | 0 | 0 | 0 | 64 | 66 | 0 | 0 | 0 | 0 | 0 | 0 | 0 | 0 | 0 | 0 | 0 | 0 | 0 | 0 | 0 | 0 | 0 | 0 |
| ASV408 | a6028017c7a | 171 | 0 | 0 | 54 | 0 | 63 | 0 | 0 | 0 | 0 | 24 | 0 | 0 | 0 | 0 | 30 | 0 | 0 | 0 | 0 | 0 | 0 | 0 | 0 | 0 | 0 | 0 | 0 | 0 | 0 | 0 | 0 | 0 | 0 | 0 | 0 |
| ASV409 | 386e761d37a | 169 | 0 | 0 | 0 | 0 | 0 | 0 | 0 | 0 | 0 | 0 | 0 | 0 | 0 | 0 | 0 | 0 | 0 | 0 | 11 | 29 | 0 | 0 | 0 | 8 | 68 | 0 | 0 | 0 | 0 | 15 | 0 | 0 | 0 | 13 | 25 |
| ASV410 | 55f9ddd0a55 | 167 | 0 | 0 | 0 | 0 | 0 | 0 | 0 | 0 | 0 | 0 | 0 | 0 | 0 | 0 | 0 | 19 | 26 | 0 | 0 | 0 | 15 | 9 | 0 | 0 | 0 | 11 | 12 | 35 | 0 | 0 | 11 | 13 | 7 | 9 | 0 |
| ASV411 | afeb7677ee9 | 166 | 0 | 0 | 0 | 0 | 0 | 0 | 0 | 0 | 0 | 0 | 0 | 0 | 0 | 0 | 0 | 0 | 0 | 12 | 0 | 0 | 0 | 0 | 13 | 14 | 0 | 0 | 0 | 65 | 57 | 5 | 0 | 0 | 0 | 0 | 0 |
| ASV412 | cf79c866027f | 165 | 21 | 20 | 54 | 32 | 7 | 15 | 0 | 0 | 0 | 0 | 0 | 0 | 0 | 5 | 0 | 0 | 0 | 0 | 0 | 5 | 0 | 0 | 0 | 6 | 0 | 0 | 0 | 0 | 0 | 0 | 0 | 0 | 0 | 0 | 0 |
| ASV413 | adbc904a96d | 164 | 0 | 0 | 33 | 29 | 22 | 16 | 0 | 0 | 0 | 0 | 0 | 0 | 0 | 15 | 10 | 0 | 0 | 13 | 0 | 0 | 0 | 0 | 0 | 0 | 0 | 0 | 0 | 17 | 0 | 9 | 0 | 0 | 0 | 0 | 0 |
| ASV414 | 997910b2ca1 | 163 | 0 | 17 | 0 | 36 | 69 | 0 | 0 | 7 | 6 | 9 | 0 | 0 | 0 | 0 | 0 | 0 | 0 | 0 | 2 | 3 | 0 | 0 | 4 | 0 | 0 | 0 | 0 | 0 | 6 | 0 | 0 | 0 | 0 | 0 | 4 |
| ASV415 | 8cc0eafcb297 | 163 | 0 | 0 | 0 | 4 | 0 | 0 | 0 | 10 | 0 | 0 | 4 | 0 | 0 | 21 | 0 | 12 | 40 | 16 | 0 | 0 | 9 | 0 | 0 | 6 | 0 | 11 | 9 | 21 | 0 | 0 | 0 | 0 | 0 | 0 | 0 |
| ASV416 | 653732bd7b3 | 161 | 0 | 0 | 0 | 0 | 0 | 0 | 0 | 0 | 0 | 0 | 0 | 0 | 0 | 0 | 0 | 61 | 66 | 0 | 0 | 0 | 0 | 0 | 0 | 0 | 0 | 0 | 0 | 34 | 0 | 0 | 0 | 0 | 0 | 0 | 0 |
| ASV417 | af935abb390 | 161 | 20 | 0 | 35 | 0 | 19 | 0 | 0 | 0 | 0 | 0 | 0 | 0 | 0 | 0 | 0 | 0 | 0 | 0 | 8 | 0 | 0 | 19 | 0 | 0 | 0 | 0 | 0 | 0 | 29 | 0 | 0 | 0 | 0 | 18 | 13 |
| ASV418 | 1fea099b799 | 160 | 0 | 0 | 0 | 0 | 0 | 0 | 0 | 0 | 0 | 0 | 0 | 0 | 0 | 0 | 0 | 32 | 79 | 0 | 0 | 0 | 0 | 0 | 0 | 0 | 0 | 0 | 0 | 0 | 0 | 0 | 0 | 0 | 0 | 49 | 0 |
| ASV419 | 2fc3478a790 | 159 | 0 | 0 | 20 | 21 | 27 | 0 | 0 | 5 | 7 | 5 | 0 | 0 | 0 | 22 | 44 | 0 | 0 | 0 | 0 | 0 | 0 | 0 | 0 | 0 | 0 | 0 | 0 | 0 | 0 | 8 | 0 | 0 | 0 | 0 | 0 |
| ASV420 | d16445c2681 | 159 | 0 | 0 | 0 | 0 | 0 | 0 | 0 | 0 | 73 | 0 | 0 | 0 | 0 | 0 | 86 | 0 | 0 | 0 | 0 | 0 | 0 | 0 | 0 | 0 | 0 | 0 | 0 | 0 | 0 | 0 | 0 | 0 | 0 | 0 | 0 |
| ASV421 | d0465cdb3fd | 158 | 0 | 0 | 0 | 0 | 0 | 0 | 0 | 10 | 30 | 12 | 0 | 0 | 0 | 0 | 27 | 0 | 0 | 0 | 3 | 13 | 0 | 0 | 0 | 0 | 32 | 0 | 0 | 0 | 4 | 15 | 0 | 0 | 0 | 0 | 12 |
| ASV422 | ebb62d3089c | 157 | 0 | 0 | 10 | 0 | 12 | 0 | 0 | 16 | 0 | 0 | 0 | 0 | 0 | 13 | 0 | 0 | 0 | 0 | 0 | 0 | 0 | 0 | 0 | 15 | 18 | 0 | 0 | 5 | 40 | 28 | 0 | 0 | 0 | 0 | 0 |
| ASV423 | b4789b7fd53 | 154 | 0 | 0 | 0 | 0 | 0 | 0 | 0 | 0 | 39 | 25 | 0 | 0 | 0 | 0 | 90 | 0 | 0 | 0 | 0 | 0 | 0 | 0 | 0 | 0 | 0 | 0 | 0 | 0 | 0 | 0 | 0 | 0 | 0 | 0 | 0 |
| ASV424 | 3d038b0c2c0 | 154 | 0 | 0 | 0 | 24 | 29 | 28 | 0 | 0 | 0 | 0 | 0 | 0 | 0 | 52 | 0 | 0 | 0 | 0 | 0 | 0 | 0 | 0 | 0 | 0 | 0 | 0 | 0 | 0 | 0 | 21 | 0 | 0 | 0 | 0 | 0 |
| ASV425 | 8cb7619b459 | 154 | 0 | 0 | 0 | 0 | 0 | 0 | 0 | 0 | 0 | 0 | 0 | 0 | 0 | 0 | 23 | 0 | 0 | 0 | 0 | 0 | 0 | 0 | 0 | 0 | 0 | 0 | 0 | 0 | 37 | 0 | 0 | 0 | 0 | 94 | 0 |
| ASV426 | aaa9c7cf85d | 154 | 0 | 0 | 0 | 0 | 0 | 0 | 0 | 0 | 0 | 0 | 0 | 0 | 0 | 0 | 0 | 0 | 0 | 0 | 0 | 0 | 0 | 0 | 0 | 0 | 0 | 17 | 0 | 0 | 0 | 0 | 82 | 16 | 15 | 24 | 0 |
| ASV427 | acf57007ceb | 152 | 0 | 0 | 0 | 0 | 0 | 0 | 0 | 0 | 0 | 0 | 0 | 0 | 0 | 0 | 0 | 50 | 83 | 13 | 0 | 0 | 0 | 0 | 0 | 0 | 0 | 6 | 0 | 0 | 0 | 0 | 0 | 0 | 0 | 0 | 0 |
| ASV428 | 8617f557db3 | 152 | 0 | 0 | 0 | 0 | 0 | 0 | 0 | 0 | 79 | 0 | 0 | 0 | 0 | 0 | 73 | 0 | 0 | 0 | 0 | 0 | 0 | 0 | 0 | 0 | 0 | 0 | 0 | 0 | 0 | 0 | 0 | 0 | 0 | 0 | 0 |
| ASV429 | 73401366214 | 152 | 0 | 0 | 0 | 0 | 32 | 25 | 0 | 0 | 0 | 0 | 0 | 0 | 0 | 0 | 66 | 0 | 0 | 0 | 0 | 0 | 0 | 0 | 13 | 0 | 0 | 0 | 0 | 0 | 16 | 0 | 0 | 0 | 0 | 0 | 0 |
| ASV430 | 1744e6c70f0 | 148 | 0 | 0 | 0 | 0 | 0 | 0 | 0 | 0 | 0 | 0 | 0 | 0 | 0 | 0 | 0 | 0 | 0 | 0 | 0 | 0 | 0 | 0 | 0 | 0 | 0 | 0 | 0 | 0 | 0 | 0 | 74 | 36 | 38 | 0 | 0 |
| ASV431 | 59095dbe42d | 148 | 0 | 0 | 16 | 0 | 15 | 0 | 0 | 0 | 33 | 0 | 0 | 0 | 0 | 0 | 26 | 0 | 0 | 0 | 15 | 0 | 0 | 0 | 0 | 0 | 0 | 0 | 0 | 0 | 0 | 0 | 43 | 0 | 0 | 0 | 0 |
| ASV432 | baf96a8b9cd | 147 | 0 | 0 | 0 | 0 | 22 | 0 | 0 | 24 | 0 | 0 | 0 | 0 | 0 | 0 | 26 | 0 | 0 | 0 | 0 | 0 | 0 | 0 | 0 | 12 | 12 | 0 | 0 | 0 | 51 | 0 | 0 | 0 | 0 | 0 | 0 |
| ASV433 | bebbae58728 | 147 | 0 | 0 | 0 | 0 | 0 | 0 | 0 | 0 | 0 | 29 | 0 | 0 | 0 | 0 | 33 | 0 | 0 | 0 | 0 | 23 | 0 | 0 | 0 | 0 | 52 | 0 | 0 | 0 | 0 | 10 | 0 | 0 | 0 | 0 | 0 |
| ASV434 | 3f3eda52b69 | 146 | 0 | 0 | 0 | 0 | 0 | 0 | 0 | 0 | 0 | 0 | 0 | 0 | 0 | 0 | 0 | 23 | 49 | 0 | 0 | 0 | 15 | 0 | 0 | 0 | 0 | 0 | 0 | 19 | 0 | 0 | 23 | 0 | 0 | 17 | 0 |
| ASV435 | 618875f9339 | 146 | 0 | 0 | 0 | 0 | 42 | 0 | 0 | 0 | 45 | 0 | 0 | 0 | 0 | 0 | 59 | 0 | 0 | 0 | 0 | 0 | 0 | 0 | 0 | 0 | 0 | 0 | 0 | 0 | 0 | 0 | 0 | 0 | 0 | 0 | 0 |
| ASV436 | 6aa1fe4a28c | 145 | 0 | 0 | 0 | 0 | 27 | 0 | 0 | 0 | 0 | 0 | 0 | 0 | 0 | 0 | 34 | 0 | 0 | 0 | 0 | 32 | 0 | 0 | 0 | 0 | 52 | 0 | 0 | 0 | 0 | 0 | 0 | 0 | 0 | 0 | 0 |
| ASV437 | 42e5e5b5960 | 145 | 0 | 0 | 4 | 0 | 34 | 0 | 0 | 0 | 27 | 33 | 0 | 0 | 0 | 0 | 38 | 0 | 0 | 0 | 0 | 0 | 0 | 0 | 0 | 0 | 0 | 0 | 0 | 0 | 0 | 0 | 0 | 0 | 0 | 0 | 9 |
| ASV438 | e8706f8a839 | 145 | 0 | 7 | 0 | 16 | 16 | 0 | 0 | 0 | 21 | 0 | 0 | 0 | 0 | 0 | 22 | 0 | 0 | 0 | 11 | 0 | 0 | 0 | 0 | 9 | 23 | 0 | 0 | 0 | 0 | 0 | 0 | 5 | 0 | 6 | 9 |
| ASV439 | 6a05b15945e | 145 | 0 | 0 | 0 | 0 | 0 | 0 | 0 | 0 | 0 | 0 | 46 | 73 | 26 | 0 | 0 | 0 | 0 | 0 | 0 | 0 | 0 | 0 | 0 | 0 | 0 | 0 | 0 | 0 | 0 | 0 | 0 | 0 | 0 | 0 | 0 |
| ASV440 | 92d82058dd0 | 143 | 0 | 0 | 0 | 0 | 0 | 0 | 0 | 0 | 0 | 0 | 0 | 0 | 0 | 0 | 0 | 0 | 0 | 0 | 0 | 0 | 0 | 0 | 0 | 7 | 0 | 0 | 0 | 12 | 58 | 16 | 0 | 0 | 0 | 40 | 10 |
| ASV441 | 4f6a759f559a | 142 | 0 | 0 | 0 | 0 | 0 | 0 | 0 | 0 | 0 | 0 | 0 | 0 | 0 | 0 | 0 | 57 | 85 | 0 | 0 | 0 | 0 | 0 | 0 | 0 | 0 | 0 | 0 | 0 | 0 | 0 | 0 | 0 | 0 | 0 | 0 |
| ASV442 | 435a4b0cafd | 141 | 0 | 0 | 0 | 0 | 17 | 0 | 0 | 0 | 0 | 26 | 0 | 0 | 0 | 0 | 32 | 0 | 0 | 0 | 0 | 13 | 0 | 0 | 0 | 0 | 0 | 0 | 0 | 0 | 12 | 18 | 0 | 0 | 0 | 0 | 23 |
| ASV443 | 27ff93acae11 | 140 | 0 | 0 | 0 | 0 | 0 | 0 | 0 | 0 | 13 | 0 | 0 | 0 | 0 | 0 | 0 | 0 | 0 | 0 | 0 | 0 | 0 | 0 | 0 | 0 | 0 | 0 | 0 | 0 | 0 | 0 | 39 | 49 | 39 | 0 | 0 |
| ASV444 | 066eac4266e | 139 | 0 | 0 | 0 | 0 | 0 | 0 | 0 | 0 | 23 | 0 | 0 | 0 | 0 | 0 | 0 | 0 | 0 | 0 | 33 | 11 | 0 | 0 | 0 | 0 | 21 | 0 | 0 | 0 | 0 | 24 | 0 | 0 | 0 | 27 | 0 |
| ASV445 | 51a13138db3 | 139 | 0 | 0 | 21 | 0 | 0 | 0 | 0 | 23 | 27 | 21 | 0 | 0 | 0 | 0 | 24 | 0 | 0 | 0 | 0 | 0 | 0 | 0 | 0 | 0 | 0 | 0 | 0 | 0 | 0 | 23 | 0 | 0 | 0 | 0 | 0 |
| ASV446 | 4ee7d437fd1 | 138 | 16 | 10 | 50 | 30 | 32 | 0 | 0 | 0 | 0 | 0 | 0 | 0 | 0 | 0 | 0 | 0 | 0 | 0 | 0 | 0 | 0 | 0 | 0 | 0 | 0 | 0 | 0 | 0 | 0 | 0 | 0 | 0 | 0 | 0 | 0 |
| ASV447 | fa74446f429 | 138 | 0 | 0 | 0 | 0 | 0 | 0 | 0 | 0 | 0 | 0 | 0 | 0 | 0 | 0 | 52 | 0 | 0 | 0 | 0 | 43 | 0 | 0 | 0 | 0 | 43 | 0 | 0 | 0 | 0 | 0 | 0 | 0 | 0 | 0 | 0 |
| ASV448 | 09bbdda11f3 | 138 | 11 | 0 | 45 | 16 | 55 | 0 | 0 | 0 | 0 | 11 | 0 | 0 | 0 | 0 | 0 | 0 | 0 | 0 | 0 | 0 | 0 | 0 | 0 | 0 | 0 | 0 | 0 | 0 | 0 | 0 | 0 | 0 | 0 | 0 | 0 |
| ASV449 | ee70504f852 | 138 | 0 | 0 | 0 | 0 | 60 | 0 | 0 | 0 | 0 | 0 | 0 | 0 | 0 | 0 | 56 | 0 | 0 | 0 | 0 | 0 | 0 | 0 | 0 | 0 | 0 | 0 | 0 | 0 | 0 | 22 | 0 | 0 | 0 | 0 | 0 |
| ASV450 | 3616465ab65 | 137 | 0 | 0 | 0 | 0 | 0 | 0 | 0 | 0 | 0 | 0 | 0 | 0 | 0 | 0 | 0 | 28 | 0 | 0 | 0 | 0 | 21 | 0 | 0 | 0 | 0 | 0 | 0 | 0 | 0 | 0 | 71 | 0 | 17 | 0 | 0 |
| ASV451 | 6e367f752ccf | 137 | 0 | 0 | 0 | 0 | 14 | 0 | 0 | 0 | 20 | 18 | 0 | 0 | 0 | 0 | 53 | 0 | 0 | 0 | 0 | 0 | 0 | 0 | 0 | 0 | 5 | 0 | 0 | 0 | 4 | 11 | 0 | 0 | 0 | 0 | 12 |
| ASV452 | b263c550540 | 136 | 0 | 0 | 0 | 0 | 0 | 0 | 0 | 0 | 0 | 0 | 0 | 14 | 11 | 13 | 0 | 0 | 0 | 0 | 0 | 0 | 0 | 0 | 0 | 0 | 0 | 0 | 0 | 0 | 0 | 0 | 39 | 32 | 27 | 0 | 0 |
| ASV453 | cf915c661e4a | 135 | 0 | 0 | 0 | 0 | 0 | 0 | 0 | 0 | 51 | 0 | 0 | 0 | 0 | 0 | 84 | 0 | 0 | 0 | 0 | 0 | 0 | 0 | 0 | 0 | 0 | 0 | 0 | 0 | 0 | 0 | 0 | 0 | 0 | 0 | 0 |
| ASV454 | 0a2463f6dd1 | 135 | 0 | 0 | 0 | 0 | 0 | 0 | 0 | 0 | 0 | 0 | 0 | 0 | 0 | 0 | 0 | 0 | 0 | 0 | 0 | 0 | 0 | 0 | 0 | 0 | 43 | 0 | 0 | 0 | 0 | 50 | 0 | 0 | 0 | 42 | 0 |
| ASV455 | f708204d5c2 | 135 | 0 | 0 | 0 | 0 | 0 | 0 | 0 | 0 | 0 | 15 | 0 | 0 | 0 | 0 | 0 | 0 | 0 | 0 | 0 | 0 | 0 | 0 | 0 | 0 | 0 | 0 | 0 | 0 | 73 | 0 | 0 | 0 | 0 | 47 | 0 |
| ASV456 | a06a4e8db20 | 135 | 33 | 26 | 15 | 15 | 1 | 16 | 29 | 0 | 0 | 0 | 0 | 0 | 0 | 0 | 0 | 0 | 0 | 0 | 0 | 0 | 0 | 0 | 0 | 0 | 0 | 0 | 0 | 0 | 0 | 0 | 0 | 0 | 0 | 0 | 0 |
| ASV457 | 0c84384e672 | 134 | 0 | 0 | 0 | 0 | 0 | 9 | 0 | 0 | 0 | 9 | 0 | 0 | 0 | 0 | 35 | 0 | 0 | 0 | 17 | 0 | 0 | 0 | 0 | 20 | 0 | 0 | 0 | 0 | 28 | 0 | 0 | 0 | 0 | 16 | 0 |
| ASV458 | d3893ebcfc1 | 134 | 13 | 0 | 46 | 24 | 51 | 0 | 0 | 0 | 0 | 0 | 0 | 0 | 0 | 0 | 0 | 0 | 0 | 0 | 0 | 0 | 0 | 0 | 0 | 0 | 0 | 0 | 0 | 0 | 0 | 0 | 0 | 0 | 0 | 0 | 0 |
| ASV459 | c33858cc31a | 133 | 0 | 0 | 0 | 0 | 0 | 0 | 0 | 0 | 0 | 0 | 0 | 0 | 0 | 0 | 0 | 0 | 0 | 0 | 0 | 0 | 0 | 0 | 0 | 0 | 0 | 0 | 0 | 0 | 0 | 0 | 83 | 0 | 50 | 0 | 0 |
| ASV460 | d444fec272c | 132 | 0 | 0 | 0 | 0 | 24 | 0 | 0 | 0 | 17 | 15 | 0 | 0 | 0 | 0 | 29 | 0 | 0 | 0 | 0 | 0 | 0 | 0 | 0 | 0 | 17 | 0 | 0 | 0 | 0 | 14 | 0 | 0 | 0 | 0 | 16 |
| ASV461 | acac251921d | 132 | 0 | 0 | 0 | 0 | 0 | 0 | 0 | 0 | 0 | 0 | 0 | 0 | 0 | 0 | 0 | 0 | 0 | 0 | 0 | 0 | 0 | 0 | 0 | 0 | 0 | 0 | 0 | 13 | 17 | 0 | 0 | 0 | 0 | 61 | 41 |
| ASV462 | a69a75d5ee0 | 131 | 0 | 0 | 0 | 0 | 0 | 0 | 0 | 0 | 0 | 0 | 0 | 0 | 0 | 0 | 0 | 47 | 40 | 0 | 0 | 0 | 0 | 0 | 0 | 0 | 0 | 0 | 0 | 0 | 0 | 0 | 44 | 0 | 0 | 0 | 0 |
| ASV463 | df64d0b5d48 | 130 | 0 | 0 | 0 | 0 | 0 | 0 | 0 | 0 | 0 | 0 | 0 | 0 | 0 | 0 | 0 | 0 | 41 | 0 | 0 | 0 | 0 | 0 | 0 | 0 | 0 | 26 | 0 | 19 | 0 | 0 | 0 | 44 | 0 | 0 | 0 |
| ASV464 | 109ca30c2be | 129 | 0 | 0 | 0 | 0 | 0 | 0 | 0 | 0 | 0 | 0 | 0 | 0 | 0 | 0 | 0 | 0 | 23 | 0 | 0 | 0 | 0 | 0 | 0 | 0 | 0 | 28 | 0 | 0 | 0 | 0 | 56 | 0 | 0 | 22 | 0 |
| ASV465 | 83c24c6a57e | 128 | 0 | 0 | 0 | 0 | 11 | 0 | 0 | 0 | 26 | 6 | 0 | 0 | 0 | 0 | 23 | 0 | 0 | 0 | 0 | 11 | 0 | 0 | 0 | 0 | 27 | 0 | 0 | 0 | 5 | 19 | 0 | 0 | 0 | 0 | 0 |
| ASV466 | edea52bf051 | 127 | 0 | 0 | 0 | 0 | 0 | 0 | 0 | 0 | 0 | 29 | 0 | 0 | 0 | 0 | 0 | 0 | 0 | 0 | 0 | 8 | 0 | 0 | 0 | 9 | 5 | 0 | 0 | 0 | 0 | 0 | 0 | 0 | 0 | 38 | 38 |
| ASV467 | e9d803cb50b | 127 | 0 | 0 | 0 | 0 | 0 | 0 | 0 | 0 | 0 | 0 | 0 | 0 | 0 | 0 | 0 | 35 | 30 | 0 | 0 | 0 | 25 | 0 | 0 | 0 | 0 | 37 | 0 | 0 | 0 | 0 | 0 | 0 | 0 | 0 | 0 |
| ASV468 | 490b60bbce7 | 126 | 44 | 0 | 4 | 0 | 0 | 0 | 0 | 0 | 0 | 0 | 0 | 0 | 0 | 0 | 0 | 0 | 0 | 0 | 0 | 0 | 0 | 0 | 0 | 0 | 0 | 41 | 0 | 37 | 0 | 0 | 0 | 0 | 0 | 0 | 0 |
| ASV469 | bfa4ab900ba | 126 | 0 | 0 | 0 | 0 | 0 | 0 | 0 | 0 | 16 | 0 | 0 | 0 | 0 | 0 | 54 | 0 | 0 | 0 | 18 | 0 | 0 | 0 | 4 | 11 | 20 | 0 | 0 | 0 | 0 | 3 | 0 | 0 | 0 | 0 | 0 |
| ASV470 | e8e59dc192a | 125 | 0 | 0 | 0 | 0 | 0 | 0 | 0 | 0 | 0 | 0 | 0 | 0 | 0 | 0 | 125 | 0 | 0 | 0 | 0 | 0 | 0 | 0 | 0 | 0 | 0 | 0 | 0 | 0 | 0 | 0 | 0 | 0 | 0 | 0 | 0 |
| ASV471 | 764f5a270af | 125 | 0 | 0 | 0 | 0 | 0 | 0 | 0 | 0 | 0 | 0 | 0 | 0 | 0 | 125 | 0 | 0 | 0 | 0 | 0 | 0 | 0 | 0 | 0 | 0 | 0 | 0 | 0 | 0 | 0 | 0 | 0 | 0 | 0 | 0 | 0 |
| ASV472 | 438927eaf1a | 124 | 0 | 0 | 0 | 12 | 0 | 0 | 0 | 6 | 0 | 0 | 0 | 10 | 4 | 0 | 0 | 26 | 20 | 9 | 0 | 0 | 0 | 8 | 0 | 0 | 0 | 9 | 8 | 0 | 0 | 0 | 0 | 0 | 5 | 7 | 0 |
| ASV473 | 4779e8ffc76 | 124 | 3 | 2 | 26 | 27 | 29 | 0 | 0 | 8 | 5 | 0 | 0 | 0 | 0 | 0 | 11 | 0 | 0 | 0 | 0 | 0 | 0 | 0 | 0 | 0 | 13 | 0 | 0 | 0 | 0 | 0 | 0 | 0 | 0 | 0 | 0 |
| ASV474 | 67b03cc55c2 | 124 | 17 | 0 | 47 | 20 | 8 | 11 | 0 | 0 | 0 | 0 | 0 | 0 | 0 | 10 | 0 | 0 | 0 | 0 | 0 | 0 | 0 | 0 | 0 | 0 | 0 | 0 | 0 | 0 | 0 | 0 | 0 | 0 | 0 | 11 | 0 |
| ASV475 | 707cea640b1 | 123 | 0 | 0 | 0 | 0 | 0 | 0 | 0 | 0 | 0 | 0 | 0 | 3 | 3 | 0 | 0 | 0 | 0 | 0 | 0 | 0 | 21 | 0 | 18 | 0 | 0 | 0 | 19 | 7 | 3 | 12 | 1 | 0 | 36 | 0 | 0 |
| ASV476 | b4c3ec66491 | 122 | 0 | 3 | 24 | 13 | 28 | 14 | 0 | 0 | 0 | 0 | 0 | 0 | 0 | 0 | 6 | 0 | 0 | 0 | 0 | 0 | 0 | 0 | 0 | 3 | 0 | 0 | 0 | 0 | 11 | 5 | 0 | 0 | 0 | 8 | 7 |
| ASV477 | 6c892d490d7 | 121 | 0 | 0 | 0 | 0 | 0 | 0 | 0 | 0 | 0 | 12 | 0 | 0 | 0 | 0 | 27 | 0 | 0 | 0 | 0 | 0 | 0 | 0 | 0 | 10 | 12 | 0 | 0 | 0 | 0 | 42 | 0 | 0 | 0 | 0 | 18 |
| ASV478 | 9e54883092a | 121 | 0 | 0 | 0 | 0 | 34 | 0 | 0 | 0 | 56 | 0 | 0 | 0 | 0 | 0 | 0 | 0 | 0 | 0 | 0 | 0 | 0 | 0 | 0 | 0 | 0 | 0 | 0 | 0 | 0 | 10 | 0 | 0 | 0 | 0 | 21 |
| ASV479 | e2a8fe5c36b | 120 | 5 | 0 | 0 | 0 | 48 | 0 | 0 | 0 | 0 | 12 | 0 | 0 | 0 | 0 | 8 | 0 | 0 | 0 | 0 | 0 | 0 | 0 | 0 | 0 | 18 | 0 | 0 | 0 | 0 | 0 | 0 | 0 | 0 | 8 | 21 |
| ASV480 | 25ace10ce6d | 118 | 0 | 0 | 0 | 0 | 22 | 0 | 0 | 0 | 0 | 8 | 0 | 0 | 0 | 0 | 57 | 0 | 0 | 0 | 0 | 0 | 0 | 0 | 0 | 0 | 0 | 0 | 0 | 0 | 0 | 11 | 0 | 0 | 0 | 0 | 20 |
| ASV481 | d295eae8eee | 117 | 0 | 0 | 0 | 46 | 43 | 0 | 6 | 0 | 0 | 0 | 0 | 0 | 14 | 0 | 0 | 0 | 0 | 0 | 0 | 0 | 0 | 0 | 8 | 0 | 0 | 0 | 0 | 0 | 0 | 0 | 0 | 0 | 0 | 0 | 0 |
| ASV482 | 6a9308a3d4b | 117 | 0 | 0 | 0 | 0 | 30 | 0 | 0 | 0 | 17 | 8 | 0 | 0 | 0 | 0 | 40 | 0 | 0 | 0 | 0 | 0 | 0 | 0 | 0 | 0 | 11 | 0 | 0 | 0 | 0 | 0 | 0 | 0 | 0 | 0 | 11 |
| ASV483 | e08b5550e72 | 116 | 0 | 0 | 0 | 0 | 0 | 0 | 0 | 0 | 74 | 0 | 0 | 0 | 0 | 0 | 0 | 0 | 0 | 0 | 0 | 42 | 0 | 0 | 0 | 0 | 0 | 0 | 0 | 0 | 0 | 0 | 0 | 0 | 0 | 0 | 0 |
| ASV484 | f0fec588b2b | 116 | 0 | 0 | 0 | 0 | 0 | 0 | 0 | 22 | 0 | 0 | 0 | 0 | 0 | 0 | 0 | 0 | 0 | 0 | 29 | 65 | 0 | 0 | 0 | 0 | 0 | 0 | 0 | 0 | 0 | 0 | 0 | 0 | 0 | 0 | 0 |
| ASV485 | e8980bb8dc2 | 115 | 0 | 0 | 0 | 0 | 0 | 0 | 0 | 0 | 0 | 0 | 15 | 38 | 0 | 0 | 0 | 62 | 0 | 0 | 0 | 0 | 0 | 0 | 0 | 0 | 0 | 0 | 0 | 0 | 0 | 0 | 0 | 0 | 0 | 0 | 0 |
| ASV486 | b0e3b77912a | 114 | 19 | 16 | 79 | 0 | 0 | 0 | 0 | 0 | 0 | 0 | 0 | 0 | 0 | 0 | 0 | 0 | 0 | 0 | 0 | 0 | 0 | 0 | 0 | 0 | 0 | 0 | 0 | 0 | 0 | 0 | 0 | 0 | 0 | 0 | 0 |
| ASV487 | 1bc1d3897b0 | 114 | 0 | 0 | 19 | 0 | 0 | 0 | 0 | 0 | 0 | 0 | 0 | 17 | 0 | 12 | 0 | 0 | 26 | 0 | 0 | 0 | 0 | 0 | 0 | 0 | 0 | 13 | 0 | 0 | 0 | 0 | 27 | 0 | 0 | 0 | 0 |
| ASV488 | 1950553995c | 113 | 0 | 0 | 0 | 0 | 0 | 0 | 0 | 0 | 0 | 0 | 0 | 0 | 0 | 0 | 0 | 0 | 27 | 0 | 0 | 0 | 0 | 0 | 0 | 0 | 0 | 30 | 0 | 0 | 0 | 0 | 0 | 56 | 0 | 0 | 0 |
| ASV489 | 2610fd16edf | 113 | 0 | 0 | 0 | 0 | 0 | 0 | 0 | 0 | 0 | 0 | 0 | 0 | 0 | 0 | 0 | 0 | 0 | 0 | 0 | 0 | 0 | 0 | 0 | 0 | 0 | 0 | 0 | 0 | 0 | 0 | 56 | 25 | 17 | 15 | 0 |
| ASV490 | 80cd4f2b322 | 113 | 0 | 0 | 53 | 0 | 0 | 26 | 0 | 15 | 0 | 0 | 0 | 0 | 0 | 0 | 0 | 0 | 0 | 0 | 0 | 0 | 0 | 0 | 0 | 19 | 0 | 0 | 0 | 0 | 0 | 0 | 0 | 0 | 0 | 0 | 0 |
| ASV491 | 741fcf923ccd | 113 | 13 | 0 | 24 | 0 | 30 | 0 | 0 | 0 | 26 | 0 | 0 | 0 | 0 | 0 | 0 | 0 | 0 | 0 | 20 | 0 | 0 | 0 | 0 | 0 | 0 | 0 | 0 | 0 | 0 | 0 | 0 | 0 | 0 | 0 | 0 |
| ASV492 | 95ee4bda57c | 112 | 0 | 0 | 0 | 0 | 0 | 0 | 0 | 0 | 0 | 24 | 0 | 0 | 0 | 0 | 0 | 0 | 0 | 0 | 18 | 22 | 0 | 0 | 0 | 12 | 0 | 0 | 0 | 0 | 18 | 18 | 0 | 0 | 0 | 0 | 0 |
| ASV493 | b5d4643929e | 111 | 0 | 0 | 0 | 0 | 0 | 0 | 0 | 0 | 0 | 0 | 0 | 0 | 0 | 0 | 0 | 0 | 0 | 0 | 0 | 0 | 74 | 37 | 0 | 0 | 0 | 0 | 0 | 0 | 0 | 0 | 0 | 0 | 0 | 0 | 0 |
| ASV494 | 583af40a1f3 | 111 | 0 | 0 | 0 | 0 | 0 | 0 | 0 | 0 | 0 | 0 | 0 | 0 | 0 | 0 | 0 | 58 | 0 | 0 | 0 | 0 | 0 | 0 | 0 | 0 | 0 | 7 | 0 | 0 | 0 | 0 | 31 | 9 | 0 | 6 | 0 |
| ASV495 | 953bf575ad5 | 111 | 0 | 0 | 0 | 0 | 0 | 0 | 0 | 0 | 0 | 0 | 0 | 0 | 0 | 0 | 0 | 48 | 63 | 0 | 0 | 0 | 0 | 0 | 0 | 0 | 0 | 0 | 0 | 0 | 0 | 0 | 0 | 0 | 0 | 0 | 0 |
| ASV496 | 1160896bd5e | 110 | 0 | 0 | 0 | 0 | 0 | 0 | 0 | 0 | 0 | 0 | 0 | 0 | 0 | 0 | 0 | 0 | 0 | 0 | 0 | 0 | 0 | 0 | 0 | 0 | 0 | 0 | 0 | 7 | 0 | 39 | 0 | 24 | 25 | 15 | 0 |
| ASV497 | 7333eff9431 | 110 | 0 | 0 | 0 | 0 | 0 | 0 | 0 | 0 | 0 | 0 | 0 | 0 | 0 | 0 | 0 | 0 | 0 | 0 | 72 | 0 | 38 | 0 | 0 | 0 | 0 | 0 | 0 | 0 | 0 | 0 | 0 | 0 | 0 | 0 | 0 |
| ASV498 | acc650be2c9 | 109 | 0 | 0 | 0 | 0 | 21 | 0 | 0 | 0 | 18 | 0 | 0 | 0 | 0 | 0 | 0 | 0 | 0 | 0 | 0 | 0 | 0 | 0 | 0 | 12 | 0 | 0 | 0 | 0 | 0 | 34 | 0 | 0 | 0 | 8 | 16 |
| ASV499 | 4ff740aace27 | 108 | 0 | 0 | 0 | 0 | 11 | 0 | 0 | 0 | 0 | 26 | 0 | 0 | 0 | 0 | 71 | 0 | 0 | 0 | 0 | 0 | 0 | 0 | 0 | 0 | 0 | 0 | 0 | 0 | 0 | 0 | 0 | 0 | 0 | 0 | 0 |
| ASV500 | f8f42cf8dc3d | 107 | 0 | 0 | 0 | 0 | 20 | 0 | 0 | 0 | 12 | 12 | 0 | 0 | 0 | 0 | 40 | 0 | 0 | 0 | 0 | 0 | 0 | 0 | 0 | 0 | 0 | 0 | 0 | 0 | 0 | 18 | 0 | 0 | 0 | 0 | 5 |
| ASV501 | dd26b396d97 | 107 | 0 | 0 | 0 | 0 | 0 | 0 | 0 | 0 | 0 | 0 | 0 | 0 | 0 | 0 | 0 | 0 | 0 | 0 | 0 | 0 | 0 | 0 | 28 | 0 | 0 | 0 | 0 | 0 | 0 | 33 | 0 | 0 | 0 | 38 | 8 |
| ASV502 | 0545daa4920 | 107 | 0 | 0 | 0 | 0 | 0 | 0 | 0 | 0 | 0 | 0 | 0 | 32 | 0 | 0 | 0 | 0 | 46 | 0 | 0 | 0 | 0 | 0 | 0 | 0 | 0 | 0 | 0 | 29 | 0 | 0 | 0 | 0 | 0 | 0 | 0 |
| ASV503 | 84b495a19bc | 107 | 0 | 0 | 0 | 8 | 12 | 0 | 0 | 0 | 15 | 26 | 0 | 0 | 0 | 0 | 22 | 0 | 0 | 0 | 0 | 0 | 0 | 0 | 0 | 0 | 4 | 0 | 0 | 0 | 0 | 13 | 0 | 0 | 0 | 0 | 7 |
| ASV504 | 1c6b4a698fd | 106 | 0 | 0 | 0 | 0 | 35 | 0 | 0 | 0 | 0 | 0 | 0 | 0 | 0 | 0 | 71 | 0 | 0 | 0 | 0 | 0 | 0 | 0 | 0 | 0 | 0 | 0 | 0 | 0 | 0 | 0 | 0 | 0 | 0 | 0 | 0 |
| ASV505 | ab6d9771255 | 105 | 0 | 0 | 0 | 0 | 0 | 0 | 0 | 0 | 0 | 0 | 0 | 0 | 0 | 0 | 0 | 0 | 57 | 12 | 0 | 0 | 0 | 0 | 0 | 0 | 0 | 0 | 0 | 21 | 0 | 0 | 15 | 0 | 0 | 0 | 0 |
| ASV506 | b193c299f6f | 105 | 0 | 0 | 0 | 0 | 0 | 0 | 0 | 0 | 0 | 60 | 0 | 0 | 0 | 0 | 45 | 0 | 0 | 0 | 0 | 0 | 0 | 0 | 0 | 0 | 0 | 0 | 0 | 0 | 0 | 0 | 0 | 0 | 0 | 0 | 0 |

**Supplementary Table S5**. Bacterioplankton 16S rRNA ASV feature table
ASVs: 1779
Seqs: 918792

| #ASV_ID | Feature_ID | total seqs | May12_0m | May12_05m | May12_25m | May12_50m | May12_75m | Jun8_0m | Jun8_05m | Jun8_25m | Jun8_50m | Jun8_75m | Jun22_0m | Jun22_05m | Jun22_25m | Jun22_50m | Jun22_75m | Jul11_0m | Jul11_05m | Jul11_25m | Jul11_50m | Jul11_75m | Jul27_0m | Jul27_05m | Jul27_25m | Jul27_50m | Jul27_75m | Aug5_0m | Aug5_05m | Aug5_25m | Aug5_50m | Aug5_75m | Sep8_0m | Sep8_05m | Sep8_25m | Sep8_50m | Sep8_75m |
|---|---|---|---|---|---|---|---|---|---|---|---|---|---|---|---|---|---|---|---|---|---|---|---|---|---|---|---|---|---|---|---|---|---|---|---|---|---|
| ASV507 | 8b96d2c683a | 104 | 0 | 0 | 4 | 0 | 10 | 0 | 0 | 0 | 7 | 9 | 0 | 0 | 0 | 0 | 16 | 0 | 0 | 0 | 0 | 5 | 0 | 0 | 0 | 0 | 15 | 0 | 0 | 0 | 3 | 12 | 0 | 0 | 0 | 0 | 23 |
| ASV508 | dbe59cce279 | 103 | 0 | 0 | 4 | 2 | 14 | 0 | 0 | 0 | 14 | 4 | 0 | 0 | 0 | 0 | 56 | 0 | 0 | 0 | 0 | 0 | 0 | 0 | 0 | 0 | 0 | 0 | 0 | 0 | 0 | 0 | 0 | 0 | 0 | 0 | 9 |
| ASV509 | c007cbfea90 | 103 | 0 | 0 | 0 | 0 | 0 | 0 | 0 | 0 | 0 | 0 | 0 | 0 | 0 | 0 | 0 | 0 | 0 | 0 | 0 | 0 | 0 | 0 | 0 | 0 | 0 | 0 | 0 | 23 | 22 | 26 | 0 | 0 | 0 | 16 | 16 |
| ASV510 | b2c416efead | 103 | 0 | 0 | 0 | 0 | 0 | 0 | 0 | 0 | 0 | 29 | 0 | 0 | 0 | 0 | 74 | 0 | 0 | 0 | 0 | 0 | 0 | 0 | 0 | 0 | 0 | 0 | 0 | 0 | 0 | 0 | 0 | 0 | 0 | 0 | 0 |
| ASV511 | c56d0651aca | 102 | 0 | 0 | 0 | 0 | 0 | 0 | 0 | 0 | 0 | 0 | 0 | 0 | 0 | 0 | 0 | 21 | 31 | 12 | 0 | 0 | 0 | 0 | 0 | 0 | 0 | 10 | 0 | 8 | 0 | 0 | 0 | 0 | 0 | 20 | 0 |
| ASV512 | c38fa7fc466f | 102 | 0 | 0 | 0 | 0 | 0 | 0 | 0 | 0 | 0 | 0 | 0 | 0 | 0 | 0 | 0 | 0 | 0 | 0 | 0 | 0 | 11 | 0 | 0 | 0 | 0 | 13 | 7 | 15 | 0 | 0 | 37 | 4 | 6 | 9 | 0 |
| ASV513 | 4b0db1d7135 | 101 | 0 | 0 | 0 | 0 | 8 | 0 | 0 | 0 | 28 | 7 | 0 | 0 | 0 | 0 | 27 | 0 | 0 | 0 | 0 | 0 | 0 | 0 | 0 | 0 | 8 | 0 | 0 | 0 | 0 | 14 | 0 | 0 | 0 | 0 | 9 |
| ASV514 | 3b660f8a4f1 | 100 | 0 | 0 | 0 | 0 | 0 | 0 | 0 | 0 | 0 | 0 | 0 | 0 | 0 | 0 | 0 | 0 | 0 | 0 | 0 | 0 | 0 | 0 | 0 | 0 | 21 | 0 | 0 | 0 | 52 | 27 | 0 | 0 | 0 | 0 | 0 |
| ASV515 | 3eac82e0aae | 98 | 8 | 0 | 9 | 0 | 27 | 0 | 0 | 0 | 0 | 0 | 0 | 0 | 0 | 35 | 0 | 0 | 0 | 0 | 0 | 0 | 0 | 0 | 0 | 0 | 0 | 0 | 0 | 0 | 15 | 4 | 0 | 0 | 0 | 0 | 0 |
| ASV516 | 8a3a7875c52 | 98 | 0 | 0 | 17 | 0 | 0 | 0 | 0 | 0 | 0 | 0 | 0 | 0 | 0 | 0 | 0 | 12 | 0 | 14 | 28 | 0 | 0 | 0 | 0 | 0 | 0 | 0 | 14 | 13 | 0 | 0 | 0 | 0 | 0 | 0 | 0 |
| ASV517 | 1eb35efbb8b | 98 | 0 | 0 | 0 | 0 | 0 | 0 | 0 | 0 | 0 | 0 | 0 | 0 | 0 | 0 | 0 | 0 | 14 | 0 | 0 | 0 | 0 | 0 | 0 | 0 | 0 | 26 | 15 | 0 | 0 | 0 | 7 | 0 | 10 | 26 | 0 |
| ASV518 | b967e649393 | 96 | 0 | 0 | 0 | 0 | 14 | 0 | 0 | 8 | 25 | 0 | 0 | 0 | 0 | 0 | 29 | 0 | 0 | 0 | 0 | 13 | 0 | 0 | 0 | 0 | 0 | 0 | 0 | 0 | 0 | 7 | 0 | 0 | 0 | 0 | 0 |
| ASV519 | 4eb74ff4aa6 | 96 | 0 | 0 | 0 | 0 | 0 | 0 | 0 | 0 | 0 | 0 | 0 | 0 | 0 | 0 | 0 | 0 | 41 | 0 | 0 | 0 | 0 | 0 | 0 | 0 | 0 | 0 | 0 | 55 | 0 | 0 | 0 | 0 | 0 | 0 | 0 |
| ASV520 | f11bc49d4ac | 96 | 0 | 0 | 0 | 0 | 0 | 0 | 0 | 0 | 0 | 0 | 0 | 0 | 0 | 0 | 0 | 0 | 0 | 0 | 0 | 0 | 0 | 0 | 0 | 0 | 0 | 0 | 0 | 0 | 0 | 0 | 58 | 25 | 13 | 0 | 0 |
| ASV521 | 65b9a59ab09 | 95 | 0 | 0 | 0 | 0 | 0 | 0 | 0 | 0 | 0 | 0 | 0 | 0 | 0 | 0 | 0 | 0 | 0 | 0 | 0 | 0 | 0 | 0 | 0 | 0 | 0 | 0 | 0 | 0 | 0 | 0 | 95 | 0 | 0 | 0 | 0 |
| ASV522 | 14f30649d0c | 95 | 0 | 0 | 0 | 0 | 0 | 0 | 0 | 0 | 0 | 0 | 0 | 0 | 0 | 0 | 0 | 14 | 0 | 0 | 0 | 0 | 12 | 0 | 0 | 0 | 0 | 23 | 13 | 0 | 0 | 0 | 22 | 11 | 0 | 0 | 0 |
| ASV523 | 6551125ac9d | 95 | 0 | 0 | 0 | 0 | 0 | 68 | 15 | 0 | 0 | 0 | 0 | 12 | 0 | 0 | 0 | 0 | 0 | 0 | 0 | 0 | 0 | 0 | 0 | 0 | 0 | 0 | 0 | 0 | 0 | 0 | 0 | 0 | 0 | 0 | 0 |
| ASV524 | 83cedb51009 | 95 | 0 | 0 | 9 | 0 | 4 | 0 | 0 | 0 | 0 | 0 | 0 | 12 | 7 | 0 | 2 | 0 | 0 | 0 | 0 | 0 | 0 | 0 | 0 | 0 | 8 | 0 | 0 | 0 | 0 | 0 | 33 | 11 | 9 | 0 | 0 |
| ASV525 | b5baf3d173d | 95 | 0 | 0 | 0 | 0 | 42 | 7 | 0 | 0 | 0 | 0 | 3 | 0 | 0 | 11 | 0 | 0 | 0 | 0 | 0 | 0 | 0 | 0 | 0 | 0 | 0 | 0 | 0 | 12 | 10 | 0 | 0 | 0 | 0 | 0 | 10 |
| ASV526 | f089527968c | 94 | 0 | 0 | 0 | 0 | 12 | 0 | 0 | 0 | 31 | 0 | 0 | 0 | 0 | 0 | 16 | 0 | 0 | 0 | 24 | 0 | 0 | 0 | 0 | 0 | 0 | 0 | 0 | 0 | 0 | 11 | 0 | 0 | 0 | 0 | 0 |
| ASV527 | 04c9d96f9a1 | 94 | 0 | 0 | 21 | 0 | 6 | 5 | 0 | 0 | 0 | 0 | 0 | 0 | 0 | 0 | 0 | 0 | 0 | 0 | 0 | 0 | 0 | 0 | 0 | 0 | 0 | 0 | 0 | 0 | 0 | 6 | 0 | 0 | 0 | 39 | 17 |
| ASV528 | fc8ab582b19 | 94 | 0 | 0 | 0 | 0 | 13 | 0 | 0 | 0 | 7 | 0 | 0 | 0 | 0 | 8 | 0 | 0 | 0 | 0 | 0 | 0 | 0 | 15 | 0 | 0 | 0 | 0 | 0 | 15 | 0 | 10 | 0 | 7 | 0 | 0 | 19 |
| ASV529 | c1be7dc0142 | 93 | 0 | 0 | 10 | 3 | 4 | 6 | 0 | 12 | 0 | 5 | 0 | 0 | 0 | 5 | 0 | 0 | 0 | 0 | 0 | 0 | 0 | 0 | 0 | 13 | 9 | 0 | 0 | 0 | 0 | 2 | 0 | 0 | 0 | 17 | 7 |
| ASV530 | e6b2e4b665d | 93 | 0 | 0 | 2 | 0 | 0 | 0 | 0 | 23 | 0 | 0 | 0 | 13 | 0 | 26 | 11 | 0 | 0 | 0 | 0 | 5 | 0 | 0 | 0 | 0 | 0 | 0 | 0 | 0 | 13 | 0 | 0 | 0 | 0 | 0 | 0 |
| ASV531 | d71e2417647 | 92 | 0 | 0 | 0 | 11 | 46 | 0 | 0 | 0 | 9 | 5 | 0 | 0 | 0 | 11 | 10 | 0 | 0 | 0 | 0 | 0 | 0 | 0 | 0 | 0 | 0 | 0 | 0 | 0 | 0 | 0 | 0 | 0 | 0 | 0 | 0 |
| ASV532 | b9e2ecb08a5 | 92 | 0 | 0 | 0 | 0 | 33 | 0 | 0 | 14 | 0 | 0 | 0 | 0 | 0 | 0 | 25 | 0 | 0 | 0 | 0 | 0 | 0 | 0 | 0 | 0 | 0 | 0 | 0 | 0 | 0 | 0 | 0 | 0 | 0 | 20 | 0 |
| ASV533 | a1a058b4498 | 92 | 0 | 0 | 0 | 0 | 0 | 0 | 0 | 0 | 46 | 46 | 0 | 0 | 0 | 0 | 0 | 0 | 0 | 0 | 0 | 0 | 0 | 0 | 0 | 0 | 0 | 0 | 0 | 0 | 0 | 0 | 0 | 0 | 0 | 0 | 0 |
| ASV534 | 90e62fbd8d5 | 91 | 0 | 0 | 4 | 0 | 0 | 0 | 0 | 0 | 0 | 0 | 0 | 1 | 1 | 25 | 0 | 7 | 0 | 0 | 0 | 0 | 0 | 0 | 0 | 9 | 0 | 19 | 0 | 0 | 0 | 0 | 0 | 0 | 0 | 14 | 11 |
| ASV535 | 3b91ffa8b3a | 91 | 0 | 0 | 0 | 0 | 0 | 58 | 0 | 0 | 0 | 0 | 0 | 0 | 0 | 0 | 0 | 0 | 0 | 0 | 0 | 0 | 0 | 0 | 0 | 33 | 0 | 0 | 0 | 0 | 0 | 0 | 0 | 0 | 0 | 0 | 0 |
| ASV536 | d0d0f02008f | 90 | 9 | 7 | 39 | 17 | 6 | 5 | 0 | 0 | 0 | 0 | 0 | 0 | 0 | 0 | 0 | 0 | 0 | 0 | 0 | 0 | 0 | 0 | 0 | 0 | 5 | 0 | 0 | 2 | 0 | 0 | 0 | 0 | 0 | 0 | 0 |
| ASV537 | 05a88fdc0f8 | 90 | 0 | 0 | 0 | 0 | 0 | 0 | 0 | 0 | 27 | 35 | 0 | 0 | 0 | 0 | 0 | 0 | 0 | 0 | 0 | 0 | 0 | 0 | 0 | 0 | 28 | 0 | 0 | 0 | 0 | 0 | 0 | 0 | 0 | 0 | 0 |
| ASV538 | 1bd6beae6c8 | 89 | 0 | 0 | 0 | 0 | 0 | 0 | 0 | 0 | 0 | 0 | 0 | 0 | 0 | 0 | 0 | 0 | 12 | 0 | 0 | 0 | 0 | 0 | 0 | 0 | 0 | 0 | 0 | 0 | 0 | 0 | 52 | 17 | 8 | 0 | 0 |
| ASV539 | c0a2c1a45a6 | 89 | 0 | 0 | 0 | 0 | 0 | 0 | 0 | 0 | 0 | 0 | 0 | 0 | 0 | 0 | 46 | 0 | 0 | 0 | 0 | 0 | 0 | 0 | 0 | 0 | 0 | 0 | 0 | 0 | 4 | 11 | 0 | 0 | 0 | 0 | 28 |
| ASV540 | 4d6a47f6ee2 | 88 | 0 | 0 | 0 | 0 | 0 | 0 | 0 | 0 | 0 | 0 | 0 | 0 | 0 | 0 | 0 | 0 | 56 | 0 | 0 | 0 | 0 | 0 | 0 | 0 | 0 | 0 | 0 | 32 | 0 | 0 | 0 | 0 | 0 | 0 | 0 |
| ASV541 | 7f959926ff2a | 88 | 0 | 0 | 0 | 0 | 0 | 0 | 0 | 0 | 0 | 0 | 0 | 0 | 0 | 0 | 0 | 41 | 35 | 0 | 0 | 0 | 0 | 12 | 0 | 0 | 0 | 0 | 0 | 0 | 0 | 0 | 0 | 0 | 0 | 0 | 0 |
| ASV542 | 0197714ed0a | 87 | 0 | 0 | 38 | 0 | 35 | 0 | 0 | 0 | 0 | 0 | 0 | 0 | 0 | 0 | 0 | 0 | 0 | 0 | 0 | 0 | 0 | 0 | 0 | 0 | 0 | 0 | 0 | 0 | 0 | 0 | 14 | 0 | 0 | 0 | 0 |
| ASV543 | 52dbb7439ee | 87 | 0 | 0 | 0 | 0 | 87 | 0 | 0 | 0 | 0 | 0 | 0 | 0 | 0 | 0 | 0 | 0 | 0 | 0 | 0 | 0 | 0 | 0 | 0 | 0 | 0 | 0 | 0 | 0 | 0 | 0 | 0 | 0 | 0 | 0 | 0 |
| ASV544 | 2a08a47582e | 87 | 0 | 0 | 0 | 0 | 0 | 0 | 0 | 0 | 0 | 0 | 0 | 0 | 0 | 0 | 0 | 0 | 0 | 0 | 87 | 0 | 0 | 0 | 0 | 0 | 0 | 0 | 0 | 0 | 0 | 0 | 0 | 0 | 0 | 0 | 0 |
| ASV545 | fb5509311f7 | 86 | 0 | 0 | 0 | 0 | 38 | 0 | 0 | 0 | 0 | 0 | 0 | 0 | 0 | 0 | 48 | 0 | 0 | 0 | 0 | 0 | 0 | 0 | 0 | 0 | 0 | 0 | 0 | 0 | 0 | 0 | 0 | 0 | 0 | 0 | 0 |
| ASV546 | 66f775d06f3 | 86 | 0 | 0 | 0 | 0 | 0 | 0 | 0 | 0 | 0 | 0 | 0 | 0 | 0 | 0 | 0 | 0 | 0 | 0 | 0 | 0 | 0 | 0 | 0 | 0 | 0 | 0 | 0 | 0 | 18 | 21 | 0 | 0 | 0 | 22 | 25 |
| ASV547 | 47354d572da | 85 | 0 | 0 | 0 | 0 | 0 | 0 | 0 | 0 | 0 | 0 | 0 | 0 | 0 | 0 | 0 | 0 | 0 | 0 | 0 | 0 | 0 | 0 | 0 | 0 | 0 | 0 | 0 | 0 | 66 | 0 | 0 | 0 | 0 | 0 | 19 |
| ASV548 | 5ae95cdb613 | 85 | 0 | 0 | 0 | 0 | 32 | 0 | 0 | 0 | 0 | 0 | 0 | 0 | 0 | 0 | 46 | 0 | 0 | 0 | 0 | 0 | 0 | 0 | 0 | 0 | 0 | 0 | 0 | 0 | 0 | 7 | 0 | 0 | 0 | 0 | 0 |
| ASV549 | c557eb0f4fa | 85 | 0 | 0 | 0 | 0 | 0 | 0 | 0 | 0 | 0 | 0 | 0 | 0 | 0 | 0 | 0 | 20 | 9 | 0 | 0 | 0 | 0 | 0 | 0 | 0 | 0 | 9 | 8 | 15 | 0 | 0 | 10 | 7 | 7 | 0 | 0 |
| ASV550 | 1711ad8be59 | 84 | 0 | 0 | 0 | 0 | 0 | 55 | 29 | 0 | 0 | 0 | 0 | 0 | 0 | 0 | 0 | 0 | 0 | 0 | 0 | 0 | 0 | 0 | 0 | 0 | 0 | 0 | 0 | 0 | 0 | 0 | 0 | 0 | 0 | 0 | 0 |
| ASV551 | f0e88f855c1 | 84 | 0 | 0 | 29 | 0 | 12 | 13 | 0 | 0 | 0 | 0 | 0 | 0 | 0 | 13 | 3 | 0 | 0 | 0 | 0 | 0 | 14 | 0 | 0 | 0 | 0 | 0 | 0 | 0 | 0 | 0 | 0 | 0 | 0 | 0 | 0 |
| ASV552 | 4e64935fcd0 | 84 | 0 | 0 | 0 | 0 | 13 | 8 | 0 | 0 | 21 | 8 | 0 | 0 | 0 | 0 | 14 | 0 | 0 | 0 | 0 | 0 | 0 | 0 | 0 | 0 | 13 | 0 | 0 | 0 | 0 | 7 | 0 | 0 | 0 | 0 | 0 |
| ASV553 | 9c609ecc137 | 83 | 0 | 0 | 83 | 0 | 0 | 0 | 0 | 0 | 0 | 0 | 0 | 0 | 0 | 0 | 0 | 0 | 0 | 0 | 0 | 0 | 0 | 0 | 0 | 0 | 0 | 0 | 0 | 0 | 0 | 0 | 0 | 0 | 0 | 0 | 0 |
| ASV554 | ab5305c1e15 | 83 | 0 | 0 | 0 | 0 | 0 | 0 | 0 | 0 | 0 | 0 | 0 | 0 | 0 | 0 | 0 | 12 | 30 | 5 | 0 | 0 | 0 | 0 | 0 | 0 | 0 | 0 | 0 | 0 | 0 | 0 | 21 | 0 | 0 | 15 | 0 |
| ASV555 | c8e0c1a3077 | 83 | 0 | 0 | 0 | 0 | 0 | 0 | 0 | 0 | 0 | 0 | 0 | 0 | 0 | 0 | 0 | 0 | 0 | 0 | 0 | 0 | 0 | 0 | 0 | 0 | 0 | 0 | 0 | 0 | 0 | 0 | 56 | 27 | 0 | 0 | 0 |
| ASV556 | 2f49cfd4834 | 83 | 0 | 0 | 0 | 0 | 0 | 0 | 0 | 0 | 0 | 0 | 0 | 0 | 0 | 0 | 0 | 0 | 0 | 0 | 0 | 0 | 0 | 0 | 0 | 0 | 0 | 0 | 0 | 0 | 0 | 32 | 0 | 0 | 0 | 0 | 51 |
| ASV557 | dafcab2d646 | 82 | 0 | 0 | 0 | 0 | 0 | 0 | 0 | 0 | 0 | 0 | 0 | 0 | 0 | 0 | 28 | 0 | 0 | 0 | 0 | 0 | 0 | 0 | 0 | 0 | 16 | 0 | 0 | 0 | 13 | 25 | 0 | 0 | 0 | 0 | 0 |
| ASV558 | d06c0c3c746 | 82 | 0 | 0 | 0 | 0 | 0 | 0 | 0 | 0 | 12 | 8 | 0 | 0 | 0 | 0 | 32 | 0 | 0 | 0 | 0 | 0 | 0 | 0 | 0 | 0 | 4 | 0 | 0 | 0 | 11 | 15 | 0 | 0 | 0 | 0 | 0 |
| ASV559 | 7c911fee228 | 82 | 0 | 0 | 0 | 0 | 0 | 0 | 0 | 0 | 0 | 7 | 0 | 0 | 0 | 0 | 39 | 0 | 0 | 0 | 0 | 0 | 0 | 0 | 0 | 0 | 17 | 0 | 0 | 0 | 0 | 0 | 0 | 0 | 0 | 0 | 19 |
| ASV560 | f19c29319a7 | 80 | 0 | 0 | 0 | 0 | 0 | 0 | 0 | 0 | 0 | 0 | 0 | 0 | 0 | 0 | 0 | 0 | 0 | 0 | 0 | 0 | 0 | 0 | 0 | 0 | 35 | 0 | 0 | 0 | 0 | 22 | 0 | 0 | 0 | 0 | 23 |
| ASV561 | ac178abd52e | 80 | 0 | 0 | 33 | 0 | 0 | 18 | 0 | 0 | 0 | 0 | 0 | 0 | 0 | 0 | 23 | 0 | 0 | 0 | 0 | 0 | 0 | 6 | 0 | 0 | 0 | 0 | 0 | 0 | 0 | 0 | 0 | 0 | 0 | 0 | 0 |
| ASV562 | 549f2f4cecba | 80 | 0 | 0 | 0 | 0 | 0 | 0 | 0 | 0 | 0 | 24 | 0 | 0 | 0 | 0 | 0 | 0 | 0 | 0 | 0 | 0 | 0 | 0 | 0 | 0 | 0 | 0 | 0 | 0 | 0 | 10 | 0 | 0 | 0 | 0 | 46 |
| ASV563 | 8ee5980ab9d | 80 | 0 | 0 | 0 | 9 | 25 | 4 | 0 | 0 | 6 | 0 | 0 | 0 | 0 | 0 | 27 | 0 | 0 | 0 | 0 | 0 | 0 | 0 | 0 | 0 | 0 | 0 | 0 | 0 | 0 | 0 | 0 | 0 | 0 | 0 | 9 |
| ASV564 | 183c25e0ca9 | 79 | 0 | 7 | 0 | 10 | 29 | 0 | 0 | 0 | 0 | 0 | 0 | 0 | 0 | 0 | 11 | 0 | 0 | 0 | 0 | 0 | 0 | 0 | 0 | 0 | 0 | 0 | 0 | 0 | 0 | 0 | 0 | 0 | 0 | 0 | 22 |
| ASV565 | 51e8dc03f87 | 79 | 21 | 0 | 34 | 0 | 17 | 0 | 0 | 0 | 0 | 0 | 0 | 0 | 0 | 0 | 0 | 0 | 0 | 0 | 0 | 0 | 0 | 0 | 0 | 0 | 0 | 0 | 0 | 0 | 0 | 0 | 0 | 0 | 0 | 7 | 0 |
| ASV566 | a291edca74e | 79 | 0 | 0 | 0 | 0 | 0 | 0 | 0 | 0 | 0 | 0 | 0 | 0 | 0 | 0 | 0 | 0 | 0 | 0 | 0 | 0 | 0 | 0 | 0 | 0 | 0 | 0 | 0 | 79 | 0 | 0 | 0 | 0 | 0 | 0 | 0 |
| ASV567 | 2ed972fdbc5 | 79 | 0 | 0 | 3 | 0 | 0 | 0 | 0 | 0 | 0 | 0 | 0 | 4 | 0 | 0 | 0 | 12 | 18 | 4 | 0 | 0 | 7 | 0 | 0 | 0 | 0 | 9 | 4 | 6 | 0 | 0 | 6 | 0 | 0 | 6 | 0 |
| ASV568 | 37bc07d963b | 79 | 0 | 0 | 0 | 0 | 0 | 0 | 0 | 0 | 79 | 0 | 0 | 0 | 0 | 0 | 0 | 0 | 0 | 0 | 0 | 0 | 0 | 0 | 0 | 0 | 0 | 0 | 0 | 0 | 0 | 0 | 0 | 0 | 0 | 0 | 0 |
| ASV569 | d4a3dd69508 | 78 | 9 | 0 | 0 | 0 | 0 | 0 | 0 | 26 | 0 | 0 | 0 | 0 | 0 | 0 | 0 | 0 | 0 | 0 | 0 | 0 | 0 | 0 | 0 | 16 | 27 | 0 | 0 | 0 | 0 | 0 | 0 | 0 | 0 | 0 | 0 |
| ASV570 | 20a7d89e310 | 77 | 0 | 0 | 0 | 0 | 11 | 0 | 0 | 3 | 0 | 5 | 0 | 0 | 0 | 0 | 4 | 0 | 0 | 0 | 0 | 5 | 0 | 0 | 0 | 0 | 5 | 0 | 0 | 0 | 19 | 25 | 0 | 0 | 0 | 0 | 0 |
| ASV571 | 545a8c7991b | 77 | 0 | 0 | 0 | 0 | 0 | 0 | 0 | 0 | 0 | 0 | 0 | 12 | 0 | 0 | 0 | 14 | 23 | 12 | 0 | 0 | 0 | 0 | 0 | 0 | 0 | 0 | 0 | 16 | 0 | 0 | 0 | 0 | 0 | 0 | 0 |
| ASV572 | 1c3a2a0910f | 77 | 0 | 0 | 23 | 0 | 25 | 0 | 0 | 0 | 0 | 0 | 0 | 0 | 0 | 0 | 19 | 0 | 0 | 0 | 0 | 0 | 0 | 0 | 0 | 0 | 0 | 0 | 0 | 0 | 0 | 10 | 0 | 0 | 0 | 0 | 0 |
| ASV573 | fb122356537 | 77 | 0 | 0 | 0 | 0 | 0 | 0 | 0 | 14 | 0 | 0 | 0 | 0 | 0 | 0 | 20 | 0 | 0 | 0 | 0 | 0 | 0 | 0 | 0 | 0 | 0 | 0 | 0 | 0 | 8 | 12 | 0 | 0 | 0 | 0 | 23 |
| ASV574 | a68a1170803 | 76 | 0 | 0 | 0 | 0 | 0 | 0 | 0 | 0 | 0 | 0 | 0 | 0 | 0 | 0 | 0 | 44 | 0 | 0 | 0 | 0 | 0 | 0 | 0 | 0 | 0 | 0 | 0 | 32 | 0 | 0 | 0 | 0 | 0 | 0 | 0 |
| ASV575 | 052ee499e58 | 76 | 0 | 0 | 0 | 0 | 0 | 0 | 0 | 0 | 0 | 0 | 0 | 0 | 0 | 9 | 67 | 0 | 0 | 0 | 0 | 0 | 0 | 0 | 0 | 0 | 0 | 0 | 0 | 0 | 0 | 0 | 0 | 0 | 0 | 0 | 0 |
| ASV576 | 8bc0014fd98 | 76 | 0 | 7 | 12 | 3 | 13 | 9 | 4 | 7 | 0 | 0 | 0 | 0 | 0 | 0 | 0 | 0 | 0 | 0 | 6 | 0 | 0 | 0 | 0 | 3 | 3 | 0 | 0 | 0 | 0 | 3 | 0 | 0 | 0 | 6 | 0 |
| ASV577 | 017b83e3cb9 | 76 | 0 | 0 | 0 | 0 | 0 | 0 | 0 | 0 | 0 | 0 | 0 | 0 | 0 | 0 | 0 | 0 | 58 | 0 | 0 | 0 | 0 | 0 | 0 | 0 | 0 | 7 | 0 | 11 | 0 | 0 | 0 | 0 | 0 | 0 | 0 |
| ASV578 | ba1f0129b1a | 76 | 0 | 0 | 0 | 28 | 0 | 0 | 0 | 0 | 0 | 0 | 0 | 0 | 0 | 0 | 0 | 12 | 0 | 22 | 0 | 0 | 0 | 0 | 0 | 0 | 0 | 0 | 0 | 0 | 14 | 0 | 0 | 0 | 0 | 0 | 0 |
| ASV579 | 88acfab92ed | 76 | 0 | 0 | 0 | 0 | 0 | 0 | 0 | 0 | 0 | 0 | 0 | 0 | 0 | 0 | 0 | 23 | 29 | 0 | 0 | 0 | 0 | 0 | 0 | 0 | 0 | 14 | 0 | 10 | 0 | 0 | 0 | 0 | 0 | 0 | 0 |
| ASV580 | 84dbc43ff05f | 76 | 0 | 0 | 4 | 0 | 0 | 0 | 0 | 9 | 0 | 3 | 0 | 0 | 0 | 0 | 0 | 0 | 0 | 0 | 2 | 0 | 0 | 0 | 0 | 20 | 12 | 0 | 0 | 0 | 13 | 13 | 0 | 0 | 0 | 0 | 0 |
| ASV581 | fd74bfe3953 | 75 | 0 | 0 | 9 | 0 | 0 | 0 | 0 | 0 | 0 | 0 | 0 | 0 | 0 | 0 | 0 | 0 | 7 | 0 | 0 | 0 | 0 | 0 | 0 | 0 | 0 | 0 | 0 | 0 | 0 | 0 | 17 | 9 | 5 | 19 | 9 |
| ASV582 | c6738714274 | 75 | 0 | 0 | 35 | 0 | 40 | 0 | 0 | 0 | 0 | 0 | 0 | 0 | 0 | 0 | 0 | 0 | 0 | 0 | 0 | 0 | 0 | 0 | 0 | 0 | 0 | 0 | 0 | 0 | 0 | 0 | 0 | 0 | 0 | 0 | 0 |
| ASV583 | ea8b0afe862 | 75 | 0 | 0 | 0 | 0 | 0 | 0 | 0 | 0 | 0 | 23 | 0 | 0 | 0 | 0 | 0 | 0 | 0 | 0 | 0 | 0 | 0 | 0 | 0 | 25 | 27 | 0 | 0 | 0 | 0 | 0 | 0 | 0 | 0 | 0 | 0 |
| ASV584 | 409bb8c4521 | 75 | 0 | 0 | 0 | 0 | 0 | 0 | 0 | 0 | 0 | 0 | 0 | 0 | 0 | 0 | 75 | 0 | 0 | 0 | 0 | 0 | 0 | 0 | 0 | 0 | 0 | 0 | 0 | 0 | 0 | 0 | 0 | 0 | 0 | 0 | 0 |
| ASV585 | c8cf89d98fc5 | 74 | 0 | 0 | 0 | 0 | 0 | 0 | 0 | 0 | 0 | 27 | 0 | 0 | 0 | 0 | 0 | 0 | 0 | 0 | 0 | 0 | 0 | 0 | 0 | 0 | 5 | 0 | 0 | 0 | 22 | 0 | 0 | 0 | 0 | 0 | 20 |
| ASV586 | 4c2378bd8d7 | 74 | 0 | 0 | 0 | 0 | 0 | 0 | 0 | 0 | 0 | 0 | 0 | 0 | 0 | 0 | 0 | 0 | 0 | 0 | 0 | 0 | 0 | 0 | 0 | 0 | 74 | 0 | 0 | 0 | 0 | 0 | 0 | 0 | 0 | 0 | 0 |
| ASV587 | e2c9ba2b677 | 73 | 0 | 0 | 0 | 0 | 12 | 0 | 0 | 0 | 22 | 10 | 0 | 0 | 0 | 0 | 18 | 0 | 0 | 0 | 3 | 0 | 0 | 0 | 0 | 0 | 4 | 0 | 0 | 0 | 0 | 4 | 0 | 0 | 0 | 0 | 0 |
| ASV588 | 742483a385e | 73 | 0 | 0 | 0 | 0 | 0 | 0 | 0 | 0 | 0 | 0 | 0 | 20 | 0 | 0 | 0 | 0 | 29 | 0 | 0 | 0 | 0 | 0 | 7 | 0 | 0 | 0 | 0 | 6 | 0 | 0 | 11 | 0 | 0 | 0 | 0 |
| ASV589 | 7b1d62f61e7 | 73 | 0 | 0 | 0 | 0 | 0 | 0 | 0 | 0 | 0 | 0 | 0 | 0 | 0 | 0 | 0 | 0 | 0 | 0 | 0 | 0 | 12 | 0 | 0 | 0 | 9 | 0 | 0 | 0 | 0 | 0 | 37 | 0 | 15 | 0 | 0 |
| ASV590 | 0d8977fdc2a | 72 | 0 | 0 | 0 | 0 | 0 | 0 | 0 | 0 | 19 | 13 | 0 | 0 | 0 | 0 | 23 | 0 | 0 | 0 | 0 | 0 | 0 | 0 | 0 | 0 | 17 | 0 | 0 | 0 | 0 | 0 | 0 | 0 | 0 | 0 | 0 |
| ASV591 | 9ac20859a7c | 72 | 0 | 0 | 0 | 0 | 7 | 0 | 0 | 0 | 13 | 0 | 0 | 0 | 0 | 0 | 46 | 0 | 0 | 0 | 0 | 0 | 0 | 0 | 0 | 0 | 0 | 0 | 0 | 0 | 0 | 6 | 0 | 0 | 0 | 0 | 0 |
| ASV592 | 874d9e136f5 | 72 | 0 | 0 | 0 | 0 | 0 | 0 | 0 | 0 | 0 | 0 | 0 | 0 | 0 | 0 | 0 | 0 | 0 | 0 | 8 | 49 | 0 | 0 | 0 | 0 | 0 | 0 | 0 | 0 | 0 | 15 | 0 | 0 | 0 | 0 | 0 |
| ASV593 | 9c44eec10b6 | 72 | 0 | 0 | 0 | 36 | 0 | 0 | 36 | 0 | 0 | 0 | 0 | 0 | 0 | 0 | 0 | 0 | 0 | 0 | 0 | 0 | 0 | 0 | 0 | 0 | 0 | 0 | 0 | 0 | 0 | 0 | 0 | 0 | 0 | 0 | 0 |
| ASV594 | c87c523c6fef | 72 | 0 | 0 | 0 | 0 | 0 | 0 | 0 | 0 | 0 | 0 | 0 | 0 | 0 | 0 | 0 | 0 | 0 | 0 | 0 | 0 | 0 | 0 | 0 | 0 | 0 | 0 | 0 | 0 | 2 | 39 | 0 | 0 | 0 | 10 | 21 |
| ASV595 | 245260e4d0f | 72 | 0 | 0 | 0 | 0 | 0 | 0 | 0 | 0 | 0 | 0 | 0 | 0 | 0 | 0 | 0 | 18 | 54 | 0 | 0 | 0 | 0 | 0 | 0 | 0 | 0 | 0 | 0 | 0 | 0 | 0 | 0 | 0 | 0 | 0 | 0 |
| ASV596 | 919cb4708a7 | 71 | 0 | 0 | 0 | 0 | 12 | 0 | 0 | 5 | 6 | 8 | 0 | 0 | 4 | 0 | 14 | 0 | 0 | 0 | 0 | 0 | 0 | 0 | 0 | 0 | 0 | 0 | 0 | 4 | 6 | 6 | 0 | 0 | 0 | 6 | 0 |
| ASV597 | fdf2a5cac1fb | 71 | 0 | 6 | 0 | 0 | 0 | 0 | 15 | 0 | 0 | 7 | 4 | 0 | 10 | 12 | 17 | 0 | 0 | 0 | 0 | 0 | 0 | 0 | 0 | 0 | 0 | 0 | 0 | 0 | 0 | 0 | 0 | 0 | 0 | 0 | 0 |
| ASV598 | 1bb16fe2b96 | 71 | 0 | 0 | 0 | 0 | 0 | 9 | 0 | 0 | 0 | 0 | 0 | 0 | 6 | 0 | 0 | 0 | 0 | 0 | 0 | 0 | 0 | 0 | 0 | 0 | 0 | 0 | 0 | 0 | 17 | 0 | 0 | 0 | 0 | 39 | 0 |
| ASV599 | 761a173d79e | 71 | 0 | 0 | 0 | 0 | 12 | 0 | 0 | 3 | 13 | 0 | 0 | 0 | 0 | 0 | 22 | 0 | 0 | 0 | 0 | 0 | 0 | 0 | 0 | 0 | 12 | 0 | 0 | 0 | 0 | 0 | 0 | 0 | 0 | 0 | 9 |
| ASV600 | cd306f4f534 | 70 | 0 | 0 | 0 | 0 | 0 | 0 | 0 | 0 | 0 | 4 | 0 | 0 | 0 | 0 | 0 | 0 | 0 | 0 | 0 | 0 | 0 | 0 | 0 | 0 | 9 | 0 | 0 | 0 | 4 | 9 | 0 | 0 | 0 | 4 | 40 |
| ASV601 | 0c65eea8e90 | 70 | 0 | 0 | 0 | 0 | 9 | 9 | 0 | 0 | 10 | 18 | 0 | 0 | 0 | 0 | 0 | 0 | 0 | 0 | 5 | 0 | 0 | 0 | 0 | 7 | 0 | 0 | 0 | 0 | 6 | 0 | 0 | 0 | 0 | 6 | 0 |
| ASV602 | ff59c48ddbe1 | 70 | 0 | 0 | 0 | 0 | 0 | 0 | 0 | 0 | 7 | 3 | 0 | 6 | 0 | 0 | 20 | 0 | 0 | 0 | 0 | 0 | 0 | 0 | 0 | 0 | 7 | 0 | 0 | 0 | 0 | 0 | 0 | 5 | 0 | 0 | 22 |
| ASV603 | 0d5094dfea1 | 70 | 0 | 0 | 9 | 8 | 4 | 23 | 0 | 0 | 0 | 0 | 0 | 16 | 10 | 0 | 0 | 0 | 0 | 0 | 0 | 0 | 0 | 0 | 0 | 0 | 0 | 0 | 0 | 0 | 0 | 0 | 0 | 0 | 0 | 0 | 0 |
| ASV604 | 6eda1a6288c | 69 | 0 | 0 | 35 | 0 | 34 | 0 | 0 | 0 | 0 | 0 | 0 | 0 | 0 | 0 | 0 | 0 | 0 | 0 | 0 | 0 | 0 | 0 | 0 | 0 | 0 | 0 | 0 | 0 | 0 | 0 | 0 | 0 | 0 | 0 | 0 |
| ASV605 | c5a0edf440b | 69 | 0 | 0 | 69 | 0 | 0 | 0 | 0 | 0 | 0 | 0 | 0 | 0 | 0 | 0 | 0 | 0 | 0 | 0 | 0 | 0 | 0 | 0 | 0 | 0 | 0 | 0 | 0 | 0 | 0 | 0 | 0 | 0 | 0 | 0 | 0 |
| ASV606 | 7ac5166f938 | 69 | 33 | 0 | 30 | 6 | 0 | 0 | 0 | 0 | 0 | 0 | 0 | 0 | 0 | 0 | 0 | 0 | 0 | 0 | 0 | 0 | 0 | 0 | 0 | 0 | 0 | 0 | 0 | 0 | 0 | 0 | 0 | 0 | 0 | 0 | 0 |
| ASV607 | 155c12b0a3a | 69 | 0 | 0 | 0 | 0 | 0 | 0 | 0 | 0 | 0 | 0 | 0 | 0 | 0 | 0 | 0 | 0 | 0 | 0 | 0 | 0 | 0 | 0 | 0 | 0 | 35 | 0 | 0 | 0 | 0 | 34 | 0 | 0 | 0 | 0 | 0 |
| ASV608 | adcb0e67dfd | 69 | 0 | 0 | 15 | 0 | 0 | 0 | 0 | 14 | 0 | 0 | 0 | 0 | 0 | 14 | 0 | 0 | 0 | 0 | 0 | 0 | 0 | 0 | 0 | 0 | 0 | 0 | 0 | 0 | 12 | 14 | 0 | 0 | 0 | 0 | 0 |
| ASV609 | 32bc678776f | 67 | 0 | 0 | 8 | 0 | 0 | 0 | 0 | 0 | 0 | 0 | 0 | 0 | 0 | 0 | 0 | 0 | 0 | 0 | 0 | 0 | 0 | 0 | 0 | 0 | 0 | 0 | 0 | 0 | 0 | 0 | 59 | 0 | 0 | 0 | 0 |
| ASV610 | 859ab5a6e6e | 67 | 2 | 0 | 5 | 8 | 4 | 0 | 2 | 0 | 0 | 0 | 0 | 0 | 0 | 0 | 0 | 0 | 0 | 0 | 0 | 0 | 0 | 0 | 0 | 2 | 0 | 0 | 0 | 0 | 0 | 0 | 8 | 0 | 0 | 28 | 8 |
| ASV611 | f7b2a213bf9a | 67 | 0 | 0 | 0 | 19 | 0 | 0 | 0 | 0 | 22 | 0 | 0 | 0 | 0 | 0 | 26 | 0 | 0 | 0 | 0 | 0 | 0 | 0 | 0 | 0 | 0 | 0 | 0 | 0 | 0 | 0 | 0 | 0 | 0 | 0 | 0 |
| ASV612 | 252e439ccbd | 67 | 0 | 0 | 0 | 0 | 0 | 0 | 0 | 0 | 0 | 0 | 0 | 0 | 0 | 0 | 0 | 20 | 0 | 32 | 0 | 0 | 0 | 0 | 0 | 0 | 0 | 0 | 15 | 0 | 0 | 0 | 0 | 0 | 0 | 0 | 0 |
| ASV613 | d61cf02c7e2 | 67 | 0 | 0 | 0 | 0 | 0 | 0 | 0 | 0 | 0 | 8 | 0 | 0 | 0 | 0 | 11 | 0 | 0 | 0 | 9 | 9 | 0 | 0 | 0 | 8 | 9 | 0 | 0 | 0 | 6 | 7 | 0 | 0 | 0 | 0 | 0 |
| ASV614 | 6b1e35bd308 | 67 | 0 | 0 | 34 | 0 | 0 | 0 | 0 | 0 | 0 | 0 | 0 | 0 | 0 | 0 | 0 | 0 | 33 | 0 | 0 | 0 | 0 | 0 | 0 | 0 | 0 | 0 | 0 | 0 | 0 | 0 | 0 | 0 | 0 | 0 | 0 |
| ASV615 | 692890348a0 | 66 | 0 | 0 | 0 | 0 | 0 | 0 | 0 | 0 | 0 | 0 | 0 | 0 | 0 | 0 | 0 | 27 | 39 | 0 | 0 | 0 | 0 | 0 | 0 | 0 | 0 | 0 | 0 | 0 | 0 | 0 | 0 | 0 | 0 | 0 | 0 |
| ASV616 | 6452c5dbadd | 66 | 0 | 0 | 0 | 0 | 0 | 0 | 0 | 0 | 0 | 0 | 0 | 0 | 0 | 0 | 0 | 0 | 0 | 0 | 0 | 0 | 0 | 0 | 0 | 0 | 0 | 0 | 0 | 0 | 0 | 0 | 50 | 16 | 0 | 0 | 0 |
| ASV617 | 8fbe91a176e | 66 | 0 | 0 | 24 | 4 | 7 | 4 | 0 | 0 | 0 | 0 | 0 | 0 | 0 | 9 | 0 | 5 | 7 | 0 | 0 | 0 | 0 | 0 | 0 | 0 | 0 | 0 | 0 | 0 | 0 | 6 | 0 | 0 | 0 | 0 | 0 |
| ASV618 | da9d4a63e9b | 66 | 0 | 0 | 11 | 0 | 0 | 8 | 0 | 0 | 0 | 0 | 0 | 0 | 0 | 4 | 0 | 0 | 0 | 0 | 0 | 0 | 13 | 0 | 11 | 0 | 0 | 0 | 0 | 0 | 0 | 0 | 0 | 0 | 0 | 19 | 0 |
| ASV619 | a774191f4bd | 66 | 0 | 0 | 0 | 0 | 10 | 0 | 0 | 0 | 0 | 0 | 0 | 0 | 0 | 0 | 0 | 0 | 0 | 0 | 0 | 0 | 0 | 0 | 0 | 0 | 0 | 0 | 0 | 0 | 43 | 13 | 0 | 0 | 0 | 0 | 0 |
| ASV620 | 32adea307c0 | 65 | 0 | 0 | 0 | 0 | 0 | 0 | 0 | 0 | 11 | 14 | 0 | 0 | 0 | 0 | 40 | 0 | 0 | 0 | 0 | 0 | 0 | 0 | 0 | 0 | 0 | 0 | 0 | 0 | 0 | 0 | 0 | 0 | 0 | 0 | 0 |
| ASV621 | b761abc16f9 | 64 | 0 | 0 | 0 | 0 | 0 | 0 | 0 | 0 | 0 | 0 | 0 | 0 | 0 | 0 | 0 | 8 | 7 | 8 | 0 | 0 | 11 | 13 | 0 | 6 | 0 | 0 | 0 | 0 | 0 | 0 | 0 | 0 | 11 | 0 | 0 |
| ASV622 | 3c9e6449d55 | 64 | 0 | 0 | 0 | 0 | 0 | 0 | 0 | 0 | 0 | 5 | 0 | 0 | 0 | 0 | 0 | 0 | 0 | 0 | 0 | 0 | 0 | 0 | 0 | 0 | 7 | 0 | 0 | 0 | 0 | 0 | 31 | 11 | 7 | 0 | 3 |
| ASV623 | d1baec9ea84 | 64 | 0 | 0 | 8 | 0 | 0 | 0 | 0 | 11 | 0 | 0 | 0 | 0 | 0 | 0 | 0 | 10 | 0 | 0 | 0 | 0 | 0 | 0 | 6 | 0 | 0 | 0 | 0 | 0 | 4 | 0 | 0 | 0 | 0 | 25 | 0 |
| ASV624 | 54f5e3b3abe | 64 | 0 | 0 | 0 | 0 | 10 | 0 | 0 | 0 | 21 | 0 | 0 | 0 | 0 | 0 | 21 | 0 | 0 | 0 | 0 | 0 | 0 | 0 | 0 | 0 | 5 | 0 | 0 | 0 | 0 | 7 | 0 | 0 | 0 | 0 | 0 |
| ASV625 | 1725f2fa7f32 | 64 | 0 | 0 | 0 | 0 | 0 | 0 | 0 | 0 | 0 | 0 | 0 | 0 | 0 | 0 | 0 | 0 | 0 | 0 | 0 | 0 | 0 | 0 | 0 | 0 | 0 | 0 | 0 | 0 | 0 | 0 | 36 | 28 | 0 | 0 | 0 |
| ASV626 | 4122b4f14ca | 62 | 0 | 0 | 0 | 0 | 0 | 0 | 0 | 0 | 0 | 0 | 0 | 0 | 0 | 0 | 0 | 0 | 0 | 0 | 0 | 0 | 0 | 0 | 0 | 0 | 20 | 0 | 0 | 0 | 0 | 19 | 0 | 0 | 0 | 0 | 23 |
| ASV627 | 500ab9e5666 | 62 | 0 | 0 | 0 | 0 | 0 | 0 | 0 | 0 | 0 | 0 | 0 | 0 | 0 | 0 | 0 | 0 | 0 | 0 | 0 | 0 | 9 | 7 | 0 | 17 | 0 | 0 | 0 | 15 | 14 | 0 | 0 | 0 | 0 | 0 | 0 |
| ASV628 | 36c205347e8 | 62 | 2 | 3 | 0 | 7 | 43 | 0 | 0 | 0 | 0 | 2 | 0 | 0 | 0 | 0 | 5 | 0 | 0 | 0 | 0 | 0 | 0 | 0 | 0 | 0 | 0 | 0 | 0 | 0 | 0 | 0 | 0 | 0 | 0 | 0 | 0 |
| ASV629 | 6d570e6a2bd | 62 | 0 | 0 | 0 | 0 | 0 | 0 | 0 | 0 | 0 | 0 | 0 | 0 | 0 | 0 | 0 | 14 | 7 | 0 | 0 | 0 | 0 | 0 | 0 | 0 | 0 | 14 | 11 | 16 | 0 | 0 | 0 | 0 | 0 | 0 | 0 |
| ASV630 | c66920b0f26 | 62 | 0 | 4 | 8 | 8 | 0 | 6 | 1 | 0 | 0 | 0 | 0 | 0 | 16 | 1 | 0 | 0 | 10 | 0 | 0 | 0 | 0 | 0 | 0 | 0 | 0 | 0 | 0 | 8 | 0 | 0 | 0 | 0 | 0 | 0 | 0 |
| ASV631 | 098cca00050 | 62 | 0 | 0 | 33 | 0 | 0 | 19 | 0 | 0 | 0 | 0 | 0 | 0 | 0 | 10 | 0 | 0 | 0 | 0 | 0 | 0 | 0 | 0 | 0 | 0 | 0 | 0 | 0 | 0 | 0 | 0 | 0 | 0 | 0 | 0 | 0 |
| ASV632 | f6502f81e1a | 62 | 0 | 0 | 0 | 0 | 0 | 0 | 0 | 0 | 0 | 0 | 0 | 0 | 0 | 0 | 0 | 3 | 0 | 5 | 0 | 0 | 19 | 29 | 0 | 0 | 0 | 0 | 0 | 0 | 6 | 0 | 0 | 0 | 0 | 0 | 0 |
| ASV633 | fd84c514cc4 | 61 | 0 | 0 | 61 | 0 | 0 | 0 | 0 | 0 | 0 | 0 | 0 | 0 | 0 | 0 | 0 | 0 | 0 | 0 | 0 | 0 | 0 | 0 | 0 | 0 | 0 | 0 | 0 | 0 | 0 | 0 | 0 | 0 | 0 | 0 | 0 |
| ASV634 | c3f089bed97 | 61 | 0 | 0 | 61 | 0 | 0 | 0 | 0 | 0 | 0 | 0 | 0 | 0 | 0 | 0 | 0 | 0 | 0 | 0 | 0 | 0 | 0 | 0 | 0 | 0 | 0 | 0 | 0 | 0 | 0 | 0 | 0 | 0 | 0 | 0 | 0 |
| ASV635 | 4cb87ce1035 | 61 | 0 | 0 | 0 | 0 | 7 | 0 | 0 | 0 | 9 | 10 | 0 | 0 | 0 | 0 | 35 | 0 | 0 | 0 | 0 | 0 | 0 | 0 | 0 | 0 | 0 | 0 | 0 | 0 | 0 | 0 | 0 | 0 | 0 | 0 | 0 |
| ASV636 | 5a83fb0df7c | 61 | 0 | 0 | 0 | 3 | 37 | 0 | 0 | 0 | 0 | 0 | 0 | 0 | 0 | 0 | 21 | 0 | 0 | 0 | 0 | 0 | 0 | 0 | 0 | 0 | 0 | 0 | 0 | 0 | 0 | 0 | 0 | 0 | 0 | 0 | 0 |
| ASV637 | 2e23cbb77c3 | 61 | 0 | 0 | 0 | 0 | 0 | 0 | 0 | 0 | 0 | 0 | 0 | 0 | 0 | 0 | 0 | 26 | 10 | 0 | 0 | 0 | 0 | 0 | 0 | 0 | 0 | 0 | 0 | 0 | 0 | 0 | 0 | 25 | 0 | 0 | 0 |
| ASV638 | 36aa2d54e71 | 61 | 0 | 0 | 0 | 0 | 0 | 0 | 0 | 0 | 0 | 20 | 0 | 0 | 0 | 0 | 26 | 0 | 0 | 0 | 0 | 0 | 0 | 0 | 0 | 0 | 0 | 0 | 0 | 0 | 0 | 0 | 0 | 0 | 0 | 0 | 15 |
| ASV639 | 96cb90db2b8 | 60 | 8 | 0 | 9 | 0 | 7 | 0 | 0 | 0 | 0 | 0 | 0 | 0 | 0 | 0 | 0 | 0 | 0 | 0 | 0 | 0 | 0 | 0 | 0 | 24 | 9 | 0 | 0 | 0 | 0 | 3 | 0 | 0 | 0 | 0 | 0 |
| ASV640 | e6f6f772608 | 60 | 0 | 0 | 0 | 0 | 0 | 40 | 0 | 0 | 0 | 0 | 0 | 0 | 0 | 0 | 0 | 0 | 0 | 0 | 0 | 0 | 0 | 0 | 0 | 0 | 0 | 0 | 0 | 0 | 20 | 0 | 0 | 0 | 0 | 0 | 0 |
| ASV641 | c418a7de7b1 | 60 | 0 | 0 | 0 | 0 | 0 | 0 | 0 | 0 | 0 | 9 | 0 | 0 | 0 | 0 | 0 | 0 | 0 | 0 | 0 | 0 | 0 | 0 | 0 | 29 | 0 | 0 | 0 | 0 | 22 | 0 | 0 | 0 | 0 | 0 | 0 |
| ASV642 | f6d026b7600 | 60 | 0 | 0 | 0 | 0 | 8 | 0 | 0 | 0 | 0 | 13 | 0 | 0 | 0 | 0 | 23 | 0 | 0 | 0 | 0 | 0 | 0 | 0 | 0 | 0 | 0 | 0 | 0 | 0 | 0 | 0 | 0 | 0 | 0 | 0 | 16 |
| ASV643 | c7524bb695b | 60 | 0 | 0 | 0 | 0 | 0 | 0 | 0 | 0 | 0 | 0 | 0 | 0 | 0 | 0 | 0 | 11 | 0 | 0 | 0 | 0 | 23 | 0 | 0 | 0 | 0 | 0 | 0 | 7 | 0 | 0 | 19 | 0 | 0 | 0 | 0 |
| ASV644 | a2d386eb6e2 | 59 | 0 | 0 | 0 | 0 | 0 | 0 | 0 | 0 | 0 | 0 | 0 | 0 | 0 | 0 | 0 | 0 | 0 | 0 | 0 | 0 | 0 | 0 | 0 | 0 | 0 | 37 | 22 | 0 | 0 | 0 | 0 | 0 | 0 | 0 | 0 |
| ASV645 | a19d7650f65 | 59 | 0 | 0 | 0 | 0 | 0 | 0 | 0 | 0 | 0 | 0 | 0 | 0 | 0 | 0 | 0 | 0 | 0 | 0 | 0 | 0 | 0 | 0 | 0 | 0 | 0 | 0 | 0 | 0 | 0 | 0 | 0 | 0 | 6 | 13 | 40 |
| ASV646 | 6d819599793 | 59 | 0 | 0 | 59 | 0 | 0 | 0 | 0 | 0 | 0 | 0 | 0 | 0 | 0 | 0 | 0 | 0 | 0 | 0 | 0 | 0 | 0 | 0 | 0 | 0 | 0 | 0 | 0 | 0 | 0 | 0 | 0 | 0 | 0 | 0 | 0 |
| ASV647 | c4380c96be5 | 59 | 0 | 0 | 9 | 21 | 17 | 0 | 0 | 6 | 0 | 0 | 0 | 0 | 0 | 6 | 0 | 0 | 0 | 0 | 0 | 0 | 0 | 0 | 0 | 0 | 0 | 0 | 0 | 0 | 0 | 0 | 0 | 0 | 0 | 0 | 0 |
| ASV648 | 938f5c6eda2 | 59 | 0 | 0 | 0 | 0 | 0 | 0 | 0 | 0 | 25 | 15 | 0 | 0 | 0 | 0 | 19 | 0 | 0 | 0 | 0 | 0 | 0 | 0 | 0 | 0 | 0 | 0 | 0 | 0 | 0 | 0 | 0 | 0 | 0 | 0 | 0 |
| ASV649 | 36a62faf56e | 59 | 0 | 2 | 8 | 3 | 9 | 0 | 0 | 0 | 0 | 0 | 0 | 0 | 0 | 0 | 7 | 0 | 0 | 0 | 0 | 0 | 0 | 0 | 0 | 11 | 3 | 0 | 0 | 0 | 5 | 8 | 0 | 0 | 0 | 3 | 0 |
| ASV650 | 1ad1f2a089b | 59 | 0 | 0 | 0 | 0 | 18 | 0 | 0 | 0 | 0 | 0 | 0 | 0 | 0 | 0 | 19 | 0 | 0 | 0 | 0 | 0 | 0 | 0 | 0 | 0 | 10 | 0 | 0 | 0 | 0 | 0 | 0 | 0 | 0 | 0 | 12 |
| ASV651 | aa24c87bfa5 | 59 | 0 | 0 | 7 | 0 | 0 | 0 | 0 | 0 | 5 | 6 | 0 | 0 | 0 | 0 | 11 | 0 | 0 | 0 | 3 | 5 | 0 | 0 | 0 | 0 | 0 | 0 | 0 | 0 | 5 | 5 | 0 | 0 | 0 | 0 | 12 |
| ASV652 | 4e7fb19f71d | 59 | 0 | 0 | 0 | 0 | 20 | 0 | 0 | 0 | 0 | 0 | 0 | 0 | 0 | 0 | 10 | 0 | 0 | 0 | 12 | 0 | 0 | 0 | 0 | 7 | 0 | 0 | 0 | 0 | 0 | 0 | 0 | 0 | 0 | 0 | 10 |
| ASV653 | 05762365530 | 58 | 0 | 0 | 18 | 0 | 0 | 0 | 0 | 0 | 0 | 0 | 0 | 0 | 0 | 0 | 0 | 0 | 0 | 0 | 0 | 0 | 0 | 0 | 0 | 0 | 0 | 0 | 0 | 11 | 29 | 0 | 0 | 0 | 0 | 0 | 0 |
| ASV654 | 7956ff617d7 | 57 | 2 | 0 | 0 | 0 | 0 | 0 | 0 | 0 | 0 | 0 | 0 | 0 | 0 | 0 | 0 | 0 | 0 | 0 | 0 | 0 | 0 | 0 | 0 | 0 | 38 | 0 | 0 | 0 | 0 | 0 | 0 | 0 | 0 | 0 | 17 |
| ASV655 | 9970e1e5f6c | 57 | 0 | 0 | 0 | 0 | 6 | 0 | 0 | 7 | 6 | 3 | 0 | 0 | 0 | 0 | 32 | 0 | 0 | 0 | 0 | 3 | 0 | 0 | 0 | 0 | 0 | 0 | 0 | 0 | 0 | 0 | 0 | 0 | 0 | 0 | 0 |
| ASV656 | 1dfb182a50f | 57 | 0 | 0 | 0 | 0 | 38 | 0 | 0 | 0 | 19 | 0 | 0 | 0 | 0 | 0 | 0 | 0 | 0 | 0 | 0 | 0 | 0 | 0 | 0 | 0 | 0 | 0 | 0 | 0 | 0 | 0 | 0 | 0 | 0 | 0 | 0 |
| ASV657 | 7f7481008cf | 57 | 0 | 0 | 0 | 0 | 0 | 0 | 0 | 0 | 0 | 0 | 0 | 0 | 0 | 0 | 0 | 0 | 0 | 0 | 0 | 0 | 0 | 0 | 0 | 0 | 57 | 0 | 0 | 0 | 0 | 0 | 0 | 0 | 0 | 0 | 0 |
| ASV658 | b582eb48adb | 57 | 0 | 0 | 0 | 14 | 0 | 0 | 15 | 0 | 0 | 0 | 0 | 0 | 0 | 5 | 0 | 0 | 0 | 0 | 0 | 0 | 0 | 0 | 0 | 0 | 11 | 0 | 0 | 0 | 12 | 0 | 0 | 0 | 0 | 0 | 0 |
| ASV659 | 1346daf56a7 | 56 | 0 | 0 | 0 | 0 | 0 | 0 | 0 | 0 | 0 | 0 | 0 | 0 | 0 | 0 | 0 | 0 | 0 | 10 | 0 | 0 | 0 | 0 | 0 | 0 | 0 | 0 | 0 | 0 | 0 | 0 | 8 | 0 | 0 | 24 | 14 |
| ASV660 | 6720d6d947a | 56 | 0 | 0 | 0 | 0 | 15 | 0 | 0 | 0 | 0 | 0 | 0 | 0 | 0 | 18 | 23 | 0 | 0 | 0 | 0 | 0 | 0 | 0 | 0 | 0 | 0 | 0 | 0 | 0 | 0 | 0 | 0 | 0 | 0 | 0 | 0 |
| ASV661 | cb134f230b9 | 55 | 0 | 0 | 0 | 0 | 0 | 0 | 0 | 0 | 0 | 0 | 0 | 0 | 0 | 0 | 0 | 22 | 0 | 0 | 0 | 0 | 0 | 0 | 0 | 0 | 0 | 0 | 0 | 0 | 0 | 0 | 2 | 31 | 0 | 0 | 0 |
| ASV662 | a077ecf7545 | 55 | 0 | 0 | 0 | 0 | 0 | 0 | 0 | 0 | 0 | 0 | 0 | 0 | 0 | 0 | 0 | 27 | 28 | 0 | 0 | 0 | 0 | 0 | 0 | 0 | 0 | 0 | 0 | 0 | 0 | 0 | 0 | 0 | 0 | 0 | 0 |
| ASV663 | e8a35fee05d | 55 | 0 | 0 | 3 | 0 | 0 | 0 | 0 | 7 | 0 | 0 | 0 | 0 | 0 | 0 | 3 | 0 | 0 | 0 | 6 | 0 | 0 | 0 | 0 | 0 | 7 | 0 | 0 | 4 | 22 | 0 | 0 | 0 | 0 | 3 | 0 |
| ASV664 | bdf6912cb04 | 55 | 0 | 0 | 10 | 0 | 0 | 0 | 0 | 10 | 0 | 0 | 0 | 0 | 0 | 0 | 18 | 0 | 0 | 0 | 0 | 0 | 0 | 0 | 0 | 0 | 6 | 0 | 0 | 0 | 0 | 11 | 0 | 0 | 0 | 0 | 0 |
| ASV665 | 4e9abbf3b24 | 55 | 0 | 0 | 0 | 0 | 0 | 0 | 0 | 0 | 0 | 18 | 0 | 0 | 0 | 0 | 37 | 0 | 0 | 0 | 0 | 0 | 0 | 0 | 0 | 0 | 0 | 0 | 0 | 0 | 0 | 0 | 0 | 0 | 0 | 0 | 0 |
| ASV666 | bd25eb9fcb0 | 55 | 0 | 0 | 0 | 0 | 0 | 0 | 0 | 0 | 0 | 0 | 0 | 0 | 0 | 0 | 0 | 16 | 20 | 0 | 0 | 0 | 0 | 0 | 0 | 0 | 0 | 3 | 0 | 16 | 0 | 0 | 0 | 0 | 0 | 0 | 0 |
| ASV667 | b33b631d66b | 54 | 0 | 0 | 0 | 0 | 0 | 0 | 0 | 0 | 0 | 0 | 0 | 0 | 0 | 0 | 0 | 0 | 0 | 0 | 0 | 0 | 19 | 35 | 0 | 0 | 0 | 0 | 0 | 0 | 0 | 0 | 0 | 0 | 0 | 0 | 0 |
| ASV668 | dd9012533ca | 54 | 0 | 0 | 0 | 0 | 0 | 0 | 0 | 14 | 12 | 7 | 0 | 0 | 0 | 10 | 0 | 0 | 0 | 0 | 0 | 0 | 0 | 0 | 0 | 0 | 0 | 0 | 0 | 0 | 0 | 11 | 0 | 0 | 0 | 0 | 0 |
| ASV669 | f85f3e981b1 | 54 | 0 | 0 | 0 | 0 | 0 | 0 | 0 | 8 | 0 | 0 | 0 | 0 | 0 | 0 | 0 | 0 | 0 | 0 | 0 | 0 | 0 | 0 | 0 | 0 | 0 | 0 | 0 | 0 | 7 | 0 | 0 | 0 | 0 | 33 | 6 |
| ASV670 | 3e89520d137 | 54 | 0 | 0 | 0 | 0 | 0 | 0 | 0 | 0 | 0 | 0 | 0 | 0 | 0 | 0 | 0 | 0 | 0 | 0 | 0 | 0 | 0 | 0 | 0 | 0 | 0 | 0 | 0 | 0 | 0 | 0 | 13 | 0 | 0 | 41 | 0 |
| ASV671 | 7bd01c648a6 | 54 | 0 | 0 | 0 | 0 | 0 | 0 | 0 | 0 | 0 | 0 | 0 | 0 | 0 | 0 | 0 | 0 | 0 | 0 | 0 | 0 | 0 | 0 | 0 | 0 | 0 | 0 | 0 | 0 | 0 | 0 | 54 | 0 | 0 | 0 | 0 |
| ASV672 | 3db31e37eb2 | 54 | 0 | 0 | 0 | 0 | 0 | 0 | 0 | 0 | 0 | 0 | 0 | 0 | 0 | 0 | 0 | 2 | 22 | 0 | 0 | 0 | 0 | 0 | 0 | 0 | 0 | 0 | 0 | 22 | 0 | 0 | 7 | 1 | 0 | 0 | 0 |
| ASV673 | 507ed460400 | 53 | 19 | 0 | 1 | 30 | 3 | 0 | 0 | 0 | 0 | 0 | 0 | 0 | 0 | 0 | 0 | 0 | 0 | 0 | 0 | 0 | 0 | 0 | 0 | 0 | 0 | 0 | 0 | 0 | 0 | 0 | 0 | 0 | 0 | 0 | 0 |
| ASV674 | 6938d6ebc15 | 53 | 0 | 0 | 0 | 0 | 24 | 14 | 0 | 0 | 0 | 0 | 0 | 0 | 0 | 15 | 0 | 0 | 0 | 0 | 0 | 0 | 0 | 0 | 0 | 0 | 0 | 0 | 0 | 0 | 0 | 0 | 0 | 0 | 0 | 0 | 0 |
| ASV675 | 69087073e43 | 53 | 0 | 0 | 0 | 0 | 0 | 0 | 0 | 0 | 0 | 0 | 0 | 0 | 0 | 0 | 0 | 23 | 30 | 0 | 0 | 0 | 0 | 0 | 0 | 0 | 0 | 0 | 0 | 0 | 0 | 0 | 0 | 0 | 0 | 0 | 0 |
| ASV676 | 41e4cb63beb | 53 | 0 | 0 | 0 | 0 | 16 | 0 | 0 | 0 | 0 | 8 | 0 | 0 | 0 | 0 | 20 | 0 | 0 | 0 | 0 | 0 | 0 | 0 | 0 | 0 | 6 | 0 | 0 | 0 | 0 | 3 | 0 | 0 | 0 | 0 | 0 |
| ASV677 | 10eab941eaa | 53 | 0 | 0 | 0 | 0 | 0 | 0 | 0 | 0 | 0 | 0 | 0 | 0 | 0 | 0 | 0 | 53 | 0 | 0 | 0 | 0 | 0 | 0 | 0 | 0 | 0 | 0 | 0 | 0 | 0 | 0 | 0 | 0 | 0 | 0 | 0 |
| ASV678 | d24db56ca95 | 52 | 0 | 0 | 0 | 0 | 0 | 0 | 0 | 0 | 0 | 0 | 0 | 0 | 0 | 0 | 0 | 0 | 0 | 0 | 18 | 16 | 0 | 0 | 6 | 12 | 0 | 0 | 0 | 0 | 0 | 0 | 0 | 0 | 0 | 0 | 0 |
| ASV679 | ce20c85816f | 52 | 0 | 0 | 18 | 7 | 13 | 0 | 0 | 0 | 0 | 0 | 0 | 0 | 0 | 0 | 14 | 0 | 0 | 0 | 0 | 0 | 0 | 0 | 0 | 0 | 0 | 0 | 0 | 0 | 0 | 0 | 0 | 0 | 0 | 0 | 0 |
| ASV680 | 1aa53435b5a | 52 | 0 | 0 | 0 | 0 | 0 | 0 | 0 | 0 | 0 | 0 | 0 | 0 | 0 | 0 | 0 | 0 | 0 | 0 | 0 | 28 | 0 | 0 | 0 | 0 | 0 | 0 | 0 | 0 | 0 | 24 | 0 | 0 | 0 | 0 | 0 |
| ASV681 | f5cf85ac34af | 52 | 0 | 0 | 3 | 7 | 12 | 0 | 0 | 0 | 0 | 7 | 0 | 0 | 0 | 0 | 12 | 0 | 0 | 0 | 0 | 4 | 0 | 0 | 0 | 0 | 0 | 0 | 0 | 0 | 0 | 0 | 0 | 0 | 0 | 0 | 7 |
| ASV682 | 5603ed8c3e5 | 52 | 0 | 0 | 0 | 0 | 0 | 0 | 0 | 0 | 0 | 0 | 0 | 0 | 0 | 0 | 0 | 0 | 0 | 0 | 0 | 52 | 0 | 0 | 0 | 0 | 0 | 0 | 0 | 0 | 0 | 0 | 0 | 0 | 0 | 0 | 0 |
| ASV683 | 4d5354dd576 | 51 | 0 | 0 | 0 | 5 | 11 | 0 | 0 | 0 | 0 | 0 | 0 | 0 | 0 | 0 | 8 | 0 | 0 | 0 | 0 | 0 | 0 | 0 | 0 | 0 | 8 | 0 | 0 | 0 | 7 | 0 | 0 | 0 | 0 | 12 | 0 |
| ASV684 | 0b048f04759 | 51 | 0 | 0 | 0 | 0 | 0 | 0 | 0 | 0 | 11 | 0 | 0 | 0 | 0 | 0 | 33 | 0 | 0 | 0 | 0 | 0 | 0 | 0 | 0 | 0 | 0 | 0 | 0 | 0 | 0 | 7 | 0 | 0 | 0 | 0 | 0 |
| ASV685 | 7284bc721fa | 51 | 0 | 0 | 0 | 0 | 3 | 0 | 0 | 0 | 0 | 0 | 0 | 0 | 0 | 0 | 0 | 9 | 4 | 0 | 0 | 0 | 0 | 13 | 0 | 0 | 13 | 2 | 7 | 0 | 0 | 0 | 0 | 0 | 0 | 0 | 0 |
| ASV686 | 23acebf2fe6 | 51 | 0 | 0 | 0 | 0 | 0 | 0 | 0 | 0 | 0 | 0 | 0 | 0 | 0 | 0 | 0 | 0 | 0 | 14 | 0 | 4 | 0 | 0 | 6 | 0 | 0 | 0 | 0 | 0 | 6 | 6 | 0 | 0 | 0 | 5 | 10 |
| ASV687 | e3347c171f1 | 51 | 0 | 0 | 0 | 0 | 0 | 0 | 0 | 0 | 0 | 0 | 0 | 0 | 0 | 0 | 0 | 0 | 0 | 0 | 13 | 0 | 0 | 0 | 0 | 0 | 0 | 0 | 0 | 0 | 0 | 0 | 0 | 0 | 0 | 38 | 0 |
| ASV688 | bfd6ebbd1a2 | 51 | 0 | 0 | 0 | 0 | 0 | 0 | 0 | 0 | 0 | 0 | 0 | 0 | 0 | 0 | 0 | 0 | 0 | 0 | 0 | 0 | 0 | 0 | 0 | 0 | 0 | 0 | 0 | 0 | 0 | 0 | 51 | 0 | 0 | 0 | 0 |
| ASV689 | 2401ffde267 | 51 | 0 | 0 | 0 | 0 | 0 | 0 | 0 | 0 | 0 | 12 | 0 | 0 | 0 | 0 | 0 | 0 | 0 | 0 | 7 | 0 | 0 | 0 | 0 | 8 | 10 | 0 | 0 | 0 | 0 | 14 | 0 | 0 | 0 | 0 | 0 |
| ASV690 | 3dd10745ff9 | 51 | 0 | 0 | 0 | 0 | 19 | 0 | 0 | 0 | 0 | 0 | 0 | 0 | 0 | 0 | 0 | 0 | 0 | 0 | 0 | 0 | 0 | 0 | 0 | 5 | 4 | 0 | 0 | 0 | 11 | 7 | 0 | 0 | 0 | 5 | 0 |
| ASV691 | cccc004d306 | 50 | 0 | 0 | 50 | 0 | 0 | 0 | 0 | 0 | 0 | 0 | 0 | 0 | 0 | 0 | 0 | 0 | 0 | 0 | 0 | 0 | 0 | 0 | 0 | 0 | 0 | 0 | 0 | 0 | 0 | 0 | 0 | 0 | 0 | 0 | 0 |
| ASV692 | b72d76feb70 | 50 | 0 | 0 | 0 | 0 | 0 | 0 | 0 | 0 | 16 | 16 | 0 | 0 | 0 | 0 | 14 | 0 | 0 | 0 | 0 | 0 | 0 | 0 | 0 | 0 | 0 | 0 | 0 | 0 | 0 | 4 | 0 | 0 | 0 | 0 | 0 |
| ASV693 | ffbd0c01dd6 | 50 | 0 | 0 | 0 | 0 | 3 | 0 | 0 | 0 | 0 | 0 | 0 | 0 | 0 | 5 | 3 | 0 | 0 | 0 | 0 | 0 | 0 | 0 | 0 | 0 | 4 | 0 | 0 | 11 | 20 | 4 | 0 | 0 | 0 | 0 | 0 |
| ASV694 | 559015c4570 | 50 | 0 | 0 | 0 | 0 | 0 | 0 | 0 | 0 | 0 | 0 | 0 | 0 | 0 | 0 | 0 | 0 | 0 | 0 | 0 | 0 | 0 | 0 | 0 | 0 | 0 | 0 | 8 | 42 | 0 | 0 | 0 | 0 | 0 | 0 | 0 |
| ASV695 | eef2ba0e710 | 50 | 0 | 0 | 0 | 0 | 0 | 0 | 0 | 0 | 0 | 0 | 0 | 0 | 0 | 0 | 0 | 0 | 0 | 0 | 0 | 0 | 0 | 0 | 0 | 0 | 0 | 0 | 16 | 0 | 0 | 0 | 34 | 0 | 0 | 0 | 0 |
| ASV696 | ffb6efb8c674 | 50 | 0 | 0 | 0 | 0 | 0 | 0 | 0 | 0 | 0 | 0 | 0 | 0 | 0 | 0 | 0 | 11 | 18 | 0 | 0 | 0 | 0 | 0 | 0 | 0 | 0 | 0 | 0 | 8 | 0 | 0 | 13 | 0 | 0 | 0 | 0 |
| ASV697 | 62e3ce37bb8 | 50 | 0 | 0 | 0 | 0 | 0 | 5 | 0 | 0 | 0 | 0 | 0 | 11 | 11 | 12 | 0 | 4 | 4 | 0 | 0 | 0 | 0 | 0 | 0 | 0 | 0 | 0 | 0 | 0 | 0 | 0 | 0 | 3 | 0 | 0 | 0 |
| ASV698 | 399bbfd7c02 | 49 | 0 | 0 | 0 | 0 | 0 | 0 | 0 | 0 | 24 | 16 | 0 | 0 | 0 | 0 | 0 | 0 | 0 | 0 | 0 | 0 | 0 | 0 | 0 | 0 | 0 | 0 | 0 | 0 | 0 | 0 | 0 | 0 | 0 | 0 | 9 |
| ASV699 | 4f0c41bbf3c | 49 | 0 | 0 | 0 | 0 | 0 | 0 | 0 | 0 | 0 | 0 | 0 | 0 | 0 | 0 | 0 | 0 | 0 | 0 | 6 | 0 | 0 | 0 | 0 | 0 | 7 | 0 | 0 | 5 | 0 | 0 | 0 | 0 | 0 | 18 | 13 |
| ASV700 | 3610432799d | 49 | 0 | 0 | 0 | 0 | 0 | 0 | 0 | 0 | 0 | 0 | 0 | 0 | 0 | 0 | 0 | 0 | 0 | 0 | 49 | 0 | 0 | 0 | 0 | 0 | 0 | 0 | 0 | 0 | 0 | 0 | 0 | 0 | 0 | 0 | 0 |
| ASV701 | 1e10d369d4b | 49 | 0 | 0 | 0 | 0 | 0 | 0 | 0 | 0 | 0 | 0 | 0 | 0 | 0 | 0 | 0 | 7 | 7 | 0 | 0 | 0 | 0 | 0 | 0 | 0 | 0 | 0 | 0 | 0 | 0 | 0 | 28 | 0 | 7 | 0 | 0 |
| ASV702 | 136e1e07b41 | 49 | 0 | 0 | 29 | 0 | 0 | 0 | 0 | 0 | 0 | 0 | 0 | 0 | 0 | 0 | 0 | 11 | 9 | 0 | 0 | 0 | 0 | 0 | 0 | 0 | 0 | 0 | 0 | 0 | 0 | 0 | 0 | 0 | 0 | 0 | 0 |
| ASV703 | 2477d6a3ed9 | 49 | 0 | 0 | 0 | 0 | 0 | 0 | 0 | 0 | 0 | 0 | 0 | 0 | 0 | 0 | 0 | 0 | 0 | 0 | 0 | 0 | 0 | 17 | 0 | 0 | 0 | 0 | 0 | 10 | 0 | 0 | 0 | 0 | 0 | 22 | 0 |
| ASV704 | ca544d3c90a | 49 | 0 | 0 | 0 | 0 | 0 | 0 | 0 | 0 | 0 | 0 | 0 | 0 | 0 | 0 | 0 | 0 | 0 | 0 | 0 | 0 | 0 | 0 | 0 | 0 | 0 | 0 | 0 | 0 | 0 | 0 | 0 | 0 | 0 | 0 | 49 |
| ASV705 | 1ddffd158d4 | 48 | 0 | 0 | 17 | 9 | 22 | 0 | 0 | 0 | 0 | 0 | 0 | 0 | 0 | 0 | 0 | 0 | 0 | 0 | 0 | 0 | 0 | 0 | 0 | 0 | 0 | 0 | 0 | 0 | 0 | 0 | 0 | 0 | 0 | 0 | 0 |
| ASV706 | 7479fa23f08 | 48 | 0 | 0 | 0 | 0 | 0 | 0 | 0 | 0 | 0 | 0 | 0 | 0 | 0 | 0 | 0 | 48 | 0 | 0 | 0 | 0 | 0 | 0 | 0 | 0 | 0 | 0 | 0 | 0 | 0 | 0 | 0 | 0 | 0 | 0 | 0 |
| ASV707 | 52bace6ee6b | 48 | 18 | 0 | 0 | 0 | 0 | 0 | 0 | 0 | 0 | 17 | 0 | 13 | 0 | 0 | 0 | 0 | 0 | 0 | 0 | 0 | 0 | 0 | 0 | 0 | 0 | 0 | 0 | 0 | 0 | 0 | 0 | 0 | 0 | 0 | 0 |
| ASV708 | 402ee8edfe9 | 48 | 0 | 0 | 0 | 0 | 0 | 0 | 0 | 7 | 0 | 0 | 0 | 0 | 0 | 0 | 41 | 0 | 0 | 0 | 0 | 0 | 0 | 0 | 0 | 0 | 0 | 0 | 0 | 0 | 0 | 0 | 0 | 0 | 0 | 0 | 0 |
| ASV709 | 9b494b22b5e | 48 | 0 | 0 | 0 | 0 | 0 | 0 | 0 | 0 | 0 | 48 | 0 | 0 | 0 | 0 | 0 | 0 | 0 | 0 | 0 | 0 | 0 | 0 | 0 | 0 | 0 | 0 | 0 | 0 | 0 | 0 | 0 | 0 | 0 | 0 | 0 |
| ASV710 | a55361bfacb | 48 | 0 | 0 | 0 | 0 | 0 | 0 | 0 | 0 | 0 | 0 | 0 | 0 | 0 | 0 | 0 | 0 | 0 | 0 | 6 | 11 | 0 | 0 | 0 | 0 | 4 | 0 | 0 | 0 | 18 | 9 | 0 | 0 | 0 | 0 | 0 |
| ASV711 | 06dfcef1d90 | 47 | 0 | 0 | 0 | 0 | 0 | 0 | 0 | 0 | 0 | 0 | 0 | 0 | 0 | 0 | 0 | 47 | 0 | 0 | 0 | 0 | 0 | 0 | 0 | 0 | 0 | 0 | 0 | 0 | 0 | 0 | 0 | 0 | 0 | 0 | 0 |
| ASV712 | d20d8dc0009 | 47 | 0 | 0 | 0 | 0 | 0 | 0 | 0 | 0 | 0 | 0 | 0 | 0 | 0 | 0 | 0 | 22 | 0 | 0 | 0 | 0 | 12 | 0 | 0 | 0 | 0 | 13 | 0 | 0 | 0 | 0 | 0 | 0 | 0 | 0 | 0 |
| ASV713 | ac5cc2d0174 | 47 | 14 | 0 | 22 | 0 | 0 | 0 | 0 | 0 | 0 | 0 | 0 | 0 | 0 | 0 | 0 | 0 | 0 | 0 | 0 | 0 | 0 | 0 | 5 | 0 | 0 | 0 | 0 | 0 | 6 | 0 | 0 | 0 | 0 | 0 | 0 |
| ASV714 | 3a579139b07 | 46 | 0 | 0 | 0 | 0 | 9 | 16 | 10 | 0 | 0 | 0 | 0 | 11 | 0 | 0 | 0 | 0 | 0 | 0 | 0 | 0 | 0 | 0 | 0 | 0 | 0 | 0 | 0 | 0 | 0 | 0 | 0 | 0 | 0 | 0 | 0 |
| ASV715 | 6a8885ddd92 | 46 | 0 | 0 | 0 | 0 | 0 | 0 | 0 | 0 | 0 | 0 | 7 | 18 | 21 | 0 | 0 | 0 | 0 | 0 | 0 | 0 | 0 | 0 | 0 | 0 | 0 | 0 | 0 | 0 | 0 | 0 | 0 | 0 | 0 | 0 | 0 |
| ASV716 | d9358f6e983 | 46 | 0 | 0 | 0 | 0 | 0 | 0 | 0 | 0 | 21 | 0 | 0 | 0 | 0 | 0 | 25 | 0 | 0 | 0 | 0 | 0 | 0 | 0 | 0 | 0 | 0 | 0 | 0 | 0 | 0 | 0 | 0 | 0 | 0 | 0 | 0 |
| ASV717 | 059ab9b4a35 | 46 | 0 | 0 | 0 | 0 | 0 | 0 | 0 | 0 | 0 | 0 | 0 | 0 | 0 | 0 | 0 | 0 | 0 | 0 | 0 | 0 | 0 | 0 | 13 | 22 | 0 | 0 | 0 | 11 | 0 | 0 | 0 | 0 | 0 | 0 | 0 |
| ASV718 | 6b9feab1259 | 46 | 0 | 0 | 0 | 0 | 0 | 0 | 0 | 0 | 0 | 0 | 0 | 0 | 0 | 0 | 0 | 0 | 0 | 0 | 0 | 0 | 0 | 0 | 0 | 0 | 0 | 0 | 0 | 0 | 0 | 46 | 0 | 0 | 0 | 0 | 0 |
| ASV719 | 36dd501e70e | 45 | 0 | 0 | 0 | 0 | 20 | 0 | 0 | 0 | 0 | 5 | 0 | 0 | 0 | 0 | 7 | 0 | 0 | 0 | 0 | 0 | 0 | 0 | 0 | 0 | 6 | 0 | 0 | 0 | 0 | 7 | 0 | 0 | 0 | 0 | 0 |
| ASV720 | 29cabdfd807 | 45 | 0 | 0 | 0 | 0 | 0 | 45 | 0 | 0 | 0 | 0 | 0 | 0 | 0 | 0 | 0 | 0 | 0 | 0 | 0 | 0 | 0 | 0 | 0 | 0 | 0 | 0 | 0 | 0 | 0 | 0 | 0 | 0 | 0 | 0 | 0 |
| ASV721 | ec4a28fe927 | 45 | 6 | 0 | 11 | 19 | 9 | 0 | 0 | 0 | 0 | 0 | 0 | 0 | 0 | 0 | 0 | 0 | 0 | 0 | 0 | 0 | 0 | 0 | 0 | 0 | 0 | 0 | 0 | 0 | 0 | 0 | 0 | 0 | 0 | 0 | 0 |
| ASV722 | e17c3bcdf29 | 45 | 0 | 0 | 0 | 0 | 45 | 0 | 0 | 0 | 0 | 0 | 0 | 0 | 0 | 0 | 0 | 0 | 0 | 0 | 0 | 0 | 0 | 0 | 0 | 0 | 0 | 0 | 0 | 0 | 0 | 0 | 0 | 0 | 0 | 0 | 0 |
| ASV723 | fea477a0ba2 | 45 | 0 | 0 | 0 | 0 | 0 | 20 | 0 | 0 | 0 | 0 | 0 | 0 | 0 | 9 | 0 | 0 | 0 | 0 | 0 | 0 | 0 | 0 | 0 | 6 | 0 | 0 | 0 | 0 | 0 | 0 | 0 | 0 | 0 | 10 | 0 |
| ASV724 | 788f21dfa7c | 45 | 0 | 0 | 16 | 12 | 17 | 0 | 0 | 0 | 0 | 0 | 0 | 0 | 0 | 0 | 0 | 0 | 0 | 0 | 0 | 0 | 0 | 0 | 0 | 0 | 0 | 0 | 0 | 0 | 0 | 0 | 0 | 0 | 0 | 0 | 0 |
| ASV725 | 1d7e23ca83a | 44 | 0 | 0 | 0 | 0 | 0 | 0 | 0 | 0 | 0 | 0 | 0 | 0 | 0 | 0 | 0 | 0 | 0 | 0 | 0 | 0 | 0 | 0 | 0 | 0 | 44 | 0 | 0 | 0 | 0 | 0 | 0 | 0 | 0 | 0 | 0 |
| ASV726 | ab48a8acbe9 | 44 | 0 | 0 | 0 | 0 | 0 | 0 | 0 | 0 | 12 | 7 | 0 | 0 | 0 | 0 | 16 | 0 | 0 | 0 | 0 | 0 | 0 | 0 | 0 | 0 | 9 | 0 | 0 | 0 | 0 | 0 | 0 | 0 | 0 | 0 | 0 |
| ASV727 | bfe56699d0d | 44 | 0 | 0 | 0 | 0 | 0 | 0 | 0 | 0 | 0 | 0 | 0 | 0 | 0 | 0 | 0 | 0 | 12 | 0 | 0 | 0 | 0 | 0 | 0 | 0 | 0 | 0 | 0 | 0 | 0 | 0 | 32 | 0 | 0 | 0 | 0 |
| ASV728 | c420d283c4b | 44 | 0 | 0 | 0 | 0 | 15 | 0 | 0 | 0 | 0 | 4 | 0 | 0 | 0 | 0 | 12 | 0 | 0 | 0 | 0 | 0 | 0 | 0 | 0 | 0 | 0 | 0 | 0 | 0 | 0 | 0 | 0 | 0 | 0 | 0 | 13 |
| ASV729 | 09f42b53f97 | 43 | 0 | 0 | 0 | 0 | 43 | 0 | 0 | 0 | 0 | 0 | 0 | 0 | 0 | 0 | 0 | 0 | 0 | 0 | 0 | 0 | 0 | 0 | 0 | 0 | 0 | 0 | 0 | 0 | 0 | 0 | 0 | 0 | 0 | 0 | 0 |
| ASV730 | 3d94f17abb6 | 43 | 0 | 0 | 0 | 0 | 0 | 11 | 0 | 0 | 0 | 0 | 12 | 0 | 15 | 0 | 0 | 0 | 0 | 0 | 0 | 0 | 0 | 0 | 0 | 5 | 0 | 0 | 0 | 0 | 0 | 0 | 0 | 0 | 0 | 0 | 0 |
| ASV731 | 0dddbe3104d | 43 | 0 | 0 | 0 | 0 | 0 | 0 | 0 | 0 | 0 | 0 | 0 | 0 | 0 | 0 | 21 | 7 | 0 | 0 | 0 | 0 | 0 | 0 | 0 | 0 | 0 | 0 | 0 | 0 | 0 | 0 | 15 | 0 | 0 | 0 | 0 |
| ASV732 | eda4911fd79 | 43 | 0 | 0 | 0 | 0 | 10 | 0 | 0 | 0 | 0 | 0 | 0 | 0 | 0 | 0 | 0 | 0 | 0 | 0 | 0 | 0 | 0 | 0 | 0 | 0 | 0 | 0 | 0 | 0 | 0 | 0 | 0 | 0 | 0 | 0 | 33 |
| ASV733 | 89ff4b670b0 | 43 | 0 | 0 | 0 | 0 | 0 | 0 | 0 | 0 | 0 | 0 | 0 | 0 | 0 | 0 | 0 | 0 | 0 | 0 | 0 | 0 | 0 | 0 | 0 | 0 | 0 | 0 | 0 | 0 | 21 | 22 | 0 | 0 | 0 | 0 | 0 |
| ASV734 | 943f12db937 | 43 | 0 | 0 | 0 | 0 | 0 | 0 | 0 | 0 | 0 | 0 | 0 | 0 | 0 | 0 | 0 | 0 | 0 | 0 | 0 | 0 | 0 | 0 | 0 | 0 | 0 | 0 | 0 | 0 | 0 | 0 | 22 | 0 | 0 | 17 | 4 |
| ASV735 | 6270e792a6a | 43 | 0 | 0 | 0 | 0 | 0 | 0 | 0 | 0 | 0 | 0 | 0 | 0 | 0 | 0 | 0 | 0 | 0 | 0 | 0 | 0 | 0 | 0 | 0 | 0 | 6 | 0 | 0 | 0 | 0 | 14 | 0 | 0 | 0 | 0 | 23 |
| ASV736 | f09630dfd03 | 42 | 0 | 0 | 0 | 0 | 0 | 0 | 0 | 0 | 0 | 0 | 0 | 0 | 0 | 0 | 0 | 0 | 42 | 0 | 0 | 0 | 0 | 0 | 0 | 0 | 0 | 0 | 0 | 0 | 0 | 0 | 0 | 0 | 0 | 0 | 0 |
| ASV737 | 6316342e2f8 | 42 | 0 | 0 | 0 | 0 | 0 | 0 | 0 | 0 | 0 | 0 | 0 | 0 | 0 | 0 | 0 | 0 | 42 | 0 | 0 | 0 | 0 | 0 | 0 | 0 | 0 | 0 | 0 | 0 | 0 | 0 | 0 | 0 | 0 | 0 | 0 |
| ASV738 | c617caf3d30 | 42 | 0 | 0 | 0 | 0 | 0 | 0 | 0 | 0 | 0 | 0 | 0 | 0 | 0 | 0 | 0 | 16 | 0 | 0 | 0 | 0 | 0 | 0 | 0 | 0 | 0 | 0 | 0 | 0 | 0 | 0 | 26 | 0 | 0 | 0 | 0 |
| ASV739 | edfb0ff2d79e | 42 | 0 | 0 | 0 | 5 | 0 | 0 | 0 | 0 | 0 | 0 | 0 | 0 | 0 | 0 | 0 | 4 | 10 | 0 | 4 | 0 | 0 | 0 | 0 | 0 | 0 | 0 | 0 | 7 | 0 | 0 | 5 | 4 | 3 | 0 | 0 |
| ASV740 | 6f9d2f842a1 | 42 | 0 | 0 | 0 | 0 | 0 | 0 | 0 | 0 | 0 | 0 | 0 | 0 | 0 | 0 | 0 | 0 | 0 | 0 | 0 | 0 | 0 | 0 | 0 | 0 | 0 | 0 | 0 | 0 | 0 | 0 | 42 | 0 | 0 | 0 | 0 |
| ASV741 | 3b15897490d | 42 | 13 | 0 | 29 | 0 | 0 | 0 | 0 | 0 | 0 | 0 | 0 | 0 | 0 | 0 | 0 | 0 | 0 | 0 | 0 | 0 | 0 | 0 | 0 | 0 | 0 | 0 | 0 | 0 | 0 | 0 | 0 | 0 | 0 | 0 | 0 |
| ASV742 | a7338f29f3c1 | 42 | 0 | 0 | 0 | 0 | 0 | 0 | 0 | 0 | 0 | 0 | 0 | 0 | 0 | 0 | 0 | 5 | 0 | 0 | 0 | 0 | 0 | 0 | 10 | 27 | 0 | 0 | 0 | 0 | 0 | 0 | 0 | 0 | 0 | 0 | 0 |
| ASV743 | a4ec7c4f7b8 | 42 | 0 | 0 | 0 | 0 | 0 | 0 | 0 | 0 | 0 | 0 | 0 | 0 | 0 | 7 | 8 | 0 | 0 | 0 | 0 | 0 | 0 | 0 | 0 | 0 | 0 | 0 | 0 | 0 | 0 | 0 | 0 | 0 | 0 | 27 | 0 |
| ASV744 | 4d5510a36e9 | 42 | 0 | 0 | 0 | 0 | 16 | 0 | 0 | 0 | 0 | 0 | 0 | 0 | 0 | 0 | 26 | 0 | 0 | 0 | 0 | 0 | 0 | 0 | 0 | 0 | 0 | 0 | 0 | 0 | 0 | 0 | 0 | 0 | 0 | 0 | 0 |
| ASV745 | add63ee675e | 42 | 0 | 0 | 0 | 0 | 0 | 0 | 0 | 0 | 0 | 0 | 0 | 0 | 0 | 0 | 0 | 0 | 0 | 0 | 0 | 0 | 0 | 0 | 0 | 0 | 0 | 10 | 0 | 6 | 0 | 0 | 3 | 10 | 13 | 0 | 0 |
| ASV746 | ef9e3487ed3 | 42 | 0 | 0 | 0 | 0 | 0 | 0 | 0 | 0 | 0 | 0 | 0 | 0 | 0 | 0 | 0 | 0 | 42 | 0 | 0 | 0 | 0 | 0 | 0 | 0 | 0 | 0 | 0 | 0 | 0 | 0 | 0 | 0 | 0 | 0 | 0 |
| ASV747 | d40abcedfd4 | 41 | 0 | 0 | 0 | 0 | 0 | 0 | 0 | 0 | 7 | 0 | 0 | 0 | 0 | 0 | 27 | 0 | 0 | 0 | 0 | 0 | 0 | 0 | 0 | 0 | 0 | 0 | 0 | 0 | 0 | 7 | 0 | 0 | 0 | 0 | 0 |
| ASV748 | 82e97640a9f | 41 | 0 | 0 | 0 | 0 | 0 | 0 | 0 | 0 | 0 | 0 | 0 | 0 | 0 | 0 | 0 | 0 | 0 | 0 | 0 | 0 | 0 | 0 | 0 | 0 | 0 | 0 | 0 | 0 | 0 | 0 | 17 | 14 | 0 | 10 | 0 |
| ASV749 | ceff91f9dd37 | 41 | 0 | 0 | 0 | 0 | 0 | 0 | 0 | 15 | 0 | 13 | 0 | 0 | 0 | 7 | 0 | 0 | 0 | 0 | 0 | 0 | 0 | 0 | 0 | 0 | 0 | 0 | 0 | 0 | 0 | 6 | 0 | 0 | 0 | 0 | 0 |
| ASV750 | 10adfc86555 | 41 | 0 | 0 | 0 | 0 | 0 | 0 | 0 | 0 | 8 | 0 | 0 | 0 | 0 | 0 | 0 | 0 | 0 | 0 | 0 | 0 | 0 | 0 | 0 | 0 | 12 | 0 | 0 | 0 | 0 | 8 | 0 | 0 | 0 | 0 | 13 |
| ASV751 | 645c030e89d | 41 | 0 | 0 | 0 | 0 | 0 | 0 | 0 | 0 | 0 | 0 | 0 | 0 | 0 | 0 | 0 | 0 | 0 | 0 | 0 | 0 | 0 | 0 | 0 | 0 | 41 | 0 | 0 | 0 | 0 | 0 | 0 | 0 | 0 | 0 | 0 |
| ASV752 | 75484a0c70a | 41 | 0 | 0 | 0 | 0 | 0 | 0 | 0 | 0 | 10 | 0 | 0 | 15 | 0 | 0 | 0 | 0 | 0 | 0 | 0 | 0 | 0 | 0 | 0 | 0 | 0 | 0 | 0 | 16 | 0 | 0 | 0 | 0 | 0 | 0 | 0 |
| ASV753 | 0f269ae1a32 | 41 | 0 | 0 | 0 | 0 | 0 | 0 | 0 | 0 | 0 | 0 | 0 | 0 | 0 | 0 | 0 | 0 | 0 | 0 | 0 | 0 | 0 | 0 | 0 | 0 | 0 | 0 | 0 | 41 | 0 | 0 | 0 | 0 | 0 | 0 | 0 |
| ASV754 | 2c360f7bc9c | 41 | 0 | 0 | 0 | 0 | 12 | 0 | 0 | 0 | 0 | 0 | 0 | 0 | 0 | 0 | 14 | 0 | 0 | 0 | 0 | 0 | 0 | 0 | 0 | 0 | 0 | 0 | 0 | 0 | 0 | 7 | 0 | 0 | 0 | 0 | 8 |
| ASV755 | 8a3f4183408 | 41 | 0 | 0 | 0 | 0 | 0 | 0 | 0 | 0 | 0 | 0 | 0 | 0 | 0 | 0 | 0 | 0 | 0 | 0 | 0 | 0 | 0 | 0 | 0 | 0 | 0 | 17 | 7 | 0 | 0 | 0 | 9 | 0 | 3 | 5 | 0 |
| ASV756 | 0998d947a55 | 41 | 0 | 0 | 0 | 0 | 0 | 10 | 0 | 7 | 0 | 0 | 0 | 0 | 0 | 7 | 0 | 0 | 0 | 0 | 0 | 0 | 0 | 0 | 0 | 0 | 4 | 0 | 0 | 0 | 5 | 0 | 0 | 0 | 0 | 8 | 0 |
| ASV757 | 2d1c8b86799 | 40 | 0 | 0 | 0 | 0 | 0 | 0 | 0 | 0 | 7 | 0 | 0 | 0 | 0 | 0 | 20 | 0 | 0 | 0 | 0 | 0 | 0 | 0 | 0 | 0 | 0 | 0 | 0 | 0 | 0 | 13 | 0 | 0 | 0 | 0 | 0 |
| ASV758 | 3a370d2e2de | 40 | 0 | 0 | 0 | 0 | 0 | 0 | 0 | 0 | 0 | 0 | 0 | 0 | 0 | 0 | 0 | 0 | 0 | 0 | 0 | 0 | 0 | 0 | 0 | 0 | 0 | 0 | 0 | 0 | 0 | 0 | 40 | 0 | 0 | 0 | 0 |
| ASV759 | 9ec07b621e5 | 40 | 0 | 0 | 7 | 0 | 20 | 0 | 0 | 0 | 0 | 0 | 0 | 0 | 0 | 0 | 11 | 0 | 0 | 0 | 2 | 0 | 0 | 0 | 0 | 0 | 0 | 0 | 0 | 0 | 0 | 0 | 0 | 0 | 0 | 0 | 0 |

**Supplementary Table S5.** Bacterioplankton 16S rRNA ASV feature table

ASVs: 1779

Seqs: 918792

| #ASV_ID | Feature_ID | total seqs | May12_0m | May12_05m | May12_25m | May12_50m | May12_75m | Jun8_0m | Jun8_05m | Jun8_25m | Jun8_50m | Jun8_75m | Jun22_0m | Jun22_05m | Jun22_25m | Jun22_50m | Jun22_75m | Jul11_0m | Jul11_05m | Jul11_25m | Jul11_50m | Jul11_75m | Jul27_0m | Jul27_05m | Jul27_25m | Jul27_50m | Jul27_75m | Aug5_0m | Aug5_05m | Aug5_25m | Aug5_50m | Aug5_75m | Sep8_0m | Sep8_05m | Sep8_25m | Sep8_50m | Sep8_75m |
|---|---|---|---|---|---|---|---|---|---|---|---|---|---|---|---|---|---|---|---|---|---|---|---|---|---|---|---|---|---|---|---|---|---|---|---|---|---|
| ASV760 | b82e2014805 | 40 | 0 | 0 | 0 | 0 | 0 | 0 | 0 | 0 | 0 | 0 | 0 | 0 | 0 | 0 | 0 | 0 | 0 | 0 | 0 | 0 | 0 | 40 | 0 | 0 | 0 | 0 | 0 | 0 | 0 | 0 | 0 | 0 | 0 | 0 | 0 |
| ASV761 | 4559531c6e4 | 40 | 0 | 0 | 0 | 0 | 40 | 0 | 0 | 0 | 0 | 0 | 0 | 0 | 0 | 0 | 0 | 0 | 0 | 0 | 0 | 0 | 0 | 0 | 0 | 0 | 0 | 0 | 0 | 0 | 0 | 0 | 0 | 0 | 0 | 0 | 0 |
| ASV762 | c5a6b7f671a | 40 | 0 | 0 | 0 | 0 | 0 | 0 | 0 | 0 | 0 | 11 | 0 | 0 | 0 | 0 | 29 | 0 | 0 | 0 | 0 | 0 | 0 | 0 | 0 | 0 | 0 | 0 | 0 | 0 | 0 | 0 | 0 | 0 | 0 | 0 | 0 |
| ASV763 | f85cac35de5 | 39 | 0 | 0 | 0 | 0 | 0 | 0 | 0 | 0 | 0 | 0 | 0 | 39 | 0 | 0 | 0 | 0 | 0 | 0 | 0 | 0 | 0 | 0 | 0 | 0 | 0 | 0 | 0 | 0 | 0 | 0 | 0 | 0 | 0 | 0 | 0 |
| ASV764 | 3ce64164af3 | 39 | 0 | 0 | 0 | 0 | 0 | 13 | 0 | 0 | 0 | 0 | 0 | 0 | 0 | 0 | 0 | 0 | 0 | 0 | 0 | 0 | 0 | 0 | 0 | 0 | 0 | 0 | 0 | 0 | 0 | 0 | 0 | 0 | 0 | 26 | 0 |
| ASV765 | eb47536874b | 39 | 0 | 0 | 0 | 0 | 5 | 0 | 0 | 0 | 4 | 6 | 0 | 0 | 0 | 0 | 5 | 0 | 0 | 0 | 0 | 0 | 0 | 0 | 0 | 0 | 0 | 0 | 0 | 0 | 14 | 5 | 0 | 0 | 0 | 0 | 0 |
| ASV766 | 06b7076c4e5 | 38 | 0 | 0 | 10 | 0 | 17 | 0 | 0 | 0 | 0 | 0 | 0 | 0 | 0 | 0 | 11 | 0 | 0 | 0 | 0 | 0 | 0 | 0 | 0 | 0 | 0 | 0 | 0 | 0 | 0 | 0 | 0 | 0 | 0 | 0 | 0 |
| ASV767 | 2c291c318a2 | 38 | 0 | 0 | 38 | 0 | 0 | 0 | 0 | 0 | 0 | 0 | 0 | 0 | 0 | 0 | 0 | 0 | 0 | 0 | 0 | 0 | 0 | 0 | 0 | 0 | 0 | 0 | 0 | 0 | 0 | 0 | 0 | 0 | 0 | 0 | 0 |
| ASV768 | ceecbd04b48 | 38 | 0 | 0 | 0 | 0 | 0 | 0 | 0 | 0 | 0 | 0 | 0 | 0 | 0 | 0 | 38 | 0 | 0 | 0 | 0 | 0 | 0 | 0 | 0 | 0 | 0 | 0 | 0 | 0 | 0 | 0 | 0 | 0 | 0 | 0 | 0 |
| ASV769 | dc4edb34d73 | 38 | 0 | 0 | 0 | 0 | 0 | 0 | 0 | 0 | 0 | 0 | 0 | 0 | 0 | 0 | 0 | 0 | 0 | 0 | 0 | 38 | 0 | 0 | 0 | 0 | 0 | 0 | 0 | 0 | 0 | 0 | 0 | 0 | 0 | 0 | 0 |
| ASV770 | cf8a13d6805 | 38 | 0 | 0 | 0 | 0 | 0 | 0 | 0 | 0 | 0 | 0 | 0 | 0 | 0 | 0 | 0 | 0 | 34 | 0 | 0 | 0 | 0 | 0 | 0 | 0 | 0 | 0 | 4 | 0 | 0 | 0 | 0 | 0 | 0 | 0 | 0 |
| ASV771 | a100291f0a1 | 37 | 0 | 0 | 0 | 0 | 0 | 0 | 0 | 0 | 0 | 0 | 0 | 0 | 0 | 0 | 0 | 0 | 0 | 0 | 0 | 0 | 0 | 0 | 0 | 0 | 0 | 0 | 0 | 17 | 20 | 0 | 0 | 0 | 0 | 0 | 0 |
| ASV772 | 8aac3aa49c8 | 37 | 0 | 0 | 0 | 0 | 10 | 0 | 0 | 0 | 0 | 0 | 0 | 0 | 0 | 0 | 27 | 0 | 0 | 0 | 0 | 0 | 0 | 0 | 0 | 0 | 0 | 0 | 0 | 0 | 0 | 0 | 0 | 0 | 0 | 0 | 0 |
| ASV773 | d9d2f8b40aa | 37 | 0 | 0 | 0 | 0 | 0 | 0 | 0 | 0 | 0 | 0 | 0 | 0 | 0 | 0 | 0 | 0 | 0 | 0 | 0 | 0 | 0 | 0 | 0 | 0 | 0 | 0 | 0 | 0 | 0 | 0 | 26 | 0 | 11 | 0 | 0 |
| ASV774 | 1842462a3ea | 37 | 0 | 0 | 0 | 0 | 0 | 0 | 0 | 0 | 0 | 0 | 0 | 0 | 0 | 0 | 0 | 11 | 19 | 0 | 0 | 0 | 0 | 0 | 0 | 0 | 0 | 0 | 0 | 0 | 0 | 0 | 7 | 0 | 0 | 0 | 0 |
| ASV775 | 906a43b16ef | 37 | 0 | 0 | 0 | 0 | 0 | 0 | 0 | 0 | 0 | 0 | 0 | 0 | 0 | 0 | 0 | 0 | 24 | 0 | 0 | 0 | 0 | 0 | 0 | 0 | 0 | 0 | 0 | 0 | 0 | 0 | 3 | 10 | 0 | 0 | 0 |
| ASV776 | ad7795885ac | 37 | 0 | 0 | 0 | 0 | 0 | 0 | 0 | 0 | 25 | 12 | 0 | 0 | 0 | 0 | 0 | 0 | 0 | 0 | 0 | 0 | 0 | 0 | 0 | 0 | 0 | 0 | 0 | 0 | 0 | 0 | 0 | 0 | 0 | 0 | 0 |
| ASV777 | efd64149119 | 37 | 0 | 0 | 0 | 0 | 0 | 10 | 13 | 0 | 0 | 0 | 0 | 4 | 0 | 0 | 0 | 3 | 7 | 0 | 0 | 0 | 0 | 0 | 0 | 0 | 0 | 0 | 0 | 0 | 0 | 0 | 0 | 0 | 0 | 0 | 0 |
| ASV778 | 33b037990c3 | 37 | 0 | 0 | 0 | 0 | 0 | 0 | 0 | 0 | 17 | 0 | 0 | 0 | 0 | 0 | 0 | 0 | 0 | 0 | 0 | 0 | 0 | 0 | 0 | 0 | 0 | 0 | 0 | 0 | 0 | 20 | 0 | 0 | 0 | 0 | 0 |
| ASV779 | 99bd4b0129e | 37 | 0 | 0 | 0 | 0 | 0 | 0 | 0 | 0 | 0 | 0 | 0 | 0 | 0 | 0 | 0 | 0 | 0 | 0 | 0 | 0 | 0 | 0 | 0 | 0 | 0 | 12 | 10 | 6 | 0 | 0 | 9 | 0 | 0 | 0 | 0 |
| ASV780 | 27513b68771 | 37 | 0 | 0 | 0 | 0 | 0 | 0 | 0 | 0 | 0 | 0 | 0 | 0 | 0 | 0 | 0 | 0 | 37 | 0 | 0 | 0 | 0 | 0 | 0 | 0 | 0 | 0 | 0 | 0 | 0 | 0 | 0 | 0 | 0 | 0 | 0 |
| ASV781 | 8a117a94fe2 | 37 | 0 | 0 | 0 | 0 | 0 | 0 | 0 | 0 | 0 | 0 | 0 | 0 | 0 | 0 | 0 | 34 | 0 | 0 | 0 | 0 | 0 | 0 | 0 | 0 | 0 | 0 | 0 | 0 | 0 | 3 | 0 | 0 | 0 | 0 | 0 |
| ASV782 | ef5bc45d1cd | 37 | 0 | 0 | 0 | 0 | 0 | 16 | 18 | 0 | 3 | 0 | 0 | 0 | 0 | 0 | 0 | 0 | 0 | 0 | 0 | 0 | 0 | 0 | 0 | 0 | 0 | 0 | 0 | 0 | 0 | 0 | 0 | 0 | 0 | 0 | 0 |
| ASV783 | 1794f8c96ca | 36 | 0 | 0 | 0 | 0 | 36 | 0 | 0 | 0 | 0 | 0 | 0 | 0 | 0 | 0 | 0 | 0 | 0 | 0 | 0 | 0 | 0 | 0 | 0 | 0 | 0 | 0 | 0 | 0 | 0 | 0 | 0 | 0 | 0 | 0 | 0 |
| ASV784 | 600d3a9b146 | 36 | 0 | 0 | 0 | 0 | 0 | 0 | 0 | 0 | 0 | 0 | 0 | 0 | 0 | 0 | 17 | 0 | 0 | 0 | 0 | 8 | 0 | 0 | 0 | 0 | 11 | 0 | 0 | 0 | 0 | 0 | 0 | 0 | 0 | 0 | 0 |
| ASV785 | 28941a14797 | 36 | 0 | 0 | 0 | 0 | 0 | 0 | 0 | 0 | 0 | 0 | 0 | 0 | 0 | 0 | 0 | 0 | 0 | 0 | 0 | 0 | 0 | 0 | 0 | 16 | 0 | 0 | 0 | 7 | 0 | 13 | 0 | 0 | 0 | 0 | 0 |
| ASV786 | fb410157f67 | 36 | 0 | 0 | 0 | 0 | 14 | 0 | 0 | 8 | 0 | 0 | 0 | 0 | 0 | 0 | 9 | 0 | 0 | 0 | 0 | 0 | 0 | 0 | 0 | 0 | 0 | 0 | 0 | 0 | 0 | 5 | 0 | 0 | 0 | 0 | 0 |
| ASV787 | 36e28bc5831 | 36 | 0 | 0 | 0 | 0 | 0 | 0 | 0 | 0 | 0 | 0 | 0 | 0 | 0 | 0 | 0 | 20 | 16 | 0 | 0 | 0 | 0 | 0 | 0 | 0 | 0 | 0 | 0 | 0 | 0 | 0 | 0 | 0 | 0 | 0 | 0 |
| ASV788 | e090cf29c31 | 35 | 0 | 0 | 0 | 0 | 0 | 0 | 0 | 0 | 0 | 0 | 0 | 0 | 0 | 0 | 0 | 13 | 14 | 0 | 0 | 0 | 0 | 0 | 0 | 0 | 0 | 0 | 0 | 0 | 0 | 0 | 8 | 0 | 0 | 0 | 0 |
| ASV789 | b13b606cb96 | 35 | 0 | 0 | 0 | 0 | 0 | 0 | 0 | 0 | 0 | 0 | 0 | 0 | 0 | 0 | 0 | 0 | 0 | 0 | 0 | 0 | 0 | 0 | 0 | 0 | 35 | 0 | 0 | 0 | 0 | 0 | 0 | 0 | 0 | 0 | 0 |
| ASV790 | df3adaa210f | 35 | 0 | 0 | 0 | 0 | 0 | 0 | 0 | 0 | 13 | 11 | 0 | 0 | 0 | 0 | 11 | 0 | 0 | 0 | 0 | 0 | 0 | 0 | 0 | 0 | 0 | 0 | 0 | 0 | 0 | 0 | 0 | 0 | 0 | 0 | 0 |
| ASV791 | 58969e6e509 | 35 | 0 | 0 | 0 | 0 | 0 | 0 | 0 | 0 | 0 | 0 | 0 | 0 | 0 | 0 | 35 | 0 | 0 | 0 | 0 | 0 | 0 | 0 | 0 | 0 | 0 | 0 | 0 | 0 | 0 | 0 | 0 | 0 | 0 | 0 | 0 |
| ASV792 | aa5493b0951 | 35 | 0 | 0 | 0 | 0 | 0 | 9 | 0 | 0 | 0 | 0 | 0 | 0 | 0 | 0 | 0 | 0 | 26 | 0 | 0 | 0 | 0 | 0 | 0 | 0 | 0 | 0 | 0 | 0 | 0 | 0 | 0 | 0 | 0 | 0 | 0 |
| ASV793 | 86172db1d4c | 35 | 0 | 0 | 0 | 0 | 0 | 0 | 0 | 0 | 0 | 0 | 0 | 0 | 0 | 0 | 35 | 0 | 0 | 0 | 0 | 0 | 0 | 0 | 0 | 0 | 0 | 0 | 0 | 0 | 0 | 0 | 0 | 0 | 0 | 0 | 0 |
| ASV794 | 674ddf111dc | 35 | 0 | 0 | 0 | 0 | 0 | 0 | 0 | 0 | 0 | 0 | 0 | 0 | 0 | 0 | 0 | 35 | 0 | 0 | 0 | 0 | 0 | 0 | 0 | 0 | 0 | 0 | 0 | 0 | 0 | 0 | 0 | 0 | 0 | 0 | 0 |
| ASV795 | ee1fbe16af5 | 34 | 0 | 0 | 0 | 0 | 0 | 0 | 0 | 0 | 0 | 0 | 0 | 0 | 0 | 0 | 0 | 0 | 2 | 0 | 0 | 0 | 13 | 0 | 0 | 0 | 0 | 0 | 0 | 0 | 0 | 0 | 12 | 7 | 0 | 0 | 0 |
| ASV796 | d8f872ff12f | 34 | 0 | 0 | 0 | 0 | 0 | 0 | 0 | 0 | 0 | 15 | 0 | 0 | 0 | 0 | 11 | 0 | 0 | 0 | 0 | 0 | 0 | 0 | 0 | 0 | 0 | 0 | 0 | 0 | 0 | 3 | 0 | 0 | 0 | 0 | 5 |
| ASV797 | 4f40256d33c | 34 | 0 | 0 | 0 | 0 | 0 | 0 | 0 | 0 | 0 | 0 | 0 | 8 | 0 | 0 | 0 | 0 | 8 | 5 | 7 | 0 | 0 | 0 | 0 | 0 | 0 | 0 | 0 | 3 | 0 | 0 | 3 | 0 | 0 | 0 | 0 |
| ASV798 | f2a5c7c217e | 34 | 0 | 0 | 0 | 11 | 23 | 0 | 0 | 0 | 0 | 0 | 0 | 0 | 0 | 0 | 0 | 0 | 0 | 0 | 0 | 0 | 0 | 0 | 0 | 0 | 0 | 0 | 0 | 0 | 0 | 0 | 0 | 0 | 0 | 0 | 0 |
| ASV799 | 58daeddf7b2 | 34 | 0 | 0 | 0 | 0 | 0 | 0 | 0 | 0 | 0 | 18 | 0 | 0 | 0 | 0 | 0 | 0 | 0 | 0 | 0 | 0 | 0 | 0 | 0 | 0 | 0 | 0 | 0 | 0 | 0 | 0 | 0 | 0 | 0 | 0 | 16 |
| ASV800 | c6d7ed89bca | 34 | 0 | 0 | 6 | 0 | 0 | 0 | 0 | 0 | 0 | 0 | 0 | 0 | 28 | 0 | 0 | 0 | 0 | 0 | 0 | 0 | 0 | 0 | 0 | 0 | 0 | 0 | 0 | 0 | 0 | 0 | 0 | 0 | 0 | 0 | 0 |
| ASV801 | 633a1a4fc54 | 34 | 0 | 0 | 0 | 21 | 0 | 0 | 0 | 0 | 0 | 0 | 0 | 0 | 0 | 0 | 0 | 0 | 0 | 0 | 0 | 0 | 0 | 0 | 0 | 0 | 0 | 0 | 0 | 0 | 0 | 0 | 0 | 0 | 0 | 13 | 0 |
| ASV802 | 183722946bc | 34 | 0 | 0 | 2 | 0 | 12 | 0 | 0 | 0 | 0 | 4 | 0 | 0 | 0 | 0 | 16 | 0 | 0 | 0 | 0 | 0 | 0 | 0 | 0 | 0 | 0 | 0 | 0 | 0 | 0 | 0 | 0 | 0 | 0 | 0 | 0 |
| ASV803 | 678cbd6ed8d | 33 | 0 | 0 | 4 | 3 | 0 | 0 | 0 | 0 | 0 | 0 | 0 | 0 | 0 | 0 | 0 | 0 | 0 | 0 | 3 | 0 | 0 | 0 | 0 | 0 | 0 | 0 | 0 | 0 | 0 | 0 | 0 | 0 | 0 | 9 | 14 |
| ASV804 | 264fb7729d1 | 33 | 0 | 0 | 0 | 0 | 0 | 0 | 0 | 0 | 0 | 33 | 0 | 0 | 0 | 0 | 0 | 0 | 0 | 0 | 0 | 0 | 0 | 0 | 0 | 0 | 0 | 0 | 0 | 0 | 0 | 0 | 0 | 0 | 0 | 0 | 0 |
| ASV805 | 76fa3adee60 | 33 | 0 | 0 | 0 | 0 | 0 | 0 | 0 | 0 | 0 | 0 | 0 | 0 | 0 | 0 | 8 | 0 | 0 | 0 | 0 | 0 | 0 | 0 | 0 | 0 | 0 | 0 | 0 | 0 | 0 | 0 | 6 | 0 | 0 | 13 | 6 |
| ASV806 | f6380181180 | 33 | 13 | 0 | 0 | 0 | 0 | 0 | 0 | 0 | 20 | 0 | 0 | 0 | 0 | 0 | 0 | 0 | 0 | 0 | 0 | 0 | 0 | 0 | 0 | 0 | 0 | 0 | 0 | 0 | 0 | 0 | 0 | 0 | 0 | 0 | 0 |
| ASV807 | b37c475b60c | 33 | 0 | 0 | 0 | 0 | 0 | 0 | 0 | 0 | 0 | 0 | 0 | 0 | 0 | 0 | 17 | 0 | 0 | 0 | 0 | 0 | 0 | 0 | 0 | 16 | 0 | 0 | 0 | 0 | 0 | 0 | 0 | 0 | 0 | 0 | 0 |
| ASV808 | 390ee06cb48 | 33 | 0 | 0 | 0 | 0 | 0 | 0 | 0 | 0 | 0 | 0 | 0 | 0 | 0 | 0 | 33 | 0 | 0 | 0 | 0 | 0 | 0 | 0 | 0 | 0 | 0 | 0 | 0 | 0 | 0 | 0 | 0 | 0 | 0 | 0 | 0 |
| ASV809 | 7200261ab7c | 33 | 0 | 0 | 0 | 0 | 0 | 0 | 0 | 0 | 0 | 0 | 0 | 0 | 0 | 0 | 0 | 33 | 0 | 0 | 0 | 0 | 0 | 0 | 0 | 0 | 0 | 0 | 0 | 0 | 0 | 0 | 0 | 0 | 0 | 0 | 0 |
| ASV810 | 5f07fc450df7 | 33 | 0 | 0 | 0 | 0 | 0 | 0 | 0 | 0 | 0 | 0 | 0 | 0 | 0 | 0 | 0 | 0 | 33 | 0 | 0 | 0 | 0 | 0 | 0 | 0 | 0 | 0 | 0 | 0 | 0 | 0 | 0 | 0 | 0 | 0 | 0 |
| ASV811 | 2893165dcb9 | 33 | 0 | 0 | 0 | 0 | 0 | 0 | 0 | 0 | 0 | 0 | 0 | 6 | 0 | 17 | 10 | 0 | 0 | 0 | 0 | 0 | 0 | 0 | 0 | 0 | 0 | 0 | 0 | 0 | 0 | 0 | 0 | 0 | 0 | 0 | 0 |
| ASV812 | 2a5c1f50e24 | 33 | 0 | 0 | 0 | 0 | 0 | 0 | 0 | 0 | 0 | 0 | 0 | 0 | 0 | 0 | 0 | 0 | 0 | 0 | 0 | 0 | 0 | 0 | 0 | 0 | 0 | 0 | 0 | 0 | 0 | 0 | 0 | 0 | 0 | 0 | 33 |
| ASV813 | 9a6775272ac | 33 | 0 | 0 | 0 | 0 | 0 | 0 | 0 | 17 | 0 | 0 | 0 | 0 | 0 | 0 | 0 | 0 | 0 | 0 | 16 | 0 | 0 | 0 | 0 | 0 | 0 | 0 | 0 | 0 | 0 | 0 | 0 | 0 | 0 | 0 | 0 |
| ASV814 | 29685b6d9e5 | 33 | 0 | 0 | 0 | 0 | 0 | 0 | 0 | 0 | 0 | 0 | 0 | 6 | 0 | 0 | 0 | 0 | 0 | 0 | 14 | 0 | 0 | 0 | 0 | 4 | 0 | 0 | 0 | 9 | 0 | 0 | 0 | 0 | 0 | 0 | 0 |
| ASV815 | ffeb4e1ae07 | 33 | 0 | 0 | 0 | 0 | 0 | 0 | 0 | 0 | 0 | 33 | 0 | 0 | 0 | 0 | 0 | 0 | 0 | 0 | 0 | 0 | 0 | 0 | 0 | 0 | 0 | 0 | 0 | 0 | 0 | 0 | 0 | 0 | 0 | 0 | 0 |
| ASV816 | 05da9fd5392 | 33 | 0 | 0 | 0 | 0 | 16 | 0 | 0 | 0 | 0 | 17 | 0 | 0 | 0 | 0 | 0 | 0 | 0 | 0 | 0 | 0 | 0 | 0 | 0 | 0 | 0 | 0 | 0 | 0 | 0 | 0 | 0 | 0 | 0 | 0 | 0 |
| ASV817 | cd8fe65bdeb | 33 | 0 | 0 | 0 | 0 | 0 | 0 | 0 | 0 | 0 | 0 | 0 | 0 | 0 | 33 | 0 | 0 | 0 | 0 | 0 | 0 | 0 | 0 | 0 | 0 | 0 | 0 | 0 | 0 | 0 | 0 | 0 | 0 | 0 | 0 | 0 |
| ASV818 | 6d2cf7f8b39 | 33 | 0 | 0 | 0 | 0 | 0 | 0 | 0 | 0 | 0 | 0 | 0 | 0 | 0 | 0 | 0 | 0 | 0 | 0 | 0 | 0 | 0 | 0 | 0 | 0 | 0 | 0 | 0 | 0 | 0 | 0 | 0 | 0 | 0 | 0 | 33 |
| ASV819 | cd7a1bb3f03 | 33 | 0 | 0 | 33 | 0 | 0 | 0 | 0 | 0 | 0 | 0 | 0 | 0 | 0 | 0 | 0 | 0 | 0 | 0 | 0 | 0 | 0 | 0 | 0 | 0 | 0 | 0 | 0 | 0 | 0 | 0 | 0 | 0 | 0 | 0 | 0 |
| ASV820 | e3d76972ac7 | 32 | 0 | 0 | 18 | 0 | 0 | 0 | 0 | 0 | 14 | 0 | 0 | 0 | 0 | 0 | 0 | 0 | 0 | 0 | 0 | 0 | 0 | 0 | 0 | 0 | 0 | 0 | 0 | 0 | 0 | 0 | 0 | 0 | 0 | 0 | 0 |
| ASV821 | 1eb1da33226 | 32 | 0 | 0 | 7 | 0 | 12 | 0 | 0 | 0 | 0 | 0 | 0 | 0 | 0 | 0 | 13 | 0 | 0 | 0 | 0 | 0 | 0 | 0 | 0 | 0 | 0 | 0 | 0 | 0 | 0 | 0 | 0 | 0 | 0 | 0 | 0 |
| ASV822 | 0fa7065c654 | 32 | 0 | 0 | 0 | 0 | 2 | 0 | 0 | 0 | 3 | 13 | 0 | 0 | 0 | 0 | 14 | 0 | 0 | 0 | 0 | 0 | 0 | 0 | 0 | 0 | 0 | 0 | 0 | 0 | 0 | 0 | 0 | 0 | 0 | 0 | 0 |
| ASV823 | c8aa00ef3ea | 32 | 0 | 0 | 5 | 0 | 0 | 0 | 0 | 0 | 0 | 0 | 2 | 0 | 0 | 0 | 0 | 0 | 0 | 0 | 0 | 0 | 0 | 0 | 0 | 0 | 12 | 0 | 0 | 0 | 5 | 8 | 0 | 0 | 0 | 0 | 0 |
| ASV824 | 0e78e0242ec | 32 | 0 | 0 | 0 | 0 | 0 | 0 | 0 | 0 | 0 | 0 | 0 | 0 | 0 | 0 | 0 | 0 | 0 | 5 | 0 | 0 | 0 | 0 | 0 | 0 | 0 | 0 | 6 | 13 | 0 | 0 | 0 | 0 | 0 | 8 | 0 |
| ASV825 | 8610ed9c52d | 32 | 0 | 0 | 0 | 0 | 0 | 0 | 0 | 0 | 0 | 0 | 0 | 0 | 0 | 0 | 0 | 32 | 0 | 0 | 0 | 0 | 0 | 0 | 0 | 0 | 0 | 0 | 0 | 0 | 0 | 0 | 0 | 0 | 0 | 0 | 0 |
| ASV826 | af7bd3e90e6 | 32 | 0 | 0 | 0 | 0 | 13 | 0 | 0 | 0 | 0 | 0 | 0 | 0 | 0 | 0 | 12 | 0 | 0 | 0 | 0 | 0 | 0 | 0 | 0 | 0 | 5 | 0 | 0 | 0 | 0 | 2 | 0 | 0 | 0 | 0 | 0 |
| ASV827 | 10006b208cc | 32 | 0 | 0 | 0 | 0 | 32 | 0 | 0 | 0 | 0 | 0 | 0 | 0 | 0 | 0 | 0 | 0 | 0 | 0 | 0 | 0 | 0 | 0 | 0 | 0 | 0 | 0 | 0 | 0 | 0 | 0 | 0 | 0 | 0 | 0 | 0 |
| ASV828 | 688001dd2fb | 32 | 0 | 0 | 0 | 0 | 0 | 0 | 0 | 0 | 0 | 0 | 18 | 0 | 0 | 0 | 0 | 0 | 0 | 0 | 0 | 0 | 0 | 0 | 0 | 0 | 14 | 0 | 0 | 0 | 0 | 0 | 0 | 0 | 0 | 0 | 0 |
| ASV829 | 6c2be4e95eb | 32 | 10 | 8 | 14 | 0 | 0 | 0 | 0 | 0 | 0 | 0 | 0 | 0 | 0 | 0 | 0 | 0 | 0 | 0 | 0 | 0 | 0 | 0 | 0 | 0 | 0 | 0 | 0 | 0 | 0 | 0 | 0 | 0 | 0 | 0 | 0 |
| ASV830 | f05b1386fdb | 32 | 0 | 0 | 0 | 0 | 0 | 0 | 0 | 0 | 0 | 0 | 0 | 0 | 0 | 0 | 0 | 0 | 0 | 0 | 0 | 0 | 0 | 0 | 0 | 0 | 0 | 0 | 0 | 0 | 0 | 0 | 0 | 32 | 0 | 0 | 0 |
| ASV831 | fd161277b49 | 32 | 0 | 0 | 0 | 7 | 0 | 0 | 0 | 0 | 9 | 8 | 0 | 0 | 0 | 0 | 0 | 0 | 0 | 0 | 8 | 0 | 0 | 0 | 0 | 0 | 0 | 0 | 0 | 0 | 0 | 0 | 0 | 0 | 0 | 0 | 0 |
| ASV832 | 63cba9f02bd | 31 | 0 | 0 | 0 | 0 | 0 | 0 | 0 | 0 | 0 | 0 | 0 | 0 | 0 | 0 | 31 | 0 | 0 | 0 | 0 | 0 | 0 | 0 | 0 | 0 | 0 | 0 | 0 | 0 | 0 | 0 | 0 | 0 | 0 | 0 | 0 |
| ASV833 | 0ea6807cfbd | 31 | 0 | 0 | 0 | 0 | 9 | 0 | 0 | 0 | 0 | 0 | 0 | 0 | 0 | 0 | 18 | 0 | 0 | 0 | 0 | 0 | 0 | 0 | 0 | 0 | 0 | 0 | 0 | 0 | 0 | 4 | 0 | 0 | 0 | 0 | 0 |
| ASV834 | 7b1c1b2c651 | 31 | 0 | 0 | 0 | 0 | 0 | 0 | 0 | 0 | 0 | 0 | 0 | 0 | 0 | 0 | 31 | 0 | 0 | 0 | 0 | 0 | 0 | 0 | 0 | 0 | 0 | 0 | 0 | 0 | 0 | 0 | 0 | 0 | 0 | 0 | 0 |
| ASV835 | 9ac94f2186f | 31 | 0 | 0 | 0 | 0 | 0 | 0 | 0 | 0 | 0 | 0 | 0 | 0 | 0 | 0 | 0 | 0 | 0 | 0 | 31 | 0 | 0 | 0 | 0 | 0 | 0 | 0 | 0 | 0 | 0 | 0 | 0 | 0 | 0 | 0 | 0 |
| ASV836 | e31241d8eb3 | 31 | 0 | 0 | 0 | 0 | 0 | 0 | 0 | 0 | 0 | 0 | 0 | 0 | 0 | 0 | 0 | 0 | 0 | 0 | 0 | 0 | 0 | 0 | 10 | 0 | 0 | 7 | 0 | 14 | 0 | 0 | 0 | 0 | 0 | 0 | 0 |
| ASV837 | 50b4087cc1b | 31 | 0 | 0 | 0 | 0 | 17 | 0 | 0 | 0 | 0 | 0 | 0 | 0 | 0 | 0 | 0 | 0 | 0 | 0 | 0 | 0 | 0 | 0 | 0 | 0 | 0 | 0 | 0 | 0 | 5 | 9 | 0 | 0 | 0 | 0 | 0 |
| ASV838 | a6f8819eb9e | 30 | 0 | 0 | 7 | 0 | 4 | 0 | 0 | 0 | 0 | 0 | 0 | 0 | 0 | 0 | 0 | 0 | 0 | 0 | 0 | 0 | 0 | 0 | 0 | 0 | 0 | 0 | 0 | 0 | 0 | 0 | 0 | 0 | 0 | 16 | 3 |
| ASV839 | dfc250acebc | 30 | 0 | 0 | 0 | 0 | 0 | 0 | 0 | 0 | 0 | 0 | 0 | 0 | 0 | 0 | 28 | 0 | 0 | 0 | 0 | 0 | 0 | 0 | 0 | 0 | 0 | 0 | 0 | 0 | 0 | 2 | 0 | 0 | 0 | 0 | 0 |
| ASV840 | 220a0f7aaf1 | 30 | 0 | 0 | 30 | 0 | 0 | 0 | 0 | 0 | 0 | 0 | 0 | 0 | 0 | 0 | 0 | 0 | 0 | 0 | 0 | 0 | 0 | 0 | 0 | 0 | 0 | 0 | 0 | 0 | 0 | 0 | 0 | 0 | 0 | 0 | 0 |
| ASV841 | 388045997bb | 30 | 0 | 0 | 0 | 0 | 0 | 0 | 0 | 0 | 0 | 0 | 0 | 0 | 0 | 0 | 0 | 0 | 0 | 0 | 0 | 0 | 0 | 0 | 0 | 0 | 0 | 0 | 0 | 0 | 0 | 0 | 30 | 0 | 0 | 0 | 0 |
| ASV842 | 1b156a88996 | 30 | 0 | 0 | 0 | 0 | 6 | 0 | 0 | 0 | 8 | 5 | 0 | 0 | 0 | 0 | 11 | 0 | 0 | 0 | 0 | 0 | 0 | 0 | 0 | 0 | 0 | 0 | 0 | 0 | 0 | 0 | 0 | 0 | 0 | 0 | 0 |
| ASV843 | 88cd7aa61ed | 30 | 0 | 0 | 0 | 0 | 0 | 0 | 0 | 0 | 0 | 0 | 0 | 0 | 0 | 0 | 0 | 0 | 0 | 0 | 0 | 0 | 0 | 0 | 0 | 0 | 0 | 0 | 0 | 0 | 0 | 0 | 0 | 0 | 0 | 8 | 22 |
| ASV844 | c32dca4586f | 30 | 0 | 0 | 0 | 0 | 0 | 0 | 0 | 0 | 0 | 0 | 0 | 0 | 0 | 0 | 0 | 0 | 0 | 0 | 0 | 0 | 0 | 0 | 0 | 0 | 0 | 0 | 0 | 30 | 0 | 0 | 0 | 0 | 0 | 0 | 0 |
| ASV845 | c8e79e64c10 | 30 | 0 | 0 | 30 | 0 | 0 | 0 | 0 | 0 | 0 | 0 | 0 | 0 | 0 | 0 | 0 | 0 | 0 | 0 | 0 | 0 | 0 | 0 | 0 | 0 | 0 | 0 | 0 | 0 | 0 | 0 | 0 | 0 | 0 | 0 | 0 |
| ASV846 | ad50aadf132 | 30 | 0 | 0 | 30 | 0 | 0 | 0 | 0 | 0 | 0 | 0 | 0 | 0 | 0 | 0 | 0 | 0 | 0 | 0 | 0 | 0 | 0 | 0 | 0 | 0 | 0 | 0 | 0 | 0 | 0 | 0 | 0 | 0 | 0 | 0 | 0 |
| ASV847 | ff9333e98d5 | 30 | 0 | 0 | 0 | 0 | 0 | 0 | 0 | 0 | 0 | 0 | 0 | 0 | 0 | 0 | 0 | 0 | 30 | 0 | 0 | 0 | 0 | 0 | 0 | 0 | 0 | 0 | 0 | 0 | 0 | 0 | 0 | 0 | 0 | 0 | 0 |
| ASV848 | c95a8f8b0c5 | 30 | 0 | 0 | 0 | 0 | 0 | 0 | 0 | 0 | 0 | 0 | 0 | 0 | 0 | 0 | 0 | 0 | 0 | 0 | 0 | 0 | 0 | 0 | 0 | 0 | 0 | 0 | 0 | 0 | 0 | 0 | 0 | 0 | 0 | 0 | 30 |
| ASV849 | e08c6823fd9 | 30 | 0 | 0 | 0 | 0 | 0 | 0 | 0 | 0 | 0 | 0 | 0 | 0 | 0 | 0 | 0 | 0 | 0 | 0 | 14 | 0 | 0 | 0 | 0 | 0 | 0 | 0 | 0 | 0 | 6 | 10 | 0 | 0 | 0 | 0 | 0 |
| ASV850 | fb05c383e03 | 30 | 0 | 0 | 0 | 0 | 0 | 0 | 0 | 0 | 0 | 0 | 0 | 0 | 0 | 0 | 0 | 0 | 0 | 0 | 0 | 0 | 0 | 0 | 0 | 0 | 0 | 0 | 0 | 0 | 0 | 0 | 30 | 0 | 0 | 0 | 0 |
| ASV851 | 9f66e6c672a | 30 | 0 | 0 | 0 | 0 | 0 | 0 | 0 | 0 | 0 | 0 | 0 | 0 | 0 | 0 | 0 | 0 | 0 | 0 | 0 | 0 | 0 | 0 | 0 | 0 | 0 | 0 | 0 | 0 | 0 | 0 | 0 | 0 | 0 | 30 | 0 |
| ASV852 | 0d7caaeae72 | 30 | 0 | 0 | 0 | 0 | 0 | 0 | 0 | 0 | 0 | 0 | 0 | 0 | 0 | 0 | 0 | 0 | 30 | 0 | 0 | 0 | 0 | 0 | 0 | 0 | 0 | 0 | 0 | 0 | 0 | 0 | 0 | 0 | 0 | 0 | 0 |
| ASV853 | b6970bf611c | 30 | 0 | 0 | 0 | 0 | 30 | 0 | 0 | 0 | 0 | 0 | 0 | 0 | 0 | 0 | 0 | 0 | 0 | 0 | 0 | 0 | 0 | 0 | 0 | 0 | 0 | 0 | 0 | 0 | 0 | 0 | 0 | 0 | 0 | 0 | 0 |
| ASV854 | a35fa0e3621 | 29 | 0 | 0 | 0 | 0 | 0 | 0 | 0 | 0 | 0 | 0 | 0 | 0 | 0 | 0 | 0 | 0 | 0 | 0 | 0 | 0 | 0 | 0 | 0 | 0 | 0 | 0 | 0 | 0 | 0 | 0 | 29 | 0 | 0 | 0 | 0 |
| ASV855 | d5f19299f9d | 29 | 18 | 0 | 0 | 0 | 0 | 2 | 9 | 0 | 0 | 0 | 0 | 0 | 0 | 0 | 0 | 0 | 0 | 0 | 0 | 0 | 0 | 0 | 0 | 0 | 0 | 0 | 0 | 0 | 0 | 0 | 0 | 0 | 0 | 0 | 0 |
| ASV856 | 239ba189230 | 29 | 0 | 0 | 0 | 0 | 9 | 0 | 0 | 0 | 0 | 0 | 0 | 0 | 0 | 0 | 0 | 0 | 0 | 0 | 0 | 0 | 0 | 0 | 0 | 0 | 0 | 0 | 0 | 0 | 8 | 5 | 0 | 0 | 0 | 0 | 7 |
| ASV857 | d59742a48c4 | 29 | 0 | 0 | 0 | 0 | 0 | 0 | 0 | 0 | 0 | 0 | 0 | 0 | 0 | 0 | 0 | 0 | 0 | 0 | 0 | 0 | 0 | 0 | 0 | 0 | 0 | 0 | 0 | 0 | 0 | 0 | 29 | 0 | 0 | 0 | 0 |
| ASV858 | 8e8dde2d3b5 | 29 | 0 | 0 | 0 | 0 | 11 | 0 | 0 | 0 | 0 | 0 | 0 | 0 | 0 | 0 | 12 | 0 | 0 | 0 | 0 | 0 | 0 | 0 | 0 | 0 | 6 | 0 | 0 | 0 | 0 | 0 | 0 | 0 | 0 | 0 | 0 |
| ASV859 | dee3cd03883 | 29 | 0 | 0 | 0 | 0 | 0 | 0 | 0 | 0 | 0 | 0 | 0 | 0 | 0 | 0 | 13 | 0 | 0 | 0 | 0 | 7 | 0 | 0 | 0 | 0 | 9 | 0 | 0 | 0 | 0 | 0 | 0 | 0 | 0 | 0 | 0 |
| ASV860 | 0bcc04aceae | 29 | 0 | 0 | 0 | 0 | 0 | 0 | 0 | 0 | 0 | 0 | 0 | 0 | 0 | 0 | 29 | 0 | 0 | 0 | 0 | 0 | 0 | 0 | 0 | 0 | 0 | 0 | 0 | 0 | 0 | 0 | 0 | 0 | 0 | 0 | 0 |
| ASV861 | 34606f52693 | 29 | 0 | 0 | 0 | 0 | 0 | 0 | 0 | 0 | 0 | 0 | 0 | 0 | 0 | 0 | 0 | 0 | 0 | 0 | 0 | 0 | 0 | 0 | 0 | 0 | 0 | 0 | 0 | 0 | 0 | 0 | 29 | 0 | 0 | 0 | 0 |
| ASV862 | a7e605d7248 | 29 | 0 | 0 | 0 | 0 | 0 | 0 | 0 | 0 | 0 | 0 | 0 | 0 | 0 | 0 | 2 | 0 | 0 | 0 | 0 | 0 | 0 | 0 | 0 | 0 | 0 | 0 | 0 | 27 | 0 | 0 | 0 | 0 | 0 | 0 | 0 |
| ASV863 | 6687d307cbf | 29 | 0 | 0 | 0 | 0 | 0 | 0 | 0 | 0 | 0 | 0 | 0 | 0 | 0 | 0 | 0 | 0 | 0 | 0 | 0 | 0 | 0 | 0 | 0 | 0 | 0 | 0 | 14 | 0 | 0 | 0 | 0 | 0 | 15 | 0 | 0 |
| ASV864 | a02578e4deb | 28 | 0 | 0 | 0 | 9 | 19 | 0 | 0 | 0 | 0 | 0 | 0 | 0 | 0 | 0 | 0 | 0 | 0 | 0 | 0 | 0 | 0 | 0 | 0 | 0 | 0 | 0 | 0 | 0 | 0 | 0 | 0 | 0 | 0 | 0 | 0 |
| ASV865 | b9abee5f6a8 | 28 | 0 | 0 | 0 | 0 | 0 | 0 | 0 | 0 | 0 | 0 | 0 | 0 | 0 | 0 | 0 | 0 | 0 | 0 | 13 | 0 | 0 | 0 | 0 | 0 | 0 | 0 | 0 | 0 | 0 | 0 | 0 | 0 | 0 | 0 | 15 |
| ASV866 | 8a2f347867e | 28 | 4 | 0 | 0 | 0 | 5 | 9 | 3 | 7 | 0 | 0 | 0 | 0 | 0 | 0 | 0 | 0 | 0 | 0 | 0 | 0 | 0 | 0 | 0 | 0 | 0 | 0 | 0 | 0 | 0 | 0 | 0 | 0 | 0 | 0 | 0 |
| ASV867 | b95b5f64e7d | 28 | 0 | 0 | 0 | 0 | 0 | 0 | 0 | 0 | 0 | 12 | 0 | 0 | 0 | 0 | 0 | 5 | 0 | 5 | 6 | 0 | 0 | 0 | 0 | 0 | 0 | 0 | 0 | 0 | 0 | 0 | 0 | 0 | 0 | 0 | 0 |
| ASV868 | 1b7981b7344 | 28 | 0 | 0 | 0 | 0 | 0 | 0 | 0 | 0 | 0 | 8 | 0 | 0 | 0 | 0 | 0 | 0 | 0 | 0 | 0 | 0 | 0 | 0 | 0 | 0 | 0 | 0 | 0 | 0 | 5 | 0 | 0 | 0 | 0 | 0 | 15 |
| ASV869 | 43dac0ed1e7 | 28 | 0 | 0 | 0 | 0 | 0 | 0 | 0 | 0 | 0 | 0 | 0 | 0 | 0 | 0 | 0 | 0 | 0 | 0 | 0 | 0 | 0 | 0 | 0 | 0 | 28 | 0 | 0 | 0 | 0 | 0 | 0 | 0 | 0 | 0 | 0 |
| ASV870 | 59f65c6f8a8 | 28 | 0 | 8 | 20 | 0 | 0 | 0 | 0 | 0 | 0 | 0 | 0 | 0 | 0 | 0 | 0 | 0 | 0 | 0 | 0 | 0 | 0 | 0 | 0 | 0 | 0 | 0 | 0 | 0 | 0 | 0 | 0 | 0 | 0 | 0 | 0 |
| ASV871 | 8957bb33867 | 28 | 3 | 0 | 13 | 7 | 0 | 0 | 0 | 0 | 0 | 0 | 0 | 0 | 0 | 0 | 0 | 0 | 0 | 0 | 0 | 0 | 0 | 0 | 0 | 0 | 5 | 0 | 0 | 0 | 0 | 0 | 0 | 0 | 0 | 0 | 0 |
| ASV872 | 0aa35bb423f | 28 | 0 | 0 | 0 | 0 | 0 | 0 | 0 | 0 | 0 | 0 | 0 | 0 | 0 | 0 | 0 | 0 | 0 | 0 | 0 | 0 | 0 | 0 | 0 | 0 | 0 | 18 | 10 | 0 | 0 | 0 | 0 | 0 | 0 | 0 | 0 |
| ASV873 | 39666b64f13 | 28 | 8 | 0 | 0 | 0 | 0 | 0 | 0 | 0 | 0 | 0 | 0 | 0 | 0 | 0 | 0 | 0 | 0 | 0 | 0 | 0 | 0 | 0 | 0 | 0 | 20 | 0 | 0 | 0 | 0 | 0 | 0 | 0 | 0 | 0 | 0 |
| ASV874 | 6bc7512f712 | 27 | 0 | 0 | 0 | 0 | 5 | 0 | 0 | 0 | 0 | 0 | 0 | 0 | 0 | 0 | 0 | 0 | 0 | 0 | 0 | 0 | 0 | 0 | 0 | 0 | 0 | 0 | 0 | 0 | 0 | 0 | 0 | 0 | 0 | 0 | 22 |
| ASV875 | 8602c264af0 | 27 | 0 | 0 | 11 | 5 | 0 | 0 | 0 | 4 | 0 | 0 | 0 | 0 | 0 | 0 | 0 | 0 | 0 | 0 | 0 | 0 | 0 | 0 | 0 | 5 | 0 | 0 | 0 | 0 | 2 | 0 | 0 | 0 | 0 | 0 | 0 |
| ASV876 | 7157ed43df2 | 27 | 0 | 10 | 17 | 0 | 0 | 0 | 0 | 0 | 0 | 0 | 0 | 0 | 0 | 0 | 0 | 0 | 0 | 0 | 0 | 0 | 0 | 0 | 0 | 0 | 0 | 0 | 0 | 0 | 0 | 0 | 0 | 0 | 0 | 0 | 0 |
| ASV877 | 0025d4e026e | 27 | 0 | 0 | 0 | 0 | 0 | 0 | 0 | 0 | 0 | 5 | 0 | 0 | 0 | 0 | 5 | 0 | 0 | 0 | 0 | 0 | 0 | 0 | 0 | 0 | 0 | 0 | 0 | 0 | 0 | 0 | 0 | 0 | 0 | 7 | 10 |
| ASV878 | ca9e427c68a | 27 | 0 | 0 | 0 | 0 | 0 | 0 | 0 | 0 | 0 | 0 | 0 | 0 | 0 | 0 | 0 | 0 | 0 | 0 | 0 | 0 | 0 | 0 | 4 | 0 | 0 | 0 | 0 | 23 | 0 | 0 | 0 | 0 | 0 | 0 | 0 |
| ASV879 | 7cf008337c9 | 27 | 0 | 0 | 0 | 0 | 0 | 0 | 0 | 0 | 0 | 12 | 0 | 0 | 0 | 0 | 15 | 0 | 0 | 0 | 0 | 0 | 0 | 0 | 0 | 0 | 0 | 0 | 0 | 0 | 0 | 0 | 0 | 0 | 0 | 0 | 0 |
| ASV880 | aaf9cfef6098 | 27 | 0 | 27 | 0 | 0 | 0 | 0 | 0 | 0 | 0 | 0 | 0 | 0 | 0 | 0 | 0 | 0 | 0 | 0 | 0 | 0 | 0 | 0 | 0 | 0 | 0 | 0 | 0 | 0 | 0 | 0 | 0 | 0 | 0 | 0 | 0 |
| ASV881 | c57b019ae5e | 27 | 0 | 0 | 10 | 0 | 8 | 0 | 0 | 0 | 0 | 0 | 0 | 0 | 0 | 0 | 0 | 0 | 0 | 0 | 0 | 0 | 0 | 0 | 0 | 0 | 7 | 0 | 0 | 0 | 2 | 0 | 0 | 0 | 0 | 0 | 0 |
| ASV882 | e0fa2fd0a5b | 26 | 0 | 0 | 0 | 0 | 0 | 0 | 0 | 0 | 0 | 0 | 0 | 0 | 0 | 0 | 26 | 0 | 0 | 0 | 0 | 0 | 0 | 0 | 0 | 0 | 0 | 0 | 0 | 0 | 0 | 0 | 0 | 0 | 0 | 0 | 0 |
| ASV883 | a4fecb717e4 | 26 | 0 | 0 | 6 | 0 | 0 | 0 | 0 | 0 | 0 | 0 | 0 | 0 | 0 | 0 | 0 | 0 | 0 | 0 | 0 | 0 | 0 | 0 | 0 | 0 | 0 | 0 | 0 | 0 | 0 | 0 | 0 | 0 | 0 | 5 | 15 |
| ASV884 | f08120a4f49 | 26 | 0 | 0 | 0 | 0 | 0 | 0 | 0 | 0 | 0 | 0 | 0 | 0 | 0 | 0 | 0 | 0 | 0 | 0 | 0 | 26 | 0 | 0 | 0 | 0 | 0 | 0 | 0 | 0 | 0 | 0 | 0 | 0 | 0 | 0 | 0 |
| ASV885 | e3e519c0159 | 26 | 0 | 0 | 0 | 0 | 0 | 0 | 0 | 0 | 0 | 0 | 0 | 0 | 0 | 0 | 0 | 26 | 0 | 0 | 0 | 0 | 0 | 0 | 0 | 0 | 0 | 0 | 0 | 0 | 0 | 0 | 0 | 0 | 0 | 0 | 0 |
| ASV886 | 580db4d4bd0 | 26 | 0 | 0 | 10 | 16 | 0 | 0 | 0 | 0 | 0 | 0 | 0 | 0 | 0 | 0 | 0 | 0 | 0 | 0 | 0 | 0 | 0 | 0 | 0 | 0 | 0 | 0 | 0 | 0 | 0 | 0 | 0 | 0 | 0 | 0 | 0 |
| ASV887 | fb6ec7a317d | 26 | 0 | 0 | 0 | 0 | 0 | 0 | 0 | 0 | 11 | 0 | 0 | 0 | 0 | 0 | 15 | 0 | 0 | 0 | 0 | 0 | 0 | 0 | 0 | 0 | 0 | 0 | 0 | 0 | 0 | 0 | 0 | 0 | 0 | 0 | 0 |
| ASV888 | dc034c38f5f3 | 26 | 0 | 0 | 0 | 0 | 0 | 0 | 0 | 0 | 0 | 0 | 0 | 0 | 0 | 0 | 0 | 26 | 0 | 0 | 0 | 0 | 0 | 0 | 0 | 0 | 0 | 0 | 0 | 0 | 0 | 0 | 0 | 0 | 0 | 0 | 0 |
| ASV889 | a0bfae3e3ba | 26 | 0 | 0 | 0 | 0 | 2 | 0 | 0 | 0 | 6 | 0 | 0 | 0 | 0 | 0 | 18 | 0 | 0 | 0 | 0 | 0 | 0 | 0 | 0 | 0 | 0 | 0 | 0 | 0 | 0 | 0 | 0 | 0 | 0 | 0 | 0 |
| ASV890 | ca46261eee6 | 26 | 0 | 0 | 0 | 0 | 0 | 0 | 0 | 0 | 0 | 0 | 0 | 0 | 0 | 0 | 0 | 0 | 0 | 0 | 0 | 0 | 0 | 0 | 0 | 0 | 0 | 0 | 0 | 0 | 0 | 0 | 0 | 0 | 0 | 13 | 13 |
| ASV891 | 9f513d43bdf | 26 | 0 | 0 | 0 | 0 | 0 | 0 | 0 | 0 | 0 | 0 | 0 | 0 | 0 | 0 | 0 | 0 | 0 | 0 | 0 | 0 | 0 | 0 | 0 | 0 | 0 | 0 | 0 | 0 | 0 | 0 | 26 | 0 | 0 | 0 | 0 |
| ASV892 | 88e8503f537 | 25 | 0 | 0 | 0 | 0 | 0 | 0 | 0 | 11 | 0 | 14 | 0 | 0 | 0 | 0 | 0 | 0 | 0 | 0 | 0 | 0 | 0 | 0 | 0 | 0 | 0 | 0 | 0 | 0 | 0 | 0 | 0 | 0 | 0 | 0 | 0 |
| ASV893 | ee763c29f15 | 25 | 0 | 0 | 0 | 0 | 25 | 0 | 0 | 0 | 0 | 0 | 0 | 0 | 0 | 0 | 0 | 0 | 0 | 0 | 0 | 0 | 0 | 0 | 0 | 0 | 0 | 0 | 0 | 0 | 0 | 0 | 0 | 0 | 0 | 0 | 0 |
| ASV894 | 771805b7556 | 25 | 0 | 0 | 0 | 0 | 0 | 0 | 0 | 0 | 0 | 0 | 0 | 0 | 0 | 0 | 25 | 0 | 0 | 0 | 0 | 0 | 0 | 0 | 0 | 0 | 0 | 0 | 0 | 0 | 0 | 0 | 0 | 0 | 0 | 0 | 0 |
| ASV895 | 6b4faa79fdf4 | 25 | 0 | 0 | 0 | 0 | 13 | 0 | 0 | 0 | 0 | 12 | 0 | 0 | 0 | 0 | 0 | 0 | 0 | 0 | 0 | 0 | 0 | 0 | 0 | 0 | 0 | 0 | 0 | 0 | 0 | 0 | 0 | 0 | 0 | 0 | 0 |
| ASV896 | af74e689e59 | 25 | 0 | 0 | 0 | 0 | 0 | 0 | 0 | 0 | 0 | 0 | 0 | 0 | 0 | 0 | 0 | 0 | 0 | 0 | 0 | 0 | 0 | 0 | 0 | 0 | 15 | 0 | 0 | 0 | 0 | 10 | 0 | 0 | 0 | 0 | 0 |
| ASV897 | b851e7a5c19 | 25 | 0 | 0 | 0 | 0 | 0 | 2 | 0 | 9 | 0 | 5 | 0 | 0 | 0 | 0 | 0 | 0 | 0 | 2 | 0 | 0 | 0 | 0 | 0 | 0 | 0 | 0 | 0 | 0 | 7 | 0 | 0 | 0 | 0 | 0 | 0 |
| ASV898 | e290d868f19 | 25 | 0 | 0 | 0 | 0 | 0 | 0 | 0 | 0 | 0 | 0 | 0 | 0 | 0 | 0 | 0 | 0 | 0 | 0 | 0 | 0 | 0 | 0 | 0 | 0 | 0 | 0 | 0 | 0 | 0 | 0 | 0 | 0 | 0 | 25 | 0 |
| ASV899 | 637d25313b4 | 25 | 0 | 0 | 0 | 0 | 0 | 0 | 0 | 0 | 0 | 0 | 0 | 0 | 0 | 0 | 0 | 15 | 10 | 0 | 0 | 0 | 0 | 0 | 0 | 0 | 0 | 0 | 0 | 0 | 0 | 0 | 0 | 0 | 0 | 0 | 0 |
| ASV900 | f80ed8ce09d | 25 | 0 | 0 | 0 | 6 | 0 | 0 | 0 | 0 | 0 | 0 | 0 | 0 | 0 | 0 | 0 | 0 | 0 | 0 | 0 | 0 | 0 | 0 | 0 | 0 | 0 | 0 | 0 | 0 | 12 | 7 | 0 | 0 | 0 | 0 | 0 |
| ASV901 | cc1c4244ac5 | 25 | 0 | 0 | 2 | 0 | 0 | 0 | 0 | 0 | 0 | 0 | 0 | 0 | 0 | 0 | 23 | 0 | 0 | 0 | 0 | 0 | 0 | 0 | 0 | 0 | 0 | 0 | 0 | 0 | 0 | 0 | 0 | 0 | 0 | 0 | 0 |
| ASV902 | d6eff63ee5d | 25 | 0 | 0 | 0 | 0 | 0 | 0 | 0 | 0 | 0 | 15 | 0 | 0 | 0 | 0 | 0 | 0 | 0 | 0 | 0 | 0 | 0 | 0 | 0 | 6 | 4 | 0 | 0 | 0 | 0 | 0 | 0 | 0 | 0 | 0 | 0 |
| ASV903 | cb4e56f878a | 24 | 0 | 0 | 0 | 0 | 0 | 0 | 0 | 0 | 0 | 0 | 0 | 0 | 0 | 0 | 0 | 0 | 0 | 0 | 0 | 0 | 0 | 0 | 0 | 0 | 0 | 0 | 0 | 24 | 0 | 0 | 0 | 0 | 0 | 0 | 0 |
| ASV904 | 31ed31c06a3 | 24 | 0 | 0 | 0 | 0 | 15 | 0 | 0 | 0 | 0 | 0 | 0 | 0 | 0 | 0 | 0 | 0 | 0 | 0 | 0 | 0 | 0 | 0 | 0 | 0 | 0 | 0 | 0 | 0 | 0 | 9 | 0 | 0 | 0 | 0 | 0 |
| ASV905 | ee1e4f82344 | 24 | 0 | 0 | 5 | 6 | 0 | 0 | 0 | 0 | 0 | 0 | 0 | 0 | 0 | 0 | 6 | 0 | 0 | 0 | 7 | 0 | 0 | 0 | 0 | 0 | 0 | 0 | 0 | 0 | 0 | 0 | 0 | 0 | 0 | 0 | 0 |
| ASV906 | 00c181aba51 | 24 | 0 | 0 | 0 | 0 | 0 | 0 | 0 | 0 | 10 | 0 | 0 | 0 | 0 | 0 | 0 | 0 | 0 | 0 | 0 | 14 | 0 | 0 | 0 | 0 | 0 | 0 | 0 | 0 | 0 | 0 | 0 | 0 | 0 | 0 | 0 |
| ASV907 | 25d7d6b9575 | 24 | 0 | 0 | 13 | 0 | 11 | 0 | 0 | 0 | 0 | 0 | 0 | 0 | 0 | 0 | 0 | 0 | 0 | 0 | 0 | 0 | 0 | 0 | 0 | 0 | 0 | 0 | 0 | 0 | 0 | 0 | 0 | 0 | 0 | 0 | 0 |
| ASV908 | 6228d1ad222 | 24 | 0 | 0 | 0 | 0 | 0 | 0 | 0 | 0 | 0 | 0 | 0 | 0 | 0 | 0 | 0 | 0 | 4 | 0 | 0 | 0 | 0 | 0 | 0 | 0 | 0 | 0 | 0 | 0 | 0 | 13 | 0 | 7 | 0 | 0 | 0 |
| ASV909 | 0f0b97057a6 | 24 | 0 | 0 | 0 | 0 | 15 | 0 | 0 | 8 | 1 | 0 | 0 | 0 | 0 | 0 | 0 | 0 | 0 | 0 | 0 | 0 | 0 | 0 | 0 | 0 | 0 | 0 | 0 | 0 | 0 | 0 | 0 | 0 | 0 | 0 | 0 |
| ASV910 | 027a873fcb6 | 24 | 0 | 0 | 3 | 0 | 0 | 0 | 0 | 0 | 0 | 0 | 0 | 12 | 0 | 0 | 0 | 0 | 9 | 0 | 0 | 0 | 0 | 0 | 0 | 0 | 0 | 0 | 0 | 0 | 0 | 0 | 0 | 0 | 0 | 0 | 0 |
| ASV911 | 6517d7c8957 | 24 | 0 | 0 | 12 | 6 | 6 | 0 | 0 | 0 | 0 | 0 | 0 | 0 | 0 | 0 | 0 | 0 | 0 | 0 | 0 | 0 | 0 | 0 | 0 | 0 | 0 | 0 | 0 | 0 | 0 | 0 | 0 | 0 | 0 | 0 | 0 |
| ASV912 | 38cb37725df | 24 | 0 | 0 | 9 | 0 | 10 | 0 | 0 | 0 | 0 | 0 | 0 | 0 | 0 | 0 | 5 | 0 | 0 | 0 | 0 | 0 | 0 | 0 | 0 | 0 | 0 | 0 | 0 | 0 | 0 | 0 | 0 | 0 | 0 | 0 | 0 |
| ASV913 | 9b4e34ffda8 | 24 | 0 | 0 | 15 | 0 | 0 | 0 | 0 | 0 | 0 | 0 | 0 | 0 | 0 | 0 | 0 | 0 | 0 | 0 | 0 | 0 | 0 | 0 | 0 | 0 | 0 | 0 | 0 | 0 | 4 | 0 | 0 | 0 | 0 | 5 | 0 |
| ASV914 | 88a94fba34b | 24 | 0 | 0 | 0 | 0 | 0 | 0 | 0 | 0 | 0 | 0 | 0 | 0 | 0 | 0 | 24 | 0 | 0 | 0 | 0 | 0 | 0 | 0 | 0 | 0 | 0 | 0 | 0 | 0 | 0 | 0 | 0 | 0 | 0 | 0 | 0 |
| ASV915 | 4b145ed5300 | 24 | 0 | 0 | 0 | 0 | 0 | 0 | 0 | 0 | 0 | 0 | 0 | 0 | 0 | 0 | 24 | 0 | 0 | 0 | 0 | 0 | 0 | 0 | 0 | 0 | 0 | 0 | 0 | 0 | 0 | 0 | 0 | 0 | 0 | 0 | 0 |
| ASV916 | 8fd69e37e3d | 23 | 11 | 0 | 0 | 0 | 0 | 0 | 0 | 0 | 0 | 0 | 0 | 0 | 0 | 12 | 0 | 0 | 0 | 0 | 0 | 0 | 0 | 0 | 0 | 0 | 0 | 0 | 0 | 0 | 0 | 0 | 0 | 0 | 0 | 0 | 0 |
| ASV917 | c2164bd99dc | 23 | 0 | 0 | 0 | 0 | 0 | 0 | 0 | 0 | 0 | 0 | 0 | 0 | 0 | 0 | 0 | 0 | 0 | 0 | 12 | 0 | 0 | 0 | 0 | 0 | 0 | 0 | 0 | 11 | 0 | 0 | 0 | 0 | 0 | 0 | 0 |
| ASV918 | 42d4a272cbf | 23 | 0 | 0 | 0 | 0 | 0 | 0 | 0 | 0 | 0 | 23 | 0 | 0 | 0 | 0 | 0 | 0 | 0 | 0 | 0 | 0 | 0 | 0 | 0 | 0 | 0 | 0 | 0 | 0 | 0 | 0 | 0 | 0 | 0 | 0 | 0 |
| ASV919 | f1878dc4551 | 23 | 0 | 0 | 0 | 0 | 0 | 0 | 0 | 0 | 0 | 0 | 0 | 0 | 0 | 0 | 12 | 0 | 0 | 0 | 0 | 0 | 0 | 0 | 0 | 0 | 11 | 0 | 0 | 0 | 0 | 0 | 0 | 0 | 0 | 0 | 0 |
| ASV920 | 620d7368c91 | 23 | 0 | 0 | 0 | 0 | 0 | 0 | 0 | 0 | 0 | 0 | 0 | 0 | 0 | 0 | 0 | 0 | 14 | 0 | 0 | 0 | 0 | 0 | 0 | 0 | 0 | 0 | 9 | 0 | 0 | 0 | 0 | 0 | 0 | 0 | 0 |
| ASV921 | c29a30bed46 | 23 | 0 | 15 | 0 | 0 | 0 | 0 | 0 | 0 | 0 | 0 | 0 | 0 | 0 | 0 | 0 | 0 | 0 | 0 | 0 | 0 | 0 | 0 | 0 | 0 | 0 | 0 | 0 | 0 | 0 | 8 | 0 | 0 | 0 | 0 | 0 |
| ASV922 | a5845fe981a | 23 | 0 | 0 | 0 | 0 | 0 | 0 | 0 | 0 | 0 | 0 | 0 | 0 | 0 | 0 | 0 | 0 | 10 | 0 | 0 | 0 | 0 | 0 | 0 | 0 | 0 | 0 | 0 | 0 | 0 | 0 | 13 | 0 | 0 | 0 | 0 |
| ASV923 | 88e5ba79e63 | 23 | 0 | 0 | 0 | 0 | 0 | 0 | 0 | 0 | 0 | 0 | 0 | 0 | 0 | 0 | 0 | 0 | 0 | 0 | 0 | 0 | 0 | 0 | 0 | 0 | 0 | 0 | 0 | 23 | 0 | 0 | 0 | 0 | 0 | 0 | 0 |
| ASV924 | e4bf9639d85 | 23 | 0 | 0 | 0 | 0 | 0 | 0 | 0 | 0 | 0 | 0 | 0 | 0 | 0 | 0 | 0 | 0 | 0 | 0 | 0 | 0 | 0 | 0 | 0 | 0 | 0 | 0 | 0 | 0 | 0 | 0 | 23 | 0 | 0 | 0 | 0 |
| ASV925 | 451eda63522 | 23 | 0 | 0 | 0 | 0 | 0 | 11 | 0 | 0 | 0 | 0 | 0 | 0 | 0 | 0 | 0 | 0 | 0 | 0 | 0 | 0 | 0 | 0 | 0 | 0 | 0 | 0 | 0 | 12 | 0 | 0 | 0 | 0 | 0 | 0 | 0 |
| ASV926 | d374259f696 | 23 | 0 | 0 | 0 | 0 | 0 | 0 | 0 | 0 | 0 | 0 | 0 | 0 | 0 | 0 | 23 | 0 | 0 | 0 | 0 | 0 | 0 | 0 | 0 | 0 | 0 | 0 | 0 | 0 | 0 | 0 | 0 | 0 | 0 | 0 | 0 |
| ASV927 | 9086635cdc2 | 23 | 0 | 0 | 0 | 0 | 0 | 0 | 0 | 0 | 0 | 8 | 0 | 0 | 0 | 0 | 15 | 0 | 0 | 0 | 0 | 0 | 0 | 0 | 0 | 0 | 0 | 0 | 0 | 0 | 0 | 0 | 0 | 0 | 0 | 0 | 0 |
| ASV928 | ae052211ae3 | 23 | 0 | 0 | 0 | 0 | 0 | 0 | 0 | 12 | 0 | 0 | 0 | 0 | 0 | 0 | 0 | 0 | 0 | 0 | 0 | 0 | 0 | 0 | 0 | 0 | 0 | 0 | 0 | 0 | 0 | 0 | 0 | 0 | 0 | 11 | 0 |
| ASV929 | 32dd08a35d3 | 23 | 0 | 0 | 0 | 0 | 0 | 0 | 0 | 0 | 0 | 0 | 0 | 0 | 0 | 0 | 23 | 0 | 0 | 0 | 0 | 0 | 0 | 0 | 0 | 0 | 0 | 0 | 0 | 0 | 0 | 0 | 0 | 0 | 0 | 0 | 0 |
| ASV930 | 6e0f61eeffdb | 23 | 0 | 0 | 0 | 0 | 0 | 0 | 0 | 0 | 0 | 0 | 0 | 0 | 0 | 0 | 0 | 0 | 7 | 12 | 0 | 0 | 0 | 0 | 0 | 0 | 0 | 0 | 0 | 0 | 0 | 0 | 0 | 0 | 0 | 4 | 0 |
| ASV931 | 1c3b99535e9 | 22 | 0 | 0 | 0 | 0 | 0 | 0 | 0 | 0 | 0 | 0 | 0 | 0 | 0 | 0 | 0 | 0 | 0 | 4 | 0 | 0 | 0 | 6 | 0 | 0 | 0 | 0 | 0 | 0 | 0 | 0 | 0 | 0 | 0 | 12 | 0 |
| ASV932 | 0dcac9c9a7c | 22 | 0 | 0 | 0 | 0 | 0 | 0 | 0 | 0 | 0 | 0 | 0 | 0 | 0 | 4 | 0 | 0 | 0 | 0 | 0 | 0 | 0 | 0 | 0 | 0 | 0 | 0 | 0 | 0 | 5 | 0 | 0 | 0 | 0 | 13 | 0 |
| ASV933 | 3fcb84cdb86 | 22 | 0 | 0 | 0 | 0 | 0 | 0 | 0 | 0 | 0 | 0 | 0 | 0 | 0 | 0 | 22 | 0 | 0 | 0 | 0 | 0 | 0 | 0 | 0 | 0 | 0 | 0 | 0 | 0 | 0 | 0 | 0 | 0 | 0 | 0 | 0 |
| ASV934 | 81df44c26a3 | 22 | 0 | 0 | 0 | 0 | 0 | 0 | 0 | 0 | 0 | 0 | 0 | 0 | 0 | 0 | 22 | 0 | 0 | 0 | 0 | 0 | 0 | 0 | 0 | 0 | 0 | 0 | 0 | 0 | 0 | 0 | 0 | 0 | 0 | 0 | 0 |
| ASV935 | b5270995850 | 22 | 0 | 0 | 22 | 0 | 0 | 0 | 0 | 0 | 0 | 0 | 0 | 0 | 0 | 0 | 0 | 0 | 0 | 0 | 0 | 0 | 0 | 0 | 0 | 0 | 0 | 0 | 0 | 0 | 0 | 0 | 0 | 0 | 0 | 0 | 0 |
| ASV936 | 9127d993e81 | 22 | 0 | 0 | 0 | 0 | 0 | 0 | 0 | 0 | 0 | 0 | 0 | 0 | 0 | 0 | 0 | 0 | 0 | 0 | 0 | 0 | 0 | 8 | 0 | 0 | 0 | 0 | 0 | 14 | 0 | 0 | 0 | 0 | 0 | 0 | 0 |
| ASV937 | d0fb0be18a4 | 22 | 0 | 0 | 0 | 0 | 0 | 0 | 0 | 0 | 0 | 0 | 0 | 0 | 0 | 0 | 0 | 0 | 4 | 0 | 10 | 0 | 0 | 0 | 0 | 0 | 0 | 0 | 0 | 8 | 0 | 0 | 0 | 0 | 0 | 0 | 0 |
| ASV938 | b76b8b9404d | 22 | 0 | 0 | 0 | 0 | 0 | 0 | 0 | 0 | 0 | 0 | 0 | 0 | 0 | 0 | 0 | 0 | 0 | 0 | 22 | 0 | 0 | 0 | 0 | 0 | 0 | 0 | 0 | 0 | 0 | 0 | 0 | 0 | 0 | 0 | 0 |
| ASV939 | 5da424385c3 | 22 | 0 | 0 | 0 | 0 | 4 | 0 | 0 | 0 | 11 | 7 | 0 | 0 | 0 | 0 | 0 | 0 | 0 | 0 | 0 | 0 | 0 | 0 | 0 | 0 | 0 | 0 | 0 | 0 | 0 | 0 | 0 | 0 | 0 | 0 | 0 |
| ASV940 | cc5a556f71fc | 22 | 0 | 0 | 0 | 0 | 0 | 0 | 0 | 0 | 0 | 0 | 0 | 2 | 0 | 0 | 0 | 0 | 9 | 0 | 0 | 0 | 0 | 0 | 0 | 0 | 0 | 7 | 0 | 0 | 0 | 4 | 0 | 0 | 0 | 0 | 0 |
| ASV941 | 6ed2144335f | 22 | 0 | 0 | 0 | 0 | 0 | 0 | 0 | 0 | 6 | 7 | 0 | 0 | 0 | 0 | 0 | 0 | 0 | 0 | 0 | 0 | 0 | 0 | 0 | 0 | 0 | 0 | 0 | 0 | 9 | 0 | 0 | 0 | 0 | 0 | 0 |
| ASV942 | 63d4c67c563 | 22 | 0 | 0 | 0 | 0 | 0 | 0 | 0 | 0 | 0 | 0 | 0 | 0 | 0 | 0 | 22 | 0 | 0 | 0 | 0 | 0 | 0 | 0 | 0 | 0 | 0 | 0 | 0 | 0 | 0 | 0 | 0 | 0 | 0 | 0 | 0 |
| ASV943 | 614bfd70d13 | 21 | 0 | 0 | 0 | 0 | 0 | 0 | 0 | 0 | 0 | 0 | 0 | 0 | 0 | 0 | 0 | 0 | 0 | 0 | 0 | 0 | 0 | 0 | 0 | 0 | 0 | 0 | 0 | 0 | 0 | 0 | 0 | 0 | 0 | 21 | 0 |
| ASV944 | 40e30ba7bf0 | 21 | 0 | 0 | 0 | 0 | 0 | 0 | 0 | 0 | 0 | 0 | 0 | 0 | 0 | 0 | 0 | 0 | 21 | 0 | 0 | 0 | 0 | 0 | 0 | 0 | 0 | 0 | 0 | 0 | 0 | 0 | 0 | 0 | 0 | 0 | 0 |
| ASV945 | d13f6e32063 | 21 | 0 | 0 | 0 | 0 | 10 | 0 | 0 | 0 | 3 | 0 | 0 | 0 | 0 | 0 | 0 | 0 | 0 | 0 | 0 | 0 | 0 | 0 | 0 | 0 | 0 | 0 | 0 | 8 | 0 | 0 | 0 | 0 | 0 | 0 | 0 |
| ASV946 | bedff13cff8c | 21 | 0 | 0 | 0 | 0 | 0 | 0 | 0 | 0 | 0 | 0 | 0 | 0 | 0 | 0 | 0 | 0 | 0 | 0 | 0 | 0 | 0 | 0 | 0 | 0 | 0 | 0 | 0 | 0 | 0 | 14 | 7 | 0 | 0 | 0 | 0 |
| ASV947 | c56cee2a8d5 | 21 | 0 | 0 | 0 | 0 | 0 | 0 | 0 | 0 | 0 | 0 | 0 | 0 | 0 | 0 | 0 | 0 | 0 | 0 | 0 | 0 | 0 | 0 | 0 | 0 | 0 | 0 | 0 | 0 | 0 | 21 | 0 | 0 | 0 | 0 | 0 |
| ASV948 | f1e4d878087 | 21 | 0 | 0 | 0 | 0 | 0 | 0 | 0 | 0 | 21 | 0 | 0 | 0 | 0 | 0 | 0 | 0 | 0 | 0 | 0 | 0 | 0 | 0 | 0 | 0 | 0 | 0 | 0 | 0 | 0 | 0 | 0 | 0 | 0 | 0 | 0 |
| ASV949 | b5179407010 | 21 | 0 | 0 | 0 | 0 | 0 | 0 | 0 | 0 | 0 | 0 | 0 | 0 | 0 | 0 | 0 | 10 | 11 | 0 | 0 | 0 | 0 | 0 | 0 | 0 | 0 | 0 | 0 | 0 | 0 | 0 | 0 | 0 | 0 | 0 | 0 |
| ASV950 | 80c777f382d | 21 | 0 | 0 | 0 | 0 | 0 | 0 | 0 | 0 | 21 | 0 | 0 | 0 | 0 | 0 | 0 | 0 | 0 | 0 | 0 | 0 | 0 | 0 | 0 | 0 | 0 | 0 | 0 | 0 | 0 | 0 | 0 | 0 | 0 | 0 | 0 |
| ASV951 | 85aaba025c8 | 21 | 21 | 0 | 0 | 0 | 0 | 0 | 0 | 0 | 0 | 0 | 0 | 0 | 0 | 0 | 0 | 0 | 0 | 0 | 0 | 0 | 0 | 0 | 0 | 0 | 0 | 0 | 0 | 0 | 0 | 0 | 0 | 0 | 0 | 0 | 0 |
| ASV952 | e7e36d3ba9a | 21 | 0 | 0 | 0 | 0 | 0 | 0 | 0 | 0 | 0 | 0 | 0 | 0 | 0 | 0 | 0 | 0 | 0 | 0 | 0 | 0 | 0 | 0 | 0 | 0 | 0 | 0 | 0 | 0 | 0 | 4 | 17 | 0 | 0 | 0 | 0 |
| ASV953 | ebba845db68 | 21 | 0 | 0 | 0 | 0 | 0 | 0 | 0 | 0 | 0 | 0 | 0 | 0 | 0 | 0 | 0 | 0 | 0 | 0 | 0 | 21 | 0 | 0 | 0 | 0 | 0 | 0 | 0 | 0 | 0 | 0 | 0 | 0 | 0 | 0 | 0 |
| ASV954 | a2934032a92 | 21 | 0 | 0 | 0 | 0 | 0 | 0 | 0 | 21 | 0 | 0 | 0 | 0 | 0 | 0 | 0 | 0 | 0 | 0 | 0 | 0 | 0 | 0 | 0 | 0 | 0 | 0 | 0 | 0 | 0 | 0 | 0 | 0 | 0 | 0 | 0 |
| ASV955 | 22eb0b7ab3d | 21 | 0 | 0 | 0 | 0 | 0 | 0 | 0 | 0 | 0 | 0 | 0 | 0 | 0 | 0 | 21 | 0 | 0 | 0 | 0 | 0 | 0 | 0 | 0 | 0 | 0 | 0 | 0 | 0 | 0 | 0 | 0 | 0 | 0 | 0 | 0 |
| ASV956 | 069529b71f2 | 21 | 0 | 0 | 0 | 0 | 0 | 0 | 0 | 0 | 0 | 0 | 0 | 0 | 0 | 0 | 0 | 0 | 0 | 0 | 0 | 0 | 0 | 0 | 0 | 0 | 0 | 0 | 0 | 0 | 0 | 0 | 0 | 0 | 0 | 21 | 0 |
| ASV957 | 506b89e9c4e | 21 | 0 | 0 | 0 | 0 | 0 | 0 | 0 | 0 | 0 | 0 | 0 | 0 | 0 | 0 | 0 | 0 | 0 | 0 | 0 | 0 | 0 | 0 | 0 | 0 | 0 | 0 | 0 | 0 | 0 | 0 | 0 | 0 | 21 | 0 | 0 |
| ASV958 | 455d144a95f | 21 | 0 | 0 | 0 | 0 | 0 | 0 | 0 | 0 | 0 | 0 | 0 | 0 | 0 | 9 | 12 | 0 | 0 | 0 | 0 | 0 | 0 | 0 | 0 | 0 | 0 | 0 | 0 | 0 | 0 | 0 | 0 | 0 | 0 | 0 | 0 |
| ASV959 | 4894ce1485e | 21 | 0 | 0 | 0 | 0 | 0 | 0 | 0 | 0 | 6 | 0 | 0 | 0 | 0 | 0 | 15 | 0 | 0 | 0 | 0 | 0 | 0 | 0 | 0 | 0 | 0 | 0 | 0 | 0 | 0 | 0 | 0 | 0 | 0 | 0 | 0 |
| ASV960 | 2266168a6e9 | 21 | 0 | 0 | 0 | 0 | 0 | 0 | 0 | 0 | 0 | 0 | 0 | 0 | 0 | 0 | 0 | 0 | 0 | 0 | 0 | 0 | 0 | 0 | 0 | 0 | 0 | 21 | 0 | 0 | 0 | 0 | 0 | 0 | 0 | 0 | 0 |
| ASV961 | 9b8029d5910 | 21 | 0 | 0 | 0 | 0 | 0 | 0 | 0 | 0 | 4 | 2 | 0 | 0 | 0 | 0 | 0 | 0 | 0 | 5 | 0 | 0 | 0 | 0 | 0 | 6 | 0 | 0 | 0 | 0 | 0 | 4 | 0 | 0 | 0 | 0 | 0 |
| ASV962 | c1603874309 | 21 | 0 | 0 | 0 | 0 | 0 | 0 | 0 | 0 | 0 | 0 | 0 | 0 | 0 | 0 | 0 | 0 | 0 | 0 | 0 | 0 | 0 | 0 | 0 | 0 | 0 | 0 | 0 | 0 | 0 | 0 | 0 | 0 | 0 | 15 | 6 |
| ASV963 | 2917e02786d | 21 | 0 | 0 | 0 | 0 | 0 | 0 | 0 | 0 | 0 | 0 | 0 | 0 | 0 | 0 | 0 | 0 | 0 | 0 | 0 | 0 | 0 | 0 | 0 | 0 | 0 | 8 | 0 | 0 | 0 | 6 | 0 | 7 | 0 | 0 | 0 |
| ASV964 | 1826547cb4f | 21 | 0 | 0 | 0 | 0 | 0 | 0 | 0 | 0 | 0 | 0 | 0 | 0 | 0 | 0 | 21 | 0 | 0 | 0 | 0 | 0 | 0 | 0 | 0 | 0 | 0 | 0 | 0 | 0 | 0 | 0 | 0 | 0 | 0 | 0 | 0 |
| ASV965 | 258ef60630a | 20 | 0 | 0 | 0 | 0 | 0 | 0 | 0 | 0 | 0 | 0 | 0 | 0 | 0 | 0 | 20 | 0 | 0 | 0 | 0 | 0 | 0 | 0 | 0 | 0 | 0 | 0 | 0 | 0 | 0 | 0 | 0 | 0 | 0 | 0 | 0 |
| ASV966 | 00fc329ff506 | 20 | 0 | 0 | 0 | 0 | 0 | 0 | 0 | 0 | 0 | 0 | 0 | 0 | 0 | 0 | 0 | 0 | 0 | 0 | 0 | 0 | 0 | 0 | 0 | 0 | 0 | 0 | 0 | 0 | 0 | 0 | 0 | 0 | 0 | 20 | 0 |
| ASV967 | 3b0a8656f57 | 20 | 0 | 0 | 0 | 0 | 14 | 0 | 0 | 0 | 0 | 0 | 0 | 0 | 0 | 0 | 6 | 0 | 0 | 0 | 0 | 0 | 0 | 0 | 0 | 0 | 0 | 0 | 0 | 0 | 0 | 0 | 0 | 0 | 0 | 0 | 0 |
| ASV968 | 13bebf1e665 | 20 | 0 | 0 | 0 | 0 | 0 | 0 | 0 | 0 | 0 | 0 | 0 | 0 | 0 | 0 | 0 | 0 | 0 | 0 | 0 | 0 | 0 | 0 | 0 | 0 | 0 | 0 | 0 | 0 | 0 | 12 | 8 | 0 | 0 | 0 | 0 |
| ASV969 | 113cf225927 | 20 | 0 | 0 | 0 | 0 | 0 | 0 | 0 | 0 | 0 | 0 | 0 | 0 | 0 | 0 | 20 | 0 | 0 | 0 | 0 | 0 | 0 | 0 | 0 | 0 | 0 | 0 | 0 | 0 | 0 | 0 | 0 | 0 | 0 | 0 | 0 |
| ASV970 | 1c1f30a08a7 | 20 | 0 | 0 | 0 | 0 | 0 | 0 | 0 | 0 | 0 | 0 | 0 | 0 | 0 | 0 | 0 | 0 | 20 | 0 | 0 | 0 | 0 | 0 | 0 | 0 | 0 | 0 | 0 | 0 | 0 | 0 | 0 | 0 | 0 | 0 | 0 |
| ASV971 | 960a44e0d9d | 20 | 0 | 0 | 0 | 0 | 3 | 0 | 0 | 0 | 0 | 0 | 0 | 0 | 0 | 0 | 0 | 0 | 0 | 0 | 0 | 0 | 0 | 0 | 0 | 0 | 0 | 0 | 0 | 0 | 0 | 3 | 0 | 0 | 0 | 0 | 14 |
| ASV972 | 9ec187f0262 | 20 | 0 | 0 | 0 | 0 | 0 | 0 | 0 | 0 | 0 | 0 | 0 | 0 | 0 | 0 | 0 | 0 | 0 | 6 | 0 | 0 | 0 | 0 | 0 | 0 | 0 | 0 | 0 | 0 | 7 | 7 | 0 | 0 | 0 | 0 | 0 |
| ASV973 | 5b41fc61b06 | 20 | 0 | 0 | 0 | 0 | 0 | 0 | 0 | 0 | 0 | 0 | 0 | 0 | 0 | 0 | 0 | 0 | 0 | 0 | 0 | 0 | 0 | 0 | 0 | 0 | 0 | 0 | 0 | 0 | 0 | 0 | 0 | 0 | 0 | 0 | 20 |
| ASV974 | 71ac71a7ad6 | 20 | 0 | 0 | 0 | 0 | 0 | 0 | 0 | 0 | 0 | 0 | 0 | 0 | 0 | 0 | 0 | 0 | 0 | 0 | 0 | 0 | 0 | 0 | 0 | 0 | 0 | 0 | 0 | 0 | 0 | 0 | 0 | 0 | 0 | 20 | 0 |
| ASV975 | 9ac2bce222b | 20 | 0 | 0 | 0 | 0 | 7 | 0 | 0 | 0 | 0 | 0 | 0 | 0 | 0 | 0 | 13 | 0 | 0 | 0 | 0 | 0 | 0 | 0 | 0 | 0 | 0 | 0 | 0 | 0 | 0 | 0 | 0 | 0 | 0 | 0 | 0 |
| ASV976 | 88968e43eec | 20 | 0 | 0 | 0 | 0 | 0 | 0 | 0 | 0 | 0 | 0 | 0 | 0 | 0 | 0 | 0 | 0 | 0 | 0 | 0 | 0 | 0 | 0 | 0 | 8 | 0 | 0 | 0 | 0 | 0 | 0 | 0 | 0 | 0 | 12 | 0 |
| ASV977 | 81d321d2a60 | 19 | 0 | 0 | 0 | 0 | 0 | 0 | 0 | 0 | 0 | 0 | 0 | 0 | 0 | 0 | 0 | 0 | 0 | 0 | 0 | 0 | 0 | 0 | 0 | 0 | 0 | 0 | 0 | 0 | 0 | 0 | 19 | 0 | 0 | 0 | 0 |
| ASV978 | e705e1eac9a | 19 | 0 | 0 | 0 | 0 | 0 | 0 | 0 | 0 | 0 | 0 | 0 | 0 | 0 | 0 | 0 | 0 | 0 | 0 | 0 | 0 | 0 | 0 | 0 | 0 | 0 | 0 | 0 | 0 | 0 | 0 | 19 | 0 | 0 | 0 | 0 |
| ASV979 | 7dd3449b4a5 | 19 | 0 | 0 | 0 | 0 | 0 | 0 | 0 | 0 | 0 | 0 | 0 | 0 | 0 | 0 | 14 | 0 | 0 | 0 | 0 | 0 | 0 | 0 | 0 | 0 | 0 | 0 | 0 | 0 | 0 | 5 | 0 | 0 | 0 | 0 | 0 |
| ASV980 | d5c306e14c9 | 19 | 0 | 0 | 0 | 0 | 0 | 0 | 0 | 0 | 19 | 0 | 0 | 0 | 0 | 0 | 0 | 0 | 0 | 0 | 0 | 0 | 0 | 0 | 0 | 0 | 0 | 0 | 0 | 0 | 0 | 0 | 0 | 0 | 0 | 0 | 0 |
| ASV981 | 76c7ab14ae6 | 19 | 0 | 0 | 0 | 0 | 0 | 0 | 0 | 0 | 0 | 0 | 0 | 0 | 0 | 0 | 19 | 0 | 0 | 0 | 0 | 0 | 0 | 0 | 0 | 0 | 0 | 0 | 0 | 0 | 0 | 0 | 0 | 0 | 0 | 0 | 0 |
| ASV982 | 2a397771272 | 19 | 0 | 0 | 0 | 0 | 0 | 0 | 0 | 0 | 0 | 0 | 0 | 0 | 0 | 0 | 0 | 0 | 0 | 0 | 0 | 0 | 0 | 0 | 0 | 0 | 7 | 0 | 0 | 0 | 0 | 12 | 0 | 0 | 0 | 0 | 0 |
| ASV983 | 6a9c0583ee9 | 19 | 0 | 0 | 0 | 0 | 19 | 0 | 0 | 0 | 0 | 0 | 0 | 0 | 0 | 0 | 0 | 0 | 0 | 0 | 0 | 0 | 0 | 0 | 0 | 0 | 0 | 0 | 0 | 0 | 0 | 0 | 0 | 0 | 0 | 0 | 0 |
| ASV984 | a048e53366d | 19 | 0 | 0 | 0 | 19 | 0 | 0 | 0 | 0 | 0 | 0 | 0 | 0 | 0 | 0 | 0 | 0 | 0 | 0 | 0 | 0 | 0 | 0 | 0 | 0 | 0 | 0 | 0 | 0 | 0 | 0 | 0 | 0 | 0 | 0 | 0 |
| ASV985 | 71ed703262a | 19 | 0 | 0 | 0 | 0 | 0 | 0 | 0 | 0 | 0 | 0 | 0 | 0 | 0 | 0 | 0 | 0 | 16 | 0 | 0 | 0 | 3 | 0 | 0 | 0 | 0 | 0 | 0 | 0 | 0 | 0 | 0 | 0 | 0 | 0 | 0 |
| ASV986 | 6abff36b05c | 19 | 0 | 0 | 0 | 0 | 0 | 0 | 0 | 0 | 0 | 0 | 0 | 0 | 0 | 0 | 0 | 0 | 0 | 0 | 0 | 0 | 0 | 0 | 0 | 0 | 0 | 0 | 0 | 0 | 0 | 0 | 9 | 10 | 0 | 0 | 0 |
| ASV987 | 90cad105041 | 19 | 0 | 0 | 0 | 0 | 0 | 0 | 0 | 0 | 0 | 0 | 0 | 0 | 0 | 0 | 0 | 0 | 0 | 0 | 0 | 19 | 0 | 0 | 0 | 0 | 0 | 0 | 0 | 0 | 0 | 0 | 0 | 0 | 0 | 0 | 0 |
| ASV988 | 0b6e5f9a4e9 | 19 | 0 | 0 | 0 | 0 | 0 | 0 | 0 | 0 | 0 | 0 | 0 | 0 | 0 | 0 | 0 | 0 | 0 | 0 | 0 | 0 | 0 | 0 | 0 | 0 | 0 | 0 | 0 | 0 | 0 | 0 | 0 | 0 | 0 | 19 | 0 |
| ASV989 | d15181c78bf | 19 | 0 | 0 | 0 | 9 | 10 | 0 | 0 | 0 | 0 | 0 | 0 | 0 | 0 | 0 | 0 | 0 | 0 | 0 | 0 | 0 | 0 | 0 | 0 | 0 | 0 | 0 | 0 | 0 | 0 | 0 | 0 | 0 | 0 | 0 | 0 |
| ASV990 | 2ee85a51dff | 19 | 0 | 0 | 9 | 10 | 0 | 0 | 0 | 0 | 0 | 0 | 0 | 0 | 0 | 0 | 0 | 0 | 0 | 0 | 0 | 0 | 0 | 0 | 0 | 0 | 0 | 0 | 0 | 0 | 0 | 0 | 0 | 0 | 0 | 0 | 0 |
| ASV991 | eaefb3e09d0 | 19 | 0 | 0 | 0 | 0 | 0 | 0 | 0 | 0 | 19 | 0 | 0 | 0 | 0 | 0 | 0 | 0 | 0 | 0 | 0 | 0 | 0 | 0 | 0 | 0 | 0 | 0 | 0 | 0 | 0 | 0 | 0 | 0 | 0 | 0 | 0 |
| ASV992 | a636832e0bc | 19 | 0 | 0 | 0 | 0 | 0 | 0 | 0 | 0 | 0 | 0 | 0 | 0 | 0 | 0 | 0 | 8 | 6 | 0 | 0 | 0 | 0 | 0 | 0 | 0 | 0 | 5 | 0 | 0 | 0 | 0 | 0 | 0 | 0 | 0 | 0 |
| ASV993 | 9724d2faa93 | 19 | 0 | 0 | 0 | 0 | 0 | 0 | 0 | 0 | 0 | 0 | 0 | 0 | 0 | 0 | 0 | 0 | 0 | 0 | 0 | 0 | 0 | 0 | 19 | 0 | 0 | 0 | 0 | 0 | 0 | 0 | 0 | 0 | 0 | 0 | 0 |
| ASV994 | 4a920fd9646 | 19 | 0 | 0 | 0 | 0 | 0 | 0 | 0 | 0 | 0 | 0 | 0 | 0 | 0 | 0 | 19 | 0 | 0 | 0 | 0 | 0 | 0 | 0 | 0 | 0 | 0 | 0 | 0 | 0 | 0 | 0 | 0 | 0 | 0 | 0 | 0 |
| ASV995 | 4128e2dd0f8 | 19 | 0 | 0 | 0 | 0 | 0 | 0 | 0 | 0 | 0 | 9 | 0 | 0 | 0 | 0 | 0 | 0 | 0 | 0 | 0 | 0 | 0 | 0 | 0 | 0 | 0 | 0 | 0 | 0 | 0 | 0 | 0 | 0 | 0 | 0 | 10 |
| ASV996 | 64cf84b7200 | 19 | 0 | 0 | 0 | 0 | 0 | 0 | 0 | 0 | 19 | 0 | 0 | 0 | 0 | 0 | 0 | 0 | 0 | 0 | 0 | 0 | 0 | 0 | 0 | 0 | 0 | 0 | 0 | 0 | 0 | 0 | 0 | 0 | 0 | 0 | 0 |
| ASV997 | 113bef0f47c | 18 | 0 | 0 | 18 | 0 | 0 | 0 | 0 | 0 | 0 | 0 | 0 | 0 | 0 | 0 | 0 | 0 | 0 | 0 | 0 | 0 | 0 | 0 | 0 | 0 | 0 | 0 | 0 | 0 | 0 | 0 | 0 | 0 | 0 | 0 | 0 |
| ASV998 | 8fcb01aba87 | 18 | 0 | 0 | 0 | 0 | 0 | 0 | 0 | 0 | 0 | 0 | 0 | 0 | 0 | 0 | 0 | 0 | 6 | 0 | 0 | 0 | 0 | 0 | 0 | 0 | 0 | 5 | 0 | 0 | 0 | 0 | 0 | 0 | 0 | 7 | 0 |
| ASV999 | 8d424edbd58 | 18 | 0 | 0 | 0 | 0 | 0 | 0 | 0 | 0 | 0 | 0 | 0 | 0 | 0 | 0 | 0 | 0 | 0 | 0 | 18 | 0 | 0 | 0 | 0 | 0 | 0 | 0 | 0 | 0 | 0 | 0 | 0 | 0 | 0 | 0 | 0 |
| ASV1000 | 6c524597d23 | 18 | 0 | 0 | 0 | 0 | 18 | 0 | 0 | 0 | 0 | 0 | 0 | 0 | 0 | 0 | 0 | 0 | 0 | 0 | 0 | 0 | 0 | 0 | 0 | 0 | 0 | 0 | 0 | 0 | 0 | 0 | 0 | 0 | 0 | 0 | 0 |
| ASV1001 | 8dd8b95a544 | 18 | 0 | 0 | 0 | 0 | 0 | 0 | 0 | 0 | 0 | 0 | 0 | 0 | 0 | 0 | 0 | 0 | 0 | 0 | 0 | 0 | 0 | 0 | 0 | 0 | 0 | 0 | 0 | 18 | 0 | 0 | 0 | 0 | 0 | 0 | 0 |
| ASV1002 | 0d510288bd3 | 18 | 0 | 0 | 0 | 0 | 0 | 0 | 0 | 0 | 0 | 0 | 0 | 0 | 0 | 0 | 0 | 0 | 0 | 0 | 0 | 0 | 0 | 0 | 0 | 0 | 18 | 0 | 0 | 0 | 0 | 0 | 0 | 0 | 0 | 0 | 0 |
| ASV1003 | 93de0b37864 | 18 | 0 | 0 | 0 | 0 | 0 | 18 | 0 | 0 | 0 | 0 | 0 | 0 | 0 | 0 | 0 | 0 | 0 | 0 | 0 | 0 | 0 | 0 | 0 | 0 | 0 | 0 | 0 | 0 | 0 | 0 | 0 | 0 | 0 | 0 | 0 |
| ASV1004 | 19522f29938 | 18 | 0 | 0 | 0 | 0 | 0 | 0 | 0 | 0 | 0 | 0 | 0 | 0 | 0 | 0 | 0 | 0 | 0 | 0 | 0 | 0 | 0 | 0 | 0 | 0 | 0 | 0 | 0 | 0 | 0 | 0 | 0 | 0 | 0 | 18 | 0 |
| ASV1005 | f19bc9d4642 | 18 | 5 | 0 | 0 | 0 | 0 | 0 | 0 | 0 | 0 | 0 | 0 | 0 | 0 | 0 | 0 | 0 | 0 | 0 | 0 | 0 | 0 | 0 | 0 | 0 | 0 | 0 | 0 | 0 | 0 | 0 | 0 | 0 | 13 | 0 | 0 |
| ASV1006 | c149a6e3943 | 18 | 0 | 0 | 0 | 0 | 0 | 0 | 0 | 0 | 0 | 18 | 0 | 0 | 0 | 0 | 0 | 0 | 0 | 0 | 0 | 0 | 0 | 0 | 0 | 0 | 0 | 0 | 0 | 0 | 0 | 0 | 0 | 0 | 0 | 0 | 0 |
| ASV1007 | a41ea6398c2 | 18 | 0 | 0 | 0 | 0 | 0 | 0 | 0 | 0 | 0 | 0 | 0 | 0 | 0 | 0 | 0 | 0 | 0 | 0 | 0 | 0 | 0 | 0 | 0 | 0 | 0 | 0 | 0 | 0 | 0 | 0 | 18 | 0 | 0 | 0 | 0 |
| ASV1008 | 466deaca70e | 18 | 0 | 0 | 0 | 0 | 0 | 0 | 0 | 0 | 0 | 0 | 0 | 0 | 0 | 0 | 0 | 0 | 18 | 0 | 0 | 0 | 0 | 0 | 0 | 0 | 0 | 0 | 0 | 0 | 0 | 0 | 0 | 0 | 0 | 0 | 0 |
| ASV1009 | e19ffd6d48b | 18 | 0 | 0 | 0 | 0 | 18 | 0 | 0 | 0 | 0 | 0 | 0 | 0 | 0 | 0 | 0 | 0 | 0 | 0 | 0 | 0 | 0 | 0 | 0 | 0 | 0 | 0 | 0 | 0 | 0 | 0 | 0 | 0 | 0 | 0 | 0 |
| ASV1010 | 9799f032f71 | 18 | 0 | 0 | 0 | 0 | 0 | 0 | 0 | 0 | 0 | 0 | 0 | 0 | 0 | 0 | 0 | 0 | 0 | 0 | 0 | 0 | 0 | 0 | 0 | 0 | 0 | 0 | 0 | 0 | 0 | 0 | 0 | 0 | 0 | 0 | 18 |
| ASV1011 | 18ba657a311 | 18 | 0 | 0 | 0 | 0 | 0 | 0 | 0 | 0 | 0 | 0 | 0 | 0 | 0 | 0 | 18 | 0 | 0 | 0 | 0 | 0 | 0 | 0 | 0 | 0 | 0 | 0 | 0 | 0 | 0 | 0 | 0 | 0 | 0 | 0 | 0 |
| ASV1012 | 15ff669914e | 18 | 0 | 0 | 2 | 0 | 0 | 0 | 0 | 0 | 0 | 0 | 0 | 0 | 0 | 0 | 5 | 0 | 0 | 0 | 0 | 0 | 0 | 0 | 0 | 6 | 0 | 0 | 0 | 0 | 0 | 5 | 0 | 0 | 0 | 0 | 0 |

**Supplementary Table S5.** Bacterioplankton 16S rRNA ASV feature table
ASVs: 1779
Seqs: 918792

| #ASV_ID | Feature_ID | total seqs | May12_0m | May12_05m | May12_25m | May12_50m | May12_75m | Jun8_0m | Jun8_05m | Jun8_25m | Jun8_50m | Jun8_75m | Jun22_0m | Jun22_05m | Jun22_25m | Jun22_50m | Jun22_75m | Jul11_0m | Jul11_05m | Jul11_25m | Jul11_50m | Jul11_75m | Jul27_0m | Jul27_05m | Jul27_25m | Jul27_50m | Jul27_75m | Aug5_0m | Aug5_05m | Aug5_25m | Aug5_50m | Aug5_75m | Sep8_0m | Sep8_05m | Sep8_25m | Sep8_50m | Sep8_75m |
|---|---|---|---|---|---|---|---|---|---|---|---|---|---|---|---|---|---|---|---|---|---|---|---|---|---|---|---|---|---|---|---|---|---|---|---|---|---|
| ASV1013 | 1649df89897 | 18 | 0 | 0 | 0 | 0 | 0 | 0 | 0 | 0 | 0 | 0 | 0 | 0 | 0 | 0 | 0 | 0 | 0 | 0 | 0 | 0 | 0 | 0 | 0 | 0 | 0 | 0 | 0 | 0 | 0 | 0 | 18 | 0 | 0 | 0 | 0 |
| ASV1014 | 1f9784a5b95 | 18 | 0 | 0 | 0 | 0 | 18 | 0 | 0 | 0 | 0 | 0 | 0 | 0 | 0 | 0 | 0 | 0 | 0 | 0 | 0 | 0 | 0 | 0 | 0 | 0 | 0 | 0 | 0 | 0 | 0 | 0 | 0 | 0 | 0 | 0 | 0 |
| ASV1015 | 9bfa918711a | 18 | 0 | 0 | 0 | 0 | 14 | 0 | 0 | 0 | 0 | 0 | 0 | 0 | 0 | 0 | 0 | 0 | 0 | 0 | 0 | 0 | 0 | 0 | 0 | 0 | 0 | 0 | 0 | 0 | 0 | 4 | 0 | 0 | 0 | 0 | 0 |
| ASV1016 | 53b1d653360 | 17 | 0 | 0 | 0 | 0 | 6 | 0 | 0 | 0 | 0 | 0 | 0 | 0 | 0 | 11 | 0 | 0 | 0 | 0 | 0 | 0 | 0 | 0 | 0 | 0 | 0 | 0 | 0 | 0 | 0 | 0 | 0 | 0 | 0 | 0 | 0 |
| ASV1017 | 10c2e08bb7c | 17 | 0 | 0 | 0 | 0 | 0 | 0 | 0 | 0 | 0 | 0 | 0 | 0 | 0 | 0 | 17 | 0 | 0 | 0 | 0 | 0 | 0 | 0 | 0 | 0 | 0 | 0 | 0 | 0 | 0 | 0 | 0 | 0 | 0 | 0 | 0 |
| ASV1018 | 242e410bd5e | 17 | 0 | 0 | 0 | 0 | 0 | 0 | 0 | 0 | 0 | 0 | 0 | 0 | 0 | 0 | 0 | 8 | 2 | 0 | 2 | 0 | 0 | 0 | 0 | 3 | 0 | 0 | 0 | 0 | 0 | 0 | 2 | 0 | 0 | 0 | 0 |
| ASV1019 | 3cf3e186111 | 17 | 0 | 0 | 0 | 0 | 10 | 0 | 0 | 0 | 0 | 0 | 0 | 0 | 0 | 0 | 7 | 0 | 0 | 0 | 0 | 0 | 0 | 0 | 0 | 0 | 0 | 0 | 0 | 0 | 0 | 0 | 0 | 0 | 0 | 0 | 0 |
| ASV1020 | f00e595d02d | 17 | 0 | 0 | 0 | 0 | 0 | 0 | 0 | 1 | 0 | 0 | 0 | 0 | 0 | 0 | 0 | 0 | 0 | 0 | 0 | 0 | 0 | 0 | 0 | 0 | 0 | 0 | 0 | 0 | 0 | 8 | 0 | 0 | 0 | 0 | 8 |
| ASV1021 | 1aebeafe2e1 | 17 | 0 | 0 | 0 | 0 | 0 | 0 | 0 | 0 | 0 | 0 | 0 | 0 | 0 | 0 | 0 | 0 | 0 | 0 | 0 | 0 | 0 | 0 | 0 | 0 | 0 | 0 | 0 | 0 | 0 | 0 | 0 | 0 | 0 | 0 | 17 |
| ASV1022 | 6b291baa9e0 | 17 | 0 | 8 | 9 | 0 | 0 | 0 | 0 | 0 | 0 | 0 | 0 | 0 | 0 | 0 | 0 | 0 | 0 | 0 | 0 | 0 | 0 | 0 | 0 | 0 | 0 | 0 | 0 | 0 | 0 | 0 | 0 | 0 | 0 | 0 | 0 |
| ASV1023 | ec5527dcd57 | 17 | 0 | 0 | 0 | 0 | 17 | 0 | 0 | 0 | 0 | 0 | 0 | 0 | 0 | 0 | 0 | 0 | 0 | 0 | 0 | 0 | 0 | 0 | 0 | 0 | 0 | 0 | 0 | 0 | 0 | 0 | 0 | 0 | 0 | 0 | 0 |
| ASV1024 | 5878eee3344 | 17 | 0 | 0 | 17 | 0 | 0 | 0 | 0 | 0 | 0 | 0 | 0 | 0 | 0 | 0 | 0 | 0 | 0 | 0 | 0 | 0 | 0 | 0 | 0 | 0 | 0 | 0 | 0 | 0 | 0 | 0 | 0 | 0 | 0 | 0 | 0 |
| ASV1025 | 8b3ee47d66e | 17 | 0 | 0 | 0 | 0 | 0 | 0 | 0 | 0 | 0 | 0 | 0 | 0 | 0 | 0 | 0 | 17 | 0 | 0 | 0 | 0 | 0 | 0 | 0 | 0 | 0 | 0 | 0 | 0 | 0 | 0 | 0 | 0 | 0 | 0 | 0 |
| ASV1026 | ef44e064fa6 | 17 | 0 | 0 | 17 | 0 | 0 | 0 | 0 | 0 | 0 | 0 | 0 | 0 | 0 | 0 | 0 | 0 | 0 | 0 | 0 | 0 | 0 | 0 | 0 | 0 | 0 | 0 | 0 | 0 | 0 | 0 | 0 | 0 | 0 | 0 | 0 |
| ASV1027 | 3b8edc60c17 | 17 | 0 | 0 | 0 | 6 | 7 | 0 | 0 | 0 | 0 | 4 | 0 | 0 | 0 | 0 | 0 | 0 | 0 | 0 | 0 | 0 | 0 | 0 | 0 | 0 | 0 | 0 | 0 | 0 | 0 | 0 | 0 | 0 | 0 | 0 | 0 |
| ASV1028 | 934d5a60529 | 17 | 0 | 0 | 0 | 0 | 0 | 0 | 0 | 0 | 0 | 0 | 0 | 0 | 0 | 0 | 9 | 0 | 0 | 0 | 0 | 0 | 0 | 0 | 0 | 0 | 0 | 0 | 0 | 0 | 0 | 0 | 0 | 0 | 0 | 0 | 8 |
| ASV1029 | 1b4b18c3282 | 17 | 0 | 0 | 0 | 0 | 0 | 0 | 0 | 0 | 0 | 0 | 0 | 0 | 0 | 0 | 0 | 0 | 0 | 0 | 0 | 0 | 0 | 0 | 0 | 0 | 17 | 0 | 0 | 0 | 0 | 0 | 0 | 0 | 0 | 0 | 0 |
| ASV1030 | f93e5daac2d | 17 | 10 | 0 | 7 | 0 | 0 | 0 | 0 | 0 | 0 | 0 | 0 | 0 | 0 | 0 | 0 | 0 | 0 | 0 | 0 | 0 | 0 | 0 | 0 | 0 | 0 | 0 | 0 | 0 | 0 | 0 | 0 | 0 | 0 | 0 | 0 |
| ASV1031 | 64cbd96da71 | 17 | 0 | 0 | 0 | 0 | 0 | 0 | 0 | 10 | 0 | 0 | 0 | 0 | 0 | 0 | 0 | 0 | 0 | 0 | 0 | 0 | 0 | 0 | 0 | 0 | 0 | 0 | 0 | 0 | 0 | 0 | 0 | 0 | 0 | 7 | 0 |
| ASV1032 | 420b912eeb1 | 16 | 0 | 0 | 0 | 0 | 0 | 0 | 0 | 0 | 0 | 0 | 0 | 0 | 0 | 0 | 16 | 0 | 0 | 0 | 0 | 0 | 0 | 0 | 0 | 0 | 0 | 0 | 0 | 0 | 0 | 0 | 0 | 0 | 0 | 0 | 0 |
| ASV1033 | 91795ad5936 | 16 | 0 | 0 | 16 | 0 | 0 | 0 | 0 | 0 | 0 | 0 | 0 | 0 | 0 | 0 | 0 | 0 | 0 | 0 | 0 | 0 | 0 | 0 | 0 | 0 | 0 | 0 | 0 | 0 | 0 | 0 | 0 | 0 | 0 | 0 | 0 |
| ASV1034 | 1c88d362891 | 16 | 0 | 0 | 0 | 0 | 0 | 0 | 0 | 0 | 0 | 0 | 0 | 0 | 0 | 0 | 0 | 0 | 0 | 0 | 0 | 0 | 0 | 0 | 0 | 0 | 0 | 0 | 0 | 0 | 0 | 0 | 16 | 0 | 0 | 0 | 0 |
| ASV1035 | c1c1f1fe56e1 | 16 | 0 | 0 | 0 | 0 | 0 | 0 | 0 | 0 | 0 | 0 | 0 | 0 | 0 | 0 | 12 | 0 | 0 | 0 | 0 | 4 | 0 | 0 | 0 | 0 | 0 | 0 | 0 | 0 | 0 | 0 | 0 | 0 | 0 | 0 | 0 |
| ASV1036 | 0b4c0501d3f | 16 | 0 | 0 | 0 | 0 | 0 | 0 | 0 | 0 | 0 | 0 | 0 | 0 | 0 | 0 | 0 | 0 | 0 | 0 | 16 | 0 | 0 | 0 | 0 | 0 | 0 | 0 | 0 | 0 | 0 | 0 | 0 | 0 | 0 | 0 | 0 |
| ASV1037 | 3fdc3646c1d | 16 | 0 | 0 | 16 | 0 | 0 | 0 | 0 | 0 | 0 | 0 | 0 | 0 | 0 | 0 | 0 | 0 | 0 | 0 | 0 | 0 | 0 | 0 | 0 | 0 | 0 | 0 | 0 | 0 | 0 | 0 | 0 | 0 | 0 | 0 | 0 |
| ASV1038 | b2f015f98dc | 16 | 0 | 0 | 0 | 9 | 0 | 0 | 0 | 0 | 0 | 0 | 0 | 0 | 0 | 0 | 0 | 0 | 0 | 0 | 0 | 0 | 0 | 0 | 0 | 0 | 7 | 0 | 0 | 0 | 0 | 0 | 0 | 0 | 0 | 0 | 0 |
| ASV1039 | 3413244e718 | 16 | 0 | 0 | 0 | 0 | 0 | 0 | 0 | 0 | 9 | 7 | 0 | 0 | 0 | 0 | 0 | 0 | 0 | 0 | 0 | 0 | 0 | 0 | 0 | 0 | 0 | 0 | 0 | 0 | 0 | 0 | 0 | 0 | 0 | 0 | 0 |
| ASV1040 | 7fb0f37f4c87 | 16 | 0 | 0 | 0 | 0 | 6 | 0 | 0 | 0 | 0 | 0 | 0 | 0 | 0 | 0 | 10 | 0 | 0 | 0 | 0 | 0 | 0 | 0 | 0 | 0 | 0 | 0 | 0 | 0 | 0 | 0 | 0 | 0 | 0 | 0 | 0 |
| ASV1041 | c83a07183f9 | 16 | 0 | 0 | 0 | 0 | 0 | 0 | 0 | 0 | 0 | 16 | 0 | 0 | 0 | 0 | 0 | 0 | 0 | 0 | 0 | 0 | 0 | 0 | 0 | 0 | 0 | 0 | 0 | 0 | 0 | 0 | 0 | 0 | 0 | 0 | 0 |
| ASV1042 | 779040b5940 | 16 | 0 | 0 | 0 | 0 | 0 | 0 | 0 | 0 | 0 | 0 | 0 | 0 | 0 | 0 | 0 | 0 | 0 | 0 | 0 | 0 | 0 | 0 | 0 | 0 | 0 | 0 | 0 | 0 | 0 | 0 | 16 | 0 | 0 | 0 | 0 |
| ASV1043 | 3fb9aab89b2 | 16 | 0 | 0 | 0 | 0 | 6 | 0 | 0 | 0 | 0 | 0 | 0 | 0 | 0 | 0 | 0 | 0 | 0 | 0 | 0 | 0 | 0 | 0 | 0 | 0 | 0 | 0 | 0 | 0 | 0 | 0 | 0 | 0 | 0 | 0 | 10 |
| ASV1044 | 6a148edba63 | 16 | 0 | 0 | 0 | 0 | 0 | 0 | 0 | 0 | 0 | 0 | 0 | 0 | 0 | 0 | 0 | 0 | 0 | 0 | 0 | 0 | 0 | 0 | 5 | 0 | 0 | 0 | 0 | 0 | 0 | 0 | 11 | 0 | 0 | 0 | 0 |
| ASV1045 | 7c2b94b80c0 | 16 | 7 | 0 | 9 | 0 | 0 | 0 | 0 | 0 | 0 | 0 | 0 | 0 | 0 | 0 | 0 | 0 | 0 | 0 | 0 | 0 | 0 | 0 | 0 | 0 | 0 | 0 | 0 | 0 | 0 | 0 | 0 | 0 | 0 | 0 | 0 |
| ASV1046 | be8e3cb9a0a | 16 | 0 | 0 | 0 | 0 | 0 | 0 | 0 | 0 | 0 | 0 | 0 | 0 | 0 | 0 | 0 | 0 | 0 | 0 | 0 | 0 | 0 | 0 | 0 | 0 | 0 | 0 | 0 | 0 | 0 | 0 | 0 | 0 | 0 | 0 | 16 |
| ASV1047 | 674c3042c46 | 16 | 0 | 0 | 0 | 0 | 0 | 0 | 0 | 0 | 0 | 0 | 0 | 0 | 0 | 0 | 0 | 0 | 0 | 0 | 16 | 0 | 0 | 0 | 0 | 0 | 0 | 0 | 0 | 0 | 0 | 0 | 0 | 0 | 0 | 0 | 0 |
| ASV1048 | 6d95c423bbd | 16 | 0 | 0 | 0 | 0 | 0 | 0 | 0 | 0 | 0 | 0 | 0 | 0 | 0 | 16 | 0 | 0 | 0 | 0 | 0 | 0 | 0 | 0 | 0 | 0 | 0 | 0 | 0 | 0 | 0 | 0 | 0 | 0 | 0 | 0 | 0 |
| ASV1049 | 2debcf34b91 | 16 | 0 | 0 | 0 | 0 | 0 | 0 | 0 | 0 | 0 | 0 | 0 | 0 | 0 | 0 | 0 | 0 | 0 | 0 | 5 | 0 | 0 | 0 | 0 | 0 | 0 | 0 | 0 | 0 | 11 | 0 | 0 | 0 | 0 | 0 | 0 |
| ASV1050 | 32cc406e27d | 16 | 0 | 0 | 0 | 0 | 0 | 0 | 0 | 0 | 0 | 0 | 0 | 0 | 0 | 0 | 0 | 0 | 0 | 0 | 0 | 0 | 0 | 0 | 0 | 0 | 0 | 0 | 0 | 0 | 0 | 0 | 3 | 0 | 0 | 13 | 0 |
| ASV1051 | 5dc579902e5 | 16 | 0 | 0 | 0 | 0 | 0 | 0 | 0 | 0 | 0 | 0 | 0 | 0 | 0 | 0 | 0 | 0 | 0 | 0 | 0 | 0 | 0 | 0 | 0 | 3 | 0 | 0 | 0 | 0 | 0 | 5 | 0 | 0 | 0 | 0 | 8 |
| ASV1052 | 9a03cc45e25 | 16 | 0 | 0 | 0 | 0 | 0 | 0 | 0 | 0 | 0 | 0 | 0 | 0 | 0 | 5 | 0 | 0 | 0 | 0 | 0 | 0 | 0 | 0 | 0 | 0 | 0 | 0 | 0 | 0 | 11 | 0 | 0 | 0 | 0 | 0 | 0 |
| ASV1053 | 01780edb0ab | 16 | 0 | 0 | 0 | 0 | 3 | 0 | 0 | 0 | 5 | 0 | 0 | 0 | 0 | 0 | 8 | 0 | 0 | 0 | 0 | 0 | 0 | 0 | 0 | 0 | 0 | 0 | 0 | 0 | 0 | 0 | 0 | 0 | 0 | 0 | 0 |
| ASV1054 | eb2d8cd6658 | 16 | 0 | 0 | 6 | 3 | 7 | 0 | 0 | 0 | 0 | 0 | 0 | 0 | 0 | 0 | 0 | 0 | 0 | 0 | 0 | 0 | 0 | 0 | 0 | 0 | 0 | 0 | 0 | 0 | 0 | 0 | 0 | 0 | 0 | 0 | 0 |
| ASV1055 | 7a9313e2d82 | 16 | 4 | 0 | 0 | 0 | 2 | 0 | 0 | 0 | 0 | 0 | 0 | 10 | 0 | 0 | 0 | 0 | 0 | 0 | 0 | 0 | 0 | 0 | 0 | 0 | 0 | 0 | 0 | 0 | 0 | 0 | 0 | 0 | 0 | 0 | 0 |
| ASV1056 | 039d12c3a75 | 16 | 0 | 0 | 0 | 0 | 0 | 0 | 0 | 0 | 0 | 0 | 0 | 0 | 0 | 0 | 0 | 0 | 0 | 0 | 0 | 0 | 0 | 0 | 0 | 0 | 0 | 0 | 0 | 0 | 0 | 0 | 16 | 0 | 0 | 0 | 0 |
| ASV1057 | 2b1673ad9db | 16 | 0 | 0 | 16 | 0 | 0 | 0 | 0 | 0 | 0 | 0 | 0 | 0 | 0 | 0 | 0 | 0 | 0 | 0 | 0 | 0 | 0 | 0 | 0 | 0 | 0 | 0 | 0 | 0 | 0 | 0 | 0 | 0 | 0 | 0 | 0 |
| ASV1058 | 8c823447f40 | 15 | 0 | 0 | 0 | 0 | 0 | 0 | 0 | 0 | 5 | 0 | 0 | 0 | 0 | 0 | 10 | 0 | 0 | 0 | 0 | 0 | 0 | 0 | 0 | 0 | 0 | 0 | 0 | 0 | 0 | 0 | 0 | 0 | 0 | 0 | 0 |
| ASV1059 | 27c3e42d173 | 15 | 0 | 0 | 0 | 0 | 0 | 0 | 0 | 0 | 0 | 0 | 0 | 0 | 0 | 0 | 15 | 0 | 0 | 0 | 0 | 0 | 0 | 0 | 0 | 0 | 0 | 0 | 0 | 0 | 0 | 0 | 0 | 0 | 0 | 0 | 0 |
| ASV1060 | 7a71b71d263 | 15 | 0 | 0 | 0 | 0 | 0 | 0 | 0 | 0 | 0 | 6 | 0 | 0 | 0 | 0 | 9 | 0 | 0 | 0 | 0 | 0 | 0 | 0 | 0 | 0 | 0 | 0 | 0 | 0 | 0 | 0 | 0 | 0 | 0 | 0 | 0 |
| ASV1061 | e6411ee5941 | 15 | 0 | 0 | 0 | 0 | 0 | 0 | 0 | 0 | 0 | 0 | 0 | 0 | 0 | 0 | 0 | 0 | 0 | 0 | 0 | 0 | 0 | 0 | 0 | 0 | 0 | 0 | 15 | 0 | 0 | 0 | 0 | 0 | 0 | 0 | 0 |
| ASV1062 | aaa9bf0307b | 15 | 0 | 0 | 0 | 0 | 0 | 0 | 0 | 0 | 0 | 0 | 0 | 0 | 0 | 0 | 10 | 0 | 0 | 0 | 0 | 0 | 0 | 0 | 0 | 0 | 0 | 0 | 0 | 0 | 0 | 5 | 0 | 0 | 0 | 0 | 0 |
| ASV1063 | b6b2f5043f1 | 15 | 0 | 0 | 0 | 0 | 0 | 0 | 0 | 0 | 0 | 0 | 0 | 0 | 0 | 0 | 0 | 0 | 0 | 0 | 0 | 0 | 0 | 0 | 0 | 0 | 0 | 0 | 0 | 15 | 0 | 0 | 0 | 0 | 0 | 0 | 0 |
| ASV1064 | dcaa50c4ae0 | 15 | 0 | 0 | 0 | 0 | 0 | 0 | 0 | 0 | 0 | 0 | 0 | 0 | 0 | 0 | 0 | 0 | 0 | 0 | 0 | 0 | 0 | 0 | 0 | 0 | 0 | 0 | 0 | 0 | 0 | 0 | 0 | 0 | 0 | 15 | 0 |
| ASV1065 | 8fd2e72fb26 | 15 | 0 | 0 | 0 | 0 | 0 | 0 | 0 | 0 | 0 | 0 | 0 | 0 | 0 | 0 | 0 | 0 | 0 | 0 | 0 | 0 | 0 | 0 | 0 | 0 | 0 | 0 | 0 | 0 | 0 | 0 | 0 | 0 | 0 | 0 | 15 |
| ASV1066 | ebce346d288 | 15 | 0 | 0 | 0 | 0 | 0 | 0 | 0 | 0 | 0 | 0 | 0 | 0 | 0 | 0 | 15 | 0 | 0 | 0 | 0 | 0 | 0 | 0 | 0 | 0 | 0 | 0 | 0 | 0 | 0 | 0 | 0 | 0 | 0 | 0 | 0 |
| ASV1067 | 5b5578251cd | 15 | 0 | 0 | 0 | 0 | 0 | 0 | 0 | 0 | 0 | 0 | 0 | 0 | 0 | 0 | 15 | 0 | 0 | 0 | 0 | 0 | 0 | 0 | 0 | 0 | 0 | 0 | 0 | 0 | 0 | 0 | 0 | 0 | 0 | 0 | 0 |
| ASV1068 | 6ec45751ec5 | 15 | 0 | 0 | 0 | 0 | 0 | 0 | 0 | 0 | 2 | 0 | 0 | 0 | 0 | 0 | 0 | 0 | 0 | 0 | 5 | 0 | 0 | 0 | 0 | 2 | 6 | 0 | 0 | 0 | 0 | 0 | 0 | 0 | 0 | 0 | 0 |
| ASV1069 | aa6ebc72e62 | 15 | 0 | 0 | 0 | 0 | 15 | 0 | 0 | 0 | 0 | 0 | 0 | 0 | 0 | 0 | 0 | 0 | 0 | 0 | 0 | 0 | 0 | 0 | 0 | 0 | 0 | 0 | 0 | 0 | 0 | 0 | 0 | 0 | 0 | 0 | 0 |
| ASV1070 | 4cd9af307d4 | 15 | 0 | 0 | 0 | 0 | 15 | 0 | 0 | 0 | 0 | 0 | 0 | 0 | 0 | 0 | 0 | 0 | 0 | 0 | 0 | 0 | 0 | 0 | 0 | 0 | 0 | 0 | 0 | 0 | 0 | 0 | 0 | 0 | 0 | 0 | 0 |
| ASV1071 | 6fc4c89b73a | 15 | 0 | 0 | 0 | 0 | 0 | 0 | 0 | 0 | 0 | 0 | 0 | 15 | 0 | 0 | 0 | 0 | 0 | 0 | 0 | 0 | 0 | 0 | 0 | 0 | 0 | 0 | 0 | 0 | 0 | 0 | 0 | 0 | 0 | 0 | 0 |
| ASV1072 | 69b79e3aab5 | 15 | 0 | 0 | 0 | 0 | 15 | 0 | 0 | 0 | 0 | 0 | 0 | 0 | 0 | 0 | 0 | 0 | 0 | 0 | 0 | 0 | 0 | 0 | 0 | 0 | 0 | 0 | 0 | 0 | 0 | 0 | 0 | 0 | 0 | 0 | 0 |
| ASV1073 | 56935a78569 | 15 | 0 | 0 | 0 | 0 | 15 | 0 | 0 | 0 | 0 | 0 | 0 | 0 | 0 | 0 | 0 | 0 | 0 | 0 | 0 | 0 | 0 | 0 | 0 | 0 | 0 | 0 | 0 | 0 | 0 | 0 | 0 | 0 | 0 | 0 | 0 |
| ASV1074 | 00a0d0f543e | 15 | 0 | 0 | 0 | 0 | 10 | 0 | 0 | 5 | 0 | 0 | 0 | 0 | 0 | 0 | 0 | 0 | 0 | 0 | 0 | 0 | 0 | 0 | 0 | 0 | 0 | 0 | 0 | 0 | 0 | 0 | 0 | 0 | 0 | 0 | 0 |
| ASV1075 | 7ccb862a0eb | 15 | 0 | 0 | 0 | 0 | 0 | 0 | 0 | 0 | 8 | 0 | 0 | 0 | 0 | 0 | 7 | 0 | 0 | 0 | 0 | 0 | 0 | 0 | 0 | 0 | 0 | 0 | 0 | 0 | 0 | 0 | 0 | 0 | 0 | 0 | 0 |
| ASV1076 | 7d026221a04 | 15 | 8 | 0 | 0 | 0 | 0 | 0 | 0 | 7 | 0 | 0 | 0 | 0 | 0 | 0 | 0 | 0 | 0 | 0 | 0 | 0 | 0 | 0 | 0 | 0 | 0 | 0 | 0 | 0 | 0 | 0 | 0 | 0 | 0 | 0 | 0 |
| ASV1077 | e4c38ff9819c | 15 | 0 | 0 | 0 | 0 | 0 | 0 | 0 | 0 | 0 | 6 | 0 | 0 | 0 | 0 | 0 | 0 | 2 | 0 | 0 | 0 | 0 | 0 | 0 | 0 | 0 | 0 | 0 | 0 | 0 | 0 | 0 | 0 | 0 | 0 | 7 |
| ASV1078 | bfce84f0234c | 15 | 0 | 0 | 0 | 0 | 0 | 0 | 0 | 0 | 15 | 0 | 0 | 0 | 0 | 0 | 0 | 0 | 0 | 0 | 0 | 0 | 0 | 0 | 0 | 0 | 0 | 0 | 0 | 0 | 0 | 0 | 0 | 0 | 0 | 0 | 0 |
| ASV1079 | 5526ae2ea4b | 15 | 0 | 0 | 9 | 0 | 0 | 0 | 0 | 0 | 6 | 0 | 0 | 0 | 0 | 0 | 0 | 0 | 0 | 0 | 0 | 0 | 0 | 0 | 0 | 0 | 0 | 0 | 0 | 0 | 0 | 0 | 0 | 0 | 0 | 0 | 0 |
| ASV1080 | 127a0c1ce82 | 15 | 0 | 0 | 0 | 0 | 15 | 0 | 0 | 0 | 0 | 0 | 0 | 0 | 0 | 0 | 0 | 0 | 0 | 0 | 0 | 0 | 0 | 0 | 0 | 0 | 0 | 0 | 0 | 0 | 0 | 0 | 0 | 0 | 0 | 0 | 0 |
| ASV1081 | 8dabe0d7869 | 14 | 0 | 0 | 0 | 0 | 0 | 0 | 0 | 0 | 0 | 0 | 0 | 0 | 0 | 0 | 0 | 0 | 14 | 0 | 0 | 0 | 0 | 0 | 0 | 0 | 0 | 0 | 0 | 0 | 0 | 0 | 0 | 0 | 0 | 0 | 0 |
| ASV1082 | 9063ece8e9f | 14 | 0 | 0 | 0 | 0 | 0 | 0 | 0 | 0 | 0 | 0 | 0 | 0 | 0 | 6 | 0 | 0 | 0 | 0 | 0 | 0 | 0 | 0 | 0 | 8 | 0 | 0 | 0 | 0 | 0 | 0 | 0 | 0 | 0 | 0 | 0 |
| ASV1083 | b61a70e0961 | 14 | 0 | 0 | 0 | 0 | 0 | 0 | 0 | 0 | 0 | 0 | 0 | 3 | 4 | 5 | 0 | 0 | 0 | 0 | 0 | 0 | 2 | 0 | 0 | 0 | 0 | 0 | 0 | 0 | 0 | 0 | 0 | 0 | 0 | 0 | 0 |
| ASV1084 | 27f0cdf9023 | 14 | 0 | 0 | 0 | 0 | 0 | 0 | 0 | 0 | 0 | 0 | 0 | 0 | 0 | 0 | 0 | 0 | 0 | 0 | 0 | 0 | 0 | 0 | 0 | 0 | 0 | 0 | 0 | 0 | 14 | 0 | 0 | 0 | 0 | 0 | 0 |
| ASV1085 | 55c0cbce327 | 14 | 0 | 0 | 0 | 0 | 0 | 0 | 0 | 0 | 0 | 0 | 0 | 0 | 0 | 0 | 0 | 0 | 0 | 0 | 0 | 0 | 0 | 0 | 0 | 0 | 0 | 0 | 0 | 0 | 0 | 0 | 0 | 0 | 0 | 14 | 0 |
| ASV1086 | e3bacd0bceb | 14 | 0 | 0 | 0 | 0 | 0 | 0 | 0 | 0 | 0 | 0 | 0 | 0 | 0 | 0 | 0 | 0 | 0 | 0 | 0 | 0 | 0 | 0 | 0 | 0 | 0 | 0 | 0 | 0 | 0 | 0 | 0 | 0 | 0 | 14 | 0 |
| ASV1087 | 0f2405a72c5 | 14 | 0 | 0 | 0 | 0 | 0 | 0 | 0 | 0 | 0 | 0 | 0 | 0 | 0 | 0 | 0 | 0 | 0 | 14 | 0 | 0 | 0 | 0 | 0 | 0 | 0 | 0 | 0 | 0 | 0 | 0 | 0 | 0 | 0 | 0 | 0 |
| ASV1088 | b4a0a8ea69e | 14 | 0 | 0 | 4 | 0 | 0 | 6 | 0 | 0 | 0 | 0 | 0 | 0 | 0 | 0 | 0 | 0 | 0 | 0 | 0 | 0 | 0 | 0 | 0 | 0 | 0 | 0 | 0 | 0 | 4 | 0 | 0 | 0 | 0 | 0 | 0 |
| ASV1089 | f34326e80f8 | 14 | 0 | 0 | 0 | 0 | 0 | 0 | 0 | 0 | 0 | 0 | 0 | 0 | 0 | 0 | 0 | 0 | 0 | 0 | 14 | 0 | 0 | 0 | 0 | 0 | 0 | 0 | 0 | 0 | 0 | 0 | 0 | 0 | 0 | 0 | 0 |
| ASV1090 | 7225bc4d50a | 14 | 0 | 0 | 0 | 0 | 0 | 0 | 0 | 0 | 4 | 0 | 0 | 0 | 0 | 0 | 10 | 0 | 0 | 0 | 0 | 0 | 0 | 0 | 0 | 0 | 0 | 0 | 0 | 0 | 0 | 0 | 0 | 0 | 0 | 0 | 0 |
| ASV1091 | 9592044e2d4 | 14 | 0 | 0 | 0 | 0 | 0 | 0 | 0 | 0 | 0 | 0 | 0 | 0 | 0 | 0 | 0 | 0 | 0 | 0 | 0 | 0 | 0 | 0 | 0 | 0 | 0 | 0 | 0 | 0 | 0 | 0 | 0 | 0 | 0 | 0 | 14 |
| ASV1092 | da6cd997737 | 14 | 0 | 0 | 0 | 0 | 0 | 0 | 0 | 0 | 0 | 0 | 0 | 0 | 0 | 0 | 0 | 0 | 14 | 0 | 0 | 0 | 0 | 0 | 0 | 0 | 0 | 0 | 0 | 0 | 0 | 0 | 0 | 0 | 0 | 0 | 0 |
| ASV1093 | 51f3434f8d9 | 14 | 0 | 0 | 0 | 0 | 0 | 0 | 0 | 0 | 0 | 0 | 0 | 0 | 0 | 0 | 0 | 0 | 0 | 0 | 0 | 0 | 0 | 4 | 10 | 0 | 0 | 0 | 0 | 0 | 0 | 0 | 0 | 0 | 0 | 0 | 0 |
| ASV1094 | e17de8149c2 | 14 | 0 | 0 | 0 | 0 | 0 | 0 | 0 | 0 | 0 | 0 | 0 | 0 | 0 | 0 | 0 | 0 | 0 | 0 | 0 | 0 | 0 | 0 | 0 | 0 | 0 | 0 | 0 | 0 | 0 | 0 | 0 | 14 | 0 | 0 | 0 |
| ASV1095 | 6ab430a4044 | 14 | 0 | 0 | 0 | 0 | 0 | 0 | 0 | 0 | 0 | 0 | 0 | 0 | 0 | 0 | 0 | 0 | 0 | 0 | 0 | 0 | 0 | 0 | 0 | 0 | 0 | 3 | 0 | 0 | 0 | 0 | 0 | 0 | 0 | 0 | 11 |
| ASV1096 | ebfd1f70b61 | 14 | 0 | 0 | 0 | 0 | 0 | 0 | 0 | 0 | 0 | 0 | 0 | 0 | 0 | 0 | 0 | 0 | 0 | 0 | 0 | 0 | 0 | 0 | 0 | 0 | 0 | 0 | 0 | 0 | 0 | 0 | 0 | 0 | 0 | 14 | 0 |
| ASV1097 | 0996ed3ed6b | 14 | 0 | 0 | 0 | 0 | 0 | 0 | 0 | 0 | 0 | 0 | 0 | 0 | 0 | 0 | 14 | 0 | 0 | 0 | 0 | 0 | 0 | 0 | 0 | 0 | 0 | 0 | 0 | 0 | 0 | 0 | 0 | 0 | 0 | 0 | 0 |
| ASV1098 | c99f9c438c3 | 14 | 0 | 0 | 0 | 0 | 0 | 0 | 0 | 0 | 0 | 0 | 0 | 0 | 0 | 0 | 0 | 14 | 0 | 0 | 0 | 0 | 0 | 0 | 0 | 0 | 0 | 0 | 0 | 0 | 0 | 0 | 0 | 0 | 0 | 0 | 0 |
| ASV1099 | 74db8edd112 | 14 | 0 | 0 | 0 | 0 | 0 | 0 | 0 | 0 | 0 | 0 | 0 | 0 | 0 | 0 | 0 | 0 | 0 | 0 | 0 | 0 | 0 | 0 | 7 | 0 | 0 | 0 | 0 | 7 | 0 | 0 | 0 | 0 | 0 | 0 | 0 |
| ASV1100 | eb9c74886ea | 14 | 0 | 0 | 0 | 0 | 0 | 0 | 0 | 0 | 0 | 0 | 0 | 0 | 0 | 0 | 0 | 0 | 8 | 0 | 0 | 0 | 0 | 0 | 0 | 0 | 0 | 0 | 0 | 0 | 0 | 0 | 6 | 0 | 0 | 0 | 0 |
| ASV1101 | b7b614066e2 | 14 | 0 | 0 | 0 | 0 | 0 | 0 | 0 | 0 | 0 | 0 | 0 | 0 | 0 | 0 | 14 | 0 | 0 | 0 | 0 | 0 | 0 | 0 | 0 | 0 | 0 | 0 | 0 | 0 | 0 | 0 | 0 | 0 | 0 | 0 | 0 |
| ASV1102 | 819c6125778 | 14 | 0 | 2 | 0 | 0 | 6 | 0 | 0 | 0 | 0 | 0 | 0 | 0 | 0 | 6 | 0 | 0 | 0 | 0 | 0 | 0 | 0 | 0 | 0 | 0 | 0 | 0 | 0 | 0 | 0 | 0 | 0 | 0 | 0 | 0 | 0 |
| ASV1103 | d87828eba5e | 14 | 0 | 0 | 0 | 0 | 14 | 0 | 0 | 0 | 0 | 0 | 0 | 0 | 0 | 0 | 0 | 0 | 0 | 0 | 0 | 0 | 0 | 0 | 0 | 0 | 0 | 0 | 0 | 0 | 0 | 0 | 0 | 0 | 0 | 0 | 0 |
| ASV1104 | 3f338bff81ad | 14 | 0 | 0 | 0 | 0 | 0 | 0 | 0 | 0 | 0 | 0 | 0 | 0 | 0 | 0 | 14 | 0 | 0 | 0 | 0 | 0 | 0 | 0 | 0 | 0 | 0 | 0 | 0 | 0 | 0 | 0 | 0 | 0 | 0 | 0 | 0 |
| ASV1105 | a4f63be1936 | 14 | 0 | 0 | 9 | 0 | 0 | 0 | 0 | 0 | 0 | 0 | 0 | 0 | 0 | 0 | 5 | 0 | 0 | 0 | 0 | 0 | 0 | 0 | 0 | 0 | 0 | 0 | 0 | 0 | 0 | 0 | 0 | 0 | 0 | 0 | 0 |
| ASV1106 | ff66de6490e | 14 | 0 | 0 | 0 | 0 | 0 | 0 | 0 | 0 | 0 | 0 | 0 | 0 | 0 | 0 | 0 | 0 | 0 | 0 | 0 | 0 | 0 | 0 | 0 | 0 | 0 | 0 | 0 | 8 | 0 | 0 | 3 | 0 | 0 | 3 | 0 |
| ASV1107 | 65dc92e561a | 14 | 0 | 0 | 0 | 0 | 0 | 0 | 0 | 0 | 0 | 0 | 0 | 0 | 0 | 0 | 0 | 0 | 14 | 0 | 0 | 0 | 0 | 0 | 0 | 0 | 0 | 0 | 0 | 0 | 0 | 0 | 0 | 0 | 0 | 0 | 0 |
| ASV1108 | 22606a3a126 | 14 | 0 | 0 | 0 | 0 | 0 | 0 | 0 | 0 | 0 | 0 | 0 | 0 | 0 | 0 | 0 | 0 | 14 | 0 | 0 | 0 | 0 | 0 | 0 | 0 | 0 | 0 | 0 | 0 | 0 | 0 | 0 | 0 | 0 | 0 | 0 |
| ASV1109 | a74512558fd | 13 | 0 | 0 | 0 | 0 | 0 | 0 | 0 | 0 | 0 | 0 | 0 | 0 | 0 | 0 | 0 | 0 | 0 | 0 | 0 | 0 | 0 | 0 | 0 | 0 | 0 | 0 | 0 | 0 | 0 | 0 | 13 | 0 | 0 | 0 | 0 |
| ASV1110 | 39ad46665d8 | 13 | 0 | 0 | 0 | 0 | 0 | 0 | 0 | 0 | 0 | 13 | 0 | 0 | 0 | 0 | 0 | 0 | 0 | 0 | 0 | 0 | 0 | 0 | 0 | 0 | 0 | 0 | 0 | 0 | 0 | 0 | 0 | 0 | 0 | 0 | 0 |
| ASV1111 | dfc288635a8 | 13 | 0 | 0 | 0 | 0 | 13 | 0 | 0 | 0 | 0 | 0 | 0 | 0 | 0 | 0 | 0 | 0 | 0 | 0 | 0 | 0 | 0 | 0 | 0 | 0 | 0 | 0 | 0 | 0 | 0 | 0 | 0 | 0 | 0 | 0 | 0 |
| ASV1112 | 52766904bbb | 13 | 0 | 0 | 0 | 0 | 0 | 0 | 0 | 0 | 0 | 13 | 0 | 0 | 0 | 0 | 0 | 0 | 0 | 0 | 0 | 0 | 0 | 0 | 0 | 0 | 0 | 0 | 0 | 0 | 0 | 0 | 0 | 0 | 0 | 0 | 0 |
| ASV1113 | 58d621afa51 | 13 | 0 | 0 | 0 | 0 | 0 | 0 | 0 | 0 | 0 | 0 | 0 | 0 | 0 | 0 | 13 | 0 | 0 | 0 | 0 | 0 | 0 | 0 | 0 | 0 | 0 | 0 | 0 | 0 | 0 | 0 | 0 | 0 | 0 | 0 | 0 |
| ASV1114 | 1a265652ef0 | 13 | 0 | 0 | 0 | 0 | 13 | 0 | 0 | 0 | 0 | 0 | 0 | 0 | 0 | 0 | 0 | 0 | 0 | 0 | 0 | 0 | 0 | 0 | 0 | 0 | 0 | 0 | 0 | 0 | 0 | 0 | 0 | 0 | 0 | 0 | 0 |
| ASV1115 | c8a9e0a31e4 | 13 | 0 | 0 | 0 | 0 | 0 | 0 | 0 | 0 | 0 | 0 | 0 | 0 | 0 | 0 | 0 | 0 | 0 | 0 | 0 | 0 | 0 | 0 | 0 | 0 | 0 | 0 | 0 | 0 | 0 | 0 | 0 | 0 | 0 | 0 | 13 |
| ASV1116 | c0a09bdf990 | 13 | 0 | 0 | 0 | 0 | 0 | 0 | 0 | 0 | 0 | 2 | 0 | 0 | 0 | 0 | 5 | 0 | 0 | 0 | 0 | 0 | 0 | 0 | 2 | 0 | 4 | 0 | 0 | 0 | 0 | 0 | 0 | 0 | 0 | 0 | 0 |
| ASV1117 | 10f0b07aad1 | 13 | 0 | 0 | 0 | 0 | 0 | 0 | 0 | 13 | 0 | 0 | 0 | 0 | 0 | 0 | 0 | 0 | 0 | 0 | 0 | 0 | 0 | 0 | 0 | 0 | 0 | 0 | 0 | 0 | 0 | 0 | 0 | 0 | 0 | 0 | 0 |
| ASV1118 | 5c337f48cfaa | 13 | 0 | 0 | 0 | 0 | 13 | 0 | 0 | 0 | 0 | 0 | 0 | 0 | 0 | 0 | 0 | 0 | 0 | 0 | 0 | 0 | 0 | 0 | 0 | 0 | 0 | 0 | 0 | 0 | 0 | 0 | 0 | 0 | 0 | 0 | 0 |
| ASV1119 | 2a45dc931eb | 13 | 13 | 0 | 0 | 0 | 0 | 0 | 0 | 0 | 0 | 0 | 0 | 0 | 0 | 0 | 0 | 0 | 0 | 0 | 0 | 0 | 0 | 0 | 0 | 0 | 0 | 0 | 0 | 0 | 0 | 0 | 0 | 0 | 0 | 0 | 0 |
| ASV1120 | ea23e76aeb2 | 13 | 0 | 0 | 0 | 0 | 0 | 0 | 0 | 0 | 0 | 0 | 0 | 0 | 0 | 0 | 13 | 0 | 0 | 0 | 0 | 0 | 0 | 0 | 0 | 0 | 0 | 0 | 0 | 0 | 0 | 0 | 0 | 0 | 0 | 0 | 0 |
| ASV1121 | a968ca64308 | 13 | 0 | 0 | 0 | 0 | 0 | 0 | 0 | 0 | 0 | 0 | 0 | 0 | 0 | 0 | 0 | 0 | 0 | 0 | 0 | 0 | 0 | 0 | 0 | 0 | 0 | 0 | 0 | 0 | 0 | 0 | 0 | 0 | 0 | 13 | 0 |
| ASV1122 | ccee959e431 | 13 | 0 | 0 | 0 | 0 | 0 | 0 | 0 | 0 | 8 | 0 | 0 | 0 | 0 | 0 | 5 | 0 | 0 | 0 | 0 | 0 | 0 | 0 | 0 | 0 | 0 | 0 | 0 | 0 | 0 | 0 | 0 | 0 | 0 | 0 | 0 |
| ASV1123 | 8efd03f85e1 | 13 | 0 | 0 | 0 | 0 | 0 | 0 | 0 | 0 | 0 | 0 | 0 | 0 | 0 | 0 | 0 | 0 | 0 | 0 | 0 | 0 | 13 | 0 | 0 | 0 | 0 | 0 | 0 | 0 | 0 | 0 | 0 | 0 | 0 | 0 | 0 |
| ASV1124 | 2942c9b812d | 13 | 0 | 0 | 0 | 0 | 0 | 0 | 0 | 0 | 0 | 0 | 0 | 0 | 0 | 0 | 0 | 0 | 0 | 0 | 0 | 0 | 0 | 0 | 0 | 0 | 13 | 0 | 0 | 0 | 0 | 0 | 0 | 0 | 0 | 0 | 0 |
| ASV1125 | 0056be93807 | 13 | 0 | 0 | 0 | 0 | 0 | 0 | 0 | 0 | 0 | 0 | 0 | 0 | 0 | 0 | 0 | 0 | 0 | 0 | 0 | 0 | 0 | 0 | 0 | 0 | 0 | 0 | 0 | 0 | 0 | 0 | 13 | 0 | 0 | 0 | 0 |
| ASV1126 | 034f7a43009 | 13 | 0 | 0 | 0 | 0 | 0 | 0 | 0 | 0 | 0 | 0 | 0 | 0 | 0 | 0 | 0 | 0 | 0 | 0 | 0 | 0 | 0 | 0 | 0 | 0 | 0 | 0 | 0 | 0 | 0 | 0 | 13 | 0 | 0 | 0 | 0 |
| ASV1127 | d5b9f90404e | 13 | 0 | 0 | 0 | 0 | 0 | 0 | 0 | 0 | 0 | 0 | 0 | 0 | 0 | 0 | 13 | 0 | 0 | 0 | 0 | 0 | 0 | 0 | 0 | 0 | 0 | 0 | 0 | 0 | 0 | 0 | 0 | 0 | 0 | 0 | 0 |
| ASV1128 | 4ffeafa7fe10 | 12 | 0 | 0 | 0 | 9 | 0 | 0 | 0 | 0 | 0 | 0 | 0 | 0 | 0 | 0 | 3 | 0 | 0 | 0 | 0 | 0 | 0 | 0 | 0 | 0 | 0 | 0 | 0 | 0 | 0 | 0 | 0 | 0 | 0 | 0 | 0 |
| ASV1129 | 837c2680dc2 | 12 | 0 | 0 | 0 | 0 | 0 | 0 | 0 | 12 | 0 | 0 | 0 | 0 | 0 | 0 | 0 | 0 | 0 | 0 | 0 | 0 | 0 | 0 | 0 | 0 | 0 | 0 | 0 | 0 | 0 | 0 | 0 | 0 | 0 | 0 | 0 |
| ASV1130 | 28033eccf64 | 12 | 0 | 0 | 8 | 4 | 0 | 0 | 0 | 0 | 0 | 0 | 0 | 0 | 0 | 0 | 0 | 0 | 0 | 0 | 0 | 0 | 0 | 0 | 0 | 0 | 0 | 0 | 0 | 0 | 0 | 0 | 0 | 0 | 0 | 0 | 0 |
| ASV1131 | 5770edaa578 | 12 | 0 | 0 | 0 | 0 | 0 | 0 | 0 | 0 | 0 | 0 | 0 | 0 | 0 | 12 | 0 | 0 | 0 | 0 | 0 | 0 | 0 | 0 | 0 | 0 | 0 | 0 | 0 | 0 | 0 | 0 | 0 | 0 | 0 | 0 | 0 |
| ASV1132 | c5589bf3a56 | 12 | 0 | 0 | 0 | 0 | 0 | 0 | 0 | 12 | 0 | 0 | 0 | 0 | 0 | 0 | 0 | 0 | 0 | 0 | 0 | 0 | 0 | 0 | 0 | 0 | 0 | 0 | 0 | 0 | 0 | 0 | 0 | 0 | 0 | 0 | 0 |
| ASV1133 | bdc7f0de378 | 12 | 0 | 0 | 0 | 0 | 0 | 12 | 0 | 0 | 0 | 0 | 0 | 0 | 0 | 0 | 0 | 0 | 0 | 0 | 0 | 0 | 0 | 0 | 0 | 0 | 0 | 0 | 0 | 0 | 0 | 0 | 0 | 0 | 0 | 0 | 0 |
| ASV1134 | c1338c53aa4 | 12 | 0 | 0 | 0 | 0 | 0 | 0 | 0 | 0 | 6 | 6 | 0 | 0 | 0 | 0 | 0 | 0 | 0 | 0 | 0 | 0 | 0 | 0 | 0 | 0 | 0 | 0 | 0 | 0 | 0 | 0 | 0 | 0 | 0 | 0 | 0 |
| ASV1135 | 29244064898 | 12 | 0 | 0 | 0 | 0 | 0 | 0 | 0 | 0 | 6 | 0 | 0 | 0 | 0 | 0 | 6 | 0 | 0 | 0 | 0 | 0 | 0 | 0 | 0 | 0 | 0 | 0 | 0 | 0 | 0 | 0 | 0 | 0 | 0 | 0 | 0 |
| ASV1136 | c48362c4d1a | 12 | 0 | 0 | 0 | 0 | 0 | 0 | 0 | 0 | 0 | 0 | 0 | 0 | 0 | 0 | 0 | 0 | 0 | 0 | 0 | 0 | 0 | 0 | 0 | 0 | 0 | 0 | 0 | 0 | 0 | 0 | 0 | 0 | 0 | 12 | 0 |
| ASV1137 | 6f4eb9151eb | 12 | 0 | 0 | 0 | 0 | 0 | 0 | 0 | 0 | 0 | 0 | 0 | 0 | 0 | 0 | 0 | 4 | 0 | 0 | 0 | 0 | 0 | 0 | 0 | 0 | 0 | 0 | 0 | 0 | 0 | 0 | 0 | 0 | 3 | 5 | 0 |
| ASV1138 | db07348ff5f0 | 12 | 0 | 0 | 0 | 0 | 0 | 0 | 0 | 0 | 0 | 0 | 0 | 0 | 0 | 0 | 0 | 0 | 0 | 0 | 0 | 0 | 0 | 0 | 0 | 0 | 0 | 0 | 0 | 0 | 0 | 0 | 0 | 0 | 0 | 12 | 0 |
| ASV1139 | 61317041500 | 12 | 0 | 0 | 0 | 0 | 0 | 0 | 0 | 0 | 0 | 0 | 0 | 0 | 0 | 0 | 0 | 12 | 0 | 0 | 0 | 0 | 0 | 0 | 0 | 0 | 0 | 0 | 0 | 0 | 0 | 0 | 0 | 0 | 0 | 0 | 0 |
| ASV1140 | 7cdff6625731 | 12 | 0 | 0 | 0 | 12 | 0 | 0 | 0 | 0 | 0 | 0 | 0 | 0 | 0 | 0 | 0 | 0 | 0 | 0 | 0 | 0 | 0 | 0 | 0 | 0 | 0 | 0 | 0 | 0 | 0 | 0 | 0 | 0 | 0 | 0 | 0 |
| ASV1141 | 7c191ba791c | 12 | 0 | 2 | 0 | 6 | 0 | 0 | 0 | 0 | 0 | 0 | 0 | 0 | 0 | 0 | 0 | 0 | 0 | 0 | 0 | 0 | 0 | 0 | 0 | 0 | 0 | 0 | 0 | 0 | 0 | 4 | 0 | 0 | 0 | 0 | 0 |
| ASV1142 | c9c77276627 | 12 | 0 | 0 | 0 | 0 | 0 | 0 | 0 | 0 | 0 | 0 | 0 | 0 | 0 | 0 | 12 | 0 | 0 | 0 | 0 | 0 | 0 | 0 | 0 | 0 | 0 | 0 | 0 | 0 | 0 | 0 | 0 | 0 | 0 | 0 | 0 |
| ASV1143 | d4b598d8ff6 | 12 | 0 | 0 | 0 | 0 | 0 | 0 | 0 | 0 | 0 | 0 | 0 | 0 | 0 | 0 | 0 | 0 | 0 | 0 | 12 | 0 | 0 | 0 | 0 | 0 | 0 | 0 | 0 | 0 | 0 | 0 | 0 | 0 | 0 | 0 | 0 |
| ASV1144 | da9d3dcbb64 | 12 | 0 | 0 | 0 | 0 | 0 | 0 | 0 | 0 | 8 | 4 | 0 | 0 | 0 | 0 | 0 | 0 | 0 | 0 | 0 | 0 | 0 | 0 | 0 | 0 | 0 | 0 | 0 | 0 | 0 | 0 | 0 | 0 | 0 | 0 | 0 |
| ASV1145 | 91f1c19d43f1 | 12 | 0 | 0 | 0 | 0 | 0 | 0 | 0 | 0 | 0 | 0 | 0 | 0 | 0 | 0 | 0 | 0 | 0 | 0 | 0 | 0 | 0 | 0 | 0 | 0 | 0 | 0 | 0 | 12 | 0 | 0 | 0 | 0 | 0 | 0 | 0 |
| ASV1146 | 87d48901d27 | 12 | 0 | 0 | 0 | 0 | 0 | 0 | 0 | 0 | 0 | 0 | 0 | 0 | 0 | 0 | 0 | 0 | 0 | 0 | 0 | 0 | 0 | 0 | 0 | 0 | 0 | 0 | 0 | 0 | 0 | 0 | 12 | 0 | 0 | 0 | 0 |
| ASV1147 | df8facef76b2 | 12 | 0 | 0 | 0 | 0 | 7 | 0 | 0 | 0 | 0 | 0 | 0 | 0 | 0 | 0 | 5 | 0 | 0 | 0 | 0 | 0 | 0 | 0 | 0 | 0 | 0 | 0 | 0 | 0 | 0 | 0 | 0 | 0 | 0 | 0 | 0 |
| ASV1148 | 1ed9ac0dbe5 | 12 | 0 | 0 | 0 | 0 | 0 | 0 | 0 | 0 | 0 | 0 | 0 | 0 | 0 | 0 | 0 | 0 | 0 | 0 | 0 | 0 | 0 | 0 | 0 | 0 | 0 | 0 | 0 | 0 | 0 | 0 | 0 | 0 | 0 | 0 | 12 |
| ASV1149 | f78737c77d6 | 12 | 0 | 0 | 0 | 0 | 12 | 0 | 0 | 0 | 0 | 0 | 0 | 0 | 0 | 0 | 0 | 0 | 0 | 0 | 0 | 0 | 0 | 0 | 0 | 0 | 0 | 0 | 0 | 0 | 0 | 0 | 0 | 0 | 0 | 0 | 0 |
| ASV1150 | b28e037a0e8 | 12 | 0 | 0 | 0 | 0 | 0 | 0 | 0 | 0 | 0 | 12 | 0 | 0 | 0 | 0 | 0 | 0 | 0 | 0 | 0 | 0 | 0 | 0 | 0 | 0 | 0 | 0 | 0 | 0 | 0 | 0 | 0 | 0 | 0 | 0 | 0 |
| ASV1151 | 928e2cdbd60 | 12 | 0 | 0 | 0 | 0 | 0 | 0 | 0 | 0 | 0 | 0 | 0 | 0 | 0 | 0 | 0 | 0 | 0 | 12 | 0 | 0 | 0 | 0 | 0 | 0 | 0 | 0 | 0 | 0 | 0 | 0 | 0 | 0 | 0 | 0 | 0 |
| ASV1152 | 8de8dbfc853 | 12 | 0 | 0 | 0 | 0 | 0 | 0 | 0 | 0 | 0 | 0 | 0 | 0 | 0 | 0 | 0 | 0 | 0 | 0 | 0 | 0 | 0 | 0 | 0 | 0 | 0 | 0 | 0 | 0 | 0 | 12 | 0 | 0 | 0 | 0 | 0 |
| ASV1153 | 91b7b5980e6 | 12 | 0 | 0 | 0 | 0 | 8 | 0 | 0 | 0 | 4 | 0 | 0 | 0 | 0 | 0 | 0 | 0 | 0 | 0 | 0 | 0 | 0 | 0 | 0 | 0 | 0 | 0 | 0 | 0 | 0 | 0 | 0 | 0 | 0 | 0 | 0 |
| ASV1154 | f19bebb06f1 | 12 | 0 | 0 | 0 | 0 | 0 | 0 | 0 | 0 | 0 | 0 | 0 | 0 | 0 | 0 | 0 | 0 | 0 | 0 | 0 | 0 | 0 | 0 | 0 | 0 | 0 | 0 | 0 | 0 | 0 | 0 | 12 | 0 | 0 | 0 | 0 |
| ASV1155 | d9a230acb01 | 12 | 0 | 0 | 0 | 0 | 0 | 0 | 0 | 0 | 0 | 0 | 0 | 0 | 0 | 0 | 0 | 0 | 0 | 0 | 0 | 0 | 0 | 0 | 0 | 0 | 0 | 0 | 0 | 0 | 0 | 0 | 0 | 0 | 0 | 0 | 12 |
| ASV1156 | 389066fa02c | 11 | 0 | 0 | 0 | 0 | 0 | 0 | 0 | 11 | 0 | 0 | 0 | 0 | 0 | 0 | 0 | 0 | 0 | 0 | 0 | 0 | 0 | 0 | 0 | 0 | 0 | 0 | 0 | 0 | 0 | 0 | 0 | 0 | 0 | 0 | 0 |
| ASV1157 | bd3b15486f3 | 11 | 0 | 0 | 0 | 0 | 0 | 0 | 0 | 0 | 0 | 0 | 0 | 11 | 0 | 0 | 0 | 0 | 0 | 0 | 0 | 0 | 0 | 0 | 0 | 0 | 0 | 0 | 0 | 0 | 0 | 0 | 0 | 0 | 0 | 0 | 0 |
| ASV1158 | c5e03ed3783 | 11 | 0 | 0 | 0 | 5 | 6 | 0 | 0 | 0 | 0 | 0 | 0 | 0 | 0 | 0 | 0 | 0 | 0 | 0 | 0 | 0 | 0 | 0 | 0 | 0 | 0 | 0 | 0 | 0 | 0 | 0 | 0 | 0 | 0 | 0 | 0 |
| ASV1159 | 164a0cba1d3 | 11 | 0 | 0 | 0 | 0 | 0 | 0 | 0 | 0 | 0 | 0 | 0 | 0 | 0 | 0 | 0 | 11 | 0 | 0 | 0 | 0 | 0 | 0 | 0 | 0 | 0 | 0 | 0 | 0 | 0 | 0 | 0 | 0 | 0 | 0 | 0 |
| ASV1160 | 1d01169d8c7 | 11 | 0 | 0 | 0 | 0 | 0 | 0 | 0 | 0 | 0 | 0 | 0 | 0 | 0 | 0 | 0 | 0 | 0 | 0 | 0 | 0 | 0 | 0 | 0 | 0 | 0 | 0 | 0 | 0 | 0 | 0 | 0 | 0 | 11 | 0 | 0 |
| ASV1161 | 243f934f46e | 11 | 0 | 0 | 0 | 0 | 0 | 0 | 0 | 0 | 0 | 0 | 0 | 11 | 0 | 0 | 0 | 0 | 0 | 0 | 0 | 0 | 0 | 0 | 0 | 0 | 0 | 0 | 0 | 0 | 0 | 0 | 0 | 0 | 0 | 0 | 0 |
| ASV1162 | b348bb98c3d | 11 | 0 | 0 | 0 | 0 | 0 | 0 | 0 | 0 | 4 | 0 | 0 | 0 | 0 | 0 | 7 | 0 | 0 | 0 | 0 | 0 | 0 | 0 | 0 | 0 | 0 | 0 | 0 | 0 | 0 | 0 | 0 | 0 | 0 | 0 | 0 |
| ASV1163 | e495b390f80 | 11 | 0 | 0 | 0 | 0 | 0 | 0 | 0 | 0 | 0 | 0 | 0 | 0 | 0 | 0 | 7 | 0 | 0 | 0 | 0 | 0 | 0 | 0 | 0 | 0 | 0 | 0 | 0 | 0 | 0 | 4 | 0 | 0 | 0 | 0 | 0 |
| ASV1164 | 3fda4837da1 | 11 | 0 | 0 | 0 | 0 | 0 | 0 | 0 | 0 | 0 | 0 | 0 | 0 | 0 | 0 | 0 | 0 | 0 | 0 | 0 | 0 | 0 | 0 | 0 | 0 | 0 | 0 | 0 | 11 | 0 | 0 | 0 | 0 | 0 | 0 | 0 |
| ASV1165 | a44dea011b7 | 11 | 0 | 0 | 0 | 0 | 0 | 0 | 0 | 0 | 0 | 11 | 0 | 0 | 0 | 0 | 0 | 0 | 0 | 0 | 0 | 0 | 0 | 0 | 0 | 0 | 0 | 0 | 0 | 0 | 0 | 0 | 0 | 0 | 0 | 0 | 0 |
| ASV1166 | fc2406b8206 | 11 | 0 | 0 | 0 | 0 | 0 | 0 | 0 | 0 | 0 | 0 | 0 | 0 | 0 | 0 | 0 | 0 | 0 | 0 | 0 | 0 | 0 | 0 | 0 | 0 | 8 | 0 | 0 | 0 | 0 | 0 | 0 | 0 | 0 | 0 | 3 |
| ASV1167 | 08e50b782be | 11 | 0 | 0 | 0 | 0 | 0 | 0 | 0 | 0 | 0 | 0 | 0 | 0 | 0 | 0 | 0 | 0 | 0 | 0 | 0 | 0 | 0 | 0 | 0 | 0 | 0 | 0 | 0 | 0 | 0 | 0 | 11 | 0 | 0 | 0 | 0 |
| ASV1168 | 174c97734e5 | 11 | 0 | 0 | 0 | 0 | 0 | 0 | 0 | 0 | 0 | 0 | 0 | 0 | 0 | 0 | 0 | 0 | 0 | 8 | 3 | 0 | 0 | 0 | 0 | 0 | 0 | 0 | 0 | 0 | 0 | 0 | 0 | 0 | 0 | 0 | 0 |
| ASV1169 | a08034cfb7b | 11 | 0 | 0 | 0 | 0 | 0 | 0 | 0 | 0 | 0 | 0 | 0 | 0 | 0 | 0 | 0 | 0 | 0 | 0 | 0 | 0 | 0 | 0 | 0 | 0 | 0 | 0 | 0 | 0 | 0 | 0 | 0 | 0 | 0 | 0 | 11 |
| ASV1170 | 78bf063117d | 11 | 0 | 0 | 0 | 0 | 0 | 0 | 0 | 0 | 0 | 0 | 0 | 0 | 0 | 0 | 0 | 0 | 0 | 0 | 0 | 0 | 0 | 0 | 0 | 0 | 0 | 0 | 0 | 0 | 11 | 0 | 0 | 0 | 0 | 0 | 0 |
| ASV1171 | 4539c99762e | 11 | 0 | 0 | 0 | 0 | 0 | 0 | 0 | 0 | 0 | 0 | 0 | 0 | 0 | 0 | 11 | 0 | 0 | 0 | 0 | 0 | 0 | 0 | 0 | 0 | 0 | 0 | 0 | 0 | 0 | 0 | 0 | 0 | 0 | 0 | 0 |
| ASV1172 | 477889c36e4 | 11 | 0 | 0 | 0 | 0 | 0 | 0 | 0 | 0 | 3 | 0 | 0 | 0 | 0 | 0 | 8 | 0 | 0 | 0 | 0 | 0 | 0 | 0 | 0 | 0 | 0 | 0 | 0 | 0 | 0 | 0 | 0 | 0 | 0 | 0 | 0 |
| ASV1173 | 15885bb1832 | 11 | 0 | 0 | 0 | 0 | 0 | 0 | 0 | 0 | 11 | 0 | 0 | 0 | 0 | 0 | 0 | 0 | 0 | 0 | 0 | 0 | 0 | 0 | 0 | 0 | 0 | 0 | 0 | 0 | 0 | 0 | 0 | 0 | 0 | 0 | 0 |
| ASV1174 | 82d8e38d3cf | 11 | 0 | 0 | 3 | 0 | 0 | 0 | 0 | 0 | 0 | 0 | 0 | 0 | 0 | 0 | 0 | 5 | 0 | 3 | 0 | 0 | 0 | 0 | 0 | 0 | 0 | 0 | 0 | 0 | 0 | 0 | 0 | 0 | 0 | 0 | 0 |
| ASV1175 | 3bf79ebc3c7 | 11 | 0 | 0 | 0 | 0 | 0 | 0 | 0 | 0 | 0 | 0 | 0 | 0 | 0 | 0 | 0 | 0 | 0 | 0 | 0 | 0 | 0 | 0 | 0 | 0 | 0 | 0 | 0 | 0 | 0 | 0 | 0 | 0 | 0 | 11 | 0 |
| ASV1176 | c4905739179 | 11 | 0 | 0 | 0 | 0 | 0 | 0 | 0 | 0 | 0 | 0 | 0 | 0 | 0 | 0 | 0 | 0 | 0 | 0 | 0 | 11 | 0 | 0 | 0 | 0 | 0 | 0 | 0 | 0 | 0 | 0 | 0 | 0 | 0 | 0 | 0 |
| ASV1177 | 6fd148441ffe | 11 | 0 | 0 | 0 | 0 | 0 | 0 | 0 | 0 | 0 | 0 | 0 | 0 | 0 | 0 | 0 | 0 | 0 | 0 | 0 | 0 | 0 | 0 | 0 | 0 | 0 | 0 | 0 | 0 | 0 | 0 | 0 | 0 | 0 | 11 | 0 |
| ASV1178 | 40dd3c7b96b | 11 | 0 | 0 | 0 | 0 | 0 | 0 | 0 | 0 | 0 | 0 | 0 | 0 | 0 | 0 | 0 | 11 | 0 | 0 | 0 | 0 | 0 | 0 | 0 | 0 | 0 | 0 | 0 | 0 | 0 | 0 | 0 | 0 | 0 | 0 | 0 |
| ASV1179 | 0242e248103 | 11 | 0 | 0 | 0 | 0 | 0 | 0 | 0 | 0 | 0 | 0 | 0 | 0 | 0 | 0 | 0 | 0 | 0 | 0 | 0 | 0 | 0 | 0 | 0 | 0 | 0 | 7 | 4 | 0 | 0 | 0 | 0 | 0 | 0 | 0 | 0 |
| ASV1180 | 95986efe6ab | 11 | 0 | 0 | 0 | 0 | 0 | 0 | 0 | 0 | 0 | 4 | 0 | 0 | 0 | 0 | 0 | 0 | 0 | 0 | 0 | 0 | 0 | 0 | 0 | 0 | 0 | 0 | 0 | 0 | 0 | 0 | 0 | 0 | 0 | 0 | 7 |
| ASV1181 | 088a9758f9e | 11 | 0 | 0 | 0 | 0 | 0 | 0 | 0 | 0 | 11 | 0 | 0 | 0 | 0 | 0 | 0 | 0 | 0 | 0 | 0 | 0 | 0 | 0 | 0 | 0 | 0 | 0 | 0 | 0 | 0 | 0 | 0 | 0 | 0 | 0 | 0 |
| ASV1182 | 8cb126c23f7 | 11 | 0 | 0 | 0 | 0 | 0 | 0 | 0 | 0 | 0 | 0 | 0 | 0 | 0 | 0 | 0 | 0 | 0 | 0 | 0 | 0 | 0 | 0 | 0 | 0 | 0 | 0 | 0 | 0 | 0 | 0 | 11 | 0 | 0 | 0 | 0 |
| ASV1183 | 505890a7b86 | 11 | 0 | 0 | 0 | 0 | 0 | 0 | 0 | 0 | 0 | 0 | 0 | 0 | 0 | 0 | 0 | 0 | 0 | 0 | 0 | 0 | 0 | 0 | 0 | 0 | 0 | 0 | 0 | 0 | 11 | 0 | 0 | 0 | 0 | 0 | 0 |
| ASV1184 | 704f4a03f69 | 11 | 0 | 0 | 0 | 0 | 11 | 0 | 0 | 0 | 0 | 0 | 0 | 0 | 0 | 0 | 0 | 0 | 0 | 0 | 0 | 0 | 0 | 0 | 0 | 0 | 0 | 0 | 0 | 0 | 0 | 0 | 0 | 0 | 0 | 0 | 0 |
| ASV1185 | 82dd528f90d | 10 | 0 | 0 | 0 | 0 | 0 | 0 | 0 | 0 | 0 | 0 | 0 | 0 | 0 | 0 | 0 | 0 | 0 | 0 | 0 | 0 | 0 | 0 | 0 | 10 | 0 | 0 | 0 | 0 | 0 | 0 | 0 | 0 | 0 | 0 | 0 |
| ASV1186 | 26841b4cd9e | 10 | 0 | 0 | 0 | 10 | 0 | 0 | 0 | 0 | 0 | 0 | 0 | 0 | 0 | 0 | 0 | 0 | 0 | 0 | 0 | 0 | 0 | 0 | 0 | 0 | 0 | 0 | 0 | 0 | 0 | 0 | 0 | 0 | 0 | 0 | 0 |
| ASV1187 | 08f7d9154e1 | 10 | 0 | 0 | 0 | 10 | 0 | 0 | 0 | 0 | 0 | 0 | 0 | 0 | 0 | 0 | 0 | 0 | 0 | 0 | 0 | 0 | 0 | 0 | 0 | 0 | 0 | 0 | 0 | 0 | 0 | 0 | 0 | 0 | 0 | 0 | 0 |
| ASV1188 | c45482a5b65 | 10 | 0 | 0 | 0 | 0 | 5 | 0 | 0 | 0 | 0 | 5 | 0 | 0 | 0 | 0 | 0 | 0 | 0 | 0 | 0 | 0 | 0 | 0 | 0 | 0 | 0 | 0 | 0 | 0 | 0 | 0 | 0 | 0 | 0 | 0 | 0 |
| ASV1189 | 3a101da2504 | 10 | 0 | 0 | 0 | 0 | 0 | 0 | 0 | 10 | 0 | 0 | 0 | 0 | 0 | 0 | 0 | 0 | 0 | 0 | 0 | 0 | 0 | 0 | 0 | 0 | 0 | 0 | 0 | 0 | 0 | 0 | 0 | 0 | 0 | 0 | 0 |
| ASV1190 | 564bf8d5970 | 10 | 0 | 0 | 0 | 0 | 0 | 0 | 0 | 0 | 0 | 0 | 0 | 0 | 0 | 0 | 0 | 0 | 0 | 0 | 0 | 0 | 0 | 0 | 0 | 0 | 0 | 0 | 0 | 10 | 0 | 0 | 0 | 0 | 0 | 0 | 0 |
| ASV1191 | eb44ac888b7 | 10 | 0 | 0 | 0 | 0 | 0 | 0 | 0 | 0 | 0 | 0 | 0 | 0 | 0 | 0 | 0 | 0 | 0 | 0 | 0 | 10 | 0 | 0 | 0 | 0 | 0 | 0 | 0 | 0 | 0 | 0 | 0 | 0 | 0 | 0 | 0 |
| ASV1192 | a8c0a170582 | 10 | 0 | 0 | 0 | 0 | 0 | 0 | 0 | 0 | 0 | 0 | 0 | 0 | 0 | 0 | 0 | 0 | 0 | 0 | 0 | 0 | 0 | 0 | 0 | 0 | 0 | 0 | 0 | 0 | 0 | 0 | 10 | 0 | 0 | 0 | 0 |
| ASV1193 | 1d2312249d6 | 10 | 0 | 0 | 0 | 0 | 0 | 0 | 0 | 0 | 0 | 0 | 0 | 0 | 0 | 0 | 0 | 0 | 0 | 0 | 0 | 0 | 0 | 0 | 0 | 0 | 0 | 0 | 0 | 0 | 0 | 0 | 10 | 0 | 0 | 0 | 0 |
| ASV1194 | 06f09aed5e3 | 10 | 0 | 0 | 0 | 0 | 3 | 0 | 0 | 0 | 0 | 0 | 0 | 0 | 0 | 0 | 5 | 0 | 0 | 0 | 0 | 0 | 0 | 0 | 0 | 0 | 0 | 0 | 0 | 0 | 0 | 2 | 0 | 0 | 0 | 0 | 0 |
| ASV1195 | 830d8ebb1ed | 10 | 0 | 0 | 0 | 0 | 0 | 0 | 0 | 0 | 0 | 0 | 0 | 0 | 0 | 0 | 0 | 0 | 0 | 0 | 0 | 0 | 0 | 0 | 0 | 0 | 0 | 0 | 0 | 0 | 0 | 0 | 0 | 10 | 0 | 0 | 0 |
| ASV1196 | 06731b15d25 | 10 | 0 | 0 | 0 | 0 | 0 | 0 | 0 | 0 | 0 | 0 | 0 | 0 | 0 | 0 | 0 | 0 | 0 | 0 | 0 | 0 | 0 | 0 | 0 | 0 | 0 | 0 | 0 | 0 | 0 | 0 | 0 | 0 | 0 | 2 | 8 |
| ASV1197 | 22caf10c09d | 10 | 0 | 0 | 0 | 0 | 0 | 0 | 0 | 10 | 0 | 0 | 0 | 0 | 0 | 0 | 0 | 0 | 0 | 0 | 0 | 0 | 0 | 0 | 0 | 0 | 0 | 0 | 0 | 0 | 0 | 0 | 0 | 0 | 0 | 0 | 0 |
| ASV1198 | 91e74e2fc1d | 10 | 0 | 0 | 0 | 0 | 0 | 10 | 0 | 0 | 0 | 0 | 0 | 0 | 0 | 0 | 0 | 0 | 0 | 0 | 0 | 0 | 0 | 0 | 0 | 0 | 0 | 0 | 0 | 0 | 0 | 0 | 0 | 0 | 0 | 0 | 0 |
| ASV1199 | 1f883e40a2e | 10 | 0 | 0 | 2 | 3 | 5 | 0 | 0 | 0 | 0 | 0 | 0 | 0 | 0 | 0 | 0 | 0 | 0 | 0 | 0 | 0 | 0 | 0 | 0 | 0 | 0 | 0 | 0 | 0 | 0 | 0 | 0 | 0 | 0 | 0 | 0 |
| ASV1200 | f820c9a3954 | 10 | 0 | 0 | 0 | 0 | 0 | 0 | 0 | 10 | 0 | 0 | 0 | 0 | 0 | 0 | 0 | 0 | 0 | 0 | 0 | 0 | 0 | 0 | 0 | 0 | 0 | 0 | 0 | 0 | 0 | 0 | 0 | 0 | 0 | 0 | 0 |
| ASV1201 | c34ac2be3bd | 10 | 0 | 0 | 0 | 0 | 2 | 0 | 0 | 0 | 0 | 0 | 0 | 0 | 0 | 0 | 8 | 0 | 0 | 0 | 0 | 0 | 0 | 0 | 0 | 0 | 0 | 0 | 0 | 0 | 0 | 0 | 0 | 0 | 0 | 0 | 0 |
| ASV1202 | 2248d33b5b4 | 10 | 0 | 0 | 0 | 0 | 0 | 0 | 5 | 0 | 0 | 0 | 0 | 0 | 0 | 0 | 0 | 0 | 0 | 0 | 0 | 0 | 0 | 0 | 0 | 5 | 0 | 0 | 0 | 0 | 0 | 0 | 0 | 0 | 0 | 0 | 0 |
| ASV1203 | d65f0c64433 | 10 | 0 | 0 | 0 | 0 | 0 | 0 | 0 | 0 | 0 | 0 | 0 | 10 | 0 | 0 | 0 | 0 | 0 | 0 | 0 | 0 | 0 | 0 | 0 | 0 | 0 | 0 | 0 | 0 | 0 | 0 | 0 | 0 | 0 | 0 | 0 |
| ASV1204 | 695676ad9e9 | 10 | 0 | 0 | 10 | 0 | 0 | 0 | 0 | 0 | 0 | 0 | 0 | 0 | 0 | 0 | 0 | 0 | 0 | 0 | 0 | 0 | 0 | 0 | 0 | 0 | 0 | 0 | 0 | 0 | 0 | 0 | 0 | 0 | 0 | 0 | 0 |
| ASV1205 | fc2c768a1b8 | 10 | 0 | 0 | 0 | 5 | 0 | 0 | 0 | 0 | 0 | 0 | 0 | 0 | 0 | 0 | 0 | 0 | 0 | 0 | 0 | 0 | 0 | 0 | 0 | 0 | 2 | 0 | 0 | 0 | 0 | 0 | 0 | 0 | 0 | 0 | 3 |
| ASV1206 | 4609b3fdee7 | 10 | 0 | 0 | 0 | 0 | 0 | 10 | 0 | 0 | 0 | 0 | 0 | 0 | 0 | 0 | 0 | 0 | 0 | 0 | 0 | 0 | 0 | 0 | 0 | 0 | 0 | 0 | 0 | 0 | 0 | 0 | 0 | 0 | 0 | 0 | 0 |
| ASV1207 | eb6c7926524 | 10 | 0 | 0 | 0 | 0 | 0 | 0 | 0 | 0 | 0 | 0 | 0 | 0 | 0 | 0 | 0 | 0 | 0 | 10 | 0 | 0 | 0 | 0 | 0 | 0 | 0 | 0 | 0 | 0 | 0 | 0 | 0 | 0 | 0 | 0 | 0 |
| ASV1208 | 1ad3a013ca3 | 10 | 0 | 0 | 0 | 0 | 3 | 0 | 0 | 7 | 0 | 0 | 0 | 0 | 0 | 0 | 0 | 0 | 0 | 0 | 0 | 0 | 0 | 0 | 0 | 0 | 0 | 0 | 0 | 0 | 0 | 0 | 0 | 0 | 0 | 0 | 0 |
| ASV1209 | 8e6e50c6ce0 | 10 | 0 | 0 | 0 | 0 | 0 | 0 | 0 | 0 | 0 | 0 | 0 | 0 | 0 | 0 | 0 | 0 | 0 | 6 | 0 | 0 | 0 | 0 | 0 | 0 | 0 | 4 | 0 | 0 | 0 | 0 | 0 | 0 | 0 | 0 | 0 |
| ASV1210 | 93811965988 | 10 | 0 | 0 | 0 | 0 | 10 | 0 | 0 | 0 | 0 | 0 | 0 | 0 | 0 | 0 | 0 | 0 | 0 | 0 | 0 | 0 | 0 | 0 | 0 | 0 | 0 | 0 | 0 | 0 | 0 | 0 | 0 | 0 | 0 | 0 | 0 |
| ASV1211 | 674ad4fd009 | 10 | 0 | 0 | 0 | 0 | 4 | 0 | 0 | 0 | 0 | 0 | 0 | 0 | 0 | 0 | 0 | 0 | 0 | 0 | 0 | 0 | 0 | 0 | 0 | 0 | 0 | 0 | 0 | 0 | 0 | 0 | 0 | 0 | 0 | 0 | 6 |
| ASV1212 | 0aeda5d66cc | 10 | 0 | 0 | 0 | 0 | 0 | 0 | 0 | 10 | 0 | 0 | 0 | 0 | 0 | 0 | 0 | 0 | 0 | 0 | 0 | 0 | 0 | 0 | 0 | 0 | 0 | 0 | 0 | 0 | 0 | 0 | 0 | 0 | 0 | 0 | 0 |
| ASV1213 | 8707c0dd066 | 10 | 0 | 0 | 0 | 0 | 0 | 0 | 0 | 10 | 0 | 0 | 0 | 0 | 0 | 0 | 0 | 0 | 0 | 0 | 0 | 0 | 0 | 0 | 0 | 0 | 0 | 0 | 0 | 0 | 0 | 0 | 0 | 0 | 0 | 0 | 0 |
| ASV1214 | e026ba5f41c | 10 | 0 | 0 | 0 | 0 | 0 | 0 | 0 | 0 | 0 | 0 | 0 | 0 | 0 | 0 | 0 | 0 | 10 | 0 | 0 | 0 | 0 | 0 | 0 | 0 | 0 | 0 | 0 | 0 | 0 | 0 | 0 | 0 | 0 | 0 | 0 |
| ASV1215 | 8fdc5ad59a2 | 10 | 0 | 0 | 0 | 0 | 10 | 0 | 0 | 0 | 0 | 0 | 0 | 0 | 0 | 0 | 0 | 0 | 0 | 0 | 0 | 0 | 0 | 0 | 0 | 0 | 0 | 0 | 0 | 0 | 0 | 0 | 0 | 0 | 0 | 0 | 0 |
| ASV1216 | 3377f176fcf2 | 10 | 0 | 0 | 0 | 0 | 0 | 0 | 0 | 0 | 0 | 0 | 0 | 0 | 0 | 0 | 0 | 0 | 0 | 0 | 0 | 0 | 0 | 0 | 0 | 10 | 0 | 0 | 0 | 0 | 0 | 0 | 0 | 0 | 0 | 0 | 0 |
| ASV1217 | 2afb47ba5ef | 10 | 0 | 0 | 0 | 0 | 0 | 0 | 0 | 0 | 0 | 0 | 0 | 0 | 0 | 0 | 0 | 0 | 0 | 0 | 0 | 0 | 0 | 0 | 0 | 0 | 0 | 0 | 0 | 0 | 0 | 10 | 0 | 0 | 0 | 0 | 0 |
| ASV1218 | fb4c659afc2f | 10 | 0 | 0 | 0 | 0 | 0 | 0 | 0 | 0 | 10 | 0 | 0 | 0 | 0 | 0 | 0 | 0 | 0 | 0 | 0 | 0 | 0 | 0 | 0 | 0 | 0 | 0 | 0 | 0 | 0 | 0 | 0 | 0 | 0 | 0 | 0 |
| ASV1219 | db495bd28b4 | 10 | 0 | 0 | 0 | 0 | 0 | 0 | 0 | 0 | 0 | 0 | 0 | 0 | 0 | 0 | 0 | 0 | 0 | 0 | 0 | 10 | 0 | 0 | 0 | 0 | 0 | 0 | 0 | 0 | 0 | 0 | 0 | 0 | 0 | 0 | 0 |
| ASV1220 | 27ba8f5586f | 10 | 0 | 0 | 0 | 0 | 0 | 0 | 0 | 0 | 0 | 0 | 0 | 0 | 0 | 0 | 10 | 0 | 0 | 0 | 0 | 0 | 0 | 0 | 0 | 0 | 0 | 0 | 0 | 0 | 0 | 0 | 0 | 0 | 0 | 0 | 0 |
| ASV1221 | c9a9f40efb9 | 10 | 0 | 0 | 0 | 0 | 0 | 0 | 0 | 0 | 0 | 0 | 0 | 0 | 0 | 0 | 0 | 0 | 0 | 0 | 0 | 0 | 0 | 0 | 0 | 0 | 0 | 10 | 0 | 0 | 0 | 0 | 0 | 0 | 0 | 0 | 0 |
| ASV1222 | 4746d060dcf | 10 | 0 | 0 | 10 | 0 | 0 | 0 | 0 | 0 | 0 | 0 | 0 | 0 | 0 | 0 | 0 | 0 | 0 | 0 | 0 | 0 | 0 | 0 | 0 | 0 | 0 | 0 | 0 | 0 | 0 | 0 | 0 | 0 | 0 | 0 | 0 |
| ASV1223 | 84a6628e19d | 10 | 0 | 0 | 0 | 0 | 0 | 0 | 0 | 0 | 0 | 0 | 0 | 0 | 0 | 0 | 10 | 0 | 0 | 0 | 0 | 0 | 0 | 0 | 0 | 0 | 0 | 0 | 0 | 0 | 0 | 0 | 0 | 0 | 0 | 0 | 0 |
| ASV1224 | c27727ce505 | 10 | 0 | 0 | 0 | 0 | 0 | 10 | 0 | 0 | 0 | 0 | 0 | 0 | 0 | 0 | 0 | 0 | 0 | 0 | 0 | 0 | 0 | 0 | 0 | 0 | 0 | 0 | 0 | 0 | 0 | 0 | 0 | 0 | 0 | 0 | 0 |
| ASV1225 | 819d506138c | 10 | 0 | 0 | 0 | 0 | 0 | 0 | 0 | 0 | 0 | 10 | 0 | 0 | 0 | 0 | 0 | 0 | 0 | 0 | 0 | 0 | 0 | 0 | 0 | 0 | 0 | 0 | 0 | 0 | 0 | 0 | 0 | 0 | 0 | 0 | 0 |
| ASV1226 | dcb98b9f9b6 | 9 | 0 | 0 | 0 | 0 | 0 | 0 | 0 | 0 | 0 | 9 | 0 | 0 | 0 | 0 | 0 | 0 | 0 | 0 | 0 | 0 | 0 | 0 | 0 | 0 | 0 | 0 | 0 | 0 | 0 | 0 | 0 | 0 | 0 | 0 | 0 |
| ASV1227 | 95af87e0bd7 | 9 | 0 | 0 | 0 | 0 | 0 | 0 | 0 | 0 | 0 | 0 | 0 | 0 | 0 | 0 | 0 | 0 | 0 | 0 | 0 | 0 | 0 | 0 | 0 | 0 | 9 | 0 | 0 | 0 | 0 | 0 | 0 | 0 | 0 | 0 | 0 |
| ASV1228 | 60d93e044da | 9 | 0 | 0 | 0 | 0 | 0 | 0 | 0 | 0 | 0 | 0 | 0 | 0 | 0 | 0 | 0 | 0 | 0 | 0 | 0 | 0 | 0 | 0 | 0 | 0 | 0 | 0 | 0 | 0 | 0 | 2 | 0 | 0 | 0 | 0 | 7 |
| ASV1229 | e34ef8095dd | 9 | 0 | 0 | 0 | 0 | 9 | 0 | 0 | 0 | 0 | 0 | 0 | 0 | 0 | 0 | 0 | 0 | 0 | 0 | 0 | 0 | 0 | 0 | 0 | 0 | 0 | 0 | 0 | 0 | 0 | 0 | 0 | 0 | 0 | 0 | 0 |
| ASV1230 | e7987a6fcb3 | 9 | 0 | 0 | 0 | 0 | 0 | 0 | 0 | 0 | 0 | 0 | 0 | 0 | 0 | 0 | 0 | 0 | 0 | 0 | 0 | 0 | 0 | 0 | 0 | 0 | 0 | 0 | 0 | 0 | 0 | 0 | 0 | 0 | 9 | 0 | 0 |
| ASV1231 | 42f32014a4d | 9 | 0 | 0 | 0 | 0 | 0 | 0 | 0 | 0 | 0 | 0 | 0 | 0 | 0 | 0 | 0 | 0 | 0 | 0 | 0 | 0 | 0 | 0 | 0 | 0 | 0 | 3 | 0 | 0 | 0 | 0 | 0 | 2 | 0 | 4 | 0 |
| ASV1232 | 3e9e854ae4f | 9 | 0 | 0 | 0 | 0 | 2 | 0 | 0 | 0 | 0 | 0 | 0 | 0 | 0 | 0 | 7 | 0 | 0 | 0 | 0 | 0 | 0 | 0 | 0 | 0 | 0 | 0 | 0 | 0 | 0 | 0 | 0 | 0 | 0 | 0 | 0 |
| ASV1233 | 1921cf951a8 | 9 | 0 | 0 | 0 | 0 | 9 | 0 | 0 | 0 | 0 | 0 | 0 | 0 | 0 | 0 | 0 | 0 | 0 | 0 | 0 | 0 | 0 | 0 | 0 | 0 | 0 | 0 | 0 | 0 | 0 | 0 | 0 | 0 | 0 | 0 | 0 |
| ASV1234 | 2ac69cc152a | 9 | 0 | 0 | 0 | 0 | 0 | 0 | 0 | 0 | 0 | 0 | 0 | 0 | 0 | 0 | 0 | 0 | 0 | 0 | 0 | 0 | 0 | 0 | 0 | 0 | 0 | 0 | 0 | 0 | 0 | 0 | 0 | 0 | 0 | 0 | 9 |
| ASV1235 | 0cfd5a997e4 | 9 | 0 | 0 | 0 | 0 | 0 | 0 | 0 | 0 | 0 | 0 | 0 | 0 | 0 | 0 | 0 | 0 | 0 | 9 | 0 | 0 | 0 | 0 | 0 | 0 | 0 | 0 | 0 | 0 | 0 | 0 | 0 | 0 | 0 | 0 | 0 |
| ASV1236 | 72478d44ccf | 9 | 0 | 0 | 0 | 0 | 9 | 0 | 0 | 0 | 0 | 0 | 0 | 0 | 0 | 0 | 0 | 0 | 0 | 0 | 0 | 0 | 0 | 0 | 0 | 0 | 0 | 0 | 0 | 0 | 0 | 0 | 0 | 0 | 0 | 0 | 0 |
| ASV1237 | bb157bfccbd | 9 | 0 | 0 | 0 | 0 | 0 | 0 | 0 | 0 | 0 | 0 | 0 | 0 | 0 | 0 | 0 | 0 | 0 | 0 | 0 | 0 | 0 | 0 | 0 | 0 | 0 | 0 | 0 | 0 | 0 | 0 | 9 | 0 | 0 | 0 | 0 |
| ASV1238 | 04aef29ff0a4 | 9 | 0 | 0 | 0 | 0 | 0 | 0 | 0 | 0 | 0 | 0 | 0 | 0 | 0 | 0 | 0 | 0 | 0 | 0 | 0 | 0 | 0 | 0 | 0 | 0 | 9 | 0 | 0 | 0 | 0 | 0 | 0 | 0 | 0 | 0 | 0 |
| ASV1239 | 17bc9a76366 | 9 | 0 | 0 | 0 | 0 | 0 | 0 | 0 | 0 | 0 | 0 | 0 | 0 | 0 | 0 | 0 | 0 | 0 | 0 | 0 | 0 | 0 | 0 | 0 | 0 | 0 | 0 | 0 | 0 | 4 | 5 | 0 | 0 | 0 | 0 | 0 |
| ASV1240 | 5cbbb3191e7 | 9 | 0 | 0 | 0 | 0 | 6 | 0 | 0 | 0 | 0 | 0 | 0 | 0 | 0 | 0 | 3 | 0 | 0 | 0 | 0 | 0 | 0 | 0 | 0 | 0 | 0 | 0 | 0 | 0 | 0 | 0 | 0 | 0 | 0 | 0 | 0 |
| ASV1241 | 580190655ef | 9 | 0 | 0 | 0 | 0 | 0 | 0 | 0 | 0 | 0 | 0 | 0 | 0 | 0 | 0 | 0 | 0 | 0 | 0 | 0 | 0 | 0 | 0 | 0 | 0 | 0 | 0 | 0 | 0 | 0 | 0 | 0 | 0 | 0 | 9 | 0 |
| ASV1242 | 3258f6ac748 | 9 | 0 | 0 | 0 | 0 | 0 | 0 | 0 | 0 | 0 | 0 | 0 | 0 | 0 | 0 | 0 | 0 | 0 | 5 | 0 | 0 | 0 | 0 | 4 | 0 | 0 | 0 | 0 | 0 | 0 | 0 | 0 | 0 | 0 | 0 | 0 |
| ASV1243 | 95e44c0cfc0 | 9 | 0 | 0 | 0 | 0 | 0 | 0 | 0 | 0 | 0 | 0 | 0 | 0 | 0 | 0 | 0 | 0 | 0 | 9 | 0 | 0 | 0 | 0 | 0 | 0 | 0 | 0 | 0 | 0 | 0 | 0 | 0 | 0 | 0 | 0 | 0 |
| ASV1244 | ce6531e8c56 | 9 | 0 | 0 | 0 | 0 | 0 | 0 | 0 | 0 | 0 | 0 | 0 | 0 | 0 | 0 | 0 | 0 | 0 | 0 | 0 | 0 | 0 | 0 | 0 | 0 | 0 | 0 | 0 | 0 | 0 | 0 | 9 | 0 | 0 | 0 | 0 |
| ASV1245 | 6598be55fe5 | 9 | 0 | 0 | 0 | 0 | 9 | 0 | 0 | 0 | 0 | 0 | 0 | 0 | 0 | 0 | 0 | 0 | 0 | 0 | 0 | 0 | 0 | 0 | 0 | 0 | 0 | 0 | 0 | 0 | 0 | 0 | 0 | 0 | 0 | 0 | 0 |
| ASV1246 | f9c91207f762 | 9 | 9 | 0 | 0 | 0 | 0 | 0 | 0 | 0 | 0 | 0 | 0 | 0 | 0 | 0 | 0 | 0 | 0 | 0 | 0 | 0 | 0 | 0 | 0 | 0 | 0 | 0 | 0 | 0 | 0 | 0 | 0 | 0 | 0 | 0 | 0 |
| ASV1247 | ebe708b42df | 9 | 0 | 0 | 0 | 9 | 0 | 0 | 0 | 0 | 0 | 0 | 0 | 0 | 0 | 0 | 0 | 0 | 0 | 0 | 0 | 0 | 0 | 0 | 0 | 0 | 0 | 0 | 0 | 0 | 0 | 0 | 0 | 0 | 0 | 0 | 0 |
| ASV1248 | 062c5c26200 | 9 | 0 | 0 | 0 | 0 | 0 | 0 | 0 | 0 | 9 | 0 | 0 | 0 | 0 | 0 | 0 | 0 | 0 | 0 | 0 | 0 | 0 | 0 | 0 | 0 | 0 | 0 | 0 | 0 | 0 | 0 | 0 | 0 | 0 | 0 | 0 |
| ASV1249 | 38f7c51a29e | 9 | 0 | 0 | 0 | 0 | 0 | 0 | 0 | 0 | 0 | 3 | 0 | 0 | 0 | 0 | 6 | 0 | 0 | 0 | 0 | 0 | 0 | 0 | 0 | 0 | 0 | 0 | 0 | 0 | 0 | 0 | 0 | 0 | 0 | 0 | 0 |
| ASV1250 | 4d56acb3808 | 9 | 0 | 0 | 0 | 0 | 0 | 0 | 0 | 0 | 0 | 0 | 0 | 0 | 0 | 0 | 0 | 0 | 0 | 0 | 0 | 0 | 9 | 0 | 0 | 0 | 0 | 0 | 0 | 0 | 0 | 0 | 0 | 0 | 0 | 0 | 0 |
| ASV1251 | ea6bc720f96 | 9 | 0 | 0 | 0 | 0 | 0 | 0 | 0 | 0 | 0 | 0 | 0 | 0 | 0 | 0 | 9 | 0 | 0 | 0 | 0 | 0 | 0 | 0 | 0 | 0 | 0 | 0 | 0 | 0 | 0 | 0 | 0 | 0 | 0 | 0 | 0 |
| ASV1252 | 8a9f2ed1917 | 9 | 0 | 0 | 0 | 0 | 0 | 0 | 0 | 0 | 0 | 0 | 0 | 0 | 0 | 0 | 0 | 0 | 0 | 0 | 0 | 9 | 0 | 0 | 0 | 0 | 0 | 0 | 0 | 0 | 0 | 0 | 0 | 0 | 0 | 0 | 0 |
| ASV1253 | b15c1b61bf9 | 9 | 0 | 0 | 0 | 0 | 0 | 3 | 0 | 0 | 0 | 0 | 0 | 0 | 6 | 0 | 0 | 0 | 0 | 0 | 0 | 0 | 0 | 0 | 0 | 0 | 0 | 0 | 0 | 0 | 0 | 0 | 0 | 0 | 0 | 0 | 0 |
| ASV1254 | 4d5c8a8090f | 9 | 0 | 0 | 0 | 0 | 0 | 0 | 0 | 0 | 0 | 0 | 0 | 0 | 0 | 0 | 0 | 0 | 0 | 0 | 0 | 9 | 0 | 0 | 0 | 0 | 0 | 0 | 0 | 0 | 0 | 0 | 0 | 0 | 0 | 0 | 0 |
| ASV1255 | e2fab2a594d | 9 | 0 | 0 | 0 | 0 | 0 | 0 | 0 | 0 | 0 | 0 | 0 | 0 | 0 | 0 | 0 | 0 | 0 | 0 | 0 | 0 | 0 | 0 | 0 | 0 | 6 | 0 | 0 | 3 | 0 | 0 | 0 | 0 | 0 | 0 | 0 |
| ASV1256 | c77a164cd24 | 9 | 0 | 0 | 0 | 0 | 9 | 0 | 0 | 0 | 0 | 0 | 0 | 0 | 0 | 0 | 0 | 0 | 0 | 0 | 0 | 0 | 0 | 0 | 0 | 0 | 0 | 0 | 0 | 0 | 0 | 0 | 0 | 0 | 0 | 0 | 0 |
| ASV1257 | 3835d0369ab | 9 | 0 | 0 | 0 | 0 | 0 | 0 | 0 | 0 | 9 | 0 | 0 | 0 | 0 | 0 | 0 | 0 | 0 | 0 | 0 | 0 | 0 | 0 | 0 | 0 | 0 | 0 | 0 | 0 | 0 | 0 | 0 | 0 | 0 | 0 | 0 |
| ASV1258 | cdfbc000e95 | 9 | 0 | 0 | 0 | 0 | 0 | 0 | 0 | 0 | 0 | 0 | 0 | 0 | 0 | 0 | 0 | 0 | 0 | 0 | 0 | 9 | 0 | 0 | 0 | 0 | 0 | 0 | 0 | 0 | 0 | 0 | 0 | 0 | 0 | 0 | 0 |
| ASV1259 | 3404f929934 | 9 | 0 | 0 | 3 | 2 | 0 | 0 | 0 | 0 | 0 | 0 | 0 | 0 | 0 | 0 | 2 | 0 | 2 | 0 | 0 | 0 | 0 | 0 | 0 | 0 | 0 | 0 | 0 | 0 | 0 | 0 | 0 | 0 | 0 | 0 | 0 |
| ASV1260 | c0cce68eb3c | 9 | 0 | 0 | 0 | 0 | 0 | 0 | 0 | 0 | 0 | 0 | 0 | 0 | 0 | 0 | 0 | 0 | 9 | 0 | 0 | 0 | 0 | 0 | 0 | 0 | 0 | 0 | 0 | 0 | 0 | 0 | 0 | 0 | 0 | 0 | 0 |
| ASV1261 | 564c3c6610a | 9 | 0 | 0 | 0 | 0 | 0 | 0 | 0 | 0 | 0 | 9 | 0 | 0 | 0 | 0 | 0 | 0 | 0 | 0 | 0 | 0 | 0 | 0 | 0 | 0 | 0 | 0 | 0 | 0 | 0 | 0 | 0 | 0 | 0 | 0 | 0 |
| ASV1262 | a5e1d9afda6 | 9 | 9 | 0 | 0 | 0 | 0 | 0 | 0 | 0 | 0 | 0 | 0 | 0 | 0 | 0 | 0 | 0 | 0 | 0 | 0 | 0 | 0 | 0 | 0 | 0 | 0 | 0 | 0 | 0 | 0 | 0 | 0 | 0 | 0 | 0 | 0 |
| ASV1263 | 95b53fb7d0f | 9 | 0 | 0 | 0 | 0 | 0 | 0 | 0 | 0 | 0 | 0 | 0 | 0 | 0 | 0 | 0 | 2 | 0 | 0 | 0 | 0 | 7 | 0 | 0 | 0 | 0 | 0 | 0 | 0 | 0 | 0 | 0 | 0 | 0 | 0 | 0 |
| ASV1264 | 1c54604c687 | 9 | 0 | 0 | 0 | 0 | 0 | 0 | 0 | 0 | 0 | 0 | 0 | 0 | 0 | 0 | 0 | 0 | 0 | 0 | 0 | 0 | 0 | 0 | 0 | 0 | 0 | 0 | 0 | 0 | 0 | 0 | 0 | 9 | 0 | 0 | 0 |
| ASV1265 | 3de83fe4c9c | 9 | 0 | 0 | 0 | 0 | 0 | 0 | 0 | 0 | 0 | 0 | 0 | 0 | 0 | 0 | 0 | 9 | 0 | 0 | 0 | 0 | 0 | 0 | 0 | 0 | 0 | 0 | 0 | 0 | 0 | 0 | 0 | 0 | 0 | 0 | 0 |

**Supplementary Table S5**. Bacterioplankton 16S rRNA ASV feature table
ASVs: 1779
Seqs: 918792

| #ASV_ID | Feature_ID | total seqs | May12_0m | May12_05m | May12_25m | May12_50m | May12_75m | Jun8_0m | Jun8_05m | Jun8_25m | Jun8_50m | Jun8_75m | Jun22_0m | Jun22_05m | Jun22_25m | Jun22_50m | Jun22_75m | Jul11_0m | Jul11_05m | Jul11_25m | Jul11_50m | Jul11_75m | Jul27_0m | Jul27_05m | Jul27_25m | Jul27_50m | Jul27_75m | Aug5_0m | Aug5_05m | Aug5_25m | Aug5_50m | Aug5_75m | Sep8_0m | Sep8_05m | Sep8_25m | Sep8_50m | Sep8_75m |
|---|---|---|---|---|---|---|---|---|---|---|---|---|---|---|---|---|---|---|---|---|---|---|---|---|---|---|---|---|---|---|---|---|---|---|---|---|---|
| ASV1266 | baa60b1c093 | 8 | 0 | 0 | 0 | 0 | 0 | 0 | 0 | 0 | 0 | 0 | 0 | 0 | 0 | 0 | 0 | 4 | 4 | 0 | 0 | 0 | 0 | 0 | 0 | 0 | 0 | 0 | 0 | 0 | 0 | 0 | 0 | 0 | 0 | 0 | 0 |
| ASV1267 | dba9f72c944 | 8 | 0 | 0 | 0 | 0 | 8 | 0 | 0 | 0 | 0 | 0 | 0 | 0 | 0 | 0 | 0 | 0 | 0 | 0 | 0 | 0 | 0 | 0 | 0 | 0 | 0 | 0 | 0 | 0 | 0 | 0 | 0 | 0 | 0 | 0 | 0 |
| ASV1268 | 87e65411867 | 8 | 0 | 0 | 0 | 4 | 4 | 0 | 0 | 0 | 0 | 0 | 0 | 0 | 0 | 0 | 0 | 0 | 0 | 0 | 0 | 0 | 0 | 0 | 0 | 0 | 0 | 0 | 0 | 0 | 0 | 0 | 0 | 0 | 0 | 0 | 0 |
| ASV1269 | f682a9827a5 | 8 | 0 | 0 | 0 | 0 | 8 | 0 | 0 | 0 | 0 | 0 | 0 | 0 | 0 | 0 | 0 | 0 | 0 | 0 | 0 | 0 | 0 | 0 | 0 | 0 | 0 | 0 | 0 | 0 | 0 | 0 | 0 | 0 | 0 | 0 | 0 |
| ASV1270 | fec41ffee3bf | 8 | 0 | 0 | 0 | 0 | 0 | 0 | 0 | 0 | 0 | 0 | 0 | 0 | 0 | 0 | 0 | 0 | 0 | 0 | 0 | 0 | 0 | 0 | 0 | 0 | 0 | 0 | 0 | 0 | 0 | 8 | 0 | 0 | 0 | 0 | 0 |
| ASV1271 | 9fdc6db7f88f | 8 | 0 | 0 | 0 | 0 | 0 | 0 | 0 | 0 | 0 | 0 | 0 | 0 | 0 | 0 | 0 | 0 | 0 | 0 | 0 | 0 | 0 | 0 | 0 | 8 | 0 | 0 | 0 | 0 | 0 | 0 | 0 | 0 | 0 | 0 | 0 |
| ASV1272 | 489760bf3d8 | 8 | 0 | 0 | 0 | 0 | 0 | 8 | 0 | 0 | 0 | 0 | 0 | 0 | 0 | 0 | 0 | 0 | 0 | 0 | 0 | 0 | 0 | 0 | 0 | 0 | 0 | 0 | 0 | 0 | 0 | 0 | 0 | 0 | 0 | 0 | 0 |
| ASV1273 | 0fae4a96139 | 8 | 0 | 0 | 0 | 0 | 0 | 0 | 0 | 0 | 0 | 0 | 0 | 0 | 0 | 0 | 8 | 0 | 0 | 0 | 0 | 0 | 0 | 0 | 0 | 0 | 0 | 0 | 0 | 0 | 0 | 0 | 0 | 0 | 0 | 0 | 0 |
| ASV1274 | f5568c69263 | 8 | 0 | 0 | 0 | 0 | 0 | 0 | 0 | 0 | 0 | 0 | 0 | 0 | 0 | 0 | 8 | 0 | 0 | 0 | 0 | 0 | 0 | 0 | 0 | 0 | 0 | 0 | 0 | 0 | 0 | 0 | 0 | 0 | 0 | 0 | 0 |
| ASV1275 | 943b298ff2d | 8 | 0 | 0 | 0 | 0 | 0 | 0 | 0 | 0 | 0 | 0 | 0 | 0 | 0 | 0 | 0 | 8 | 0 | 0 | 0 | 0 | 0 | 0 | 0 | 0 | 0 | 0 | 0 | 0 | 0 | 0 | 0 | 0 | 0 | 0 | 0 |
| ASV1276 | b7c31f68ffdb | 8 | 0 | 0 | 0 | 0 | 0 | 0 | 0 | 0 | 0 | 0 | 0 | 8 | 0 | 0 | 0 | 0 | 0 | 0 | 0 | 0 | 0 | 0 | 0 | 0 | 0 | 0 | 0 | 0 | 0 | 0 | 0 | 0 | 0 | 0 | 0 |
| ASV1277 | 2ba659a32f3 | 8 | 0 | 0 | 0 | 8 | 0 | 0 | 0 | 0 | 0 | 0 | 0 | 0 | 0 | 0 | 0 | 0 | 0 | 0 | 0 | 0 | 0 | 0 | 0 | 0 | 0 | 0 | 0 | 0 | 0 | 0 | 0 | 0 | 0 | 0 | 0 |
| ASV1278 | c70043f9bb0 | 8 | 0 | 0 | 0 | 0 | 0 | 8 | 0 | 0 | 0 | 0 | 0 | 0 | 0 | 0 | 0 | 0 | 0 | 0 | 0 | 0 | 0 | 0 | 0 | 0 | 0 | 0 | 0 | 0 | 0 | 0 | 0 | 0 | 0 | 0 | 0 |
| ASV1279 | 59cdf625fe5 | 8 | 0 | 0 | 0 | 0 | 0 | 0 | 0 | 0 | 0 | 0 | 0 | 0 | 0 | 0 | 0 | 0 | 0 | 0 | 0 | 8 | 0 | 0 | 0 | 0 | 0 | 0 | 0 | 0 | 0 | 0 | 0 | 0 | 0 | 0 | 0 |
| ASV1280 | b7b973f5199 | 8 | 0 | 0 | 0 | 0 | 0 | 0 | 0 | 0 | 0 | 0 | 0 | 0 | 8 | 0 | 0 | 0 | 0 | 0 | 0 | 0 | 0 | 0 | 0 | 0 | 0 | 0 | 0 | 0 | 0 | 0 | 0 | 0 | 0 | 0 | 0 |
| ASV1281 | 8e8214b1c5b | 8 | 0 | 0 | 0 | 0 | 0 | 0 | 0 | 0 | 8 | 0 | 0 | 0 | 0 | 0 | 0 | 0 | 0 | 0 | 0 | 0 | 0 | 0 | 0 | 0 | 0 | 0 | 0 | 0 | 0 | 0 | 0 | 0 | 0 | 0 | 0 |
| ASV1282 | 72aa622a680 | 8 | 0 | 0 | 0 | 0 | 0 | 0 | 0 | 0 | 0 | 0 | 0 | 0 | 0 | 0 | 0 | 0 | 0 | 0 | 0 | 0 | 0 | 0 | 0 | 0 | 0 | 0 | 0 | 0 | 0 | 8 | 0 | 0 | 0 | 0 | 0 |
| ASV1283 | 922f28646c7 | 8 | 0 | 0 | 0 | 0 | 0 | 0 | 0 | 0 | 0 | 0 | 0 | 0 | 0 | 0 | 0 | 0 | 0 | 0 | 0 | 0 | 0 | 0 | 0 | 0 | 0 | 0 | 0 | 0 | 0 | 0 | 8 | 0 | 0 | 0 | 0 |
| ASV1284 | 6e9fb661da4 | 8 | 0 | 0 | 0 | 0 | 0 | 0 | 0 | 0 | 0 | 0 | 0 | 0 | 0 | 0 | 0 | 0 | 0 | 0 | 0 | 0 | 0 | 0 | 0 | 8 | 0 | 0 | 0 | 0 | 0 | 0 | 0 | 0 | 0 | 0 | 0 |
| ASV1285 | 66f5120c0c6 | 8 | 0 | 0 | 0 | 0 | 0 | 8 | 0 | 0 | 0 | 0 | 0 | 0 | 0 | 0 | 0 | 0 | 0 | 0 | 0 | 0 | 0 | 0 | 0 | 0 | 0 | 0 | 0 | 0 | 0 | 0 | 0 | 0 | 0 | 0 | 0 |
| ASV1286 | 97e80f21afbf | 8 | 8 | 0 | 0 | 0 | 0 | 0 | 0 | 0 | 0 | 0 | 0 | 0 | 0 | 0 | 0 | 0 | 0 | 0 | 0 | 0 | 0 | 0 | 0 | 0 | 0 | 0 | 0 | 0 | 0 | 0 | 0 | 0 | 0 | 0 | 0 |
| ASV1287 | 94a0ddc0e96 | 8 | 0 | 0 | 8 | 0 | 0 | 0 | 0 | 0 | 0 | 0 | 0 | 0 | 0 | 0 | 0 | 0 | 0 | 0 | 0 | 0 | 0 | 0 | 0 | 0 | 0 | 0 | 0 | 0 | 0 | 0 | 0 | 0 | 0 | 0 | 0 |
| ASV1288 | a3b2da88ac8 | 8 | 0 | 0 | 0 | 0 | 4 | 0 | 0 | 0 | 0 | 0 | 0 | 0 | 0 | 0 | 4 | 0 | 0 | 0 | 0 | 0 | 0 | 0 | 0 | 0 | 0 | 0 | 0 | 0 | 0 | 0 | 0 | 0 | 0 | 0 | 0 |
| ASV1289 | 53d4e655ba7 | 8 | 0 | 0 | 0 | 0 | 8 | 0 | 0 | 0 | 0 | 0 | 0 | 0 | 0 | 0 | 0 | 0 | 0 | 0 | 0 | 0 | 0 | 0 | 0 | 0 | 0 | 0 | 0 | 0 | 0 | 0 | 0 | 0 | 0 | 0 | 0 |
| ASV1290 | 509d18c6dbf | 8 | 0 | 0 | 0 | 0 | 0 | 0 | 0 | 0 | 0 | 0 | 0 | 0 | 0 | 0 | 8 | 0 | 0 | 0 | 0 | 0 | 0 | 0 | 0 | 0 | 0 | 0 | 0 | 0 | 0 | 0 | 0 | 0 | 0 | 0 | 0 |
| ASV1291 | ab20d637985 | 8 | 0 | 0 | 0 | 0 | 8 | 0 | 0 | 0 | 0 | 0 | 0 | 0 | 0 | 0 | 0 | 0 | 0 | 0 | 0 | 0 | 0 | 0 | 0 | 0 | 0 | 0 | 0 | 0 | 0 | 0 | 0 | 0 | 0 | 0 | 0 |
| ASV1292 | 726cf59156c | 8 | 0 | 0 | 0 | 0 | 0 | 0 | 0 | 0 | 0 | 0 | 0 | 0 | 0 | 0 | 0 | 0 | 0 | 0 | 0 | 0 | 0 | 0 | 0 | 0 | 0 | 0 | 0 | 0 | 0 | 4 | 0 | 0 | 0 | 4 | 0 |
| ASV1293 | 6e4f053a510 | 8 | 0 | 0 | 0 | 0 | 0 | 0 | 0 | 0 | 0 | 0 | 0 | 0 | 0 | 0 | 0 | 0 | 0 | 0 | 0 | 0 | 0 | 0 | 0 | 0 | 0 | 0 | 0 | 6 | 2 | 0 | 0 | 0 | 0 | 0 | 0 |
| ASV1294 | ecbe8a8ccb1 | 8 | 0 | 0 | 0 | 0 | 0 | 0 | 0 | 0 | 0 | 0 | 0 | 0 | 0 | 0 | 0 | 0 | 8 | 0 | 0 | 0 | 0 | 0 | 0 | 0 | 0 | 0 | 0 | 0 | 0 | 0 | 0 | 0 | 0 | 0 | 0 |
| ASV1295 | d26cd7f6bdb | 8 | 0 | 0 | 0 | 0 | 0 | 0 | 0 | 0 | 0 | 0 | 0 | 0 | 0 | 0 | 0 | 0 | 0 | 0 | 0 | 0 | 0 | 0 | 0 | 0 | 0 | 0 | 0 | 0 | 8 | 0 | 0 | 0 | 0 | 0 | 0 |
| ASV1296 | 3ab2f61acc2 | 8 | 0 | 0 | 0 | 0 | 0 | 0 | 0 | 0 | 0 | 0 | 0 | 0 | 0 | 0 | 0 | 0 | 0 | 8 | 0 | 0 | 0 | 0 | 0 | 0 | 0 | 0 | 0 | 0 | 0 | 0 | 0 | 0 | 0 | 0 | 0 |
| ASV1297 | f333eb1f71a | 8 | 0 | 0 | 0 | 0 | 0 | 0 | 0 | 0 | 0 | 0 | 0 | 0 | 0 | 0 | 0 | 0 | 8 | 0 | 0 | 0 | 0 | 0 | 0 | 0 | 0 | 0 | 0 | 0 | 0 | 0 | 0 | 0 | 0 | 0 | 0 |
| ASV1298 | d070a05a7ab | 8 | 4 | 0 | 0 | 0 | 4 | 0 | 0 | 0 | 0 | 0 | 0 | 0 | 0 | 0 | 0 | 0 | 0 | 0 | 0 | 0 | 0 | 0 | 0 | 0 | 0 | 0 | 0 | 0 | 0 | 0 | 0 | 0 | 0 | 0 | 0 |
| ASV1299 | 15f5ea541d3 | 8 | 0 | 0 | 0 | 0 | 0 | 0 | 0 | 0 | 0 | 0 | 0 | 0 | 0 | 0 | 0 | 0 | 0 | 0 | 0 | 0 | 8 | 0 | 0 | 0 | 0 | 0 | 0 | 0 | 0 | 0 | 0 | 0 | 0 | 0 | 0 |
| ASV1300 | 9edef7d5515 | 8 | 0 | 0 | 4 | 4 | 0 | 0 | 0 | 0 | 0 | 0 | 0 | 0 | 0 | 0 | 0 | 0 | 0 | 0 | 0 | 0 | 0 | 0 | 0 | 0 | 0 | 0 | 0 | 0 | 0 | 0 | 0 | 0 | 0 | 0 | 0 |
| ASV1301 | 33e705ea1eb | 8 | 0 | 0 | 0 | 0 | 0 | 0 | 0 | 0 | 5 | 3 | 0 | 0 | 0 | 0 | 0 | 0 | 0 | 0 | 0 | 0 | 0 | 0 | 0 | 0 | 0 | 0 | 0 | 0 | 0 | 0 | 0 | 0 | 0 | 0 | 0 |
| ASV1302 | 2d56d9d44ff | 8 | 0 | 0 | 0 | 0 | 0 | 0 | 0 | 0 | 0 | 0 | 0 | 0 | 0 | 0 | 0 | 0 | 0 | 0 | 0 | 0 | 0 | 0 | 0 | 0 | 0 | 0 | 0 | 8 | 0 | 0 | 0 | 0 | 0 | 0 | 0 |
| ASV1303 | 8a78b2a83c2 | 8 | 0 | 0 | 0 | 0 | 0 | 0 | 0 | 0 | 0 | 0 | 0 | 0 | 0 | 0 | 0 | 8 | 0 | 0 | 0 | 0 | 0 | 0 | 0 | 0 | 0 | 0 | 0 | 0 | 0 | 0 | 0 | 0 | 0 | 0 | 0 |
| ASV1304 | 2e4be662f07 | 8 | 0 | 0 | 0 | 8 | 0 | 0 | 0 | 0 | 0 | 0 | 0 | 0 | 0 | 0 | 0 | 0 | 0 | 0 | 0 | 0 | 0 | 0 | 0 | 0 | 0 | 0 | 0 | 0 | 0 | 0 | 0 | 0 | 0 | 0 | 0 |
| ASV1305 | 98f11dae43b | 8 | 0 | 0 | 0 | 0 | 0 | 0 | 0 | 0 | 0 | 0 | 0 | 0 | 0 | 0 | 0 | 0 | 0 | 0 | 0 | 0 | 0 | 0 | 0 | 8 | 0 | 0 | 0 | 0 | 0 | 0 | 0 | 0 | 0 | 0 | 0 |
| ASV1306 | 1393f76bb10 | 8 | 0 | 0 | 0 | 0 | 8 | 0 | 0 | 0 | 0 | 0 | 0 | 0 | 0 | 0 | 0 | 0 | 0 | 0 | 0 | 0 | 0 | 0 | 0 | 0 | 0 | 0 | 0 | 0 | 0 | 0 | 0 | 0 | 0 | 0 | 0 |
| ASV1307 | a808d351235 | 8 | 0 | 0 | 0 | 8 | 0 | 0 | 0 | 0 | 0 | 0 | 0 | 0 | 0 | 0 | 0 | 0 | 0 | 0 | 0 | 0 | 0 | 0 | 0 | 0 | 0 | 0 | 0 | 0 | 0 | 0 | 0 | 0 | 0 | 0 | 0 |
| ASV1308 | 7f0420aa9f2 | 8 | 0 | 0 | 0 | 0 | 0 | 0 | 0 | 0 | 0 | 0 | 0 | 0 | 0 | 0 | 0 | 0 | 0 | 0 | 0 | 0 | 0 | 0 | 0 | 0 | 0 | 0 | 0 | 0 | 0 | 0 | 0 | 8 | 0 | 0 | 0 |
| ASV1309 | 108581dd41a | 8 | 0 | 0 | 0 | 0 | 0 | 0 | 0 | 0 | 0 | 0 | 0 | 0 | 0 | 0 | 0 | 0 | 0 | 0 | 0 | 0 | 0 | 0 | 0 | 0 | 0 | 8 | 0 | 0 | 0 | 0 | 0 | 0 | 0 | 0 | 0 |
| ASV1310 | 3a5cd96f54d | 8 | 0 | 0 | 0 | 0 | 0 | 0 | 0 | 0 | 0 | 0 | 0 | 0 | 0 | 0 | 8 | 0 | 0 | 0 | 0 | 0 | 0 | 0 | 0 | 0 | 0 | 0 | 0 | 0 | 0 | 0 | 0 | 0 | 0 | 0 | 0 |
| ASV1311 | ac3694e776b | 8 | 0 | 0 | 0 | 0 | 0 | 0 | 0 | 0 | 0 | 0 | 0 | 0 | 0 | 0 | 0 | 0 | 0 | 0 | 0 | 0 | 0 | 0 | 0 | 0 | 0 | 0 | 0 | 8 | 0 | 0 | 0 | 0 | 0 | 0 | 0 |
| ASV1312 | 9a40dc9e0f4 | 8 | 0 | 0 | 0 | 0 | 0 | 0 | 0 | 0 | 0 | 0 | 0 | 0 | 0 | 0 | 8 | 0 | 0 | 0 | 0 | 0 | 0 | 0 | 0 | 0 | 0 | 0 | 0 | 0 | 0 | 0 | 0 | 0 | 0 | 0 | 0 |
| ASV1313 | 3554e46b91e | 8 | 0 | 0 | 0 | 0 | 0 | 0 | 0 | 0 | 0 | 0 | 0 | 0 | 0 | 0 | 0 | 0 | 0 | 0 | 0 | 0 | 0 | 0 | 0 | 0 | 0 | 0 | 0 | 0 | 8 | 0 | 0 | 0 | 0 | 0 | 0 |
| ASV1314 | 246b7de821c | 8 | 0 | 0 | 0 | 0 | 0 | 2 | 0 | 0 | 0 | 0 | 0 | 0 | 0 | 0 | 0 | 0 | 0 | 0 | 0 | 0 | 0 | 0 | 0 | 0 | 6 | 0 | 0 | 0 | 0 | 0 | 0 | 0 | 0 | 0 | 0 |
| ASV1315 | 0e9b8b829db | 8 | 0 | 0 | 0 | 0 | 0 | 0 | 0 | 0 | 0 | 8 | 0 | 0 | 0 | 0 | 0 | 0 | 0 | 0 | 0 | 0 | 0 | 0 | 0 | 0 | 0 | 0 | 0 | 0 | 0 | 0 | 0 | 0 | 0 | 0 | 0 |
| ASV1316 | 23cd2b04141 | 8 | 0 | 0 | 0 | 0 | 0 | 0 | 0 | 0 | 0 | 0 | 0 | 0 | 0 | 0 | 0 | 0 | 0 | 0 | 0 | 0 | 0 | 0 | 0 | 0 | 0 | 0 | 0 | 0 | 0 | 0 | 0 | 0 | 8 | 0 | 0 |
| ASV1317 | 62d822227d0 | 8 | 0 | 0 | 0 | 0 | 8 | 0 | 0 | 0 | 0 | 0 | 0 | 0 | 0 | 0 | 0 | 0 | 0 | 0 | 0 | 0 | 0 | 0 | 0 | 0 | 0 | 0 | 0 | 0 | 0 | 0 | 0 | 0 | 0 | 0 | 0 |
| ASV1318 | ac0f3d11539 | 8 | 0 | 0 | 0 | 0 | 8 | 0 | 0 | 0 | 0 | 0 | 0 | 0 | 0 | 0 | 0 | 0 | 0 | 0 | 0 | 0 | 0 | 0 | 0 | 0 | 0 | 0 | 0 | 0 | 0 | 0 | 0 | 0 | 0 | 0 | 0 |
| ASV1319 | 6e54cec1bab | 8 | 0 | 0 | 0 | 0 | 0 | 0 | 0 | 0 | 0 | 0 | 0 | 0 | 0 | 8 | 0 | 0 | 0 | 0 | 0 | 0 | 0 | 0 | 0 | 0 | 0 | 0 | 0 | 0 | 0 | 0 | 0 | 0 | 0 | 0 | 0 |
| ASV1320 | a651f5af1c2 | 8 | 0 | 0 | 0 | 0 | 0 | 0 | 0 | 0 | 0 | 8 | 0 | 0 | 0 | 0 | 0 | 0 | 0 | 0 | 0 | 0 | 0 | 0 | 0 | 0 | 0 | 0 | 0 | 0 | 0 | 0 | 0 | 0 | 0 | 0 | 0 |
| ASV1321 | f7cba05fcaee | 8 | 0 | 0 | 0 | 0 | 0 | 0 | 0 | 0 | 0 | 0 | 0 | 0 | 0 | 0 | 0 | 0 | 0 | 0 | 0 | 0 | 0 | 0 | 0 | 0 | 0 | 0 | 0 | 0 | 0 | 8 | 0 | 0 | 0 | 0 | 0 |
| ASV1322 | 263c7a8bb12 | 8 | 0 | 0 | 0 | 0 | 0 | 0 | 0 | 0 | 0 | 0 | 0 | 0 | 0 | 0 | 0 | 8 | 0 | 0 | 0 | 0 | 0 | 0 | 0 | 0 | 0 | 0 | 0 | 0 | 0 | 0 | 0 | 0 | 0 | 0 | 0 |
| ASV1323 | af42e6e4fd9 | 7 | 0 | 0 | 0 | 0 | 0 | 4 | 3 | 0 | 0 | 0 | 0 | 0 | 0 | 0 | 0 | 0 | 0 | 0 | 0 | 0 | 0 | 0 | 0 | 0 | 0 | 0 | 0 | 0 | 0 | 0 | 0 | 0 | 0 | 0 | 0 |
| ASV1324 | 9621167ddea | 7 | 0 | 0 | 0 | 0 | 0 | 0 | 0 | 0 | 0 | 0 | 0 | 0 | 0 | 0 | 0 | 0 | 0 | 0 | 0 | 0 | 0 | 0 | 0 | 0 | 7 | 0 | 0 | 0 | 0 | 0 | 0 | 0 | 0 | 0 | 0 |
| ASV1325 | 89a258b55e5 | 7 | 0 | 0 | 0 | 0 | 0 | 0 | 0 | 0 | 0 | 0 | 0 | 0 | 0 | 0 | 0 | 0 | 0 | 0 | 0 | 0 | 0 | 0 | 0 | 0 | 0 | 0 | 0 | 0 | 0 | 0 | 0 | 0 | 0 | 0 | 7 |
| ASV1326 | b0dc5f42017 | 7 | 0 | 0 | 0 | 0 | 0 | 0 | 0 | 0 | 0 | 0 | 0 | 0 | 0 | 0 | 0 | 0 | 0 | 0 | 3 | 0 | 0 | 0 | 0 | 0 | 0 | 0 | 0 | 0 | 0 | 0 | 0 | 4 | 0 | 0 | 0 |
| ASV1327 | 7d3e1f71754 | 7 | 0 | 0 | 0 | 0 | 0 | 0 | 0 | 0 | 0 | 0 | 0 | 0 | 0 | 0 | 0 | 0 | 5 | 2 | 0 | 0 | 0 | 0 | 0 | 0 | 0 | 0 | 0 | 0 | 0 | 0 | 0 | 0 | 0 | 0 | 0 |
| ASV1328 | 49fd7322a61 | 7 | 0 | 0 | 0 | 0 | 0 | 0 | 0 | 0 | 0 | 0 | 0 | 0 | 0 | 7 | 0 | 0 | 0 | 0 | 0 | 0 | 0 | 0 | 0 | 0 | 0 | 0 | 0 | 0 | 0 | 0 | 0 | 0 | 0 | 0 | 0 |
| ASV1329 | 42f24da52fcf | 7 | 0 | 0 | 0 | 0 | 0 | 0 | 0 | 0 | 0 | 0 | 0 | 0 | 0 | 0 | 7 | 0 | 0 | 0 | 0 | 0 | 0 | 0 | 0 | 0 | 0 | 0 | 0 | 0 | 0 | 0 | 0 | 0 | 0 | 0 | 0 |
| ASV1330 | 0e4dc486dd0 | 7 | 0 | 0 | 0 | 0 | 0 | 0 | 0 | 0 | 0 | 0 | 0 | 0 | 0 | 0 | 7 | 0 | 0 | 0 | 0 | 0 | 0 | 0 | 0 | 0 | 0 | 0 | 0 | 0 | 0 | 0 | 0 | 0 | 0 | 0 | 0 |
| ASV1331 | 0eb26f6155f | 7 | 0 | 0 | 0 | 0 | 0 | 0 | 0 | 0 | 0 | 0 | 0 | 0 | 0 | 0 | 0 | 0 | 0 | 0 | 0 | 0 | 7 | 0 | 0 | 0 | 0 | 0 | 0 | 0 | 0 | 0 | 0 | 0 | 0 | 0 | 0 |
| ASV1332 | d4e873350e9 | 7 | 0 | 0 | 0 | 0 | 0 | 0 | 0 | 0 | 0 | 0 | 0 | 0 | 0 | 0 | 0 | 0 | 0 | 0 | 0 | 0 | 0 | 0 | 0 | 0 | 0 | 0 | 0 | 0 | 0 | 0 | 7 | 0 | 0 | 0 | 0 |
| ASV1333 | 3f08bb4aa92 | 7 | 0 | 0 | 0 | 0 | 0 | 0 | 0 | 0 | 0 | 0 | 0 | 0 | 0 | 0 | 0 | 0 | 0 | 0 | 0 | 0 | 0 | 0 | 0 | 0 | 0 | 0 | 0 | 0 | 0 | 0 | 7 | 0 | 0 | 0 | 0 |
| ASV1334 | 71ef1116b46 | 7 | 0 | 0 | 0 | 0 | 0 | 0 | 0 | 0 | 0 | 7 | 0 | 0 | 0 | 0 | 0 | 0 | 0 | 0 | 0 | 0 | 0 | 0 | 0 | 0 | 0 | 0 | 0 | 0 | 0 | 0 | 0 | 0 | 0 | 0 | 0 |
| ASV1335 | 93843b3fbf0 | 7 | 0 | 0 | 0 | 2 | 5 | 0 | 0 | 0 | 0 | 0 | 0 | 0 | 0 | 0 | 0 | 0 | 0 | 0 | 0 | 0 | 0 | 0 | 0 | 0 | 0 | 0 | 0 | 0 | 0 | 0 | 0 | 0 | 0 | 0 | 0 |
| ASV1336 | 8d59216f6a8 | 7 | 0 | 0 | 7 | 0 | 0 | 0 | 0 | 0 | 0 | 0 | 0 | 0 | 0 | 0 | 0 | 0 | 0 | 0 | 0 | 0 | 0 | 0 | 0 | 0 | 0 | 0 | 0 | 0 | 0 | 0 | 0 | 0 | 0 | 0 | 0 |
| ASV1337 | f9ca9b47174 | 7 | 0 | 0 | 0 | 0 | 0 | 0 | 0 | 0 | 0 | 0 | 0 | 0 | 0 | 0 | 0 | 0 | 0 | 0 | 0 | 0 | 0 | 7 | 0 | 0 | 0 | 0 | 0 | 0 | 0 | 0 | 0 | 0 | 0 | 0 | 0 |
| ASV1338 | 49a3ac4f569 | 7 | 0 | 0 | 3 | 0 | 0 | 4 | 0 | 0 | 0 | 0 | 0 | 0 | 0 | 0 | 0 | 0 | 0 | 0 | 0 | 0 | 0 | 0 | 0 | 0 | 0 | 0 | 0 | 0 | 0 | 0 | 0 | 0 | 0 | 0 | 0 |
| ASV1339 | 3a2337d2829 | 7 | 0 | 0 | 0 | 0 | 0 | 0 | 0 | 0 | 0 | 0 | 0 | 0 | 0 | 0 | 7 | 0 | 0 | 0 | 0 | 0 | 0 | 0 | 0 | 0 | 0 | 0 | 0 | 0 | 0 | 0 | 0 | 0 | 0 | 0 | 0 |
| ASV1340 | e4298f09f6a | 7 | 0 | 0 | 0 | 0 | 0 | 0 | 0 | 0 | 0 | 0 | 0 | 0 | 0 | 0 | 0 | 0 | 0 | 0 | 7 | 0 | 0 | 0 | 0 | 0 | 0 | 0 | 0 | 0 | 0 | 0 | 0 | 0 | 0 | 0 | 0 |
| ASV1341 | a44e518aa92 | 7 | 0 | 0 | 0 | 0 | 0 | 0 | 0 | 0 | 0 | 0 | 0 | 0 | 0 | 0 | 0 | 7 | 0 | 0 | 0 | 0 | 0 | 0 | 0 | 0 | 0 | 0 | 0 | 0 | 0 | 0 | 0 | 0 | 0 | 0 | 0 |
| ASV1342 | 2785a702537 | 7 | 0 | 0 | 0 | 0 | 7 | 0 | 0 | 0 | 0 | 0 | 0 | 0 | 0 | 0 | 0 | 0 | 0 | 0 | 0 | 0 | 0 | 0 | 0 | 0 | 0 | 0 | 0 | 0 | 0 | 0 | 0 | 0 | 0 | 0 | 0 |
| ASV1343 | 6f3554b8e49 | 7 | 0 | 0 | 0 | 0 | 0 | 0 | 0 | 0 | 0 | 0 | 0 | 0 | 0 | 0 | 0 | 0 | 0 | 0 | 0 | 0 | 0 | 0 | 0 | 0 | 0 | 0 | 0 | 0 | 0 | 0 | 0 | 0 | 7 | 0 | 0 |
| ASV1344 | ca0727f74f0 | 7 | 0 | 0 | 0 | 0 | 0 | 0 | 0 | 0 | 0 | 0 | 0 | 0 | 0 | 0 | 0 | 0 | 7 | 0 | 0 | 0 | 0 | 0 | 0 | 0 | 0 | 0 | 0 | 0 | 0 | 0 | 0 | 0 | 0 | 0 | 0 |
| ASV1345 | d770a3bc8c4 | 7 | 0 | 0 | 0 | 0 | 0 | 0 | 0 | 0 | 0 | 7 | 0 | 0 | 0 | 0 | 0 | 0 | 0 | 0 | 0 | 0 | 0 | 0 | 0 | 0 | 0 | 0 | 0 | 0 | 0 | 0 | 0 | 0 | 0 | 0 | 0 |
| ASV1346 | 6cc38a14c3d | 7 | 0 | 0 | 0 | 0 | 0 | 0 | 0 | 0 | 0 | 7 | 0 | 0 | 0 | 0 | 0 | 0 | 0 | 0 | 0 | 0 | 0 | 0 | 0 | 0 | 0 | 0 | 0 | 0 | 0 | 0 | 0 | 0 | 0 | 0 | 0 |
| ASV1347 | 9e7e8df738a | 7 | 0 | 0 | 0 | 0 | 0 | 0 | 0 | 0 | 0 | 0 | 0 | 0 | 0 | 0 | 0 | 0 | 0 | 0 | 0 | 0 | 0 | 0 | 0 | 0 | 0 | 0 | 0 | 0 | 0 | 2 | 0 | 0 | 0 | 5 | 0 |
| ASV1348 | 793d5df6499 | 7 | 0 | 0 | 0 | 0 | 0 | 0 | 0 | 0 | 0 | 0 | 0 | 0 | 0 | 0 | 0 | 0 | 0 | 0 | 0 | 0 | 0 | 0 | 0 | 0 | 0 | 0 | 0 | 0 | 7 | 0 | 0 | 0 | 0 | 0 | 0 |
| ASV1349 | 6150f78b7fa | 7 | 0 | 0 | 0 | 0 | 0 | 0 | 0 | 0 | 0 | 0 | 0 | 0 | 0 | 0 | 0 | 0 | 0 | 0 | 0 | 0 | 0 | 0 | 0 | 0 | 0 | 0 | 0 | 0 | 0 | 0 | 0 | 0 | 0 | 7 | 0 |
| ASV1350 | fe2e9c776a5 | 7 | 0 | 0 | 0 | 0 | 0 | 0 | 0 | 0 | 0 | 0 | 0 | 0 | 0 | 0 | 0 | 0 | 0 | 0 | 0 | 0 | 0 | 0 | 0 | 0 | 7 | 0 | 0 | 0 | 0 | 0 | 0 | 0 | 0 | 0 | 0 |
| ASV1351 | 23e5352a7b4 | 7 | 0 | 7 | 0 | 0 | 0 | 0 | 0 | 0 | 0 | 0 | 0 | 0 | 0 | 0 | 0 | 0 | 0 | 0 | 0 | 0 | 0 | 0 | 0 | 0 | 0 | 0 | 0 | 0 | 0 | 0 | 0 | 0 | 0 | 0 | 0 |
| ASV1352 | c20d4ad50c0 | 7 | 0 | 0 | 0 | 0 | 7 | 0 | 0 | 0 | 0 | 0 | 0 | 0 | 0 | 0 | 0 | 0 | 0 | 0 | 0 | 0 | 0 | 0 | 0 | 0 | 0 | 0 | 0 | 0 | 0 | 0 | 0 | 0 | 0 | 0 | 0 |
| ASV1353 | c8eed146433 | 7 | 0 | 0 | 0 | 0 | 0 | 0 | 0 | 0 | 0 | 0 | 0 | 0 | 0 | 0 | 0 | 0 | 0 | 0 | 0 | 0 | 0 | 0 | 0 | 0 | 0 | 0 | 0 | 0 | 0 | 0 | 0 | 0 | 0 | 4 | 3 |
| ASV1354 | 3036c113bed | 7 | 0 | 0 | 0 | 0 | 0 | 0 | 0 | 0 | 0 | 0 | 0 | 0 | 0 | 0 | 7 | 0 | 0 | 0 | 0 | 0 | 0 | 0 | 0 | 0 | 0 | 0 | 0 | 0 | 0 | 0 | 0 | 0 | 0 | 0 | 0 |
| ASV1355 | 66e1ef49396 | 7 | 0 | 0 | 0 | 0 | 0 | 0 | 0 | 0 | 0 | 0 | 0 | 0 | 0 | 0 | 0 | 0 | 0 | 7 | 0 | 0 | 0 | 0 | 0 | 0 | 0 | 0 | 0 | 0 | 0 | 0 | 0 | 0 | 0 | 0 | 0 |
| ASV1356 | a88539f7763 | 7 | 0 | 0 | 0 | 0 | 0 | 0 | 0 | 0 | 0 | 0 | 0 | 0 | 0 | 0 | 7 | 0 | 0 | 0 | 0 | 0 | 0 | 0 | 0 | 0 | 0 | 0 | 0 | 0 | 0 | 0 | 0 | 0 | 0 | 0 | 0 |
| ASV1357 | 3d651a342f8 | 7 | 0 | 0 | 0 | 0 | 0 | 0 | 0 | 0 | 0 | 0 | 0 | 0 | 0 | 0 | 0 | 0 | 0 | 0 | 0 | 0 | 0 | 0 | 0 | 0 | 0 | 0 | 0 | 0 | 0 | 7 | 0 | 0 | 0 | 0 | 0 |
| ASV1358 | bdcf1f0f9d60 | 7 | 0 | 0 | 0 | 0 | 0 | 0 | 0 | 0 | 0 | 3 | 0 | 0 | 0 | 0 | 4 | 0 | 0 | 0 | 0 | 0 | 0 | 0 | 0 | 0 | 0 | 0 | 0 | 0 | 0 | 0 | 0 | 0 | 0 | 0 | 0 |
| ASV1359 | 5caa316ccb1 | 7 | 0 | 0 | 0 | 0 | 0 | 0 | 0 | 0 | 0 | 0 | 0 | 0 | 0 | 0 | 0 | 0 | 0 | 0 | 0 | 0 | 7 | 0 | 0 | 0 | 0 | 0 | 0 | 0 | 0 | 0 | 0 | 0 | 0 | 0 | 0 |
| ASV1360 | 1176e7dc97e | 7 | 0 | 0 | 0 | 0 | 0 | 0 | 0 | 0 | 0 | 0 | 0 | 0 | 0 | 0 | 0 | 0 | 0 | 0 | 0 | 0 | 0 | 0 | 0 | 0 | 0 | 0 | 0 | 0 | 0 | 0 | 0 | 0 | 0 | 0 | 7 |
| ASV1361 | 8cd7b7a8c14 | 7 | 0 | 0 | 0 | 0 | 0 | 0 | 0 | 0 | 0 | 0 | 0 | 0 | 0 | 0 | 0 | 0 | 0 | 0 | 0 | 0 | 0 | 0 | 0 | 3 | 0 | 0 | 0 | 0 | 0 | 4 | 0 | 0 | 0 | 0 | 0 |
| ASV1362 | 0bcab4ff927 | 7 | 0 | 0 | 0 | 0 | 0 | 0 | 0 | 0 | 0 | 0 | 0 | 0 | 0 | 0 | 0 | 0 | 0 | 0 | 0 | 0 | 0 | 0 | 0 | 0 | 0 | 0 | 0 | 0 | 0 | 0 | 7 | 0 | 0 | 0 | 0 |
| ASV1363 | d4929e92dd0 | 7 | 0 | 0 | 0 | 0 | 0 | 0 | 0 | 0 | 7 | 0 | 0 | 0 | 0 | 0 | 0 | 0 | 0 | 0 | 0 | 0 | 0 | 0 | 0 | 0 | 0 | 0 | 0 | 0 | 0 | 0 | 0 | 0 | 0 | 0 | 0 |
| ASV1364 | 200462cdae2 | 7 | 0 | 0 | 0 | 0 | 0 | 0 | 0 | 0 | 0 | 0 | 0 | 0 | 0 | 0 | 0 | 0 | 0 | 0 | 7 | 0 | 0 | 0 | 0 | 0 | 0 | 0 | 0 | 0 | 0 | 0 | 0 | 0 | 0 | 0 | 0 |
| ASV1365 | afe5bed304b | 7 | 0 | 0 | 0 | 0 | 7 | 0 | 0 | 0 | 0 | 0 | 0 | 0 | 0 | 0 | 0 | 0 | 0 | 0 | 0 | 0 | 0 | 0 | 0 | 0 | 0 | 0 | 0 | 0 | 0 | 0 | 0 | 0 | 0 | 0 | 0 |
| ASV1366 | 2276e0df2a2 | 7 | 0 | 0 | 3 | 0 | 0 | 0 | 0 | 0 | 0 | 0 | 0 | 0 | 0 | 0 | 0 | 0 | 0 | 0 | 4 | 0 | 0 | 0 | 0 | 0 | 0 | 0 | 0 | 0 | 0 | 0 | 0 | 0 | 0 | 0 | 0 |
| ASV1367 | 4d94478a371 | 7 | 0 | 0 | 0 | 0 | 0 | 0 | 0 | 0 | 0 | 0 | 0 | 0 | 0 | 0 | 0 | 0 | 7 | 0 | 0 | 0 | 0 | 0 | 0 | 0 | 0 | 0 | 0 | 0 | 0 | 0 | 0 | 0 | 0 | 0 | 0 |
| ASV1368 | d949487c027 | 7 | 0 | 0 | 0 | 0 | 0 | 0 | 0 | 0 | 0 | 0 | 0 | 0 | 0 | 0 | 0 | 0 | 0 | 0 | 0 | 0 | 0 | 0 | 5 | 0 | 0 | 0 | 0 | 2 | 0 | 0 | 0 | 0 | 0 | 0 | 0 |
| ASV1369 | 8fc4f917919 | 7 | 0 | 0 | 0 | 0 | 0 | 0 | 0 | 0 | 0 | 0 | 0 | 0 | 0 | 0 | 0 | 0 | 0 | 0 | 0 | 0 | 0 | 0 | 0 | 0 | 0 | 0 | 0 | 0 | 0 | 0 | 0 | 0 | 0 | 7 | 0 |
| ASV1370 | 2193f1a58af | 7 | 0 | 0 | 0 | 0 | 0 | 0 | 0 | 0 | 0 | 0 | 0 | 0 | 0 | 0 | 0 | 0 | 0 | 0 | 0 | 0 | 0 | 0 | 0 | 0 | 0 | 0 | 0 | 0 | 0 | 0 | 0 | 0 | 0 | 0 | 7 |
| ASV1371 | 5c1723f82a8 | 7 | 0 | 0 | 0 | 0 | 0 | 0 | 0 | 0 | 0 | 0 | 0 | 0 | 0 | 0 | 0 | 0 | 0 | 0 | 0 | 0 | 0 | 0 | 0 | 0 | 0 | 0 | 0 | 0 | 0 | 0 | 0 | 0 | 0 | 7 | 0 |
| ASV1372 | 48b1d0d4a44 | 7 | 0 | 0 | 0 | 0 | 0 | 0 | 0 | 0 | 0 | 0 | 0 | 0 | 0 | 0 | 0 | 0 | 0 | 0 | 0 | 0 | 0 | 0 | 0 | 0 | 0 | 0 | 0 | 0 | 0 | 0 | 0 | 7 | 0 | 0 | 0 |
| ASV1373 | 74edab1156c | 7 | 0 | 0 | 0 | 0 | 0 | 0 | 0 | 0 | 0 | 0 | 0 | 0 | 0 | 0 | 7 | 0 | 0 | 0 | 0 | 0 | 0 | 0 | 0 | 0 | 0 | 0 | 0 | 0 | 0 | 0 | 0 | 0 | 0 | 0 | 0 |
| ASV1374 | 001de51c1b9 | 7 | 0 | 0 | 0 | 0 | 0 | 0 | 0 | 0 | 0 | 0 | 0 | 0 | 0 | 0 | 0 | 0 | 0 | 0 | 0 | 0 | 0 | 0 | 0 | 0 | 0 | 0 | 0 | 0 | 0 | 7 | 0 | 0 | 0 | 0 | 0 |
| ASV1375 | 398b7a80dc2 | 7 | 0 | 0 | 0 | 0 | 0 | 0 | 0 | 0 | 0 | 0 | 0 | 0 | 0 | 0 | 0 | 0 | 0 | 0 | 0 | 0 | 0 | 0 | 0 | 0 | 0 | 0 | 0 | 0 | 0 | 0 | 7 | 0 | 0 | 0 | 0 |
| ASV1376 | c9f02f1cb79 | 7 | 0 | 0 | 0 | 0 | 0 | 0 | 0 | 0 | 0 | 0 | 0 | 0 | 0 | 0 | 7 | 0 | 0 | 0 | 0 | 0 | 0 | 0 | 0 | 0 | 0 | 0 | 0 | 0 | 0 | 0 | 0 | 0 | 0 | 0 | 0 |
| ASV1377 | 65b974e2d7f | 6 | 0 | 0 | 0 | 0 | 0 | 0 | 0 | 0 | 0 | 0 | 0 | 0 | 0 | 0 | 0 | 0 | 0 | 0 | 0 | 0 | 0 | 0 | 0 | 6 | 0 | 0 | 0 | 0 | 0 | 0 | 0 | 0 | 0 | 0 | 0 |
| ASV1378 | 3354179ec2f | 6 | 0 | 0 | 0 | 0 | 6 | 0 | 0 | 0 | 0 | 0 | 0 | 0 | 0 | 0 | 0 | 0 | 0 | 0 | 0 | 0 | 0 | 0 | 0 | 0 | 0 | 0 | 0 | 0 | 0 | 0 | 0 | 0 | 0 | 0 | 0 |
| ASV1379 | 1b9bd4a7185 | 6 | 0 | 0 | 0 | 0 | 0 | 0 | 0 | 0 | 0 | 0 | 0 | 0 | 0 | 0 | 0 | 0 | 0 | 0 | 0 | 0 | 0 | 0 | 0 | 0 | 0 | 0 | 0 | 0 | 0 | 6 | 0 | 0 | 0 | 0 | 0 |
| ASV1380 | 62e1e6a5e25 | 6 | 0 | 0 | 0 | 0 | 0 | 0 | 0 | 0 | 0 | 0 | 0 | 0 | 0 | 0 | 0 | 0 | 0 | 0 | 0 | 0 | 0 | 0 | 0 | 0 | 6 | 0 | 0 | 0 | 0 | 0 | 0 | 0 | 0 | 0 | 0 |
| ASV1381 | cd8c09391fa | 6 | 0 | 0 | 0 | 0 | 0 | 0 | 0 | 0 | 0 | 0 | 0 | 0 | 0 | 0 | 0 | 0 | 0 | 0 | 0 | 0 | 0 | 0 | 0 | 0 | 0 | 0 | 0 | 0 | 6 | 0 | 0 | 0 | 0 | 0 | 0 |
| ASV1382 | 8e9e1c3f472 | 6 | 0 | 0 | 0 | 0 | 2 | 0 | 0 | 0 | 0 | 0 | 0 | 0 | 0 | 0 | 4 | 0 | 0 | 0 | 0 | 0 | 0 | 0 | 0 | 0 | 0 | 0 | 0 | 0 | 0 | 0 | 0 | 0 | 0 | 0 | 0 |
| ASV1383 | aa225ae35df | 6 | 0 | 0 | 0 | 0 | 0 | 0 | 0 | 0 | 0 | 0 | 0 | 0 | 0 | 0 | 0 | 0 | 4 | 0 | 0 | 0 | 0 | 0 | 0 | 0 | 0 | 0 | 0 | 0 | 0 | 0 | 0 | 0 | 0 | 2 | 0 |
| ASV1384 | 10390c2c16e | 6 | 6 | 0 | 0 | 0 | 0 | 0 | 0 | 0 | 0 | 0 | 0 | 0 | 0 | 0 | 0 | 0 | 0 | 0 | 0 | 0 | 0 | 0 | 0 | 0 | 0 | 0 | 0 | 0 | 0 | 0 | 0 | 0 | 0 | 0 | 0 |
| ASV1385 | 105102e9216 | 6 | 0 | 0 | 0 | 0 | 0 | 0 | 0 | 0 | 0 | 0 | 6 | 0 | 0 | 0 | 0 | 0 | 0 | 0 | 0 | 0 | 0 | 0 | 0 | 0 | 0 | 0 | 0 | 0 | 0 | 0 | 0 | 0 | 0 | 0 | 0 |
| ASV1386 | 0e6ac77416b | 6 | 0 | 0 | 0 | 0 | 0 | 0 | 0 | 0 | 0 | 0 | 0 | 0 | 0 | 0 | 0 | 0 | 0 | 0 | 0 | 0 | 0 | 0 | 0 | 0 | 0 | 0 | 0 | 6 | 0 | 0 | 0 | 0 | 0 | 0 | 0 |
| ASV1387 | 3d0e8c71d8d | 6 | 0 | 0 | 6 | 0 | 0 | 0 | 0 | 0 | 0 | 0 | 0 | 0 | 0 | 0 | 0 | 0 | 0 | 0 | 0 | 0 | 0 | 0 | 0 | 0 | 0 | 0 | 0 | 0 | 0 | 0 | 0 | 0 | 0 | 0 | 0 |
| ASV1388 | 8e45207056e | 6 | 0 | 0 | 0 | 0 | 0 | 0 | 0 | 0 | 0 | 0 | 0 | 0 | 0 | 0 | 0 | 0 | 0 | 0 | 0 | 0 | 0 | 2 | 0 | 0 | 4 | 0 | 0 | 0 | 0 | 0 | 0 | 0 | 0 | 0 | 0 |
| ASV1389 | 96a513a3e8b | 6 | 0 | 0 | 0 | 0 | 0 | 2 | 0 | 4 | 0 | 0 | 0 | 0 | 0 | 0 | 0 | 0 | 0 | 0 | 0 | 0 | 0 | 0 | 0 | 0 | 0 | 0 | 0 | 0 | 0 | 0 | 0 | 0 | 0 | 0 | 0 |
| ASV1390 | c79df9f8a10 | 6 | 0 | 0 | 0 | 0 | 6 | 0 | 0 | 0 | 0 | 0 | 0 | 0 | 0 | 0 | 0 | 0 | 0 | 0 | 0 | 0 | 0 | 0 | 0 | 0 | 0 | 0 | 0 | 0 | 0 | 0 | 0 | 0 | 0 | 0 | 0 |
| ASV1391 | 9a0cc027d73 | 6 | 0 | 0 | 0 | 0 | 0 | 0 | 0 | 0 | 0 | 0 | 0 | 0 | 0 | 0 | 0 | 0 | 0 | 6 | 0 | 0 | 0 | 0 | 0 | 0 | 0 | 0 | 0 | 0 | 0 | 0 | 0 | 0 | 0 | 0 | 0 |
| ASV1392 | 24f27c51e96 | 6 | 0 | 0 | 0 | 0 | 0 | 0 | 0 | 0 | 0 | 0 | 0 | 0 | 0 | 0 | 0 | 0 | 0 | 0 | 0 | 0 | 0 | 0 | 6 | 0 | 0 | 0 | 0 | 0 | 0 | 0 | 0 | 0 | 0 | 0 | 0 |
| ASV1393 | 04e1a13c375 | 6 | 0 | 0 | 0 | 6 | 0 | 0 | 0 | 0 | 0 | 0 | 0 | 0 | 0 | 0 | 0 | 0 | 0 | 0 | 0 | 0 | 0 | 0 | 0 | 0 | 0 | 0 | 0 | 0 | 0 | 0 | 0 | 0 | 0 | 0 | 0 |
| ASV1394 | d03872c3a7f | 6 | 0 | 0 | 0 | 0 | 6 | 0 | 0 | 0 | 0 | 0 | 0 | 0 | 0 | 0 | 0 | 0 | 0 | 0 | 0 | 0 | 0 | 0 | 0 | 0 | 0 | 0 | 0 | 0 | 0 | 0 | 0 | 0 | 0 | 0 | 0 |
| ASV1395 | 15928f46e09 | 6 | 0 | 0 | 0 | 0 | 0 | 0 | 0 | 0 | 0 | 0 | 0 | 0 | 0 | 0 | 0 | 0 | 0 | 0 | 0 | 0 | 6 | 0 | 0 | 0 | 0 | 0 | 0 | 0 | 0 | 0 | 0 | 0 | 0 | 0 | 0 |
| ASV1396 | fb6d8e6ae6a | 6 | 0 | 3 | 0 | 0 | 0 | 0 | 0 | 0 | 0 | 0 | 0 | 0 | 0 | 0 | 0 | 0 | 0 | 0 | 0 | 0 | 0 | 0 | 0 | 3 | 0 | 0 | 0 | 0 | 0 | 0 | 0 | 0 | 0 | 0 | 0 |
| ASV1397 | c252e2ba6c3 | 6 | 0 | 0 | 0 | 0 | 0 | 6 | 0 | 0 | 0 | 0 | 0 | 0 | 0 | 0 | 0 | 0 | 0 | 0 | 0 | 0 | 0 | 0 | 0 | 0 | 0 | 0 | 0 | 0 | 0 | 0 | 0 | 0 | 0 | 0 | 0 |
| ASV1398 | 5e34faab597 | 6 | 0 | 0 | 6 | 0 | 0 | 0 | 0 | 0 | 0 | 0 | 0 | 0 | 0 | 0 | 0 | 0 | 0 | 0 | 0 | 0 | 0 | 0 | 0 | 0 | 0 | 0 | 0 | 0 | 0 | 0 | 0 | 0 | 0 | 0 | 0 |
| ASV1399 | 43b5d7c5fea | 6 | 0 | 0 | 0 | 0 | 0 | 0 | 0 | 0 | 0 | 0 | 0 | 0 | 0 | 0 | 6 | 0 | 0 | 0 | 0 | 0 | 0 | 0 | 0 | 0 | 0 | 0 | 0 | 0 | 0 | 0 | 0 | 0 | 0 | 0 | 0 |
| ASV1400 | 0db1f515050 | 6 | 0 | 0 | 0 | 0 | 0 | 0 | 0 | 0 | 0 | 0 | 0 | 0 | 0 | 0 | 6 | 0 | 0 | 0 | 0 | 0 | 0 | 0 | 0 | 0 | 0 | 0 | 0 | 0 | 0 | 0 | 0 | 0 | 0 | 0 | 0 |
| ASV1401 | bd39bf6933a | 6 | 0 | 0 | 0 | 0 | 0 | 0 | 0 | 6 | 0 | 0 | 0 | 0 | 0 | 0 | 0 | 0 | 0 | 0 | 0 | 0 | 0 | 0 | 0 | 0 | 0 | 0 | 0 | 0 | 0 | 0 | 0 | 0 | 0 | 0 | 0 |
| ASV1402 | 75c45a45ff4 | 6 | 0 | 0 | 0 | 0 | 0 | 0 | 0 | 0 | 0 | 0 | 0 | 0 | 0 | 0 | 0 | 0 | 0 | 0 | 0 | 0 | 0 | 0 | 0 | 0 | 6 | 0 | 0 | 0 | 0 | 0 | 0 | 0 | 0 | 0 | 0 |
| ASV1403 | 6f52ad7b849 | 6 | 0 | 0 | 0 | 0 | 0 | 0 | 0 | 0 | 0 | 6 | 0 | 0 | 0 | 0 | 0 | 0 | 0 | 0 | 0 | 0 | 0 | 0 | 0 | 0 | 0 | 0 | 0 | 0 | 0 | 0 | 0 | 0 | 0 | 0 | 0 |
| ASV1404 | 520391183ab | 6 | 0 | 0 | 0 | 0 | 0 | 0 | 0 | 0 | 0 | 0 | 0 | 0 | 0 | 0 | 0 | 0 | 0 | 0 | 0 | 0 | 0 | 0 | 0 | 6 | 0 | 0 | 0 | 0 | 0 | 0 | 0 | 0 | 0 | 0 | 0 |
| ASV1405 | e2c9a0e5e62 | 6 | 0 | 0 | 0 | 0 | 0 | 0 | 0 | 0 | 0 | 0 | 0 | 0 | 0 | 0 | 1 | 0 | 0 | 0 | 0 | 0 | 0 | 0 | 0 | 0 | 0 | 0 | 0 | 0 | 0 | 5 | 0 | 0 | 0 | 0 | 0 |
| ASV1406 | e9df451896a | 6 | 0 | 0 | 0 | 0 | 0 | 0 | 0 | 0 | 0 | 0 | 0 | 0 | 0 | 0 | 0 | 0 | 0 | 0 | 0 | 0 | 0 | 6 | 0 | 0 | 0 | 0 | 0 | 0 | 0 | 0 | 0 | 0 | 0 | 0 | 0 |
| ASV1407 | 9a08cb6c81b | 6 | 0 | 0 | 6 | 0 | 0 | 0 | 0 | 0 | 0 | 0 | 0 | 0 | 0 | 0 | 0 | 0 | 0 | 0 | 0 | 0 | 0 | 0 | 0 | 0 | 0 | 0 | 0 | 0 | 0 | 0 | 0 | 0 | 0 | 0 | 0 |
| ASV1408 | 4103ac66d2a | 6 | 0 | 0 | 0 | 0 | 0 | 0 | 0 | 0 | 0 | 0 | 0 | 0 | 0 | 0 | 0 | 0 | 0 | 0 | 0 | 0 | 0 | 0 | 0 | 0 | 6 | 0 | 0 | 0 | 0 | 0 | 0 | 0 | 0 | 0 | 0 |
| ASV1409 | 7ebe7391dbd | 6 | 0 | 0 | 0 | 0 | 0 | 0 | 0 | 0 | 0 | 0 | 0 | 0 | 0 | 0 | 6 | 0 | 0 | 0 | 0 | 0 | 0 | 0 | 0 | 0 | 0 | 0 | 0 | 0 | 0 | 0 | 0 | 0 | 0 | 0 | 0 |
| ASV1410 | de1c7559fd0 | 6 | 0 | 0 | 0 | 0 | 6 | 0 | 0 | 0 | 0 | 0 | 0 | 0 | 0 | 0 | 0 | 0 | 0 | 0 | 0 | 0 | 0 | 0 | 0 | 0 | 0 | 0 | 0 | 0 | 0 | 0 | 0 | 0 | 0 | 0 | 0 |
| ASV1411 | d9b94b24f2f | 6 | 0 | 0 | 0 | 0 | 0 | 0 | 0 | 0 | 0 | 0 | 0 | 0 | 0 | 0 | 0 | 0 | 0 | 0 | 6 | 0 | 0 | 0 | 0 | 0 | 0 | 0 | 0 | 0 | 0 | 0 | 0 | 0 | 0 | 0 | 0 |
| ASV1412 | 2d8cdefe87d | 6 | 0 | 0 | 0 | 0 | 0 | 0 | 0 | 0 | 0 | 0 | 0 | 0 | 0 | 0 | 0 | 0 | 0 | 0 | 0 | 0 | 0 | 0 | 0 | 0 | 0 | 0 | 0 | 0 | 0 | 0 | 0 | 0 | 0 | 0 | 6 |
| ASV1413 | 5397d90c9cd | 6 | 0 | 0 | 0 | 0 | 0 | 6 | 0 | 0 | 0 | 0 | 0 | 0 | 0 | 0 | 0 | 0 | 0 | 0 | 0 | 0 | 0 | 0 | 0 | 0 | 0 | 0 | 0 | 0 | 0 | 0 | 0 | 0 | 0 | 0 | 0 |
| ASV1414 | 901f7d86a89 | 6 | 0 | 0 | 0 | 0 | 0 | 0 | 0 | 0 | 0 | 0 | 0 | 0 | 0 | 0 | 0 | 0 | 0 | 0 | 0 | 0 | 0 | 0 | 0 | 0 | 0 | 0 | 0 | 0 | 0 | 0 | 6 | 0 | 0 | 0 | 0 |
| ASV1415 | 35b3ee3c85c | 6 | 0 | 0 | 0 | 0 | 6 | 0 | 0 | 0 | 0 | 0 | 0 | 0 | 0 | 0 | 0 | 0 | 0 | 0 | 0 | 0 | 0 | 0 | 0 | 0 | 0 | 0 | 0 | 0 | 0 | 0 | 0 | 0 | 0 | 0 | 0 |
| ASV1416 | 24dcc3462a9 | 6 | 0 | 0 | 0 | 0 | 0 | 0 | 0 | 0 | 0 | 0 | 0 | 0 | 0 | 0 | 0 | 0 | 0 | 0 | 6 | 0 | 0 | 0 | 0 | 0 | 0 | 0 | 0 | 0 | 0 | 0 | 0 | 0 | 0 | 0 | 0 |
| ASV1417 | 6e30705e019 | 6 | 0 | 0 | 0 | 0 | 0 | 0 | 0 | 0 | 0 | 0 | 0 | 0 | 0 | 0 | 0 | 0 | 0 | 0 | 0 | 0 | 0 | 0 | 0 | 0 | 6 | 0 | 0 | 0 | 0 | 0 | 0 | 0 | 0 | 0 | 0 |
| ASV1418 | 6313cb7de0b | 6 | 0 | 0 | 0 | 0 | 0 | 0 | 0 | 0 | 0 | 0 | 0 | 0 | 0 | 0 | 0 | 0 | 0 | 6 | 0 | 0 | 0 | 0 | 0 | 0 | 0 | 0 | 0 | 0 | 0 | 0 | 0 | 0 | 0 | 0 | 0 |
| ASV1419 | 7b67152335d | 6 | 0 | 0 | 0 | 0 | 0 | 0 | 0 | 0 | 0 | 6 | 0 | 0 | 0 | 0 | 0 | 0 | 0 | 0 | 0 | 0 | 0 | 0 | 0 | 0 | 0 | 0 | 0 | 0 | 0 | 0 | 0 | 0 | 0 | 0 | 0 |
| ASV1420 | b30ec1714cf | 6 | 0 | 0 | 0 | 0 | 6 | 0 | 0 | 0 | 0 | 0 | 0 | 0 | 0 | 0 | 0 | 0 | 0 | 0 | 0 | 0 | 0 | 0 | 0 | 0 | 0 | 0 | 0 | 0 | 0 | 0 | 0 | 0 | 0 | 0 | 0 |
| ASV1421 | 8fda521c823 | 6 | 0 | 0 | 0 | 0 | 0 | 0 | 0 | 0 | 0 | 0 | 0 | 0 | 0 | 0 | 0 | 0 | 0 | 0 | 0 | 0 | 0 | 0 | 0 | 0 | 6 | 0 | 0 | 0 | 0 | 0 | 0 | 0 | 0 | 0 | 0 |
| ASV1422 | 1eec9b16acc | 6 | 0 | 0 | 0 | 0 | 0 | 0 | 0 | 0 | 0 | 0 | 0 | 0 | 0 | 0 | 0 | 0 | 0 | 0 | 0 | 6 | 0 | 0 | 0 | 0 | 0 | 0 | 0 | 0 | 0 | 0 | 0 | 0 | 0 | 0 | 0 |
| ASV1423 | 689ec83d9bd | 6 | 0 | 0 | 0 | 0 | 6 | 0 | 0 | 0 | 0 | 0 | 0 | 0 | 0 | 0 | 0 | 0 | 0 | 0 | 0 | 0 | 0 | 0 | 0 | 0 | 0 | 0 | 0 | 0 | 0 | 0 | 0 | 0 | 0 | 0 | 0 |
| ASV1424 | 34d3b687c4d | 6 | 0 | 0 | 0 | 6 | 0 | 0 | 0 | 0 | 0 | 0 | 0 | 0 | 0 | 0 | 0 | 0 | 0 | 0 | 0 | 0 | 0 | 0 | 0 | 0 | 0 | 0 | 0 | 0 | 0 | 0 | 0 | 0 | 0 | 0 | 0 |
| ASV1425 | cece916144b | 6 | 0 | 0 | 0 | 0 | 0 | 0 | 0 | 0 | 0 | 0 | 0 | 0 | 0 | 0 | 0 | 0 | 0 | 0 | 0 | 0 | 0 | 0 | 0 | 0 | 0 | 0 | 0 | 0 | 0 | 0 | 6 | 0 | 0 | 0 | 0 |
| ASV1426 | 528e6043f6c | 6 | 0 | 0 | 0 | 0 | 0 | 0 | 0 | 0 | 0 | 6 | 0 | 0 | 0 | 0 | 0 | 0 | 0 | 0 | 0 | 0 | 0 | 0 | 0 | 0 | 0 | 0 | 0 | 0 | 0 | 0 | 0 | 0 | 0 | 0 | 0 |
| ASV1427 | 90669f55d64 | 6 | 0 | 0 | 0 | 6 | 0 | 0 | 0 | 0 | 0 | 0 | 0 | 0 | 0 | 0 | 0 | 0 | 0 | 0 | 0 | 0 | 0 | 0 | 0 | 0 | 0 | 0 | 0 | 0 | 0 | 0 | 0 | 0 | 0 | 0 | 0 |
| ASV1428 | 171bbc6a554 | 6 | 0 | 0 | 0 | 0 | 0 | 0 | 0 | 0 | 0 | 0 | 0 | 0 | 0 | 0 | 0 | 0 | 0 | 0 | 0 | 0 | 0 | 0 | 0 | 0 | 0 | 0 | 0 | 0 | 6 | 0 | 0 | 0 | 0 | 0 | 0 |
| ASV1429 | 4eefa79728b | 6 | 0 | 0 | 0 | 0 | 0 | 0 | 0 | 0 | 6 | 0 | 0 | 0 | 0 | 0 | 0 | 0 | 0 | 0 | 0 | 0 | 0 | 0 | 0 | 0 | 0 | 0 | 0 | 0 | 0 | 0 | 0 | 0 | 0 | 0 | 0 |
| ASV1430 | 243337808ed | 6 | 0 | 0 | 6 | 0 | 0 | 0 | 0 | 0 | 0 | 0 | 0 | 0 | 0 | 0 | 0 | 0 | 0 | 0 | 0 | 0 | 0 | 0 | 0 | 0 | 0 | 0 | 0 | 0 | 0 | 0 | 0 | 0 | 0 | 0 | 0 |
| ASV1431 | 278156bf3df | 6 | 0 | 0 | 0 | 0 | 0 | 0 | 0 | 0 | 0 | 0 | 0 | 0 | 0 | 0 | 0 | 0 | 0 | 0 | 0 | 0 | 0 | 0 | 0 | 0 | 6 | 0 | 0 | 0 | 0 | 0 | 0 | 0 | 0 | 0 | 0 |
| ASV1432 | 8dfd63f862fa | 6 | 0 | 0 | 0 | 0 | 0 | 0 | 0 | 0 | 0 | 0 | 6 | 0 | 0 | 0 | 0 | 0 | 0 | 0 | 0 | 0 | 0 | 0 | 0 | 0 | 0 | 0 | 0 | 0 | 0 | 0 | 0 | 0 | 0 | 0 | 0 |
| ASV1433 | b8a0949c76b | 6 | 0 | 0 | 0 | 0 | 0 | 0 | 0 | 0 | 0 | 0 | 0 | 6 | 0 | 0 | 0 | 0 | 0 | 0 | 0 | 0 | 0 | 0 | 0 | 0 | 0 | 0 | 0 | 0 | 0 | 0 | 0 | 0 | 0 | 0 | 0 |
| ASV1434 | c6b4b1189dc | 6 | 0 | 0 | 0 | 0 | 0 | 0 | 0 | 0 | 0 | 0 | 0 | 0 | 0 | 0 | 0 | 0 | 0 | 0 | 6 | 0 | 0 | 0 | 0 | 0 | 0 | 0 | 0 | 0 | 0 | 0 | 0 | 0 | 0 | 0 | 0 |
| ASV1435 | de3f928b57f | 6 | 0 | 0 | 0 | 0 | 0 | 0 | 0 | 0 | 0 | 0 | 0 | 0 | 0 | 0 | 0 | 0 | 0 | 0 | 0 | 0 | 0 | 0 | 0 | 0 | 0 | 0 | 0 | 0 | 0 | 0 | 6 | 0 | 0 | 0 | 0 |
| ASV1436 | 526515e3063 | 6 | 0 | 0 | 0 | 0 | 6 | 0 | 0 | 0 | 0 | 0 | 0 | 0 | 0 | 0 | 0 | 0 | 0 | 0 | 0 | 0 | 0 | 0 | 0 | 0 | 0 | 0 | 0 | 0 | 0 | 0 | 0 | 0 | 0 | 0 | 0 |
| ASV1437 | aa215cef5ee | 6 | 0 | 0 | 0 | 0 | 0 | 0 | 0 | 0 | 0 | 0 | 0 | 0 | 0 | 0 | 0 | 0 | 0 | 0 | 0 | 0 | 0 | 0 | 0 | 0 | 0 | 0 | 0 | 6 | 0 | 0 | 0 | 0 | 0 | 0 | 0 |
| ASV1438 | 76ddac6b0bb | 6 | 0 | 0 | 0 | 0 | 0 | 0 | 0 | 0 | 0 | 6 | 0 | 0 | 0 | 0 | 0 | 0 | 0 | 0 | 0 | 0 | 0 | 0 | 0 | 0 | 0 | 0 | 0 | 0 | 0 | 0 | 0 | 0 | 0 | 0 | 0 |
| ASV1439 | 325ac32109f | 6 | 0 | 0 | 0 | 0 | 0 | 0 | 0 | 0 | 0 | 0 | 0 | 0 | 0 | 0 | 6 | 0 | 0 | 0 | 0 | 0 | 0 | 0 | 0 | 0 | 0 | 0 | 0 | 0 | 0 | 0 | 0 | 0 | 0 | 0 | 0 |
| ASV1440 | 09efea11e0d | 6 | 0 | 0 | 0 | 0 | 0 | 0 | 0 | 0 | 0 | 0 | 0 | 0 | 0 | 0 | 0 | 0 | 0 | 0 | 0 | 0 | 0 | 0 | 0 | 0 | 0 | 0 | 0 | 0 | 0 | 0 | 6 | 0 | 0 | 0 | 0 |
| ASV1441 | 4914d60eb34 | 5 | 0 | 0 | 0 | 0 | 0 | 0 | 0 | 0 | 0 | 0 | 0 | 0 | 0 | 0 | 0 | 0 | 0 | 0 | 0 | 0 | 0 | 0 | 0 | 0 | 0 | 0 | 0 | 0 | 0 | 0 | 0 | 0 | 5 | 0 | 0 |
| ASV1442 | df431af4616 | 5 | 0 | 0 | 0 | 0 | 0 | 0 | 0 | 0 | 5 | 0 | 0 | 0 | 0 | 0 | 0 | 0 | 0 | 0 | 0 | 0 | 0 | 0 | 0 | 0 | 0 | 0 | 0 | 0 | 0 | 0 | 0 | 0 | 0 | 0 | 0 |
| ASV1443 | 7128721f0a3 | 5 | 0 | 0 | 0 | 0 | 0 | 0 | 0 | 0 | 0 | 0 | 0 | 0 | 0 | 0 | 0 | 0 | 0 | 0 | 0 | 0 | 0 | 0 | 0 | 0 | 0 | 0 | 0 | 0 | 0 | 0 | 0 | 0 | 0 | 5 | 0 |
| ASV1444 | 20e4f3f6724 | 5 | 0 | 0 | 0 | 0 | 0 | 0 | 0 | 0 | 0 | 5 | 0 | 0 | 0 | 0 | 0 | 0 | 0 | 0 | 0 | 0 | 0 | 0 | 0 | 0 | 0 | 0 | 0 | 0 | 0 | 0 | 0 | 0 | 0 | 0 | 0 |
| ASV1445 | 44fa2b32fe4 | 5 | 0 | 3 | 0 | 0 | 0 | 0 | 0 | 0 | 0 | 0 | 0 | 0 | 2 | 0 | 0 | 0 | 0 | 0 | 0 | 0 | 0 | 0 | 0 | 0 | 0 | 0 | 0 | 0 | 0 | 0 | 0 | 0 | 0 | 0 | 0 |
| ASV1446 | 007609a1092 | 5 | 0 | 0 | 0 | 0 | 0 | 0 | 0 | 0 | 0 | 5 | 0 | 0 | 0 | 0 | 0 | 0 | 0 | 0 | 0 | 0 | 0 | 0 | 0 | 0 | 0 | 0 | 0 | 0 | 0 | 0 | 0 | 0 | 0 | 0 | 0 |
| ASV1447 | f05c96b76bb | 5 | 0 | 0 | 0 | 0 | 0 | 0 | 0 | 0 | 0 | 0 | 0 | 0 | 0 | 0 | 0 | 0 | 0 | 0 | 0 | 0 | 5 | 0 | 0 | 0 | 0 | 0 | 0 | 0 | 0 | 0 | 0 | 0 | 0 | 0 | 0 |
| ASV1448 | fc4f9cc518e1 | 5 | 0 | 0 | 0 | 0 | 0 | 0 | 0 | 0 | 0 | 0 | 0 | 0 | 5 | 0 | 0 | 0 | 0 | 0 | 0 | 0 | 0 | 0 | 0 | 0 | 0 | 0 | 0 | 0 | 0 | 0 | 0 | 0 | 0 | 0 | 0 |
| ASV1449 | 587bb680282 | 5 | 0 | 0 | 0 | 0 | 0 | 0 | 0 | 0 | 0 | 0 | 0 | 0 | 0 | 0 | 5 | 0 | 0 | 0 | 0 | 0 | 0 | 0 | 0 | 0 | 0 | 0 | 0 | 0 | 0 | 0 | 0 | 0 | 0 | 0 | 0 |
| ASV1450 | 440e78141b1 | 5 | 0 | 0 | 0 | 0 | 0 | 5 | 0 | 0 | 0 | 0 | 0 | 0 | 0 | 0 | 0 | 0 | 0 | 0 | 0 | 0 | 0 | 0 | 0 | 0 | 0 | 0 | 0 | 0 | 0 | 0 | 0 | 0 | 0 | 0 | 0 |
| ASV1451 | f231f9253aa | 5 | 0 | 0 | 0 | 0 | 0 | 0 | 0 | 0 | 0 | 0 | 0 | 0 | 0 | 0 | 0 | 0 | 0 | 0 | 0 | 0 | 0 | 0 | 0 | 0 | 0 | 0 | 5 | 0 | 0 | 0 | 0 | 0 | 0 | 0 | 0 |
| ASV1452 | 9be3e682703 | 5 | 0 | 0 | 0 | 0 | 0 | 0 | 0 | 0 | 0 | 0 | 0 | 0 | 0 | 0 | 0 | 0 | 0 | 0 | 0 | 0 | 0 | 0 | 0 | 5 | 0 | 0 | 0 | 0 | 0 | 0 | 0 | 0 | 0 | 0 | 0 |
| ASV1453 | e1274bc2a7b | 5 | 0 | 0 | 0 | 0 | 5 | 0 | 0 | 0 | 0 | 0 | 0 | 0 | 0 | 0 | 0 | 0 | 0 | 0 | 0 | 0 | 0 | 0 | 0 | 0 | 0 | 0 | 0 | 0 | 0 | 0 | 0 | 0 | 0 | 0 | 0 |
| ASV1454 | da1fcf4c87e0 | 5 | 0 | 0 | 0 | 0 | 0 | 0 | 0 | 0 | 5 | 0 | 0 | 0 | 0 | 0 | 0 | 0 | 0 | 0 | 0 | 0 | 0 | 0 | 0 | 0 | 0 | 0 | 0 | 0 | 0 | 0 | 0 | 0 | 0 | 0 | 0 |
| ASV1455 | a9f80c705a5 | 5 | 0 | 0 | 0 | 0 | 0 | 0 | 0 | 0 | 0 | 0 | 0 | 0 | 0 | 0 | 0 | 0 | 0 | 0 | 0 | 0 | 0 | 0 | 0 | 0 | 0 | 0 | 0 | 0 | 0 | 0 | 5 | 0 | 0 | 0 | 0 |
| ASV1456 | c37ce5ba819 | 5 | 0 | 0 | 0 | 0 | 0 | 0 | 0 | 0 | 0 | 5 | 0 | 0 | 0 | 0 | 0 | 0 | 0 | 0 | 0 | 0 | 0 | 0 | 0 | 0 | 0 | 0 | 0 | 0 | 0 | 0 | 0 | 0 | 0 | 0 | 0 |
| ASV1457 | 199e7e3b09d | 5 | 0 | 0 | 0 | 0 | 0 | 0 | 0 | 0 | 0 | 5 | 0 | 0 | 0 | 0 | 0 | 0 | 0 | 0 | 0 | 0 | 0 | 0 | 0 | 0 | 0 | 0 | 0 | 0 | 0 | 0 | 0 | 0 | 0 | 0 | 0 |
| ASV1458 | 5db00182d76 | 5 | 0 | 0 | 0 | 0 | 0 | 0 | 0 | 0 | 0 | 0 | 0 | 0 | 0 | 0 | 0 | 0 | 0 | 0 | 0 | 0 | 0 | 0 | 0 | 0 | 0 | 0 | 0 | 5 | 0 | 0 | 0 | 0 | 0 | 0 | 0 |
| ASV1459 | fbd60c0f882 | 5 | 0 | 0 | 0 | 0 | 0 | 0 | 0 | 0 | 0 | 0 | 0 | 0 | 0 | 0 | 5 | 0 | 0 | 0 | 0 | 0 | 0 | 0 | 0 | 0 | 0 | 0 | 0 | 0 | 0 | 0 | 0 | 0 | 0 | 0 | 0 |
| ASV1460 | 84f4194acd7 | 5 | 0 | 0 | 0 | 0 | 0 | 0 | 0 | 0 | 0 | 0 | 0 | 0 | 0 | 0 | 0 | 0 | 0 | 0 | 0 | 0 | 0 | 0 | 0 | 0 | 0 | 0 | 0 | 5 | 0 | 0 | 0 | 0 | 0 | 0 | 0 |
| ASV1461 | cee750c9da0 | 5 | 0 | 0 | 0 | 0 | 0 | 0 | 0 | 0 | 0 | 0 | 0 | 0 | 0 | 0 | 0 | 0 | 0 | 0 | 0 | 0 | 0 | 0 | 0 | 5 | 0 | 0 | 0 | 0 | 0 | 0 | 0 | 0 | 0 | 0 | 0 |
| ASV1462 | 2c96e1f2b66 | 5 | 0 | 0 | 0 | 0 | 0 | 0 | 0 | 0 | 0 | 5 | 0 | 0 | 0 | 0 | 0 | 0 | 0 | 0 | 0 | 0 | 0 | 0 | 0 | 0 | 0 | 0 | 0 | 0 | 0 | 0 | 0 | 0 | 0 | 0 | 0 |
| ASV1463 | 17e3fe1edd7 | 5 | 0 | 0 | 0 | 0 | 0 | 0 | 0 | 0 | 0 | 0 | 0 | 0 | 0 | 0 | 0 | 0 | 0 | 0 | 0 | 0 | 0 | 0 | 0 | 5 | 0 | 0 | 0 | 0 | 0 | 0 | 0 | 0 | 0 | 0 | 0 |
| ASV1464 | da78a9d089f | 5 | 0 | 0 | 0 | 0 | 0 | 0 | 5 | 0 | 0 | 0 | 0 | 0 | 0 | 0 | 0 | 0 | 0 | 0 | 0 | 0 | 0 | 0 | 0 | 0 | 0 | 0 | 0 | 0 | 0 | 0 | 0 | 0 | 0 | 0 | 0 |
| ASV1465 | b9565621e58 | 5 | 0 | 0 | 0 | 0 | 0 | 0 | 0 | 0 | 0 | 0 | 0 | 0 | 0 | 0 | 0 | 0 | 0 | 0 | 0 | 0 | 0 | 0 | 0 | 0 | 0 | 0 | 0 | 0 | 0 | 0 | 0 | 0 | 5 | 0 | 0 |
| ASV1466 | 12350db43e3 | 5 | 0 | 0 | 0 | 0 | 5 | 0 | 0 | 0 | 0 | 0 | 0 | 0 | 0 | 0 | 0 | 0 | 0 | 0 | 0 | 0 | 0 | 0 | 0 | 0 | 0 | 0 | 0 | 0 | 0 | 0 | 0 | 0 | 0 | 0 | 0 |
| ASV1467 | 8334ac344c0 | 5 | 0 | 0 | 0 | 0 | 0 | 0 | 0 | 0 | 0 | 0 | 0 | 0 | 0 | 0 | 5 | 0 | 0 | 0 | 0 | 0 | 0 | 0 | 0 | 0 | 0 | 0 | 0 | 0 | 0 | 0 | 0 | 0 | 0 | 0 | 0 |
| ASV1468 | 672c650be25 | 5 | 0 | 0 | 0 | 0 | 0 | 0 | 0 | 0 | 0 | 0 | 0 | 0 | 0 | 0 | 0 | 0 | 0 | 0 | 0 | 0 | 0 | 0 | 0 | 5 | 0 | 0 | 0 | 0 | 0 | 0 | 0 | 0 | 0 | 0 | 0 |
| ASV1469 | eddf0245aed | 5 | 0 | 0 | 0 | 0 | 0 | 0 | 5 | 0 | 0 | 0 | 0 | 0 | 0 | 0 | 0 | 0 | 0 | 0 | 0 | 0 | 0 | 0 | 0 | 0 | 0 | 0 | 0 | 0 | 0 | 0 | 0 | 0 | 0 | 0 | 0 |
| ASV1470 | 81c6ed55e02 | 5 | 0 | 0 | 0 | 0 | 0 | 0 | 0 | 0 | 0 | 0 | 0 | 0 | 0 | 0 | 0 | 0 | 0 | 0 | 0 | 0 | 0 | 0 | 0 | 0 | 0 | 0 | 0 | 0 | 0 | 0 | 0 | 0 | 5 | 0 | 0 |
| ASV1471 | a54c2dd068c | 5 | 0 | 5 | 0 | 0 | 0 | 0 | 0 | 0 | 0 | 0 | 0 | 0 | 0 | 0 | 0 | 0 | 0 | 0 | 0 | 0 | 0 | 0 | 0 | 0 | 0 | 0 | 0 | 0 | 0 | 0 | 0 | 0 | 0 | 0 | 0 |
| ASV1472 | ab04c03d9ef | 5 | 0 | 0 | 0 | 0 | 5 | 0 | 0 | 0 | 0 | 0 | 0 | 0 | 0 | 0 | 0 | 0 | 0 | 0 | 0 | 0 | 0 | 0 | 0 | 0 | 0 | 0 | 0 | 0 | 0 | 0 | 0 | 0 | 0 | 0 | 0 |
| ASV1473 | a5968078fe6 | 5 | 0 | 0 | 0 | 0 | 0 | 0 | 0 | 0 | 0 | 0 | 0 | 0 | 0 | 0 | 0 | 0 | 0 | 0 | 0 | 0 | 0 | 0 | 0 | 0 | 0 | 0 | 0 | 0 | 5 | 0 | 0 | 0 | 0 | 0 | 0 |
| ASV1474 | ab3630d2ff5 | 5 | 0 | 0 | 0 | 0 | 0 | 0 | 0 | 0 | 0 | 0 | 0 | 0 | 0 | 0 | 5 | 0 | 0 | 0 | 0 | 0 | 0 | 0 | 0 | 0 | 0 | 0 | 0 | 0 | 0 | 0 | 0 | 0 | 0 | 0 | 0 |
| ASV1475 | 9b2582585c8 | 5 | 0 | 0 | 0 | 0 | 5 | 0 | 0 | 0 | 0 | 0 | 0 | 0 | 0 | 0 | 0 | 0 | 0 | 0 | 0 | 0 | 0 | 0 | 0 | 0 | 0 | 0 | 0 | 0 | 0 | 0 | 0 | 0 | 0 | 0 | 0 |
| ASV1476 | b5102527585 | 5 | 0 | 0 | 0 | 0 | 0 | 0 | 0 | 0 | 0 | 0 | 0 | 0 | 0 | 0 | 0 | 0 | 0 | 0 | 0 | 0 | 0 | 0 | 0 | 0 | 0 | 0 | 0 | 0 | 0 | 0 | 0 | 0 | 0 | 5 | 0 |
| ASV1477 | 9eee8f818ee | 5 | 0 | 0 | 0 | 0 | 0 | 0 | 0 | 0 | 0 | 0 | 0 | 0 | 0 | 0 | 0 | 0 | 0 | 0 | 5 | 0 | 0 | 0 | 0 | 0 | 0 | 0 | 0 | 0 | 0 | 0 | 0 | 0 | 0 | 0 | 0 |
| ASV1478 | 0bc9da7ebfa | 5 | 0 | 0 | 0 | 0 | 0 | 0 | 0 | 0 | 0 | 0 | 0 | 0 | 0 | 0 | 0 | 0 | 0 | 0 | 0 | 5 | 0 | 0 | 0 | 0 | 0 | 0 | 0 | 0 | 0 | 0 | 0 | 0 | 0 | 0 | 0 |
| ASV1479 | 7be9301973d | 5 | 0 | 0 | 0 | 0 | 0 | 0 | 0 | 0 | 0 | 0 | 0 | 0 | 0 | 0 | 0 | 0 | 0 | 0 | 0 | 0 | 0 | 0 | 0 | 0 | 0 | 0 | 0 | 0 | 0 | 5 | 0 | 0 | 0 | 0 | 0 |
| ASV1480 | 178639bcfd4 | 5 | 0 | 0 | 0 | 0 | 0 | 0 | 0 | 0 | 0 | 0 | 0 | 0 | 0 | 0 | 0 | 0 | 0 | 0 | 0 | 0 | 5 | 0 | 0 | 0 | 0 | 0 | 0 | 0 | 0 | 0 | 0 | 0 | 0 | 0 | 0 |
| ASV1481 | 6fa2d5478f3 | 5 | 0 | 0 | 0 | 0 | 0 | 0 | 0 | 0 | 5 | 0 | 0 | 0 | 0 | 0 | 0 | 0 | 0 | 0 | 0 | 0 | 0 | 0 | 0 | 0 | 0 | 0 | 0 | 0 | 0 | 0 | 0 | 0 | 0 | 0 | 0 |
| ASV1482 | b4f8d588bab | 5 | 0 | 0 | 0 | 0 | 0 | 0 | 0 | 0 | 0 | 0 | 0 | 0 | 0 | 0 | 0 | 0 | 0 | 5 | 0 | 0 | 0 | 0 | 0 | 0 | 0 | 0 | 0 | 0 | 0 | 0 | 0 | 0 | 0 | 0 | 0 |
| ASV1483 | 53f24703416 | 5 | 0 | 0 | 0 | 0 | 0 | 0 | 0 | 0 | 0 | 0 | 0 | 0 | 0 | 0 | 0 | 0 | 5 | 0 | 0 | 0 | 0 | 0 | 0 | 0 | 0 | 0 | 0 | 0 | 0 | 0 | 0 | 0 | 0 | 0 | 0 |
| ASV1484 | 8bcbb796a33 | 5 | 0 | 0 | 0 | 0 | 0 | 0 | 0 | 0 | 0 | 0 | 0 | 0 | 0 | 0 | 0 | 0 | 0 | 0 | 0 | 0 | 0 | 0 | 0 | 0 | 0 | 0 | 0 | 5 | 0 | 0 | 0 | 0 | 0 | 0 | 0 |
| ASV1485 | 173014114c7 | 5 | 0 | 0 | 0 | 0 | 0 | 0 | 0 | 0 | 0 | 0 | 0 | 0 | 0 | 0 | 0 | 0 | 0 | 0 | 0 | 0 | 0 | 0 | 0 | 0 | 0 | 0 | 0 | 0 | 0 | 0 | 0 | 0 | 0 | 0 | 5 |
| ASV1486 | 8dc8e2bcd7c | 5 | 0 | 0 | 5 | 0 | 0 | 0 | 0 | 0 | 0 | 0 | 0 | 0 | 0 | 0 | 0 | 0 | 0 | 0 | 0 | 0 | 0 | 0 | 0 | 0 | 0 | 0 | 0 | 0 | 0 | 0 | 0 | 0 | 0 | 0 | 0 |
| ASV1487 | 6a2c29d16a6 | 5 | 0 | 0 | 0 | 0 | 5 | 0 | 0 | 0 | 0 | 0 | 0 | 0 | 0 | 0 | 0 | 0 | 0 | 0 | 0 | 0 | 0 | 0 | 0 | 0 | 0 | 0 | 0 | 0 | 0 | 0 | 0 | 0 | 0 | 0 | 0 |
| ASV1488 | 49a86b3a708 | 5 | 0 | 0 | 0 | 0 | 0 | 0 | 0 | 0 | 0 | 0 | 0 | 0 | 0 | 0 | 5 | 0 | 0 | 0 | 0 | 0 | 0 | 0 | 0 | 0 | 0 | 0 | 0 | 0 | 0 | 0 | 0 | 0 | 0 | 0 | 0 |
| ASV1489 | 0fabb1c309a | 5 | 0 | 0 | 0 | 0 | 0 | 0 | 0 | 5 | 0 | 0 | 0 | 0 | 0 | 0 | 0 | 0 | 0 | 0 | 0 | 0 | 0 | 0 | 0 | 0 | 0 | 0 | 0 | 0 | 0 | 0 | 0 | 0 | 0 | 0 | 0 |
| ASV1490 | 8b2414e92a8 | 5 | 0 | 0 | 0 | 0 | 0 | 0 | 0 | 0 | 0 | 0 | 0 | 0 | 0 | 0 | 0 | 0 | 0 | 0 | 0 | 0 | 5 | 0 | 0 | 0 | 0 | 0 | 0 | 0 | 0 | 0 | 0 | 0 | 0 | 0 | 0 |
| ASV1491 | 5b8092bd1e7 | 5 | 0 | 0 | 0 | 0 | 5 | 0 | 0 | 0 | 0 | 0 | 0 | 0 | 0 | 0 | 0 | 0 | 0 | 0 | 0 | 0 | 0 | 0 | 0 | 0 | 0 | 0 | 0 | 0 | 0 | 0 | 0 | 0 | 0 | 0 | 0 |
| ASV1492 | 1f83d9e9221 | 5 | 0 | 0 | 0 | 0 | 0 | 0 | 0 | 0 | 0 | 0 | 0 | 0 | 0 | 0 | 0 | 0 | 0 | 0 | 0 | 0 | 0 | 0 | 0 | 0 | 0 | 0 | 0 | 0 | 0 | 0 | 0 | 0 | 0 | 5 | 0 |
| ASV1493 | 1445692ac0f | 5 | 0 | 0 | 0 | 0 | 0 | 0 | 0 | 0 | 0 | 0 | 0 | 0 | 0 | 0 | 0 | 0 | 0 | 0 | 0 | 0 | 0 | 0 | 0 | 0 | 0 | 0 | 0 | 5 | 0 | 0 | 0 | 0 | 0 | 0 | 0 |
| ASV1494 | 73f9c4cf86ef | 5 | 0 | 0 | 0 | 0 | 0 | 0 | 0 | 0 | 0 | 0 | 0 | 0 | 0 | 0 | 0 | 0 | 0 | 0 | 0 | 0 | 0 | 0 | 0 | 0 | 0 | 0 | 0 | 0 | 0 | 0 | 0 | 0 | 0 | 0 | 5 |
| ASV1495 | 9a442e6daf4 | 5 | 0 | 0 | 0 | 0 | 0 | 0 | 0 | 0 | 0 | 0 | 0 | 0 | 0 | 0 | 0 | 0 | 0 | 0 | 0 | 0 | 0 | 0 | 0 | 0 | 0 | 0 | 0 | 0 | 0 | 0 | 0 | 0 | 0 | 0 | 5 |
| ASV1496 | 14408a7f003 | 5 | 0 | 0 | 0 | 0 | 0 | 0 | 0 | 0 | 0 | 0 | 0 | 0 | 0 | 0 | 0 | 0 | 0 | 0 | 0 | 2 | 0 | 0 | 0 | 0 | 0 | 0 | 0 | 0 | 0 | 0 | 0 | 0 | 0 | 0 | 3 |
| ASV1497 | 3615f1a7df5 | 5 | 0 | 0 | 0 | 0 | 0 | 0 | 0 | 0 | 0 | 0 | 0 | 0 | 0 | 0 | 5 | 0 | 0 | 0 | 0 | 0 | 0 | 0 | 0 | 0 | 0 | 0 | 0 | 0 | 0 | 0 | 0 | 0 | 0 | 0 | 0 |
| ASV1498 | 818898f8bffc | 5 | 0 | 0 | 0 | 5 | 0 | 0 | 0 | 0 | 0 | 0 | 0 | 0 | 0 | 0 | 0 | 0 | 0 | 0 | 0 | 0 | 0 | 0 | 0 | 0 | 0 | 0 | 0 | 0 | 0 | 0 | 0 | 0 | 0 | 0 | 0 |
| ASV1499 | 6953c7da1a5 | 5 | 0 | 0 | 0 | 0 | 0 | 5 | 0 | 0 | 0 | 0 | 0 | 0 | 0 | 0 | 0 | 0 | 0 | 0 | 0 | 0 | 0 | 0 | 0 | 0 | 0 | 0 | 0 | 0 | 0 | 0 | 0 | 0 | 0 | 0 | 0 |
| ASV1500 | a1d78803716 | 5 | 0 | 5 | 0 | 0 | 0 | 0 | 0 | 0 | 0 | 0 | 0 | 0 | 0 | 0 | 0 | 0 | 0 | 0 | 0 | 0 | 0 | 0 | 0 | 0 | 0 | 0 | 0 | 0 | 0 | 0 | 0 | 0 | 0 | 0 | 0 |
| ASV1501 | d60c0fa2a9b | 5 | 0 | 0 | 0 | 0 | 0 | 0 | 0 | 5 | 0 | 0 | 0 | 0 | 0 | 0 | 0 | 0 | 0 | 0 | 0 | 0 | 0 | 0 | 0 | 0 | 0 | 0 | 0 | 0 | 0 | 0 | 0 | 0 | 0 | 0 | 0 |
| ASV1502 | 2f1416e0cb2 | 5 | 0 | 0 | 0 | 0 | 0 | 5 | 0 | 0 | 0 | 0 | 0 | 0 | 0 | 0 | 0 | 0 | 0 | 0 | 0 | 0 | 0 | 0 | 0 | 0 | 0 | 0 | 0 | 0 | 0 | 0 | 0 | 0 | 0 | 0 | 0 |
| ASV1503 | 80da5f0b030 | 5 | 0 | 0 | 0 | 0 | 0 | 0 | 0 | 0 | 5 | 0 | 0 | 0 | 0 | 0 | 0 | 0 | 0 | 0 | 0 | 0 | 0 | 0 | 0 | 0 | 0 | 0 | 0 | 0 | 0 | 0 | 0 | 0 | 0 | 0 | 0 |
| ASV1504 | eda8a677029 | 5 | 0 | 0 | 0 | 0 | 0 | 0 | 0 | 0 | 0 | 0 | 0 | 0 | 0 | 0 | 0 | 0 | 0 | 0 | 0 | 0 | 0 | 0 | 0 | 0 | 0 | 0 | 0 | 0 | 0 | 0 | 0 | 0 | 0 | 0 | 5 |
| ASV1505 | 0207e404bea | 5 | 0 | 0 | 0 | 0 | 0 | 0 | 0 | 0 | 0 | 0 | 0 | 0 | 0 | 0 | 0 | 0 | 0 | 0 | 0 | 0 | 0 | 0 | 0 | 0 | 0 | 0 | 0 | 0 | 0 | 0 | 0 | 0 | 0 | 0 | 5 |
| ASV1506 | b457abff0c0 | 5 | 0 | 0 | 0 | 0 | 0 | 0 | 0 | 0 | 0 | 0 | 0 | 0 | 0 | 0 | 5 | 0 | 0 | 0 | 0 | 0 | 0 | 0 | 0 | 0 | 0 | 0 | 0 | 0 | 0 | 0 | 0 | 0 | 0 | 0 | 0 |
| ASV1507 | a0d3971a66c | 4 | 0 | 0 | 0 | 0 | 0 | 0 | 0 | 0 | 4 | 0 | 0 | 0 | 0 | 0 | 0 | 0 | 0 | 0 | 0 | 0 | 0 | 0 | 0 | 0 | 0 | 0 | 0 | 0 | 0 | 0 | 0 | 0 | 0 | 0 | 0 |
| ASV1508 | 34bdb4d45e8 | 4 | 0 | 0 | 0 | 0 | 0 | 0 | 0 | 0 | 0 | 4 | 0 | 0 | 0 | 0 | 0 | 0 | 0 | 0 | 0 | 0 | 0 | 0 | 0 | 0 | 0 | 0 | 0 | 0 | 0 | 0 | 0 | 0 | 0 | 0 | 0 |
| ASV1509 | e8483dc5ea4 | 4 | 0 | 0 | 0 | 4 | 0 | 0 | 0 | 0 | 0 | 0 | 0 | 0 | 0 | 0 | 0 | 0 | 0 | 0 | 0 | 0 | 0 | 0 | 0 | 0 | 0 | 0 | 0 | 0 | 0 | 0 | 0 | 0 | 0 | 0 | 0 |
| ASV1510 | c0535dfac73 | 4 | 0 | 0 | 0 | 0 | 0 | 0 | 4 | 0 | 0 | 0 | 0 | 0 | 0 | 0 | 0 | 0 | 0 | 0 | 0 | 0 | 0 | 0 | 0 | 0 | 0 | 0 | 0 | 0 | 0 | 0 | 0 | 0 | 0 | 0 | 0 |
| ASV1511 | 372b0e58df1 | 4 | 0 | 0 | 0 | 0 | 0 | 4 | 0 | 0 | 0 | 0 | 0 | 0 | 0 | 0 | 0 | 0 | 0 | 0 | 0 | 0 | 0 | 0 | 0 | 0 | 0 | 0 | 0 | 0 | 0 | 0 | 0 | 0 | 0 | 0 | 0 |
| ASV1512 | f1cf3687e81 | 4 | 0 | 0 | 0 | 0 | 0 | 0 | 0 | 0 | 0 | 0 | 0 | 0 | 0 | 0 | 0 | 0 | 0 | 0 | 0 | 0 | 0 | 0 | 0 | 4 | 0 | 0 | 0 | 0 | 0 | 0 | 0 | 0 | 0 | 0 | 0 |
| ASV1513 | d8a292b392a | 4 | 0 | 0 | 0 | 0 | 0 | 0 | 0 | 0 | 0 | 0 | 0 | 0 | 0 | 0 | 0 | 0 | 0 | 0 | 0 | 0 | 0 | 0 | 0 | 0 | 0 | 0 | 0 | 0 | 0 | 0 | 0 | 0 | 0 | 0 | 4 |
| ASV1514 | ab2163ba359 | 4 | 0 | 0 | 0 | 0 | 0 | 0 | 0 | 4 | 0 | 0 | 0 | 0 | 0 | 0 | 0 | 0 | 0 | 0 | 0 | 0 | 0 | 0 | 0 | 0 | 0 | 0 | 0 | 0 | 0 | 0 | 0 | 0 | 0 | 0 | 0 |
| ASV1515 | ea9ae875c76 | 4 | 0 | 0 | 0 | 0 | 0 | 0 | 0 | 0 | 0 | 0 | 0 | 0 | 0 | 0 | 0 | 0 | 0 | 0 | 0 | 0 | 0 | 0 | 0 | 0 | 0 | 0 | 0 | 0 | 0 | 0 | 0 | 0 | 0 | 0 | 4 |
| ASV1516 | e657989a25a | 4 | 0 | 0 | 0 | 0 | 0 | 0 | 0 | 0 | 0 | 0 | 0 | 0 | 0 | 0 | 0 | 0 | 0 | 0 | 0 | 0 | 0 | 0 | 0 | 0 | 0 | 0 | 0 | 0 | 0 | 0 | 0 | 0 | 0 | 4 | 0 |
| ASV1517 | 2bea4e2bd82 | 4 | 0 | 0 | 0 | 0 | 0 | 0 | 0 | 0 | 0 | 0 | 0 | 0 | 0 | 0 | 4 | 0 | 0 | 0 | 0 | 0 | 0 | 0 | 0 | 0 | 0 | 0 | 0 | 0 | 0 | 0 | 0 | 0 | 0 | 0 | 0 |
| ASV1518 | 3646a09ac8f | 4 | 0 | 0 | 0 | 0 | 0 | 0 | 0 | 0 | 0 | 0 | 0 | 0 | 0 | 0 | 0 | 0 | 0 | 0 | 0 | 0 | 0 | 0 | 0 | 0 | 0 | 0 | 0 | 0 | 4 | 0 | 0 | 0 | 0 | 0 | 0 |

**Supplementary Table S5.** Bacterioplankton 16S rRNA ASV feature table
ASVs: 1779
Seqs: 918792

| #ASV_ID | Feature_ID | total seqs | May12_0m | May12_05m | May12_25m | May12_50m | May12_75m | Jun8_0m | Jun8_05m | Jun8_25m | Jun8_50m | Jun8_75m | Jun22_0m | Jun22_05m | Jun22_25m | Jun22_50m | Jun22_75m | Jul11_0m | Jul11_05m | Jul11_25m | Jul11_50m | Jul11_75m | Jul27_0m | Jul27_05m | Jul27_25m | Jul27_50m | Jul27_75m | Aug5_0m | Aug5_05m | Aug5_25m | Aug5_50m | Aug5_75m | Sep8_0m | Sep8_05m | Sep8_25m | Sep8_50m | Sep8_75m |
|---|---|---|---|---|---|---|---|---|---|---|---|---|---|---|---|---|---|---|---|---|---|---|---|---|---|---|---|---|---|---|---|---|---|---|---|---|---|
| ASV1519 | f1d297fd71ae | 4 | 0 | 0 | 0 | 0 | 0 | 0 | 0 | 0 | 0 | 0 | 0 | 0 | 0 | 0 | 0 | 0 | 0 | 0 | 0 | 0 | 0 | 0 | 0 | 0 | 0 | 0 | 0 | 0 | 0 | 0 | 0 | 4 | 0 | 0 | 0 |
| ASV1520 | 92a5cf6d0f6a | 4 | 0 | 0 | 0 | 0 | 0 | 0 | 0 | 0 | 0 | 0 | 0 | 0 | 0 | 0 | 0 | 0 | 0 | 0 | 0 | 0 | 0 | 0 | 0 | 0 | 0 | 0 | 0 | 0 | 0 | 0 | 0 | 0 | 0 | 4 | 0 |
| ASV1521 | 4d0ece734c5 | 4 | 0 | 4 | 0 | 0 | 0 | 0 | 0 | 0 | 0 | 0 | 0 | 0 | 0 | 0 | 0 | 0 | 0 | 0 | 0 | 0 | 0 | 0 | 0 | 0 | 0 | 0 | 0 | 0 | 0 | 0 | 0 | 0 | 0 | 0 | 0 |
| ASV1522 | 55e87a00a72 | 4 | 0 | 0 | 0 | 0 | 0 | 0 | 0 | 0 | 0 | 4 | 0 | 0 | 0 | 0 | 0 | 0 | 0 | 0 | 0 | 0 | 0 | 0 | 0 | 0 | 0 | 0 | 0 | 0 | 0 | 0 | 0 | 0 | 0 | 0 | 0 |
| ASV1523 | 5242c543602 | 4 | 0 | 2 | 0 | 2 | 0 | 0 | 0 | 0 | 0 | 0 | 0 | 0 | 0 | 0 | 0 | 0 | 0 | 0 | 0 | 0 | 0 | 0 | 0 | 0 | 0 | 0 | 0 | 0 | 0 | 0 | 0 | 0 | 0 | 0 | 0 |
| ASV1524 | b0cc16ef0aa | 4 | 0 | 0 | 0 | 0 | 0 | 0 | 0 | 0 | 0 | 0 | 0 | 0 | 0 | 0 | 4 | 0 | 0 | 0 | 0 | 0 | 0 | 0 | 0 | 0 | 0 | 0 | 0 | 0 | 0 | 0 | 0 | 0 | 0 | 0 | 0 |
| ASV1525 | 7b9ecdcf8513 | 4 | 0 | 0 | 0 | 0 | 0 | 0 | 0 | 0 | 0 | 0 | 0 | 0 | 0 | 0 | 0 | 0 | 0 | 0 | 0 | 0 | 0 | 0 | 0 | 0 | 0 | 0 | 0 | 0 | 0 | 0 | 0 | 0 | 0 | 0 | 4 |
| ASV1526 | f328237b633 | 4 | 0 | 0 | 0 | 0 | 0 | 0 | 0 | 0 | 0 | 0 | 0 | 0 | 0 | 0 | 0 | 0 | 0 | 0 | 0 | 0 | 0 | 0 | 0 | 0 | 0 | 0 | 0 | 4 | 0 | 0 | 0 | 0 | 0 | 0 | 0 |
| ASV1527 | ab9a06144b6 | 4 | 0 | 0 | 0 | 0 | 0 | 0 | 0 | 0 | 0 | 0 | 0 | 0 | 0 | 0 | 0 | 0 | 0 | 0 | 0 | 0 | 0 | 0 | 0 | 0 | 0 | 0 | 0 | 0 | 0 | 0 | 0 | 0 | 0 | 0 | 4 |
| ASV1528 | 8fe6daf570f8 | 4 | 0 | 0 | 0 | 0 | 0 | 0 | 0 | 0 | 0 | 0 | 0 | 0 | 0 | 0 | 0 | 0 | 4 | 0 | 0 | 0 | 0 | 0 | 0 | 0 | 0 | 0 | 0 | 0 | 0 | 0 | 0 | 0 | 0 | 0 | 0 |
| ASV1529 | df1e0a4f564f | 4 | 0 | 0 | 0 | 0 | 0 | 0 | 0 | 0 | 0 | 4 | 0 | 0 | 0 | 0 | 0 | 0 | 0 | 0 | 0 | 0 | 0 | 0 | 0 | 0 | 0 | 0 | 0 | 0 | 0 | 0 | 0 | 0 | 0 | 0 | 0 |
| ASV1530 | 39604659583 | 4 | 0 | 0 | 0 | 0 | 0 | 0 | 0 | 0 | 0 | 0 | 0 | 0 | 0 | 0 | 0 | 0 | 0 | 0 | 0 | 0 | 0 | 0 | 0 | 0 | 0 | 0 | 0 | 0 | 0 | 0 | 4 | 0 | 0 | 0 | 0 |
| ASV1531 | 20e2bc59d79 | 4 | 0 | 0 | 0 | 0 | 0 | 0 | 0 | 0 | 0 | 0 | 0 | 0 | 4 | 0 | 0 | 0 | 0 | 0 | 0 | 0 | 0 | 0 | 0 | 0 | 0 | 0 | 0 | 0 | 0 | 0 | 0 | 0 | 0 | 0 | 0 |
| ASV1532 | 90316d25d47 | 4 | 0 | 0 | 0 | 0 | 0 | 0 | 0 | 0 | 0 | 0 | 0 | 0 | 0 | 0 | 0 | 4 | 0 | 0 | 0 | 0 | 0 | 0 | 0 | 0 | 0 | 0 | 0 | 0 | 0 | 0 | 0 | 0 | 0 | 0 | 0 |
| ASV1533 | 6bf2daf6e15 | 4 | 0 | 0 | 0 | 0 | 0 | 0 | 0 | 0 | 0 | 0 | 0 | 0 | 0 | 0 | 4 | 0 | 0 | 0 | 0 | 0 | 0 | 0 | 0 | 0 | 0 | 0 | 0 | 0 | 0 | 0 | 0 | 0 | 0 | 0 | 0 |
| ASV1534 | 18b5a613386 | 4 | 0 | 0 | 0 | 0 | 0 | 0 | 0 | 0 | 0 | 0 | 0 | 0 | 0 | 0 | 0 | 0 | 0 | 0 | 0 | 0 | 0 | 0 | 0 | 0 | 0 | 0 | 0 | 0 | 0 | 0 | 0 | 4 | 0 | 0 | 0 |
| ASV1535 | d6698271c81 | 4 | 0 | 4 | 0 | 0 | 0 | 0 | 0 | 0 | 0 | 0 | 0 | 0 | 0 | 0 | 0 | 0 | 0 | 0 | 0 | 0 | 0 | 0 | 0 | 0 | 0 | 0 | 0 | 0 | 0 | 0 | 0 | 0 | 0 | 0 | 0 |
| ASV1536 | 3939bbb1803 | 4 | 0 | 0 | 0 | 0 | 0 | 4 | 0 | 0 | 0 | 0 | 0 | 0 | 0 | 0 | 0 | 0 | 0 | 0 | 0 | 0 | 0 | 0 | 0 | 0 | 0 | 0 | 0 | 0 | 0 | 0 | 0 | 0 | 0 | 0 | 0 |
| ASV1537 | cf50ac71972 | 4 | 0 | 0 | 0 | 0 | 0 | 0 | 0 | 0 | 0 | 0 | 0 | 0 | 0 | 0 | 0 | 0 | 0 | 0 | 4 | 0 | 0 | 0 | 0 | 0 | 0 | 0 | 0 | 0 | 0 | 0 | 0 | 0 | 0 | 0 | 0 |
| ASV1538 | d4c9e63d6b4 | 4 | 0 | 0 | 0 | 0 | 4 | 0 | 0 | 0 | 0 | 0 | 0 | 0 | 0 | 0 | 0 | 0 | 0 | 0 | 0 | 0 | 0 | 0 | 0 | 0 | 0 | 0 | 0 | 0 | 0 | 0 | 0 | 0 | 0 | 0 | 0 |
| ASV1539 | 865d95bfe65 | 4 | 0 | 0 | 0 | 0 | 0 | 0 | 0 | 0 | 0 | 0 | 0 | 0 | 0 | 0 | 0 | 4 | 0 | 0 | 0 | 0 | 0 | 0 | 0 | 0 | 0 | 0 | 0 | 0 | 0 | 0 | 0 | 0 | 0 | 0 | 0 |
| ASV1540 | c5ee1fcd89e | 4 | 0 | 0 | 0 | 0 | 4 | 0 | 0 | 0 | 0 | 0 | 0 | 0 | 0 | 0 | 0 | 0 | 0 | 0 | 0 | 0 | 0 | 0 | 0 | 0 | 0 | 0 | 0 | 0 | 0 | 0 | 0 | 0 | 0 | 0 | 0 |
| ASV1541 | c82305cfe51 | 4 | 0 | 0 | 0 | 0 | 0 | 0 | 0 | 0 | 0 | 0 | 0 | 0 | 0 | 0 | 0 | 0 | 0 | 0 | 0 | 0 | 4 | 0 | 0 | 0 | 0 | 0 | 0 | 0 | 0 | 0 | 0 | 0 | 0 | 0 | 0 |
| ASV1542 | df52e9427f7 | 4 | 0 | 0 | 0 | 0 | 0 | 0 | 0 | 0 | 0 | 0 | 0 | 0 | 0 | 0 | 0 | 0 | 0 | 0 | 0 | 0 | 0 | 0 | 0 | 0 | 0 | 0 | 0 | 0 | 0 | 0 | 4 | 0 | 0 | 0 | 0 |
| ASV1543 | 3ca6b8c53d1 | 4 | 0 | 0 | 0 | 0 | 0 | 0 | 0 | 0 | 0 | 2 | 0 | 0 | 0 | 0 | 0 | 0 | 0 | 0 | 0 | 0 | 0 | 0 | 0 | 0 | 0 | 0 | 0 | 0 | 0 | 2 | 0 | 0 | 0 | 0 | 0 |
| ASV1544 | 0ff69bf5a7ef | 4 | 0 | 0 | 0 | 0 | 0 | 0 | 0 | 0 | 0 | 0 | 0 | 0 | 0 | 0 | 0 | 0 | 0 | 0 | 0 | 0 | 0 | 0 | 0 | 0 | 0 | 0 | 0 | 0 | 0 | 0 | 4 | 0 | 0 | 0 | 0 |
| ASV1545 | c98092e9ee7 | 4 | 0 | 0 | 0 | 0 | 0 | 0 | 0 | 0 | 0 | 0 | 0 | 0 | 0 | 0 | 0 | 0 | 0 | 0 | 0 | 0 | 0 | 0 | 0 | 0 | 0 | 0 | 0 | 0 | 0 | 0 | 0 | 0 | 0 | 4 | 0 |
| ASV1546 | 06b802ac5ad | 4 | 0 | 0 | 0 | 0 | 0 | 0 | 0 | 0 | 0 | 0 | 0 | 0 | 0 | 0 | 0 | 0 | 0 | 0 | 0 | 0 | 0 | 0 | 0 | 0 | 0 | 0 | 4 | 0 | 0 | 0 | 0 | 0 | 0 | 0 | 0 |
| ASV1547 | 441919abe88 | 4 | 0 | 0 | 0 | 0 | 4 | 0 | 0 | 0 | 0 | 0 | 0 | 0 | 0 | 0 | 0 | 0 | 0 | 0 | 0 | 0 | 0 | 0 | 0 | 0 | 0 | 0 | 0 | 0 | 0 | 0 | 0 | 0 | 0 | 0 | 0 |
| ASV1548 | 4a79d27b813 | 4 | 0 | 0 | 0 | 0 | 4 | 0 | 0 | 0 | 0 | 0 | 0 | 0 | 0 | 0 | 0 | 0 | 0 | 0 | 0 | 0 | 0 | 0 | 0 | 0 | 0 | 0 | 0 | 0 | 0 | 0 | 0 | 0 | 0 | 0 | 0 |
| ASV1549 | 99a2b7b9b67 | 4 | 0 | 0 | 0 | 0 | 0 | 0 | 0 | 0 | 0 | 0 | 0 | 0 | 0 | 0 | 4 | 0 | 0 | 0 | 0 | 0 | 0 | 0 | 0 | 0 | 0 | 0 | 0 | 0 | 0 | 0 | 0 | 0 | 0 | 0 | 0 |
| ASV1550 | ffba9d1c736 | 4 | 0 | 0 | 0 | 0 | 0 | 0 | 0 | 0 | 0 | 0 | 0 | 0 | 0 | 0 | 0 | 0 | 0 | 0 | 0 | 0 | 0 | 0 | 0 | 0 | 0 | 0 | 0 | 0 | 0 | 0 | 0 | 0 | 0 | 0 | 4 |
| ASV1551 | 13110171537 | 4 | 0 | 0 | 0 | 4 | 0 | 0 | 0 | 0 | 0 | 0 | 0 | 0 | 0 | 0 | 0 | 0 | 0 | 0 | 0 | 0 | 0 | 0 | 0 | 0 | 0 | 0 | 0 | 0 | 0 | 0 | 0 | 0 | 0 | 0 | 0 |
| ASV1552 | 8b749642527 | 4 | 0 | 0 | 0 | 0 | 0 | 0 | 0 | 0 | 0 | 0 | 0 | 0 | 0 | 0 | 0 | 0 | 0 | 0 | 0 | 0 | 0 | 0 | 0 | 0 | 0 | 0 | 0 | 0 | 0 | 0 | 0 | 0 | 0 | 0 | 4 |
| ASV1553 | 4209435f807 | 4 | 0 | 0 | 0 | 0 | 0 | 0 | 0 | 0 | 0 | 0 | 0 | 0 | 0 | 0 | 0 | 0 | 0 | 0 | 0 | 0 | 0 | 0 | 0 | 0 | 0 | 0 | 0 | 0 | 0 | 0 | 4 | 0 | 0 | 0 | 0 |
| ASV1554 | 889187307fa | 4 | 0 | 0 | 0 | 0 | 0 | 0 | 0 | 0 | 4 | 0 | 0 | 0 | 0 | 0 | 0 | 0 | 0 | 0 | 0 | 0 | 0 | 0 | 0 | 0 | 0 | 0 | 0 | 0 | 0 | 0 | 0 | 0 | 0 | 0 | 0 |
| ASV1555 | 3659fa8e677 | 4 | 0 | 0 | 0 | 0 | 0 | 0 | 0 | 0 | 0 | 0 | 0 | 0 | 0 | 4 | 0 | 0 | 0 | 0 | 0 | 0 | 0 | 0 | 0 | 0 | 0 | 0 | 0 | 0 | 0 | 0 | 0 | 0 | 0 | 0 | 0 |
| ASV1556 | f6f24174dbc | 4 | 0 | 0 | 0 | 0 | 0 | 0 | 0 | 0 | 0 | 0 | 0 | 0 | 0 | 0 | 0 | 0 | 0 | 0 | 0 | 0 | 0 | 0 | 0 | 0 | 0 | 0 | 0 | 0 | 0 | 0 | 0 | 0 | 0 | 0 | 4 |
| ASV1557 | 243344b04b8 | 4 | 0 | 0 | 0 | 0 | 0 | 0 | 0 | 0 | 0 | 0 | 0 | 0 | 0 | 0 | 0 | 0 | 0 | 0 | 0 | 0 | 0 | 0 | 0 | 0 | 0 | 0 | 0 | 0 | 0 | 4 | 0 | 0 | 0 | 0 | 0 |
| ASV1558 | 2454fdcb79a | 4 | 0 | 0 | 4 | 0 | 0 | 0 | 0 | 0 | 0 | 0 | 0 | 0 | 0 | 0 | 0 | 0 | 0 | 0 | 0 | 0 | 0 | 0 | 0 | 0 | 0 | 0 | 0 | 0 | 0 | 0 | 0 | 0 | 0 | 0 | 0 |
| ASV1559 | 2a860a97764 | 4 | 0 | 0 | 0 | 0 | 0 | 0 | 0 | 0 | 4 | 0 | 0 | 0 | 0 | 0 | 0 | 0 | 0 | 0 | 0 | 0 | 0 | 0 | 0 | 0 | 0 | 0 | 0 | 0 | 0 | 0 | 0 | 0 | 0 | 0 | 0 |
| ASV1560 | 71d30286b6f | 4 | 0 | 4 | 0 | 0 | 0 | 0 | 0 | 0 | 0 | 0 | 0 | 0 | 0 | 0 | 0 | 0 | 0 | 0 | 0 | 0 | 0 | 0 | 0 | 0 | 0 | 0 | 0 | 0 | 0 | 0 | 0 | 0 | 0 | 0 | 0 |
| ASV1561 | 9e84a367e99 | 4 | 0 | 0 | 0 | 0 | 0 | 0 | 0 | 0 | 0 | 4 | 0 | 0 | 0 | 0 | 0 | 0 | 0 | 0 | 0 | 0 | 0 | 0 | 0 | 0 | 0 | 0 | 0 | 0 | 0 | 0 | 0 | 0 | 0 | 0 | 0 |
| ASV1562 | c48f41767b1 | 4 | 0 | 0 | 0 | 0 | 0 | 0 | 0 | 0 | 0 | 0 | 0 | 0 | 0 | 0 | 0 | 0 | 0 | 0 | 0 | 0 | 0 | 0 | 0 | 0 | 0 | 0 | 0 | 0 | 0 | 0 | 0 | 0 | 0 | 0 | 4 |
| ASV1563 | b76d5b55446 | 4 | 0 | 0 | 0 | 0 | 0 | 0 | 0 | 0 | 0 | 0 | 0 | 0 | 0 | 0 | 0 | 0 | 0 | 0 | 0 | 0 | 0 | 0 | 0 | 0 | 0 | 0 | 0 | 0 | 0 | 0 | 0 | 0 | 0 | 4 | 0 |
| ASV1564 | d7ea75c97d8 | 4 | 0 | 0 | 0 | 0 | 0 | 0 | 0 | 0 | 0 | 0 | 0 | 0 | 0 | 4 | 0 | 0 | 0 | 0 | 0 | 0 | 0 | 0 | 0 | 0 | 0 | 0 | 0 | 0 | 0 | 0 | 0 | 0 | 0 | 0 | 0 |
| ASV1565 | d4e3121ddb0 | 4 | 0 | 0 | 0 | 0 | 0 | 4 | 0 | 0 | 0 | 0 | 0 | 0 | 0 | 0 | 0 | 0 | 0 | 0 | 0 | 0 | 0 | 0 | 0 | 0 | 0 | 0 | 0 | 0 | 0 | 0 | 0 | 0 | 0 | 0 | 0 |
| ASV1566 | e79caaf336d | 4 | 0 | 0 | 0 | 4 | 0 | 0 | 0 | 0 | 0 | 0 | 0 | 0 | 0 | 0 | 0 | 0 | 0 | 0 | 0 | 0 | 0 | 0 | 0 | 0 | 0 | 0 | 0 | 0 | 0 | 0 | 0 | 0 | 0 | 0 | 0 |
| ASV1567 | 973207b51ad | 4 | 0 | 0 | 0 | 0 | 0 | 0 | 0 | 0 | 0 | 0 | 0 | 0 | 0 | 0 | 0 | 4 | 0 | 0 | 0 | 0 | 0 | 0 | 0 | 0 | 0 | 0 | 0 | 0 | 0 | 0 | 0 | 0 | 0 | 0 | 0 |
| ASV1568 | 3e617551815 | 4 | 0 | 0 | 0 | 0 | 0 | 0 | 0 | 0 | 0 | 0 | 0 | 0 | 0 | 4 | 0 | 0 | 0 | 0 | 0 | 0 | 0 | 0 | 0 | 0 | 0 | 0 | 0 | 0 | 0 | 0 | 0 | 0 | 0 | 0 | 0 |
| ASV1569 | 885b34a8326 | 4 | 0 | 0 | 0 | 0 | 0 | 0 | 0 | 0 | 0 | 0 | 0 | 0 | 0 | 0 | 4 | 0 | 0 | 0 | 0 | 0 | 0 | 0 | 0 | 0 | 0 | 0 | 0 | 0 | 0 | 0 | 0 | 0 | 0 | 0 | 0 |
| ASV1570 | bf8c7e778f80 | 4 | 0 | 0 | 0 | 0 | 0 | 0 | 0 | 0 | 0 | 4 | 0 | 0 | 0 | 0 | 0 | 0 | 0 | 0 | 0 | 0 | 0 | 0 | 0 | 0 | 0 | 0 | 0 | 0 | 0 | 0 | 0 | 0 | 0 | 0 | 0 |
| ASV1571 | 060be68394e | 3 | 0 | 0 | 0 | 0 | 0 | 0 | 0 | 0 | 3 | 0 | 0 | 0 | 0 | 0 | 0 | 0 | 0 | 0 | 0 | 0 | 0 | 0 | 0 | 0 | 0 | 0 | 0 | 0 | 0 | 0 | 0 | 0 | 0 | 0 | 0 |
| ASV1572 | 1df81a0fc27a | 3 | 0 | 0 | 0 | 0 | 0 | 0 | 0 | 0 | 0 | 0 | 0 | 0 | 0 | 0 | 0 | 0 | 0 | 0 | 0 | 0 | 0 | 0 | 0 | 0 | 0 | 0 | 0 | 3 | 0 | 0 | 0 | 0 | 0 | 0 | 0 |
| ASV1573 | 2c6075d66e6 | 3 | 0 | 0 | 0 | 0 | 3 | 0 | 0 | 0 | 0 | 0 | 0 | 0 | 0 | 0 | 0 | 0 | 0 | 0 | 0 | 0 | 0 | 0 | 0 | 0 | 0 | 0 | 0 | 0 | 0 | 0 | 0 | 0 | 0 | 0 | 0 |
| ASV1574 | ceff7e2f6423 | 3 | 0 | 0 | 0 | 0 | 0 | 0 | 0 | 0 | 3 | 0 | 0 | 0 | 0 | 0 | 0 | 0 | 0 | 0 | 0 | 0 | 0 | 0 | 0 | 0 | 0 | 0 | 0 | 0 | 0 | 0 | 0 | 0 | 0 | 0 | 0 |
| ASV1575 | 74a382c0aa3 | 3 | 0 | 0 | 0 | 0 | 0 | 0 | 0 | 0 | 0 | 0 | 0 | 0 | 0 | 0 | 0 | 0 | 0 | 0 | 0 | 0 | 0 | 0 | 0 | 3 | 0 | 0 | 0 | 0 | 0 | 0 | 0 | 0 | 0 | 0 | 0 |
| ASV1576 | 79b0cf973f2f | 3 | 0 | 0 | 3 | 0 | 0 | 0 | 0 | 0 | 0 | 0 | 0 | 0 | 0 | 0 | 0 | 0 | 0 | 0 | 0 | 0 | 0 | 0 | 0 | 0 | 0 | 0 | 0 | 0 | 0 | 0 | 0 | 0 | 0 | 0 | 0 |
| ASV1577 | ab89265ceb6 | 3 | 0 | 0 | 0 | 0 | 3 | 0 | 0 | 0 | 0 | 0 | 0 | 0 | 0 | 0 | 0 | 0 | 0 | 0 | 0 | 0 | 0 | 0 | 0 | 0 | 0 | 0 | 0 | 0 | 0 | 0 | 0 | 0 | 0 | 0 | 0 |
| ASV1578 | b64938b7e62 | 3 | 0 | 0 | 0 | 0 | 0 | 0 | 0 | 0 | 0 | 0 | 0 | 0 | 0 | 0 | 0 | 0 | 0 | 0 | 0 | 0 | 0 | 0 | 0 | 0 | 0 | 0 | 0 | 0 | 3 | 0 | 0 | 0 | 0 | 0 | 0 |
| ASV1579 | 4691037dc93 | 3 | 0 | 0 | 0 | 0 | 0 | 0 | 0 | 0 | 0 | 0 | 0 | 0 | 0 | 0 | 0 | 0 | 0 | 0 | 0 | 0 | 0 | 0 | 0 | 0 | 0 | 0 | 0 | 3 | 0 | 0 | 0 | 0 | 0 | 0 | 0 |
| ASV1580 | 1f1b47cc9d4 | 3 | 0 | 0 | 0 | 0 | 0 | 0 | 0 | 0 | 0 | 0 | 0 | 0 | 0 | 0 | 0 | 3 | 0 | 0 | 0 | 0 | 0 | 0 | 0 | 0 | 0 | 0 | 0 | 0 | 0 | 0 | 0 | 0 | 0 | 0 | 0 |
| ASV1581 | 2cfca4b4278 | 3 | 0 | 0 | 0 | 0 | 0 | 0 | 0 | 0 | 0 | 0 | 0 | 0 | 0 | 0 | 0 | 0 | 0 | 0 | 3 | 0 | 0 | 0 | 0 | 0 | 0 | 0 | 0 | 0 | 0 | 0 | 0 | 0 | 0 | 0 | 0 |
| ASV1582 | 95b582d7871 | 3 | 0 | 3 | 0 | 0 | 0 | 0 | 0 | 0 | 0 | 0 | 0 | 0 | 0 | 0 | 0 | 0 | 0 | 0 | 0 | 0 | 0 | 0 | 0 | 0 | 0 | 0 | 0 | 0 | 0 | 0 | 0 | 0 | 0 | 0 | 0 |
| ASV1583 | 063f45b1609 | 3 | 0 | 0 | 0 | 0 | 0 | 0 | 0 | 0 | 0 | 0 | 0 | 0 | 0 | 0 | 0 | 0 | 0 | 0 | 0 | 0 | 0 | 0 | 0 | 0 | 0 | 0 | 0 | 0 | 0 | 0 | 0 | 0 | 0 | 0 | 3 |
| ASV1584 | 7b631e625a5 | 3 | 0 | 0 | 0 | 0 | 0 | 0 | 0 | 0 | 0 | 0 | 0 | 0 | 0 | 0 | 0 | 0 | 0 | 0 | 0 | 0 | 0 | 0 | 3 | 0 | 0 | 0 | 0 | 0 | 0 | 0 | 0 | 0 | 0 | 0 | 0 |
| ASV1585 | 1e58ed425f1 | 3 | 0 | 0 | 0 | 0 | 0 | 0 | 0 | 3 | 0 | 0 | 0 | 0 | 0 | 0 | 0 | 0 | 0 | 0 | 0 | 0 | 0 | 0 | 0 | 0 | 0 | 0 | 0 | 0 | 0 | 0 | 0 | 0 | 0 | 0 | 0 |
| ASV1586 | 640ebce32e6 | 3 | 0 | 0 | 0 | 0 | 0 | 3 | 0 | 0 | 0 | 0 | 0 | 0 | 0 | 0 | 0 | 0 | 0 | 0 | 0 | 0 | 0 | 0 | 0 | 0 | 0 | 0 | 0 | 0 | 0 | 0 | 0 | 0 | 0 | 0 | 0 |
| ASV1587 | 1b2ea70c0f6 | 3 | 0 | 0 | 0 | 0 | 0 | 0 | 0 | 0 | 0 | 0 | 0 | 0 | 0 | 0 | 0 | 0 | 0 | 0 | 0 | 0 | 0 | 0 | 0 | 0 | 0 | 0 | 0 | 0 | 0 | 0 | 0 | 0 | 0 | 3 | 0 |
| ASV1588 | b567ad23661 | 3 | 0 | 0 | 0 | 0 | 0 | 0 | 0 | 0 | 0 | 0 | 0 | 0 | 0 | 0 | 0 | 0 | 0 | 0 | 0 | 0 | 0 | 0 | 0 | 0 | 0 | 0 | 0 | 0 | 0 | 0 | 3 | 0 | 0 | 0 | 0 |
| ASV1589 | 3953c46d669 | 3 | 0 | 0 | 0 | 0 | 0 | 0 | 3 | 0 | 0 | 0 | 0 | 0 | 0 | 0 | 0 | 0 | 0 | 0 | 0 | 0 | 0 | 0 | 0 | 0 | 0 | 0 | 0 | 0 | 0 | 0 | 0 | 0 | 0 | 0 | 0 |
| ASV1590 | f0ecd50f752a | 3 | 0 | 0 | 0 | 0 | 0 | 0 | 0 | 0 | 0 | 0 | 0 | 0 | 0 | 0 | 0 | 0 | 0 | 0 | 0 | 0 | 0 | 0 | 0 | 0 | 0 | 0 | 0 | 3 | 0 | 0 | 0 | 0 | 0 | 0 | 0 |
| ASV1591 | f555a90112b | 3 | 0 | 0 | 3 | 0 | 0 | 0 | 0 | 0 | 0 | 0 | 0 | 0 | 0 | 0 | 0 | 0 | 0 | 0 | 0 | 0 | 0 | 0 | 0 | 0 | 0 | 0 | 0 | 0 | 0 | 0 | 0 | 0 | 0 | 0 | 0 |
| ASV1592 | 48cbc0483fe | 3 | 0 | 0 | 0 | 0 | 0 | 0 | 0 | 0 | 0 | 0 | 0 | 0 | 0 | 3 | 0 | 0 | 0 | 0 | 0 | 0 | 0 | 0 | 0 | 0 | 0 | 0 | 0 | 0 | 0 | 0 | 0 | 0 | 0 | 0 | 0 |
| ASV1593 | 41c5a697151 | 3 | 0 | 0 | 0 | 0 | 0 | 0 | 0 | 3 | 0 | 0 | 0 | 0 | 0 | 0 | 0 | 0 | 0 | 0 | 0 | 0 | 0 | 0 | 0 | 0 | 0 | 0 | 0 | 0 | 0 | 0 | 0 | 0 | 0 | 0 | 0 |
| ASV1594 | 502ff238dd7 | 3 | 0 | 0 | 0 | 0 | 0 | 0 | 0 | 0 | 0 | 0 | 0 | 0 | 0 | 0 | 0 | 0 | 0 | 0 | 0 | 0 | 0 | 0 | 0 | 0 | 0 | 0 | 0 | 0 | 0 | 0 | 0 | 0 | 3 | 0 | 0 |
| ASV1595 | 467f23910e6 | 3 | 0 | 0 | 0 | 0 | 0 | 0 | 0 | 3 | 0 | 0 | 0 | 0 | 0 | 0 | 0 | 0 | 0 | 0 | 0 | 0 | 0 | 0 | 0 | 0 | 0 | 0 | 0 | 0 | 0 | 0 | 0 | 0 | 0 | 0 | 0 |
| ASV1596 | 1660941eec3 | 3 | 0 | 0 | 0 | 0 | 0 | 0 | 0 | 0 | 0 | 3 | 0 | 0 | 0 | 0 | 0 | 0 | 0 | 0 | 0 | 0 | 0 | 0 | 0 | 0 | 0 | 0 | 0 | 0 | 0 | 0 | 0 | 0 | 0 | 0 | 0 |
| ASV1597 | 0e8a2ce7a56 | 3 | 0 | 0 | 0 | 0 | 0 | 0 | 0 | 0 | 0 | 0 | 0 | 0 | 0 | 0 | 3 | 0 | 0 | 0 | 0 | 0 | 0 | 0 | 0 | 0 | 0 | 0 | 0 | 0 | 0 | 0 | 0 | 0 | 0 | 0 | 0 |
| ASV1598 | 06e4826650d | 3 | 0 | 0 | 0 | 0 | 3 | 0 | 0 | 0 | 0 | 0 | 0 | 0 | 0 | 0 | 0 | 0 | 0 | 0 | 0 | 0 | 0 | 0 | 0 | 0 | 0 | 0 | 0 | 0 | 0 | 0 | 0 | 0 | 0 | 0 | 0 |
| ASV1599 | 9b45962bf31 | 3 | 0 | 0 | 0 | 0 | 0 | 0 | 0 | 0 | 0 | 0 | 3 | 0 | 0 | 0 | 0 | 0 | 0 | 0 | 0 | 0 | 0 | 0 | 0 | 0 | 0 | 0 | 0 | 0 | 0 | 0 | 0 | 0 | 0 | 0 | 0 |
| ASV1600 | db8a20878bd | 3 | 0 | 0 | 0 | 0 | 0 | 0 | 0 | 0 | 0 | 0 | 0 | 0 | 0 | 0 | 3 | 0 | 0 | 0 | 0 | 0 | 0 | 0 | 0 | 0 | 0 | 0 | 0 | 0 | 0 | 0 | 0 | 0 | 0 | 0 | 0 |
| ASV1601 | d30a36e9fcfe | 3 | 0 | 0 | 0 | 0 | 0 | 0 | 0 | 0 | 0 | 0 | 0 | 0 | 3 | 0 | 0 | 0 | 0 | 0 | 0 | 0 | 0 | 0 | 0 | 0 | 0 | 0 | 0 | 0 | 0 | 0 | 0 | 0 | 0 | 0 | 0 |
| ASV1602 | 3bee117d5da | 3 | 0 | 0 | 0 | 0 | 0 | 0 | 0 | 0 | 0 | 0 | 0 | 0 | 0 | 0 | 0 | 0 | 3 | 0 | 0 | 0 | 0 | 0 | 0 | 0 | 0 | 0 | 0 | 0 | 0 | 0 | 0 | 0 | 0 | 0 | 0 |
| ASV1603 | b3ff176f9e9d | 3 | 0 | 0 | 0 | 3 | 0 | 0 | 0 | 0 | 0 | 0 | 0 | 0 | 0 | 0 | 0 | 0 | 0 | 0 | 0 | 0 | 0 | 0 | 0 | 0 | 0 | 0 | 0 | 0 | 0 | 0 | 0 | 0 | 0 | 0 | 0 |
| ASV1604 | bbcb32b8ed4 | 3 | 0 | 0 | 0 | 3 | 0 | 0 | 0 | 0 | 0 | 0 | 0 | 0 | 0 | 0 | 0 | 0 | 0 | 0 | 0 | 0 | 0 | 0 | 0 | 0 | 0 | 0 | 0 | 0 | 0 | 0 | 0 | 0 | 0 | 0 | 0 |
| ASV1605 | 83144d5ea68 | 3 | 0 | 0 | 0 | 0 | 0 | 0 | 0 | 0 | 0 | 0 | 0 | 0 | 0 | 0 | 0 | 0 | 0 | 0 | 0 | 0 | 0 | 0 | 0 | 0 | 0 | 0 | 0 | 0 | 0 | 0 | 0 | 0 | 3 | 0 | 0 |
| ASV1606 | 20c2a3a2c1c | 3 | 0 | 0 | 0 | 0 | 0 | 0 | 0 | 0 | 0 | 0 | 0 | 0 | 0 | 0 | 0 | 0 | 0 | 0 | 0 | 0 | 0 | 0 | 0 | 3 | 0 | 0 | 0 | 0 | 0 | 0 | 0 | 0 | 0 | 0 | 0 |
| ASV1607 | 271ae02863a | 3 | 0 | 0 | 0 | 0 | 0 | 0 | 0 | 0 | 0 | 0 | 0 | 0 | 0 | 0 | 0 | 0 | 0 | 0 | 0 | 0 | 0 | 0 | 3 | 0 | 0 | 0 | 0 | 0 | 0 | 0 | 0 | 0 | 0 | 0 | 0 |
| ASV1608 | 1ca133cbfe1 | 3 | 0 | 0 | 0 | 0 | 0 | 3 | 0 | 0 | 0 | 0 | 0 | 0 | 0 | 0 | 0 | 0 | 0 | 0 | 0 | 0 | 0 | 0 | 0 | 0 | 0 | 0 | 0 | 0 | 0 | 0 | 0 | 0 | 0 | 0 | 0 |
| ASV1609 | 9f7e6f2877fa | 3 | 0 | 0 | 0 | 0 | 0 | 0 | 0 | 0 | 0 | 0 | 0 | 0 | 0 | 0 | 0 | 0 | 0 | 0 | 0 | 0 | 0 | 0 | 0 | 0 | 0 | 0 | 0 | 0 | 0 | 0 | 0 | 0 | 0 | 0 | 3 |
| ASV1610 | 4e88736f7eb | 3 | 0 | 0 | 0 | 0 | 0 | 0 | 0 | 0 | 0 | 0 | 0 | 0 | 0 | 0 | 0 | 0 | 0 | 0 | 0 | 0 | 0 | 0 | 0 | 0 | 0 | 0 | 0 | 0 | 0 | 0 | 0 | 0 | 0 | 3 | 0 |
| ASV1611 | c94e14eead2 | 3 | 0 | 0 | 0 | 0 | 0 | 0 | 0 | 0 | 0 | 0 | 0 | 0 | 0 | 0 | 3 | 0 | 0 | 0 | 0 | 0 | 0 | 0 | 0 | 0 | 0 | 0 | 0 | 0 | 0 | 0 | 0 | 0 | 0 | 0 | 0 |
| ASV1612 | e45f610b2ea | 3 | 0 | 0 | 0 | 0 | 0 | 0 | 0 | 0 | 0 | 0 | 0 | 0 | 0 | 0 | 0 | 0 | 0 | 0 | 0 | 0 | 0 | 0 | 0 | 0 | 0 | 0 | 0 | 0 | 0 | 3 | 0 | 0 | 0 | 0 | 0 |
| ASV1613 | 8bbaec5dba9 | 3 | 0 | 0 | 0 | 0 | 0 | 0 | 3 | 0 | 0 | 0 | 0 | 0 | 0 | 0 | 0 | 0 | 0 | 0 | 0 | 0 | 0 | 0 | 0 | 0 | 0 | 0 | 0 | 0 | 0 | 0 | 0 | 0 | 0 | 0 | 0 |
| ASV1614 | 8d26e8a6c10 | 3 | 0 | 0 | 0 | 0 | 0 | 0 | 0 | 0 | 0 | 0 | 0 | 0 | 0 | 0 | 0 | 0 | 0 | 0 | 0 | 0 | 0 | 0 | 0 | 0 | 0 | 0 | 0 | 0 | 0 | 0 | 0 | 0 | 0 | 0 | 3 |
| ASV1615 | 795565ed022 | 3 | 0 | 0 | 0 | 0 | 0 | 0 | 0 | 0 | 0 | 0 | 0 | 0 | 0 | 0 | 0 | 0 | 0 | 0 | 0 | 0 | 0 | 0 | 0 | 0 | 3 | 0 | 0 | 0 | 0 | 0 | 0 | 0 | 0 | 0 | 0 |
| ASV1616 | 46739cf0e79 | 3 | 0 | 0 | 0 | 0 | 3 | 0 | 0 | 0 | 0 | 0 | 0 | 0 | 0 | 0 | 0 | 0 | 0 | 0 | 0 | 0 | 0 | 0 | 0 | 0 | 0 | 0 | 0 | 0 | 0 | 0 | 0 | 0 | 0 | 0 | 0 |
| ASV1617 | 987c9175dc6 | 3 | 0 | 0 | 0 | 0 | 0 | 0 | 0 | 0 | 0 | 0 | 0 | 0 | 0 | 0 | 0 | 0 | 0 | 0 | 0 | 0 | 0 | 0 | 0 | 0 | 0 | 0 | 0 | 0 | 0 | 0 | 0 | 3 | 0 | 0 | 0 |
| ASV1618 | f8817219b8e | 3 | 0 | 0 | 0 | 0 | 3 | 0 | 0 | 0 | 0 | 0 | 0 | 0 | 0 | 0 | 0 | 0 | 0 | 0 | 0 | 0 | 0 | 0 | 0 | 0 | 0 | 0 | 0 | 0 | 0 | 0 | 0 | 0 | 0 | 0 | 0 |
| ASV1619 | 5eec1b46e05 | 3 | 0 | 0 | 3 | 0 | 0 | 0 | 0 | 0 | 0 | 0 | 0 | 0 | 0 | 0 | 0 | 0 | 0 | 0 | 0 | 0 | 0 | 0 | 0 | 0 | 0 | 0 | 0 | 0 | 0 | 0 | 0 | 0 | 0 | 0 | 0 |
| ASV1620 | 72d6d1a9083 | 3 | 0 | 0 | 0 | 0 | 0 | 0 | 0 | 0 | 0 | 0 | 0 | 0 | 0 | 0 | 0 | 0 | 0 | 0 | 0 | 0 | 0 | 0 | 0 | 0 | 0 | 0 | 0 | 0 | 0 | 0 | 3 | 0 | 0 | 0 | 0 |
| ASV1621 | 527dec942f3 | 3 | 0 | 0 | 0 | 0 | 0 | 0 | 0 | 0 | 0 | 0 | 0 | 0 | 0 | 0 | 0 | 0 | 0 | 0 | 0 | 0 | 0 | 0 | 0 | 0 | 0 | 0 | 0 | 0 | 0 | 0 | 0 | 0 | 0 | 0 | 3 |
| ASV1622 | 7b6cccf3c22c | 3 | 0 | 0 | 0 | 0 | 0 | 0 | 0 | 0 | 0 | 0 | 0 | 0 | 0 | 0 | 0 | 0 | 0 | 0 | 0 | 0 | 0 | 0 | 0 | 3 | 0 | 0 | 0 | 0 | 0 | 0 | 0 | 0 | 0 | 0 | 0 |
| ASV1623 | ad5d7acebf4 | 3 | 0 | 0 | 0 | 0 | 0 | 0 | 0 | 0 | 0 | 0 | 0 | 0 | 0 | 3 | 0 | 0 | 0 | 0 | 0 | 0 | 0 | 0 | 0 | 0 | 0 | 0 | 0 | 0 | 0 | 0 | 0 | 0 | 0 | 0 | 0 |
| ASV1624 | f11bd7b1f30f | 3 | 0 | 0 | 0 | 0 | 0 | 0 | 0 | 0 | 0 | 0 | 0 | 0 | 0 | 0 | 0 | 0 | 0 | 0 | 0 | 0 | 0 | 3 | 0 | 0 | 0 | 0 | 0 | 0 | 0 | 0 | 0 | 0 | 0 | 0 | 0 |
| ASV1625 | 041e39e9319 | 3 | 0 | 0 | 0 | 0 | 3 | 0 | 0 | 0 | 0 | 0 | 0 | 0 | 0 | 0 | 0 | 0 | 0 | 0 | 0 | 0 | 0 | 0 | 0 | 0 | 0 | 0 | 0 | 0 | 0 | 0 | 0 | 0 | 0 | 0 | 0 |
| ASV1626 | 0dccdbb119b | 3 | 0 | 0 | 0 | 0 | 0 | 0 | 0 | 0 | 0 | 0 | 0 | 0 | 0 | 0 | 0 | 0 | 0 | 0 | 0 | 0 | 0 | 0 | 0 | 0 | 0 | 0 | 0 | 3 | 0 | 0 | 0 | 0 | 0 | 0 | 0 |
| ASV1627 | b65d8b1e41b | 3 | 0 | 0 | 0 | 0 | 0 | 0 | 0 | 0 | 0 | 0 | 0 | 0 | 0 | 0 | 3 | 0 | 0 | 0 | 0 | 0 | 0 | 0 | 0 | 0 | 0 | 0 | 0 | 0 | 0 | 0 | 0 | 0 | 0 | 0 | 0 |
| ASV1628 | cabbe95cc39 | 3 | 0 | 0 | 0 | 0 | 3 | 0 | 0 | 0 | 0 | 0 | 0 | 0 | 0 | 0 | 0 | 0 | 0 | 0 | 0 | 0 | 0 | 0 | 0 | 0 | 0 | 0 | 0 | 0 | 0 | 0 | 0 | 0 | 0 | 0 | 0 |
| ASV1629 | 862114615c3 | 3 | 0 | 0 | 0 | 0 | 0 | 0 | 0 | 0 | 0 | 0 | 0 | 0 | 0 | 0 | 0 | 0 | 0 | 0 | 0 | 0 | 0 | 0 | 0 | 0 | 0 | 0 | 0 | 0 | 0 | 3 | 0 | 0 | 0 | 0 | 0 |
| ASV1630 | 211a4522cb1 | 3 | 0 | 0 | 3 | 0 | 0 | 0 | 0 | 0 | 0 | 0 | 0 | 0 | 0 | 0 | 0 | 0 | 0 | 0 | 0 | 0 | 0 | 0 | 0 | 0 | 0 | 0 | 0 | 0 | 0 | 0 | 0 | 0 | 0 | 0 | 0 |
| ASV1631 | 720a4420b88 | 3 | 0 | 0 | 0 | 0 | 0 | 0 | 0 | 0 | 0 | 0 | 0 | 0 | 0 | 0 | 0 | 0 | 0 | 0 | 0 | 0 | 3 | 0 | 0 | 0 | 0 | 0 | 0 | 0 | 0 | 0 | 0 | 0 | 0 | 0 | 0 |
| ASV1632 | dc630db72ca | 3 | 0 | 0 | 0 | 0 | 0 | 0 | 0 | 0 | 0 | 0 | 0 | 0 | 0 | 0 | 0 | 0 | 3 | 0 | 0 | 0 | 0 | 0 | 0 | 0 | 0 | 0 | 0 | 0 | 0 | 0 | 0 | 0 | 0 | 0 | 0 |
| ASV1633 | 8efd07f86b0 | 3 | 0 | 0 | 0 | 0 | 0 | 0 | 0 | 0 | 0 | 3 | 0 | 0 | 0 | 0 | 0 | 0 | 0 | 0 | 0 | 0 | 0 | 0 | 0 | 0 | 0 | 0 | 0 | 0 | 0 | 0 | 0 | 0 | 0 | 0 | 0 |
| ASV1634 | 3797c84fb3a | 3 | 0 | 0 | 0 | 0 | 0 | 0 | 0 | 0 | 0 | 0 | 0 | 0 | 0 | 0 | 0 | 0 | 3 | 0 | 0 | 0 | 0 | 0 | 0 | 0 | 0 | 0 | 0 | 0 | 0 | 0 | 0 | 0 | 0 | 0 | 0 |
| ASV1635 | cbb82d9f44c | 3 | 0 | 0 | 0 | 0 | 0 | 0 | 0 | 0 | 0 | 0 | 0 | 0 | 0 | 0 | 3 | 0 | 0 | 0 | 0 | 0 | 0 | 0 | 0 | 0 | 0 | 0 | 0 | 0 | 0 | 0 | 0 | 0 | 0 | 0 | 0 |
| ASV1636 | 194c9098513 | 3 | 0 | 0 | 0 | 0 | 0 | 0 | 0 | 0 | 0 | 0 | 0 | 0 | 0 | 0 | 0 | 0 | 0 | 0 | 0 | 0 | 0 | 0 | 0 | 0 | 0 | 0 | 0 | 0 | 0 | 0 | 3 | 0 | 0 | 0 | 0 |
| ASV1637 | 6f1e4320d15 | 3 | 0 | 0 | 0 | 0 | 0 | 0 | 0 | 0 | 0 | 3 | 0 | 0 | 0 | 0 | 0 | 0 | 0 | 0 | 0 | 0 | 0 | 0 | 0 | 0 | 0 | 0 | 0 | 0 | 0 | 0 | 0 | 0 | 0 | 0 | 0 |
| ASV1638 | 700766015f6 | 3 | 0 | 0 | 0 | 0 | 0 | 0 | 0 | 0 | 0 | 0 | 0 | 0 | 0 | 0 | 0 | 0 | 0 | 0 | 0 | 0 | 0 | 0 | 0 | 0 | 0 | 3 | 0 | 0 | 0 | 0 | 0 | 0 | 0 | 0 | 0 |
| ASV1639 | eaba607e554 | 3 | 0 | 0 | 0 | 0 | 0 | 0 | 0 | 0 | 0 | 0 | 0 | 0 | 0 | 0 | 0 | 0 | 0 | 0 | 0 | 0 | 0 | 0 | 0 | 0 | 0 | 0 | 0 | 0 | 0 | 0 | 3 | 0 | 0 | 0 | 0 |
| ASV1640 | 55ddb1d5456 | 3 | 0 | 0 | 0 | 0 | 0 | 0 | 0 | 0 | 0 | 0 | 0 | 0 | 0 | 0 | 0 | 0 | 0 | 0 | 0 | 0 | 0 | 0 | 0 | 0 | 0 | 0 | 0 | 0 | 0 | 3 | 0 | 0 | 0 | 0 | 0 |
| ASV1641 | de4b641533d | 3 | 0 | 0 | 0 | 0 | 0 | 0 | 0 | 0 | 0 | 0 | 0 | 0 | 0 | 0 | 0 | 0 | 0 | 0 | 0 | 0 | 0 | 0 | 0 | 3 | 0 | 0 | 0 | 0 | 0 | 0 | 0 | 0 | 0 | 0 | 0 |
| ASV1642 | 57fe7211471 | 3 | 0 | 0 | 0 | 0 | 0 | 3 | 0 | 0 | 0 | 0 | 0 | 0 | 0 | 0 | 0 | 0 | 0 | 0 | 0 | 0 | 0 | 0 | 0 | 0 | 0 | 0 | 0 | 0 | 0 | 0 | 0 | 0 | 0 | 0 | 0 |
| ASV1643 | 22d1c875fe6 | 3 | 0 | 0 | 0 | 0 | 0 | 0 | 0 | 0 | 0 | 0 | 0 | 0 | 0 | 0 | 0 | 0 | 0 | 0 | 0 | 0 | 0 | 0 | 0 | 0 | 3 | 0 | 0 | 0 | 0 | 0 | 0 | 0 | 0 | 0 | 0 |
| ASV1644 | 0d1771f0238 | 3 | 0 | 0 | 0 | 0 | 0 | 0 | 0 | 3 | 0 | 0 | 0 | 0 | 0 | 0 | 0 | 0 | 0 | 0 | 0 | 0 | 0 | 0 | 0 | 0 | 0 | 0 | 0 | 0 | 0 | 0 | 0 | 0 | 0 | 0 | 0 |
| ASV1645 | 4fb3beb75df | 3 | 0 | 0 | 3 | 0 | 0 | 0 | 0 | 0 | 0 | 0 | 0 | 0 | 0 | 0 | 0 | 0 | 0 | 0 | 0 | 0 | 0 | 0 | 0 | 0 | 0 | 0 | 0 | 0 | 0 | 0 | 0 | 0 | 0 | 0 | 0 |
| ASV1646 | eb1f76bfcd72 | 3 | 0 | 0 | 0 | 0 | 0 | 0 | 0 | 0 | 3 | 0 | 0 | 0 | 0 | 0 | 0 | 0 | 0 | 0 | 0 | 0 | 0 | 0 | 0 | 0 | 0 | 0 | 0 | 0 | 0 | 0 | 0 | 0 | 0 | 0 | 0 |
| ASV1647 | b994cbc19e9 | 3 | 0 | 0 | 0 | 0 | 0 | 0 | 0 | 0 | 0 | 0 | 0 | 0 | 0 | 0 | 0 | 0 | 0 | 3 | 0 | 0 | 0 | 0 | 0 | 0 | 0 | 0 | 0 | 0 | 0 | 0 | 0 | 0 | 0 | 0 | 0 |
| ASV1648 | 52cf327326d | 3 | 0 | 0 | 0 | 0 | 0 | 0 | 0 | 0 | 0 | 0 | 0 | 0 | 0 | 0 | 0 | 0 | 0 | 0 | 0 | 0 | 0 | 0 | 0 | 0 | 0 | 0 | 0 | 0 | 0 | 0 | 3 | 0 | 0 | 0 | 0 |
| ASV1649 | 2d7f0eab395 | 3 | 0 | 0 | 0 | 0 | 0 | 0 | 0 | 0 | 0 | 0 | 0 | 0 | 0 | 0 | 0 | 0 | 0 | 0 | 0 | 0 | 0 | 0 | 0 | 0 | 3 | 0 | 0 | 0 | 0 | 0 | 0 | 0 | 0 | 0 | 0 |
| ASV1650 | ee64c77c947 | 3 | 0 | 0 | 0 | 0 | 0 | 0 | 0 | 0 | 0 | 3 | 0 | 0 | 0 | 0 | 0 | 0 | 0 | 0 | 0 | 0 | 0 | 0 | 0 | 0 | 0 | 0 | 0 | 0 | 0 | 0 | 0 | 0 | 0 | 0 | 0 |
| ASV1651 | ac214674600 | 3 | 0 | 0 | 3 | 0 | 0 | 0 | 0 | 0 | 0 | 0 | 0 | 0 | 0 | 0 | 0 | 0 | 0 | 0 | 0 | 0 | 0 | 0 | 0 | 0 | 0 | 0 | 0 | 0 | 0 | 0 | 0 | 0 | 0 | 0 | 0 |
| ASV1652 | 44b08a0a937 | 3 | 0 | 0 | 0 | 0 | 0 | 0 | 0 | 0 | 0 | 0 | 0 | 0 | 0 | 0 | 0 | 0 | 0 | 0 | 0 | 0 | 0 | 0 | 0 | 0 | 0 | 0 | 0 | 0 | 0 | 3 | 0 | 0 | 0 | 0 | 0 |
| ASV1653 | 71d2cac4c42 | 3 | 0 | 0 | 0 | 0 | 0 | 0 | 0 | 0 | 0 | 0 | 0 | 0 | 0 | 0 | 0 | 0 | 0 | 0 | 3 | 0 | 0 | 0 | 0 | 0 | 0 | 0 | 0 | 0 | 0 | 0 | 0 | 0 | 0 | 0 | 0 |
| ASV1654 | ba7bb52bc70 | 3 | 0 | 0 | 0 | 0 | 0 | 0 | 0 | 0 | 0 | 0 | 0 | 0 | 0 | 0 | 0 | 0 | 0 | 0 | 0 | 0 | 0 | 0 | 0 | 0 | 0 | 0 | 3 | 0 | 0 | 0 | 0 | 0 | 0 | 0 | 0 |
| ASV1655 | 4d456631155 | 3 | 0 | 0 | 0 | 0 | 0 | 0 | 0 | 0 | 0 | 0 | 0 | 0 | 0 | 0 | 0 | 0 | 0 | 0 | 0 | 0 | 0 | 0 | 0 | 0 | 0 | 0 | 0 | 0 | 3 | 0 | 0 | 0 | 0 | 0 | 0 |
| ASV1656 | a328ebe3a24 | 3 | 0 | 0 | 0 | 0 | 0 | 0 | 0 | 0 | 0 | 0 | 0 | 0 | 0 | 0 | 0 | 3 | 0 | 0 | 0 | 0 | 0 | 0 | 0 | 0 | 0 | 0 | 0 | 0 | 0 | 0 | 0 | 0 | 0 | 0 | 0 |
| ASV1657 | ea2ffaf32d4e | 2 | 0 | 0 | 0 | 0 | 2 | 0 | 0 | 0 | 0 | 0 | 0 | 0 | 0 | 0 | 0 | 0 | 0 | 0 | 0 | 0 | 0 | 0 | 0 | 0 | 0 | 0 | 0 | 0 | 0 | 0 | 0 | 0 | 0 | 0 | 0 |
| ASV1658 | 500e1a71fab | 2 | 0 | 0 | 0 | 0 | 0 | 2 | 0 | 0 | 0 | 0 | 0 | 0 | 0 | 0 | 0 | 0 | 0 | 0 | 0 | 0 | 0 | 0 | 0 | 0 | 0 | 0 | 0 | 0 | 0 | 0 | 0 | 0 | 0 | 0 | 0 |
| ASV1659 | 4d07490bc41 | 2 | 0 | 0 | 0 | 0 | 0 | 0 | 0 | 0 | 0 | 0 | 0 | 0 | 0 | 0 | 0 | 0 | 0 | 0 | 0 | 0 | 0 | 0 | 0 | 0 | 0 | 0 | 0 | 0 | 0 | 2 | 0 | 0 | 0 | 0 | 0 |
| ASV1660 | c7752b2f352 | 2 | 0 | 0 | 0 | 0 | 2 | 0 | 0 | 0 | 0 | 0 | 0 | 0 | 0 | 0 | 0 | 0 | 0 | 0 | 0 | 0 | 0 | 0 | 0 | 0 | 0 | 0 | 0 | 0 | 0 | 0 | 0 | 0 | 0 | 0 | 0 |
| ASV1661 | eab6d4a131c | 2 | 0 | 0 | 0 | 0 | 0 | 0 | 0 | 0 | 0 | 2 | 0 | 0 | 0 | 0 | 0 | 0 | 0 | 0 | 0 | 0 | 0 | 0 | 0 | 0 | 0 | 0 | 0 | 0 | 0 | 0 | 0 | 0 | 0 | 0 | 0 |
| ASV1662 | 656ea0e46f3 | 2 | 0 | 0 | 0 | 0 | 0 | 0 | 0 | 0 | 0 | 0 | 0 | 0 | 2 | 0 | 0 | 0 | 0 | 0 | 0 | 0 | 0 | 0 | 0 | 0 | 0 | 0 | 0 | 0 | 0 | 0 | 0 | 0 | 0 | 0 | 0 |
| ASV1663 | b2b8a8c9475 | 2 | 0 | 0 | 0 | 0 | 0 | 0 | 0 | 0 | 0 | 0 | 0 | 0 | 0 | 0 | 0 | 0 | 0 | 0 | 0 | 0 | 0 | 2 | 0 | 0 | 0 | 0 | 0 | 0 | 0 | 0 | 0 | 0 | 0 | 0 | 0 |
| ASV1664 | 0293224acd7 | 2 | 0 | 0 | 0 | 0 | 0 | 0 | 0 | 0 | 0 | 2 | 0 | 0 | 0 | 0 | 0 | 0 | 0 | 0 | 0 | 0 | 0 | 0 | 0 | 0 | 0 | 0 | 0 | 0 | 0 | 0 | 0 | 0 | 0 | 0 | 0 |
| ASV1665 | 06e730d3fa6 | 2 | 0 | 0 | 0 | 0 | 0 | 0 | 0 | 0 | 0 | 0 | 0 | 0 | 0 | 0 | 0 | 0 | 0 | 0 | 0 | 0 | 0 | 0 | 0 | 0 | 0 | 0 | 0 | 0 | 2 | 0 | 0 | 0 | 0 | 0 | 0 |
| ASV1666 | a71f6d2e333 | 2 | 0 | 0 | 0 | 0 | 0 | 0 | 0 | 0 | 0 | 0 | 0 | 0 | 0 | 0 | 0 | 0 | 0 | 0 | 0 | 0 | 0 | 0 | 0 | 0 | 0 | 0 | 0 | 0 | 0 | 0 | 0 | 0 | 2 | 0 | 0 |
| ASV1667 | 319e508ce31 | 2 | 0 | 0 | 0 | 0 | 0 | 0 | 0 | 0 | 0 | 0 | 0 | 0 | 0 | 0 | 0 | 0 | 0 | 0 | 0 | 0 | 0 | 0 | 0 | 0 | 0 | 0 | 0 | 0 | 0 | 0 | 2 | 0 | 0 | 0 | 0 |
| ASV1668 | b4d2191952e | 2 | 0 | 0 | 0 | 0 | 2 | 0 | 0 | 0 | 0 | 0 | 0 | 0 | 0 | 0 | 0 | 0 | 0 | 0 | 0 | 0 | 0 | 0 | 0 | 0 | 0 | 0 | 0 | 0 | 0 | 0 | 0 | 0 | 0 | 0 | 0 |
| ASV1669 | 63ec8b0c3dd | 2 | 0 | 0 | 0 | 0 | 0 | 0 | 0 | 0 | 0 | 0 | 0 | 0 | 0 | 0 | 0 | 0 | 0 | 0 | 0 | 0 | 0 | 0 | 0 | 0 | 0 | 0 | 0 | 0 | 0 | 2 | 0 | 0 | 0 | 0 | 0 |
| ASV1670 | 376f506f232 | 2 | 2 | 0 | 0 | 0 | 0 | 0 | 0 | 0 | 0 | 0 | 0 | 0 | 0 | 0 | 0 | 0 | 0 | 0 | 0 | 0 | 0 | 0 | 0 | 0 | 0 | 0 | 0 | 0 | 0 | 0 | 0 | 0 | 0 | 0 | 0 |
| ASV1671 | ea1b1a98376 | 2 | 0 | 0 | 0 | 0 | 0 | 0 | 0 | 0 | 0 | 0 | 0 | 0 | 0 | 0 | 0 | 0 | 0 | 0 | 0 | 0 | 0 | 0 | 0 | 0 | 0 | 2 | 0 | 0 | 0 | 0 | 0 | 0 | 0 | 0 | 0 |
| ASV1672 | 3827820c615 | 2 | 0 | 0 | 0 | 0 | 0 | 0 | 0 | 0 | 0 | 0 | 0 | 0 | 0 | 0 | 0 | 0 | 0 | 0 | 0 | 0 | 0 | 0 | 0 | 0 | 0 | 0 | 0 | 0 | 0 | 0 | 0 | 0 | 0 | 2 | 0 |
| ASV1673 | fa787dac5fb | 2 | 0 | 0 | 0 | 0 | 0 | 0 | 0 | 0 | 0 | 0 | 0 | 0 | 0 | 0 | 0 | 0 | 0 | 0 | 0 | 0 | 0 | 0 | 0 | 0 | 0 | 0 | 0 | 0 | 0 | 0 | 0 | 2 | 0 | 0 | 0 |
| ASV1674 | 85c7bc111a3 | 2 | 0 | 0 | 0 | 0 | 2 | 0 | 0 | 0 | 0 | 0 | 0 | 0 | 0 | 0 | 0 | 0 | 0 | 0 | 0 | 0 | 0 | 0 | 0 | 0 | 0 | 0 | 0 | 0 | 0 | 0 | 0 | 0 | 0 | 0 | 0 |
| ASV1675 | cea09f9f364c | 2 | 0 | 0 | 0 | 0 | 0 | 0 | 0 | 0 | 0 | 0 | 0 | 0 | 0 | 0 | 0 | 0 | 0 | 0 | 0 | 0 | 0 | 0 | 0 | 0 | 0 | 0 | 0 | 0 | 0 | 0 | 0 | 0 | 0 | 0 | 2 |
| ASV1676 | 757ff749295 | 2 | 0 | 0 | 0 | 0 | 0 | 0 | 0 | 0 | 0 | 0 | 0 | 0 | 0 | 0 | 0 | 0 | 0 | 0 | 0 | 0 | 0 | 0 | 0 | 0 | 0 | 0 | 0 | 0 | 0 | 0 | 2 | 0 | 0 | 0 | 0 |
| ASV1677 | daf3e8d6a1d | 2 | 0 | 0 | 0 | 0 | 0 | 0 | 0 | 0 | 0 | 0 | 0 | 0 | 0 | 0 | 0 | 0 | 0 | 0 | 0 | 0 | 0 | 0 | 0 | 0 | 0 | 0 | 0 | 0 | 0 | 2 | 0 | 0 | 0 | 0 | 0 |
| ASV1678 | de721b10771 | 2 | 0 | 0 | 0 | 0 | 0 | 0 | 0 | 0 | 2 | 0 | 0 | 0 | 0 | 0 | 0 | 0 | 0 | 0 | 0 | 0 | 0 | 0 | 0 | 0 | 0 | 0 | 0 | 0 | 0 | 0 | 0 | 0 | 0 | 0 | 0 |
| ASV1679 | e5dc7b37b16 | 2 | 0 | 0 | 0 | 0 | 0 | 0 | 0 | 0 | 0 | 0 | 0 | 0 | 0 | 0 | 0 | 0 | 0 | 0 | 0 | 0 | 0 | 0 | 0 | 0 | 0 | 0 | 0 | 0 | 0 | 0 | 0 | 0 | 2 | 0 | 0 |
| ASV1680 | 2037353a449 | 2 | 0 | 0 | 0 | 0 | 0 | 0 | 0 | 0 | 0 | 0 | 0 | 0 | 0 | 0 | 0 | 0 | 0 | 0 | 0 | 0 | 0 | 0 | 0 | 0 | 0 | 0 | 0 | 0 | 0 | 0 | 0 | 0 | 0 | 2 | 0 |
| ASV1681 | 88a322662a8 | 2 | 0 | 0 | 0 | 0 | 0 | 0 | 0 | 0 | 0 | 0 | 0 | 0 | 0 | 0 | 0 | 0 | 0 | 0 | 0 | 0 | 0 | 0 | 0 | 0 | 0 | 0 | 0 | 0 | 0 | 0 | 2 | 0 | 0 | 0 | 0 |
| ASV1682 | be557549dcc | 2 | 0 | 0 | 0 | 0 | 0 | 0 | 0 | 0 | 0 | 0 | 0 | 0 | 0 | 0 | 0 | 0 | 2 | 0 | 0 | 0 | 0 | 0 | 0 | 0 | 0 | 0 | 0 | 0 | 0 | 0 | 0 | 0 | 0 | 0 | 0 |
| ASV1683 | 2075c3dcd08 | 2 | 0 | 0 | 0 | 0 | 0 | 0 | 0 | 0 | 0 | 0 | 0 | 0 | 0 | 0 | 0 | 0 | 0 | 0 | 0 | 2 | 0 | 0 | 0 | 0 | 0 | 0 | 0 | 0 | 0 | 0 | 0 | 0 | 0 | 0 | 0 |
| ASV1684 | 1caab32ff200 | 2 | 0 | 0 | 0 | 0 | 2 | 0 | 0 | 0 | 0 | 0 | 0 | 0 | 0 | 0 | 0 | 0 | 0 | 0 | 0 | 0 | 0 | 0 | 0 | 0 | 0 | 0 | 0 | 0 | 0 | 0 | 0 | 0 | 0 | 0 | 0 |
| ASV1685 | 89c861363bf | 2 | 0 | 2 | 0 | 0 | 0 | 0 | 0 | 0 | 0 | 0 | 0 | 0 | 0 | 0 | 0 | 0 | 0 | 0 | 0 | 0 | 0 | 0 | 0 | 0 | 0 | 0 | 0 | 0 | 0 | 0 | 0 | 0 | 0 | 0 | 0 |
| ASV1686 | ce294127f9a | 2 | 0 | 0 | 0 | 0 | 0 | 0 | 0 | 0 | 0 | 0 | 0 | 0 | 0 | 0 | 0 | 0 | 0 | 0 | 0 | 0 | 2 | 0 | 0 | 0 | 0 | 0 | 0 | 0 | 0 | 0 | 0 | 0 | 0 | 0 | 0 |
| ASV1687 | a62a9e8d592 | 2 | 2 | 0 | 0 | 0 | 0 | 0 | 0 | 0 | 0 | 0 | 0 | 0 | 0 | 0 | 0 | 0 | 0 | 0 | 0 | 0 | 0 | 0 | 0 | 0 | 0 | 0 | 0 | 0 | 0 | 0 | 0 | 0 | 0 | 0 | 0 |
| ASV1688 | d9caf276795 | 2 | 0 | 0 | 0 | 0 | 0 | 0 | 0 | 0 | 0 | 0 | 0 | 0 | 0 | 0 | 0 | 0 | 0 | 0 | 0 | 0 | 0 | 0 | 0 | 0 | 0 | 0 | 0 | 0 | 0 | 0 | 2 | 0 | 0 | 0 | 0 |
| ASV1689 | e4dc9940e2d | 2 | 0 | 0 | 0 | 0 | 0 | 0 | 0 | 0 | 2 | 0 | 0 | 0 | 0 | 0 | 0 | 0 | 0 | 0 | 0 | 0 | 0 | 0 | 0 | 0 | 0 | 0 | 0 | 0 | 0 | 0 | 0 | 0 | 0 | 0 | 0 |
| ASV1690 | e01085ec4f0 | 2 | 0 | 0 | 0 | 0 | 0 | 0 | 0 | 0 | 0 | 2 | 0 | 0 | 0 | 0 | 0 | 0 | 0 | 0 | 0 | 0 | 0 | 0 | 0 | 0 | 0 | 0 | 0 | 0 | 0 | 0 | 0 | 0 | 0 | 0 | 0 |
| ASV1691 | bc70c55e3fa | 2 | 0 | 2 | 0 | 0 | 0 | 0 | 0 | 0 | 0 | 0 | 0 | 0 | 0 | 0 | 0 | 0 | 0 | 0 | 0 | 0 | 0 | 0 | 0 | 0 | 0 | 0 | 0 | 0 | 0 | 0 | 0 | 0 | 0 | 0 | 0 |
| ASV1692 | 55793d1d78c | 2 | 0 | 0 | 0 | 0 | 0 | 0 | 0 | 0 | 0 | 0 | 0 | 0 | 0 | 0 | 0 | 0 | 0 | 0 | 0 | 0 | 0 | 0 | 0 | 0 | 0 | 0 | 0 | 0 | 0 | 2 | 0 | 0 | 0 | 0 | 0 |
| ASV1693 | 92ad9e74972 | 2 | 0 | 0 | 0 | 0 | 0 | 0 | 0 | 0 | 0 | 0 | 0 | 0 | 0 | 0 | 0 | 0 | 0 | 0 | 0 | 0 | 0 | 0 | 0 | 0 | 0 | 0 | 0 | 0 | 0 | 0 | 0 | 0 | 0 | 2 | 0 |
| ASV1694 | aa5a62173f2 | 2 | 0 | 0 | 0 | 0 | 0 | 0 | 0 | 0 | 0 | 0 | 0 | 0 | 0 | 0 | 0 | 0 | 0 | 0 | 0 | 0 | 0 | 0 | 0 | 0 | 0 | 0 | 0 | 0 | 0 | 0 | 0 | 0 | 0 | 2 | 0 |
| ASV1695 | c0c27820c72 | 2 | 0 | 0 | 0 | 2 | 0 | 0 | 0 | 0 | 0 | 0 | 0 | 0 | 0 | 0 | 0 | 0 | 0 | 0 | 0 | 0 | 0 | 0 | 0 | 0 | 0 | 0 | 0 | 0 | 0 | 0 | 0 | 0 | 0 | 0 | 0 |
| ASV1696 | 461e5f49467 | 2 | 0 | 0 | 0 | 0 | 0 | 0 | 0 | 0 | 0 | 0 | 0 | 0 | 0 | 0 | 0 | 0 | 0 | 0 | 0 | 0 | 0 | 0 | 0 | 0 | 0 | 0 | 0 | 0 | 2 | 0 | 0 | 0 | 0 | 0 | 0 |
| ASV1697 | de9a2905a64 | 2 | 0 | 0 | 0 | 0 | 0 | 0 | 0 | 0 | 0 | 0 | 0 | 0 | 0 | 0 | 0 | 0 | 0 | 0 | 0 | 0 | 0 | 0 | 0 | 0 | 0 | 0 | 0 | 0 | 0 | 0 | 0 | 0 | 0 | 0 | 2 |
| ASV1698 | f97dbc45358 | 2 | 0 | 0 | 0 | 0 | 0 | 0 | 0 | 0 | 2 | 0 | 0 | 0 | 0 | 0 | 0 | 0 | 0 | 0 | 0 | 0 | 0 | 0 | 0 | 0 | 0 | 0 | 0 | 0 | 0 | 0 | 0 | 0 | 0 | 0 | 0 |
| ASV1699 | 8a33676217d | 2 | 0 | 0 | 2 | 0 | 0 | 0 | 0 | 0 | 0 | 0 | 0 | 0 | 0 | 0 | 0 | 0 | 0 | 0 | 0 | 0 | 0 | 0 | 0 | 0 | 0 | 0 | 0 | 0 | 0 | 0 | 0 | 0 | 0 | 0 | 0 |
| ASV1700 | 50a31f6b196 | 2 | 0 | 0 | 0 | 0 | 0 | 0 | 0 | 0 | 0 | 0 | 0 | 0 | 0 | 0 | 0 | 0 | 0 | 0 | 0 | 0 | 0 | 0 | 0 | 0 | 0 | 0 | 2 | 0 | 0 | 0 | 0 | 0 | 0 | 0 | 0 |
| ASV1701 | f5e881da9a0 | 2 | 0 | 0 | 0 | 0 | 0 | 0 | 0 | 0 | 0 | 0 | 0 | 0 | 0 | 0 | 0 | 0 | 2 | 0 | 0 | 0 | 0 | 0 | 0 | 0 | 0 | 0 | 0 | 0 | 0 | 0 | 0 | 0 | 0 | 0 | 0 |
| ASV1702 | d80b58c5235 | 2 | 0 | 0 | 0 | 0 | 0 | 0 | 0 | 0 | 0 | 0 | 0 | 0 | 0 | 0 | 0 | 0 | 0 | 0 | 0 | 0 | 0 | 0 | 0 | 0 | 0 | 0 | 0 | 0 | 0 | 0 | 0 | 2 | 0 | 0 | 0 |
| ASV1703 | 573115e964a | 2 | 0 | 0 | 0 | 0 | 0 | 0 | 0 | 0 | 0 | 0 | 0 | 0 | 0 | 0 | 0 | 0 | 0 | 0 | 0 | 0 | 0 | 2 | 0 | 0 | 0 | 0 | 0 | 0 | 0 | 0 | 0 | 0 | 0 | 0 | 0 |
| ASV1704 | d7d214b1152 | 2 | 0 | 0 | 0 | 0 | 0 | 0 | 0 | 0 | 0 | 0 | 0 | 0 | 0 | 0 | 0 | 0 | 0 | 0 | 0 | 0 | 0 | 0 | 0 | 0 | 2 | 0 | 0 | 0 | 0 | 0 | 0 | 0 | 0 | 0 | 0 |
| ASV1705 | 7d5294ecf2c | 2 | 0 | 0 | 0 | 0 | 0 | 0 | 0 | 0 | 0 | 0 | 0 | 0 | 0 | 2 | 0 | 0 | 0 | 0 | 0 | 0 | 0 | 0 | 0 | 0 | 0 | 0 | 0 | 0 | 0 | 0 | 0 | 0 | 0 | 0 | 0 |
| ASV1706 | 8463537d406 | 2 | 0 | 0 | 0 | 0 | 0 | 0 | 0 | 0 | 0 | 0 | 0 | 0 | 0 | 0 | 2 | 0 | 0 | 0 | 0 | 0 | 0 | 0 | 0 | 0 | 0 | 0 | 0 | 0 | 0 | 0 | 0 | 0 | 0 | 0 | 0 |
| ASV1707 | e67f3df98f4b | 2 | 0 | 0 | 0 | 0 | 0 | 0 | 0 | 0 | 0 | 0 | 0 | 0 | 0 | 0 | 0 | 0 | 0 | 0 | 0 | 0 | 0 | 0 | 0 | 0 | 0 | 0 | 0 | 0 | 0 | 0 | 0 | 0 | 0 | 0 | 2 |
| ASV1708 | b41f4d87061 | 2 | 0 | 0 | 0 | 0 | 0 | 0 | 0 | 0 | 0 | 0 | 0 | 0 | 0 | 0 | 2 | 0 | 0 | 0 | 0 | 0 | 0 | 0 | 0 | 0 | 0 | 0 | 0 | 0 | 0 | 0 | 0 | 0 | 0 | 0 | 0 |
| ASV1709 | dc1e524db5b | 2 | 0 | 0 | 0 | 0 | 0 | 0 | 0 | 0 | 0 | 0 | 0 | 0 | 0 | 0 | 0 | 0 | 0 | 0 | 0 | 0 | 0 | 0 | 0 | 0 | 0 | 0 | 0 | 0 | 0 | 2 | 0 | 0 | 0 | 0 | 0 |
| ASV1710 | b3f1eebb5a6 | 2 | 0 | 0 | 0 | 0 | 0 | 0 | 0 | 0 | 0 | 0 | 0 | 0 | 0 | 0 | 0 | 0 | 0 | 0 | 0 | 0 | 0 | 0 | 0 | 0 | 0 | 0 | 0 | 0 | 0 | 0 | 0 | 0 | 2 | 0 | 0 |
| ASV1711 | c067a201755 | 2 | 0 | 2 | 0 | 0 | 0 | 0 | 0 | 0 | 0 | 0 | 0 | 0 | 0 | 0 | 0 | 0 | 0 | 0 | 0 | 0 | 0 | 0 | 0 | 0 | 0 | 0 | 0 | 0 | 0 | 0 | 0 | 0 | 0 | 0 | 0 |
| ASV1712 | 0a3b2ecf7f7 | 2 | 0 | 0 | 0 | 0 | 0 | 0 | 0 | 0 | 0 | 0 | 0 | 2 | 0 | 0 | 0 | 0 | 0 | 0 | 0 | 0 | 0 | 0 | 0 | 0 | 0 | 0 | 0 | 0 | 0 | 0 | 0 | 0 | 0 | 0 | 0 |
| ASV1713 | af2dc8ee24b | 2 | 0 | 0 | 0 | 0 | 0 | 0 | 0 | 0 | 0 | 0 | 0 | 0 | 0 | 0 | 0 | 0 | 2 | 0 | 0 | 0 | 0 | 0 | 0 | 0 | 0 | 0 | 0 | 0 | 0 | 0 | 0 | 0 | 0 | 0 | 0 |
| ASV1714 | b8348d8601c | 2 | 0 | 0 | 0 | 0 | 0 | 2 | 0 | 0 | 0 | 0 | 0 | 0 | 0 | 0 | 0 | 0 | 0 | 0 | 0 | 0 | 0 | 0 | 0 | 0 | 0 | 0 | 0 | 0 | 0 | 0 | 0 | 0 | 0 | 0 | 0 |
| ASV1715 | 936d27e23f8 | 2 | 0 | 0 | 0 | 0 | 0 | 0 | 0 | 0 | 0 | 0 | 0 | 0 | 0 | 0 | 0 | 0 | 0 | 0 | 0 | 0 | 0 | 0 | 0 | 0 | 2 | 0 | 0 | 0 | 0 | 0 | 0 | 0 | 0 | 0 | 0 |
| ASV1716 | afdf3a5808a | 2 | 0 | 0 | 0 | 0 | 0 | 0 | 0 | 0 | 0 | 0 | 0 | 0 | 0 | 0 | 0 | 0 | 0 | 0 | 0 | 2 | 0 | 0 | 0 | 0 | 0 | 0 | 0 | 0 | 0 | 0 | 0 | 0 | 0 | 0 | 0 |
| ASV1717 | 99178e8d301 | 2 | 0 | 0 | 0 | 0 | 0 | 0 | 0 | 0 | 0 | 0 | 0 | 0 | 0 | 0 | 0 | 0 | 0 | 0 | 0 | 0 | 0 | 0 | 0 | 0 | 0 | 0 | 0 | 0 | 0 | 0 | 0 | 0 | 0 | 2 | 0 |
| ASV1718 | 193e863ef3d | 2 | 0 | 0 | 0 | 0 | 0 | 0 | 0 | 0 | 0 | 0 | 0 | 0 | 0 | 0 | 0 | 0 | 0 | 0 | 0 | 0 | 0 | 0 | 0 | 0 | 0 | 0 | 2 | 0 | 0 | 0 | 0 | 0 | 0 | 0 | 0 |
| ASV1719 | ef6514c3180 | 2 | 2 | 0 | 0 | 0 | 0 | 0 | 0 | 0 | 0 | 0 | 0 | 0 | 0 | 0 | 0 | 0 | 0 | 0 | 0 | 0 | 0 | 0 | 0 | 0 | 0 | 0 | 0 | 0 | 0 | 0 | 0 | 0 | 0 | 0 | 0 |
| ASV1720 | fc026e2c89a | 2 | 0 | 0 | 0 | 0 | 0 | 0 | 0 | 0 | 0 | 0 | 0 | 0 | 0 | 0 | 0 | 0 | 0 | 0 | 0 | 0 | 0 | 0 | 0 | 2 | 0 | 0 | 0 | 0 | 0 | 0 | 0 | 0 | 0 | 0 | 0 |
| ASV1721 | d16a1386b99 | 2 | 0 | 0 | 0 | 0 | 0 | 0 | 0 | 0 | 0 | 0 | 2 | 0 | 0 | 0 | 0 | 0 | 0 | 0 | 0 | 0 | 0 | 0 | 0 | 0 | 0 | 0 | 0 | 0 | 0 | 0 | 0 | 0 | 0 | 0 | 0 |
| ASV1722 | 02d9429323e | 2 | 0 | 0 | 0 | 2 | 0 | 0 | 0 | 0 | 0 | 0 | 0 | 0 | 0 | 0 | 0 | 0 | 0 | 0 | 0 | 0 | 0 | 0 | 0 | 0 | 0 | 0 | 0 | 0 | 0 | 0 | 0 | 0 | 0 | 0 | 0 |
| ASV1723 | d500d56d74f | 2 | 0 | 0 | 0 | 0 | 0 | 0 | 0 | 0 | 0 | 0 | 0 | 0 | 0 | 0 | 0 | 0 | 0 | 0 | 0 | 0 | 0 | 0 | 0 | 0 | 2 | 0 | 0 | 0 | 0 | 0 | 0 | 0 | 0 | 0 | 0 |
| ASV1724 | 155d7c348f3 | 2 | 0 | 0 | 2 | 0 | 0 | 0 | 0 | 0 | 0 | 0 | 0 | 0 | 0 | 0 | 0 | 0 | 0 | 0 | 0 | 0 | 0 | 0 | 0 | 0 | 0 | 0 | 0 | 0 | 0 | 0 | 0 | 0 | 0 | 0 | 0 |
| ASV1725 | cbfd06baa4e | 2 | 0 | 0 | 0 | 0 | 0 | 0 | 0 | 0 | 0 | 0 | 0 | 0 | 0 | 0 | 0 | 0 | 0 | 0 | 0 | 0 | 0 | 0 | 0 | 0 | 0 | 0 | 0 | 0 | 0 | 2 | 0 | 0 | 0 | 0 | 0 |
| ASV1726 | 27a816f6ed9 | 2 | 0 | 0 | 0 | 0 | 0 | 0 | 0 | 0 | 0 | 0 | 0 | 0 | 0 | 0 | 0 | 0 | 0 | 0 | 0 | 0 | 0 | 0 | 0 | 0 | 0 | 0 | 0 | 0 | 0 | 0 | 0 | 0 | 0 | 0 | 2 |
| ASV1727 | 18dfde1fcfd0 | 2 | 0 | 0 | 0 | 0 | 0 | 0 | 0 | 0 | 0 | 0 | 0 | 0 | 0 | 0 | 0 | 0 | 0 | 0 | 0 | 0 | 0 | 0 | 0 | 0 | 0 | 0 | 0 | 0 | 0 | 0 | 0 | 2 | 0 | 0 | 0 |
| ASV1728 | 1ecf1353a5d | 2 | 0 | 0 | 0 | 0 | 0 | 0 | 0 | 0 | 0 | 0 | 0 | 0 | 0 | 0 | 0 | 0 | 0 | 0 | 0 | 0 | 0 | 0 | 0 | 0 | 0 | 0 | 0 | 0 | 0 | 0 | 2 | 0 | 0 | 0 | 0 |
| ASV1729 | 6c59936d3fa | 2 | 0 | 0 | 0 | 0 | 0 | 0 | 0 | 0 | 0 | 0 | 0 | 0 | 0 | 2 | 0 | 0 | 0 | 0 | 0 | 0 | 0 | 0 | 0 | 0 | 0 | 0 | 0 | 0 | 0 | 0 | 0 | 0 | 0 | 0 | 0 |
| ASV1730 | fbacaf25ec02 | 2 | 0 | 0 | 0 | 0 | 0 | 0 | 0 | 0 | 0 | 2 | 0 | 0 | 0 | 0 | 0 | 0 | 0 | 0 | 0 | 0 | 0 | 0 | 0 | 0 | 0 | 0 | 0 | 0 | 0 | 0 | 0 | 0 | 0 | 0 | 0 |
| ASV1731 | f38ddf60368f | 2 | 0 | 0 | 0 | 0 | 0 | 0 | 0 | 0 | 0 | 0 | 0 | 0 | 0 | 0 | 0 | 0 | 0 | 0 | 0 | 0 | 0 | 0 | 0 | 0 | 0 | 0 | 2 | 0 | 0 | 0 | 0 | 0 | 0 | 0 | 0 |
| ASV1732 | 3286a679a2c | 2 | 0 | 0 | 0 | 0 | 0 | 0 | 0 | 0 | 0 | 0 | 0 | 0 | 0 | 0 | 0 | 0 | 2 | 0 | 0 | 0 | 0 | 0 | 0 | 0 | 0 | 0 | 0 | 0 | 0 | 0 | 0 | 0 | 0 | 0 | 0 |
| ASV1733 | fe5dfe6da0d | 2 | 0 | 0 | 0 | 0 | 0 | 0 | 0 | 0 | 0 | 0 | 0 | 0 | 0 | 0 | 0 | 2 | 0 | 0 | 0 | 0 | 0 | 0 | 0 | 0 | 0 | 0 | 0 | 0 | 0 | 0 | 0 | 0 | 0 | 0 | 0 |
| ASV1734 | 351ff60066fc | 2 | 0 | 0 | 0 | 0 | 2 | 0 | 0 | 0 | 0 | 0 | 0 | 0 | 0 | 0 | 0 | 0 | 0 | 0 | 0 | 0 | 0 | 0 | 0 | 0 | 0 | 0 | 0 | 0 | 0 | 0 | 0 | 0 | 0 | 0 | 0 |
| ASV1735 | e1e5809eb35 | 2 | 2 | 0 | 0 | 0 | 0 | 0 | 0 | 0 | 0 | 0 | 0 | 0 | 0 | 0 | 0 | 0 | 0 | 0 | 0 | 0 | 0 | 0 | 0 | 0 | 0 | 0 | 0 | 0 | 0 | 0 | 0 | 0 | 0 | 0 | 0 |
| ASV1736 | 2b2dad8ddae | 2 | 0 | 0 | 0 | 0 | 0 | 0 | 0 | 0 | 0 | 0 | 0 | 0 | 0 | 0 | 0 | 0 | 0 | 0 | 0 | 0 | 0 | 0 | 0 | 0 | 0 | 0 | 0 | 0 | 0 | 2 | 0 | 0 | 0 | 0 | 0 |
| ASV1737 | 47b8a40cbc5 | 2 | 0 | 0 | 0 | 0 | 0 | 0 | 0 | 0 | 0 | 0 | 0 | 0 | 0 | 0 | 0 | 0 | 0 | 0 | 0 | 0 | 0 | 0 | 0 | 0 | 0 | 0 | 0 | 0 | 0 | 0 | 0 | 0 | 2 | 0 | 0 |
| ASV1738 | ca25f1b88c3 | 2 | 0 | 0 | 2 | 0 | 0 | 0 | 0 | 0 | 0 | 0 | 0 | 0 | 0 | 0 | 0 | 0 | 0 | 0 | 0 | 0 | 0 | 0 | 0 | 0 | 0 | 0 | 0 | 0 | 0 | 0 | 0 | 0 | 0 | 0 | 0 |
| ASV1739 | a35eff4dfbc0 | 2 | 0 | 0 | 0 | 0 | 0 | 0 | 0 | 0 | 0 | 0 | 0 | 0 | 0 | 0 | 0 | 0 | 0 | 0 | 0 | 0 | 0 | 0 | 0 | 2 | 0 | 0 | 0 | 0 | 0 | 0 | 0 | 0 | 0 | 0 | 0 |
| ASV1740 | 0b6c7cb6533 | 2 | 0 | 0 | 0 | 0 | 0 | 0 | 0 | 0 | 0 | 0 | 0 | 0 | 0 | 0 | 0 | 0 | 0 | 0 | 0 | 0 | 0 | 0 | 0 | 2 | 0 | 0 | 0 | 0 | 0 | 0 | 0 | 0 | 0 | 0 | 0 |
| ASV1741 | 4e85fdab57a | 2 | 0 | 0 | 0 | 0 | 0 | 0 | 0 | 0 | 0 | 0 | 2 | 0 | 0 | 0 | 0 | 0 | 0 | 0 | 0 | 0 | 0 | 0 | 0 | 0 | 0 | 0 | 0 | 0 | 0 | 0 | 0 | 0 | 0 | 0 | 0 |
| ASV1742 | db122da0233 | 2 | 0 | 0 | 0 | 0 | 0 | 0 | 0 | 0 | 0 | 0 | 0 | 0 | 0 | 0 | 0 | 0 | 0 | 0 | 0 | 0 | 0 | 2 | 0 | 0 | 0 | 0 | 0 | 0 | 0 | 0 | 0 | 0 | 0 | 0 | 0 |
| ASV1743 | a7fa870a72f | 2 | 0 | 0 | 0 | 0 | 0 | 0 | 0 | 0 | 0 | 0 | 0 | 0 | 0 | 0 | 0 | 0 | 0 | 0 | 0 | 0 | 0 | 2 | 0 | 0 | 0 | 0 | 0 | 0 | 0 | 0 | 0 | 0 | 0 | 0 | 0 |
| ASV1744 | 02553af81ed | 2 | 0 | 0 | 0 | 0 | 0 | 0 | 0 | 0 | 0 | 0 | 0 | 0 | 0 | 0 | 0 | 0 | 0 | 0 | 0 | 0 | 0 | 0 | 0 | 0 | 2 | 0 | 0 | 0 | 0 | 0 | 0 | 0 | 0 | 0 | 0 |
| ASV1745 | 7dbe029ba4e | 2 | 0 | 0 | 0 | 0 | 0 | 0 | 0 | 0 | 0 | 0 | 0 | 0 | 0 | 0 | 0 | 0 | 0 | 0 | 0 | 0 | 0 | 2 | 0 | 0 | 0 | 0 | 0 | 0 | 0 | 0 | 0 | 0 | 0 | 0 | 0 |
| ASV1746 | 563fce8b936 | 2 | 0 | 0 | 0 | 2 | 0 | 0 | 0 | 0 | 0 | 0 | 0 | 0 | 0 | 0 | 0 | 0 | 0 | 0 | 0 | 0 | 0 | 0 | 0 | 0 | 0 | 0 | 0 | 0 | 0 | 0 | 0 | 0 | 0 | 0 | 0 |
| ASV1747 | 4a30345adaf | 2 | 0 | 0 | 2 | 0 | 0 | 0 | 0 | 0 | 0 | 0 | 0 | 0 | 0 | 0 | 0 | 0 | 0 | 0 | 0 | 0 | 0 | 0 | 0 | 0 | 0 | 0 | 0 | 0 | 0 | 0 | 0 | 0 | 0 | 0 | 0 |
| ASV1748 | 0890ca596fb | 2 | 0 | 0 | 0 | 0 | 0 | 0 | 0 | 0 | 0 | 0 | 0 | 0 | 0 | 0 | 0 | 0 | 0 | 0 | 0 | 0 | 0 | 2 | 0 | 0 | 0 | 0 | 0 | 0 | 0 | 0 | 0 | 0 | 0 | 0 | 0 |
| ASV1749 | a03f77553da | 2 | 0 | 0 | 0 | 0 | 0 | 0 | 0 | 0 | 0 | 0 | 0 | 0 | 0 | 0 | 0 | 0 | 0 | 0 | 0 | 0 | 0 | 0 | 0 | 0 | 0 | 0 | 0 | 0 | 0 | 0 | 0 | 0 | 0 | 0 | 2 |
| ASV1750 | 2630ab298a7 | 2 | 0 | 0 | 0 | 0 | 0 | 0 | 0 | 0 | 0 | 0 | 0 | 0 | 0 | 0 | 0 | 0 | 0 | 0 | 0 | 0 | 0 | 0 | 0 | 0 | 0 | 0 | 2 | 0 | 0 | 0 | 0 | 0 | 0 | 0 | 0 |
| ASV1751 | 2d8b0db2bd0 | 2 | 0 | 0 | 0 | 0 | 0 | 0 | 0 | 0 | 2 | 0 | 0 | 0 | 0 | 0 | 0 | 0 | 0 | 0 | 0 | 0 | 0 | 0 | 0 | 0 | 0 | 0 | 0 | 0 | 0 | 0 | 0 | 0 | 0 | 0 | 0 |
| ASV1752 | 5af725f5754 | 2 | 0 | 0 | 0 | 0 | 0 | 0 | 0 | 0 | 0 | 0 | 0 | 0 | 0 | 0 | 0 | 0 | 0 | 0 | 0 | 0 | 0 | 0 | 0 | 0 | 0 | 0 | 0 | 0 | 0 | 0 | 0 | 0 | 0 | 2 | 0 |
| ASV1753 | d95bdb90872 | 2 | 0 | 0 | 0 | 0 | 0 | 0 | 0 | 0 | 0 | 0 | 0 | 0 | 0 | 0 | 0 | 0 | 2 | 0 | 0 | 0 | 0 | 0 | 0 | 0 | 0 | 0 | 0 | 0 | 0 | 0 | 0 | 0 | 0 | 0 | 0 |
| ASV1754 | 316cc31f486 | 2 | 0 | 0 | 0 | 0 | 0 | 0 | 0 | 0 | 0 | 2 | 0 | 0 | 0 | 0 | 0 | 0 | 0 | 0 | 0 | 0 | 0 | 0 | 0 | 0 | 0 | 0 | 0 | 0 | 0 | 0 | 0 | 0 | 0 | 0 | 0 |
| ASV1755 | 06f31d9b89d | 2 | 0 | 0 | 0 | 0 | 0 | 0 | 0 | 0 | 0 | 0 | 0 | 0 | 0 | 0 | 0 | 0 | 0 | 0 | 0 | 0 | 0 | 0 | 0 | 0 | 0 | 0 | 0 | 0 | 0 | 0 | 0 | 0 | 0 | 0 | 2 |
| ASV1756 | 28d11587b73 | 2 | 0 | 0 | 0 | 0 | 0 | 0 | 0 | 0 | 2 | 0 | 0 | 0 | 0 | 0 | 0 | 0 | 0 | 0 | 0 | 0 | 0 | 0 | 0 | 0 | 0 | 0 | 0 | 0 | 0 | 0 | 0 | 0 | 0 | 0 | 0 |
| ASV1757 | 41ca5ce8d28 | 2 | 0 | 0 | 0 | 0 | 0 | 0 | 0 | 0 | 0 | 0 | 0 | 0 | 0 | 0 | 2 | 0 | 0 | 0 | 0 | 0 | 0 | 0 | 0 | 0 | 0 | 0 | 0 | 0 | 0 | 0 | 0 | 0 | 0 | 0 | 0 |
| ASV1758 | 1a555dba755 | 2 | 0 | 0 | 0 | 0 | 0 | 2 | 0 | 0 | 0 | 0 | 0 | 0 | 0 | 0 | 0 | 0 | 0 | 0 | 0 | 0 | 0 | 0 | 0 | 0 | 0 | 0 | 0 | 0 | 0 | 0 | 0 | 0 | 0 | 0 | 0 |
| ASV1759 | be55a4adbb1 | 2 | 0 | 0 | 0 | 0 | 0 | 0 | 0 | 0 | 0 | 0 | 0 | 0 | 0 | 0 | 0 | 0 | 0 | 0 | 0 | 0 | 0 | 0 | 0 | 0 | 0 | 0 | 0 | 0 | 0 | 0 | 0 | 0 | 2 | 0 | 0 |
| ASV1760 | 5c6c50dd5bb | 2 | 0 | 0 | 0 | 0 | 0 | 2 | 0 | 0 | 0 | 0 | 0 | 0 | 0 | 0 | 0 | 0 | 0 | 0 | 0 | 0 | 0 | 0 | 0 | 0 | 0 | 0 | 0 | 0 | 0 | 0 | 0 | 0 | 0 | 0 | 0 |
| ASV1761 | 9e47775fe27 | 2 | 0 | 0 | 0 | 0 | 0 | 0 | 0 | 0 | 0 | 0 | 0 | 0 | 0 | 0 | 2 | 0 | 0 | 0 | 0 | 0 | 0 | 0 | 0 | 0 | 0 | 0 | 0 | 0 | 0 | 0 | 0 | 0 | 0 | 0 | 0 |
| ASV1762 | 6aa70ce7d90 | 2 | 0 | 2 | 0 | 0 | 0 | 0 | 0 | 0 | 0 | 0 | 0 | 0 | 0 | 0 | 0 | 0 | 0 | 0 | 0 | 0 | 0 | 0 | 0 | 0 | 0 | 0 | 0 | 0 | 0 | 0 | 0 | 0 | 0 | 0 | 0 |
| ASV1763 | ac0250c03da | 2 | 0 | 0 | 0 | 0 | 0 | 0 | 0 | 0 | 0 | 0 | 0 | 0 | 0 | 0 | 0 | 0 | 0 | 0 | 0 | 0 | 0 | 2 | 0 | 0 | 0 | 0 | 0 | 0 | 0 | 0 | 0 | 0 | 0 | 0 | 0 |
| ASV1764 | aad97b97b6d | 2 | 0 | 0 | 0 | 0 | 0 | 0 | 0 | 0 | 0 | 2 | 0 | 0 | 0 | 0 | 0 | 0 | 0 | 0 | 0 | 0 | 0 | 0 | 0 | 0 | 0 | 0 | 0 | 0 | 0 | 0 | 0 | 0 | 0 | 0 | 0 |
| ASV1765 | ed5bf1ff620e | 2 | 0 | 0 | 0 | 0 | 0 | 0 | 0 | 0 | 0 | 0 | 0 | 0 | 0 | 0 | 0 | 0 | 0 | 0 | 0 | 0 | 0 | 0 | 0 | 0 | 2 | 0 | 0 | 0 | 0 | 0 | 0 | 0 | 0 | 0 | 0 |
| ASV1766 | 818f93a7f90 | 2 | 0 | 0 | 0 | 0 | 0 | 2 | 0 | 0 | 0 | 0 | 0 | 0 | 0 | 0 | 0 | 0 | 0 | 0 | 0 | 0 | 0 | 0 | 0 | 0 | 0 | 0 | 0 | 0 | 0 | 0 | 0 | 0 | 0 | 0 | 0 |
| ASV1767 | f95ef8dc3b9 | 2 | 0 | 0 | 0 | 0 | 0 | 2 | 0 | 0 | 0 | 0 | 0 | 0 | 0 | 0 | 0 | 0 | 0 | 0 | 0 | 0 | 0 | 0 | 0 | 0 | 0 | 0 | 0 | 0 | 0 | 0 | 0 | 0 | 0 | 0 | 0 |
| ASV1768 | 00836362a8f | 2 | 0 | 0 | 0 | 0 | 0 | 0 | 0 | 0 | 0 | 0 | 0 | 0 | 0 | 0 | 0 | 2 | 0 | 0 | 0 | 0 | 0 | 0 | 0 | 0 | 0 | 0 | 0 | 0 | 0 | 0 | 0 | 0 | 0 | 0 | 0 |
| ASV1769 | 377ba6e5a43 | 2 | 0 | 0 | 0 | 0 | 0 | 0 | 0 | 0 | 0 | 0 | 0 | 0 | 0 | 0 | 0 | 0 | 0 | 0 | 0 | 2 | 0 | 0 | 0 | 0 | 0 | 0 | 0 | 0 | 0 | 0 | 0 | 0 | 0 | 0 | 0 |
| ASV1770 | 8445de4328e | 2 | 0 | 0 | 0 | 0 | 0 | 0 | 0 | 0 | 0 | 0 | 0 | 0 | 0 | 0 | 0 | 0 | 0 | 0 | 0 | 0 | 0 | 0 | 0 | 0 | 0 | 0 | 0 | 0 | 0 | 0 | 0 | 0 | 2 | 0 | 0 |
| ASV1771 | 69da0c771e9 | 2 | 0 | 0 | 0 | 0 | 0 | 0 | 0 | 0 | 0 | 0 | 0 | 0 | 0 | 0 | 0 | 0 | 0 | 0 | 0 | 0 | 0 | 0 | 0 | 0 | 0 | 0 | 0 | 2 | 0 | 0 | 0 | 0 | 0 | 0 | 0 |

**Supplementary Table S5**. Bacterioplankton 16S rRNA ASV feature table
ASVs: 1779
Seqs: 918792

| #ASV_ID | Feature_ID | total seqs | May12_0m | May12_05m | May12_25m | May12_50m | May12_75m | Jun8_0m | Jun8_05m | Jun8_25m | Jun8_50m | Jun8_75m | Jun22_0m | Jun22_05m | Jun22_25m | Jun22_50m | Jun22_75m |
|---|---|---|---|---|---|---|---|---|---|---|---|---|---|---|---|---|---|
| ASV1772 | 985b3ce3095 | 2 | 2 | 0 | 0 | 0 | 0 | 0 | 0 | 0 | 0 | 0 | 0 | 0 | 0 | 0 | 0 |
| ASV1773 | 2bcdb0f74ff8 | 2 | 0 | 0 | 2 | 0 | 0 | 0 | 0 | 0 | 0 | 0 | 0 | 0 | 0 | 0 | 0 |
| ASV1774 | a094d9f9868 | 2 | 2 | 0 | 0 | 0 | 0 | 0 | 0 | 0 | 0 | 0 | 0 | 0 | 0 | 0 | 0 |
| ASV1775 | 5a576b5f62a | 2 | 0 | 0 | 0 | 0 | 0 | 0 | 0 | 0 | 0 | 0 | 0 | 0 | 0 | 0 | 0 |
| ASV1776 | be50a2ea634 | 1 | 0 | 0 | 0 | 1 | 0 | 0 | 0 | 0 | 0 | 0 | 0 | 0 | 0 | 0 | 0 |
| ASV1777 | 077ec792baa | 1 | 0 | 0 | 0 | 0 | 0 | 0 | 0 | 0 | 0 | 0 | 0 | 0 | 0 | 0 | 0 |
| ASV1778 | 7556c38714d | 1 | 0 | 0 | 0 | 0 | 0 | 0 | 0 | 0 | 1 | 0 | 0 | 0 | 0 | 0 | 0 |
| ASV1779 | b1c785e8fd5 | 1 | 0 | 0 | 0 | 0 | 0 | 0 | 0 | 0 | 0 | 1 | 0 | 0 | 0 | 0 | 0 |

| #ASV_ID | Jul11_0m | Jul11_05m | Jul11_25m | Jul11_50m | Jul11_75m | Jul27_0m | Jul27_05m | Jul27_25m | Jul27_50m | Jul27_75m | Aug5_0m | Aug5_05m | Aug5_25m | Aug5_50m | Aug5_75m | Sep8_0m | Sep8_05m | Sep8_25m | Sep8_50m | Sep8_75m |
|---|---|---|---|---|---|---|---|---|---|---|---|---|---|---|---|---|---|---|---|---|
| ASV1772 | 0 | 0 | 0 | 0 | 0 | 0 | 0 | 0 | 0 | 0 | 0 | 0 | 0 | 0 | 0 | 0 | 0 | 0 | 0 | 0 |
| ASV1773 | 0 | 0 | 0 | 0 | 0 | 0 | 0 | 0 | 0 | 0 | 0 | 0 | 0 | 0 | 0 | 0 | 0 | 0 | 0 | 0 |
| ASV1774 | 0 | 0 | 0 | 0 | 0 | 0 | 0 | 0 | 0 | 0 | 0 | 0 | 0 | 0 | 0 | 0 | 0 | 0 | 0 | 0 |
| ASV1775 | 0 | 0 | 0 | 0 | 0 | 0 | 0 | 0 | 0 | 2 | 0 | 0 | 0 | 0 | 0 | 0 | 0 | 0 | 0 | 0 |
| ASV1776 | 0 | 0 | 0 | 0 | 0 | 0 | 0 | 0 | 0 | 0 | 0 | 0 | 0 | 0 | 0 | 0 | 0 | 0 | 0 | 0 |
| ASV1777 | 0 | 0 | 0 | 0 | 0 | 0 | 0 | 0 | 0 | 0 | 1 | 0 | 0 | 0 | 0 | 0 | 0 | 0 | 0 | 0 |
| ASV1778 | 0 | 0 | 0 | 0 | 0 | 0 | 0 | 0 | 0 | 0 | 0 | 0 | 0 | 0 | 0 | 0 | 0 | 0 | 0 | 0 |
| ASV1779 | 0 | 0 | 0 | 0 | 0 | 0 | 0 | 0 | 0 | 0 | 0 | 0 | 0 | 0 | 0 | 0 | 0 | 0 | 0 | 0 |

**Supplementary Table S6**. 16S ASVs in 29 samples with RTPR amplicons, with total count >100, and with at least 10 counts in at least 10 samples

Total nuc seqs: 680781

| #OTU ID | May12_05m | May12_25m | May12_50m | May12_75m | Jun8_25m | Jun8_50m | Jun8_75m | Jun22_0m | Jun22_05m | Jun22_50m | Jun22_75m | Jul11_0m | Jul11_05m | Jul11_25m | Jul11_50m | Jul11_75m | Jul27_05m | Jul27_25m | Jul27_50m | Jul27_75m | Aug5_0m | Aug5_05m | Aug5_25m | Aug5_50m | Aug5_75m | Sep8_0m | Sep8_05m | Sep8_25m | Sep8_50m |
|---|---|---|---|---|---|---|---|---|---|---|---|---|---|---|---|---|---|---|---|---|---|---|---|---|---|---|---|---|---|
| e5cad0f83c5da3a8e4f138aac9423460 | 832 | 1593 | 1024 | 671 | 1078 | 209 | 542 | 1126 | 2924 | 363 | 208 | 5720 | 8003 | 3002 | 951 | 74 | 2024 | 1872 | 2203 | 423 | 3306 | 2156 | 6601 | 4155 | 394 | 3140 | 1602 | 1321 | 1227 |
| 36b651dfb1ecf5381fe35ea3c4a1061f | 646 | 2439 | 1325 | 2190 | 1557 | 647 | 448 | 594 | 2486 | 1481 | 1183 | 2081 | 2012 | 1025 | 643 | 294 | 1082 | 769 | 533 | 719 | 927 | 547 | 1325 | 978 | 908 | 375 | 315 | 190 | 526 |
| 1d72f3f992bcd714b2a0c1b37d614dd2 | 1378 | 2375 | 1395 | 1116 | 1559 | 248 | 410 | 417 | 943 | 2791 | 366 | 254 | 404 | 54 | 39 | 19 | 108 | 93 | 176 | 155 | 217 | 102 | 868 | 1151 | 322 | 116 | 0 | 21 | 110 |
| 936be975fb97fd3b6b5829e1c83be6bd | 299 | 768 | 465 | 306 | 263 | 76 | 113 | 315 | 1035 | 127 | 129 | 1679 | 2895 | 474 | 213 | 63 | 440 | 174 | 179 | 117 | 1385 | 624 | 829 | 271 | 123 | 1709 | 759 | 496 | 181 |
| a6a79b8a486f3cdaf899162d34b8035a | 298 | 861 | 481 | 758 | 395 | 311 | 176 | 99 | 492 | 163 | 389 | 1480 | 1603 | 423 | 304 | 213 | 686 | 404 | 181 | 202 | 1357 | 804 | 987 | 328 | 224 | 1030 | 697 | 465 | 385 |
| 1122e6b7ea8ece945016e945302ee1ad | 98 | 549 | 284 | 402 | 343 | 394 | 114 | 179 | 750 | 323 | 532 | 1044 | 1064 | 657 | 410 | 212 | 413 | 345 | 312 | 385 | 562 | 413 | 499 | 331 | 325 | 1680 | 1160 | 883 | 369 |
| a3d4f90910b762e33dd4d344f63357f8 | 336 | 1063 | 609 | 997 | 674 | 699 | 419 | 121 | 511 | 286 | 767 | 618 | 691 | 440 | 511 | 395 | 389 | 356 | 391 | 717 | 647 | 395 | 601 | 419 | 439 | 420 | 271 | 231 | 467 |
| a756a146af3f113055fcd3b5ebebe18f | 387 | 1027 | 602 | 1100 | 494 | 454 | 306 | 115 | 507 | 236 | 515 | 664 | 827 | 413 | 423 | 245 | 402 | 328 | 288 | 363 | 713 | 484 | 661 | 408 | 337 | 491 | 360 | 247 | 364 |
| b77bf87ad1c17314baa9d7c9acc10a3b | 282 | 676 | 399 | 317 | 168 | 40 | 66 | 370 | 1077 | 220 | 98 | 1335 | 1894 | 596 | 190 | 32 | 491 | 373 | 323 | 170 | 778 | 489 | 1263 | 804 | 310 | 147 | 46 | 45 | 155 |
| 2d4bbf1e16d0492f979b8c5510decc4c | 114 | 398 | 191 | 377 | 328 | 244 | 185 | 172 | 604 | 227 | 424 | 759 | 898 | 523 | 292 | 135 | 338 | 251 | 247 | 421 | 458 | 294 | 430 | 304 | 231 | 1832 | 1294 | 884 | 239 |
| 0194fcb71fb1f171d26db7ddaeb20811 | 90 | 394 | 236 | 236 | 271 | 74 | 106 | 134 | 408 | 98 | 148 | 1133 | 1995 | 197 | 117 | 56 | 279 | 116 | 79 | 46 | 740 | 384 | 834 | 192 | 114 | 2099 | 811 | 468 | 249 |
| 2b7c6eeca2efe4e826759346836efc05 | 126 | 382 | 279 | 157 | 200 | 61 | 111 | 250 | 634 | 82 | 0 | 1000 | 1534 | 627 | 172 | 0 | 311 | 416 | 517 | 104 | 526 | 284 | 1368 | 987 | 0 | 630 | 213 | 232 | 198 |
| c1b1851dde5aaca57cc59dc350c37be7 | 356 | 792 | 530 | 610 | 228 | 121 | 147 | 153 | 561 | 61 | 200 | 719 | 1009 | 407 | 298 | 92 | 226 | 265 | 269 | 144 | 648 | 397 | 973 | 465 | 162 | 308 | 147 | 112 | 374 |
| 8b0192de06c3b10f85bc482236c92366 | 148 | 263 | 122 | 79 | 126 | 0 | 66 | 101 | 110 | 123 | 0 | 307 | 421 | 71 | 0 | 0 | 204 | 118 | 107 | 0 | 422 | 237 | 857 | 471 | 88 | 3400 | 1113 | 976 | 695 |
| c7d8e6d1fd5ae641fff2b96fa1112d7b | 325 | 1387 | 837 | 1416 | 601 | 320 | 308 | 34 | 193 | 315 | 751 | 35 | 58 | 133 | 259 | 151 | 84 | 137 | 277 | 401 | 87 | 57 | 531 | 894 | 445 | 13 | 0 | 12 | 174 |
| 81092899f1c0c2db02ec2ab6e65d86f2 | 189 | 651 | 389 | 646 | 428 | 365 | 195 | 78 | 324 | 167 | 436 | 545 | 627 | 296 | 300 | 217 | 236 | 192 | 209 | 316 | 462 | 290 | 414 | 257 | 218 | 350 | 254 | 200 | 260 |
| 1c54ed87c1705610cad2896671526efe | 0 | 52 | 38 | 51 | 620 | 945 | 1950 | 0 | 0 | 53 | 491 | 0 | 0 | 0 | 634 | 416 | 0 | 89 | 1126 | 1011 | 0 | 0 | 232 | 1034 | 479 | 0 | 0 | 0 | 243 |
| a5e6de786770e07687674cc32d097319 | 105 | 245 | 189 | 183 | 201 | 48 | 48 | 190 | 538 | 105 | 86 | 817 | 1273 | 537 | 219 | 64 | 198 | 203 | 220 | 135 | 420 | 318 | 839 | 461 | 99 | 340 | 226 | 135 | 155 |
| b4aaa30f8f64f72ba6cfd139f2023a8d | 182 | 581 | 427 | 630 | 367 | 264 | 224 | 136 | 369 | 345 | 693 | 207 | 381 | 147 | 169 | 129 | 200 | 170 | 245 | 320 | 239 | 139 | 602 | 683 | 384 | 78 | 53 | 54 | 155 |
| e43703ac625ad9fda149e97e827b0553 | 103 | 501 | 249 | 397 | 225 | 205 | 219 | 20 | 141 | 107 | 262 | 166 | 209 | 192 | 277 | 161 | 236 | 175 | 215 | 250 | 326 | 195 | 1120 | 637 | 312 | 373 | 214 | 200 | 281 |
| a35da92fe92a502bc206941a91a633bc | 204 | 605 | 393 | 535 | 440 | 213 | 325 | 71 | 236 | 368 | 348 | 127 | 218 | 81 | 190 | 116 | 117 | 117 | 315 | 320 | 158 | 81 | 490 | 513 | 249 | 318 | 122 | 129 | 151 |
| 91f687fdb62026d040baa83be4ad4556 | 0 | 0 | 0 | 0 | 252 | 445 | 1060 | 0 | 0 | 0 | 293 | 0 | 0 | 51 | 627 | 434 | 0 | 100 | 1175 | 1059 | 0 | 0 | 131 | 889 | 375 | 0 | 0 | 0 | 267 |
| 5234f65d442af2c647f49e161bb35ffa | 244 | 1484 | 637 | 459 | 103 | 15 | 34 | 26 | 51 | 64 | 111 | 59 | 257 | 18 | 15 | 0 | 54 | 13 | 28 | 21 | 205 | 165 | 684 | 395 | 104 | 857 | 366 | 260 | 333 |
| 8941b912fff03d55d958a89a8510b9c2 | 84 | 369 | 213 | 140 | 128 | 20 | 28 | 76 | 257 | 57 | 64 | 384 | 641 | 108 | 85 | 18 | 187 | 95 | 41 | 29 | 573 | 354 | 664 | 140 | 36 | 1028 | 535 | 388 | 231 |
| 3ab38fc647166c931ad18a473daa0560 | 186 | 587 | 321 | 164 | 122 | 0 | 0 | 281 | 907 | 174 | 29 | 331 | 789 | 186 | 20 | 0 | 132 | 61 | 33 | 9 | 330 | 179 | 415 | 118 | 18 | 558 | 188 | 173 | 172 |
| 46eb9145a13d564c00f01e905866be62 | 179 | 441 | 290 | 397 | 201 | 130 | 282 | 40 | 118 | 411 | 339 | 58 | 190 | 62 | 105 | 65 | 111 | 63 | 337 | 357 | 32 | 38 | 324 | 793 | 701 | 59 | 0 | 0 | 250 |
| 1b980624ac540cbc5036300a8919d6da | 102 | 310 | 162 | 159 | 226 | 27 | 67 | 161 | 573 | 378 | 169 | 481 | 857 | 206 | 64 | 29 | 194 | 68 | 120 | 121 | 271 | 173 | 437 | 264 | 180 | 121 | 64 | 27 | 131 |
| 05932a0c91ddcdbca8bcd2c6ad0849b7 | 121 | 377 | 286 | 416 | 191 | 97 | 101 | 108 | 354 | 225 | 379 | 261 | 451 | 143 | 75 | 57 | 185 | 125 | 137 | 176 | 243 | 120 | 555 | 438 | 207 | 64 | 29 | 27 | 111 |
| e057f66ccacf6c694b1b69b11f19533f | 75 | 253 | 166 | 110 | 62 | 53 | 84 | 35 | 231 | 171 | 64 | 329 | 529 | 124 | 70 | 35 | 335 | 184 | 172 | 87 | 213 | 169 | 867 | 424 | 172 | 291 | 132 | 103 | 119 |
| 2c6e619dbaab11dccebfaa0c0bc5d1d3 | 281 | 712 | 434 | 471 | 161 | 11 | 103 | 113 | 242 | 106 | 156 | 166 | 238 | 142 | 164 | 54 | 122 | 107 | 198 | 108 | 172 | 77 | 370 | 311 | 128 | 99 | 42 | 51 | 256 |
| de6ce53e1a0ea2d20425f04a7b4e733f | 158 | 504 | 357 | 659 | 205 | 186 | 246 | 0 | 177 | 393 | 439 | 0 | 0 | 86 | 156 | 92 | 40 | 103 | 261 | 241 | 0 | 0 | 290 | 511 | 319 | 0 | 0 | 0 | 51 |
| 08223140d04d57bdcd77795646bbb117 | 76 | 192 | 108 | 134 | 124 | 0 | 0 | 147 | 409 | 303 | 100 | 408 | 745 | 198 | 67 | 0 | 202 | 74 | 152 | 115 | 330 | 135 | 438 | 235 | 211 | 231 | 69 | 72 | 151 |
| 539970532548c674f88ff4566f27590b | 264 | 765 | 427 | 279 | 118 | 0 | 0 | 183 | 593 | 80 | 51 | 417 | 762 | 211 | 68 | 0 | 119 | 70 | 110 | 47 | 180 | 92 | 249 | 136 | 0 | 0 | 0 | 0 | 48 |
| 7db24e7c57b22d168f4d3b7bba68a3ea | 28 | 182 | 123 | 123 | 302 | 66 | 36 | 16 | 137 | 368 | 91 | 95 | 120 | 89 | 206 | 61 | 169 | 116 | 186 | 202 | 89 | 85 | 313 | 375 | 314 | 288 | 260 | 225 | 479 |
| bb26c60c17351a7aef92a8c89d3b4dd9 | 329 | 735 | 472 | 564 | 194 | 39 | 74 | 111 | 306 | 355 | 167 | 99 | 268 | 86 | 57 | 8 | 70 | 52 | 79 | 91 | 77 | 64 | 262 | 272 | 135 | 26 | 0 | 16 | 68 |
| 9eb3f4b8a7fb033b51fb84fdc95e4775 | 128 | 425 | 225 | 404 | 155 | 71 | 114 | 71 | 295 | 184 | 166 | 350 | 595 | 210 | 336 | 77 | 36 | 74 | 166 | 150 | 85 | 48 | 154 | 172 | 141 | 41 | 0 | 51 | 145 |
| b85ed7a908f42de3eea96eea6301f7f4 | 162 | 340 | 332 | 583 | 271 | 228 | 258 | 7 | 0 | 98 | 568 | 45 | 36 | 69 | 240 | 85 | 47 | 76 | 312 | 328 | 31 | 35 | 281 | 342 | 134 | 39 | 16 | 0 | 72 |
| c95be457c741775ab35521676639b910 | 29 | 189 | 107 | 208 | 204 | 103 | 278 | 8 | 37 | 96 | 245 | 0 | 126 | 42 | 124 | 103 | 92 | 54 | 316 | 313 | 42 | 32 | 352 | 632 | 492 | 166 | 60 | 17 | 300 |
| a6a7d35754f59891dba1d67c6c7144bf | 292 | 1272 | 636 | 356 | 187 | 40 | 44 | 98 | 354 | 280 | 128 | 134 | 209 | 140 | 58 | 0 | 14 | 24 | 65 | 31 | 19 | 16 | 86 | 113 | 24 | 10 | 12 | 0 | 47 |
| 7290b3cdf5690aa2bd532a5dc67d77ad | 224 | 854 | 496 | 810 | 178 | 92 | 151 | 0 | 126 | 446 | 519 | 0 | 0 | 0 | 76 | 31 | 0 | 0 | 143 | 181 | 0 | 0 | 0 | 209 | 118 | 0 | 0 | 0 | 28 |
| 5472553891fee5a529081d99d11dea18 | 59 | 244 | 123 | 173 | 117 | 74 | 167 | 15 | 52 | 52 | 92 | 74 | 309 | 44 | 145 | 57 | 146 | 89 | 177 | 136 | 77 | 75 | 489 | 426 | 290 | 278 | 134 | 92 | 388 |
| 477382dd44cb51c8fe5d476693824054 | 57 | 295 | 133 | 28 | 171 | 0 | 14 | 98 | 325 | 28 | 16 | 430 | 669 | 146 | 61 | 0 | 69 | 43 | 36 | 10 | 330 | 204 | 246 | 88 | 20 | 491 | 205 | 154 | 136 |
| 584ade95e9d1b271b83a5f45e85a4152 | 46 | 248 | 129 | 263 | 200 | 284 | 155 | 0 | 158 | 125 | 377 | 86 | 75 | 135 | 244 | 183 | 71 | 91 | 111 | 253 | 76 | 62 | 88 | 120 | 201 | 252 | 145 | 161 | 162 |
| b1c4d3c70082054ddc4a26eb17563fbc | 132 | 292 | 257 | 451 | 222 | 128 | 203 | 30 | 111 | 285 | 384 | 0 | 0 | 50 | 109 | 81 | 0 | 70 | 238 | 239 | 47 | 6 | 225 | 565 | 274 | 0 | 0 | 0 | 56 |
| 5d51ef8a47229f789328b35ea0b5b39a | 0 | 75 | 58 | 39 | 30 | 0 | 0 | 134 | 422 | 95 | 33 | 605 | 984 | 242 | 39 | 0 | 141 | 74 | 26 | 0 | 271 | 218 | 414 | 105 | 22 | 70 | 0 | 0 | 68 |
| 191d7e7da61a2dc4cbe8c2f898d070c4 | 154 | 588 | 346 | 408 | 120 | 59 | 61 | 67 | 195 | 185 | 136 | 179 | 280 | 41 | 79 | 19 | 72 | 18 | 25 | 28 | 104 | 72 | 224 | 70 | 44 | 288 | 107 | 90 | 43 |
| 0b3451bfacccfe3f45e24b486da99a9c | 153 | 977 | 408 | 636 | 183 | 72 | 95 | 52 | 145 | 334 | 149 | 99 | 228 | 51 | 10 | 16 | 39 | 14 | 9 | 31 | 51 | 20 | 121 | 70 | 74 | 0 | 14 | 0 | 49 |
| dbd6a30965f3370841c1850f121f6c4e | 158 | 546 | 370 | 430 | 67 | 34 | 48 | 69 | 239 | 114 | 133 | 510 | 687 | 91 | 31 | 16 | 33 | 25 | 21 | 49 | 76 | 36 | 114 | 78 | 47 | 0 | 0 | 4 | 18 |
| bbaa4707c6e1d69a212df3828254e70f | 36 | 115 | 48 | 68 | 142 | 101 | 86 | 41 | 113 | 89 | 90 | 179 | 306 | 67 | 115 | 70 | 14 | 110 | 208 | 146 | 132 | 71 | 168 | 205 | 133 | 459 | 260 | 213 | 105 |
| b3ede366cfbb90e778be76dcbd29aa11 | 248 | 746 | 435 | 435 | 59 | 13 | 11 | 99 | 212 | 61 | 27 | 132 | 302 | 85 | 7 | 0 | 33 | 32 | 22 | 0 | 203 | 93 | 221 | 47 | 27 | 41 | 13 | 20 | 181 |
| 35b220edde7a107dfd6e8c88f1e36ec5 | 0 | 0 | 0 | 0 | 218 | 321 | 816 | 0 | 0 | 0 | 311 | 0 | 0 | 0 | 184 | 246 | 0 | 0 | 452 | 415 | 0 | 0 | 119 | 390 | 136 | 0 | 0 | 0 | 122 |
| f67951137456a41df3608df7675ae032 | 0 | 156 | 90 | 147 | 148 | 136 | 52 | 0 | 164 | 66 | 197 | 150 | 206 | 115 | 123 | 85 | 72 | 89 | 60 | 138 | 93 | 70 | 112 | 109 | 79 | 389 | 230 | 226 | 99 |
| 802c4a4234381e27c4ea1730da37a783 | 95 | 337 | 187 | 109 | 37 | 0 | 0 | 153 | 373 | 206 | 33 | 113 | 218 | 116 | 70 | 0 | 130 | 106 | 187 | 42 | 48 | 52 | 410 | 348 | 148 | 0 | 0 | 0 | 43 |
| 9f4e97a31dd378385a259d3f043220af | 80 | 132 | 83 | 125 | 114 | 91 | 69 | 27 | 113 | 71 | 128 | 227 | 286 | 166 | 110 | 73 | 72 | 66 | 110 | 85 | 79 | 85 | 72 | 55 | 45 | 341 | 278 | 246 | 124 |
| 44c88775a796caf1fc5f29bf7bc73033 | 0 | 238 | 120 | 281 | 121 | 77 | 0 | 0 | 151 | 0 | 83 | 296 | 347 | 175 | 115 | 67 | 102 | 142 | 0 | 47 | 160 | 71 | 183 | 75 | 49 | 241 | 143 | 106 | 131 |
| b97409614c1176bec58b078d86169ffb | 47 | 164 | 178 | 515 | 190 | 238 | 212 | 7 | 43 | 210 | 713 | 0 | 0 | 5 | 45 | 84 | 0 | 25 | 107 | 170 | 10 | 11 | 81 | 268 | 173 | 0 | 0 | 0 | 20 |
| 10884bc8cb1cbbbd39749627ef5e323f | 61 | 174 | 124 | 126 | 61 | 39 | 56 | 14 | 71 | 48 | 49 | 182 | 314 | 65 | 43 | 18 | 69 | 33 | 49 | 36 | 123 | 49 | 248 | 108 | 47 | 657 | 225 | 224 | 82 |
| 9e7dfa8ee4c2408adfd9f088ed34ecd5 | 0 | 31 | 15 | 0 | 53 | 0 | 28 | 0 | 33 | 0 | 0 | 171 | 487 | 37 | 90 | 33 | 153 | 51 | 69 | 0 | 155 | 137 | 479 | 172 | 76 | 393 | 136 | 151 | 370 |
| d077a35cd9abcb1074362008c48f0cd3 | 0 | 0 | 0 | 0 | 0 | 0 | 0 | 0 | 166 | 0 | 0 | 343 | 613 | 220 | 0 | 0 | 102 | 129 | 147 | 0 | 177 | 0 | 586 | 387 | 0 | 231 | 95 | 0 | 0 |
| 3582e1beb44986ccf8fed703ecd0a314 | 169 | 344 | 251 | 213 | 68 | 41 | 43 | 86 | 220 | 147 | 80 | 136 | 239 | 74 | 97 | 0 | 83 | 40 | 97 | 69 | 71 | 49 | 206 | 147 | 57 | 46 | 0 | 0 | 101 |
| a8f894d55f7034b905cb45a15df31bc8 | 133 | 227 | 91 | 111 | 93 | 147 | 109 | 49 | 78 | 118 | 162 | 174 | 179 | 136 | 119 | 104 | 47 | 87 | 116 | 79 | 102 | 68 | 74 | 174 | 91 | 81 | 71 | 45 | 108 |
| f4c50f1ab46aaf3860b962135d8ac239 | 16 | 96 | 55 | 91 | 387 | 242 | 279 | 0 | 29 | 138 | 617 | 0 | 0 | 10 | 57 | 85 | 0 | 18 | 99 | 260 | 0 | 0 | 75 | 192 | 291 | 0 | 0 | 0 | 51 |
| daac7046e78b2c355f0518e9d7cc5692 | 11 | 40 | 38 | 184 | 117 | 320 | 334 | 0 | 0 | 24 | 817 | 0 | 0 | 0 | 118 | 184 | 0 | 24 | 178 | 360 | 0 | 0 | 19 | 115 | 141 | 0 | 0 | 0 | 42 |
| 47ff44a42565e4d0db3a806b626dd11f | 39 | 187 | 104 | 164 | 80 | 0 | 0 | 79 | 147 | 41 | 0 | 249 | 366 | 96 | 0 | 0 | 50 | 59 | 0 | 0 | 208 | 147 | 253 | 102 | 0 | 244 | 142 | 117 | 86 |
| c582f5e706adebf9fa8aa9dba825e821 | 0 | 112 | 41 | 79 | 31 | 0 | 0 | 0 | 79 | 30 | 35 | 199 | 284 | 37 | 67 | 45 | 62 | 39 | 0 | 28 | 190 | 110 | 266 | 60 | 23 | 654 | 196 | 223 | 60 |
| fe8f740f4d7f9ae55b4eda61d15dc311 | 89 | 248 | 88 | 75 | 197 | 92 | 0 | 103 | 137 | 74 | 96 | 118 | 243 | 108 | 112 | 0 | 187 | 150 | 222 | 105 | 0 | 0 | 129 | 208 | 50 | 0 | 0 | 0 | 0 |
| 6d99895665fb90d94a10f0204333500d | 103 | 271 | 238 | 235 | 40 | 19 | 34 | 55 | 154 | 67 | 77 | 149 | 230 | 68 | 61 | 13 | 41 | 29 | 68 | 46 | 79 | 80 | 179 | 85 | 44 | 93 | 20 | 42 | 145 |
| a10d9d5d48bef4fab665520e8981884d | 110 | 795 | 282 | 63 | 65 | 9 | 0 | 33 | 178 | 8 | 0 | 246 | 99 | 42 | 32 | 0 | 8 | 18 | 5 | 0 | 160 | 90 | 71 | 9 | 0 | 135 | 152 | 74 | 49 |
| b516660d69b8590d4917b26e8a493dcf | 0 | 123 | 47 | 32 | 31 | 0 | 0 | 34 | 161 | 24 | 13 | 262 | 396 | 108 | 57 | 0 | 64 | 37 | 49 | 15 | 203 | 128 | 191 | 73 | 24 | 277 | 136 | 122 | 115 |
| 3c59436521bef1ebffc31d2c98c50dd8 | 0 | 0 | 0 | 0 | 0 | 5 | 0 | 28 | 73 | 0 | 0 | 461 | 583 | 148 | 74 | 12 | 101 | 111 | 69 | 5 | 95 | 59 | 233 | 94 | 9 | 221 | 116 | 90 | 59 |
| 04a389e30cb0917b65770980426f47e9 | 68 | 371 | 182 | 184 | 132 | 20 | 18 | 14 | 36 | 76 | 41 | 149 | 176 | 58 | 63 | 22 | 34 | 31 | 78 | 56 | 28 | 0 | 102 | 113 | 87 | 78 | 53 | 32 | 251 |
| 992fe20d2957d584c944da77375e4496 | 0 | 133 | 46 | 0 | 53 | 0 | 9 | 205 | 796 | 111 | 0 | 119 | 327 | 92 | 0 | 0 | 0 | 0 | 13 | 0 | 123 | 53 | 138 | 31 | 0 | 126 | 33 | 0 | 116 |
| 26746171a93079b0b9d8fad0b5468038 | 187 | 191 | 142 | 168 | 210 | 61 | 130 | 0 | 76 | 436 | 140 | 0 | 0 | 0 | 30 | 0 | 0 | 0 | 37 | 25 | 0 | 0 | 242 | 314 | 47 | 0 | 0 | 0 | 86 |
| 9ed82d76e0e6c7b93614e35870c76787 | 28 | 91 | 78 | 341 | 80 | 238 | 230 | 0 | 0 | 58 | 330 | 0 | 0 | 0 | 61 | 126 | 0 | 0 | 111 | 188 | 0 | 0 | 64 | 151 | 197 | 0 | 0 | 0 | 61 |
| 6cf2f2a5b633ef318ec491e8379f02fe | 58 | 262 | 130 | 49 | 114 | 16 | 22 | 30 | 144 | 52 | 33 | 157 | 311 | 84 | 139 | 35 | 66 | 47 | 72 | 56 | 82 | 70 | 146 | 83 | 32 | 35 | 9 | 0 | 47 |
| d054169d4bf17f483a2af0da2aa2d686 | 116 | 552 | 222 | 272 | 207 | 43 | 36 | 20 | 74 | 265 | 62 | 0 | 60 | 41 | 52 | 0 | 0 | 25 | 0 | 41 | 0 | 0 | 26 | 99 | 67 | 0 | 0 | 0 | 25 |
| 6b31940a8beebbbf26dc9f013108f1d5 | 46 | 99 | 89 | 180 | 102 | 78 | 147 | 0 | 50 | 75 | 96 | 57 | 103 | 48 | 87 | 33 | 60 | 34 | 45 | 71 | 8 | 37 | 164 | 193 | 101 | 117 | 56 | 36 | 70 |
| 252554d217904f1bd746191daf86ff98 | 45 | 180 | 81 | 57 | 121 | 0 | 0 | 42 | 154 | 156 | 80 | 64 | 152 | 77 | 207 | 28 | 54 | 58 | 123 | 110 | 7 | 0 | 70 | 139 | 95 | 53 | 0 | 0 | 0 |
| 5ca72a029c2aff7c4ee9e6d7b0787e24 | 0 | 0 | 0 | 0 | 37 | 0 | 53 | 0 | 70 | 0 | 87 | 228 | 323 | 65 | 34 | 0 | 105 | 0 | 0 | 0 | 105 | 103 | 107 | 31 | 0 | 424 | 201 | 117 | 41 |
| 77643ca079d7ae633ff198b7b17d75cf | 106 | 158 | 159 | 219 | 60 | 52 | 68 | 20 | 63 | 75 | 209 | 3 | 76 | 59 | 54 | 29 | 22 | 47 | 61 | 69 | 0 | 39 | 154 | 145 | 74 | 40 | 5 | 0 | 59 |
| 4a1f5b9c4d1ae591ffaa0b264dd24d60 | 41 | 120 | 89 | 82 | 99 | 42 | 35 | 53 | 142 | 96 | 44 | 97 | 200 | 56 | 37 | 26 | 65 | 33 | 39 | 69 | 98 | 43 | 164 | 135 | 69 | 54 | 34 | 34 | 28 |
| db13c5b283352a7f288dd617c0adb847 | 9 | 84 | 31 | 73 | 30 | 4 | 39 | 0 | 0 | 0 | 37 | 42 | 91 | 10 | 74 | 29 | 28 | 39 | 197 | 129 | 68 | 58 | 272 | 428 | 214 | 0 | 0 | 0 | 134 |
| b17f25f6a3ce44c2802197794cc61846 | 107 | 332 | 229 | 286 | 125 | 105 | 119 | 13 | 57 | 62 | 222 | 0 | 22 | 24 | 95 | 24 | 0 | 18 | 33 | 44 | 0 | 0 | 47 | 52 | 36 | 14 | 0 | 0 | 37 |
| 88c221ea1bba372ca003d01321272911 | 19 | 73 | 92 | 411 | 72 | 141 | 142 | 0 | 0 | 26 | 360 | 0 | 0 | 0 | 26 | 85 | 0 | 0 | 53 | 141 | 5 | 0 | 33 | 168 | 219 | 0 | 0 | 0 | 31 |
| 781bf210724ebca9606340c2ce9fa190 | 37 | 76 | 45 | 70 | 43 | 13 | 17 | 15 | 51 | 7 | 46 | 93 | 171 | 68 | 73 | 17 | 35 | 52 | 98 | 62 | 188 | 97 | 233 | 138 | 34 | 135 | 66 | 43 | 67 |
| 289dd5fdf6016e756a0ace95d394acd8 | 19 | 116 | 89 | 176 | 53 | 0 | 0 | 17 | 73 | 0 | 85 | 101 | 122 | 0 | 34 | 0 | 53 | 48 | 21 | 0 | 119 | 92 | 186 | 134 | 69 | 230 | 98 | 57 | 98 |
| bde0f7cede6a0f51d7c7e78513f6cca2 | 0 | 105 | 62 | 97 | 75 | 94 | 95 | 0 | 66 | 33 | 132 | 70 | 55 | 62 | 83 | 70 | 0 | 44 | 28 | 85 | 51 | 40 | 35 | 42 | 56 | 206 | 129 | 111 | 99 |
| 3cc0470e5ae9eecee24011e0f9708b45 | 68 | 231 | 116 | 93 | 37 | 26 | 0 | 44 | 151 | 60 | 42 | 152 | 282 | 101 | 28 | 0 | 67 | 37 | 0 | 0 | 89 | 63 | 185 | 46 | 0 | 34 | 0 | 0 | 36 |
| 931b5b8dcc30783ebb945f1bf7008850 | 104 | 182 | 121 | 93 | 76 | 29 | 31 | 41 | 82 | 65 | 63 | 102 | 156 | 80 | 39 | 19 | 53 | 30 | 87 | 46 | 47 | 24 | 129 | 111 | 41 | 24 | 17 | 0 | 57 |
| 95aac9dadcf8629bed8e50619d7bf87c | 14 | 64 | 29 | 19 | 15 | 0 | 0 | 22 | 49 | 6 | 0 | 170 | 244 | 74 | 19 | 0 | 73 | 53 | 18 | 3 | 246 | 149 | 302 | 90 | 24 | 59 | 41 | 21 | 129 |
| d4025c348719cedb2ea9c148e1d8837f | 0 | 0 | 0 | 0 | 0 | 0 | 0 | 0 | 28 | 0 | 0 | 129 | 343 | 51 | 0 | 0 | 97 | 22 | 0 | 7 | 101 | 80 | 408 | 94 | 31 | 255 | 86 | 76 | 45 |
| ed85ffc35c0865b3bc572cdf5fda97b8 | 35 | 252 | 100 | 163 | 163 | 32 | 28 | 24 | 59 | 499 | 91 | 8 | 10 | 10 | 32 | 0 | 8 | 8 | 54 | 58 | 3 | 0 | 16 | 97 | 78 | 0 | 0 | 0 | 20 |
| ce5f5fefa9a0156716cc33690a335dcb | 31 | 134 | 77 | 146 | 56 | 65 | 49 | 19 | 95 | 59 | 151 | 49 | 86 | 37 | 36 | 31 | 26 | 36 | 54 | 62 | 54 | 32 | 144 | 150 | 71 | 20 | 12 | 0 | 23 |
| f8b4478984e7ecd3a28c667cc9d29100 | 34 | 256 | 102 | 112 | 35 | 0 | 15 | 26 | 138 | 56 | 25 | 117 | 213 | 56 | 31 | 0 | 51 | 29 | 0 | 0 | 89 | 36 | 157 | 61 | 0 | 80 | 28 | 0 | 42 |
| 4e1167fbbbc5e772fad1530d2b3d2b3d | 69 | 270 | 90 | 45 | 63 | 0 | 0 | 26 | 109 | 37 | 6 | 172 | 261 | 37 | 28 | 0 | 0 | 16 | 22 | 10 | 107 | 47 | 119 | 19 | 0 | 69 | 27 | 18 | 80 |
| 291deb67818a7f99faadc6ddaacb2842 | 0 | 54 | 54 | 205 | 49 | 146 | 227 | 0 | 0 | 12 | 252 | 0 | 0 | 0 | 110 | 147 | 0 | 0 | 81 | 175 | 0 | 0 | 38 | 83 | 92 | 0 | 0 | 0 | 22 |
| 7dcb42dd3b53586e69d0d52a42ad9519 | 0 | 206 | 111 | 237 | 98 | 95 | 62 | 0 | 86 | 110 | 94 | 64 | 58 | 53 | 71 | 76 | 0 | 0 | 0 | 70 | 73 | 0 | 58 | 57 | 65 | 0 | 0 | 0 | 0 |
| e0a62ac838662cb1b9fa1caba62aad66 | 24 | 261 | 111 | 177 | 158 | 39 | 13 | 11 | 47 | 154 | 103 | 0 | 0 | 0 | 69 | 13 | 0 | 28 | 58 | 58 | 0 | 15 | 0 | 91 | 117 | 0 | 0 | 0 | 174 |
| 05794f49d61313bcf77f6880ae90cfa0 | 117 | 492 | 253 | 230 | 48 | 0 | 9 | 0 | 0 | 21 | 79 | 0 | 0 | 0 | 18 | 0 | 0 | 0 | 54 | 14 | 5 | 0 | 26 | 161 | 75 | 0 | 0 | 4 | 67 |
| f4dfaada866a3f67086adc50567f5065 | 0 | 0 | 0 | 0 | 0 | 0 | 0 | 34 | 127 | 0 | 0 | 435 | 515 | 162 | 0 | 0 | 35 | 0 | 0 | 0 | 0 | 42 | 33 | 38 | 83 | 42 | 0 | 0 | 0 |
| 63b47a5c24b84d4bcce61a8e4d500bd7 | 54 | 461 | 187 | 102 | 113 | 0 | 10 | 4 | 43 | 77 | 17 | 41 | 19 | 33 | 17 | 0 | 11 | 24 | 27 | 18 | 41 | 30 | 66 | 37 | 10 | 0 | 12 | 10 | 70 |
| e46b4d859dc6df8f7c07cd3cf053284f | 32 | 294 | 119 | 83 | 0 | 11 | 15 | 71 | 379 | 130 | 25 | 55 | 124 | 0 | 44 | 0 | 0 | 0 | 0 | 0 | 35 | 0 | 44 | 0 | 0 | 0 | 0 | 0 | 51 |
| c45b037c35d97b0afa741dbad837674f | 45 | 154 | 112 | 133 | 49 | 43 | 89 | 7 | 38 | 13 | 96 | 25 | 39 | 18 | 83 | 41 | 16 | 27 | 65 | 42 | 8 | 10 | 58 | 82 | 54 | 38 | 17 | 21 | 86 |
| 1b7e6b1555ef9b7c5aba76214e21bfff | 0 | 147 | 94 | 160 | 112 | 119 | 79 | 0 | 0 | 161 | 165 | 0 | 0 | 0 | 59 | 56 | 0 | 0 | 127 | 78 | 0 | 0 | 0 | 0 | 82 | 0 | 0 | 0 | 0 |
| 71f6bf6f8f5090b414c3ea8dd9eb713b | 32 | 74 | 85 | 167 | 84 | 124 | 111 | 0 | 5 | 70 | 173 | 0 | 0 | 0 | 46 | 44 | 0 | 16 | 47 | 94 | 7 | 0 | 11 | 109 | 109 | 0 | 0 | 0 | 20 |
| b9ec28fefa62f2e28a7189169b9c5734 | 13 | 28 | 27 | 214 | 34 | 33 | 70 | 0 | 0 | 12 | 230 | 0 | 0 | 0 | 23 | 26 | 0 | 13 | 92 | 77 | 11 | 6 | 51 | 284 | 138 | 0 | 0 | 0 | 23 |
| 5cbccd30f9dd319ee8f0f6e2b8dfec37 | 14 | 32 | 22 | 0 | 0 | 12 | 0 | 0 | 0 | 0 | 0 | 33 | 45 | 13 | 0 | 0 | 49 | 29 | 0 | 0 | 22 | 0 | 128 | 42 | 0 | 429 | 203 | 164 | 114 |
| 9d240954c03c9e53333e3a4d2f10d6e6 | 14 | 88 | 57 | 98 | 51 | 0 | 26 | 0 | 6 | 33 | 93 | 0 | 0 | 0 | 35 | 0 | 0 | 0 | 88 | 70 | 0 | 0 | 33 | 265 | 190 | 0 | 0 | 0 | 166 |
| dfc67b35046703839f39ea644d857bbd | 0 | 0 | 0 | 0 | 0 | 0 | 0 | 0 | 0 | 0 | 0 | 166 | 143 | 89 | 0 | 0 | 49 | 0 | 0 | 0 | 67 | 39 | 50 | 0 | 0 | 332 | 210 | 162 | 0 |
| 85631eb6b9bc924e5ed6053447181c3e | 16 | 159 | 63 | 24 | 18 | 0 | 6 | 10 | 0 | 23 | 0 | 94 | 302 | 0 | 0 | 0 | 106 | 37 | 13 | 0 | 88 | 62 | 146 | 85 | 40 | 0 | 0 | 0 | 12 |
| 15c5581ecf3c9229c2c206379d488ce6 | 0 | 16 | 0 | 0 | 16 | 0 | 0 | 4 | 27 | 0 | 0 | 110 | 94 | 27 | 39 | 0 | 34 | 26 | 0 | 0 | 90 | 65 | 119 | 16 | 6 | 157 | 166 | 134 | 149 |
| db06db4029f8cb3b6f899ffbe76c03aa | 57 | 279 | 196 | 176 | 38 | 24 | 0 | 0 | 31 | 21 | 54 | 0 | 12 | 0 | 13 | 0 | 0 | 34 | 14 | 23 | 0 | 0 | 96 | 114 | 60 | 0 | 0 | 0 | 29 |
| f4a6e50e16e7c6f7a8d4ebb1229e1326 | 0 | 180 | 66 | 96 | 98 | 0 | 0 | 9 | 53 | 162 | 64 | 29 | 0 | 0 | 0 | 0 | 16 | 0 | 75 | 55 | 0 | 0 | 75 | 101 | 116 | 0 | 0 | 0 | 73 |
| f9730793fd8ff31c44cb686796bb6188 | 85 | 235 | 152 | 133 | 31 | 0 | 0 | 21 | 73 | 56 | 50 | 28 | 46 | 28 | 34 | 0 | 19 | 6 | 0 | 47 | 24 | 25 | 58 | 44 | 25 | 16 | 0 | 0 | 20 |
| 4bf887a6d1b93c8b0b13ca9a677800a1 | 0 | 0 | 0 | 0 | 0 | 0 | 0 | 0 | 73 | 0 | 0 | 188 | 346 | 68 | 0 | 0 | 0 | 27 | 0 | 0 | 87 | 107 | 98 | 0 | 0 | 112 | 41 | 0 | 81 |
| 1a6128b13c51f1f35cd10c0456a6b8e2 | 50 | 261 | 100 | 85 | 24 | 0 | 0 | 17 | 65 | 52 | 0 | 0 | 76 | 24 | 6 | 0 | 20 | 13 | 0 | 0 | 41 | 35 | 96 | 9 | 0 | 65 | 31 | 19 | 119 |
| 6efc1f46a7713ccbf6592587f88c5d9e | 55 | 84 | 24 | 35 | 67 | 64 | 65 | 0 | 0 | 297 | 150 | 25 | 39 | 0 | 0 | 0 | 0 | 0 | 0 | 40 | 25 | 0 | 30 | 87 | 27 | 39 | 0 | 0 | 42 |
| 5afd2e8cb777b78f3999b91d97dd305a | 45 | 92 | 80 | 211 | 73 | 47 | 38 | 0 | 10 | 97 | 160 | 0 | 0 | 7 | 21 | 0 | 0 | 0 | 28 | 58 | 0 | 0 | 19 | 64 | 111 | 0 | 0 | 0 | 18 |
| 843859b79828f1386a1f9ae0f630774b | 0 | 69 | 87 | 89 | 62 | 41 | 25 | 9 | 73 | 48 | 71 | 0 | 27 | 45 | 77 | 37 | 0 | 21 | 58 | 78 | 22 | 0 | 32 | 107 | 66 | 10 | 0 | 0 | 21 |
| 08f5c4a8d1100d6ed3b234abbc6ef0b1 | 0 | 88 | 44 | 126 | 52 | 0 | 42 | 0 | 0 | 72 | 46 | 0 | 38 | 0 | 0 | 0 | 0 | 0 | 72 | 53 | 0 | 0 | 101 | 195 | 167 | 0 | 11 | 11 | 44 |
| efe10567db1cbd5febc8ffb808575e98 | 15 | 54 | 18 | 0 | 40 | 11 | 0 | 8 | 62 | 16 | 0 | 75 | 104 | 38 | 48 | 0 | 73 | 0 | 0 | 0 | 48 | 30 | 95 | 24 | 0 | 178 | 122 | 75 | 25 |

**Supplementary Table S6**. 16S ASVs in 29 samples with RTPR amplicons, with total count >100, and with at least 10 counts in at least 10 samples

Total nuc seqs: 680781

| #OTU ID | May12_05m | May12_25m | May12_50m | May12_75m | Jun8_25m | Jun8_50m | Jun8_75m | Jun22_0m | Jun22_05m | Jun22_50m | Jun22_75m | Jul11_0m | Jul11_05m | Jul11_25m | Jul11_50m | Jul11_75m | Jul27_05m | Jul27_25m | Jul27_50m | Jul27_75m | Aug5_0m | Aug5_05m | Aug5_25m | Aug5_50m | Aug5_75m | Sep8_0m | Sep8_05m | Sep8_25m | Sep8_50m |
|---|---|---|---|---|---|---|---|---|---|---|---|---|---|---|---|---|---|---|---|---|---|---|---|---|---|---|---|---|---|
| 07f203cb593fa3e98a355dc445235d40 | 29 | 227 | 99 | 170 | 122 | 16 | 0 | 0 | 36 | 163 | 84 | 0 | 0 | 0 | 0 | 0 | 0 | 0 | 22 | 27 | 0 | 0 | 29 | 38 | 41 | 0 | 0 | 0 | 44 |
| 0d04c71dcf004a6321b5a41cb75f12fa | 0 | 32 | 0 | 0 | 78 | 0 | 21 | 57 | 152 | 66 | 0 | 94 | 148 | 43 | 32 | 0 | 48 | 0 | 0 | 28 | 0 | 0 | 101 | 31 | 0 | 107 | 57 | 0 | 0 |
| e71e2cdbc8f65de93e08721c92b89c90 | 28 | 142 | 89 | 204 | 22 | 24 | 46 | 0 | 4 | 9 | 119 | 0 | 0 | 0 | 7 | 9 | 0 | 2 | 29 | 32 | 0 | 0 | 10 | 80 | 95 | 4 | 0 | 3 | 121 |
| 664af6a633547b3dd5b07372f47ee95b | 16 | 51 | 42 | 95 | 28 | 30 | 34 | 3 | 10 | 26 | 110 | 0 | 0 | 0 | 15 | 17 | 0 | 20 | 50 | 48 | 17 | 12 | 84 | 148 | 130 | 0 | 0 | 0 | 73 |
| ef58a82c6abb005e535d37100f1ca82a | 0 | 8 | 0 | 0 | 14 | 32 | 69 | 0 | 22 | 0 | 117 | 6 | 17 | 49 | 95 | 38 | 12 | 29 | 85 | 46 | 10 | 0 | 62 | 91 | 27 | 87 | 44 | 31 | 48 |
| 85809ab6d10133261b503338a1f1cd60 | 32 | 70 | 41 | 64 | 24 | 0 | 0 | 31 | 82 | 53 | 106 | 106 | 129 | 43 | 20 | 0 | 37 | 13 | 6 | 11 | 40 | 25 | 64 | 26 | 12 | 0 | 0 | 0 | 0 |
| aa4e24fcdfd89ec545f1d14d1d3e9300 | 6 | 13 | 11 | 113 | 31 | 85 | 54 | 0 | 0 | 0 | 290 | 0 | 0 | 0 | 11 | 36 | 0 | 20 | 30 | 94 | 4 | 0 | 35 | 92 | 89 | 0 | 0 | 0 | 18 |
| bcbbb314a8fa5891dae2d2ac6cfe667e | 0 | 0 | 0 | 0 | 0 | 0 | 0 | 0 | 0 | 0 | 0 | 93 | 222 | 19 | 0 | 0 | 38 | 0 | 0 | 0 | 60 | 31 | 117 | 0 | 0 | 266 | 87 | 95 | 0 |
| 82d8d63aec8f3d3860586defded5be32 | 31 | 77 | 83 | 151 | 48 | 32 | 40 | 0 | 4 | 10 | 120 | 0 | 0 | 8 | 58 | 35 | 0 | 14 | 28 | 52 | 0 | 10 | 54 | 77 | 51 | 0 | 0 | 0 | 37 |
| 605559dba01eee0ccfdfb68e4dd35cb8 | 0 | 82 | 55 | 86 | 30 | 100 | 93 | 0 | 24 | 0 | 159 | 0 | 0 | 0 | 60 | 66 | 0 | 38 | 0 | 75 | 14 | 0 | 17 | 0 | 87 | 18 | 8 | 8 | 0 |
| de392404ac3ef9a07e3629ed62484d85 | 0 | 0 | 0 | 0 | 0 | 0 | 4 | 0 | 44 | 0 | 11 | 248 | 324 | 57 | 38 | 0 | 16 | 23 | 0 | 0 | 0 | 13 | 77 | 0 | 0 | 101 | 35 | 28 | 0 |
| b6e8e35011fbccc0d4c70b5870f9961b | 93 | 92 | 73 | 36 | 100 | 0 | 0 | 0 | 46 | 109 | 0 | 0 | 0 | 0 | 0 | 0 | 0 | 0 | 31 | 0 | 0 | 0 | 40 | 108 | 39 | 0 | 0 | 0 | 238 |
| e30d19fe5b0ebf9554c052840c4d956a | 0 | 92 | 60 | 72 | 64 | 0 | 0 | 0 | 45 | 0 | 77 | 88 | 104 | 58 | 0 | 0 | 0 | 0 | 0 | 63 | 57 | 47 | 0 | 0 | 0 | 69 | 55 | 41 | 0 |
| 9e893d8a87f8fb7eade73e5095b731c9 | 90 | 238 | 130 | 146 | 150 | 0 | 0 | 0 | 55 | 102 | 25 | 0 | 0 | 0 | 0 | 0 | 0 | 0 | 0 | 24 | 0 | 0 | 0 | 30 | 0 | 0 | 0 | 0 | 0 |
| 82636c13aeb820d1aebf7dedaecf8cd2 | 27 | 118 | 73 | 78 | 85 | 16 | 18 | 8 | 18 | 64 | 52 | 0 | 0 | 0 | 31 | 0 | 0 | 8 | 21 | 11 | 18 | 8 | 83 | 115 | 41 | 15 | 11 | 7 | 57 |
| 3e8607ee4c179ad7f18201b67f76863a | 0 | 0 | 0 | 0 | 0 | 0 | 0 | 0 | 0 | 0 | 0 | 94 | 210 | 46 | 0 | 0 | 43 | 21 | 8 | 0 | 134 | 121 | 149 | 15 | 0 | 54 | 0 | 32 | 46 |
| 55fba4512510e83ea3af0871e3cbd291 | 19 | 133 | 69 | 119 | 22 | 6 | 0 | 11 | 57 | 118 | 15 | 47 | 76 | 49 | 24 | 0 | 15 | 17 | 8 | 21 | 7 | 12 | 64 | 33 | 17 | 0 | 0 | 0 | 13 |
| 8a13fa187cbfd57eeba9e8067576048a | 0 | 61 | 31 | 63 | 38 | 41 | 42 | 0 | 39 | 0 | 53 | 0 | 0 | 0 | 51 | 43 | 0 | 25 | 0 | 34 | 36 | 0 | 33 | 0 | 34 | 156 | 96 | 83 | 0 |
| 7c5a0fd8a87be2cb862f42d207bfbcb0 | 0 | 44 | 59 | 147 | 62 | 64 | 66 | 0 | 0 | 53 | 200 | 0 | 0 | 0 | 0 | 0 | 0 | 0 | 32 | 75 | 0 | 0 | 16 | 82 | 50 | 0 | 0 | 0 | 0 |
| 98c93109ee2ee98c7b1fabbf0075f0e5 | 0 | 0 | 0 | 0 | 0 | 0 | 0 | 15 | 31 | 0 | 0 | 190 | 279 | 75 | 0 | 0 | 0 | 0 | 0 | 0 | 75 | 21 | 88 | 0 | 0 | 94 | 35 | 0 | 45 |
| 753197e4f87bd09a6ffc97956daa03a8 | 0 | 33 | 11 | 8 | 53 | 0 | 18 | 0 | 12 | 56 | 18 | 58 | 78 | 19 | 30 | 12 | 21 | 17 | 51 | 59 | 25 | 9 | 64 | 89 | 60 | 28 | 15 | 18 | 83 |
| e78765602973483890504954a9bdc9b0 | 15 | 70 | 39 | 41 | 50 | 0 | 0 | 0 | 12 | 105 | 0 | 14 | 28 | 0 | 0 | 0 | 31 | 0 | 48 | 37 | 23 | 0 | 88 | 146 | 103 | 0 | 0 | 0 | 92 |
| 73bb08decc3b8de17a42365a3e530dad | 0 | 0 | 0 | 0 | 7 | 0 | 0 | 5 | 27 | 8 | 0 | 79 | 88 | 36 | 0 | 6 | 25 | 25 | 6 | 9 | 53 | 26 | 45 | 2 | 0 | 246 | 124 | 88 | 8 |
| d2f397f5b7eaa4e05f0f54e9e9b19611 | 0 | 0 | 0 | 0 | 0 | 0 | 0 | 63 | 102 | 0 | 0 | 151 | 286 | 22 | 18 | 0 | 0 | 0 | 0 | 0 | 53 | 38 | 53 | 7 | 0 | 60 | 21 | 16 | 14 |
| cc092684f2a7f76179a466e844d29f23 | 16 | 39 | 70 | 254 | 26 | 58 | 35 | 0 | 0 | 0 | 187 | 0 | 0 | 0 | 0 | 32 | 0 | 0 | 0 | 61 | 0 | 0 | 19 | 39 | 37 | 0 | 0 | 0 | 0 |
| c01b1b54d3ec820b9861c26638955c30 | 0 | 0 | 0 | 120 | 19 | 79 | 30 | 0 | 0 | 45 | 364 | 0 | 0 | 0 | 0 | 11 | 0 | 0 | 0 | 38 | 0 | 0 | 0 | 36 | 110 | 0 | 0 | 0 | 0 |
| 599689144411ce9c897d565b1439ed0f | 0 | 39 | 29 | 0 | 0 | 0 | 0 | 0 | 93 | 29 | 0 | 68 | 156 | 23 | 39 | 0 | 40 | 15 | 0 | 0 | 26 | 50 | 53 | 108 | 0 | 42 | 0 | 30 | 0 |
| 03a52768936e3d07ef55f97972de1bc7 | 0 | 31 | 19 | 0 | 0 | 0 | 0 | 22 | 40 | 0 | 0 | 107 | 234 | 0 | 0 | 0 | 0 | 0 | 0 | 0 | 78 | 54 | 129 | 45 | 0 | 31 | 0 | 34 | 0 |
| 71362904ece78f66227f6d6a23402ecf | 0 | 0 | 0 | 0 | 0 | 0 | 0 | 0 | 0 | 0 | 0 | 71 | 191 | 0 | 0 | 0 | 29 | 0 | 0 | 0 | 20 | 20 | 106 | 0 | 0 | 195 | 77 | 51 | 36 |
| c5034d5ed01af74802cdae21bda00b7a | 0 | 96 | 48 | 0 | 59 | 14 | 28 | 0 | 65 | 39 | 0 | 24 | 82 | 33 | 82 | 0 | 29 | 0 | 0 | 0 | 16 | 17 | 63 | 24 | 37 | 18 | 10 | 0 | 9 |
| c5dbf2ea44cadec702ee97926cfe8184 | 0 | 50 | 0 | 31 | 0 | 0 | 0 | 0 | 28 | 0 | 0 | 69 | 138 | 23 | 0 | 0 | 30 | 24 | 0 | 0 | 50 | 29 | 65 | 0 | 15 | 64 | 76 | 42 | 46 |
| 4d072cfea4d0d51d223a1de459017a90 | 0 | 19 | 0 | 0 | 0 | 0 | 0 | 19 | 20 | 0 | 0 | 78 | 163 | 23 | 15 | 0 | 31 | 28 | 16 | 0 | 71 | 48 | 52 | 15 | 5 | 102 | 26 | 20 | 11 |
| b6b738a3d99d9e95203548f7dfba8e32 | 0 | 56 | 18 | 0 | 60 | 0 | 11 | 42 | 70 | 0 | 0 | 86 | 198 | 25 | 17 | 0 | 0 | 8 | 0 | 0 | 49 | 34 | 52 | 33 | 0 | 0 | 0 | 0 | 0 |
| bf18173ef06b0e8cd0a55ce86358c75f | 5 | 18 | 22 | 62 | 26 | 0 | 18 | 0 | 7 | 0 | 71 | 0 | 7 | 16 | 56 | 33 | 0 | 51 | 52 | 0 | 10 | 18 | 111 | 104 | 0 | 31 | 3 | 10 | 0 |
| c53dad5262ba5e488c5284f6a083e9b3 | 0 | 42 | 0 | 17 | 0 | 0 | 0 | 7 | 32 | 0 | 0 | 86 | 195 | 36 | 24 | 0 | 0 | 0 | 11 | 0 | 35 | 27 | 34 | 0 | 0 | 44 | 38 | 34 | 60 |
| 6b3d10862ba35a1b4736215214f4e0ae | 0 | 0 | 0 | 22 | 71 | 36 | 60 | 0 | 0 | 18 | 69 | 0 | 0 | 0 | 0 | 19 | 0 | 0 | 72 | 78 | 0 | 0 | 26 | 62 | 112 | 0 | 0 | 0 | 54 |
| d3cdda9df90471774e94fad3fa66968a | 0 | 14 | 10 | 0 | 5 | 4 | 0 | 0 | 0 | 28 | 11 | 0 | 0 | 0 | 9 | 0 | 0 | 7 | 41 | 21 | 15 | 5 | 97 | 223 | 110 | 5 | 5 | 8 | 79 |
| a8d6e8aaa8fdfc9e88461c8f4c5a694f | 3 | 12 | 11 | 33 | 35 | 39 | 56 | 0 | 0 | 0 | 67 | 0 | 0 | 0 | 27 | 12 | 0 | 0 | 62 | 70 | 0 | 0 | 0 | 102 | 122 | 0 | 0 | 0 | 42 |
| fe37c03cee5f1b60dc84416bbfec8213 | 6 | 45 | 24 | 23 | 9 | 0 | 0 | 27 | 45 | 21 | 10 | 44 | 109 | 13 | 19 | 0 | 18 | 11 | 7 | 0 | 40 | 21 | 75 | 29 | 15 | 34 | 0 | 8 | 31 |
| a5555d4fb3c468646e2ccbc60debf2eb | 0 | 20 | 25 | 18 | 19 | 9 | 0 | 0 | 40 | 0 | 21 | 95 | 150 | 21 | 0 | 0 | 21 | 0 | 0 | 0 | 69 | 41 | 49 | 13 | 0 | 24 | 8 | 8 | 0 |
| b6c0d89b177b3dd41142025330da6cba | 15 | 56 | 47 | 75 | 31 | 56 | 61 | 0 | 9 | 17 | 76 | 0 | 0 | 10 | 65 | 44 | 0 | 4 | 12 | 38 | 0 | 0 | 5 | 22 | 0 | 0 | 0 | 0 | 0 |
| 07915a1137f699de14f72c85f5bbd2a6 | 6 | 35 | 21 | 30 | 24 | 26 | 28 | 0 | 13 | 0 | 0 | 18 | 37 | 18 | 31 | 19 | 11 | 17 | 46 | 0 | 14 | 13 | 55 | 19 | 32 | 29 | 22 | 18 | 60 |
| a67d31cadaa0705b60077cea1b01eb3c | 0 | 21 | 6 | 0 | 36 | 0 | 0 | 11 | 41 | 21 | 10 | 64 | 101 | 50 | 11 | 0 | 25 | 5 | 20 | 8 | 16 | 14 | 20 | 19 | 23 | 35 | 17 | 0 | 59 |
| 62131078606f33fa2aee4fb1e24d1325 | 0 | 44 | 32 | 54 | 74 | 45 | 56 | 3 | 0 | 42 | 48 | 0 | 0 | 0 | 22 | 22 | 0 | 12 | 0 | 24 | 0 | 0 | 0 | 48 | 65 | 0 | 0 | 0 | 21 |
| 269c56d6f6d510fca9e695dd4ef3aded | 33 | 154 | 88 | 85 | 22 | 0 | 0 | 0 | 0 | 8 | 23 | 0 | 0 | 0 | 0 | 0 | 0 | 0 | 15 | 7 | 0 | 0 | 12 | 55 | 27 | 0 | 0 | 0 | 77 |
| 635b862162c7602ce467943c39341ec6 | 34 | 32 | 34 | 62 | 35 | 23 | 50 | 12 | 22 | 0 | 27 | 19 | 50 | 22 | 49 | 0 | 0 | 14 | 0 | 32 | 0 | 0 | 14 | 48 | 0 | 12 | 13 | 0 | 0 |
| 0c74eae4dbf73f8e229295b0106c102a | 0 | 64 | 67 | 67 | 22 | 0 | 17 | 0 | 28 | 15 | 31 | 0 | 22 | 0 | 26 | 0 | 0 | 0 | 46 | 17 | 18 | 0 | 27 | 0 | 31 | 0 | 8 | 0 | 93 |
| 46dcd9314a0ff27108e1825c812d2eb0 | 0 | 78 | 53 | 138 | 44 | 60 | 47 | 0 | 0 | 0 | 51 | 0 | 0 | 0 | 44 | 0 | 0 | 0 | 0 | 0 | 0 | 0 | 0 | 35 | 0 | 0 | 0 | 0 | 26 |
| 4756f9f656811df6fd09118ba685b062 | 10 | 35 | 31 | 135 | 25 | 48 | 25 | 0 | 8 | 32 | 135 | 0 | 0 | 0 | 0 | 0 | 0 | 0 | 0 | 16 | 0 | 0 | 0 | 39 | 15 | 0 | 0 | 0 | 0 |
| cd803d1bccf1637b1b70fd30b8e8fd1c | 0 | 65 | 33 | 108 | 20 | 34 | 58 | 0 | 0 | 105 | 63 | 0 | 0 | 0 | 0 | 0 | 0 | 0 | 0 | 29 | 0 | 0 | 0 | 19 | 11 | 0 | 0 | 0 | 0 |
| bdc0a8e3e90a5e0985147bf5144416be | 0 | 18 | 6 | 19 | 30 | 0 | 0 | 0 | 10 | 43 | 22 | 7 | 0 | 16 | 12 | 0 | 12 | 27 | 11 | 0 | 9 | 8 | 54 | 66 | 30 | 40 | 26 | 21 | 55 |
| ce330d07615fe7d3ec50c3016a5633f4 | 0 | 0 | 0 | 5 | 0 | 0 | 0 | 0 | 0 | 0 | 0 | 61 | 98 | 31 | 19 | 0 | 51 | 0 | 0 | 0 | 49 | 61 | 52 | 0 | 0 | 48 | 0 | 0 | 65 |
| 314b54066917a95d63f9354217b50cb5 | 0 | 17 | 16 | 80 | 16 | 21 | 11 | 0 | 0 | 7 | 201 | 0 | 0 | 0 | 2 | 11 | 0 | 5 | 7 | 42 | 0 | 0 | 24 | 40 | 34 | 0 | 0 | 0 | 6 |
| dcb5ea7fb45ddcc0b93680167a8b7c44 | 40 | 55 | 49 | 0 | 0 | 0 | 0 | 27 | 31 | 0 | 10 | 42 | 134 | 0 | 0 | 0 | 11 | 0 | 0 | 0 | 60 | 11 | 51 | 0 | 0 | 0 | 0 | 0 | 18 |
| c2ffe73bcebceeeb92743ada0e34d4fb | 0 | 0 | 0 | 0 | 0 | 0 | 0 | 0 | 33 | 0 | 0 | 58 | 28 | 62 | 0 | 0 | 15 | 0 | 0 | 0 | 31 | 0 | 50 | 0 | 0 | 56 | 42 | 59 | 96 |
| 2219cd2896936f3ef4554239084b8846 | 27 | 52 | 45 | 72 | 5 | 0 | 0 | 0 | 14 | 0 | 20 | 19 | 50 | 21 | 0 | 0 | 0 | 7 | 11 | 0 | 44 | 27 | 57 | 22 | 4 | 14 | 0 | 0 | 15 |
| 09e533a38f7cea3dbdc6ddc568385026 | 26 | 131 | 69 | 23 | 4 | 0 | 0 | 21 | 62 | 9 | 0 | 33 | 49 | 15 | 3 | 0 | 7 | 12 | 0 | 0 | 27 | 0 | 19 | 7 | 3 | 0 | 0 | 0 | 0 |
| ff02653b16980833c506a2b99a8820c8 | 8 | 77 | 35 | 52 | 20 | 0 | 0 | 22 | 95 | 108 | 22 | 0 | 0 | 0 | 0 | 0 | 0 | 0 | 0 | 0 | 0 | 0 | 52 | 26 | 0 | 0 | 0 | 0 | 0 |
| b014d868ab6ea2b0ef47d3a7c8c8ae90 | 0 | 0 | 0 | 0 | 0 | 0 | 0 | 0 | 8 | 0 | 0 | 91 | 103 | 21 | 4 | 0 | 10 | 13 | 0 | 0 | 66 | 28 | 40 | 3 | 0 | 62 | 28 | 27 | 8 |
| 24c0daead3c6eb74983e67a111a6d5b7 | 4 | 23 | 0 | 17 | 0 | 5 | 0 | 4 | 19 | 0 | 9 | 71 | 140 | 28 | 17 | 0 | 13 | 0 | 0 | 0 | 42 | 35 | 13 | 0 | 0 | 54 | 0 | 0 | 14 |
| e57d8fdef0d6b6588e27684430cfa57c | 5 | 51 | 30 | 115 | 38 | 33 | 6 | 0 | 0 | 20 | 131 | 0 | 0 | 0 | 0 | 5 | 0 | 0 | 8 | 14 | 0 | 0 | 0 | 10 | 28 | 0 | 0 | 0 | 10 |
| 75bd04ce5abd401ad49f6d6a41635e1b | 0 | 14 | 0 | 5 | 0 | 0 | 0 | 0 | 16 | 0 | 0 | 68 | 115 | 23 | 8 | 0 | 23 | 0 | 7 | 0 | 49 | 15 | 25 | 7 | 0 | 51 | 43 | 14 | 19 |
| 3c618a9049bc88d4ac4eff7acb13eda4 | 12 | 24 | 24 | 85 | 18 | 41 | 18 | 0 | 0 | 25 | 131 | 0 | 0 | 0 | 0 | 13 | 0 | 0 | 14 | 49 | 0 | 0 | 0 | 23 | 24 | 0 | 0 | 0 | 0 |
| 5a8040f87477bb12b890b5887c52c2fb | 6 | 45 | 53 | 58 | 45 | 30 | 31 | 0 | 17 | 63 | 49 | 0 | 0 | 12 | 0 | 0 | 0 | 15 | 13 | 17 | 0 | 0 | 14 | 23 | 0 | 0 | 0 | 0 | 6 |
| 38e67f409674a5d8f3df37ddbbb771a0 | 0 | 49 | 36 | 66 | 45 | 26 | 37 | 0 | 0 | 16 | 82 | 0 | 0 | 0 | 0 | 0 | 0 | 0 | 37 | 37 | 0 | 0 | 17 | 46 | 0 | 0 | 0 | 0 | 0 |
| e08fbb4b67723c99b3d2d43e56045915 | 12 | 60 | 26 | 25 | 31 | 0 | 19 | 2 | 4 | 20 | 14 | 0 | 0 | 0 | 25 | 0 | 0 | 8 | 39 | 11 | 0 | 4 | 30 | 63 | 22 | 0 | 0 | 0 | 72 |
| 868e7facae9a1b8f62b269fa7bfaa730 | 0 | 53 | 0 | 0 | 27 | 0 | 0 | 23 | 54 | 0 | 0 | 31 | 73 | 63 | 0 | 0 | 0 | 0 | 17 | 4 | 0 | 23 | 0 | 0 | 7 | 76 | 12 | 19 | 0 |
| f4a251958883c58397ad4ba8175c930d | 0 | 0 | 0 | 0 | 26 | 30 | 61 | 0 | 0 | 0 | 41 | 0 | 0 | 0 | 48 | 40 | 0 | 0 | 48 | 80 | 0 | 0 | 0 | 27 | 54 | 0 | 0 | 0 | 16 |
| c7763753d3f0e889277fcdae148eb371 | 0 | 0 | 0 | 0 | 0 | 0 | 0 | 0 | 0 | 0 | 0 | 55 | 96 | 47 | 0 | 0 | 18 | 14 | 0 | 0 | 51 | 14 | 21 | 0 | 0 | 105 | 18 | 19 | 11 |
| a4f1edf6207afee5cede60c8a78ab396 | 54 | 78 | 43 | 33 | 40 | 0 | 0 | 0 | 21 | 0 | 0 | 25 | 28 | 0 | 0 | 0 | 0 | 0 | 0 | 0 | 10 | 0 | 39 | 32 | 0 | 18 | 0 | 0 | 32 |
| 0774d9e02be877dc80ca29d200c55c4b | 14 | 62 | 53 | 110 | 18 | 16 | 12 | 0 | 0 | 6 | 18 | 0 | 0 | 0 | 9 | 17 | 0 | 8 | 13 | 9 | 0 | 0 | 15 | 29 | 24 | 0 | 0 | 0 | 18 |
| e7ba9027d00b4757a5c590c5a3854b5e | 9 | 47 | 27 | 56 | 38 | 37 | 28 | 0 | 0 | 26 | 40 | 0 | 0 | 0 | 18 | 0 | 0 | 13 | 0 | 0 | 0 | 0 | 19 | 51 | 33 | 0 | 0 | 0 | 0 |
| a80ce521498d598368b58e246b4202ae | 0 | 0 | 0 | 25 | 0 | 0 | 9 | 0 | 0 | 21 | 238 | 33 | 0 | 0 | 15 | 25 | 0 | 0 | 6 | 13 | 10 | 0 | 0 | 3 | 18 | 0 | 0 | 5 | 10 |
| 50c80c9baee4fa5da977674a08447452 | 34 | 107 | 72 | 42 | 26 | 5 | 7 | 0 | 0 | 0 | 6 | 0 | 0 | 0 | 33 | 6 | 0 | 0 | 17 | 3 | 0 | 0 | 18 | 17 | 0 | 14 | 11 | 0 | 11 |
| c2de020fd6003f4489484e3e0d69db71 | 17 | 27 | 20 | 73 | 21 | 13 | 7 | 0 | 0 | 39 | 83 | 0 | 0 | 0 | 0 | 8 | 0 | 0 | 7 | 15 | 0 | 0 | 10 | 34 | 37 | 0 | 0 | 0 | 16 |
| 77ccdc9de7f2053dadebbb04c863c012 | 0 | 60 | 25 | 19 | 0 | 0 | 0 | 0 | 14 | 0 | 0 | 51 | 15 | 8 | 0 | 0 | 19 | 0 | 0 | 0 | 24 | 29 | 38 | 0 | 0 | 9 | 13 | 19 | 57 |
| a5b834320abae36afa162bf9ea064f9a | 27 | 81 | 48 | 102 | 26 | 0 | 11 | 0 | 0 | 19 | 31 | 0 | 0 | 0 | 0 | 0 | 0 | 0 | 11 | 16 | 0 | 0 | 0 | 21 | 0 | 0 | 0 | 0 | 0 |
| 9fa3553e1f858086b01c3e1277682a7b | 15 | 50 | 18 | 47 | 28 | 21 | 22 | 0 | 0 | 15 | 20 | 0 | 0 | 0 | 0 | 7 | 0 | 0 | 12 | 20 | 5 | 4 | 4 | 30 | 62 | 0 | 0 | 0 | 6 |
| 5da7c5389fac9c8aa4f254d7864439e5 | 16 | 31 | 14 | 19 | 18 | 0 | 0 | 9 | 21 | 10 | 0 | 34 | 26 | 23 | 8 | 0 | 12 | 0 | 0 | 0 | 21 | 21 | 23 | 0 | 0 | 19 | 21 | 12 | 20 |
| 4a1c9b400c293e6639ee864ef90732df | 9 | 70 | 14 | 58 | 22 | 20 | 0 | 0 | 22 | 60 | 19 | 0 | 0 | 0 | 0 | 0 | 0 | 0 | 0 | 10 | 6 | 0 | 0 | 21 | 27 | 0 | 0 | 0 | 17 |
| d7716b0e4b28cf56e18343f29efe53b3 | 0 | 0 | 0 | 0 | 0 | 0 | 0 | 0 | 0 | 0 | 0 | 57 | 29 | 32 | 15 | 0 | 0 | 0 | 20 | 0 | 39 | 38 | 52 | 28 | 16 | 20 | 28 | 0 | 0 |
| 00e9343d8fc364bc3c93a300efef122a | 0 | 17 | 11 | 10 | 13 | 18 | 35 | 0 | 8 | 6 | 33 | 0 | 0 | 13 | 33 | 18 | 0 | 0 | 28 | 32 | 0 | 0 | 0 | 16 | 18 | 13 | 17 | 10 | 16 |
| c074ffe7bde46fb34a5fb3d12b1d16f1 | 0 | 34 | 8 | 8 | 0 | 0 | 0 | 0 | 21 | 37 | 0 | 32 | 35 | 16 | 0 | 0 | 31 | 14 | 7 | 0 | 38 | 13 | 46 | 12 | 0 | 0 | 0 | 0 | 7 |
| 16cf0460d6cd711c823a97642f23c7c8 | 0 | 31 | 43 | 96 | 32 | 7 | 33 | 0 | 0 | 14 | 20 | 0 | 0 | 0 | 0 | 17 | 0 | 14 | 14 | 0 | 0 | 0 | 0 | 26 | 12 | 0 | 0 | 0 | 0 |
| f06900dc9acb0a6894ff5db2bb7ec67e | 0 | 0 | 0 | 0 | 0 | 0 | 0 | 0 | 0 | 7 | 0 | 23 | 56 | 22 | 0 | 0 | 55 | 12 | 0 | 0 | 28 | 15 | 90 | 13 | 7 | 5 | 0 | 0 | 26 |
| 3b521bfe4081d4e6a5f952bc58deee87 | 0 | 0 | 0 | 15 | 11 | 18 | 28 | 0 | 0 | 0 | 43 | 0 | 0 | 0 | 15 | 37 | 0 | 0 | 13 | 80 | 0 | 0 | 0 | 17 | 51 | 0 | 0 | 0 | 19 |
| d772fc4bf76190ab1f52ab252da575a2 | 17 | 95 | 46 | 82 | 21 | 0 | 14 | 0 | 10 | 9 | 0 | 0 | 15 | 0 | 0 | 0 | 0 | 0 | 0 | 0 | 15 | 21 | 0 | 0 | 0 | 0 | 0 | 0 | 0 |
| f07714adb6189b375fa21a85940c5592 | 6 | 22 | 0 | 43 | 16 | 9 | 15 | 0 | 32 | 21 | 23 | 0 | 7 | 17 | 20 | 0 | 19 | 11 | 21 | 22 | 0 | 0 | 15 | 24 | 0 | 0 | 0 | 0 | 0 |
| 8cebec5a31a2bad840a6f58d37a6dc16 | 0 | 0 | 0 | 0 | 0 | 0 | 0 | 0 | 20 | 0 | 0 | 2 | 39 | 26 | 0 | 0 | 15 | 0 | 0 | 0 | 22 | 20 | 61 | 0 | 0 | 71 | 20 | 21 | 21 |
| ef08dbff58adabf9fb8e78614c9c9d83 | 0 | 19 | 0 | 42 | 0 | 0 | 27 | 0 | 0 | 0 | 53 | 0 | 0 | 0 | 47 | 14 | 0 | 0 | 21 | 31 | 0 | 0 | 6 | 41 | 8 | 0 | 0 | 0 | 10 |
| 63d8db74a2f9118f15242f0f04cace91 | 31 | 61 | 26 | 0 | 0 | 0 | 0 | 11 | 29 | 10 | 0 | 31 | 63 | 6 | 0 | 6 | 0 | 0 | 0 | 0 | 19 | 5 | 13 | 0 | 0 | 0 | 0 | 3 | 0 |
| f8c92c975389e2b8865c4c9491f10ea5 | 0 | 15 | 31 | 48 | 15 | 37 | 32 | 0 | 0 | 0 | 54 | 0 | 0 | 0 | 0 | 0 | 0 | 0 | 0 | 33 | 0 | 0 | 10 | 0 | 17 | 0 | 0 | 0 | 0 |
| dba6da18688461248a5e43144412a1be | 0 | 0 | 0 | 27 | 20 | 0 | 16 | 0 | 0 | 27 | 70 | 0 | 0 | 0 | 0 | 5 | 0 | 0 | 14 | 23 | 0 | 0 | 0 | 23 | 50 | 0 | 0 | 0 | 16 |
| 204e689938a2d8b443b249866bd369b6 | 11 | 84 | 31 | 23 | 8 | 0 | 0 | 11 | 17 | 23 | 0 | 0 | 0 | 0 | 0 | 0 | 0 | 0 | 0 | 0 | 22 | 17 | 25 | 0 | 0 | 0 | 0 | 6 | 0 |
| d1af6b6bc0d37351821bc3652777aaa4 | 4 | 0 | 0 | 0 | 18 | 0 | 0 | 17 | 0 | 0 | 0 | 18 | 0 | 0 | 17 | 0 | 23 | 0 | 0 | 13 | 31 | 14 | 12 | 0 | 10 | 45 | 25 | 23 | 0 |
| ff40d8496e342401901cc4330e3118bc | 0 | 0 | 0 | 0 | 26 | 15 | 0 | 0 | 0 | 0 | 27 | 0 | 41 | 0 | 13 | 0 | 0 | 0 | 16 | 36 | 0 | 0 | 11 | 0 | 0 | 34 | 9 | 0 | 16 |
| b3a4e7f7a8e676839126c78d23367ce9 | 5 | 24 | 10 | 13 | 18 | 0 | 0 | 0 | 0 | 22 | 11 | 0 | 0 | 0 | 0 | 0 | 0 | 0 | 13 | 18 | 0 | 0 | 0 | 43 | 43 | 0 | 0 | 0 | 0 |
| 8381bfa1787dd0d5fbfa1a5108465af9 | 0 | 9 | 18 | 33 | 14 | 13 | 10 | 0 | 0 | 0 | 41 | 0 | 0 | 0 | 16 | 0 | 0 | 0 | 14 | 16 | 0 | 0 | 5 | 0 | 20 | 0 | 0 | 0 | 0 |
| 5b467f21a97309d3b35f9cd90691a62e | 0 | 0 | 0 | 0 | 0 | 0 | 0 | 0 | 0 | 0 | 0 | 22 | 13 | 21 | 0 | 0 | 15 | 24 | 0 | 0 | 0 | 11 | 9 | 15 | 0 | 24 | 11 | 24 | 0 |
| 3e9b7bc1b40e8d02d12d76820c7dbd27 | 0 | 10 | 7 | 14 | 0 | 12 | 21 | 0 | 0 | 0 | 0 | 0 | 12 | 5 | 23 | 8 | 0 | 0 | 0 | 12 | 0 | 0 | 13 | 7 | 7 | 18 | 13 | 0 | 3 |

**Supplementary Table S7**. Common Bacterio- and virioplankton RTPR 98% OTUs in 27 samples with RTPR amplicons in both fractions

Total seqs: 47997; Subtotal seqs in retained

OTUs: 41046

| OTUs with >=10 seqs: 254 | OTU subtotal | May12_05m bacterioplankton | May12_05m virioplankton | May12_25m bacterioplankton | May12_25m virioplankton | May12_50m bacterioplankton | May12_50m virioplankton | May12_75m bacterioplankton | May12_75m virioplankton | Jun8_25m bacterioplankton | Jun8_25m virioplankton | Jun8_50m bacterioplankton | Jun8_50m virioplankton | Jun8_75m bacterioplankton | Jun8_75m virioplankton | Jun22_0m bacterioplankton | Jun22_0m virioplankton | Jun22_50m bacterioplankton | Jun22_50m virioplankton | Jun22_75m bacterioplankton | Jun22_75m virioplankton | Jul11_0m bacterioplankton | Jul11_0m virioplankton | Jul11_05m bacterioplankton | Jul11_05m virioplankton | Jul11_25m bacterioplankton | Jul11_25m virioplankton | Jul11_50m bacterioplankton | Jul11_50m virioplankton | Jul11_75m bacterioplankton | Jul11_75m virioplankton | Jul27_05m bacterioplankton | Jul27_05m virioplankton | Jul27_25m bacterioplankton | Jul27_25m virioplankton | Jul27_75m bacterioplankton | Jul27_75m virioplankton | Aug5_0m bacterioplankton | Aug5_0m virioplankton | Aug5_05m bacterioplankton | Aug5_05m virioplankton | Aug5_25m bacterioplankton | Aug5_25m virioplankton | Aug5_50m bacterioplankton | Aug5_50m virioplankton | Aug5_75m bacterioplankton | Aug5_75m virioplankton | Sep8_0m bacterioplankton | Sep8_0m virioplankton | Sep8_05m bacterioplankton | Sep8_05m virioplankton | Sep8_25m bacterioplankton | Sep8_25m virioplankton | Sep8_50m bacterioplankton | Sep8_50m virioplankton |
|---|---|---|---|---|---|---|---|---|---|---|---|---|---|---|---|---|---|---|---|---|---|---|---|---|---|---|---|---|---|---|---|---|---|---|---|---|---|---|---|---|---|---|---|---|---|---|---|---|---|---|---|---|---|---|---|
| bacteria_Aug5d25b_1039 | 4761 | 6 | 108 | 120 | 130 | 85 | 261 | 13 | 432 | 329 | 84 | 254 | 340 | 31 | 118 | 31 | 18 | 43 | 261 | 168 | 290 | 8 | 9 | 3 | 28 | 9 | 56 | 29 | 139 | 97 | 112 | 38 | 16 | 39 | 30 | 341 | 22 | 4 | 23 | 3 | 12 | 44 | 22 | 64 | 22 | 75 | 55 | 4 | 7 | 23 | 48 | 8 | 199 | 21 | 29 |
| bacteria_Jul11d0b_35947 | 3280 | 0 | 2 | 2 | 4 | 7 | 9 | 0 | 3 | 33 | 10 | 0 | 14 | 1 | 0 | 43 | 19 | 19 | 235 | 13 | 22 | 36 | 12 | 9 | 90 | 41 | 112 | 33 | 38 | 33 | 9 | 186 | 195 | 182 | 200 | 131 | 24 | 64 | 206 | 6 | 102 | 267 | 160 | 76 | 154 | 10 | 78 | 10 | 39 | 78 | 3 | 3 | 28 | 9 | 220 |
| bacteria_Jul11d0b_349 | 2880 | 0 | 1 | 0 | 0 | 3 | 3 | 2 | 4 | 0 | 0 | 1 | 0 | 0 | 0 | 3 | 6 | 0 | 11 | 13 | 1 | 37 | 4 | 15 | 13 | 10 | 0 | 9 | 0 | 24 | 1 | 531 | 170 | 259 | 65 | 0 | 1 | 87 | 362 | 20 | 92 | 489 | 306 | 47 | 26 | 3 | 10 | 7 | 58 | 70 | 0 | 12 | 1 | 5 | 98 |
| bacteria_Jul11d0b_5406 | 2347 | 1 | 38 | 9 | 47 | 7 | 140 | 0 | 37 | 32 | 20 | 4 | 57 | 2 | 4 | 20 | 56 | 5 | 76 | 6 | 39 | 64 | 41 | 27 | 63 | 48 | 66 | 15 | 34 | 19 | 23 | 79 | 93 | 32 | 41 | 14 | 7 | 34 | 221 | 11 | 103 | 59 | 124 | 16 | 24 | 2 | 19 | 14 | 45 | 213 | 20 | 10 | 7 | 1 | 158 |
| bacteria_Aug5d25b_42861 | 2111 | 5 | 58 | 233 | 37 | 69 | 93 | 11 | 56 | 280 | 20 | 48 | 63 | 45 | 7 | 8 | 26 | 267 | 191 | 38 | 21 | 2 | 1 | 1 | 11 | 3 | 40 | 9 | 22 | 16 | 53 | 11 | 7 | 17 | 9 | 77 | 8 | 1 | 12 | 0 | 11 | 16 | 19 | 62 | 29 | 23 | 24 | 0 | 6 | 4 | 18 | 0 | 5 | 3 | 15 |
| bacteria_Jul11d0b_35378 | 1523 | 1 | 69 | 40 | 81 | 37 | 146 | 4 | 66 | 3 | 21 | 8 | 28 | 2 | 4 | 28 | 79 | 8 | 121 | 10 | 29 | 35 | 77 | 9 | 49 | 9 | 22 | 2 | 18 | 14 | 10 | 24 | 44 | 18 | 34 | 16 | 3 | 16 | 62 | 3 | 33 | 38 | 21 | 4 | 15 | 1 | 22 | 2 | 43 | 22 | 37 | 2 | 6 | 1 | 26 |
| bacteria_Jul11d0b_26490 | 1501 | 1 | 32 | 6 | 31 | 3 | 34 | 4 | 26 | 38 | 14 | 2 | 29 | 4 | 8 | 15 | 26 | 0 | 31 | 8 | 20 | 38 | 37 | 14 | 63 | 29 | 56 | 5 | 17 | 10 | 19 | 86 | 66 | 24 | 33 | 9 | 5 | 41 | 126 | 14 | 67 | 50 | 67 | 4 | 11 | 0 | 12 | 10 | 56 | 127 | 6 | 2 | 4 | 2 | 59 |
| bacteria_Aug5d5b_22963 | 1500 | 2 | 96 | 4 | 96 | 29 | 100 | 2 | 175 | 55 | 19 | 31 | 147 | 1 | 5 | 9 | 21 | 6 | 33 | 16 | 120 | 6 | 14 | 2 | 17 | 7 | 11 | 11 | 28 | 39 | 67 | 14 | 16 | 17 | 21 | 45 | 29 | 1 | 11 | 2 | 16 | 15 | 10 | 5 | 4 | 7 | 48 | 0 | 4 | 6 | 30 | 5 | 11 | 3 | 11 |
| bacteria_Jul11d0b_24544 | 1377 | 1 | 36 | 6 | 29 | 10 | 73 | 3 | 36 | 32 | 10 | 3 | 14 | 1 | 4 | 11 | 27 | 2 | 30 | 7 | 32 | 50 | 40 | 18 | 40 | 14 | 43 | 4 | 29 | 11 | 26 | 52 | 58 | 22 | 28 | 7 | 2 | 31 | 112 | 7 | 50 | 45 | 69 | 11 | 15 | 0 | 10 | 7 | 33 | 102 | 7 | 8 | 1 | 1 | 57 |
| bacteria_Jul11d0b_624 | 920 | 1 | 31 | 5 | 35 | 7 | 27 | 0 | 25 | 4 | 9 | 2 | 63 | 0 | 6 | 12 | 13 | 0 | 34 | 3 | 62 | 9 | 12 | 2 | 20 | 4 | 11 | 1 | 8 | 9 | 31 | 19 | 25 | 16 | 13 | 12 | 4 | 16 | 52 | 5 | 25 | 27 | 13 | 2 | 6 | 2 | 14 | 6 | 16 | 38 | 33 | 1 | 2 | 1 | 126 |
| bacteria_Jul11d0b_26699 | 867 | 1 | 39 | 5 | 31 | 36 | 11 | 2 | 77 | 108 | 22 | 21 | 69 | 1 | 44 | 11 | 3 | 1 | 25 | 27 | 59 | 6 | 2 | 3 | 17 | 4 | 27 | 3 | 18 | 10 | 17 | 5 | 5 | 1 | 2 | 15 | 3 | 0 | 2 | 0 | 2 | 1 | 0 | 2 | 3 | 4 | 5 | 0 | 1 | 3 | 100 | 3 | 4 | 3 | 3 |
| bacteria_Aug5d25b_2112 | 853 | 4 | 67 | 42 | 70 | 8 | 84 | 1 | 23 | 5 | 10 | 1 | 9 | 0 | 6 | 18 | 15 | 2 | 49 | 1 | 19 | 19 | 15 | 4 | 52 | 5 | 18 | 0 | 4 | 6 | 10 | 24 | 39 | 15 | 8 | 7 | 1 | 5 | 35 | 2 | 18 | 21 | 12 | 1 | 7 | 0 | 15 | 3 | 7 | 2 | 47 | 2 | 2 | 1 | 12 |
| bacteria_May12d50b_252 | 721 | 0 | 13 | 6 | 21 | 11 | 39 | 0 | 18 | 12 | 10 | 6 | 78 | 5 | 12 | 6 | 11 | 1 | 20 | 8 | 26 | 6 | 7 | 1 | 20 | 2 | 20 | 6 | 27 | 9 | 36 | 16 | 18 | 21 | 16 | 15 | 11 | 5 | 11 | 1 | 11 | 23 | 13 | 2 | 10 | 1 | 9 | 1 | 5 | 21 | 31 | 3 | 4 | 2 | 34 |
| virus_Jul11.25v_8557 | 601 | 0 | 23 | 4 | 18 | 10 | 30 | 3 | 12 | 18 | 14 | 3 | 75 | 1 | 8 | 6 | 11 | 1 | 21 | 12 | 29 | 9 | 2 | 2 | 18 | 3 | 7 | 5 | 15 | 9 | 45 | 9 | 16 | 20 | 10 | 14 | 3 | 0 | 4 | 0 | 9 | 14 | 6 | 5 | 6 | 3 | 7 | 4 | 1 | 13 | 12 | 1 | 4 | 2 | 24 |
| virus_Aug5.5v_32649 | 537 | 0 | 0 | 0 | 0 | 1 | 0 | 0 | 0 | 0 | 0 | 0 | 0 | 0 | 0 | 0 | 1 | 1 | 0 | 1 | 0 | 10 | 24 | 18 | 25 | 4 | 8 | 1 | 0 | 2 | 0 | 11 | 28 | 7 | 35 | 1 | 0 | 7 | 170 | 8 | 30 | 11 | 67 | 7 | 8 | 0 | 7 | 0 | 12 | 7 | 0 | 2 | 1 | 1 | 21 |
| bacteria_Aug5d50b_36848 | 494 | 6 | 0 | 229 | 0 | 40 | 0 | 2 | 0 | 87 | 0 | 23 | 0 | 19 | 0 | 0 | 0 | 0 | 0 | 21 | 0 | 0 | 0 | 0 | 0 | 0 | 0 | 6 | 0 | 1 | 0 | 1 | 0 | 1 | 0 | 52 | 0 | 0 | 0 | 0 | 0 | 4 | 0 | 2 | 0 | 0 | 0 | 0 | 0 | 0 | 0 | 0 | 0 | 0 | 0 |
| virus_Jul11.25v_14520 | 480 | 0 | 15 | 6 | 5 | 18 | 21 | 5 | 20 | 8 | 10 | 5 | 52 | 4 | 10 | 2 | 2 | 2 | 21 | 14 | 29 | 1 | 2 | 3 | 11 | 3 | 9 | 4 | 11 | 12 | 21 | 8 | 11 | 18 | 7 | 14 | 6 | 1 | 6 | 0 | 8 | 13 | 4 | 5 | 8 | 0 | 9 | 0 | 3 | 5 | 11 | 0 | 0 | 2 | 15 |
| bacteria_Aug5d25b_1458 | 474 | 2 | 7 | 1 | 5 | 1 | 9 | 0 | 10 | 31 | 1 | 2 | 3 | 0 | 0 | 2 | 7 | 8 | 35 | 18 | 18 | 3 | 5 | 3 | 5 | 10 | 5 | 9 | 20 | 8 | 7 | 11 | 3 | 9 | 8 | 21 | 4 | 6 | 32 | 7 | 8 | 10 | 24 | 6 | 17 | 0 | 25 | 1 | 6 | 9 | 1 | 3 | 2 | 1 | 25 |
| bacteria_Jul27d5b_14425 | 419 | 0 | 8 | 1 | 11 | 1 | 10 | 1 | 5 | 5 | 3 | 1 | 10 | 1 | 0 | 0 | 9 | 1 | 10 | 4 | 7 | 1 | 7 | 2 | 6 | 4 | 6 | 1 | 7 | 8 | 5 | 17 | 17 | 18 | 10 | 5 | 1 | 4 | 37 | 2 | 20 | 17 | 21 | 4 | 11 | 0 | 17 | 0 | 7 | 4 | 3 | 1 | 4 | 2 | 62 |
| bacteria_Aug5d5b_43576 | 360 | 0 | 6 | 1 | 4 | 1 | 9 | 0 | 9 | 6 | 4 | 0 | 2 | 1 | 0 | 4 | 6 | 1 | 4 | 3 | 8 | 12 | 10 | 2 | 16 | 7 | 9 | 2 | 4 | 3 | 3 | 25 | 21 | 10 | 12 | 3 | 0 | 11 | 42 | 5 | 20 | 11 | 9 | 0 | 2 | 0 | 3 | 0 | 8 | 19 | 2 | 2 | 0 | 0 | 18 |
| virus_Aug5.25v_30214 | 359 | 0 | 1 | 0 | 1 | 1 | 0 | 0 | 1 | 0 | 0 | 0 | 0 | 0 | 0 | 0 | 2 | 2 | 0 | 2 | 0 | 9 | 14 | 12 | 18 | 3 | 6 | 0 | 1 | 7 | 0 | 15 | 19 | 7 | 21 | 0 | 0 | 5 | 107 | 5 | 14 | 6 | 48 | 3 | 3 | 0 | 8 | 0 | 5 | 1 | 0 | 0 | 0 | 0 | 12 |
| bacteria_Jul11d0b_19157 | 341 | 1 | 0 | 1 | 4 | 0 | 5 | 0 | 8 | 5 | 0 | 2 | 4 | 0 | 1 | 0 | 1 | 0 | 2 | 2 | 2 | 9 | 3 | 5 | 10 | 3 | 4 | 1 | 1 | 3 | 0 | 28 | 20 | 20 | 10 | 2 | 0 | 10 | 34 | 4 | 16 | 48 | 23 | 2 | 3 | 0 | 1 | 0 | 13 | 17 | 3 | 2 | 0 | 1 | 7 |

**Supplementary Table S7**. Common Bacterio- and virioplankton RTPR 98% OTUs in 27 samples with RTPR amplicons in both fractions

Total seqs: 47997; Subtotal seqs in retained

| OTUs: 41046 | | May12_05m | | May12_25m | | May12_50m | | May12_75m | | Jun8_25m | | Jun8_50m | | Jun8_75m | | Jun22_0m | | Jun22_50m | | Jun22_75m | | Jul11_0m | | Jul11_05m | | Jul11_25m | | Jul11_50m | | Jul11_75m | | Jul27_05m | | Jul27_25m | | Jul27_75m | | Aug5_0m | | Aug5_05m | | Aug5_25m | | Aug5_50m | | Aug5_75m | | Sep8_0m | | Sep8_05m | | Sep8_25m | | Sep8_50m | |
|---|---|---|---|---|---|---|---|---|---|---|---|---|---|---|---|---|---|---|---|---|---|---|---|---|---|---|---|---|---|---|---|---|---|---|---|---|---|---|---|---|---|---|---|---|---|---|---|---|---|---|---|---|---|---|---|
| OTUs with >=10 seqs: 254 | OTU subtotal | bacterioplankton | virioplankton | bacterioplankton | virioplankton | bacterioplankton | virioplankton | bacterioplankton | virioplankton | bacterioplankton | virioplankton | bacterioplankton | virioplankton | bacterioplankton | virioplankton | bacterioplankton | virioplankton | bacterioplankton | virioplankton | bacterioplankton | virioplankton | bacterioplankton | virioplankton | bacterioplankton | virioplankton | bacterioplankton | virioplankton | bacterioplankton | virioplankton | bacterioplankton | virioplankton | bacterioplankton | virioplankton | bacterioplankton | virioplankton | bacterioplankton | virioplankton | bacterioplankton | virioplankton | bacterioplankton | virioplankton | bacterioplankton | virioplankton | bacterioplankton | virioplankton | bacterioplankton | virioplankton | bacterioplankton | virioplankton | bacterioplankton | virioplankton | bacterioplankton | virioplankton | bacterioplankton | virioplankton |
| bacteria_Aug5d25b_1149 | 8 | 0 | 0 | 0 | 0 | 0 | 0 | 0 | 0 | 0 | 0 | 0 | 0 | 0 | 0 | 0 | 0 | 0 | 1 | 0 | 0 | 0 | 0 | 0 | 0 | 0 | 1 | 0 | 0 | 0 | 0 | 0 | 0 | 1 | 1 | 0 | 0 | 1 | 0 | 0 | 0 | 1 | 2 | 0 | 0 | 0 | 0 | 0 | 0 | 0 | 0 | 0 | 0 | 0 | 0 |
| bacteria_Jul27d5b_23117 | 8 | 0 | 0 | 0 | 0 | 0 | 0 | 0 | 1 | 0 | 0 | 0 | 0 | 0 | 0 | 0 | 0 | 0 | 1 | 0 | 0 | 0 | 0 | 0 | 0 | 0 | 0 | 0 | 0 | 0 | 0 | 1 | 0 | 0 | 1 | 0 | 0 | 0 | 0 | 0 | 0 | 0 | 1 | 0 | 0 | 0 | 0 | 0 | 1 | 2 | 0 | 0 | 0 | 0 | 0 |
| bacteria_Jul27d75b_2754 | 8 | 0 | 0 | 0 | 0 | 0 | 0 | 1 | 0 | 0 | 0 | 0 | 0 | 1 | 0 | 0 | 0 | 0 | 0 | 0 | 0 | 0 | 0 | 0 | 0 | 0 | 0 | 0 | 0 | 0 | 0 | 0 | 0 | 1 | 0 | 3 | 0 | 0 | 0 | 0 | 0 | 0 | 0 | 0 | 0 | 0 | 0 | 0 | 0 | 2 | 0 | 0 | 0 | 0 | 0 |
| bacteria_Jun8d25b_1057 | 8 | 0 | 0 | 0 | 0 | 0 | 0 | 0 | 0 | 6 | 0 | 0 | 0 | 0 | 0 | 0 | 0 | 0 | 0 | 0 | 1 | 0 | 0 | 0 | 0 | 0 | 0 | 0 | 0 | 0 | 0 | 0 | 0 | 0 | 0 | 1 | 0 | 0 | 0 | 0 | 0 | 0 | 0 | 0 | 0 | 0 | 0 | 0 | 0 | 0 | 0 | 0 | 0 | 0 | 0 |
| bacteria_Jun8d25b_12904 | 8 | 0 | 0 | 1 | 0 | 0 | 0 | 0 | 0 | 6 | 0 | 0 | 0 | 1 | 0 | 0 | 0 | 0 | 0 | 0 | 0 | 0 | 0 | 0 | 0 | 0 | 0 | 0 | 0 | 0 | 0 | 0 | 0 | 0 | 0 | 0 | 0 | 0 | 0 | 0 | 0 | 0 | 0 | 0 | 0 | 0 | 0 | 0 | 0 | 0 | 0 | 0 | 0 | 0 | 0 |
| bacteria_Jun22d0b_69465 | 8 | 0 | 0 | 0 | 0 | 0 | 0 | 0 | 0 | 3 | 0 | 0 | 0 | 0 | 0 | 1 | 0 | 0 | 0 | 0 | 0 | 1 | 0 | 0 | 0 | 0 | 0 | 0 | 0 | 0 | 0 | 0 | 0 | 1 | 0 | 0 | 0 | 0 | 0 | 0 | 0 | 0 | 0 | 0 | 0 | 0 | 0 | 0 | 0 | 2 | 0 | 0 | 0 | 0 | 0 |
| bacteria_Jun22d75b_7265 | 8 | 0 | 0 | 0 | 0 | 0 | 1 | 0 | 1 | 0 | 0 | 0 | 1 | 0 | 0 | 0 | 0 | 0 | 1 | 1 | 0 | 0 | 0 | 0 | 0 | 0 | 0 | 0 | 0 | 0 | 1 | 0 | 0 | 0 | 0 | 0 | 2 | 0 | 0 | 0 | 0 | 0 | 0 | 0 | 0 | 0 | 0 | 0 | 0 | 0 | 0 | 0 | 0 | 0 | 0 |
| virus_Aug5.5v_19438 | 8 | 0 | 0 | 0 | 0 | 0 | 0 | 0 | 0 | 0 | 0 | 0 | 0 | 0 | 0 | 0 | 0 | 0 | 0 | 0 | 0 | 0 | 0 | 0 | 0 | 0 | 0 | 0 | 0 | 1 | 1 | 0 | 1 | 0 | 0 | 0 | 0 | 0 | 1 | 0 | 1 | 0 | 1 | 0 | 0 | 0 | 0 | 0 | 1 | 0 | 0 | 0 | 1 | 0 | 0 |
| virus_Aug5.25v_25126 | 8 | 0 | 0 | 0 | 0 | 0 | 1 | 0 | 1 | 0 | 0 | 0 | 0 | 0 | 0 | 0 | 0 | 0 | 0 | 1 | 1 | 0 | 0 | 0 | 0 | 0 | 0 | 0 | 0 | 0 | 1 | 0 | 0 | 0 | 0 | 0 | 0 | 0 | 0 | 0 | 0 | 0 | 2 | 1 | 0 | 0 | 0 | 0 | 0 | 0 | 0 | 0 | 0 | 0 | 0 |
| virus_Aug5.25v_31556 | 8 | 0 | 0 | 0 | 0 | 0 | 0 | 0 | 0 | 0 | 0 | 0 | 0 | 0 | 0 | 1 | 0 | 0 | 0 | 0 | 0 | 0 | 0 | 0 | 0 | 0 | 0 | 0 | 0 | 0 | 0 | 0 | 2 | 0 | 0 | 1 | 0 | 0 | 0 | 0 | 0 | 2 | 1 | 0 | 1 | 0 | 0 | 0 | 0 | 0 | 0 | 0 | 0 | 0 | 0 |
| virus_Jul27.5v_16280 | 8 | 0 | 0 | 0 | 0 | 0 | 0 | 0 | 0 | 0 | 0 | 0 | 0 | 0 | 0 | 0 | 0 | 0 | 0 | 0 | 0 | 0 | 1 | 1 | 0 | 0 | 1 | 0 | 0 | 0 | 0 | 0 | 2 | 0 | 0 | 0 | 1 | 0 | 0 | 0 | 0 | 1 | 0 | 0 | 0 | 0 | 0 | 0 | 1 | 0 | 0 | 0 | 0 | 0 | 0 |
| virus_Jul27.25v_10497 | 8 | 0 | 0 | 0 | 0 | 0 | 0 | 0 | 0 | 0 | 0 | 0 | 1 | 0 | 0 | 0 | 0 | 0 | 0 | 0 | 0 | 1 | 0 | 0 | 0 | 0 | 0 | 0 | 0 | 0 | 0 | 1 | 1 | 0 | 2 | 2 | 0 | 0 | 0 | 0 | 0 | 0 | 0 | 0 | 0 | 0 | 0 | 0 | 0 | 0 | 0 | 0 | 0 | 0 | 0 |
| virus_Jun22.0v_3245 | 8 | 0 | 0 | 0 | 0 | 0 | 1 | 0 | 1 | 0 | 0 | 0 | 0 | 0 | 0 | 0 | 1 | 0 | 0 | 0 | 0 | 0 | 0 | 0 | 0 | 0 | 0 | 0 | 0 | 0 | 1 | 0 | 0 | 0 | 0 | 0 | 0 | 0 | 0 | 0 | 0 | 0 | 0 | 0 | 0 | 0 | 0 | 0 | 0 | 0 | 2 | 0 | 2 | 0 | 0 |
| virus_Sep8.50v_3182 | 8 | 0 | 0 | 0 | 0 | 0 | 1 | 0 | 0 | 0 | 0 | 0 | 1 | 0 | 0 | 0 | 0 | 0 | 0 | 0 | 0 | 0 | 0 | 0 | 0 | 0 | 0 | 0 | 0 | 0 | 1 | 0 | 1 | 0 | 0 | 0 | 0 | 0 | 1 | 0 | 0 | 0 | 0 | 0 | 0 | 0 | 0 | 0 | 0 | 0 | 1 | 1 | 0 | 0 | 1 |
| bacteria_Aug5d25b_1001 | 8 | 0 | 0 | 0 | 0 | 0 | 0 | 0 | 0 | 0 | 1 | 0 | 0 | 0 | 0 | 0 | 0 | 0 | 0 | 1 | 0 | 0 | 0 | 0 | 1 | 0 | 0 | 0 | 0 | 0 | 0 | 0 | 2 | 0 | 0 | 1 | 0 | 0 | 0 | 0 | 0 | 2 | 0 | 0 | 0 | 0 | 0 | 0 | 0 | 0 | 0 | 0 | 0 | 0 | 0 |
| bacteria_Jul27d5b_985 | 8 | 0 | 0 | 0 | 0 | 0 | 0 | 0 | 0 | 1 | 0 | 0 | 0 | 0 | 0 | 1 | 0 | 0 | 0 | 0 | 0 | 0 | 0 | 0 | 0 | 0 | 0 | 0 | 0 | 0 | 0 | 1 | 1 | 0 | 0 | 0 | 0 | 0 | 0 | 0 | 0 | 1 | 1 | 0 | 0 | 0 | 0 | 0 | 0 | 0 | 0 | 0 | 0 | 0 | 2 |
| bacteria_Jun8d25b_57524 | 8 | 0 | 0 | 0 | 0 | 0 | 1 | 0 | 1 | 2 | 0 | 0 | 0 | 0 | 0 | 0 | 0 | 0 | 0 | 0 | 0 | 0 | 0 | 0 | 1 | 0 | 0 | 0 | 0 | 0 | 0 | 1 | 0 | 0 | 0 | 1 | 0 | 0 | 0 | 0 | 0 | 0 | 0 | 0 | 0 | 0 | 0 | 0 | 0 | 0 | 0 | 0 | 0 | 1 | 0 |
| bacteria_May12d50b_399 | 8 | 0 | 1 | 0 | 1 | 2 | 0 | 0 | 1 | 0 | 0 | 0 | 2 | 0 | 0 | 0 | 1 | 0 | 0 | 0 | 0 | 0 | 0 | 0 | 0 | 0 | 0 | 0 | 0 | 0 | 0 | 0 | 0 | 0 | 0 | 0 | 0 | 0 | 0 | 0 | 0 | 0 | 0 | 0 | 0 | 0 | 0 | 0 | 0 | 0 | 0 | 0 | 0 | 0 | 0 |
| virus_Aug5.0v_4580 | 8 | 0 | 0 | 0 | 0 | 0 | 0 | 0 | 0 | 0 | 0 | 0 | 0 | 0 | 0 | 0 | 0 | 0 | 0 | 0 | 0 | 0 | 0 | 0 | 0 | 0 | 0 | 0 | 0 | 0 | 0 | 0 | 1 | 0 | 0 | 0 | 0 | 0 | 5 | 1 | 1 | 0 | 0 | 0 | 0 | 0 | 0 | 0 | 0 | 0 | 0 | 0 | 0 | 0 | 0 |
| virus_Jul11.25v_7068 | 8 | 0 | 0 | 0 | 0 | 0 | 0 | 0 | 0 | 0 | 0 | 0 | 0 | 0 | 1 | 0 | 0 | 0 | 1 | 0 | 1 | 0 | 0 | 0 | 2 | 0 | 1 | 0 | 0 | 0 | 1 | 0 | 0 | 0 | 0 | 0 | 0 | 0 | 0 | 0 | 0 | 0 | 0 | 0 | 0 | 0 | 0 | 0 | 0 | 0 | 1 | 0 | 0 | 0 | 0 |
| virus_Jul27.5v_13813 | 8 | 0 | 0 | 0 | 0 | 0 | 0 | 0 | 0 | 0 | 0 | 0 | 0 | 0 | 0 | 0 | 0 | 0 | 1 | 0 | 0 | 0 | 0 | 0 | 0 | 0 | 0 | 0 | 0 | 0 | 0 | 0 | 2 | 2 | 0 | 0 | 0 | 0 | 0 | 0 | 1 | 0 | 1 | 0 | 0 | 1 | 0 | 0 | 0 | 0 | 0 | 0 | 0 | 0 | 0 |
| virus_May12.75v_10506 | 8 | 0 | 0 | 0 | 0 | 0 | 0 | 0 | 3 | 0 | 1 | 1 | 0 | 0 | 0 | 0 | 0 | 0 | 1 | 0 | 1 | 0 | 0 | 0 | 1 | 0 | 0 | 0 | 0 | 0 | 0 | 0 | 0 | 0 | 0 | 0 | 0 | 0 | 0 | 0 | 0 | 0 | 0 | 0 | 0 | 0 | 0 | 0 | 0 | 0 | 0 | 0 | 0 | 0 | 0 |
| virus_Jul11.5v_5239 | 8 | 0 | 0 | 0 | 0 | 0 | 0 | 0 | 0 | 0 | 0 | 0 | 2 | 0 | 0 | 0 | 0 | 0 | 0 | 0 | 0 | 0 | 0 | 0 | 3 | 0 | 0 | 0 | 1 | 0 | 0 | 0 | 0 | 0 | 0 | 0 | 0 | 0 | 0 | 0 | 0 | 0 | 1 | 0 | 0 | 0 | 1 | 0 | 0 | 0 | 0 | 0 | 0 | 0 | 0 |
| bacteria_Aug5d25b_1585 | 8 | 0 | 0 | 0 | 1 | 0 | 0 | 0 | 0 | 0 | 0 | 0 | 0 | 0 | 0 | 0 | 0 | 0 | 1 | 0 | 0 | 1 | 0 | 0 | 0 | 1 | 0 | 0 | 0 | 1 | 0 | 0 | 0 | 0 | 0 | 1 | 0 | 0 | 0 | 0 | 0 | 1 | 0 | 0 | 1 | 0 | 0 | 0 | 0 | 0 | 0 | 0 | 0 | 0 | 0 |
| virus_Aug5.75v_30379 | 8 | 0 | 1 | 0 | 0 | 0 | 1 | 0 | 0 | 0 | 0 | 0 | 0 | 0 | 0 | 0 | 0 | 0 | 0 | 0 | 0 | 0 | 0 | 0 | 0 | 0 | 1 | 0 | 0 | 0 | 0 | 0 | 2 | 0 | 0 | 0 | 0 | 0 | 0 | 0 | 0 | 0 | 0 | 0 | 0 | 0 | 1 | 0 | 1 | 0 | 1 | 0 | 0 | 0 | 0 |
| virus_May12.75v_8282 | 8 | 0 | 0 | 0 | 1 | 0 | 0 | 0 | 2 | 0 | 0 | 1 | 1 | 0 | 0 | 0 | 0 | 0 | 0 | 0 | 0 | 0 | 0 | 0 | 0 | 0 | 0 | 0 | 0 | 0 | 0 | 0 | 0 | 0 | 0 | 0 | 0 | 0 | 0 | 0 | 0 | 0 | 0 | 0 | 0 | 0 | 0 | 0 | 0 | 0 | 0 | 0 | 0 | 1 | 2 |
| virus_Aug5.0v_5090 | 8 | 0 | 0 | 0 | 1 | 0 | 0 | 0 | 0 | 0 | 0 | 0 | 0 | 0 | 0 | 0 | 0 | 0 | 1 | 0 | 0 | 0 | 0 | 0 | 0 | 0 | 0 | 0 | 0 | 0 | 1 | 0 | 2 | 0 | 0 | 0 | 0 | 0 | 2 | 0 | 0 | 0 | 0 | 0 | 0 | 0 | 0 | 0 | 0 | 1 | 0 | 0 | 0 | 0 | 0 |
| virus_Aug5.75v_24035 | 8 | 0 | 0 | 0 | 1 | 0 | 0 | 0 | 0 | 0 | 1 | 0 | 0 | 0 | 0 | 0 | 0 | 0 | 0 | 0 | 1 | 0 | 0 | 0 | 0 | 0 | 0 | 0 | 0 | 0 | 0 | 0 | 0 | 0 | 0 | 0 | 1 | 1 | 1 | 0 | 0 | 0 | 0 | 0 | 0 | 0 | 1 | 0 | 0 | 0 | 0 | 0 | 1 | 0 | 0 |
| bacteria_Aug5d25b_3120 | 7 | 0 | 1 | 0 | 0 | 0 | 0 | 0 | 0 | 0 | 0 | 0 | 0 | 0 | 0 | 0 | 1 | 0 | 0 | 0 | 0 | 0 | 0 | 0 | 0 | 1 | 0 | 0 | 0 | 0 | 0 | 0 | 0 | 0 | 0 | 0 | 0 | 0 | 0 | 0 | 0 | 1 | 0 | 1 | 0 | 0 | 0 | 0 | 0 | 1 | 0 | 0 | 1 | 0 | 0 |
| bacteria_Aug5d25b_6891 | 7 | 0 | 0 | 0 | 0 | 0 | 0 | 0 | 0 | 0 | 0 | 0 | 0 | 0 | 0 | 0 | 0 | 0 | 0 | 0 | 0 | 0 | 0 | 0 | 0 | 0 | 0 | 0 | 0 | 0 | 0 | 2 | 0 | 1 | 0 | 0 | 0 | 0 | 0 | 0 | 0 | 3 | 0 | 0 | 0 | 0 | 0 | 0 | 0 | 1 | 0 | 0 | 0 | 0 | 0 |
| bacteria_Jul11d25b_195 | 7 | 0 | 0 | 0 | 0 | 0 | 0 | 0 | 0 | 4 | 0 | 0 | 0 | 0 | 0 | 0 | 0 | 0 | 0 | 0 | 0 | 0 | 0 | 0 | 0 | 1 | 0 | 0 | 0 | 0 | 0 | 0 | 0 | 0 | 0 | 0 | 0 | 0 | 0 | 0 | 0 | 1 | 0 | 0 | 0 | 0 | 0 | 0 | 0 | 1 | 0 | 0 | 0 | 0 | 0 |
| bacteria_Jul11d25b_7263 | 7 | 0 | 0 | 0 | 0 | 1 | 0 | 0 | 1 | 0 | 0 | 0 | 0 | 0 | 0 | 0 | 0 | 0 | 0 | 0 | 0 | 0 | 0 | 0 | 0 | 2 | 0 | 0 | 0 | 0 | 0 | 0 | 0 | 0 | 0 | 0 | 0 | 0 | 0 | 0 | 0 | 1 | 0 | 0 | 0 | 0 | 0 | 0 | 1 | 0 | 0 | 0 | 0 | 0 | 1 |
| bacteria_Jul11d75b_1743 | 7 | 0 | 0 | 0 | 1 | 1 | 0 | 0 | 0 | 1 | 0 | 0 | 1 | 0 | 0 | 0 | 0 | 0 | 0 | 0 | 0 | 0 | 1 | 0 | 0 | 0 | 0 | 0 | 0 | 1 | 0 | 0 | 0 | 0 | 0 | 0 | 1 | 0 | 0 | 0 | 0 | 0 | 0 | 0 | 0 | 0 | 0 | 0 | 0 | 0 | 0 | 0 | 0 | 0 | 0 |
| bacteria_Jul11d75b_2479 | 7 | 0 | 0 | 0 | 0 | 0 | 0 | 0 | 1 | 0 | 0 | 0 | 0 | 0 | 0 | 0 | 0 | 0 | 0 | 0 | 0 | 0 | 0 | 0 | 0 | 0 | 1 | 0 | 1 | 2 | 0 | 0 | 0 | 1 | 0 | 0 | 0 | 0 | 0 | 0 | 0 | 0 | 0 | 0 | 1 | 0 | 0 | 0 | 0 | 0 | 0 | 0 | 0 | 0 | 0 |
| bacteria_Jun8d50b_14625 | 7 | 0 | 0 | 0 | 0 | 0 | 0 | 0 | 0 | 0 | 0 | 4 | 0 | 0 | 0 | 0 | 0 | 0 | 0 | 0 | 0 | 0 | 0 | 1 | 0 | 0 | 0 | 0 | 0 | 1 | 0 | 1 | 0 | 0 | 0 | 0 | 0 | 0 | 0 | 0 | 0 | 0 | 0 | 0 | 0 | 0 | 0 | 0 | 0 | 0 | 0 | 0 | 0 | 0 | 0 |
| bacteria_Jun22d75b_5443 | 7 | 0 | 0 | 0 | 0 | 0 | 0 | 0 | 0 | 0 | 0 | 0 | 0 | 0 | 0 | 0 | 0 | 0 | 1 | 1 | 0 | 0 | 0 | 0 | 0 | 0 | 0 | 0 | 0 | 0 | 0 | 0 | 1 | 0 | 0 | 0 | 0 | 1 | 1 | 0 | 0 | 0 | 2 | 0 | 0 | 0 | 0 | 0 | 0 | 0 | 0 | 0 | 0 | 0 | 0 |
| bacteria_May12d25b_152 | 7 | 0 | 0 | 3 | 0 | 0 | 0 | 0 | 0 | 2 | 0 | 0 | 0 | 0 | 0 | 0 | 0 | 0 | 0 | 0 | 0 | 0 | 0 | 0 | 0 | 0 | 0 | 0 | 0 | 0 | 0 | 0 | 0 | 1 | 0 | 1 | 0 | 0 | 0 | 0 | 0 | 0 | 0 | 0 | 0 | 0 | 0 | 0 | 0 | 0 | 0 | 0 | 0 | 0 | 0 |

[illegible]

**Supplementary Table S7**. Common Bacterio- and virioplankton RTPR 98% OTUs in 27 samples with RTPR amplicons in both fractions

Total seqs: 47997; Subtotal seqs in retained

OTUs: 41046

| OTUs with >=10 seqs: 254 | OTU subtotal | May12_05m | | May12_25m | | May12_50m | | May12_75m | | Jun8_25m | | Jun8_50m | | Jun8_75m | | Jun22_0m | | Jun22_50m | | Jun22_75m | | Jul11_0m | | Jul11_05m | | Jul11_25m | | Jul11_50m | | Jul11_75m | | Jul27_05m | | Jul27_25m | | Jul27_75m | | Aug5_0m | | Aug5_05m | | Aug5_25m | | Aug5_50m | | Aug5_75m | | Sep8_0m | | Sep8_05m | | Sep8_25m | | Sep8_50m | |
|---|---|---|---|---|---|---|---|---|---|---|---|---|---|---|---|---|---|---|---|---|---|---|---|---|---|---|---|---|---|---|---|---|---|---|---|---|---|---|---|---|---|---|---|---|---|---|---|---|---|---|---|---|---|---|---|
| | | bacterioplankton | virioplankton | bacterioplankton | virioplankton | bacterioplankton | virioplankton | bacterioplankton | virioplankton | bacterioplankton | virioplankton | bacterioplankton | virioplankton | bacterioplankton | virioplankton | bacterioplankton | virioplankton | bacterioplankton | virioplankton | bacterioplankton | virioplankton | bacterioplankton | virioplankton | bacterioplankton | virioplankton | bacterioplankton | virioplankton | bacterioplankton | virioplankton | bacterioplankton | virioplankton | bacterioplankton | virioplankton | bacterioplankton | virioplankton | bacterioplankton | virioplankton | bacterioplankton | virioplankton | bacterioplankton | virioplankton | bacterioplankton | virioplankton | bacterioplankton | virioplankton | bacterioplankton | virioplankton | bacterioplankton | virioplankton | bacterioplankton | virioplankton | bacterioplankton | virioplankton | bacterioplankton | virioplankton |
| bacteria_Jun8d25b_37071 | 3 | 0 | 0 | 0 | 0 | 0 | 0 | 0 | 0 | 1 | 0 | 1 | 0 | 0 | 0 | 0 | 0 | 0 | 0 | 0 | 0 | 0 | 0 | 1 | 0 | 0 | 0 | [illegible] | [illegible] | [illegible] | [illegible] | [illegible] | [illegible] | [illegible] | [illegible] | [illegible] | [illegible] | [illegible] | [illegible] | [illegible] | [illegible] | [illegible] | [illegible] | [illegible] | [illegible] | [illegible] | [illegible] | [illegible] | [illegible] | [illegible] | [illegible] | [illegible] | [illegible] | [illegible] | [illegible] |
| bacteria_Jun8d25b_49086 | 3 | 0 | 0 | 0 | 0 | 0 | 0 | 0 | 0 | 1 | 0 | 0 | 0 | 0 | 0 | 0 | 0 | 0 | 0 | 0 | 0 | 0 | 0 | 0 | 0 | 0 | 0 | [illegible] | [illegible] | [illegible] | [illegible] | [illegible] | [illegible] | [illegible] | [illegible] | [illegible] | [illegible] | [illegible] | [illegible] | [illegible] | [illegible] | [illegible] | [illegible] | [illegible] | [illegible] | [illegible] | [illegible] | [illegible] | [illegible] | [illegible] | [illegible] | [illegible] | [illegible] | [illegible] | [illegible] |
| bacteria_Jun8d75b_42996 | 3 | 0 | 0 | 0 | 0 | 0 | 0 | 0 | 0 | 0 | 0 | 0 | 0 | 2 | 0 | 0 | 0 | 0 | 0 | 0 | 0 | 0 | 0 | 0 | 0 | 0 | 0 | [illegible] | [illegible] | [illegible] | [illegible] | [illegible] | [illegible] | [illegible] | [illegible] | [illegible] | [illegible] | [illegible] | [illegible] | [illegible] | [illegible] | [illegible] | [illegible] | [illegible] | [illegible] | [illegible] | [illegible] | [illegible] | [illegible] | [illegible] | [illegible] | [illegible] | [illegible] | [illegible] | [illegible] |
| bacteria_Jun8d75b_64247 | 3 | 0 | 0 | 0 | 0 | 0 | 0 | 0 | 1 | 0 | 0 | 0 | 0 | 1 | 0 | 0 | 0 | 0 | 0 | 0 | 0 | 0 | 0 | 0 | 0 | 0 | 0 | [illegible] | [illegible] | [illegible] | [illegible] | [illegible] | [illegible] | [illegible] | [illegible] | [illegible] | [illegible] | [illegible] | [illegible] | [illegible] | [illegible] | [illegible] | [illegible] | [illegible] | [illegible] | [illegible] | [illegible] | [illegible] | [illegible] | [illegible] | [illegible] | [illegible] | [illegible] | [illegible] | [illegible] |
| bacteria_Jun22d0b_15273 | 3 | 0 | 0 | 1 | 0 | 0 | 0 | 0 | 0 | 0 | 0 | 0 | 0 | 0 | 0 | 1 | 0 | 0 | 0 | 0 | 0 | 0 | 0 | 0 | 0 | 0 | 0 | [illegible] | [illegible] | [illegible] | [illegible] | [illegible] | [illegible] | [illegible] | [illegible] | [illegible] | [illegible] | [illegible] | [illegible] | [illegible] | [illegible] | [illegible] | [illegible] | [illegible] | [illegible] | [illegible] | [illegible] | [illegible] | [illegible] | [illegible] | [illegible] | [illegible] | [illegible] | [illegible] | [illegible] |
| bacteria_May12d25b_622 | 3 | 0 | 0 | 1 | 0 | 0 | 0 | 0 | 0 | 2 | 0 | 0 | 0 | 0 | 0 | 0 | 0 | 0 | 0 | 0 | 0 | 0 | 0 | 0 | 0 | 0 | 0 | [illegible] | [illegible] | [illegible] | [illegible] | [illegible] | [illegible] | [illegible] | [illegible] | [illegible] | [illegible] | [illegible] | [illegible] | [illegible] | [illegible] | [illegible] | [illegible] | [illegible] | [illegible] | [illegible] | [illegible] | [illegible] | [illegible] | [illegible] | [illegible] | [illegible] | [illegible] | [illegible] | [illegible] |
| bacteria_May12d50b_228 | 3 | 0 | 0 | 0 | 0 | 3 | 0 | 0 | 0 | 0 | 0 | 0 | 0 | 0 | 0 | 0 | 0 | 0 | 0 | 0 | 0 | 0 | 0 | 0 | 0 | 0 | 0 | [illegible] | [illegible] | [illegible] | [illegible] | [illegible] | [illegible] | [illegible] | [illegible] | [illegible] | [illegible] | [illegible] | [illegible] | [illegible] | [illegible] | [illegible] | [illegible] | [illegible] | [illegible] | [illegible] | [illegible] | [illegible] | [illegible] | [illegible] | [illegible] | [illegible] | [illegible] | [illegible] | [illegible] |
| bacteria_Sep8d0b_37415 | 3 | 0 | 0 | 0 | 0 | 0 | 0 | 0 | 0 | 0 | 0 | 0 | 0 | 1 | 0 | 0 | 0 | 0 | 0 | 0 | 0 | 0 | 0 | 0 | 0 | 0 | 0 | [illegible] | [illegible] | [illegible] | [illegible] | [illegible] | [illegible] | [illegible] | [illegible] | [illegible] | [illegible] | [illegible] | [illegible] | [illegible] | [illegible] | [illegible] | [illegible] | [illegible] | [illegible] | [illegible] | [illegible] | [illegible] | [illegible] | [illegible] | [illegible] | [illegible] | [illegible] | [illegible] | [illegible] |
| virus_Aug5.0v_1120 | 3 | 0 | 0 | 0 | 0 | 0 | 0 | 0 | 0 | 0 | 0 | 0 | 0 | 0 | 0 | 0 | 0 | 0 | 1 | 0 | 0 | 0 | 0 | 0 | 0 | 0 | 0 | [illegible] | [illegible] | [illegible] | [illegible] | [illegible] | [illegible] | [illegible] | [illegible] | [illegible] | [illegible] | [illegible] | [illegible] | [illegible] | [illegible] | [illegible] | [illegible] | [illegible] | [illegible] | [illegible] | [illegible] | [illegible] | [illegible] | [illegible] | [illegible] | [illegible] | [illegible] | [illegible] | [illegible] |
| virus_Aug5.0v_5038 | 3 | 0 | 0 | 0 | 0 | 0 | 0 | 0 | 0 | 0 | 0 | 0 | 0 | 0 | 0 | 0 | 0 | 0 | 0 | 0 | 0 | 0 | 0 | 0 | 0 | 0 | 0 | [illegible] | [illegible] | [illegible] | [illegible] | [illegible] | [illegible] | [illegible] | [illegible] | [illegible] | [illegible] | [illegible] | [illegible] | [illegible] | [illegible] | [illegible] | [illegible] | [illegible] | [illegible] | [illegible] | [illegible] | [illegible] | [illegible] | [illegible] | [illegible] | [illegible] | [illegible] | [illegible] | [illegible] |
| virus_Aug5.0v_5057 | 3 | 0 | 0 | 0 | 0 | 0 | 0 | 0 | 0 | 0 | 0 | 0 | 0 | 0 | 0 | 0 | 0 | 0 | 0 | 0 | 0 | 0 | 0 | 0 | 0 | 0 | 0 | [illegible] | [illegible] | [illegible] | [illegible] | [illegible] | [illegible] | [illegible] | [illegible] | [illegible] | [illegible] | [illegible] | [illegible] | [illegible] | [illegible] | [illegible] | [illegible] | [illegible] | [illegible] | [illegible] | [illegible] | [illegible] | [illegible] | [illegible] | [illegible] | [illegible] | [illegible] | [illegible] | [illegible] |
| virus_Aug5.25v_36661 | 3 | 0 | 0 | 0 | 0 | 0 | 0 | 0 | 0 | 0 | 0 | 0 | 0 | 0 | 0 | 0 | 0 | 0 | 0 | 0 | 0 | 0 | 0 | 0 | 0 | 1 | 0 | [illegible] | [illegible] | [illegible] | [illegible] | [illegible] | [illegible] | [illegible] | [illegible] | [illegible] | [illegible] | [illegible] | [illegible] | [illegible] | [illegible] | [illegible] | [illegible] | [illegible] | [illegible] | [illegible] | [illegible] | [illegible] | [illegible] | [illegible] | [illegible] | [illegible] | [illegible] | [illegible] | [illegible] |
| virus_Aug5.50v_37809 | 3 | 0 | 0 | 0 | 0 | 0 | 0 | 0 | 0 | 0 | 0 | 0 | 0 | 0 | 0 | 0 | 0 | 0 | 0 | 0 | 0 | 0 | 0 | 0 | 0 | 0 | 0 | [illegible] | [illegible] | [illegible] | [illegible] | [illegible] | [illegible] | [illegible] | [illegible] | [illegible] | [illegible] | [illegible] | [illegible] | [illegible] | [illegible] | [illegible] | [illegible] | [illegible] | [illegible] | [illegible] | [illegible] | [illegible] | [illegible] | [illegible] | [illegible] | [illegible] | [illegible] | [illegible] | [illegible] |
| virus_Aug5.75v_17455 | 3 | 0 | 0 | 0 | 0 | 0 | 0 | 0 | 1 | 0 | 0 | 0 | 0 | 0 | 0 | 0 | 0 | 0 | 0 | 0 | 0 | 0 | 0 | 0 | 0 | 0 | 0 | [illegible] | [illegible] | [illegible] | [illegible] | [illegible] | [illegible] | [illegible] | [illegible] | [illegible] | [illegible] | [illegible] | [illegible] | [illegible] | [illegible] | [illegible] | [illegible] | [illegible] | [illegible] | [illegible] | [illegible] | [illegible] | [illegible] | [illegible] | [illegible] | [illegible] | [illegible] | [illegible] | [illegible] |
| virus_Jul11.5v_1149 | 3 | 0 | 0 | 0 | 0 | 0 | 0 | 0 | 0 | 0 | 0 | 0 | 0 | 0 | 0 | 0 | 0 | 0 | 0 | 0 | 0 | 0 | 0 | 0 | 1 | 0 | 0 | [illegible] | [illegible] | [illegible] | [illegible] | [illegible] | [illegible] | [illegible] | [illegible] | [illegible] | [illegible] | [illegible] | [illegible] | [illegible] | [illegible] | [illegible] | [illegible] | [illegible] | [illegible] | [illegible] | [illegible] | [illegible] | [illegible] | [illegible] | [illegible] | [illegible] | [illegible] | [illegible] | [illegible] |
| virus_Jul11.5v_3354 | 3 | 0 | 0 | 0 | 0 | 0 | 1 | 0 | 0 | 0 | 0 | 0 | 0 | 0 | 0 | 0 | 0 | 0 | 0 | 0 | 0 | 0 | 0 | 0 | 2 | 0 | 0 | [illegible] | [illegible] | [illegible] | [illegible] | [illegible] | [illegible] | [illegible] | [illegible] | [illegible] | [illegible] | [illegible] | [illegible] | [illegible] | [illegible] | [illegible] | [illegible] | [illegible] | [illegible] | [illegible] | [illegible] | [illegible] | [illegible] | [illegible] | [illegible] | [illegible] | [illegible] | [illegible] | [illegible] |
| virus_Jul11.25v_13997 | 3 | 0 | 0 | 0 | 0 | 0 | 0 | 0 | 1 | 0 | 0 | 0 | 0 | 0 | 0 | 0 | 0 | 0 | 1 | 0 | 0 | 0 | 0 | 0 | 0 | 0 | 1 | [illegible] | [illegible] | [illegible] | [illegible] | [illegible] | [illegible] | [illegible] | [illegible] | [illegible] | [illegible] | [illegible] | [illegible] | [illegible] | [illegible] | [illegible] | [illegible] | [illegible] | [illegible] | [illegible] | [illegible] | [illegible] | [illegible] | [illegible] | [illegible] | [illegible] | [illegible] | [illegible] | [illegible] |
| virus_Jul11.75v_32525 | 3 | 0 | 0 | 0 | 1 | 0 | 0 | 0 | 0 | 0 | 1 | 0 | 0 | 0 | 0 | 0 | 0 | 0 | 0 | 0 | 0 | 0 | 0 | 0 | 0 | 0 | 0 | [illegible] | [illegible] | [illegible] | [illegible] | [illegible] | [illegible] | [illegible] | [illegible] | [illegible] | [illegible] | [illegible] | [illegible] | [illegible] | [illegible] | [illegible] | [illegible] | [illegible] | [illegible] | [illegible] | [illegible] | [illegible] | [illegible] | [illegible] | [illegible] | [illegible] | [illegible] | [illegible] | [illegible] |
| virus_Jul27.5v_2368 | 3 | 0 | 0 | 0 | 0 | 0 | 0 | 0 | 0 | 0 | 0 | 0 | 0 | 0 | 0 | 0 | 0 | 0 | 1 | 0 | 0 | 0 | 0 | 0 | 0 | 0 | 0 | [illegible] | [illegible] | [illegible] | [illegible] | [illegible] | [illegible] | [illegible] | [illegible] | [illegible] | [illegible] | [illegible] | [illegible] | [illegible] | [illegible] | [illegible] | [illegible] | [illegible] | [illegible] | [illegible] | [illegible] | [illegible] | [illegible] | [illegible] | [illegible] | [illegible] | [illegible] | [illegible] | [illegible] |
| virus_Jul27.25v_13490 | 3 | 0 | 0 | 0 | 0 | 0 | 0 | 0 | 0 | 0 | 0 | 0 | 0 | 0 | 0 | 1 | 0 | 0 | 0 | 0 | 0 | 0 | 0 | 0 | 0 | 0 | 0 | [illegible] | [illegible] | [illegible] | [illegible] | [illegible] | [illegible] | [illegible] | [illegible] | [illegible] | [illegible] | [illegible] | [illegible] | [illegible] | [illegible] | [illegible] | [illegible] | [illegible] | [illegible] | [illegible] | [illegible] | [illegible] | [illegible] | [illegible] | [illegible] | [illegible] | [illegible] | [illegible] | [illegible] |
| virus_Jul27.25v_27552 | 3 | 0 | 0 | 0 | 0 | 0 | 0 | 0 | 0 | 0 | 0 | 0 | 0 | 0 | 0 | 0 | 0 | 0 | 0 | 0 | 0 | 0 | 0 | 0 | 0 | 0 | 0 | [illegible] | [illegible] | [illegible] | [illegible] | [illegible] | [illegible] | [illegible] | [illegible] | [illegible] | [illegible] | [illegible] | [illegible] | [illegible] | [illegible] | [illegible] | [illegible] | [illegible] | [illegible] | [illegible] | [illegible] | [illegible] | [illegible] | [illegible] | [illegible] | [illegible] | [illegible] | [illegible] | [illegible] |
| virus_Jun22.0v_1288 | 3 | 0 | 0 | 0 | 0 | 0 | 0 | 0 | 0 | 0 | 0 | 0 | 0 | 0 | 0 | 0 | 1 | 0 | 1 | 0 | 0 | 0 | 0 | 1 | 0 | 0 | 0 | [illegible] | [illegible] | [illegible] | [illegible] | [illegible] | [illegible] | [illegible] | [illegible] | [illegible] | [illegible] | [illegible] | [illegible] | [illegible] | [illegible] | [illegible] | [illegible] | [illegible] | [illegible] | [illegible] | [illegible] | [illegible] | [illegible] | [illegible] | [illegible] | [illegible] | [illegible] | [illegible] | [illegible] |
| virus_Jun22.50v_2361 | 3 | 0 | 0 | 0 | 0 | 0 | 0 | 0 | 0 | 0 | 0 | 0 | 0 | 0 | 0 | 0 | 0 | 0 | 1 | 0 | 0 | 0 | 0 | 0 | 0 | 0 | 0 | [illegible] | [illegible] | [illegible] | [illegible] | [illegible] | [illegible] | [illegible] | [illegible] | [illegible] | [illegible] | [illegible] | [illegible] | [illegible] | [illegible] | [illegible] | [illegible] | [illegible] | [illegible] | [illegible] | [illegible] | [illegible] | [illegible] | [illegible] | [illegible] | [illegible] | [illegible] | [illegible] | [illegible] |
| virus_May12.50v_8935 | 3 | 0 | 0 | 0 | 0 | 0 | 1 | 0 | 0 | 0 | 0 | 0 | 0 | 0 | 0 | 0 | 0 | 0 | 0 | 0 | 0 | 0 | 0 | 0 | 0 | 0 | 0 | [illegible] | [illegible] | [illegible] | [illegible] | [illegible] | [illegible] | [illegible] | [illegible] | [illegible] | [illegible] | [illegible] | [illegible] | [illegible] | [illegible] | [illegible] | [illegible] | [illegible] | [illegible] | [illegible] | [illegible] | [illegible] | [illegible] | [illegible] | [illegible] | [illegible] | [illegible] | [illegible] | [illegible] |
| virus_May12.50v_26155 | 3 | 0 | 0 | 0 | 0 | 0 | 1 | 0 | 2 | 0 | 0 | 0 | 0 | 0 | 0 | 0 | 0 | 0 | 0 | 0 | 0 | 0 | 0 | 0 | 0 | 0 | 0 | [illegible] | [illegible] | [illegible] | [illegible] | [illegible] | [illegible] | [illegible] | [illegible] | [illegible] | [illegible] | [illegible] | [illegible] | [illegible] | [illegible] | [illegible] | [illegible] | [illegible] | [illegible] | [illegible] | [illegible] | [illegible] | [illegible] | [illegible] | [illegible] | [illegible] | [illegible] | [illegible] | [illegible] |
| virus_May12.75v_24463 | 3 | 0 | 1 | 0 | 0 | 0 | 0 | 0 | 1 | 0 | 0 | 0 | 0 | 0 | 0 | 0 | 0 | 0 | 0 | 0 | 0 | 0 | 0 | 0 | 0 | 0 | 1 | [illegible] | [illegible] | [illegible] | [illegible] | [illegible] | [illegible] | [illegible] | [illegible] | [illegible] | [illegible] | [illegible] | [illegible] | [illegible] | [illegible] | [illegible] | [illegible] | [illegible] | [illegible] | [illegible] | [illegible] | [illegible] | [illegible] | [illegible] | [illegible] | [illegible] | [illegible] | [illegible] | [illegible] |
| virus_Sep8.0v_1170 | 3 | 0 | 0 | 0 | 0 | 0 | 0 | 0 | 0 | 0 | 0 | 0 | 0 | 0 | 0 | 0 | 0 | 0 | 0 | 0 | 0 | 0 | 0 | 1 | 0 | 0 | 0 | [illegible] | [illegible] | [illegible] | [illegible] | [illegible] | [illegible] | [illegible] | [illegible] | [illegible] | [illegible] | [illegible] | [illegible] | [illegible] | [illegible] | [illegible] | [illegible] | [illegible] | [illegible] | [illegible] | [illegible] | [illegible] | [illegible] | [illegible] | [illegible] | [illegible] | [illegible] | [illegible] | [illegible] |
| virus_Sep8.0v_18231 | 3 | 0 | 0 | 0 | 0 | 0 | 0 | 0 | 0 | 0 | 0 | 0 | 0 | 0 | 0 | 0 | 0 | 0 | 0 | 0 | 0 | 0 | 0 | 0 | 0 | 0 | 0 | [illegible] | [illegible] | [illegible] | [illegible] | [illegible] | [illegible] | [illegible] | [illegible] | [illegible] | [illegible] | [illegible] | [illegible] | [illegible] | [illegible] | [illegible] | [illegible] | [illegible] | [illegible] | [illegible] | [illegible] | [illegible] | [illegible] | [illegible] | [illegible] | [illegible] | [illegible] | [illegible] | [illegible] |
| virus_Sep8.0v_20528 | 3 | 0 | 0 | 0 | 0 | 0 | 1 | 0 | 0 | 0 | 1 | 0 | 0 | 0 | 0 | 0 | 0 | 0 | 0 | 0 | 0 | 0 | 0 | 0 | 0 | 0 | 0 | [illegible] | [illegible] | [illegible] | [illegible] | [illegible] | [illegible] | [illegible] | [illegible] | [illegible] | [illegible] | [illegible] | [illegible] | [illegible] | [illegible] | [illegible] | [illegible] | [illegible] | [illegible] | [illegible] | [illegible] | [illegible] | [illegible] | [illegible] | [illegible] | [illegible] | [illegible] | [illegible] | [illegible] |
| virus_Sep8.50v_32593 | 3 | 0 | 0 | 0 | 0 | 0 | 0 | 0 | 0 | 0 | 0 | 0 | 0 | 0 | 0 | 0 | 0 | 0 | 0 | 0 | 0 | 0 | 0 | 0 | 1 | 0 | 0 | [illegible] | [illegible] | [illegible] | [illegible] | [illegible] | [illegible] | [illegible] | [illegible] | [illegible] | [illegible] | [illegible] | [illegible] | [illegible] | [illegible] | [illegible] | [illegible] | [illegible] | [illegible] | [illegible] | [illegible] | [illegible] | [illegible] | [illegible] | [illegible] | [illegible] | [illegible] | [illegible] | [illegible] |
| bacteria_Jul11d75b_3897 | 3 | 0 | 0 | 0 | 0 | 0 | 0 | 0 | 0 | 0 | 0 | 0 | 0 | 0 | 0 | 0 | 0 | 0 | 0 | 0 | 0 | 0 | 0 | 0 | 0 | 0 | 0 | [illegible] | [illegible] | [illegible] | [illegible] | [illegible] | [illegible] | [illegible] | [illegible] | [illegible] | [illegible] | [illegible] | [illegible] | [illegible] | [illegible] | [illegible] | [illegible] | [illegible] | [illegible] | [illegible] | [illegible] | [illegible] | [illegible] | [illegible] | [illegible] | [illegible] | [illegible] | [illegible] | [illegible] |
| bacteria_Jul27d75b_3855 | 3 | 0 | 0 | 0 | 0 | 0 | 0 | 0 | 0 | 0 | 0 | 0 | 0 | 0 | 0 | 0 | 0 | 0 | 0 | 0 | 0 | 0 | 0 | 0 | 0 | 0 | 0 | [illegible] | [illegible] | [illegible] | [illegible] | [illegible] | [illegible] | [illegible] | [illegible] | [illegible] | [illegible] | [illegible] | [illegible] | [illegible] | [illegible] | [illegible] | [illegible] | [illegible] | [illegible] | [illegible] | [illegible] | [illegible] | [illegible] | [illegible] | [illegible] | [illegible] | [illegible] | [illegible] | [illegible] |
| bacteria_Jul27d75b_7174 | 3 | 0 | 0 | 1 | 0 | 1 | 0 | 0 | 0 | 0 | 0 | 0 | 0 | 0 | 0 | 0 | 0 | 0 | 0 | 0 | 0 | 0 | 0 | 0 | 0 | 0 | 0 | [illegible] | [illegible] | [illegible] | [illegible] | [illegible] | [illegible] | [illegible] | [illegible] | [illegible] | [illegible] | [illegible] | [illegible] | [illegible] | [illegible] | [illegible] | [illegible] | [illegible] | [illegible] | [illegible] | [illegible] | [illegible] | [illegible] | [illegible] | [illegible] | [illegible] | [illegible] | [illegible] | [illegible] |
| bacteria_Jun8d25b_64582 | 3 | 0 | 0 | 0 | 0 | 0 | 0 | 0 | 0 | 1 | 0 | 0 | 0 | 0 | 0 | 0 | 0 | 1 | 0 | 1 | 0 | 0 | 0 | 0 | 0 | 0 | 0 | [illegible] | [illegible] | [illegible] | [illegible] | [illegible] | [illegible] | [illegible] | [illegible] | [illegible] | [illegible] | [illegible] | [illegible] | [illegible] | [illegible] | [illegible] | [illegible] | [illegible] | [illegible] | [illegible] | [illegible] | [illegible] | [illegible] | [illegible] | [illegible] | [illegible] | [illegible] | [illegible] | [illegible] |
| bacteria_Jun22d0b_73272 | 3 | 0 | 0 | 0 | 0 | 0 | 0 | 0 | 0 | 0 | 0 | 0 | 0 | 0 | 0 | 1 | 0 | 0 | 0 | 0 | 0 | 0 | 0 | 1 | 0 | 0 | 0 | [illegible] | [illegible] | [illegible] | [illegible] | [illegible] | [illegible] | [illegible] | [illegible] | [illegible] | [illegible] | [illegible] | [illegible] | [illegible] | [illegible] | [illegible] | [illegible] | [illegible] | [illegible] | [illegible] | [illegible] | [illegible] | [illegible] | [illegible] | [illegible] | [illegible] | [illegible] | [illegible] | [illegible] |
| bacteria_Jun22d50b_3319 | 3 | 0 | 0 | 0 | 0 | 0 | 0 | 0 | 0 | 0 | 0 | 0 | 0 | 0 | 0 | 0 | 0 | 1 | 0 | 0 | 0 | 0 | 0 | 0 | 0 | 0 | 0 | [illegible] | [illegible] | [illegible] | [illegible] | [illegible] | [illegible] | [illegible] | [illegible] | [illegible] | [illegible] | [illegible] | [illegible] | [illegible] | [illegible] | [illegible] | [illegible] | [illegible] | [illegible] | [illegible] | [illegible] | [illegible] | [illegible] | [illegible] | [illegible] | [illegible] | [illegible] | [illegible] | [illegible] |
| bacteria_May12d25b_163 | 3 | 0 | 0 | 2 | 0 | 0 | 0 | 0 | 0 | 1 | 0 | 0 | 0 | 0 | 0 | 0 | 0 | 0 | 0 | 0 | 0 | 0 | 0 | 0 | 0 | 0 | 0 | [illegible] | [illegible] | [illegible] | [illegible] | [illegible] | [illegible] | [illegible] | [illegible] | [illegible] | [illegible] | [illegible] | [illegible] | [illegible] | [illegible] | [illegible] | [illegible] | [illegible] | [illegible] | [illegible] | [illegible] | [illegible] | [illegible] | [illegible] | [illegible] | [illegible] | [illegible] | [illegible] | [illegible] |
| bacteria_May12d50b_350 | 3 | 0 | 0 | 0 | 0 | 3 | 0 | 0 | 0 | 0 | 0 | 0 | 0 | 0 | 0 | 0 | 0 | 0 | 0 | 0 | 0 | 0 | 0 | 0 | 0 | 0 | 0 | [illegible] | [illegible] | [illegible] | [illegible] | [illegible] | [illegible] | [illegible] | [illegible] | [illegible] | [illegible] | [illegible] | [illegible] | [illegible] | [illegible] | [illegible] | [illegible] | [illegible] | [illegible] | [illegible] | [illegible] | [illegible] | [illegible] | [illegible] | [illegible] | [illegible] | [illegible] | [illegible] | [illegible] |
| virus_Aug5.0v_1060 | 3 | 0 | 0 | 0 | 0 | 0 | 0 | 0 | 0 | 0 | 0 | 0 | 1 | 0 | 0 | 0 | 0 | 0 | 0 | 0 | 0 | 0 | 0 | 0 | 0 | 0 | 0 | [illegible] | [illegible] | [illegible] | [illegible] | [illegible] | [illegible] | [illegible] | [illegible] | [illegible] | [illegible] | [illegible] | [illegible] | [illegible] | [illegible] | [illegible] | [illegible] | [illegible] | [illegible] | [illegible] | [illegible] | [illegible] | [illegible] | [illegible] | [illegible] | [illegible] | [illegible] | [illegible] | [illegible] |
| virus_Aug5.50v_3350 | 3 | 0 | 0 | 0 | 0 | 0 | 0 | 0 | 0 | 0 | 0 | 0 | 0 | 0 | 0 | 0 | 0 | 0 | 0 | 0 | 0 | 0 | 0 | 0 | 0 | 0 | 1 | [illegible] | [illegible] | [illegible] | [illegible] | [illegible] | [illegible] | [illegible] | [illegible] | [illegible] | [illegible] | [illegible] | [illegible] | [illegible] | [illegible] | [illegible] | [illegible] | [illegible] | [illegible] | [illegible] | [illegible] | [illegible] | [illegible] | [illegible] | [illegible] | [illegible] | [illegible] | [illegible] | [illegible] |
| virus_Jul11.25v_16470 | 3 | 0 | 0 | 0 | 0 | 0 | 2 | 0 | 0 | 0 | 0 | 0 | 0 | 0 | 0 | 0 | 0 | 0 | 0 | 0 | 0 | 0 | 0 | 0 | 0 | 0 | 1 | [illegible] | [illegible] | [illegible] | [illegible] | [illegible] | [illegible] | [illegible] | [illegible] | [illegible] | [illegible] | [illegible] | [illegible] | [illegible] | [illegible] | [illegible] | [illegible] | [illegible] | [illegible] | [illegible] | [illegible] | [illegible] | [illegible] | [illegible] | [illegible] | [illegible] | [illegible] | [illegible] | [illegible] |
| virus_Jul11.50v_17700 | 3 | 0 | 0 | 0 | 0 | 0 | 0 | 0 | 0 | 0 | 0 | 0 | 0 | 0 | 0 | 0 | 0 | 0 | 0 | 0 | 0 | 0 | 1 | 0 | 0 | 0 | 0 | [illegible] | [illegible] | [illegible] | [illegible] | [illegible] | [illegible] | [illegible] | [illegible] | [illegible] | [illegible] | [illegible] | [illegible] | [illegible] | [illegible] | [illegible] | [illegible] | [illegible] | [illegible] | [illegible] | [illegible] | [illegible] | [illegible] | [illegible] | [illegible] | [illegible] | [illegible] | [illegible] | [illegible] |
| virus_Jul11.75v_36830 | 3 | 0 | 0 | 0 | 0 | 0 | 0 | 0 | 1 | 0 | 0 | 0 | 0 | 0 | 0 | 0 | 0 | 0 | 0 | 0 | 0 | 0 | 0 | 0 | 0 | 0 | 0 | [illegible] | [illegible] | [illegible] | [illegible] | [illegible] | [illegible] | [illegible] | [illegible] | [illegible] | [illegible] | [illegible] | [illegible] | [illegible] | [illegible] | [illegible] | [illegible] | [illegible] | [illegible] | [illegible] | [illegible] | [illegible] | [illegible] | [illegible] | [illegible] | [illegible] | [illegible] | [illegible] | [illegible] |
| virus_Jul27.5v_8684 | 3 | 0 | 0 | 0 | 0 | 0 | 0 | 0 | 0 | 0 | 0 | 0 | 0 | 0 | 0 | 0 | 0 | 0 | 0 | 0 | 0 | 0 | 0 | 0 | 0 | 0 | 0 | [illegible] | [illegible] | [illegible] | [illegible] | [illegible] | [illegible] | [illegible] | [illegible] | [illegible] | [illegible] | [illegible] | [illegible] | [illegible] | [illegible] | [illegible] | [illegible] | [illegible] | [illegible] | [illegible] | [illegible] | [illegible] | [illegible] | [illegible] | [illegible] | [illegible] | [illegible] | [illegible] | [illegible] |
| virus_Jul27.5v_12484 | 3 | 0 | 0 | 0 | 0 | 0 | 0 | 0 | 0 | 1 | 0 | 0 | 0 | 0 | 0 | 0 | 0 | 0 | 0 | 0 | 0 | 0 | 0 | 0 | 0 | 0 | 0 | [illegible] | [illegible] | [illegible] | [illegible] | [illegible] | [illegible] | [illegible] | [illegible] | [illegible] | [illegible] | [illegible] | [illegible] | [illegible] | [illegible] | [illegible] | [illegible] | [illegible] | [illegible] | [illegible] | [illegible] | [illegible] | [illegible] | [illegible] | [illegible] | [illegible] | [illegible] | [illegible] | [illegible] |
| virus_Jul27.25v_2238 | 3 | 0 | 0 | 0 | 0 | 0 | 0 | 0 | 0 | 0 | 0 | 0 | 0 | 0 | 0 | 0 | 0 | 0 | 0 | 0 | 0 | 0 | 0 | 0 | 0 | 0 | 0 | [illegible] | [illegible] | [illegible] | [illegible] | [illegible] | [illegible] | [illegible] | [illegible] | [illegible] | [illegible] | [illegible] | [illegible] | [illegible] | [illegible] | [illegible] | [illegible] | [illegible] | [illegible] | [illegible] | [illegible] | [illegible] | [illegible] | [illegible] | [illegible] | [illegible] | [illegible] | [illegible] | [illegible] |
| virus_Jun8.50v_20040 | 3 | 0 | 1 | 0 | 0 | 1 | 0 | 0 | 0 | 0 | 0 | 0 | 1 | 0 | 0 | 0 | 0 | 0 | 0 | 0 | 0 | 0 | 0 | 0 | 0 | 0 | 0 | [illegible] | [illegible] | [illegible] | [illegible] | [illegible] | [illegible] | [illegible] | [illegible] | [illegible] | [illegible] | [illegible] | [illegible] | [illegible] | [illegible] | [illegible] | [illegible] | [illegible] | [illegible] | [illegible] | [illegible] | [illegible] | [illegible] | [illegible] | [illegible] | [illegible] | [illegible] | [illegible] | [illegible] |
| virus_Jun8.75v_6822 | 3 | 0 | 0 | 0 | 0 | 0 | 0 | 0 | 0 | 0 | 0 | 0 | 0 | 0 | 1 | 0 | 1 | 0 | 0 | 0 | 0 | 0 | 0 | 0 | 0 | 0 | 0 | [illegible] | [illegible] | [illegible] | [illegible] | [illegible] | [illegible] | [illegible] | [illegible] | [illegible] | [illegible] | [illegible] | [illegible] | [illegible] | [illegible] | [illegible] | [illegible] | [illegible] | [illegible] | [illegible] | [illegible] | [illegible] | [illegible] | [illegible] | [illegible] | [illegible] | [illegible] | [illegible] | [illegible] |
| virus_Jun22.50v_10904 | 3 | 0 | 0 | 0 | 0 | 0 | 0 | 0 | 0 | 0 | 0 | 0 | 0 | 0 | 0 | 0 | 0 | 0 | 1 | 0 | 0 | 0 | 0 | 0 | 0 | 0 | 0 | [illegible] | [illegible] | [illegible] | [illegible] | [illegible] | [illegible] | [illegible] | [illegible] | [illegible] | [illegible] | [illegible] | [illegible] | [illegible] | [illegible] | [illegible] | [illegible] | [illegible] | [illegible] | [illegible] | [illegible] | [illegible] | [illegible] | [illegible] | [illegible] | [illegible] | [illegible] | [illegible] | [illegible] |

**Supplementary Table S7**. Common Bacterio- and virioplankton RTPR 98% OTUs in 27 samples with RTPR amplicons in both fractions

Total seqs: 47997; Subtotal seqs in retained

OTUs: 41046

| | | May12_05m | | May12_25m | | May12_50m | | May12_75m | | Jun8_25m | | Jun8_50m | | Jun8_75m | | Jun22_0m | | Jun22_50m | | Jun22_75m | | Jul11_0m | | Jul11_05m | | Jul11_25m | | Jul11_50m | | Jul11_75m | | Jul27_05m | | Jul27_25m | | Jul27_75m | | Aug5_0m | | Aug5_05m | | Aug5_25m | | Aug5_50m | | Aug5_75m | | Sep8_0m | | Sep8_05m | | Sep8_25m | | Sep8_50m | |
|---|---|---|---|---|---|---|---|---|---|---|---|---|---|---|---|---|---|---|---|---|---|---|---|---|---|---|---|---|---|---|---|---|---|---|---|---|---|---|---|---|---|---|---|---|---|---|---|---|---|---|---|---|---|---|---|
| OTUs with >=10 seqs: 254 OTU | subtotal | bacterioplankton | virioplankton | bacterioplankton | virioplankton | bacterioplankton | virioplankton | bacterioplankton | virioplankton | bacterioplankton | virioplankton | bacterioplankton | virioplankton | bacterioplankton | virioplankton | bacterioplankton | virioplankton | bacterioplankton | virioplankton | bacterioplankton | virioplankton | bacterioplankton | virioplankton | bacterioplankton | virioplankton | bacterioplankton | virioplankton | bacterioplankton | virioplankton | bacterioplankton | virioplankton | bacterioplankton | virioplankton | bacterioplankton | virioplankton | bacterioplankton | virioplankton | bacterioplankton | virioplankton | bacterioplankton | virioplankton | bacterioplankton | virioplankton | bacterioplankton | virioplankton | bacterioplankton | virioplankton | bacterioplankton | virioplankton | bacterioplankton | virioplankton | bacterioplankton | virioplankton | bacterioplankton | virioplankton |
| bacteria_May12d50b_402 | 2 | 0 | 0 | 0 | 0 | 2 | 0 | 0 | 0 | 0 | 0 | 0 | 0 | 0 | 0 | 0 | 0 | 0 | 0 | 0 | 0 | 0 | 0 | 0 | 0 | 0 | 0 | 0 | 0 | 0 | 0 | 0 | 0 | 0 | 0 | 0 | 0 | 0 | 0 | 0 | 0 | 0 | 0 | 0 | 0 | 0 | 0 | 0 | 0 | 0 | 0 | 0 | 0 | 0 | 0 |
| bacteria_Sep8d0b_69954 | 2 | 0 | 0 | 0 | 0 | 0 | 0 | 0 | 0 | 0 | 0 | 0 | 0 | 0 | 0 | 0 | 0 | 0 | 0 | 0 | 0 | 0 | 0 | 0 | 0 | 0 | 0 | 0 | 0 | 0 | 0 | 0 | 0 | 0 | 0 | 0 | 0 | 0 | 0 | 0 | 0 | 0 | 0 | 0 | 0 | 0 | 0 | 2 | 0 | 0 | 0 | 0 | 0 | 0 | 0 |
| bacteria_Sep8d5b_26646 | 2 | 0 | 0 | 0 | 0 | 0 | 0 | 0 | 0 | 0 | 0 | 0 | 0 | 0 | 0 | 0 | 0 | 0 | 0 | 0 | 0 | 0 | 0 | 0 | 0 | 0 | 0 | 0 | 0 | 0 | 0 | 0 | 0 | 0 | 0 | 0 | 0 | 0 | 0 | 0 | 0 | 0 | 0 | 0 | 0 | 0 | 0 | 0 | 0 | 2 | 0 | 0 | 0 | 0 | 0 |
| bacteria_Sep8d5b_33456 | 2 | 0 | 0 | 0 | 0 | 0 | 0 | 0 | 0 | 0 | 0 | 0 | 0 | 0 | 0 | 0 | 0 | 0 | 0 | 0 | 0 | 0 | 0 | 0 | 0 | 0 | 0 | 0 | 0 | 0 | 0 | 0 | 0 | 0 | 0 | 0 | 0 | 0 | 0 | 0 | 0 | 0 | 0 | 0 | 0 | 0 | 0 | 0 | 0 | 1 | 0 | 0 | 0 | 0 | 1 |
| bacteria_Sep8d5b_35871 | 2 | 0 | 0 | 0 | 0 | 0 | 0 | 0 | 0 | 0 | 0 | 0 | 0 | 0 | 0 | 0 | 0 | 0 | 0 | 0 | 0 | 1 | 0 | 0 | 0 | 0 | 0 | 0 | 0 | 0 | 0 | 0 | 0 | 0 | 0 | 0 | 0 | 0 | 0 | 0 | 0 | 0 | 0 | 0 | 0 | 0 | 0 | 0 | 0 | 1 | 0 | 0 | 0 | 0 | 0 |
| bacteria_Sep8d25b_4177 | 2 | 0 | 0 | 0 | 0 | 0 | 0 | 0 | 0 | 0 | 0 | 0 | 0 | 0 | 0 | 0 | 0 | 0 | 1 | 0 | 0 | 0 | 0 | 0 | 0 | 0 | 0 | 0 | 0 | 0 | 0 | 0 | 0 | 0 | 0 | 0 | 0 | 0 | 0 | 0 | 0 | 0 | 0 | 0 | 0 | 0 | 0 | 0 | 0 | 0 | 0 | 1 | 0 | 0 | 0 |
| virus_Aug5.0v_1816 | 2 | 0 | 0 | 0 | 0 | 0 | 0 | 0 | 0 | 0 | 0 | 0 | 0 | 0 | 0 | 0 | 0 | 0 | 0 | 0 | 0 | 0 | 0 | 0 | 0 | 0 | 0 | 0 | 0 | 0 | 0 | 0 | 0 | 0 | 0 | 0 | 0 | 0 | 1 | 0 | 0 | 0 | 1 | 0 | 0 | 0 | 0 | 0 | 0 | 0 | 0 | 0 | 0 | 0 | 0 |
| virus_Aug5.5v_18066 | 2 | 0 | 0 | 0 | 0 | 0 | 0 | 0 | 0 | 0 | 0 | 0 | 0 | 0 | 0 | 0 | 0 | 0 | 0 | 0 | 0 | 0 | 1 | 0 | 0 | 0 | 0 | 0 | 0 | 0 | 0 | 0 | 0 | 0 | 0 | 0 | 0 | 0 | 0 | 0 | 1 | 0 | 0 | 0 | 0 | 0 | 0 | 0 | 0 | 0 | 0 | 0 | 0 | 0 | 0 |
| virus_Aug5.5v_23135 | 2 | 0 | 0 | 0 | 0 | 0 | 0 | 0 | 0 | 0 | 0 | 0 | 0 | 0 | 0 | 0 | 0 | 0 | 0 | 0 | 0 | 0 | 0 | 0 | 0 | 0 | 0 | 0 | 1 | 0 | 0 | 0 | 0 | 0 | 0 | 0 | 0 | 0 | 0 | 0 | 1 | 0 | 0 | 0 | 0 | 0 | 0 | 0 | 0 | 0 | 0 | 0 | 0 | 0 | 0 |
| virus_Aug5.5v_35539 | 2 | 0 | 0 | 0 | 0 | 0 | 0 | 0 | 0 | 1 | 0 | 0 | 0 | 0 | 0 | 0 | 0 | 0 | 0 | 0 | 0 | 0 | 0 | 0 | 0 | 0 | 0 | 0 | 0 | 0 | 0 | 0 | 0 | 0 | 0 | 0 | 0 | 0 | 0 | 0 | 1 | 0 | 0 | 0 | 0 | 0 | 0 | 0 | 0 | 0 | 0 | 0 | 0 | 0 | 0 |
| virus_Jul11.5v_20187 | 2 | 0 | 0 | 0 | 0 | 0 | 1 | 0 | 0 | 0 | 0 | 0 | 0 | 0 | 0 | 0 | 0 | 0 | 0 | 0 | 0 | 0 | 0 | 0 | 1 | 0 | 0 | 0 | 0 | 0 | 0 | 0 | 0 | 0 | 0 | 0 | 0 | 0 | 0 | 0 | 0 | 0 | 0 | 0 | 0 | 0 | 0 | 0 | 0 | 0 | 0 | 0 | 0 | 0 | 0 |
| virus_Jul11.25v_19575 | 2 | 0 | 0 | 0 | 0 | 0 | 0 | 0 | 0 | 0 | 0 | 0 | 0 | 0 | 0 | 0 | 0 | 0 | 0 | 0 | 0 | 0 | 0 | 0 | 0 | 0 | 1 | 0 | 0 | 0 | 0 | 0 | 0 | 0 | 0 | 0 | 0 | 0 | 0 | 0 | 0 | 0 | 1 | 0 | 0 | 0 | 0 | 0 | 0 | 0 | 0 | 0 | 0 | 0 | 0 |
| virus_Jul27.5v_3275 | 2 | 0 | 0 | 0 | 0 | 0 | 0 | 0 | 0 | 0 | 0 | 0 | 0 | 0 | 0 | 0 | 0 | 0 | 0 | 0 | 0 | 0 | 0 | 0 | 0 | 0 | 0 | 0 | 0 | 0 | 0 | 0 | 1 | 0 | 0 | 0 | 0 | 0 | 0 | 0 | 0 | 0 | 0 | 0 | 0 | 0 | 0 | 0 | 0 | 0 | 0 | 0 | 0 | 0 | 1 |
| virus_Jul27.5v_6689 | 2 | 0 | 0 | 0 | 0 | 0 | 0 | 0 | 0 | 0 | 0 | 0 | 0 | 0 | 0 | 0 | 0 | 0 | 0 | 1 | 0 | 0 | 0 | 0 | 0 | 0 | 0 | 0 | 0 | 0 | 0 | 0 | 1 | 0 | 0 | 0 | 0 | 0 | 0 | 0 | 0 | 0 | 0 | 0 | 0 | 0 | 0 | 0 | 0 | 0 | 0 | 0 | 0 | 0 | 0 |
| virus_Jul27.5v_13132 | 2 | 0 | 0 | 0 | 0 | 0 | 0 | 0 | 0 | 0 | 0 | 0 | 0 | 0 | 0 | 1 | 0 | 0 | 0 | 0 | 0 | 0 | 0 | 0 | 0 | 0 | 0 | 0 | 0 | 0 | 0 | 0 | 1 | 0 | 0 | 0 | 0 | 0 | 0 | 0 | 0 | 0 | 0 | 0 | 0 | 0 | 0 | 0 | 0 | 0 | 0 | 0 | 0 | 0 | 0 |
| virus_Jul27.5v_14080 | 2 | 0 | 0 | 0 | 0 | 0 | 0 | 0 | 0 | 0 | 0 | 0 | 0 | 0 | 0 | 0 | 0 | 0 | 0 | 0 | 0 | 0 | 0 | 0 | 0 | 0 | 1 | 0 | 0 | 0 | 0 | 0 | 1 | 0 | 0 | 0 | 0 | 0 | 0 | 0 | 0 | 0 | 0 | 0 | 0 | 0 | 0 | 0 | 0 | 0 | 0 | 0 | 0 | 0 | 0 |
| virus_Jun8.50v_13174 | 2 | 0 | 0 | 0 | 0 | 0 | 0 | 0 | 1 | 0 | 0 | 0 | 1 | 0 | 0 | 0 | 0 | 0 | 0 | 0 | 0 | 0 | 0 | 0 | 0 | 0 | 0 | 0 | 0 | 0 | 0 | 0 | 0 | 0 | 0 | 0 | 0 | 0 | 0 | 0 | 0 | 0 | 0 | 0 | 0 | 0 | 0 | 0 | 0 | 0 | 0 | 0 | 0 | 0 | 0 |
| virus_Jun8.50v_39939 | 2 | 0 | 0 | 0 | 0 | 0 | 0 | 0 | 0 | 0 | 0 | 0 | 2 | 0 | 0 | 0 | 0 | 0 | 0 | 0 | 0 | 0 | 0 | 0 | 0 | 0 | 0 | 0 | 0 | 0 | 0 | 0 | 0 | 0 | 0 | 0 | 0 | 0 | 0 | 0 | 0 | 0 | 0 | 0 | 0 | 0 | 0 | 0 | 0 | 0 | 0 | 0 | 0 | 0 | 0 |
| virus_Jun22.50v_15762 | 2 | 0 | 0 | 0 | 0 | 0 | 0 | 0 | 0 | 0 | 0 | 0 | 0 | 0 | 0 | 0 | 0 | 0 | 1 | 0 | 0 | 0 | 0 | 0 | 0 | 0 | 0 | 0 | 0 | 0 | 0 | 0 | 1 | 0 | 0 | 0 | 0 | 0 | 0 | 0 | 0 | 0 | 0 | 0 | 0 | 0 | 0 | 0 | 0 | 0 | 0 | 0 | 0 | 0 | 0 |
| virus_Jun22.50v_24832 | 2 | 0 | 0 | 0 | 0 | 0 | 0 | 0 | 0 | 0 | 0 | 0 | 0 | 0 | 0 | 0 | 0 | 0 | 1 | 0 | 0 | 0 | 0 | 0 | 0 | 0 | 0 | 0 | 0 | 0 | 0 | 0 | 0 | 0 | 0 | 0 | 0 | 0 | 0 | 0 | 1 | 0 | 0 | 0 | 0 | 0 | 0 | 0 | 0 | 0 | 0 | 0 | 0 | 0 | 0 |
| virus_Jun22.50v_38646 | 2 | 0 | 0 | 0 | 0 | 0 | 0 | 0 | 0 | 0 | 0 | 0 | 0 | 0 | 0 | 0 | 0 | 0 | 1 | 0 | 0 | 0 | 0 | 0 | 0 | 0 | 0 | 0 | 0 | 0 | 1 | 0 | 0 | 0 | 0 | 0 | 0 | 0 | 0 | 0 | 0 | 0 | 0 | 0 | 0 | 0 | 0 | 0 | 0 | 0 | 0 | 0 | 0 | 0 | 0 |
| virus_Jun22.75v_36044 | 2 | 0 | 0 | 0 | 0 | 0 | 0 | 0 | 0 | 0 | 0 | 0 | 0 | 0 | 0 | 0 | 0 | 0 | 0 | 0 | 1 | 0 | 0 | 0 | 0 | 0 | 0 | 0 | 0 | 0 | 0 | 0 | 0 | 0 | 0 | 0 | 0 | 0 | 1 | 0 | 0 | 0 | 0 | 0 | 0 | 0 | 0 | 0 | 0 | 0 | 0 | 0 | 0 | 0 | 0 |
| virus_May12.50v_12297 | 2 | 0 | 0 | 0 | 0 | 0 | 1 | 0 | 1 | 0 | 0 | 0 | 0 | 0 | 0 | 0 | 0 | 0 | 0 | 0 | 0 | 0 | 0 | 0 | 0 | 0 | 0 | 0 | 0 | 0 | 0 | 0 | 0 | 0 | 0 | 0 | 0 | 0 | 0 | 0 | 0 | 0 | 0 | 0 | 0 | 0 | 0 | 0 | 0 | 0 | 0 | 0 | 0 | 0 | 0 |

**Supplementary Table S7**. Common Bacterio- and virioplankton RTPR 98% OTUs in 27 samples with RTPR amplicons in both fractions

Total seqs: 47997; Subtotal seqs in retained

| OTUs: 41046 | | May12_05m | | May12_25m | | May12_50m | | May12_75m | | Jun8_25m | | Jun8_50m | | Jun8_75m | | Jun22_0m | | Jun22_50m | | Jun22_75m | | Jul11_0m | | Jul11_05m | | Jul11_25m | | Jul11_50m | | Jul11_75m | | Jul27_05m | | Jul27_25m | | Jul27_75m | | Aug5_0m | | Aug5_05m | | Aug5_25m | | Aug5_50m | | Aug5_75m | | Sep8_0m | | Sep8_05m | | Sep8_25m | | Sep8_50m | |
|---|---|---|---|---|---|---|---|---|---|---|---|---|---|---|---|---|---|---|---|---|---|---|---|---|---|---|---|---|---|---|---|---|---|---|---|---|---|---|---|---|---|---|---|---|---|---|---|---|---|---|---|---|---|---|---|
| OTUs with >=10 seqs: 254 OTU subtotal | | bacterioplankton | virioplankton | bacterioplankton | virioplankton | bacterioplankton | virioplankton | bacterioplankton | virioplankton | bacterioplankton | virioplankton | bacterioplankton | virioplankton | bacterioplankton | virioplankton | bacterioplankton | virioplankton | bacterioplankton | virioplankton | bacterioplankton | virioplankton | bacterioplankton | virioplankton | bacterioplankton | virioplankton | bacterioplankton | virioplankton | bacterioplankton | virioplankton | bacterioplankton | virioplankton | bacterioplankton | virioplankton | bacterioplankton | virioplankton | bacterioplankton | virioplankton | bacterioplankton | virioplankton | bacterioplankton | virioplankton | bacterioplankton | virioplankton | bacterioplankton | virioplankton | bacterioplankton | virioplankton | bacterioplankton | virioplankton | bacterioplankton | virioplankton | bacterioplankton | virioplankton | bacterioplankton | virioplankton |
| virus_Aug5.75v_25496 | 1 | 0 | 0 | 0 | 0 | 0 | 0 | 0 | 0 | 0 | 0 | 0 | 0 | 0 | 0 | 0 | 0 | 0 | 0 | 0 | 0 | 0 | 0 | 0 | 0 | 0 | 0 | 0 | 0 | 0 | 0 | 0 | 0 | 0 | 0 | 0 | 0 | 0 | 0 | 0 | 0 | 0 | 0 | 0 | 0 | 0 | 1 | 0 | 0 | 0 | 0 | 0 | 0 | 0 | 0 |
| virus_Aug5.75v_34716 | 1 | 0 | 0 | 0 | 0 | 0 | 0 | 0 | 0 | 0 | 0 | 0 | 0 | 0 | 0 | 0 | 0 | 0 | 0 | 0 | 0 | 0 | 0 | 0 | 0 | 0 | 0 | 0 | 0 | 0 | 0 | 0 | 0 | 0 | 0 | 0 | 0 | 0 | 0 | 0 | 0 | 0 | 0 | 0 | 0 | 0 | 1 | 0 | 0 | 0 | 0 | 0 | 0 | 0 | 0 |
| virus_Aug5.75v_36446 | 1 | 0 | 0 | 0 | 0 | 0 | 0 | 0 | 0 | 0 | 0 | 0 | 0 | 0 | 0 | 0 | 0 | 0 | 0 | 0 | 0 | 0 | 0 | 0 | 0 | 0 | 0 | 0 | 0 | 0 | 0 | 0 | 0 | 0 | 0 | 0 | 0 | 0 | 0 | 0 | 0 | 0 | 0 | 0 | 0 | 0 | 1 | 0 | 0 | 0 | 0 | 0 | 0 | 0 | 0 |
| virus_Jul11.0v_1306 | 1 | 0 | 0 | 0 | 0 | 0 | 0 | 0 | 0 | 0 | 0 | 0 | 0 | 0 | 0 | 0 | 0 | 0 | 0 | 0 | 0 | 0 | 1 | 0 | 0 | 0 | 0 | 0 | 0 | 0 | 0 | 0 | 0 | 0 | 0 | 0 | 0 | 0 | 0 | 0 | 0 | 0 | 0 | 0 | 0 | 0 | 0 | 0 | 0 | 0 | 0 | 0 | 0 | 0 | 0 |
| virus_Jul11.0v_9287 | 1 | 0 | 0 | 0 | 0 | 0 | 0 | 0 | 0 | 0 | 0 | 0 | 0 | 0 | 0 | 0 | 0 | 0 | 0 | 0 | 0 | 0 | 1 | 0 | 0 | 0 | 0 | 0 | 0 | 0 | 0 | 0 | 0 | 0 | 0 | 0 | 0 | 0 | 0 | 0 | 0 | 0 | 0 | 0 | 0 | 0 | 0 | 0 | 0 | 0 | 0 | 0 | 0 | 0 | 0 |
| virus_Jul11.0v_11308 | 1 | 0 | 0 | 0 | 0 | 0 | 0 | 0 | 0 | 0 | 0 | 0 | 0 | 0 | 0 | 0 | 0 | 0 | 0 | 0 | 0 | 0 | 1 | 0 | 0 | 0 | 0 | 0 | 0 | 0 | 0 | 0 | 0 | 0 | 0 | 0 | 0 | 0 | 0 | 0 | 0 | 0 | 0 | 0 | 0 | 0 | 0 | 0 | 0 | 0 | 0 | 0 | 0 | 0 | 0 |
| virus_Jul11.0v_11656 | 1 | 0 | 0 | 0 | 0 | 0 | 0 | 0 | 0 | 0 | 0 | 0 | 0 | 0 | 0 | 0 | 0 | 0 | 0 | 0 | 0 | 0 | 1 | 0 | 0 | 0 | 0 | 0 | 0 | 0 | 0 | 0 | 0 | 0 | 0 | 0 | 0 | 0 | 0 | 0 | 0 | 0 | 0 | 0 | 0 | 0 | 0 | 0 | 0 | 0 | 0 | 0 | 0 | 0 | 0 |
| virus_Jul11.0v_16432 | 1 | 0 | 0 | 0 | 0 | 0 | 0 | 0 | 0 | 0 | 0 | 0 | 0 | 0 | 0 | 0 | 0 | 0 | 0 | 0 | 0 | 0 | 1 | 0 | 0 | 0 | 0 | 0 | 0 | 0 | 0 | 0 | 0 | 0 | 0 | 0 | 0 | 0 | 0 | 0 | 0 | 0 | 0 | 0 | 0 | 0 | 0 | 0 | 0 | 0 | 0 | 0 | 0 | 0 | 0 |
| virus_Jul11.0v_19603 | 1 | 0 | 0 | 0 | 0 | 0 | 0 | 0 | 0 | 0 | 0 | 0 | 0 | 0 | 0 | 0 | 0 | 0 | 0 | 0 | 0 | 0 | 1 | 0 | 0 | 0 | 0 | 0 | 0 | 0 | 0 | 0 | 0 | 0 | 0 | 0 | 0 | 0 | 0 | 0 | 0 | 0 | 0 | 0 | 0 | 0 | 0 | 0 | 0 | 0 | 0 | 0 | 0 | 0 | 0 |
| virus_Jul11.5v_4614 | 1 | 0 | 0 | 0 | 0 | 0 | 0 | 0 | 0 | 0 | 0 | 0 | 0 | 0 | 0 | 0 | 0 | 0 | 0 | 0 | 0 | 0 | 0 | 0 | 1 | 0 | 0 | 0 | 0 | 0 | 0 | 0 | 0 | 0 | 0 | 0 | 0 | 0 | 0 | 0 | 0 | 0 | 0 | 0 | 0 | 0 | 0 | 0 | 0 | 0 | 0 | 0 | 0 | 0 | 0 |
| virus_Jul11.5v_11265 | 1 | 0 | 0 | 0 | 0 | 0 | 0 | 0 | 0 | 0 | 0 | 0 | 0 | 0 | 0 | 0 | 0 | 0 | 0 | 0 | 0 | 0 | 0 | 0 | 1 | 0 | 0 | 0 | 0 | 0 | 0 | 0 | 0 | 0 | 0 | 0 | 0 | 0 | 0 | 0 | 0 | 0 | 0 | 0 | 0 | 0 | 0 | 0 | 0 | 0 | 0 | 0 | 0 | 0 | 0 |
| virus_Jul11.5v_12327 | 1 | 0 | 0 | 0 | 0 | 0 | 0 | 0 | 0 | 0 | 0 | 0 | 0 | 0 | 0 | 0 | 0 | 0 | 0 | 0 | 0 | 0 | 0 | 0 | 1 | 0 | 0 | 0 | 0 | 0 | 0 | 0 | 0 | 0 | 0 | 0 | 0 | 0 | 0 | 0 | 0 | 0 | 0 | 0 | 0 | 0 | 0 | 0 | 0 | 0 | 0 | 0 | 0 | 0 | 0 |
| virus_Jul11.5v_12620 | 1 | 0 | 0 | 0 | 0 | 0 | 0 | 0 | 0 | 0 | 0 | 0 | 0 | 0 | 0 | 0 | 0 | 0 | 0 | 0 | 0 | 0 | 0 | 0 | 1 | 0 | 0 | 0 | 0 | 0 | 0 | 0 | 0 | 0 | 0 | 0 | 0 | 0 | 0 | 0 | 0 | 0 | 0 | 0 | 0 | 0 | 0 | 0 | 0 | 0 | 0 | 0 | 0 | 0 | 0 |
| virus_Jul11.5v_13628 | 1 | 0 | 0 | 0 | 0 | 0 | 0 | 0 | 0 | 0 | 0 | 0 | 0 | 0 | 0 | 0 | 0 | 0 | 0 | 0 | 0 | 0 | 0 | 0 | 1 | 0 | 0 | 0 | 0 | 0 | 0 | 0 | 0 | 0 | 0 | 0 | 0 | 0 | 0 | 0 | 0 | 0 | 0 | 0 | 0 | 0 | 0 | 0 | 0 | 0 | 0 | 0 | 0 | 0 | 0 |
| virus_Jul11.5v_15098 | 1 | 0 | 0 | 0 | 0 | 0 | 0 | 0 | 0 | 0 | 0 | 0 | 0 | 0 | 0 | 0 | 0 | 0 | 0 | 0 | 0 | 0 | 0 | 0 | 1 | 0 | 0 | 0 | 0 | 0 | 0 | 0 | 0 | 0 | 0 | 0 | 0 | 0 | 0 | 0 | 0 | 0 | 0 | 0 | 0 | 0 | 0 | 0 | 0 | 0 | 0 | 0 | 0 | 0 | 0 |
| virus_Jul11.5v_15679 | 1 | 0 | 0 | 0 | 0 | 0 | 0 | 0 | 0 | 0 | 0 | 0 | 0 | 0 | 0 | 0 | 0 | 0 | 0 | 0 | 0 | 0 | 0 | 0 | 1 | 0 | 0 | 0 | 0 | 0 | 0 | 0 | 0 | 0 | 0 | 0 | 0 | 0 | 0 | 0 | 0 | 0 | 0 | 0 | 0 | 0 | 0 | 0 | 0 | 0 | 0 | 0 | 0 | 0 | 0 |
| virus_Jul11.25v_5561 | 1 | 0 | 0 | 0 | 0 | 0 | 0 | 0 | 0 | 0 | 0 | 0 | 0 | 0 | 0 | 0 | 0 | 0 | 0 | 0 | 0 | 0 | 0 | 0 | 0 | 0 | 1 | 0 | 0 | 0 | 0 | 0 | 0 | 0 | 0 | 0 | 0 | 0 | 0 | 0 | 0 | 0 | 0 | 0 | 0 | 0 | 0 | 0 | 0 | 0 | 0 | 0 | 0 | 0 | 0 |
| virus_Jul11.25v_10897 | 1 | 0 | 0 | 0 | 0 | 0 | 0 | 0 | 0 | 0 | 0 | 0 | 0 | 0 | 0 | 0 | 0 | 0 | 0 | 0 | 0 | 0 | 0 | 0 | 0 | 0 | 1 | 0 | 0 | 0 | 0 | 0 | 0 | 0 | 0 | 0 | 0 | 0 | 0 | 0 | 0 | 0 | 0 | 0 | 0 | 0 | 0 | 0 | 0 | 0 | 0 | 0 | 0 | 0 | 0 |
| virus_Jul11.25v_16320 | 1 | 0 | 0 | 0 | 0 | 0 | 0 | 0 | 0 | 0 | 0 | 0 | 0 | 0 | 0 | 0 | 0 | 0 | 0 | 0 | 0 | 0 | 0 | 0 | 0 | 0 | 1 | 0 | 0 | 0 | 0 | 0 | 0 | 0 | 0 | 0 | 0 | 0 | 0 | 0 | 0 | 0 | 0 | 0 | 0 | 0 | 0 | 0 | 0 | 0 | 0 | 0 | 0 | 0 | 0 |
| virus_Jul11.25v_16748 | 1 | 0 | 0 | 0 | 0 | 0 | 0 | 0 | 0 | 0 | 0 | 0 | 0 | 0 | 0 | 0 | 0 | 0 | 0 | 0 | 0 | 0 | 0 | 0 | 0 | 0 | 1 | 0 | 0 | 0 | 0 | 0 | 0 | 0 | 0 | 0 | 0 | 0 | 0 | 0 | 0 | 0 | 0 | 0 | 0 | 0 | 0 | 0 | 0 | 0 | 0 | 0 | 0 | 0 | 0 |
| virus_Jul11.25v_17316 | 1 | 0 | 0 | 0 | 0 | 0 | 0 | 0 | 0 | 0 | 0 | 0 | 0 | 0 | 0 | 0 | 0 | 0 | 0 | 0 | 0 | 0 | 0 | 0 | 0 | 0 | 1 | 0 | 0 | 0 | 0 | 0 | 0 | 0 | 0 | 0 | 0 | 0 | 0 | 0 | 0 | 0 | 0 | 0 | 0 | 0 | 0 | 0 | 0 | 0 | 0 | 0 | 0 | 0 | 0 |
| virus_Jul11.50v_1249 | 1 | 0 | 0 | 0 | 0 | 0 | 0 | 0 | 0 | 0 | 0 | 0 | 0 | 0 | 0 | 0 | 0 | 0 | 0 | 0 | 0 | 0 | 0 | 0 | 0 | 0 | 0 | 0 | 1 | 0 | 0 | 0 | 0 | 0 | 0 | 0 | 0 | 0 | 0 | 0 | 0 | 0 | 0 | 0 | 0 | 0 | 0 | 0 | 0 | 0 | 0 | 0 | 0 | 0 | 0 |
| virus_Jul11.50v_5024 | 1 | 0 | 0 | 0 | 0 | 0 | 0 | 0 | 0 | 0 | 0 | 0 | 0 | 0 | 0 | 0 | 0 | 0 | 0 | 0 | 0 | 0 | 0 | 0 | 0 | 0 | 0 | 0 | 1 | 0 | 0 | 0 | 0 | 0 | 0 | 0 | 0 | 0 | 0 | 0 | 0 | 0 | 0 | 0 | 0 | 0 | 0 | 0 | 0 | 0 | 0 | 0 | 0 | 0 | 0 |
| virus_Jul11.50v_9111 | 1 | 0 | 0 | 0 | 0 | 0 | 0 | 0 | 0 | 0 | 0 | 0 | 0 | 0 | 0 | 0 | 0 | 0 | 0 | 0 | 0 | 0 | 0 | 0 | 0 | 0 | 0 | 0 | 1 | 0 | 0 | 0 | 0 | 0 | 0 | 0 | 0 | 0 | 0 | 0 | 0 | 0 | 0 | 0 | 0 | 0 | 0 | 0 | 0 | 0 | 0 | 0 | 0 | 0 | 0 |
| virus_Jul11.50v_13741 | 1 | 0 | 0 | 0 | 0 | 0 | 0 | 0 | 0 | 0 | 0 | 0 | 0 | 0 | 0 | 0 | 0 | 0 | 0 | 0 | 0 | 0 | 0 | 0 | 0 | 0 | 0 | 0 | 1 | 0 | 0 | 0 | 0 | 0 | 0 | 0 | 0 | 0 | 0 | 0 | 0 | 0 | 0 | 0 | 0 | 0 | 0 | 0 | 0 | 0 | 0 | 0 | 0 | 0 | 0 |
| virus_Jul11.50v_14684 | 1 | 0 | 0 | 0 | 0 | 0 | 0 | 0 | 0 | 0 | 0 | 0 | 0 | 0 | 0 | 0 | 0 | 0 | 0 | 0 | 0 | 0 | 0 | 0 | 0 | 0 | 0 | 0 | 1 | 0 | 0 | 0 | 0 | 0 | 0 | 0 | 0 | 0 | 0 | 0 | 0 | 0 | 0 | 0 | 0 | 0 | 0 | 0 | 0 | 0 | 0 | 0 | 0 | 0 | 0 |
| virus_Jul11.50v_16904 | 1 | 0 | 0 | 0 | 0 | 0 | 0 | 0 | 0 | 0 | 0 | 0 | 0 | 0 | 0 | 0 | 0 | 0 | 0 | 0 | 0 | 0 | 0 | 0 | 0 | 0 | 0 | 0 | 1 | 0 | 0 | 0 | 0 | 0 | 0 | 0 | 0 | 0 | 0 | 0 | 0 | 0 | 0 | 0 | 0 | 0 | 0 | 0 | 0 | 0 | 0 | 0 | 0 | 0 | 0 |
| virus_Jul11.50v_20336 | 1 | 0 | 0 | 0 | 0 | 0 | 0 | 0 | 0 | 0 | 0 | 0 | 0 | 0 | 0 | 0 | 0 | 0 | 0 | 0 | 0 | 0 | 0 | 0 | 0 | 0 | 0 | 0 | 1 | 0 | 0 | 0 | 0 | 0 | 0 | 0 | 0 | 0 | 0 | 0 | 0 | 0 | 0 | 0 | 0 | 0 | 0 | 0 | 0 | 0 | 0 | 0 | 0 | 0 | 0 |
| virus_Jul11.75v_179 | 1 | 0 | 0 | 0 | 0 | 0 | 0 | 0 | 0 | 0 | 0 | 0 | 0 | 0 | 0 | 0 | 0 | 0 | 0 | 0 | 0 | 0 | 0 | 0 | 0 | 0 | 0 | 0 | 0 | 0 | 1 | 0 | 0 | 0 | 0 | 0 | 0 | 0 | 0 | 0 | 0 | 0 | 0 | 0 | 0 | 0 | 0 | 0 | 0 | 0 | 0 | 0 | 0 | 0 | 0 |
| virus_Jul11.75v_5240 | 1 | 0 | 0 | 0 | 0 | 0 | 0 | 0 | 0 | 0 | 0 | 0 | 0 | 0 | 0 | 0 | 0 | 0 | 0 | 0 | 0 | 0 | 0 | 0 | 0 | 0 | 0 | 0 | 0 | 0 | 1 | 0 | 0 | 0 | 0 | 0 | 0 | 0 | 0 | 0 | 0 | 0 | 0 | 0 | 0 | 0 | 0 | 0 | 0 | 0 | 0 | 0 | 0 | 0 | 0 |
| virus_Jul11.75v_19905 | 1 | 0 | 0 | 0 | 0 | 0 | 0 | 0 | 0 | 0 | 0 | 0 | 0 | 0 | 0 | 0 | 0 | 0 | 0 | 0 | 0 | 0 | 0 | 0 | 0 | 0 | 0 | 0 | 0 | 0 | 1 | 0 | 0 | 0 | 0 | 0 | 0 | 0 | 0 | 0 | 0 | 0 | 0 | 0 | 0 | 0 | 0 | 0 | 0 | 0 | 0 | 0 | 0 | 0 | 0 |
| virus_Jul11.75v_23605 | 1 | 0 | 0 | 0 | 0 | 0 | 0 | 0 | 0 | 0 | 0 | 0 | 0 | 0 | 0 | 0 | 0 | 0 | 0 | 0 | 0 | 0 | 0 | 0 | 0 | 0 | 0 | 0 | 0 | 0 | 1 | 0 | 0 | 0 | 0 | 0 | 0 | 0 | 0 | 0 | 0 | 0 | 0 | 0 | 0 | 0 | 0 | 0 | 0 | 0 | 0 | 0 | 0 | 0 | 0 |
| virus_Jul11.75v_35941 | 1 | 0 | 0 | 0 | 0 | 0 | 0 | 0 | 0 | 0 | 0 | 0 | 0 | 0 | 0 | 0 | 0 | 0 | 0 | 0 | 0 | 0 | 0 | 0 | 0 | 0 | 0 | 0 | 0 | 0 | 1 | 0 | 0 | 0 | 0 | 0 | 0 | 0 | 0 | 0 | 0 | 0 | 0 | 0 | 0 | 0 | 0 | 0 | 0 | 0 | 0 | 0 | 0 | 0 | 0 |
| virus_Jul11.75v_40569 | 1 | 0 | 0 | 0 | 0 | 0 | 0 | 0 | 0 | 0 | 0 | 0 | 0 | 0 | 0 | 0 | 0 | 0 | 0 | 0 | 0 | 0 | 0 | 0 | 0 | 0 | 0 | 0 | 0 | 0 | 1 | 0 | 0 | 0 | 0 | 0 | 0 | 0 | 0 | 0 | 0 | 0 | 0 | 0 | 0 | 0 | 0 | 0 | 0 | 0 | 0 | 0 | 0 | 0 | 0 |
| virus_Jul27.5v_1426 | 1 | 0 | 0 | 0 | 0 | 0 | 0 | 0 | 0 | 0 | 0 | 0 | 0 | 0 | 0 | 0 | 0 | 0 | 0 | 0 | 0 | 0 | 0 | 0 | 0 | 0 | 0 | 0 | 0 | 0 | 0 | 0 | 1 | 0 | 0 | 0 | 0 | 0 | 0 | 0 | 0 | 0 | 0 | 0 | 0 | 0 | 0 | 0 | 0 | 0 | 0 | 0 | 0 | 0 | 0 |
| virus_Jul27.5v_5032 | 1 | 0 | 0 | 0 | 0 | 0 | 0 | 0 | 0 | 0 | 0 | 0 | 0 | 0 | 0 | 0 | 0 | 0 | 0 | 0 | 0 | 0 | 0 | 0 | 0 | 0 | 0 | 0 | 0 | 0 | 0 | 0 | 1 | 0 | 0 | 0 | 0 | 0 | 0 | 0 | 0 | 0 | 0 | 0 | 0 | 0 | 0 | 0 | 0 | 0 | 0 | 0 | 0 | 0 | 0 |
| virus_Jul27.5v_5739 | 1 | 0 | 0 | 0 | 0 | 0 | 0 | 0 | 0 | 0 | 0 | 0 | 0 | 0 | 0 | 0 | 0 | 0 | 0 | 0 | 0 | 0 | 0 | 0 | 0 | 0 | 0 | 0 | 0 | 0 | 0 | 0 | 1 | 0 | 0 | 0 | 0 | 0 | 0 | 0 | 0 | 0 | 0 | 0 | 0 | 0 | 0 | 0 | 0 | 0 | 0 | 0 | 0 | 0 | 0 |
| virus_Jul27.5v_5916 | 1 | 0 | 0 | 0 | 0 | 0 | 0 | 0 | 0 | 0 | 0 | 0 | 0 | 0 | 0 | 0 | 0 | 0 | 0 | 0 | 0 | 0 | 0 | 0 | 0 | 0 | 0 | 0 | 0 | 0 | 0 | 0 | 1 | 0 | 0 | 0 | 0 | 0 | 0 | 0 | 0 | 0 | 0 | 0 | 0 | 0 | 0 | 0 | 0 | 0 | 0 | 0 | 0 | 0 | 0 |
| virus_Jul27.5v_7114 | 1 | 0 | 0 | 0 | 0 | 0 | 0 | 0 | 0 | 0 | 0 | 0 | 0 | 0 | 0 | 0 | 0 | 0 | 0 | 0 | 0 | 0 | 0 | 0 | 0 | 0 | 0 | 0 | 0 | 0 | 0 | 0 | 1 | 0 | 0 | 0 | 0 | 0 | 0 | 0 | 0 | 0 | 0 | 0 | 0 | 0 | 0 | 0 | 0 | 0 | 0 | 0 | 0 | 0 | 0 |
| virus_Jul27.5v_7801 | 1 | 0 | 0 | 0 | 0 | 0 | 0 | 0 | 0 | 0 | 0 | 0 | 0 | 0 | 0 | 0 | 0 | 0 | 0 | 0 | 0 | 0 | 0 | 0 | 0 | 0 | 0 | 0 | 0 | 0 | 0 | 0 | 1 | 0 | 0 | 0 | 0 | 0 | 0 | 0 | 0 | 0 | 0 | 0 | 0 | 0 | 0 | 0 | 0 | 0 | 0 | 0 | 0 | 0 | 0 |
| virus_Jul27.5v_9493 | 1 | 0 | 0 | 0 | 0 | 0 | 0 | 0 | 0 | 0 | 0 | 0 | 0 | 0 | 0 | 0 | 0 | 0 | 0 | 0 | 0 | 0 | 0 | 0 | 0 | 0 | 0 | 0 | 0 | 0 | 0 | 0 | 1 | 0 | 0 | 0 | 0 | 0 | 0 | 0 | 0 | 0 | 0 | 0 | 0 | 0 | 0 | 0 | 0 | 0 | 0 | 0 | 0 | 0 | 0 |
| virus_Jul27.5v_9941 | 1 | 0 | 0 | 0 | 0 | 0 | 0 | 0 | 0 | 0 | 0 | 0 | 0 | 0 | 0 | 0 | 0 | 0 | 0 | 0 | 0 | 0 | 0 | 0 | 0 | 0 | 0 | 0 | 0 | 0 | 0 | 0 | 1 | 0 | 0 | 0 | 0 | 0 | 0 | 0 | 0 | 0 | 0 | 0 | 0 | 0 | 0 | 0 | 0 | 0 | 0 | 0 | 0 | 0 | 0 |
| virus_Jul27.5v_10641 | 1 | 0 | 0 | 0 | 0 | 0 | 0 | 0 | 0 | 0 | 0 | 0 | 0 | 0 | 0 | 0 | 0 | 0 | 0 | 0 | 0 | 0 | 0 | 0 | 0 | 0 | 0 | 0 | 0 | 0 | 0 | 0 | 1 | 0 | 0 | 0 | 0 | 0 | 0 | 0 | 0 | 0 | 0 | 0 | 0 | 0 | 0 | 0 | 0 | 0 | 0 | 0 | 0 | 0 | 0 |
| virus_Jul27.5v_11478 | 1 | 0 | 0 | 0 | 0 | 0 | 0 | 0 | 0 | 0 | 0 | 0 | 0 | 0 | 0 | 0 | 0 | 0 | 0 | 0 | 0 | 0 | 0 | 0 | 0 | 0 | 0 | 0 | 0 | 0 | 0 | 0 | 1 | 0 | 0 | 0 | 0 | 0 | 0 | 0 | 0 | 0 | 0 | 0 | 0 | 0 | 0 | 0 | 0 | 0 | 0 | 0 | 0 | 0 | 0 |
| virus_Jul27.5v_13960 | 1 | 0 | 0 | 0 | 0 | 0 | 0 | 0 | 0 | 0 | 0 | 0 | 0 | 0 | 0 | 0 | 0 | 0 | 0 | 0 | 0 | 0 | 0 | 0 | 0 | 0 | 0 | 0 | 0 | 0 | 0 | 0 | 1 | 0 | 0 | 0 | 0 | 0 | 0 | 0 | 0 | 0 | 0 | 0 | 0 | 0 | 0 | 0 | 0 | 0 | 0 | 0 | 0 | 0 | 0 |
| virus_Jul27.5v_14499 | 1 | 0 | 0 | 0 | 0 | 0 | 0 | 0 | 0 | 0 | 0 | 0 | 0 | 0 | 0 | 0 | 0 | 0 | 0 | 0 | 0 | 0 | 0 | 0 | 0 | 0 | 0 | 0 | 0 | 0 | 0 | 0 | 1 | 0 | 0 | 0 | 0 | 0 | 0 | 0 | 0 | 0 | 0 | 0 | 0 | 0 | 0 | 0 | 0 | 0 | 0 | 0 | 0 | 0 | 0 |
| virus_Jul27.5v_15916 | 1 | 0 | 0 | 0 | 0 | 0 | 0 | 0 | 0 | 0 | 0 | 0 | 0 | 0 | 0 | 0 | 0 | 0 | 0 | 0 | 0 | 0 | 0 | 0 | 0 | 0 | 0 | 0 | 0 | 0 | 0 | 0 | 1 | 0 | 0 | 0 | 0 | 0 | 0 | 0 | 0 | 0 | 0 | 0 | 0 | 0 | 0 | 0 | 0 | 0 | 0 | 0 | 0 | 0 | 0 |
| virus_Jul27.5v_19523 | 1 | 0 | 0 | 0 | 0 | 0 | 0 | 0 | 0 | 0 | 0 | 0 | 0 | 0 | 0 | 0 | 0 | 0 | 0 | 0 | 0 | 0 | 0 | 0 | 0 | 0 | 0 | 0 | 0 | 0 | 0 | 0 | 1 | 0 | 0 | 0 | 0 | 0 | 0 | 0 | 0 | 0 | 0 | 0 | 0 | 0 | 0 | 0 | 0 | 0 | 0 | 0 | 0 | 0 | 0 |
| virus_Jul27.5v_20148 | 1 | 0 | 0 | 0 | 0 | 0 | 0 | 0 | 0 | 0 | 0 | 0 | 0 | 0 | 0 | 0 | 0 | 0 | 0 | 0 | 0 | 0 | 0 | 0 | 0 | 0 | 0 | 0 | 0 | 0 | 0 | 0 | 1 | 0 | 0 | 0 | 0 | 0 | 0 | 0 | 0 | 0 | 0 | 0 | 0 | 0 | 0 | 0 | 0 | 0 | 0 | 0 | 0 | 0 | 0 |
| virus_Jul27.5v_20233 | 1 | 0 | 0 | 0 | 0 | 0 | 0 | 0 | 0 | 0 | 0 | 0 | 0 | 0 | 0 | 0 | 0 | 0 | 0 | 0 | 0 | 0 | 0 | 0 | 0 | 0 | 0 | 0 | 0 | 0 | 0 | 0 | 1 | 0 | 0 | 0 | 0 | 0 | 0 | 0 | 0 | 0 | 0 | 0 | 0 | 0 | 0 | 0 | 0 | 0 | 0 | 0 | 0 | 0 | 0 |
| virus_Jul27.25v_1517 | 1 | 0 | 0 | 0 | 0 | 0 | 0 | 0 | 0 | 0 | 0 | 0 | 0 | 0 | 0 | 0 | 0 | 0 | 0 | 0 | 0 | 0 | 0 | 0 | 0 | 0 | 0 | 0 | 0 | 0 | 0 | 0 | 0 | 0 | 1 | 0 | 0 | 0 | 0 | 0 | 0 | 0 | 0 | 0 | 0 | 0 | 0 | 0 | 0 | 0 | 0 | 0 | 0 | 0 | 0 |
| virus_Jul27.25v_6981 | 1 | 0 | 0 | 0 | 0 | 0 | 0 | 0 | 0 | 0 | 0 | 0 | 0 | 0 | 0 | 0 | 0 | 0 | 0 | 0 | 0 | 0 | 0 | 0 | 0 | 0 | 0 | 0 | 0 | 0 | 0 | 0 | 0 | 0 | 1 | 0 | 0 | 0 | 0 | 0 | 0 | 0 | 0 | 0 | 0 | 0 | 0 | 0 | 0 | 0 | 0 | 0 | 0 | 0 | 0 |
| virus_Jul27.25v_14448 | 1 | 0 | 0 | 0 | 0 | 0 | 0 | 0 | 0 | 0 | 0 | 0 | 0 | 0 | 0 | 0 | 0 | 0 | 0 | 0 | 0 | 0 | 0 | 0 | 0 | 0 | 0 | 0 | 0 | 0 | 0 | 0 | 0 | 0 | 1 | 0 | 0 | 0 | 0 | 0 | 0 | 0 | 0 | 0 | 0 | 0 | 0 | 0 | 0 | 0 | 0 | 0 | 0 | 0 | 0 |
| virus_Jul27.25v_15557 | 1 | 0 | 0 | 0 | 0 | 0 | 0 | 0 | 0 | 0 | 0 | 0 | 0 | 0 | 0 | 0 | 0 | 0 | 0 | 0 | 0 | 0 | 0 | 0 | 0 | 0 | 0 | 0 | 0 | 0 | 0 | 0 | 0 | 0 | 1 | 0 | 0 | 0 | 0 | 0 | 0 | 0 | 0 | 0 | 0 | 0 | 0 | 0 | 0 | 0 | 0 | 0 | 0 | 0 | 0 |
| virus_Jul27.25v_24394 | 1 | 0 | 0 | 0 | 0 | 0 | 0 | 0 | 0 | 0 | 0 | 0 | 0 | 0 | 0 | 0 | 0 | 0 | 0 | 0 | 0 | 0 | 0 | 0 | 0 | 0 | 0 | 0 | 0 | 0 | 0 | 0 | 0 | 0 | 1 | 0 | 0 | 0 | 0 | 0 | 0 | 0 | 0 | 0 | 0 | 0 | 0 | 0 | 0 | 0 | 0 | 0 | 0 | 0 | 0 |
| virus_Jul27.25v_27779 | 1 | 0 | 0 | 0 | 0 | 0 | 0 | 0 | 0 | 0 | 0 | 0 | 0 | 0 | 0 | 0 | 0 | 0 | 0 | 0 | 0 | 0 | 0 | 0 | 0 | 0 | 0 | 0 | 0 | 0 | 0 | 0 | 0 | 0 | 1 | 0 | 0 | 0 | 0 | 0 | 0 | 0 | 0 | 0 | 0 | 0 | 0 | 0 | 0 | 0 | 0 | 0 | 0 | 0 | 0 |
| virus_Jul27.25v_28837 | 1 | 0 | 0 | 0 | 0 | 0 | 0 | 0 | 0 | 0 | 0 | 0 | 0 | 0 | 0 | 0 | 0 | 0 | 0 | 0 | 0 | 0 | 0 | 0 | 0 | 0 | 0 | 0 | 0 | 0 | 0 | 0 | 0 | 0 | 1 | 0 | 0 | 0 | 0 | 0 | 0 | 0 | 0 | 0 | 0 | 0 | 0 | 0 | 0 | 0 | 0 | 0 | 0 | 0 | 0 |
| virus_Jul27.25v_31066 | 1 | 0 | 0 | 0 | 0 | 0 | 0 | 0 | 0 | 0 | 0 | 0 | 0 | 0 | 0 | 0 | 0 | 0 | 0 | 0 | 0 | 0 | 0 | 0 | 0 | 0 | 0 | 0 | 0 | 0 | 0 | 0 | 0 | 0 | 1 | 0 | 0 | 0 | 0 | 0 | 0 | 0 | 0 | 0 | 0 | 0 | 0 | 0 | 0 | 0 | 0 | 0 | 0 | 0 | 0 |
| virus_Jul27.25v_32372 | 1 | 0 | 0 | 0 | 0 | 0 | 0 | 0 | 0 | 0 | 0 | 0 | 0 | 0 | 0 | 0 | 0 | 0 | 0 | 0 | 0 | 0 | 0 | 0 | 0 | 0 | 0 | 0 | 0 | 0 | 0 | 0 | 0 | 0 | 1 | 0 | 0 | 0 | 0 | 0 | 0 | 0 | 0 | 0 | 0 | 0 | 0 | 0 | 0 | 0 | 0 | 0 | 0 | 0 | 0 |
| virus_Jul27.25v_34410 | 1 | 0 | 0 | 0 | 0 | 0 | 0 | 0 | 0 | 0 | 0 | 0 | 0 | 0 | 0 | 0 | 0 | 0 | 0 | 0 | 0 | 0 | 0 | 0 | 0 | 0 | 0 | 0 | 0 | 0 | 0 | 0 | 0 | 0 | 1 | 0 | 0 | 0 | 0 | 0 | 0 | 0 | 0 | 0 | 0 | 0 | 0 | 0 | 0 | 0 | 0 | 0 | 0 | 0 | 0 |
| virus_Jul27.25v_35759 | 1 | 0 | 0 | 0 | 0 | 0 | 0 | 0 | 0 | 0 | 0 | 0 | 0 | 0 | 0 | 0 | 0 | 0 | 0 | 0 | 0 | 0 | 0 | 0 | 0 | 0 | 0 | 0 | 0 | 0 | 0 | 0 | 0 | 0 | 1 | 0 | 0 | 0 | 0 | 0 | 0 | 0 | 0 | 0 | 0 | 0 | 0 | 0 | 0 | 0 | 0 | 0 | 0 | 0 | 0 |
| virus_Jul27.25v_39770 | 1 | 0 | 0 | 0 | 0 | 0 | 0 | 0 | 0 | 0 | 0 | 0 | 0 | 0 | 0 | 0 | 0 | 0 | 0 | 0 | 0 | 0 | 0 | 0 | 0 | 0 | 0 | 0 | 0 | 0 | 0 | 0 | 0 | 0 | 1 | 0 | 0 | 0 | 0 | 0 | 0 | 0 | 0 | 0 | 0 | 0 | 0 | 0 | 0 | 0 | 0 | 0 | 0 | 0 | 0 |
| virus_Jul27.75v_1947 | 1 | 0 | 0 | 0 | 0 | 0 | 0 | 0 | 0 | 0 | 0 | 0 | 0 | 0 | 0 | 0 | 0 | 0 | 0 | 0 | 0 | 0 | 0 | 0 | 0 | 0 | 0 | 0 | 0 | 0 | 0 | 0 | 0 | 0 | 0 | 0 | 1 | 0 | 0 | 0 | 0 | 0 | 0 | 0 | 0 | 0 | 0 | 0 | 0 | 0 | 0 | 0 | 0 | 0 | 0 |
| virus_Jul27.75v_3620 | 1 | 0 | 0 | 0 | 0 | 0 | 0 | 0 | 0 | 0 | 0 | 0 | 0 | 0 | 0 | 0 | 0 | 0 | 0 | 0 | 0 | 0 | 0 | 0 | 0 | 0 | 0 | 0 | 0 | 0 | 0 | 0 | 0 | 0 | 0 | 0 | 1 | 0 | 0 | 0 | 0 | 0 | 0 | 0 | 0 | 0 | 0 | 0 | 0 | 0 | 0 | 0 | 0 | 0 | 0 |
| virus_Jul27.75v_18162 | 1 | 0 | 0 | 0 | 0 | 0 | 0 | 0 | 0 | 0 | 0 | 0 | 0 | 0 | 0 | 0 | 0 | 0 | 0 | 0 | 0 | 0 | 0 | 0 | 0 | 0 | 0 | 0 | 0 | 0 | 0 | 0 | 0 | 0 | 0 | 0 | 1 | 0 | 0 | 0 | 0 | 0 | 0 | 0 | 0 | 0 | 0 | 0 | 0 | 0 | 0 | 0 | 0 | 0 | 0 |
| virus_Jul27.75v_18179 | 1 | 0 | 0 | 0 | 0 | 0 | 0 | 0 | 0 | 0 | 0 | 0 | 0 | 0 | 0 | 0 | 0 | 0 | 0 | 0 | 0 | 0 | 0 | 0 | 0 | 0 | 0 | 0 | 0 | 0 | 0 | 0 | 0 | 0 | 0 | 0 | 1 | 0 | 0 | 0 | 0 | 0 | 0 | 0 | 0 | 0 | 0 | 0 | 0 | 0 | 0 | 0 | 0 | 0 | 0 |
| virus_Jun8.25v_23058 | 1 | 0 | 0 | 0 | 0 | 0 | 0 | 0 | 0 | 0 | 1 | 0 | 0 | 0 | 0 | 0 | 0 | 0 | 0 | 0 | 0 | 0 | 0 | 0 | 0 | 0 | 0 | 0 | 0 | 0 | 0 | 0 | 0 | 0 | 0 | 0 | 0 | 0 | 0 | 0 | 0 | 0 | 0 | 0 | 0 | 0 | 0 | 0 | 0 | 0 | 0 | 0 | 0 | 0 | 0 |
| virus_Jun8.50v_3281 | 1 | 0 | 0 | 0 | 0 | 0 | 0 | 0 | 0 | 0 | 0 | 0 | 1 | 0 | 0 | 0 | 0 | 0 | 0 | 0 | 0 | 0 | 0 | 0 | 0 | 0 | 0 | 0 | 0 | 0 | 0 | 0 | 0 | 0 | 0 | 0 | 0 | 0 | 0 | 0 | 0 | 0 | 0 | 0 | 0 | 0 | 0 | 0 | 0 | 0 | 0 | 0 | 0 | 0 | 0 |
| virus_Jun8.50v_8532 | 1 | 0 | 0 | 0 | 0 | 0 | 0 | 0 | 0 | 0 | 0 | 0 | 1 | 0 | 0 | 0 | 0 | 0 | 0 | 0 | 0 | 0 | 0 | 0 | 0 | 0 | 0 | 0 | 0 | 0 | 0 | 0 | 0 | 0 | 0 | 0 | 0 | 0 | 0 | 0 | 0 | 0 | 0 | 0 | 0 | 0 | 0 | 0 | 0 | 0 | 0 | 0 | 0 | 0 | 0 |
| virus_Jun8.50v_24602 | 1 | 0 | 0 | 0 | 0 | 0 | 0 | 0 | 0 | 0 | 0 | 0 | 1 | 0 | 0 | 0 | 0 | 0 | 0 | 0 | 0 | 0 | 0 | 0 | 0 | 0 | 0 | 0 | 0 | 0 | 0 | 0 | 0 | 0 | 0 | 0 | 0 | 0 | 0 | 0 | 0 | 0 | 0 | 0 | 0 | 0 | 0 | 0 | 0 | 0 | 0 | 0 | 0 | 0 | 0 |
| virus_Jun8.50v_28378 | 1 | 0 | 0 | 0 | 0 | 0 | 0 | 0 | 0 | 0 | 0 | 0 | 1 | 0 | 0 | 0 | 0 | 0 | 0 | 0 | 0 | 0 | 0 | 0 | 0 | 0 | 0 | 0 | 0 | 0 | 0 | 0 | 0 | 0 | 0 | 0 | 0 | 0 | 0 | 0 | 0 | 0 | 0 | 0 | 0 | 0 | 0 | 0 | 0 | 0 | 0 | 0 | 0 | 0 | 0 |
| virus_Jun8.50v_36988 | 1 | 0 | 0 | 0 | 0 | 0 | 0 | 0 | 0 | 0 | 0 | 0 | 1 | 0 | 0 | 0 | 0 | 0 | 0 | 0 | 0 | 0 | 0 | 0 | 0 | 0 | 0 | 0 | 0 | 0 | 0 | 0 | 0 | 0 | 0 | 0 | 0 | 0 | 0 | 0 | 0 | 0 | 0 | 0 | 0 | 0 | 0 | 0 | 0 | 0 | 0 | 0 | 0 | 0 | 0 |
| virus_Jun8.75v_1576 | 1 | 0 | 0 | 0 | 0 | 0 | 0 | 0 | 0 | 0 | 0 | 0 | 0 | 0 | 1 | 0 | 0 | 0 | 0 | 0 | 0 | 0 | 0 | 0 | 0 | 0 | 0 | 0 | 0 | 0 | 0 | 0 | 0 | 0 | 0 | 0 | 0 | 0 | 0 | 0 | 0 | 0 | 0 | 0 | 0 | 0 | 0 | 0 | 0 | 0 | 0 | 0 | 0 | 0 | 0 |
| virus_Jun8.75v_3585 | 1 | 0 | 0 | 0 | 0 | 0 | 0 | 0 | 0 | 0 | 0 | 0 | 0 | 0 | 1 | 0 | 0 | 0 | 0 | 0 | 0 | 0 | 0 | 0 | 0 | 0 | 0 | 0 | 0 | 0 | 0 | 0 | 0 | 0 | 0 | 0 | 0 | 0 | 0 | 0 | 0 | 0 | 0 | 0 | 0 | 0 | 0 | 0 | 0 | 0 | 0 | 0 | 0 | 0 | 0 |
| virus_Jun22.0v_2504 | 1 | 0 | 0 | 0 | 0 | 0 | 0 | 0 | 0 | 0 | 0 | 0 | 0 | 0 | 0 | 0 | 1 | 0 | 0 | 0 | 0 | 0 | 0 | 0 | 0 | 0 | 0 | 0 | 0 | 0 | 0 | 0 | 0 | 0 | 0 | 0 | 0 | 0 | 0 | 0 | 0 | 0 | 0 | 0 | 0 | 0 | 0 | 0 | 0 | 0 | 0 | 0 | 0 | 0 | 0 |
| virus_Jun22.0v_3466 | 1 | 0 | 0 | 0 | 0 | 0 | 0 | 0 | 0 | 0 | 0 | 0 | 0 | 0 | 0 | 0 | 1 | 0 | 0 | 0 | 0 | 0 | 0 | 0 | 0 | 0 | 0 | 0 | 0 | 0 | 0 | 0 | 0 | 0 | 0 | 0 | 0 | 0 | 0 | 0 | 0 | 0 | 0 | 0 | 0 | 0 | 0 | 0 | 0 | 0 | 0 | 0 | 0 | 0 | 0 |
| virus_Jun22.0v_6617 | 1 | 0 | 0 | 0 | 0 | 0 | 0 | 0 | 0 | 0 | 0 | 0 | 0 | 0 | 0 | 0 | 1 | 0 | 0 | 0 | 0 | 0 | 0 | 0 | 0 | 0 | 0 | 0 | 0 | 0 | 0 | 0 | 0 | 0 | 0 | 0 | 0 | 0 | 0 | 0 | 0 | 0 | 0 | 0 | 0 | 0 | 0 | 0 | 0 | 0 | 0 | 0 | 0 | 0 | 0 |
| virus_Jun22.0v_6931 | 1 | 0 | 0 | 0 | 0 | 0 | 0 | 0 | 0 | 0 | 0 | 0 | 0 | 0 | 0 | 0 | 1 | 0 | 0 | 0 | 0 | 0 | 0 | 0 | 0 | 0 | 0 | 0 | 0 | 0 | 0 | 0 | 0 | 0 | 0 | 0 | 0 | 0 | 0 | 0 | 0 | 0 | 0 | 0 | 0 | 0 | 0 | 0 | 0 | 0 | 0 | 0 | 0 | 0 | 0 |
| virus_Jun22.0v_17823 | 1 | 0 | 0 | 0 | 0 | 0 | 0 | 0 | 0 | 0 | 0 | 0 | 0 | 0 | 0 | 0 | 1 | 0 | 0 | 0 | 0 | 0 | 0 | 0 | 0 | 0 | 0 | 0 | 0 | 0 | 0 | 0 | 0 | 0 | 0 | 0 | 0 | 0 | 0 | 0 | 0 | 0 | 0 | 0 | 0 | 0 | 0 | 0 | 0 | 0 | 0 | 0 | 0 | 0 | 0 |
| virus_Jun22.0v_17853 | 1 | 0 | 0 | 0 | 0 | 0 | 0 | 0 | 0 | 0 | 0 | 0 | 0 | 0 | 0 | 0 | 1 | 0 | 0 | 0 | 0 | 0 | 0 | 0 | 0 | 0 | 0 | 0 | 0 | 0 | 0 | 0 | 0 | 0 | 0 | 0 | 0 | 0 | 0 | 0 | 0 | 0 | 0 | 0 | 0 | 0 | 0 | 0 | 0 | 0 | 0 | 0 | 0 | 0 | 0 |
| virus_Jun22.50v_13859 | 1 | 0 | 0 | 0 | 0 | 0 | 0 | 0 | 0 | 0 | 0 | 0 | 0 | 0 | 0 | 0 | 0 | 0 | 1 | 0 | 0 | 0 | 0 | 0 | 0 | 0 | 0 | 0 | 0 | 0 | 0 | 0 | 0 | 0 | 0 | 0 | 0 | 0 | 0 | 0 | 0 | 0 | 0 | 0 | 0 | 0 | 0 | 0 | 0 | 0 | 0 | 0 | 0 | 0 | 0 |
| virus_Jun22.50v_15780 | 1 | 0 | 0 | 0 | 0 | 0 | 0 | 0 | 0 | 0 | 0 | 0 | 0 | 0 | 0 | 0 | 0 | 0 | 1 | 0 | 0 | 0 | 0 | 0 | 0 | 0 | 0 | 0 | 0 | 0 | 0 | 0 | 0 | 0 | 0 | 0 | 0 | 0 | 0 | 0 | 0 | 0 | 0 | 0 | 0 | 0 | 0 | 0 | 0 | 0 | 0 | 0 | 0 | 0 | 0 |
| virus_Jun22.50v_15993 | 1 | 0 | 0 | 0 | 0 | 0 | 0 | 0 | 0 | 0 | 0 | 0 | 0 | 0 | 0 | 0 | 0 | 0 | 1 | 0 | 0 | 0 | 0 | 0 | 0 | 0 | 0 | 0 | 0 | 0 | 0 | 0 | 0 | 0 | 0 | 0 | 0 | 0 | 0 | 0 | 0 | 0 | 0 | 0 | 0 | 0 | 0 | 0 | 0 | 0 | 0 | 0 | 0 | 0 | 0 |
| virus_Jun22.50v_16105 | 1 | 0 | 0 | 0 | 0 | 0 | 0 | 0 | 0 | 0 | 0 | 0 | 0 | 0 | 0 | 0 | 0 | 0 | 1 | 0 | 0 | 0 | 0 | 0 | 0 | 0 | 0 | 0 | 0 | 0 | 0 | 0 | 0 | 0 | 0 | 0 | 0 | 0 | 0 | 0 | 0 | 0 | 0 | 0 | 0 | 0 | 0 | 0 | 0 | 0 | 0 | 0 | 0 | 0 | 0 |
| virus_Jun22.50v_17216 | 1 | 0 | 0 | 0 | 0 | 0 | 0 | 0 | 0 | 0 | 0 | 0 | 0 | 0 | 0 | 0 | 0 | 0 | 1 | 0 | 0 | 0 | 0 | 0 | 0 | 0 | 0 | 0 | 0 | 0 | 0 | 0 | 0 | 0 | 0 | 0 | 0 | 0 | 0 | 0 | 0 | 0 | 0 | 0 | 0 | 0 | 0 | 0 | 0 | 0 | 0 | 0 | 0 | 0 | 0 |
| virus_Jun22.50v_18262 | 1 | 0 | 0 | 0 | 0 | 0 | 0 | 0 | 0 | 0 | 0 | 0 | 0 | 0 | 0 | 0 | 0 | 0 | 1 | 0 | 0 | 0 | 0 | 0 | 0 | 0 | 0 | 0 | 0 | 0 | 0 | 0 | 0 | 0 | 0 | 0 | 0 | 0 | 0 | 0 | 0 | 0 | 0 | 0 | 0 | 0 | 0 | 0 | 0 | 0 | 0 | 0 | 0 | 0 | 0 |
| virus_Jun22.50v_27550 | 1 | 0 | 0 | 0 | 0 | 0 | 0 | 0 | 0 | 0 | 0 | 0 | 0 | 0 | 0 | 0 | 0 | 0 | 1 | 0 | 0 | 0 | 0 | 0 | 0 | 0 | 0 | 0 | 0 | 0 | 0 | 0 | 0 | 0 | 0 | 0 | 0 | 0 | 0 | 0 | 0 | 0 | 0 | 0 | 0 | 0 | 0 | 0 | 0 | 0 | 0 | 0 | 0 | 0 | 0 |
| virus_Jun22.50v_30159 | 1 | 0 | 0 | 0 | 0 | 0 | 0 | 0 | 0 | 0 | 0 | 0 | 0 | 0 | 0 | 0 | 0 | 0 | 1 | 0 | 0 | 0 | 0 | 0 | 0 | 0 | 0 | 0 | 0 | 0 | 0 | 0 | 0 | 0 | 0 | 0 | 0 | 0 | 0 | 0 | 0 | 0 | 0 | 0 | 0 | 0 | 0 | 0 | 0 | 0 | 0 | 0 | 0 | 0 | 0 |
| virus_Jun22.50v_31867 | 1 | 0 | 0 | 0 | 0 | 0 | 0 | 0 | 0 | 0 | 0 | 0 | 0 | 0 | 0 | 0 | 0 | 0 | 1 | 0 | 0 | 0 | 0 | 0 | 0 | 0 | 0 | 0 | 0 | 0 | 0 | 0 | 0 | 0 | 0 | 0 | 0 | 0 | 0 | 0 | 0 | 0 | 0 | 0 | 0 | 0 | 0 | 0 | 0 | 0 | 0 | 0 | 0 | 0 | 0 |
| virus_Jun22.50v_34570 | 1 | 0 | 0 | 0 | 0 | 0 | 0 | 0 | 0 | 0 | 0 | 0 | 0 | 0 | 0 | 0 | 0 | 0 | 1 | 0 | 0 | 0 | 0 | 0 | 0 | 0 | 0 | 0 | 0 | 0 | 0 | 0 | 0 | 0 | 0 | 0 | 0 | 0 | 0 | 0 | 0 | 0 | 0 | 0 | 0 | 0 | 0 | 0 | 0 | 0 | 0 | 0 | 0 | 0 | 0 |
| virus_Jun22.50v_38582 | 1 | 0 | 0 | 0 | 0 | 0 | 0 | 0 | 0 | 0 | 0 | 0 | 0 | 0 | 0 | 0 | 0 | 0 | 1 | 0 | 0 | 0 | 0 | 0 | 0 | 0 | 0 | 0 | 0 | 0 | 0 | 0 | 0 | 0 | 0 | 0 | 0 | 0 | 0 | 0 | 0 | 0 | 0 | 0 | 0 | 0 | 0 | 0 | 0 | 0 | 0 | 0 | 0 | 0 | 0 |
| virus_Jun22.50v_38796 | 1 | 0 | 0 | 0 | 0 | 0 | 0 | 0 | 0 | 0 | 0 | 0 | 0 | 0 | 0 | 0 | 0 | 0 | 1 | 0 | 0 | 0 | 0 | 0 | 0 | 0 | 0 | 0 | 0 | 0 | 0 | 0 | 0 | 0 | 0 | 0 | 0 | 0 | 0 | 0 | 0 | 0 | 0 | 0 | 0 | 0 | 0 | 0 | 0 | 0 | 0 | 0 | 0 | 0 | 0 |
| virus_Jun22.50v_40223 | 1 | 0 | 0 | 0 | 0 | 0 | 0 | 0 | 0 | 0 | 0 | 0 | 0 | 0 | 0 | 0 | 0 | 0 | 1 | 0 | 0 | 0 | 0 | 0 | 0 | 0 | 0 | 0 | 0 | 0 | 0 | 0 | 0 | 0 | 0 | 0 | 0 | 0 | 0 | 0 | 0 | 0 | 0 | 0 | 0 | 0 | 0 | 0 | 0 | 0 | 0 | 0 | 0 | 0 | 0 |
| virus_Jun22.75v_2628 | 1 | 0 | 0 | 0 | 0 | 0 | 0 | 0 | 0 | 0 | 0 | 0 | 0 | 0 | 0 | 0 | 0 | 0 | 0 | 0 | 1 | 0 | 0 | 0 | 0 | 0 | 0 | 0 | 0 | 0 | 0 | 0 | 0 | 0 | 0 | 0 | 0 | 0 | 0 | 0 | 0 | 0 | 0 | 0 | 0 | 0 | 0 | 0 | 0 | 0 | 0 | 0 | 0 | 0 | 0 |
| virus_Jun22.75v_4010 | 1 | 0 | 0 | 0 | 0 | 0 | 0 | 0 | 0 | 0 | 0 | 0 | 0 | 0 | 0 | 0 | 0 | 0 | 0 | 0 | 1 | 0 | 0 | 0 | 0 | 0 | 0 | 0 | 0 | 0 | 0 | 0 | 0 | 0 | 0 | 0 | 0 | 0 | 0 | 0 | 0 | 0 | 0 | 0 | 0 | 0 | 0 | 0 | 0 | 0 | 0 | 0 | 0 | 0 | 0 |
| virus_Jun22.75v_5319 | 1 | 0 | 0 | 0 | 0 | 0 | 0 | 0 | 0 | 0 | 0 | 0 | 0 | 0 | 0 | 0 | 0 | 0 | 0 | 0 | 1 | 0 | 0 | 0 | 0 | 0 | 0 | 0 | 0 | 0 | 0 | 0 | 0 | 0 | 0 | 0 | 0 | 0 | 0 | 0 | 0 | 0 | 0 | 0 | 0 | 0 | 0 | 0 | 0 | 0 | 0 | 0 | 0 | 0 | 0 |
| virus_Jun22.75v_9284 | 1 | 0 | 0 | 0 | 0 | 0 | 0 | 0 | 0 | 0 | 0 | 0 | 0 | 0 | 0 | 0 | 0 | 0 | 0 | 0 | 1 | 0 | 0 | 0 | 0 | 0 | 0 | 0 | 0 | 0 | 0 | 0 | 0 | 0 | 0 | 0 | 0 | 0 | 0 | 0 | 0 | 0 | 0 | 0 | 0 | 0 | 0 | 0 | 0 | 0 | 0 | 0 | 0 | 0 | 0 |
| virus_Jun22.75v_13651 | 1 | 0 | 0 | 0 | 0 | 0 | 0 | 0 | 0 | 0 | 0 | 0 | 0 | 0 | 0 | 0 | 0 | 0 | 0 | 0 | 1 | 0 | 0 | 0 | 0 | 0 | 0 | 0 | 0 | 0 | 0 | 0 | 0 | 0 | 0 | 0 | 0 | 0 | 0 | 0 | 0 | 0 | 0 | 0 | 0 | 0 | 0 | 0 | 0 | 0 | 0 | 0 | 0 | 0 | 0 |
| virus_Jun22.75v_19461 | 1 | 0 | 0 | 0 | 0 | 0 | 0 | 0 | 0 | 0 | 0 | 0 | 0 | 0 | 0 | 0 | 0 | 0 | 0 | 0 | 1 | 0 | 0 | 0 | 0 | 0 | 0 | 0 | 0 | 0 | 0 | 0 | 0 | 0 | 0 | 0 | 0 | 0 | 0 | 0 | 0 | 0 | 0 | 0 | 0 | 0 | 0 | 0 | 0 | 0 | 0 | 0 | 0 | 0 | 0 |
| virus_Jun22.75v_21291 | 1 | 0 | 0 | 0 | 0 | 0 | 0 | 0 | 0 | 0 | 0 | 0 | 0 | 0 | 0 | 0 | 0 | 0 | 0 | 0 | 1 | 0 | 0 | 0 | 0 | 0 | 0 | 0 | 0 | 0 | 0 | 0 | 0 | 0 | 0 | 0 | 0 | 0 | 0 | 0 | 0 | 0 | 0 | 0 | 0 | 0 | 0 | 0 | 0 | 0 | 0 | 0 | 0 | 0 | 0 |
| virus_Jun22.75v_32291 | 1 | 0 | 0 | 0 | 0 | 0 | 0 | 0 | 0 | 0 | 0 | 0 | 0 | 0 | 0 | 0 | 0 | 0 | 0 | 0 | 1 | 0 | 0 | 0 | 0 | 0 | 0 | 0 | 0 | 0 | 0 | 0 | 0 | 0 | 0 | 0 | 0 | 0 | 0 | 0 | 0 | 0 | 0 | 0 | 0 | 0 | 0 | 0 | 0 | 0 | 0 | 0 | 0 | 0 | 0 |
| virus_Jun22.75v_34153 | 1 | 0 | 0 | 0 | 0 | 0 | 0 | 0 | 0 | 0 | 0 | 0 | 0 | 0 | 0 | 0 | 0 | 0 | 0 | 0 | 1 | 0 | 0 | 0 | 0 | 0 | 0 | 0 | 0 | 0 | 0 | 0 | 0 | 0 | 0 | 0 | 0 | 0 | 0 | 0 | 0 | 0 | 0 | 0 | 0 | 0 | 0 | 0 | 0 | 0 | 0 | 0 | 0 | 0 | 0 |
| virus_Jun22.75v_41066 | 1 | 0 | 0 | 0 | 0 | 0 | 0 | 0 | 0 | 0 | 0 | 0 | 0 | 0 | 0 | 0 | 0 | 0 | 0 | 0 | 1 | 0 | 0 | 0 | 0 | 0 | 0 | 0 | 0 | 0 | 0 | 0 | 0 | 0 | 0 | 0 | 0 | 0 | 0 | 0 | 0 | 0 | 0 | 0 | 0 | 0 | 0 | 0 | 0 | 0 | 0 | 0 | 0 | 0 | 0 |
| virus_May12.5v_21702 | 1 | 0 | 1 | 0 | 0 | 0 | 0 | 0 | 0 | 0 | 0 | 0 | 0 | 0 | 0 | 0 | 0 | 0 | 0 | 0 | 0 | 0 | 0 | 0 | 0 | 0 | 0 | 0 | 0 | 0 | 0 | 0 | 0 | 0 | 0 | 0 | 0 | 0 | 0 | 0 | 0 | 0 | 0 | 0 | 0 | 0 | 0 | 0 | 0 | 0 | 0 | 0 | 0 | 0 | 0 |
| virus_May12.25v_2423 | 1 | 0 | 0 | 0 | 1 | 0 | 0 | 0 | 0 | 0 | 0 | 0 | 0 | 0 | 0 | 0 | 0 | 0 | 0 | 0 | 0 | 0 | 0 | 0 | 0 | 0 | 0 | 0 | 0 | 0 | 0 | 0 | 0 | 0 | 0 | 0 | 0 | 0 | 0 | 0 | 0 | 0 | 0 | 0 | 0 | 0 | 0 | 0 | 0 | 0 | 0 | 0 | 0 | 0 | 0 |
| virus_May12.25v_5937 | 1 | 0 | 0 | 0 | 1 | 0 | 0 | 0 | 0 | 0 | 0 | 0 | 0 | 0 | 0 | 0 | 0 | 0 | 0 | 0 | 0 | 0 | 0 | 0 | 0 | 0 | 0 | 0 | 0 | 0 | 0 | 0 | 0 | 0 | 0 | 0 | 0 | 0 | 0 | 0 | 0 | 0 | 0 | 0 | 0 | 0 | 0 | 0 | 0 | 0 | 0 | 0 | 0 | 0 | 0 |
| virus_May12.25v_13721 | 1 | 0 | 0 | 0 | 1 | 0 | 0 | 0 | 0 | 0 | 0 | 0 | 0 | 0 | 0 | 0 | 0 | 0 | 0 | 0 | 0 | 0 | 0 | 0 | 0 | 0 | 0 | 0 | 0 | 0 | 0 | 0 | 0 | 0 | 0 | 0 | 0 | 0 | 0 | 0 | 0 | 0 | 0 | 0 | 0 | 0 | 0 | 0 | 0 | 0 | 0 | 0 | 0 | 0 | 0 |
| virus_May12.25v_29362 | 1 | 0 | 0 | 0 | 1 | 0 | 0 | 0 | 0 | 0 | 0 | 0 | 0 | 0 | 0 | 0 | 0 | 0 | 0 | 0 | 0 | 0 | 0 | 0 | 0 | 0 | 0 | 0 | 0 | 0 | 0 | 0 | 0 | 0 | 0 | 0 | 0 | 0 | 0 | 0 | 0 | 0 | 0 | 0 | 0 | 0 | 0 | 0 | 0 | 0 | 0 | 0 | 0 | 0 | 0 |
| virus_May12.25v_31200 | 1 | 0 | 0 | 0 | 1 | 0 | 0 | 0 | 0 | 0 | 0 | 0 | 0 | 0 | 0 | 0 | 0 | 0 | 0 | 0 | 0 | 0 | 0 | 0 | 0 | 0 | 0 | 0 | 0 | 0 | 0 | 0 | 0 | 0 | 0 | 0 | 0 | 0 | 0 | 0 | 0 | 0 | 0 | 0 | 0 | 0 | 0 | 0 | 0 | 0 | 0 | 0 | 0 | 0 | 0 |
| virus_May12.25v_33418 | 1 | 0 | 0 | 0 | 1 | 0 | 0 | 0 | 0 | 0 | 0 | 0 | 0 | 0 | 0 | 0 | 0 | 0 | 0 | 0 | 0 | 0 | 0 | 0 | 0 | 0 | 0 | 0 | 0 | 0 | 0 | 0 | 0 | 0 | 0 | 0 | 0 | 0 | 0 | 0 | 0 | 0 | 0 | 0 | 0 | 0 | 0 | 0 | 0 | 0 | 0 | 0 | 0 | 0 | 0 |
| virus_May12.25v_35142 | 1 | 0 | 0 | 0 | 1 | 0 | 0 | 0 | 0 | 0 | 0 | 0 | 0 | 0 | 0 | 0 | 0 | 0 | 0 | 0 | 0 | 0 | 0 | 0 | 0 | 0 | 0 | 0 | 0 | 0 | 0 | 0 | 0 | 0 | 0 | 0 | 0 | 0 | 0 | 0 | 0 | 0 | 0 | 0 | 0 | 0 | 0 | 0 | 0 | 0 | 0 | 0 | 0 | 0 | 0 |
| virus_May12.25v_35881 | 1 | 0 | 0 | 0 | 1 | 0 | 0 | 0 | 0 | 0 | 0 | 0 | 0 | 0 | 0 | 0 | 0 | 0 | 0 | 0 | 0 | 0 | 0 | 0 | 0 | 0 | 0 | 0 | 0 | 0 | 0 | 0 | 0 | 0 | 0 | 0 | 0 | 0 | 0 | 0 | 0 | 0 | 0 | 0 | 0 | 0 | 0 | 0 | 0 | 0 | 0 | 0 | 0 | 0 | 0 |
| virus_May12.50v_9895 | 1 | 0 | 0 | 0 | 0 | 0 | 1 | 0 | 0 | 0 | 0 | 0 | 0 | 0 | 0 | 0 | 0 | 0 | 0 | 0 | 0 | 0 | 0 | 0 | 0 | 0 | 0 | 0 | 0 | 0 | 0 | 0 | 0 | 0 | 0 | 0 | 0 | 0 | 0 | 0 | 0 | 0 | 0 | 0 | 0 | 0 | 0 | 0 | 0 | 0 | 0 | 0 | 0 | 0 | 0 |
| virus_May12.50v_9925 | 1 | 0 | 0 | 0 | 0 | 0 | 1 | 0 | 0 | 0 | 0 | 0 | 0 | 0 | 0 | 0 | 0 | 0 | 0 | 0 | 0 | 0 | 0 | 0 | 0 | 0 | 0 | 0 | 0 | 0 | 0 | 0 | 0 | 0 | 0 | 0 | 0 | 0 | 0 | 0 | 0 | 0 | 0 | 0 | 0 | 0 | 0 | 0 | 0 | 0 | 0 | 0 | 0 | 0 | 0 |
| virus_May12.50v_26510 | 1 | 0 | 0 | 0 | 0 | 0 | 1 | 0 | 0 | 0 | 0 | 0 | 0 | 0 | 0 | 0 | 0 | 0 | 0 | 0 | 0 | 0 | 0 | 0 | 0 | 0 | 0 | 0 | 0 | 0 | 0 | 0 | 0 | 0 | 0 | 0 | 0 | 0 | 0 | 0 | 0 | 0 | 0 | 0 | 0 | 0 | 0 | 0 | 0 | 0 | 0 | 0 | 0 | 0 | 0 |
| virus_May12.50v_29669 | 1 | 0 | 0 | 0 | 0 | 0 | 1 | 0 | 0 | 0 | 0 | 0 | 0 | 0 | 0 | 0 | 0 | 0 | 0 | 0 | 0 | 0 | 0 | 0 | 0 | 0 | 0 | 0 | 0 | 0 | 0 | 0 | 0 | 0 | 0 | 0 | 0 | 0 | 0 | 0 | 0 | 0 | 0 | 0 | 0 | 0 | 0 | 0 | 0 | 0 | 0 | 0 | 0 | 0 | 0 |
| virus_May12.50v_35559 | 1 | 0 | 0 | 0 | 0 | 0 | 1 | 0 | 0 | 0 | 0 | 0 | 0 | 0 | 0 | 0 | 0 | 0 | 0 | 0 | 0 | 0 | 0 | 0 | 0 | 0 | 0 | 0 | 0 | 0 | 0 | 0 | 0 | 0 | 0 | 0 | 0 | 0 | 0 | 0 | 0 | 0 | 0 | 0 | 0 | 0 | 0 | 0 | 0 | 0 | 0 | 0 | 0 | 0 | 0 |
| virus_May12.50v_35624 | 1 | 0 | 0 | 0 | 0 | 0 | 1 | 0 | 0 | 0 | 0 | 0 | 0 | 0 | 0 | 0 | 0 | 0 | 0 | 0 | 0 | 0 | 0 | 0 | 0 | 0 | 0 | 0 | 0 | 0 | 0 | 0 | 0 | 0 | 0 | 0 | 0 | 0 | 0 | 0 | 0 | 0 | 0 | 0 | 0 | 0 | 0 | 0 | 0 | 0 | 0 | 0 | 0 | 0 | 0 |
| virus_May12.50v_36443 | 1 | 0 | 0 | 0 | 0 | 0 | 1 | 0 | 0 | 0 | 0 | 0 | 0 | 0 | 0 | 0 | 0 | 0 | 0 | 0 | 0 | 0 | 0 | 0 | 0 | 0 | 0 | 0 | 0 | 0 | 0 | 0 | 0 | 0 | 0 | 0 | 0 | 0 | 0 | 0 | 0 | 0 | 0 | 0 | 0 | 0 | 0 | 0 | 0 | 0 | 0 | 0 | 0 | 0 | 0 |
| virus_May12.50v_36683 | 1 | 0 | 0 | 0 | 0 | 0 | 1 | 0 | 0 | 0 | 0 | 0 | 0 | 0 | 0 | 0 | 0 | 0 | 0 | 0 | 0 | 0 | 0 | 0 | 0 | 0 | 0 | 0 | 0 | 0 | 0 | 0 | 0 | 0 | 0 | 0 | 0 | 0 | 0 | 0 | 0 | 0 | 0 | 0 | 0 | 0 | 0 | 0 | 0 | 0 | 0 | 0 | 0 | 0 | 0 |
| virus_May12.50v_36914 | 1 | 0 | 0 | 0 | 0 | 0 | 1 | 0 | 0 | 0 | 0 | 0 | 0 | 0 | 0 | 0 | 0 | 0 | 0 | 0 | 0 | 0 | 0 | 0 | 0 | 0 | 0 | 0 | 0 | 0 | 0 | 0 | 0 | 0 | 0 | 0 | 0 | 0 | 0 | 0 | 0 | 0 | 0 | 0 | 0 | 0 | 0 | 0 | 0 | 0 | 0 | 0 | 0 | 0 | 0 |
| virus_May12.50v_39522 | 1 | 0 | 0 | 0 | 0 | 0 | 1 | 0 | 0 | 0 | 0 | 0 | 0 | 0 | 0 | 0 | 0 | 0 | 0 | 0 | 0 | 0 | 0 | 0 | 0 | 0 | 0 | 0 | 0 | 0 | 0 | 0 | 0 | 0 | 0 | 0 | 0 | 0 | 0 | 0 | 0 | 0 | 0 | 0 | 0 | 0 | 0 | 0 | 0 | 0 | 0 | 0 | 0 | 0 | 0 |
| virus_May12.75v_1135 | 1 | 0 | 0 | 0 | 0 | 0 | 0 | 0 | 1 | 0 | 0 | 0 | 0 | 0 | 0 | 0 | 0 | 0 | 0 | 0 | 0 | 0 | 0 | 0 | 0 | 0 | 0 | 0 | 0 | 0 | 0 | 0 | 0 | 0 | 0 | 0 | 0 | 0 | 0 | 0 | 0 | 0 | 0 | 0 | 0 | 0 | 0 | 0 | 0 | 0 | 0 | 0 | 0 | 0 | 0 |
| virus_May12.75v_6881 | 1 | 0 | 0 | 0 | 0 | 0 | 0 | 0 | 1 | 0 | 0 | 0 | 0 | 0 | 0 | 0 | 0 | 0 | 0 | 0 | 0 | 0 | 0 | 0 | 0 | 0 | 0 | 0 | 0 | 0 | 0 | 0 | 0 | 0 | 0 | 0 | 0 | 0 | 0 | 0 | 0 | 0 | 0 | 0 | 0 | 0 | 0 | 0 | 0 | 0 | 0 | 0 | 0 | 0 | 0 |
| virus_May12.75v_11710 | 1 | 0 | 0 | 0 | 0 | 0 | 0 | 0 | 1 | 0 | 0 | 0 | 0 | 0 | 0 | 0 | 0 | 0 | 0 | 0 | 0 | 0 | 0 | 0 | 0 | 0 | 0 | 0 | 0 | 0 | 0 | 0 | 0 | 0 | 0 | 0 | 0 | 0 | 0 | 0 | 0 | 0 | 0 | 0 | 0 | 0 | 0 | 0 | 0 | 0 | 0 | 0 | 0 | 0 | 0 |
| virus_May12.75v_11814 | 1 | 0 | 0 | 0 | 0 | 0 | 0 | 0 | 1 | 0 | 0 | 0 | 0 | 0 | 0 | 0 | 0 | 0 | 0 | 0 | 0 | 0 | 0 | 0 | 0 | 0 | 0 | 0 | 0 | 0 | 0 | 0 | 0 | 0 | 0 | 0 | 0 | 0 | 0 | 0 | 0 | 0 | 0 | 0 | 0 | 0 | 0 | 0 | 0 | 0 | 0 | 0 | 0 | 0 | 0 |
| virus_May12.75v_14086 | 1 | 0 | 0 | 0 | 0 | 0 | 0 | 0 | 1 | 0 | 0 | 0 | 0 | 0 | 0 | 0 | 0 | 0 | 0 | 0 | 0 | 0 | 0 | 0 | 0 | 0 | 0 | 0 | 0 | 0 | 0 | 0 | 0 | 0 | 0 | 0 | 0 | 0 | 0 | 0 | 0 | 0 | 0 | 0 | 0 | 0 | 0 | 0 | 0 | 0 | 0 | 0 | 0 | 0 | 0 |
| virus_May12.75v_16053 | 1 | 0 | 0 | 0 | 0 | 0 | 0 | 0 | 1 | 0 | 0 | 0 | 0 | 0 | 0 | 0 | 0 | 0 | 0 | 0 | 0 | 0 | 0 | 0 | 0 | 0 | 0 | 0 | 0 | 0 | 0 | 0 | 0 | 0 | 0 | 0 | 0 | 0 | 0 | 0 | 0 | 0 | 0 | 0 | 0 | 0 | 0 | 0 | 0 | 0 | 0 | 0 | 0 | 0 | 0 |
| virus_May12.75v_16591 | 1 | 0 | 0 | 0 | 0 | 0 | 0 | 0 | 1 | 0 | 0 | 0 | 0 | 0 | 0 | 0 | 0 | 0 | 0 | 0 | 0 | 0 | 0 | 0 | 0 | 0 | 0 | 0 | 0 | 0 | 0 | 0 | 0 | 0 | 0 | 0 | 0 | 0 | 0 | 0 | 0 | 0 | 0 | 0 | 0 | 0 | 0 | 0 | 0 | 0 | 0 | 0 | 0 | 0 | 0 |
| virus_May12.75v_18089 | 1 | 0 | 0 | 0 | 0 | 0 | 0 | 0 | 1 | 0 | 0 | 0 | 0 | 0 | 0 | 0 | 0 | 0 | 0 | 0 | 0 | 0 | 0 | 0 | 0 | 0 | 0 | 0 | 0 | 0 | 0 | 0 | 0 | 0 | 0 | 0 | 0 | 0 | 0 | 0 | 0 | 0 | 0 | 0 | 0 | 0 | 0 | 0 | 0 | 0 | 0 | 0 | 0 | 0 | 0 |
| virus_May12.75v_26070 | 1 | 0 | 0 | 0 | 0 | 0 | 0 | 0 | 1 | 0 | 0 | 0 | 0 | 0 | 0 | 0 | 0 | 0 | 0 | 0 | 0 | 0 | 0 | 0 | 0 | 0 | 0 | 0 | 0 | 0 | 0 | 0 | 0 | 0 | 0 | 0 | 0 | 0 | 0 | 0 | 0 | 0 | 0 | 0 | 0 | 0 | 0 | 0 | 0 | 0 | 0 | 0 | 0 | 0 | 0 |
| virus_Sep8.0v_20500 | 1 | 0 | 0 | 0 | 0 | 0 | 0 | 0 | 0 | 0 | 0 | 0 | 0 | 0 | 0 | 0 | 0 | 0 | 0 | 0 | 0 | 0 | 0 | 0 | 0 | 0 | 0 | 0 | 0 | 0 | 0 | 0 | 0 | 0 | 0 | 0 | 0 | 0 | 0 | 0 | 0 | 0 | 0 | 0 | 0 | 0 | 0 | 0 | 1 | 0 | 0 | 0 | 0 | 0 | 0 |
| virus_Sep8.5v_498 | 1 | 0 | 0 | 0 | 0 | 0 | 0 | 0 | 0 | 0 | 0 | 0 | 0 | 0 | 0 | 0 | 0 | 0 | 0 | 0 | 0 | 0 | 0 | 0 | 0 | 0 | 0 | 0 | 0 | 0 | 0 | 0 | 0 | 0 | 0 | 0 | 0 | 0 | 0 | 0 | 0 | 0 | 0 | 0 | 0 | 0 | 0 | 0 | 0 | 0 | 1 | 0 | 0 | 0 | 0 |
| virus_Sep8.5v_2658 | 1 | 0 | 0 | 0 | 0 | 0 | 0 | 0 | 0 | 0 | 0 | 0 | 0 | 0 | 0 | 0 | 0 | 0 | 0 | 0 | 0 | 0 | 0 | 0 | 0 | 0 | 0 | 0 | 0 | 0 | 0 | 0 | 0 | 0 | 0 | 0 | 0 | 0 | 0 | 0 | 0 | 0 | 0 | 0 | 0 | 0 | 0 | 0 | 0 | 0 | 1 | 0 | 0 | 0 | 0 |
| virus_Sep8.5v_3819 | 1 | 0 | 0 | 0 | 0 | 0 | 0 | 0 | 0 | 0 | 0 | 0 | 0 | 0 | 0 | 0 | 0 | 0 | 0 | 0 | 0 | 0 | 0 | 0 | 0 | 0 | 0 | 0 | 0 | 0 | 0 | 0 | 0 | 0 | 0 | 0 | 0 | 0 | 0 | 0 | 0 | 0 | 0 | 0 | 0 | 0 | 0 | 0 | 0 | 0 | 1 | 0 | 0 | 0 | 0 |
| virus_Sep8.5v_5681 | 1 | 0 | 0 | 0 | 0 | 0 | 0 | 0 | 0 | 0 | 0 | 0 | 0 | 0 | 0 | 0 | 0 | 0 | 0 | 0 | 0 | 0 | 0 | 0 | 0 | 0 | 0 | 0 | 0 | 0 | 0 | 0 | 0 | 0 | 0 | 0 | 0 | 0 | 0 | 0 | 0 | 0 | 0 | 0 | 0 | 0 | 0 | 0 | 0 | 0 | 1 | 0 | 0 | 0 | 0 |
| virus_Sep8.5v_10931 | 1 | 0 | 0 | 0 | 0 | 0 | 0 | 0 | 0 | 0 | 0 | 0 | 0 | 0 | 0 | 0 | 0 | 0 | 0 | 0 | 0 | 0 | 0 | 0 | 0 | 0 | 0 | 0 | 0 | 0 | 0 | 0 | 0 | 0 | 0 | 0 | 0 | 0 | 0 | 0 | 0 | 0 | 0 | 0 | 0 | 0 | 0 | 0 | 0 | 0 | 1 | 0 | 0 | 0 | 0 |
| virus_Sep8.5v_11546 | 1 | 0 | 0 | 0 | 0 | 0 | 0 | 0 | 0 | 0 | 0 | 0 | 0 | 0 | 0 | 0 | 0 | 0 | 0 | 0 | 0 | 0 | 0 | 0 | 0 | 0 | 0 | 0 | 0 | 0 | 0 | 0 | 0 | 0 | 0 | 0 | 0 | 0 | 0 | 0 | 0 | 0 | 0 | 0 | 0 | 0 | 0 | 0 | 0 | 0 | 1 | 0 | 0 | 0 | 0 |
| virus_Sep8.5v_12583 | 1 | 0 | 0 | 0 | 0 | 0 | 0 | 0 | 0 | 0 | 0 | 0 | 0 | 0 | 0 | 0 | 0 | 0 | 0 | 0 | 0 | 0 | 0 | 0 | 0 | 0 | 0 | 0 | 0 | 0 | 0 | 0 | 0 | 0 | 0 | 0 | 0 | 0 | 0 | 0 | 0 | 0 | 0 | 0 | 0 | 0 | 0 | 0 | 0 | 0 | 1 | 0 | 0 | 0 | 0 |
| virus_Sep8.25v_293 | 1 | 0 | 0 | 0 | 0 | 0 | 0 | 0 | 0 | 0 | 0 | 0 | 0 | 0 | 0 | 0 | 0 | 0 | 0 | 0 | 0 | 0 | 0 | 0 | 0 | 0 | 0 | 0 | 0 | 0 | 0 | 0 | 0 | 0 | 0 | 0 | 0 | 0 | 0 | 0 | 0 | 0 | 0 | 0 | 0 | 0 | 0 | 0 | 0 | 0 | 0 | 0 | 1 | 0 | 0 |
| virus_Sep8.25v_1130 | 1 | 0 | 0 | 0 | 0 | 0 | 0 | 0 | 0 | 0 | 0 | 0 | 0 | 0 | 0 | 0 | 0 | 0 | 0 | 0 | 0 | 0 | 0 | 0 | 0 | 0 | 0 | 0 | 0 | 0 | 0 | 0 | 0 | 0 | 0 | 0 | 0 | 0 | 0 | 0 | 0 | 0 | 0 | 0 | 0 | 0 | 0 | 0 | 0 | 0 | 0 | 0 | 1 | 0 | 0 |
| virus_Sep8.25v_6070 | 1 | 0 | 0 | 0 | 0 | 0 | 0 | 0 | 0 | 0 | 0 | 0 | 0 | 0 | 0 | 0 | 0 | 0 | 0 | 0 | 0 | 0 | 0 | 0 | 0 | 0 | 0 | 0 | 0 | 0 | 0 | 0 | 0 | 0 | 0 | 0 | 0 | 0 | 0 | 0 | 0 | 0 | 0 | 0 | 0 | 0 | 0 | 0 | 0 | 0 | 0 | 0 | 1 | 0 | 0 |
| virus_Sep8.25v_11003 | 1 | 0 | 0 | 0 | 0 | 0 | 0 | 0 | 0 | 0 | 0 | 0 | 0 | 0 | 0 | 0 | 0 | 0 | 0 | 0 | 0 | 0 | 0 | 0 | 0 | 0 | 0 | 0 | 0 | 0 | 0 | 0 | 0 | 0 | 0 | 0 | 0 | 0 | 0 | 0 | 0 | 0 | 0 | 0 | 0 | 0 | 0 | 0 | 0 | 0 | 0 | 0 | 1 | 0 | 0 |
| virus_Sep8.50v_906 | 1 | 0 | 0 | 0 | 0 | 0 | 0 | 0 | 0 | 0 | 0 | 0 | 0 | 0 | 0 | 0 | 0 | 0 | 0 | 0 | 0 | 0 | 0 | 0 | 0 | 0 | 0 | 0 | 0 | 0 | 0 | 0 | 0 | 0 | 0 | 0 | 0 | 0 | 0 | 0 | 0 | 0 | 0 | 0 | 0 | 0 | 0 | 0 | 0 | 0 | 0 | 0 | 0 | 0 | 1 |
| virus_Sep8.50v_5030 | 1 | 0 | 0 | 0 | 0 | 0 | 0 | 0 | 0 | 0 | 0 | 0 | 0 | 0 | 0 | 0 | 0 | 0 | 0 | 0 | 0 | 0 | 0 | 0 | 0 | 0 | 0 | 0 | 0 | 0 | 0 | 0 | 0 | 0 | 0 | 0 | 0 | 0 | 0 | 0 | 0 | 0 | 0 | 0 | 0 | 0 | 0 | 0 | 0 | 0 | 0 | 0 | 0 | 0 | 1 |
| virus_Sep8.50v_8612 | 1 | 0 | 0 | 0 | 0 | 0 | 0 | 0 | 0 | 0 | 0 | 0 | 0 | 0 | 0 | 0 | 0 | 0 | 0 | 0 | 0 | 0 | 0 | 0 | 0 | 0 | 0 | 0 | 0 | 0 | 0 | 0 | 0 | 0 | 0 | 0 | 0 | 0 | 0 | 0 | 0 | 0 | 0 | 0 | 0 | 0 | 0 | 0 | 0 | 0 | 0 | 0 | 0 | 0 | 1 |
| virus_Sep8.50v_9119 | 1 | 0 | 0 | 0 | 0 | 0 | 0 | 0 | 0 | 0 | 0 | 0 | 0 | 0 | 0 | 0 | 0 | 0 | 0 | 0 | 0 | 0 | 0 | 0 | 0 | 0 | 0 | 0 | 0 | 0 | 0 | 0 | 0 | 0 | 0 | 0 | 0 | 0 | 0 | 0 | 0 | 0 | 0 | 0 | 0 | 0 | 0 | 0 | 0 | 0 | 0 | 0 | 0 | 0 | 1 |
| virus_Sep8.50v_11061 | 1 | 0 | 0 | 0 | 0 | 0 | 0 | 0 | 0 | 0 | 0 | 0 | 0 | 0 | 0 | 0 | 0 | 0 | 0 | 0 | 0 | 0 | 0 | 0 | 0 | 0 | 0 | 0 | 0 | 0 | 0 | 0 | 0 | 0 | 0 | 0 | 0 | 0 | 0 | 0 | 0 | 0 | 0 | 0 | 0 | 0 | 0 | 0 | 0 | 0 | 0 | 0 | 0 | 0 | 1 |
| virus_Sep8.50v_11476 | 1 | 0 | 0 | 0 | 0 | 0 | 0 | 0 | 0 | 0 | 0 | 0 | 0 | 0 | 0 | 0 | 0 | 0 | 0 | 0 | 0 | 0 | 0 | 0 | 0 | 0 | 0 | 0 | 0 | 0 | 0 | 0 | 0 | 0 | 0 | 0 | 0 | 0 | 0 | 0 | 0 | 0 | 0 | 0 | 0 | 0 | 0 | 0 | 0 | 0 | 0 | 0 | 0 | 0 | 1 |
| virus_Sep8.50v_16464 | 1 | 0 | 0 | 0 | 0 | 0 | 0 | 0 | 0 | 0 | 0 | 0 | 0 | 0 | 0 | 0 | 0 | 0 | 0 | 0 | 0 | 0 | 0 | 0 | 0 | 0 | 0 | 0 | 0 | 0 | 0 | 0 | 0 | 0 | 0 | 0 | 0 | 0 | 0 | 0 | 0 | 0 | 0 | 0 | 0 | 0 | 0 | 0 | 0 | 0 | 0 | 0 | 0 | 0 | 1 |
| virus_Sep8.50v_19346 | 1 | 0 | 0 | 0 | 0 | 0 | 0 | 0 | 0 | 0 | 0 | 0 | 0 | 0 | 0 | 0 | 0 | 0 | 0 | 0 | 0 | 0 | 0 | 0 | 0 | 0 | 0 | 0 | 0 | 0 | 0 | 0 | 0 | 0 | 0 | 0 | 0 | 0 | 0 | 0 | 0 | 0 | 0 | 0 | 0 | 0 | 0 | 0 | 0 | 0 | 0 | 0 | 0 | 0 | 1 |
| virus_Sep8.50v_20095 | 1 | 0 | 0 | 0 | 0 | 0 | 0 | 0 | 0 | 0 | 0 | 0 | 0 | 0 | 0 | 0 | 0 | 0 | 0 | 0 | 0 | 0 | 0 | 0 | 0 | 0 | 0 | 0 | 0 | 0 | 0 | 0 | 0 | 0 | 0 | 0 | 0 | 0 | 0 | 0 | 0 | 0 | 0 | 0 | 0 | 0 | 0 | 0 | 0 | 0 | 0 | 0 | 0 | 0 | 1 |
| virus_Sep8.50v_25302 | 1 | 0 | 0 | 0 | 0 | 0 | 0 | 0 | 0 | 0 | 0 | 0 | 0 | 0 | 0 | 0 | 0 | 0 | 0 | 0 | 0 | 0 | 0 | 0 | 0 | 0 | 0 | 0 | 0 | 0 | 0 | 0 | 0 | 0 | 0 | 0 | 0 | 0 | 0 | 0 | 0 | 0 | 0 | 0 | 0 | 0 | 0 | 0 | 0 | 0 | 0 | 0 | 0 | 0 | 1 |
| virus_Sep8.50v_29766 | 1 | 0 | 0 | 0 | 0 | 0 | 0 | 0 | 0 | 0 | 0 | 0 | 0 | 0 | 0 | 0 | 0 | 0 | 0 | 0 | 0 | 0 | 0 | 0 | 0 | 0 | 0 | 0 | 0 | 0 | 0 | 0 | 0 | 0 | 0 | 0 | 0 | 0 | 0 | 0 | 0 | 0 | 0 | 0 | 0 | 0 | 0 | 0 | 0 | 0 | 0 | 0 | 0 | 0 | 1 |
| virus_Sep8.50v_34884 | 1 | 0 | 0 | 0 | 0 | 0 | 0 | 0 | 0 | 0 | 0 | 0 | 0 | 0 | 0 | 0 | 0 | 0 | 0 | 0 | 0 | 0 | 0 | 0 | 0 | 0 | 0 | 0 | 0 | 0 | 0 | 0 | 0 | 0 | 0 | 0 | 0 | 0 | 0 | 0 | 0 | 0 | 0 | 0 | 0 | 0 | 0 | 0 | 0 | 0 | 0 | 0 | 0 | 0 | 1 |
| virus_Sep8.50v_40247 | 1 | 0 | 0 | 0 | 0 | 0 | 0 | 0 | 0 | 0 | 0 | 0 | 0 | 0 | 0 | 0 | 0 | 0 | 0 | 0 | 0 | 0 | 0 | 0 | 0 | 0 | 0 | 0 | 0 | 0 | 0 | 0 | 0 | 0 | 0 | 0 | 0 | 0 | 0 | 0 | 0 | 0 | 0 | 0 | 0 | 0 | 0 | 0 | 0 | 0 | 0 | 0 | 0 | 0 | 1 |
| bacteria_Jul11d0b_57039 | 1 | 0 | 0 | 0 | 0 | 0 | 0 | 0 | 0 | 0 | 0 | 0 | 0 | 0 | 0 | 0 | 0 | 0 | 0 | 0 | 0 | 1 | 0 | 0 | 0 | 0 | 0 | 0 | 0 | 0 | 0 | 0 | 0 | 0 | 0 | 0 | 0 | 0 | 0 | 0 | 0 | 0 | 0 | 0 | 0 | 0 | 0 | 0 | 0 | 0 | 0 | 0 | 0 | 0 | 0 |
| bacteria_Jul11d0b_57590 | 1 | 0 | 0 | 0 | 0 | 0 | 0 | 0 | 0 | 0 | 0 | 0 | 0 | 0 | 0 | 0 | 0 | 0 | 0 | 0 | 0 | 1 | 0 | 0 | 0 | 0 | 0 | 0 | 0 | 0 | 0 | 0 | 0 | 0 | 0 | 0 | 0 | 0 | 0 | 0 | 0 | 0 | 0 | 0 | 0 | 0 | 0 | 0 | 0 | 0 | 0 | 0 | 0 | 0 | 0 |
| bacteria_Aug5d0b_3944 | 1 | 0 | 0 | 0 | 0 | 0 | 0 | 0 | 0 | 0 | 0 | 0 | 0 | 0 | 0 | 0 | 0 | 0 | 0 | 0 | 0 | 0 | 0 | 0 | 0 | 0 | 0 | 0 | 0 | 0 | 0 | 0 | 0 | 0 | 0 | 0 | 0 | 1 | 0 | 0 | 0 | 0 | 0 | 0 | 0 | 0 | 0 | 0 | 0 | 0 | 0 | 0 | 0 | 0 | 0 |
| bacteria_Aug5d0b_10700 | 1 | 0 | 0 | 0 | 0 | 0 | 0 | 0 | 0 | 0 | 0 | 0 | 0 | 0 | 0 | 0 | 0 | 0 | 0 | 0 | 0 | 0 | 0 | 0 | 0 | 0 | 0 | 0 | 0 | 0 | 0 | 0 | 0 | 0 | 0 | 0 | 0 | 1 | 0 | 0 | 0 | 0 | 0 | 0 | 0 | 0 | 0 | 0 | 0 | 0 | 0 | 0 | 0 | 0 | 0 |
| bacteria_Aug5d0b_21195 | 1 | 0 | 0 | 0 | 0 | 0 | 0 | 0 | 0 | 0 | 0 | 0 | 0 | 0 | 0 | 0 | 0 | 0 | 0 | 0 | 0 | 0 | 0 | 0 | 0 | 0 | 0 | 0 | 0 | 0 | 0 | 0 | 0 | 0 | 0 | 0 | 0 | 1 | 0 | 0 | 0 | 0 | 0 | 0 | 0 | 0 | 0 | 0 | 0 | 0 | 0 | 0 | 0 | 0 | 0 |
| bacteria_Aug5d0b_27442 | 1 | 0 | 0 | 0 | 0 | 0 | 0 | 0 | 0 | 0 | 0 | 0 | 0 | 0 | 0 | 0 | 0 | 0 | 0 | 0 | 0 | 0 | 0 | 0 | 0 | 0 | 0 | 0 | 0 | 0 | 0 | 0 | 0 | 0 | 0 | 0 | 0 | 1 | 0 | 0 | 0 | 0 | 0 | 0 | 0 | 0 | 0 | 0 | 0 | 0 | 0 | 0 | 0 | 0 | 0 |
| bacteria_Aug5d0b_38140 | 1 | 0 | 0 | 0 | 0 | 0 | 0 | 0 | 0 | 0 | 0 | 0 | 0 | 0 | 0 | 0 | 0 | 0 | 0 | 0 | 0 | 0 | 0 | 0 | 0 | 0 | 0 | 0 | 0 | 0 | 0 | 0 | 0 | 0 | 0 | 0 | 0 | 1 | 0 | 0 | 0 | 0 | 0 | 0 | 0 | 0 | 0 | 0 | 0 | 0 | 0 | 0 | 0 | 0 | 0 |
| bacteria_Aug5d0b_45479 | 1 | 0 | 0 | 0 | 0 | 0 | 0 | 0 | 0 | 0 | 0 | 0 | 0 | 0 | 0 | 0 | 0 | 0 | 0 | 0 | 0 | 0 | 0 | 0 | 0 | 0 | 0 | 0 | 0 | 0 | 0 | 0 | 0 | 0 | 0 | 0 | 0 | 1 | 0 | 0 | 0 | 0 | 0 | 0 | 0 | 0 | 0 | 0 | 0 | 0 | 0 | 0 | 0 | 0 | 0 |
| bacteria_Aug5d0b_49411 | 1 | 0 | 0 | 0 | 0 | 0 | 0 | 0 | 0 | 0 | 0 | 0 | 0 | 0 | 0 | 0 | 0 | 0 | 0 | 0 | 0 | 0 | 0 | 0 | 0 | 0 | 0 | 0 | 0 | 0 | 0 | 0 | 0 | 0 | 0 | 0 | 0 | 1 | 0 | 0 | 0 | 0 | 0 | 0 | 0 | 0 | 0 | 0 | 0 | 0 | 0 | 0 | 0 | 0 | 0 |
| bacteria_Aug5d5b_28863 | 1 | 0 | 0 | 0 | 0 | 0 | 0 | 0 | 0 | 0 | 0 | 0 | 0 | 0 | 0 | 0 | 0 | 0 | 0 | 0 | 0 | 0 | 0 | 0 | 0 | 0 | 0 | 0 | 0 | 0 | 0 | 0 | 0 | 0 | 0 | 0 | 0 | 0 | 0 | 1 | 0 | 0 | 0 | 0 | 0 | 0 | 0 | 0 | 0 | 0 | 0 | 0 | 0 | 0 | 0 |
| bacteria_Aug5d5b_68029 | 1 | 0 | 0 | 0 | 0 | 0 | 0 | 0 | 0 | 0 | 0 | 0 | 0 | 0 | 0 | 0 | 0 | 0 | 0 | 0 | 0 | 0 | 0 | 0 | 0 | 0 | 0 | 0 | 0 | 0 | 0 | 0 | 0 | 0 | 0 | 0 | 0 | 0 | 0 | 1 | 0 | 0 | 0 | 0 | 0 | 0 | 0 | 0 | 0 | 0 | 0 | 0 | 0 | 0 | 0 |
| bacteria_Aug5d25b_5118 | 1 | 0 | 0 | 0 | 0 | 0 | 0 | 0 | 0 | 0 | 0 | 0 | 0 | 0 | 0 | 0 | 0 | 0 | 0 | 0 | 0 | 0 | 0 | 0 | 0 | 0 | 0 | 0 | 0 | 0 | 0 | 0 | 0 | 0 | 0 | 0 | 0 | 0 | 0 | 0 | 0 | 1 | 0 | 0 | 0 | 0 | 0 | 0 | 0 | 0 | 0 | 0 | 0 | 0 | 0 |
| bacteria_Aug5d25b_9853 | 1 | 0 | 0 | 0 | 0 | 0 | 0 | 0 | 0 | 0 | 0 | 0 | 0 | 0 | 0 | 0 | 0 | 0 | 0 | 0 | 0 | 0 | 0 | 0 | 0 | 0 | 0 | 0 | 0 | 0 | 0 | 0 | 0 | 0 | 0 | 0 | 0 | 0 | 0 | 0 | 0 | 1 | 0 | 0 | 0 | 0 | 0 | 0 | 0 | 0 | 0 | 0 | 0 | 0 | 0 |
| bacteria_Aug5d25b_9970 | 1 | 0 | 0 | 0 | 0 | 0 | 0 | 0 | 0 | 0 | 0 | 0 | 0 | 0 | 0 | 0 | 0 | 0 | 0 | 0 | 0 | 0 | 0 | 0 | 0 | 0 | 0 | 0 | 0 | 0 | 0 | 0 | 0 | 0 | 0 | 0 | 0 | 0 | 0 | 0 | 0 | 1 | 0 | 0 | 0 | 0 | 0 | 0 | 0 | 0 | 0 | 0 | 0 | 0 | 0 |
| bacteria_Aug5d25b_11149 | 1 | 0 | 0 | 0 | 0 | 0 | 0 | 0 | 0 | 0 | 0 | 0 | 0 | 0 | 0 | 0 | 0 | 0 | 0 | 0 | 0 | 0 | 0 | 0 | 0 | 0 | 0 | 0 | 0 | 0 | 0 | 0 | 0 | 0 | 0 | 0 | 0 | 0 | 0 | 0 | 0 | 1 | 0 | 0 | 0 | 0 | 0 | 0 | 0 | 0 | 0 | 0 | 0 | 0 | 0 |
| bacteria_Aug5d25b_11719 | 1 | 0 | 0 | 0 | 0 | 0 | 0 | 0 | 0 | 0 | 0 | 0 | 0 | 0 | 0 | 0 | 0 | 0 | 0 | 0 | 0 | 0 | 0 | 0 | 0 | 0 | 0 | 0 | 0 | 0 | 0 | 0 | 0 | 0 | 0 | 0 | 0 | 0 | 0 | 0 | 0 | 1 | 0 | 0 | 0 | 0 | 0 | 0 | 0 | 0 | 0 | 0 | 0 | 0 | 0 |
| bacteria_Aug5d25b_17410 | 1 | 0 | 0 | 0 | 0 | 0 | 0 | 0 | 0 | 0 | 0 | 0 | 0 | 0 | 0 | 0 | 0 | 0 | 0 | 0 | 0 | 0 | 0 | 0 | 0 | 0 | 0 | 0 | 0 | 0 | 0 | 0 | 0 | 0 | 0 | 0 | 0 | 0 | 0 | 0 | 0 | 1 | 0 | 0 | 0 | 0 | 0 | 0 | 0 | 0 | 0 | 0 | 0 | 0 | 0 |
| bacteria_Aug5d25b_27681 | 1 | 0 | 0 | 0 | 0 | 0 | 0 | 0 | 0 | 0 | 0 | 0 | 0 | 0 | 0 | 0 | 0 | 0 | 0 | 0 | 0 | 0 | 0 | 0 | 0 | 0 | 0 | 0 | 0 | 0 | 0 | 0 | 0 | 0 | 0 | 0 | 0 | 0 | 0 | 0 | 0 | 1 | 0 | 0 | 0 | 0 | 0 | 0 | 0 | 0 | 0 | 0 | 0 | 0 | 0 |
| bacteria_Aug5d25b_36633 | 1 | 0 | 0 | 0 | 0 | 0 | 0 | 0 | 0 | 0 | 0 | 0 | 0 | 0 | 0 | 0 | 0 | 0 | 0 | 0 | 0 | 0 | 0 | 0 | 0 | 0 | 0 | 0 | 0 | 0 | 0 | 0 | 0 | 0 | 0 | 0 | 0 | 0 | 0 | 0 | 0 | 1 | 0 | 0 | 0 | 0 | 0 | 0 | 0 | 0 | 0 | 0 | 0 | 0 | 0 |
| bacteria_Aug5d25b_41414 | 1 | 0 | 0 | 0 | 0 | 0 | 0 | 0 | 0 | 0 | 0 | 0 | 0 | 0 | 0 | 0 | 0 | 0 | 0 | 0 | 0 | 0 | 0 | 0 | 0 | 0 | 0 | 0 | 0 | 0 | 0 | 0 | 0 | 0 | 0 | 0 | 0 | 0 | 0 | 0 | 0 | 1 | 0 | 0 | 0 | 0 | 0 | 0 | 0 | 0 | 0 | 0 | 0 | 0 | 0 |
| bacteria_Aug5d25b_42877 | 1 | 0 | 0 | 0 | 0 | 0 | 0 | 0 | 0 | 0 | 0 | 0 | 0 | 0 | 0 | 0 | 0 | 0 | 0 | 0 | 0 | 0 | 0 | 0 | 0 | 0 | 0 | 0 | 0 | 0 | 0 | 0 | 0 | 0 | 0 | 0 | 0 | 0 | 0 | 0 | 0 | 1 | 0 | 0 | 0 | 0 | 0 | 0 | 0 | 0 | 0 | 0 | 0 | 0 | 0 |
| bacteria_Aug5d25b_44898 | 1 | 0 | 0 | 0 | 0 | 0 | 0 | 0 | 0 | 0 | 0 | 0 | 0 | 0 | 0 | 0 | 0 | 0 | 0 | 0 | 0 | 0 | 0 | 0 | 0 | 0 | 0 | 0 | 0 | 0 | 0 | 0 | 0 | 0 | 0 | 0 | 0 | 0 | 0 | 0 | 0 | 1 | 0 | 0 | 0 | 0 | 0 | 0 | 0 | 0 | 0 | 0 | 0 | 0 | 0 |
| bacteria_Aug5d25b_46724 | 1 | 0 | 0 | 0 | 0 | 0 | 0 | 0 | 0 | 0 | 0 | 0 | 0 | 0 | 0 | 0 | 0 | 0 | 0 | 0 | 0 | 0 | 0 | 0 | 0 | 0 | 0 | 0 | 0 | 0 | 0 | 0 | 0 | 0 | 0 | 0 | 0 | 0 | 0 | 0 | 0 | 1 | 0 | 0 | 0 | 0 | 0 | 0 | 0 | 0 | 0 | 0 | 0 | 0 | 0 |
| bacteria_Aug5d25b_46892 | 1 | 0 | 0 | 0 | 0 | 0 | 0 | 0 | 0 | 0 | 0 | 0 | 0 | 0 | 0 | 0 | 0 | 0 | 0 | 0 | 0 | 0 | 0 | 0 | 0 | 0 | 0 | 0 | 0 | 0 | 0 | 0 | 0 | 0 | 0 | 0 | 0 | 0 | 0 | 0 | 0 | 1 | 0 | 0 | 0 | 0 | 0 | 0 | 0 | 0 | 0 | 0 | 0 | 0 | 0 |
| bacteria_Aug5d25b_52110 | 1 | 0 | 0 | 0 | 0 | 0 | 0 | 0 | 0 | 0 | 0 | 0 | 0 | 0 | 0 | 0 | 0 | 0 | 0 | 0 | 0 | 0 | 0 | 0 | 0 | 0 | 0 | 0 | 0 | 0 | 0 | 0 | 0 | 0 | 0 | 0 | 0 | 0 | 0 | 0 | 0 | 1 | 0 | 0 | 0 | 0 | 0 | 0 | 0 | 0 | 0 | 0 | 0 | 0 | 0 |
| bacteria_Aug5d25b_52686 | 1 | 0 | 0 | 0 | 0 | 0 | 0 | 0 | 0 | 0 | 0 | 0 | 0 | 0 | 0 | 0 | 0 | 0 | 0 | 0 | 0 | 0 | 0 | 0 | 0 | 0 | 0 | 0 | 0 | 0 | 0 | 0 | 0 | 0 | 0 | 0 | 0 | 0 | 0 | 0 | 0 | 1 | 0 | 0 | 0 | 0 | 0 | 0 | 0 | 0 | 0 | 0 | 0 | 0 | 0 |
| bacteria_Aug5d25b_55450 | 1 | 0 | 0 | 0 | 0 | 0 | 0 | 0 | 0 | 0 | 0 | 0 | 0 | 0 | 0 | 0 | 0 | 0 | 0 | 0 | 0 | 0 | 0 | 0 | 0 | 0 | 0 | 0 | 0 | 0 | 0 | 0 | 0 | 0 | 0 | 0 | 0 | 0 | 0 | 0 | 0 | 1 | 0 | 0 | 0 | 0 | 0 | 0 | 0 | 0 | 0 | 0 | 0 | 0 | 0 |
| bacteria_Aug5d25b_58274 | 1 | 0 | 0 | 0 | 0 | 0 | 0 | 0 | 0 | 0 | 0 | 0 | 0 | 0 | 0 | 0 | 0 | 0 | 0 | 0 | 0 | 0 | 0 | 0 | 0 | 0 | 0 | 0 | 0 | 0 | 0 | 0 | 0 | 0 | 0 | 0 | 0 | 0 | 0 | 0 | 0 | 1 | 0 | 0 | 0 | 0 | 0 | 0 | 0 | 0 | 0 | 0 | 0 | 0 | 0 |
| bacteria_Aug5d25b_65829 | 1 | 0 | 0 | 0 | 0 | 0 | 0 | 0 | 0 | 0 | 0 | 0 | 0 | 0 | 0 | 0 | 0 | 0 | 0 | 0 | 0 | 0 | 0 | 0 | 0 | 0 | 0 | 0 | 0 | 0 | 0 | 0 | 0 | 0 | 0 | 0 | 0 | 0 | 0 | 0 | 0 | 1 | 0 | 0 | 0 | 0 | 0 | 0 | 0 | 0 | 0 | 0 | 0 | 0 | 0 |
| bacteria_Aug5d50b_39196 | 1 | 0 | 0 | 0 | 0 | 0 | 0 | 0 | 0 | 0 | 0 | 0 | 0 | 0 | 0 | 0 | 0 | 0 | 0 | 0 | 0 | 0 | 0 | 0 | 0 | 0 | 0 | 0 | 0 | 0 | 0 | 0 | 0 | 0 | 0 | 0 | 0 | 0 | 0 | 0 | 0 | 0 | 0 | 1 | 0 | 0 | 0 | 0 | 0 | 0 | 0 | 0 | 0 | 0 | 0 |
| bacteria_Jul11d5b_16587 | 1 | 0 | 0 | 0 | 0 | 0 | 0 | 0 | 0 | 0 | 0 | 0 | 0 | 0 | 0 | 0 | 0 | 0 | 0 | 0 | 0 | 0 | 0 | 1 | 0 | 0 | 0 | 0 | 0 | 0 | 0 | 0 | 0 | 0 | 0 | 0 | 0 | 0 | 0 | 0 | 0 | 0 | 0 | 0 | 0 | 0 | 0 | 0 | 0 | 0 | 0 | 0 | 0 | 0 | 0 |
| bacteria_Jul11d25b_23949 | 1 | 0 | 0 | 0 | 0 | 0 | 0 | 0 | 0 | 0 | 0 | 0 | 0 | 0 | 0 | 0 | 0 | 0 | 0 | 0 | 0 | 0 | 0 | 0 | 0 | 1 | 0 | 0 | 0 | 0 | 0 | 0 | 0 | 0 | 0 | 0 | 0 | 0 | 0 | 0 | 0 | 0 | 0 | 0 | 0 | 0 | 0 | 0 | 0 | 0 | 0 | 0 | 0 | 0 | 0 |
| bacteria_Jul11d25b_27699 | 1 | 0 | 0 | 0 | 0 | 0 | 0 | 0 | 0 | 0 | 0 | 0 | 0 | 0 | 0 | 0 | 0 | 0 | 0 | 0 | 0 | 0 | 0 | 0 | 0 | 1 | 0 | 0 | 0 | 0 | 0 | 0 | 0 | 0 | 0 | 0 | 0 | 0 | 0 | 0 | 0 | 0 | 0 | 0 | 0 | 0 | 0 | 0 | 0 | 0 | 0 | 0 | 0 | 0 | 0 |
| bacteria_Jul11d25b_41160 | 1 | 0 | 0 | 0 | 0 | 0 | 0 | 0 | 0 | 0 | 0 | 0 | 0 | 0 | 0 | 0 | 0 | 0 | 0 | 0 | 0 | 0 | 0 | 0 | 0 | 1 | 0 | 0 | 0 | 0 | 0 | 0 | 0 | 0 | 0 | 0 | 0 | 0 | 0 | 0 | 0 | 0 | 0 | 0 | 0 | 0 | 0 | 0 | 0 | 0 | 0 | 0 | 0 | 0 | 0 |
| bacteria_Jul11d50b_4516 | 1 | 0 | 0 | 0 | 0 | 0 | 0 | 0 | 0 | 0 | 0 | 0 | 0 | 0 | 0 | 0 | 0 | 0 | 0 | 0 | 0 | 0 | 0 | 0 | 0 | 0 | 0 | 1 | 0 | 0 | 0 | 0 | 0 | 0 | 0 | 0 | 0 | 0 | 0 | 0 | 0 | 0 | 0 | 0 | 0 | 0 | 0 | 0 | 0 | 0 | 0 | 0 | 0 | 0 | 0 |
| bacteria_Jul11d50b_23069 | 1 | 0 | 0 | 0 | 0 | 0 | 0 | 0 | 0 | 0 | 0 | 0 | 0 | 0 | 0 | 0 | 0 | 0 | 0 | 0 | 0 | 0 | 0 | 0 | 0 | 0 | 0 | 1 | 0 | 0 | 0 | 0 | 0 | 0 | 0 | 0 | 0 | 0 | 0 | 0 | 0 | 0 | 0 | 0 | 0 | 0 | 0 | 0 | 0 | 0 | 0 | 0 | 0 | 0 | 0 |
| bacteria_Jul11d50b_48239 | 1 | 0 | 0 | 0 | 0 | 0 | 0 | 0 | 0 | 0 | 0 | 0 | 0 | 0 | 0 | 0 | 0 | 0 | 0 | 0 | 0 | 0 | 0 | 0 | 0 | 0 | 0 | 1 | 0 | 0 | 0 | 0 | 0 | 0 | 0 | 0 | 0 | 0 | 0 | 0 | 0 | 0 | 0 | 0 | 0 | 0 | 0 | 0 | 0 | 0 | 0 | 0 | 0 | 0 | 0 |
| bacteria_Jul11d75b_3634 | 1 | 0 | 0 | 0 | 0 | 0 | 0 | 0 | 0 | 0 | 0 | 0 | 0 | 0 | 0 | 0 | 0 | 0 | 0 | 0 | 0 | 0 | 0 | 0 | 0 | 0 | 0 | 0 | 0 | 1 | 0 | 0 | 0 | 0 | 0 | 0 | 0 | 0 | 0 | 0 | 0 | 0 | 0 | 0 | 0 | 0 | 0 | 0 | 0 | 0 | 0 | 0 | 0 | 0 | 0 |
| bacteria_Jul11d75b_26807 | 1 | 0 | 0 | 0 | 0 | 0 | 0 | 0 | 0 | 0 | 0 | 0 | 0 | 0 | 0 | 0 | 0 | 0 | 0 | 0 | 0 | 0 | 0 | 0 | 0 | 0 | 0 | 0 | 0 | 1 | 0 | 0 | 0 | 0 | 0 | 0 | 0 | 0 | 0 | 0 | 0 | 0 | 0 | 0 | 0 | 0 | 0 | 0 | 0 | 0 | 0 | 0 | 0 | 0 | 0 |
| bacteria_Jul11d75b_30582 | 1 | 0 | 0 | 0 | 0 | 0 | 0 | 0 | 0 | 0 | 0 | 0 | 0 | 0 | 0 | 0 | 0 | 0 | 0 | 0 | 0 | 0 | 0 | 0 | 0 | 0 | 0 | 0 | 0 | 1 | 0 | 0 | 0 | 0 | 0 | 0 | 0 | 0 | 0 | 0 | 0 | 0 | 0 | 0 | 0 | 0 | 0 | 0 | 0 | 0 | 0 | 0 | 0 | 0 | 0 |
| bacteria_Jul11d75b_32512 | 1 | 0 | 0 | 0 | 0 | 0 | 0 | 0 | 0 | 0 | 0 | 0 | 0 | 0 | 0 | 0 | 0 | 0 | 0 | 0 | 0 | 0 | 0 | 0 | 0 | 0 | 0 | 0 | 0 | 1 | 0 | 0 | 0 | 0 | 0 | 0 | 0 | 0 | 0 | 0 | 0 | 0 | 0 | 0 | 0 | 0 | 0 | 0 | 0 | 0 | 0 | 0 | 0 | 0 | 0 |
| bacteria_Jul11d75b_41369 | 1 | 0 | 0 | 0 | 0 | 0 | 0 | 0 | 0 | 0 | 0 | 0 | 0 | 0 | 0 | 0 | 0 | 0 | 0 | 0 | 0 | 0 | 0 | 0 | 0 | 0 | 0 | 0 | 0 | 1 | 0 | 0 | 0 | 0 | 0 | 0 | 0 | 0 | 0 | 0 | 0 | 0 | 0 | 0 | 0 | 0 | 0 | 0 | 0 | 0 | 0 | 0 | 0 | 0 | 0 |
| bacteria_Jul11d75b_41847 | 1 | 0 | 0 | 0 | 0 | 0 | 0 | 0 | 0 | 0 | 0 | 0 | 0 | 0 | 0 | 0 | 0 | 0 | 0 | 0 | 0 | 0 | 0 | 0 | 0 | 0 | 0 | 0 | 0 | 1 | 0 | 0 | 0 | 0 | 0 | 0 | 0 | 0 | 0 | 0 | 0 | 0 | 0 | 0 | 0 | 0 | 0 | 0 | 0 | 0 | 0 | 0 | 0 | 0 | 0 |
| bacteria_Jul11d75b_48633 | 1 | 0 | 0 | 0 | 0 | 0 | 0 | 0 | 0 | 0 | 0 | 0 | 0 | 0 | 0 | 0 | 0 | 0 | 0 | 0 | 0 | 0 | 0 | 0 | 0 | 0 | 0 | 0 | 0 | 1 | 0 | 0 | 0 | 0 | 0 | 0 | 0 | 0 | 0 | 0 | 0 | 0 | 0 | 0 | 0 | 0 | 0 | 0 | 0 | 0 | 0 | 0 | 0 | 0 | 0 |
| bacteria_Jul11d75b_49870 | 1 | 0 | 0 | 0 | 0 | 0 | 0 | 0 | 0 | 0 | 0 | 0 | 0 | 0 | 0 | 0 | 0 | 0 | 0 | 0 | 0 | 0 | 0 | 0 | 0 | 0 | 0 | 0 | 0 | 1 | 0 | 0 | 0 | 0 | 0 | 0 | 0 | 0 | 0 | 0 | 0 | 0 | 0 | 0 | 0 | 0 | 0 | 0 | 0 | 0 | 0 | 0 | 0 | 0 | 0 |
| bacteria_Jul11d75b_50049 | 1 | 0 | 0 | 0 | 0 | 0 | 0 | 0 | 0 | 0 | 0 | 0 | 0 | 0 | 0 | 0 | 0 | 0 | 0 | 0 | 0 | 0 | 0 | 0 | 0 | 0 | 0 | 0 | 0 | 1 | 0 | 0 | 0 | 0 | 0 | 0 | 0 | 0 | 0 | 0 | 0 | 0 | 0 | 0 | 0 | 0 | 0 | 0 | 0 | 0 | 0 | 0 | 0 | 0 | 0 |
| bacteria_Jul27d5b_266 | 1 | 0 | 0 | 0 | 0 | 0 | 0 | 0 | 0 | 0 | 0 | 0 | 0 | 0 | 0 | 0 | 0 | 0 | 0 | 0 | 0 | 0 | 0 | 0 | 0 | 0 | 0 | 0 | 0 | 0 | 0 | 1 | 0 | 0 | 0 | 0 | 0 | 0 | 0 | 0 | 0 | 0 | 0 | 0 | 0 | 0 | 0 | 0 | 0 | 0 | 0 | 0 | 0 | 0 | 0 |
| bacteria_Jul27d5b_1964 | 1 | 0 | 0 | 0 | 0 | 0 | 0 | 0 | 0 | 0 | 0 | 0 | 0 | 0 | 0 | 0 | 0 | 0 | 0 | 0 | 0 | 0 | 0 | 0 | 0 | 0 | 0 | 0 | 0 | 0 | 0 | 1 | 0 | 0 | 0 | 0 | 0 | 0 | 0 | 0 | 0 | 0 | 0 | 0 | 0 | 0 | 0 | 0 | 0 | 0 | 0 | 0 | 0 | 0 | 0 |
| bacteria_Jul27d5b_16291 | 1 | 0 | 0 | 0 | 0 | 0 | 0 | 0 | 0 | 0 | 0 | 0 | 0 | 0 | 0 | 0 | 0 | 0 | 0 | 0 | 0 | 0 | 0 | 0 | 0 | 0 | 0 | 0 | 0 | 0 | 0 | 1 | 0 | 0 | 0 | 0 | 0 | 0 | 0 | 0 | 0 | 0 | 0 | 0 | 0 | 0 | 0 | 0 | 0 | 0 | 0 | 0 | 0 | 0 | 0 |
| bacteria_Jul27d5b_20185 | 1 | 0 | 0 | 0 | 0 | 0 | 0 | 0 | 0 | 0 | 0 | 0 | 0 | 0 | 0 | 0 | 0 | 0 | 0 | 0 | 0 | 0 | 0 | 0 | 0 | 0 | 0 | 0 | 0 | 0 | 0 | 1 | 0 | 0 | 0 | 0 | 0 | 0 | 0 | 0 | 0 | 0 | 0 | 0 | 0 | 0 | 0 | 0 | 0 | 0 | 0 | 0 | 0 | 0 | 0 |
| bacteria_Jul27d5b_20306 | 1 | 0 | 0 | 0 | 0 | 0 | 0 | 0 | 0 | 0 | 0 | 0 | 0 | 0 | 0 | 0 | 0 | 0 | 0 | 0 | 0 | 0 | 0 | 0 | 0 | 0 | 0 | 0 | 0 | 0 | 0 | 1 | 0 | 0 | 0 | 0 | 0 | 0 | 0 | 0 | 0 | 0 | 0 | 0 | 0 | 0 | 0 | 0 | 0 | 0 | 0 | 0 | 0 | 0 | 0 |
| bacteria_Jul27d5b_23686 | 1 | 0 | 0 | 0 | 0 | 0 | 0 | 0 | 0 | 0 | 0 | 0 | 0 | 0 | 0 | 0 | 0 | 0 | 0 | 0 | 0 | 0 | 0 | 0 | 0 | 0 | 0 | 0 | 0 | 0 | 0 | 1 | 0 | 0 | 0 | 0 | 0 | 0 | 0 | 0 | 0 | 0 | 0 | 0 | 0 | 0 | 0 | 0 | 0 | 0 | 0 | 0 | 0 | 0 | 0 |
| bacteria_Jul27d5b_27118 | 1 | 0 | 0 | 0 | 0 | 0 | 0 | 0 | 0 | 0 | 0 | 0 | 0 | 0 | 0 | 0 | 0 | 0 | 0 | 0 | 0 | 0 | 0 | 0 | 0 | 0 | 0 | 0 | 0 | 0 | 0 | 1 | 0 | 0 | 0 | 0 | 0 | 0 | 0 | 0 | 0 | 0 | 0 | 0 | 0 | 0 | 0 | 0 | 0 | 0 | 0 | 0 | 0 | 0 | 0 |
| bacteria_Jul27d5b_29462 | 1 | 0 | 0 | 0 | 0 | 0 | 0 | 0 | 0 | 0 | 0 | 0 | 0 | 0 | 0 | 0 | 0 | 0 | 0 | 0 | 0 | 0 | 0 | 0 | 0 | 0 | 0 | 0 | 0 | 0 | 0 | 1 | 0 | 0 | 0 | 0 | 0 | 0 | 0 | 0 | 0 | 0 | 0 | 0 | 0 | 0 | 0 | 0 | 0 | 0 | 0 | 0 | 0 | 0 | 0 |
| bacteria_Jul27d5b_31136 | 1 | 0 | 0 | 0 | 0 | 0 | 0 | 0 | 0 | 0 | 0 | 0 | 0 | 0 | 0 | 0 | 0 | 0 | 0 | 0 | 0 | 0 | 0 | 0 | 0 | 0 | 0 | 0 | 0 | 0 | 0 | 1 | 0 | 0 | 0 | 0 | 0 | 0 | 0 | 0 | 0 | 0 | 0 | 0 | 0 | 0 | 0 | 0 | 0 | 0 | 0 | 0 | 0 | 0 | 0 |
| bacteria_Jul27d5b_39640 | 1 | 0 | 0 | 0 | 0 | 0 | 0 | 0 | 0 | 0 | 0 | 0 | 0 | 0 | 0 | 0 | 0 | 0 | 0 | 0 | 0 | 0 | 0 | 0 | 0 | 0 | 0 | 0 | 0 | 0 | 0 | 1 | 0 | 0 | 0 | 0 | 0 | 0 | 0 | 0 | 0 | 0 | 0 | 0 | 0 | 0 | 0 | 0 | 0 | 0 | 0 | 0 | 0 | 0 | 0 |
| bacteria_Jul27d5b_39687 | 1 | 0 | 0 | 0 | 0 | 0 | 0 | 0 | 0 | 0 | 0 | 0 | 0 | 0 | 0 | 0 | 0 | 0 | 0 | 0 | 0 | 0 | 0 | 0 | 0 | 0 | 0 | 0 | 0 | 0 | 0 | 1 | 0 | 0 | 0 | 0 | 0 | 0 | 0 | 0 | 0 | 0 | 0 | 0 | 0 | 0 | 0 | 0 | 0 | 0 | 0 | 0 | 0 | 0 | 0 |
| bacteria_Jul27d5b_44472 | 1 | 0 | 0 | 0 | 0 | 0 | 0 | 0 | 0 | 0 | 0 | 0 | 0 | 0 | 0 | 0 | 0 | 0 | 0 | 0 | 0 | 0 | 0 | 0 | 0 | 0 | 0 | 0 | 0 | 0 | 0 | 1 | 0 | 0 | 0 | 0 | 0 | 0 | 0 | 0 | 0 | 0 | 0 | 0 | 0 | 0 | 0 | 0 | 0 | 0 | 0 | 0 | 0 | 0 | 0 |
| bacteria_Jul27d25b_19148 | 1 | 0 | 0 | 0 | 0 | 0 | 0 | 0 | 0 | 0 | 0 | 0 | 0 | 0 | 0 | 0 | 0 | 0 | 0 | 0 | 0 | 0 | 0 | 0 | 0 | 0 | 0 | 0 | 0 | 0 | 0 | 0 | 0 | 1 | 0 | 0 | 0 | 0 | 0 | 0 | 0 | 0 | 0 | 0 | 0 | 0 | 0 | 0 | 0 | 0 | 0 | 0 | 0 | 0 | 0 |
| bacteria_Jul27d25b_27671 | 1 | 0 | 0 | 0 | 0 | 0 | 0 | 0 | 0 | 0 | 0 | 0 | 0 | 0 | 0 | 0 | 0 | 0 | 0 | 0 | 0 | 0 | 0 | 0 | 0 | 0 | 0 | 0 | 0 | 0 | 0 | 0 | 0 | 1 | 0 | 0 | 0 | 0 | 0 | 0 | 0 | 0 | 0 | 0 | 0 | 0 | 0 | 0 | 0 | 0 | 0 | 0 | 0 | 0 | 0 |
| bacteria_Jul27d25b_31851 | 1 | 0 | 0 | 0 | 0 | 0 | 0 | 0 | 0 | 0 | 0 | 0 | 0 | 0 | 0 | 0 | 0 | 0 | 0 | 0 | 0 | 0 | 0 | 0 | 0 | 0 | 0 | 0 | 0 | 0 | 0 | 0 | 0 | 1 | 0 | 0 | 0 | 0 | 0 | 0 | 0 | 0 | 0 | 0 | 0 | 0 | 0 | 0 | 0 | 0 | 0 | 0 | 0 | 0 | 0 |
| bacteria_Jul27d25b_31859 | 1 | 0 | 0 | 0 | 0 | 0 | 0 | 0 | 0 | 0 | 0 | 0 | 0 | 0 | 0 | 0 | 0 | 0 | 0 | 0 | 0 | 0 | 0 | 0 | 0 | 0 | 0 | 0 | 0 | 0 | 0 | 0 | 0 | 1 | 0 | 0 | 0 | 0 | 0 | 0 | 0 | 0 | 0 | 0 | 0 | 0 | 0 | 0 | 0 | 0 | 0 | 0 | 0 | 0 | 0 |
| bacteria_Jul27d25b_31871 | 1 | 0 | 0 | 0 | 0 | 0 | 0 | 0 | 0 | 0 | 0 | 0 | 0 | 0 | 0 | 0 | 0 | 0 | 0 | 0 | 0 | 0 | 0 | 0 | 0 | 0 | 0 | 0 | 0 | 0 | 0 | 0 | 0 | 1 | 0 | 0 | 0 | 0 | 0 | 0 | 0 | 0 | 0 | 0 | 0 | 0 | 0 | 0 | 0 | 0 | 0 | 0 | 0 | 0 | 0 |
| bacteria_Jul27d25b_32000 | 1 | 0 | 0 | 0 | 0 | 0 | 0 | 0 | 0 | 0 | 0 | 0 | 0 | 0 | 0 | 0 | 0 | 0 | 0 | 0 | 0 | 0 | 0 | 0 | 0 | 0 | 0 | 0 | 0 | 0 | 0 | 0 | 0 | 1 | 0 | 0 | 0 | 0 | 0 | 0 | 0 | 0 | 0 | 0 | 0 | 0 | 0 | 0 | 0 | 0 | 0 | 0 | 0 | 0 | 0 |
| bacteria_Jul27d25b_36843 | 1 | 0 | 0 | 0 | 0 | 0 | 0 | 0 | 0 | 0 | 0 | 0 | 0 | 0 | 0 | 0 | 0 | 0 | 0 | 0 | 0 | 0 | 0 | 0 | 0 | 0 | 0 | 0 | 0 | 0 | 0 | 0 | 0 | 1 | 0 | 0 | 0 | 0 | 0 | 0 | 0 | 0 | 0 | 0 | 0 | 0 | 0 | 0 | 0 | 0 | 0 | 0 | 0 | 0 | 0 |
| bacteria_Jul27d25b_47179 | 1 | 0 | 0 | 0 | 0 | 0 | 0 | 0 | 0 | 0 | 0 | 0 | 0 | 0 | 0 | 0 | 0 | 0 | 0 | 0 | 0 | 0 | 0 | 0 | 0 | 0 | 0 | 0 | 0 | 0 | 0 | 0 | 0 | 1 | 0 | 0 | 0 | 0 | 0 | 0 | 0 | 0 | 0 | 0 | 0 | 0 | 0 | 0 | 0 | 0 | 0 | 0 | 0 | 0 | 0 |
| bacteria_Jul27d25b_55213 | 1 | 0 | 0 | 0 | 0 | 0 | 0 | 0 | 0 | 0 | 0 | 0 | 0 | 0 | 0 | 0 | 0 | 0 | 0 | 0 | 0 | 0 | 0 | 0 | 0 | 0 | 0 | 0 | 0 | 0 | 0 | 0 | 0 | 1 | 0 | 0 | 0 | 0 | 0 | 0 | 0 | 0 | 0 | 0 | 0 | 0 | 0 | 0 | 0 | 0 | 0 | 0 | 0 | 0 | 0 |
| bacteria_Jul27d25b_73669 | 1 | 0 | 0 | 0 | 0 | 0 | 0 | 0 | 0 | 0 | 0 | 0 | 0 | 0 | 0 | 0 | 0 | 0 | 0 | 0 | 0 | 0 | 0 | 0 | 0 | 0 | 0 | 0 | 0 | 0 | 0 | 0 | 0 | 1 | 0 | 0 | 0 | 0 | 0 | 0 | 0 | 0 | 0 | 0 | 0 | 0 | 0 | 0 | 0 | 0 | 0 | 0 | 0 | 0 | 0 |
| bacteria_Jul27d25b_75133 | 1 | 0 | 0 | 0 | 0 | 0 | 0 | 0 | 0 | 0 | 0 | 0 | 0 | 0 | 0 | 0 | 0 | 0 | 0 | 0 | 0 | 0 | 0 | 0 | 0 | 0 | 0 | 0 | 0 | 0 | 0 | 0 | 0 | 1 | 0 | 0 | 0 | 0 | 0 | 0 | 0 | 0 | 0 | 0 | 0 | 0 | 0 | 0 | 0 | 0 | 0 | 0 | 0 | 0 | 0 |
| bacteria_Jul27d75b_1937 | 1 | 0 | 0 | 0 | 0 | 0 | 0 | 0 | 0 | 0 | 0 | 0 | 0 | 0 | 0 | 0 | 0 | 0 | 0 | 0 | 0 | 0 | 0 | 0 | 0 | 0 | 0 | 0 | 0 | 0 | 0 | 0 | 0 | 0 | 0 | 1 | 0 | 0 | 0 | 0 | 0 | 0 | 0 | 0 | 0 | 0 | 0 | 0 | 0 | 0 | 0 | 0 | 0 | 0 | 0 |
| bacteria_Jul27d75b_12967 | 1 | 0 | 0 | 0 | 0 | 0 | 0 | 0 | 0 | 0 | 0 | 0 | 0 | 0 | 0 | 0 | 0 | 0 | 0 | 0 | 0 | 0 | 0 | 0 | 0 | 0 | 0 | 0 | 0 | 0 | 0 | 0 | 0 | 0 | 0 | 1 | 0 | 0 | 0 | 0 | 0 | 0 | 0 | 0 | 0 | 0 | 0 | 0 | 0 | 0 | 0 | 0 | 0 | 0 | 0 |
| bacteria_Jul27d75b_35743 | 1 | 0 | 0 | 0 | 0 | 0 | 0 | 0 | 0 | 0 | 0 | 0 | 0 | 0 | 0 | 0 | 0 | 0 | 0 | 0 | 0 | 0 | 0 | 0 | 0 | 0 | 0 | 0 | 0 | 0 | 0 | 0 | 0 | 0 | 0 | 1 | 0 | 0 | 0 | 0 | 0 | 0 | 0 | 0 | 0 | 0 | 0 | 0 | 0 | 0 | 0 | 0 | 0 | 0 | 0 |
| bacteria_Jul27d75b_42798 | 1 | 0 | 0 | 0 | 0 | 0 | 0 | 0 | 0 | 0 | 0 | 0 | 0 | 0 | 0 | 0 | 0 | 0 | 0 | 0 | 0 | 0 | 0 | 0 | 0 | 0 | 0 | 0 | 0 | 0 | 0 | 0 | 0 | 0 | 0 | 1 | 0 | 0 | 0 | 0 | 0 | 0 | 0 | 0 | 0 | 0 | 0 | 0 | 0 | 0 | 0 | 0 | 0 | 0 | 0 |
| bacteria_Jul27d75b_67019 | 1 | 0 | 0 | 0 | 0 | 0 | 0 | 0 | 0 | 0 | 0 | 0 | 0 | 0 | 0 | 0 | 0 | 0 | 0 | 0 | 0 | 0 | 0 | 0 | 0 | 0 | 0 | 0 | 0 | 0 | 0 | 0 | 0 | 0 | 0 | 1 | 0 | 0 | 0 | 0 | 0 | 0 | 0 | 0 | 0 | 0 | 0 | 0 | 0 | 0 | 0 | 0 | 0 | 0 | 0 |
| bacteria_Jun8d25b_7303 | 1 | 0 | 0 | 0 | 0 | 0 | 0 | 0 | 0 | 1 | 0 | 0 | 0 | 0 | 0 | 0 | 0 | 0 | 0 | 0 | 0 | 0 | 0 | 0 | 0 | 0 | 0 | 0 | 0 | 0 | 0 | 0 | 0 | 0 | 0 | 0 | 0 | 0 | 0 | 0 | 0 | 0 | 0 | 0 | 0 | 0 | 0 | 0 | 0 | 0 | 0 | 0 | 0 | 0 | 0 |
| bacteria_Jun8d25b_7643 | 1 | 0 | 0 | 0 | 0 | 0 | 0 | 0 | 0 | 1 | 0 | 0 | 0 | 0 | 0 | 0 | 0 | 0 | 0 | 0 | 0 | 0 | 0 | 0 | 0 | 0 | 0 | 0 | 0 | 0 | 0 | 0 | 0 | 0 | 0 | 0 | 0 | 0 | 0 | 0 | 0 | 0 | 0 | 0 | 0 | 0 | 0 | 0 | 0 | 0 | 0 | 0 | 0 | 0 | 0 |
| bacteria_Jun8d25b_11474 | 1 | 0 | 0 | 0 | 0 | 0 | 0 | 0 | 0 | 1 | 0 | 0 | 0 | 0 | 0 | 0 | 0 | 0 | 0 | 0 | 0 | 0 | 0 | 0 | 0 | 0 | 0 | 0 | 0 | 0 | 0 | 0 | 0 | 0 | 0 | 0 | 0 | 0 | 0 | 0 | 0 | 0 | 0 | 0 | 0 | 0 | 0 | 0 | 0 | 0 | 0 | 0 | 0 | 0 | 0 |
| bacteria_Jun8d25b_15489 | 1 | 0 | 0 | 0 | 0 | 0 | 0 | 0 | 0 | 1 | 0 | 0 | 0 | 0 | 0 | 0 | 0 | 0 | 0 | 0 | 0 | 0 | 0 | 0 | 0 | 0 | 0 | 0 | 0 | 0 | 0 | 0 | 0 | 0 | 0 | 0 | 0 | 0 | 0 | 0 | 0 | 0 | 0 | 0 | 0 | 0 | 0 | 0 | 0 | 0 | 0 | 0 | 0 | 0 | 0 |
| bacteria_Jun8d25b_16198 | 1 | 0 | 0 | 0 | 0 | 0 | 0 | 0 | 0 | 1 | 0 | 0 | 0 | 0 | 0 | 0 | 0 | 0 | 0 | 0 | 0 | 0 | 0 | 0 | 0 | 0 | 0 | 0 | 0 | 0 | 0 | 0 | 0 | 0 | 0 | 0 | 0 | 0 | 0 | 0 | 0 | 0 | 0 | 0 | 0 | 0 | 0 | 0 | 0 | 0 | 0 | 0 | 0 | 0 | 0 |
| bacteria_Jun8d25b_58090 | 1 | 0 | 0 | 0 | 0 | 0 | 0 | 0 | 0 | 1 | 0 | 0 | 0 | 0 | 0 | 0 | 0 | 0 | 0 | 0 | 0 | 0 | 0 | 0 | 0 | 0 | 0 | 0 | 0 | 0 | 0 | 0 | 0 | 0 | 0 | 0 | 0 | 0 | 0 | 0 | 0 | 0 | 0 | 0 | 0 | 0 | 0 | 0 | 0 | 0 | 0 | 0 | 0 | 0 | 0 |
| bacteria_Jun8d25b_59223 | 1 | 0 | 0 | 0 | 0 | 0 | 0 | 0 | 0 | 1 | 0 | 0 | 0 | 0 | 0 | 0 | 0 | 0 | 0 | 0 | 0 | 0 | 0 | 0 | 0 | 0 | 0 | 0 | 0 | 0 | 0 | 0 | 0 | 0 | 0 | 0 | 0 | 0 | 0 | 0 | 0 | 0 | 0 | 0 | 0 | 0 | 0 | 0 | 0 | 0 | 0 | 0 | 0 | 0 | 0 |
| bacteria_Jun8d25b_60249 | 1 | 0 | 0 | 0 | 0 | 0 | 0 | 0 | 0 | 1 | 0 | 0 | 0 | 0 | 0 | 0 | 0 | 0 | 0 | 0 | 0 | 0 | 0 | 0 | 0 | 0 | 0 | 0 | 0 | 0 | 0 | 0 | 0 | 0 | 0 | 0 | 0 | 0 | 0 | 0 | 0 | 0 | 0 | 0 | 0 | 0 | 0 | 0 | 0 | 0 | 0 | 0 | 0 | 0 | 0 |
| bacteria_Jun8d25b_70783 | 1 | 0 | 0 | 0 | 0 | 0 | 0 | 0 | 0 | 1 | 0 | 0 | 0 | 0 | 0 | 0 | 0 | 0 | 0 | 0 | 0 | 0 | 0 | 0 | 0 | 0 | 0 | 0 | 0 | 0 | 0 | 0 | 0 | 0 | 0 | 0 | 0 | 0 | 0 | 0 | 0 | 0 | 0 | 0 | 0 | 0 | 0 | 0 | 0 | 0 | 0 | 0 | 0 | 0 | 0 |
| bacteria_Jun8d50b_14277 | 1 | 0 | 0 | 0 | 0 | 0 | 0 | 0 | 0 | 0 | 0 | 1 | 0 | 0 | 0 | 0 | 0 | 0 | 0 | 0 | 0 | 0 | 0 | 0 | 0 | 0 | 0 | 0 | 0 | 0 | 0 | 0 | 0 | 0 | 0 | 0 | 0 | 0 | 0 | 0 | 0 | 0 | 0 | 0 | 0 | 0 | 0 | 0 | 0 | 0 | 0 | 0 | 0 | 0 | 0 |
| bacteria_Jun8d50b_26056 | 1 | 0 | 0 | 0 | 0 | 0 | 0 | 0 | 0 | 0 | 0 | 1 | 0 | 0 | 0 | 0 | 0 | 0 | 0 | 0 | 0 | 0 | 0 | 0 | 0 | 0 | 0 | 0 | 0 | 0 | 0 | 0 | 0 | 0 | 0 | 0 | 0 | 0 | 0 | 0 | 0 | 0 | 0 | 0 | 0 | 0 | 0 | 0 | 0 | 0 | 0 | 0 | 0 | 0 | 0 |
| bacteria_Jun8d50b_41294 | 1 | 0 | 0 | 0 | 0 | 0 | 0 | 0 | 0 | 0 | 0 | 1 | 0 | 0 | 0 | 0 | 0 | 0 | 0 | 0 | 0 | 0 | 0 | 0 | 0 | 0 | 0 | 0 | 0 | 0 | 0 | 0 | 0 | 0 | 0 | 0 | 0 | 0 | 0 | 0 | 0 | 0 | 0 | 0 | 0 | 0 | 0 | 0 | 0 | 0 | 0 | 0 | 0 | 0 | 0 |
| bacteria_Jun8d50b_47497 | 1 | 0 | 0 | 0 | 0 | 0 | 0 | 0 | 0 | 0 | 0 | 1 | 0 | 0 | 0 | 0 | 0 | 0 | 0 | 0 | 0 | 0 | 0 | 0 | 0 | 0 | 0 | 0 | 0 | 0 | 0 | 0 | 0 | 0 | 0 | 0 | 0 | 0 | 0 | 0 | 0 | 0 | 0 | 0 | 0 | 0 | 0 | 0 | 0 | 0 | 0 | 0 | 0 | 0 | 0 |
| bacteria_Jun8d75b_3886 | 1 | 0 | 0 | 0 | 0 | 0 | 0 | 0 | 0 | 0 | 0 | 0 | 0 | 1 | 0 | 0 | 0 | 0 | 0 | 0 | 0 | 0 | 0 | 0 | 0 | 0 | 0 | 0 | 0 | 0 | 0 | 0 | 0 | 0 | 0 | 0 | 0 | 0 | 0 | 0 | 0 | 0 | 0 | 0 | 0 | 0 | 0 | 0 | 0 | 0 | 0 | 0 | 0 | 0 | 0 |
| bacteria_Jun22d0b_21445 | 1 | 0 | 0 | 0 | 0 | 0 | 0 | 0 | 0 | 0 | 0 | 0 | 0 | 0 | 0 | 1 | 0 | 0 | 0 | 0 | 0 | 0 | 0 | 0 | 0 | 0 | 0 | 0 | 0 | 0 | 0 | 0 | 0 | 0 | 0 | 0 | 0 | 0 | 0 | 0 | 0 | 0 | 0 | 0 | 0 | 0 | 0 | 0 | 0 | 0 | 0 | 0 | 0 | 0 | 0 |
| bacteria_Jun22d0b_21911 | 1 | 0 | 0 | 0 | 0 | 0 | 0 | 0 | 0 | 0 | 0 | 0 | 0 | 0 | 0 | 1 | 0 | 0 | 0 | 0 | 0 | 0 | 0 | 0 | 0 | 0 | 0 | 0 | 0 | 0 | 0 | 0 | 0 | 0 | 0 | 0 | 0 | 0 | 0 | 0 | 0 | 0 | 0 | 0 | 0 | 0 | 0 | 0 | 0 | 0 | 0 | 0 | 0 | 0 | 0 |
| bacteria_Jun22d50b_1620 | 1 | 0 | 0 | 0 | 0 | 0 | 0 | 0 | 0 | 0 | 0 | 0 | 0 | 0 | 0 | 0 | 0 | 1 | 0 | 0 | 0 | 0 | 0 | 0 | 0 | 0 | 0 | 0 | 0 | 0 | 0 | 0 | 0 | 0 | 0 | 0 | 0 | 0 | 0 | 0 | 0 | 0 | 0 | 0 | 0 | 0 | 0 | 0 | 0 | 0 | 0 | 0 | 0 | 0 | 0 |
| bacteria_Jun22d50b_3167 | 1 | 0 | 0 | 0 | 0 | 0 | 0 | 0 | 0 | 0 | 0 | 0 | 0 | 0 | 0 | 0 | 0 | 1 | 0 | 0 | 0 | 0 | 0 | 0 | 0 | 0 | 0 | 0 | 0 | 0 | 0 | 0 | 0 | 0 | 0 | 0 | 0 | 0 | 0 | 0 | 0 | 0 | 0 | 0 | 0 | 0 | 0 | 0 | 0 | 0 | 0 | 0 | 0 | 0 | 0 |
| bacteria_Jun22d50b_4234 | 1 | 0 | 0 | 0 | 0 | 0 | 0 | 0 | 0 | 0 | 0 | 0 | 0 | 0 | 0 | 0 | 0 | 1 | 0 | 0 | 0 | 0 | 0 | 0 | 0 | 0 | 0 | 0 | 0 | 0 | 0 | 0 | 0 | 0 | 0 | 0 | 0 | 0 | 0 | 0 | 0 | 0 | 0 | 0 | 0 | 0 | 0 | 0 | 0 | 0 | 0 | 0 | 0 | 0 | 0 |
| bacteria_Jun22d75b_1642 | 1 | 0 | 0 | 0 | 0 | 0 | 0 | 0 | 0 | 0 | 0 | 0 | 0 | 0 | 0 | 0 | 0 | 0 | 0 | 1 | 0 | 0 | 0 | 0 | 0 | 0 | 0 | 0 | 0 | 0 | 0 | 0 | 0 | 0 | 0 | 0 | 0 | 0 | 0 | 0 | 0 | 0 | 0 | 0 | 0 | 0 | 0 | 0 | 0 | 0 | 0 | 0 | 0 | 0 | 0 |
| bacteria_Jun22d75b_5102 | 1 | 0 | 0 | 0 | 0 | 0 | 0 | 0 | 0 | 0 | 0 | 0 | 0 | 0 | 0 | 0 | 0 | 0 | 0 | 1 | 0 | 0 | 0 | 0 | 0 | 0 | 0 | 0 | 0 | 0 | 0 | 0 | 0 | 0 | 0 | 0 | 0 | 0 | 0 | 0 | 0 | 0 | 0 | 0 | 0 | 0 | 0 | 0 | 0 | 0 | 0 | 0 | 0 | 0 | 0 |
| bacteria_Jun22d75b_5810 | 1 | 0 | 0 | 0 | 0 | 0 | 0 | 0 | 0 | 0 | 0 | 0 | 0 | 0 | 0 | 0 | 0 | 0 | 0 | 1 | 0 | 0 | 0 | 0 | 0 | 0 | 0 | 0 | 0 | 0 | 0 | 0 | 0 | 0 | 0 | 0 | 0 | 0 | 0 | 0 | 0 | 0 | 0 | 0 | 0 | 0 | 0 | 0 | 0 | 0 | 0 | 0 | 0 | 0 | 0 |
| bacteria_May12d25b_101 | 1 | 0 | 0 | 1 | 0 | 0 | 0 | 0 | 0 | 0 | 0 | 0 | 0 | 0 | 0 | 0 | 0 | 0 | 0 | 0 | 0 | 0 | 0 | 0 | 0 | 0 | 0 | 0 | 0 | 0 | 0 | 0 | 0 | 0 | 0 | 0 | 0 | 0 | 0 | 0 | 0 | 0 | 0 | 0 | 0 | 0 | 0 | 0 | 0 | 0 | 0 | 0 | 0 | 0 | 0 |
| bacteria_May12d25b_130 | 1 | 0 | 0 | 1 | 0 | 0 | 0 | 0 | 0 | 0 | 0 | 0 | 0 | 0 | 0 | 0 | 0 | 0 | 0 | 0 | 0 | 0 | 0 | 0 | 0 | 0 | 0 | 0 | 0 | 0 | 0 | 0 | 0 | 0 | 0 | 0 | 0 | 0 | 0 | 0 | 0 | 0 | 0 | 0 | 0 | 0 | 0 | 0 | 0 | 0 | 0 | 0 | 0 | 0 | 0 |
| bacteria_May12d25b_530 | 1 | 0 | 0 | 1 | 0 | 0 | 0 | 0 | 0 | 0 | 0 | 0 | 0 | 0 | 0 | 0 | 0 | 0 | 0 | 0 | 0 | 0 | 0 | 0 | 0 | 0 | 0 | 0 | 0 | 0 | 0 | 0 | 0 | 0 | 0 | 0 | 0 | 0 | 0 | 0 | 0 | 0 | 0 | 0 | 0 | 0 | 0 | 0 | 0 | 0 | 0 | 0 | 0 | 0 | 0 |
| bacteria_May12d25b_622 | 1 | 0 | 0 | 1 | 0 | 0 | 0 | 0 | 0 | 0 | 0 | 0 | 0 | 0 | 0 | 0 | 0 | 0 | 0 | 0 | 0 | 0 | 0 | 0 | 0 | 0 | 0 | 0 | 0 | 0 | 0 | 0 | 0 | 0 | 0 | 0 | 0 | 0 | 0 | 0 | 0 | 0 | 0 | 0 | 0 | 0 | 0 | 0 | 0 | 0 | 0 | 0 | 0 | 0 | 0 |
| bacteria_May12d25b_624 | 1 | 0 | 0 | 1 | 0 | 0 | 0 | 0 | 0 | 0 | 0 | 0 | 0 | 0 | 0 | 0 | 0 | 0 | 0 | 0 | 0 | 0 | 0 | 0 | 0 | 0 | 0 | 0 | 0 | 0 | 0 | 0 | 0 | 0 | 0 | 0 | 0 | 0 | 0 | 0 | 0 | 0 | 0 | 0 | 0 | 0 | 0 | 0 | 0 | 0 | 0 | 0 | 0 | 0 | 0 |
| bacteria_May12d25b_637 | 1 | 0 | 0 | 1 | 0 | 0 | 0 | 0 | 0 | 0 | 0 | 0 | 0 | 0 | 0 | 0 | 0 | 0 | 0 | 0 | 0 | 0 | 0 | 0 | 0 | 0 | 0 | 0 | 0 | 0 | 0 | 0 | 0 | 0 | 0 | 0 | 0 | 0 | 0 | 0 | 0 | 0 | 0 | 0 | 0 | 0 | 0 | 0 | 0 | 0 | 0 | 0 | 0 | 0 | 0 |
| bacteria_May12d25b_683 | 1 | 0 | 0 | 1 | 0 | 0 | 0 | 0 | 0 | 0 | 0 | 0 | 0 | 0 | 0 | 0 | 0 | 0 | 0 | 0 | 0 | 0 | 0 | 0 | 0 | 0 | 0 | 0 | 0 | 0 | 0 | 0 | 0 | 0 | 0 | 0 | 0 | 0 | 0 | 0 | 0 | 0 | 0 | 0 | 0 | 0 | 0 | 0 | 0 | 0 | 0 | 0 | 0 | 0 | 0 |
| bacteria_May12d50b_179 | 1 | 0 | 0 | 0 | 0 | 1 | 0 | 0 | 0 | 0 | 0 | 0 | 0 | 0 | 0 | 0 | 0 | 0 | 0 | 0 | 0 | 0 | 0 | 0 | 0 | 0 | 0 | 0 | 0 | 0 | 0 | 0 | 0 | 0 | 0 | 0 | 0 | 0 | 0 | 0 | 0 | 0 | 0 | 0 | 0 | 0 | 0 | 0 | 0 | 0 | 0 | 0 | 0 | 0 | 0 |
| bacteria_May12d50b_113 | 1 | 0 | 0 | 0 | 0 | 1 | 0 | 0 | 0 | 0 | 0 | 0 | 0 | 0 | 0 | 0 | 0 | 0 | 0 | 0 | 0 | 0 | 0 | 0 | 0 | 0 | 0 | 0 | 0 | 0 | 0 | 0 | 0 | 0 | 0 | 0 | 0 | 0 | 0 | 0 | 0 | 0 | 0 | 0 | 0 | 0 | 0 | 0 | 0 | 0 | 0 | 0 | 0 | 0 | 0 |
| bacteria_May12d50b_146 | 1 | 0 | 0 | 0 | 0 | 1 | 0 | 0 | 0 | 0 | 0 | 0 | 0 | 0 | 0 | 0 | 0 | 0 | 0 | 0 | 0 | 0 | 0 | 0 | 0 | 0 | 0 | 0 | 0 | 0 | 0 | 0 | 0 | 0 | 0 | 0 | 0 | 0 | 0 | 0 | 0 | 0 | 0 | 0 | 0 | 0 | 0 | 0 | 0 | 0 | 0 | 0 | 0 | 0 | 0 |
| bacteria_May12d50b_152 | 1 | 0 | 0 | 0 | 0 | 1 | 0 | 0 | 0 | 0 | 0 | 0 | 0 | 0 | 0 | 0 | 0 | 0 | 0 | 0 | 0 | 0 | 0 | 0 | 0 | 0 | 0 | 0 | 0 | 0 | 0 | 0 | 0 | 0 | 0 | 0 | 0 | 0 | 0 | 0 | 0 | 0 | 0 | 0 | 0 | 0 | 0 | 0 | 0 | 0 | 0 | 0 | 0 | 0 | 0 |
| bacteria_May12d50b_184 | 1 | 0 | 0 | 0 | 0 | 1 | 0 | 0 | 0 | 0 | 0 | 0 | 0 | 0 | 0 | 0 | 0 | 0 | 0 | 0 | 0 | 0 | 0 | 0 | 0 | 0 | 0 | 0 | 0 | 0 | 0 | 0 | 0 | 0 | 0 | 0 | 0 | 0 | 0 | 0 | 0 | 0 | 0 | 0 | 0 | 0 | 0 | 0 | 0 | 0 | 0 | 0 | 0 | 0 | 0 |
| bacteria_May12d50b_185 | 1 | 0 | 0 | 0 | 0 | 1 | 0 | 0 | 0 | 0 | 0 | 0 | 0 | 0 | 0 | 0 | 0 | 0 | 0 | 0 | 0 | 0 | 0 | 0 | 0 | 0 | 0 | 0 | 0 | 0 | 0 | 0 | 0 | 0 | 0 | 0 | 0 | 0 | 0 | 0 | 0 | 0 | 0 | 0 | 0 | 0 | 0 | 0 | 0 | 0 | 0 | 0 | 0 | 0 | 0 |
| bacteria_May12d50b_191 | 1 | 0 | 0 | 0 | 0 | 1 | 0 | 0 | 0 | 0 | 0 | 0 | 0 | 0 | 0 | 0 | 0 | 0 | 0 | 0 | 0 | 0 | 0 | 0 | 0 | 0 | 0 | 0 | 0 | 0 | 0 | 0 | 0 | 0 | 0 | 0 | 0 | 0 | 0 | 0 | 0 | 0 | 0 | 0 | 0 | 0 | 0 | 0 | 0 | 0 | 0 | 0 | 0 | 0 | 0 |
| bacteria_May12d50b_462 | 1 | 0 | 0 | 0 | 0 | 1 | 0 | 0 | 0 | 0 | 0 | 0 | 0 | 0 | 0 | 0 | 0 | 0 | 0 | 0 | 0 | 0 | 0 | 0 | 0 | 0 | 0 | 0 | 0 | 0 | 0 | 0 | 0 | 0 | 0 | 0 | 0 | 0 | 0 | 0 | 0 | 0 | 0 | 0 | 0 | 0 | 0 | 0 | 0 | 0 | 0 | 0 | 0 | 0 | 0 |
| bacteria_Sep8d0b_44016 | 1 | 0 | 0 | 0 | 0 | 0 | 0 | 0 | 0 | 0 | 0 | 0 | 0 | 0 | 0 | 0 | 0 | 0 | 0 | 0 | 0 | 0 | 0 | 0 | 0 | 0 | 0 | 0 | 0 | 0 | 0 | 0 | 0 | 0 | 0 | 0 | 0 | 0 | 0 | 0 | 0 | 0 | 0 | 0 | 0 | 0 | 0 | 1 | 0 | 0 | 0 | 0 | 0 | 0 | 0 |
| bacteria_Sep8d5b_19474 | 1 | 0 | 0 | 0 | 0 | 0 | 0 | 0 | 0 | 0 | 0 | 0 | 0 | 0 | 0 | 0 | 0 | 0 | 0 | 0 | 0 | 0 | 0 | 0 | 0 | 0 | 0 | 0 | 0 | 0 | 0 | 0 | 0 | 0 | 0 | 0 | 0 | 0 | 0 | 0 | 0 | 0 | 0 | 0 | 0 | 0 | 0 | 0 | 0 | 1 | 0 | 0 | 0 | 0 | 0 |
| bacteria_Sep8d5b_44398 | 1 | 0 | 0 | 0 | 0 | 0 | 0 | 0 | 0 | 0 | 0 | 0 | 0 | 0 | 0 | 0 | 0 | 0 | 0 | 0 | 0 | 0 | 0 | 0 | 0 | 0 | 0 | 0 | 0 | 0 | 0 | 0 | 0 | 0 | 0 | 0 | 0 | 0 | 0 | 0 | 0 | 0 | 0 | 0 | 0 | 0 | 0 | 0 | 0 | 1 | 0 | 0 | 0 | 0 | 0 |
| bacteria_Sep8d5b_44813 | 1 | 0 | 0 | 0 | 0 | 0 | 0 | 0 | 0 | 0 | 0 | 0 | 0 | 0 | 0 | 0 | 0 | 0 | 0 | 0 | 0 | 0 | 0 | 0 | 0 | 0 | 0 | 0 | 0 | 0 | 0 | 0 | 0 | 0 | 0 | 0 | 0 | 0 | 0 | 0 | 0 | 0 | 0 | 0 | 0 | 0 | 0 | 0 | 0 | 1 | 0 | 0 | 0 | 0 | 0 |
| bacteria_Sep8d5b_51298 | 1 | 0 | 0 | 0 | 0 | 0 | 0 | 0 | 0 | 0 | 0 | 0 | 0 | 0 | 0 | 0 | 0 | 0 | 0 | 0 | 0 | 0 | 0 | 0 | 0 | 0 | 0 | 0 | 0 | 0 | 0 | 0 | 0 | 0 | 0 | 0 | 0 | 0 | 0 | 0 | 0 | 0 | 0 | 0 | 0 | 0 | 0 | 0 | 0 | 1 | 0 | 0 | 0 | 0 | 0 |

**Supplementary Table S7**. Common Bacterio- and virioplankton RTPR 98% OTUs in 27 samples with RTPR amplicons in both fractions

Total seqs: 47997; Subtotal seqs in retained

OTUs: 41046

| OTUs with >=10 seqs: 254 | OTU subtotal | May12_05m | | May12_25m | | May12_50m | | May12_75m | | Jun8_25m | | Jun8_50m | | Jun8_75m | | Jun22_0m | | Jun22_50m | | Jun22_75m | | Jul11_0m | | Jul11_05m | | Jul11_25m | | Jul11_50m | | Jul11_75m | | Jul27_05m | | Jul27_25m | | Jul27_75m | | Aug5_0m | | Aug5_05m | | Aug5_25m | | Aug5_50m | | Aug5_75m | | Sep8_0m | | Sep8_05m | | Sep8_25m | | Sep8_50m | |
|---|---|---|---|---|---|---|---|---|---|---|---|---|---|---|---|---|---|---|---|---|---|---|---|---|---|---|---|---|---|---|---|---|---|---|---|---|---|---|---|---|---|---|---|---|---|---|---|---|---|---|---|---|---|---|---|
| | | bacterioplankton | virioplankton | bacterioplankton | virioplankton | bacterioplankton | virioplankton | bacterioplankton | virioplankton | bacterioplankton | virioplankton | bacterioplankton | virioplankton | bacterioplankton | virioplankton | bacterioplankton | virioplankton | bacterioplankton | virioplankton | bacterioplankton | virioplankton | bacterioplankton | virioplankton | bacterioplankton | virioplankton | bacterioplankton | virioplankton | bacterioplankton | virioplankton | bacterioplankton | virioplankton | bacterioplankton | virioplankton | bacterioplankton | virioplankton | bacterioplankton | virioplankton | bacterioplankton | virioplankton | bacterioplankton | virioplankton | bacterioplankton | virioplankton | bacterioplankton | virioplankton | bacterioplankton | virioplankton | bacterioplankton | virioplankton | bacterioplankton | virioplankton | bacterioplankton | virioplankton | bacterioplankton | virioplankton |
| bacteria_Sep8d5b_53048 | 1 | 0 | 0 | 0 | 0 | 0 | 0 | 0 | 0 | 0 | 0 | 0 | 0 | 0 | 0 | 0 | 0 | 0 | 0 | 0 | 0 | 0 | 0 | 0 | 0 | 0 | 0 | 0 | 0 | 0 | 0 | 0 | 0 | 0 | 0 | 0 | 0 | 0 | 0 | 0 | 0 | 0 | 0 | 0 | 0 | 0 | 0 | 0 | 0 | 1 | 0 | 0 | 0 | 0 | 0 |
| bacteria_Sep8d5b_69339 | 1 | 0 | 0 | 0 | 0 | 0 | 0 | 0 | 0 | 0 | 0 | 0 | 0 | 0 | 0 | 0 | 0 | 0 | 0 | 0 | 0 | 0 | 0 | 0 | 0 | 0 | 0 | 0 | 0 | 0 | 0 | 0 | 0 | 0 | 0 | 0 | 0 | 0 | 0 | 0 | 0 | 0 | 0 | 0 | 0 | 0 | 0 | 0 | 0 | 1 | 0 | 0 | 0 | 0 | 0 |
| bacteria_Sep8d25b_12041 | 1 | 0 | 0 | 0 | 0 | 0 | 0 | 0 | 0 | 0 | 0 | 0 | 0 | 0 | 0 | 0 | 0 | 0 | 0 | 0 | 0 | 0 | 0 | 0 | 0 | 0 | 0 | 0 | 0 | 0 | 0 | 0 | 0 | 0 | 0 | 0 | 0 | 0 | 0 | 0 | 0 | 0 | 0 | 0 | 0 | 0 | 0 | 0 | 0 | 0 | 0 | 1 | 0 | 0 | 0 |
| bacteria_Sep8d25b_24038 | 1 | 0 | 0 | 0 | 0 | 0 | 0 | 0 | 0 | 0 | 0 | 0 | 0 | 0 | 0 | 0 | 0 | 0 | 0 | 0 | 0 | 0 | 0 | 0 | 0 | 0 | 0 | 0 | 0 | 0 | 0 | 0 | 0 | 0 | 0 | 0 | 0 | 0 | 0 | 0 | 0 | 0 | 0 | 0 | 0 | 0 | 0 | 0 | 0 | 0 | 0 | 1 | 0 | 0 | 0 |
| bacteria_Sep8d25b_45808 | 1 | 0 | 0 | 0 | 0 | 0 | 0 | 0 | 0 | 0 | 0 | 0 | 0 | 0 | 0 | 0 | 0 | 0 | 0 | 0 | 0 | 0 | 0 | 0 | 0 | 0 | 0 | 0 | 0 | 0 | 0 | 0 | 0 | 0 | 0 | 0 | 0 | 0 | 0 | 0 | 0 | 0 | 0 | 0 | 0 | 0 | 0 | 0 | 0 | 0 | 0 | 1 | 0 | 0 | 0 |
| bacteria_Sep8d25b_51194 | 1 | 0 | 0 | 0 | 0 | 0 | 0 | 0 | 0 | 0 | 0 | 0 | 0 | 0 | 0 | 0 | 0 | 0 | 0 | 0 | 0 | 0 | 0 | 0 | 0 | 0 | 0 | 0 | 0 | 0 | 0 | 0 | 0 | 0 | 0 | 0 | 0 | 0 | 0 | 0 | 0 | 0 | 0 | 0 | 0 | 0 | 0 | 0 | 0 | 0 | 0 | 1 | 0 | 0 | 0 |
| bacteria_Sep8d50b_15224 | 1 | 0 | 0 | 0 | 0 | 0 | 0 | 0 | 0 | 0 | 0 | 0 | 0 | 0 | 0 | 0 | 0 | 0 | 0 | 0 | 0 | 0 | 0 | 0 | 0 | 0 | 0 | 0 | 0 | 0 | 0 | 0 | 0 | 0 | 0 | 0 | 0 | 0 | 0 | 0 | 0 | 0 | 0 | 0 | 0 | 0 | 0 | 0 | 0 | 0 | 0 | 0 | 0 | 1 | 0 |
| bacteria_Sep8d50b_37428 | 1 | 0 | 0 | 0 | 0 | 0 | 0 | 0 | 0 | 0 | 0 | 0 | 0 | 0 | 0 | 0 | 0 | 0 | 0 | 0 | 0 | 0 | 0 | 0 | 0 | 0 | 0 | 0 | 0 | 0 | 0 | 0 | 0 | 0 | 0 | 0 | 0 | 0 | 0 | 0 | 0 | 0 | 0 | 0 | 0 | 0 | 0 | 0 | 0 | 0 | 0 | 0 | 0 | 1 | 0 |
| bacteria_Sep8d50b_38337 | 1 | 0 | 0 | 0 | 0 | 0 | 0 | 0 | 0 | 0 | 0 | 0 | 0 | 0 | 0 | 0 | 0 | 0 | 0 | 0 | 0 | 0 | 0 | 0 | 0 | 0 | 0 | 0 | 0 | 0 | 0 | 0 | 0 | 0 | 0 | 0 | 0 | 0 | 0 | 0 | 0 | 0 | 0 | 0 | 0 | 0 | 0 | 0 | 0 | 0 | 0 | 0 | 0 | 1 | 0 |
| virus_Aug5.0v_202 | 1 | 0 | 0 | 0 | 0 | 0 | 0 | 0 | 0 | 0 | 0 | 0 | 0 | 0 | 0 | 0 | 0 | 0 | 0 | 0 | 0 | 0 | 0 | 0 | 0 | 0 | 0 | 0 | 0 | 0 | 0 | 0 | 0 | 0 | 0 | 0 | 0 | 0 | 1 | 0 | 0 | 0 | 0 | 0 | 0 | 0 | 0 | 0 | 0 | 0 | 0 | 0 | 0 | 0 | 0 |
| virus_Aug5.0v_793 | 1 | 0 | 0 | 0 | 0 | 0 | 0 | 0 | 0 | 0 | 0 | 0 | 0 | 0 | 0 | 0 | 0 | 0 | 0 | 0 | 0 | 0 | 0 | 0 | 0 | 0 | 0 | 0 | 0 | 0 | 0 | 0 | 0 | 0 | 0 | 0 | 0 | 0 | 1 | 0 | 0 | 0 | 0 | 0 | 0 | 0 | 0 | 0 | 0 | 0 | 0 | 0 | 0 | 0 | 0 |
| virus_Aug5.0v_1193 | 1 | 0 | 0 | 0 | 0 | 0 | 0 | 0 | 0 | 0 | 0 | 0 | 0 | 0 | 0 | 0 | 0 | 0 | 0 | 0 | 0 | 0 | 0 | 0 | 0 | 0 | 0 | 0 | 0 | 0 | 0 | 0 | 0 | 0 | 0 | 0 | 0 | 0 | 1 | 0 | 0 | 0 | 0 | 0 | 0 | 0 | 0 | 0 | 0 | 0 | 0 | 0 | 0 | 0 | 0 |
| virus_Aug5.0v_1519 | 1 | 0 | 0 | 0 | 0 | 0 | 0 | 0 | 0 | 0 | 0 | 0 | 0 | 0 | 0 | 0 | 0 | 0 | 0 | 0 | 0 | 0 | 0 | 0 | 0 | 0 | 0 | 0 | 0 | 0 | 0 | 0 | 0 | 0 | 0 | 0 | 0 | 0 | 1 | 0 | 0 | 0 | 0 | 0 | 0 | 0 | 0 | 0 | 0 | 0 | 0 | 0 | 0 | 0 | 0 |
| virus_Aug5.0v_1537 | 1 | 0 | 0 | 0 | 0 | 0 | 0 | 0 | 0 | 0 | 0 | 0 | 0 | 0 | 0 | 0 | 0 | 0 | 0 | 0 | 0 | 0 | 0 | 0 | 0 | 0 | 0 | 0 | 0 | 0 | 0 | 0 | 0 | 0 | 0 | 0 | 0 | 0 | 1 | 0 | 0 | 0 | 0 | 0 | 0 | 0 | 0 | 0 | 0 | 0 | 0 | 0 | 0 | 0 | 0 |
| virus_Aug5.0v_1609 | 1 | 0 | 0 | 0 | 0 | 0 | 0 | 0 | 0 | 0 | 0 | 0 | 0 | 0 | 0 | 0 | 0 | 0 | 0 | 0 | 0 | 0 | 0 | 0 | 0 | 0 | 0 | 0 | 0 | 0 | 0 | 0 | 0 | 0 | 0 | 0 | 0 | 0 | 1 | 0 | 0 | 0 | 0 | 0 | 0 | 0 | 0 | 0 | 0 | 0 | 0 | 0 | 0 | 0 | 0 |
| virus_Aug5.0v_2246 | 1 | 0 | 0 | 0 | 0 | 0 | 0 | 0 | 0 | 0 | 0 | 0 | 0 | 0 | 0 | 0 | 0 | 0 | 0 | 0 | 0 | 0 | 0 | 0 | 0 | 0 | 0 | 0 | 0 | 0 | 0 | 0 | 0 | 0 | 0 | 0 | 0 | 0 | 1 | 0 | 0 | 0 | 0 | 0 | 0 | 0 | 0 | 0 | 0 | 0 | 0 | 0 | 0 | 0 | 0 |
| virus_Aug5.0v_2268 | 1 | 0 | 0 | 0 | 0 | 0 | 0 | 0 | 0 | 0 | 0 | 0 | 0 | 0 | 0 | 0 | 0 | 0 | 0 | 0 | 0 | 0 | 0 | 0 | 0 | 0 | 0 | 0 | 0 | 0 | 0 | 0 | 0 | 0 | 0 | 0 | 0 | 0 | 1 | 0 | 0 | 0 | 0 | 0 | 0 | 0 | 0 | 0 | 0 | 0 | 0 | 0 | 0 | 0 | 0 |
| virus_Aug5.0v_2799 | 1 | 0 | 0 | 0 | 0 | 0 | 0 | 0 | 0 | 0 | 0 | 0 | 0 | 0 | 0 | 0 | 0 | 0 | 0 | 0 | 0 | 0 | 0 | 0 | 0 | 0 | 0 | 0 | 0 | 0 | 0 | 0 | 0 | 0 | 0 | 0 | 0 | 0 | 1 | 0 | 0 | 0 | 0 | 0 | 0 | 0 | 0 | 0 | 0 | 0 | 0 | 0 | 0 | 0 | 0 |
| virus_Aug5.0v_3301 | 1 | 0 | 0 | 0 | 0 | 0 | 0 | 0 | 0 | 0 | 0 | 0 | 0 | 0 | 0 | 0 | 0 | 0 | 0 | 0 | 0 | 0 | 0 | 0 | 0 | 0 | 0 | 0 | 0 | 0 | 0 | 0 | 0 | 0 | 0 | 0 | 0 | 0 | 1 | 0 | 0 | 0 | 0 | 0 | 0 | 0 | 0 | 0 | 0 | 0 | 0 | 0 | 0 | 0 | 0 |
| virus_Aug5.0v_3389 | 1 | 0 | 0 | 0 | 0 | 0 | 0 | 0 | 0 | 0 | 0 | 0 | 0 | 0 | 0 | 0 | 0 | 0 | 0 | 0 | 0 | 0 | 0 | 0 | 0 | 0 | 0 | 0 | 0 | 0 | 0 | 0 | 0 | 0 | 0 | 0 | 0 | 0 | 1 | 0 | 0 | 0 | 0 | 0 | 0 | 0 | 0 | 0 | 0 | 0 | 0 | 0 | 0 | 0 | 0 |
| virus_Aug5.0v_3520 | 1 | 0 | 0 | 0 | 0 | 0 | 0 | 0 | 0 | 0 | 0 | 0 | 0 | 0 | 0 | 0 | 0 | 0 | 0 | 0 | 0 | 0 | 0 | 0 | 0 | 0 | 0 | 0 | 0 | 0 | 0 | 0 | 0 | 0 | 0 | 0 | 0 | 0 | 1 | 0 | 0 | 0 | 0 | 0 | 0 | 0 | 0 | 0 | 0 | 0 | 0 | 0 | 0 | 0 | 0 |
| virus_Aug5.0v_4562 | 1 | 0 | 0 | 0 | 0 | 0 | 0 | 0 | 0 | 0 | 0 | 0 | 0 | 0 | 0 | 0 | 0 | 0 | 0 | 0 | 0 | 0 | 0 | 0 | 0 | 0 | 0 | 0 | 0 | 0 | 0 | 0 | 0 | 0 | 0 | 0 | 0 | 0 | 1 | 0 | 0 | 0 | 0 | 0 | 0 | 0 | 0 | 0 | 0 | 0 | 0 | 0 | 0 | 0 | 0 |
| virus_Aug5.0v_4614 | 1 | 0 | 0 | 0 | 0 | 0 | 0 | 0 | 0 | 0 | 0 | 0 | 0 | 0 | 0 | 0 | 0 | 0 | 0 | 0 | 0 | 0 | 0 | 0 | 0 | 0 | 0 | 0 | 0 | 0 | 0 | 0 | 0 | 0 | 0 | 0 | 0 | 0 | 1 | 0 | 0 | 0 | 0 | 0 | 0 | 0 | 0 | 0 | 0 | 0 | 0 | 0 | 0 | 0 | 0 |
| virus_Aug5.0v_5281 | 1 | 0 | 0 | 0 | 0 | 0 | 0 | 0 | 0 | 0 | 0 | 0 | 0 | 0 | 0 | 0 | 0 | 0 | 0 | 0 | 0 | 0 | 0 | 0 | 0 | 0 | 0 | 0 | 0 | 0 | 0 | 0 | 0 | 0 | 0 | 0 | 0 | 0 | 1 | 0 | 0 | 0 | 0 | 0 | 0 | 0 | 0 | 0 | 0 | 0 | 0 | 0 | 0 | 0 | 0 |
| virus_Aug5.0v_5312 | 1 | 0 | 0 | 0 | 0 | 0 | 0 | 0 | 0 | 0 | 0 | 0 | 0 | 0 | 0 | 0 | 0 | 0 | 0 | 0 | 0 | 0 | 0 | 0 | 0 | 0 | 0 | 0 | 0 | 0 | 0 | 0 | 0 | 0 | 0 | 0 | 0 | 0 | 1 | 0 | 0 | 0 | 0 | 0 | 0 | 0 | 0 | 0 | 0 | 0 | 0 | 0 | 0 | 0 | 0 |
| virus_Aug5.0v_5323 | 1 | 0 | 0 | 0 | 0 | 0 | 0 | 0 | 0 | 0 | 0 | 0 | 0 | 0 | 0 | 0 | 0 | 0 | 0 | 0 | 0 | 0 | 0 | 0 | 0 | 0 | 0 | 0 | 0 | 0 | 0 | 0 | 0 | 0 | 0 | 0 | 0 | 0 | 1 | 0 | 0 | 0 | 0 | 0 | 0 | 0 | 0 | 0 | 0 | 0 | 0 | 0 | 0 | 0 | 0 |
| virus_Aug5.0v_5341 | 1 | 0 | 0 | 0 | 0 | 0 | 0 | 0 | 0 | 0 | 0 | 0 | 0 | 0 | 0 | 0 | 0 | 0 | 0 | 0 | 0 | 0 | 0 | 0 | 0 | 0 | 0 | 0 | 0 | 0 | 0 | 0 | 0 | 0 | 0 | 0 | 0 | 0 | 1 | 0 | 0 | 0 | 0 | 0 | 0 | 0 | 0 | 0 | 0 | 0 | 0 | 0 | 0 | 0 | 0 |
| virus_Aug5.5v_3959 | 1 | 0 | 0 | 0 | 0 | 0 | 0 | 0 | 0 | 0 | 0 | 0 | 0 | 0 | 0 | 0 | 0 | 0 | 0 | 0 | 0 | 0 | 0 | 0 | 0 | 0 | 0 | 0 | 0 | 0 | 0 | 0 | 0 | 0 | 0 | 0 | 0 | 0 | 0 | 0 | 1 | 0 | 0 | 0 | 0 | 0 | 0 | 0 | 0 | 0 | 0 | 0 | 0 | 0 | 0 |
| virus_Aug5.5v_4974 | 1 | 0 | 0 | 0 | 0 | 0 | 0 | 0 | 0 | 0 | 0 | 0 | 0 | 0 | 0 | 0 | 0 | 0 | 0 | 0 | 0 | 0 | 0 | 0 | 0 | 0 | 0 | 0 | 0 | 0 | 0 | 0 | 0 | 0 | 0 | 0 | 0 | 0 | 0 | 0 | 1 | 0 | 0 | 0 | 0 | 0 | 0 | 0 | 0 | 0 | 0 | 0 | 0 | 0 | 0 |
| virus_Aug5.5v_11623 | 1 | 0 | 0 | 0 | 0 | 0 | 0 | 0 | 0 | 0 | 0 | 0 | 0 | 0 | 0 | 0 | 0 | 0 | 0 | 0 | 0 | 0 | 0 | 0 | 0 | 0 | 0 | 0 | 0 | 0 | 0 | 0 | 0 | 0 | 0 | 0 | 0 | 0 | 0 | 0 | 1 | 0 | 0 | 0 | 0 | 0 | 0 | 0 | 0 | 0 | 0 | 0 | 0 | 0 | 0 |
| virus_Aug5.5v_15186 | 1 | 0 | 0 | 0 | 0 | 0 | 0 | 0 | 0 | 0 | 0 | 0 | 0 | 0 | 0 | 0 | 0 | 0 | 0 | 0 | 0 | 0 | 0 | 0 | 0 | 0 | 0 | 0 | 0 | 0 | 0 | 0 | 0 | 0 | 0 | 0 | 0 | 0 | 0 | 0 | 1 | 0 | 0 | 0 | 0 | 0 | 0 | 0 | 0 | 0 | 0 | 0 | 0 | 0 | 0 |
| virus_Aug5.5v_17227 | 1 | 0 | 0 | 0 | 0 | 0 | 0 | 0 | 0 | 0 | 0 | 0 | 0 | 0 | 0 | 0 | 0 | 0 | 0 | 0 | 0 | 0 | 0 | 0 | 0 | 0 | 0 | 0 | 0 | 0 | 0 | 0 | 0 | 0 | 0 | 0 | 0 | 0 | 0 | 0 | 1 | 0 | 0 | 0 | 0 | 0 | 0 | 0 | 0 | 0 | 0 | 0 | 0 | 0 | 0 |
| virus_Aug5.5v_18550 | 1 | 0 | 0 | 0 | 0 | 0 | 0 | 0 | 0 | 0 | 0 | 0 | 0 | 0 | 0 | 0 | 0 | 0 | 0 | 0 | 0 | 0 | 0 | 0 | 0 | 0 | 0 | 0 | 0 | 0 | 0 | 0 | 0 | 0 | 0 | 0 | 0 | 0 | 0 | 0 | 1 | 0 | 0 | 0 | 0 | 0 | 0 | 0 | 0 | 0 | 0 | 0 | 0 | 0 | 0 |
| virus_Aug5.5v_33657 | 1 | 0 | 0 | 0 | 0 | 0 | 0 | 0 | 0 | 0 | 0 | 0 | 0 | 0 | 0 | 0 | 0 | 0 | 0 | 0 | 0 | 0 | 0 | 0 | 0 | 0 | 0 | 0 | 0 | 0 | 0 | 0 | 0 | 0 | 0 | 0 | 0 | 0 | 0 | 0 | 1 | 0 | 0 | 0 | 0 | 0 | 0 | 0 | 0 | 0 | 0 | 0 | 0 | 0 | 0 |
| virus_Aug5.5v_35015 | 1 | 0 | 0 | 0 | 0 | 0 | 0 | 0 | 0 | 0 | 0 | 0 | 0 | 0 | 0 | 0 | 0 | 0 | 0 | 0 | 0 | 0 | 0 | 0 | 0 | 0 | 0 | 0 | 0 | 0 | 0 | 0 | 0 | 0 | 0 | 0 | 0 | 0 | 0 | 0 | 1 | 0 | 0 | 0 | 0 | 0 | 0 | 0 | 0 | 0 | 0 | 0 | 0 | 0 | 0 |
| virus_Aug5.25v_86 | 1 | 0 | 0 | 0 | 0 | 0 | 0 | 0 | 0 | 0 | 0 | 0 | 0 | 0 | 0 | 0 | 0 | 0 | 0 | 0 | 0 | 0 | 0 | 0 | 0 | 0 | 0 | 0 | 0 | 0 | 0 | 0 | 0 | 0 | 0 | 0 | 0 | 0 | 0 | 0 | 0 | 0 | 1 | 0 | 0 | 0 | 0 | 0 | 0 | 0 | 0 | 0 | 0 | 0 | 0 |
| virus_Aug5.25v_552 | 1 | 0 | 0 | 0 | 0 | 0 | 0 | 0 | 0 | 0 | 0 | 0 | 0 | 0 | 0 | 0 | 0 | 0 | 0 | 0 | 0 | 0 | 0 | 0 | 0 | 0 | 0 | 0 | 0 | 0 | 0 | 0 | 0 | 0 | 0 | 0 | 0 | 0 | 0 | 0 | 0 | 0 | 1 | 0 | 0 | 0 | 0 | 0 | 0 | 0 | 0 | 0 | 0 | 0 | 0 |
| virus_Aug5.25v_2583 | 1 | 0 | 0 | 0 | 0 | 0 | 0 | 0 | 0 | 0 | 0 | 0 | 0 | 0 | 0 | 0 | 0 | 0 | 0 | 0 | 0 | 0 | 0 | 0 | 0 | 0 | 0 | 0 | 0 | 0 | 0 | 0 | 0 | 0 | 0 | 0 | 0 | 0 | 0 | 0 | 0 | 0 | 1 | 0 | 0 | 0 | 0 | 0 | 0 | 0 | 0 | 0 | 0 | 0 | 0 |
| virus_Aug5.25v_3738 | 1 | 0 | 0 | 0 | 0 | 0 | 0 | 0 | 0 | 0 | 0 | 0 | 0 | 0 | 0 | 0 | 0 | 0 | 0 | 0 | 0 | 0 | 0 | 0 | 0 | 0 | 0 | 0 | 0 | 0 | 0 | 0 | 0 | 0 | 0 | 0 | 0 | 0 | 0 | 0 | 0 | 0 | 1 | 0 | 0 | 0 | 0 | 0 | 0 | 0 | 0 | 0 | 0 | 0 | 0 |
| virus_Aug5.25v_4597 | 1 | 0 | 0 | 0 | 0 | 0 | 0 | 0 | 0 | 0 | 0 | 0 | 0 | 0 | 0 | 0 | 0 | 0 | 0 | 0 | 0 | 0 | 0 | 0 | 0 | 0 | 0 | 0 | 0 | 0 | 0 | 0 | 0 | 0 | 0 | 0 | 0 | 0 | 0 | 0 | 0 | 0 | 1 | 0 | 0 | 0 | 0 | 0 | 0 | 0 | 0 | 0 | 0 | 0 | 0 |
| virus_Aug5.25v_4730 | 1 | 0 | 0 | 0 | 0 | 0 | 0 | 0 | 0 | 0 | 0 | 0 | 0 | 0 | 0 | 0 | 0 | 0 | 0 | 0 | 0 | 0 | 0 | 0 | 0 | 0 | 0 | 0 | 0 | 0 | 0 | 0 | 0 | 0 | 0 | 0 | 0 | 0 | 0 | 0 | 0 | 0 | 1 | 0 | 0 | 0 | 0 | 0 | 0 | 0 | 0 | 0 | 0 | 0 | 0 |
| virus_Aug5.25v_8581 | 1 | 0 | 0 | 0 | 0 | 0 | 0 | 0 | 0 | 0 | 0 | 0 | 0 | 0 | 0 | 0 | 0 | 0 | 0 | 0 | 0 | 0 | 0 | 0 | 0 | 0 | 0 | 0 | 0 | 0 | 0 | 0 | 0 | 0 | 0 | 0 | 0 | 0 | 0 | 0 | 0 | 0 | 1 | 0 | 0 | 0 | 0 | 0 | 0 | 0 | 0 | 0 | 0 | 0 | 0 |
| virus_Aug5.25v_10776 | 1 | 0 | 0 | 0 | 0 | 0 | 0 | 0 | 0 | 0 | 0 | 0 | 0 | 0 | 0 | 0 | 0 | 0 | 0 | 0 | 0 | 0 | 0 | 0 | 0 | 0 | 0 | 0 | 0 | 0 | 0 | 0 | 0 | 0 | 0 | 0 | 0 | 0 | 0 | 0 | 0 | 0 | 1 | 0 | 0 | 0 | 0 | 0 | 0 | 0 | 0 | 0 | 0 | 0 | 0 |
| virus_Aug5.25v_20427 | 1 | 0 | 0 | 0 | 0 | 0 | 0 | 0 | 0 | 0 | 0 | 0 | 0 | 0 | 0 | 0 | 0 | 0 | 0 | 0 | 0 | 0 | 0 | 0 | 0 | 0 | 0 | 0 | 0 | 0 | 0 | 0 | 0 | 0 | 0 | 0 | 0 | 0 | 0 | 0 | 0 | 0 | 1 | 0 | 0 | 0 | 0 | 0 | 0 | 0 | 0 | 0 | 0 | 0 | 0 |
| virus_Aug5.25v_20700 | 1 | 0 | 0 | 0 | 0 | 0 | 0 | 0 | 0 | 0 | 0 | 0 | 0 | 0 | 0 | 0 | 0 | 0 | 0 | 0 | 0 | 0 | 0 | 0 | 0 | 0 | 0 | 0 | 0 | 0 | 0 | 0 | 0 | 0 | 0 | 0 | 0 | 0 | 0 | 0 | 0 | 0 | 1 | 0 | 0 | 0 | 0 | 0 | 0 | 0 | 0 | 0 | 0 | 0 | 0 |
| virus_Aug5.25v_21512 | 1 | 0 | 0 | 0 | 0 | 0 | 0 | 0 | 0 | 0 | 0 | 0 | 0 | 0 | 0 | 0 | 0 | 0 | 0 | 0 | 0 | 0 | 0 | 0 | 0 | 0 | 0 | 0 | 0 | 0 | 0 | 0 | 0 | 0 | 0 | 0 | 0 | 0 | 0 | 0 | 0 | 0 | 1 | 0 | 0 | 0 | 0 | 0 | 0 | 0 | 0 | 0 | 0 | 0 | 0 |
| virus_Aug5.25v_24273 | 1 | 0 | 0 | 0 | 0 | 0 | 0 | 0 | 0 | 0 | 0 | 0 | 0 | 0 | 0 | 0 | 0 | 0 | 0 | 0 | 0 | 0 | 0 | 0 | 0 | 0 | 0 | 0 | 0 | 0 | 0 | 0 | 0 | 0 | 0 | 0 | 0 | 0 | 0 | 0 | 0 | 0 | 1 | 0 | 0 | 0 | 0 | 0 | 0 | 0 | 0 | 0 | 0 | 0 | 0 |
| virus_Aug5.25v_25788 | 1 | 0 | 0 | 0 | 0 | 0 | 0 | 0 | 0 | 0 | 0 | 0 | 0 | 0 | 0 | 0 | 0 | 0 | 0 | 0 | 0 | 0 | 0 | 0 | 0 | 0 | 0 | 0 | 0 | 0 | 0 | 0 | 0 | 0 | 0 | 0 | 0 | 0 | 0 | 0 | 0 | 0 | 1 | 0 | 0 | 0 | 0 | 0 | 0 | 0 | 0 | 0 | 0 | 0 | 0 |
| virus_Aug5.25v_25920 | 1 | 0 | 0 | 0 | 0 | 0 | 0 | 0 | 0 | 0 | 0 | 0 | 0 | 0 | 0 | 0 | 0 | 0 | 0 | 0 | 0 | 0 | 0 | 0 | 0 | 0 | 0 | 0 | 0 | 0 | 0 | 0 | 0 | 0 | 0 | 0 | 0 | 0 | 0 | 0 | 0 | 0 | 1 | 0 | 0 | 0 | 0 | 0 | 0 | 0 | 0 | 0 | 0 | 0 | 0 |
| virus_Aug5.25v_28677 | 1 | 0 | 0 | 0 | 0 | 0 | 0 | 0 | 0 | 0 | 0 | 0 | 0 | 0 | 0 | 0 | 0 | 0 | 0 | 0 | 0 | 0 | 0 | 0 | 0 | 0 | 0 | 0 | 0 | 0 | 0 | 0 | 0 | 0 | 0 | 0 | 0 | 0 | 0 | 0 | 0 | 0 | 1 | 0 | 0 | 0 | 0 | 0 | 0 | 0 | 0 | 0 | 0 | 0 | 0 |
| virus_Aug5.25v_33587 | 1 | 0 | 0 | 0 | 0 | 0 | 0 | 0 | 0 | 0 | 0 | 0 | 0 | 0 | 0 | 0 | 0 | 0 | 0 | 0 | 0 | 0 | 0 | 0 | 0 | 0 | 0 | 0 | 0 | 0 | 0 | 0 | 0 | 0 | 0 | 0 | 0 | 0 | 0 | 0 | 0 | 0 | 1 | 0 | 0 | 0 | 0 | 0 | 0 | 0 | 0 | 0 | 0 | 0 | 0 |
| virus_Aug5.25v_34030 | 1 | 0 | 0 | 0 | 0 | 0 | 0 | 0 | 0 | 0 | 0 | 0 | 0 | 0 | 0 | 0 | 0 | 0 | 0 | 0 | 0 | 0 | 0 | 0 | 0 | 0 | 0 | 0 | 0 | 0 | 0 | 0 | 0 | 0 | 0 | 0 | 0 | 0 | 0 | 0 | 0 | 0 | 1 | 0 | 0 | 0 | 0 | 0 | 0 | 0 | 0 | 0 | 0 | 0 | 0 |
| virus_Aug5.25v_40869 | 1 | 0 | 0 | 0 | 0 | 0 | 0 | 0 | 0 | 0 | 0 | 0 | 0 | 0 | 0 | 0 | 0 | 0 | 0 | 0 | 0 | 0 | 0 | 0 | 0 | 0 | 0 | 0 | 0 | 0 | 0 | 0 | 0 | 0 | 0 | 0 | 0 | 0 | 0 | 0 | 0 | 0 | 1 | 0 | 0 | 0 | 0 | 0 | 0 | 0 | 0 | 0 | 0 | 0 | 0 |
| virus_Aug5.50v_4473 | 1 | 0 | 0 | 0 | 0 | 0 | 0 | 0 | 0 | 0 | 0 | 0 | 0 | 0 | 0 | 0 | 0 | 0 | 0 | 0 | 0 | 0 | 0 | 0 | 0 | 0 | 0 | 0 | 0 | 0 | 0 | 0 | 0 | 0 | 0 | 0 | 0 | 0 | 0 | 0 | 0 | 0 | 0 | 0 | 1 | 0 | 0 | 0 | 0 | 0 | 0 | 0 | 0 | 0 | 0 |
| virus_Aug5.50v_9849 | 1 | 0 | 0 | 0 | 0 | 0 | 0 | 0 | 0 | 0 | 0 | 0 | 0 | 0 | 0 | 0 | 0 | 0 | 0 | 0 | 0 | 0 | 0 | 0 | 0 | 0 | 0 | 0 | 0 | 0 | 0 | 0 | 0 | 0 | 0 | 0 | 0 | 0 | 0 | 0 | 0 | 0 | 0 | 0 | 1 | 0 | 0 | 0 | 0 | 0 | 0 | 0 | 0 | 0 | 0 |
| virus_Aug5.50v_16031 | 1 | 0 | 0 | 0 | 0 | 0 | 0 | 0 | 0 | 0 | 0 | 0 | 0 | 0 | 0 | 0 | 0 | 0 | 0 | 0 | 0 | 0 | 0 | 0 | 0 | 0 | 0 | 0 | 0 | 0 | 0 | 0 | 0 | 0 | 0 | 0 | 0 | 0 | 0 | 0 | 0 | 0 | 0 | 0 | 1 | 0 | 0 | 0 | 0 | 0 | 0 | 0 | 0 | 0 | 0 |
| virus_Aug5.50v_30098 | 1 | 0 | 0 | 0 | 0 | 0 | 0 | 0 | 0 | 0 | 0 | 0 | 0 | 0 | 0 | 0 | 0 | 0 | 0 | 0 | 0 | 0 | 0 | 0 | 0 | 0 | 0 | 0 | 0 | 0 | 0 | 0 | 0 | 0 | 0 | 0 | 0 | 0 | 0 | 0 | 0 | 0 | 0 | 0 | 1 | 0 | 0 | 0 | 0 | 0 | 0 | 0 | 0 | 0 | 0 |
| virus_Aug5.75v_865 | 1 | 0 | 0 | 0 | 0 | 0 | 0 | 0 | 0 | 0 | 0 | 0 | 0 | 0 | 0 | 0 | 0 | 0 | 0 | 0 | 0 | 0 | 0 | 0 | 0 | 0 | 0 | 0 | 0 | 0 | 0 | 0 | 0 | 0 | 0 | 0 | 0 | 0 | 0 | 0 | 0 | 0 | 0 | 0 | 0 | 0 | 1 | 0 | 0 | 0 | 0 | 0 | 0 | 0 | 0 |
| virus_Aug5.75v_2824 | 1 | 0 | 0 | 0 | 0 | 0 | 0 | 0 | 0 | 0 | 0 | 0 | 0 | 0 | 0 | 0 | 0 | 0 | 0 | 0 | 0 | 0 | 0 | 0 | 0 | 0 | 0 | 0 | 0 | 0 | 0 | 0 | 0 | 0 | 0 | 0 | 0 | 0 | 0 | 0 | 0 | 0 | 0 | 0 | 0 | 0 | 1 | 0 | 0 | 0 | 0 | 0 | 0 | 0 | 0 |
| virus_Aug5.75v_17539 | 1 | 0 | 0 | 0 | 0 | 0 | 0 | 0 | 0 | 0 | 0 | 0 | 0 | 0 | 0 | 0 | 0 | 0 | 0 | 0 | 0 | 0 | 0 | 0 | 0 | 0 | 0 | 0 | 0 | 0 | 0 | 0 | 0 | 0 | 0 | 0 | 0 | 0 | 0 | 0 | 0 | 0 | 0 | 0 | 0 | 0 | 1 | 0 | 0 | 0 | 0 | 0 | 0 | 0 | 0 |
| virus_Aug5.75v_28329 | 1 | 0 | 0 | 0 | 0 | 0 | 0 | 0 | 0 | 0 | 0 | 0 | 0 | 0 | 0 | 0 | 0 | 0 | 0 | 0 | 0 | 0 | 0 | 0 | 0 | 0 | 0 | 0 | 0 | 0 | 0 | 0 | 0 | 0 | 0 | 0 | 0 | 0 | 0 | 0 | 0 | 0 | 0 | 0 | 0 | 0 | 1 | 0 | 0 | 0 | 0 | 0 | 0 | 0 | 0 |
| virus_Aug5.75v_34066 | 1 | 0 | 0 | 0 | 0 | 0 | 0 | 0 | 0 | 0 | 0 | 0 | 0 | 0 | 0 | 0 | 0 | 0 | 0 | 0 | 0 | 0 | 0 | 0 | 0 | 0 | 0 | 0 | 0 | 0 | 0 | 0 | 0 | 0 | 0 | 0 | 0 | 0 | 0 | 0 | 0 | 0 | 0 | 0 | 0 | 0 | 1 | 0 | 0 | 0 | 0 | 0 | 0 | 0 | 0 |
| virus_Jul11.0v_16073 | 1 | 0 | 0 | 0 | 0 | 0 | 0 | 0 | 0 | 0 | 0 | 0 | 0 | 0 | 0 | 0 | 0 | 0 | 0 | 0 | 0 | 0 | 1 | 0 | 0 | 0 | 0 | 0 | 0 | 0 | 0 | 0 | 0 | 0 | 0 | 0 | 0 | 0 | 0 | 0 | 0 | 0 | 0 | 0 | 0 | 0 | 0 | 0 | 0 | 0 | 0 | 0 | 0 | 0 | 0 |
| virus_Jul11.0v_17606 | 1 | 0 | 0 | 0 | 0 | 0 | 0 | 0 | 0 | 0 | 0 | 0 | 0 | 0 | 0 | 0 | 0 | 0 | 0 | 0 | 0 | 0 | 1 | 0 | 0 | 0 | 0 | 0 | 0 | 0 | 0 | 0 | 0 | 0 | 0 | 0 | 0 | 0 | 0 | 0 | 0 | 0 | 0 | 0 | 0 | 0 | 0 | 0 | 0 | 0 | 0 | 0 | 0 | 0 | 0 |
| virus_Jul11.0v_20942 | 1 | 0 | 0 | 0 | 0 | 0 | 0 | 0 | 0 | 0 | 0 | 0 | 0 | 0 | 0 | 0 | 0 | 0 | 0 | 0 | 0 | 0 | 1 | 0 | 0 | 0 | 0 | 0 | 0 | 0 | 0 | 0 | 0 | 0 | 0 | 0 | 0 | 0 | 0 | 0 | 0 | 0 | 0 | 0 | 0 | 0 | 0 | 0 | 0 | 0 | 0 | 0 | 0 | 0 | 0 |
| virus_Jul11.5v_2015 | 1 | 0 | 0 | 0 | 0 | 0 | 0 | 0 | 0 | 0 | 0 | 0 | 0 | 0 | 0 | 0 | 0 | 0 | 0 | 0 | 0 | 0 | 0 | 0 | 1 | 0 | 0 | 0 | 0 | 0 | 0 | 0 | 0 | 0 | 0 | 0 | 0 | 0 | 0 | 0 | 0 | 0 | 0 | 0 | 0 | 0 | 0 | 0 | 0 | 0 | 0 | 0 | 0 | 0 | 0 |
| virus_Jul11.5v_11287 | 1 | 0 | 0 | 0 | 0 | 0 | 0 | 0 | 0 | 0 | 0 | 0 | 0 | 0 | 0 | 0 | 0 | 0 | 0 | 0 | 0 | 0 | 0 | 0 | 1 | 0 | 0 | 0 | 0 | 0 | 0 | 0 | 0 | 0 | 0 | 0 | 0 | 0 | 0 | 0 | 0 | 0 | 0 | 0 | 0 | 0 | 0 | 0 | 0 | 0 | 0 | 0 | 0 | 0 | 0 |
| virus_Jul11.5v_11372 | 1 | 0 | 0 | 0 | 0 | 0 | 0 | 0 | 0 | 0 | 0 | 0 | 0 | 0 | 0 | 0 | 0 | 0 | 0 | 0 | 0 | 0 | 0 | 0 | 1 | 0 | 0 | 0 | 0 | 0 | 0 | 0 | 0 | 0 | 0 | 0 | 0 | 0 | 0 | 0 | 0 | 0 | 0 | 0 | 0 | 0 | 0 | 0 | 0 | 0 | 0 | 0 | 0 | 0 | 0 |
| virus_Jul11.5v_13541 | 1 | 0 | 0 | 0 | 0 | 0 | 0 | 0 | 0 | 0 | 0 | 0 | 0 | 0 | 0 | 0 | 0 | 0 | 0 | 0 | 0 | 0 | 0 | 0 | 1 | 0 | 0 | 0 | 0 | 0 | 0 | 0 | 0 | 0 | 0 | 0 | 0 | 0 | 0 | 0 | 0 | 0 | 0 | 0 | 0 | 0 | 0 | 0 | 0 | 0 | 0 | 0 | 0 | 0 | 0 |
| virus_Jul11.5v_13921 | 1 | 0 | 0 | 0 | 0 | 0 | 0 | 0 | 0 | 0 | 0 | 0 | 0 | 0 | 0 | 0 | 0 | 0 | 0 | 0 | 0 | 0 | 0 | 0 | 1 | 0 | 0 | 0 | 0 | 0 | 0 | 0 | 0 | 0 | 0 | 0 | 0 | 0 | 0 | 0 | 0 | 0 | 0 | 0 | 0 | 0 | 0 | 0 | 0 | 0 | 0 | 0 | 0 | 0 | 0 |
| virus_Jul11.5v_15683 | 1 | 0 | 0 | 0 | 0 | 0 | 0 | 0 | 0 | 0 | 0 | 0 | 0 | 0 | 0 | 0 | 0 | 0 | 0 | 0 | 0 | 0 | 0 | 0 | 1 | 0 | 0 | 0 | 0 | 0 | 0 | 0 | 0 | 0 | 0 | 0 | 0 | 0 | 0 | 0 | 0 | 0 | 0 | 0 | 0 | 0 | 0 | 0 | 0 | 0 | 0 | 0 | 0 | 0 | 0 |
| virus_Jul11.5v_18298 | 1 | 0 | 0 | 0 | 0 | 0 | 0 | 0 | 0 | 0 | 0 | 0 | 0 | 0 | 0 | 0 | 0 | 0 | 0 | 0 | 0 | 0 | 0 | 0 | 1 | 0 | 0 | 0 | 0 | 0 | 0 | 0 | 0 | 0 | 0 | 0 | 0 | 0 | 0 | 0 | 0 | 0 | 0 | 0 | 0 | 0 | 0 | 0 | 0 | 0 | 0 | 0 | 0 | 0 | 0 |
| virus_Jul11.5v_18843 | 1 | 0 | 0 | 0 | 0 | 0 | 0 | 0 | 0 | 0 | 0 | 0 | 0 | 0 | 0 | 0 | 0 | 0 | 0 | 0 | 0 | 0 | 0 | 0 | 1 | 0 | 0 | 0 | 0 | 0 | 0 | 0 | 0 | 0 | 0 | 0 | 0 | 0 | 0 | 0 | 0 | 0 | 0 | 0 | 0 | 0 | 0 | 0 | 0 | 0 | 0 | 0 | 0 | 0 | 0 |
| virus_Jul11.25v_400 | 1 | 0 | 0 | 0 | 0 | 0 | 0 | 0 | 0 | 0 | 0 | 0 | 0 | 0 | 0 | 0 | 0 | 0 | 0 | 0 | 0 | 0 | 0 | 0 | 0 | 0 | 1 | 0 | 0 | 0 | 0 | 0 | 0 | 0 | 0 | 0 | 0 | 0 | 0 | 0 | 0 | 0 | 0 | 0 | 0 | 0 | 0 | 0 | 0 | 0 | 0 | 0 | 0 | 0 | 0 |
| virus_Jul11.25v_914 | 1 | 0 | 0 | 0 | 0 | 0 | 0 | 0 | 0 | 0 | 0 | 0 | 0 | 0 | 0 | 0 | 0 | 0 | 0 | 0 | 0 | 0 | 0 | 0 | 0 | 0 | 1 | 0 | 0 | 0 | 0 | 0 | 0 | 0 | 0 | 0 | 0 | 0 | 0 | 0 | 0 | 0 | 0 | 0 | 0 | 0 | 0 | 0 | 0 | 0 | 0 | 0 | 0 | 0 | 0 |
| virus_Jul11.25v_2745 | 1 | 0 | 0 | 0 | 0 | 0 | 0 | 0 | 0 | 0 | 0 | 0 | 0 | 0 | 0 | 0 | 0 | 0 | 0 | 0 | 0 | 0 | 0 | 0 | 0 | 0 | 1 | 0 | 0 | 0 | 0 | 0 | 0 | 0 | 0 | 0 | 0 | 0 | 0 | 0 | 0 | 0 | 0 | 0 | 0 | 0 | 0 | 0 | 0 | 0 | 0 | 0 | 0 | 0 | 0 |
| virus_Jul11.25v_2818 | 1 | 0 | 0 | 0 | 0 | 0 | 0 | 0 | 0 | 0 | 0 | 0 | 0 | 0 | 0 | 0 | 0 | 0 | 0 | 0 | 0 | 0 | 0 | 0 | 0 | 0 | 1 | 0 | 0 | 0 | 0 | 0 | 0 | 0 | 0 | 0 | 0 | 0 | 0 | 0 | 0 | 0 | 0 | 0 | 0 | 0 | 0 | 0 | 0 | 0 | 0 | 0 | 0 | 0 | 0 |
| virus_Jul11.25v_3200 | 1 | 0 | 0 | 0 | 0 | 0 | 0 | 0 | 0 | 0 | 0 | 0 | 0 | 0 | 0 | 0 | 0 | 0 | 0 | 0 | 0 | 0 | 0 | 0 | 0 | 0 | 1 | 0 | 0 | 0 | 0 | 0 | 0 | 0 | 0 | 0 | 0 | 0 | 0 | 0 | 0 | 0 | 0 | 0 | 0 | 0 | 0 | 0 | 0 | 0 | 0 | 0 | 0 | 0 | 0 |
| virus_Jul11.25v_5072 | 1 | 0 | 0 | 0 | 0 | 0 | 0 | 0 | 0 | 0 | 0 | 0 | 0 | 0 | 0 | 0 | 0 | 0 | 0 | 0 | 0 | 0 | 0 | 0 | 0 | 0 | 1 | 0 | 0 | 0 | 0 | 0 | 0 | 0 | 0 | 0 | 0 | 0 | 0 | 0 | 0 | 0 | 0 | 0 | 0 | 0 | 0 | 0 | 0 | 0 | 0 | 0 | 0 | 0 | 0 |
| virus_Jul11.25v_5616 | 1 | 0 | 0 | 0 | 0 | 0 | 0 | 0 | 0 | 0 | 0 | 0 | 0 | 0 | 0 | 0 | 0 | 0 | 0 | 0 | 0 | 0 | 0 | 0 | 0 | 0 | 1 | 0 | 0 | 0 | 0 | 0 | 0 | 0 | 0 | 0 | 0 | 0 | 0 | 0 | 0 | 0 | 0 | 0 | 0 | 0 | 0 | 0 | 0 | 0 | 0 | 0 | 0 | 0 | 0 |
| virus_Jul11.25v_6604 | 1 | 0 | 0 | 0 | 0 | 0 | 0 | 0 | 0 | 0 | 0 | 0 | 0 | 0 | 0 | 0 | 0 | 0 | 0 | 0 | 0 | 0 | 0 | 0 | 0 | 0 | 1 | 0 | 0 | 0 | 0 | 0 | 0 | 0 | 0 | 0 | 0 | 0 | 0 | 0 | 0 | 0 | 0 | 0 | 0 | 0 | 0 | 0 | 0 | 0 | 0 | 0 | 0 | 0 | 0 |
| virus_Jul11.25v_7261 | 1 | 0 | 0 | 0 | 0 | 0 | 0 | 0 | 0 | 0 | 0 | 0 | 0 | 0 | 0 | 0 | 0 | 0 | 0 | 0 | 0 | 0 | 0 | 0 | 0 | 0 | 1 | 0 | 0 | 0 | 0 | 0 | 0 | 0 | 0 | 0 | 0 | 0 | 0 | 0 | 0 | 0 | 0 | 0 | 0 | 0 | 0 | 0 | 0 | 0 | 0 | 0 | 0 | 0 | 0 |
| virus_Jul11.25v_9154 | 1 | 0 | 0 | 0 | 0 | 0 | 0 | 0 | 0 | 0 | 0 | 0 | 0 | 0 | 0 | 0 | 0 | 0 | 0 | 0 | 0 | 0 | 0 | 0 | 0 | 0 | 1 | 0 | 0 | 0 | 0 | 0 | 0 | 0 | 0 | 0 | 0 | 0 | 0 | 0 | 0 | 0 | 0 | 0 | 0 | 0 | 0 | 0 | 0 | 0 | 0 | 0 | 0 | 0 | 0 |
| virus_Jul11.25v_11137 | 1 | 0 | 0 | 0 | 0 | 0 | 0 | 0 | 0 | 0 | 0 | 0 | 0 | 0 | 0 | 0 | 0 | 0 | 0 | 0 | 0 | 0 | 0 | 0 | 0 | 0 | 1 | 0 | 0 | 0 | 0 | 0 | 0 | 0 | 0 | 0 | 0 | 0 | 0 | 0 | 0 | 0 | 0 | 0 | 0 | 0 | 0 | 0 | 0 | 0 | 0 | 0 | 0 | 0 | 0 |
| virus_Jul11.25v_11902 | 1 | 0 | 0 | 0 | 0 | 0 | 0 | 0 | 0 | 0 | 0 | 0 | 0 | 0 | 0 | 0 | 0 | 0 | 0 | 0 | 0 | 0 | 0 | 0 | 0 | 0 | 1 | 0 | 0 | 0 | 0 | 0 | 0 | 0 | 0 | 0 | 0 | 0 | 0 | 0 | 0 | 0 | 0 | 0 | 0 | 0 | 0 | 0 | 0 | 0 | 0 | 0 | 0 | 0 | 0 |
| virus_Jul11.25v_12432 | 1 | 0 | 0 | 0 | 0 | 0 | 0 | 0 | 0 | 0 | 0 | 0 | 0 | 0 | 0 | 0 | 0 | 0 | 0 | 0 | 0 | 0 | 0 | 0 | 0 | 0 | 1 | 0 | 0 | 0 | 0 | 0 | 0 | 0 | 0 | 0 | 0 | 0 | 0 | 0 | 0 | 0 | 0 | 0 | 0 | 0 | 0 | 0 | 0 | 0 | 0 | 0 | 0 | 0 | 0 |
| virus_Jul11.25v_12881 | 1 | 0 | 0 | 0 | 0 | 0 | 0 | 0 | 0 | 0 | 0 | 0 | 0 | 0 | 0 | 0 | 0 | 0 | 0 | 0 | 0 | 0 | 0 | 0 | 0 | 0 | 1 | 0 | 0 | 0 | 0 | 0 | 0 | 0 | 0 | 0 | 0 | 0 | 0 | 0 | 0 | 0 | 0 | 0 | 0 | 0 | 0 | 0 | 0 | 0 | 0 | 0 | 0 | 0 | 0 |
| virus_Jul11.25v_12958 | 1 | 0 | 0 | 0 | 0 | 0 | 0 | 0 | 0 | 0 | 0 | 0 | 0 | 0 | 0 | 0 | 0 | 0 | 0 | 0 | 0 | 0 | 0 | 0 | 0 | 0 | 1 | 0 | 0 | 0 | 0 | 0 | 0 | 0 | 0 | 0 | 0 | 0 | 0 | 0 | 0 | 0 | 0 | 0 | 0 | 0 | 0 | 0 | 0 | 0 | 0 | 0 | 0 | 0 | 0 |
| virus_Jul11.25v_13332 | 1 | 0 | 0 | 0 | 0 | 0 | 0 | 0 | 0 | 0 | 0 | 0 | 0 | 0 | 0 | 0 | 0 | 0 | 0 | 0 | 0 | 0 | 0 | 0 | 0 | 0 | 1 | 0 | 0 | 0 | 0 | 0 | 0 | 0 | 0 | 0 | 0 | 0 | 0 | 0 | 0 | 0 | 0 | 0 | 0 | 0 | 0 | 0 | 0 | 0 | 0 | 0 | 0 | 0 | 0 |
| virus_Jul11.25v_14963 | 1 | 0 | 0 | 0 | 0 | 0 | 0 | 0 | 0 | 0 | 0 | 0 | 0 | 0 | 0 | 0 | 0 | 0 | 0 | 0 | 0 | 0 | 0 | 0 | 0 | 0 | 1 | 0 | 0 | 0 | 0 | 0 | 0 | 0 | 0 | 0 | 0 | 0 | 0 | 0 | 0 | 0 | 0 | 0 | 0 | 0 | 0 | 0 | 0 | 0 | 0 | 0 | 0 | 0 | 0 |
| virus_Jul11.25v_15112 | 1 | 0 | 0 | 0 | 0 | 0 | 0 | 0 | 0 | 0 | 0 | 0 | 0 | 0 | 0 | 0 | 0 | 0 | 0 | 0 | 0 | 0 | 0 | 0 | 0 | 0 | 1 | 0 | 0 | 0 | 0 | 0 | 0 | 0 | 0 | 0 | 0 | 0 | 0 | 0 | 0 | 0 | 0 | 0 | 0 | 0 | 0 | 0 | 0 | 0 | 0 | 0 | 0 | 0 | 0 |
| virus_Jul11.25v_18292 | 1 | 0 | 0 | 0 | 0 | 0 | 0 | 0 | 0 | 0 | 0 | 0 | 0 | 0 | 0 | 0 | 0 | 0 | 0 | 0 | 0 | 0 | 0 | 0 | 0 | 0 | 1 | 0 | 0 | 0 | 0 | 0 | 0 | 0 | 0 | 0 | 0 | 0 | 0 | 0 | 0 | 0 | 0 | 0 | 0 | 0 | 0 | 0 | 0 | 0 | 0 | 0 | 0 | 0 | 0 |
| virus_Jul11.25v_20780 | 1 | 0 | 0 | 0 | 0 | 0 | 0 | 0 | 0 | 0 | 0 | 0 | 0 | 0 | 0 | 0 | 0 | 0 | 0 | 0 | 0 | 0 | 0 | 0 | 0 | 0 | 1 | 0 | 0 | 0 | 0 | 0 | 0 | 0 | 0 | 0 | 0 | 0 | 0 | 0 | 0 | 0 | 0 | 0 | 0 | 0 | 0 | 0 | 0 | 0 | 0 | 0 | 0 | 0 | 0 |
| virus_Jul11.50v_1408 | 1 | 0 | 0 | 0 | 0 | 0 | 0 | 0 | 0 | 0 | 0 | 0 | 0 | 0 | 0 | 0 | 0 | 0 | 0 | 0 | 0 | 0 | 0 | 0 | 0 | 0 | 0 | 0 | 1 | 0 | 0 | 0 | 0 | 0 | 0 | 0 | 0 | 0 | 0 | 0 | 0 | 0 | 0 | 0 | 0 | 0 | 0 | 0 | 0 | 0 | 0 | 0 | 0 | 0 | 0 |
| virus_Jul11.50v_4166 | 1 | 0 | 0 | 0 | 0 | 0 | 0 | 0 | 0 | 0 | 0 | 0 | 0 | 0 | 0 | 0 | 0 | 0 | 0 | 0 | 0 | 0 | 0 | 0 | 0 | 0 | 0 | 0 | 1 | 0 | 0 | 0 | 0 | 0 | 0 | 0 | 0 | 0 | 0 | 0 | 0 | 0 | 0 | 0 | 0 | 0 | 0 | 0 | 0 | 0 | 0 | 0 | 0 | 0 | 0 |
| virus_Jul11.50v_4788 | 1 | 0 | 0 | 0 | 0 | 0 | 0 | 0 | 0 | 0 | 0 | 0 | 0 | 0 | 0 | 0 | 0 | 0 | 0 | 0 | 0 | 0 | 0 | 0 | 0 | 0 | 0 | 0 | 1 | 0 | 0 | 0 | 0 | 0 | 0 | 0 | 0 | 0 | 0 | 0 | 0 | 0 | 0 | 0 | 0 | 0 | 0 | 0 | 0 | 0 | 0 | 0 | 0 | 0 | 0 |
| virus_Jul11.50v_6014 | 1 | 0 | 0 | 0 | 0 | 0 | 0 | 0 | 0 | 0 | 0 | 0 | 0 | 0 | 0 | 0 | 0 | 0 | 0 | 0 | 0 | 0 | 0 | 0 | 0 | 0 | 0 | 0 | 1 | 0 | 0 | 0 | 0 | 0 | 0 | 0 | 0 | 0 | 0 | 0 | 0 | 0 | 0 | 0 | 0 | 0 | 0 | 0 | 0 | 0 | 0 | 0 | 0 | 0 | 0 |
| virus_Jul11.50v_9390 | 1 | 0 | 0 | 0 | 0 | 0 | 0 | 0 | 0 | 0 | 0 | 0 | 0 | 0 | 0 | 0 | 0 | 0 | 0 | 0 | 0 | 0 | 0 | 0 | 0 | 0 | 0 | 0 | 1 | 0 | 0 | 0 | 0 | 0 | 0 | 0 | 0 | 0 | 0 | 0 | 0 | 0 | 0 | 0 | 0 | 0 | 0 | 0 | 0 | 0 | 0 | 0 | 0 | 0 | 0 |
| virus_Jul11.50v_9468 | 1 | 0 | 0 | 0 | 0 | 0 | 0 | 0 | 0 | 0 | 0 | 0 | 0 | 0 | 0 | 0 | 0 | 0 | 0 | 0 | 0 | 0 | 0 | 0 | 0 | 0 | 0 | 0 | 1 | 0 | 0 | 0 | 0 | 0 | 0 | 0 | 0 | 0 | 0 | 0 | 0 | 0 | 0 | 0 | 0 | 0 | 0 | 0 | 0 | 0 | 0 | 0 | 0 | 0 | 0 |
| virus_Jul11.50v_10110 | 1 | 0 | 0 | 0 | 0 | 0 | 0 | 0 | 0 | 0 | 0 | 0 | 0 | 0 | 0 | 0 | 0 | 0 | 0 | 0 | 0 | 0 | 0 | 0 | 0 | 0 | 0 | 0 | 1 | 0 | 0 | 0 | 0 | 0 | 0 | 0 | 0 | 0 | 0 | 0 | 0 | 0 | 0 | 0 | 0 | 0 | 0 | 0 | 0 | 0 | 0 | 0 | 0 | 0 | 0 |
| virus_Jul11.50v_11281 | 1 | 0 | 0 | 0 | 0 | 0 | 0 | 0 | 0 | 0 | 0 | 0 | 0 | 0 | 0 | 0 | 0 | 0 | 0 | 0 | 0 | 0 | 0 | 0 | 0 | 0 | 0 | 0 | 1 | 0 | 0 | 0 | 0 | 0 | 0 | 0 | 0 | 0 | 0 | 0 | 0 | 0 | 0 | 0 | 0 | 0 | 0 | 0 | 0 | 0 | 0 | 0 | 0 | 0 | 0 |
| virus_Jul11.50v_16973 | 1 | 0 | 0 | 0 | 0 | 0 | 0 | 0 | 0 | 0 | 0 | 0 | 0 | 0 | 0 | 0 | 0 | 0 | 0 | 0 | 0 | 0 | 0 | 0 | 0 | 0 | 0 | 0 | 1 | 0 | 0 | 0 | 0 | 0 | 0 | 0 | 0 | 0 | 0 | 0 | 0 | 0 | 0 | 0 | 0 | 0 | 0 | 0 | 0 | 0 | 0 | 0 | 0 | 0 | 0 |
| virus_Jul11.75v_3084 | 1 | 0 | 0 | 0 | 0 | 0 | 0 | 0 | 0 | 0 | 0 | 0 | 0 | 0 | 0 | 0 | 0 | 0 | 0 | 0 | 0 | 0 | 0 | 0 | 0 | 0 | 0 | 0 | 0 | 0 | 1 | 0 | 0 | 0 | 0 | 0 | 0 | 0 | 0 | 0 | 0 | 0 | 0 | 0 | 0 | 0 | 0 | 0 | 0 | 0 | 0 | 0 | 0 | 0 | 0 |
| virus_Jul11.75v_4190 | 1 | 0 | 0 | 0 | 0 | 0 | 0 | 0 | 0 | 0 | 0 | 0 | 0 | 0 | 0 | 0 | 0 | 0 | 0 | 0 | 0 | 0 | 0 | 0 | 0 | 0 | 0 | 0 | 0 | 0 | 1 | 0 | 0 | 0 | 0 | 0 | 0 | 0 | 0 | 0 | 0 | 0 | 0 | 0 | 0 | 0 | 0 | 0 | 0 | 0 | 0 | 0 | 0 | 0 | 0 |
| virus_Jul11.75v_17025 | 1 | 0 | 0 | 0 | 0 | 0 | 0 | 0 | 0 | 0 | 0 | 0 | 0 | 0 | 0 | 0 | 0 | 0 | 0 | 0 | 0 | 0 | 0 | 0 | 0 | 0 | 0 | 0 | 0 | 0 | 1 | 0 | 0 | 0 | 0 | 0 | 0 | 0 | 0 | 0 | 0 | 0 | 0 | 0 | 0 | 0 | 0 | 0 | 0 | 0 | 0 | 0 | 0 | 0 | 0 |
| virus_Jul11.75v_17405 | 1 | 0 | 0 | 0 | 0 | 0 | 0 | 0 | 0 | 0 | 0 | 0 | 0 | 0 | 0 | 0 | 0 | 0 | 0 | 0 | 0 | 0 | 0 | 0 | 0 | 0 | 0 | 0 | 0 | 0 | 1 | 0 | 0 | 0 | 0 | 0 | 0 | 0 | 0 | 0 | 0 | 0 | 0 | 0 | 0 | 0 | 0 | 0 | 0 | 0 | 0 | 0 | 0 | 0 | 0 |
| virus_Jul11.75v_23737 | 1 | 0 | 0 | 0 | 0 | 0 | 0 | 0 | 0 | 0 | 0 | 0 | 0 | 0 | 0 | 0 | 0 | 0 | 0 | 0 | 0 | 0 | 0 | 0 | 0 | 0 | 0 | 0 | 0 | 0 | 1 | 0 | 0 | 0 | 0 | 0 | 0 | 0 | 0 | 0 | 0 | 0 | 0 | 0 | 0 | 0 | 0 | 0 | 0 | 0 | 0 | 0 | 0 | 0 | 0 |
| virus_Jul11.75v_37018 | 1 | 0 | 0 | 0 | 0 | 0 | 0 | 0 | 0 | 0 | 0 | 0 | 0 | 0 | 0 | 0 | 0 | 0 | 0 | 0 | 0 | 0 | 0 | 0 | 0 | 0 | 0 | 0 | 0 | 0 | 1 | 0 | 0 | 0 | 0 | 0 | 0 | 0 | 0 | 0 | 0 | 0 | 0 | 0 | 0 | 0 | 0 | 0 | 0 | 0 | 0 | 0 | 0 | 0 | 0 |
| virus_Jul11.75v_39594 | 1 | 0 | 0 | 0 | 0 | 0 | 0 | 0 | 0 | 0 | 0 | 0 | 0 | 0 | 0 | 0 | 0 | 0 | 0 | 0 | 0 | 0 | 0 | 0 | 0 | 0 | 0 | 0 | 0 | 0 | 1 | 0 | 0 | 0 | 0 | 0 | 0 | 0 | 0 | 0 | 0 | 0 | 0 | 0 | 0 | 0 | 0 | 0 | 0 | 0 | 0 | 0 | 0 | 0 | 0 |
| virus_Jul11.75v_40073 | 1 | 0 | 0 | 0 | 0 | 0 | 0 | 0 | 0 | 0 | 0 | 0 | 0 | 0 | 0 | 0 | 0 | 0 | 0 | 0 | 0 | 0 | 0 | 0 | 0 | 0 | 0 | 0 | 0 | 0 | 1 | 0 | 0 | 0 | 0 | 0 | 0 | 0 | 0 | 0 | 0 | 0 | 0 | 0 | 0 | 0 | 0 | 0 | 0 | 0 | 0 | 0 | 0 | 0 | 0 |
| virus_Jul27.5v_43 | 1 | 0 | 0 | 0 | 0 | 0 | 0 | 0 | 0 | 0 | 0 | 0 | 0 | 0 | 0 | 0 | 0 | 0 | 0 | 0 | 0 | 0 | 0 | 0 | 0 | 0 | 0 | 0 | 0 | 0 | 0 | 0 | 1 | 0 | 0 | 0 | 0 | 0 | 0 | 0 | 0 | 0 | 0 | 0 | 0 | 0 | 0 | 0 | 0 | 0 | 0 | 0 | 0 | 0 | 0 |
| virus_Jul27.5v_380 | 1 | 0 | 0 | 0 | 0 | 0 | 0 | 0 | 0 | 0 | 0 | 0 | 0 | 0 | 0 | 0 | 0 | 0 | 0 | 0 | 0 | 0 | 0 | 0 | 0 | 0 | 0 | 0 | 0 | 0 | 0 | 0 | 1 | 0 | 0 | 0 | 0 | 0 | 0 | 0 | 0 | 0 | 0 | 0 | 0 | 0 | 0 | 0 | 0 | 0 | 0 | 0 | 0 | 0 | 0 |
| virus_Jul27.5v_1937 | 1 | 0 | 0 | 0 | 0 | 0 | 0 | 0 | 0 | 0 | 0 | 0 | 0 | 0 | 0 | 0 | 0 | 0 | 0 | 0 | 0 | 0 | 0 | 0 | 0 | 0 | 0 | 0 | 0 | 0 | 0 | 0 | 1 | 0 | 0 | 0 | 0 | 0 | 0 | 0 | 0 | 0 | 0 | 0 | 0 | 0 | 0 | 0 | 0 | 0 | 0 | 0 | 0 | 0 | 0 |
| virus_Jul27.5v_5152 | 1 | 0 | 0 | 0 | 0 | 0 | 0 | 0 | 0 | 0 | 0 | 0 | 0 | 0 | 0 | 0 | 0 | 0 | 0 | 0 | 0 | 0 | 0 | 0 | 0 | 0 | 0 | 0 | 0 | 0 | 0 | 0 | 1 | 0 | 0 | 0 | 0 | 0 | 0 | 0 | 0 | 0 | 0 | 0 | 0 | 0 | 0 | 0 | 0 | 0 | 0 | 0 | 0 | 0 | 0 |
| virus_Jul27.5v_7463 | 1 | 0 | 0 | 0 | 0 | 0 | 0 | 0 | 0 | 0 | 0 | 0 | 0 | 0 | 0 | 0 | 0 | 0 | 0 | 0 | 0 | 0 | 0 | 0 | 0 | 0 | 0 | 0 | 0 | 0 | 0 | 0 | 1 | 0 | 0 | 0 | 0 | 0 | 0 | 0 | 0 | 0 | 0 | 0 | 0 | 0 | 0 | 0 | 0 | 0 | 0 | 0 | 0 | 0 | 0 |
| virus_Jul27.5v_7704 | 1 | 0 | 0 | 0 | 0 | 0 | 0 | 0 | 0 | 0 | 0 | 0 | 0 | 0 | 0 | 0 | 0 | 0 | 0 | 0 | 0 | 0 | 0 | 0 | 0 | 0 | 0 | 0 | 0 | 0 | 0 | 0 | 1 | 0 | 0 | 0 | 0 | 0 | 0 | 0 | 0 | 0 | 0 | 0 | 0 | 0 | 0 | 0 | 0 | 0 | 0 | 0 | 0 | 0 | 0 |
| virus_Jul27.5v_8055 | 1 | 0 | 0 | 0 | 0 | 0 | 0 | 0 | 0 | 0 | 0 | 0 | 0 | 0 | 0 | 0 | 0 | 0 | 0 | 0 | 0 | 0 | 0 | 0 | 0 | 0 | 0 | 0 | 0 | 0 | 0 | 0 | 1 | 0 | 0 | 0 | 0 | 0 | 0 | 0 | 0 | 0 | 0 | 0 | 0 | 0 | 0 | 0 | 0 | 0 | 0 | 0 | 0 | 0 | 0 |
| virus_Jul27.5v_8443 | 1 | 0 | 0 | 0 | 0 | 0 | 0 | 0 | 0 | 0 | 0 | 0 | 0 | 0 | 0 | 0 | 0 | 0 | 0 | 0 | 0 | 0 | 0 | 0 | 0 | 0 | 0 | 0 | 0 | 0 | 0 | 0 | 1 | 0 | 0 | 0 | 0 | 0 | 0 | 0 | 0 | 0 | 0 | 0 | 0 | 0 | 0 | 0 | 0 | 0 | 0 | 0 | 0 | 0 | 0 |
| virus_Jul27.5v_12688 | 1 | 0 | 0 | 0 | 0 | 0 | 0 | 0 | 0 | 0 | 0 | 0 | 0 | 0 | 0 | 0 | 0 | 0 | 0 | 0 | 0 | 0 | 0 | 0 | 0 | 0 | 0 | 0 | 0 | 0 | 0 | 0 | 1 | 0 | 0 | 0 | 0 | 0 | 0 | 0 | 0 | 0 | 0 | 0 | 0 | 0 | 0 | 0 | 0 | 0 | 0 | 0 | 0 | 0 | 0 |
| virus_Jul27.5v_13580 | 1 | 0 | 0 | 0 | 0 | 0 | 0 | 0 | 0 | 0 | 0 | 0 | 0 | 0 | 0 | 0 | 0 | 0 | 0 | 0 | 0 | 0 | 0 | 0 | 0 | 0 | 0 | 0 | 0 | 0 | 0 | 0 | 1 | 0 | 0 | 0 | 0 | 0 | 0 | 0 | 0 | 0 | 0 | 0 | 0 | 0 | 0 | 0 | 0 | 0 | 0 | 0 | 0 | 0 | 0 |
| virus_Jul27.5v_13776 | 1 | 0 | 0 | 0 | 0 | 0 | 0 | 0 | 0 | 0 | 0 | 0 | 0 | 0 | 0 | 0 | 0 | 0 | 0 | 0 | 0 | 0 | 0 | 0 | 0 | 0 | 0 | 0 | 0 | 0 | 0 | 0 | 1 | 0 | 0 | 0 | 0 | 0 | 0 | 0 | 0 | 0 | 0 | 0 | 0 | 0 | 0 | 0 | 0 | 0 | 0 | 0 | 0 | 0 | 0 |
| virus_Jul27.5v_14695 | 1 | 0 | 0 | 0 | 0 | 0 | 0 | 0 | 0 | 0 | 0 | 0 | 0 | 0 | 0 | 0 | 0 | 0 | 0 | 0 | 0 | 0 | 0 | 0 | 0 | 0 | 0 | 0 | 0 | 0 | 0 | 0 | 1 | 0 | 0 | 0 | 0 | 0 | 0 | 0 | 0 | 0 | 0 | 0 | 0 | 0 | 0 | 0 | 0 | 0 | 0 | 0 | 0 | 0 | 0 |
| virus_Jul27.5v_16922 | 1 | 0 | 0 | 0 | 0 | 0 | 0 | 0 | 0 | 0 | 0 | 0 | 0 | 0 | 0 | 0 | 0 | 0 | 0 | 0 | 0 | 0 | 0 | 0 | 0 | 0 | 0 | 0 | 0 | 0 | 0 | 0 | 1 | 0 | 0 | 0 | 0 | 0 | 0 | 0 | 0 | 0 | 0 | 0 | 0 | 0 | 0 | 0 | 0 | 0 | 0 | 0 | 0 | 0 | 0 |
| virus_Jul27.5v_18729 | 1 | 0 | 0 | 0 | 0 | 0 | 0 | 0 | 0 | 0 | 0 | 0 | 0 | 0 | 0 | 0 | 0 | 0 | 0 | 0 | 0 | 0 | 0 | 0 | 0 | 0 | 0 | 0 | 0 | 0 | 0 | 0 | 1 | 0 | 0 | 0 | 0 | 0 | 0 | 0 | 0 | 0 | 0 | 0 | 0 | 0 | 0 | 0 | 0 | 0 | 0 | 0 | 0 | 0 | 0 |
| virus_Jul27.5v_19719 | 1 | 0 | 0 | 0 | 0 | 0 | 0 | 0 | 0 | 0 | 0 | 0 | 0 | 0 | 0 | 0 | 0 | 0 | 0 | 0 | 0 | 0 | 0 | 0 | 0 | 0 | 0 | 0 | 0 | 0 | 0 | 0 | 1 | 0 | 0 | 0 | 0 | 0 | 0 | 0 | 0 | 0 | 0 | 0 | 0 | 0 | 0 | 0 | 0 | 0 | 0 | 0 | 0 | 0 | 0 |
| virus_Jul27.5v_19974 | 1 | 0 | 0 | 0 | 0 | 0 | 0 | 0 | 0 | 0 | 0 | 0 | 0 | 0 | 0 | 0 | 0 | 0 | 0 | 0 | 0 | 0 | 0 | 0 | 0 | 0 | 0 | 0 | 0 | 0 | 0 | 0 | 1 | 0 | 0 | 0 | 0 | 0 | 0 | 0 | 0 | 0 | 0 | 0 | 0 | 0 | 0 | 0 | 0 | 0 | 0 | 0 | 0 | 0 | 0 |
| virus_Jul27.25v_5795 | 1 | 0 | 0 | 0 | 0 | 0 | 0 | 0 | 0 | 0 | 0 | 0 | 0 | 0 | 0 | 0 | 0 | 0 | 0 | 0 | 0 | 0 | 0 | 0 | 0 | 0 | 0 | 0 | 0 | 0 | 0 | 0 | 0 | 0 | 1 | 0 | 0 | 0 | 0 | 0 | 0 | 0 | 0 | 0 | 0 | 0 | 0 | 0 | 0 | 0 | 0 | 0 | 0 | 0 | 0 |
| virus_Jul27.25v_7668 | 1 | 0 | 0 | 0 | 0 | 0 | 0 | 0 | 0 | 0 | 0 | 0 | 0 | 0 | 0 | 0 | 0 | 0 | 0 | 0 | 0 | 0 | 0 | 0 | 0 | 0 | 0 | 0 | 0 | 0 | 0 | 0 | 0 | 0 | 1 | 0 | 0 | 0 | 0 | 0 | 0 | 0 | 0 | 0 | 0 | 0 | 0 | 0 | 0 | 0 | 0 | 0 | 0 | 0 | 0 |
| virus_Jul27.25v_8840 | 1 | 0 | 0 | 0 | 0 | 0 | 0 | 0 | 0 | 0 | 0 | 0 | 0 | 0 | 0 | 0 | 0 | 0 | 0 | 0 | 0 | 0 | 0 | 0 | 0 | 0 | 0 | 0 | 0 | 0 | 0 | 0 | 0 | 0 | 1 | 0 | 0 | 0 | 0 | 0 | 0 | 0 | 0 | 0 | 0 | 0 | 0 | 0 | 0 | 0 | 0 | 0 | 0 | 0 | 0 |
| virus_Jul27.25v_9816 | 1 | 0 | 0 | 0 | 0 | 0 | 0 | 0 | 0 | 0 | 0 | 0 | 0 | 0 | 0 | 0 | 0 | 0 | 0 | 0 | 0 | 0 | 0 | 0 | 0 | 0 | 0 | 0 | 0 | 0 | 0 | 0 | 0 | 0 | 1 | 0 | 0 | 0 | 0 | 0 | 0 | 0 | 0 | 0 | 0 | 0 | 0 | 0 | 0 | 0 | 0 | 0 | 0 | 0 | 0 |
| virus_Jul27.25v_10141 | 1 | 0 | 0 | 0 | 0 | 0 | 0 | 0 | 0 | 0 | 0 | 0 | 0 | 0 | 0 | 0 | 0 | 0 | 0 | 0 | 0 | 0 | 0 | 0 | 0 | 0 | 0 | 0 | 0 | 0 | 0 | 0 | 0 | 0 | 1 | 0 | 0 | 0 | 0 | 0 | 0 | 0 | 0 | 0 | 0 | 0 | 0 | 0 | 0 | 0 | 0 | 0 | 0 | 0 | 0 |
| virus_Jul27.25v_15898 | 1 | 0 | 0 | 0 | 0 | 0 | 0 | 0 | 0 | 0 | 0 | 0 | 0 | 0 | 0 | 0 | 0 | 0 | 0 | 0 | 0 | 0 | 0 | 0 | 0 | 0 | 0 | 0 | 0 | 0 | 0 | 0 | 0 | 0 | 1 | 0 | 0 | 0 | 0 | 0 | 0 | 0 | 0 | 0 | 0 | 0 | 0 | 0 | 0 | 0 | 0 | 0 | 0 | 0 | 0 |
| virus_Jul27.25v_17052 | 1 | 0 | 0 | 0 | 0 | 0 | 0 | 0 | 0 | 0 | 0 | 0 | 0 | 0 | 0 | 0 | 0 | 0 | 0 | 0 | 0 | 0 | 0 | 0 | 0 | 0 | 0 | 0 | 0 | 0 | 0 | 0 | 0 | 0 | 1 | 0 | 0 | 0 | 0 | 0 | 0 | 0 | 0 | 0 | 0 | 0 | 0 | 0 | 0 | 0 | 0 | 0 | 0 | 0 | 0 |
| virus_Jul27.25v_20364 | 1 | 0 | 0 | 0 | 0 | 0 | 0 | 0 | 0 | 0 | 0 | 0 | 0 | 0 | 0 | 0 | 0 | 0 | 0 | 0 | 0 | 0 | 0 | 0 | 0 | 0 | 0 | 0 | 0 | 0 | 0 | 0 | 0 | 0 | 1 | 0 | 0 | 0 | 0 | 0 | 0 | 0 | 0 | 0 | 0 | 0 | 0 | 0 | 0 | 0 | 0 | 0 | 0 | 0 | 0 |
| virus_Jul27.25v_24189 | 1 | 0 | 0 | 0 | 0 | 0 | 0 | 0 | 0 | 0 | 0 | 0 | 0 | 0 | 0 | 0 | 0 | 0 | 0 | 0 | 0 | 0 | 0 | 0 | 0 | 0 | 0 | 0 | 0 | 0 | 0 | 0 | 0 | 0 | 1 | 0 | 0 | 0 | 0 | 0 | 0 | 0 | 0 | 0 | 0 | 0 | 0 | 0 | 0 | 0 | 0 | 0 | 0 | 0 | 0 |
| virus_Jul27.25v_25363 | 1 | 0 | 0 | 0 | 0 | 0 | 0 | 0 | 0 | 0 | 0 | 0 | 0 | 0 | 0 | 0 | 0 | 0 | 0 | 0 | 0 | 0 | 0 | 0 | 0 | 0 | 0 | 0 | 0 | 0 | 0 | 0 | 0 | 0 | 1 | 0 | 0 | 0 | 0 | 0 | 0 | 0 | 0 | 0 | 0 | 0 | 0 | 0 | 0 | 0 | 0 | 0 | 0 | 0 | 0 |
| virus_Jul27.25v_34099 | 1 | 0 | 0 | 0 | 0 | 0 | 0 | 0 | 0 | 0 | 0 | 0 | 0 | 0 | 0 | 0 | 0 | 0 | 0 | 0 | 0 | 0 | 0 | 0 | 0 | 0 | 0 | 0 | 0 | 0 | 0 | 0 | 0 | 0 | 1 | 0 | 0 | 0 | 0 | 0 | 0 | 0 | 0 | 0 | 0 | 0 | 0 | 0 | 0 | 0 | 0 | 0 | 0 | 0 | 0 |
| virus_Jul27.25v_34387 | 1 | 0 | 0 | 0 | 0 | 0 | 0 | 0 | 0 | 0 | 0 | 0 | 0 | 0 | 0 | 0 | 0 | 0 | 0 | 0 | 0 | 0 | 0 | 0 | 0 | 0 | 0 | 0 | 0 | 0 | 0 | 0 | 0 | 0 | 1 | 0 | 0 | 0 | 0 | 0 | 0 | 0 | 0 | 0 | 0 | 0 | 0 | 0 | 0 | 0 | 0 | 0 | 0 | 0 | 0 |
| virus_Jul27.25v_38302 | 1 | 0 | 0 | 0 | 0 | 0 | 0 | 0 | 0 | 0 | 0 | 0 | 0 | 0 | 0 | 0 | 0 | 0 | 0 | 0 | 0 | 0 | 0 | 0 | 0 | 0 | 0 | 0 | 0 | 0 | 0 | 0 | 0 | 0 | 1 | 0 | 0 | 0 | 0 | 0 | 0 | 0 | 0 | 0 | 0 | 0 | 0 | 0 | 0 | 0 | 0 | 0 | 0 | 0 | 0 |
| virus_Jul27.75v_1714 | 1 | 0 | 0 | 0 | 0 | 0 | 0 | 0 | 0 | 0 | 0 | 0 | 0 | 0 | 0 | 0 | 0 | 0 | 0 | 0 | 0 | 0 | 0 | 0 | 0 | 0 | 0 | 0 | 0 | 0 | 0 | 0 | 0 | 0 | 0 | 0 | 1 | 0 | 0 | 0 | 0 | 0 | 0 | 0 | 0 | 0 | 0 | 0 | 0 | 0 | 0 | 0 | 0 | 0 | 0 |
| virus_Jul27.75v_2711 | 1 | 0 | 0 | 0 | 0 | 0 | 0 | 0 | 0 | 0 | 0 | 0 | 0 | 0 | 0 | 0 | 0 | 0 | 0 | 0 | 0 | 0 | 0 | 0 | 0 | 0 | 0 | 0 | 0 | 0 | 0 | 0 | 0 | 0 | 0 | 0 | 1 | 0 | 0 | 0 | 0 | 0 | 0 | 0 | 0 | 0 | 0 | 0 | 0 | 0 | 0 | 0 | 0 | 0 | 0 |
| virus_Jul27.75v_10732 | 1 | 0 | 0 | 0 | 0 | 0 | 0 | 0 | 0 | 0 | 0 | 0 | 0 | 0 | 0 | 0 | 0 | 0 | 0 | 0 | 0 | 0 | 0 | 0 | 0 | 0 | 0 | 0 | 0 | 0 | 0 | 0 | 0 | 0 | 0 | 0 | 1 | 0 | 0 | 0 | 0 | 0 | 0 | 0 | 0 | 0 | 0 | 0 | 0 | 0 | 0 | 0 | 0 | 0 | 0 |
| virus_Jun8.25v_9630 | 1 | 0 | 0 | 0 | 0 | 0 | 0 | 0 | 0 | 0 | 1 | 0 | 0 | 0 | 0 | 0 | 0 | 0 | 0 | 0 | 0 | 0 | 0 | 0 | 0 | 0 | 0 | 0 | 0 | 0 | 0 | 0 | 0 | 0 | 0 | 0 | 0 | 0 | 0 | 0 | 0 | 0 | 0 | 0 | 0 | 0 | 0 | 0 | 0 | 0 | 0 | 0 | 0 | 0 | 0 |
| virus_Jun8.25v_16511 | 1 | 0 | 0 | 0 | 0 | 0 | 0 | 0 | 0 | 0 | 1 | 0 | 0 | 0 | 0 | 0 | 0 | 0 | 0 | 0 | 0 | 0 | 0 | 0 | 0 | 0 | 0 | 0 | 0 | 0 | 0 | 0 | 0 | 0 | 0 | 0 | 0 | 0 | 0 | 0 | 0 | 0 | 0 | 0 | 0 | 0 | 0 | 0 | 0 | 0 | 0 | 0 | 0 | 0 | 0 |
| virus_Jun8.25v_20975 | 1 | 0 | 0 | 0 | 0 | 0 | 0 | 0 | 0 | 0 | 1 | 0 | 0 | 0 | 0 | 0 | 0 | 0 | 0 | 0 | 0 | 0 | 0 | 0 | 0 | 0 | 0 | 0 | 0 | 0 | 0 | 0 | 0 | 0 | 0 | 0 | 0 | 0 | 0 | 0 | 0 | 0 | 0 | 0 | 0 | 0 | 0 | 0 | 0 | 0 | 0 | 0 | 0 | 0 | 0 |
| virus_Jun8.50v_7739 | 1 | 0 | 0 | 0 | 0 | 0 | 0 | 0 | 0 | 0 | 0 | 0 | 1 | 0 | 0 | 0 | 0 | 0 | 0 | 0 | 0 | 0 | 0 | 0 | 0 | 0 | 0 | 0 | 0 | 0 | 0 | 0 | 0 | 0 | 0 | 0 | 0 | 0 | 0 | 0 | 0 | 0 | 0 | 0 | 0 | 0 | 0 | 0 | 0 | 0 | 0 | 0 | 0 | 0 | 0 |
| virus_Jun8.50v_16449 | 1 | 0 | 0 | 0 | 0 | 0 | 0 | 0 | 0 | 0 | 0 | 0 | 1 | 0 | 0 | 0 | 0 | 0 | 0 | 0 | 0 | 0 | 0 | 0 | 0 | 0 | 0 | 0 | 0 | 0 | 0 | 0 | 0 | 0 | 0 | 0 | 0 | 0 | 0 | 0 | 0 | 0 | 0 | 0 | 0 | 0 | 0 | 0 | 0 | 0 | 0 | 0 | 0 | 0 | 0 |
| virus_Jun8.50v_19047 | 1 | 0 | 0 | 0 | 0 | 0 | 0 | 0 | 0 | 0 | 0 | 0 | 1 | 0 | 0 | 0 | 0 | 0 | 0 | 0 | 0 | 0 | 0 | 0 | 0 | 0 | 0 | 0 | 0 | 0 | 0 | 0 | 0 | 0 | 0 | 0 | 0 | 0 | 0 | 0 | 0 | 0 | 0 | 0 | 0 | 0 | 0 | 0 | 0 | 0 | 0 | 0 | 0 | 0 | 0 |
| virus_Jun8.50v_20410 | 1 | 0 | 0 | 0 | 0 | 0 | 0 | 0 | 0 | 0 | 0 | 0 | 1 | 0 | 0 | 0 | 0 | 0 | 0 | 0 | 0 | 0 | 0 | 0 | 0 | 0 | 0 | 0 | 0 | 0 | 0 | 0 | 0 | 0 | 0 | 0 | 0 | 0 | 0 | 0 | 0 | 0 | 0 | 0 | 0 | 0 | 0 | 0 | 0 | 0 | 0 | 0 | 0 | 0 | 0 |
| virus_Jun8.50v_30737 | 1 | 0 | 0 | 0 | 0 | 0 | 0 | 0 | 0 | 0 | 0 | 0 | 1 | 0 | 0 | 0 | 0 | 0 | 0 | 0 | 0 | 0 | 0 | 0 | 0 | 0 | 0 | 0 | 0 | 0 | 0 | 0 | 0 | 0 | 0 | 0 | 0 | 0 | 0 | 0 | 0 | 0 | 0 | 0 | 0 | 0 | 0 | 0 | 0 | 0 | 0 | 0 | 0 | 0 | 0 |
| virus_Jun8.50v_32582 | 1 | 0 | 0 | 0 | 0 | 0 | 0 | 0 | 0 | 0 | 0 | 0 | 1 | 0 | 0 | 0 | 0 | 0 | 0 | 0 | 0 | 0 | 0 | 0 | 0 | 0 | 0 | 0 | 0 | 0 | 0 | 0 | 0 | 0 | 0 | 0 | 0 | 0 | 0 | 0 | 0 | 0 | 0 | 0 | 0 | 0 | 0 | 0 | 0 | 0 | 0 | 0 | 0 | 0 | 0 |
| virus_Jun8.50v_40686 | 1 | 0 | 0 | 0 | 0 | 0 | 0 | 0 | 0 | 0 | 0 | 0 | 1 | 0 | 0 | 0 | 0 | 0 | 0 | 0 | 0 | 0 | 0 | 0 | 0 | 0 | 0 | 0 | 0 | 0 | 0 | 0 | 0 | 0 | 0 | 0 | 0 | 0 | 0 | 0 | 0 | 0 | 0 | 0 | 0 | 0 | 0 | 0 | 0 | 0 | 0 | 0 | 0 | 0 | 0 |
| virus_Jun8.75v_561 | 1 | 0 | 0 | 0 | 0 | 0 | 0 | 0 | 0 | 0 | 0 | 0 | 0 | 0 | 1 | 0 | 0 | 0 | 0 | 0 | 0 | 0 | 0 | 0 | 0 | 0 | 0 | 0 | 0 | 0 | 0 | 0 | 0 | 0 | 0 | 0 | 0 | 0 | 0 | 0 | 0 | 0 | 0 | 0 | 0 | 0 | 0 | 0 | 0 | 0 | 0 | 0 | 0 | 0 | 0 |
| virus_Jun8.75v_2728 | 1 | 0 | 0 | 0 | 0 | 0 | 0 | 0 | 0 | 0 | 0 | 0 | 0 | 0 | 1 | 0 | 0 | 0 | 0 | 0 | 0 | 0 | 0 | 0 | 0 | 0 | 0 | 0 | 0 | 0 | 0 | 0 | 0 | 0 | 0 | 0 | 0 | 0 | 0 | 0 | 0 | 0 | 0 | 0 | 0 | 0 | 0 | 0 | 0 | 0 | 0 | 0 | 0 | 0 | 0 |
| virus_Jun8.75v_8412 | 1 | 0 | 0 | 0 | 0 | 0 | 0 | 0 | 0 | 0 | 0 | 0 | 0 | 0 | 1 | 0 | 0 | 0 | 0 | 0 | 0 | 0 | 0 | 0 | 0 | 0 | 0 | 0 | 0 | 0 | 0 | 0 | 0 | 0 | 0 | 0 | 0 | 0 | 0 | 0 | 0 | 0 | 0 | 0 | 0 | 0 | 0 | 0 | 0 | 0 | 0 | 0 | 0 | 0 | 0 |
| virus_Jun8.75v_15985 | 1 | 0 | 0 | 0 | 0 | 0 | 0 | 0 | 0 | 0 | 0 | 0 | 0 | 0 | 1 | 0 | 0 | 0 | 0 | 0 | 0 | 0 | 0 | 0 | 0 | 0 | 0 | 0 | 0 | 0 | 0 | 0 | 0 | 0 | 0 | 0 | 0 | 0 | 0 | 0 | 0 | 0 | 0 | 0 | 0 | 0 | 0 | 0 | 0 | 0 | 0 | 0 | 0 | 0 | 0 |
| virus_Jun8.75v_18738 | 1 | 0 | 0 | 0 | 0 | 0 | 0 | 0 | 0 | 0 | 0 | 0 | 0 | 0 | 1 | 0 | 0 | 0 | 0 | 0 | 0 | 0 | 0 | 0 | 0 | 0 | 0 | 0 | 0 | 0 | 0 | 0 | 0 | 0 | 0 | 0 | 0 | 0 | 0 | 0 | 0 | 0 | 0 | 0 | 0 | 0 | 0 | 0 | 0 | 0 | 0 | 0 | 0 | 0 | 0 |
| virus_Jun8.75v_19828 | 1 | 0 | 0 | 0 | 0 | 0 | 0 | 0 | 0 | 0 | 0 | 0 | 0 | 0 | 1 | 0 | 0 | 0 | 0 | 0 | 0 | 0 | 0 | 0 | 0 | 0 | 0 | 0 | 0 | 0 | 0 | 0 | 0 | 0 | 0 | 0 | 0 | 0 | 0 | 0 | 0 | 0 | 0 | 0 | 0 | 0 | 0 | 0 | 0 | 0 | 0 | 0 | 0 | 0 | 0 |
| virus_Jun22.0v_1661 | 1 | 0 | 0 | 0 | 0 | 0 | 0 | 0 | 0 | 0 | 0 | 0 | 0 | 0 | 0 | 0 | 1 | 0 | 0 | 0 | 0 | 0 | 0 | 0 | 0 | 0 | 0 | 0 | 0 | 0 | 0 | 0 | 0 | 0 | 0 | 0 | 0 | 0 | 0 | 0 | 0 | 0 | 0 | 0 | 0 | 0 | 0 | 0 | 0 | 0 | 0 | 0 | 0 | 0 | 0 |
| virus_Jun22.0v_2351 | 1 | 0 | 0 | 0 | 0 | 0 | 0 | 0 | 0 | 0 | 0 | 0 | 0 | 0 | 0 | 0 | 1 | 0 | 0 | 0 | 0 | 0 | 0 | 0 | 0 | 0 | 0 | 0 | 0 | 0 | 0 | 0 | 0 | 0 | 0 | 0 | 0 | 0 | 0 | 0 | 0 | 0 | 0 | 0 | 0 | 0 | 0 | 0 | 0 | 0 | 0 | 0 | 0 | 0 | 0 |
| virus_Jun22.0v_2549 | 1 | 0 | 0 | 0 | 0 | 0 | 0 | 0 | 0 | 0 | 0 | 0 | 0 | 0 | 0 | 0 | 1 | 0 | 0 | 0 | 0 | 0 | 0 | 0 | 0 | 0 | 0 | 0 | 0 | 0 | 0 | 0 | 0 | 0 | 0 | 0 | 0 | 0 | 0 | 0 | 0 | 0 | 0 | 0 | 0 | 0 | 0 | 0 | 0 | 0 | 0 | 0 | 0 | 0 | 0 |
| virus_Jun22.0v_8194 | 1 | 0 | 0 | 0 | 0 | 0 | 0 | 0 | 0 | 0 | 0 | 0 | 0 | 0 | 0 | 0 | 1 | 0 | 0 | 0 | 0 | 0 | 0 | 0 | 0 | 0 | 0 | 0 | 0 | 0 | 0 | 0 | 0 | 0 | 0 | 0 | 0 | 0 | 0 | 0 | 0 | 0 | 0 | 0 | 0 | 0 | 0 | 0 | 0 | 0 | 0 | 0 | 0 | 0 | 0 |
| virus_Jun22.0v_12992 | 1 | 0 | 0 | 0 | 0 | 0 | 0 | 0 | 0 | 0 | 0 | 0 | 0 | 0 | 0 | 0 | 1 | 0 | 0 | 0 | 0 | 0 | 0 | 0 | 0 | 0 | 0 | 0 | 0 | 0 | 0 | 0 | 0 | 0 | 0 | 0 | 0 | 0 | 0 | 0 | 0 | 0 | 0 | 0 | 0 | 0 | 0 | 0 | 0 | 0 | 0 | 0 | 0 | 0 | 0 |
| virus_Jun22.0v_17047 | 1 | 0 | 0 | 0 | 0 | 0 | 0 | 0 | 0 | 0 | 0 | 0 | 0 | 0 | 0 | 0 | 1 | 0 | 0 | 0 | 0 | 0 | 0 | 0 | 0 | 0 | 0 | 0 | 0 | 0 | 0 | 0 | 0 | 0 | 0 | 0 | 0 | 0 | 0 | 0 | 0 | 0 | 0 | 0 | 0 | 0 | 0 | 0 | 0 | 0 | 0 | 0 | 0 | 0 | 0 |
| virus_Jun22.0v_18187 | 1 | 0 | 0 | 0 | 0 | 0 | 0 | 0 | 0 | 0 | 0 | 0 | 0 | 0 | 0 | 0 | 1 | 0 | 0 | 0 | 0 | 0 | 0 | 0 | 0 | 0 | 0 | 0 | 0 | 0 | 0 | 0 | 0 | 0 | 0 | 0 | 0 | 0 | 0 | 0 | 0 | 0 | 0 | 0 | 0 | 0 | 0 | 0 | 0 | 0 | 0 | 0 | 0 | 0 | 0 |
| virus_Jun22.0v_18741 | 1 | 0 | 0 | 0 | 0 | 0 | 0 | 0 | 0 | 0 | 0 | 0 | 0 | 0 | 0 | 0 | 1 | 0 | 0 | 0 | 0 | 0 | 0 | 0 | 0 | 0 | 0 | 0 | 0 | 0 | 0 | 0 | 0 | 0 | 0 | 0 | 0 | 0 | 0 | 0 | 0 | 0 | 0 | 0 | 0 | 0 | 0 | 0 | 0 | 0 | 0 | 0 | 0 | 0 | 0 |
| virus_Jun22.0v_20897 | 1 | 0 | 0 | 0 | 0 | 0 | 0 | 0 | 0 | 0 | 0 | 0 | 0 | 0 | 0 | 0 | 1 | 0 | 0 | 0 | 0 | 0 | 0 | 0 | 0 | 0 | 0 | 0 | 0 | 0 | 0 | 0 | 0 | 0 | 0 | 0 | 0 | 0 | 0 | 0 | 0 | 0 | 0 | 0 | 0 | 0 | 0 | 0 | 0 | 0 | 0 | 0 | 0 | 0 | 0 |
| virus_Jun22.50v_6516 | 1 | 0 | 0 | 0 | 0 | 0 | 0 | 0 | 0 | 0 | 0 | 0 | 0 | 0 | 0 | 0 | 0 | 0 | 1 | 0 | 0 | 0 | 0 | 0 | 0 | 0 | 0 | 0 | 0 | 0 | 0 | 0 | 0 | 0 | 0 | 0 | 0 | 0 | 0 | 0 | 0 | 0 | 0 | 0 | 0 | 0 | 0 | 0 | 0 | 0 | 0 | 0 | 0 | 0 | 0 |
| virus_Jun22.50v_8936 | 1 | 0 | 0 | 0 | 0 | 0 | 0 | 0 | 0 | 0 | 0 | 0 | 0 | 0 | 0 | 0 | 0 | 0 | 1 | 0 | 0 | 0 | 0 | 0 | 0 | 0 | 0 | 0 | 0 | 0 | 0 | 0 | 0 | 0 | 0 | 0 | 0 | 0 | 0 | 0 | 0 | 0 | 0 | 0 | 0 | 0 | 0 | 0 | 0 | 0 | 0 | 0 | 0 | 0 | 0 |
| virus_Jun22.50v_9120 | 1 | 0 | 0 | 0 | 0 | 0 | 0 | 0 | 0 | 0 | 0 | 0 | 0 | 0 | 0 | 0 | 0 | 0 | 1 | 0 | 0 | 0 | 0 | 0 | 0 | 0 | 0 | 0 | 0 | 0 | 0 | 0 | 0 | 0 | 0 | 0 | 0 | 0 | 0 | 0 | 0 | 0 | 0 | 0 | 0 | 0 | 0 | 0 | 0 | 0 | 0 | 0 | 0 | 0 | 0 |
| virus_Jun22.50v_9291 | 1 | 0 | 0 | 0 | 0 | 0 | 0 | 0 | 0 | 0 | 0 | 0 | 0 | 0 | 0 | 0 | 0 | 0 | 1 | 0 | 0 | 0 | 0 | 0 | 0 | 0 | 0 | 0 | 0 | 0 | 0 | 0 | 0 | 0 | 0 | 0 | 0 | 0 | 0 | 0 | 0 | 0 | 0 | 0 | 0 | 0 | 0 | 0 | 0 | 0 | 0 | 0 | 0 | 0 | 0 |
| virus_Jun22.50v_14807 | 1 | 0 | 0 | 0 | 0 | 0 | 0 | 0 | 0 | 0 | 0 | 0 | 0 | 0 | 0 | 0 | 0 | 0 | 1 | 0 | 0 | 0 | 0 | 0 | 0 | 0 | 0 | 0 | 0 | 0 | 0 | 0 | 0 | 0 | 0 | 0 | 0 | 0 | 0 | 0 | 0 | 0 | 0 | 0 | 0 | 0 | 0 | 0 | 0 | 0 | 0 | 0 | 0 | 0 | 0 |
| virus_Jun22.50v_17696 | 1 | 0 | 0 | 0 | 0 | 0 | 0 | 0 | 0 | 0 | 0 | 0 | 0 | 0 | 0 | 0 | 0 | 0 | 1 | 0 | 0 | 0 | 0 | 0 | 0 | 0 | 0 | 0 | 0 | 0 | 0 | 0 | 0 | 0 | 0 | 0 | 0 | 0 | 0 | 0 | 0 | 0 | 0 | 0 | 0 | 0 | 0 | 0 | 0 | 0 | 0 | 0 | 0 | 0 | 0 |
| virus_Jun22.50v_19008 | 1 | 0 | 0 | 0 | 0 | 0 | 0 | 0 | 0 | 0 | 0 | 0 | 0 | 0 | 0 | 0 | 0 | 0 | 1 | 0 | 0 | 0 | 0 | 0 | 0 | 0 | 0 | 0 | 0 | 0 | 0 | 0 | 0 | 0 | 0 | 0 | 0 | 0 | 0 | 0 | 0 | 0 | 0 | 0 | 0 | 0 | 0 | 0 | 0 | 0 | 0 | 0 | 0 | 0 | 0 |
| virus_Jun22.50v_20514 | 1 | 0 | 0 | 0 | 0 | 0 | 0 | 0 | 0 | 0 | 0 | 0 | 0 | 0 | 0 | 0 | 0 | 0 | 1 | 0 | 0 | 0 | 0 | 0 | 0 | 0 | 0 | 0 | 0 | 0 | 0 | 0 | 0 | 0 | 0 | 0 | 0 | 0 | 0 | 0 | 0 | 0 | 0 | 0 | 0 | 0 | 0 | 0 | 0 | 0 | 0 | 0 | 0 | 0 | 0 |
| virus_Jun22.50v_24843 | 1 | 0 | 0 | 0 | 0 | 0 | 0 | 0 | 0 | 0 | 0 | 0 | 0 | 0 | 0 | 0 | 0 | 0 | 1 | 0 | 0 | 0 | 0 | 0 | 0 | 0 | 0 | 0 | 0 | 0 | 0 | 0 | 0 | 0 | 0 | 0 | 0 | 0 | 0 | 0 | 0 | 0 | 0 | 0 | 0 | 0 | 0 | 0 | 0 | 0 | 0 | 0 | 0 | 0 | 0 |
| virus_Jun22.50v_26935 | 1 | 0 | 0 | 0 | 0 | 0 | 0 | 0 | 0 | 0 | 0 | 0 | 0 | 0 | 0 | 0 | 0 | 0 | 1 | 0 | 0 | 0 | 0 | 0 | 0 | 0 | 0 | 0 | 0 | 0 | 0 | 0 | 0 | 0 | 0 | 0 | 0 | 0 | 0 | 0 | 0 | 0 | 0 | 0 | 0 | 0 | 0 | 0 | 0 | 0 | 0 | 0 | 0 | 0 | 0 |
| virus_Jun22.50v_30078 | 1 | 0 | 0 | 0 | 0 | 0 | 0 | 0 | 0 | 0 | 0 | 0 | 0 | 0 | 0 | 0 | 0 | 0 | 1 | 0 | 0 | 0 | 0 | 0 | 0 | 0 | 0 | 0 | 0 | 0 | 0 | 0 | 0 | 0 | 0 | 0 | 0 | 0 | 0 | 0 | 0 | 0 | 0 | 0 | 0 | 0 | 0 | 0 | 0 | 0 | 0 | 0 | 0 | 0 | 0 |
| virus_Jun22.50v_31467 | 1 | 0 | 0 | 0 | 0 | 0 | 0 | 0 | 0 | 0 | 0 | 0 | 0 | 0 | 0 | 0 | 0 | 0 | 1 | 0 | 0 | 0 | 0 | 0 | 0 | 0 | 0 | 0 | 0 | 0 | 0 | 0 | 0 | 0 | 0 | 0 | 0 | 0 | 0 | 0 | 0 | 0 | 0 | 0 | 0 | 0 | 0 | 0 | 0 | 0 | 0 | 0 | 0 | 0 | 0 |
| virus_Jun22.50v_33089 | 1 | 0 | 0 | 0 | 0 | 0 | 0 | 0 | 0 | 0 | 0 | 0 | 0 | 0 | 0 | 0 | 0 | 0 | 1 | 0 | 0 | 0 | 0 | 0 | 0 | 0 | 0 | 0 | 0 | 0 | 0 | 0 | 0 | 0 | 0 | 0 | 0 | 0 | 0 | 0 | 0 | 0 | 0 | 0 | 0 | 0 | 0 | 0 | 0 | 0 | 0 | 0 | 0 | 0 | 0 |
| virus_Jun22.50v_33799 | 1 | 0 | 0 | 0 | 0 | 0 | 0 | 0 | 0 | 0 | 0 | 0 | 0 | 0 | 0 | 0 | 0 | 0 | 1 | 0 | 0 | 0 | 0 | 0 | 0 | 0 | 0 | 0 | 0 | 0 | 0 | 0 | 0 | 0 | 0 | 0 | 0 | 0 | 0 | 0 | 0 | 0 | 0 | 0 | 0 | 0 | 0 | 0 | 0 | 0 | 0 | 0 | 0 | 0 | 0 |
| virus_Jun22.50v_39505 | 1 | 0 | 0 | 0 | 0 | 0 | 0 | 0 | 0 | 0 | 0 | 0 | 0 | 0 | 0 | 0 | 0 | 0 | 1 | 0 | 0 | 0 | 0 | 0 | 0 | 0 | 0 | 0 | 0 | 0 | 0 | 0 | 0 | 0 | 0 | 0 | 0 | 0 | 0 | 0 | 0 | 0 | 0 | 0 | 0 | 0 | 0 | 0 | 0 | 0 | 0 | 0 | 0 | 0 | 0 |
| virus_Jun22.50v_40677 | 1 | 0 | 0 | 0 | 0 | 0 | 0 | 0 | 0 | 0 | 0 | 0 | 0 | 0 | 0 | 0 | 0 | 0 | 1 | 0 | 0 | 0 | 0 | 0 | 0 | 0 | 0 | 0 | 0 | 0 | 0 | 0 | 0 | 0 | 0 | 0 | 0 | 0 | 0 | 0 | 0 | 0 | 0 | 0 | 0 | 0 | 0 | 0 | 0 | 0 | 0 | 0 | 0 | 0 | 0 |
| virus_Jun22.75v_12338 | 1 | 0 | 0 | 0 | 0 | 0 | 0 | 0 | 0 | 0 | 0 | 0 | 0 | 0 | 0 | 0 | 0 | 0 | 0 | 0 | 1 | 0 | 0 | 0 | 0 | 0 | 0 | 0 | 0 | 0 | 0 | 0 | 0 | 0 | 0 | 0 | 0 | 0 | 0 | 0 | 0 | 0 | 0 | 0 | 0 | 0 | 0 | 0 | 0 | 0 | 0 | 0 | 0 | 0 | 0 |
| virus_Jun22.75v_12846 | 1 | 0 | 0 | 0 | 0 | 0 | 0 | 0 | 0 | 0 | 0 | 0 | 0 | 0 | 0 | 0 | 0 | 0 | 0 | 0 | 1 | 0 | 0 | 0 | 0 | 0 | 0 | 0 | 0 | 0 | 0 | 0 | 0 | 0 | 0 | 0 | 0 | 0 | 0 | 0 | 0 | 0 | 0 | 0 | 0 | 0 | 0 | 0 | 0 | 0 | 0 | 0 | 0 | 0 | 0 |
| virus_Jun22.75v_18459 | 1 | 0 | 0 | 0 | 0 | 0 | 0 | 0 | 0 | 0 | 0 | 0 | 0 | 0 | 0 | 0 | 0 | 0 | 0 | 0 | 1 | 0 | 0 | 0 | 0 | 0 | 0 | 0 | 0 | 0 | 0 | 0 | 0 | 0 | 0 | 0 | 0 | 0 | 0 | 0 | 0 | 0 | 0 | 0 | 0 | 0 | 0 | 0 | 0 | 0 | 0 | 0 | 0 | 0 | 0 |
| virus_Jun22.75v_29454 | 1 | 0 | 0 | 0 | 0 | 0 | 0 | 0 | 0 | 0 | 0 | 0 | 0 | 0 | 0 | 0 | 0 | 0 | 0 | 0 | 1 | 0 | 0 | 0 | 0 | 0 | 0 | 0 | 0 | 0 | 0 | 0 | 0 | 0 | 0 | 0 | 0 | 0 | 0 | 0 | 0 | 0 | 0 | 0 | 0 | 0 | 0 | 0 | 0 | 0 | 0 | 0 | 0 | 0 | 0 |
| virus_Jun22.75v_36613 | 1 | 0 | 0 | 0 | 0 | 0 | 0 | 0 | 0 | 0 | 0 | 0 | 0 | 0 | 0 | 0 | 0 | 0 | 0 | 0 | 1 | 0 | 0 | 0 | 0 | 0 | 0 | 0 | 0 | 0 | 0 | 0 | 0 | 0 | 0 | 0 | 0 | 0 | 0 | 0 | 0 | 0 | 0 | 0 | 0 | 0 | 0 | 0 | 0 | 0 | 0 | 0 | 0 | 0 | 0 |
| virus_Jun22.75v_40825 | 1 | 0 | 0 | 0 | 0 | 0 | 0 | 0 | 0 | 0 | 0 | 0 | 0 | 0 | 0 | 0 | 0 | 0 | 0 | 0 | 1 | 0 | 0 | 0 | 0 | 0 | 0 | 0 | 0 | 0 | 0 | 0 | 0 | 0 | 0 | 0 | 0 | 0 | 0 | 0 | 0 | 0 | 0 | 0 | 0 | 0 | 0 | 0 | 0 | 0 | 0 | 0 | 0 | 0 | 0 |
| virus_May12.5v_2938 | 1 | 0 | 1 | 0 | 0 | 0 | 0 | 0 | 0 | 0 | 0 | 0 | 0 | 0 | 0 | 0 | 0 | 0 | 0 | 0 | 0 | 0 | 0 | 0 | 0 | 0 | 0 | 0 | 0 | 0 | 0 | 0 | 0 | 0 | 0 | 0 | 0 | 0 | 0 | 0 | 0 | 0 | 0 | 0 | 0 | 0 | 0 | 0 | 0 | 0 | 0 | 0 | 0 | 0 | 0 |
| virus_May12.5v_13330 | 1 | 0 | 1 | 0 | 0 | 0 | 0 | 0 | 0 | 0 | 0 | 0 | 0 | 0 | 0 | 0 | 0 | 0 | 0 | 0 | 0 | 0 | 0 | 0 | 0 | 0 | 0 | 0 | 0 | 0 | 0 | 0 | 0 | 0 | 0 | 0 | 0 | 0 | 0 | 0 | 0 | 0 | 0 | 0 | 0 | 0 | 0 | 0 | 0 | 0 | 0 | 0 | 0 | 0 | 0 |
| virus_May12.5v_23168 | 1 | 0 | 1 | 0 | 0 | 0 | 0 | 0 | 0 | 0 | 0 | 0 | 0 | 0 | 0 | 0 | 0 | 0 | 0 | 0 | 0 | 0 | 0 | 0 | 0 | 0 | 0 | 0 | 0 | 0 | 0 | 0 | 0 | 0 | 0 | 0 | 0 | 0 | 0 | 0 | 0 | 0 | 0 | 0 | 0 | 0 | 0 | 0 | 0 | 0 | 0 | 0 | 0 | 0 | 0 |
| virus_May12.5v_27624 | 1 | 0 | 1 | 0 | 0 | 0 | 0 | 0 | 0 | 0 | 0 | 0 | 0 | 0 | 0 | 0 | 0 | 0 | 0 | 0 | 0 | 0 | 0 | 0 | 0 | 0 | 0 | 0 | 0 | 0 | 0 | 0 | 0 | 0 | 0 | 0 | 0 | 0 | 0 | 0 | 0 | 0 | 0 | 0 | 0 | 0 | 0 | 0 | 0 | 0 | 0 | 0 | 0 | 0 | 0 |
| virus_May12.5v_29202 | 1 | 0 | 1 | 0 | 0 | 0 | 0 | 0 | 0 | 0 | 0 | 0 | 0 | 0 | 0 | 0 | 0 | 0 | 0 | 0 | 0 | 0 | 0 | 0 | 0 | 0 | 0 | 0 | 0 | 0 | 0 | 0 | 0 | 0 | 0 | 0 | 0 | 0 | 0 | 0 | 0 | 0 | 0 | 0 | 0 | 0 | 0 | 0 | 0 | 0 | 0 | 0 | 0 | 0 | 0 |
| virus_May12.5v_33029 | 1 | 0 | 1 | 0 | 0 | 0 | 0 | 0 | 0 | 0 | 0 | 0 | 0 | 0 | 0 | 0 | 0 | 0 | 0 | 0 | 0 | 0 | 0 | 0 | 0 | 0 | 0 | 0 | 0 | 0 | 0 | 0 | 0 | 0 | 0 | 0 | 0 | 0 | 0 | 0 | 0 | 0 | 0 | 0 | 0 | 0 | 0 | 0 | 0 | 0 | 0 | 0 | 0 | 0 | 0 |
| virus_May12.5v_33832 | 1 | 0 | 1 | 0 | 0 | 0 | 0 | 0 | 0 | 0 | 0 | 0 | 0 | 0 | 0 | 0 | 0 | 0 | 0 | 0 | 0 | 0 | 0 | 0 | 0 | 0 | 0 | 0 | 0 | 0 | 0 | 0 | 0 | 0 | 0 | 0 | 0 | 0 | 0 | 0 | 0 | 0 | 0 | 0 | 0 | 0 | 0 | 0 | 0 | 0 | 0 | 0 | 0 | 0 | 0 |
| virus_May12.5v_37184 | 1 | 0 | 1 | 0 | 0 | 0 | 0 | 0 | 0 | 0 | 0 | 0 | 0 | 0 | 0 | 0 | 0 | 0 | 0 | 0 | 0 | 0 | 0 | 0 | 0 | 0 | 0 | 0 | 0 | 0 | 0 | 0 | 0 | 0 | 0 | 0 | 0 | 0 | 0 | 0 | 0 | 0 | 0 | 0 | 0 | 0 | 0 | 0 | 0 | 0 | 0 | 0 | 0 | 0 | 0 |
| virus_May12.5v_41032 | 1 | 0 | 1 | 0 | 0 | 0 | 0 | 0 | 0 | 0 | 0 | 0 | 0 | 0 | 0 | 0 | 0 | 0 | 0 | 0 | 0 | 0 | 0 | 0 | 0 | 0 | 0 | 0 | 0 | 0 | 0 | 0 | 0 | 0 | 0 | 0 | 0 | 0 | 0 | 0 | 0 | 0 | 0 | 0 | 0 | 0 | 0 | 0 | 0 | 0 | 0 | 0 | 0 | 0 | 0 |
| virus_May12.25v_288 | 1 | 0 | 0 | 0 | 1 | 0 | 0 | 0 | 0 | 0 | 0 | 0 | 0 | 0 | 0 | 0 | 0 | 0 | 0 | 0 | 0 | 0 | 0 | 0 | 0 | 0 | 0 | 0 | 0 | 0 | 0 | 0 | 0 | 0 | 0 | 0 | 0 | 0 | 0 | 0 | 0 | 0 | 0 | 0 | 0 | 0 | 0 | 0 | 0 | 0 | 0 | 0 | 0 | 0 | 0 |
| virus_May12.25v_3047 | 1 | 0 | 0 | 0 | 1 | 0 | 0 | 0 | 0 | 0 | 0 | 0 | 0 | 0 | 0 | 0 | 0 | 0 | 0 | 0 | 0 | 0 | 0 | 0 | 0 | 0 | 0 | 0 | 0 | 0 | 0 | 0 | 0 | 0 | 0 | 0 | 0 | 0 | 0 | 0 | 0 | 0 | 0 | 0 | 0 | 0 | 0 | 0 | 0 | 0 | 0 | 0 | 0 | 0 | 0 |
| virus_May12.25v_8994 | 1 | 0 | 0 | 0 | 1 | 0 | 0 | 0 | 0 | 0 | 0 | 0 | 0 | 0 | 0 | 0 | 0 | 0 | 0 | 0 | 0 | 0 | 0 | 0 | 0 | 0 | 0 | 0 | 0 | 0 | 0 | 0 | 0 | 0 | 0 | 0 | 0 | 0 | 0 | 0 | 0 | 0 | 0 | 0 | 0 | 0 | 0 | 0 | 0 | 0 | 0 | 0 | 0 | 0 | 0 |
| virus_May12.25v_13023 | 1 | 0 | 0 | 0 | 1 | 0 | 0 | 0 | 0 | 0 | 0 | 0 | 0 | 0 | 0 | 0 | 0 | 0 | 0 | 0 | 0 | 0 | 0 | 0 | 0 | 0 | 0 | 0 | 0 | 0 | 0 | 0 | 0 | 0 | 0 | 0 | 0 | 0 | 0 | 0 | 0 | 0 | 0 | 0 | 0 | 0 | 0 | 0 | 0 | 0 | 0 | 0 | 0 | 0 | 0 |
| virus_May12.25v_16726 | 1 | 0 | 0 | 0 | 1 | 0 | 0 | 0 | 0 | 0 | 0 | 0 | 0 | 0 | 0 | 0 | 0 | 0 | 0 | 0 | 0 | 0 | 0 | 0 | 0 | 0 | 0 | 0 | 0 | 0 | 0 | 0 | 0 | 0 | 0 | 0 | 0 | 0 | 0 | 0 | 0 | 0 | 0 | 0 | 0 | 0 | 0 | 0 | 0 | 0 | 0 | 0 | 0 | 0 | 0 |
| virus_May12.25v_26714 | 1 | 0 | 0 | 0 | 1 | 0 | 0 | 0 | 0 | 0 | 0 | 0 | 0 | 0 | 0 | 0 | 0 | 0 | 0 | 0 | 0 | 0 | 0 | 0 | 0 | 0 | 0 | 0 | 0 | 0 | 0 | 0 | 0 | 0 | 0 | 0 | 0 | 0 | 0 | 0 | 0 | 0 | 0 | 0 | 0 | 0 | 0 | 0 | 0 | 0 | 0 | 0 | 0 | 0 | 0 |
| virus_May12.25v_28165 | 1 | 0 | 0 | 0 | 1 | 0 | 0 | 0 | 0 | 0 | 0 | 0 | 0 | 0 | 0 | 0 | 0 | 0 | 0 | 0 | 0 | 0 | 0 | 0 | 0 | 0 | 0 | 0 | 0 | 0 | 0 | 0 | 0 | 0 | 0 | 0 | 0 | 0 | 0 | 0 | 0 | 0 | 0 | 0 | 0 | 0 | 0 | 0 | 0 | 0 | 0 | 0 | 0 | 0 | 0 |
| virus_May12.25v_35885 | 1 | 0 | 0 | 0 | 1 | 0 | 0 | 0 | 0 | 0 | 0 | 0 | 0 | 0 | 0 | 0 | 0 | 0 | 0 | 0 | 0 | 0 | 0 | 0 | 0 | 0 | 0 | 0 | 0 | 0 | 0 | 0 | 0 | 0 | 0 | 0 | 0 | 0 | 0 | 0 | 0 | 0 | 0 | 0 | 0 | 0 | 0 | 0 | 0 | 0 | 0 | 0 | 0 | 0 | 0 |
| virus_May12.50v_7190 | 1 | 0 | 0 | 0 | 0 | 0 | 1 | 0 | 0 | 0 | 0 | 0 | 0 | 0 | 0 | 0 | 0 | 0 | 0 | 0 | 0 | 0 | 0 | 0 | 0 | 0 | 0 | 0 | 0 | 0 | 0 | 0 | 0 | 0 | 0 | 0 | 0 | 0 | 0 | 0 | 0 | 0 | 0 | 0 | 0 | 0 | 0 | 0 | 0 | 0 | 0 | 0 | 0 | 0 | 0 |
| virus_May12.50v_9400 | 1 | 0 | 0 | 0 | 0 | 0 | 1 | 0 | 0 | 0 | 0 | 0 | 0 | 0 | 0 | 0 | 0 | 0 | 0 | 0 | 0 | 0 | 0 | 0 | 0 | 0 | 0 | 0 | 0 | 0 | 0 | 0 | 0 | 0 | 0 | 0 | 0 | 0 | 0 | 0 | 0 | 0 | 0 | 0 | 0 | 0 | 0 | 0 | 0 | 0 | 0 | 0 | 0 | 0 | 0 |
| virus_May12.50v_9880 | 1 | 0 | 0 | 0 | 0 | 0 | 1 | 0 | 0 | 0 | 0 | 0 | 0 | 0 | 0 | 0 | 0 | 0 | 0 | 0 | 0 | 0 | 0 | 0 | 0 | 0 | 0 | 0 | 0 | 0 | 0 | 0 | 0 | 0 | 0 | 0 | 0 | 0 | 0 | 0 | 0 | 0 | 0 | 0 | 0 | 0 | 0 | 0 | 0 | 0 | 0 | 0 | 0 | 0 | 0 |
| virus_May12.50v_11531 | 1 | 0 | 0 | 0 | 0 | 0 | 1 | 0 | 0 | 0 | 0 | 0 | 0 | 0 | 0 | 0 | 0 | 0 | 0 | 0 | 0 | 0 | 0 | 0 | 0 | 0 | 0 | 0 | 0 | 0 | 0 | 0 | 0 | 0 | 0 | 0 | 0 | 0 | 0 | 0 | 0 | 0 | 0 | 0 | 0 | 0 | 0 | 0 | 0 | 0 | 0 | 0 | 0 | 0 | 0 |
| virus_May12.50v_11953 | 1 | 0 | 0 | 0 | 0 | 0 | 1 | 0 | 0 | 0 | 0 | 0 | 0 | 0 | 0 | 0 | 0 | 0 | 0 | 0 | 0 | 0 | 0 | 0 | 0 | 0 | 0 | 0 | 0 | 0 | 0 | 0 | 0 | 0 | 0 | 0 | 0 | 0 | 0 | 0 | 0 | 0 | 0 | 0 | 0 | 0 | 0 | 0 | 0 | 0 | 0 | 0 | 0 | 0 | 0 |
| virus_May12.50v_19340 | 1 | 0 | 0 | 0 | 0 | 0 | 1 | 0 | 0 | 0 | 0 | 0 | 0 | 0 | 0 | 0 | 0 | 0 | 0 | 0 | 0 | 0 | 0 | 0 | 0 | 0 | 0 | 0 | 0 | 0 | 0 | 0 | 0 | 0 | 0 | 0 | 0 | 0 | 0 | 0 | 0 | 0 | 0 | 0 | 0 | 0 | 0 | 0 | 0 | 0 | 0 | 0 | 0 | 0 | 0 |
| virus_May12.50v_22672 | 1 | 0 | 0 | 0 | 0 | 0 | 1 | 0 | 0 | 0 | 0 | 0 | 0 | 0 | 0 | 0 | 0 | 0 | 0 | 0 | 0 | 0 | 0 | 0 | 0 | 0 | 0 | 0 | 0 | 0 | 0 | 0 | 0 | 0 | 0 | 0 | 0 | 0 | 0 | 0 | 0 | 0 | 0 | 0 | 0 | 0 | 0 | 0 | 0 | 0 | 0 | 0 | 0 | 0 | 0 |
| virus_May12.50v_22804 | 1 | 0 | 0 | 0 | 0 | 0 | 1 | 0 | 0 | 0 | 0 | 0 | 0 | 0 | 0 | 0 | 0 | 0 | 0 | 0 | 0 | 0 | 0 | 0 | 0 | 0 | 0 | 0 | 0 | 0 | 0 | 0 | 0 | 0 | 0 | 0 | 0 | 0 | 0 | 0 | 0 | 0 | 0 | 0 | 0 | 0 | 0 | 0 | 0 | 0 | 0 | 0 | 0 | 0 | 0 |
| virus_May12.50v_25185 | 1 | 0 | 0 | 0 | 0 | 0 | 1 | 0 | 0 | 0 | 0 | 0 | 0 | 0 | 0 | 0 | 0 | 0 | 0 | 0 | 0 | 0 | 0 | 0 | 0 | 0 | 0 | 0 | 0 | 0 | 0 | 0 | 0 | 0 | 0 | 0 | 0 | 0 | 0 | 0 | 0 | 0 | 0 | 0 | 0 | 0 | 0 | 0 | 0 | 0 | 0 | 0 | 0 | 0 | 0 |
| virus_May12.50v_27974 | 1 | 0 | 0 | 0 | 0 | 0 | 1 | 0 | 0 | 0 | 0 | 0 | 0 | 0 | 0 | 0 | 0 | 0 | 0 | 0 | 0 | 0 | 0 | 0 | 0 | 0 | 0 | 0 | 0 | 0 | 0 | 0 | 0 | 0 | 0 | 0 | 0 | 0 | 0 | 0 | 0 | 0 | 0 | 0 | 0 | 0 | 0 | 0 | 0 | 0 | 0 | 0 | 0 | 0 | 0 |
| virus_May12.50v_32397 | 1 | 0 | 0 | 0 | 0 | 0 | 1 | 0 | 0 | 0 | 0 | 0 | 0 | 0 | 0 | 0 | 0 | 0 | 0 | 0 | 0 | 0 | 0 | 0 | 0 | 0 | 0 | 0 | 0 | 0 | 0 | 0 | 0 | 0 | 0 | 0 | 0 | 0 | 0 | 0 | 0 | 0 | 0 | 0 | 0 | 0 | 0 | 0 | 0 | 0 | 0 | 0 | 0 | 0 | 0 |
| virus_May12.50v_38567 | 1 | 0 | 0 | 0 | 0 | 0 | 1 | 0 | 0 | 0 | 0 | 0 | 0 | 0 | 0 | 0 | 0 | 0 | 0 | 0 | 0 | 0 | 0 | 0 | 0 | 0 | 0 | 0 | 0 | 0 | 0 | 0 | 0 | 0 | 0 | 0 | 0 | 0 | 0 | 0 | 0 | 0 | 0 | 0 | 0 | 0 | 0 | 0 | 0 | 0 | 0 | 0 | 0 | 0 | 0 |
| virus_May12.50v_40245 | 1 | 0 | 0 | 0 | 0 | 0 | 1 | 0 | 0 | 0 | 0 | 0 | 0 | 0 | 0 | 0 | 0 | 0 | 0 | 0 | 0 | 0 | 0 | 0 | 0 | 0 | 0 | 0 | 0 | 0 | 0 | 0 | 0 | 0 | 0 | 0 | 0 | 0 | 0 | 0 | 0 | 0 | 0 | 0 | 0 | 0 | 0 | 0 | 0 | 0 | 0 | 0 | 0 | 0 | 0 |
| virus_May12.50v_40416 | 1 | 0 | 0 | 0 | 0 | 0 | 1 | 0 | 0 | 0 | 0 | 0 | 0 | 0 | 0 | 0 | 0 | 0 | 0 | 0 | 0 | 0 | 0 | 0 | 0 | 0 | 0 | 0 | 0 | 0 | 0 | 0 | 0 | 0 | 0 | 0 | 0 | 0 | 0 | 0 | 0 | 0 | 0 | 0 | 0 | 0 | 0 | 0 | 0 | 0 | 0 | 0 | 0 | 0 | 0 |
| virus_May12.75v_8355 | 1 | 0 | 0 | 0 | 0 | 0 | 0 | 0 | 1 | 0 | 0 | 0 | 0 | 0 | 0 | 0 | 0 | 0 | 0 | 0 | 0 | 0 | 0 | 0 | 0 | 0 | 0 | 0 | 0 | 0 | 0 | 0 | 0 | 0 | 0 | 0 | 0 | 0 | 0 | 0 | 0 | 0 | 0 | 0 | 0 | 0 | 0 | 0 | 0 | 0 | 0 | 0 | 0 | 0 | 0 |
| virus_May12.75v_13516 | 1 | 0 | 0 | 0 | 0 | 0 | 0 | 0 | 1 | 0 | 0 | 0 | 0 | 0 | 0 | 0 | 0 | 0 | 0 | 0 | 0 | 0 | 0 | 0 | 0 | 0 | 0 | 0 | 0 | 0 | 0 | 0 | 0 | 0 | 0 | 0 | 0 | 0 | 0 | 0 | 0 | 0 | 0 | 0 | 0 | 0 | 0 | 0 | 0 | 0 | 0 | 0 | 0 | 0 | 0 |
| virus_May12.75v_13829 | 1 | 0 | 0 | 0 | 0 | 0 | 0 | 0 | 1 | 0 | 0 | 0 | 0 | 0 | 0 | 0 | 0 | 0 | 0 | 0 | 0 | 0 | 0 | 0 | 0 | 0 | 0 | 0 | 0 | 0 | 0 | 0 | 0 | 0 | 0 | 0 | 0 | 0 | 0 | 0 | 0 | 0 | 0 | 0 | 0 | 0 | 0 | 0 | 0 | 0 | 0 | 0 | 0 | 0 | 0 |
| virus_May12.75v_27240 | 1 | 0 | 0 | 0 | 0 | 0 | 0 | 0 | 1 | 0 | 0 | 0 | 0 | 0 | 0 | 0 | 0 | 0 | 0 | 0 | 0 | 0 | 0 | 0 | 0 | 0 | 0 | 0 | 0 | 0 | 0 | 0 | 0 | 0 | 0 | 0 | 0 | 0 | 0 | 0 | 0 | 0 | 0 | 0 | 0 | 0 | 0 | 0 | 0 | 0 | 0 | 0 | 0 | 0 | 0 |
| virus_May12.75v_30830 | 1 | 0 | 0 | 0 | 0 | 0 | 0 | 0 | 1 | 0 | 0 | 0 | 0 | 0 | 0 | 0 | 0 | 0 | 0 | 0 | 0 | 0 | 0 | 0 | 0 | 0 | 0 | 0 | 0 | 0 | 0 | 0 | 0 | 0 | 0 | 0 | 0 | 0 | 0 | 0 | 0 | 0 | 0 | 0 | 0 | 0 | 0 | 0 | 0 | 0 | 0 | 0 | 0 | 0 | 0 |
| virus_May12.75v_31618 | 1 | 0 | 0 | 0 | 0 | 0 | 0 | 0 | 1 | 0 | 0 | 0 | 0 | 0 | 0 | 0 | 0 | 0 | 0 | 0 | 0 | 0 | 0 | 0 | 0 | 0 | 0 | 0 | 0 | 0 | 0 | 0 | 0 | 0 | 0 | 0 | 0 | 0 | 0 | 0 | 0 | 0 | 0 | 0 | 0 | 0 | 0 | 0 | 0 | 0 | 0 | 0 | 0 | 0 | 0 |
| virus_May12.75v_39250 | 1 | 0 | 0 | 0 | 0 | 0 | 0 | 0 | 1 | 0 | 0 | 0 | 0 | 0 | 0 | 0 | 0 | 0 | 0 | 0 | 0 | 0 | 0 | 0 | 0 | 0 | 0 | 0 | 0 | 0 | 0 | 0 | 0 | 0 | 0 | 0 | 0 | 0 | 0 | 0 | 0 | 0 | 0 | 0 | 0 | 0 | 0 | 0 | 0 | 0 | 0 | 0 | 0 | 0 | 0 |
| virus_May12.75v_39704 | 1 | 0 | 0 | 0 | 0 | 0 | 0 | 0 | 1 | 0 | 0 | 0 | 0 | 0 | 0 | 0 | 0 | 0 | 0 | 0 | 0 | 0 | 0 | 0 | 0 | 0 | 0 | 0 | 0 | 0 | 0 | 0 | 0 | 0 | 0 | 0 | 0 | 0 | 0 | 0 | 0 | 0 | 0 | 0 | 0 | 0 | 0 | 0 | 0 | 0 | 0 | 0 | 0 | 0 | 0 |
| virus_Sep8.0v_2419 | 1 | 0 | 0 | 0 | 0 | 0 | 0 | 0 | 0 | 0 | 0 | 0 | 0 | 0 | 0 | 0 | 0 | 0 | 0 | 0 | 0 | 0 | 0 | 0 | 0 | 0 | 0 | 0 | 0 | 0 | 0 | 0 | 0 | 0 | 0 | 0 | 0 | 0 | 0 | 0 | 0 | 0 | 0 | 0 | 0 | 0 | 0 | 0 | 1 | 0 | 0 | 0 | 0 | 0 | 0 |
| virus_Sep8.0v_5444 | 1 | 0 | 0 | 0 | 0 | 0 | 0 | 0 | 0 | 0 | 0 | 0 | 0 | 0 | 0 | 0 | 0 | 0 | 0 | 0 | 0 | 0 | 0 | 0 | 0 | 0 | 0 | 0 | 0 | 0 | 0 | 0 | 0 | 0 | 0 | 0 | 0 | 0 | 0 | 0 | 0 | 0 | 0 | 0 | 0 | 0 | 0 | 0 | 1 | 0 | 0 | 0 | 0 | 0 | 0 |
| virus_Sep8.0v_6319 | 1 | 0 | 0 | 0 | 0 | 0 | 0 | 0 | 0 | 0 | 0 | 0 | 0 | 0 | 0 | 0 | 0 | 0 | 0 | 0 | 0 | 0 | 0 | 0 | 0 | 0 | 0 | 0 | 0 | 0 | 0 | 0 | 0 | 0 | 0 | 0 | 0 | 0 | 0 | 0 | 0 | 0 | 0 | 0 | 0 | 0 | 0 | 0 | 1 | 0 | 0 | 0 | 0 | 0 | 0 |
| virus_Sep8.0v_7000 | 1 | 0 | 0 | 0 | 0 | 0 | 0 | 0 | 0 | 0 | 0 | 0 | 0 | 0 | 0 | 0 | 0 | 0 | 0 | 0 | 0 | 0 | 0 | 0 | 0 | 0 | 0 | 0 | 0 | 0 | 0 | 0 | 0 | 0 | 0 | 0 | 0 | 0 | 0 | 0 | 0 | 0 | 0 | 0 | 0 | 0 | 0 | 0 | 1 | 0 | 0 | 0 | 0 | 0 | 0 |
| virus_Sep8.0v_10367 | 1 | 0 | 0 | 0 | 0 | 0 | 0 | 0 | 0 | 0 | 0 | 0 | 0 | 0 | 0 | 0 | 0 | 0 | 0 | 0 | 0 | 0 | 0 | 0 | 0 | 0 | 0 | 0 | 0 | 0 | 0 | 0 | 0 | 0 | 0 | 0 | 0 | 0 | 0 | 0 | 0 | 0 | 0 | 0 | 0 | 0 | 0 | 0 | 1 | 0 | 0 | 0 | 0 | 0 | 0 |
| virus_Sep8.0v_15316 | 1 | 0 | 0 | 0 | 0 | 0 | 0 | 0 | 0 | 0 | 0 | 0 | 0 | 0 | 0 | 0 | 0 | 0 | 0 | 0 | 0 | 0 | 0 | 0 | 0 | 0 | 0 | 0 | 0 | 0 | 0 | 0 | 0 | 0 | 0 | 0 | 0 | 0 | 0 | 0 | 0 | 0 | 0 | 0 | 0 | 0 | 0 | 0 | 1 | 0 | 0 | 0 | 0 | 0 | 0 |
| virus_Sep8.0v_18705 | 1 | 0 | 0 | 0 | 0 | 0 | 0 | 0 | 0 | 0 | 0 | 0 | 0 | 0 | 0 | 0 | 0 | 0 | 0 | 0 | 0 | 0 | 0 | 0 | 0 | 0 | 0 | 0 | 0 | 0 | 0 | 0 | 0 | 0 | 0 | 0 | 0 | 0 | 0 | 0 | 0 | 0 | 0 | 0 | 0 | 0 | 0 | 0 | 1 | 0 | 0 | 0 | 0 | 0 | 0 |
| virus_Sep8.5v_5564 | 1 | 0 | 0 | 0 | 0 | 0 | 0 | 0 | 0 | 0 | 0 | 0 | 0 | 0 | 0 | 0 | 0 | 0 | 0 | 0 | 0 | 0 | 0 | 0 | 0 | 0 | 0 | 0 | 0 | 0 | 0 | 0 | 0 | 0 | 0 | 0 | 0 | 0 | 0 | 0 | 0 | 0 | 0 | 0 | 0 | 0 | 0 | 0 | 0 | 0 | 1 | 0 | 0 | 0 | 0 |
| virus_Sep8.5v_9257 | 1 | 0 | 0 | 0 | 0 | 0 | 0 | 0 | 0 | 0 | 0 | 0 | 0 | 0 | 0 | 0 | 0 | 0 | 0 | 0 | 0 | 0 | 0 | 0 | 0 | 0 | 0 | 0 | 0 | 0 | 0 | 0 | 0 | 0 | 0 | 0 | 0 | 0 | 0 | 0 | 0 | 0 | 0 | 0 | 0 | 0 | 0 | 0 | 0 | 0 | 1 | 0 | 0 | 0 | 0 |
| virus_Sep8.5v_10759 | 1 | 0 | 0 | 0 | 0 | 0 | 0 | 0 | 0 | 0 | 0 | 0 | 0 | 0 | 0 | 0 | 0 | 0 | 0 | 0 | 0 | 0 | 0 | 0 | 0 | 0 | 0 | 0 | 0 | 0 | 0 | 0 | 0 | 0 | 0 | 0 | 0 | 0 | 0 | 0 | 0 | 0 | 0 | 0 | 0 | 0 | 0 | 0 | 0 | 0 | 1 | 0 | 0 | 0 | 0 |
| virus_Sep8.5v_11860 | 1 | 0 | 0 | 0 | 0 | 0 | 0 | 0 | 0 | 0 | 0 | 0 | 0 | 0 | 0 | 0 | 0 | 0 | 0 | 0 | 0 | 0 | 0 | 0 | 0 | 0 | 0 | 0 | 0 | 0 | 0 | 0 | 0 | 0 | 0 | 0 | 0 | 0 | 0 | 0 | 0 | 0 | 0 | 0 | 0 | 0 | 0 | 0 | 0 | 0 | 1 | 0 | 0 | 0 | 0 |
| virus_Sep8.5v_13376 | 1 | 0 | 0 | 0 | 0 | 0 | 0 | 0 | 0 | 0 | 0 | 0 | 0 | 0 | 0 | 0 | 0 | 0 | 0 | 0 | 0 | 0 | 0 | 0 | 0 | 0 | 0 | 0 | 0 | 0 | 0 | 0 | 0 | 0 | 0 | 0 | 0 | 0 | 0 | 0 | 0 | 0 | 0 | 0 | 0 | 0 | 0 | 0 | 0 | 0 | 1 | 0 | 0 | 0 | 0 |
| virus_Sep8.5v_15067 | 1 | 0 | 0 | 0 | 0 | 0 | 0 | 0 | 0 | 0 | 0 | 0 | 0 | 0 | 0 | 0 | 0 | 0 | 0 | 0 | 0 | 0 | 0 | 0 | 0 | 0 | 0 | 0 | 0 | 0 | 0 | 0 | 0 | 0 | 0 | 0 | 0 | 0 | 0 | 0 | 0 | 0 | 0 | 0 | 0 | 0 | 0 | 0 | 0 | 0 | 1 | 0 | 0 | 0 | 0 |
| virus_Sep8.25v_7208 | 1 | 0 | 0 | 0 | 0 | 0 | 0 | 0 | 0 | 0 | 0 | 0 | 0 | 0 | 0 | 0 | 0 | 0 | 0 | 0 | 0 | 0 | 0 | 0 | 0 | 0 | 0 | 0 | 0 | 0 | 0 | 0 | 0 | 0 | 0 | 0 | 0 | 0 | 0 | 0 | 0 | 0 | 0 | 0 | 0 | 0 | 0 | 0 | 0 | 0 | 0 | 0 | 1 | 0 | 0 |
| virus_Sep8.25v_9123 | 1 | 0 | 0 | 0 | 0 | 0 | 0 | 0 | 0 | 0 | 0 | 0 | 0 | 0 | 0 | 0 | 0 | 0 | 0 | 0 | 0 | 0 | 0 | 0 | 0 | 0 | 0 | 0 | 0 | 0 | 0 | 0 | 0 | 0 | 0 | 0 | 0 | 0 | 0 | 0 | 0 | 0 | 0 | 0 | 0 | 0 | 0 | 0 | 0 | 0 | 0 | 0 | 1 | 0 | 0 |
| virus_Sep8.25v_12326 | 1 | 0 | 0 | 0 | 0 | 0 | 0 | 0 | 0 | 0 | 0 | 0 | 0 | 0 | 0 | 0 | 0 | 0 | 0 | 0 | 0 | 0 | 0 | 0 | 0 | 0 | 0 | 0 | 0 | 0 | 0 | 0 | 0 | 0 | 0 | 0 | 0 | 0 | 0 | 0 | 0 | 0 | 0 | 0 | 0 | 0 | 0 | 0 | 0 | 0 | 0 | 0 | 1 | 0 | 0 |
| virus_Sep8.25v_18163 | 1 | 0 | 0 | 0 | 0 | 0 | 0 | 0 | 0 | 0 | 0 | 0 | 0 | 0 | 0 | 0 | 0 | 0 | 0 | 0 | 0 | 0 | 0 | 0 | 0 | 0 | 0 | 0 | 0 | 0 | 0 | 0 | 0 | 0 | 0 | 0 | 0 | 0 | 0 | 0 | 0 | 0 | 0 | 0 | 0 | 0 | 0 | 0 | 0 | 0 | 0 | 0 | 1 | 0 | 0 |
| virus_Sep8.25v_21004 | 1 | 0 | 0 | 0 | 0 | 0 | 0 | 0 | 0 | 0 | 0 | 0 | 0 | 0 | 0 | 0 | 0 | 0 | 0 | 0 | 0 | 0 | 0 | 0 | 0 | 0 | 0 | 0 | 0 | 0 | 0 | 0 | 0 | 0 | 0 | 0 | 0 | 0 | 0 | 0 | 0 | 0 | 0 | 0 | 0 | 0 | 0 | 0 | 0 | 0 | 0 | 0 | 1 | 0 | 0 |
| virus_Sep8.50v_3217 | 1 | 0 | 0 | 0 | 0 | 0 | 0 | 0 | 0 | 0 | 0 | 0 | 0 | 0 | 0 | 0 | 0 | 0 | 0 | 0 | 0 | 0 | 0 | 0 | 0 | 0 | 0 | 0 | 0 | 0 | 0 | 0 | 0 | 0 | 0 | 0 | 0 | 0 | 0 | 0 | 0 | 0 | 0 | 0 | 0 | 0 | 0 | 0 | 0 | 0 | 0 | 0 | 0 | 0 | 1 |
| virus_Sep8.50v_5179 | 1 | 0 | 0 | 0 | 0 | 0 | 0 | 0 | 0 | 0 | 0 | 0 | 0 | 0 | 0 | 0 | 0 | 0 | 0 | 0 | 0 | 0 | 0 | 0 | 0 | 0 | 0 | 0 | 0 | 0 | 0 | 0 | 0 | 0 | 0 | 0 | 0 | 0 | 0 | 0 | 0 | 0 | 0 | 0 | 0 | 0 | 0 | 0 | 0 | 0 | 0 | 0 | 0 | 0 | 1 |
| virus_Sep8.50v_12252 | 1 | 0 | 0 | 0 | 0 | 0 | 0 | 0 | 0 | 0 | 0 | 0 | 0 | 0 | 0 | 0 | 0 | 0 | 0 | 0 | 0 | 0 | 0 | 0 | 0 | 0 | 0 | 0 | 0 | 0 | 0 | 0 | 0 | 0 | 0 | 0 | 0 | 0 | 0 | 0 | 0 | 0 | 0 | 0 | 0 | 0 | 0 | 0 | 0 | 0 | 0 | 0 | 0 | 0 | 1 |
| virus_Sep8.50v_12628 | 1 | 0 | 0 | 0 | 0 | 0 | 0 | 0 | 0 | 0 | 0 | 0 | 0 | 0 | 0 | 0 | 0 | 0 | 0 | 0 | 0 | 0 | 0 | 0 | 0 | 0 | 0 | 0 | 0 | 0 | 0 | 0 | 0 | 0 | 0 | 0 | 0 | 0 | 0 | 0 | 0 | 0 | 0 | 0 | 0 | 0 | 0 | 0 | 0 | 0 | 0 | 0 | 0 | 0 | 1 |
| virus_Sep8.50v_15779 | 1 | 0 | 0 | 0 | 0 | 0 | 0 | 0 | 0 | 0 | 0 | 0 | 0 | 0 | 0 | 0 | 0 | 0 | 0 | 0 | 0 | 0 | 0 | 0 | 0 | 0 | 0 | 0 | 0 | 0 | 0 | 0 | 0 | 0 | 0 | 0 | 0 | 0 | 0 | 0 | 0 | 0 | 0 | 0 | 0 | 0 | 0 | 0 | 0 | 0 | 0 | 0 | 0 | 0 | 1 |
| virus_Sep8.50v_18102 | 1 | 0 | 0 | 0 | 0 | 0 | 0 | 0 | 0 | 0 | 0 | 0 | 0 | 0 | 0 | 0 | 0 | 0 | 0 | 0 | 0 | 0 | 0 | 0 | 0 | 0 | 0 | 0 | 0 | 0 | 0 | 0 | 0 | 0 | 0 | 0 | 0 | 0 | 0 | 0 | 0 | 0 | 0 | 0 | 0 | 0 | 0 | 0 | 0 | 0 | 0 | 0 | 0 | 0 | 1 |
| virus_Sep8.50v_27082 | 1 | 0 | 0 | 0 | 0 | 0 | 0 | 0 | 0 | 0 | 0 | 0 | 0 | 0 | 0 | 0 | 0 | 0 | 0 | 0 | 0 | 0 | 0 | 0 | 0 | 0 | 0 | 0 | 0 | 0 | 0 | 0 | 0 | 0 | 0 | 0 | 0 | 0 | 0 | 0 | 0 | 0 | 0 | 0 | 0 | 0 | 0 | 0 | 0 | 0 | 0 | 0 | 0 | 0 | 1 |
| virus_Sep8.50v_28279 | 1 | 0 | 0 | 0 | 0 | 0 | 0 | 0 | 0 | 0 | 0 | 0 | 0 | 0 | 0 | 0 | 0 | 0 | 0 | 0 | 0 | 0 | 0 | 0 | 0 | 0 | 0 | 0 | 0 | 0 | 0 | 0 | 0 | 0 | 0 | 0 | 0 | 0 | 0 | 0 | 0 | 0 | 0 | 0 | 0 | 0 | 0 | 0 | 0 | 0 | 0 | 0 | 0 | 0 | 1 |
| virus_Sep8.50v_30979 | 1 | 0 | 0 | 0 | 0 | 0 | 0 | 0 | 0 | 0 | 0 | 0 | 0 | 0 | 0 | 0 | 0 | 0 | 0 | 0 | 0 | 0 | 0 | 0 | 0 | 0 | 0 | 0 | 0 | 0 | 0 | 0 | 0 | 0 | 0 | 0 | 0 | 0 | 0 | 0 | 0 | 0 | 0 | 0 | 0 | 0 | 0 | 0 | 0 | 0 | 0 | 0 | 0 | 0 | 1 |
| virus_Sep8.50v_31633 | 1 | 0 | 0 | 0 | 0 | 0 | 0 | 0 | 0 | 0 | 0 | 0 | 0 | 0 | 0 | 0 | 0 | 0 | 0 | 0 | 0 | 0 | 0 | 0 | 0 | 0 | 0 | 0 | 0 | 0 | 0 | 0 | 0 | 0 | 0 | 0 | 0 | 0 | 0 | 0 | 0 | 0 | 0 | 0 | 0 | 0 | 0 | 0 | 0 | 0 | 0 | 0 | 0 | 0 | 1 |
| virus_Sep8.50v_34011 | 1 | 0 | 0 | 0 | 0 | 0 | 0 | 0 | 0 | 0 | 0 | 0 | 0 | 0 | 0 | 0 | 0 | 0 | 0 | 0 | 0 | 0 | 0 | 0 | 0 | 0 | 0 | 0 | 0 | 0 | 0 | 0 | 0 | 0 | 0 | 0 | 0 | 0 | 0 | 0 | 0 | 0 | 0 | 0 | 0 | 0 | 0 | 0 | 0 | 0 | 0 | 0 | 0 | 0 | 1 |
| virus_Sep8.50v_37801 | 1 | 0 | 0 | 0 | 0 | 0 | 0 | 0 | 0 | 0 | 0 | 0 | 0 | 0 | 0 | 0 | 0 | 0 | 0 | 0 | 0 | 0 | 0 | 0 | 0 | 0 | 0 | 0 | 0 | 0 | 0 | 0 | 0 | 0 | 0 | 0 | 0 | 0 | 0 | 0 | 0 | 0 | 0 | 0 | 0 | 0 | 0 | 0 | 0 | 0 | 0 | 0 | 0 | 0 | 1 |
| bacteria_Jul11d0b_10479 | 1 | 0 | 0 | 0 | 0 | 0 | 0 | 0 | 0 | 0 | 0 | 0 | 0 | 0 | 0 | 0 | 0 | 0 | 0 | 0 | 0 | 1 | 0 | 0 | 0 | 0 | 0 | 0 | 0 | 0 | 0 | 0 | 0 | 0 | 0 | 0 | 0 | 0 | 0 | 0 | 0 | 0 | 0 | 0 | 0 | 0 | 0 | 0 | 0 | 0 | 0 | 0 | 0 | 0 | 0 |
| bacteria_Jul11d0b_16778 | 1 | 0 | 0 | 0 | 0 | 0 | 0 | 0 | 0 | 0 | 0 | 0 | 0 | 0 | 0 | 0 | 0 | 0 | 0 | 0 | 0 | 1 | 0 | 0 | 0 | 0 | 0 | 0 | 0 | 0 | 0 | 0 | 0 | 0 | 0 | 0 | 0 | 0 | 0 | 0 | 0 | 0 | 0 | 0 | 0 | 0 | 0 | 0 | 0 | 0 | 0 | 0 | 0 | 0 | 0 |
| bacteria_Jul11d0b_29309 | 1 | 0 | 0 | 0 | 0 | 0 | 0 | 0 | 0 | 0 | 0 | 0 | 0 | 0 | 0 | 0 | 0 | 0 | 0 | 0 | 0 | 1 | 0 | 0 | 0 | 0 | 0 | 0 | 0 | 0 | 0 | 0 | 0 | 0 | 0 | 0 | 0 | 0 | 0 | 0 | 0 | 0 | 0 | 0 | 0 | 0 | 0 | 0 | 0 | 0 | 0 | 0 | 0 | 0 | 0 |
| bacteria_Jul11d0b_39221 | 1 | 0 | 0 | 0 | 0 | 0 | 0 | 0 | 0 | 0 | 0 | 0 | 0 | 0 | 0 | 0 | 0 | 0 | 0 | 0 | 0 | 1 | 0 | 0 | 0 | 0 | 0 | 0 | 0 | 0 | 0 | 0 | 0 | 0 | 0 | 0 | 0 | 0 | 0 | 0 | 0 | 0 | 0 | 0 | 0 | 0 | 0 | 0 | 0 | 0 | 0 | 0 | 0 | 0 | 0 |
| bacteria_Jul11d0b_50644 | 1 | 0 | 0 | 0 | 0 | 0 | 0 | 0 | 0 | 0 | 0 | 0 | 0 | 0 | 0 | 0 | 0 | 0 | 0 | 0 | 0 | 1 | 0 | 0 | 0 | 0 | 0 | 0 | 0 | 0 | 0 | 0 | 0 | 0 | 0 | 0 | 0 | 0 | 0 | 0 | 0 | 0 | 0 | 0 | 0 | 0 | 0 | 0 | 0 | 0 | 0 | 0 | 0 | 0 | 0 |
| bacteria_Aug5d0b_29542 | 1 | 0 | 0 | 0 | 0 | 0 | 0 | 0 | 0 | 0 | 0 | 0 | 0 | 0 | 0 | 0 | 0 | 0 | 0 | 0 | 0 | 0 | 0 | 0 | 0 | 0 | 0 | 0 | 0 | 0 | 0 | 0 | 0 | 0 | 0 | 0 | 0 | 1 | 0 | 0 | 0 | 0 | 0 | 0 | 0 | 0 | 0 | 0 | 0 | 0 | 0 | 0 | 0 | 0 | 0 |
| bacteria_Aug5d0b_36096 | 1 | 0 | 0 | 0 | 0 | 0 | 0 | 0 | 0 | 0 | 0 | 0 | 0 | 0 | 0 | 0 | 0 | 0 | 0 | 0 | 0 | 0 | 0 | 0 | 0 | 0 | 0 | 0 | 0 | 0 | 0 | 0 | 0 | 0 | 0 | 0 | 0 | 1 | 0 | 0 | 0 | 0 | 0 | 0 | 0 | 0 | 0 | 0 | 0 | 0 | 0 | 0 | 0 | 0 | 0 |
| bacteria_Aug5d0b_57371 | 1 | 0 | 0 | 0 | 0 | 0 | 0 | 0 | 0 | 0 | 0 | 0 | 0 | 0 | 0 | 0 | 0 | 0 | 0 | 0 | 0 | 0 | 0 | 0 | 0 | 0 | 0 | 0 | 0 | 0 | 0 | 0 | 0 | 0 | 0 | 0 | 0 | 1 | 0 | 0 | 0 | 0 | 0 | 0 | 0 | 0 | 0 | 0 | 0 | 0 | 0 | 0 | 0 | 0 | 0 |
| bacteria_Aug5d0b_61259 | 1 | 0 | 0 | 0 | 0 | 0 | 0 | 0 | 0 | 0 | 0 | 0 | 0 | 0 | 0 | 0 | 0 | 0 | 0 | 0 | 0 | 0 | 0 | 0 | 0 | 0 | 0 | 0 | 0 | 0 | 0 | 0 | 0 | 0 | 0 | 0 | 0 | 1 | 0 | 0 | 0 | 0 | 0 | 0 | 0 | 0 | 0 | 0 | 0 | 0 | 0 | 0 | 0 | 0 | 0 |
| bacteria_Aug5d5b_2271 | 1 | 0 | 0 | 0 | 0 | 0 | 0 | 0 | 0 | 0 | 0 | 0 | 0 | 0 | 0 | 0 | 0 | 0 | 0 | 0 | 0 | 0 | 0 | 0 | 0 | 0 | 0 | 0 | 0 | 0 | 0 | 0 | 0 | 0 | 0 | 0 | 0 | 0 | 0 | 1 | 0 | 0 | 0 | 0 | 0 | 0 | 0 | 0 | 0 | 0 | 0 | 0 | 0 | 0 | 0 |
| bacteria_Aug5d25b_3118 | 1 | 0 | 0 | 0 | 0 | 0 | 0 | 0 | 0 | 0 | 0 | 0 | 0 | 0 | 0 | 0 | 0 | 0 | 0 | 0 | 0 | 0 | 0 | 0 | 0 | 0 | 0 | 0 | 0 | 0 | 0 | 0 | 0 | 0 | 0 | 0 | 0 | 0 | 0 | 0 | 0 | 1 | 0 | 0 | 0 | 0 | 0 | 0 | 0 | 0 | 0 | 0 | 0 | 0 | 0 |
| bacteria_Aug5d25b_16314 | 1 | 0 | 0 | 0 | 0 | 0 | 0 | 0 | 0 | 0 | 0 | 0 | 0 | 0 | 0 | 0 | 0 | 0 | 0 | 0 | 0 | 0 | 0 | 0 | 0 | 0 | 0 | 0 | 0 | 0 | 0 | 0 | 0 | 0 | 0 | 0 | 0 | 0 | 0 | 0 | 0 | 1 | 0 | 0 | 0 | 0 | 0 | 0 | 0 | 0 | 0 | 0 | 0 | 0 | 0 |
| bacteria_Aug5d25b_23204 | 1 | 0 | 0 | 0 | 0 | 0 | 0 | 0 | 0 | 0 | 0 | 0 | 0 | 0 | 0 | 0 | 0 | 0 | 0 | 0 | 0 | 0 | 0 | 0 | 0 | 0 | 0 | 0 | 0 | 0 | 0 | 0 | 0 | 0 | 0 | 0 | 0 | 0 | 0 | 0 | 0 | 1 | 0 | 0 | 0 | 0 | 0 | 0 | 0 | 0 | 0 | 0 | 0 | 0 | 0 |
| bacteria_Aug5d25b_25404 | 1 | 0 | 0 | 0 | 0 | 0 | 0 | 0 | 0 | 0 | 0 | 0 | 0 | 0 | 0 | 0 | 0 | 0 | 0 | 0 | 0 | 0 | 0 | 0 | 0 | 0 | 0 | 0 | 0 | 0 | 0 | 0 | 0 | 0 | 0 | 0 | 0 | 0 | 0 | 0 | 0 | 1 | 0 | 0 | 0 | 0 | 0 | 0 | 0 | 0 | 0 | 0 | 0 | 0 | 0 |

**Supplementary Table S7**. Common Bacterio- and virioplankton RTPR 98% OTUs in 27 samples with RTPR amplicons in both fractions

Total seqs: 47997; Subtotal seqs in retained

| OTUs: 41046 | | May12_05m | | May12_25m | | May12_50m | | May12_75m | | Jun8_25m | | Jun8_50m | | Jun8_75m | | Jun22_0m | | Jun22_50m | | Jun22_75m | | Jul11_0m | | Jul11_05m | | Jul11_25m | | Jul11_50m | | Jul11_75m | | Jul27_05m | | Jul27_25m | | Jul27_75m | | Aug5_0m | | Aug5_05m | | Aug5_25m | | Aug5_50m | | Aug5_75m | | Sep8_0m | | Sep8_05m | | Sep8_25m | | Sep8_50m | |
|---|---|---|---|---|---|---|---|---|---|---|---|---|---|---|---|---|---|---|---|---|---|---|---|---|---|---|---|---|---|---|---|---|---|---|---|---|---|---|---|---|---|---|---|---|---|---|---|---|---|---|---|---|---|---|---|
| OTUs with >=10 seqs: 254 | OTU subtotal | bacterioplankton | virioplankton | bacterioplankton | virioplankton | bacterioplankton | virioplankton | bacterioplankton | virioplankton | bacterioplankton | virioplankton | bacterioplankton | virioplankton | bacterioplankton | virioplankton | bacterioplankton | virioplankton | bacterioplankton | virioplankton | bacterioplankton | virioplankton | bacterioplankton | virioplankton | bacterioplankton | virioplankton | bacterioplankton | virioplankton | bacterioplankton | virioplankton | bacterioplankton | virioplankton | bacterioplankton | virioplankton | bacterioplankton | virioplankton | bacterioplankton | virioplankton | bacterioplankton | virioplankton | bacterioplankton | virioplankton | bacterioplankton | virioplankton | bacterioplankton | virioplankton | bacterioplankton | virioplankton | bacterioplankton | virioplankton | bacterioplankton | virioplankton | bacterioplankton | virioplankton | bacterioplankton | virioplankton |
|bacteria_Aug5d25b_2847|1|0|0|0|0|0|0|0|0|0|0|0|0|0|0|0|0|0|0|0|0|0|0|0|0|0|0|0|0|0|0|0|0|0|0|0|0|0|0|0|0|1|0|0|0|0|0|0|0|0|0|0|0|0|0|
|bacteria_Aug5d25b_2986|1|0|0|0|0|0|0|0|0|0|0|0|0|0|0|0|0|0|0|0|0|0|0|0|0|0|0|0|0|0|0|0|0|0|0|0|0|0|0|0|0|1|0|0|0|0|0|0|0|0|0|0|0|0|0|
|bacteria_Aug5d25b_3976|1|0|0|0|0|0|0|0|0|0|0|0|0|0|0|0|0|0|0|0|0|0|0|0|0|0|0|0|0|0|0|0|0|0|0|0|0|0|0|0|0|1|0|0|0|0|0|0|0|0|0|0|0|0|0|
|bacteria_Aug5d25b_4000|1|0|0|0|0|0|0|0|0|0|0|0|0|0|0|0|0|0|0|0|0|0|0|0|0|0|0|0|0|0|0|0|0|0|0|0|0|0|0|0|0|1|0|0|0|0|0|0|0|0|0|0|0|0|0|
|bacteria_Aug5d25b_5754|1|0|0|0|0|0|0|0|0|0|0|0|0|0|0|0|0|0|0|0|0|0|0|0|0|0|0|0|0|0|0|0|0|0|0|0|0|0|0|0|0|1|0|0|0|0|0|0|0|0|0|0|0|0|0|
|bacteria_Aug5d25b_6405|1|0|0|0|0|0|0|0|0|0|0|0|0|0|0|0|0|0|0|0|0|0|0|0|0|0|0|0|0|0|0|0|0|0|0|0|0|0|0|0|0|1|0|0|0|0|0|0|0|0|0|0|0|0|0|
|bacteria_Aug5d25b_7014|1|0|0|0|0|0|0|0|0|0|0|0|0|0|0|0|0|0|0|0|0|0|0|0|0|0|0|0|0|0|0|0|0|0|0|0|0|0|0|0|0|1|0|0|0|0|0|0|0|0|0|0|0|0|0|
|bacteria_Aug5d25b_7082|1|0|0|0|0|0|0|0|0|0|0|0|0|0|0|0|0|0|0|0|0|0|0|0|0|0|0|0|0|0|0|0|0|0|0|0|0|0|0|0|0|1|0|0|0|0|0|0|0|0|0|0|0|0|0|
|bacteria_Aug5d50b_2535|1|0|0|0|0|0|0|0|0|0|0|0|0|0|0|0|0|0|0|0|0|0|0|0|0|0|0|0|0|0|0|0|0|0|0|0|0|0|0|0|0|0|0|1|0|0|0|0|0|0|0|0|0|0|0|
|bacteria_Aug5d50b_1050|1|0|0|0|0|0|0|0|0|0|0|0|0|0|0|0|0|0|0|0|0|0|0|0|0|0|0|0|0|0|0|0|0|0|0|0|0|0|0|0|0|0|0|1|0|0|0|0|0|0|0|0|0|0|0|
|bacteria_Aug5d50b_2392|1|0|0|0|0|0|0|0|0|0|0|0|0|0|0|0|0|0|0|0|0|0|0|0|0|0|0|0|0|0|0|0|0|0|0|0|0|0|0|0|0|0|0|1|0|0|0|0|0|0|0|0|0|0|0|
|bacteria_Aug5d50b_3465|1|0|0|0|0|0|0|0|0|0|0|0|0|0|0|0|0|0|0|0|0|0|0|0|0|0|0|0|0|0|0|0|0|0|0|0|0|0|0|0|0|0|0|1|0|0|0|0|0|0|0|0|0|0|0|
|bacteria_Aug5d50b_4162|1|0|0|0|0|0|0|0|0|0|0|0|0|0|0|0|0|0|0|0|0|0|0|0|0|0|0|0|0|0|0|0|0|0|0|0|0|0|0|0|0|0|0|1|0|0|0|0|0|0|0|0|0|0|0|
|bacteria_Aug5d50b_4703|1|0|0|0|0|0|0|0|0|0|0|0|0|0|0|0|0|0|0|0|0|0|0|0|0|0|0|0|0|0|0|0|0|0|0|0|0|0|0|0|0|0|0|1|0|0|0|0|0|0|0|0|0|0|0|
|bacteria_Aug5d50b_4762|1|0|0|0|0|0|0|0|0|0|0|0|0|0|0|0|0|0|0|0|0|0|0|0|0|0|0|0|0|0|0|0|0|0|0|0|0|0|0|0|0|0|0|1|0|0|0|0|0|0|0|0|0|0|0|
|bacteria_Aug5d50b_6207|1|0|0|0|0|0|0|0|0|0|0|0|0|0|0|0|0|0|0|0|0|0|0|0|0|0|0|0|0|0|0|0|0|0|0|0|0|0|0|0|0|0|0|1|0|0|0|0|0|0|0|0|0|0|0|
|bacteria_Aug5d50b_7518|1|0|0|0|0|0|0|0|0|0|0|0|0|0|0|0|0|0|0|0|0|0|0|0|0|0|0|0|0|0|0|0|0|0|0|0|0|0|0|0|0|0|0|1|0|0|0|0|0|0|0|0|0|0|0|
|bacteria_Aug5d75b_2964|1|0|0|0|0|0|0|0|0|0|0|0|0|0|0|0|0|0|0|0|0|0|0|0|0|0|0|0|0|0|0|0|0|0|0|0|0|0|0|0|0|0|0|0|0|1|0|0|0|0|0|0|0|0|0|
|bacteria_Jul11d5b_3608|1|0|0|0|0|0|0|0|0|0|0|0|0|0|0|0|0|0|0|0|0|0|0|1|0|0|0|0|0|0|0|0|0|0|0|0|0|0|0|0|0|0|0|0|0|0|0|0|0|0|0|0|0|0|0|
|bacteria_Jul11d5b_8474|1|0|0|0|0|0|0|0|0|0|0|0|0|0|0|0|0|0|0|0|0|0|0|1|0|0|0|0|0|0|0|0|0|0|0|0|0|0|0|0|0|0|0|0|0|0|0|0|0|0|0|0|0|0|0|
|bacteria_Jul11d5b_10588|1|0|0|0|0|0|0|0|0|0|0|0|0|0|0|0|0|0|0|0|0|0|0|1|0|0|0|0|0|0|0|0|0|0|0|0|0|0|0|0|0|0|0|0|0|0|0|0|0|0|0|0|0|0|0|
|bacteria_Jul11d5b_23901|1|0|0|0|0|0|0|0|0|0|0|0|0|0|0|0|0|0|0|0|0|0|0|1|0|0|0|0|0|0|0|0|0|0|0|0|0|0|0|0|0|0|0|0|0|0|0|0|0|0|0|0|0|0|0|
|bacteria_Jul11d5b_41355|1|0|0|0|0|0|0|0|0|0|0|0|0|0|0|0|0|0|0|0|0|0|0|1|0|0|0|0|0|0|0|0|0|0|0|0|0|0|0|0|0|0|0|0|0|0|0|0|0|0|0|0|0|0|0|
|bacteria_Jul11d25b_5650|1|0|0|0|0|0|0|0|0|0|0|0|0|0|0|0|0|0|0|0|0|0|0|0|0|1|0|0|0|0|0|0|0|0|0|0|0|0|0|0|0|0|0|0|0|0|0|0|0|0|0|0|0|0|0|
|bacteria_Jul11d25b_1167|1|0|0|0|0|0|0|0|0|0|0|0|0|0|0|0|0|0|0|0|0|0|0|0|0|1|0|0|0|0|0|0|0|0|0|0|0|0|0|0|0|0|0|0|0|0|0|0|0|0|0|0|0|0|0|
|bacteria_Jul11d25b_1579|1|0|0|0|0|0|0|0|0|0|0|0|0|0|0|0|0|0|0|0|0|0|0|0|0|1|0|0|0|0|0|0|0|0|0|0|0|0|0|0|0|0|0|0|0|0|0|0|0|0|0|0|0|0|0|
|bacteria_Jul11d25b_2052|1|0|0|0|0|0|0|0|0|0|0|0|0|0|0|0|0|0|0|0|0|0|0|0|0|1|0|0|0|0|0|0|0|0|0|0|0|0|0|0|0|0|0|0|0|0|0|0|0|0|0|0|0|0|0|
|bacteria_Jul11d25b_2505|1|0|0|0|0|0|0|0|0|0|0|0|0|0|0|0|0|0|0|0|0|0|0|0|0|1|0|0|0|0|0|0|0|0|0|0|0|0|0|0|0|0|0|0|0|0|0|0|0|0|0|0|0|0|0|
|bacteria_Jul11d25b_3993|1|0|0|0|0|0|0|0|0|0|0|0|0|0|0|0|0|0|0|0|0|0|0|0|0|1|0|0|0|0|0|0|0|0|0|0|0|0|0|0|0|0|0|0|0|0|0|0|0|0|0|0|0|0|0|
|bacteria_Jul11d25b_4055|1|0|0|0|0|0|0|0|0|0|0|0|0|0|0|0|0|0|0|0|0|0|0|0|0|1|0|0|0|0|0|0|0|0|0|0|0|0|0|0|0|0|0|0|0|0|0|0|0|0|0|0|0|0|0|
|bacteria_Jul11d25b_4379|1|0|0|0|0|0|0|0|0|0|0|0|0|0|0|0|0|0|0|0|0|0|0|0|0|1|0|0|0|0|0|0|0|0|0|0|0|0|0|0|0|0|0|0|0|0|0|0|0|0|0|0|0|0|0|
|bacteria_Jul11d50b_3236|1|0|0|0|0|0|0|0|0|0|0|0|0|0|0|0|0|0|0|0|0|0|0|0|0|0|0|1|0|0|0|0|0|0|0|0|0|0|0|0|0|0|0|0|0|0|0|0|0|0|0|0|0|0|0|
|bacteria_Jul11d75b_826|1|0|0|0|0|0|0|0|0|0|0|0|0|0|0|0|0|0|0|0|0|0|0|0|0|0|0|0|0|1|0|0|0|0|0|0|0|0|0|0|0|0|0|0|0|0|0|0|0|0|0|0|0|0|0|
|bacteria_Jul11d75b_1469|1|0|0|0|0|0|0|0|0|0|0|0|0|0|0|0|0|0|0|0|0|0|0|0|0|0|0|0|0|1|0|0|0|0|0|0|0|0|0|0|0|0|0|0|0|0|0|0|0|0|0|0|0|0|0|
|bacteria_Jul11d75b_2405|1|0|0|0|0|0|0|0|0|0|0|0|0|0|0|0|0|0|0|0|0|0|0|0|0|0|0|0|0|1|0|0|0|0|0|0|0|0|0|0|0|0|0|0|0|0|0|0|0|0|0|0|0|0|0|
|bacteria_Jul11d75b_1479|1|0|0|0|0|0|0|0|0|0|0|0|0|0|0|0|0|0|0|0|0|0|0|0|0|0|0|0|0|1|0|0|0|0|0|0|0|0|0|0|0|0|0|0|0|0|0|0|0|0|0|0|0|0|0|
|bacteria_Jul11d75b_1914|1|0|0|0|0|0|0|0|0|0|0|0|0|0|0|0|0|0|0|0|0|0|0|0|0|0|0|0|0|1|0|0|0|0|0|0|0|0|0|0|0|0|0|0|0|0|0|0|0|0|0|0|0|0|0|
|bacteria_Jul11d75b_2391|1|0|0|0|0|0|0|0|0|0|0|0|0|0|0|0|0|0|0|0|0|0|0|0|0|0|0|0|0|1|0|0|0|0|0|0|0|0|0|0|0|0|0|0|0|0|0|0|0|0|0|0|0|0|0|
|bacteria_Jul11d75b_3104|1|0|0|0|0|0|0|0|0|0|0|0|0|0|0|0|0|0|0|0|0|0|0|0|0|0|0|0|0|1|0|0|0|0|0|0|0|0|0|0|0|0|0|0|0|0|0|0|0|0|0|0|0|0|0|
|bacteria_Jul11d75b_3365|1|0|0|0|0|0|0|0|0|0|0|0|0|0|0|0|0|0|0|0|0|0|0|0|0|0|0|0|0|1|0|0|0|0|0|0|0|0|0|0|0|0|0|0|0|0|0|0|0|0|0|0|0|0|0|
|bacteria_Jul11d75b_3831|1|0|0|0|0|0|0|0|0|0|0|0|0|0|0|0|0|0|0|0|0|0|0|0|0|0|0|0|0|1|0|0|0|0|0|0|0|0|0|0|0|0|0|0|0|0|0|0|0|0|0|0|0|0|0|
|bacteria_Jul11d75b_3891|1|0|0|0|0|0|0|0|0|0|0|0|0|0|0|0|0|0|0|0|0|0|0|0|0|0|0|0|0|1|0|0|0|0|0|0|0|0|0|0|0|0|0|0|0|0|0|0|0|0|0|0|0|0|0|
|bacteria_Jul11d75b_3932|1|0|0|0|0|0|0|0|0|0|0|0|0|0|0|0|0|0|0|0|0|0|0|0|0|0|0|0|0|1|0|0|0|0|0|0|0|0|0|0|0|0|0|0|0|0|0|0|0|0|0|0|0|0|0|
|bacteria_Jul11d75b_4317|1|0|0|0|0|0|0|0|0|0|0|0|0|0|0|0|0|0|0|0|0|0|0|0|0|0|0|0|0|1|0|0|0|0|0|0|0|0|0|0|0|0|0|0|0|0|0|0|0|0|0|0|0|0|0|
|bacteria_Jul11d75b_4543|1|0|0|0|0|0|0|0|0|0|0|0|0|0|0|0|0|0|0|0|0|0|0|0|0|0|0|0|0|1|0|0|0|0|0|0|0|0|0|0|0|0|0|0|0|0|0|0|0|0|0|0|0|0|0|
|bacteria_Jul27d5b_6962|1|0|0|0|0|0|0|0|0|0|0|0|0|0|0|0|0|0|0|0|0|0|0|0|0|0|0|0|0|0|0|1|0|0|0|0|0|0|0|0|0|0|0|0|0|0|0|0|0|0|0|0|0|0|0|
|bacteria_Jul27d5b_8807|1|0|0|0|0|0|0|0|0|0|0|0|0|0|0|0|0|0|0|0|0|0|0|0|0|0|0|0|0|0|0|1|0|0|0|0|0|0|0|0|0|0|0|0|0|0|0|0|0|0|0|0|0|0|0|
|bacteria_Jul27d5b_20737|1|0|0|0|0|0|0|0|0|0|0|0|0|0|0|0|0|0|0|0|0|0|0|0|0|0|0|0|0|0|0|1|0|0|0|0|0|0|0|0|0|0|0|0|0|0|0|0|0|0|0|0|0|0|0|
|bacteria_Jul27d5b_22999|1|0|0|0|0|0|0|0|0|0|0|0|0|0|0|0|0|0|0|0|0|0|0|0|0|0|0|0|0|0|0|1|0|0|0|0|0|0|0|0|0|0|0|0|0|0|0|0|0|0|0|0|0|0|0|
|bacteria_Jul27d5b_25455|1|0|0|0|0|0|0|0|0|0|0|0|0|0|0|0|0|0|0|0|0|0|0|0|0|0|0|0|0|0|0|1|0|0|0|0|0|0|0|0|0|0|0|0|0|0|0|0|0|0|0|0|0|0|0|
|bacteria_Jul27d5b_27004|1|0|0|0|0|0|0|0|0|0|0|0|0|0|0|0|0|0|0|0|0|0|0|0|0|0|0|0|0|0|0|1|0|0|0|0|0|0|0|0|0|0|0|0|0|0|0|0|0|0|0|0|0|0|0|
|bacteria_Jul27d5b_33034|1|0|0|0|0|0|0|0|0|0|0|0|0|0|0|0|0|0|0|0|0|0|0|0|0|0|0|0|0|0|0|1|0|0|0|0|0|0|0|0|0|0|0|0|0|0|0|0|0|0|0|0|0|0|0|
|bacteria_Jul27d5b_40933|1|0|0|0|0|0|0|0|0|0|0|0|0|0|0|0|0|0|0|0|0|0|0|0|0|0|0|0|0|0|0|1|0|0|0|0|0|0|0|0|0|0|0|0|0|0|0|0|0|0|0|0|0|0|0|
|bacteria_Jul27d5b_43272|1|0|0|0|0|0|0|0|0|0|0|0|0|0|0|0|0|0|0|0|0|0|0|0|0|0|0|0|0|0|0|1|0|0|0|0|0|0|0|0|0|0|0|0|0|0|0|0|0|0|0|0|0|0|0|
|bacteria_Jul27d5b_44170|1|0|0|0|0|0|0|0|0|0|0|0|0|0|0|0|0|0|0|0|0|0|0|0|0|0|0|0|0|0|0|1|0|0|0|0|0|0|0|0|0|0|0|0|0|0|0|0|0|0|0|0|0|0|0|
|bacteria_Jul27d5b_49546|1|0|0|0|0|0|0|0|0|0|0|0|0|0|0|0|0|0|0|0|0|0|0|0|0|0|0|0|0|0|0|1|0|0|0|0|0|0|0|0|0|0|0|0|0|0|0|0|0|0|0|0|0|0|0|
|bacteria_Jul27d5b_51156|1|0|0|0|0|0|0|0|0|0|0|0|0|0|0|0|0|0|0|0|0|0|0|0|0|0|0|0|0|0|0|1|0|0|0|0|0|0|0|0|0|0|0|0|0|0|0|0|0|0|0|0|0|0|0|
|bacteria_Jul27d25b_6046|1|0|0|0|0|0|0|0|0|0|0|0|0|0|0|0|0|0|0|0|0|0|0|0|0|0|0|0|0|0|0|0|0|1|0|0|0|0|0|0|0|0|0|0|0|0|0|0|0|0|0|0|0|0|0|
|bacteria_Jul27d25b_2495|1|0|0|0|0|0|0|0|0|0|0|0|0|0|0|0|0|0|0|0|0|0|0|0|0|0|0|0|0|0|0|0|0|1|0|0|0|0|0|0|0|0|0|0|0|0|0|0|0|0|0|0|0|0|0|
|bacteria_Jul27d25b_2536|1|0|0|0|0|0|0|0|0|0|0|0|0|0|0|0|0|0|0|0|0|0|0|0|0|0|0|0|0|0|0|0|0|1|0|0|0|0|0|0|0|0|0|0|0|0|0|0|0|0|0|0|0|0|0|
|bacteria_Jul27d25b_2621|1|0|0|0|0|0|0|0|0|0|0|0|0|0|0|0|0|0|0|0|0|0|0|0|0|0|0|0|0|0|0|0|0|1|0|0|0|0|0|0|0|0|0|0|0|0|0|0|0|0|0|0|0|0|0|
|bacteria_Jul27d25b_2746|1|0|0|0|0|0|0|0|0|0|0|0|0|0|0|0|0|0|0|0|0|0|0|0|0|0|0|0|0|0|0|0|0|1|0|0|0|0|0|0|0|0|0|0|0|0|0|0|0|0|0|0|0|0|0|
|bacteria_Jul27d25b_2931|1|0|0|0|0|0|0|0|0|0|0|0|0|0|0|0|0|0|0|0|0|0|0|0|0|0|0|0|0|0|0|0|0|1|0|0|0|0|0|0|0|0|0|0|0|0|0|0|0|0|0|0|0|0|0|
|bacteria_Jul27d25b_3107|1|0|0|0|0|0|0|0|0|0|0|0|0|0|0|0|0|0|0|0|0|0|0|0|0|0|0|0|0|0|0|0|0|1|0|0|0|0|0|0|0|0|0|0|0|0|0|0|0|0|0|0|0|0|0|
|bacteria_Jul27d25b_4853|1|0|0|0|0|0|0|0|0|0|0|0|0|0|0|0|0|0|0|0|0|0|0|0|0|0|0|0|0|0|0|0|0|1|0|0|0|0|0|0|0|0|0|0|0|0|0|0|0|0|0|0|0|0|0|
|bacteria_Jul27d25b_5430|1|0|0|0|0|0|0|0|0|0|0|0|0|0|0|0|0|0|0|0|0|0|0|0|0|0|0|0|0|0|0|0|0|1|0|0|0|0|0|0|0|0|0|0|0|0|0|0|0|0|0|0|0|0|0|
|bacteria_Jul27d25b_6167|1|0|0|0|0|0|0|0|0|0|0|0|0|0|0|0|0|0|0|0|0|0|0|0|0|0|0|0|0|0|0|0|0|1|0|0|0|0|0|0|0|0|0|0|0|0|0|0|0|0|0|0|0|0|0|
|bacteria_Jul27d25b_6593|1|0|0|0|0|0|0|0|0|0|0|0|0|0|0|0|0|0|0|0|0|0|0|0|0|0|0|0|0|0|0|0|0|1|0|0|0|0|0|0|0|0|0|0|0|0|0|0|0|0|0|0|0|0|0|
|bacteria_Jul27d25b_6692|1|0|0|0|0|0|0|0|0|0|0|0|0|0|0|0|0|0|0|0|0|0|0|0|0|0|0|0|0|0|0|0|0|1|0|0|0|0|0|0|0|0|0|0|0|0|0|0|0|0|0|0|0|0|0|
|bacteria_Jul27d25b_6965|1|0|0|0|0|0|0|0|0|0|0|0|0|0|0|0|0|0|0|0|0|0|0|0|0|0|0|0|0|0|0|0|0|1|0|0|0|0|0|0|0|0|0|0|0|0|0|0|0|0|0|0|0|0|0|
|bacteria_Jul27d75b_5032|1|0|0|0|0|0|0|0|0|0|0|0|0|0|0|0|0|0|0|0|0|0|0|0|0|0|0|0|0|0|0|0|0|0|0|1|0|0|0|0|0|0|0|0|0|0|0|0|0|0|0|0|0|0|0|
|bacteria_Jul27d75b_1277|1|0|0|0|0|0|0|0|0|0|0|0|0|0|0|0|0|0|0|0|0|0|0|0|0|0|0|0|0|0|0|0|0|0|0|1|0|0|0|0|0|0|0|0|0|0|0|0|0|0|0|0|0|0|0|
|bacteria_Jul27d75b_1369|1|0|0|0|0|0|0|0|0|0|0|0|0|0|0|0|0|0|0|0|0|0|0|0|0|0|0|0|0|0|0|0|0|0|0|1|0|0|0|0|0|0|0|0|0|0|0|0|0|0|0|0|0|0|0|
|bacteria_Jul27d75b_2223|1|0|0|0|0|0|0|0|0|0|0|0|0|0|0|0|0|0|0|0|0|0|0|0|0|0|0|0|0|0|0|0|0|0|0|1|0|0|0|0|0|0|0|0|0|0|0|0|0|0|0|0|0|0|0|
|bacteria_Jul27d75b_3048|1|0|0|0|0|0|0|0|0|0|0|0|0|0|0|0|0|0|0|0|0|0|0|0|0|0|0|0|0|0|0|0|0|0|0|1|0|0|0|0|0|0|0|0|0|0|0|0|0|0|0|0|0|0|0|
|bacteria_Jul27d75b_3433|1|0|0|0|0|0|0|0|0|0|0|0|0|0|0|0|0|0|0|0|0|0|0|0|0|0|0|0|0|0|0|0|0|0|0|1|0|0|0|0|0|0|0|0|0|0|0|0|0|0|0|0|0|0|0|
|bacteria_Jul27d75b_5152|1|0|0|0|0|0|0|0|0|0|0|0|0|0|0|0|0|0|0|0|0|0|0|0|0|0|0|0|0|0|0|0|0|0|0|1|0|0|0|0|0|0|0|0|0|0|0|0|0|0|0|0|0|0|0|
|bacteria_Jul27d75b_5195|1|0|0|0|0|0|0|0|0|0|0|0|0|0|0|0|0|0|0|0|0|0|0|0|0|0|0|0|0|0|0|0|0|0|0|1|0|0|0|0|0|0|0|0|0|0|0|0|0|0|0|0|0|0|0|
|bacteria_Jul27d75b_5782|1|0|0|0|0|0|0|0|0|0|0|0|0|0|0|0|0|0|0|0|0|0|0|0|0|0|0|0|0|0|0|0|0|0|0|1|0|0|0|0|0|0|0|0|0|0|0|0|0|0|0|0|0|0|0|
|bacteria_Jul27d75b_5918|1|0|0|0|0|0|0|0|0|0|0|0|0|0|0|0|0|0|0|0|0|0|0|0|0|0|0|0|0|0|0|0|0|0|0|1|0|0|0|0|0|0|0|0|0|0|0|0|0|0|0|0|0|0|0|
|bacteria_Jul27d75b_6152|1|0|0|0|0|0|0|0|0|0|0|0|0|0|0|0|0|0|0|0|0|0|0|0|0|0|0|0|0|0|0|0|0|0|0|1|0|0|0|0|0|0|0|0|0|0|0|0|0|0|0|0|0|0|0|
|bacteria_Jul27d75b_6903|1|0|0|0|0|0|0|0|0|0|0|0|0|0|0|0|0|0|0|0|0|0|0|0|0|0|0|0|0|0|0|0|0|0|0|1|0|0|0|0|0|0|0|0|0|0|0|0|0|0|0|0|0|0|0|
|bacteria_Jul27d75b_7401|1|0|0|0|0|0|0|0|0|0|0|0|0|0|0|0|0|0|0|0|0|0|0|0|0|0|0|0|0|0|0|0|0|0|0|1|0|0|0|0|0|0|0|0|0|0|0|0|0|0|0|0|0|0|0|
|bacteria_Jun8d25b_12666|1|0|0|0|0|0|0|0|0|1|0|0|0|0|0|0|0|0|0|0|0|0|0|0|0|0|0|0|0|0|0|0|0|0|0|0|0|0|0|0|0|0|0|0|0|0|0|0|0|0|0|0|0|0|0|
|bacteria_Jun8d25b_31827|1|0|0|0|0|0|0|0|0|1|0|0|0|0|0|0|0|0|0|0|0|0|0|0|0|0|0|0|0|0|0|0|0|0|0|0|0|0|0|0|0|0|0|0|0|0|0|0|0|0|0|0|0|0|0|
|bacteria_Jun8d25b_34108|1|0|0|0|0|0|0|0|0|1|0|0|0|0|0|0|0|0|0|0|0|0|0|0|0|0|0|0|0|0|0|0|0|0|0|0|0|0|0|0|0|0|0|0|0|0|0|0|0|0|0|0|0|0|0|
|bacteria_Jun8d25b_39401|1|0|0|0|0|0|0|0|0|1|0|0|0|0|0|0|0|0|0|0|0|0|0|0|0|0|0|0|0|0|0|0|0|0|0|0|0|0|0|0|0|0|0|0|0|0|0|0|0|0|0|0|0|0|0|
|bacteria_Jun8d25b_41365|1|0|0|0|0|0|0|0|0|1|0|0|0|0|0|0|0|0|0|0|0|0|0|0|0|0|0|0|0|0|0|0|0|0|0|0|0|0|0|0|0|0|0|0|0|0|0|0|0|0|0|0|0|0|0|
|bacteria_Jun8d25b_46307|1|0|0|0|0|0|0|0|0|1|0|0|0|0|0|0|0|0|0|0|0|0|0|0|0|0|0|0|0|0|0|0|0|0|0|0|0|0|0|0|0|0|0|0|0|0|0|0|0|0|0|0|0|0|0|
|bacteria_Jun8d25b_49754|1|0|0|0|0|0|0|0|0|1|0|0|0|0|0|0|0|0|0|0|0|0|0|0|0|0|0|0|0|0|0|0|0|0|0|0|0|0|0|0|0|0|0|0|0|0|0|0|0|0|0|0|0|0|0|
|bacteria_Jun8d25b_54041|1|0|0|0|0|0|0|0|0|1|0|0|0|0|0|0|0|0|0|0|0|0|0|0|0|0|0|0|0|0|0|0|0|0|0|0|0|0|0|0|0|0|0|0|0|0|0|0|0|0|0|0|0|0|0|
|bacteria_Jun8d25b_68066|1|0|0|0|0|0|0|0|0|1|0|0|0|0|0|0|0|0|0|0|0|0|0|0|0|0|0|0|0|0|0|0|0|0|0|0|0|0|0|0|0|0|0|0|0|0|0|0|0|0|0|0|0|0|0|
|bacteria_Jun8d25b_73990|1|0|0|0|0|0|0|0|0|1|0|0|0|0|0|0|0|0|0|0|0|0|0|0|0|0|0|0|0|0|0|0|0|0|0|0|0|0|0|0|0|0|0|0|0|0|0|0|0|0|0|0|0|0|0|
|bacteria_Jun8d50b_4413|1|0|0|0|0|0|0|0|0|0|0|1|0|0|0|0|0|0|0|0|0|0|0|0|0|0|0|0|0|0|0|0|0|0|0|0|0|0|0|0|0|0|0|0|0|0|0|0|0|0|0|0|0|0|0|
|bacteria_Jun8d50b_16298|1|0|0|0|0|0|0|0|0|0|0|1|0|0|0|0|0|0|0|0|0|0|0|0|0|0|0|0|0|0|0|0|0|0|0|0|0|0|0|0|0|0|0|0|0|0|0|0|0|0|0|0|0|0|0|
|bacteria_Jun8d50b_30780|1|0|0|0|0|0|0|0|0|0|0|1|0|0|0|0|0|0|0|0|0|0|0|0|0|0|0|0|0|0|0|0|0|0|0|0|0|0|0|0|0|0|0|0|0|0|0|0|0|0|0|0|0|0|0|
|bacteria_Jun8d75b_53038|1|0|0|0|0|0|0|0|0|0|0|0|0|1|0|0|0|0|0|0|0|0|0|0|0|0|0|0|0|0|0|0|0|0|0|0|0|0|0|0|0|0|0|0|0|0|0|0|0|0|0|0|0|0|0|
|bacteria_Jun22d0b_5193|1|0|0|0|0|0|0|0|0|0|0|0|0|0|0|1|0|0|0|0|0|0|0|0|0|0|0|0|0|0|0|0|0|0|0|0|0|0|0|0|0|0|0|0|0|0|0|0|0|0|0|0|0|0|0|
|bacteria_Jun22d0b_14922|1|0|0|0|0|0|0|0|0|0|0|0|0|0|0|1|0|0|0|0|0|0|0|0|0|0|0|0|0|0|0|0|0|0|0|0|0|0|0|0|0|0|0|0|0|0|0|0|0|0|0|0|0|0|0|
|bacteria_Jun22d0b_34569|1|0|0|0|0|0|0|0|0|0|0|0|0|0|0|1|0|0|0|0|0|0|0|0|0|0|0|0|0|0|0|0|0|0|0|0|0|0|0|0|0|0|0|0|0|0|0|0|0|0|0|0|0|0|0|
|bacteria_Jun22d50b_2567|1|0|0|0|0|0|0|0|0|0|0|0|0|0|0|0|0|1|0|0|0|0|0|0|0|0|0|0|0|0|0|0|0|0|0|0|0|0|0|0|0|0|0|0|0|0|0|0|0|0|0|0|0|0|0|
|bacteria_Jun22d50b_1345|1|0|0|0|0|0|0|0|0|0|0|0|0|0|0|0|0|1|0|0|0|0|0|0|0|0|0|0|0|0|0|0|0|0|0|0|0|0|0|0|0|0|0|0|0|0|0|0|0|0|0|0|0|0|0|
|bacteria_Jun22d50b_1537|1|0|0|0|0|0|0|0|0|0|0|0|0|0|0|0|0|1|0|0|0|0|0|0|0|0|0|0|0|0|0|0|0|0|0|0|0|0|0|0|0|0|0|0|0|0|0|0|0|0|0|0|0|0|0|
|bacteria_Jun22d50b_2693|1|0|0|0|0|0|0|0|0|0|0|0|0|0|0|0|0|1|0|0|0|0|0|0|0|0|0|0|0|0|0|0|0|0|0|0|0|0|0|0|0|0|0|0|0|0|0|0|0|0|0|0|0|0|0|
|bacteria_Jun22d50b_3665|1|0|0|0|0|0|0|0|0|0|0|0|0|0|0|0|0|1|0|0|0|0|0|0|0|0|0|0|0|0|0|0|0|0|0|0|0|0|0|0|0|0|0|0|0|0|0|0|0|0|0|0|0|0|0|
|bacteria_Jun22d75b_4712|1|0|0|0|0|0|0|0|0|0|0|0|0|0|0|0|0|0|0|1|0|0|0|0|0|0|0|0|0|0|0|0|0|0|0|0|0|0|0|0|0|0|0|0|0|0|0|0|0|0|0|0|0|0|0|
|bacteria_Jun22d75b_4086|1|0|0|0|0|0|0|0|0|0|0|0|0|0|0|0|0|0|0|1|0|0|0|0|0|0|0|0|0|0|0|0|0|0|0|0|0|0|0|0|0|0|0|0|0|0|0|0|0|0|0|0|0|0|0|
|bacteria_Jun22d75b_4256|1|0|0|0|0|0|0|0|0|0|0|0|0|0|0|0|0|0|0|1|0|0|0|0|0|0|0|0|0|0|0|0|0|0|0|0|0|0|0|0|0|0|0|0|0|0|0|0|0|0|0|0|0|0|0|
|bacteria_Jun22d75b_4572|1|0|0|0|0|0|0|0|0|0|0|0|0|0|0|0|0|0|0|1|0|0|0|0|0|0|0|0|0|0|0|0|0|0|0|0|0|0|0|0|0|0|0|0|0|0|0|0|0|0|0|0|0|0|0|
|bacteria_Jun22d75b_5134|1|0|0|0|0|0|0|0|0|0|0|0|0|0|0|0|0|0|0|1|0|0|0|0|0|0|0|0|0|0|0|0|0|0|0|0|0|0|0|0|0|0|0|0|0|0|0|0|0|0|0|0|0|0|0|
|bacteria_Jun22d75b_6256|1|0|0|0|0|0|0|0|0|0|0|0|0|0|0|0|0|0|0|1|0|0|0|0|0|0|0|0|0|0|0|0|0|0|0|0|0|0|0|0|0|0|0|0|0|0|0|0|0|0|0|0|0|0|0|
|bacteria_May12d5b_1486|1|1|0|0|0|0|0|0|0|0|0|0|0|0|0|0|0|0|0|0|0|0|0|0|0|0|0|0|0|0|0|0|0|0|0|0|0|0|0|0|0|0|0|0|0|0|0|0|0|0|0|0|0|0|0|
|bacteria_May12d5b_4758|1|1|0|0|0|0|0|0|0|0|0|0|0|0|0|0|0|0|0|0|0|0|0|0|0|0|0|0|0|0|0|0|0|0|0|0|0|0|0|0|0|0|0|0|0|0|0|0|0|0|0|0|0|0|0|
|bacteria_May12d25b_147|1|0|0|1|0|0|0|0|0|0|0|0|0|0|0|0|0|0|0|0|0|0|0|0|0|0|0|0|0|0|0|0|0|0|0|0|0|0|0|0|0|0|0|0|0|0|0|0|0|0|0|0|0|0|0|
|bacteria_May12d25b_357|1|0|0|1|0|0|0|0|0|0|0|0|0|0|0|0|0|0|0|0|0|0|0|0|0|0|0|0|0|0|0|0|0|0|0|0|0|0|0|0|0|0|0|0|0|0|0|0|0|0|0|0|0|0|0|
|bacteria_May12d25b_951|1|0|0|1|0|0|0|0|0|0|0|0|0|0|0|0|0|0|0|0|0|0|0|0|0|0|0|0|0|0|0|0|0|0|0|0|0|0|0|0|0|0|0|0|0|0|0|0|0|0|0|0|0|0|0|
|bacteria_May12d25b_502|1|0|0|1|0|0|0|0|0|0|0|0|0|0|0|0|0|0|0|0|0|0|0|0|0|0|0|0|0|0|0|0|0|0|0|0|0|0|0|0|0|0|0|0|0|0|0|0|0|0|0|0|0|0|0|
|bacteria_May12d25b_564|1|0|0|1|0|0|0|0|0|0|0|0|0|0|0|0|0|0|0|0|0|0|0|0|0|0|0|0|0|0|0|0|0|0|0|0|0|0|0|0|0|0|0|0|0|0|0|0|0|0|0|0|0|0|0|
|bacteria_May12d25b_598|1|0|0|1|0|0|0|0|0|0|0|0|0|0|0|0|0|0|0|0|0|0|0|0|0|0|0|0|0|0|0|0|0|0|0|0|0|0|0|0|0|0|0|0|0|0|0|0|0|0|0|0|0|0|0|
|bacteria_May12d25b_684|1|0|0|1|0|0|0|0|0|0|0|0|0|0|0|0|0|0|0|0|0|0|0|0|0|0|0|0|0|0|0|0|0|0|0|0|0|0|0|0|0|0|0|0|0|0|0|0|0|0|0|0|0|0|0|
|bacteria_May12d50b_801|1|0|0|0|0|1|0|0|0|0|0|0|0|0|0|0|0|0|0|0|0|0|0|0|0|0|0|0|0|0|0|0|0|0|0|0|0|0|0|0|0|0|0|0|0|0|0|0|0|0|0|0|0|0|0|
|bacteria_May12d50b_112|1|0|0|0|0|1|0|0|0|0|0|0|0|0|0|0|0|0|0|0|0|0|0|0|0|0|0|0|0|0|0|0|0|0|0|0|0|0|0|0|0|0|0|0|0|0|0|0|0|0|0|0|0|0|0|
|bacteria_May12d50b_211|1|0|0|0|0|1|0|0|0|0|0|0|0|0|0|0|0|0|0|0|0|0|0|0|0|0|0|0|0|0|0|0|0|0|0|0|0|0|0|0|0|0|0|0|0|0|0|0|0|0|0|0|0|0|0|
|bacteria_May12d50b_217|1|0|0|0|0|1|0|0|0|0|0|0|0|0|0|0|0|0|0|0|0|0|0|0|0|0|0|0|0|0|0|0|0|0|0|0|0|0|0|0|0|0|0|0|0|0|0|0|0|0|0|0|0|0|0|
|bacteria_May12d50b_238|1|0|0|0|0|1|0|0|0|0|0|0|0|0|0|0|0|0|0|0|0|0|0|0|0|0|0|0|0|0|0|0|0|0|0|0|0|0|0|0|0|0|0|0|0|0|0|0|0|0|0|0|0|0|0|
|bacteria_May12d50b_472|1|0|0|0|0|1|0|0|0|0|0|0|0|0|0|0|0|0|0|0|0|0|0|0|0|0|0|0|0|0|0|0|0|0|0|0|0|0|0|0|0|0|0|0|0|0|0|0|0|0|0|0|0|0|0|
|bacteria_May12d50b_510|1|0|0|0|0|1|0|0|0|0|0|0|0|0|0|0|0|0|0|0|0|0|0|0|0|0|0|0|0|0|0|0|0|0|0|0|0|0|0|0|0|0|0|0|0|0|0|0|0|0|0|0|0|0|0|
|bacteria_May12d75b_388|1|0|0|0|0|0|0|1|0|0|0|0|0|0|0|0|0|0|0|0|0|0|0|0|0|0|0|0|0|0|0|0|0|0|0|0|0|0|0|0|0|0|0|0|0|0|0|0|0|0|0|0|0|0|0|
|bacteria_Sep8d0b_25219|1|0|0|0|0|0|0|0|0|0|0|0|0|0|0|0|0|0|0|0|0|0|0|0|0|0|0|0|0|0|0|0|0|0|0|0|0|0|0|0|0|0|0|0|0|0|0|1|0|0|0|0|0|0|0|
|bacteria_Sep8d0b_26669|1|0|0|0|0|0|0|0|0|0|0|0|0|0|0|0|0|0|0|0|0|0|0|0|0|0|0|0|0|0|0|0|0|0|0|0|0|0|0|0|0|0|0|0|0|0|0|1|0|0|0|0|0|0|0|
|bacteria_Sep8d0b_36515|1|0|0|0|0|0|0|0|0|0|0|0|0|0|0|0|0|0|0|0|0|0|0|0|0|0|0|0|0|0|0|0|0|0|0|0|0|0|0|0|0|0|0|0|0|0|0|1|0|0|0|0|0|0|0|
|bacteria_Sep8d5b_3542|1|0|0|0|0|0|0|0|0|0|0|0|0|0|0|0|0|0|0|0|0|0|0|0|0|0|0|0|0|0|0|0|0|0|0|0|0|0|0|0|0|0|0|0|0|0|0|0|0|1|0|0|0|0|0|
|bacteria_Sep8d5b_10378|1|0|0|0|0|0|0|0|0|0|0|0|0|0|0|0|0|0|0|0|0|0|0|0|0|0|0|0|0|0|0|0|0|0|0|0|0|0|0|0|0|0|0|0|0|0|0|0|0|1|0|0|0|0|0|
|bacteria_Sep8d5b_16031|1|0|0|0|0|0|0|0|0|0|0|0|0|0|0|0|0|0|0|0|0|0|0|0|0|0|0|0|0|0|0|0|0|0|0|0|0|0|0|0|0|0|0|0|0|0|0|0|0|1|0|0|0|0|0|
|bacteria_Sep8d5b_24502|1|0|0|0|0|0|0|0|0|0|0|0|0|0|0|0|0|0|0|0|0|0|0|0|0|0|0|0|0|0|0|0|0|0|0|0|0|0|0|0|0|0|0|0|0|0|0|0|0|1|0|0|0|0|0|
|bacteria_Sep8d5b_39511|1|0|0|0|0|0|0|0|0|0|0|0|0|0|0|0|0|0|0|0|0|0|0|0|0|0|0|0|0|0|0|0|0|0|0|0|0|0|0|0|0|0|0|0|0|0|0|0|0|1|0|0|0|0|0|
|bacteria_Sep8d5b_42190|1|0|0|0|0|0|0|0|0|0|0|0|0|0|0|0|0|0|0|0|0|0|0|0|0|0|0|0|0|0|0|0|0|0|0|0|0|0|0|0|0|0|0|0|0|0|0|0|0|1|0|0|0|0|0|
|bacteria_Sep8d5b_43629|1|0|0|0|0|0|0|0|0|0|0|0|0|0|0|0|0|0|0|0|0|0|0|0|0|0|0|0|0|0|0|0|0|0|0|0|0|0|0|0|0|0|0|0|0|0|0|0|0|1|0|0|0|0|0|
|bacteria_Sep8d5b_43979|1|0|0|0|0|0|0|0|0|0|0|0|0|0|0|0|0|0|0|0|0|0|0|0|0|0|0|0|0|0|0|0|0|0|0|0|0|0|0|0|0|0|0|0|0|0|0|0|0|1|0|0|0|0|0|
|bacteria_Sep8d5b_45292|1|0|0|0|0|0|0|0|0|0|0|0|0|0|0|0|0|0|0|0|0|0|0|0|0|0|0|0|0|0|0|0|0|0|0|0|0|0|0|0|0|0|0|0|0|0|0|0|0|1|0|0|0|0|0|
|bacteria_Sep8d5b_50208|1|0|0|0|0|0|0|0|0|0|0|0|0|0|0|0|0|0|0|0|0|0|0|0|0|0|0|0|0|0|0|0|0|0|0|0|0|0|0|0|0|0|0|0|0|0|0|0|0|1|0|0|0|0|0|
|bacteria_Sep8d5b_70638|1|0|0|0|0|0|0|0|0|0|0|0|0|0|0|0|0|0|0|0|0|0|0|0|0|0|0|0|0|0|0|0|0|0|0|0|0|0|0|0|0|0|0|0|0|0|0|0|0|1|0|0|0|0|0|
|bacteria_Sep8d5b_70779|1|0|0|0|0|0|0|0|0|0|0|0|0|0|0|0|0|0|0|0|0|0|0|0|0|0|0|0|0|0|0|0|0|0|0|0|0|0|0|0|0|0|0|0|0|0|0|0|0|1|0|0|0|0|0|
|bacteria_Sep8d25b_1033|1|0|0|0|0|0|0|0|0|0|0|0|0|0|0|0|0|0|0|0|0|0|0|0|0|0|0|0|0|0|0|0|0|0|0|0|0|0|0|0|0|0|0|0|0|0|0|0|0|0|0|1|0|0|0|
|bacteria_Sep8d25b_1691|1|0|0|0|0|0|0|0|0|0|0|0|0|0|0|0|0|0|0|0|0|0|0|0|0|0|0|0|0|0|0|0|0|0|0|0|0|0|0|0|0|0|0|0|0|0|0|0|0|0|0|1|0|0|0|
|bacteria_Sep8d25b_2409|1|0|0|0|0|0|0|0|0|0|0|0|0|0|0|0|0|0|0|0|0|0|0|0|0|0|0|0|0|0|0|0|0|0|0|0|0|0|0|0|0|0|0|0|0|0|0|0|0|0|0|1|0|0|0|
|bacteria_Sep8d25b_3603|1|0|0|0|0|0|0|0|0|0|0|0|0|0|0|0|0|0|0|0|0|0|0|0|0|0|0|0|0|0|0|0|0|0|0|0|0|0|0|0|0|0|0|0|0|0|0|0|0|0|0|1|0|0|0|
|bacteria_Sep8d50b_3989|1|0|0|0|0|0|0|0|0|0|0|0|0|0|0|0|0|0|0|0|0|0|0|0|0|0|0|0|0|0|0|0|0|0|0|0|0|0|0|0|0|0|0|0|0|0|0|0|0|0|0|0|0|1|0|
|bacteria_Sep8d50b_5041|1|0|0|0|0|0|0|0|0|0|0|0|0|0|0|0|0|0|0|0|0|0|0|0|0|0|0|0|0|0|0|0|0|0|0|0|0|0|0|0|0|0|0|0|0|0|0|0|0|0|0|0|0|1|0|
|virus_Aug5.0v_789|1|0|0|0|0|0|0|0|0|0|0|0|0|0|0|0|0|0|0|0|0|0|0|0|0|0|0|0|0|0|0|0|0|0|0|0|0|0|1|0|0|0|0|0|0|0|0|0|0|0|0|0|0|0|0|
|virus_Aug5.0v_1297|1|0|0|0|0|0|0|0|0|0|0|0|0|0|0|0|0|0|0|0|0|0|0|0|0|0|0|0|0|0|0|0|0|0|0|0|0|0|1|0|0|0|0|0|0|0|0|0|0|0|0|0|0|0|0|
|virus_Aug5.0v_1593|1|0|0|0|0|0|0|0|0|0|0|0|0|0|0|0|0|0|0|0|0|0|0|0|0|0|0|0|0|0|0|0|0|0|0|0|0|0|1|0|0|0|0|0|0|0|0|0|0|0|0|0|0|0|0|
|virus_Aug5.0v_1658|1|0|0|0|0|0|0|0|0|0|0|0|0|0|0|0|0|0|0|0|0|0|0|0|0|0|0|0|0|0|0|0|0|0|0|0|0|0|1|0|0|0|0|0|0|0|0|0|0|0|0|0|0|0|0|
|virus_Aug5.0v_2120|1|0|0|0|0|0|0|0|0|0|0|0|0|0|0|0|0|0|0|0|0|0|0|0|0|0|0|0|0|0|0|0|0|0|0|0|0|0|1|0|0|0|0|0|0|0|0|0|0|0|0|0|0|0|0|
|virus_Aug5.0v_2155|1|0|0|0|0|0|0|0|0|0|0|0|0|0|0|0|0|0|0|0|0|0|0|0|0|0|0|0|0|0|0|0|0|0|0|0|0|0|1|0|0|0|0|0|0|0|0|0|0|0|0|0|0|0|0|
|virus_Aug5.0v_2189|1|0|0|0|0|0|0|0|0|0|0|0|0|0|0|0|0|0|0|0|0|0|0|0|0|0|0|0|0|0|0|0|0|0|0|0|0|0|1|0|0|0|0|0|0|0|0|0|0|0|0|0|0|0|0|
|virus_Aug5.0v_3190|1|0|0|0|0|0|0|0|0|0|0|0|0|0|0|0|0|0|0|0|0|0|0|0|0|0|0|0|0|0|0|0|0|0|0|0|0|0|1|0|0|0|0|0|0|0|0|0|0|0|0|0|0|0|0|
|virus_Aug5.0v_3240|1|0|0|0|0|0|0|0|0|0|0|0|0|0|0|0|0|0|0|0|0|0|0|0|0|0|0|0|0|0|0|0|0|0|0|0|0|0|1|0|0|0|0|0|0|0|0|0|0|0|0|0|0|0|0|
|virus_Aug5.0v_3609|1|0|0|0|0|0|0|0|0|0|0|0|0|0|0|0|0|0|0|0|0|0|0|0|0|0|0|0|0|0|0|0|0|0|0|0|0|0|1|0|0|0|0|0|0|0|0|0|0|0|0|0|0|0|0|
|virus_Aug5.0v_3795|1|0|0|0|0|0|0|0|0|0|0|0|0|0|0|0|0|0|0|0|0|0|0|0|0|0|0|0|0|0|0|0|0|0|0|0|0|0|1|0|0|0|0|0|0|0|0|0|0|0|0|0|0|0|0|
|virus_Aug5.0v_4066|1|0|0|0|0|0|0|0|0|0|0|0|0|0|0|0|0|0|0|0|0|0|0|0|0|0|0|0|0|0|0|0|0|0|0|0|0|0|1|0|0|0|0|0|0|0|0|0|0|0|0|0|0|0|0|
|virus_Aug5.0v_4136|1|0|0|0|0|0|0|0|0|0|0|0|0|0|0|0|0|0|0|0|0|0|0|0|0|0|0|0|0|0|0|0|0|0|0|0|0|0|1|0|0|0|0|0|0|0|0|0|0|0|0|0|0|0|0|
|virus_Aug5.0v_4370|1|0|0|0|0|0|0|0|0|0|0|0|0|0|0|0|0|0|0|0|0|0|0|0|0|0|0|0|0|0|0|0|0|0|0|0|0|0|1|0|0|0|0|0|0|0|0|0|0|0|0|0|0|0|0|
|virus_Aug5.0v_4499|1|0|0|0|0|0|0|0|0|0|0|0|0|0|0|0|0|0|0|0|0|0|0|0|0|0|0|0|0|0|0|0|0|0|0|0|0|0|1|0|0|0|0|0|0|0|0|0|0|0|0|0|0|0|0|
|virus_Aug5.0v_4554|1|0|0|0|0|0|0|0|0|0|0|0|0|0|0|0|0|0|0|0|0|0|0|0|0|0|0|0|0|0|0|0|0|0|0|0|0|0|1|0|0|0|0|0|0|0|0|0|0|0|0|0|0|0|0|
|virus_Aug5.0v_5152|1|0|0|0|0|0|0|0|0|0|0|0|0|0|0|0|0|0|0|0|0|0|0|0|0|0|0|0|0|0|0|0|0|0|0|0|0|0|1|0|0|0|0|0|0|0|0|0|0|0|0|0|0|0|0|
|virus_Aug5.0v_5172|1|0|0|0|0|0|0|0|0|0|0|0|0|0|0|0|0|0|0|0|0|0|0|0|0|0|0|0|0|0|0|0|0|0|0|0|0|0|1|0|0|0|0|0|0|0|0|0|0|0|0|0|0|0|0|
|virus_Aug5.0v_5204|1|0|0|0|0|0|0|0|0|0|0|0|0|0|0|0|0|0|0|0|0|0|0|0|0|0|0|0|0|0|0|0|0|0|0|0|0|0|1|0|0|0|0|0|0|0|0|0|0|0|0|0|0|0|0|
|virus_Aug5.0v_5230|1|0|0|0|0|0|0|0|0|0|0|0|0|0|0|0|0|0|0|0|0|0|0|0|0|0|0|0|0|0|0|0|0|0|0|0|0|0|1|0|0|0|0|0|0|0|0|0|0|0|0|0|0|0|0|
|virus_Aug5.0v_5247|1|0|0|0|0|0|0|0|0|0|0|0|0|0|0|0|0|0|0|0|0|0|0|0|0|0|0|0|0|0|0|0|0|0|0|0|0|0|1|0|0|0|0|0|0|0|0|0|0|0|0|0|0|0|0|
|virus_Aug5.0v_5393|1|0|0|0|0|0|0|0|0|0|0|0|0|0|0|0|0|0|0|0|0|0|0|0|0|0|0|0|0|0|0|0|0|0|0|0|0|0|1|0|0|0|0|0|0|0|0|0|0|0|0|0|0|0|0|
|virus_Aug5.5v_2571|1|0|0|0|0|0|0|0|0|0|0|0|0|0|0|0|0|0|0|0|0|0|0|0|0|0|0|0|0|0|0|0|0|0|0|0|0|0|0|0|1|0|0|0|0|0|0|0|0|0|0|0|0|0|0|
|virus_Aug5.5v_3543|1|0|0|0|0|0|0|0|0|0|0|0|0|0|0|0|0|0|0|0|0|0|0|0|0|0|0|0|0|0|0|0|0|0|0|0|0|0|0|0|1|0|0|0|0|0|0|0|0|0|0|0|0|0|0|
|virus_Aug5.5v_26014|1|0|0|0|0|0|0|0|0|0|0|0|0|0|0|0|0|0|0|0|0|0|0|0|0|0|0|0|0|0|0|0|0|0|0|0|0|0|0|0|1|0|0|0|0|0|0|0|0|0|0|0|0|0|0|
|virus_Aug5.5v_34765|1|0|0|0|0|0|0|0|0|0|0|0|0|0|0|0|0|0|0|0|0|0|0|0|0|0|0|0|0|0|0|0|0|0|0|0|0|0|0|0|1|0|0|0|0|0|0|0|0|0|0|0|0|0|0|
|virus_Aug5.5v_40500|1|0|0|0|0|0|0|0|0|0|0|0|0|0|0|0|0|0|0|0|0|0|0|0|0|0|0|0|0|0|0|0|0|0|0|0|0|0|0|0|1|0|0|0|0|0|0|0|0|0|0|0|0|0|0|
|virus_Aug5.5v_40891|1|0|0|0|0|0|0|0|0|0|0|0|0|0|0|0|0|0|0|0|0|0|0|0|0|0|0|0|0|0|0|0|0|0|0|0|0|0|0|0|1|0|0|0|0|0|0|0|0|0|0|0|0|0|0|
|virus_Aug5.25v_5930|1|0|0|0|0|0|0|0|0|0|0|0|0|0|0|0|0|0|0|0|0|0|0|0|0|0|0|0|0|0|0|0|0|0|0|0|0|0|0|0|0|0|1|0|0|0|0|0|0|0|0|0|0|0|0|
|virus_Aug5.25v_6480|1|0|0|0|0|0|0|0|0|0|0|0|0|0|0|0|0|0|0|0|0|0|0|0|0|0|0|0|0|0|0|0|0|0|0|0|0|0|0|0|0|0|1|0|0|0|0|0|0|0|0|0|0|0|0|
|virus_Aug5.25v_7470|1|0|0|0|0|0|0|0|0|0|0|0|0|0|0|0|0|0|0|0|0|0|0|0|0|0|0|0|0|0|0|0|0|0|0|0|0|0|0|0|0|0|1|0|0|0|0|0|0|0|0|0|0|0|0|
|virus_Aug5.25v_11359|1|0|0|0|0|0|0|0|0|0|0|0|0|0|0|0|0|0|0|0|0|0|0|0|0|0|0|0|0|0|0|0|0|0|0|0|0|0|0|0|0|0|1|0|0|0|0|0|0|0|0|0|0|0|0|
|virus_Aug5.25v_13128|1|0|0|0|0|0|0|0|0|0|0|0|0|0|0|0|0|0|0|0|0|0|0|0|0|0|0|0|0|0|0|0|0|0|0|0|0|0|0|0|0|0|1|0|0|0|0|0|0|0|0|0|0|0|0|
|virus_Aug5.25v_13924|1|0|0|0|0|0|0|0|0|0|0|0|0|0|0|0|0|0|0|0|0|0|0|0|0|0|0|0|0|0|0|0|0|0|0|0|0|0|0|0|0|0|1|0|0|0|0|0|0|0|0|0|0|0|0|
|virus_Aug5.25v_14876|1|0|0|0|0|0|0|0|0|0|0|0|0|0|0|0|0|0|0|0|0|0|0|0|0|0|0|0|0|0|0|0|0|0|0|0|0|0|0|0|0|0|1|0|0|0|0|0|0|0|0|0|0|0|0|
|virus_Aug5.25v_18357|1|0|0|0|0|0|0|0|0|0|0|0|0|0|0|0|0|0|0|0|0|0|0|0|0|0|0|0|0|0|0|0|0|0|0|0|0|0|0|0|0|0|1|0|0|0|0|0|0|0|0|0|0|0|0|
|virus_Aug5.25v_18580|1|0|0|0|0|0|0|0|0|0|0|0|0|0|0|0|0|0|0|0|0|0|0|0|0|0|0|0|0|0|0|0|0|0|0|0|0|0|0|0|0|0|1|0|0|0|0|0|0|0|0|0|0|0|0|
|virus_Aug5.25v_18767|1|0|0|0|0|0|0|0|0|0|0|0|0|0|0|0|0|0|0|0|0|0|0|0|0|0|0|0|0|0|0|0|0|0|0|0|0|0|0|0|0|0|1|0|0|0|0|0|0|0|0|0|0|0|0|
|virus_Aug5.25v_19083|1|0|0|0|0|0|0|0|0|0|0|0|0|0|0|0|0|0|0|0|0|0|0|0|0|0|0|0|0|0|0|0|0|0|0|0|0|0|0|0|0|0|1|0|0|0|0|0|0|0|0|0|0|0|0|
|virus_Aug5.25v_22167|1|0|0|0|0|0|0|0|0|0|0|0|0|0|0|0|0|0|0|0|0|0|0|0|0|0|0|0|0|0|0|0|0|0|0|0|0|0|0|0|0|0|1|0|0|0|0|0|0|0|0|0|0|0|0|
|virus_Aug5.25v_23454|1|0|0|0|0|0|0|0|0|0|0|0|0|0|0|0|0|0|0|0|0|0|0|0|0|0|0|0|0|0|0|0|0|0|0|0|0|0|0|0|0|0|1|0|0|0|0|0|0|0|0|0|0|0|0|
|virus_Aug5.25v_27923|1|0|0|0|0|0|0|0|0|0|0|0|0|0|0|0|0|0|0|0|0|0|0|0|0|0|0|0|0|0|0|0|0|0|0|0|0|0|0|0|0|0|1|0|0|0|0|0|0|0|0|0|0|0|0|
|virus_Aug5.25v_35124|1|0|0|0|0|0|0|0|0|0|0|0|0|0|0|0|0|0|0|0|0|0|0|0|0|0|0|0|0|0|0|0|0|0|0|0|0|0|0|0|0|0|1|0|0|0|0|0|0|0|0|0|0|0|0|
|virus_Aug5.25v_35953|1|0|0|0|0|0|0|0|0|0|0|0|0|0|0|0|0|0|0|0|0|0|0|0|0|0|0|0|0|0|0|0|0|0|0|0|0|0|0|0|0|0|1|0|0|0|0|0|0|0|0|0|0|0|0|
|virus_Aug5.25v_37357|1|0|0|0|0|0|0|0|0|0|0|0|0|0|0|0|0|0|0|0|0|0|0|0|0|0|0|0|0|0|0|0|0|0|0|0|0|0|0|0|0|0|1|0|0|0|0|0|0|0|0|0|0|0|0|
|virus_Aug5.25v_38436|1|0|0|0|0|0|0|0|0|0|0|0|0|0|0|0|0|0|0|0|0|0|0|0|0|0|0|0|0|0|0|0|0|0|0|0|0|0|0|0|0|0|1|0|0|0|0|0|0|0|0|0|0|0|0|
|virus_Aug5.25v_38579|1|0|0|0|0|0|0|0|0|0|0|0|0|0|0|0|0|0|0|0|0|0|0|0|0|0|0|0|0|0|0|0|0|0|0|0|0|0|0|0|0|0|1|0|0|0|0|0|0|0|0|0|0|0|0|
|virus_Aug5.50v_6581|1|0|0|0|0|0|0|0|0|0|0|0|0|0|0|0|0|0|0|0|0|0|0|0|0|0|0|0|0|0|0|0|0|0|0|0|0|0|0|0|0|0|0|0|1|0|0|0|0|0|0|0|0|0|0|
|virus_Aug5.50v_8883|1|0|0|0|0|0|0|0|0|0|0|0|0|0|0|0|0|0|0|0|0|0|0|0|0|0|0|0|0|0|0|0|0|0|0|0|0|0|0|0|0|0|0|0|1|0|0|0|0|0|0|0|0|0|0|
|virus_Aug5.50v_12710|1|0|0|0|0|0|0|0|0|0|0|0|0|0|0|0|0|0|0|0|0|0|0|0|0|0|0|0|0|0|0|0|0|0|0|0|0|0|0|0|0|0|0|0|1|0|0|0|0|0|0|0|0|0|0|
|virus_Aug5.50v_24607|1|0|0|0|0|0|0|0|0|0|0|0|0|0|0|0|0|0|0|0|0|0|0|0|0|0|0|0|0|0|0|0|0|0|0|0|0|0|0|0|0|0|0|0|1|0|0|0|0|0|0|0|0|0|0|
|virus_Aug5.50v_38847|1|0|0|0|0|0|0|0|0|0|0|0|0|0|0|0|0|0|0|0|0|0|0|0|0|0|0|0|0|0|0|0|0|0|0|0|0|0|0|0|0|0|0|0|1|0|0|0|0|0|0|0|0|0|0|
|virus_Aug5.75v_13162|1|0|0|0|0|0|0|0|0|0|0|0|0|0|0|0|0|0|0|0|0|0|0|0|0|0|0|0|0|0|0|0|0|0|0|0|0|0|0|0|0|0|0|0|0|0|1|0|0|0|0|0|0|0|0|
|virus_Aug5.75v_13845|1|0|0|0|0|0|0|0|0|0|0|0|0|0|0|0|0|0|0|0|0|0|0|0|0|0|0|0|0|0|0|0|0|0|0|0|0|0|0|0|0|0|0|0|0|0|1|0|0|0|0|0|0|0|0|
|virus_Aug5.75v_21894|1|0|0|0|0|0|0|0|0|0|0|0|0|0|0|0|0|0|0|0|0|0|0|0|0|0|0|0|0|0|0|0|0|0|0|0|0|0|0|0|0|0|0|0|0|0|1|0|0|0|0|0|0|0|0|
|virus_Aug5.75v_39888|1|0|0|0|0|0|0|0|0|0|0|0|0|0|0|0|0|0|0|0|0|0|0|0|0|0|0|0|0|0|0|0|0|0|0|0|0|0|0|0|0|0|0|0|0|0|1|0|0|0|0|0|0|0|0|
|virus_Aug5.75v_41101|1|0|0|0|0|0|0|0|0|0|0|0|0|0|0|0|0|0|0|0|0|0|0|0|0|0|0|0|0|0|0|0|0|0|0|0|0|0|0|0|0|0|0|0|0|0|1|0|0|0|0|0|0|0|0|
|virus_Jul11.0v_5281|1|0|0|0|0|0|0|0|0|0|0|0|0|0|0|0|0|0|0|0|0|0|1|0|0|0|0|0|0|0|0|0|0|0|0|0|0|0|0|0|0|0|0|0|0|0|0|0|0|0|0|0|0|0|0|
|virus_Jul11.0v_14736|1|0|0|0|0|0|0|0|0|0|0|0|0|0|0|0|0|0|0|0|0|0|1|0|0|0|0|0|0|0|0|0|0|0|0|0|0|0|0|0|0|0|0|0|0|0|0|0|0|0|0|0|0|0|0|
|virus_Jul11.0v_14923|1|0|0|0|0|0|0|0|0|0|0|0|0|0|0|0|0|0|0|0|0|0|1|0|0|0|0|0|0|0|0|0|0|0|0|0|0|0|0|0|0|0|0|0|0|0|0|0|0|0|0|0|0|0|0|
|virus_Jul11.0v_17430|1|0|0|0|0|0|0|0|0|0|0|0|0|0|0|0|0|0|0|0|0|0|1|0|0|0|0|0|0|0|0|0|0|0|0|0|0|0|0|0|0|0|0|0|0|0|0|0|0|0|0|0|0|0|0|
|virus_Jul11.0v_19171|1|0|0|0|0|0|0|0|0|0|0|0|0|0|0|0|0|0|0|0|0|0|1|0|0|0|0|0|0|0|0|0|0|0|0|0|0|0|0|0|0|0|0|0|0|0|0|0|0|0|0|0|0|0|0|
|virus_Jul11.5v_1317|1|0|0|0|0|0|0|0|0|0|0|0|0|0|0|0|0|0|0|0|0|0|0|0|1|0|0|0|0|0|0|0|0|0|0|0|0|0|0|0|0|0|0|0|0|0|0|0|0|0|0|0|0|0|0|
|virus_Jul11.5v_1898|1|0|0|0|0|0|0|0|0|0|0|0|0|0|0|0|0|0|0|0|0|0|0|0|1|0|0|0|0|0|0|0|0|0|0|0|0|0|0|0|0|0|0|0|0|0|0|0|0|0|0|0|0|0|0|
|virus_Jul11.5v_2359|1|0|0|0|0|0|0|0|0|0|0|0|0|0|0|0|0|0|0|0|0|0|0|0|1|0|0|0|0|0|0|0|0|0|0|0|0|0|0|0|0|0|0|0|0|0|0|0|0|0|0|0|0|0|0|
|virus_Jul11.5v_3203|1|0|0|0|0|0|0|0|0|0|0|0|0|0|0|0|0|0|0|0|0|0|0|0|1|0|0|0|0|0|0|0|0|0|0|0|0|0|0|0|0|0|0|0|0|0|0|0|0|0|0|0|0|0|0|
|virus_Jul11.5v_4175|1|0|0|0|0|0|0|0|0|0|0|0|0|0|0|0|0|0|0|0|0|0|0|0|1|0|0|0|0|0|0|0|0|0|0|0|0|0|0|0|0|0|0|0|0|0|0|0|0|0|0|0|0|0|0|
|virus_Jul11.5v_4210|1|0|0|0|0|0|0|0|0|0|0|0|0|0|0|0|0|0|0|0|0|0|0|0|1|0|0|0|0|0|0|0|0|0|0|0|0|0|0|0|0|0|0|0|0|0|0|0|0|0|0|0|0|0|0|
|virus_Jul11.5v_9168|1|0|0|0|0|0|0|0|0|0|0|0|0|0|0|0|0|0|0|0|0|0|0|0|1|0|0|0|0|0|0|0|0|0|0|0|0|0|0|0|0|0|0|0|0|0|0|0|0|0|0|0|0|0|0|
|virus_Jul11.5v_9652|1|0|0|0|0|0|0|0|0|0|0|0|0|0|0|0|0|0|0|0|0|0|0|0|1|0|0|0|0|0|0|0|0|0|0|0|0|0|0|0|0|0|0|0|0|0|0|0|0|0|0|0|0|0|0|
|virus_Jul11.5v_9989|1|0|0|0|0|0|0|0|0|0|0|0|0|0|0|0|0|0|0|0|0|0|0|0|1|0|0|0|0|0|0|0|0|0|0|0|0|0|0|0|0|0|0|0|0|0|0|0|0|0|0|0|0|0|0|
|virus_Jul11.5v_11436|1|0|0|0|0|0|0|0|0|0|0|0|0|0|0|0|0|0|0|0|0|0|0|0|1|0|0|0|0|0|0|0|0|0|0|0|0|0|0|0|0|0|0|0|0|0|0|0|0|0|0|0|0|0|0|
|virus_Jul11.5v_11629|1|0|0|0|0|0|0|0|0|0|0|0|0|0|0|0|0|0|0|0|0|0|0|0|1|0|0|0|0|0|0|0|0|0|0|0|0|0|0|0|0|0|0|0|0|0|0|0|0|0|0|0|0|0|0|
|virus_Jul11.5v_12625|1|0|0|0|0|0|0|0|0|0|0|0|0|0|0|0|0|0|0|0|0|0|0|0|1|0|0|0|0|0|0|0|0|0|0|0|0|0|0|0|0|0|0|0|0|0|0|0|0|0|0|0|0|0|0|
|virus_Jul11.5v_15222|1|0|0|0|0|0|0|0|0|0|0|0|0|0|0|0|0|0|0|0|0|0|0|0|1|0|0|0|0|0|0|0|0|0|0|0|0|0|0|0|0|0|0|0|0|0|0|0|0|0|0|0|0|0|0|
|virus_Jul11.5v_17996|1|0|0|0|0|0|0|0|0|0|0|0|0|0|0|0|0|0|0|0|0|0|0|0|1|0|0|0|0|0|0|0|0|0|0|0|0|0|0|0|0|0|0|0|0|0|0|0|0|0|0|0|0|0|0|
|virus_Jul11.5v_19242|1|0|0|0|0|0|0|0|0|0|0|0|0|0|0|0|0|0|0|0|0|0|0|0|1|0|0|0|0|0|0|0|0|0|0|0|0|0|0|0|0|0|0|0|0|0|0|0|0|0|0|0|0|0|0|
|virus_Jul11.5v_20935|1|0|0|0|0|0|0|0|0|0|0|0|0|0|0|0|0|0|0|0|0|0|0|0|1|0|0|0|0|0|0|0|0|0|0|0|0|0|0|0|0|0|0|0|0|0|0|0|0|0|0|0|0|0|0|
|virus_Jul11.25v_976|1|0|0|0|0|0|0|0|0|0|0|0|0|0|0|0|0|0|0|0|0|0|0|0|0|0|1|0|0|0|0|0|0|0|0|0|0|0|0|0|0|0|0|0|0|0|0|0|0|0|0|0|0|0|0|
|virus_Jul11.25v_1782|1|0|0|0|0|0|0|0|0|0|0|0|0|0|0|0|0|0|0|0|0|0|0|0|0|0|1|0|0|0|0|0|0|0|0|0|0|0|0|0|0|0|0|0|0|0|0|0|0|0|0|0|0|0|0|
|virus_Jul11.25v_8380|1|0|0|0|0|0|0|0|0|0|0|0|0|0|0|0|0|0|0|0|0|0|0|0|0|0|1|0|0|0|0|0|0|0|0|0|0|0|0|0|0|0|0|0|0|0|0|0|0|0|0|0|0|0|0|
|virus_Jul11.25v_10088|1|0|0|0|0|0|0|0|0|0|0|0|0|0|0|0|0|0|0|0|0|0|0|0|0|0|1|0|0|0|0|0|0|0|0|0|0|0|0|0|0|0|0|0|0|0|0|0|0|0|0|0|0|0|0|
|virus_Jul11.25v_11920|1|0|0|0|0|0|0|0|0|0|0|0|0|0|0|0|0|0|0|0|0|0|0|0|0|0|1|0|0|0|0|0|0|0|0|0|0|0|0|0|0|0|0|0|0|0|0|0|0|0|0|0|0|0|0|
|virus_Jul11.25v_14119|1|0|0|0|0|0|0|0|0|0|0|0|0|0|0|0|0|0|0|0|0|0|0|0|0|0|1|0|0|0|0|0|0|0|0|0|0|0|0|0|0|0|0|0|0|0|0|0|0|0|0|0|0|0|0|
|virus_Jul11.25v_19522|1|0|0|0|0|0|0|0|0|0|0|0|0|0|0|0|0|0|0|0|0|0|0|0|0|0|1|0|0|0|0|0|0|0|0|0|0|0|0|0|0|0|0|0|0|0|0|0|0|0|0|0|0|0|0|
|virus_Jul11.25v_20568|1|0|0|0|0|0|0|0|0|0|0|0|0|0|0|0|0|0|0|0|0|0|0|0|0|0|1|0|0|0|0|0|0|0|0|0|0|0|0|0|0|0|0|0|0|0|0|0|0|0|0|0|0|0|0|
|virus_Jul11.50v_1396|1|0|0|0|0|0|0|0|0|0|0|0|0|0|0|0|0|0|0|0|0|0|0|0|0|0|0|0|1|0|0|0|0|0|0|0|0|0|0|0|0|0|0|0|0|0|0|0|0|0|0|0|0|0|0|
|virus_Jul11.50v_1579|1|0|0|0|0|0|0|0|0|0|0|0|0|0|0|0|0|0|0|0|0|0|0|0|0|0|0|0|1|0|0|0|0|0|0|0|0|0|0|0|0|0|0|0|0|0|0|0|0|0|0|0|0|0|0|
|virus_Jul11.50v_2116|1|0|0|0|0|0|0|0|0|0|0|0|0|0|0|0|0|0|0|0|0|0|0|0|0|0|0|0|1|0|0|0|0|0|0|0|0|0|0|0|0|0|0|0|0|0|0|0|0|0|0|0|0|0|0|
|virus_Jul11.50v_6507|1|0|0|0|0|0|0|0|0|0|0|0|0|0|0|0|0|0|0|0|0|0|0|0|0|0|0|0|1|0|0|0|0|0|0|0|0|0|0|0|0|0|0|0|0|0|0|0|0|0|0|0|0|0|0|
|virus_Jul11.50v_9863|1|0|0|0|0|0|0|0|0|0|0|0|0|0|0|0|0|0|0|0|0|0|0|0|0|0|0|0|1|0|0|0|0|0|0|0|0|0|0|0|0|0|0|0|0|0|0|0|0|0|0|0|0|0|0|
|virus_Jul11.50v_12927|1|0|0|0|0|0|0|0|0|0|0|0|0|0|0|0|0|0|0|0|0|0|0|0|0|0|0|0|1|0|0|0|0|0|0|0|0|0|0|0|0|0|0|0|0|0|0|0|0|0|0|0|0|0|0|
|virus_Jul11.50v_13993|1|0|0|0|0|0|0|0|0|0|0|0|0|0|0|0|0|0|0|0|0|0|0|0|0|0|0|0|1|0|0|0|0|0|0|0|0|0|0|0|0|0|0|0|0|0|0|0|0|0|0|0|0|0|0|
|virus_Jul11.50v_14675|1|0|0|0|0|0|0|0|0|0|0|0|0|0|0|0|0|0|0|0|0|0|0|0|0|0|0|0|1|0|0|0|0|0|0|0|0|0|0|0|0|0|0|0|0|0|0|0|0|0|0|0|0|0|0|
|virus_Jul11.50v_15423|1|0|0|0|0|0|0|0|0|0|0|0|0|0|0|0|0|0|0|0|0|0|0|0|0|0|0|0|1|0|0|0|0|0|0|0|0|0|0|0|0|0|0|0|0|0|0|0|0|0|0|0|0|0|0|
|virus_Jul11.50v_17503|1|0|0|0|0|0|0|0|0|0|0|0|0|0|0|0|0|0|0|0|0|0|0|0|0|0|0|0|1|0|0|0|0|0|0|0|0|0|0|0|0|0|0|0|0|0|0|0|0|0|0|0|0|0|0|
|virus_Jul11.50v_17530|1|0|0|0|0|0|0|0|0|0|0|0|0|0|0|0|0|0|0|0|0|0|0|0|0|0|0|0|1|0|0|0|0|0|0|0|0|0|0|0|0|0|0|0|0|0|0|0|0|0|0|0|0|0|0|
|virus_Jul11.50v_18986|1|0|0|0|0|0|0|0|0|0|0|0|0|0|0|0|0|0|0|0|0|0|0|0|0|0|0|0|1|0|0|0|0|0|0|0|0|0|0|0|0|0|0|0|0|0|0|0|0|0|0|0|0|0|0|
|virus_Jul11.75v_6438|1|0|0|0|0|0|0|0|0|0|0|0|0|0|0|0|0|0|0|0|0|0|0|0|0|0|0|0|0|0|1|0|0|0|0|0|0|0|0|0|0|0|0|0|0|0|0|0|0|0|0|0|0|0|0|
|virus_Jul11.75v_8152|1|0|0|0|0|0|0|0|0|0|0|0|0|0|0|0|0|0|0|0|0|0|0|0|0|0|0|0|0|0|1|0|0|0|0|0|0|0|0|0|0|0|0|0|0|0|0|0|0|0|0|0|0|0|0|
|virus_Jul11.75v_10703|1|0|0|0|0|0|0|0|0|0|0|0|0|0|0|0|0|0|0|0|0|0|0|0|0|0|0|0|0|0|1|0|0|0|0|0|0|0|0|0|0|0|0|0|0|0|0|0|0|0|0|0|0|0|0|
|virus_Jul11.75v_12927|1|0|0|0|0|0|0|0|0|0|0|0|0|0|0|0|0|0|0|0|0|0|0|0|0|0|0|0|0|0|1|0|0|0|0|0|0|0|0|0|0|0|0|0|0|0|0|0|0|0|0|0|0|0|0|
|virus_Jul11.75v_23390|1|0|0|0|0|0|0|0|0|0|0|0|0|0|0|0|0|0|0|0|0|0|0|0|0|0|0|0|0|0|1|0|0|0|0|0|0|0|0|0|0|0|0|0|0|0|0|0|0|0|0|0|0|0|0|
|virus_Jul11.75v_28208|1|0|0|0|0|0|0|0|0|0|0|0|0|0|0|0|0|0|0|0|0|0|0|0|0|0|0|0|0|0|1|0|0|0|0|0|0|0|0|0|0|0|0|0|0|0|0|0|0|0|0|0|0|0|0|
|virus_Jul11.75v_38785|1|0|0|0|0|0|0|0|0|0|0|0|0|0|0|0|0|0|0|0|0|0|0|0|0|0|0|0|0|0|1|0|0|0|0|0|0|0|0|0|0|0|0|0|0|0|0|0|0|0|0|0|0|0|0|
|virus_Jul11.75v_38853|1|0|0|0|0|0|0|0|0|0|0|0|0|0|0|0|0|0|0|0|0|0|0|0|0|0|0|0|0|0|1|0|0|0|0|0|0|0|0|0|0|0|0|0|0|0|0|0|0|0|0|0|0|0|0|
|virus_Jul27.5v_878|1|0|0|0|0|0|0|0|0|0|0|0|0|0|0|0|0|0|0|0|0|0|0|0|0|0|0|0|0|0|0|0|1|0|0|0|0|0|0|0|0|0|0|0|0|0|0|0|0|0|0|0|0|0|0|
|virus_Jul27.5v_2522|1|0|0|0|0|0|0|0|0|0|0|0|0|0|0|0|0|0|0|0|0|0|0|0|0|0|0|0|0|0|0|0|1|0|0|0|0|0|0|0|0|0|0|0|0|0|0|0|0|0|0|0|0|0|0|
|virus_Jul27.5v_7214|1|0|0|0|0|0|0|0|0|0|0|0|0|0|0|0|0|0|0|0|0|0|0|0|0|0|0|0|0|0|0|0|1|0|0|0|0|0|0|0|0|0|0|0|0|0|0|0|0|0|0|0|0|0|0|
|virus_Jul27.5v_11959|1|0|0|0|0|0|0|0|0|0|0|0|0|0|0|0|0|0|0|0|0|0|0|0|0|0|0|0|0|0|0|0|1|0|0|0|0|0|0|0|0|0|0|0|0|0|0|0|0|0|0|0|0|0|0|
|virus_Jul27.5v_15103|1|0|0|0|0|0|0|0|0|0|0|0|0|0|0|0|0|0|0|0|0|0|0|0|0|0|0|0|0|0|0|0|1|0|0|0|0|0|0|0|0|0|0|0|0|0|0|0|0|0|0|0|0|0|0|
|virus_Jul27.5v_15110|1|0|0|0|0|0|0|0|0|0|0|0|0|0|0|0|0|0|0|0|0|0|0|0|0|0|0|0|0|0|0|0|1|0|0|0|0|0|0|0|0|0|0|0|0|0|0|0|0|0|0|0|0|0|0|
|virus_Jul27.5v_15470|1|0|0|0|0|0|0|0|0|0|0|0|0|0|0|0|0|0|0|0|0|0|0|0|0|0|0|0|0|0|0|0|1|0|0|0|0|0|0|0|0|0|0|0|0|0|0|0|0|0|0|0|0|0|0|
|virus_Jul27.5v_17149|1|0|0|0|0|0|0|0|0|0|0|0|0|0|0|0|0|0|0|0|0|0|0|0|0|0|0|0|0|0|0|0|1|0|0|0|0|0|0|0|0|0|0|0|0|0|0|0|0|0|0|0|0|0|0|
|virus_Jul27.5v_20206|1|0|0|0|0|0|0|0|0|0|0|0|0|0|0|0|0|0|0|0|0|0|0|0|0|0|0|0|0|0|0|0|1|0|0|0|0|0|0|0|0|0|0|0|0|0|0|0|0|0|0|0|0|0|0|
|virus_Jul27.25v_1425|1|0|0|0|0|0|0|0|0|0|0|0|0|0|0|0|0|0|0|0|0|0|0|0|0|0|0|0|0|0|0|0|0|0|1|0|0|0|0|0|0|0|0|0|0|0|0|0|0|0|0|0|0|0|0|
|virus_Jul27.25v_1689|1|0|0|0|0|0|0|0|0|0|0|0|0|0|0|0|0|0|0|0|0|0|0|0|0|0|0|0|0|0|0|0|0|0|1|0|0|0|0|0|0|0|0|0|0|0|0|0|0|0|0|0|0|0|0|
|virus_Jul27.25v_2009|1|0|0|0|0|0|0|0|0|0|0|0|0|0|0|0|0|0|0|0|0|0|0|0|0|0|0|0|0|0|0|0|0|0|1|0|0|0|0|0|0|0|0|0|0|0|0|0|0|0|0|0|0|0|0|
|virus_Jul27.25v_6038|1|0|0|0|0|0|0|0|0|0|0|0|0|0|0|0|0|0|0|0|0|0|0|0|0|0|0|0|0|0|0|0|0|0|1|0|0|0|0|0|0|0|0|0|0|0|0|0|0|0|0|0|0|0|0|
|virus_Jul27.25v_11119|1|0|0|0|0|0|0|0|0|0|0|0|0|0|0|0|0|0|0|0|0|0|0|0|0|0|0|0|0|0|0|0|0|0|1|0|0|0|0|0|0|0|0|0|0|0|0|0|0|0|0|0|0|0|0|
|virus_Jul27.25v_13759|1|0|0|0|0|0|0|0|0|0|0|0|0|0|0|0|0|0|0|0|0|0|0|0|0|0|0|0|0|0|0|0|0|0|1|0|0|0|0|0|0|0|0|0|0|0|0|0|0|0|0|0|0|0|0|
|virus_Jul27.25v_20090|1|0|0|0|0|0|0|0|0|0|0|0|0|0|0|0|0|0|0|0|0|0|0|0|0|0|0|0|0|0|0|0|0|0|1|0|0|0|0|0|0|0|0|0|0|0|0|0|0|0|0|0|0|0|0|
|virus_Jul27.25v_32053|1|0|0|0|0|0|0|0|0|0|0|0|0|0|0|0|0|0|0|0|0|0|0|0|0|0|0|0|0|0|0|0|0|0|1|0|0|0|0|0|0|0|0|0|0|0|0|0|0|0|0|0|0|0|0|

**Supplementary Table S7.** Common Bacterio- and virioplankton RTPR 98% OTUs in 27 samples with RTPR amplicons in both fractions

Total seqs: 47997; Subtotal seqs in retained

| OTUs: 41046 | | May12_05m | | May12_25m | | May12_50m | | May12_75m | | Jun8_25m | | Jun8_50m | | Jun8_75m | | Jun22_0m | | Jun22_50m | | Jun22_75m | | Jul11_0m | | Jul11_05m | | Jul11_25m | | Jul11_50m | | Jul11_75m | | Jul27_05m | | Jul27_25m | | Jul27_75m | | Aug5_0m | | Aug5_05m | | Aug5_25m | | Aug5_50m | | Aug5_75m | | Sep8_0m | | Sep8_05m | | Sep8_25m | | Sep8_50m | |
|---|---|---|---|---|---|---|---|---|---|---|---|---|---|---|---|---|---|---|---|---|---|---|---|---|---|---|---|---|---|---|---|---|---|---|---|---|---|---|---|---|---|---|---|---|---|---|---|---|---|---|---|---|---|---|---|
|OTUs with >=10 seqs: 254|OTU subtotal|bacterioplankton|virioplankton|bacterioplankton|virioplankton|bacterioplankton|virioplankton|bacterioplankton|virioplankton|bacterioplankton|virioplankton|bacterioplankton|virioplankton|bacterioplankton|virioplankton|bacterioplankton|virioplankton|bacterioplankton|virioplankton|bacterioplankton|virioplankton|bacterioplankton|virioplankton|bacterioplankton|virioplankton|bacterioplankton|virioplankton|bacterioplankton|virioplankton|bacterioplankton|virioplankton|bacterioplankton|virioplankton|bacterioplankton|virioplankton|bacterioplankton|virioplankton|bacterioplankton|virioplankton|bacterioplankton|virioplankton|bacterioplankton|virioplankton|bacterioplankton|virioplankton|bacterioplankton|virioplankton|bacterioplankton|virioplankton|bacterioplankton|virioplankton|bacterioplankton|virioplankton|bacterioplankton|virioplankton|
|virus_Jul27.25v_39795|1|0|0|0|0|0|0|0|0|0|0|0|0|0|0|0|0|0|0|0|0|0|0|0|0|0|0|0|0|0|0|0|0|0|1|0|0|0|0|0|0|0|0|0|0|0|0|0|0|0|0|0|0|0|0|
|virus_Jul27.75v_735|1|0|0|0|0|0|0|0|0|0|0|0|0|0|0|0|0|0|0|0|0|0|0|0|0|0|0|0|0|0|0|0|0|0|0|0|1|0|0|0|0|0|0|0|0|0|0|0|0|0|0|0|0|0|0|
|virus_Jul27.75v_975|1|0|0|0|0|0|0|0|0|0|0|0|0|0|0|0|0|0|0|0|0|0|0|0|0|0|0|0|0|0|0|0|0|0|0|0|1|0|0|0|0|0|0|0|0|0|0|0|0|0|0|0|0|0|0|
|virus_Jul27.75v_3642|1|0|0|0|0|0|0|0|0|0|0|0|0|0|0|0|0|0|0|0|0|0|0|0|0|0|0|0|0|0|0|0|0|0|0|0|1|0|0|0|0|0|0|0|0|0|0|0|0|0|0|0|0|0|0|
|virus_Jul27.75v_8888|1|0|0|0|0|0|0|0|0|0|0|0|0|0|0|0|0|0|0|0|0|0|0|0|0|0|0|0|0|0|0|0|0|0|0|0|1|0|0|0|0|0|0|0|0|0|0|0|0|0|0|0|0|0|0|
|virus_Jul27.75v_17542|1|0|0|0|0|0|0|0|0|0|0|0|0|0|0|0|0|0|0|0|0|0|0|0|0|0|0|0|0|0|0|0|0|0|0|0|1|0|0|0|0|0|0|0|0|0|0|0|0|0|0|0|0|0|0|
|virus_Jun8.25v_10081|1|0|0|0|0|0|0|0|0|0|1|0|0|0|0|0|0|0|0|0|0|0|0|0|0|0|0|0|0|0|0|0|0|0|0|0|0|0|0|0|0|0|0|0|0|0|0|0|0|0|0|0|0|0|0|
|virus_Jun8.25v_11433|1|0|0|0|0|0|0|0|0|0|1|0|0|0|0|0|0|0|0|0|0|0|0|0|0|0|0|0|0|0|0|0|0|0|0|0|0|0|0|0|0|0|0|0|0|0|0|0|0|0|0|0|0|0|0|
|virus_Jun8.25v_35482|1|0|0|0|0|0|0|0|0|0|1|0|0|0|0|0|0|0|0|0|0|0|0|0|0|0|0|0|0|0|0|0|0|0|0|0|0|0|0|0|0|0|0|0|0|0|0|0|0|0|0|0|0|0|0|
|virus_Jun8.50v_3611|1|0|0|0|0|0|0|0|0|0|0|0|1|0|0|0|0|0|0|0|0|0|0|0|0|0|0|0|0|0|0|0|0|0|0|0|0|0|0|0|0|0|0|0|0|0|0|0|0|0|0|0|0|0|0|
|virus_Jun8.50v_3759|1|0|0|0|0|0|0|0|0|0|0|0|1|0|0|0|0|0|0|0|0|0|0|0|0|0|0|0|0|0|0|0|0|0|0|0|0|0|0|0|0|0|0|0|0|0|0|0|0|0|0|0|0|0|0|
|virus_Jun8.50v_8834|1|0|0|0|0|0|0|0|0|0|0|0|1|0|0|0|0|0|0|0|0|0|0|0|0|0|0|0|0|0|0|0|0|0|0|0|0|0|0|0|0|0|0|0|0|0|0|0|0|0|0|0|0|0|0|
|virus_Jun8.50v_12310|1|0|0|0|0|0|0|0|0|0|0|0|1|0|0|0|0|0|0|0|0|0|0|0|0|0|0|0|0|0|0|0|0|0|0|0|0|0|0|0|0|0|0|0|0|0|0|0|0|0|0|0|0|0|0|
|virus_Jun8.50v_13238|1|0|0|0|0|0|0|0|0|0|0|0|1|0|0|0|0|0|0|0|0|0|0|0|0|0|0|0|0|0|0|0|0|0|0|0|0|0|0|0|0|0|0|0|0|0|0|0|0|0|0|0|0|0|0|
|virus_Jun8.50v_17308|1|0|0|0|0|0|0|0|0|0|0|0|1|0|0|0|0|0|0|0|0|0|0|0|0|0|0|0|0|0|0|0|0|0|0|0|0|0|0|0|0|0|0|0|0|0|0|0|0|0|0|0|0|0|0|
|virus_Jun8.50v_22104|1|0|0|0|0|0|0|0|0|0|0|0|1|0|0|0|0|0|0|0|0|0|0|0|0|0|0|0|0|0|0|0|0|0|0|0|0|0|0|0|0|0|0|0|0|0|0|0|0|0|0|0|0|0|0|
|virus_Jun8.50v_22877|1|0|0|0|0|0|0|0|0|0|0|0|1|0|0|0|0|0|0|0|0|0|0|0|0|0|0|0|0|0|0|0|0|0|0|0|0|0|0|0|0|0|0|0|0|0|0|0|0|0|0|0|0|0|0|
|virus_Jun8.50v_25735|1|0|0|0|0|0|0|0|0|0|0|0|1|0|0|0|0|0|0|0|0|0|0|0|0|0|0|0|0|0|0|0|0|0|0|0|0|0|0|0|0|0|0|0|0|0|0|0|0|0|0|0|0|0|0|
|virus_Jun8.50v_34579|1|0|0|0|0|0|0|0|0|0|0|0|1|0|0|0|0|0|0|0|0|0|0|0|0|0|0|0|0|0|0|0|0|0|0|0|0|0|0|0|0|0|0|0|0|0|0|0|0|0|0|0|0|0|0|
|virus_Jun8.50v_35343|1|0|0|0|0|0|0|0|0|0|0|0|1|0|0|0|0|0|0|0|0|0|0|0|0|0|0|0|0|0|0|0|0|0|0|0|0|0|0|0|0|0|0|0|0|0|0|0|0|0|0|0|0|0|0|
|virus_Jun8.50v_40458|1|0|0|0|0|0|0|0|0|0|0|0|1|0|0|0|0|0|0|0|0|0|0|0|0|0|0|0|0|0|0|0|0|0|0|0|0|0|0|0|0|0|0|0|0|0|0|0|0|0|0|0|0|0|0|
|virus_Jun8.75v_12607|1|0|0|0|0|0|0|0|0|0|0|0|0|0|1|0|0|0|0|0|0|0|0|0|0|0|0|0|0|0|0|0|0|0|0|0|0|0|0|0|0|0|0|0|0|0|0|0|0|0|0|0|0|0|0|
|virus_Jun8.75v_20644|1|0|0|0|0|0|0|0|0|0|0|0|0|0|1|0|0|0|0|0|0|0|0|0|0|0|0|0|0|0|0|0|0|0|0|0|0|0|0|0|0|0|0|0|0|0|0|0|0|0|0|0|0|0|0|
|virus_Jun22.0v_7656|1|0|0|0|0|0|0|0|0|0|0|0|0|0|0|0|1|0|0|0|0|0|0|0|0|0|0|0|0|0|0|0|0|0|0|0|0|0|0|0|0|0|0|0|0|0|0|0|0|0|0|0|0|0|0|
|virus_Jun22.0v_9662|1|0|0|0|0|0|0|0|0|0|0|0|0|0|0|0|1|0|0|0|0|0|0|0|0|0|0|0|0|0|0|0|0|0|0|0|0|0|0|0|0|0|0|0|0|0|0|0|0|0|0|0|0|0|0|
|virus_Jun22.0v_10421|1|0|0|0|0|0|0|0|0|0|0|0|0|0|0|0|1|0|0|0|0|0|0|0|0|0|0|0|0|0|0|0|0|0|0|0|0|0|0|0|0|0|0|0|0|0|0|0|0|0|0|0|0|0|0|
|virus_Jun22.0v_11636|1|0|0|0|0|0|0|0|0|0|0|0|0|0|0|0|1|0|0|0|0|0|0|0|0|0|0|0|0|0|0|0|0|0|0|0|0|0|0|0|0|0|0|0|0|0|0|0|0|0|0|0|0|0|0|
|virus_Jun22.0v_12525|1|0|0|0|0|0|0|0|0|0|0|0|0|0|0|0|1|0|0|0|0|0|0|0|0|0|0|0|0|0|0|0|0|0|0|0|0|0|0|0|0|0|0|0|0|0|0|0|0|0|0|0|0|0|0|
|virus_Jun22.0v_12960|1|0|0|0|0|0|0|0|0|0|0|0|0|0|0|0|1|0|0|0|0|0|0|0|0|0|0|0|0|0|0|0|0|0|0|0|0|0|0|0|0|0|0|0|0|0|0|0|0|0|0|0|0|0|0|
|virus_Jun22.0v_13049|1|0|0|0|0|0|0|0|0|0|0|0|0|0|0|0|1|0|0|0|0|0|0|0|0|0|0|0|0|0|0|0|0|0|0|0|0|0|0|0|0|0|0|0|0|0|0|0|0|0|0|0|0|0|0|
|virus_Jun22.0v_13574|1|0|0|0|0|0|0|0|0|0|0|0|0|0|0|0|1|0|0|0|0|0|0|0|0|0|0|0|0|0|0|0|0|0|0|0|0|0|0|0|0|0|0|0|0|0|0|0|0|0|0|0|0|0|0|
|virus_Jun22.0v_20743|1|0|0|0|0|0|0|0|0|0|0|0|0|0|0|0|1|0|0|0|0|0|0|0|0|0|0|0|0|0|0|0|0|0|0|0|0|0|0|0|0|0|0|0|0|0|0|0|0|0|0|0|0|0|0|
|virus_Jun22.0v_20826|1|0|0|0|0|0|0|0|0|0|0|0|0|0|0|0|1|0|0|0|0|0|0|0|0|0|0|0|0|0|0|0|0|0|0|0|0|0|0|0|0|0|0|0|0|0|0|0|0|0|0|0|0|0|0|
|virus_Jun22.50v_2786|1|0|0|0|0|0|0|0|0|0|0|0|0|0|0|0|0|0|1|0|0|0|0|0|0|0|0|0|0|0|0|0|0|0|0|0|0|0|0|0|0|0|0|0|0|0|0|0|0|0|0|0|0|0|0|
|virus_Jun22.50v_3075|1|0|0|0|0|0|0|0|0|0|0|0|0|0|0|0|0|0|1|0|0|0|0|0|0|0|0|0|0|0|0|0|0|0|0|0|0|0|0|0|0|0|0|0|0|0|0|0|0|0|0|0|0|0|0|
|virus_Jun22.50v_3567|1|0|0|0|0|0|0|0|0|0|0|0|0|0|0|0|0|0|1|0|0|0|0|0|0|0|0|0|0|0|0|0|0|0|0|0|0|0|0|0|0|0|0|0|0|0|0|0|0|0|0|0|0|0|0|
|virus_Jun22.50v_3841|1|0|0|0|0|0|0|0|0|0|0|0|0|0|0|0|0|0|1|0|0|0|0|0|0|0|0|0|0|0|0|0|0|0|0|0|0|0|0|0|0|0|0|0|0|0|0|0|0|0|0|0|0|0|0|
|virus_Jun22.50v_6918|1|0|0|0|0|0|0|0|0|0|0|0|0|0|0|0|0|0|1|0|0|0|0|0|0|0|0|0|0|0|0|0|0|0|0|0|0|0|0|0|0|0|0|0|0|0|0|0|0|0|0|0|0|0|0|
|virus_Jun22.50v_9386|1|0|0|0|0|0|0|0|0|0|0|0|0|0|0|0|0|0|1|0|0|0|0|0|0|0|0|0|0|0|0|0|0|0|0|0|0|0|0|0|0|0|0|0|0|0|0|0|0|0|0|0|0|0|0|
|virus_Jun22.50v_11727|1|0|0|0|0|0|0|0|0|0|0|0|0|0|0|0|0|0|1|0|0|0|0|0|0|0|0|0|0|0|0|0|0|0|0|0|0|0|0|0|0|0|0|0|0|0|0|0|0|0|0|0|0|0|0|
|virus_Jun22.50v_16876|1|0|0|0|0|0|0|0|0|0|0|0|0|0|0|0|0|0|1|0|0|0|0|0|0|0|0|0|0|0|0|0|0|0|0|0|0|0|0|0|0|0|0|0|0|0|0|0|0|0|0|0|0|0|0|
|virus_Jun22.50v_17698|1|0|0|0|0|0|0|0|0|0|0|0|0|0|0|0|0|0|1|0|0|0|0|0|0|0|0|0|0|0|0|0|0|0|0|0|0|0|0|0|0|0|0|0|0|0|0|0|0|0|0|0|0|0|0|
|virus_Jun22.50v_18819|1|0|0|0|0|0|0|0|0|0|0|0|0|0|0|0|0|0|1|0|0|0|0|0|0|0|0|0|0|0|0|0|0|0|0|0|0|0|0|0|0|0|0|0|0|0|0|0|0|0|0|0|0|0|0|
|virus_Jun22.50v_23542|1|0|0|0|0|0|0|0|0|0|0|0|0|0|0|0|0|0|1|0|0|0|0|0|0|0|0|0|0|0|0|0|0|0|0|0|0|0|0|0|0|0|0|0|0|0|0|0|0|0|0|0|0|0|0|
|virus_Jun22.50v_27048|1|0|0|0|0|0|0|0|0|0|0|0|0|0|0|0|0|0|1|0|0|0|0|0|0|0|0|0|0|0|0|0|0|0|0|0|0|0|0|0|0|0|0|0|0|0|0|0|0|0|0|0|0|0|0|
|virus_Jun22.50v_31296|1|0|0|0|0|0|0|0|0|0|0|0|0|0|0|0|0|0|1|0|0|0|0|0|0|0|0|0|0|0|0|0|0|0|0|0|0|0|0|0|0|0|0|0|0|0|0|0|0|0|0|0|0|0|0|
|virus_Jun22.50v_33174|1|0|0|0|0|0|0|0|0|0|0|0|0|0|0|0|0|0|1|0|0|0|0|0|0|0|0|0|0|0|0|0|0|0|0|0|0|0|0|0|0|0|0|0|0|0|0|0|0|0|0|0|0|0|0|
|virus_Jun22.50v_39821|1|0|0|0|0|0|0|0|0|0|0|0|0|0|0|0|0|0|1|0|0|0|0|0|0|0|0|0|0|0|0|0|0|0|0|0|0|0|0|0|0|0|0|0|0|0|0|0|0|0|0|0|0|0|0|
|virus_Jun22.75v_731|1|0|0|0|0|0|0|0|0|0|0|0|0|0|0|0|0|0|0|0|1|0|0|0|0|0|0|0|0|0|0|0|0|0|0|0|0|0|0|0|0|0|0|0|0|0|0|0|0|0|0|0|0|0|0|
|virus_Jun22.75v_3480|1|0|0|0|0|0|0|0|0|0|0|0|0|0|0|0|0|0|0|0|1|0|0|0|0|0|0|0|0|0|0|0|0|0|0|0|0|0|0|0|0|0|0|0|0|0|0|0|0|0|0|0|0|0|0|
|virus_Jun22.75v_5791|1|0|0|0|0|0|0|0|0|0|0|0|0|0|0|0|0|0|0|0|1|0|0|0|0|0|0|0|0|0|0|0|0|0|0|0|0|0|0|0|0|0|0|0|0|0|0|0|0|0|0|0|0|0|0|
|virus_Jun22.75v_7618|1|0|0|0|0|0|0|0|0|0|0|0|0|0|0|0|0|0|0|0|1|0|0|0|0|0|0|0|0|0|0|0|0|0|0|0|0|0|0|0|0|0|0|0|0|0|0|0|0|0|0|0|0|0|0|
|virus_Jun22.75v_24388|1|0|0|0|0|0|0|0|0|0|0|0|0|0|0|0|0|0|0|0|1|0|0|0|0|0|0|0|0|0|0|0|0|0|0|0|0|0|0|0|0|0|0|0|0|0|0|0|0|0|0|0|0|0|0|
|virus_Jun22.75v_25721|1|0|0|0|0|0|0|0|0|0|0|0|0|0|0|0|0|0|0|0|1|0|0|0|0|0|0|0|0|0|0|0|0|0|0|0|0|0|0|0|0|0|0|0|0|0|0|0|0|0|0|0|0|0|0|
|virus_Jun22.75v_35587|1|0|0|0|0|0|0|0|0|0|0|0|0|0|0|0|0|0|0|0|1|0|0|0|0|0|0|0|0|0|0|0|0|0|0|0|0|0|0|0|0|0|0|0|0|0|0|0|0|0|0|0|0|0|0|
|virus_Jun22.75v_39233|1|0|0|0|0|0|0|0|0|0|0|0|0|0|0|0|0|0|0|0|1|0|0|0|0|0|0|0|0|0|0|0|0|0|0|0|0|0|0|0|0|0|0|0|0|0|0|0|0|0|0|0|0|0|0|
|virus_May12.5v_10392|1|0|1|0|0|0|0|0|0|0|0|0|0|0|0|0|0|0|0|0|0|0|0|0|0|0|0|0|0|0|0|0|0|0|0|0|0|0|0|0|0|0|0|0|0|0|0|0|0|0|0|0|0|0|0|
|virus_May12.5v_16119|1|0|1|0|0|0|0|0|0|0|0|0|0|0|0|0|0|0|0|0|0|0|0|0|0|0|0|0|0|0|0|0|0|0|0|0|0|0|0|0|0|0|0|0|0|0|0|0|0|0|0|0|0|0|0|
|virus_May12.5v_30925|1|0|1|0|0|0|0|0|0|0|0|0|0|0|0|0|0|0|0|0|0|0|0|0|0|0|0|0|0|0|0|0|0|0|0|0|0|0|0|0|0|0|0|0|0|0|0|0|0|0|0|0|0|0|0|
|virus_May12.5v_38399|1|0|1|0|0|0|0|0|0|0|0|0|0|0|0|0|0|0|0|0|0|0|0|0|0|0|0|0|0|0|0|0|0|0|0|0|0|0|0|0|0|0|0|0|0|0|0|0|0|0|0|0|0|0|0|
|virus_May12.25v_1198|1|0|0|0|1|0|0|0|0|0|0|0|0|0|0|0|0|0|0|0|0|0|0|0|0|0|0|0|0|0|0|0|0|0|0|0|0|0|0|0|0|0|0|0|0|0|0|0|0|0|0|0|0|0|0|
|virus_May12.25v_1851|1|0|0|0|1|0|0|0|0|0|0|0|0|0|0|0|0|0|0|0|0|0|0|0|0|0|0|0|0|0|0|0|0|0|0|0|0|0|0|0|0|0|0|0|0|0|0|0|0|0|0|0|0|0|0|
|virus_May12.25v_2853|1|0|0|0|1|0|0|0|0|0|0|0|0|0|0|0|0|0|0|0|0|0|0|0|0|0|0|0|0|0|0|0|0|0|0|0|0|0|0|0|0|0|0|0|0|0|0|0|0|0|0|0|0|0|0|
|virus_May12.25v_6517|1|0|0|0|1|0|0|0|0|0|0|0|0|0|0|0|0|0|0|0|0|0|0|0|0|0|0|0|0|0|0|0|0|0|0|0|0|0|0|0|0|0|0|0|0|0|0|0|0|0|0|0|0|0|0|
|virus_May12.25v_6591|1|0|0|0|1|0|0|0|0|0|0|0|0|0|0|0|0|0|0|0|0|0|0|0|0|0|0|0|0|0|0|0|0|0|0|0|0|0|0|0|0|0|0|0|0|0|0|0|0|0|0|0|0|0|0|
|virus_May12.25v_13968|1|0|0|0|1|0|0|0|0|0|0|0|0|0|0|0|0|0|0|0|0|0|0|0|0|0|0|0|0|0|0|0|0|0|0|0|0|0|0|0|0|0|0|0|0|0|0|0|0|0|0|0|0|0|0|
|virus_May12.25v_21905|1|0|0|0|1|0|0|0|0|0|0|0|0|0|0|0|0|0|0|0|0|0|0|0|0|0|0|0|0|0|0|0|0|0|0|0|0|0|0|0|0|0|0|0|0|0|0|0|0|0|0|0|0|0|0|
|virus_May12.25v_24101|1|0|0|0|1|0|0|0|0|0|0|0|0|0|0|0|0|0|0|0|0|0|0|0|0|0|0|0|0|0|0|0|0|0|0|0|0|0|0|0|0|0|0|0|0|0|0|0|0|0|0|0|0|0|0|
|virus_May12.25v_26384|1|0|0|0|1|0|0|0|0|0|0|0|0|0|0|0|0|0|0|0|0|0|0|0|0|0|0|0|0|0|0|0|0|0|0|0|0|0|0|0|0|0|0|0|0|0|0|0|0|0|0|0|0|0|0|
|virus_May12.25v_33564|1|0|0|0|1|0|0|0|0|0|0|0|0|0|0|0|0|0|0|0|0|0|0|0|0|0|0|0|0|0|0|0|0|0|0|0|0|0|0|0|0|0|0|0|0|0|0|0|0|0|0|0|0|0|0|
|virus_May12.25v_34745|1|0|0|0|1|0|0|0|0|0|0|0|0|0|0|0|0|0|0|0|0|0|0|0|0|0|0|0|0|0|0|0|0|0|0|0|0|0|0|0|0|0|0|0|0|0|0|0|0|0|0|0|0|0|0|
|virus_May12.50v_254|1|0|0|0|0|0|1|0|0|0|0|0|0|0|0|0|0|0|0|0|0|0|0|0|0|0|0|0|0|0|0|0|0|0|0|0|0|0|0|0|0|0|0|0|0|0|0|0|0|0|0|0|0|0|0|
|virus_May12.50v_2606|1|0|0|0|0|0|1|0|0|0|0|0|0|0|0|0|0|0|0|0|0|0|0|0|0|0|0|0|0|0|0|0|0|0|0|0|0|0|0|0|0|0|0|0|0|0|0|0|0|0|0|0|0|0|0|
|virus_May12.50v_5666|1|0|0|0|0|0|1|0|0|0|0|0|0|0|0|0|0|0|0|0|0|0|0|0|0|0|0|0|0|0|0|0|0|0|0|0|0|0|0|0|0|0|0|0|0|0|0|0|0|0|0|0|0|0|0|
|virus_May12.50v_6376|1|0|0|0|0|0|1|0|0|0|0|0|0|0|0|0|0|0|0|0|0|0|0|0|0|0|0|0|0|0|0|0|0|0|0|0|0|0|0|0|0|0|0|0|0|0|0|0|0|0|0|0|0|0|0|
|virus_May12.50v_13126|1|0|0|0|0|0|1|0|0|0|0|0|0|0|0|0|0|0|0|0|0|0|0|0|0|0|0|0|0|0|0|0|0|0|0|0|0|0|0|0|0|0|0|0|0|0|0|0|0|0|0|0|0|0|0|
|virus_May12.50v_14452|1|0|0|0|0|0|1|0|0|0|0|0|0|0|0|0|0|0|0|0|0|0|0|0|0|0|0|0|0|0|0|0|0|0|0|0|0|0|0|0|0|0|0|0|0|0|0|0|0|0|0|0|0|0|0|
|virus_May12.50v_29396|1|0|0|0|0|0|1|0|0|0|0|0|0|0|0|0|0|0|0|0|0|0|0|0|0|0|0|0|0|0|0|0|0|0|0|0|0|0|0|0|0|0|0|0|0|0|0|0|0|0|0|0|0|0|0|
|virus_May12.50v_29767|1|0|0|0|0|0|1|0|0|0|0|0|0|0|0|0|0|0|0|0|0|0|0|0|0|0|0|0|0|0|0|0|0|0|0|0|0|0|0|0|0|0|0|0|0|0|0|0|0|0|0|0|0|0|0|
|virus_May12.50v_31451|1|0|0|0|0|0|1|0|0|0|0|0|0|0|0|0|0|0|0|0|0|0|0|0|0|0|0|0|0|0|0|0|0|0|0|0|0|0|0|0|0|0|0|0|0|0|0|0|0|0|0|0|0|0|0|
|virus_May12.50v_32115|1|0|0|0|0|0|1|0|0|0|0|0|0|0|0|0|0|0|0|0|0|0|0|0|0|0|0|0|0|0|0|0|0|0|0|0|0|0|0|0|0|0|0|0|0|0|0|0|0|0|0|0|0|0|0|
|virus_May12.50v_33445|1|0|0|0|0|0|1|0|0|0|0|0|0|0|0|0|0|0|0|0|0|0|0|0|0|0|0|0|0|0|0|0|0|0|0|0|0|0|0|0|0|0|0|0|0|0|0|0|0|0|0|0|0|0|0|
|virus_May12.50v_35971|1|0|0|0|0|0|1|0|0|0|0|0|0|0|0|0|0|0|0|0|0|0|0|0|0|0|0|0|0|0|0|0|0|0|0|0|0|0|0|0|0|0|0|0|0|0|0|0|0|0|0|0|0|0|0|
|virus_May12.50v_39645|1|0|0|0|0|0|1|0|0|0|0|0|0|0|0|0|0|0|0|0|0|0|0|0|0|0|0|0|0|0|0|0|0|0|0|0|0|0|0|0|0|0|0|0|0|0|0|0|0|0|0|0|0|0|0|
|virus_May12.50v_40192|1|0|0|0|0|0|1|0|0|0|0|0|0|0|0|0|0|0|0|0|0|0|0|0|0|0|0|0|0|0|0|0|0|0|0|0|0|0|0|0|0|0|0|0|0|0|0|0|0|0|0|0|0|0|0|
|virus_May12.75v_113|1|0|0|0|0|0|0|0|1|0|0|0|0|0|0|0|0|0|0|0|0|0|0|0|0|0|0|0|0|0|0|0|0|0|0|0|0|0|0|0|0|0|0|0|0|0|0|0|0|0|0|0|0|0|0|
|virus_May12.75v_6261|1|0|0|0|0|0|0|0|1|0|0|0|0|0|0|0|0|0|0|0|0|0|0|0|0|0|0|0|0|0|0|0|0|0|0|0|0|0|0|0|0|0|0|0|0|0|0|0|0|0|0|0|0|0|0|
|virus_May12.75v_9003|1|0|0|0|0|0|0|0|1|0|0|0|0|0|0|0|0|0|0|0|0|0|0|0|0|0|0|0|0|0|0|0|0|0|0|0|0|0|0|0|0|0|0|0|0|0|0|0|0|0|0|0|0|0|0|
|virus_May12.75v_15769|1|0|0|0|0|0|0|0|1|0|0|0|0|0|0|0|0|0|0|0|0|0|0|0|0|0|0|0|0|0|0|0|0|0|0|0|0|0|0|0|0|0|0|0|0|0|0|0|0|0|0|0|0|0|0|
|virus_May12.75v_20408|1|0|0|0|0|0|0|0|1|0|0|0|0|0|0|0|0|0|0|0|0|0|0|0|0|0|0|0|0|0|0|0|0|0|0|0|0|0|0|0|0|0|0|0|0|0|0|0|0|0|0|0|0|0|0|
|virus_May12.75v_20647|1|0|0|0|0|0|0|0|1|0|0|0|0|0|0|0|0|0|0|0|0|0|0|0|0|0|0|0|0|0|0|0|0|0|0|0|0|0|0|0|0|0|0|0|0|0|0|0|0|0|0|0|0|0|0|
|virus_May12.75v_20654|1|0|0|0|0|0|0|0|1|0|0|0|0|0|0|0|0|0|0|0|0|0|0|0|0|0|0|0|0|0|0|0|0|0|0|0|0|0|0|0|0|0|0|0|0|0|0|0|0|0|0|0|0|0|0|
|virus_May12.75v_22749|1|0|0|0|0|0|0|0|1|0|0|0|0|0|0|0|0|0|0|0|0|0|0|0|0|0|0|0|0|0|0|0|0|0|0|0|0|0|0|0|0|0|0|0|0|0|0|0|0|0|0|0|0|0|0|
|virus_May12.75v_29897|1|0|0|0|0|0|0|0|1|0|0|0|0|0|0|0|0|0|0|0|0|0|0|0|0|0|0|0|0|0|0|0|0|0|0|0|0|0|0|0|0|0|0|0|0|0|0|0|0|0|0|0|0|0|0|
|virus_May12.75v_31871|1|0|0|0|0|0|0|0|1|0|0|0|0|0|0|0|0|0|0|0|0|0|0|0|0|0|0|0|0|0|0|0|0|0|0|0|0|0|0|0|0|0|0|0|0|0|0|0|0|0|0|0|0|0|0|
|virus_May12.75v_37898|1|0|0|0|0|0|0|0|1|0|0|0|0|0|0|0|0|0|0|0|0|0|0|0|0|0|0|0|0|0|0|0|0|0|0|0|0|0|0|0|0|0|0|0|0|0|0|0|0|0|0|0|0|0|0|
|virus_Sep8.0v_3905|1|0|0|0|0|0|0|0|0|0|0|0|0|0|0|0|0|0|0|0|0|0|0|0|0|0|0|0|0|0|0|0|0|0|0|0|0|0|0|0|0|0|0|0|0|0|0|0|1|0|0|0|0|0|0|
|virus_Sep8.0v_6064|1|0|0|0|0|0|0|0|0|0|0|0|0|0|0|0|0|0|0|0|0|0|0|0|0|0|0|0|0|0|0|0|0|0|0|0|0|0|0|0|0|0|0|0|0|0|0|0|1|0|0|0|0|0|0|
|virus_Sep8.0v_6839|1|0|0|0|0|0|0|0|0|0|0|0|0|0|0|0|0|0|0|0|0|0|0|0|0|0|0|0|0|0|0|0|0|0|0|0|0|0|0|0|0|0|0|0|0|0|0|0|1|0|0|0|0|0|0|
|virus_Sep8.0v_9317|1|0|0|0|0|0|0|0|0|0|0|0|0|0|0|0|0|0|0|0|0|0|0|0|0|0|0|0|0|0|0|0|0|0|0|0|0|0|0|0|0|0|0|0|0|0|0|0|1|0|0|0|0|0|0|
|virus_Sep8.0v_11663|1|0|0|0|0|0|0|0|0|0|0|0|0|0|0|0|0|0|0|0|0|0|0|0|0|0|0|0|0|0|0|0|0|0|0|0|0|0|0|0|0|0|0|0|0|0|0|0|1|0|0|0|0|0|0|
|virus_Sep8.0v_14253|1|0|0|0|0|0|0|0|0|0|0|0|0|0|0|0|0|0|0|0|0|0|0|0|0|0|0|0|0|0|0|0|0|0|0|0|0|0|0|0|0|0|0|0|0|0|0|0|1|0|0|0|0|0|0|
|virus_Sep8.0v_15334|1|0|0|0|0|0|0|0|0|0|0|0|0|0|0|0|0|0|0|0|0|0|0|0|0|0|0|0|0|0|0|0|0|0|0|0|0|0|0|0|0|0|0|0|0|0|0|0|1|0|0|0|0|0|0|
|virus_Sep8.0v_16261|1|0|0|0|0|0|0|0|0|0|0|0|0|0|0|0|0|0|0|0|0|0|0|0|0|0|0|0|0|0|0|0|0|0|0|0|0|0|0|0|0|0|0|0|0|0|0|0|1|0|0|0|0|0|0|
|virus_Sep8.0v_17426|1|0|0|0|0|0|0|0|0|0|0|0|0|0|0|0|0|0|0|0|0|0|0|0|0|0|0|0|0|0|0|0|0|0|0|0|0|0|0|0|0|0|0|0|0|0|0|0|1|0|0|0|0|0|0|
|virus_Sep8.0v_18634|1|0|0|0|0|0|0|0|0|0|0|0|0|0|0|0|0|0|0|0|0|0|0|0|0|0|0|0|0|0|0|0|0|0|0|0|0|0|0|0|0|0|0|0|0|0|0|0|1|0|0|0|0|0|0|
|virus_Sep8.0v_20158|1|0|0|0|0|0|0|0|0|0|0|0|0|0|0|0|0|0|0|0|0|0|0|0|0|0|0|0|0|0|0|0|0|0|0|0|0|0|0|0|0|0|0|0|0|0|0|0|1|0|0|0|0|0|0|
|virus_Sep8.5v_15643|1|0|0|0|0|0|0|0|0|0|0|0|0|0|0|0|0|0|0|0|0|0|0|0|0|0|0|0|0|0|0|0|0|0|0|0|0|0|0|0|0|0|0|0|0|0|0|0|0|0|1|0|0|0|0|
|virus_Sep8.25v_4126|1|0|0|0|0|0|0|0|0|0|0|0|0|0|0|0|0|0|0|0|0|0|0|0|0|0|0|0|0|0|0|0|0|0|0|0|0|0|0|0|0|0|0|0|0|0|0|0|0|0|0|0|1|0|0|
|virus_Sep8.25v_13893|1|0|0|0|0|0|0|0|0|0|0|0|0|0|0|0|0|0|0|0|0|0|0|0|0|0|0|0|0|0|0|0|0|0|0|0|0|0|0|0|0|0|0|0|0|0|0|0|0|0|0|0|1|0|0|
|virus_Sep8.25v_16511|1|0|0|0|0|0|0|0|0|0|0|0|0|0|0|0|0|0|0|0|0|0|0|0|0|0|0|0|0|0|0|0|0|0|0|0|0|0|0|0|0|0|0|0|0|0|0|0|0|0|0|0|1|0|0|
|virus_Sep8.25v_20096|1|0|0|0|0|0|0|0|0|0|0|0|0|0|0|0|0|0|0|0|0|0|0|0|0|0|0|0|0|0|0|0|0|0|0|0|0|0|0|0|0|0|0|0|0|0|0|0|0|0|0|0|1|0|0|
|virus_Sep8.50v_1340|1|0|0|0|0|0|0|0|0|0|0|0|0|0|0|0|0|0|0|0|0|0|0|0|0|0|0|0|0|0|0|0|0|0|0|0|0|0|0|0|0|0|0|0|0|0|0|0|0|0|0|0|0|0|1|
|virus_Sep8.50v_7825|1|0|0|0|0|0|0|0|0|0|0|0|0|0|0|0|0|0|0|0|0|0|0|0|0|0|0|0|0|0|0|0|0|0|0|0|0|0|0|0|0|0|0|0|0|0|0|0|0|0|0|0|0|0|1|
|virus_Sep8.50v_15806|1|0|0|0|0|0|0|0|0|0|0|0|0|0|0|0|0|0|0|0|0|0|0|0|0|0|0|0|0|0|0|0|0|0|0|0|0|0|0|0|0|0|0|0|0|0|0|0|0|0|0|0|0|0|1|
|virus_Sep8.50v_16868|1|0|0|0|0|0|0|0|0|0|0|0|0|0|0|0|0|0|0|0|0|0|0|0|0|0|0|0|0|0|0|0|0|0|0|0|0|0|0|0|0|0|0|0|0|0|0|0|0|0|0|0|0|0|1|
|virus_Sep8.50v_20345|1|0|0|0|0|0|0|0|0|0|0|0|0|0|0|0|0|0|0|0|0|0|0|0|0|0|0|0|0|0|0|0|0|0|0|0|0|0|0|0|0|0|0|0|0|0|0|0|0|0|0|0|0|0|1|
|virus_Sep8.50v_27827|1|0|0|0|0|0|0|0|0|0|0|0|0|0|0|0|0|0|0|0|0|0|0|0|0|0|0|0|0|0|0|0|0|0|0|0|0|0|0|0|0|0|0|0|0|0|0|0|0|0|0|0|0|0|1|
|virus_Sep8.50v_30163|1|0|0|0|0|0|0|0|0|0|0|0|0|0|0|0|0|0|0|0|0|0|0|0|0|0|0|0|0|0|0|0|0|0|0|0|0|0|0|0|0|0|0|0|0|0|0|0|0|0|0|0|0|0|1|
|virus_Sep8.50v_33206|1|0|0|0|0|0|0|0|0|0|0|0|0|0|0|0|0|0|0|0|0|0|0|0|0|0|0|0|0|0|0|0|0|0|0|0|0|0|0|0|0|0|0|0|0|0|0|0|0|0|0|0|0|0|1|
|virus_Sep8.50v_36820|1|0|0|0|0|0|0|0|0|0|0|0|0|0|0|0|0|0|0|0|0|0|0|0|0|0|0|0|0|0|0|0|0|0|0|0|0|0|0|0|0|0|0|0|0|0|0|0|0|0|0|0|0|0|1|
|virus_Sep8.50v_37938|1|0|0|0|0|0|0|0|0|0|0|0|0|0|0|0|0|0|0|0|0|0|0|0|0|0|0|0|0|0|0|0|0|0|0|0|0|0|0|0|0|0|0|0|0|0|0|0|0|0|0|0|0|0|1|
|virus_Sep8.50v_39605|1|0|0|0|0|0|0|0|0|0|0|0|0|0|0|0|0|0|0|0|0|0|0|0|0|0|0|0|0|0|0|0|0|0|0|0|0|0|0|0|0|0|0|0|0|0|0|0|0|0|0|0|0|0|1|
|bacteria_Jul11d0b_5361|1|0|0|0|0|0|0|0|0|0|0|0|0|0|0|0|0|0|0|0|0|1|0|0|0|0|0|0|0|0|0|0|0|0|0|0|0|0|0|0|0|0|0|0|0|0|0|0|0|0|0|0|0|0|0|
|bacteria_Jul11d0b_30174|1|0|0|0|0|0|0|0|0|0|0|0|0|0|0|0|0|0|0|0|0|1|0|0|0|0|0|0|0|0|0|0|0|0|0|0|0|0|0|0|0|0|0|0|0|0|0|0|0|0|0|0|0|0|0|
|bacteria_Jul11d0b_37224|1|0|0|0|0|0|0|0|0|0|0|0|0|0|0|0|0|0|0|0|0|1|0|0|0|0|0|0|0|0|0|0|0|0|0|0|0|0|0|0|0|0|0|0|0|0|0|0|0|0|0|0|0|0|0|
|bacteria_Jul11d0b_38022|1|0|0|0|0|0|0|0|0|0|0|0|0|0|0|0|0|0|0|0|0|1|0|0|0|0|0|0|0|0|0|0|0|0|0|0|0|0|0|0|0|0|0|0|0|0|0|0|0|0|0|0|0|0|0|
|bacteria_Jul11d0b_43588|1|0|0|0|0|0|0|0|0|0|0|0|0|0|0|0|0|0|0|0|0|1|0|0|0|0|0|0|0|0|0|0|0|0|0|0|0|0|0|0|0|0|0|0|0|0|0|0|0|0|0|0|0|0|0|
|bacteria_Aug5d0b_18242|1|0|0|0|0|0|0|0|0|0|0|0|0|0|0|0|0|0|0|0|0|0|0|0|0|0|0|0|0|0|0|0|0|0|0|0|0|1|0|0|0|0|0|0|0|0|0|0|0|0|0|0|0|0|0|
|bacteria_Aug5d5b_65398|1|0|0|0|0|0|0|0|0|0|0|0|0|0|0|0|0|0|0|0|0|0|0|0|0|0|0|0|0|0|0|0|0|0|0|0|0|0|0|1|0|0|0|0|0|0|0|0|0|0|0|0|0|0|0|
|bacteria_Aug5d25b_73|1|0|0|0|0|0|0|0|0|0|0|0|0|0|0|0|0|0|0|0|0|0|0|0|0|0|0|0|0|0|0|0|0|0|0|0|0|0|0|0|0|1|0|0|0|0|0|0|0|0|0|0|0|0|0|
|bacteria_Aug5d25b_11334|1|0|0|0|0|0|0|0|0|0|0|0|0|0|0|0|0|0|0|0|0|0|0|0|0|0|0|0|0|0|0|0|0|0|0|0|0|0|0|0|0|1|0|0|0|0|0|0|0|0|0|0|0|0|0|
|bacteria_Aug5d25b_21156|1|0|0|0|0|0|0|0|0|0|0|0|0|0|0|0|0|0|0|0|0|0|0|0|0|0|0|0|0|0|0|0|0|0|0|0|0|0|0|0|0|1|0|0|0|0|0|0|0|0|0|0|0|0|0|
|bacteria_Aug5d25b_21262|1|0|0|0|0|0|0|0|0|0|0|0|0|0|0|0|0|0|0|0|0|0|0|0|0|0|0|0|0|0|0|0|0|0|0|0|0|0|0|0|0|1|0|0|0|0|0|0|0|0|0|0|0|0|0|
|bacteria_Aug5d25b_23361|1|0|0|0|0|0|0|0|0|0|0|0|0|0|0|0|0|0|0|0|0|0|0|0|0|0|0|0|0|0|0|0|0|0|0|0|0|0|0|0|0|1|0|0|0|0|0|0|0|0|0|0|0|0|0|
|bacteria_Aug5d25b_33607|1|0|0|0|0|0|0|0|0|0|0|0|0|0|0|0|0|0|0|0|0|0|0|0|0|0|0|0|0|0|0|0|0|0|0|0|0|0|0|0|0|1|0|0|0|0|0|0|0|0|0|0|0|0|0|
|bacteria_Aug5d25b_33619|1|0|0|0|0|0|0|0|0|0|0|0|0|0|0|0|0|0|0|0|0|0|0|0|0|0|0|0|0|0|0|0|0|0|0|0|0|0|0|0|0|1|0|0|0|0|0|0|0|0|0|0|0|0|0|
|bacteria_Aug5d25b_37158|1|0|0|0|0|0|0|0|0|0|0|0|0|0|0|0|0|0|0|0|0|0|0|0|0|0|0|0|0|0|0|0|0|0|0|0|0|0|0|0|0|1|0|0|0|0|0|0|0|0|0|0|0|0|0|
|bacteria_Aug5d25b_39379|1|0|0|0|0|0|0|0|0|0|0|0|0|0|0|0|0|0|0|0|0|0|0|0|0|0|0|0|0|0|0|0|0|0|0|0|0|0|0|0|0|1|0|0|0|0|0|0|0|0|0|0|0|0|0|
|bacteria_Aug5d25b_46040|1|0|0|0|0|0|0|0|0|0|0|0|0|0|0|0|0|0|0|0|0|0|0|0|0|0|0|0|0|0|0|0|0|0|0|0|0|0|0|0|0|1|0|0|0|0|0|0|0|0|0|0|0|0|0|
|bacteria_Aug5d25b_50009|1|0|0|0|0|0|0|0|0|0|0|0|0|0|0|0|0|0|0|0|0|0|0|0|0|0|0|0|0|0|0|0|0|0|0|0|0|0|0|0|0|1|0|0|0|0|0|0|0|0|0|0|0|0|0|
|bacteria_Aug5d25b_51012|1|0|0|0|0|0|0|0|0|0|0|0|0|0|0|0|0|0|0|0|0|0|0|0|0|0|0|0|0|0|0|0|0|0|0|0|0|0|0|0|0|1|0|0|0|0|0|0|0|0|0|0|0|0|0|
|bacteria_Aug5d25b_67053|1|0|0|0|0|0|0|0|0|0|0|0|0|0|0|0|0|0|0|0|0|0|0|0|0|0|0|0|0|0|0|0|0|0|0|0|0|0|0|0|0|1|0|0|0|0|0|0|0|0|0|0|0|0|0|
|bacteria_Aug5d25b_67151|1|0|0|0|0|0|0|0|0|0|0|0|0|0|0|0|0|0|0|0|0|0|0|0|0|0|0|0|0|0|0|0|0|0|0|0|0|0|0|0|0|1|0|0|0|0|0|0|0|0|0|0|0|0|0|
|bacteria_Aug5d25b_68097|1|0|0|0|0|0|0|0|0|0|0|0|0|0|0|0|0|0|0|0|0|0|0|0|0|0|0|0|0|0|0|0|0|0|0|0|0|0|0|0|0|1|0|0|0|0|0|0|0|0|0|0|0|0|0|
|bacteria_Aug5d50b_2721|1|0|0|0|0|0|0|0|0|0|0|0|0|0|0|0|0|0|0|0|0|0|0|0|0|0|0|0|0|0|0|0|0|0|0|0|0|0|0|0|0|0|0|1|0|0|0|0|0|0|0|0|0|0|0|
|bacteria_Aug5d50b_3371|1|0|0|0|0|0|0|0|0|0|0|0|0|0|0|0|0|0|0|0|0|0|0|0|0|0|0|0|0|0|0|0|0|0|0|0|0|0|0|0|0|0|0|1|0|0|0|0|0|0|0|0|0|0|0|
|bacteria_Aug5d50b_27643|1|0|0|0|0|0|0|0|0|0|0|0|0|0|0|0|0|0|0|0|0|0|0|0|0|0|0|0|0|0|0|0|0|0|0|0|0|0|0|0|0|0|0|1|0|0|0|0|0|0|0|0|0|0|0|
|bacteria_Aug5d50b_52726|1|0|0|0|0|0|0|0|0|0|0|0|0|0|0|0|0|0|0|0|0|0|0|0|0|0|0|0|0|0|0|0|0|0|0|0|0|0|0|0|0|0|0|1|0|0|0|0|0|0|0|0|0|0|0|
|bacteria_Aug5d50b_70364|1|0|0|0|0|0|0|0|0|0|0|0|0|0|0|0|0|0|0|0|0|0|0|0|0|0|0|0|0|0|0|0|0|0|0|0|0|0|0|0|0|0|0|1|0|0|0|0|0|0|0|0|0|0|0|
|bacteria_Jul11d5b_1349|1|0|0|0|0|0|0|0|0|0|0|0|0|0|0|0|0|0|0|0|0|0|0|1|0|0|0|0|0|0|0|0|0|0|0|0|0|0|0|0|0|0|0|0|0|0|0|0|0|0|0|0|0|0|0|
|bacteria_Jul11d5b_23323|1|0|0|0|0|0|0|0|0|0|0|0|0|0|0|0|0|0|0|0|0|0|0|1|0|0|0|0|0|0|0|0|0|0|0|0|0|0|0|0|0|0|0|0|0|0|0|0|0|0|0|0|0|0|0|
|bacteria_Jul11d5b_23680|1|0|0|0|0|0|0|0|0|0|0|0|0|0|0|0|0|0|0|0|0|0|0|1|0|0|0|0|0|0|0|0|0|0|0|0|0|0|0|0|0|0|0|0|0|0|0|0|0|0|0|0|0|0|0|
|bacteria_Jul11d5b_27296|1|0|0|0|0|0|0|0|0|0|0|0|0|0|0|0|0|0|0|0|0|0|0|1|0|0|0|0|0|0|0|0|0|0|0|0|0|0|0|0|0|0|0|0|0|0|0|0|0|0|0|0|0|0|0|
|bacteria_Jul11d5b_38272|1|0|0|0|0|0|0|0|0|0|0|0|0|0|0|0|0|0|0|0|0|0|0|1|0|0|0|0|0|0|0|0|0|0|0|0|0|0|0|0|0|0|0|0|0|0|0|0|0|0|0|0|0|0|0|
|bacteria_Jul11d5b_45099|1|0|0|0|0|0|0|0|0|0|0|0|0|0|0|0|0|0|0|0|0|0|0|1|0|0|0|0|0|0|0|0|0|0|0|0|0|0|0|0|0|0|0|0|0|0|0|0|0|0|0|0|0|0|0|
|bacteria_Jul11d25b_10267|1|0|0|0|0|0|0|0|0|0|0|0|0|0|0|0|0|0|0|0|0|0|0|0|0|1|0|0|0|0|0|0|0|0|0|0|0|0|0|0|0|0|0|0|0|0|0|0|0|0|0|0|0|0|0|
|bacteria_Jul11d25b_14302|1|0|0|0|0|0|0|0|0|0|0|0|0|0|0|0|0|0|0|0|0|0|0|0|0|1|0|0|0|0|0|0|0|0|0|0|0|0|0|0|0|0|0|0|0|0|0|0|0|0|0|0|0|0|0|
|bacteria_Jul11d25b_19241|1|0|0|0|0|0|0|0|0|0|0|0|0|0|0|0|0|0|0|0|0|0|0|0|0|1|0|0|0|0|0|0|0|0|0|0|0|0|0|0|0|0|0|0|0|0|0|0|0|0|0|0|0|0|0|
|bacteria_Jul11d25b_35754|1|0|0|0|0|0|0|0|0|0|0|0|0|0|0|0|0|0|0|0|0|0|0|0|0|1|0|0|0|0|0|0|0|0|0|0|0|0|0|0|0|0|0|0|0|0|0|0|0|0|0|0|0|0|0|
|bacteria_Jul11d25b_50924|1|0|0|0|0|0|0|0|0|0|0|0|0|0|0|0|0|0|0|0|0|0|0|0|0|1|0|0|0|0|0|0|0|0|0|0|0|0|0|0|0|0|0|0|0|0|0|0|0|0|0|0|0|0|0|
|bacteria_Jul11d50b_29403|1|0|0|0|0|0|0|0|0|0|0|0|0|0|0|0|0|0|0|0|0|0|0|0|0|0|0|1|0|0|0|0|0|0|0|0|0|0|0|0|0|0|0|0|0|0|0|0|0|0|0|0|0|0|0|
|bacteria_Jul11d50b_44736|1|0|0|0|0|0|0|0|0|0|0|0|0|0|0|0|0|0|0|0|0|0|0|0|0|0|0|1|0|0|0|0|0|0|0|0|0|0|0|0|0|0|0|0|0|0|0|0|0|0|0|0|0|0|0|
|bacteria_Jul11d50b_47362|1|0|0|0|0|0|0|0|0|0|0|0|0|0|0|0|0|0|0|0|0|0|0|0|0|0|0|1|0|0|0|0|0|0|0|0|0|0|0|0|0|0|0|0|0|0|0|0|0|0|0|0|0|0|0|
|bacteria_Jul11d50b_52033|1|0|0|0|0|0|0|0|0|0|0|0|0|0|0|0|0|0|0|0|0|0|0|0|0|0|0|1|0|0|0|0|0|0|0|0|0|0|0|0|0|0|0|0|0|0|0|0|0|0|0|0|0|0|0|
|bacteria_Jul11d75b_8184|1|0|0|0|0|0|0|0|0|0|0|0|0|0|0|0|0|0|0|0|0|0|0|0|0|0|0|0|0|1|0|0|0|0|0|0|0|0|0|0|0|0|0|0|0|0|0|0|0|0|0|0|0|0|0|
|bacteria_Jul11d75b_11391|1|0|0|0|0|0|0|0|0|0|0|0|0|0|0|0|0|0|0|0|0|0|0|0|0|0|0|0|0|1|0|0|0|0|0|0|0|0|0|0|0|0|0|0|0|0|0|0|0|0|0|0|0|0|0|
|bacteria_Jul11d75b_16431|1|0|0|0|0|0|0|0|0|0|0|0|0|0|0|0|0|0|0|0|0|0|0|0|0|0|0|0|0|1|0|0|0|0|0|0|0|0|0|0|0|0|0|0|0|0|0|0|0|0|0|0|0|0|0|
|bacteria_Jul11d75b_20357|1|0|0|0|0|0|0|0|0|0|0|0|0|0|0|0|0|0|0|0|0|0|0|0|0|0|0|0|0|1|0|0|0|0|0|0|0|0|0|0|0|0|0|0|0|0|0|0|0|0|0|0|0|0|0|
|bacteria_Jul11d75b_23171|1|0|0|0|0|0|0|0|0|0|0|0|0|0|0|0|0|0|0|0|0|0|0|0|0|0|0|0|0|1|0|0|0|0|0|0|0|0|0|0|0|0|0|0|0|0|0|0|0|0|0|0|0|0|0|
|bacteria_Jul11d75b_25120|1|0|0|0|0|0|0|0|0|0|0|0|0|0|0|0|0|0|0|0|0|0|0|0|0|0|0|0|0|1|0|0|0|0|0|0|0|0|0|0|0|0|0|0|0|0|0|0|0|0|0|0|0|0|0|
|bacteria_Jul11d75b_28029|1|0|0|0|0|0|0|0|0|0|0|0|0|0|0|0|0|0|0|0|0|0|0|0|0|0|0|0|0|1|0|0|0|0|0|0|0|0|0|0|0|0|0|0|0|0|0|0|0|0|0|0|0|0|0|
|bacteria_Jul11d75b_29883|1|0|0|0|0|0|0|0|0|0|0|0|0|0|0|0|0|0|0|0|0|0|0|0|0|0|0|0|0|1|0|0|0|0|0|0|0|0|0|0|0|0|0|0|0|0|0|0|0|0|0|0|0|0|0|
|bacteria_Jul11d75b_32862|1|0|0|0|0|0|0|0|0|0|0|0|0|0|0|0|0|0|0|0|0|0|0|0|0|0|0|0|0|1|0|0|0|0|0|0|0|0|0|0|0|0|0|0|0|0|0|0|0|0|0|0|0|0|0|
|bacteria_Jul11d75b_33219|1|0|0|0|0|0|0|0|0|0|0|0|0|0|0|0|0|0|0|0|0|0|0|0|0|0|0|0|0|1|0|0|0|0|0|0|0|0|0|0|0|0|0|0|0|0|0|0|0|0|0|0|0|0|0|
|bacteria_Jul11d75b_43477|1|0|0|0|0|0|0|0|0|0|0|0|0|0|0|0|0|0|0|0|0|0|0|0|0|0|0|0|0|1|0|0|0|0|0|0|0|0|0|0|0|0|0|0|0|0|0|0|0|0|0|0|0|0|0|
|bacteria_Jul11d75b_46998|1|0|0|0|0|0|0|0|0|0|0|0|0|0|0|0|0|0|0|0|0|0|0|0|0|0|0|0|0|1|0|0|0|0|0|0|0|0|0|0|0|0|0|0|0|0|0|0|0|0|0|0|0|0|0|
|bacteria_Jul11d75b_50166|1|0|0|0|0|0|0|0|0|0|0|0|0|0|0|0|0|0|0|0|0|0|0|0|0|0|0|0|0|1|0|0|0|0|0|0|0|0|0|0|0|0|0|0|0|0|0|0|0|0|0|0|0|0|0|
|bacteria_Jul27d5b_3120|1|0|0|0|0|0|0|0|0|0|0|0|0|0|0|0|0|0|0|0|0|0|0|0|0|0|0|0|0|0|0|1|0|0|0|0|0|0|0|0|0|0|0|0|0|0|0|0|0|0|0|0|0|0|0|
|bacteria_Jul27d5b_5428|1|0|0|0|0|0|0|0|0|0|0|0|0|0|0|0|0|0|0|0|0|0|0|0|0|0|0|0|0|0|0|1|0|0|0|0|0|0|0|0|0|0|0|0|0|0|0|0|0|0|0|0|0|0|0|
|bacteria_Jul27d5b_10360|1|0|0|0|0|0|0|0|0|0|0|0|0|0|0|0|0|0|0|0|0|0|0|0|0|0|0|0|0|0|0|1|0|0|0|0|0|0|0|0|0|0|0|0|0|0|0|0|0|0|0|0|0|0|0|
|bacteria_Jul27d5b_12140|1|0|0|0|0|0|0|0|0|0|0|0|0|0|0|0|0|0|0|0|0|0|0|0|0|0|0|0|0|0|0|1|0|0|0|0|0|0|0|0|0|0|0|0|0|0|0|0|0|0|0|0|0|0|0|
|bacteria_Jul27d5b_15953|1|0|0|0|0|0|0|0|0|0|0|0|0|0|0|0|0|0|0|0|0|0|0|0|0|0|0|0|0|0|0|1|0|0|0|0|0|0|0|0|0|0|0|0|0|0|0|0|0|0|0|0|0|0|0|
|bacteria_Jul27d5b_26185|1|0|0|0|0|0|0|0|0|0|0|0|0|0|0|0|0|0|0|0|0|0|0|0|0|0|0|0|0|0|0|1|0|0|0|0|0|0|0|0|0|0|0|0|0|0|0|0|0|0|0|0|0|0|0|
|bacteria_Jul27d5b_26651|1|0|0|0|0|0|0|0|0|0|0|0|0|0|0|0|0|0|0|0|0|0|0|0|0|0|0|0|0|0|0|1|0|0|0|0|0|0|0|0|0|0|0|0|0|0|0|0|0|0|0|0|0|0|0|
|bacteria_Jul27d5b_33096|1|0|0|0|0|0|0|0|0|0|0|0|0|0|0|0|0|0|0|0|0|0|0|0|0|0|0|0|0|0|0|1|0|0|0|0|0|0|0|0|0|0|0|0|0|0|0|0|0|0|0|0|0|0|0|
|bacteria_Jul27d5b_33101|1|0|0|0|0|0|0|0|0|0|0|0|0|0|0|0|0|0|0|0|0|0|0|0|0|0|0|0|0|0|0|1|0|0|0|0|0|0|0|0|0|0|0|0|0|0|0|0|0|0|0|0|0|0|0|
|bacteria_Jul27d5b_35075|1|0|0|0|0|0|0|0|0|0|0|0|0|0|0|0|0|0|0|0|0|0|0|0|0|0|0|0|0|0|0|1|0|0|0|0|0|0|0|0|0|0|0|0|0|0|0|0|0|0|0|0|0|0|0|
|bacteria_Jul27d5b_35909|1|0|0|0|0|0|0|0|0|0|0|0|0|0|0|0|0|0|0|0|0|0|0|0|0|0|0|0|0|0|0|1|0|0|0|0|0|0|0|0|0|0|0|0|0|0|0|0|0|0|0|0|0|0|0|
|bacteria_Jul27d5b_48854|1|0|0|0|0|0|0|0|0|0|0|0|0|0|0|0|0|0|0|0|0|0|0|0|0|0|0|0|0|0|0|1|0|0|0|0|0|0|0|0|0|0|0|0|0|0|0|0|0|0|0|0|0|0|0|
|bacteria_Jul27d5b_49859|1|0|0|0|0|0|0|0|0|0|0|0|0|0|0|0|0|0|0|0|0|0|0|0|0|0|0|0|0|0|0|1|0|0|0|0|0|0|0|0|0|0|0|0|0|0|0|0|0|0|0|0|0|0|0|
|bacteria_Jul27d25b_7972|1|0|0|0|0|0|0|0|0|0|0|0|0|0|0|0|0|0|0|0|0|0|0|0|0|0|0|0|0|0|0|0|0|1|0|0|0|0|0|0|0|0|0|0|0|0|0|0|0|0|0|0|0|0|0|
|bacteria_Jul27d25b_15951|1|0|0|0|0|0|0|0|0|0|0|0|0|0|0|0|0|0|0|0|0|0|0|0|0|0|0|0|0|0|0|0|0|1|0|0|0|0|0|0|0|0|0|0|0|0|0|0|0|0|0|0|0|0|0|
|bacteria_Jul27d25b_20487|1|0|0|0|0|0|0|0|0|0|0|0|0|0|0|0|0|0|0|0|0|0|0|0|0|0|0|0|0|0|0|0|0|1|0|0|0|0|0|0|0|0|0|0|0|0|0|0|0|0|0|0|0|0|0|
|bacteria_Jul27d25b_22563|1|0|0|0|0|0|0|0|0|0|0|0|0|0|0|0|0|0|0|0|0|0|0|0|0|0|0|0|0|0|0|0|0|1|0|0|0|0|0|0|0|0|0|0|0|0|0|0|0|0|0|0|0|0|0|
|bacteria_Jul27d25b_25276|1|0|0|0|0|0|0|0|0|0|0|0|0|0|0|0|0|0|0|0|0|0|0|0|0|0|0|0|0|0|0|0|0|1|0|0|0|0|0|0|0|0|0|0|0|0|0|0|0|0|0|0|0|0|0|
|bacteria_Jul27d25b_25284|1|0|0|0|0|0|0|0|0|0|0|0|0|0|0|0|0|0|0|0|0|0|0|0|0|0|0|0|0|0|0|0|0|1|0|0|0|0|0|0|0|0|0|0|0|0|0|0|0|0|0|0|0|0|0|
|bacteria_Jul27d25b_26744|1|0|0|0|0|0|0|0|0|0|0|0|0|0|0|0|0|0|0|0|0|0|0|0|0|0|0|0|0|0|0|0|0|1|0|0|0|0|0|0|0|0|0|0|0|0|0|0|0|0|0|0|0|0|0|
|bacteria_Jul27d25b_30574|1|0|0|0|0|0|0|0|0|0|0|0|0|0|0|0|0|0|0|0|0|0|0|0|0|0|0|0|0|0|0|0|0|1|0|0|0|0|0|0|0|0|0|0|0|0|0|0|0|0|0|0|0|0|0|
|bacteria_Jul27d25b_31099|1|0|0|0|0|0|0|0|0|0|0|0|0|0|0|0|0|0|0|0|0|0|0|0|0|0|0|0|0|0|0|0|0|1|0|0|0|0|0|0|0|0|0|0|0|0|0|0|0|0|0|0|0|0|0|
|bacteria_Jul27d25b_51427|1|0|0|0|0|0|0|0|0|0|0|0|0|0|0|0|0|0|0|0|0|0|0|0|0|0|0|0|0|0|0|0|0|1|0|0|0|0|0|0|0|0|0|0|0|0|0|0|0|0|0|0|0|0|0|
|bacteria_Jul27d25b_51956|1|0|0|0|0|0|0|0|0|0|0|0|0|0|0|0|0|0|0|0|0|0|0|0|0|0|0|0|0|0|0|0|0|1|0|0|0|0|0|0|0|0|0|0|0|0|0|0|0|0|0|0|0|0|0|
|bacteria_Jul27d25b_60261|1|0|0|0|0|0|0|0|0|0|0|0|0|0|0|0|0|0|0|0|0|0|0|0|0|0|0|0|0|0|0|0|0|1|0|0|0|0|0|0|0|0|0|0|0|0|0|0|0|0|0|0|0|0|0|
|bacteria_Jul27d25b_62967|1|0|0|0|0|0|0|0|0|0|0|0|0|0|0|0|0|0|0|0|0|0|0|0|0|0|0|0|0|0|0|0|0|1|0|0|0|0|0|0|0|0|0|0|0|0|0|0|0|0|0|0|0|0|0|
|bacteria_Jul27d25b_64491|1|0|0|0|0|0|0|0|0|0|0|0|0|0|0|0|0|0|0|0|0|0|0|0|0|0|0|0|0|0|0|0|0|1|0|0|0|0|0|0|0|0|0|0|0|0|0|0|0|0|0|0|0|0|0|
|bacteria_Jul27d25b_73563|1|0|0|0|0|0|0|0|0|0|0|0|0|0|0|0|0|0|0|0|0|0|0|0|0|0|0|0|0|0|0|0|0|1|0|0|0|0|0|0|0|0|0|0|0|0|0|0|0|0|0|0|0|0|0|
|bacteria_Jul27d75b_16064|1|0|0|0|0|0|0|0|0|0|0|0|0|0|0|0|0|0|0|0|0|0|0|0|0|0|0|0|0|0|0|0|0|0|0|1|0|0|0|0|0|0|0|0|0|0|0|0|0|0|0|0|0|0|0|
|bacteria_Jul27d75b_28594|1|0|0|0|0|0|0|0|0|0|0|0|0|0|0|0|0|0|0|0|0|0|0|0|0|0|0|0|0|0|0|0|0|0|0|1|0|0|0|0|0|0|0|0|0|0|0|0|0|0|0|0|0|0|0|
|bacteria_Jul27d75b_52028|1|0|0|0|0|0|0|0|0|0|0|0|0|0|0|0|0|0|0|0|0|0|0|0|0|0|0|0|0|0|0|0|0|0|0|1|0|0|0|0|0|0|0|0|0|0|0|0|0|0|0|0|0|0|0|
|bacteria_Jul27d75b_53903|1|0|0|0|0|0|0|0|0|0|0|0|0|0|0|0|0|0|0|0|0|0|0|0|0|0|0|0|0|0|0|0|0|0|0|1|0|0|0|0|0|0|0|0|0|0|0|0|0|0|0|0|0|0|0|
|bacteria_Jul27d75b_55566|1|0|0|0|0|0|0|0|0|0|0|0|0|0|0|0|0|0|0|0|0|0|0|0|0|0|0|0|0|0|0|0|0|0|0|1|0|0|0|0|0|0|0|0|0|0|0|0|0|0|0|0|0|0|0|
|bacteria_Jul27d75b_58121|1|0|0|0|0|0|0|0|0|0|0|0|0|0|0|0|0|0|0|0|0|0|0|0|0|0|0|0|0|0|0|0|0|0|0|1|0|0|0|0|0|0|0|0|0|0|0|0|0|0|0|0|0|0|0|
|bacteria_Jun8d25b_7145|1|0|0|0|0|0|0|0|0|1|0|0|0|0|0|0|0|0|0|0|0|0|0|0|0|0|0|0|0|0|0|0|0|0|0|0|0|0|0|0|0|0|0|0|0|0|0|0|0|0|0|0|0|0|0|
|bacteria_Jun8d25b_20564|1|0|0|0|0|0|0|0|0|1|0|0|0|0|0|0|0|0|0|0|0|0|0|0|0|0|0|0|0|0|0|0|0|0|0|0|0|0|0|0|0|0|0|0|0|0|0|0|0|0|0|0|0|0|0|
|bacteria_Jun8d25b_21602|1|0|0|0|0|0|0|0|0|1|0|0|0|0|0|0|0|0|0|0|0|0|0|0|0|0|0|0|0|0|0|0|0|0|0|0|0|0|0|0|0|0|0|0|0|0|0|0|0|0|0|0|0|0|0|
|bacteria_Jun8d25b_34360|1|0|0|0|0|0|0|0|0|1|0|0|0|0|0|0|0|0|0|0|0|0|0|0|0|0|0|0|0|0|0|0|0|0|0|0|0|0|0|0|0|0|0|0|0|0|0|0|0|0|0|0|0|0|0|
|bacteria_Jun8d25b_34461|1|0|0|0|0|0|0|0|0|1|0|0|0|0|0|0|0|0|0|0|0|0|0|0|0|0|0|0|0|0|0|0|0|0|0|0|0|0|0|0|0|0|0|0|0|0|0|0|0|0|0|0|0|0|0|
|bacteria_Jun8d25b_41983|1|0|0|0|0|0|0|0|0|1|0|0|0|0|0|0|0|0|0|0|0|0|0|0|0|0|0|0|0|0|0|0|0|0|0|0|0|0|0|0|0|0|0|0|0|0|0|0|0|0|0|0|0|0|0|
|bacteria_Jun8d25b_45038|1|0|0|0|0|0|0|0|0|1|0|0|0|0|0|0|0|0|0|0|0|0|0|0|0|0|0|0|0|0|0|0|0|0|0|0|0|0|0|0|0|0|0|0|0|0|0|0|0|0|0|0|0|0|0|
|bacteria_Jun8d25b_63896|1|0|0|0|0|0|0|0|0|1|0|0|0|0|0|0|0|0|0|0|0|0|0|0|0|0|0|0|0|0|0|0|0|0|0|0|0|0|0|0|0|0|0|0|0|0|0|0|0|0|0|0|0|0|0|
|bacteria_Jun8d25b_72308|1|0|0|0|0|0|0|0|0|1|0|0|0|0|0|0|0|0|0|0|0|0|0|0|0|0|0|0|0|0|0|0|0|0|0|0|0|0|0|0|0|0|0|0|0|0|0|0|0|0|0|0|0|0|0|
|bacteria_Jun8d25b_73418|1|0|0|0|0|0|0|0|0|1|0|0|0|0|0|0|0|0|0|0|0|0|0|0|0|0|0|0|0|0|0|0|0|0|0|0|0|0|0|0|0|0|0|0|0|0|0|0|0|0|0|0|0|0|0|
|bacteria_Jun8d50b_20442|1|0|0|0|0|0|0|0|0|0|0|1|0|0|0|0|0|0|0|0|0|0|0|0|0|0|0|0|0|0|0|0|0|0|0|0|0|0|0|0|0|0|0|0|0|0|0|0|0|0|0|0|0|0|0|
|bacteria_Jun8d50b_21639|1|0|0|0|0|0|0|0|0|0|0|1|0|0|0|0|0|0|0|0|0|0|0|0|0|0|0|0|0|0|0|0|0|0|0|0|0|0|0|0|0|0|0|0|0|0|0|0|0|0|0|0|0|0|0|
|bacteria_Jun8d50b_34821|1|0|0|0|0|0|0|0|0|0|0|1|0|0|0|0|0|0|0|0|0|0|0|0|0|0|0|0|0|0|0|0|0|0|0|0|0|0|0|0|0|0|0|0|0|0|0|0|0|0|0|0|0|0|0|
|bacteria_Jun8d50b_46464|1|0|0|0|0|0|0|0|0|0|0|1|0|0|0|0|0|0|0|0|0|0|0|0|0|0|0|0|0|0|0|0|0|0|0|0|0|0|0|0|0|0|0|0|0|0|0|0|0|0|0|0|0|0|0|
|bacteria_Jun8d50b_50881|1|0|0|0|0|0|0|0|0|0|0|1|0|0|0|0|0|0|0|0|0|0|0|0|0|0|0|0|0|0|0|0|0|0|0|0|0|0|0|0|0|0|0|0|0|0|0|0|0|0|0|0|0|0|0|
|bacteria_Jun8d75b_13815|1|0|0|0|0|0|0|0|0|0|0|0|0|1|0|0|0|0|0|0|0|0|0|0|0|0|0|0|0|0|0|0|0|0|0|0|0|0|0|0|0|0|0|0|0|0|0|0|0|0|0|0|0|0|0|
|bacteria_Jun22d75b_545|1|0|0|0|0|0|0|0|0|0|0|0|0|0|0|0|0|0|0|1|0|0|0|0|0|0|0|0|0|0|0|0|0|0|0|0|0|0|0|0|0|0|0|0|0|0|0|0|0|0|0|0|0|0|0|
|bacteria_Jun22d75b_2717|1|0|0|0|0|0|0|0|0|0|0|0|0|0|0|0|0|0|0|1|0|0|0|0|0|0|0|0|0|0|0|0|0|0|0|0|0|0|0|0|0|0|0|0|0|0|0|0|0|0|0|0|0|0|0|
|bacteria_Jun22d75b_3369|1|0|0|0|0|0|0|0|0|0|0|0|0|0|0|0|0|0|0|1|0|0|0|0|0|0|0|0|0|0|0|0|0|0|0|0|0|0|0|0|0|0|0|0|0|0|0|0|0|0|0|0|0|0|0|
|bacteria_Jun22d75b_6917|1|0|0|0|0|0|0|0|0|0|0|0|0|0|0|0|0|0|0|1|0|0|0|0|0|0|0|0|0|0|0|0|0|0|0|0|0|0|0|0|0|0|0|0|0|0|0|0|0|0|0|0|0|0|0|
|bacteria_May12d25b_513|1|0|0|1|0|0|0|0|0|0|0|0|0|0|0|0|0|0|0|0|0|0|0|0|0|0|0|0|0|0|0|0|0|0|0|0|0|0|0|0|0|0|0|0|0|0|0|0|0|0|0|0|0|0|0|
|bacteria_May12d25b_192|1|0|0|1|0|0|0|0|0|0|0|0|0|0|0|0|0|0|0|0|0|0|0|0|0|0|0|0|0|0|0|0|0|0|0|0|0|0|0|0|0|0|0|0|0|0|0|0|0|0|0|0|0|0|0|
|bacteria_May12d25b_238|1|0|0|1|0|0|0|0|0|0|0|0|0|0|0|0|0|0|0|0|0|0|0|0|0|0|0|0|0|0|0|0|0|0|0|0|0|0|0|0|0|0|0|0|0|0|0|0|0|0|0|0|0|0|0|
|bacteria_May12d25b_270|1|0|0|1|0|0|0|0|0|0|0|0|0|0|0|0|0|0|0|0|0|0|0|0|0|0|0|0|0|0|0|0|0|0|0|0|0|0|0|0|0|0|0|0|0|0|0|0|0|0|0|0|0|0|0|
|bacteria_May12d25b_283|1|0|0|1|0|0|0|0|0|0|0|0|0|0|0|0|0|0|0|0|0|0|0|0|0|0|0|0|0|0|0|0|0|0|0|0|0|0|0|0|0|0|0|0|0|0|0|0|0|0|0|0|0|0|0|
|bacteria_May12d25b_403|1|0|0|1|0|0|0|0|0|0|0|0|0|0|0|0|0|0|0|0|0|0|0|0|0|0|0|0|0|0|0|0|0|0|0|0|0|0|0|0|0|0|0|0|0|0|0|0|0|0|0|0|0|0|0|
|bacteria_May12d25b_431|1|0|0|1|0|0|0|0|0|0|0|0|0|0|0|0|0|0|0|0|0|0|0|0|0|0|0|0|0|0|0|0|0|0|0|0|0|0|0|0|0|0|0|0|0|0|0|0|0|0|0|0|0|0|0|
|bacteria_May12d25b_478|1|0|0|1|0|0|0|0|0|0|0|0|0|0|0|0|0|0|0|0|0|0|0|0|0|0|0|0|0|0|0|0|0|0|0|0|0|0|0|0|0|0|0|0|0|0|0|0|0|0|0|0|0|0|0|
|bacteria_May12d25b_478|1|0|0|1|0|0|0|0|0|0|0|0|0|0|0|0|0|0|0|0|0|0|0|0|0|0|0|0|0|0|0|0|0|0|0|0|0|0|0|0|0|0|0|0|0|0|0|0|0|0|0|0|0|0|0|
|bacteria_May12d25b_511|1|0|0|1|0|0|0|0|0|0|0|0|0|0|0|0|0|0|0|0|0|0|0|0|0|0|0|0|0|0|0|0|0|0|0|0|0|0|0|0|0|0|0|0|0|0|0|0|0|0|0|0|0|0|0|
|bacteria_May12d25b_527|1|0|0|1|0|0|0|0|0|0|0|0|0|0|0|0|0|0|0|0|0|0|0|0|0|0|0|0|0|0|0|0|0|0|0|0|0|0|0|0|0|0|0|0|0|0|0|0|0|0|0|0|0|0|0|
|bacteria_May12d50b_692|1|0|0|0|0|1|0|0|0|0|0|0|0|0|0|0|0|0|0|0|0|0|0|0|0|0|0|0|0|0|0|0|0|0|0|0|0|0|0|0|0|0|0|0|0|0|0|0|0|0|0|0|0|0|0|
|bacteria_May12d50b_232|1|0|0|0|0|1|0|0|0|0|0|0|0|0|0|0|0|0|0|0|0|0|0|0|0|0|0|0|0|0|0|0|0|0|0|0|0|0|0|0|0|0|0|0|0|0|0|0|0|0|0|0|0|0|0|
|bacteria_May12d50b_242|1|0|0|0|0|1|0|0|0|0|0|0|0|0|0|0|0|0|0|0|0|0|0|0|0|0|0|0|0|0|0|0|0|0|0|0|0|0|0|0|0|0|0|0|0|0|0|0|0|0|0|0|0|0|0|
|bacteria_May12d50b_277|1|0|0|0|0|1|0|0|0|0|0|0|0|0|0|0|0|0|0|0|0|0|0|0|0|0|0|0|0|0|0|0|0|0|0|0|0|0|0|0|0|0|0|0|0|0|0|0|0|0|0|0|0|0|0|
|bacteria_May12d50b_419|1|0|0|0|0|1|0|0|0|0|0|0|0|0|0|0|0|0|0|0|0|0|0|0|0|0|0|0|0|0|0|0|0|0|0|0|0|0|0|0|0|0|0|0|0|0|0|0|0|0|0|0|0|0|0|
|bacteria_May12d50b_471|1|0|0|0|0|1|0|0|0|0|0|0|0|0|0|0|0|0|0|0|0|0|0|0|0|0|0|0|0|0|0|0|0|0|0|0|0|0|0|0|0|0|0|0|0|0|0|0|0|0|0|0|0|0|0|
|bacteria_May12d50b_479|1|0|0|0|0|1|0|0|0|0|0|0|0|0|0|0|0|0|0|0|0|0|0|0|0|0|0|0|0|0|0|0|0|0|0|0|0|0|0|0|0|0|0|0|0|0|0|0|0|0|0|0|0|0|0|
|bacteria_Sep8d0b_352|1|0|0|0|0|0|0|0|0|0|0|0|0|0|0|0|0|0|0|0|0|0|0|0|0|0|0|0|0|0|0|0|0|0|0|0|0|0|0|0|0|0|0|0|0|0|0|1|0|0|0|0|0|0|0|
|bacteria_Sep8d5b_3363|1|0|0|0|0|0|0|0|0|0|0|0|0|0|0|0|0|0|0|0|0|0|0|0|0|0|0|0|0|0|0|0|0|0|0|0|0|0|0|0|0|0|0|0|0|0|0|0|0|1|0|0|0|0|0|
|bacteria_Sep8d5b_12958|1|0|0|0|0|0|0|0|0|0|0|0|0|0|0|0|0|0|0|0|0|0|0|0|0|0|0|0|0|0|0|0|0|0|0|0|0|0|0|0|0|0|0|0|0|0|0|0|0|1|0|0|0|0|0|
|bacteria_Sep8d5b_26342|1|0|0|0|0|0|0|0|0|0|0|0|0|0|0|0|0|0|0|0|0|0|0|0|0|0|0|0|0|0|0|0|0|0|0|0|0|0|0|0|0|0|0|0|0|0|0|0|0|1|0|0|0|0|0|
|bacteria_Sep8d5b_26406|1|0|0|0|0|0|0|0|0|0|0|0|0|0|0|0|0|0|0|0|0|0|0|0|0|0|0|0|0|0|0|0|0|0|0|0|0|0|0|0|0|0|0|0|0|0|0|0|0|1|0|0|0|0|0|
|bacteria_Sep8d5b_49612|1|0|0|0|0|0|0|0|0|0|0|0|0|0|0|0|0|0|0|0|0|0|0|0|0|0|0|0|0|0|0|0|0|0|0|0|0|0|0|0|0|0|0|0|0|0|0|0|0|1|0|0|0|0|0|
|bacteria_Sep8d5b_53005|1|0|0|0|0|0|0|0|0|0|0|0|0|0|0|0|0|0|0|0|0|0|0|0|0|0|0|0|0|0|0|0|0|0|0|0|0|0|0|0|0|0|0|0|0|0|0|0|0|1|0|0|0|0|0|
|bacteria_Sep8d5b_54964|1|0|0|0|0|0|0|0|0|0|0|0|0|0|0|0|0|0|0|0|0|0|0|0|0|0|0|0|0|0|0|0|0|0|0|0|0|0|0|0|0|0|0|0|0|0|0|0|0|1|0|0|0|0|0|
|bacteria_Sep8d5b_58266|1|0|0|0|0|0|0|0|0|0|0|0|0|0|0|0|0|0|0|0|0|0|0|0|0|0|0|0|0|0|0|0|0|0|0|0|0|0|0|0|0|0|0|0|0|0|0|0|0|1|0|0|0|0|0|
|bacteria_Sep8d5b_63780|1|0|0|0|0|0|0|0|0|0|0|0|0|0|0|0|0|0|0|0|0|0|0|0|0|0|0|0|0|0|0|0|0|0|0|0|0|0|0|0|0|0|0|0|0|0|0|0|0|1|0|0|0|0|0|
|bacteria_Sep8d5b_66338|1|0|0|0|0|0|0|0|0|0|0|0|0|0|0|0|0|0|0|0|0|0|0|0|0|0|0|0|0|0|0|0|0|0|0|0|0|0|0|0|0|0|0|0|0|0|0|0|0|1|0|0|0|0|0|
|bacteria_Sep8d5b_66689|1|0|0|0|0|0|0|0|0|0|0|0|0|0|0|0|0|0|0|0|0|0|0|0|0|0|0|0|0|0|0|0|0|0|0|0|0|0|0|0|0|0|0|0|0|0|0|0|0|1|0|0|0|0|0|
|bacteria_Sep8d25b_899|1|0|0|0|0|0|0|0|0|0|0|0|0|0|0|0|0|0|0|0|0|0|0|0|0|0|0|0|0|0|0|0|0|0|0|0|0|0|0|0|0|0|0|0|0|0|0|0|0|0|0|1|0|0|0|
|bacteria_Sep8d25b_7083|1|0|0|0|0|0|0|0|0|0|0|0|0|0|0|0|0|0|0|0|0|0|0|0|0|0|0|0|0|0|0|0|0|0|0|0|0|0|0|0|0|0|0|0|0|0|0|0|0|0|0|1|0|0|0|
|bacteria_Sep8d25b_17783|1|0|0|0|0|0|0|0|0|0|0|0|0|0|0|0|0|0|0|0|0|0|0|0|0|0|0|0|0|0|0|0|0|0|0|0|0|0|0|0|0|0|0|0|0|0|0|0|0|0|0|1|0|0|0|
|bacteria_Sep8d25b_38783|1|0|0|0|0|0|0|0|0|0|0|0|0|0|0|0|0|0|0|0|0|0|0|0|0|0|0|0|0|0|0|0|0|0|0|0|0|0|0|0|0|0|0|0|0|0|0|0|0|0|0|1|0|0|0|
|bacteria_Sep8d50b_12570|1|0|0|0|0|0|0|0|0|0|0|0|0|0|0|0|0|0|0|0|0|0|0|0|0|0|0|0|0|0|0|0|0|0|0|0|0|0|0|0|0|0|0|0|0|0|0|0|0|0|0|0|0|1|0|
|virus_Aug5.0v_14|1|0|0|0|0|0|0|0|0|0|0|0|0|0|0|0|0|0|0|0|0|0|0|0|0|0|0|0|0|0|0|0|0|0|0|0|0|0|1|0|0|0|0|0|0|0|0|0|0|0|0|0|0|0|0|
|virus_Aug5.0v_212|1|0|0|0|0|0|0|0|0|0|0|0|0|0|0|0|0|0|0|0|0|0|0|0|0|0|0|0|0|0|0|0|0|0|0|0|0|0|1|0|0|0|0|0|0|0|0|0|0|0|0|0|0|0|0|
|virus_Aug5.0v_329|1|0|0|0|0|0|0|0|0|0|0|0|0|0|0|0|0|0|0|0|0|0|0|0|0|0|0|0|0|0|0|0|0|0|0|0|0|0|1|0|0|0|0|0|0|0|0|0|0|0|0|0|0|0|0|
|virus_Aug5.0v_422|1|0|0|0|0|0|0|0|0|0|0|0|0|0|0|0|0|0|0|0|0|0|0|0|0|0|0|0|0|0|0|0|0|0|0|0|0|0|1|0|0|0|0|0|0|0|0|0|0|0|0|0|0|0|0|
|virus_Aug5.0v_1810|1|0|0|0|0|0|0|0|0|0|0|0|0|0|0|0|0|0|0|0|0|0|0|0|0|0|0|0|0|0|0|0|0|0|0|0|0|0|1|0|0|0|0|0|0|0|0|0|0|0|0|0|0|0|0|

**Supplementary Table S7**. Common Bacterio- and virioplankton RTPR 98% OTUs in 27 samples with RTPR amplicons in both fractions

Total seqs: 47997; Subtotal seqs in retained

| OTUs: 41046 | | May12_05m | | May12_25m | | May12_50m | | May12_75m | | Jun8_25m | | Jun8_50m | | Jun8_75m | | Jun22_0m | | Jun22_50m | | Jun22_75m | | Jul11_0m | | Jul11_05m | | Jul11_25m | | Jul11_50m | | Jul11_75m | | Jul27_05m | | Jul27_25m | | Jul27_75m | | Aug5_0m | | Aug5_05m | | Aug5_25m | | Aug5_50m | | Aug5_75m | | Sep8_0m | | Sep8_05m | | Sep8_25m | | Sep8_50m | |
|---|---|---|---|---|---|---|---|---|---|---|---|---|---|---|---|---|---|---|---|---|---|---|---|---|---|---|---|---|---|---|---|---|---|---|---|---|---|---|---|---|---|---|---|---|---|---|---|---|---|---|---|---|---|---|---|
| OTUs with >=10 seqs: 254 | OTU subtotal | bacterioplankton | virioplankton | bacterioplankton | virioplankton | bacterioplankton | virioplankton | bacterioplankton | virioplankton | bacterioplankton | virioplankton | bacterioplankton | virioplankton | bacterioplankton | virioplankton | bacterioplankton | virioplankton | bacterioplankton | virioplankton | bacterioplankton | virioplankton | bacterioplankton | virioplankton | bacterioplankton | virioplankton | bacterioplankton | virioplankton | bacterioplankton | virioplankton | bacterioplankton | virioplankton | bacterioplankton | virioplankton | bacterioplankton | virioplankton | bacterioplankton | virioplankton | bacterioplankton | virioplankton | bacterioplankton | virioplankton | bacterioplankton | virioplankton | bacterioplankton | virioplankton | bacterioplankton | virioplankton | bacterioplankton | virioplankton | bacterioplankton | virioplankton | bacterioplankton | virioplankton | bacterioplankton | virioplankton |
| virus_Aug5.0v_1821 | 1 | 0 | 0 | 0 | 0 | 0 | 0 | 0 | 0 | 0 | 0 | 0 | 0 | 0 | 0 | 0 | 0 | 0 | 0 | 0 | 0 | 0 | 0 | 0 | 0 | 0 | 0 | 0 | 0 | 0 | 0 | 0 | 0 | 0 | 0 | 0 | 0 | 0 | 1 | 0 | 0 | 0 | 0 | 0 | 0 | 0 | 0 | 0 | 0 | 0 | 0 | 0 | 0 | 0 | 0 |
| virus_Aug5.0v_2129 | 1 | 0 | 0 | 0 | 0 | 0 | 0 | 0 | 0 | 0 | 0 | 0 | 0 | 0 | 0 | 0 | 0 | 0 | 0 | 0 | 0 | 0 | 0 | 0 | 0 | 0 | 0 | 0 | 0 | 0 | 0 | 0 | 0 | 0 | 0 | 0 | 0 | 0 | 1 | 0 | 0 | 0 | 0 | 0 | 0 | 0 | 0 | 0 | 0 | 0 | 0 | 0 | 0 | 0 | 0 |
| virus_Aug5.0v_2186 | 1 | 0 | 0 | 0 | 0 | 0 | 0 | 0 | 0 | 0 | 0 | 0 | 0 | 0 | 0 | 0 | 0 | 0 | 0 | 0 | 0 | 0 | 0 | 0 | 0 | 0 | 0 | 0 | 0 | 0 | 0 | 0 | 0 | 0 | 0 | 0 | 0 | 0 | 1 | 0 | 0 | 0 | 0 | 0 | 0 | 0 | 0 | 0 | 0 | 0 | 0 | 0 | 0 | 0 | 0 |
| virus_Aug5.0v_2365 | 1 | 0 | 0 | 0 | 0 | 0 | 0 | 0 | 0 | 0 | 0 | 0 | 0 | 0 | 0 | 0 | 0 | 0 | 0 | 0 | 0 | 0 | 0 | 0 | 0 | 0 | 0 | 0 | 0 | 0 | 0 | 0 | 0 | 0 | 0 | 0 | 0 | 0 | 1 | 0 | 0 | 0 | 0 | 0 | 0 | 0 | 0 | 0 | 0 | 0 | 0 | 0 | 0 | 0 | 0 |
| virus_Aug5.0v_2471 | 1 | 0 | 0 | 0 | 0 | 0 | 0 | 0 | 0 | 0 | 0 | 0 | 0 | 0 | 0 | 0 | 0 | 0 | 0 | 0 | 0 | 0 | 0 | 0 | 0 | 0 | 0 | 0 | 0 | 0 | 0 | 0 | 0 | 0 | 0 | 0 | 0 | 0 | 1 | 0 | 0 | 0 | 0 | 0 | 0 | 0 | 0 | 0 | 0 | 0 | 0 | 0 | 0 | 0 | 0 |
| virus_Aug5.0v_2521 | 1 | 0 | 0 | 0 | 0 | 0 | 0 | 0 | 0 | 0 | 0 | 0 | 0 | 0 | 0 | 0 | 0 | 0 | 0 | 0 | 0 | 0 | 0 | 0 | 0 | 0 | 0 | 0 | 0 | 0 | 0 | 0 | 0 | 0 | 0 | 0 | 0 | 0 | 1 | 0 | 0 | 0 | 0 | 0 | 0 | 0 | 0 | 0 | 0 | 0 | 0 | 0 | 0 | 0 | 0 |
| virus_Aug5.0v_2618 | 1 | 0 | 0 | 0 | 0 | 0 | 0 | 0 | 0 | 0 | 0 | 0 | 0 | 0 | 0 | 0 | 0 | 0 | 0 | 0 | 0 | 0 | 0 | 0 | 0 | 0 | 0 | 0 | 0 | 0 | 0 | 0 | 0 | 0 | 0 | 0 | 0 | 0 | 1 | 0 | 0 | 0 | 0 | 0 | 0 | 0 | 0 | 0 | 0 | 0 | 0 | 0 | 0 | 0 | 0 |
| virus_Aug5.0v_2756 | 1 | 0 | 0 | 0 | 0 | 0 | 0 | 0 | 0 | 0 | 0 | 0 | 0 | 0 | 0 | 0 | 0 | 0 | 0 | 0 | 0 | 0 | 0 | 0 | 0 | 0 | 0 | 0 | 0 | 0 | 0 | 0 | 0 | 0 | 0 | 0 | 0 | 0 | 1 | 0 | 0 | 0 | 0 | 0 | 0 | 0 | 0 | 0 | 0 | 0 | 0 | 0 | 0 | 0 | 0 |
| virus_Aug5.0v_2789 | 1 | 0 | 0 | 0 | 0 | 0 | 0 | 0 | 0 | 0 | 0 | 0 | 0 | 0 | 0 | 0 | 0 | 0 | 0 | 0 | 0 | 0 | 0 | 0 | 0 | 0 | 0 | 0 | 0 | 0 | 0 | 0 | 0 | 0 | 0 | 0 | 0 | 0 | 1 | 0 | 0 | 0 | 0 | 0 | 0 | 0 | 0 | 0 | 0 | 0 | 0 | 0 | 0 | 0 | 0 |
| virus_Aug5.0v_2905 | 1 | 0 | 0 | 0 | 0 | 0 | 0 | 0 | 0 | 0 | 0 | 0 | 0 | 0 | 0 | 0 | 0 | 0 | 0 | 0 | 0 | 0 | 0 | 0 | 0 | 0 | 0 | 0 | 0 | 0 | 0 | 0 | 0 | 0 | 0 | 0 | 0 | 0 | 1 | 0 | 0 | 0 | 0 | 0 | 0 | 0 | 0 | 0 | 0 | 0 | 0 | 0 | 0 | 0 | 0 |
| virus_Aug5.0v_2957 | 1 | 0 | 0 | 0 | 0 | 0 | 0 | 0 | 0 | 0 | 0 | 0 | 0 | 0 | 0 | 0 | 0 | 0 | 0 | 0 | 0 | 0 | 0 | 0 | 0 | 0 | 0 | 0 | 0 | 0 | 0 | 0 | 0 | 0 | 0 | 0 | 0 | 0 | 1 | 0 | 0 | 0 | 0 | 0 | 0 | 0 | 0 | 0 | 0 | 0 | 0 | 0 | 0 | 0 | 0 |
| virus_Aug5.0v_3560 | 1 | 0 | 0 | 0 | 0 | 0 | 0 | 0 | 0 | 0 | 0 | 0 | 0 | 0 | 0 | 0 | 0 | 0 | 0 | 0 | 0 | 0 | 0 | 0 | 0 | 0 | 0 | 0 | 0 | 0 | 0 | 0 | 0 | 0 | 0 | 0 | 0 | 0 | 1 | 0 | 0 | 0 | 0 | 0 | 0 | 0 | 0 | 0 | 0 | 0 | 0 | 0 | 0 | 0 | 0 |
| virus_Aug5.0v_3981 | 1 | 0 | 0 | 0 | 0 | 0 | 0 | 0 | 0 | 0 | 0 | 0 | 0 | 0 | 0 | 0 | 0 | 0 | 0 | 0 | 0 | 0 | 0 | 0 | 0 | 0 | 0 | 0 | 0 | 0 | 0 | 0 | 0 | 0 | 0 | 0 | 0 | 0 | 1 | 0 | 0 | 0 | 0 | 0 | 0 | 0 | 0 | 0 | 0 | 0 | 0 | 0 | 0 | 0 | 0 |
| virus_Aug5.0v_4043 | 1 | 0 | 0 | 0 | 0 | 0 | 0 | 0 | 0 | 0 | 0 | 0 | 0 | 0 | 0 | 0 | 0 | 0 | 0 | 0 | 0 | 0 | 0 | 0 | 0 | 0 | 0 | 0 | 0 | 0 | 0 | 0 | 0 | 0 | 0 | 0 | 0 | 0 | 1 | 0 | 0 | 0 | 0 | 0 | 0 | 0 | 0 | 0 | 0 | 0 | 0 | 0 | 0 | 0 | 0 |
| virus_Aug5.0v_4310 | 1 | 0 | 0 | 0 | 0 | 0 | 0 | 0 | 0 | 0 | 0 | 0 | 0 | 0 | 0 | 0 | 0 | 0 | 0 | 0 | 0 | 0 | 0 | 0 | 0 | 0 | 0 | 0 | 0 | 0 | 0 | 0 | 0 | 0 | 0 | 0 | 0 | 0 | 1 | 0 | 0 | 0 | 0 | 0 | 0 | 0 | 0 | 0 | 0 | 0 | 0 | 0 | 0 | 0 | 0 |
| virus_Aug5.0v_4450 | 1 | 0 | 0 | 0 | 0 | 0 | 0 | 0 | 0 | 0 | 0 | 0 | 0 | 0 | 0 | 0 | 0 | 0 | 0 | 0 | 0 | 0 | 0 | 0 | 0 | 0 | 0 | 0 | 0 | 0 | 0 | 0 | 0 | 0 | 0 | 0 | 0 | 0 | 1 | 0 | 0 | 0 | 0 | 0 | 0 | 0 | 0 | 0 | 0 | 0 | 0 | 0 | 0 | 0 | 0 |
| virus_Aug5.0v_4926 | 1 | 0 | 0 | 0 | 0 | 0 | 0 | 0 | 0 | 0 | 0 | 0 | 0 | 0 | 0 | 0 | 0 | 0 | 0 | 0 | 0 | 0 | 0 | 0 | 0 | 0 | 0 | 0 | 0 | 0 | 0 | 0 | 0 | 0 | 0 | 0 | 0 | 0 | 1 | 0 | 0 | 0 | 0 | 0 | 0 | 0 | 0 | 0 | 0 | 0 | 0 | 0 | 0 | 0 | 0 |
| virus_Aug5.0v_4936 | 1 | 0 | 0 | 0 | 0 | 0 | 0 | 0 | 0 | 0 | 0 | 0 | 0 | 0 | 0 | 0 | 0 | 0 | 0 | 0 | 0 | 0 | 0 | 0 | 0 | 0 | 0 | 0 | 0 | 0 | 0 | 0 | 0 | 0 | 0 | 0 | 0 | 0 | 1 | 0 | 0 | 0 | 0 | 0 | 0 | 0 | 0 | 0 | 0 | 0 | 0 | 0 | 0 | 0 | 0 |
| virus_Aug5.0v_4945 | 1 | 0 | 0 | 0 | 0 | 0 | 0 | 0 | 0 | 0 | 0 | 0 | 0 | 0 | 0 | 0 | 0 | 0 | 0 | 0 | 0 | 0 | 0 | 0 | 0 | 0 | 0 | 0 | 0 | 0 | 0 | 0 | 0 | 0 | 0 | 0 | 0 | 0 | 1 | 0 | 0 | 0 | 0 | 0 | 0 | 0 | 0 | 0 | 0 | 0 | 0 | 0 | 0 | 0 | 0 |
| virus_Aug5.0v_4969 | 1 | 0 | 0 | 0 | 0 | 0 | 0 | 0 | 0 | 0 | 0 | 0 | 0 | 0 | 0 | 0 | 0 | 0 | 0 | 0 | 0 | 0 | 0 | 0 | 0 | 0 | 0 | 0 | 0 | 0 | 0 | 0 | 0 | 0 | 0 | 0 | 0 | 0 | 1 | 0 | 0 | 0 | 0 | 0 | 0 | 0 | 0 | 0 | 0 | 0 | 0 | 0 | 0 | 0 | 0 |
| virus_Aug5.0v_5286 | 1 | 0 | 0 | 0 | 0 | 0 | 0 | 0 | 0 | 0 | 0 | 0 | 0 | 0 | 0 | 0 | 0 | 0 | 0 | 0 | 0 | 0 | 0 | 0 | 0 | 0 | 0 | 0 | 0 | 0 | 0 | 0 | 0 | 0 | 0 | 0 | 0 | 0 | 1 | 0 | 0 | 0 | 0 | 0 | 0 | 0 | 0 | 0 | 0 | 0 | 0 | 0 | 0 | 0 | 0 |
| virus_Aug5.0v_5428 | 1 | 0 | 0 | 0 | 0 | 0 | 0 | 0 | 0 | 0 | 0 | 0 | 0 | 0 | 0 | 0 | 0 | 0 | 0 | 0 | 0 | 0 | 0 | 0 | 0 | 0 | 0 | 0 | 0 | 0 | 0 | 0 | 0 | 0 | 0 | 0 | 0 | 0 | 1 | 0 | 0 | 0 | 0 | 0 | 0 | 0 | 0 | 0 | 0 | 0 | 0 | 0 | 0 | 0 | 0 |
| virus_Aug5.5v_357 | 1 | 0 | 0 | 0 | 0 | 0 | 0 | 0 | 0 | 0 | 0 | 0 | 0 | 0 | 0 | 0 | 0 | 0 | 0 | 0 | 0 | 0 | 0 | 0 | 0 | 0 | 0 | 0 | 0 | 0 | 0 | 0 | 0 | 0 | 0 | 0 | 0 | 0 | 0 | 0 | 1 | 0 | 0 | 0 | 0 | 0 | 0 | 0 | 0 | 0 | 0 | 0 | 0 | 0 | 0 |
| virus_Aug5.5v_7957 | 1 | 0 | 0 | 0 | 0 | 0 | 0 | 0 | 0 | 0 | 0 | 0 | 0 | 0 | 0 | 0 | 0 | 0 | 0 | 0 | 0 | 0 | 0 | 0 | 0 | 0 | 0 | 0 | 0 | 0 | 0 | 0 | 0 | 0 | 0 | 0 | 0 | 0 | 0 | 0 | 1 | 0 | 0 | 0 | 0 | 0 | 0 | 0 | 0 | 0 | 0 | 0 | 0 | 0 | 0 |
| virus_Aug5.5v_13568 | 1 | 0 | 0 | 0 | 0 | 0 | 0 | 0 | 0 | 0 | 0 | 0 | 0 | 0 | 0 | 0 | 0 | 0 | 0 | 0 | 0 | 0 | 0 | 0 | 0 | 0 | 0 | 0 | 0 | 0 | 0 | 0 | 0 | 0 | 0 | 0 | 0 | 0 | 0 | 0 | 1 | 0 | 0 | 0 | 0 | 0 | 0 | 0 | 0 | 0 | 0 | 0 | 0 | 0 | 0 |
| virus_Aug5.5v_16378 | 1 | 0 | 0 | 0 | 0 | 0 | 0 | 0 | 0 | 0 | 0 | 0 | 0 | 0 | 0 | 0 | 0 | 0 | 0 | 0 | 0 | 0 | 0 | 0 | 0 | 0 | 0 | 0 | 0 | 0 | 0 | 0 | 0 | 0 | 0 | 0 | 0 | 0 | 0 | 0 | 1 | 0 | 0 | 0 | 0 | 0 | 0 | 0 | 0 | 0 | 0 | 0 | 0 | 0 | 0 |
| virus_Aug5.5v_18078 | 1 | 0 | 0 | 0 | 0 | 0 | 0 | 0 | 0 | 0 | 0 | 0 | 0 | 0 | 0 | 0 | 0 | 0 | 0 | 0 | 0 | 0 | 0 | 0 | 0 | 0 | 0 | 0 | 0 | 0 | 0 | 0 | 0 | 0 | 0 | 0 | 0 | 0 | 0 | 0 | 1 | 0 | 0 | 0 | 0 | 0 | 0 | 0 | 0 | 0 | 0 | 0 | 0 | 0 | 0 |
| virus_Aug5.5v_20966 | 1 | 0 | 0 | 0 | 0 | 0 | 0 | 0 | 0 | 0 | 0 | 0 | 0 | 0 | 0 | 0 | 0 | 0 | 0 | 0 | 0 | 0 | 0 | 0 | 0 | 0 | 0 | 0 | 0 | 0 | 0 | 0 | 0 | 0 | 0 | 0 | 0 | 0 | 0 | 0 | 1 | 0 | 0 | 0 | 0 | 0 | 0 | 0 | 0 | 0 | 0 | 0 | 0 | 0 | 0 |
| virus_Aug5.5v_21274 | 1 | 0 | 0 | 0 | 0 | 0 | 0 | 0 | 0 | 0 | 0 | 0 | 0 | 0 | 0 | 0 | 0 | 0 | 0 | 0 | 0 | 0 | 0 | 0 | 0 | 0 | 0 | 0 | 0 | 0 | 0 | 0 | 0 | 0 | 0 | 0 | 0 | 0 | 0 | 0 | 1 | 0 | 0 | 0 | 0 | 0 | 0 | 0 | 0 | 0 | 0 | 0 | 0 | 0 | 0 |
| virus_Aug5.5v_36929 | 1 | 0 | 0 | 0 | 0 | 0 | 0 | 0 | 0 | 0 | 0 | 0 | 0 | 0 | 0 | 0 | 0 | 0 | 0 | 0 | 0 | 0 | 0 | 0 | 0 | 0 | 0 | 0 | 0 | 0 | 0 | 0 | 0 | 0 | 0 | 0 | 0 | 0 | 0 | 0 | 1 | 0 | 0 | 0 | 0 | 0 | 0 | 0 | 0 | 0 | 0 | 0 | 0 | 0 | 0 |
| virus_Aug5.5v_38127 | 1 | 0 | 0 | 0 | 0 | 0 | 0 | 0 | 0 | 0 | 0 | 0 | 0 | 0 | 0 | 0 | 0 | 0 | 0 | 0 | 0 | 0 | 0 | 0 | 0 | 0 | 0 | 0 | 0 | 0 | 0 | 0 | 0 | 0 | 0 | 0 | 0 | 0 | 0 | 0 | 1 | 0 | 0 | 0 | 0 | 0 | 0 | 0 | 0 | 0 | 0 | 0 | 0 | 0 | 0 |
| virus_Aug5.25v_2 | 1 | 0 | 0 | 0 | 0 | 0 | 0 | 0 | 0 | 0 | 0 | 0 | 0 | 0 | 0 | 0 | 0 | 0 | 0 | 0 | 0 | 0 | 0 | 0 | 0 | 0 | 0 | 0 | 0 | 0 | 0 | 0 | 0 | 0 | 0 | 0 | 0 | 0 | 0 | 0 | 0 | 0 | 1 | 0 | 0 | 0 | 0 | 0 | 0 | 0 | 0 | 0 | 0 | 0 | 0 |
| virus_Aug5.25v_3142 | 1 | 0 | 0 | 0 | 0 | 0 | 0 | 0 | 0 | 0 | 0 | 0 | 0 | 0 | 0 | 0 | 0 | 0 | 0 | 0 | 0 | 0 | 0 | 0 | 0 | 0 | 0 | 0 | 0 | 0 | 0 | 0 | 0 | 0 | 0 | 0 | 0 | 0 | 0 | 0 | 0 | 0 | 1 | 0 | 0 | 0 | 0 | 0 | 0 | 0 | 0 | 0 | 0 | 0 | 0 |
| virus_Aug5.25v_4313 | 1 | 0 | 0 | 0 | 0 | 0 | 0 | 0 | 0 | 0 | 0 | 0 | 0 | 0 | 0 | 0 | 0 | 0 | 0 | 0 | 0 | 0 | 0 | 0 | 0 | 0 | 0 | 0 | 0 | 0 | 0 | 0 | 0 | 0 | 0 | 0 | 0 | 0 | 0 | 0 | 0 | 0 | 1 | 0 | 0 | 0 | 0 | 0 | 0 | 0 | 0 | 0 | 0 | 0 | 0 |
| virus_Aug5.25v_7175 | 1 | 0 | 0 | 0 | 0 | 0 | 0 | 0 | 0 | 0 | 0 | 0 | 0 | 0 | 0 | 0 | 0 | 0 | 0 | 0 | 0 | 0 | 0 | 0 | 0 | 0 | 0 | 0 | 0 | 0 | 0 | 0 | 0 | 0 | 0 | 0 | 0 | 0 | 0 | 0 | 0 | 0 | 1 | 0 | 0 | 0 | 0 | 0 | 0 | 0 | 0 | 0 | 0 | 0 | 0 |
| virus_Aug5.25v_8171 | 1 | 0 | 0 | 0 | 0 | 0 | 0 | 0 | 0 | 0 | 0 | 0 | 0 | 0 | 0 | 0 | 0 | 0 | 0 | 0 | 0 | 0 | 0 | 0 | 0 | 0 | 0 | 0 | 0 | 0 | 0 | 0 | 0 | 0 | 0 | 0 | 0 | 0 | 0 | 0 | 0 | 0 | 1 | 0 | 0 | 0 | 0 | 0 | 0 | 0 | 0 | 0 | 0 | 0 | 0 |
| virus_Aug5.25v_9437 | 1 | 0 | 0 | 0 | 0 | 0 | 0 | 0 | 0 | 0 | 0 | 0 | 0 | 0 | 0 | 0 | 0 | 0 | 0 | 0 | 0 | 0 | 0 | 0 | 0 | 0 | 0 | 0 | 0 | 0 | 0 | 0 | 0 | 0 | 0 | 0 | 0 | 0 | 0 | 0 | 0 | 0 | 1 | 0 | 0 | 0 | 0 | 0 | 0 | 0 | 0 | 0 | 0 | 0 | 0 |
| virus_Aug5.25v_9868 | 1 | 0 | 0 | 0 | 0 | 0 | 0 | 0 | 0 | 0 | 0 | 0 | 0 | 0 | 0 | 0 | 0 | 0 | 0 | 0 | 0 | 0 | 0 | 0 | 0 | 0 | 0 | 0 | 0 | 0 | 0 | 0 | 0 | 0 | 0 | 0 | 0 | 0 | 0 | 0 | 0 | 0 | 1 | 0 | 0 | 0 | 0 | 0 | 0 | 0 | 0 | 0 | 0 | 0 | 0 |
| virus_Aug5.25v_11199 | 1 | 0 | 0 | 0 | 0 | 0 | 0 | 0 | 0 | 0 | 0 | 0 | 0 | 0 | 0 | 0 | 0 | 0 | 0 | 0 | 0 | 0 | 0 | 0 | 0 | 0 | 0 | 0 | 0 | 0 | 0 | 0 | 0 | 0 | 0 | 0 | 0 | 0 | 0 | 0 | 0 | 0 | 1 | 0 | 0 | 0 | 0 | 0 | 0 | 0 | 0 | 0 | 0 | 0 | 0 |
| virus_Aug5.25v_11227 | 1 | 0 | 0 | 0 | 0 | 0 | 0 | 0 | 0 | 0 | 0 | 0 | 0 | 0 | 0 | 0 | 0 | 0 | 0 | 0 | 0 | 0 | 0 | 0 | 0 | 0 | 0 | 0 | 0 | 0 | 0 | 0 | 0 | 0 | 0 | 0 | 0 | 0 | 0 | 0 | 0 | 0 | 1 | 0 | 0 | 0 | 0 | 0 | 0 | 0 | 0 | 0 | 0 | 0 | 0 |
| virus_Aug5.25v_17104 | 1 | 0 | 0 | 0 | 0 | 0 | 0 | 0 | 0 | 0 | 0 | 0 | 0 | 0 | 0 | 0 | 0 | 0 | 0 | 0 | 0 | 0 | 0 | 0 | 0 | 0 | 0 | 0 | 0 | 0 | 0 | 0 | 0 | 0 | 0 | 0 | 0 | 0 | 0 | 0 | 0 | 0 | 1 | 0 | 0 | 0 | 0 | 0 | 0 | 0 | 0 | 0 | 0 | 0 | 0 |
| virus_Aug5.25v_19259 | 1 | 0 | 0 | 0 | 0 | 0 | 0 | 0 | 0 | 0 | 0 | 0 | 0 | 0 | 0 | 0 | 0 | 0 | 0 | 0 | 0 | 0 | 0 | 0 | 0 | 0 | 0 | 0 | 0 | 0 | 0 | 0 | 0 | 0 | 0 | 0 | 0 | 0 | 0 | 0 | 0 | 0 | 1 | 0 | 0 | 0 | 0 | 0 | 0 | 0 | 0 | 0 | 0 | 0 | 0 |
| virus_Aug5.25v_20583 | 1 | 0 | 0 | 0 | 0 | 0 | 0 | 0 | 0 | 0 | 0 | 0 | 0 | 0 | 0 | 0 | 0 | 0 | 0 | 0 | 0 | 0 | 0 | 0 | 0 | 0 | 0 | 0 | 0 | 0 | 0 | 0 | 0 | 0 | 0 | 0 | 0 | 0 | 0 | 0 | 0 | 0 | 1 | 0 | 0 | 0 | 0 | 0 | 0 | 0 | 0 | 0 | 0 | 0 | 0 |
| virus_Aug5.25v_21003 | 1 | 0 | 0 | 0 | 0 | 0 | 0 | 0 | 0 | 0 | 0 | 0 | 0 | 0 | 0 | 0 | 0 | 0 | 0 | 0 | 0 | 0 | 0 | 0 | 0 | 0 | 0 | 0 | 0 | 0 | 0 | 0 | 0 | 0 | 0 | 0 | 0 | 0 | 0 | 0 | 0 | 0 | 1 | 0 | 0 | 0 | 0 | 0 | 0 | 0 | 0 | 0 | 0 | 0 | 0 |
| virus_Aug5.25v_21305 | 1 | 0 | 0 | 0 | 0 | 0 | 0 | 0 | 0 | 0 | 0 | 0 | 0 | 0 | 0 | 0 | 0 | 0 | 0 | 0 | 0 | 0 | 0 | 0 | 0 | 0 | 0 | 0 | 0 | 0 | 0 | 0 | 0 | 0 | 0 | 0 | 0 | 0 | 0 | 0 | 0 | 0 | 1 | 0 | 0 | 0 | 0 | 0 | 0 | 0 | 0 | 0 | 0 | 0 | 0 |
| virus_Aug5.25v_24050 | 1 | 0 | 0 | 0 | 0 | 0 | 0 | 0 | 0 | 0 | 0 | 0 | 0 | 0 | 0 | 0 | 0 | 0 | 0 | 0 | 0 | 0 | 0 | 0 | 0 | 0 | 0 | 0 | 0 | 0 | 0 | 0 | 0 | 0 | 0 | 0 | 0 | 0 | 0 | 0 | 0 | 0 | 1 | 0 | 0 | 0 | 0 | 0 | 0 | 0 | 0 | 0 | 0 | 0 | 0 |
| virus_Aug5.25v_28937 | 1 | 0 | 0 | 0 | 0 | 0 | 0 | 0 | 0 | 0 | 0 | 0 | 0 | 0 | 0 | 0 | 0 | 0 | 0 | 0 | 0 | 0 | 0 | 0 | 0 | 0 | 0 | 0 | 0 | 0 | 0 | 0 | 0 | 0 | 0 | 0 | 0 | 0 | 0 | 0 | 0 | 0 | 1 | 0 | 0 | 0 | 0 | 0 | 0 | 0 | 0 | 0 | 0 | 0 | 0 |
| virus_Aug5.25v_36597 | 1 | 0 | 0 | 0 | 0 | 0 | 0 | 0 | 0 | 0 | 0 | 0 | 0 | 0 | 0 | 0 | 0 | 0 | 0 | 0 | 0 | 0 | 0 | 0 | 0 | 0 | 0 | 0 | 0 | 0 | 0 | 0 | 0 | 0 | 0 | 0 | 0 | 0 | 0 | 0 | 0 | 0 | 1 | 0 | 0 | 0 | 0 | 0 | 0 | 0 | 0 | 0 | 0 | 0 | 0 |
| virus_Aug5.25v_40304 | 1 | 0 | 0 | 0 | 0 | 0 | 0 | 0 | 0 | 0 | 0 | 0 | 0 | 0 | 0 | 0 | 0 | 0 | 0 | 0 | 0 | 0 | 0 | 0 | 0 | 0 | 0 | 0 | 0 | 0 | 0 | 0 | 0 | 0 | 0 | 0 | 0 | 0 | 0 | 0 | 0 | 0 | 1 | 0 | 0 | 0 | 0 | 0 | 0 | 0 | 0 | 0 | 0 | 0 | 0 |
| virus_Aug5.50v_4328 | 1 | 0 | 0 | 0 | 0 | 0 | 0 | 0 | 0 | 0 | 0 | 0 | 0 | 0 | 0 | 0 | 0 | 0 | 0 | 0 | 0 | 0 | 0 | 0 | 0 | 0 | 0 | 0 | 0 | 0 | 0 | 0 | 0 | 0 | 0 | 0 | 0 | 0 | 0 | 0 | 0 | 0 | 0 | 0 | 1 | 0 | 0 | 0 | 0 | 0 | 0 | 0 | 0 | 0 | 0 |
| virus_Aug5.50v_7119 | 1 | 0 | 0 | 0 | 0 | 0 | 0 | 0 | 0 | 0 | 0 | 0 | 0 | 0 | 0 | 0 | 0 | 0 | 0 | 0 | 0 | 0 | 0 | 0 | 0 | 0 | 0 | 0 | 0 | 0 | 0 | 0 | 0 | 0 | 0 | 0 | 0 | 0 | 0 | 0 | 0 | 0 | 0 | 0 | 1 | 0 | 0 | 0 | 0 | 0 | 0 | 0 | 0 | 0 | 0 |
| virus_Aug5.50v_11588 | 1 | 0 | 0 | 0 | 0 | 0 | 0 | 0 | 0 | 0 | 0 | 0 | 0 | 0 | 0 | 0 | 0 | 0 | 0 | 0 | 0 | 0 | 0 | 0 | 0 | 0 | 0 | 0 | 0 | 0 | 0 | 0 | 0 | 0 | 0 | 0 | 0 | 0 | 0 | 0 | 0 | 0 | 0 | 0 | 1 | 0 | 0 | 0 | 0 | 0 | 0 | 0 | 0 | 0 | 0 |
| virus_Aug5.50v_13084 | 1 | 0 | 0 | 0 | 0 | 0 | 0 | 0 | 0 | 0 | 0 | 0 | 0 | 0 | 0 | 0 | 0 | 0 | 0 | 0 | 0 | 0 | 0 | 0 | 0 | 0 | 0 | 0 | 0 | 0 | 0 | 0 | 0 | 0 | 0 | 0 | 0 | 0 | 0 | 0 | 0 | 0 | 0 | 0 | 1 | 0 | 0 | 0 | 0 | 0 | 0 | 0 | 0 | 0 | 0 |
| virus_Aug5.50v_13508 | 1 | 0 | 0 | 0 | 0 | 0 | 0 | 0 | 0 | 0 | 0 | 0 | 0 | 0 | 0 | 0 | 0 | 0 | 0 | 0 | 0 | 0 | 0 | 0 | 0 | 0 | 0 | 0 | 0 | 0 | 0 | 0 | 0 | 0 | 0 | 0 | 0 | 0 | 0 | 0 | 0 | 0 | 0 | 0 | 1 | 0 | 0 | 0 | 0 | 0 | 0 | 0 | 0 | 0 | 0 |
| virus_Aug5.50v_21345 | 1 | 0 | 0 | 0 | 0 | 0 | 0 | 0 | 0 | 0 | 0 | 0 | 0 | 0 | 0 | 0 | 0 | 0 | 0 | 0 | 0 | 0 | 0 | 0 | 0 | 0 | 0 | 0 | 0 | 0 | 0 | 0 | 0 | 0 | 0 | 0 | 0 | 0 | 0 | 0 | 0 | 0 | 0 | 0 | 1 | 0 | 0 | 0 | 0 | 0 | 0 | 0 | 0 | 0 | 0 |
| virus_Aug5.50v_22866 | 1 | 0 | 0 | 0 | 0 | 0 | 0 | 0 | 0 | 0 | 0 | 0 | 0 | 0 | 0 | 0 | 0 | 0 | 0 | 0 | 0 | 0 | 0 | 0 | 0 | 0 | 0 | 0 | 0 | 0 | 0 | 0 | 0 | 0 | 0 | 0 | 0 | 0 | 0 | 0 | 0 | 0 | 0 | 0 | 1 | 0 | 0 | 0 | 0 | 0 | 0 | 0 | 0 | 0 | 0 |
| virus_Aug5.50v_38232 | 1 | 0 | 0 | 0 | 0 | 0 | 0 | 0 | 0 | 0 | 0 | 0 | 0 | 0 | 0 | 0 | 0 | 0 | 0 | 0 | 0 | 0 | 0 | 0 | 0 | 0 | 0 | 0 | 0 | 0 | 0 | 0 | 0 | 0 | 0 | 0 | 0 | 0 | 0 | 0 | 0 | 0 | 0 | 0 | 1 | 0 | 0 | 0 | 0 | 0 | 0 | 0 | 0 | 0 | 0 |
| virus_Aug5.75v_1000 | 1 | 0 | 0 | 0 | 0 | 0 | 0 | 0 | 0 | 0 | 0 | 0 | 0 | 0 | 0 | 0 | 0 | 0 | 0 | 0 | 0 | 0 | 0 | 0 | 0 | 0 | 0 | 0 | 0 | 0 | 0 | 0 | 0 | 0 | 0 | 0 | 0 | 0 | 0 | 0 | 0 | 0 | 0 | 0 | 0 | 0 | 1 | 0 | 0 | 0 | 0 | 0 | 0 | 0 | 0 |
| virus_Aug5.75v_4421 | 1 | 0 | 0 | 0 | 0 | 0 | 0 | 0 | 0 | 0 | 0 | 0 | 0 | 0 | 0 | 0 | 0 | 0 | 0 | 0 | 0 | 0 | 0 | 0 | 0 | 0 | 0 | 0 | 0 | 0 | 0 | 0 | 0 | 0 | 0 | 0 | 0 | 0 | 0 | 0 | 0 | 0 | 0 | 0 | 0 | 0 | 1 | 0 | 0 | 0 | 0 | 0 | 0 | 0 | 0 |
| virus_Aug5.75v_11084 | 1 | 0 | 0 | 0 | 0 | 0 | 0 | 0 | 0 | 0 | 0 | 0 | 0 | 0 | 0 | 0 | 0 | 0 | 0 | 0 | 0 | 0 | 0 | 0 | 0 | 0 | 0 | 0 | 0 | 0 | 0 | 0 | 0 | 0 | 0 | 0 | 0 | 0 | 0 | 0 | 0 | 0 | 0 | 0 | 0 | 0 | 1 | 0 | 0 | 0 | 0 | 0 | 0 | 0 | 0 |
| virus_Aug5.75v_18383 | 1 | 0 | 0 | 0 | 0 | 0 | 0 | 0 | 0 | 0 | 0 | 0 | 0 | 0 | 0 | 0 | 0 | 0 | 0 | 0 | 0 | 0 | 0 | 0 | 0 | 0 | 0 | 0 | 0 | 0 | 0 | 0 | 0 | 0 | 0 | 0 | 0 | 0 | 0 | 0 | 0 | 0 | 0 | 0 | 0 | 0 | 1 | 0 | 0 | 0 | 0 | 0 | 0 | 0 | 0 |
| virus_Aug5.75v_27104 | 1 | 0 | 0 | 0 | 0 | 0 | 0 | 0 | 0 | 0 | 0 | 0 | 0 | 0 | 0 | 0 | 0 | 0 | 0 | 0 | 0 | 0 | 0 | 0 | 0 | 0 | 0 | 0 | 0 | 0 | 0 | 0 | 0 | 0 | 0 | 0 | 0 | 0 | 0 | 0 | 0 | 0 | 0 | 0 | 0 | 0 | 1 | 0 | 0 | 0 | 0 | 0 | 0 | 0 | 0 |
| virus_Aug5.75v_28751 | 1 | 0 | 0 | 0 | 0 | 0 | 0 | 0 | 0 | 0 | 0 | 0 | 0 | 0 | 0 | 0 | 0 | 0 | 0 | 0 | 0 | 0 | 0 | 0 | 0 | 0 | 0 | 0 | 0 | 0 | 0 | 0 | 0 | 0 | 0 | 0 | 0 | 0 | 0 | 0 | 0 | 0 | 0 | 0 | 0 | 0 | 1 | 0 | 0 | 0 | 0 | 0 | 0 | 0 | 0 |
| virus_Aug5.75v_30233 | 1 | 0 | 0 | 0 | 0 | 0 | 0 | 0 | 0 | 0 | 0 | 0 | 0 | 0 | 0 | 0 | 0 | 0 | 0 | 0 | 0 | 0 | 0 | 0 | 0 | 0 | 0 | 0 | 0 | 0 | 0 | 0 | 0 | 0 | 0 | 0 | 0 | 0 | 0 | 0 | 0 | 0 | 0 | 0 | 0 | 0 | 1 | 0 | 0 | 0 | 0 | 0 | 0 | 0 | 0 |
| virus_Aug5.75v_31313 | 1 | 0 | 0 | 0 | 0 | 0 | 0 | 0 | 0 | 0 | 0 | 0 | 0 | 0 | 0 | 0 | 0 | 0 | 0 | 0 | 0 | 0 | 0 | 0 | 0 | 0 | 0 | 0 | 0 | 0 | 0 | 0 | 0 | 0 | 0 | 0 | 0 | 0 | 0 | 0 | 0 | 0 | 0 | 0 | 0 | 0 | 1 | 0 | 0 | 0 | 0 | 0 | 0 | 0 | 0 |
| virus_Aug5.75v_32334 | 1 | 0 | 0 | 0 | 0 | 0 | 0 | 0 | 0 | 0 | 0 | 0 | 0 | 0 | 0 | 0 | 0 | 0 | 0 | 0 | 0 | 0 | 0 | 0 | 0 | 0 | 0 | 0 | 0 | 0 | 0 | 0 | 0 | 0 | 0 | 0 | 0 | 0 | 0 | 0 | 0 | 0 | 0 | 0 | 0 | 0 | 1 | 0 | 0 | 0 | 0 | 0 | 0 | 0 | 0 |
| virus_Jul11.0v_1841 | 1 | 0 | 0 | 0 | 0 | 0 | 0 | 0 | 0 | 0 | 0 | 0 | 0 | 0 | 0 | 0 | 0 | 0 | 0 | 0 | 0 | 0 | 1 | 0 | 0 | 0 | 0 | 0 | 0 | 0 | 0 | 0 | 0 | 0 | 0 | 0 | 0 | 0 | 0 | 0 | 0 | 0 | 0 | 0 | 0 | 0 | 0 | 0 | 0 | 0 | 0 | 0 | 0 | 0 | 0 |
| virus_Jul11.0v_5433 | 1 | 0 | 0 | 0 | 0 | 0 | 0 | 0 | 0 | 0 | 0 | 0 | 0 | 0 | 0 | 0 | 0 | 0 | 0 | 0 | 0 | 0 | 1 | 0 | 0 | 0 | 0 | 0 | 0 | 0 | 0 | 0 | 0 | 0 | 0 | 0 | 0 | 0 | 0 | 0 | 0 | 0 | 0 | 0 | 0 | 0 | 0 | 0 | 0 | 0 | 0 | 0 | 0 | 0 | 0 |
| virus_Jul11.0v_5545 | 1 | 0 | 0 | 0 | 0 | 0 | 0 | 0 | 0 | 0 | 0 | 0 | 0 | 0 | 0 | 0 | 0 | 0 | 0 | 0 | 0 | 0 | 1 | 0 | 0 | 0 | 0 | 0 | 0 | 0 | 0 | 0 | 0 | 0 | 0 | 0 | 0 | 0 | 0 | 0 | 0 | 0 | 0 | 0 | 0 | 0 | 0 | 0 | 0 | 0 | 0 | 0 | 0 | 0 | 0 |
| virus_Jul11.0v_12161 | 1 | 0 | 0 | 0 | 0 | 0 | 0 | 0 | 0 | 0 | 0 | 0 | 0 | 0 | 0 | 0 | 0 | 0 | 0 | 0 | 0 | 0 | 1 | 0 | 0 | 0 | 0 | 0 | 0 | 0 | 0 | 0 | 0 | 0 | 0 | 0 | 0 | 0 | 0 | 0 | 0 | 0 | 0 | 0 | 0 | 0 | 0 | 0 | 0 | 0 | 0 | 0 | 0 | 0 | 0 |
| virus_Jul11.0v_16249 | 1 | 0 | 0 | 0 | 0 | 0 | 0 | 0 | 0 | 0 | 0 | 0 | 0 | 0 | 0 | 0 | 0 | 0 | 0 | 0 | 0 | 0 | 1 | 0 | 0 | 0 | 0 | 0 | 0 | 0 | 0 | 0 | 0 | 0 | 0 | 0 | 0 | 0 | 0 | 0 | 0 | 0 | 0 | 0 | 0 | 0 | 0 | 0 | 0 | 0 | 0 | 0 | 0 | 0 | 0 |
| virus_Jul11.0v_17580 | 1 | 0 | 0 | 0 | 0 | 0 | 0 | 0 | 0 | 0 | 0 | 0 | 0 | 0 | 0 | 0 | 0 | 0 | 0 | 0 | 0 | 0 | 1 | 0 | 0 | 0 | 0 | 0 | 0 | 0 | 0 | 0 | 0 | 0 | 0 | 0 | 0 | 0 | 0 | 0 | 0 | 0 | 0 | 0 | 0 | 0 | 0 | 0 | 0 | 0 | 0 | 0 | 0 | 0 | 0 |
| virus_Jul11.0v_18060 | 1 | 0 | 0 | 0 | 0 | 0 | 0 | 0 | 0 | 0 | 0 | 0 | 0 | 0 | 0 | 0 | 0 | 0 | 0 | 0 | 0 | 0 | 1 | 0 | 0 | 0 | 0 | 0 | 0 | 0 | 0 | 0 | 0 | 0 | 0 | 0 | 0 | 0 | 0 | 0 | 0 | 0 | 0 | 0 | 0 | 0 | 0 | 0 | 0 | 0 | 0 | 0 | 0 | 0 | 0 |
| virus_Jul11.0v_18988 | 1 | 0 | 0 | 0 | 0 | 0 | 0 | 0 | 0 | 0 | 0 | 0 | 0 | 0 | 0 | 0 | 0 | 0 | 0 | 0 | 0 | 0 | 1 | 0 | 0 | 0 | 0 | 0 | 0 | 0 | 0 | 0 | 0 | 0 | 0 | 0 | 0 | 0 | 0 | 0 | 0 | 0 | 0 | 0 | 0 | 0 | 0 | 0 | 0 | 0 | 0 | 0 | 0 | 0 | 0 |
| virus_Jul11.5v_505 | 1 | 0 | 0 | 0 | 0 | 0 | 0 | 0 | 0 | 0 | 0 | 0 | 0 | 0 | 0 | 0 | 0 | 0 | 0 | 0 | 0 | 0 | 0 | 0 | 1 | 0 | 0 | 0 | 0 | 0 | 0 | 0 | 0 | 0 | 0 | 0 | 0 | 0 | 0 | 0 | 0 | 0 | 0 | 0 | 0 | 0 | 0 | 0 | 0 | 0 | 0 | 0 | 0 | 0 | 0 |
| virus_Jul11.5v_1015 | 1 | 0 | 0 | 0 | 0 | 0 | 0 | 0 | 0 | 0 | 0 | 0 | 0 | 0 | 0 | 0 | 0 | 0 | 0 | 0 | 0 | 0 | 0 | 0 | 1 | 0 | 0 | 0 | 0 | 0 | 0 | 0 | 0 | 0 | 0 | 0 | 0 | 0 | 0 | 0 | 0 | 0 | 0 | 0 | 0 | 0 | 0 | 0 | 0 | 0 | 0 | 0 | 0 | 0 | 0 |
| virus_Jul11.5v_2601 | 1 | 0 | 0 | 0 | 0 | 0 | 0 | 0 | 0 | 0 | 0 | 0 | 0 | 0 | 0 | 0 | 0 | 0 | 0 | 0 | 0 | 0 | 0 | 0 | 1 | 0 | 0 | 0 | 0 | 0 | 0 | 0 | 0 | 0 | 0 | 0 | 0 | 0 | 0 | 0 | 0 | 0 | 0 | 0 | 0 | 0 | 0 | 0 | 0 | 0 | 0 | 0 | 0 | 0 | 0 |
| virus_Jul11.5v_3734 | 1 | 0 | 0 | 0 | 0 | 0 | 0 | 0 | 0 | 0 | 0 | 0 | 0 | 0 | 0 | 0 | 0 | 0 | 0 | 0 | 0 | 0 | 0 | 0 | 1 | 0 | 0 | 0 | 0 | 0 | 0 | 0 | 0 | 0 | 0 | 0 | 0 | 0 | 0 | 0 | 0 | 0 | 0 | 0 | 0 | 0 | 0 | 0 | 0 | 0 | 0 | 0 | 0 | 0 | 0 |
| virus_Jul11.5v_4525 | 1 | 0 | 0 | 0 | 0 | 0 | 0 | 0 | 0 | 0 | 0 | 0 | 0 | 0 | 0 | 0 | 0 | 0 | 0 | 0 | 0 | 0 | 0 | 0 | 1 | 0 | 0 | 0 | 0 | 0 | 0 | 0 | 0 | 0 | 0 | 0 | 0 | 0 | 0 | 0 | 0 | 0 | 0 | 0 | 0 | 0 | 0 | 0 | 0 | 0 | 0 | 0 | 0 | 0 | 0 |
| virus_Jul11.5v_4595 | 1 | 0 | 0 | 0 | 0 | 0 | 0 | 0 | 0 | 0 | 0 | 0 | 0 | 0 | 0 | 0 | 0 | 0 | 0 | 0 | 0 | 0 | 0 | 0 | 1 | 0 | 0 | 0 | 0 | 0 | 0 | 0 | 0 | 0 | 0 | 0 | 0 | 0 | 0 | 0 | 0 | 0 | 0 | 0 | 0 | 0 | 0 | 0 | 0 | 0 | 0 | 0 | 0 | 0 | 0 |
| virus_Jul11.5v_4718 | 1 | 0 | 0 | 0 | 0 | 0 | 0 | 0 | 0 | 0 | 0 | 0 | 0 | 0 | 0 | 0 | 0 | 0 | 0 | 0 | 0 | 0 | 0 | 0 | 1 | 0 | 0 | 0 | 0 | 0 | 0 | 0 | 0 | 0 | 0 | 0 | 0 | 0 | 0 | 0 | 0 | 0 | 0 | 0 | 0 | 0 | 0 | 0 | 0 | 0 | 0 | 0 | 0 | 0 | 0 |
| virus_Jul11.5v_8294 | 1 | 0 | 0 | 0 | 0 | 0 | 0 | 0 | 0 | 0 | 0 | 0 | 0 | 0 | 0 | 0 | 0 | 0 | 0 | 0 | 0 | 0 | 0 | 0 | 1 | 0 | 0 | 0 | 0 | 0 | 0 | 0 | 0 | 0 | 0 | 0 | 0 | 0 | 0 | 0 | 0 | 0 | 0 | 0 | 0 | 0 | 0 | 0 | 0 | 0 | 0 | 0 | 0 | 0 | 0 |
| virus_Jul11.5v_10738 | 1 | 0 | 0 | 0 | 0 | 0 | 0 | 0 | 0 | 0 | 0 | 0 | 0 | 0 | 0 | 0 | 0 | 0 | 0 | 0 | 0 | 0 | 0 | 0 | 1 | 0 | 0 | 0 | 0 | 0 | 0 | 0 | 0 | 0 | 0 | 0 | 0 | 0 | 0 | 0 | 0 | 0 | 0 | 0 | 0 | 0 | 0 | 0 | 0 | 0 | 0 | 0 | 0 | 0 | 0 |
| virus_Jul11.5v_13320 | 1 | 0 | 0 | 0 | 0 | 0 | 0 | 0 | 0 | 0 | 0 | 0 | 0 | 0 | 0 | 0 | 0 | 0 | 0 | 0 | 0 | 0 | 0 | 0 | 1 | 0 | 0 | 0 | 0 | 0 | 0 | 0 | 0 | 0 | 0 | 0 | 0 | 0 | 0 | 0 | 0 | 0 | 0 | 0 | 0 | 0 | 0 | 0 | 0 | 0 | 0 | 0 | 0 | 0 | 0 |
| virus_Jul11.5v_14609 | 1 | 0 | 0 | 0 | 0 | 0 | 0 | 0 | 0 | 0 | 0 | 0 | 0 | 0 | 0 | 0 | 0 | 0 | 0 | 0 | 0 | 0 | 0 | 0 | 1 | 0 | 0 | 0 | 0 | 0 | 0 | 0 | 0 | 0 | 0 | 0 | 0 | 0 | 0 | 0 | 0 | 0 | 0 | 0 | 0 | 0 | 0 | 0 | 0 | 0 | 0 | 0 | 0 | 0 | 0 |
| virus_Jul11.5v_15561 | 1 | 0 | 0 | 0 | 0 | 0 | 0 | 0 | 0 | 0 | 0 | 0 | 0 | 0 | 0 | 0 | 0 | 0 | 0 | 0 | 0 | 0 | 0 | 0 | 1 | 0 | 0 | 0 | 0 | 0 | 0 | 0 | 0 | 0 | 0 | 0 | 0 | 0 | 0 | 0 | 0 | 0 | 0 | 0 | 0 | 0 | 0 | 0 | 0 | 0 | 0 | 0 | 0 | 0 | 0 |
| virus_Jul11.5v_16835 | 1 | 0 | 0 | 0 | 0 | 0 | 0 | 0 | 0 | 0 | 0 | 0 | 0 | 0 | 0 | 0 | 0 | 0 | 0 | 0 | 0 | 0 | 0 | 0 | 1 | 0 | 0 | 0 | 0 | 0 | 0 | 0 | 0 | 0 | 0 | 0 | 0 | 0 | 0 | 0 | 0 | 0 | 0 | 0 | 0 | 0 | 0 | 0 | 0 | 0 | 0 | 0 | 0 | 0 | 0 |
| virus_Jul11.5v_17515 | 1 | 0 | 0 | 0 | 0 | 0 | 0 | 0 | 0 | 0 | 0 | 0 | 0 | 0 | 0 | 0 | 0 | 0 | 0 | 0 | 0 | 0 | 0 | 0 | 1 | 0 | 0 | 0 | 0 | 0 | 0 | 0 | 0 | 0 | 0 | 0 | 0 | 0 | 0 | 0 | 0 | 0 | 0 | 0 | 0 | 0 | 0 | 0 | 0 | 0 | 0 | 0 | 0 | 0 | 0 |
| virus_Jul11.25v_348 | 1 | 0 | 0 | 0 | 0 | 0 | 0 | 0 | 0 | 0 | 0 | 0 | 0 | 0 | 0 | 0 | 0 | 0 | 0 | 0 | 0 | 0 | 0 | 0 | 0 | 0 | 1 | 0 | 0 | 0 | 0 | 0 | 0 | 0 | 0 | 0 | 0 | 0 | 0 | 0 | 0 | 0 | 0 | 0 | 0 | 0 | 0 | 0 | 0 | 0 | 0 | 0 | 0 | 0 | 0 |
| virus_Jul11.25v_3684 | 1 | 0 | 0 | 0 | 0 | 0 | 0 | 0 | 0 | 0 | 0 | 0 | 0 | 0 | 0 | 0 | 0 | 0 | 0 | 0 | 0 | 0 | 0 | 0 | 0 | 0 | 1 | 0 | 0 | 0 | 0 | 0 | 0 | 0 | 0 | 0 | 0 | 0 | 0 | 0 | 0 | 0 | 0 | 0 | 0 | 0 | 0 | 0 | 0 | 0 | 0 | 0 | 0 | 0 | 0 |
| virus_Jul11.25v_3735 | 1 | 0 | 0 | 0 | 0 | 0 | 0 | 0 | 0 | 0 | 0 | 0 | 0 | 0 | 0 | 0 | 0 | 0 | 0 | 0 | 0 | 0 | 0 | 0 | 0 | 0 | 1 | 0 | 0 | 0 | 0 | 0 | 0 | 0 | 0 | 0 | 0 | 0 | 0 | 0 | 0 | 0 | 0 | 0 | 0 | 0 | 0 | 0 | 0 | 0 | 0 | 0 | 0 | 0 | 0 |
| virus_Jul11.25v_4792 | 1 | 0 | 0 | 0 | 0 | 0 | 0 | 0 | 0 | 0 | 0 | 0 | 0 | 0 | 0 | 0 | 0 | 0 | 0 | 0 | 0 | 0 | 0 | 0 | 0 | 0 | 1 | 0 | 0 | 0 | 0 | 0 | 0 | 0 | 0 | 0 | 0 | 0 | 0 | 0 | 0 | 0 | 0 | 0 | 0 | 0 | 0 | 0 | 0 | 0 | 0 | 0 | 0 | 0 | 0 |
| virus_Jul11.25v_6271 | 1 | 0 | 0 | 0 | 0 | 0 | 0 | 0 | 0 | 0 | 0 | 0 | 0 | 0 | 0 | 0 | 0 | 0 | 0 | 0 | 0 | 0 | 0 | 0 | 0 | 0 | 1 | 0 | 0 | 0 | 0 | 0 | 0 | 0 | 0 | 0 | 0 | 0 | 0 | 0 | 0 | 0 | 0 | 0 | 0 | 0 | 0 | 0 | 0 | 0 | 0 | 0 | 0 | 0 | 0 |
| virus_Jul11.25v_6769 | 1 | 0 | 0 | 0 | 0 | 0 | 0 | 0 | 0 | 0 | 0 | 0 | 0 | 0 | 0 | 0 | 0 | 0 | 0 | 0 | 0 | 0 | 0 | 0 | 0 | 0 | 1 | 0 | 0 | 0 | 0 | 0 | 0 | 0 | 0 | 0 | 0 | 0 | 0 | 0 | 0 | 0 | 0 | 0 | 0 | 0 | 0 | 0 | 0 | 0 | 0 | 0 | 0 | 0 | 0 |
| virus_Jul11.25v_9543 | 1 | 0 | 0 | 0 | 0 | 0 | 0 | 0 | 0 | 0 | 0 | 0 | 0 | 0 | 0 | 0 | 0 | 0 | 0 | 0 | 0 | 0 | 0 | 0 | 0 | 0 | 1 | 0 | 0 | 0 | 0 | 0 | 0 | 0 | 0 | 0 | 0 | 0 | 0 | 0 | 0 | 0 | 0 | 0 | 0 | 0 | 0 | 0 | 0 | 0 | 0 | 0 | 0 | 0 | 0 |
| virus_Jul11.25v_11244 | 1 | 0 | 0 | 0 | 0 | 0 | 0 | 0 | 0 | 0 | 0 | 0 | 0 | 0 | 0 | 0 | 0 | 0 | 0 | 0 | 0 | 0 | 0 | 0 | 0 | 0 | 1 | 0 | 0 | 0 | 0 | 0 | 0 | 0 | 0 | 0 | 0 | 0 | 0 | 0 | 0 | 0 | 0 | 0 | 0 | 0 | 0 | 0 | 0 | 0 | 0 | 0 | 0 | 0 | 0 |
| virus_Jul11.25v_14845 | 1 | 0 | 0 | 0 | 0 | 0 | 0 | 0 | 0 | 0 | 0 | 0 | 0 | 0 | 0 | 0 | 0 | 0 | 0 | 0 | 0 | 0 | 0 | 0 | 0 | 0 | 1 | 0 | 0 | 0 | 0 | 0 | 0 | 0 | 0 | 0 | 0 | 0 | 0 | 0 | 0 | 0 | 0 | 0 | 0 | 0 | 0 | 0 | 0 | 0 | 0 | 0 | 0 | 0 | 0 |
| virus_Jul11.25v_15608 | 1 | 0 | 0 | 0 | 0 | 0 | 0 | 0 | 0 | 0 | 0 | 0 | 0 | 0 | 0 | 0 | 0 | 0 | 0 | 0 | 0 | 0 | 0 | 0 | 0 | 0 | 1 | 0 | 0 | 0 | 0 | 0 | 0 | 0 | 0 | 0 | 0 | 0 | 0 | 0 | 0 | 0 | 0 | 0 | 0 | 0 | 0 | 0 | 0 | 0 | 0 | 0 | 0 | 0 | 0 |
| virus_Jul11.25v_18324 | 1 | 0 | 0 | 0 | 0 | 0 | 0 | 0 | 0 | 0 | 0 | 0 | 0 | 0 | 0 | 0 | 0 | 0 | 0 | 0 | 0 | 0 | 0 | 0 | 0 | 0 | 1 | 0 | 0 | 0 | 0 | 0 | 0 | 0 | 0 | 0 | 0 | 0 | 0 | 0 | 0 | 0 | 0 | 0 | 0 | 0 | 0 | 0 | 0 | 0 | 0 | 0 | 0 | 0 | 0 |
| virus_Jul11.25v_19146 | 1 | 0 | 0 | 0 | 0 | 0 | 0 | 0 | 0 | 0 | 0 | 0 | 0 | 0 | 0 | 0 | 0 | 0 | 0 | 0 | 0 | 0 | 0 | 0 | 0 | 0 | 1 | 0 | 0 | 0 | 0 | 0 | 0 | 0 | 0 | 0 | 0 | 0 | 0 | 0 | 0 | 0 | 0 | 0 | 0 | 0 | 0 | 0 | 0 | 0 | 0 | 0 | 0 | 0 | 0 |
| virus_Jul11.50v_1478 | 1 | 0 | 0 | 0 | 0 | 0 | 0 | 0 | 0 | 0 | 0 | 0 | 0 | 0 | 0 | 0 | 0 | 0 | 0 | 0 | 0 | 0 | 0 | 0 | 0 | 0 | 0 | 0 | 1 | 0 | 0 | 0 | 0 | 0 | 0 | 0 | 0 | 0 | 0 | 0 | 0 | 0 | 0 | 0 | 0 | 0 | 0 | 0 | 0 | 0 | 0 | 0 | 0 | 0 | 0 |
| virus_Jul11.50v_1692 | 1 | 0 | 0 | 0 | 0 | 0 | 0 | 0 | 0 | 0 | 0 | 0 | 0 | 0 | 0 | 0 | 0 | 0 | 0 | 0 | 0 | 0 | 0 | 0 | 0 | 0 | 0 | 0 | 1 | 0 | 0 | 0 | 0 | 0 | 0 | 0 | 0 | 0 | 0 | 0 | 0 | 0 | 0 | 0 | 0 | 0 | 0 | 0 | 0 | 0 | 0 | 0 | 0 | 0 | 0 |
| virus_Jul11.50v_2377 | 1 | 0 | 0 | 0 | 0 | 0 | 0 | 0 | 0 | 0 | 0 | 0 | 0 | 0 | 0 | 0 | 0 | 0 | 0 | 0 | 0 | 0 | 0 | 0 | 0 | 0 | 0 | 0 | 1 | 0 | 0 | 0 | 0 | 0 | 0 | 0 | 0 | 0 | 0 | 0 | 0 | 0 | 0 | 0 | 0 | 0 | 0 | 0 | 0 | 0 | 0 | 0 | 0 | 0 | 0 |
| virus_Jul11.50v_8236 | 1 | 0 | 0 | 0 | 0 | 0 | 0 | 0 | 0 | 0 | 0 | 0 | 0 | 0 | 0 | 0 | 0 | 0 | 0 | 0 | 0 | 0 | 0 | 0 | 0 | 0 | 0 | 0 | 1 | 0 | 0 | 0 | 0 | 0 | 0 | 0 | 0 | 0 | 0 | 0 | 0 | 0 | 0 | 0 | 0 | 0 | 0 | 0 | 0 | 0 | 0 | 0 | 0 | 0 | 0 |
| virus_Jul11.50v_9034 | 1 | 0 | 0 | 0 | 0 | 0 | 0 | 0 | 0 | 0 | 0 | 0 | 0 | 0 | 0 | 0 | 0 | 0 | 0 | 0 | 0 | 0 | 0 | 0 | 0 | 0 | 0 | 0 | 1 | 0 | 0 | 0 | 0 | 0 | 0 | 0 | 0 | 0 | 0 | 0 | 0 | 0 | 0 | 0 | 0 | 0 | 0 | 0 | 0 | 0 | 0 | 0 | 0 | 0 | 0 |
| virus_Jul11.50v_9634 | 1 | 0 | 0 | 0 | 0 | 0 | 0 | 0 | 0 | 0 | 0 | 0 | 0 | 0 | 0 | 0 | 0 | 0 | 0 | 0 | 0 | 0 | 0 | 0 | 0 | 0 | 0 | 0 | 1 | 0 | 0 | 0 | 0 | 0 | 0 | 0 | 0 | 0 | 0 | 0 | 0 | 0 | 0 | 0 | 0 | 0 | 0 | 0 | 0 | 0 | 0 | 0 | 0 | 0 | 0 |
| virus_Jul11.50v_10452 | 1 | 0 | 0 | 0 | 0 | 0 | 0 | 0 | 0 | 0 | 0 | 0 | 0 | 0 | 0 | 0 | 0 | 0 | 0 | 0 | 0 | 0 | 0 | 0 | 0 | 0 | 0 | 0 | 1 | 0 | 0 | 0 | 0 | 0 | 0 | 0 | 0 | 0 | 0 | 0 | 0 | 0 | 0 | 0 | 0 | 0 | 0 | 0 | 0 | 0 | 0 | 0 | 0 | 0 | 0 |
| virus_Jul11.50v_10799 | 1 | 0 | 0 | 0 | 0 | 0 | 0 | 0 | 0 | 0 | 0 | 0 | 0 | 0 | 0 | 0 | 0 | 0 | 0 | 0 | 0 | 0 | 0 | 0 | 0 | 0 | 0 | 0 | 1 | 0 | 0 | 0 | 0 | 0 | 0 | 0 | 0 | 0 | 0 | 0 | 0 | 0 | 0 | 0 | 0 | 0 | 0 | 0 | 0 | 0 | 0 | 0 | 0 | 0 | 0 |
| virus_Jul11.50v_10873 | 1 | 0 | 0 | 0 | 0 | 0 | 0 | 0 | 0 | 0 | 0 | 0 | 0 | 0 | 0 | 0 | 0 | 0 | 0 | 0 | 0 | 0 | 0 | 0 | 0 | 0 | 0 | 0 | 1 | 0 | 0 | 0 | 0 | 0 | 0 | 0 | 0 | 0 | 0 | 0 | 0 | 0 | 0 | 0 | 0 | 0 | 0 | 0 | 0 | 0 | 0 | 0 | 0 | 0 | 0 |
| virus_Jul11.50v_11059 | 1 | 0 | 0 | 0 | 0 | 0 | 0 | 0 | 0 | 0 | 0 | 0 | 0 | 0 | 0 | 0 | 0 | 0 | 0 | 0 | 0 | 0 | 0 | 0 | 0 | 0 | 0 | 0 | 1 | 0 | 0 | 0 | 0 | 0 | 0 | 0 | 0 | 0 | 0 | 0 | 0 | 0 | 0 | 0 | 0 | 0 | 0 | 0 | 0 | 0 | 0 | 0 | 0 | 0 | 0 |
| virus_Jul11.50v_11565 | 1 | 0 | 0 | 0 | 0 | 0 | 0 | 0 | 0 | 0 | 0 | 0 | 0 | 0 | 0 | 0 | 0 | 0 | 0 | 0 | 0 | 0 | 0 | 0 | 0 | 0 | 0 | 0 | 1 | 0 | 0 | 0 | 0 | 0 | 0 | 0 | 0 | 0 | 0 | 0 | 0 | 0 | 0 | 0 | 0 | 0 | 0 | 0 | 0 | 0 | 0 | 0 | 0 | 0 | 0 |
| virus_Jul11.50v_13821 | 1 | 0 | 0 | 0 | 0 | 0 | 0 | 0 | 0 | 0 | 0 | 0 | 0 | 0 | 0 | 0 | 0 | 0 | 0 | 0 | 0 | 0 | 0 | 0 | 0 | 0 | 0 | 0 | 1 | 0 | 0 | 0 | 0 | 0 | 0 | 0 | 0 | 0 | 0 | 0 | 0 | 0 | 0 | 0 | 0 | 0 | 0 | 0 | 0 | 0 | 0 | 0 | 0 | 0 | 0 |
| virus_Jul11.50v_14199 | 1 | 0 | 0 | 0 | 0 | 0 | 0 | 0 | 0 | 0 | 0 | 0 | 0 | 0 | 0 | 0 | 0 | 0 | 0 | 0 | 0 | 0 | 0 | 0 | 0 | 0 | 0 | 0 | 1 | 0 | 0 | 0 | 0 | 0 | 0 | 0 | 0 | 0 | 0 | 0 | 0 | 0 | 0 | 0 | 0 | 0 | 0 | 0 | 0 | 0 | 0 | 0 | 0 | 0 | 0 |
| virus_Jul11.50v_18621 | 1 | 0 | 0 | 0 | 0 | 0 | 0 | 0 | 0 | 0 | 0 | 0 | 0 | 0 | 0 | 0 | 0 | 0 | 0 | 0 | 0 | 0 | 0 | 0 | 0 | 0 | 0 | 0 | 1 | 0 | 0 | 0 | 0 | 0 | 0 | 0 | 0 | 0 | 0 | 0 | 0 | 0 | 0 | 0 | 0 | 0 | 0 | 0 | 0 | 0 | 0 | 0 | 0 | 0 | 0 |
| virus_Jul11.50v_19785 | 1 | 0 | 0 | 0 | 0 | 0 | 0 | 0 | 0 | 0 | 0 | 0 | 0 | 0 | 0 | 0 | 0 | 0 | 0 | 0 | 0 | 0 | 0 | 0 | 0 | 0 | 0 | 0 | 1 | 0 | 0 | 0 | 0 | 0 | 0 | 0 | 0 | 0 | 0 | 0 | 0 | 0 | 0 | 0 | 0 | 0 | 0 | 0 | 0 | 0 | 0 | 0 | 0 | 0 | 0 |
| virus_Jul11.50v_19926 | 1 | 0 | 0 | 0 | 0 | 0 | 0 | 0 | 0 | 0 | 0 | 0 | 0 | 0 | 0 | 0 | 0 | 0 | 0 | 0 | 0 | 0 | 0 | 0 | 0 | 0 | 0 | 0 | 1 | 0 | 0 | 0 | 0 | 0 | 0 | 0 | 0 | 0 | 0 | 0 | 0 | 0 | 0 | 0 | 0 | 0 | 0 | 0 | 0 | 0 | 0 | 0 | 0 | 0 | 0 |
| virus_Jul11.75v_3391 | 1 | 0 | 0 | 0 | 0 | 0 | 0 | 0 | 0 | 0 | 0 | 0 | 0 | 0 | 0 | 0 | 0 | 0 | 0 | 0 | 0 | 0 | 0 | 0 | 0 | 0 | 0 | 0 | 0 | 0 | 1 | 0 | 0 | 0 | 0 | 0 | 0 | 0 | 0 | 0 | 0 | 0 | 0 | 0 | 0 | 0 | 0 | 0 | 0 | 0 | 0 | 0 | 0 | 0 | 0 |
| virus_Jul11.75v_24568 | 1 | 0 | 0 | 0 | 0 | 0 | 0 | 0 | 0 | 0 | 0 | 0 | 0 | 0 | 0 | 0 | 0 | 0 | 0 | 0 | 0 | 0 | 0 | 0 | 0 | 0 | 0 | 0 | 0 | 0 | 1 | 0 | 0 | 0 | 0 | 0 | 0 | 0 | 0 | 0 | 0 | 0 | 0 | 0 | 0 | 0 | 0 | 0 | 0 | 0 | 0 | 0 | 0 | 0 | 0 |
| virus_Jul11.75v_34355 | 1 | 0 | 0 | 0 | 0 | 0 | 0 | 0 | 0 | 0 | 0 | 0 | 0 | 0 | 0 | 0 | 0 | 0 | 0 | 0 | 0 | 0 | 0 | 0 | 0 | 0 | 0 | 0 | 0 | 0 | 1 | 0 | 0 | 0 | 0 | 0 | 0 | 0 | 0 | 0 | 0 | 0 | 0 | 0 | 0 | 0 | 0 | 0 | 0 | 0 | 0 | 0 | 0 | 0 | 0 |
| virus_Jul11.75v_37277 | 1 | 0 | 0 | 0 | 0 | 0 | 0 | 0 | 0 | 0 | 0 | 0 | 0 | 0 | 0 | 0 | 0 | 0 | 0 | 0 | 0 | 0 | 0 | 0 | 0 | 0 | 0 | 0 | 0 | 0 | 1 | 0 | 0 | 0 | 0 | 0 | 0 | 0 | 0 | 0 | 0 | 0 | 0 | 0 | 0 | 0 | 0 | 0 | 0 | 0 | 0 | 0 | 0 | 0 | 0 |
| virus_Jul27.5v_843 | 1 | 0 | 0 | 0 | 0 | 0 | 0 | 0 | 0 | 0 | 0 | 0 | 0 | 0 | 0 | 0 | 0 | 0 | 0 | 0 | 0 | 0 | 0 | 0 | 0 | 0 | 0 | 0 | 0 | 0 | 0 | 0 | 1 | 0 | 0 | 0 | 0 | 0 | 0 | 0 | 0 | 0 | 0 | 0 | 0 | 0 | 0 | 0 | 0 | 0 | 0 | 0 | 0 | 0 | 0 |
| virus_Jul27.5v_950 | 1 | 0 | 0 | 0 | 0 | 0 | 0 | 0 | 0 | 0 | 0 | 0 | 0 | 0 | 0 | 0 | 0 | 0 | 0 | 0 | 0 | 0 | 0 | 0 | 0 | 0 | 0 | 0 | 0 | 0 | 0 | 0 | 1 | 0 | 0 | 0 | 0 | 0 | 0 | 0 | 0 | 0 | 0 | 0 | 0 | 0 | 0 | 0 | 0 | 0 | 0 | 0 | 0 | 0 | 0 |
| virus_Jul27.5v_1171 | 1 | 0 | 0 | 0 | 0 | 0 | 0 | 0 | 0 | 0 | 0 | 0 | 0 | 0 | 0 | 0 | 0 | 0 | 0 | 0 | 0 | 0 | 0 | 0 | 0 | 0 | 0 | 0 | 0 | 0 | 0 | 0 | 1 | 0 | 0 | 0 | 0 | 0 | 0 | 0 | 0 | 0 | 0 | 0 | 0 | 0 | 0 | 0 | 0 | 0 | 0 | 0 | 0 | 0 | 0 |
| virus_Jul27.5v_1182 | 1 | 0 | 0 | 0 | 0 | 0 | 0 | 0 | 0 | 0 | 0 | 0 | 0 | 0 | 0 | 0 | 0 | 0 | 0 | 0 | 0 | 0 | 0 | 0 | 0 | 0 | 0 | 0 | 0 | 0 | 0 | 0 | 1 | 0 | 0 | 0 | 0 | 0 | 0 | 0 | 0 | 0 | 0 | 0 | 0 | 0 | 0 | 0 | 0 | 0 | 0 | 0 | 0 | 0 | 0 |
| virus_Jul27.5v_2857 | 1 | 0 | 0 | 0 | 0 | 0 | 0 | 0 | 0 | 0 | 0 | 0 | 0 | 0 | 0 | 0 | 0 | 0 | 0 | 0 | 0 | 0 | 0 | 0 | 0 | 0 | 0 | 0 | 0 | 0 | 0 | 0 | 1 | 0 | 0 | 0 | 0 | 0 | 0 | 0 | 0 | 0 | 0 | 0 | 0 | 0 | 0 | 0 | 0 | 0 | 0 | 0 | 0 | 0 | 0 |
| virus_Jul27.5v_3540 | 1 | 0 | 0 | 0 | 0 | 0 | 0 | 0 | 0 | 0 | 0 | 0 | 0 | 0 | 0 | 0 | 0 | 0 | 0 | 0 | 0 | 0 | 0 | 0 | 0 | 0 | 0 | 0 | 0 | 0 | 0 | 0 | 1 | 0 | 0 | 0 | 0 | 0 | 0 | 0 | 0 | 0 | 0 | 0 | 0 | 0 | 0 | 0 | 0 | 0 | 0 | 0 | 0 | 0 | 0 |
| virus_Jul27.5v_3810 | 1 | 0 | 0 | 0 | 0 | 0 | 0 | 0 | 0 | 0 | 0 | 0 | 0 | 0 | 0 | 0 | 0 | 0 | 0 | 0 | 0 | 0 | 0 | 0 | 0 | 0 | 0 | 0 | 0 | 0 | 0 | 0 | 1 | 0 | 0 | 0 | 0 | 0 | 0 | 0 | 0 | 0 | 0 | 0 | 0 | 0 | 0 | 0 | 0 | 0 | 0 | 0 | 0 | 0 | 0 |
| virus_Jul27.5v_4018 | 1 | 0 | 0 | 0 | 0 | 0 | 0 | 0 | 0 | 0 | 0 | 0 | 0 | 0 | 0 | 0 | 0 | 0 | 0 | 0 | 0 | 0 | 0 | 0 | 0 | 0 | 0 | 0 | 0 | 0 | 0 | 0 | 1 | 0 | 0 | 0 | 0 | 0 | 0 | 0 | 0 | 0 | 0 | 0 | 0 | 0 | 0 | 0 | 0 | 0 | 0 | 0 | 0 | 0 | 0 |
| virus_Jul27.5v_8906 | 1 | 0 | 0 | 0 | 0 | 0 | 0 | 0 | 0 | 0 | 0 | 0 | 0 | 0 | 0 | 0 | 0 | 0 | 0 | 0 | 0 | 0 | 0 | 0 | 0 | 0 | 0 | 0 | 0 | 0 | 0 | 0 | 1 | 0 | 0 | 0 | 0 | 0 | 0 | 0 | 0 | 0 | 0 | 0 | 0 | 0 | 0 | 0 | 0 | 0 | 0 | 0 | 0 | 0 | 0 |
| virus_Jul27.5v_9686 | 1 | 0 | 0 | 0 | 0 | 0 | 0 | 0 | 0 | 0 | 0 | 0 | 0 | 0 | 0 | 0 | 0 | 0 | 0 | 0 | 0 | 0 | 0 | 0 | 0 | 0 | 0 | 0 | 0 | 0 | 0 | 0 | 1 | 0 | 0 | 0 | 0 | 0 | 0 | 0 | 0 | 0 | 0 | 0 | 0 | 0 | 0 | 0 | 0 | 0 | 0 | 0 | 0 | 0 | 0 |
| virus_Jul27.5v_10371 | 1 | 0 | 0 | 0 | 0 | 0 | 0 | 0 | 0 | 0 | 0 | 0 | 0 | 0 | 0 | 0 | 0 | 0 | 0 | 0 | 0 | 0 | 0 | 0 | 0 | 0 | 0 | 0 | 0 | 0 | 0 | 0 | 1 | 0 | 0 | 0 | 0 | 0 | 0 | 0 | 0 | 0 | 0 | 0 | 0 | 0 | 0 | 0 | 0 | 0 | 0 | 0 | 0 | 0 | 0 |
| virus_Jul27.5v_10375 | 1 | 0 | 0 | 0 | 0 | 0 | 0 | 0 | 0 | 0 | 0 | 0 | 0 | 0 | 0 | 0 | 0 | 0 | 0 | 0 | 0 | 0 | 0 | 0 | 0 | 0 | 0 | 0 | 0 | 0 | 0 | 0 | 1 | 0 | 0 | 0 | 0 | 0 | 0 | 0 | 0 | 0 | 0 | 0 | 0 | 0 | 0 | 0 | 0 | 0 | 0 | 0 | 0 | 0 | 0 |
| virus_Jul27.5v_10681 | 1 | 0 | 0 | 0 | 0 | 0 | 0 | 0 | 0 | 0 | 0 | 0 | 0 | 0 | 0 | 0 | 0 | 0 | 0 | 0 | 0 | 0 | 0 | 0 | 0 | 0 | 0 | 0 | 0 | 0 | 0 | 0 | 1 | 0 | 0 | 0 | 0 | 0 | 0 | 0 | 0 | 0 | 0 | 0 | 0 | 0 | 0 | 0 | 0 | 0 | 0 | 0 | 0 | 0 | 0 |
| virus_Jul27.5v_10725 | 1 | 0 | 0 | 0 | 0 | 0 | 0 | 0 | 0 | 0 | 0 | 0 | 0 | 0 | 0 | 0 | 0 | 0 | 0 | 0 | 0 | 0 | 0 | 0 | 0 | 0 | 0 | 0 | 0 | 0 | 0 | 0 | 1 | 0 | 0 | 0 | 0 | 0 | 0 | 0 | 0 | 0 | 0 | 0 | 0 | 0 | 0 | 0 | 0 | 0 | 0 | 0 | 0 | 0 | 0 |
| virus_Jul27.5v_11919 | 1 | 0 | 0 | 0 | 0 | 0 | 0 | 0 | 0 | 0 | 0 | 0 | 0 | 0 | 0 | 0 | 0 | 0 | 0 | 0 | 0 | 0 | 0 | 0 | 0 | 0 | 0 | 0 | 0 | 0 | 0 | 0 | 1 | 0 | 0 | 0 | 0 | 0 | 0 | 0 | 0 | 0 | 0 | 0 | 0 | 0 | 0 | 0 | 0 | 0 | 0 | 0 | 0 | 0 | 0 |
| virus_Jul27.5v_12166 | 1 | 0 | 0 | 0 | 0 | 0 | 0 | 0 | 0 | 0 | 0 | 0 | 0 | 0 | 0 | 0 | 0 | 0 | 0 | 0 | 0 | 0 | 0 | 0 | 0 | 0 | 0 | 0 | 0 | 0 | 0 | 0 | 1 | 0 | 0 | 0 | 0 | 0 | 0 | 0 | 0 | 0 | 0 | 0 | 0 | 0 | 0 | 0 | 0 | 0 | 0 | 0 | 0 | 0 | 0 |
| virus_Jul27.5v_13408 | 1 | 0 | 0 | 0 | 0 | 0 | 0 | 0 | 0 | 0 | 0 | 0 | 0 | 0 | 0 | 0 | 0 | 0 | 0 | 0 | 0 | 0 | 0 | 0 | 0 | 0 | 0 | 0 | 0 | 0 | 0 | 0 | 1 | 0 | 0 | 0 | 0 | 0 | 0 | 0 | 0 | 0 | 0 | 0 | 0 | 0 | 0 | 0 | 0 | 0 | 0 | 0 | 0 | 0 | 0 |
| virus_Jul27.5v_14291 | 1 | 0 | 0 | 0 | 0 | 0 | 0 | 0 | 0 | 0 | 0 | 0 | 0 | 0 | 0 | 0 | 0 | 0 | 0 | 0 | 0 | 0 | 0 | 0 | 0 | 0 | 0 | 0 | 0 | 0 | 0 | 0 | 1 | 0 | 0 | 0 | 0 | 0 | 0 | 0 | 0 | 0 | 0 | 0 | 0 | 0 | 0 | 0 | 0 | 0 | 0 | 0 | 0 | 0 | 0 |
| virus_Jul27.5v_15997 | 1 | 0 | 0 | 0 | 0 | 0 | 0 | 0 | 0 | 0 | 0 | 0 | 0 | 0 | 0 | 0 | 0 | 0 | 0 | 0 | 0 | 0 | 0 | 0 | 0 | 0 | 0 | 0 | 0 | 0 | 0 | 0 | 1 | 0 | 0 | 0 | 0 | 0 | 0 | 0 | 0 | 0 | 0 | 0 | 0 | 0 | 0 | 0 | 0 | 0 | 0 | 0 | 0 | 0 | 0 |
| virus_Jul27.5v_16519 | 1 | 0 | 0 | 0 | 0 | 0 | 0 | 0 | 0 | 0 | 0 | 0 | 0 | 0 | 0 | 0 | 0 | 0 | 0 | 0 | 0 | 0 | 0 | 0 | 0 | 0 | 0 | 0 | 0 | 0 | 0 | 0 | 1 | 0 | 0 | 0 | 0 | 0 | 0 | 0 | 0 | 0 | 0 | 0 | 0 | 0 | 0 | 0 | 0 | 0 | 0 | 0 | 0 | 0 | 0 |
| virus_Jul27.5v_16814 | 1 | 0 | 0 | 0 | 0 | 0 | 0 | 0 | 0 | 0 | 0 | 0 | 0 | 0 | 0 | 0 | 0 | 0 | 0 | 0 | 0 | 0 | 0 | 0 | 0 | 0 | 0 | 0 | 0 | 0 | 0 | 0 | 1 | 0 | 0 | 0 | 0 | 0 | 0 | 0 | 0 | 0 | 0 | 0 | 0 | 0 | 0 | 0 | 0 | 0 | 0 | 0 | 0 | 0 | 0 |
| virus_Jul27.5v_18724 | 1 | 0 | 0 | 0 | 0 | 0 | 0 | 0 | 0 | 0 | 0 | 0 | 0 | 0 | 0 | 0 | 0 | 0 | 0 | 0 | 0 | 0 | 0 | 0 | 0 | 0 | 0 | 0 | 0 | 0 | 0 | 0 | 1 | 0 | 0 | 0 | 0 | 0 | 0 | 0 | 0 | 0 | 0 | 0 | 0 | 0 | 0 | 0 | 0 | 0 | 0 | 0 | 0 | 0 | 0 |
| virus_Jul27.5v_18780 | 1 | 0 | 0 | 0 | 0 | 0 | 0 | 0 | 0 | 0 | 0 | 0 | 0 | 0 | 0 | 0 | 0 | 0 | 0 | 0 | 0 | 0 | 0 | 0 | 0 | 0 | 0 | 0 | 0 | 0 | 0 | 0 | 1 | 0 | 0 | 0 | 0 | 0 | 0 | 0 | 0 | 0 | 0 | 0 | 0 | 0 | 0 | 0 | 0 | 0 | 0 | 0 | 0 | 0 | 0 |
| virus_Jul27.5v_20008 | 1 | 0 | 0 | 0 | 0 | 0 | 0 | 0 | 0 | 0 | 0 | 0 | 0 | 0 | 0 | 0 | 0 | 0 | 0 | 0 | 0 | 0 | 0 | 0 | 0 | 0 | 0 | 0 | 0 | 0 | 0 | 0 | 1 | 0 | 0 | 0 | 0 | 0 | 0 | 0 | 0 | 0 | 0 | 0 | 0 | 0 | 0 | 0 | 0 | 0 | 0 | 0 | 0 | 0 | 0 |
| virus_Jul27.5v_20962 | 1 | 0 | 0 | 0 | 0 | 0 | 0 | 0 | 0 | 0 | 0 | 0 | 0 | 0 | 0 | 0 | 0 | 0 | 0 | 0 | 0 | 0 | 0 | 0 | 0 | 0 | 0 | 0 | 0 | 0 | 0 | 0 | 1 | 0 | 0 | 0 | 0 | 0 | 0 | 0 | 0 | 0 | 0 | 0 | 0 | 0 | 0 | 0 | 0 | 0 | 0 | 0 | 0 | 0 | 0 |
| virus_Jul27.5v_21101 | 1 | 0 | 0 | 0 | 0 | 0 | 0 | 0 | 0 | 0 | 0 | 0 | 0 | 0 | 0 | 0 | 0 | 0 | 0 | 0 | 0 | 0 | 0 | 0 | 0 | 0 | 0 | 0 | 0 | 0 | 0 | 0 | 1 | 0 | 0 | 0 | 0 | 0 | 0 | 0 | 0 | 0 | 0 | 0 | 0 | 0 | 0 | 0 | 0 | 0 | 0 | 0 | 0 | 0 | 0 |
| virus_Jul27.25v_469 | 1 | 0 | 0 | 0 | 0 | 0 | 0 | 0 | 0 | 0 | 0 | 0 | 0 | 0 | 0 | 0 | 0 | 0 | 0 | 0 | 0 | 0 | 0 | 0 | 0 | 0 | 0 | 0 | 0 | 0 | 0 | 0 | 0 | 0 | 1 | 0 | 0 | 0 | 0 | 0 | 0 | 0 | 0 | 0 | 0 | 0 | 0 | 0 | 0 | 0 | 0 | 0 | 0 | 0 | 0 |
| virus_Jul27.25v_2097 | 1 | 0 | 0 | 0 | 0 | 0 | 0 | 0 | 0 | 0 | 0 | 0 | 0 | 0 | 0 | 0 | 0 | 0 | 0 | 0 | 0 | 0 | 0 | 0 | 0 | 0 | 0 | 0 | 0 | 0 | 0 | 0 | 0 | 0 | 1 | 0 | 0 | 0 | 0 | 0 | 0 | 0 | 0 | 0 | 0 | 0 | 0 | 0 | 0 | 0 | 0 | 0 | 0 | 0 | 0 |
| virus_Jul27.25v_3424 | 1 | 0 | 0 | 0 | 0 | 0 | 0 | 0 | 0 | 0 | 0 | 0 | 0 | 0 | 0 | 0 | 0 | 0 | 0 | 0 | 0 | 0 | 0 | 0 | 0 | 0 | 0 | 0 | 0 | 0 | 0 | 0 | 0 | 0 | 1 | 0 | 0 | 0 | 0 | 0 | 0 | 0 | 0 | 0 | 0 | 0 | 0 | 0 | 0 | 0 | 0 | 0 | 0 | 0 | 0 |
| virus_Jul27.25v_6615 | 1 | 0 | 0 | 0 | 0 | 0 | 0 | 0 | 0 | 0 | 0 | 0 | 0 | 0 | 0 | 0 | 0 | 0 | 0 | 0 | 0 | 0 | 0 | 0 | 0 | 0 | 0 | 0 | 0 | 0 | 0 | 0 | 0 | 0 | 1 | 0 | 0 | 0 | 0 | 0 | 0 | 0 | 0 | 0 | 0 | 0 | 0 | 0 | 0 | 0 | 0 | 0 | 0 | 0 | 0 |
| virus_Jul27.25v_6801 | 1 | 0 | 0 | 0 | 0 | 0 | 0 | 0 | 0 | 0 | 0 | 0 | 0 | 0 | 0 | 0 | 0 | 0 | 0 | 0 | 0 | 0 | 0 | 0 | 0 | 0 | 0 | 0 | 0 | 0 | 0 | 0 | 0 | 0 | 1 | 0 | 0 | 0 | 0 | 0 | 0 | 0 | 0 | 0 | 0 | 0 | 0 | 0 | 0 | 0 | 0 | 0 | 0 | 0 | 0 |
| virus_Jul27.25v_10565 | 1 | 0 | 0 | 0 | 0 | 0 | 0 | 0 | 0 | 0 | 0 | 0 | 0 | 0 | 0 | 0 | 0 | 0 | 0 | 0 | 0 | 0 | 0 | 0 | 0 | 0 | 0 | 0 | 0 | 0 | 0 | 0 | 0 | 0 | 1 | 0 | 0 | 0 | 0 | 0 | 0 | 0 | 0 | 0 | 0 | 0 | 0 | 0 | 0 | 0 | 0 | 0 | 0 | 0 | 0 |
| virus_Jul27.25v_12240 | 1 | 0 | 0 | 0 | 0 | 0 | 0 | 0 | 0 | 0 | 0 | 0 | 0 | 0 | 0 | 0 | 0 | 0 | 0 | 0 | 0 | 0 | 0 | 0 | 0 | 0 | 0 | 0 | 0 | 0 | 0 | 0 | 0 | 0 | 1 | 0 | 0 | 0 | 0 | 0 | 0 | 0 | 0 | 0 | 0 | 0 | 0 | 0 | 0 | 0 | 0 | 0 | 0 | 0 | 0 |
| virus_Jul27.25v_13264 | 1 | 0 | 0 | 0 | 0 | 0 | 0 | 0 | 0 | 0 | 0 | 0 | 0 | 0 | 0 | 0 | 0 | 0 | 0 | 0 | 0 | 0 | 0 | 0 | 0 | 0 | 0 | 0 | 0 | 0 | 0 | 0 | 0 | 0 | 1 | 0 | 0 | 0 | 0 | 0 | 0 | 0 | 0 | 0 | 0 | 0 | 0 | 0 | 0 | 0 | 0 | 0 | 0 | 0 | 0 |
| virus_Jul27.25v_14754 | 1 | 0 | 0 | 0 | 0 | 0 | 0 | 0 | 0 | 0 | 0 | 0 | 0 | 0 | 0 | 0 | 0 | 0 | 0 | 0 | 0 | 0 | 0 | 0 | 0 | 0 | 0 | 0 | 0 | 0 | 0 | 0 | 0 | 0 | 1 | 0 | 0 | 0 | 0 | 0 | 0 | 0 | 0 | 0 | 0 | 0 | 0 | 0 | 0 | 0 | 0 | 0 | 0 | 0 | 0 |
| virus_Jul27.25v_22688 | 1 | 0 | 0 | 0 | 0 | 0 | 0 | 0 | 0 | 0 | 0 | 0 | 0 | 0 | 0 | 0 | 0 | 0 | 0 | 0 | 0 | 0 | 0 | 0 | 0 | 0 | 0 | 0 | 0 | 0 | 0 | 0 | 0 | 0 | 1 | 0 | 0 | 0 | 0 | 0 | 0 | 0 | 0 | 0 | 0 | 0 | 0 | 0 | 0 | 0 | 0 | 0 | 0 | 0 | 0 |
| virus_Jul27.25v_25885 | 1 | 0 | 0 | 0 | 0 | 0 | 0 | 0 | 0 | 0 | 0 | 0 | 0 | 0 | 0 | 0 | 0 | 0 | 0 | 0 | 0 | 0 | 0 | 0 | 0 | 0 | 0 | 0 | 0 | 0 | 0 | 0 | 0 | 0 | 1 | 0 | 0 | 0 | 0 | 0 | 0 | 0 | 0 | 0 | 0 | 0 | 0 | 0 | 0 | 0 | 0 | 0 | 0 | 0 | 0 |
| virus_Jul27.25v_32275 | 1 | 0 | 0 | 0 | 0 | 0 | 0 | 0 | 0 | 0 | 0 | 0 | 0 | 0 | 0 | 0 | 0 | 0 | 0 | 0 | 0 | 0 | 0 | 0 | 0 | 0 | 0 | 0 | 0 | 0 | 0 | 0 | 0 | 0 | 1 | 0 | 0 | 0 | 0 | 0 | 0 | 0 | 0 | 0 | 0 | 0 | 0 | 0 | 0 | 0 | 0 | 0 | 0 | 0 | 0 |
| virus_Jul27.25v_32554 | 1 | 0 | 0 | 0 | 0 | 0 | 0 | 0 | 0 | 0 | 0 | 0 | 0 | 0 | 0 | 0 | 0 | 0 | 0 | 0 | 0 | 0 | 0 | 0 | 0 | 0 | 0 | 0 | 0 | 0 | 0 | 0 | 0 | 0 | 1 | 0 | 0 | 0 | 0 | 0 | 0 | 0 | 0 | 0 | 0 | 0 | 0 | 0 | 0 | 0 | 0 | 0 | 0 | 0 | 0 |
| virus_Jul27.25v_35277 | 1 | 0 | 0 | 0 | 0 | 0 | 0 | 0 | 0 | 0 | 0 | 0 | 0 | 0 | 0 | 0 | 0 | 0 | 0 | 0 | 0 | 0 | 0 | 0 | 0 | 0 | 0 | 0 | 0 | 0 | 0 | 0 | 0 | 0 | 1 | 0 | 0 | 0 | 0 | 0 | 0 | 0 | 0 | 0 | 0 | 0 | 0 | 0 | 0 | 0 | 0 | 0 | 0 | 0 | 0 |
| virus_Jul27.75v_3174 | 1 | 0 | 0 | 0 | 0 | 0 | 0 | 0 | 0 | 0 | 0 | 0 | 0 | 0 | 0 | 0 | 0 | 0 | 0 | 0 | 0 | 0 | 0 | 0 | 0 | 0 | 0 | 0 | 0 | 0 | 0 | 0 | 0 | 0 | 0 | 0 | 1 | 0 | 0 | 0 | 0 | 0 | 0 | 0 | 0 | 0 | 0 | 0 | 0 | 0 | 0 | 0 | 0 | 0 | 0 |
| virus_Jul27.75v_11058 | 1 | 0 | 0 | 0 | 0 | 0 | 0 | 0 | 0 | 0 | 0 | 0 | 0 | 0 | 0 | 0 | 0 | 0 | 0 | 0 | 0 | 0 | 0 | 0 | 0 | 0 | 0 | 0 | 0 | 0 | 0 | 0 | 0 | 0 | 0 | 0 | 1 | 0 | 0 | 0 | 0 | 0 | 0 | 0 | 0 | 0 | 0 | 0 | 0 | 0 | 0 | 0 | 0 | 0 | 0 |
| virus_Jun8.25v_4189 | 1 | 0 | 0 | 0 | 0 | 0 | 0 | 0 | 0 | 0 | 1 | 0 | 0 | 0 | 0 | 0 | 0 | 0 | 0 | 0 | 0 | 0 | 0 | 0 | 0 | 0 | 0 | 0 | 0 | 0 | 0 | 0 | 0 | 0 | 0 | 0 | 0 | 0 | 0 | 0 | 0 | 0 | 0 | 0 | 0 | 0 | 0 | 0 | 0 | 0 | 0 | 0 | 0 | 0 | 0 |
| virus_Jun8.25v_4654 | 1 | 0 | 0 | 0 | 0 | 0 | 0 | 0 | 0 | 0 | 1 | 0 | 0 | 0 | 0 | 0 | 0 | 0 | 0 | 0 | 0 | 0 | 0 | 0 | 0 | 0 | 0 | 0 | 0 | 0 | 0 | 0 | 0 | 0 | 0 | 0 | 0 | 0 | 0 | 0 | 0 | 0 | 0 | 0 | 0 | 0 | 0 | 0 | 0 | 0 | 0 | 0 | 0 | 0 | 0 |
| virus_Jun8.25v_5811 | 1 | 0 | 0 | 0 | 0 | 0 | 0 | 0 | 0 | 0 | 1 | 0 | 0 | 0 | 0 | 0 | 0 | 0 | 0 | 0 | 0 | 0 | 0 | 0 | 0 | 0 | 0 | 0 | 0 | 0 | 0 | 0 | 0 | 0 | 0 | 0 | 0 | 0 | 0 | 0 | 0 | 0 | 0 | 0 | 0 | 0 | 0 | 0 | 0 | 0 | 0 | 0 | 0 | 0 | 0 |
| virus_Jun8.25v_37889 | 1 | 0 | 0 | 0 | 0 | 0 | 0 | 0 | 0 | 0 | 1 | 0 | 0 | 0 | 0 | 0 | 0 | 0 | 0 | 0 | 0 | 0 | 0 | 0 | 0 | 0 | 0 | 0 | 0 | 0 | 0 | 0 | 0 | 0 | 0 | 0 | 0 | 0 | 0 | 0 | 0 | 0 | 0 | 0 | 0 | 0 | 0 | 0 | 0 | 0 | 0 | 0 | 0 | 0 | 0 |
| virus_Jun8.50v_502 | 1 | 0 | 0 | 0 | 0 | 0 | 0 | 0 | 0 | 0 | 0 | 0 | 1 | 0 | 0 | 0 | 0 | 0 | 0 | 0 | 0 | 0 | 0 | 0 | 0 | 0 | 0 | 0 | 0 | 0 | 0 | 0 | 0 | 0 | 0 | 0 | 0 | 0 | 0 | 0 | 0 | 0 | 0 | 0 | 0 | 0 | 0 | 0 | 0 | 0 | 0 | 0 | 0 | 0 | 0 |
| virus_Jun8.50v_5068 | 1 | 0 | 0 | 0 | 0 | 0 | 0 | 0 | 0 | 0 | 0 | 0 | 1 | 0 | 0 | 0 | 0 | 0 | 0 | 0 | 0 | 0 | 0 | 0 | 0 | 0 | 0 | 0 | 0 | 0 | 0 | 0 | 0 | 0 | 0 | 0 | 0 | 0 | 0 | 0 | 0 | 0 | 0 | 0 | 0 | 0 | 0 | 0 | 0 | 0 | 0 | 0 | 0 | 0 | 0 |
| virus_Jun8.50v_7917 | 1 | 0 | 0 | 0 | 0 | 0 | 0 | 0 | 0 | 0 | 0 | 0 | 1 | 0 | 0 | 0 | 0 | 0 | 0 | 0 | 0 | 0 | 0 | 0 | 0 | 0 | 0 | 0 | 0 | 0 | 0 | 0 | 0 | 0 | 0 | 0 | 0 | 0 | 0 | 0 | 0 | 0 | 0 | 0 | 0 | 0 | 0 | 0 | 0 | 0 | 0 | 0 | 0 | 0 | 0 |
| virus_Jun8.50v_17672 | 1 | 0 | 0 | 0 | 0 | 0 | 0 | 0 | 0 | 0 | 0 | 0 | 1 | 0 | 0 | 0 | 0 | 0 | 0 | 0 | 0 | 0 | 0 | 0 | 0 | 0 | 0 | 0 | 0 | 0 | 0 | 0 | 0 | 0 | 0 | 0 | 0 | 0 | 0 | 0 | 0 | 0 | 0 | 0 | 0 | 0 | 0 | 0 | 0 | 0 | 0 | 0 | 0 | 0 | 0 |
| virus_Jun8.50v_20182 | 1 | 0 | 0 | 0 | 0 | 0 | 0 | 0 | 0 | 0 | 0 | 0 | 1 | 0 | 0 | 0 | 0 | 0 | 0 | 0 | 0 | 0 | 0 | 0 | 0 | 0 | 0 | 0 | 0 | 0 | 0 | 0 | 0 | 0 | 0 | 0 | 0 | 0 | 0 | 0 | 0 | 0 | 0 | 0 | 0 | 0 | 0 | 0 | 0 | 0 | 0 | 0 | 0 | 0 | 0 |
| virus_Jun8.50v_24700 | 1 | 0 | 0 | 0 | 0 | 0 | 0 | 0 | 0 | 0 | 0 | 0 | 1 | 0 | 0 | 0 | 0 | 0 | 0 | 0 | 0 | 0 | 0 | 0 | 0 | 0 | 0 | 0 | 0 | 0 | 0 | 0 | 0 | 0 | 0 | 0 | 0 | 0 | 0 | 0 | 0 | 0 | 0 | 0 | 0 | 0 | 0 | 0 | 0 | 0 | 0 | 0 | 0 | 0 | 0 |
| virus_Jun8.50v_27475 | 1 | 0 | 0 | 0 | 0 | 0 | 0 | 0 | 0 | 0 | 0 | 0 | 1 | 0 | 0 | 0 | 0 | 0 | 0 | 0 | 0 | 0 | 0 | 0 | 0 | 0 | 0 | 0 | 0 | 0 | 0 | 0 | 0 | 0 | 0 | 0 | 0 | 0 | 0 | 0 | 0 | 0 | 0 | 0 | 0 | 0 | 0 | 0 | 0 | 0 | 0 | 0 | 0 | 0 | 0 |
| virus_Jun8.75v_115 | 1 | 0 | 0 | 0 | 0 | 0 | 0 | 0 | 0 | 0 | 0 | 0 | 0 | 0 | 1 | 0 | 0 | 0 | 0 | 0 | 0 | 0 | 0 | 0 | 0 | 0 | 0 | 0 | 0 | 0 | 0 | 0 | 0 | 0 | 0 | 0 | 0 | 0 | 0 | 0 | 0 | 0 | 0 | 0 | 0 | 0 | 0 | 0 | 0 | 0 | 0 | 0 | 0 | 0 | 0 |
| virus_Jun8.75v_10966 | 1 | 0 | 0 | 0 | 0 | 0 | 0 | 0 | 0 | 0 | 0 | 0 | 0 | 0 | 1 | 0 | 0 | 0 | 0 | 0 | 0 | 0 | 0 | 0 | 0 | 0 | 0 | 0 | 0 | 0 | 0 | 0 | 0 | 0 | 0 | 0 | 0 | 0 | 0 | 0 | 0 | 0 | 0 | 0 | 0 | 0 | 0 | 0 | 0 | 0 | 0 | 0 | 0 | 0 | 0 |
| virus_Jun8.75v_12670 | 1 | 0 | 0 | 0 | 0 | 0 | 0 | 0 | 0 | 0 | 0 | 0 | 0 | 0 | 1 | 0 | 0 | 0 | 0 | 0 | 0 | 0 | 0 | 0 | 0 | 0 | 0 | 0 | 0 | 0 | 0 | 0 | 0 | 0 | 0 | 0 | 0 | 0 | 0 | 0 | 0 | 0 | 0 | 0 | 0 | 0 | 0 | 0 | 0 | 0 | 0 | 0 | 0 | 0 | 0 |
| virus_Jun8.75v_14242 | 1 | 0 | 0 | 0 | 0 | 0 | 0 | 0 | 0 | 0 | 0 | 0 | 0 | 0 | 1 | 0 | 0 | 0 | 0 | 0 | 0 | 0 | 0 | 0 | 0 | 0 | 0 | 0 | 0 | 0 | 0 | 0 | 0 | 0 | 0 | 0 | 0 | 0 | 0 | 0 | 0 | 0 | 0 | 0 | 0 | 0 | 0 | 0 | 0 | 0 | 0 | 0 | 0 | 0 | 0 |
| virus_Jun8.75v_17613 | 1 | 0 | 0 | 0 | 0 | 0 | 0 | 0 | 0 | 0 | 0 | 0 | 0 | 0 | 1 | 0 | 0 | 0 | 0 | 0 | 0 | 0 | 0 | 0 | 0 | 0 | 0 | 0 | 0 | 0 | 0 | 0 | 0 | 0 | 0 | 0 | 0 | 0 | 0 | 0 | 0 | 0 | 0 | 0 | 0 | 0 | 0 | 0 | 0 | 0 | 0 | 0 | 0 | 0 | 0 |
| virus_Jun22.0v_3590 | 1 | 0 | 0 | 0 | 0 | 0 | 0 | 0 | 0 | 0 | 0 | 0 | 0 | 0 | 0 | 0 | 1 | 0 | 0 | 0 | 0 | 0 | 0 | 0 | 0 | 0 | 0 | 0 | 0 | 0 | 0 | 0 | 0 | 0 | 0 | 0 | 0 | 0 | 0 | 0 | 0 | 0 | 0 | 0 | 0 | 0 | 0 | 0 | 0 | 0 | 0 | 0 | 0 | 0 | 0 |
| virus_Jun22.0v_5725 | 1 | 0 | 0 | 0 | 0 | 0 | 0 | 0 | 0 | 0 | 0 | 0 | 0 | 0 | 0 | 0 | 1 | 0 | 0 | 0 | 0 | 0 | 0 | 0 | 0 | 0 | 0 | 0 | 0 | 0 | 0 | 0 | 0 | 0 | 0 | 0 | 0 | 0 | 0 | 0 | 0 | 0 | 0 | 0 | 0 | 0 | 0 | 0 | 0 | 0 | 0 | 0 | 0 | 0 | 0 |
| virus_Jun22.0v_6413 | 1 | 0 | 0 | 0 | 0 | 0 | 0 | 0 | 0 | 0 | 0 | 0 | 0 | 0 | 0 | 0 | 1 | 0 | 0 | 0 | 0 | 0 | 0 | 0 | 0 | 0 | 0 | 0 | 0 | 0 | 0 | 0 | 0 | 0 | 0 | 0 | 0 | 0 | 0 | 0 | 0 | 0 | 0 | 0 | 0 | 0 | 0 | 0 | 0 | 0 | 0 | 0 | 0 | 0 | 0 |
| virus_Jun22.0v_8397 | 1 | 0 | 0 | 0 | 0 | 0 | 0 | 0 | 0 | 0 | 0 | 0 | 0 | 0 | 0 | 0 | 1 | 0 | 0 | 0 | 0 | 0 | 0 | 0 | 0 | 0 | 0 | 0 | 0 | 0 | 0 | 0 | 0 | 0 | 0 | 0 | 0 | 0 | 0 | 0 | 0 | 0 | 0 | 0 | 0 | 0 | 0 | 0 | 0 | 0 | 0 | 0 | 0 | 0 | 0 |
| virus_Jun22.0v_9242 | 1 | 0 | 0 | 0 | 0 | 0 | 0 | 0 | 0 | 0 | 0 | 0 | 0 | 0 | 0 | 0 | 1 | 0 | 0 | 0 | 0 | 0 | 0 | 0 | 0 | 0 | 0 | 0 | 0 | 0 | 0 | 0 | 0 | 0 | 0 | 0 | 0 | 0 | 0 | 0 | 0 | 0 | 0 | 0 | 0 | 0 | 0 | 0 | 0 | 0 | 0 | 0 | 0 | 0 | 0 |
| virus_Jun22.0v_9832 | 1 | 0 | 0 | 0 | 0 | 0 | 0 | 0 | 0 | 0 | 0 | 0 | 0 | 0 | 0 | 0 | 1 | 0 | 0 | 0 | 0 | 0 | 0 | 0 | 0 | 0 | 0 | 0 | 0 | 0 | 0 | 0 | 0 | 0 | 0 | 0 | 0 | 0 | 0 | 0 | 0 | 0 | 0 | 0 | 0 | 0 | 0 | 0 | 0 | 0 | 0 | 0 | 0 | 0 | 0 |
| virus_Jun22.0v_10533 | 1 | 0 | 0 | 0 | 0 | 0 | 0 | 0 | 0 | 0 | 0 | 0 | 0 | 0 | 0 | 0 | 1 | 0 | 0 | 0 | 0 | 0 | 0 | 0 | 0 | 0 | 0 | 0 | 0 | 0 | 0 | 0 | 0 | 0 | 0 | 0 | 0 | 0 | 0 | 0 | 0 | 0 | 0 | 0 | 0 | 0 | 0 | 0 | 0 | 0 | 0 | 0 | 0 | 0 | 0 |
| virus_Jun22.0v_10752 | 1 | 0 | 0 | 0 | 0 | 0 | 0 | 0 | 0 | 0 | 0 | 0 | 0 | 0 | 0 | 0 | 1 | 0 | 0 | 0 | 0 | 0 | 0 | 0 | 0 | 0 | 0 | 0 | 0 | 0 | 0 | 0 | 0 | 0 | 0 | 0 | 0 | 0 | 0 | 0 | 0 | 0 | 0 | 0 | 0 | 0 | 0 | 0 | 0 | 0 | 0 | 0 | 0 | 0 | 0 |
| virus_Jun22.0v_13380 | 1 | 0 | 0 | 0 | 0 | 0 | 0 | 0 | 0 | 0 | 0 | 0 | 0 | 0 | 0 | 0 | 1 | 0 | 0 | 0 | 0 | 0 | 0 | 0 | 0 | 0 | 0 | 0 | 0 | 0 | 0 | 0 | 0 | 0 | 0 | 0 | 0 | 0 | 0 | 0 | 0 | 0 | 0 | 0 | 0 | 0 | 0 | 0 | 0 | 0 | 0 | 0 | 0 | 0 | 0 |
| virus_Jun22.0v_15281 | 1 | 0 | 0 | 0 | 0 | 0 | 0 | 0 | 0 | 0 | 0 | 0 | 0 | 0 | 0 | 0 | 1 | 0 | 0 | 0 | 0 | 0 | 0 | 0 | 0 | 0 | 0 | 0 | 0 | 0 | 0 | 0 | 0 | 0 | 0 | 0 | 0 | 0 | 0 | 0 | 0 | 0 | 0 | 0 | 0 | 0 | 0 | 0 | 0 | 0 | 0 | 0 | 0 | 0 | 0 |
| virus_Jun22.0v_15498 | 1 | 0 | 0 | 0 | 0 | 0 | 0 | 0 | 0 | 0 | 0 | 0 | 0 | 0 | 0 | 0 | 1 | 0 | 0 | 0 | 0 | 0 | 0 | 0 | 0 | 0 | 0 | 0 | 0 | 0 | 0 | 0 | 0 | 0 | 0 | 0 | 0 | 0 | 0 | 0 | 0 | 0 | 0 | 0 | 0 | 0 | 0 | 0 | 0 | 0 | 0 | 0 | 0 | 0 | 0 |
| virus_Jun22.0v_19381 | 1 | 0 | 0 | 0 | 0 | 0 | 0 | 0 | 0 | 0 | 0 | 0 | 0 | 0 | 0 | 0 | 1 | 0 | 0 | 0 | 0 | 0 | 0 | 0 | 0 | 0 | 0 | 0 | 0 | 0 | 0 | 0 | 0 | 0 | 0 | 0 | 0 | 0 | 0 | 0 | 0 | 0 | 0 | 0 | 0 | 0 | 0 | 0 | 0 | 0 | 0 | 0 | 0 | 0 | 0 |
| virus_Jun22.0v_20016 | 1 | 0 | 0 | 0 | 0 | 0 | 0 | 0 | 0 | 0 | 0 | 0 | 0 | 0 | 0 | 0 | 1 | 0 | 0 | 0 | 0 | 0 | 0 | 0 | 0 | 0 | 0 | 0 | 0 | 0 | 0 | 0 | 0 | 0 | 0 | 0 | 0 | 0 | 0 | 0 | 0 | 0 | 0 | 0 | 0 | 0 | 0 | 0 | 0 | 0 | 0 | 0 | 0 | 0 | 0 |
| virus_Jun22.50v_62 | 1 | 0 | 0 | 0 | 0 | 0 | 0 | 0 | 0 | 0 | 0 | 0 | 0 | 0 | 0 | 0 | 0 | 0 | 1 | 0 | 0 | 0 | 0 | 0 | 0 | 0 | 0 | 0 | 0 | 0 | 0 | 0 | 0 | 0 | 0 | 0 | 0 | 0 | 0 | 0 | 0 | 0 | 0 | 0 | 0 | 0 | 0 | 0 | 0 | 0 | 0 | 0 | 0 | 0 | 0 |
| virus_Jun22.50v_441 | 1 | 0 | 0 | 0 | 0 | 0 | 0 | 0 | 0 | 0 | 0 | 0 | 0 | 0 | 0 | 0 | 0 | 0 | 1 | 0 | 0 | 0 | 0 | 0 | 0 | 0 | 0 | 0 | 0 | 0 | 0 | 0 | 0 | 0 | 0 | 0 | 0 | 0 | 0 | 0 | 0 | 0 | 0 | 0 | 0 | 0 | 0 | 0 | 0 | 0 | 0 | 0 | 0 | 0 | 0 |
| virus_Jun22.50v_3420 | 1 | 0 | 0 | 0 | 0 | 0 | 0 | 0 | 0 | 0 | 0 | 0 | 0 | 0 | 0 | 0 | 0 | 0 | 1 | 0 | 0 | 0 | 0 | 0 | 0 | 0 | 0 | 0 | 0 | 0 | 0 | 0 | 0 | 0 | 0 | 0 | 0 | 0 | 0 | 0 | 0 | 0 | 0 | 0 | 0 | 0 | 0 | 0 | 0 | 0 | 0 | 0 | 0 | 0 | 0 |
| virus_Jun22.50v_4277 | 1 | 0 | 0 | 0 | 0 | 0 | 0 | 0 | 0 | 0 | 0 | 0 | 0 | 0 | 0 | 0 | 0 | 0 | 1 | 0 | 0 | 0 | 0 | 0 | 0 | 0 | 0 | 0 | 0 | 0 | 0 | 0 | 0 | 0 | 0 | 0 | 0 | 0 | 0 | 0 | 0 | 0 | 0 | 0 | 0 | 0 | 0 | 0 | 0 | 0 | 0 | 0 | 0 | 0 | 0 |
| virus_Jun22.50v_6586 | 1 | 0 | 0 | 0 | 0 | 0 | 0 | 0 | 0 | 0 | 0 | 0 | 0 | 0 | 0 | 0 | 0 | 0 | 1 | 0 | 0 | 0 | 0 | 0 | 0 | 0 | 0 | 0 | 0 | 0 | 0 | 0 | 0 | 0 | 0 | 0 | 0 | 0 | 0 | 0 | 0 | 0 | 0 | 0 | 0 | 0 | 0 | 0 | 0 | 0 | 0 | 0 | 0 | 0 | 0 |
| virus_Jun22.50v_9933 | 1 | 0 | 0 | 0 | 0 | 0 | 0 | 0 | 0 | 0 | 0 | 0 | 0 | 0 | 0 | 0 | 0 | 0 | 1 | 0 | 0 | 0 | 0 | 0 | 0 | 0 | 0 | 0 | 0 | 0 | 0 | 0 | 0 | 0 | 0 | 0 | 0 | 0 | 0 | 0 | 0 | 0 | 0 | 0 | 0 | 0 | 0 | 0 | 0 | 0 | 0 | 0 | 0 | 0 | 0 |
| virus_Jun22.50v_12043 | 1 | 0 | 0 | 0 | 0 | 0 | 0 | 0 | 0 | 0 | 0 | 0 | 0 | 0 | 0 | 0 | 0 | 0 | 1 | 0 | 0 | 0 | 0 | 0 | 0 | 0 | 0 | 0 | 0 | 0 | 0 | 0 | 0 | 0 | 0 | 0 | 0 | 0 | 0 | 0 | 0 | 0 | 0 | 0 | 0 | 0 | 0 | 0 | 0 | 0 | 0 | 0 | 0 | 0 | 0 |
| virus_Jun22.50v_14067 | 1 | 0 | 0 | 0 | 0 | 0 | 0 | 0 | 0 | 0 | 0 | 0 | 0 | 0 | 0 | 0 | 0 | 0 | 1 | 0 | 0 | 0 | 0 | 0 | 0 | 0 | 0 | 0 | 0 | 0 | 0 | 0 | 0 | 0 | 0 | 0 | 0 | 0 | 0 | 0 | 0 | 0 | 0 | 0 | 0 | 0 | 0 | 0 | 0 | 0 | 0 | 0 | 0 | 0 | 0 |
| virus_Jun22.50v_16786 | 1 | 0 | 0 | 0 | 0 | 0 | 0 | 0 | 0 | 0 | 0 | 0 | 0 | 0 | 0 | 0 | 0 | 0 | 1 | 0 | 0 | 0 | 0 | 0 | 0 | 0 | 0 | 0 | 0 | 0 | 0 | 0 | 0 | 0 | 0 | 0 | 0 | 0 | 0 | 0 | 0 | 0 | 0 | 0 | 0 | 0 | 0 | 0 | 0 | 0 | 0 | 0 | 0 | 0 | 0 |
| virus_Jun22.50v_18709 | 1 | 0 | 0 | 0 | 0 | 0 | 0 | 0 | 0 | 0 | 0 | 0 | 0 | 0 | 0 | 0 | 0 | 0 | 1 | 0 | 0 | 0 | 0 | 0 | 0 | 0 | 0 | 0 | 0 | 0 | 0 | 0 | 0 | 0 | 0 | 0 | 0 | 0 | 0 | 0 | 0 | 0 | 0 | 0 | 0 | 0 | 0 | 0 | 0 | 0 | 0 | 0 | 0 | 0 | 0 |
| virus_Jun22.50v_19423 | 1 | 0 | 0 | 0 | 0 | 0 | 0 | 0 | 0 | 0 | 0 | 0 | 0 | 0 | 0 | 0 | 0 | 0 | 1 | 0 | 0 | 0 | 0 | 0 | 0 | 0 | 0 | 0 | 0 | 0 | 0 | 0 | 0 | 0 | 0 | 0 | 0 | 0 | 0 | 0 | 0 | 0 | 0 | 0 | 0 | 0 | 0 | 0 | 0 | 0 | 0 | 0 | 0 | 0 | 0 |
| virus_Jun22.50v_26207 | 1 | 0 | 0 | 0 | 0 | 0 | 0 | 0 | 0 | 0 | 0 | 0 | 0 | 0 | 0 | 0 | 0 | 0 | 1 | 0 | 0 | 0 | 0 | 0 | 0 | 0 | 0 | 0 | 0 | 0 | 0 | 0 | 0 | 0 | 0 | 0 | 0 | 0 | 0 | 0 | 0 | 0 | 0 | 0 | 0 | 0 | 0 | 0 | 0 | 0 | 0 | 0 | 0 | 0 | 0 |
| virus_Jun22.50v_28387 | 1 | 0 | 0 | 0 | 0 | 0 | 0 | 0 | 0 | 0 | 0 | 0 | 0 | 0 | 0 | 0 | 0 | 0 | 1 | 0 | 0 | 0 | 0 | 0 | 0 | 0 | 0 | 0 | 0 | 0 | 0 | 0 | 0 | 0 | 0 | 0 | 0 | 0 | 0 | 0 | 0 | 0 | 0 | 0 | 0 | 0 | 0 | 0 | 0 | 0 | 0 | 0 | 0 | 0 | 0 |
| virus_Jun22.50v_31049 | 1 | 0 | 0 | 0 | 0 | 0 | 0 | 0 | 0 | 0 | 0 | 0 | 0 | 0 | 0 | 0 | 0 | 0 | 1 | 0 | 0 | 0 | 0 | 0 | 0 | 0 | 0 | 0 | 0 | 0 | 0 | 0 | 0 | 0 | 0 | 0 | 0 | 0 | 0 | 0 | 0 | 0 | 0 | 0 | 0 | 0 | 0 | 0 | 0 | 0 | 0 | 0 | 0 | 0 | 0 |
| virus_Jun22.50v_33443 | 1 | 0 | 0 | 0 | 0 | 0 | 0 | 0 | 0 | 0 | 0 | 0 | 0 | 0 | 0 | 0 | 0 | 0 | 1 | 0 | 0 | 0 | 0 | 0 | 0 | 0 | 0 | 0 | 0 | 0 | 0 | 0 | 0 | 0 | 0 | 0 | 0 | 0 | 0 | 0 | 0 | 0 | 0 | 0 | 0 | 0 | 0 | 0 | 0 | 0 | 0 | 0 | 0 | 0 | 0 |
| virus_Jun22.50v_34947 | 1 | 0 | 0 | 0 | 0 | 0 | 0 | 0 | 0 | 0 | 0 | 0 | 0 | 0 | 0 | 0 | 0 | 0 | 1 | 0 | 0 | 0 | 0 | 0 | 0 | 0 | 0 | 0 | 0 | 0 | 0 | 0 | 0 | 0 | 0 | 0 | 0 | 0 | 0 | 0 | 0 | 0 | 0 | 0 | 0 | 0 | 0 | 0 | 0 | 0 | 0 | 0 | 0 | 0 | 0 |
| virus_Jun22.50v_39947 | 1 | 0 | 0 | 0 | 0 | 0 | 0 | 0 | 0 | 0 | 0 | 0 | 0 | 0 | 0 | 0 | 0 | 0 | 1 | 0 | 0 | 0 | 0 | 0 | 0 | 0 | 0 | 0 | 0 | 0 | 0 | 0 | 0 | 0 | 0 | 0 | 0 | 0 | 0 | 0 | 0 | 0 | 0 | 0 | 0 | 0 | 0 | 0 | 0 | 0 | 0 | 0 | 0 | 0 | 0 |
| virus_Jun22.50v_40701 | 1 | 0 | 0 | 0 | 0 | 0 | 0 | 0 | 0 | 0 | 0 | 0 | 0 | 0 | 0 | 0 | 0 | 0 | 1 | 0 | 0 | 0 | 0 | 0 | 0 | 0 | 0 | 0 | 0 | 0 | 0 | 0 | 0 | 0 | 0 | 0 | 0 | 0 | 0 | 0 | 0 | 0 | 0 | 0 | 0 | 0 | 0 | 0 | 0 | 0 | 0 | 0 | 0 | 0 | 0 |
| virus_Jun22.75v_8444 | 1 | 0 | 0 | 0 | 0 | 0 | 0 | 0 | 0 | 0 | 0 | 0 | 0 | 0 | 0 | 0 | 0 | 0 | 0 | 0 | 1 | 0 | 0 | 0 | 0 | 0 | 0 | 0 | 0 | 0 | 0 | 0 | 0 | 0 | 0 | 0 | 0 | 0 | 0 | 0 | 0 | 0 | 0 | 0 | 0 | 0 | 0 | 0 | 0 | 0 | 0 | 0 | 0 | 0 | 0 |
| virus_Jun22.75v_21281 | 1 | 0 | 0 | 0 | 0 | 0 | 0 | 0 | 0 | 0 | 0 | 0 | 0 | 0 | 0 | 0 | 0 | 0 | 0 | 0 | 1 | 0 | 0 | 0 | 0 | 0 | 0 | 0 | 0 | 0 | 0 | 0 | 0 | 0 | 0 | 0 | 0 | 0 | 0 | 0 | 0 | 0 | 0 | 0 | 0 | 0 | 0 | 0 | 0 | 0 | 0 | 0 | 0 | 0 | 0 |
| virus_Jun22.75v_23329 | 1 | 0 | 0 | 0 | 0 | 0 | 0 | 0 | 0 | 0 | 0 | 0 | 0 | 0 | 0 | 0 | 0 | 0 | 0 | 0 | 1 | 0 | 0 | 0 | 0 | 0 | 0 | 0 | 0 | 0 | 0 | 0 | 0 | 0 | 0 | 0 | 0 | 0 | 0 | 0 | 0 | 0 | 0 | 0 | 0 | 0 | 0 | 0 | 0 | 0 | 0 | 0 | 0 | 0 | 0 |
| virus_Jun22.75v_30527 | 1 | 0 | 0 | 0 | 0 | 0 | 0 | 0 | 0 | 0 | 0 | 0 | 0 | 0 | 0 | 0 | 0 | 0 | 0 | 0 | 1 | 0 | 0 | 0 | 0 | 0 | 0 | 0 | 0 | 0 | 0 | 0 | 0 | 0 | 0 | 0 | 0 | 0 | 0 | 0 | 0 | 0 | 0 | 0 | 0 | 0 | 0 | 0 | 0 | 0 | 0 | 0 | 0 | 0 | 0 |
| virus_Jun22.75v_34611 | 1 | 0 | 0 | 0 | 0 | 0 | 0 | 0 | 0 | 0 | 0 | 0 | 0 | 0 | 0 | 0 | 0 | 0 | 0 | 0 | 1 | 0 | 0 | 0 | 0 | 0 | 0 | 0 | 0 | 0 | 0 | 0 | 0 | 0 | 0 | 0 | 0 | 0 | 0 | 0 | 0 | 0 | 0 | 0 | 0 | 0 | 0 | 0 | 0 | 0 | 0 | 0 | 0 | 0 | 0 |
| virus_Jun22.75v_36530 | 1 | 0 | 0 | 0 | 0 | 0 | 0 | 0 | 0 | 0 | 0 | 0 | 0 | 0 | 0 | 0 | 0 | 0 | 0 | 0 | 1 | 0 | 0 | 0 | 0 | 0 | 0 | 0 | 0 | 0 | 0 | 0 | 0 | 0 | 0 | 0 | 0 | 0 | 0 | 0 | 0 | 0 | 0 | 0 | 0 | 0 | 0 | 0 | 0 | 0 | 0 | 0 | 0 | 0 | 0 |
| virus_May12.5v_10726 | 1 | 0 | 1 | 0 | 0 | 0 | 0 | 0 | 0 | 0 | 0 | 0 | 0 | 0 | 0 | 0 | 0 | 0 | 0 | 0 | 0 | 0 | 0 | 0 | 0 | 0 | 0 | 0 | 0 | 0 | 0 | 0 | 0 | 0 | 0 | 0 | 0 | 0 | 0 | 0 | 0 | 0 | 0 | 0 | 0 | 0 | 0 | 0 | 0 | 0 | 0 | 0 | 0 | 0 | 0 |
| virus_May12.5v_13906 | 1 | 0 | 1 | 0 | 0 | 0 | 0 | 0 | 0 | 0 | 0 | 0 | 0 | 0 | 0 | 0 | 0 | 0 | 0 | 0 | 0 | 0 | 0 | 0 | 0 | 0 | 0 | 0 | 0 | 0 | 0 | 0 | 0 | 0 | 0 | 0 | 0 | 0 | 0 | 0 | 0 | 0 | 0 | 0 | 0 | 0 | 0 | 0 | 0 | 0 | 0 | 0 | 0 | 0 | 0 |
| virus_May12.5v_20244 | 1 | 0 | 1 | 0 | 0 | 0 | 0 | 0 | 0 | 0 | 0 | 0 | 0 | 0 | 0 | 0 | 0 | 0 | 0 | 0 | 0 | 0 | 0 | 0 | 0 | 0 | 0 | 0 | 0 | 0 | 0 | 0 | 0 | 0 | 0 | 0 | 0 | 0 | 0 | 0 | 0 | 0 | 0 | 0 | 0 | 0 | 0 | 0 | 0 | 0 | 0 | 0 | 0 | 0 | 0 |
| virus_May12.5v_26333 | 1 | 0 | 1 | 0 | 0 | 0 | 0 | 0 | 0 | 0 | 0 | 0 | 0 | 0 | 0 | 0 | 0 | 0 | 0 | 0 | 0 | 0 | 0 | 0 | 0 | 0 | 0 | 0 | 0 | 0 | 0 | 0 | 0 | 0 | 0 | 0 | 0 | 0 | 0 | 0 | 0 | 0 | 0 | 0 | 0 | 0 | 0 | 0 | 0 | 0 | 0 | 0 | 0 | 0 | 0 |
| virus_May12.5v_29920 | 1 | 0 | 1 | 0 | 0 | 0 | 0 | 0 | 0 | 0 | 0 | 0 | 0 | 0 | 0 | 0 | 0 | 0 | 0 | 0 | 0 | 0 | 0 | 0 | 0 | 0 | 0 | 0 | 0 | 0 | 0 | 0 | 0 | 0 | 0 | 0 | 0 | 0 | 0 | 0 | 0 | 0 | 0 | 0 | 0 | 0 | 0 | 0 | 0 | 0 | 0 | 0 | 0 | 0 | 0 |
| virus_May12.5v_30341 | 1 | 0 | 1 | 0 | 0 | 0 | 0 | 0 | 0 | 0 | 0 | 0 | 0 | 0 | 0 | 0 | 0 | 0 | 0 | 0 | 0 | 0 | 0 | 0 | 0 | 0 | 0 | 0 | 0 | 0 | 0 | 0 | 0 | 0 | 0 | 0 | 0 | 0 | 0 | 0 | 0 | 0 | 0 | 0 | 0 | 0 | 0 | 0 | 0 | 0 | 0 | 0 | 0 | 0 | 0 |
| virus_May12.25v_2024 | 1 | 0 | 0 | 0 | 1 | 0 | 0 | 0 | 0 | 0 | 0 | 0 | 0 | 0 | 0 | 0 | 0 | 0 | 0 | 0 | 0 | 0 | 0 | 0 | 0 | 0 | 0 | 0 | 0 | 0 | 0 | 0 | 0 | 0 | 0 | 0 | 0 | 0 | 0 | 0 | 0 | 0 | 0 | 0 | 0 | 0 | 0 | 0 | 0 | 0 | 0 | 0 | 0 | 0 | 0 |
| virus_May12.25v_8993 | 1 | 0 | 0 | 0 | 1 | 0 | 0 | 0 | 0 | 0 | 0 | 0 | 0 | 0 | 0 | 0 | 0 | 0 | 0 | 0 | 0 | 0 | 0 | 0 | 0 | 0 | 0 | 0 | 0 | 0 | 0 | 0 | 0 | 0 | 0 | 0 | 0 | 0 | 0 | 0 | 0 | 0 | 0 | 0 | 0 | 0 | 0 | 0 | 0 | 0 | 0 | 0 | 0 | 0 | 0 |
| virus_May12.25v_10234 | 1 | 0 | 0 | 0 | 1 | 0 | 0 | 0 | 0 | 0 | 0 | 0 | 0 | 0 | 0 | 0 | 0 | 0 | 0 | 0 | 0 | 0 | 0 | 0 | 0 | 0 | 0 | 0 | 0 | 0 | 0 | 0 | 0 | 0 | 0 | 0 | 0 | 0 | 0 | 0 | 0 | 0 | 0 | 0 | 0 | 0 | 0 | 0 | 0 | 0 | 0 | 0 | 0 | 0 | 0 |
| virus_May12.25v_18546 | 1 | 0 | 0 | 0 | 1 | 0 | 0 | 0 | 0 | 0 | 0 | 0 | 0 | 0 | 0 | 0 | 0 | 0 | 0 | 0 | 0 | 0 | 0 | 0 | 0 | 0 | 0 | 0 | 0 | 0 | 0 | 0 | 0 | 0 | 0 | 0 | 0 | 0 | 0 | 0 | 0 | 0 | 0 | 0 | 0 | 0 | 0 | 0 | 0 | 0 | 0 | 0 | 0 | 0 | 0 |
| virus_May12.25v_19002 | 1 | 0 | 0 | 0 | 1 | 0 | 0 | 0 | 0 | 0 | 0 | 0 | 0 | 0 | 0 | 0 | 0 | 0 | 0 | 0 | 0 | 0 | 0 | 0 | 0 | 0 | 0 | 0 | 0 | 0 | 0 | 0 | 0 | 0 | 0 | 0 | 0 | 0 | 0 | 0 | 0 | 0 | 0 | 0 | 0 | 0 | 0 | 0 | 0 | 0 | 0 | 0 | 0 | 0 | 0 |
| virus_May12.25v_32287 | 1 | 0 | 0 | 0 | 1 | 0 | 0 | 0 | 0 | 0 | 0 | 0 | 0 | 0 | 0 | 0 | 0 | 0 | 0 | 0 | 0 | 0 | 0 | 0 | 0 | 0 | 0 | 0 | 0 | 0 | 0 | 0 | 0 | 0 | 0 | 0 | 0 | 0 | 0 | 0 | 0 | 0 | 0 | 0 | 0 | 0 | 0 | 0 | 0 | 0 | 0 | 0 | 0 | 0 | 0 |
| virus_May12.50v_2546 | 1 | 0 | 0 | 0 | 0 | 0 | 1 | 0 | 0 | 0 | 0 | 0 | 0 | 0 | 0 | 0 | 0 | 0 | 0 | 0 | 0 | 0 | 0 | 0 | 0 | 0 | 0 | 0 | 0 | 0 | 0 | 0 | 0 | 0 | 0 | 0 | 0 | 0 | 0 | 0 | 0 | 0 | 0 | 0 | 0 | 0 | 0 | 0 | 0 | 0 | 0 | 0 | 0 | 0 | 0 |
| virus_May12.50v_3821 | 1 | 0 | 0 | 0 | 0 | 0 | 1 | 0 | 0 | 0 | 0 | 0 | 0 | 0 | 0 | 0 | 0 | 0 | 0 | 0 | 0 | 0 | 0 | 0 | 0 | 0 | 0 | 0 | 0 | 0 | 0 | 0 | 0 | 0 | 0 | 0 | 0 | 0 | 0 | 0 | 0 | 0 | 0 | 0 | 0 | 0 | 0 | 0 | 0 | 0 | 0 | 0 | 0 | 0 | 0 |
| virus_May12.50v_5278 | 1 | 0 | 0 | 0 | 0 | 0 | 1 | 0 | 0 | 0 | 0 | 0 | 0 | 0 | 0 | 0 | 0 | 0 | 0 | 0 | 0 | 0 | 0 | 0 | 0 | 0 | 0 | 0 | 0 | 0 | 0 | 0 | 0 | 0 | 0 | 0 | 0 | 0 | 0 | 0 | 0 | 0 | 0 | 0 | 0 | 0 | 0 | 0 | 0 | 0 | 0 | 0 | 0 | 0 | 0 |
| virus_May12.50v_7981 | 1 | 0 | 0 | 0 | 0 | 0 | 1 | 0 | 0 | 0 | 0 | 0 | 0 | 0 | 0 | 0 | 0 | 0 | 0 | 0 | 0 | 0 | 0 | 0 | 0 | 0 | 0 | 0 | 0 | 0 | 0 | 0 | 0 | 0 | 0 | 0 | 0 | 0 | 0 | 0 | 0 | 0 | 0 | 0 | 0 | 0 | 0 | 0 | 0 | 0 | 0 | 0 | 0 | 0 | 0 |
| virus_May12.50v_8873 | 1 | 0 | 0 | 0 | 0 | 0 | 1 | 0 | 0 | 0 | 0 | 0 | 0 | 0 | 0 | 0 | 0 | 0 | 0 | 0 | 0 | 0 | 0 | 0 | 0 | 0 | 0 | 0 | 0 | 0 | 0 | 0 | 0 | 0 | 0 | 0 | 0 | 0 | 0 | 0 | 0 | 0 | 0 | 0 | 0 | 0 | 0 | 0 | 0 | 0 | 0 | 0 | 0 | 0 | 0 |
| virus_May12.50v_10361 | 1 | 0 | 0 | 0 | 0 | 0 | 1 | 0 | 0 | 0 | 0 | 0 | 0 | 0 | 0 | 0 | 0 | 0 | 0 | 0 | 0 | 0 | 0 | 0 | 0 | 0 | 0 | 0 | 0 | 0 | 0 | 0 | 0 | 0 | 0 | 0 | 0 | 0 | 0 | 0 | 0 | 0 | 0 | 0 | 0 | 0 | 0 | 0 | 0 | 0 | 0 | 0 | 0 | 0 | 0 |
| virus_May12.50v_11921 | 1 | 0 | 0 | 0 | 0 | 0 | 1 | 0 | 0 | 0 | 0 | 0 | 0 | 0 | 0 | 0 | 0 | 0 | 0 | 0 | 0 | 0 | 0 | 0 | 0 | 0 | 0 | 0 | 0 | 0 | 0 | 0 | 0 | 0 | 0 | 0 | 0 | 0 | 0 | 0 | 0 | 0 | 0 | 0 | 0 | 0 | 0 | 0 | 0 | 0 | 0 | 0 | 0 | 0 | 0 |
| virus_May12.50v_13239 | 1 | 0 | 0 | 0 | 0 | 0 | 1 | 0 | 0 | 0 | 0 | 0 | 0 | 0 | 0 | 0 | 0 | 0 | 0 | 0 | 0 | 0 | 0 | 0 | 0 | 0 | 0 | 0 | 0 | 0 | 0 | 0 | 0 | 0 | 0 | 0 | 0 | 0 | 0 | 0 | 0 | 0 | 0 | 0 | 0 | 0 | 0 | 0 | 0 | 0 | 0 | 0 | 0 | 0 | 0 |
| virus_May12.50v_18083 | 1 | 0 | 0 | 0 | 0 | 0 | 1 | 0 | 0 | 0 | 0 | 0 | 0 | 0 | 0 | 0 | 0 | 0 | 0 | 0 | 0 | 0 | 0 | 0 | 0 | 0 | 0 | 0 | 0 | 0 | 0 | 0 | 0 | 0 | 0 | 0 | 0 | 0 | 0 | 0 | 0 | 0 | 0 | 0 | 0 | 0 | 0 | 0 | 0 | 0 | 0 | 0 | 0 | 0 | 0 |
| virus_May12.50v_19435 | 1 | 0 | 0 | 0 | 0 | 0 | 1 | 0 | 0 | 0 | 0 | 0 | 0 | 0 | 0 | 0 | 0 | 0 | 0 | 0 | 0 | 0 | 0 | 0 | 0 | 0 | 0 | 0 | 0 | 0 | 0 | 0 | 0 | 0 | 0 | 0 | 0 | 0 | 0 | 0 | 0 | 0 | 0 | 0 | 0 | 0 | 0 | 0 | 0 | 0 | 0 | 0 | 0 | 0 | 0 |
| virus_May12.50v_26459 | 1 | 0 | 0 | 0 | 0 | 0 | 1 | 0 | 0 | 0 | 0 | 0 | 0 | 0 | 0 | 0 | 0 | 0 | 0 | 0 | 0 | 0 | 0 | 0 | 0 | 0 | 0 | 0 | 0 | 0 | 0 | 0 | 0 | 0 | 0 | 0 | 0 | 0 | 0 | 0 | 0 | 0 | 0 | 0 | 0 | 0 | 0 | 0 | 0 | 0 | 0 | 0 | 0 | 0 | 0 |
| virus_May12.50v_26716 | 1 | 0 | 0 | 0 | 0 | 0 | 1 | 0 | 0 | 0 | 0 | 0 | 0 | 0 | 0 | 0 | 0 | 0 | 0 | 0 | 0 | 0 | 0 | 0 | 0 | 0 | 0 | 0 | 0 | 0 | 0 | 0 | 0 | 0 | 0 | 0 | 0 | 0 | 0 | 0 | 0 | 0 | 0 | 0 | 0 | 0 | 0 | 0 | 0 | 0 | 0 | 0 | 0 | 0 | 0 |
| virus_May12.50v_26768 | 1 | 0 | 0 | 0 | 0 | 0 | 1 | 0 | 0 | 0 | 0 | 0 | 0 | 0 | 0 | 0 | 0 | 0 | 0 | 0 | 0 | 0 | 0 | 0 | 0 | 0 | 0 | 0 | 0 | 0 | 0 | 0 | 0 | 0 | 0 | 0 | 0 | 0 | 0 | 0 | 0 | 0 | 0 | 0 | 0 | 0 | 0 | 0 | 0 | 0 | 0 | 0 | 0 | 0 | 0 |
| virus_May12.50v_30347 | 1 | 0 | 0 | 0 | 0 | 0 | 1 | 0 | 0 | 0 | 0 | 0 | 0 | 0 | 0 | 0 | 0 | 0 | 0 | 0 | 0 | 0 | 0 | 0 | 0 | 0 | 0 | 0 | 0 | 0 | 0 | 0 | 0 | 0 | 0 | 0 | 0 | 0 | 0 | 0 | 0 | 0 | 0 | 0 | 0 | 0 | 0 | 0 | 0 | 0 | 0 | 0 | 0 | 0 | 0 |
| virus_May12.50v_30643 | 1 | 0 | 0 | 0 | 0 | 0 | 1 | 0 | 0 | 0 | 0 | 0 | 0 | 0 | 0 | 0 | 0 | 0 | 0 | 0 | 0 | 0 | 0 | 0 | 0 | 0 | 0 | 0 | 0 | 0 | 0 | 0 | 0 | 0 | 0 | 0 | 0 | 0 | 0 | 0 | 0 | 0 | 0 | 0 | 0 | 0 | 0 | 0 | 0 | 0 | 0 | 0 | 0 | 0 | 0 |
| virus_May12.50v_30988 | 1 | 0 | 0 | 0 | 0 | 0 | 1 | 0 | 0 | 0 | 0 | 0 | 0 | 0 | 0 | 0 | 0 | 0 | 0 | 0 | 0 | 0 | 0 | 0 | 0 | 0 | 0 | 0 | 0 | 0 | 0 | 0 | 0 | 0 | 0 | 0 | 0 | 0 | 0 | 0 | 0 | 0 | 0 | 0 | 0 | 0 | 0 | 0 | 0 | 0 | 0 | 0 | 0 | 0 | 0 |
| virus_May12.50v_34468 | 1 | 0 | 0 | 0 | 0 | 0 | 1 | 0 | 0 | 0 | 0 | 0 | 0 | 0 | 0 | 0 | 0 | 0 | 0 | 0 | 0 | 0 | 0 | 0 | 0 | 0 | 0 | 0 | 0 | 0 | 0 | 0 | 0 | 0 | 0 | 0 | 0 | 0 | 0 | 0 | 0 | 0 | 0 | 0 | 0 | 0 | 0 | 0 | 0 | 0 | 0 | 0 | 0 | 0 | 0 |
| virus_May12.50v_41150 | 1 | 0 | 0 | 0 | 0 | 0 | 1 | 0 | 0 | 0 | 0 | 0 | 0 | 0 | 0 | 0 | 0 | 0 | 0 | 0 | 0 | 0 | 0 | 0 | 0 | 0 | 0 | 0 | 0 | 0 | 0 | 0 | 0 | 0 | 0 | 0 | 0 | 0 | 0 | 0 | 0 | 0 | 0 | 0 | 0 | 0 | 0 | 0 | 0 | 0 | 0 | 0 | 0 | 0 | 0 |
| virus_May12.75v_1619 | 1 | 0 | 0 | 0 | 0 | 0 | 0 | 0 | 1 | 0 | 0 | 0 | 0 | 0 | 0 | 0 | 0 | 0 | 0 | 0 | 0 | 0 | 0 | 0 | 0 | 0 | 0 | 0 | 0 | 0 | 0 | 0 | 0 | 0 | 0 | 0 | 0 | 0 | 0 | 0 | 0 | 0 | 0 | 0 | 0 | 0 | 0 | 0 | 0 | 0 | 0 | 0 | 0 | 0 | 0 |
| virus_May12.75v_5902 | 1 | 0 | 0 | 0 | 0 | 0 | 0 | 0 | 1 | 0 | 0 | 0 | 0 | 0 | 0 | 0 | 0 | 0 | 0 | 0 | 0 | 0 | 0 | 0 | 0 | 0 | 0 | 0 | 0 | 0 | 0 | 0 | 0 | 0 | 0 | 0 | 0 | 0 | 0 | 0 | 0 | 0 | 0 | 0 | 0 | 0 | 0 | 0 | 0 | 0 | 0 | 0 | 0 | 0 | 0 |
| virus_May12.75v_7263 | 1 | 0 | 0 | 0 | 0 | 0 | 0 | 0 | 1 | 0 | 0 | 0 | 0 | 0 | 0 | 0 | 0 | 0 | 0 | 0 | 0 | 0 | 0 | 0 | 0 | 0 | 0 | 0 | 0 | 0 | 0 | 0 | 0 | 0 | 0 | 0 | 0 | 0 | 0 | 0 | 0 | 0 | 0 | 0 | 0 | 0 | 0 | 0 | 0 | 0 | 0 | 0 | 0 | 0 | 0 |
| virus_May12.75v_12328 | 1 | 0 | 0 | 0 | 0 | 0 | 0 | 0 | 1 | 0 | 0 | 0 | 0 | 0 | 0 | 0 | 0 | 0 | 0 | 0 | 0 | 0 | 0 | 0 | 0 | 0 | 0 | 0 | 0 | 0 | 0 | 0 | 0 | 0 | 0 | 0 | 0 | 0 | 0 | 0 | 0 | 0 | 0 | 0 | 0 | 0 | 0 | 0 | 0 | 0 | 0 | 0 | 0 | 0 | 0 |
| virus_May12.75v_13736 | 1 | 0 | 0 | 0 | 0 | 0 | 0 | 0 | 1 | 0 | 0 | 0 | 0 | 0 | 0 | 0 | 0 | 0 | 0 | 0 | 0 | 0 | 0 | 0 | 0 | 0 | 0 | 0 | 0 | 0 | 0 | 0 | 0 | 0 | 0 | 0 | 0 | 0 | 0 | 0 | 0 | 0 | 0 | 0 | 0 | 0 | 0 | 0 | 0 | 0 | 0 | 0 | 0 | 0 | 0 |
| virus_May12.75v_15365 | 1 | 0 | 0 | 0 | 0 | 0 | 0 | 0 | 1 | 0 | 0 | 0 | 0 | 0 | 0 | 0 | 0 | 0 | 0 | 0 | 0 | 0 | 0 | 0 | 0 | 0 | 0 | 0 | 0 | 0 | 0 | 0 | 0 | 0 | 0 | 0 | 0 | 0 | 0 | 0 | 0 | 0 | 0 | 0 | 0 | 0 | 0 | 0 | 0 | 0 | 0 | 0 | 0 | 0 | 0 |
| virus_May12.75v_16696 | 1 | 0 | 0 | 0 | 0 | 0 | 0 | 0 | 1 | 0 | 0 | 0 | 0 | 0 | 0 | 0 | 0 | 0 | 0 | 0 | 0 | 0 | 0 | 0 | 0 | 0 | 0 | 0 | 0 | 0 | 0 | 0 | 0 | 0 | 0 | 0 | 0 | 0 | 0 | 0 | 0 | 0 | 0 | 0 | 0 | 0 | 0 | 0 | 0 | 0 | 0 | 0 | 0 | 0 | 0 |
| virus_May12.75v_16871 | 1 | 0 | 0 | 0 | 0 | 0 | 0 | 0 | 1 | 0 | 0 | 0 | 0 | 0 | 0 | 0 | 0 | 0 | 0 | 0 | 0 | 0 | 0 | 0 | 0 | 0 | 0 | 0 | 0 | 0 | 0 | 0 | 0 | 0 | 0 | 0 | 0 | 0 | 0 | 0 | 0 | 0 | 0 | 0 | 0 | 0 | 0 | 0 | 0 | 0 | 0 | 0 | 0 | 0 | 0 |
| virus_May12.75v_23165 | 1 | 0 | 0 | 0 | 0 | 0 | 0 | 0 | 1 | 0 | 0 | 0 | 0 | 0 | 0 | 0 | 0 | 0 | 0 | 0 | 0 | 0 | 0 | 0 | 0 | 0 | 0 | 0 | 0 | 0 | 0 | 0 | 0 | 0 | 0 | 0 | 0 | 0 | 0 | 0 | 0 | 0 | 0 | 0 | 0 | 0 | 0 | 0 | 0 | 0 | 0 | 0 | 0 | 0 | 0 |
| virus_May12.75v_23945 | 1 | 0 | 0 | 0 | 0 | 0 | 0 | 0 | 1 | 0 | 0 | 0 | 0 | 0 | 0 | 0 | 0 | 0 | 0 | 0 | 0 | 0 | 0 | 0 | 0 | 0 | 0 | 0 | 0 | 0 | 0 | 0 | 0 | 0 | 0 | 0 | 0 | 0 | 0 | 0 | 0 | 0 | 0 | 0 | 0 | 0 | 0 | 0 | 0 | 0 | 0 | 0 | 0 | 0 | 0 |
| virus_May12.75v_26613 | 1 | 0 | 0 | 0 | 0 | 0 | 0 | 0 | 1 | 0 | 0 | 0 | 0 | 0 | 0 | 0 | 0 | 0 | 0 | 0 | 0 | 0 | 0 | 0 | 0 | 0 | 0 | 0 | 0 | 0 | 0 | 0 | 0 | 0 | 0 | 0 | 0 | 0 | 0 | 0 | 0 | 0 | 0 | 0 | 0 | 0 | 0 | 0 | 0 | 0 | 0 | 0 | 0 | 0 | 0 |
| virus_May12.75v_31817 | 1 | 0 | 0 | 0 | 0 | 0 | 0 | 0 | 1 | 0 | 0 | 0 | 0 | 0 | 0 | 0 | 0 | 0 | 0 | 0 | 0 | 0 | 0 | 0 | 0 | 0 | 0 | 0 | 0 | 0 | 0 | 0 | 0 | 0 | 0 | 0 | 0 | 0 | 0 | 0 | 0 | 0 | 0 | 0 | 0 | 0 | 0 | 0 | 0 | 0 | 0 | 0 | 0 | 0 | 0 |
| virus_May12.75v_35955 | 1 | 0 | 0 | 0 | 0 | 0 | 0 | 0 | 1 | 0 | 0 | 0 | 0 | 0 | 0 | 0 | 0 | 0 | 0 | 0 | 0 | 0 | 0 | 0 | 0 | 0 | 0 | 0 | 0 | 0 | 0 | 0 | 0 | 0 | 0 | 0 | 0 | 0 | 0 | 0 | 0 | 0 | 0 | 0 | 0 | 0 | 0 | 0 | 0 | 0 | 0 | 0 | 0 | 0 | 0 |
| virus_Sep8.0v_2763 | 1 | 0 | 0 | 0 | 0 | 0 | 0 | 0 | 0 | 0 | 0 | 0 | 0 | 0 | 0 | 0 | 0 | 0 | 0 | 0 | 0 | 0 | 0 | 0 | 0 | 0 | 0 | 0 | 0 | 0 | 0 | 0 | 0 | 0 | 0 | 0 | 0 | 0 | 0 | 0 | 0 | 0 | 0 | 0 | 0 | 0 | 0 | 0 | 1 | 0 | 0 | 0 | 0 | 0 | 0 |
| virus_Sep8.0v_2914 | 1 | 0 | 0 | 0 | 0 | 0 | 0 | 0 | 0 | 0 | 0 | 0 | 0 | 0 | 0 | 0 | 0 | 0 | 0 | 0 | 0 | 0 | 0 | 0 | 0 | 0 | 0 | 0 | 0 | 0 | 0 | 0 | 0 | 0 | 0 | 0 | 0 | 0 | 0 | 0 | 0 | 0 | 0 | 0 | 0 | 0 | 0 | 0 | 1 | 0 | 0 | 0 | 0 | 0 | 0 |
| virus_Sep8.0v_3366 | 1 | 0 | 0 | 0 | 0 | 0 | 0 | 0 | 0 | 0 | 0 | 0 | 0 | 0 | 0 | 0 | 0 | 0 | 0 | 0 | 0 | 0 | 0 | 0 | 0 | 0 | 0 | 0 | 0 | 0 | 0 | 0 | 0 | 0 | 0 | 0 | 0 | 0 | 0 | 0 | 0 | 0 | 0 | 0 | 0 | 0 | 0 | 0 | 1 | 0 | 0 | 0 | 0 | 0 | 0 |
| virus_Sep8.0v_6021 | 1 | 0 | 0 | 0 | 0 | 0 | 0 | 0 | 0 | 0 | 0 | 0 | 0 | 0 | 0 | 0 | 0 | 0 | 0 | 0 | 0 | 0 | 0 | 0 | 0 | 0 | 0 | 0 | 0 | 0 | 0 | 0 | 0 | 0 | 0 | 0 | 0 | 0 | 0 | 0 | 0 | 0 | 0 | 0 | 0 | 0 | 0 | 0 | 1 | 0 | 0 | 0 | 0 | 0 | 0 |
| virus_Sep8.0v_7698 | 1 | 0 | 0 | 0 | 0 | 0 | 0 | 0 | 0 | 0 | 0 | 0 | 0 | 0 | 0 | 0 | 0 | 0 | 0 | 0 | 0 | 0 | 0 | 0 | 0 | 0 | 0 | 0 | 0 | 0 | 0 | 0 | 0 | 0 | 0 | 0 | 0 | 0 | 0 | 0 | 0 | 0 | 0 | 0 | 0 | 0 | 0 | 0 | 1 | 0 | 0 | 0 | 0 | 0 | 0 |
| virus_Sep8.0v_12226 | 1 | 0 | 0 | 0 | 0 | 0 | 0 | 0 | 0 | 0 | 0 | 0 | 0 | 0 | 0 | 0 | 0 | 0 | 0 | 0 | 0 | 0 | 0 | 0 | 0 | 0 | 0 | 0 | 0 | 0 | 0 | 0 | 0 | 0 | 0 | 0 | 0 | 0 | 0 | 0 | 0 | 0 | 0 | 0 | 0 | 0 | 0 | 0 | 1 | 0 | 0 | 0 | 0 | 0 | 0 |
| virus_Sep8.0v_14852 | 1 | 0 | 0 | 0 | 0 | 0 | 0 | 0 | 0 | 0 | 0 | 0 | 0 | 0 | 0 | 0 | 0 | 0 | 0 | 0 | 0 | 0 | 0 | 0 | 0 | 0 | 0 | 0 | 0 | 0 | 0 | 0 | 0 | 0 | 0 | 0 | 0 | 0 | 0 | 0 | 0 | 0 | 0 | 0 | 0 | 0 | 0 | 0 | 1 | 0 | 0 | 0 | 0 | 0 | 0 |
| virus_Sep8.0v_16804 | 1 | 0 | 0 | 0 | 0 | 0 | 0 | 0 | 0 | 0 | 0 | 0 | 0 | 0 | 0 | 0 | 0 | 0 | 0 | 0 | 0 | 0 | 0 | 0 | 0 | 0 | 0 | 0 | 0 | 0 | 0 | 0 | 0 | 0 | 0 | 0 | 0 | 0 | 0 | 0 | 0 | 0 | 0 | 0 | 0 | 0 | 0 | 0 | 1 | 0 | 0 | 0 | 0 | 0 | 0 |
| virus_Sep8.5v_4344 | 1 | 0 | 0 | 0 | 0 | 0 | 0 | 0 | 0 | 0 | 0 | 0 | 0 | 0 | 0 | 0 | 0 | 0 | 0 | 0 | 0 | 0 | 0 | 0 | 0 | 0 | 0 | 0 | 0 | 0 | 0 | 0 | 0 | 0 | 0 | 0 | 0 | 0 | 0 | 0 | 0 | 0 | 0 | 0 | 0 | 0 | 0 | 0 | 0 | 0 | 1 | 0 | 0 | 0 | 0 |
| virus_Sep8.5v_4658 | 1 | 0 | 0 | 0 | 0 | 0 | 0 | 0 | 0 | 0 | 0 | 0 | 0 | 0 | 0 | 0 | 0 | 0 | 0 | 0 | 0 | 0 | 0 | 0 | 0 | 0 | 0 | 0 | 0 | 0 | 0 | 0 | 0 | 0 | 0 | 0 | 0 | 0 | 0 | 0 | 0 | 0 | 0 | 0 | 0 | 0 | 0 | 0 | 0 | 0 | 1 | 0 | 0 | 0 | 0 |
| virus_Sep8.5v_6795 | 1 | 0 | 0 | 0 | 0 | 0 | 0 | 0 | 0 | 0 | 0 | 0 | 0 | 0 | 0 | 0 | 0 | 0 | 0 | 0 | 0 | 0 | 0 | 0 | 0 | 0 | 0 | 0 | 0 | 0 | 0 | 0 | 0 | 0 | 0 | 0 | 0 | 0 | 0 | 0 | 0 | 0 | 0 | 0 | 0 | 0 | 0 | 0 | 0 | 0 | 1 | 0 | 0 | 0 | 0 |
| virus_Sep8.5v_7952 | 1 | 0 | 0 | 0 | 0 | 0 | 0 | 0 | 0 | 0 | 0 | 0 | 0 | 0 | 0 | 0 | 0 | 0 | 0 | 0 | 0 | 0 | 0 | 0 | 0 | 0 | 0 | 0 | 0 | 0 | 0 | 0 | 0 | 0 | 0 | 0 | 0 | 0 | 0 | 0 | 0 | 0 | 0 | 0 | 0 | 0 | 0 | 0 | 0 | 0 | 1 | 0 | 0 | 0 | 0 |
| virus_Sep8.5v_9010 | 1 | 0 | 0 | 0 | 0 | 0 | 0 | 0 | 0 | 0 | 0 | 0 | 0 | 0 | 0 | 0 | 0 | 0 | 0 | 0 | 0 | 0 | 0 | 0 | 0 | 0 | 0 | 0 | 0 | 0 | 0 | 0 | 0 | 0 | 0 | 0 | 0 | 0 | 0 | 0 | 0 | 0 | 0 | 0 | 0 | 0 | 0 | 0 | 0 | 0 | 1 | 0 | 0 | 0 | 0 |
| virus_Sep8.5v_11686 | 1 | 0 | 0 | 0 | 0 | 0 | 0 | 0 | 0 | 0 | 0 | 0 | 0 | 0 | 0 | 0 | 0 | 0 | 0 | 0 | 0 | 0 | 0 | 0 | 0 | 0 | 0 | 0 | 0 | 0 | 0 | 0 | 0 | 0 | 0 | 0 | 0 | 0 | 0 | 0 | 0 | 0 | 0 | 0 | 0 | 0 | 0 | 0 | 0 | 0 | 1 | 0 | 0 | 0 | 0 |

**Supplementary Table S7**. Common Bacterio- and virioplankton RTPR 98% OTUs in 27 samples with RTPR amplicons in both fractions

Total seqs: 47997; Subtotal seqs in retained

| OTUs: 41046 | | May12_05m | | May12_25m | | May12_50m | | May12_75m | | Jun8_25m | | Jun8_50m | | Jun8_75m | | Jun22_0m | | Jun22_50m | | Jun22_75m | | Jul11_0m | | Jul11_05m | | Jul11_25m | | Jul11_50m | | Jul11_75m | | Jul27_05m | | Jul27_25m | | Jul27_75m | | Aug5_0m | | Aug5_05m | | Aug5_25m | | Aug5_50m | | Aug5_75m | | Sep8_0m | | Sep8_05m | | Sep8_25m | | Sep8_50m | |
|---|---|---|---|---|---|---|---|---|---|---|---|---|---|---|---|---|---|---|---|---|---|---|---|---|---|---|---|---|---|---|---|---|---|---|---|---|---|---|---|---|---|---|---|---|---|---|---|---|---|---|---|---|---|---|---|
| OTUs with >=10 seqs: 254 | OTU subtotal | bacterioplankton | virioplankton | bacterioplankton | virioplankton | bacterioplankton | virioplankton | bacterioplankton | virioplankton | bacterioplankton | virioplankton | bacterioplankton | virioplankton | bacterioplankton | virioplankton | bacterioplankton | virioplankton | bacterioplankton | virioplankton | bacterioplankton | virioplankton | bacterioplankton | virioplankton | bacterioplankton | virioplankton | bacterioplankton | virioplankton | bacterioplankton | virioplankton | bacterioplankton | virioplankton | bacterioplankton | virioplankton | bacterioplankton | virioplankton | bacterioplankton | virioplankton | bacterioplankton | virioplankton | bacterioplankton | virioplankton | bacterioplankton | virioplankton | bacterioplankton | virioplankton | bacterioplankton | virioplankton | bacterioplankton | virioplankton | bacterioplankton | virioplankton | bacterioplankton | virioplankton | bacterioplankton | virioplankton |
| virus_Sep8.5v_14931 | 1 | 0 | 0 | 0 | 0 | 0 | 0 | 0 | 0 | 0 | 0 | 0 | 0 | 0 | 0 | 0 | 0 | 0 | 0 | 0 | 0 | 0 | 0 | 0 | 0 | 0 | 0 | 0 | 0 | 0 | 0 | 0 | 0 | 0 | 0 | 0 | 0 | 0 | 0 | 0 | 0 | 0 | 0 | 0 | 0 | 0 | 0 | 0 | 0 | 0 | 1 | 0 | 0 | 0 | 0 |
| virus_Sep8.5v_16321 | 1 | 0 | 0 | 0 | 0 | 0 | 0 | 0 | 0 | 0 | 0 | 0 | 0 | 0 | 0 | 0 | 0 | 0 | 0 | 0 | 0 | 0 | 0 | 0 | 0 | 0 | 0 | 0 | 0 | 0 | 0 | 0 | 0 | 0 | 0 | 0 | 0 | 0 | 0 | 0 | 0 | 0 | 0 | 0 | 0 | 0 | 0 | 0 | 0 | 0 | 1 | 0 | 0 | 0 | 0 |
| virus_Sep8.25v_10438 | 1 | 0 | 0 | 0 | 0 | 0 | 0 | 0 | 0 | 0 | 0 | 0 | 0 | 0 | 0 | 0 | 0 | 0 | 0 | 0 | 0 | 0 | 0 | 0 | 0 | 0 | 0 | 0 | 0 | 0 | 0 | 0 | 0 | 0 | 0 | 0 | 0 | 0 | 0 | 0 | 0 | 0 | 0 | 0 | 0 | 0 | 0 | 0 | 0 | 0 | 0 | 0 | 1 | 0 | 0 |
| virus_Sep8.25v_20688 | 1 | 0 | 0 | 0 | 0 | 0 | 0 | 0 | 0 | 0 | 0 | 0 | 0 | 0 | 0 | 0 | 0 | 0 | 0 | 0 | 0 | 0 | 0 | 0 | 0 | 0 | 0 | 0 | 0 | 0 | 0 | 0 | 0 | 0 | 0 | 0 | 0 | 0 | 0 | 0 | 0 | 0 | 0 | 0 | 0 | 0 | 0 | 0 | 0 | 0 | 0 | 0 | 1 | 0 | 0 |
| virus_Sep8.50v_4677 | 1 | 0 | 0 | 0 | 0 | 0 | 0 | 0 | 0 | 0 | 0 | 0 | 0 | 0 | 0 | 0 | 0 | 0 | 0 | 0 | 0 | 0 | 0 | 0 | 0 | 0 | 0 | 0 | 0 | 0 | 0 | 0 | 0 | 0 | 0 | 0 | 0 | 0 | 0 | 0 | 0 | 0 | 0 | 0 | 0 | 0 | 0 | 0 | 0 | 0 | 0 | 0 | 0 | 0 | 1 |
| virus_Sep8.50v_5365 | 1 | 0 | 0 | 0 | 0 | 0 | 0 | 0 | 0 | 0 | 0 | 0 | 0 | 0 | 0 | 0 | 0 | 0 | 0 | 0 | 0 | 0 | 0 | 0 | 0 | 0 | 0 | 0 | 0 | 0 | 0 | 0 | 0 | 0 | 0 | 0 | 0 | 0 | 0 | 0 | 0 | 0 | 0 | 0 | 0 | 0 | 0 | 0 | 0 | 0 | 0 | 0 | 0 | 0 | 1 |
| virus_Sep8.50v_13663 | 1 | 0 | 0 | 0 | 0 | 0 | 0 | 0 | 0 | 0 | 0 | 0 | 0 | 0 | 0 | 0 | 0 | 0 | 0 | 0 | 0 | 0 | 0 | 0 | 0 | 0 | 0 | 0 | 0 | 0 | 0 | 0 | 0 | 0 | 0 | 0 | 0 | 0 | 0 | 0 | 0 | 0 | 0 | 0 | 0 | 0 | 0 | 0 | 0 | 0 | 0 | 0 | 0 | 0 | 1 |
| virus_Sep8.50v_17016 | 1 | 0 | 0 | 0 | 0 | 0 | 0 | 0 | 0 | 0 | 0 | 0 | 0 | 0 | 0 | 0 | 0 | 0 | 0 | 0 | 0 | 0 | 0 | 0 | 0 | 0 | 0 | 0 | 0 | 0 | 0 | 0 | 0 | 0 | 0 | 0 | 0 | 0 | 0 | 0 | 0 | 0 | 0 | 0 | 0 | 0 | 0 | 0 | 0 | 0 | 0 | 0 | 0 | 0 | 1 |
| virus_Sep8.50v_17084 | 1 | 0 | 0 | 0 | 0 | 0 | 0 | 0 | 0 | 0 | 0 | 0 | 0 | 0 | 0 | 0 | 0 | 0 | 0 | 0 | 0 | 0 | 0 | 0 | 0 | 0 | 0 | 0 | 0 | 0 | 0 | 0 | 0 | 0 | 0 | 0 | 0 | 0 | 0 | 0 | 0 | 0 | 0 | 0 | 0 | 0 | 0 | 0 | 0 | 0 | 0 | 0 | 0 | 0 | 1 |
| virus_Sep8.50v_17827 | 1 | 0 | 0 | 0 | 0 | 0 | 0 | 0 | 0 | 0 | 0 | 0 | 0 | 0 | 0 | 0 | 0 | 0 | 0 | 0 | 0 | 0 | 0 | 0 | 0 | 0 | 0 | 0 | 0 | 0 | 0 | 0 | 0 | 0 | 0 | 0 | 0 | 0 | 0 | 0 | 0 | 0 | 0 | 0 | 0 | 0 | 0 | 0 | 0 | 0 | 0 | 0 | 0 | 0 | 1 |
| virus_Sep8.50v_30838 | 1 | 0 | 0 | 0 | 0 | 0 | 0 | 0 | 0 | 0 | 0 | 0 | 0 | 0 | 0 | 0 | 0 | 0 | 0 | 0 | 0 | 0 | 0 | 0 | 0 | 0 | 0 | 0 | 0 | 0 | 0 | 0 | 0 | 0 | 0 | 0 | 0 | 0 | 0 | 0 | 0 | 0 | 0 | 0 | 0 | 0 | 0 | 0 | 0 | 0 | 0 | 0 | 0 | 0 | 1 |
| virus_Sep8.50v_32867 | 1 | 0 | 0 | 0 | 0 | 0 | 0 | 0 | 0 | 0 | 0 | 0 | 0 | 0 | 0 | 0 | 0 | 0 | 0 | 0 | 0 | 0 | 0 | 0 | 0 | 0 | 0 | 0 | 0 | 0 | 0 | 0 | 0 | 0 | 0 | 0 | 0 | 0 | 0 | 0 | 0 | 0 | 0 | 0 | 0 | 0 | 0 | 0 | 0 | 0 | 0 | 0 | 0 | 0 | 1 |
| virus_Sep8.50v_33302 | 1 | 0 | 0 | 0 | 0 | 0 | 0 | 0 | 0 | 0 | 0 | 0 | 0 | 0 | 0 | 0 | 0 | 0 | 0 | 0 | 0 | 0 | 0 | 0 | 0 | 0 | 0 | 0 | 0 | 0 | 0 | 0 | 0 | 0 | 0 | 0 | 0 | 0 | 0 | 0 | 0 | 0 | 0 | 0 | 0 | 0 | 0 | 0 | 0 | 0 | 0 | 0 | 0 | 0 | 1 |
| virus_Sep8.50v_36214 | 1 | 0 | 0 | 0 | 0 | 0 | 0 | 0 | 0 | 0 | 0 | 0 | 0 | 0 | 0 | 0 | 0 | 0 | 0 | 0 | 0 | 0 | 0 | 0 | 0 | 0 | 0 | 0 | 0 | 0 | 0 | 0 | 0 | 0 | 0 | 0 | 0 | 0 | 0 | 0 | 0 | 0 | 0 | 0 | 0 | 0 | 0 | 0 | 0 | 0 | 0 | 0 | 0 | 0 | 1 |
| virus_Sep8.50v_36971 | 1 | 0 | 0 | 0 | 0 | 0 | 0 | 0 | 0 | 0 | 0 | 0 | 0 | 0 | 0 | 0 | 0 | 0 | 0 | 0 | 0 | 0 | 0 | 0 | 0 | 0 | 0 | 0 | 0 | 0 | 0 | 0 | 0 | 0 | 0 | 0 | 0 | 0 | 0 | 0 | 0 | 0 | 0 | 0 | 0 | 0 | 0 | 0 | 0 | 0 | 0 | 0 | 0 | 0 | 1 |
| virus_Sep8.50v_38954 | 1 | 0 | 0 | 0 | 0 | 0 | 0 | 0 | 0 | 0 | 0 | 0 | 0 | 0 | 0 | 0 | 0 | 0 | 0 | 0 | 0 | 0 | 0 | 0 | 0 | 0 | 0 | 0 | 0 | 0 | 0 | 0 | 0 | 0 | 0 | 0 | 0 | 0 | 0 | 0 | 0 | 0 | 0 | 0 | 0 | 0 | 0 | 0 | 0 | 0 | 0 | 0 | 0 | 0 | 1 |
| virus_Sep8.50v_40927 | 1 | 0 | 0 | 0 | 0 | 0 | 0 | 0 | 0 | 0 | 0 | 0 | 0 | 0 | 0 | 0 | 0 | 0 | 0 | 0 | 0 | 0 | 0 | 0 | 0 | 0 | 0 | 0 | 0 | 0 | 0 | 0 | 0 | 0 | 0 | 0 | 0 | 0 | 0 | 0 | 0 | 0 | 0 | 0 | 0 | 0 | 0 | 0 | 0 | 0 | 0 | 0 | 0 | 0 | 1 |
| bacteria_Jul11d0b_24044 | 1 | 0 | 0 | 0 | 0 | 0 | 0 | 0 | 0 | 0 | 0 | 0 | 0 | 0 | 0 | 0 | 0 | 0 | 0 | 0 | 0 | 1 | 0 | 0 | 0 | 0 | 0 | 0 | 0 | 0 | 0 | 0 | 0 | 0 | 0 | 0 | 0 | 0 | 0 | 0 | 0 | 0 | 0 | 0 | 0 | 0 | 0 | 0 | 0 | 0 | 0 | 0 | 0 | 0 | 0 |
| bacteria_Jul11d0b_30788 | 1 | 0 | 0 | 0 | 0 | 0 | 0 | 0 | 0 | 0 | 0 | 0 | 0 | 0 | 0 | 0 | 0 | 0 | 0 | 0 | 0 | 1 | 0 | 0 | 0 | 0 | 0 | 0 | 0 | 0 | 0 | 0 | 0 | 0 | 0 | 0 | 0 | 0 | 0 | 0 | 0 | 0 | 0 | 0 | 0 | 0 | 0 | 0 | 0 | 0 | 0 | 0 | 0 | 0 | 0 |
| bacteria_Jul11d0b_34563 | 1 | 0 | 0 | 0 | 0 | 0 | 0 | 0 | 0 | 0 | 0 | 0 | 0 | 0 | 0 | 0 | 0 | 0 | 0 | 0 | 0 | 1 | 0 | 0 | 0 | 0 | 0 | 0 | 0 | 0 | 0 | 0 | 0 | 0 | 0 | 0 | 0 | 0 | 0 | 0 | 0 | 0 | 0 | 0 | 0 | 0 | 0 | 0 | 0 | 0 | 0 | 0 | 0 | 0 | 0 |
| bacteria_Jul11d0b_43429 | 1 | 0 | 0 | 0 | 0 | 0 | 0 | 0 | 0 | 0 | 0 | 0 | 0 | 0 | 0 | 0 | 0 | 0 | 0 | 0 | 0 | 1 | 0 | 0 | 0 | 0 | 0 | 0 | 0 | 0 | 0 | 0 | 0 | 0 | 0 | 0 | 0 | 0 | 0 | 0 | 0 | 0 | 0 | 0 | 0 | 0 | 0 | 0 | 0 | 0 | 0 | 0 | 0 | 0 | 0 |
| bacteria_Jul11d0b_50464 | 1 | 0 | 0 | 0 | 0 | 0 | 0 | 0 | 0 | 0 | 0 | 0 | 0 | 0 | 0 | 0 | 0 | 0 | 0 | 0 | 0 | 1 | 0 | 0 | 0 | 0 | 0 | 0 | 0 | 0 | 0 | 0 | 0 | 0 | 0 | 0 | 0 | 0 | 0 | 0 | 0 | 0 | 0 | 0 | 0 | 0 | 0 | 0 | 0 | 0 | 0 | 0 | 0 | 0 | 0 |
| bacteria_Jul11d0b_53423 | 1 | 0 | 0 | 0 | 0 | 0 | 0 | 0 | 0 | 0 | 0 | 0 | 0 | 0 | 0 | 0 | 0 | 0 | 0 | 0 | 0 | 1 | 0 | 0 | 0 | 0 | 0 | 0 | 0 | 0 | 0 | 0 | 0 | 0 | 0 | 0 | 0 | 0 | 0 | 0 | 0 | 0 | 0 | 0 | 0 | 0 | 0 | 0 | 0 | 0 | 0 | 0 | 0 | 0 | 0 |
| bacteria_Jul11d0b_54836 | 1 | 0 | 0 | 0 | 0 | 0 | 0 | 0 | 0 | 0 | 0 | 0 | 0 | 0 | 0 | 0 | 0 | 0 | 0 | 0 | 0 | 1 | 0 | 0 | 0 | 0 | 0 | 0 | 0 | 0 | 0 | 0 | 0 | 0 | 0 | 0 | 0 | 0 | 0 | 0 | 0 | 0 | 0 | 0 | 0 | 0 | 0 | 0 | 0 | 0 | 0 | 0 | 0 | 0 | 0 |
| bacteria_Jul11d0b_65078 | 1 | 0 | 0 | 0 | 0 | 0 | 0 | 0 | 0 | 0 | 0 | 0 | 0 | 0 | 0 | 0 | 0 | 0 | 0 | 0 | 0 | 1 | 0 | 0 | 0 | 0 | 0 | 0 | 0 | 0 | 0 | 0 | 0 | 0 | 0 | 0 | 0 | 0 | 0 | 0 | 0 | 0 | 0 | 0 | 0 | 0 | 0 | 0 | 0 | 0 | 0 | 0 | 0 | 0 | 0 |
| bacteria_Jul11d0b_71634 | 1 | 0 | 0 | 0 | 0 | 0 | 0 | 0 | 0 | 0 | 0 | 0 | 0 | 0 | 0 | 0 | 0 | 0 | 0 | 0 | 0 | 1 | 0 | 0 | 0 | 0 | 0 | 0 | 0 | 0 | 0 | 0 | 0 | 0 | 0 | 0 | 0 | 0 | 0 | 0 | 0 | 0 | 0 | 0 | 0 | 0 | 0 | 0 | 0 | 0 | 0 | 0 | 0 | 0 | 0 |
| bacteria_Aug5d0b_7396 | 1 | 0 | 0 | 0 | 0 | 0 | 0 | 0 | 0 | 0 | 0 | 0 | 0 | 0 | 0 | 0 | 0 | 0 | 0 | 0 | 0 | 0 | 0 | 0 | 0 | 0 | 0 | 0 | 0 | 0 | 0 | 0 | 0 | 0 | 0 | 0 | 0 | 1 | 0 | 0 | 0 | 0 | 0 | 0 | 0 | 0 | 0 | 0 | 0 | 0 | 0 | 0 | 0 | 0 | 0 |
| bacteria_Aug5d0b_11150 | 1 | 0 | 0 | 0 | 0 | 0 | 0 | 0 | 0 | 0 | 0 | 0 | 0 | 0 | 0 | 0 | 0 | 0 | 0 | 0 | 0 | 0 | 0 | 0 | 0 | 0 | 0 | 0 | 0 | 0 | 0 | 0 | 0 | 0 | 0 | 0 | 0 | 1 | 0 | 0 | 0 | 0 | 0 | 0 | 0 | 0 | 0 | 0 | 0 | 0 | 0 | 0 | 0 | 0 | 0 |
| bacteria_Aug5d0b_33754 | 1 | 0 | 0 | 0 | 0 | 0 | 0 | 0 | 0 | 0 | 0 | 0 | 0 | 0 | 0 | 0 | 0 | 0 | 0 | 0 | 0 | 0 | 0 | 0 | 0 | 0 | 0 | 0 | 0 | 0 | 0 | 0 | 0 | 0 | 0 | 0 | 0 | 1 | 0 | 0 | 0 | 0 | 0 | 0 | 0 | 0 | 0 | 0 | 0 | 0 | 0 | 0 | 0 | 0 | 0 |
| bacteria_Aug5d0b_64213 | 1 | 0 | 0 | 0 | 0 | 0 | 0 | 0 | 0 | 0 | 0 | 0 | 0 | 0 | 0 | 0 | 0 | 0 | 0 | 0 | 0 | 0 | 0 | 0 | 0 | 0 | 0 | 0 | 0 | 0 | 0 | 0 | 0 | 0 | 0 | 0 | 0 | 1 | 0 | 0 | 0 | 0 | 0 | 0 | 0 | 0 | 0 | 0 | 0 | 0 | 0 | 0 | 0 | 0 | 0 |
| bacteria_Aug5d5b_2148 | 1 | 0 | 0 | 0 | 0 | 0 | 0 | 0 | 0 | 0 | 0 | 0 | 0 | 0 | 0 | 0 | 0 | 0 | 0 | 0 | 0 | 0 | 0 | 0 | 0 | 0 | 0 | 0 | 0 | 0 | 0 | 0 | 0 | 0 | 0 | 0 | 0 | 0 | 0 | 1 | 0 | 0 | 0 | 0 | 0 | 0 | 0 | 0 | 0 | 0 | 0 | 0 | 0 | 0 | 0 |
| bacteria_Aug5d5b_8186 | 1 | 0 | 0 | 0 | 0 | 0 | 0 | 0 | 0 | 0 | 0 | 0 | 0 | 0 | 0 | 0 | 0 | 0 | 0 | 0 | 0 | 0 | 0 | 0 | 0 | 0 | 0 | 0 | 0 | 0 | 0 | 0 | 0 | 0 | 0 | 0 | 0 | 0 | 0 | 1 | 0 | 0 | 0 | 0 | 0 | 0 | 0 | 0 | 0 | 0 | 0 | 0 | 0 | 0 | 0 |
| bacteria_Aug5d5b_26848 | 1 | 0 | 0 | 0 | 0 | 0 | 0 | 0 | 0 | 0 | 0 | 0 | 0 | 0 | 0 | 0 | 0 | 0 | 0 | 0 | 0 | 0 | 0 | 0 | 0 | 0 | 0 | 0 | 0 | 0 | 0 | 0 | 0 | 0 | 0 | 0 | 0 | 0 | 0 | 1 | 0 | 0 | 0 | 0 | 0 | 0 | 0 | 0 | 0 | 0 | 0 | 0 | 0 | 0 | 0 |
| bacteria_Aug5d5b_28101 | 1 | 0 | 0 | 0 | 0 | 0 | 0 | 0 | 0 | 0 | 0 | 0 | 0 | 0 | 0 | 0 | 0 | 0 | 0 | 0 | 0 | 0 | 0 | 0 | 0 | 0 | 0 | 0 | 0 | 0 | 0 | 0 | 0 | 0 | 0 | 0 | 0 | 0 | 0 | 1 | 0 | 0 | 0 | 0 | 0 | 0 | 0 | 0 | 0 | 0 | 0 | 0 | 0 | 0 | 0 |
| bacteria_Aug5d5b_55827 | 1 | 0 | 0 | 0 | 0 | 0 | 0 | 0 | 0 | 0 | 0 | 0 | 0 | 0 | 0 | 0 | 0 | 0 | 0 | 0 | 0 | 0 | 0 | 0 | 0 | 0 | 0 | 0 | 0 | 0 | 0 | 0 | 0 | 0 | 0 | 0 | 0 | 0 | 0 | 1 | 0 | 0 | 0 | 0 | 0 | 0 | 0 | 0 | 0 | 0 | 0 | 0 | 0 | 0 | 0 |
| bacteria_Aug5d5b_58626 | 1 | 0 | 0 | 0 | 0 | 0 | 0 | 0 | 0 | 0 | 0 | 0 | 0 | 0 | 0 | 0 | 0 | 0 | 0 | 0 | 0 | 0 | 0 | 0 | 0 | 0 | 0 | 0 | 0 | 0 | 0 | 0 | 0 | 0 | 0 | 0 | 0 | 0 | 0 | 1 | 0 | 0 | 0 | 0 | 0 | 0 | 0 | 0 | 0 | 0 | 0 | 0 | 0 | 0 | 0 |
| bacteria_Aug5d25b_6197 | 1 | 0 | 0 | 0 | 0 | 0 | 0 | 0 | 0 | 0 | 0 | 0 | 0 | 0 | 0 | 0 | 0 | 0 | 0 | 0 | 0 | 0 | 0 | 0 | 0 | 0 | 0 | 0 | 0 | 0 | 0 | 0 | 0 | 0 | 0 | 0 | 0 | 0 | 0 | 0 | 0 | 1 | 0 | 0 | 0 | 0 | 0 | 0 | 0 | 0 | 0 | 0 | 0 | 0 | 0 |
| bacteria_Aug5d25b_12313 | 1 | 0 | 0 | 0 | 0 | 0 | 0 | 0 | 0 | 0 | 0 | 0 | 0 | 0 | 0 | 0 | 0 | 0 | 0 | 0 | 0 | 0 | 0 | 0 | 0 | 0 | 0 | 0 | 0 | 0 | 0 | 0 | 0 | 0 | 0 | 0 | 0 | 0 | 0 | 0 | 0 | 1 | 0 | 0 | 0 | 0 | 0 | 0 | 0 | 0 | 0 | 0 | 0 | 0 | 0 |
| bacteria_Aug5d25b_13559 | 1 | 0 | 0 | 0 | 0 | 0 | 0 | 0 | 0 | 0 | 0 | 0 | 0 | 0 | 0 | 0 | 0 | 0 | 0 | 0 | 0 | 0 | 0 | 0 | 0 | 0 | 0 | 0 | 0 | 0 | 0 | 0 | 0 | 0 | 0 | 0 | 0 | 0 | 0 | 0 | 0 | 1 | 0 | 0 | 0 | 0 | 0 | 0 | 0 | 0 | 0 | 0 | 0 | 0 | 0 |
| bacteria_Aug5d25b_14783 | 1 | 0 | 0 | 0 | 0 | 0 | 0 | 0 | 0 | 0 | 0 | 0 | 0 | 0 | 0 | 0 | 0 | 0 | 0 | 0 | 0 | 0 | 0 | 0 | 0 | 0 | 0 | 0 | 0 | 0 | 0 | 0 | 0 | 0 | 0 | 0 | 0 | 0 | 0 | 0 | 0 | 1 | 0 | 0 | 0 | 0 | 0 | 0 | 0 | 0 | 0 | 0 | 0 | 0 | 0 |
| bacteria_Aug5d25b_16579 | 1 | 0 | 0 | 0 | 0 | 0 | 0 | 0 | 0 | 0 | 0 | 0 | 0 | 0 | 0 | 0 | 0 | 0 | 0 | 0 | 0 | 0 | 0 | 0 | 0 | 0 | 0 | 0 | 0 | 0 | 0 | 0 | 0 | 0 | 0 | 0 | 0 | 0 | 0 | 0 | 0 | 1 | 0 | 0 | 0 | 0 | 0 | 0 | 0 | 0 | 0 | 0 | 0 | 0 | 0 |
| bacteria_Aug5d25b_21783 | 1 | 0 | 0 | 0 | 0 | 0 | 0 | 0 | 0 | 0 | 0 | 0 | 0 | 0 | 0 | 0 | 0 | 0 | 0 | 0 | 0 | 0 | 0 | 0 | 0 | 0 | 0 | 0 | 0 | 0 | 0 | 0 | 0 | 0 | 0 | 0 | 0 | 0 | 0 | 0 | 0 | 1 | 0 | 0 | 0 | 0 | 0 | 0 | 0 | 0 | 0 | 0 | 0 | 0 | 0 |
| bacteria_Aug5d25b_26381 | 1 | 0 | 0 | 0 | 0 | 0 | 0 | 0 | 0 | 0 | 0 | 0 | 0 | 0 | 0 | 0 | 0 | 0 | 0 | 0 | 0 | 0 | 0 | 0 | 0 | 0 | 0 | 0 | 0 | 0 | 0 | 0 | 0 | 0 | 0 | 0 | 0 | 0 | 0 | 0 | 0 | 1 | 0 | 0 | 0 | 0 | 0 | 0 | 0 | 0 | 0 | 0 | 0 | 0 | 0 |
| bacteria_Aug5d25b_30172 | 1 | 0 | 0 | 0 | 0 | 0 | 0 | 0 | 0 | 0 | 0 | 0 | 0 | 0 | 0 | 0 | 0 | 0 | 0 | 0 | 0 | 0 | 0 | 0 | 0 | 0 | 0 | 0 | 0 | 0 | 0 | 0 | 0 | 0 | 0 | 0 | 0 | 0 | 0 | 0 | 0 | 1 | 0 | 0 | 0 | 0 | 0 | 0 | 0 | 0 | 0 | 0 | 0 | 0 | 0 |
| bacteria_Aug5d25b_30549 | 1 | 0 | 0 | 0 | 0 | 0 | 0 | 0 | 0 | 0 | 0 | 0 | 0 | 0 | 0 | 0 | 0 | 0 | 0 | 0 | 0 | 0 | 0 | 0 | 0 | 0 | 0 | 0 | 0 | 0 | 0 | 0 | 0 | 0 | 0 | 0 | 0 | 0 | 0 | 0 | 0 | 1 | 0 | 0 | 0 | 0 | 0 | 0 | 0 | 0 | 0 | 0 | 0 | 0 | 0 |
| bacteria_Aug5d25b_44387 | 1 | 0 | 0 | 0 | 0 | 0 | 0 | 0 | 0 | 0 | 0 | 0 | 0 | 0 | 0 | 0 | 0 | 0 | 0 | 0 | 0 | 0 | 0 | 0 | 0 | 0 | 0 | 0 | 0 | 0 | 0 | 0 | 0 | 0 | 0 | 0 | 0 | 0 | 0 | 0 | 0 | 1 | 0 | 0 | 0 | 0 | 0 | 0 | 0 | 0 | 0 | 0 | 0 | 0 | 0 |
| bacteria_Aug5d25b_57302 | 1 | 0 | 0 | 0 | 0 | 0 | 0 | 0 | 0 | 0 | 0 | 0 | 0 | 0 | 0 | 0 | 0 | 0 | 0 | 0 | 0 | 0 | 0 | 0 | 0 | 0 | 0 | 0 | 0 | 0 | 0 | 0 | 0 | 0 | 0 | 0 | 0 | 0 | 0 | 0 | 0 | 1 | 0 | 0 | 0 | 0 | 0 | 0 | 0 | 0 | 0 | 0 | 0 | 0 | 0 |
| bacteria_Aug5d25b_72088 | 1 | 0 | 0 | 0 | 0 | 0 | 0 | 0 | 0 | 0 | 0 | 0 | 0 | 0 | 0 | 0 | 0 | 0 | 0 | 0 | 0 | 0 | 0 | 0 | 0 | 0 | 0 | 0 | 0 | 0 | 0 | 0 | 0 | 0 | 0 | 0 | 0 | 0 | 0 | 0 | 0 | 1 | 0 | 0 | 0 | 0 | 0 | 0 | 0 | 0 | 0 | 0 | 0 | 0 | 0 |
| bacteria_Aug5d25b_72849 | 1 | 0 | 0 | 0 | 0 | 0 | 0 | 0 | 0 | 0 | 0 | 0 | 0 | 0 | 0 | 0 | 0 | 0 | 0 | 0 | 0 | 0 | 0 | 0 | 0 | 0 | 0 | 0 | 0 | 0 | 0 | 0 | 0 | 0 | 0 | 0 | 0 | 0 | 0 | 0 | 0 | 1 | 0 | 0 | 0 | 0 | 0 | 0 | 0 | 0 | 0 | 0 | 0 | 0 | 0 |
| bacteria_Aug5d25b_73102 | 1 | 0 | 0 | 0 | 0 | 0 | 0 | 0 | 0 | 0 | 0 | 0 | 0 | 0 | 0 | 0 | 0 | 0 | 0 | 0 | 0 | 0 | 0 | 0 | 0 | 0 | 0 | 0 | 0 | 0 | 0 | 0 | 0 | 0 | 0 | 0 | 0 | 0 | 0 | 0 | 0 | 1 | 0 | 0 | 0 | 0 | 0 | 0 | 0 | 0 | 0 | 0 | 0 | 0 | 0 |
| bacteria_Aug5d25b_73860 | 1 | 0 | 0 | 0 | 0 | 0 | 0 | 0 | 0 | 0 | 0 | 0 | 0 | 0 | 0 | 0 | 0 | 0 | 0 | 0 | 0 | 0 | 0 | 0 | 0 | 0 | 0 | 0 | 0 | 0 | 0 | 0 | 0 | 0 | 0 | 0 | 0 | 0 | 0 | 0 | 0 | 1 | 0 | 0 | 0 | 0 | 0 | 0 | 0 | 0 | 0 | 0 | 0 | 0 | 0 |
| bacteria_Aug5d50b_37663 | 1 | 0 | 0 | 0 | 0 | 0 | 0 | 0 | 0 | 0 | 0 | 0 | 0 | 0 | 0 | 0 | 0 | 0 | 0 | 0 | 0 | 0 | 0 | 0 | 0 | 0 | 0 | 0 | 0 | 0 | 0 | 0 | 0 | 0 | 0 | 0 | 0 | 0 | 0 | 0 | 0 | 0 | 0 | 1 | 0 | 0 | 0 | 0 | 0 | 0 | 0 | 0 | 0 | 0 | 0 |
| bacteria_Aug5d50b_48721 | 1 | 0 | 0 | 0 | 0 | 0 | 0 | 0 | 0 | 0 | 0 | 0 | 0 | 0 | 0 | 0 | 0 | 0 | 0 | 0 | 0 | 0 | 0 | 0 | 0 | 0 | 0 | 0 | 0 | 0 | 0 | 0 | 0 | 0 | 0 | 0 | 0 | 0 | 0 | 0 | 0 | 0 | 0 | 1 | 0 | 0 | 0 | 0 | 0 | 0 | 0 | 0 | 0 | 0 | 0 |
| bacteria_Aug5d50b_64610 | 1 | 0 | 0 | 0 | 0 | 0 | 0 | 0 | 0 | 0 | 0 | 0 | 0 | 0 | 0 | 0 | 0 | 0 | 0 | 0 | 0 | 0 | 0 | 0 | 0 | 0 | 0 | 0 | 0 | 0 | 0 | 0 | 0 | 0 | 0 | 0 | 0 | 0 | 0 | 0 | 0 | 0 | 0 | 1 | 0 | 0 | 0 | 0 | 0 | 0 | 0 | 0 | 0 | 0 | 0 |
| bacteria_Aug5d50b_65293 | 1 | 0 | 0 | 0 | 0 | 0 | 0 | 0 | 0 | 0 | 0 | 0 | 0 | 0 | 0 | 0 | 0 | 0 | 0 | 0 | 0 | 0 | 0 | 0 | 0 | 0 | 0 | 0 | 0 | 0 | 0 | 0 | 0 | 0 | 0 | 0 | 0 | 0 | 0 | 0 | 0 | 0 | 0 | 1 | 0 | 0 | 0 | 0 | 0 | 0 | 0 | 0 | 0 | 0 | 0 |
| bacteria_Jul11d5b_48719 | 1 | 0 | 0 | 0 | 0 | 0 | 0 | 0 | 0 | 0 | 0 | 0 | 0 | 0 | 0 | 0 | 0 | 0 | 0 | 0 | 0 | 0 | 0 | 1 | 0 | 0 | 0 | 0 | 0 | 0 | 0 | 0 | 0 | 0 | 0 | 0 | 0 | 0 | 0 | 0 | 0 | 0 | 0 | 0 | 0 | 0 | 0 | 0 | 0 | 0 | 0 | 0 | 0 | 0 | 0 |
| bacteria_Jul11d25b_1024 | 1 | 0 | 0 | 0 | 0 | 0 | 0 | 0 | 0 | 0 | 0 | 0 | 0 | 0 | 0 | 0 | 0 | 0 | 0 | 0 | 0 | 0 | 0 | 0 | 0 | 1 | 0 | 0 | 0 | 0 | 0 | 0 | 0 | 0 | 0 | 0 | 0 | 0 | 0 | 0 | 0 | 0 | 0 | 0 | 0 | 0 | 0 | 0 | 0 | 0 | 0 | 0 | 0 | 0 | 0 |
| bacteria_Jul11d25b_6161 | 1 | 0 | 0 | 0 | 0 | 0 | 0 | 0 | 0 | 0 | 0 | 0 | 0 | 0 | 0 | 0 | 0 | 0 | 0 | 0 | 0 | 0 | 0 | 0 | 0 | 1 | 0 | 0 | 0 | 0 | 0 | 0 | 0 | 0 | 0 | 0 | 0 | 0 | 0 | 0 | 0 | 0 | 0 | 0 | 0 | 0 | 0 | 0 | 0 | 0 | 0 | 0 | 0 | 0 | 0 |
| bacteria_Jul11d25b_10720 | 1 | 0 | 0 | 0 | 0 | 0 | 0 | 0 | 0 | 0 | 0 | 0 | 0 | 0 | 0 | 0 | 0 | 0 | 0 | 0 | 0 | 0 | 0 | 0 | 0 | 1 | 0 | 0 | 0 | 0 | 0 | 0 | 0 | 0 | 0 | 0 | 0 | 0 | 0 | 0 | 0 | 0 | 0 | 0 | 0 | 0 | 0 | 0 | 0 | 0 | 0 | 0 | 0 | 0 | 0 |
| bacteria_Jul11d25b_12879 | 1 | 0 | 0 | 0 | 0 | 0 | 0 | 0 | 0 | 0 | 0 | 0 | 0 | 0 | 0 | 0 | 0 | 0 | 0 | 0 | 0 | 0 | 0 | 0 | 0 | 1 | 0 | 0 | 0 | 0 | 0 | 0 | 0 | 0 | 0 | 0 | 0 | 0 | 0 | 0 | 0 | 0 | 0 | 0 | 0 | 0 | 0 | 0 | 0 | 0 | 0 | 0 | 0 | 0 | 0 |
| bacteria_Jul11d25b_13173 | 1 | 0 | 0 | 0 | 0 | 0 | 0 | 0 | 0 | 0 | 0 | 0 | 0 | 0 | 0 | 0 | 0 | 0 | 0 | 0 | 0 | 0 | 0 | 0 | 0 | 1 | 0 | 0 | 0 | 0 | 0 | 0 | 0 | 0 | 0 | 0 | 0 | 0 | 0 | 0 | 0 | 0 | 0 | 0 | 0 | 0 | 0 | 0 | 0 | 0 | 0 | 0 | 0 | 0 | 0 |
| bacteria_Jul11d25b_15054 | 1 | 0 | 0 | 0 | 0 | 0 | 0 | 0 | 0 | 0 | 0 | 0 | 0 | 0 | 0 | 0 | 0 | 0 | 0 | 0 | 0 | 0 | 0 | 0 | 0 | 1 | 0 | 0 | 0 | 0 | 0 | 0 | 0 | 0 | 0 | 0 | 0 | 0 | 0 | 0 | 0 | 0 | 0 | 0 | 0 | 0 | 0 | 0 | 0 | 0 | 0 | 0 | 0 | 0 | 0 |
| bacteria_Jul11d25b_15678 | 1 | 0 | 0 | 0 | 0 | 0 | 0 | 0 | 0 | 0 | 0 | 0 | 0 | 0 | 0 | 0 | 0 | 0 | 0 | 0 | 0 | 0 | 0 | 0 | 0 | 1 | 0 | 0 | 0 | 0 | 0 | 0 | 0 | 0 | 0 | 0 | 0 | 0 | 0 | 0 | 0 | 0 | 0 | 0 | 0 | 0 | 0 | 0 | 0 | 0 | 0 | 0 | 0 | 0 | 0 |
| bacteria_Jul11d25b_51857 | 1 | 0 | 0 | 0 | 0 | 0 | 0 | 0 | 0 | 0 | 0 | 0 | 0 | 0 | 0 | 0 | 0 | 0 | 0 | 0 | 0 | 0 | 0 | 0 | 0 | 1 | 0 | 0 | 0 | 0 | 0 | 0 | 0 | 0 | 0 | 0 | 0 | 0 | 0 | 0 | 0 | 0 | 0 | 0 | 0 | 0 | 0 | 0 | 0 | 0 | 0 | 0 | 0 | 0 | 0 |
| bacteria_Jul11d50b_3464 | 1 | 0 | 0 | 0 | 0 | 0 | 0 | 0 | 0 | 0 | 0 | 0 | 0 | 0 | 0 | 0 | 0 | 0 | 0 | 0 | 0 | 0 | 0 | 0 | 0 | 0 | 0 | 1 | 0 | 0 | 0 | 0 | 0 | 0 | 0 | 0 | 0 | 0 | 0 | 0 | 0 | 0 | 0 | 0 | 0 | 0 | 0 | 0 | 0 | 0 | 0 | 0 | 0 | 0 | 0 |
| bacteria_Jul11d50b_16219 | 1 | 0 | 0 | 0 | 0 | 0 | 0 | 0 | 0 | 0 | 0 | 0 | 0 | 0 | 0 | 0 | 0 | 0 | 0 | 0 | 0 | 0 | 0 | 0 | 0 | 0 | 0 | 1 | 0 | 0 | 0 | 0 | 0 | 0 | 0 | 0 | 0 | 0 | 0 | 0 | 0 | 0 | 0 | 0 | 0 | 0 | 0 | 0 | 0 | 0 | 0 | 0 | 0 | 0 | 0 |
| bacteria_Jul11d50b_32894 | 1 | 0 | 0 | 0 | 0 | 0 | 0 | 0 | 0 | 0 | 0 | 0 | 0 | 0 | 0 | 0 | 0 | 0 | 0 | 0 | 0 | 0 | 0 | 0 | 0 | 0 | 0 | 1 | 0 | 0 | 0 | 0 | 0 | 0 | 0 | 0 | 0 | 0 | 0 | 0 | 0 | 0 | 0 | 0 | 0 | 0 | 0 | 0 | 0 | 0 | 0 | 0 | 0 | 0 | 0 |
| bacteria_Jul11d50b_41228 | 1 | 0 | 0 | 0 | 0 | 0 | 0 | 0 | 0 | 0 | 0 | 0 | 0 | 0 | 0 | 0 | 0 | 0 | 0 | 0 | 0 | 0 | 0 | 0 | 0 | 0 | 0 | 1 | 0 | 0 | 0 | 0 | 0 | 0 | 0 | 0 | 0 | 0 | 0 | 0 | 0 | 0 | 0 | 0 | 0 | 0 | 0 | 0 | 0 | 0 | 0 | 0 | 0 | 0 | 0 |
| bacteria_Jul11d50b_42406 | 1 | 0 | 0 | 0 | 0 | 0 | 0 | 0 | 0 | 0 | 0 | 0 | 0 | 0 | 0 | 0 | 0 | 0 | 0 | 0 | 0 | 0 | 0 | 0 | 0 | 0 | 0 | 1 | 0 | 0 | 0 | 0 | 0 | 0 | 0 | 0 | 0 | 0 | 0 | 0 | 0 | 0 | 0 | 0 | 0 | 0 | 0 | 0 | 0 | 0 | 0 | 0 | 0 | 0 | 0 |
| bacteria_Jul11d50b_47170 | 1 | 0 | 0 | 0 | 0 | 0 | 0 | 0 | 0 | 0 | 0 | 0 | 0 | 0 | 0 | 0 | 0 | 0 | 0 | 0 | 0 | 0 | 0 | 0 | 0 | 0 | 0 | 1 | 0 | 0 | 0 | 0 | 0 | 0 | 0 | 0 | 0 | 0 | 0 | 0 | 0 | 0 | 0 | 0 | 0 | 0 | 0 | 0 | 0 | 0 | 0 | 0 | 0 | 0 | 0 |
| bacteria_Jul11d75b_2126 | 1 | 0 | 0 | 0 | 0 | 0 | 0 | 0 | 0 | 0 | 0 | 0 | 0 | 0 | 0 | 0 | 0 | 0 | 0 | 0 | 0 | 0 | 0 | 0 | 0 | 0 | 0 | 0 | 0 | 1 | 0 | 0 | 0 | 0 | 0 | 0 | 0 | 0 | 0 | 0 | 0 | 0 | 0 | 0 | 0 | 0 | 0 | 0 | 0 | 0 | 0 | 0 | 0 | 0 | 0 |
| bacteria_Jul11d75b_5378 | 1 | 0 | 0 | 0 | 0 | 0 | 0 | 0 | 0 | 0 | 0 | 0 | 0 | 0 | 0 | 0 | 0 | 0 | 0 | 0 | 0 | 0 | 0 | 0 | 0 | 0 | 0 | 0 | 0 | 1 | 0 | 0 | 0 | 0 | 0 | 0 | 0 | 0 | 0 | 0 | 0 | 0 | 0 | 0 | 0 | 0 | 0 | 0 | 0 | 0 | 0 | 0 | 0 | 0 | 0 |
| bacteria_Jul11d75b_7061 | 1 | 0 | 0 | 0 | 0 | 0 | 0 | 0 | 0 | 0 | 0 | 0 | 0 | 0 | 0 | 0 | 0 | 0 | 0 | 0 | 0 | 0 | 0 | 0 | 0 | 0 | 0 | 0 | 0 | 1 | 0 | 0 | 0 | 0 | 0 | 0 | 0 | 0 | 0 | 0 | 0 | 0 | 0 | 0 | 0 | 0 | 0 | 0 | 0 | 0 | 0 | 0 | 0 | 0 | 0 |
| bacteria_Jul11d75b_20424 | 1 | 0 | 0 | 0 | 0 | 0 | 0 | 0 | 0 | 0 | 0 | 0 | 0 | 0 | 0 | 0 | 0 | 0 | 0 | 0 | 0 | 0 | 0 | 0 | 0 | 0 | 0 | 0 | 0 | 1 | 0 | 0 | 0 | 0 | 0 | 0 | 0 | 0 | 0 | 0 | 0 | 0 | 0 | 0 | 0 | 0 | 0 | 0 | 0 | 0 | 0 | 0 | 0 | 0 | 0 |
| bacteria_Jul11d75b_22001 | 1 | 0 | 0 | 0 | 0 | 0 | 0 | 0 | 0 | 0 | 0 | 0 | 0 | 0 | 0 | 0 | 0 | 0 | 0 | 0 | 0 | 0 | 0 | 0 | 0 | 0 | 0 | 0 | 0 | 1 | 0 | 0 | 0 | 0 | 0 | 0 | 0 | 0 | 0 | 0 | 0 | 0 | 0 | 0 | 0 | 0 | 0 | 0 | 0 | 0 | 0 | 0 | 0 | 0 | 0 |
| bacteria_Jul11d75b_22382 | 1 | 0 | 0 | 0 | 0 | 0 | 0 | 0 | 0 | 0 | 0 | 0 | 0 | 0 | 0 | 0 | 0 | 0 | 0 | 0 | 0 | 0 | 0 | 0 | 0 | 0 | 0 | 0 | 0 | 1 | 0 | 0 | 0 | 0 | 0 | 0 | 0 | 0 | 0 | 0 | 0 | 0 | 0 | 0 | 0 | 0 | 0 | 0 | 0 | 0 | 0 | 0 | 0 | 0 | 0 |
| bacteria_Jul11d75b_40531 | 1 | 0 | 0 | 0 | 0 | 0 | 0 | 0 | 0 | 0 | 0 | 0 | 0 | 0 | 0 | 0 | 0 | 0 | 0 | 0 | 0 | 0 | 0 | 0 | 0 | 0 | 0 | 0 | 0 | 1 | 0 | 0 | 0 | 0 | 0 | 0 | 0 | 0 | 0 | 0 | 0 | 0 | 0 | 0 | 0 | 0 | 0 | 0 | 0 | 0 | 0 | 0 | 0 | 0 | 0 |
| bacteria_Jul11d75b_44131 | 1 | 0 | 0 | 0 | 0 | 0 | 0 | 0 | 0 | 0 | 0 | 0 | 0 | 0 | 0 | 0 | 0 | 0 | 0 | 0 | 0 | 0 | 0 | 0 | 0 | 0 | 0 | 0 | 0 | 1 | 0 | 0 | 0 | 0 | 0 | 0 | 0 | 0 | 0 | 0 | 0 | 0 | 0 | 0 | 0 | 0 | 0 | 0 | 0 | 0 | 0 | 0 | 0 | 0 | 0 |
| bacteria_Jul11d75b_44469 | 1 | 0 | 0 | 0 | 0 | 0 | 0 | 0 | 0 | 0 | 0 | 0 | 0 | 0 | 0 | 0 | 0 | 0 | 0 | 0 | 0 | 0 | 0 | 0 | 0 | 0 | 0 | 0 | 0 | 1 | 0 | 0 | 0 | 0 | 0 | 0 | 0 | 0 | 0 | 0 | 0 | 0 | 0 | 0 | 0 | 0 | 0 | 0 | 0 | 0 | 0 | 0 | 0 | 0 | 0 |
| bacteria_Jul11d75b_47026 | 1 | 0 | 0 | 0 | 0 | 0 | 0 | 0 | 0 | 0 | 0 | 0 | 0 | 0 | 0 | 0 | 0 | 0 | 0 | 0 | 0 | 0 | 0 | 0 | 0 | 0 | 0 | 0 | 0 | 1 | 0 | 0 | 0 | 0 | 0 | 0 | 0 | 0 | 0 | 0 | 0 | 0 | 0 | 0 | 0 | 0 | 0 | 0 | 0 | 0 | 0 | 0 | 0 | 0 | 0 |
| bacteria_Jul11d75b_48668 | 1 | 0 | 0 | 0 | 0 | 0 | 0 | 0 | 0 | 0 | 0 | 0 | 0 | 0 | 0 | 0 | 0 | 0 | 0 | 0 | 0 | 0 | 0 | 0 | 0 | 0 | 0 | 0 | 0 | 1 | 0 | 0 | 0 | 0 | 0 | 0 | 0 | 0 | 0 | 0 | 0 | 0 | 0 | 0 | 0 | 0 | 0 | 0 | 0 | 0 | 0 | 0 | 0 | 0 | 0 |
| bacteria_Jul11d75b_51199 | 1 | 0 | 0 | 0 | 0 | 0 | 0 | 0 | 0 | 0 | 0 | 0 | 0 | 0 | 0 | 0 | 0 | 0 | 0 | 0 | 0 | 0 | 0 | 0 | 0 | 0 | 0 | 0 | 0 | 1 | 0 | 0 | 0 | 0 | 0 | 0 | 0 | 0 | 0 | 0 | 0 | 0 | 0 | 0 | 0 | 0 | 0 | 0 | 0 | 0 | 0 | 0 | 0 | 0 | 0 |
| bacteria_Jul27d5b_371 | 1 | 0 | 0 | 0 | 0 | 0 | 0 | 0 | 0 | 0 | 0 | 0 | 0 | 0 | 0 | 0 | 0 | 0 | 0 | 0 | 0 | 0 | 0 | 0 | 0 | 0 | 0 | 0 | 0 | 0 | 0 | 1 | 0 | 0 | 0 | 0 | 0 | 0 | 0 | 0 | 0 | 0 | 0 | 0 | 0 | 0 | 0 | 0 | 0 | 0 | 0 | 0 | 0 | 0 | 0 |
| bacteria_Jul27d5b_3836 | 1 | 0 | 0 | 0 | 0 | 0 | 0 | 0 | 0 | 0 | 0 | 0 | 0 | 0 | 0 | 0 | 0 | 0 | 0 | 0 | 0 | 0 | 0 | 0 | 0 | 0 | 0 | 0 | 0 | 0 | 0 | 1 | 0 | 0 | 0 | 0 | 0 | 0 | 0 | 0 | 0 | 0 | 0 | 0 | 0 | 0 | 0 | 0 | 0 | 0 | 0 | 0 | 0 | 0 | 0 |
| bacteria_Jul27d5b_4533 | 1 | 0 | 0 | 0 | 0 | 0 | 0 | 0 | 0 | 0 | 0 | 0 | 0 | 0 | 0 | 0 | 0 | 0 | 0 | 0 | 0 | 0 | 0 | 0 | 0 | 0 | 0 | 0 | 0 | 0 | 0 | 1 | 0 | 0 | 0 | 0 | 0 | 0 | 0 | 0 | 0 | 0 | 0 | 0 | 0 | 0 | 0 | 0 | 0 | 0 | 0 | 0 | 0 | 0 | 0 |
| bacteria_Jul27d5b_5095 | 1 | 0 | 0 | 0 | 0 | 0 | 0 | 0 | 0 | 0 | 0 | 0 | 0 | 0 | 0 | 0 | 0 | 0 | 0 | 0 | 0 | 0 | 0 | 0 | 0 | 0 | 0 | 0 | 0 | 0 | 0 | 1 | 0 | 0 | 0 | 0 | 0 | 0 | 0 | 0 | 0 | 0 | 0 | 0 | 0 | 0 | 0 | 0 | 0 | 0 | 0 | 0 | 0 | 0 | 0 |
| bacteria_Jul27d5b_8610 | 1 | 0 | 0 | 0 | 0 | 0 | 0 | 0 | 0 | 0 | 0 | 0 | 0 | 0 | 0 | 0 | 0 | 0 | 0 | 0 | 0 | 0 | 0 | 0 | 0 | 0 | 0 | 0 | 0 | 0 | 0 | 1 | 0 | 0 | 0 | 0 | 0 | 0 | 0 | 0 | 0 | 0 | 0 | 0 | 0 | 0 | 0 | 0 | 0 | 0 | 0 | 0 | 0 | 0 | 0 |
| bacteria_Jul27d5b_8809 | 1 | 0 | 0 | 0 | 0 | 0 | 0 | 0 | 0 | 0 | 0 | 0 | 0 | 0 | 0 | 0 | 0 | 0 | 0 | 0 | 0 | 0 | 0 | 0 | 0 | 0 | 0 | 0 | 0 | 0 | 0 | 1 | 0 | 0 | 0 | 0 | 0 | 0 | 0 | 0 | 0 | 0 | 0 | 0 | 0 | 0 | 0 | 0 | 0 | 0 | 0 | 0 | 0 | 0 | 0 |
| bacteria_Jul27d5b_11294 | 1 | 0 | 0 | 0 | 0 | 0 | 0 | 0 | 0 | 0 | 0 | 0 | 0 | 0 | 0 | 0 | 0 | 0 | 0 | 0 | 0 | 0 | 0 | 0 | 0 | 0 | 0 | 0 | 0 | 0 | 0 | 1 | 0 | 0 | 0 | 0 | 0 | 0 | 0 | 0 | 0 | 0 | 0 | 0 | 0 | 0 | 0 | 0 | 0 | 0 | 0 | 0 | 0 | 0 | 0 |
| bacteria_Jul27d5b_14307 | 1 | 0 | 0 | 0 | 0 | 0 | 0 | 0 | 0 | 0 | 0 | 0 | 0 | 0 | 0 | 0 | 0 | 0 | 0 | 0 | 0 | 0 | 0 | 0 | 0 | 0 | 0 | 0 | 0 | 0 | 0 | 1 | 0 | 0 | 0 | 0 | 0 | 0 | 0 | 0 | 0 | 0 | 0 | 0 | 0 | 0 | 0 | 0 | 0 | 0 | 0 | 0 | 0 | 0 | 0 |
| bacteria_Jul27d5b_16666 | 1 | 0 | 0 | 0 | 0 | 0 | 0 | 0 | 0 | 0 | 0 | 0 | 0 | 0 | 0 | 0 | 0 | 0 | 0 | 0 | 0 | 0 | 0 | 0 | 0 | 0 | 0 | 0 | 0 | 0 | 0 | 1 | 0 | 0 | 0 | 0 | 0 | 0 | 0 | 0 | 0 | 0 | 0 | 0 | 0 | 0 | 0 | 0 | 0 | 0 | 0 | 0 | 0 | 0 | 0 |
| bacteria_Jul27d5b_19792 | 1 | 0 | 0 | 0 | 0 | 0 | 0 | 0 | 0 | 0 | 0 | 0 | 0 | 0 | 0 | 0 | 0 | 0 | 0 | 0 | 0 | 0 | 0 | 0 | 0 | 0 | 0 | 0 | 0 | 0 | 0 | 1 | 0 | 0 | 0 | 0 | 0 | 0 | 0 | 0 | 0 | 0 | 0 | 0 | 0 | 0 | 0 | 0 | 0 | 0 | 0 | 0 | 0 | 0 | 0 |
| bacteria_Jul27d5b_20733 | 1 | 0 | 0 | 0 | 0 | 0 | 0 | 0 | 0 | 0 | 0 | 0 | 0 | 0 | 0 | 0 | 0 | 0 | 0 | 0 | 0 | 0 | 0 | 0 | 0 | 0 | 0 | 0 | 0 | 0 | 0 | 1 | 0 | 0 | 0 | 0 | 0 | 0 | 0 | 0 | 0 | 0 | 0 | 0 | 0 | 0 | 0 | 0 | 0 | 0 | 0 | 0 | 0 | 0 | 0 |
| bacteria_Jul27d5b_22807 | 1 | 0 | 0 | 0 | 0 | 0 | 0 | 0 | 0 | 0 | 0 | 0 | 0 | 0 | 0 | 0 | 0 | 0 | 0 | 0 | 0 | 0 | 0 | 0 | 0 | 0 | 0 | 0 | 0 | 0 | 0 | 1 | 0 | 0 | 0 | 0 | 0 | 0 | 0 | 0 | 0 | 0 | 0 | 0 | 0 | 0 | 0 | 0 | 0 | 0 | 0 | 0 | 0 | 0 | 0 |
| bacteria_Jul27d5b_27485 | 1 | 0 | 0 | 0 | 0 | 0 | 0 | 0 | 0 | 0 | 0 | 0 | 0 | 0 | 0 | 0 | 0 | 0 | 0 | 0 | 0 | 0 | 0 | 0 | 0 | 0 | 0 | 0 | 0 | 0 | 0 | 1 | 0 | 0 | 0 | 0 | 0 | 0 | 0 | 0 | 0 | 0 | 0 | 0 | 0 | 0 | 0 | 0 | 0 | 0 | 0 | 0 | 0 | 0 | 0 |
| bacteria_Jul27d5b_30402 | 1 | 0 | 0 | 0 | 0 | 0 | 0 | 0 | 0 | 0 | 0 | 0 | 0 | 0 | 0 | 0 | 0 | 0 | 0 | 0 | 0 | 0 | 0 | 0 | 0 | 0 | 0 | 0 | 0 | 0 | 0 | 1 | 0 | 0 | 0 | 0 | 0 | 0 | 0 | 0 | 0 | 0 | 0 | 0 | 0 | 0 | 0 | 0 | 0 | 0 | 0 | 0 | 0 | 0 | 0 |
| bacteria_Jul27d5b_33079 | 1 | 0 | 0 | 0 | 0 | 0 | 0 | 0 | 0 | 0 | 0 | 0 | 0 | 0 | 0 | 0 | 0 | 0 | 0 | 0 | 0 | 0 | 0 | 0 | 0 | 0 | 0 | 0 | 0 | 0 | 0 | 1 | 0 | 0 | 0 | 0 | 0 | 0 | 0 | 0 | 0 | 0 | 0 | 0 | 0 | 0 | 0 | 0 | 0 | 0 | 0 | 0 | 0 | 0 | 0 |
| bacteria_Jul27d5b_34794 | 1 | 0 | 0 | 0 | 0 | 0 | 0 | 0 | 0 | 0 | 0 | 0 | 0 | 0 | 0 | 0 | 0 | 0 | 0 | 0 | 0 | 0 | 0 | 0 | 0 | 0 | 0 | 0 | 0 | 0 | 0 | 1 | 0 | 0 | 0 | 0 | 0 | 0 | 0 | 0 | 0 | 0 | 0 | 0 | 0 | 0 | 0 | 0 | 0 | 0 | 0 | 0 | 0 | 0 | 0 |
| bacteria_Jul27d5b_34881 | 1 | 0 | 0 | 0 | 0 | 0 | 0 | 0 | 0 | 0 | 0 | 0 | 0 | 0 | 0 | 0 | 0 | 0 | 0 | 0 | 0 | 0 | 0 | 0 | 0 | 0 | 0 | 0 | 0 | 0 | 0 | 1 | 0 | 0 | 0 | 0 | 0 | 0 | 0 | 0 | 0 | 0 | 0 | 0 | 0 | 0 | 0 | 0 | 0 | 0 | 0 | 0 | 0 | 0 | 0 |
| bacteria_Jul27d5b_35273 | 1 | 0 | 0 | 0 | 0 | 0 | 0 | 0 | 0 | 0 | 0 | 0 | 0 | 0 | 0 | 0 | 0 | 0 | 0 | 0 | 0 | 0 | 0 | 0 | 0 | 0 | 0 | 0 | 0 | 0 | 0 | 1 | 0 | 0 | 0 | 0 | 0 | 0 | 0 | 0 | 0 | 0 | 0 | 0 | 0 | 0 | 0 | 0 | 0 | 0 | 0 | 0 | 0 | 0 | 0 |
| bacteria_Jul27d5b_39022 | 1 | 0 | 0 | 0 | 0 | 0 | 0 | 0 | 0 | 0 | 0 | 0 | 0 | 0 | 0 | 0 | 0 | 0 | 0 | 0 | 0 | 0 | 0 | 0 | 0 | 0 | 0 | 0 | 0 | 0 | 0 | 1 | 0 | 0 | 0 | 0 | 0 | 0 | 0 | 0 | 0 | 0 | 0 | 0 | 0 | 0 | 0 | 0 | 0 | 0 | 0 | 0 | 0 | 0 | 0 |
| bacteria_Jul27d5b_39135 | 1 | 0 | 0 | 0 | 0 | 0 | 0 | 0 | 0 | 0 | 0 | 0 | 0 | 0 | 0 | 0 | 0 | 0 | 0 | 0 | 0 | 0 | 0 | 0 | 0 | 0 | 0 | 0 | 0 | 0 | 0 | 1 | 0 | 0 | 0 | 0 | 0 | 0 | 0 | 0 | 0 | 0 | 0 | 0 | 0 | 0 | 0 | 0 | 0 | 0 | 0 | 0 | 0 | 0 | 0 |
| bacteria_Jul27d5b_43532 | 1 | 0 | 0 | 0 | 0 | 0 | 0 | 0 | 0 | 0 | 0 | 0 | 0 | 0 | 0 | 0 | 0 | 0 | 0 | 0 | 0 | 0 | 0 | 0 | 0 | 0 | 0 | 0 | 0 | 0 | 0 | 1 | 0 | 0 | 0 | 0 | 0 | 0 | 0 | 0 | 0 | 0 | 0 | 0 | 0 | 0 | 0 | 0 | 0 | 0 | 0 | 0 | 0 | 0 | 0 |
| bacteria_Jul27d5b_49001 | 1 | 0 | 0 | 0 | 0 | 0 | 0 | 0 | 0 | 0 | 0 | 0 | 0 | 0 | 0 | 0 | 0 | 0 | 0 | 0 | 0 | 0 | 0 | 0 | 0 | 0 | 0 | 0 | 0 | 0 | 0 | 1 | 0 | 0 | 0 | 0 | 0 | 0 | 0 | 0 | 0 | 0 | 0 | 0 | 0 | 0 | 0 | 0 | 0 | 0 | 0 | 0 | 0 | 0 | 0 |
| bacteria_Jul27d5b_50003 | 1 | 0 | 0 | 0 | 0 | 0 | 0 | 0 | 0 | 0 | 0 | 0 | 0 | 0 | 0 | 0 | 0 | 0 | 0 | 0 | 0 | 0 | 0 | 0 | 0 | 0 | 0 | 0 | 0 | 0 | 0 | 1 | 0 | 0 | 0 | 0 | 0 | 0 | 0 | 0 | 0 | 0 | 0 | 0 | 0 | 0 | 0 | 0 | 0 | 0 | 0 | 0 | 0 | 0 | 0 |
| bacteria_Jul27d5b_51746 | 1 | 0 | 0 | 0 | 0 | 0 | 0 | 0 | 0 | 0 | 0 | 0 | 0 | 0 | 0 | 0 | 0 | 0 | 0 | 0 | 0 | 0 | 0 | 0 | 0 | 0 | 0 | 0 | 0 | 0 | 0 | 1 | 0 | 0 | 0 | 0 | 0 | 0 | 0 | 0 | 0 | 0 | 0 | 0 | 0 | 0 | 0 | 0 | 0 | 0 | 0 | 0 | 0 | 0 | 0 |
| bacteria_Jul27d5b_52090 | 1 | 0 | 0 | 0 | 0 | 0 | 0 | 0 | 0 | 0 | 0 | 0 | 0 | 0 | 0 | 0 | 0 | 0 | 0 | 0 | 0 | 0 | 0 | 0 | 0 | 0 | 0 | 0 | 0 | 0 | 0 | 1 | 0 | 0 | 0 | 0 | 0 | 0 | 0 | 0 | 0 | 0 | 0 | 0 | 0 | 0 | 0 | 0 | 0 | 0 | 0 | 0 | 0 | 0 | 0 |
| bacteria_Jul27d25b_1530 | 1 | 0 | 0 | 0 | 0 | 0 | 0 | 0 | 0 | 0 | 0 | 0 | 0 | 0 | 0 | 0 | 0 | 0 | 0 | 0 | 0 | 0 | 0 | 0 | 0 | 0 | 0 | 0 | 0 | 0 | 0 | 0 | 0 | 1 | 0 | 0 | 0 | 0 | 0 | 0 | 0 | 0 | 0 | 0 | 0 | 0 | 0 | 0 | 0 | 0 | 0 | 0 | 0 | 0 | 0 |
| bacteria_Jul27d25b_8485 | 1 | 0 | 0 | 0 | 0 | 0 | 0 | 0 | 0 | 0 | 0 | 0 | 0 | 0 | 0 | 0 | 0 | 0 | 0 | 0 | 0 | 0 | 0 | 0 | 0 | 0 | 0 | 0 | 0 | 0 | 0 | 0 | 0 | 1 | 0 | 0 | 0 | 0 | 0 | 0 | 0 | 0 | 0 | 0 | 0 | 0 | 0 | 0 | 0 | 0 | 0 | 0 | 0 | 0 | 0 |
| bacteria_Jul27d25b_19749 | 1 | 0 | 0 | 0 | 0 | 0 | 0 | 0 | 0 | 0 | 0 | 0 | 0 | 0 | 0 | 0 | 0 | 0 | 0 | 0 | 0 | 0 | 0 | 0 | 0 | 0 | 0 | 0 | 0 | 0 | 0 | 0 | 0 | 1 | 0 | 0 | 0 | 0 | 0 | 0 | 0 | 0 | 0 | 0 | 0 | 0 | 0 | 0 | 0 | 0 | 0 | 0 | 0 | 0 | 0 |
| bacteria_Jul27d25b_25726 | 1 | 0 | 0 | 0 | 0 | 0 | 0 | 0 | 0 | 0 | 0 | 0 | 0 | 0 | 0 | 0 | 0 | 0 | 0 | 0 | 0 | 0 | 0 | 0 | 0 | 0 | 0 | 0 | 0 | 0 | 0 | 0 | 0 | 1 | 0 | 0 | 0 | 0 | 0 | 0 | 0 | 0 | 0 | 0 | 0 | 0 | 0 | 0 | 0 | 0 | 0 | 0 | 0 | 0 | 0 |
| bacteria_Jul27d25b_26121 | 1 | 0 | 0 | 0 | 0 | 0 | 0 | 0 | 0 | 0 | 0 | 0 | 0 | 0 | 0 | 0 | 0 | 0 | 0 | 0 | 0 | 0 | 0 | 0 | 0 | 0 | 0 | 0 | 0 | 0 | 0 | 0 | 0 | 1 | 0 | 0 | 0 | 0 | 0 | 0 | 0 | 0 | 0 | 0 | 0 | 0 | 0 | 0 | 0 | 0 | 0 | 0 | 0 | 0 | 0 |
| bacteria_Jul27d25b_28840 | 1 | 0 | 0 | 0 | 0 | 0 | 0 | 0 | 0 | 0 | 0 | 0 | 0 | 0 | 0 | 0 | 0 | 0 | 0 | 0 | 0 | 0 | 0 | 0 | 0 | 0 | 0 | 0 | 0 | 0 | 0 | 0 | 0 | 1 | 0 | 0 | 0 | 0 | 0 | 0 | 0 | 0 | 0 | 0 | 0 | 0 | 0 | 0 | 0 | 0 | 0 | 0 | 0 | 0 | 0 |
| bacteria_Jul27d25b_29152 | 1 | 0 | 0 | 0 | 0 | 0 | 0 | 0 | 0 | 0 | 0 | 0 | 0 | 0 | 0 | 0 | 0 | 0 | 0 | 0 | 0 | 0 | 0 | 0 | 0 | 0 | 0 | 0 | 0 | 0 | 0 | 0 | 0 | 1 | 0 | 0 | 0 | 0 | 0 | 0 | 0 | 0 | 0 | 0 | 0 | 0 | 0 | 0 | 0 | 0 | 0 | 0 | 0 | 0 | 0 |
| bacteria_Jul27d25b_30149 | 1 | 0 | 0 | 0 | 0 | 0 | 0 | 0 | 0 | 0 | 0 | 0 | 0 | 0 | 0 | 0 | 0 | 0 | 0 | 0 | 0 | 0 | 0 | 0 | 0 | 0 | 0 | 0 | 0 | 0 | 0 | 0 | 0 | 1 | 0 | 0 | 0 | 0 | 0 | 0 | 0 | 0 | 0 | 0 | 0 | 0 | 0 | 0 | 0 | 0 | 0 | 0 | 0 | 0 | 0 |
| bacteria_Jul27d25b_39334 | 1 | 0 | 0 | 0 | 0 | 0 | 0 | 0 | 0 | 0 | 0 | 0 | 0 | 0 | 0 | 0 | 0 | 0 | 0 | 0 | 0 | 0 | 0 | 0 | 0 | 0 | 0 | 0 | 0 | 0 | 0 | 0 | 0 | 1 | 0 | 0 | 0 | 0 | 0 | 0 | 0 | 0 | 0 | 0 | 0 | 0 | 0 | 0 | 0 | 0 | 0 | 0 | 0 | 0 | 0 |
| bacteria_Jul27d25b_40619 | 1 | 0 | 0 | 0 | 0 | 0 | 0 | 0 | 0 | 0 | 0 | 0 | 0 | 0 | 0 | 0 | 0 | 0 | 0 | 0 | 0 | 0 | 0 | 0 | 0 | 0 | 0 | 0 | 0 | 0 | 0 | 0 | 0 | 1 | 0 | 0 | 0 | 0 | 0 | 0 | 0 | 0 | 0 | 0 | 0 | 0 | 0 | 0 | 0 | 0 | 0 | 0 | 0 | 0 | 0 |
| bacteria_Jul27d25b_47389 | 1 | 0 | 0 | 0 | 0 | 0 | 0 | 0 | 0 | 0 | 0 | 0 | 0 | 0 | 0 | 0 | 0 | 0 | 0 | 0 | 0 | 0 | 0 | 0 | 0 | 0 | 0 | 0 | 0 | 0 | 0 | 0 | 0 | 1 | 0 | 0 | 0 | 0 | 0 | 0 | 0 | 0 | 0 | 0 | 0 | 0 | 0 | 0 | 0 | 0 | 0 | 0 | 0 | 0 | 0 |
| bacteria_Jul27d25b_55853 | 1 | 0 | 0 | 0 | 0 | 0 | 0 | 0 | 0 | 0 | 0 | 0 | 0 | 0 | 0 | 0 | 0 | 0 | 0 | 0 | 0 | 0 | 0 | 0 | 0 | 0 | 0 | 0 | 0 | 0 | 0 | 0 | 0 | 1 | 0 | 0 | 0 | 0 | 0 | 0 | 0 | 0 | 0 | 0 | 0 | 0 | 0 | 0 | 0 | 0 | 0 | 0 | 0 | 0 | 0 |
| bacteria_Jul27d25b_57018 | 1 | 0 | 0 | 0 | 0 | 0 | 0 | 0 | 0 | 0 | 0 | 0 | 0 | 0 | 0 | 0 | 0 | 0 | 0 | 0 | 0 | 0 | 0 | 0 | 0 | 0 | 0 | 0 | 0 | 0 | 0 | 0 | 0 | 1 | 0 | 0 | 0 | 0 | 0 | 0 | 0 | 0 | 0 | 0 | 0 | 0 | 0 | 0 | 0 | 0 | 0 | 0 | 0 | 0 | 0 |
| bacteria_Jul27d25b_58438 | 1 | 0 | 0 | 0 | 0 | 0 | 0 | 0 | 0 | 0 | 0 | 0 | 0 | 0 | 0 | 0 | 0 | 0 | 0 | 0 | 0 | 0 | 0 | 0 | 0 | 0 | 0 | 0 | 0 | 0 | 0 | 0 | 0 | 1 | 0 | 0 | 0 | 0 | 0 | 0 | 0 | 0 | 0 | 0 | 0 | 0 | 0 | 0 | 0 | 0 | 0 | 0 | 0 | 0 | 0 |
| bacteria_Jul27d25b_63781 | 1 | 0 | 0 | 0 | 0 | 0 | 0 | 0 | 0 | 0 | 0 | 0 | 0 | 0 | 0 | 0 | 0 | 0 | 0 | 0 | 0 | 0 | 0 | 0 | 0 | 0 | 0 | 0 | 0 | 0 | 0 | 0 | 0 | 1 | 0 | 0 | 0 | 0 | 0 | 0 | 0 | 0 | 0 | 0 | 0 | 0 | 0 | 0 | 0 | 0 | 0 | 0 | 0 | 0 | 0 |
| bacteria_Jul27d25b_71708 | 1 | 0 | 0 | 0 | 0 | 0 | 0 | 0 | 0 | 0 | 0 | 0 | 0 | 0 | 0 | 0 | 0 | 0 | 0 | 0 | 0 | 0 | 0 | 0 | 0 | 0 | 0 | 0 | 0 | 0 | 0 | 0 | 0 | 1 | 0 | 0 | 0 | 0 | 0 | 0 | 0 | 0 | 0 | 0 | 0 | 0 | 0 | 0 | 0 | 0 | 0 | 0 | 0 | 0 | 0 |
| bacteria_Jul27d25b_73132 | 1 | 0 | 0 | 0 | 0 | 0 | 0 | 0 | 0 | 0 | 0 | 0 | 0 | 0 | 0 | 0 | 0 | 0 | 0 | 0 | 0 | 0 | 0 | 0 | 0 | 0 | 0 | 0 | 0 | 0 | 0 | 0 | 0 | 1 | 0 | 0 | 0 | 0 | 0 | 0 | 0 | 0 | 0 | 0 | 0 | 0 | 0 | 0 | 0 | 0 | 0 | 0 | 0 | 0 | 0 |
| bacteria_Jul27d75b_18089 | 1 | 0 | 0 | 0 | 0 | 0 | 0 | 0 | 0 | 0 | 0 | 0 | 0 | 0 | 0 | 0 | 0 | 0 | 0 | 0 | 0 | 0 | 0 | 0 | 0 | 0 | 0 | 0 | 0 | 0 | 0 | 0 | 0 | 0 | 0 | 1 | 0 | 0 | 0 | 0 | 0 | 0 | 0 | 0 | 0 | 0 | 0 | 0 | 0 | 0 | 0 | 0 | 0 | 0 | 0 |
| bacteria_Jul27d75b_21158 | 1 | 0 | 0 | 0 | 0 | 0 | 0 | 0 | 0 | 0 | 0 | 0 | 0 | 0 | 0 | 0 | 0 | 0 | 0 | 0 | 0 | 0 | 0 | 0 | 0 | 0 | 0 | 0 | 0 | 0 | 0 | 0 | 0 | 0 | 0 | 1 | 0 | 0 | 0 | 0 | 0 | 0 | 0 | 0 | 0 | 0 | 0 | 0 | 0 | 0 | 0 | 0 | 0 | 0 | 0 |
| bacteria_Jul27d75b_22650 | 1 | 0 | 0 | 0 | 0 | 0 | 0 | 0 | 0 | 0 | 0 | 0 | 0 | 0 | 0 | 0 | 0 | 0 | 0 | 0 | 0 | 0 | 0 | 0 | 0 | 0 | 0 | 0 | 0 | 0 | 0 | 0 | 0 | 0 | 0 | 1 | 0 | 0 | 0 | 0 | 0 | 0 | 0 | 0 | 0 | 0 | 0 | 0 | 0 | 0 | 0 | 0 | 0 | 0 | 0 |
| bacteria_Jul27d75b_26096 | 1 | 0 | 0 | 0 | 0 | 0 | 0 | 0 | 0 | 0 | 0 | 0 | 0 | 0 | 0 | 0 | 0 | 0 | 0 | 0 | 0 | 0 | 0 | 0 | 0 | 0 | 0 | 0 | 0 | 0 | 0 | 0 | 0 | 0 | 0 | 1 | 0 | 0 | 0 | 0 | 0 | 0 | 0 | 0 | 0 | 0 | 0 | 0 | 0 | 0 | 0 | 0 | 0 | 0 | 0 |
| bacteria_Jul27d75b_31732 | 1 | 0 | 0 | 0 | 0 | 0 | 0 | 0 | 0 | 0 | 0 | 0 | 0 | 0 | 0 | 0 | 0 | 0 | 0 | 0 | 0 | 0 | 0 | 0 | 0 | 0 | 0 | 0 | 0 | 0 | 0 | 0 | 0 | 0 | 0 | 1 | 0 | 0 | 0 | 0 | 0 | 0 | 0 | 0 | 0 | 0 | 0 | 0 | 0 | 0 | 0 | 0 | 0 | 0 | 0 |
| bacteria_Jul27d75b_33579 | 1 | 0 | 0 | 0 | 0 | 0 | 0 | 0 | 0 | 0 | 0 | 0 | 0 | 0 | 0 | 0 | 0 | 0 | 0 | 0 | 0 | 0 | 0 | 0 | 0 | 0 | 0 | 0 | 0 | 0 | 0 | 0 | 0 | 0 | 0 | 1 | 0 | 0 | 0 | 0 | 0 | 0 | 0 | 0 | 0 | 0 | 0 | 0 | 0 | 0 | 0 | 0 | 0 | 0 | 0 |
| bacteria_Jul27d75b_48580 | 1 | 0 | 0 | 0 | 0 | 0 | 0 | 0 | 0 | 0 | 0 | 0 | 0 | 0 | 0 | 0 | 0 | 0 | 0 | 0 | 0 | 0 | 0 | 0 | 0 | 0 | 0 | 0 | 0 | 0 | 0 | 0 | 0 | 0 | 0 | 1 | 0 | 0 | 0 | 0 | 0 | 0 | 0 | 0 | 0 | 0 | 0 | 0 | 0 | 0 | 0 | 0 | 0 | 0 | 0 |
| bacteria_Jul27d75b_53502 | 1 | 0 | 0 | 0 | 0 | 0 | 0 | 0 | 0 | 0 | 0 | 0 | 0 | 0 | 0 | 0 | 0 | 0 | 0 | 0 | 0 | 0 | 0 | 0 | 0 | 0 | 0 | 0 | 0 | 0 | 0 | 0 | 0 | 0 | 0 | 1 | 0 | 0 | 0 | 0 | 0 | 0 | 0 | 0 | 0 | 0 | 0 | 0 | 0 | 0 | 0 | 0 | 0 | 0 | 0 |
| bacteria_Jul27d75b_56542 | 1 | 0 | 0 | 0 | 0 | 0 | 0 | 0 | 0 | 0 | 0 | 0 | 0 | 0 | 0 | 0 | 0 | 0 | 0 | 0 | 0 | 0 | 0 | 0 | 0 | 0 | 0 | 0 | 0 | 0 | 0 | 0 | 0 | 0 | 0 | 1 | 0 | 0 | 0 | 0 | 0 | 0 | 0 | 0 | 0 | 0 | 0 | 0 | 0 | 0 | 0 | 0 | 0 | 0 | 0 |
| bacteria_Jul27d75b_63474 | 1 | 0 | 0 | 0 | 0 | 0 | 0 | 0 | 0 | 0 | 0 | 0 | 0 | 0 | 0 | 0 | 0 | 0 | 0 | 0 | 0 | 0 | 0 | 0 | 0 | 0 | 0 | 0 | 0 | 0 | 0 | 0 | 0 | 0 | 0 | 1 | 0 | 0 | 0 | 0 | 0 | 0 | 0 | 0 | 0 | 0 | 0 | 0 | 0 | 0 | 0 | 0 | 0 | 0 | 0 |
| bacteria_Jul27d75b_67767 | 1 | 0 | 0 | 0 | 0 | 0 | 0 | 0 | 0 | 0 | 0 | 0 | 0 | 0 | 0 | 0 | 0 | 0 | 0 | 0 | 0 | 0 | 0 | 0 | 0 | 0 | 0 | 0 | 0 | 0 | 0 | 0 | 0 | 0 | 0 | 1 | 0 | 0 | 0 | 0 | 0 | 0 | 0 | 0 | 0 | 0 | 0 | 0 | 0 | 0 | 0 | 0 | 0 | 0 | 0 |
| bacteria_Jul27d75b_68680 | 1 | 0 | 0 | 0 | 0 | 0 | 0 | 0 | 0 | 0 | 0 | 0 | 0 | 0 | 0 | 0 | 0 | 0 | 0 | 0 | 0 | 0 | 0 | 0 | 0 | 0 | 0 | 0 | 0 | 0 | 0 | 0 | 0 | 0 | 0 | 1 | 0 | 0 | 0 | 0 | 0 | 0 | 0 | 0 | 0 | 0 | 0 | 0 | 0 | 0 | 0 | 0 | 0 | 0 | 0 |
| bacteria_Jul27d75b_68855 | 1 | 0 | 0 | 0 | 0 | 0 | 0 | 0 | 0 | 0 | 0 | 0 | 0 | 0 | 0 | 0 | 0 | 0 | 0 | 0 | 0 | 0 | 0 | 0 | 0 | 0 | 0 | 0 | 0 | 0 | 0 | 0 | 0 | 0 | 0 | 1 | 0 | 0 | 0 | 0 | 0 | 0 | 0 | 0 | 0 | 0 | 0 | 0 | 0 | 0 | 0 | 0 | 0 | 0 | 0 |
| bacteria_Jul27d75b_72237 | 1 | 0 | 0 | 0 | 0 | 0 | 0 | 0 | 0 | 0 | 0 | 0 | 0 | 0 | 0 | 0 | 0 | 0 | 0 | 0 | 0 | 0 | 0 | 0 | 0 | 0 | 0 | 0 | 0 | 0 | 0 | 0 | 0 | 0 | 0 | 1 | 0 | 0 | 0 | 0 | 0 | 0 | 0 | 0 | 0 | 0 | 0 | 0 | 0 | 0 | 0 | 0 | 0 | 0 | 0 |
| bacteria_Jun8d25b_3103 | 1 | 0 | 0 | 0 | 0 | 0 | 0 | 0 | 0 | 1 | 0 | 0 | 0 | 0 | 0 | 0 | 0 | 0 | 0 | 0 | 0 | 0 | 0 | 0 | 0 | 0 | 0 | 0 | 0 | 0 | 0 | 0 | 0 | 0 | 0 | 0 | 0 | 0 | 0 | 0 | 0 | 0 | 0 | 0 | 0 | 0 | 0 | 0 | 0 | 0 | 0 | 0 | 0 | 0 | 0 |
| bacteria_Jun8d25b_34399 | 1 | 0 | 0 | 0 | 0 | 0 | 0 | 0 | 0 | 1 | 0 | 0 | 0 | 0 | 0 | 0 | 0 | 0 | 0 | 0 | 0 | 0 | 0 | 0 | 0 | 0 | 0 | 0 | 0 | 0 | 0 | 0 | 0 | 0 | 0 | 0 | 0 | 0 | 0 | 0 | 0 | 0 | 0 | 0 | 0 | 0 | 0 | 0 | 0 | 0 | 0 | 0 | 0 | 0 | 0 |
| bacteria_Jun8d25b_37545 | 1 | 0 | 0 | 0 | 0 | 0 | 0 | 0 | 0 | 1 | 0 | 0 | 0 | 0 | 0 | 0 | 0 | 0 | 0 | 0 | 0 | 0 | 0 | 0 | 0 | 0 | 0 | 0 | 0 | 0 | 0 | 0 | 0 | 0 | 0 | 0 | 0 | 0 | 0 | 0 | 0 | 0 | 0 | 0 | 0 | 0 | 0 | 0 | 0 | 0 | 0 | 0 | 0 | 0 | 0 |
| bacteria_Jun8d25b_39385 | 1 | 0 | 0 | 0 | 0 | 0 | 0 | 0 | 0 | 1 | 0 | 0 | 0 | 0 | 0 | 0 | 0 | 0 | 0 | 0 | 0 | 0 | 0 | 0 | 0 | 0 | 0 | 0 | 0 | 0 | 0 | 0 | 0 | 0 | 0 | 0 | 0 | 0 | 0 | 0 | 0 | 0 | 0 | 0 | 0 | 0 | 0 | 0 | 0 | 0 | 0 | 0 | 0 | 0 | 0 |
| bacteria_Jun8d25b_44318 | 1 | 0 | 0 | 0 | 0 | 0 | 0 | 0 | 0 | 1 | 0 | 0 | 0 | 0 | 0 | 0 | 0 | 0 | 0 | 0 | 0 | 0 | 0 | 0 | 0 | 0 | 0 | 0 | 0 | 0 | 0 | 0 | 0 | 0 | 0 | 0 | 0 | 0 | 0 | 0 | 0 | 0 | 0 | 0 | 0 | 0 | 0 | 0 | 0 | 0 | 0 | 0 | 0 | 0 | 0 |
| bacteria_Jun8d25b_52047 | 1 | 0 | 0 | 0 | 0 | 0 | 0 | 0 | 0 | 1 | 0 | 0 | 0 | 0 | 0 | 0 | 0 | 0 | 0 | 0 | 0 | 0 | 0 | 0 | 0 | 0 | 0 | 0 | 0 | 0 | 0 | 0 | 0 | 0 | 0 | 0 | 0 | 0 | 0 | 0 | 0 | 0 | 0 | 0 | 0 | 0 | 0 | 0 | 0 | 0 | 0 | 0 | 0 | 0 | 0 |
| bacteria_Jun8d25b_57777 | 1 | 0 | 0 | 0 | 0 | 0 | 0 | 0 | 0 | 1 | 0 | 0 | 0 | 0 | 0 | 0 | 0 | 0 | 0 | 0 | 0 | 0 | 0 | 0 | 0 | 0 | 0 | 0 | 0 | 0 | 0 | 0 | 0 | 0 | 0 | 0 | 0 | 0 | 0 | 0 | 0 | 0 | 0 | 0 | 0 | 0 | 0 | 0 | 0 | 0 | 0 | 0 | 0 | 0 | 0 |
| bacteria_Jun8d25b_62757 | 1 | 0 | 0 | 0 | 0 | 0 | 0 | 0 | 0 | 1 | 0 | 0 | 0 | 0 | 0 | 0 | 0 | 0 | 0 | 0 | 0 | 0 | 0 | 0 | 0 | 0 | 0 | 0 | 0 | 0 | 0 | 0 | 0 | 0 | 0 | 0 | 0 | 0 | 0 | 0 | 0 | 0 | 0 | 0 | 0 | 0 | 0 | 0 | 0 | 0 | 0 | 0 | 0 | 0 | 0 |
| bacteria_Jun8d25b_71380 | 1 | 0 | 0 | 0 | 0 | 0 | 0 | 0 | 0 | 1 | 0 | 0 | 0 | 0 | 0 | 0 | 0 | 0 | 0 | 0 | 0 | 0 | 0 | 0 | 0 | 0 | 0 | 0 | 0 | 0 | 0 | 0 | 0 | 0 | 0 | 0 | 0 | 0 | 0 | 0 | 0 | 0 | 0 | 0 | 0 | 0 | 0 | 0 | 0 | 0 | 0 | 0 | 0 | 0 | 0 |
| bacteria_Jun8d50b_341 | 1 | 0 | 0 | 0 | 0 | 0 | 0 | 0 | 0 | 0 | 0 | 1 | 0 | 0 | 0 | 0 | 0 | 0 | 0 | 0 | 0 | 0 | 0 | 0 | 0 | 0 | 0 | 0 | 0 | 0 | 0 | 0 | 0 | 0 | 0 | 0 | 0 | 0 | 0 | 0 | 0 | 0 | 0 | 0 | 0 | 0 | 0 | 0 | 0 | 0 | 0 | 0 | 0 | 0 | 0 |
| bacteria_Jun8d50b_4841 | 1 | 0 | 0 | 0 | 0 | 0 | 0 | 0 | 0 | 0 | 0 | 1 | 0 | 0 | 0 | 0 | 0 | 0 | 0 | 0 | 0 | 0 | 0 | 0 | 0 | 0 | 0 | 0 | 0 | 0 | 0 | 0 | 0 | 0 | 0 | 0 | 0 | 0 | 0 | 0 | 0 | 0 | 0 | 0 | 0 | 0 | 0 | 0 | 0 | 0 | 0 | 0 | 0 | 0 | 0 |
| bacteria_Jun8d50b_24396 | 1 | 0 | 0 | 0 | 0 | 0 | 0 | 0 | 0 | 0 | 0 | 1 | 0 | 0 | 0 | 0 | 0 | 0 | 0 | 0 | 0 | 0 | 0 | 0 | 0 | 0 | 0 | 0 | 0 | 0 | 0 | 0 | 0 | 0 | 0 | 0 | 0 | 0 | 0 | 0 | 0 | 0 | 0 | 0 | 0 | 0 | 0 | 0 | 0 | 0 | 0 | 0 | 0 | 0 | 0 |
| bacteria_Jun8d50b_36238 | 1 | 0 | 0 | 0 | 0 | 0 | 0 | 0 | 0 | 0 | 0 | 1 | 0 | 0 | 0 | 0 | 0 | 0 | 0 | 0 | 0 | 0 | 0 | 0 | 0 | 0 | 0 | 0 | 0 | 0 | 0 | 0 | 0 | 0 | 0 | 0 | 0 | 0 | 0 | 0 | 0 | 0 | 0 | 0 | 0 | 0 | 0 | 0 | 0 | 0 | 0 | 0 | 0 | 0 | 0 |
| bacteria_Jun8d50b_45077 | 1 | 0 | 0 | 0 | 0 | 0 | 0 | 0 | 0 | 0 | 0 | 1 | 0 | 0 | 0 | 0 | 0 | 0 | 0 | 0 | 0 | 0 | 0 | 0 | 0 | 0 | 0 | 0 | 0 | 0 | 0 | 0 | 0 | 0 | 0 | 0 | 0 | 0 | 0 | 0 | 0 | 0 | 0 | 0 | 0 | 0 | 0 | 0 | 0 | 0 | 0 | 0 | 0 | 0 | 0 |
| bacteria_Jun8d50b_46634 | 1 | 0 | 0 | 0 | 0 | 0 | 0 | 0 | 0 | 0 | 0 | 1 | 0 | 0 | 0 | 0 | 0 | 0 | 0 | 0 | 0 | 0 | 0 | 0 | 0 | 0 | 0 | 0 | 0 | 0 | 0 | 0 | 0 | 0 | 0 | 0 | 0 | 0 | 0 | 0 | 0 | 0 | 0 | 0 | 0 | 0 | 0 | 0 | 0 | 0 | 0 | 0 | 0 | 0 | 0 |
| bacteria_Jun8d75b_24040 | 1 | 0 | 0 | 0 | 0 | 0 | 0 | 0 | 0 | 0 | 0 | 0 | 0 | 1 | 0 | 0 | 0 | 0 | 0 | 0 | 0 | 0 | 0 | 0 | 0 | 0 | 0 | 0 | 0 | 0 | 0 | 0 | 0 | 0 | 0 | 0 | 0 | 0 | 0 | 0 | 0 | 0 | 0 | 0 | 0 | 0 | 0 | 0 | 0 | 0 | 0 | 0 | 0 | 0 | 0 |
| bacteria_Jun22d0b_1872 | 1 | 0 | 0 | 0 | 0 | 0 | 0 | 0 | 0 | 0 | 0 | 0 | 0 | 0 | 0 | 1 | 0 | 0 | 0 | 0 | 0 | 0 | 0 | 0 | 0 | 0 | 0 | 0 | 0 | 0 | 0 | 0 | 0 | 0 | 0 | 0 | 0 | 0 | 0 | 0 | 0 | 0 | 0 | 0 | 0 | 0 | 0 | 0 | 0 | 0 | 0 | 0 | 0 | 0 | 0 |
| bacteria_Jun22d0b_13561 | 1 | 0 | 0 | 0 | 0 | 0 | 0 | 0 | 0 | 0 | 0 | 0 | 0 | 0 | 0 | 1 | 0 | 0 | 0 | 0 | 0 | 0 | 0 | 0 | 0 | 0 | 0 | 0 | 0 | 0 | 0 | 0 | 0 | 0 | 0 | 0 | 0 | 0 | 0 | 0 | 0 | 0 | 0 | 0 | 0 | 0 | 0 | 0 | 0 | 0 | 0 | 0 | 0 | 0 | 0 |
| bacteria_Jun22d50b_1117 | 1 | 0 | 0 | 0 | 0 | 0 | 0 | 0 | 0 | 0 | 0 | 0 | 0 | 0 | 0 | 0 | 0 | 1 | 0 | 0 | 0 | 0 | 0 | 0 | 0 | 0 | 0 | 0 | 0 | 0 | 0 | 0 | 0 | 0 | 0 | 0 | 0 | 0 | 0 | 0 | 0 | 0 | 0 | 0 | 0 | 0 | 0 | 0 | 0 | 0 | 0 | 0 | 0 | 0 | 0 |
| bacteria_Jun22d50b_3703 | 1 | 0 | 0 | 0 | 0 | 0 | 0 | 0 | 0 | 0 | 0 | 0 | 0 | 0 | 0 | 0 | 0 | 1 | 0 | 0 | 0 | 0 | 0 | 0 | 0 | 0 | 0 | 0 | 0 | 0 | 0 | 0 | 0 | 0 | 0 | 0 | 0 | 0 | 0 | 0 | 0 | 0 | 0 | 0 | 0 | 0 | 0 | 0 | 0 | 0 | 0 | 0 | 0 | 0 | 0 |
| bacteria_Jun22d50b_4484 | 1 | 0 | 0 | 0 | 0 | 0 | 0 | 0 | 0 | 0 | 0 | 0 | 0 | 0 | 0 | 0 | 0 | 1 | 0 | 0 | 0 | 0 | 0 | 0 | 0 | 0 | 0 | 0 | 0 | 0 | 0 | 0 | 0 | 0 | 0 | 0 | 0 | 0 | 0 | 0 | 0 | 0 | 0 | 0 | 0 | 0 | 0 | 0 | 0 | 0 | 0 | 0 | 0 | 0 | 0 |
| bacteria_Jun22d75b_1705 | 1 | 0 | 0 | 0 | 0 | 0 | 0 | 0 | 0 | 0 | 0 | 0 | 0 | 0 | 0 | 0 | 0 | 0 | 0 | 1 | 0 | 0 | 0 | 0 | 0 | 0 | 0 | 0 | 0 | 0 | 0 | 0 | 0 | 0 | 0 | 0 | 0 | 0 | 0 | 0 | 0 | 0 | 0 | 0 | 0 | 0 | 0 | 0 | 0 | 0 | 0 | 0 | 0 | 0 | 0 |
| bacteria_Jun22d75b_4105 | 1 | 0 | 0 | 0 | 0 | 0 | 0 | 0 | 0 | 0 | 0 | 0 | 0 | 0 | 0 | 0 | 0 | 0 | 0 | 1 | 0 | 0 | 0 | 0 | 0 | 0 | 0 | 0 | 0 | 0 | 0 | 0 | 0 | 0 | 0 | 0 | 0 | 0 | 0 | 0 | 0 | 0 | 0 | 0 | 0 | 0 | 0 | 0 | 0 | 0 | 0 | 0 | 0 | 0 | 0 |
| bacteria_Jun22d75b_6146 | 1 | 0 | 0 | 0 | 0 | 0 | 0 | 0 | 0 | 0 | 0 | 0 | 0 | 0 | 0 | 0 | 0 | 0 | 0 | 1 | 0 | 0 | 0 | 0 | 0 | 0 | 0 | 0 | 0 | 0 | 0 | 0 | 0 | 0 | 0 | 0 | 0 | 0 | 0 | 0 | 0 | 0 | 0 | 0 | 0 | 0 | 0 | 0 | 0 | 0 | 0 | 0 | 0 | 0 | 0 |
| bacteria_Jun22d75b_6543 | 1 | 0 | 0 | 0 | 0 | 0 | 0 | 0 | 0 | 0 | 0 | 0 | 0 | 0 | 0 | 0 | 0 | 0 | 0 | 1 | 0 | 0 | 0 | 0 | 0 | 0 | 0 | 0 | 0 | 0 | 0 | 0 | 0 | 0 | 0 | 0 | 0 | 0 | 0 | 0 | 0 | 0 | 0 | 0 | 0 | 0 | 0 | 0 | 0 | 0 | 0 | 0 | 0 | 0 | 0 |
| bacteria_Jun22d75b_7289 | 1 | 0 | 0 | 0 | 0 | 0 | 0 | 0 | 0 | 0 | 0 | 0 | 0 | 0 | 0 | 0 | 0 | 0 | 0 | 1 | 0 | 0 | 0 | 0 | 0 | 0 | 0 | 0 | 0 | 0 | 0 | 0 | 0 | 0 | 0 | 0 | 0 | 0 | 0 | 0 | 0 | 0 | 0 | 0 | 0 | 0 | 0 | 0 | 0 | 0 | 0 | 0 | 0 | 0 | 0 |
| bacteria_May12d25b_997 | 1 | 0 | 0 | 1 | 0 | 0 | 0 | 0 | 0 | 0 | 0 | 0 | 0 | 0 | 0 | 0 | 0 | 0 | 0 | 0 | 0 | 0 | 0 | 0 | 0 | 0 | 0 | 0 | 0 | 0 | 0 | 0 | 0 | 0 | 0 | 0 | 0 | 0 | 0 | 0 | 0 | 0 | 0 | 0 | 0 | 0 | 0 | 0 | 0 | 0 | 0 | 0 | 0 | 0 | 0 |
| bacteria_May12d25b_131 | 1 | 0 | 0 | 1 | 0 | 0 | 0 | 0 | 0 | 0 | 0 | 0 | 0 | 0 | 0 | 0 | 0 | 0 | 0 | 0 | 0 | 0 | 0 | 0 | 0 | 0 | 0 | 0 | 0 | 0 | 0 | 0 | 0 | 0 | 0 | 0 | 0 | 0 | 0 | 0 | 0 | 0 | 0 | 0 | 0 | 0 | 0 | 0 | 0 | 0 | 0 | 0 | 0 | 0 | 0 |
| bacteria_May12d25b_131 | 1 | 0 | 0 | 1 | 0 | 0 | 0 | 0 | 0 | 0 | 0 | 0 | 0 | 0 | 0 | 0 | 0 | 0 | 0 | 0 | 0 | 0 | 0 | 0 | 0 | 0 | 0 | 0 | 0 | 0 | 0 | 0 | 0 | 0 | 0 | 0 | 0 | 0 | 0 | 0 | 0 | 0 | 0 | 0 | 0 | 0 | 0 | 0 | 0 | 0 | 0 | 0 | 0 | 0 | 0 |
| bacteria_May12d25b_432 | 1 | 0 | 0 | 1 | 0 | 0 | 0 | 0 | 0 | 0 | 0 | 0 | 0 | 0 | 0 | 0 | 0 | 0 | 0 | 0 | 0 | 0 | 0 | 0 | 0 | 0 | 0 | 0 | 0 | 0 | 0 | 0 | 0 | 0 | 0 | 0 | 0 | 0 | 0 | 0 | 0 | 0 | 0 | 0 | 0 | 0 | 0 | 0 | 0 | 0 | 0 | 0 | 0 | 0 | 0 |
| bacteria_May12d25b_541 | 1 | 0 | 0 | 1 | 0 | 0 | 0 | 0 | 0 | 0 | 0 | 0 | 0 | 0 | 0 | 0 | 0 | 0 | 0 | 0 | 0 | 0 | 0 | 0 | 0 | 0 | 0 | 0 | 0 | 0 | 0 | 0 | 0 | 0 | 0 | 0 | 0 | 0 | 0 | 0 | 0 | 0 | 0 | 0 | 0 | 0 | 0 | 0 | 0 | 0 | 0 | 0 | 0 | 0 | 0 |
| bacteria_May12d25b_676 | 1 | 0 | 0 | 1 | 0 | 0 | 0 | 0 | 0 | 0 | 0 | 0 | 0 | 0 | 0 | 0 | 0 | 0 | 0 | 0 | 0 | 0 | 0 | 0 | 0 | 0 | 0 | 0 | 0 | 0 | 0 | 0 | 0 | 0 | 0 | 0 | 0 | 0 | 0 | 0 | 0 | 0 | 0 | 0 | 0 | 0 | 0 | 0 | 0 | 0 | 0 | 0 | 0 | 0 | 0 |
| bacteria_May12d50b_505 | 1 | 0 | 0 | 0 | 0 | 1 | 0 | 0 | 0 | 0 | 0 | 0 | 0 | 0 | 0 | 0 | 0 | 0 | 0 | 0 | 0 | 0 | 0 | 0 | 0 | 0 | 0 | 0 | 0 | 0 | 0 | 0 | 0 | 0 | 0 | 0 | 0 | 0 | 0 | 0 | 0 | 0 | 0 | 0 | 0 | 0 | 0 | 0 | 0 | 0 | 0 | 0 | 0 | 0 | 0 |
| bacteria_May12d50b_843 | 1 | 0 | 0 | 0 | 0 | 1 | 0 | 0 | 0 | 0 | 0 | 0 | 0 | 0 | 0 | 0 | 0 | 0 | 0 | 0 | 0 | 0 | 0 | 0 | 0 | 0 | 0 | 0 | 0 | 0 | 0 | 0 | 0 | 0 | 0 | 0 | 0 | 0 | 0 | 0 | 0 | 0 | 0 | 0 | 0 | 0 | 0 | 0 | 0 | 0 | 0 | 0 | 0 | 0 | 0 |
| bacteria_May12d50b_206 | 1 | 0 | 0 | 0 | 0 | 1 | 0 | 0 | 0 | 0 | 0 | 0 | 0 | 0 | 0 | 0 | 0 | 0 | 0 | 0 | 0 | 0 | 0 | 0 | 0 | 0 | 0 | 0 | 0 | 0 | 0 | 0 | 0 | 0 | 0 | 0 | 0 | 0 | 0 | 0 | 0 | 0 | 0 | 0 | 0 | 0 | 0 | 0 | 0 | 0 | 0 | 0 | 0 | 0 | 0 |
| bacteria_May12d50b_272 | 1 | 0 | 0 | 0 | 0 | 1 | 0 | 0 | 0 | 0 | 0 | 0 | 0 | 0 | 0 | 0 | 0 | 0 | 0 | 0 | 0 | 0 | 0 | 0 | 0 | 0 | 0 | 0 | 0 | 0 | 0 | 0 | 0 | 0 | 0 | 0 | 0 | 0 | 0 | 0 | 0 | 0 | 0 | 0 | 0 | 0 | 0 | 0 | 0 | 0 | 0 | 0 | 0 | 0 | 0 |
| bacteria_May12d50b_444 | 1 | 0 | 0 | 0 | 0 | 1 | 0 | 0 | 0 | 0 | 0 | 0 | 0 | 0 | 0 | 0 | 0 | 0 | 0 | 0 | 0 | 0 | 0 | 0 | 0 | 0 | 0 | 0 | 0 | 0 | 0 | 0 | 0 | 0 | 0 | 0 | 0 | 0 | 0 | 0 | 0 | 0 | 0 | 0 | 0 | 0 | 0 | 0 | 0 | 0 | 0 | 0 | 0 | 0 | 0 |
| bacteria_May12d50b_447 | 1 | 0 | 0 | 0 | 0 | 1 | 0 | 0 | 0 | 0 | 0 | 0 | 0 | 0 | 0 | 0 | 0 | 0 | 0 | 0 | 0 | 0 | 0 | 0 | 0 | 0 | 0 | 0 | 0 | 0 | 0 | 0 | 0 | 0 | 0 | 0 | 0 | 0 | 0 | 0 | 0 | 0 | 0 | 0 | 0 | 0 | 0 | 0 | 0 | 0 | 0 | 0 | 0 | 0 | 0 |
| bacteria_May12d50b_464 | 1 | 0 | 0 | 0 | 0 | 1 | 0 | 0 | 0 | 0 | 0 | 0 | 0 | 0 | 0 | 0 | 0 | 0 | 0 | 0 | 0 | 0 | 0 | 0 | 0 | 0 | 0 | 0 | 0 | 0 | 0 | 0 | 0 | 0 | 0 | 0 | 0 | 0 | 0 | 0 | 0 | 0 | 0 | 0 | 0 | 0 | 0 | 0 | 0 | 0 | 0 | 0 | 0 | 0 | 0 |
| bacteria_May12d50b_489 | 1 | 0 | 0 | 0 | 0 | 1 | 0 | 0 | 0 | 0 | 0 | 0 | 0 | 0 | 0 | 0 | 0 | 0 | 0 | 0 | 0 | 0 | 0 | 0 | 0 | 0 | 0 | 0 | 0 | 0 | 0 | 0 | 0 | 0 | 0 | 0 | 0 | 0 | 0 | 0 | 0 | 0 | 0 | 0 | 0 | 0 | 0 | 0 | 0 | 0 | 0 | 0 | 0 | 0 | 0 |
| bacteria_May12d50b_500 | 1 | 0 | 0 | 0 | 0 | 1 | 0 | 0 | 0 | 0 | 0 | 0 | 0 | 0 | 0 | 0 | 0 | 0 | 0 | 0 | 0 | 0 | 0 | 0 | 0 | 0 | 0 | 0 | 0 | 0 | 0 | 0 | 0 | 0 | 0 | 0 | 0 | 0 | 0 | 0 | 0 | 0 | 0 | 0 | 0 | 0 | 0 | 0 | 0 | 0 | 0 | 0 | 0 | 0 | 0 |
| bacteria_May12d50b_510 | 1 | 0 | 0 | 0 | 0 | 1 | 0 | 0 | 0 | 0 | 0 | 0 | 0 | 0 | 0 | 0 | 0 | 0 | 0 | 0 | 0 | 0 | 0 | 0 | 0 | 0 | 0 | 0 | 0 | 0 | 0 | 0 | 0 | 0 | 0 | 0 | 0 | 0 | 0 | 0 | 0 | 0 | 0 | 0 | 0 | 0 | 0 | 0 | 0 | 0 | 0 | 0 | 0 | 0 | 0 |
| bacteria_Sep8d5b_2772 | 1 | 0 | 0 | 0 | 0 | 0 | 0 | 0 | 0 | 0 | 0 | 0 | 0 | 0 | 0 | 0 | 0 | 0 | 0 | 0 | 0 | 0 | 0 | 0 | 0 | 0 | 0 | 0 | 0 | 0 | 0 | 0 | 0 | 0 | 0 | 0 | 0 | 0 | 0 | 0 | 0 | 0 | 0 | 0 | 0 | 0 | 0 | 0 | 0 | 1 | 0 | 0 | 0 | 0 | 0 |
| bacteria_Sep8d5b_4362 | 1 | 0 | 0 | 0 | 0 | 0 | 0 | 0 | 0 | 0 | 0 | 0 | 0 | 0 | 0 | 0 | 0 | 0 | 0 | 0 | 0 | 0 | 0 | 0 | 0 | 0 | 0 | 0 | 0 | 0 | 0 | 0 | 0 | 0 | 0 | 0 | 0 | 0 | 0 | 0 | 0 | 0 | 0 | 0 | 0 | 0 | 0 | 0 | 0 | 1 | 0 | 0 | 0 | 0 | 0 |
| bacteria_Sep8d5b_4996 | 1 | 0 | 0 | 0 | 0 | 0 | 0 | 0 | 0 | 0 | 0 | 0 | 0 | 0 | 0 | 0 | 0 | 0 | 0 | 0 | 0 | 0 | 0 | 0 | 0 | 0 | 0 | 0 | 0 | 0 | 0 | 0 | 0 | 0 | 0 | 0 | 0 | 0 | 0 | 0 | 0 | 0 | 0 | 0 | 0 | 0 | 0 | 0 | 0 | 1 | 0 | 0 | 0 | 0 | 0 |
| bacteria_Sep8d5b_5046 | 1 | 0 | 0 | 0 | 0 | 0 | 0 | 0 | 0 | 0 | 0 | 0 | 0 | 0 | 0 | 0 | 0 | 0 | 0 | 0 | 0 | 0 | 0 | 0 | 0 | 0 | 0 | 0 | 0 | 0 | 0 | 0 | 0 | 0 | 0 | 0 | 0 | 0 | 0 | 0 | 0 | 0 | 0 | 0 | 0 | 0 | 0 | 0 | 0 | 1 | 0 | 0 | 0 | 0 | 0 |
| bacteria_Sep8d5b_5518 | 1 | 0 | 0 | 0 | 0 | 0 | 0 | 0 | 0 | 0 | 0 | 0 | 0 | 0 | 0 | 0 | 0 | 0 | 0 | 0 | 0 | 0 | 0 | 0 | 0 | 0 | 0 | 0 | 0 | 0 | 0 | 0 | 0 | 0 | 0 | 0 | 0 | 0 | 0 | 0 | 0 | 0 | 0 | 0 | 0 | 0 | 0 | 0 | 0 | 1 | 0 | 0 | 0 | 0 | 0 |
| bacteria_Sep8d5b_10019 | 1 | 0 | 0 | 0 | 0 | 0 | 0 | 0 | 0 | 0 | 0 | 0 | 0 | 0 | 0 | 0 | 0 | 0 | 0 | 0 | 0 | 0 | 0 | 0 | 0 | 0 | 0 | 0 | 0 | 0 | 0 | 0 | 0 | 0 | 0 | 0 | 0 | 0 | 0 | 0 | 0 | 0 | 0 | 0 | 0 | 0 | 0 | 0 | 0 | 1 | 0 | 0 | 0 | 0 | 0 |
| bacteria_Sep8d5b_10717 | 1 | 0 | 0 | 0 | 0 | 0 | 0 | 0 | 0 | 0 | 0 | 0 | 0 | 0 | 0 | 0 | 0 | 0 | 0 | 0 | 0 | 0 | 0 | 0 | 0 | 0 | 0 | 0 | 0 | 0 | 0 | 0 | 0 | 0 | 0 | 0 | 0 | 0 | 0 | 0 | 0 | 0 | 0 | 0 | 0 | 0 | 0 | 0 | 0 | 1 | 0 | 0 | 0 | 0 | 0 |
| bacteria_Sep8d5b_12119 | 1 | 0 | 0 | 0 | 0 | 0 | 0 | 0 | 0 | 0 | 0 | 0 | 0 | 0 | 0 | 0 | 0 | 0 | 0 | 0 | 0 | 0 | 0 | 0 | 0 | 0 | 0 | 0 | 0 | 0 | 0 | 0 | 0 | 0 | 0 | 0 | 0 | 0 | 0 | 0 | 0 | 0 | 0 | 0 | 0 | 0 | 0 | 0 | 0 | 1 | 0 | 0 | 0 | 0 | 0 |
| bacteria_Sep8d5b_18164 | 1 | 0 | 0 | 0 | 0 | 0 | 0 | 0 | 0 | 0 | 0 | 0 | 0 | 0 | 0 | 0 | 0 | 0 | 0 | 0 | 0 | 0 | 0 | 0 | 0 | 0 | 0 | 0 | 0 | 0 | 0 | 0 | 0 | 0 | 0 | 0 | 0 | 0 | 0 | 0 | 0 | 0 | 0 | 0 | 0 | 0 | 0 | 0 | 0 | 1 | 0 | 0 | 0 | 0 | 0 |
| bacteria_Sep8d5b_19593 | 1 | 0 | 0 | 0 | 0 | 0 | 0 | 0 | 0 | 0 | 0 | 0 | 0 | 0 | 0 | 0 | 0 | 0 | 0 | 0 | 0 | 0 | 0 | 0 | 0 | 0 | 0 | 0 | 0 | 0 | 0 | 0 | 0 | 0 | 0 | 0 | 0 | 0 | 0 | 0 | 0 | 0 | 0 | 0 | 0 | 0 | 0 | 0 | 0 | 1 | 0 | 0 | 0 | 0 | 0 |
| bacteria_Sep8d5b_24411 | 1 | 0 | 0 | 0 | 0 | 0 | 0 | 0 | 0 | 0 | 0 | 0 | 0 | 0 | 0 | 0 | 0 | 0 | 0 | 0 | 0 | 0 | 0 | 0 | 0 | 0 | 0 | 0 | 0 | 0 | 0 | 0 | 0 | 0 | 0 | 0 | 0 | 0 | 0 | 0 | 0 | 0 | 0 | 0 | 0 | 0 | 0 | 0 | 0 | 1 | 0 | 0 | 0 | 0 | 0 |
| bacteria_Sep8d5b_28614 | 1 | 0 | 0 | 0 | 0 | 0 | 0 | 0 | 0 | 0 | 0 | 0 | 0 | 0 | 0 | 0 | 0 | 0 | 0 | 0 | 0 | 0 | 0 | 0 | 0 | 0 | 0 | 0 | 0 | 0 | 0 | 0 | 0 | 0 | 0 | 0 | 0 | 0 | 0 | 0 | 0 | 0 | 0 | 0 | 0 | 0 | 0 | 0 | 0 | 1 | 0 | 0 | 0 | 0 | 0 |
| bacteria_Sep8d5b_30494 | 1 | 0 | 0 | 0 | 0 | 0 | 0 | 0 | 0 | 0 | 0 | 0 | 0 | 0 | 0 | 0 | 0 | 0 | 0 | 0 | 0 | 0 | 0 | 0 | 0 | 0 | 0 | 0 | 0 | 0 | 0 | 0 | 0 | 0 | 0 | 0 | 0 | 0 | 0 | 0 | 0 | 0 | 0 | 0 | 0 | 0 | 0 | 0 | 0 | 1 | 0 | 0 | 0 | 0 | 0 |
| bacteria_Sep8d5b_62465 | 1 | 0 | 0 | 0 | 0 | 0 | 0 | 0 | 0 | 0 | 0 | 0 | 0 | 0 | 0 | 0 | 0 | 0 | 0 | 0 | 0 | 0 | 0 | 0 | 0 | 0 | 0 | 0 | 0 | 0 | 0 | 0 | 0 | 0 | 0 | 0 | 0 | 0 | 0 | 0 | 0 | 0 | 0 | 0 | 0 | 0 | 0 | 0 | 0 | 1 | 0 | 0 | 0 | 0 | 0 |
| bacteria_Sep8d5b_64306 | 1 | 0 | 0 | 0 | 0 | 0 | 0 | 0 | 0 | 0 | 0 | 0 | 0 | 0 | 0 | 0 | 0 | 0 | 0 | 0 | 0 | 0 | 0 | 0 | 0 | 0 | 0 | 0 | 0 | 0 | 0 | 0 | 0 | 0 | 0 | 0 | 0 | 0 | 0 | 0 | 0 | 0 | 0 | 0 | 0 | 0 | 0 | 0 | 0 | 1 | 0 | 0 | 0 | 0 | 0 |
| bacteria_Sep8d25b_35008 | 1 | 0 | 0 | 0 | 0 | 0 | 0 | 0 | 0 | 0 | 0 | 0 | 0 | 0 | 0 | 0 | 0 | 0 | 0 | 0 | 0 | 0 | 0 | 0 | 0 | 0 | 0 | 0 | 0 | 0 | 0 | 0 | 0 | 0 | 0 | 0 | 0 | 0 | 0 | 0 | 0 | 0 | 0 | 0 | 0 | 0 | 0 | 0 | 0 | 0 | 0 | 1 | 0 | 0 | 0 |
| bacteria_Sep8d25b_45121 | 1 | 0 | 0 | 0 | 0 | 0 | 0 | 0 | 0 | 0 | 0 | 0 | 0 | 0 | 0 | 0 | 0 | 0 | 0 | 0 | 0 | 0 | 0 | 0 | 0 | 0 | 0 | 0 | 0 | 0 | 0 | 0 | 0 | 0 | 0 | 0 | 0 | 0 | 0 | 0 | 0 | 0 | 0 | 0 | 0 | 0 | 0 | 0 | 0 | 0 | 0 | 1 | 0 | 0 | 0 |
| bacteria_Sep8d50b_5207 | 1 | 0 | 0 | 0 | 0 | 0 | 0 | 0 | 0 | 0 | 0 | 0 | 0 | 0 | 0 | 0 | 0 | 0 | 0 | 0 | 0 | 0 | 0 | 0 | 0 | 0 | 0 | 0 | 0 | 0 | 0 | 0 | 0 | 0 | 0 | 0 | 0 | 0 | 0 | 0 | 0 | 0 | 0 | 0 | 0 | 0 | 0 | 0 | 0 | 0 | 0 | 0 | 0 | 1 | 0 |
| bacteria_Sep8d50b_51343 | 1 | 0 | 0 | 0 | 0 | 0 | 0 | 0 | 0 | 0 | 0 | 0 | 0 | 0 | 0 | 0 | 0 | 0 | 0 | 0 | 0 | 0 | 0 | 0 | 0 | 0 | 0 | 0 | 0 | 0 | 0 | 0 | 0 | 0 | 0 | 0 | 0 | 0 | 0 | 0 | 0 | 0 | 0 | 0 | 0 | 0 | 0 | 0 | 0 | 0 | 0 | 0 | 0 | 1 | 0 |
| virus_Aug5.0v_393 | 1 | 0 | 0 | 0 | 0 | 0 | 0 | 0 | 0 | 0 | 0 | 0 | 0 | 0 | 0 | 0 | 0 | 0 | 0 | 0 | 0 | 0 | 0 | 0 | 0 | 0 | 0 | 0 | 0 | 0 | 0 | 0 | 0 | 0 | 0 | 0 | 0 | 0 | 1 | 0 | 0 | 0 | 0 | 0 | 0 | 0 | 0 | 0 | 0 | 0 | 0 | 0 | 0 | 0 | 0 |
| virus_Aug5.0v_397 | 1 | 0 | 0 | 0 | 0 | 0 | 0 | 0 | 0 | 0 | 0 | 0 | 0 | 0 | 0 | 0 | 0 | 0 | 0 | 0 | 0 | 0 | 0 | 0 | 0 | 0 | 0 | 0 | 0 | 0 | 0 | 0 | 0 | 0 | 0 | 0 | 0 | 0 | 1 | 0 | 0 | 0 | 0 | 0 | 0 | 0 | 0 | 0 | 0 | 0 | 0 | 0 | 0 | 0 | 0 |
| virus_Aug5.0v_705 | 1 | 0 | 0 | 0 | 0 | 0 | 0 | 0 | 0 | 0 | 0 | 0 | 0 | 0 | 0 | 0 | 0 | 0 | 0 | 0 | 0 | 0 | 0 | 0 | 0 | 0 | 0 | 0 | 0 | 0 | 0 | 0 | 0 | 0 | 0 | 0 | 0 | 0 | 1 | 0 | 0 | 0 | 0 | 0 | 0 | 0 | 0 | 0 | 0 | 0 | 0 | 0 | 0 | 0 | 0 |
| virus_Aug5.0v_722 | 1 | 0 | 0 | 0 | 0 | 0 | 0 | 0 | 0 | 0 | 0 | 0 | 0 | 0 | 0 | 0 | 0 | 0 | 0 | 0 | 0 | 0 | 0 | 0 | 0 | 0 | 0 | 0 | 0 | 0 | 0 | 0 | 0 | 0 | 0 | 0 | 0 | 0 | 1 | 0 | 0 | 0 | 0 | 0 | 0 | 0 | 0 | 0 | 0 | 0 | 0 | 0 | 0 | 0 | 0 |
| virus_Aug5.0v_1135 | 1 | 0 | 0 | 0 | 0 | 0 | 0 | 0 | 0 | 0 | 0 | 0 | 0 | 0 | 0 | 0 | 0 | 0 | 0 | 0 | 0 | 0 | 0 | 0 | 0 | 0 | 0 | 0 | 0 | 0 | 0 | 0 | 0 | 0 | 0 | 0 | 0 | 0 | 1 | 0 | 0 | 0 | 0 | 0 | 0 | 0 | 0 | 0 | 0 | 0 | 0 | 0 | 0 | 0 | 0 |
| virus_Aug5.0v_1409 | 1 | 0 | 0 | 0 | 0 | 0 | 0 | 0 | 0 | 0 | 0 | 0 | 0 | 0 | 0 | 0 | 0 | 0 | 0 | 0 | 0 | 0 | 0 | 0 | 0 | 0 | 0 | 0 | 0 | 0 | 0 | 0 | 0 | 0 | 0 | 0 | 0 | 0 | 1 | 0 | 0 | 0 | 0 | 0 | 0 | 0 | 0 | 0 | 0 | 0 | 0 | 0 | 0 | 0 | 0 |
| virus_Aug5.0v_1855 | 1 | 0 | 0 | 0 | 0 | 0 | 0 | 0 | 0 | 0 | 0 | 0 | 0 | 0 | 0 | 0 | 0 | 0 | 0 | 0 | 0 | 0 | 0 | 0 | 0 | 0 | 0 | 0 | 0 | 0 | 0 | 0 | 0 | 0 | 0 | 0 | 0 | 0 | 1 | 0 | 0 | 0 | 0 | 0 | 0 | 0 | 0 | 0 | 0 | 0 | 0 | 0 | 0 | 0 | 0 |
| virus_Aug5.0v_1978 | 1 | 0 | 0 | 0 | 0 | 0 | 0 | 0 | 0 | 0 | 0 | 0 | 0 | 0 | 0 | 0 | 0 | 0 | 0 | 0 | 0 | 0 | 0 | 0 | 0 | 0 | 0 | 0 | 0 | 0 | 0 | 0 | 0 | 0 | 0 | 0 | 0 | 0 | 1 | 0 | 0 | 0 | 0 | 0 | 0 | 0 | 0 | 0 | 0 | 0 | 0 | 0 | 0 | 0 | 0 |
| virus_Aug5.0v_2018 | 1 | 0 | 0 | 0 | 0 | 0 | 0 | 0 | 0 | 0 | 0 | 0 | 0 | 0 | 0 | 0 | 0 | 0 | 0 | 0 | 0 | 0 | 0 | 0 | 0 | 0 | 0 | 0 | 0 | 0 | 0 | 0 | 0 | 0 | 0 | 0 | 0 | 0 | 1 | 0 | 0 | 0 | 0 | 0 | 0 | 0 | 0 | 0 | 0 | 0 | 0 | 0 | 0 | 0 | 0 |
| virus_Aug5.0v_2075 | 1 | 0 | 0 | 0 | 0 | 0 | 0 | 0 | 0 | 0 | 0 | 0 | 0 | 0 | 0 | 0 | 0 | 0 | 0 | 0 | 0 | 0 | 0 | 0 | 0 | 0 | 0 | 0 | 0 | 0 | 0 | 0 | 0 | 0 | 0 | 0 | 0 | 0 | 1 | 0 | 0 | 0 | 0 | 0 | 0 | 0 | 0 | 0 | 0 | 0 | 0 | 0 | 0 | 0 | 0 |
| virus_Aug5.0v_2760 | 1 | 0 | 0 | 0 | 0 | 0 | 0 | 0 | 0 | 0 | 0 | 0 | 0 | 0 | 0 | 0 | 0 | 0 | 0 | 0 | 0 | 0 | 0 | 0 | 0 | 0 | 0 | 0 | 0 | 0 | 0 | 0 | 0 | 0 | 0 | 0 | 0 | 0 | 1 | 0 | 0 | 0 | 0 | 0 | 0 | 0 | 0 | 0 | 0 | 0 | 0 | 0 | 0 | 0 | 0 |
| virus_Aug5.0v_3014 | 1 | 0 | 0 | 0 | 0 | 0 | 0 | 0 | 0 | 0 | 0 | 0 | 0 | 0 | 0 | 0 | 0 | 0 | 0 | 0 | 0 | 0 | 0 | 0 | 0 | 0 | 0 | 0 | 0 | 0 | 0 | 0 | 0 | 0 | 0 | 0 | 0 | 0 | 1 | 0 | 0 | 0 | 0 | 0 | 0 | 0 | 0 | 0 | 0 | 0 | 0 | 0 | 0 | 0 | 0 |
| virus_Aug5.0v_3711 | 1 | 0 | 0 | 0 | 0 | 0 | 0 | 0 | 0 | 0 | 0 | 0 | 0 | 0 | 0 | 0 | 0 | 0 | 0 | 0 | 0 | 0 | 0 | 0 | 0 | 0 | 0 | 0 | 0 | 0 | 0 | 0 | 0 | 0 | 0 | 0 | 0 | 0 | 1 | 0 | 0 | 0 | 0 | 0 | 0 | 0 | 0 | 0 | 0 | 0 | 0 | 0 | 0 | 0 | 0 |
| virus_Aug5.0v_3725 | 1 | 0 | 0 | 0 | 0 | 0 | 0 | 0 | 0 | 0 | 0 | 0 | 0 | 0 | 0 | 0 | 0 | 0 | 0 | 0 | 0 | 0 | 0 | 0 | 0 | 0 | 0 | 0 | 0 | 0 | 0 | 0 | 0 | 0 | 0 | 0 | 0 | 0 | 1 | 0 | 0 | 0 | 0 | 0 | 0 | 0 | 0 | 0 | 0 | 0 | 0 | 0 | 0 | 0 | 0 |
| virus_Aug5.0v_3810 | 1 | 0 | 0 | 0 | 0 | 0 | 0 | 0 | 0 | 0 | 0 | 0 | 0 | 0 | 0 | 0 | 0 | 0 | 0 | 0 | 0 | 0 | 0 | 0 | 0 | 0 | 0 | 0 | 0 | 0 | 0 | 0 | 0 | 0 | 0 | 0 | 0 | 0 | 1 | 0 | 0 | 0 | 0 | 0 | 0 | 0 | 0 | 0 | 0 | 0 | 0 | 0 | 0 | 0 | 0 |
| virus_Aug5.0v_3893 | 1 | 0 | 0 | 0 | 0 | 0 | 0 | 0 | 0 | 0 | 0 | 0 | 0 | 0 | 0 | 0 | 0 | 0 | 0 | 0 | 0 | 0 | 0 | 0 | 0 | 0 | 0 | 0 | 0 | 0 | 0 | 0 | 0 | 0 | 0 | 0 | 0 | 0 | 1 | 0 | 0 | 0 | 0 | 0 | 0 | 0 | 0 | 0 | 0 | 0 | 0 | 0 | 0 | 0 | 0 |
| virus_Aug5.0v_4361 | 1 | 0 | 0 | 0 | 0 | 0 | 0 | 0 | 0 | 0 | 0 | 0 | 0 | 0 | 0 | 0 | 0 | 0 | 0 | 0 | 0 | 0 | 0 | 0 | 0 | 0 | 0 | 0 | 0 | 0 | 0 | 0 | 0 | 0 | 0 | 0 | 0 | 0 | 1 | 0 | 0 | 0 | 0 | 0 | 0 | 0 | 0 | 0 | 0 | 0 | 0 | 0 | 0 | 0 | 0 |
| virus_Aug5.0v_4464 | 1 | 0 | 0 | 0 | 0 | 0 | 0 | 0 | 0 | 0 | 0 | 0 | 0 | 0 | 0 | 0 | 0 | 0 | 0 | 0 | 0 | 0 | 0 | 0 | 0 | 0 | 0 | 0 | 0 | 0 | 0 | 0 | 0 | 0 | 0 | 0 | 0 | 0 | 1 | 0 | 0 | 0 | 0 | 0 | 0 | 0 | 0 | 0 | 0 | 0 | 0 | 0 | 0 | 0 | 0 |
| virus_Aug5.0v_4490 | 1 | 0 | 0 | 0 | 0 | 0 | 0 | 0 | 0 | 0 | 0 | 0 | 0 | 0 | 0 | 0 | 0 | 0 | 0 | 0 | 0 | 0 | 0 | 0 | 0 | 0 | 0 | 0 | 0 | 0 | 0 | 0 | 0 | 0 | 0 | 0 | 0 | 0 | 1 | 0 | 0 | 0 | 0 | 0 | 0 | 0 | 0 | 0 | 0 | 0 | 0 | 0 | 0 | 0 | 0 |
| virus_Aug5.0v_4573 | 1 | 0 | 0 | 0 | 0 | 0 | 0 | 0 | 0 | 0 | 0 | 0 | 0 | 0 | 0 | 0 | 0 | 0 | 0 | 0 | 0 | 0 | 0 | 0 | 0 | 0 | 0 | 0 | 0 | 0 | 0 | 0 | 0 | 0 | 0 | 0 | 0 | 0 | 1 | 0 | 0 | 0 | 0 | 0 | 0 | 0 | 0 | 0 | 0 | 0 | 0 | 0 | 0 | 0 | 0 |
| virus_Aug5.0v_4613 | 1 | 0 | 0 | 0 | 0 | 0 | 0 | 0 | 0 | 0 | 0 | 0 | 0 | 0 | 0 | 0 | 0 | 0 | 0 | 0 | 0 | 0 | 0 | 0 | 0 | 0 | 0 | 0 | 0 | 0 | 0 | 0 | 0 | 0 | 0 | 0 | 0 | 0 | 1 | 0 | 0 | 0 | 0 | 0 | 0 | 0 | 0 | 0 | 0 | 0 | 0 | 0 | 0 | 0 | 0 |
| virus_Aug5.0v_4629 | 1 | 0 | 0 | 0 | 0 | 0 | 0 | 0 | 0 | 0 | 0 | 0 | 0 | 0 | 0 | 0 | 0 | 0 | 0 | 0 | 0 | 0 | 0 | 0 | 0 | 0 | 0 | 0 | 0 | 0 | 0 | 0 | 0 | 0 | 0 | 0 | 0 | 0 | 1 | 0 | 0 | 0 | 0 | 0 | 0 | 0 | 0 | 0 | 0 | 0 | 0 | 0 | 0 | 0 | 0 |
| virus_Aug5.0v_4761 | 1 | 0 | 0 | 0 | 0 | 0 | 0 | 0 | 0 | 0 | 0 | 0 | 0 | 0 | 0 | 0 | 0 | 0 | 0 | 0 | 0 | 0 | 0 | 0 | 0 | 0 | 0 | 0 | 0 | 0 | 0 | 0 | 0 | 0 | 0 | 0 | 0 | 0 | 1 | 0 | 0 | 0 | 0 | 0 | 0 | 0 | 0 | 0 | 0 | 0 | 0 | 0 | 0 | 0 | 0 |
| virus_Aug5.0v_4774 | 1 | 0 | 0 | 0 | 0 | 0 | 0 | 0 | 0 | 0 | 0 | 0 | 0 | 0 | 0 | 0 | 0 | 0 | 0 | 0 | 0 | 0 | 0 | 0 | 0 | 0 | 0 | 0 | 0 | 0 | 0 | 0 | 0 | 0 | 0 | 0 | 0 | 0 | 1 | 0 | 0 | 0 | 0 | 0 | 0 | 0 | 0 | 0 | 0 | 0 | 0 | 0 | 0 | 0 | 0 |
| virus_Aug5.0v_5373 | 1 | 0 | 0 | 0 | 0 | 0 | 0 | 0 | 0 | 0 | 0 | 0 | 0 | 0 | 0 | 0 | 0 | 0 | 0 | 0 | 0 | 0 | 0 | 0 | 0 | 0 | 0 | 0 | 0 | 0 | 0 | 0 | 0 | 0 | 0 | 0 | 0 | 0 | 1 | 0 | 0 | 0 | 0 | 0 | 0 | 0 | 0 | 0 | 0 | 0 | 0 | 0 | 0 | 0 | 0 |
| virus_Aug5.5v_7503 | 1 | 0 | 0 | 0 | 0 | 0 | 0 | 0 | 0 | 0 | 0 | 0 | 0 | 0 | 0 | 0 | 0 | 0 | 0 | 0 | 0 | 0 | 0 | 0 | 0 | 0 | 0 | 0 | 0 | 0 | 0 | 0 | 0 | 0 | 0 | 0 | 0 | 0 | 0 | 0 | 1 | 0 | 0 | 0 | 0 | 0 | 0 | 0 | 0 | 0 | 0 | 0 | 0 | 0 | 0 |
| virus_Aug5.5v_10513 | 1 | 0 | 0 | 0 | 0 | 0 | 0 | 0 | 0 | 0 | 0 | 0 | 0 | 0 | 0 | 0 | 0 | 0 | 0 | 0 | 0 | 0 | 0 | 0 | 0 | 0 | 0 | 0 | 0 | 0 | 0 | 0 | 0 | 0 | 0 | 0 | 0 | 0 | 0 | 0 | 1 | 0 | 0 | 0 | 0 | 0 | 0 | 0 | 0 | 0 | 0 | 0 | 0 | 0 | 0 |
| virus_Aug5.5v_15316 | 1 | 0 | 0 | 0 | 0 | 0 | 0 | 0 | 0 | 0 | 0 | 0 | 0 | 0 | 0 | 0 | 0 | 0 | 0 | 0 | 0 | 0 | 0 | 0 | 0 | 0 | 0 | 0 | 0 | 0 | 0 | 0 | 0 | 0 | 0 | 0 | 0 | 0 | 0 | 0 | 1 | 0 | 0 | 0 | 0 | 0 | 0 | 0 | 0 | 0 | 0 | 0 | 0 | 0 | 0 |
| virus_Aug5.5v_17722 | 1 | 0 | 0 | 0 | 0 | 0 | 0 | 0 | 0 | 0 | 0 | 0 | 0 | 0 | 0 | 0 | 0 | 0 | 0 | 0 | 0 | 0 | 0 | 0 | 0 | 0 | 0 | 0 | 0 | 0 | 0 | 0 | 0 | 0 | 0 | 0 | 0 | 0 | 0 | 0 | 1 | 0 | 0 | 0 | 0 | 0 | 0 | 0 | 0 | 0 | 0 | 0 | 0 | 0 | 0 |
| virus_Aug5.5v_22369 | 1 | 0 | 0 | 0 | 0 | 0 | 0 | 0 | 0 | 0 | 0 | 0 | 0 | 0 | 0 | 0 | 0 | 0 | 0 | 0 | 0 | 0 | 0 | 0 | 0 | 0 | 0 | 0 | 0 | 0 | 0 | 0 | 0 | 0 | 0 | 0 | 0 | 0 | 0 | 0 | 1 | 0 | 0 | 0 | 0 | 0 | 0 | 0 | 0 | 0 | 0 | 0 | 0 | 0 | 0 |
| virus_Aug5.5v_28614 | 1 | 0 | 0 | 0 | 0 | 0 | 0 | 0 | 0 | 0 | 0 | 0 | 0 | 0 | 0 | 0 | 0 | 0 | 0 | 0 | 0 | 0 | 0 | 0 | 0 | 0 | 0 | 0 | 0 | 0 | 0 | 0 | 0 | 0 | 0 | 0 | 0 | 0 | 0 | 0 | 1 | 0 | 0 | 0 | 0 | 0 | 0 | 0 | 0 | 0 | 0 | 0 | 0 | 0 | 0 |
| virus_Aug5.5v_28796 | 1 | 0 | 0 | 0 | 0 | 0 | 0 | 0 | 0 | 0 | 0 | 0 | 0 | 0 | 0 | 0 | 0 | 0 | 0 | 0 | 0 | 0 | 0 | 0 | 0 | 0 | 0 | 0 | 0 | 0 | 0 | 0 | 0 | 0 | 0 | 0 | 0 | 0 | 0 | 0 | 1 | 0 | 0 | 0 | 0 | 0 | 0 | 0 | 0 | 0 | 0 | 0 | 0 | 0 | 0 |
| virus_Aug5.5v_35563 | 1 | 0 | 0 | 0 | 0 | 0 | 0 | 0 | 0 | 0 | 0 | 0 | 0 | 0 | 0 | 0 | 0 | 0 | 0 | 0 | 0 | 0 | 0 | 0 | 0 | 0 | 0 | 0 | 0 | 0 | 0 | 0 | 0 | 0 | 0 | 0 | 0 | 0 | 0 | 0 | 1 | 0 | 0 | 0 | 0 | 0 | 0 | 0 | 0 | 0 | 0 | 0 | 0 | 0 | 0 |
| virus_Aug5.5v_39677 | 1 | 0 | 0 | 0 | 0 | 0 | 0 | 0 | 0 | 0 | 0 | 0 | 0 | 0 | 0 | 0 | 0 | 0 | 0 | 0 | 0 | 0 | 0 | 0 | 0 | 0 | 0 | 0 | 0 | 0 | 0 | 0 | 0 | 0 | 0 | 0 | 0 | 0 | 0 | 0 | 1 | 0 | 0 | 0 | 0 | 0 | 0 | 0 | 0 | 0 | 0 | 0 | 0 | 0 | 0 |
| virus_Aug5.5v_41029 | 1 | 0 | 0 | 0 | 0 | 0 | 0 | 0 | 0 | 0 | 0 | 0 | 0 | 0 | 0 | 0 | 0 | 0 | 0 | 0 | 0 | 0 | 0 | 0 | 0 | 0 | 0 | 0 | 0 | 0 | 0 | 0 | 0 | 0 | 0 | 0 | 0 | 0 | 0 | 0 | 1 | 0 | 0 | 0 | 0 | 0 | 0 | 0 | 0 | 0 | 0 | 0 | 0 | 0 | 0 |
| virus_Aug5.25v_1114 | 1 | 0 | 0 | 0 | 0 | 0 | 0 | 0 | 0 | 0 | 0 | 0 | 0 | 0 | 0 | 0 | 0 | 0 | 0 | 0 | 0 | 0 | 0 | 0 | 0 | 0 | 0 | 0 | 0 | 0 | 0 | 0 | 0 | 0 | 0 | 0 | 0 | 0 | 0 | 0 | 0 | 0 | 1 | 0 | 0 | 0 | 0 | 0 | 0 | 0 | 0 | 0 | 0 | 0 | 0 |
| virus_Aug5.25v_1472 | 1 | 0 | 0 | 0 | 0 | 0 | 0 | 0 | 0 | 0 | 0 | 0 | 0 | 0 | 0 | 0 | 0 | 0 | 0 | 0 | 0 | 0 | 0 | 0 | 0 | 0 | 0 | 0 | 0 | 0 | 0 | 0 | 0 | 0 | 0 | 0 | 0 | 0 | 0 | 0 | 0 | 0 | 1 | 0 | 0 | 0 | 0 | 0 | 0 | 0 | 0 | 0 | 0 | 0 | 0 |
| virus_Aug5.25v_2209 | 1 | 0 | 0 | 0 | 0 | 0 | 0 | 0 | 0 | 0 | 0 | 0 | 0 | 0 | 0 | 0 | 0 | 0 | 0 | 0 | 0 | 0 | 0 | 0 | 0 | 0 | 0 | 0 | 0 | 0 | 0 | 0 | 0 | 0 | 0 | 0 | 0 | 0 | 0 | 0 | 0 | 0 | 1 | 0 | 0 | 0 | 0 | 0 | 0 | 0 | 0 | 0 | 0 | 0 | 0 |
| virus_Aug5.25v_3366 | 1 | 0 | 0 | 0 | 0 | 0 | 0 | 0 | 0 | 0 | 0 | 0 | 0 | 0 | 0 | 0 | 0 | 0 | 0 | 0 | 0 | 0 | 0 | 0 | 0 | 0 | 0 | 0 | 0 | 0 | 0 | 0 | 0 | 0 | 0 | 0 | 0 | 0 | 0 | 0 | 0 | 0 | 1 | 0 | 0 | 0 | 0 | 0 | 0 | 0 | 0 | 0 | 0 | 0 | 0 |
| virus_Aug5.25v_6601 | 1 | 0 | 0 | 0 | 0 | 0 | 0 | 0 | 0 | 0 | 0 | 0 | 0 | 0 | 0 | 0 | 0 | 0 | 0 | 0 | 0 | 0 | 0 | 0 | 0 | 0 | 0 | 0 | 0 | 0 | 0 | 0 | 0 | 0 | 0 | 0 | 0 | 0 | 0 | 0 | 0 | 0 | 1 | 0 | 0 | 0 | 0 | 0 | 0 | 0 | 0 | 0 | 0 | 0 | 0 |
| virus_Aug5.25v_8939 | 1 | 0 | 0 | 0 | 0 | 0 | 0 | 0 | 0 | 0 | 0 | 0 | 0 | 0 | 0 | 0 | 0 | 0 | 0 | 0 | 0 | 0 | 0 | 0 | 0 | 0 | 0 | 0 | 0 | 0 | 0 | 0 | 0 | 0 | 0 | 0 | 0 | 0 | 0 | 0 | 0 | 0 | 1 | 0 | 0 | 0 | 0 | 0 | 0 | 0 | 0 | 0 | 0 | 0 | 0 |
| virus_Aug5.25v_9357 | 1 | 0 | 0 | 0 | 0 | 0 | 0 | 0 | 0 | 0 | 0 | 0 | 0 | 0 | 0 | 0 | 0 | 0 | 0 | 0 | 0 | 0 | 0 | 0 | 0 | 0 | 0 | 0 | 0 | 0 | 0 | 0 | 0 | 0 | 0 | 0 | 0 | 0 | 0 | 0 | 0 | 0 | 1 | 0 | 0 | 0 | 0 | 0 | 0 | 0 | 0 | 0 | 0 | 0 | 0 |
| virus_Aug5.25v_10907 | 1 | 0 | 0 | 0 | 0 | 0 | 0 | 0 | 0 | 0 | 0 | 0 | 0 | 0 | 0 | 0 | 0 | 0 | 0 | 0 | 0 | 0 | 0 | 0 | 0 | 0 | 0 | 0 | 0 | 0 | 0 | 0 | 0 | 0 | 0 | 0 | 0 | 0 | 0 | 0 | 0 | 0 | 1 | 0 | 0 | 0 | 0 | 0 | 0 | 0 | 0 | 0 | 0 | 0 | 0 |
| virus_Aug5.25v_12783 | 1 | 0 | 0 | 0 | 0 | 0 | 0 | 0 | 0 | 0 | 0 | 0 | 0 | 0 | 0 | 0 | 0 | 0 | 0 | 0 | 0 | 0 | 0 | 0 | 0 | 0 | 0 | 0 | 0 | 0 | 0 | 0 | 0 | 0 | 0 | 0 | 0 | 0 | 0 | 0 | 0 | 0 | 1 | 0 | 0 | 0 | 0 | 0 | 0 | 0 | 0 | 0 | 0 | 0 | 0 |
| virus_Aug5.25v_12787 | 1 | 0 | 0 | 0 | 0 | 0 | 0 | 0 | 0 | 0 | 0 | 0 | 0 | 0 | 0 | 0 | 0 | 0 | 0 | 0 | 0 | 0 | 0 | 0 | 0 | 0 | 0 | 0 | 0 | 0 | 0 | 0 | 0 | 0 | 0 | 0 | 0 | 0 | 0 | 0 | 0 | 0 | 1 | 0 | 0 | 0 | 0 | 0 | 0 | 0 | 0 | 0 | 0 | 0 | 0 |
| virus_Aug5.25v_15452 | 1 | 0 | 0 | 0 | 0 | 0 | 0 | 0 | 0 | 0 | 0 | 0 | 0 | 0 | 0 | 0 | 0 | 0 | 0 | 0 | 0 | 0 | 0 | 0 | 0 | 0 | 0 | 0 | 0 | 0 | 0 | 0 | 0 | 0 | 0 | 0 | 0 | 0 | 0 | 0 | 0 | 0 | 1 | 0 | 0 | 0 | 0 | 0 | 0 | 0 | 0 | 0 | 0 | 0 | 0 |
| virus_Aug5.25v_19510 | 1 | 0 | 0 | 0 | 0 | 0 | 0 | 0 | 0 | 0 | 0 | 0 | 0 | 0 | 0 | 0 | 0 | 0 | 0 | 0 | 0 | 0 | 0 | 0 | 0 | 0 | 0 | 0 | 0 | 0 | 0 | 0 | 0 | 0 | 0 | 0 | 0 | 0 | 0 | 0 | 0 | 0 | 1 | 0 | 0 | 0 | 0 | 0 | 0 | 0 | 0 | 0 | 0 | 0 | 0 |
| virus_Aug5.25v_20101 | 1 | 0 | 0 | 0 | 0 | 0 | 0 | 0 | 0 | 0 | 0 | 0 | 0 | 0 | 0 | 0 | 0 | 0 | 0 | 0 | 0 | 0 | 0 | 0 | 0 | 0 | 0 | 0 | 0 | 0 | 0 | 0 | 0 | 0 | 0 | 0 | 0 | 0 | 0 | 0 | 0 | 0 | 1 | 0 | 0 | 0 | 0 | 0 | 0 | 0 | 0 | 0 | 0 | 0 | 0 |
| virus_Aug5.25v_21421 | 1 | 0 | 0 | 0 | 0 | 0 | 0 | 0 | 0 | 0 | 0 | 0 | 0 | 0 | 0 | 0 | 0 | 0 | 0 | 0 | 0 | 0 | 0 | 0 | 0 | 0 | 0 | 0 | 0 | 0 | 0 | 0 | 0 | 0 | 0 | 0 | 0 | 0 | 0 | 0 | 0 | 0 | 1 | 0 | 0 | 0 | 0 | 0 | 0 | 0 | 0 | 0 | 0 | 0 | 0 |
| virus_Aug5.25v_26529 | 1 | 0 | 0 | 0 | 0 | 0 | 0 | 0 | 0 | 0 | 0 | 0 | 0 | 0 | 0 | 0 | 0 | 0 | 0 | 0 | 0 | 0 | 0 | 0 | 0 | 0 | 0 | 0 | 0 | 0 | 0 | 0 | 0 | 0 | 0 | 0 | 0 | 0 | 0 | 0 | 0 | 0 | 1 | 0 | 0 | 0 | 0 | 0 | 0 | 0 | 0 | 0 | 0 | 0 | 0 |
| virus_Aug5.25v_27987 | 1 | 0 | 0 | 0 | 0 | 0 | 0 | 0 | 0 | 0 | 0 | 0 | 0 | 0 | 0 | 0 | 0 | 0 | 0 | 0 | 0 | 0 | 0 | 0 | 0 | 0 | 0 | 0 | 0 | 0 | 0 | 0 | 0 | 0 | 0 | 0 | 0 | 0 | 0 | 0 | 0 | 0 | 1 | 0 | 0 | 0 | 0 | 0 | 0 | 0 | 0 | 0 | 0 | 0 | 0 |
| virus_Aug5.25v_33871 | 1 | 0 | 0 | 0 | 0 | 0 | 0 | 0 | 0 | 0 | 0 | 0 | 0 | 0 | 0 | 0 | 0 | 0 | 0 | 0 | 0 | 0 | 0 | 0 | 0 | 0 | 0 | 0 | 0 | 0 | 0 | 0 | 0 | 0 | 0 | 0 | 0 | 0 | 0 | 0 | 0 | 0 | 1 | 0 | 0 | 0 | 0 | 0 | 0 | 0 | 0 | 0 | 0 | 0 | 0 |
| virus_Aug5.25v_33981 | 1 | 0 | 0 | 0 | 0 | 0 | 0 | 0 | 0 | 0 | 0 | 0 | 0 | 0 | 0 | 0 | 0 | 0 | 0 | 0 | 0 | 0 | 0 | 0 | 0 | 0 | 0 | 0 | 0 | 0 | 0 | 0 | 0 | 0 | 0 | 0 | 0 | 0 | 0 | 0 | 0 | 0 | 1 | 0 | 0 | 0 | 0 | 0 | 0 | 0 | 0 | 0 | 0 | 0 | 0 |
| virus_Aug5.25v_34476 | 1 | 0 | 0 | 0 | 0 | 0 | 0 | 0 | 0 | 0 | 0 | 0 | 0 | 0 | 0 | 0 | 0 | 0 | 0 | 0 | 0 | 0 | 0 | 0 | 0 | 0 | 0 | 0 | 0 | 0 | 0 | 0 | 0 | 0 | 0 | 0 | 0 | 0 | 0 | 0 | 0 | 0 | 1 | 0 | 0 | 0 | 0 | 0 | 0 | 0 | 0 | 0 | 0 | 0 | 0 |
| virus_Aug5.25v_34563 | 1 | 0 | 0 | 0 | 0 | 0 | 0 | 0 | 0 | 0 | 0 | 0 | 0 | 0 | 0 | 0 | 0 | 0 | 0 | 0 | 0 | 0 | 0 | 0 | 0 | 0 | 0 | 0 | 0 | 0 | 0 | 0 | 0 | 0 | 0 | 0 | 0 | 0 | 0 | 0 | 0 | 0 | 1 | 0 | 0 | 0 | 0 | 0 | 0 | 0 | 0 | 0 | 0 | 0 | 0 |
| virus_Aug5.50v_4994 | 1 | 0 | 0 | 0 | 0 | 0 | 0 | 0 | 0 | 0 | 0 | 0 | 0 | 0 | 0 | 0 | 0 | 0 | 0 | 0 | 0 | 0 | 0 | 0 | 0 | 0 | 0 | 0 | 0 | 0 | 0 | 0 | 0 | 0 | 0 | 0 | 0 | 0 | 0 | 0 | 0 | 0 | 0 | 0 | 1 | 0 | 0 | 0 | 0 | 0 | 0 | 0 | 0 | 0 | 0 |
| virus_Aug5.50v_15274 | 1 | 0 | 0 | 0 | 0 | 0 | 0 | 0 | 0 | 0 | 0 | 0 | 0 | 0 | 0 | 0 | 0 | 0 | 0 | 0 | 0 | 0 | 0 | 0 | 0 | 0 | 0 | 0 | 0 | 0 | 0 | 0 | 0 | 0 | 0 | 0 | 0 | 0 | 0 | 0 | 0 | 0 | 0 | 0 | 1 | 0 | 0 | 0 | 0 | 0 | 0 | 0 | 0 | 0 | 0 |
| virus_Aug5.50v_23067 | 1 | 0 | 0 | 0 | 0 | 0 | 0 | 0 | 0 | 0 | 0 | 0 | 0 | 0 | 0 | 0 | 0 | 0 | 0 | 0 | 0 | 0 | 0 | 0 | 0 | 0 | 0 | 0 | 0 | 0 | 0 | 0 | 0 | 0 | 0 | 0 | 0 | 0 | 0 | 0 | 0 | 0 | 0 | 0 | 1 | 0 | 0 | 0 | 0 | 0 | 0 | 0 | 0 | 0 | 0 |
| virus_Aug5.50v_25883 | 1 | 0 | 0 | 0 | 0 | 0 | 0 | 0 | 0 | 0 | 0 | 0 | 0 | 0 | 0 | 0 | 0 | 0 | 0 | 0 | 0 | 0 | 0 | 0 | 0 | 0 | 0 | 0 | 0 | 0 | 0 | 0 | 0 | 0 | 0 | 0 | 0 | 0 | 0 | 0 | 0 | 0 | 0 | 0 | 1 | 0 | 0 | 0 | 0 | 0 | 0 | 0 | 0 | 0 | 0 |
| virus_Aug5.50v_37660 | 1 | 0 | 0 | 0 | 0 | 0 | 0 | 0 | 0 | 0 | 0 | 0 | 0 | 0 | 0 | 0 | 0 | 0 | 0 | 0 | 0 | 0 | 0 | 0 | 0 | 0 | 0 | 0 | 0 | 0 | 0 | 0 | 0 | 0 | 0 | 0 | 0 | 0 | 0 | 0 | 0 | 0 | 0 | 0 | 1 | 0 | 0 | 0 | 0 | 0 | 0 | 0 | 0 | 0 | 0 |
| virus_Aug5.50v_40182 | 1 | 0 | 0 | 0 | 0 | 0 | 0 | 0 | 0 | 0 | 0 | 0 | 0 | 0 | 0 | 0 | 0 | 0 | 0 | 0 | 0 | 0 | 0 | 0 | 0 | 0 | 0 | 0 | 0 | 0 | 0 | 0 | 0 | 0 | 0 | 0 | 0 | 0 | 0 | 0 | 0 | 0 | 0 | 0 | 1 | 0 | 0 | 0 | 0 | 0 | 0 | 0 | 0 | 0 | 0 |
| virus_Aug5.75v_14261 | 1 | 0 | 0 | 0 | 0 | 0 | 0 | 0 | 0 | 0 | 0 | 0 | 0 | 0 | 0 | 0 | 0 | 0 | 0 | 0 | 0 | 0 | 0 | 0 | 0 | 0 | 0 | 0 | 0 | 0 | 0 | 0 | 0 | 0 | 0 | 0 | 0 | 0 | 0 | 0 | 0 | 0 | 0 | 0 | 0 | 0 | 1 | 0 | 0 | 0 | 0 | 0 | 0 | 0 | 0 |
| virus_Aug5.75v_14842 | 1 | 0 | 0 | 0 | 0 | 0 | 0 | 0 | 0 | 0 | 0 | 0 | 0 | 0 | 0 | 0 | 0 | 0 | 0 | 0 | 0 | 0 | 0 | 0 | 0 | 0 | 0 | 0 | 0 | 0 | 0 | 0 | 0 | 0 | 0 | 0 | 0 | 0 | 0 | 0 | 0 | 0 | 0 | 0 | 0 | 0 | 1 | 0 | 0 | 0 | 0 | 0 | 0 | 0 | 0 |
| virus_Aug5.75v_19722 | 1 | 0 | 0 | 0 | 0 | 0 | 0 | 0 | 0 | 0 | 0 | 0 | 0 | 0 | 0 | 0 | 0 | 0 | 0 | 0 | 0 | 0 | 0 | 0 | 0 | 0 | 0 | 0 | 0 | 0 | 0 | 0 | 0 | 0 | 0 | 0 | 0 | 0 | 0 | 0 | 0 | 0 | 0 | 0 | 0 | 0 | 1 | 0 | 0 | 0 | 0 | 0 | 0 | 0 | 0 |
| virus_Aug5.75v_26312 | 1 | 0 | 0 | 0 | 0 | 0 | 0 | 0 | 0 | 0 | 0 | 0 | 0 | 0 | 0 | 0 | 0 | 0 | 0 | 0 | 0 | 0 | 0 | 0 | 0 | 0 | 0 | 0 | 0 | 0 | 0 | 0 | 0 | 0 | 0 | 0 | 0 | 0 | 0 | 0 | 0 | 0 | 0 | 0 | 0 | 0 | 1 | 0 | 0 | 0 | 0 | 0 | 0 | 0 | 0 |
| virus_Aug5.75v_33388 | 1 | 0 | 0 | 0 | 0 | 0 | 0 | 0 | 0 | 0 | 0 | 0 | 0 | 0 | 0 | 0 | 0 | 0 | 0 | 0 | 0 | 0 | 0 | 0 | 0 | 0 | 0 | 0 | 0 | 0 | 0 | 0 | 0 | 0 | 0 | 0 | 0 | 0 | 0 | 0 | 0 | 0 | 0 | 0 | 0 | 0 | 1 | 0 | 0 | 0 | 0 | 0 | 0 | 0 | 0 |
| virus_Aug5.75v_36458 | 1 | 0 | 0 | 0 | 0 | 0 | 0 | 0 | 0 | 0 | 0 | 0 | 0 | 0 | 0 | 0 | 0 | 0 | 0 | 0 | 0 | 0 | 0 | 0 | 0 | 0 | 0 | 0 | 0 | 0 | 0 | 0 | 0 | 0 | 0 | 0 | 0 | 0 | 0 | 0 | 0 | 0 | 0 | 0 | 0 | 0 | 1 | 0 | 0 | 0 | 0 | 0 | 0 | 0 | 0 |
| virus_Aug5.75v_37349 | 1 | 0 | 0 | 0 | 0 | 0 | 0 | 0 | 0 | 0 | 0 | 0 | 0 | 0 | 0 | 0 | 0 | 0 | 0 | 0 | 0 | 0 | 0 | 0 | 0 | 0 | 0 | 0 | 0 | 0 | 0 | 0 | 0 | 0 | 0 | 0 | 0 | 0 | 0 | 0 | 0 | 0 | 0 | 0 | 0 | 0 | 1 | 0 | 0 | 0 | 0 | 0 | 0 | 0 | 0 |
| virus_Aug5.75v_38675 | 1 | 0 | 0 | 0 | 0 | 0 | 0 | 0 | 0 | 0 | 0 | 0 | 0 | 0 | 0 | 0 | 0 | 0 | 0 | 0 | 0 | 0 | 0 | 0 | 0 | 0 | 0 | 0 | 0 | 0 | 0 | 0 | 0 | 0 | 0 | 0 | 0 | 0 | 0 | 0 | 0 | 0 | 0 | 0 | 0 | 0 | 1 | 0 | 0 | 0 | 0 | 0 | 0 | 0 | 0 |
| virus_Jul11.0v_130 | 1 | 0 | 0 | 0 | 0 | 0 | 0 | 0 | 0 | 0 | 0 | 0 | 0 | 0 | 0 | 0 | 0 | 0 | 0 | 0 | 0 | 0 | 1 | 0 | 0 | 0 | 0 | 0 | 0 | 0 | 0 | 0 | 0 | 0 | 0 | 0 | 0 | 0 | 0 | 0 | 0 | 0 | 0 | 0 | 0 | 0 | 0 | 0 | 0 | 0 | 0 | 0 | 0 | 0 | 0 |
| virus_Jul11.0v_11429 | 1 | 0 | 0 | 0 | 0 | 0 | 0 | 0 | 0 | 0 | 0 | 0 | 0 | 0 | 0 | 0 | 0 | 0 | 0 | 0 | 0 | 0 | 1 | 0 | 0 | 0 | 0 | 0 | 0 | 0 | 0 | 0 | 0 | 0 | 0 | 0 | 0 | 0 | 0 | 0 | 0 | 0 | 0 | 0 | 0 | 0 | 0 | 0 | 0 | 0 | 0 | 0 | 0 | 0 | 0 |
| virus_Jul11.0v_12253 | 1 | 0 | 0 | 0 | 0 | 0 | 0 | 0 | 0 | 0 | 0 | 0 | 0 | 0 | 0 | 0 | 0 | 0 | 0 | 0 | 0 | 0 | 1 | 0 | 0 | 0 | 0 | 0 | 0 | 0 | 0 | 0 | 0 | 0 | 0 | 0 | 0 | 0 | 0 | 0 | 0 | 0 | 0 | 0 | 0 | 0 | 0 | 0 | 0 | 0 | 0 | 0 | 0 | 0 | 0 |
| virus_Jul11.0v_12503 | 1 | 0 | 0 | 0 | 0 | 0 | 0 | 0 | 0 | 0 | 0 | 0 | 0 | 0 | 0 | 0 | 0 | 0 | 0 | 0 | 0 | 0 | 1 | 0 | 0 | 0 | 0 | 0 | 0 | 0 | 0 | 0 | 0 | 0 | 0 | 0 | 0 | 0 | 0 | 0 | 0 | 0 | 0 | 0 | 0 | 0 | 0 | 0 | 0 | 0 | 0 | 0 | 0 | 0 | 0 |

**Supplementary Table S7**. Common Bacterio- and virioplankton RTPR 98% OTUs in 27 samples with RTPR amplicons in both fractions

Total seqs: 47997; Subtotal seqs in retained

| OTUs: 41046 | | May12_05m | | May12_25m | | May12_50m | | May12_75m | | Jun8_25m | | Jun8_50m | | Jun8_75m | | Jun22_0m | | Jun22_50m | | Jun22_75m | | Jul11_0m | | Jul11_05m | | Jul11_25m | | Jul11_50m | | Jul11_75m | | Jul27_05m | | Jul27_25m | | Jul27_75m | | Aug5_0m | | Aug5_05m | | Aug5_25m | | Aug5_50m | | Aug5_75m | | Sep8_0m | | Sep8_05m | | Sep8_25m | | Sep8_50m | |
|---|---|---|---|---|---|---|---|---|---|---|---|---|---|---|---|---|---|---|---|---|---|---|---|---|---|---|---|---|---|---|---|---|---|---|---|---|---|---|---|---|---|---|---|---|---|---|---|---|---|---|---|---|---|---|---|
| OTUs with >=10 seqs: 254 | OTU subtotal | bacterioplankton | virioplankton | bacterioplankton | virioplankton | bacterioplankton | virioplankton | bacterioplankton | virioplankton | bacterioplankton | virioplankton | bacterioplankton | virioplankton | bacterioplankton | virioplankton | bacterioplankton | virioplankton | bacterioplankton | virioplankton | bacterioplankton | virioplankton | bacterioplankton | virioplankton | bacterioplankton | virioplankton | bacterioplankton | virioplankton | bacterioplankton | virioplankton | bacterioplankton | virioplankton | bacterioplankton | virioplankton | bacterioplankton | virioplankton | bacterioplankton | virioplankton | bacterioplankton | virioplankton | bacterioplankton | virioplankton | bacterioplankton | virioplankton | bacterioplankton | virioplankton | bacterioplankton | virioplankton | bacterioplankton | virioplankton | bacterioplankton | virioplankton | bacterioplankton | virioplankton | bacterioplankton | virioplankton |
|virus_Jul11.0v_19984|1|0|0|0|0|0|0|0|0|0|0|0|0|0|0|0|0|0|0|0|0|0|1|0|0|0|0|0|0|0|0|0|0|0|0|0|0|0|0|0|0|0|0|0|0|0|0|0|0|0|0|0|0|0|0|
|virus_Jul11.0v_20214|1|0|0|0|0|0|0|0|0|0|0|0|0|0|0|0|0|0|0|0|0|0|1|0|0|0|0|0|0|0|0|0|0|0|0|0|0|0|0|0|0|0|0|0|0|0|0|0|0|0|0|0|0|0|0|
|virus_Jul11.5v_381|1|0|0|0|0|0|0|0|0|0|0|0|0|0|0|0|0|0|0|0|0|0|0|0|1|0|0|0|0|0|0|0|0|0|0|0|0|0|0|0|0|0|0|0|0|0|0|0|0|0|0|0|0|0|0|
|virus_Jul11.5v_673|1|0|0|0|0|0|0|0|0|0|0|0|0|0|0|0|0|0|0|0|0|0|0|0|1|0|0|0|0|0|0|0|0|0|0|0|0|0|0|0|0|0|0|0|0|0|0|0|0|0|0|0|0|0|0|
|virus_Jul11.5v_1191|1|0|0|0|0|0|0|0|0|0|0|0|0|0|0|0|0|0|0|0|0|0|0|0|1|0|0|0|0|0|0|0|0|0|0|0|0|0|0|0|0|0|0|0|0|0|0|0|0|0|0|0|0|0|0|
|virus_Jul11.5v_2794|1|0|0|0|0|0|0|0|0|0|0|0|0|0|0|0|0|0|0|0|0|0|0|0|1|0|0|0|0|0|0|0|0|0|0|0|0|0|0|0|0|0|0|0|0|0|0|0|0|0|0|0|0|0|0|
|virus_Jul11.5v_5256|1|0|0|0|0|0|0|0|0|0|0|0|0|0|0|0|0|0|0|0|0|0|0|0|1|0|0|0|0|0|0|0|0|0|0|0|0|0|0|0|0|0|0|0|0|0|0|0|0|0|0|0|0|0|0|
|virus_Jul11.5v_6463|1|0|0|0|0|0|0|0|0|0|0|0|0|0|0|0|0|0|0|0|0|0|0|0|1|0|0|0|0|0|0|0|0|0|0|0|0|0|0|0|0|0|0|0|0|0|0|0|0|0|0|0|0|0|0|
|virus_Jul11.5v_7391|1|0|0|0|0|0|0|0|0|0|0|0|0|0|0|0|0|0|0|0|0|0|0|0|1|0|0|0|0|0|0|0|0|0|0|0|0|0|0|0|0|0|0|0|0|0|0|0|0|0|0|0|0|0|0|
|virus_Jul11.5v_7560|1|0|0|0|0|0|0|0|0|0|0|0|0|0|0|0|0|0|0|0|0|0|0|0|1|0|0|0|0|0|0|0|0|0|0|0|0|0|0|0|0|0|0|0|0|0|0|0|0|0|0|0|0|0|0|
|virus_Jul11.5v_8070|1|0|0|0|0|0|0|0|0|0|0|0|0|0|0|0|0|0|0|0|0|0|0|0|1|0|0|0|0|0|0|0|0|0|0|0|0|0|0|0|0|0|0|0|0|0|0|0|0|0|0|0|0|0|0|
|virus_Jul11.5v_8177|1|0|0|0|0|0|0|0|0|0|0|0|0|0|0|0|0|0|0|0|0|0|0|0|1|0|0|0|0|0|0|0|0|0|0|0|0|0|0|0|0|0|0|0|0|0|0|0|0|0|0|0|0|0|0|
|virus_Jul11.5v_11526|1|0|0|0|0|0|0|0|0|0|0|0|0|0|0|0|0|0|0|0|0|0|0|0|1|0|0|0|0|0|0|0|0|0|0|0|0|0|0|0|0|0|0|0|0|0|0|0|0|0|0|0|0|0|0|
|virus_Jul11.5v_12569|1|0|0|0|0|0|0|0|0|0|0|0|0|0|0|0|0|0|0|0|0|0|0|0|1|0|0|0|0|0|0|0|0|0|0|0|0|0|0|0|0|0|0|0|0|0|0|0|0|0|0|0|0|0|0|
|virus_Jul11.5v_13808|1|0|0|0|0|0|0|0|0|0|0|0|0|0|0|0|0|0|0|0|0|0|0|0|1|0|0|0|0|0|0|0|0|0|0|0|0|0|0|0|0|0|0|0|0|0|0|0|0|0|0|0|0|0|0|
|virus_Jul11.5v_14896|1|0|0|0|0|0|0|0|0|0|0|0|0|0|0|0|0|0|0|0|0|0|0|0|1|0|0|0|0|0|0|0|0|0|0|0|0|0|0|0|0|0|0|0|0|0|0|0|0|0|0|0|0|0|0|
|virus_Jul11.5v_15011|1|0|0|0|0|0|0|0|0|0|0|0|0|0|0|0|0|0|0|0|0|0|0|0|1|0|0|0|0|0|0|0|0|0|0|0|0|0|0|0|0|0|0|0|0|0|0|0|0|0|0|0|0|0|0|
|virus_Jul11.5v_15540|1|0|0|0|0|0|0|0|0|0|0|0|0|0|0|0|0|0|0|0|0|0|0|0|1|0|0|0|0|0|0|0|0|0|0|0|0|0|0|0|0|0|0|0|0|0|0|0|0|0|0|0|0|0|0|
|virus_Jul11.5v_18487|1|0|0|0|0|0|0|0|0|0|0|0|0|0|0|0|0|0|0|0|0|0|0|0|1|0|0|0|0|0|0|0|0|0|0|0|0|0|0|0|0|0|0|0|0|0|0|0|0|0|0|0|0|0|0|
|virus_Jul11.5v_19652|1|0|0|0|0|0|0|0|0|0|0|0|0|0|0|0|0|0|0|0|0|0|0|0|1|0|0|0|0|0|0|0|0|0|0|0|0|0|0|0|0|0|0|0|0|0|0|0|0|0|0|0|0|0|0|
|virus_Jul11.25v_543|1|0|0|0|0|0|0|0|0|0|0|0|0|0|0|0|0|0|0|0|0|0|0|0|0|0|1|0|0|0|0|0|0|0|0|0|0|0|0|0|0|0|0|0|0|0|0|0|0|0|0|0|0|0|0|
|virus_Jul11.25v_3211|1|0|0|0|0|0|0|0|0|0|0|0|0|0|0|0|0|0|0|0|0|0|0|0|0|0|1|0|0|0|0|0|0|0|0|0|0|0|0|0|0|0|0|0|0|0|0|0|0|0|0|0|0|0|0|
|virus_Jul11.25v_7446|1|0|0|0|0|0|0|0|0|0|0|0|0|0|0|0|0|0|0|0|0|0|0|0|0|0|1|0|0|0|0|0|0|0|0|0|0|0|0|0|0|0|0|0|0|0|0|0|0|0|0|0|0|0|0|
|virus_Jul11.25v_8637|1|0|0|0|0|0|0|0|0|0|0|0|0|0|0|0|0|0|0|0|0|0|0|0|0|0|1|0|0|0|0|0|0|0|0|0|0|0|0|0|0|0|0|0|0|0|0|0|0|0|0|0|0|0|0|
|virus_Jul11.25v_12490|1|0|0|0|0|0|0|0|0|0|0|0|0|0|0|0|0|0|0|0|0|0|0|0|0|0|1|0|0|0|0|0|0|0|0|0|0|0|0|0|0|0|0|0|0|0|0|0|0|0|0|0|0|0|0|
|virus_Jul11.25v_16201|1|0|0|0|0|0|0|0|0|0|0|0|0|0|0|0|0|0|0|0|0|0|0|0|0|0|1|0|0|0|0|0|0|0|0|0|0|0|0|0|0|0|0|0|0|0|0|0|0|0|0|0|0|0|0|
|virus_Jul11.25v_17500|1|0|0|0|0|0|0|0|0|0|0|0|0|0|0|0|0|0|0|0|0|0|0|0|0|0|1|0|0|0|0|0|0|0|0|0|0|0|0|0|0|0|0|0|0|0|0|0|0|0|0|0|0|0|0|
|virus_Jul11.50v_5842|1|0|0|0|0|0|0|0|0|0|0|0|0|0|0|0|0|0|0|0|0|0|0|0|0|0|0|0|1|0|0|0|0|0|0|0|0|0|0|0|0|0|0|0|0|0|0|0|0|0|0|0|0|0|0|
|virus_Jul11.50v_7997|1|0|0|0|0|0|0|0|0|0|0|0|0|0|0|0|0|0|0|0|0|0|0|0|0|0|0|0|1|0|0|0|0|0|0|0|0|0|0|0|0|0|0|0|0|0|0|0|0|0|0|0|0|0|0|
|virus_Jul11.50v_9849|1|0|0|0|0|0|0|0|0|0|0|0|0|0|0|0|0|0|0|0|0|0|0|0|0|0|0|0|1|0|0|0|0|0|0|0|0|0|0|0|0|0|0|0|0|0|0|0|0|0|0|0|0|0|0|
|virus_Jul11.50v_9853|1|0|0|0|0|0|0|0|0|0|0|0|0|0|0|0|0|0|0|0|0|0|0|0|0|0|0|0|1|0|0|0|0|0|0|0|0|0|0|0|0|0|0|0|0|0|0|0|0|0|0|0|0|0|0|
|virus_Jul11.50v_11789|1|0|0|0|0|0|0|0|0|0|0|0|0|0|0|0|0|0|0|0|0|0|0|0|0|0|0|0|1|0|0|0|0|0|0|0|0|0|0|0|0|0|0|0|0|0|0|0|0|0|0|0|0|0|0|
|virus_Jul11.50v_11809|1|0|0|0|0|0|0|0|0|0|0|0|0|0|0|0|0|0|0|0|0|0|0|0|0|0|0|0|1|0|0|0|0|0|0|0|0|0|0|0|0|0|0|0|0|0|0|0|0|0|0|0|0|0|0|
|virus_Jul11.50v_13214|1|0|0|0|0|0|0|0|0|0|0|0|0|0|0|0|0|0|0|0|0|0|0|0|0|0|0|0|1|0|0|0|0|0|0|0|0|0|0|0|0|0|0|0|0|0|0|0|0|0|0|0|0|0|0|
|virus_Jul11.50v_14411|1|0|0|0|0|0|0|0|0|0|0|0|0|0|0|0|0|0|0|0|0|0|0|0|0|0|0|0|1|0|0|0|0|0|0|0|0|0|0|0|0|0|0|0|0|0|0|0|0|0|0|0|0|0|0|
|virus_Jul11.50v_16194|1|0|0|0|0|0|0|0|0|0|0|0|0|0|0|0|0|0|0|0|0|0|0|0|0|0|0|0|1|0|0|0|0|0|0|0|0|0|0|0|0|0|0|0|0|0|0|0|0|0|0|0|0|0|0|
|virus_Jul11.50v_18630|1|0|0|0|0|0|0|0|0|0|0|0|0|0|0|0|0|0|0|0|0|0|0|0|0|0|0|0|1|0|0|0|0|0|0|0|0|0|0|0|0|0|0|0|0|0|0|0|0|0|0|0|0|0|0|
|virus_Jul11.75v_2531|1|0|0|0|0|0|0|0|0|0|0|0|0|0|0|0|0|0|0|0|0|0|0|0|0|0|0|0|0|0|1|0|0|0|0|0|0|0|0|0|0|0|0|0|0|0|0|0|0|0|0|0|0|0|0|
|virus_Jul11.75v_12470|1|0|0|0|0|0|0|0|0|0|0|0|0|0|0|0|0|0|0|0|0|0|0|0|0|0|0|0|0|0|1|0|0|0|0|0|0|0|0|0|0|0|0|0|0|0|0|0|0|0|0|0|0|0|0|
|virus_Jul11.75v_23511|1|0|0|0|0|0|0|0|0|0|0|0|0|0|0|0|0|0|0|0|0|0|0|0|0|0|0|0|0|0|1|0|0|0|0|0|0|0|0|0|0|0|0|0|0|0|0|0|0|0|0|0|0|0|0|
|virus_Jul11.75v_27674|1|0|0|0|0|0|0|0|0|0|0|0|0|0|0|0|0|0|0|0|0|0|0|0|0|0|0|0|0|0|1|0|0|0|0|0|0|0|0|0|0|0|0|0|0|0|0|0|0|0|0|0|0|0|0|
|virus_Jul11.75v_32918|1|0|0|0|0|0|0|0|0|0|0|0|0|0|0|0|0|0|0|0|0|0|0|0|0|0|0|0|0|0|1|0|0|0|0|0|0|0|0|0|0|0|0|0|0|0|0|0|0|0|0|0|0|0|0|
|virus_Jul11.75v_34246|1|0|0|0|0|0|0|0|0|0|0|0|0|0|0|0|0|0|0|0|0|0|0|0|0|0|0|0|0|0|1|0|0|0|0|0|0|0|0|0|0|0|0|0|0|0|0|0|0|0|0|0|0|0|0|
|virus_Jul11.75v_38552|1|0|0|0|0|0|0|0|0|0|0|0|0|0|0|0|0|0|0|0|0|0|0|0|0|0|0|0|0|0|1|0|0|0|0|0|0|0|0|0|0|0|0|0|0|0|0|0|0|0|0|0|0|0|0|
|virus_Jul27.5v_370|1|0|0|0|0|0|0|0|0|0|0|0|0|0|0|0|0|0|0|0|0|0|0|0|0|0|0|0|0|0|0|0|1|0|0|0|0|0|0|0|0|0|0|0|0|0|0|0|0|0|0|0|0|0|0|
|virus_Jul27.5v_1138|1|0|0|0|0|0|0|0|0|0|0|0|0|0|0|0|0|0|0|0|0|0|0|0|0|0|0|0|0|0|0|0|1|0|0|0|0|0|0|0|0|0|0|0|0|0|0|0|0|0|0|0|0|0|0|
|virus_Jul27.5v_3543|1|0|0|0|0|0|0|0|0|0|0|0|0|0|0|0|0|0|0|0|0|0|0|0|0|0|0|0|0|0|0|0|1|0|0|0|0|0|0|0|0|0|0|0|0|0|0|0|0|0|0|0|0|0|0|
|virus_Jul27.5v_3646|1|0|0|0|0|0|0|0|0|0|0|0|0|0|0|0|0|0|0|0|0|0|0|0|0|0|0|0|0|0|0|0|1|0|0|0|0|0|0|0|0|0|0|0|0|0|0|0|0|0|0|0|0|0|0|
|virus_Jul27.5v_6013|1|0|0|0|0|0|0|0|0|0|0|0|0|0|0|0|0|0|0|0|0|0|0|0|0|0|0|0|0|0|0|0|1|0|0|0|0|0|0|0|0|0|0|0|0|0|0|0|0|0|0|0|0|0|0|
|virus_Jul27.5v_6125|1|0|0|0|0|0|0|0|0|0|0|0|0|0|0|0|0|0|0|0|0|0|0|0|0|0|0|0|0|0|0|0|1|0|0|0|0|0|0|0|0|0|0|0|0|0|0|0|0|0|0|0|0|0|0|
|virus_Jul27.5v_8885|1|0|0|0|0|0|0|0|0|0|0|0|0|0|0|0|0|0|0|0|0|0|0|0|0|0|0|0|0|0|0|0|1|0|0|0|0|0|0|0|0|0|0|0|0|0|0|0|0|0|0|0|0|0|0|
|virus_Jul27.5v_8922|1|0|0|0|0|0|0|0|0|0|0|0|0|0|0|0|0|0|0|0|0|0|0|0|0|0|0|0|0|0|0|0|1|0|0|0|0|0|0|0|0|0|0|0|0|0|0|0|0|0|0|0|0|0|0|
|virus_Jul27.5v_9777|1|0|0|0|0|0|0|0|0|0|0|0|0|0|0|0|0|0|0|0|0|0|0|0|0|0|0|0|0|0|0|0|1|0|0|0|0|0|0|0|0|0|0|0|0|0|0|0|0|0|0|0|0|0|0|
|virus_Jul27.5v_9857|1|0|0|0|0|0|0|0|0|0|0|0|0|0|0|0|0|0|0|0|0|0|0|0|0|0|0|0|0|0|0|0|1|0|0|0|0|0|0|0|0|0|0|0|0|0|0|0|0|0|0|0|0|0|0|
|virus_Jul27.5v_10389|1|0|0|0|0|0|0|0|0|0|0|0|0|0|0|0|0|0|0|0|0|0|0|0|0|0|0|0|0|0|0|0|1|0|0|0|0|0|0|0|0|0|0|0|0|0|0|0|0|0|0|0|0|0|0|
|virus_Jul27.5v_11463|1|0|0|0|0|0|0|0|0|0|0|0|0|0|0|0|0|0|0|0|0|0|0|0|0|0|0|0|0|0|0|0|1|0|0|0|0|0|0|0|0|0|0|0|0|0|0|0|0|0|0|0|0|0|0|
|virus_Jul27.5v_12791|1|0|0|0|0|0|0|0|0|0|0|0|0|0|0|0|0|0|0|0|0|0|0|0|0|0|0|0|0|0|0|0|1|0|0|0|0|0|0|0|0|0|0|0|0|0|0|0|0|0|0|0|0|0|0|
|virus_Jul27.5v_14925|1|0|0|0|0|0|0|0|0|0|0|0|0|0|0|0|0|0|0|0|0|0|0|0|0|0|0|0|0|0|0|0|1|0|0|0|0|0|0|0|0|0|0|0|0|0|0|0|0|0|0|0|0|0|0|
|virus_Jul27.5v_16264|1|0|0|0|0|0|0|0|0|0|0|0|0|0|0|0|0|0|0|0|0|0|0|0|0|0|0|0|0|0|0|0|1|0|0|0|0|0|0|0|0|0|0|0|0|0|0|0|0|0|0|0|0|0|0|
|virus_Jul27.5v_16298|1|0|0|0|0|0|0|0|0|0|0|0|0|0|0|0|0|0|0|0|0|0|0|0|0|0|0|0|0|0|0|0|1|0|0|0|0|0|0|0|0|0|0|0|0|0|0|0|0|0|0|0|0|0|0|
|virus_Jul27.5v_17000|1|0|0|0|0|0|0|0|0|0|0|0|0|0|0|0|0|0|0|0|0|0|0|0|0|0|0|0|0|0|0|0|1|0|0|0|0|0|0|0|0|0|0|0|0|0|0|0|0|0|0|0|0|0|0|
|virus_Jul27.5v_17644|1|0|0|0|0|0|0|0|0|0|0|0|0|0|0|0|0|0|0|0|0|0|0|0|0|0|0|0|0|0|0|0|1|0|0|0|0|0|0|0|0|0|0|0|0|0|0|0|0|0|0|0|0|0|0|
|virus_Jul27.5v_18255|1|0|0|0|0|0|0|0|0|0|0|0|0|0|0|0|0|0|0|0|0|0|0|0|0|0|0|0|0|0|0|0|1|0|0|0|0|0|0|0|0|0|0|0|0|0|0|0|0|0|0|0|0|0|0|
|virus_Jul27.5v_18965|1|0|0|0|0|0|0|0|0|0|0|0|0|0|0|0|0|0|0|0|0|0|0|0|0|0|0|0|0|0|0|0|1|0|0|0|0|0|0|0|0|0|0|0|0|0|0|0|0|0|0|0|0|0|0|
|virus_Jul27.5v_19824|1|0|0|0|0|0|0|0|0|0|0|0|0|0|0|0|0|0|0|0|0|0|0|0|0|0|0|0|0|0|0|0|1|0|0|0|0|0|0|0|0|0|0|0|0|0|0|0|0|0|0|0|0|0|0|
|virus_Jul27.5v_20005|1|0|0|0|0|0|0|0|0|0|0|0|0|0|0|0|0|0|0|0|0|0|0|0|0|0|0|0|0|0|0|0|1|0|0|0|0|0|0|0|0|0|0|0|0|0|0|0|0|0|0|0|0|0|0|
|virus_Jul27.5v_20218|1|0|0|0|0|0|0|0|0|0|0|0|0|0|0|0|0|0|0|0|0|0|0|0|0|0|0|0|0|0|0|0|1|0|0|0|0|0|0|0|0|0|0|0|0|0|0|0|0|0|0|0|0|0|0|
|virus_Jul27.25v_193|1|0|0|0|0|0|0|0|0|0|0|0|0|0|0|0|0|0|0|0|0|0|0|0|0|0|0|0|0|0|0|0|0|0|1|0|0|0|0|0|0|0|0|0|0|0|0|0|0|0|0|0|0|0|0|
|virus_Jul27.25v_412|1|0|0|0|0|0|0|0|0|0|0|0|0|0|0|0|0|0|0|0|0|0|0|0|0|0|0|0|0|0|0|0|0|0|1|0|0|0|0|0|0|0|0|0|0|0|0|0|0|0|0|0|0|0|0|
|virus_Jul27.25v_861|1|0|0|0|0|0|0|0|0|0|0|0|0|0|0|0|0|0|0|0|0|0|0|0|0|0|0|0|0|0|0|0|0|0|1|0|0|0|0|0|0|0|0|0|0|0|0|0|0|0|0|0|0|0|0|
|virus_Jul27.25v_9152|1|0|0|0|0|0|0|0|0|0|0|0|0|0|0|0|0|0|0|0|0|0|0|0|0|0|0|0|0|0|0|0|0|0|1|0|0|0|0|0|0|0|0|0|0|0|0|0|0|0|0|0|0|0|0|
|virus_Jul27.25v_9384|1|0|0|0|0|0|0|0|0|0|0|0|0|0|0|0|0|0|0|0|0|0|0|0|0|0|0|0|0|0|0|0|0|0|1|0|0|0|0|0|0|0|0|0|0|0|0|0|0|0|0|0|0|0|0|
|virus_Jul27.25v_10021|1|0|0|0|0|0|0|0|0|0|0|0|0|0|0|0|0|0|0|0|0|0|0|0|0|0|0|0|0|0|0|0|0|0|1|0|0|0|0|0|0|0|0|0|0|0|0|0|0|0|0|0|0|0|0|
|virus_Jul27.25v_12717|1|0|0|0|0|0|0|0|0|0|0|0|0|0|0|0|0|0|0|0|0|0|0|0|0|0|0|0|0|0|0|0|0|0|1|0|0|0|0|0|0|0|0|0|0|0|0|0|0|0|0|0|0|0|0|
|virus_Jul27.25v_16514|1|0|0|0|0|0|0|0|0|0|0|0|0|0|0|0|0|0|0|0|0|0|0|0|0|0|0|0|0|0|0|0|0|0|1|0|0|0|0|0|0|0|0|0|0|0|0|0|0|0|0|0|0|0|0|
|virus_Jul27.25v_17182|1|0|0|0|0|0|0|0|0|0|0|0|0|0|0|0|0|0|0|0|0|0|0|0|0|0|0|0|0|0|0|0|0|0|1|0|0|0|0|0|0|0|0|0|0|0|0|0|0|0|0|0|0|0|0|
|virus_Jul27.25v_19006|1|0|0|0|0|0|0|0|0|0|0|0|0|0|0|0|0|0|0|0|0|0|0|0|0|0|0|0|0|0|0|0|0|0|1|0|0|0|0|0|0|0|0|0|0|0|0|0|0|0|0|0|0|0|0|
|virus_Jul27.25v_19311|1|0|0|0|0|0|0|0|0|0|0|0|0|0|0|0|0|0|0|0|0|0|0|0|0|0|0|0|0|0|0|0|0|0|1|0|0|0|0|0|0|0|0|0|0|0|0|0|0|0|0|0|0|0|0|
|virus_Jul27.25v_25018|1|0|0|0|0|0|0|0|0|0|0|0|0|0|0|0|0|0|0|0|0|0|0|0|0|0|0|0|0|0|0|0|0|0|1|0|0|0|0|0|0|0|0|0|0|0|0|0|0|0|0|0|0|0|0|
|virus_Jul27.25v_39273|1|0|0|0|0|0|0|0|0|0|0|0|0|0|0|0|0|0|0|0|0|0|0|0|0|0|0|0|0|0|0|0|0|0|1|0|0|0|0|0|0|0|0|0|0|0|0|0|0|0|0|0|0|0|0|
|virus_Jul27.25v_39279|1|0|0|0|0|0|0|0|0|0|0|0|0|0|0|0|0|0|0|0|0|0|0|0|0|0|0|0|0|0|0|0|0|0|1|0|0|0|0|0|0|0|0|0|0|0|0|0|0|0|0|0|0|0|0|
|virus_Jul27.75v_3872|1|0|0|0|0|0|0|0|0|0|0|0|0|0|0|0|0|0|0|0|0|0|0|0|0|0|0|0|0|0|0|0|0|0|0|0|1|0|0|0|0|0|0|0|0|0|0|0|0|0|0|0|0|0|0|
|virus_Jun8.25v_468|1|0|0|0|0|0|0|0|0|0|1|0|0|0|0|0|0|0|0|0|0|0|0|0|0|0|0|0|0|0|0|0|0|0|0|0|0|0|0|0|0|0|0|0|0|0|0|0|0|0|0|0|0|0|0|
|virus_Jun8.25v_2122|1|0|0|0|0|0|0|0|0|0|1|0|0|0|0|0|0|0|0|0|0|0|0|0|0|0|0|0|0|0|0|0|0|0|0|0|0|0|0|0|0|0|0|0|0|0|0|0|0|0|0|0|0|0|0|
|virus_Jun8.25v_2418|1|0|0|0|0|0|0|0|0|0|1|0|0|0|0|0|0|0|0|0|0|0|0|0|0|0|0|0|0|0|0|0|0|0|0|0|0|0|0|0|0|0|0|0|0|0|0|0|0|0|0|0|0|0|0|
|virus_Jun8.25v_36017|1|0|0|0|0|0|0|0|0|0|1|0|0|0|0|0|0|0|0|0|0|0|0|0|0|0|0|0|0|0|0|0|0|0|0|0|0|0|0|0|0|0|0|0|0|0|0|0|0|0|0|0|0|0|0|
|virus_Jun8.50v_686|1|0|0|0|0|0|0|0|0|0|0|0|1|0|0|0|0|0|0|0|0|0|0|0|0|0|0|0|0|0|0|0|0|0|0|0|0|0|0|0|0|0|0|0|0|0|0|0|0|0|0|0|0|0|0|
|virus_Jun8.50v_2793|1|0|0|0|0|0|0|0|0|0|0|0|1|0|0|0|0|0|0|0|0|0|0|0|0|0|0|0|0|0|0|0|0|0|0|0|0|0|0|0|0|0|0|0|0|0|0|0|0|0|0|0|0|0|0|
|virus_Jun8.50v_4633|1|0|0|0|0|0|0|0|0|0|0|0|1|0|0|0|0|0|0|0|0|0|0|0|0|0|0|0|0|0|0|0|0|0|0|0|0|0|0|0|0|0|0|0|0|0|0|0|0|0|0|0|0|0|0|
|virus_Jun8.50v_9271|1|0|0|0|0|0|0|0|0|0|0|0|1|0|0|0|0|0|0|0|0|0|0|0|0|0|0|0|0|0|0|0|0|0|0|0|0|0|0|0|0|0|0|0|0|0|0|0|0|0|0|0|0|0|0|
|virus_Jun8.50v_11912|1|0|0|0|0|0|0|0|0|0|0|0|1|0|0|0|0|0|0|0|0|0|0|0|0|0|0|0|0|0|0|0|0|0|0|0|0|0|0|0|0|0|0|0|0|0|0|0|0|0|0|0|0|0|0|
|virus_Jun8.50v_13861|1|0|0|0|0|0|0|0|0|0|0|0|1|0|0|0|0|0|0|0|0|0|0|0|0|0|0|0|0|0|0|0|0|0|0|0|0|0|0|0|0|0|0|0|0|0|0|0|0|0|0|0|0|0|0|
|virus_Jun8.50v_18944|1|0|0|0|0|0|0|0|0|0|0|0|1|0|0|0|0|0|0|0|0|0|0|0|0|0|0|0|0|0|0|0|0|0|0|0|0|0|0|0|0|0|0|0|0|0|0|0|0|0|0|0|0|0|0|
|virus_Jun8.50v_19908|1|0|0|0|0|0|0|0|0|0|0|0|1|0|0|0|0|0|0|0|0|0|0|0|0|0|0|0|0|0|0|0|0|0|0|0|0|0|0|0|0|0|0|0|0|0|0|0|0|0|0|0|0|0|0|
|virus_Jun8.50v_23587|1|0|0|0|0|0|0|0|0|0|0|0|1|0|0|0|0|0|0|0|0|0|0|0|0|0|0|0|0|0|0|0|0|0|0|0|0|0|0|0|0|0|0|0|0|0|0|0|0|0|0|0|0|0|0|
|virus_Jun8.50v_32285|1|0|0|0|0|0|0|0|0|0|0|0|1|0|0|0|0|0|0|0|0|0|0|0|0|0|0|0|0|0|0|0|0|0|0|0|0|0|0|0|0|0|0|0|0|0|0|0|0|0|0|0|0|0|0|
|virus_Jun8.50v_32983|1|0|0|0|0|0|0|0|0|0|0|0|1|0|0|0|0|0|0|0|0|0|0|0|0|0|0|0|0|0|0|0|0|0|0|0|0|0|0|0|0|0|0|0|0|0|0|0|0|0|0|0|0|0|0|
|virus_Jun8.50v_37130|1|0|0|0|0|0|0|0|0|0|0|0|1|0|0|0|0|0|0|0|0|0|0|0|0|0|0|0|0|0|0|0|0|0|0|0|0|0|0|0|0|0|0|0|0|0|0|0|0|0|0|0|0|0|0|
|virus_Jun8.75v_6003|1|0|0|0|0|0|0|0|0|0|0|0|0|0|1|0|0|0|0|0|0|0|0|0|0|0|0|0|0|0|0|0|0|0|0|0|0|0|0|0|0|0|0|0|0|0|0|0|0|0|0|0|0|0|0|
|virus_Jun8.75v_8467|1|0|0|0|0|0|0|0|0|0|0|0|0|0|1|0|0|0|0|0|0|0|0|0|0|0|0|0|0|0|0|0|0|0|0|0|0|0|0|0|0|0|0|0|0|0|0|0|0|0|0|0|0|0|0|
|virus_Jun22.0v_839|1|0|0|0|0|0|0|0|0|0|0|0|0|0|0|0|1|0|0|0|0|0|0|0|0|0|0|0|0|0|0|0|0|0|0|0|0|0|0|0|0|0|0|0|0|0|0|0|0|0|0|0|0|0|0|
|virus_Jun22.0v_2092|1|0|0|0|0|0|0|0|0|0|0|0|0|0|0|0|1|0|0|0|0|0|0|0|0|0|0|0|0|0|0|0|0|0|0|0|0|0|0|0|0|0|0|0|0|0|0|0|0|0|0|0|0|0|0|
|virus_Jun22.0v_13800|1|0|0|0|0|0|0|0|0|0|0|0|0|0|0|0|1|0|0|0|0|0|0|0|0|0|0|0|0|0|0|0|0|0|0|0|0|0|0|0|0|0|0|0|0|0|0|0|0|0|0|0|0|0|0|
|virus_Jun22.0v_17340|1|0|0|0|0|0|0|0|0|0|0|0|0|0|0|0|1|0|0|0|0|0|0|0|0|0|0|0|0|0|0|0|0|0|0|0|0|0|0|0|0|0|0|0|0|0|0|0|0|0|0|0|0|0|0|
|virus_Jun22.0v_19526|1|0|0|0|0|0|0|0|0|0|0|0|0|0|0|0|1|0|0|0|0|0|0|0|0|0|0|0|0|0|0|0|0|0|0|0|0|0|0|0|0|0|0|0|0|0|0|0|0|0|0|0|0|0|0|
|virus_Jun22.50v_1067|1|0|0|0|0|0|0|0|0|0|0|0|0|0|0|0|0|0|1|0|0|0|0|0|0|0|0|0|0|0|0|0|0|0|0|0|0|0|0|0|0|0|0|0|0|0|0|0|0|0|0|0|0|0|0|
|virus_Jun22.50v_7192|1|0|0|0|0|0|0|0|0|0|0|0|0|0|0|0|0|0|1|0|0|0|0|0|0|0|0|0|0|0|0|0|0|0|0|0|0|0|0|0|0|0|0|0|0|0|0|0|0|0|0|0|0|0|0|
|virus_Jun22.50v_9394|1|0|0|0|0|0|0|0|0|0|0|0|0|0|0|0|0|0|1|0|0|0|0|0|0|0|0|0|0|0|0|0|0|0|0|0|0|0|0|0|0|0|0|0|0|0|0|0|0|0|0|0|0|0|0|
|virus_Jun22.50v_11217|1|0|0|0|0|0|0|0|0|0|0|0|0|0|0|0|0|0|1|0|0|0|0|0|0|0|0|0|0|0|0|0|0|0|0|0|0|0|0|0|0|0|0|0|0|0|0|0|0|0|0|0|0|0|0|
|virus_Jun22.50v_13684|1|0|0|0|0|0|0|0|0|0|0|0|0|0|0|0|0|0|1|0|0|0|0|0|0|0|0|0|0|0|0|0|0|0|0|0|0|0|0|0|0|0|0|0|0|0|0|0|0|0|0|0|0|0|0|
|virus_Jun22.50v_15849|1|0|0|0|0|0|0|0|0|0|0|0|0|0|0|0|0|0|1|0|0|0|0|0|0|0|0|0|0|0|0|0|0|0|0|0|0|0|0|0|0|0|0|0|0|0|0|0|0|0|0|0|0|0|0|
|virus_Jun22.50v_16076|1|0|0|0|0|0|0|0|0|0|0|0|0|0|0|0|0|0|1|0|0|0|0|0|0|0|0|0|0|0|0|0|0|0|0|0|0|0|0|0|0|0|0|0|0|0|0|0|0|0|0|0|0|0|0|
|virus_Jun22.50v_16395|1|0|0|0|0|0|0|0|0|0|0|0|0|0|0|0|0|0|1|0|0|0|0|0|0|0|0|0|0|0|0|0|0|0|0|0|0|0|0|0|0|0|0|0|0|0|0|0|0|0|0|0|0|0|0|
|virus_Jun22.50v_19149|1|0|0|0|0|0|0|0|0|0|0|0|0|0|0|0|0|0|1|0|0|0|0|0|0|0|0|0|0|0|0|0|0|0|0|0|0|0|0|0|0|0|0|0|0|0|0|0|0|0|0|0|0|0|0|
|virus_Jun22.50v_23024|1|0|0|0|0|0|0|0|0|0|0|0|0|0|0|0|0|0|1|0|0|0|0|0|0|0|0|0|0|0|0|0|0|0|0|0|0|0|0|0|0|0|0|0|0|0|0|0|0|0|0|0|0|0|0|
|virus_Jun22.50v_25119|1|0|0|0|0|0|0|0|0|0|0|0|0|0|0|0|0|0|1|0|0|0|0|0|0|0|0|0|0|0|0|0|0|0|0|0|0|0|0|0|0|0|0|0|0|0|0|0|0|0|0|0|0|0|0|
|virus_Jun22.50v_25905|1|0|0|0|0|0|0|0|0|0|0|0|0|0|0|0|0|0|1|0|0|0|0|0|0|0|0|0|0|0|0|0|0|0|0|0|0|0|0|0|0|0|0|0|0|0|0|0|0|0|0|0|0|0|0|
|virus_Jun22.50v_31945|1|0|0|0|0|0|0|0|0|0|0|0|0|0|0|0|0|0|1|0|0|0|0|0|0|0|0|0|0|0|0|0|0|0|0|0|0|0|0|0|0|0|0|0|0|0|0|0|0|0|0|0|0|0|0|
|virus_Jun22.50v_32309|1|0|0|0|0|0|0|0|0|0|0|0|0|0|0|0|0|0|1|0|0|0|0|0|0|0|0|0|0|0|0|0|0|0|0|0|0|0|0|0|0|0|0|0|0|0|0|0|0|0|0|0|0|0|0|
|virus_Jun22.50v_33242|1|0|0|0|0|0|0|0|0|0|0|0|0|0|0|0|0|0|1|0|0|0|0|0|0|0|0|0|0|0|0|0|0|0|0|0|0|0|0|0|0|0|0|0|0|0|0|0|0|0|0|0|0|0|0|
|virus_Jun22.50v_33321|1|0|0|0|0|0|0|0|0|0|0|0|0|0|0|0|0|0|1|0|0|0|0|0|0|0|0|0|0|0|0|0|0|0|0|0|0|0|0|0|0|0|0|0|0|0|0|0|0|0|0|0|0|0|0|
|virus_Jun22.50v_35089|1|0|0|0|0|0|0|0|0|0|0|0|0|0|0|0|0|0|1|0|0|0|0|0|0|0|0|0|0|0|0|0|0|0|0|0|0|0|0|0|0|0|0|0|0|0|0|0|0|0|0|0|0|0|0|
|virus_Jun22.50v_38892|1|0|0|0|0|0|0|0|0|0|0|0|0|0|0|0|0|0|1|0|0|0|0|0|0|0|0|0|0|0|0|0|0|0|0|0|0|0|0|0|0|0|0|0|0|0|0|0|0|0|0|0|0|0|0|
|virus_Jun22.50v_39859|1|0|0|0|0|0|0|0|0|0|0|0|0|0|0|0|0|0|1|0|0|0|0|0|0|0|0|0|0|0|0|0|0|0|0|0|0|0|0|0|0|0|0|0|0|0|0|0|0|0|0|0|0|0|0|
|virus_Jun22.50v_40139|1|0|0|0|0|0|0|0|0|0|0|0|0|0|0|0|0|0|1|0|0|0|0|0|0|0|0|0|0|0|0|0|0|0|0|0|0|0|0|0|0|0|0|0|0|0|0|0|0|0|0|0|0|0|0|
|virus_Jun22.50v_40868|1|0|0|0|0|0|0|0|0|0|0|0|0|0|0|0|0|0|1|0|0|0|0|0|0|0|0|0|0|0|0|0|0|0|0|0|0|0|0|0|0|0|0|0|0|0|0|0|0|0|0|0|0|0|0|
|virus_Jun22.75v_2594|1|0|0|0|0|0|0|0|0|0|0|0|0|0|0|0|0|0|0|0|1|0|0|0|0|0|0|0|0|0|0|0|0|0|0|0|0|0|0|0|0|0|0|0|0|0|0|0|0|0|0|0|0|0|0|
|virus_Jun22.75v_7335|1|0|0|0|0|0|0|0|0|0|0|0|0|0|0|0|0|0|0|0|1|0|0|0|0|0|0|0|0|0|0|0|0|0|0|0|0|0|0|0|0|0|0|0|0|0|0|0|0|0|0|0|0|0|0|
|virus_Jun22.75v_13569|1|0|0|0|0|0|0|0|0|0|0|0|0|0|0|0|0|0|0|0|1|0|0|0|0|0|0|0|0|0|0|0|0|0|0|0|0|0|0|0|0|0|0|0|0|0|0|0|0|0|0|0|0|0|0|
|virus_Jun22.75v_20936|1|0|0|0|0|0|0|0|0|0|0|0|0|0|0|0|0|0|0|0|1|0|0|0|0|0|0|0|0|0|0|0|0|0|0|0|0|0|0|0|0|0|0|0|0|0|0|0|0|0|0|0|0|0|0|
|virus_Jun22.75v_28096|1|0|0|0|0|0|0|0|0|0|0|0|0|0|0|0|0|0|0|0|1|0|0|0|0|0|0|0|0|0|0|0|0|0|0|0|0|0|0|0|0|0|0|0|0|0|0|0|0|0|0|0|0|0|0|
|virus_May12.5v_4693|1|0|1|0|0|0|0|0|0|0|0|0|0|0|0|0|0|0|0|0|0|0|0|0|0|0|0|0|0|0|0|0|0|0|0|0|0|0|0|0|0|0|0|0|0|0|0|0|0|0|0|0|0|0|0|
|virus_May12.5v_7454|1|0|1|0|0|0|0|0|0|0|0|0|0|0|0|0|0|0|0|0|0|0|0|0|0|0|0|0|0|0|0|0|0|0|0|0|0|0|0|0|0|0|0|0|0|0|0|0|0|0|0|0|0|0|0|
|virus_May12.5v_10900|1|0|1|0|0|0|0|0|0|0|0|0|0|0|0|0|0|0|0|0|0|0|0|0|0|0|0|0|0|0|0|0|0|0|0|0|0|0|0|0|0|0|0|0|0|0|0|0|0|0|0|0|0|0|0|
|virus_May12.5v_11516|1|0|1|0|0|0|0|0|0|0|0|0|0|0|0|0|0|0|0|0|0|0|0|0|0|0|0|0|0|0|0|0|0|0|0|0|0|0|0|0|0|0|0|0|0|0|0|0|0|0|0|0|0|0|0|
|virus_May12.5v_19292|1|0|1|0|0|0|0|0|0|0|0|0|0|0|0|0|0|0|0|0|0|0|0|0|0|0|0|0|0|0|0|0|0|0|0|0|0|0|0|0|0|0|0|0|0|0|0|0|0|0|0|0|0|0|0|
|virus_May12.5v_29532|1|0|1|0|0|0|0|0|0|0|0|0|0|0|0|0|0|0|0|0|0|0|0|0|0|0|0|0|0|0|0|0|0|0|0|0|0|0|0|0|0|0|0|0|0|0|0|0|0|0|0|0|0|0|0|
|virus_May12.5v_38052|1|0|1|0|0|0|0|0|0|0|0|0|0|0|0|0|0|0|0|0|0|0|0|0|0|0|0|0|0|0|0|0|0|0|0|0|0|0|0|0|0|0|0|0|0|0|0|0|0|0|0|0|0|0|0|
|virus_May12.25v_9039|1|0|0|0|1|0|0|0|0|0|0|0|0|0|0|0|0|0|0|0|0|0|0|0|0|0|0|0|0|0|0|0|0|0|0|0|0|0|0|0|0|0|0|0|0|0|0|0|0|0|0|0|0|0|0|
|virus_May12.25v_9204|1|0|0|0|1|0|0|0|0|0|0|0|0|0|0|0|0|0|0|0|0|0|0|0|0|0|0|0|0|0|0|0|0|0|0|0|0|0|0|0|0|0|0|0|0|0|0|0|0|0|0|0|0|0|0|
|virus_May12.25v_17390|1|0|0|0|1|0|0|0|0|0|0|0|0|0|0|0|0|0|0|0|0|0|0|0|0|0|0|0|0|0|0|0|0|0|0|0|0|0|0|0|0|0|0|0|0|0|0|0|0|0|0|0|0|0|0|
|virus_May12.25v_22625|1|0|0|0|1|0|0|0|0|0|0|0|0|0|0|0|0|0|0|0|0|0|0|0|0|0|0|0|0|0|0|0|0|0|0|0|0|0|0|0|0|0|0|0|0|0|0|0|0|0|0|0|0|0|0|
|virus_May12.25v_22808|1|0|0|0|1|0|0|0|0|0|0|0|0|0|0|0|0|0|0|0|0|0|0|0|0|0|0|0|0|0|0|0|0|0|0|0|0|0|0|0|0|0|0|0|0|0|0|0|0|0|0|0|0|0|0|
|virus_May12.25v_31397|1|0|0|0|1|0|0|0|0|0|0|0|0|0|0|0|0|0|0|0|0|0|0|0|0|0|0|0|0|0|0|0|0|0|0|0|0|0|0|0|0|0|0|0|0|0|0|0|0|0|0|0|0|0|0|
|virus_May12.25v_39988|1|0|0|0|1|0|0|0|0|0|0|0|0|0|0|0|0|0|0|0|0|0|0|0|0|0|0|0|0|0|0|0|0|0|0|0|0|0|0|0|0|0|0|0|0|0|0|0|0|0|0|0|0|0|0|
|virus_May12.50v_1250|1|0|0|0|0|0|1|0|0|0|0|0|0|0|0|0|0|0|0|0|0|0|0|0|0|0|0|0|0|0|0|0|0|0|0|0|0|0|0|0|0|0|0|0|0|0|0|0|0|0|0|0|0|0|0|
|virus_May12.50v_10400|1|0|0|0|0|0|1|0|0|0|0|0|0|0|0|0|0|0|0|0|0|0|0|0|0|0|0|0|0|0|0|0|0|0|0|0|0|0|0|0|0|0|0|0|0|0|0|0|0|0|0|0|0|0|0|
|virus_May12.50v_12063|1|0|0|0|0|0|1|0|0|0|0|0|0|0|0|0|0|0|0|0|0|0|0|0|0|0|0|0|0|0|0|0|0|0|0|0|0|0|0|0|0|0|0|0|0|0|0|0|0|0|0|0|0|0|0|
|virus_May12.50v_17053|1|0|0|0|0|0|1|0|0|0|0|0|0|0|0|0|0|0|0|0|0|0|0|0|0|0|0|0|0|0|0|0|0|0|0|0|0|0|0|0|0|0|0|0|0|0|0|0|0|0|0|0|0|0|0|
|virus_May12.50v_20096|1|0|0|0|0|0|1|0|0|0|0|0|0|0|0|0|0|0|0|0|0|0|0|0|0|0|0|0|0|0|0|0|0|0|0|0|0|0|0|0|0|0|0|0|0|0|0|0|0|0|0|0|0|0|0|
|virus_May12.50v_20803|1|0|0|0|0|0|1|0|0|0|0|0|0|0|0|0|0|0|0|0|0|0|0|0|0|0|0|0|0|0|0|0|0|0|0|0|0|0|0|0|0|0|0|0|0|0|0|0|0|0|0|0|0|0|0|
|virus_May12.50v_23630|1|0|0|0|0|0|1|0|0|0|0|0|0|0|0|0|0|0|0|0|0|0|0|0|0|0|0|0|0|0|0|0|0|0|0|0|0|0|0|0|0|0|0|0|0|0|0|0|0|0|0|0|0|0|0|
|virus_May12.50v_24948|1|0|0|0|0|0|1|0|0|0|0|0|0|0|0|0|0|0|0|0|0|0|0|0|0|0|0|0|0|0|0|0|0|0|0|0|0|0|0|0|0|0|0|0|0|0|0|0|0|0|0|0|0|0|0|
|virus_May12.50v_27008|1|0|0|0|0|0|1|0|0|0|0|0|0|0|0|0|0|0|0|0|0|0|0|0|0|0|0|0|0|0|0|0|0|0|0|0|0|0|0|0|0|0|0|0|0|0|0|0|0|0|0|0|0|0|0|
|virus_May12.50v_29488|1|0|0|0|0|0|1|0|0|0|0|0|0|0|0|0|0|0|0|0|0|0|0|0|0|0|0|0|0|0|0|0|0|0|0|0|0|0|0|0|0|0|0|0|0|0|0|0|0|0|0|0|0|0|0|
|virus_May12.50v_30715|1|0|0|0|0|0|1|0|0|0|0|0|0|0|0|0|0|0|0|0|0|0|0|0|0|0|0|0|0|0|0|0|0|0|0|0|0|0|0|0|0|0|0|0|0|0|0|0|0|0|0|0|0|0|0|
|virus_May12.50v_32540|1|0|0|0|0|0|1|0|0|0|0|0|0|0|0|0|0|0|0|0|0|0|0|0|0|0|0|0|0|0|0|0|0|0|0|0|0|0|0|0|0|0|0|0|0|0|0|0|0|0|0|0|0|0|0|
|virus_May12.50v_39712|1|0|0|0|0|0|1|0|0|0|0|0|0|0|0|0|0|0|0|0|0|0|0|0|0|0|0|0|0|0|0|0|0|0|0|0|0|0|0|0|0|0|0|0|0|0|0|0|0|0|0|0|0|0|0|
|virus_May12.50v_39758|1|0|0|0|0|0|1|0|0|0|0|0|0|0|0|0|0|0|0|0|0|0|0|0|0|0|0|0|0|0|0|0|0|0|0|0|0|0|0|0|0|0|0|0|0|0|0|0|0|0|0|0|0|0|0|
|virus_May12.75v_14789|1|0|0|0|0|0|0|0|1|0|0|0|0|0|0|0|0|0|0|0|0|0|0|0|0|0|0|0|0|0|0|0|0|0|0|0|0|0|0|0|0|0|0|0|0|0|0|0|0|0|0|0|0|0|0|
|virus_May12.75v_20699|1|0|0|0|0|0|0|0|1|0|0|0|0|0|0|0|0|0|0|0|0|0|0|0|0|0|0|0|0|0|0|0|0|0|0|0|0|0|0|0|0|0|0|0|0|0|0|0|0|0|0|0|0|0|0|
|virus_May12.75v_22093|1|0|0|0|0|0|0|0|1|0|0|0|0|0|0|0|0|0|0|0|0|0|0|0|0|0|0|0|0|0|0|0|0|0|0|0|0|0|0|0|0|0|0|0|0|0|0|0|0|0|0|0|0|0|0|
|virus_May12.75v_26017|1|0|0|0|0|0|0|0|1|0|0|0|0|0|0|0|0|0|0|0|0|0|0|0|0|0|0|0|0|0|0|0|0|0|0|0|0|0|0|0|0|0|0|0|0|0|0|0|0|0|0|0|0|0|0|
|virus_May12.75v_29072|1|0|0|0|0|0|0|0|1|0|0|0|0|0|0|0|0|0|0|0|0|0|0|0|0|0|0|0|0|0|0|0|0|0|0|0|0|0|0|0|0|0|0|0|0|0|0|0|0|0|0|0|0|0|0|
|virus_May12.75v_32293|1|0|0|0|0|0|0|0|1|0|0|0|0|0|0|0|0|0|0|0|0|0|0|0|0|0|0|0|0|0|0|0|0|0|0|0|0|0|0|0|0|0|0|0|0|0|0|0|0|0|0|0|0|0|0|
|virus_May12.75v_33047|1|0|0|0|0|0|0|0|1|0|0|0|0|0|0|0|0|0|0|0|0|0|0|0|0|0|0|0|0|0|0|0|0|0|0|0|0|0|0|0|0|0|0|0|0|0|0|0|0|0|0|0|0|0|0|
|virus_Sep8.0v_6322|1|0|0|0|0|0|0|0|0|0|0|0|0|0|0|0|0|0|0|0|0|0|0|0|0|0|0|0|0|0|0|0|0|0|0|0|0|0|0|0|0|0|0|0|0|0|0|0|1|0|0|0|0|0|0|
|virus_Sep8.0v_7899|1|0|0|0|0|0|0|0|0|0|0|0|0|0|0|0|0|0|0|0|0|0|0|0|0|0|0|0|0|0|0|0|0|0|0|0|0|0|0|0|0|0|0|0|0|0|0|0|1|0|0|0|0|0|0|
|virus_Sep8.0v_8244|1|0|0|0|0|0|0|0|0|0|0|0|0|0|0|0|0|0|0|0|0|0|0|0|0|0|0|0|0|0|0|0|0|0|0|0|0|0|0|0|0|0|0|0|0|0|0|0|1|0|0|0|0|0|0|
|virus_Sep8.0v_9273|1|0|0|0|0|0|0|0|0|0|0|0|0|0|0|0|0|0|0|0|0|0|0|0|0|0|0|0|0|0|0|0|0|0|0|0|0|0|0|0|0|0|0|0|0|0|0|0|1|0|0|0|0|0|0|
|virus_Sep8.0v_10379|1|0|0|0|0|0|0|0|0|0|0|0|0|0|0|0|0|0|0|0|0|0|0|0|0|0|0|0|0|0|0|0|0|0|0|0|0|0|0|0|0|0|0|0|0|0|0|0|1|0|0|0|0|0|0|
|virus_Sep8.0v_13182|1|0|0|0|0|0|0|0|0|0|0|0|0|0|0|0|0|0|0|0|0|0|0|0|0|0|0|0|0|0|0|0|0|0|0|0|0|0|0|0|0|0|0|0|0|0|0|0|1|0|0|0|0|0|0|
|virus_Sep8.0v_13245|1|0|0|0|0|0|0|0|0|0|0|0|0|0|0|0|0|0|0|0|0|0|0|0|0|0|0|0|0|0|0|0|0|0|0|0|0|0|0|0|0|0|0|0|0|0|0|0|1|0|0|0|0|0|0|
|virus_Sep8.0v_14474|1|0|0|0|0|0|0|0|0|0|0|0|0|0|0|0|0|0|0|0|0|0|0|0|0|0|0|0|0|0|0|0|0|0|0|0|0|0|0|0|0|0|0|0|0|0|0|0|1|0|0|0|0|0|0|
|virus_Sep8.0v_15743|1|0|0|0|0|0|0|0|0|0|0|0|0|0|0|0|0|0|0|0|0|0|0|0|0|0|0|0|0|0|0|0|0|0|0|0|0|0|0|0|0|0|0|0|0|0|0|0|1|0|0|0|0|0|0|
|virus_Sep8.0v_18564|1|0|0|0|0|0|0|0|0|0|0|0|0|0|0|0|0|0|0|0|0|0|0|0|0|0|0|0|0|0|0|0|0|0|0|0|0|0|0|0|0|0|0|0|0|0|0|0|1|0|0|0|0|0|0|
|virus_Sep8.5v_616|1|0|0|0|0|0|0|0|0|0|0|0|0|0|0|0|0|0|0|0|0|0|0|0|0|0|0|0|0|0|0|0|0|0|0|0|0|0|0|0|0|0|0|0|0|0|0|0|0|0|1|0|0|0|0|
|virus_Sep8.5v_5247|1|0|0|0|0|0|0|0|0|0|0|0|0|0|0|0|0|0|0|0|0|0|0|0|0|0|0|0|0|0|0|0|0|0|0|0|0|0|0|0|0|0|0|0|0|0|0|0|0|0|1|0|0|0|0|
|virus_Sep8.5v_16613|1|0|0|0|0|0|0|0|0|0|0|0|0|0|0|0|0|0|0|0|0|0|0|0|0|0|0|0|0|0|0|0|0|0|0|0|0|0|0|0|0|0|0|0|0|0|0|0|0|0|1|0|0|0|0|
|virus_Sep8.5v_17091|1|0|0|0|0|0|0|0|0|0|0|0|0|0|0|0|0|0|0|0|0|0|0|0|0|0|0|0|0|0|0|0|0|0|0|0|0|0|0|0|0|0|0|0|0|0|0|0|0|0|1|0|0|0|0|
|virus_Sep8.25v_2227|1|0|0|0|0|0|0|0|0|0|0|0|0|0|0|0|0|0|0|0|0|0|0|0|0|0|0|0|0|0|0|0|0|0|0|0|0|0|0|0|0|0|0|0|0|0|0|0|0|0|0|0|1|0|0|
|virus_Sep8.25v_11905|1|0|0|0|0|0|0|0|0|0|0|0|0|0|0|0|0|0|0|0|0|0|0|0|0|0|0|0|0|0|0|0|0|0|0|0|0|0|0|0|0|0|0|0|0|0|0|0|0|0|0|0|1|0|0|
|virus_Sep8.25v_16285|1|0|0|0|0|0|0|0|0|0|0|0|0|0|0|0|0|0|0|0|0|0|0|0|0|0|0|0|0|0|0|0|0|0|0|0|0|0|0|0|0|0|0|0|0|0|0|0|0|0|0|0|1|0|0|
|virus_Sep8.25v_19115|1|0|0|0|0|0|0|0|0|0|0|0|0|0|0|0|0|0|0|0|0|0|0|0|0|0|0|0|0|0|0|0|0|0|0|0|0|0|0|0|0|0|0|0|0|0|0|0|0|0|0|0|1|0|0|
|virus_Sep8.25v_19723|1|0|0|0|0|0|0|0|0|0|0|0|0|0|0|0|0|0|0|0|0|0|0|0|0|0|0|0|0|0|0|0|0|0|0|0|0|0|0|0|0|0|0|0|0|0|0|0|0|0|0|0|1|0|0|
|virus_Sep8.25v_19816|1|0|0|0|0|0|0|0|0|0|0|0|0|0|0|0|0|0|0|0|0|0|0|0|0|0|0|0|0|0|0|0|0|0|0|0|0|0|0|0|0|0|0|0|0|0|0|0|0|0|0|0|1|0|0|
|virus_Sep8.50v_128|1|0|0|0|0|0|0|0|0|0|0|0|0|0|0|0|0|0|0|0|0|0|0|0|0|0|0|0|0|0|0|0|0|0|0|0|0|0|0|0|0|0|0|0|0|0|0|0|0|0|0|0|0|0|1|
|virus_Sep8.50v_1458|1|0|0|0|0|0|0|0|0|0|0|0|0|0|0|0|0|0|0|0|0|0|0|0|0|0|0|0|0|0|0|0|0|0|0|0|0|0|0|0|0|0|0|0|0|0|0|0|0|0|0|0|0|0|1|
|virus_Sep8.50v_10422|1|0|0|0|0|0|0|0|0|0|0|0|0|0|0|0|0|0|0|0|0|0|0|0|0|0|0|0|0|0|0|0|0|0|0|0|0|0|0|0|0|0|0|0|0|0|0|0|0|0|0|0|0|0|1|
|virus_Sep8.50v_13225|1|0|0|0|0|0|0|0|0|0|0|0|0|0|0|0|0|0|0|0|0|0|0|0|0|0|0|0|0|0|0|0|0|0|0|0|0|0|0|0|0|0|0|0|0|0|0|0|0|0|0|0|0|0|1|
|virus_Sep8.50v_16320|1|0|0|0|0|0|0|0|0|0|0|0|0|0|0|0|0|0|0|0|0|0|0|0|0|0|0|0|0|0|0|0|0|0|0|0|0|0|0|0|0|0|0|0|0|0|0|0|0|0|0|0|0|0|1|
|virus_Sep8.50v_21840|1|0|0|0|0|0|0|0|0|0|0|0|0|0|0|0|0|0|0|0|0|0|0|0|0|0|0|0|0|0|0|0|0|0|0|0|0|0|0|0|0|0|0|0|0|0|0|0|0|0|0|0|0|0|1|
|virus_Sep8.50v_24019|1|0|0|0|0|0|0|0|0|0|0|0|0|0|0|0|0|0|0|0|0|0|0|0|0|0|0|0|0|0|0|0|0|0|0|0|0|0|0|0|0|0|0|0|0|0|0|0|0|0|0|0|0|0|1|
|virus_Sep8.50v_27465|1|0|0|0|0|0|0|0|0|0|0|0|0|0|0|0|0|0|0|0|0|0|0|0|0|0|0|0|0|0|0|0|0|0|0|0|0|0|0|0|0|0|0|0|0|0|0|0|0|0|0|0|0|0|1|
|virus_Sep8.50v_32091|1|0|0|0|0|0|0|0|0|0|0|0|0|0|0|0|0|0|0|0|0|0|0|0|0|0|0|0|0|0|0|0|0|0|0|0|0|0|0|0|0|0|0|0|0|0|0|0|0|0|0|0|0|0|1|
|virus_Sep8.50v_32558|1|0|0|0|0|0|0|0|0|0|0|0|0|0|0|0|0|0|0|0|0|0|0|0|0|0|0|0|0|0|0|0|0|0|0|0|0|0|0|0|0|0|0|0|0|0|0|0|0|0|0|0|0|0|1|
|virus_Sep8.50v_38153|1|0|0|0|0|0|0|0|0|0|0|0|0|0|0|0|0|0|0|0|0|0|0|0|0|0|0|0|0|0|0|0|0|0|0|0|0|0|0|0|0|0|0|0|0|0|0|0|0|0|0|0|0|0|1|
|virus_Sep8.50v_39493|1|0|0|0|0|0|0|0|0|0|0|0|0|0|0|0|0|0|0|0|0|0|0|0|0|0|0|0|0|0|0|0|0|0|0|0|0|0|0|0|0|0|0|0|0|0|0|0|0|0|0|0|0|0|1|
|virus_Sep8.50v_40089|1|0|0|0|0|0|0|0|0|0|0|0|0|0|0|0|0|0|0|0|0|0|0|0|0|0|0|0|0|0|0|0|0|0|0|0|0|0|0|0|0|0|0|0|0|0|0|0|0|0|0|0|0|0|1|
|bacteria_Jul11d0b_11488|1|0|0|0|0|0|0|0|0|0|0|0|0|0|0|0|0|0|0|0|0|1|0|0|0|0|0|0|0|0|0|0|0|0|0|0|0|0|0|0|0|0|0|0|0|0|0|0|0|0|0|0|0|0|0|
|bacteria_Jul11d0b_13870|1|0|0|0|0|0|0|0|0|0|0|0|0|0|0|0|0|0|0|0|0|1|0|0|0|0|0|0|0|0|0|0|0|0|0|0|0|0|0|0|0|0|0|0|0|0|0|0|0|0|0|0|0|0|0|
|bacteria_Jul11d0b_18094|1|0|0|0|0|0|0|0|0|0|0|0|0|0|0|0|0|0|0|0|0|1|0|0|0|0|0|0|0|0|0|0|0|0|0|0|0|0|0|0|0|0|0|0|0|0|0|0|0|0|0|0|0|0|0|
|bacteria_Jul11d0b_24169|1|0|0|0|0|0|0|0|0|0|0|0|0|0|0|0|0|0|0|0|0|1|0|0|0|0|0|0|0|0|0|0|0|0|0|0|0|0|0|0|0|0|0|0|0|0|0|0|0|0|0|0|0|0|0|
|bacteria_Jul11d0b_28117|1|0|0|0|0|0|0|0|0|0|0|0|0|0|0|0|0|0|0|0|0|1|0|0|0|0|0|0|0|0|0|0|0|0|0|0|0|0|0|0|0|0|0|0|0|0|0|0|0|0|0|0|0|0|0|
|bacteria_Jul11d0b_38701|1|0|0|0|0|0|0|0|0|0|0|0|0|0|0|0|0|0|0|0|0|1|0|0|0|0|0|0|0|0|0|0|0|0|0|0|0|0|0|0|0|0|0|0|0|0|0|0|0|0|0|0|0|0|0|
|bacteria_Jul11d0b_39470|1|0|0|0|0|0|0|0|0|0|0|0|0|0|0|0|0|0|0|0|0|1|0|0|0|0|0|0|0|0|0|0|0|0|0|0|0|0|0|0|0|0|0|0|0|0|0|0|0|0|0|0|0|0|0|
|bacteria_Jul11d0b_39953|1|0|0|0|0|0|0|0|0|0|0|0|0|0|0|0|0|0|0|0|0|1|0|0|0|0|0|0|0|0|0|0|0|0|0|0|0|0|0|0|0|0|0|0|0|0|0|0|0|0|0|0|0|0|0|
|bacteria_Jul11d0b_50942|1|0|0|0|0|0|0|0|0|0|0|0|0|0|0|0|0|0|0|0|0|1|0|0|0|0|0|0|0|0|0|0|0|0|0|0|0|0|0|0|0|0|0|0|0|0|0|0|0|0|0|0|0|0|0|
|bacteria_Jul11d0b_64156|1|0|0|0|0|0|0|0|0|0|0|0|0|0|0|0|0|0|0|0|0|1|0|0|0|0|0|0|0|0|0|0|0|0|0|0|0|0|0|0|0|0|0|0|0|0|0|0|0|0|0|0|0|0|0|
|bacteria_Aug5d0b_32446|1|0|0|0|0|0|0|0|0|0|0|0|0|0|0|0|0|0|0|0|0|0|0|0|0|0|0|0|0|0|0|0|0|0|0|0|0|1|0|0|0|0|0|0|0|0|0|0|0|0|0|0|0|0|0|
|bacteria_Aug5d0b_72428|1|0|0|0|0|0|0|0|0|0|0|0|0|0|0|0|0|0|0|0|0|0|0|0|0|0|0|0|0|0|0|0|0|0|0|0|0|1|0|0|0|0|0|0|0|0|0|0|0|0|0|0|0|0|0|
|bacteria_Aug5d5b_10045|1|0|0|0|0|0|0|0|0|0|0|0|0|0|0|0|0|0|0|0|0|0|0|0|0|0|0|0|0|0|0|0|0|0|0|0|0|0|0|1|0|0|0|0|0|0|0|0|0|0|0|0|0|0|0|
|bacteria_Aug5d5b_30134|1|0|0|0|0|0|0|0|0|0|0|0|0|0|0|0|0|0|0|0|0|0|0|0|0|0|0|0|0|0|0|0|0|0|0|0|0|0|0|1|0|0|0|0|0|0|0|0|0|0|0|0|0|0|0|
|bacteria_Aug5d25b_9580|1|0|0|0|0|0|0|0|0|0|0|0|0|0|0|0|0|0|0|0|0|0|0|0|0|0|0|0|0|0|0|0|0|0|0|0|0|0|0|0|0|1|0|0|0|0|0|0|0|0|0|0|0|0|0|
|bacteria_Aug5d25b_10628|1|0|0|0|0|0|0|0|0|0|0|0|0|0|0|0|0|0|0|0|0|0|0|0|0|0|0|0|0|0|0|0|0|0|0|0|0|0|0|0|0|1|0|0|0|0|0|0|0|0|0|0|0|0|0|
|bacteria_Aug5d25b_13592|1|0|0|0|0|0|0|0|0|0|0|0|0|0|0|0|0|0|0|0|0|0|0|0|0|0|0|0|0|0|0|0|0|0|0|0|0|0|0|0|0|1|0|0|0|0|0|0|0|0|0|0|0|0|0|
|bacteria_Aug5d25b_16026|1|0|0|0|0|0|0|0|0|0|0|0|0|0|0|0|0|0|0|0|0|0|0|0|0|0|0|0|0|0|0|0|0|0|0|0|0|0|0|0|0|1|0|0|0|0|0|0|0|0|0|0|0|0|0|
|bacteria_Aug5d25b_18342|1|0|0|0|0|0|0|0|0|0|0|0|0|0|0|0|0|0|0|0|0|0|0|0|0|0|0|0|0|0|0|0|0|0|0|0|0|0|0|0|0|1|0|0|0|0|0|0|0|0|0|0|0|0|0|
|bacteria_Aug5d25b_18729|1|0|0|0|0|0|0|0|0|0|0|0|0|0|0|0|0|0|0|0|0|0|0|0|0|0|0|0|0|0|0|0|0|0|0|0|0|0|0|0|0|1|0|0|0|0|0|0|0|0|0|0|0|0|0|
|bacteria_Aug5d25b_20128|1|0|0|0|0|0|0|0|0|0|0|0|0|0|0|0|0|0|0|0|0|0|0|0|0|0|0|0|0|0|0|0|0|0|0|0|0|0|0|0|0|1|0|0|0|0|0|0|0|0|0|0|0|0|0|
|bacteria_Aug5d25b_22027|1|0|0|0|0|0|0|0|0|0|0|0|0|0|0|0|0|0|0|0|0|0|0|0|0|0|0|0|0|0|0|0|0|0|0|0|0|0|0|0|0|1|0|0|0|0|0|0|0|0|0|0|0|0|0|
|bacteria_Aug5d25b_22709|1|0|0|0|0|0|0|0|0|0|0|0|0|0|0|0|0|0|0|0|0|0|0|0|0|0|0|0|0|0|0|0|0|0|0|0|0|0|0|0|0|1|0|0|0|0|0|0|0|0|0|0|0|0|0|
|bacteria_Aug5d25b_31402|1|0|0|0|0|0|0|0|0|0|0|0|0|0|0|0|0|0|0|0|0|0|0|0|0|0|0|0|0|0|0|0|0|0|0|0|0|0|0|0|0|1|0|0|0|0|0|0|0|0|0|0|0|0|0|
|bacteria_Aug5d25b_46283|1|0|0|0|0|0|0|0|0|0|0|0|0|0|0|0|0|0|0|0|0|0|0|0|0|0|0|0|0|0|0|0|0|0|0|0|0|0|0|0|0|1|0|0|0|0|0|0|0|0|0|0|0|0|0|
|bacteria_Aug5d25b_48257|1|0|0|0|0|0|0|0|0|0|0|0|0|0|0|0|0|0|0|0|0|0|0|0|0|0|0|0|0|0|0|0|0|0|0|0|0|0|0|0|0|1|0|0|0|0|0|0|0|0|0|0|0|0|0|
|bacteria_Aug5d50b_4093|1|0|0|0|0|0|0|0|0|0|0|0|0|0|0|0|0|0|0|0|0|0|0|0|0|0|0|0|0|0|0|0|0|0|0|0|0|0|0|0|0|0|0|1|0|0|0|0|0|0|0|0|0|0|0|
|bacteria_Aug5d50b_11288|1|0|0|0|0|0|0|0|0|0|0|0|0|0|0|0|0|0|0|0|0|0|0|0|0|0|0|0|0|0|0|0|0|0|0|0|0|0|0|0|0|0|0|1|0|0|0|0|0|0|0|0|0|0|0|
|bacteria_Aug5d50b_63029|1|0|0|0|0|0|0|0|0|0|0|0|0|0|0|0|0|0|0|0|0|0|0|0|0|0|0|0|0|0|0|0|0|0|0|0|0|0|0|0|0|0|0|1|0|0|0|0|0|0|0|0|0|0|0|
|bacteria_Aug5d50b_70446|1|0|0|0|0|0|0|0|0|0|0|0|0|0|0|0|0|0|0|0|0|0|0|0|0|0|0|0|0|0|0|0|0|0|0|0|0|0|0|0|0|0|0|1|0|0|0|0|0|0|0|0|0|0|0|
|bacteria_Jul11d5b_6987|1|0|0|0|0|0|0|0|0|0|0|0|0|0|0|0|0|0|0|0|0|0|0|1|0|0|0|0|0|0|0|0|0|0|0|0|0|0|0|0|0|0|0|0|0|0|0|0|0|0|0|0|0|0|0|
|bacteria_Jul11d5b_13001|1|0|0|0|0|0|0|0|0|0|0|0|0|0|0|0|0|0|0|0|0|0|0|1|0|0|0|0|0|0|0|0|0|0|0|0|0|0|0|0|0|0|0|0|0|0|0|0|0|0|0|0|0|0|0|
|bacteria_Jul11d5b_25213|1|0|0|0|0|0|0|0|0|0|0|0|0|0|0|0|0|0|0|0|0|0|0|1|0|0|0|0|0|0|0|0|0|0|0|0|0|0|0|0|0|0|0|0|0|0|0|0|0|0|0|0|0|0|0|
|bacteria_Jul11d5b_37105|1|0|0|0|0|0|0|0|0|0|0|0|0|0|0|0|0|0|0|0|0|0|0|1|0|0|0|0|0|0|0|0|0|0|0|0|0|0|0|0|0|0|0|0|0|0|0|0|0|0|0|0|0|0|0|
|bacteria_Jul11d5b_42893|1|0|0|0|0|0|0|0|0|0|0|0|0|0|0|0|0|0|0|0|0|0|0|1|0|0|0|0|0|0|0|0|0|0|0|0|0|0|0|0|0|0|0|0|0|0|0|0|0|0|0|0|0|0|0|
|bacteria_Jul11d25b_3094|1|0|0|0|0|0|0|0|0|0|0|0|0|0|0|0|0|0|0|0|0|0|0|0|0|1|0|0|0|0|0|0|0|0|0|0|0|0|0|0|0|0|0|0|0|0|0|0|0|0|0|0|0|0|0|
|bacteria_Jul11d25b_11944|1|0|0|0|0|0|0|0|0|0|0|0|0|0|0|0|0|0|0|0|0|0|0|0|0|1|0|0|0|0|0|0|0|0|0|0|0|0|0|0|0|0|0|0|0|0|0|0|0|0|0|0|0|0|0|
|bacteria_Jul11d25b_13107|1|0|0|0|0|0|0|0|0|0|0|0|0|0|0|0|0|0|0|0|0|0|0|0|0|1|0|0|0|0|0|0|0|0|0|0|0|0|0|0|0|0|0|0|0|0|0|0|0|0|0|0|0|0|0|
|bacteria_Jul11d25b_14040|1|0|0|0|0|0|0|0|0|0|0|0|0|0|0|0|0|0|0|0|0|0|0|0|0|1|0|0|0|0|0|0|0|0|0|0|0|0|0|0|0|0|0|0|0|0|0|0|0|0|0|0|0|0|0|
|bacteria_Jul11d25b_20003|1|0|0|0|0|0|0|0|0|0|0|0|0|0|0|0|0|0|0|0|0|0|0|0|0|1|0|0|0|0|0|0|0|0|0|0|0|0|0|0|0|0|0|0|0|0|0|0|0|0|0|0|0|0|0|
|bacteria_Jul11d25b_38601|1|0|0|0|0|0|0|0|0|0|0|0|0|0|0|0|0|0|0|0|0|0|0|0|0|1|0|0|0|0|0|0|0|0|0|0|0|0|0|0|0|0|0|0|0|0|0|0|0|0|0|0|0|0|0|
|bacteria_Jul11d25b_51429|1|0|0|0|0|0|0|0|0|0|0|0|0|0|0|0|0|0|0|0|0|0|0|0|0|1|0|0|0|0|0|0|0|0|0|0|0|0|0|0|0|0|0|0|0|0|0|0|0|0|0|0|0|0|0|
|bacteria_Jul11d25b_51486|1|0|0|0|0|0|0|0|0|0|0|0|0|0|0|0|0|0|0|0|0|0|0|0|0|1|0|0|0|0|0|0|0|0|0|0|0|0|0|0|0|0|0|0|0|0|0|0|0|0|0|0|0|0|0|
|bacteria_Jul11d25b_51509|1|0|0|0|0|0|0|0|0|0|0|0|0|0|0|0|0|0|0|0|0|0|0|0|0|1|0|0|0|0|0|0|0|0|0|0|0|0|0|0|0|0|0|0|0|0|0|0|0|0|0|0|0|0|0|
|bacteria_Jul11d50b_1675|1|0|0|0|0|0|0|0|0|0|0|0|0|0|0|0|0|0|0|0|0|0|0|0|0|0|0|1|0|0|0|0|0|0|0|0|0|0|0|0|0|0|0|0|0|0|0|0|0|0|0|0|0|0|0|
|bacteria_Jul11d50b_15432|1|0|0|0|0|0|0|0|0|0|0|0|0|0|0|0|0|0|0|0|0|0|0|0|0|0|0|1|0|0|0|0|0|0|0|0|0|0|0|0|0|0|0|0|0|0|0|0|0|0|0|0|0|0|0|
|bacteria_Jul11d50b_46901|1|0|0|0|0|0|0|0|0|0|0|0|0|0|0|0|0|0|0|0|0|0|0|0|0|0|0|1|0|0|0|0|0|0|0|0|0|0|0|0|0|0|0|0|0|0|0|0|0|0|0|0|0|0|0|
|bacteria_Jul11d50b_51108|1|0|0|0|0|0|0|0|0|0|0|0|0|0|0|0|0|0|0|0|0|0|0|0|0|0|0|1|0|0|0|0|0|0|0|0|0|0|0|0|0|0|0|0|0|0|0|0|0|0|0|0|0|0|0|
|bacteria_Jul11d75b_3746|1|0|0|0|0|0|0|0|0|0|0|0|0|0|0|0|0|0|0|0|0|0|0|0|0|0|0|0|0|1|0|0|0|0|0|0|0|0|0|0|0|0|0|0|0|0|0|0|0|0|0|0|0|0|0|
|bacteria_Jul11d75b_10346|1|0|0|0|0|0|0|0|0|0|0|0|0|0|0|0|0|0|0|0|0|0|0|0|0|0|0|0|0|1|0|0|0|0|0|0|0|0|0|0|0|0|0|0|0|0|0|0|0|0|0|0|0|0|0|
|bacteria_Jul11d75b_13970|1|0|0|0|0|0|0|0|0|0|0|0|0|0|0|0|0|0|0|0|0|0|0|0|0|0|0|0|0|1|0|0|0|0|0|0|0|0|0|0|0|0|0|0|0|0|0|0|0|0|0|0|0|0|0|
|bacteria_Jul11d75b_17694|1|0|0|0|0|0|0|0|0|0|0|0|0|0|0|0|0|0|0|0|0|0|0|0|0|0|0|0|0|1|0|0|0|0|0|0|0|0|0|0|0|0|0|0|0|0|0|0|0|0|0|0|0|0|0|
|bacteria_Jul11d75b_17803|1|0|0|0|0|0|0|0|0|0|0|0|0|0|0|0|0|0|0|0|0|0|0|0|0|0|0|0|0|1|0|0|0|0|0|0|0|0|0|0|0|0|0|0|0|0|0|0|0|0|0|0|0|0|0|
|bacteria_Jul11d75b_38527|1|0|0|0|0|0|0|0|0|0|0|0|0|0|0|0|0|0|0|0|0|0|0|0|0|0|0|0|0|1|0|0|0|0|0|0|0|0|0|0|0|0|0|0|0|0|0|0|0|0|0|0|0|0|0|
|bacteria_Jul11d75b_46826|1|0|0|0|0|0|0|0|0|0|0|0|0|0|0|0|0|0|0|0|0|0|0|0|0|0|0|0|0|1|0|0|0|0|0|0|0|0|0|0|0|0|0|0|0|0|0|0|0|0|0|0|0|0|0|
|bacteria_Jul11d75b_49356|1|0|0|0|0|0|0|0|0|0|0|0|0|0|0|0|0|0|0|0|0|0|0|0|0|0|0|0|0|1|0|0|0|0|0|0|0|0|0|0|0|0|0|0|0|0|0|0|0|0|0|0|0|0|0|
|bacteria_Jul11d75b_50938|1|0|0|0|0|0|0|0|0|0|0|0|0|0|0|0|0|0|0|0|0|0|0|0|0|0|0|0|0|1|0|0|0|0|0|0|0|0|0|0|0|0|0|0|0|0|0|0|0|0|0|0|0|0|0|
|bacteria_Jul27d5b_2731|1|0|0|0|0|0|0|0|0|0|0|0|0|0|0|0|0|0|0|0|0|0|0|0|0|0|0|0|0|0|0|1|0|0|0|0|0|0|0|0|0|0|0|0|0|0|0|0|0|0|0|0|0|0|0|
|bacteria_Jul27d5b_4398|1|0|0|0|0|0|0|0|0|0|0|0|0|0|0|0|0|0|0|0|0|0|0|0|0|0|0|0|0|0|0|1|0|0|0|0|0|0|0|0|0|0|0|0|0|0|0|0|0|0|0|0|0|0|0|
|bacteria_Jul27d5b_12258|1|0|0|0|0|0|0|0|0|0|0|0|0|0|0|0|0|0|0|0|0|0|0|0|0|0|0|0|0|0|0|1|0|0|0|0|0|0|0|0|0|0|0|0|0|0|0|0|0|0|0|0|0|0|0|
|bacteria_Jul27d5b_12885|1|0|0|0|0|0|0|0|0|0|0|0|0|0|0|0|0|0|0|0|0|0|0|0|0|0|0|0|0|0|0|1|0|0|0|0|0|0|0|0|0|0|0|0|0|0|0|0|0|0|0|0|0|0|0|
|bacteria_Jul27d5b_15744|1|0|0|0|0|0|0|0|0|0|0|0|0|0|0|0|0|0|0|0|0|0|0|0|0|0|0|0|0|0|0|1|0|0|0|0|0|0|0|0|0|0|0|0|0|0|0|0|0|0|0|0|0|0|0|
|bacteria_Jul27d5b_16149|1|0|0|0|0|0|0|0|0|0|0|0|0|0|0|0|0|0|0|0|0|0|0|0|0|0|0|0|0|0|0|1|0|0|0|0|0|0|0|0|0|0|0|0|0|0|0|0|0|0|0|0|0|0|0|
|bacteria_Jul27d5b_24390|1|0|0|0|0|0|0|0|0|0|0|0|0|0|0|0|0|0|0|0|0|0|0|0|0|0|0|0|0|0|0|1|0|0|0|0|0|0|0|0|0|0|0|0|0|0|0|0|0|0|0|0|0|0|0|
|bacteria_Jul27d5b_24460|1|0|0|0|0|0|0|0|0|0|0|0|0|0|0|0|0|0|0|0|0|0|0|0|0|0|0|0|0|0|0|1|0|0|0|0|0|0|0|0|0|0|0|0|0|0|0|0|0|0|0|0|0|0|0|
|bacteria_Jul27d5b_27471|1|0|0|0|0|0|0|0|0|0|0|0|0|0|0|0|0|0|0|0|0|0|0|0|0|0|0|0|0|0|0|1|0|0|0|0|0|0|0|0|0|0|0|0|0|0|0|0|0|0|0|0|0|0|0|
|bacteria_Jul27d5b_28308|1|0|0|0|0|0|0|0|0|0|0|0|0|0|0|0|0|0|0|0|0|0|0|0|0|0|0|0|0|0|0|1|0|0|0|0|0|0|0|0|0|0|0|0|0|0|0|0|0|0|0|0|0|0|0|
|bacteria_Jul27d5b_29681|1|0|0|0|0|0|0|0|0|0|0|0|0|0|0|0|0|0|0|0|0|0|0|0|0|0|0|0|0|0|0|1|0|0|0|0|0|0|0|0|0|0|0|0|0|0|0|0|0|0|0|0|0|0|0|
|bacteria_Jul27d5b_29806|1|0|0|0|0|0|0|0|0|0|0|0|0|0|0|0|0|0|0|0|0|0|0|0|0|0|0|0|0|0|0|1|0|0|0|0|0|0|0|0|0|0|0|0|0|0|0|0|0|0|0|0|0|0|0|
|bacteria_Jul27d5b_37915|1|0|0|0|0|0|0|0|0|0|0|0|0|0|0|0|0|0|0|0|0|0|0|0|0|0|0|0|0|0|0|1|0|0|0|0|0|0|0|0|0|0|0|0|0|0|0|0|0|0|0|0|0|0|0|
|bacteria_Jul27d5b_45730|1|0|0|0|0|0|0|0|0|0|0|0|0|0|0|0|0|0|0|0|0|0|0|0|0|0|0|0|0|0|0|1|0|0|0|0|0|0|0|0|0|0|0|0|0|0|0|0|0|0|0|0|0|0|0|
|bacteria_Jul27d5b_45881|1|0|0|0|0|0|0|0|0|0|0|0|0|0|0|0|0|0|0|0|0|0|0|0|0|0|0|0|0|0|0|1|0|0|0|0|0|0|0|0|0|0|0|0|0|0|0|0|0|0|0|0|0|0|0|
|bacteria_Jul27d5b_47242|1|0|0|0|0|0|0|0|0|0|0|0|0|0|0|0|0|0|0|0|0|0|0|0|0|0|0|0|0|0|0|1|0|0|0|0|0|0|0|0|0|0|0|0|0|0|0|0|0|0|0|0|0|0|0|

**Supplementary Table S7**. Common Bacterio- and virioplankton RTPR 98% OTUs in 27 samples with RTPR amplicons in both fractions

Total seqs: 47997; Subtotal seqs in retained

| OTUs: 41046 | | May12_05m | | May12_25m | | May12_50m | | May12_75m | | Jun8_25m | | Jun8_50m | | Jun8_75m | | Jun22_0m | | Jun22_50m | | Jun22_75m | | Jul11_0m | | Jul11_05m | | Jul11_25m | | Jul11_50m | | Jul11_75m | | Jul27_05m | | Jul27_25m | | Jul27_75m | | Aug5_0m | | Aug5_05m | | Aug5_25m | | Aug5_50m | | Aug5_75m | | Sep8_0m | | Sep8_05m | | Sep8_25m | | Sep8_50m | |
|---|---|---|---|---|---|---|---|---|---|---|---|---|---|---|---|---|---|---|---|---|---|---|---|---|---|---|---|---|---|---|---|---|---|---|---|---|---|---|---|---|---|---|---|---|---|---|---|---|---|---|---|---|---|---|---|
| OTUs with >=10 seqs: 254 | OTU subtotal | bacterioplankton | virioplankton | bacterioplankton | virioplankton | bacterioplankton | virioplankton | bacterioplankton | virioplankton | bacterioplankton | virioplankton | bacterioplankton | virioplankton | bacterioplankton | virioplankton | bacterioplankton | virioplankton | bacterioplankton | virioplankton | bacterioplankton | virioplankton | bacterioplankton | virioplankton | bacterioplankton | virioplankton | bacterioplankton | virioplankton | bacterioplankton | virioplankton | bacterioplankton | virioplankton | bacterioplankton | virioplankton | bacterioplankton | virioplankton | bacterioplankton | virioplankton | bacterioplankton | virioplankton | bacterioplankton | virioplankton | bacterioplankton | virioplankton | bacterioplankton | virioplankton | bacterioplankton | virioplankton | bacterioplankton | virioplankton | bacterioplankton | virioplankton | bacterioplankton | virioplankton | bacterioplankton | virioplankton |
| bacteria_Jul27d5b_50670 | 1 | 0 | 0 | 0 | 0 | 0 | 0 | 0 | 0 | 0 | 0 | 0 | 0 | 0 | 0 | 0 | 0 | 0 | 0 | 0 | 0 | 0 | 0 | 0 | 0 | 0 | 0 | 0 | 0 | 0 | 0 | 1 | 0 | 0 | 0 | 0 | 0 | 0 | 0 | 0 | 0 | 0 | 0 | 0 | 0 | 0 | 0 | 0 | 0 | 0 | 0 | 0 | 0 | 0 | 0 |
| bacteria_Jul27d5b_52174 | 1 | 0 | 0 | 0 | 0 | 0 | 0 | 0 | 0 | 0 | 0 | 0 | 0 | 0 | 0 | 0 | 0 | 0 | 0 | 0 | 0 | 0 | 0 | 0 | 0 | 0 | 0 | 0 | 0 | 0 | 0 | 1 | 0 | 0 | 0 | 0 | 0 | 0 | 0 | 0 | 0 | 0 | 0 | 0 | 0 | 0 | 0 | 0 | 0 | 0 | 0 | 0 | 0 | 0 | 0 |
| bacteria_Jul27d25b_1138 | 1 | 0 | 0 | 0 | 0 | 0 | 0 | 0 | 0 | 0 | 0 | 0 | 0 | 0 | 0 | 0 | 0 | 0 | 0 | 0 | 0 | 0 | 0 | 0 | 0 | 0 | 0 | 0 | 0 | 0 | 0 | 0 | 0 | 1 | 0 | 0 | 0 | 0 | 0 | 0 | 0 | 0 | 0 | 0 | 0 | 0 | 0 | 0 | 0 | 0 | 0 | 0 | 0 | 0 | 0 |
| bacteria_Jul27d25b_1792 | 1 | 0 | 0 | 0 | 0 | 0 | 0 | 0 | 0 | 0 | 0 | 0 | 0 | 0 | 0 | 0 | 0 | 0 | 0 | 0 | 0 | 0 | 0 | 0 | 0 | 0 | 0 | 0 | 0 | 0 | 0 | 0 | 0 | 1 | 0 | 0 | 0 | 0 | 0 | 0 | 0 | 0 | 0 | 0 | 0 | 0 | 0 | 0 | 0 | 0 | 0 | 0 | 0 | 0 | 0 |
| bacteria_Jul27d25b_2054 | 1 | 0 | 0 | 0 | 0 | 0 | 0 | 0 | 0 | 0 | 0 | 0 | 0 | 0 | 0 | 0 | 0 | 0 | 0 | 0 | 0 | 0 | 0 | 0 | 0 | 0 | 0 | 0 | 0 | 0 | 0 | 0 | 0 | 1 | 0 | 0 | 0 | 0 | 0 | 0 | 0 | 0 | 0 | 0 | 0 | 0 | 0 | 0 | 0 | 0 | 0 | 0 | 0 | 0 | 0 |
| bacteria_Jul27d25b_2238 | 1 | 0 | 0 | 0 | 0 | 0 | 0 | 0 | 0 | 0 | 0 | 0 | 0 | 0 | 0 | 0 | 0 | 0 | 0 | 0 | 0 | 0 | 0 | 0 | 0 | 0 | 0 | 0 | 0 | 0 | 0 | 0 | 0 | 1 | 0 | 0 | 0 | 0 | 0 | 0 | 0 | 0 | 0 | 0 | 0 | 0 | 0 | 0 | 0 | 0 | 0 | 0 | 0 | 0 | 0 |
| bacteria_Jul27d25b_2657 | 1 | 0 | 0 | 0 | 0 | 0 | 0 | 0 | 0 | 0 | 0 | 0 | 0 | 0 | 0 | 0 | 0 | 0 | 0 | 0 | 0 | 0 | 0 | 0 | 0 | 0 | 0 | 0 | 0 | 0 | 0 | 0 | 0 | 1 | 0 | 0 | 0 | 0 | 0 | 0 | 0 | 0 | 0 | 0 | 0 | 0 | 0 | 0 | 0 | 0 | 0 | 0 | 0 | 0 | 0 |
| bacteria_Jul27d25b_2874 | 1 | 0 | 0 | 0 | 0 | 0 | 0 | 0 | 0 | 0 | 0 | 0 | 0 | 0 | 0 | 0 | 0 | 0 | 0 | 0 | 0 | 0 | 0 | 0 | 0 | 0 | 0 | 0 | 0 | 0 | 0 | 0 | 0 | 1 | 0 | 0 | 0 | 0 | 0 | 0 | 0 | 0 | 0 | 0 | 0 | 0 | 0 | 0 | 0 | 0 | 0 | 0 | 0 | 0 | 0 |
| bacteria_Jul27d25b_3060 | 1 | 0 | 0 | 0 | 0 | 0 | 0 | 0 | 0 | 0 | 0 | 0 | 0 | 0 | 0 | 0 | 0 | 0 | 0 | 0 | 0 | 0 | 0 | 0 | 0 | 0 | 0 | 0 | 0 | 0 | 0 | 0 | 0 | 1 | 0 | 0 | 0 | 0 | 0 | 0 | 0 | 0 | 0 | 0 | 0 | 0 | 0 | 0 | 0 | 0 | 0 | 0 | 0 | 0 | 0 |
| bacteria_Jul27d25b_3426 | 1 | 0 | 0 | 0 | 0 | 0 | 0 | 0 | 0 | 0 | 0 | 0 | 0 | 0 | 0 | 0 | 0 | 0 | 0 | 0 | 0 | 0 | 0 | 0 | 0 | 0 | 0 | 0 | 0 | 0 | 0 | 0 | 0 | 1 | 0 | 0 | 0 | 0 | 0 | 0 | 0 | 0 | 0 | 0 | 0 | 0 | 0 | 0 | 0 | 0 | 0 | 0 | 0 | 0 | 0 |
| bacteria_Jul27d25b_3482 | 1 | 0 | 0 | 0 | 0 | 0 | 0 | 0 | 0 | 0 | 0 | 0 | 0 | 0 | 0 | 0 | 0 | 0 | 0 | 0 | 0 | 0 | 0 | 0 | 0 | 0 | 0 | 0 | 0 | 0 | 0 | 0 | 0 | 1 | 0 | 0 | 0 | 0 | 0 | 0 | 0 | 0 | 0 | 0 | 0 | 0 | 0 | 0 | 0 | 0 | 0 | 0 | 0 | 0 | 0 |
| bacteria_Jul27d25b_4122 | 1 | 0 | 0 | 0 | 0 | 0 | 0 | 0 | 0 | 0 | 0 | 0 | 0 | 0 | 0 | 0 | 0 | 0 | 0 | 0 | 0 | 0 | 0 | 0 | 0 | 0 | 0 | 0 | 0 | 0 | 0 | 0 | 0 | 1 | 0 | 0 | 0 | 0 | 0 | 0 | 0 | 0 | 0 | 0 | 0 | 0 | 0 | 0 | 0 | 0 | 0 | 0 | 0 | 0 | 0 |
| bacteria_Jul27d25b_4515 | 1 | 0 | 0 | 0 | 0 | 0 | 0 | 0 | 0 | 0 | 0 | 0 | 0 | 0 | 0 | 0 | 0 | 0 | 0 | 0 | 0 | 0 | 0 | 0 | 0 | 0 | 0 | 0 | 0 | 0 | 0 | 0 | 0 | 1 | 0 | 0 | 0 | 0 | 0 | 0 | 0 | 0 | 0 | 0 | 0 | 0 | 0 | 0 | 0 | 0 | 0 | 0 | 0 | 0 | 0 |
| bacteria_Jul27d25b_5003 | 1 | 0 | 0 | 0 | 0 | 0 | 0 | 0 | 0 | 0 | 0 | 0 | 0 | 0 | 0 | 0 | 0 | 0 | 0 | 0 | 0 | 0 | 0 | 0 | 0 | 0 | 0 | 0 | 0 | 0 | 0 | 0 | 0 | 1 | 0 | 0 | 0 | 0 | 0 | 0 | 0 | 0 | 0 | 0 | 0 | 0 | 0 | 0 | 0 | 0 | 0 | 0 | 0 | 0 | 0 |
| bacteria_Jul27d25b_5044 | 1 | 0 | 0 | 0 | 0 | 0 | 0 | 0 | 0 | 0 | 0 | 0 | 0 | 0 | 0 | 0 | 0 | 0 | 0 | 0 | 0 | 0 | 0 | 0 | 0 | 0 | 0 | 0 | 0 | 0 | 0 | 0 | 0 | 1 | 0 | 0 | 0 | 0 | 0 | 0 | 0 | 0 | 0 | 0 | 0 | 0 | 0 | 0 | 0 | 0 | 0 | 0 | 0 | 0 | 0 |
| bacteria_Jul27d25b_5139 | 1 | 0 | 0 | 0 | 0 | 0 | 0 | 0 | 0 | 0 | 0 | 0 | 0 | 0 | 0 | 0 | 0 | 0 | 0 | 0 | 0 | 0 | 0 | 0 | 0 | 0 | 0 | 0 | 0 | 0 | 0 | 0 | 0 | 1 | 0 | 0 | 0 | 0 | 0 | 0 | 0 | 0 | 0 | 0 | 0 | 0 | 0 | 0 | 0 | 0 | 0 | 0 | 0 | 0 | 0 |
| bacteria_Jul27d25b_6715 | 1 | 0 | 0 | 0 | 0 | 0 | 0 | 0 | 0 | 0 | 0 | 0 | 0 | 0 | 0 | 0 | 0 | 0 | 0 | 0 | 0 | 0 | 0 | 0 | 0 | 0 | 0 | 0 | 0 | 0 | 0 | 0 | 0 | 1 | 0 | 0 | 0 | 0 | 0 | 0 | 0 | 0 | 0 | 0 | 0 | 0 | 0 | 0 | 0 | 0 | 0 | 0 | 0 | 0 | 0 |
| bacteria_Jul27d25b_6827 | 1 | 0 | 0 | 0 | 0 | 0 | 0 | 0 | 0 | 0 | 0 | 0 | 0 | 0 | 0 | 0 | 0 | 0 | 0 | 0 | 0 | 0 | 0 | 0 | 0 | 0 | 0 | 0 | 0 | 0 | 0 | 0 | 0 | 1 | 0 | 0 | 0 | 0 | 0 | 0 | 0 | 0 | 0 | 0 | 0 | 0 | 0 | 0 | 0 | 0 | 0 | 0 | 0 | 0 | 0 |
| bacteria_Jul27d25b_7127 | 1 | 0 | 0 | 0 | 0 | 0 | 0 | 0 | 0 | 0 | 0 | 0 | 0 | 0 | 0 | 0 | 0 | 0 | 0 | 0 | 0 | 0 | 0 | 0 | 0 | 0 | 0 | 0 | 0 | 0 | 0 | 0 | 0 | 1 | 0 | 0 | 0 | 0 | 0 | 0 | 0 | 0 | 0 | 0 | 0 | 0 | 0 | 0 | 0 | 0 | 0 | 0 | 0 | 0 | 0 |
| bacteria_Jul27d25b_7254 | 1 | 0 | 0 | 0 | 0 | 0 | 0 | 0 | 0 | 0 | 0 | 0 | 0 | 0 | 0 | 0 | 0 | 0 | 0 | 0 | 0 | 0 | 0 | 0 | 0 | 0 | 0 | 0 | 0 | 0 | 0 | 0 | 0 | 1 | 0 | 0 | 0 | 0 | 0 | 0 | 0 | 0 | 0 | 0 | 0 | 0 | 0 | 0 | 0 | 0 | 0 | 0 | 0 | 0 | 0 |
| bacteria_Jul27d25b_7281 | 1 | 0 | 0 | 0 | 0 | 0 | 0 | 0 | 0 | 0 | 0 | 0 | 0 | 0 | 0 | 0 | 0 | 0 | 0 | 0 | 0 | 0 | 0 | 0 | 0 | 0 | 0 | 0 | 0 | 0 | 0 | 0 | 0 | 1 | 0 | 0 | 0 | 0 | 0 | 0 | 0 | 0 | 0 | 0 | 0 | 0 | 0 | 0 | 0 | 0 | 0 | 0 | 0 | 0 | 0 |
| bacteria_Jul27d75b_1935 | 1 | 0 | 0 | 0 | 0 | 0 | 0 | 0 | 0 | 0 | 0 | 0 | 0 | 0 | 0 | 0 | 0 | 0 | 0 | 0 | 0 | 0 | 0 | 0 | 0 | 0 | 0 | 0 | 0 | 0 | 0 | 0 | 0 | 0 | 0 | 1 | 0 | 0 | 0 | 0 | 0 | 0 | 0 | 0 | 0 | 0 | 0 | 0 | 0 | 0 | 0 | 0 | 0 | 0 | 0 |
| bacteria_Jul27d75b_2931 | 1 | 0 | 0 | 0 | 0 | 0 | 0 | 0 | 0 | 0 | 0 | 0 | 0 | 0 | 0 | 0 | 0 | 0 | 0 | 0 | 0 | 0 | 0 | 0 | 0 | 0 | 0 | 0 | 0 | 0 | 0 | 0 | 0 | 0 | 0 | 1 | 0 | 0 | 0 | 0 | 0 | 0 | 0 | 0 | 0 | 0 | 0 | 0 | 0 | 0 | 0 | 0 | 0 | 0 | 0 |
| bacteria_Jul27d75b_6865 | 1 | 0 | 0 | 0 | 0 | 0 | 0 | 0 | 0 | 0 | 0 | 0 | 0 | 0 | 0 | 0 | 0 | 0 | 0 | 0 | 0 | 0 | 0 | 0 | 0 | 0 | 0 | 0 | 0 | 0 | 0 | 0 | 0 | 0 | 0 | 1 | 0 | 0 | 0 | 0 | 0 | 0 | 0 | 0 | 0 | 0 | 0 | 0 | 0 | 0 | 0 | 0 | 0 | 0 | 0 |
| bacteria_Jul27d75b_9384 | 1 | 0 | 0 | 0 | 0 | 0 | 0 | 0 | 0 | 0 | 0 | 0 | 0 | 0 | 0 | 0 | 0 | 0 | 0 | 0 | 0 | 0 | 0 | 0 | 0 | 0 | 0 | 0 | 0 | 0 | 0 | 0 | 0 | 0 | 0 | 1 | 0 | 0 | 0 | 0 | 0 | 0 | 0 | 0 | 0 | 0 | 0 | 0 | 0 | 0 | 0 | 0 | 0 | 0 | 0 |
| bacteria_Jul27d75b_9429 | 1 | 0 | 0 | 0 | 0 | 0 | 0 | 0 | 0 | 0 | 0 | 0 | 0 | 0 | 0 | 0 | 0 | 0 | 0 | 0 | 0 | 0 | 0 | 0 | 0 | 0 | 0 | 0 | 0 | 0 | 0 | 0 | 0 | 0 | 0 | 1 | 0 | 0 | 0 | 0 | 0 | 0 | 0 | 0 | 0 | 0 | 0 | 0 | 0 | 0 | 0 | 0 | 0 | 0 | 0 |
| bacteria_Jul27d75b_1146 | 1 | 0 | 0 | 0 | 0 | 0 | 0 | 0 | 0 | 0 | 0 | 0 | 0 | 0 | 0 | 0 | 0 | 0 | 0 | 0 | 0 | 0 | 0 | 0 | 0 | 0 | 0 | 0 | 0 | 0 | 0 | 0 | 0 | 0 | 0 | 1 | 0 | 0 | 0 | 0 | 0 | 0 | 0 | 0 | 0 | 0 | 0 | 0 | 0 | 0 | 0 | 0 | 0 | 0 | 0 |
| bacteria_Jul27d75b_1257 | 1 | 0 | 0 | 0 | 0 | 0 | 0 | 0 | 0 | 0 | 0 | 0 | 0 | 0 | 0 | 0 | 0 | 0 | 0 | 0 | 0 | 0 | 0 | 0 | 0 | 0 | 0 | 0 | 0 | 0 | 0 | 0 | 0 | 0 | 0 | 1 | 0 | 0 | 0 | 0 | 0 | 0 | 0 | 0 | 0 | 0 | 0 | 0 | 0 | 0 | 0 | 0 | 0 | 0 | 0 |
| bacteria_Jul27d75b_1264 | 1 | 0 | 0 | 0 | 0 | 0 | 0 | 0 | 0 | 0 | 0 | 0 | 0 | 0 | 0 | 0 | 0 | 0 | 0 | 0 | 0 | 0 | 0 | 0 | 0 | 0 | 0 | 0 | 0 | 0 | 0 | 0 | 0 | 0 | 0 | 1 | 0 | 0 | 0 | 0 | 0 | 0 | 0 | 0 | 0 | 0 | 0 | 0 | 0 | 0 | 0 | 0 | 0 | 0 | 0 |
| bacteria_Jul27d75b_1277 | 1 | 0 | 0 | 0 | 0 | 0 | 0 | 0 | 0 | 0 | 0 | 0 | 0 | 0 | 0 | 0 | 0 | 0 | 0 | 0 | 0 | 0 | 0 | 0 | 0 | 0 | 0 | 0 | 0 | 0 | 0 | 0 | 0 | 0 | 0 | 1 | 0 | 0 | 0 | 0 | 0 | 0 | 0 | 0 | 0 | 0 | 0 | 0 | 0 | 0 | 0 | 0 | 0 | 0 | 0 |
| bacteria_Jul27d75b_1352 | 1 | 0 | 0 | 0 | 0 | 0 | 0 | 0 | 0 | 0 | 0 | 0 | 0 | 0 | 0 | 0 | 0 | 0 | 0 | 0 | 0 | 0 | 0 | 0 | 0 | 0 | 0 | 0 | 0 | 0 | 0 | 0 | 0 | 0 | 0 | 1 | 0 | 0 | 0 | 0 | 0 | 0 | 0 | 0 | 0 | 0 | 0 | 0 | 0 | 0 | 0 | 0 | 0 | 0 | 0 |
| bacteria_Jul27d75b_1574 | 1 | 0 | 0 | 0 | 0 | 0 | 0 | 0 | 0 | 0 | 0 | 0 | 0 | 0 | 0 | 0 | 0 | 0 | 0 | 0 | 0 | 0 | 0 | 0 | 0 | 0 | 0 | 0 | 0 | 0 | 0 | 0 | 0 | 0 | 0 | 1 | 0 | 0 | 0 | 0 | 0 | 0 | 0 | 0 | 0 | 0 | 0 | 0 | 0 | 0 | 0 | 0 | 0 | 0 | 0 |
| bacteria_Jul27d75b_2060 | 1 | 0 | 0 | 0 | 0 | 0 | 0 | 0 | 0 | 0 | 0 | 0 | 0 | 0 | 0 | 0 | 0 | 0 | 0 | 0 | 0 | 0 | 0 | 0 | 0 | 0 | 0 | 0 | 0 | 0 | 0 | 0 | 0 | 0 | 0 | 1 | 0 | 0 | 0 | 0 | 0 | 0 | 0 | 0 | 0 | 0 | 0 | 0 | 0 | 0 | 0 | 0 | 0 | 0 | 0 |
| bacteria_Jul27d75b_2203 | 1 | 0 | 0 | 0 | 0 | 0 | 0 | 0 | 0 | 0 | 0 | 0 | 0 | 0 | 0 | 0 | 0 | 0 | 0 | 0 | 0 | 0 | 0 | 0 | 0 | 0 | 0 | 0 | 0 | 0 | 0 | 0 | 0 | 0 | 0 | 1 | 0 | 0 | 0 | 0 | 0 | 0 | 0 | 0 | 0 | 0 | 0 | 0 | 0 | 0 | 0 | 0 | 0 | 0 | 0 |
| bacteria_Jul27d75b_2971 | 1 | 0 | 0 | 0 | 0 | 0 | 0 | 0 | 0 | 0 | 0 | 0 | 0 | 0 | 0 | 0 | 0 | 0 | 0 | 0 | 0 | 0 | 0 | 0 | 0 | 0 | 0 | 0 | 0 | 0 | 0 | 0 | 0 | 0 | 0 | 1 | 0 | 0 | 0 | 0 | 0 | 0 | 0 | 0 | 0 | 0 | 0 | 0 | 0 | 0 | 0 | 0 | 0 | 0 | 0 |
| bacteria_Jul27d75b_3408 | 1 | 0 | 0 | 0 | 0 | 0 | 0 | 0 | 0 | 0 | 0 | 0 | 0 | 0 | 0 | 0 | 0 | 0 | 0 | 0 | 0 | 0 | 0 | 0 | 0 | 0 | 0 | 0 | 0 | 0 | 0 | 0 | 0 | 0 | 0 | 1 | 0 | 0 | 0 | 0 | 0 | 0 | 0 | 0 | 0 | 0 | 0 | 0 | 0 | 0 | 0 | 0 | 0 | 0 | 0 |
| bacteria_Jul27d75b_6077 | 1 | 0 | 0 | 0 | 0 | 0 | 0 | 0 | 0 | 0 | 0 | 0 | 0 | 0 | 0 | 0 | 0 | 0 | 0 | 0 | 0 | 0 | 0 | 0 | 0 | 0 | 0 | 0 | 0 | 0 | 0 | 0 | 0 | 0 | 0 | 1 | 0 | 0 | 0 | 0 | 0 | 0 | 0 | 0 | 0 | 0 | 0 | 0 | 0 | 0 | 0 | 0 | 0 | 0 | 0 |
| bacteria_Jul27d75b_6691 | 1 | 0 | 0 | 0 | 0 | 0 | 0 | 0 | 0 | 0 | 0 | 0 | 0 | 0 | 0 | 0 | 0 | 0 | 0 | 0 | 0 | 0 | 0 | 0 | 0 | 0 | 0 | 0 | 0 | 0 | 0 | 0 | 0 | 0 | 0 | 1 | 0 | 0 | 0 | 0 | 0 | 0 | 0 | 0 | 0 | 0 | 0 | 0 | 0 | 0 | 0 | 0 | 0 | 0 | 0 |
| bacteria_Jun8d25b_30135 | 1 | 0 | 0 | 0 | 0 | 0 | 0 | 0 | 0 | 1 | 0 | 0 | 0 | 0 | 0 | 0 | 0 | 0 | 0 | 0 | 0 | 0 | 0 | 0 | 0 | 0 | 0 | 0 | 0 | 0 | 0 | 0 | 0 | 0 | 0 | 0 | 0 | 0 | 0 | 0 | 0 | 0 | 0 | 0 | 0 | 0 | 0 | 0 | 0 | 0 | 0 | 0 | 0 | 0 | 0 |
| bacteria_Jun8d25b_38642 | 1 | 0 | 0 | 0 | 0 | 0 | 0 | 0 | 0 | 1 | 0 | 0 | 0 | 0 | 0 | 0 | 0 | 0 | 0 | 0 | 0 | 0 | 0 | 0 | 0 | 0 | 0 | 0 | 0 | 0 | 0 | 0 | 0 | 0 | 0 | 0 | 0 | 0 | 0 | 0 | 0 | 0 | 0 | 0 | 0 | 0 | 0 | 0 | 0 | 0 | 0 | 0 | 0 | 0 | 0 |
| bacteria_Jun8d25b_44065 | 1 | 0 | 0 | 0 | 0 | 0 | 0 | 0 | 0 | 1 | 0 | 0 | 0 | 0 | 0 | 0 | 0 | 0 | 0 | 0 | 0 | 0 | 0 | 0 | 0 | 0 | 0 | 0 | 0 | 0 | 0 | 0 | 0 | 0 | 0 | 0 | 0 | 0 | 0 | 0 | 0 | 0 | 0 | 0 | 0 | 0 | 0 | 0 | 0 | 0 | 0 | 0 | 0 | 0 | 0 |
| bacteria_Jun8d25b_48717 | 1 | 0 | 0 | 0 | 0 | 0 | 0 | 0 | 0 | 1 | 0 | 0 | 0 | 0 | 0 | 0 | 0 | 0 | 0 | 0 | 0 | 0 | 0 | 0 | 0 | 0 | 0 | 0 | 0 | 0 | 0 | 0 | 0 | 0 | 0 | 0 | 0 | 0 | 0 | 0 | 0 | 0 | 0 | 0 | 0 | 0 | 0 | 0 | 0 | 0 | 0 | 0 | 0 | 0 | 0 |
| bacteria_Jun8d25b_51731 | 1 | 0 | 0 | 0 | 0 | 0 | 0 | 0 | 0 | 1 | 0 | 0 | 0 | 0 | 0 | 0 | 0 | 0 | 0 | 0 | 0 | 0 | 0 | 0 | 0 | 0 | 0 | 0 | 0 | 0 | 0 | 0 | 0 | 0 | 0 | 0 | 0 | 0 | 0 | 0 | 0 | 0 | 0 | 0 | 0 | 0 | 0 | 0 | 0 | 0 | 0 | 0 | 0 | 0 | 0 |
| bacteria_Jun8d25b_57047 | 1 | 0 | 0 | 0 | 0 | 0 | 0 | 0 | 0 | 1 | 0 | 0 | 0 | 0 | 0 | 0 | 0 | 0 | 0 | 0 | 0 | 0 | 0 | 0 | 0 | 0 | 0 | 0 | 0 | 0 | 0 | 0 | 0 | 0 | 0 | 0 | 0 | 0 | 0 | 0 | 0 | 0 | 0 | 0 | 0 | 0 | 0 | 0 | 0 | 0 | 0 | 0 | 0 | 0 | 0 |
| bacteria_Jun8d25b_65983 | 1 | 0 | 0 | 0 | 0 | 0 | 0 | 0 | 0 | 1 | 0 | 0 | 0 | 0 | 0 | 0 | 0 | 0 | 0 | 0 | 0 | 0 | 0 | 0 | 0 | 0 | 0 | 0 | 0 | 0 | 0 | 0 | 0 | 0 | 0 | 0 | 0 | 0 | 0 | 0 | 0 | 0 | 0 | 0 | 0 | 0 | 0 | 0 | 0 | 0 | 0 | 0 | 0 | 0 | 0 |
| bacteria_Jun8d25b_73142 | 1 | 0 | 0 | 0 | 0 | 0 | 0 | 0 | 0 | 1 | 0 | 0 | 0 | 0 | 0 | 0 | 0 | 0 | 0 | 0 | 0 | 0 | 0 | 0 | 0 | 0 | 0 | 0 | 0 | 0 | 0 | 0 | 0 | 0 | 0 | 0 | 0 | 0 | 0 | 0 | 0 | 0 | 0 | 0 | 0 | 0 | 0 | 0 | 0 | 0 | 0 | 0 | 0 | 0 | 0 |
| bacteria_Jun8d50b_9073 | 1 | 0 | 0 | 0 | 0 | 0 | 0 | 0 | 0 | 0 | 0 | 1 | 0 | 0 | 0 | 0 | 0 | 0 | 0 | 0 | 0 | 0 | 0 | 0 | 0 | 0 | 0 | 0 | 0 | 0 | 0 | 0 | 0 | 0 | 0 | 0 | 0 | 0 | 0 | 0 | 0 | 0 | 0 | 0 | 0 | 0 | 0 | 0 | 0 | 0 | 0 | 0 | 0 | 0 | 0 |
| bacteria_Jun8d50b_23697 | 1 | 0 | 0 | 0 | 0 | 0 | 0 | 0 | 0 | 0 | 0 | 1 | 0 | 0 | 0 | 0 | 0 | 0 | 0 | 0 | 0 | 0 | 0 | 0 | 0 | 0 | 0 | 0 | 0 | 0 | 0 | 0 | 0 | 0 | 0 | 0 | 0 | 0 | 0 | 0 | 0 | 0 | 0 | 0 | 0 | 0 | 0 | 0 | 0 | 0 | 0 | 0 | 0 | 0 | 0 |
| bacteria_Jun8d50b_23918 | 1 | 0 | 0 | 0 | 0 | 0 | 0 | 0 | 0 | 0 | 0 | 1 | 0 | 0 | 0 | 0 | 0 | 0 | 0 | 0 | 0 | 0 | 0 | 0 | 0 | 0 | 0 | 0 | 0 | 0 | 0 | 0 | 0 | 0 | 0 | 0 | 0 | 0 | 0 | 0 | 0 | 0 | 0 | 0 | 0 | 0 | 0 | 0 | 0 | 0 | 0 | 0 | 0 | 0 | 0 |
| bacteria_Jun8d50b_37342 | 1 | 0 | 0 | 0 | 0 | 0 | 0 | 0 | 0 | 0 | 0 | 1 | 0 | 0 | 0 | 0 | 0 | 0 | 0 | 0 | 0 | 0 | 0 | 0 | 0 | 0 | 0 | 0 | 0 | 0 | 0 | 0 | 0 | 0 | 0 | 0 | 0 | 0 | 0 | 0 | 0 | 0 | 0 | 0 | 0 | 0 | 0 | 0 | 0 | 0 | 0 | 0 | 0 | 0 | 0 |
| bacteria_Jun8d50b_38853 | 1 | 0 | 0 | 0 | 0 | 0 | 0 | 0 | 0 | 0 | 0 | 1 | 0 | 0 | 0 | 0 | 0 | 0 | 0 | 0 | 0 | 0 | 0 | 0 | 0 | 0 | 0 | 0 | 0 | 0 | 0 | 0 | 0 | 0 | 0 | 0 | 0 | 0 | 0 | 0 | 0 | 0 | 0 | 0 | 0 | 0 | 0 | 0 | 0 | 0 | 0 | 0 | 0 | 0 | 0 |
| bacteria_Jun8d50b_40026 | 1 | 0 | 0 | 0 | 0 | 0 | 0 | 0 | 0 | 0 | 0 | 1 | 0 | 0 | 0 | 0 | 0 | 0 | 0 | 0 | 0 | 0 | 0 | 0 | 0 | 0 | 0 | 0 | 0 | 0 | 0 | 0 | 0 | 0 | 0 | 0 | 0 | 0 | 0 | 0 | 0 | 0 | 0 | 0 | 0 | 0 | 0 | 0 | 0 | 0 | 0 | 0 | 0 | 0 | 0 |
| bacteria_Jun8d50b_44799 | 1 | 0 | 0 | 0 | 0 | 0 | 0 | 0 | 0 | 0 | 0 | 1 | 0 | 0 | 0 | 0 | 0 | 0 | 0 | 0 | 0 | 0 | 0 | 0 | 0 | 0 | 0 | 0 | 0 | 0 | 0 | 0 | 0 | 0 | 0 | 0 | 0 | 0 | 0 | 0 | 0 | 0 | 0 | 0 | 0 | 0 | 0 | 0 | 0 | 0 | 0 | 0 | 0 | 0 | 0 |
| bacteria_Jun8d75b_16458 | 1 | 0 | 0 | 0 | 0 | 0 | 0 | 0 | 0 | 0 | 0 | 0 | 0 | 1 | 0 | 0 | 0 | 0 | 0 | 0 | 0 | 0 | 0 | 0 | 0 | 0 | 0 | 0 | 0 | 0 | 0 | 0 | 0 | 0 | 0 | 0 | 0 | 0 | 0 | 0 | 0 | 0 | 0 | 0 | 0 | 0 | 0 | 0 | 0 | 0 | 0 | 0 | 0 | 0 | 0 |
| bacteria_Jun8d75b_27066 | 1 | 0 | 0 | 0 | 0 | 0 | 0 | 0 | 0 | 0 | 0 | 0 | 0 | 1 | 0 | 0 | 0 | 0 | 0 | 0 | 0 | 0 | 0 | 0 | 0 | 0 | 0 | 0 | 0 | 0 | 0 | 0 | 0 | 0 | 0 | 0 | 0 | 0 | 0 | 0 | 0 | 0 | 0 | 0 | 0 | 0 | 0 | 0 | 0 | 0 | 0 | 0 | 0 | 0 | 0 |
| bacteria_Jun8d75b_72425 | 1 | 0 | 0 | 0 | 0 | 0 | 0 | 0 | 0 | 0 | 0 | 0 | 0 | 1 | 0 | 0 | 0 | 0 | 0 | 0 | 0 | 0 | 0 | 0 | 0 | 0 | 0 | 0 | 0 | 0 | 0 | 0 | 0 | 0 | 0 | 0 | 0 | 0 | 0 | 0 | 0 | 0 | 0 | 0 | 0 | 0 | 0 | 0 | 0 | 0 | 0 | 0 | 0 | 0 | 0 |
| bacteria_Jun22d0b_30541 | 1 | 0 | 0 | 0 | 0 | 0 | 0 | 0 | 0 | 0 | 0 | 0 | 0 | 0 | 0 | 1 | 0 | 0 | 0 | 0 | 0 | 0 | 0 | 0 | 0 | 0 | 0 | 0 | 0 | 0 | 0 | 0 | 0 | 0 | 0 | 0 | 0 | 0 | 0 | 0 | 0 | 0 | 0 | 0 | 0 | 0 | 0 | 0 | 0 | 0 | 0 | 0 | 0 | 0 | 0 |
| bacteria_Jun22d0b_55410 | 1 | 0 | 0 | 0 | 0 | 0 | 0 | 0 | 0 | 0 | 0 | 0 | 0 | 0 | 0 | 1 | 0 | 0 | 0 | 0 | 0 | 0 | 0 | 0 | 0 | 0 | 0 | 0 | 0 | 0 | 0 | 0 | 0 | 0 | 0 | 0 | 0 | 0 | 0 | 0 | 0 | 0 | 0 | 0 | 0 | 0 | 0 | 0 | 0 | 0 | 0 | 0 | 0 | 0 | 0 |
| bacteria_Jun22d0b_66924 | 1 | 0 | 0 | 0 | 0 | 0 | 0 | 0 | 0 | 0 | 0 | 0 | 0 | 0 | 0 | 1 | 0 | 0 | 0 | 0 | 0 | 0 | 0 | 0 | 0 | 0 | 0 | 0 | 0 | 0 | 0 | 0 | 0 | 0 | 0 | 0 | 0 | 0 | 0 | 0 | 0 | 0 | 0 | 0 | 0 | 0 | 0 | 0 | 0 | 0 | 0 | 0 | 0 | 0 | 0 |
| bacteria_Jun22d75b_3931 | 1 | 0 | 0 | 0 | 0 | 0 | 0 | 0 | 0 | 0 | 0 | 0 | 0 | 0 | 0 | 0 | 0 | 0 | 0 | 1 | 0 | 0 | 0 | 0 | 0 | 0 | 0 | 0 | 0 | 0 | 0 | 0 | 0 | 0 | 0 | 0 | 0 | 0 | 0 | 0 | 0 | 0 | 0 | 0 | 0 | 0 | 0 | 0 | 0 | 0 | 0 | 0 | 0 | 0 | 0 |
| bacteria_Jun22d75b_5194 | 1 | 0 | 0 | 0 | 0 | 0 | 0 | 0 | 0 | 0 | 0 | 0 | 0 | 0 | 0 | 0 | 0 | 0 | 0 | 1 | 0 | 0 | 0 | 0 | 0 | 0 | 0 | 0 | 0 | 0 | 0 | 0 | 0 | 0 | 0 | 0 | 0 | 0 | 0 | 0 | 0 | 0 | 0 | 0 | 0 | 0 | 0 | 0 | 0 | 0 | 0 | 0 | 0 | 0 | 0 |
| bacteria_Jun22d75b_5463 | 1 | 0 | 0 | 0 | 0 | 0 | 0 | 0 | 0 | 0 | 0 | 0 | 0 | 0 | 0 | 0 | 0 | 0 | 0 | 1 | 0 | 0 | 0 | 0 | 0 | 0 | 0 | 0 | 0 | 0 | 0 | 0 | 0 | 0 | 0 | 0 | 0 | 0 | 0 | 0 | 0 | 0 | 0 | 0 | 0 | 0 | 0 | 0 | 0 | 0 | 0 | 0 | 0 | 0 | 0 |
| bacteria_Jun22d75b_5572 | 1 | 0 | 0 | 0 | 0 | 0 | 0 | 0 | 0 | 0 | 0 | 0 | 0 | 0 | 0 | 0 | 0 | 0 | 0 | 1 | 0 | 0 | 0 | 0 | 0 | 0 | 0 | 0 | 0 | 0 | 0 | 0 | 0 | 0 | 0 | 0 | 0 | 0 | 0 | 0 | 0 | 0 | 0 | 0 | 0 | 0 | 0 | 0 | 0 | 0 | 0 | 0 | 0 | 0 | 0 |
| bacteria_Jun22d75b_6065 | 1 | 0 | 0 | 0 | 0 | 0 | 0 | 0 | 0 | 0 | 0 | 0 | 0 | 0 | 0 | 0 | 0 | 0 | 0 | 1 | 0 | 0 | 0 | 0 | 0 | 0 | 0 | 0 | 0 | 0 | 0 | 0 | 0 | 0 | 0 | 0 | 0 | 0 | 0 | 0 | 0 | 0 | 0 | 0 | 0 | 0 | 0 | 0 | 0 | 0 | 0 | 0 | 0 | 0 | 0 |
| bacteria_Jun22d75b_6400 | 1 | 0 | 0 | 0 | 0 | 0 | 0 | 0 | 0 | 0 | 0 | 0 | 0 | 0 | 0 | 0 | 0 | 0 | 0 | 1 | 0 | 0 | 0 | 0 | 0 | 0 | 0 | 0 | 0 | 0 | 0 | 0 | 0 | 0 | 0 | 0 | 0 | 0 | 0 | 0 | 0 | 0 | 0 | 0 | 0 | 0 | 0 | 0 | 0 | 0 | 0 | 0 | 0 | 0 | 0 |
| bacteria_Jun22d75b_6747 | 1 | 0 | 0 | 0 | 0 | 0 | 0 | 0 | 0 | 0 | 0 | 0 | 0 | 0 | 0 | 0 | 0 | 0 | 0 | 1 | 0 | 0 | 0 | 0 | 0 | 0 | 0 | 0 | 0 | 0 | 0 | 0 | 0 | 0 | 0 | 0 | 0 | 0 | 0 | 0 | 0 | 0 | 0 | 0 | 0 | 0 | 0 | 0 | 0 | 0 | 0 | 0 | 0 | 0 | 0 |
| bacteria_May12d5b_6614 | 1 | 1 | 0 | 0 | 0 | 0 | 0 | 0 | 0 | 0 | 0 | 0 | 0 | 0 | 0 | 0 | 0 | 0 | 0 | 0 | 0 | 0 | 0 | 0 | 0 | 0 | 0 | 0 | 0 | 0 | 0 | 0 | 0 | 0 | 0 | 0 | 0 | 0 | 0 | 0 | 0 | 0 | 0 | 0 | 0 | 0 | 0 | 0 | 0 | 0 | 0 | 0 | 0 | 0 | 0 |
| bacteria_May12d25b_443 | 1 | 0 | 0 | 1 | 0 | 0 | 0 | 0 | 0 | 0 | 0 | 0 | 0 | 0 | 0 | 0 | 0 | 0 | 0 | 0 | 0 | 0 | 0 | 0 | 0 | 0 | 0 | 0 | 0 | 0 | 0 | 0 | 0 | 0 | 0 | 0 | 0 | 0 | 0 | 0 | 0 | 0 | 0 | 0 | 0 | 0 | 0 | 0 | 0 | 0 | 0 | 0 | 0 | 0 | 0 |
| bacteria_May12d25b_213 | 1 | 0 | 0 | 1 | 0 | 0 | 0 | 0 | 0 | 0 | 0 | 0 | 0 | 0 | 0 | 0 | 0 | 0 | 0 | 0 | 0 | 0 | 0 | 0 | 0 | 0 | 0 | 0 | 0 | 0 | 0 | 0 | 0 | 0 | 0 | 0 | 0 | 0 | 0 | 0 | 0 | 0 | 0 | 0 | 0 | 0 | 0 | 0 | 0 | 0 | 0 | 0 | 0 | 0 | 0 |
| bacteria_May12d25b_371 | 1 | 0 | 0 | 1 | 0 | 0 | 0 | 0 | 0 | 0 | 0 | 0 | 0 | 0 | 0 | 0 | 0 | 0 | 0 | 0 | 0 | 0 | 0 | 0 | 0 | 0 | 0 | 0 | 0 | 0 | 0 | 0 | 0 | 0 | 0 | 0 | 0 | 0 | 0 | 0 | 0 | 0 | 0 | 0 | 0 | 0 | 0 | 0 | 0 | 0 | 0 | 0 | 0 | 0 | 0 |
| bacteria_May12d25b_395 | 1 | 0 | 0 | 1 | 0 | 0 | 0 | 0 | 0 | 0 | 0 | 0 | 0 | 0 | 0 | 0 | 0 | 0 | 0 | 0 | 0 | 0 | 0 | 0 | 0 | 0 | 0 | 0 | 0 | 0 | 0 | 0 | 0 | 0 | 0 | 0 | 0 | 0 | 0 | 0 | 0 | 0 | 0 | 0 | 0 | 0 | 0 | 0 | 0 | 0 | 0 | 0 | 0 | 0 | 0 |
| bacteria_May12d25b_613 | 1 | 0 | 0 | 1 | 0 | 0 | 0 | 0 | 0 | 0 | 0 | 0 | 0 | 0 | 0 | 0 | 0 | 0 | 0 | 0 | 0 | 0 | 0 | 0 | 0 | 0 | 0 | 0 | 0 | 0 | 0 | 0 | 0 | 0 | 0 | 0 | 0 | 0 | 0 | 0 | 0 | 0 | 0 | 0 | 0 | 0 | 0 | 0 | 0 | 0 | 0 | 0 | 0 | 0 | 0 |
| bacteria_May12d25b_640 | 1 | 0 | 0 | 1 | 0 | 0 | 0 | 0 | 0 | 0 | 0 | 0 | 0 | 0 | 0 | 0 | 0 | 0 | 0 | 0 | 0 | 0 | 0 | 0 | 0 | 0 | 0 | 0 | 0 | 0 | 0 | 0 | 0 | 0 | 0 | 0 | 0 | 0 | 0 | 0 | 0 | 0 | 0 | 0 | 0 | 0 | 0 | 0 | 0 | 0 | 0 | 0 | 0 | 0 | 0 |
| bacteria_May12d25b_722 | 1 | 0 | 0 | 1 | 0 | 0 | 0 | 0 | 0 | 0 | 0 | 0 | 0 | 0 | 0 | 0 | 0 | 0 | 0 | 0 | 0 | 0 | 0 | 0 | 0 | 0 | 0 | 0 | 0 | 0 | 0 | 0 | 0 | 0 | 0 | 0 | 0 | 0 | 0 | 0 | 0 | 0 | 0 | 0 | 0 | 0 | 0 | 0 | 0 | 0 | 0 | 0 | 0 | 0 | 0 |
| bacteria_May12d50b_133 | 1 | 0 | 0 | 0 | 0 | 1 | 0 | 0 | 0 | 0 | 0 | 0 | 0 | 0 | 0 | 0 | 0 | 0 | 0 | 0 | 0 | 0 | 0 | 0 | 0 | 0 | 0 | 0 | 0 | 0 | 0 | 0 | 0 | 0 | 0 | 0 | 0 | 0 | 0 | 0 | 0 | 0 | 0 | 0 | 0 | 0 | 0 | 0 | 0 | 0 | 0 | 0 | 0 | 0 | 0 |
| bacteria_May12d50b_507 | 1 | 0 | 0 | 0 | 0 | 1 | 0 | 0 | 0 | 0 | 0 | 0 | 0 | 0 | 0 | 0 | 0 | 0 | 0 | 0 | 0 | 0 | 0 | 0 | 0 | 0 | 0 | 0 | 0 | 0 | 0 | 0 | 0 | 0 | 0 | 0 | 0 | 0 | 0 | 0 | 0 | 0 | 0 | 0 | 0 | 0 | 0 | 0 | 0 | 0 | 0 | 0 | 0 | 0 | 0 |
| bacteria_May12d50b_589 | 1 | 0 | 0 | 0 | 0 | 1 | 0 | 0 | 0 | 0 | 0 | 0 | 0 | 0 | 0 | 0 | 0 | 0 | 0 | 0 | 0 | 0 | 0 | 0 | 0 | 0 | 0 | 0 | 0 | 0 | 0 | 0 | 0 | 0 | 0 | 0 | 0 | 0 | 0 | 0 | 0 | 0 | 0 | 0 | 0 | 0 | 0 | 0 | 0 | 0 | 0 | 0 | 0 | 0 | 0 |
| bacteria_May12d50b_112 | 1 | 0 | 0 | 0 | 0 | 1 | 0 | 0 | 0 | 0 | 0 | 0 | 0 | 0 | 0 | 0 | 0 | 0 | 0 | 0 | 0 | 0 | 0 | 0 | 0 | 0 | 0 | 0 | 0 | 0 | 0 | 0 | 0 | 0 | 0 | 0 | 0 | 0 | 0 | 0 | 0 | 0 | 0 | 0 | 0 | 0 | 0 | 0 | 0 | 0 | 0 | 0 | 0 | 0 | 0 |
| bacteria_May12d50b_118 | 1 | 0 | 0 | 0 | 0 | 1 | 0 | 0 | 0 | 0 | 0 | 0 | 0 | 0 | 0 | 0 | 0 | 0 | 0 | 0 | 0 | 0 | 0 | 0 | 0 | 0 | 0 | 0 | 0 | 0 | 0 | 0 | 0 | 0 | 0 | 0 | 0 | 0 | 0 | 0 | 0 | 0 | 0 | 0 | 0 | 0 | 0 | 0 | 0 | 0 | 0 | 0 | 0 | 0 | 0 |
| bacteria_May12d50b_133 | 1 | 0 | 0 | 0 | 0 | 1 | 0 | 0 | 0 | 0 | 0 | 0 | 0 | 0 | 0 | 0 | 0 | 0 | 0 | 0 | 0 | 0 | 0 | 0 | 0 | 0 | 0 | 0 | 0 | 0 | 0 | 0 | 0 | 0 | 0 | 0 | 0 | 0 | 0 | 0 | 0 | 0 | 0 | 0 | 0 | 0 | 0 | 0 | 0 | 0 | 0 | 0 | 0 | 0 | 0 |
| bacteria_May12d50b_149 | 1 | 0 | 0 | 0 | 0 | 1 | 0 | 0 | 0 | 0 | 0 | 0 | 0 | 0 | 0 | 0 | 0 | 0 | 0 | 0 | 0 | 0 | 0 | 0 | 0 | 0 | 0 | 0 | 0 | 0 | 0 | 0 | 0 | 0 | 0 | 0 | 0 | 0 | 0 | 0 | 0 | 0 | 0 | 0 | 0 | 0 | 0 | 0 | 0 | 0 | 0 | 0 | 0 | 0 | 0 |
| bacteria_May12d50b_312 | 1 | 0 | 0 | 0 | 0 | 1 | 0 | 0 | 0 | 0 | 0 | 0 | 0 | 0 | 0 | 0 | 0 | 0 | 0 | 0 | 0 | 0 | 0 | 0 | 0 | 0 | 0 | 0 | 0 | 0 | 0 | 0 | 0 | 0 | 0 | 0 | 0 | 0 | 0 | 0 | 0 | 0 | 0 | 0 | 0 | 0 | 0 | 0 | 0 | 0 | 0 | 0 | 0 | 0 | 0 |
| bacteria_May12d50b_400 | 1 | 0 | 0 | 0 | 0 | 1 | 0 | 0 | 0 | 0 | 0 | 0 | 0 | 0 | 0 | 0 | 0 | 0 | 0 | 0 | 0 | 0 | 0 | 0 | 0 | 0 | 0 | 0 | 0 | 0 | 0 | 0 | 0 | 0 | 0 | 0 | 0 | 0 | 0 | 0 | 0 | 0 | 0 | 0 | 0 | 0 | 0 | 0 | 0 | 0 | 0 | 0 | 0 | 0 | 0 |
| bacteria_May12d50b_473 | 1 | 0 | 0 | 0 | 0 | 1 | 0 | 0 | 0 | 0 | 0 | 0 | 0 | 0 | 0 | 0 | 0 | 0 | 0 | 0 | 0 | 0 | 0 | 0 | 0 | 0 | 0 | 0 | 0 | 0 | 0 | 0 | 0 | 0 | 0 | 0 | 0 | 0 | 0 | 0 | 0 | 0 | 0 | 0 | 0 | 0 | 0 | 0 | 0 | 0 | 0 | 0 | 0 | 0 | 0 |
| bacteria_May12d75b_227 | 1 | 0 | 0 | 0 | 0 | 0 | 0 | 1 | 0 | 0 | 0 | 0 | 0 | 0 | 0 | 0 | 0 | 0 | 0 | 0 | 0 | 0 | 0 | 0 | 0 | 0 | 0 | 0 | 0 | 0 | 0 | 0 | 0 | 0 | 0 | 0 | 0 | 0 | 0 | 0 | 0 | 0 | 0 | 0 | 0 | 0 | 0 | 0 | 0 | 0 | 0 | 0 | 0 | 0 | 0 |
| bacteria_Sep8d0b_54399 | 1 | 0 | 0 | 0 | 0 | 0 | 0 | 0 | 0 | 0 | 0 | 0 | 0 | 0 | 0 | 0 | 0 | 0 | 0 | 0 | 0 | 0 | 0 | 0 | 0 | 0 | 0 | 0 | 0 | 0 | 0 | 0 | 0 | 0 | 0 | 0 | 0 | 0 | 0 | 0 | 0 | 0 | 0 | 0 | 0 | 0 | 0 | 1 | 0 | 0 | 0 | 0 | 0 | 0 | 0 |
| bacteria_Sep8d5b_843 | 1 | 0 | 0 | 0 | 0 | 0 | 0 | 0 | 0 | 0 | 0 | 0 | 0 | 0 | 0 | 0 | 0 | 0 | 0 | 0 | 0 | 0 | 0 | 0 | 0 | 0 | 0 | 0 | 0 | 0 | 0 | 0 | 0 | 0 | 0 | 0 | 0 | 0 | 0 | 0 | 0 | 0 | 0 | 0 | 0 | 0 | 0 | 0 | 0 | 1 | 0 | 0 | 0 | 0 | 0 |
| bacteria_Sep8d5b_1310 | 1 | 0 | 0 | 0 | 0 | 0 | 0 | 0 | 0 | 0 | 0 | 0 | 0 | 0 | 0 | 0 | 0 | 0 | 0 | 0 | 0 | 0 | 0 | 0 | 0 | 0 | 0 | 0 | 0 | 0 | 0 | 0 | 0 | 0 | 0 | 0 | 0 | 0 | 0 | 0 | 0 | 0 | 0 | 0 | 0 | 0 | 0 | 0 | 0 | 1 | 0 | 0 | 0 | 0 | 0 |
| bacteria_Sep8d5b_12186 | 1 | 0 | 0 | 0 | 0 | 0 | 0 | 0 | 0 | 0 | 0 | 0 | 0 | 0 | 0 | 0 | 0 | 0 | 0 | 0 | 0 | 0 | 0 | 0 | 0 | 0 | 0 | 0 | 0 | 0 | 0 | 0 | 0 | 0 | 0 | 0 | 0 | 0 | 0 | 0 | 0 | 0 | 0 | 0 | 0 | 0 | 0 | 0 | 0 | 1 | 0 | 0 | 0 | 0 | 0 |
| bacteria_Sep8d5b_12323 | 1 | 0 | 0 | 0 | 0 | 0 | 0 | 0 | 0 | 0 | 0 | 0 | 0 | 0 | 0 | 0 | 0 | 0 | 0 | 0 | 0 | 0 | 0 | 0 | 0 | 0 | 0 | 0 | 0 | 0 | 0 | 0 | 0 | 0 | 0 | 0 | 0 | 0 | 0 | 0 | 0 | 0 | 0 | 0 | 0 | 0 | 0 | 0 | 0 | 1 | 0 | 0 | 0 | 0 | 0 |
| bacteria_Sep8d5b_16248 | 1 | 0 | 0 | 0 | 0 | 0 | 0 | 0 | 0 | 0 | 0 | 0 | 0 | 0 | 0 | 0 | 0 | 0 | 0 | 0 | 0 | 0 | 0 | 0 | 0 | 0 | 0 | 0 | 0 | 0 | 0 | 0 | 0 | 0 | 0 | 0 | 0 | 0 | 0 | 0 | 0 | 0 | 0 | 0 | 0 | 0 | 0 | 0 | 0 | 1 | 0 | 0 | 0 | 0 | 0 |
| bacteria_Sep8d5b_22301 | 1 | 0 | 0 | 0 | 0 | 0 | 0 | 0 | 0 | 0 | 0 | 0 | 0 | 0 | 0 | 0 | 0 | 0 | 0 | 0 | 0 | 0 | 0 | 0 | 0 | 0 | 0 | 0 | 0 | 0 | 0 | 0 | 0 | 0 | 0 | 0 | 0 | 0 | 0 | 0 | 0 | 0 | 0 | 0 | 0 | 0 | 0 | 0 | 0 | 1 | 0 | 0 | 0 | 0 | 0 |
| bacteria_Sep8d5b_25121 | 1 | 0 | 0 | 0 | 0 | 0 | 0 | 0 | 0 | 0 | 0 | 0 | 0 | 0 | 0 | 0 | 0 | 0 | 0 | 0 | 0 | 0 | 0 | 0 | 0 | 0 | 0 | 0 | 0 | 0 | 0 | 0 | 0 | 0 | 0 | 0 | 0 | 0 | 0 | 0 | 0 | 0 | 0 | 0 | 0 | 0 | 0 | 0 | 0 | 1 | 0 | 0 | 0 | 0 | 0 |
| bacteria_Sep8d5b_27115 | 1 | 0 | 0 | 0 | 0 | 0 | 0 | 0 | 0 | 0 | 0 | 0 | 0 | 0 | 0 | 0 | 0 | 0 | 0 | 0 | 0 | 0 | 0 | 0 | 0 | 0 | 0 | 0 | 0 | 0 | 0 | 0 | 0 | 0 | 0 | 0 | 0 | 0 | 0 | 0 | 0 | 0 | 0 | 0 | 0 | 0 | 0 | 0 | 0 | 1 | 0 | 0 | 0 | 0 | 0 |
| bacteria_Sep8d5b_27802 | 1 | 0 | 0 | 0 | 0 | 0 | 0 | 0 | 0 | 0 | 0 | 0 | 0 | 0 | 0 | 0 | 0 | 0 | 0 | 0 | 0 | 0 | 0 | 0 | 0 | 0 | 0 | 0 | 0 | 0 | 0 | 0 | 0 | 0 | 0 | 0 | 0 | 0 | 0 | 0 | 0 | 0 | 0 | 0 | 0 | 0 | 0 | 0 | 0 | 1 | 0 | 0 | 0 | 0 | 0 |
| bacteria_Sep8d5b_28161 | 1 | 0 | 0 | 0 | 0 | 0 | 0 | 0 | 0 | 0 | 0 | 0 | 0 | 0 | 0 | 0 | 0 | 0 | 0 | 0 | 0 | 0 | 0 | 0 | 0 | 0 | 0 | 0 | 0 | 0 | 0 | 0 | 0 | 0 | 0 | 0 | 0 | 0 | 0 | 0 | 0 | 0 | 0 | 0 | 0 | 0 | 0 | 0 | 0 | 1 | 0 | 0 | 0 | 0 | 0 |
| bacteria_Sep8d5b_29068 | 1 | 0 | 0 | 0 | 0 | 0 | 0 | 0 | 0 | 0 | 0 | 0 | 0 | 0 | 0 | 0 | 0 | 0 | 0 | 0 | 0 | 0 | 0 | 0 | 0 | 0 | 0 | 0 | 0 | 0 | 0 | 0 | 0 | 0 | 0 | 0 | 0 | 0 | 0 | 0 | 0 | 0 | 0 | 0 | 0 | 0 | 0 | 0 | 0 | 1 | 0 | 0 | 0 | 0 | 0 |
| bacteria_Sep8d5b_39467 | 1 | 0 | 0 | 0 | 0 | 0 | 0 | 0 | 0 | 0 | 0 | 0 | 0 | 0 | 0 | 0 | 0 | 0 | 0 | 0 | 0 | 0 | 0 | 0 | 0 | 0 | 0 | 0 | 0 | 0 | 0 | 0 | 0 | 0 | 0 | 0 | 0 | 0 | 0 | 0 | 0 | 0 | 0 | 0 | 0 | 0 | 0 | 0 | 0 | 1 | 0 | 0 | 0 | 0 | 0 |
| bacteria_Sep8d5b_40166 | 1 | 0 | 0 | 0 | 0 | 0 | 0 | 0 | 0 | 0 | 0 | 0 | 0 | 0 | 0 | 0 | 0 | 0 | 0 | 0 | 0 | 0 | 0 | 0 | 0 | 0 | 0 | 0 | 0 | 0 | 0 | 0 | 0 | 0 | 0 | 0 | 0 | 0 | 0 | 0 | 0 | 0 | 0 | 0 | 0 | 0 | 0 | 0 | 0 | 1 | 0 | 0 | 0 | 0 | 0 |
| bacteria_Sep8d5b_42221 | 1 | 0 | 0 | 0 | 0 | 0 | 0 | 0 | 0 | 0 | 0 | 0 | 0 | 0 | 0 | 0 | 0 | 0 | 0 | 0 | 0 | 0 | 0 | 0 | 0 | 0 | 0 | 0 | 0 | 0 | 0 | 0 | 0 | 0 | 0 | 0 | 0 | 0 | 0 | 0 | 0 | 0 | 0 | 0 | 0 | 0 | 0 | 0 | 0 | 1 | 0 | 0 | 0 | 0 | 0 |
| bacteria_Sep8d5b_44029 | 1 | 0 | 0 | 0 | 0 | 0 | 0 | 0 | 0 | 0 | 0 | 0 | 0 | 0 | 0 | 0 | 0 | 0 | 0 | 0 | 0 | 0 | 0 | 0 | 0 | 0 | 0 | 0 | 0 | 0 | 0 | 0 | 0 | 0 | 0 | 0 | 0 | 0 | 0 | 0 | 0 | 0 | 0 | 0 | 0 | 0 | 0 | 0 | 0 | 1 | 0 | 0 | 0 | 0 | 0 |
| bacteria_Sep8d5b_44060 | 1 | 0 | 0 | 0 | 0 | 0 | 0 | 0 | 0 | 0 | 0 | 0 | 0 | 0 | 0 | 0 | 0 | 0 | 0 | 0 | 0 | 0 | 0 | 0 | 0 | 0 | 0 | 0 | 0 | 0 | 0 | 0 | 0 | 0 | 0 | 0 | 0 | 0 | 0 | 0 | 0 | 0 | 0 | 0 | 0 | 0 | 0 | 0 | 0 | 1 | 0 | 0 | 0 | 0 | 0 |
| bacteria_Sep8d5b_51717 | 1 | 0 | 0 | 0 | 0 | 0 | 0 | 0 | 0 | 0 | 0 | 0 | 0 | 0 | 0 | 0 | 0 | 0 | 0 | 0 | 0 | 0 | 0 | 0 | 0 | 0 | 0 | 0 | 0 | 0 | 0 | 0 | 0 | 0 | 0 | 0 | 0 | 0 | 0 | 0 | 0 | 0 | 0 | 0 | 0 | 0 | 0 | 0 | 0 | 1 | 0 | 0 | 0 | 0 | 0 |
| bacteria_Sep8d5b_57398 | 1 | 0 | 0 | 0 | 0 | 0 | 0 | 0 | 0 | 0 | 0 | 0 | 0 | 0 | 0 | 0 | 0 | 0 | 0 | 0 | 0 | 0 | 0 | 0 | 0 | 0 | 0 | 0 | 0 | 0 | 0 | 0 | 0 | 0 | 0 | 0 | 0 | 0 | 0 | 0 | 0 | 0 | 0 | 0 | 0 | 0 | 0 | 0 | 0 | 1 | 0 | 0 | 0 | 0 | 0 |
| bacteria_Sep8d5b_58396 | 1 | 0 | 0 | 0 | 0 | 0 | 0 | 0 | 0 | 0 | 0 | 0 | 0 | 0 | 0 | 0 | 0 | 0 | 0 | 0 | 0 | 0 | 0 | 0 | 0 | 0 | 0 | 0 | 0 | 0 | 0 | 0 | 0 | 0 | 0 | 0 | 0 | 0 | 0 | 0 | 0 | 0 | 0 | 0 | 0 | 0 | 0 | 0 | 0 | 1 | 0 | 0 | 0 | 0 | 0 |
| bacteria_Sep8d5b_62351 | 1 | 0 | 0 | 0 | 0 | 0 | 0 | 0 | 0 | 0 | 0 | 0 | 0 | 0 | 0 | 0 | 0 | 0 | 0 | 0 | 0 | 0 | 0 | 0 | 0 | 0 | 0 | 0 | 0 | 0 | 0 | 0 | 0 | 0 | 0 | 0 | 0 | 0 | 0 | 0 | 0 | 0 | 0 | 0 | 0 | 0 | 0 | 0 | 0 | 1 | 0 | 0 | 0 | 0 | 0 |
| bacteria_Sep8d5b_71080 | 1 | 0 | 0 | 0 | 0 | 0 | 0 | 0 | 0 | 0 | 0 | 0 | 0 | 0 | 0 | 0 | 0 | 0 | 0 | 0 | 0 | 0 | 0 | 0 | 0 | 0 | 0 | 0 | 0 | 0 | 0 | 0 | 0 | 0 | 0 | 0 | 0 | 0 | 0 | 0 | 0 | 0 | 0 | 0 | 0 | 0 | 0 | 0 | 0 | 1 | 0 | 0 | 0 | 0 | 0 |
| bacteria_Sep8d25b_2747 | 1 | 0 | 0 | 0 | 0 | 0 | 0 | 0 | 0 | 0 | 0 | 0 | 0 | 0 | 0 | 0 | 0 | 0 | 0 | 0 | 0 | 0 | 0 | 0 | 0 | 0 | 0 | 0 | 0 | 0 | 0 | 0 | 0 | 0 | 0 | 0 | 0 | 0 | 0 | 0 | 0 | 0 | 0 | 0 | 0 | 0 | 0 | 0 | 0 | 0 | 0 | 1 | 0 | 0 | 0 |
| bacteria_Sep8d25b_2183 | 1 | 0 | 0 | 0 | 0 | 0 | 0 | 0 | 0 | 0 | 0 | 0 | 0 | 0 | 0 | 0 | 0 | 0 | 0 | 0 | 0 | 0 | 0 | 0 | 0 | 0 | 0 | 0 | 0 | 0 | 0 | 0 | 0 | 0 | 0 | 0 | 0 | 0 | 0 | 0 | 0 | 0 | 0 | 0 | 0 | 0 | 0 | 0 | 0 | 0 | 0 | 1 | 0 | 0 | 0 |
| bacteria_Sep8d25b_4024 | 1 | 0 | 0 | 0 | 0 | 0 | 0 | 0 | 0 | 0 | 0 | 0 | 0 | 0 | 0 | 0 | 0 | 0 | 0 | 0 | 0 | 0 | 0 | 0 | 0 | 0 | 0 | 0 | 0 | 0 | 0 | 0 | 0 | 0 | 0 | 0 | 0 | 0 | 0 | 0 | 0 | 0 | 0 | 0 | 0 | 0 | 0 | 0 | 0 | 0 | 0 | 1 | 0 | 0 | 0 |
| bacteria_Sep8d25b_4292 | 1 | 0 | 0 | 0 | 0 | 0 | 0 | 0 | 0 | 0 | 0 | 0 | 0 | 0 | 0 | 0 | 0 | 0 | 0 | 0 | 0 | 0 | 0 | 0 | 0 | 0 | 0 | 0 | 0 | 0 | 0 | 0 | 0 | 0 | 0 | 0 | 0 | 0 | 0 | 0 | 0 | 0 | 0 | 0 | 0 | 0 | 0 | 0 | 0 | 0 | 0 | 1 | 0 | 0 | 0 |
| bacteria_Sep8d25b_4527 | 1 | 0 | 0 | 0 | 0 | 0 | 0 | 0 | 0 | 0 | 0 | 0 | 0 | 0 | 0 | 0 | 0 | 0 | 0 | 0 | 0 | 0 | 0 | 0 | 0 | 0 | 0 | 0 | 0 | 0 | 0 | 0 | 0 | 0 | 0 | 0 | 0 | 0 | 0 | 0 | 0 | 0 | 0 | 0 | 0 | 0 | 0 | 0 | 0 | 0 | 0 | 1 | 0 | 0 | 0 |
| bacteria_Sep8d50b_922 | 1 | 0 | 0 | 0 | 0 | 0 | 0 | 0 | 0 | 0 | 0 | 0 | 0 | 0 | 0 | 0 | 0 | 0 | 0 | 0 | 0 | 0 | 0 | 0 | 0 | 0 | 0 | 0 | 0 | 0 | 0 | 0 | 0 | 0 | 0 | 0 | 0 | 0 | 0 | 0 | 0 | 0 | 0 | 0 | 0 | 0 | 0 | 0 | 0 | 0 | 0 | 0 | 0 | 1 | 0 |
| bacteria_Sep8d50b_8892 | 1 | 0 | 0 | 0 | 0 | 0 | 0 | 0 | 0 | 0 | 0 | 0 | 0 | 0 | 0 | 0 | 0 | 0 | 0 | 0 | 0 | 0 | 0 | 0 | 0 | 0 | 0 | 0 | 0 | 0 | 0 | 0 | 0 | 0 | 0 | 0 | 0 | 0 | 0 | 0 | 0 | 0 | 0 | 0 | 0 | 0 | 0 | 0 | 0 | 0 | 0 | 0 | 0 | 1 | 0 |
| bacteria_Sep8d50b_2068 | 1 | 0 | 0 | 0 | 0 | 0 | 0 | 0 | 0 | 0 | 0 | 0 | 0 | 0 | 0 | 0 | 0 | 0 | 0 | 0 | 0 | 0 | 0 | 0 | 0 | 0 | 0 | 0 | 0 | 0 | 0 | 0 | 0 | 0 | 0 | 0 | 0 | 0 | 0 | 0 | 0 | 0 | 0 | 0 | 0 | 0 | 0 | 0 | 0 | 0 | 0 | 0 | 0 | 1 | 0 |
| bacteria_Sep8d50b_4189 | 1 | 0 | 0 | 0 | 0 | 0 | 0 | 0 | 0 | 0 | 0 | 0 | 0 | 0 | 0 | 0 | 0 | 0 | 0 | 0 | 0 | 0 | 0 | 0 | 0 | 0 | 0 | 0 | 0 | 0 | 0 | 0 | 0 | 0 | 0 | 0 | 0 | 0 | 0 | 0 | 0 | 0 | 0 | 0 | 0 | 0 | 0 | 0 | 0 | 0 | 0 | 0 | 0 | 1 | 0 |
| virus_Aug5.0v_144 | 1 | 0 | 0 | 0 | 0 | 0 | 0 | 0 | 0 | 0 | 0 | 0 | 0 | 0 | 0 | 0 | 0 | 0 | 0 | 0 | 0 | 0 | 0 | 0 | 0 | 0 | 0 | 0 | 0 | 0 | 0 | 0 | 0 | 0 | 0 | 0 | 0 | 0 | 1 | 0 | 0 | 0 | 0 | 0 | 0 | 0 | 0 | 0 | 0 | 0 | 0 | 0 | 0 | 0 | 0 |
| virus_Aug5.0v_357 | 1 | 0 | 0 | 0 | 0 | 0 | 0 | 0 | 0 | 0 | 0 | 0 | 0 | 0 | 0 | 0 | 0 | 0 | 0 | 0 | 0 | 0 | 0 | 0 | 0 | 0 | 0 | 0 | 0 | 0 | 0 | 0 | 0 | 0 | 0 | 0 | 0 | 0 | 1 | 0 | 0 | 0 | 0 | 0 | 0 | 0 | 0 | 0 | 0 | 0 | 0 | 0 | 0 | 0 | 0 |
| virus_Aug5.0v_508 | 1 | 0 | 0 | 0 | 0 | 0 | 0 | 0 | 0 | 0 | 0 | 0 | 0 | 0 | 0 | 0 | 0 | 0 | 0 | 0 | 0 | 0 | 0 | 0 | 0 | 0 | 0 | 0 | 0 | 0 | 0 | 0 | 0 | 0 | 0 | 0 | 0 | 0 | 1 | 0 | 0 | 0 | 0 | 0 | 0 | 0 | 0 | 0 | 0 | 0 | 0 | 0 | 0 | 0 | 0 |
| virus_Aug5.0v_648 | 1 | 0 | 0 | 0 | 0 | 0 | 0 | 0 | 0 | 0 | 0 | 0 | 0 | 0 | 0 | 0 | 0 | 0 | 0 | 0 | 0 | 0 | 0 | 0 | 0 | 0 | 0 | 0 | 0 | 0 | 0 | 0 | 0 | 0 | 0 | 0 | 0 | 0 | 1 | 0 | 0 | 0 | 0 | 0 | 0 | 0 | 0 | 0 | 0 | 0 | 0 | 0 | 0 | 0 | 0 |
| virus_Aug5.0v_809 | 1 | 0 | 0 | 0 | 0 | 0 | 0 | 0 | 0 | 0 | 0 | 0 | 0 | 0 | 0 | 0 | 0 | 0 | 0 | 0 | 0 | 0 | 0 | 0 | 0 | 0 | 0 | 0 | 0 | 0 | 0 | 0 | 0 | 0 | 0 | 0 | 0 | 0 | 1 | 0 | 0 | 0 | 0 | 0 | 0 | 0 | 0 | 0 | 0 | 0 | 0 | 0 | 0 | 0 | 0 |
| virus_Aug5.0v_1201 | 1 | 0 | 0 | 0 | 0 | 0 | 0 | 0 | 0 | 0 | 0 | 0 | 0 | 0 | 0 | 0 | 0 | 0 | 0 | 0 | 0 | 0 | 0 | 0 | 0 | 0 | 0 | 0 | 0 | 0 | 0 | 0 | 0 | 0 | 0 | 0 | 0 | 0 | 1 | 0 | 0 | 0 | 0 | 0 | 0 | 0 | 0 | 0 | 0 | 0 | 0 | 0 | 0 | 0 | 0 |
| virus_Aug5.0v_1242 | 1 | 0 | 0 | 0 | 0 | 0 | 0 | 0 | 0 | 0 | 0 | 0 | 0 | 0 | 0 | 0 | 0 | 0 | 0 | 0 | 0 | 0 | 0 | 0 | 0 | 0 | 0 | 0 | 0 | 0 | 0 | 0 | 0 | 0 | 0 | 0 | 0 | 0 | 1 | 0 | 0 | 0 | 0 | 0 | 0 | 0 | 0 | 0 | 0 | 0 | 0 | 0 | 0 | 0 | 0 |
| virus_Aug5.0v_1502 | 1 | 0 | 0 | 0 | 0 | 0 | 0 | 0 | 0 | 0 | 0 | 0 | 0 | 0 | 0 | 0 | 0 | 0 | 0 | 0 | 0 | 0 | 0 | 0 | 0 | 0 | 0 | 0 | 0 | 0 | 0 | 0 | 0 | 0 | 0 | 0 | 0 | 0 | 1 | 0 | 0 | 0 | 0 | 0 | 0 | 0 | 0 | 0 | 0 | 0 | 0 | 0 | 0 | 0 | 0 |
| virus_Aug5.0v_1877 | 1 | 0 | 0 | 0 | 0 | 0 | 0 | 0 | 0 | 0 | 0 | 0 | 0 | 0 | 0 | 0 | 0 | 0 | 0 | 0 | 0 | 0 | 0 | 0 | 0 | 0 | 0 | 0 | 0 | 0 | 0 | 0 | 0 | 0 | 0 | 0 | 0 | 0 | 1 | 0 | 0 | 0 | 0 | 0 | 0 | 0 | 0 | 0 | 0 | 0 | 0 | 0 | 0 | 0 | 0 |
| virus_Aug5.0v_2151 | 1 | 0 | 0 | 0 | 0 | 0 | 0 | 0 | 0 | 0 | 0 | 0 | 0 | 0 | 0 | 0 | 0 | 0 | 0 | 0 | 0 | 0 | 0 | 0 | 0 | 0 | 0 | 0 | 0 | 0 | 0 | 0 | 0 | 0 | 0 | 0 | 0 | 0 | 1 | 0 | 0 | 0 | 0 | 0 | 0 | 0 | 0 | 0 | 0 | 0 | 0 | 0 | 0 | 0 | 0 |
| virus_Aug5.0v_2249 | 1 | 0 | 0 | 0 | 0 | 0 | 0 | 0 | 0 | 0 | 0 | 0 | 0 | 0 | 0 | 0 | 0 | 0 | 0 | 0 | 0 | 0 | 0 | 0 | 0 | 0 | 0 | 0 | 0 | 0 | 0 | 0 | 0 | 0 | 0 | 0 | 0 | 0 | 1 | 0 | 0 | 0 | 0 | 0 | 0 | 0 | 0 | 0 | 0 | 0 | 0 | 0 | 0 | 0 | 0 |
| virus_Aug5.0v_2548 | 1 | 0 | 0 | 0 | 0 | 0 | 0 | 0 | 0 | 0 | 0 | 0 | 0 | 0 | 0 | 0 | 0 | 0 | 0 | 0 | 0 | 0 | 0 | 0 | 0 | 0 | 0 | 0 | 0 | 0 | 0 | 0 | 0 | 0 | 0 | 0 | 0 | 0 | 1 | 0 | 0 | 0 | 0 | 0 | 0 | 0 | 0 | 0 | 0 | 0 | 0 | 0 | 0 | 0 | 0 |
| virus_Aug5.0v_2599 | 1 | 0 | 0 | 0 | 0 | 0 | 0 | 0 | 0 | 0 | 0 | 0 | 0 | 0 | 0 | 0 | 0 | 0 | 0 | 0 | 0 | 0 | 0 | 0 | 0 | 0 | 0 | 0 | 0 | 0 | 0 | 0 | 0 | 0 | 0 | 0 | 0 | 0 | 1 | 0 | 0 | 0 | 0 | 0 | 0 | 0 | 0 | 0 | 0 | 0 | 0 | 0 | 0 | 0 | 0 |
| virus_Aug5.0v_2941 | 1 | 0 | 0 | 0 | 0 | 0 | 0 | 0 | 0 | 0 | 0 | 0 | 0 | 0 | 0 | 0 | 0 | 0 | 0 | 0 | 0 | 0 | 0 | 0 | 0 | 0 | 0 | 0 | 0 | 0 | 0 | 0 | 0 | 0 | 0 | 0 | 0 | 0 | 1 | 0 | 0 | 0 | 0 | 0 | 0 | 0 | 0 | 0 | 0 | 0 | 0 | 0 | 0 | 0 | 0 |
| virus_Aug5.0v_3054 | 1 | 0 | 0 | 0 | 0 | 0 | 0 | 0 | 0 | 0 | 0 | 0 | 0 | 0 | 0 | 0 | 0 | 0 | 0 | 0 | 0 | 0 | 0 | 0 | 0 | 0 | 0 | 0 | 0 | 0 | 0 | 0 | 0 | 0 | 0 | 0 | 0 | 0 | 1 | 0 | 0 | 0 | 0 | 0 | 0 | 0 | 0 | 0 | 0 | 0 | 0 | 0 | 0 | 0 | 0 |
| virus_Aug5.0v_3215 | 1 | 0 | 0 | 0 | 0 | 0 | 0 | 0 | 0 | 0 | 0 | 0 | 0 | 0 | 0 | 0 | 0 | 0 | 0 | 0 | 0 | 0 | 0 | 0 | 0 | 0 | 0 | 0 | 0 | 0 | 0 | 0 | 0 | 0 | 0 | 0 | 0 | 0 | 1 | 0 | 0 | 0 | 0 | 0 | 0 | 0 | 0 | 0 | 0 | 0 | 0 | 0 | 0 | 0 | 0 |
| virus_Aug5.0v_3596 | 1 | 0 | 0 | 0 | 0 | 0 | 0 | 0 | 0 | 0 | 0 | 0 | 0 | 0 | 0 | 0 | 0 | 0 | 0 | 0 | 0 | 0 | 0 | 0 | 0 | 0 | 0 | 0 | 0 | 0 | 0 | 0 | 0 | 0 | 0 | 0 | 0 | 0 | 1 | 0 | 0 | 0 | 0 | 0 | 0 | 0 | 0 | 0 | 0 | 0 | 0 | 0 | 0 | 0 | 0 |
| virus_Aug5.0v_3749 | 1 | 0 | 0 | 0 | 0 | 0 | 0 | 0 | 0 | 0 | 0 | 0 | 0 | 0 | 0 | 0 | 0 | 0 | 0 | 0 | 0 | 0 | 0 | 0 | 0 | 0 | 0 | 0 | 0 | 0 | 0 | 0 | 0 | 0 | 0 | 0 | 0 | 0 | 1 | 0 | 0 | 0 | 0 | 0 | 0 | 0 | 0 | 0 | 0 | 0 | 0 | 0 | 0 | 0 | 0 |
| virus_Aug5.0v_4286 | 1 | 0 | 0 | 0 | 0 | 0 | 0 | 0 | 0 | 0 | 0 | 0 | 0 | 0 | 0 | 0 | 0 | 0 | 0 | 0 | 0 | 0 | 0 | 0 | 0 | 0 | 0 | 0 | 0 | 0 | 0 | 0 | 0 | 0 | 0 | 0 | 0 | 0 | 1 | 0 | 0 | 0 | 0 | 0 | 0 | 0 | 0 | 0 | 0 | 0 | 0 | 0 | 0 | 0 | 0 |
| virus_Aug5.0v_4436 | 1 | 0 | 0 | 0 | 0 | 0 | 0 | 0 | 0 | 0 | 0 | 0 | 0 | 0 | 0 | 0 | 0 | 0 | 0 | 0 | 0 | 0 | 0 | 0 | 0 | 0 | 0 | 0 | 0 | 0 | 0 | 0 | 0 | 0 | 0 | 0 | 0 | 0 | 1 | 0 | 0 | 0 | 0 | 0 | 0 | 0 | 0 | 0 | 0 | 0 | 0 | 0 | 0 | 0 | 0 |
| virus_Aug5.0v_5325 | 1 | 0 | 0 | 0 | 0 | 0 | 0 | 0 | 0 | 0 | 0 | 0 | 0 | 0 | 0 | 0 | 0 | 0 | 0 | 0 | 0 | 0 | 0 | 0 | 0 | 0 | 0 | 0 | 0 | 0 | 0 | 0 | 0 | 0 | 0 | 0 | 0 | 0 | 1 | 0 | 0 | 0 | 0 | 0 | 0 | 0 | 0 | 0 | 0 | 0 | 0 | 0 | 0 | 0 | 0 |
| virus_Aug5.0v_5390 | 1 | 0 | 0 | 0 | 0 | 0 | 0 | 0 | 0 | 0 | 0 | 0 | 0 | 0 | 0 | 0 | 0 | 0 | 0 | 0 | 0 | 0 | 0 | 0 | 0 | 0 | 0 | 0 | 0 | 0 | 0 | 0 | 0 | 0 | 0 | 0 | 0 | 0 | 1 | 0 | 0 | 0 | 0 | 0 | 0 | 0 | 0 | 0 | 0 | 0 | 0 | 0 | 0 | 0 | 0 |
| virus_Aug5.5v_1196 | 1 | 0 | 0 | 0 | 0 | 0 | 0 | 0 | 0 | 0 | 0 | 0 | 0 | 0 | 0 | 0 | 0 | 0 | 0 | 0 | 0 | 0 | 0 | 0 | 0 | 0 | 0 | 0 | 0 | 0 | 0 | 0 | 0 | 0 | 0 | 0 | 0 | 0 | 0 | 0 | 1 | 0 | 0 | 0 | 0 | 0 | 0 | 0 | 0 | 0 | 0 | 0 | 0 | 0 | 0 |
| virus_Aug5.5v_14907 | 1 | 0 | 0 | 0 | 0 | 0 | 0 | 0 | 0 | 0 | 0 | 0 | 0 | 0 | 0 | 0 | 0 | 0 | 0 | 0 | 0 | 0 | 0 | 0 | 0 | 0 | 0 | 0 | 0 | 0 | 0 | 0 | 0 | 0 | 0 | 0 | 0 | 0 | 0 | 0 | 1 | 0 | 0 | 0 | 0 | 0 | 0 | 0 | 0 | 0 | 0 | 0 | 0 | 0 | 0 |
| virus_Aug5.5v_16686 | 1 | 0 | 0 | 0 | 0 | 0 | 0 | 0 | 0 | 0 | 0 | 0 | 0 | 0 | 0 | 0 | 0 | 0 | 0 | 0 | 0 | 0 | 0 | 0 | 0 | 0 | 0 | 0 | 0 | 0 | 0 | 0 | 0 | 0 | 0 | 0 | 0 | 0 | 0 | 0 | 1 | 0 | 0 | 0 | 0 | 0 | 0 | 0 | 0 | 0 | 0 | 0 | 0 | 0 | 0 |
| virus_Aug5.5v_17871 | 1 | 0 | 0 | 0 | 0 | 0 | 0 | 0 | 0 | 0 | 0 | 0 | 0 | 0 | 0 | 0 | 0 | 0 | 0 | 0 | 0 | 0 | 0 | 0 | 0 | 0 | 0 | 0 | 0 | 0 | 0 | 0 | 0 | 0 | 0 | 0 | 0 | 0 | 0 | 0 | 1 | 0 | 0 | 0 | 0 | 0 | 0 | 0 | 0 | 0 | 0 | 0 | 0 | 0 | 0 |
| virus_Aug5.5v_20459 | 1 | 0 | 0 | 0 | 0 | 0 | 0 | 0 | 0 | 0 | 0 | 0 | 0 | 0 | 0 | 0 | 0 | 0 | 0 | 0 | 0 | 0 | 0 | 0 | 0 | 0 | 0 | 0 | 0 | 0 | 0 | 0 | 0 | 0 | 0 | 0 | 0 | 0 | 0 | 0 | 1 | 0 | 0 | 0 | 0 | 0 | 0 | 0 | 0 | 0 | 0 | 0 | 0 | 0 | 0 |
| virus_Aug5.5v_27451 | 1 | 0 | 0 | 0 | 0 | 0 | 0 | 0 | 0 | 0 | 0 | 0 | 0 | 0 | 0 | 0 | 0 | 0 | 0 | 0 | 0 | 0 | 0 | 0 | 0 | 0 | 0 | 0 | 0 | 0 | 0 | 0 | 0 | 0 | 0 | 0 | 0 | 0 | 0 | 0 | 1 | 0 | 0 | 0 | 0 | 0 | 0 | 0 | 0 | 0 | 0 | 0 | 0 | 0 | 0 |
| virus_Aug5.5v_34422 | 1 | 0 | 0 | 0 | 0 | 0 | 0 | 0 | 0 | 0 | 0 | 0 | 0 | 0 | 0 | 0 | 0 | 0 | 0 | 0 | 0 | 0 | 0 | 0 | 0 | 0 | 0 | 0 | 0 | 0 | 0 | 0 | 0 | 0 | 0 | 0 | 0 | 0 | 0 | 0 | 1 | 0 | 0 | 0 | 0 | 0 | 0 | 0 | 0 | 0 | 0 | 0 | 0 | 0 | 0 |
| virus_Aug5.5v_36555 | 1 | 0 | 0 | 0 | 0 | 0 | 0 | 0 | 0 | 0 | 0 | 0 | 0 | 0 | 0 | 0 | 0 | 0 | 0 | 0 | 0 | 0 | 0 | 0 | 0 | 0 | 0 | 0 | 0 | 0 | 0 | 0 | 0 | 0 | 0 | 0 | 0 | 0 | 0 | 0 | 1 | 0 | 0 | 0 | 0 | 0 | 0 | 0 | 0 | 0 | 0 | 0 | 0 | 0 | 0 |
| virus_Aug5.5v_37292 | 1 | 0 | 0 | 0 | 0 | 0 | 0 | 0 | 0 | 0 | 0 | 0 | 0 | 0 | 0 | 0 | 0 | 0 | 0 | 0 | 0 | 0 | 0 | 0 | 0 | 0 | 0 | 0 | 0 | 0 | 0 | 0 | 0 | 0 | 0 | 0 | 0 | 0 | 0 | 0 | 1 | 0 | 0 | 0 | 0 | 0 | 0 | 0 | 0 | 0 | 0 | 0 | 0 | 0 | 0 |
| virus_Aug5.5v_37490 | 1 | 0 | 0 | 0 | 0 | 0 | 0 | 0 | 0 | 0 | 0 | 0 | 0 | 0 | 0 | 0 | 0 | 0 | 0 | 0 | 0 | 0 | 0 | 0 | 0 | 0 | 0 | 0 | 0 | 0 | 0 | 0 | 0 | 0 | 0 | 0 | 0 | 0 | 0 | 0 | 1 | 0 | 0 | 0 | 0 | 0 | 0 | 0 | 0 | 0 | 0 | 0 | 0 | 0 | 0 |
| virus_Aug5.5v_38688 | 1 | 0 | 0 | 0 | 0 | 0 | 0 | 0 | 0 | 0 | 0 | 0 | 0 | 0 | 0 | 0 | 0 | 0 | 0 | 0 | 0 | 0 | 0 | 0 | 0 | 0 | 0 | 0 | 0 | 0 | 0 | 0 | 0 | 0 | 0 | 0 | 0 | 0 | 0 | 0 | 1 | 0 | 0 | 0 | 0 | 0 | 0 | 0 | 0 | 0 | 0 | 0 | 0 | 0 | 0 |
| virus_Aug5.25v_7723 | 1 | 0 | 0 | 0 | 0 | 0 | 0 | 0 | 0 | 0 | 0 | 0 | 0 | 0 | 0 | 0 | 0 | 0 | 0 | 0 | 0 | 0 | 0 | 0 | 0 | 0 | 0 | 0 | 0 | 0 | 0 | 0 | 0 | 0 | 0 | 0 | 0 | 0 | 0 | 0 | 0 | 0 | 1 | 0 | 0 | 0 | 0 | 0 | 0 | 0 | 0 | 0 | 0 | 0 | 0 |
| virus_Aug5.25v_10034 | 1 | 0 | 0 | 0 | 0 | 0 | 0 | 0 | 0 | 0 | 0 | 0 | 0 | 0 | 0 | 0 | 0 | 0 | 0 | 0 | 0 | 0 | 0 | 0 | 0 | 0 | 0 | 0 | 0 | 0 | 0 | 0 | 0 | 0 | 0 | 0 | 0 | 0 | 0 | 0 | 0 | 0 | 1 | 0 | 0 | 0 | 0 | 0 | 0 | 0 | 0 | 0 | 0 | 0 | 0 |
| virus_Aug5.25v_11212 | 1 | 0 | 0 | 0 | 0 | 0 | 0 | 0 | 0 | 0 | 0 | 0 | 0 | 0 | 0 | 0 | 0 | 0 | 0 | 0 | 0 | 0 | 0 | 0 | 0 | 0 | 0 | 0 | 0 | 0 | 0 | 0 | 0 | 0 | 0 | 0 | 0 | 0 | 0 | 0 | 0 | 0 | 1 | 0 | 0 | 0 | 0 | 0 | 0 | 0 | 0 | 0 | 0 | 0 | 0 |
| virus_Aug5.25v_14747 | 1 | 0 | 0 | 0 | 0 | 0 | 0 | 0 | 0 | 0 | 0 | 0 | 0 | 0 | 0 | 0 | 0 | 0 | 0 | 0 | 0 | 0 | 0 | 0 | 0 | 0 | 0 | 0 | 0 | 0 | 0 | 0 | 0 | 0 | 0 | 0 | 0 | 0 | 0 | 0 | 0 | 0 | 1 | 0 | 0 | 0 | 0 | 0 | 0 | 0 | 0 | 0 | 0 | 0 | 0 |
| virus_Aug5.25v_16298 | 1 | 0 | 0 | 0 | 0 | 0 | 0 | 0 | 0 | 0 | 0 | 0 | 0 | 0 | 0 | 0 | 0 | 0 | 0 | 0 | 0 | 0 | 0 | 0 | 0 | 0 | 0 | 0 | 0 | 0 | 0 | 0 | 0 | 0 | 0 | 0 | 0 | 0 | 0 | 0 | 0 | 0 | 1 | 0 | 0 | 0 | 0 | 0 | 0 | 0 | 0 | 0 | 0 | 0 | 0 |
| virus_Aug5.25v_17243 | 1 | 0 | 0 | 0 | 0 | 0 | 0 | 0 | 0 | 0 | 0 | 0 | 0 | 0 | 0 | 0 | 0 | 0 | 0 | 0 | 0 | 0 | 0 | 0 | 0 | 0 | 0 | 0 | 0 | 0 | 0 | 0 | 0 | 0 | 0 | 0 | 0 | 0 | 0 | 0 | 0 | 0 | 1 | 0 | 0 | 0 | 0 | 0 | 0 | 0 | 0 | 0 | 0 | 0 | 0 |
| virus_Aug5.25v_17930 | 1 | 0 | 0 | 0 | 0 | 0 | 0 | 0 | 0 | 0 | 0 | 0 | 0 | 0 | 0 | 0 | 0 | 0 | 0 | 0 | 0 | 0 | 0 | 0 | 0 | 0 | 0 | 0 | 0 | 0 | 0 | 0 | 0 | 0 | 0 | 0 | 0 | 0 | 0 | 0 | 0 | 0 | 1 | 0 | 0 | 0 | 0 | 0 | 0 | 0 | 0 | 0 | 0 | 0 | 0 |
| virus_Aug5.25v_19585 | 1 | 0 | 0 | 0 | 0 | 0 | 0 | 0 | 0 | 0 | 0 | 0 | 0 | 0 | 0 | 0 | 0 | 0 | 0 | 0 | 0 | 0 | 0 | 0 | 0 | 0 | 0 | 0 | 0 | 0 | 0 | 0 | 0 | 0 | 0 | 0 | 0 | 0 | 0 | 0 | 0 | 0 | 1 | 0 | 0 | 0 | 0 | 0 | 0 | 0 | 0 | 0 | 0 | 0 | 0 |
| virus_Aug5.25v_21808 | 1 | 0 | 0 | 0 | 0 | 0 | 0 | 0 | 0 | 0 | 0 | 0 | 0 | 0 | 0 | 0 | 0 | 0 | 0 | 0 | 0 | 0 | 0 | 0 | 0 | 0 | 0 | 0 | 0 | 0 | 0 | 0 | 0 | 0 | 0 | 0 | 0 | 0 | 0 | 0 | 0 | 0 | 1 | 0 | 0 | 0 | 0 | 0 | 0 | 0 | 0 | 0 | 0 | 0 | 0 |
| virus_Aug5.25v_21957 | 1 | 0 | 0 | 0 | 0 | 0 | 0 | 0 | 0 | 0 | 0 | 0 | 0 | 0 | 0 | 0 | 0 | 0 | 0 | 0 | 0 | 0 | 0 | 0 | 0 | 0 | 0 | 0 | 0 | 0 | 0 | 0 | 0 | 0 | 0 | 0 | 0 | 0 | 0 | 0 | 0 | 0 | 1 | 0 | 0 | 0 | 0 | 0 | 0 | 0 | 0 | 0 | 0 | 0 | 0 |
| virus_Aug5.25v_22285 | 1 | 0 | 0 | 0 | 0 | 0 | 0 | 0 | 0 | 0 | 0 | 0 | 0 | 0 | 0 | 0 | 0 | 0 | 0 | 0 | 0 | 0 | 0 | 0 | 0 | 0 | 0 | 0 | 0 | 0 | 0 | 0 | 0 | 0 | 0 | 0 | 0 | 0 | 0 | 0 | 0 | 0 | 1 | 0 | 0 | 0 | 0 | 0 | 0 | 0 | 0 | 0 | 0 | 0 | 0 |
| virus_Aug5.25v_23714 | 1 | 0 | 0 | 0 | 0 | 0 | 0 | 0 | 0 | 0 | 0 | 0 | 0 | 0 | 0 | 0 | 0 | 0 | 0 | 0 | 0 | 0 | 0 | 0 | 0 | 0 | 0 | 0 | 0 | 0 | 0 | 0 | 0 | 0 | 0 | 0 | 0 | 0 | 0 | 0 | 0 | 0 | 1 | 0 | 0 | 0 | 0 | 0 | 0 | 0 | 0 | 0 | 0 | 0 | 0 |
| virus_Aug5.25v_26919 | 1 | 0 | 0 | 0 | 0 | 0 | 0 | 0 | 0 | 0 | 0 | 0 | 0 | 0 | 0 | 0 | 0 | 0 | 0 | 0 | 0 | 0 | 0 | 0 | 0 | 0 | 0 | 0 | 0 | 0 | 0 | 0 | 0 | 0 | 0 | 0 | 0 | 0 | 0 | 0 | 0 | 0 | 1 | 0 | 0 | 0 | 0 | 0 | 0 | 0 | 0 | 0 | 0 | 0 | 0 |
| virus_Aug5.25v_30444 | 1 | 0 | 0 | 0 | 0 | 0 | 0 | 0 | 0 | 0 | 0 | 0 | 0 | 0 | 0 | 0 | 0 | 0 | 0 | 0 | 0 | 0 | 0 | 0 | 0 | 0 | 0 | 0 | 0 | 0 | 0 | 0 | 0 | 0 | 0 | 0 | 0 | 0 | 0 | 0 | 0 | 0 | 1 | 0 | 0 | 0 | 0 | 0 | 0 | 0 | 0 | 0 | 0 | 0 | 0 |
| virus_Aug5.25v_31255 | 1 | 0 | 0 | 0 | 0 | 0 | 0 | 0 | 0 | 0 | 0 | 0 | 0 | 0 | 0 | 0 | 0 | 0 | 0 | 0 | 0 | 0 | 0 | 0 | 0 | 0 | 0 | 0 | 0 | 0 | 0 | 0 | 0 | 0 | 0 | 0 | 0 | 0 | 0 | 0 | 0 | 0 | 1 | 0 | 0 | 0 | 0 | 0 | 0 | 0 | 0 | 0 | 0 | 0 | 0 |
| virus_Aug5.25v_32541 | 1 | 0 | 0 | 0 | 0 | 0 | 0 | 0 | 0 | 0 | 0 | 0 | 0 | 0 | 0 | 0 | 0 | 0 | 0 | 0 | 0 | 0 | 0 | 0 | 0 | 0 | 0 | 0 | 0 | 0 | 0 | 0 | 0 | 0 | 0 | 0 | 0 | 0 | 0 | 0 | 0 | 0 | 1 | 0 | 0 | 0 | 0 | 0 | 0 | 0 | 0 | 0 | 0 | 0 | 0 |
| virus_Aug5.25v_36318 | 1 | 0 | 0 | 0 | 0 | 0 | 0 | 0 | 0 | 0 | 0 | 0 | 0 | 0 | 0 | 0 | 0 | 0 | 0 | 0 | 0 | 0 | 0 | 0 | 0 | 0 | 0 | 0 | 0 | 0 | 0 | 0 | 0 | 0 | 0 | 0 | 0 | 0 | 0 | 0 | 0 | 0 | 1 | 0 | 0 | 0 | 0 | 0 | 0 | 0 | 0 | 0 | 0 | 0 | 0 |
| virus_Aug5.25v_36353 | 1 | 0 | 0 | 0 | 0 | 0 | 0 | 0 | 0 | 0 | 0 | 0 | 0 | 0 | 0 | 0 | 0 | 0 | 0 | 0 | 0 | 0 | 0 | 0 | 0 | 0 | 0 | 0 | 0 | 0 | 0 | 0 | 0 | 0 | 0 | 0 | 0 | 0 | 0 | 0 | 0 | 0 | 1 | 0 | 0 | 0 | 0 | 0 | 0 | 0 | 0 | 0 | 0 | 0 | 0 |
| virus_Aug5.25v_38368 | 1 | 0 | 0 | 0 | 0 | 0 | 0 | 0 | 0 | 0 | 0 | 0 | 0 | 0 | 0 | 0 | 0 | 0 | 0 | 0 | 0 | 0 | 0 | 0 | 0 | 0 | 0 | 0 | 0 | 0 | 0 | 0 | 0 | 0 | 0 | 0 | 0 | 0 | 0 | 0 | 0 | 0 | 1 | 0 | 0 | 0 | 0 | 0 | 0 | 0 | 0 | 0 | 0 | 0 | 0 |
| virus_Aug5.25v_38987 | 1 | 0 | 0 | 0 | 0 | 0 | 0 | 0 | 0 | 0 | 0 | 0 | 0 | 0 | 0 | 0 | 0 | 0 | 0 | 0 | 0 | 0 | 0 | 0 | 0 | 0 | 0 | 0 | 0 | 0 | 0 | 0 | 0 | 0 | 0 | 0 | 0 | 0 | 0 | 0 | 0 | 0 | 1 | 0 | 0 | 0 | 0 | 0 | 0 | 0 | 0 | 0 | 0 | 0 | 0 |
| virus_Aug5.25v_40327 | 1 | 0 | 0 | 0 | 0 | 0 | 0 | 0 | 0 | 0 | 0 | 0 | 0 | 0 | 0 | 0 | 0 | 0 | 0 | 0 | 0 | 0 | 0 | 0 | 0 | 0 | 0 | 0 | 0 | 0 | 0 | 0 | 0 | 0 | 0 | 0 | 0 | 0 | 0 | 0 | 0 | 0 | 1 | 0 | 0 | 0 | 0 | 0 | 0 | 0 | 0 | 0 | 0 | 0 | 0 |
| virus_Aug5.25v_40792 | 1 | 0 | 0 | 0 | 0 | 0 | 0 | 0 | 0 | 0 | 0 | 0 | 0 | 0 | 0 | 0 | 0 | 0 | 0 | 0 | 0 | 0 | 0 | 0 | 0 | 0 | 0 | 0 | 0 | 0 | 0 | 0 | 0 | 0 | 0 | 0 | 0 | 0 | 0 | 0 | 0 | 0 | 1 | 0 | 0 | 0 | 0 | 0 | 0 | 0 | 0 | 0 | 0 | 0 | 0 |
| virus_Aug5.25v_41255 | 1 | 0 | 0 | 0 | 0 | 0 | 0 | 0 | 0 | 0 | 0 | 0 | 0 | 0 | 0 | 0 | 0 | 0 | 0 | 0 | 0 | 0 | 0 | 0 | 0 | 0 | 0 | 0 | 0 | 0 | 0 | 0 | 0 | 0 | 0 | 0 | 0 | 0 | 0 | 0 | 0 | 0 | 1 | 0 | 0 | 0 | 0 | 0 | 0 | 0 | 0 | 0 | 0 | 0 | 0 |
| virus_Aug5.50v_5862 | 1 | 0 | 0 | 0 | 0 | 0 | 0 | 0 | 0 | 0 | 0 | 0 | 0 | 0 | 0 | 0 | 0 | 0 | 0 | 0 | 0 | 0 | 0 | 0 | 0 | 0 | 0 | 0 | 0 | 0 | 0 | 0 | 0 | 0 | 0 | 0 | 0 | 0 | 0 | 0 | 0 | 0 | 0 | 0 | 1 | 0 | 0 | 0 | 0 | 0 | 0 | 0 | 0 | 0 | 0 |
| virus_Aug5.50v_25014 | 1 | 0 | 0 | 0 | 0 | 0 | 0 | 0 | 0 | 0 | 0 | 0 | 0 | 0 | 0 | 0 | 0 | 0 | 0 | 0 | 0 | 0 | 0 | 0 | 0 | 0 | 0 | 0 | 0 | 0 | 0 | 0 | 0 | 0 | 0 | 0 | 0 | 0 | 0 | 0 | 0 | 0 | 0 | 0 | 1 | 0 | 0 | 0 | 0 | 0 | 0 | 0 | 0 | 0 | 0 |
| virus_Aug5.50v_27756 | 1 | 0 | 0 | 0 | 0 | 0 | 0 | 0 | 0 | 0 | 0 | 0 | 0 | 0 | 0 | 0 | 0 | 0 | 0 | 0 | 0 | 0 | 0 | 0 | 0 | 0 | 0 | 0 | 0 | 0 | 0 | 0 | 0 | 0 | 0 | 0 | 0 | 0 | 0 | 0 | 0 | 0 | 0 | 0 | 1 | 0 | 0 | 0 | 0 | 0 | 0 | 0 | 0 | 0 | 0 |
| virus_Aug5.50v_40106 | 1 | 0 | 0 | 0 | 0 | 0 | 0 | 0 | 0 | 0 | 0 | 0 | 0 | 0 | 0 | 0 | 0 | 0 | 0 | 0 | 0 | 0 | 0 | 0 | 0 | 0 | 0 | 0 | 0 | 0 | 0 | 0 | 0 | 0 | 0 | 0 | 0 | 0 | 0 | 0 | 0 | 0 | 0 | 0 | 1 | 0 | 0 | 0 | 0 | 0 | 0 | 0 | 0 | 0 | 0 |
| virus_Aug5.75v_3968 | 1 | 0 | 0 | 0 | 0 | 0 | 0 | 0 | 0 | 0 | 0 | 0 | 0 | 0 | 0 | 0 | 0 | 0 | 0 | 0 | 0 | 0 | 0 | 0 | 0 | 0 | 0 | 0 | 0 | 0 | 0 | 0 | 0 | 0 | 0 | 0 | 0 | 0 | 0 | 0 | 0 | 0 | 0 | 0 | 0 | 0 | 1 | 0 | 0 | 0 | 0 | 0 | 0 | 0 | 0 |
| virus_Aug5.75v_5047 | 1 | 0 | 0 | 0 | 0 | 0 | 0 | 0 | 0 | 0 | 0 | 0 | 0 | 0 | 0 | 0 | 0 | 0 | 0 | 0 | 0 | 0 | 0 | 0 | 0 | 0 | 0 | 0 | 0 | 0 | 0 | 0 | 0 | 0 | 0 | 0 | 0 | 0 | 0 | 0 | 0 | 0 | 0 | 0 | 0 | 0 | 1 | 0 | 0 | 0 | 0 | 0 | 0 | 0 | 0 |
| virus_Jul11.0v_1613 | 1 | 0 | 0 | 0 | 0 | 0 | 0 | 0 | 0 | 0 | 0 | 0 | 0 | 0 | 0 | 0 | 0 | 0 | 0 | 0 | 0 | 0 | 1 | 0 | 0 | 0 | 0 | 0 | 0 | 0 | 0 | 0 | 0 | 0 | 0 | 0 | 0 | 0 | 0 | 0 | 0 | 0 | 0 | 0 | 0 | 0 | 0 | 0 | 0 | 0 | 0 | 0 | 0 | 0 | 0 |
| virus_Jul11.0v_3874 | 1 | 0 | 0 | 0 | 0 | 0 | 0 | 0 | 0 | 0 | 0 | 0 | 0 | 0 | 0 | 0 | 0 | 0 | 0 | 0 | 0 | 0 | 1 | 0 | 0 | 0 | 0 | 0 | 0 | 0 | 0 | 0 | 0 | 0 | 0 | 0 | 0 | 0 | 0 | 0 | 0 | 0 | 0 | 0 | 0 | 0 | 0 | 0 | 0 | 0 | 0 | 0 | 0 | 0 | 0 |
| virus_Jul11.0v_7815 | 1 | 0 | 0 | 0 | 0 | 0 | 0 | 0 | 0 | 0 | 0 | 0 | 0 | 0 | 0 | 0 | 0 | 0 | 0 | 0 | 0 | 0 | 1 | 0 | 0 | 0 | 0 | 0 | 0 | 0 | 0 | 0 | 0 | 0 | 0 | 0 | 0 | 0 | 0 | 0 | 0 | 0 | 0 | 0 | 0 | 0 | 0 | 0 | 0 | 0 | 0 | 0 | 0 | 0 | 0 |
| virus_Jul11.0v_14307 | 1 | 0 | 0 | 0 | 0 | 0 | 0 | 0 | 0 | 0 | 0 | 0 | 0 | 0 | 0 | 0 | 0 | 0 | 0 | 0 | 0 | 0 | 1 | 0 | 0 | 0 | 0 | 0 | 0 | 0 | 0 | 0 | 0 | 0 | 0 | 0 | 0 | 0 | 0 | 0 | 0 | 0 | 0 | 0 | 0 | 0 | 0 | 0 | 0 | 0 | 0 | 0 | 0 | 0 | 0 |
| virus_Jul11.0v_14910 | 1 | 0 | 0 | 0 | 0 | 0 | 0 | 0 | 0 | 0 | 0 | 0 | 0 | 0 | 0 | 0 | 0 | 0 | 0 | 0 | 0 | 0 | 1 | 0 | 0 | 0 | 0 | 0 | 0 | 0 | 0 | 0 | 0 | 0 | 0 | 0 | 0 | 0 | 0 | 0 | 0 | 0 | 0 | 0 | 0 | 0 | 0 | 0 | 0 | 0 | 0 | 0 | 0 | 0 | 0 |
| virus_Jul11.0v_18936 | 1 | 0 | 0 | 0 | 0 | 0 | 0 | 0 | 0 | 0 | 0 | 0 | 0 | 0 | 0 | 0 | 0 | 0 | 0 | 0 | 0 | 0 | 1 | 0 | 0 | 0 | 0 | 0 | 0 | 0 | 0 | 0 | 0 | 0 | 0 | 0 | 0 | 0 | 0 | 0 | 0 | 0 | 0 | 0 | 0 | 0 | 0 | 0 | 0 | 0 | 0 | 0 | 0 | 0 | 0 |
| virus_Jul11.0v_20475 | 1 | 0 | 0 | 0 | 0 | 0 | 0 | 0 | 0 | 0 | 0 | 0 | 0 | 0 | 0 | 0 | 0 | 0 | 0 | 0 | 0 | 0 | 1 | 0 | 0 | 0 | 0 | 0 | 0 | 0 | 0 | 0 | 0 | 0 | 0 | 0 | 0 | 0 | 0 | 0 | 0 | 0 | 0 | 0 | 0 | 0 | 0 | 0 | 0 | 0 | 0 | 0 | 0 | 0 | 0 |
| virus_Jul11.5v_948 | 1 | 0 | 0 | 0 | 0 | 0 | 0 | 0 | 0 | 0 | 0 | 0 | 0 | 0 | 0 | 0 | 0 | 0 | 0 | 0 | 0 | 0 | 0 | 0 | 1 | 0 | 0 | 0 | 0 | 0 | 0 | 0 | 0 | 0 | 0 | 0 | 0 | 0 | 0 | 0 | 0 | 0 | 0 | 0 | 0 | 0 | 0 | 0 | 0 | 0 | 0 | 0 | 0 | 0 | 0 |
| virus_Jul11.5v_1952 | 1 | 0 | 0 | 0 | 0 | 0 | 0 | 0 | 0 | 0 | 0 | 0 | 0 | 0 | 0 | 0 | 0 | 0 | 0 | 0 | 0 | 0 | 0 | 0 | 1 | 0 | 0 | 0 | 0 | 0 | 0 | 0 | 0 | 0 | 0 | 0 | 0 | 0 | 0 | 0 | 0 | 0 | 0 | 0 | 0 | 0 | 0 | 0 | 0 | 0 | 0 | 0 | 0 | 0 | 0 |
| virus_Jul11.5v_3073 | 1 | 0 | 0 | 0 | 0 | 0 | 0 | 0 | 0 | 0 | 0 | 0 | 0 | 0 | 0 | 0 | 0 | 0 | 0 | 0 | 0 | 0 | 0 | 0 | 1 | 0 | 0 | 0 | 0 | 0 | 0 | 0 | 0 | 0 | 0 | 0 | 0 | 0 | 0 | 0 | 0 | 0 | 0 | 0 | 0 | 0 | 0 | 0 | 0 | 0 | 0 | 0 | 0 | 0 | 0 |
| virus_Jul11.5v_3138 | 1 | 0 | 0 | 0 | 0 | 0 | 0 | 0 | 0 | 0 | 0 | 0 | 0 | 0 | 0 | 0 | 0 | 0 | 0 | 0 | 0 | 0 | 0 | 0 | 1 | 0 | 0 | 0 | 0 | 0 | 0 | 0 | 0 | 0 | 0 | 0 | 0 | 0 | 0 | 0 | 0 | 0 | 0 | 0 | 0 | 0 | 0 | 0 | 0 | 0 | 0 | 0 | 0 | 0 | 0 |
| virus_Jul11.5v_4412 | 1 | 0 | 0 | 0 | 0 | 0 | 0 | 0 | 0 | 0 | 0 | 0 | 0 | 0 | 0 | 0 | 0 | 0 | 0 | 0 | 0 | 0 | 0 | 0 | 1 | 0 | 0 | 0 | 0 | 0 | 0 | 0 | 0 | 0 | 0 | 0 | 0 | 0 | 0 | 0 | 0 | 0 | 0 | 0 | 0 | 0 | 0 | 0 | 0 | 0 | 0 | 0 | 0 | 0 | 0 |
| virus_Jul11.5v_6076 | 1 | 0 | 0 | 0 | 0 | 0 | 0 | 0 | 0 | 0 | 0 | 0 | 0 | 0 | 0 | 0 | 0 | 0 | 0 | 0 | 0 | 0 | 0 | 0 | 1 | 0 | 0 | 0 | 0 | 0 | 0 | 0 | 0 | 0 | 0 | 0 | 0 | 0 | 0 | 0 | 0 | 0 | 0 | 0 | 0 | 0 | 0 | 0 | 0 | 0 | 0 | 0 | 0 | 0 | 0 |
| virus_Jul11.5v_6230 | 1 | 0 | 0 | 0 | 0 | 0 | 0 | 0 | 0 | 0 | 0 | 0 | 0 | 0 | 0 | 0 | 0 | 0 | 0 | 0 | 0 | 0 | 0 | 0 | 1 | 0 | 0 | 0 | 0 | 0 | 0 | 0 | 0 | 0 | 0 | 0 | 0 | 0 | 0 | 0 | 0 | 0 | 0 | 0 | 0 | 0 | 0 | 0 | 0 | 0 | 0 | 0 | 0 | 0 | 0 |
| virus_Jul11.5v_6464 | 1 | 0 | 0 | 0 | 0 | 0 | 0 | 0 | 0 | 0 | 0 | 0 | 0 | 0 | 0 | 0 | 0 | 0 | 0 | 0 | 0 | 0 | 0 | 0 | 1 | 0 | 0 | 0 | 0 | 0 | 0 | 0 | 0 | 0 | 0 | 0 | 0 | 0 | 0 | 0 | 0 | 0 | 0 | 0 | 0 | 0 | 0 | 0 | 0 | 0 | 0 | 0 | 0 | 0 | 0 |
| virus_Jul11.5v_6977 | 1 | 0 | 0 | 0 | 0 | 0 | 0 | 0 | 0 | 0 | 0 | 0 | 0 | 0 | 0 | 0 | 0 | 0 | 0 | 0 | 0 | 0 | 0 | 0 | 1 | 0 | 0 | 0 | 0 | 0 | 0 | 0 | 0 | 0 | 0 | 0 | 0 | 0 | 0 | 0 | 0 | 0 | 0 | 0 | 0 | 0 | 0 | 0 | 0 | 0 | 0 | 0 | 0 | 0 | 0 |
| virus_Jul11.5v_9419 | 1 | 0 | 0 | 0 | 0 | 0 | 0 | 0 | 0 | 0 | 0 | 0 | 0 | 0 | 0 | 0 | 0 | 0 | 0 | 0 | 0 | 0 | 0 | 0 | 1 | 0 | 0 | 0 | 0 | 0 | 0 | 0 | 0 | 0 | 0 | 0 | 0 | 0 | 0 | 0 | 0 | 0 | 0 | 0 | 0 | 0 | 0 | 0 | 0 | 0 | 0 | 0 | 0 | 0 | 0 |
| virus_Jul11.5v_9901 | 1 | 0 | 0 | 0 | 0 | 0 | 0 | 0 | 0 | 0 | 0 | 0 | 0 | 0 | 0 | 0 | 0 | 0 | 0 | 0 | 0 | 0 | 0 | 0 | 1 | 0 | 0 | 0 | 0 | 0 | 0 | 0 | 0 | 0 | 0 | 0 | 0 | 0 | 0 | 0 | 0 | 0 | 0 | 0 | 0 | 0 | 0 | 0 | 0 | 0 | 0 | 0 | 0 | 0 | 0 |
| virus_Jul11.5v_13471 | 1 | 0 | 0 | 0 | 0 | 0 | 0 | 0 | 0 | 0 | 0 | 0 | 0 | 0 | 0 | 0 | 0 | 0 | 0 | 0 | 0 | 0 | 0 | 0 | 1 | 0 | 0 | 0 | 0 | 0 | 0 | 0 | 0 | 0 | 0 | 0 | 0 | 0 | 0 | 0 | 0 | 0 | 0 | 0 | 0 | 0 | 0 | 0 | 0 | 0 | 0 | 0 | 0 | 0 | 0 |
| virus_Jul11.5v_15182 | 1 | 0 | 0 | 0 | 0 | 0 | 0 | 0 | 0 | 0 | 0 | 0 | 0 | 0 | 0 | 0 | 0 | 0 | 0 | 0 | 0 | 0 | 0 | 0 | 1 | 0 | 0 | 0 | 0 | 0 | 0 | 0 | 0 | 0 | 0 | 0 | 0 | 0 | 0 | 0 | 0 | 0 | 0 | 0 | 0 | 0 | 0 | 0 | 0 | 0 | 0 | 0 | 0 | 0 | 0 |
| virus_Jul11.5v_17185 | 1 | 0 | 0 | 0 | 0 | 0 | 0 | 0 | 0 | 0 | 0 | 0 | 0 | 0 | 0 | 0 | 0 | 0 | 0 | 0 | 0 | 0 | 0 | 0 | 1 | 0 | 0 | 0 | 0 | 0 | 0 | 0 | 0 | 0 | 0 | 0 | 0 | 0 | 0 | 0 | 0 | 0 | 0 | 0 | 0 | 0 | 0 | 0 | 0 | 0 | 0 | 0 | 0 | 0 | 0 |
| virus_Jul11.5v_18128 | 1 | 0 | 0 | 0 | 0 | 0 | 0 | 0 | 0 | 0 | 0 | 0 | 0 | 0 | 0 | 0 | 0 | 0 | 0 | 0 | 0 | 0 | 0 | 0 | 1 | 0 | 0 | 0 | 0 | 0 | 0 | 0 | 0 | 0 | 0 | 0 | 0 | 0 | 0 | 0 | 0 | 0 | 0 | 0 | 0 | 0 | 0 | 0 | 0 | 0 | 0 | 0 | 0 | 0 | 0 |
| virus_Jul11.5v_18322 | 1 | 0 | 0 | 0 | 0 | 0 | 0 | 0 | 0 | 0 | 0 | 0 | 0 | 0 | 0 | 0 | 0 | 0 | 0 | 0 | 0 | 0 | 0 | 0 | 1 | 0 | 0 | 0 | 0 | 0 | 0 | 0 | 0 | 0 | 0 | 0 | 0 | 0 | 0 | 0 | 0 | 0 | 0 | 0 | 0 | 0 | 0 | 0 | 0 | 0 | 0 | 0 | 0 | 0 | 0 |
| virus_Jul11.25v_1151 | 1 | 0 | 0 | 0 | 0 | 0 | 0 | 0 | 0 | 0 | 0 | 0 | 0 | 0 | 0 | 0 | 0 | 0 | 0 | 0 | 0 | 0 | 0 | 0 | 0 | 0 | 1 | 0 | 0 | 0 | 0 | 0 | 0 | 0 | 0 | 0 | 0 | 0 | 0 | 0 | 0 | 0 | 0 | 0 | 0 | 0 | 0 | 0 | 0 | 0 | 0 | 0 | 0 | 0 | 0 |
| virus_Jul11.25v_1165 | 1 | 0 | 0 | 0 | 0 | 0 | 0 | 0 | 0 | 0 | 0 | 0 | 0 | 0 | 0 | 0 | 0 | 0 | 0 | 0 | 0 | 0 | 0 | 0 | 0 | 0 | 1 | 0 | 0 | 0 | 0 | 0 | 0 | 0 | 0 | 0 | 0 | 0 | 0 | 0 | 0 | 0 | 0 | 0 | 0 | 0 | 0 | 0 | 0 | 0 | 0 | 0 | 0 | 0 | 0 |
| virus_Jul11.25v_1628 | 1 | 0 | 0 | 0 | 0 | 0 | 0 | 0 | 0 | 0 | 0 | 0 | 0 | 0 | 0 | 0 | 0 | 0 | 0 | 0 | 0 | 0 | 0 | 0 | 0 | 0 | 1 | 0 | 0 | 0 | 0 | 0 | 0 | 0 | 0 | 0 | 0 | 0 | 0 | 0 | 0 | 0 | 0 | 0 | 0 | 0 | 0 | 0 | 0 | 0 | 0 | 0 | 0 | 0 | 0 |
| virus_Jul11.25v_3496 | 1 | 0 | 0 | 0 | 0 | 0 | 0 | 0 | 0 | 0 | 0 | 0 | 0 | 0 | 0 | 0 | 0 | 0 | 0 | 0 | 0 | 0 | 0 | 0 | 0 | 0 | 1 | 0 | 0 | 0 | 0 | 0 | 0 | 0 | 0 | 0 | 0 | 0 | 0 | 0 | 0 | 0 | 0 | 0 | 0 | 0 | 0 | 0 | 0 | 0 | 0 | 0 | 0 | 0 | 0 |
| virus_Jul11.25v_3823 | 1 | 0 | 0 | 0 | 0 | 0 | 0 | 0 | 0 | 0 | 0 | 0 | 0 | 0 | 0 | 0 | 0 | 0 | 0 | 0 | 0 | 0 | 0 | 0 | 0 | 0 | 1 | 0 | 0 | 0 | 0 | 0 | 0 | 0 | 0 | 0 | 0 | 0 | 0 | 0 | 0 | 0 | 0 | 0 | 0 | 0 | 0 | 0 | 0 | 0 | 0 | 0 | 0 | 0 | 0 |
| virus_Jul11.25v_4548 | 1 | 0 | 0 | 0 | 0 | 0 | 0 | 0 | 0 | 0 | 0 | 0 | 0 | 0 | 0 | 0 | 0 | 0 | 0 | 0 | 0 | 0 | 0 | 0 | 0 | 0 | 1 | 0 | 0 | 0 | 0 | 0 | 0 | 0 | 0 | 0 | 0 | 0 | 0 | 0 | 0 | 0 | 0 | 0 | 0 | 0 | 0 | 0 | 0 | 0 | 0 | 0 | 0 | 0 | 0 |
| virus_Jul11.25v_5123 | 1 | 0 | 0 | 0 | 0 | 0 | 0 | 0 | 0 | 0 | 0 | 0 | 0 | 0 | 0 | 0 | 0 | 0 | 0 | 0 | 0 | 0 | 0 | 0 | 0 | 0 | 1 | 0 | 0 | 0 | 0 | 0 | 0 | 0 | 0 | 0 | 0 | 0 | 0 | 0 | 0 | 0 | 0 | 0 | 0 | 0 | 0 | 0 | 0 | 0 | 0 | 0 | 0 | 0 | 0 |
| virus_Jul11.25v_5315 | 1 | 0 | 0 | 0 | 0 | 0 | 0 | 0 | 0 | 0 | 0 | 0 | 0 | 0 | 0 | 0 | 0 | 0 | 0 | 0 | 0 | 0 | 0 | 0 | 0 | 0 | 1 | 0 | 0 | 0 | 0 | 0 | 0 | 0 | 0 | 0 | 0 | 0 | 0 | 0 | 0 | 0 | 0 | 0 | 0 | 0 | 0 | 0 | 0 | 0 | 0 | 0 | 0 | 0 | 0 |
| virus_Jul11.25v_11011 | 1 | 0 | 0 | 0 | 0 | 0 | 0 | 0 | 0 | 0 | 0 | 0 | 0 | 0 | 0 | 0 | 0 | 0 | 0 | 0 | 0 | 0 | 0 | 0 | 0 | 0 | 1 | 0 | 0 | 0 | 0 | 0 | 0 | 0 | 0 | 0 | 0 | 0 | 0 | 0 | 0 | 0 | 0 | 0 | 0 | 0 | 0 | 0 | 0 | 0 | 0 | 0 | 0 | 0 | 0 |
| virus_Jul11.25v_11317 | 1 | 0 | 0 | 0 | 0 | 0 | 0 | 0 | 0 | 0 | 0 | 0 | 0 | 0 | 0 | 0 | 0 | 0 | 0 | 0 | 0 | 0 | 0 | 0 | 0 | 0 | 1 | 0 | 0 | 0 | 0 | 0 | 0 | 0 | 0 | 0 | 0 | 0 | 0 | 0 | 0 | 0 | 0 | 0 | 0 | 0 | 0 | 0 | 0 | 0 | 0 | 0 | 0 | 0 | 0 |
| virus_Jul11.25v_11801 | 1 | 0 | 0 | 0 | 0 | 0 | 0 | 0 | 0 | 0 | 0 | 0 | 0 | 0 | 0 | 0 | 0 | 0 | 0 | 0 | 0 | 0 | 0 | 0 | 0 | 0 | 1 | 0 | 0 | 0 | 0 | 0 | 0 | 0 | 0 | 0 | 0 | 0 | 0 | 0 | 0 | 0 | 0 | 0 | 0 | 0 | 0 | 0 | 0 | 0 | 0 | 0 | 0 | 0 | 0 |
| virus_Jul11.25v_13785 | 1 | 0 | 0 | 0 | 0 | 0 | 0 | 0 | 0 | 0 | 0 | 0 | 0 | 0 | 0 | 0 | 0 | 0 | 0 | 0 | 0 | 0 | 0 | 0 | 0 | 0 | 1 | 0 | 0 | 0 | 0 | 0 | 0 | 0 | 0 | 0 | 0 | 0 | 0 | 0 | 0 | 0 | 0 | 0 | 0 | 0 | 0 | 0 | 0 | 0 | 0 | 0 | 0 | 0 | 0 |
| virus_Jul11.25v_16828 | 1 | 0 | 0 | 0 | 0 | 0 | 0 | 0 | 0 | 0 | 0 | 0 | 0 | 0 | 0 | 0 | 0 | 0 | 0 | 0 | 0 | 0 | 0 | 0 | 0 | 0 | 1 | 0 | 0 | 0 | 0 | 0 | 0 | 0 | 0 | 0 | 0 | 0 | 0 | 0 | 0 | 0 | 0 | 0 | 0 | 0 | 0 | 0 | 0 | 0 | 0 | 0 | 0 | 0 | 0 |
| virus_Jul11.25v_17692 | 1 | 0 | 0 | 0 | 0 | 0 | 0 | 0 | 0 | 0 | 0 | 0 | 0 | 0 | 0 | 0 | 0 | 0 | 0 | 0 | 0 | 0 | 0 | 0 | 0 | 0 | 1 | 0 | 0 | 0 | 0 | 0 | 0 | 0 | 0 | 0 | 0 | 0 | 0 | 0 | 0 | 0 | 0 | 0 | 0 | 0 | 0 | 0 | 0 | 0 | 0 | 0 | 0 | 0 | 0 |
| virus_Jul11.25v_20956 | 1 | 0 | 0 | 0 | 0 | 0 | 0 | 0 | 0 | 0 | 0 | 0 | 0 | 0 | 0 | 0 | 0 | 0 | 0 | 0 | 0 | 0 | 0 | 0 | 0 | 0 | 1 | 0 | 0 | 0 | 0 | 0 | 0 | 0 | 0 | 0 | 0 | 0 | 0 | 0 | 0 | 0 | 0 | 0 | 0 | 0 | 0 | 0 | 0 | 0 | 0 | 0 | 0 | 0 | 0 |
| virus_Jul11.50v_1658 | 1 | 0 | 0 | 0 | 0 | 0 | 0 | 0 | 0 | 0 | 0 | 0 | 0 | 0 | 0 | 0 | 0 | 0 | 0 | 0 | 0 | 0 | 0 | 0 | 0 | 0 | 0 | 0 | 1 | 0 | 0 | 0 | 0 | 0 | 0 | 0 | 0 | 0 | 0 | 0 | 0 | 0 | 0 | 0 | 0 | 0 | 0 | 0 | 0 | 0 | 0 | 0 | 0 | 0 | 0 |
| virus_Jul11.50v_3134 | 1 | 0 | 0 | 0 | 0 | 0 | 0 | 0 | 0 | 0 | 0 | 0 | 0 | 0 | 0 | 0 | 0 | 0 | 0 | 0 | 0 | 0 | 0 | 0 | 0 | 0 | 0 | 0 | 1 | 0 | 0 | 0 | 0 | 0 | 0 | 0 | 0 | 0 | 0 | 0 | 0 | 0 | 0 | 0 | 0 | 0 | 0 | 0 | 0 | 0 | 0 | 0 | 0 | 0 | 0 |
| virus_Jul11.50v_3570 | 1 | 0 | 0 | 0 | 0 | 0 | 0 | 0 | 0 | 0 | 0 | 0 | 0 | 0 | 0 | 0 | 0 | 0 | 0 | 0 | 0 | 0 | 0 | 0 | 0 | 0 | 0 | 0 | 1 | 0 | 0 | 0 | 0 | 0 | 0 | 0 | 0 | 0 | 0 | 0 | 0 | 0 | 0 | 0 | 0 | 0 | 0 | 0 | 0 | 0 | 0 | 0 | 0 | 0 | 0 |
| virus_Jul11.50v_5386 | 1 | 0 | 0 | 0 | 0 | 0 | 0 | 0 | 0 | 0 | 0 | 0 | 0 | 0 | 0 | 0 | 0 | 0 | 0 | 0 | 0 | 0 | 0 | 0 | 0 | 0 | 0 | 0 | 1 | 0 | 0 | 0 | 0 | 0 | 0 | 0 | 0 | 0 | 0 | 0 | 0 | 0 | 0 | 0 | 0 | 0 | 0 | 0 | 0 | 0 | 0 | 0 | 0 | 0 | 0 |
| virus_Jul11.50v_8132 | 1 | 0 | 0 | 0 | 0 | 0 | 0 | 0 | 0 | 0 | 0 | 0 | 0 | 0 | 0 | 0 | 0 | 0 | 0 | 0 | 0 | 0 | 0 | 0 | 0 | 0 | 0 | 0 | 1 | 0 | 0 | 0 | 0 | 0 | 0 | 0 | 0 | 0 | 0 | 0 | 0 | 0 | 0 | 0 | 0 | 0 | 0 | 0 | 0 | 0 | 0 | 0 | 0 | 0 | 0 |
| virus_Jul11.50v_9013 | 1 | 0 | 0 | 0 | 0 | 0 | 0 | 0 | 0 | 0 | 0 | 0 | 0 | 0 | 0 | 0 | 0 | 0 | 0 | 0 | 0 | 0 | 0 | 0 | 0 | 0 | 0 | 0 | 1 | 0 | 0 | 0 | 0 | 0 | 0 | 0 | 0 | 0 | 0 | 0 | 0 | 0 | 0 | 0 | 0 | 0 | 0 | 0 | 0 | 0 | 0 | 0 | 0 | 0 | 0 |
| virus_Jul11.50v_9145 | 1 | 0 | 0 | 0 | 0 | 0 | 0 | 0 | 0 | 0 | 0 | 0 | 0 | 0 | 0 | 0 | 0 | 0 | 0 | 0 | 0 | 0 | 0 | 0 | 0 | 0 | 0 | 0 | 1 | 0 | 0 | 0 | 0 | 0 | 0 | 0 | 0 | 0 | 0 | 0 | 0 | 0 | 0 | 0 | 0 | 0 | 0 | 0 | 0 | 0 | 0 | 0 | 0 | 0 | 0 |
| virus_Jul11.50v_14073 | 1 | 0 | 0 | 0 | 0 | 0 | 0 | 0 | 0 | 0 | 0 | 0 | 0 | 0 | 0 | 0 | 0 | 0 | 0 | 0 | 0 | 0 | 0 | 0 | 0 | 0 | 0 | 0 | 1 | 0 | 0 | 0 | 0 | 0 | 0 | 0 | 0 | 0 | 0 | 0 | 0 | 0 | 0 | 0 | 0 | 0 | 0 | 0 | 0 | 0 | 0 | 0 | 0 | 0 | 0 |
| virus_Jul11.50v_14817 | 1 | 0 | 0 | 0 | 0 | 0 | 0 | 0 | 0 | 0 | 0 | 0 | 0 | 0 | 0 | 0 | 0 | 0 | 0 | 0 | 0 | 0 | 0 | 0 | 0 | 0 | 0 | 0 | 1 | 0 | 0 | 0 | 0 | 0 | 0 | 0 | 0 | 0 | 0 | 0 | 0 | 0 | 0 | 0 | 0 | 0 | 0 | 0 | 0 | 0 | 0 | 0 | 0 | 0 | 0 |
| virus_Jul11.50v_15596 | 1 | 0 | 0 | 0 | 0 | 0 | 0 | 0 | 0 | 0 | 0 | 0 | 0 | 0 | 0 | 0 | 0 | 0 | 0 | 0 | 0 | 0 | 0 | 0 | 0 | 0 | 0 | 0 | 1 | 0 | 0 | 0 | 0 | 0 | 0 | 0 | 0 | 0 | 0 | 0 | 0 | 0 | 0 | 0 | 0 | 0 | 0 | 0 | 0 | 0 | 0 | 0 | 0 | 0 | 0 |
| virus_Jul11.50v_15921 | 1 | 0 | 0 | 0 | 0 | 0 | 0 | 0 | 0 | 0 | 0 | 0 | 0 | 0 | 0 | 0 | 0 | 0 | 0 | 0 | 0 | 0 | 0 | 0 | 0 | 0 | 0 | 0 | 1 | 0 | 0 | 0 | 0 | 0 | 0 | 0 | 0 | 0 | 0 | 0 | 0 | 0 | 0 | 0 | 0 | 0 | 0 | 0 | 0 | 0 | 0 | 0 | 0 | 0 | 0 |
| virus_Jul11.50v_16057 | 1 | 0 | 0 | 0 | 0 | 0 | 0 | 0 | 0 | 0 | 0 | 0 | 0 | 0 | 0 | 0 | 0 | 0 | 0 | 0 | 0 | 0 | 0 | 0 | 0 | 0 | 0 | 0 | 1 | 0 | 0 | 0 | 0 | 0 | 0 | 0 | 0 | 0 | 0 | 0 | 0 | 0 | 0 | 0 | 0 | 0 | 0 | 0 | 0 | 0 | 0 | 0 | 0 | 0 | 0 |
| virus_Jul11.50v_17405 | 1 | 0 | 0 | 0 | 0 | 0 | 0 | 0 | 0 | 0 | 0 | 0 | 0 | 0 | 0 | 0 | 0 | 0 | 0 | 0 | 0 | 0 | 0 | 0 | 0 | 0 | 0 | 0 | 1 | 0 | 0 | 0 | 0 | 0 | 0 | 0 | 0 | 0 | 0 | 0 | 0 | 0 | 0 | 0 | 0 | 0 | 0 | 0 | 0 | 0 | 0 | 0 | 0 | 0 | 0 |
| virus_Jul11.50v_18812 | 1 | 0 | 0 | 0 | 0 | 0 | 0 | 0 | 0 | 0 | 0 | 0 | 0 | 0 | 0 | 0 | 0 | 0 | 0 | 0 | 0 | 0 | 0 | 0 | 0 | 0 | 0 | 0 | 1 | 0 | 0 | 0 | 0 | 0 | 0 | 0 | 0 | 0 | 0 | 0 | 0 | 0 | 0 | 0 | 0 | 0 | 0 | 0 | 0 | 0 | 0 | 0 | 0 | 0 | 0 |
| virus_Jul11.50v_19569 | 1 | 0 | 0 | 0 | 0 | 0 | 0 | 0 | 0 | 0 | 0 | 0 | 0 | 0 | 0 | 0 | 0 | 0 | 0 | 0 | 0 | 0 | 0 | 0 | 0 | 0 | 0 | 0 | 1 | 0 | 0 | 0 | 0 | 0 | 0 | 0 | 0 | 0 | 0 | 0 | 0 | 0 | 0 | 0 | 0 | 0 | 0 | 0 | 0 | 0 | 0 | 0 | 0 | 0 | 0 |
| virus_Jul11.75v_22505 | 1 | 0 | 0 | 0 | 0 | 0 | 0 | 0 | 0 | 0 | 0 | 0 | 0 | 0 | 0 | 0 | 0 | 0 | 0 | 0 | 0 | 0 | 0 | 0 | 0 | 0 | 0 | 0 | 0 | 0 | 1 | 0 | 0 | 0 | 0 | 0 | 0 | 0 | 0 | 0 | 0 | 0 | 0 | 0 | 0 | 0 | 0 | 0 | 0 | 0 | 0 | 0 | 0 | 0 | 0 |
| virus_Jul11.75v_23409 | 1 | 0 | 0 | 0 | 0 | 0 | 0 | 0 | 0 | 0 | 0 | 0 | 0 | 0 | 0 | 0 | 0 | 0 | 0 | 0 | 0 | 0 | 0 | 0 | 0 | 0 | 0 | 0 | 0 | 0 | 1 | 0 | 0 | 0 | 0 | 0 | 0 | 0 | 0 | 0 | 0 | 0 | 0 | 0 | 0 | 0 | 0 | 0 | 0 | 0 | 0 | 0 | 0 | 0 | 0 |
| virus_Jul11.75v_26829 | 1 | 0 | 0 | 0 | 0 | 0 | 0 | 0 | 0 | 0 | 0 | 0 | 0 | 0 | 0 | 0 | 0 | 0 | 0 | 0 | 0 | 0 | 0 | 0 | 0 | 0 | 0 | 0 | 0 | 0 | 1 | 0 | 0 | 0 | 0 | 0 | 0 | 0 | 0 | 0 | 0 | 0 | 0 | 0 | 0 | 0 | 0 | 0 | 0 | 0 | 0 | 0 | 0 | 0 | 0 |
| virus_Jul11.75v_27124 | 1 | 0 | 0 | 0 | 0 | 0 | 0 | 0 | 0 | 0 | 0 | 0 | 0 | 0 | 0 | 0 | 0 | 0 | 0 | 0 | 0 | 0 | 0 | 0 | 0 | 0 | 0 | 0 | 0 | 0 | 1 | 0 | 0 | 0 | 0 | 0 | 0 | 0 | 0 | 0 | 0 | 0 | 0 | 0 | 0 | 0 | 0 | 0 | 0 | 0 | 0 | 0 | 0 | 0 | 0 |
| virus_Jul11.75v_28798 | 1 | 0 | 0 | 0 | 0 | 0 | 0 | 0 | 0 | 0 | 0 | 0 | 0 | 0 | 0 | 0 | 0 | 0 | 0 | 0 | 0 | 0 | 0 | 0 | 0 | 0 | 0 | 0 | 0 | 0 | 1 | 0 | 0 | 0 | 0 | 0 | 0 | 0 | 0 | 0 | 0 | 0 | 0 | 0 | 0 | 0 | 0 | 0 | 0 | 0 | 0 | 0 | 0 | 0 | 0 |
| virus_Jul11.75v_31337 | 1 | 0 | 0 | 0 | 0 | 0 | 0 | 0 | 0 | 0 | 0 | 0 | 0 | 0 | 0 | 0 | 0 | 0 | 0 | 0 | 0 | 0 | 0 | 0 | 0 | 0 | 0 | 0 | 0 | 0 | 1 | 0 | 0 | 0 | 0 | 0 | 0 | 0 | 0 | 0 | 0 | 0 | 0 | 0 | 0 | 0 | 0 | 0 | 0 | 0 | 0 | 0 | 0 | 0 | 0 |
| virus_Jul11.75v_36835 | 1 | 0 | 0 | 0 | 0 | 0 | 0 | 0 | 0 | 0 | 0 | 0 | 0 | 0 | 0 | 0 | 0 | 0 | 0 | 0 | 0 | 0 | 0 | 0 | 0 | 0 | 0 | 0 | 0 | 0 | 1 | 0 | 0 | 0 | 0 | 0 | 0 | 0 | 0 | 0 | 0 | 0 | 0 | 0 | 0 | 0 | 0 | 0 | 0 | 0 | 0 | 0 | 0 | 0 | 0 |
| virus_Jul11.75v_37165 | 1 | 0 | 0 | 0 | 0 | 0 | 0 | 0 | 0 | 0 | 0 | 0 | 0 | 0 | 0 | 0 | 0 | 0 | 0 | 0 | 0 | 0 | 0 | 0 | 0 | 0 | 0 | 0 | 0 | 0 | 1 | 0 | 0 | 0 | 0 | 0 | 0 | 0 | 0 | 0 | 0 | 0 | 0 | 0 | 0 | 0 | 0 | 0 | 0 | 0 | 0 | 0 | 0 | 0 | 0 |
| virus_Jul27.5v_213 | 1 | 0 | 0 | 0 | 0 | 0 | 0 | 0 | 0 | 0 | 0 | 0 | 0 | 0 | 0 | 0 | 0 | 0 | 0 | 0 | 0 | 0 | 0 | 0 | 0 | 0 | 0 | 0 | 0 | 0 | 0 | 0 | 1 | 0 | 0 | 0 | 0 | 0 | 0 | 0 | 0 | 0 | 0 | 0 | 0 | 0 | 0 | 0 | 0 | 0 | 0 | 0 | 0 | 0 | 0 |
| virus_Jul27.5v_354 | 1 | 0 | 0 | 0 | 0 | 0 | 0 | 0 | 0 | 0 | 0 | 0 | 0 | 0 | 0 | 0 | 0 | 0 | 0 | 0 | 0 | 0 | 0 | 0 | 0 | 0 | 0 | 0 | 0 | 0 | 0 | 0 | 1 | 0 | 0 | 0 | 0 | 0 | 0 | 0 | 0 | 0 | 0 | 0 | 0 | 0 | 0 | 0 | 0 | 0 | 0 | 0 | 0 | 0 | 0 |
| virus_Jul27.5v_650 | 1 | 0 | 0 | 0 | 0 | 0 | 0 | 0 | 0 | 0 | 0 | 0 | 0 | 0 | 0 | 0 | 0 | 0 | 0 | 0 | 0 | 0 | 0 | 0 | 0 | 0 | 0 | 0 | 0 | 0 | 0 | 0 | 1 | 0 | 0 | 0 | 0 | 0 | 0 | 0 | 0 | 0 | 0 | 0 | 0 | 0 | 0 | 0 | 0 | 0 | 0 | 0 | 0 | 0 | 0 |
| virus_Jul27.5v_1734 | 1 | 0 | 0 | 0 | 0 | 0 | 0 | 0 | 0 | 0 | 0 | 0 | 0 | 0 | 0 | 0 | 0 | 0 | 0 | 0 | 0 | 0 | 0 | 0 | 0 | 0 | 0 | 0 | 0 | 0 | 0 | 0 | 1 | 0 | 0 | 0 | 0 | 0 | 0 | 0 | 0 | 0 | 0 | 0 | 0 | 0 | 0 | 0 | 0 | 0 | 0 | 0 | 0 | 0 | 0 |
| virus_Jul27.5v_2154 | 1 | 0 | 0 | 0 | 0 | 0 | 0 | 0 | 0 | 0 | 0 | 0 | 0 | 0 | 0 | 0 | 0 | 0 | 0 | 0 | 0 | 0 | 0 | 0 | 0 | 0 | 0 | 0 | 0 | 0 | 0 | 0 | 1 | 0 | 0 | 0 | 0 | 0 | 0 | 0 | 0 | 0 | 0 | 0 | 0 | 0 | 0 | 0 | 0 | 0 | 0 | 0 | 0 | 0 | 0 |
| virus_Jul27.5v_2593 | 1 | 0 | 0 | 0 | 0 | 0 | 0 | 0 | 0 | 0 | 0 | 0 | 0 | 0 | 0 | 0 | 0 | 0 | 0 | 0 | 0 | 0 | 0 | 0 | 0 | 0 | 0 | 0 | 0 | 0 | 0 | 0 | 1 | 0 | 0 | 0 | 0 | 0 | 0 | 0 | 0 | 0 | 0 | 0 | 0 | 0 | 0 | 0 | 0 | 0 | 0 | 0 | 0 | 0 | 0 |
| virus_Jul27.5v_3707 | 1 | 0 | 0 | 0 | 0 | 0 | 0 | 0 | 0 | 0 | 0 | 0 | 0 | 0 | 0 | 0 | 0 | 0 | 0 | 0 | 0 | 0 | 0 | 0 | 0 | 0 | 0 | 0 | 0 | 0 | 0 | 0 | 1 | 0 | 0 | 0 | 0 | 0 | 0 | 0 | 0 | 0 | 0 | 0 | 0 | 0 | 0 | 0 | 0 | 0 | 0 | 0 | 0 | 0 | 0 |
| virus_Jul27.5v_4775 | 1 | 0 | 0 | 0 | 0 | 0 | 0 | 0 | 0 | 0 | 0 | 0 | 0 | 0 | 0 | 0 | 0 | 0 | 0 | 0 | 0 | 0 | 0 | 0 | 0 | 0 | 0 | 0 | 0 | 0 | 0 | 0 | 1 | 0 | 0 | 0 | 0 | 0 | 0 | 0 | 0 | 0 | 0 | 0 | 0 | 0 | 0 | 0 | 0 | 0 | 0 | 0 | 0 | 0 | 0 |
| virus_Jul27.5v_5298 | 1 | 0 | 0 | 0 | 0 | 0 | 0 | 0 | 0 | 0 | 0 | 0 | 0 | 0 | 0 | 0 | 0 | 0 | 0 | 0 | 0 | 0 | 0 | 0 | 0 | 0 | 0 | 0 | 0 | 0 | 0 | 0 | 1 | 0 | 0 | 0 | 0 | 0 | 0 | 0 | 0 | 0 | 0 | 0 | 0 | 0 | 0 | 0 | 0 | 0 | 0 | 0 | 0 | 0 | 0 |
| virus_Jul27.5v_6476 | 1 | 0 | 0 | 0 | 0 | 0 | 0 | 0 | 0 | 0 | 0 | 0 | 0 | 0 | 0 | 0 | 0 | 0 | 0 | 0 | 0 | 0 | 0 | 0 | 0 | 0 | 0 | 0 | 0 | 0 | 0 | 0 | 1 | 0 | 0 | 0 | 0 | 0 | 0 | 0 | 0 | 0 | 0 | 0 | 0 | 0 | 0 | 0 | 0 | 0 | 0 | 0 | 0 | 0 | 0 |
| virus_Jul27.5v_6482 | 1 | 0 | 0 | 0 | 0 | 0 | 0 | 0 | 0 | 0 | 0 | 0 | 0 | 0 | 0 | 0 | 0 | 0 | 0 | 0 | 0 | 0 | 0 | 0 | 0 | 0 | 0 | 0 | 0 | 0 | 0 | 0 | 1 | 0 | 0 | 0 | 0 | 0 | 0 | 0 | 0 | 0 | 0 | 0 | 0 | 0 | 0 | 0 | 0 | 0 | 0 | 0 | 0 | 0 | 0 |
| virus_Jul27.5v_8541 | 1 | 0 | 0 | 0 | 0 | 0 | 0 | 0 | 0 | 0 | 0 | 0 | 0 | 0 | 0 | 0 | 0 | 0 | 0 | 0 | 0 | 0 | 0 | 0 | 0 | 0 | 0 | 0 | 0 | 0 | 0 | 0 | 1 | 0 | 0 | 0 | 0 | 0 | 0 | 0 | 0 | 0 | 0 | 0 | 0 | 0 | 0 | 0 | 0 | 0 | 0 | 0 | 0 | 0 | 0 |
| virus_Jul27.5v_8588 | 1 | 0 | 0 | 0 | 0 | 0 | 0 | 0 | 0 | 0 | 0 | 0 | 0 | 0 | 0 | 0 | 0 | 0 | 0 | 0 | 0 | 0 | 0 | 0 | 0 | 0 | 0 | 0 | 0 | 0 | 0 | 0 | 1 | 0 | 0 | 0 | 0 | 0 | 0 | 0 | 0 | 0 | 0 | 0 | 0 | 0 | 0 | 0 | 0 | 0 | 0 | 0 | 0 | 0 | 0 |
| virus_Jul27.5v_9251 | 1 | 0 | 0 | 0 | 0 | 0 | 0 | 0 | 0 | 0 | 0 | 0 | 0 | 0 | 0 | 0 | 0 | 0 | 0 | 0 | 0 | 0 | 0 | 0 | 0 | 0 | 0 | 0 | 0 | 0 | 0 | 0 | 1 | 0 | 0 | 0 | 0 | 0 | 0 | 0 | 0 | 0 | 0 | 0 | 0 | 0 | 0 | 0 | 0 | 0 | 0 | 0 | 0 | 0 | 0 |
| virus_Jul27.5v_9916 | 1 | 0 | 0 | 0 | 0 | 0 | 0 | 0 | 0 | 0 | 0 | 0 | 0 | 0 | 0 | 0 | 0 | 0 | 0 | 0 | 0 | 0 | 0 | 0 | 0 | 0 | 0 | 0 | 0 | 0 | 0 | 0 | 1 | 0 | 0 | 0 | 0 | 0 | 0 | 0 | 0 | 0 | 0 | 0 | 0 | 0 | 0 | 0 | 0 | 0 | 0 | 0 | 0 | 0 | 0 |
| virus_Jul27.5v_10429 | 1 | 0 | 0 | 0 | 0 | 0 | 0 | 0 | 0 | 0 | 0 | 0 | 0 | 0 | 0 | 0 | 0 | 0 | 0 | 0 | 0 | 0 | 0 | 0 | 0 | 0 | 0 | 0 | 0 | 0 | 0 | 0 | 1 | 0 | 0 | 0 | 0 | 0 | 0 | 0 | 0 | 0 | 0 | 0 | 0 | 0 | 0 | 0 | 0 | 0 | 0 | 0 | 0 | 0 | 0 |
| virus_Jul27.5v_10453 | 1 | 0 | 0 | 0 | 0 | 0 | 0 | 0 | 0 | 0 | 0 | 0 | 0 | 0 | 0 | 0 | 0 | 0 | 0 | 0 | 0 | 0 | 0 | 0 | 0 | 0 | 0 | 0 | 0 | 0 | 0 | 0 | 1 | 0 | 0 | 0 | 0 | 0 | 0 | 0 | 0 | 0 | 0 | 0 | 0 | 0 | 0 | 0 | 0 | 0 | 0 | 0 | 0 | 0 | 0 |
| virus_Jul27.5v_11706 | 1 | 0 | 0 | 0 | 0 | 0 | 0 | 0 | 0 | 0 | 0 | 0 | 0 | 0 | 0 | 0 | 0 | 0 | 0 | 0 | 0 | 0 | 0 | 0 | 0 | 0 | 0 | 0 | 0 | 0 | 0 | 0 | 1 | 0 | 0 | 0 | 0 | 0 | 0 | 0 | 0 | 0 | 0 | 0 | 0 | 0 | 0 | 0 | 0 | 0 | 0 | 0 | 0 | 0 | 0 |
| virus_Jul27.5v_12269 | 1 | 0 | 0 | 0 | 0 | 0 | 0 | 0 | 0 | 0 | 0 | 0 | 0 | 0 | 0 | 0 | 0 | 0 | 0 | 0 | 0 | 0 | 0 | 0 | 0 | 0 | 0 | 0 | 0 | 0 | 0 | 0 | 1 | 0 | 0 | 0 | 0 | 0 | 0 | 0 | 0 | 0 | 0 | 0 | 0 | 0 | 0 | 0 | 0 | 0 | 0 | 0 | 0 | 0 | 0 |
| virus_Jul27.5v_12825 | 1 | 0 | 0 | 0 | 0 | 0 | 0 | 0 | 0 | 0 | 0 | 0 | 0 | 0 | 0 | 0 | 0 | 0 | 0 | 0 | 0 | 0 | 0 | 0 | 0 | 0 | 0 | 0 | 0 | 0 | 0 | 0 | 1 | 0 | 0 | 0 | 0 | 0 | 0 | 0 | 0 | 0 | 0 | 0 | 0 | 0 | 0 | 0 | 0 | 0 | 0 | 0 | 0 | 0 | 0 |
| virus_Jul27.5v_12858 | 1 | 0 | 0 | 0 | 0 | 0 | 0 | 0 | 0 | 0 | 0 | 0 | 0 | 0 | 0 | 0 | 0 | 0 | 0 | 0 | 0 | 0 | 0 | 0 | 0 | 0 | 0 | 0 | 0 | 0 | 0 | 0 | 1 | 0 | 0 | 0 | 0 | 0 | 0 | 0 | 0 | 0 | 0 | 0 | 0 | 0 | 0 | 0 | 0 | 0 | 0 | 0 | 0 | 0 | 0 |
| virus_Jul27.5v_13352 | 1 | 0 | 0 | 0 | 0 | 0 | 0 | 0 | 0 | 0 | 0 | 0 | 0 | 0 | 0 | 0 | 0 | 0 | 0 | 0 | 0 | 0 | 0 | 0 | 0 | 0 | 0 | 0 | 0 | 0 | 0 | 0 | 1 | 0 | 0 | 0 | 0 | 0 | 0 | 0 | 0 | 0 | 0 | 0 | 0 | 0 | 0 | 0 | 0 | 0 | 0 | 0 | 0 | 0 | 0 |
| virus_Jul27.5v_15851 | 1 | 0 | 0 | 0 | 0 | 0 | 0 | 0 | 0 | 0 | 0 | 0 | 0 | 0 | 0 | 0 | 0 | 0 | 0 | 0 | 0 | 0 | 0 | 0 | 0 | 0 | 0 | 0 | 0 | 0 | 0 | 0 | 1 | 0 | 0 | 0 | 0 | 0 | 0 | 0 | 0 | 0 | 0 | 0 | 0 | 0 | 0 | 0 | 0 | 0 | 0 | 0 | 0 | 0 | 0 |
| virus_Jul27.5v_16211 | 1 | 0 | 0 | 0 | 0 | 0 | 0 | 0 | 0 | 0 | 0 | 0 | 0 | 0 | 0 | 0 | 0 | 0 | 0 | 0 | 0 | 0 | 0 | 0 | 0 | 0 | 0 | 0 | 0 | 0 | 0 | 0 | 1 | 0 | 0 | 0 | 0 | 0 | 0 | 0 | 0 | 0 | 0 | 0 | 0 | 0 | 0 | 0 | 0 | 0 | 0 | 0 | 0 | 0 | 0 |
| virus_Jul27.5v_17045 | 1 | 0 | 0 | 0 | 0 | 0 | 0 | 0 | 0 | 0 | 0 | 0 | 0 | 0 | 0 | 0 | 0 | 0 | 0 | 0 | 0 | 0 | 0 | 0 | 0 | 0 | 0 | 0 | 0 | 0 | 0 | 0 | 1 | 0 | 0 | 0 | 0 | 0 | 0 | 0 | 0 | 0 | 0 | 0 | 0 | 0 | 0 | 0 | 0 | 0 | 0 | 0 | 0 | 0 | 0 |
| virus_Jul27.5v_17195 | 1 | 0 | 0 | 0 | 0 | 0 | 0 | 0 | 0 | 0 | 0 | 0 | 0 | 0 | 0 | 0 | 0 | 0 | 0 | 0 | 0 | 0 | 0 | 0 | 0 | 0 | 0 | 0 | 0 | 0 | 0 | 0 | 1 | 0 | 0 | 0 | 0 | 0 | 0 | 0 | 0 | 0 | 0 | 0 | 0 | 0 | 0 | 0 | 0 | 0 | 0 | 0 | 0 | 0 | 0 |
| virus_Jul27.5v_18023 | 1 | 0 | 0 | 0 | 0 | 0 | 0 | 0 | 0 | 0 | 0 | 0 | 0 | 0 | 0 | 0 | 0 | 0 | 0 | 0 | 0 | 0 | 0 | 0 | 0 | 0 | 0 | 0 | 0 | 0 | 0 | 0 | 1 | 0 | 0 | 0 | 0 | 0 | 0 | 0 | 0 | 0 | 0 | 0 | 0 | 0 | 0 | 0 | 0 | 0 | 0 | 0 | 0 | 0 | 0 |
| virus_Jul27.5v_18282 | 1 | 0 | 0 | 0 | 0 | 0 | 0 | 0 | 0 | 0 | 0 | 0 | 0 | 0 | 0 | 0 | 0 | 0 | 0 | 0 | 0 | 0 | 0 | 0 | 0 | 0 | 0 | 0 | 0 | 0 | 0 | 0 | 1 | 0 | 0 | 0 | 0 | 0 | 0 | 0 | 0 | 0 | 0 | 0 | 0 | 0 | 0 | 0 | 0 | 0 | 0 | 0 | 0 | 0 | 0 |
| virus_Jul27.5v_18332 | 1 | 0 | 0 | 0 | 0 | 0 | 0 | 0 | 0 | 0 | 0 | 0 | 0 | 0 | 0 | 0 | 0 | 0 | 0 | 0 | 0 | 0 | 0 | 0 | 0 | 0 | 0 | 0 | 0 | 0 | 0 | 0 | 1 | 0 | 0 | 0 | 0 | 0 | 0 | 0 | 0 | 0 | 0 | 0 | 0 | 0 | 0 | 0 | 0 | 0 | 0 | 0 | 0 | 0 | 0 |
| virus_Jul27.5v_18638 | 1 | 0 | 0 | 0 | 0 | 0 | 0 | 0 | 0 | 0 | 0 | 0 | 0 | 0 | 0 | 0 | 0 | 0 | 0 | 0 | 0 | 0 | 0 | 0 | 0 | 0 | 0 | 0 | 0 | 0 | 0 | 0 | 1 | 0 | 0 | 0 | 0 | 0 | 0 | 0 | 0 | 0 | 0 | 0 | 0 | 0 | 0 | 0 | 0 | 0 | 0 | 0 | 0 | 0 | 0 |
| virus_Jul27.5v_18651 | 1 | 0 | 0 | 0 | 0 | 0 | 0 | 0 | 0 | 0 | 0 | 0 | 0 | 0 | 0 | 0 | 0 | 0 | 0 | 0 | 0 | 0 | 0 | 0 | 0 | 0 | 0 | 0 | 0 | 0 | 0 | 0 | 1 | 0 | 0 | 0 | 0 | 0 | 0 | 0 | 0 | 0 | 0 | 0 | 0 | 0 | 0 | 0 | 0 | 0 | 0 | 0 | 0 | 0 | 0 |
| virus_Jul27.5v_19951 | 1 | 0 | 0 | 0 | 0 | 0 | 0 | 0 | 0 | 0 | 0 | 0 | 0 | 0 | 0 | 0 | 0 | 0 | 0 | 0 | 0 | 0 | 0 | 0 | 0 | 0 | 0 | 0 | 0 | 0 | 0 | 0 | 1 | 0 | 0 | 0 | 0 | 0 | 0 | 0 | 0 | 0 | 0 | 0 | 0 | 0 | 0 | 0 | 0 | 0 | 0 | 0 | 0 | 0 | 0 |
| virus_Jul27.5v_20723 | 1 | 0 | 0 | 0 | 0 | 0 | 0 | 0 | 0 | 0 | 0 | 0 | 0 | 0 | 0 | 0 | 0 | 0 | 0 | 0 | 0 | 0 | 0 | 0 | 0 | 0 | 0 | 0 | 0 | 0 | 0 | 0 | 1 | 0 | 0 | 0 | 0 | 0 | 0 | 0 | 0 | 0 | 0 | 0 | 0 | 0 | 0 | 0 | 0 | 0 | 0 | 0 | 0 | 0 | 0 |

**Supplementary Table S7.** Common Bacterio- and virioplankton RTPR 98% OTUs in 27 samples with RTPR amplicons in both fractions

Total seqs: 47997; Subtotal seqs in retained

| OTUs: 41046 | | May12_05m | | May12_25m | | May12_50m | | May12_75m | | Jun8_25m | | Jun8_50m | | Jun8_75m | | Jun22_0m | | Jun22_50m | | Jun22_75m | | Jul11_0m | | Jul11_05m | | Jul11_25m | | Jul11_50m | | Jul11_75m | | Jul27_05m | | Jul27_25m | | Jul27_75m | | Aug5_0m | | Aug5_05m | | Aug5_25m | | Aug5_50m | | Aug5_75m | | Sep8_0m | | Sep8_05m | | Sep8_25m | | Sep8_50m | |
|---|---|---|---|---|---|---|---|---|---|---|---|---|---|---|---|---|---|---|---|---|---|---|---|---|---|---|---|---|---|---|---|---|---|---|---|---|---|---|---|---|---|---|---|---|---|---|---|---|---|---|---|---|---|---|---|
| OTUs with >=10 seqs: 254 | OTU subtotal | bacterioplankton | virioplankton | bacterioplankton | virioplankton | bacterioplankton | virioplankton | bacterioplankton | virioplankton | bacterioplankton | virioplankton | bacterioplankton | virioplankton | bacterioplankton | virioplankton | bacterioplankton | virioplankton | bacterioplankton | virioplankton | bacterioplankton | virioplankton | bacterioplankton | virioplankton | bacterioplankton | virioplankton | bacterioplankton | virioplankton | bacterioplankton | virioplankton | bacterioplankton | virioplankton | bacterioplankton | virioplankton | bacterioplankton | virioplankton | bacterioplankton | virioplankton | bacterioplankton | virioplankton | bacterioplankton | virioplankton | bacterioplankton | virioplankton | bacterioplankton | virioplankton | bacterioplankton | virioplankton | bacterioplankton | virioplankton | bacterioplankton | virioplankton | bacterioplankton | virioplankton | bacterioplankton | virioplankton |
|virus_Jul27.5v_20949|1| 0|0|0|0|0|0|0|0|0|0| 0|0|0|0|0|0|0|0|0|0| 0|0|0|0|0|0|0|0|0|0| 0|1| 0|0|0|0|0|0|0|0|0|0| 0|0|0|0|0|0|0|0|0|0| 0|0|
|virus_Jul27.25v_4130|1| 0|0|0|0|0|0|0|0|0|0| 0|0|0|0|0|0|0|0|0|0| 0|0|0|0|0|0|0|0|0|0| 0|0|0|1| 0|0|0|0|0|0|0|0|0|0| 0|0|0|0|0|0|0|0|0|0|
|virus_Jul27.25v_6746|1| 0|0|0|0|0|0|0|0|0|0| 0|0|0|0|0|0|0|0|0|0| 0|0|0|0|0|0|0|0|0|0| 0|0|0|1| 0|0|0|0|0|0|0|0|0|0| 0|0|0|0|0|0|0|0|0|0|
|virus_Jul27.25v_9489|1| 0|0|0|0|0|0|0|0|0|0| 0|0|0|0|0|0|0|0|0|0| 0|0|0|0|0|0|0|0|0|0| 0|0|0|1| 0|0|0|0|0|0|0|0|0|0| 0|0|0|0|0|0|0|0|0|0|
|virus_Jul27.25v_12888|1| 0|0|0|0|0|0|0|0|0|0| 0|0|0|0|0|0|0|0|0|0| 0|0|0|0|0|0|0|0|0|0| 0|0|0|1| 0|0|0|0|0|0|0|0|0|0| 0|0|0|0|0|0|0|0|0|0|
|virus_Jul27.25v_13260|1| 0|0|0|0|0|0|0|0|0|0| 0|0|0|0|0|0|0|0|0|0| 0|0|0|0|0|0|0|0|0|0| 0|0|0|1| 0|0|0|0|0|0|0|0|0|0| 0|0|0|0|0|0|0|0|0|0|
|virus_Jul27.25v_18132|1| 0|0|0|0|0|0|0|0|0|0| 0|0|0|0|0|0|0|0|0|0| 0|0|0|0|0|0|0|0|0|0| 0|0|0|1| 0|0|0|0|0|0|0|0|0|0| 0|0|0|0|0|0|0|0|0|0|
|virus_Jul27.25v_19173|1| 0|0|0|0|0|0|0|0|0|0| 0|0|0|0|0|0|0|0|0|0| 0|0|0|0|0|0|0|0|0|0| 0|0|0|1| 0|0|0|0|0|0|0|0|0|0| 0|0|0|0|0|0|0|0|0|0|
|virus_Jul27.25v_21996|1| 0|0|0|0|0|0|0|0|0|0| 0|0|0|0|0|0|0|0|0|0| 0|0|0|0|0|0|0|0|0|0| 0|0|0|1| 0|0|0|0|0|0|0|0|0|0| 0|0|0|0|0|0|0|0|0|0|
|virus_Jul27.25v_22871|1| 0|0|0|0|0|0|0|0|0|0| 0|0|0|0|0|0|0|0|0|0| 0|0|0|0|0|0|0|0|0|0| 0|0|0|1| 0|0|0|0|0|0|0|0|0|0| 0|0|0|0|0|0|0|0|0|0|
|virus_Jul27.25v_29083|1| 0|0|0|0|0|0|0|0|0|0| 0|0|0|0|0|0|0|0|0|0| 0|0|0|0|0|0|0|0|0|0| 0|0|0|1| 0|0|0|0|0|0|0|0|0|0| 0|0|0|0|0|0|0|0|0|0|
|virus_Jul27.25v_29742|1| 0|0|0|0|0|0|0|0|0|0| 0|0|0|0|0|0|0|0|0|0| 0|0|0|0|0|0|0|0|0|0| 0|0|0|1| 0|0|0|0|0|0|0|0|0|0| 0|0|0|0|0|0|0|0|0|0|
|virus_Jul27.25v_32099|1| 0|0|0|0|0|0|0|0|0|0| 0|0|0|0|0|0|0|0|0|0| 0|0|0|0|0|0|0|0|0|0| 0|0|0|1| 0|0|0|0|0|0|0|0|0|0| 0|0|0|0|0|0|0|0|0|0|
|virus_Jul27.75v_15687|1| 0|0|0|0|0|0|0|0|0|0| 0|0|0|0|0|0|0|0|0|0| 0|0|0|0|0|0|0|0|0|0| 0|0|0|0|0|1| 0|0|0|0|0|0|0|0|0|0| 0|0|0|0|0|0|0|0|
|virus_Jun8.25v_8775|1| 0|0|0|0|0|0|0|0|0|1| 0|0|0|0|0|0|0|0|0|0| 0|0|0|0|0|0|0|0|0|0| 0|0|0|0|0|0|0|0|0|0| 0|0|0|0|0|0|0|0|0|0| 0|0|0|0|
|virus_Jun8.25v_13954|1| 0|0|0|0|0|0|0|0|0|1| 0|0|0|0|0|0|0|0|0|0| 0|0|0|0|0|0|0|0|0|0| 0|0|0|0|0|0|0|0|0|0| 0|0|0|0|0|0|0|0|0|0| 0|0|0|0|
|virus_Jun8.25v_21043|1| 0|0|0|0|0|0|0|0|0|1| 0|0|0|0|0|0|0|0|0|0| 0|0|0|0|0|0|0|0|0|0| 0|0|0|0|0|0|0|0|0|0| 0|0|0|0|0|0|0|0|0|0| 0|0|0|0|
|virus_Jun8.25v_24974|1| 0|0|0|0|0|0|0|0|0|1| 0|0|0|0|0|0|0|0|0|0| 0|0|0|0|0|0|0|0|0|0| 0|0|0|0|0|0|0|0|0|0| 0|0|0|0|0|0|0|0|0|0| 0|0|0|0|
|virus_Jun8.25v_38200|1| 0|0|0|0|0|0|0|0|0|1| 0|0|0|0|0|0|0|0|0|0| 0|0|0|0|0|0|0|0|0|0| 0|0|0|0|0|0|0|0|0|0| 0|0|0|0|0|0|0|0|0|0| 0|0|0|0|
|virus_Jun8.25v_39198|1| 0|0|0|0|0|0|0|0|0|1| 0|0|0|0|0|0|0|0|0|0| 0|0|0|0|0|0|0|0|0|0| 0|0|0|0|0|0|0|0|0|0| 0|0|0|0|0|0|0|0|0|0| 0|0|0|0|
|virus_Jun8.50v_2049|1| 0|0|0|0|0|0|0|0|0|0| 0|1| 0|0|0|0|0|0|0|0|0|0| 0|0|0|0|0|0|0|0|0|0| 0|0|0|0|0|0|0|0|0|0| 0|0|0|0|0|0|0|0|0|0| 0|0|
|virus_Jun8.50v_4763|1| 0|0|0|0|0|0|0|0|0|0| 0|1| 0|0|0|0|0|0|0|0|0|0| 0|0|0|0|0|0|0|0|0|0| 0|0|0|0|0|0|0|0|0|0| 0|0|0|0|0|0|0|0|0|0| 0|0|
|virus_Jun8.50v_5089|1| 0|0|0|0|0|0|0|0|0|0| 0|1| 0|0|0|0|0|0|0|0|0|0| 0|0|0|0|0|0|0|0|0|0| 0|0|0|0|0|0|0|0|0|0| 0|0|0|0|0|0|0|0|0|0| 0|0|
|virus_Jun8.50v_6995|1| 0|0|0|0|0|0|0|0|0|0| 0|1| 0|0|0|0|0|0|0|0|0|0| 0|0|0|0|0|0|0|0|0|0| 0|0|0|0|0|0|0|0|0|0| 0|0|0|0|0|0|0|0|0|0| 0|0|
|virus_Jun8.50v_8716|1| 0|0|0|0|0|0|0|0|0|0| 0|1| 0|0|0|0|0|0|0|0|0|0| 0|0|0|0|0|0|0|0|0|0| 0|0|0|0|0|0|0|0|0|0| 0|0|0|0|0|0|0|0|0|0| 0|0|
|virus_Jun8.50v_13937|1| 0|0|0|0|0|0|0|0|0|0| 0|1| 0|0|0|0|0|0|0|0|0|0| 0|0|0|0|0|0|0|0|0|0| 0|0|0|0|0|0|0|0|0|0| 0|0|0|0|0|0|0|0|0|0| 0|0|
|virus_Jun8.50v_16334|1| 0|0|0|0|0|0|0|0|0|0| 0|1| 0|0|0|0|0|0|0|0|0|0| 0|0|0|0|0|0|0|0|0|0| 0|0|0|0|0|0|0|0|0|0| 0|0|0|0|0|0|0|0|0|0| 0|0|
|virus_Jun8.50v_17015|1| 0|0|0|0|0|0|0|0|0|0| 0|1| 0|0|0|0|0|0|0|0|0|0| 0|0|0|0|0|0|0|0|0|0| 0|0|0|0|0|0|0|0|0|0| 0|0|0|0|0|0|0|0|0|0| 0|0|
|virus_Jun8.50v_17419|1| 0|0|0|0|0|0|0|0|0|0| 0|1| 0|0|0|0|0|0|0|0|0|0| 0|0|0|0|0|0|0|0|0|0| 0|0|0|0|0|0|0|0|0|0| 0|0|0|0|0|0|0|0|0|0| 0|0|
|virus_Jun8.50v_17880|1| 0|0|0|0|0|0|0|0|0|0| 0|1| 0|0|0|0|0|0|0|0|0|0| 0|0|0|0|0|0|0|0|0|0| 0|0|0|0|0|0|0|0|0|0| 0|0|0|0|0|0|0|0|0|0| 0|0|
|virus_Jun8.50v_19106|1| 0|0|0|0|0|0|0|0|0|0| 0|1| 0|0|0|0|0|0|0|0|0|0| 0|0|0|0|0|0|0|0|0|0| 0|0|0|0|0|0|0|0|0|0| 0|0|0|0|0|0|0|0|0|0| 0|0|
|virus_Jun8.50v_19367|1| 0|0|0|0|0|0|0|0|0|0| 0|1| 0|0|0|0|0|0|0|0|0|0| 0|0|0|0|0|0|0|0|0|0| 0|0|0|0|0|0|0|0|0|0| 0|0|0|0|0|0|0|0|0|0| 0|0|
|virus_Jun8.50v_20991|1| 0|0|0|0|0|0|0|0|0|0| 0|1| 0|0|0|0|0|0|0|0|0|0| 0|0|0|0|0|0|0|0|0|0| 0|0|0|0|0|0|0|0|0|0| 0|0|0|0|0|0|0|0|0|0| 0|0|
|virus_Jun8.50v_26042|1| 0|0|0|0|0|0|0|0|0|0| 0|1| 0|0|0|0|0|0|0|0|0|0| 0|0|0|0|0|0|0|0|0|0| 0|0|0|0|0|0|0|0|0|0| 0|0|0|0|0|0|0|0|0|0| 0|0|
|virus_Jun8.50v_28327|1| 0|0|0|0|0|0|0|0|0|0| 0|1| 0|0|0|0|0|0|0|0|0|0| 0|0|0|0|0|0|0|0|0|0| 0|0|0|0|0|0|0|0|0|0| 0|0|0|0|0|0|0|0|0|0| 0|0|
|virus_Jun8.50v_28675|1| 0|0|0|0|0|0|0|0|0|0| 0|1| 0|0|0|0|0|0|0|0|0|0| 0|0|0|0|0|0|0|0|0|0| 0|0|0|0|0|0|0|0|0|0| 0|0|0|0|0|0|0|0|0|0| 0|0|
|virus_Jun8.50v_30966|1| 0|0|0|0|0|0|0|0|0|0| 0|1| 0|0|0|0|0|0|0|0|0|0| 0|0|0|0|0|0|0|0|0|0| 0|0|0|0|0|0|0|0|0|0| 0|0|0|0|0|0|0|0|0|0| 0|0|
|virus_Jun8.50v_36205|1| 0|0|0|0|0|0|0|0|0|0| 0|1| 0|0|0|0|0|0|0|0|0|0| 0|0|0|0|0|0|0|0|0|0| 0|0|0|0|0|0|0|0|0|0| 0|0|0|0|0|0|0|0|0|0| 0|0|
|virus_Jun8.50v_40474|1| 0|0|0|0|0|0|0|0|0|0| 0|1| 0|0|0|0|0|0|0|0|0|0| 0|0|0|0|0|0|0|0|0|0| 0|0|0|0|0|0|0|0|0|0| 0|0|0|0|0|0|0|0|0|0| 0|0|
|virus_Jun8.75v_451|1| 0|0|0|0|0|0|0|0|0|0| 0|0|0|1| 0|0|0|0|0|0|0|0|0|0| 0|0|0|0|0|0|0|0|0|0| 0|0|0|0|0|0|0|0|0|0| 0|0|0|0|0|0|0|0|0|0|
|virus_Jun8.75v_6517|1| 0|0|0|0|0|0|0|0|0|0| 0|0|0|1| 0|0|0|0|0|0|0|0|0|0| 0|0|0|0|0|0|0|0|0|0| 0|0|0|0|0|0|0|0|0|0| 0|0|0|0|0|0|0|0|0|0|
|virus_Jun22.0v_1189|1| 0|0|0|0|0|0|0|0|0|0| 0|0|0|0|0|1| 0|0|0|0|0|0|0|0|0|0| 0|0|0|0|0|0|0|0|0|0| 0|0|0|0|0|0|0|0|0|0| 0|0|0|0|0|0|0|0|
|virus_Jun22.0v_7061|1| 0|0|0|0|0|0|0|0|0|0| 0|0|0|0|0|1| 0|0|0|0|0|0|0|0|0|0| 0|0|0|0|0|0|0|0|0|0| 0|0|0|0|0|0|0|0|0|0| 0|0|0|0|0|0|0|0|
|virus_Jun22.0v_7637|1| 0|0|0|0|0|0|0|0|0|0| 0|0|0|0|0|1| 0|0|0|0|0|0|0|0|0|0| 0|0|0|0|0|0|0|0|0|0| 0|0|0|0|0|0|0|0|0|0| 0|0|0|0|0|0|0|0|
|virus_Jun22.0v_8025|1| 0|0|0|0|0|0|0|0|0|0| 0|0|0|0|0|1| 0|0|0|0|0|0|0|0|0|0| 0|0|0|0|0|0|0|0|0|0| 0|0|0|0|0|0|0|0|0|0| 0|0|0|0|0|0|0|0|
|virus_Jun22.0v_9080|1| 0|0|0|0|0|0|0|0|0|0| 0|0|0|0|0|1| 0|0|0|0|0|0|0|0|0|0| 0|0|0|0|0|0|0|0|0|0| 0|0|0|0|0|0|0|0|0|0| 0|0|0|0|0|0|0|0|
|virus_Jun22.0v_11812|1| 0|0|0|0|0|0|0|0|0|0| 0|0|0|0|0|1| 0|0|0|0|0|0|0|0|0|0| 0|0|0|0|0|0|0|0|0|0| 0|0|0|0|0|0|0|0|0|0| 0|0|0|0|0|0|0|0|
|virus_Jun22.0v_12447|1| 0|0|0|0|0|0|0|0|0|0| 0|0|0|0|0|1| 0|0|0|0|0|0|0|0|0|0| 0|0|0|0|0|0|0|0|0|0| 0|0|0|0|0|0|0|0|0|0| 0|0|0|0|0|0|0|0|
|virus_Jun22.0v_15566|1| 0|0|0|0|0|0|0|0|0|0| 0|0|0|0|0|1| 0|0|0|0|0|0|0|0|0|0| 0|0|0|0|0|0|0|0|0|0| 0|0|0|0|0|0|0|0|0|0| 0|0|0|0|0|0|0|0|
|virus_Jun22.0v_19488|1| 0|0|0|0|0|0|0|0|0|0| 0|0|0|0|0|1| 0|0|0|0|0|0|0|0|0|0| 0|0|0|0|0|0|0|0|0|0| 0|0|0|0|0|0|0|0|0|0| 0|0|0|0|0|0|0|0|
|virus_Jun22.0v_19850|1| 0|0|0|0|0|0|0|0|0|0| 0|0|0|0|0|1| 0|0|0|0|0|0|0|0|0|0| 0|0|0|0|0|0|0|0|0|0| 0|0|0|0|0|0|0|0|0|0| 0|0|0|0|0|0|0|0|
|virus_Jun22.50v_1367|1| 0|0|0|0|0|0|0|0|0|0| 0|0|0|0|0|0|0|1| 0|0|0|0|0|0|0|0|0|0| 0|0|0|0|0|0|0|0|0|0| 0|0|0|0|0|0|0|0|0|0| 0|0|0|0|0|0|
|virus_Jun22.50v_1537|1| 0|0|0|0|0|0|0|0|0|0| 0|0|0|0|0|0|0|1| 0|0|0|0|0|0|0|0|0|0| 0|0|0|0|0|0|0|0|0|0| 0|0|0|0|0|0|0|0|0|0| 0|0|0|0|0|0|
|virus_Jun22.50v_1966|1| 0|0|0|0|0|0|0|0|0|0| 0|0|0|0|0|0|0|1| 0|0|0|0|0|0|0|0|0|0| 0|0|0|0|0|0|0|0|0|0| 0|0|0|0|0|0|0|0|0|0| 0|0|0|0|0|0|
|virus_Jun22.50v_2620|1| 0|0|0|0|0|0|0|0|0|0| 0|0|0|0|0|0|0|1| 0|0|0|0|0|0|0|0|0|0| 0|0|0|0|0|0|0|0|0|0| 0|0|0|0|0|0|0|0|0|0| 0|0|0|0|0|0|
|virus_Jun22.50v_3703|1| 0|0|0|0|0|0|0|0|0|0| 0|0|0|0|0|0|0|1| 0|0|0|0|0|0|0|0|0|0| 0|0|0|0|0|0|0|0|0|0| 0|0|0|0|0|0|0|0|0|0| 0|0|0|0|0|0|
|virus_Jun22.50v_4782|1| 0|0|0|0|0|0|0|0|0|0| 0|0|0|0|0|0|0|1| 0|0|0|0|0|0|0|0|0|0| 0|0|0|0|0|0|0|0|0|0| 0|0|0|0|0|0|0|0|0|0| 0|0|0|0|0|0|
|virus_Jun22.50v_8144|1| 0|0|0|0|0|0|0|0|0|0| 0|0|0|0|0|0|0|1| 0|0|0|0|0|0|0|0|0|0| 0|0|0|0|0|0|0|0|0|0| 0|0|0|0|0|0|0|0|0|0| 0|0|0|0|0|0|
|virus_Jun22.50v_9466|1| 0|0|0|0|0|0|0|0|0|0| 0|0|0|0|0|0|0|1| 0|0|0|0|0|0|0|0|0|0| 0|0|0|0|0|0|0|0|0|0| 0|0|0|0|0|0|0|0|0|0| 0|0|0|0|0|0|
|virus_Jun22.50v_9692|1| 0|0|0|0|0|0|0|0|0|0| 0|0|0|0|0|0|0|1| 0|0|0|0|0|0|0|0|0|0| 0|0|0|0|0|0|0|0|0|0| 0|0|0|0|0|0|0|0|0|0| 0|0|0|0|0|0|
|virus_Jun22.50v_13919|1| 0|0|0|0|0|0|0|0|0|0| 0|0|0|0|0|0|0|1| 0|0|0|0|0|0|0|0|0|0| 0|0|0|0|0|0|0|0|0|0| 0|0|0|0|0|0|0|0|0|0| 0|0|0|0|0|0|
|virus_Jun22.50v_15738|1| 0|0|0|0|0|0|0|0|0|0| 0|0|0|0|0|0|0|1| 0|0|0|0|0|0|0|0|0|0| 0|0|0|0|0|0|0|0|0|0| 0|0|0|0|0|0|0|0|0|0| 0|0|0|0|0|0|
|virus_Jun22.50v_16087|1| 0|0|0|0|0|0|0|0|0|0| 0|0|0|0|0|0|0|1| 0|0|0|0|0|0|0|0|0|0| 0|0|0|0|0|0|0|0|0|0| 0|0|0|0|0|0|0|0|0|0| 0|0|0|0|0|0|
|virus_Jun22.50v_19633|1| 0|0|0|0|0|0|0|0|0|0| 0|0|0|0|0|0|0|1| 0|0|0|0|0|0|0|0|0|0| 0|0|0|0|0|0|0|0|0|0| 0|0|0|0|0|0|0|0|0|0| 0|0|0|0|0|0|
|virus_Jun22.50v_20126|1| 0|0|0|0|0|0|0|0|0|0| 0|0|0|0|0|0|0|1| 0|0|0|0|0|0|0|0|0|0| 0|0|0|0|0|0|0|0|0|0| 0|0|0|0|0|0|0|0|0|0| 0|0|0|0|0|0|
|virus_Jun22.50v_25683|1| 0|0|0|0|0|0|0|0|0|0| 0|0|0|0|0|0|0|1| 0|0|0|0|0|0|0|0|0|0| 0|0|0|0|0|0|0|0|0|0| 0|0|0|0|0|0|0|0|0|0| 0|0|0|0|0|0|
|virus_Jun22.50v_27917|1| 0|0|0|0|0|0|0|0|0|0| 0|0|0|0|0|0|0|1| 0|0|0|0|0|0|0|0|0|0| 0|0|0|0|0|0|0|0|0|0| 0|0|0|0|0|0|0|0|0|0| 0|0|0|0|0|0|
|virus_Jun22.50v_34629|1| 0|0|0|0|0|0|0|0|0|0| 0|0|0|0|0|0|0|1| 0|0|0|0|0|0|0|0|0|0| 0|0|0|0|0|0|0|0|0|0| 0|0|0|0|0|0|0|0|0|0| 0|0|0|0|0|0|
|virus_Jun22.50v_35523|1| 0|0|0|0|0|0|0|0|0|0| 0|0|0|0|0|0|0|1| 0|0|0|0|0|0|0|0|0|0| 0|0|0|0|0|0|0|0|0|0| 0|0|0|0|0|0|0|0|0|0| 0|0|0|0|0|0|
|virus_Jun22.50v_36714|1| 0|0|0|0|0|0|0|0|0|0| 0|0|0|0|0|0|0|1| 0|0|0|0|0|0|0|0|0|0| 0|0|0|0|0|0|0|0|0|0| 0|0|0|0|0|0|0|0|0|0| 0|0|0|0|0|0|
|virus_Jun22.50v_38051|1| 0|0|0|0|0|0|0|0|0|0| 0|0|0|0|0|0|0|1| 0|0|0|0|0|0|0|0|0|0| 0|0|0|0|0|0|0|0|0|0| 0|0|0|0|0|0|0|0|0|0| 0|0|0|0|0|0|
|virus_Jun22.75v_223|1| 0|0|0|0|0|0|0|0|0|0| 0|0|0|0|0|0|0|0|0|1| 0|0|0|0|0|0|0|0|0|0| 0|0|0|0|0|0|0|0|0|0| 0|0|0|0|0|0|0|0|0|0| 0|0|0|0|
|virus_Jun22.75v_12929|1| 0|0|0|0|0|0|0|0|0|0| 0|0|0|0|0|0|0|0|0|1| 0|0|0|0|0|0|0|0|0|0| 0|0|0|0|0|0|0|0|0|0| 0|0|0|0|0|0|0|0|0|0| 0|0|0|0|
|virus_Jun22.75v_20691|1| 0|0|0|0|0|0|0|0|0|0| 0|0|0|0|0|0|0|0|0|1| 0|0|0|0|0|0|0|0|0|0| 0|0|0|0|0|0|0|0|0|0| 0|0|0|0|0|0|0|0|0|0| 0|0|0|0|
|virus_Jun22.75v_31125|1| 0|0|0|0|0|0|0|0|0|0| 0|0|0|0|0|0|0|0|0|1| 0|0|0|0|0|0|0|0|0|0| 0|0|0|0|0|0|0|0|0|0| 0|0|0|0|0|0|0|0|0|0| 0|0|0|0|
|virus_Jun22.75v_31544|1| 0|0|0|0|0|0|0|0|0|0| 0|0|0|0|0|0|0|0|0|1| 0|0|0|0|0|0|0|0|0|0| 0|0|0|0|0|0|0|0|0|0| 0|0|0|0|0|0|0|0|0|0| 0|0|0|0|
|virus_Jun22.75v_33788|1| 0|0|0|0|0|0|0|0|0|0| 0|0|0|0|0|0|0|0|0|1| 0|0|0|0|0|0|0|0|0|0| 0|0|0|0|0|0|0|0|0|0| 0|0|0|0|0|0|0|0|0|0| 0|0|0|0|
|virus_Jun22.75v_35460|1| 0|0|0|0|0|0|0|0|0|0| 0|0|0|0|0|0|0|0|0|1| 0|0|0|0|0|0|0|0|0|0| 0|0|0|0|0|0|0|0|0|0| 0|0|0|0|0|0|0|0|0|0| 0|0|0|0|
|virus_Jun22.75v_36894|1| 0|0|0|0|0|0|0|0|0|0| 0|0|0|0|0|0|0|0|0|1| 0|0|0|0|0|0|0|0|0|0| 0|0|0|0|0|0|0|0|0|0| 0|0|0|0|0|0|0|0|0|0| 0|0|0|0|
|virus_Jun22.75v_39617|1| 0|0|0|0|0|0|0|0|0|0| 0|0|0|0|0|0|0|0|0|1| 0|0|0|0|0|0|0|0|0|0| 0|0|0|0|0|0|0|0|0|0| 0|0|0|0|0|0|0|0|0|0| 0|0|0|0|
|virus_May12.5v_7184|1| 0|1| 0|0|0|0|0|0|0|0|0|0| 0|0|0|0|0|0|0|0|0|0| 0|0|0|0|0|0|0|0|0|0| 0|0|0|0|0|0|0|0|0|0| 0|0|0|0|0|0|0|0|0|0| 0|0|
|virus_May12.5v_8804|1| 0|1| 0|0|0|0|0|0|0|0|0|0| 0|0|0|0|0|0|0|0|0|0| 0|0|0|0|0|0|0|0|0|0| 0|0|0|0|0|0|0|0|0|0| 0|0|0|0|0|0|0|0|0|0| 0|0|
|virus_May12.5v_21886|1| 0|1| 0|0|0|0|0|0|0|0|0|0| 0|0|0|0|0|0|0|0|0|0| 0|0|0|0|0|0|0|0|0|0| 0|0|0|0|0|0|0|0|0|0| 0|0|0|0|0|0|0|0|0|0| 0|0|
|virus_May12.5v_34493|1| 0|1| 0|0|0|0|0|0|0|0|0|0| 0|0|0|0|0|0|0|0|0|0| 0|0|0|0|0|0|0|0|0|0| 0|0|0|0|0|0|0|0|0|0| 0|0|0|0|0|0|0|0|0|0| 0|0|
|virus_May12.5v_39318|1| 0|1| 0|0|0|0|0|0|0|0|0|0| 0|0|0|0|0|0|0|0|0|0| 0|0|0|0|0|0|0|0|0|0| 0|0|0|0|0|0|0|0|0|0| 0|0|0|0|0|0|0|0|0|0| 0|0|
|virus_May12.5v_40344|1| 0|1| 0|0|0|0|0|0|0|0|0|0| 0|0|0|0|0|0|0|0|0|0| 0|0|0|0|0|0|0|0|0|0| 0|0|0|0|0|0|0|0|0|0| 0|0|0|0|0|0|0|0|0|0| 0|0|
|virus_May12.25v_6074|1| 0|0|0|1| 0|0|0|0|0|0|0|0|0|0| 0|0|0|0|0|0|0|0|0|0| 0|0|0|0|0|0|0|0|0|0| 0|0|0|0|0|0|0|0|0|0| 0|0|0|0|0|0|0|0|0|0|
|virus_May12.25v_13980|1| 0|0|0|1| 0|0|0|0|0|0|0|0|0|0| 0|0|0|0|0|0|0|0|0|0| 0|0|0|0|0|0|0|0|0|0| 0|0|0|0|0|0|0|0|0|0| 0|0|0|0|0|0|0|0|0|0|
|virus_May12.25v_18305|1| 0|0|0|1| 0|0|0|0|0|0|0|0|0|0| 0|0|0|0|0|0|0|0|0|0| 0|0|0|0|0|0|0|0|0|0| 0|0|0|0|0|0|0|0|0|0| 0|0|0|0|0|0|0|0|0|0|
|virus_May12.25v_20845|1| 0|0|0|1| 0|0|0|0|0|0|0|0|0|0| 0|0|0|0|0|0|0|0|0|0| 0|0|0|0|0|0|0|0|0|0| 0|0|0|0|0|0|0|0|0|0| 0|0|0|0|0|0|0|0|0|0|
|virus_May12.25v_22797|1| 0|0|0|1| 0|0|0|0|0|0|0|0|0|0| 0|0|0|0|0|0|0|0|0|0| 0|0|0|0|0|0|0|0|0|0| 0|0|0|0|0|0|0|0|0|0| 0|0|0|0|0|0|0|0|0|0|
|virus_May12.25v_33105|1| 0|0|0|1| 0|0|0|0|0|0|0|0|0|0| 0|0|0|0|0|0|0|0|0|0| 0|0|0|0|0|0|0|0|0|0| 0|0|0|0|0|0|0|0|0|0| 0|0|0|0|0|0|0|0|0|0|
|virus_May12.50v_11507|1| 0|0|0|0|0|1| 0|0|0|0|0|0|0|0|0|0| 0|0|0|0|0|0|0|0|0|0| 0|0|0|0|0|0|0|0|0|0| 0|0|0|0|0|0|0|0|0|0| 0|0|0|0|0|0|0|0|
|virus_May12.50v_13686|1| 0|0|0|0|0|1| 0|0|0|0|0|0|0|0|0|0| 0|0|0|0|0|0|0|0|0|0| 0|0|0|0|0|0|0|0|0|0| 0|0|0|0|0|0|0|0|0|0| 0|0|0|0|0|0|0|0|
|virus_May12.50v_14520|1| 0|0|0|0|0|1| 0|0|0|0|0|0|0|0|0|0| 0|0|0|0|0|0|0|0|0|0| 0|0|0|0|0|0|0|0|0|0| 0|0|0|0|0|0|0|0|0|0| 0|0|0|0|0|0|0|0|
|virus_May12.50v_16193|1| 0|0|0|0|0|1| 0|0|0|0|0|0|0|0|0|0| 0|0|0|0|0|0|0|0|0|0| 0|0|0|0|0|0|0|0|0|0| 0|0|0|0|0|0|0|0|0|0| 0|0|0|0|0|0|0|0|
|virus_May12.50v_18124|1| 0|0|0|0|0|1| 0|0|0|0|0|0|0|0|0|0| 0|0|0|0|0|0|0|0|0|0| 0|0|0|0|0|0|0|0|0|0| 0|0|0|0|0|0|0|0|0|0| 0|0|0|0|0|0|0|0|
|virus_May12.50v_20309|1| 0|0|0|0|0|1| 0|0|0|0|0|0|0|0|0|0| 0|0|0|0|0|0|0|0|0|0| 0|0|0|0|0|0|0|0|0|0| 0|0|0|0|0|0|0|0|0|0| 0|0|0|0|0|0|0|0|
|virus_May12.50v_21848|1| 0|0|0|0|0|1| 0|0|0|0|0|0|0|0|0|0| 0|0|0|0|0|0|0|0|0|0| 0|0|0|0|0|0|0|0|0|0| 0|0|0|0|0|0|0|0|0|0| 0|0|0|0|0|0|0|0|
|virus_May12.50v_22108|1| 0|0|0|0|0|1| 0|0|0|0|0|0|0|0|0|0| 0|0|0|0|0|0|0|0|0|0| 0|0|0|0|0|0|0|0|0|0| 0|0|0|0|0|0|0|0|0|0| 0|0|0|0|0|0|0|0|
|virus_May12.50v_22484|1| 0|0|0|0|0|1| 0|0|0|0|0|0|0|0|0|0| 0|0|0|0|0|0|0|0|0|0| 0|0|0|0|0|0|0|0|0|0| 0|0|0|0|0|0|0|0|0|0| 0|0|0|0|0|0|0|0|
|virus_May12.50v_23050|1| 0|0|0|0|0|1| 0|0|0|0|0|0|0|0|0|0| 0|0|0|0|0|0|0|0|0|0| 0|0|0|0|0|0|0|0|0|0| 0|0|0|0|0|0|0|0|0|0| 0|0|0|0|0|0|0|0|
|virus_May12.50v_24689|1| 0|0|0|0|0|1| 0|0|0|0|0|0|0|0|0|0| 0|0|0|0|0|0|0|0|0|0| 0|0|0|0|0|0|0|0|0|0| 0|0|0|0|0|0|0|0|0|0| 0|0|0|0|0|0|0|0|
|virus_May12.50v_24998|1| 0|0|0|0|0|1| 0|0|0|0|0|0|0|0|0|0| 0|0|0|0|0|0|0|0|0|0| 0|0|0|0|0|0|0|0|0|0| 0|0|0|0|0|0|0|0|0|0| 0|0|0|0|0|0|0|0|
|virus_May12.50v_28989|1| 0|0|0|0|0|1| 0|0|0|0|0|0|0|0|0|0| 0|0|0|0|0|0|0|0|0|0| 0|0|0|0|0|0|0|0|0|0| 0|0|0|0|0|0|0|0|0|0| 0|0|0|0|0|0|0|0|
|virus_May12.50v_37033|1| 0|0|0|0|0|1| 0|0|0|0|0|0|0|0|0|0| 0|0|0|0|0|0|0|0|0|0| 0|0|0|0|0|0|0|0|0|0| 0|0|0|0|0|0|0|0|0|0| 0|0|0|0|0|0|0|0|
|virus_May12.50v_38219|1| 0|0|0|0|0|1| 0|0|0|0|0|0|0|0|0|0| 0|0|0|0|0|0|0|0|0|0| 0|0|0|0|0|0|0|0|0|0| 0|0|0|0|0|0|0|0|0|0| 0|0|0|0|0|0|0|0|
|virus_May12.50v_40959|1| 0|0|0|0|0|1| 0|0|0|0|0|0|0|0|0|0| 0|0|0|0|0|0|0|0|0|0| 0|0|0|0|0|0|0|0|0|0| 0|0|0|0|0|0|0|0|0|0| 0|0|0|0|0|0|0|0|
|virus_May12.75v_5824|1| 0|0|0|0|0|0|0|1| 0|0|0|0|0|0|0|0|0|0| 0|0|0|0|0|0|0|0|0|0| 0|0|0|0|0|0|0|0|0|0| 0|0|0|0|0|0|0|0|0|0| 0|0|0|0|0|0|
|virus_May12.75v_7621|1| 0|0|0|0|0|0|0|1| 0|0|0|0|0|0|0|0|0|0| 0|0|0|0|0|0|0|0|0|0| 0|0|0|0|0|0|0|0|0|0| 0|0|0|0|0|0|0|0|0|0| 0|0|0|0|0|0|
|virus_May12.75v_8932|1| 0|0|0|0|0|0|0|1| 0|0|0|0|0|0|0|0|0|0| 0|0|0|0|0|0|0|0|0|0| 0|0|0|0|0|0|0|0|0|0| 0|0|0|0|0|0|0|0|0|0| 0|0|0|0|0|0|
|virus_May12.75v_17207|1| 0|0|0|0|0|0|0|1| 0|0|0|0|0|0|0|0|0|0| 0|0|0|0|0|0|0|0|0|0| 0|0|0|0|0|0|0|0|0|0| 0|0|0|0|0|0|0|0|0|0| 0|0|0|0|0|0|
|virus_May12.75v_19288|1| 0|0|0|0|0|0|0|1| 0|0|0|0|0|0|0|0|0|0| 0|0|0|0|0|0|0|0|0|0| 0|0|0|0|0|0|0|0|0|0| 0|0|0|0|0|0|0|0|0|0| 0|0|0|0|0|0|
|virus_May12.75v_20779|1| 0|0|0|0|0|0|0|1| 0|0|0|0|0|0|0|0|0|0| 0|0|0|0|0|0|0|0|0|0| 0|0|0|0|0|0|0|0|0|0| 0|0|0|0|0|0|0|0|0|0| 0|0|0|0|0|0|
|virus_May12.75v_26924|1| 0|0|0|0|0|0|0|1| 0|0|0|0|0|0|0|0|0|0| 0|0|0|0|0|0|0|0|0|0| 0|0|0|0|0|0|0|0|0|0| 0|0|0|0|0|0|0|0|0|0| 0|0|0|0|0|0|
|virus_Sep8.0v_344|1| 0|0|0|0|0|0|0|0|0|0| 0|0|0|0|0|0|0|0|0|0| 0|0|0|0|0|0|0|0|0|0| 0|0|0|0|0|0|0|0|0|0| 0|0|0|0|0|0|0|1| 0|0|0|0|0|0|
|virus_Sep8.0v_4929|1| 0|0|0|0|0|0|0|0|0|0| 0|0|0|0|0|0|0|0|0|0| 0|0|0|0|0|0|0|0|0|0| 0|0|0|0|0|0|0|0|0|0| 0|0|0|0|0|0|0|1| 0|0|0|0|0|0|
|virus_Sep8.0v_6668|1| 0|0|0|0|0|0|0|0|0|0| 0|0|0|0|0|0|0|0|0|0| 0|0|0|0|0|0|0|0|0|0| 0|0|0|0|0|0|0|0|0|0| 0|0|0|0|0|0|0|1| 0|0|0|0|0|0|
|virus_Sep8.0v_8381|1| 0|0|0|0|0|0|0|0|0|0| 0|0|0|0|0|0|0|0|0|0| 0|0|0|0|0|0|0|0|0|0| 0|0|0|0|0|0|0|0|0|0| 0|0|0|0|0|0|0|1| 0|0|0|0|0|0|
|virus_Sep8.0v_14734|1| 0|0|0|0|0|0|0|0|0|0| 0|0|0|0|0|0|0|0|0|0| 0|0|0|0|0|0|0|0|0|0| 0|0|0|0|0|0|0|0|0|0| 0|0|0|0|0|0|0|1| 0|0|0|0|0|0|
|virus_Sep8.0v_15059|1| 0|0|0|0|0|0|0|0|0|0| 0|0|0|0|0|0|0|0|0|0| 0|0|0|0|0|0|0|0|0|0| 0|0|0|0|0|0|0|0|0|0| 0|0|0|0|0|0|0|1| 0|0|0|0|0|0|
|virus_Sep8.0v_15617|1| 0|0|0|0|0|0|0|0|0|0| 0|0|0|0|0|0|0|0|0|0| 0|0|0|0|0|0|0|0|0|0| 0|0|0|0|0|0|0|0|0|0| 0|0|0|0|0|0|0|1| 0|0|0|0|0|0|
|virus_Sep8.0v_16579|1| 0|0|0|0|0|0|0|0|0|0| 0|0|0|0|0|0|0|0|0|0| 0|0|0|0|0|0|0|0|0|0| 0|0|0|0|0|0|0|0|0|0| 0|0|0|0|0|0|0|1| 0|0|0|0|0|0|
|virus_Sep8.0v_17533|1| 0|0|0|0|0|0|0|0|0|0| 0|0|0|0|0|0|0|0|0|0| 0|0|0|0|0|0|0|0|0|0| 0|0|0|0|0|0|0|0|0|0| 0|0|0|0|0|0|0|1| 0|0|0|0|0|0|
|virus_Sep8.5v_4513|1| 0|0|0|0|0|0|0|0|0|0| 0|0|0|0|0|0|0|0|0|0| 0|0|0|0|0|0|0|0|0|0| 0|0|0|0|0|0|0|0|0|0| 0|0|0|0|0|0|0|0|0|1| 0|0|0|0|
|virus_Sep8.5v_15177|1| 0|0|0|0|0|0|0|0|0|0| 0|0|0|0|0|0|0|0|0|0| 0|0|0|0|0|0|0|0|0|0| 0|0|0|0|0|0|0|0|0|0| 0|0|0|0|0|0|0|0|0|1| 0|0|0|0|
|virus_Sep8.5v_19322|1| 0|0|0|0|0|0|0|0|0|0| 0|0|0|0|0|0|0|0|0|0| 0|0|0|0|0|0|0|0|0|0| 0|0|0|0|0|0|0|0|0|0| 0|0|0|0|0|0|0|0|0|1| 0|0|0|0|
|virus_Sep8.5v_19493|1| 0|0|0|0|0|0|0|0|0|0| 0|0|0|0|0|0|0|0|0|0| 0|0|0|0|0|0|0|0|0|0| 0|0|0|0|0|0|0|0|0|0| 0|0|0|0|0|0|0|0|0|1| 0|0|0|0|
|virus_Sep8.25v_3482|1| 0|0|0|0|0|0|0|0|0|0| 0|0|0|0|0|0|0|0|0|0| 0|0|0|0|0|0|0|0|0|0| 0|0|0|0|0|0|0|0|0|0| 0|0|0|0|0|0|0|0|0|0| 0|1| 0|0|
|virus_Sep8.25v_8066|1| 0|0|0|0|0|0|0|0|0|0| 0|0|0|0|0|0|0|0|0|0| 0|0|0|0|0|0|0|0|0|0| 0|0|0|0|0|0|0|0|0|0| 0|0|0|0|0|0|0|0|0|0| 0|1| 0|0|
|virus_Sep8.25v_10270|1| 0|0|0|0|0|0|0|0|0|0| 0|0|0|0|0|0|0|0|0|0| 0|0|0|0|0|0|0|0|0|0| 0|0|0|0|0|0|0|0|0|0| 0|0|0|0|0|0|0|0|0|0| 0|1| 0|0|
|virus_Sep8.25v_13433|1| 0|0|0|0|0|0|0|0|0|0| 0|0|0|0|0|0|0|0|0|0| 0|0|0|0|0|0|0|0|0|0| 0|0|0|0|0|0|0|0|0|0| 0|0|0|0|0|0|0|0|0|0| 0|1| 0|0|
|virus_Sep8.25v_15394|1| 0|0|0|0|0|0|0|0|0|0| 0|0|0|0|0|0|0|0|0|0| 0|0|0|0|0|0|0|0|0|0| 0|0|0|0|0|0|0|0|0|0| 0|0|0|0|0|0|0|0|0|0| 0|1| 0|0|
|virus_Sep8.50v_1937|1| 0|0|0|0|0|0|0|0|0|0| 0|0|0|0|0|0|0|0|0|0| 0|0|0|0|0|0|0|0|0|0| 0|0|0|0|0|0|0|0|0|0| 0|0|0|0|0|0|0|0|0|0| 0|0|0|1|
|virus_Sep8.50v_4736|1| 0|0|0|0|0|0|0|0|0|0| 0|0|0|0|0|0|0|0|0|0| 0|0|0|0|0|0|0|0|0|0| 0|0|0|0|0|0|0|0|0|0| 0|0|0|0|0|0|0|0|0|0| 0|0|0|1|
|virus_Sep8.50v_8381|1| 0|0|0|0|0|0|0|0|0|0| 0|0|0|0|0|0|0|0|0|0| 0|0|0|0|0|0|0|0|0|0| 0|0|0|0|0|0|0|0|0|0| 0|0|0|0|0|0|0|0|0|0| 0|0|0|1|
|virus_Sep8.50v_11308|1| 0|0|0|0|0|0|0|0|0|0| 0|0|0|0|0|0|0|0|0|0| 0|0|0|0|0|0|0|0|0|0| 0|0|0|0|0|0|0|0|0|0| 0|0|0|0|0|0|0|0|0|0| 0|0|0|1|
|virus_Sep8.50v_12759|1| 0|0|0|0|0|0|0|0|0|0| 0|0|0|0|0|0|0|0|0|0| 0|0|0|0|0|0|0|0|0|0| 0|0|0|0|0|0|0|0|0|0| 0|0|0|0|0|0|0|0|0|0| 0|0|0|1|
|virus_Sep8.50v_17838|1| 0|0|0|0|0|0|0|0|0|0| 0|0|0|0|0|0|0|0|0|0| 0|0|0|0|0|0|0|0|0|0| 0|0|0|0|0|0|0|0|0|0| 0|0|0|0|0|0|0|0|0|0| 0|0|0|1|
|virus_Sep8.50v_18193|1| 0|0|0|0|0|0|0|0|0|0| 0|0|0|0|0|0|0|0|0|0| 0|0|0|0|0|0|0|0|0|0| 0|0|0|0|0|0|0|0|0|0| 0|0|0|0|0|0|0|0|0|0| 0|0|0|1|
|virus_Sep8.50v_19579|1| 0|0|0|0|0|0|0|0|0|0| 0|0|0|0|0|0|0|0|0|0| 0|0|0|0|0|0|0|0|0|0| 0|0|0|0|0|0|0|0|0|0| 0|0|0|0|0|0|0|0|0|0| 0|0|0|1|
|virus_Sep8.50v_22677|1| 0|0|0|0|0|0|0|0|0|0| 0|0|0|0|0|0|0|0|0|0| 0|0|0|0|0|0|0|0|0|0| 0|0|0|0|0|0|0|0|0|0| 0|0|0|0|0|0|0|0|0|0| 0|0|0|1|
|virus_Sep8.50v_23280|1| 0|0|0|0|0|0|0|0|0|0| 0|0|0|0|0|0|0|0|0|0| 0|0|0|0|0|0|0|0|0|0| 0|0|0|0|0|0|0|0|0|0| 0|0|0|0|0|0|0|0|0|0| 0|0|0|1|
|virus_Sep8.50v_24505|1| 0|0|0|0|0|0|0|0|0|0| 0|0|0|0|0|0|0|0|0|0| 0|0|0|0|0|0|0|0|0|0| 0|0|0|0|0|0|0|0|0|0| 0|0|0|0|0|0|0|0|0|0| 0|0|0|1|
|virus_Sep8.50v_25933|1| 0|0|0|0|0|0|0|0|0|0| 0|0|0|0|0|0|0|0|0|0| 0|0|0|0|0|0|0|0|0|0| 0|0|0|0|0|0|0|0|0|0| 0|0|0|0|0|0|0|0|0|0| 0|0|0|1|
|virus_Sep8.50v_27680|1| 0|0|0|0|0|0|0|0|0|0| 0|0|0|0|0|0|0|0|0|0| 0|0|0|0|0|0|0|0|0|0| 0|0|0|0|0|0|0|0|0|0| 0|0|0|0|0|0|0|0|0|0| 0|0|0|1|
|virus_Sep8.50v_28215|1| 0|0|0|0|0|0|0|0|0|0| 0|0|0|0|0|0|0|0|0|0| 0|0|0|0|0|0|0|0|0|0| 0|0|0|0|0|0|0|0|0|0| 0|0|0|0|0|0|0|0|0|0| 0|0|0|1|
|virus_Sep8.50v_36519|1| 0|0|0|0|0|0|0|0|0|0| 0|0|0|0|0|0|0|0|0|0| 0|0|0|0|0|0|0|0|0|0| 0|0|0|0|0|0|0|0|0|0| 0|0|0|0|0|0|0|0|0|0| 0|0|0|1|
|virus_Sep8.50v_39051|1| 0|0|0|0|0|0|0|0|0|0| 0|0|0|0|0|0|0|0|0|0| 0|0|0|0|0|0|0|0|0|0| 0|0|0|0|0|0|0|0|0|0| 0|0|0|0|0|0|0|0|0|0| 0|0|0|1|
|bacteria_Jul11d0b_8428|1| 0|0|0|0|0|0|0|0|0|0| 0|0|0|0|0|0|0|0|0|0| 1| 0|0|0|0|0|0|0|0|0|0| 0|0|0|0|0|0|0|0|0|0| 0|0|0|0|0|0|0|0|0|0| 0|0|0|
|bacteria_Jul11d0b_10918|1| 0|0|0|0|0|0|0|0|0|0| 0|0|0|0|0|0|0|0|0|0| 1| 0|0|0|0|0|0|0|0|0|0| 0|0|0|0|0|0|0|0|0|0| 0|0|0|0|0|0|0|0|0|0| 0|0|0|
|bacteria_Jul11d0b_12494|1| 0|0|0|0|0|0|0|0|0|0| 0|0|0|0|0|0|0|0|0|0| 1| 0|0|0|0|0|0|0|0|0|0| 0|0|0|0|0|0|0|0|0|0| 0|0|0|0|0|0|0|0|0|0| 0|0|0|
|bacteria_Jul11d0b_22352|1| 0|0|0|0|0|0|0|0|0|0| 0|0|0|0|0|0|0|0|0|0| 1| 0|0|0|0|0|0|0|0|0|0| 0|0|0|0|0|0|0|0|0|0| 0|0|0|0|0|0|0|0|0|0| 0|0|0|
|bacteria_Jul11d0b_24258|1| 0|0|0|0|0|0|0|0|0|0| 0|0|0|0|0|0|0|0|0|0| 1| 0|0|0|0|0|0|0|0|0|0| 0|0|0|0|0|0|0|0|0|0| 0|0|0|0|0|0|0|0|0|0| 0|0|0|
|bacteria_Jul11d0b_24428|1| 0|0|0|0|0|0|0|0|0|0| 0|0|0|0|0|0|0|0|0|0| 1| 0|0|0|0|0|0|0|0|0|0| 0|0|0|0|0|0|0|0|0|0| 0|0|0|0|0|0|0|0|0|0| 0|0|0|
|bacteria_Jul11d0b_38423|1| 0|0|0|0|0|0|0|0|0|0| 0|0|0|0|0|0|0|0|0|0| 1| 0|0|0|0|0|0|0|0|0|0| 0|0|0|0|0|0|0|0|0|0| 0|0|0|0|0|0|0|0|0|0| 0|0|0|
|bacteria_Jul11d0b_41022|1| 0|0|0|0|0|0|0|0|0|0| 0|0|0|0|0|0|0|0|0|0| 1| 0|0|0|0|0|0|0|0|0|0| 0|0|0|0|0|0|0|0|0|0| 0|0|0|0|0|0|0|0|0|0| 0|0|0|
|bacteria_Jul11d0b_41903|1| 0|0|0|0|0|0|0|0|0|0| 0|0|0|0|0|0|0|0|0|0| 1| 0|0|0|0|0|0|0|0|0|0| 0|0|0|0|0|0|0|0|0|0| 0|0|0|0|0|0|0|0|0|0| 0|0|0|
|bacteria_Jul11d0b_54474|1| 0|0|0|0|0|0|0|0|0|0| 0|0|0|0|0|0|0|0|0|0| 1| 0|0|0|0|0|0|0|0|0|0| 0|0|0|0|0|0|0|0|0|0| 0|0|0|0|0|0|0|0|0|0| 0|0|0|
|bacteria_Jul11d0b_58257|1| 0|0|0|0|0|0|0|0|0|0| 0|0|0|0|0|0|0|0|0|0| 1| 0|0|0|0|0|0|0|0|0|0| 0|0|0|0|0|0|0|0|0|0| 0|0|0|0|0|0|0|0|0|0| 0|0|0|
|bacteria_Jul11d0b_59524|1| 0|0|0|0|0|0|0|0|0|0| 0|0|0|0|0|0|0|0|0|0| 1| 0|0|0|0|0|0|0|0|0|0| 0|0|0|0|0|0|0|0|0|0| 0|0|0|0|0|0|0|0|0|0| 0|0|0|
|bacteria_Jul11d0b_61002|1| 0|0|0|0|0|0|0|0|0|0| 0|0|0|0|0|0|0|0|0|0| 1| 0|0|0|0|0|0|0|0|0|0| 0|0|0|0|0|0|0|0|0|0| 0|0|0|0|0|0|0|0|0|0| 0|0|0|
|bacteria_Jul11d0b_72150|1| 0|0|0|0|0|0|0|0|0|0| 0|0|0|0|0|0|0|0|0|0| 1| 0|0|0|0|0|0|0|0|0|0| 0|0|0|0|0|0|0|0|0|0| 0|0|0|0|0|0|0|0|0|0| 0|0|0|
|bacteria_Aug5d0b_48375|1| 0|0|0|0|0|0|0|0|0|0| 0|0|0|0|0|0|0|0|0|0| 0|0|0|0|0|0|0|0|0|0| 0|0|0|0|0|0|1| 0|0|0|0|0|0|0|0|0|0| 0|0|0|0|0|0|0|
|bacteria_Aug5d5b_5022|1| 0|0|0|0|0|0|0|0|0|0| 0|0|0|0|0|0|0|0|0|0| 0|0|0|0|0|0|0|0|0|0| 0|0|0|0|0|0|0|0|1| 0|0|0|0|0|0|0|0|0|0| 0|0|0|0|0|
|bacteria_Aug5d5b_7350|1| 0|0|0|0|0|0|0|0|0|0| 0|0|0|0|0|0|0|0|0|0| 0|0|0|0|0|0|0|0|0|0| 0|0|0|0|0|0|0|0|1| 0|0|0|0|0|0|0|0|0|0| 0|0|0|0|0|
|bacteria_Aug5d5b_36130|1| 0|0|0|0|0|0|0|0|0|0| 0|0|0|0|0|0|0|0|0|0| 0|0|0|0|0|0|0|0|0|0| 0|0|0|0|0|0|0|0|1| 0|0|0|0|0|0|0|0|0|0| 0|0|0|0|0|
|bacteria_Aug5d25b_1978|1| 0|0|0|0|0|0|0|0|0|0| 0|0|0|0|0|0|0|0|0|0| 0|0|0|0|0|0|0|0|0|0| 0|0|0|0|0|0|0|0|0|0| 1| 0|0|0|0|0|0|0|0|0|0| 0|0|0|
|bacteria_Aug5d25b_3474|1| 0|0|0|0|0|0|0|0|0|0| 0|0|0|0|0|0|0|0|0|0| 0|0|0|0|0|0|0|0|0|0| 0|0|0|0|0|0|0|0|0|0| 1| 0|0|0|0|0|0|0|0|0|0| 0|0|0|
|bacteria_Aug5d25b_6289|1| 0|0|0|0|0|0|0|0|0|0| 0|0|0|0|0|0|0|0|0|0| 0|0|0|0|0|0|0|0|0|0| 0|0|0|0|0|0|0|0|0|0| 1| 0|0|0|0|0|0|0|0|0|0| 0|0|0|
|bacteria_Aug5d25b_20212|1| 0|0|0|0|0|0|0|0|0|0| 0|0|0|0|0|0|0|0|0|0| 0|0|0|0|0|0|0|0|0|0| 0|0|0|0|0|0|0|0|0|0| 1| 0|0|0|0|0|0|0|0|0|0| 0|0|0|
|bacteria_Aug5d25b_22280|1| 0|0|0|0|0|0|0|0|0|0| 0|0|0|0|0|0|0|0|0|0| 0|0|0|0|0|0|0|0|0|0| 0|0|0|0|0|0|0|0|0|0| 1| 0|0|0|0|0|0|0|0|0|0| 0|0|0|
|bacteria_Aug5d25b_25496|1| 0|0|0|0|0|0|0|0|0|0| 0|0|0|0|0|0|0|0|0|0| 0|0|0|0|0|0|0|0|0|0| 0|0|0|0|0|0|0|0|0|0| 1| 0|0|0|0|0|0|0|0|0|0| 0|0|0|
|bacteria_Aug5d25b_31739|1| 0|0|0|0|0|0|0|0|0|0| 0|0|0|0|0|0|0|0|0|0| 0|0|0|0|0|0|0|0|0|0| 0|0|0|0|0|0|0|0|0|0| 1| 0|0|0|0|0|0|0|0|0|0| 0|0|0|
|bacteria_Aug5d25b_33783|1| 0|0|0|0|0|0|0|0|0|0| 0|0|0|0|0|0|0|0|0|0| 0|0|0|0|0|0|0|0|0|0| 0|0|0|0|0|0|0|0|0|0| 1| 0|0|0|0|0|0|0|0|0|0| 0|0|0|
|bacteria_Aug5d25b_40949|1| 0|0|0|0|0|0|0|0|0|0| 0|0|0|0|0|0|0|0|0|0| 0|0|0|0|0|0|0|0|0|0| 0|0|0|0|0|0|0|0|0|0| 1| 0|0|0|0|0|0|0|0|0|0| 0|0|0|
|bacteria_Aug5d25b_42824|1| 0|0|0|0|0|0|0|0|0|0| 0|0|0|0|0|0|0|0|0|0| 0|0|0|0|0|0|0|0|0|0| 0|0|0|0|0|0|0|0|0|0| 1| 0|0|0|0|0|0|0|0|0|0| 0|0|0|
|bacteria_Aug5d25b_46191|1| 0|0|0|0|0|0|0|0|0|0| 0|0|0|0|0|0|0|0|0|0| 0|0|0|0|0|0|0|0|0|0| 0|0|0|0|0|0|0|0|0|0| 1| 0|0|0|0|0|0|0|0|0|0| 0|0|0|
|bacteria_Aug5d25b_47332|1| 0|0|0|0|0|0|0|0|0|0| 0|0|0|0|0|0|0|0|0|0| 0|0|0|0|0|0|0|0|0|0| 0|0|0|0|0|0|0|0|0|0| 1| 0|0|0|0|0|0|0|0|0|0| 0|0|0|
|bacteria_Aug5d25b_47879|1| 0|0|0|0|0|0|0|0|0|0| 0|0|0|0|0|0|0|0|0|0| 0|0|0|0|0|0|0|0|0|0| 0|0|0|0|0|0|0|0|0|0| 1| 0|0|0|0|0|0|0|0|0|0| 0|0|0|
|bacteria_Aug5d25b_48501|1| 0|0|0|0|0|0|0|0|0|0| 0|0|0|0|0|0|0|0|0|0| 0|0|0|0|0|0|0|0|0|0| 0|0|0|0|0|0|0|0|0|0| 1| 0|0|0|0|0|0|0|0|0|0| 0|0|0|
|bacteria_Aug5d25b_49248|1| 0|0|0|0|0|0|0|0|0|0| 0|0|0|0|0|0|0|0|0|0| 0|0|0|0|0|0|0|0|0|0| 0|0|0|0|0|0|0|0|0|0| 1| 0|0|0|0|0|0|0|0|0|0| 0|0|0|
|bacteria_Aug5d25b_56458|1| 0|0|0|0|0|0|0|0|0|0| 0|0|0|0|0|0|0|0|0|0| 0|0|0|0|0|0|0|0|0|0| 0|0|0|0|0|0|0|0|0|0| 1| 0|0|0|0|0|0|0|0|0|0| 0|0|0|
|bacteria_Aug5d25b_59654|1| 0|0|0|0|0|0|0|0|0|0| 0|0|0|0|0|0|0|0|0|0| 0|0|0|0|0|0|0|0|0|0| 0|0|0|0|0|0|0|0|0|0| 1| 0|0|0|0|0|0|0|0|0|0| 0|0|0|
|bacteria_Aug5d25b_60503|1| 0|0|0|0|0|0|0|0|0|0| 0|0|0|0|0|0|0|0|0|0| 0|0|0|0|0|0|0|0|0|0| 0|0|0|0|0|0|0|0|0|0| 1| 0|0|0|0|0|0|0|0|0|0| 0|0|0|
|bacteria_Aug5d25b_62399|1| 0|0|0|0|0|0|0|0|0|0| 0|0|0|0|0|0|0|0|0|0| 0|0|0|0|0|0|0|0|0|0| 0|0|0|0|0|0|0|0|0|0| 1| 0|0|0|0|0|0|0|0|0|0| 0|0|0|
|bacteria_Aug5d25b_63089|1| 0|0|0|0|0|0|0|0|0|0| 0|0|0|0|0|0|0|0|0|0| 0|0|0|0|0|0|0|0|0|0| 0|0|0|0|0|0|0|0|0|0| 1| 0|0|0|0|0|0|0|0|0|0| 0|0|0|
|bacteria_Aug5d25b_66662|1| 0|0|0|0|0|0|0|0|0|0| 0|0|0|0|0|0|0|0|0|0| 0|0|0|0|0|0|0|0|0|0| 0|0|0|0|0|0|0|0|0|0| 1| 0|0|0|0|0|0|0|0|0|0| 0|0|0|
|bacteria_Aug5d25b_67028|1| 0|0|0|0|0|0|0|0|0|0| 0|0|0|0|0|0|0|0|0|0| 0|0|0|0|0|0|0|0|0|0| 0|0|0|0|0|0|0|0|0|0| 1| 0|0|0|0|0|0|0|0|0|0| 0|0|0|
|bacteria_Aug5d25b_72871|1| 0|0|0|0|0|0|0|0|0|0| 0|0|0|0|0|0|0|0|0|0| 0|0|0|0|0|0|0|0|0|0| 0|0|0|0|0|0|0|0|0|0| 1| 0|0|0|0|0|0|0|0|0|0| 0|0|0|
|bacteria_Aug5d50b_4420|1| 0|0|0|0|0|0|0|0|0|0| 0|0|0|0|0|0|0|0|0|0| 0|0|0|0|0|0|0|0|0|0| 0|0|0|0|0|0|0|0|0|0| 0|0|1| 0|0|0|0|0|0|0|0|0|0| 0|
|bacteria_Aug5d50b_6225|1| 0|0|0|0|0|0|0|0|0|0| 0|0|0|0|0|0|0|0|0|0| 0|0|0|0|0|0|0|0|0|0| 0|0|0|0|0|0|0|0|0|0| 0|0|1| 0|0|0|0|0|0|0|0|0|0| 0|
|bacteria_Aug5d50b_16661|1| 0|0|0|0|0|0|0|0|0|0| 0|0|0|0|0|0|0|0|0|0| 0|0|0|0|0|0|0|0|0|0| 0|0|0|0|0|0|0|0|0|0| 0|0|1| 0|0|0|0|0|0|0|0|0|0| 0|
|bacteria_Aug5d50b_33163|1| 0|0|0|0|0|0|0|0|0|0| 0|0|0|0|0|0|0|0|0|0| 0|0|0|0|0|0|0|0|0|0| 0|0|0|0|0|0|0|0|0|0| 0|0|1| 0|0|0|0|0|0|0|0|0|0| 0|
|bacteria_Aug5d50b_65429|1| 0|0|0|0|0|0|0|0|0|0| 0|0|0|0|0|0|0|0|0|0| 0|0|0|0|0|0|0|0|0|0| 0|0|0|0|0|0|0|0|0|0| 0|0|1| 0|0|0|0|0|0|0|0|0|0| 0|
|bacteria_Aug5d50b_70461|1| 0|0|0|0|0|0|0|0|0|0| 0|0|0|0|0|0|0|0|0|0| 0|0|0|0|0|0|0|0|0|0| 0|0|0|0|0|0|0|0|0|0| 0|0|1| 0|0|0|0|0|0|0|0|0|0| 0|
|bacteria_Aug5d75b_33872|1| 0|0|0|0|0|0|0|0|0|0| 0|0|0|0|0|0|0|0|0|0| 0|0|0|0|0|0|0|0|0|0| 0|0|0|0|0|0|0|0|0|0| 0|0|0|0|1| 0|0|0|0|0|0|0|0|0|
|bacteria_Aug5d75b_43197|1| 0|0|0|0|0|0|0|0|0|0| 0|0|0|0|0|0|0|0|0|0| 0|0|0|0|0|0|0|0|0|0| 0|0|0|0|0|0|0|0|0|0| 0|0|0|0|1| 0|0|0|0|0|0|0|0|0|
|bacteria_Jul11d5b_5193|1| 0|0|0|0|0|0|0|0|0|0| 0|0|0|0|0|0|0|0|0|0| 0|0|1| 0|0|0|0|0|0|0|0|0|0| 0|0|0|0|0|0|0|0|0|0| 0|0|0|0|0|0|0|0|0|0| 0|
|bacteria_Jul11d5b_11014|1| 0|0|0|0|0|0|0|0|0|0| 0|0|0|0|0|0|0|0|0|0| 0|0|1| 0|0|0|0|0|0|0|0|0|0| 0|0|0|0|0|0|0|0|0|0| 0|0|0|0|0|0|0|0|0|0| 0|
|bacteria_Jul11d5b_11355|1| 0|0|0|0|0|0|0|0|0|0| 0|0|0|0|0|0|0|0|0|0| 0|0|1| 0|0|0|0|0|0|0|0|0|0| 0|0|0|0|0|0|0|0|0|0| 0|0|0|0|0|0|0|0|0|0| 0|
|bacteria_Jul11d5b_12692|1| 0|0|0|0|0|0|0|0|0|0| 0|0|0|0|0|0|0|0|0|0| 0|0|1| 0|0|0|0|0|0|0|0|0|0| 0|0|0|0|0|0|0|0|0|0| 0|0|0|0|0|0|0|0|0|0| 0|
|bacteria_Jul11d5b_22577|1| 0|0|0|0|0|0|0|0|0|0| 0|0|0|0|0|0|0|0|0|0| 0|0|1| 0|0|0|0|0|0|0|0|0|0| 0|0|0|0|0|0|0|0|0|0| 0|0|0|0|0|0|0|0|0|0| 0|
|bacteria_Jul11d5b_24110|1| 0|0|0|0|0|0|0|0|0|0| 0|0|0|0|0|0|0|0|0|0| 0|0|1| 0|0|0|0|0|0|0|0|0|0| 0|0|0|0|0|0|0|0|0|0| 0|0|0|0|0|0|0|0|0|0| 0|
|bacteria_Jul11d5b_30495|1| 0|0|0|0|0|0|0|0|0|0| 0|0|0|0|0|0|0|0|0|0| 0|0|1| 0|0|0|0|0|0|0|0|0|0| 0|0|0|0|0|0|0|0|0|0| 0|0|0|0|0|0|0|0|0|0| 0|
|bacteria_Jul11d5b_30935|1| 0|0|0|0|0|0|0|0|0|0| 0|0|0|0|0|0|0|0|0|0| 0|0|1| 0|0|0|0|0|0|0|0|0|0| 0|0|0|0|0|0|0|0|0|0| 0|0|0|0|0|0|0|0|0|0| 0|
|bacteria_Jul11d5b_43132|1| 0|0|0|0|0|0|0|0|0|0| 0|0|0|0|0|0|0|0|0|0| 0|0|1| 0|0|0|0|0|0|0|0|0|0| 0|0|0|0|0|0|0|0|0|0| 0|0|0|0|0|0|0|0|0|0| 0|
|bacteria_Jul11d5b_44630|1| 0|0|0|0|0|0|0|0|0|0| 0|0|0|0|0|0|0|0|0|0| 0|0|1| 0|0|0|0|0|0|0|0|0|0| 0|0|0|0|0|0|0|0|0|0| 0|0|0|0|0|0|0|0|0|0| 0|
|bacteria_Jul11d5b_46404|1| 0|0|0|0|0|0|0|0|0|0| 0|0|0|0|0|0|0|0|0|0| 0|0|1| 0|0|0|0|0|0|0|0|0|0| 0|0|0|0|0|0|0|0|0|0| 0|0|0|0|0|0|0|0|0|0| 0|
|bacteria_Jul11d5b_48870|1| 0|0|0|0|0|0|0|0|0|0| 0|0|0|0|0|0|0|0|0|0| 0|0|1| 0|0|0|0|0|0|0|0|0|0| 0|0|0|0|0|0|0|0|0|0| 0|0|0|0|0|0|0|0|0|0| 0|
|bacteria_Jul11d25b_2700|1| 0|0|0|0|0|0|0|0|0|0| 0|0|0|0|0|0|0|0|0|0| 0|0|0|0|1| 0|0|0|0|0|0|0|0|0|0| 0|0|0|0|0|0|0|0|0|0| 0|0|0|0|0|0|0|0|0|
|bacteria_Jul11d25b_15213|1| 0|0|0|0|0|0|0|0|0|0| 0|0|0|0|0|0|0|0|0|0| 0|0|0|0|1| 0|0|0|0|0|0|0|0|0|0| 0|0|0|0|0|0|0|0|0|0| 0|0|0|0|0|0|0|0|0|
|bacteria_Jul11d25b_18916|1| 0|0|0|0|0|0|0|0|0|0| 0|0|0|0|0|0|0|0|0|0| 0|0|0|0|1| 0|0|0|0|0|0|0|0|0|0| 0|0|0|0|0|0|0|0|0|0| 0|0|0|0|0|0|0|0|0|
|bacteria_Jul11d25b_22182|1| 0|0|0|0|0|0|0|0|0|0| 0|0|0|0|0|0|0|0|0|0| 0|0|0|0|1| 0|0|0|0|0|0|0|0|0|0| 0|0|0|0|0|0|0|0|0|0| 0|0|0|0|0|0|0|0|0|
|bacteria_Jul11d25b_34247|1| 0|0|0|0|0|0|0|0|0|0| 0|0|0|0|0|0|0|0|0|0| 0|0|0|0|1| 0|0|0|0|0|0|0|0|0|0| 0|0|0|0|0|0|0|0|0|0| 0|0|0|0|0|0|0|0|0|
|bacteria_Jul11d25b_34379|1| 0|0|0|0|0|0|0|0|0|0| 0|0|0|0|0|0|0|0|0|0| 0|0|0|0|1| 0|0|0|0|0|0|0|0|0|0| 0|0|0|0|0|0|0|0|0|0| 0|0|0|0|0|0|0|0|0|
|bacteria_Jul11d50b_16078|1| 0|0|0|0|0|0|0|0|0|0| 0|0|0|0|0|0|0|0|0|0| 0|0|0|0|0|0|1| 0|0|0|0|0|0|0|0|0|0| 0|0|0|0|0|0|0|0|0|0| 0|0|0|0|0|0|0|
|bacteria_Jul11d50b_23104|1| 0|0|0|0|0|0|0|0|0|0| 0|0|0|0|0|0|0|0|0|0| 0|0|0|0|0|0|1| 0|0|0|0|0|0|0|0|0|0| 0|0|0|0|0|0|0|0|0|0| 0|0|0|0|0|0|0|
|bacteria_Jul11d50b_40497|1| 0|0|0|0|0|0|0|0|0|0| 0|0|0|0|0|0|0|0|0|0| 0|0|0|0|0|0|1| 0|0|0|0|0|0|0|0|0|0| 0|0|0|0|0|0|0|0|0|0| 0|0|0|0|0|0|0|
|bacteria_Jul11d50b_47920|1| 0|0|0|0|0|0|0|0|0|0| 0|0|0|0|0|0|0|0|0|0| 0|0|0|0|0|0|1| 0|0|0|0|0|0|0|0|0|0| 0|0|0|0|0|0|0|0|0|0| 0|0|0|0|0|0|0|
|bacteria_Jul11d50b_47967|1| 0|0|0|0|0|0|0|0|0|0| 0|0|0|0|0|0|0|0|0|0| 0|0|0|0|0|0|1| 0|0|0|0|0|0|0|0|0|0| 0|0|0|0|0|0|0|0|0|0| 0|0|0|0|0|0|0|
|bacteria_Jul11d50b_48240|1| 0|0|0|0|0|0|0|0|0|0| 0|0|0|0|0|0|0|0|0|0| 0|0|0|0|0|0|1| 0|0|0|0|0|0|0|0|0|0| 0|0|0|0|0|0|0|0|0|0| 0|0|0|0|0|0|0|
|bacteria_Jul11d75b_1312|1| 0|0|0|0|0|0|0|0|0|0| 0|0|0|0|0|0|0|0|0|0| 0|0|0|0|0|0|0|0|1| 0|0|0|0|0|0|0|0|0|0| 0|0|0|0|0|0|0|0|0|0| 0|0|0|0|0|
|bacteria_Jul11d75b_10964|1| 0|0|0|0|0|0|0|0|0|0| 0|0|0|0|0|0|0|0|0|0| 0|0|0|0|0|0|0|0|1| 0|0|0|0|0|0|0|0|0|0| 0|0|0|0|0|0|0|0|0|0| 0|0|0|0|0|
|bacteria_Jul11d75b_11531|1| 0|0|0|0|0|0|0|0|0|0| 0|0|0|0|0|0|0|0|0|0| 0|0|0|0|0|0|0|0|1| 0|0|0|0|0|0|0|0|0|0| 0|0|0|0|0|0|0|0|0|0| 0|0|0|0|0|
|bacteria_Jul11d75b_16422|1| 0|0|0|0|0|0|0|0|0|0| 0|0|0|0|0|0|0|0|0|0| 0|0|0|0|0|0|0|0|1| 0|0|0|0|0|0|0|0|0|0| 0|0|0|0|0|0|0|0|0|0| 0|0|0|0|0|
|bacteria_Jul11d75b_16739|1| 0|0|0|0|0|0|0|0|0|0| 0|0|0|0|0|0|0|0|0|0| 0|0|0|0|0|0|0|0|1| 0|0|0|0|0|0|0|0|0|0| 0|0|0|0|0|0|0|0|0|0| 0|0|0|0|0|
|bacteria_Jul11d75b_20620|1| 0|0|0|0|0|0|0|0|0|0| 0|0|0|0|0|0|0|0|0|0| 0|0|0|0|0|0|0|0|1| 0|0|0|0|0|0|0|0|0|0| 0|0|0|0|0|0|0|0|0|0| 0|0|0|0|0|
|bacteria_Jul11d75b_36557|1| 0|0|0|0|0|0|0|0|0|0| 0|0|0|0|0|0|0|0|0|0| 0|0|0|0|0|0|0|0|1| 0|0|0|0|0|0|0|0|0|0| 0|0|0|0|0|0|0|0|0|0| 0|0|0|0|0|
|bacteria_Jul11d75b_38056|1| 0|0|0|0|0|0|0|0|0|0| 0|0|0|0|0|0|0|0|0|0| 0|0|0|0|0|0|0|0|1| 0|0|0|0|0|0|0|0|0|0| 0|0|0|0|0|0|0|0|0|0| 0|0|0|0|0|
|bacteria_Jul11d75b_38237|1| 0|0|0|0|0|0|0|0|0|0| 0|0|0|0|0|0|0|0|0|0| 0|0|0|0|0|0|0|0|1| 0|0|0|0|0|0|0|0|0|0| 0|0|0|0|0|0|0|0|0|0| 0|0|0|0|0|
|bacteria_Jul11d75b_43467|1| 0|0|0|0|0|0|0|0|0|0| 0|0|0|0|0|0|0|0|0|0| 0|0|0|0|0|0|0|0|1| 0|0|0|0|0|0|0|0|0|0| 0|0|0|0|0|0|0|0|0|0| 0|0|0|0|0|
|bacteria_Jul11d75b_47780|1| 0|0|0|0|0|0|0|0|0|0| 0|0|0|0|0|0|0|0|0|0| 0|0|0|0|0|0|0|0|1| 0|0|0|0|0|0|0|0|0|0| 0|0|0|0|0|0|0|0|0|0| 0|0|0|0|0|
|bacteria_Jul11d75b_48402|1| 0|0|0|0|0|0|0|0|0|0| 0|0|0|0|0|0|0|0|0|0| 0|0|0|0|0|0|0|0|1| 0|0|0|0|0|0|0|0|0|0| 0|0|0|0|0|0|0|0|0|0| 0|0|0|0|0|
|bacteria_Jul11d75b_51001|1| 0|0|0|0|0|0|0|0|0|0| 0|0|0|0|0|0|0|0|0|0| 0|0|0|0|0|0|0|0|1| 0|0|0|0|0|0|0|0|0|0| 0|0|0|0|0|0|0|0|0|0| 0|0|0|0|0|
|bacteria_Jul27d5b_686|1| 0|0|0|0|0|0|0|0|0|0| 0|0|0|0|0|0|0|0|0|0| 0|0|0|0|0|0|0|0|0|0| 1| 0|0|0|0|0|0|0|0|0|0| 0|0|0|0|0|0|0|0|0|0| 0|0|0|
|bacteria_Jul27d5b_12734|1| 0|0|0|0|0|0|0|0|0|0| 0|0|0|0|0|0|0|0|0|0| 0|0|0|0|0|0|0|0|0|0| 1| 0|0|0|0|0|0|0|0|0|0| 0|0|0|0|0|0|0|0|0|0| 0|0|0|
|bacteria_Jul27d5b_14326|1| 0|0|0|0|0|0|0|0|0|0| 0|0|0|0|0|0|0|0|0|0| 0|0|0|0|0|0|0|0|0|0| 1| 0|0|0|0|0|0|0|0|0|0| 0|0|0|0|0|0|0|0|0|0| 0|0|0|
|bacteria_Jul27d5b_16153|1| 0|0|0|0|0|0|0|0|0|0| 0|0|0|0|0|0|0|0|0|0| 0|0|0|0|0|0|0|0|0|0| 1| 0|0|0|0|0|0|0|0|0|0| 0|0|0|0|0|0|0|0|0|0| 0|0|0|
|bacteria_Jul27d5b_19267|1| 0|0|0|0|0|0|0|0|0|0| 0|0|0|0|0|0|0|0|0|0| 0|0|0|0|0|0|0|0|0|0| 1| 0|0|0|0|0|0|0|0|0|0| 0|0|0|0|0|0|0|0|0|0| 0|0|0|
|bacteria_Jul27d5b_19286|1| 0|0|0|0|0|0|0|0|0|0| 0|0|0|0|0|0|0|0|0|0| 0|0|0|0|0|0|0|0|0|0| 1| 0|0|0|0|0|0|0|0|0|0| 0|0|0|0|0|0|0|0|0|0| 0|0|0|
|bacteria_Jul27d5b_21977|1| 0|0|0|0|0|0|0|0|0|0| 0|0|0|0|0|0|0|0|0|0| 0|0|0|0|0|0|0|0|0|0| 1| 0|0|0|0|0|0|0|0|0|0| 0|0|0|0|0|0|0|0|0|0| 0|0|0|
|bacteria_Jul27d5b_27108|1| 0|0|0|0|0|0|0|0|0|0| 0|0|0|0|0|0|0|0|0|0| 0|0|0|0|0|0|0|0|0|0| 1| 0|0|0|0|0|0|0|0|0|0| 0|0|0|0|0|0|0|0|0|0| 0|0|0|
|bacteria_Jul27d5b_28753|1| 0|0|0|0|0|0|0|0|0|0| 0|0|0|0|0|0|0|0|0|0| 0|0|0|0|0|0|0|0|0|0| 1| 0|0|0|0|0|0|0|0|0|0| 0|0|0|0|0|0|0|0|0|0| 0|0|0|
|bacteria_Jul27d5b_29495|1| 0|0|0|0|0|0|0|0|0|0| 0|0|0|0|0|0|0|0|0|0| 0|0|0|0|0|0|0|0|0|0| 1| 0|0|0|0|0|0|0|0|0|0| 0|0|0|0|0|0|0|0|0|0| 0|0|0|
|bacteria_Jul27d5b_32759|1| 0|0|0|0|0|0|0|0|0|0| 0|0|0|0|0|0|0|0|0|0| 0|0|0|0|0|0|0|0|0|0| 1| 0|0|0|0|0|0|0|0|0|0| 0|0|0|0|0|0|0|0|0|0| 0|0|0|
|bacteria_Jul27d5b_35645|1| 0|0|0|0|0|0|0|0|0|0| 0|0|0|0|0|0|0|0|0|0| 0|0|0|0|0|0|0|0|0|0| 1| 0|0|0|0|0|0|0|0|0|0| 0|0|0|0|0|0|0|0|0|0| 0|0|0|
|bacteria_Jul27d5b_45172|1| 0|0|0|0|0|0|0|0|0|0| 0|0|0|0|0|0|0|0|0|0| 0|0|0|0|0|0|0|0|0|0| 1| 0|0|0|0|0|0|0|0|0|0| 0|0|0|0|0|0|0|0|0|0| 0|0|0|
|bacteria_Jul27d5b_50606|1| 0|0|0|0|0|0|0|0|0|0| 0|0|0|0|0|0|0|0|0|0| 0|0|0|0|0|0|0|0|0|0| 1| 0|0|0|0|0|0|0|0|0|0| 0|0|0|0|0|0|0|0|0|0| 0|0|0|
|bacteria_Jul27d5b_50718|1| 0|0|0|0|0|0|0|0|0|0| 0|0|0|0|0|0|0|0|0|0| 0|0|0|0|0|0|0|0|0|0| 1| 0|0|0|0|0|0|0|0|0|0| 0|0|0|0|0|0|0|0|0|0| 0|0|0|
|bacteria_Jul27d25b_502|1| 0|0|0|0|0|0|0|0|0|0| 0|0|0|0|0|0|0|0|0|0| 0|0|0|0|0|0|0|0|0|0| 0|0|1| 0|0|0|0|0|0|0|0|0|0| 0|0|0|0|0|0|0|0|0|0| 0|
|bacteria_Jul27d25b_5613|1| 0|0|0|0|0|0|0|0|0|0| 0|0|0|0|0|0|0|0|0|0| 0|0|0|0|0|0|0|0|0|0| 0|0|1| 0|0|0|0|0|0|0|0|0|0| 0|0|0|0|0|0|0|0|0|0| 0|
|bacteria_Jul27d25b_7793|1| 0|0|0|0|0|0|0|0|0|0| 0|0|0|0|0|0|0|0|0|0| 0|0|0|0|0|0|0|0|0|0| 0|0|1| 0|0|0|0|0|0|0|0|0|0| 0|0|0|0|0|0|0|0|0|0| 0|
|bacteria_Jul27d25b_10581|1| 0|0|0|0|0|0|0|0|0|0| 0|0|0|0|0|0|0|0|0|0| 0|0|0|0|0|0|0|0|0|0| 0|0|1| 0|0|0|0|0|0|0|0|0|0| 0|0|0|0|0|0|0|0|0|0| 0|
|bacteria_Jul27d25b_11446|1| 0|0|0|0|0|0|0|0|0|0| 0|0|0|0|0|0|0|0|0|0| 0|0|0|0|0|0|0|0|0|0| 0|0|1| 0|0|0|0|0|0|0|0|0|0| 0|0|0|0|0|0|0|0|0|0| 0|
|bacteria_Jul27d25b_18520|1| 0|0|0|0|0|0|0|0|0|0| 0|0|0|0|0|0|0|0|0|0| 0|0|0|0|0|0|0|0|0|0| 0|0|1| 0|0|0|0|0|0|0|0|0|0| 0|0|0|0|0|0|0|0|0|0| 0|
|bacteria_Jul27d25b_28904|1| 0|0|0|0|0|0|0|0|0|0| 0|0|0|0|0|0|0|0|0|0| 0|0|0|0|0|0|0|0|0|0| 0|0|1| 0|0|0|0|0|0|0|0|0|0| 0|0|0|0|0|0|0|0|0|0| 0|
|bacteria_Jul27d25b_29322|1| 0|0|0|0|0|0|0|0|0|0| 0|0|0|0|0|0|0|0|0|0| 0|0|0|0|0|0|0|0|0|0| 0|0|1| 0|0|0|0|0|0|0|0|0|0| 0|0|0|0|0|0|0|0|0|0| 0|
|bacteria_Jul27d25b_30209|1| 0|0|0|0|0|0|0|0|0|0| 0|0|0|0|0|0|0|0|0|0| 0|0|0|0|0|0|0|0|0|0| 0|0|1| 0|0|0|0|0|0|0|0|0|0| 0|0|0|0|0|0|0|0|0|0| 0|
|bacteria_Jul27d25b_42959|1| 0|0|0|0|0|0|0|0|0|0| 0|0|0|0|0|0|0|0|0|0| 0|0|0|0|0|0|0|0|0|0| 0|0|1| 0|0|0|0|0|0|0|0|0|0| 0|0|0|0|0|0|0|0|0|0| 0|
|bacteria_Jul27d25b_47334|1| 0|0|0|0|0|0|0|0|0|0| 0|0|0|0|0|0|0|0|0|0| 0|0|0|0|0|0|0|0|0|0| 0|0|1| 0|0|0|0|0|0|0|0|0|0| 0|0|0|0|0|0|0|0|0|0| 0|
|bacteria_Jul27d25b_47345|1| 0|0|0|0|0|0|0|0|0|0| 0|0|0|0|0|0|0|0|0|0| 0|0|0|0|0|0|0|0|0|0| 0|0|1| 0|0|0|0|0|0|0|0|0|0| 0|0|0|0|0|0|0|0|0|0| 0|
|bacteria_Jul27d25b_53264|1| 0|0|0|0|0|0|0|0|0|0| 0|0|0|0|0|0|0|0|0|0| 0|0|0|0|0|0|0|0|0|0| 0|0|1| 0|0|0|0|0|0|0|0|0|0| 0|0|0|0|0|0|0|0|0|0| 0|
|bacteria_Jul27d25b_58082|1| 0|0|0|0|0|0|0|0|0|0| 0|0|0|0|0|0|0|0|0|0| 0|0|0|0|0|0|0|0|0|0| 0|0|1| 0|0|0|0|0|0|0|0|0|0| 0|0|0|0|0|0|0|0|0|0| 0|
|bacteria_Jul27d25b_74608|1| 0|0|0|0|0|0|0|0|0|0| 0|0|0|0|0|0|0|0|0|0| 0|0|0|0|0|0|0|0|0|0| 0|0|1| 0|0|0|0|0|0|0|0|0|0| 0|0|0|0|0|0|0|0|0|0| 0|
|bacteria_Jul27d75b_8199|1| 0|0|0|0|0|0|0|0|0|0| 0|0|0|0|0|0|0|0|0|0| 0|0|0|0|0|0|0|0|0|0| 0|0|0|0|1| 0|0|0|0|0|0|0|0|0|0| 0|0|0|0|0|0|0|0|0|
|bacteria_Jul27d75b_15254|1| 0|0|0|0|0|0|0|0|0|0| 0|0|0|0|0|0|0|0|0|0| 0|0|0|0|0|0|0|0|0|0| 0|0|0|0|1| 0|0|0|0|0|0|0|0|0|0| 0|0|0|0|0|0|0|0|0|
|bacteria_Jul27d75b_16713|1| 0|0|0|0|0|0|0|0|0|0| 0|0|0|0|0|0|0|0|0|0| 0|0|0|0|0|0|0|0|0|0| 0|0|0|0|1| 0|0|0|0|0|0|0|0|0|0| 0|0|0|0|0|0|0|0|0|
|bacteria_Jul27d75b_25740|1| 0|0|0|0|0|0|0|0|0|0| 0|0|0|0|0|0|0|0|0|0| 0|0|0|0|0|0|0|0|0|0| 0|0|0|0|1| 0|0|0|0|0|0|0|0|0|0| 0|0|0|0|0|0|0|0|0|
|bacteria_Jul27d75b_32002|1| 0|0|0|0|0|0|0|0|0|0| 0|0|0|0|0|0|0|0|0|0| 0|0|0|0|0|0|0|0|0|0| 0|0|0|0|1| 0|0|0|0|0|0|0|0|0|0| 0|0|0|0|0|0|0|0|0|
|bacteria_Jul27d75b_35408|1| 0|0|0|0|0|0|0|0|0|0| 0|0|0|0|0|0|0|0|0|0| 0|0|0|0|0|0|0|0|0|0| 0|0|0|0|1| 0|0|0|0|0|0|0|0|0|0| 0|0|0|0|0|0|0|0|0|
|bacteria_Jul27d75b_42054|1| 0|0|0|0|0|0|0|0|0|0| 0|0|0|0|0|0|0|0|0|0| 0|0|0|0|0|0|0|0|0|0| 0|0|0|0|1| 0|0|0|0|0|0|0|0|0|0| 0|0|0|0|0|0|0|0|0|

**Supplementary Table S7**. Common Bacterio- and virioplankton RTPR 98% OTUs in 27 samples with RTPR amplicons in both fractions

Total seqs: 47997; Subtotal seqs in retained

| OTUs: 41046 | | May12_05m | | May12_25m | | May12_50m | | May12_75m | | Jun8_25m | | Jun8_50m | | Jun8_75m | | Jun22_0m | | Jun22_50m | | Jun22_75m | | Jul11_0m | | Jul11_05m | | Jul11_25m | | Jul11_50m | | Jul11_75m | | Jul27_05m | | Jul27_25m | | Jul27_75m | | Aug5_0m | | Aug5_05m | | Aug5_25m | | Aug5_50m | | Aug5_75m | | Sep8_0m | | Sep8_05m | | Sep8_25m | | Sep8_50m | |
|---|---|---|---|---|---|---|---|---|---|---|---|---|---|---|---|---|---|---|---|---|---|---|---|---|---|---|---|---|---|---|---|---|---|---|---|---|---|---|---|---|---|---|---|---|---|---|---|---|---|---|---|---|---|---|---|
| OTUs with >=10 seqs: 254 | OTU subtotal | bacterioplankton | virioplankton | bacterioplankton | virioplankton | bacterioplankton | virioplankton | bacterioplankton | virioplankton | bacterioplankton | virioplankton | bacterioplankton | virioplankton | bacterioplankton | virioplankton | bacterioplankton | virioplankton | bacterioplankton | virioplankton | bacterioplankton | virioplankton | bacterioplankton | virioplankton | bacterioplankton | virioplankton | bacterioplankton | virioplankton | bacterioplankton | virioplankton | bacterioplankton | virioplankton | bacterioplankton | virioplankton | bacterioplankton | virioplankton | bacterioplankton | virioplankton | bacterioplankton | virioplankton | bacterioplankton | virioplankton | bacterioplankton | virioplankton | bacterioplankton | virioplankton | bacterioplankton | virioplankton | bacterioplankton | virioplankton | bacterioplankton | virioplankton | bacterioplankton | virioplankton | bacterioplankton | virioplankton |
| bacteria_Jul27d75b_4807 | 1 | 0 | 0 | 0 | 0 | 0 | 0 | 0 | 0 | 0 | 0 | 0 | 0 | 0 | 0 | 0 | 0 | 0 | 0 | 0 | 0 | 0 | 0 | 0 | 0 | 0 | 0 | 0 | 0 | 0 | 0 | 0 | 0 | 0 | 0 | 1 | 0 | 0 | 0 | 0 | 0 | 0 | 0 | 0 | 0 | 0 | 0 | 0 | 0 | 0 | 0 | 0 | 0 | 0 | 0 |
| bacteria_Jul27d75b_5533 | 1 | 0 | 0 | 0 | 0 | 0 | 0 | 0 | 0 | 0 | 0 | 0 | 0 | 0 | 0 | 0 | 0 | 0 | 0 | 0 | 0 | 0 | 0 | 0 | 0 | 0 | 0 | 0 | 0 | 0 | 0 | 0 | 0 | 0 | 0 | 1 | 0 | 0 | 0 | 0 | 0 | 0 | 0 | 0 | 0 | 0 | 0 | 0 | 0 | 0 | 0 | 0 | 0 | 0 | 0 |
| bacteria_Jul27d75b_7325 | 1 | 0 | 0 | 0 | 0 | 0 | 0 | 0 | 0 | 0 | 0 | 0 | 0 | 0 | 0 | 0 | 0 | 0 | 0 | 0 | 0 | 0 | 0 | 0 | 0 | 0 | 0 | 0 | 0 | 0 | 0 | 0 | 0 | 0 | 0 | 1 | 0 | 0 | 0 | 0 | 0 | 0 | 0 | 0 | 0 | 0 | 0 | 0 | 0 | 0 | 0 | 0 | 0 | 0 | 0 |
| bacteria_Jun8d25b_13733 | 1 | 0 | 0 | 0 | 0 | 0 | 0 | 0 | 0 | 1 | 0 | 0 | 0 | 0 | 0 | 0 | 0 | 0 | 0 | 0 | 0 | 0 | 0 | 0 | 0 | 0 | 0 | 0 | 0 | 0 | 0 | 0 | 0 | 0 | 0 | 0 | 0 | 0 | 0 | 0 | 0 | 0 | 0 | 0 | 0 | 0 | 0 | 0 | 0 | 0 | 0 | 0 | 0 | 0 | 0 |
| bacteria_Jun8d25b_19595 | 1 | 0 | 0 | 0 | 0 | 0 | 0 | 0 | 0 | 1 | 0 | 0 | 0 | 0 | 0 | 0 | 0 | 0 | 0 | 0 | 0 | 0 | 0 | 0 | 0 | 0 | 0 | 0 | 0 | 0 | 0 | 0 | 0 | 0 | 0 | 0 | 0 | 0 | 0 | 0 | 0 | 0 | 0 | 0 | 0 | 0 | 0 | 0 | 0 | 0 | 0 | 0 | 0 | 0 | 0 |
| bacteria_Jun8d25b_34124 | 1 | 0 | 0 | 0 | 0 | 0 | 0 | 0 | 0 | 1 | 0 | 0 | 0 | 0 | 0 | 0 | 0 | 0 | 0 | 0 | 0 | 0 | 0 | 0 | 0 | 0 | 0 | 0 | 0 | 0 | 0 | 0 | 0 | 0 | 0 | 0 | 0 | 0 | 0 | 0 | 0 | 0 | 0 | 0 | 0 | 0 | 0 | 0 | 0 | 0 | 0 | 0 | 0 | 0 | 0 |
| bacteria_Jun8d25b_34838 | 1 | 0 | 0 | 0 | 0 | 0 | 0 | 0 | 0 | 1 | 0 | 0 | 0 | 0 | 0 | 0 | 0 | 0 | 0 | 0 | 0 | 0 | 0 | 0 | 0 | 0 | 0 | 0 | 0 | 0 | 0 | 0 | 0 | 0 | 0 | 0 | 0 | 0 | 0 | 0 | 0 | 0 | 0 | 0 | 0 | 0 | 0 | 0 | 0 | 0 | 0 | 0 | 0 | 0 | 0 |
| bacteria_Jun8d25b_36928 | 1 | 0 | 0 | 0 | 0 | 0 | 0 | 0 | 0 | 1 | 0 | 0 | 0 | 0 | 0 | 0 | 0 | 0 | 0 | 0 | 0 | 0 | 0 | 0 | 0 | 0 | 0 | 0 | 0 | 0 | 0 | 0 | 0 | 0 | 0 | 0 | 0 | 0 | 0 | 0 | 0 | 0 | 0 | 0 | 0 | 0 | 0 | 0 | 0 | 0 | 0 | 0 | 0 | 0 | 0 |
| bacteria_Jun8d25b_43589 | 1 | 0 | 0 | 0 | 0 | 0 | 0 | 0 | 0 | 1 | 0 | 0 | 0 | 0 | 0 | 0 | 0 | 0 | 0 | 0 | 0 | 0 | 0 | 0 | 0 | 0 | 0 | 0 | 0 | 0 | 0 | 0 | 0 | 0 | 0 | 0 | 0 | 0 | 0 | 0 | 0 | 0 | 0 | 0 | 0 | 0 | 0 | 0 | 0 | 0 | 0 | 0 | 0 | 0 | 0 |
| bacteria_Jun8d25b_48941 | 1 | 0 | 0 | 0 | 0 | 0 | 0 | 0 | 0 | 1 | 0 | 0 | 0 | 0 | 0 | 0 | 0 | 0 | 0 | 0 | 0 | 0 | 0 | 0 | 0 | 0 | 0 | 0 | 0 | 0 | 0 | 0 | 0 | 0 | 0 | 0 | 0 | 0 | 0 | 0 | 0 | 0 | 0 | 0 | 0 | 0 | 0 | 0 | 0 | 0 | 0 | 0 | 0 | 0 | 0 |
| bacteria_Jun8d25b_58158 | 1 | 0 | 0 | 0 | 0 | 0 | 0 | 0 | 0 | 1 | 0 | 0 | 0 | 0 | 0 | 0 | 0 | 0 | 0 | 0 | 0 | 0 | 0 | 0 | 0 | 0 | 0 | 0 | 0 | 0 | 0 | 0 | 0 | 0 | 0 | 0 | 0 | 0 | 0 | 0 | 0 | 0 | 0 | 0 | 0 | 0 | 0 | 0 | 0 | 0 | 0 | 0 | 0 | 0 | 0 |
| bacteria_Jun8d25b_60913 | 1 | 0 | 0 | 0 | 0 | 0 | 0 | 0 | 0 | 1 | 0 | 0 | 0 | 0 | 0 | 0 | 0 | 0 | 0 | 0 | 0 | 0 | 0 | 0 | 0 | 0 | 0 | 0 | 0 | 0 | 0 | 0 | 0 | 0 | 0 | 0 | 0 | 0 | 0 | 0 | 0 | 0 | 0 | 0 | 0 | 0 | 0 | 0 | 0 | 0 | 0 | 0 | 0 | 0 | 0 |
| bacteria_Jun8d25b_64834 | 1 | 0 | 0 | 0 | 0 | 0 | 0 | 0 | 0 | 1 | 0 | 0 | 0 | 0 | 0 | 0 | 0 | 0 | 0 | 0 | 0 | 0 | 0 | 0 | 0 | 0 | 0 | 0 | 0 | 0 | 0 | 0 | 0 | 0 | 0 | 0 | 0 | 0 | 0 | 0 | 0 | 0 | 0 | 0 | 0 | 0 | 0 | 0 | 0 | 0 | 0 | 0 | 0 | 0 | 0 |
| bacteria_Jun8d25b_69873 | 1 | 0 | 0 | 0 | 0 | 0 | 0 | 0 | 0 | 1 | 0 | 0 | 0 | 0 | 0 | 0 | 0 | 0 | 0 | 0 | 0 | 0 | 0 | 0 | 0 | 0 | 0 | 0 | 0 | 0 | 0 | 0 | 0 | 0 | 0 | 0 | 0 | 0 | 0 | 0 | 0 | 0 | 0 | 0 | 0 | 0 | 0 | 0 | 0 | 0 | 0 | 0 | 0 | 0 | 0 |
| bacteria_Jun8d50b_775 | 1 | 0 | 0 | 0 | 0 | 0 | 0 | 0 | 0 | 0 | 0 | 1 | 0 | 0 | 0 | 0 | 0 | 0 | 0 | 0 | 0 | 0 | 0 | 0 | 0 | 0 | 0 | 0 | 0 | 0 | 0 | 0 | 0 | 0 | 0 | 0 | 0 | 0 | 0 | 0 | 0 | 0 | 0 | 0 | 0 | 0 | 0 | 0 | 0 | 0 | 0 | 0 | 0 | 0 | 0 |
| bacteria_Jun8d50b_1368 | 1 | 0 | 0 | 0 | 0 | 0 | 0 | 0 | 0 | 0 | 0 | 1 | 0 | 0 | 0 | 0 | 0 | 0 | 0 | 0 | 0 | 0 | 0 | 0 | 0 | 0 | 0 | 0 | 0 | 0 | 0 | 0 | 0 | 0 | 0 | 0 | 0 | 0 | 0 | 0 | 0 | 0 | 0 | 0 | 0 | 0 | 0 | 0 | 0 | 0 | 0 | 0 | 0 | 0 | 0 |
| bacteria_Jun8d50b_2490 | 1 | 0 | 0 | 0 | 0 | 0 | 0 | 0 | 0 | 0 | 0 | 1 | 0 | 0 | 0 | 0 | 0 | 0 | 0 | 0 | 0 | 0 | 0 | 0 | 0 | 0 | 0 | 0 | 0 | 0 | 0 | 0 | 0 | 0 | 0 | 0 | 0 | 0 | 0 | 0 | 0 | 0 | 0 | 0 | 0 | 0 | 0 | 0 | 0 | 0 | 0 | 0 | 0 | 0 | 0 |
| bacteria_Jun8d50b_4374 | 1 | 0 | 0 | 0 | 0 | 0 | 0 | 0 | 0 | 0 | 0 | 1 | 0 | 0 | 0 | 0 | 0 | 0 | 0 | 0 | 0 | 0 | 0 | 0 | 0 | 0 | 0 | 0 | 0 | 0 | 0 | 0 | 0 | 0 | 0 | 0 | 0 | 0 | 0 | 0 | 0 | 0 | 0 | 0 | 0 | 0 | 0 | 0 | 0 | 0 | 0 | 0 | 0 | 0 | 0 |
| bacteria_Jun8d50b_18591 | 1 | 0 | 0 | 0 | 0 | 0 | 0 | 0 | 0 | 0 | 0 | 1 | 0 | 0 | 0 | 0 | 0 | 0 | 0 | 0 | 0 | 0 | 0 | 0 | 0 | 0 | 0 | 0 | 0 | 0 | 0 | 0 | 0 | 0 | 0 | 0 | 0 | 0 | 0 | 0 | 0 | 0 | 0 | 0 | 0 | 0 | 0 | 0 | 0 | 0 | 0 | 0 | 0 | 0 | 0 |
| bacteria_Jun8d50b_48805 | 1 | 0 | 0 | 0 | 0 | 0 | 0 | 0 | 0 | 0 | 0 | 1 | 0 | 0 | 0 | 0 | 0 | 0 | 0 | 0 | 0 | 0 | 0 | 0 | 0 | 0 | 0 | 0 | 0 | 0 | 0 | 0 | 0 | 0 | 0 | 0 | 0 | 0 | 0 | 0 | 0 | 0 | 0 | 0 | 0 | 0 | 0 | 0 | 0 | 0 | 0 | 0 | 0 | 0 | 0 |
| bacteria_Jun8d50b_50538 | 1 | 0 | 0 | 0 | 0 | 0 | 0 | 0 | 0 | 0 | 0 | 1 | 0 | 0 | 0 | 0 | 0 | 0 | 0 | 0 | 0 | 0 | 0 | 0 | 0 | 0 | 0 | 0 | 0 | 0 | 0 | 0 | 0 | 0 | 0 | 0 | 0 | 0 | 0 | 0 | 0 | 0 | 0 | 0 | 0 | 0 | 0 | 0 | 0 | 0 | 0 | 0 | 0 | 0 | 0 |
| bacteria_Jun8d75b_57768 | 1 | 0 | 0 | 0 | 0 | 0 | 0 | 0 | 0 | 0 | 0 | 0 | 0 | 1 | 0 | 0 | 0 | 0 | 0 | 0 | 0 | 0 | 0 | 0 | 0 | 0 | 0 | 0 | 0 | 0 | 0 | 0 | 0 | 0 | 0 | 0 | 0 | 0 | 0 | 0 | 0 | 0 | 0 | 0 | 0 | 0 | 0 | 0 | 0 | 0 | 0 | 0 | 0 | 0 | 0 |
| bacteria_Jun8d75b_64161 | 1 | 0 | 0 | 0 | 0 | 0 | 0 | 0 | 0 | 0 | 0 | 0 | 0 | 1 | 0 | 0 | 0 | 0 | 0 | 0 | 0 | 0 | 0 | 0 | 0 | 0 | 0 | 0 | 0 | 0 | 0 | 0 | 0 | 0 | 0 | 0 | 0 | 0 | 0 | 0 | 0 | 0 | 0 | 0 | 0 | 0 | 0 | 0 | 0 | 0 | 0 | 0 | 0 | 0 | 0 |
| bacteria_Jun8d75b_73646 | 1 | 0 | 0 | 0 | 0 | 0 | 0 | 0 | 0 | 0 | 0 | 0 | 0 | 1 | 0 | 0 | 0 | 0 | 0 | 0 | 0 | 0 | 0 | 0 | 0 | 0 | 0 | 0 | 0 | 0 | 0 | 0 | 0 | 0 | 0 | 0 | 0 | 0 | 0 | 0 | 0 | 0 | 0 | 0 | 0 | 0 | 0 | 0 | 0 | 0 | 0 | 0 | 0 | 0 | 0 |
| bacteria_Jun22d0b_14396 | 1 | 0 | 0 | 0 | 0 | 0 | 0 | 0 | 0 | 0 | 0 | 0 | 0 | 0 | 0 | 1 | 0 | 0 | 0 | 0 | 0 | 0 | 0 | 0 | 0 | 0 | 0 | 0 | 0 | 0 | 0 | 0 | 0 | 0 | 0 | 0 | 0 | 0 | 0 | 0 | 0 | 0 | 0 | 0 | 0 | 0 | 0 | 0 | 0 | 0 | 0 | 0 | 0 | 0 | 0 |
| bacteria_Jun22d50b_4723 | 1 | 0 | 0 | 0 | 0 | 0 | 0 | 0 | 0 | 0 | 0 | 0 | 0 | 0 | 0 | 0 | 0 | 1 | 0 | 0 | 0 | 0 | 0 | 0 | 0 | 0 | 0 | 0 | 0 | 0 | 0 | 0 | 0 | 0 | 0 | 0 | 0 | 0 | 0 | 0 | 0 | 0 | 0 | 0 | 0 | 0 | 0 | 0 | 0 | 0 | 0 | 0 | 0 | 0 | 0 |
| bacteria_Jun22d50b_1452 | 1 | 0 | 0 | 0 | 0 | 0 | 0 | 0 | 0 | 0 | 0 | 0 | 0 | 0 | 0 | 0 | 0 | 1 | 0 | 0 | 0 | 0 | 0 | 0 | 0 | 0 | 0 | 0 | 0 | 0 | 0 | 0 | 0 | 0 | 0 | 0 | 0 | 0 | 0 | 0 | 0 | 0 | 0 | 0 | 0 | 0 | 0 | 0 | 0 | 0 | 0 | 0 | 0 | 0 | 0 |
| bacteria_Jun22d50b_1717 | 1 | 0 | 0 | 0 | 0 | 0 | 0 | 0 | 0 | 0 | 0 | 0 | 0 | 0 | 0 | 0 | 0 | 1 | 0 | 0 | 0 | 0 | 0 | 0 | 0 | 0 | 0 | 0 | 0 | 0 | 0 | 0 | 0 | 0 | 0 | 0 | 0 | 0 | 0 | 0 | 0 | 0 | 0 | 0 | 0 | 0 | 0 | 0 | 0 | 0 | 0 | 0 | 0 | 0 | 0 |
| bacteria_Jun22d50b_1717 | 1 | 0 | 0 | 0 | 0 | 0 | 0 | 0 | 0 | 0 | 0 | 0 | 0 | 0 | 0 | 0 | 0 | 1 | 0 | 0 | 0 | 0 | 0 | 0 | 0 | 0 | 0 | 0 | 0 | 0 | 0 | 0 | 0 | 0 | 0 | 0 | 0 | 0 | 0 | 0 | 0 | 0 | 0 | 0 | 0 | 0 | 0 | 0 | 0 | 0 | 0 | 0 | 0 | 0 | 0 |
| bacteria_Jun22d50b_3123 | 1 | 0 | 0 | 0 | 0 | 0 | 0 | 0 | 0 | 0 | 0 | 0 | 0 | 0 | 0 | 0 | 0 | 1 | 0 | 0 | 0 | 0 | 0 | 0 | 0 | 0 | 0 | 0 | 0 | 0 | 0 | 0 | 0 | 0 | 0 | 0 | 0 | 0 | 0 | 0 | 0 | 0 | 0 | 0 | 0 | 0 | 0 | 0 | 0 | 0 | 0 | 0 | 0 | 0 | 0 |
| bacteria_Jun22d75b_9016 | 1 | 0 | 0 | 0 | 0 | 0 | 0 | 0 | 0 | 0 | 0 | 0 | 0 | 0 | 0 | 0 | 0 | 0 | 0 | 1 | 0 | 0 | 0 | 0 | 0 | 0 | 0 | 0 | 0 | 0 | 0 | 0 | 0 | 0 | 0 | 0 | 0 | 0 | 0 | 0 | 0 | 0 | 0 | 0 | 0 | 0 | 0 | 0 | 0 | 0 | 0 | 0 | 0 | 0 | 0 |
| bacteria_Jun22d75b_2295 | 1 | 0 | 0 | 0 | 0 | 0 | 0 | 0 | 0 | 0 | 0 | 0 | 0 | 0 | 0 | 0 | 0 | 0 | 0 | 1 | 0 | 0 | 0 | 0 | 0 | 0 | 0 | 0 | 0 | 0 | 0 | 0 | 0 | 0 | 0 | 0 | 0 | 0 | 0 | 0 | 0 | 0 | 0 | 0 | 0 | 0 | 0 | 0 | 0 | 0 | 0 | 0 | 0 | 0 | 0 |
| bacteria_Jun22d75b_2506 | 1 | 0 | 0 | 0 | 0 | 0 | 0 | 0 | 0 | 0 | 0 | 0 | 0 | 0 | 0 | 0 | 0 | 0 | 0 | 1 | 0 | 0 | 0 | 0 | 0 | 0 | 0 | 0 | 0 | 0 | 0 | 0 | 0 | 0 | 0 | 0 | 0 | 0 | 0 | 0 | 0 | 0 | 0 | 0 | 0 | 0 | 0 | 0 | 0 | 0 | 0 | 0 | 0 | 0 | 0 |
| bacteria_Jun22d75b_3053 | 1 | 0 | 0 | 0 | 0 | 0 | 0 | 0 | 0 | 0 | 0 | 0 | 0 | 0 | 0 | 0 | 0 | 0 | 0 | 1 | 0 | 0 | 0 | 0 | 0 | 0 | 0 | 0 | 0 | 0 | 0 | 0 | 0 | 0 | 0 | 0 | 0 | 0 | 0 | 0 | 0 | 0 | 0 | 0 | 0 | 0 | 0 | 0 | 0 | 0 | 0 | 0 | 0 | 0 | 0 |
| bacteria_Jun22d75b_3382 | 1 | 0 | 0 | 0 | 0 | 0 | 0 | 0 | 0 | 0 | 0 | 0 | 0 | 0 | 0 | 0 | 0 | 0 | 0 | 1 | 0 | 0 | 0 | 0 | 0 | 0 | 0 | 0 | 0 | 0 | 0 | 0 | 0 | 0 | 0 | 0 | 0 | 0 | 0 | 0 | 0 | 0 | 0 | 0 | 0 | 0 | 0 | 0 | 0 | 0 | 0 | 0 | 0 | 0 | 0 |
| bacteria_Jun22d75b_3543 | 1 | 0 | 0 | 0 | 0 | 0 | 0 | 0 | 0 | 0 | 0 | 0 | 0 | 0 | 0 | 0 | 0 | 0 | 0 | 1 | 0 | 0 | 0 | 0 | 0 | 0 | 0 | 0 | 0 | 0 | 0 | 0 | 0 | 0 | 0 | 0 | 0 | 0 | 0 | 0 | 0 | 0 | 0 | 0 | 0 | 0 | 0 | 0 | 0 | 0 | 0 | 0 | 0 | 0 | 0 |
| bacteria_Jun22d75b_3919 | 1 | 0 | 0 | 0 | 0 | 0 | 0 | 0 | 0 | 0 | 0 | 0 | 0 | 0 | 0 | 0 | 0 | 0 | 0 | 1 | 0 | 0 | 0 | 0 | 0 | 0 | 0 | 0 | 0 | 0 | 0 | 0 | 0 | 0 | 0 | 0 | 0 | 0 | 0 | 0 | 0 | 0 | 0 | 0 | 0 | 0 | 0 | 0 | 0 | 0 | 0 | 0 | 0 | 0 | 0 |
| bacteria_Jun22d75b_4791 | 1 | 0 | 0 | 0 | 0 | 0 | 0 | 0 | 0 | 0 | 0 | 0 | 0 | 0 | 0 | 0 | 0 | 0 | 0 | 1 | 0 | 0 | 0 | 0 | 0 | 0 | 0 | 0 | 0 | 0 | 0 | 0 | 0 | 0 | 0 | 0 | 0 | 0 | 0 | 0 | 0 | 0 | 0 | 0 | 0 | 0 | 0 | 0 | 0 | 0 | 0 | 0 | 0 | 0 | 0 |
| bacteria_Jun22d75b_5687 | 1 | 0 | 0 | 0 | 0 | 0 | 0 | 0 | 0 | 0 | 0 | 0 | 0 | 0 | 0 | 0 | 0 | 0 | 0 | 1 | 0 | 0 | 0 | 0 | 0 | 0 | 0 | 0 | 0 | 0 | 0 | 0 | 0 | 0 | 0 | 0 | 0 | 0 | 0 | 0 | 0 | 0 | 0 | 0 | 0 | 0 | 0 | 0 | 0 | 0 | 0 | 0 | 0 | 0 | 0 |
| bacteria_Jun22d75b_7202 | 1 | 0 | 0 | 0 | 0 | 0 | 0 | 0 | 0 | 0 | 0 | 0 | 0 | 0 | 0 | 0 | 0 | 0 | 0 | 1 | 0 | 0 | 0 | 0 | 0 | 0 | 0 | 0 | 0 | 0 | 0 | 0 | 0 | 0 | 0 | 0 | 0 | 0 | 0 | 0 | 0 | 0 | 0 | 0 | 0 | 0 | 0 | 0 | 0 | 0 | 0 | 0 | 0 | 0 | 0 |
| bacteria_May12d5b_3683 | 1 | 1 | 0 | 0 | 0 | 0 | 0 | 0 | 0 | 0 | 0 | 0 | 0 | 0 | 0 | 0 | 0 | 0 | 0 | 0 | 0 | 0 | 0 | 0 | 0 | 0 | 0 | 0 | 0 | 0 | 0 | 0 | 0 | 0 | 0 | 0 | 0 | 0 | 0 | 0 | 0 | 0 | 0 | 0 | 0 | 0 | 0 | 0 | 0 | 0 | 0 | 0 | 0 | 0 | 0 |
| bacteria_May12d25b_335 | 1 | 0 | 0 | 1 | 0 | 0 | 0 | 0 | 0 | 0 | 0 | 0 | 0 | 0 | 0 | 0 | 0 | 0 | 0 | 0 | 0 | 0 | 0 | 0 | 0 | 0 | 0 | 0 | 0 | 0 | 0 | 0 | 0 | 0 | 0 | 0 | 0 | 0 | 0 | 0 | 0 | 0 | 0 | 0 | 0 | 0 | 0 | 0 | 0 | 0 | 0 | 0 | 0 | 0 | 0 |
| bacteria_May12d25b_337 | 1 | 0 | 0 | 1 | 0 | 0 | 0 | 0 | 0 | 0 | 0 | 0 | 0 | 0 | 0 | 0 | 0 | 0 | 0 | 0 | 0 | 0 | 0 | 0 | 0 | 0 | 0 | 0 | 0 | 0 | 0 | 0 | 0 | 0 | 0 | 0 | 0 | 0 | 0 | 0 | 0 | 0 | 0 | 0 | 0 | 0 | 0 | 0 | 0 | 0 | 0 | 0 | 0 | 0 | 0 |
| bacteria_May12d25b_629 | 1 | 0 | 0 | 1 | 0 | 0 | 0 | 0 | 0 | 0 | 0 | 0 | 0 | 0 | 0 | 0 | 0 | 0 | 0 | 0 | 0 | 0 | 0 | 0 | 0 | 0 | 0 | 0 | 0 | 0 | 0 | 0 | 0 | 0 | 0 | 0 | 0 | 0 | 0 | 0 | 0 | 0 | 0 | 0 | 0 | 0 | 0 | 0 | 0 | 0 | 0 | 0 | 0 | 0 | 0 |
| bacteria_May12d25b_118 | 1 | 0 | 0 | 1 | 0 | 0 | 0 | 0 | 0 | 0 | 0 | 0 | 0 | 0 | 0 | 0 | 0 | 0 | 0 | 0 | 0 | 0 | 0 | 0 | 0 | 0 | 0 | 0 | 0 | 0 | 0 | 0 | 0 | 0 | 0 | 0 | 0 | 0 | 0 | 0 | 0 | 0 | 0 | 0 | 0 | 0 | 0 | 0 | 0 | 0 | 0 | 0 | 0 | 0 | 0 |
| bacteria_May12d25b_214 | 1 | 0 | 0 | 1 | 0 | 0 | 0 | 0 | 0 | 0 | 0 | 0 | 0 | 0 | 0 | 0 | 0 | 0 | 0 | 0 | 0 | 0 | 0 | 0 | 0 | 0 | 0 | 0 | 0 | 0 | 0 | 0 | 0 | 0 | 0 | 0 | 0 | 0 | 0 | 0 | 0 | 0 | 0 | 0 | 0 | 0 | 0 | 0 | 0 | 0 | 0 | 0 | 0 | 0 | 0 |
| bacteria_May12d25b_245 | 1 | 0 | 0 | 1 | 0 | 0 | 0 | 0 | 0 | 0 | 0 | 0 | 0 | 0 | 0 | 0 | 0 | 0 | 0 | 0 | 0 | 0 | 0 | 0 | 0 | 0 | 0 | 0 | 0 | 0 | 0 | 0 | 0 | 0 | 0 | 0 | 0 | 0 | 0 | 0 | 0 | 0 | 0 | 0 | 0 | 0 | 0 | 0 | 0 | 0 | 0 | 0 | 0 | 0 | 0 |
| bacteria_May12d25b_254 | 1 | 0 | 0 | 1 | 0 | 0 | 0 | 0 | 0 | 0 | 0 | 0 | 0 | 0 | 0 | 0 | 0 | 0 | 0 | 0 | 0 | 0 | 0 | 0 | 0 | 0 | 0 | 0 | 0 | 0 | 0 | 0 | 0 | 0 | 0 | 0 | 0 | 0 | 0 | 0 | 0 | 0 | 0 | 0 | 0 | 0 | 0 | 0 | 0 | 0 | 0 | 0 | 0 | 0 | 0 |
| bacteria_May12d25b_483 | 1 | 0 | 0 | 1 | 0 | 0 | 0 | 0 | 0 | 0 | 0 | 0 | 0 | 0 | 0 | 0 | 0 | 0 | 0 | 0 | 0 | 0 | 0 | 0 | 0 | 0 | 0 | 0 | 0 | 0 | 0 | 0 | 0 | 0 | 0 | 0 | 0 | 0 | 0 | 0 | 0 | 0 | 0 | 0 | 0 | 0 | 0 | 0 | 0 | 0 | 0 | 0 | 0 | 0 | 0 |
| bacteria_May12d25b_688 | 1 | 0 | 0 | 1 | 0 | 0 | 0 | 0 | 0 | 0 | 0 | 0 | 0 | 0 | 0 | 0 | 0 | 0 | 0 | 0 | 0 | 0 | 0 | 0 | 0 | 0 | 0 | 0 | 0 | 0 | 0 | 0 | 0 | 0 | 0 | 0 | 0 | 0 | 0 | 0 | 0 | 0 | 0 | 0 | 0 | 0 | 0 | 0 | 0 | 0 | 0 | 0 | 0 | 0 | 0 |
| bacteria_May12d25b_732 | 1 | 0 | 0 | 1 | 0 | 0 | 0 | 0 | 0 | 0 | 0 | 0 | 0 | 0 | 0 | 0 | 0 | 0 | 0 | 0 | 0 | 0 | 0 | 0 | 0 | 0 | 0 | 0 | 0 | 0 | 0 | 0 | 0 | 0 | 0 | 0 | 0 | 0 | 0 | 0 | 0 | 0 | 0 | 0 | 0 | 0 | 0 | 0 | 0 | 0 | 0 | 0 | 0 | 0 | 0 |
| bacteria_May12d25b_752 | 1 | 0 | 0 | 1 | 0 | 0 | 0 | 0 | 0 | 0 | 0 | 0 | 0 | 0 | 0 | 0 | 0 | 0 | 0 | 0 | 0 | 0 | 0 | 0 | 0 | 0 | 0 | 0 | 0 | 0 | 0 | 0 | 0 | 0 | 0 | 0 | 0 | 0 | 0 | 0 | 0 | 0 | 0 | 0 | 0 | 0 | 0 | 0 | 0 | 0 | 0 | 0 | 0 | 0 | 0 |
| bacteria_May12d50b_425 | 1 | 0 | 0 | 0 | 0 | 1 | 0 | 0 | 0 | 0 | 0 | 0 | 0 | 0 | 0 | 0 | 0 | 0 | 0 | 0 | 0 | 0 | 0 | 0 | 0 | 0 | 0 | 0 | 0 | 0 | 0 | 0 | 0 | 0 | 0 | 0 | 0 | 0 | 0 | 0 | 0 | 0 | 0 | 0 | 0 | 0 | 0 | 0 | 0 | 0 | 0 | 0 | 0 | 0 | 0 |
| bacteria_May12d50b_486 | 1 | 0 | 0 | 0 | 0 | 1 | 0 | 0 | 0 | 0 | 0 | 0 | 0 | 0 | 0 | 0 | 0 | 0 | 0 | 0 | 0 | 0 | 0 | 0 | 0 | 0 | 0 | 0 | 0 | 0 | 0 | 0 | 0 | 0 | 0 | 0 | 0 | 0 | 0 | 0 | 0 | 0 | 0 | 0 | 0 | 0 | 0 | 0 | 0 | 0 | 0 | 0 | 0 | 0 | 0 |
| bacteria_May12d50b_539 | 1 | 0 | 0 | 0 | 0 | 1 | 0 | 0 | 0 | 0 | 0 | 0 | 0 | 0 | 0 | 0 | 0 | 0 | 0 | 0 | 0 | 0 | 0 | 0 | 0 | 0 | 0 | 0 | 0 | 0 | 0 | 0 | 0 | 0 | 0 | 0 | 0 | 0 | 0 | 0 | 0 | 0 | 0 | 0 | 0 | 0 | 0 | 0 | 0 | 0 | 0 | 0 | 0 | 0 | 0 |
| bacteria_May12d50b_107 | 1 | 0 | 0 | 0 | 0 | 1 | 0 | 0 | 0 | 0 | 0 | 0 | 0 | 0 | 0 | 0 | 0 | 0 | 0 | 0 | 0 | 0 | 0 | 0 | 0 | 0 | 0 | 0 | 0 | 0 | 0 | 0 | 0 | 0 | 0 | 0 | 0 | 0 | 0 | 0 | 0 | 0 | 0 | 0 | 0 | 0 | 0 | 0 | 0 | 0 | 0 | 0 | 0 | 0 | 0 |
| bacteria_May12d50b_280 | 1 | 0 | 0 | 0 | 0 | 1 | 0 | 0 | 0 | 0 | 0 | 0 | 0 | 0 | 0 | 0 | 0 | 0 | 0 | 0 | 0 | 0 | 0 | 0 | 0 | 0 | 0 | 0 | 0 | 0 | 0 | 0 | 0 | 0 | 0 | 0 | 0 | 0 | 0 | 0 | 0 | 0 | 0 | 0 | 0 | 0 | 0 | 0 | 0 | 0 | 0 | 0 | 0 | 0 | 0 |
| bacteria_May12d50b_294 | 1 | 0 | 0 | 0 | 0 | 1 | 0 | 0 | 0 | 0 | 0 | 0 | 0 | 0 | 0 | 0 | 0 | 0 | 0 | 0 | 0 | 0 | 0 | 0 | 0 | 0 | 0 | 0 | 0 | 0 | 0 | 0 | 0 | 0 | 0 | 0 | 0 | 0 | 0 | 0 | 0 | 0 | 0 | 0 | 0 | 0 | 0 | 0 | 0 | 0 | 0 | 0 | 0 | 0 | 0 |
| bacteria_May12d50b_304 | 1 | 0 | 0 | 0 | 0 | 1 | 0 | 0 | 0 | 0 | 0 | 0 | 0 | 0 | 0 | 0 | 0 | 0 | 0 | 0 | 0 | 0 | 0 | 0 | 0 | 0 | 0 | 0 | 0 | 0 | 0 | 0 | 0 | 0 | 0 | 0 | 0 | 0 | 0 | 0 | 0 | 0 | 0 | 0 | 0 | 0 | 0 | 0 | 0 | 0 | 0 | 0 | 0 | 0 | 0 |
| bacteria_May12d50b_349 | 1 | 0 | 0 | 0 | 0 | 1 | 0 | 0 | 0 | 0 | 0 | 0 | 0 | 0 | 0 | 0 | 0 | 0 | 0 | 0 | 0 | 0 | 0 | 0 | 0 | 0 | 0 | 0 | 0 | 0 | 0 | 0 | 0 | 0 | 0 | 0 | 0 | 0 | 0 | 0 | 0 | 0 | 0 | 0 | 0 | 0 | 0 | 0 | 0 | 0 | 0 | 0 | 0 | 0 | 0 |
| bacteria_May12d50b_436 | 1 | 0 | 0 | 0 | 0 | 1 | 0 | 0 | 0 | 0 | 0 | 0 | 0 | 0 | 0 | 0 | 0 | 0 | 0 | 0 | 0 | 0 | 0 | 0 | 0 | 0 | 0 | 0 | 0 | 0 | 0 | 0 | 0 | 0 | 0 | 0 | 0 | 0 | 0 | 0 | 0 | 0 | 0 | 0 | 0 | 0 | 0 | 0 | 0 | 0 | 0 | 0 | 0 | 0 | 0 |
| bacteria_May12d50b_439 | 1 | 0 | 0 | 0 | 0 | 1 | 0 | 0 | 0 | 0 | 0 | 0 | 0 | 0 | 0 | 0 | 0 | 0 | 0 | 0 | 0 | 0 | 0 | 0 | 0 | 0 | 0 | 0 | 0 | 0 | 0 | 0 | 0 | 0 | 0 | 0 | 0 | 0 | 0 | 0 | 0 | 0 | 0 | 0 | 0 | 0 | 0 | 0 | 0 | 0 | 0 | 0 | 0 | 0 | 0 |
| bacteria_May12d50b_448 | 1 | 0 | 0 | 0 | 0 | 1 | 0 | 0 | 0 | 0 | 0 | 0 | 0 | 0 | 0 | 0 | 0 | 0 | 0 | 0 | 0 | 0 | 0 | 0 | 0 | 0 | 0 | 0 | 0 | 0 | 0 | 0 | 0 | 0 | 0 | 0 | 0 | 0 | 0 | 0 | 0 | 0 | 0 | 0 | 0 | 0 | 0 | 0 | 0 | 0 | 0 | 0 | 0 | 0 | 0 |
| bacteria_May12d50b_452 | 1 | 0 | 0 | 0 | 0 | 1 | 0 | 0 | 0 | 0 | 0 | 0 | 0 | 0 | 0 | 0 | 0 | 0 | 0 | 0 | 0 | 0 | 0 | 0 | 0 | 0 | 0 | 0 | 0 | 0 | 0 | 0 | 0 | 0 | 0 | 0 | 0 | 0 | 0 | 0 | 0 | 0 | 0 | 0 | 0 | 0 | 0 | 0 | 0 | 0 | 0 | 0 | 0 | 0 | 0 |
| bacteria_Sep8d0b_64274 | 1 | 0 | 0 | 0 | 0 | 0 | 0 | 0 | 0 | 0 | 0 | 0 | 0 | 0 | 0 | 0 | 0 | 0 | 0 | 0 | 0 | 0 | 0 | 0 | 0 | 0 | 0 | 0 | 0 | 0 | 0 | 0 | 0 | 0 | 0 | 0 | 0 | 0 | 0 | 0 | 0 | 0 | 0 | 0 | 0 | 0 | 0 | 1 | 0 | 0 | 0 | 0 | 0 | 0 | 0 |
| bacteria_Sep8d5b_2421 | 1 | 0 | 0 | 0 | 0 | 0 | 0 | 0 | 0 | 0 | 0 | 0 | 0 | 0 | 0 | 0 | 0 | 0 | 0 | 0 | 0 | 0 | 0 | 0 | 0 | 0 | 0 | 0 | 0 | 0 | 0 | 0 | 0 | 0 | 0 | 0 | 0 | 0 | 0 | 0 | 0 | 0 | 0 | 0 | 0 | 0 | 0 | 0 | 0 | 1 | 0 | 0 | 0 | 0 | 0 |
| bacteria_Sep8d5b_7498 | 1 | 0 | 0 | 0 | 0 | 0 | 0 | 0 | 0 | 0 | 0 | 0 | 0 | 0 | 0 | 0 | 0 | 0 | 0 | 0 | 0 | 0 | 0 | 0 | 0 | 0 | 0 | 0 | 0 | 0 | 0 | 0 | 0 | 0 | 0 | 0 | 0 | 0 | 0 | 0 | 0 | 0 | 0 | 0 | 0 | 0 | 0 | 0 | 0 | 1 | 0 | 0 | 0 | 0 | 0 |
| bacteria_Sep8d5b_9113 | 1 | 0 | 0 | 0 | 0 | 0 | 0 | 0 | 0 | 0 | 0 | 0 | 0 | 0 | 0 | 0 | 0 | 0 | 0 | 0 | 0 | 0 | 0 | 0 | 0 | 0 | 0 | 0 | 0 | 0 | 0 | 0 | 0 | 0 | 0 | 0 | 0 | 0 | 0 | 0 | 0 | 0 | 0 | 0 | 0 | 0 | 0 | 0 | 0 | 1 | 0 | 0 | 0 | 0 | 0 |
| bacteria_Sep8d5b_13510 | 1 | 0 | 0 | 0 | 0 | 0 | 0 | 0 | 0 | 0 | 0 | 0 | 0 | 0 | 0 | 0 | 0 | 0 | 0 | 0 | 0 | 0 | 0 | 0 | 0 | 0 | 0 | 0 | 0 | 0 | 0 | 0 | 0 | 0 | 0 | 0 | 0 | 0 | 0 | 0 | 0 | 0 | 0 | 0 | 0 | 0 | 0 | 0 | 0 | 1 | 0 | 0 | 0 | 0 | 0 |
| bacteria_Sep8d5b_18132 | 1 | 0 | 0 | 0 | 0 | 0 | 0 | 0 | 0 | 0 | 0 | 0 | 0 | 0 | 0 | 0 | 0 | 0 | 0 | 0 | 0 | 0 | 0 | 0 | 0 | 0 | 0 | 0 | 0 | 0 | 0 | 0 | 0 | 0 | 0 | 0 | 0 | 0 | 0 | 0 | 0 | 0 | 0 | 0 | 0 | 0 | 0 | 0 | 0 | 1 | 0 | 0 | 0 | 0 | 0 |
| bacteria_Sep8d5b_23650 | 1 | 0 | 0 | 0 | 0 | 0 | 0 | 0 | 0 | 0 | 0 | 0 | 0 | 0 | 0 | 0 | 0 | 0 | 0 | 0 | 0 | 0 | 0 | 0 | 0 | 0 | 0 | 0 | 0 | 0 | 0 | 0 | 0 | 0 | 0 | 0 | 0 | 0 | 0 | 0 | 0 | 0 | 0 | 0 | 0 | 0 | 0 | 0 | 0 | 1 | 0 | 0 | 0 | 0 | 0 |
| bacteria_Sep8d5b_30965 | 1 | 0 | 0 | 0 | 0 | 0 | 0 | 0 | 0 | 0 | 0 | 0 | 0 | 0 | 0 | 0 | 0 | 0 | 0 | 0 | 0 | 0 | 0 | 0 | 0 | 0 | 0 | 0 | 0 | 0 | 0 | 0 | 0 | 0 | 0 | 0 | 0 | 0 | 0 | 0 | 0 | 0 | 0 | 0 | 0 | 0 | 0 | 0 | 0 | 1 | 0 | 0 | 0 | 0 | 0 |
| bacteria_Sep8d5b_40204 | 1 | 0 | 0 | 0 | 0 | 0 | 0 | 0 | 0 | 0 | 0 | 0 | 0 | 0 | 0 | 0 | 0 | 0 | 0 | 0 | 0 | 0 | 0 | 0 | 0 | 0 | 0 | 0 | 0 | 0 | 0 | 0 | 0 | 0 | 0 | 0 | 0 | 0 | 0 | 0 | 0 | 0 | 0 | 0 | 0 | 0 | 0 | 0 | 0 | 1 | 0 | 0 | 0 | 0 | 0 |
| bacteria_Sep8d5b_49247 | 1 | 0 | 0 | 0 | 0 | 0 | 0 | 0 | 0 | 0 | 0 | 0 | 0 | 0 | 0 | 0 | 0 | 0 | 0 | 0 | 0 | 0 | 0 | 0 | 0 | 0 | 0 | 0 | 0 | 0 | 0 | 0 | 0 | 0 | 0 | 0 | 0 | 0 | 0 | 0 | 0 | 0 | 0 | 0 | 0 | 0 | 0 | 0 | 0 | 1 | 0 | 0 | 0 | 0 | 0 |
| bacteria_Sep8d5b_54102 | 1 | 0 | 0 | 0 | 0 | 0 | 0 | 0 | 0 | 0 | 0 | 0 | 0 | 0 | 0 | 0 | 0 | 0 | 0 | 0 | 0 | 0 | 0 | 0 | 0 | 0 | 0 | 0 | 0 | 0 | 0 | 0 | 0 | 0 | 0 | 0 | 0 | 0 | 0 | 0 | 0 | 0 | 0 | 0 | 0 | 0 | 0 | 0 | 0 | 1 | 0 | 0 | 0 | 0 | 0 |
| bacteria_Sep8d5b_61998 | 1 | 0 | 0 | 0 | 0 | 0 | 0 | 0 | 0 | 0 | 0 | 0 | 0 | 0 | 0 | 0 | 0 | 0 | 0 | 0 | 0 | 0 | 0 | 0 | 0 | 0 | 0 | 0 | 0 | 0 | 0 | 0 | 0 | 0 | 0 | 0 | 0 | 0 | 0 | 0 | 0 | 0 | 0 | 0 | 0 | 0 | 0 | 0 | 0 | 1 | 0 | 0 | 0 | 0 | 0 |
| bacteria_Sep8d25b_5510 | 1 | 0 | 0 | 0 | 0 | 0 | 0 | 0 | 0 | 0 | 0 | 0 | 0 | 0 | 0 | 0 | 0 | 0 | 0 | 0 | 0 | 0 | 0 | 0 | 0 | 0 | 0 | 0 | 0 | 0 | 0 | 0 | 0 | 0 | 0 | 0 | 0 | 0 | 0 | 0 | 0 | 0 | 0 | 0 | 0 | 0 | 0 | 0 | 0 | 0 | 0 | 1 | 0 | 0 | 0 |
| bacteria_Sep8d25b_2344 | 1 | 0 | 0 | 0 | 0 | 0 | 0 | 0 | 0 | 0 | 0 | 0 | 0 | 0 | 0 | 0 | 0 | 0 | 0 | 0 | 0 | 0 | 0 | 0 | 0 | 0 | 0 | 0 | 0 | 0 | 0 | 0 | 0 | 0 | 0 | 0 | 0 | 0 | 0 | 0 | 0 | 0 | 0 | 0 | 0 | 0 | 0 | 0 | 0 | 0 | 0 | 1 | 0 | 0 | 0 |
| bacteria_Sep8d25b_27187 | 1 | 0 | 0 | 0 | 0 | 0 | 0 | 0 | 0 | 0 | 0 | 0 | 0 | 0 | 0 | 0 | 0 | 0 | 0 | 0 | 0 | 0 | 0 | 0 | 0 | 0 | 0 | 0 | 0 | 0 | 0 | 0 | 0 | 0 | 0 | 0 | 0 | 0 | 0 | 0 | 0 | 0 | 0 | 0 | 0 | 0 | 0 | 0 | 0 | 0 | 0 | 1 | 0 | 0 | 0 |
| bacteria_Sep8d25b_36989 | 1 | 0 | 0 | 0 | 0 | 0 | 0 | 0 | 0 | 0 | 0 | 0 | 0 | 0 | 0 | 0 | 0 | 0 | 0 | 0 | 0 | 0 | 0 | 0 | 0 | 0 | 0 | 0 | 0 | 0 | 0 | 0 | 0 | 0 | 0 | 0 | 0 | 0 | 0 | 0 | 0 | 0 | 0 | 0 | 0 | 0 | 0 | 0 | 0 | 0 | 0 | 1 | 0 | 0 | 0 |
| bacteria_Sep8d50b_37121 | 1 | 0 | 0 | 0 | 0 | 0 | 0 | 0 | 0 | 0 | 0 | 0 | 0 | 0 | 0 | 0 | 0 | 0 | 0 | 0 | 0 | 0 | 0 | 0 | 0 | 0 | 0 | 0 | 0 | 0 | 0 | 0 | 0 | 0 | 0 | 0 | 0 | 0 | 0 | 0 | 0 | 0 | 0 | 0 | 0 | 0 | 0 | 0 | 0 | 0 | 0 | 0 | 0 | 1 | 0 |
| virus_Aug5.0v_306 | 1 | 0 | 0 | 0 | 0 | 0 | 0 | 0 | 0 | 0 | 0 | 0 | 0 | 0 | 0 | 0 | 0 | 0 | 0 | 0 | 0 | 0 | 0 | 0 | 0 | 0 | 0 | 0 | 0 | 0 | 0 | 0 | 0 | 0 | 0 | 0 | 0 | 0 | 1 | 0 | 0 | 0 | 0 | 0 | 0 | 0 | 0 | 0 | 0 | 0 | 0 | 0 | 0 | 0 | 0 |
| virus_Aug5.0v_366 | 1 | 0 | 0 | 0 | 0 | 0 | 0 | 0 | 0 | 0 | 0 | 0 | 0 | 0 | 0 | 0 | 0 | 0 | 0 | 0 | 0 | 0 | 0 | 0 | 0 | 0 | 0 | 0 | 0 | 0 | 0 | 0 | 0 | 0 | 0 | 0 | 0 | 0 | 1 | 0 | 0 | 0 | 0 | 0 | 0 | 0 | 0 | 0 | 0 | 0 | 0 | 0 | 0 | 0 | 0 |
| virus_Aug5.0v_386 | 1 | 0 | 0 | 0 | 0 | 0 | 0 | 0 | 0 | 0 | 0 | 0 | 0 | 0 | 0 | 0 | 0 | 0 | 0 | 0 | 0 | 0 | 0 | 0 | 0 | 0 | 0 | 0 | 0 | 0 | 0 | 0 | 0 | 0 | 0 | 0 | 0 | 0 | 1 | 0 | 0 | 0 | 0 | 0 | 0 | 0 | 0 | 0 | 0 | 0 | 0 | 0 | 0 | 0 | 0 |
| virus_Aug5.0v_549 | 1 | 0 | 0 | 0 | 0 | 0 | 0 | 0 | 0 | 0 | 0 | 0 | 0 | 0 | 0 | 0 | 0 | 0 | 0 | 0 | 0 | 0 | 0 | 0 | 0 | 0 | 0 | 0 | 0 | 0 | 0 | 0 | 0 | 0 | 0 | 0 | 0 | 0 | 1 | 0 | 0 | 0 | 0 | 0 | 0 | 0 | 0 | 0 | 0 | 0 | 0 | 0 | 0 | 0 | 0 |
| virus_Aug5.0v_624 | 1 | 0 | 0 | 0 | 0 | 0 | 0 | 0 | 0 | 0 | 0 | 0 | 0 | 0 | 0 | 0 | 0 | 0 | 0 | 0 | 0 | 0 | 0 | 0 | 0 | 0 | 0 | 0 | 0 | 0 | 0 | 0 | 0 | 0 | 0 | 0 | 0 | 0 | 1 | 0 | 0 | 0 | 0 | 0 | 0 | 0 | 0 | 0 | 0 | 0 | 0 | 0 | 0 | 0 | 0 |
| virus_Aug5.0v_658 | 1 | 0 | 0 | 0 | 0 | 0 | 0 | 0 | 0 | 0 | 0 | 0 | 0 | 0 | 0 | 0 | 0 | 0 | 0 | 0 | 0 | 0 | 0 | 0 | 0 | 0 | 0 | 0 | 0 | 0 | 0 | 0 | 0 | 0 | 0 | 0 | 0 | 0 | 1 | 0 | 0 | 0 | 0 | 0 | 0 | 0 | 0 | 0 | 0 | 0 | 0 | 0 | 0 | 0 | 0 |
| virus_Aug5.0v_708 | 1 | 0 | 0 | 0 | 0 | 0 | 0 | 0 | 0 | 0 | 0 | 0 | 0 | 0 | 0 | 0 | 0 | 0 | 0 | 0 | 0 | 0 | 0 | 0 | 0 | 0 | 0 | 0 | 0 | 0 | 0 | 0 | 0 | 0 | 0 | 0 | 0 | 0 | 1 | 0 | 0 | 0 | 0 | 0 | 0 | 0 | 0 | 0 | 0 | 0 | 0 | 0 | 0 | 0 | 0 |
| virus_Aug5.0v_879 | 1 | 0 | 0 | 0 | 0 | 0 | 0 | 0 | 0 | 0 | 0 | 0 | 0 | 0 | 0 | 0 | 0 | 0 | 0 | 0 | 0 | 0 | 0 | 0 | 0 | 0 | 0 | 0 | 0 | 0 | 0 | 0 | 0 | 0 | 0 | 0 | 0 | 0 | 1 | 0 | 0 | 0 | 0 | 0 | 0 | 0 | 0 | 0 | 0 | 0 | 0 | 0 | 0 | 0 | 0 |
| virus_Aug5.0v_987 | 1 | 0 | 0 | 0 | 0 | 0 | 0 | 0 | 0 | 0 | 0 | 0 | 0 | 0 | 0 | 0 | 0 | 0 | 0 | 0 | 0 | 0 | 0 | 0 | 0 | 0 | 0 | 0 | 0 | 0 | 0 | 0 | 0 | 0 | 0 | 0 | 0 | 0 | 1 | 0 | 0 | 0 | 0 | 0 | 0 | 0 | 0 | 0 | 0 | 0 | 0 | 0 | 0 | 0 | 0 |
| virus_Aug5.0v_1092 | 1 | 0 | 0 | 0 | 0 | 0 | 0 | 0 | 0 | 0 | 0 | 0 | 0 | 0 | 0 | 0 | 0 | 0 | 0 | 0 | 0 | 0 | 0 | 0 | 0 | 0 | 0 | 0 | 0 | 0 | 0 | 0 | 0 | 0 | 0 | 0 | 0 | 0 | 1 | 0 | 0 | 0 | 0 | 0 | 0 | 0 | 0 | 0 | 0 | 0 | 0 | 0 | 0 | 0 | 0 |
| virus_Aug5.0v_1251 | 1 | 0 | 0 | 0 | 0 | 0 | 0 | 0 | 0 | 0 | 0 | 0 | 0 | 0 | 0 | 0 | 0 | 0 | 0 | 0 | 0 | 0 | 0 | 0 | 0 | 0 | 0 | 0 | 0 | 0 | 0 | 0 | 0 | 0 | 0 | 0 | 0 | 0 | 1 | 0 | 0 | 0 | 0 | 0 | 0 | 0 | 0 | 0 | 0 | 0 | 0 | 0 | 0 | 0 | 0 |
| virus_Aug5.0v_1426 | 1 | 0 | 0 | 0 | 0 | 0 | 0 | 0 | 0 | 0 | 0 | 0 | 0 | 0 | 0 | 0 | 0 | 0 | 0 | 0 | 0 | 0 | 0 | 0 | 0 | 0 | 0 | 0 | 0 | 0 | 0 | 0 | 0 | 0 | 0 | 0 | 0 | 0 | 1 | 0 | 0 | 0 | 0 | 0 | 0 | 0 | 0 | 0 | 0 | 0 | 0 | 0 | 0 | 0 | 0 |
| virus_Aug5.0v_1512 | 1 | 0 | 0 | 0 | 0 | 0 | 0 | 0 | 0 | 0 | 0 | 0 | 0 | 0 | 0 | 0 | 0 | 0 | 0 | 0 | 0 | 0 | 0 | 0 | 0 | 0 | 0 | 0 | 0 | 0 | 0 | 0 | 0 | 0 | 0 | 0 | 0 | 0 | 1 | 0 | 0 | 0 | 0 | 0 | 0 | 0 | 0 | 0 | 0 | 0 | 0 | 0 | 0 | 0 | 0 |
| virus_Aug5.0v_1608 | 1 | 0 | 0 | 0 | 0 | 0 | 0 | 0 | 0 | 0 | 0 | 0 | 0 | 0 | 0 | 0 | 0 | 0 | 0 | 0 | 0 | 0 | 0 | 0 | 0 | 0 | 0 | 0 | 0 | 0 | 0 | 0 | 0 | 0 | 0 | 0 | 0 | 0 | 1 | 0 | 0 | 0 | 0 | 0 | 0 | 0 | 0 | 0 | 0 | 0 | 0 | 0 | 0 | 0 | 0 |
| virus_Aug5.0v_1667 | 1 | 0 | 0 | 0 | 0 | 0 | 0 | 0 | 0 | 0 | 0 | 0 | 0 | 0 | 0 | 0 | 0 | 0 | 0 | 0 | 0 | 0 | 0 | 0 | 0 | 0 | 0 | 0 | 0 | 0 | 0 | 0 | 0 | 0 | 0 | 0 | 0 | 0 | 1 | 0 | 0 | 0 | 0 | 0 | 0 | 0 | 0 | 0 | 0 | 0 | 0 | 0 | 0 | 0 | 0 |
| virus_Aug5.0v_1708 | 1 | 0 | 0 | 0 | 0 | 0 | 0 | 0 | 0 | 0 | 0 | 0 | 0 | 0 | 0 | 0 | 0 | 0 | 0 | 0 | 0 | 0 | 0 | 0 | 0 | 0 | 0 | 0 | 0 | 0 | 0 | 0 | 0 | 0 | 0 | 0 | 0 | 0 | 1 | 0 | 0 | 0 | 0 | 0 | 0 | 0 | 0 | 0 | 0 | 0 | 0 | 0 | 0 | 0 | 0 |
| virus_Aug5.0v_1835 | 1 | 0 | 0 | 0 | 0 | 0 | 0 | 0 | 0 | 0 | 0 | 0 | 0 | 0 | 0 | 0 | 0 | 0 | 0 | 0 | 0 | 0 | 0 | 0 | 0 | 0 | 0 | 0 | 0 | 0 | 0 | 0 | 0 | 0 | 0 | 0 | 0 | 0 | 1 | 0 | 0 | 0 | 0 | 0 | 0 | 0 | 0 | 0 | 0 | 0 | 0 | 0 | 0 | 0 | 0 |
| virus_Aug5.0v_2441 | 1 | 0 | 0 | 0 | 0 | 0 | 0 | 0 | 0 | 0 | 0 | 0 | 0 | 0 | 0 | 0 | 0 | 0 | 0 | 0 | 0 | 0 | 0 | 0 | 0 | 0 | 0 | 0 | 0 | 0 | 0 | 0 | 0 | 0 | 0 | 0 | 0 | 0 | 1 | 0 | 0 | 0 | 0 | 0 | 0 | 0 | 0 | 0 | 0 | 0 | 0 | 0 | 0 | 0 | 0 |
| virus_Aug5.0v_3602 | 1 | 0 | 0 | 0 | 0 | 0 | 0 | 0 | 0 | 0 | 0 | 0 | 0 | 0 | 0 | 0 | 0 | 0 | 0 | 0 | 0 | 0 | 0 | 0 | 0 | 0 | 0 | 0 | 0 | 0 | 0 | 0 | 0 | 0 | 0 | 0 | 0 | 0 | 1 | 0 | 0 | 0 | 0 | 0 | 0 | 0 | 0 | 0 | 0 | 0 | 0 | 0 | 0 | 0 | 0 |
| virus_Aug5.0v_4028 | 1 | 0 | 0 | 0 | 0 | 0 | 0 | 0 | 0 | 0 | 0 | 0 | 0 | 0 | 0 | 0 | 0 | 0 | 0 | 0 | 0 | 0 | 0 | 0 | 0 | 0 | 0 | 0 | 0 | 0 | 0 | 0 | 0 | 0 | 0 | 0 | 0 | 0 | 1 | 0 | 0 | 0 | 0 | 0 | 0 | 0 | 0 | 0 | 0 | 0 | 0 | 0 | 0 | 0 | 0 |
| virus_Aug5.0v_4122 | 1 | 0 | 0 | 0 | 0 | 0 | 0 | 0 | 0 | 0 | 0 | 0 | 0 | 0 | 0 | 0 | 0 | 0 | 0 | 0 | 0 | 0 | 0 | 0 | 0 | 0 | 0 | 0 | 0 | 0 | 0 | 0 | 0 | 0 | 0 | 0 | 0 | 0 | 1 | 0 | 0 | 0 | 0 | 0 | 0 | 0 | 0 | 0 | 0 | 0 | 0 | 0 | 0 | 0 | 0 |
| virus_Aug5.0v_4191 | 1 | 0 | 0 | 0 | 0 | 0 | 0 | 0 | 0 | 0 | 0 | 0 | 0 | 0 | 0 | 0 | 0 | 0 | 0 | 0 | 0 | 0 | 0 | 0 | 0 | 0 | 0 | 0 | 0 | 0 | 0 | 0 | 0 | 0 | 0 | 0 | 0 | 0 | 1 | 0 | 0 | 0 | 0 | 0 | 0 | 0 | 0 | 0 | 0 | 0 | 0 | 0 | 0 | 0 | 0 |
| virus_Aug5.0v_4641 | 1 | 0 | 0 | 0 | 0 | 0 | 0 | 0 | 0 | 0 | 0 | 0 | 0 | 0 | 0 | 0 | 0 | 0 | 0 | 0 | 0 | 0 | 0 | 0 | 0 | 0 | 0 | 0 | 0 | 0 | 0 | 0 | 0 | 0 | 0 | 0 | 0 | 0 | 1 | 0 | 0 | 0 | 0 | 0 | 0 | 0 | 0 | 0 | 0 | 0 | 0 | 0 | 0 | 0 | 0 |
| virus_Aug5.0v_4912 | 1 | 0 | 0 | 0 | 0 | 0 | 0 | 0 | 0 | 0 | 0 | 0 | 0 | 0 | 0 | 0 | 0 | 0 | 0 | 0 | 0 | 0 | 0 | 0 | 0 | 0 | 0 | 0 | 0 | 0 | 0 | 0 | 0 | 0 | 0 | 0 | 0 | 0 | 1 | 0 | 0 | 0 | 0 | 0 | 0 | 0 | 0 | 0 | 0 | 0 | 0 | 0 | 0 | 0 | 0 |
| virus_Aug5.0v_5136 | 1 | 0 | 0 | 0 | 0 | 0 | 0 | 0 | 0 | 0 | 0 | 0 | 0 | 0 | 0 | 0 | 0 | 0 | 0 | 0 | 0 | 0 | 0 | 0 | 0 | 0 | 0 | 0 | 0 | 0 | 0 | 0 | 0 | 0 | 0 | 0 | 0 | 0 | 1 | 0 | 0 | 0 | 0 | 0 | 0 | 0 | 0 | 0 | 0 | 0 | 0 | 0 | 0 | 0 | 0 |
| virus_Aug5.0v_5212 | 1 | 0 | 0 | 0 | 0 | 0 | 0 | 0 | 0 | 0 | 0 | 0 | 0 | 0 | 0 | 0 | 0 | 0 | 0 | 0 | 0 | 0 | 0 | 0 | 0 | 0 | 0 | 0 | 0 | 0 | 0 | 0 | 0 | 0 | 0 | 0 | 0 | 0 | 1 | 0 | 0 | 0 | 0 | 0 | 0 | 0 | 0 | 0 | 0 | 0 | 0 | 0 | 0 | 0 | 0 |
| virus_Aug5.0v_5430 | 1 | 0 | 0 | 0 | 0 | 0 | 0 | 0 | 0 | 0 | 0 | 0 | 0 | 0 | 0 | 0 | 0 | 0 | 0 | 0 | 0 | 0 | 0 | 0 | 0 | 0 | 0 | 0 | 0 | 0 | 0 | 0 | 0 | 0 | 0 | 0 | 0 | 0 | 1 | 0 | 0 | 0 | 0 | 0 | 0 | 0 | 0 | 0 | 0 | 0 | 0 | 0 | 0 | 0 | 0 |
| virus_Aug5.5v_427 | 1 | 0 | 0 | 0 | 0 | 0 | 0 | 0 | 0 | 0 | 0 | 0 | 0 | 0 | 0 | 0 | 0 | 0 | 0 | 0 | 0 | 0 | 0 | 0 | 0 | 0 | 0 | 0 | 0 | 0 | 0 | 0 | 0 | 0 | 0 | 0 | 0 | 0 | 0 | 0 | 1 | 0 | 0 | 0 | 0 | 0 | 0 | 0 | 0 | 0 | 0 | 0 | 0 | 0 | 0 |
| virus_Aug5.5v_8232 | 1 | 0 | 0 | 0 | 0 | 0 | 0 | 0 | 0 | 0 | 0 | 0 | 0 | 0 | 0 | 0 | 0 | 0 | 0 | 0 | 0 | 0 | 0 | 0 | 0 | 0 | 0 | 0 | 0 | 0 | 0 | 0 | 0 | 0 | 0 | 0 | 0 | 0 | 0 | 0 | 1 | 0 | 0 | 0 | 0 | 0 | 0 | 0 | 0 | 0 | 0 | 0 | 0 | 0 | 0 |
| virus_Aug5.5v_9607 | 1 | 0 | 0 | 0 | 0 | 0 | 0 | 0 | 0 | 0 | 0 | 0 | 0 | 0 | 0 | 0 | 0 | 0 | 0 | 0 | 0 | 0 | 0 | 0 | 0 | 0 | 0 | 0 | 0 | 0 | 0 | 0 | 0 | 0 | 0 | 0 | 0 | 0 | 0 | 0 | 1 | 0 | 0 | 0 | 0 | 0 | 0 | 0 | 0 | 0 | 0 | 0 | 0 | 0 | 0 |
| virus_Aug5.5v_11130 | 1 | 0 | 0 | 0 | 0 | 0 | 0 | 0 | 0 | 0 | 0 | 0 | 0 | 0 | 0 | 0 | 0 | 0 | 0 | 0 | 0 | 0 | 0 | 0 | 0 | 0 | 0 | 0 | 0 | 0 | 0 | 0 | 0 | 0 | 0 | 0 | 0 | 0 | 0 | 0 | 1 | 0 | 0 | 0 | 0 | 0 | 0 | 0 | 0 | 0 | 0 | 0 | 0 | 0 | 0 |
| virus_Aug5.5v_13719 | 1 | 0 | 0 | 0 | 0 | 0 | 0 | 0 | 0 | 0 | 0 | 0 | 0 | 0 | 0 | 0 | 0 | 0 | 0 | 0 | 0 | 0 | 0 | 0 | 0 | 0 | 0 | 0 | 0 | 0 | 0 | 0 | 0 | 0 | 0 | 0 | 0 | 0 | 0 | 0 | 1 | 0 | 0 | 0 | 0 | 0 | 0 | 0 | 0 | 0 | 0 | 0 | 0 | 0 | 0 |
| virus_Aug5.5v_15451 | 1 | 0 | 0 | 0 | 0 | 0 | 0 | 0 | 0 | 0 | 0 | 0 | 0 | 0 | 0 | 0 | 0 | 0 | 0 | 0 | 0 | 0 | 0 | 0 | 0 | 0 | 0 | 0 | 0 | 0 | 0 | 0 | 0 | 0 | 0 | 0 | 0 | 0 | 0 | 0 | 1 | 0 | 0 | 0 | 0 | 0 | 0 | 0 | 0 | 0 | 0 | 0 | 0 | 0 | 0 |
| virus_Aug5.5v_19338 | 1 | 0 | 0 | 0 | 0 | 0 | 0 | 0 | 0 | 0 | 0 | 0 | 0 | 0 | 0 | 0 | 0 | 0 | 0 | 0 | 0 | 0 | 0 | 0 | 0 | 0 | 0 | 0 | 0 | 0 | 0 | 0 | 0 | 0 | 0 | 0 | 0 | 0 | 0 | 0 | 1 | 0 | 0 | 0 | 0 | 0 | 0 | 0 | 0 | 0 | 0 | 0 | 0 | 0 | 0 |
| virus_Aug5.5v_24009 | 1 | 0 | 0 | 0 | 0 | 0 | 0 | 0 | 0 | 0 | 0 | 0 | 0 | 0 | 0 | 0 | 0 | 0 | 0 | 0 | 0 | 0 | 0 | 0 | 0 | 0 | 0 | 0 | 0 | 0 | 0 | 0 | 0 | 0 | 0 | 0 | 0 | 0 | 0 | 0 | 1 | 0 | 0 | 0 | 0 | 0 | 0 | 0 | 0 | 0 | 0 | 0 | 0 | 0 | 0 |
| virus_Aug5.5v_24892 | 1 | 0 | 0 | 0 | 0 | 0 | 0 | 0 | 0 | 0 | 0 | 0 | 0 | 0 | 0 | 0 | 0 | 0 | 0 | 0 | 0 | 0 | 0 | 0 | 0 | 0 | 0 | 0 | 0 | 0 | 0 | 0 | 0 | 0 | 0 | 0 | 0 | 0 | 0 | 0 | 1 | 0 | 0 | 0 | 0 | 0 | 0 | 0 | 0 | 0 | 0 | 0 | 0 | 0 | 0 |
| virus_Aug5.5v_24908 | 1 | 0 | 0 | 0 | 0 | 0 | 0 | 0 | 0 | 0 | 0 | 0 | 0 | 0 | 0 | 0 | 0 | 0 | 0 | 0 | 0 | 0 | 0 | 0 | 0 | 0 | 0 | 0 | 0 | 0 | 0 | 0 | 0 | 0 | 0 | 0 | 0 | 0 | 0 | 0 | 1 | 0 | 0 | 0 | 0 | 0 | 0 | 0 | 0 | 0 | 0 | 0 | 0 | 0 | 0 |
| virus_Aug5.5v_25676 | 1 | 0 | 0 | 0 | 0 | 0 | 0 | 0 | 0 | 0 | 0 | 0 | 0 | 0 | 0 | 0 | 0 | 0 | 0 | 0 | 0 | 0 | 0 | 0 | 0 | 0 | 0 | 0 | 0 | 0 | 0 | 0 | 0 | 0 | 0 | 0 | 0 | 0 | 0 | 0 | 1 | 0 | 0 | 0 | 0 | 0 | 0 | 0 | 0 | 0 | 0 | 0 | 0 | 0 | 0 |
| virus_Aug5.5v_27007 | 1 | 0 | 0 | 0 | 0 | 0 | 0 | 0 | 0 | 0 | 0 | 0 | 0 | 0 | 0 | 0 | 0 | 0 | 0 | 0 | 0 | 0 | 0 | 0 | 0 | 0 | 0 | 0 | 0 | 0 | 0 | 0 | 0 | 0 | 0 | 0 | 0 | 0 | 0 | 0 | 1 | 0 | 0 | 0 | 0 | 0 | 0 | 0 | 0 | 0 | 0 | 0 | 0 | 0 | 0 |
| virus_Aug5.5v_30894 | 1 | 0 | 0 | 0 | 0 | 0 | 0 | 0 | 0 | 0 | 0 | 0 | 0 | 0 | 0 | 0 | 0 | 0 | 0 | 0 | 0 | 0 | 0 | 0 | 0 | 0 | 0 | 0 | 0 | 0 | 0 | 0 | 0 | 0 | 0 | 0 | 0 | 0 | 0 | 0 | 1 | 0 | 0 | 0 | 0 | 0 | 0 | 0 | 0 | 0 | 0 | 0 | 0 | 0 | 0 |
| virus_Aug5.5v_31730 | 1 | 0 | 0 | 0 | 0 | 0 | 0 | 0 | 0 | 0 | 0 | 0 | 0 | 0 | 0 | 0 | 0 | 0 | 0 | 0 | 0 | 0 | 0 | 0 | 0 | 0 | 0 | 0 | 0 | 0 | 0 | 0 | 0 | 0 | 0 | 0 | 0 | 0 | 0 | 0 | 1 | 0 | 0 | 0 | 0 | 0 | 0 | 0 | 0 | 0 | 0 | 0 | 0 | 0 | 0 |
| virus_Aug5.5v_31869 | 1 | 0 | 0 | 0 | 0 | 0 | 0 | 0 | 0 | 0 | 0 | 0 | 0 | 0 | 0 | 0 | 0 | 0 | 0 | 0 | 0 | 0 | 0 | 0 | 0 | 0 | 0 | 0 | 0 | 0 | 0 | 0 | 0 | 0 | 0 | 0 | 0 | 0 | 0 | 0 | 1 | 0 | 0 | 0 | 0 | 0 | 0 | 0 | 0 | 0 | 0 | 0 | 0 | 0 | 0 |
| virus_Aug5.5v_33528 | 1 | 0 | 0 | 0 | 0 | 0 | 0 | 0 | 0 | 0 | 0 | 0 | 0 | 0 | 0 | 0 | 0 | 0 | 0 | 0 | 0 | 0 | 0 | 0 | 0 | 0 | 0 | 0 | 0 | 0 | 0 | 0 | 0 | 0 | 0 | 0 | 0 | 0 | 0 | 0 | 1 | 0 | 0 | 0 | 0 | 0 | 0 | 0 | 0 | 0 | 0 | 0 | 0 | 0 | 0 |
| virus_Aug5.25v_7919 | 1 | 0 | 0 | 0 | 0 | 0 | 0 | 0 | 0 | 0 | 0 | 0 | 0 | 0 | 0 | 0 | 0 | 0 | 0 | 0 | 0 | 0 | 0 | 0 | 0 | 0 | 0 | 0 | 0 | 0 | 0 | 0 | 0 | 0 | 0 | 0 | 0 | 0 | 0 | 0 | 0 | 0 | 1 | 0 | 0 | 0 | 0 | 0 | 0 | 0 | 0 | 0 | 0 | 0 | 0 |
| virus_Aug5.25v_18348 | 1 | 0 | 0 | 0 | 0 | 0 | 0 | 0 | 0 | 0 | 0 | 0 | 0 | 0 | 0 | 0 | 0 | 0 | 0 | 0 | 0 | 0 | 0 | 0 | 0 | 0 | 0 | 0 | 0 | 0 | 0 | 0 | 0 | 0 | 0 | 0 | 0 | 0 | 0 | 0 | 0 | 0 | 1 | 0 | 0 | 0 | 0 | 0 | 0 | 0 | 0 | 0 | 0 | 0 | 0 |
| virus_Aug5.25v_19141 | 1 | 0 | 0 | 0 | 0 | 0 | 0 | 0 | 0 | 0 | 0 | 0 | 0 | 0 | 0 | 0 | 0 | 0 | 0 | 0 | 0 | 0 | 0 | 0 | 0 | 0 | 0 | 0 | 0 | 0 | 0 | 0 | 0 | 0 | 0 | 0 | 0 | 0 | 0 | 0 | 0 | 0 | 1 | 0 | 0 | 0 | 0 | 0 | 0 | 0 | 0 | 0 | 0 | 0 | 0 |
| virus_Aug5.25v_21303 | 1 | 0 | 0 | 0 | 0 | 0 | 0 | 0 | 0 | 0 | 0 | 0 | 0 | 0 | 0 | 0 | 0 | 0 | 0 | 0 | 0 | 0 | 0 | 0 | 0 | 0 | 0 | 0 | 0 | 0 | 0 | 0 | 0 | 0 | 0 | 0 | 0 | 0 | 0 | 0 | 0 | 0 | 1 | 0 | 0 | 0 | 0 | 0 | 0 | 0 | 0 | 0 | 0 | 0 | 0 |
| virus_Aug5.25v_22287 | 1 | 0 | 0 | 0 | 0 | 0 | 0 | 0 | 0 | 0 | 0 | 0 | 0 | 0 | 0 | 0 | 0 | 0 | 0 | 0 | 0 | 0 | 0 | 0 | 0 | 0 | 0 | 0 | 0 | 0 | 0 | 0 | 0 | 0 | 0 | 0 | 0 | 0 | 0 | 0 | 0 | 0 | 1 | 0 | 0 | 0 | 0 | 0 | 0 | 0 | 0 | 0 | 0 | 0 | 0 |
| virus_Aug5.25v_22964 | 1 | 0 | 0 | 0 | 0 | 0 | 0 | 0 | 0 | 0 | 0 | 0 | 0 | 0 | 0 | 0 | 0 | 0 | 0 | 0 | 0 | 0 | 0 | 0 | 0 | 0 | 0 | 0 | 0 | 0 | 0 | 0 | 0 | 0 | 0 | 0 | 0 | 0 | 0 | 0 | 0 | 0 | 1 | 0 | 0 | 0 | 0 | 0 | 0 | 0 | 0 | 0 | 0 | 0 | 0 |
| virus_Aug5.25v_24202 | 1 | 0 | 0 | 0 | 0 | 0 | 0 | 0 | 0 | 0 | 0 | 0 | 0 | 0 | 0 | 0 | 0 | 0 | 0 | 0 | 0 | 0 | 0 | 0 | 0 | 0 | 0 | 0 | 0 | 0 | 0 | 0 | 0 | 0 | 0 | 0 | 0 | 0 | 0 | 0 | 0 | 0 | 1 | 0 | 0 | 0 | 0 | 0 | 0 | 0 | 0 | 0 | 0 | 0 | 0 |
| virus_Aug5.25v_28461 | 1 | 0 | 0 | 0 | 0 | 0 | 0 | 0 | 0 | 0 | 0 | 0 | 0 | 0 | 0 | 0 | 0 | 0 | 0 | 0 | 0 | 0 | 0 | 0 | 0 | 0 | 0 | 0 | 0 | 0 | 0 | 0 | 0 | 0 | 0 | 0 | 0 | 0 | 0 | 0 | 0 | 0 | 1 | 0 | 0 | 0 | 0 | 0 | 0 | 0 | 0 | 0 | 0 | 0 | 0 |
| virus_Aug5.25v_30080 | 1 | 0 | 0 | 0 | 0 | 0 | 0 | 0 | 0 | 0 | 0 | 0 | 0 | 0 | 0 | 0 | 0 | 0 | 0 | 0 | 0 | 0 | 0 | 0 | 0 | 0 | 0 | 0 | 0 | 0 | 0 | 0 | 0 | 0 | 0 | 0 | 0 | 0 | 0 | 0 | 0 | 0 | 1 | 0 | 0 | 0 | 0 | 0 | 0 | 0 | 0 | 0 | 0 | 0 | 0 |
| virus_Aug5.25v_30997 | 1 | 0 | 0 | 0 | 0 | 0 | 0 | 0 | 0 | 0 | 0 | 0 | 0 | 0 | 0 | 0 | 0 | 0 | 0 | 0 | 0 | 0 | 0 | 0 | 0 | 0 | 0 | 0 | 0 | 0 | 0 | 0 | 0 | 0 | 0 | 0 | 0 | 0 | 0 | 0 | 0 | 0 | 1 | 0 | 0 | 0 | 0 | 0 | 0 | 0 | 0 | 0 | 0 | 0 | 0 |
| virus_Aug5.25v_35003 | 1 | 0 | 0 | 0 | 0 | 0 | 0 | 0 | 0 | 0 | 0 | 0 | 0 | 0 | 0 | 0 | 0 | 0 | 0 | 0 | 0 | 0 | 0 | 0 | 0 | 0 | 0 | 0 | 0 | 0 | 0 | 0 | 0 | 0 | 0 | 0 | 0 | 0 | 0 | 0 | 0 | 0 | 1 | 0 | 0 | 0 | 0 | 0 | 0 | 0 | 0 | 0 | 0 | 0 | 0 |
| virus_Aug5.25v_35301 | 1 | 0 | 0 | 0 | 0 | 0 | 0 | 0 | 0 | 0 | 0 | 0 | 0 | 0 | 0 | 0 | 0 | 0 | 0 | 0 | 0 | 0 | 0 | 0 | 0 | 0 | 0 | 0 | 0 | 0 | 0 | 0 | 0 | 0 | 0 | 0 | 0 | 0 | 0 | 0 | 0 | 0 | 1 | 0 | 0 | 0 | 0 | 0 | 0 | 0 | 0 | 0 | 0 | 0 | 0 |
| virus_Aug5.25v_38816 | 1 | 0 | 0 | 0 | 0 | 0 | 0 | 0 | 0 | 0 | 0 | 0 | 0 | 0 | 0 | 0 | 0 | 0 | 0 | 0 | 0 | 0 | 0 | 0 | 0 | 0 | 0 | 0 | 0 | 0 | 0 | 0 | 0 | 0 | 0 | 0 | 0 | 0 | 0 | 0 | 0 | 0 | 1 | 0 | 0 | 0 | 0 | 0 | 0 | 0 | 0 | 0 | 0 | 0 | 0 |
| virus_Aug5.25v_39418 | 1 | 0 | 0 | 0 | 0 | 0 | 0 | 0 | 0 | 0 | 0 | 0 | 0 | 0 | 0 | 0 | 0 | 0 | 0 | 0 | 0 | 0 | 0 | 0 | 0 | 0 | 0 | 0 | 0 | 0 | 0 | 0 | 0 | 0 | 0 | 0 | 0 | 0 | 0 | 0 | 0 | 0 | 1 | 0 | 0 | 0 | 0 | 0 | 0 | 0 | 0 | 0 | 0 | 0 | 0 |
| virus_Aug5.50v_7594 | 1 | 0 | 0 | 0 | 0 | 0 | 0 | 0 | 0 | 0 | 0 | 0 | 0 | 0 | 0 | 0 | 0 | 0 | 0 | 0 | 0 | 0 | 0 | 0 | 0 | 0 | 0 | 0 | 0 | 0 | 0 | 0 | 0 | 0 | 0 | 0 | 0 | 0 | 0 | 0 | 0 | 0 | 0 | 0 | 1 | 0 | 0 | 0 | 0 | 0 | 0 | 0 | 0 | 0 | 0 |
| virus_Aug5.50v_7694 | 1 | 0 | 0 | 0 | 0 | 0 | 0 | 0 | 0 | 0 | 0 | 0 | 0 | 0 | 0 | 0 | 0 | 0 | 0 | 0 | 0 | 0 | 0 | 0 | 0 | 0 | 0 | 0 | 0 | 0 | 0 | 0 | 0 | 0 | 0 | 0 | 0 | 0 | 0 | 0 | 0 | 0 | 0 | 0 | 1 | 0 | 0 | 0 | 0 | 0 | 0 | 0 | 0 | 0 | 0 |
| virus_Aug5.50v_18854 | 1 | 0 | 0 | 0 | 0 | 0 | 0 | 0 | 0 | 0 | 0 | 0 | 0 | 0 | 0 | 0 | 0 | 0 | 0 | 0 | 0 | 0 | 0 | 0 | 0 | 0 | 0 | 0 | 0 | 0 | 0 | 0 | 0 | 0 | 0 | 0 | 0 | 0 | 0 | 0 | 0 | 0 | 0 | 0 | 1 | 0 | 0 | 0 | 0 | 0 | 0 | 0 | 0 | 0 | 0 |
| virus_Aug5.50v_22967 | 1 | 0 | 0 | 0 | 0 | 0 | 0 | 0 | 0 | 0 | 0 | 0 | 0 | 0 | 0 | 0 | 0 | 0 | 0 | 0 | 0 | 0 | 0 | 0 | 0 | 0 | 0 | 0 | 0 | 0 | 0 | 0 | 0 | 0 | 0 | 0 | 0 | 0 | 0 | 0 | 0 | 0 | 0 | 0 | 1 | 0 | 0 | 0 | 0 | 0 | 0 | 0 | 0 | 0 | 0 |
| virus_Aug5.50v_26854 | 1 | 0 | 0 | 0 | 0 | 0 | 0 | 0 | 0 | 0 | 0 | 0 | 0 | 0 | 0 | 0 | 0 | 0 | 0 | 0 | 0 | 0 | 0 | 0 | 0 | 0 | 0 | 0 | 0 | 0 | 0 | 0 | 0 | 0 | 0 | 0 | 0 | 0 | 0 | 0 | 0 | 0 | 0 | 0 | 1 | 0 | 0 | 0 | 0 | 0 | 0 | 0 | 0 | 0 | 0 |
| virus_Aug5.50v_31053 | 1 | 0 | 0 | 0 | 0 | 0 | 0 | 0 | 0 | 0 | 0 | 0 | 0 | 0 | 0 | 0 | 0 | 0 | 0 | 0 | 0 | 0 | 0 | 0 | 0 | 0 | 0 | 0 | 0 | 0 | 0 | 0 | 0 | 0 | 0 | 0 | 0 | 0 | 0 | 0 | 0 | 0 | 0 | 0 | 1 | 0 | 0 | 0 | 0 | 0 | 0 | 0 | 0 | 0 | 0 |
| virus_Aug5.50v_31512 | 1 | 0 | 0 | 0 | 0 | 0 | 0 | 0 | 0 | 0 | 0 | 0 | 0 | 0 | 0 | 0 | 0 | 0 | 0 | 0 | 0 | 0 | 0 | 0 | 0 | 0 | 0 | 0 | 0 | 0 | 0 | 0 | 0 | 0 | 0 | 0 | 0 | 0 | 0 | 0 | 0 | 0 | 0 | 0 | 1 | 0 | 0 | 0 | 0 | 0 | 0 | 0 | 0 | 0 | 0 |
| virus_Aug5.50v_33561 | 1 | 0 | 0 | 0 | 0 | 0 | 0 | 0 | 0 | 0 | 0 | 0 | 0 | 0 | 0 | 0 | 0 | 0 | 0 | 0 | 0 | 0 | 0 | 0 | 0 | 0 | 0 | 0 | 0 | 0 | 0 | 0 | 0 | 0 | 0 | 0 | 0 | 0 | 0 | 0 | 0 | 0 | 0 | 0 | 1 | 0 | 0 | 0 | 0 | 0 | 0 | 0 | 0 | 0 | 0 |
| virus_Aug5.50v_36307 | 1 | 0 | 0 | 0 | 0 | 0 | 0 | 0 | 0 | 0 | 0 | 0 | 0 | 0 | 0 | 0 | 0 | 0 | 0 | 0 | 0 | 0 | 0 | 0 | 0 | 0 | 0 | 0 | 0 | 0 | 0 | 0 | 0 | 0 | 0 | 0 | 0 | 0 | 0 | 0 | 0 | 0 | 0 | 0 | 1 | 0 | 0 | 0 | 0 | 0 | 0 | 0 | 0 | 0 | 0 |
| virus_Aug5.75v_11363 | 1 | 0 | 0 | 0 | 0 | 0 | 0 | 0 | 0 | 0 | 0 | 0 | 0 | 0 | 0 | 0 | 0 | 0 | 0 | 0 | 0 | 0 | 0 | 0 | 0 | 0 | 0 | 0 | 0 | 0 | 0 | 0 | 0 | 0 | 0 | 0 | 0 | 0 | 0 | 0 | 0 | 0 | 0 | 0 | 0 | 0 | 1 | 0 | 0 | 0 | 0 | 0 | 0 | 0 | 0 |
| virus_Aug5.75v_24045 | 1 | 0 | 0 | 0 | 0 | 0 | 0 | 0 | 0 | 0 | 0 | 0 | 0 | 0 | 0 | 0 | 0 | 0 | 0 | 0 | 0 | 0 | 0 | 0 | 0 | 0 | 0 | 0 | 0 | 0 | 0 | 0 | 0 | 0 | 0 | 0 | 0 | 0 | 0 | 0 | 0 | 0 | 0 | 0 | 0 | 0 | 1 | 0 | 0 | 0 | 0 | 0 | 0 | 0 | 0 |
| virus_Aug5.75v_27115 | 1 | 0 | 0 | 0 | 0 | 0 | 0 | 0 | 0 | 0 | 0 | 0 | 0 | 0 | 0 | 0 | 0 | 0 | 0 | 0 | 0 | 0 | 0 | 0 | 0 | 0 | 0 | 0 | 0 | 0 | 0 | 0 | 0 | 0 | 0 | 0 | 0 | 0 | 0 | 0 | 0 | 0 | 0 | 0 | 0 | 0 | 1 | 0 | 0 | 0 | 0 | 0 | 0 | 0 | 0 |
| virus_Aug5.75v_27904 | 1 | 0 | 0 | 0 | 0 | 0 | 0 | 0 | 0 | 0 | 0 | 0 | 0 | 0 | 0 | 0 | 0 | 0 | 0 | 0 | 0 | 0 | 0 | 0 | 0 | 0 | 0 | 0 | 0 | 0 | 0 | 0 | 0 | 0 | 0 | 0 | 0 | 0 | 0 | 0 | 0 | 0 | 0 | 0 | 0 | 0 | 1 | 0 | 0 | 0 | 0 | 0 | 0 | 0 | 0 |
| virus_Aug5.75v_28382 | 1 | 0 | 0 | 0 | 0 | 0 | 0 | 0 | 0 | 0 | 0 | 0 | 0 | 0 | 0 | 0 | 0 | 0 | 0 | 0 | 0 | 0 | 0 | 0 | 0 | 0 | 0 | 0 | 0 | 0 | 0 | 0 | 0 | 0 | 0 | 0 | 0 | 0 | 0 | 0 | 0 | 0 | 0 | 0 | 0 | 0 | 1 | 0 | 0 | 0 | 0 | 0 | 0 | 0 | 0 |
| virus_Aug5.75v_32948 | 1 | 0 | 0 | 0 | 0 | 0 | 0 | 0 | 0 | 0 | 0 | 0 | 0 | 0 | 0 | 0 | 0 | 0 | 0 | 0 | 0 | 0 | 0 | 0 | 0 | 0 | 0 | 0 | 0 | 0 | 0 | 0 | 0 | 0 | 0 | 0 | 0 | 0 | 0 | 0 | 0 | 0 | 0 | 0 | 0 | 0 | 1 | 0 | 0 | 0 | 0 | 0 | 0 | 0 | 0 |
| virus_Aug5.75v_37632 | 1 | 0 | 0 | 0 | 0 | 0 | 0 | 0 | 0 | 0 | 0 | 0 | 0 | 0 | 0 | 0 | 0 | 0 | 0 | 0 | 0 | 0 | 0 | 0 | 0 | 0 | 0 | 0 | 0 | 0 | 0 | 0 | 0 | 0 | 0 | 0 | 0 | 0 | 0 | 0 | 0 | 0 | 0 | 0 | 0 | 0 | 1 | 0 | 0 | 0 | 0 | 0 | 0 | 0 | 0 |
| virus_Aug5.75v_39058 | 1 | 0 | 0 | 0 | 0 | 0 | 0 | 0 | 0 | 0 | 0 | 0 | 0 | 0 | 0 | 0 | 0 | 0 | 0 | 0 | 0 | 0 | 0 | 0 | 0 | 0 | 0 | 0 | 0 | 0 | 0 | 0 | 0 | 0 | 0 | 0 | 0 | 0 | 0 | 0 | 0 | 0 | 0 | 0 | 0 | 0 | 1 | 0 | 0 | 0 | 0 | 0 | 0 | 0 | 0 |
| virus_Jul11.0v_542 | 1 | 0 | 0 | 0 | 0 | 0 | 0 | 0 | 0 | 0 | 0 | 0 | 0 | 0 | 0 | 0 | 0 | 0 | 0 | 0 | 0 | 0 | 1 | 0 | 0 | 0 | 0 | 0 | 0 | 0 | 0 | 0 | 0 | 0 | 0 | 0 | 0 | 0 | 0 | 0 | 0 | 0 | 0 | 0 | 0 | 0 | 0 | 0 | 0 | 0 | 0 | 0 | 0 | 0 | 0 |
| virus_Jul11.0v_3981 | 1 | 0 | 0 | 0 | 0 | 0 | 0 | 0 | 0 | 0 | 0 | 0 | 0 | 0 | 0 | 0 | 0 | 0 | 0 | 0 | 0 | 0 | 1 | 0 | 0 | 0 | 0 | 0 | 0 | 0 | 0 | 0 | 0 | 0 | 0 | 0 | 0 | 0 | 0 | 0 | 0 | 0 | 0 | 0 | 0 | 0 | 0 | 0 | 0 | 0 | 0 | 0 | 0 | 0 | 0 |
| virus_Jul11.0v_5391 | 1 | 0 | 0 | 0 | 0 | 0 | 0 | 0 | 0 | 0 | 0 | 0 | 0 | 0 | 0 | 0 | 0 | 0 | 0 | 0 | 0 | 0 | 1 | 0 | 0 | 0 | 0 | 0 | 0 | 0 | 0 | 0 | 0 | 0 | 0 | 0 | 0 | 0 | 0 | 0 | 0 | 0 | 0 | 0 | 0 | 0 | 0 | 0 | 0 | 0 | 0 | 0 | 0 | 0 | 0 |
| virus_Jul11.0v_8193 | 1 | 0 | 0 | 0 | 0 | 0 | 0 | 0 | 0 | 0 | 0 | 0 | 0 | 0 | 0 | 0 | 0 | 0 | 0 | 0 | 0 | 0 | 1 | 0 | 0 | 0 | 0 | 0 | 0 | 0 | 0 | 0 | 0 | 0 | 0 | 0 | 0 | 0 | 0 | 0 | 0 | 0 | 0 | 0 | 0 | 0 | 0 | 0 | 0 | 0 | 0 | 0 | 0 | 0 | 0 |
| virus_Jul11.0v_9887 | 1 | 0 | 0 | 0 | 0 | 0 | 0 | 0 | 0 | 0 | 0 | 0 | 0 | 0 | 0 | 0 | 0 | 0 | 0 | 0 | 0 | 0 | 1 | 0 | 0 | 0 | 0 | 0 | 0 | 0 | 0 | 0 | 0 | 0 | 0 | 0 | 0 | 0 | 0 | 0 | 0 | 0 | 0 | 0 | 0 | 0 | 0 | 0 | 0 | 0 | 0 | 0 | 0 | 0 | 0 |
| virus_Jul11.0v_13563 | 1 | 0 | 0 | 0 | 0 | 0 | 0 | 0 | 0 | 0 | 0 | 0 | 0 | 0 | 0 | 0 | 0 | 0 | 0 | 0 | 0 | 0 | 1 | 0 | 0 | 0 | 0 | 0 | 0 | 0 | 0 | 0 | 0 | 0 | 0 | 0 | 0 | 0 | 0 | 0 | 0 | 0 | 0 | 0 | 0 | 0 | 0 | 0 | 0 | 0 | 0 | 0 | 0 | 0 | 0 |
| virus_Jul11.0v_19224 | 1 | 0 | 0 | 0 | 0 | 0 | 0 | 0 | 0 | 0 | 0 | 0 | 0 | 0 | 0 | 0 | 0 | 0 | 0 | 0 | 0 | 0 | 1 | 0 | 0 | 0 | 0 | 0 | 0 | 0 | 0 | 0 | 0 | 0 | 0 | 0 | 0 | 0 | 0 | 0 | 0 | 0 | 0 | 0 | 0 | 0 | 0 | 0 | 0 | 0 | 0 | 0 | 0 | 0 | 0 |
| virus_Jul11.5v_1837 | 1 | 0 | 0 | 0 | 0 | 0 | 0 | 0 | 0 | 0 | 0 | 0 | 0 | 0 | 0 | 0 | 0 | 0 | 0 | 0 | 0 | 0 | 0 | 0 | 1 | 0 | 0 | 0 | 0 | 0 | 0 | 0 | 0 | 0 | 0 | 0 | 0 | 0 | 0 | 0 | 0 | 0 | 0 | 0 | 0 | 0 | 0 | 0 | 0 | 0 | 0 | 0 | 0 | 0 | 0 |
| virus_Jul11.5v_2027 | 1 | 0 | 0 | 0 | 0 | 0 | 0 | 0 | 0 | 0 | 0 | 0 | 0 | 0 | 0 | 0 | 0 | 0 | 0 | 0 | 0 | 0 | 0 | 0 | 1 | 0 | 0 | 0 | 0 | 0 | 0 | 0 | 0 | 0 | 0 | 0 | 0 | 0 | 0 | 0 | 0 | 0 | 0 | 0 | 0 | 0 | 0 | 0 | 0 | 0 | 0 | 0 | 0 | 0 | 0 |
| virus_Jul11.5v_2104 | 1 | 0 | 0 | 0 | 0 | 0 | 0 | 0 | 0 | 0 | 0 | 0 | 0 | 0 | 0 | 0 | 0 | 0 | 0 | 0 | 0 | 0 | 0 | 0 | 1 | 0 | 0 | 0 | 0 | 0 | 0 | 0 | 0 | 0 | 0 | 0 | 0 | 0 | 0 | 0 | 0 | 0 | 0 | 0 | 0 | 0 | 0 | 0 | 0 | 0 | 0 | 0 | 0 | 0 | 0 |
| virus_Jul11.5v_2369 | 1 | 0 | 0 | 0 | 0 | 0 | 0 | 0 | 0 | 0 | 0 | 0 | 0 | 0 | 0 | 0 | 0 | 0 | 0 | 0 | 0 | 0 | 0 | 0 | 1 | 0 | 0 | 0 | 0 | 0 | 0 | 0 | 0 | 0 | 0 | 0 | 0 | 0 | 0 | 0 | 0 | 0 | 0 | 0 | 0 | 0 | 0 | 0 | 0 | 0 | 0 | 0 | 0 | 0 | 0 |
| virus_Jul11.5v_3342 | 1 | 0 | 0 | 0 | 0 | 0 | 0 | 0 | 0 | 0 | 0 | 0 | 0 | 0 | 0 | 0 | 0 | 0 | 0 | 0 | 0 | 0 | 0 | 0 | 1 | 0 | 0 | 0 | 0 | 0 | 0 | 0 | 0 | 0 | 0 | 0 | 0 | 0 | 0 | 0 | 0 | 0 | 0 | 0 | 0 | 0 | 0 | 0 | 0 | 0 | 0 | 0 | 0 | 0 | 0 |
| virus_Jul11.5v_3420 | 1 | 0 | 0 | 0 | 0 | 0 | 0 | 0 | 0 | 0 | 0 | 0 | 0 | 0 | 0 | 0 | 0 | 0 | 0 | 0 | 0 | 0 | 0 | 0 | 1 | 0 | 0 | 0 | 0 | 0 | 0 | 0 | 0 | 0 | 0 | 0 | 0 | 0 | 0 | 0 | 0 | 0 | 0 | 0 | 0 | 0 | 0 | 0 | 0 | 0 | 0 | 0 | 0 | 0 | 0 |
| virus_Jul11.5v_3575 | 1 | 0 | 0 | 0 | 0 | 0 | 0 | 0 | 0 | 0 | 0 | 0 | 0 | 0 | 0 | 0 | 0 | 0 | 0 | 0 | 0 | 0 | 0 | 0 | 1 | 0 | 0 | 0 | 0 | 0 | 0 | 0 | 0 | 0 | 0 | 0 | 0 | 0 | 0 | 0 | 0 | 0 | 0 | 0 | 0 | 0 | 0 | 0 | 0 | 0 | 0 | 0 | 0 | 0 | 0 |
| virus_Jul11.5v_5632 | 1 | 0 | 0 | 0 | 0 | 0 | 0 | 0 | 0 | 0 | 0 | 0 | 0 | 0 | 0 | 0 | 0 | 0 | 0 | 0 | 0 | 0 | 0 | 0 | 1 | 0 | 0 | 0 | 0 | 0 | 0 | 0 | 0 | 0 | 0 | 0 | 0 | 0 | 0 | 0 | 0 | 0 | 0 | 0 | 0 | 0 | 0 | 0 | 0 | 0 | 0 | 0 | 0 | 0 | 0 |
| virus_Jul11.5v_9729 | 1 | 0 | 0 | 0 | 0 | 0 | 0 | 0 | 0 | 0 | 0 | 0 | 0 | 0 | 0 | 0 | 0 | 0 | 0 | 0 | 0 | 0 | 0 | 0 | 1 | 0 | 0 | 0 | 0 | 0 | 0 | 0 | 0 | 0 | 0 | 0 | 0 | 0 | 0 | 0 | 0 | 0 | 0 | 0 | 0 | 0 | 0 | 0 | 0 | 0 | 0 | 0 | 0 | 0 | 0 |
| virus_Jul11.5v_12061 | 1 | 0 | 0 | 0 | 0 | 0 | 0 | 0 | 0 | 0 | 0 | 0 | 0 | 0 | 0 | 0 | 0 | 0 | 0 | 0 | 0 | 0 | 0 | 0 | 1 | 0 | 0 | 0 | 0 | 0 | 0 | 0 | 0 | 0 | 0 | 0 | 0 | 0 | 0 | 0 | 0 | 0 | 0 | 0 | 0 | 0 | 0 | 0 | 0 | 0 | 0 | 0 | 0 | 0 | 0 |
| virus_Jul11.5v_17463 | 1 | 0 | 0 | 0 | 0 | 0 | 0 | 0 | 0 | 0 | 0 | 0 | 0 | 0 | 0 | 0 | 0 | 0 | 0 | 0 | 0 | 0 | 0 | 0 | 1 | 0 | 0 | 0 | 0 | 0 | 0 | 0 | 0 | 0 | 0 | 0 | 0 | 0 | 0 | 0 | 0 | 0 | 0 | 0 | 0 | 0 | 0 | 0 | 0 | 0 | 0 | 0 | 0 | 0 | 0 |
| virus_Jul11.5v_18582 | 1 | 0 | 0 | 0 | 0 | 0 | 0 | 0 | 0 | 0 | 0 | 0 | 0 | 0 | 0 | 0 | 0 | 0 | 0 | 0 | 0 | 0 | 0 | 0 | 1 | 0 | 0 | 0 | 0 | 0 | 0 | 0 | 0 | 0 | 0 | 0 | 0 | 0 | 0 | 0 | 0 | 0 | 0 | 0 | 0 | 0 | 0 | 0 | 0 | 0 | 0 | 0 | 0 | 0 | 0 |
| virus_Jul11.5v_19033 | 1 | 0 | 0 | 0 | 0 | 0 | 0 | 0 | 0 | 0 | 0 | 0 | 0 | 0 | 0 | 0 | 0 | 0 | 0 | 0 | 0 | 0 | 0 | 0 | 1 | 0 | 0 | 0 | 0 | 0 | 0 | 0 | 0 | 0 | 0 | 0 | 0 | 0 | 0 | 0 | 0 | 0 | 0 | 0 | 0 | 0 | 0 | 0 | 0 | 0 | 0 | 0 | 0 | 0 | 0 |
| virus_Jul11.5v_19624 | 1 | 0 | 0 | 0 | 0 | 0 | 0 | 0 | 0 | 0 | 0 | 0 | 0 | 0 | 0 | 0 | 0 | 0 | 0 | 0 | 0 | 0 | 0 | 0 | 1 | 0 | 0 | 0 | 0 | 0 | 0 | 0 | 0 | 0 | 0 | 0 | 0 | 0 | 0 | 0 | 0 | 0 | 0 | 0 | 0 | 0 | 0 | 0 | 0 | 0 | 0 | 0 | 0 | 0 | 0 |
| virus_Jul11.5v_20599 | 1 | 0 | 0 | 0 | 0 | 0 | 0 | 0 | 0 | 0 | 0 | 0 | 0 | 0 | 0 | 0 | 0 | 0 | 0 | 0 | 0 | 0 | 0 | 0 | 1 | 0 | 0 | 0 | 0 | 0 | 0 | 0 | 0 | 0 | 0 | 0 | 0 | 0 | 0 | 0 | 0 | 0 | 0 | 0 | 0 | 0 | 0 | 0 | 0 | 0 | 0 | 0 | 0 | 0 | 0 |
| virus_Jul11.5v_21037 | 1 | 0 | 0 | 0 | 0 | 0 | 0 | 0 | 0 | 0 | 0 | 0 | 0 | 0 | 0 | 0 | 0 | 0 | 0 | 0 | 0 | 0 | 0 | 0 | 1 | 0 | 0 | 0 | 0 | 0 | 0 | 0 | 0 | 0 | 0 | 0 | 0 | 0 | 0 | 0 | 0 | 0 | 0 | 0 | 0 | 0 | 0 | 0 | 0 | 0 | 0 | 0 | 0 | 0 | 0 |
| virus_Jul11.25v_125 | 1 | 0 | 0 | 0 | 0 | 0 | 0 | 0 | 0 | 0 | 0 | 0 | 0 | 0 | 0 | 0 | 0 | 0 | 0 | 0 | 0 | 0 | 0 | 0 | 0 | 0 | 1 | 0 | 0 | 0 | 0 | 0 | 0 | 0 | 0 | 0 | 0 | 0 | 0 | 0 | 0 | 0 | 0 | 0 | 0 | 0 | 0 | 0 | 0 | 0 | 0 | 0 | 0 | 0 | 0 |
| virus_Jul11.25v_1142 | 1 | 0 | 0 | 0 | 0 | 0 | 0 | 0 | 0 | 0 | 0 | 0 | 0 | 0 | 0 | 0 | 0 | 0 | 0 | 0 | 0 | 0 | 0 | 0 | 0 | 0 | 1 | 0 | 0 | 0 | 0 | 0 | 0 | 0 | 0 | 0 | 0 | 0 | 0 | 0 | 0 | 0 | 0 | 0 | 0 | 0 | 0 | 0 | 0 | 0 | 0 | 0 | 0 | 0 | 0 |
| virus_Jul11.25v_1365 | 1 | 0 | 0 | 0 | 0 | 0 | 0 | 0 | 0 | 0 | 0 | 0 | 0 | 0 | 0 | 0 | 0 | 0 | 0 | 0 | 0 | 0 | 0 | 0 | 0 | 0 | 1 | 0 | 0 | 0 | 0 | 0 | 0 | 0 | 0 | 0 | 0 | 0 | 0 | 0 | 0 | 0 | 0 | 0 | 0 | 0 | 0 | 0 | 0 | 0 | 0 | 0 | 0 | 0 | 0 |
| virus_Jul11.25v_7826 | 1 | 0 | 0 | 0 | 0 | 0 | 0 | 0 | 0 | 0 | 0 | 0 | 0 | 0 | 0 | 0 | 0 | 0 | 0 | 0 | 0 | 0 | 0 | 0 | 0 | 0 | 1 | 0 | 0 | 0 | 0 | 0 | 0 | 0 | 0 | 0 | 0 | 0 | 0 | 0 | 0 | 0 | 0 | 0 | 0 | 0 | 0 | 0 | 0 | 0 | 0 | 0 | 0 | 0 | 0 |
| virus_Jul11.25v_9234 | 1 | 0 | 0 | 0 | 0 | 0 | 0 | 0 | 0 | 0 | 0 | 0 | 0 | 0 | 0 | 0 | 0 | 0 | 0 | 0 | 0 | 0 | 0 | 0 | 0 | 0 | 1 | 0 | 0 | 0 | 0 | 0 | 0 | 0 | 0 | 0 | 0 | 0 | 0 | 0 | 0 | 0 | 0 | 0 | 0 | 0 | 0 | 0 | 0 | 0 | 0 | 0 | 0 | 0 | 0 |
| virus_Jul11.25v_13802 | 1 | 0 | 0 | 0 | 0 | 0 | 0 | 0 | 0 | 0 | 0 | 0 | 0 | 0 | 0 | 0 | 0 | 0 | 0 | 0 | 0 | 0 | 0 | 0 | 0 | 0 | 1 | 0 | 0 | 0 | 0 | 0 | 0 | 0 | 0 | 0 | 0 | 0 | 0 | 0 | 0 | 0 | 0 | 0 | 0 | 0 | 0 | 0 | 0 | 0 | 0 | 0 | 0 | 0 | 0 |
| virus_Jul11.25v_16091 | 1 | 0 | 0 | 0 | 0 | 0 | 0 | 0 | 0 | 0 | 0 | 0 | 0 | 0 | 0 | 0 | 0 | 0 | 0 | 0 | 0 | 0 | 0 | 0 | 0 | 0 | 1 | 0 | 0 | 0 | 0 | 0 | 0 | 0 | 0 | 0 | 0 | 0 | 0 | 0 | 0 | 0 | 0 | 0 | 0 | 0 | 0 | 0 | 0 | 0 | 0 | 0 | 0 | 0 | 0 |
| virus_Jul11.25v_16658 | 1 | 0 | 0 | 0 | 0 | 0 | 0 | 0 | 0 | 0 | 0 | 0 | 0 | 0 | 0 | 0 | 0 | 0 | 0 | 0 | 0 | 0 | 0 | 0 | 0 | 0 | 1 | 0 | 0 | 0 | 0 | 0 | 0 | 0 | 0 | 0 | 0 | 0 | 0 | 0 | 0 | 0 | 0 | 0 | 0 | 0 | 0 | 0 | 0 | 0 | 0 | 0 | 0 | 0 | 0 |
| virus_Jul11.25v_16824 | 1 | 0 | 0 | 0 | 0 | 0 | 0 | 0 | 0 | 0 | 0 | 0 | 0 | 0 | 0 | 0 | 0 | 0 | 0 | 0 | 0 | 0 | 0 | 0 | 0 | 0 | 1 | 0 | 0 | 0 | 0 | 0 | 0 | 0 | 0 | 0 | 0 | 0 | 0 | 0 | 0 | 0 | 0 | 0 | 0 | 0 | 0 | 0 | 0 | 0 | 0 | 0 | 0 | 0 | 0 |
| virus_Jul11.25v_17732 | 1 | 0 | 0 | 0 | 0 | 0 | 0 | 0 | 0 | 0 | 0 | 0 | 0 | 0 | 0 | 0 | 0 | 0 | 0 | 0 | 0 | 0 | 0 | 0 | 0 | 0 | 1 | 0 | 0 | 0 | 0 | 0 | 0 | 0 | 0 | 0 | 0 | 0 | 0 | 0 | 0 | 0 | 0 | 0 | 0 | 0 | 0 | 0 | 0 | 0 | 0 | 0 | 0 | 0 | 0 |
| virus_Jul11.25v_19545 | 1 | 0 | 0 | 0 | 0 | 0 | 0 | 0 | 0 | 0 | 0 | 0 | 0 | 0 | 0 | 0 | 0 | 0 | 0 | 0 | 0 | 0 | 0 | 0 | 0 | 0 | 1 | 0 | 0 | 0 | 0 | 0 | 0 | 0 | 0 | 0 | 0 | 0 | 0 | 0 | 0 | 0 | 0 | 0 | 0 | 0 | 0 | 0 | 0 | 0 | 0 | 0 | 0 | 0 | 0 |
| virus_Jul11.50v_32 | 1 | 0 | 0 | 0 | 0 | 0 | 0 | 0 | 0 | 0 | 0 | 0 | 0 | 0 | 0 | 0 | 0 | 0 | 0 | 0 | 0 | 0 | 0 | 0 | 0 | 0 | 0 | 0 | 1 | 0 | 0 | 0 | 0 | 0 | 0 | 0 | 0 | 0 | 0 | 0 | 0 | 0 | 0 | 0 | 0 | 0 | 0 | 0 | 0 | 0 | 0 | 0 | 0 | 0 | 0 |
| virus_Jul11.50v_391 | 1 | 0 | 0 | 0 | 0 | 0 | 0 | 0 | 0 | 0 | 0 | 0 | 0 | 0 | 0 | 0 | 0 | 0 | 0 | 0 | 0 | 0 | 0 | 0 | 0 | 0 | 0 | 0 | 1 | 0 | 0 | 0 | 0 | 0 | 0 | 0 | 0 | 0 | 0 | 0 | 0 | 0 | 0 | 0 | 0 | 0 | 0 | 0 | 0 | 0 | 0 | 0 | 0 | 0 | 0 |
| virus_Jul11.50v_5094 | 1 | 0 | 0 | 0 | 0 | 0 | 0 | 0 | 0 | 0 | 0 | 0 | 0 | 0 | 0 | 0 | 0 | 0 | 0 | 0 | 0 | 0 | 0 | 0 | 0 | 0 | 0 | 0 | 1 | 0 | 0 | 0 | 0 | 0 | 0 | 0 | 0 | 0 | 0 | 0 | 0 | 0 | 0 | 0 | 0 | 0 | 0 | 0 | 0 | 0 | 0 | 0 | 0 | 0 | 0 |
| virus_Jul11.50v_6504 | 1 | 0 | 0 | 0 | 0 | 0 | 0 | 0 | 0 | 0 | 0 | 0 | 0 | 0 | 0 | 0 | 0 | 0 | 0 | 0 | 0 | 0 | 0 | 0 | 0 | 0 | 0 | 0 | 1 | 0 | 0 | 0 | 0 | 0 | 0 | 0 | 0 | 0 | 0 | 0 | 0 | 0 | 0 | 0 | 0 | 0 | 0 | 0 | 0 | 0 | 0 | 0 | 0 | 0 | 0 |
| virus_Jul11.50v_9430 | 1 | 0 | 0 | 0 | 0 | 0 | 0 | 0 | 0 | 0 | 0 | 0 | 0 | 0 | 0 | 0 | 0 | 0 | 0 | 0 | 0 | 0 | 0 | 0 | 0 | 0 | 0 | 0 | 1 | 0 | 0 | 0 | 0 | 0 | 0 | 0 | 0 | 0 | 0 | 0 | 0 | 0 | 0 | 0 | 0 | 0 | 0 | 0 | 0 | 0 | 0 | 0 | 0 | 0 | 0 |
| virus_Jul11.50v_9436 | 1 | 0 | 0 | 0 | 0 | 0 | 0 | 0 | 0 | 0 | 0 | 0 | 0 | 0 | 0 | 0 | 0 | 0 | 0 | 0 | 0 | 0 | 0 | 0 | 0 | 0 | 0 | 0 | 1 | 0 | 0 | 0 | 0 | 0 | 0 | 0 | 0 | 0 | 0 | 0 | 0 | 0 | 0 | 0 | 0 | 0 | 0 | 0 | 0 | 0 | 0 | 0 | 0 | 0 | 0 |
| virus_Jul11.50v_9556 | 1 | 0 | 0 | 0 | 0 | 0 | 0 | 0 | 0 | 0 | 0 | 0 | 0 | 0 | 0 | 0 | 0 | 0 | 0 | 0 | 0 | 0 | 0 | 0 | 0 | 0 | 0 | 0 | 1 | 0 | 0 | 0 | 0 | 0 | 0 | 0 | 0 | 0 | 0 | 0 | 0 | 0 | 0 | 0 | 0 | 0 | 0 | 0 | 0 | 0 | 0 | 0 | 0 | 0 | 0 |
| virus_Jul11.50v_14336 | 1 | 0 | 0 | 0 | 0 | 0 | 0 | 0 | 0 | 0 | 0 | 0 | 0 | 0 | 0 | 0 | 0 | 0 | 0 | 0 | 0 | 0 | 0 | 0 | 0 | 0 | 0 | 0 | 1 | 0 | 0 | 0 | 0 | 0 | 0 | 0 | 0 | 0 | 0 | 0 | 0 | 0 | 0 | 0 | 0 | 0 | 0 | 0 | 0 | 0 | 0 | 0 | 0 | 0 | 0 |
| virus_Jul11.50v_14475 | 1 | 0 | 0 | 0 | 0 | 0 | 0 | 0 | 0 | 0 | 0 | 0 | 0 | 0 | 0 | 0 | 0 | 0 | 0 | 0 | 0 | 0 | 0 | 0 | 0 | 0 | 0 | 0 | 1 | 0 | 0 | 0 | 0 | 0 | 0 | 0 | 0 | 0 | 0 | 0 | 0 | 0 | 0 | 0 | 0 | 0 | 0 | 0 | 0 | 0 | 0 | 0 | 0 | 0 | 0 |
| virus_Jul11.50v_14820 | 1 | 0 | 0 | 0 | 0 | 0 | 0 | 0 | 0 | 0 | 0 | 0 | 0 | 0 | 0 | 0 | 0 | 0 | 0 | 0 | 0 | 0 | 0 | 0 | 0 | 0 | 0 | 0 | 1 | 0 | 0 | 0 | 0 | 0 | 0 | 0 | 0 | 0 | 0 | 0 | 0 | 0 | 0 | 0 | 0 | 0 | 0 | 0 | 0 | 0 | 0 | 0 | 0 | 0 | 0 |
| virus_Jul11.50v_15682 | 1 | 0 | 0 | 0 | 0 | 0 | 0 | 0 | 0 | 0 | 0 | 0 | 0 | 0 | 0 | 0 | 0 | 0 | 0 | 0 | 0 | 0 | 0 | 0 | 0 | 0 | 0 | 0 | 1 | 0 | 0 | 0 | 0 | 0 | 0 | 0 | 0 | 0 | 0 | 0 | 0 | 0 | 0 | 0 | 0 | 0 | 0 | 0 | 0 | 0 | 0 | 0 | 0 | 0 | 0 |
| virus_Jul11.50v_15968 | 1 | 0 | 0 | 0 | 0 | 0 | 0 | 0 | 0 | 0 | 0 | 0 | 0 | 0 | 0 | 0 | 0 | 0 | 0 | 0 | 0 | 0 | 0 | 0 | 0 | 0 | 0 | 0 | 1 | 0 | 0 | 0 | 0 | 0 | 0 | 0 | 0 | 0 | 0 | 0 | 0 | 0 | 0 | 0 | 0 | 0 | 0 | 0 | 0 | 0 | 0 | 0 | 0 | 0 | 0 |
| virus_Jul11.50v_16447 | 1 | 0 | 0 | 0 | 0 | 0 | 0 | 0 | 0 | 0 | 0 | 0 | 0 | 0 | 0 | 0 | 0 | 0 | 0 | 0 | 0 | 0 | 0 | 0 | 0 | 0 | 0 | 0 | 1 | 0 | 0 | 0 | 0 | 0 | 0 | 0 | 0 | 0 | 0 | 0 | 0 | 0 | 0 | 0 | 0 | 0 | 0 | 0 | 0 | 0 | 0 | 0 | 0 | 0 | 0 |
| virus_Jul11.50v_17919 | 1 | 0 | 0 | 0 | 0 | 0 | 0 | 0 | 0 | 0 | 0 | 0 | 0 | 0 | 0 | 0 | 0 | 0 | 0 | 0 | 0 | 0 | 0 | 0 | 0 | 0 | 0 | 0 | 1 | 0 | 0 | 0 | 0 | 0 | 0 | 0 | 0 | 0 | 0 | 0 | 0 | 0 | 0 | 0 | 0 | 0 | 0 | 0 | 0 | 0 | 0 | 0 | 0 | 0 | 0 |
| virus_Jul11.75v_20354 | 1 | 0 | 0 | 0 | 0 | 0 | 0 | 0 | 0 | 0 | 0 | 0 | 0 | 0 | 0 | 0 | 0 | 0 | 0 | 0 | 0 | 0 | 0 | 0 | 0 | 0 | 0 | 0 | 0 | 0 | 1 | 0 | 0 | 0 | 0 | 0 | 0 | 0 | 0 | 0 | 0 | 0 | 0 | 0 | 0 | 0 | 0 | 0 | 0 | 0 | 0 | 0 | 0 | 0 | 0 |
| virus_Jul11.75v_23749 | 1 | 0 | 0 | 0 | 0 | 0 | 0 | 0 | 0 | 0 | 0 | 0 | 0 | 0 | 0 | 0 | 0 | 0 | 0 | 0 | 0 | 0 | 0 | 0 | 0 | 0 | 0 | 0 | 0 | 0 | 1 | 0 | 0 | 0 | 0 | 0 | 0 | 0 | 0 | 0 | 0 | 0 | 0 | 0 | 0 | 0 | 0 | 0 | 0 | 0 | 0 | 0 | 0 | 0 | 0 |
| virus_Jul11.75v_33809 | 1 | 0 | 0 | 0 | 0 | 0 | 0 | 0 | 0 | 0 | 0 | 0 | 0 | 0 | 0 | 0 | 0 | 0 | 0 | 0 | 0 | 0 | 0 | 0 | 0 | 0 | 0 | 0 | 0 | 0 | 1 | 0 | 0 | 0 | 0 | 0 | 0 | 0 | 0 | 0 | 0 | 0 | 0 | 0 | 0 | 0 | 0 | 0 | 0 | 0 | 0 | 0 | 0 | 0 | 0 |
| virus_Jul27.5v_170 | 1 | 0 | 0 | 0 | 0 | 0 | 0 | 0 | 0 | 0 | 0 | 0 | 0 | 0 | 0 | 0 | 0 | 0 | 0 | 0 | 0 | 0 | 0 | 0 | 0 | 0 | 0 | 0 | 0 | 0 | 0 | 0 | 1 | 0 | 0 | 0 | 0 | 0 | 0 | 0 | 0 | 0 | 0 | 0 | 0 | 0 | 0 | 0 | 0 | 0 | 0 | 0 | 0 | 0 | 0 |
| virus_Jul27.5v_179 | 1 | 0 | 0 | 0 | 0 | 0 | 0 | 0 | 0 | 0 | 0 | 0 | 0 | 0 | 0 | 0 | 0 | 0 | 0 | 0 | 0 | 0 | 0 | 0 | 0 | 0 | 0 | 0 | 0 | 0 | 0 | 0 | 1 | 0 | 0 | 0 | 0 | 0 | 0 | 0 | 0 | 0 | 0 | 0 | 0 | 0 | 0 | 0 | 0 | 0 | 0 | 0 | 0 | 0 | 0 |
| virus_Jul27.5v_364 | 1 | 0 | 0 | 0 | 0 | 0 | 0 | 0 | 0 | 0 | 0 | 0 | 0 | 0 | 0 | 0 | 0 | 0 | 0 | 0 | 0 | 0 | 0 | 0 | 0 | 0 | 0 | 0 | 0 | 0 | 0 | 0 | 1 | 0 | 0 | 0 | 0 | 0 | 0 | 0 | 0 | 0 | 0 | 0 | 0 | 0 | 0 | 0 | 0 | 0 | 0 | 0 | 0 | 0 | 0 |
| virus_Jul27.5v_519 | 1 | 0 | 0 | 0 | 0 | 0 | 0 | 0 | 0 | 0 | 0 | 0 | 0 | 0 | 0 | 0 | 0 | 0 | 0 | 0 | 0 | 0 | 0 | 0 | 0 | 0 | 0 | 0 | 0 | 0 | 0 | 0 | 1 | 0 | 0 | 0 | 0 | 0 | 0 | 0 | 0 | 0 | 0 | 0 | 0 | 0 | 0 | 0 | 0 | 0 | 0 | 0 | 0 | 0 | 0 |
| virus_Jul27.5v_3024 | 1 | 0 | 0 | 0 | 0 | 0 | 0 | 0 | 0 | 0 | 0 | 0 | 0 | 0 | 0 | 0 | 0 | 0 | 0 | 0 | 0 | 0 | 0 | 0 | 0 | 0 | 0 | 0 | 0 | 0 | 0 | 0 | 1 | 0 | 0 | 0 | 0 | 0 | 0 | 0 | 0 | 0 | 0 | 0 | 0 | 0 | 0 | 0 | 0 | 0 | 0 | 0 | 0 | 0 | 0 |
| virus_Jul27.5v_3767 | 1 | 0 | 0 | 0 | 0 | 0 | 0 | 0 | 0 | 0 | 0 | 0 | 0 | 0 | 0 | 0 | 0 | 0 | 0 | 0 | 0 | 0 | 0 | 0 | 0 | 0 | 0 | 0 | 0 | 0 | 0 | 0 | 1 | 0 | 0 | 0 | 0 | 0 | 0 | 0 | 0 | 0 | 0 | 0 | 0 | 0 | 0 | 0 | 0 | 0 | 0 | 0 | 0 | 0 | 0 |
| virus_Jul27.5v_5539 | 1 | 0 | 0 | 0 | 0 | 0 | 0 | 0 | 0 | 0 | 0 | 0 | 0 | 0 | 0 | 0 | 0 | 0 | 0 | 0 | 0 | 0 | 0 | 0 | 0 | 0 | 0 | 0 | 0 | 0 | 0 | 0 | 1 | 0 | 0 | 0 | 0 | 0 | 0 | 0 | 0 | 0 | 0 | 0 | 0 | 0 | 0 | 0 | 0 | 0 | 0 | 0 | 0 | 0 | 0 |
| virus_Jul27.5v_6121 | 1 | 0 | 0 | 0 | 0 | 0 | 0 | 0 | 0 | 0 | 0 | 0 | 0 | 0 | 0 | 0 | 0 | 0 | 0 | 0 | 0 | 0 | 0 | 0 | 0 | 0 | 0 | 0 | 0 | 0 | 0 | 0 | 1 | 0 | 0 | 0 | 0 | 0 | 0 | 0 | 0 | 0 | 0 | 0 | 0 | 0 | 0 | 0 | 0 | 0 | 0 | 0 | 0 | 0 | 0 |
| virus_Jul27.5v_7323 | 1 | 0 | 0 | 0 | 0 | 0 | 0 | 0 | 0 | 0 | 0 | 0 | 0 | 0 | 0 | 0 | 0 | 0 | 0 | 0 | 0 | 0 | 0 | 0 | 0 | 0 | 0 | 0 | 0 | 0 | 0 | 0 | 1 | 0 | 0 | 0 | 0 | 0 | 0 | 0 | 0 | 0 | 0 | 0 | 0 | 0 | 0 | 0 | 0 | 0 | 0 | 0 | 0 | 0 | 0 |
| virus_Jul27.5v_8652 | 1 | 0 | 0 | 0 | 0 | 0 | 0 | 0 | 0 | 0 | 0 | 0 | 0 | 0 | 0 | 0 | 0 | 0 | 0 | 0 | 0 | 0 | 0 | 0 | 0 | 0 | 0 | 0 | 0 | 0 | 0 | 0 | 1 | 0 | 0 | 0 | 0 | 0 | 0 | 0 | 0 | 0 | 0 | 0 | 0 | 0 | 0 | 0 | 0 | 0 | 0 | 0 | 0 | 0 | 0 |
| virus_Jul27.5v_8945 | 1 | 0 | 0 | 0 | 0 | 0 | 0 | 0 | 0 | 0 | 0 | 0 | 0 | 0 | 0 | 0 | 0 | 0 | 0 | 0 | 0 | 0 | 0 | 0 | 0 | 0 | 0 | 0 | 0 | 0 | 0 | 0 | 1 | 0 | 0 | 0 | 0 | 0 | 0 | 0 | 0 | 0 | 0 | 0 | 0 | 0 | 0 | 0 | 0 | 0 | 0 | 0 | 0 | 0 | 0 |
| virus_Jul27.5v_9148 | 1 | 0 | 0 | 0 | 0 | 0 | 0 | 0 | 0 | 0 | 0 | 0 | 0 | 0 | 0 | 0 | 0 | 0 | 0 | 0 | 0 | 0 | 0 | 0 | 0 | 0 | 0 | 0 | 0 | 0 | 0 | 0 | 1 | 0 | 0 | 0 | 0 | 0 | 0 | 0 | 0 | 0 | 0 | 0 | 0 | 0 | 0 | 0 | 0 | 0 | 0 | 0 | 0 | 0 | 0 |
| virus_Jul27.5v_11613 | 1 | 0 | 0 | 0 | 0 | 0 | 0 | 0 | 0 | 0 | 0 | 0 | 0 | 0 | 0 | 0 | 0 | 0 | 0 | 0 | 0 | 0 | 0 | 0 | 0 | 0 | 0 | 0 | 0 | 0 | 0 | 0 | 1 | 0 | 0 | 0 | 0 | 0 | 0 | 0 | 0 | 0 | 0 | 0 | 0 | 0 | 0 | 0 | 0 | 0 | 0 | 0 | 0 | 0 | 0 |
| virus_Jul27.5v_13084 | 1 | 0 | 0 | 0 | 0 | 0 | 0 | 0 | 0 | 0 | 0 | 0 | 0 | 0 | 0 | 0 | 0 | 0 | 0 | 0 | 0 | 0 | 0 | 0 | 0 | 0 | 0 | 0 | 0 | 0 | 0 | 0 | 1 | 0 | 0 | 0 | 0 | 0 | 0 | 0 | 0 | 0 | 0 | 0 | 0 | 0 | 0 | 0 | 0 | 0 | 0 | 0 | 0 | 0 | 0 |
| virus_Jul27.5v_13706 | 1 | 0 | 0 | 0 | 0 | 0 | 0 | 0 | 0 | 0 | 0 | 0 | 0 | 0 | 0 | 0 | 0 | 0 | 0 | 0 | 0 | 0 | 0 | 0 | 0 | 0 | 0 | 0 | 0 | 0 | 0 | 0 | 1 | 0 | 0 | 0 | 0 | 0 | 0 | 0 | 0 | 0 | 0 | 0 | 0 | 0 | 0 | 0 | 0 | 0 | 0 | 0 | 0 | 0 | 0 |
| virus_Jul27.5v_13910 | 1 | 0 | 0 | 0 | 0 | 0 | 0 | 0 | 0 | 0 | 0 | 0 | 0 | 0 | 0 | 0 | 0 | 0 | 0 | 0 | 0 | 0 | 0 | 0 | 0 | 0 | 0 | 0 | 0 | 0 | 0 | 0 | 1 | 0 | 0 | 0 | 0 | 0 | 0 | 0 | 0 | 0 | 0 | 0 | 0 | 0 | 0 | 0 | 0 | 0 | 0 | 0 | 0 | 0 | 0 |
| virus_Jul27.5v_15771 | 1 | 0 | 0 | 0 | 0 | 0 | 0 | 0 | 0 | 0 | 0 | 0 | 0 | 0 | 0 | 0 | 0 | 0 | 0 | 0 | 0 | 0 | 0 | 0 | 0 | 0 | 0 | 0 | 0 | 0 | 0 | 0 | 1 | 0 | 0 | 0 | 0 | 0 | 0 | 0 | 0 | 0 | 0 | 0 | 0 | 0 | 0 | 0 | 0 | 0 | 0 | 0 | 0 | 0 | 0 |
| virus_Jul27.5v_16350 | 1 | 0 | 0 | 0 | 0 | 0 | 0 | 0 | 0 | 0 | 0 | 0 | 0 | 0 | 0 | 0 | 0 | 0 | 0 | 0 | 0 | 0 | 0 | 0 | 0 | 0 | 0 | 0 | 0 | 0 | 0 | 0 | 1 | 0 | 0 | 0 | 0 | 0 | 0 | 0 | 0 | 0 | 0 | 0 | 0 | 0 | 0 | 0 | 0 | 0 | 0 | 0 | 0 | 0 | 0 |
| virus_Jul27.5v_16972 | 1 | 0 | 0 | 0 | 0 | 0 | 0 | 0 | 0 | 0 | 0 | 0 | 0 | 0 | 0 | 0 | 0 | 0 | 0 | 0 | 0 | 0 | 0 | 0 | 0 | 0 | 0 | 0 | 0 | 0 | 0 | 0 | 1 | 0 | 0 | 0 | 0 | 0 | 0 | 0 | 0 | 0 | 0 | 0 | 0 | 0 | 0 | 0 | 0 | 0 | 0 | 0 | 0 | 0 | 0 |
| virus_Jul27.5v_17153 | 1 | 0 | 0 | 0 | 0 | 0 | 0 | 0 | 0 | 0 | 0 | 0 | 0 | 0 | 0 | 0 | 0 | 0 | 0 | 0 | 0 | 0 | 0 | 0 | 0 | 0 | 0 | 0 | 0 | 0 | 0 | 0 | 1 | 0 | 0 | 0 | 0 | 0 | 0 | 0 | 0 | 0 | 0 | 0 | 0 | 0 | 0 | 0 | 0 | 0 | 0 | 0 | 0 | 0 | 0 |
| virus_Jul27.5v_17304 | 1 | 0 | 0 | 0 | 0 | 0 | 0 | 0 | 0 | 0 | 0 | 0 | 0 | 0 | 0 | 0 | 0 | 0 | 0 | 0 | 0 | 0 | 0 | 0 | 0 | 0 | 0 | 0 | 0 | 0 | 0 | 0 | 1 | 0 | 0 | 0 | 0 | 0 | 0 | 0 | 0 | 0 | 0 | 0 | 0 | 0 | 0 | 0 | 0 | 0 | 0 | 0 | 0 | 0 | 0 |
| virus_Jul27.5v_17778 | 1 | 0 | 0 | 0 | 0 | 0 | 0 | 0 | 0 | 0 | 0 | 0 | 0 | 0 | 0 | 0 | 0 | 0 | 0 | 0 | 0 | 0 | 0 | 0 | 0 | 0 | 0 | 0 | 0 | 0 | 0 | 0 | 1 | 0 | 0 | 0 | 0 | 0 | 0 | 0 | 0 | 0 | 0 | 0 | 0 | 0 | 0 | 0 | 0 | 0 | 0 | 0 | 0 | 0 | 0 |
| virus_Jul27.5v_17934 | 1 | 0 | 0 | 0 | 0 | 0 | 0 | 0 | 0 | 0 | 0 | 0 | 0 | 0 | 0 | 0 | 0 | 0 | 0 | 0 | 0 | 0 | 0 | 0 | 0 | 0 | 0 | 0 | 0 | 0 | 0 | 0 | 1 | 0 | 0 | 0 | 0 | 0 | 0 | 0 | 0 | 0 | 0 | 0 | 0 | 0 | 0 | 0 | 0 | 0 | 0 | 0 | 0 | 0 | 0 |
| virus_Jul27.5v_18031 | 1 | 0 | 0 | 0 | 0 | 0 | 0 | 0 | 0 | 0 | 0 | 0 | 0 | 0 | 0 | 0 | 0 | 0 | 0 | 0 | 0 | 0 | 0 | 0 | 0 | 0 | 0 | 0 | 0 | 0 | 0 | 0 | 1 | 0 | 0 | 0 | 0 | 0 | 0 | 0 | 0 | 0 | 0 | 0 | 0 | 0 | 0 | 0 | 0 | 0 | 0 | 0 | 0 | 0 | 0 |
| virus_Jul27.5v_18355 | 1 | 0 | 0 | 0 | 0 | 0 | 0 | 0 | 0 | 0 | 0 | 0 | 0 | 0 | 0 | 0 | 0 | 0 | 0 | 0 | 0 | 0 | 0 | 0 | 0 | 0 | 0 | 0 | 0 | 0 | 0 | 0 | 1 | 0 | 0 | 0 | 0 | 0 | 0 | 0 | 0 | 0 | 0 | 0 | 0 | 0 | 0 | 0 | 0 | 0 | 0 | 0 | 0 | 0 | 0 |
| virus_Jul27.5v_18500 | 1 | 0 | 0 | 0 | 0 | 0 | 0 | 0 | 0 | 0 | 0 | 0 | 0 | 0 | 0 | 0 | 0 | 0 | 0 | 0 | 0 | 0 | 0 | 0 | 0 | 0 | 0 | 0 | 0 | 0 | 0 | 0 | 1 | 0 | 0 | 0 | 0 | 0 | 0 | 0 | 0 | 0 | 0 | 0 | 0 | 0 | 0 | 0 | 0 | 0 | 0 | 0 | 0 | 0 | 0 |
| virus_Jul27.5v_19079 | 1 | 0 | 0 | 0 | 0 | 0 | 0 | 0 | 0 | 0 | 0 | 0 | 0 | 0 | 0 | 0 | 0 | 0 | 0 | 0 | 0 | 0 | 0 | 0 | 0 | 0 | 0 | 0 | 0 | 0 | 0 | 0 | 1 | 0 | 0 | 0 | 0 | 0 | 0 | 0 | 0 | 0 | 0 | 0 | 0 | 0 | 0 | 0 | 0 | 0 | 0 | 0 | 0 | 0 | 0 |
| virus_Jul27.5v_19245 | 1 | 0 | 0 | 0 | 0 | 0 | 0 | 0 | 0 | 0 | 0 | 0 | 0 | 0 | 0 | 0 | 0 | 0 | 0 | 0 | 0 | 0 | 0 | 0 | 0 | 0 | 0 | 0 | 0 | 0 | 0 | 0 | 1 | 0 | 0 | 0 | 0 | 0 | 0 | 0 | 0 | 0 | 0 | 0 | 0 | 0 | 0 | 0 | 0 | 0 | 0 | 0 | 0 | 0 | 0 |
| virus_Jul27.25v_626 | 1 | 0 | 0 | 0 | 0 | 0 | 0 | 0 | 0 | 0 | 0 | 0 | 0 | 0 | 0 | 0 | 0 | 0 | 0 | 0 | 0 | 0 | 0 | 0 | 0 | 0 | 0 | 0 | 0 | 0 | 0 | 0 | 0 | 0 | 1 | 0 | 0 | 0 | 0 | 0 | 0 | 0 | 0 | 0 | 0 | 0 | 0 | 0 | 0 | 0 | 0 | 0 | 0 | 0 | 0 |
| virus_Jul27.25v_1516 | 1 | 0 | 0 | 0 | 0 | 0 | 0 | 0 | 0 | 0 | 0 | 0 | 0 | 0 | 0 | 0 | 0 | 0 | 0 | 0 | 0 | 0 | 0 | 0 | 0 | 0 | 0 | 0 | 0 | 0 | 0 | 0 | 0 | 0 | 1 | 0 | 0 | 0 | 0 | 0 | 0 | 0 | 0 | 0 | 0 | 0 | 0 | 0 | 0 | 0 | 0 | 0 | 0 | 0 | 0 |
| virus_Jul27.25v_3062 | 1 | 0 | 0 | 0 | 0 | 0 | 0 | 0 | 0 | 0 | 0 | 0 | 0 | 0 | 0 | 0 | 0 | 0 | 0 | 0 | 0 | 0 | 0 | 0 | 0 | 0 | 0 | 0 | 0 | 0 | 0 | 0 | 0 | 0 | 1 | 0 | 0 | 0 | 0 | 0 | 0 | 0 | 0 | 0 | 0 | 0 | 0 | 0 | 0 | 0 | 0 | 0 | 0 | 0 | 0 |
| virus_Jul27.25v_3573 | 1 | 0 | 0 | 0 | 0 | 0 | 0 | 0 | 0 | 0 | 0 | 0 | 0 | 0 | 0 | 0 | 0 | 0 | 0 | 0 | 0 | 0 | 0 | 0 | 0 | 0 | 0 | 0 | 0 | 0 | 0 | 0 | 0 | 0 | 1 | 0 | 0 | 0 | 0 | 0 | 0 | 0 | 0 | 0 | 0 | 0 | 0 | 0 | 0 | 0 | 0 | 0 | 0 | 0 | 0 |
| virus_Jul27.25v_3941 | 1 | 0 | 0 | 0 | 0 | 0 | 0 | 0 | 0 | 0 | 0 | 0 | 0 | 0 | 0 | 0 | 0 | 0 | 0 | 0 | 0 | 0 | 0 | 0 | 0 | 0 | 0 | 0 | 0 | 0 | 0 | 0 | 0 | 0 | 1 | 0 | 0 | 0 | 0 | 0 | 0 | 0 | 0 | 0 | 0 | 0 | 0 | 0 | 0 | 0 | 0 | 0 | 0 | 0 | 0 |
| virus_Jul27.25v_10421 | 1 | 0 | 0 | 0 | 0 | 0 | 0 | 0 | 0 | 0 | 0 | 0 | 0 | 0 | 0 | 0 | 0 | 0 | 0 | 0 | 0 | 0 | 0 | 0 | 0 | 0 | 0 | 0 | 0 | 0 | 0 | 0 | 0 | 0 | 1 | 0 | 0 | 0 | 0 | 0 | 0 | 0 | 0 | 0 | 0 | 0 | 0 | 0 | 0 | 0 | 0 | 0 | 0 | 0 | 0 |
| virus_Jul27.25v_17730 | 1 | 0 | 0 | 0 | 0 | 0 | 0 | 0 | 0 | 0 | 0 | 0 | 0 | 0 | 0 | 0 | 0 | 0 | 0 | 0 | 0 | 0 | 0 | 0 | 0 | 0 | 0 | 0 | 0 | 0 | 0 | 0 | 0 | 0 | 1 | 0 | 0 | 0 | 0 | 0 | 0 | 0 | 0 | 0 | 0 | 0 | 0 | 0 | 0 | 0 | 0 | 0 | 0 | 0 | 0 |
| virus_Jul27.25v_18805 | 1 | 0 | 0 | 0 | 0 | 0 | 0 | 0 | 0 | 0 | 0 | 0 | 0 | 0 | 0 | 0 | 0 | 0 | 0 | 0 | 0 | 0 | 0 | 0 | 0 | 0 | 0 | 0 | 0 | 0 | 0 | 0 | 0 | 0 | 1 | 0 | 0 | 0 | 0 | 0 | 0 | 0 | 0 | 0 | 0 | 0 | 0 | 0 | 0 | 0 | 0 | 0 | 0 | 0 | 0 |
| virus_Jul27.25v_29797 | 1 | 0 | 0 | 0 | 0 | 0 | 0 | 0 | 0 | 0 | 0 | 0 | 0 | 0 | 0 | 0 | 0 | 0 | 0 | 0 | 0 | 0 | 0 | 0 | 0 | 0 | 0 | 0 | 0 | 0 | 0 | 0 | 0 | 0 | 1 | 0 | 0 | 0 | 0 | 0 | 0 | 0 | 0 | 0 | 0 | 0 | 0 | 0 | 0 | 0 | 0 | 0 | 0 | 0 | 0 |
| virus_Jul27.25v_32502 | 1 | 0 | 0 | 0 | 0 | 0 | 0 | 0 | 0 | 0 | 0 | 0 | 0 | 0 | 0 | 0 | 0 | 0 | 0 | 0 | 0 | 0 | 0 | 0 | 0 | 0 | 0 | 0 | 0 | 0 | 0 | 0 | 0 | 0 | 1 | 0 | 0 | 0 | 0 | 0 | 0 | 0 | 0 | 0 | 0 | 0 | 0 | 0 | 0 | 0 | 0 | 0 | 0 | 0 | 0 |
| virus_Jul27.25v_35958 | 1 | 0 | 0 | 0 | 0 | 0 | 0 | 0 | 0 | 0 | 0 | 0 | 0 | 0 | 0 | 0 | 0 | 0 | 0 | 0 | 0 | 0 | 0 | 0 | 0 | 0 | 0 | 0 | 0 | 0 | 0 | 0 | 0 | 0 | 1 | 0 | 0 | 0 | 0 | 0 | 0 | 0 | 0 | 0 | 0 | 0 | 0 | 0 | 0 | 0 | 0 | 0 | 0 | 0 | 0 |
| virus_Jul27.25v_36115 | 1 | 0 | 0 | 0 | 0 | 0 | 0 | 0 | 0 | 0 | 0 | 0 | 0 | 0 | 0 | 0 | 0 | 0 | 0 | 0 | 0 | 0 | 0 | 0 | 0 | 0 | 0 | 0 | 0 | 0 | 0 | 0 | 0 | 0 | 1 | 0 | 0 | 0 | 0 | 0 | 0 | 0 | 0 | 0 | 0 | 0 | 0 | 0 | 0 | 0 | 0 | 0 | 0 | 0 | 0 |
| virus_Jul27.25v_37529 | 1 | 0 | 0 | 0 | 0 | 0 | 0 | 0 | 0 | 0 | 0 | 0 | 0 | 0 | 0 | 0 | 0 | 0 | 0 | 0 | 0 | 0 | 0 | 0 | 0 | 0 | 0 | 0 | 0 | 0 | 0 | 0 | 0 | 0 | 1 | 0 | 0 | 0 | 0 | 0 | 0 | 0 | 0 | 0 | 0 | 0 | 0 | 0 | 0 | 0 | 0 | 0 | 0 | 0 | 0 |
| virus_Jul27.25v_41111 | 1 | 0 | 0 | 0 | 0 | 0 | 0 | 0 | 0 | 0 | 0 | 0 | 0 | 0 | 0 | 0 | 0 | 0 | 0 | 0 | 0 | 0 | 0 | 0 | 0 | 0 | 0 | 0 | 0 | 0 | 0 | 0 | 0 | 0 | 1 | 0 | 0 | 0 | 0 | 0 | 0 | 0 | 0 | 0 | 0 | 0 | 0 | 0 | 0 | 0 | 0 | 0 | 0 | 0 | 0 |
| virus_Jul27.75v_5232 | 1 | 0 | 0 | 0 | 0 | 0 | 0 | 0 | 0 | 0 | 0 | 0 | 0 | 0 | 0 | 0 | 0 | 0 | 0 | 0 | 0 | 0 | 0 | 0 | 0 | 0 | 0 | 0 | 0 | 0 | 0 | 0 | 0 | 0 | 0 | 0 | 1 | 0 | 0 | 0 | 0 | 0 | 0 | 0 | 0 | 0 | 0 | 0 | 0 | 0 | 0 | 0 | 0 | 0 | 0 |
| virus_Jul27.75v_6139 | 1 | 0 | 0 | 0 | 0 | 0 | 0 | 0 | 0 | 0 | 0 | 0 | 0 | 0 | 0 | 0 | 0 | 0 | 0 | 0 | 0 | 0 | 0 | 0 | 0 | 0 | 0 | 0 | 0 | 0 | 0 | 0 | 0 | 0 | 0 | 0 | 1 | 0 | 0 | 0 | 0 | 0 | 0 | 0 | 0 | 0 | 0 | 0 | 0 | 0 | 0 | 0 | 0 | 0 | 0 |
| virus_Jul27.75v_18436 | 1 | 0 | 0 | 0 | 0 | 0 | 0 | 0 | 0 | 0 | 0 | 0 | 0 | 0 | 0 | 0 | 0 | 0 | 0 | 0 | 0 | 0 | 0 | 0 | 0 | 0 | 0 | 0 | 0 | 0 | 0 | 0 | 0 | 0 | 0 | 0 | 1 | 0 | 0 | 0 | 0 | 0 | 0 | 0 | 0 | 0 | 0 | 0 | 0 | 0 | 0 | 0 | 0 | 0 | 0 |
| virus_Jun8.25v_36706 | 1 | 0 | 0 | 0 | 0 | 0 | 0 | 0 | 0 | 0 | 1 | 0 | 0 | 0 | 0 | 0 | 0 | 0 | 0 | 0 | 0 | 0 | 0 | 0 | 0 | 0 | 0 | 0 | 0 | 0 | 0 | 0 | 0 | 0 | 0 | 0 | 0 | 0 | 0 | 0 | 0 | 0 | 0 | 0 | 0 | 0 | 0 | 0 | 0 | 0 | 0 | 0 | 0 | 0 | 0 |
| virus_Jun8.50v_12566 | 1 | 0 | 0 | 0 | 0 | 0 | 0 | 0 | 0 | 0 | 0 | 0 | 1 | 0 | 0 | 0 | 0 | 0 | 0 | 0 | 0 | 0 | 0 | 0 | 0 | 0 | 0 | 0 | 0 | 0 | 0 | 0 | 0 | 0 | 0 | 0 | 0 | 0 | 0 | 0 | 0 | 0 | 0 | 0 | 0 | 0 | 0 | 0 | 0 | 0 | 0 | 0 | 0 | 0 | 0 |
| virus_Jun8.50v_15301 | 1 | 0 | 0 | 0 | 0 | 0 | 0 | 0 | 0 | 0 | 0 | 0 | 1 | 0 | 0 | 0 | 0 | 0 | 0 | 0 | 0 | 0 | 0 | 0 | 0 | 0 | 0 | 0 | 0 | 0 | 0 | 0 | 0 | 0 | 0 | 0 | 0 | 0 | 0 | 0 | 0 | 0 | 0 | 0 | 0 | 0 | 0 | 0 | 0 | 0 | 0 | 0 | 0 | 0 | 0 |
| virus_Jun8.50v_25689 | 1 | 0 | 0 | 0 | 0 | 0 | 0 | 0 | 0 | 0 | 0 | 0 | 1 | 0 | 0 | 0 | 0 | 0 | 0 | 0 | 0 | 0 | 0 | 0 | 0 | 0 | 0 | 0 | 0 | 0 | 0 | 0 | 0 | 0 | 0 | 0 | 0 | 0 | 0 | 0 | 0 | 0 | 0 | 0 | 0 | 0 | 0 | 0 | 0 | 0 | 0 | 0 | 0 | 0 | 0 |
| virus_Jun8.50v_27280 | 1 | 0 | 0 | 0 | 0 | 0 | 0 | 0 | 0 | 0 | 0 | 0 | 1 | 0 | 0 | 0 | 0 | 0 | 0 | 0 | 0 | 0 | 0 | 0 | 0 | 0 | 0 | 0 | 0 | 0 | 0 | 0 | 0 | 0 | 0 | 0 | 0 | 0 | 0 | 0 | 0 | 0 | 0 | 0 | 0 | 0 | 0 | 0 | 0 | 0 | 0 | 0 | 0 | 0 | 0 |
| virus_Jun8.50v_29234 | 1 | 0 | 0 | 0 | 0 | 0 | 0 | 0 | 0 | 0 | 0 | 0 | 1 | 0 | 0 | 0 | 0 | 0 | 0 | 0 | 0 | 0 | 0 | 0 | 0 | 0 | 0 | 0 | 0 | 0 | 0 | 0 | 0 | 0 | 0 | 0 | 0 | 0 | 0 | 0 | 0 | 0 | 0 | 0 | 0 | 0 | 0 | 0 | 0 | 0 | 0 | 0 | 0 | 0 | 0 |
| virus_Jun8.50v_29243 | 1 | 0 | 0 | 0 | 0 | 0 | 0 | 0 | 0 | 0 | 0 | 0 | 1 | 0 | 0 | 0 | 0 | 0 | 0 | 0 | 0 | 0 | 0 | 0 | 0 | 0 | 0 | 0 | 0 | 0 | 0 | 0 | 0 | 0 | 0 | 0 | 0 | 0 | 0 | 0 | 0 | 0 | 0 | 0 | 0 | 0 | 0 | 0 | 0 | 0 | 0 | 0 | 0 | 0 | 0 |
| virus_Jun8.50v_30352 | 1 | 0 | 0 | 0 | 0 | 0 | 0 | 0 | 0 | 0 | 0 | 0 | 1 | 0 | 0 | 0 | 0 | 0 | 0 | 0 | 0 | 0 | 0 | 0 | 0 | 0 | 0 | 0 | 0 | 0 | 0 | 0 | 0 | 0 | 0 | 0 | 0 | 0 | 0 | 0 | 0 | 0 | 0 | 0 | 0 | 0 | 0 | 0 | 0 | 0 | 0 | 0 | 0 | 0 | 0 |
| virus_Jun8.50v_31585 | 1 | 0 | 0 | 0 | 0 | 0 | 0 | 0 | 0 | 0 | 0 | 0 | 1 | 0 | 0 | 0 | 0 | 0 | 0 | 0 | 0 | 0 | 0 | 0 | 0 | 0 | 0 | 0 | 0 | 0 | 0 | 0 | 0 | 0 | 0 | 0 | 0 | 0 | 0 | 0 | 0 | 0 | 0 | 0 | 0 | 0 | 0 | 0 | 0 | 0 | 0 | 0 | 0 | 0 | 0 |
| virus_Jun8.75v_6905 | 1 | 0 | 0 | 0 | 0 | 0 | 0 | 0 | 0 | 0 | 0 | 0 | 0 | 0 | 1 | 0 | 0 | 0 | 0 | 0 | 0 | 0 | 0 | 0 | 0 | 0 | 0 | 0 | 0 | 0 | 0 | 0 | 0 | 0 | 0 | 0 | 0 | 0 | 0 | 0 | 0 | 0 | 0 | 0 | 0 | 0 | 0 | 0 | 0 | 0 | 0 | 0 | 0 | 0 | 0 |
| virus_Jun8.75v_17411 | 1 | 0 | 0 | 0 | 0 | 0 | 0 | 0 | 0 | 0 | 0 | 0 | 0 | 0 | 1 | 0 | 0 | 0 | 0 | 0 | 0 | 0 | 0 | 0 | 0 | 0 | 0 | 0 | 0 | 0 | 0 | 0 | 0 | 0 | 0 | 0 | 0 | 0 | 0 | 0 | 0 | 0 | 0 | 0 | 0 | 0 | 0 | 0 | 0 | 0 | 0 | 0 | 0 | 0 | 0 |
| virus_Jun22.0v_2695 | 1 | 0 | 0 | 0 | 0 | 0 | 0 | 0 | 0 | 0 | 0 | 0 | 0 | 0 | 0 | 0 | 1 | 0 | 0 | 0 | 0 | 0 | 0 | 0 | 0 | 0 | 0 | 0 | 0 | 0 | 0 | 0 | 0 | 0 | 0 | 0 | 0 | 0 | 0 | 0 | 0 | 0 | 0 | 0 | 0 | 0 | 0 | 0 | 0 | 0 | 0 | 0 | 0 | 0 | 0 |
| virus_Jun22.0v_2799 | 1 | 0 | 0 | 0 | 0 | 0 | 0 | 0 | 0 | 0 | 0 | 0 | 0 | 0 | 0 | 0 | 1 | 0 | 0 | 0 | 0 | 0 | 0 | 0 | 0 | 0 | 0 | 0 | 0 | 0 | 0 | 0 | 0 | 0 | 0 | 0 | 0 | 0 | 0 | 0 | 0 | 0 | 0 | 0 | 0 | 0 | 0 | 0 | 0 | 0 | 0 | 0 | 0 | 0 | 0 |
| virus_Jun22.0v_5371 | 1 | 0 | 0 | 0 | 0 | 0 | 0 | 0 | 0 | 0 | 0 | 0 | 0 | 0 | 0 | 0 | 1 | 0 | 0 | 0 | 0 | 0 | 0 | 0 | 0 | 0 | 0 | 0 | 0 | 0 | 0 | 0 | 0 | 0 | 0 | 0 | 0 | 0 | 0 | 0 | 0 | 0 | 0 | 0 | 0 | 0 | 0 | 0 | 0 | 0 | 0 | 0 | 0 | 0 | 0 |
| virus_Jun22.0v_10868 | 1 | 0 | 0 | 0 | 0 | 0 | 0 | 0 | 0 | 0 | 0 | 0 | 0 | 0 | 0 | 0 | 1 | 0 | 0 | 0 | 0 | 0 | 0 | 0 | 0 | 0 | 0 | 0 | 0 | 0 | 0 | 0 | 0 | 0 | 0 | 0 | 0 | 0 | 0 | 0 | 0 | 0 | 0 | 0 | 0 | 0 | 0 | 0 | 0 | 0 | 0 | 0 | 0 | 0 | 0 |
| virus_Jun22.0v_12841 | 1 | 0 | 0 | 0 | 0 | 0 | 0 | 0 | 0 | 0 | 0 | 0 | 0 | 0 | 0 | 0 | 1 | 0 | 0 | 0 | 0 | 0 | 0 | 0 | 0 | 0 | 0 | 0 | 0 | 0 | 0 | 0 | 0 | 0 | 0 | 0 | 0 | 0 | 0 | 0 | 0 | 0 | 0 | 0 | 0 | 0 | 0 | 0 | 0 | 0 | 0 | 0 | 0 | 0 | 0 |
| virus_Jun22.0v_15014 | 1 | 0 | 0 | 0 | 0 | 0 | 0 | 0 | 0 | 0 | 0 | 0 | 0 | 0 | 0 | 0 | 1 | 0 | 0 | 0 | 0 | 0 | 0 | 0 | 0 | 0 | 0 | 0 | 0 | 0 | 0 | 0 | 0 | 0 | 0 | 0 | 0 | 0 | 0 | 0 | 0 | 0 | 0 | 0 | 0 | 0 | 0 | 0 | 0 | 0 | 0 | 0 | 0 | 0 | 0 |
| virus_Jun22.0v_16126 | 1 | 0 | 0 | 0 | 0 | 0 | 0 | 0 | 0 | 0 | 0 | 0 | 0 | 0 | 0 | 0 | 1 | 0 | 0 | 0 | 0 | 0 | 0 | 0 | 0 | 0 | 0 | 0 | 0 | 0 | 0 | 0 | 0 | 0 | 0 | 0 | 0 | 0 | 0 | 0 | 0 | 0 | 0 | 0 | 0 | 0 | 0 | 0 | 0 | 0 | 0 | 0 | 0 | 0 | 0 |
| virus_Jun22.0v_17581 | 1 | 0 | 0 | 0 | 0 | 0 | 0 | 0 | 0 | 0 | 0 | 0 | 0 | 0 | 0 | 0 | 1 | 0 | 0 | 0 | 0 | 0 | 0 | 0 | 0 | 0 | 0 | 0 | 0 | 0 | 0 | 0 | 0 | 0 | 0 | 0 | 0 | 0 | 0 | 0 | 0 | 0 | 0 | 0 | 0 | 0 | 0 | 0 | 0 | 0 | 0 | 0 | 0 | 0 | 0 |
| virus_Jun22.0v_18752 | 1 | 0 | 0 | 0 | 0 | 0 | 0 | 0 | 0 | 0 | 0 | 0 | 0 | 0 | 0 | 0 | 1 | 0 | 0 | 0 | 0 | 0 | 0 | 0 | 0 | 0 | 0 | 0 | 0 | 0 | 0 | 0 | 0 | 0 | 0 | 0 | 0 | 0 | 0 | 0 | 0 | 0 | 0 | 0 | 0 | 0 | 0 | 0 | 0 | 0 | 0 | 0 | 0 | 0 | 0 |
| virus_Jun22.0v_19176 | 1 | 0 | 0 | 0 | 0 | 0 | 0 | 0 | 0 | 0 | 0 | 0 | 0 | 0 | 0 | 0 | 1 | 0 | 0 | 0 | 0 | 0 | 0 | 0 | 0 | 0 | 0 | 0 | 0 | 0 | 0 | 0 | 0 | 0 | 0 | 0 | 0 | 0 | 0 | 0 | 0 | 0 | 0 | 0 | 0 | 0 | 0 | 0 | 0 | 0 | 0 | 0 | 0 | 0 | 0 |

**Supplementary Table S7**. Common Bacterio- and virioplankton RTPR 98% OTUs in 27 samples with RTPR amplicons in both fractions
Total seqs: 47997; Subtotal seqs in retained

| OTUs: 41046 | | May12_05m | | May12_25m | | May12_50m | | May12_75m | | Jun8_25m | | Jun8_50m | | Jun8_75m | | Jun22_0m | | Jun22_50m | | Jun22_75m | | Jul11_0m | | Jul11_05m | | Jul11_25m | | Jul11_50m | | Jul11_75m | | Jul27_05m | | Jul27_25m | | Jul27_75m | | Aug5_0m | | Aug5_05m | | Aug5_25m | | Aug5_50m | | Aug5_75m | | Sep8_0m | | Sep8_05m | | Sep8_25m | | Sep8_50m | |
|---|---|---|---|---|---|---|---|---|---|---|---|---|---|---|---|---|---|---|---|---|---|---|---|---|---|---|---|---|---|---|---|---|---|---|---|---|---|---|---|---|---|---|---|---|---|---|---|---|---|---|---|---|---|---|---|
| OTUs with >=10 seqs: 254 OTU subtotal | | bacterioplankton | virioplankton | bacterioplankton | virioplankton | bacterioplankton | virioplankton | bacterioplankton | virioplankton | bacterioplankton | virioplankton | bacterioplankton | virioplankton | bacterioplankton | virioplankton | bacterioplankton | virioplankton | bacterioplankton | virioplankton | bacterioplankton | virioplankton | bacterioplankton | virioplankton | bacterioplankton | virioplankton | bacterioplankton | virioplankton | bacterioplankton | virioplankton | bacterioplankton | virioplankton | bacterioplankton | virioplankton | bacterioplankton | virioplankton | bacterioplankton | virioplankton | bacterioplankton | virioplankton | bacterioplankton | virioplankton | bacterioplankton | virioplankton | bacterioplankton | virioplankton | bacterioplankton | virioplankton | bacterioplankton | virioplankton | bacterioplankton | virioplankton | bacterioplankton | virioplankton | bacterioplankton | virioplankton |
| virus_Jun22.50v_1479 | 1 | 0 | 0 | 0 | 0 | 0 | 0 | 0 | 0 | 0 | 0 | 0 | 0 | 0 | 0 | 0 | 0 | 0 | 1 | 0 | 0 | 0 | 0 | 0 | 0 | 0 | 0 | 0 | 0 | 0 | 0 | 0 | 0 | 0 | 0 | 0 | 0 | 0 | 0 | 0 | 0 | 0 | 0 | 0 | 0 | 0 | 0 | 0 | 0 | 0 | 0 | 0 | 0 | 0 | 0 |
| virus_Jun22.50v_4224 | 1 | 0 | 0 | 0 | 0 | 0 | 0 | 0 | 0 | 0 | 0 | 0 | 0 | 0 | 0 | 0 | 0 | 0 | 1 | 0 | 0 | 0 | 0 | 0 | 0 | 0 | 0 | 0 | 0 | 0 | 0 | 0 | 0 | 0 | 0 | 0 | 0 | 0 | 0 | 0 | 0 | 0 | 0 | 0 | 0 | 0 | 0 | 0 | 0 | 0 | 0 | 0 | 0 | 0 | 0 |
| virus_Jun22.50v_4841 | 1 | 0 | 0 | 0 | 0 | 0 | 0 | 0 | 0 | 0 | 0 | 0 | 0 | 0 | 0 | 0 | 0 | 0 | 1 | 0 | 0 | 0 | 0 | 0 | 0 | 0 | 0 | 0 | 0 | 0 | 0 | 0 | 0 | 0 | 0 | 0 | 0 | 0 | 0 | 0 | 0 | 0 | 0 | 0 | 0 | 0 | 0 | 0 | 0 | 0 | 0 | 0 | 0 | 0 | 0 |
| virus_Jun22.50v_5419 | 1 | 0 | 0 | 0 | 0 | 0 | 0 | 0 | 0 | 0 | 0 | 0 | 0 | 0 | 0 | 0 | 0 | 0 | 1 | 0 | 0 | 0 | 0 | 0 | 0 | 0 | 0 | 0 | 0 | 0 | 0 | 0 | 0 | 0 | 0 | 0 | 0 | 0 | 0 | 0 | 0 | 0 | 0 | 0 | 0 | 0 | 0 | 0 | 0 | 0 | 0 | 0 | 0 | 0 | 0 |
| virus_Jun22.50v_7359 | 1 | 0 | 0 | 0 | 0 | 0 | 0 | 0 | 0 | 0 | 0 | 0 | 0 | 0 | 0 | 0 | 0 | 0 | 1 | 0 | 0 | 0 | 0 | 0 | 0 | 0 | 0 | 0 | 0 | 0 | 0 | 0 | 0 | 0 | 0 | 0 | 0 | 0 | 0 | 0 | 0 | 0 | 0 | 0 | 0 | 0 | 0 | 0 | 0 | 0 | 0 | 0 | 0 | 0 | 0 |
| virus_Jun22.50v_7769 | 1 | 0 | 0 | 0 | 0 | 0 | 0 | 0 | 0 | 0 | 0 | 0 | 0 | 0 | 0 | 0 | 0 | 0 | 1 | 0 | 0 | 0 | 0 | 0 | 0 | 0 | 0 | 0 | 0 | 0 | 0 | 0 | 0 | 0 | 0 | 0 | 0 | 0 | 0 | 0 | 0 | 0 | 0 | 0 | 0 | 0 | 0 | 0 | 0 | 0 | 0 | 0 | 0 | 0 | 0 |
| virus_Jun22.50v_8292 | 1 | 0 | 0 | 0 | 0 | 0 | 0 | 0 | 0 | 0 | 0 | 0 | 0 | 0 | 0 | 0 | 0 | 0 | 1 | 0 | 0 | 0 | 0 | 0 | 0 | 0 | 0 | 0 | 0 | 0 | 0 | 0 | 0 | 0 | 0 | 0 | 0 | 0 | 0 | 0 | 0 | 0 | 0 | 0 | 0 | 0 | 0 | 0 | 0 | 0 | 0 | 0 | 0 | 0 | 0 |
| virus_Jun22.50v_8773 | 1 | 0 | 0 | 0 | 0 | 0 | 0 | 0 | 0 | 0 | 0 | 0 | 0 | 0 | 0 | 0 | 0 | 0 | 1 | 0 | 0 | 0 | 0 | 0 | 0 | 0 | 0 | 0 | 0 | 0 | 0 | 0 | 0 | 0 | 0 | 0 | 0 | 0 | 0 | 0 | 0 | 0 | 0 | 0 | 0 | 0 | 0 | 0 | 0 | 0 | 0 | 0 | 0 | 0 | 0 |
| virus_Jun22.50v_9376 | 1 | 0 | 0 | 0 | 0 | 0 | 0 | 0 | 0 | 0 | 0 | 0 | 0 | 0 | 0 | 0 | 0 | 0 | 1 | 0 | 0 | 0 | 0 | 0 | 0 | 0 | 0 | 0 | 0 | 0 | 0 | 0 | 0 | 0 | 0 | 0 | 0 | 0 | 0 | 0 | 0 | 0 | 0 | 0 | 0 | 0 | 0 | 0 | 0 | 0 | 0 | 0 | 0 | 0 | 0 |
| virus_Jun22.50v_11549 | 1 | 0 | 0 | 0 | 0 | 0 | 0 | 0 | 0 | 0 | 0 | 0 | 0 | 0 | 0 | 0 | 0 | 0 | 1 | 0 | 0 | 0 | 0 | 0 | 0 | 0 | 0 | 0 | 0 | 0 | 0 | 0 | 0 | 0 | 0 | 0 | 0 | 0 | 0 | 0 | 0 | 0 | 0 | 0 | 0 | 0 | 0 | 0 | 0 | 0 | 0 | 0 | 0 | 0 | 0 |
| virus_Jun22.50v_12093 | 1 | 0 | 0 | 0 | 0 | 0 | 0 | 0 | 0 | 0 | 0 | 0 | 0 | 0 | 0 | 0 | 0 | 0 | 1 | 0 | 0 | 0 | 0 | 0 | 0 | 0 | 0 | 0 | 0 | 0 | 0 | 0 | 0 | 0 | 0 | 0 | 0 | 0 | 0 | 0 | 0 | 0 | 0 | 0 | 0 | 0 | 0 | 0 | 0 | 0 | 0 | 0 | 0 | 0 | 0 |
| virus_Jun22.50v_13394 | 1 | 0 | 0 | 0 | 0 | 0 | 0 | 0 | 0 | 0 | 0 | 0 | 0 | 0 | 0 | 0 | 0 | 0 | 1 | 0 | 0 | 0 | 0 | 0 | 0 | 0 | 0 | 0 | 0 | 0 | 0 | 0 | 0 | 0 | 0 | 0 | 0 | 0 | 0 | 0 | 0 | 0 | 0 | 0 | 0 | 0 | 0 | 0 | 0 | 0 | 0 | 0 | 0 | 0 | 0 |
| virus_Jun22.50v_15153 | 1 | 0 | 0 | 0 | 0 | 0 | 0 | 0 | 0 | 0 | 0 | 0 | 0 | 0 | 0 | 0 | 0 | 0 | 1 | 0 | 0 | 0 | 0 | 0 | 0 | 0 | 0 | 0 | 0 | 0 | 0 | 0 | 0 | 0 | 0 | 0 | 0 | 0 | 0 | 0 | 0 | 0 | 0 | 0 | 0 | 0 | 0 | 0 | 0 | 0 | 0 | 0 | 0 | 0 | 0 |
| virus_Jun22.50v_15625 | 1 | 0 | 0 | 0 | 0 | 0 | 0 | 0 | 0 | 0 | 0 | 0 | 0 | 0 | 0 | 0 | 0 | 0 | 1 | 0 | 0 | 0 | 0 | 0 | 0 | 0 | 0 | 0 | 0 | 0 | 0 | 0 | 0 | 0 | 0 | 0 | 0 | 0 | 0 | 0 | 0 | 0 | 0 | 0 | 0 | 0 | 0 | 0 | 0 | 0 | 0 | 0 | 0 | 0 | 0 |
| virus_Jun22.50v_18588 | 1 | 0 | 0 | 0 | 0 | 0 | 0 | 0 | 0 | 0 | 0 | 0 | 0 | 0 | 0 | 0 | 0 | 0 | 1 | 0 | 0 | 0 | 0 | 0 | 0 | 0 | 0 | 0 | 0 | 0 | 0 | 0 | 0 | 0 | 0 | 0 | 0 | 0 | 0 | 0 | 0 | 0 | 0 | 0 | 0 | 0 | 0 | 0 | 0 | 0 | 0 | 0 | 0 | 0 | 0 |
| virus_Jun22.50v_23531 | 1 | 0 | 0 | 0 | 0 | 0 | 0 | 0 | 0 | 0 | 0 | 0 | 0 | 0 | 0 | 0 | 0 | 0 | 1 | 0 | 0 | 0 | 0 | 0 | 0 | 0 | 0 | 0 | 0 | 0 | 0 | 0 | 0 | 0 | 0 | 0 | 0 | 0 | 0 | 0 | 0 | 0 | 0 | 0 | 0 | 0 | 0 | 0 | 0 | 0 | 0 | 0 | 0 | 0 | 0 |
| virus_Jun22.50v_28668 | 1 | 0 | 0 | 0 | 0 | 0 | 0 | 0 | 0 | 0 | 0 | 0 | 0 | 0 | 0 | 0 | 0 | 0 | 1 | 0 | 0 | 0 | 0 | 0 | 0 | 0 | 0 | 0 | 0 | 0 | 0 | 0 | 0 | 0 | 0 | 0 | 0 | 0 | 0 | 0 | 0 | 0 | 0 | 0 | 0 | 0 | 0 | 0 | 0 | 0 | 0 | 0 | 0 | 0 | 0 |
| virus_Jun22.50v_29227 | 1 | 0 | 0 | 0 | 0 | 0 | 0 | 0 | 0 | 0 | 0 | 0 | 0 | 0 | 0 | 0 | 0 | 0 | 1 | 0 | 0 | 0 | 0 | 0 | 0 | 0 | 0 | 0 | 0 | 0 | 0 | 0 | 0 | 0 | 0 | 0 | 0 | 0 | 0 | 0 | 0 | 0 | 0 | 0 | 0 | 0 | 0 | 0 | 0 | 0 | 0 | 0 | 0 | 0 | 0 |
| virus_Jun22.50v_31012 | 1 | 0 | 0 | 0 | 0 | 0 | 0 | 0 | 0 | 0 | 0 | 0 | 0 | 0 | 0 | 0 | 0 | 0 | 1 | 0 | 0 | 0 | 0 | 0 | 0 | 0 | 0 | 0 | 0 | 0 | 0 | 0 | 0 | 0 | 0 | 0 | 0 | 0 | 0 | 0 | 0 | 0 | 0 | 0 | 0 | 0 | 0 | 0 | 0 | 0 | 0 | 0 | 0 | 0 | 0 |
| virus_Jun22.50v_31086 | 1 | 0 | 0 | 0 | 0 | 0 | 0 | 0 | 0 | 0 | 0 | 0 | 0 | 0 | 0 | 0 | 0 | 0 | 1 | 0 | 0 | 0 | 0 | 0 | 0 | 0 | 0 | 0 | 0 | 0 | 0 | 0 | 0 | 0 | 0 | 0 | 0 | 0 | 0 | 0 | 0 | 0 | 0 | 0 | 0 | 0 | 0 | 0 | 0 | 0 | 0 | 0 | 0 | 0 | 0 |
| virus_Jun22.50v_31215 | 1 | 0 | 0 | 0 | 0 | 0 | 0 | 0 | 0 | 0 | 0 | 0 | 0 | 0 | 0 | 0 | 0 | 0 | 1 | 0 | 0 | 0 | 0 | 0 | 0 | 0 | 0 | 0 | 0 | 0 | 0 | 0 | 0 | 0 | 0 | 0 | 0 | 0 | 0 | 0 | 0 | 0 | 0 | 0 | 0 | 0 | 0 | 0 | 0 | 0 | 0 | 0 | 0 | 0 | 0 |
| virus_Jun22.50v_32786 | 1 | 0 | 0 | 0 | 0 | 0 | 0 | 0 | 0 | 0 | 0 | 0 | 0 | 0 | 0 | 0 | 0 | 0 | 1 | 0 | 0 | 0 | 0 | 0 | 0 | 0 | 0 | 0 | 0 | 0 | 0 | 0 | 0 | 0 | 0 | 0 | 0 | 0 | 0 | 0 | 0 | 0 | 0 | 0 | 0 | 0 | 0 | 0 | 0 | 0 | 0 | 0 | 0 | 0 | 0 |
| virus_Jun22.50v_33173 | 1 | 0 | 0 | 0 | 0 | 0 | 0 | 0 | 0 | 0 | 0 | 0 | 0 | 0 | 0 | 0 | 0 | 0 | 1 | 0 | 0 | 0 | 0 | 0 | 0 | 0 | 0 | 0 | 0 | 0 | 0 | 0 | 0 | 0 | 0 | 0 | 0 | 0 | 0 | 0 | 0 | 0 | 0 | 0 | 0 | 0 | 0 | 0 | 0 | 0 | 0 | 0 | 0 | 0 | 0 |
| virus_Jun22.50v_37659 | 1 | 0 | 0 | 0 | 0 | 0 | 0 | 0 | 0 | 0 | 0 | 0 | 0 | 0 | 0 | 0 | 0 | 0 | 1 | 0 | 0 | 0 | 0 | 0 | 0 | 0 | 0 | 0 | 0 | 0 | 0 | 0 | 0 | 0 | 0 | 0 | 0 | 0 | 0 | 0 | 0 | 0 | 0 | 0 | 0 | 0 | 0 | 0 | 0 | 0 | 0 | 0 | 0 | 0 | 0 |
| virus_Jun22.50v_40415 | 1 | 0 | 0 | 0 | 0 | 0 | 0 | 0 | 0 | 0 | 0 | 0 | 0 | 0 | 0 | 0 | 0 | 0 | 1 | 0 | 0 | 0 | 0 | 0 | 0 | 0 | 0 | 0 | 0 | 0 | 0 | 0 | 0 | 0 | 0 | 0 | 0 | 0 | 0 | 0 | 0 | 0 | 0 | 0 | 0 | 0 | 0 | 0 | 0 | 0 | 0 | 0 | 0 | 0 | 0 |
| virus_Jun22.75v_3347 | 1 | 0 | 0 | 0 | 0 | 0 | 0 | 0 | 0 | 0 | 0 | 0 | 0 | 0 | 0 | 0 | 0 | 0 | 0 | 0 | 1 | 0 | 0 | 0 | 0 | 0 | 0 | 0 | 0 | 0 | 0 | 0 | 0 | 0 | 0 | 0 | 0 | 0 | 0 | 0 | 0 | 0 | 0 | 0 | 0 | 0 | 0 | 0 | 0 | 0 | 0 | 0 | 0 | 0 | 0 |
| virus_Jun22.75v_9026 | 1 | 0 | 0 | 0 | 0 | 0 | 0 | 0 | 0 | 0 | 0 | 0 | 0 | 0 | 0 | 0 | 0 | 0 | 0 | 0 | 1 | 0 | 0 | 0 | 0 | 0 | 0 | 0 | 0 | 0 | 0 | 0 | 0 | 0 | 0 | 0 | 0 | 0 | 0 | 0 | 0 | 0 | 0 | 0 | 0 | 0 | 0 | 0 | 0 | 0 | 0 | 0 | 0 | 0 | 0 |
| virus_Jun22.75v_11856 | 1 | 0 | 0 | 0 | 0 | 0 | 0 | 0 | 0 | 0 | 0 | 0 | 0 | 0 | 0 | 0 | 0 | 0 | 0 | 0 | 1 | 0 | 0 | 0 | 0 | 0 | 0 | 0 | 0 | 0 | 0 | 0 | 0 | 0 | 0 | 0 | 0 | 0 | 0 | 0 | 0 | 0 | 0 | 0 | 0 | 0 | 0 | 0 | 0 | 0 | 0 | 0 | 0 | 0 | 0 |
| virus_Jun22.75v_28013 | 1 | 0 | 0 | 0 | 0 | 0 | 0 | 0 | 0 | 0 | 0 | 0 | 0 | 0 | 0 | 0 | 0 | 0 | 0 | 0 | 1 | 0 | 0 | 0 | 0 | 0 | 0 | 0 | 0 | 0 | 0 | 0 | 0 | 0 | 0 | 0 | 0 | 0 | 0 | 0 | 0 | 0 | 0 | 0 | 0 | 0 | 0 | 0 | 0 | 0 | 0 | 0 | 0 | 0 | 0 |
| virus_Jun22.75v_28044 | 1 | 0 | 0 | 0 | 0 | 0 | 0 | 0 | 0 | 0 | 0 | 0 | 0 | 0 | 0 | 0 | 0 | 0 | 0 | 0 | 1 | 0 | 0 | 0 | 0 | 0 | 0 | 0 | 0 | 0 | 0 | 0 | 0 | 0 | 0 | 0 | 0 | 0 | 0 | 0 | 0 | 0 | 0 | 0 | 0 | 0 | 0 | 0 | 0 | 0 | 0 | 0 | 0 | 0 | 0 |
| virus_Jun22.75v_35672 | 1 | 0 | 0 | 0 | 0 | 0 | 0 | 0 | 0 | 0 | 0 | 0 | 0 | 0 | 0 | 0 | 0 | 0 | 0 | 0 | 1 | 0 | 0 | 0 | 0 | 0 | 0 | 0 | 0 | 0 | 0 | 0 | 0 | 0 | 0 | 0 | 0 | 0 | 0 | 0 | 0 | 0 | 0 | 0 | 0 | 0 | 0 | 0 | 0 | 0 | 0 | 0 | 0 | 0 | 0 |
| virus_Jun22.75v_36577 | 1 | 0 | 0 | 0 | 0 | 0 | 0 | 0 | 0 | 0 | 0 | 0 | 0 | 0 | 0 | 0 | 0 | 0 | 0 | 0 | 1 | 0 | 0 | 0 | 0 | 0 | 0 | 0 | 0 | 0 | 0 | 0 | 0 | 0 | 0 | 0 | 0 | 0 | 0 | 0 | 0 | 0 | 0 | 0 | 0 | 0 | 0 | 0 | 0 | 0 | 0 | 0 | 0 | 0 | 0 |
| virus_Jun22.75v_37712 | 1 | 0 | 0 | 0 | 0 | 0 | 0 | 0 | 0 | 0 | 0 | 0 | 0 | 0 | 0 | 0 | 0 | 0 | 0 | 0 | 1 | 0 | 0 | 0 | 0 | 0 | 0 | 0 | 0 | 0 | 0 | 0 | 0 | 0 | 0 | 0 | 0 | 0 | 0 | 0 | 0 | 0 | 0 | 0 | 0 | 0 | 0 | 0 | 0 | 0 | 0 | 0 | 0 | 0 | 0 |
| virus_May12.5v_229 | 1 | 0 | 1 | 0 | 0 | 0 | 0 | 0 | 0 | 0 | 0 | 0 | 0 | 0 | 0 | 0 | 0 | 0 | 0 | 0 | 0 | 0 | 0 | 0 | 0 | 0 | 0 | 0 | 0 | 0 | 0 | 0 | 0 | 0 | 0 | 0 | 0 | 0 | 0 | 0 | 0 | 0 | 0 | 0 | 0 | 0 | 0 | 0 | 0 | 0 | 0 | 0 | 0 | 0 | 0 |
| virus_May12.5v_776 | 1 | 0 | 1 | 0 | 0 | 0 | 0 | 0 | 0 | 0 | 0 | 0 | 0 | 0 | 0 | 0 | 0 | 0 | 0 | 0 | 0 | 0 | 0 | 0 | 0 | 0 | 0 | 0 | 0 | 0 | 0 | 0 | 0 | 0 | 0 | 0 | 0 | 0 | 0 | 0 | 0 | 0 | 0 | 0 | 0 | 0 | 0 | 0 | 0 | 0 | 0 | 0 | 0 | 0 | 0 |
| virus_May12.5v_3677 | 1 | 0 | 1 | 0 | 0 | 0 | 0 | 0 | 0 | 0 | 0 | 0 | 0 | 0 | 0 | 0 | 0 | 0 | 0 | 0 | 0 | 0 | 0 | 0 | 0 | 0 | 0 | 0 | 0 | 0 | 0 | 0 | 0 | 0 | 0 | 0 | 0 | 0 | 0 | 0 | 0 | 0 | 0 | 0 | 0 | 0 | 0 | 0 | 0 | 0 | 0 | 0 | 0 | 0 | 0 |
| virus_May12.5v_5978 | 1 | 0 | 1 | 0 | 0 | 0 | 0 | 0 | 0 | 0 | 0 | 0 | 0 | 0 | 0 | 0 | 0 | 0 | 0 | 0 | 0 | 0 | 0 | 0 | 0 | 0 | 0 | 0 | 0 | 0 | 0 | 0 | 0 | 0 | 0 | 0 | 0 | 0 | 0 | 0 | 0 | 0 | 0 | 0 | 0 | 0 | 0 | 0 | 0 | 0 | 0 | 0 | 0 | 0 | 0 |
| virus_May12.5v_14498 | 1 | 0 | 1 | 0 | 0 | 0 | 0 | 0 | 0 | 0 | 0 | 0 | 0 | 0 | 0 | 0 | 0 | 0 | 0 | 0 | 0 | 0 | 0 | 0 | 0 | 0 | 0 | 0 | 0 | 0 | 0 | 0 | 0 | 0 | 0 | 0 | 0 | 0 | 0 | 0 | 0 | 0 | 0 | 0 | 0 | 0 | 0 | 0 | 0 | 0 | 0 | 0 | 0 | 0 | 0 |
| virus_May12.5v_20478 | 1 | 0 | 1 | 0 | 0 | 0 | 0 | 0 | 0 | 0 | 0 | 0 | 0 | 0 | 0 | 0 | 0 | 0 | 0 | 0 | 0 | 0 | 0 | 0 | 0 | 0 | 0 | 0 | 0 | 0 | 0 | 0 | 0 | 0 | 0 | 0 | 0 | 0 | 0 | 0 | 0 | 0 | 0 | 0 | 0 | 0 | 0 | 0 | 0 | 0 | 0 | 0 | 0 | 0 | 0 |
| virus_May12.5v_26165 | 1 | 0 | 1 | 0 | 0 | 0 | 0 | 0 | 0 | 0 | 0 | 0 | 0 | 0 | 0 | 0 | 0 | 0 | 0 | 0 | 0 | 0 | 0 | 0 | 0 | 0 | 0 | 0 | 0 | 0 | 0 | 0 | 0 | 0 | 0 | 0 | 0 | 0 | 0 | 0 | 0 | 0 | 0 | 0 | 0 | 0 | 0 | 0 | 0 | 0 | 0 | 0 | 0 | 0 | 0 |
| virus_May12.5v_30061 | 1 | 0 | 1 | 0 | 0 | 0 | 0 | 0 | 0 | 0 | 0 | 0 | 0 | 0 | 0 | 0 | 0 | 0 | 0 | 0 | 0 | 0 | 0 | 0 | 0 | 0 | 0 | 0 | 0 | 0 | 0 | 0 | 0 | 0 | 0 | 0 | 0 | 0 | 0 | 0 | 0 | 0 | 0 | 0 | 0 | 0 | 0 | 0 | 0 | 0 | 0 | 0 | 0 | 0 | 0 |
| virus_May12.5v_30272 | 1 | 0 | 1 | 0 | 0 | 0 | 0 | 0 | 0 | 0 | 0 | 0 | 0 | 0 | 0 | 0 | 0 | 0 | 0 | 0 | 0 | 0 | 0 | 0 | 0 | 0 | 0 | 0 | 0 | 0 | 0 | 0 | 0 | 0 | 0 | 0 | 0 | 0 | 0 | 0 | 0 | 0 | 0 | 0 | 0 | 0 | 0 | 0 | 0 | 0 | 0 | 0 | 0 | 0 | 0 |
| virus_May12.5v_34438 | 1 | 0 | 1 | 0 | 0 | 0 | 0 | 0 | 0 | 0 | 0 | 0 | 0 | 0 | 0 | 0 | 0 | 0 | 0 | 0 | 0 | 0 | 0 | 0 | 0 | 0 | 0 | 0 | 0 | 0 | 0 | 0 | 0 | 0 | 0 | 0 | 0 | 0 | 0 | 0 | 0 | 0 | 0 | 0 | 0 | 0 | 0 | 0 | 0 | 0 | 0 | 0 | 0 | 0 | 0 |
| virus_May12.5v_38517 | 1 | 0 | 1 | 0 | 0 | 0 | 0 | 0 | 0 | 0 | 0 | 0 | 0 | 0 | 0 | 0 | 0 | 0 | 0 | 0 | 0 | 0 | 0 | 0 | 0 | 0 | 0 | 0 | 0 | 0 | 0 | 0 | 0 | 0 | 0 | 0 | 0 | 0 | 0 | 0 | 0 | 0 | 0 | 0 | 0 | 0 | 0 | 0 | 0 | 0 | 0 | 0 | 0 | 0 | 0 |
| virus_May12.25v_6734 | 1 | 0 | 0 | 0 | 1 | 0 | 0 | 0 | 0 | 0 | 0 | 0 | 0 | 0 | 0 | 0 | 0 | 0 | 0 | 0 | 0 | 0 | 0 | 0 | 0 | 0 | 0 | 0 | 0 | 0 | 0 | 0 | 0 | 0 | 0 | 0 | 0 | 0 | 0 | 0 | 0 | 0 | 0 | 0 | 0 | 0 | 0 | 0 | 0 | 0 | 0 | 0 | 0 | 0 | 0 |
| virus_May12.25v_13063 | 1 | 0 | 0 | 0 | 1 | 0 | 0 | 0 | 0 | 0 | 0 | 0 | 0 | 0 | 0 | 0 | 0 | 0 | 0 | 0 | 0 | 0 | 0 | 0 | 0 | 0 | 0 | 0 | 0 | 0 | 0 | 0 | 0 | 0 | 0 | 0 | 0 | 0 | 0 | 0 | 0 | 0 | 0 | 0 | 0 | 0 | 0 | 0 | 0 | 0 | 0 | 0 | 0 | 0 | 0 |
| virus_May12.25v_18789 | 1 | 0 | 0 | 0 | 1 | 0 | 0 | 0 | 0 | 0 | 0 | 0 | 0 | 0 | 0 | 0 | 0 | 0 | 0 | 0 | 0 | 0 | 0 | 0 | 0 | 0 | 0 | 0 | 0 | 0 | 0 | 0 | 0 | 0 | 0 | 0 | 0 | 0 | 0 | 0 | 0 | 0 | 0 | 0 | 0 | 0 | 0 | 0 | 0 | 0 | 0 | 0 | 0 | 0 | 0 |
| virus_May12.25v_20561 | 1 | 0 | 0 | 0 | 1 | 0 | 0 | 0 | 0 | 0 | 0 | 0 | 0 | 0 | 0 | 0 | 0 | 0 | 0 | 0 | 0 | 0 | 0 | 0 | 0 | 0 | 0 | 0 | 0 | 0 | 0 | 0 | 0 | 0 | 0 | 0 | 0 | 0 | 0 | 0 | 0 | 0 | 0 | 0 | 0 | 0 | 0 | 0 | 0 | 0 | 0 | 0 | 0 | 0 | 0 |
| virus_May12.25v_22028 | 1 | 0 | 0 | 0 | 1 | 0 | 0 | 0 | 0 | 0 | 0 | 0 | 0 | 0 | 0 | 0 | 0 | 0 | 0 | 0 | 0 | 0 | 0 | 0 | 0 | 0 | 0 | 0 | 0 | 0 | 0 | 0 | 0 | 0 | 0 | 0 | 0 | 0 | 0 | 0 | 0 | 0 | 0 | 0 | 0 | 0 | 0 | 0 | 0 | 0 | 0 | 0 | 0 | 0 | 0 |
| virus_May12.25v_26376 | 1 | 0 | 0 | 0 | 1 | 0 | 0 | 0 | 0 | 0 | 0 | 0 | 0 | 0 | 0 | 0 | 0 | 0 | 0 | 0 | 0 | 0 | 0 | 0 | 0 | 0 | 0 | 0 | 0 | 0 | 0 | 0 | 0 | 0 | 0 | 0 | 0 | 0 | 0 | 0 | 0 | 0 | 0 | 0 | 0 | 0 | 0 | 0 | 0 | 0 | 0 | 0 | 0 | 0 | 0 |
| virus_May12.25v_37667 | 1 | 0 | 0 | 0 | 1 | 0 | 0 | 0 | 0 | 0 | 0 | 0 | 0 | 0 | 0 | 0 | 0 | 0 | 0 | 0 | 0 | 0 | 0 | 0 | 0 | 0 | 0 | 0 | 0 | 0 | 0 | 0 | 0 | 0 | 0 | 0 | 0 | 0 | 0 | 0 | 0 | 0 | 0 | 0 | 0 | 0 | 0 | 0 | 0 | 0 | 0 | 0 | 0 | 0 | 0 |
| virus_May12.25v_40561 | 1 | 0 | 0 | 0 | 1 | 0 | 0 | 0 | 0 | 0 | 0 | 0 | 0 | 0 | 0 | 0 | 0 | 0 | 0 | 0 | 0 | 0 | 0 | 0 | 0 | 0 | 0 | 0 | 0 | 0 | 0 | 0 | 0 | 0 | 0 | 0 | 0 | 0 | 0 | 0 | 0 | 0 | 0 | 0 | 0 | 0 | 0 | 0 | 0 | 0 | 0 | 0 | 0 | 0 | 0 |
| virus_May12.50v_517 | 1 | 0 | 0 | 0 | 0 | 0 | 1 | 0 | 0 | 0 | 0 | 0 | 0 | 0 | 0 | 0 | 0 | 0 | 0 | 0 | 0 | 0 | 0 | 0 | 0 | 0 | 0 | 0 | 0 | 0 | 0 | 0 | 0 | 0 | 0 | 0 | 0 | 0 | 0 | 0 | 0 | 0 | 0 | 0 | 0 | 0 | 0 | 0 | 0 | 0 | 0 | 0 | 0 | 0 | 0 |
| virus_May12.50v_1873 | 1 | 0 | 0 | 0 | 0 | 0 | 1 | 0 | 0 | 0 | 0 | 0 | 0 | 0 | 0 | 0 | 0 | 0 | 0 | 0 | 0 | 0 | 0 | 0 | 0 | 0 | 0 | 0 | 0 | 0 | 0 | 0 | 0 | 0 | 0 | 0 | 0 | 0 | 0 | 0 | 0 | 0 | 0 | 0 | 0 | 0 | 0 | 0 | 0 | 0 | 0 | 0 | 0 | 0 | 0 |
| virus_May12.50v_4359 | 1 | 0 | 0 | 0 | 0 | 0 | 1 | 0 | 0 | 0 | 0 | 0 | 0 | 0 | 0 | 0 | 0 | 0 | 0 | 0 | 0 | 0 | 0 | 0 | 0 | 0 | 0 | 0 | 0 | 0 | 0 | 0 | 0 | 0 | 0 | 0 | 0 | 0 | 0 | 0 | 0 | 0 | 0 | 0 | 0 | 0 | 0 | 0 | 0 | 0 | 0 | 0 | 0 | 0 | 0 |
| virus_May12.50v_6436 | 1 | 0 | 0 | 0 | 0 | 0 | 1 | 0 | 0 | 0 | 0 | 0 | 0 | 0 | 0 | 0 | 0 | 0 | 0 | 0 | 0 | 0 | 0 | 0 | 0 | 0 | 0 | 0 | 0 | 0 | 0 | 0 | 0 | 0 | 0 | 0 | 0 | 0 | 0 | 0 | 0 | 0 | 0 | 0 | 0 | 0 | 0 | 0 | 0 | 0 | 0 | 0 | 0 | 0 | 0 |
| virus_May12.50v_7012 | 1 | 0 | 0 | 0 | 0 | 0 | 1 | 0 | 0 | 0 | 0 | 0 | 0 | 0 | 0 | 0 | 0 | 0 | 0 | 0 | 0 | 0 | 0 | 0 | 0 | 0 | 0 | 0 | 0 | 0 | 0 | 0 | 0 | 0 | 0 | 0 | 0 | 0 | 0 | 0 | 0 | 0 | 0 | 0 | 0 | 0 | 0 | 0 | 0 | 0 | 0 | 0 | 0 | 0 | 0 |
| virus_May12.50v_7840 | 1 | 0 | 0 | 0 | 0 | 0 | 1 | 0 | 0 | 0 | 0 | 0 | 0 | 0 | 0 | 0 | 0 | 0 | 0 | 0 | 0 | 0 | 0 | 0 | 0 | 0 | 0 | 0 | 0 | 0 | 0 | 0 | 0 | 0 | 0 | 0 | 0 | 0 | 0 | 0 | 0 | 0 | 0 | 0 | 0 | 0 | 0 | 0 | 0 | 0 | 0 | 0 | 0 | 0 | 0 |
| virus_May12.50v_12855 | 1 | 0 | 0 | 0 | 0 | 0 | 1 | 0 | 0 | 0 | 0 | 0 | 0 | 0 | 0 | 0 | 0 | 0 | 0 | 0 | 0 | 0 | 0 | 0 | 0 | 0 | 0 | 0 | 0 | 0 | 0 | 0 | 0 | 0 | 0 | 0 | 0 | 0 | 0 | 0 | 0 | 0 | 0 | 0 | 0 | 0 | 0 | 0 | 0 | 0 | 0 | 0 | 0 | 0 | 0 |
| virus_May12.50v_14301 | 1 | 0 | 0 | 0 | 0 | 0 | 1 | 0 | 0 | 0 | 0 | 0 | 0 | 0 | 0 | 0 | 0 | 0 | 0 | 0 | 0 | 0 | 0 | 0 | 0 | 0 | 0 | 0 | 0 | 0 | 0 | 0 | 0 | 0 | 0 | 0 | 0 | 0 | 0 | 0 | 0 | 0 | 0 | 0 | 0 | 0 | 0 | 0 | 0 | 0 | 0 | 0 | 0 | 0 | 0 |
| virus_May12.50v_16267 | 1 | 0 | 0 | 0 | 0 | 0 | 1 | 0 | 0 | 0 | 0 | 0 | 0 | 0 | 0 | 0 | 0 | 0 | 0 | 0 | 0 | 0 | 0 | 0 | 0 | 0 | 0 | 0 | 0 | 0 | 0 | 0 | 0 | 0 | 0 | 0 | 0 | 0 | 0 | 0 | 0 | 0 | 0 | 0 | 0 | 0 | 0 | 0 | 0 | 0 | 0 | 0 | 0 | 0 | 0 |
| virus_May12.50v_19591 | 1 | 0 | 0 | 0 | 0 | 0 | 1 | 0 | 0 | 0 | 0 | 0 | 0 | 0 | 0 | 0 | 0 | 0 | 0 | 0 | 0 | 0 | 0 | 0 | 0 | 0 | 0 | 0 | 0 | 0 | 0 | 0 | 0 | 0 | 0 | 0 | 0 | 0 | 0 | 0 | 0 | 0 | 0 | 0 | 0 | 0 | 0 | 0 | 0 | 0 | 0 | 0 | 0 | 0 | 0 |
| virus_May12.50v_21730 | 1 | 0 | 0 | 0 | 0 | 0 | 1 | 0 | 0 | 0 | 0 | 0 | 0 | 0 | 0 | 0 | 0 | 0 | 0 | 0 | 0 | 0 | 0 | 0 | 0 | 0 | 0 | 0 | 0 | 0 | 0 | 0 | 0 | 0 | 0 | 0 | 0 | 0 | 0 | 0 | 0 | 0 | 0 | 0 | 0 | 0 | 0 | 0 | 0 | 0 | 0 | 0 | 0 | 0 | 0 |
| virus_May12.50v_22241 | 1 | 0 | 0 | 0 | 0 | 0 | 1 | 0 | 0 | 0 | 0 | 0 | 0 | 0 | 0 | 0 | 0 | 0 | 0 | 0 | 0 | 0 | 0 | 0 | 0 | 0 | 0 | 0 | 0 | 0 | 0 | 0 | 0 | 0 | 0 | 0 | 0 | 0 | 0 | 0 | 0 | 0 | 0 | 0 | 0 | 0 | 0 | 0 | 0 | 0 | 0 | 0 | 0 | 0 | 0 |
| virus_May12.50v_24086 | 1 | 0 | 0 | 0 | 0 | 0 | 1 | 0 | 0 | 0 | 0 | 0 | 0 | 0 | 0 | 0 | 0 | 0 | 0 | 0 | 0 | 0 | 0 | 0 | 0 | 0 | 0 | 0 | 0 | 0 | 0 | 0 | 0 | 0 | 0 | 0 | 0 | 0 | 0 | 0 | 0 | 0 | 0 | 0 | 0 | 0 | 0 | 0 | 0 | 0 | 0 | 0 | 0 | 0 | 0 |
| virus_May12.50v_24282 | 1 | 0 | 0 | 0 | 0 | 0 | 1 | 0 | 0 | 0 | 0 | 0 | 0 | 0 | 0 | 0 | 0 | 0 | 0 | 0 | 0 | 0 | 0 | 0 | 0 | 0 | 0 | 0 | 0 | 0 | 0 | 0 | 0 | 0 | 0 | 0 | 0 | 0 | 0 | 0 | 0 | 0 | 0 | 0 | 0 | 0 | 0 | 0 | 0 | 0 | 0 | 0 | 0 | 0 | 0 |
| virus_May12.50v_34656 | 1 | 0 | 0 | 0 | 0 | 0 | 1 | 0 | 0 | 0 | 0 | 0 | 0 | 0 | 0 | 0 | 0 | 0 | 0 | 0 | 0 | 0 | 0 | 0 | 0 | 0 | 0 | 0 | 0 | 0 | 0 | 0 | 0 | 0 | 0 | 0 | 0 | 0 | 0 | 0 | 0 | 0 | 0 | 0 | 0 | 0 | 0 | 0 | 0 | 0 | 0 | 0 | 0 | 0 | 0 |
| virus_May12.50v_37498 | 1 | 0 | 0 | 0 | 0 | 0 | 1 | 0 | 0 | 0 | 0 | 0 | 0 | 0 | 0 | 0 | 0 | 0 | 0 | 0 | 0 | 0 | 0 | 0 | 0 | 0 | 0 | 0 | 0 | 0 | 0 | 0 | 0 | 0 | 0 | 0 | 0 | 0 | 0 | 0 | 0 | 0 | 0 | 0 | 0 | 0 | 0 | 0 | 0 | 0 | 0 | 0 | 0 | 0 | 0 |
| virus_May12.50v_39241 | 1 | 0 | 0 | 0 | 0 | 0 | 1 | 0 | 0 | 0 | 0 | 0 | 0 | 0 | 0 | 0 | 0 | 0 | 0 | 0 | 0 | 0 | 0 | 0 | 0 | 0 | 0 | 0 | 0 | 0 | 0 | 0 | 0 | 0 | 0 | 0 | 0 | 0 | 0 | 0 | 0 | 0 | 0 | 0 | 0 | 0 | 0 | 0 | 0 | 0 | 0 | 0 | 0 | 0 | 0 |
| virus_May12.50v_40642 | 1 | 0 | 0 | 0 | 0 | 0 | 1 | 0 | 0 | 0 | 0 | 0 | 0 | 0 | 0 | 0 | 0 | 0 | 0 | 0 | 0 | 0 | 0 | 0 | 0 | 0 | 0 | 0 | 0 | 0 | 0 | 0 | 0 | 0 | 0 | 0 | 0 | 0 | 0 | 0 | 0 | 0 | 0 | 0 | 0 | 0 | 0 | 0 | 0 | 0 | 0 | 0 | 0 | 0 | 0 |
| virus_May12.75v_5955 | 1 | 0 | 0 | 0 | 0 | 0 | 0 | 0 | 1 | 0 | 0 | 0 | 0 | 0 | 0 | 0 | 0 | 0 | 0 | 0 | 0 | 0 | 0 | 0 | 0 | 0 | 0 | 0 | 0 | 0 | 0 | 0 | 0 | 0 | 0 | 0 | 0 | 0 | 0 | 0 | 0 | 0 | 0 | 0 | 0 | 0 | 0 | 0 | 0 | 0 | 0 | 0 | 0 | 0 | 0 |
| virus_May12.75v_6315 | 1 | 0 | 0 | 0 | 0 | 0 | 0 | 0 | 1 | 0 | 0 | 0 | 0 | 0 | 0 | 0 | 0 | 0 | 0 | 0 | 0 | 0 | 0 | 0 | 0 | 0 | 0 | 0 | 0 | 0 | 0 | 0 | 0 | 0 | 0 | 0 | 0 | 0 | 0 | 0 | 0 | 0 | 0 | 0 | 0 | 0 | 0 | 0 | 0 | 0 | 0 | 0 | 0 | 0 | 0 |
| virus_May12.75v_12526 | 1 | 0 | 0 | 0 | 0 | 0 | 0 | 0 | 1 | 0 | 0 | 0 | 0 | 0 | 0 | 0 | 0 | 0 | 0 | 0 | 0 | 0 | 0 | 0 | 0 | 0 | 0 | 0 | 0 | 0 | 0 | 0 | 0 | 0 | 0 | 0 | 0 | 0 | 0 | 0 | 0 | 0 | 0 | 0 | 0 | 0 | 0 | 0 | 0 | 0 | 0 | 0 | 0 | 0 | 0 |
| virus_May12.75v_13232 | 1 | 0 | 0 | 0 | 0 | 0 | 0 | 0 | 1 | 0 | 0 | 0 | 0 | 0 | 0 | 0 | 0 | 0 | 0 | 0 | 0 | 0 | 0 | 0 | 0 | 0 | 0 | 0 | 0 | 0 | 0 | 0 | 0 | 0 | 0 | 0 | 0 | 0 | 0 | 0 | 0 | 0 | 0 | 0 | 0 | 0 | 0 | 0 | 0 | 0 | 0 | 0 | 0 | 0 | 0 |
| virus_May12.75v_17992 | 1 | 0 | 0 | 0 | 0 | 0 | 0 | 0 | 1 | 0 | 0 | 0 | 0 | 0 | 0 | 0 | 0 | 0 | 0 | 0 | 0 | 0 | 0 | 0 | 0 | 0 | 0 | 0 | 0 | 0 | 0 | 0 | 0 | 0 | 0 | 0 | 0 | 0 | 0 | 0 | 0 | 0 | 0 | 0 | 0 | 0 | 0 | 0 | 0 | 0 | 0 | 0 | 0 | 0 | 0 |
| virus_May12.75v_19401 | 1 | 0 | 0 | 0 | 0 | 0 | 0 | 0 | 1 | 0 | 0 | 0 | 0 | 0 | 0 | 0 | 0 | 0 | 0 | 0 | 0 | 0 | 0 | 0 | 0 | 0 | 0 | 0 | 0 | 0 | 0 | 0 | 0 | 0 | 0 | 0 | 0 | 0 | 0 | 0 | 0 | 0 | 0 | 0 | 0 | 0 | 0 | 0 | 0 | 0 | 0 | 0 | 0 | 0 | 0 |
| virus_May12.75v_22279 | 1 | 0 | 0 | 0 | 0 | 0 | 0 | 0 | 1 | 0 | 0 | 0 | 0 | 0 | 0 | 0 | 0 | 0 | 0 | 0 | 0 | 0 | 0 | 0 | 0 | 0 | 0 | 0 | 0 | 0 | 0 | 0 | 0 | 0 | 0 | 0 | 0 | 0 | 0 | 0 | 0 | 0 | 0 | 0 | 0 | 0 | 0 | 0 | 0 | 0 | 0 | 0 | 0 | 0 | 0 |
| virus_May12.75v_23095 | 1 | 0 | 0 | 0 | 0 | 0 | 0 | 0 | 1 | 0 | 0 | 0 | 0 | 0 | 0 | 0 | 0 | 0 | 0 | 0 | 0 | 0 | 0 | 0 | 0 | 0 | 0 | 0 | 0 | 0 | 0 | 0 | 0 | 0 | 0 | 0 | 0 | 0 | 0 | 0 | 0 | 0 | 0 | 0 | 0 | 0 | 0 | 0 | 0 | 0 | 0 | 0 | 0 | 0 | 0 |
| virus_May12.75v_23746 | 1 | 0 | 0 | 0 | 0 | 0 | 0 | 0 | 1 | 0 | 0 | 0 | 0 | 0 | 0 | 0 | 0 | 0 | 0 | 0 | 0 | 0 | 0 | 0 | 0 | 0 | 0 | 0 | 0 | 0 | 0 | 0 | 0 | 0 | 0 | 0 | 0 | 0 | 0 | 0 | 0 | 0 | 0 | 0 | 0 | 0 | 0 | 0 | 0 | 0 | 0 | 0 | 0 | 0 | 0 |
| virus_May12.75v_26200 | 1 | 0 | 0 | 0 | 0 | 0 | 0 | 0 | 1 | 0 | 0 | 0 | 0 | 0 | 0 | 0 | 0 | 0 | 0 | 0 | 0 | 0 | 0 | 0 | 0 | 0 | 0 | 0 | 0 | 0 | 0 | 0 | 0 | 0 | 0 | 0 | 0 | 0 | 0 | 0 | 0 | 0 | 0 | 0 | 0 | 0 | 0 | 0 | 0 | 0 | 0 | 0 | 0 | 0 | 0 |
| virus_May12.75v_28437 | 1 | 0 | 0 | 0 | 0 | 0 | 0 | 0 | 1 | 0 | 0 | 0 | 0 | 0 | 0 | 0 | 0 | 0 | 0 | 0 | 0 | 0 | 0 | 0 | 0 | 0 | 0 | 0 | 0 | 0 | 0 | 0 | 0 | 0 | 0 | 0 | 0 | 0 | 0 | 0 | 0 | 0 | 0 | 0 | 0 | 0 | 0 | 0 | 0 | 0 | 0 | 0 | 0 | 0 | 0 |
| virus_May12.75v_33213 | 1 | 0 | 0 | 0 | 0 | 0 | 0 | 0 | 1 | 0 | 0 | 0 | 0 | 0 | 0 | 0 | 0 | 0 | 0 | 0 | 0 | 0 | 0 | 0 | 0 | 0 | 0 | 0 | 0 | 0 | 0 | 0 | 0 | 0 | 0 | 0 | 0 | 0 | 0 | 0 | 0 | 0 | 0 | 0 | 0 | 0 | 0 | 0 | 0 | 0 | 0 | 0 | 0 | 0 | 0 |
| virus_May12.75v_38720 | 1 | 0 | 0 | 0 | 0 | 0 | 0 | 0 | 1 | 0 | 0 | 0 | 0 | 0 | 0 | 0 | 0 | 0 | 0 | 0 | 0 | 0 | 0 | 0 | 0 | 0 | 0 | 0 | 0 | 0 | 0 | 0 | 0 | 0 | 0 | 0 | 0 | 0 | 0 | 0 | 0 | 0 | 0 | 0 | 0 | 0 | 0 | 0 | 0 | 0 | 0 | 0 | 0 | 0 | 0 |
| virus_Sep8.0v_1101 | 1 | 0 | 0 | 0 | 0 | 0 | 0 | 0 | 0 | 0 | 0 | 0 | 0 | 0 | 0 | 0 | 0 | 0 | 0 | 0 | 0 | 0 | 0 | 0 | 0 | 0 | 0 | 0 | 0 | 0 | 0 | 0 | 0 | 0 | 0 | 0 | 0 | 0 | 0 | 0 | 0 | 0 | 0 | 0 | 0 | 0 | 0 | 0 | 1 | 0 | 0 | 0 | 0 | 0 | 0 |
| virus_Sep8.0v_1275 | 1 | 0 | 0 | 0 | 0 | 0 | 0 | 0 | 0 | 0 | 0 | 0 | 0 | 0 | 0 | 0 | 0 | 0 | 0 | 0 | 0 | 0 | 0 | 0 | 0 | 0 | 0 | 0 | 0 | 0 | 0 | 0 | 0 | 0 | 0 | 0 | 0 | 0 | 0 | 0 | 0 | 0 | 0 | 0 | 0 | 0 | 0 | 0 | 1 | 0 | 0 | 0 | 0 | 0 | 0 |
| virus_Sep8.0v_10736 | 1 | 0 | 0 | 0 | 0 | 0 | 0 | 0 | 0 | 0 | 0 | 0 | 0 | 0 | 0 | 0 | 0 | 0 | 0 | 0 | 0 | 0 | 0 | 0 | 0 | 0 | 0 | 0 | 0 | 0 | 0 | 0 | 0 | 0 | 0 | 0 | 0 | 0 | 0 | 0 | 0 | 0 | 0 | 0 | 0 | 0 | 0 | 0 | 1 | 0 | 0 | 0 | 0 | 0 | 0 |
| virus_Sep8.0v_13128 | 1 | 0 | 0 | 0 | 0 | 0 | 0 | 0 | 0 | 0 | 0 | 0 | 0 | 0 | 0 | 0 | 0 | 0 | 0 | 0 | 0 | 0 | 0 | 0 | 0 | 0 | 0 | 0 | 0 | 0 | 0 | 0 | 0 | 0 | 0 | 0 | 0 | 0 | 0 | 0 | 0 | 0 | 0 | 0 | 0 | 0 | 0 | 0 | 1 | 0 | 0 | 0 | 0 | 0 | 0 |
| virus_Sep8.0v_14349 | 1 | 0 | 0 | 0 | 0 | 0 | 0 | 0 | 0 | 0 | 0 | 0 | 0 | 0 | 0 | 0 | 0 | 0 | 0 | 0 | 0 | 0 | 0 | 0 | 0 | 0 | 0 | 0 | 0 | 0 | 0 | 0 | 0 | 0 | 0 | 0 | 0 | 0 | 0 | 0 | 0 | 0 | 0 | 0 | 0 | 0 | 0 | 0 | 1 | 0 | 0 | 0 | 0 | 0 | 0 |
| virus_Sep8.0v_15372 | 1 | 0 | 0 | 0 | 0 | 0 | 0 | 0 | 0 | 0 | 0 | 0 | 0 | 0 | 0 | 0 | 0 | 0 | 0 | 0 | 0 | 0 | 0 | 0 | 0 | 0 | 0 | 0 | 0 | 0 | 0 | 0 | 0 | 0 | 0 | 0 | 0 | 0 | 0 | 0 | 0 | 0 | 0 | 0 | 0 | 0 | 0 | 0 | 1 | 0 | 0 | 0 | 0 | 0 | 0 |
| virus_Sep8.0v_16220 | 1 | 0 | 0 | 0 | 0 | 0 | 0 | 0 | 0 | 0 | 0 | 0 | 0 | 0 | 0 | 0 | 0 | 0 | 0 | 0 | 0 | 0 | 0 | 0 | 0 | 0 | 0 | 0 | 0 | 0 | 0 | 0 | 0 | 0 | 0 | 0 | 0 | 0 | 0 | 0 | 0 | 0 | 0 | 0 | 0 | 0 | 0 | 0 | 1 | 0 | 0 | 0 | 0 | 0 | 0 |
| virus_Sep8.0v_17221 | 1 | 0 | 0 | 0 | 0 | 0 | 0 | 0 | 0 | 0 | 0 | 0 | 0 | 0 | 0 | 0 | 0 | 0 | 0 | 0 | 0 | 0 | 0 | 0 | 0 | 0 | 0 | 0 | 0 | 0 | 0 | 0 | 0 | 0 | 0 | 0 | 0 | 0 | 0 | 0 | 0 | 0 | 0 | 0 | 0 | 0 | 0 | 0 | 1 | 0 | 0 | 0 | 0 | 0 | 0 |
| virus_Sep8.0v_18019 | 1 | 0 | 0 | 0 | 0 | 0 | 0 | 0 | 0 | 0 | 0 | 0 | 0 | 0 | 0 | 0 | 0 | 0 | 0 | 0 | 0 | 0 | 0 | 0 | 0 | 0 | 0 | 0 | 0 | 0 | 0 | 0 | 0 | 0 | 0 | 0 | 0 | 0 | 0 | 0 | 0 | 0 | 0 | 0 | 0 | 0 | 0 | 0 | 1 | 0 | 0 | 0 | 0 | 0 | 0 |
| virus_Sep8.0v_19152 | 1 | 0 | 0 | 0 | 0 | 0 | 0 | 0 | 0 | 0 | 0 | 0 | 0 | 0 | 0 | 0 | 0 | 0 | 0 | 0 | 0 | 0 | 0 | 0 | 0 | 0 | 0 | 0 | 0 | 0 | 0 | 0 | 0 | 0 | 0 | 0 | 0 | 0 | 0 | 0 | 0 | 0 | 0 | 0 | 0 | 0 | 0 | 0 | 1 | 0 | 0 | 0 | 0 | 0 | 0 |
| virus_Sep8.0v_19482 | 1 | 0 | 0 | 0 | 0 | 0 | 0 | 0 | 0 | 0 | 0 | 0 | 0 | 0 | 0 | 0 | 0 | 0 | 0 | 0 | 0 | 0 | 0 | 0 | 0 | 0 | 0 | 0 | 0 | 0 | 0 | 0 | 0 | 0 | 0 | 0 | 0 | 0 | 0 | 0 | 0 | 0 | 0 | 0 | 0 | 0 | 0 | 0 | 1 | 0 | 0 | 0 | 0 | 0 | 0 |
| virus_Sep8.5v_1832 | 1 | 0 | 0 | 0 | 0 | 0 | 0 | 0 | 0 | 0 | 0 | 0 | 0 | 0 | 0 | 0 | 0 | 0 | 0 | 0 | 0 | 0 | 0 | 0 | 0 | 0 | 0 | 0 | 0 | 0 | 0 | 0 | 0 | 0 | 0 | 0 | 0 | 0 | 0 | 0 | 0 | 0 | 0 | 0 | 0 | 0 | 0 | 0 | 0 | 0 | 1 | 0 | 0 | 0 | 0 |
| virus_Sep8.5v_6144 | 1 | 0 | 0 | 0 | 0 | 0 | 0 | 0 | 0 | 0 | 0 | 0 | 0 | 0 | 0 | 0 | 0 | 0 | 0 | 0 | 0 | 0 | 0 | 0 | 0 | 0 | 0 | 0 | 0 | 0 | 0 | 0 | 0 | 0 | 0 | 0 | 0 | 0 | 0 | 0 | 0 | 0 | 0 | 0 | 0 | 0 | 0 | 0 | 0 | 0 | 1 | 0 | 0 | 0 | 0 |
| virus_Sep8.5v_8500 | 1 | 0 | 0 | 0 | 0 | 0 | 0 | 0 | 0 | 0 | 0 | 0 | 0 | 0 | 0 | 0 | 0 | 0 | 0 | 0 | 0 | 0 | 0 | 0 | 0 | 0 | 0 | 0 | 0 | 0 | 0 | 0 | 0 | 0 | 0 | 0 | 0 | 0 | 0 | 0 | 0 | 0 | 0 | 0 | 0 | 0 | 0 | 0 | 0 | 0 | 1 | 0 | 0 | 0 | 0 |
| virus_Sep8.5v_8644 | 1 | 0 | 0 | 0 | 0 | 0 | 0 | 0 | 0 | 0 | 0 | 0 | 0 | 0 | 0 | 0 | 0 | 0 | 0 | 0 | 0 | 0 | 0 | 0 | 0 | 0 | 0 | 0 | 0 | 0 | 0 | 0 | 0 | 0 | 0 | 0 | 0 | 0 | 0 | 0 | 0 | 0 | 0 | 0 | 0 | 0 | 0 | 0 | 0 | 0 | 1 | 0 | 0 | 0 | 0 |
| virus_Sep8.5v_17440 | 1 | 0 | 0 | 0 | 0 | 0 | 0 | 0 | 0 | 0 | 0 | 0 | 0 | 0 | 0 | 0 | 0 | 0 | 0 | 0 | 0 | 0 | 0 | 0 | 0 | 0 | 0 | 0 | 0 | 0 | 0 | 0 | 0 | 0 | 0 | 0 | 0 | 0 | 0 | 0 | 0 | 0 | 0 | 0 | 0 | 0 | 0 | 0 | 0 | 0 | 1 | 0 | 0 | 0 | 0 |
| virus_Sep8.5v_19205 | 1 | 0 | 0 | 0 | 0 | 0 | 0 | 0 | 0 | 0 | 0 | 0 | 0 | 0 | 0 | 0 | 0 | 0 | 0 | 0 | 0 | 0 | 0 | 0 | 0 | 0 | 0 | 0 | 0 | 0 | 0 | 0 | 0 | 0 | 0 | 0 | 0 | 0 | 0 | 0 | 0 | 0 | 0 | 0 | 0 | 0 | 0 | 0 | 0 | 0 | 1 | 0 | 0 | 0 | 0 |
| virus_Sep8.25v_1879 | 1 | 0 | 0 | 0 | 0 | 0 | 0 | 0 | 0 | 0 | 0 | 0 | 0 | 0 | 0 | 0 | 0 | 0 | 0 | 0 | 0 | 0 | 0 | 0 | 0 | 0 | 0 | 0 | 0 | 0 | 0 | 0 | 0 | 0 | 0 | 0 | 0 | 0 | 0 | 0 | 0 | 0 | 0 | 0 | 0 | 0 | 0 | 0 | 0 | 0 | 0 | 0 | 1 | 0 | 0 |
| virus_Sep8.25v_2068 | 1 | 0 | 0 | 0 | 0 | 0 | 0 | 0 | 0 | 0 | 0 | 0 | 0 | 0 | 0 | 0 | 0 | 0 | 0 | 0 | 0 | 0 | 0 | 0 | 0 | 0 | 0 | 0 | 0 | 0 | 0 | 0 | 0 | 0 | 0 | 0 | 0 | 0 | 0 | 0 | 0 | 0 | 0 | 0 | 0 | 0 | 0 | 0 | 0 | 0 | 0 | 0 | 1 | 0 | 0 |
| virus_Sep8.25v_5364 | 1 | 0 | 0 | 0 | 0 | 0 | 0 | 0 | 0 | 0 | 0 | 0 | 0 | 0 | 0 | 0 | 0 | 0 | 0 | 0 | 0 | 0 | 0 | 0 | 0 | 0 | 0 | 0 | 0 | 0 | 0 | 0 | 0 | 0 | 0 | 0 | 0 | 0 | 0 | 0 | 0 | 0 | 0 | 0 | 0 | 0 | 0 | 0 | 0 | 0 | 0 | 0 | 1 | 0 | 0 |
| virus_Sep8.25v_8232 | 1 | 0 | 0 | 0 | 0 | 0 | 0 | 0 | 0 | 0 | 0 | 0 | 0 | 0 | 0 | 0 | 0 | 0 | 0 | 0 | 0 | 0 | 0 | 0 | 0 | 0 | 0 | 0 | 0 | 0 | 0 | 0 | 0 | 0 | 0 | 0 | 0 | 0 | 0 | 0 | 0 | 0 | 0 | 0 | 0 | 0 | 0 | 0 | 0 | 0 | 0 | 0 | 1 | 0 | 0 |
| virus_Sep8.25v_9026 | 1 | 0 | 0 | 0 | 0 | 0 | 0 | 0 | 0 | 0 | 0 | 0 | 0 | 0 | 0 | 0 | 0 | 0 | 0 | 0 | 0 | 0 | 0 | 0 | 0 | 0 | 0 | 0 | 0 | 0 | 0 | 0 | 0 | 0 | 0 | 0 | 0 | 0 | 0 | 0 | 0 | 0 | 0 | 0 | 0 | 0 | 0 | 0 | 0 | 0 | 0 | 0 | 1 | 0 | 0 |
| virus_Sep8.25v_14057 | 1 | 0 | 0 | 0 | 0 | 0 | 0 | 0 | 0 | 0 | 0 | 0 | 0 | 0 | 0 | 0 | 0 | 0 | 0 | 0 | 0 | 0 | 0 | 0 | 0 | 0 | 0 | 0 | 0 | 0 | 0 | 0 | 0 | 0 | 0 | 0 | 0 | 0 | 0 | 0 | 0 | 0 | 0 | 0 | 0 | 0 | 0 | 0 | 0 | 0 | 0 | 0 | 1 | 0 | 0 |
| virus_Sep8.25v_17217 | 1 | 0 | 0 | 0 | 0 | 0 | 0 | 0 | 0 | 0 | 0 | 0 | 0 | 0 | 0 | 0 | 0 | 0 | 0 | 0 | 0 | 0 | 0 | 0 | 0 | 0 | 0 | 0 | 0 | 0 | 0 | 0 | 0 | 0 | 0 | 0 | 0 | 0 | 0 | 0 | 0 | 0 | 0 | 0 | 0 | 0 | 0 | 0 | 0 | 0 | 0 | 0 | 1 | 0 | 0 |
| virus_Sep8.25v_17295 | 1 | 0 | 0 | 0 | 0 | 0 | 0 | 0 | 0 | 0 | 0 | 0 | 0 | 0 | 0 | 0 | 0 | 0 | 0 | 0 | 0 | 0 | 0 | 0 | 0 | 0 | 0 | 0 | 0 | 0 | 0 | 0 | 0 | 0 | 0 | 0 | 0 | 0 | 0 | 0 | 0 | 0 | 0 | 0 | 0 | 0 | 0 | 0 | 0 | 0 | 0 | 0 | 1 | 0 | 0 |
| virus_Sep8.25v_18280 | 1 | 0 | 0 | 0 | 0 | 0 | 0 | 0 | 0 | 0 | 0 | 0 | 0 | 0 | 0 | 0 | 0 | 0 | 0 | 0 | 0 | 0 | 0 | 0 | 0 | 0 | 0 | 0 | 0 | 0 | 0 | 0 | 0 | 0 | 0 | 0 | 0 | 0 | 0 | 0 | 0 | 0 | 0 | 0 | 0 | 0 | 0 | 0 | 0 | 0 | 0 | 0 | 1 | 0 | 0 |
| virus_Sep8.50v_1433 | 1 | 0 | 0 | 0 | 0 | 0 | 0 | 0 | 0 | 0 | 0 | 0 | 0 | 0 | 0 | 0 | 0 | 0 | 0 | 0 | 0 | 0 | 0 | 0 | 0 | 0 | 0 | 0 | 0 | 0 | 0 | 0 | 0 | 0 | 0 | 0 | 0 | 0 | 0 | 0 | 0 | 0 | 0 | 0 | 0 | 0 | 0 | 0 | 0 | 0 | 0 | 0 | 0 | 0 | 1 |
| virus_Sep8.50v_1984 | 1 | 0 | 0 | 0 | 0 | 0 | 0 | 0 | 0 | 0 | 0 | 0 | 0 | 0 | 0 | 0 | 0 | 0 | 0 | 0 | 0 | 0 | 0 | 0 | 0 | 0 | 0 | 0 | 0 | 0 | 0 | 0 | 0 | 0 | 0 | 0 | 0 | 0 | 0 | 0 | 0 | 0 | 0 | 0 | 0 | 0 | 0 | 0 | 0 | 0 | 0 | 0 | 0 | 0 | 1 |
| virus_Sep8.50v_7969 | 1 | 0 | 0 | 0 | 0 | 0 | 0 | 0 | 0 | 0 | 0 | 0 | 0 | 0 | 0 | 0 | 0 | 0 | 0 | 0 | 0 | 0 | 0 | 0 | 0 | 0 | 0 | 0 | 0 | 0 | 0 | 0 | 0 | 0 | 0 | 0 | 0 | 0 | 0 | 0 | 0 | 0 | 0 | 0 | 0 | 0 | 0 | 0 | 0 | 0 | 0 | 0 | 0 | 0 | 1 |
| virus_Sep8.50v_13391 | 1 | 0 | 0 | 0 | 0 | 0 | 0 | 0 | 0 | 0 | 0 | 0 | 0 | 0 | 0 | 0 | 0 | 0 | 0 | 0 | 0 | 0 | 0 | 0 | 0 | 0 | 0 | 0 | 0 | 0 | 0 | 0 | 0 | 0 | 0 | 0 | 0 | 0 | 0 | 0 | 0 | 0 | 0 | 0 | 0 | 0 | 0 | 0 | 0 | 0 | 0 | 0 | 0 | 0 | 1 |
| virus_Sep8.50v_20658 | 1 | 0 | 0 | 0 | 0 | 0 | 0 | 0 | 0 | 0 | 0 | 0 | 0 | 0 | 0 | 0 | 0 | 0 | 0 | 0 | 0 | 0 | 0 | 0 | 0 | 0 | 0 | 0 | 0 | 0 | 0 | 0 | 0 | 0 | 0 | 0 | 0 | 0 | 0 | 0 | 0 | 0 | 0 | 0 | 0 | 0 | 0 | 0 | 0 | 0 | 0 | 0 | 0 | 0 | 1 |
| virus_Sep8.50v_22833 | 1 | 0 | 0 | 0 | 0 | 0 | 0 | 0 | 0 | 0 | 0 | 0 | 0 | 0 | 0 | 0 | 0 | 0 | 0 | 0 | 0 | 0 | 0 | 0 | 0 | 0 | 0 | 0 | 0 | 0 | 0 | 0 | 0 | 0 | 0 | 0 | 0 | 0 | 0 | 0 | 0 | 0 | 0 | 0 | 0 | 0 | 0 | 0 | 0 | 0 | 0 | 0 | 0 | 0 | 1 |
| virus_Sep8.50v_25890 | 1 | 0 | 0 | 0 | 0 | 0 | 0 | 0 | 0 | 0 | 0 | 0 | 0 | 0 | 0 | 0 | 0 | 0 | 0 | 0 | 0 | 0 | 0 | 0 | 0 | 0 | 0 | 0 | 0 | 0 | 0 | 0 | 0 | 0 | 0 | 0 | 0 | 0 | 0 | 0 | 0 | 0 | 0 | 0 | 0 | 0 | 0 | 0 | 0 | 0 | 0 | 0 | 0 | 0 | 1 |
| virus_Sep8.50v_32743 | 1 | 0 | 0 | 0 | 0 | 0 | 0 | 0 | 0 | 0 | 0 | 0 | 0 | 0 | 0 | 0 | 0 | 0 | 0 | 0 | 0 | 0 | 0 | 0 | 0 | 0 | 0 | 0 | 0 | 0 | 0 | 0 | 0 | 0 | 0 | 0 | 0 | 0 | 0 | 0 | 0 | 0 | 0 | 0 | 0 | 0 | 0 | 0 | 0 | 0 | 0 | 0 | 0 | 0 | 1 |
| virus_Sep8.50v_36946 | 1 | 0 | 0 | 0 | 0 | 0 | 0 | 0 | 0 | 0 | 0 | 0 | 0 | 0 | 0 | 0 | 0 | 0 | 0 | 0 | 0 | 0 | 0 | 0 | 0 | 0 | 0 | 0 | 0 | 0 | 0 | 0 | 0 | 0 | 0 | 0 | 0 | 0 | 0 | 0 | 0 | 0 | 0 | 0 | 0 | 0 | 0 | 0 | 0 | 0 | 0 | 0 | 0 | 0 | 1 |
| virus_Sep8.50v_39897 | 1 | 0 | 0 | 0 | 0 | 0 | 0 | 0 | 0 | 0 | 0 | 0 | 0 | 0 | 0 | 0 | 0 | 0 | 0 | 0 | 0 | 0 | 0 | 0 | 0 | 0 | 0 | 0 | 0 | 0 | 0 | 0 | 0 | 0 | 0 | 0 | 0 | 0 | 0 | 0 | 0 | 0 | 0 | 0 | 0 | 0 | 0 | 0 | 0 | 0 | 0 | 0 | 0 | 0 | 1 |
| bacteria_Jul11d0b_3604 | 1 | 0 | 0 | 0 | 0 | 0 | 0 | 0 | 0 | 0 | 0 | 0 | 0 | 0 | 0 | 0 | 0 | 0 | 0 | 0 | 0 | 1 | 0 | 0 | 0 | 0 | 0 | 0 | 0 | 0 | 0 | 0 | 0 | 0 | 0 | 0 | 0 | 0 | 0 | 0 | 0 | 0 | 0 | 0 | 0 | 0 | 0 | 0 | 0 | 0 | 0 | 0 | 0 | 0 | 0 |
| bacteria_Jul11d0b_5048 | 1 | 0 | 0 | 0 | 0 | 0 | 0 | 0 | 0 | 0 | 0 | 0 | 0 | 0 | 0 | 0 | 0 | 0 | 0 | 0 | 0 | 1 | 0 | 0 | 0 | 0 | 0 | 0 | 0 | 0 | 0 | 0 | 0 | 0 | 0 | 0 | 0 | 0 | 0 | 0 | 0 | 0 | 0 | 0 | 0 | 0 | 0 | 0 | 0 | 0 | 0 | 0 | 0 | 0 | 0 |
| bacteria_Jul11d0b_15193 | 1 | 0 | 0 | 0 | 0 | 0 | 0 | 0 | 0 | 0 | 0 | 0 | 0 | 0 | 0 | 0 | 0 | 0 | 0 | 0 | 0 | 1 | 0 | 0 | 0 | 0 | 0 | 0 | 0 | 0 | 0 | 0 | 0 | 0 | 0 | 0 | 0 | 0 | 0 | 0 | 0 | 0 | 0 | 0 | 0 | 0 | 0 | 0 | 0 | 0 | 0 | 0 | 0 | 0 | 0 |
| bacteria_Jul11d0b_19957 | 1 | 0 | 0 | 0 | 0 | 0 | 0 | 0 | 0 | 0 | 0 | 0 | 0 | 0 | 0 | 0 | 0 | 0 | 0 | 0 | 0 | 1 | 0 | 0 | 0 | 0 | 0 | 0 | 0 | 0 | 0 | 0 | 0 | 0 | 0 | 0 | 0 | 0 | 0 | 0 | 0 | 0 | 0 | 0 | 0 | 0 | 0 | 0 | 0 | 0 | 0 | 0 | 0 | 0 | 0 |
| bacteria_Jul11d0b_32605 | 1 | 0 | 0 | 0 | 0 | 0 | 0 | 0 | 0 | 0 | 0 | 0 | 0 | 0 | 0 | 0 | 0 | 0 | 0 | 0 | 0 | 1 | 0 | 0 | 0 | 0 | 0 | 0 | 0 | 0 | 0 | 0 | 0 | 0 | 0 | 0 | 0 | 0 | 0 | 0 | 0 | 0 | 0 | 0 | 0 | 0 | 0 | 0 | 0 | 0 | 0 | 0 | 0 | 0 | 0 |
| bacteria_Jul11d0b_34466 | 1 | 0 | 0 | 0 | 0 | 0 | 0 | 0 | 0 | 0 | 0 | 0 | 0 | 0 | 0 | 0 | 0 | 0 | 0 | 0 | 0 | 1 | 0 | 0 | 0 | 0 | 0 | 0 | 0 | 0 | 0 | 0 | 0 | 0 | 0 | 0 | 0 | 0 | 0 | 0 | 0 | 0 | 0 | 0 | 0 | 0 | 0 | 0 | 0 | 0 | 0 | 0 | 0 | 0 | 0 |
| bacteria_Jul11d0b_47693 | 1 | 0 | 0 | 0 | 0 | 0 | 0 | 0 | 0 | 0 | 0 | 0 | 0 | 0 | 0 | 0 | 0 | 0 | 0 | 0 | 0 | 1 | 0 | 0 | 0 | 0 | 0 | 0 | 0 | 0 | 0 | 0 | 0 | 0 | 0 | 0 | 0 | 0 | 0 | 0 | 0 | 0 | 0 | 0 | 0 | 0 | 0 | 0 | 0 | 0 | 0 | 0 | 0 | 0 | 0 |
| bacteria_Jul11d0b_59949 | 1 | 0 | 0 | 0 | 0 | 0 | 0 | 0 | 0 | 0 | 0 | 0 | 0 | 0 | 0 | 0 | 0 | 0 | 0 | 0 | 0 | 1 | 0 | 0 | 0 | 0 | 0 | 0 | 0 | 0 | 0 | 0 | 0 | 0 | 0 | 0 | 0 | 0 | 0 | 0 | 0 | 0 | 0 | 0 | 0 | 0 | 0 | 0 | 0 | 0 | 0 | 0 | 0 | 0 | 0 |
| bacteria_Jul11d0b_66100 | 1 | 0 | 0 | 0 | 0 | 0 | 0 | 0 | 0 | 0 | 0 | 0 | 0 | 0 | 0 | 0 | 0 | 0 | 0 | 0 | 0 | 1 | 0 | 0 | 0 | 0 | 0 | 0 | 0 | 0 | 0 | 0 | 0 | 0 | 0 | 0 | 0 | 0 | 0 | 0 | 0 | 0 | 0 | 0 | 0 | 0 | 0 | 0 | 0 | 0 | 0 | 0 | 0 | 0 | 0 |
| bacteria_Jul11d0b_74640 | 1 | 0 | 0 | 0 | 0 | 0 | 0 | 0 | 0 | 0 | 0 | 0 | 0 | 0 | 0 | 0 | 0 | 0 | 0 | 0 | 0 | 1 | 0 | 0 | 0 | 0 | 0 | 0 | 0 | 0 | 0 | 0 | 0 | 0 | 0 | 0 | 0 | 0 | 0 | 0 | 0 | 0 | 0 | 0 | 0 | 0 | 0 | 0 | 0 | 0 | 0 | 0 | 0 | 0 | 0 |
| bacteria_Aug5d0b_10414 | 1 | 0 | 0 | 0 | 0 | 0 | 0 | 0 | 0 | 0 | 0 | 0 | 0 | 0 | 0 | 0 | 0 | 0 | 0 | 0 | 0 | 0 | 0 | 0 | 0 | 0 | 0 | 0 | 0 | 0 | 0 | 0 | 0 | 0 | 0 | 0 | 0 | 1 | 0 | 0 | 0 | 0 | 0 | 0 | 0 | 0 | 0 | 0 | 0 | 0 | 0 | 0 | 0 | 0 | 0 |
| bacteria_Aug5d0b_15290 | 1 | 0 | 0 | 0 | 0 | 0 | 0 | 0 | 0 | 0 | 0 | 0 | 0 | 0 | 0 | 0 | 0 | 0 | 0 | 0 | 0 | 0 | 0 | 0 | 0 | 0 | 0 | 0 | 0 | 0 | 0 | 0 | 0 | 0 | 0 | 0 | 0 | 1 | 0 | 0 | 0 | 0 | 0 | 0 | 0 | 0 | 0 | 0 | 0 | 0 | 0 | 0 | 0 | 0 | 0 |
| bacteria_Aug5d0b_20121 | 1 | 0 | 0 | 0 | 0 | 0 | 0 | 0 | 0 | 0 | 0 | 0 | 0 | 0 | 0 | 0 | 0 | 0 | 0 | 0 | 0 | 0 | 0 | 0 | 0 | 0 | 0 | 0 | 0 | 0 | 0 | 0 | 0 | 0 | 0 | 0 | 0 | 1 | 0 | 0 | 0 | 0 | 0 | 0 | 0 | 0 | 0 | 0 | 0 | 0 | 0 | 0 | 0 | 0 | 0 |
| bacteria_Aug5d0b_25710 | 1 | 0 | 0 | 0 | 0 | 0 | 0 | 0 | 0 | 0 | 0 | 0 | 0 | 0 | 0 | 0 | 0 | 0 | 0 | 0 | 0 | 0 | 0 | 0 | 0 | 0 | 0 | 0 | 0 | 0 | 0 | 0 | 0 | 0 | 0 | 0 | 0 | 1 | 0 | 0 | 0 | 0 | 0 | 0 | 0 | 0 | 0 | 0 | 0 | 0 | 0 | 0 | 0 | 0 | 0 |
| bacteria_Aug5d5b_14816 | 1 | 0 | 0 | 0 | 0 | 0 | 0 | 0 | 0 | 0 | 0 | 0 | 0 | 0 | 0 | 0 | 0 | 0 | 0 | 0 | 0 | 0 | 0 | 0 | 0 | 0 | 0 | 0 | 0 | 0 | 0 | 0 | 0 | 0 | 0 | 0 | 0 | 0 | 0 | 1 | 0 | 0 | 0 | 0 | 0 | 0 | 0 | 0 | 0 | 0 | 0 | 0 | 0 | 0 | 0 |
| bacteria_Aug5d5b_52619 | 1 | 0 | 0 | 0 | 0 | 0 | 0 | 0 | 0 | 0 | 0 | 0 | 0 | 0 | 0 | 0 | 0 | 0 | 0 | 0 | 0 | 0 | 0 | 0 | 0 | 0 | 0 | 0 | 0 | 0 | 0 | 0 | 0 | 0 | 0 | 0 | 0 | 0 | 0 | 1 | 0 | 0 | 0 | 0 | 0 | 0 | 0 | 0 | 0 | 0 | 0 | 0 | 0 | 0 | 0 |
| bacteria_Aug5d5b_60951 | 1 | 0 | 0 | 0 | 0 | 0 | 0 | 0 | 0 | 0 | 0 | 0 | 0 | 0 | 0 | 0 | 0 | 0 | 0 | 0 | 0 | 0 | 0 | 0 | 0 | 0 | 0 | 0 | 0 | 0 | 0 | 0 | 0 | 0 | 0 | 0 | 0 | 0 | 0 | 1 | 0 | 0 | 0 | 0 | 0 | 0 | 0 | 0 | 0 | 0 | 0 | 0 | 0 | 0 | 0 |
| bacteria_Aug5d25b_49 | 1 | 0 | 0 | 0 | 0 | 0 | 0 | 0 | 0 | 0 | 0 | 0 | 0 | 0 | 0 | 0 | 0 | 0 | 0 | 0 | 0 | 0 | 0 | 0 | 0 | 0 | 0 | 0 | 0 | 0 | 0 | 0 | 0 | 0 | 0 | 0 | 0 | 0 | 0 | 0 | 0 | 1 | 0 | 0 | 0 | 0 | 0 | 0 | 0 | 0 | 0 | 0 | 0 | 0 | 0 |
| bacteria_Aug5d25b_4558 | 1 | 0 | 0 | 0 | 0 | 0 | 0 | 0 | 0 | 0 | 0 | 0 | 0 | 0 | 0 | 0 | 0 | 0 | 0 | 0 | 0 | 0 | 0 | 0 | 0 | 0 | 0 | 0 | 0 | 0 | 0 | 0 | 0 | 0 | 0 | 0 | 0 | 0 | 0 | 0 | 0 | 1 | 0 | 0 | 0 | 0 | 0 | 0 | 0 | 0 | 0 | 0 | 0 | 0 | 0 |
| bacteria_Aug5d25b_5385 | 1 | 0 | 0 | 0 | 0 | 0 | 0 | 0 | 0 | 0 | 0 | 0 | 0 | 0 | 0 | 0 | 0 | 0 | 0 | 0 | 0 | 0 | 0 | 0 | 0 | 0 | 0 | 0 | 0 | 0 | 0 | 0 | 0 | 0 | 0 | 0 | 0 | 0 | 0 | 0 | 0 | 1 | 0 | 0 | 0 | 0 | 0 | 0 | 0 | 0 | 0 | 0 | 0 | 0 | 0 |
| bacteria_Aug5d25b_7514 | 1 | 0 | 0 | 0 | 0 | 0 | 0 | 0 | 0 | 0 | 0 | 0 | 0 | 0 | 0 | 0 | 0 | 0 | 0 | 0 | 0 | 0 | 0 | 0 | 0 | 0 | 0 | 0 | 0 | 0 | 0 | 0 | 0 | 0 | 0 | 0 | 0 | 0 | 0 | 0 | 0 | 1 | 0 | 0 | 0 | 0 | 0 | 0 | 0 | 0 | 0 | 0 | 0 | 0 | 0 |
| bacteria_Aug5d25b_7927 | 1 | 0 | 0 | 0 | 0 | 0 | 0 | 0 | 0 | 0 | 0 | 0 | 0 | 0 | 0 | 0 | 0 | 0 | 0 | 0 | 0 | 0 | 0 | 0 | 0 | 0 | 0 | 0 | 0 | 0 | 0 | 0 | 0 | 0 | 0 | 0 | 0 | 0 | 0 | 0 | 0 | 1 | 0 | 0 | 0 | 0 | 0 | 0 | 0 | 0 | 0 | 0 | 0 | 0 | 0 |
| bacteria_Aug5d25b_2157 | 1 | 0 | 0 | 0 | 0 | 0 | 0 | 0 | 0 | 0 | 0 | 0 | 0 | 0 | 0 | 0 | 0 | 0 | 0 | 0 | 0 | 0 | 0 | 0 | 0 | 0 | 0 | 0 | 0 | 0 | 0 | 0 | 0 | 0 | 0 | 0 | 0 | 0 | 0 | 0 | 0 | 1 | 0 | 0 | 0 | 0 | 0 | 0 | 0 | 0 | 0 | 0 | 0 | 0 | 0 |
| bacteria_Aug5d25b_2735 | 1 | 0 | 0 | 0 | 0 | 0 | 0 | 0 | 0 | 0 | 0 | 0 | 0 | 0 | 0 | 0 | 0 | 0 | 0 | 0 | 0 | 0 | 0 | 0 | 0 | 0 | 0 | 0 | 0 | 0 | 0 | 0 | 0 | 0 | 0 | 0 | 0 | 0 | 0 | 0 | 0 | 1 | 0 | 0 | 0 | 0 | 0 | 0 | 0 | 0 | 0 | 0 | 0 | 0 | 0 |
| bacteria_Aug5d25b_3170 | 1 | 0 | 0 | 0 | 0 | 0 | 0 | 0 | 0 | 0 | 0 | 0 | 0 | 0 | 0 | 0 | 0 | 0 | 0 | 0 | 0 | 0 | 0 | 0 | 0 | 0 | 0 | 0 | 0 | 0 | 0 | 0 | 0 | 0 | 0 | 0 | 0 | 0 | 0 | 0 | 0 | 1 | 0 | 0 | 0 | 0 | 0 | 0 | 0 | 0 | 0 | 0 | 0 | 0 | 0 |
| bacteria_Aug5d25b_3381 | 1 | 0 | 0 | 0 | 0 | 0 | 0 | 0 | 0 | 0 | 0 | 0 | 0 | 0 | 0 | 0 | 0 | 0 | 0 | 0 | 0 | 0 | 0 | 0 | 0 | 0 | 0 | 0 | 0 | 0 | 0 | 0 | 0 | 0 | 0 | 0 | 0 | 0 | 0 | 0 | 0 | 1 | 0 | 0 | 0 | 0 | 0 | 0 | 0 | 0 | 0 | 0 | 0 | 0 | 0 |
| bacteria_Aug5d25b_4169 | 1 | 0 | 0 | 0 | 0 | 0 | 0 | 0 | 0 | 0 | 0 | 0 | 0 | 0 | 0 | 0 | 0 | 0 | 0 | 0 | 0 | 0 | 0 | 0 | 0 | 0 | 0 | 0 | 0 | 0 | 0 | 0 | 0 | 0 | 0 | 0 | 0 | 0 | 0 | 0 | 0 | 1 | 0 | 0 | 0 | 0 | 0 | 0 | 0 | 0 | 0 | 0 | 0 | 0 | 0 |
| bacteria_Aug5d25b_4691 | 1 | 0 | 0 | 0 | 0 | 0 | 0 | 0 | 0 | 0 | 0 | 0 | 0 | 0 | 0 | 0 | 0 | 0 | 0 | 0 | 0 | 0 | 0 | 0 | 0 | 0 | 0 | 0 | 0 | 0 | 0 | 0 | 0 | 0 | 0 | 0 | 0 | 0 | 0 | 0 | 0 | 1 | 0 | 0 | 0 | 0 | 0 | 0 | 0 | 0 | 0 | 0 | 0 | 0 | 0 |
| bacteria_Aug5d25b_5281 | 1 | 0 | 0 | 0 | 0 | 0 | 0 | 0 | 0 | 0 | 0 | 0 | 0 | 0 | 0 | 0 | 0 | 0 | 0 | 0 | 0 | 0 | 0 | 0 | 0 | 0 | 0 | 0 | 0 | 0 | 0 | 0 | 0 | 0 | 0 | 0 | 0 | 0 | 0 | 0 | 0 | 1 | 0 | 0 | 0 | 0 | 0 | 0 | 0 | 0 | 0 | 0 | 0 | 0 | 0 |
| bacteria_Aug5d25b_5648 | 1 | 0 | 0 | 0 | 0 | 0 | 0 | 0 | 0 | 0 | 0 | 0 | 0 | 0 | 0 | 0 | 0 | 0 | 0 | 0 | 0 | 0 | 0 | 0 | 0 | 0 | 0 | 0 | 0 | 0 | 0 | 0 | 0 | 0 | 0 | 0 | 0 | 0 | 0 | 0 | 0 | 1 | 0 | 0 | 0 | 0 | 0 | 0 | 0 | 0 | 0 | 0 | 0 | 0 | 0 |
| bacteria_Aug5d25b_5755 | 1 | 0 | 0 | 0 | 0 | 0 | 0 | 0 | 0 | 0 | 0 | 0 | 0 | 0 | 0 | 0 | 0 | 0 | 0 | 0 | 0 | 0 | 0 | 0 | 0 | 0 | 0 | 0 | 0 | 0 | 0 | 0 | 0 | 0 | 0 | 0 | 0 | 0 | 0 | 0 | 0 | 1 | 0 | 0 | 0 | 0 | 0 | 0 | 0 | 0 | 0 | 0 | 0 | 0 | 0 |
| bacteria_Aug5d25b_5966 | 1 | 0 | 0 | 0 | 0 | 0 | 0 | 0 | 0 | 0 | 0 | 0 | 0 | 0 | 0 | 0 | 0 | 0 | 0 | 0 | 0 | 0 | 0 | 0 | 0 | 0 | 0 | 0 | 0 | 0 | 0 | 0 | 0 | 0 | 0 | 0 | 0 | 0 | 0 | 0 | 0 | 1 | 0 | 0 | 0 | 0 | 0 | 0 | 0 | 0 | 0 | 0 | 0 | 0 | 0 |
| bacteria_Aug5d25b_7064 | 1 | 0 | 0 | 0 | 0 | 0 | 0 | 0 | 0 | 0 | 0 | 0 | 0 | 0 | 0 | 0 | 0 | 0 | 0 | 0 | 0 | 0 | 0 | 0 | 0 | 0 | 0 | 0 | 0 | 0 | 0 | 0 | 0 | 0 | 0 | 0 | 0 | 0 | 0 | 0 | 0 | 1 | 0 | 0 | 0 | 0 | 0 | 0 | 0 | 0 | 0 | 0 | 0 | 0 | 0 |
| bacteria_Aug5d25b_7503 | 1 | 0 | 0 | 0 | 0 | 0 | 0 | 0 | 0 | 0 | 0 | 0 | 0 | 0 | 0 | 0 | 0 | 0 | 0 | 0 | 0 | 0 | 0 | 0 | 0 | 0 | 0 | 0 | 0 | 0 | 0 | 0 | 0 | 0 | 0 | 0 | 0 | 0 | 0 | 0 | 0 | 1 | 0 | 0 | 0 | 0 | 0 | 0 | 0 | 0 | 0 | 0 | 0 | 0 | 0 |
| bacteria_Aug5d50b_8022 | 1 | 0 | 0 | 0 | 0 | 0 | 0 | 0 | 0 | 0 | 0 | 0 | 0 | 0 | 0 | 0 | 0 | 0 | 0 | 0 | 0 | 0 | 0 | 0 | 0 | 0 | 0 | 0 | 0 | 0 | 0 | 0 | 0 | 0 | 0 | 0 | 0 | 0 | 0 | 0 | 0 | 0 | 0 | 1 | 0 | 0 | 0 | 0 | 0 | 0 | 0 | 0 | 0 | 0 | 0 |
| bacteria_Aug5d50b_2506 | 1 | 0 | 0 | 0 | 0 | 0 | 0 | 0 | 0 | 0 | 0 | 0 | 0 | 0 | 0 | 0 | 0 | 0 | 0 | 0 | 0 | 0 | 0 | 0 | 0 | 0 | 0 | 0 | 0 | 0 | 0 | 0 | 0 | 0 | 0 | 0 | 0 | 0 | 0 | 0 | 0 | 0 | 0 | 1 | 0 | 0 | 0 | 0 | 0 | 0 | 0 | 0 | 0 | 0 | 0 |
| bacteria_Aug5d50b_3926 | 1 | 0 | 0 | 0 | 0 | 0 | 0 | 0 | 0 | 0 | 0 | 0 | 0 | 0 | 0 | 0 | 0 | 0 | 0 | 0 | 0 | 0 | 0 | 0 | 0 | 0 | 0 | 0 | 0 | 0 | 0 | 0 | 0 | 0 | 0 | 0 | 0 | 0 | 0 | 0 | 0 | 0 | 0 | 1 | 0 | 0 | 0 | 0 | 0 | 0 | 0 | 0 | 0 | 0 | 0 |
| bacteria_Aug5d50b_4922 | 1 | 0 | 0 | 0 | 0 | 0 | 0 | 0 | 0 | 0 | 0 | 0 | 0 | 0 | 0 | 0 | 0 | 0 | 0 | 0 | 0 | 0 | 0 | 0 | 0 | 0 | 0 | 0 | 0 | 0 | 0 | 0 | 0 | 0 | 0 | 0 | 0 | 0 | 0 | 0 | 0 | 0 | 0 | 1 | 0 | 0 | 0 | 0 | 0 | 0 | 0 | 0 | 0 | 0 | 0 |
| bacteria_Aug5d50b_5121 | 1 | 0 | 0 | 0 | 0 | 0 | 0 | 0 | 0 | 0 | 0 | 0 | 0 | 0 | 0 | 0 | 0 | 0 | 0 | 0 | 0 | 0 | 0 | 0 | 0 | 0 | 0 | 0 | 0 | 0 | 0 | 0 | 0 | 0 | 0 | 0 | 0 | 0 | 0 | 0 | 0 | 0 | 0 | 1 | 0 | 0 | 0 | 0 | 0 | 0 | 0 | 0 | 0 | 0 | 0 |
| bacteria_Aug5d50b_5148 | 1 | 0 | 0 | 0 | 0 | 0 | 0 | 0 | 0 | 0 | 0 | 0 | 0 | 0 | 0 | 0 | 0 | 0 | 0 | 0 | 0 | 0 | 0 | 0 | 0 | 0 | 0 | 0 | 0 | 0 | 0 | 0 | 0 | 0 | 0 | 0 | 0 | 0 | 0 | 0 | 0 | 0 | 0 | 1 | 0 | 0 | 0 | 0 | 0 | 0 | 0 | 0 | 0 | 0 | 0 |
| bacteria_Aug5d50b_6945 | 1 | 0 | 0 | 0 | 0 | 0 | 0 | 0 | 0 | 0 | 0 | 0 | 0 | 0 | 0 | 0 | 0 | 0 | 0 | 0 | 0 | 0 | 0 | 0 | 0 | 0 | 0 | 0 | 0 | 0 | 0 | 0 | 0 | 0 | 0 | 0 | 0 | 0 | 0 | 0 | 0 | 0 | 0 | 1 | 0 | 0 | 0 | 0 | 0 | 0 | 0 | 0 | 0 | 0 | 0 |
| bacteria_Aug5d50b_7071 | 1 | 0 | 0 | 0 | 0 | 0 | 0 | 0 | 0 | 0 | 0 | 0 | 0 | 0 | 0 | 0 | 0 | 0 | 0 | 0 | 0 | 0 | 0 | 0 | 0 | 0 | 0 | 0 | 0 | 0 | 0 | 0 | 0 | 0 | 0 | 0 | 0 | 0 | 0 | 0 | 0 | 0 | 0 | 1 | 0 | 0 | 0 | 0 | 0 | 0 | 0 | 0 | 0 | 0 | 0 |
| bacteria_Aug5d75b_3359 | 1 | 0 | 0 | 0 | 0 | 0 | 0 | 0 | 0 | 0 | 0 | 0 | 0 | 0 | 0 | 0 | 0 | 0 | 0 | 0 | 0 | 0 | 0 | 0 | 0 | 0 | 0 | 0 | 0 | 0 | 0 | 0 | 0 | 0 | 0 | 0 | 0 | 0 | 0 | 0 | 0 | 0 | 0 | 0 | 0 | 1 | 0 | 0 | 0 | 0 | 0 | 0 | 0 | 0 | 0 |
| bacteria_Jul11d5b_18268 | 1 | 0 | 0 | 0 | 0 | 0 | 0 | 0 | 0 | 0 | 0 | 0 | 0 | 0 | 0 | 0 | 0 | 0 | 0 | 0 | 0 | 0 | 0 | 1 | 0 | 0 | 0 | 0 | 0 | 0 | 0 | 0 | 0 | 0 | 0 | 0 | 0 | 0 | 0 | 0 | 0 | 0 | 0 | 0 | 0 | 0 | 0 | 0 | 0 | 0 | 0 | 0 | 0 | 0 | 0 |
| bacteria_Jul11d5b_23101 | 1 | 0 | 0 | 0 | 0 | 0 | 0 | 0 | 0 | 0 | 0 | 0 | 0 | 0 | 0 | 0 | 0 | 0 | 0 | 0 | 0 | 0 | 0 | 1 | 0 | 0 | 0 | 0 | 0 | 0 | 0 | 0 | 0 | 0 | 0 | 0 | 0 | 0 | 0 | 0 | 0 | 0 | 0 | 0 | 0 | 0 | 0 | 0 | 0 | 0 | 0 | 0 | 0 | 0 | 0 |
| bacteria_Jul11d25b_4548 | 1 | 0 | 0 | 0 | 0 | 0 | 0 | 0 | 0 | 0 | 0 | 0 | 0 | 0 | 0 | 0 | 0 | 0 | 0 | 0 | 0 | 0 | 0 | 0 | 0 | 1 | 0 | 0 | 0 | 0 | 0 | 0 | 0 | 0 | 0 | 0 | 0 | 0 | 0 | 0 | 0 | 0 | 0 | 0 | 0 | 0 | 0 | 0 | 0 | 0 | 0 | 0 | 0 | 0 | 0 |
| bacteria_Jul11d25b_1131 | 1 | 0 | 0 | 0 | 0 | 0 | 0 | 0 | 0 | 0 | 0 | 0 | 0 | 0 | 0 | 0 | 0 | 0 | 0 | 0 | 0 | 0 | 0 | 0 | 0 | 1 | 0 | 0 | 0 | 0 | 0 | 0 | 0 | 0 | 0 | 0 | 0 | 0 | 0 | 0 | 0 | 0 | 0 | 0 | 0 | 0 | 0 | 0 | 0 | 0 | 0 | 0 | 0 | 0 | 0 |
| bacteria_Jul11d25b_1895 | 1 | 0 | 0 | 0 | 0 | 0 | 0 | 0 | 0 | 0 | 0 | 0 | 0 | 0 | 0 | 0 | 0 | 0 | 0 | 0 | 0 | 0 | 0 | 0 | 0 | 1 | 0 | 0 | 0 | 0 | 0 | 0 | 0 | 0 | 0 | 0 | 0 | 0 | 0 | 0 | 0 | 0 | 0 | 0 | 0 | 0 | 0 | 0 | 0 | 0 | 0 | 0 | 0 | 0 | 0 |
| bacteria_Jul11d25b_2145 | 1 | 0 | 0 | 0 | 0 | 0 | 0 | 0 | 0 | 0 | 0 | 0 | 0 | 0 | 0 | 0 | 0 | 0 | 0 | 0 | 0 | 0 | 0 | 0 | 0 | 1 | 0 | 0 | 0 | 0 | 0 | 0 | 0 | 0 | 0 | 0 | 0 | 0 | 0 | 0 | 0 | 0 | 0 | 0 | 0 | 0 | 0 | 0 | 0 | 0 | 0 | 0 | 0 | 0 | 0 |
| bacteria_Jul11d25b_2575 | 1 | 0 | 0 | 0 | 0 | 0 | 0 | 0 | 0 | 0 | 0 | 0 | 0 | 0 | 0 | 0 | 0 | 0 | 0 | 0 | 0 | 0 | 0 | 0 | 0 | 1 | 0 | 0 | 0 | 0 | 0 | 0 | 0 | 0 | 0 | 0 | 0 | 0 | 0 | 0 | 0 | 0 | 0 | 0 | 0 | 0 | 0 | 0 | 0 | 0 | 0 | 0 | 0 | 0 | 0 |
| bacteria_Jul11d25b_3172 | 1 | 0 | 0 | 0 | 0 | 0 | 0 | 0 | 0 | 0 | 0 | 0 | 0 | 0 | 0 | 0 | 0 | 0 | 0 | 0 | 0 | 0 | 0 | 0 | 0 | 1 | 0 | 0 | 0 | 0 | 0 | 0 | 0 | 0 | 0 | 0 | 0 | 0 | 0 | 0 | 0 | 0 | 0 | 0 | 0 | 0 | 0 | 0 | 0 | 0 | 0 | 0 | 0 | 0 | 0 |
| bacteria_Jul11d25b_3200 | 1 | 0 | 0 | 0 | 0 | 0 | 0 | 0 | 0 | 0 | 0 | 0 | 0 | 0 | 0 | 0 | 0 | 0 | 0 | 0 | 0 | 0 | 0 | 0 | 0 | 1 | 0 | 0 | 0 | 0 | 0 | 0 | 0 | 0 | 0 | 0 | 0 | 0 | 0 | 0 | 0 | 0 | 0 | 0 | 0 | 0 | 0 | 0 | 0 | 0 | 0 | 0 | 0 | 0 | 0 |
| bacteria_Jul11d25b_3707 | 1 | 0 | 0 | 0 | 0 | 0 | 0 | 0 | 0 | 0 | 0 | 0 | 0 | 0 | 0 | 0 | 0 | 0 | 0 | 0 | 0 | 0 | 0 | 0 | 0 | 1 | 0 | 0 | 0 | 0 | 0 | 0 | 0 | 0 | 0 | 0 | 0 | 0 | 0 | 0 | 0 | 0 | 0 | 0 | 0 | 0 | 0 | 0 | 0 | 0 | 0 | 0 | 0 | 0 | 0 |
| bacteria_Jul11d25b_4291 | 1 | 0 | 0 | 0 | 0 | 0 | 0 | 0 | 0 | 0 | 0 | 0 | 0 | 0 | 0 | 0 | 0 | 0 | 0 | 0 | 0 | 0 | 0 | 0 | 0 | 1 | 0 | 0 | 0 | 0 | 0 | 0 | 0 | 0 | 0 | 0 | 0 | 0 | 0 | 0 | 0 | 0 | 0 | 0 | 0 | 0 | 0 | 0 | 0 | 0 | 0 | 0 | 0 | 0 | 0 |
| bacteria_Jul11d25b_4477 | 1 | 0 | 0 | 0 | 0 | 0 | 0 | 0 | 0 | 0 | 0 | 0 | 0 | 0 | 0 | 0 | 0 | 0 | 0 | 0 | 0 | 0 | 0 | 0 | 0 | 1 | 0 | 0 | 0 | 0 | 0 | 0 | 0 | 0 | 0 | 0 | 0 | 0 | 0 | 0 | 0 | 0 | 0 | 0 | 0 | 0 | 0 | 0 | 0 | 0 | 0 | 0 | 0 | 0 | 0 |
| bacteria_Jul11d25b_4772 | 1 | 0 | 0 | 0 | 0 | 0 | 0 | 0 | 0 | 0 | 0 | 0 | 0 | 0 | 0 | 0 | 0 | 0 | 0 | 0 | 0 | 0 | 0 | 0 | 0 | 1 | 0 | 0 | 0 | 0 | 0 | 0 | 0 | 0 | 0 | 0 | 0 | 0 | 0 | 0 | 0 | 0 | 0 | 0 | 0 | 0 | 0 | 0 | 0 | 0 | 0 | 0 | 0 | 0 | 0 |
| bacteria_Jul11d50b_924 | 1 | 0 | 0 | 0 | 0 | 0 | 0 | 0 | 0 | 0 | 0 | 0 | 0 | 0 | 0 | 0 | 0 | 0 | 0 | 0 | 0 | 0 | 0 | 0 | 0 | 0 | 0 | 1 | 0 | 0 | 0 | 0 | 0 | 0 | 0 | 0 | 0 | 0 | 0 | 0 | 0 | 0 | 0 | 0 | 0 | 0 | 0 | 0 | 0 | 0 | 0 | 0 | 0 | 0 | 0 |
| bacteria_Jul11d50b_3387 | 1 | 0 | 0 | 0 | 0 | 0 | 0 | 0 | 0 | 0 | 0 | 0 | 0 | 0 | 0 | 0 | 0 | 0 | 0 | 0 | 0 | 0 | 0 | 0 | 0 | 0 | 0 | 1 | 0 | 0 | 0 | 0 | 0 | 0 | 0 | 0 | 0 | 0 | 0 | 0 | 0 | 0 | 0 | 0 | 0 | 0 | 0 | 0 | 0 | 0 | 0 | 0 | 0 | 0 | 0 |
| bacteria_Jul11d50b_7734 | 1 | 0 | 0 | 0 | 0 | 0 | 0 | 0 | 0 | 0 | 0 | 0 | 0 | 0 | 0 | 0 | 0 | 0 | 0 | 0 | 0 | 0 | 0 | 0 | 0 | 0 | 0 | 1 | 0 | 0 | 0 | 0 | 0 | 0 | 0 | 0 | 0 | 0 | 0 | 0 | 0 | 0 | 0 | 0 | 0 | 0 | 0 | 0 | 0 | 0 | 0 | 0 | 0 | 0 | 0 |
| bacteria_Jul11d50b_3479 | 1 | 0 | 0 | 0 | 0 | 0 | 0 | 0 | 0 | 0 | 0 | 0 | 0 | 0 | 0 | 0 | 0 | 0 | 0 | 0 | 0 | 0 | 0 | 0 | 0 | 0 | 0 | 1 | 0 | 0 | 0 | 0 | 0 | 0 | 0 | 0 | 0 | 0 | 0 | 0 | 0 | 0 | 0 | 0 | 0 | 0 | 0 | 0 | 0 | 0 | 0 | 0 | 0 | 0 | 0 |
| bacteria_Jul11d50b_3833 | 1 | 0 | 0 | 0 | 0 | 0 | 0 | 0 | 0 | 0 | 0 | 0 | 0 | 0 | 0 | 0 | 0 | 0 | 0 | 0 | 0 | 0 | 0 | 0 | 0 | 0 | 0 | 1 | 0 | 0 | 0 | 0 | 0 | 0 | 0 | 0 | 0 | 0 | 0 | 0 | 0 | 0 | 0 | 0 | 0 | 0 | 0 | 0 | 0 | 0 | 0 | 0 | 0 | 0 | 0 |
| bacteria_Jul11d50b_4848 | 1 | 0 | 0 | 0 | 0 | 0 | 0 | 0 | 0 | 0 | 0 | 0 | 0 | 0 | 0 | 0 | 0 | 0 | 0 | 0 | 0 | 0 | 0 | 0 | 0 | 0 | 0 | 1 | 0 | 0 | 0 | 0 | 0 | 0 | 0 | 0 | 0 | 0 | 0 | 0 | 0 | 0 | 0 | 0 | 0 | 0 | 0 | 0 | 0 | 0 | 0 | 0 | 0 | 0 | 0 |
| bacteria_Jul11d50b_5211 | 1 | 0 | 0 | 0 | 0 | 0 | 0 | 0 | 0 | 0 | 0 | 0 | 0 | 0 | 0 | 0 | 0 | 0 | 0 | 0 | 0 | 0 | 0 | 0 | 0 | 0 | 0 | 1 | 0 | 0 | 0 | 0 | 0 | 0 | 0 | 0 | 0 | 0 | 0 | 0 | 0 | 0 | 0 | 0 | 0 | 0 | 0 | 0 | 0 | 0 | 0 | 0 | 0 | 0 | 0 |
| bacteria_Jul11d75b_5601 | 1 | 0 | 0 | 0 | 0 | 0 | 0 | 0 | 0 | 0 | 0 | 0 | 0 | 0 | 0 | 0 | 0 | 0 | 0 | 0 | 0 | 0 | 0 | 0 | 0 | 0 | 0 | 0 | 0 | 1 | 0 | 0 | 0 | 0 | 0 | 0 | 0 | 0 | 0 | 0 | 0 | 0 | 0 | 0 | 0 | 0 | 0 | 0 | 0 | 0 | 0 | 0 | 0 | 0 | 0 |
| bacteria_Jul11d75b_1115 | 1 | 0 | 0 | 0 | 0 | 0 | 0 | 0 | 0 | 0 | 0 | 0 | 0 | 0 | 0 | 0 | 0 | 0 | 0 | 0 | 0 | 0 | 0 | 0 | 0 | 0 | 0 | 0 | 0 | 1 | 0 | 0 | 0 | 0 | 0 | 0 | 0 | 0 | 0 | 0 | 0 | 0 | 0 | 0 | 0 | 0 | 0 | 0 | 0 | 0 | 0 | 0 | 0 | 0 | 0 |
| bacteria_Jul11d75b_1342 | 1 | 0 | 0 | 0 | 0 | 0 | 0 | 0 | 0 | 0 | 0 | 0 | 0 | 0 | 0 | 0 | 0 | 0 | 0 | 0 | 0 | 0 | 0 | 0 | 0 | 0 | 0 | 0 | 0 | 1 | 0 | 0 | 0 | 0 | 0 | 0 | 0 | 0 | 0 | 0 | 0 | 0 | 0 | 0 | 0 | 0 | 0 | 0 | 0 | 0 | 0 | 0 | 0 | 0 | 0 |
| bacteria_Jul11d75b_2118 | 1 | 0 | 0 | 0 | 0 | 0 | 0 | 0 | 0 | 0 | 0 | 0 | 0 | 0 | 0 | 0 | 0 | 0 | 0 | 0 | 0 | 0 | 0 | 0 | 0 | 0 | 0 | 0 | 0 | 1 | 0 | 0 | 0 | 0 | 0 | 0 | 0 | 0 | 0 | 0 | 0 | 0 | 0 | 0 | 0 | 0 | 0 | 0 | 0 | 0 | 0 | 0 | 0 | 0 | 0 |
| bacteria_Jul11d75b_2601 | 1 | 0 | 0 | 0 | 0 | 0 | 0 | 0 | 0 | 0 | 0 | 0 | 0 | 0 | 0 | 0 | 0 | 0 | 0 | 0 | 0 | 0 | 0 | 0 | 0 | 0 | 0 | 0 | 0 | 1 | 0 | 0 | 0 | 0 | 0 | 0 | 0 | 0 | 0 | 0 | 0 | 0 | 0 | 0 | 0 | 0 | 0 | 0 | 0 | 0 | 0 | 0 | 0 | 0 | 0 |
| bacteria_Jul11d75b_2616 | 1 | 0 | 0 | 0 | 0 | 0 | 0 | 0 | 0 | 0 | 0 | 0 | 0 | 0 | 0 | 0 | 0 | 0 | 0 | 0 | 0 | 0 | 0 | 0 | 0 | 0 | 0 | 0 | 0 | 1 | 0 | 0 | 0 | 0 | 0 | 0 | 0 | 0 | 0 | 0 | 0 | 0 | 0 | 0 | 0 | 0 | 0 | 0 | 0 | 0 | 0 | 0 | 0 | 0 | 0 |
| bacteria_Jul11d75b_2892 | 1 | 0 | 0 | 0 | 0 | 0 | 0 | 0 | 0 | 0 | 0 | 0 | 0 | 0 | 0 | 0 | 0 | 0 | 0 | 0 | 0 | 0 | 0 | 0 | 0 | 0 | 0 | 0 | 0 | 1 | 0 | 0 | 0 | 0 | 0 | 0 | 0 | 0 | 0 | 0 | 0 | 0 | 0 | 0 | 0 | 0 | 0 | 0 | 0 | 0 | 0 | 0 | 0 | 0 | 0 |
| bacteria_Jul11d75b_3000 | 1 | 0 | 0 | 0 | 0 | 0 | 0 | 0 | 0 | 0 | 0 | 0 | 0 | 0 | 0 | 0 | 0 | 0 | 0 | 0 | 0 | 0 | 0 | 0 | 0 | 0 | 0 | 0 | 0 | 1 | 0 | 0 | 0 | 0 | 0 | 0 | 0 | 0 | 0 | 0 | 0 | 0 | 0 | 0 | 0 | 0 | 0 | 0 | 0 | 0 | 0 | 0 | 0 | 0 | 0 |
| bacteria_Jul11d75b_3047 | 1 | 0 | 0 | 0 | 0 | 0 | 0 | 0 | 0 | 0 | 0 | 0 | 0 | 0 | 0 | 0 | 0 | 0 | 0 | 0 | 0 | 0 | 0 | 0 | 0 | 0 | 0 | 0 | 0 | 1 | 0 | 0 | 0 | 0 | 0 | 0 | 0 | 0 | 0 | 0 | 0 | 0 | 0 | 0 | 0 | 0 | 0 | 0 | 0 | 0 | 0 | 0 | 0 | 0 | 0 |
| bacteria_Jul11d75b_3054 | 1 | 0 | 0 | 0 | 0 | 0 | 0 | 0 | 0 | 0 | 0 | 0 | 0 | 0 | 0 | 0 | 0 | 0 | 0 | 0 | 0 | 0 | 0 | 0 | 0 | 0 | 0 | 0 | 0 | 1 | 0 | 0 | 0 | 0 | 0 | 0 | 0 | 0 | 0 | 0 | 0 | 0 | 0 | 0 | 0 | 0 | 0 | 0 | 0 | 0 | 0 | 0 | 0 | 0 | 0 |
| bacteria_Jul11d75b_3790 | 1 | 0 | 0 | 0 | 0 | 0 | 0 | 0 | 0 | 0 | 0 | 0 | 0 | 0 | 0 | 0 | 0 | 0 | 0 | 0 | 0 | 0 | 0 | 0 | 0 | 0 | 0 | 0 | 0 | 1 | 0 | 0 | 0 | 0 | 0 | 0 | 0 | 0 | 0 | 0 | 0 | 0 | 0 | 0 | 0 | 0 | 0 | 0 | 0 | 0 | 0 | 0 | 0 | 0 | 0 |
| bacteria_Jul11d75b_4444 | 1 | 0 | 0 | 0 | 0 | 0 | 0 | 0 | 0 | 0 | 0 | 0 | 0 | 0 | 0 | 0 | 0 | 0 | 0 | 0 | 0 | 0 | 0 | 0 | 0 | 0 | 0 | 0 | 0 | 1 | 0 | 0 | 0 | 0 | 0 | 0 | 0 | 0 | 0 | 0 | 0 | 0 | 0 | 0 | 0 | 0 | 0 | 0 | 0 | 0 | 0 | 0 | 0 | 0 | 0 |
| bacteria_Jul11d75b_4979 | 1 | 0 | 0 | 0 | 0 | 0 | 0 | 0 | 0 | 0 | 0 | 0 | 0 | 0 | 0 | 0 | 0 | 0 | 0 | 0 | 0 | 0 | 0 | 0 | 0 | 0 | 0 | 0 | 0 | 1 | 0 | 0 | 0 | 0 | 0 | 0 | 0 | 0 | 0 | 0 | 0 | 0 | 0 | 0 | 0 | 0 | 0 | 0 | 0 | 0 | 0 | 0 | 0 | 0 | 0 |
| bacteria_Jul27d5b_39 | 1 | 0 | 0 | 0 | 0 | 0 | 0 | 0 | 0 | 0 | 0 | 0 | 0 | 0 | 0 | 0 | 0 | 0 | 0 | 0 | 0 | 0 | 0 | 0 | 0 | 0 | 0 | 0 | 0 | 0 | 0 | 1 | 0 | 0 | 0 | 0 | 0 | 0 | 0 | 0 | 0 | 0 | 0 | 0 | 0 | 0 | 0 | 0 | 0 | 0 | 0 | 0 | 0 | 0 | 0 |
| bacteria_Jul27d5b_6063 | 1 | 0 | 0 | 0 | 0 | 0 | 0 | 0 | 0 | 0 | 0 | 0 | 0 | 0 | 0 | 0 | 0 | 0 | 0 | 0 | 0 | 0 | 0 | 0 | 0 | 0 | 0 | 0 | 0 | 0 | 0 | 1 | 0 | 0 | 0 | 0 | 0 | 0 | 0 | 0 | 0 | 0 | 0 | 0 | 0 | 0 | 0 | 0 | 0 | 0 | 0 | 0 | 0 | 0 | 0 |
| bacteria_Jul27d5b_8869 | 1 | 0 | 0 | 0 | 0 | 0 | 0 | 0 | 0 | 0 | 0 | 0 | 0 | 0 | 0 | 0 | 0 | 0 | 0 | 0 | 0 | 0 | 0 | 0 | 0 | 0 | 0 | 0 | 0 | 0 | 0 | 1 | 0 | 0 | 0 | 0 | 0 | 0 | 0 | 0 | 0 | 0 | 0 | 0 | 0 | 0 | 0 | 0 | 0 | 0 | 0 | 0 | 0 | 0 | 0 |
| bacteria_Jul27d5b_8981 | 1 | 0 | 0 | 0 | 0 | 0 | 0 | 0 | 0 | 0 | 0 | 0 | 0 | 0 | 0 | 0 | 0 | 0 | 0 | 0 | 0 | 0 | 0 | 0 | 0 | 0 | 0 | 0 | 0 | 0 | 0 | 1 | 0 | 0 | 0 | 0 | 0 | 0 | 0 | 0 | 0 | 0 | 0 | 0 | 0 | 0 | 0 | 0 | 0 | 0 | 0 | 0 | 0 | 0 | 0 |
| bacteria_Jul27d5b_9683 | 1 | 0 | 0 | 0 | 0 | 0 | 0 | 0 | 0 | 0 | 0 | 0 | 0 | 0 | 0 | 0 | 0 | 0 | 0 | 0 | 0 | 0 | 0 | 0 | 0 | 0 | 0 | 0 | 0 | 0 | 0 | 1 | 0 | 0 | 0 | 0 | 0 | 0 | 0 | 0 | 0 | 0 | 0 | 0 | 0 | 0 | 0 | 0 | 0 | 0 | 0 | 0 | 0 | 0 | 0 |
| bacteria_Jul27d5b_11900 | 1 | 0 | 0 | 0 | 0 | 0 | 0 | 0 | 0 | 0 | 0 | 0 | 0 | 0 | 0 | 0 | 0 | 0 | 0 | 0 | 0 | 0 | 0 | 0 | 0 | 0 | 0 | 0 | 0 | 0 | 0 | 1 | 0 | 0 | 0 | 0 | 0 | 0 | 0 | 0 | 0 | 0 | 0 | 0 | 0 | 0 | 0 | 0 | 0 | 0 | 0 | 0 | 0 | 0 | 0 |
| bacteria_Jul27d5b_12737 | 1 | 0 | 0 | 0 | 0 | 0 | 0 | 0 | 0 | 0 | 0 | 0 | 0 | 0 | 0 | 0 | 0 | 0 | 0 | 0 | 0 | 0 | 0 | 0 | 0 | 0 | 0 | 0 | 0 | 0 | 0 | 1 | 0 | 0 | 0 | 0 | 0 | 0 | 0 | 0 | 0 | 0 | 0 | 0 | 0 | 0 | 0 | 0 | 0 | 0 | 0 | 0 | 0 | 0 | 0 |
| bacteria_Jul27d5b_17374 | 1 | 0 | 0 | 0 | 0 | 0 | 0 | 0 | 0 | 0 | 0 | 0 | 0 | 0 | 0 | 0 | 0 | 0 | 0 | 0 | 0 | 0 | 0 | 0 | 0 | 0 | 0 | 0 | 0 | 0 | 0 | 1 | 0 | 0 | 0 | 0 | 0 | 0 | 0 | 0 | 0 | 0 | 0 | 0 | 0 | 0 | 0 | 0 | 0 | 0 | 0 | 0 | 0 | 0 | 0 |
| bacteria_Jul27d5b_22266 | 1 | 0 | 0 | 0 | 0 | 0 | 0 | 0 | 0 | 0 | 0 | 0 | 0 | 0 | 0 | 0 | 0 | 0 | 0 | 0 | 0 | 0 | 0 | 0 | 0 | 0 | 0 | 0 | 0 | 0 | 0 | 1 | 0 | 0 | 0 | 0 | 0 | 0 | 0 | 0 | 0 | 0 | 0 | 0 | 0 | 0 | 0 | 0 | 0 | 0 | 0 | 0 | 0 | 0 | 0 |
| bacteria_Jul27d5b_25331 | 1 | 0 | 0 | 0 | 0 | 0 | 0 | 0 | 0 | 0 | 0 | 0 | 0 | 0 | 0 | 0 | 0 | 0 | 0 | 0 | 0 | 0 | 0 | 0 | 0 | 0 | 0 | 0 | 0 | 0 | 0 | 1 | 0 | 0 | 0 | 0 | 0 | 0 | 0 | 0 | 0 | 0 | 0 | 0 | 0 | 0 | 0 | 0 | 0 | 0 | 0 | 0 | 0 | 0 | 0 |
| bacteria_Jul27d5b_26540 | 1 | 0 | 0 | 0 | 0 | 0 | 0 | 0 | 0 | 0 | 0 | 0 | 0 | 0 | 0 | 0 | 0 | 0 | 0 | 0 | 0 | 0 | 0 | 0 | 0 | 0 | 0 | 0 | 0 | 0 | 0 | 1 | 0 | 0 | 0 | 0 | 0 | 0 | 0 | 0 | 0 | 0 | 0 | 0 | 0 | 0 | 0 | 0 | 0 | 0 | 0 | 0 | 0 | 0 | 0 |
| bacteria_Jul27d5b_29220 | 1 | 0 | 0 | 0 | 0 | 0 | 0 | 0 | 0 | 0 | 0 | 0 | 0 | 0 | 0 | 0 | 0 | 0 | 0 | 0 | 0 | 0 | 0 | 0 | 0 | 0 | 0 | 0 | 0 | 0 | 0 | 1 | 0 | 0 | 0 | 0 | 0 | 0 | 0 | 0 | 0 | 0 | 0 | 0 | 0 | 0 | 0 | 0 | 0 | 0 | 0 | 0 | 0 | 0 | 0 |
| bacteria_Jul27d5b_32879 | 1 | 0 | 0 | 0 | 0 | 0 | 0 | 0 | 0 | 0 | 0 | 0 | 0 | 0 | 0 | 0 | 0 | 0 | 0 | 0 | 0 | 0 | 0 | 0 | 0 | 0 | 0 | 0 | 0 | 0 | 0 | 1 | 0 | 0 | 0 | 0 | 0 | 0 | 0 | 0 | 0 | 0 | 0 | 0 | 0 | 0 | 0 | 0 | 0 | 0 | 0 | 0 | 0 | 0 | 0 |
| bacteria_Jul27d5b_34322 | 1 | 0 | 0 | 0 | 0 | 0 | 0 | 0 | 0 | 0 | 0 | 0 | 0 | 0 | 0 | 0 | 0 | 0 | 0 | 0 | 0 | 0 | 0 | 0 | 0 | 0 | 0 | 0 | 0 | 0 | 0 | 1 | 0 | 0 | 0 | 0 | 0 | 0 | 0 | 0 | 0 | 0 | 0 | 0 | 0 | 0 | 0 | 0 | 0 | 0 | 0 | 0 | 0 | 0 | 0 |
| bacteria_Jul27d5b_35460 | 1 | 0 | 0 | 0 | 0 | 0 | 0 | 0 | 0 | 0 | 0 | 0 | 0 | 0 | 0 | 0 | 0 | 0 | 0 | 0 | 0 | 0 | 0 | 0 | 0 | 0 | 0 | 0 | 0 | 0 | 0 | 1 | 0 | 0 | 0 | 0 | 0 | 0 | 0 | 0 | 0 | 0 | 0 | 0 | 0 | 0 | 0 | 0 | 0 | 0 | 0 | 0 | 0 | 0 | 0 |
| bacteria_Jul27d5b_40006 | 1 | 0 | 0 | 0 | 0 | 0 | 0 | 0 | 0 | 0 | 0 | 0 | 0 | 0 | 0 | 0 | 0 | 0 | 0 | 0 | 0 | 0 | 0 | 0 | 0 | 0 | 0 | 0 | 0 | 0 | 0 | 1 | 0 | 0 | 0 | 0 | 0 | 0 | 0 | 0 | 0 | 0 | 0 | 0 | 0 | 0 | 0 | 0 | 0 | 0 | 0 | 0 | 0 | 0 | 0 |
| bacteria_Jul27d5b_42270 | 1 | 0 | 0 | 0 | 0 | 0 | 0 | 0 | 0 | 0 | 0 | 0 | 0 | 0 | 0 | 0 | 0 | 0 | 0 | 0 | 0 | 0 | 0 | 0 | 0 | 0 | 0 | 0 | 0 | 0 | 0 | 1 | 0 | 0 | 0 | 0 | 0 | 0 | 0 | 0 | 0 | 0 | 0 | 0 | 0 | 0 | 0 | 0 | 0 | 0 | 0 | 0 | 0 | 0 | 0 |
| bacteria_Jul27d5b_42482 | 1 | 0 | 0 | 0 | 0 | 0 | 0 | 0 | 0 | 0 | 0 | 0 | 0 | 0 | 0 | 0 | 0 | 0 | 0 | 0 | 0 | 0 | 0 | 0 | 0 | 0 | 0 | 0 | 0 | 0 | 0 | 1 | 0 | 0 | 0 | 0 | 0 | 0 | 0 | 0 | 0 | 0 | 0 | 0 | 0 | 0 | 0 | 0 | 0 | 0 | 0 | 0 | 0 | 0 | 0 |
| bacteria_Jul27d5b_49113 | 1 | 0 | 0 | 0 | 0 | 0 | 0 | 0 | 0 | 0 | 0 | 0 | 0 | 0 | 0 | 0 | 0 | 0 | 0 | 0 | 0 | 0 | 0 | 0 | 0 | 0 | 0 | 0 | 0 | 0 | 0 | 1 | 0 | 0 | 0 | 0 | 0 | 0 | 0 | 0 | 0 | 0 | 0 | 0 | 0 | 0 | 0 | 0 | 0 | 0 | 0 | 0 | 0 | 0 | 0 |
| bacteria_Jul27d5b_49118 | 1 | 0 | 0 | 0 | 0 | 0 | 0 | 0 | 0 | 0 | 0 | 0 | 0 | 0 | 0 | 0 | 0 | 0 | 0 | 0 | 0 | 0 | 0 | 0 | 0 | 0 | 0 | 0 | 0 | 0 | 0 | 1 | 0 | 0 | 0 | 0 | 0 | 0 | 0 | 0 | 0 | 0 | 0 | 0 | 0 | 0 | 0 | 0 | 0 | 0 | 0 | 0 | 0 | 0 | 0 |
| bacteria_Jul27d25b_1325 | 1 | 0 | 0 | 0 | 0 | 0 | 0 | 0 | 0 | 0 | 0 | 0 | 0 | 0 | 0 | 0 | 0 | 0 | 0 | 0 | 0 | 0 | 0 | 0 | 0 | 0 | 0 | 0 | 0 | 0 | 0 | 0 | 0 | 1 | 0 | 0 | 0 | 0 | 0 | 0 | 0 | 0 | 0 | 0 | 0 | 0 | 0 | 0 | 0 | 0 | 0 | 0 | 0 | 0 | 0 |
| bacteria_Jul27d25b_6232 | 1 | 0 | 0 | 0 | 0 | 0 | 0 | 0 | 0 | 0 | 0 | 0 | 0 | 0 | 0 | 0 | 0 | 0 | 0 | 0 | 0 | 0 | 0 | 0 | 0 | 0 | 0 | 0 | 0 | 0 | 0 | 0 | 0 | 1 | 0 | 0 | 0 | 0 | 0 | 0 | 0 | 0 | 0 | 0 | 0 | 0 | 0 | 0 | 0 | 0 | 0 | 0 | 0 | 0 | 0 |
| bacteria_Jul27d25b_1060 | 1 | 0 | 0 | 0 | 0 | 0 | 0 | 0 | 0 | 0 | 0 | 0 | 0 | 0 | 0 | 0 | 0 | 0 | 0 | 0 | 0 | 0 | 0 | 0 | 0 | 0 | 0 | 0 | 0 | 0 | 0 | 0 | 0 | 1 | 0 | 0 | 0 | 0 | 0 | 0 | 0 | 0 | 0 | 0 | 0 | 0 | 0 | 0 | 0 | 0 | 0 | 0 | 0 | 0 | 0 |
| bacteria_Jul27d25b_1119 | 1 | 0 | 0 | 0 | 0 | 0 | 0 | 0 | 0 | 0 | 0 | 0 | 0 | 0 | 0 | 0 | 0 | 0 | 0 | 0 | 0 | 0 | 0 | 0 | 0 | 0 | 0 | 0 | 0 | 0 | 0 | 0 | 0 | 1 | 0 | 0 | 0 | 0 | 0 | 0 | 0 | 0 | 0 | 0 | 0 | 0 | 0 | 0 | 0 | 0 | 0 | 0 | 0 | 0 | 0 |
| bacteria_Jul27d25b_1517 | 1 | 0 | 0 | 0 | 0 | 0 | 0 | 0 | 0 | 0 | 0 | 0 | 0 | 0 | 0 | 0 | 0 | 0 | 0 | 0 | 0 | 0 | 0 | 0 | 0 | 0 | 0 | 0 | 0 | 0 | 0 | 0 | 0 | 1 | 0 | 0 | 0 | 0 | 0 | 0 | 0 | 0 | 0 | 0 | 0 | 0 | 0 | 0 | 0 | 0 | 0 | 0 | 0 | 0 | 0 |
| bacteria_Jul27d25b_1523 | 1 | 0 | 0 | 0 | 0 | 0 | 0 | 0 | 0 | 0 | 0 | 0 | 0 | 0 | 0 | 0 | 0 | 0 | 0 | 0 | 0 | 0 | 0 | 0 | 0 | 0 | 0 | 0 | 0 | 0 | 0 | 0 | 0 | 1 | 0 | 0 | 0 | 0 | 0 | 0 | 0 | 0 | 0 | 0 | 0 | 0 | 0 | 0 | 0 | 0 | 0 | 0 | 0 | 0 | 0 |
| bacteria_Jul27d25b_1948 | 1 | 0 | 0 | 0 | 0 | 0 | 0 | 0 | 0 | 0 | 0 | 0 | 0 | 0 | 0 | 0 | 0 | 0 | 0 | 0 | 0 | 0 | 0 | 0 | 0 | 0 | 0 | 0 | 0 | 0 | 0 | 0 | 0 | 1 | 0 | 0 | 0 | 0 | 0 | 0 | 0 | 0 | 0 | 0 | 0 | 0 | 0 | 0 | 0 | 0 | 0 | 0 | 0 | 0 | 0 |
| bacteria_Jul27d25b_2556 | 1 | 0 | 0 | 0 | 0 | 0 | 0 | 0 | 0 | 0 | 0 | 0 | 0 | 0 | 0 | 0 | 0 | 0 | 0 | 0 | 0 | 0 | 0 | 0 | 0 | 0 | 0 | 0 | 0 | 0 | 0 | 0 | 0 | 1 | 0 | 0 | 0 | 0 | 0 | 0 | 0 | 0 | 0 | 0 | 0 | 0 | 0 | 0 | 0 | 0 | 0 | 0 | 0 | 0 | 0 |
| bacteria_Jul27d25b_2763 | 1 | 0 | 0 | 0 | 0 | 0 | 0 | 0 | 0 | 0 | 0 | 0 | 0 | 0 | 0 | 0 | 0 | 0 | 0 | 0 | 0 | 0 | 0 | 0 | 0 | 0 | 0 | 0 | 0 | 0 | 0 | 0 | 0 | 1 | 0 | 0 | 0 | 0 | 0 | 0 | 0 | 0 | 0 | 0 | 0 | 0 | 0 | 0 | 0 | 0 | 0 | 0 | 0 | 0 | 0 |
| bacteria_Jul27d25b_3314 | 1 | 0 | 0 | 0 | 0 | 0 | 0 | 0 | 0 | 0 | 0 | 0 | 0 | 0 | 0 | 0 | 0 | 0 | 0 | 0 | 0 | 0 | 0 | 0 | 0 | 0 | 0 | 0 | 0 | 0 | 0 | 0 | 0 | 1 | 0 | 0 | 0 | 0 | 0 | 0 | 0 | 0 | 0 | 0 | 0 | 0 | 0 | 0 | 0 | 0 | 0 | 0 | 0 | 0 | 0 |
| bacteria_Jul27d25b_3427 | 1 | 0 | 0 | 0 | 0 | 0 | 0 | 0 | 0 | 0 | 0 | 0 | 0 | 0 | 0 | 0 | 0 | 0 | 0 | 0 | 0 | 0 | 0 | 0 | 0 | 0 | 0 | 0 | 0 | 0 | 0 | 0 | 0 | 1 | 0 | 0 | 0 | 0 | 0 | 0 | 0 | 0 | 0 | 0 | 0 | 0 | 0 | 0 | 0 | 0 | 0 | 0 | 0 | 0 | 0 |
| bacteria_Jul27d25b_3774 | 1 | 0 | 0 | 0 | 0 | 0 | 0 | 0 | 0 | 0 | 0 | 0 | 0 | 0 | 0 | 0 | 0 | 0 | 0 | 0 | 0 | 0 | 0 | 0 | 0 | 0 | 0 | 0 | 0 | 0 | 0 | 0 | 0 | 1 | 0 | 0 | 0 | 0 | 0 | 0 | 0 | 0 | 0 | 0 | 0 | 0 | 0 | 0 | 0 | 0 | 0 | 0 | 0 | 0 | 0 |
| bacteria_Jul27d25b_3844 | 1 | 0 | 0 | 0 | 0 | 0 | 0 | 0 | 0 | 0 | 0 | 0 | 0 | 0 | 0 | 0 | 0 | 0 | 0 | 0 | 0 | 0 | 0 | 0 | 0 | 0 | 0 | 0 | 0 | 0 | 0 | 0 | 0 | 1 | 0 | 0 | 0 | 0 | 0 | 0 | 0 | 0 | 0 | 0 | 0 | 0 | 0 | 0 | 0 | 0 | 0 | 0 | 0 | 0 | 0 |
| bacteria_Jul27d25b_4377 | 1 | 0 | 0 | 0 | 0 | 0 | 0 | 0 | 0 | 0 | 0 | 0 | 0 | 0 | 0 | 0 | 0 | 0 | 0 | 0 | 0 | 0 | 0 | 0 | 0 | 0 | 0 | 0 | 0 | 0 | 0 | 0 | 0 | 1 | 0 | 0 | 0 | 0 | 0 | 0 | 0 | 0 | 0 | 0 | 0 | 0 | 0 | 0 | 0 | 0 | 0 | 0 | 0 | 0 | 0 |
| bacteria_Jul27d25b_4802 | 1 | 0 | 0 | 0 | 0 | 0 | 0 | 0 | 0 | 0 | 0 | 0 | 0 | 0 | 0 | 0 | 0 | 0 | 0 | 0 | 0 | 0 | 0 | 0 | 0 | 0 | 0 | 0 | 0 | 0 | 0 | 0 | 0 | 1 | 0 | 0 | 0 | 0 | 0 | 0 | 0 | 0 | 0 | 0 | 0 | 0 | 0 | 0 | 0 | 0 | 0 | 0 | 0 | 0 | 0 |
| bacteria_Jul27d25b_5227 | 1 | 0 | 0 | 0 | 0 | 0 | 0 | 0 | 0 | 0 | 0 | 0 | 0 | 0 | 0 | 0 | 0 | 0 | 0 | 0 | 0 | 0 | 0 | 0 | 0 | 0 | 0 | 0 | 0 | 0 | 0 | 0 | 0 | 1 | 0 | 0 | 0 | 0 | 0 | 0 | 0 | 0 | 0 | 0 | 0 | 0 | 0 | 0 | 0 | 0 | 0 | 0 | 0 | 0 | 0 |
| bacteria_Jul27d25b_5287 | 1 | 0 | 0 | 0 | 0 | 0 | 0 | 0 | 0 | 0 | 0 | 0 | 0 | 0 | 0 | 0 | 0 | 0 | 0 | 0 | 0 | 0 | 0 | 0 | 0 | 0 | 0 | 0 | 0 | 0 | 0 | 0 | 0 | 1 | 0 | 0 | 0 | 0 | 0 | 0 | 0 | 0 | 0 | 0 | 0 | 0 | 0 | 0 | 0 | 0 | 0 | 0 | 0 | 0 | 0 |
| bacteria_Jul27d25b_5707 | 1 | 0 | 0 | 0 | 0 | 0 | 0 | 0 | 0 | 0 | 0 | 0 | 0 | 0 | 0 | 0 | 0 | 0 | 0 | 0 | 0 | 0 | 0 | 0 | 0 | 0 | 0 | 0 | 0 | 0 | 0 | 0 | 0 | 1 | 0 | 0 | 0 | 0 | 0 | 0 | 0 | 0 | 0 | 0 | 0 | 0 | 0 | 0 | 0 | 0 | 0 | 0 | 0 | 0 | 0 |
| bacteria_Jul27d25b_5876 | 1 | 0 | 0 | 0 | 0 | 0 | 0 | 0 | 0 | 0 | 0 | 0 | 0 | 0 | 0 | 0 | 0 | 0 | 0 | 0 | 0 | 0 | 0 | 0 | 0 | 0 | 0 | 0 | 0 | 0 | 0 | 0 | 0 | 1 | 0 | 0 | 0 | 0 | 0 | 0 | 0 | 0 | 0 | 0 | 0 | 0 | 0 | 0 | 0 | 0 | 0 | 0 | 0 | 0 | 0 |
| bacteria_Jul27d25b_6675 | 1 | 0 | 0 | 0 | 0 | 0 | 0 | 0 | 0 | 0 | 0 | 0 | 0 | 0 | 0 | 0 | 0 | 0 | 0 | 0 | 0 | 0 | 0 | 0 | 0 | 0 | 0 | 0 | 0 | 0 | 0 | 0 | 0 | 1 | 0 | 0 | 0 | 0 | 0 | 0 | 0 | 0 | 0 | 0 | 0 | 0 | 0 | 0 | 0 | 0 | 0 | 0 | 0 | 0 | 0 |
| bacteria_Jul27d25b_6836 | 1 | 0 | 0 | 0 | 0 | 0 | 0 | 0 | 0 | 0 | 0 | 0 | 0 | 0 | 0 | 0 | 0 | 0 | 0 | 0 | 0 | 0 | 0 | 0 | 0 | 0 | 0 | 0 | 0 | 0 | 0 | 0 | 0 | 1 | 0 | 0 | 0 | 0 | 0 | 0 | 0 | 0 | 0 | 0 | 0 | 0 | 0 | 0 | 0 | 0 | 0 | 0 | 0 | 0 | 0 |
| bacteria_Jul27d25b_6909 | 1 | 0 | 0 | 0 | 0 | 0 | 0 | 0 | 0 | 0 | 0 | 0 | 0 | 0 | 0 | 0 | 0 | 0 | 0 | 0 | 0 | 0 | 0 | 0 | 0 | 0 | 0 | 0 | 0 | 0 | 0 | 0 | 0 | 1 | 0 | 0 | 0 | 0 | 0 | 0 | 0 | 0 | 0 | 0 | 0 | 0 | 0 | 0 | 0 | 0 | 0 | 0 | 0 | 0 | 0 |
| bacteria_Jul27d25b_7275 | 1 | 0 | 0 | 0 | 0 | 0 | 0 | 0 | 0 | 0 | 0 | 0 | 0 | 0 | 0 | 0 | 0 | 0 | 0 | 0 | 0 | 0 | 0 | 0 | 0 | 0 | 0 | 0 | 0 | 0 | 0 | 0 | 0 | 1 | 0 | 0 | 0 | 0 | 0 | 0 | 0 | 0 | 0 | 0 | 0 | 0 | 0 | 0 | 0 | 0 | 0 | 0 | 0 | 0 | 0 |
| bacteria_Jul27d75b_533 | 1 | 0 | 0 | 0 | 0 | 0 | 0 | 0 | 0 | 0 | 0 | 0 | 0 | 0 | 0 | 0 | 0 | 0 | 0 | 0 | 0 | 0 | 0 | 0 | 0 | 0 | 0 | 0 | 0 | 0 | 0 | 0 | 0 | 0 | 0 | 1 | 0 | 0 | 0 | 0 | 0 | 0 | 0 | 0 | 0 | 0 | 0 | 0 | 0 | 0 | 0 | 0 | 0 | 0 | 0 |
| bacteria_Jul27d75b_1878 | 1 | 0 | 0 | 0 | 0 | 0 | 0 | 0 | 0 | 0 | 0 | 0 | 0 | 0 | 0 | 0 | 0 | 0 | 0 | 0 | 0 | 0 | 0 | 0 | 0 | 0 | 0 | 0 | 0 | 0 | 0 | 0 | 0 | 0 | 0 | 1 | 0 | 0 | 0 | 0 | 0 | 0 | 0 | 0 | 0 | 0 | 0 | 0 | 0 | 0 | 0 | 0 | 0 | 0 | 0 |
| bacteria_Jul27d75b_8694 | 1 | 0 | 0 | 0 | 0 | 0 | 0 | 0 | 0 | 0 | 0 | 0 | 0 | 0 | 0 | 0 | 0 | 0 | 0 | 0 | 0 | 0 | 0 | 0 | 0 | 0 | 0 | 0 | 0 | 0 | 0 | 0 | 0 | 0 | 0 | 1 | 0 | 0 | 0 | 0 | 0 | 0 | 0 | 0 | 0 | 0 | 0 | 0 | 0 | 0 | 0 | 0 | 0 | 0 | 0 |
| bacteria_Jul27d75b_1493 | 1 | 0 | 0 | 0 | 0 | 0 | 0 | 0 | 0 | 0 | 0 | 0 | 0 | 0 | 0 | 0 | 0 | 0 | 0 | 0 | 0 | 0 | 0 | 0 | 0 | 0 | 0 | 0 | 0 | 0 | 0 | 0 | 0 | 0 | 0 | 1 | 0 | 0 | 0 | 0 | 0 | 0 | 0 | 0 | 0 | 0 | 0 | 0 | 0 | 0 | 0 | 0 | 0 | 0 | 0 |
| bacteria_Jul27d75b_1689 | 1 | 0 | 0 | 0 | 0 | 0 | 0 | 0 | 0 | 0 | 0 | 0 | 0 | 0 | 0 | 0 | 0 | 0 | 0 | 0 | 0 | 0 | 0 | 0 | 0 | 0 | 0 | 0 | 0 | 0 | 0 | 0 | 0 | 0 | 0 | 1 | 0 | 0 | 0 | 0 | 0 | 0 | 0 | 0 | 0 | 0 | 0 | 0 | 0 | 0 | 0 | 0 | 0 | 0 | 0 |
| bacteria_Jul27d75b_1736 | 1 | 0 | 0 | 0 | 0 | 0 | 0 | 0 | 0 | 0 | 0 | 0 | 0 | 0 | 0 | 0 | 0 | 0 | 0 | 0 | 0 | 0 | 0 | 0 | 0 | 0 | 0 | 0 | 0 | 0 | 0 | 0 | 0 | 0 | 0 | 1 | 0 | 0 | 0 | 0 | 0 | 0 | 0 | 0 | 0 | 0 | 0 | 0 | 0 | 0 | 0 | 0 | 0 | 0 | 0 |
| bacteria_Jul27d75b_2823 | 1 | 0 | 0 | 0 | 0 | 0 | 0 | 0 | 0 | 0 | 0 | 0 | 0 | 0 | 0 | 0 | 0 | 0 | 0 | 0 | 0 | 0 | 0 | 0 | 0 | 0 | 0 | 0 | 0 | 0 | 0 | 0 | 0 | 0 | 0 | 1 | 0 | 0 | 0 | 0 | 0 | 0 | 0 | 0 | 0 | 0 | 0 | 0 | 0 | 0 | 0 | 0 | 0 | 0 | 0 |
| bacteria_Jul27d75b_2940 | 1 | 0 | 0 | 0 | 0 | 0 | 0 | 0 | 0 | 0 | 0 | 0 | 0 | 0 | 0 | 0 | 0 | 0 | 0 | 0 | 0 | 0 | 0 | 0 | 0 | 0 | 0 | 0 | 0 | 0 | 0 | 0 | 0 | 0 | 0 | 1 | 0 | 0 | 0 | 0 | 0 | 0 | 0 | 0 | 0 | 0 | 0 | 0 | 0 | 0 | 0 | 0 | 0 | 0 | 0 |
| bacteria_Jul27d75b_4590 | 1 | 0 | 0 | 0 | 0 | 0 | 0 | 0 | 0 | 0 | 0 | 0 | 0 | 0 | 0 | 0 | 0 | 0 | 0 | 0 | 0 | 0 | 0 | 0 | 0 | 0 | 0 | 0 | 0 | 0 | 0 | 0 | 0 | 0 | 0 | 1 | 0 | 0 | 0 | 0 | 0 | 0 | 0 | 0 | 0 | 0 | 0 | 0 | 0 | 0 | 0 | 0 | 0 | 0 | 0 |
| bacteria_Jul27d75b_4769 | 1 | 0 | 0 | 0 | 0 | 0 | 0 | 0 | 0 | 0 | 0 | 0 | 0 | 0 | 0 | 0 | 0 | 0 | 0 | 0 | 0 | 0 | 0 | 0 | 0 | 0 | 0 | 0 | 0 | 0 | 0 | 0 | 0 | 0 | 0 | 1 | 0 | 0 | 0 | 0 | 0 | 0 | 0 | 0 | 0 | 0 | 0 | 0 | 0 | 0 | 0 | 0 | 0 | 0 | 0 |
| bacteria_Jul27d75b_5539 | 1 | 0 | 0 | 0 | 0 | 0 | 0 | 0 | 0 | 0 | 0 | 0 | 0 | 0 | 0 | 0 | 0 | 0 | 0 | 0 | 0 | 0 | 0 | 0 | 0 | 0 | 0 | 0 | 0 | 0 | 0 | 0 | 0 | 0 | 0 | 1 | 0 | 0 | 0 | 0 | 0 | 0 | 0 | 0 | 0 | 0 | 0 | 0 | 0 | 0 | 0 | 0 | 0 | 0 | 0 |
| bacteria_Jul27d75b_5722 | 1 | 0 | 0 | 0 | 0 | 0 | 0 | 0 | 0 | 0 | 0 | 0 | 0 | 0 | 0 | 0 | 0 | 0 | 0 | 0 | 0 | 0 | 0 | 0 | 0 | 0 | 0 | 0 | 0 | 0 | 0 | 0 | 0 | 0 | 0 | 1 | 0 | 0 | 0 | 0 | 0 | 0 | 0 | 0 | 0 | 0 | 0 | 0 | 0 | 0 | 0 | 0 | 0 | 0 | 0 |
| bacteria_Jul27d75b_5886 | 1 | 0 | 0 | 0 | 0 | 0 | 0 | 0 | 0 | 0 | 0 | 0 | 0 | 0 | 0 | 0 | 0 | 0 | 0 | 0 | 0 | 0 | 0 | 0 | 0 | 0 | 0 | 0 | 0 | 0 | 0 | 0 | 0 | 0 | 0 | 1 | 0 | 0 | 0 | 0 | 0 | 0 | 0 | 0 | 0 | 0 | 0 | 0 | 0 | 0 | 0 | 0 | 0 | 0 | 0 |
| bacteria_Jul27d75b_6429 | 1 | 0 | 0 | 0 | 0 | 0 | 0 | 0 | 0 | 0 | 0 | 0 | 0 | 0 | 0 | 0 | 0 | 0 | 0 | 0 | 0 | 0 | 0 | 0 | 0 | 0 | 0 | 0 | 0 | 0 | 0 | 0 | 0 | 0 | 0 | 1 | 0 | 0 | 0 | 0 | 0 | 0 | 0 | 0 | 0 | 0 | 0 | 0 | 0 | 0 | 0 | 0 | 0 | 0 | 0 |
| bacteria_Jul27d75b_6571 | 1 | 0 | 0 | 0 | 0 | 0 | 0 | 0 | 0 | 0 | 0 | 0 | 0 | 0 | 0 | 0 | 0 | 0 | 0 | 0 | 0 | 0 | 0 | 0 | 0 | 0 | 0 | 0 | 0 | 0 | 0 | 0 | 0 | 0 | 0 | 1 | 0 | 0 | 0 | 0 | 0 | 0 | 0 | 0 | 0 | 0 | 0 | 0 | 0 | 0 | 0 | 0 | 0 | 0 | 0 |
| bacteria_Jul27d75b_6835 | 1 | 0 | 0 | 0 | 0 | 0 | 0 | 0 | 0 | 0 | 0 | 0 | 0 | 0 | 0 | 0 | 0 | 0 | 0 | 0 | 0 | 0 | 0 | 0 | 0 | 0 | 0 | 0 | 0 | 0 | 0 | 0 | 0 | 0 | 0 | 1 | 0 | 0 | 0 | 0 | 0 | 0 | 0 | 0 | 0 | 0 | 0 | 0 | 0 | 0 | 0 | 0 | 0 | 0 | 0 |
| bacteria_Jul27d75b_6884 | 1 | 0 | 0 | 0 | 0 | 0 | 0 | 0 | 0 | 0 | 0 | 0 | 0 | 0 | 0 | 0 | 0 | 0 | 0 | 0 | 0 | 0 | 0 | 0 | 0 | 0 | 0 | 0 | 0 | 0 | 0 | 0 | 0 | 0 | 0 | 1 | 0 | 0 | 0 | 0 | 0 | 0 | 0 | 0 | 0 | 0 | 0 | 0 | 0 | 0 | 0 | 0 | 0 | 0 | 0 |
| bacteria_Jun8d25b_1031 | 1 | 0 | 0 | 0 | 0 | 0 | 0 | 0 | 0 | 1 | 0 | 0 | 0 | 0 | 0 | 0 | 0 | 0 | 0 | 0 | 0 | 0 | 0 | 0 | 0 | 0 | 0 | 0 | 0 | 0 | 0 | 0 | 0 | 0 | 0 | 0 | 0 | 0 | 0 | 0 | 0 | 0 | 0 | 0 | 0 | 0 | 0 | 0 | 0 | 0 | 0 | 0 | 0 | 0 | 0 |
| bacteria_Jun8d25b_21844 | 1 | 0 | 0 | 0 | 0 | 0 | 0 | 0 | 0 | 1 | 0 | 0 | 0 | 0 | 0 | 0 | 0 | 0 | 0 | 0 | 0 | 0 | 0 | 0 | 0 | 0 | 0 | 0 | 0 | 0 | 0 | 0 | 0 | 0 | 0 | 0 | 0 | 0 | 0 | 0 | 0 | 0 | 0 | 0 | 0 | 0 | 0 | 0 | 0 | 0 | 0 | 0 | 0 | 0 | 0 |
| bacteria_Jun8d25b_27181 | 1 | 0 | 0 | 0 | 0 | 0 | 0 | 0 | 0 | 1 | 0 | 0 | 0 | 0 | 0 | 0 | 0 | 0 | 0 | 0 | 0 | 0 | 0 | 0 | 0 | 0 | 0 | 0 | 0 | 0 | 0 | 0 | 0 | 0 | 0 | 0 | 0 | 0 | 0 | 0 | 0 | 0 | 0 | 0 | 0 | 0 | 0 | 0 | 0 | 0 | 0 | 0 | 0 | 0 | 0 |
| bacteria_Jun8d25b_31268 | 1 | 0 | 0 | 0 | 0 | 0 | 0 | 0 | 0 | 1 | 0 | 0 | 0 | 0 | 0 | 0 | 0 | 0 | 0 | 0 | 0 | 0 | 0 | 0 | 0 | 0 | 0 | 0 | 0 | 0 | 0 | 0 | 0 | 0 | 0 | 0 | 0 | 0 | 0 | 0 | 0 | 0 | 0 | 0 | 0 | 0 | 0 | 0 | 0 | 0 | 0 | 0 | 0 | 0 | 0 |
| bacteria_Jun8d25b_45582 | 1 | 0 | 0 | 0 | 0 | 0 | 0 | 0 | 0 | 1 | 0 | 0 | 0 | 0 | 0 | 0 | 0 | 0 | 0 | 0 | 0 | 0 | 0 | 0 | 0 | 0 | 0 | 0 | 0 | 0 | 0 | 0 | 0 | 0 | 0 | 0 | 0 | 0 | 0 | 0 | 0 | 0 | 0 | 0 | 0 | 0 | 0 | 0 | 0 | 0 | 0 | 0 | 0 | 0 | 0 |
| bacteria_Jun8d25b_53591 | 1 | 0 | 0 | 0 | 0 | 0 | 0 | 0 | 0 | 1 | 0 | 0 | 0 | 0 | 0 | 0 | 0 | 0 | 0 | 0 | 0 | 0 | 0 | 0 | 0 | 0 | 0 | 0 | 0 | 0 | 0 | 0 | 0 | 0 | 0 | 0 | 0 | 0 | 0 | 0 | 0 | 0 | 0 | 0 | 0 | 0 | 0 | 0 | 0 | 0 | 0 | 0 | 0 | 0 | 0 |
| bacteria_Jun8d25b_55128 | 1 | 0 | 0 | 0 | 0 | 0 | 0 | 0 | 0 | 1 | 0 | 0 | 0 | 0 | 0 | 0 | 0 | 0 | 0 | 0 | 0 | 0 | 0 | 0 | 0 | 0 | 0 | 0 | 0 | 0 | 0 | 0 | 0 | 0 | 0 | 0 | 0 | 0 | 0 | 0 | 0 | 0 | 0 | 0 | 0 | 0 | 0 | 0 | 0 | 0 | 0 | 0 | 0 | 0 | 0 |
| bacteria_Jun8d25b_59492 | 1 | 0 | 0 | 0 | 0 | 0 | 0 | 0 | 0 | 1 | 0 | 0 | 0 | 0 | 0 | 0 | 0 | 0 | 0 | 0 | 0 | 0 | 0 | 0 | 0 | 0 | 0 | 0 | 0 | 0 | 0 | 0 | 0 | 0 | 0 | 0 | 0 | 0 | 0 | 0 | 0 | 0 | 0 | 0 | 0 | 0 | 0 | 0 | 0 | 0 | 0 | 0 | 0 | 0 | 0 |
| bacteria_Jun8d25b_63591 | 1 | 0 | 0 | 0 | 0 | 0 | 0 | 0 | 0 | 1 | 0 | 0 | 0 | 0 | 0 | 0 | 0 | 0 | 0 | 0 | 0 | 0 | 0 | 0 | 0 | 0 | 0 | 0 | 0 | 0 | 0 | 0 | 0 | 0 | 0 | 0 | 0 | 0 | 0 | 0 | 0 | 0 | 0 | 0 | 0 | 0 | 0 | 0 | 0 | 0 | 0 | 0 | 0 | 0 | 0 |
| bacteria_Jun8d50b_9577 | 1 | 0 | 0 | 0 | 0 | 0 | 0 | 0 | 0 | 0 | 0 | 1 | 0 | 0 | 0 | 0 | 0 | 0 | 0 | 0 | 0 | 0 | 0 | 0 | 0 | 0 | 0 | 0 | 0 | 0 | 0 | 0 | 0 | 0 | 0 | 0 | 0 | 0 | 0 | 0 | 0 | 0 | 0 | 0 | 0 | 0 | 0 | 0 | 0 | 0 | 0 | 0 | 0 | 0 | 0 |
| bacteria_Jun8d50b_13763 | 1 | 0 | 0 | 0 | 0 | 0 | 0 | 0 | 0 | 0 | 0 | 1 | 0 | 0 | 0 | 0 | 0 | 0 | 0 | 0 | 0 | 0 | 0 | 0 | 0 | 0 | 0 | 0 | 0 | 0 | 0 | 0 | 0 | 0 | 0 | 0 | 0 | 0 | 0 | 0 | 0 | 0 | 0 | 0 | 0 | 0 | 0 | 0 | 0 | 0 | 0 | 0 | 0 | 0 | 0 |
| bacteria_Jun8d50b_40756 | 1 | 0 | 0 | 0 | 0 | 0 | 0 | 0 | 0 | 0 | 0 | 1 | 0 | 0 | 0 | 0 | 0 | 0 | 0 | 0 | 0 | 0 | 0 | 0 | 0 | 0 | 0 | 0 | 0 | 0 | 0 | 0 | 0 | 0 | 0 | 0 | 0 | 0 | 0 | 0 | 0 | 0 | 0 | 0 | 0 | 0 | 0 | 0 | 0 | 0 | 0 | 0 | 0 | 0 | 0 |
| bacteria_Jun8d75b_24465 | 1 | 0 | 0 | 0 | 0 | 0 | 0 | 0 | 0 | 0 | 0 | 0 | 0 | 1 | 0 | 0 | 0 | 0 | 0 | 0 | 0 | 0 | 0 | 0 | 0 | 0 | 0 | 0 | 0 | 0 | 0 | 0 | 0 | 0 | 0 | 0 | 0 | 0 | 0 | 0 | 0 | 0 | 0 | 0 | 0 | 0 | 0 | 0 | 0 | 0 | 0 | 0 | 0 | 0 | 0 |
| bacteria_Jun22d0b_29245 | 1 | 0 | 0 | 0 | 0 | 0 | 0 | 0 | 0 | 0 | 0 | 0 | 0 | 0 | 0 | 1 | 0 | 0 | 0 | 0 | 0 | 0 | 0 | 0 | 0 | 0 | 0 | 0 | 0 | 0 | 0 | 0 | 0 | 0 | 0 | 0 | 0 | 0 | 0 | 0 | 0 | 0 | 0 | 0 | 0 | 0 | 0 | 0 | 0 | 0 | 0 | 0 | 0 | 0 | 0 |
| bacteria_Jun22d0b_43496 | 1 | 0 | 0 | 0 | 0 | 0 | 0 | 0 | 0 | 0 | 0 | 0 | 0 | 0 | 0 | 1 | 0 | 0 | 0 | 0 | 0 | 0 | 0 | 0 | 0 | 0 | 0 | 0 | 0 | 0 | 0 | 0 | 0 | 0 | 0 | 0 | 0 | 0 | 0 | 0 | 0 | 0 | 0 | 0 | 0 | 0 | 0 | 0 | 0 | 0 | 0 | 0 | 0 | 0 | 0 |
| bacteria_Jun22d0b_66483 | 1 | 0 | 0 | 0 | 0 | 0 | 0 | 0 | 0 | 0 | 0 | 0 | 0 | 0 | 0 | 1 | 0 | 0 | 0 | 0 | 0 | 0 | 0 | 0 | 0 | 0 | 0 | 0 | 0 | 0 | 0 | 0 | 0 | 0 | 0 | 0 | 0 | 0 | 0 | 0 | 0 | 0 | 0 | 0 | 0 | 0 | 0 | 0 | 0 | 0 | 0 | 0 | 0 | 0 | 0 |
| bacteria_Jun22d0b_68415 | 1 | 0 | 0 | 0 | 0 | 0 | 0 | 0 | 0 | 0 | 0 | 0 | 0 | 0 | 0 | 1 | 0 | 0 | 0 | 0 | 0 | 0 | 0 | 0 | 0 | 0 | 0 | 0 | 0 | 0 | 0 | 0 | 0 | 0 | 0 | 0 | 0 | 0 | 0 | 0 | 0 | 0 | 0 | 0 | 0 | 0 | 0 | 0 | 0 | 0 | 0 | 0 | 0 | 0 | 0 |

**Supplementary Table S7**. Common Bacterio- and virioplankton RTPR 98% OTUs in 27 samples with RTPR amplicons in both fractions

Total seqs: 47997; Subtotal seqs in retained

| OTUs: 41046 | | May12_05m | | May12_25m | | May12_50m | | May12_75m | | Jun8_25m | | Jun8_50m | | Jun8_75m | | Jun22_0m | | Jun22_50m | | Jun22_75m | | Jul11_0m | | Jul11_05m | | Jul11_25m | | Jul11_50m | | Jul11_75m | | Jul27_05m | | Jul27_25m | | Jul27_75m | | Aug5_0m | | Aug5_05m | | Aug5_25m | | Aug5_50m | | Aug5_75m | | Sep8_0m | | Sep8_05m | | Sep8_25m | | Sep8_50m | |
|---|---|---|---|---|---|---|---|---|---|---|---|---|---|---|---|---|---|---|---|---|---|---|---|---|---|---|---|---|---|---|---|---|---|---|---|---|---|---|---|---|---|---|---|---|---|---|---|---|---|---|---|---|---|---|---|
| OTUs with >=10 seqs: 254 | OTU subtotal | bacterioplankton | virioplankton | bacterioplankton | virioplankton | bacterioplankton | virioplankton | bacterioplankton | virioplankton | bacterioplankton | virioplankton | bacterioplankton | virioplankton | bacterioplankton | virioplankton | bacterioplankton | virioplankton | bacterioplankton | virioplankton | bacterioplankton | virioplankton | bacterioplankton | virioplankton | bacterioplankton | virioplankton | bacterioplankton | virioplankton | bacterioplankton | virioplankton | bacterioplankton | virioplankton | bacterioplankton | virioplankton | bacterioplankton | virioplankton | bacterioplankton | virioplankton | bacterioplankton | virioplankton | bacterioplankton | virioplankton | bacterioplankton | virioplankton | bacterioplankton | virioplankton | bacterioplankton | virioplankton | bacterioplankton | virioplankton | bacterioplankton | virioplankton | bacterioplankton | virioplankton | bacterioplankton | virioplankton |
| bacteria_Jun22d50b_4976 | 1 | 0 | 0 | 0 | 0 | 0 | 0 | 0 | 0 | 0 | 0 | 0 | 0 | 0 | 0 | 0 | 0 | 1 | 0 | 0 | 0 | 0 | 0 | 0 | 0 | 0 | 0 | 0 | 0 | 0 | 0 | 0 | 0 | 0 | 0 | 0 | 0 | 0 | 0 | 0 | 0 | 0 | 0 | 0 | 0 | 0 | 0 | 0 | 0 | 0 | 0 | 0 | 0 | 0 | 0 |
| bacteria_Jun22d50b_1481 | 1 | 0 | 0 | 0 | 0 | 0 | 0 | 0 | 0 | 0 | 0 | 0 | 0 | 0 | 0 | 0 | 0 | 1 | 0 | 0 | 0 | 0 | 0 | 0 | 0 | 0 | 0 | 0 | 0 | 0 | 0 | 0 | 0 | 0 | 0 | 0 | 0 | 0 | 0 | 0 | 0 | 0 | 0 | 0 | 0 | 0 | 0 | 0 | 0 | 0 | 0 | 0 | 0 | 0 | 0 |
| bacteria_Jun22d50b_2401 | 1 | 0 | 0 | 0 | 0 | 0 | 0 | 0 | 0 | 0 | 0 | 0 | 0 | 0 | 0 | 0 | 0 | 1 | 0 | 0 | 0 | 0 | 0 | 0 | 0 | 0 | 0 | 0 | 0 | 0 | 0 | 0 | 0 | 0 | 0 | 0 | 0 | 0 | 0 | 0 | 0 | 0 | 0 | 0 | 0 | 0 | 0 | 0 | 0 | 0 | 0 | 0 | 0 | 0 | 0 |
| bacteria_Jun22d50b_4653 | 1 | 0 | 0 | 0 | 0 | 0 | 0 | 0 | 0 | 0 | 0 | 0 | 0 | 0 | 0 | 0 | 0 | 1 | 0 | 0 | 0 | 0 | 0 | 0 | 0 | 0 | 0 | 0 | 0 | 0 | 0 | 0 | 0 | 0 | 0 | 0 | 0 | 0 | 0 | 0 | 0 | 0 | 0 | 0 | 0 | 0 | 0 | 0 | 0 | 0 | 0 | 0 | 0 | 0 | 0 |
| bacteria_Jun22d75b_1740 | 1 | 0 | 0 | 0 | 0 | 0 | 0 | 0 | 0 | 0 | 0 | 0 | 0 | 0 | 0 | 0 | 0 | 0 | 0 | 1 | 0 | 0 | 0 | 0 | 0 | 0 | 0 | 0 | 0 | 0 | 0 | 0 | 0 | 0 | 0 | 0 | 0 | 0 | 0 | 0 | 0 | 0 | 0 | 0 | 0 | 0 | 0 | 0 | 0 | 0 | 0 | 0 | 0 | 0 | 0 |
| bacteria_Jun22d75b_1265 | 1 | 0 | 0 | 0 | 0 | 0 | 0 | 0 | 0 | 0 | 0 | 0 | 0 | 0 | 0 | 0 | 0 | 0 | 0 | 1 | 0 | 0 | 0 | 0 | 0 | 0 | 0 | 0 | 0 | 0 | 0 | 0 | 0 | 0 | 0 | 0 | 0 | 0 | 0 | 0 | 0 | 0 | 0 | 0 | 0 | 0 | 0 | 0 | 0 | 0 | 0 | 0 | 0 | 0 | 0 |
| bacteria_Jun22d75b_1335 | 1 | 0 | 0 | 0 | 0 | 0 | 0 | 0 | 0 | 0 | 0 | 0 | 0 | 0 | 0 | 0 | 0 | 0 | 0 | 1 | 0 | 0 | 0 | 0 | 0 | 0 | 0 | 0 | 0 | 0 | 0 | 0 | 0 | 0 | 0 | 0 | 0 | 0 | 0 | 0 | 0 | 0 | 0 | 0 | 0 | 0 | 0 | 0 | 0 | 0 | 0 | 0 | 0 | 0 | 0 |
| bacteria_Jun22d75b_1984 | 1 | 0 | 0 | 0 | 0 | 0 | 0 | 0 | 0 | 0 | 0 | 0 | 0 | 0 | 0 | 0 | 0 | 0 | 0 | 1 | 0 | 0 | 0 | 0 | 0 | 0 | 0 | 0 | 0 | 0 | 0 | 0 | 0 | 0 | 0 | 0 | 0 | 0 | 0 | 0 | 0 | 0 | 0 | 0 | 0 | 0 | 0 | 0 | 0 | 0 | 0 | 0 | 0 | 0 | 0 |
| bacteria_Jun22d75b_2850 | 1 | 0 | 0 | 0 | 0 | 0 | 0 | 0 | 0 | 0 | 0 | 0 | 0 | 0 | 0 | 0 | 0 | 0 | 0 | 1 | 0 | 0 | 0 | 0 | 0 | 0 | 0 | 0 | 0 | 0 | 0 | 0 | 0 | 0 | 0 | 0 | 0 | 0 | 0 | 0 | 0 | 0 | 0 | 0 | 0 | 0 | 0 | 0 | 0 | 0 | 0 | 0 | 0 | 0 | 0 |
| bacteria_Jun22d75b_3492 | 1 | 0 | 0 | 0 | 0 | 0 | 0 | 0 | 0 | 0 | 0 | 0 | 0 | 0 | 0 | 0 | 0 | 0 | 0 | 1 | 0 | 0 | 0 | 0 | 0 | 0 | 0 | 0 | 0 | 0 | 0 | 0 | 0 | 0 | 0 | 0 | 0 | 0 | 0 | 0 | 0 | 0 | 0 | 0 | 0 | 0 | 0 | 0 | 0 | 0 | 0 | 0 | 0 | 0 | 0 |
| bacteria_Jun22d75b_7152 | 1 | 0 | 0 | 0 | 0 | 0 | 0 | 0 | 0 | 0 | 0 | 0 | 0 | 0 | 0 | 0 | 0 | 0 | 0 | 1 | 0 | 0 | 0 | 0 | 0 | 0 | 0 | 0 | 0 | 0 | 0 | 0 | 0 | 0 | 0 | 0 | 0 | 0 | 0 | 0 | 0 | 0 | 0 | 0 | 0 | 0 | 0 | 0 | 0 | 0 | 0 | 0 | 0 | 0 | 0 |
| bacteria_May12d25b_189 | 1 | 0 | 0 | 1 | 0 | 0 | 0 | 0 | 0 | 0 | 0 | 0 | 0 | 0 | 0 | 0 | 0 | 0 | 0 | 0 | 0 | 0 | 0 | 0 | 0 | 0 | 0 | 0 | 0 | 0 | 0 | 0 | 0 | 0 | 0 | 0 | 0 | 0 | 0 | 0 | 0 | 0 | 0 | 0 | 0 | 0 | 0 | 0 | 0 | 0 | 0 | 0 | 0 | 0 | 0 |
| bacteria_May12d25b_224 | 1 | 0 | 0 | 1 | 0 | 0 | 0 | 0 | 0 | 0 | 0 | 0 | 0 | 0 | 0 | 0 | 0 | 0 | 0 | 0 | 0 | 0 | 0 | 0 | 0 | 0 | 0 | 0 | 0 | 0 | 0 | 0 | 0 | 0 | 0 | 0 | 0 | 0 | 0 | 0 | 0 | 0 | 0 | 0 | 0 | 0 | 0 | 0 | 0 | 0 | 0 | 0 | 0 | 0 | 0 |
| bacteria_May12d25b_273 | 1 | 0 | 0 | 1 | 0 | 0 | 0 | 0 | 0 | 0 | 0 | 0 | 0 | 0 | 0 | 0 | 0 | 0 | 0 | 0 | 0 | 0 | 0 | 0 | 0 | 0 | 0 | 0 | 0 | 0 | 0 | 0 | 0 | 0 | 0 | 0 | 0 | 0 | 0 | 0 | 0 | 0 | 0 | 0 | 0 | 0 | 0 | 0 | 0 | 0 | 0 | 0 | 0 | 0 | 0 |
| bacteria_May12d25b_325 | 1 | 0 | 0 | 1 | 0 | 0 | 0 | 0 | 0 | 0 | 0 | 0 | 0 | 0 | 0 | 0 | 0 | 0 | 0 | 0 | 0 | 0 | 0 | 0 | 0 | 0 | 0 | 0 | 0 | 0 | 0 | 0 | 0 | 0 | 0 | 0 | 0 | 0 | 0 | 0 | 0 | 0 | 0 | 0 | 0 | 0 | 0 | 0 | 0 | 0 | 0 | 0 | 0 | 0 | 0 |
| bacteria_May12d25b_435 | 1 | 0 | 0 | 1 | 0 | 0 | 0 | 0 | 0 | 0 | 0 | 0 | 0 | 0 | 0 | 0 | 0 | 0 | 0 | 0 | 0 | 0 | 0 | 0 | 0 | 0 | 0 | 0 | 0 | 0 | 0 | 0 | 0 | 0 | 0 | 0 | 0 | 0 | 0 | 0 | 0 | 0 | 0 | 0 | 0 | 0 | 0 | 0 | 0 | 0 | 0 | 0 | 0 | 0 | 0 |
| bacteria_May12d25b_442 | 1 | 0 | 0 | 1 | 0 | 0 | 0 | 0 | 0 | 0 | 0 | 0 | 0 | 0 | 0 | 0 | 0 | 0 | 0 | 0 | 0 | 0 | 0 | 0 | 0 | 0 | 0 | 0 | 0 | 0 | 0 | 0 | 0 | 0 | 0 | 0 | 0 | 0 | 0 | 0 | 0 | 0 | 0 | 0 | 0 | 0 | 0 | 0 | 0 | 0 | 0 | 0 | 0 | 0 | 0 |
| bacteria_May12d25b_447 | 1 | 0 | 0 | 1 | 0 | 0 | 0 | 0 | 0 | 0 | 0 | 0 | 0 | 0 | 0 | 0 | 0 | 0 | 0 | 0 | 0 | 0 | 0 | 0 | 0 | 0 | 0 | 0 | 0 | 0 | 0 | 0 | 0 | 0 | 0 | 0 | 0 | 0 | 0 | 0 | 0 | 0 | 0 | 0 | 0 | 0 | 0 | 0 | 0 | 0 | 0 | 0 | 0 | 0 | 0 |
| bacteria_May12d25b_499 | 1 | 0 | 0 | 1 | 0 | 0 | 0 | 0 | 0 | 0 | 0 | 0 | 0 | 0 | 0 | 0 | 0 | 0 | 0 | 0 | 0 | 0 | 0 | 0 | 0 | 0 | 0 | 0 | 0 | 0 | 0 | 0 | 0 | 0 | 0 | 0 | 0 | 0 | 0 | 0 | 0 | 0 | 0 | 0 | 0 | 0 | 0 | 0 | 0 | 0 | 0 | 0 | 0 | 0 | 0 |
| bacteria_May12d25b_581 | 1 | 0 | 0 | 1 | 0 | 0 | 0 | 0 | 0 | 0 | 0 | 0 | 0 | 0 | 0 | 0 | 0 | 0 | 0 | 0 | 0 | 0 | 0 | 0 | 0 | 0 | 0 | 0 | 0 | 0 | 0 | 0 | 0 | 0 | 0 | 0 | 0 | 0 | 0 | 0 | 0 | 0 | 0 | 0 | 0 | 0 | 0 | 0 | 0 | 0 | 0 | 0 | 0 | 0 | 0 |
| bacteria_May12d25b_607 | 1 | 0 | 0 | 1 | 0 | 0 | 0 | 0 | 0 | 0 | 0 | 0 | 0 | 0 | 0 | 0 | 0 | 0 | 0 | 0 | 0 | 0 | 0 | 0 | 0 | 0 | 0 | 0 | 0 | 0 | 0 | 0 | 0 | 0 | 0 | 0 | 0 | 0 | 0 | 0 | 0 | 0 | 0 | 0 | 0 | 0 | 0 | 0 | 0 | 0 | 0 | 0 | 0 | 0 | 0 |
| bacteria_May12d25b_683 | 1 | 0 | 0 | 1 | 0 | 0 | 0 | 0 | 0 | 0 | 0 | 0 | 0 | 0 | 0 | 0 | 0 | 0 | 0 | 0 | 0 | 0 | 0 | 0 | 0 | 0 | 0 | 0 | 0 | 0 | 0 | 0 | 0 | 0 | 0 | 0 | 0 | 0 | 0 | 0 | 0 | 0 | 0 | 0 | 0 | 0 | 0 | 0 | 0 | 0 | 0 | 0 | 0 | 0 | 0 |
| bacteria_May12d50b_135 | 1 | 0 | 0 | 0 | 0 | 1 | 0 | 0 | 0 | 0 | 0 | 0 | 0 | 0 | 0 | 0 | 0 | 0 | 0 | 0 | 0 | 0 | 0 | 0 | 0 | 0 | 0 | 0 | 0 | 0 | 0 | 0 | 0 | 0 | 0 | 0 | 0 | 0 | 0 | 0 | 0 | 0 | 0 | 0 | 0 | 0 | 0 | 0 | 0 | 0 | 0 | 0 | 0 | 0 | 0 |
| bacteria_May12d50b_146 | 1 | 0 | 0 | 0 | 0 | 1 | 0 | 0 | 0 | 0 | 0 | 0 | 0 | 0 | 0 | 0 | 0 | 0 | 0 | 0 | 0 | 0 | 0 | 0 | 0 | 0 | 0 | 0 | 0 | 0 | 0 | 0 | 0 | 0 | 0 | 0 | 0 | 0 | 0 | 0 | 0 | 0 | 0 | 0 | 0 | 0 | 0 | 0 | 0 | 0 | 0 | 0 | 0 | 0 | 0 |
| bacteria_May12d50b_273 | 1 | 0 | 0 | 0 | 0 | 1 | 0 | 0 | 0 | 0 | 0 | 0 | 0 | 0 | 0 | 0 | 0 | 0 | 0 | 0 | 0 | 0 | 0 | 0 | 0 | 0 | 0 | 0 | 0 | 0 | 0 | 0 | 0 | 0 | 0 | 0 | 0 | 0 | 0 | 0 | 0 | 0 | 0 | 0 | 0 | 0 | 0 | 0 | 0 | 0 | 0 | 0 | 0 | 0 | 0 |
| bacteria_May12d50b_300 | 1 | 0 | 0 | 0 | 0 | 1 | 0 | 0 | 0 | 0 | 0 | 0 | 0 | 0 | 0 | 0 | 0 | 0 | 0 | 0 | 0 | 0 | 0 | 0 | 0 | 0 | 0 | 0 | 0 | 0 | 0 | 0 | 0 | 0 | 0 | 0 | 0 | 0 | 0 | 0 | 0 | 0 | 0 | 0 | 0 | 0 | 0 | 0 | 0 | 0 | 0 | 0 | 0 | 0 | 0 |
| bacteria_May12d50b_308 | 1 | 0 | 0 | 0 | 0 | 1 | 0 | 0 | 0 | 0 | 0 | 0 | 0 | 0 | 0 | 0 | 0 | 0 | 0 | 0 | 0 | 0 | 0 | 0 | 0 | 0 | 0 | 0 | 0 | 0 | 0 | 0 | 0 | 0 | 0 | 0 | 0 | 0 | 0 | 0 | 0 | 0 | 0 | 0 | 0 | 0 | 0 | 0 | 0 | 0 | 0 | 0 | 0 | 0 | 0 |
| bacteria_May12d50b_396 | 1 | 0 | 0 | 0 | 0 | 1 | 0 | 0 | 0 | 0 | 0 | 0 | 0 | 0 | 0 | 0 | 0 | 0 | 0 | 0 | 0 | 0 | 0 | 0 | 0 | 0 | 0 | 0 | 0 | 0 | 0 | 0 | 0 | 0 | 0 | 0 | 0 | 0 | 0 | 0 | 0 | 0 | 0 | 0 | 0 | 0 | 0 | 0 | 0 | 0 | 0 | 0 | 0 | 0 | 0 |
| bacteria_May12d50b_483 | 1 | 0 | 0 | 0 | 0 | 1 | 0 | 0 | 0 | 0 | 0 | 0 | 0 | 0 | 0 | 0 | 0 | 0 | 0 | 0 | 0 | 0 | 0 | 0 | 0 | 0 | 0 | 0 | 0 | 0 | 0 | 0 | 0 | 0 | 0 | 0 | 0 | 0 | 0 | 0 | 0 | 0 | 0 | 0 | 0 | 0 | 0 | 0 | 0 | 0 | 0 | 0 | 0 | 0 | 0 |
| bacteria_May12d75b_538 | 1 | 0 | 0 | 0 | 0 | 0 | 0 | 1 | 0 | 0 | 0 | 0 | 0 | 0 | 0 | 0 | 0 | 0 | 0 | 0 | 0 | 0 | 0 | 0 | 0 | 0 | 0 | 0 | 0 | 0 | 0 | 0 | 0 | 0 | 0 | 0 | 0 | 0 | 0 | 0 | 0 | 0 | 0 | 0 | 0 | 0 | 0 | 0 | 0 | 0 | 0 | 0 | 0 | 0 | 0 |
| bacteria_Sep8d0b_30961 | 1 | 0 | 0 | 0 | 0 | 0 | 0 | 0 | 0 | 0 | 0 | 0 | 0 | 0 | 0 | 0 | 0 | 0 | 0 | 0 | 0 | 0 | 0 | 0 | 0 | 0 | 0 | 0 | 0 | 0 | 0 | 0 | 0 | 0 | 0 | 0 | 0 | 0 | 0 | 0 | 0 | 0 | 0 | 0 | 0 | 0 | 0 | 1 | 0 | 0 | 0 | 0 | 0 | 0 | 0 |
| bacteria_Sep8d0b_35968 | 1 | 0 | 0 | 0 | 0 | 0 | 0 | 0 | 0 | 0 | 0 | 0 | 0 | 0 | 0 | 0 | 0 | 0 | 0 | 0 | 0 | 0 | 0 | 0 | 0 | 0 | 0 | 0 | 0 | 0 | 0 | 0 | 0 | 0 | 0 | 0 | 0 | 0 | 0 | 0 | 0 | 0 | 0 | 0 | 0 | 0 | 0 | 1 | 0 | 0 | 0 | 0 | 0 | 0 | 0 |
| bacteria_Sep8d5b_9448 | 1 | 0 | 0 | 0 | 0 | 0 | 0 | 0 | 0 | 0 | 0 | 0 | 0 | 0 | 0 | 0 | 0 | 0 | 0 | 0 | 0 | 0 | 0 | 0 | 0 | 0 | 0 | 0 | 0 | 0 | 0 | 0 | 0 | 0 | 0 | 0 | 0 | 0 | 0 | 0 | 0 | 0 | 0 | 0 | 0 | 0 | 0 | 0 | 0 | 1 | 0 | 0 | 0 | 0 | 0 |
| bacteria_Sep8d5b_12068 | 1 | 0 | 0 | 0 | 0 | 0 | 0 | 0 | 0 | 0 | 0 | 0 | 0 | 0 | 0 | 0 | 0 | 0 | 0 | 0 | 0 | 0 | 0 | 0 | 0 | 0 | 0 | 0 | 0 | 0 | 0 | 0 | 0 | 0 | 0 | 0 | 0 | 0 | 0 | 0 | 0 | 0 | 0 | 0 | 0 | 0 | 0 | 0 | 0 | 1 | 0 | 0 | 0 | 0 | 0 |
| bacteria_Sep8d5b_14374 | 1 | 0 | 0 | 0 | 0 | 0 | 0 | 0 | 0 | 0 | 0 | 0 | 0 | 0 | 0 | 0 | 0 | 0 | 0 | 0 | 0 | 0 | 0 | 0 | 0 | 0 | 0 | 0 | 0 | 0 | 0 | 0 | 0 | 0 | 0 | 0 | 0 | 0 | 0 | 0 | 0 | 0 | 0 | 0 | 0 | 0 | 0 | 0 | 0 | 1 | 0 | 0 | 0 | 0 | 0 |
| bacteria_Sep8d5b_19536 | 1 | 0 | 0 | 0 | 0 | 0 | 0 | 0 | 0 | 0 | 0 | 0 | 0 | 0 | 0 | 0 | 0 | 0 | 0 | 0 | 0 | 0 | 0 | 0 | 0 | 0 | 0 | 0 | 0 | 0 | 0 | 0 | 0 | 0 | 0 | 0 | 0 | 0 | 0 | 0 | 0 | 0 | 0 | 0 | 0 | 0 | 0 | 0 | 0 | 1 | 0 | 0 | 0 | 0 | 0 |
| bacteria_Sep8d5b_26480 | 1 | 0 | 0 | 0 | 0 | 0 | 0 | 0 | 0 | 0 | 0 | 0 | 0 | 0 | 0 | 0 | 0 | 0 | 0 | 0 | 0 | 0 | 0 | 0 | 0 | 0 | 0 | 0 | 0 | 0 | 0 | 0 | 0 | 0 | 0 | 0 | 0 | 0 | 0 | 0 | 0 | 0 | 0 | 0 | 0 | 0 | 0 | 0 | 0 | 1 | 0 | 0 | 0 | 0 | 0 |
| bacteria_Sep8d5b_29126 | 1 | 0 | 0 | 0 | 0 | 0 | 0 | 0 | 0 | 0 | 0 | 0 | 0 | 0 | 0 | 0 | 0 | 0 | 0 | 0 | 0 | 0 | 0 | 0 | 0 | 0 | 0 | 0 | 0 | 0 | 0 | 0 | 0 | 0 | 0 | 0 | 0 | 0 | 0 | 0 | 0 | 0 | 0 | 0 | 0 | 0 | 0 | 0 | 0 | 1 | 0 | 0 | 0 | 0 | 0 |
| bacteria_Sep8d5b_35637 | 1 | 0 | 0 | 0 | 0 | 0 | 0 | 0 | 0 | 0 | 0 | 0 | 0 | 0 | 0 | 0 | 0 | 0 | 0 | 0 | 0 | 0 | 0 | 0 | 0 | 0 | 0 | 0 | 0 | 0 | 0 | 0 | 0 | 0 | 0 | 0 | 0 | 0 | 0 | 0 | 0 | 0 | 0 | 0 | 0 | 0 | 0 | 0 | 0 | 1 | 0 | 0 | 0 | 0 | 0 |
| bacteria_Sep8d5b_36022 | 1 | 0 | 0 | 0 | 0 | 0 | 0 | 0 | 0 | 0 | 0 | 0 | 0 | 0 | 0 | 0 | 0 | 0 | 0 | 0 | 0 | 0 | 0 | 0 | 0 | 0 | 0 | 0 | 0 | 0 | 0 | 0 | 0 | 0 | 0 | 0 | 0 | 0 | 0 | 0 | 0 | 0 | 0 | 0 | 0 | 0 | 0 | 0 | 0 | 1 | 0 | 0 | 0 | 0 | 0 |
| bacteria_Sep8d5b_38265 | 1 | 0 | 0 | 0 | 0 | 0 | 0 | 0 | 0 | 0 | 0 | 0 | 0 | 0 | 0 | 0 | 0 | 0 | 0 | 0 | 0 | 0 | 0 | 0 | 0 | 0 | 0 | 0 | 0 | 0 | 0 | 0 | 0 | 0 | 0 | 0 | 0 | 0 | 0 | 0 | 0 | 0 | 0 | 0 | 0 | 0 | 0 | 0 | 0 | 1 | 0 | 0 | 0 | 0 | 0 |
| bacteria_Sep8d5b_41465 | 1 | 0 | 0 | 0 | 0 | 0 | 0 | 0 | 0 | 0 | 0 | 0 | 0 | 0 | 0 | 0 | 0 | 0 | 0 | 0 | 0 | 0 | 0 | 0 | 0 | 0 | 0 | 0 | 0 | 0 | 0 | 0 | 0 | 0 | 0 | 0 | 0 | 0 | 0 | 0 | 0 | 0 | 0 | 0 | 0 | 0 | 0 | 0 | 0 | 1 | 0 | 0 | 0 | 0 | 0 |
| bacteria_Sep8d5b_50190 | 1 | 0 | 0 | 0 | 0 | 0 | 0 | 0 | 0 | 0 | 0 | 0 | 0 | 0 | 0 | 0 | 0 | 0 | 0 | 0 | 0 | 0 | 0 | 0 | 0 | 0 | 0 | 0 | 0 | 0 | 0 | 0 | 0 | 0 | 0 | 0 | 0 | 0 | 0 | 0 | 0 | 0 | 0 | 0 | 0 | 0 | 0 | 0 | 0 | 1 | 0 | 0 | 0 | 0 | 0 |
| bacteria_Sep8d5b_52952 | 1 | 0 | 0 | 0 | 0 | 0 | 0 | 0 | 0 | 0 | 0 | 0 | 0 | 0 | 0 | 0 | 0 | 0 | 0 | 0 | 0 | 0 | 0 | 0 | 0 | 0 | 0 | 0 | 0 | 0 | 0 | 0 | 0 | 0 | 0 | 0 | 0 | 0 | 0 | 0 | 0 | 0 | 0 | 0 | 0 | 0 | 0 | 0 | 0 | 1 | 0 | 0 | 0 | 0 | 0 |
| bacteria_Sep8d5b_53435 | 1 | 0 | 0 | 0 | 0 | 0 | 0 | 0 | 0 | 0 | 0 | 0 | 0 | 0 | 0 | 0 | 0 | 0 | 0 | 0 | 0 | 0 | 0 | 0 | 0 | 0 | 0 | 0 | 0 | 0 | 0 | 0 | 0 | 0 | 0 | 0 | 0 | 0 | 0 | 0 | 0 | 0 | 0 | 0 | 0 | 0 | 0 | 0 | 0 | 1 | 0 | 0 | 0 | 0 | 0 |
| bacteria_Sep8d5b_55699 | 1 | 0 | 0 | 0 | 0 | 0 | 0 | 0 | 0 | 0 | 0 | 0 | 0 | 0 | 0 | 0 | 0 | 0 | 0 | 0 | 0 | 0 | 0 | 0 | 0 | 0 | 0 | 0 | 0 | 0 | 0 | 0 | 0 | 0 | 0 | 0 | 0 | 0 | 0 | 0 | 0 | 0 | 0 | 0 | 0 | 0 | 0 | 0 | 0 | 1 | 0 | 0 | 0 | 0 | 0 |
| bacteria_Sep8d5b_57191 | 1 | 0 | 0 | 0 | 0 | 0 | 0 | 0 | 0 | 0 | 0 | 0 | 0 | 0 | 0 | 0 | 0 | 0 | 0 | 0 | 0 | 0 | 0 | 0 | 0 | 0 | 0 | 0 | 0 | 0 | 0 | 0 | 0 | 0 | 0 | 0 | 0 | 0 | 0 | 0 | 0 | 0 | 0 | 0 | 0 | 0 | 0 | 0 | 0 | 1 | 0 | 0 | 0 | 0 | 0 |
| bacteria_Sep8d5b_59121 | 1 | 0 | 0 | 0 | 0 | 0 | 0 | 0 | 0 | 0 | 0 | 0 | 0 | 0 | 0 | 0 | 0 | 0 | 0 | 0 | 0 | 0 | 0 | 0 | 0 | 0 | 0 | 0 | 0 | 0 | 0 | 0 | 0 | 0 | 0 | 0 | 0 | 0 | 0 | 0 | 0 | 0 | 0 | 0 | 0 | 0 | 0 | 0 | 0 | 1 | 0 | 0 | 0 | 0 | 0 |
| bacteria_Sep8d5b_59920 | 1 | 0 | 0 | 0 | 0 | 0 | 0 | 0 | 0 | 0 | 0 | 0 | 0 | 0 | 0 | 0 | 0 | 0 | 0 | 0 | 0 | 0 | 0 | 0 | 0 | 0 | 0 | 0 | 0 | 0 | 0 | 0 | 0 | 0 | 0 | 0 | 0 | 0 | 0 | 0 | 0 | 0 | 0 | 0 | 0 | 0 | 0 | 0 | 0 | 1 | 0 | 0 | 0 | 0 | 0 |
| bacteria_Sep8d5b_60883 | 1 | 0 | 0 | 0 | 0 | 0 | 0 | 0 | 0 | 0 | 0 | 0 | 0 | 0 | 0 | 0 | 0 | 0 | 0 | 0 | 0 | 0 | 0 | 0 | 0 | 0 | 0 | 0 | 0 | 0 | 0 | 0 | 0 | 0 | 0 | 0 | 0 | 0 | 0 | 0 | 0 | 0 | 0 | 0 | 0 | 0 | 0 | 0 | 0 | 1 | 0 | 0 | 0 | 0 | 0 |
| bacteria_Sep8d5b_61021 | 1 | 0 | 0 | 0 | 0 | 0 | 0 | 0 | 0 | 0 | 0 | 0 | 0 | 0 | 0 | 0 | 0 | 0 | 0 | 0 | 0 | 0 | 0 | 0 | 0 | 0 | 0 | 0 | 0 | 0 | 0 | 0 | 0 | 0 | 0 | 0 | 0 | 0 | 0 | 0 | 0 | 0 | 0 | 0 | 0 | 0 | 0 | 0 | 0 | 1 | 0 | 0 | 0 | 0 | 0 |
| bacteria_Sep8d5b_63599 | 1 | 0 | 0 | 0 | 0 | 0 | 0 | 0 | 0 | 0 | 0 | 0 | 0 | 0 | 0 | 0 | 0 | 0 | 0 | 0 | 0 | 0 | 0 | 0 | 0 | 0 | 0 | 0 | 0 | 0 | 0 | 0 | 0 | 0 | 0 | 0 | 0 | 0 | 0 | 0 | 0 | 0 | 0 | 0 | 0 | 0 | 0 | 0 | 0 | 1 | 0 | 0 | 0 | 0 | 0 |
| bacteria_Sep8d5b_67463 | 1 | 0 | 0 | 0 | 0 | 0 | 0 | 0 | 0 | 0 | 0 | 0 | 0 | 0 | 0 | 0 | 0 | 0 | 0 | 0 | 0 | 0 | 0 | 0 | 0 | 0 | 0 | 0 | 0 | 0 | 0 | 0 | 0 | 0 | 0 | 0 | 0 | 0 | 0 | 0 | 0 | 0 | 0 | 0 | 0 | 0 | 0 | 0 | 0 | 1 | 0 | 0 | 0 | 0 | 0 |
| bacteria_Sep8d5b_68026 | 1 | 0 | 0 | 0 | 0 | 0 | 0 | 0 | 0 | 0 | 0 | 0 | 0 | 0 | 0 | 0 | 0 | 0 | 0 | 0 | 0 | 0 | 0 | 0 | 0 | 0 | 0 | 0 | 0 | 0 | 0 | 0 | 0 | 0 | 0 | 0 | 0 | 0 | 0 | 0 | 0 | 0 | 0 | 0 | 0 | 0 | 0 | 0 | 0 | 1 | 0 | 0 | 0 | 0 | 0 |
| bacteria_Sep8d5b_70353 | 1 | 0 | 0 | 0 | 0 | 0 | 0 | 0 | 0 | 0 | 0 | 0 | 0 | 0 | 0 | 0 | 0 | 0 | 0 | 0 | 0 | 0 | 0 | 0 | 0 | 0 | 0 | 0 | 0 | 0 | 0 | 0 | 0 | 0 | 0 | 0 | 0 | 0 | 0 | 0 | 0 | 0 | 0 | 0 | 0 | 0 | 0 | 0 | 0 | 1 | 0 | 0 | 0 | 0 | 0 |
| bacteria_Sep8d5b_74335 | 1 | 0 | 0 | 0 | 0 | 0 | 0 | 0 | 0 | 0 | 0 | 0 | 0 | 0 | 0 | 0 | 0 | 0 | 0 | 0 | 0 | 0 | 0 | 0 | 0 | 0 | 0 | 0 | 0 | 0 | 0 | 0 | 0 | 0 | 0 | 0 | 0 | 0 | 0 | 0 | 0 | 0 | 0 | 0 | 0 | 0 | 0 | 0 | 0 | 1 | 0 | 0 | 0 | 0 | 0 |
| bacteria_Sep8d25b_31062 | 1 | 0 | 0 | 0 | 0 | 0 | 0 | 0 | 0 | 0 | 0 | 0 | 0 | 0 | 0 | 0 | 0 | 0 | 0 | 0 | 0 | 0 | 0 | 0 | 0 | 0 | 0 | 0 | 0 | 0 | 0 | 0 | 0 | 0 | 0 | 0 | 0 | 0 | 0 | 0 | 0 | 0 | 0 | 0 | 0 | 0 | 0 | 0 | 0 | 0 | 0 | 1 | 0 | 0 | 0 |
| bacteria_Sep8d25b_43934 | 1 | 0 | 0 | 0 | 0 | 0 | 0 | 0 | 0 | 0 | 0 | 0 | 0 | 0 | 0 | 0 | 0 | 0 | 0 | 0 | 0 | 0 | 0 | 0 | 0 | 0 | 0 | 0 | 0 | 0 | 0 | 0 | 0 | 0 | 0 | 0 | 0 | 0 | 0 | 0 | 0 | 0 | 0 | 0 | 0 | 0 | 0 | 0 | 0 | 0 | 0 | 1 | 0 | 0 | 0 |
| bacteria_Sep8d50b_464 | 1 | 0 | 0 | 0 | 0 | 0 | 0 | 0 | 0 | 0 | 0 | 0 | 0 | 0 | 0 | 0 | 0 | 0 | 0 | 0 | 0 | 0 | 0 | 0 | 0 | 0 | 0 | 0 | 0 | 0 | 0 | 0 | 0 | 0 | 0 | 0 | 0 | 0 | 0 | 0 | 0 | 0 | 0 | 0 | 0 | 0 | 0 | 0 | 0 | 0 | 0 | 0 | 0 | 1 | 0 |
| virus_Aug5.0v_17 | 1 | 0 | 0 | 0 | 0 | 0 | 0 | 0 | 0 | 0 | 0 | 0 | 0 | 0 | 0 | 0 | 0 | 0 | 0 | 0 | 0 | 0 | 0 | 0 | 0 | 0 | 0 | 0 | 0 | 0 | 0 | 0 | 0 | 0 | 0 | 0 | 0 | 0 | 1 | 0 | 0 | 0 | 0 | 0 | 0 | 0 | 0 | 0 | 0 | 0 | 0 | 0 | 0 | 0 | 0 |
| virus_Aug5.0v_188 | 1 | 0 | 0 | 0 | 0 | 0 | 0 | 0 | 0 | 0 | 0 | 0 | 0 | 0 | 0 | 0 | 0 | 0 | 0 | 0 | 0 | 0 | 0 | 0 | 0 | 0 | 0 | 0 | 0 | 0 | 0 | 0 | 0 | 0 | 0 | 0 | 0 | 0 | 1 | 0 | 0 | 0 | 0 | 0 | 0 | 0 | 0 | 0 | 0 | 0 | 0 | 0 | 0 | 0 | 0 |
| virus_Aug5.0v_468 | 1 | 0 | 0 | 0 | 0 | 0 | 0 | 0 | 0 | 0 | 0 | 0 | 0 | 0 | 0 | 0 | 0 | 0 | 0 | 0 | 0 | 0 | 0 | 0 | 0 | 0 | 0 | 0 | 0 | 0 | 0 | 0 | 0 | 0 | 0 | 0 | 0 | 0 | 1 | 0 | 0 | 0 | 0 | 0 | 0 | 0 | 0 | 0 | 0 | 0 | 0 | 0 | 0 | 0 | 0 |
| virus_Aug5.0v_1188 | 1 | 0 | 0 | 0 | 0 | 0 | 0 | 0 | 0 | 0 | 0 | 0 | 0 | 0 | 0 | 0 | 0 | 0 | 0 | 0 | 0 | 0 | 0 | 0 | 0 | 0 | 0 | 0 | 0 | 0 | 0 | 0 | 0 | 0 | 0 | 0 | 0 | 0 | 1 | 0 | 0 | 0 | 0 | 0 | 0 | 0 | 0 | 0 | 0 | 0 | 0 | 0 | 0 | 0 | 0 |
| virus_Aug5.0v_1224 | 1 | 0 | 0 | 0 | 0 | 0 | 0 | 0 | 0 | 0 | 0 | 0 | 0 | 0 | 0 | 0 | 0 | 0 | 0 | 0 | 0 | 0 | 0 | 0 | 0 | 0 | 0 | 0 | 0 | 0 | 0 | 0 | 0 | 0 | 0 | 0 | 0 | 0 | 1 | 0 | 0 | 0 | 0 | 0 | 0 | 0 | 0 | 0 | 0 | 0 | 0 | 0 | 0 | 0 | 0 |
| virus_Aug5.0v_1237 | 1 | 0 | 0 | 0 | 0 | 0 | 0 | 0 | 0 | 0 | 0 | 0 | 0 | 0 | 0 | 0 | 0 | 0 | 0 | 0 | 0 | 0 | 0 | 0 | 0 | 0 | 0 | 0 | 0 | 0 | 0 | 0 | 0 | 0 | 0 | 0 | 0 | 0 | 1 | 0 | 0 | 0 | 0 | 0 | 0 | 0 | 0 | 0 | 0 | 0 | 0 | 0 | 0 | 0 | 0 |
| virus_Aug5.0v_1632 | 1 | 0 | 0 | 0 | 0 | 0 | 0 | 0 | 0 | 0 | 0 | 0 | 0 | 0 | 0 | 0 | 0 | 0 | 0 | 0 | 0 | 0 | 0 | 0 | 0 | 0 | 0 | 0 | 0 | 0 | 0 | 0 | 0 | 0 | 0 | 0 | 0 | 0 | 1 | 0 | 0 | 0 | 0 | 0 | 0 | 0 | 0 | 0 | 0 | 0 | 0 | 0 | 0 | 0 | 0 |
| virus_Aug5.0v_1656 | 1 | 0 | 0 | 0 | 0 | 0 | 0 | 0 | 0 | 0 | 0 | 0 | 0 | 0 | 0 | 0 | 0 | 0 | 0 | 0 | 0 | 0 | 0 | 0 | 0 | 0 | 0 | 0 | 0 | 0 | 0 | 0 | 0 | 0 | 0 | 0 | 0 | 0 | 1 | 0 | 0 | 0 | 0 | 0 | 0 | 0 | 0 | 0 | 0 | 0 | 0 | 0 | 0 | 0 | 0 |
| virus_Aug5.0v_2181 | 1 | 0 | 0 | 0 | 0 | 0 | 0 | 0 | 0 | 0 | 0 | 0 | 0 | 0 | 0 | 0 | 0 | 0 | 0 | 0 | 0 | 0 | 0 | 0 | 0 | 0 | 0 | 0 | 0 | 0 | 0 | 0 | 0 | 0 | 0 | 0 | 0 | 0 | 1 | 0 | 0 | 0 | 0 | 0 | 0 | 0 | 0 | 0 | 0 | 0 | 0 | 0 | 0 | 0 | 0 |
| virus_Aug5.0v_2555 | 1 | 0 | 0 | 0 | 0 | 0 | 0 | 0 | 0 | 0 | 0 | 0 | 0 | 0 | 0 | 0 | 0 | 0 | 0 | 0 | 0 | 0 | 0 | 0 | 0 | 0 | 0 | 0 | 0 | 0 | 0 | 0 | 0 | 0 | 0 | 0 | 0 | 0 | 1 | 0 | 0 | 0 | 0 | 0 | 0 | 0 | 0 | 0 | 0 | 0 | 0 | 0 | 0 | 0 | 0 |
| virus_Aug5.0v_2672 | 1 | 0 | 0 | 0 | 0 | 0 | 0 | 0 | 0 | 0 | 0 | 0 | 0 | 0 | 0 | 0 | 0 | 0 | 0 | 0 | 0 | 0 | 0 | 0 | 0 | 0 | 0 | 0 | 0 | 0 | 0 | 0 | 0 | 0 | 0 | 0 | 0 | 0 | 1 | 0 | 0 | 0 | 0 | 0 | 0 | 0 | 0 | 0 | 0 | 0 | 0 | 0 | 0 | 0 | 0 |
| virus_Aug5.0v_2718 | 1 | 0 | 0 | 0 | 0 | 0 | 0 | 0 | 0 | 0 | 0 | 0 | 0 | 0 | 0 | 0 | 0 | 0 | 0 | 0 | 0 | 0 | 0 | 0 | 0 | 0 | 0 | 0 | 0 | 0 | 0 | 0 | 0 | 0 | 0 | 0 | 0 | 0 | 1 | 0 | 0 | 0 | 0 | 0 | 0 | 0 | 0 | 0 | 0 | 0 | 0 | 0 | 0 | 0 | 0 |
| virus_Aug5.0v_2990 | 1 | 0 | 0 | 0 | 0 | 0 | 0 | 0 | 0 | 0 | 0 | 0 | 0 | 0 | 0 | 0 | 0 | 0 | 0 | 0 | 0 | 0 | 0 | 0 | 0 | 0 | 0 | 0 | 0 | 0 | 0 | 0 | 0 | 0 | 0 | 0 | 0 | 0 | 1 | 0 | 0 | 0 | 0 | 0 | 0 | 0 | 0 | 0 | 0 | 0 | 0 | 0 | 0 | 0 | 0 |
| virus_Aug5.0v_3220 | 1 | 0 | 0 | 0 | 0 | 0 | 0 | 0 | 0 | 0 | 0 | 0 | 0 | 0 | 0 | 0 | 0 | 0 | 0 | 0 | 0 | 0 | 0 | 0 | 0 | 0 | 0 | 0 | 0 | 0 | 0 | 0 | 0 | 0 | 0 | 0 | 0 | 0 | 1 | 0 | 0 | 0 | 0 | 0 | 0 | 0 | 0 | 0 | 0 | 0 | 0 | 0 | 0 | 0 | 0 |
| virus_Aug5.0v_3323 | 1 | 0 | 0 | 0 | 0 | 0 | 0 | 0 | 0 | 0 | 0 | 0 | 0 | 0 | 0 | 0 | 0 | 0 | 0 | 0 | 0 | 0 | 0 | 0 | 0 | 0 | 0 | 0 | 0 | 0 | 0 | 0 | 0 | 0 | 0 | 0 | 0 | 0 | 1 | 0 | 0 | 0 | 0 | 0 | 0 | 0 | 0 | 0 | 0 | 0 | 0 | 0 | 0 | 0 | 0 |
| virus_Aug5.0v_3567 | 1 | 0 | 0 | 0 | 0 | 0 | 0 | 0 | 0 | 0 | 0 | 0 | 0 | 0 | 0 | 0 | 0 | 0 | 0 | 0 | 0 | 0 | 0 | 0 | 0 | 0 | 0 | 0 | 0 | 0 | 0 | 0 | 0 | 0 | 0 | 0 | 0 | 0 | 1 | 0 | 0 | 0 | 0 | 0 | 0 | 0 | 0 | 0 | 0 | 0 | 0 | 0 | 0 | 0 | 0 |
| virus_Aug5.0v_3574 | 1 | 0 | 0 | 0 | 0 | 0 | 0 | 0 | 0 | 0 | 0 | 0 | 0 | 0 | 0 | 0 | 0 | 0 | 0 | 0 | 0 | 0 | 0 | 0 | 0 | 0 | 0 | 0 | 0 | 0 | 0 | 0 | 0 | 0 | 0 | 0 | 0 | 0 | 1 | 0 | 0 | 0 | 0 | 0 | 0 | 0 | 0 | 0 | 0 | 0 | 0 | 0 | 0 | 0 | 0 |
| virus_Aug5.0v_4127 | 1 | 0 | 0 | 0 | 0 | 0 | 0 | 0 | 0 | 0 | 0 | 0 | 0 | 0 | 0 | 0 | 0 | 0 | 0 | 0 | 0 | 0 | 0 | 0 | 0 | 0 | 0 | 0 | 0 | 0 | 0 | 0 | 0 | 0 | 0 | 0 | 0 | 0 | 1 | 0 | 0 | 0 | 0 | 0 | 0 | 0 | 0 | 0 | 0 | 0 | 0 | 0 | 0 | 0 | 0 |
| virus_Aug5.0v_4322 | 1 | 0 | 0 | 0 | 0 | 0 | 0 | 0 | 0 | 0 | 0 | 0 | 0 | 0 | 0 | 0 | 0 | 0 | 0 | 0 | 0 | 0 | 0 | 0 | 0 | 0 | 0 | 0 | 0 | 0 | 0 | 0 | 0 | 0 | 0 | 0 | 0 | 0 | 1 | 0 | 0 | 0 | 0 | 0 | 0 | 0 | 0 | 0 | 0 | 0 | 0 | 0 | 0 | 0 | 0 |
| virus_Aug5.0v_4410 | 1 | 0 | 0 | 0 | 0 | 0 | 0 | 0 | 0 | 0 | 0 | 0 | 0 | 0 | 0 | 0 | 0 | 0 | 0 | 0 | 0 | 0 | 0 | 0 | 0 | 0 | 0 | 0 | 0 | 0 | 0 | 0 | 0 | 0 | 0 | 0 | 0 | 0 | 1 | 0 | 0 | 0 | 0 | 0 | 0 | 0 | 0 | 0 | 0 | 0 | 0 | 0 | 0 | 0 | 0 |
| virus_Aug5.0v_4542 | 1 | 0 | 0 | 0 | 0 | 0 | 0 | 0 | 0 | 0 | 0 | 0 | 0 | 0 | 0 | 0 | 0 | 0 | 0 | 0 | 0 | 0 | 0 | 0 | 0 | 0 | 0 | 0 | 0 | 0 | 0 | 0 | 0 | 0 | 0 | 0 | 0 | 0 | 1 | 0 | 0 | 0 | 0 | 0 | 0 | 0 | 0 | 0 | 0 | 0 | 0 | 0 | 0 | 0 | 0 |
| virus_Aug5.0v_4731 | 1 | 0 | 0 | 0 | 0 | 0 | 0 | 0 | 0 | 0 | 0 | 0 | 0 | 0 | 0 | 0 | 0 | 0 | 0 | 0 | 0 | 0 | 0 | 0 | 0 | 0 | 0 | 0 | 0 | 0 | 0 | 0 | 0 | 0 | 0 | 0 | 0 | 0 | 1 | 0 | 0 | 0 | 0 | 0 | 0 | 0 | 0 | 0 | 0 | 0 | 0 | 0 | 0 | 0 | 0 |
| virus_Aug5.0v_4824 | 1 | 0 | 0 | 0 | 0 | 0 | 0 | 0 | 0 | 0 | 0 | 0 | 0 | 0 | 0 | 0 | 0 | 0 | 0 | 0 | 0 | 0 | 0 | 0 | 0 | 0 | 0 | 0 | 0 | 0 | 0 | 0 | 0 | 0 | 0 | 0 | 0 | 0 | 1 | 0 | 0 | 0 | 0 | 0 | 0 | 0 | 0 | 0 | 0 | 0 | 0 | 0 | 0 | 0 | 0 |
| virus_Aug5.0v_4956 | 1 | 0 | 0 | 0 | 0 | 0 | 0 | 0 | 0 | 0 | 0 | 0 | 0 | 0 | 0 | 0 | 0 | 0 | 0 | 0 | 0 | 0 | 0 | 0 | 0 | 0 | 0 | 0 | 0 | 0 | 0 | 0 | 0 | 0 | 0 | 0 | 0 | 0 | 1 | 0 | 0 | 0 | 0 | 0 | 0 | 0 | 0 | 0 | 0 | 0 | 0 | 0 | 0 | 0 | 0 |
| virus_Aug5.0v_5074 | 1 | 0 | 0 | 0 | 0 | 0 | 0 | 0 | 0 | 0 | 0 | 0 | 0 | 0 | 0 | 0 | 0 | 0 | 0 | 0 | 0 | 0 | 0 | 0 | 0 | 0 | 0 | 0 | 0 | 0 | 0 | 0 | 0 | 0 | 0 | 0 | 0 | 0 | 1 | 0 | 0 | 0 | 0 | 0 | 0 | 0 | 0 | 0 | 0 | 0 | 0 | 0 | 0 | 0 | 0 |
| virus_Aug5.0v_5383 | 1 | 0 | 0 | 0 | 0 | 0 | 0 | 0 | 0 | 0 | 0 | 0 | 0 | 0 | 0 | 0 | 0 | 0 | 0 | 0 | 0 | 0 | 0 | 0 | 0 | 0 | 0 | 0 | 0 | 0 | 0 | 0 | 0 | 0 | 0 | 0 | 0 | 0 | 1 | 0 | 0 | 0 | 0 | 0 | 0 | 0 | 0 | 0 | 0 | 0 | 0 | 0 | 0 | 0 | 0 |
| virus_Aug5.5v_449 | 1 | 0 | 0 | 0 | 0 | 0 | 0 | 0 | 0 | 0 | 0 | 0 | 0 | 0 | 0 | 0 | 0 | 0 | 0 | 0 | 0 | 0 | 0 | 0 | 0 | 0 | 0 | 0 | 0 | 0 | 0 | 0 | 0 | 0 | 0 | 0 | 0 | 0 | 0 | 0 | 1 | 0 | 0 | 0 | 0 | 0 | 0 | 0 | 0 | 0 | 0 | 0 | 0 | 0 | 0 |
| virus_Aug5.5v_10746 | 1 | 0 | 0 | 0 | 0 | 0 | 0 | 0 | 0 | 0 | 0 | 0 | 0 | 0 | 0 | 0 | 0 | 0 | 0 | 0 | 0 | 0 | 0 | 0 | 0 | 0 | 0 | 0 | 0 | 0 | 0 | 0 | 0 | 0 | 0 | 0 | 0 | 0 | 0 | 0 | 1 | 0 | 0 | 0 | 0 | 0 | 0 | 0 | 0 | 0 | 0 | 0 | 0 | 0 | 0 |
| virus_Aug5.5v_12704 | 1 | 0 | 0 | 0 | 0 | 0 | 0 | 0 | 0 | 0 | 0 | 0 | 0 | 0 | 0 | 0 | 0 | 0 | 0 | 0 | 0 | 0 | 0 | 0 | 0 | 0 | 0 | 0 | 0 | 0 | 0 | 0 | 0 | 0 | 0 | 0 | 0 | 0 | 0 | 0 | 1 | 0 | 0 | 0 | 0 | 0 | 0 | 0 | 0 | 0 | 0 | 0 | 0 | 0 | 0 |
| virus_Aug5.5v_14009 | 1 | 0 | 0 | 0 | 0 | 0 | 0 | 0 | 0 | 0 | 0 | 0 | 0 | 0 | 0 | 0 | 0 | 0 | 0 | 0 | 0 | 0 | 0 | 0 | 0 | 0 | 0 | 0 | 0 | 0 | 0 | 0 | 0 | 0 | 0 | 0 | 0 | 0 | 0 | 0 | 1 | 0 | 0 | 0 | 0 | 0 | 0 | 0 | 0 | 0 | 0 | 0 | 0 | 0 | 0 |
| virus_Aug5.5v_23090 | 1 | 0 | 0 | 0 | 0 | 0 | 0 | 0 | 0 | 0 | 0 | 0 | 0 | 0 | 0 | 0 | 0 | 0 | 0 | 0 | 0 | 0 | 0 | 0 | 0 | 0 | 0 | 0 | 0 | 0 | 0 | 0 | 0 | 0 | 0 | 0 | 0 | 0 | 0 | 0 | 1 | 0 | 0 | 0 | 0 | 0 | 0 | 0 | 0 | 0 | 0 | 0 | 0 | 0 | 0 |
| virus_Aug5.5v_26095 | 1 | 0 | 0 | 0 | 0 | 0 | 0 | 0 | 0 | 0 | 0 | 0 | 0 | 0 | 0 | 0 | 0 | 0 | 0 | 0 | 0 | 0 | 0 | 0 | 0 | 0 | 0 | 0 | 0 | 0 | 0 | 0 | 0 | 0 | 0 | 0 | 0 | 0 | 0 | 0 | 1 | 0 | 0 | 0 | 0 | 0 | 0 | 0 | 0 | 0 | 0 | 0 | 0 | 0 | 0 |
| virus_Aug5.5v_26523 | 1 | 0 | 0 | 0 | 0 | 0 | 0 | 0 | 0 | 0 | 0 | 0 | 0 | 0 | 0 | 0 | 0 | 0 | 0 | 0 | 0 | 0 | 0 | 0 | 0 | 0 | 0 | 0 | 0 | 0 | 0 | 0 | 0 | 0 | 0 | 0 | 0 | 0 | 0 | 0 | 1 | 0 | 0 | 0 | 0 | 0 | 0 | 0 | 0 | 0 | 0 | 0 | 0 | 0 | 0 |
| virus_Aug5.5v_27701 | 1 | 0 | 0 | 0 | 0 | 0 | 0 | 0 | 0 | 0 | 0 | 0 | 0 | 0 | 0 | 0 | 0 | 0 | 0 | 0 | 0 | 0 | 0 | 0 | 0 | 0 | 0 | 0 | 0 | 0 | 0 | 0 | 0 | 0 | 0 | 0 | 0 | 0 | 0 | 0 | 1 | 0 | 0 | 0 | 0 | 0 | 0 | 0 | 0 | 0 | 0 | 0 | 0 | 0 | 0 |
| virus_Aug5.5v_28182 | 1 | 0 | 0 | 0 | 0 | 0 | 0 | 0 | 0 | 0 | 0 | 0 | 0 | 0 | 0 | 0 | 0 | 0 | 0 | 0 | 0 | 0 | 0 | 0 | 0 | 0 | 0 | 0 | 0 | 0 | 0 | 0 | 0 | 0 | 0 | 0 | 0 | 0 | 0 | 0 | 1 | 0 | 0 | 0 | 0 | 0 | 0 | 0 | 0 | 0 | 0 | 0 | 0 | 0 | 0 |
| virus_Aug5.5v_31422 | 1 | 0 | 0 | 0 | 0 | 0 | 0 | 0 | 0 | 0 | 0 | 0 | 0 | 0 | 0 | 0 | 0 | 0 | 0 | 0 | 0 | 0 | 0 | 0 | 0 | 0 | 0 | 0 | 0 | 0 | 0 | 0 | 0 | 0 | 0 | 0 | 0 | 0 | 0 | 0 | 1 | 0 | 0 | 0 | 0 | 0 | 0 | 0 | 0 | 0 | 0 | 0 | 0 | 0 | 0 |
| virus_Aug5.5v_35456 | 1 | 0 | 0 | 0 | 0 | 0 | 0 | 0 | 0 | 0 | 0 | 0 | 0 | 0 | 0 | 0 | 0 | 0 | 0 | 0 | 0 | 0 | 0 | 0 | 0 | 0 | 0 | 0 | 0 | 0 | 0 | 0 | 0 | 0 | 0 | 0 | 0 | 0 | 0 | 0 | 1 | 0 | 0 | 0 | 0 | 0 | 0 | 0 | 0 | 0 | 0 | 0 | 0 | 0 | 0 |
| virus_Aug5.5v_38006 | 1 | 0 | 0 | 0 | 0 | 0 | 0 | 0 | 0 | 0 | 0 | 0 | 0 | 0 | 0 | 0 | 0 | 0 | 0 | 0 | 0 | 0 | 0 | 0 | 0 | 0 | 0 | 0 | 0 | 0 | 0 | 0 | 0 | 0 | 0 | 0 | 0 | 0 | 0 | 0 | 1 | 0 | 0 | 0 | 0 | 0 | 0 | 0 | 0 | 0 | 0 | 0 | 0 | 0 | 0 |
| virus_Aug5.5v_40425 | 1 | 0 | 0 | 0 | 0 | 0 | 0 | 0 | 0 | 0 | 0 | 0 | 0 | 0 | 0 | 0 | 0 | 0 | 0 | 0 | 0 | 0 | 0 | 0 | 0 | 0 | 0 | 0 | 0 | 0 | 0 | 0 | 0 | 0 | 0 | 0 | 0 | 0 | 0 | 0 | 1 | 0 | 0 | 0 | 0 | 0 | 0 | 0 | 0 | 0 | 0 | 0 | 0 | 0 | 0 |
| virus_Aug5.25v_4744 | 1 | 0 | 0 | 0 | 0 | 0 | 0 | 0 | 0 | 0 | 0 | 0 | 0 | 0 | 0 | 0 | 0 | 0 | 0 | 0 | 0 | 0 | 0 | 0 | 0 | 0 | 0 | 0 | 0 | 0 | 0 | 0 | 0 | 0 | 0 | 0 | 0 | 0 | 0 | 0 | 0 | 0 | 1 | 0 | 0 | 0 | 0 | 0 | 0 | 0 | 0 | 0 | 0 | 0 | 0 |
| virus_Aug5.25v_6835 | 1 | 0 | 0 | 0 | 0 | 0 | 0 | 0 | 0 | 0 | 0 | 0 | 0 | 0 | 0 | 0 | 0 | 0 | 0 | 0 | 0 | 0 | 0 | 0 | 0 | 0 | 0 | 0 | 0 | 0 | 0 | 0 | 0 | 0 | 0 | 0 | 0 | 0 | 0 | 0 | 0 | 0 | 1 | 0 | 0 | 0 | 0 | 0 | 0 | 0 | 0 | 0 | 0 | 0 | 0 |
| virus_Aug5.25v_8038 | 1 | 0 | 0 | 0 | 0 | 0 | 0 | 0 | 0 | 0 | 0 | 0 | 0 | 0 | 0 | 0 | 0 | 0 | 0 | 0 | 0 | 0 | 0 | 0 | 0 | 0 | 0 | 0 | 0 | 0 | 0 | 0 | 0 | 0 | 0 | 0 | 0 | 0 | 0 | 0 | 0 | 0 | 1 | 0 | 0 | 0 | 0 | 0 | 0 | 0 | 0 | 0 | 0 | 0 | 0 |
| virus_Aug5.25v_8334 | 1 | 0 | 0 | 0 | 0 | 0 | 0 | 0 | 0 | 0 | 0 | 0 | 0 | 0 | 0 | 0 | 0 | 0 | 0 | 0 | 0 | 0 | 0 | 0 | 0 | 0 | 0 | 0 | 0 | 0 | 0 | 0 | 0 | 0 | 0 | 0 | 0 | 0 | 0 | 0 | 0 | 0 | 1 | 0 | 0 | 0 | 0 | 0 | 0 | 0 | 0 | 0 | 0 | 0 | 0 |
| virus_Aug5.25v_10634 | 1 | 0 | 0 | 0 | 0 | 0 | 0 | 0 | 0 | 0 | 0 | 0 | 0 | 0 | 0 | 0 | 0 | 0 | 0 | 0 | 0 | 0 | 0 | 0 | 0 | 0 | 0 | 0 | 0 | 0 | 0 | 0 | 0 | 0 | 0 | 0 | 0 | 0 | 0 | 0 | 0 | 0 | 1 | 0 | 0 | 0 | 0 | 0 | 0 | 0 | 0 | 0 | 0 | 0 | 0 |
| virus_Aug5.25v_12087 | 1 | 0 | 0 | 0 | 0 | 0 | 0 | 0 | 0 | 0 | 0 | 0 | 0 | 0 | 0 | 0 | 0 | 0 | 0 | 0 | 0 | 0 | 0 | 0 | 0 | 0 | 0 | 0 | 0 | 0 | 0 | 0 | 0 | 0 | 0 | 0 | 0 | 0 | 0 | 0 | 0 | 0 | 1 | 0 | 0 | 0 | 0 | 0 | 0 | 0 | 0 | 0 | 0 | 0 | 0 |
| virus_Aug5.25v_12350 | 1 | 0 | 0 | 0 | 0 | 0 | 0 | 0 | 0 | 0 | 0 | 0 | 0 | 0 | 0 | 0 | 0 | 0 | 0 | 0 | 0 | 0 | 0 | 0 | 0 | 0 | 0 | 0 | 0 | 0 | 0 | 0 | 0 | 0 | 0 | 0 | 0 | 0 | 0 | 0 | 0 | 0 | 1 | 0 | 0 | 0 | 0 | 0 | 0 | 0 | 0 | 0 | 0 | 0 | 0 |
| virus_Aug5.25v_14138 | 1 | 0 | 0 | 0 | 0 | 0 | 0 | 0 | 0 | 0 | 0 | 0 | 0 | 0 | 0 | 0 | 0 | 0 | 0 | 0 | 0 | 0 | 0 | 0 | 0 | 0 | 0 | 0 | 0 | 0 | 0 | 0 | 0 | 0 | 0 | 0 | 0 | 0 | 0 | 0 | 0 | 0 | 1 | 0 | 0 | 0 | 0 | 0 | 0 | 0 | 0 | 0 | 0 | 0 | 0 |
| virus_Aug5.25v_18338 | 1 | 0 | 0 | 0 | 0 | 0 | 0 | 0 | 0 | 0 | 0 | 0 | 0 | 0 | 0 | 0 | 0 | 0 | 0 | 0 | 0 | 0 | 0 | 0 | 0 | 0 | 0 | 0 | 0 | 0 | 0 | 0 | 0 | 0 | 0 | 0 | 0 | 0 | 0 | 0 | 0 | 0 | 1 | 0 | 0 | 0 | 0 | 0 | 0 | 0 | 0 | 0 | 0 | 0 | 0 |
| virus_Aug5.25v_20746 | 1 | 0 | 0 | 0 | 0 | 0 | 0 | 0 | 0 | 0 | 0 | 0 | 0 | 0 | 0 | 0 | 0 | 0 | 0 | 0 | 0 | 0 | 0 | 0 | 0 | 0 | 0 | 0 | 0 | 0 | 0 | 0 | 0 | 0 | 0 | 0 | 0 | 0 | 0 | 0 | 0 | 0 | 1 | 0 | 0 | 0 | 0 | 0 | 0 | 0 | 0 | 0 | 0 | 0 | 0 |
| virus_Aug5.25v_20934 | 1 | 0 | 0 | 0 | 0 | 0 | 0 | 0 | 0 | 0 | 0 | 0 | 0 | 0 | 0 | 0 | 0 | 0 | 0 | 0 | 0 | 0 | 0 | 0 | 0 | 0 | 0 | 0 | 0 | 0 | 0 | 0 | 0 | 0 | 0 | 0 | 0 | 0 | 0 | 0 | 0 | 0 | 1 | 0 | 0 | 0 | 0 | 0 | 0 | 0 | 0 | 0 | 0 | 0 | 0 |
| virus_Aug5.25v_22083 | 1 | 0 | 0 | 0 | 0 | 0 | 0 | 0 | 0 | 0 | 0 | 0 | 0 | 0 | 0 | 0 | 0 | 0 | 0 | 0 | 0 | 0 | 0 | 0 | 0 | 0 | 0 | 0 | 0 | 0 | 0 | 0 | 0 | 0 | 0 | 0 | 0 | 0 | 0 | 0 | 0 | 0 | 1 | 0 | 0 | 0 | 0 | 0 | 0 | 0 | 0 | 0 | 0 | 0 | 0 |
| virus_Aug5.25v_24775 | 1 | 0 | 0 | 0 | 0 | 0 | 0 | 0 | 0 | 0 | 0 | 0 | 0 | 0 | 0 | 0 | 0 | 0 | 0 | 0 | 0 | 0 | 0 | 0 | 0 | 0 | 0 | 0 | 0 | 0 | 0 | 0 | 0 | 0 | 0 | 0 | 0 | 0 | 0 | 0 | 0 | 0 | 1 | 0 | 0 | 0 | 0 | 0 | 0 | 0 | 0 | 0 | 0 | 0 | 0 |
| virus_Aug5.25v_27289 | 1 | 0 | 0 | 0 | 0 | 0 | 0 | 0 | 0 | 0 | 0 | 0 | 0 | 0 | 0 | 0 | 0 | 0 | 0 | 0 | 0 | 0 | 0 | 0 | 0 | 0 | 0 | 0 | 0 | 0 | 0 | 0 | 0 | 0 | 0 | 0 | 0 | 0 | 0 | 0 | 0 | 0 | 1 | 0 | 0 | 0 | 0 | 0 | 0 | 0 | 0 | 0 | 0 | 0 | 0 |
| virus_Aug5.25v_31155 | 1 | 0 | 0 | 0 | 0 | 0 | 0 | 0 | 0 | 0 | 0 | 0 | 0 | 0 | 0 | 0 | 0 | 0 | 0 | 0 | 0 | 0 | 0 | 0 | 0 | 0 | 0 | 0 | 0 | 0 | 0 | 0 | 0 | 0 | 0 | 0 | 0 | 0 | 0 | 0 | 0 | 0 | 1 | 0 | 0 | 0 | 0 | 0 | 0 | 0 | 0 | 0 | 0 | 0 | 0 |
| virus_Aug5.25v_32749 | 1 | 0 | 0 | 0 | 0 | 0 | 0 | 0 | 0 | 0 | 0 | 0 | 0 | 0 | 0 | 0 | 0 | 0 | 0 | 0 | 0 | 0 | 0 | 0 | 0 | 0 | 0 | 0 | 0 | 0 | 0 | 0 | 0 | 0 | 0 | 0 | 0 | 0 | 0 | 0 | 0 | 0 | 1 | 0 | 0 | 0 | 0 | 0 | 0 | 0 | 0 | 0 | 0 | 0 | 0 |
| virus_Aug5.25v_32847 | 1 | 0 | 0 | 0 | 0 | 0 | 0 | 0 | 0 | 0 | 0 | 0 | 0 | 0 | 0 | 0 | 0 | 0 | 0 | 0 | 0 | 0 | 0 | 0 | 0 | 0 | 0 | 0 | 0 | 0 | 0 | 0 | 0 | 0 | 0 | 0 | 0 | 0 | 0 | 0 | 0 | 0 | 1 | 0 | 0 | 0 | 0 | 0 | 0 | 0 | 0 | 0 | 0 | 0 | 0 |
| virus_Aug5.25v_32890 | 1 | 0 | 0 | 0 | 0 | 0 | 0 | 0 | 0 | 0 | 0 | 0 | 0 | 0 | 0 | 0 | 0 | 0 | 0 | 0 | 0 | 0 | 0 | 0 | 0 | 0 | 0 | 0 | 0 | 0 | 0 | 0 | 0 | 0 | 0 | 0 | 0 | 0 | 0 | 0 | 0 | 0 | 1 | 0 | 0 | 0 | 0 | 0 | 0 | 0 | 0 | 0 | 0 | 0 | 0 |
| virus_Aug5.25v_33426 | 1 | 0 | 0 | 0 | 0 | 0 | 0 | 0 | 0 | 0 | 0 | 0 | 0 | 0 | 0 | 0 | 0 | 0 | 0 | 0 | 0 | 0 | 0 | 0 | 0 | 0 | 0 | 0 | 0 | 0 | 0 | 0 | 0 | 0 | 0 | 0 | 0 | 0 | 0 | 0 | 0 | 0 | 1 | 0 | 0 | 0 | 0 | 0 | 0 | 0 | 0 | 0 | 0 | 0 | 0 |
| virus_Aug5.25v_35473 | 1 | 0 | 0 | 0 | 0 | 0 | 0 | 0 | 0 | 0 | 0 | 0 | 0 | 0 | 0 | 0 | 0 | 0 | 0 | 0 | 0 | 0 | 0 | 0 | 0 | 0 | 0 | 0 | 0 | 0 | 0 | 0 | 0 | 0 | 0 | 0 | 0 | 0 | 0 | 0 | 0 | 0 | 1 | 0 | 0 | 0 | 0 | 0 | 0 | 0 | 0 | 0 | 0 | 0 | 0 |
| virus_Aug5.25v_35775 | 1 | 0 | 0 | 0 | 0 | 0 | 0 | 0 | 0 | 0 | 0 | 0 | 0 | 0 | 0 | 0 | 0 | 0 | 0 | 0 | 0 | 0 | 0 | 0 | 0 | 0 | 0 | 0 | 0 | 0 | 0 | 0 | 0 | 0 | 0 | 0 | 0 | 0 | 0 | 0 | 0 | 0 | 1 | 0 | 0 | 0 | 0 | 0 | 0 | 0 | 0 | 0 | 0 | 0 | 0 |
| virus_Aug5.50v_78 | 1 | 0 | 0 | 0 | 0 | 0 | 0 | 0 | 0 | 0 | 0 | 0 | 0 | 0 | 0 | 0 | 0 | 0 | 0 | 0 | 0 | 0 | 0 | 0 | 0 | 0 | 0 | 0 | 0 | 0 | 0 | 0 | 0 | 0 | 0 | 0 | 0 | 0 | 0 | 0 | 0 | 0 | 0 | 0 | 1 | 0 | 0 | 0 | 0 | 0 | 0 | 0 | 0 | 0 | 0 |
| virus_Aug5.50v_7705 | 1 | 0 | 0 | 0 | 0 | 0 | 0 | 0 | 0 | 0 | 0 | 0 | 0 | 0 | 0 | 0 | 0 | 0 | 0 | 0 | 0 | 0 | 0 | 0 | 0 | 0 | 0 | 0 | 0 | 0 | 0 | 0 | 0 | 0 | 0 | 0 | 0 | 0 | 0 | 0 | 0 | 0 | 0 | 0 | 1 | 0 | 0 | 0 | 0 | 0 | 0 | 0 | 0 | 0 | 0 |
| virus_Aug5.50v_21180 | 1 | 0 | 0 | 0 | 0 | 0 | 0 | 0 | 0 | 0 | 0 | 0 | 0 | 0 | 0 | 0 | 0 | 0 | 0 | 0 | 0 | 0 | 0 | 0 | 0 | 0 | 0 | 0 | 0 | 0 | 0 | 0 | 0 | 0 | 0 | 0 | 0 | 0 | 0 | 0 | 0 | 0 | 0 | 0 | 1 | 0 | 0 | 0 | 0 | 0 | 0 | 0 | 0 | 0 | 0 |
| virus_Aug5.50v_22303 | 1 | 0 | 0 | 0 | 0 | 0 | 0 | 0 | 0 | 0 | 0 | 0 | 0 | 0 | 0 | 0 | 0 | 0 | 0 | 0 | 0 | 0 | 0 | 0 | 0 | 0 | 0 | 0 | 0 | 0 | 0 | 0 | 0 | 0 | 0 | 0 | 0 | 0 | 0 | 0 | 0 | 0 | 0 | 0 | 1 | 0 | 0 | 0 | 0 | 0 | 0 | 0 | 0 | 0 | 0 |
| virus_Aug5.50v_28901 | 1 | 0 | 0 | 0 | 0 | 0 | 0 | 0 | 0 | 0 | 0 | 0 | 0 | 0 | 0 | 0 | 0 | 0 | 0 | 0 | 0 | 0 | 0 | 0 | 0 | 0 | 0 | 0 | 0 | 0 | 0 | 0 | 0 | 0 | 0 | 0 | 0 | 0 | 0 | 0 | 0 | 0 | 0 | 0 | 1 | 0 | 0 | 0 | 0 | 0 | 0 | 0 | 0 | 0 | 0 |
| virus_Aug5.50v_31718 | 1 | 0 | 0 | 0 | 0 | 0 | 0 | 0 | 0 | 0 | 0 | 0 | 0 | 0 | 0 | 0 | 0 | 0 | 0 | 0 | 0 | 0 | 0 | 0 | 0 | 0 | 0 | 0 | 0 | 0 | 0 | 0 | 0 | 0 | 0 | 0 | 0 | 0 | 0 | 0 | 0 | 0 | 0 | 0 | 1 | 0 | 0 | 0 | 0 | 0 | 0 | 0 | 0 | 0 | 0 |
| virus_Aug5.50v_35175 | 1 | 0 | 0 | 0 | 0 | 0 | 0 | 0 | 0 | 0 | 0 | 0 | 0 | 0 | 0 | 0 | 0 | 0 | 0 | 0 | 0 | 0 | 0 | 0 | 0 | 0 | 0 | 0 | 0 | 0 | 0 | 0 | 0 | 0 | 0 | 0 | 0 | 0 | 0 | 0 | 0 | 0 | 0 | 0 | 1 | 0 | 0 | 0 | 0 | 0 | 0 | 0 | 0 | 0 | 0 |
| virus_Aug5.75v_7742 | 1 | 0 | 0 | 0 | 0 | 0 | 0 | 0 | 0 | 0 | 0 | 0 | 0 | 0 | 0 | 0 | 0 | 0 | 0 | 0 | 0 | 0 | 0 | 0 | 0 | 0 | 0 | 0 | 0 | 0 | 0 | 0 | 0 | 0 | 0 | 0 | 0 | 0 | 0 | 0 | 0 | 0 | 0 | 0 | 0 | 0 | 1 | 0 | 0 | 0 | 0 | 0 | 0 | 0 | 0 |
| virus_Aug5.75v_15194 | 1 | 0 | 0 | 0 | 0 | 0 | 0 | 0 | 0 | 0 | 0 | 0 | 0 | 0 | 0 | 0 | 0 | 0 | 0 | 0 | 0 | 0 | 0 | 0 | 0 | 0 | 0 | 0 | 0 | 0 | 0 | 0 | 0 | 0 | 0 | 0 | 0 | 0 | 0 | 0 | 0 | 0 | 0 | 0 | 0 | 0 | 1 | 0 | 0 | 0 | 0 | 0 | 0 | 0 | 0 |
| virus_Aug5.75v_16458 | 1 | 0 | 0 | 0 | 0 | 0 | 0 | 0 | 0 | 0 | 0 | 0 | 0 | 0 | 0 | 0 | 0 | 0 | 0 | 0 | 0 | 0 | 0 | 0 | 0 | 0 | 0 | 0 | 0 | 0 | 0 | 0 | 0 | 0 | 0 | 0 | 0 | 0 | 0 | 0 | 0 | 0 | 0 | 0 | 0 | 0 | 1 | 0 | 0 | 0 | 0 | 0 | 0 | 0 | 0 |
| virus_Aug5.75v_26667 | 1 | 0 | 0 | 0 | 0 | 0 | 0 | 0 | 0 | 0 | 0 | 0 | 0 | 0 | 0 | 0 | 0 | 0 | 0 | 0 | 0 | 0 | 0 | 0 | 0 | 0 | 0 | 0 | 0 | 0 | 0 | 0 | 0 | 0 | 0 | 0 | 0 | 0 | 0 | 0 | 0 | 0 | 0 | 0 | 0 | 0 | 1 | 0 | 0 | 0 | 0 | 0 | 0 | 0 | 0 |
| virus_Aug5.75v_31974 | 1 | 0 | 0 | 0 | 0 | 0 | 0 | 0 | 0 | 0 | 0 | 0 | 0 | 0 | 0 | 0 | 0 | 0 | 0 | 0 | 0 | 0 | 0 | 0 | 0 | 0 | 0 | 0 | 0 | 0 | 0 | 0 | 0 | 0 | 0 | 0 | 0 | 0 | 0 | 0 | 0 | 0 | 0 | 0 | 0 | 0 | 1 | 0 | 0 | 0 | 0 | 0 | 0 | 0 | 0 |
| virus_Aug5.75v_32037 | 1 | 0 | 0 | 0 | 0 | 0 | 0 | 0 | 0 | 0 | 0 | 0 | 0 | 0 | 0 | 0 | 0 | 0 | 0 | 0 | 0 | 0 | 0 | 0 | 0 | 0 | 0 | 0 | 0 | 0 | 0 | 0 | 0 | 0 | 0 | 0 | 0 | 0 | 0 | 0 | 0 | 0 | 0 | 0 | 0 | 0 | 1 | 0 | 0 | 0 | 0 | 0 | 0 | 0 | 0 |
| virus_Jul11.0v_1929 | 1 | 0 | 0 | 0 | 0 | 0 | 0 | 0 | 0 | 0 | 0 | 0 | 0 | 0 | 0 | 0 | 0 | 0 | 0 | 0 | 0 | 0 | 1 | 0 | 0 | 0 | 0 | 0 | 0 | 0 | 0 | 0 | 0 | 0 | 0 | 0 | 0 | 0 | 0 | 0 | 0 | 0 | 0 | 0 | 0 | 0 | 0 | 0 | 0 | 0 | 0 | 0 | 0 | 0 | 0 |
| virus_Jul11.0v_3574 | 1 | 0 | 0 | 0 | 0 | 0 | 0 | 0 | 0 | 0 | 0 | 0 | 0 | 0 | 0 | 0 | 0 | 0 | 0 | 0 | 0 | 0 | 1 | 0 | 0 | 0 | 0 | 0 | 0 | 0 | 0 | 0 | 0 | 0 | 0 | 0 | 0 | 0 | 0 | 0 | 0 | 0 | 0 | 0 | 0 | 0 | 0 | 0 | 0 | 0 | 0 | 0 | 0 | 0 | 0 |
| virus_Jul11.0v_3581 | 1 | 0 | 0 | 0 | 0 | 0 | 0 | 0 | 0 | 0 | 0 | 0 | 0 | 0 | 0 | 0 | 0 | 0 | 0 | 0 | 0 | 0 | 1 | 0 | 0 | 0 | 0 | 0 | 0 | 0 | 0 | 0 | 0 | 0 | 0 | 0 | 0 | 0 | 0 | 0 | 0 | 0 | 0 | 0 | 0 | 0 | 0 | 0 | 0 | 0 | 0 | 0 | 0 | 0 | 0 |
| virus_Jul11.0v_10567 | 1 | 0 | 0 | 0 | 0 | 0 | 0 | 0 | 0 | 0 | 0 | 0 | 0 | 0 | 0 | 0 | 0 | 0 | 0 | 0 | 0 | 0 | 1 | 0 | 0 | 0 | 0 | 0 | 0 | 0 | 0 | 0 | 0 | 0 | 0 | 0 | 0 | 0 | 0 | 0 | 0 | 0 | 0 | 0 | 0 | 0 | 0 | 0 | 0 | 0 | 0 | 0 | 0 | 0 | 0 |
| virus_Jul11.0v_14863 | 1 | 0 | 0 | 0 | 0 | 0 | 0 | 0 | 0 | 0 | 0 | 0 | 0 | 0 | 0 | 0 | 0 | 0 | 0 | 0 | 0 | 0 | 1 | 0 | 0 | 0 | 0 | 0 | 0 | 0 | 0 | 0 | 0 | 0 | 0 | 0 | 0 | 0 | 0 | 0 | 0 | 0 | 0 | 0 | 0 | 0 | 0 | 0 | 0 | 0 | 0 | 0 | 0 | 0 | 0 |
| virus_Jul11.0v_19251 | 1 | 0 | 0 | 0 | 0 | 0 | 0 | 0 | 0 | 0 | 0 | 0 | 0 | 0 | 0 | 0 | 0 | 0 | 0 | 0 | 0 | 0 | 1 | 0 | 0 | 0 | 0 | 0 | 0 | 0 | 0 | 0 | 0 | 0 | 0 | 0 | 0 | 0 | 0 | 0 | 0 | 0 | 0 | 0 | 0 | 0 | 0 | 0 | 0 | 0 | 0 | 0 | 0 | 0 | 0 |
| virus_Jul11.0v_20973 | 1 | 0 | 0 | 0 | 0 | 0 | 0 | 0 | 0 | 0 | 0 | 0 | 0 | 0 | 0 | 0 | 0 | 0 | 0 | 0 | 0 | 0 | 1 | 0 | 0 | 0 | 0 | 0 | 0 | 0 | 0 | 0 | 0 | 0 | 0 | 0 | 0 | 0 | 0 | 0 | 0 | 0 | 0 | 0 | 0 | 0 | 0 | 0 | 0 | 0 | 0 | 0 | 0 | 0 | 0 |
| virus_Jul11.5v_3195 | 1 | 0 | 0 | 0 | 0 | 0 | 0 | 0 | 0 | 0 | 0 | 0 | 0 | 0 | 0 | 0 | 0 | 0 | 0 | 0 | 0 | 0 | 0 | 0 | 1 | 0 | 0 | 0 | 0 | 0 | 0 | 0 | 0 | 0 | 0 | 0 | 0 | 0 | 0 | 0 | 0 | 0 | 0 | 0 | 0 | 0 | 0 | 0 | 0 | 0 | 0 | 0 | 0 | 0 | 0 |
| virus_Jul11.5v_6366 | 1 | 0 | 0 | 0 | 0 | 0 | 0 | 0 | 0 | 0 | 0 | 0 | 0 | 0 | 0 | 0 | 0 | 0 | 0 | 0 | 0 | 0 | 0 | 0 | 1 | 0 | 0 | 0 | 0 | 0 | 0 | 0 | 0 | 0 | 0 | 0 | 0 | 0 | 0 | 0 | 0 | 0 | 0 | 0 | 0 | 0 | 0 | 0 | 0 | 0 | 0 | 0 | 0 | 0 | 0 |
| virus_Jul11.5v_8622 | 1 | 0 | 0 | 0 | 0 | 0 | 0 | 0 | 0 | 0 | 0 | 0 | 0 | 0 | 0 | 0 | 0 | 0 | 0 | 0 | 0 | 0 | 0 | 0 | 1 | 0 | 0 | 0 | 0 | 0 | 0 | 0 | 0 | 0 | 0 | 0 | 0 | 0 | 0 | 0 | 0 | 0 | 0 | 0 | 0 | 0 | 0 | 0 | 0 | 0 | 0 | 0 | 0 | 0 | 0 |
| virus_Jul11.5v_9693 | 1 | 0 | 0 | 0 | 0 | 0 | 0 | 0 | 0 | 0 | 0 | 0 | 0 | 0 | 0 | 0 | 0 | 0 | 0 | 0 | 0 | 0 | 0 | 0 | 1 | 0 | 0 | 0 | 0 | 0 | 0 | 0 | 0 | 0 | 0 | 0 | 0 | 0 | 0 | 0 | 0 | 0 | 0 | 0 | 0 | 0 | 0 | 0 | 0 | 0 | 0 | 0 | 0 | 0 | 0 |
| virus_Jul11.5v_10872 | 1 | 0 | 0 | 0 | 0 | 0 | 0 | 0 | 0 | 0 | 0 | 0 | 0 | 0 | 0 | 0 | 0 | 0 | 0 | 0 | 0 | 0 | 0 | 0 | 1 | 0 | 0 | 0 | 0 | 0 | 0 | 0 | 0 | 0 | 0 | 0 | 0 | 0 | 0 | 0 | 0 | 0 | 0 | 0 | 0 | 0 | 0 | 0 | 0 | 0 | 0 | 0 | 0 | 0 | 0 |
| virus_Jul11.5v_12056 | 1 | 0 | 0 | 0 | 0 | 0 | 0 | 0 | 0 | 0 | 0 | 0 | 0 | 0 | 0 | 0 | 0 | 0 | 0 | 0 | 0 | 0 | 0 | 0 | 1 | 0 | 0 | 0 | 0 | 0 | 0 | 0 | 0 | 0 | 0 | 0 | 0 | 0 | 0 | 0 | 0 | 0 | 0 | 0 | 0 | 0 | 0 | 0 | 0 | 0 | 0 | 0 | 0 | 0 | 0 |
| virus_Jul11.5v_12999 | 1 | 0 | 0 | 0 | 0 | 0 | 0 | 0 | 0 | 0 | 0 | 0 | 0 | 0 | 0 | 0 | 0 | 0 | 0 | 0 | 0 | 0 | 0 | 0 | 1 | 0 | 0 | 0 | 0 | 0 | 0 | 0 | 0 | 0 | 0 | 0 | 0 | 0 | 0 | 0 | 0 | 0 | 0 | 0 | 0 | 0 | 0 | 0 | 0 | 0 | 0 | 0 | 0 | 0 | 0 |
| virus_Jul11.5v_15655 | 1 | 0 | 0 | 0 | 0 | 0 | 0 | 0 | 0 | 0 | 0 | 0 | 0 | 0 | 0 | 0 | 0 | 0 | 0 | 0 | 0 | 0 | 0 | 0 | 1 | 0 | 0 | 0 | 0 | 0 | 0 | 0 | 0 | 0 | 0 | 0 | 0 | 0 | 0 | 0 | 0 | 0 | 0 | 0 | 0 | 0 | 0 | 0 | 0 | 0 | 0 | 0 | 0 | 0 | 0 |
| virus_Jul11.5v_18759 | 1 | 0 | 0 | 0 | 0 | 0 | 0 | 0 | 0 | 0 | 0 | 0 | 0 | 0 | 0 | 0 | 0 | 0 | 0 | 0 | 0 | 0 | 0 | 0 | 1 | 0 | 0 | 0 | 0 | 0 | 0 | 0 | 0 | 0 | 0 | 0 | 0 | 0 | 0 | 0 | 0 | 0 | 0 | 0 | 0 | 0 | 0 | 0 | 0 | 0 | 0 | 0 | 0 | 0 | 0 |
| virus_Jul11.5v_19132 | 1 | 0 | 0 | 0 | 0 | 0 | 0 | 0 | 0 | 0 | 0 | 0 | 0 | 0 | 0 | 0 | 0 | 0 | 0 | 0 | 0 | 0 | 0 | 0 | 1 | 0 | 0 | 0 | 0 | 0 | 0 | 0 | 0 | 0 | 0 | 0 | 0 | 0 | 0 | 0 | 0 | 0 | 0 | 0 | 0 | 0 | 0 | 0 | 0 | 0 | 0 | 0 | 0 | 0 | 0 |
| virus_Jul11.25v_858 | 1 | 0 | 0 | 0 | 0 | 0 | 0 | 0 | 0 | 0 | 0 | 0 | 0 | 0 | 0 | 0 | 0 | 0 | 0 | 0 | 0 | 0 | 0 | 0 | 0 | 0 | 1 | 0 | 0 | 0 | 0 | 0 | 0 | 0 | 0 | 0 | 0 | 0 | 0 | 0 | 0 | 0 | 0 | 0 | 0 | 0 | 0 | 0 | 0 | 0 | 0 | 0 | 0 | 0 | 0 |
| virus_Jul11.25v_4155 | 1 | 0 | 0 | 0 | 0 | 0 | 0 | 0 | 0 | 0 | 0 | 0 | 0 | 0 | 0 | 0 | 0 | 0 | 0 | 0 | 0 | 0 | 0 | 0 | 0 | 0 | 1 | 0 | 0 | 0 | 0 | 0 | 0 | 0 | 0 | 0 | 0 | 0 | 0 | 0 | 0 | 0 | 0 | 0 | 0 | 0 | 0 | 0 | 0 | 0 | 0 | 0 | 0 | 0 | 0 |
| virus_Jul11.25v_7357 | 1 | 0 | 0 | 0 | 0 | 0 | 0 | 0 | 0 | 0 | 0 | 0 | 0 | 0 | 0 | 0 | 0 | 0 | 0 | 0 | 0 | 0 | 0 | 0 | 0 | 0 | 1 | 0 | 0 | 0 | 0 | 0 | 0 | 0 | 0 | 0 | 0 | 0 | 0 | 0 | 0 | 0 | 0 | 0 | 0 | 0 | 0 | 0 | 0 | 0 | 0 | 0 | 0 | 0 | 0 |
| virus_Jul11.25v_9931 | 1 | 0 | 0 | 0 | 0 | 0 | 0 | 0 | 0 | 0 | 0 | 0 | 0 | 0 | 0 | 0 | 0 | 0 | 0 | 0 | 0 | 0 | 0 | 0 | 0 | 0 | 1 | 0 | 0 | 0 | 0 | 0 | 0 | 0 | 0 | 0 | 0 | 0 | 0 | 0 | 0 | 0 | 0 | 0 | 0 | 0 | 0 | 0 | 0 | 0 | 0 | 0 | 0 | 0 | 0 |
| virus_Jul11.25v_14451 | 1 | 0 | 0 | 0 | 0 | 0 | 0 | 0 | 0 | 0 | 0 | 0 | 0 | 0 | 0 | 0 | 0 | 0 | 0 | 0 | 0 | 0 | 0 | 0 | 0 | 0 | 1 | 0 | 0 | 0 | 0 | 0 | 0 | 0 | 0 | 0 | 0 | 0 | 0 | 0 | 0 | 0 | 0 | 0 | 0 | 0 | 0 | 0 | 0 | 0 | 0 | 0 | 0 | 0 | 0 |
| virus_Jul11.25v_15347 | 1 | 0 | 0 | 0 | 0 | 0 | 0 | 0 | 0 | 0 | 0 | 0 | 0 | 0 | 0 | 0 | 0 | 0 | 0 | 0 | 0 | 0 | 0 | 0 | 0 | 0 | 1 | 0 | 0 | 0 | 0 | 0 | 0 | 0 | 0 | 0 | 0 | 0 | 0 | 0 | 0 | 0 | 0 | 0 | 0 | 0 | 0 | 0 | 0 | 0 | 0 | 0 | 0 | 0 | 0 |
| virus_Jul11.25v_17097 | 1 | 0 | 0 | 0 | 0 | 0 | 0 | 0 | 0 | 0 | 0 | 0 | 0 | 0 | 0 | 0 | 0 | 0 | 0 | 0 | 0 | 0 | 0 | 0 | 0 | 0 | 1 | 0 | 0 | 0 | 0 | 0 | 0 | 0 | 0 | 0 | 0 | 0 | 0 | 0 | 0 | 0 | 0 | 0 | 0 | 0 | 0 | 0 | 0 | 0 | 0 | 0 | 0 | 0 | 0 |
| virus_Jul11.25v_18940 | 1 | 0 | 0 | 0 | 0 | 0 | 0 | 0 | 0 | 0 | 0 | 0 | 0 | 0 | 0 | 0 | 0 | 0 | 0 | 0 | 0 | 0 | 0 | 0 | 0 | 0 | 1 | 0 | 0 | 0 | 0 | 0 | 0 | 0 | 0 | 0 | 0 | 0 | 0 | 0 | 0 | 0 | 0 | 0 | 0 | 0 | 0 | 0 | 0 | 0 | 0 | 0 | 0 | 0 | 0 |
| virus_Jul11.25v_19468 | 1 | 0 | 0 | 0 | 0 | 0 | 0 | 0 | 0 | 0 | 0 | 0 | 0 | 0 | 0 | 0 | 0 | 0 | 0 | 0 | 0 | 0 | 0 | 0 | 0 | 0 | 1 | 0 | 0 | 0 | 0 | 0 | 0 | 0 | 0 | 0 | 0 | 0 | 0 | 0 | 0 | 0 | 0 | 0 | 0 | 0 | 0 | 0 | 0 | 0 | 0 | 0 | 0 | 0 | 0 |
| virus_Jul11.50v_514 | 1 | 0 | 0 | 0 | 0 | 0 | 0 | 0 | 0 | 0 | 0 | 0 | 0 | 0 | 0 | 0 | 0 | 0 | 0 | 0 | 0 | 0 | 0 | 0 | 0 | 0 | 0 | 0 | 1 | 0 | 0 | 0 | 0 | 0 | 0 | 0 | 0 | 0 | 0 | 0 | 0 | 0 | 0 | 0 | 0 | 0 | 0 | 0 | 0 | 0 | 0 | 0 | 0 | 0 | 0 |
| virus_Jul11.50v_4248 | 1 | 0 | 0 | 0 | 0 | 0 | 0 | 0 | 0 | 0 | 0 | 0 | 0 | 0 | 0 | 0 | 0 | 0 | 0 | 0 | 0 | 0 | 0 | 0 | 0 | 0 | 0 | 0 | 1 | 0 | 0 | 0 | 0 | 0 | 0 | 0 | 0 | 0 | 0 | 0 | 0 | 0 | 0 | 0 | 0 | 0 | 0 | 0 | 0 | 0 | 0 | 0 | 0 | 0 | 0 |
| virus_Jul11.50v_4694 | 1 | 0 | 0 | 0 | 0 | 0 | 0 | 0 | 0 | 0 | 0 | 0 | 0 | 0 | 0 | 0 | 0 | 0 | 0 | 0 | 0 | 0 | 0 | 0 | 0 | 0 | 0 | 0 | 1 | 0 | 0 | 0 | 0 | 0 | 0 | 0 | 0 | 0 | 0 | 0 | 0 | 0 | 0 | 0 | 0 | 0 | 0 | 0 | 0 | 0 | 0 | 0 | 0 | 0 | 0 |
| virus_Jul11.50v_6169 | 1 | 0 | 0 | 0 | 0 | 0 | 0 | 0 | 0 | 0 | 0 | 0 | 0 | 0 | 0 | 0 | 0 | 0 | 0 | 0 | 0 | 0 | 0 | 0 | 0 | 0 | 0 | 0 | 1 | 0 | 0 | 0 | 0 | 0 | 0 | 0 | 0 | 0 | 0 | 0 | 0 | 0 | 0 | 0 | 0 | 0 | 0 | 0 | 0 | 0 | 0 | 0 | 0 | 0 | 0 |
| virus_Jul11.50v_6448 | 1 | 0 | 0 | 0 | 0 | 0 | 0 | 0 | 0 | 0 | 0 | 0 | 0 | 0 | 0 | 0 | 0 | 0 | 0 | 0 | 0 | 0 | 0 | 0 | 0 | 0 | 0 | 0 | 1 | 0 | 0 | 0 | 0 | 0 | 0 | 0 | 0 | 0 | 0 | 0 | 0 | 0 | 0 | 0 | 0 | 0 | 0 | 0 | 0 | 0 | 0 | 0 | 0 | 0 | 0 |
| virus_Jul11.50v_7910 | 1 | 0 | 0 | 0 | 0 | 0 | 0 | 0 | 0 | 0 | 0 | 0 | 0 | 0 | 0 | 0 | 0 | 0 | 0 | 0 | 0 | 0 | 0 | 0 | 0 | 0 | 0 | 0 | 1 | 0 | 0 | 0 | 0 | 0 | 0 | 0 | 0 | 0 | 0 | 0 | 0 | 0 | 0 | 0 | 0 | 0 | 0 | 0 | 0 | 0 | 0 | 0 | 0 | 0 | 0 |
| virus_Jul11.50v_8038 | 1 | 0 | 0 | 0 | 0 | 0 | 0 | 0 | 0 | 0 | 0 | 0 | 0 | 0 | 0 | 0 | 0 | 0 | 0 | 0 | 0 | 0 | 0 | 0 | 0 | 0 | 0 | 0 | 1 | 0 | 0 | 0 | 0 | 0 | 0 | 0 | 0 | 0 | 0 | 0 | 0 | 0 | 0 | 0 | 0 | 0 | 0 | 0 | 0 | 0 | 0 | 0 | 0 | 0 | 0 |
| virus_Jul11.50v_8950 | 1 | 0 | 0 | 0 | 0 | 0 | 0 | 0 | 0 | 0 | 0 | 0 | 0 | 0 | 0 | 0 | 0 | 0 | 0 | 0 | 0 | 0 | 0 | 0 | 0 | 0 | 0 | 0 | 1 | 0 | 0 | 0 | 0 | 0 | 0 | 0 | 0 | 0 | 0 | 0 | 0 | 0 | 0 | 0 | 0 | 0 | 0 | 0 | 0 | 0 | 0 | 0 | 0 | 0 | 0 |
| virus_Jul11.50v_10028 | 1 | 0 | 0 | 0 | 0 | 0 | 0 | 0 | 0 | 0 | 0 | 0 | 0 | 0 | 0 | 0 | 0 | 0 | 0 | 0 | 0 | 0 | 0 | 0 | 0 | 0 | 0 | 0 | 1 | 0 | 0 | 0 | 0 | 0 | 0 | 0 | 0 | 0 | 0 | 0 | 0 | 0 | 0 | 0 | 0 | 0 | 0 | 0 | 0 | 0 | 0 | 0 | 0 | 0 | 0 |
| virus_Jul11.50v_11322 | 1 | 0 | 0 | 0 | 0 | 0 | 0 | 0 | 0 | 0 | 0 | 0 | 0 | 0 | 0 | 0 | 0 | 0 | 0 | 0 | 0 | 0 | 0 | 0 | 0 | 0 | 0 | 0 | 1 | 0 | 0 | 0 | 0 | 0 | 0 | 0 | 0 | 0 | 0 | 0 | 0 | 0 | 0 | 0 | 0 | 0 | 0 | 0 | 0 | 0 | 0 | 0 | 0 | 0 | 0 |
| virus_Jul11.50v_12821 | 1 | 0 | 0 | 0 | 0 | 0 | 0 | 0 | 0 | 0 | 0 | 0 | 0 | 0 | 0 | 0 | 0 | 0 | 0 | 0 | 0 | 0 | 0 | 0 | 0 | 0 | 0 | 0 | 1 | 0 | 0 | 0 | 0 | 0 | 0 | 0 | 0 | 0 | 0 | 0 | 0 | 0 | 0 | 0 | 0 | 0 | 0 | 0 | 0 | 0 | 0 | 0 | 0 | 0 | 0 |
| virus_Jul11.50v_14197 | 1 | 0 | 0 | 0 | 0 | 0 | 0 | 0 | 0 | 0 | 0 | 0 | 0 | 0 | 0 | 0 | 0 | 0 | 0 | 0 | 0 | 0 | 0 | 0 | 0 | 0 | 0 | 0 | 1 | 0 | 0 | 0 | 0 | 0 | 0 | 0 | 0 | 0 | 0 | 0 | 0 | 0 | 0 | 0 | 0 | 0 | 0 | 0 | 0 | 0 | 0 | 0 | 0 | 0 | 0 |
| virus_Jul11.50v_14384 | 1 | 0 | 0 | 0 | 0 | 0 | 0 | 0 | 0 | 0 | 0 | 0 | 0 | 0 | 0 | 0 | 0 | 0 | 0 | 0 | 0 | 0 | 0 | 0 | 0 | 0 | 0 | 0 | 1 | 0 | 0 | 0 | 0 | 0 | 0 | 0 | 0 | 0 | 0 | 0 | 0 | 0 | 0 | 0 | 0 | 0 | 0 | 0 | 0 | 0 | 0 | 0 | 0 | 0 | 0 |
| virus_Jul11.50v_18584 | 1 | 0 | 0 | 0 | 0 | 0 | 0 | 0 | 0 | 0 | 0 | 0 | 0 | 0 | 0 | 0 | 0 | 0 | 0 | 0 | 0 | 0 | 0 | 0 | 0 | 0 | 0 | 0 | 1 | 0 | 0 | 0 | 0 | 0 | 0 | 0 | 0 | 0 | 0 | 0 | 0 | 0 | 0 | 0 | 0 | 0 | 0 | 0 | 0 | 0 | 0 | 0 | 0 | 0 | 0 |
| virus_Jul11.50v_18898 | 1 | 0 | 0 | 0 | 0 | 0 | 0 | 0 | 0 | 0 | 0 | 0 | 0 | 0 | 0 | 0 | 0 | 0 | 0 | 0 | 0 | 0 | 0 | 0 | 0 | 0 | 0 | 0 | 1 | 0 | 0 | 0 | 0 | 0 | 0 | 0 | 0 | 0 | 0 | 0 | 0 | 0 | 0 | 0 | 0 | 0 | 0 | 0 | 0 | 0 | 0 | 0 | 0 | 0 | 0 |
| virus_Jul11.75v_11827 | 1 | 0 | 0 | 0 | 0 | 0 | 0 | 0 | 0 | 0 | 0 | 0 | 0 | 0 | 0 | 0 | 0 | 0 | 0 | 0 | 0 | 0 | 0 | 0 | 0 | 0 | 0 | 0 | 0 | 0 | 1 | 0 | 0 | 0 | 0 | 0 | 0 | 0 | 0 | 0 | 0 | 0 | 0 | 0 | 0 | 0 | 0 | 0 | 0 | 0 | 0 | 0 | 0 | 0 | 0 |
| virus_Jul11.75v_18794 | 1 | 0 | 0 | 0 | 0 | 0 | 0 | 0 | 0 | 0 | 0 | 0 | 0 | 0 | 0 | 0 | 0 | 0 | 0 | 0 | 0 | 0 | 0 | 0 | 0 | 0 | 0 | 0 | 0 | 0 | 1 | 0 | 0 | 0 | 0 | 0 | 0 | 0 | 0 | 0 | 0 | 0 | 0 | 0 | 0 | 0 | 0 | 0 | 0 | 0 | 0 | 0 | 0 | 0 | 0 |
| virus_Jul11.75v_25365 | 1 | 0 | 0 | 0 | 0 | 0 | 0 | 0 | 0 | 0 | 0 | 0 | 0 | 0 | 0 | 0 | 0 | 0 | 0 | 0 | 0 | 0 | 0 | 0 | 0 | 0 | 0 | 0 | 0 | 0 | 1 | 0 | 0 | 0 | 0 | 0 | 0 | 0 | 0 | 0 | 0 | 0 | 0 | 0 | 0 | 0 | 0 | 0 | 0 | 0 | 0 | 0 | 0 | 0 | 0 |
| virus_Jul11.75v_26150 | 1 | 0 | 0 | 0 | 0 | 0 | 0 | 0 | 0 | 0 | 0 | 0 | 0 | 0 | 0 | 0 | 0 | 0 | 0 | 0 | 0 | 0 | 0 | 0 | 0 | 0 | 0 | 0 | 0 | 0 | 1 | 0 | 0 | 0 | 0 | 0 | 0 | 0 | 0 | 0 | 0 | 0 | 0 | 0 | 0 | 0 | 0 | 0 | 0 | 0 | 0 | 0 | 0 | 0 | 0 |
| virus_Jul11.75v_26801 | 1 | 0 | 0 | 0 | 0 | 0 | 0 | 0 | 0 | 0 | 0 | 0 | 0 | 0 | 0 | 0 | 0 | 0 | 0 | 0 | 0 | 0 | 0 | 0 | 0 | 0 | 0 | 0 | 0 | 0 | 1 | 0 | 0 | 0 | 0 | 0 | 0 | 0 | 0 | 0 | 0 | 0 | 0 | 0 | 0 | 0 | 0 | 0 | 0 | 0 | 0 | 0 | 0 | 0 | 0 |
| virus_Jul11.75v_36647 | 1 | 0 | 0 | 0 | 0 | 0 | 0 | 0 | 0 | 0 | 0 | 0 | 0 | 0 | 0 | 0 | 0 | 0 | 0 | 0 | 0 | 0 | 0 | 0 | 0 | 0 | 0 | 0 | 0 | 0 | 1 | 0 | 0 | 0 | 0 | 0 | 0 | 0 | 0 | 0 | 0 | 0 | 0 | 0 | 0 | 0 | 0 | 0 | 0 | 0 | 0 | 0 | 0 | 0 | 0 |
| virus_Jul27.5v_37 | 1 | 0 | 0 | 0 | 0 | 0 | 0 | 0 | 0 | 0 | 0 | 0 | 0 | 0 | 0 | 0 | 0 | 0 | 0 | 0 | 0 | 0 | 0 | 0 | 0 | 0 | 0 | 0 | 0 | 0 | 0 | 0 | 1 | 0 | 0 | 0 | 0 | 0 | 0 | 0 | 0 | 0 | 0 | 0 | 0 | 0 | 0 | 0 | 0 | 0 | 0 | 0 | 0 | 0 | 0 |
| virus_Jul27.5v_186 | 1 | 0 | 0 | 0 | 0 | 0 | 0 | 0 | 0 | 0 | 0 | 0 | 0 | 0 | 0 | 0 | 0 | 0 | 0 | 0 | 0 | 0 | 0 | 0 | 0 | 0 | 0 | 0 | 0 | 0 | 0 | 0 | 1 | 0 | 0 | 0 | 0 | 0 | 0 | 0 | 0 | 0 | 0 | 0 | 0 | 0 | 0 | 0 | 0 | 0 | 0 | 0 | 0 | 0 | 0 |
| virus_Jul27.5v_1242 | 1 | 0 | 0 | 0 | 0 | 0 | 0 | 0 | 0 | 0 | 0 | 0 | 0 | 0 | 0 | 0 | 0 | 0 | 0 | 0 | 0 | 0 | 0 | 0 | 0 | 0 | 0 | 0 | 0 | 0 | 0 | 0 | 1 | 0 | 0 | 0 | 0 | 0 | 0 | 0 | 0 | 0 | 0 | 0 | 0 | 0 | 0 | 0 | 0 | 0 | 0 | 0 | 0 | 0 | 0 |
| virus_Jul27.5v_2891 | 1 | 0 | 0 | 0 | 0 | 0 | 0 | 0 | 0 | 0 | 0 | 0 | 0 | 0 | 0 | 0 | 0 | 0 | 0 | 0 | 0 | 0 | 0 | 0 | 0 | 0 | 0 | 0 | 0 | 0 | 0 | 0 | 1 | 0 | 0 | 0 | 0 | 0 | 0 | 0 | 0 | 0 | 0 | 0 | 0 | 0 | 0 | 0 | 0 | 0 | 0 | 0 | 0 | 0 | 0 |
| virus_Jul27.5v_3224 | 1 | 0 | 0 | 0 | 0 | 0 | 0 | 0 | 0 | 0 | 0 | 0 | 0 | 0 | 0 | 0 | 0 | 0 | 0 | 0 | 0 | 0 | 0 | 0 | 0 | 0 | 0 | 0 | 0 | 0 | 0 | 0 | 1 | 0 | 0 | 0 | 0 | 0 | 0 | 0 | 0 | 0 | 0 | 0 | 0 | 0 | 0 | 0 | 0 | 0 | 0 | 0 | 0 | 0 | 0 |
| virus_Jul27.5v_3497 | 1 | 0 | 0 | 0 | 0 | 0 | 0 | 0 | 0 | 0 | 0 | 0 | 0 | 0 | 0 | 0 | 0 | 0 | 0 | 0 | 0 | 0 | 0 | 0 | 0 | 0 | 0 | 0 | 0 | 0 | 0 | 0 | 1 | 0 | 0 | 0 | 0 | 0 | 0 | 0 | 0 | 0 | 0 | 0 | 0 | 0 | 0 | 0 | 0 | 0 | 0 | 0 | 0 | 0 | 0 |
| virus_Jul27.5v_3834 | 1 | 0 | 0 | 0 | 0 | 0 | 0 | 0 | 0 | 0 | 0 | 0 | 0 | 0 | 0 | 0 | 0 | 0 | 0 | 0 | 0 | 0 | 0 | 0 | 0 | 0 | 0 | 0 | 0 | 0 | 0 | 0 | 1 | 0 | 0 | 0 | 0 | 0 | 0 | 0 | 0 | 0 | 0 | 0 | 0 | 0 | 0 | 0 | 0 | 0 | 0 | 0 | 0 | 0 | 0 |
| virus_Jul27.5v_5273 | 1 | 0 | 0 | 0 | 0 | 0 | 0 | 0 | 0 | 0 | 0 | 0 | 0 | 0 | 0 | 0 | 0 | 0 | 0 | 0 | 0 | 0 | 0 | 0 | 0 | 0 | 0 | 0 | 0 | 0 | 0 | 0 | 1 | 0 | 0 | 0 | 0 | 0 | 0 | 0 | 0 | 0 | 0 | 0 | 0 | 0 | 0 | 0 | 0 | 0 | 0 | 0 | 0 | 0 | 0 |
| virus_Jul27.5v_6098 | 1 | 0 | 0 | 0 | 0 | 0 | 0 | 0 | 0 | 0 | 0 | 0 | 0 | 0 | 0 | 0 | 0 | 0 | 0 | 0 | 0 | 0 | 0 | 0 | 0 | 0 | 0 | 0 | 0 | 0 | 0 | 0 | 1 | 0 | 0 | 0 | 0 | 0 | 0 | 0 | 0 | 0 | 0 | 0 | 0 | 0 | 0 | 0 | 0 | 0 | 0 | 0 | 0 | 0 | 0 |
| virus_Jul27.5v_9291 | 1 | 0 | 0 | 0 | 0 | 0 | 0 | 0 | 0 | 0 | 0 | 0 | 0 | 0 | 0 | 0 | 0 | 0 | 0 | 0 | 0 | 0 | 0 | 0 | 0 | 0 | 0 | 0 | 0 | 0 | 0 | 0 | 1 | 0 | 0 | 0 | 0 | 0 | 0 | 0 | 0 | 0 | 0 | 0 | 0 | 0 | 0 | 0 | 0 | 0 | 0 | 0 | 0 | 0 | 0 |
| virus_Jul27.5v_10821 | 1 | 0 | 0 | 0 | 0 | 0 | 0 | 0 | 0 | 0 | 0 | 0 | 0 | 0 | 0 | 0 | 0 | 0 | 0 | 0 | 0 | 0 | 0 | 0 | 0 | 0 | 0 | 0 | 0 | 0 | 0 | 0 | 1 | 0 | 0 | 0 | 0 | 0 | 0 | 0 | 0 | 0 | 0 | 0 | 0 | 0 | 0 | 0 | 0 | 0 | 0 | 0 | 0 | 0 | 0 |
| virus_Jul27.5v_11134 | 1 | 0 | 0 | 0 | 0 | 0 | 0 | 0 | 0 | 0 | 0 | 0 | 0 | 0 | 0 | 0 | 0 | 0 | 0 | 0 | 0 | 0 | 0 | 0 | 0 | 0 | 0 | 0 | 0 | 0 | 0 | 0 | 1 | 0 | 0 | 0 | 0 | 0 | 0 | 0 | 0 | 0 | 0 | 0 | 0 | 0 | 0 | 0 | 0 | 0 | 0 | 0 | 0 | 0 | 0 |
| virus_Jul27.5v_11482 | 1 | 0 | 0 | 0 | 0 | 0 | 0 | 0 | 0 | 0 | 0 | 0 | 0 | 0 | 0 | 0 | 0 | 0 | 0 | 0 | 0 | 0 | 0 | 0 | 0 | 0 | 0 | 0 | 0 | 0 | 0 | 0 | 1 | 0 | 0 | 0 | 0 | 0 | 0 | 0 | 0 | 0 | 0 | 0 | 0 | 0 | 0 | 0 | 0 | 0 | 0 | 0 | 0 | 0 | 0 |
| virus_Jul27.5v_12822 | 1 | 0 | 0 | 0 | 0 | 0 | 0 | 0 | 0 | 0 | 0 | 0 | 0 | 0 | 0 | 0 | 0 | 0 | 0 | 0 | 0 | 0 | 0 | 0 | 0 | 0 | 0 | 0 | 0 | 0 | 0 | 0 | 1 | 0 | 0 | 0 | 0 | 0 | 0 | 0 | 0 | 0 | 0 | 0 | 0 | 0 | 0 | 0 | 0 | 0 | 0 | 0 | 0 | 0 | 0 |
| virus_Jul27.5v_12864 | 1 | 0 | 0 | 0 | 0 | 0 | 0 | 0 | 0 | 0 | 0 | 0 | 0 | 0 | 0 | 0 | 0 | 0 | 0 | 0 | 0 | 0 | 0 | 0 | 0 | 0 | 0 | 0 | 0 | 0 | 0 | 0 | 1 | 0 | 0 | 0 | 0 | 0 | 0 | 0 | 0 | 0 | 0 | 0 | 0 | 0 | 0 | 0 | 0 | 0 | 0 | 0 | 0 | 0 | 0 |
| virus_Jul27.5v_13223 | 1 | 0 | 0 | 0 | 0 | 0 | 0 | 0 | 0 | 0 | 0 | 0 | 0 | 0 | 0 | 0 | 0 | 0 | 0 | 0 | 0 | 0 | 0 | 0 | 0 | 0 | 0 | 0 | 0 | 0 | 0 | 0 | 1 | 0 | 0 | 0 | 0 | 0 | 0 | 0 | 0 | 0 | 0 | 0 | 0 | 0 | 0 | 0 | 0 | 0 | 0 | 0 | 0 | 0 | 0 |
| virus_Jul27.5v_13230 | 1 | 0 | 0 | 0 | 0 | 0 | 0 | 0 | 0 | 0 | 0 | 0 | 0 | 0 | 0 | 0 | 0 | 0 | 0 | 0 | 0 | 0 | 0 | 0 | 0 | 0 | 0 | 0 | 0 | 0 | 0 | 0 | 1 | 0 | 0 | 0 | 0 | 0 | 0 | 0 | 0 | 0 | 0 | 0 | 0 | 0 | 0 | 0 | 0 | 0 | 0 | 0 | 0 | 0 | 0 |
| virus_Jul27.5v_14174 | 1 | 0 | 0 | 0 | 0 | 0 | 0 | 0 | 0 | 0 | 0 | 0 | 0 | 0 | 0 | 0 | 0 | 0 | 0 | 0 | 0 | 0 | 0 | 0 | 0 | 0 | 0 | 0 | 0 | 0 | 0 | 0 | 1 | 0 | 0 | 0 | 0 | 0 | 0 | 0 | 0 | 0 | 0 | 0 | 0 | 0 | 0 | 0 | 0 | 0 | 0 | 0 | 0 | 0 | 0 |
| virus_Jul27.5v_17154 | 1 | 0 | 0 | 0 | 0 | 0 | 0 | 0 | 0 | 0 | 0 | 0 | 0 | 0 | 0 | 0 | 0 | 0 | 0 | 0 | 0 | 0 | 0 | 0 | 0 | 0 | 0 | 0 | 0 | 0 | 0 | 0 | 1 | 0 | 0 | 0 | 0 | 0 | 0 | 0 | 0 | 0 | 0 | 0 | 0 | 0 | 0 | 0 | 0 | 0 | 0 | 0 | 0 | 0 | 0 |
| virus_Jul27.5v_20216 | 1 | 0 | 0 | 0 | 0 | 0 | 0 | 0 | 0 | 0 | 0 | 0 | 0 | 0 | 0 | 0 | 0 | 0 | 0 | 0 | 0 | 0 | 0 | 0 | 0 | 0 | 0 | 0 | 0 | 0 | 0 | 0 | 1 | 0 | 0 | 0 | 0 | 0 | 0 | 0 | 0 | 0 | 0 | 0 | 0 | 0 | 0 | 0 | 0 | 0 | 0 | 0 | 0 | 0 | 0 |
| virus_Jul27.5v_21090 | 1 | 0 | 0 | 0 | 0 | 0 | 0 | 0 | 0 | 0 | 0 | 0 | 0 | 0 | 0 | 0 | 0 | 0 | 0 | 0 | 0 | 0 | 0 | 0 | 0 | 0 | 0 | 0 | 0 | 0 | 0 | 0 | 1 | 0 | 0 | 0 | 0 | 0 | 0 | 0 | 0 | 0 | 0 | 0 | 0 | 0 | 0 | 0 | 0 | 0 | 0 | 0 | 0 | 0 | 0 |
| virus_Jul27.25v_2860 | 1 | 0 | 0 | 0 | 0 | 0 | 0 | 0 | 0 | 0 | 0 | 0 | 0 | 0 | 0 | 0 | 0 | 0 | 0 | 0 | 0 | 0 | 0 | 0 | 0 | 0 | 0 | 0 | 0 | 0 | 0 | 0 | 0 | 0 | 1 | 0 | 0 | 0 | 0 | 0 | 0 | 0 | 0 | 0 | 0 | 0 | 0 | 0 | 0 | 0 | 0 | 0 | 0 | 0 | 0 |
| virus_Jul27.25v_5587 | 1 | 0 | 0 | 0 | 0 | 0 | 0 | 0 | 0 | 0 | 0 | 0 | 0 | 0 | 0 | 0 | 0 | 0 | 0 | 0 | 0 | 0 | 0 | 0 | 0 | 0 | 0 | 0 | 0 | 0 | 0 | 0 | 0 | 0 | 1 | 0 | 0 | 0 | 0 | 0 | 0 | 0 | 0 | 0 | 0 | 0 | 0 | 0 | 0 | 0 | 0 | 0 | 0 | 0 | 0 |
| virus_Jul27.25v_13389 | 1 | 0 | 0 | 0 | 0 | 0 | 0 | 0 | 0 | 0 | 0 | 0 | 0 | 0 | 0 | 0 | 0 | 0 | 0 | 0 | 0 | 0 | 0 | 0 | 0 | 0 | 0 | 0 | 0 | 0 | 0 | 0 | 0 | 0 | 1 | 0 | 0 | 0 | 0 | 0 | 0 | 0 | 0 | 0 | 0 | 0 | 0 | 0 | 0 | 0 | 0 | 0 | 0 | 0 | 0 |
| virus_Jul27.25v_14334 | 1 | 0 | 0 | 0 | 0 | 0 | 0 | 0 | 0 | 0 | 0 | 0 | 0 | 0 | 0 | 0 | 0 | 0 | 0 | 0 | 0 | 0 | 0 | 0 | 0 | 0 | 0 | 0 | 0 | 0 | 0 | 0 | 0 | 0 | 1 | 0 | 0 | 0 | 0 | 0 | 0 | 0 | 0 | 0 | 0 | 0 | 0 | 0 | 0 | 0 | 0 | 0 | 0 | 0 | 0 |
| virus_Jul27.25v_21311 | 1 | 0 | 0 | 0 | 0 | 0 | 0 | 0 | 0 | 0 | 0 | 0 | 0 | 0 | 0 | 0 | 0 | 0 | 0 | 0 | 0 | 0 | 0 | 0 | 0 | 0 | 0 | 0 | 0 | 0 | 0 | 0 | 0 | 0 | 1 | 0 | 0 | 0 | 0 | 0 | 0 | 0 | 0 | 0 | 0 | 0 | 0 | 0 | 0 | 0 | 0 | 0 | 0 | 0 | 0 |
| virus_Jul27.25v_21411 | 1 | 0 | 0 | 0 | 0 | 0 | 0 | 0 | 0 | 0 | 0 | 0 | 0 | 0 | 0 | 0 | 0 | 0 | 0 | 0 | 0 | 0 | 0 | 0 | 0 | 0 | 0 | 0 | 0 | 0 | 0 | 0 | 0 | 0 | 1 | 0 | 0 | 0 | 0 | 0 | 0 | 0 | 0 | 0 | 0 | 0 | 0 | 0 | 0 | 0 | 0 | 0 | 0 | 0 | 0 |
| virus_Jul27.25v_22225 | 1 | 0 | 0 | 0 | 0 | 0 | 0 | 0 | 0 | 0 | 0 | 0 | 0 | 0 | 0 | 0 | 0 | 0 | 0 | 0 | 0 | 0 | 0 | 0 | 0 | 0 | 0 | 0 | 0 | 0 | 0 | 0 | 0 | 0 | 1 | 0 | 0 | 0 | 0 | 0 | 0 | 0 | 0 | 0 | 0 | 0 | 0 | 0 | 0 | 0 | 0 | 0 | 0 | 0 | 0 |
| virus_Jul27.25v_26383 | 1 | 0 | 0 | 0 | 0 | 0 | 0 | 0 | 0 | 0 | 0 | 0 | 0 | 0 | 0 | 0 | 0 | 0 | 0 | 0 | 0 | 0 | 0 | 0 | 0 | 0 | 0 | 0 | 0 | 0 | 0 | 0 | 0 | 0 | 1 | 0 | 0 | 0 | 0 | 0 | 0 | 0 | 0 | 0 | 0 | 0 | 0 | 0 | 0 | 0 | 0 | 0 | 0 | 0 | 0 |
| virus_Jul27.25v_28250 | 1 | 0 | 0 | 0 | 0 | 0 | 0 | 0 | 0 | 0 | 0 | 0 | 0 | 0 | 0 | 0 | 0 | 0 | 0 | 0 | 0 | 0 | 0 | 0 | 0 | 0 | 0 | 0 | 0 | 0 | 0 | 0 | 0 | 0 | 1 | 0 | 0 | 0 | 0 | 0 | 0 | 0 | 0 | 0 | 0 | 0 | 0 | 0 | 0 | 0 | 0 | 0 | 0 | 0 | 0 |
| virus_Jul27.25v_29812 | 1 | 0 | 0 | 0 | 0 | 0 | 0 | 0 | 0 | 0 | 0 | 0 | 0 | 0 | 0 | 0 | 0 | 0 | 0 | 0 | 0 | 0 | 0 | 0 | 0 | 0 | 0 | 0 | 0 | 0 | 0 | 0 | 0 | 0 | 1 | 0 | 0 | 0 | 0 | 0 | 0 | 0 | 0 | 0 | 0 | 0 | 0 | 0 | 0 | 0 | 0 | 0 | 0 | 0 | 0 |
| virus_Jul27.25v_34443 | 1 | 0 | 0 | 0 | 0 | 0 | 0 | 0 | 0 | 0 | 0 | 0 | 0 | 0 | 0 | 0 | 0 | 0 | 0 | 0 | 0 | 0 | 0 | 0 | 0 | 0 | 0 | 0 | 0 | 0 | 0 | 0 | 0 | 0 | 1 | 0 | 0 | 0 | 0 | 0 | 0 | 0 | 0 | 0 | 0 | 0 | 0 | 0 | 0 | 0 | 0 | 0 | 0 | 0 | 0 |
| virus_Jul27.75v_898 | 1 | 0 | 0 | 0 | 0 | 0 | 0 | 0 | 0 | 0 | 0 | 0 | 0 | 0 | 0 | 0 | 0 | 0 | 0 | 0 | 0 | 0 | 0 | 0 | 0 | 0 | 0 | 0 | 0 | 0 | 0 | 0 | 0 | 0 | 0 | 0 | 1 | 0 | 0 | 0 | 0 | 0 | 0 | 0 | 0 | 0 | 0 | 0 | 0 | 0 | 0 | 0 | 0 | 0 | 0 |
| virus_Jul27.75v_8322 | 1 | 0 | 0 | 0 | 0 | 0 | 0 | 0 | 0 | 0 | 0 | 0 | 0 | 0 | 0 | 0 | 0 | 0 | 0 | 0 | 0 | 0 | 0 | 0 | 0 | 0 | 0 | 0 | 0 | 0 | 0 | 0 | 0 | 0 | 0 | 0 | 1 | 0 | 0 | 0 | 0 | 0 | 0 | 0 | 0 | 0 | 0 | 0 | 0 | 0 | 0 | 0 | 0 | 0 | 0 |
| virus_Jul27.75v_10122 | 1 | 0 | 0 | 0 | 0 | 0 | 0 | 0 | 0 | 0 | 0 | 0 | 0 | 0 | 0 | 0 | 0 | 0 | 0 | 0 | 0 | 0 | 0 | 0 | 0 | 0 | 0 | 0 | 0 | 0 | 0 | 0 | 0 | 0 | 0 | 0 | 1 | 0 | 0 | 0 | 0 | 0 | 0 | 0 | 0 | 0 | 0 | 0 | 0 | 0 | 0 | 0 | 0 | 0 | 0 |
| virus_Jul27.75v_10776 | 1 | 0 | 0 | 0 | 0 | 0 | 0 | 0 | 0 | 0 | 0 | 0 | 0 | 0 | 0 | 0 | 0 | 0 | 0 | 0 | 0 | 0 | 0 | 0 | 0 | 0 | 0 | 0 | 0 | 0 | 0 | 0 | 0 | 0 | 0 | 0 | 1 | 0 | 0 | 0 | 0 | 0 | 0 | 0 | 0 | 0 | 0 | 0 | 0 | 0 | 0 | 0 | 0 | 0 | 0 |
| virus_Jul27.75v_14092 | 1 | 0 | 0 | 0 | 0 | 0 | 0 | 0 | 0 | 0 | 0 | 0 | 0 | 0 | 0 | 0 | 0 | 0 | 0 | 0 | 0 | 0 | 0 | 0 | 0 | 0 | 0 | 0 | 0 | 0 | 0 | 0 | 0 | 0 | 0 | 0 | 1 | 0 | 0 | 0 | 0 | 0 | 0 | 0 | 0 | 0 | 0 | 0 | 0 | 0 | 0 | 0 | 0 | 0 | 0 |
| virus_Jul27.75v_18316 | 1 | 0 | 0 | 0 | 0 | 0 | 0 | 0 | 0 | 0 | 0 | 0 | 0 | 0 | 0 | 0 | 0 | 0 | 0 | 0 | 0 | 0 | 0 | 0 | 0 | 0 | 0 | 0 | 0 | 0 | 0 | 0 | 0 | 0 | 0 | 0 | 1 | 0 | 0 | 0 | 0 | 0 | 0 | 0 | 0 | 0 | 0 | 0 | 0 | 0 | 0 | 0 | 0 | 0 | 0 |
| virus_Jul27.75v_19148 | 1 | 0 | 0 | 0 | 0 | 0 | 0 | 0 | 0 | 0 | 0 | 0 | 0 | 0 | 0 | 0 | 0 | 0 | 0 | 0 | 0 | 0 | 0 | 0 | 0 | 0 | 0 | 0 | 0 | 0 | 0 | 0 | 0 | 0 | 0 | 0 | 1 | 0 | 0 | 0 | 0 | 0 | 0 | 0 | 0 | 0 | 0 | 0 | 0 | 0 | 0 | 0 | 0 | 0 | 0 |
| virus_Jun8.25v_16698 | 1 | 0 | 0 | 0 | 0 | 0 | 0 | 0 | 0 | 0 | 1 | 0 | 0 | 0 | 0 | 0 | 0 | 0 | 0 | 0 | 0 | 0 | 0 | 0 | 0 | 0 | 0 | 0 | 0 | 0 | 0 | 0 | 0 | 0 | 0 | 0 | 0 | 0 | 0 | 0 | 0 | 0 | 0 | 0 | 0 | 0 | 0 | 0 | 0 | 0 | 0 | 0 | 0 | 0 | 0 |
| virus_Jun8.25v_26408 | 1 | 0 | 0 | 0 | 0 | 0 | 0 | 0 | 0 | 0 | 1 | 0 | 0 | 0 | 0 | 0 | 0 | 0 | 0 | 0 | 0 | 0 | 0 | 0 | 0 | 0 | 0 | 0 | 0 | 0 | 0 | 0 | 0 | 0 | 0 | 0 | 0 | 0 | 0 | 0 | 0 | 0 | 0 | 0 | 0 | 0 | 0 | 0 | 0 | 0 | 0 | 0 | 0 | 0 | 0 |
| virus_Jun8.25v_27945 | 1 | 0 | 0 | 0 | 0 | 0 | 0 | 0 | 0 | 0 | 1 | 0 | 0 | 0 | 0 | 0 | 0 | 0 | 0 | 0 | 0 | 0 | 0 | 0 | 0 | 0 | 0 | 0 | 0 | 0 | 0 | 0 | 0 | 0 | 0 | 0 | 0 | 0 | 0 | 0 | 0 | 0 | 0 | 0 | 0 | 0 | 0 | 0 | 0 | 0 | 0 | 0 | 0 | 0 | 0 |
| virus_Jun8.25v_29954 | 1 | 0 | 0 | 0 | 0 | 0 | 0 | 0 | 0 | 0 | 1 | 0 | 0 | 0 | 0 | 0 | 0 | 0 | 0 | 0 | 0 | 0 | 0 | 0 | 0 | 0 | 0 | 0 | 0 | 0 | 0 | 0 | 0 | 0 | 0 | 0 | 0 | 0 | 0 | 0 | 0 | 0 | 0 | 0 | 0 | 0 | 0 | 0 | 0 | 0 | 0 | 0 | 0 | 0 | 0 |
| virus_Jun8.50v_8853 | 1 | 0 | 0 | 0 | 0 | 0 | 0 | 0 | 0 | 0 | 0 | 0 | 1 | 0 | 0 | 0 | 0 | 0 | 0 | 0 | 0 | 0 | 0 | 0 | 0 | 0 | 0 | 0 | 0 | 0 | 0 | 0 | 0 | 0 | 0 | 0 | 0 | 0 | 0 | 0 | 0 | 0 | 0 | 0 | 0 | 0 | 0 | 0 | 0 | 0 | 0 | 0 | 0 | 0 | 0 |
| virus_Jun8.50v_13689 | 1 | 0 | 0 | 0 | 0 | 0 | 0 | 0 | 0 | 0 | 0 | 0 | 1 | 0 | 0 | 0 | 0 | 0 | 0 | 0 | 0 | 0 | 0 | 0 | 0 | 0 | 0 | 0 | 0 | 0 | 0 | 0 | 0 | 0 | 0 | 0 | 0 | 0 | 0 | 0 | 0 | 0 | 0 | 0 | 0 | 0 | 0 | 0 | 0 | 0 | 0 | 0 | 0 | 0 | 0 |
| virus_Jun8.50v_17712 | 1 | 0 | 0 | 0 | 0 | 0 | 0 | 0 | 0 | 0 | 0 | 0 | 1 | 0 | 0 | 0 | 0 | 0 | 0 | 0 | 0 | 0 | 0 | 0 | 0 | 0 | 0 | 0 | 0 | 0 | 0 | 0 | 0 | 0 | 0 | 0 | 0 | 0 | 0 | 0 | 0 | 0 | 0 | 0 | 0 | 0 | 0 | 0 | 0 | 0 | 0 | 0 | 0 | 0 | 0 |
| virus_Jun8.50v_20481 | 1 | 0 | 0 | 0 | 0 | 0 | 0 | 0 | 0 | 0 | 0 | 0 | 1 | 0 | 0 | 0 | 0 | 0 | 0 | 0 | 0 | 0 | 0 | 0 | 0 | 0 | 0 | 0 | 0 | 0 | 0 | 0 | 0 | 0 | 0 | 0 | 0 | 0 | 0 | 0 | 0 | 0 | 0 | 0 | 0 | 0 | 0 | 0 | 0 | 0 | 0 | 0 | 0 | 0 | 0 |
| virus_Jun8.50v_27147 | 1 | 0 | 0 | 0 | 0 | 0 | 0 | 0 | 0 | 0 | 0 | 0 | 1 | 0 | 0 | 0 | 0 | 0 | 0 | 0 | 0 | 0 | 0 | 0 | 0 | 0 | 0 | 0 | 0 | 0 | 0 | 0 | 0 | 0 | 0 | 0 | 0 | 0 | 0 | 0 | 0 | 0 | 0 | 0 | 0 | 0 | 0 | 0 | 0 | 0 | 0 | 0 | 0 | 0 | 0 |
| virus_Jun8.50v_29052 | 1 | 0 | 0 | 0 | 0 | 0 | 0 | 0 | 0 | 0 | 0 | 0 | 1 | 0 | 0 | 0 | 0 | 0 | 0 | 0 | 0 | 0 | 0 | 0 | 0 | 0 | 0 | 0 | 0 | 0 | 0 | 0 | 0 | 0 | 0 | 0 | 0 | 0 | 0 | 0 | 0 | 0 | 0 | 0 | 0 | 0 | 0 | 0 | 0 | 0 | 0 | 0 | 0 | 0 | 0 |
| virus_Jun8.50v_29123 | 1 | 0 | 0 | 0 | 0 | 0 | 0 | 0 | 0 | 0 | 0 | 0 | 1 | 0 | 0 | 0 | 0 | 0 | 0 | 0 | 0 | 0 | 0 | 0 | 0 | 0 | 0 | 0 | 0 | 0 | 0 | 0 | 0 | 0 | 0 | 0 | 0 | 0 | 0 | 0 | 0 | 0 | 0 | 0 | 0 | 0 | 0 | 0 | 0 | 0 | 0 | 0 | 0 | 0 | 0 |
| virus_Jun8.50v_32615 | 1 | 0 | 0 | 0 | 0 | 0 | 0 | 0 | 0 | 0 | 0 | 0 | 1 | 0 | 0 | 0 | 0 | 0 | 0 | 0 | 0 | 0 | 0 | 0 | 0 | 0 | 0 | 0 | 0 | 0 | 0 | 0 | 0 | 0 | 0 | 0 | 0 | 0 | 0 | 0 | 0 | 0 | 0 | 0 | 0 | 0 | 0 | 0 | 0 | 0 | 0 | 0 | 0 | 0 | 0 |
| virus_Jun8.50v_35056 | 1 | 0 | 0 | 0 | 0 | 0 | 0 | 0 | 0 | 0 | 0 | 0 | 1 | 0 | 0 | 0 | 0 | 0 | 0 | 0 | 0 | 0 | 0 | 0 | 0 | 0 | 0 | 0 | 0 | 0 | 0 | 0 | 0 | 0 | 0 | 0 | 0 | 0 | 0 | 0 | 0 | 0 | 0 | 0 | 0 | 0 | 0 | 0 | 0 | 0 | 0 | 0 | 0 | 0 | 0 |
| virus_Jun8.50v_35836 | 1 | 0 | 0 | 0 | 0 | 0 | 0 | 0 | 0 | 0 | 0 | 0 | 1 | 0 | 0 | 0 | 0 | 0 | 0 | 0 | 0 | 0 | 0 | 0 | 0 | 0 | 0 | 0 | 0 | 0 | 0 | 0 | 0 | 0 | 0 | 0 | 0 | 0 | 0 | 0 | 0 | 0 | 0 | 0 | 0 | 0 | 0 | 0 | 0 | 0 | 0 | 0 | 0 | 0 | 0 |
| virus_Jun8.50v_36178 | 1 | 0 | 0 | 0 | 0 | 0 | 0 | 0 | 0 | 0 | 0 | 0 | 1 | 0 | 0 | 0 | 0 | 0 | 0 | 0 | 0 | 0 | 0 | 0 | 0 | 0 | 0 | 0 | 0 | 0 | 0 | 0 | 0 | 0 | 0 | 0 | 0 | 0 | 0 | 0 | 0 | 0 | 0 | 0 | 0 | 0 | 0 | 0 | 0 | 0 | 0 | 0 | 0 | 0 | 0 |
| virus_Jun8.75v_596 | 1 | 0 | 0 | 0 | 0 | 0 | 0 | 0 | 0 | 0 | 0 | 0 | 0 | 0 | 1 | 0 | 0 | 0 | 0 | 0 | 0 | 0 | 0 | 0 | 0 | 0 | 0 | 0 | 0 | 0 | 0 | 0 | 0 | 0 | 0 | 0 | 0 | 0 | 0 | 0 | 0 | 0 | 0 | 0 | 0 | 0 | 0 | 0 | 0 | 0 | 0 | 0 | 0 | 0 | 0 |
| virus_Jun8.75v_2130 | 1 | 0 | 0 | 0 | 0 | 0 | 0 | 0 | 0 | 0 | 0 | 0 | 0 | 0 | 1 | 0 | 0 | 0 | 0 | 0 | 0 | 0 | 0 | 0 | 0 | 0 | 0 | 0 | 0 | 0 | 0 | 0 | 0 | 0 | 0 | 0 | 0 | 0 | 0 | 0 | 0 | 0 | 0 | 0 | 0 | 0 | 0 | 0 | 0 | 0 | 0 | 0 | 0 | 0 | 0 |
| virus_Jun8.75v_20642 | 1 | 0 | 0 | 0 | 0 | 0 | 0 | 0 | 0 | 0 | 0 | 0 | 0 | 0 | 1 | 0 | 0 | 0 | 0 | 0 | 0 | 0 | 0 | 0 | 0 | 0 | 0 | 0 | 0 | 0 | 0 | 0 | 0 | 0 | 0 | 0 | 0 | 0 | 0 | 0 | 0 | 0 | 0 | 0 | 0 | 0 | 0 | 0 | 0 | 0 | 0 | 0 | 0 | 0 | 0 |
| virus_Jun22.0v_397 | 1 | 0 | 0 | 0 | 0 | 0 | 0 | 0 | 0 | 0 | 0 | 0 | 0 | 0 | 0 | 0 | 1 | 0 | 0 | 0 | 0 | 0 | 0 | 0 | 0 | 0 | 0 | 0 | 0 | 0 | 0 | 0 | 0 | 0 | 0 | 0 | 0 | 0 | 0 | 0 | 0 | 0 | 0 | 0 | 0 | 0 | 0 | 0 | 0 | 0 | 0 | 0 | 0 | 0 | 0 |
| virus_Jun22.0v_2785 | 1 | 0 | 0 | 0 | 0 | 0 | 0 | 0 | 0 | 0 | 0 | 0 | 0 | 0 | 0 | 0 | 1 | 0 | 0 | 0 | 0 | 0 | 0 | 0 | 0 | 0 | 0 | 0 | 0 | 0 | 0 | 0 | 0 | 0 | 0 | 0 | 0 | 0 | 0 | 0 | 0 | 0 | 0 | 0 | 0 | 0 | 0 | 0 | 0 | 0 | 0 | 0 | 0 | 0 | 0 |
| virus_Jun22.0v_4319 | 1 | 0 | 0 | 0 | 0 | 0 | 0 | 0 | 0 | 0 | 0 | 0 | 0 | 0 | 0 | 0 | 1 | 0 | 0 | 0 | 0 | 0 | 0 | 0 | 0 | 0 | 0 | 0 | 0 | 0 | 0 | 0 | 0 | 0 | 0 | 0 | 0 | 0 | 0 | 0 | 0 | 0 | 0 | 0 | 0 | 0 | 0 | 0 | 0 | 0 | 0 | 0 | 0 | 0 | 0 |
| virus_Jun22.0v_5997 | 1 | 0 | 0 | 0 | 0 | 0 | 0 | 0 | 0 | 0 | 0 | 0 | 0 | 0 | 0 | 0 | 1 | 0 | 0 | 0 | 0 | 0 | 0 | 0 | 0 | 0 | 0 | 0 | 0 | 0 | 0 | 0 | 0 | 0 | 0 | 0 | 0 | 0 | 0 | 0 | 0 | 0 | 0 | 0 | 0 | 0 | 0 | 0 | 0 | 0 | 0 | 0 | 0 | 0 | 0 |
| virus_Jun22.0v_6860 | 1 | 0 | 0 | 0 | 0 | 0 | 0 | 0 | 0 | 0 | 0 | 0 | 0 | 0 | 0 | 0 | 1 | 0 | 0 | 0 | 0 | 0 | 0 | 0 | 0 | 0 | 0 | 0 | 0 | 0 | 0 | 0 | 0 | 0 | 0 | 0 | 0 | 0 | 0 | 0 | 0 | 0 | 0 | 0 | 0 | 0 | 0 | 0 | 0 | 0 | 0 | 0 | 0 | 0 | 0 |
| virus_Jun22.0v_7074 | 1 | 0 | 0 | 0 | 0 | 0 | 0 | 0 | 0 | 0 | 0 | 0 | 0 | 0 | 0 | 0 | 1 | 0 | 0 | 0 | 0 | 0 | 0 | 0 | 0 | 0 | 0 | 0 | 0 | 0 | 0 | 0 | 0 | 0 | 0 | 0 | 0 | 0 | 0 | 0 | 0 | 0 | 0 | 0 | 0 | 0 | 0 | 0 | 0 | 0 | 0 | 0 | 0 | 0 | 0 |
| virus_Jun22.0v_7652 | 1 | 0 | 0 | 0 | 0 | 0 | 0 | 0 | 0 | 0 | 0 | 0 | 0 | 0 | 0 | 0 | 1 | 0 | 0 | 0 | 0 | 0 | 0 | 0 | 0 | 0 | 0 | 0 | 0 | 0 | 0 | 0 | 0 | 0 | 0 | 0 | 0 | 0 | 0 | 0 | 0 | 0 | 0 | 0 | 0 | 0 | 0 | 0 | 0 | 0 | 0 | 0 | 0 | 0 | 0 |
| virus_Jun22.0v_12106 | 1 | 0 | 0 | 0 | 0 | 0 | 0 | 0 | 0 | 0 | 0 | 0 | 0 | 0 | 0 | 0 | 1 | 0 | 0 | 0 | 0 | 0 | 0 | 0 | 0 | 0 | 0 | 0 | 0 | 0 | 0 | 0 | 0 | 0 | 0 | 0 | 0 | 0 | 0 | 0 | 0 | 0 | 0 | 0 | 0 | 0 | 0 | 0 | 0 | 0 | 0 | 0 | 0 | 0 | 0 |
| virus_Jun22.0v_16886 | 1 | 0 | 0 | 0 | 0 | 0 | 0 | 0 | 0 | 0 | 0 | 0 | 0 | 0 | 0 | 0 | 1 | 0 | 0 | 0 | 0 | 0 | 0 | 0 | 0 | 0 | 0 | 0 | 0 | 0 | 0 | 0 | 0 | 0 | 0 | 0 | 0 | 0 | 0 | 0 | 0 | 0 | 0 | 0 | 0 | 0 | 0 | 0 | 0 | 0 | 0 | 0 | 0 | 0 | 0 |
| virus_Jun22.0v_16908 | 1 | 0 | 0 | 0 | 0 | 0 | 0 | 0 | 0 | 0 | 0 | 0 | 0 | 0 | 0 | 0 | 1 | 0 | 0 | 0 | 0 | 0 | 0 | 0 | 0 | 0 | 0 | 0 | 0 | 0 | 0 | 0 | 0 | 0 | 0 | 0 | 0 | 0 | 0 | 0 | 0 | 0 | 0 | 0 | 0 | 0 | 0 | 0 | 0 | 0 | 0 | 0 | 0 | 0 | 0 |
| virus_Jun22.0v_17757 | 1 | 0 | 0 | 0 | 0 | 0 | 0 | 0 | 0 | 0 | 0 | 0 | 0 | 0 | 0 | 0 | 1 | 0 | 0 | 0 | 0 | 0 | 0 | 0 | 0 | 0 | 0 | 0 | 0 | 0 | 0 | 0 | 0 | 0 | 0 | 0 | 0 | 0 | 0 | 0 | 0 | 0 | 0 | 0 | 0 | 0 | 0 | 0 | 0 | 0 | 0 | 0 | 0 | 0 | 0 |
| virus_Jun22.0v_17979 | 1 | 0 | 0 | 0 | 0 | 0 | 0 | 0 | 0 | 0 | 0 | 0 | 0 | 0 | 0 | 0 | 1 | 0 | 0 | 0 | 0 | 0 | 0 | 0 | 0 | 0 | 0 | 0 | 0 | 0 | 0 | 0 | 0 | 0 | 0 | 0 | 0 | 0 | 0 | 0 | 0 | 0 | 0 | 0 | 0 | 0 | 0 | 0 | 0 | 0 | 0 | 0 | 0 | 0 | 0 |
| virus_Jun22.0v_19010 | 1 | 0 | 0 | 0 | 0 | 0 | 0 | 0 | 0 | 0 | 0 | 0 | 0 | 0 | 0 | 0 | 1 | 0 | 0 | 0 | 0 | 0 | 0 | 0 | 0 | 0 | 0 | 0 | 0 | 0 | 0 | 0 | 0 | 0 | 0 | 0 | 0 | 0 | 0 | 0 | 0 | 0 | 0 | 0 | 0 | 0 | 0 | 0 | 0 | 0 | 0 | 0 | 0 | 0 | 0 |
| virus_Jun22.0v_19116 | 1 | 0 | 0 | 0 | 0 | 0 | 0 | 0 | 0 | 0 | 0 | 0 | 0 | 0 | 0 | 0 | 1 | 0 | 0 | 0 | 0 | 0 | 0 | 0 | 0 | 0 | 0 | 0 | 0 | 0 | 0 | 0 | 0 | 0 | 0 | 0 | 0 | 0 | 0 | 0 | 0 | 0 | 0 | 0 | 0 | 0 | 0 | 0 | 0 | 0 | 0 | 0 | 0 | 0 | 0 |
| virus_Jun22.0v_19212 | 1 | 0 | 0 | 0 | 0 | 0 | 0 | 0 | 0 | 0 | 0 | 0 | 0 | 0 | 0 | 0 | 1 | 0 | 0 | 0 | 0 | 0 | 0 | 0 | 0 | 0 | 0 | 0 | 0 | 0 | 0 | 0 | 0 | 0 | 0 | 0 | 0 | 0 | 0 | 0 | 0 | 0 | 0 | 0 | 0 | 0 | 0 | 0 | 0 | 0 | 0 | 0 | 0 | 0 | 0 |
| virus_Jun22.50v_170 | 1 | 0 | 0 | 0 | 0 | 0 | 0 | 0 | 0 | 0 | 0 | 0 | 0 | 0 | 0 | 0 | 0 | 0 | 1 | 0 | 0 | 0 | 0 | 0 | 0 | 0 | 0 | 0 | 0 | 0 | 0 | 0 | 0 | 0 | 0 | 0 | 0 | 0 | 0 | 0 | 0 | 0 | 0 | 0 | 0 | 0 | 0 | 0 | 0 | 0 | 0 | 0 | 0 | 0 | 0 |
| virus_Jun22.50v_754 | 1 | 0 | 0 | 0 | 0 | 0 | 0 | 0 | 0 | 0 | 0 | 0 | 0 | 0 | 0 | 0 | 0 | 0 | 1 | 0 | 0 | 0 | 0 | 0 | 0 | 0 | 0 | 0 | 0 | 0 | 0 | 0 | 0 | 0 | 0 | 0 | 0 | 0 | 0 | 0 | 0 | 0 | 0 | 0 | 0 | 0 | 0 | 0 | 0 | 0 | 0 | 0 | 0 | 0 | 0 |
| virus_Jun22.50v_9417 | 1 | 0 | 0 | 0 | 0 | 0 | 0 | 0 | 0 | 0 | 0 | 0 | 0 | 0 | 0 | 0 | 0 | 0 | 1 | 0 | 0 | 0 | 0 | 0 | 0 | 0 | 0 | 0 | 0 | 0 | 0 | 0 | 0 | 0 | 0 | 0 | 0 | 0 | 0 | 0 | 0 | 0 | 0 | 0 | 0 | 0 | 0 | 0 | 0 | 0 | 0 | 0 | 0 | 0 | 0 |
| virus_Jun22.50v_16142 | 1 | 0 | 0 | 0 | 0 | 0 | 0 | 0 | 0 | 0 | 0 | 0 | 0 | 0 | 0 | 0 | 0 | 0 | 1 | 0 | 0 | 0 | 0 | 0 | 0 | 0 | 0 | 0 | 0 | 0 | 0 | 0 | 0 | 0 | 0 | 0 | 0 | 0 | 0 | 0 | 0 | 0 | 0 | 0 | 0 | 0 | 0 | 0 | 0 | 0 | 0 | 0 | 0 | 0 | 0 |
| virus_Jun22.50v_16177 | 1 | 0 | 0 | 0 | 0 | 0 | 0 | 0 | 0 | 0 | 0 | 0 | 0 | 0 | 0 | 0 | 0 | 0 | 1 | 0 | 0 | 0 | 0 | 0 | 0 | 0 | 0 | 0 | 0 | 0 | 0 | 0 | 0 | 0 | 0 | 0 | 0 | 0 | 0 | 0 | 0 | 0 | 0 | 0 | 0 | 0 | 0 | 0 | 0 | 0 | 0 | 0 | 0 | 0 | 0 |
| virus_Jun22.50v_17142 | 1 | 0 | 0 | 0 | 0 | 0 | 0 | 0 | 0 | 0 | 0 | 0 | 0 | 0 | 0 | 0 | 0 | 0 | 1 | 0 | 0 | 0 | 0 | 0 | 0 | 0 | 0 | 0 | 0 | 0 | 0 | 0 | 0 | 0 | 0 | 0 | 0 | 0 | 0 | 0 | 0 | 0 | 0 | 0 | 0 | 0 | 0 | 0 | 0 | 0 | 0 | 0 | 0 | 0 | 0 |
| virus_Jun22.50v_18075 | 1 | 0 | 0 | 0 | 0 | 0 | 0 | 0 | 0 | 0 | 0 | 0 | 0 | 0 | 0 | 0 | 0 | 0 | 1 | 0 | 0 | 0 | 0 | 0 | 0 | 0 | 0 | 0 | 0 | 0 | 0 | 0 | 0 | 0 | 0 | 0 | 0 | 0 | 0 | 0 | 0 | 0 | 0 | 0 | 0 | 0 | 0 | 0 | 0 | 0 | 0 | 0 | 0 | 0 | 0 |
| virus_Jun22.50v_21417 | 1 | 0 | 0 | 0 | 0 | 0 | 0 | 0 | 0 | 0 | 0 | 0 | 0 | 0 | 0 | 0 | 0 | 0 | 1 | 0 | 0 | 0 | 0 | 0 | 0 | 0 | 0 | 0 | 0 | 0 | 0 | 0 | 0 | 0 | 0 | 0 | 0 | 0 | 0 | 0 | 0 | 0 | 0 | 0 | 0 | 0 | 0 | 0 | 0 | 0 | 0 | 0 | 0 | 0 | 0 |
| virus_Jun22.50v_22685 | 1 | 0 | 0 | 0 | 0 | 0 | 0 | 0 | 0 | 0 | 0 | 0 | 0 | 0 | 0 | 0 | 0 | 0 | 1 | 0 | 0 | 0 | 0 | 0 | 0 | 0 | 0 | 0 | 0 | 0 | 0 | 0 | 0 | 0 | 0 | 0 | 0 | 0 | 0 | 0 | 0 | 0 | 0 | 0 | 0 | 0 | 0 | 0 | 0 | 0 | 0 | 0 | 0 | 0 | 0 |
| virus_Jun22.50v_23272 | 1 | 0 | 0 | 0 | 0 | 0 | 0 | 0 | 0 | 0 | 0 | 0 | 0 | 0 | 0 | 0 | 0 | 0 | 1 | 0 | 0 | 0 | 0 | 0 | 0 | 0 | 0 | 0 | 0 | 0 | 0 | 0 | 0 | 0 | 0 | 0 | 0 | 0 | 0 | 0 | 0 | 0 | 0 | 0 | 0 | 0 | 0 | 0 | 0 | 0 | 0 | 0 | 0 | 0 | 0 |
| virus_Jun22.50v_27084 | 1 | 0 | 0 | 0 | 0 | 0 | 0 | 0 | 0 | 0 | 0 | 0 | 0 | 0 | 0 | 0 | 0 | 0 | 1 | 0 | 0 | 0 | 0 | 0 | 0 | 0 | 0 | 0 | 0 | 0 | 0 | 0 | 0 | 0 | 0 | 0 | 0 | 0 | 0 | 0 | 0 | 0 | 0 | 0 | 0 | 0 | 0 | 0 | 0 | 0 | 0 | 0 | 0 | 0 | 0 |
| virus_Jun22.50v_27535 | 1 | 0 | 0 | 0 | 0 | 0 | 0 | 0 | 0 | 0 | 0 | 0 | 0 | 0 | 0 | 0 | 0 | 0 | 1 | 0 | 0 | 0 | 0 | 0 | 0 | 0 | 0 | 0 | 0 | 0 | 0 | 0 | 0 | 0 | 0 | 0 | 0 | 0 | 0 | 0 | 0 | 0 | 0 | 0 | 0 | 0 | 0 | 0 | 0 | 0 | 0 | 0 | 0 | 0 | 0 |
| virus_Jun22.50v_29457 | 1 | 0 | 0 | 0 | 0 | 0 | 0 | 0 | 0 | 0 | 0 | 0 | 0 | 0 | 0 | 0 | 0 | 0 | 1 | 0 | 0 | 0 | 0 | 0 | 0 | 0 | 0 | 0 | 0 | 0 | 0 | 0 | 0 | 0 | 0 | 0 | 0 | 0 | 0 | 0 | 0 | 0 | 0 | 0 | 0 | 0 | 0 | 0 | 0 | 0 | 0 | 0 | 0 | 0 | 0 |
| virus_Jun22.50v_29617 | 1 | 0 | 0 | 0 | 0 | 0 | 0 | 0 | 0 | 0 | 0 | 0 | 0 | 0 | 0 | 0 | 0 | 0 | 1 | 0 | 0 | 0 | 0 | 0 | 0 | 0 | 0 | 0 | 0 | 0 | 0 | 0 | 0 | 0 | 0 | 0 | 0 | 0 | 0 | 0 | 0 | 0 | 0 | 0 | 0 | 0 | 0 | 0 | 0 | 0 | 0 | 0 | 0 | 0 | 0 |
| virus_Jun22.50v_32780 | 1 | 0 | 0 | 0 | 0 | 0 | 0 | 0 | 0 | 0 | 0 | 0 | 0 | 0 | 0 | 0 | 0 | 0 | 1 | 0 | 0 | 0 | 0 | 0 | 0 | 0 | 0 | 0 | 0 | 0 | 0 | 0 | 0 | 0 | 0 | 0 | 0 | 0 | 0 | 0 | 0 | 0 | 0 | 0 | 0 | 0 | 0 | 0 | 0 | 0 | 0 | 0 | 0 | 0 | 0 |
| virus_Jun22.75v_2016 | 1 | 0 | 0 | 0 | 0 | 0 | 0 | 0 | 0 | 0 | 0 | 0 | 0 | 0 | 0 | 0 | 0 | 0 | 0 | 0 | 1 | 0 | 0 | 0 | 0 | 0 | 0 | 0 | 0 | 0 | 0 | 0 | 0 | 0 | 0 | 0 | 0 | 0 | 0 | 0 | 0 | 0 | 0 | 0 | 0 | 0 | 0 | 0 | 0 | 0 | 0 | 0 | 0 | 0 | 0 |
| virus_Jun22.75v_5295 | 1 | 0 | 0 | 0 | 0 | 0 | 0 | 0 | 0 | 0 | 0 | 0 | 0 | 0 | 0 | 0 | 0 | 0 | 0 | 0 | 1 | 0 | 0 | 0 | 0 | 0 | 0 | 0 | 0 | 0 | 0 | 0 | 0 | 0 | 0 | 0 | 0 | 0 | 0 | 0 | 0 | 0 | 0 | 0 | 0 | 0 | 0 | 0 | 0 | 0 | 0 | 0 | 0 | 0 | 0 |
| virus_Jun22.75v_13437 | 1 | 0 | 0 | 0 | 0 | 0 | 0 | 0 | 0 | 0 | 0 | 0 | 0 | 0 | 0 | 0 | 0 | 0 | 0 | 0 | 1 | 0 | 0 | 0 | 0 | 0 | 0 | 0 | 0 | 0 | 0 | 0 | 0 | 0 | 0 | 0 | 0 | 0 | 0 | 0 | 0 | 0 | 0 | 0 | 0 | 0 | 0 | 0 | 0 | 0 | 0 | 0 | 0 | 0 | 0 |
| virus_Jun22.75v_21990 | 1 | 0 | 0 | 0 | 0 | 0 | 0 | 0 | 0 | 0 | 0 | 0 | 0 | 0 | 0 | 0 | 0 | 0 | 0 | 0 | 1 | 0 | 0 | 0 | 0 | 0 | 0 | 0 | 0 | 0 | 0 | 0 | 0 | 0 | 0 | 0 | 0 | 0 | 0 | 0 | 0 | 0 | 0 | 0 | 0 | 0 | 0 | 0 | 0 | 0 | 0 | 0 | 0 | 0 | 0 |
| virus_Jun22.75v_22208 | 1 | 0 | 0 | 0 | 0 | 0 | 0 | 0 | 0 | 0 | 0 | 0 | 0 | 0 | 0 | 0 | 0 | 0 | 0 | 0 | 1 | 0 | 0 | 0 | 0 | 0 | 0 | 0 | 0 | 0 | 0 | 0 | 0 | 0 | 0 | 0 | 0 | 0 | 0 | 0 | 0 | 0 | 0 | 0 | 0 | 0 | 0 | 0 | 0 | 0 | 0 | 0 | 0 | 0 | 0 |
| virus_Jun22.75v_23111 | 1 | 0 | 0 | 0 | 0 | 0 | 0 | 0 | 0 | 0 | 0 | 0 | 0 | 0 | 0 | 0 | 0 | 0 | 0 | 0 | 1 | 0 | 0 | 0 | 0 | 0 | 0 | 0 | 0 | 0 | 0 | 0 | 0 | 0 | 0 | 0 | 0 | 0 | 0 | 0 | 0 | 0 | 0 | 0 | 0 | 0 | 0 | 0 | 0 | 0 | 0 | 0 | 0 | 0 | 0 |

**Supplementary Table S7**. Common Bacterio- and virioplankton RTPR 98% OTUs in 27 samples with RTPR amplicons in both fractions

Total seqs: 47997; Subtotal seqs in retained

| OTUs: 41046 | | May12_05m | | May12_25m | | May12_50m | | May12_75m | | Jun8_25m | | Jun8_50m | | Jun8_75m | | Jun22_0m | | Jun22_50m | | Jun22_75m | | Jul11_0m | | Jul11_05m | | Jul11_25m | | Jul11_50m | | Jul11_75m | | Jul27_05m | | Jul27_25m | | Jul27_75m | | Aug5_0m | | Aug5_05m | | Aug5_25m | | Aug5_50m | | Aug5_75m | | Sep8_0m | | Sep8_05m | | Sep8_25m | | Sep8_50m | |
|---|---|---|---|---|---|---|---|---|---|---|---|---|---|---|---|---|---|---|---|---|---|---|---|---|---|---|---|---|---|---|---|---|---|---|---|---|---|---|---|---|---|---|---|---|---|---|---|---|---|---|---|---|---|---|---|
| OTUs with >=10 seqs: 254 | OTU subtotal | bacterioplankton | virioplankton | bacterioplankton | virioplankton | bacterioplankton | virioplankton | bacterioplankton | virioplankton | bacterioplankton | virioplankton | bacterioplankton | virioplankton | bacterioplankton | virioplankton | bacterioplankton | virioplankton | bacterioplankton | virioplankton | bacterioplankton | virioplankton | bacterioplankton | virioplankton | bacterioplankton | virioplankton | bacterioplankton | virioplankton | bacterioplankton | virioplankton | bacterioplankton | virioplankton | bacterioplankton | virioplankton | bacterioplankton | virioplankton | bacterioplankton | virioplankton | bacterioplankton | virioplankton | bacterioplankton | virioplankton | bacterioplankton | virioplankton | bacterioplankton | virioplankton | bacterioplankton | virioplankton | bacterioplankton | virioplankton | bacterioplankton | virioplankton | bacterioplankton | virioplankton | bacterioplankton | virioplankton |
| virus_Jun22.75v_37444 | 1 | 0 | 0 | 0 | 0 | 0 | 0 | 0 | 0 | 0 | 0 | 0 | 0 | 0 | 0 | 0 | 0 | 0 | 0 | 0 | 1 | 0 | 0 | 0 | 0 | 0 | 0 | 0 | 0 | 0 | 0 | 0 | 0 | 0 | 0 | 0 | 0 | 0 | 0 | 0 | 0 | 0 | 0 | 0 | 0 | 0 | 0 | 0 | 0 | 0 | 0 | 0 | 0 | 0 | 0 |
| virus_Jun22.75v_37735 | 1 | 0 | 0 | 0 | 0 | 0 | 0 | 0 | 0 | 0 | 0 | 0 | 0 | 0 | 0 | 0 | 0 | 0 | 0 | 0 | 1 | 0 | 0 | 0 | 0 | 0 | 0 | 0 | 0 | 0 | 0 | 0 | 0 | 0 | 0 | 0 | 0 | 0 | 0 | 0 | 0 | 0 | 0 | 0 | 0 | 0 | 0 | 0 | 0 | 0 | 0 | 0 | 0 | 0 | 0 |
| virus_May12.5v_10352 | 1 | 0 | 1 | 0 | 0 | 0 | 0 | 0 | 0 | 0 | 0 | 0 | 0 | 0 | 0 | 0 | 0 | 0 | 0 | 0 | 0 | 0 | 0 | 0 | 0 | 0 | 0 | 0 | 0 | 0 | 0 | 0 | 0 | 0 | 0 | 0 | 0 | 0 | 0 | 0 | 0 | 0 | 0 | 0 | 0 | 0 | 0 | 0 | 0 | 0 | 0 | 0 | 0 | 0 | 0 |
| virus_May12.5v_12219 | 1 | 0 | 1 | 0 | 0 | 0 | 0 | 0 | 0 | 0 | 0 | 0 | 0 | 0 | 0 | 0 | 0 | 0 | 0 | 0 | 0 | 0 | 0 | 0 | 0 | 0 | 0 | 0 | 0 | 0 | 0 | 0 | 0 | 0 | 0 | 0 | 0 | 0 | 0 | 0 | 0 | 0 | 0 | 0 | 0 | 0 | 0 | 0 | 0 | 0 | 0 | 0 | 0 | 0 | 0 |
| virus_May12.5v_20341 | 1 | 0 | 1 | 0 | 0 | 0 | 0 | 0 | 0 | 0 | 0 | 0 | 0 | 0 | 0 | 0 | 0 | 0 | 0 | 0 | 0 | 0 | 0 | 0 | 0 | 0 | 0 | 0 | 0 | 0 | 0 | 0 | 0 | 0 | 0 | 0 | 0 | 0 | 0 | 0 | 0 | 0 | 0 | 0 | 0 | 0 | 0 | 0 | 0 | 0 | 0 | 0 | 0 | 0 | 0 |
| virus_May12.5v_24933 | 1 | 0 | 1 | 0 | 0 | 0 | 0 | 0 | 0 | 0 | 0 | 0 | 0 | 0 | 0 | 0 | 0 | 0 | 0 | 0 | 0 | 0 | 0 | 0 | 0 | 0 | 0 | 0 | 0 | 0 | 0 | 0 | 0 | 0 | 0 | 0 | 0 | 0 | 0 | 0 | 0 | 0 | 0 | 0 | 0 | 0 | 0 | 0 | 0 | 0 | 0 | 0 | 0 | 0 | 0 |
| virus_May12.5v_29781 | 1 | 0 | 1 | 0 | 0 | 0 | 0 | 0 | 0 | 0 | 0 | 0 | 0 | 0 | 0 | 0 | 0 | 0 | 0 | 0 | 0 | 0 | 0 | 0 | 0 | 0 | 0 | 0 | 0 | 0 | 0 | 0 | 0 | 0 | 0 | 0 | 0 | 0 | 0 | 0 | 0 | 0 | 0 | 0 | 0 | 0 | 0 | 0 | 0 | 0 | 0 | 0 | 0 | 0 | 0 |
| virus_May12.5v_33866 | 1 | 0 | 1 | 0 | 0 | 0 | 0 | 0 | 0 | 0 | 0 | 0 | 0 | 0 | 0 | 0 | 0 | 0 | 0 | 0 | 0 | 0 | 0 | 0 | 0 | 0 | 0 | 0 | 0 | 0 | 0 | 0 | 0 | 0 | 0 | 0 | 0 | 0 | 0 | 0 | 0 | 0 | 0 | 0 | 0 | 0 | 0 | 0 | 0 | 0 | 0 | 0 | 0 | 0 | 0 |
| virus_May12.5v_35782 | 1 | 0 | 1 | 0 | 0 | 0 | 0 | 0 | 0 | 0 | 0 | 0 | 0 | 0 | 0 | 0 | 0 | 0 | 0 | 0 | 0 | 0 | 0 | 0 | 0 | 0 | 0 | 0 | 0 | 0 | 0 | 0 | 0 | 0 | 0 | 0 | 0 | 0 | 0 | 0 | 0 | 0 | 0 | 0 | 0 | 0 | 0 | 0 | 0 | 0 | 0 | 0 | 0 | 0 | 0 |
| virus_May12.25v_3512 | 1 | 0 | 0 | 0 | 1 | 0 | 0 | 0 | 0 | 0 | 0 | 0 | 0 | 0 | 0 | 0 | 0 | 0 | 0 | 0 | 0 | 0 | 0 | 0 | 0 | 0 | 0 | 0 | 0 | 0 | 0 | 0 | 0 | 0 | 0 | 0 | 0 | 0 | 0 | 0 | 0 | 0 | 0 | 0 | 0 | 0 | 0 | 0 | 0 | 0 | 0 | 0 | 0 | 0 | 0 |
| virus_May12.25v_9922 | 1 | 0 | 0 | 0 | 1 | 0 | 0 | 0 | 0 | 0 | 0 | 0 | 0 | 0 | 0 | 0 | 0 | 0 | 0 | 0 | 0 | 0 | 0 | 0 | 0 | 0 | 0 | 0 | 0 | 0 | 0 | 0 | 0 | 0 | 0 | 0 | 0 | 0 | 0 | 0 | 0 | 0 | 0 | 0 | 0 | 0 | 0 | 0 | 0 | 0 | 0 | 0 | 0 | 0 | 0 |
| virus_May12.25v_13925 | 1 | 0 | 0 | 0 | 1 | 0 | 0 | 0 | 0 | 0 | 0 | 0 | 0 | 0 | 0 | 0 | 0 | 0 | 0 | 0 | 0 | 0 | 0 | 0 | 0 | 0 | 0 | 0 | 0 | 0 | 0 | 0 | 0 | 0 | 0 | 0 | 0 | 0 | 0 | 0 | 0 | 0 | 0 | 0 | 0 | 0 | 0 | 0 | 0 | 0 | 0 | 0 | 0 | 0 | 0 |
| virus_May12.25v_23099 | 1 | 0 | 0 | 0 | 1 | 0 | 0 | 0 | 0 | 0 | 0 | 0 | 0 | 0 | 0 | 0 | 0 | 0 | 0 | 0 | 0 | 0 | 0 | 0 | 0 | 0 | 0 | 0 | 0 | 0 | 0 | 0 | 0 | 0 | 0 | 0 | 0 | 0 | 0 | 0 | 0 | 0 | 0 | 0 | 0 | 0 | 0 | 0 | 0 | 0 | 0 | 0 | 0 | 0 | 0 |
| virus_May12.25v_23907 | 1 | 0 | 0 | 0 | 1 | 0 | 0 | 0 | 0 | 0 | 0 | 0 | 0 | 0 | 0 | 0 | 0 | 0 | 0 | 0 | 0 | 0 | 0 | 0 | 0 | 0 | 0 | 0 | 0 | 0 | 0 | 0 | 0 | 0 | 0 | 0 | 0 | 0 | 0 | 0 | 0 | 0 | 0 | 0 | 0 | 0 | 0 | 0 | 0 | 0 | 0 | 0 | 0 | 0 | 0 |
| virus_May12.25v_25260 | 1 | 0 | 0 | 0 | 1 | 0 | 0 | 0 | 0 | 0 | 0 | 0 | 0 | 0 | 0 | 0 | 0 | 0 | 0 | 0 | 0 | 0 | 0 | 0 | 0 | 0 | 0 | 0 | 0 | 0 | 0 | 0 | 0 | 0 | 0 | 0 | 0 | 0 | 0 | 0 | 0 | 0 | 0 | 0 | 0 | 0 | 0 | 0 | 0 | 0 | 0 | 0 | 0 | 0 | 0 |
| virus_May12.25v_26035 | 1 | 0 | 0 | 0 | 1 | 0 | 0 | 0 | 0 | 0 | 0 | 0 | 0 | 0 | 0 | 0 | 0 | 0 | 0 | 0 | 0 | 0 | 0 | 0 | 0 | 0 | 0 | 0 | 0 | 0 | 0 | 0 | 0 | 0 | 0 | 0 | 0 | 0 | 0 | 0 | 0 | 0 | 0 | 0 | 0 | 0 | 0 | 0 | 0 | 0 | 0 | 0 | 0 | 0 | 0 |
| virus_May12.50v_2829 | 1 | 0 | 0 | 0 | 0 | 0 | 1 | 0 | 0 | 0 | 0 | 0 | 0 | 0 | 0 | 0 | 0 | 0 | 0 | 0 | 0 | 0 | 0 | 0 | 0 | 0 | 0 | 0 | 0 | 0 | 0 | 0 | 0 | 0 | 0 | 0 | 0 | 0 | 0 | 0 | 0 | 0 | 0 | 0 | 0 | 0 | 0 | 0 | 0 | 0 | 0 | 0 | 0 | 0 | 0 |
| virus_May12.50v_3419 | 1 | 0 | 0 | 0 | 0 | 0 | 1 | 0 | 0 | 0 | 0 | 0 | 0 | 0 | 0 | 0 | 0 | 0 | 0 | 0 | 0 | 0 | 0 | 0 | 0 | 0 | 0 | 0 | 0 | 0 | 0 | 0 | 0 | 0 | 0 | 0 | 0 | 0 | 0 | 0 | 0 | 0 | 0 | 0 | 0 | 0 | 0 | 0 | 0 | 0 | 0 | 0 | 0 | 0 | 0 |
| virus_May12.50v_5753 | 1 | 0 | 0 | 0 | 0 | 0 | 1 | 0 | 0 | 0 | 0 | 0 | 0 | 0 | 0 | 0 | 0 | [illegible] | [illegible] | [illegible] | [illegible] | [illegible] | [illegible] | [illegible] | [illegible] | [illegible] | [illegible] | [illegible] | [illegible] | [illegible] | [illegible] | [illegible] | [illegible] | [illegible] | [illegible] | [illegible] | [illegible] | [illegible] | [illegible] | [illegible] | [illegible] | [illegible] | [illegible] | [illegible] | [illegible] | [illegible] | [illegible] | [illegible] | [illegible] | [illegible] | [illegible] | [illegible] | [illegible] | [illegible] | [illegible] |
| virus_May12.50v_6335 | 1 | 0 | 0 | 0 | 0 | 0 | 1 | 0 | 0 | 0 | 0 | 0 | 0 | 0 | 0 | 0 | 0 | [illegible] | [illegible] | [illegible] | [illegible] | [illegible] | [illegible] | [illegible] | [illegible] | [illegible] | [illegible] | [illegible] | [illegible] | [illegible] | [illegible] | [illegible] | [illegible] | [illegible] | [illegible] | [illegible] | [illegible] | [illegible] | [illegible] | [illegible] | [illegible] | [illegible] | [illegible] | [illegible] | [illegible] | [illegible] | [illegible] | [illegible] | [illegible] | [illegible] | [illegible] | [illegible] | [illegible] | [illegible] | [illegible] |
| virus_May12.50v_6736 | 1 | 0 | 0 | 0 | 0 | 0 | 1 | 0 | 0 | 0 | 0 | 0 | 0 | 0 | 0 | 0 | 0 | [illegible] | [illegible] | [illegible] | [illegible] | [illegible] | [illegible] | [illegible] | [illegible] | [illegible] | [illegible] | [illegible] | [illegible] | [illegible] | [illegible] | [illegible] | [illegible] | [illegible] | [illegible] | [illegible] | [illegible] | [illegible] | [illegible] | [illegible] | [illegible] | [illegible] | [illegible] | [illegible] | [illegible] | [illegible] | [illegible] | [illegible] | [illegible] | [illegible] | [illegible] | [illegible] | [illegible] | [illegible] | [illegible] |
| virus_May12.50v_10133 | 1 | 0 | 0 | 0 | 0 | 0 | 1 | 0 | 0 | 0 | 0 | 0 | 0 | 0 | 0 | 0 | 0 | [illegible] | [illegible] | [illegible] | [illegible] | [illegible] | [illegible] | [illegible] | [illegible] | [illegible] | [illegible] | [illegible] | [illegible] | [illegible] | [illegible] | [illegible] | [illegible] | [illegible] | [illegible] | [illegible] | [illegible] | [illegible] | [illegible] | [illegible] | [illegible] | [illegible] | [illegible] | [illegible] | [illegible] | [illegible] | [illegible] | [illegible] | [illegible] | [illegible] | [illegible] | [illegible] | [illegible] | [illegible] | [illegible] |
| [illegible] | | | | | | | | | | | | | | | | | | | | | | | | | | | | | | | | | | | | | | | | | | | | | | | | | | | | | | | |

**Supplementary Table S7**. Common Bacterio- and virioplankton RTPR 98% OTUs in 27 samples with RTPR amplicons in both fractions

Total seqs: 47997; Subtotal seqs in retained

| OTUs: 41046 | | May12_05m | | May12_25m | | May12_50m | | May12_75m | | Jun8_25m | | Jun8_50m | | Jun8_75m | | Jun22_0m | | Jun22_50m | | Jun22_75m | | Jul11_0m | | Jul11_05m | | Jul11_25m | | Jul11_50m | | Jul11_75m | | Jul27_05m | | Jul27_25m | | Jul27_75m | | Aug5_0m | | Aug5_05m | | Aug5_25m | | Aug5_50m | | Aug5_75m | | Sep8_0m | | Sep8_05m | | Sep8_25m | | Sep8_50m | |
|---|---|---|---|---|---|---|---|---|---|---|---|---|---|---|---|---|---|---|---|---|---|---|---|---|---|---|---|---|---|---|---|---|---|---|---|---|---|---|---|---|---|---|---|---|---|---|---|---|---|---|---|---|---|---|---|
| OTUs with >=10 seqs: 254 | OTU subtotal | bacterioplankton | virioplankton | bacterioplankton | virioplankton | bacterioplankton | virioplankton | bacterioplankton | virioplankton | bacterioplankton | virioplankton | bacterioplankton | virioplankton | bacterioplankton | virioplankton | bacterioplankton | virioplankton | bacterioplankton | virioplankton | bacterioplankton | virioplankton | bacterioplankton | virioplankton | bacterioplankton | virioplankton | bacterioplankton | virioplankton | bacterioplankton | virioplankton | bacterioplankton | virioplankton | bacterioplankton | virioplankton | bacterioplankton | virioplankton | bacterioplankton | virioplankton | bacterioplankton | virioplankton | bacterioplankton | virioplankton | bacterioplankton | virioplankton | bacterioplankton | virioplankton | bacterioplankton | virioplankton | bacterioplankton | virioplankton | bacterioplankton | virioplankton | bacterioplankton | virioplankton | bacterioplankton | virioplankton |
| virus_Aug5.0v_4007 | 1 | 0 | 0 | 0 | 0 | 0 | 0 | 0 | 0 | 0 | 0 | 0 | 0 | 0 | 0 | 0 | 0 | 0 | 0 | 0 | 0 | 0 | 0 | 0 | 0 | 0 | 0 | 0 | 0 | 0 | 0 | 0 | 0 | 0 | 0 | 0 | 0 | 0 | 1 | 0 | 0 | 0 | 0 | 0 | 0 | 0 | 0 | 0 | 0 | 0 | 0 | 0 | 0 | 0 | 0 |
| virus_Aug5.0v_4045 | 1 | 0 | 0 | 0 | 0 | 0 | 0 | 0 | 0 | 0 | 0 | 0 | 0 | 0 | 0 | 0 | 0 | 0 | 0 | 0 | 0 | 0 | 0 | 0 | 0 | 0 | 0 | 0 | 0 | 0 | 0 | 0 | 0 | 0 | 0 | 0 | 0 | 0 | 1 | 0 | 0 | 0 | 0 | 0 | 0 | 0 | 0 | 0 | 0 | 0 | 0 | 0 | 0 | 0 | 0 |
| virus_Aug5.0v_4098 | 1 | 0 | 0 | 0 | 0 | 0 | 0 | 0 | 0 | 0 | 0 | 0 | 0 | 0 | 0 | 0 | 0 | 0 | 0 | 0 | 0 | 0 | 0 | 0 | 0 | 0 | 0 | 0 | 0 | 0 | 0 | 0 | 0 | 0 | 0 | 0 | 0 | 0 | 1 | 0 | 0 | 0 | 0 | 0 | 0 | 0 | 0 | 0 | 0 | 0 | 0 | 0 | 0 | 0 | 0 |
| virus_Aug5.0v_4228 | 1 | 0 | 0 | 0 | 0 | 0 | 0 | 0 | 0 | 0 | 0 | 0 | 0 | 0 | 0 | 0 | 0 | 0 | 0 | 0 | 0 | 0 | 0 | 0 | 0 | 0 | 0 | 0 | 0 | 0 | 0 | 0 | 0 | 0 | 0 | 0 | 0 | 0 | 1 | 0 | 0 | 0 | 0 | 0 | 0 | 0 | 0 | 0 | 0 | 0 | 0 | 0 | 0 | 0 | 0 |
| virus_Aug5.0v_4777 | 1 | 0 | 0 | 0 | 0 | 0 | 0 | 0 | 0 | 0 | 0 | 0 | 0 | 0 | 0 | 0 | 0 | 0 | 0 | 0 | 0 | 0 | 0 | 0 | 0 | 0 | 0 | 0 | 0 | 0 | 0 | 0 | 0 | 0 | 0 | 0 | 0 | 0 | 1 | 0 | 0 | 0 | 0 | 0 | 0 | 0 | 0 | 0 | 0 | 0 | 0 | 0 | 0 | 0 | 0 |
| virus_Aug5.0v_4813 | 1 | 0 | 0 | 0 | 0 | 0 | 0 | 0 | 0 | 0 | 0 | 0 | 0 | 0 | 0 | 0 | 0 | 0 | 0 | 0 | 0 | 0 | 0 | 0 | 0 | 0 | 0 | 0 | 0 | 0 | 0 | 0 | 0 | 0 | 0 | 0 | 0 | 0 | 1 | 0 | 0 | 0 | 0 | 0 | 0 | 0 | 0 | 0 | 0 | 0 | 0 | 0 | 0 | 0 | 0 |
| virus_Aug5.0v_5214 | 1 | 0 | 0 | 0 | 0 | 0 | 0 | 0 | 0 | 0 | 0 | 0 | 0 | 0 | 0 | 0 | 0 | 0 | 0 | 0 | 0 | 0 | 0 | 0 | 0 | 0 | 0 | 0 | 0 | 0 | 0 | 0 | 0 | 0 | 0 | 0 | 0 | 0 | 1 | 0 | 0 | 0 | 0 | 0 | 0 | 0 | 0 | 0 | 0 | 0 | 0 | 0 | 0 | 0 | 0 |
| virus_Aug5.5v_1027 | 1 | 0 | 0 | 0 | 0 | 0 | 0 | 0 | 0 | 0 | 0 | 0 | 0 | 0 | 0 | 0 | 0 | 0 | 0 | 0 | 0 | 0 | 0 | 0 | 0 | 0 | 0 | 0 | 0 | 0 | 0 | 0 | 0 | 0 | 0 | 0 | 0 | 0 | 0 | 0 | 1 | 0 | 0 | 0 | 0 | 0 | 0 | 0 | 0 | 0 | 0 | 0 | 0 | 0 | 0 |
| virus_Aug5.5v_11011 | 1 | 0 | 0 | 0 | 0 | 0 | 0 | 0 | 0 | 0 | 0 | 0 | 0 | 0 | 0 | 0 | 0 | 0 | 0 | 0 | 0 | 0 | 0 | 0 | 0 | 0 | 0 | 0 | 0 | 0 | 0 | 0 | 0 | 0 | 0 | 0 | 0 | 0 | 0 | 0 | 1 | 0 | 0 | 0 | 0 | 0 | 0 | 0 | 0 | 0 | 0 | 0 | 0 | 0 | 0 |
| virus_Aug5.5v_19223 | 1 | 0 | 0 | 0 | 0 | 0 | 0 | 0 | 0 | 0 | 0 | 0 | 0 | 0 | 0 | 0 | 0 | 0 | 0 | 0 | 0 | 0 | 0 | 0 | 0 | 0 | 0 | 0 | 0 | 0 | 0 | 0 | 0 | 0 | 0 | 0 | 0 | 0 | 0 | 0 | 1 | 0 | 0 | 0 | 0 | 0 | 0 | 0 | 0 | 0 | 0 | 0 | 0 | 0 | 0 |
| virus_Aug5.5v_19590 | 1 | 0 | 0 | 0 | 0 | 0 | 0 | 0 | 0 | 0 | 0 | 0 | 0 | 0 | 0 | 0 | 0 | 0 | 0 | 0 | 0 | 0 | 0 | 0 | 0 | 0 | 0 | 0 | 0 | 0 | 0 | 0 | 0 | 0 | 0 | 0 | 0 | 0 | 0 | 0 | 1 | 0 | 0 | 0 | 0 | 0 | 0 | 0 | 0 | 0 | 0 | 0 | 0 | 0 | 0 |
| virus_Aug5.5v_21555 | 1 | 0 | 0 | 0 | 0 | 0 | 0 | 0 | 0 | 0 | 0 | 0 | 0 | 0 | 0 | 0 | 0 | 0 | 0 | 0 | 0 | 0 | 0 | 0 | 0 | 0 | 0 | 0 | 0 | 0 | 0 | 0 | 0 | 0 | 0 | 0 | 0 | 0 | 0 | 0 | 1 | 0 | 0 | 0 | 0 | 0 | 0 | 0 | 0 | 0 | 0 | 0 | 0 | 0 | 0 |
| virus_Aug5.5v_27059 | 1 | 0 | 0 | 0 | 0 | 0 | 0 | 0 | 0 | 0 | 0 | 0 | 0 | 0 | 0 | 0 | 0 | 0 | 0 | 0 | 0 | 0 | 0 | 0 | 0 | 0 | 0 | 0 | 0 | 0 | 0 | 0 | 0 | 0 | 0 | 0 | 0 | 0 | 0 | 0 | 1 | 0 | 0 | 0 | 0 | 0 | 0 | 0 | 0 | 0 | 0 | 0 | 0 | 0 | 0 |
| virus_Aug5.5v_34126 | 1 | 0 | 0 | 0 | 0 | 0 | 0 | 0 | 0 | 0 | 0 | 0 | 0 | 0 | 0 | 0 | 0 | 0 | 0 | 0 | 0 | 0 | 0 | 0 | 0 | 0 | 0 | 0 | 0 | 0 | 0 | 0 | 0 | 0 | 0 | 0 | 0 | 0 | 0 | 0 | 1 | 0 | 0 | 0 | 0 | 0 | 0 | 0 | 0 | 0 | 0 | 0 | 0 | 0 | 0 |
| virus_Aug5.5v_36038 | 1 | 0 | 0 | 0 | 0 | 0 | 0 | 0 | 0 | 0 | 0 | 0 | 0 | 0 | 0 | 0 | 0 | 0 | 0 | 0 | 0 | 0 | 0 | 0 | 0 | 0 | 0 | 0 | 0 | 0 | 0 | 0 | 0 | 0 | 0 | 0 | 0 | 0 | 0 | 0 | 1 | 0 | 0 | 0 | 0 | 0 | 0 | 0 | 0 | 0 | 0 | 0 | 0 | 0 | 0 |
| virus_Aug5.5v_36607 | 1 | 0 | 0 | 0 | 0 | 0 | 0 | 0 | 0 | 0 | 0 | 0 | 0 | 0 | 0 | 0 | 0 | 0 | 0 | 0 | 0 | 0 | 0 | 0 | 0 | 0 | 0 | 0 | 0 | 0 | 0 | 0 | 0 | 0 | 0 | 0 | 0 | 0 | 0 | 0 | 1 | 0 | 0 | 0 | 0 | 0 | 0 | 0 | 0 | 0 | 0 | 0 | 0 | 0 | 0 |
| virus_Aug5.5v_39651 | 1 | 0 | 0 | 0 | 0 | 0 | 0 | 0 | 0 | 0 | 0 | 0 | 0 | 0 | 0 | 0 | 0 | 0 | 0 | 0 | 0 | 0 | 0 | 0 | 0 | 0 | 0 | 0 | 0 | 0 | 0 | 0 | 0 | 0 | 0 | 0 | 0 | 0 | 0 | 0 | 1 | 0 | 0 | 0 | 0 | 0 | 0 | 0 | 0 | 0 | 0 | 0 | 0 | 0 | 0 |
| virus_Aug5.25v_5018 | 1 | 0 | 0 | 0 | 0 | 0 | 0 | 0 | 0 | 0 | 0 | 0 | 0 | 0 | 0 | 0 | 0 | 0 | 0 | 0 | 0 | 0 | 0 | 0 | 0 | 0 | 0 | 0 | 0 | 0 | 0 | 0 | 0 | 0 | 0 | 0 | 0 | 0 | 0 | 0 | 0 | 0 | 1 | 0 | 0 | 0 | 0 | 0 | 0 | 0 | 0 | 0 | 0 | 0 | 0 |
| virus_Aug5.25v_5611 | 1 | 0 | 0 | 0 | 0 | 0 | 0 | 0 | 0 | 0 | 0 | 0 | 0 | 0 | 0 | 0 | 0 | 0 | 0 | 0 | 0 | 0 | 0 | 0 | 0 | 0 | 0 | 0 | 0 | 0 | 0 | 0 | 0 | 0 | 0 | 0 | 0 | 0 | 0 | 0 | 0 | 0 | 1 | 0 | 0 | 0 | 0 | 0 | 0 | 0 | 0 | 0 | 0 | 0 | 0 |
| virus_Aug5.25v_6389 | 1 | 0 | 0 | 0 | 0 | 0 | 0 | 0 | 0 | 0 | 0 | 0 | 0 | 0 | 0 | 0 | 0 | 0 | 0 | 0 | 0 | 0 | 0 | 0 | 0 | 0 | 0 | 0 | 0 | 0 | 0 | 0 | 0 | 0 | 0 | 0 | 0 | 0 | 0 | 0 | 0 | 0 | 1 | 0 | 0 | 0 | 0 | 0 | 0 | 0 | 0 | 0 | 0 | 0 | 0 |
| virus_Aug5.25v_8358 | 1 | 0 | 0 | 0 | 0 | 0 | 0 | 0 | 0 | 0 | 0 | 0 | 0 | 0 | 0 | 0 | 0 | 0 | 0 | 0 | 0 | 0 | 0 | 0 | 0 | 0 | 0 | 0 | 0 | 0 | 0 | 0 | 0 | 0 | 0 | 0 | 0 | 0 | 0 | 0 | 0 | 0 | 1 | 0 | 0 | 0 | 0 | 0 | 0 | 0 | 0 | 0 | 0 | 0 | 0 |
| virus_Aug5.25v_9617 | 1 | 0 | 0 | 0 | 0 | 0 | 0 | 0 | 0 | 0 | 0 | 0 | 0 | 0 | 0 | 0 | 0 | 0 | 0 | 0 | 0 | 0 | 0 | 0 | 0 | 0 | 0 | 0 | 0 | 0 | 0 | 0 | 0 | 0 | 0 | 0 | 0 | 0 | 0 | 0 | 0 | 0 | 1 | 0 | 0 | 0 | 0 | 0 | 0 | 0 | 0 | 0 | 0 | 0 | 0 |
| virus_Aug5.25v_13579 | 1 | 0 | 0 | 0 | 0 | 0 | 0 | 0 | 0 | 0 | 0 | 0 | 0 | 0 | 0 | 0 | 0 | 0 | 0 | 0 | 0 | 0 | 0 | 0 | 0 | 0 | 0 | 0 | 0 | 0 | 0 | 0 | 0 | 0 | 0 | 0 | 0 | 0 | 0 | 0 | 0 | 0 | 1 | 0 | 0 | 0 | 0 | 0 | 0 | 0 | 0 | 0 | 0 | 0 | 0 |
| virus_Aug5.25v_15035 | 1 | 0 | 0 | 0 | 0 | 0 | 0 | 0 | 0 | 0 | 0 | 0 | 0 | 0 | 0 | 0 | 0 | 0 | 0 | 0 | 0 | 0 | 0 | 0 | 0 | 0 | 0 | 0 | 0 | 0 | 0 | 0 | 0 | 0 | 0 | 0 | 0 | 0 | 0 | 0 | 0 | 0 | 1 | 0 | 0 | 0 | 0 | 0 | 0 | 0 | 0 | 0 | 0 | 0 | 0 |
| virus_Aug5.25v_15566 | 1 | 0 | 0 | 0 | 0 | 0 | 0 | 0 | 0 | 0 | 0 | 0 | 0 | 0 | 0 | 0 | 0 | 0 | 0 | 0 | 0 | 0 | 0 | 0 | 0 | 0 | 0 | 0 | 0 | 0 | 0 | 0 | 0 | 0 | 0 | 0 | 0 | 0 | 0 | 0 | 0 | 0 | 1 | 0 | 0 | 0 | 0 | 0 | 0 | 0 | 0 | 0 | 0 | 0 | 0 |
| virus_Aug5.25v_16100 | 1 | 0 | 0 | 0 | 0 | 0 | 0 | 0 | 0 | 0 | 0 | 0 | 0 | 0 | 0 | 0 | 0 | 0 | 0 | 0 | 0 | 0 | 0 | 0 | 0 | 0 | 0 | 0 | 0 | 0 | 0 | 0 | 0 | 0 | 0 | 0 | 0 | 0 | 0 | 0 | 0 | 0 | 1 | 0 | 0 | 0 | 0 | 0 | 0 | 0 | 0 | 0 | 0 | 0 | 0 |
| virus_Aug5.25v_19477 | 1 | 0 | 0 | 0 | 0 | 0 | 0 | 0 | 0 | 0 | 0 | 0 | 0 | 0 | 0 | 0 | 0 | 0 | 0 | 0 | 0 | 0 | 0 | 0 | 0 | 0 | 0 | 0 | 0 | 0 | 0 | 0 | 0 | 0 | 0 | 0 | 0 | 0 | 0 | 0 | 0 | 0 | 1 | 0 | 0 | 0 | 0 | 0 | 0 | 0 | 0 | 0 | 0 | 0 | 0 |
| virus_Aug5.25v_22020 | 1 | 0 | 0 | 0 | 0 | 0 | 0 | 0 | 0 | 0 | 0 | 0 | 0 | 0 | 0 | 0 | 0 | 0 | 0 | 0 | 0 | 0 | 0 | 0 | 0 | 0 | 0 | 0 | 0 | 0 | 0 | 0 | 0 | 0 | 0 | 0 | 0 | 0 | 0 | 0 | 0 | 0 | 1 | 0 | 0 | 0 | 0 | 0 | 0 | 0 | 0 | 0 | 0 | 0 | 0 |
| virus_Aug5.25v_22567 | 1 | 0 | 0 | 0 | 0 | 0 | 0 | 0 | 0 | 0 | 0 | 0 | 0 | 0 | 0 | 0 | 0 | 0 | 0 | 0 | 0 | 0 | 0 | 0 | 0 | 0 | 0 | 0 | 0 | 0 | 0 | 0 | 0 | 0 | 0 | 0 | 0 | 0 | 0 | 0 | 0 | 0 | 1 | 0 | 0 | 0 | 0 | 0 | 0 | 0 | 0 | 0 | 0 | 0 | 0 |
| virus_Aug5.25v_23323 | 1 | 0 | 0 | 0 | 0 | 0 | 0 | 0 | 0 | 0 | 0 | 0 | 0 | 0 | 0 | 0 | 0 | 0 | 0 | 0 | 0 | 0 | 0 | 0 | 0 | 0 | 0 | 0 | 0 | 0 | 0 | 0 | 0 | 0 | 0 | 0 | 0 | 0 | 0 | 0 | 0 | 0 | 1 | 0 | 0 | 0 | 0 | 0 | 0 | 0 | 0 | 0 | 0 | 0 | 0 |
| virus_Aug5.25v_23484 | 1 | 0 | 0 | 0 | 0 | 0 | 0 | 0 | 0 | 0 | 0 | 0 | 0 | 0 | 0 | 0 | 0 | 0 | 0 | 0 | 0 | 0 | 0 | 0 | 0 | 0 | 0 | 0 | 0 | 0 | 0 | 0 | 0 | 0 | 0 | 0 | 0 | 0 | 0 | 0 | 0 | 0 | 1 | 0 | 0 | 0 | 0 | 0 | 0 | 0 | 0 | 0 | 0 | 0 | 0 |
| virus_Aug5.25v_23672 | 1 | 0 | 0 | 0 | 0 | 0 | 0 | 0 | 0 | 0 | 0 | 0 | 0 | 0 | 0 | 0 | 0 | 0 | 0 | 0 | 0 | 0 | 0 | 0 | 0 | 0 | 0 | 0 | 0 | 0 | 0 | 0 | 0 | 0 | 0 | 0 | 0 | 0 | 0 | 0 | 0 | 0 | 1 | 0 | 0 | 0 | 0 | 0 | 0 | 0 | 0 | 0 | 0 | 0 | 0 |
| virus_Aug5.25v_25630 | 1 | 0 | 0 | 0 | 0 | 0 | 0 | 0 | 0 | 0 | 0 | 0 | 0 | 0 | 0 | 0 | 0 | 0 | 0 | 0 | 0 | 0 | 0 | 0 | 0 | 0 | 0 | 0 | 0 | 0 | 0 | 0 | 0 | 0 | 0 | 0 | 0 | 0 | 0 | 0 | 0 | 0 | 1 | 0 | 0 | 0 | 0 | 0 | 0 | 0 | 0 | 0 | 0 | 0 | 0 |
| virus_Aug5.25v_26039 | 1 | 0 | 0 | 0 | 0 | 0 | 0 | 0 | 0 | 0 | 0 | 0 | 0 | 0 | 0 | 0 | 0 | 0 | 0 | 0 | 0 | 0 | 0 | 0 | 0 | 0 | 0 | 0 | 0 | 0 | 0 | 0 | 0 | 0 | 0 | 0 | 0 | 0 | 0 | 0 | 0 | 0 | 1 | 0 | 0 | 0 | 0 | 0 | 0 | 0 | 0 | 0 | 0 | 0 | 0 |
| virus_Aug5.25v_27480 | 1 | 0 | 0 | 0 | 0 | 0 | 0 | 0 | 0 | 0 | 0 | 0 | 0 | 0 | 0 | 0 | 0 | 0 | 0 | 0 | 0 | 0 | 0 | 0 | 0 | 0 | 0 | 0 | 0 | 0 | 0 | 0 | 0 | 0 | 0 | 0 | 0 | 0 | 0 | 0 | 0 | 0 | 1 | 0 | 0 | 0 | 0 | 0 | 0 | 0 | 0 | 0 | 0 | 0 | 0 |
| virus_Aug5.25v_27726 | 1 | 0 | 0 | 0 | 0 | 0 | 0 | 0 | 0 | 0 | 0 | 0 | 0 | 0 | 0 | 0 | 0 | 0 | 0 | 0 | 0 | 0 | 0 | 0 | 0 | 0 | 0 | 0 | 0 | 0 | 0 | 0 | 0 | 0 | 0 | 0 | 0 | 0 | 0 | 0 | 0 | 0 | 1 | 0 | 0 | 0 | 0 | 0 | 0 | 0 | 0 | 0 | 0 | 0 | 0 |
| virus_Aug5.25v_28987 | 1 | 0 | 0 | 0 | 0 | 0 | 0 | 0 | 0 | 0 | 0 | 0 | 0 | 0 | 0 | 0 | 0 | 0 | 0 | 0 | 0 | 0 | 0 | 0 | 0 | 0 | 0 | 0 | 0 | 0 | 0 | 0 | 0 | 0 | 0 | 0 | 0 | 0 | 0 | 0 | 0 | 0 | 1 | 0 | 0 | 0 | 0 | 0 | 0 | 0 | 0 | 0 | 0 | 0 | 0 |
| virus_Aug5.25v_29117 | 1 | 0 | 0 | 0 | 0 | 0 | 0 | 0 | 0 | 0 | 0 | 0 | 0 | 0 | 0 | 0 | 0 | 0 | 0 | 0 | 0 | 0 | 0 | 0 | 0 | 0 | 0 | 0 | 0 | 0 | 0 | 0 | 0 | 0 | 0 | 0 | 0 | 0 | 0 | 0 | 0 | 0 | 1 | 0 | 0 | 0 | 0 | 0 | 0 | 0 | 0 | 0 | 0 | 0 | 0 |
| virus_Aug5.25v_33610 | 1 | 0 | 0 | 0 | 0 | 0 | 0 | 0 | 0 | 0 | 0 | 0 | 0 | 0 | 0 | 0 | 0 | 0 | 0 | 0 | 0 | 0 | 0 | 0 | 0 | 0 | 0 | 0 | 0 | 0 | 0 | 0 | 0 | 0 | 0 | 0 | 0 | 0 | 0 | 0 | 0 | 0 | 1 | 0 | 0 | 0 | 0 | 0 | 0 | 0 | 0 | 0 | 0 | 0 | 0 |
| virus_Aug5.25v_34955 | 1 | 0 | 0 | 0 | 0 | 0 | 0 | 0 | 0 | 0 | 0 | 0 | 0 | 0 | 0 | 0 | 0 | 0 | 0 | 0 | 0 | 0 | 0 | 0 | 0 | 0 | 0 | 0 | 0 | 0 | 0 | 0 | 0 | 0 | 0 | 0 | 0 | 0 | 0 | 0 | 0 | 0 | 1 | 0 | 0 | 0 | 0 | 0 | 0 | 0 | 0 | 0 | 0 | 0 | 0 |
| virus_Aug5.25v_39753 | 1 | 0 | 0 | 0 | 0 | 0 | 0 | 0 | 0 | 0 | 0 | 0 | 0 | 0 | 0 | 0 | 0 | 0 | 0 | 0 | 0 | 0 | 0 | 0 | 0 | 0 | 0 | 0 | 0 | 0 | 0 | 0 | 0 | 0 | 0 | 0 | 0 | 0 | 0 | 0 | 0 | 0 | 1 | 0 | 0 | 0 | 0 | 0 | 0 | 0 | 0 | 0 | 0 | 0 | 0 |
| virus_Aug5.25v_40751 | 1 | 0 | 0 | 0 | 0 | 0 | 0 | 0 | 0 | 0 | 0 | 0 | 0 | 0 | 0 | 0 | 0 | 0 | 0 | 0 | 0 | 0 | 0 | 0 | 0 | 0 | 0 | 0 | 0 | 0 | 0 | 0 | 0 | 0 | 0 | 0 | 0 | 0 | 0 | 0 | 0 | 0 | 1 | 0 | 0 | 0 | 0 | 0 | 0 | 0 | 0 | 0 | 0 | 0 | 0 |
| virus_Aug5.50v_720 | 1 | 0 | 0 | 0 | 0 | 0 | 0 | 0 | 0 | 0 | 0 | 0 | 0 | 0 | 0 | 0 | 0 | 0 | 0 | 0 | 0 | 0 | 0 | 0 | 0 | 0 | 0 | 0 | 0 | 0 | 0 | 0 | 0 | 0 | 0 | 0 | 0 | 0 | 0 | 0 | 0 | 0 | 0 | 0 | 1 | 0 | 0 | 0 | 0 | 0 | 0 | 0 | 0 | 0 | 0 |
| virus_Aug5.50v_14187 | 1 | 0 | 0 | 0 | 0 | 0 | 0 | 0 | 0 | 0 | 0 | 0 | 0 | 0 | 0 | 0 | 0 | 0 | 0 | 0 | 0 | 0 | 0 | 0 | 0 | 0 | 0 | 0 | 0 | 0 | 0 | 0 | 0 | 0 | 0 | 0 | 0 | 0 | 0 | 0 | 0 | 0 | 0 | 0 | 1 | 0 | 0 | 0 | 0 | 0 | 0 | 0 | 0 | 0 | 0 |
| virus_Aug5.50v_15857 | 1 | 0 | 0 | 0 | 0 | 0 | 0 | 0 | 0 | 0 | 0 | 0 | 0 | 0 | 0 | 0 | 0 | 0 | 0 | 0 | 0 | 0 | 0 | 0 | 0 | 0 | 0 | 0 | 0 | 0 | 0 | 0 | 0 | 0 | 0 | 0 | 0 | 0 | 0 | 0 | 0 | 0 | 0 | 0 | 1 | 0 | 0 | 0 | 0 | 0 | 0 | 0 | 0 | 0 | 0 |
| virus_Aug5.50v_18806 | 1 | 0 | 0 | 0 | 0 | 0 | 0 | 0 | 0 | 0 | 0 | 0 | 0 | 0 | 0 | 0 | 0 | 0 | 0 | 0 | 0 | 0 | 0 | 0 | 0 | 0 | 0 | 0 | 0 | 0 | 0 | 0 | 0 | 0 | 0 | 0 | 0 | 0 | 0 | 0 | 0 | 0 | 0 | 0 | 1 | 0 | 0 | 0 | 0 | 0 | 0 | 0 | 0 | 0 | 0 |
| virus_Aug5.50v_25476 | 1 | 0 | 0 | 0 | 0 | 0 | 0 | 0 | 0 | 0 | 0 | 0 | 0 | 0 | 0 | 0 | 0 | 0 | 0 | 0 | 0 | 0 | 0 | 0 | 0 | 0 | 0 | 0 | 0 | 0 | 0 | 0 | 0 | 0 | 0 | 0 | 0 | 0 | 0 | 0 | 0 | 0 | 0 | 0 | 1 | 0 | 0 | 0 | 0 | 0 | 0 | 0 | 0 | 0 | 0 |
| virus_Aug5.50v_26588 | 1 | 0 | 0 | 0 | 0 | 0 | 0 | 0 | 0 | 0 | 0 | 0 | 0 | 0 | 0 | 0 | 0 | 0 | 0 | 0 | 0 | 0 | 0 | 0 | 0 | 0 | 0 | 0 | 0 | 0 | 0 | 0 | 0 | 0 | 0 | 0 | 0 | 0 | 0 | 0 | 0 | 0 | 0 | 0 | 1 | 0 | 0 | 0 | 0 | 0 | 0 | 0 | 0 | 0 | 0 |
| virus_Aug5.50v_32577 | 1 | 0 | 0 | 0 | 0 | 0 | 0 | 0 | 0 | 0 | 0 | 0 | 0 | 0 | 0 | 0 | 0 | 0 | 0 | 0 | 0 | 0 | 0 | 0 | 0 | 0 | 0 | 0 | 0 | 0 | 0 | 0 | 0 | 0 | 0 | 0 | 0 | 0 | 0 | 0 | 0 | 0 | 0 | 0 | 1 | 0 | 0 | 0 | 0 | 0 | 0 | 0 | 0 | 0 | 0 |
| virus_Aug5.50v_32970 | 1 | 0 | 0 | 0 | 0 | 0 | 0 | 0 | 0 | 0 | 0 | 0 | 0 | 0 | 0 | 0 | 0 | 0 | 0 | 0 | 0 | 0 | 0 | 0 | 0 | 0 | 0 | 0 | 0 | 0 | 0 | 0 | 0 | 0 | 0 | 0 | 0 | 0 | 0 | 0 | 0 | 0 | 0 | 0 | 1 | 0 | 0 | 0 | 0 | 0 | 0 | 0 | 0 | 0 | 0 |
| virus_Aug5.50v_37664 | 1 | 0 | 0 | 0 | 0 | 0 | 0 | 0 | 0 | 0 | 0 | 0 | 0 | 0 | 0 | 0 | 0 | 0 | 0 | 0 | 0 | 0 | 0 | 0 | 0 | 0 | 0 | 0 | 0 | 0 | 0 | 0 | 0 | 0 | 0 | 0 | 0 | 0 | 0 | 0 | 0 | 0 | 0 | 0 | 1 | 0 | 0 | 0 | 0 | 0 | 0 | 0 | 0 | 0 | 0 |
| virus_Aug5.50v_41091 | 1 | 0 | 0 | 0 | 0 | 0 | 0 | 0 | 0 | 0 | 0 | 0 | 0 | 0 | 0 | 0 | 0 | 0 | 0 | 0 | 0 | 0 | 0 | 0 | 0 | 0 | 0 | 0 | 0 | 0 | 0 | 0 | 0 | 0 | 0 | 0 | 0 | 0 | 0 | 0 | 0 | 0 | 0 | 0 | 1 | 0 | 0 | 0 | 0 | 0 | 0 | 0 | 0 | 0 | 0 |
| virus_Aug5.75v_9587 | 1 | 0 | 0 | 0 | 0 | 0 | 0 | 0 | 0 | 0 | 0 | 0 | 0 | 0 | 0 | 0 | 0 | 0 | 0 | 0 | 0 | 0 | 0 | 0 | 0 | 0 | 0 | 0 | 0 | 0 | 0 | 0 | 0 | 0 | 0 | 0 | 0 | 0 | 0 | 0 | 0 | 0 | 0 | 0 | 0 | 0 | 1 | 0 | 0 | 0 | 0 | 0 | 0 | 0 | 0 |
| virus_Aug5.75v_16676 | 1 | 0 | 0 | 0 | 0 | 0 | 0 | 0 | 0 | 0 | 0 | 0 | 0 | 0 | 0 | 0 | 0 | 0 | 0 | 0 | 0 | 0 | 0 | 0 | 0 | 0 | 0 | 0 | 0 | 0 | 0 | 0 | 0 | 0 | 0 | 0 | 0 | 0 | 0 | 0 | 0 | 0 | 0 | 0 | 0 | 0 | 1 | 0 | 0 | 0 | 0 | 0 | 0 | 0 | 0 |
| virus_Aug5.75v_17768 | 1 | 0 | 0 | 0 | 0 | 0 | 0 | 0 | 0 | 0 | 0 | 0 | 0 | 0 | 0 | 0 | 0 | 0 | 0 | 0 | 0 | 0 | 0 | 0 | 0 | 0 | 0 | 0 | 0 | 0 | 0 | 0 | 0 | 0 | 0 | 0 | 0 | 0 | 0 | 0 | 0 | 0 | 0 | 0 | 0 | 0 | 1 | 0 | 0 | 0 | 0 | 0 | 0 | 0 | 0 |
| virus_Aug5.75v_19491 | 1 | 0 | 0 | 0 | 0 | 0 | 0 | 0 | 0 | 0 | 0 | 0 | 0 | 0 | 0 | 0 | 0 | 0 | 0 | 0 | 0 | 0 | 0 | 0 | 0 | 0 | 0 | 0 | 0 | 0 | 0 | 0 | 0 | 0 | 0 | 0 | 0 | 0 | 0 | 0 | 0 | 0 | 0 | 0 | 0 | 0 | 1 | 0 | 0 | 0 | 0 | 0 | 0 | 0 | 0 |
| virus_Jul11.0v_442 | 1 | 0 | 0 | 0 | 0 | 0 | 0 | 0 | 0 | 0 | 0 | 0 | 0 | 0 | 0 | 0 | 0 | 0 | 0 | 0 | 0 | 0 | 1 | 0 | 0 | 0 | 0 | 0 | 0 | 0 | 0 | 0 | 0 | 0 | 0 | 0 | 0 | 0 | 0 | 0 | 0 | 0 | 0 | 0 | 0 | 0 | 0 | 0 | 0 | 0 | 0 | 0 | 0 | 0 | 0 |
| virus_Jul11.0v_4604 | 1 | 0 | 0 | 0 | 0 | 0 | 0 | 0 | 0 | 0 | 0 | 0 | 0 | 0 | 0 | 0 | 0 | 0 | 0 | 0 | 0 | 0 | 1 | 0 | 0 | 0 | 0 | 0 | 0 | 0 | 0 | 0 | 0 | 0 | 0 | 0 | 0 | 0 | 0 | 0 | 0 | 0 | 0 | 0 | 0 | 0 | 0 | 0 | 0 | 0 | 0 | 0 | 0 | 0 | 0 |
| virus_Jul11.0v_5551 | 1 | 0 | 0 | 0 | 0 | 0 | 0 | 0 | 0 | 0 | 0 | 0 | 0 | 0 | 0 | 0 | 0 | 0 | 0 | 0 | 0 | 0 | 1 | 0 | 0 | 0 | 0 | 0 | 0 | 0 | 0 | 0 | 0 | 0 | 0 | 0 | 0 | 0 | 0 | 0 | 0 | 0 | 0 | 0 | 0 | 0 | 0 | 0 | 0 | 0 | 0 | 0 | 0 | 0 | 0 |
| virus_Jul11.0v_7048 | 1 | 0 | 0 | 0 | 0 | 0 | 0 | 0 | 0 | 0 | 0 | 0 | 0 | 0 | 0 | 0 | 0 | 0 | 0 | 0 | 0 | 0 | 1 | 0 | 0 | 0 | 0 | 0 | 0 | 0 | 0 | 0 | 0 | 0 | 0 | 0 | 0 | 0 | 0 | 0 | 0 | 0 | 0 | 0 | 0 | 0 | 0 | 0 | 0 | 0 | 0 | 0 | 0 | 0 | 0 |
| virus_Jul11.0v_8126 | 1 | 0 | 0 | 0 | 0 | 0 | 0 | 0 | 0 | 0 | 0 | 0 | 0 | 0 | 0 | 0 | 0 | 0 | 0 | 0 | 0 | 0 | 1 | 0 | 0 | 0 | 0 | 0 | 0 | 0 | 0 | 0 | 0 | 0 | 0 | 0 | 0 | 0 | 0 | 0 | 0 | 0 | 0 | 0 | 0 | 0 | 0 | 0 | 0 | 0 | 0 | 0 | 0 | 0 | 0 |
| virus_Jul11.0v_9962 | 1 | 0 | 0 | 0 | 0 | 0 | 0 | 0 | 0 | 0 | 0 | 0 | 0 | 0 | 0 | 0 | 0 | 0 | 0 | 0 | 0 | 0 | 1 | 0 | 0 | 0 | 0 | 0 | 0 | 0 | 0 | 0 | 0 | 0 | 0 | 0 | 0 | 0 | 0 | 0 | 0 | 0 | 0 | 0 | 0 | 0 | 0 | 0 | 0 | 0 | 0 | 0 | 0 | 0 | 0 |
| virus_Jul11.0v_10133 | 1 | 0 | 0 | 0 | 0 | 0 | 0 | 0 | 0 | 0 | 0 | 0 | 0 | 0 | 0 | 0 | 0 | 0 | 0 | 0 | 0 | 0 | 1 | 0 | 0 | 0 | 0 | 0 | 0 | 0 | 0 | 0 | 0 | 0 | 0 | 0 | 0 | 0 | 0 | 0 | 0 | 0 | 0 | 0 | 0 | 0 | 0 | 0 | 0 | 0 | 0 | 0 | 0 | 0 | 0 |
| virus_Jul11.0v_14097 | 1 | 0 | 0 | 0 | 0 | 0 | 0 | 0 | 0 | 0 | 0 | 0 | 0 | 0 | 0 | 0 | 0 | 0 | 0 | 0 | 0 | 0 | 1 | 0 | 0 | 0 | 0 | 0 | 0 | 0 | 0 | 0 | 0 | 0 | 0 | 0 | 0 | 0 | 0 | 0 | 0 | 0 | 0 | 0 | 0 | 0 | 0 | 0 | 0 | 0 | 0 | 0 | 0 | 0 | 0 |
| virus_Jul11.5v_96 | 1 | 0 | 0 | 0 | 0 | 0 | 0 | 0 | 0 | 0 | 0 | 0 | 0 | 0 | 0 | 0 | 0 | 0 | 0 | 0 | 0 | 0 | 0 | 0 | 1 | 0 | 0 | 0 | 0 | 0 | 0 | 0 | 0 | 0 | 0 | 0 | 0 | 0 | 0 | 0 | 0 | 0 | 0 | 0 | 0 | 0 | 0 | 0 | 0 | 0 | 0 | 0 | 0 | 0 | 0 |
| virus_Jul11.5v_2249 | 1 | 0 | 0 | 0 | 0 | 0 | 0 | 0 | 0 | 0 | 0 | 0 | 0 | 0 | 0 | 0 | 0 | 0 | 0 | 0 | 0 | 0 | 0 | 0 | 1 | 0 | 0 | 0 | 0 | 0 | 0 | 0 | 0 | 0 | 0 | 0 | 0 | 0 | 0 | 0 | 0 | 0 | 0 | 0 | 0 | 0 | 0 | 0 | 0 | 0 | 0 | 0 | 0 | 0 | 0 |
| virus_Jul11.5v_2297 | 1 | 0 | 0 | 0 | 0 | 0 | 0 | 0 | 0 | 0 | 0 | 0 | 0 | 0 | 0 | 0 | 0 | 0 | 0 | 0 | 0 | 0 | 0 | 0 | 1 | 0 | 0 | 0 | 0 | 0 | 0 | 0 | 0 | 0 | 0 | 0 | 0 | 0 | 0 | 0 | 0 | 0 | 0 | 0 | 0 | 0 | 0 | 0 | 0 | 0 | 0 | 0 | 0 | 0 | 0 |
| virus_Jul11.5v_3433 | 1 | 0 | 0 | 0 | 0 | 0 | 0 | 0 | 0 | 0 | 0 | 0 | 0 | 0 | 0 | 0 | 0 | 0 | 0 | 0 | 0 | 0 | 0 | 0 | 1 | 0 | 0 | 0 | 0 | 0 | 0 | 0 | 0 | 0 | 0 | 0 | 0 | 0 | 0 | 0 | 0 | 0 | 0 | 0 | 0 | 0 | 0 | 0 | 0 | 0 | 0 | 0 | 0 | 0 | 0 |
| virus_Jul11.5v_4199 | 1 | 0 | 0 | 0 | 0 | 0 | 0 | 0 | 0 | 0 | 0 | 0 | 0 | 0 | 0 | 0 | 0 | 0 | 0 | 0 | 0 | 0 | 0 | 0 | 1 | 0 | 0 | 0 | 0 | 0 | 0 | 0 | 0 | 0 | 0 | 0 | 0 | 0 | 0 | 0 | 0 | 0 | 0 | 0 | 0 | 0 | 0 | 0 | 0 | 0 | 0 | 0 | 0 | 0 | 0 |
| virus_Jul11.5v_5665 | 1 | 0 | 0 | 0 | 0 | 0 | 0 | 0 | 0 | 0 | 0 | 0 | 0 | 0 | 0 | 0 | 0 | 0 | 0 | 0 | 0 | 0 | 0 | 0 | 1 | 0 | 0 | 0 | 0 | 0 | 0 | 0 | 0 | 0 | 0 | 0 | 0 | 0 | 0 | 0 | 0 | 0 | 0 | 0 | 0 | 0 | 0 | 0 | 0 | 0 | 0 | 0 | 0 | 0 | 0 |
| virus_Jul11.5v_8581 | 1 | 0 | 0 | 0 | 0 | 0 | 0 | 0 | 0 | 0 | 0 | 0 | 0 | 0 | 0 | 0 | 0 | 0 | 0 | 0 | 0 | 0 | 0 | 0 | 1 | 0 | 0 | 0 | 0 | 0 | 0 | 0 | 0 | 0 | 0 | 0 | 0 | 0 | 0 | 0 | 0 | 0 | 0 | 0 | 0 | 0 | 0 | 0 | 0 | 0 | 0 | 0 | 0 | 0 | 0 |
| virus_Jul11.5v_9486 | 1 | 0 | 0 | 0 | 0 | 0 | 0 | 0 | 0 | 0 | 0 | 0 | 0 | 0 | 0 | 0 | 0 | 0 | 0 | 0 | 0 | 0 | 0 | 0 | 1 | 0 | 0 | 0 | 0 | 0 | 0 | 0 | 0 | 0 | 0 | 0 | 0 | 0 | 0 | 0 | 0 | 0 | 0 | 0 | 0 | 0 | 0 | 0 | 0 | 0 | 0 | 0 | 0 | 0 | 0 |
| virus_Jul11.5v_12100 | 1 | 0 | 0 | 0 | 0 | 0 | 0 | 0 | 0 | 0 | 0 | 0 | 0 | 0 | 0 | 0 | 0 | 0 | 0 | 0 | 0 | 0 | 0 | 0 | 1 | 0 | 0 | 0 | 0 | 0 | 0 | 0 | 0 | 0 | 0 | 0 | 0 | 0 | 0 | 0 | 0 | 0 | 0 | 0 | 0 | 0 | 0 | 0 | 0 | 0 | 0 | 0 | 0 | 0 | 0 |
| virus_Jul11.5v_12724 | 1 | 0 | 0 | 0 | 0 | 0 | 0 | 0 | 0 | 0 | 0 | 0 | 0 | 0 | 0 | 0 | 0 | 0 | 0 | 0 | 0 | 0 | 0 | 0 | 1 | 0 | 0 | 0 | 0 | 0 | 0 | 0 | 0 | 0 | 0 | 0 | 0 | 0 | 0 | 0 | 0 | 0 | 0 | 0 | 0 | 0 | 0 | 0 | 0 | 0 | 0 | 0 | 0 | 0 | 0 |
| virus_Jul11.5v_13492 | 1 | 0 | 0 | 0 | 0 | 0 | 0 | 0 | 0 | 0 | 0 | 0 | 0 | 0 | 0 | 0 | 0 | 0 | 0 | 0 | 0 | 0 | 0 | 0 | 1 | 0 | 0 | 0 | 0 | 0 | 0 | 0 | 0 | 0 | 0 | 0 | 0 | 0 | 0 | 0 | 0 | 0 | 0 | 0 | 0 | 0 | 0 | 0 | 0 | 0 | 0 | 0 | 0 | 0 | 0 |
| virus_Jul11.5v_14178 | 1 | 0 | 0 | 0 | 0 | 0 | 0 | 0 | 0 | 0 | 0 | 0 | 0 | 0 | 0 | 0 | 0 | 0 | 0 | 0 | 0 | 0 | 0 | 0 | 1 | 0 | 0 | 0 | 0 | 0 | 0 | 0 | 0 | 0 | 0 | 0 | 0 | 0 | 0 | 0 | 0 | 0 | 0 | 0 | 0 | 0 | 0 | 0 | 0 | 0 | 0 | 0 | 0 | 0 | 0 |
| virus_Jul11.5v_14940 | 1 | 0 | 0 | 0 | 0 | 0 | 0 | 0 | 0 | 0 | 0 | 0 | 0 | 0 | 0 | 0 | 0 | 0 | 0 | 0 | 0 | 0 | 0 | 0 | 1 | 0 | 0 | 0 | 0 | 0 | 0 | 0 | 0 | 0 | 0 | 0 | 0 | 0 | 0 | 0 | 0 | 0 | 0 | 0 | 0 | 0 | 0 | 0 | 0 | 0 | 0 | 0 | 0 | 0 | 0 |
| virus_Jul11.5v_15366 | 1 | 0 | 0 | 0 | 0 | 0 | 0 | 0 | 0 | 0 | 0 | 0 | 0 | 0 | 0 | 0 | 0 | 0 | 0 | 0 | 0 | 0 | 0 | 0 | 1 | 0 | 0 | 0 | 0 | 0 | 0 | 0 | 0 | 0 | 0 | 0 | 0 | 0 | 0 | 0 | 0 | 0 | 0 | 0 | 0 | 0 | 0 | 0 | 0 | 0 | 0 | 0 | 0 | 0 | 0 |
| virus_Jul11.5v_15420 | 1 | 0 | 0 | 0 | 0 | 0 | 0 | 0 | 0 | 0 | 0 | 0 | 0 | 0 | 0 | 0 | 0 | 0 | 0 | 0 | 0 | 0 | 0 | 0 | 1 | 0 | 0 | 0 | 0 | 0 | 0 | 0 | 0 | 0 | 0 | 0 | 0 | 0 | 0 | 0 | 0 | 0 | 0 | 0 | 0 | 0 | 0 | 0 | 0 | 0 | 0 | 0 | 0 | 0 | 0 |
| virus_Jul11.5v_17790 | 1 | 0 | 0 | 0 | 0 | 0 | 0 | 0 | 0 | 0 | 0 | 0 | 0 | 0 | 0 | 0 | 0 | 0 | 0 | 0 | 0 | 0 | 0 | 0 | 1 | 0 | 0 | 0 | 0 | 0 | 0 | 0 | 0 | 0 | 0 | 0 | 0 | 0 | 0 | 0 | 0 | 0 | 0 | 0 | 0 | 0 | 0 | 0 | 0 | 0 | 0 | 0 | 0 | 0 | 0 |
| virus_Jul11.5v_18748 | 1 | 0 | 0 | 0 | 0 | 0 | 0 | 0 | 0 | 0 | 0 | 0 | 0 | 0 | 0 | 0 | 0 | 0 | 0 | 0 | 0 | 0 | 0 | 0 | 1 | 0 | 0 | 0 | 0 | 0 | 0 | 0 | 0 | 0 | 0 | 0 | 0 | 0 | 0 | 0 | 0 | 0 | 0 | 0 | 0 | 0 | 0 | 0 | 0 | 0 | 0 | 0 | 0 | 0 | 0 |
| virus_Jul11.5v_20090 | 1 | 0 | 0 | 0 | 0 | 0 | 0 | 0 | 0 | 0 | 0 | 0 | 0 | 0 | 0 | 0 | 0 | 0 | 0 | 0 | 0 | 0 | 0 | 0 | 1 | 0 | 0 | 0 | 0 | 0 | 0 | 0 | 0 | 0 | 0 | 0 | 0 | 0 | 0 | 0 | 0 | 0 | 0 | 0 | 0 | 0 | 0 | 0 | 0 | 0 | 0 | 0 | 0 | 0 | 0 |
| virus_Jul11.5v_20509 | 1 | 0 | 0 | 0 | 0 | 0 | 0 | 0 | 0 | 0 | 0 | 0 | 0 | 0 | 0 | 0 | 0 | 0 | 0 | 0 | 0 | 0 | 0 | 0 | 1 | 0 | 0 | 0 | 0 | 0 | 0 | 0 | 0 | 0 | 0 | 0 | 0 | 0 | 0 | 0 | 0 | 0 | 0 | 0 | 0 | 0 | 0 | 0 | 0 | 0 | 0 | 0 | 0 | 0 | 0 |
| virus_Jul11.25v_1083 | 1 | 0 | 0 | 0 | 0 | 0 | 0 | 0 | 0 | 0 | 0 | 0 | 0 | 0 | 0 | 0 | 0 | 0 | 0 | 0 | 0 | 0 | 0 | 0 | 0 | 0 | 1 | 0 | 0 | 0 | 0 | 0 | 0 | 0 | 0 | 0 | 0 | 0 | 0 | 0 | 0 | 0 | 0 | 0 | 0 | 0 | 0 | 0 | 0 | 0 | 0 | 0 | 0 | 0 | 0 |
| virus_Jul11.25v_3761 | 1 | 0 | 0 | 0 | 0 | 0 | 0 | 0 | 0 | 0 | 0 | 0 | 0 | 0 | 0 | 0 | 0 | 0 | 0 | 0 | 0 | 0 | 0 | 0 | 0 | 0 | 1 | 0 | 0 | 0 | 0 | 0 | 0 | 0 | 0 | 0 | 0 | 0 | 0 | 0 | 0 | 0 | 0 | 0 | 0 | 0 | 0 | 0 | 0 | 0 | 0 | 0 | 0 | 0 | 0 |
| virus_Jul11.25v_4477 | 1 | 0 | 0 | 0 | 0 | 0 | 0 | 0 | 0 | 0 | 0 | 0 | 0 | 0 | 0 | 0 | 0 | 0 | 0 | 0 | 0 | 0 | 0 | 0 | 0 | 0 | 1 | 0 | 0 | 0 | 0 | 0 | 0 | 0 | 0 | 0 | 0 | 0 | 0 | 0 | 0 | 0 | 0 | 0 | 0 | 0 | 0 | 0 | 0 | 0 | 0 | 0 | 0 | 0 | 0 |
| virus_Jul11.25v_6383 | 1 | 0 | 0 | 0 | 0 | 0 | 0 | 0 | 0 | 0 | 0 | 0 | 0 | 0 | 0 | 0 | 0 | 0 | 0 | 0 | 0 | 0 | 0 | 0 | 0 | 0 | 1 | 0 | 0 | 0 | 0 | 0 | 0 | 0 | 0 | 0 | 0 | 0 | 0 | 0 | 0 | 0 | 0 | 0 | 0 | 0 | 0 | 0 | 0 | 0 | 0 | 0 | 0 | 0 | 0 |
| virus_Jul11.25v_8611 | 1 | 0 | 0 | 0 | 0 | 0 | 0 | 0 | 0 | 0 | 0 | 0 | 0 | 0 | 0 | 0 | 0 | 0 | 0 | 0 | 0 | 0 | 0 | 0 | 0 | 0 | 1 | 0 | 0 | 0 | 0 | 0 | 0 | 0 | 0 | 0 | 0 | 0 | 0 | 0 | 0 | 0 | 0 | 0 | 0 | 0 | 0 | 0 | 0 | 0 | 0 | 0 | 0 | 0 | 0 |
| virus_Jul11.25v_13188 | 1 | 0 | 0 | 0 | 0 | 0 | 0 | 0 | 0 | 0 | 0 | 0 | 0 | 0 | 0 | 0 | 0 | 0 | 0 | 0 | 0 | 0 | 0 | 0 | 0 | 0 | 1 | 0 | 0 | 0 | 0 | 0 | 0 | 0 | 0 | 0 | 0 | 0 | 0 | 0 | 0 | 0 | 0 | 0 | 0 | 0 | 0 | 0 | 0 | 0 | 0 | 0 | 0 | 0 | 0 |
| virus_Jul11.25v_15692 | 1 | 0 | 0 | 0 | 0 | 0 | 0 | 0 | 0 | 0 | 0 | 0 | 0 | 0 | 0 | 0 | 0 | 0 | 0 | 0 | 0 | 0 | 0 | 0 | 0 | 0 | 1 | 0 | 0 | 0 | 0 | 0 | 0 | 0 | 0 | 0 | 0 | 0 | 0 | 0 | 0 | 0 | 0 | 0 | 0 | 0 | 0 | 0 | 0 | 0 | 0 | 0 | 0 | 0 | 0 |
| virus_Jul11.25v_17401 | 1 | 0 | 0 | 0 | 0 | 0 | 0 | 0 | 0 | 0 | 0 | 0 | 0 | 0 | 0 | 0 | 0 | 0 | 0 | 0 | 0 | 0 | 0 | 0 | 0 | 0 | 1 | 0 | 0 | 0 | 0 | 0 | 0 | 0 | 0 | 0 | 0 | 0 | 0 | 0 | 0 | 0 | 0 | 0 | 0 | 0 | 0 | 0 | 0 | 0 | 0 | 0 | 0 | 0 | 0 |
| virus_Jul11.25v_18199 | 1 | 0 | 0 | 0 | 0 | 0 | 0 | 0 | 0 | 0 | 0 | 0 | 0 | 0 | 0 | 0 | 0 | 0 | 0 | 0 | 0 | 0 | 0 | 0 | 0 | 0 | 1 | 0 | 0 | 0 | 0 | 0 | 0 | 0 | 0 | 0 | 0 | 0 | 0 | 0 | 0 | 0 | 0 | 0 | 0 | 0 | 0 | 0 | 0 | 0 | 0 | 0 | 0 | 0 | 0 |
| virus_Jul11.25v_19730 | 1 | 0 | 0 | 0 | 0 | 0 | 0 | 0 | 0 | 0 | 0 | 0 | 0 | 0 | 0 | 0 | 0 | 0 | 0 | 0 | 0 | 0 | 0 | 0 | 0 | 0 | 1 | 0 | 0 | 0 | 0 | 0 | 0 | 0 | 0 | 0 | 0 | 0 | 0 | 0 | 0 | 0 | 0 | 0 | 0 | 0 | 0 | 0 | 0 | 0 | 0 | 0 | 0 | 0 | 0 |
| virus_Jul11.50v_760 | 1 | 0 | 0 | 0 | 0 | 0 | 0 | 0 | 0 | 0 | 0 | 0 | 0 | 0 | 0 | 0 | 0 | 0 | 0 | 0 | 0 | 0 | 0 | 0 | 0 | 0 | 0 | 0 | 1 | 0 | 0 | 0 | 0 | 0 | 0 | 0 | 0 | 0 | 0 | 0 | 0 | 0 | 0 | 0 | 0 | 0 | 0 | 0 | 0 | 0 | 0 | 0 | 0 | 0 | 0 |
| virus_Jul11.50v_1622 | 1 | 0 | 0 | 0 | 0 | 0 | 0 | 0 | 0 | 0 | 0 | 0 | 0 | 0 | 0 | 0 | 0 | 0 | 0 | 0 | 0 | 0 | 0 | 0 | 0 | 0 | 0 | 0 | 1 | 0 | 0 | 0 | 0 | 0 | 0 | 0 | 0 | 0 | 0 | 0 | 0 | 0 | 0 | 0 | 0 | 0 | 0 | 0 | 0 | 0 | 0 | 0 | 0 | 0 | 0 |
| virus_Jul11.50v_2621 | 1 | 0 | 0 | 0 | 0 | 0 | 0 | 0 | 0 | 0 | 0 | 0 | 0 | 0 | 0 | 0 | 0 | 0 | 0 | 0 | 0 | 0 | 0 | 0 | 0 | 0 | 0 | 0 | 1 | 0 | 0 | 0 | 0 | 0 | 0 | 0 | 0 | 0 | 0 | 0 | 0 | 0 | 0 | 0 | 0 | 0 | 0 | 0 | 0 | 0 | 0 | 0 | 0 | 0 | 0 |
| virus_Jul11.50v_3228 | 1 | 0 | 0 | 0 | 0 | 0 | 0 | 0 | 0 | 0 | 0 | 0 | 0 | 0 | 0 | 0 | 0 | 0 | 0 | 0 | 0 | 0 | 0 | 0 | 0 | 0 | 0 | 0 | 1 | 0 | 0 | 0 | 0 | 0 | 0 | 0 | 0 | 0 | 0 | 0 | 0 | 0 | 0 | 0 | 0 | 0 | 0 | 0 | 0 | 0 | 0 | 0 | 0 | 0 | 0 |
| virus_Jul11.50v_9019 | 1 | 0 | 0 | 0 | 0 | 0 | 0 | 0 | 0 | 0 | 0 | 0 | 0 | 0 | 0 | 0 | 0 | 0 | 0 | 0 | 0 | 0 | 0 | 0 | 0 | 0 | 0 | 0 | 1 | 0 | 0 | 0 | 0 | 0 | 0 | 0 | 0 | 0 | 0 | 0 | 0 | 0 | 0 | 0 | 0 | 0 | 0 | 0 | 0 | 0 | 0 | 0 | 0 | 0 | 0 |
| virus_Jul11.50v_9942 | 1 | 0 | 0 | 0 | 0 | 0 | 0 | 0 | 0 | 0 | 0 | 0 | 0 | 0 | 0 | 0 | 0 | 0 | 0 | 0 | 0 | 0 | 0 | 0 | 0 | 0 | 0 | 0 | 1 | 0 | 0 | 0 | 0 | 0 | 0 | 0 | 0 | 0 | 0 | 0 | 0 | 0 | 0 | 0 | 0 | 0 | 0 | 0 | 0 | 0 | 0 | 0 | 0 | 0 | 0 |
| virus_Jul11.50v_10932 | 1 | 0 | 0 | 0 | 0 | 0 | 0 | 0 | 0 | 0 | 0 | 0 | 0 | 0 | 0 | 0 | 0 | 0 | 0 | 0 | 0 | 0 | 0 | 0 | 0 | 0 | 0 | 0 | 1 | 0 | 0 | 0 | 0 | 0 | 0 | 0 | 0 | 0 | 0 | 0 | 0 | 0 | 0 | 0 | 0 | 0 | 0 | 0 | 0 | 0 | 0 | 0 | 0 | 0 | 0 |
| virus_Jul11.50v_12102 | 1 | 0 | 0 | 0 | 0 | 0 | 0 | 0 | 0 | 0 | 0 | 0 | 0 | 0 | 0 | 0 | 0 | 0 | 0 | 0 | 0 | 0 | 0 | 0 | 0 | 0 | 0 | 0 | 1 | 0 | 0 | 0 | 0 | 0 | 0 | 0 | 0 | 0 | 0 | 0 | 0 | 0 | 0 | 0 | 0 | 0 | 0 | 0 | 0 | 0 | 0 | 0 | 0 | 0 | 0 |
| virus_Jul11.50v_12580 | 1 | 0 | 0 | 0 | 0 | 0 | 0 | 0 | 0 | 0 | 0 | 0 | 0 | 0 | 0 | 0 | 0 | 0 | 0 | 0 | 0 | 0 | 0 | 0 | 0 | 0 | 0 | 0 | 1 | 0 | 0 | 0 | 0 | 0 | 0 | 0 | 0 | 0 | 0 | 0 | 0 | 0 | 0 | 0 | 0 | 0 | 0 | 0 | 0 | 0 | 0 | 0 | 0 | 0 | 0 |
| virus_Jul11.50v_12998 | 1 | 0 | 0 | 0 | 0 | 0 | 0 | 0 | 0 | 0 | 0 | 0 | 0 | 0 | 0 | 0 | 0 | 0 | 0 | 0 | 0 | 0 | 0 | 0 | 0 | 0 | 0 | 0 | 1 | 0 | 0 | 0 | 0 | 0 | 0 | 0 | 0 | 0 | 0 | 0 | 0 | 0 | 0 | 0 | 0 | 0 | 0 | 0 | 0 | 0 | 0 | 0 | 0 | 0 | 0 |
| virus_Jul11.50v_14281 | 1 | 0 | 0 | 0 | 0 | 0 | 0 | 0 | 0 | 0 | 0 | 0 | 0 | 0 | 0 | 0 | 0 | 0 | 0 | 0 | 0 | 0 | 0 | 0 | 0 | 0 | 0 | 0 | 1 | 0 | 0 | 0 | 0 | 0 | 0 | 0 | 0 | 0 | 0 | 0 | 0 | 0 | 0 | 0 | 0 | 0 | 0 | 0 | 0 | 0 | 0 | 0 | 0 | 0 | 0 |
| virus_Jul11.50v_19238 | 1 | 0 | 0 | 0 | 0 | 0 | 0 | 0 | 0 | 0 | 0 | 0 | 0 | 0 | 0 | 0 | 0 | 0 | 0 | 0 | 0 | 0 | 0 | 0 | 0 | 0 | 0 | 0 | 1 | 0 | 0 | 0 | 0 | 0 | 0 | 0 | 0 | 0 | 0 | 0 | 0 | 0 | 0 | 0 | 0 | 0 | 0 | 0 | 0 | 0 | 0 | 0 | 0 | 0 | 0 |
| virus_Jul11.75v_310 | 1 | 0 | 0 | 0 | 0 | 0 | 0 | 0 | 0 | 0 | 0 | 0 | 0 | 0 | 0 | 0 | 0 | 0 | 0 | 0 | 0 | 0 | 0 | 0 | 0 | 0 | 0 | 0 | 0 | 0 | 1 | 0 | 0 | 0 | 0 | 0 | 0 | 0 | 0 | 0 | 0 | 0 | 0 | 0 | 0 | 0 | 0 | 0 | 0 | 0 | 0 | 0 | 0 | 0 | 0 |
| virus_Jul11.75v_5099 | 1 | 0 | 0 | 0 | 0 | 0 | 0 | 0 | 0 | 0 | 0 | 0 | 0 | 0 | 0 | 0 | 0 | 0 | 0 | 0 | 0 | 0 | 0 | 0 | 0 | 0 | 0 | 0 | 0 | 0 | 1 | 0 | 0 | 0 | 0 | 0 | 0 | 0 | 0 | 0 | 0 | 0 | 0 | 0 | 0 | 0 | 0 | 0 | 0 | 0 | 0 | 0 | 0 | 0 | 0 |
| virus_Jul11.75v_20708 | 1 | 0 | 0 | 0 | 0 | 0 | 0 | 0 | 0 | 0 | 0 | 0 | 0 | 0 | 0 | 0 | 0 | 0 | 0 | 0 | 0 | 0 | 0 | 0 | 0 | 0 | 0 | 0 | 0 | 0 | 1 | 0 | 0 | 0 | 0 | 0 | 0 | 0 | 0 | 0 | 0 | 0 | 0 | 0 | 0 | 0 | 0 | 0 | 0 | 0 | 0 | 0 | 0 | 0 | 0 |
| virus_Jul11.75v_22172 | 1 | 0 | 0 | 0 | 0 | 0 | 0 | 0 | 0 | 0 | 0 | 0 | 0 | 0 | 0 | 0 | 0 | 0 | 0 | 0 | 0 | 0 | 0 | 0 | 0 | 0 | 0 | 0 | 0 | 0 | 1 | 0 | 0 | 0 | 0 | 0 | 0 | 0 | 0 | 0 | 0 | 0 | 0 | 0 | 0 | 0 | 0 | 0 | 0 | 0 | 0 | 0 | 0 | 0 | 0 |
| virus_Jul11.75v_28690 | 1 | 0 | 0 | 0 | 0 | 0 | 0 | 0 | 0 | 0 | 0 | 0 | 0 | 0 | 0 | 0 | 0 | 0 | 0 | 0 | 0 | 0 | 0 | 0 | 0 | 0 | 0 | 0 | 0 | 0 | 1 | 0 | 0 | 0 | 0 | 0 | 0 | 0 | 0 | 0 | 0 | 0 | 0 | 0 | 0 | 0 | 0 | 0 | 0 | 0 | 0 | 0 | 0 | 0 | 0 |
| virus_Jul11.75v_30859 | 1 | 0 | 0 | 0 | 0 | 0 | 0 | 0 | 0 | 0 | 0 | 0 | 0 | 0 | 0 | 0 | 0 | 0 | 0 | 0 | 0 | 0 | 0 | 0 | 0 | 0 | 0 | 0 | 0 | 0 | 1 | 0 | 0 | 0 | 0 | 0 | 0 | 0 | 0 | 0 | 0 | 0 | 0 | 0 | 0 | 0 | 0 | 0 | 0 | 0 | 0 | 0 | 0 | 0 | 0 |
| virus_Jul11.75v_32281 | 1 | 0 | 0 | 0 | 0 | 0 | 0 | 0 | 0 | 0 | 0 | 0 | 0 | 0 | 0 | 0 | 0 | 0 | 0 | 0 | 0 | 0 | 0 | 0 | 0 | 0 | 0 | 0 | 0 | 0 | 1 | 0 | 0 | 0 | 0 | 0 | 0 | 0 | 0 | 0 | 0 | 0 | 0 | 0 | 0 | 0 | 0 | 0 | 0 | 0 | 0 | 0 | 0 | 0 | 0 |
| virus_Jul11.75v_32550 | 1 | 0 | 0 | 0 | 0 | 0 | 0 | 0 | 0 | 0 | 0 | 0 | 0 | 0 | 0 | 0 | 0 | 0 | 0 | 0 | 0 | 0 | 0 | 0 | 0 | 0 | 0 | 0 | 0 | 0 | 1 | 0 | 0 | 0 | 0 | 0 | 0 | 0 | 0 | 0 | 0 | 0 | 0 | 0 | 0 | 0 | 0 | 0 | 0 | 0 | 0 | 0 | 0 | 0 | 0 |
| virus_Jul11.75v_35753 | 1 | 0 | 0 | 0 | 0 | 0 | 0 | 0 | 0 | 0 | 0 | 0 | 0 | 0 | 0 | 0 | 0 | 0 | 0 | 0 | 0 | 0 | 0 | 0 | 0 | 0 | 0 | 0 | 0 | 0 | 1 | 0 | 0 | 0 | 0 | 0 | 0 | 0 | 0 | 0 | 0 | 0 | 0 | 0 | 0 | 0 | 0 | 0 | 0 | 0 | 0 | 0 | 0 | 0 | 0 |
| virus_Jul11.75v_38733 | 1 | 0 | 0 | 0 | 0 | 0 | 0 | 0 | 0 | 0 | 0 | 0 | 0 | 0 | 0 | 0 | 0 | 0 | 0 | 0 | 0 | 0 | 0 | 0 | 0 | 0 | 0 | 0 | 0 | 0 | 1 | 0 | 0 | 0 | 0 | 0 | 0 | 0 | 0 | 0 | 0 | 0 | 0 | 0 | 0 | 0 | 0 | 0 | 0 | 0 | 0 | 0 | 0 | 0 | 0 |
| virus_Jul11.75v_40626 | 1 | 0 | 0 | 0 | 0 | 0 | 0 | 0 | 0 | 0 | 0 | 0 | 0 | 0 | 0 | 0 | 0 | 0 | 0 | 0 | 0 | 0 | 0 | 0 | 0 | 0 | 0 | 0 | 0 | 0 | 1 | 0 | 0 | 0 | 0 | 0 | 0 | 0 | 0 | 0 | 0 | 0 | 0 | 0 | 0 | 0 | 0 | 0 | 0 | 0 | 0 | 0 | 0 | 0 | 0 |
| virus_Jul27.5v_612 | 1 | 0 | 0 | 0 | 0 | 0 | 0 | 0 | 0 | 0 | 0 | 0 | 0 | 0 | 0 | 0 | 0 | 0 | 0 | 0 | 0 | 0 | 0 | 0 | 0 | 0 | 0 | 0 | 0 | 0 | 0 | 0 | 1 | 0 | 0 | 0 | 0 | 0 | 0 | 0 | 0 | 0 | 0 | 0 | 0 | 0 | 0 | 0 | 0 | 0 | 0 | 0 | 0 | 0 | 0 |
| virus_Jul27.5v_1625 | 1 | 0 | 0 | 0 | 0 | 0 | 0 | 0 | 0 | 0 | 0 | 0 | 0 | 0 | 0 | 0 | 0 | 0 | 0 | 0 | 0 | 0 | 0 | 0 | 0 | 0 | 0 | 0 | 0 | 0 | 0 | 0 | 1 | 0 | 0 | 0 | 0 | 0 | 0 | 0 | 0 | 0 | 0 | 0 | 0 | 0 | 0 | 0 | 0 | 0 | 0 | 0 | 0 | 0 | 0 |
| virus_Jul27.5v_1963 | 1 | 0 | 0 | 0 | 0 | 0 | 0 | 0 | 0 | 0 | 0 | 0 | 0 | 0 | 0 | 0 | 0 | 0 | 0 | 0 | 0 | 0 | 0 | 0 | 0 | 0 | 0 | 0 | 0 | 0 | 0 | 0 | 1 | 0 | 0 | 0 | 0 | 0 | 0 | 0 | 0 | 0 | 0 | 0 | 0 | 0 | 0 | 0 | 0 | 0 | 0 | 0 | 0 | 0 | 0 |
| virus_Jul27.5v_2044 | 1 | 0 | 0 | 0 | 0 | 0 | 0 | 0 | 0 | 0 | 0 | 0 | 0 | 0 | 0 | 0 | 0 | 0 | 0 | 0 | 0 | 0 | 0 | 0 | 0 | 0 | 0 | 0 | 0 | 0 | 0 | 0 | 1 | 0 | 0 | 0 | 0 | 0 | 0 | 0 | 0 | 0 | 0 | 0 | 0 | 0 | 0 | 0 | 0 | 0 | 0 | 0 | 0 | 0 | 0 |
| virus_Jul27.5v_2291 | 1 | 0 | 0 | 0 | 0 | 0 | 0 | 0 | 0 | 0 | 0 | 0 | 0 | 0 | 0 | 0 | 0 | 0 | 0 | 0 | 0 | 0 | 0 | 0 | 0 | 0 | 0 | 0 | 0 | 0 | 0 | 0 | 1 | 0 | 0 | 0 | 0 | 0 | 0 | 0 | 0 | 0 | 0 | 0 | 0 | 0 | 0 | 0 | 0 | 0 | 0 | 0 | 0 | 0 | 0 |
| virus_Jul27.5v_2985 | 1 | 0 | 0 | 0 | 0 | 0 | 0 | 0 | 0 | 0 | 0 | 0 | 0 | 0 | 0 | 0 | 0 | 0 | 0 | 0 | 0 | 0 | 0 | 0 | 0 | 0 | 0 | 0 | 0 | 0 | 0 | 0 | 1 | 0 | 0 | 0 | 0 | 0 | 0 | 0 | 0 | 0 | 0 | 0 | 0 | 0 | 0 | 0 | 0 | 0 | 0 | 0 | 0 | 0 | 0 |
| virus_Jul27.5v_4301 | 1 | 0 | 0 | 0 | 0 | 0 | 0 | 0 | 0 | 0 | 0 | 0 | 0 | 0 | 0 | 0 | 0 | 0 | 0 | 0 | 0 | 0 | 0 | 0 | 0 | 0 | 0 | 0 | 0 | 0 | 0 | 0 | 1 | 0 | 0 | 0 | 0 | 0 | 0 | 0 | 0 | 0 | 0 | 0 | 0 | 0 | 0 | 0 | 0 | 0 | 0 | 0 | 0 | 0 | 0 |
| virus_Jul27.5v_5003 | 1 | 0 | 0 | 0 | 0 | 0 | 0 | 0 | 0 | 0 | 0 | 0 | 0 | 0 | 0 | 0 | 0 | 0 | 0 | 0 | 0 | 0 | 0 | 0 | 0 | 0 | 0 | 0 | 0 | 0 | 0 | 0 | 1 | 0 | 0 | 0 | 0 | 0 | 0 | 0 | 0 | 0 | 0 | 0 | 0 | 0 | 0 | 0 | 0 | 0 | 0 | 0 | 0 | 0 | 0 |
| virus_Jul27.5v_5430 | 1 | 0 | 0 | 0 | 0 | 0 | 0 | 0 | 0 | 0 | 0 | 0 | 0 | 0 | 0 | 0 | 0 | 0 | 0 | 0 | 0 | 0 | 0 | 0 | 0 | 0 | 0 | 0 | 0 | 0 | 0 | 0 | 1 | 0 | 0 | 0 | 0 | 0 | 0 | 0 | 0 | 0 | 0 | 0 | 0 | 0 | 0 | 0 | 0 | 0 | 0 | 0 | 0 | 0 | 0 |
| virus_Jul27.5v_5955 | 1 | 0 | 0 | 0 | 0 | 0 | 0 | 0 | 0 | 0 | 0 | 0 | 0 | 0 | 0 | 0 | 0 | 0 | 0 | 0 | 0 | 0 | 0 | 0 | 0 | 0 | 0 | 0 | 0 | 0 | 0 | 0 | 1 | 0 | 0 | 0 | 0 | 0 | 0 | 0 | 0 | 0 | 0 | 0 | 0 | 0 | 0 | 0 | 0 | 0 | 0 | 0 | 0 | 0 | 0 |
| virus_Jul27.5v_7634 | 1 | 0 | 0 | 0 | 0 | 0 | 0 | 0 | 0 | 0 | 0 | 0 | 0 | 0 | 0 | 0 | 0 | 0 | 0 | 0 | 0 | 0 | 0 | 0 | 0 | 0 | 0 | 0 | 0 | 0 | 0 | 0 | 1 | 0 | 0 | 0 | 0 | 0 | 0 | 0 | 0 | 0 | 0 | 0 | 0 | 0 | 0 | 0 | 0 | 0 | 0 | 0 | 0 | 0 | 0 |
| virus_Jul27.5v_8937 | 1 | 0 | 0 | 0 | 0 | 0 | 0 | 0 | 0 | 0 | 0 | 0 | 0 | 0 | 0 | 0 | 0 | 0 | 0 | 0 | 0 | 0 | 0 | 0 | 0 | 0 | 0 | 0 | 0 | 0 | 0 | 0 | 1 | 0 | 0 | 0 | 0 | 0 | 0 | 0 | 0 | 0 | 0 | 0 | 0 | 0 | 0 | 0 | 0 | 0 | 0 | 0 | 0 | 0 | 0 |
| virus_Jul27.5v_10211 | 1 | 0 | 0 | 0 | 0 | 0 | 0 | 0 | 0 | 0 | 0 | 0 | 0 | 0 | 0 | 0 | 0 | 0 | 0 | 0 | 0 | 0 | 0 | 0 | 0 | 0 | 0 | 0 | 0 | 0 | 0 | 0 | 1 | 0 | 0 | 0 | 0 | 0 | 0 | 0 | 0 | 0 | 0 | 0 | 0 | 0 | 0 | 0 | 0 | 0 | 0 | 0 | 0 | 0 | 0 |
| virus_Jul27.5v_11104 | 1 | 0 | 0 | 0 | 0 | 0 | 0 | 0 | 0 | 0 | 0 | 0 | 0 | 0 | 0 | 0 | 0 | 0 | 0 | 0 | 0 | 0 | 0 | 0 | 0 | 0 | 0 | 0 | 0 | 0 | 0 | 0 | 1 | 0 | 0 | 0 | 0 | 0 | 0 | 0 | 0 | 0 | 0 | 0 | 0 | 0 | 0 | 0 | 0 | 0 | 0 | 0 | 0 | 0 | 0 |
| virus_Jul27.5v_11617 | 1 | 0 | 0 | 0 | 0 | 0 | 0 | 0 | 0 | 0 | 0 | 0 | 0 | 0 | 0 | 0 | 0 | 0 | 0 | 0 | 0 | 0 | 0 | 0 | 0 | 0 | 0 | 0 | 0 | 0 | 0 | 0 | 1 | 0 | 0 | 0 | 0 | 0 | 0 | 0 | 0 | 0 | 0 | 0 | 0 | 0 | 0 | 0 | 0 | 0 | 0 | 0 | 0 | 0 | 0 |
| virus_Jul27.5v_11739 | 1 | 0 | 0 | 0 | 0 | 0 | 0 | 0 | 0 | 0 | 0 | 0 | 0 | 0 | 0 | 0 | 0 | 0 | 0 | 0 | 0 | 0 | 0 | 0 | 0 | 0 | 0 | 0 | 0 | 0 | 0 | 0 | 1 | 0 | 0 | 0 | 0 | 0 | 0 | 0 | 0 | 0 | 0 | 0 | 0 | 0 | 0 | 0 | 0 | 0 | 0 | 0 | 0 | 0 | 0 |
| virus_Jul27.5v_12604 | 1 | 0 | 0 | 0 | 0 | 0 | 0 | 0 | 0 | 0 | 0 | 0 | 0 | 0 | 0 | 0 | 0 | 0 | 0 | 0 | 0 | 0 | 0 | 0 | 0 | 0 | 0 | 0 | 0 | 0 | 0 | 0 | 1 | 0 | 0 | 0 | 0 | 0 | 0 | 0 | 0 | 0 | 0 | 0 | 0 | 0 | 0 | 0 | 0 | 0 | 0 | 0 | 0 | 0 | 0 |
| virus_Jul27.5v_13178 | 1 | 0 | 0 | 0 | 0 | 0 | 0 | 0 | 0 | 0 | 0 | 0 | 0 | 0 | 0 | 0 | 0 | 0 | 0 | 0 | 0 | 0 | 0 | 0 | 0 | 0 | 0 | 0 | 0 | 0 | 0 | 0 | 1 | 0 | 0 | 0 | 0 | 0 | 0 | 0 | 0 | 0 | 0 | 0 | 0 | 0 | 0 | 0 | 0 | 0 | 0 | 0 | 0 | 0 | 0 |
| virus_Jul27.5v_13640 | 1 | 0 | 0 | 0 | 0 | 0 | 0 | 0 | 0 | 0 | 0 | 0 | 0 | 0 | 0 | 0 | 0 | 0 | 0 | 0 | 0 | 0 | 0 | 0 | 0 | 0 | 0 | 0 | 0 | 0 | 0 | 0 | 1 | 0 | 0 | 0 | 0 | 0 | 0 | 0 | 0 | 0 | 0 | 0 | 0 | 0 | 0 | 0 | 0 | 0 | 0 | 0 | 0 | 0 | 0 |
| virus_Jul27.5v_14443 | 1 | 0 | 0 | 0 | 0 | 0 | 0 | 0 | 0 | 0 | 0 | 0 | 0 | 0 | 0 | 0 | 0 | 0 | 0 | 0 | 0 | 0 | 0 | 0 | 0 | 0 | 0 | 0 | 0 | 0 | 0 | 0 | 1 | 0 | 0 | 0 | 0 | 0 | 0 | 0 | 0 | 0 | 0 | 0 | 0 | 0 | 0 | 0 | 0 | 0 | 0 | 0 | 0 | 0 | 0 |
| virus_Jul27.5v_15185 | 1 | 0 | 0 | 0 | 0 | 0 | 0 | 0 | 0 | 0 | 0 | 0 | 0 | 0 | 0 | 0 | 0 | 0 | 0 | 0 | 0 | 0 | 0 | 0 | 0 | 0 | 0 | 0 | 0 | 0 | 0 | 0 | 1 | 0 | 0 | 0 | 0 | 0 | 0 | 0 | 0 | 0 | 0 | 0 | 0 | 0 | 0 | 0 | 0 | 0 | 0 | 0 | 0 | 0 | 0 |
| virus_Jul27.5v_15390 | 1 | 0 | 0 | 0 | 0 | 0 | 0 | 0 | 0 | 0 | 0 | 0 | 0 | 0 | 0 | 0 | 0 | 0 | 0 | 0 | 0 | 0 | 0 | 0 | 0 | 0 | 0 | 0 | 0 | 0 | 0 | 0 | 1 | 0 | 0 | 0 | 0 | 0 | 0 | 0 | 0 | 0 | 0 | 0 | 0 | 0 | 0 | 0 | 0 | 0 | 0 | 0 | 0 | 0 | 0 |
| virus_Jul27.5v_15609 | 1 | 0 | 0 | 0 | 0 | 0 | 0 | 0 | 0 | 0 | 0 | 0 | 0 | 0 | 0 | 0 | 0 | 0 | 0 | 0 | 0 | 0 | 0 | 0 | 0 | 0 | 0 | 0 | 0 | 0 | 0 | 0 | 1 | 0 | 0 | 0 | 0 | 0 | 0 | 0 | 0 | 0 | 0 | 0 | 0 | 0 | 0 | 0 | 0 | 0 | 0 | 0 | 0 | 0 | 0 |
| virus_Jul27.5v_15809 | 1 | 0 | 0 | 0 | 0 | 0 | 0 | 0 | 0 | 0 | 0 | 0 | 0 | 0 | 0 | 0 | 0 | 0 | 0 | 0 | 0 | 0 | 0 | 0 | 0 | 0 | 0 | 0 | 0 | 0 | 0 | 0 | 1 | 0 | 0 | 0 | 0 | 0 | 0 | 0 | 0 | 0 | 0 | 0 | 0 | 0 | 0 | 0 | 0 | 0 | 0 | 0 | 0 | 0 | 0 |
| virus_Jul27.5v_16884 | 1 | 0 | 0 | 0 | 0 | 0 | 0 | 0 | 0 | 0 | 0 | 0 | 0 | 0 | 0 | 0 | 0 | 0 | 0 | 0 | 0 | 0 | 0 | 0 | 0 | 0 | 0 | 0 | 0 | 0 | 0 | 0 | 1 | 0 | 0 | 0 | 0 | 0 | 0 | 0 | 0 | 0 | 0 | 0 | 0 | 0 | 0 | 0 | 0 | 0 | 0 | 0 | 0 | 0 | 0 |
| virus_Jul27.5v_16984 | 1 | 0 | 0 | 0 | 0 | 0 | 0 | 0 | 0 | 0 | 0 | 0 | 0 | 0 | 0 | 0 | 0 | 0 | 0 | 0 | 0 | 0 | 0 | 0 | 0 | 0 | 0 | 0 | 0 | 0 | 0 | 0 | 1 | 0 | 0 | 0 | 0 | 0 | 0 | 0 | 0 | 0 | 0 | 0 | 0 | 0 | 0 | 0 | 0 | 0 | 0 | 0 | 0 | 0 | 0 |
| virus_Jul27.5v_19180 | 1 | 0 | 0 | 0 | 0 | 0 | 0 | 0 | 0 | 0 | 0 | 0 | 0 | 0 | 0 | 0 | 0 | 0 | 0 | 0 | 0 | 0 | 0 | 0 | 0 | 0 | 0 | 0 | 0 | 0 | 0 | 0 | 1 | 0 | 0 | 0 | 0 | 0 | 0 | 0 | 0 | 0 | 0 | 0 | 0 | 0 | 0 | 0 | 0 | 0 | 0 | 0 | 0 | 0 | 0 |
| virus_Jul27.5v_19187 | 1 | 0 | 0 | 0 | 0 | 0 | 0 | 0 | 0 | 0 | 0 | 0 | 0 | 0 | 0 | 0 | 0 | 0 | 0 | 0 | 0 | 0 | 0 | 0 | 0 | 0 | 0 | 0 | 0 | 0 | 0 | 0 | 1 | 0 | 0 | 0 | 0 | 0 | 0 | 0 | 0 | 0 | 0 | 0 | 0 | 0 | 0 | 0 | 0 | 0 | 0 | 0 | 0 | 0 | 0 |
| virus_Jul27.5v_20806 | 1 | 0 | 0 | 0 | 0 | 0 | 0 | 0 | 0 | 0 | 0 | 0 | 0 | 0 | 0 | 0 | 0 | 0 | 0 | 0 | 0 | 0 | 0 | 0 | 0 | 0 | 0 | 0 | 0 | 0 | 0 | 0 | 1 | 0 | 0 | 0 | 0 | 0 | 0 | 0 | 0 | 0 | 0 | 0 | 0 | 0 | 0 | 0 | 0 | 0 | 0 | 0 | 0 | 0 | 0 |
| virus_Jul27.5v_21031 | 1 | 0 | 0 | 0 | 0 | 0 | 0 | 0 | 0 | 0 | 0 | 0 | 0 | 0 | 0 | 0 | 0 | 0 | 0 | 0 | 0 | 0 | 0 | 0 | 0 | 0 | 0 | 0 | 0 | 0 | 0 | 0 | 1 | 0 | 0 | 0 | 0 | 0 | 0 | 0 | 0 | 0 | 0 | 0 | 0 | 0 | 0 | 0 | 0 | 0 | 0 | 0 | 0 | 0 | 0 |
| virus_Jul27.25v_578 | 1 | 0 | 0 | 0 | 0 | 0 | 0 | 0 | 0 | 0 | 0 | 0 | 0 | 0 | 0 | 0 | 0 | 0 | 0 | 0 | 0 | 0 | 0 | 0 | 0 | 0 | 0 | 0 | 0 | 0 | 0 | 0 | 0 | 0 | 1 | 0 | 0 | 0 | 0 | 0 | 0 | 0 | 0 | 0 | 0 | 0 | 0 | 0 | 0 | 0 | 0 | 0 | 0 | 0 | 0 |
| virus_Jul27.25v_765 | 1 | 0 | 0 | 0 | 0 | 0 | 0 | 0 | 0 | 0 | 0 | 0 | 0 | 0 | 0 | 0 | 0 | 0 | 0 | 0 | 0 | 0 | 0 | 0 | 0 | 0 | 0 | 0 | 0 | 0 | 0 | 0 | 0 | 0 | 1 | 0 | 0 | 0 | 0 | 0 | 0 | 0 | 0 | 0 | 0 | 0 | 0 | 0 | 0 | 0 | 0 | 0 | 0 | 0 | 0 |
| virus_Jul27.25v_3153 | 1 | 0 | 0 | 0 | 0 | 0 | 0 | 0 | 0 | 0 | 0 | 0 | 0 | 0 | 0 | 0 | 0 | 0 | 0 | 0 | 0 | 0 | 0 | 0 | 0 | 0 | 0 | 0 | 0 | 0 | 0 | 0 | 0 | 0 | 1 | 0 | 0 | 0 | 0 | 0 | 0 | 0 | 0 | 0 | 0 | 0 | 0 | 0 | 0 | 0 | 0 | 0 | 0 | 0 | 0 |
| virus_Jul27.25v_4579 | 1 | 0 | 0 | 0 | 0 | 0 | 0 | 0 | 0 | 0 | 0 | 0 | 0 | 0 | 0 | 0 | 0 | 0 | 0 | 0 | 0 | 0 | 0 | 0 | 0 | 0 | 0 | 0 | 0 | 0 | 0 | 0 | 0 | 0 | 1 | 0 | 0 | 0 | 0 | 0 | 0 | 0 | 0 | 0 | 0 | 0 | 0 | 0 | 0 | 0 | 0 | 0 | 0 | 0 | 0 |
| virus_Jul27.25v_17282 | 1 | 0 | 0 | 0 | 0 | 0 | 0 | 0 | 0 | 0 | 0 | 0 | 0 | 0 | 0 | 0 | 0 | 0 | 0 | 0 | 0 | 0 | 0 | 0 | 0 | 0 | 0 | 0 | 0 | 0 | 0 | 0 | 0 | 0 | 1 | 0 | 0 | 0 | 0 | 0 | 0 | 0 | 0 | 0 | 0 | 0 | 0 | 0 | 0 | 0 | 0 | 0 | 0 | 0 | 0 |
| virus_Jul27.25v_18466 | 1 | 0 | 0 | 0 | 0 | 0 | 0 | 0 | 0 | 0 | 0 | 0 | 0 | 0 | 0 | 0 | 0 | 0 | 0 | 0 | 0 | 0 | 0 | 0 | 0 | 0 | 0 | 0 | 0 | 0 | 0 | 0 | 0 | 0 | 1 | 0 | 0 | 0 | 0 | 0 | 0 | 0 | 0 | 0 | 0 | 0 | 0 | 0 | 0 | 0 | 0 | 0 | 0 | 0 | 0 |
| virus_Jul27.25v_21341 | 1 | 0 | 0 | 0 | 0 | 0 | 0 | 0 | 0 | 0 | 0 | 0 | 0 | 0 | 0 | 0 | 0 | 0 | 0 | 0 | 0 | 0 | 0 | 0 | 0 | 0 | 0 | 0 | 0 | 0 | 0 | 0 | 0 | 0 | 1 | 0 | 0 | 0 | 0 | 0 | 0 | 0 | 0 | 0 | 0 | 0 | 0 | 0 | 0 | 0 | 0 | 0 | 0 | 0 | 0 |
| virus_Jul27.25v_28362 | 1 | 0 | 0 | 0 | 0 | 0 | 0 | 0 | 0 | 0 | 0 | 0 | 0 | 0 | 0 | 0 | 0 | 0 | 0 | 0 | 0 | 0 | 0 | 0 | 0 | 0 | 0 | 0 | 0 | 0 | 0 | 0 | 0 | 0 | 1 | 0 | 0 | 0 | 0 | 0 | 0 | 0 | 0 | 0 | 0 | 0 | 0 | 0 | 0 | 0 | 0 | 0 | 0 | 0 | 0 |
| virus_Jul27.25v_31892 | 1 | 0 | 0 | 0 | 0 | 0 | 0 | 0 | 0 | 0 | 0 | 0 | 0 | 0 | 0 | 0 | 0 | 0 | 0 | 0 | 0 | 0 | 0 | 0 | 0 | 0 | 0 | 0 | 0 | 0 | 0 | 0 | 0 | 0 | 1 | 0 | 0 | 0 | 0 | 0 | 0 | 0 | 0 | 0 | 0 | 0 | 0 | 0 | 0 | 0 | 0 | 0 | 0 | 0 | 0 |
| virus_Jul27.25v_32817 | 1 | 0 | 0 | 0 | 0 | 0 | 0 | 0 | 0 | 0 | 0 | 0 | 0 | 0 | 0 | 0 | 0 | 0 | 0 | 0 | 0 | 0 | 0 | 0 | 0 | 0 | 0 | 0 | 0 | 0 | 0 | 0 | 0 | 0 | 1 | 0 | 0 | 0 | 0 | 0 | 0 | 0 | 0 | 0 | 0 | 0 | 0 | 0 | 0 | 0 | 0 | 0 | 0 | 0 | 0 |
| virus_Jul27.25v_33044 | 1 | 0 | 0 | 0 | 0 | 0 | 0 | 0 | 0 | 0 | 0 | 0 | 0 | 0 | 0 | 0 | 0 | 0 | 0 | 0 | 0 | 0 | 0 | 0 | 0 | 0 | 0 | 0 | 0 | 0 | 0 | 0 | 0 | 0 | 1 | 0 | 0 | 0 | 0 | 0 | 0 | 0 | 0 | 0 | 0 | 0 | 0 | 0 | 0 | 0 | 0 | 0 | 0 | 0 | 0 |
| virus_Jul27.25v_33364 | 1 | 0 | 0 | 0 | 0 | 0 | 0 | 0 | 0 | 0 | 0 | 0 | 0 | 0 | 0 | 0 | 0 | 0 | 0 | 0 | 0 | 0 | 0 | 0 | 0 | 0 | 0 | 0 | 0 | 0 | 0 | 0 | 0 | 0 | 1 | 0 | 0 | 0 | 0 | 0 | 0 | 0 | 0 | 0 | 0 | 0 | 0 | 0 | 0 | 0 | 0 | 0 | 0 | 0 | 0 |
| virus_Jul27.25v_34509 | 1 | 0 | 0 | 0 | 0 | 0 | 0 | 0 | 0 | 0 | 0 | 0 | 0 | 0 | 0 | 0 | 0 | 0 | 0 | 0 | 0 | 0 | 0 | 0 | 0 | 0 | 0 | 0 | 0 | 0 | 0 | 0 | 0 | 0 | 1 | 0 | 0 | 0 | 0 | 0 | 0 | 0 | 0 | 0 | 0 | 0 | 0 | 0 | 0 | 0 | 0 | 0 | 0 | 0 | 0 |
| virus_Jul27.25v_36312 | 1 | 0 | 0 | 0 | 0 | 0 | 0 | 0 | 0 | 0 | 0 | 0 | 0 | 0 | 0 | 0 | 0 | 0 | 0 | 0 | 0 | 0 | 0 | 0 | 0 | 0 | 0 | 0 | 0 | 0 | 0 | 0 | 0 | 0 | 1 | 0 | 0 | 0 | 0 | 0 | 0 | 0 | 0 | 0 | 0 | 0 | 0 | 0 | 0 | 0 | 0 | 0 | 0 | 0 | 0 |
| virus_Jul27.25v_37678 | 1 | 0 | 0 | 0 | 0 | 0 | 0 | 0 | 0 | 0 | 0 | 0 | 0 | 0 | 0 | 0 | 0 | 0 | 0 | 0 | 0 | 0 | 0 | 0 | 0 | 0 | 0 | 0 | 0 | 0 | 0 | 0 | 0 | 0 | 1 | 0 | 0 | 0 | 0 | 0 | 0 | 0 | 0 | 0 | 0 | 0 | 0 | 0 | 0 | 0 | 0 | 0 | 0 | 0 | 0 |
| virus_Jul27.25v_39511 | 1 | 0 | 0 | 0 | 0 | 0 | 0 | 0 | 0 | 0 | 0 | 0 | 0 | 0 | 0 | 0 | 0 | 0 | 0 | 0 | 0 | 0 | 0 | 0 | 0 | 0 | 0 | 0 | 0 | 0 | 0 | 0 | 0 | 0 | 1 | 0 | 0 | 0 | 0 | 0 | 0 | 0 | 0 | 0 | 0 | 0 | 0 | 0 | 0 | 0 | 0 | 0 | 0 | 0 | 0 |
| virus_Jul27.75v_1676 | 1 | 0 | 0 | 0 | 0 | 0 | 0 | 0 | 0 | 0 | 0 | 0 | 0 | 0 | 0 | 0 | 0 | 0 | 0 | 0 | 0 | 0 | 0 | 0 | 0 | 0 | 0 | 0 | 0 | 0 | 0 | 0 | 0 | 0 | 0 | 0 | 1 | 0 | 0 | 0 | 0 | 0 | 0 | 0 | 0 | 0 | 0 | 0 | 0 | 0 | 0 | 0 | 0 | 0 | 0 |
| virus_Jul27.75v_12255 | 1 | 0 | 0 | 0 | 0 | 0 | 0 | 0 | 0 | 0 | 0 | 0 | 0 | 0 | 0 | 0 | 0 | 0 | 0 | 0 | 0 | 0 | 0 | 0 | 0 | 0 | 0 | 0 | 0 | 0 | 0 | 0 | 0 | 0 | 0 | 0 | 1 | 0 | 0 | 0 | 0 | 0 | 0 | 0 | 0 | 0 | 0 | 0 | 0 | 0 | 0 | 0 | 0 | 0 | 0 |
| virus_Jun8.25v_4802 | 1 | 0 | 0 | 0 | 0 | 0 | 0 | 0 | 0 | 0 | 1 | 0 | 0 | 0 | 0 | 0 | 0 | 0 | 0 | 0 | 0 | 0 | 0 | 0 | 0 | 0 | 0 | 0 | 0 | 0 | 0 | 0 | 0 | 0 | 0 | 0 | 0 | 0 | 0 | 0 | 0 | 0 | 0 | 0 | 0 | 0 | 0 | 0 | 0 | 0 | 0 | 0 | 0 | 0 | 0 |
| virus_Jun8.25v_5246 | 1 | 0 | 0 | 0 | 0 | 0 | 0 | 0 | 0 | 0 | 1 | 0 | 0 | 0 | 0 | 0 | 0 | 0 | 0 | 0 | 0 | 0 | 0 | 0 | 0 | 0 | 0 | 0 | 0 | 0 | 0 | 0 | 0 | 0 | 0 | 0 | 0 | 0 | 0 | 0 | 0 | 0 | 0 | 0 | 0 | 0 | 0 | 0 | 0 | 0 | 0 | 0 | 0 | 0 | 0 |
| virus_Jun8.25v_23093 | 1 | 0 | 0 | 0 | 0 | 0 | 0 | 0 | 0 | 0 | 1 | 0 | 0 | 0 | 0 | 0 | 0 | 0 | 0 | 0 | 0 | 0 | 0 | 0 | 0 | 0 | 0 | 0 | 0 | 0 | 0 | 0 | 0 | 0 | 0 | 0 | 0 | 0 | 0 | 0 | 0 | 0 | 0 | 0 | 0 | 0 | 0 | 0 | 0 | 0 | 0 | 0 | 0 | 0 | 0 |
| virus_Jun8.25v_27165 | 1 | 0 | 0 | 0 | 0 | 0 | 0 | 0 | 0 | 0 | 1 | 0 | 0 | 0 | 0 | 0 | 0 | 0 | 0 | 0 | 0 | 0 | 0 | 0 | 0 | 0 | 0 | 0 | 0 | 0 | 0 | 0 | 0 | 0 | 0 | 0 | 0 | 0 | 0 | 0 | 0 | 0 | 0 | 0 | 0 | 0 | 0 | 0 | 0 | 0 | 0 | 0 | 0 | 0 | 0 |
| virus_Jun8.50v_4437 | 1 | 0 | 0 | 0 | 0 | 0 | 0 | 0 | 0 | 0 | 0 | 0 | 1 | 0 | 0 | 0 | 0 | 0 | 0 | 0 | 0 | 0 | 0 | 0 | 0 | 0 | 0 | 0 | 0 | 0 | 0 | 0 | 0 | 0 | 0 | 0 | 0 | 0 | 0 | 0 | 0 | 0 | 0 | 0 | 0 | 0 | 0 | 0 | 0 | 0 | 0 | 0 | 0 | 0 | 0 |
| virus_Jun8.50v_5126 | 1 | 0 | 0 | 0 | 0 | 0 | 0 | 0 | 0 | 0 | 0 | 0 | 1 | 0 | 0 | 0 | 0 | 0 | 0 | 0 | 0 | 0 | 0 | 0 | 0 | 0 | 0 | 0 | 0 | 0 | 0 | 0 | 0 | 0 | 0 | 0 | 0 | 0 | 0 | 0 | 0 | 0 | 0 | 0 | 0 | 0 | 0 | 0 | 0 | 0 | 0 | 0 | 0 | 0 | 0 |
| virus_Jun8.50v_13375 | 1 | 0 | 0 | 0 | 0 | 0 | 0 | 0 | 0 | 0 | 0 | 0 | 1 | 0 | 0 | 0 | 0 | 0 | 0 | 0 | 0 | 0 | 0 | 0 | 0 | 0 | 0 | 0 | 0 | 0 | 0 | 0 | 0 | 0 | 0 | 0 | 0 | 0 | 0 | 0 | 0 | 0 | 0 | 0 | 0 | 0 | 0 | 0 | 0 | 0 | 0 | 0 | 0 | 0 | 0 |
| virus_Jun8.50v_16842 | 1 | 0 | 0 | 0 | 0 | 0 | 0 | 0 | 0 | 0 | 0 | 0 | 1 | 0 | 0 | 0 | 0 | 0 | 0 | 0 | 0 | 0 | 0 | 0 | 0 | 0 | 0 | 0 | 0 | 0 | 0 | 0 | 0 | 0 | 0 | 0 | 0 | 0 | 0 | 0 | 0 | 0 | 0 | 0 | 0 | 0 | 0 | 0 | 0 | 0 | 0 | 0 | 0 | 0 | 0 |
| virus_Jun8.50v_17840 | 1 | 0 | 0 | 0 | 0 | 0 | 0 | 0 | 0 | 0 | 0 | 0 | 1 | 0 | 0 | 0 | 0 | 0 | 0 | 0 | 0 | 0 | 0 | 0 | 0 | 0 | 0 | 0 | 0 | 0 | 0 | 0 | 0 | 0 | 0 | 0 | 0 | 0 | 0 | 0 | 0 | 0 | 0 | 0 | 0 | 0 | 0 | 0 | 0 | 0 | 0 | 0 | 0 | 0 | 0 |
| virus_Jun8.50v_20729 | 1 | 0 | 0 | 0 | 0 | 0 | 0 | 0 | 0 | 0 | 0 | 0 | 1 | 0 | 0 | 0 | 0 | 0 | 0 | 0 | 0 | 0 | 0 | 0 | 0 | 0 | 0 | 0 | 0 | 0 | 0 | 0 | 0 | 0 | 0 | 0 | 0 | 0 | 0 | 0 | 0 | 0 | 0 | 0 | 0 | 0 | 0 | 0 | 0 | 0 | 0 | 0 | 0 | 0 | 0 |
| virus_Jun8.50v_30353 | 1 | 0 | 0 | 0 | 0 | 0 | 0 | 0 | 0 | 0 | 0 | 0 | 1 | 0 | 0 | 0 | 0 | 0 | 0 | 0 | 0 | 0 | 0 | 0 | 0 | 0 | 0 | 0 | 0 | 0 | 0 | 0 | 0 | 0 | 0 | 0 | 0 | 0 | 0 | 0 | 0 | 0 | 0 | 0 | 0 | 0 | 0 | 0 | 0 | 0 | 0 | 0 | 0 | 0 | 0 |
| virus_Jun8.50v_37590 | 1 | 0 | 0 | 0 | 0 | 0 | 0 | 0 | 0 | 0 | 0 | 0 | 1 | 0 | 0 | 0 | 0 | 0 | 0 | 0 | 0 | 0 | 0 | 0 | 0 | 0 | 0 | 0 | 0 | 0 | 0 | 0 | 0 | 0 | 0 | 0 | 0 | 0 | 0 | 0 | 0 | 0 | 0 | 0 | 0 | 0 | 0 | 0 | 0 | 0 | 0 | 0 | 0 | 0 | 0 |
| virus_Jun8.50v_37882 | 1 | 0 | 0 | 0 | 0 | 0 | 0 | 0 | 0 | 0 | 0 | 0 | 1 | 0 | 0 | 0 | 0 | 0 | 0 | 0 | 0 | 0 | 0 | 0 | 0 | 0 | 0 | 0 | 0 | 0 | 0 | 0 | 0 | 0 | 0 | 0 | 0 | 0 | 0 | 0 | 0 | 0 | 0 | 0 | 0 | 0 | 0 | 0 | 0 | 0 | 0 | 0 | 0 | 0 | 0 |
| virus_Jun8.75v_3226 | 1 | 0 | 0 | 0 | 0 | 0 | 0 | 0 | 0 | 0 | 0 | 0 | 0 | 0 | 1 | 0 | 0 | 0 | 0 | 0 | 0 | 0 | 0 | 0 | 0 | 0 | 0 | 0 | 0 | 0 | 0 | 0 | 0 | 0 | 0 | 0 | 0 | 0 | 0 | 0 | 0 | 0 | 0 | 0 | 0 | 0 | 0 | 0 | 0 | 0 | 0 | 0 | 0 | 0 | 0 |
| virus_Jun8.75v_18911 | 1 | 0 | 0 | 0 | 0 | 0 | 0 | 0 | 0 | 0 | 0 | 0 | 0 | 0 | 1 | 0 | 0 | 0 | 0 | 0 | 0 | 0 | 0 | 0 | 0 | 0 | 0 | 0 | 0 | 0 | 0 | 0 | 0 | 0 | 0 | 0 | 0 | 0 | 0 | 0 | 0 | 0 | 0 | 0 | 0 | 0 | 0 | 0 | 0 | 0 | 0 | 0 | 0 | 0 | 0 |
| virus_Jun22.0v_556 | 1 | 0 | 0 | 0 | 0 | 0 | 0 | 0 | 0 | 0 | 0 | 0 | 0 | 0 | 0 | 0 | 1 | 0 | 0 | 0 | 0 | 0 | 0 | 0 | 0 | 0 | 0 | 0 | 0 | 0 | 0 | 0 | 0 | 0 | 0 | 0 | 0 | 0 | 0 | 0 | 0 | 0 | 0 | 0 | 0 | 0 | 0 | 0 | 0 | 0 | 0 | 0 | 0 | 0 | 0 |
| virus_Jun22.0v_861 | 1 | 0 | 0 | 0 | 0 | 0 | 0 | 0 | 0 | 0 | 0 | 0 | 0 | 0 | 0 | 0 | 1 | 0 | 0 | 0 | 0 | 0 | 0 | 0 | 0 | 0 | 0 | 0 | 0 | 0 | 0 | 0 | 0 | 0 | 0 | 0 | 0 | 0 | 0 | 0 | 0 | 0 | 0 | 0 | 0 | 0 | 0 | 0 | 0 | 0 | 0 | 0 | 0 | 0 | 0 |
| virus_Jun22.0v_1274 | 1 | 0 | 0 | 0 | 0 | 0 | 0 | 0 | 0 | 0 | 0 | 0 | 0 | 0 | 0 | 0 | 1 | 0 | 0 | 0 | 0 | 0 | 0 | 0 | 0 | 0 | 0 | 0 | 0 | 0 | 0 | 0 | 0 | 0 | 0 | 0 | 0 | 0 | 0 | 0 | 0 | 0 | 0 | 0 | 0 | 0 | 0 | 0 | 0 | 0 | 0 | 0 | 0 | 0 | 0 |
| virus_Jun22.0v_3429 | 1 | 0 | 0 | 0 | 0 | 0 | 0 | 0 | 0 | 0 | 0 | 0 | 0 | 0 | 0 | 0 | 1 | 0 | 0 | 0 | 0 | 0 | 0 | 0 | 0 | 0 | 0 | 0 | 0 | 0 | 0 | 0 | 0 | 0 | 0 | 0 | 0 | 0 | 0 | 0 | 0 | 0 | 0 | 0 | 0 | 0 | 0 | 0 | 0 | 0 | 0 | 0 | 0 | 0 | 0 |
| virus_Jun22.0v_4953 | 1 | 0 | 0 | 0 | 0 | 0 | 0 | 0 | 0 | 0 | 0 | 0 | 0 | 0 | 0 | 0 | 1 | 0 | 0 | 0 | 0 | 0 | 0 | 0 | 0 | 0 | 0 | 0 | 0 | 0 | 0 | 0 | 0 | 0 | 0 | 0 | 0 | 0 | 0 | 0 | 0 | 0 | 0 | 0 | 0 | 0 | 0 | 0 | 0 | 0 | 0 | 0 | 0 | 0 | 0 |
| virus_Jun22.0v_8658 | 1 | 0 | 0 | 0 | 0 | 0 | 0 | 0 | 0 | 0 | 0 | 0 | 0 | 0 | 0 | 0 | 1 | 0 | 0 | 0 | 0 | 0 | 0 | 0 | 0 | 0 | 0 | 0 | 0 | 0 | 0 | 0 | 0 | 0 | 0 | 0 | 0 | 0 | 0 | 0 | 0 | 0 | 0 | 0 | 0 | 0 | 0 | 0 | 0 | 0 | 0 | 0 | 0 | 0 | 0 |
| virus_Jun22.0v_11126 | 1 | 0 | 0 | 0 | 0 | 0 | 0 | 0 | 0 | 0 | 0 | 0 | 0 | 0 | 0 | 0 | 1 | 0 | 0 | 0 | 0 | 0 | 0 | 0 | 0 | 0 | 0 | 0 | 0 | 0 | 0 | 0 | 0 | 0 | 0 | 0 | 0 | 0 | 0 | 0 | 0 | 0 | 0 | 0 | 0 | 0 | 0 | 0 | 0 | 0 | 0 | 0 | 0 | 0 | 0 |
| virus_Jun22.0v_11947 | 1 | 0 | 0 | 0 | 0 | 0 | 0 | 0 | 0 | 0 | 0 | 0 | 0 | 0 | 0 | 0 | 1 | 0 | 0 | 0 | 0 | 0 | 0 | 0 | 0 | 0 | 0 | 0 | 0 | 0 | 0 | 0 | 0 | 0 | 0 | 0 | 0 | 0 | 0 | 0 | 0 | 0 | 0 | 0 | 0 | 0 | 0 | 0 | 0 | 0 | 0 | 0 | 0 | 0 | 0 |
| virus_Jun22.0v_15447 | 1 | 0 | 0 | 0 | 0 | 0 | 0 | 0 | 0 | 0 | 0 | 0 | 0 | 0 | 0 | 0 | 1 | 0 | 0 | 0 | 0 | 0 | 0 | 0 | 0 | 0 | 0 | 0 | 0 | 0 | 0 | 0 | 0 | 0 | 0 | 0 | 0 | 0 | 0 | 0 | 0 | 0 | 0 | 0 | 0 | 0 | 0 | 0 | 0 | 0 | 0 | 0 | 0 | 0 | 0 |
| virus_Jun22.0v_19577 | 1 | 0 | 0 | 0 | 0 | 0 | 0 | 0 | 0 | 0 | 0 | 0 | 0 | 0 | 0 | 0 | 1 | 0 | 0 | 0 | 0 | 0 | 0 | 0 | 0 | 0 | 0 | 0 | 0 | 0 | 0 | 0 | 0 | 0 | 0 | 0 | 0 | 0 | 0 | 0 | 0 | 0 | 0 | 0 | 0 | 0 | 0 | 0 | 0 | 0 | 0 | 0 | 0 | 0 | 0 |
| virus_Jun22.50v_4968 | 1 | 0 | 0 | 0 | 0 | 0 | 0 | 0 | 0 | 0 | 0 | 0 | 0 | 0 | 0 | 0 | 0 | 0 | 1 | 0 | 0 | 0 | 0 | 0 | 0 | 0 | 0 | 0 | 0 | 0 | 0 | 0 | 0 | 0 | 0 | 0 | 0 | 0 | 0 | 0 | 0 | 0 | 0 | 0 | 0 | 0 | 0 | 0 | 0 | 0 | 0 | 0 | 0 | 0 | 0 |
| virus_Jun22.50v_8234 | 1 | 0 | 0 | 0 | 0 | 0 | 0 | 0 | 0 | 0 | 0 | 0 | 0 | 0 | 0 | 0 | 0 | 0 | 1 | 0 | 0 | 0 | 0 | 0 | 0 | 0 | 0 | 0 | 0 | 0 | 0 | 0 | 0 | 0 | 0 | 0 | 0 | 0 | 0 | 0 | 0 | 0 | 0 | 0 | 0 | 0 | 0 | 0 | 0 | 0 | 0 | 0 | 0 | 0 | 0 |
| virus_Jun22.50v_8302 | 1 | 0 | 0 | 0 | 0 | 0 | 0 | 0 | 0 | 0 | 0 | 0 | 0 | 0 | 0 | 0 | 0 | 0 | 1 | 0 | 0 | 0 | 0 | 0 | 0 | 0 | 0 | 0 | 0 | 0 | 0 | 0 | 0 | 0 | 0 | 0 | 0 | 0 | 0 | 0 | 0 | 0 | 0 | 0 | 0 | 0 | 0 | 0 | 0 | 0 | 0 | 0 | 0 | 0 | 0 |
| virus_Jun22.50v_11598 | 1 | 0 | 0 | 0 | 0 | 0 | 0 | 0 | 0 | 0 | 0 | 0 | 0 | 0 | 0 | 0 | 0 | 0 | 1 | 0 | 0 | 0 | 0 | 0 | 0 | 0 | 0 | 0 | 0 | 0 | 0 | 0 | 0 | 0 | 0 | 0 | 0 | 0 | 0 | 0 | 0 | 0 | 0 | 0 | 0 | 0 | 0 | 0 | 0 | 0 | 0 | 0 | 0 | 0 | 0 |
| virus_Jun22.50v_15338 | 1 | 0 | 0 | 0 | 0 | 0 | 0 | 0 | 0 | 0 | 0 | 0 | 0 | 0 | 0 | 0 | 0 | 0 | 1 | 0 | 0 | 0 | 0 | 0 | 0 | 0 | 0 | 0 | 0 | 0 | 0 | 0 | 0 | 0 | 0 | 0 | 0 | 0 | 0 | 0 | 0 | 0 | 0 | 0 | 0 | 0 | 0 | 0 | 0 | 0 | 0 | 0 | 0 | 0 | 0 |
| virus_Jun22.50v_15599 | 1 | 0 | 0 | 0 | 0 | 0 | 0 | 0 | 0 | 0 | 0 | 0 | 0 | 0 | 0 | 0 | 0 | 0 | 1 | 0 | 0 | 0 | 0 | 0 | 0 | 0 | 0 | 0 | 0 | 0 | 0 | 0 | 0 | 0 | 0 | 0 | 0 | 0 | 0 | 0 | 0 | 0 | 0 | 0 | 0 | 0 | 0 | 0 | 0 | 0 | 0 | 0 | 0 | 0 | 0 |
| virus_Jun22.50v_19630 | 1 | 0 | 0 | 0 | 0 | 0 | 0 | 0 | 0 | 0 | 0 | 0 | 0 | 0 | 0 | 0 | 0 | 0 | 1 | 0 | 0 | 0 | 0 | 0 | 0 | 0 | 0 | 0 | 0 | 0 | 0 | 0 | 0 | 0 | 0 | 0 | 0 | 0 | 0 | 0 | 0 | 0 | 0 | 0 | 0 | 0 | 0 | 0 | 0 | 0 | 0 | 0 | 0 | 0 | 0 |
| virus_Jun22.50v_19641 | 1 | 0 | 0 | 0 | 0 | 0 | 0 | 0 | 0 | 0 | 0 | 0 | 0 | 0 | 0 | 0 | 0 | 0 | 1 | 0 | 0 | 0 | 0 | 0 | 0 | 0 | 0 | 0 | 0 | 0 | 0 | 0 | 0 | 0 | 0 | 0 | 0 | 0 | 0 | 0 | 0 | 0 | 0 | 0 | 0 | 0 | 0 | 0 | 0 | 0 | 0 | 0 | 0 | 0 | 0 |
| virus_Jun22.50v_20864 | 1 | 0 | 0 | 0 | 0 | 0 | 0 | 0 | 0 | 0 | 0 | 0 | 0 | 0 | 0 | 0 | 0 | 0 | 1 | 0 | 0 | 0 | 0 | 0 | 0 | 0 | 0 | 0 | 0 | 0 | 0 | 0 | 0 | 0 | 0 | 0 | 0 | 0 | 0 | 0 | 0 | 0 | 0 | 0 | 0 | 0 | 0 | 0 | 0 | 0 | 0 | 0 | 0 | 0 | 0 |
| virus_Jun22.50v_23741 | 1 | 0 | 0 | 0 | 0 | 0 | 0 | 0 | 0 | 0 | 0 | 0 | 0 | 0 | 0 | 0 | 0 | 0 | 1 | 0 | 0 | 0 | 0 | 0 | 0 | 0 | 0 | 0 | 0 | 0 | 0 | 0 | 0 | 0 | 0 | 0 | 0 | 0 | 0 | 0 | 0 | 0 | 0 | 0 | 0 | 0 | 0 | 0 | 0 | 0 | 0 | 0 | 0 | 0 | 0 |
| virus_Jun22.50v_25125 | 1 | 0 | 0 | 0 | 0 | 0 | 0 | 0 | 0 | 0 | 0 | 0 | 0 | 0 | 0 | 0 | 0 | 0 | 1 | 0 | 0 | 0 | 0 | 0 | 0 | 0 | 0 | 0 | 0 | 0 | 0 | 0 | 0 | 0 | 0 | 0 | 0 | 0 | 0 | 0 | 0 | 0 | 0 | 0 | 0 | 0 | 0 | 0 | 0 | 0 | 0 | 0 | 0 | 0 | 0 |
| virus_Jun22.50v_27762 | 1 | 0 | 0 | 0 | 0 | 0 | 0 | 0 | 0 | 0 | 0 | 0 | 0 | 0 | 0 | 0 | 0 | 0 | 1 | 0 | 0 | 0 | 0 | 0 | 0 | 0 | 0 | 0 | 0 | 0 | 0 | 0 | 0 | 0 | 0 | 0 | 0 | 0 | 0 | 0 | 0 | 0 | 0 | 0 | 0 | 0 | 0 | 0 | 0 | 0 | 0 | 0 | 0 | 0 | 0 |
| virus_Jun22.50v_29776 | 1 | 0 | 0 | 0 | 0 | 0 | 0 | 0 | 0 | 0 | 0 | 0 | 0 | 0 | 0 | 0 | 0 | 0 | 1 | 0 | 0 | 0 | 0 | 0 | 0 | 0 | 0 | 0 | 0 | 0 | 0 | 0 | 0 | 0 | 0 | 0 | 0 | 0 | 0 | 0 | 0 | 0 | 0 | 0 | 0 | 0 | 0 | 0 | 0 | 0 | 0 | 0 | 0 | 0 | 0 |
| virus_Jun22.50v_32459 | 1 | 0 | 0 | 0 | 0 | 0 | 0 | 0 | 0 | 0 | 0 | 0 | 0 | 0 | 0 | 0 | 0 | 0 | 1 | 0 | 0 | 0 | 0 | 0 | 0 | 0 | 0 | 0 | 0 | 0 | 0 | 0 | 0 | 0 | 0 | 0 | 0 | 0 | 0 | 0 | 0 | 0 | 0 | 0 | 0 | 0 | 0 | 0 | 0 | 0 | 0 | 0 | 0 | 0 | 0 |
| virus_Jun22.50v_32722 | 1 | 0 | 0 | 0 | 0 | 0 | 0 | 0 | 0 | 0 | 0 | 0 | 0 | 0 | 0 | 0 | 0 | 0 | 1 | 0 | 0 | 0 | 0 | 0 | 0 | 0 | 0 | 0 | 0 | 0 | 0 | 0 | 0 | 0 | 0 | 0 | 0 | 0 | 0 | 0 | 0 | 0 | 0 | 0 | 0 | 0 | 0 | 0 | 0 | 0 | 0 | 0 | 0 | 0 | 0 |
| virus_Jun22.50v_38736 | 1 | 0 | 0 | 0 | 0 | 0 | 0 | 0 | 0 | 0 | 0 | 0 | 0 | 0 | 0 | 0 | 0 | 0 | 1 | 0 | 0 | 0 | 0 | 0 | 0 | 0 | 0 | 0 | 0 | 0 | 0 | 0 | 0 | 0 | 0 | 0 | 0 | 0 | 0 | 0 | 0 | 0 | 0 | 0 | 0 | 0 | 0 | 0 | 0 | 0 | 0 | 0 | 0 | 0 | 0 |
| virus_Jun22.75v_2967 | 1 | 0 | 0 | 0 | 0 | 0 | 0 | 0 | 0 | 0 | 0 | 0 | 0 | 0 | 0 | 0 | 0 | 0 | 0 | 0 | 1 | 0 | 0 | 0 | 0 | 0 | 0 | 0 | 0 | 0 | 0 | 0 | 0 | 0 | 0 | 0 | 0 | 0 | 0 | 0 | 0 | 0 | 0 | 0 | 0 | 0 | 0 | 0 | 0 | 0 | 0 | 0 | 0 | 0 | 0 |
| virus_Jun22.75v_3511 | 1 | 0 | 0 | 0 | 0 | 0 | 0 | 0 | 0 | 0 | 0 | 0 | 0 | 0 | 0 | 0 | 0 | 0 | 0 | 0 | 1 | 0 | 0 | 0 | 0 | 0 | 0 | 0 | 0 | 0 | 0 | 0 | 0 | 0 | 0 | 0 | 0 | 0 | 0 | 0 | 0 | 0 | 0 | 0 | 0 | 0 | 0 | 0 | 0 | 0 | 0 | 0 | 0 | 0 | 0 |
| virus_Jun22.75v_4608 | 1 | 0 | 0 | 0 | 0 | 0 | 0 | 0 | 0 | 0 | 0 | 0 | 0 | 0 | 0 | 0 | 0 | 0 | 0 | 0 | 1 | 0 | 0 | 0 | 0 | 0 | 0 | 0 | 0 | 0 | 0 | 0 | 0 | 0 | 0 | 0 | 0 | 0 | 0 | 0 | 0 | 0 | 0 | 0 | 0 | 0 | 0 | 0 | 0 | 0 | 0 | 0 | 0 | 0 | 0 |
| virus_Jun22.75v_9197 | 1 | 0 | 0 | 0 | 0 | 0 | 0 | 0 | 0 | 0 | 0 | 0 | 0 | 0 | 0 | 0 | 0 | 0 | 0 | 0 | 1 | 0 | 0 | 0 | 0 | 0 | 0 | 0 | 0 | 0 | 0 | 0 | 0 | 0 | 0 | 0 | 0 | 0 | 0 | 0 | 0 | 0 | 0 | 0 | 0 | 0 | 0 | 0 | 0 | 0 | 0 | 0 | 0 | 0 | 0 |
| virus_Jun22.75v_11180 | 1 | 0 | 0 | 0 | 0 | 0 | 0 | 0 | 0 | 0 | 0 | 0 | 0 | 0 | 0 | 0 | 0 | 0 | 0 | 0 | 1 | 0 | 0 | 0 | 0 | 0 | 0 | 0 | 0 | 0 | 0 | 0 | 0 | 0 | 0 | 0 | 0 | 0 | 0 | 0 | 0 | 0 | 0 | 0 | 0 | 0 | 0 | 0 | 0 | 0 | 0 | 0 | 0 | 0 | 0 |
| virus_Jun22.75v_15887 | 1 | 0 | 0 | 0 | 0 | 0 | 0 | 0 | 0 | 0 | 0 | 0 | 0 | 0 | 0 | 0 | 0 | 0 | 0 | 0 | 1 | 0 | 0 | 0 | 0 | 0 | 0 | 0 | 0 | 0 | 0 | 0 | 0 | 0 | 0 | 0 | 0 | 0 | 0 | 0 | 0 | 0 | 0 | 0 | 0 | 0 | 0 | 0 | 0 | 0 | 0 | 0 | 0 | 0 | 0 |
| virus_Jun22.75v_17235 | 1 | 0 | 0 | 0 | 0 | 0 | 0 | 0 | 0 | 0 | 0 | 0 | 0 | 0 | 0 | 0 | 0 | 0 | 0 | 0 | 1 | 0 | 0 | 0 | 0 | 0 | 0 | 0 | 0 | 0 | 0 | 0 | 0 | 0 | 0 | 0 | 0 | 0 | 0 | 0 | 0 | 0 | 0 | 0 | 0 | 0 | 0 | 0 | 0 | 0 | 0 | 0 | 0 | 0 | 0 |
| virus_Jun22.75v_17750 | 1 | 0 | 0 | 0 | 0 | 0 | 0 | 0 | 0 | 0 | 0 | 0 | 0 | 0 | 0 | 0 | 0 | 0 | 0 | 0 | 1 | 0 | 0 | 0 | 0 | 0 | 0 | 0 | 0 | 0 | 0 | 0 | 0 | 0 | 0 | 0 | 0 | 0 | 0 | 0 | 0 | 0 | 0 | 0 | 0 | 0 | 0 | 0 | 0 | 0 | 0 | 0 | 0 | 0 | 0 |
| virus_Jun22.75v_19113 | 1 | 0 | 0 | 0 | 0 | 0 | 0 | 0 | 0 | 0 | 0 | 0 | 0 | 0 | 0 | 0 | 0 | 0 | 0 | 0 | 1 | 0 | 0 | 0 | 0 | 0 | 0 | 0 | 0 | 0 | 0 | 0 | 0 | 0 | 0 | 0 | 0 | 0 | 0 | 0 | 0 | 0 | 0 | 0 | 0 | 0 | 0 | 0 | 0 | 0 | 0 | 0 | 0 | 0 | 0 |
| virus_Jun22.75v_21972 | 1 | 0 | 0 | 0 | 0 | 0 | 0 | 0 | 0 | 0 | 0 | 0 | 0 | 0 | 0 | 0 | 0 | 0 | 0 | 0 | 1 | 0 | 0 | 0 | 0 | 0 | 0 | 0 | 0 | 0 | 0 | 0 | 0 | 0 | 0 | 0 | 0 | 0 | 0 | 0 | 0 | 0 | 0 | 0 | 0 | 0 | 0 | 0 | 0 | 0 | 0 | 0 | 0 | 0 | 0 |
| virus_Jun22.75v_25000 | 1 | 0 | 0 | 0 | 0 | 0 | 0 | 0 | 0 | 0 | 0 | 0 | 0 | 0 | 0 | 0 | 0 | 0 | 0 | 0 | 1 | 0 | 0 | 0 | 0 | 0 | 0 | 0 | 0 | 0 | 0 | 0 | 0 | 0 | 0 | 0 | 0 | 0 | 0 | 0 | 0 | 0 | 0 | 0 | 0 | 0 | 0 | 0 | 0 | 0 | 0 | 0 | 0 | 0 | 0 |
| virus_Jun22.75v_27225 | 1 | 0 | 0 | 0 | 0 | 0 | 0 | 0 | 0 | 0 | 0 | 0 | 0 | 0 | 0 | 0 | 0 | 0 | 0 | 0 | 1 | 0 | 0 | 0 | 0 | 0 | 0 | 0 | 0 | 0 | 0 | 0 | 0 | 0 | 0 | 0 | 0 | 0 | 0 | 0 | 0 | 0 | 0 | 0 | 0 | 0 | 0 | 0 | 0 | 0 | 0 | 0 | 0 | 0 | 0 |
| virus_Jun22.75v_30843 | 1 | 0 | 0 | 0 | 0 | 0 | 0 | 0 | 0 | 0 | 0 | 0 | 0 | 0 | 0 | 0 | 0 | 0 | 0 | 0 | 1 | 0 | 0 | 0 | 0 | 0 | 0 | 0 | 0 | 0 | 0 | 0 | 0 | 0 | 0 | 0 | 0 | 0 | 0 | 0 | 0 | 0 | 0 | 0 | 0 | 0 | 0 | 0 | 0 | 0 | 0 | 0 | 0 | 0 | 0 |
| virus_Jun22.75v_30907 | 1 | 0 | 0 | 0 | 0 | 0 | 0 | 0 | 0 | 0 | 0 | 0 | 0 | 0 | 0 | 0 | 0 | 0 | 0 | 0 | 1 | 0 | 0 | 0 | 0 | 0 | 0 | 0 | 0 | 0 | 0 | 0 | 0 | 0 | 0 | 0 | 0 | 0 | 0 | 0 | 0 | 0 | 0 | 0 | 0 | 0 | 0 | 0 | 0 | 0 | 0 | 0 | 0 | 0 | 0 |
| virus_Jun22.75v_32988 | 1 | 0 | 0 | 0 | 0 | 0 | 0 | 0 | 0 | 0 | 0 | 0 | 0 | 0 | 0 | 0 | 0 | 0 | 0 | 0 | 1 | 0 | 0 | 0 | 0 | 0 | 0 | 0 | 0 | 0 | 0 | 0 | 0 | 0 | 0 | 0 | 0 | 0 | 0 | 0 | 0 | 0 | 0 | 0 | 0 | 0 | 0 | 0 | 0 | 0 | 0 | 0 | 0 | 0 | 0 |
| virus_Jun22.75v_39688 | 1 | 0 | 0 | 0 | 0 | 0 | 0 | 0 | 0 | 0 | 0 | 0 | 0 | 0 | 0 | 0 | 0 | 0 | 0 | 0 | 1 | 0 | 0 | 0 | 0 | 0 | 0 | 0 | 0 | 0 | 0 | 0 | 0 | 0 | 0 | 0 | 0 | 0 | 0 | 0 | 0 | 0 | 0 | 0 | 0 | 0 | 0 | 0 | 0 | 0 | 0 | 0 | 0 | 0 | 0 |
| virus_Jun22.75v_40555 | 1 | 0 | 0 | 0 | 0 | 0 | 0 | 0 | 0 | 0 | 0 | 0 | 0 | 0 | 0 | 0 | 0 | 0 | 0 | 0 | 1 | 0 | 0 | 0 | 0 | 0 | 0 | 0 | 0 | 0 | 0 | 0 | 0 | 0 | 0 | 0 | 0 | 0 | 0 | 0 | 0 | 0 | 0 | 0 | 0 | 0 | 0 | 0 | 0 | 0 | 0 | 0 | 0 | 0 | 0 |
| virus_May12.5v_19120 | 1 | 0 | 1 | 0 | 0 | 0 | 0 | 0 | 0 | 0 | 0 | 0 | 0 | 0 | 0 | 0 | 0 | 0 | 0 | 0 | 0 | 0 | 0 | 0 | 0 | 0 | 0 | 0 | 0 | 0 | 0 | 0 | 0 | 0 | 0 | 0 | 0 | 0 | 0 | 0 | 0 | 0 | 0 | 0 | 0 | 0 | 0 | 0 | 0 | 0 | 0 | 0 | 0 | 0 | 0 |
| virus_May12.5v_19860 | 1 | 0 | 1 | 0 | 0 | 0 | 0 | 0 | 0 | 0 | 0 | 0 | 0 | 0 | 0 | 0 | 0 | 0 | 0 | 0 | 0 | 0 | 0 | 0 | 0 | 0 | 0 | 0 | 0 | 0 | 0 | 0 | 0 | 0 | 0 | 0 | 0 | 0 | 0 | 0 | 0 | 0 | 0 | 0 | 0 | 0 | 0 | 0 | 0 | 0 | 0 | 0 | 0 | 0 | 0 |
| virus_May12.5v_19934 | 1 | 0 | 1 | 0 | 0 | 0 | 0 | 0 | 0 | 0 | 0 | 0 | 0 | 0 | 0 | 0 | 0 | 0 | 0 | 0 | 0 | 0 | 0 | 0 | 0 | 0 | 0 | 0 | 0 | 0 | 0 | 0 | 0 | 0 | 0 | 0 | 0 | 0 | 0 | 0 | 0 | 0 | 0 | 0 | 0 | 0 | 0 | 0 | 0 | 0 | 0 | 0 | 0 | 0 | 0 |
| virus_May12.5v_25221 | 1 | 0 | 1 | 0 | 0 | 0 | 0 | 0 | 0 | 0 | 0 | 0 | 0 | 0 | 0 | 0 | 0 | 0 | 0 | 0 | 0 | 0 | 0 | 0 | 0 | 0 | 0 | 0 | 0 | 0 | 0 | 0 | 0 | 0 | 0 | 0 | 0 | 0 | 0 | 0 | 0 | 0 | 0 | 0 | 0 | 0 | 0 | 0 | 0 | 0 | 0 | 0 | 0 | 0 | 0 |
| virus_May12.5v_26224 | 1 | 0 | 1 | 0 | 0 | 0 | 0 | 0 | 0 | 0 | 0 | 0 | 0 | 0 | 0 | 0 | 0 | 0 | 0 | 0 | 0 | 0 | 0 | 0 | 0 | 0 | 0 | 0 | 0 | 0 | 0 | 0 | 0 | 0 | 0 | 0 | 0 | 0 | 0 | 0 | 0 | 0 | 0 | 0 | 0 | 0 | 0 | 0 | 0 | 0 | 0 | 0 | 0 | 0 | 0 |
| virus_May12.5v_31990 | 1 | 0 | 1 | 0 | 0 | 0 | 0 | 0 | 0 | 0 | 0 | 0 | 0 | 0 | 0 | 0 | 0 | 0 | 0 | 0 | 0 | 0 | 0 | 0 | 0 | 0 | 0 | 0 | 0 | 0 | 0 | 0 | 0 | 0 | 0 | 0 | 0 | 0 | 0 | 0 | 0 | 0 | 0 | 0 | 0 | 0 | 0 | 0 | 0 | 0 | 0 | 0 | 0 | 0 | 0 |
| virus_May12.5v_39748 | 1 | 0 | 1 | 0 | 0 | 0 | 0 | 0 | 0 | 0 | 0 | 0 | 0 | 0 | 0 | 0 | 0 | 0 | 0 | 0 | 0 | 0 | 0 | 0 | 0 | 0 | 0 | 0 | 0 | 0 | 0 | 0 | 0 | 0 | 0 | 0 | 0 | 0 | 0 | 0 | 0 | 0 | 0 | 0 | 0 | 0 | 0 | 0 | 0 | 0 | 0 | 0 | 0 | 0 | 0 |
| virus_May12.5v_40204 | 1 | 0 | 1 | 0 | 0 | 0 | 0 | 0 | 0 | 0 | 0 | 0 | 0 | 0 | 0 | 0 | 0 | 0 | 0 | 0 | 0 | 0 | 0 | 0 | 0 | 0 | 0 | 0 | 0 | 0 | 0 | 0 | 0 | 0 | 0 | 0 | 0 | 0 | 0 | 0 | 0 | 0 | 0 | 0 | 0 | 0 | 0 | 0 | 0 | 0 | 0 | 0 | 0 | 0 | 0 |
| virus_May12.5v_40693 | 1 | 0 | 1 | 0 | 0 | 0 | 0 | 0 | 0 | 0 | 0 | 0 | 0 | 0 | 0 | 0 | 0 | 0 | 0 | 0 | 0 | 0 | 0 | 0 | 0 | 0 | 0 | 0 | 0 | 0 | 0 | 0 | 0 | 0 | 0 | 0 | 0 | 0 | 0 | 0 | 0 | 0 | 0 | 0 | 0 | 0 | 0 | 0 | 0 | 0 | 0 | 0 | 0 | 0 | 0 |
| virus_May12.25v_3798 | 1 | 0 | 0 | 0 | 1 | 0 | 0 | 0 | 0 | 0 | 0 | 0 | 0 | 0 | 0 | 0 | 0 | 0 | 0 | 0 | 0 | 0 | 0 | 0 | 0 | 0 | 0 | 0 | 0 | 0 | 0 | 0 | 0 | 0 | 0 | 0 | 0 | 0 | 0 | 0 | 0 | 0 | 0 | 0 | 0 | 0 | 0 | 0 | 0 | 0 | 0 | 0 | 0 | 0 | 0 |
| virus_May12.25v_8321 | 1 | 0 | 0 | 0 | 1 | 0 | 0 | 0 | 0 | 0 | 0 | 0 | 0 | 0 | 0 | 0 | 0 | 0 | 0 | 0 | 0 | 0 | 0 | 0 | 0 | 0 | 0 | 0 | 0 | 0 | 0 | 0 | 0 | 0 | 0 | 0 | 0 | 0 | 0 | 0 | 0 | 0 | 0 | 0 | 0 | 0 | 0 | 0 | 0 | 0 | 0 | 0 | 0 | 0 | 0 |
| virus_May12.25v_9339 | 1 | 0 | 0 | 0 | 1 | 0 | 0 | 0 | 0 | 0 | 0 | 0 | 0 | 0 | 0 | 0 | 0 | 0 | 0 | 0 | 0 | 0 | 0 | 0 | 0 | 0 | 0 | 0 | 0 | 0 | 0 | 0 | 0 | 0 | 0 | 0 | 0 | 0 | 0 | 0 | 0 | 0 | 0 | 0 | 0 | 0 | 0 | 0 | 0 | 0 | 0 | 0 | 0 | 0 | 0 |
| virus_May12.25v_17687 | 1 | 0 | 0 | 0 | 1 | 0 | 0 | 0 | 0 | 0 | 0 | 0 | 0 | 0 | 0 | 0 | 0 | 0 | 0 | 0 | 0 | 0 | 0 | 0 | 0 | 0 | 0 | 0 | 0 | 0 | 0 | 0 | 0 | 0 | 0 | 0 | 0 | 0 | 0 | 0 | 0 | 0 | 0 | 0 | 0 | 0 | 0 | 0 | 0 | 0 | 0 | 0 | 0 | 0 | 0 |
| virus_May12.25v_18938 | 1 | 0 | 0 | 0 | 1 | 0 | 0 | 0 | 0 | 0 | 0 | 0 | 0 | 0 | 0 | 0 | 0 | 0 | 0 | 0 | 0 | 0 | 0 | 0 | 0 | 0 | 0 | 0 | 0 | 0 | 0 | 0 | 0 | 0 | 0 | 0 | 0 | 0 | 0 | 0 | 0 | 0 | 0 | 0 | 0 | 0 | 0 | 0 | 0 | 0 | 0 | 0 | 0 | 0 | 0 |
| virus_May12.25v_24167 | 1 | 0 | 0 | 0 | 1 | 0 | 0 | 0 | 0 | 0 | 0 | 0 | 0 | 0 | 0 | 0 | 0 | 0 | 0 | 0 | 0 | 0 | 0 | 0 | 0 | 0 | 0 | 0 | 0 | 0 | 0 | 0 | 0 | 0 | 0 | 0 | 0 | 0 | 0 | 0 | 0 | 0 | 0 | 0 | 0 | 0 | 0 | 0 | 0 | 0 | 0 | 0 | 0 | 0 | 0 |
| virus_May12.25v_31660 | 1 | 0 | 0 | 0 | 1 | 0 | 0 | 0 | 0 | 0 | 0 | 0 | 0 | 0 | 0 | 0 | 0 | 0 | 0 | 0 | 0 | 0 | 0 | 0 | 0 | 0 | 0 | 0 | 0 | 0 | 0 | 0 | 0 | 0 | 0 | 0 | 0 | 0 | 0 | 0 | 0 | 0 | 0 | 0 | 0 | 0 | 0 | 0 | 0 | 0 | 0 | 0 | 0 | 0 | 0 |
| virus_May12.25v_31949 | 1 | 0 | 0 | 0 | 1 | 0 | 0 | 0 | 0 | 0 | 0 | 0 | 0 | 0 | 0 | 0 | 0 | 0 | 0 | 0 | 0 | 0 | 0 | 0 | 0 | 0 | 0 | 0 | 0 | 0 | 0 | 0 | 0 | 0 | 0 | 0 | 0 | 0 | 0 | 0 | 0 | 0 | 0 | 0 | 0 | 0 | 0 | 0 | 0 | 0 | 0 | 0 | 0 | 0 | 0 |
| virus_May12.25v_33909 | 1 | 0 | 0 | 0 | 1 | 0 | 0 | 0 | 0 | 0 | 0 | 0 | 0 | 0 | 0 | 0 | 0 | 0 | 0 | 0 | 0 | 0 | 0 | 0 | 0 | 0 | 0 | 0 | 0 | 0 | 0 | 0 | 0 | 0 | 0 | 0 | 0 | 0 | 0 | 0 | 0 | 0 | 0 | 0 | 0 | 0 | 0 | 0 | 0 | 0 | 0 | 0 | 0 | 0 | 0 |
| virus_May12.50v_5143 | 1 | 0 | 0 | 0 | 0 | 0 | 1 | 0 | 0 | 0 | 0 | 0 | 0 | 0 | 0 | 0 | 0 | 0 | 0 | 0 | 0 | 0 | 0 | 0 | 0 | 0 | 0 | 0 | 0 | 0 | 0 | 0 | 0 | 0 | 0 | 0 | 0 | 0 | 0 | 0 | 0 | 0 | 0 | 0 | 0 | 0 | 0 | 0 | 0 | 0 | 0 | 0 | 0 | 0 | 0 |
| virus_May12.50v_11557 | 1 | 0 | 0 | 0 | 0 | 0 | 1 | 0 | 0 | 0 | 0 | 0 | 0 | 0 | 0 | 0 | 0 | 0 | 0 | 0 | 0 | 0 | 0 | 0 | 0 | 0 | 0 | 0 | 0 | 0 | 0 | 0 | 0 | 0 | 0 | 0 | 0 | 0 | 0 | 0 | 0 | 0 | 0 | 0 | 0 | 0 | 0 | 0 | 0 | 0 | 0 | 0 | 0 | 0 | 0 |
| virus_May12.50v_11845 | 1 | 0 | 0 | 0 | 0 | 0 | 1 | 0 | 0 | 0 | 0 | 0 | 0 | 0 | 0 | 0 | 0 | 0 | 0 | 0 | 0 | 0 | 0 | 0 | 0 | 0 | 0 | 0 | 0 | 0 | 0 | 0 | 0 | 0 | 0 | 0 | 0 | 0 | 0 | 0 | 0 | 0 | 0 | 0 | 0 | 0 | 0 | 0 | 0 | 0 | 0 | 0 | 0 | 0 | 0 |
| virus_May12.50v_12016 | 1 | 0 | 0 | 0 | 0 | 0 | 1 | 0 | 0 | 0 | 0 | 0 | 0 | 0 | 0 | 0 | 0 | 0 | 0 | 0 | 0 | 0 | 0 | 0 | 0 | 0 | 0 | 0 | 0 | 0 | 0 | 0 | 0 | 0 | 0 | 0 | 0 | 0 | 0 | 0 | 0 | 0 | 0 | 0 | 0 | 0 | 0 | 0 | 0 | 0 | 0 | 0 | 0 | 0 | 0 |
| virus_May12.50v_12239 | 1 | 0 | 0 | 0 | 0 | 0 | 1 | 0 | 0 | 0 | 0 | 0 | 0 | 0 | 0 | 0 | 0 | 0 | 0 | 0 | 0 | 0 | 0 | 0 | 0 | 0 | 0 | 0 | 0 | 0 | 0 | 0 | 0 | 0 | 0 | 0 | 0 | 0 | 0 | 0 | 0 | 0 | 0 | 0 | 0 | 0 | 0 | 0 | 0 | 0 | 0 | 0 | 0 | 0 | 0 |
| virus_May12.50v_12696 | 1 | 0 | 0 | 0 | 0 | 0 | 1 | 0 | 0 | 0 | 0 | 0 | 0 | 0 | 0 | 0 | 0 | 0 | 0 | 0 | 0 | 0 | 0 | 0 | 0 | 0 | 0 | 0 | 0 | 0 | 0 | 0 | 0 | 0 | 0 | 0 | 0 | 0 | 0 | 0 | 0 | 0 | 0 | 0 | 0 | 0 | 0 | 0 | 0 | 0 | 0 | 0 | 0 | 0 | 0 |
| virus_May12.50v_19565 | 1 | 0 | 0 | 0 | 0 | 0 | 1 | 0 | 0 | 0 | 0 | 0 | 0 | 0 | 0 | 0 | 0 | 0 | 0 | 0 | 0 | 0 | 0 | 0 | 0 | 0 | 0 | 0 | 0 | 0 | 0 | 0 | 0 | 0 | 0 | 0 | 0 | 0 | 0 | 0 | 0 | 0 | 0 | 0 | 0 | 0 | 0 | 0 | 0 | 0 | 0 | 0 | 0 | 0 | 0 |
| virus_May12.50v_19765 | 1 | 0 | 0 | 0 | 0 | 0 | 1 | 0 | 0 | 0 | 0 | 0 | 0 | 0 | 0 | 0 | 0 | 0 | 0 | 0 | 0 | 0 | 0 | 0 | 0 | 0 | 0 | 0 | 0 | 0 | 0 | 0 | 0 | 0 | 0 | 0 | 0 | 0 | 0 | 0 | 0 | 0 | 0 | 0 | 0 | 0 | 0 | 0 | 0 | 0 | 0 | 0 | 0 | 0 | 0 |
| virus_May12.50v_33817 | 1 | 0 | 0 | 0 | 0 | 0 | 1 | 0 | 0 | 0 | 0 | 0 | 0 | 0 | 0 | 0 | 0 | 0 | 0 | 0 | 0 | 0 | 0 | 0 | 0 | 0 | 0 | 0 | 0 | 0 | 0 | 0 | 0 | 0 | 0 | 0 | 0 | 0 | 0 | 0 | 0 | 0 | 0 | 0 | 0 | 0 | 0 | 0 | 0 | 0 | 0 | 0 | 0 | 0 | 0 |
| virus_May12.50v_36826 | 1 | 0 | 0 | 0 | 0 | 0 | 1 | 0 | 0 | 0 | 0 | 0 | 0 | 0 | 0 | 0 | 0 | 0 | 0 | 0 | 0 | 0 | 0 | 0 | 0 | 0 | 0 | 0 | 0 | 0 | 0 | 0 | 0 | 0 | 0 | 0 | 0 | 0 | 0 | 0 | 0 | 0 | 0 | 0 | 0 | 0 | 0 | 0 | 0 | 0 | 0 | 0 | 0 | 0 | 0 |
| virus_May12.75v_19610 | 1 | 0 | 0 | 0 | 0 | 0 | 0 | 0 | 1 | 0 | 0 | 0 | 0 | 0 | 0 | 0 | 0 | 0 | 0 | 0 | 0 | 0 | 0 | 0 | 0 | 0 | 0 | 0 | 0 | 0 | 0 | 0 | 0 | 0 | 0 | 0 | 0 | 0 | 0 | 0 | 0 | 0 | 0 | 0 | 0 | 0 | 0 | 0 | 0 | 0 | 0 | 0 | 0 | 0 | 0 |
| virus_May12.75v_21318 | 1 | 0 | 0 | 0 | 0 | 0 | 0 | 0 | 1 | 0 | 0 | 0 | 0 | 0 | 0 | 0 | 0 | 0 | 0 | 0 | 0 | 0 | 0 | 0 | 0 | 0 | 0 | 0 | 0 | 0 | 0 | 0 | 0 | 0 | 0 | 0 | 0 | 0 | 0 | 0 | 0 | 0 | 0 | 0 | 0 | 0 | 0 | 0 | 0 | 0 | 0 | 0 | 0 | 0 | 0 |
| virus_May12.75v_22671 | 1 | 0 | 0 | 0 | 0 | 0 | 0 | 0 | 1 | 0 | 0 | 0 | 0 | 0 | 0 | 0 | 0 | 0 | 0 | 0 | 0 | 0 | 0 | 0 | 0 | 0 | 0 | 0 | 0 | 0 | 0 | 0 | 0 | 0 | 0 | 0 | 0 | 0 | 0 | 0 | 0 | 0 | 0 | 0 | 0 | 0 | 0 | 0 | 0 | 0 | 0 | 0 | 0 | 0 | 0 |
| virus_May12.75v_23551 | 1 | 0 | 0 | 0 | 0 | 0 | 0 | 0 | 1 | 0 | 0 | 0 | 0 | 0 | 0 | 0 | 0 | 0 | 0 | 0 | 0 | 0 | 0 | 0 | 0 | 0 | 0 | 0 | 0 | 0 | 0 | 0 | 0 | 0 | 0 | 0 | 0 | 0 | 0 | 0 | 0 | 0 | 0 | 0 | 0 | 0 | 0 | 0 | 0 | 0 | 0 | 0 | 0 | 0 | 0 |
| virus_May12.75v_33699 | 1 | 0 | 0 | 0 | 0 | 0 | 0 | 0 | 1 | 0 | 0 | 0 | 0 | 0 | 0 | 0 | 0 | 0 | 0 | 0 | 0 | 0 | 0 | 0 | 0 | 0 | 0 | 0 | 0 | 0 | 0 | 0 | 0 | 0 | 0 | 0 | 0 | 0 | 0 | 0 | 0 | 0 | 0 | 0 | 0 | 0 | 0 | 0 | 0 | 0 | 0 | 0 | 0 | 0 | 0 |
| virus_May12.75v_35375 | 1 | 0 | 0 | 0 | 0 | 0 | 0 | 0 | 1 | 0 | 0 | 0 | 0 | 0 | 0 | 0 | 0 | 0 | 0 | 0 | 0 | 0 | 0 | 0 | 0 | 0 | 0 | 0 | 0 | 0 | 0 | 0 | 0 | 0 | 0 | 0 | 0 | 0 | 0 | 0 | 0 | 0 | 0 | 0 | 0 | 0 | 0 | 0 | 0 | 0 | 0 | 0 | 0 | 0 | 0 |
| virus_Sep8.0v_1296 | 1 | 0 | 0 | 0 | 0 | 0 | 0 | 0 | 0 | 0 | 0 | 0 | 0 | 0 | 0 | 0 | 0 | 0 | 0 | 0 | 0 | 0 | 0 | 0 | 0 | 0 | 0 | 0 | 0 | 0 | 0 | 0 | 0 | 0 | 0 | 0 | 0 | 0 | 0 | 0 | 0 | 0 | 0 | 0 | 0 | 0 | 0 | 0 | 1 | 0 | 0 | 0 | 0 | 0 | 0 |
| virus_Sep8.0v_5587 | 1 | 0 | 0 | 0 | 0 | 0 | 0 | 0 | 0 | 0 | 0 | 0 | 0 | 0 | 0 | 0 | 0 | 0 | 0 | 0 | 0 | 0 | 0 | 0 | 0 | 0 | 0 | 0 | 0 | 0 | 0 | 0 | 0 | 0 | 0 | 0 | 0 | 0 | 0 | 0 | 0 | 0 | 0 | 0 | 0 | 0 | 0 | 0 | 1 | 0 | 0 | 0 | 0 | 0 | 0 |
| virus_Sep8.0v_6401 | 1 | 0 | 0 | 0 | 0 | 0 | 0 | 0 | 0 | 0 | 0 | 0 | 0 | 0 | 0 | 0 | 0 | 0 | 0 | 0 | 0 | 0 | 0 | 0 | 0 | 0 | 0 | 0 | 0 | 0 | 0 | 0 | 0 | 0 | 0 | 0 | 0 | 0 | 0 | 0 | 0 | 0 | 0 | 0 | 0 | 0 | 0 | 0 | 1 | 0 | 0 | 0 | 0 | 0 | 0 |
| virus_Sep8.0v_7485 | 1 | 0 | 0 | 0 | 0 | 0 | 0 | 0 | 0 | 0 | 0 | 0 | 0 | 0 | 0 | 0 | 0 | 0 | 0 | 0 | 0 | 0 | 0 | 0 | 0 | 0 | 0 | 0 | 0 | 0 | 0 | 0 | 0 | 0 | 0 | 0 | 0 | 0 | 0 | 0 | 0 | 0 | 0 | 0 | 0 | 0 | 0 | 0 | 1 | 0 | 0 | 0 | 0 | 0 | 0 |
| virus_Sep8.0v_8165 | 1 | 0 | 0 | 0 | 0 | 0 | 0 | 0 | 0 | 0 | 0 | 0 | 0 | 0 | 0 | 0 | 0 | 0 | 0 | 0 | 0 | 0 | 0 | 0 | 0 | 0 | 0 | 0 | 0 | 0 | 0 | 0 | 0 | 0 | 0 | 0 | 0 | 0 | 0 | 0 | 0 | 0 | 0 | 0 | 0 | 0 | 0 | 0 | 1 | 0 | 0 | 0 | 0 | 0 | 0 |
| virus_Sep8.0v_9852 | 1 | 0 | 0 | 0 | 0 | 0 | 0 | 0 | 0 | 0 | 0 | 0 | 0 | 0 | 0 | 0 | 0 | 0 | 0 | 0 | 0 | 0 | 0 | 0 | 0 | 0 | 0 | 0 | 0 | 0 | 0 | 0 | 0 | 0 | 0 | 0 | 0 | 0 | 0 | 0 | 0 | 0 | 0 | 0 | 0 | 0 | 0 | 0 | 1 | 0 | 0 | 0 | 0 | 0 | 0 |
| virus_Sep8.0v_12375 | 1 | 0 | 0 | 0 | 0 | 0 | 0 | 0 | 0 | 0 | 0 | 0 | 0 | 0 | 0 | 0 | 0 | 0 | 0 | 0 | 0 | 0 | 0 | 0 | 0 | 0 | 0 | 0 | 0 | 0 | 0 | 0 | 0 | 0 | 0 | 0 | 0 | 0 | 0 | 0 | 0 | 0 | 0 | 0 | 0 | 0 | 0 | 0 | 1 | 0 | 0 | 0 | 0 | 0 | 0 |
| virus_Sep8.0v_12732 | 1 | 0 | 0 | 0 | 0 | 0 | 0 | 0 | 0 | 0 | 0 | 0 | 0 | 0 | 0 | 0 | 0 | 0 | 0 | 0 | 0 | 0 | 0 | 0 | 0 | 0 | 0 | 0 | 0 | 0 | 0 | 0 | 0 | 0 | 0 | 0 | 0 | 0 | 0 | 0 | 0 | 0 | 0 | 0 | 0 | 0 | 0 | 0 | 1 | 0 | 0 | 0 | 0 | 0 | 0 |
| virus_Sep8.0v_14290 | 1 | 0 | 0 | 0 | 0 | 0 | 0 | 0 | 0 | 0 | 0 | 0 | 0 | 0 | 0 | 0 | 0 | 0 | 0 | 0 | 0 | 0 | 0 | 0 | 0 | 0 | 0 | 0 | 0 | 0 | 0 | 0 | 0 | 0 | 0 | 0 | 0 | 0 | 0 | 0 | 0 | 0 | 0 | 0 | 0 | 0 | 0 | 0 | 1 | 0 | 0 | 0 | 0 | 0 | 0 |
| virus_Sep8.0v_15345 | 1 | 0 | 0 | 0 | 0 | 0 | 0 | 0 | 0 | 0 | 0 | 0 | 0 | 0 | 0 | 0 | 0 | 0 | 0 | 0 | 0 | 0 | 0 | 0 | 0 | 0 | 0 | 0 | 0 | 0 | 0 | 0 | 0 | 0 | 0 | 0 | 0 | 0 | 0 | 0 | 0 | 0 | 0 | 0 | 0 | 0 | 0 | 0 | 1 | 0 | 0 | 0 | 0 | 0 | 0 |
| virus_Sep8.0v_15453 | 1 | 0 | 0 | 0 | 0 | 0 | 0 | 0 | 0 | 0 | 0 | 0 | 0 | 0 | 0 | 0 | 0 | 0 | 0 | 0 | 0 | 0 | 0 | 0 | 0 | 0 | 0 | 0 | 0 | 0 | 0 | 0 | 0 | 0 | 0 | 0 | 0 | 0 | 0 | 0 | 0 | 0 | 0 | 0 | 0 | 0 | 0 | 0 | 1 | 0 | 0 | 0 | 0 | 0 | 0 |
| virus_Sep8.0v_16923 | 1 | 0 | 0 | 0 | 0 | 0 | 0 | 0 | 0 | 0 | 0 | 0 | 0 | 0 | 0 | 0 | 0 | 0 | 0 | 0 | 0 | 0 | 0 | 0 | 0 | 0 | 0 | 0 | 0 | 0 | 0 | 0 | 0 | 0 | 0 | 0 | 0 | 0 | 0 | 0 | 0 | 0 | 0 | 0 | 0 | 0 | 0 | 0 | 1 | 0 | 0 | 0 | 0 | 0 | 0 |
| virus_Sep8.5v_1908 | 1 | 0 | 0 | 0 | 0 | 0 | 0 | 0 | 0 | 0 | 0 | 0 | 0 | 0 | 0 | 0 | 0 | 0 | 0 | 0 | 0 | 0 | 0 | 0 | 0 | 0 | 0 | 0 | 0 | 0 | 0 | 0 | 0 | 0 | 0 | 0 | 0 | 0 | 0 | 0 | 0 | 0 | 0 | 0 | 0 | 0 | 0 | 0 | 0 | 0 | 1 | 0 | 0 | 0 | 0 |
| virus_Sep8.5v_3741 | 1 | 0 | 0 | 0 | 0 | 0 | 0 | 0 | 0 | 0 | 0 | 0 | 0 | 0 | 0 | 0 | 0 | 0 | 0 | 0 | 0 | 0 | 0 | 0 | 0 | 0 | 0 | 0 | 0 | 0 | 0 | 0 | 0 | 0 | 0 | 0 | 0 | 0 | 0 | 0 | 0 | 0 | 0 | 0 | 0 | 0 | 0 | 0 | 0 | 0 | 1 | 0 | 0 | 0 | 0 |
| virus_Sep8.5v_8003 | 1 | 0 | 0 | 0 | 0 | 0 | 0 | 0 | 0 | 0 | 0 | 0 | 0 | 0 | 0 | 0 | 0 | 0 | 0 | 0 | 0 | 0 | 0 | 0 | 0 | 0 | 0 | 0 | 0 | 0 | 0 | 0 | 0 | 0 | 0 | 0 | 0 | 0 | 0 | 0 | 0 | 0 | 0 | 0 | 0 | 0 | 0 | 0 | 0 | 0 | 1 | 0 | 0 | 0 | 0 |
| virus_Sep8.5v_8909 | 1 | 0 | 0 | 0 | 0 | 0 | 0 | 0 | 0 | 0 | 0 | 0 | 0 | 0 | 0 | 0 | 0 | 0 | 0 | 0 | 0 | 0 | 0 | 0 | 0 | 0 | 0 | 0 | 0 | 0 | 0 | 0 | 0 | 0 | 0 | 0 | 0 | 0 | 0 | 0 | 0 | 0 | 0 | 0 | 0 | 0 | 0 | 0 | 0 | 0 | 1 | 0 | 0 | 0 | 0 |

**Supplementary Table S7**. Common Bacterio- and virioplankton RTPR 98% OTUs in 27 samples with RTPR amplicons in both fractions

Total seqs: 47997; Subtotal seqs in retained

| OTUs: 41046 | | May12_05m | | May12_25m | | May12_50m | | May12_75m | | Jun8_25m | | Jun8_50m | | Jun8_75m | | Jun22_0m | | Jun22_50m | | Jun22_75m | | Jul11_0m | | Jul11_05m | | Jul11_25m | | Jul11_50m | | Jul11_75m | | Jul27_05m | | Jul27_25m | | Jul27_75m | | Aug5_0m | | Aug5_05m | | Aug5_25m | | Aug5_50m | | Aug5_75m | | Sep8_0m | | Sep8_05m | | Sep8_25m | | Sep8_50m | |
|---|---|---|---|---|---|---|---|---|---|---|---|---|---|---|---|---|---|---|---|---|---|---|---|---|---|---|---|---|---|---|---|---|---|---|---|---|---|---|---|---|---|---|---|---|---|---|---|---|---|---|---|---|---|---|---|
| OTUs with >=10 seqs: 254 OTU subtotal | | bacterioplankton | virioplankton | bacterioplankton | virioplankton | bacterioplankton | virioplankton | bacterioplankton | virioplankton | bacterioplankton | virioplankton | bacterioplankton | virioplankton | bacterioplankton | virioplankton | bacterioplankton | virioplankton | bacterioplankton | virioplankton | bacterioplankton | virioplankton | bacterioplankton | virioplankton | bacterioplankton | virioplankton | bacterioplankton | virioplankton | bacterioplankton | virioplankton | bacterioplankton | virioplankton | bacterioplankton | virioplankton | bacterioplankton | virioplankton | bacterioplankton | virioplankton | bacterioplankton | virioplankton | bacterioplankton | virioplankton | bacterioplankton | virioplankton | bacterioplankton | virioplankton | bacterioplankton | virioplankton | bacterioplankton | virioplankton | bacterioplankton | virioplankton | bacterioplankton | virioplankton | bacterioplankton | virioplankton |
| virus_Sep8.5v_12833 | 1 | 0 | 0 | 0 | 0 | 0 | 0 | 0 | 0 | 0 | 0 | 0 | 0 | 0 | 0 | 0 | 0 | 0 | 0 | 0 | 0 | 0 | 0 | 0 | 0 | 0 | 0 | 0 | 0 | 0 | 0 | 0 | 0 | 0 | 0 | 0 | 0 | 0 | 0 | 0 | 0 | 0 | 0 | 0 | 0 | 0 | 0 | 0 | 0 | 0 | 1 | 0 | 0 | 0 | 0 |
| virus_Sep8.5v_13874 | 1 | 0 | 0 | 0 | 0 | 0 | 0 | 0 | 0 | 0 | 0 | 0 | 0 | 0 | 0 | 0 | 0 | 0 | 0 | 0 | 0 | 0 | 0 | 0 | 0 | 0 | 0 | 0 | 0 | 0 | 0 | 0 | 0 | 0 | 0 | 0 | 0 | 0 | 0 | 0 | 0 | 0 | 0 | 0 | 0 | 0 | 0 | 0 | 0 | 0 | 1 | 0 | 0 | 0 | 0 |
| virus_Sep8.5v_13987 | 1 | 0 | 0 | 0 | 0 | 0 | 0 | 0 | 0 | 0 | 0 | 0 | 0 | 0 | 0 | 0 | 0 | 0 | 0 | 0 | 0 | 0 | 0 | 0 | 0 | 0 | 0 | 0 | 0 | 0 | 0 | 0 | 0 | 0 | 0 | 0 | 0 | 0 | 0 | 0 | 0 | 0 | 0 | 0 | 0 | 0 | 0 | 0 | 0 | 0 | 1 | 0 | 0 | 0 | 0 |
| virus_Sep8.5v_15731 | 1 | 0 | 0 | 0 | 0 | 0 | 0 | 0 | 0 | 0 | 0 | 0 | 0 | 0 | 0 | 0 | 0 | 0 | 0 | 0 | 0 | 0 | 0 | 0 | 0 | 0 | 0 | 0 | 0 | 0 | 0 | 0 | 0 | 0 | 0 | 0 | 0 | 0 | 0 | 0 | 0 | 0 | 0 | 0 | 0 | 0 | 0 | 0 | 0 | 0 | 1 | 0 | 0 | 0 | 0 |
| virus_Sep8.25v_3817 | 1 | 0 | 0 | 0 | 0 | 0 | 0 | 0 | 0 | 0 | 0 | 0 | 0 | 0 | 0 | 0 | 0 | 0 | 0 | 0 | 0 | 0 | 0 | 0 | 0 | 0 | 0 | 0 | 0 | 0 | 0 | 0 | 0 | 0 | 0 | 0 | 0 | 0 | 0 | 0 | 0 | 0 | 0 | 0 | 0 | 0 | 0 | 0 | 0 | 0 | 0 | 0 | 1 | 0 | 0 |
| virus_Sep8.25v_4686 | 1 | 0 | 0 | 0 | 0 | 0 | 0 | 0 | 0 | 0 | 0 | 0 | 0 | 0 | 0 | 0 | 0 | 0 | 0 | 0 | 0 | 0 | 0 | 0 | 0 | 0 | 0 | 0 | 0 | 0 | 0 | 0 | 0 | 0 | 0 | 0 | 0 | 0 | 0 | 0 | 0 | 0 | 0 | 0 | 0 | 0 | 0 | 0 | 0 | 0 | 0 | 0 | 1 | 0 | 0 |
| virus_Sep8.25v_9252 | 1 | 0 | 0 | 0 | 0 | 0 | 0 | 0 | 0 | 0 | 0 | 0 | 0 | 0 | 0 | 0 | 0 | 0 | 0 | 0 | 0 | 0 | 0 | 0 | 0 | 0 | 0 | 0 | 0 | 0 | 0 | 0 | 0 | 0 | 0 | 0 | 0 | 0 | 0 | 0 | 0 | 0 | 0 | 0 | 0 | 0 | 0 | 0 | 0 | 0 | 0 | 0 | 1 | 0 | 0 |
| virus_Sep8.25v_11089 | 1 | 0 | 0 | 0 | 0 | 0 | 0 | 0 | 0 | 0 | 0 | 0 | 0 | 0 | 0 | 0 | 0 | 0 | 0 | 0 | 0 | 0 | 0 | 0 | 0 | 0 | 0 | 0 | 0 | 0 | 0 | 0 | 0 | 0 | 0 | 0 | 0 | 0 | 0 | 0 | 0 | 0 | 0 | 0 | 0 | 0 | 0 | 0 | 0 | 0 | 0 | 0 | 1 | 0 | 0 |
| virus_Sep8.25v_11794 | 1 | 0 | 0 | 0 | 0 | 0 | 0 | 0 | 0 | 0 | 0 | 0 | 0 | 0 | 0 | 0 | 0 | 0 | 0 | 0 | 0 | 0 | 0 | 0 | 0 | 0 | 0 | 0 | 0 | 0 | 0 | 0 | 0 | 0 | 0 | 0 | 0 | 0 | 0 | 0 | 0 | 0 | 0 | 0 | 0 | 0 | 0 | 0 | 0 | 0 | 0 | 0 | 1 | 0 | 0 |
| virus_Sep8.25v_13890 | 1 | 0 | 0 | 0 | 0 | 0 | 0 | 0 | 0 | 0 | 0 | 0 | 0 | 0 | 0 | 0 | 0 | 0 | 0 | 0 | 0 | 0 | 0 | 0 | 0 | 0 | 0 | 0 | 0 | 0 | 0 | 0 | 0 | 0 | 0 | 0 | 0 | 0 | 0 | 0 | 0 | 0 | 0 | 0 | 0 | 0 | 0 | 0 | 0 | 0 | 0 | 0 | 1 | 0 | 0 |
| virus_Sep8.25v_15359 | 1 | 0 | 0 | 0 | 0 | 0 | 0 | 0 | 0 | 0 | 0 | 0 | 0 | 0 | 0 | 0 | 0 | 0 | 0 | 0 | 0 | 0 | 0 | 0 | 0 | 0 | 0 | 0 | 0 | 0 | 0 | 0 | 0 | 0 | 0 | 0 | 0 | 0 | 0 | 0 | 0 | 0 | 0 | 0 | 0 | 0 | 0 | 0 | 0 | 0 | 0 | 0 | 1 | 0 | 0 |
| virus_Sep8.25v_17635 | 1 | 0 | 0 | 0 | 0 | 0 | 0 | 0 | 0 | 0 | 0 | 0 | 0 | 0 | 0 | 0 | 0 | 0 | 0 | 0 | 0 | 0 | 0 | 0 | 0 | 0 | 0 | 0 | 0 | 0 | 0 | 0 | 0 | 0 | 0 | 0 | 0 | 0 | 0 | 0 | 0 | 0 | 0 | 0 | 0 | 0 | 0 | 0 | 0 | 0 | 0 | 0 | 1 | 0 | 0 |
| virus_Sep8.50v_1245 | 1 | 0 | 0 | 0 | 0 | 0 | 0 | 0 | 0 | 0 | 0 | 0 | 0 | 0 | 0 | 0 | 0 | 0 | 0 | 0 | 0 | 0 | 0 | 0 | 0 | 0 | 0 | 0 | 0 | 0 | 0 | 0 | 0 | 0 | 0 | 0 | 0 | 0 | 0 | 0 | 0 | 0 | 0 | 0 | 0 | 0 | 0 | 0 | 0 | 0 | 0 | 0 | 0 | 0 | 1 |
| virus_Sep8.50v_2515 | 1 | 0 | 0 | 0 | 0 | 0 | 0 | 0 | 0 | 0 | 0 | 0 | 0 | 0 | 0 | 0 | 0 | 0 | 0 | 0 | 0 | 0 | 0 | 0 | 0 | 0 | 0 | 0 | 0 | 0 | 0 | 0 | 0 | 0 | 0 | 0 | 0 | 0 | 0 | 0 | 0 | 0 | 0 | 0 | 0 | 0 | 0 | 0 | 0 | 0 | 0 | 0 | 0 | 0 | 1 |
| virus_Sep8.50v_6130 | 1 | 0 | 0 | 0 | 0 | 0 | 0 | 0 | 0 | 0 | 0 | 0 | 0 | 0 | 0 | 0 | 0 | 0 | 0 | 0 | 0 | 0 | 0 | 0 | 0 | 0 | 0 | 0 | 0 | 0 | 0 | 0 | 0 | 0 | 0 | 0 | 0 | 0 | 0 | 0 | 0 | 0 | 0 | 0 | 0 | 0 | 0 | 0 | 0 | 0 | 0 | 0 | 0 | 0 | 1 |
| virus_Sep8.50v_6763 | 1 | 0 | 0 | 0 | 0 | 0 | 0 | 0 | 0 | 0 | 0 | 0 | 0 | 0 | 0 | 0 | 0 | 0 | 0 | 0 | 0 | 0 | 0 | 0 | 0 | 0 | 0 | 0 | 0 | 0 | 0 | 0 | 0 | 0 | 0 | 0 | 0 | 0 | 0 | 0 | 0 | 0 | 0 | 0 | 0 | 0 | 0 | 0 | 0 | 0 | 0 | 0 | 0 | 0 | 1 |
| virus_Sep8.50v_8608 | 1 | 0 | 0 | 0 | 0 | 0 | 0 | 0 | 0 | 0 | 0 | 0 | 0 | 0 | 0 | 0 | 0 | 0 | 0 | 0 | 0 | 0 | 0 | 0 | 0 | 0 | 0 | 0 | 0 | 0 | 0 | 0 | 0 | 0 | 0 | 0 | 0 | 0 | 0 | 0 | 0 | 0 | 0 | 0 | 0 | 0 | 0 | 0 | 0 | 0 | 0 | 0 | 0 | 0 | 1 |
| virus_Sep8.50v_9662 | 1 | 0 | 0 | 0 | 0 | 0 | 0 | 0 | 0 | 0 | 0 | 0 | 0 | 0 | 0 | 0 | 0 | 0 | 0 | 0 | 0 | 0 | 0 | 0 | 0 | 0 | 0 | 0 | 0 | 0 | 0 | 0 | 0 | 0 | 0 | 0 | 0 | 0 | 0 | 0 | 0 | 0 | 0 | 0 | 0 | 0 | 0 | 0 | 0 | 0 | 0 | 0 | 0 | 0 | 1 |
| virus_Sep8.50v_10971 | 1 | 0 | 0 | 0 | 0 | 0 | 0 | 0 | 0 | 0 | 0 | 0 | 0 | 0 | 0 | 0 | 0 | 0 | 0 | 0 | 0 | 0 | 0 | 0 | 0 | 0 | 0 | 0 | 0 | 0 | 0 | 0 | 0 | 0 | 0 | 0 | 0 | 0 | 0 | 0 | 0 | 0 | 0 | 0 | 0 | 0 | 0 | 0 | 0 | 0 | 0 | 0 | 0 | 0 | 1 |
| virus_Sep8.50v_11571 | 1 | 0 | 0 | 0 | 0 | 0 | 0 | 0 | 0 | 0 | 0 | 0 | 0 | 0 | 0 | 0 | 0 | 0 | 0 | 0 | 0 | 0 | 0 | 0 | 0 | 0 | 0 | 0 | 0 | 0 | 0 | 0 | 0 | 0 | 0 | 0 | 0 | 0 | 0 | 0 | 0 | 0 | 0 | 0 | 0 | 0 | 0 | 0 | 0 | 0 | 0 | 0 | 0 | 0 | 1 |
| virus_Sep8.50v_14560 | 1 | 0 | 0 | 0 | 0 | 0 | 0 | 0 | 0 | 0 | 0 | 0 | 0 | 0 | 0 | 0 | 0 | 0 | 0 | 0 | 0 | 0 | 0 | 0 | 0 | 0 | 0 | 0 | 0 | 0 | 0 | 0 | 0 | 0 | 0 | 0 | 0 | 0 | 0 | 0 | 0 | 0 | 0 | 0 | 0 | 0 | 0 | 0 | 0 | 0 | 0 | 0 | 0 | 0 | 1 |
| virus_Sep8.50v_14612 | 1 | 0 | 0 | 0 | 0 | 0 | 0 | 0 | 0 | 0 | 0 | 0 | 0 | 0 | 0 | 0 | 0 | 0 | 0 | 0 | 0 | 0 | 0 | 0 | 0 | 0 | 0 | 0 | 0 | 0 | 0 | 0 | 0 | 0 | 0 | 0 | 0 | 0 | 0 | 0 | 0 | 0 | 0 | 0 | 0 | 0 | 0 | 0 | 0 | 0 | 0 | 0 | 0 | 0 | 1 |
| virus_Sep8.50v_18336 | 1 | 0 | 0 | 0 | 0 | 0 | 0 | 0 | 0 | 0 | 0 | 0 | 0 | 0 | 0 | 0 | 0 | 0 | 0 | 0 | 0 | 0 | 0 | 0 | 0 | 0 | 0 | 0 | 0 | 0 | 0 | 0 | 0 | 0 | 0 | 0 | 0 | 0 | 0 | 0 | 0 | 0 | 0 | 0 | 0 | 0 | 0 | 0 | 0 | 0 | 0 | 0 | 0 | 0 | 1 |
| virus_Sep8.50v_19636 | 1 | 0 | 0 | 0 | 0 | 0 | 0 | 0 | 0 | 0 | 0 | 0 | 0 | 0 | 0 | 0 | 0 | 0 | 0 | 0 | 0 | 0 | 0 | 0 | 0 | 0 | 0 | 0 | 0 | 0 | 0 | 0 | 0 | 0 | 0 | 0 | 0 | 0 | 0 | 0 | 0 | 0 | 0 | 0 | 0 | 0 | 0 | 0 | 0 | 0 | 0 | 0 | 0 | 0 | 1 |
| virus_Sep8.50v_21263 | 1 | 0 | 0 | 0 | 0 | 0 | 0 | 0 | 0 | 0 | 0 | 0 | 0 | 0 | 0 | 0 | 0 | 0 | 0 | 0 | 0 | 0 | 0 | 0 | 0 | 0 | 0 | 0 | 0 | 0 | 0 | 0 | 0 | 0 | 0 | 0 | 0 | 0 | 0 | 0 | 0 | 0 | 0 | 0 | 0 | 0 | 0 | 0 | 0 | 0 | 0 | 0 | 0 | 0 | 1 |
| virus_Sep8.50v_28086 | 1 | 0 | 0 | 0 | 0 | 0 | 0 | 0 | 0 | 0 | 0 | 0 | 0 | 0 | 0 | 0 | 0 | 0 | 0 | 0 | 0 | 0 | 0 | 0 | 0 | 0 | 0 | 0 | 0 | 0 | 0 | 0 | 0 | 0 | 0 | 0 | 0 | 0 | 0 | 0 | 0 | 0 | 0 | 0 | 0 | 0 | 0 | 0 | 0 | 0 | 0 | 0 | 0 | 0 | 1 |
| virus_Sep8.50v_37486 | 1 | 0 | 0 | 0 | 0 | 0 | 0 | 0 | 0 | 0 | 0 | 0 | 0 | 0 | 0 | 0 | 0 | 0 | 0 | 0 | 0 | 0 | 0 | 0 | 0 | 0 | 0 | 0 | 0 | 0 | 0 | 0 | 0 | 0 | 0 | 0 | 0 | 0 | 0 | 0 | 0 | 0 | 0 | 0 | 0 | 0 | 0 | 0 | 0 | 0 | 0 | 0 | 0 | 0 | 1 |
| virus_Sep8.50v_37785 | 1 | 0 | 0 | 0 | 0 | 0 | 0 | 0 | 0 | 0 | 0 | 0 | 0 | 0 | 0 | 0 | 0 | 0 | 0 | 0 | 0 | 0 | 0 | 0 | 0 | 0 | 0 | 0 | 0 | 0 | 0 | 0 | 0 | 0 | 0 | 0 | 0 | 0 | 0 | 0 | 0 | 0 | 0 | 0 | 0 | 0 | 0 | 0 | 0 | 0 | 0 | 0 | 0 | 0 | 1 |
| bacteria_Jul11d0b_4064 | 1 | 0 | 0 | 0 | 0 | 0 | 0 | 0 | 0 | 0 | 0 | 0 | 0 | 0 | 0 | 0 | 0 | 0 | 0 | 0 | 0 | 1 | 0 | 0 | 0 | 0 | 0 | 0 | 0 | 0 | 0 | 0 | 0 | 0 | 0 | 0 | 0 | 0 | 0 | 0 | 0 | 0 | 0 | 0 | 0 | 0 | 0 | 0 | 0 | 0 | 0 | 0 | 0 | 0 | 0 |
| bacteria_Jul11d0b_7450 | 1 | 0 | 0 | 0 | 0 | 0 | 0 | 0 | 0 | 0 | 0 | 0 | 0 | 0 | 0 | 0 | 0 | 0 | 0 | 0 | 0 | 1 | 0 | 0 | 0 | 0 | 0 | 0 | 0 | 0 | 0 | 0 | 0 | 0 | 0 | 0 | 0 | 0 | 0 | 0 | 0 | 0 | 0 | 0 | 0 | 0 | 0 | 0 | 0 | 0 | 0 | 0 | 0 | 0 | 0 |
| bacteria_Jul11d0b_10933 | 1 | 0 | 0 | 0 | 0 | 0 | 0 | 0 | 0 | 0 | 0 | 0 | 0 | 0 | 0 | 0 | 0 | 0 | 0 | 0 | 0 | 1 | 0 | 0 | 0 | 0 | 0 | 0 | 0 | 0 | 0 | 0 | 0 | 0 | 0 | 0 | 0 | 0 | 0 | 0 | 0 | 0 | 0 | 0 | 0 | 0 | 0 | 0 | 0 | 0 | 0 | 0 | 0 | 0 | 0 |
| bacteria_Jul11d0b_25686 | 1 | 0 | 0 | 0 | 0 | 0 | 0 | 0 | 0 | 0 | 0 | 0 | 0 | 0 | 0 | 0 | 0 | 0 | 0 | 0 | 0 | 1 | 0 | 0 | 0 | 0 | 0 | 0 | 0 | 0 | 0 | 0 | 0 | 0 | 0 | 0 | 0 | 0 | 0 | 0 | 0 | 0 | 0 | 0 | 0 | 0 | 0 | 0 | 0 | 0 | 0 | 0 | 0 | 0 | 0 |
| bacteria_Jul11d0b_33463 | 1 | 0 | 0 | 0 | 0 | 0 | 0 | 0 | 0 | 0 | 0 | 0 | 0 | 0 | 0 | 0 | 0 | 0 | 0 | 0 | 0 | 1 | 0 | 0 | 0 | 0 | 0 | 0 | 0 | 0 | 0 | 0 | 0 | 0 | 0 | 0 | 0 | 0 | 0 | 0 | 0 | 0 | 0 | 0 | 0 | 0 | 0 | 0 | 0 | 0 | 0 | 0 | 0 | 0 | 0 |
| bacteria_Jul11d0b_33558 | 1 | 0 | 0 | 0 | 0 | 0 | 0 | 0 | 0 | 0 | 0 | 0 | 0 | 0 | 0 | 0 | 0 | 0 | 0 | 0 | 0 | 1 | 0 | 0 | 0 | 0 | 0 | 0 | 0 | 0 | 0 | 0 | 0 | 0 | 0 | 0 | 0 | 0 | 0 | 0 | 0 | 0 | 0 | 0 | 0 | 0 | 0 | 0 | 0 | 0 | 0 | 0 | 0 | 0 | 0 |
| bacteria_Jul11d0b_54020 | 1 | 0 | 0 | 0 | 0 | 0 | 0 | 0 | 0 | 0 | 0 | 0 | 0 | 0 | 0 | 0 | 0 | 0 | 0 | 0 | 0 | 1 | 0 | 0 | 0 | 0 | 0 | 0 | 0 | 0 | 0 | 0 | 0 | 0 | 0 | 0 | 0 | 0 | 0 | 0 | 0 | 0 | 0 | 0 | 0 | 0 | 0 | 0 | 0 | 0 | 0 | 0 | 0 | 0 | 0 |
| bacteria_Aug5d0b_33601 | 1 | 0 | 0 | 0 | 0 | 0 | 0 | 0 | 0 | 0 | 0 | 0 | 0 | 0 | 0 | 0 | 0 | 0 | 0 | 0 | 0 | 0 | 0 | 0 | 0 | 0 | 0 | 0 | 0 | 0 | 0 | 0 | 0 | 0 | 0 | 0 | 0 | 1 | 0 | 0 | 0 | 0 | 0 | 0 | 0 | 0 | 0 | 0 | 0 | 0 | 0 | 0 | 0 | 0 | 0 |
| bacteria_Aug5d0b_54816 | 1 | 0 | 0 | 0 | 0 | 0 | 0 | 0 | 0 | 0 | 0 | 0 | 0 | 0 | 0 | 0 | 0 | 0 | 0 | 0 | 0 | 0 | 0 | 0 | 0 | 0 | 0 | 0 | 0 | 0 | 0 | 0 | 0 | 0 | 0 | 0 | 0 | 1 | 0 | 0 | 0 | 0 | 0 | 0 | 0 | 0 | 0 | 0 | 0 | 0 | 0 | 0 | 0 | 0 | 0 |
| bacteria_Aug5d0b_60065 | 1 | 0 | 0 | 0 | 0 | 0 | 0 | 0 | 0 | 0 | 0 | 0 | 0 | 0 | 0 | 0 | 0 | 0 | 0 | 0 | 0 | 0 | 0 | 0 | 0 | 0 | 0 | 0 | 0 | 0 | 0 | 0 | 0 | 0 | 0 | 0 | 0 | 1 | 0 | 0 | 0 | 0 | 0 | 0 | 0 | 0 | 0 | 0 | 0 | 0 | 0 | 0 | 0 | 0 | 0 |
| bacteria_Aug5d0b_60239 | 1 | 0 | 0 | 0 | 0 | 0 | 0 | 0 | 0 | 0 | 0 | 0 | 0 | 0 | 0 | 0 | 0 | 0 | 0 | 0 | 0 | 0 | 0 | 0 | 0 | 0 | 0 | 0 | 0 | 0 | 0 | 0 | 0 | 0 | 0 | 0 | 0 | 1 | 0 | 0 | 0 | 0 | 0 | 0 | 0 | 0 | 0 | 0 | 0 | 0 | 0 | 0 | 0 | 0 | 0 |
| bacteria_Aug5d0b_60568 | 1 | 0 | 0 | 0 | 0 | 0 | 0 | 0 | 0 | 0 | 0 | 0 | 0 | 0 | 0 | 0 | 0 | 0 | 0 | 0 | 0 | 0 | 0 | 0 | 0 | 0 | 0 | 0 | 0 | 0 | 0 | 0 | 0 | 0 | 0 | 0 | 0 | 1 | 0 | 0 | 0 | 0 | 0 | 0 | 0 | 0 | 0 | 0 | 0 | 0 | 0 | 0 | 0 | 0 | 0 |
| bacteria_Aug5d5b_5724 | 1 | 0 | 0 | 0 | 0 | 0 | 0 | 0 | 0 | 0 | 0 | 0 | 0 | 0 | 0 | 0 | 0 | 0 | 0 | 0 | 0 | 0 | 0 | 0 | 0 | 0 | 0 | 0 | 0 | 0 | 0 | 0 | 0 | 0 | 0 | 0 | 0 | 0 | 0 | 1 | 0 | 0 | 0 | 0 | 0 | 0 | 0 | 0 | 0 | 0 | 0 | 0 | 0 | 0 | 0 |
| bacteria_Aug5d5b_19122 | 1 | 0 | 0 | 0 | 0 | 0 | 0 | 0 | 0 | 0 | 0 | 0 | 0 | 0 | 0 | 0 | 0 | 0 | 0 | 0 | 0 | 0 | 0 | 0 | 0 | 0 | 0 | 0 | 0 | 0 | 0 | 0 | 0 | 0 | 0 | 0 | 0 | 0 | 0 | 1 | 0 | 0 | 0 | 0 | 0 | 0 | 0 | 0 | 0 | 0 | 0 | 0 | 0 | 0 | 0 |
| bacteria_Aug5d5b_61659 | 1 | 0 | 0 | 0 | 0 | 0 | 0 | 0 | 0 | 0 | 0 | 0 | 0 | 0 | 0 | 0 | 0 | 0 | 0 | 0 | 0 | 0 | 0 | 0 | 0 | 0 | 0 | 0 | 0 | 0 | 0 | 0 | 0 | 0 | 0 | 0 | 0 | 0 | 0 | 1 | 0 | 0 | 0 | 0 | 0 | 0 | 0 | 0 | 0 | 0 | 0 | 0 | 0 | 0 | 0 |
| bacteria_Aug5d25b_791 | 1 | 0 | 0 | 0 | 0 | 0 | 0 | 0 | 0 | 0 | 0 | 0 | 0 | 0 | 0 | 0 | 0 | 0 | 0 | 0 | 0 | 0 | 0 | 0 | 0 | 0 | 0 | 0 | 0 | 0 | 0 | 0 | 0 | 0 | 0 | 0 | 0 | 0 | 0 | 0 | 0 | 1 | 0 | 0 | 0 | 0 | 0 | 0 | 0 | 0 | 0 | 0 | 0 | 0 | 0 |
| bacteria_Aug5d25b_2222 | 1 | 0 | 0 | 0 | 0 | 0 | 0 | 0 | 0 | 0 | 0 | 0 | 0 | 0 | 0 | 0 | 0 | 0 | 0 | 0 | 0 | 0 | 0 | 0 | 0 | 0 | 0 | 0 | 0 | 0 | 0 | 0 | 0 | 0 | 0 | 0 | 0 | 0 | 0 | 0 | 0 | 1 | 0 | 0 | 0 | 0 | 0 | 0 | 0 | 0 | 0 | 0 | 0 | 0 | 0 |
| bacteria_Aug5d25b_2981 | 1 | 0 | 0 | 0 | 0 | 0 | 0 | 0 | 0 | 0 | 0 | 0 | 0 | 0 | 0 | 0 | 0 | 0 | 0 | 0 | 0 | 0 | 0 | 0 | 0 | 0 | 0 | 0 | 0 | 0 | 0 | 0 | 0 | 0 | 0 | 0 | 0 | 0 | 0 | 0 | 0 | 1 | 0 | 0 | 0 | 0 | 0 | 0 | 0 | 0 | 0 | 0 | 0 | 0 | 0 |
| bacteria_Aug5d25b_7759 | 1 | 0 | 0 | 0 | 0 | 0 | 0 | 0 | 0 | 0 | 0 | 0 | 0 | 0 | 0 | 0 | 0 | 0 | 0 | 0 | 0 | 0 | 0 | 0 | 0 | 0 | 0 | 0 | 0 | 0 | 0 | 0 | 0 | 0 | 0 | 0 | 0 | 0 | 0 | 0 | 0 | 1 | 0 | 0 | 0 | 0 | 0 | 0 | 0 | 0 | 0 | 0 | 0 | 0 | 0 |
| bacteria_Aug5d25b_12071 | 1 | 0 | 0 | 0 | 0 | 0 | 0 | 0 | 0 | 0 | 0 | 0 | 0 | 0 | 0 | 0 | 0 | 0 | 0 | 0 | 0 | 0 | 0 | 0 | 0 | 0 | 0 | 0 | 0 | 0 | 0 | 0 | 0 | 0 | 0 | 0 | 0 | 0 | 0 | 0 | 0 | 1 | 0 | 0 | 0 | 0 | 0 | 0 | 0 | 0 | 0 | 0 | 0 | 0 | 0 |
| bacteria_Aug5d25b_23554 | 1 | 0 | 0 | 0 | 0 | 0 | 0 | 0 | 0 | 0 | 0 | 0 | 0 | 0 | 0 | 0 | 0 | 0 | 0 | 0 | 0 | 0 | 0 | 0 | 0 | 0 | 0 | 0 | 0 | 0 | 0 | 0 | 0 | 0 | 0 | 0 | 0 | 0 | 0 | 0 | 0 | 1 | 0 | 0 | 0 | 0 | 0 | 0 | 0 | 0 | 0 | 0 | 0 | 0 | 0 |
| bacteria_Aug5d25b_25050 | 1 | 0 | 0 | 0 | 0 | 0 | 0 | 0 | 0 | 0 | 0 | 0 | 0 | 0 | 0 | 0 | 0 | 0 | 0 | 0 | 0 | 0 | 0 | 0 | 0 | 0 | 0 | 0 | 0 | 0 | 0 | 0 | 0 | 0 | 0 | 0 | 0 | 0 | 0 | 0 | 0 | 1 | 0 | 0 | 0 | 0 | 0 | 0 | 0 | 0 | 0 | 0 | 0 | 0 | 0 |
| bacteria_Aug5d25b_25665 | 1 | 0 | 0 | 0 | 0 | 0 | 0 | 0 | 0 | 0 | 0 | 0 | 0 | 0 | 0 | 0 | 0 | 0 | 0 | 0 | 0 | 0 | 0 | 0 | 0 | 0 | 0 | 0 | 0 | 0 | 0 | 0 | 0 | 0 | 0 | 0 | 0 | 0 | 0 | 0 | 0 | 1 | 0 | 0 | 0 | 0 | 0 | 0 | 0 | 0 | 0 | 0 | 0 | 0 | 0 |
| bacteria_Aug5d25b_26824 | 1 | 0 | 0 | 0 | 0 | 0 | 0 | 0 | 0 | 0 | 0 | 0 | 0 | 0 | 0 | 0 | 0 | 0 | 0 | 0 | 0 | 0 | 0 | 0 | 0 | 0 | 0 | 0 | 0 | 0 | 0 | 0 | 0 | 0 | 0 | 0 | 0 | 0 | 0 | 0 | 0 | 1 | 0 | 0 | 0 | 0 | 0 | 0 | 0 | 0 | 0 | 0 | 0 | 0 | 0 |
| bacteria_Aug5d25b_28046 | 1 | 0 | 0 | 0 | 0 | 0 | 0 | 0 | 0 | 0 | 0 | 0 | 0 | 0 | 0 | 0 | 0 | 0 | 0 | 0 | 0 | 0 | 0 | 0 | 0 | 0 | 0 | 0 | 0 | 0 | 0 | 0 | 0 | 0 | 0 | 0 | 0 | 0 | 0 | 0 | 0 | 1 | 0 | 0 | 0 | 0 | 0 | 0 | 0 | 0 | 0 | 0 | 0 | 0 | 0 |
| bacteria_Aug5d25b_31025 | 1 | 0 | 0 | 0 | 0 | 0 | 0 | 0 | 0 | 0 | 0 | 0 | 0 | 0 | 0 | 0 | 0 | 0 | 0 | 0 | 0 | 0 | 0 | 0 | 0 | 0 | 0 | 0 | 0 | 0 | 0 | 0 | 0 | 0 | 0 | 0 | 0 | 0 | 0 | 0 | 0 | 1 | 0 | 0 | 0 | 0 | 0 | 0 | 0 | 0 | 0 | 0 | 0 | 0 | 0 |
| bacteria_Aug5d25b_35349 | 1 | 0 | 0 | 0 | 0 | 0 | 0 | 0 | 0 | 0 | 0 | 0 | 0 | 0 | 0 | 0 | 0 | 0 | 0 | 0 | 0 | 0 | 0 | 0 | 0 | 0 | 0 | 0 | 0 | 0 | 0 | 0 | 0 | 0 | 0 | 0 | 0 | 0 | 0 | 0 | 0 | 1 | 0 | 0 | 0 | 0 | 0 | 0 | 0 | 0 | 0 | 0 | 0 | 0 | 0 |
| bacteria_Aug5d25b_35959 | 1 | 0 | 0 | 0 | 0 | 0 | 0 | 0 | 0 | 0 | 0 | 0 | 0 | 0 | 0 | 0 | 0 | 0 | 0 | 0 | 0 | 0 | 0 | 0 | 0 | 0 | 0 | 0 | 0 | 0 | 0 | 0 | 0 | 0 | 0 | 0 | 0 | 0 | 0 | 0 | 0 | 1 | 0 | 0 | 0 | 0 | 0 | 0 | 0 | 0 | 0 | 0 | 0 | 0 | 0 |
| bacteria_Aug5d25b_39343 | 1 | 0 | 0 | 0 | 0 | 0 | 0 | 0 | 0 | 0 | 0 | 0 | 0 | 0 | 0 | 0 | 0 | 0 | 0 | 0 | 0 | 0 | 0 | 0 | 0 | 0 | 0 | 0 | 0 | 0 | 0 | 0 | 0 | 0 | 0 | 0 | 0 | 0 | 0 | 0 | 0 | 1 | 0 | 0 | 0 | 0 | 0 | 0 | 0 | 0 | 0 | 0 | 0 | 0 | 0 |
| bacteria_Aug5d25b_46921 | 1 | 0 | 0 | 0 | 0 | 0 | 0 | 0 | 0 | 0 | 0 | 0 | 0 | 0 | 0 | 0 | 0 | 0 | 0 | 0 | 0 | 0 | 0 | 0 | 0 | 0 | 0 | 0 | 0 | 0 | 0 | 0 | 0 | 0 | 0 | 0 | 0 | 0 | 0 | 0 | 0 | 1 | 0 | 0 | 0 | 0 | 0 | 0 | 0 | 0 | 0 | 0 | 0 | 0 | 0 |
| bacteria_Aug5d25b_48644 | 1 | 0 | 0 | 0 | 0 | 0 | 0 | 0 | 0 | 0 | 0 | 0 | 0 | 0 | 0 | 0 | 0 | 0 | 0 | 0 | 0 | 0 | 0 | 0 | 0 | 0 | 0 | 0 | 0 | 0 | 0 | 0 | 0 | 0 | 0 | 0 | 0 | 0 | 0 | 0 | 0 | 1 | 0 | 0 | 0 | 0 | 0 | 0 | 0 | 0 | 0 | 0 | 0 | 0 | 0 |
| bacteria_Aug5d25b_60264 | 1 | 0 | 0 | 0 | 0 | 0 | 0 | 0 | 0 | 0 | 0 | 0 | 0 | 0 | 0 | 0 | 0 | 0 | 0 | 0 | 0 | 0 | 0 | 0 | 0 | 0 | 0 | 0 | 0 | 0 | 0 | 0 | 0 | 0 | 0 | 0 | 0 | 0 | 0 | 0 | 0 | 1 | 0 | 0 | 0 | 0 | 0 | 0 | 0 | 0 | 0 | 0 | 0 | 0 | 0 |
| bacteria_Aug5d25b_61264 | 1 | 0 | 0 | 0 | 0 | 0 | 0 | 0 | 0 | 0 | 0 | 0 | 0 | 0 | 0 | 0 | 0 | 0 | 0 | 0 | 0 | 0 | 0 | 0 | 0 | 0 | 0 | 0 | 0 | 0 | 0 | 0 | 0 | 0 | 0 | 0 | 0 | 0 | 0 | 0 | 0 | 1 | 0 | 0 | 0 | 0 | 0 | 0 | 0 | 0 | 0 | 0 | 0 | 0 | 0 |
| bacteria_Aug5d25b_61912 | 1 | 0 | 0 | 0 | 0 | 0 | 0 | 0 | 0 | 0 | 0 | 0 | 0 | 0 | 0 | 0 | 0 | 0 | 0 | 0 | 0 | 0 | 0 | 0 | 0 | 0 | 0 | 0 | 0 | 0 | 0 | 0 | 0 | 0 | 0 | 0 | 0 | 0 | 0 | 0 | 0 | 1 | 0 | 0 | 0 | 0 | 0 | 0 | 0 | 0 | 0 | 0 | 0 | 0 | 0 |
| bacteria_Aug5d25b_64549 | 1 | 0 | 0 | 0 | 0 | 0 | 0 | 0 | 0 | 0 | 0 | 0 | 0 | 0 | 0 | 0 | 0 | 0 | 0 | 0 | 0 | 0 | 0 | 0 | 0 | 0 | 0 | 0 | 0 | 0 | 0 | 0 | 0 | 0 | 0 | 0 | 0 | 0 | 0 | 0 | 0 | 1 | 0 | 0 | 0 | 0 | 0 | 0 | 0 | 0 | 0 | 0 | 0 | 0 | 0 |
| bacteria_Aug5d25b_70162 | 1 | 0 | 0 | 0 | 0 | 0 | 0 | 0 | 0 | 0 | 0 | 0 | 0 | 0 | 0 | 0 | 0 | 0 | 0 | 0 | 0 | 0 | 0 | 0 | 0 | 0 | 0 | 0 | 0 | 0 | 0 | 0 | 0 | 0 | 0 | 0 | 0 | 0 | 0 | 0 | 0 | 1 | 0 | 0 | 0 | 0 | 0 | 0 | 0 | 0 | 0 | 0 | 0 | 0 | 0 |
| bacteria_Aug5d50b_8785 | 1 | 0 | 0 | 0 | 0 | 0 | 0 | 0 | 0 | 0 | 0 | 0 | 0 | 0 | 0 | 0 | 0 | 0 | 0 | 0 | 0 | 0 | 0 | 0 | 0 | 0 | 0 | 0 | 0 | 0 | 0 | 0 | 0 | 0 | 0 | 0 | 0 | 0 | 0 | 0 | 0 | 0 | 0 | 1 | 0 | 0 | 0 | 0 | 0 | 0 | 0 | 0 | 0 | 0 | 0 |
| bacteria_Aug5d50b_26982 | 1 | 0 | 0 | 0 | 0 | 0 | 0 | 0 | 0 | 0 | 0 | 0 | 0 | 0 | 0 | 0 | 0 | 0 | 0 | 0 | 0 | 0 | 0 | 0 | 0 | 0 | 0 | 0 | 0 | 0 | 0 | 0 | 0 | 0 | 0 | 0 | 0 | 0 | 0 | 0 | 0 | 0 | 0 | 1 | 0 | 0 | 0 | 0 | 0 | 0 | 0 | 0 | 0 | 0 | 0 |
| bacteria_Aug5d50b_41700 | 1 | 0 | 0 | 0 | 0 | 0 | 0 | 0 | 0 | 0 | 0 | 0 | 0 | 0 | 0 | 0 | 0 | 0 | 0 | 0 | 0 | 0 | 0 | 0 | 0 | 0 | 0 | 0 | 0 | 0 | 0 | 0 | 0 | 0 | 0 | 0 | 0 | 0 | 0 | 0 | 0 | 0 | 0 | 1 | 0 | 0 | 0 | 0 | 0 | 0 | 0 | 0 | 0 | 0 | 0 |
| bacteria_Aug5d50b_62509 | 1 | 0 | 0 | 0 | 0 | 0 | 0 | 0 | 0 | 0 | 0 | 0 | 0 | 0 | 0 | 0 | 0 | 0 | 0 | 0 | 0 | 0 | 0 | 0 | 0 | 0 | 0 | 0 | 0 | 0 | 0 | 0 | 0 | 0 | 0 | 0 | 0 | 0 | 0 | 0 | 0 | 0 | 0 | 1 | 0 | 0 | 0 | 0 | 0 | 0 | 0 | 0 | 0 | 0 | 0 |
| bacteria_Aug5d75b_33727 | 1 | 0 | 0 | 0 | 0 | 0 | 0 | 0 | 0 | 0 | 0 | 0 | 0 | 0 | 0 | 0 | 0 | 0 | 0 | 0 | 0 | 0 | 0 | 0 | 0 | 0 | 0 | 0 | 0 | 0 | 0 | 0 | 0 | 0 | 0 | 0 | 0 | 0 | 0 | 0 | 0 | 0 | 0 | 0 | 0 | 1 | 0 | 0 | 0 | 0 | 0 | 0 | 0 | 0 | 0 |
| bacteria_Jul11d5b_31800 | 1 | 0 | 0 | 0 | 0 | 0 | 0 | 0 | 0 | 0 | 0 | 0 | 0 | 0 | 0 | 0 | 0 | 0 | 0 | 0 | 0 | 0 | 0 | 1 | 0 | 0 | 0 | 0 | 0 | 0 | 0 | 0 | 0 | 0 | 0 | 0 | 0 | 0 | 0 | 0 | 0 | 0 | 0 | 0 | 0 | 0 | 0 | 0 | 0 | 0 | 0 | 0 | 0 | 0 | 0 |
| bacteria_Jul11d5b_36453 | 1 | 0 | 0 | 0 | 0 | 0 | 0 | 0 | 0 | 0 | 0 | 0 | 0 | 0 | 0 | 0 | 0 | 0 | 0 | 0 | 0 | 0 | 0 | 1 | 0 | 0 | 0 | 0 | 0 | 0 | 0 | 0 | 0 | 0 | 0 | 0 | 0 | 0 | 0 | 0 | 0 | 0 | 0 | 0 | 0 | 0 | 0 | 0 | 0 | 0 | 0 | 0 | 0 | 0 | 0 |
| bacteria_Jul11d5b_38728 | 1 | 0 | 0 | 0 | 0 | 0 | 0 | 0 | 0 | 0 | 0 | 0 | 0 | 0 | 0 | 0 | 0 | 0 | 0 | 0 | 0 | 0 | 0 | 1 | 0 | 0 | 0 | 0 | 0 | 0 | 0 | 0 | 0 | 0 | 0 | 0 | 0 | 0 | 0 | 0 | 0 | 0 | 0 | 0 | 0 | 0 | 0 | 0 | 0 | 0 | 0 | 0 | 0 | 0 | 0 |
| bacteria_Jul11d5b_42764 | 1 | 0 | 0 | 0 | 0 | 0 | 0 | 0 | 0 | 0 | 0 | 0 | 0 | 0 | 0 | 0 | 0 | 0 | 0 | 0 | 0 | 0 | 0 | 1 | 0 | 0 | 0 | 0 | 0 | 0 | 0 | 0 | 0 | 0 | 0 | 0 | 0 | 0 | 0 | 0 | 0 | 0 | 0 | 0 | 0 | 0 | 0 | 0 | 0 | 0 | 0 | 0 | 0 | 0 | 0 |
| bacteria_Jul11d5b_47970 | 1 | 0 | 0 | 0 | 0 | 0 | 0 | 0 | 0 | 0 | 0 | 0 | 0 | 0 | 0 | 0 | 0 | 0 | 0 | 0 | 0 | 0 | 0 | 1 | 0 | 0 | 0 | 0 | 0 | 0 | 0 | 0 | 0 | 0 | 0 | 0 | 0 | 0 | 0 | 0 | 0 | 0 | 0 | 0 | 0 | 0 | 0 | 0 | 0 | 0 | 0 | 0 | 0 | 0 | 0 |
| bacteria_Jul11d5b_51347 | 1 | 0 | 0 | 0 | 0 | 0 | 0 | 0 | 0 | 0 | 0 | 0 | 0 | 0 | 0 | 0 | 0 | 0 | 0 | 0 | 0 | 0 | 0 | 1 | 0 | 0 | 0 | 0 | 0 | 0 | 0 | 0 | 0 | 0 | 0 | 0 | 0 | 0 | 0 | 0 | 0 | 0 | 0 | 0 | 0 | 0 | 0 | 0 | 0 | 0 | 0 | 0 | 0 | 0 | 0 |
| bacteria_Jul11d25b_5809 | 1 | 0 | 0 | 0 | 0 | 0 | 0 | 0 | 0 | 0 | 0 | 0 | 0 | 0 | 0 | 0 | 0 | 0 | 0 | 0 | 0 | 0 | 0 | 0 | 0 | 1 | 0 | 0 | 0 | 0 | 0 | 0 | 0 | 0 | 0 | 0 | 0 | 0 | 0 | 0 | 0 | 0 | 0 | 0 | 0 | 0 | 0 | 0 | 0 | 0 | 0 | 0 | 0 | 0 | 0 |
| bacteria_Jul11d25b_12907 | 1 | 0 | 0 | 0 | 0 | 0 | 0 | 0 | 0 | 0 | 0 | 0 | 0 | 0 | 0 | 0 | 0 | 0 | 0 | 0 | 0 | 0 | 0 | 0 | 0 | 1 | 0 | 0 | 0 | 0 | 0 | 0 | 0 | 0 | 0 | 0 | 0 | 0 | 0 | 0 | 0 | 0 | 0 | 0 | 0 | 0 | 0 | 0 | 0 | 0 | 0 | 0 | 0 | 0 | 0 |
| bacteria_Jul11d25b_13423 | 1 | 0 | 0 | 0 | 0 | 0 | 0 | 0 | 0 | 0 | 0 | 0 | 0 | 0 | 0 | 0 | 0 | 0 | 0 | 0 | 0 | 0 | 0 | 0 | 0 | 1 | 0 | 0 | 0 | 0 | 0 | 0 | 0 | 0 | 0 | 0 | 0 | 0 | 0 | 0 | 0 | 0 | 0 | 0 | 0 | 0 | 0 | 0 | 0 | 0 | 0 | 0 | 0 | 0 | 0 |
| bacteria_Jul11d25b_14189 | 1 | 0 | 0 | 0 | 0 | 0 | 0 | 0 | 0 | 0 | 0 | 0 | 0 | 0 | 0 | 0 | 0 | 0 | 0 | 0 | 0 | 0 | 0 | 0 | 0 | 1 | 0 | 0 | 0 | 0 | 0 | 0 | 0 | 0 | 0 | 0 | 0 | 0 | 0 | 0 | 0 | 0 | 0 | 0 | 0 | 0 | 0 | 0 | 0 | 0 | 0 | 0 | 0 | 0 | 0 |
| bacteria_Jul11d50b_10287 | 1 | 0 | 0 | 0 | 0 | 0 | 0 | 0 | 0 | 0 | 0 | 0 | 0 | 0 | 0 | 0 | 0 | 0 | 0 | 0 | 0 | 0 | 0 | 0 | 0 | 0 | 0 | 1 | 0 | 0 | 0 | 0 | 0 | 0 | 0 | 0 | 0 | 0 | 0 | 0 | 0 | 0 | 0 | 0 | 0 | 0 | 0 | 0 | 0 | 0 | 0 | 0 | 0 | 0 | 0 |
| bacteria_Jul11d50b_15301 | 1 | 0 | 0 | 0 | 0 | 0 | 0 | 0 | 0 | 0 | 0 | 0 | 0 | 0 | 0 | 0 | 0 | 0 | 0 | 0 | 0 | 0 | 0 | 0 | 0 | 0 | 0 | 1 | 0 | 0 | 0 | 0 | 0 | 0 | 0 | 0 | 0 | 0 | 0 | 0 | 0 | 0 | 0 | 0 | 0 | 0 | 0 | 0 | 0 | 0 | 0 | 0 | 0 | 0 | 0 |
| bacteria_Jul11d50b_16716 | 1 | 0 | 0 | 0 | 0 | 0 | 0 | 0 | 0 | 0 | 0 | 0 | 0 | 0 | 0 | 0 | 0 | 0 | 0 | 0 | 0 | 0 | 0 | 0 | 0 | 0 | 0 | 1 | 0 | 0 | 0 | 0 | 0 | 0 | 0 | 0 | 0 | 0 | 0 | 0 | 0 | 0 | 0 | 0 | 0 | 0 | 0 | 0 | 0 | 0 | 0 | 0 | 0 | 0 | 0 |
| bacteria_Jul11d50b_20983 | 1 | 0 | 0 | 0 | 0 | 0 | 0 | 0 | 0 | 0 | 0 | 0 | 0 | 0 | 0 | 0 | 0 | 0 | 0 | 0 | 0 | 0 | 0 | 0 | 0 | 0 | 0 | 1 | 0 | 0 | 0 | 0 | 0 | 0 | 0 | 0 | 0 | 0 | 0 | 0 | 0 | 0 | 0 | 0 | 0 | 0 | 0 | 0 | 0 | 0 | 0 | 0 | 0 | 0 | 0 |
| bacteria_Jul11d50b_40527 | 1 | 0 | 0 | 0 | 0 | 0 | 0 | 0 | 0 | 0 | 0 | 0 | 0 | 0 | 0 | 0 | 0 | 0 | 0 | 0 | 0 | 0 | 0 | 0 | 0 | 0 | 0 | 1 | 0 | 0 | 0 | 0 | 0 | 0 | 0 | 0 | 0 | 0 | 0 | 0 | 0 | 0 | 0 | 0 | 0 | 0 | 0 | 0 | 0 | 0 | 0 | 0 | 0 | 0 | 0 |
| bacteria_Jul11d50b_41113 | 1 | 0 | 0 | 0 | 0 | 0 | 0 | 0 | 0 | 0 | 0 | 0 | 0 | 0 | 0 | 0 | 0 | 0 | 0 | 0 | 0 | 0 | 0 | 0 | 0 | 0 | 0 | 1 | 0 | 0 | 0 | 0 | 0 | 0 | 0 | 0 | 0 | 0 | 0 | 0 | 0 | 0 | 0 | 0 | 0 | 0 | 0 | 0 | 0 | 0 | 0 | 0 | 0 | 0 | 0 |
| bacteria_Jul11d50b_51144 | 1 | 0 | 0 | 0 | 0 | 0 | 0 | 0 | 0 | 0 | 0 | 0 | 0 | 0 | 0 | 0 | 0 | 0 | 0 | 0 | 0 | 0 | 0 | 0 | 0 | 0 | 0 | 1 | 0 | 0 | 0 | 0 | 0 | 0 | 0 | 0 | 0 | 0 | 0 | 0 | 0 | 0 | 0 | 0 | 0 | 0 | 0 | 0 | 0 | 0 | 0 | 0 | 0 | 0 | 0 |
| bacteria_Jul11d75b_6043 | 1 | 0 | 0 | 0 | 0 | 0 | 0 | 0 | 0 | 0 | 0 | 0 | 0 | 0 | 0 | 0 | 0 | 0 | 0 | 0 | 0 | 0 | 0 | 0 | 0 | 0 | 0 | 0 | 0 | 1 | 0 | 0 | 0 | 0 | 0 | 0 | 0 | 0 | 0 | 0 | 0 | 0 | 0 | 0 | 0 | 0 | 0 | 0 | 0 | 0 | 0 | 0 | 0 | 0 | 0 |
| bacteria_Jul11d75b_15586 | 1 | 0 | 0 | 0 | 0 | 0 | 0 | 0 | 0 | 0 | 0 | 0 | 0 | 0 | 0 | 0 | 0 | 0 | 0 | 0 | 0 | 0 | 0 | 0 | 0 | 0 | 0 | 0 | 0 | 1 | 0 | 0 | 0 | 0 | 0 | 0 | 0 | 0 | 0 | 0 | 0 | 0 | 0 | 0 | 0 | 0 | 0 | 0 | 0 | 0 | 0 | 0 | 0 | 0 | 0 |
| bacteria_Jul11d75b_19273 | 1 | 0 | 0 | 0 | 0 | 0 | 0 | 0 | 0 | 0 | 0 | 0 | 0 | 0 | 0 | 0 | 0 | 0 | 0 | 0 | 0 | 0 | 0 | 0 | 0 | 0 | 0 | 0 | 0 | 1 | 0 | 0 | 0 | 0 | 0 | 0 | 0 | 0 | 0 | 0 | 0 | 0 | 0 | 0 | 0 | 0 | 0 | 0 | 0 | 0 | 0 | 0 | 0 | 0 | 0 |
| bacteria_Jul11d75b_22809 | 1 | 0 | 0 | 0 | 0 | 0 | 0 | 0 | 0 | 0 | 0 | 0 | 0 | 0 | 0 | 0 | 0 | 0 | 0 | 0 | 0 | 0 | 0 | 0 | 0 | 0 | 0 | 0 | 0 | 1 | 0 | 0 | 0 | 0 | 0 | 0 | 0 | 0 | 0 | 0 | 0 | 0 | 0 | 0 | 0 | 0 | 0 | 0 | 0 | 0 | 0 | 0 | 0 | 0 | 0 |
| bacteria_Jul11d75b_22856 | 1 | 0 | 0 | 0 | 0 | 0 | 0 | 0 | 0 | 0 | 0 | 0 | 0 | 0 | 0 | 0 | 0 | 0 | 0 | 0 | 0 | 0 | 0 | 0 | 0 | 0 | 0 | 0 | 0 | 1 | 0 | 0 | 0 | 0 | 0 | 0 | 0 | 0 | 0 | 0 | 0 | 0 | 0 | 0 | 0 | 0 | 0 | 0 | 0 | 0 | 0 | 0 | 0 | 0 | 0 |
| bacteria_Jul11d75b_23815 | 1 | 0 | 0 | 0 | 0 | 0 | 0 | 0 | 0 | 0 | 0 | 0 | 0 | 0 | 0 | 0 | 0 | 0 | 0 | 0 | 0 | 0 | 0 | 0 | 0 | 0 | 0 | 0 | 0 | 1 | 0 | 0 | 0 | 0 | 0 | 0 | 0 | 0 | 0 | 0 | 0 | 0 | 0 | 0 | 0 | 0 | 0 | 0 | 0 | 0 | 0 | 0 | 0 | 0 | 0 |
| bacteria_Jul11d75b_25694 | 1 | 0 | 0 | 0 | 0 | 0 | 0 | 0 | 0 | 0 | 0 | 0 | 0 | 0 | 0 | 0 | 0 | 0 | 0 | 0 | 0 | 0 | 0 | 0 | 0 | 0 | 0 | 0 | 0 | 1 | 0 | 0 | 0 | 0 | 0 | 0 | 0 | 0 | 0 | 0 | 0 | 0 | 0 | 0 | 0 | 0 | 0 | 0 | 0 | 0 | 0 | 0 | 0 | 0 | 0 |
| bacteria_Jul11d75b_35410 | 1 | 0 | 0 | 0 | 0 | 0 | 0 | 0 | 0 | 0 | 0 | 0 | 0 | 0 | 0 | 0 | 0 | 0 | 0 | 0 | 0 | 0 | 0 | 0 | 0 | 0 | 0 | 0 | 0 | 1 | 0 | 0 | 0 | 0 | 0 | 0 | 0 | 0 | 0 | 0 | 0 | 0 | 0 | 0 | 0 | 0 | 0 | 0 | 0 | 0 | 0 | 0 | 0 | 0 | 0 |
| bacteria_Jul11d75b_43579 | 1 | 0 | 0 | 0 | 0 | 0 | 0 | 0 | 0 | 0 | 0 | 0 | 0 | 0 | 0 | 0 | 0 | 0 | 0 | 0 | 0 | 0 | 0 | 0 | 0 | 0 | 0 | 0 | 0 | 1 | 0 | 0 | 0 | 0 | 0 | 0 | 0 | 0 | 0 | 0 | 0 | 0 | 0 | 0 | 0 | 0 | 0 | 0 | 0 | 0 | 0 | 0 | 0 | 0 | 0 |
| bacteria_Jul11d75b_47361 | 1 | 0 | 0 | 0 | 0 | 0 | 0 | 0 | 0 | 0 | 0 | 0 | 0 | 0 | 0 | 0 | 0 | 0 | 0 | 0 | 0 | 0 | 0 | 0 | 0 | 0 | 0 | 0 | 0 | 1 | 0 | 0 | 0 | 0 | 0 | 0 | 0 | 0 | 0 | 0 | 0 | 0 | 0 | 0 | 0 | 0 | 0 | 0 | 0 | 0 | 0 | 0 | 0 | 0 | 0 |
| bacteria_Jul11d75b_50279 | 1 | 0 | 0 | 0 | 0 | 0 | 0 | 0 | 0 | 0 | 0 | 0 | 0 | 0 | 0 | 0 | 0 | 0 | 0 | 0 | 0 | 0 | 0 | 0 | 0 | 0 | 0 | 0 | 0 | 1 | 0 | 0 | 0 | 0 | 0 | 0 | 0 | 0 | 0 | 0 | 0 | 0 | 0 | 0 | 0 | 0 | 0 | 0 | 0 | 0 | 0 | 0 | 0 | 0 | 0 |
| bacteria_Jul11d75b_50788 | 1 | 0 | 0 | 0 | 0 | 0 | 0 | 0 | 0 | 0 | 0 | 0 | 0 | 0 | 0 | 0 | 0 | 0 | 0 | 0 | 0 | 0 | 0 | 0 | 0 | 0 | 0 | 0 | 0 | 1 | 0 | 0 | 0 | 0 | 0 | 0 | 0 | 0 | 0 | 0 | 0 | 0 | 0 | 0 | 0 | 0 | 0 | 0 | 0 | 0 | 0 | 0 | 0 | 0 | 0 |
| bacteria_Jul27d5b_1828 | 1 | 0 | 0 | 0 | 0 | 0 | 0 | 0 | 0 | 0 | 0 | 0 | 0 | 0 | 0 | 0 | 0 | 0 | 0 | 0 | 0 | 0 | 0 | 0 | 0 | 0 | 0 | 0 | 0 | 0 | 0 | 1 | 0 | 0 | 0 | 0 | 0 | 0 | 0 | 0 | 0 | 0 | 0 | 0 | 0 | 0 | 0 | 0 | 0 | 0 | 0 | 0 | 0 | 0 | 0 |
| bacteria_Jul27d5b_2410 | 1 | 0 | 0 | 0 | 0 | 0 | 0 | 0 | 0 | 0 | 0 | 0 | 0 | 0 | 0 | 0 | 0 | 0 | 0 | 0 | 0 | 0 | 0 | 0 | 0 | 0 | 0 | 0 | 0 | 0 | 0 | 1 | 0 | 0 | 0 | 0 | 0 | 0 | 0 | 0 | 0 | 0 | 0 | 0 | 0 | 0 | 0 | 0 | 0 | 0 | 0 | 0 | 0 | 0 | 0 |
| bacteria_Jul27d5b_5023 | 1 | 0 | 0 | 0 | 0 | 0 | 0 | 0 | 0 | 0 | 0 | 0 | 0 | 0 | 0 | 0 | 0 | 0 | 0 | 0 | 0 | 0 | 0 | 0 | 0 | 0 | 0 | 0 | 0 | 0 | 0 | 1 | 0 | 0 | 0 | 0 | 0 | 0 | 0 | 0 | 0 | 0 | 0 | 0 | 0 | 0 | 0 | 0 | 0 | 0 | 0 | 0 | 0 | 0 | 0 |
| bacteria_Jul27d5b_5198 | 1 | 0 | 0 | 0 | 0 | 0 | 0 | 0 | 0 | 0 | 0 | 0 | 0 | 0 | 0 | 0 | 0 | 0 | 0 | 0 | 0 | 0 | 0 | 0 | 0 | 0 | 0 | 0 | 0 | 0 | 0 | 1 | 0 | 0 | 0 | 0 | 0 | 0 | 0 | 0 | 0 | 0 | 0 | 0 | 0 | 0 | 0 | 0 | 0 | 0 | 0 | 0 | 0 | 0 | 0 |
| bacteria_Jul27d5b_9568 | 1 | 0 | 0 | 0 | 0 | 0 | 0 | 0 | 0 | 0 | 0 | 0 | 0 | 0 | 0 | 0 | 0 | 0 | 0 | 0 | 0 | 0 | 0 | 0 | 0 | 0 | 0 | 0 | 0 | 0 | 0 | 1 | 0 | 0 | 0 | 0 | 0 | 0 | 0 | 0 | 0 | 0 | 0 | 0 | 0 | 0 | 0 | 0 | 0 | 0 | 0 | 0 | 0 | 0 | 0 |
| bacteria_Jul27d5b_10242 | 1 | 0 | 0 | 0 | 0 | 0 | 0 | 0 | 0 | 0 | 0 | 0 | 0 | 0 | 0 | 0 | 0 | 0 | 0 | 0 | 0 | 0 | 0 | 0 | 0 | 0 | 0 | 0 | 0 | 0 | 0 | 1 | 0 | 0 | 0 | 0 | 0 | 0 | 0 | 0 | 0 | 0 | 0 | 0 | 0 | 0 | 0 | 0 | 0 | 0 | 0 | 0 | 0 | 0 | 0 |
| bacteria_Jul27d5b_16318 | 1 | 0 | 0 | 0 | 0 | 0 | 0 | 0 | 0 | 0 | 0 | 0 | 0 | 0 | 0 | 0 | 0 | 0 | 0 | 0 | 0 | 0 | 0 | 0 | 0 | 0 | 0 | 0 | 0 | 0 | 0 | 1 | 0 | 0 | 0 | 0 | 0 | 0 | 0 | 0 | 0 | 0 | 0 | 0 | 0 | 0 | 0 | 0 | 0 | 0 | 0 | 0 | 0 | 0 | 0 |
| bacteria_Jul27d5b_16574 | 1 | 0 | 0 | 0 | 0 | 0 | 0 | 0 | 0 | 0 | 0 | 0 | 0 | 0 | 0 | 0 | 0 | 0 | 0 | 0 | 0 | 0 | 0 | 0 | 0 | 0 | 0 | 0 | 0 | 0 | 0 | 1 | 0 | 0 | 0 | 0 | 0 | 0 | 0 | 0 | 0 | 0 | 0 | 0 | 0 | 0 | 0 | 0 | 0 | 0 | 0 | 0 | 0 | 0 | 0 |
| bacteria_Jul27d5b_22465 | 1 | 0 | 0 | 0 | 0 | 0 | 0 | 0 | 0 | 0 | 0 | 0 | 0 | 0 | 0 | 0 | 0 | 0 | 0 | 0 | 0 | 0 | 0 | 0 | 0 | 0 | 0 | 0 | 0 | 0 | 0 | 1 | 0 | 0 | 0 | 0 | 0 | 0 | 0 | 0 | 0 | 0 | 0 | 0 | 0 | 0 | 0 | 0 | 0 | 0 | 0 | 0 | 0 | 0 | 0 |
| bacteria_Jul27d5b_27923 | 1 | 0 | 0 | 0 | 0 | 0 | 0 | 0 | 0 | 0 | 0 | 0 | 0 | 0 | 0 | 0 | 0 | 0 | 0 | 0 | 0 | 0 | 0 | 0 | 0 | 0 | 0 | 0 | 0 | 0 | 0 | 1 | 0 | 0 | 0 | 0 | 0 | 0 | 0 | 0 | 0 | 0 | 0 | 0 | 0 | 0 | 0 | 0 | 0 | 0 | 0 | 0 | 0 | 0 | 0 |
| bacteria_Jul27d5b_29555 | 1 | 0 | 0 | 0 | 0 | 0 | 0 | 0 | 0 | 0 | 0 | 0 | 0 | 0 | 0 | 0 | 0 | 0 | 0 | 0 | 0 | 0 | 0 | 0 | 0 | 0 | 0 | 0 | 0 | 0 | 0 | 1 | 0 | 0 | 0 | 0 | 0 | 0 | 0 | 0 | 0 | 0 | 0 | 0 | 0 | 0 | 0 | 0 | 0 | 0 | 0 | 0 | 0 | 0 | 0 |
| bacteria_Jul27d5b_33466 | 1 | 0 | 0 | 0 | 0 | 0 | 0 | 0 | 0 | 0 | 0 | 0 | 0 | 0 | 0 | 0 | 0 | 0 | 0 | 0 | 0 | 0 | 0 | 0 | 0 | 0 | 0 | 0 | 0 | 0 | 0 | 1 | 0 | 0 | 0 | 0 | 0 | 0 | 0 | 0 | 0 | 0 | 0 | 0 | 0 | 0 | 0 | 0 | 0 | 0 | 0 | 0 | 0 | 0 | 0 |
| bacteria_Jul27d5b_34460 | 1 | 0 | 0 | 0 | 0 | 0 | 0 | 0 | 0 | 0 | 0 | 0 | 0 | 0 | 0 | 0 | 0 | 0 | 0 | 0 | 0 | 0 | 0 | 0 | 0 | 0 | 0 | 0 | 0 | 0 | 0 | 1 | 0 | 0 | 0 | 0 | 0 | 0 | 0 | 0 | 0 | 0 | 0 | 0 | 0 | 0 | 0 | 0 | 0 | 0 | 0 | 0 | 0 | 0 | 0 |
| bacteria_Jul27d5b_35314 | 1 | 0 | 0 | 0 | 0 | 0 | 0 | 0 | 0 | 0 | 0 | 0 | 0 | 0 | 0 | 0 | 0 | 0 | 0 | 0 | 0 | 0 | 0 | 0 | 0 | 0 | 0 | 0 | 0 | 0 | 0 | 1 | 0 | 0 | 0 | 0 | 0 | 0 | 0 | 0 | 0 | 0 | 0 | 0 | 0 | 0 | 0 | 0 | 0 | 0 | 0 | 0 | 0 | 0 | 0 |
| bacteria_Jul27d5b_36998 | 1 | 0 | 0 | 0 | 0 | 0 | 0 | 0 | 0 | 0 | 0 | 0 | 0 | 0 | 0 | 0 | 0 | 0 | 0 | 0 | 0 | 0 | 0 | 0 | 0 | 0 | 0 | 0 | 0 | 0 | 0 | 1 | 0 | 0 | 0 | 0 | 0 | 0 | 0 | 0 | 0 | 0 | 0 | 0 | 0 | 0 | 0 | 0 | 0 | 0 | 0 | 0 | 0 | 0 | 0 |
| bacteria_Jul27d5b_39938 | 1 | 0 | 0 | 0 | 0 | 0 | 0 | 0 | 0 | 0 | 0 | 0 | 0 | 0 | 0 | 0 | 0 | 0 | 0 | 0 | 0 | 0 | 0 | 0 | 0 | 0 | 0 | 0 | 0 | 0 | 0 | 1 | 0 | 0 | 0 | 0 | 0 | 0 | 0 | 0 | 0 | 0 | 0 | 0 | 0 | 0 | 0 | 0 | 0 | 0 | 0 | 0 | 0 | 0 | 0 |
| bacteria_Jul27d5b_39975 | 1 | 0 | 0 | 0 | 0 | 0 | 0 | 0 | 0 | 0 | 0 | 0 | 0 | 0 | 0 | 0 | 0 | 0 | 0 | 0 | 0 | 0 | 0 | 0 | 0 | 0 | 0 | 0 | 0 | 0 | 0 | 1 | 0 | 0 | 0 | 0 | 0 | 0 | 0 | 0 | 0 | 0 | 0 | 0 | 0 | 0 | 0 | 0 | 0 | 0 | 0 | 0 | 0 | 0 | 0 |
| bacteria_Jul27d5b_40256 | 1 | 0 | 0 | 0 | 0 | 0 | 0 | 0 | 0 | 0 | 0 | 0 | 0 | 0 | 0 | 0 | 0 | 0 | 0 | 0 | 0 | 0 | 0 | 0 | 0 | 0 | 0 | 0 | 0 | 0 | 0 | 1 | 0 | 0 | 0 | 0 | 0 | 0 | 0 | 0 | 0 | 0 | 0 | 0 | 0 | 0 | 0 | 0 | 0 | 0 | 0 | 0 | 0 | 0 | 0 |
| bacteria_Jul27d5b_41588 | 1 | 0 | 0 | 0 | 0 | 0 | 0 | 0 | 0 | 0 | 0 | 0 | 0 | 0 | 0 | 0 | 0 | 0 | 0 | 0 | 0 | 0 | 0 | 0 | 0 | 0 | 0 | 0 | 0 | 0 | 0 | 1 | 0 | 0 | 0 | 0 | 0 | 0 | 0 | 0 | 0 | 0 | 0 | 0 | 0 | 0 | 0 | 0 | 0 | 0 | 0 | 0 | 0 | 0 | 0 |
| bacteria_Jul27d5b_42493 | 1 | 0 | 0 | 0 | 0 | 0 | 0 | 0 | 0 | 0 | 0 | 0 | 0 | 0 | 0 | 0 | 0 | 0 | 0 | 0 | 0 | 0 | 0 | 0 | 0 | 0 | 0 | 0 | 0 | 0 | 0 | 1 | 0 | 0 | 0 | 0 | 0 | 0 | 0 | 0 | 0 | 0 | 0 | 0 | 0 | 0 | 0 | 0 | 0 | 0 | 0 | 0 | 0 | 0 | 0 |
| bacteria_Jul27d5b_43443 | 1 | 0 | 0 | 0 | 0 | 0 | 0 | 0 | 0 | 0 | 0 | 0 | 0 | 0 | 0 | 0 | 0 | 0 | 0 | 0 | 0 | 0 | 0 | 0 | 0 | 0 | 0 | 0 | 0 | 0 | 0 | 1 | 0 | 0 | 0 | 0 | 0 | 0 | 0 | 0 | 0 | 0 | 0 | 0 | 0 | 0 | 0 | 0 | 0 | 0 | 0 | 0 | 0 | 0 | 0 |
| bacteria_Jul27d5b_44981 | 1 | 0 | 0 | 0 | 0 | 0 | 0 | 0 | 0 | 0 | 0 | 0 | 0 | 0 | 0 | 0 | 0 | 0 | 0 | 0 | 0 | 0 | 0 | 0 | 0 | 0 | 0 | 0 | 0 | 0 | 0 | 1 | 0 | 0 | 0 | 0 | 0 | 0 | 0 | 0 | 0 | 0 | 0 | 0 | 0 | 0 | 0 | 0 | 0 | 0 | 0 | 0 | 0 | 0 | 0 |
| bacteria_Jul27d5b_50313 | 1 | 0 | 0 | 0 | 0 | 0 | 0 | 0 | 0 | 0 | 0 | 0 | 0 | 0 | 0 | 0 | 0 | 0 | 0 | 0 | 0 | 0 | 0 | 0 | 0 | 0 | 0 | 0 | 0 | 0 | 0 | 1 | 0 | 0 | 0 | 0 | 0 | 0 | 0 | 0 | 0 | 0 | 0 | 0 | 0 | 0 | 0 | 0 | 0 | 0 | 0 | 0 | 0 | 0 | 0 |
| bacteria_Jul27d25b_2841 | 1 | 0 | 0 | 0 | 0 | 0 | 0 | 0 | 0 | 0 | 0 | 0 | 0 | 0 | 0 | 0 | 0 | 0 | 0 | 0 | 0 | 0 | 0 | 0 | 0 | 0 | 0 | 0 | 0 | 0 | 0 | 0 | 0 | 1 | 0 | 0 | 0 | 0 | 0 | 0 | 0 | 0 | 0 | 0 | 0 | 0 | 0 | 0 | 0 | 0 | 0 | 0 | 0 | 0 | 0 |
| bacteria_Jul27d25b_8443 | 1 | 0 | 0 | 0 | 0 | 0 | 0 | 0 | 0 | 0 | 0 | 0 | 0 | 0 | 0 | 0 | 0 | 0 | 0 | 0 | 0 | 0 | 0 | 0 | 0 | 0 | 0 | 0 | 0 | 0 | 0 | 0 | 0 | 1 | 0 | 0 | 0 | 0 | 0 | 0 | 0 | 0 | 0 | 0 | 0 | 0 | 0 | 0 | 0 | 0 | 0 | 0 | 0 | 0 | 0 |
| bacteria_Jul27d25b_8988 | 1 | 0 | 0 | 0 | 0 | 0 | 0 | 0 | 0 | 0 | 0 | 0 | 0 | 0 | 0 | 0 | 0 | 0 | 0 | 0 | 0 | 0 | 0 | 0 | 0 | 0 | 0 | 0 | 0 | 0 | 0 | 0 | 0 | 1 | 0 | 0 | 0 | 0 | 0 | 0 | 0 | 0 | 0 | 0 | 0 | 0 | 0 | 0 | 0 | 0 | 0 | 0 | 0 | 0 | 0 |
| bacteria_Jul27d25b_9583 | 1 | 0 | 0 | 0 | 0 | 0 | 0 | 0 | 0 | 0 | 0 | 0 | 0 | 0 | 0 | 0 | 0 | 0 | 0 | 0 | 0 | 0 | 0 | 0 | 0 | 0 | 0 | 0 | 0 | 0 | 0 | 0 | 0 | 1 | 0 | 0 | 0 | 0 | 0 | 0 | 0 | 0 | 0 | 0 | 0 | 0 | 0 | 0 | 0 | 0 | 0 | 0 | 0 | 0 | 0 |
| bacteria_Jul27d25b_9926 | 1 | 0 | 0 | 0 | 0 | 0 | 0 | 0 | 0 | 0 | 0 | 0 | 0 | 0 | 0 | 0 | 0 | 0 | 0 | 0 | 0 | 0 | 0 | 0 | 0 | 0 | 0 | 0 | 0 | 0 | 0 | 0 | 0 | 1 | 0 | 0 | 0 | 0 | 0 | 0 | 0 | 0 | 0 | 0 | 0 | 0 | 0 | 0 | 0 | 0 | 0 | 0 | 0 | 0 | 0 |
| bacteria_Jul27d25b_16877 | 1 | 0 | 0 | 0 | 0 | 0 | 0 | 0 | 0 | 0 | 0 | 0 | 0 | 0 | 0 | 0 | 0 | 0 | 0 | 0 | 0 | 0 | 0 | 0 | 0 | 0 | 0 | 0 | 0 | 0 | 0 | 0 | 0 | 1 | 0 | 0 | 0 | 0 | 0 | 0 | 0 | 0 | 0 | 0 | 0 | 0 | 0 | 0 | 0 | 0 | 0 | 0 | 0 | 0 | 0 |
| bacteria_Jul27d25b_21227 | 1 | 0 | 0 | 0 | 0 | 0 | 0 | 0 | 0 | 0 | 0 | 0 | 0 | 0 | 0 | 0 | 0 | 0 | 0 | 0 | 0 | 0 | 0 | 0 | 0 | 0 | 0 | 0 | 0 | 0 | 0 | 0 | 0 | 1 | 0 | 0 | 0 | 0 | 0 | 0 | 0 | 0 | 0 | 0 | 0 | 0 | 0 | 0 | 0 | 0 | 0 | 0 | 0 | 0 | 0 |
| bacteria_Jul27d25b_28632 | 1 | 0 | 0 | 0 | 0 | 0 | 0 | 0 | 0 | 0 | 0 | 0 | 0 | 0 | 0 | 0 | 0 | 0 | 0 | 0 | 0 | 0 | 0 | 0 | 0 | 0 | 0 | 0 | 0 | 0 | 0 | 0 | 0 | 1 | 0 | 0 | 0 | 0 | 0 | 0 | 0 | 0 | 0 | 0 | 0 | 0 | 0 | 0 | 0 | 0 | 0 | 0 | 0 | 0 | 0 |
| bacteria_Jul27d25b_31910 | 1 | 0 | 0 | 0 | 0 | 0 | 0 | 0 | 0 | 0 | 0 | 0 | 0 | 0 | 0 | 0 | 0 | 0 | 0 | 0 | 0 | 0 | 0 | 0 | 0 | 0 | 0 | 0 | 0 | 0 | 0 | 0 | 0 | 1 | 0 | 0 | 0 | 0 | 0 | 0 | 0 | 0 | 0 | 0 | 0 | 0 | 0 | 0 | 0 | 0 | 0 | 0 | 0 | 0 | 0 |
| bacteria_Jul27d25b_31926 | 1 | 0 | 0 | 0 | 0 | 0 | 0 | 0 | 0 | 0 | 0 | 0 | 0 | 0 | 0 | 0 | 0 | 0 | 0 | 0 | 0 | 0 | 0 | 0 | 0 | 0 | 0 | 0 | 0 | 0 | 0 | 0 | 0 | 1 | 0 | 0 | 0 | 0 | 0 | 0 | 0 | 0 | 0 | 0 | 0 | 0 | 0 | 0 | 0 | 0 | 0 | 0 | 0 | 0 | 0 |
| bacteria_Jul27d25b_37673 | 1 | 0 | 0 | 0 | 0 | 0 | 0 | 0 | 0 | 0 | 0 | 0 | 0 | 0 | 0 | 0 | 0 | 0 | 0 | 0 | 0 | 0 | 0 | 0 | 0 | 0 | 0 | 0 | 0 | 0 | 0 | 0 | 0 | 1 | 0 | 0 | 0 | 0 | 0 | 0 | 0 | 0 | 0 | 0 | 0 | 0 | 0 | 0 | 0 | 0 | 0 | 0 | 0 | 0 | 0 |
| bacteria_Jul27d25b_40649 | 1 | 0 | 0 | 0 | 0 | 0 | 0 | 0 | 0 | 0 | 0 | 0 | 0 | 0 | 0 | 0 | 0 | 0 | 0 | 0 | 0 | 0 | 0 | 0 | 0 | 0 | 0 | 0 | 0 | 0 | 0 | 0 | 0 | 1 | 0 | 0 | 0 | 0 | 0 | 0 | 0 | 0 | 0 | 0 | 0 | 0 | 0 | 0 | 0 | 0 | 0 | 0 | 0 | 0 | 0 |
| bacteria_Jul27d25b_42449 | 1 | 0 | 0 | 0 | 0 | 0 | 0 | 0 | 0 | 0 | 0 | 0 | 0 | 0 | 0 | 0 | 0 | 0 | 0 | 0 | 0 | 0 | 0 | 0 | 0 | 0 | 0 | 0 | 0 | 0 | 0 | 0 | 0 | 1 | 0 | 0 | 0 | 0 | 0 | 0 | 0 | 0 | 0 | 0 | 0 | 0 | 0 | 0 | 0 | 0 | 0 | 0 | 0 | 0 | 0 |
| bacteria_Jul27d25b_53679 | 1 | 0 | 0 | 0 | 0 | 0 | 0 | 0 | 0 | 0 | 0 | 0 | 0 | 0 | 0 | 0 | 0 | 0 | 0 | 0 | 0 | 0 | 0 | 0 | 0 | 0 | 0 | 0 | 0 | 0 | 0 | 0 | 0 | 1 | 0 | 0 | 0 | 0 | 0 | 0 | 0 | 0 | 0 | 0 | 0 | 0 | 0 | 0 | 0 | 0 | 0 | 0 | 0 | 0 | 0 |
| bacteria_Jul27d25b_59509 | 1 | 0 | 0 | 0 | 0 | 0 | 0 | 0 | 0 | 0 | 0 | 0 | 0 | 0 | 0 | 0 | 0 | 0 | 0 | 0 | 0 | 0 | 0 | 0 | 0 | 0 | 0 | 0 | 0 | 0 | 0 | 0 | 0 | 1 | 0 | 0 | 0 | 0 | 0 | 0 | 0 | 0 | 0 | 0 | 0 | 0 | 0 | 0 | 0 | 0 | 0 | 0 | 0 | 0 | 0 |
| bacteria_Jul27d25b_61017 | 1 | 0 | 0 | 0 | 0 | 0 | 0 | 0 | 0 | 0 | 0 | 0 | 0 | 0 | 0 | 0 | 0 | 0 | 0 | 0 | 0 | 0 | 0 | 0 | 0 | 0 | 0 | 0 | 0 | 0 | 0 | 0 | 0 | 1 | 0 | 0 | 0 | 0 | 0 | 0 | 0 | 0 | 0 | 0 | 0 | 0 | 0 | 0 | 0 | 0 | 0 | 0 | 0 | 0 | 0 |
| bacteria_Jul27d25b_64458 | 1 | 0 | 0 | 0 | 0 | 0 | 0 | 0 | 0 | 0 | 0 | 0 | 0 | 0 | 0 | 0 | 0 | 0 | 0 | 0 | 0 | 0 | 0 | 0 | 0 | 0 | 0 | 0 | 0 | 0 | 0 | 0 | 0 | 1 | 0 | 0 | 0 | 0 | 0 | 0 | 0 | 0 | 0 | 0 | 0 | 0 | 0 | 0 | 0 | 0 | 0 | 0 | 0 | 0 | 0 |
| bacteria_Jul27d25b_64687 | 1 | 0 | 0 | 0 | 0 | 0 | 0 | 0 | 0 | 0 | 0 | 0 | 0 | 0 | 0 | 0 | 0 | 0 | 0 | 0 | 0 | 0 | 0 | 0 | 0 | 0 | 0 | 0 | 0 | 0 | 0 | 0 | 0 | 1 | 0 | 0 | 0 | 0 | 0 | 0 | 0 | 0 | 0 | 0 | 0 | 0 | 0 | 0 | 0 | 0 | 0 | 0 | 0 | 0 | 0 |
| bacteria_Jul27d25b_67249 | 1 | 0 | 0 | 0 | 0 | 0 | 0 | 0 | 0 | 0 | 0 | 0 | 0 | 0 | 0 | 0 | 0 | 0 | 0 | 0 | 0 | 0 | 0 | 0 | 0 | 0 | 0 | 0 | 0 | 0 | 0 | 0 | 0 | 1 | 0 | 0 | 0 | 0 | 0 | 0 | 0 | 0 | 0 | 0 | 0 | 0 | 0 | 0 | 0 | 0 | 0 | 0 | 0 | 0 | 0 |
| bacteria_Jul27d25b_70877 | 1 | 0 | 0 | 0 | 0 | 0 | 0 | 0 | 0 | 0 | 0 | 0 | 0 | 0 | 0 | 0 | 0 | 0 | 0 | 0 | 0 | 0 | 0 | 0 | 0 | 0 | 0 | 0 | 0 | 0 | 0 | 0 | 0 | 1 | 0 | 0 | 0 | 0 | 0 | 0 | 0 | 0 | 0 | 0 | 0 | 0 | 0 | 0 | 0 | 0 | 0 | 0 | 0 | 0 | 0 |
| bacteria_Jul27d75b_5075 | 1 | 0 | 0 | 0 | 0 | 0 | 0 | 0 | 0 | 0 | 0 | 0 | 0 | 0 | 0 | 0 | 0 | 0 | 0 | 0 | 0 | 0 | 0 | 0 | 0 | 0 | 0 | 0 | 0 | 0 | 0 | 0 | 0 | 0 | 0 | 1 | 0 | 0 | 0 | 0 | 0 | 0 | 0 | 0 | 0 | 0 | 0 | 0 | 0 | 0 | 0 | 0 | 0 | 0 | 0 |
| bacteria_Jul27d75b_9102 | 1 | 0 | 0 | 0 | 0 | 0 | 0 | 0 | 0 | 0 | 0 | 0 | 0 | 0 | 0 | 0 | 0 | 0 | 0 | 0 | 0 | 0 | 0 | 0 | 0 | 0 | 0 | 0 | 0 | 0 | 0 | 0 | 0 | 0 | 0 | 1 | 0 | 0 | 0 | 0 | 0 | 0 | 0 | 0 | 0 | 0 | 0 | 0 | 0 | 0 | 0 | 0 | 0 | 0 | 0 |
| bacteria_Jul27d75b_11137 | 1 | 0 | 0 | 0 | 0 | 0 | 0 | 0 | 0 | 0 | 0 | 0 | 0 | 0 | 0 | 0 | 0 | 0 | 0 | 0 | 0 | 0 | 0 | 0 | 0 | 0 | 0 | 0 | 0 | 0 | 0 | 0 | 0 | 0 | 0 | 1 | 0 | 0 | 0 | 0 | 0 | 0 | 0 | 0 | 0 | 0 | 0 | 0 | 0 | 0 | 0 | 0 | 0 | 0 | 0 |
| bacteria_Jul27d75b_20773 | 1 | 0 | 0 | 0 | 0 | 0 | 0 | 0 | 0 | 0 | 0 | 0 | 0 | 0 | 0 | 0 | 0 | 0 | 0 | 0 | 0 | 0 | 0 | 0 | 0 | 0 | 0 | 0 | 0 | 0 | 0 | 0 | 0 | 0 | 0 | 1 | 0 | 0 | 0 | 0 | 0 | 0 | 0 | 0 | 0 | 0 | 0 | 0 | 0 | 0 | 0 | 0 | 0 | 0 | 0 |
| bacteria_Jul27d75b_21908 | 1 | 0 | 0 | 0 | 0 | 0 | 0 | 0 | 0 | 0 | 0 | 0 | 0 | 0 | 0 | 0 | 0 | 0 | 0 | 0 | 0 | 0 | 0 | 0 | 0 | 0 | 0 | 0 | 0 | 0 | 0 | 0 | 0 | 0 | 0 | 1 | 0 | 0 | 0 | 0 | 0 | 0 | 0 | 0 | 0 | 0 | 0 | 0 | 0 | 0 | 0 | 0 | 0 | 0 | 0 |
| bacteria_Jul27d75b_26709 | 1 | 0 | 0 | 0 | 0 | 0 | 0 | 0 | 0 | 0 | 0 | 0 | 0 | 0 | 0 | 0 | 0 | 0 | 0 | 0 | 0 | 0 | 0 | 0 | 0 | 0 | 0 | 0 | 0 | 0 | 0 | 0 | 0 | 0 | 0 | 1 | 0 | 0 | 0 | 0 | 0 | 0 | 0 | 0 | 0 | 0 | 0 | 0 | 0 | 0 | 0 | 0 | 0 | 0 | 0 |
| bacteria_Jul27d75b_27684 | 1 | 0 | 0 | 0 | 0 | 0 | 0 | 0 | 0 | 0 | 0 | 0 | 0 | 0 | 0 | 0 | 0 | 0 | 0 | 0 | 0 | 0 | 0 | 0 | 0 | 0 | 0 | 0 | 0 | 0 | 0 | 0 | 0 | 0 | 0 | 1 | 0 | 0 | 0 | 0 | 0 | 0 | 0 | 0 | 0 | 0 | 0 | 0 | 0 | 0 | 0 | 0 | 0 | 0 | 0 |
| bacteria_Jul27d75b_30548 | 1 | 0 | 0 | 0 | 0 | 0 | 0 | 0 | 0 | 0 | 0 | 0 | 0 | 0 | 0 | 0 | 0 | 0 | 0 | 0 | 0 | 0 | 0 | 0 | 0 | 0 | 0 | 0 | 0 | 0 | 0 | 0 | 0 | 0 | 0 | 1 | 0 | 0 | 0 | 0 | 0 | 0 | 0 | 0 | 0 | 0 | 0 | 0 | 0 | 0 | 0 | 0 | 0 | 0 | 0 |
| bacteria_Jul27d75b_35681 | 1 | 0 | 0 | 0 | 0 | 0 | 0 | 0 | 0 | 0 | 0 | 0 | 0 | 0 | 0 | 0 | 0 | 0 | 0 | 0 | 0 | 0 | 0 | 0 | 0 | 0 | 0 | 0 | 0 | 0 | 0 | 0 | 0 | 0 | 0 | 1 | 0 | 0 | 0 | 0 | 0 | 0 | 0 | 0 | 0 | 0 | 0 | 0 | 0 | 0 | 0 | 0 | 0 | 0 | 0 |
| bacteria_Jul27d75b_42367 | 1 | 0 | 0 | 0 | 0 | 0 | 0 | 0 | 0 | 0 | 0 | 0 | 0 | 0 | 0 | 0 | 0 | 0 | 0 | 0 | 0 | 0 | 0 | 0 | 0 | 0 | 0 | 0 | 0 | 0 | 0 | 0 | 0 | 0 | 0 | 1 | 0 | 0 | 0 | 0 | 0 | 0 | 0 | 0 | 0 | 0 | 0 | 0 | 0 | 0 | 0 | 0 | 0 | 0 | 0 |
| bacteria_Jul27d75b_67952 | 1 | 0 | 0 | 0 | 0 | 0 | 0 | 0 | 0 | 0 | 0 | 0 | 0 | 0 | 0 | 0 | 0 | 0 | 0 | 0 | 0 | 0 | 0 | 0 | 0 | 0 | 0 | 0 | 0 | 0 | 0 | 0 | 0 | 0 | 0 | 1 | 0 | 0 | 0 | 0 | 0 | 0 | 0 | 0 | 0 | 0 | 0 | 0 | 0 | 0 | 0 | 0 | 0 | 0 | 0 |
| bacteria_Jul27d75b_70493 | 1 | 0 | 0 | 0 | 0 | 0 | 0 | 0 | 0 | 0 | 0 | 0 | 0 | 0 | 0 | 0 | 0 | 0 | 0 | 0 | 0 | 0 | 0 | 0 | 0 | 0 | 0 | 0 | 0 | 0 | 0 | 0 | 0 | 0 | 0 | 1 | 0 | 0 | 0 | 0 | 0 | 0 | 0 | 0 | 0 | 0 | 0 | 0 | 0 | 0 | 0 | 0 | 0 | 0 | 0 |
| bacteria_Jun8d25b_2004 | 1 | 0 | 0 | 0 | 0 | 0 | 0 | 0 | 0 | 1 | 0 | 0 | 0 | 0 | 0 | 0 | 0 | 0 | 0 | 0 | 0 | 0 | 0 | 0 | 0 | 0 | 0 | 0 | 0 | 0 | 0 | 0 | 0 | 0 | 0 | 0 | 0 | 0 | 0 | 0 | 0 | 0 | 0 | 0 | 0 | 0 | 0 | 0 | 0 | 0 | 0 | 0 | 0 | 0 | 0 |
| bacteria_Jun8d25b_3182 | 1 | 0 | 0 | 0 | 0 | 0 | 0 | 0 | 0 | 1 | 0 | 0 | 0 | 0 | 0 | 0 | 0 | 0 | 0 | 0 | 0 | 0 | 0 | 0 | 0 | 0 | 0 | 0 | 0 | 0 | 0 | 0 | 0 | 0 | 0 | 0 | 0 | 0 | 0 | 0 | 0 | 0 | 0 | 0 | 0 | 0 | 0 | 0 | 0 | 0 | 0 | 0 | 0 | 0 | 0 |
| bacteria_Jun8d25b_3988 | 1 | 0 | 0 | 0 | 0 | 0 | 0 | 0 | 0 | 1 | 0 | 0 | 0 | 0 | 0 | 0 | 0 | 0 | 0 | 0 | 0 | 0 | 0 | 0 | 0 | 0 | 0 | 0 | 0 | 0 | 0 | 0 | 0 | 0 | 0 | 0 | 0 | 0 | 0 | 0 | 0 | 0 | 0 | 0 | 0 | 0 | 0 | 0 | 0 | 0 | 0 | 0 | 0 | 0 | 0 |
| bacteria_Jun8d25b_4019 | 1 | 0 | 0 | 0 | 0 | 0 | 0 | 0 | 0 | 1 | 0 | 0 | 0 | 0 | 0 | 0 | 0 | 0 | 0 | 0 | 0 | 0 | 0 | 0 | 0 | 0 | 0 | 0 | 0 | 0 | 0 | 0 | 0 | 0 | 0 | 0 | 0 | 0 | 0 | 0 | 0 | 0 | 0 | 0 | 0 | 0 | 0 | 0 | 0 | 0 | 0 | 0 | 0 | 0 | 0 |
| bacteria_Jun8d25b_5230 | 1 | 0 | 0 | 0 | 0 | 0 | 0 | 0 | 0 | 1 | 0 | 0 | 0 | 0 | 0 | 0 | 0 | 0 | 0 | 0 | 0 | 0 | 0 | 0 | 0 | 0 | 0 | 0 | 0 | 0 | 0 | 0 | 0 | 0 | 0 | 0 | 0 | 0 | 0 | 0 | 0 | 0 | 0 | 0 | 0 | 0 | 0 | 0 | 0 | 0 | 0 | 0 | 0 | 0 | 0 |
| bacteria_Jun8d25b_10147 | 1 | 0 | 0 | 0 | 0 | 0 | 0 | 0 | 0 | 1 | 0 | 0 | 0 | 0 | 0 | 0 | 0 | 0 | 0 | 0 | 0 | 0 | 0 | 0 | 0 | 0 | 0 | 0 | 0 | 0 | 0 | 0 | 0 | 0 | 0 | 0 | 0 | 0 | 0 | 0 | 0 | 0 | 0 | 0 | 0 | 0 | 0 | 0 | 0 | 0 | 0 | 0 | 0 | 0 | 0 |
| bacteria_Jun8d25b_12595 | 1 | 0 | 0 | 0 | 0 | 0 | 0 | 0 | 0 | 1 | 0 | 0 | 0 | 0 | 0 | 0 | 0 | 0 | 0 | 0 | 0 | 0 | 0 | 0 | 0 | 0 | 0 | 0 | 0 | 0 | 0 | 0 | 0 | 0 | 0 | 0 | 0 | 0 | 0 | 0 | 0 | 0 | 0 | 0 | 0 | 0 | 0 | 0 | 0 | 0 | 0 | 0 | 0 | 0 | 0 |
| bacteria_Jun8d25b_30233 | 1 | 0 | 0 | 0 | 0 | 0 | 0 | 0 | 0 | 1 | 0 | 0 | 0 | 0 | 0 | 0 | 0 | 0 | 0 | 0 | 0 | 0 | 0 | 0 | 0 | 0 | 0 | 0 | 0 | 0 | 0 | 0 | 0 | 0 | 0 | 0 | 0 | 0 | 0 | 0 | 0 | 0 | 0 | 0 | 0 | 0 | 0 | 0 | 0 | 0 | 0 | 0 | 0 | 0 | 0 |
| bacteria_Jun8d25b_41464 | 1 | 0 | 0 | 0 | 0 | 0 | 0 | 0 | 0 | 1 | 0 | 0 | 0 | 0 | 0 | 0 | 0 | 0 | 0 | 0 | 0 | 0 | 0 | 0 | 0 | 0 | 0 | 0 | 0 | 0 | 0 | 0 | 0 | 0 | 0 | 0 | 0 | 0 | 0 | 0 | 0 | 0 | 0 | 0 | 0 | 0 | 0 | 0 | 0 | 0 | 0 | 0 | 0 | 0 | 0 |
| bacteria_Jun8d25b_45795 | 1 | 0 | 0 | 0 | 0 | 0 | 0 | 0 | 0 | 1 | 0 | 0 | 0 | 0 | 0 | 0 | 0 | 0 | 0 | 0 | 0 | 0 | 0 | 0 | 0 | 0 | 0 | 0 | 0 | 0 | 0 | 0 | 0 | 0 | 0 | 0 | 0 | 0 | 0 | 0 | 0 | 0 | 0 | 0 | 0 | 0 | 0 | 0 | 0 | 0 | 0 | 0 | 0 | 0 | 0 |
| bacteria_Jun8d25b_46747 | 1 | 0 | 0 | 0 | 0 | 0 | 0 | 0 | 0 | 1 | 0 | 0 | 0 | 0 | 0 | 0 | 0 | 0 | 0 | 0 | 0 | 0 | 0 | 0 | 0 | 0 | 0 | 0 | 0 | 0 | 0 | 0 | 0 | 0 | 0 | 0 | 0 | 0 | 0 | 0 | 0 | 0 | 0 | 0 | 0 | 0 | 0 | 0 | 0 | 0 | 0 | 0 | 0 | 0 | 0 |
| bacteria_Jun8d25b_70968 | 1 | 0 | 0 | 0 | 0 | 0 | 0 | 0 | 0 | 1 | 0 | 0 | 0 | 0 | 0 | 0 | 0 | 0 | 0 | 0 | 0 | 0 | 0 | 0 | 0 | 0 | 0 | 0 | 0 | 0 | 0 | 0 | 0 | 0 | 0 | 0 | 0 | 0 | 0 | 0 | 0 | 0 | 0 | 0 | 0 | 0 | 0 | 0 | 0 | 0 | 0 | 0 | 0 | 0 | 0 |
| bacteria_Jun8d50b_6659 | 1 | 0 | 0 | 0 | 0 | 0 | 0 | 0 | 0 | 0 | 0 | 1 | 0 | 0 | 0 | 0 | 0 | 0 | 0 | 0 | 0 | 0 | 0 | 0 | 0 | 0 | 0 | 0 | 0 | 0 | 0 | 0 | 0 | 0 | 0 | 0 | 0 | 0 | 0 | 0 | 0 | 0 | 0 | 0 | 0 | 0 | 0 | 0 | 0 | 0 | 0 | 0 | 0 | 0 | 0 |
| bacteria_Jun8d75b_66505 | 1 | 0 | 0 | 0 | 0 | 0 | 0 | 0 | 0 | 0 | 0 | 0 | 0 | 1 | 0 | 0 | 0 | 0 | 0 | 0 | 0 | 0 | 0 | 0 | 0 | 0 | 0 | 0 | 0 | 0 | 0 | 0 | 0 | 0 | 0 | 0 | 0 | 0 | 0 | 0 | 0 | 0 | 0 | 0 | 0 | 0 | 0 | 0 | 0 | 0 | 0 | 0 | 0 | 0 | 0 |
| bacteria_Jun22d0b_13028 | 1 | 0 | 0 | 0 | 0 | 0 | 0 | 0 | 0 | 0 | 0 | 0 | 0 | 0 | 0 | 1 | 0 | 0 | 0 | 0 | 0 | 0 | 0 | 0 | 0 | 0 | 0 | 0 | 0 | 0 | 0 | 0 | 0 | 0 | 0 | 0 | 0 | 0 | 0 | 0 | 0 | 0 | 0 | 0 | 0 | 0 | 0 | 0 | 0 | 0 | 0 | 0 | 0 | 0 | 0 |
| bacteria_Jun22d0b_21865 | 1 | 0 | 0 | 0 | 0 | 0 | 0 | 0 | 0 | 0 | 0 | 0 | 0 | 0 | 0 | 1 | 0 | 0 | 0 | 0 | 0 | 0 | 0 | 0 | 0 | 0 | 0 | 0 | 0 | 0 | 0 | 0 | 0 | 0 | 0 | 0 | 0 | 0 | 0 | 0 | 0 | 0 | 0 | 0 | 0 | 0 | 0 | 0 | 0 | 0 | 0 | 0 | 0 | 0 | 0 |
| bacteria_Jun22d0b_23038 | 1 | 0 | 0 | 0 | 0 | 0 | 0 | 0 | 0 | 0 | 0 | 0 | 0 | 0 | 0 | 1 | 0 | 0 | 0 | 0 | 0 | 0 | 0 | 0 | 0 | 0 | 0 | 0 | 0 | 0 | 0 | 0 | 0 | 0 | 0 | 0 | 0 | 0 | 0 | 0 | 0 | 0 | 0 | 0 | 0 | 0 | 0 | 0 | 0 | 0 | 0 | 0 | 0 | 0 | 0 |
| bacteria_Jun22d0b_42214 | 1 | 0 | 0 | 0 | 0 | 0 | 0 | 0 | 0 | 0 | 0 | 0 | 0 | 0 | 0 | 1 | 0 | 0 | 0 | 0 | 0 | 0 | 0 | 0 | 0 | 0 | 0 | 0 | 0 | 0 | 0 | 0 | 0 | 0 | 0 | 0 | 0 | 0 | 0 | 0 | 0 | 0 | 0 | 0 | 0 | 0 | 0 | 0 | 0 | 0 | 0 | 0 | 0 | 0 | 0 |
| bacteria_Jun22d50b_3291 | 1 | 0 | 0 | 0 | 0 | 0 | 0 | 0 | 0 | 0 | 0 | 0 | 0 | 0 | 0 | 0 | 0 | 1 | 0 | 0 | 0 | 0 | 0 | 0 | 0 | 0 | 0 | 0 | 0 | 0 | 0 | 0 | 0 | 0 | 0 | 0 | 0 | 0 | 0 | 0 | 0 | 0 | 0 | 0 | 0 | 0 | 0 | 0 | 0 | 0 | 0 | 0 | 0 | 0 | 0 |
| bacteria_Jun22d50b_1156 | 1 | 0 | 0 | 0 | 0 | 0 | 0 | 0 | 0 | 0 | 0 | 0 | 0 | 0 | 0 | 0 | 0 | 1 | 0 | 0 | 0 | 0 | 0 | 0 | 0 | 0 | 0 | 0 | 0 | 0 | 0 | 0 | 0 | 0 | 0 | 0 | 0 | 0 | 0 | 0 | 0 | 0 | 0 | 0 | 0 | 0 | 0 | 0 | 0 | 0 | 0 | 0 | 0 | 0 | 0 |
| bacteria_Jun22d50b_2111 | 1 | 0 | 0 | 0 | 0 | 0 | 0 | 0 | 0 | 0 | 0 | 0 | 0 | 0 | 0 | 0 | 0 | 1 | 0 | 0 | 0 | 0 | 0 | 0 | 0 | 0 | 0 | 0 | 0 | 0 | 0 | 0 | 0 | 0 | 0 | 0 | 0 | 0 | 0 | 0 | 0 | 0 | 0 | 0 | 0 | 0 | 0 | 0 | 0 | 0 | 0 | 0 | 0 | 0 | 0 |
| bacteria_Jun22d50b_3249 | 1 | 0 | 0 | 0 | 0 | 0 | 0 | 0 | 0 | 0 | 0 | 0 | 0 | 0 | 0 | 0 | 0 | 1 | 0 | 0 | 0 | 0 | 0 | 0 | 0 | 0 | 0 | 0 | 0 | 0 | 0 | 0 | 0 | 0 | 0 | 0 | 0 | 0 | 0 | 0 | 0 | 0 | 0 | 0 | 0 | 0 | 0 | 0 | 0 | 0 | 0 | 0 | 0 | 0 | 0 |
| bacteria_Jun22d50b_3415 | 1 | 0 | 0 | 0 | 0 | 0 | 0 | 0 | 0 | 0 | 0 | 0 | 0 | 0 | 0 | 0 | 0 | 1 | 0 | 0 | 0 | 0 | 0 | 0 | 0 | 0 | 0 | 0 | 0 | 0 | 0 | 0 | 0 | 0 | 0 | 0 | 0 | 0 | 0 | 0 | 0 | 0 | 0 | 0 | 0 | 0 | 0 | 0 | 0 | 0 | 0 | 0 | 0 | 0 | 0 |
| bacteria_Jun22d75b_2954 | 1 | 0 | 0 | 0 | 0 | 0 | 0 | 0 | 0 | 0 | 0 | 0 | 0 | 0 | 0 | 0 | 0 | 0 | 0 | 1 | 0 | 0 | 0 | 0 | 0 | 0 | 0 | 0 | 0 | 0 | 0 | 0 | 0 | 0 | 0 | 0 | 0 | 0 | 0 | 0 | 0 | 0 | 0 | 0 | 0 | 0 | 0 | 0 | 0 | 0 | 0 | 0 | 0 | 0 | 0 |
| bacteria_Jun22d75b_3312 | 1 | 0 | 0 | 0 | 0 | 0 | 0 | 0 | 0 | 0 | 0 | 0 | 0 | 0 | 0 | 0 | 0 | 0 | 0 | 1 | 0 | 0 | 0 | 0 | 0 | 0 | 0 | 0 | 0 | 0 | 0 | 0 | 0 | 0 | 0 | 0 | 0 | 0 | 0 | 0 | 0 | 0 | 0 | 0 | 0 | 0 | 0 | 0 | 0 | 0 | 0 | 0 | 0 | 0 | 0 |
| bacteria_Jun22d75b_8601 | 1 | 0 | 0 | 0 | 0 | 0 | 0 | 0 | 0 | 0 | 0 | 0 | 0 | 0 | 0 | 0 | 0 | 0 | 0 | 1 | 0 | 0 | 0 | 0 | 0 | 0 | 0 | 0 | 0 | 0 | 0 | 0 | 0 | 0 | 0 | 0 | 0 | 0 | 0 | 0 | 0 | 0 | 0 | 0 | 0 | 0 | 0 | 0 | 0 | 0 | 0 | 0 | 0 | 0 | 0 |
| bacteria_Jun22d75b_1278 | 1 | 0 | 0 | 0 | 0 | 0 | 0 | 0 | 0 | 0 | 0 | 0 | 0 | 0 | 0 | 0 | 0 | 0 | 0 | 1 | 0 | 0 | 0 | 0 | 0 | 0 | 0 | 0 | 0 | 0 | 0 | 0 | 0 | 0 | 0 | 0 | 0 | 0 | 0 | 0 | 0 | 0 | 0 | 0 | 0 | 0 | 0 | 0 | 0 | 0 | 0 | 0 | 0 | 0 | 0 |
| bacteria_Jun22d75b_1526 | 1 | 0 | 0 | 0 | 0 | 0 | 0 | 0 | 0 | 0 | 0 | 0 | 0 | 0 | 0 | 0 | 0 | 0 | 0 | 1 | 0 | 0 | 0 | 0 | 0 | 0 | 0 | 0 | 0 | 0 | 0 | 0 | 0 | 0 | 0 | 0 | 0 | 0 | 0 | 0 | 0 | 0 | 0 | 0 | 0 | 0 | 0 | 0 | 0 | 0 | 0 | 0 | 0 | 0 | 0 |
| bacteria_Jun22d75b_2272 | 1 | 0 | 0 | 0 | 0 | 0 | 0 | 0 | 0 | 0 | 0 | 0 | 0 | 0 | 0 | 0 | 0 | 0 | 0 | 1 | 0 | 0 | 0 | 0 | 0 | 0 | 0 | 0 | 0 | 0 | 0 | 0 | 0 | 0 | 0 | 0 | 0 | 0 | 0 | 0 | 0 | 0 | 0 | 0 | 0 | 0 | 0 | 0 | 0 | 0 | 0 | 0 | 0 | 0 | 0 |
| bacteria_Jun22d75b_2979 | 1 | 0 | 0 | 0 | 0 | 0 | 0 | 0 | 0 | 0 | 0 | 0 | 0 | 0 | 0 | 0 | 0 | 0 | 0 | 1 | 0 | 0 | 0 | 0 | 0 | 0 | 0 | 0 | 0 | 0 | 0 | 0 | 0 | 0 | 0 | 0 | 0 | 0 | 0 | 0 | 0 | 0 | 0 | 0 | 0 | 0 | 0 | 0 | 0 | 0 | 0 | 0 | 0 | 0 | 0 |
| bacteria_Jun22d75b_3525 | 1 | 0 | 0 | 0 | 0 | 0 | 0 | 0 | 0 | 0 | 0 | 0 | 0 | 0 | 0 | 0 | 0 | 0 | 0 | 1 | 0 | 0 | 0 | 0 | 0 | 0 | 0 | 0 | 0 | 0 | 0 | 0 | 0 | 0 | 0 | 0 | 0 | 0 | 0 | 0 | 0 | 0 | 0 | 0 | 0 | 0 | 0 | 0 | 0 | 0 | 0 | 0 | 0 | 0 | 0 |
| bacteria_Jun22d75b_4262 | 1 | 0 | 0 | 0 | 0 | 0 | 0 | 0 | 0 | 0 | 0 | 0 | 0 | 0 | 0 | 0 | 0 | 0 | 0 | 1 | 0 | 0 | 0 | 0 | 0 | 0 | 0 | 0 | 0 | 0 | 0 | 0 | 0 | 0 | 0 | 0 | 0 | 0 | 0 | 0 | 0 | 0 | 0 | 0 | 0 | 0 | 0 | 0 | 0 | 0 | 0 | 0 | 0 | 0 | 0 |
| bacteria_Jun22d75b_4726 | 1 | 0 | 0 | 0 | 0 | 0 | 0 | 0 | 0 | 0 | 0 | 0 | 0 | 0 | 0 | 0 | 0 | 0 | 0 | 1 | 0 | 0 | 0 | 0 | 0 | 0 | 0 | 0 | 0 | 0 | 0 | 0 | 0 | 0 | 0 | 0 | 0 | 0 | 0 | 0 | 0 | 0 | 0 | 0 | 0 | 0 | 0 | 0 | 0 | 0 | 0 | 0 | 0 | 0 | 0 |
| bacteria_Jun22d75b_5584 | 1 | 0 | 0 | 0 | 0 | 0 | 0 | 0 | 0 | 0 | 0 | 0 | 0 | 0 | 0 | 0 | 0 | 0 | 0 | 1 | 0 | 0 | 0 | 0 | 0 | 0 | 0 | 0 | 0 | 0 | 0 | 0 | 0 | 0 | 0 | 0 | 0 | 0 | 0 | 0 | 0 | 0 | 0 | 0 | 0 | 0 | 0 | 0 | 0 | 0 | 0 | 0 | 0 | 0 | 0 |
| bacteria_Jun22d75b_5889 | 1 | 0 | 0 | 0 | 0 | 0 | 0 | 0 | 0 | 0 | 0 | 0 | 0 | 0 | 0 | 0 | 0 | 0 | 0 | 1 | 0 | 0 | 0 | 0 | 0 | 0 | 0 | 0 | 0 | 0 | 0 | 0 | 0 | 0 | 0 | 0 | 0 | 0 | 0 | 0 | 0 | 0 | 0 | 0 | 0 | 0 | 0 | 0 | 0 | 0 | 0 | 0 | 0 | 0 | 0 |
| bacteria_May12d5b_6907 | 1 | 1 | 0 | 0 | 0 | 0 | 0 | 0 | 0 | 0 | 0 | 0 | 0 | 0 | 0 | 0 | 0 | 0 | 0 | 0 | 0 | 0 | 0 | 0 | 0 | 0 | 0 | 0 | 0 | 0 | 0 | 0 | 0 | 0 | 0 | 0 | 0 | 0 | 0 | 0 | 0 | 0 | 0 | 0 | 0 | 0 | 0 | 0 | 0 | 0 | 0 | 0 | 0 | 0 | 0 |
| bacteria_May12d5b_4683 | 1 | 1 | 0 | 0 | 0 | 0 | 0 | 0 | 0 | 0 | 0 | 0 | 0 | 0 | 0 | 0 | 0 | 0 | 0 | 0 | 0 | 0 | 0 | 0 | 0 | 0 | 0 | 0 | 0 | 0 | 0 | 0 | 0 | 0 | 0 | 0 | 0 | 0 | 0 | 0 | 0 | 0 | 0 | 0 | 0 | 0 | 0 | 0 | 0 | 0 | 0 | 0 | 0 | 0 | 0 |
| bacteria_May12d25b_207 | 1 | 0 | 0 | 1 | 0 | 0 | 0 | 0 | 0 | 0 | 0 | 0 | 0 | 0 | 0 | 0 | 0 | 0 | 0 | 0 | 0 | 0 | 0 | 0 | 0 | 0 | 0 | 0 | 0 | 0 | 0 | 0 | 0 | 0 | 0 | 0 | 0 | 0 | 0 | 0 | 0 | 0 | 0 | 0 | 0 | 0 | 0 | 0 | 0 | 0 | 0 | 0 | 0 | 0 | 0 |
| bacteria_May12d25b_608 | 1 | 0 | 0 | 1 | 0 | 0 | 0 | 0 | 0 | 0 | 0 | 0 | 0 | 0 | 0 | 0 | 0 | 0 | 0 | 0 | 0 | 0 | 0 | 0 | 0 | 0 | 0 | 0 | 0 | 0 | 0 | 0 | 0 | 0 | 0 | 0 | 0 | 0 | 0 | 0 | 0 | 0 | 0 | 0 | 0 | 0 | 0 | 0 | 0 | 0 | 0 | 0 | 0 | 0 | 0 |
| bacteria_May12d25b_712 | 1 | 0 | 0 | 1 | 0 | 0 | 0 | 0 | 0 | 0 | 0 | 0 | 0 | 0 | 0 | 0 | 0 | 0 | 0 | 0 | 0 | 0 | 0 | 0 | 0 | 0 | 0 | 0 | 0 | 0 | 0 | 0 | 0 | 0 | 0 | 0 | 0 | 0 | 0 | 0 | 0 | 0 | 0 | 0 | 0 | 0 | 0 | 0 | 0 | 0 | 0 | 0 | 0 | 0 | 0 |
| bacteria_May12d25b_121 | 1 | 0 | 0 | 1 | 0 | 0 | 0 | 0 | 0 | 0 | 0 | 0 | 0 | 0 | 0 | 0 | 0 | 0 | 0 | 0 | 0 | 0 | 0 | 0 | 0 | 0 | 0 | 0 | 0 | 0 | 0 | 0 | 0 | 0 | 0 | 0 | 0 | 0 | 0 | 0 | 0 | 0 | 0 | 0 | 0 | 0 | 0 | 0 | 0 | 0 | 0 | 0 | 0 | 0 | 0 |
| bacteria_May12d25b_124 | 1 | 0 | 0 | 1 | 0 | 0 | 0 | 0 | 0 | 0 | 0 | 0 | 0 | 0 | 0 | 0 | 0 | 0 | 0 | 0 | 0 | 0 | 0 | 0 | 0 | 0 | 0 | 0 | 0 | 0 | 0 | 0 | 0 | 0 | 0 | 0 | 0 | 0 | 0 | 0 | 0 | 0 | 0 | 0 | 0 | 0 | 0 | 0 | 0 | 0 | 0 | 0 | 0 | 0 | 0 |
| bacteria_May12d25b_317 | 1 | 0 | 0 | 1 | 0 | 0 | 0 | 0 | 0 | 0 | 0 | 0 | 0 | 0 | 0 | 0 | 0 | 0 | 0 | 0 | 0 | 0 | 0 | 0 | 0 | 0 | 0 | 0 | 0 | 0 | 0 | 0 | 0 | 0 | 0 | 0 | 0 | 0 | 0 | 0 | 0 | 0 | 0 | 0 | 0 | 0 | 0 | 0 | 0 | 0 | 0 | 0 | 0 | 0 | 0 |
| bacteria_May12d25b_322 | 1 | 0 | 0 | 1 | 0 | 0 | 0 | 0 | 0 | 0 | 0 | 0 | 0 | 0 | 0 | 0 | 0 | 0 | 0 | 0 | 0 | 0 | 0 | 0 | 0 | 0 | 0 | 0 | 0 | 0 | 0 | 0 | 0 | 0 | 0 | 0 | 0 | 0 | 0 | 0 | 0 | 0 | 0 | 0 | 0 | 0 | 0 | 0 | 0 | 0 | 0 | 0 | 0 | 0 | 0 |
| bacteria_May12d25b_382 | 1 | 0 | 0 | 1 | 0 | 0 | 0 | 0 | 0 | 0 | 0 | 0 | 0 | 0 | 0 | 0 | 0 | 0 | 0 | 0 | 0 | 0 | 0 | 0 | 0 | 0 | 0 | 0 | 0 | 0 | 0 | 0 | 0 | 0 | 0 | 0 | 0 | 0 | 0 | 0 | 0 | 0 | 0 | 0 | 0 | 0 | 0 | 0 | 0 | 0 | 0 | 0 | 0 | 0 | 0 |
| bacteria_May12d25b_408 | 1 | 0 | 0 | 1 | 0 | 0 | 0 | 0 | 0 | 0 | 0 | 0 | 0 | 0 | 0 | 0 | 0 | 0 | 0 | 0 | 0 | 0 | 0 | 0 | 0 | 0 | 0 | 0 | 0 | 0 | 0 | 0 | 0 | 0 | 0 | 0 | 0 | 0 | 0 | 0 | 0 | 0 | 0 | 0 | 0 | 0 | 0 | 0 | 0 | 0 | 0 | 0 | 0 | 0 | 0 |
| bacteria_May12d25b_460 | 1 | 0 | 0 | 1 | 0 | 0 | 0 | 0 | 0 | 0 | 0 | 0 | 0 | 0 | 0 | 0 | 0 | 0 | 0 | 0 | 0 | 0 | 0 | 0 | 0 | 0 | 0 | 0 | 0 | 0 | 0 | 0 | 0 | 0 | 0 | 0 | 0 | 0 | 0 | 0 | 0 | 0 | 0 | 0 | 0 | 0 | 0 | 0 | 0 | 0 | 0 | 0 | 0 | 0 | 0 |
| bacteria_May12d50b_158 | 1 | 0 | 0 | 0 | 0 | 1 | 0 | 0 | 0 | 0 | 0 | 0 | 0 | 0 | 0 | 0 | 0 | 0 | 0 | 0 | 0 | 0 | 0 | 0 | 0 | 0 | 0 | 0 | 0 | 0 | 0 | 0 | 0 | 0 | 0 | 0 | 0 | 0 | 0 | 0 | 0 | 0 | 0 | 0 | 0 | 0 | 0 | 0 | 0 | 0 | 0 | 0 | 0 | 0 | 0 |
| bacteria_May12d50b_239 | 1 | 0 | 0 | 0 | 0 | 1 | 0 | 0 | 0 | 0 | 0 | 0 | 0 | 0 | 0 | 0 | 0 | 0 | 0 | 0 | 0 | 0 | 0 | 0 | 0 | 0 | 0 | 0 | 0 | 0 | 0 | 0 | 0 | 0 | 0 | 0 | 0 | 0 | 0 | 0 | 0 | 0 | 0 | 0 | 0 | 0 | 0 | 0 | 0 | 0 | 0 | 0 | 0 | 0 | 0 |
| bacteria_May12d50b_256 | 1 | 0 | 0 | 0 | 0 | 1 | 0 | 0 | 0 | 0 | 0 | 0 | 0 | 0 | 0 | 0 | 0 | 0 | 0 | 0 | 0 | 0 | 0 | 0 | 0 | 0 | 0 | 0 | 0 | 0 | 0 | 0 | 0 | 0 | 0 | 0 | 0 | 0 | 0 | 0 | 0 | 0 | 0 | 0 | 0 | 0 | 0 | 0 | 0 | 0 | 0 | 0 | 0 | 0 | 0 |
| bacteria_May12d50b_459 | 1 | 0 | 0 | 0 | 0 | 1 | 0 | 0 | 0 | 0 | 0 | 0 | 0 | 0 | 0 | 0 | 0 | 0 | 0 | 0 | 0 | 0 | 0 | 0 | 0 | 0 | 0 | 0 | 0 | 0 | 0 | 0 | 0 | 0 | 0 | 0 | 0 | 0 | 0 | 0 | 0 | 0 | 0 | 0 | 0 | 0 | 0 | 0 | 0 | 0 | 0 | 0 | 0 | 0 | 0 |
| bacteria_May12d50b_511 | 1 | 0 | 0 | 0 | 0 | 1 | 0 | 0 | 0 | 0 | 0 | 0 | 0 | 0 | 0 | 0 | 0 | 0 | 0 | 0 | 0 | 0 | 0 | 0 | 0 | 0 | 0 | 0 | 0 | 0 | 0 | 0 | 0 | 0 | 0 | 0 | 0 | 0 | 0 | 0 | 0 | 0 | 0 | 0 | 0 | 0 | 0 | 0 | 0 | 0 | 0 | 0 | 0 | 0 | 0 |
| bacteria_May12d75b_601 | 1 | 0 | 0 | 0 | 0 | 0 | 0 | 1 | 0 | 0 | 0 | 0 | 0 | 0 | 0 | 0 | 0 | 0 | 0 | 0 | 0 | 0 | 0 | 0 | 0 | 0 | 0 | 0 | 0 | 0 | 0 | 0 | 0 | 0 | 0 | 0 | 0 | 0 | 0 | 0 | 0 | 0 | 0 | 0 | 0 | 0 | 0 | 0 | 0 | 0 | 0 | 0 | 0 | 0 | 0 |
| bacteria_Sep8d5b_2294 | 1 | 0 | 0 | 0 | 0 | 0 | 0 | 0 | 0 | 0 | 0 | 0 | 0 | 0 | 0 | 0 | 0 | 0 | 0 | 0 | 0 | 0 | 0 | 0 | 0 | 0 | 0 | 0 | 0 | 0 | 0 | 0 | 0 | 0 | 0 | 0 | 0 | 0 | 0 | 0 | 0 | 0 | 0 | 0 | 0 | 0 | 0 | 0 | 0 | 1 | 0 | 0 | 0 | 0 | 0 |
| bacteria_Sep8d5b_12590 | 1 | 0 | 0 | 0 | 0 | 0 | 0 | 0 | 0 | 0 | 0 | 0 | 0 | 0 | 0 | 0 | 0 | 0 | 0 | 0 | 0 | 0 | 0 | 0 | 0 | 0 | 0 | 0 | 0 | 0 | 0 | 0 | 0 | 0 | 0 | 0 | 0 | 0 | 0 | 0 | 0 | 0 | 0 | 0 | 0 | 0 | 0 | 0 | 0 | 1 | 0 | 0 | 0 | 0 | 0 |
| bacteria_Sep8d5b_14868 | 1 | 0 | 0 | 0 | 0 | 0 | 0 | 0 | 0 | 0 | 0 | 0 | 0 | 0 | 0 | 0 | 0 | 0 | 0 | 0 | 0 | 0 | 0 | 0 | 0 | 0 | 0 | 0 | 0 | 0 | 0 | 0 | 0 | 0 | 0 | 0 | 0 | 0 | 0 | 0 | 0 | 0 | 0 | 0 | 0 | 0 | 0 | 0 | 0 | 1 | 0 | 0 | 0 | 0 | 0 |
| bacteria_Sep8d5b_17067 | 1 | 0 | 0 | 0 | 0 | 0 | 0 | 0 | 0 | 0 | 0 | 0 | 0 | 0 | 0 | 0 | 0 | 0 | 0 | 0 | 0 | 0 | 0 | 0 | 0 | 0 | 0 | 0 | 0 | 0 | 0 | 0 | 0 | 0 | 0 | 0 | 0 | 0 | 0 | 0 | 0 | 0 | 0 | 0 | 0 | 0 | 0 | 0 | 0 | 1 | 0 | 0 | 0 | 0 | 0 |
| bacteria_Sep8d5b_18883 | 1 | 0 | 0 | 0 | 0 | 0 | 0 | 0 | 0 | 0 | 0 | 0 | 0 | 0 | 0 | 0 | 0 | 0 | 0 | 0 | 0 | 0 | 0 | 0 | 0 | 0 | 0 | 0 | 0 | 0 | 0 | 0 | 0 | 0 | 0 | 0 | 0 | 0 | 0 | 0 | 0 | 0 | 0 | 0 | 0 | 0 | 0 | 0 | 0 | 1 | 0 | 0 | 0 | 0 | 0 |
| bacteria_Sep8d5b_23224 | 1 | 0 | 0 | 0 | 0 | 0 | 0 | 0 | 0 | 0 | 0 | 0 | 0 | 0 | 0 | 0 | 0 | 0 | 0 | 0 | 0 | 0 | 0 | 0 | 0 | 0 | 0 | 0 | 0 | 0 | 0 | 0 | 0 | 0 | 0 | 0 | 0 | 0 | 0 | 0 | 0 | 0 | 0 | 0 | 0 | 0 | 0 | 0 | 0 | 1 | 0 | 0 | 0 | 0 | 0 |
| bacteria_Sep8d5b_27106 | 1 | 0 | 0 | 0 | 0 | 0 | 0 | 0 | 0 | 0 | 0 | 0 | 0 | 0 | 0 | 0 | 0 | 0 | 0 | 0 | 0 | 0 | 0 | 0 | 0 | 0 | 0 | 0 | 0 | 0 | 0 | 0 | 0 | 0 | 0 | 0 | 0 | 0 | 0 | 0 | 0 | 0 | 0 | 0 | 0 | 0 | 0 | 0 | 0 | 1 | 0 | 0 | 0 | 0 | 0 |
| bacteria_Sep8d5b_30260 | 1 | 0 | 0 | 0 | 0 | 0 | 0 | 0 | 0 | 0 | 0 | 0 | 0 | 0 | 0 | 0 | 0 | 0 | 0 | 0 | 0 | 0 | 0 | 0 | 0 | 0 | 0 | 0 | 0 | 0 | 0 | 0 | 0 | 0 | 0 | 0 | 0 | 0 | 0 | 0 | 0 | 0 | 0 | 0 | 0 | 0 | 0 | 0 | 0 | 1 | 0 | 0 | 0 | 0 | 0 |
| bacteria_Sep8d5b_32539 | 1 | 0 | 0 | 0 | 0 | 0 | 0 | 0 | 0 | 0 | 0 | 0 | 0 | 0 | 0 | 0 | 0 | 0 | 0 | 0 | 0 | 0 | 0 | 0 | 0 | 0 | 0 | 0 | 0 | 0 | 0 | 0 | 0 | 0 | 0 | 0 | 0 | 0 | 0 | 0 | 0 | 0 | 0 | 0 | 0 | 0 | 0 | 0 | 0 | 1 | 0 | 0 | 0 | 0 | 0 |
| bacteria_Sep8d5b_33226 | 1 | 0 | 0 | 0 | 0 | 0 | 0 | 0 | 0 | 0 | 0 | 0 | 0 | 0 | 0 | 0 | 0 | 0 | 0 | 0 | 0 | 0 | 0 | 0 | 0 | 0 | 0 | 0 | 0 | 0 | 0 | 0 | 0 | 0 | 0 | 0 | 0 | 0 | 0 | 0 | 0 | 0 | 0 | 0 | 0 | 0 | 0 | 0 | 0 | 1 | 0 | 0 | 0 | 0 | 0 |
| bacteria_Sep8d5b_39706 | 1 | 0 | 0 | 0 | 0 | 0 | 0 | 0 | 0 | 0 | 0 | 0 | 0 | 0 | 0 | 0 | 0 | 0 | 0 | 0 | 0 | 0 | 0 | 0 | 0 | 0 | 0 | 0 | 0 | 0 | 0 | 0 | 0 | 0 | 0 | 0 | 0 | 0 | 0 | 0 | 0 | 0 | 0 | 0 | 0 | 0 | 0 | 0 | 0 | 1 | 0 | 0 | 0 | 0 | 0 |
| bacteria_Sep8d5b_43802 | 1 | 0 | 0 | 0 | 0 | 0 | 0 | 0 | 0 | 0 | 0 | 0 | 0 | 0 | 0 | 0 | 0 | 0 | 0 | 0 | 0 | 0 | 0 | 0 | 0 | 0 | 0 | 0 | 0 | 0 | 0 | 0 | 0 | 0 | 0 | 0 | 0 | 0 | 0 | 0 | 0 | 0 | 0 | 0 | 0 | 0 | 0 | 0 | 0 | 1 | 0 | 0 | 0 | 0 | 0 |
| bacteria_Sep8d5b_45766 | 1 | 0 | 0 | 0 | 0 | 0 | 0 | 0 | 0 | 0 | 0 | 0 | 0 | 0 | 0 | 0 | 0 | 0 | 0 | 0 | 0 | 0 | 0 | 0 | 0 | 0 | 0 | 0 | 0 | 0 | 0 | 0 | 0 | 0 | 0 | 0 | 0 | 0 | 0 | 0 | 0 | 0 | 0 | 0 | 0 | 0 | 0 | 0 | 0 | 1 | 0 | 0 | 0 | 0 | 0 |
| bacteria_Sep8d5b_46599 | 1 | 0 | 0 | 0 | 0 | 0 | 0 | 0 | 0 | 0 | 0 | 0 | 0 | 0 | 0 | 0 | 0 | 0 | 0 | 0 | 0 | 0 | 0 | 0 | 0 | 0 | 0 | 0 | 0 | 0 | 0 | 0 | 0 | 0 | 0 | 0 | 0 | 0 | 0 | 0 | 0 | 0 | 0 | 0 | 0 | 0 | 0 | 0 | 0 | 1 | 0 | 0 | 0 | 0 | 0 |
| bacteria_Sep8d5b_53057 | 1 | 0 | 0 | 0 | 0 | 0 | 0 | 0 | 0 | 0 | 0 | 0 | 0 | 0 | 0 | 0 | 0 | 0 | 0 | 0 | 0 | 0 | 0 | 0 | 0 | 0 | 0 | 0 | 0 | 0 | 0 | 0 | 0 | 0 | 0 | 0 | 0 | 0 | 0 | 0 | 0 | 0 | 0 | 0 | 0 | 0 | 0 | 0 | 0 | 1 | 0 | 0 | 0 | 0 | 0 |
| bacteria_Sep8d5b_57581 | 1 | 0 | 0 | 0 | 0 | 0 | 0 | 0 | 0 | 0 | 0 | 0 | 0 | 0 | 0 | 0 | 0 | 0 | 0 | 0 | 0 | 0 | 0 | 0 | 0 | 0 | 0 | 0 | 0 | 0 | 0 | 0 | 0 | 0 | 0 | 0 | 0 | 0 | 0 | 0 | 0 | 0 | 0 | 0 | 0 | 0 | 0 | 0 | 0 | 1 | 0 | 0 | 0 | 0 | 0 |
| bacteria_Sep8d5b_62136 | 1 | 0 | 0 | 0 | 0 | 0 | 0 | 0 | 0 | 0 | 0 | 0 | 0 | 0 | 0 | 0 | 0 | 0 | 0 | 0 | 0 | 0 | 0 | 0 | 0 | 0 | 0 | 0 | 0 | 0 | 0 | 0 | 0 | 0 | 0 | 0 | 0 | 0 | 0 | 0 | 0 | 0 | 0 | 0 | 0 | 0 | 0 | 0 | 0 | 1 | 0 | 0 | 0 | 0 | 0 |
| bacteria_Sep8d5b_62790 | 1 | 0 | 0 | 0 | 0 | 0 | 0 | 0 | 0 | 0 | 0 | 0 | 0 | 0 | 0 | 0 | 0 | 0 | 0 | 0 | 0 | 0 | 0 | 0 | 0 | 0 | 0 | 0 | 0 | 0 | 0 | 0 | 0 | 0 | 0 | 0 | 0 | 0 | 0 | 0 | 0 | 0 | 0 | 0 | 0 | 0 | 0 | 0 | 0 | 1 | 0 | 0 | 0 | 0 | 0 |
| bacteria_Sep8d5b_65244 | 1 | 0 | 0 | 0 | 0 | 0 | 0 | 0 | 0 | 0 | 0 | 0 | 0 | 0 | 0 | 0 | 0 | 0 | 0 | 0 | 0 | 0 | 0 | 0 | 0 | 0 | 0 | 0 | 0 | 0 | 0 | 0 | 0 | 0 | 0 | 0 | 0 | 0 | 0 | 0 | 0 | 0 | 0 | 0 | 0 | 0 | 0 | 0 | 0 | 1 | 0 | 0 | 0 | 0 | 0 |
| bacteria_Sep8d5b_67161 | 1 | 0 | 0 | 0 | 0 | 0 | 0 | 0 | 0 | 0 | 0 | 0 | 0 | 0 | 0 | 0 | 0 | 0 | 0 | 0 | 0 | 0 | 0 | 0 | 0 | 0 | 0 | 0 | 0 | 0 | 0 | 0 | 0 | 0 | 0 | 0 | 0 | 0 | 0 | 0 | 0 | 0 | 0 | 0 | 0 | 0 | 0 | 0 | 0 | 1 | 0 | 0 | 0 | 0 | 0 |
| bacteria_Sep8d5b_68717 | 1 | 0 | 0 | 0 | 0 | 0 | 0 | 0 | 0 | 0 | 0 | 0 | 0 | 0 | 0 | 0 | 0 | 0 | 0 | 0 | 0 | 0 | 0 | 0 | 0 | 0 | 0 | 0 | 0 | 0 | 0 | 0 | 0 | 0 | 0 | 0 | 0 | 0 | 0 | 0 | 0 | 0 | 0 | 0 | 0 | 0 | 0 | 0 | 0 | 1 | 0 | 0 | 0 | 0 | 0 |
| bacteria_Sep8d5b_69240 | 1 | 0 | 0 | 0 | 0 | 0 | 0 | 0 | 0 | 0 | 0 | 0 | 0 | 0 | 0 | 0 | 0 | 0 | 0 | 0 | 0 | 0 | 0 | 0 | 0 | 0 | 0 | 0 | 0 | 0 | 0 | 0 | 0 | 0 | 0 | 0 | 0 | 0 | 0 | 0 | 0 | 0 | 0 | 0 | 0 | 0 | 0 | 0 | 0 | 1 | 0 | 0 | 0 | 0 | 0 |
| bacteria_Sep8d25b_13259 | 1 | 0 | 0 | 0 | 0 | 0 | 0 | 0 | 0 | 0 | 0 | 0 | 0 | 0 | 0 | 0 | 0 | 0 | 0 | 0 | 0 | 0 | 0 | 0 | 0 | 0 | 0 | 0 | 0 | 0 | 0 | 0 | 0 | 0 | 0 | 0 | 0 | 0 | 0 | 0 | 0 | 0 | 0 | 0 | 0 | 0 | 0 | 0 | 0 | 0 | 0 | 1 | 0 | 0 | 0 |
| bacteria_Sep8d25b_16890 | 1 | 0 | 0 | 0 | 0 | 0 | 0 | 0 | 0 | 0 | 0 | 0 | 0 | 0 | 0 | 0 | 0 | 0 | 0 | 0 | 0 | 0 | 0 | 0 | 0 | 0 | 0 | 0 | 0 | 0 | 0 | 0 | 0 | 0 | 0 | 0 | 0 | 0 | 0 | 0 | 0 | 0 | 0 | 0 | 0 | 0 | 0 | 0 | 0 | 0 | 0 | 1 | 0 | 0 | 0 |
| bacteria_Sep8d25b_47178 | 1 | 0 | 0 | 0 | 0 | 0 | 0 | 0 | 0 | 0 | 0 | 0 | 0 | 0 | 0 | 0 | 0 | 0 | 0 | 0 | 0 | 0 | 0 | 0 | 0 | 0 | 0 | 0 | 0 | 0 | 0 | 0 | 0 | 0 | 0 | 0 | 0 | 0 | 0 | 0 | 0 | 0 | 0 | 0 | 0 | 0 | 0 | 0 | 0 | 0 | 0 | 1 | 0 | 0 | 0 |
| bacteria_Sep8d50b_29588 | 1 | 0 | 0 | 0 | 0 | 0 | 0 | 0 | 0 | 0 | 0 | 0 | 0 | 0 | 0 | 0 | 0 | 0 | 0 | 0 | 0 | 0 | 0 | 0 | 0 | 0 | 0 | 0 | 0 | 0 | 0 | 0 | 0 | 0 | 0 | 0 | 0 | 0 | 0 | 0 | 0 | 0 | 0 | 0 | 0 | 0 | 0 | 0 | 0 | 0 | 0 | 0 | 0 | 1 | 0 |
| virus_Aug5.0v_290 | 1 | 0 | 0 | 0 | 0 | 0 | 0 | 0 | 0 | 0 | 0 | 0 | 0 | 0 | 0 | 0 | 0 | 0 | 0 | 0 | 0 | 0 | 0 | 0 | 0 | 0 | 0 | 0 | 0 | 0 | 0 | 0 | 0 | 0 | 0 | 0 | 0 | 0 | 1 | 0 | 0 | 0 | 0 | 0 | 0 | 0 | 0 | 0 | 0 | 0 | 0 | 0 | 0 | 0 | 0 |
| virus_Aug5.0v_348 | 1 | 0 | 0 | 0 | 0 | 0 | 0 | 0 | 0 | 0 | 0 | 0 | 0 | 0 | 0 | 0 | 0 | 0 | 0 | 0 | 0 | 0 | 0 | 0 | 0 | 0 | 0 | 0 | 0 | 0 | 0 | 0 | 0 | 0 | 0 | 0 | 0 | 0 | 1 | 0 | 0 | 0 | 0 | 0 | 0 | 0 | 0 | 0 | 0 | 0 | 0 | 0 | 0 | 0 | 0 |
| virus_Aug5.0v_390 | 1 | 0 | 0 | 0 | 0 | 0 | 0 | 0 | 0 | 0 | 0 | 0 | 0 | 0 | 0 | 0 | 0 | 0 | 0 | 0 | 0 | 0 | 0 | 0 | 0 | 0 | 0 | 0 | 0 | 0 | 0 | 0 | 0 | 0 | 0 | 0 | 0 | 0 | 1 | 0 | 0 | 0 | 0 | 0 | 0 | 0 | 0 | 0 | 0 | 0 | 0 | 0 | 0 | 0 | 0 |
| virus_Aug5.0v_626 | 1 | 0 | 0 | 0 | 0 | 0 | 0 | 0 | 0 | 0 | 0 | 0 | 0 | 0 | 0 | 0 | 0 | 0 | 0 | 0 | 0 | 0 | 0 | 0 | 0 | 0 | 0 | 0 | 0 | 0 | 0 | 0 | 0 | 0 | 0 | 0 | 0 | 0 | 1 | 0 | 0 | 0 | 0 | 0 | 0 | 0 | 0 | 0 | 0 | 0 | 0 | 0 | 0 | 0 | 0 |
| virus_Aug5.0v_695 | 1 | 0 | 0 | 0 | 0 | 0 | 0 | 0 | 0 | 0 | 0 | 0 | 0 | 0 | 0 | 0 | 0 | 0 | 0 | 0 | 0 | 0 | 0 | 0 | 0 | 0 | 0 | 0 | 0 | 0 | 0 | 0 | 0 | 0 | 0 | 0 | 0 | 0 | 1 | 0 | 0 | 0 | 0 | 0 | 0 | 0 | 0 | 0 | 0 | 0 | 0 | 0 | 0 | 0 | 0 |
| virus_Aug5.0v_845 | 1 | 0 | 0 | 0 | 0 | 0 | 0 | 0 | 0 | 0 | 0 | 0 | 0 | 0 | 0 | 0 | 0 | 0 | 0 | 0 | 0 | 0 | 0 | 0 | 0 | 0 | 0 | 0 | 0 | 0 | 0 | 0 | 0 | 0 | 0 | 0 | 0 | 0 | 1 | 0 | 0 | 0 | 0 | 0 | 0 | 0 | 0 | 0 | 0 | 0 | 0 | 0 | 0 | 0 | 0 |
| virus_Aug5.0v_1048 | 1 | 0 | 0 | 0 | 0 | 0 | 0 | 0 | 0 | 0 | 0 | 0 | 0 | 0 | 0 | 0 | 0 | 0 | 0 | 0 | 0 | 0 | 0 | 0 | 0 | 0 | 0 | 0 | 0 | 0 | 0 | 0 | 0 | 0 | 0 | 0 | 0 | 0 | 1 | 0 | 0 | 0 | 0 | 0 | 0 | 0 | 0 | 0 | 0 | 0 | 0 | 0 | 0 | 0 | 0 |
| virus_Aug5.0v_1451 | 1 | 0 | 0 | 0 | 0 | 0 | 0 | 0 | 0 | 0 | 0 | 0 | 0 | 0 | 0 | 0 | 0 | 0 | 0 | 0 | 0 | 0 | 0 | 0 | 0 | 0 | 0 | 0 | 0 | 0 | 0 | 0 | 0 | 0 | 0 | 0 | 0 | 0 | 1 | 0 | 0 | 0 | 0 | 0 | 0 | 0 | 0 | 0 | 0 | 0 | 0 | 0 | 0 | 0 | 0 |
| virus_Aug5.0v_1478 | 1 | 0 | 0 | 0 | 0 | 0 | 0 | 0 | 0 | 0 | 0 | 0 | 0 | 0 | 0 | 0 | 0 | 0 | 0 | 0 | 0 | 0 | 0 | 0 | 0 | 0 | 0 | 0 | 0 | 0 | 0 | 0 | 0 | 0 | 0 | 0 | 0 | 0 | 1 | 0 | 0 | 0 | 0 | 0 | 0 | 0 | 0 | 0 | 0 | 0 | 0 | 0 | 0 | 0 | 0 |
| virus_Aug5.0v_1688 | 1 | 0 | 0 | 0 | 0 | 0 | 0 | 0 | 0 | 0 | 0 | 0 | 0 | 0 | 0 | 0 | 0 | 0 | 0 | 0 | 0 | 0 | 0 | 0 | 0 | 0 | 0 | 0 | 0 | 0 | 0 | 0 | 0 | 0 | 0 | 0 | 0 | 0 | 1 | 0 | 0 | 0 | 0 | 0 | 0 | 0 | 0 | 0 | 0 | 0 | 0 | 0 | 0 | 0 | 0 |
| virus_Aug5.0v_2536 | 1 | 0 | 0 | 0 | 0 | 0 | 0 | 0 | 0 | 0 | 0 | 0 | 0 | 0 | 0 | 0 | 0 | 0 | 0 | 0 | 0 | 0 | 0 | 0 | 0 | 0 | 0 | 0 | 0 | 0 | 0 | 0 | 0 | 0 | 0 | 0 | 0 | 0 | 1 | 0 | 0 | 0 | 0 | 0 | 0 | 0 | 0 | 0 | 0 | 0 | 0 | 0 | 0 | 0 | 0 |
| virus_Aug5.0v_3006 | 1 | 0 | 0 | 0 | 0 | 0 | 0 | 0 | 0 | 0 | 0 | 0 | 0 | 0 | 0 | 0 | 0 | 0 | 0 | 0 | 0 | 0 | 0 | 0 | 0 | 0 | 0 | 0 | 0 | 0 | 0 | 0 | 0 | 0 | 0 | 0 | 0 | 0 | 1 | 0 | 0 | 0 | 0 | 0 | 0 | 0 | 0 | 0 | 0 | 0 | 0 | 0 | 0 | 0 | 0 |
| virus_Aug5.0v_3052 | 1 | 0 | 0 | 0 | 0 | 0 | 0 | 0 | 0 | 0 | 0 | 0 | 0 | 0 | 0 | 0 | 0 | 0 | 0 | 0 | 0 | 0 | 0 | 0 | 0 | 0 | 0 | 0 | 0 | 0 | 0 | 0 | 0 | 0 | 0 | 0 | 0 | 0 | 1 | 0 | 0 | 0 | 0 | 0 | 0 | 0 | 0 | 0 | 0 | 0 | 0 | 0 | 0 | 0 | 0 |
| virus_Aug5.0v_3248 | 1 | 0 | 0 | 0 | 0 | 0 | 0 | 0 | 0 | 0 | 0 | 0 | 0 | 0 | 0 | 0 | 0 | 0 | 0 | 0 | 0 | 0 | 0 | 0 | 0 | 0 | 0 | 0 | 0 | 0 | 0 | 0 | 0 | 0 | 0 | 0 | 0 | 0 | 1 | 0 | 0 | 0 | 0 | 0 | 0 | 0 | 0 | 0 | 0 | 0 | 0 | 0 | 0 | 0 | 0 |
| virus_Aug5.0v_3442 | 1 | 0 | 0 | 0 | 0 | 0 | 0 | 0 | 0 | 0 | 0 | 0 | 0 | 0 | 0 | 0 | 0 | 0 | 0 | 0 | 0 | 0 | 0 | 0 | 0 | 0 | 0 | 0 | 0 | 0 | 0 | 0 | 0 | 0 | 0 | 0 | 0 | 0 | 1 | 0 | 0 | 0 | 0 | 0 | 0 | 0 | 0 | 0 | 0 | 0 | 0 | 0 | 0 | 0 | 0 |
| virus_Aug5.0v_3538 | 1 | 0 | 0 | 0 | 0 | 0 | 0 | 0 | 0 | 0 | 0 | 0 | 0 | 0 | 0 | 0 | 0 | 0 | 0 | 0 | 0 | 0 | 0 | 0 | 0 | 0 | 0 | 0 | 0 | 0 | 0 | 0 | 0 | 0 | 0 | 0 | 0 | 0 | 1 | 0 | 0 | 0 | 0 | 0 | 0 | 0 | 0 | 0 | 0 | 0 | 0 | 0 | 0 | 0 | 0 |
| virus_Aug5.0v_3549 | 1 | 0 | 0 | 0 | 0 | 0 | 0 | 0 | 0 | 0 | 0 | 0 | 0 | 0 | 0 | 0 | 0 | 0 | 0 | 0 | 0 | 0 | 0 | 0 | 0 | 0 | 0 | 0 | 0 | 0 | 0 | 0 | 0 | 0 | 0 | 0 | 0 | 0 | 1 | 0 | 0 | 0 | 0 | 0 | 0 | 0 | 0 | 0 | 0 | 0 | 0 | 0 | 0 | 0 | 0 |
| virus_Aug5.0v_3564 | 1 | 0 | 0 | 0 | 0 | 0 | 0 | 0 | 0 | 0 | 0 | 0 | 0 | 0 | 0 | 0 | 0 | 0 | 0 | 0 | 0 | 0 | 0 | 0 | 0 | 0 | 0 | 0 | 0 | 0 | 0 | 0 | 0 | 0 | 0 | 0 | 0 | 0 | 1 | 0 | 0 | 0 | 0 | 0 | 0 | 0 | 0 | 0 | 0 | 0 | 0 | 0 | 0 | 0 | 0 |
| virus_Aug5.0v_3625 | 1 | 0 | 0 | 0 | 0 | 0 | 0 | 0 | 0 | 0 | 0 | 0 | 0 | 0 | 0 | 0 | 0 | 0 | 0 | 0 | 0 | 0 | 0 | 0 | 0 | 0 | 0 | 0 | 0 | 0 | 0 | 0 | 0 | 0 | 0 | 0 | 0 | 0 | 1 | 0 | 0 | 0 | 0 | 0 | 0 | 0 | 0 | 0 | 0 | 0 | 0 | 0 | 0 | 0 | 0 |
| virus_Aug5.0v_3985 | 1 | 0 | 0 | 0 | 0 | 0 | 0 | 0 | 0 | 0 | 0 | 0 | 0 | 0 | 0 | 0 | 0 | 0 | 0 | 0 | 0 | 0 | 0 | 0 | 0 | 0 | 0 | 0 | 0 | 0 | 0 | 0 | 0 | 0 | 0 | 0 | 0 | 0 | 1 | 0 | 0 | 0 | 0 | 0 | 0 | 0 | 0 | 0 | 0 | 0 | 0 | 0 | 0 | 0 | 0 |
| virus_Aug5.0v_4021 | 1 | 0 | 0 | 0 | 0 | 0 | 0 | 0 | 0 | 0 | 0 | 0 | 0 | 0 | 0 | 0 | 0 | 0 | 0 | 0 | 0 | 0 | 0 | 0 | 0 | 0 | 0 | 0 | 0 | 0 | 0 | 0 | 0 | 0 | 0 | 0 | 0 | 0 | 1 | 0 | 0 | 0 | 0 | 0 | 0 | 0 | 0 | 0 | 0 | 0 | 0 | 0 | 0 | 0 | 0 |
| virus_Aug5.0v_4206 | 1 | 0 | 0 | 0 | 0 | 0 | 0 | 0 | 0 | 0 | 0 | 0 | 0 | 0 | 0 | 0 | 0 | 0 | 0 | 0 | 0 | 0 | 0 | 0 | 0 | 0 | 0 | 0 | 0 | 0 | 0 | 0 | 0 | 0 | 0 | 0 | 0 | 0 | 1 | 0 | 0 | 0 | 0 | 0 | 0 | 0 | 0 | 0 | 0 | 0 | 0 | 0 | 0 | 0 | 0 |
| virus_Aug5.0v_4337 | 1 | 0 | 0 | 0 | 0 | 0 | 0 | 0 | 0 | 0 | 0 | 0 | 0 | 0 | 0 | 0 | 0 | 0 | 0 | 0 | 0 | 0 | 0 | 0 | 0 | 0 | 0 | 0 | 0 | 0 | 0 | 0 | 0 | 0 | 0 | 0 | 0 | 0 | 1 | 0 | 0 | 0 | 0 | 0 | 0 | 0 | 0 | 0 | 0 | 0 | 0 | 0 | 0 | 0 | 0 |
| virus_Aug5.0v_4514 | 1 | 0 | 0 | 0 | 0 | 0 | 0 | 0 | 0 | 0 | 0 | 0 | 0 | 0 | 0 | 0 | 0 | 0 | 0 | 0 | 0 | 0 | 0 | 0 | 0 | 0 | 0 | 0 | 0 | 0 | 0 | 0 | 0 | 0 | 0 | 0 | 0 | 0 | 1 | 0 | 0 | 0 | 0 | 0 | 0 | 0 | 0 | 0 | 0 | 0 | 0 | 0 | 0 | 0 | 0 |
| virus_Aug5.0v_4642 | 1 | 0 | 0 | 0 | 0 | 0 | 0 | 0 | 0 | 0 | 0 | 0 | 0 | 0 | 0 | 0 | 0 | 0 | 0 | 0 | 0 | 0 | 0 | 0 | 0 | 0 | 0 | 0 | 0 | 0 | 0 | 0 | 0 | 0 | 0 | 0 | 0 | 0 | 1 | 0 | 0 | 0 | 0 | 0 | 0 | 0 | 0 | 0 | 0 | 0 | 0 | 0 | 0 | 0 | 0 |
| virus_Aug5.0v_4674 | 1 | 0 | 0 | 0 | 0 | 0 | 0 | 0 | 0 | 0 | 0 | 0 | 0 | 0 | 0 | 0 | 0 | 0 | 0 | 0 | 0 | 0 | 0 | 0 | 0 | 0 | 0 | 0 | 0 | 0 | 0 | 0 | 0 | 0 | 0 | 0 | 0 | 0 | 1 | 0 | 0 | 0 | 0 | 0 | 0 | 0 | 0 | 0 | 0 | 0 | 0 | 0 | 0 | 0 | 0 |
| virus_Aug5.0v_4814 | 1 | 0 | 0 | 0 | 0 | 0 | 0 | 0 | 0 | 0 | 0 | 0 | 0 | 0 | 0 | 0 | 0 | 0 | 0 | 0 | 0 | 0 | 0 | 0 | 0 | 0 | 0 | 0 | 0 | 0 | 0 | 0 | 0 | 0 | 0 | 0 | 0 | 0 | 1 | 0 | 0 | 0 | 0 | 0 | 0 | 0 | 0 | 0 | 0 | 0 | 0 | 0 | 0 | 0 | 0 |
| virus_Aug5.0v_5066 | 1 | 0 | 0 | 0 | 0 | 0 | 0 | 0 | 0 | 0 | 0 | 0 | 0 | 0 | 0 | 0 | 0 | 0 | 0 | 0 | 0 | 0 | 0 | 0 | 0 | 0 | 0 | 0 | 0 | 0 | 0 | 0 | 0 | 0 | 0 | 0 | 0 | 0 | 1 | 0 | 0 | 0 | 0 | 0 | 0 | 0 | 0 | 0 | 0 | 0 | 0 | 0 | 0 | 0 | 0 |
| virus_Aug5.0v_5222 | 1 | 0 | 0 | 0 | 0 | 0 | 0 | 0 | 0 | 0 | 0 | 0 | 0 | 0 | 0 | 0 | 0 | 0 | 0 | 0 | 0 | 0 | 0 | 0 | 0 | 0 | 0 | 0 | 0 | 0 | 0 | 0 | 0 | 0 | 0 | 0 | 0 | 0 | 1 | 0 | 0 | 0 | 0 | 0 | 0 | 0 | 0 | 0 | 0 | 0 | 0 | 0 | 0 | 0 | 0 |
| virus_Aug5.0v_5270 | 1 | 0 | 0 | 0 | 0 | 0 | 0 | 0 | 0 | 0 | 0 | 0 | 0 | 0 | 0 | 0 | 0 | 0 | 0 | 0 | 0 | 0 | 0 | 0 | 0 | 0 | 0 | 0 | 0 | 0 | 0 | 0 | 0 | 0 | 0 | 0 | 0 | 0 | 1 | 0 | 0 | 0 | 0 | 0 | 0 | 0 | 0 | 0 | 0 | 0 | 0 | 0 | 0 | 0 | 0 |
| virus_Aug5.5v_4964 | 1 | 0 | 0 | 0 | 0 | 0 | 0 | 0 | 0 | 0 | 0 | 0 | 0 | 0 | 0 | 0 | 0 | 0 | 0 | 0 | 0 | 0 | 0 | 0 | 0 | 0 | 0 | 0 | 0 | 0 | 0 | 0 | 0 | 0 | 0 | 0 | 0 | 0 | 0 | 0 | 1 | 0 | 0 | 0 | 0 | 0 | 0 | 0 | 0 | 0 | 0 | 0 | 0 | 0 | 0 |
| virus_Aug5.5v_4987 | 1 | 0 | 0 | 0 | 0 | 0 | 0 | 0 | 0 | 0 | 0 | 0 | 0 | 0 | 0 | 0 | 0 | 0 | 0 | 0 | 0 | 0 | 0 | 0 | 0 | 0 | 0 | 0 | 0 | 0 | 0 | 0 | 0 | 0 | 0 | 0 | 0 | 0 | 0 | 0 | 1 | 0 | 0 | 0 | 0 | 0 | 0 | 0 | 0 | 0 | 0 | 0 | 0 | 0 | 0 |
| virus_Aug5.5v_5667 | 1 | 0 | 0 | 0 | 0 | 0 | 0 | 0 | 0 | 0 | 0 | 0 | 0 | 0 | 0 | 0 | 0 | 0 | 0 | 0 | 0 | 0 | 0 | 0 | 0 | 0 | 0 | 0 | 0 | 0 | 0 | 0 | 0 | 0 | 0 | 0 | 0 | 0 | 0 | 0 | 1 | 0 | 0 | 0 | 0 | 0 | 0 | 0 | 0 | 0 | 0 | 0 | 0 | 0 | 0 |
| virus_Aug5.5v_6323 | 1 | 0 | 0 | 0 | 0 | 0 | 0 | 0 | 0 | 0 | 0 | 0 | 0 | 0 | 0 | 0 | 0 | 0 | 0 | 0 | 0 | 0 | 0 | 0 | 0 | 0 | 0 | 0 | 0 | 0 | 0 | 0 | 0 | 0 | 0 | 0 | 0 | 0 | 0 | 0 | 1 | 0 | 0 | 0 | 0 | 0 | 0 | 0 | 0 | 0 | 0 | 0 | 0 | 0 | 0 |
| virus_Aug5.5v_6613 | 1 | 0 | 0 | 0 | 0 | 0 | 0 | 0 | 0 | 0 | 0 | 0 | 0 | 0 | 0 | 0 | 0 | 0 | 0 | 0 | 0 | 0 | 0 | 0 | 0 | 0 | 0 | 0 | 0 | 0 | 0 | 0 | 0 | 0 | 0 | 0 | 0 | 0 | 0 | 0 | 1 | 0 | 0 | 0 | 0 | 0 | 0 | 0 | 0 | 0 | 0 | 0 | 0 | 0 | 0 |
| virus_Aug5.5v_7763 | 1 | 0 | 0 | 0 | 0 | 0 | 0 | 0 | 0 | 0 | 0 | 0 | 0 | 0 | 0 | 0 | 0 | 0 | 0 | 0 | 0 | 0 | 0 | 0 | 0 | 0 | 0 | 0 | 0 | 0 | 0 | 0 | 0 | 0 | 0 | 0 | 0 | 0 | 0 | 0 | 1 | 0 | 0 | 0 | 0 | 0 | 0 | 0 | 0 | 0 | 0 | 0 | 0 | 0 | 0 |
| virus_Aug5.5v_21254 | 1 | 0 | 0 | 0 | 0 | 0 | 0 | 0 | 0 | 0 | 0 | 0 | 0 | 0 | 0 | 0 | 0 | 0 | 0 | 0 | 0 | 0 | 0 | 0 | 0 | 0 | 0 | 0 | 0 | 0 | 0 | 0 | 0 | 0 | 0 | 0 | 0 | 0 | 0 | 0 | 1 | 0 | 0 | 0 | 0 | 0 | 0 | 0 | 0 | 0 | 0 | 0 | 0 | 0 | 0 |
| virus_Aug5.5v_23129 | 1 | 0 | 0 | 0 | 0 | 0 | 0 | 0 | 0 | 0 | 0 | 0 | 0 | 0 | 0 | 0 | 0 | 0 | 0 | 0 | 0 | 0 | 0 | 0 | 0 | 0 | 0 | 0 | 0 | 0 | 0 | 0 | 0 | 0 | 0 | 0 | 0 | 0 | 0 | 0 | 1 | 0 | 0 | 0 | 0 | 0 | 0 | 0 | 0 | 0 | 0 | 0 | 0 | 0 | 0 |
| virus_Aug5.5v_33291 | 1 | 0 | 0 | 0 | 0 | 0 | 0 | 0 | 0 | 0 | 0 | 0 | 0 | 0 | 0 | 0 | 0 | 0 | 0 | 0 | 0 | 0 | 0 | 0 | 0 | 0 | 0 | 0 | 0 | 0 | 0 | 0 | 0 | 0 | 0 | 0 | 0 | 0 | 0 | 0 | 1 | 0 | 0 | 0 | 0 | 0 | 0 | 0 | 0 | 0 | 0 | 0 | 0 | 0 | 0 |
| virus_Aug5.5v_36082 | 1 | 0 | 0 | 0 | 0 | 0 | 0 | 0 | 0 | 0 | 0 | 0 | 0 | 0 | 0 | 0 | 0 | 0 | 0 | 0 | 0 | 0 | 0 | 0 | 0 | 0 | 0 | 0 | 0 | 0 | 0 | 0 | 0 | 0 | 0 | 0 | 0 | 0 | 0 | 0 | 1 | 0 | 0 | 0 | 0 | 0 | 0 | 0 | 0 | 0 | 0 | 0 | 0 | 0 | 0 |

**Supplementary Table S7**. Common Bacterio- and virioplankton RTPR 98% OTUs in 27 samples with RTPR amplicons in both fractions

Total seqs: 47997; Subtotal seqs in retained

| OTUs: 41046 | | May12_05m | | May12_25m | | May12_50m | | May12_75m | | Jun8_25m | | Jun8_50m | | Jun8_75m | | Jun22_0m | | Jun22_50m | | Jun22_75m | | Jul11_0m | | Jul11_05m | | Jul11_25m | | Jul11_50m | | Jul11_75m | | Jul27_05m | | Jul27_25m | | Jul27_75m | | Aug5_0m | | Aug5_05m | | Aug5_25m | | Aug5_50m | | Aug5_75m | | Sep8_0m | | Sep8_05m | | Sep8_25m | | Sep8_50m | |
|---|---|---|---|---|---|---|---|---|---|---|---|---|---|---|---|---|---|---|---|---|---|---|---|---|---|---|---|---|---|---|---|---|---|---|---|---|---|---|---|---|---|---|---|---|---|---|---|---|---|---|---|---|---|---|---|
| OTUs with >=10 seqs: 254 | OTU subtotal | bacterioplankton | virioplankton | bacterioplankton | virioplankton | bacterioplankton | virioplankton | bacterioplankton | virioplankton | bacterioplankton | virioplankton | bacterioplankton | virioplankton | bacterioplankton | virioplankton | bacterioplankton | virioplankton | bacterioplankton | virioplankton | bacterioplankton | virioplankton | bacterioplankton | virioplankton | bacterioplankton | virioplankton | bacterioplankton | virioplankton | bacterioplankton | virioplankton | bacterioplankton | virioplankton | bacterioplankton | virioplankton | bacterioplankton | virioplankton | bacterioplankton | virioplankton | bacterioplankton | virioplankton | bacterioplankton | virioplankton | bacterioplankton | virioplankton | bacterioplankton | virioplankton | bacterioplankton | virioplankton | bacterioplankton | virioplankton | bacterioplankton | virioplankton | bacterioplankton | virioplankton | bacterioplankton | virioplankton |
| virus_Aug5.5v_36926 | 1 | 0 | 0 | 0 | 0 | 0 | 0 | 0 | 0 | 0 | 0 | 0 | 0 | 0 | 0 | 0 | 0 | 0 | 0 | 0 | 0 | 0 | 0 | 0 | 0 | 0 | 0 | 0 | 0 | 0 | 0 | 0 | 0 | 0 | 0 | 0 | 0 | 0 | 0 | 0 | 1 | 0 | 0 | 0 | 0 | 0 | 0 | 0 | 0 | 0 | 0 | 0 | 0 | 0 | 0 |
| virus_Aug5.5v_36992 | 1 | 0 | 0 | 0 | 0 | 0 | 0 | 0 | 0 | 0 | 0 | 0 | 0 | 0 | 0 | 0 | 0 | 0 | 0 | 0 | 0 | 0 | 0 | 0 | 0 | 0 | 0 | 0 | 0 | 0 | 0 | 0 | 0 | 0 | 0 | 0 | 0 | 0 | 0 | 0 | 1 | 0 | 0 | 0 | 0 | 0 | 0 | 0 | 0 | 0 | 0 | 0 | 0 | 0 | 0 |
| virus_Aug5.25v_652 | 1 | 0 | 0 | 0 | 0 | 0 | 0 | 0 | 0 | 0 | 0 | 0 | 0 | 0 | 0 | 0 | 0 | 0 | 0 | 0 | 0 | 0 | 0 | 0 | 0 | 0 | 0 | 0 | 0 | 0 | 0 | 0 | 0 | 0 | 0 | 0 | 0 | 0 | 0 | 0 | 0 | 0 | 1 | 0 | 0 | 0 | 0 | 0 | 0 | 0 | 0 | 0 | 0 | 0 | 0 |
| virus_Aug5.25v_1084 | 1 | 0 | 0 | 0 | 0 | 0 | 0 | 0 | 0 | 0 | 0 | 0 | 0 | 0 | 0 | 0 | 0 | 0 | 0 | 0 | 0 | 0 | 0 | 0 | 0 | 0 | 0 | 0 | 0 | 0 | 0 | 0 | 0 | 0 | 0 | 0 | 0 | 0 | 0 | 0 | 0 | 0 | 1 | 0 | 0 | 0 | 0 | 0 | 0 | 0 | 0 | 0 | 0 | 0 | 0 |
| virus_Aug5.25v_1465 | 1 | 0 | 0 | 0 | 0 | 0 | 0 | 0 | 0 | 0 | 0 | 0 | 0 | 0 | 0 | 0 | 0 | 0 | 0 | 0 | 0 | 0 | 0 | 0 | 0 | 0 | 0 | 0 | 0 | 0 | 0 | 0 | 0 | 0 | 0 | 0 | 0 | 0 | 0 | 0 | 0 | 0 | 1 | 0 | 0 | 0 | 0 | 0 | 0 | 0 | 0 | 0 | 0 | 0 | 0 |
| virus_Aug5.25v_5891 | 1 | 0 | 0 | 0 | 0 | 0 | 0 | 0 | 0 | 0 | 0 | 0 | 0 | 0 | 0 | 0 | 0 | 0 | 0 | 0 | 0 | 0 | 0 | 0 | 0 | 0 | 0 | 0 | 0 | 0 | 0 | 0 | 0 | 0 | 0 | 0 | 0 | 0 | 0 | 0 | 0 | 0 | 1 | 0 | 0 | 0 | 0 | 0 | 0 | 0 | 0 | 0 | 0 | 0 | 0 |
| virus_Aug5.25v_6260 | 1 | 0 | 0 | 0 | 0 | 0 | 0 | 0 | 0 | 0 | 0 | 0 | 0 | 0 | 0 | 0 | 0 | 0 | 0 | 0 | 0 | 0 | 0 | 0 | 0 | 0 | 0 | 0 | 0 | 0 | 0 | 0 | 0 | 0 | 0 | 0 | 0 | 0 | 0 | 0 | 0 | 0 | 1 | 0 | 0 | 0 | 0 | 0 | 0 | 0 | 0 | 0 | 0 | 0 | 0 |
| virus_Aug5.25v_12315 | 1 | 0 | 0 | 0 | 0 | 0 | 0 | 0 | 0 | 0 | 0 | 0 | 0 | 0 | 0 | 0 | 0 | 0 | 0 | 0 | 0 | 0 | 0 | 0 | 0 | 0 | 0 | 0 | 0 | 0 | 0 | 0 | 0 | 0 | 0 | 0 | 0 | 0 | 0 | 0 | 0 | 0 | 1 | 0 | 0 | 0 | 0 | 0 | 0 | 0 | 0 | 0 | 0 | 0 | 0 |
| virus_Aug5.25v_13513 | 1 | 0 | 0 | 0 | 0 | 0 | 0 | 0 | 0 | 0 | 0 | 0 | 0 | 0 | 0 | 0 | 0 | 0 | 0 | 0 | 0 | 0 | 0 | 0 | 0 | 0 | 0 | 0 | 0 | 0 | 0 | 0 | 0 | 0 | 0 | 0 | 0 | 0 | 0 | 0 | 0 | 0 | 1 | 0 | 0 | 0 | 0 | 0 | 0 | 0 | 0 | 0 | 0 | 0 | 0 |
| virus_Aug5.25v_17057 | 1 | 0 | 0 | 0 | 0 | 0 | 0 | 0 | 0 | 0 | 0 | 0 | 0 | 0 | 0 | 0 | 0 | 0 | 0 | 0 | 0 | 0 | 0 | 0 | 0 | 0 | 0 | 0 | 0 | 0 | 0 | 0 | 0 | 0 | 0 | 0 | 0 | 0 | 0 | 0 | 0 | 0 | 1 | 0 | 0 | 0 | 0 | 0 | 0 | 0 | 0 | 0 | 0 | 0 | 0 |
| virus_Aug5.25v_18271 | 1 | 0 | 0 | 0 | 0 | 0 | 0 | 0 | 0 | 0 | 0 | 0 | 0 | 0 | 0 | 0 | 0 | 0 | 0 | 0 | 0 | 0 | 0 | 0 | 0 | 0 | 0 | 0 | 0 | 0 | 0 | 0 | 0 | 0 | 0 | 0 | 0 | 0 | 0 | 0 | 0 | 0 | 1 | 0 | 0 | 0 | 0 | 0 | 0 | 0 | 0 | 0 | 0 | 0 | 0 |
| virus_Aug5.25v_18561 | 1 | 0 | 0 | 0 | 0 | 0 | 0 | 0 | 0 | 0 | 0 | 0 | 0 | 0 | 0 | 0 | 0 | 0 | 0 | 0 | 0 | 0 | 0 | 0 | 0 | 0 | 0 | 0 | 0 | 0 | 0 | 0 | 0 | 0 | 0 | 0 | 0 | 0 | 0 | 0 | 0 | 0 | 1 | 0 | 0 | 0 | 0 | 0 | 0 | 0 | 0 | 0 | 0 | 0 | 0 |
| virus_Aug5.25v_18955 | 1 | 0 | 0 | 0 | 0 | 0 | 0 | 0 | 0 | 0 | 0 | 0 | 0 | 0 | 0 | 0 | 0 | 0 | 0 | 0 | 0 | 0 | 0 | 0 | 0 | 0 | 0 | 0 | 0 | 0 | 0 | 0 | 0 | 0 | 0 | 0 | 0 | 0 | 0 | 0 | 0 | 0 | 1 | 0 | 0 | 0 | 0 | 0 | 0 | 0 | 0 | 0 | 0 | 0 | 0 |
| virus_Aug5.25v_21604 | 1 | 0 | 0 | 0 | 0 | 0 | 0 | 0 | 0 | 0 | 0 | 0 | 0 | 0 | 0 | 0 | 0 | 0 | 0 | 0 | 0 | 0 | 0 | 0 | 0 | 0 | 0 | 0 | 0 | 0 | 0 | 0 | 0 | 0 | 0 | 0 | 0 | 0 | 0 | 0 | 0 | 0 | 1 | 0 | 0 | 0 | 0 | 0 | 0 | 0 | 0 | 0 | 0 | 0 | 0 |
| virus_Aug5.25v_22660 | 1 | 0 | 0 | 0 | 0 | 0 | 0 | 0 | 0 | 0 | 0 | 0 | 0 | 0 | 0 | 0 | 0 | 0 | 0 | 0 | 0 | 0 | 0 | 0 | 0 | 0 | 0 | 0 | 0 | 0 | 0 | 0 | 0 | 0 | 0 | 0 | 0 | 0 | 0 | 0 | 0 | 0 | 1 | 0 | 0 | 0 | 0 | 0 | 0 | 0 | 0 | 0 | 0 | 0 | 0 |
| virus_Aug5.25v_22819 | 1 | 0 | 0 | 0 | 0 | 0 | 0 | 0 | 0 | 0 | 0 | 0 | 0 | 0 | 0 | 0 | 0 | 0 | 0 | 0 | 0 | 0 | 0 | 0 | 0 | 0 | 0 | 0 | 0 | 0 | 0 | 0 | 0 | 0 | 0 | 0 | 0 | 0 | 0 | 0 | 0 | 0 | 1 | 0 | 0 | 0 | 0 | 0 | 0 | 0 | 0 | 0 | 0 | 0 | 0 |
| virus_Aug5.25v_24413 | 1 | 0 | 0 | 0 | 0 | 0 | 0 | 0 | 0 | 0 | 0 | 0 | 0 | 0 | 0 | 0 | 0 | 0 | 0 | 0 | 0 | 0 | 0 | 0 | 0 | 0 | 0 | 0 | 0 | 0 | 0 | 0 | 0 | 0 | 0 | 0 | 0 | 0 | 0 | 0 | 0 | 0 | 1 | 0 | 0 | 0 | 0 | 0 | 0 | 0 | 0 | 0 | 0 | 0 | 0 |
| virus_Aug5.25v_24767 | 1 | 0 | 0 | 0 | 0 | 0 | 0 | 0 | 0 | 0 | 0 | 0 | 0 | 0 | 0 | 0 | 0 | 0 | 0 | 0 | 0 | 0 | 0 | 0 | 0 | 0 | 0 | 0 | 0 | 0 | 0 | 0 | 0 | 0 | 0 | 0 | 0 | 0 | 0 | 0 | 0 | 0 | 1 | 0 | 0 | 0 | 0 | 0 | 0 | 0 | 0 | 0 | 0 | 0 | 0 |
| virus_Aug5.25v_27579 | 1 | 0 | 0 | 0 | 0 | 0 | 0 | 0 | 0 | 0 | 0 | 0 | 0 | 0 | 0 | 0 | 0 | 0 | 0 | 0 | 0 | 0 | 0 | 0 | 0 | 0 | 0 | 0 | 0 | 0 | 0 | 0 | 0 | 0 | 0 | 0 | 0 | 0 | 0 | 0 | 0 | 0 | 1 | 0 | 0 | 0 | 0 | 0 | 0 | 0 | 0 | 0 | 0 | 0 | 0 |
| virus_Aug5.25v_29883 | 1 | 0 | 0 | 0 | 0 | 0 | 0 | 0 | 0 | 0 | 0 | 0 | 0 | 0 | 0 | 0 | 0 | 0 | 0 | 0 | 0 | 0 | 0 | 0 | 0 | 0 | 0 | 0 | 0 | 0 | 0 | 0 | 0 | 0 | 0 | 0 | 0 | 0 | 0 | 0 | 0 | 0 | 1 | 0 | 0 | 0 | 0 | 0 | 0 | 0 | 0 | 0 | 0 | 0 | 0 |
| virus_Aug5.25v_32838 | 1 | 0 | 0 | 0 | 0 | 0 | 0 | 0 | 0 | 0 | 0 | 0 | 0 | 0 | 0 | 0 | 0 | 0 | 0 | 0 | 0 | 0 | 0 | 0 | 0 | 0 | 0 | 0 | 0 | 0 | 0 | 0 | 0 | 0 | 0 | 0 | 0 | 0 | 0 | 0 | 0 | 0 | 1 | 0 | 0 | 0 | 0 | 0 | 0 | 0 | 0 | 0 | 0 | 0 | 0 |
| virus_Aug5.25v_33658 | 1 | 0 | 0 | 0 | 0 | 0 | 0 | 0 | 0 | 0 | 0 | 0 | 0 | 0 | 0 | 0 | 0 | 0 | 0 | 0 | 0 | 0 | 0 | 0 | 0 | 0 | 0 | 0 | 0 | 0 | 0 | 0 | 0 | 0 | 0 | 0 | 0 | 0 | 0 | 0 | 0 | 0 | 1 | 0 | 0 | 0 | 0 | 0 | 0 | 0 | 0 | 0 | 0 | 0 | 0 |
| virus_Aug5.25v_33983 | 1 | 0 | 0 | 0 | 0 | 0 | 0 | 0 | 0 | 0 | 0 | 0 | 0 | 0 | 0 | 0 | 0 | 0 | 0 | 0 | 0 | 0 | 0 | 0 | 0 | 0 | 0 | 0 | 0 | 0 | 0 | 0 | 0 | 0 | 0 | 0 | 0 | 0 | 0 | 0 | 0 | 0 | 1 | 0 | 0 | 0 | 0 | 0 | 0 | 0 | 0 | 0 | 0 | 0 | 0 |
| virus_Aug5.25v_35907 | 1 | 0 | 0 | 0 | 0 | 0 | 0 | 0 | 0 | 0 | 0 | 0 | 0 | 0 | 0 | 0 | 0 | 0 | 0 | 0 | 0 | 0 | 0 | 0 | 0 | 0 | 0 | 0 | 0 | 0 | 0 | 0 | 0 | 0 | 0 | 0 | 0 | 0 | 0 | 0 | 0 | 0 | 1 | 0 | 0 | 0 | 0 | 0 | 0 | 0 | 0 | 0 | 0 | 0 | 0 |
| virus_Aug5.25v_37846 | 1 | 0 | 0 | 0 | 0 | 0 | 0 | 0 | 0 | 0 | 0 | 0 | 0 | 0 | 0 | 0 | 0 | 0 | 0 | 0 | 0 | 0 | 0 | 0 | 0 | 0 | 0 | 0 | 0 | 0 | 0 | 0 | 0 | 0 | 0 | 0 | 0 | 0 | 0 | 0 | 0 | 0 | 1 | 0 | 0 | 0 | 0 | 0 | 0 | 0 | 0 | 0 | 0 | 0 | 0 |
| virus_Aug5.25v_37860 | 1 | 0 | 0 | 0 | 0 | 0 | 0 | 0 | 0 | 0 | 0 | 0 | 0 | 0 | 0 | 0 | 0 | 0 | 0 | 0 | 0 | 0 | 0 | 0 | 0 | 0 | 0 | 0 | 0 | 0 | 0 | 0 | 0 | 0 | 0 | 0 | 0 | 0 | 0 | 0 | 0 | 0 | 1 | 0 | 0 | 0 | 0 | 0 | 0 | 0 | 0 | 0 | 0 | 0 | 0 |
| virus_Aug5.25v_38021 | 1 | 0 | 0 | 0 | 0 | 0 | 0 | 0 | 0 | 0 | 0 | 0 | 0 | 0 | 0 | 0 | 0 | 0 | 0 | 0 | 0 | 0 | 0 | 0 | 0 | 0 | 0 | 0 | 0 | 0 | 0 | 0 | 0 | 0 | 0 | 0 | 0 | 0 | 0 | 0 | 0 | 0 | 1 | 0 | 0 | 0 | 0 | 0 | 0 | 0 | 0 | 0 | 0 | 0 | 0 |
| virus_Aug5.50v_28836 | 1 | 0 | 0 | 0 | 0 | 0 | 0 | 0 | 0 | 0 | 0 | 0 | 0 | 0 | 0 | 0 | 0 | 0 | 0 | 0 | 0 | 0 | 0 | 0 | 0 | 0 | 0 | 0 | 0 | 0 | 0 | 0 | 0 | 0 | 0 | 0 | 0 | 0 | 0 | 0 | 0 | 0 | 0 | 0 | 1 | 0 | 0 | 0 | 0 | 0 | 0 | 0 | 0 | 0 | 0 |
| virus_Aug5.50v_31030 | 1 | 0 | 0 | 0 | 0 | 0 | 0 | 0 | 0 | 0 | 0 | 0 | 0 | 0 | 0 | 0 | 0 | 0 | 0 | 0 | 0 | 0 | 0 | 0 | 0 | 0 | 0 | 0 | 0 | 0 | 0 | 0 | 0 | 0 | 0 | 0 | 0 | 0 | 0 | 0 | 0 | 0 | 0 | 0 | 1 | 0 | 0 | 0 | 0 | 0 | 0 | 0 | 0 | 0 | 0 |
| virus_Aug5.50v_31317 | 1 | 0 | 0 | 0 | 0 | 0 | 0 | 0 | 0 | 0 | 0 | 0 | 0 | 0 | 0 | 0 | 0 | 0 | 0 | 0 | 0 | 0 | 0 | 0 | 0 | 0 | 0 | 0 | 0 | 0 | 0 | 0 | 0 | 0 | 0 | 0 | 0 | 0 | 0 | 0 | 0 | 0 | 0 | 0 | 1 | 0 | 0 | 0 | 0 | 0 | 0 | 0 | 0 | 0 | 0 |
| virus_Aug5.50v_32846 | 1 | 0 | 0 | 0 | 0 | 0 | 0 | 0 | 0 | 0 | 0 | 0 | 0 | 0 | 0 | 0 | 0 | 0 | 0 | 0 | 0 | 0 | 0 | 0 | 0 | 0 | 0 | 0 | 0 | 0 | 0 | 0 | 0 | 0 | 0 | 0 | 0 | 0 | 0 | 0 | 0 | 0 | 0 | 0 | 1 | 0 | 0 | 0 | 0 | 0 | 0 | 0 | 0 | 0 | 0 |
| virus_Aug5.50v_39142 | 1 | 0 | 0 | 0 | 0 | 0 | 0 | 0 | 0 | 0 | 0 | 0 | 0 | 0 | 0 | 0 | 0 | 0 | 0 | 0 | 0 | 0 | 0 | 0 | 0 | 0 | 0 | 0 | 0 | 0 | 0 | 0 | 0 | 0 | 0 | 0 | 0 | 0 | 0 | 0 | 0 | 0 | 0 | 0 | 1 | 0 | 0 | 0 | 0 | 0 | 0 | 0 | 0 | 0 | 0 |
| virus_Aug5.75v_499 | 1 | 0 | 0 | 0 | 0 | 0 | 0 | 0 | 0 | 0 | 0 | 0 | 0 | 0 | 0 | 0 | 0 | 0 | 0 | 0 | 0 | 0 | 0 | 0 | 0 | 0 | 0 | 0 | 0 | 0 | 0 | 0 | 0 | 0 | 0 | 0 | 0 | 0 | 0 | 0 | 0 | 0 | 0 | 0 | 0 | 0 | 1 | 0 | 0 | 0 | 0 | 0 | 0 | 0 | 0 |
| virus_Aug5.75v_2997 | 1 | 0 | 0 | 0 | 0 | 0 | 0 | 0 | 0 | 0 | 0 | 0 | 0 | 0 | 0 | 0 | 0 | 0 | 0 | 0 | 0 | 0 | 0 | 0 | 0 | 0 | 0 | 0 | 0 | 0 | 0 | 0 | 0 | 0 | 0 | 0 | 0 | 0 | 0 | 0 | 0 | 0 | 0 | 0 | 0 | 0 | 1 | 0 | 0 | 0 | 0 | 0 | 0 | 0 | 0 |
| virus_Aug5.75v_11712 | 1 | 0 | 0 | 0 | 0 | 0 | 0 | 0 | 0 | 0 | 0 | 0 | 0 | 0 | 0 | 0 | 0 | 0 | 0 | 0 | 0 | 0 | 0 | 0 | 0 | 0 | 0 | 0 | 0 | 0 | 0 | 0 | 0 | 0 | 0 | 0 | 0 | 0 | 0 | 0 | 0 | 0 | 0 | 0 | 0 | 0 | 1 | 0 | 0 | 0 | 0 | 0 | 0 | 0 | 0 |
| virus_Aug5.75v_16093 | 1 | 0 | 0 | 0 | 0 | 0 | 0 | 0 | 0 | 0 | 0 | 0 | 0 | 0 | 0 | 0 | 0 | 0 | 0 | 0 | 0 | 0 | 0 | 0 | 0 | 0 | 0 | 0 | 0 | 0 | 0 | 0 | 0 | 0 | 0 | 0 | 0 | 0 | 0 | 0 | 0 | 0 | 0 | 0 | 0 | 0 | 1 | 0 | 0 | 0 | 0 | 0 | 0 | 0 | 0 |
| virus_Aug5.75v_19284 | 1 | 0 | 0 | 0 | 0 | 0 | 0 | 0 | 0 | 0 | 0 | 0 | 0 | 0 | 0 | 0 | 0 | 0 | 0 | 0 | 0 | 0 | 0 | 0 | 0 | 0 | 0 | 0 | 0 | 0 | 0 | 0 | 0 | 0 | 0 | 0 | 0 | 0 | 0 | 0 | 0 | 0 | 0 | 0 | 0 | 0 | 1 | 0 | 0 | 0 | 0 | 0 | 0 | 0 | 0 |
| virus_Aug5.75v_26254 | 1 | 0 | 0 | 0 | 0 | 0 | 0 | 0 | 0 | 0 | 0 | 0 | 0 | 0 | 0 | 0 | 0 | 0 | 0 | 0 | 0 | 0 | 0 | 0 | 0 | 0 | 0 | 0 | 0 | 0 | 0 | 0 | 0 | 0 | 0 | 0 | 0 | 0 | 0 | 0 | 0 | 0 | 0 | 0 | 0 | 0 | 1 | 0 | 0 | 0 | 0 | 0 | 0 | 0 | 0 |
| virus_Aug5.75v_26509 | 1 | 0 | 0 | 0 | 0 | 0 | 0 | 0 | 0 | 0 | 0 | 0 | 0 | 0 | 0 | 0 | 0 | 0 | 0 | 0 | 0 | 0 | 0 | 0 | 0 | 0 | 0 | 0 | 0 | 0 | 0 | 0 | 0 | 0 | 0 | 0 | 0 | 0 | 0 | 0 | 0 | 0 | 0 | 0 | 0 | 0 | 1 | 0 | 0 | 0 | 0 | 0 | 0 | 0 | 0 |
| virus_Aug5.75v_26937 | 1 | 0 | 0 | 0 | 0 | 0 | 0 | 0 | 0 | 0 | 0 | 0 | 0 | 0 | 0 | 0 | 0 | 0 | 0 | 0 | 0 | 0 | 0 | 0 | 0 | 0 | 0 | 0 | 0 | 0 | 0 | 0 | 0 | 0 | 0 | 0 | 0 | 0 | 0 | 0 | 0 | 0 | 0 | 0 | 0 | 0 | 1 | 0 | 0 | 0 | 0 | 0 | 0 | 0 | 0 |
| virus_Aug5.75v_40920 | 1 | 0 | 0 | 0 | 0 | 0 | 0 | 0 | 0 | 0 | 0 | 0 | 0 | 0 | 0 | 0 | 0 | 0 | 0 | 0 | 0 | 0 | 0 | 0 | 0 | 0 | 0 | 0 | 0 | 0 | 0 | 0 | 0 | 0 | 0 | 0 | 0 | 0 | 0 | 0 | 0 | 0 | 0 | 0 | 0 | 0 | 1 | 0 | 0 | 0 | 0 | 0 | 0 | 0 | 0 |
| virus_Jul11.0v_583 | 1 | 0 | 0 | 0 | 0 | 0 | 0 | 0 | 0 | 0 | 0 | 0 | 0 | 0 | 0 | 0 | 0 | 0 | 0 | 0 | 0 | 0 | 1 | 0 | 0 | 0 | 0 | 0 | 0 | 0 | 0 | 0 | 0 | 0 | 0 | 0 | 0 | 0 | 0 | 0 | 0 | 0 | 0 | 0 | 0 | 0 | 0 | 0 | 0 | 0 | 0 | 0 | 0 | 0 | 0 |
| virus_Jul11.0v_2963 | 1 | 0 | 0 | 0 | 0 | 0 | 0 | 0 | 0 | 0 | 0 | 0 | 0 | 0 | 0 | 0 | 0 | 0 | 0 | 0 | 0 | 0 | 1 | 0 | 0 | 0 | 0 | 0 | 0 | 0 | 0 | 0 | 0 | 0 | 0 | 0 | 0 | 0 | 0 | 0 | 0 | 0 | 0 | 0 | 0 | 0 | 0 | 0 | 0 | 0 | 0 | 0 | 0 | 0 | 0 |
| virus_Jul11.0v_5945 | 1 | 0 | 0 | 0 | 0 | 0 | 0 | 0 | 0 | 0 | 0 | 0 | 0 | 0 | 0 | 0 | 0 | 0 | 0 | 0 | 0 | 0 | 1 | 0 | 0 | 0 | 0 | 0 | 0 | 0 | 0 | 0 | 0 | 0 | 0 | 0 | 0 | 0 | 0 | 0 | 0 | 0 | 0 | 0 | 0 | 0 | 0 | 0 | 0 | 0 | 0 | 0 | 0 | 0 | 0 |
| virus_Jul11.0v_7008 | 1 | 0 | 0 | 0 | 0 | 0 | 0 | 0 | 0 | 0 | 0 | 0 | 0 | 0 | 0 | 0 | 0 | 0 | 0 | 0 | 0 | 0 | 1 | 0 | 0 | 0 | 0 | 0 | 0 | 0 | 0 | 0 | 0 | 0 | 0 | 0 | 0 | 0 | 0 | 0 | 0 | 0 | 0 | 0 | 0 | 0 | 0 | 0 | 0 | 0 | 0 | 0 | 0 | 0 | 0 |
| virus_Jul11.0v_8246 | 1 | 0 | 0 | 0 | 0 | 0 | 0 | 0 | 0 | 0 | 0 | 0 | 0 | 0 | 0 | 0 | 0 | 0 | 0 | 0 | 0 | 0 | 1 | 0 | 0 | 0 | 0 | 0 | 0 | 0 | 0 | 0 | 0 | 0 | 0 | 0 | 0 | 0 | 0 | 0 | 0 | 0 | 0 | 0 | 0 | 0 | 0 | 0 | 0 | 0 | 0 | 0 | 0 | 0 | 0 |
| virus_Jul11.0v_9478 | 1 | 0 | 0 | 0 | 0 | 0 | 0 | 0 | 0 | 0 | 0 | 0 | 0 | 0 | 0 | 0 | 0 | 0 | 0 | 0 | 0 | 0 | 1 | 0 | 0 | 0 | 0 | 0 | 0 | 0 | 0 | 0 | 0 | 0 | 0 | 0 | 0 | 0 | 0 | 0 | 0 | 0 | 0 | 0 | 0 | 0 | 0 | 0 | 0 | 0 | 0 | 0 | 0 | 0 | 0 |
| virus_Jul11.0v_9971 | 1 | 0 | 0 | 0 | 0 | 0 | 0 | 0 | 0 | 0 | 0 | 0 | 0 | 0 | 0 | 0 | 0 | 0 | 0 | 0 | 0 | 0 | 1 | 0 | 0 | 0 | 0 | 0 | 0 | 0 | 0 | 0 | 0 | 0 | 0 | 0 | 0 | 0 | 0 | 0 | 0 | 0 | 0 | 0 | 0 | 0 | 0 | 0 | 0 | 0 | 0 | 0 | 0 | 0 | 0 |
| virus_Jul11.0v_10217 | 1 | 0 | 0 | 0 | 0 | 0 | 0 | 0 | 0 | 0 | 0 | 0 | 0 | 0 | 0 | 0 | 0 | 0 | 0 | 0 | 0 | 0 | 1 | 0 | 0 | 0 | 0 | 0 | 0 | 0 | 0 | 0 | 0 | 0 | 0 | 0 | 0 | 0 | 0 | 0 | 0 | 0 | 0 | 0 | 0 | 0 | 0 | 0 | 0 | 0 | 0 | 0 | 0 | 0 | 0 |
| virus_Jul11.0v_13147 | 1 | 0 | 0 | 0 | 0 | 0 | 0 | 0 | 0 | 0 | 0 | 0 | 0 | 0 | 0 | 0 | 0 | 0 | 0 | 0 | 0 | 0 | 1 | 0 | 0 | 0 | 0 | 0 | 0 | 0 | 0 | 0 | 0 | 0 | 0 | 0 | 0 | 0 | 0 | 0 | 0 | 0 | 0 | 0 | 0 | 0 | 0 | 0 | 0 | 0 | 0 | 0 | 0 | 0 | 0 |
| virus_Jul11.0v_17969 | 1 | 0 | 0 | 0 | 0 | 0 | 0 | 0 | 0 | 0 | 0 | 0 | 0 | 0 | 0 | 0 | 0 | 0 | 0 | 0 | 0 | 0 | 1 | 0 | 0 | 0 | 0 | 0 | 0 | 0 | 0 | 0 | 0 | 0 | 0 | 0 | 0 | 0 | 0 | 0 | 0 | 0 | 0 | 0 | 0 | 0 | 0 | 0 | 0 | 0 | 0 | 0 | 0 | 0 | 0 |
| virus_Jul11.5v_412 | 1 | 0 | 0 | 0 | 0 | 0 | 0 | 0 | 0 | 0 | 0 | 0 | 0 | 0 | 0 | 0 | 0 | 0 | 0 | 0 | 0 | 0 | 0 | 0 | 1 | 0 | 0 | 0 | 0 | 0 | 0 | 0 | 0 | 0 | 0 | 0 | 0 | 0 | 0 | 0 | 0 | 0 | 0 | 0 | 0 | 0 | 0 | 0 | 0 | 0 | 0 | 0 | 0 | 0 | 0 |
| virus_Jul11.5v_3527 | 1 | 0 | 0 | 0 | 0 | 0 | 0 | 0 | 0 | 0 | 0 | 0 | 0 | 0 | 0 | 0 | 0 | 0 | 0 | 0 | 0 | 0 | 0 | 0 | 1 | 0 | 0 | 0 | 0 | 0 | 0 | 0 | 0 | 0 | 0 | 0 | 0 | 0 | 0 | 0 | 0 | 0 | 0 | 0 | 0 | 0 | 0 | 0 | 0 | 0 | 0 | 0 | 0 | 0 | 0 |
| virus_Jul11.5v_4015 | 1 | 0 | 0 | 0 | 0 | 0 | 0 | 0 | 0 | 0 | 0 | 0 | 0 | 0 | 0 | 0 | 0 | 0 | 0 | 0 | 0 | 0 | 0 | 0 | 1 | 0 | 0 | 0 | 0 | 0 | 0 | 0 | 0 | 0 | 0 | 0 | 0 | 0 | 0 | 0 | 0 | 0 | 0 | 0 | 0 | 0 | 0 | 0 | 0 | 0 | 0 | 0 | 0 | 0 | 0 |
| virus_Jul11.5v_4145 | 1 | 0 | 0 | 0 | 0 | 0 | 0 | 0 | 0 | 0 | 0 | 0 | 0 | 0 | 0 | 0 | 0 | 0 | 0 | 0 | 0 | 0 | 0 | 0 | 1 | 0 | 0 | 0 | 0 | 0 | 0 | 0 | 0 | 0 | 0 | 0 | 0 | 0 | 0 | 0 | 0 | 0 | 0 | 0 | 0 | 0 | 0 | 0 | 0 | 0 | 0 | 0 | 0 | 0 | 0 |
| virus_Jul11.5v_9420 | 1 | 0 | 0 | 0 | 0 | 0 | 0 | 0 | 0 | 0 | 0 | 0 | 0 | 0 | 0 | 0 | 0 | 0 | 0 | 0 | 0 | 0 | 0 | 0 | 1 | 0 | 0 | 0 | 0 | 0 | 0 | 0 | 0 | 0 | 0 | 0 | 0 | 0 | 0 | 0 | 0 | 0 | 0 | 0 | 0 | 0 | 0 | 0 | 0 | 0 | 0 | 0 | 0 | 0 | 0 |
| virus_Jul11.5v_11138 | 1 | 0 | 0 | 0 | 0 | 0 | 0 | 0 | 0 | 0 | 0 | 0 | 0 | 0 | 0 | 0 | 0 | 0 | 0 | 0 | 0 | 0 | 0 | 0 | 1 | 0 | 0 | 0 | 0 | 0 | 0 | 0 | 0 | 0 | 0 | 0 | 0 | 0 | 0 | 0 | 0 | 0 | 0 | 0 | 0 | 0 | 0 | 0 | 0 | 0 | 0 | 0 | 0 | 0 | 0 |
| virus_Jul11.5v_14679 | 1 | 0 | 0 | 0 | 0 | 0 | 0 | 0 | 0 | 0 | 0 | 0 | 0 | 0 | 0 | 0 | 0 | 0 | 0 | 0 | 0 | 0 | 0 | 0 | 1 | 0 | 0 | 0 | 0 | 0 | 0 | 0 | 0 | 0 | 0 | 0 | 0 | 0 | 0 | 0 | 0 | 0 | 0 | 0 | 0 | 0 | 0 | 0 | 0 | 0 | 0 | 0 | 0 | 0 | 0 |
| virus_Jul11.5v_20285 | 1 | 0 | 0 | 0 | 0 | 0 | 0 | 0 | 0 | 0 | 0 | 0 | 0 | 0 | 0 | 0 | 0 | 0 | 0 | 0 | 0 | 0 | 0 | 0 | 1 | 0 | 0 | 0 | 0 | 0 | 0 | 0 | 0 | 0 | 0 | 0 | 0 | 0 | 0 | 0 | 0 | 0 | 0 | 0 | 0 | 0 | 0 | 0 | 0 | 0 | 0 | 0 | 0 | 0 | 0 |
| virus_Jul11.25v_89 | 1 | 0 | 0 | 0 | 0 | 0 | 0 | 0 | 0 | 0 | 0 | 0 | 0 | 0 | 0 | 0 | 0 | 0 | 0 | 0 | 0 | 0 | 0 | 0 | 0 | 0 | 1 | 0 | 0 | 0 | 0 | 0 | 0 | 0 | 0 | 0 | 0 | 0 | 0 | 0 | 0 | 0 | 0 | 0 | 0 | 0 | 0 | 0 | 0 | 0 | 0 | 0 | 0 | 0 | 0 |
| virus_Jul11.25v_670 | 1 | 0 | 0 | 0 | 0 | 0 | 0 | 0 | 0 | 0 | 0 | 0 | 0 | 0 | 0 | 0 | 0 | 0 | 0 | 0 | 0 | 0 | 0 | 0 | 0 | 0 | 1 | 0 | 0 | 0 | 0 | 0 | 0 | 0 | 0 | 0 | 0 | 0 | 0 | 0 | 0 | 0 | 0 | 0 | 0 | 0 | 0 | 0 | 0 | 0 | 0 | 0 | 0 | 0 | 0 |
| virus_Jul11.25v_1359 | 1 | 0 | 0 | 0 | 0 | 0 | 0 | 0 | 0 | 0 | 0 | 0 | 0 | 0 | 0 | 0 | 0 | 0 | 0 | 0 | 0 | 0 | 0 | 0 | 0 | 0 | 1 | 0 | 0 | 0 | 0 | 0 | 0 | 0 | 0 | 0 | 0 | 0 | 0 | 0 | 0 | 0 | 0 | 0 | 0 | 0 | 0 | 0 | 0 | 0 | 0 | 0 | 0 | 0 | 0 |
| virus_Jul11.25v_2817 | 1 | 0 | 0 | 0 | 0 | 0 | 0 | 0 | 0 | 0 | 0 | 0 | 0 | 0 | 0 | 0 | 0 | 0 | 0 | 0 | 0 | 0 | 0 | 0 | 0 | 0 | 1 | 0 | 0 | 0 | 0 | 0 | 0 | 0 | 0 | 0 | 0 | 0 | 0 | 0 | 0 | 0 | 0 | 0 | 0 | 0 | 0 | 0 | 0 | 0 | 0 | 0 | 0 | 0 | 0 |
| virus_Jul11.25v_8754 | 1 | 0 | 0 | 0 | 0 | 0 | 0 | 0 | 0 | 0 | 0 | 0 | 0 | 0 | 0 | 0 | 0 | 0 | 0 | 0 | 0 | 0 | 0 | 0 | 0 | 0 | 1 | 0 | 0 | 0 | 0 | 0 | 0 | 0 | 0 | 0 | 0 | 0 | 0 | 0 | 0 | 0 | 0 | 0 | 0 | 0 | 0 | 0 | 0 | 0 | 0 | 0 | 0 | 0 | 0 |
| virus_Jul11.25v_10130 | 1 | 0 | 0 | 0 | 0 | 0 | 0 | 0 | 0 | 0 | 0 | 0 | 0 | 0 | 0 | 0 | 0 | 0 | 0 | 0 | 0 | 0 | 0 | 0 | 0 | 0 | 1 | 0 | 0 | 0 | 0 | 0 | 0 | 0 | 0 | 0 | 0 | 0 | 0 | 0 | 0 | 0 | 0 | 0 | 0 | 0 | 0 | 0 | 0 | 0 | 0 | 0 | 0 | 0 | 0 |
| virus_Jul11.25v_12389 | 1 | 0 | 0 | 0 | 0 | 0 | 0 | 0 | 0 | 0 | 0 | 0 | 0 | 0 | 0 | 0 | 0 | 0 | 0 | 0 | 0 | 0 | 0 | 0 | 0 | 0 | 1 | 0 | 0 | 0 | 0 | 0 | 0 | 0 | 0 | 0 | 0 | 0 | 0 | 0 | 0 | 0 | 0 | 0 | 0 | 0 | 0 | 0 | 0 | 0 | 0 | 0 | 0 | 0 | 0 |
| virus_Jul11.25v_14332 | 1 | 0 | 0 | 0 | 0 | 0 | 0 | 0 | 0 | 0 | 0 | 0 | 0 | 0 | 0 | 0 | 0 | 0 | 0 | 0 | 0 | 0 | 0 | 0 | 0 | 0 | 1 | 0 | 0 | 0 | 0 | 0 | 0 | 0 | 0 | 0 | 0 | 0 | 0 | 0 | 0 | 0 | 0 | 0 | 0 | 0 | 0 | 0 | 0 | 0 | 0 | 0 | 0 | 0 | 0 |
| virus_Jul11.25v_14388 | 1 | 0 | 0 | 0 | 0 | 0 | 0 | 0 | 0 | 0 | 0 | 0 | 0 | 0 | 0 | 0 | 0 | 0 | 0 | 0 | 0 | 0 | 0 | 0 | 0 | 0 | 1 | 0 | 0 | 0 | 0 | 0 | 0 | 0 | 0 | 0 | 0 | 0 | 0 | 0 | 0 | 0 | 0 | 0 | 0 | 0 | 0 | 0 | 0 | 0 | 0 | 0 | 0 | 0 | 0 |
| virus_Jul11.25v_17868 | 1 | 0 | 0 | 0 | 0 | 0 | 0 | 0 | 0 | 0 | 0 | 0 | 0 | 0 | 0 | 0 | 0 | 0 | 0 | 0 | 0 | 0 | 0 | 0 | 0 | 0 | 1 | 0 | 0 | 0 | 0 | 0 | 0 | 0 | 0 | 0 | 0 | 0 | 0 | 0 | 0 | 0 | 0 | 0 | 0 | 0 | 0 | 0 | 0 | 0 | 0 | 0 | 0 | 0 | 0 |
| virus_Jul11.50v_1076 | 1 | 0 | 0 | 0 | 0 | 0 | 0 | 0 | 0 | 0 | 0 | 0 | 0 | 0 | 0 | 0 | 0 | 0 | 0 | 0 | 0 | 0 | 0 | 0 | 0 | 0 | 0 | 0 | 1 | 0 | 0 | 0 | 0 | 0 | 0 | 0 | 0 | 0 | 0 | 0 | 0 | 0 | 0 | 0 | 0 | 0 | 0 | 0 | 0 | 0 | 0 | 0 | 0 | 0 | 0 |
| virus_Jul11.50v_1938 | 1 | 0 | 0 | 0 | 0 | 0 | 0 | 0 | 0 | 0 | 0 | 0 | 0 | 0 | 0 | 0 | 0 | 0 | 0 | 0 | 0 | 0 | 0 | 0 | 0 | 0 | 0 | 0 | 1 | 0 | 0 | 0 | 0 | 0 | 0 | 0 | 0 | 0 | 0 | 0 | 0 | 0 | 0 | 0 | 0 | 0 | 0 | 0 | 0 | 0 | 0 | 0 | 0 | 0 | 0 |
| virus_Jul11.50v_3146 | 1 | 0 | 0 | 0 | 0 | 0 | 0 | 0 | 0 | 0 | 0 | 0 | 0 | 0 | 0 | 0 | 0 | 0 | 0 | 0 | 0 | 0 | 0 | 0 | 0 | 0 | 0 | 0 | 1 | 0 | 0 | 0 | 0 | 0 | 0 | 0 | 0 | 0 | 0 | 0 | 0 | 0 | 0 | 0 | 0 | 0 | 0 | 0 | 0 | 0 | 0 | 0 | 0 | 0 | 0 |
| virus_Jul11.50v_7126 | 1 | 0 | 0 | 0 | 0 | 0 | 0 | 0 | 0 | 0 | 0 | 0 | 0 | 0 | 0 | 0 | 0 | 0 | 0 | 0 | 0 | 0 | 0 | 0 | 0 | 0 | 0 | 0 | 1 | 0 | 0 | 0 | 0 | 0 | 0 | 0 | 0 | 0 | 0 | 0 | 0 | 0 | 0 | 0 | 0 | 0 | 0 | 0 | 0 | 0 | 0 | 0 | 0 | 0 | 0 |
| virus_Jul11.50v_8009 | 1 | 0 | 0 | 0 | 0 | 0 | 0 | 0 | 0 | 0 | 0 | 0 | 0 | 0 | 0 | 0 | 0 | 0 | 0 | 0 | 0 | 0 | 0 | 0 | 0 | 0 | 0 | 0 | 1 | 0 | 0 | 0 | 0 | 0 | 0 | 0 | 0 | 0 | 0 | 0 | 0 | 0 | 0 | 0 | 0 | 0 | 0 | 0 | 0 | 0 | 0 | 0 | 0 | 0 | 0 |
| virus_Jul11.50v_8700 | 1 | 0 | 0 | 0 | 0 | 0 | 0 | 0 | 0 | 0 | 0 | 0 | 0 | 0 | 0 | 0 | 0 | 0 | 0 | 0 | 0 | 0 | 0 | 0 | 0 | 0 | 0 | 0 | 1 | 0 | 0 | 0 | 0 | 0 | 0 | 0 | 0 | 0 | 0 | 0 | 0 | 0 | 0 | 0 | 0 | 0 | 0 | 0 | 0 | 0 | 0 | 0 | 0 | 0 | 0 |
| virus_Jul11.50v_11036 | 1 | 0 | 0 | 0 | 0 | 0 | 0 | 0 | 0 | 0 | 0 | 0 | 0 | 0 | 0 | 0 | 0 | 0 | 0 | 0 | 0 | 0 | 0 | 0 | 0 | 0 | 0 | 0 | 1 | 0 | 0 | 0 | 0 | 0 | 0 | 0 | 0 | 0 | 0 | 0 | 0 | 0 | 0 | 0 | 0 | 0 | 0 | 0 | 0 | 0 | 0 | 0 | 0 | 0 | 0 |
| virus_Jul11.50v_11517 | 1 | 0 | 0 | 0 | 0 | 0 | 0 | 0 | 0 | 0 | 0 | 0 | 0 | 0 | 0 | 0 | 0 | 0 | 0 | 0 | 0 | 0 | 0 | 0 | 0 | 0 | 0 | 0 | 1 | 0 | 0 | 0 | 0 | 0 | 0 | 0 | 0 | 0 | 0 | 0 | 0 | 0 | 0 | 0 | 0 | 0 | 0 | 0 | 0 | 0 | 0 | 0 | 0 | 0 | 0 |
| virus_Jul11.50v_18833 | 1 | 0 | 0 | 0 | 0 | 0 | 0 | 0 | 0 | 0 | 0 | 0 | 0 | 0 | 0 | 0 | 0 | 0 | 0 | 0 | 0 | 0 | 0 | 0 | 0 | 0 | 0 | 0 | 1 | 0 | 0 | 0 | 0 | 0 | 0 | 0 | 0 | 0 | 0 | 0 | 0 | 0 | 0 | 0 | 0 | 0 | 0 | 0 | 0 | 0 | 0 | 0 | 0 | 0 | 0 |
| virus_Jul11.50v_20177 | 1 | 0 | 0 | 0 | 0 | 0 | 0 | 0 | 0 | 0 | 0 | 0 | 0 | 0 | 0 | 0 | 0 | 0 | 0 | 0 | 0 | 0 | 0 | 0 | 0 | 0 | 0 | 0 | 1 | 0 | 0 | 0 | 0 | 0 | 0 | 0 | 0 | 0 | 0 | 0 | 0 | 0 | 0 | 0 | 0 | 0 | 0 | 0 | 0 | 0 | 0 | 0 | 0 | 0 | 0 |
| virus_Jul11.50v_20537 | 1 | 0 | 0 | 0 | 0 | 0 | 0 | 0 | 0 | 0 | 0 | 0 | 0 | 0 | 0 | 0 | 0 | 0 | 0 | 0 | 0 | 0 | 0 | 0 | 0 | 0 | 0 | 0 | 1 | 0 | 0 | 0 | 0 | 0 | 0 | 0 | 0 | 0 | 0 | 0 | 0 | 0 | 0 | 0 | 0 | 0 | 0 | 0 | 0 | 0 | 0 | 0 | 0 | 0 | 0 |
| virus_Jul11.75v_3327 | 1 | 0 | 0 | 0 | 0 | 0 | 0 | 0 | 0 | 0 | 0 | 0 | 0 | 0 | 0 | 0 | 0 | 0 | 0 | 0 | 0 | 0 | 0 | 0 | 0 | 0 | 0 | 0 | 0 | 0 | 1 | 0 | 0 | 0 | 0 | 0 | 0 | 0 | 0 | 0 | 0 | 0 | 0 | 0 | 0 | 0 | 0 | 0 | 0 | 0 | 0 | 0 | 0 | 0 | 0 |
| virus_Jul11.75v_6584 | 1 | 0 | 0 | 0 | 0 | 0 | 0 | 0 | 0 | 0 | 0 | 0 | 0 | 0 | 0 | 0 | 0 | 0 | 0 | 0 | 0 | 0 | 0 | 0 | 0 | 0 | 0 | 0 | 0 | 0 | 1 | 0 | 0 | 0 | 0 | 0 | 0 | 0 | 0 | 0 | 0 | 0 | 0 | 0 | 0 | 0 | 0 | 0 | 0 | 0 | 0 | 0 | 0 | 0 | 0 |
| virus_Jul11.75v_11755 | 1 | 0 | 0 | 0 | 0 | 0 | 0 | 0 | 0 | 0 | 0 | 0 | 0 | 0 | 0 | 0 | 0 | 0 | 0 | 0 | 0 | 0 | 0 | 0 | 0 | 0 | 0 | 0 | 0 | 0 | 1 | 0 | 0 | 0 | 0 | 0 | 0 | 0 | 0 | 0 | 0 | 0 | 0 | 0 | 0 | 0 | 0 | 0 | 0 | 0 | 0 | 0 | 0 | 0 | 0 |
| virus_Jul11.75v_20801 | 1 | 0 | 0 | 0 | 0 | 0 | 0 | 0 | 0 | 0 | 0 | 0 | 0 | 0 | 0 | 0 | 0 | 0 | 0 | 0 | 0 | 0 | 0 | 0 | 0 | 0 | 0 | 0 | 0 | 0 | 1 | 0 | 0 | 0 | 0 | 0 | 0 | 0 | 0 | 0 | 0 | 0 | 0 | 0 | 0 | 0 | 0 | 0 | 0 | 0 | 0 | 0 | 0 | 0 | 0 |
| virus_Jul11.75v_24996 | 1 | 0 | 0 | 0 | 0 | 0 | 0 | 0 | 0 | 0 | 0 | 0 | 0 | 0 | 0 | 0 | 0 | 0 | 0 | 0 | 0 | 0 | 0 | 0 | 0 | 0 | 0 | 0 | 0 | 0 | 1 | 0 | 0 | 0 | 0 | 0 | 0 | 0 | 0 | 0 | 0 | 0 | 0 | 0 | 0 | 0 | 0 | 0 | 0 | 0 | 0 | 0 | 0 | 0 | 0 |
| virus_Jul11.75v_29943 | 1 | 0 | 0 | 0 | 0 | 0 | 0 | 0 | 0 | 0 | 0 | 0 | 0 | 0 | 0 | 0 | 0 | 0 | 0 | 0 | 0 | 0 | 0 | 0 | 0 | 0 | 0 | 0 | 0 | 0 | 1 | 0 | 0 | 0 | 0 | 0 | 0 | 0 | 0 | 0 | 0 | 0 | 0 | 0 | 0 | 0 | 0 | 0 | 0 | 0 | 0 | 0 | 0 | 0 | 0 |
| virus_Jul11.75v_35859 | 1 | 0 | 0 | 0 | 0 | 0 | 0 | 0 | 0 | 0 | 0 | 0 | 0 | 0 | 0 | 0 | 0 | 0 | 0 | 0 | 0 | 0 | 0 | 0 | 0 | 0 | 0 | 0 | 0 | 0 | 1 | 0 | 0 | 0 | 0 | 0 | 0 | 0 | 0 | 0 | 0 | 0 | 0 | 0 | 0 | 0 | 0 | 0 | 0 | 0 | 0 | 0 | 0 | 0 | 0 |
| virus_Jul27.5v_1048 | 1 | 0 | 0 | 0 | 0 | 0 | 0 | 0 | 0 | 0 | 0 | 0 | 0 | 0 | 0 | 0 | 0 | 0 | 0 | 0 | 0 | 0 | 0 | 0 | 0 | 0 | 0 | 0 | 0 | 0 | 0 | 0 | 1 | 0 | 0 | 0 | 0 | 0 | 0 | 0 | 0 | 0 | 0 | 0 | 0 | 0 | 0 | 0 | 0 | 0 | 0 | 0 | 0 | 0 | 0 |
| virus_Jul27.5v_1642 | 1 | 0 | 0 | 0 | 0 | 0 | 0 | 0 | 0 | 0 | 0 | 0 | 0 | 0 | 0 | 0 | 0 | 0 | 0 | 0 | 0 | 0 | 0 | 0 | 0 | 0 | 0 | 0 | 0 | 0 | 0 | 0 | 1 | 0 | 0 | 0 | 0 | 0 | 0 | 0 | 0 | 0 | 0 | 0 | 0 | 0 | 0 | 0 | 0 | 0 | 0 | 0 | 0 | 0 | 0 |
| virus_Jul27.5v_2167 | 1 | 0 | 0 | 0 | 0 | 0 | 0 | 0 | 0 | 0 | 0 | 0 | 0 | 0 | 0 | 0 | 0 | 0 | 0 | 0 | 0 | 0 | 0 | 0 | 0 | 0 | 0 | 0 | 0 | 0 | 0 | 0 | 1 | 0 | 0 | 0 | 0 | 0 | 0 | 0 | 0 | 0 | 0 | 0 | 0 | 0 | 0 | 0 | 0 | 0 | 0 | 0 | 0 | 0 | 0 |
| virus_Jul27.5v_2735 | 1 | 0 | 0 | 0 | 0 | 0 | 0 | 0 | 0 | 0 | 0 | 0 | 0 | 0 | 0 | 0 | 0 | 0 | 0 | 0 | 0 | 0 | 0 | 0 | 0 | 0 | 0 | 0 | 0 | 0 | 0 | 0 | 1 | 0 | 0 | 0 | 0 | 0 | 0 | 0 | 0 | 0 | 0 | 0 | 0 | 0 | 0 | 0 | 0 | 0 | 0 | 0 | 0 | 0 | 0 |
| virus_Jul27.5v_4455 | 1 | 0 | 0 | 0 | 0 | 0 | 0 | 0 | 0 | 0 | 0 | 0 | 0 | 0 | 0 | 0 | 0 | 0 | 0 | 0 | 0 | 0 | 0 | 0 | 0 | 0 | 0 | 0 | 0 | 0 | 0 | 0 | 1 | 0 | 0 | 0 | 0 | 0 | 0 | 0 | 0 | 0 | 0 | 0 | 0 | 0 | 0 | 0 | 0 | 0 | 0 | 0 | 0 | 0 | 0 |
| virus_Jul27.5v_7648 | 1 | 0 | 0 | 0 | 0 | 0 | 0 | 0 | 0 | 0 | 0 | 0 | 0 | 0 | 0 | 0 | 0 | 0 | 0 | 0 | 0 | 0 | 0 | 0 | 0 | 0 | 0 | 0 | 0 | 0 | 0 | 0 | 1 | 0 | 0 | 0 | 0 | 0 | 0 | 0 | 0 | 0 | 0 | 0 | 0 | 0 | 0 | 0 | 0 | 0 | 0 | 0 | 0 | 0 | 0 |
| virus_Jul27.5v_8002 | 1 | 0 | 0 | 0 | 0 | 0 | 0 | 0 | 0 | 0 | 0 | 0 | 0 | 0 | 0 | 0 | 0 | 0 | 0 | 0 | 0 | 0 | 0 | 0 | 0 | 0 | 0 | 0 | 0 | 0 | 0 | 0 | 1 | 0 | 0 | 0 | 0 | 0 | 0 | 0 | 0 | 0 | 0 | 0 | 0 | 0 | 0 | 0 | 0 | 0 | 0 | 0 | 0 | 0 | 0 |
| virus_Jul27.5v_9780 | 1 | 0 | 0 | 0 | 0 | 0 | 0 | 0 | 0 | 0 | 0 | 0 | 0 | 0 | 0 | 0 | 0 | 0 | 0 | 0 | 0 | 0 | 0 | 0 | 0 | 0 | 0 | 0 | 0 | 0 | 0 | 0 | 1 | 0 | 0 | 0 | 0 | 0 | 0 | 0 | 0 | 0 | 0 | 0 | 0 | 0 | 0 | 0 | 0 | 0 | 0 | 0 | 0 | 0 | 0 |
| virus_Jul27.5v_12543 | 1 | 0 | 0 | 0 | 0 | 0 | 0 | 0 | 0 | 0 | 0 | 0 | 0 | 0 | 0 | 0 | 0 | 0 | 0 | 0 | 0 | 0 | 0 | 0 | 0 | 0 | 0 | 0 | 0 | 0 | 0 | 0 | 1 | 0 | 0 | 0 | 0 | 0 | 0 | 0 | 0 | 0 | 0 | 0 | 0 | 0 | 0 | 0 | 0 | 0 | 0 | 0 | 0 | 0 | 0 |
| virus_Jul27.5v_13117 | 1 | 0 | 0 | 0 | 0 | 0 | 0 | 0 | 0 | 0 | 0 | 0 | 0 | 0 | 0 | 0 | 0 | 0 | 0 | 0 | 0 | 0 | 0 | 0 | 0 | 0 | 0 | 0 | 0 | 0 | 0 | 0 | 1 | 0 | 0 | 0 | 0 | 0 | 0 | 0 | 0 | 0 | 0 | 0 | 0 | 0 | 0 | 0 | 0 | 0 | 0 | 0 | 0 | 0 | 0 |
| virus_Jul27.5v_13619 | 1 | 0 | 0 | 0 | 0 | 0 | 0 | 0 | 0 | 0 | 0 | 0 | 0 | 0 | 0 | 0 | 0 | 0 | 0 | 0 | 0 | 0 | 0 | 0 | 0 | 0 | 0 | 0 | 0 | 0 | 0 | 0 | 1 | 0 | 0 | 0 | 0 | 0 | 0 | 0 | 0 | 0 | 0 | 0 | 0 | 0 | 0 | 0 | 0 | 0 | 0 | 0 | 0 | 0 | 0 |
| virus_Jul27.5v_14056 | 1 | 0 | 0 | 0 | 0 | 0 | 0 | 0 | 0 | 0 | 0 | 0 | 0 | 0 | 0 | 0 | 0 | 0 | 0 | 0 | 0 | 0 | 0 | 0 | 0 | 0 | 0 | 0 | 0 | 0 | 0 | 0 | 1 | 0 | 0 | 0 | 0 | 0 | 0 | 0 | 0 | 0 | 0 | 0 | 0 | 0 | 0 | 0 | 0 | 0 | 0 | 0 | 0 | 0 | 0 |
| virus_Jul27.5v_14572 | 1 | 0 | 0 | 0 | 0 | 0 | 0 | 0 | 0 | 0 | 0 | 0 | 0 | 0 | 0 | 0 | 0 | 0 | 0 | 0 | 0 | 0 | 0 | 0 | 0 | 0 | 0 | 0 | 0 | 0 | 0 | 0 | 1 | 0 | 0 | 0 | 0 | 0 | 0 | 0 | 0 | 0 | 0 | 0 | 0 | 0 | 0 | 0 | 0 | 0 | 0 | 0 | 0 | 0 | 0 |
| virus_Jul27.5v_14825 | 1 | 0 | 0 | 0 | 0 | 0 | 0 | 0 | 0 | 0 | 0 | 0 | 0 | 0 | 0 | 0 | 0 | 0 | 0 | 0 | 0 | 0 | 0 | 0 | 0 | 0 | 0 | 0 | 0 | 0 | 0 | 0 | 1 | 0 | 0 | 0 | 0 | 0 | 0 | 0 | 0 | 0 | 0 | 0 | 0 | 0 | 0 | 0 | 0 | 0 | 0 | 0 | 0 | 0 | 0 |
| virus_Jul27.5v_17228 | 1 | 0 | 0 | 0 | 0 | 0 | 0 | 0 | 0 | 0 | 0 | 0 | 0 | 0 | 0 | 0 | 0 | 0 | 0 | 0 | 0 | 0 | 0 | 0 | 0 | 0 | 0 | 0 | 0 | 0 | 0 | 0 | 1 | 0 | 0 | 0 | 0 | 0 | 0 | 0 | 0 | 0 | 0 | 0 | 0 | 0 | 0 | 0 | 0 | 0 | 0 | 0 | 0 | 0 | 0 |
| virus_Jul27.5v_18212 | 1 | 0 | 0 | 0 | 0 | 0 | 0 | 0 | 0 | 0 | 0 | 0 | 0 | 0 | 0 | 0 | 0 | 0 | 0 | 0 | 0 | 0 | 0 | 0 | 0 | 0 | 0 | 0 | 0 | 0 | 0 | 0 | 1 | 0 | 0 | 0 | 0 | 0 | 0 | 0 | 0 | 0 | 0 | 0 | 0 | 0 | 0 | 0 | 0 | 0 | 0 | 0 | 0 | 0 | 0 |
| virus_Jul27.5v_19511 | 1 | 0 | 0 | 0 | 0 | 0 | 0 | 0 | 0 | 0 | 0 | 0 | 0 | 0 | 0 | 0 | 0 | 0 | 0 | 0 | 0 | 0 | 0 | 0 | 0 | 0 | 0 | 0 | 0 | 0 | 0 | 0 | 1 | 0 | 0 | 0 | 0 | 0 | 0 | 0 | 0 | 0 | 0 | 0 | 0 | 0 | 0 | 0 | 0 | 0 | 0 | 0 | 0 | 0 | 0 |
| virus_Jul27.5v_20574 | 1 | 0 | 0 | 0 | 0 | 0 | 0 | 0 | 0 | 0 | 0 | 0 | 0 | 0 | 0 | 0 | 0 | 0 | 0 | 0 | 0 | 0 | 0 | 0 | 0 | 0 | 0 | 0 | 0 | 0 | 0 | 0 | 1 | 0 | 0 | 0 | 0 | 0 | 0 | 0 | 0 | 0 | 0 | 0 | 0 | 0 | 0 | 0 | 0 | 0 | 0 | 0 | 0 | 0 | 0 |
| virus_Jul27.25v_5603 | 1 | 0 | 0 | 0 | 0 | 0 | 0 | 0 | 0 | 0 | 0 | 0 | 0 | 0 | 0 | 0 | 0 | 0 | 0 | 0 | 0 | 0 | 0 | 0 | 0 | 0 | 0 | 0 | 0 | 0 | 0 | 0 | 0 | 0 | 1 | 0 | 0 | 0 | 0 | 0 | 0 | 0 | 0 | 0 | 0 | 0 | 0 | 0 | 0 | 0 | 0 | 0 | 0 | 0 | 0 |
| virus_Jul27.25v_11841 | 1 | 0 | 0 | 0 | 0 | 0 | 0 | 0 | 0 | 0 | 0 | 0 | 0 | 0 | 0 | 0 | 0 | 0 | 0 | 0 | 0 | 0 | 0 | 0 | 0 | 0 | 0 | 0 | 0 | 0 | 0 | 0 | 0 | 0 | 1 | 0 | 0 | 0 | 0 | 0 | 0 | 0 | 0 | 0 | 0 | 0 | 0 | 0 | 0 | 0 | 0 | 0 | 0 | 0 | 0 |
| virus_Jul27.25v_13112 | 1 | 0 | 0 | 0 | 0 | 0 | 0 | 0 | 0 | 0 | 0 | 0 | 0 | 0 | 0 | 0 | 0 | 0 | 0 | 0 | 0 | 0 | 0 | 0 | 0 | 0 | 0 | 0 | 0 | 0 | 0 | 0 | 0 | 0 | 1 | 0 | 0 | 0 | 0 | 0 | 0 | 0 | 0 | 0 | 0 | 0 | 0 | 0 | 0 | 0 | 0 | 0 | 0 | 0 | 0 |
| virus_Jul27.25v_16854 | 1 | 0 | 0 | 0 | 0 | 0 | 0 | 0 | 0 | 0 | 0 | 0 | 0 | 0 | 0 | 0 | 0 | 0 | 0 | 0 | 0 | 0 | 0 | 0 | 0 | 0 | 0 | 0 | 0 | 0 | 0 | 0 | 0 | 0 | 1 | 0 | 0 | 0 | 0 | 0 | 0 | 0 | 0 | 0 | 0 | 0 | 0 | 0 | 0 | 0 | 0 | 0 | 0 | 0 | 0 |
| virus_Jul27.25v_19127 | 1 | 0 | 0 | 0 | 0 | 0 | 0 | 0 | 0 | 0 | 0 | 0 | 0 | 0 | 0 | 0 | 0 | 0 | 0 | 0 | 0 | 0 | 0 | 0 | 0 | 0 | 0 | 0 | 0 | 0 | 0 | 0 | 0 | 0 | 1 | 0 | 0 | 0 | 0 | 0 | 0 | 0 | 0 | 0 | 0 | 0 | 0 | 0 | 0 | 0 | 0 | 0 | 0 | 0 | 0 |
| virus_Jul27.25v_27739 | 1 | 0 | 0 | 0 | 0 | 0 | 0 | 0 | 0 | 0 | 0 | 0 | 0 | 0 | 0 | 0 | 0 | 0 | 0 | 0 | 0 | 0 | 0 | 0 | 0 | 0 | 0 | 0 | 0 | 0 | 0 | 0 | 0 | 0 | 1 | 0 | 0 | 0 | 0 | 0 | 0 | 0 | 0 | 0 | 0 | 0 | 0 | 0 | 0 | 0 | 0 | 0 | 0 | 0 | 0 |
| virus_Jul27.25v_28745 | 1 | 0 | 0 | 0 | 0 | 0 | 0 | 0 | 0 | 0 | 0 | 0 | 0 | 0 | 0 | 0 | 0 | 0 | 0 | 0 | 0 | 0 | 0 | 0 | 0 | 0 | 0 | 0 | 0 | 0 | 0 | 0 | 0 | 0 | 1 | 0 | 0 | 0 | 0 | 0 | 0 | 0 | 0 | 0 | 0 | 0 | 0 | 0 | 0 | 0 | 0 | 0 | 0 | 0 | 0 |
| virus_Jul27.25v_32390 | 1 | 0 | 0 | 0 | 0 | 0 | 0 | 0 | 0 | 0 | 0 | 0 | 0 | 0 | 0 | 0 | 0 | 0 | 0 | 0 | 0 | 0 | 0 | 0 | 0 | 0 | 0 | 0 | 0 | 0 | 0 | 0 | 0 | 0 | 1 | 0 | 0 | 0 | 0 | 0 | 0 | 0 | 0 | 0 | 0 | 0 | 0 | 0 | 0 | 0 | 0 | 0 | 0 | 0 | 0 |
| virus_Jul27.25v_32724 | 1 | 0 | 0 | 0 | 0 | 0 | 0 | 0 | 0 | 0 | 0 | 0 | 0 | 0 | 0 | 0 | 0 | 0 | 0 | 0 | 0 | 0 | 0 | 0 | 0 | 0 | 0 | 0 | 0 | 0 | 0 | 0 | 0 | 0 | 1 | 0 | 0 | 0 | 0 | 0 | 0 | 0 | 0 | 0 | 0 | 0 | 0 | 0 | 0 | 0 | 0 | 0 | 0 | 0 | 0 |
| virus_Jul27.25v_33018 | 1 | 0 | 0 | 0 | 0 | 0 | 0 | 0 | 0 | 0 | 0 | 0 | 0 | 0 | 0 | 0 | 0 | 0 | 0 | 0 | 0 | 0 | 0 | 0 | 0 | 0 | 0 | 0 | 0 | 0 | 0 | 0 | 0 | 0 | 1 | 0 | 0 | 0 | 0 | 0 | 0 | 0 | 0 | 0 | 0 | 0 | 0 | 0 | 0 | 0 | 0 | 0 | 0 | 0 | 0 |
| virus_Jul27.25v_35410 | 1 | 0 | 0 | 0 | 0 | 0 | 0 | 0 | 0 | 0 | 0 | 0 | 0 | 0 | 0 | 0 | 0 | 0 | 0 | 0 | 0 | 0 | 0 | 0 | 0 | 0 | 0 | 0 | 0 | 0 | 0 | 0 | 0 | 0 | 1 | 0 | 0 | 0 | 0 | 0 | 0 | 0 | 0 | 0 | 0 | 0 | 0 | 0 | 0 | 0 | 0 | 0 | 0 | 0 | 0 |
| virus_Jul27.25v_37043 | 1 | 0 | 0 | 0 | 0 | 0 | 0 | 0 | 0 | 0 | 0 | 0 | 0 | 0 | 0 | 0 | 0 | 0 | 0 | 0 | 0 | 0 | 0 | 0 | 0 | 0 | 0 | 0 | 0 | 0 | 0 | 0 | 0 | 0 | 1 | 0 | 0 | 0 | 0 | 0 | 0 | 0 | 0 | 0 | 0 | 0 | 0 | 0 | 0 | 0 | 0 | 0 | 0 | 0 | 0 |
| virus_Jul27.25v_39824 | 1 | 0 | 0 | 0 | 0 | 0 | 0 | 0 | 0 | 0 | 0 | 0 | 0 | 0 | 0 | 0 | 0 | 0 | 0 | 0 | 0 | 0 | 0 | 0 | 0 | 0 | 0 | 0 | 0 | 0 | 0 | 0 | 0 | 0 | 1 | 0 | 0 | 0 | 0 | 0 | 0 | 0 | 0 | 0 | 0 | 0 | 0 | 0 | 0 | 0 | 0 | 0 | 0 | 0 | 0 |
| virus_Jul27.75v_2284 | 1 | 0 | 0 | 0 | 0 | 0 | 0 | 0 | 0 | 0 | 0 | 0 | 0 | 0 | 0 | 0 | 0 | 0 | 0 | 0 | 0 | 0 | 0 | 0 | 0 | 0 | 0 | 0 | 0 | 0 | 0 | 0 | 0 | 0 | 0 | 0 | 1 | 0 | 0 | 0 | 0 | 0 | 0 | 0 | 0 | 0 | 0 | 0 | 0 | 0 | 0 | 0 | 0 | 0 | 0 |
| virus_Jul27.75v_2820 | 1 | 0 | 0 | 0 | 0 | 0 | 0 | 0 | 0 | 0 | 0 | 0 | 0 | 0 | 0 | 0 | 0 | 0 | 0 | 0 | 0 | 0 | 0 | 0 | 0 | 0 | 0 | 0 | 0 | 0 | 0 | 0 | 0 | 0 | 0 | 0 | 1 | 0 | 0 | 0 | 0 | 0 | 0 | 0 | 0 | 0 | 0 | 0 | 0 | 0 | 0 | 0 | 0 | 0 | 0 |
| virus_Jul27.75v_4233 | 1 | 0 | 0 | 0 | 0 | 0 | 0 | 0 | 0 | 0 | 0 | 0 | 0 | 0 | 0 | 0 | 0 | 0 | 0 | 0 | 0 | 0 | 0 | 0 | 0 | 0 | 0 | 0 | 0 | 0 | 0 | 0 | 0 | 0 | 0 | 0 | 1 | 0 | 0 | 0 | 0 | 0 | 0 | 0 | 0 | 0 | 0 | 0 | 0 | 0 | 0 | 0 | 0 | 0 | 0 |
| virus_Jul27.75v_6963 | 1 | 0 | 0 | 0 | 0 | 0 | 0 | 0 | 0 | 0 | 0 | 0 | 0 | 0 | 0 | 0 | 0 | 0 | 0 | 0 | 0 | 0 | 0 | 0 | 0 | 0 | 0 | 0 | 0 | 0 | 0 | 0 | 0 | 0 | 0 | 0 | 1 | 0 | 0 | 0 | 0 | 0 | 0 | 0 | 0 | 0 | 0 | 0 | 0 | 0 | 0 | 0 | 0 | 0 | 0 |
| virus_Jul27.75v_18485 | 1 | 0 | 0 | 0 | 0 | 0 | 0 | 0 | 0 | 0 | 0 | 0 | 0 | 0 | 0 | 0 | 0 | 0 | 0 | 0 | 0 | 0 | 0 | 0 | 0 | 0 | 0 | 0 | 0 | 0 | 0 | 0 | 0 | 0 | 0 | 0 | 1 | 0 | 0 | 0 | 0 | 0 | 0 | 0 | 0 | 0 | 0 | 0 | 0 | 0 | 0 | 0 | 0 | 0 | 0 |
| virus_Jun8.25v_21223 | 1 | 0 | 0 | 0 | 0 | 0 | 0 | 0 | 0 | 0 | 1 | 0 | 0 | 0 | 0 | 0 | 0 | 0 | 0 | 0 | 0 | 0 | 0 | 0 | 0 | 0 | 0 | 0 | 0 | 0 | 0 | 0 | 0 | 0 | 0 | 0 | 0 | 0 | 0 | 0 | 0 | 0 | 0 | 0 | 0 | 0 | 0 | 0 | 0 | 0 | 0 | 0 | 0 | 0 | 0 |
| virus_Jun8.25v_27543 | 1 | 0 | 0 | 0 | 0 | 0 | 0 | 0 | 0 | 0 | 1 | 0 | 0 | 0 | 0 | 0 | 0 | 0 | 0 | 0 | 0 | 0 | 0 | 0 | 0 | 0 | 0 | 0 | 0 | 0 | 0 | 0 | 0 | 0 | 0 | 0 | 0 | 0 | 0 | 0 | 0 | 0 | 0 | 0 | 0 | 0 | 0 | 0 | 0 | 0 | 0 | 0 | 0 | 0 | 0 |
| virus_Jun8.50v_610 | 1 | 0 | 0 | 0 | 0 | 0 | 0 | 0 | 0 | 0 | 0 | 0 | 1 | 0 | 0 | 0 | 0 | 0 | 0 | 0 | 0 | 0 | 0 | 0 | 0 | 0 | 0 | 0 | 0 | 0 | 0 | 0 | 0 | 0 | 0 | 0 | 0 | 0 | 0 | 0 | 0 | 0 | 0 | 0 | 0 | 0 | 0 | 0 | 0 | 0 | 0 | 0 | 0 | 0 | 0 |
| virus_Jun8.50v_663 | 1 | 0 | 0 | 0 | 0 | 0 | 0 | 0 | 0 | 0 | 0 | 0 | 1 | 0 | 0 | 0 | 0 | 0 | 0 | 0 | 0 | 0 | 0 | 0 | 0 | 0 | 0 | 0 | 0 | 0 | 0 | 0 | 0 | 0 | 0 | 0 | 0 | 0 | 0 | 0 | 0 | 0 | 0 | 0 | 0 | 0 | 0 | 0 | 0 | 0 | 0 | 0 | 0 | 0 | 0 |
| virus_Jun8.50v_5606 | 1 | 0 | 0 | 0 | 0 | 0 | 0 | 0 | 0 | 0 | 0 | 0 | 1 | 0 | 0 | 0 | 0 | 0 | 0 | 0 | 0 | 0 | 0 | 0 | 0 | 0 | 0 | 0 | 0 | 0 | 0 | 0 | 0 | 0 | 0 | 0 | 0 | 0 | 0 | 0 | 0 | 0 | 0 | 0 | 0 | 0 | 0 | 0 | 0 | 0 | 0 | 0 | 0 | 0 | 0 |
| virus_Jun8.50v_9744 | 1 | 0 | 0 | 0 | 0 | 0 | 0 | 0 | 0 | 0 | 0 | 0 | 1 | 0 | 0 | 0 | 0 | 0 | 0 | 0 | 0 | 0 | 0 | 0 | 0 | 0 | 0 | 0 | 0 | 0 | 0 | 0 | 0 | 0 | 0 | 0 | 0 | 0 | 0 | 0 | 0 | 0 | 0 | 0 | 0 | 0 | 0 | 0 | 0 | 0 | 0 | 0 | 0 | 0 | 0 |
| virus_Jun8.50v_12412 | 1 | 0 | 0 | 0 | 0 | 0 | 0 | 0 | 0 | 0 | 0 | 0 | 1 | 0 | 0 | 0 | 0 | 0 | 0 | 0 | 0 | 0 | 0 | 0 | 0 | 0 | 0 | 0 | 0 | 0 | 0 | 0 | 0 | 0 | 0 | 0 | 0 | 0 | 0 | 0 | 0 | 0 | 0 | 0 | 0 | 0 | 0 | 0 | 0 | 0 | 0 | 0 | 0 | 0 | 0 |
| virus_Jun8.50v_16697 | 1 | 0 | 0 | 0 | 0 | 0 | 0 | 0 | 0 | 0 | 0 | 0 | 1 | 0 | 0 | 0 | 0 | 0 | 0 | 0 | 0 | 0 | 0 | 0 | 0 | 0 | 0 | 0 | 0 | 0 | 0 | 0 | 0 | 0 | 0 | 0 | 0 | 0 | 0 | 0 | 0 | 0 | 0 | 0 | 0 | 0 | 0 | 0 | 0 | 0 | 0 | 0 | 0 | 0 | 0 |
| virus_Jun8.50v_17108 | 1 | 0 | 0 | 0 | 0 | 0 | 0 | 0 | 0 | 0 | 0 | 0 | 1 | 0 | 0 | 0 | 0 | 0 | 0 | 0 | 0 | 0 | 0 | 0 | 0 | 0 | 0 | 0 | 0 | 0 | 0 | 0 | 0 | 0 | 0 | 0 | 0 | 0 | 0 | 0 | 0 | 0 | 0 | 0 | 0 | 0 | 0 | 0 | 0 | 0 | 0 | 0 | 0 | 0 | 0 |
| virus_Jun8.50v_19258 | 1 | 0 | 0 | 0 | 0 | 0 | 0 | 0 | 0 | 0 | 0 | 0 | 1 | 0 | 0 | 0 | 0 | 0 | 0 | 0 | 0 | 0 | 0 | 0 | 0 | 0 | 0 | 0 | 0 | 0 | 0 | 0 | 0 | 0 | 0 | 0 | 0 | 0 | 0 | 0 | 0 | 0 | 0 | 0 | 0 | 0 | 0 | 0 | 0 | 0 | 0 | 0 | 0 | 0 | 0 |
| virus_Jun8.50v_21260 | 1 | 0 | 0 | 0 | 0 | 0 | 0 | 0 | 0 | 0 | 0 | 0 | 1 | 0 | 0 | 0 | 0 | 0 | 0 | 0 | 0 | 0 | 0 | 0 | 0 | 0 | 0 | 0 | 0 | 0 | 0 | 0 | 0 | 0 | 0 | 0 | 0 | 0 | 0 | 0 | 0 | 0 | 0 | 0 | 0 | 0 | 0 | 0 | 0 | 0 | 0 | 0 | 0 | 0 | 0 |
| virus_Jun8.50v_23094 | 1 | 0 | 0 | 0 | 0 | 0 | 0 | 0 | 0 | 0 | 0 | 0 | 1 | 0 | 0 | 0 | 0 | 0 | 0 | 0 | 0 | 0 | 0 | 0 | 0 | 0 | 0 | 0 | 0 | 0 | 0 | 0 | 0 | 0 | 0 | 0 | 0 | 0 | 0 | 0 | 0 | 0 | 0 | 0 | 0 | 0 | 0 | 0 | 0 | 0 | 0 | 0 | 0 | 0 | 0 |
| virus_Jun8.50v_32175 | 1 | 0 | 0 | 0 | 0 | 0 | 0 | 0 | 0 | 0 | 0 | 0 | 1 | 0 | 0 | 0 | 0 | 0 | 0 | 0 | 0 | 0 | 0 | 0 | 0 | 0 | 0 | 0 | 0 | 0 | 0 | 0 | 0 | 0 | 0 | 0 | 0 | 0 | 0 | 0 | 0 | 0 | 0 | 0 | 0 | 0 | 0 | 0 | 0 | 0 | 0 | 0 | 0 | 0 | 0 |
| virus_Jun8.50v_37894 | 1 | 0 | 0 | 0 | 0 | 0 | 0 | 0 | 0 | 0 | 0 | 0 | 1 | 0 | 0 | 0 | 0 | 0 | 0 | 0 | 0 | 0 | 0 | 0 | 0 | 0 | 0 | 0 | 0 | 0 | 0 | 0 | 0 | 0 | 0 | 0 | 0 | 0 | 0 | 0 | 0 | 0 | 0 | 0 | 0 | 0 | 0 | 0 | 0 | 0 | 0 | 0 | 0 | 0 | 0 |
| virus_Jun8.75v_14531 | 1 | 0 | 0 | 0 | 0 | 0 | 0 | 0 | 0 | 0 | 0 | 0 | 0 | 0 | 1 | 0 | 0 | 0 | 0 | 0 | 0 | 0 | 0 | 0 | 0 | 0 | 0 | 0 | 0 | 0 | 0 | 0 | 0 | 0 | 0 | 0 | 0 | 0 | 0 | 0 | 0 | 0 | 0 | 0 | 0 | 0 | 0 | 0 | 0 | 0 | 0 | 0 | 0 | 0 | 0 |
| virus_Jun8.75v_16443 | 1 | 0 | 0 | 0 | 0 | 0 | 0 | 0 | 0 | 0 | 0 | 0 | 0 | 0 | 1 | 0 | 0 | 0 | 0 | 0 | 0 | 0 | 0 | 0 | 0 | 0 | 0 | 0 | 0 | 0 | 0 | 0 | 0 | 0 | 0 | 0 | 0 | 0 | 0 | 0 | 0 | 0 | 0 | 0 | 0 | 0 | 0 | 0 | 0 | 0 | 0 | 0 | 0 | 0 | 0 |
| virus_Jun8.75v_17650 | 1 | 0 | 0 | 0 | 0 | 0 | 0 | 0 | 0 | 0 | 0 | 0 | 0 | 0 | 1 | 0 | 0 | 0 | 0 | 0 | 0 | 0 | 0 | 0 | 0 | 0 | 0 | 0 | 0 | 0 | 0 | 0 | 0 | 0 | 0 | 0 | 0 | 0 | 0 | 0 | 0 | 0 | 0 | 0 | 0 | 0 | 0 | 0 | 0 | 0 | 0 | 0 | 0 | 0 | 0 |
| virus_Jun22.0v_628 | 1 | 0 | 0 | 0 | 0 | 0 | 0 | 0 | 0 | 0 | 0 | 0 | 0 | 0 | 0 | 0 | 1 | 0 | 0 | 0 | 0 | 0 | 0 | 0 | 0 | 0 | 0 | 0 | 0 | 0 | 0 | 0 | 0 | 0 | 0 | 0 | 0 | 0 | 0 | 0 | 0 | 0 | 0 | 0 | 0 | 0 | 0 | 0 | 0 | 0 | 0 | 0 | 0 | 0 | 0 |
| virus_Jun22.0v_6661 | 1 | 0 | 0 | 0 | 0 | 0 | 0 | 0 | 0 | 0 | 0 | 0 | 0 | 0 | 0 | 0 | 1 | 0 | 0 | 0 | 0 | 0 | 0 | 0 | 0 | 0 | 0 | 0 | 0 | 0 | 0 | 0 | 0 | 0 | 0 | 0 | 0 | 0 | 0 | 0 | 0 | 0 | 0 | 0 | 0 | 0 | 0 | 0 | 0 | 0 | 0 | 0 | 0 | 0 | 0 |
| virus_Jun22.0v_8077 | 1 | 0 | 0 | 0 | 0 | 0 | 0 | 0 | 0 | 0 | 0 | 0 | 0 | 0 | 0 | 0 | 1 | 0 | 0 | 0 | 0 | 0 | 0 | 0 | 0 | 0 | 0 | 0 | 0 | 0 | 0 | 0 | 0 | 0 | 0 | 0 | 0 | 0 | 0 | 0 | 0 | 0 | 0 | 0 | 0 | 0 | 0 | 0 | 0 | 0 | 0 | 0 | 0 | 0 | 0 |
| virus_Jun22.0v_10341 | 1 | 0 | 0 | 0 | 0 | 0 | 0 | 0 | 0 | 0 | 0 | 0 | 0 | 0 | 0 | 0 | 1 | 0 | 0 | 0 | 0 | 0 | 0 | 0 | 0 | 0 | 0 | 0 | 0 | 0 | 0 | 0 | 0 | 0 | 0 | 0 | 0 | 0 | 0 | 0 | 0 | 0 | 0 | 0 | 0 | 0 | 0 | 0 | 0 | 0 | 0 | 0 | 0 | 0 | 0 |
| virus_Jun22.0v_11443 | 1 | 0 | 0 | 0 | 0 | 0 | 0 | 0 | 0 | 0 | 0 | 0 | 0 | 0 | 0 | 0 | 1 | 0 | 0 | 0 | 0 | 0 | 0 | 0 | 0 | 0 | 0 | 0 | 0 | 0 | 0 | 0 | 0 | 0 | 0 | 0 | 0 | 0 | 0 | 0 | 0 | 0 | 0 | 0 | 0 | 0 | 0 | 0 | 0 | 0 | 0 | 0 | 0 | 0 | 0 |
| virus_Jun22.0v_13622 | 1 | 0 | 0 | 0 | 0 | 0 | 0 | 0 | 0 | 0 | 0 | 0 | 0 | 0 | 0 | 0 | 1 | 0 | 0 | 0 | 0 | 0 | 0 | 0 | 0 | 0 | 0 | 0 | 0 | 0 | 0 | 0 | 0 | 0 | 0 | 0 | 0 | 0 | 0 | 0 | 0 | 0 | 0 | 0 | 0 | 0 | 0 | 0 | 0 | 0 | 0 | 0 | 0 | 0 | 0 |
| virus_Jun22.0v_14147 | 1 | 0 | 0 | 0 | 0 | 0 | 0 | 0 | 0 | 0 | 0 | 0 | 0 | 0 | 0 | 0 | 1 | 0 | 0 | 0 | 0 | 0 | 0 | 0 | 0 | 0 | 0 | 0 | 0 | 0 | 0 | 0 | 0 | 0 | 0 | 0 | 0 | 0 | 0 | 0 | 0 | 0 | 0 | 0 | 0 | 0 | 0 | 0 | 0 | 0 | 0 | 0 | 0 | 0 | 0 |
| virus_Jun22.0v_15267 | 1 | 0 | 0 | 0 | 0 | 0 | 0 | 0 | 0 | 0 | 0 | 0 | 0 | 0 | 0 | 0 | 1 | 0 | 0 | 0 | 0 | 0 | 0 | 0 | 0 | 0 | 0 | 0 | 0 | 0 | 0 | 0 | 0 | 0 | 0 | 0 | 0 | 0 | 0 | 0 | 0 | 0 | 0 | 0 | 0 | 0 | 0 | 0 | 0 | 0 | 0 | 0 | 0 | 0 | 0 |
| virus_Jun22.0v_15322 | 1 | 0 | 0 | 0 | 0 | 0 | 0 | 0 | 0 | 0 | 0 | 0 | 0 | 0 | 0 | 0 | 1 | 0 | 0 | 0 | 0 | 0 | 0 | 0 | 0 | 0 | 0 | 0 | 0 | 0 | 0 | 0 | 0 | 0 | 0 | 0 | 0 | 0 | 0 | 0 | 0 | 0 | 0 | 0 | 0 | 0 | 0 | 0 | 0 | 0 | 0 | 0 | 0 | 0 | 0 |
| virus_Jun22.0v_17184 | 1 | 0 | 0 | 0 | 0 | 0 | 0 | 0 | 0 | 0 | 0 | 0 | 0 | 0 | 0 | 0 | 1 | 0 | 0 | 0 | 0 | 0 | 0 | 0 | 0 | 0 | 0 | 0 | 0 | 0 | 0 | 0 | 0 | 0 | 0 | 0 | 0 | 0 | 0 | 0 | 0 | 0 | 0 | 0 | 0 | 0 | 0 | 0 | 0 | 0 | 0 | 0 | 0 | 0 | 0 |
| virus_Jun22.0v_18743 | 1 | 0 | 0 | 0 | 0 | 0 | 0 | 0 | 0 | 0 | 0 | 0 | 0 | 0 | 0 | 0 | 1 | 0 | 0 | 0 | 0 | 0 | 0 | 0 | 0 | 0 | 0 | 0 | 0 | 0 | 0 | 0 | 0 | 0 | 0 | 0 | 0 | 0 | 0 | 0 | 0 | 0 | 0 | 0 | 0 | 0 | 0 | 0 | 0 | 0 | 0 | 0 | 0 | 0 | 0 |
| virus_Jun22.0v_19420 | 1 | 0 | 0 | 0 | 0 | 0 | 0 | 0 | 0 | 0 | 0 | 0 | 0 | 0 | 0 | 0 | 1 | 0 | 0 | 0 | 0 | 0 | 0 | 0 | 0 | 0 | 0 | 0 | 0 | 0 | 0 | 0 | 0 | 0 | 0 | 0 | 0 | 0 | 0 | 0 | 0 | 0 | 0 | 0 | 0 | 0 | 0 | 0 | 0 | 0 | 0 | 0 | 0 | 0 | 0 |
| virus_Jun22.0v_19636 | 1 | 0 | 0 | 0 | 0 | 0 | 0 | 0 | 0 | 0 | 0 | 0 | 0 | 0 | 0 | 0 | 1 | 0 | 0 | 0 | 0 | 0 | 0 | 0 | 0 | 0 | 0 | 0 | 0 | 0 | 0 | 0 | 0 | 0 | 0 | 0 | 0 | 0 | 0 | 0 | 0 | 0 | 0 | 0 | 0 | 0 | 0 | 0 | 0 | 0 | 0 | 0 | 0 | 0 | 0 |
| virus_Jun22.50v_1003 | 1 | 0 | 0 | 0 | 0 | 0 | 0 | 0 | 0 | 0 | 0 | 0 | 0 | 0 | 0 | 0 | 0 | 0 | 1 | 0 | 0 | 0 | 0 | 0 | 0 | 0 | 0 | 0 | 0 | 0 | 0 | 0 | 0 | 0 | 0 | 0 | 0 | 0 | 0 | 0 | 0 | 0 | 0 | 0 | 0 | 0 | 0 | 0 | 0 | 0 | 0 | 0 | 0 | 0 | 0 |
| virus_Jun22.50v_3776 | 1 | 0 | 0 | 0 | 0 | 0 | 0 | 0 | 0 | 0 | 0 | 0 | 0 | 0 | 0 | 0 | 0 | 0 | 1 | 0 | 0 | 0 | 0 | 0 | 0 | 0 | 0 | 0 | 0 | 0 | 0 | 0 | 0 | 0 | 0 | 0 | 0 | 0 | 0 | 0 | 0 | 0 | 0 | 0 | 0 | 0 | 0 | 0 | 0 | 0 | 0 | 0 | 0 | 0 | 0 |
| virus_Jun22.50v_5212 | 1 | 0 | 0 | 0 | 0 | 0 | 0 | 0 | 0 | 0 | 0 | 0 | 0 | 0 | 0 | 0 | 0 | 0 | 1 | 0 | 0 | 0 | 0 | 0 | 0 | 0 | 0 | 0 | 0 | 0 | 0 | 0 | 0 | 0 | 0 | 0 | 0 | 0 | 0 | 0 | 0 | 0 | 0 | 0 | 0 | 0 | 0 | 0 | 0 | 0 | 0 | 0 | 0 | 0 | 0 |
| virus_Jun22.50v_5520 | 1 | 0 | 0 | 0 | 0 | 0 | 0 | 0 | 0 | 0 | 0 | 0 | 0 | 0 | 0 | 0 | 0 | 0 | 1 | 0 | 0 | 0 | 0 | 0 | 0 | 0 | 0 | 0 | 0 | 0 | 0 | 0 | 0 | 0 | 0 | 0 | 0 | 0 | 0 | 0 | 0 | 0 | 0 | 0 | 0 | 0 | 0 | 0 | 0 | 0 | 0 | 0 | 0 | 0 | 0 |
| virus_Jun22.50v_10823 | 1 | 0 | 0 | 0 | 0 | 0 | 0 | 0 | 0 | 0 | 0 | 0 | 0 | 0 | 0 | 0 | 0 | 0 | 1 | 0 | 0 | 0 | 0 | 0 | 0 | 0 | 0 | 0 | 0 | 0 | 0 | 0 | 0 | 0 | 0 | 0 | 0 | 0 | 0 | 0 | 0 | 0 | 0 | 0 | 0 | 0 | 0 | 0 | 0 | 0 | 0 | 0 | 0 | 0 | 0 |
| virus_Jun22.50v_16755 | 1 | 0 | 0 | 0 | 0 | 0 | 0 | 0 | 0 | 0 | 0 | 0 | 0 | 0 | 0 | 0 | 0 | 0 | 1 | 0 | 0 | 0 | 0 | 0 | 0 | 0 | 0 | 0 | 0 | 0 | 0 | 0 | 0 | 0 | 0 | 0 | 0 | 0 | 0 | 0 | 0 | 0 | 0 | 0 | 0 | 0 | 0 | 0 | 0 | 0 | 0 | 0 | 0 | 0 | 0 |
| virus_Jun22.50v_20221 | 1 | 0 | 0 | 0 | 0 | 0 | 0 | 0 | 0 | 0 | 0 | 0 | 0 | 0 | 0 | 0 | 0 | 0 | 1 | 0 | 0 | 0 | 0 | 0 | 0 | 0 | 0 | 0 | 0 | 0 | 0 | 0 | 0 | 0 | 0 | 0 | 0 | 0 | 0 | 0 | 0 | 0 | 0 | 0 | 0 | 0 | 0 | 0 | 0 | 0 | 0 | 0 | 0 | 0 | 0 |
| virus_Jun22.50v_24243 | 1 | 0 | 0 | 0 | 0 | 0 | 0 | 0 | 0 | 0 | 0 | 0 | 0 | 0 | 0 | 0 | 0 | 0 | 1 | 0 | 0 | 0 | 0 | 0 | 0 | 0 | 0 | 0 | 0 | 0 | 0 | 0 | 0 | 0 | 0 | 0 | 0 | 0 | 0 | 0 | 0 | 0 | 0 | 0 | 0 | 0 | 0 | 0 | 0 | 0 | 0 | 0 | 0 | 0 | 0 |
| virus_Jun22.50v_29480 | 1 | 0 | 0 | 0 | 0 | 0 | 0 | 0 | 0 | 0 | 0 | 0 | 0 | 0 | 0 | 0 | 0 | 0 | 1 | 0 | 0 | 0 | 0 | 0 | 0 | 0 | 0 | 0 | 0 | 0 | 0 | 0 | 0 | 0 | 0 | 0 | 0 | 0 | 0 | 0 | 0 | 0 | 0 | 0 | 0 | 0 | 0 | 0 | 0 | 0 | 0 | 0 | 0 | 0 | 0 |
| virus_Jun22.50v_36324 | 1 | 0 | 0 | 0 | 0 | 0 | 0 | 0 | 0 | 0 | 0 | 0 | 0 | 0 | 0 | 0 | 0 | 0 | 1 | 0 | 0 | 0 | 0 | 0 | 0 | 0 | 0 | 0 | 0 | 0 | 0 | 0 | 0 | 0 | 0 | 0 | 0 | 0 | 0 | 0 | 0 | 0 | 0 | 0 | 0 | 0 | 0 | 0 | 0 | 0 | 0 | 0 | 0 | 0 | 0 |
| virus_Jun22.50v_36973 | 1 | 0 | 0 | 0 | 0 | 0 | 0 | 0 | 0 | 0 | 0 | 0 | 0 | 0 | 0 | 0 | 0 | 0 | 1 | 0 | 0 | 0 | 0 | 0 | 0 | 0 | 0 | 0 | 0 | 0 | 0 | 0 | 0 | 0 | 0 | 0 | 0 | 0 | 0 | 0 | 0 | 0 | 0 | 0 | 0 | 0 | 0 | 0 | 0 | 0 | 0 | 0 | 0 | 0 | 0 |
| virus_Jun22.50v_39809 | 1 | 0 | 0 | 0 | 0 | 0 | 0 | 0 | 0 | 0 | 0 | 0 | 0 | 0 | 0 | 0 | 0 | 0 | 1 | 0 | 0 | 0 | 0 | 0 | 0 | 0 | 0 | 0 | 0 | 0 | 0 | 0 | 0 | 0 | 0 | 0 | 0 | 0 | 0 | 0 | 0 | 0 | 0 | 0 | 0 | 0 | 0 | 0 | 0 | 0 | 0 | 0 | 0 | 0 | 0 |
| virus_Jun22.50v_41054 | 1 | 0 | 0 | 0 | 0 | 0 | 0 | 0 | 0 | 0 | 0 | 0 | 0 | 0 | 0 | 0 | 0 | 0 | 1 | 0 | 0 | 0 | 0 | 0 | 0 | 0 | 0 | 0 | 0 | 0 | 0 | 0 | 0 | 0 | 0 | 0 | 0 | 0 | 0 | 0 | 0 | 0 | 0 | 0 | 0 | 0 | 0 | 0 | 0 | 0 | 0 | 0 | 0 | 0 | 0 |
| virus_Jun22.75v_705 | 1 | 0 | 0 | 0 | 0 | 0 | 0 | 0 | 0 | 0 | 0 | 0 | 0 | 0 | 0 | 0 | 0 | 0 | 0 | 0 | 1 | 0 | 0 | 0 | 0 | 0 | 0 | 0 | 0 | 0 | 0 | 0 | 0 | 0 | 0 | 0 | 0 | 0 | 0 | 0 | 0 | 0 | 0 | 0 | 0 | 0 | 0 | 0 | 0 | 0 | 0 | 0 | 0 | 0 | 0 |
| virus_Jun22.75v_1417 | 1 | 0 | 0 | 0 | 0 | 0 | 0 | 0 | 0 | 0 | 0 | 0 | 0 | 0 | 0 | 0 | 0 | 0 | 0 | 0 | 1 | 0 | 0 | 0 | 0 | 0 | 0 | 0 | 0 | 0 | 0 | 0 | 0 | 0 | 0 | 0 | 0 | 0 | 0 | 0 | 0 | 0 | 0 | 0 | 0 | 0 | 0 | 0 | 0 | 0 | 0 | 0 | 0 | 0 | 0 |
| virus_Jun22.75v_3936 | 1 | 0 | 0 | 0 | 0 | 0 | 0 | 0 | 0 | 0 | 0 | 0 | 0 | 0 | 0 | 0 | 0 | 0 | 0 | 0 | 1 | 0 | 0 | 0 | 0 | 0 | 0 | 0 | 0 | 0 | 0 | 0 | 0 | 0 | 0 | 0 | 0 | 0 | 0 | 0 | 0 | 0 | 0 | 0 | 0 | 0 | 0 | 0 | 0 | 0 | 0 | 0 | 0 | 0 | 0 |
| virus_Jun22.75v_4550 | 1 | 0 | 0 | 0 | 0 | 0 | 0 | 0 | 0 | 0 | 0 | 0 | 0 | 0 | 0 | 0 | 0 | 0 | 0 | 0 | 1 | 0 | 0 | 0 | 0 | 0 | 0 | 0 | 0 | 0 | 0 | 0 | 0 | 0 | 0 | 0 | 0 | 0 | 0 | 0 | 0 | 0 | 0 | 0 | 0 | 0 | 0 | 0 | 0 | 0 | 0 | 0 | 0 | 0 | 0 |
| virus_Jun22.75v_20711 | 1 | 0 | 0 | 0 | 0 | 0 | 0 | 0 | 0 | 0 | 0 | 0 | 0 | 0 | 0 | 0 | 0 | 0 | 0 | 0 | 1 | 0 | 0 | 0 | 0 | 0 | 0 | 0 | 0 | 0 | 0 | 0 | 0 | 0 | 0 | 0 | 0 | 0 | 0 | 0 | 0 | 0 | 0 | 0 | 0 | 0 | 0 | 0 | 0 | 0 | 0 | 0 | 0 | 0 | 0 |
| virus_Jun22.75v_26391 | 1 | 0 | 0 | 0 | 0 | 0 | 0 | 0 | 0 | 0 | 0 | 0 | 0 | 0 | 0 | 0 | 0 | 0 | 0 | 0 | 1 | 0 | 0 | 0 | 0 | 0 | 0 | 0 | 0 | 0 | 0 | 0 | 0 | 0 | 0 | 0 | 0 | 0 | 0 | 0 | 0 | 0 | 0 | 0 | 0 | 0 | 0 | 0 | 0 | 0 | 0 | 0 | 0 | 0 | 0 |
| virus_Jun22.75v_28430 | 1 | 0 | 0 | 0 | 0 | 0 | 0 | 0 | 0 | 0 | 0 | 0 | 0 | 0 | 0 | 0 | 0 | 0 | 0 | 0 | 1 | 0 | 0 | 0 | 0 | 0 | 0 | 0 | 0 | 0 | 0 | 0 | 0 | 0 | 0 | 0 | 0 | 0 | 0 | 0 | 0 | 0 | 0 | 0 | 0 | 0 | 0 | 0 | 0 | 0 | 0 | 0 | 0 | 0 | 0 |
| virus_Jun22.75v_36860 | 1 | 0 | 0 | 0 | 0 | 0 | 0 | 0 | 0 | 0 | 0 | 0 | 0 | 0 | 0 | 0 | 0 | 0 | 0 | 0 | 1 | 0 | 0 | 0 | 0 | 0 | 0 | 0 | 0 | 0 | 0 | 0 | 0 | 0 | 0 | 0 | 0 | 0 | 0 | 0 | 0 | 0 | 0 | 0 | 0 | 0 | 0 | 0 | 0 | 0 | 0 | 0 | 0 | 0 | 0 |
| virus_Jun22.75v_38535 | 1 | 0 | 0 | 0 | 0 | 0 | 0 | 0 | 0 | 0 | 0 | 0 | 0 | 0 | 0 | 0 | 0 | 0 | 0 | 0 | 1 | 0 | 0 | 0 | 0 | 0 | 0 | 0 | 0 | 0 | 0 | 0 | 0 | 0 | 0 | 0 | 0 | 0 | 0 | 0 | 0 | 0 | 0 | 0 | 0 | 0 | 0 | 0 | 0 | 0 | 0 | 0 | 0 | 0 | 0 |
| virus_May12.5v_5981 | 1 | 0 | 1 | 0 | 0 | 0 | 0 | 0 | 0 | 0 | 0 | 0 | 0 | 0 | 0 | 0 | 0 | 0 | 0 | 0 | 0 | 0 | 0 | 0 | 0 | 0 | 0 | 0 | 0 | 0 | 0 | 0 | 0 | 0 | 0 | 0 | 0 | 0 | 0 | 0 | 0 | 0 | 0 | 0 | 0 | 0 | 0 | 0 | 0 | 0 | 0 | 0 | 0 | 0 | 0 |
| virus_May12.5v_9250 | 1 | 0 | 1 | 0 | 0 | 0 | 0 | 0 | 0 | 0 | 0 | 0 | 0 | 0 | 0 | 0 | 0 | 0 | 0 | 0 | 0 | 0 | 0 | 0 | 0 | 0 | 0 | 0 | 0 | 0 | 0 | 0 | 0 | 0 | 0 | 0 | 0 | 0 | 0 | 0 | 0 | 0 | 0 | 0 | 0 | 0 | 0 | 0 | 0 | 0 | 0 | 0 | 0 | 0 | 0 |
| virus_May12.5v_14281 | 1 | 0 | 1 | 0 | 0 | 0 | 0 | 0 | 0 | 0 | 0 | 0 | 0 | 0 | 0 | 0 | 0 | 0 | 0 | 0 | 0 | 0 | 0 | 0 | 0 | 0 | 0 | 0 | 0 | 0 | 0 | 0 | 0 | 0 | 0 | 0 | 0 | 0 | 0 | 0 | 0 | 0 | 0 | 0 | 0 | 0 | 0 | 0 | 0 | 0 | 0 | 0 | 0 | 0 | 0 |
| virus_May12.5v_16513 | 1 | 0 | 1 | 0 | 0 | 0 | 0 | 0 | 0 | 0 | 0 | 0 | 0 | 0 | 0 | 0 | 0 | 0 | 0 | 0 | 0 | 0 | 0 | 0 | 0 | 0 | 0 | 0 | 0 | 0 | 0 | 0 | 0 | 0 | 0 | 0 | 0 | 0 | 0 | 0 | 0 | 0 | 0 | 0 | 0 | 0 | 0 | 0 | 0 | 0 | 0 | 0 | 0 | 0 | 0 |
| virus_May12.5v_20899 | 1 | 0 | 1 | 0 | 0 | 0 | 0 | 0 | 0 | 0 | 0 | 0 | 0 | 0 | 0 | 0 | 0 | 0 | 0 | 0 | 0 | 0 | 0 | 0 | 0 | 0 | 0 | 0 | 0 | 0 | 0 | 0 | 0 | 0 | 0 | 0 | 0 | 0 | 0 | 0 | 0 | 0 | 0 | 0 | 0 | 0 | 0 | 0 | 0 | 0 | 0 | 0 | 0 | 0 | 0 |
| virus_May12.5v_35100 | 1 | 0 | 1 | 0 | 0 | 0 | 0 | 0 | 0 | 0 | 0 | 0 | 0 | 0 | 0 | 0 | 0 | 0 | 0 | 0 | 0 | 0 | 0 | 0 | 0 | 0 | 0 | 0 | 0 | 0 | 0 | 0 | 0 | 0 | 0 | 0 | 0 | 0 | 0 | 0 | 0 | 0 | 0 | 0 | 0 | 0 | 0 | 0 | 0 | 0 | 0 | 0 | 0 | 0 | 0 |
| virus_May12.5v_35952 | 1 | 0 | 1 | 0 | 0 | 0 | 0 | 0 | 0 | 0 | 0 | 0 | 0 | 0 | 0 | 0 | 0 | 0 | 0 | 0 | 0 | 0 | 0 | 0 | 0 | 0 | 0 | 0 | 0 | 0 | 0 | 0 | 0 | 0 | 0 | 0 | 0 | 0 | 0 | 0 | 0 | 0 | 0 | 0 | 0 | 0 | 0 | 0 | 0 | 0 | 0 | 0 | 0 | 0 | 0 |
| virus_May12.25v_3321 | 1 | 0 | 0 | 0 | 1 | 0 | 0 | 0 | 0 | 0 | 0 | 0 | 0 | 0 | 0 | 0 | 0 | 0 | 0 | 0 | 0 | 0 | 0 | 0 | 0 | 0 | 0 | 0 | 0 | 0 | 0 | 0 | 0 | 0 | 0 | 0 | 0 | 0 | 0 | 0 | 0 | 0 | 0 | 0 | 0 | 0 | 0 | 0 | 0 | 0 | 0 | 0 | 0 | 0 | 0 |
| virus_May12.25v_6494 | 1 | 0 | 0 | 0 | 1 | 0 | 0 | 0 | 0 | 0 | 0 | 0 | 0 | 0 | 0 | 0 | 0 | 0 | 0 | 0 | 0 | 0 | 0 | 0 | 0 | 0 | 0 | 0 | 0 | 0 | 0 | 0 | 0 | 0 | 0 | 0 | 0 | 0 | 0 | 0 | 0 | 0 | 0 | 0 | 0 | 0 | 0 | 0 | 0 | 0 | 0 | 0 | 0 | 0 | 0 |
| virus_May12.25v_12960 | 1 | 0 | 0 | 0 | 1 | 0 | 0 | 0 | 0 | 0 | 0 | 0 | 0 | 0 | 0 | 0 | 0 | 0 | 0 | 0 | 0 | 0 | 0 | 0 | 0 | 0 | 0 | 0 | 0 | 0 | 0 | 0 | 0 | 0 | 0 | 0 | 0 | 0 | 0 | 0 | 0 | 0 | 0 | 0 | 0 | 0 | 0 | 0 | 0 | 0 | 0 | 0 | 0 | 0 | 0 |
| virus_May12.25v_13959 | 1 | 0 | 0 | 0 | 1 | 0 | 0 | 0 | 0 | 0 | 0 | 0 | 0 | 0 | 0 | 0 | 0 | 0 | 0 | 0 | 0 | 0 | 0 | 0 | 0 | 0 | 0 | 0 | 0 | 0 | 0 | 0 | 0 | 0 | 0 | 0 | 0 | 0 | 0 | 0 | 0 | 0 | 0 | 0 | 0 | 0 | 0 | 0 | 0 | 0 | 0 | 0 | 0 | 0 | 0 |
| virus_May12.25v_18041 | 1 | 0 | 0 | 0 | 1 | 0 | 0 | 0 | 0 | 0 | 0 | 0 | 0 | 0 | 0 | 0 | 0 | 0 | 0 | 0 | 0 | 0 | 0 | 0 | 0 | 0 | 0 | 0 | 0 | 0 | 0 | 0 | 0 | 0 | 0 | 0 | 0 | 0 | 0 | 0 | 0 | 0 | 0 | 0 | 0 | 0 | 0 | 0 | 0 | 0 | 0 | 0 | 0 | 0 | 0 |
| virus_May12.25v_24338 | 1 | 0 | 0 | 0 | 1 | 0 | 0 | 0 | 0 | 0 | 0 | 0 | 0 | 0 | 0 | 0 | 0 | 0 | 0 | 0 | 0 | 0 | 0 | 0 | 0 | 0 | 0 | 0 | 0 | 0 | 0 | 0 | 0 | 0 | 0 | 0 | 0 | 0 | 0 | 0 | 0 | 0 | 0 | 0 | 0 | 0 | 0 | 0 | 0 | 0 | 0 | 0 | 0 | 0 | 0 |
| virus_May12.25v_32267 | 1 | 0 | 0 | 0 | 1 | 0 | 0 | 0 | 0 | 0 | 0 | 0 | 0 | 0 | 0 | 0 | 0 | 0 | 0 | 0 | 0 | 0 | 0 | 0 | 0 | 0 | 0 | 0 | 0 | 0 | 0 | 0 | 0 | 0 | 0 | 0 | 0 | 0 | 0 | 0 | 0 | 0 | 0 | 0 | 0 | 0 | 0 | 0 | 0 | 0 | 0 | 0 | 0 | 0 | 0 |
| virus_May12.25v_32412 | 1 | 0 | 0 | 0 | 1 | 0 | 0 | 0 | 0 | 0 | 0 | 0 | 0 | 0 | 0 | 0 | 0 | 0 | 0 | 0 | 0 | 0 | 0 | 0 | 0 | 0 | 0 | 0 | 0 | 0 | 0 | 0 | 0 | 0 | 0 | 0 | 0 | 0 | 0 | 0 | 0 | 0 | 0 | 0 | 0 | 0 | 0 | 0 | 0 | 0 | 0 | 0 | 0 | 0 | 0 |
| virus_May12.50v_2088 | 1 | 0 | 0 | 0 | 0 | 0 | 1 | 0 | 0 | 0 | 0 | 0 | 0 | 0 | 0 | 0 | 0 | 0 | 0 | 0 | 0 | 0 | 0 | 0 | 0 | 0 | 0 | 0 | 0 | 0 | 0 | 0 | 0 | 0 | 0 | 0 | 0 | 0 | 0 | 0 | 0 | 0 | 0 | 0 | 0 | 0 | 0 | 0 | 0 | 0 | 0 | 0 | 0 | 0 | 0 |
| virus_May12.50v_2141 | 1 | 0 | 0 | 0 | 0 | 0 | 1 | 0 | 0 | 0 | 0 | 0 | 0 | 0 | 0 | 0 | 0 | 0 | 0 | 0 | 0 | 0 | 0 | 0 | 0 | 0 | 0 | 0 | 0 | 0 | 0 | 0 | 0 | 0 | 0 | 0 | 0 | 0 | 0 | 0 | 0 | 0 | 0 | 0 | 0 | 0 | 0 | 0 | 0 | 0 | 0 | 0 | 0 | 0 | 0 |
| virus_May12.50v_3367 | 1 | 0 | 0 | 0 | 0 | 0 | 1 | 0 | 0 | 0 | 0 | 0 | 0 | 0 | 0 | 0 | 0 | 0 | 0 | 0 | 0 | 0 | 0 | 0 | 0 | 0 | 0 | 0 | 0 | 0 | 0 | 0 | 0 | 0 | 0 | 0 | 0 | 0 | 0 | 0 | 0 | 0 | 0 | 0 | 0 | 0 | 0 | 0 | 0 | 0 | 0 | 0 | 0 | 0 | 0 |
| virus_May12.50v_4477 | 1 | 0 | 0 | 0 | 0 | 0 | 1 | 0 | 0 | 0 | 0 | 0 | 0 | 0 | 0 | 0 | 0 | 0 | 0 | 0 | 0 | 0 | 0 | 0 | 0 | 0 | 0 | 0 | 0 | 0 | 0 | 0 | 0 | 0 | 0 | 0 | 0 | 0 | 0 | 0 | 0 | 0 | 0 | 0 | 0 | 0 | 0 | 0 | 0 | 0 | 0 | 0 | 0 | 0 | 0 |
| virus_May12.50v_5150 | 1 | 0 | 0 | 0 | 0 | 0 | 1 | 0 | 0 | 0 | 0 | 0 | 0 | 0 | 0 | 0 | 0 | 0 | 0 | 0 | 0 | 0 | 0 | 0 | 0 | 0 | 0 | 0 | 0 | 0 | 0 | 0 | 0 | 0 | 0 | 0 | 0 | 0 | 0 | 0 | 0 | 0 | 0 | 0 | 0 | 0 | 0 | 0 | 0 | 0 | 0 | 0 | 0 | 0 | 0 |
| virus_May12.50v_5341 | 1 | 0 | 0 | 0 | 0 | 0 | 1 | 0 | 0 | 0 | 0 | 0 | 0 | 0 | 0 | 0 | 0 | 0 | 0 | 0 | 0 | 0 | 0 | 0 | 0 | 0 | 0 | 0 | 0 | 0 | 0 | 0 | 0 | 0 | 0 | 0 | 0 | 0 | 0 | 0 | 0 | 0 | 0 | 0 | 0 | 0 | 0 | 0 | 0 | 0 | 0 | 0 | 0 | 0 | 0 |
| virus_May12.50v_9190 | 1 | 0 | 0 | 0 | 0 | 0 | 1 | 0 | 0 | 0 | 0 | 0 | 0 | 0 | 0 | 0 | 0 | 0 | 0 | 0 | 0 | 0 | 0 | 0 | 0 | 0 | 0 | 0 | 0 | 0 | 0 | 0 | 0 | 0 | 0 | 0 | 0 | 0 | 0 | 0 | 0 | 0 | 0 | 0 | 0 | 0 | 0 | 0 | 0 | 0 | 0 | 0 | 0 | 0 | 0 |
| virus_May12.50v_10911 | 1 | 0 | 0 | 0 | 0 | 0 | 1 | 0 | 0 | 0 | 0 | 0 | 0 | 0 | 0 | 0 | 0 | 0 | 0 | 0 | 0 | 0 | 0 | 0 | 0 | 0 | 0 | 0 | 0 | 0 | 0 | 0 | 0 | 0 | 0 | 0 | 0 | 0 | 0 | 0 | 0 | 0 | 0 | 0 | 0 | 0 | 0 | 0 | 0 | 0 | 0 | 0 | 0 | 0 | 0 |
| virus_May12.50v_14924 | 1 | 0 | 0 | 0 | 0 | 0 | 1 | 0 | 0 | 0 | 0 | 0 | 0 | 0 | 0 | 0 | 0 | 0 | 0 | 0 | 0 | 0 | 0 | 0 | 0 | 0 | 0 | 0 | 0 | 0 | 0 | 0 | 0 | 0 | 0 | 0 | 0 | 0 | 0 | 0 | 0 | 0 | 0 | 0 | 0 | 0 | 0 | 0 | 0 | 0 | 0 | 0 | 0 | 0 | 0 |
| virus_May12.50v_16415 | 1 | 0 | 0 | 0 | 0 | 0 | 1 | 0 | 0 | 0 | 0 | 0 | 0 | 0 | 0 | 0 | 0 | 0 | 0 | 0 | 0 | 0 | 0 | 0 | 0 | 0 | 0 | 0 | 0 | 0 | 0 | 0 | 0 | 0 | 0 | 0 | 0 | 0 | 0 | 0 | 0 | 0 | 0 | 0 | 0 | 0 | 0 | 0 | 0 | 0 | 0 | 0 | 0 | 0 | 0 |
| virus_May12.50v_16743 | 1 | 0 | 0 | 0 | 0 | 0 | 1 | 0 | 0 | 0 | 0 | 0 | 0 | 0 | 0 | 0 | 0 | 0 | 0 | 0 | 0 | 0 | 0 | 0 | 0 | 0 | 0 | 0 | 0 | 0 | 0 | 0 | 0 | 0 | 0 | 0 | 0 | 0 | 0 | 0 | 0 | 0 | 0 | 0 | 0 | 0 | 0 | 0 | 0 | 0 | 0 | 0 | 0 | 0 | 0 |
| virus_May12.50v_23648 | 1 | 0 | 0 | 0 | 0 | 0 | 1 | 0 | 0 | 0 | 0 | 0 | 0 | 0 | 0 | 0 | 0 | 0 | 0 | 0 | 0 | 0 | 0 | 0 | 0 | 0 | 0 | 0 | 0 | 0 | 0 | 0 | 0 | 0 | 0 | 0 | 0 | 0 | 0 | 0 | 0 | 0 | 0 | 0 | 0 | 0 | 0 | 0 | 0 | 0 | 0 | 0 | 0 | 0 | 0 |
| virus_May12.50v_35285 | 1 | 0 | 0 | 0 | 0 | 0 | 1 | 0 | 0 | 0 | 0 | 0 | 0 | 0 | 0 | 0 | 0 | 0 | 0 | 0 | 0 | 0 | 0 | 0 | 0 | 0 | 0 | 0 | 0 | 0 | 0 | 0 | 0 | 0 | 0 | 0 | 0 | 0 | 0 | 0 | 0 | 0 | 0 | 0 | 0 | 0 | 0 | 0 | 0 | 0 | 0 | 0 | 0 | 0 | 0 |
| virus_May12.75v_5202 | 1 | 0 | 0 | 0 | 0 | 0 | 0 | 0 | 1 | 0 | 0 | 0 | 0 | 0 | 0 | 0 | 0 | 0 | 0 | 0 | 0 | 0 | 0 | 0 | 0 | 0 | 0 | 0 | 0 | 0 | 0 | 0 | 0 | 0 | 0 | 0 | 0 | 0 | 0 | 0 | 0 | 0 | 0 | 0 | 0 | 0 | 0 | 0 | 0 | 0 | 0 | 0 | 0 | 0 | 0 |
| virus_May12.75v_7025 | 1 | 0 | 0 | 0 | 0 | 0 | 0 | 0 | 1 | 0 | 0 | 0 | 0 | 0 | 0 | 0 | 0 | 0 | 0 | 0 | 0 | 0 | 0 | 0 | 0 | 0 | 0 | 0 | 0 | 0 | 0 | 0 | 0 | 0 | 0 | 0 | 0 | 0 | 0 | 0 | 0 | 0 | 0 | 0 | 0 | 0 | 0 | 0 | 0 | 0 | 0 | 0 | 0 | 0 | 0 |
| virus_May12.75v_7338 | 1 | 0 | 0 | 0 | 0 | 0 | 0 | 0 | 1 | 0 | 0 | 0 | 0 | 0 | 0 | 0 | 0 | 0 | 0 | 0 | 0 | 0 | 0 | 0 | 0 | 0 | 0 | 0 | 0 | 0 | 0 | 0 | 0 | 0 | 0 | 0 | 0 | 0 | 0 | 0 | 0 | 0 | 0 | 0 | 0 | 0 | 0 | 0 | 0 | 0 | 0 | 0 | 0 | 0 | 0 |
| virus_May12.75v_7417 | 1 | 0 | 0 | 0 | 0 | 0 | 0 | 0 | 1 | 0 | 0 | 0 | 0 | 0 | 0 | 0 | 0 | 0 | 0 | 0 | 0 | 0 | 0 | 0 | 0 | 0 | 0 | 0 | 0 | 0 | 0 | 0 | 0 | 0 | 0 | 0 | 0 | 0 | 0 | 0 | 0 | 0 | 0 | 0 | 0 | 0 | 0 | 0 | 0 | 0 | 0 | 0 | 0 | 0 | 0 |
| virus_May12.75v_13775 | 1 | 0 | 0 | 0 | 0 | 0 | 0 | 0 | 1 | 0 | 0 | 0 | 0 | 0 | 0 | 0 | 0 | 0 | 0 | 0 | 0 | 0 | 0 | 0 | 0 | 0 | 0 | 0 | 0 | 0 | 0 | 0 | 0 | 0 | 0 | 0 | 0 | 0 | 0 | 0 | 0 | 0 | 0 | 0 | 0 | 0 | 0 | 0 | 0 | 0 | 0 | 0 | 0 | 0 | 0 |
| virus_May12.75v_13877 | 1 | 0 | 0 | 0 | 0 | 0 | 0 | 0 | 1 | 0 | 0 | 0 | 0 | 0 | 0 | 0 | 0 | 0 | 0 | 0 | 0 | 0 | 0 | 0 | 0 | 0 | 0 | 0 | 0 | 0 | 0 | 0 | 0 | 0 | 0 | 0 | 0 | 0 | 0 | 0 | 0 | 0 | 0 | 0 | 0 | 0 | 0 | 0 | 0 | 0 | 0 | 0 | 0 | 0 | 0 |
| virus_May12.75v_14700 | 1 | 0 | 0 | 0 | 0 | 0 | 0 | 0 | 1 | 0 | 0 | 0 | 0 | 0 | 0 | 0 | 0 | 0 | 0 | 0 | 0 | 0 | 0 | 0 | 0 | 0 | 0 | 0 | 0 | 0 | 0 | 0 | 0 | 0 | 0 | 0 | 0 | 0 | 0 | 0 | 0 | 0 | 0 | 0 | 0 | 0 | 0 | 0 | 0 | 0 | 0 | 0 | 0 | 0 | 0 |
| virus_May12.75v_21511 | 1 | 0 | 0 | 0 | 0 | 0 | 0 | 0 | 1 | 0 | 0 | 0 | 0 | 0 | 0 | 0 | 0 | 0 | 0 | 0 | 0 | 0 | 0 | 0 | 0 | 0 | 0 | 0 | 0 | 0 | 0 | 0 | 0 | 0 | 0 | 0 | 0 | 0 | 0 | 0 | 0 | 0 | 0 | 0 | 0 | 0 | 0 | 0 | 0 | 0 | 0 | 0 | 0 | 0 | 0 |
| virus_May12.75v_24328 | 1 | 0 | 0 | 0 | 0 | 0 | 0 | 0 | 1 | 0 | 0 | 0 | 0 | 0 | 0 | 0 | 0 | 0 | 0 | 0 | 0 | 0 | 0 | 0 | 0 | 0 | 0 | 0 | 0 | 0 | 0 | 0 | 0 | 0 | 0 | 0 | 0 | 0 | 0 | 0 | 0 | 0 | 0 | 0 | 0 | 0 | 0 | 0 | 0 | 0 | 0 | 0 | 0 | 0 | 0 |
| virus_May12.75v_26799 | 1 | 0 | 0 | 0 | 0 | 0 | 0 | 0 | 1 | 0 | 0 | 0 | 0 | 0 | 0 | 0 | 0 | 0 | 0 | 0 | 0 | 0 | 0 | 0 | 0 | 0 | 0 | 0 | 0 | 0 | 0 | 0 | 0 | 0 | 0 | 0 | 0 | 0 | 0 | 0 | 0 | 0 | 0 | 0 | 0 | 0 | 0 | 0 | 0 | 0 | 0 | 0 | 0 | 0 | 0 |
| virus_May12.75v_26962 | 1 | 0 | 0 | 0 | 0 | 0 | 0 | 0 | 1 | 0 | 0 | 0 | 0 | 0 | 0 | 0 | 0 | 0 | 0 | 0 | 0 | 0 | 0 | 0 | 0 | 0 | 0 | 0 | 0 | 0 | 0 | 0 | 0 | 0 | 0 | 0 | 0 | 0 | 0 | 0 | 0 | 0 | 0 | 0 | 0 | 0 | 0 | 0 | 0 | 0 | 0 | 0 | 0 | 0 | 0 |
| virus_May12.75v_38312 | 1 | 0 | 0 | 0 | 0 | 0 | 0 | 0 | 1 | 0 | 0 | 0 | 0 | 0 | 0 | 0 | 0 | 0 | 0 | 0 | 0 | 0 | 0 | 0 | 0 | 0 | 0 | 0 | 0 | 0 | 0 | 0 | 0 | 0 | 0 | 0 | 0 | 0 | 0 | 0 | 0 | 0 | 0 | 0 | 0 | 0 | 0 | 0 | 0 | 0 | 0 | 0 | 0 | 0 | 0 |
| virus_Sep8.0v_205 | 1 | 0 | 0 | 0 | 0 | 0 | 0 | 0 | 0 | 0 | 0 | 0 | 0 | 0 | 0 | 0 | 0 | 0 | 0 | 0 | 0 | 0 | 0 | 0 | 0 | 0 | 0 | 0 | 0 | 0 | 0 | 0 | 0 | 0 | 0 | 0 | 0 | 0 | 0 | 0 | 0 | 0 | 0 | 0 | 0 | 0 | 0 | 0 | 1 | 0 | 0 | 0 | 0 | 0 | 0 |
| virus_Sep8.0v_2148 | 1 | 0 | 0 | 0 | 0 | 0 | 0 | 0 | 0 | 0 | 0 | 0 | 0 | 0 | 0 | 0 | 0 | 0 | 0 | 0 | 0 | 0 | 0 | 0 | 0 | 0 | 0 | 0 | 0 | 0 | 0 | 0 | 0 | 0 | 0 | 0 | 0 | 0 | 0 | 0 | 0 | 0 | 0 | 0 | 0 | 0 | 0 | 0 | 1 | 0 | 0 | 0 | 0 | 0 | 0 |
| virus_Sep8.0v_3627 | 1 | 0 | 0 | 0 | 0 | 0 | 0 | 0 | 0 | 0 | 0 | 0 | 0 | 0 | 0 | 0 | 0 | 0 | 0 | 0 | 0 | 0 | 0 | 0 | 0 | 0 | 0 | 0 | 0 | 0 | 0 | 0 | 0 | 0 | 0 | 0 | 0 | 0 | 0 | 0 | 0 | 0 | 0 | 0 | 0 | 0 | 0 | 0 | 1 | 0 | 0 | 0 | 0 | 0 | 0 |
| virus_Sep8.0v_3910 | 1 | 0 | 0 | 0 | 0 | 0 | 0 | 0 | 0 | 0 | 0 | 0 | 0 | 0 | 0 | 0 | 0 | 0 | 0 | 0 | 0 | 0 | 0 | 0 | 0 | 0 | 0 | 0 | 0 | 0 | 0 | 0 | 0 | 0 | 0 | 0 | 0 | 0 | 0 | 0 | 0 | 0 | 0 | 0 | 0 | 0 | 0 | 0 | 1 | 0 | 0 | 0 | 0 | 0 | 0 |
| virus_Sep8.0v_4218 | 1 | 0 | 0 | 0 | 0 | 0 | 0 | 0 | 0 | 0 | 0 | 0 | 0 | 0 | 0 | 0 | 0 | 0 | 0 | 0 | 0 | 0 | 0 | 0 | 0 | 0 | 0 | 0 | 0 | 0 | 0 | 0 | 0 | 0 | 0 | 0 | 0 | 0 | 0 | 0 | 0 | 0 | 0 | 0 | 0 | 0 | 0 | 0 | 1 | 0 | 0 | 0 | 0 | 0 | 0 |
| virus_Sep8.0v_4467 | 1 | 0 | 0 | 0 | 0 | 0 | 0 | 0 | 0 | 0 | 0 | 0 | 0 | 0 | 0 | 0 | 0 | 0 | 0 | 0 | 0 | 0 | 0 | 0 | 0 | 0 | 0 | 0 | 0 | 0 | 0 | 0 | 0 | 0 | 0 | 0 | 0 | 0 | 0 | 0 | 0 | 0 | 0 | 0 | 0 | 0 | 0 | 0 | 1 | 0 | 0 | 0 | 0 | 0 | 0 |
| virus_Sep8.0v_6813 | 1 | 0 | 0 | 0 | 0 | 0 | 0 | 0 | 0 | 0 | 0 | 0 | 0 | 0 | 0 | 0 | 0 | 0 | 0 | 0 | 0 | 0 | 0 | 0 | 0 | 0 | 0 | 0 | 0 | 0 | 0 | 0 | 0 | 0 | 0 | 0 | 0 | 0 | 0 | 0 | 0 | 0 | 0 | 0 | 0 | 0 | 0 | 0 | 1 | 0 | 0 | 0 | 0 | 0 | 0 |
| virus_Sep8.0v_11375 | 1 | 0 | 0 | 0 | 0 | 0 | 0 | 0 | 0 | 0 | 0 | 0 | 0 | 0 | 0 | 0 | 0 | 0 | 0 | 0 | 0 | 0 | 0 | 0 | 0 | 0 | 0 | 0 | 0 | 0 | 0 | 0 | 0 | 0 | 0 | 0 | 0 | 0 | 0 | 0 | 0 | 0 | 0 | 0 | 0 | 0 | 0 | 0 | 1 | 0 | 0 | 0 | 0 | 0 | 0 |
| virus_Sep8.0v_11931 | 1 | 0 | 0 | 0 | 0 | 0 | 0 | 0 | 0 | 0 | 0 | 0 | 0 | 0 | 0 | 0 | 0 | 0 | 0 | 0 | 0 | 0 | 0 | 0 | 0 | 0 | 0 | 0 | 0 | 0 | 0 | 0 | 0 | 0 | 0 | 0 | 0 | 0 | 0 | 0 | 0 | 0 | 0 | 0 | 0 | 0 | 0 | 0 | 1 | 0 | 0 | 0 | 0 | 0 | 0 |
| virus_Sep8.0v_15950 | 1 | 0 | 0 | 0 | 0 | 0 | 0 | 0 | 0 | 0 | 0 | 0 | 0 | 0 | 0 | 0 | 0 | 0 | 0 | 0 | 0 | 0 | 0 | 0 | 0 | 0 | 0 | 0 | 0 | 0 | 0 | 0 | 0 | 0 | 0 | 0 | 0 | 0 | 0 | 0 | 0 | 0 | 0 | 0 | 0 | 0 | 0 | 0 | 1 | 0 | 0 | 0 | 0 | 0 | 0 |
| virus_Sep8.0v_17805 | 1 | 0 | 0 | 0 | 0 | 0 | 0 | 0 | 0 | 0 | 0 | 0 | 0 | 0 | 0 | 0 | 0 | 0 | 0 | 0 | 0 | 0 | 0 | 0 | 0 | 0 | 0 | 0 | 0 | 0 | 0 | 0 | 0 | 0 | 0 | 0 | 0 | 0 | 0 | 0 | 0 | 0 | 0 | 0 | 0 | 0 | 0 | 0 | 1 | 0 | 0 | 0 | 0 | 0 | 0 |
| virus_Sep8.5v_460 | 1 | 0 | 0 | 0 | 0 | 0 | 0 | 0 | 0 | 0 | 0 | 0 | 0 | 0 | 0 | 0 | 0 | 0 | 0 | 0 | 0 | 0 | 0 | 0 | 0 | 0 | 0 | 0 | 0 | 0 | 0 | 0 | 0 | 0 | 0 | 0 | 0 | 0 | 0 | 0 | 0 | 0 | 0 | 0 | 0 | 0 | 0 | 0 | 0 | 0 | 1 | 0 | 0 | 0 | 0 |
| virus_Sep8.5v_4351 | 1 | 0 | 0 | 0 | 0 | 0 | 0 | 0 | 0 | 0 | 0 | 0 | 0 | 0 | 0 | 0 | 0 | 0 | 0 | 0 | 0 | 0 | 0 | 0 | 0 | 0 | 0 | 0 | 0 | 0 | 0 | 0 | 0 | 0 | 0 | 0 | 0 | 0 | 0 | 0 | 0 | 0 | 0 | 0 | 0 | 0 | 0 | 0 | 0 | 0 | 1 | 0 | 0 | 0 | 0 |
| virus_Sep8.5v_15891 | 1 | 0 | 0 | 0 | 0 | 0 | 0 | 0 | 0 | 0 | 0 | 0 | 0 | 0 | 0 | 0 | 0 | 0 | 0 | 0 | 0 | 0 | 0 | 0 | 0 | 0 | 0 | 0 | 0 | 0 | 0 | 0 | 0 | 0 | 0 | 0 | 0 | 0 | 0 | 0 | 0 | 0 | 0 | 0 | 0 | 0 | 0 | 0 | 0 | 0 | 1 | 0 | 0 | 0 | 0 |
| virus_Sep8.5v_20490 | 1 | 0 | 0 | 0 | 0 | 0 | 0 | 0 | 0 | 0 | 0 | 0 | 0 | 0 | 0 | 0 | 0 | 0 | 0 | 0 | 0 | 0 | 0 | 0 | 0 | 0 | 0 | 0 | 0 | 0 | 0 | 0 | 0 | 0 | 0 | 0 | 0 | 0 | 0 | 0 | 0 | 0 | 0 | 0 | 0 | 0 | 0 | 0 | 0 | 0 | 1 | 0 | 0 | 0 | 0 |
| virus_Sep8.25v_19340 | 1 | 0 | 0 | 0 | 0 | 0 | 0 | 0 | 0 | 0 | 0 | 0 | 0 | 0 | 0 | 0 | 0 | 0 | 0 | 0 | 0 | 0 | 0 | 0 | 0 | 0 | 0 | 0 | 0 | 0 | 0 | 0 | 0 | 0 | 0 | 0 | 0 | 0 | 0 | 0 | 0 | 0 | 0 | 0 | 0 | 0 | 0 | 0 | 0 | 0 | 0 | 0 | 1 | 0 | 0 |
| virus_Sep8.50v_1272 | 1 | 0 | 0 | 0 | 0 | 0 | 0 | 0 | 0 | 0 | 0 | 0 | 0 | 0 | 0 | 0 | 0 | 0 | 0 | 0 | 0 | 0 | 0 | 0 | 0 | 0 | 0 | 0 | 0 | 0 | 0 | 0 | 0 | 0 | 0 | 0 | 0 | 0 | 0 | 0 | 0 | 0 | 0 | 0 | 0 | 0 | 0 | 0 | 0 | 0 | 0 | 0 | 0 | 0 | 1 |
| virus_Sep8.50v_5442 | 1 | 0 | 0 | 0 | 0 | 0 | 0 | 0 | 0 | 0 | 0 | 0 | 0 | 0 | 0 | 0 | 0 | 0 | 0 | 0 | 0 | 0 | 0 | 0 | 0 | 0 | 0 | 0 | 0 | 0 | 0 | 0 | 0 | 0 | 0 | 0 | 0 | 0 | 0 | 0 | 0 | 0 | 0 | 0 | 0 | 0 | 0 | 0 | 0 | 0 | 0 | 0 | 0 | 0 | 1 |
| virus_Sep8.50v_7321 | 1 | 0 | 0 | 0 | 0 | 0 | 0 | 0 | 0 | 0 | 0 | 0 | 0 | 0 | 0 | 0 | 0 | 0 | 0 | 0 | 0 | 0 | 0 | 0 | 0 | 0 | 0 | 0 | 0 | 0 | 0 | 0 | 0 | 0 | 0 | 0 | 0 | 0 | 0 | 0 | 0 | 0 | 0 | 0 | 0 | 0 | 0 | 0 | 0 | 0 | 0 | 0 | 0 | 0 | 1 |
| virus_Sep8.50v_9303 | 1 | 0 | 0 | 0 | 0 | 0 | 0 | 0 | 0 | 0 | 0 | 0 | 0 | 0 | 0 | 0 | 0 | 0 | 0 | 0 | 0 | 0 | 0 | 0 | 0 | 0 | 0 | 0 | 0 | 0 | 0 | 0 | 0 | 0 | 0 | 0 | 0 | 0 | 0 | 0 | 0 | 0 | 0 | 0 | 0 | 0 | 0 | 0 | 0 | 0 | 0 | 0 | 0 | 0 | 1 |
| virus_Sep8.50v_11296 | 1 | 0 | 0 | 0 | 0 | 0 | 0 | 0 | 0 | 0 | 0 | 0 | 0 | 0 | 0 | 0 | 0 | 0 | 0 | 0 | 0 | 0 | 0 | 0 | 0 | 0 | 0 | 0 | 0 | 0 | 0 | 0 | 0 | 0 | 0 | 0 | 0 | 0 | 0 | 0 | 0 | 0 | 0 | 0 | 0 | 0 | 0 | 0 | 0 | 0 | 0 | 0 | 0 | 0 | 1 |
| virus_Sep8.50v_17089 | 1 | 0 | 0 | 0 | 0 | 0 | 0 | 0 | 0 | 0 | 0 | 0 | 0 | 0 | 0 | 0 | 0 | 0 | 0 | 0 | 0 | 0 | 0 | 0 | 0 | 0 | 0 | 0 | 0 | 0 | 0 | 0 | 0 | 0 | 0 | 0 | 0 | 0 | 0 | 0 | 0 | 0 | 0 | 0 | 0 | 0 | 0 | 0 | 0 | 0 | 0 | 0 | 0 | 0 | 1 |
| virus_Sep8.50v_21946 | 1 | 0 | 0 | 0 | 0 | 0 | 0 | 0 | 0 | 0 | 0 | 0 | 0 | 0 | 0 | 0 | 0 | 0 | 0 | 0 | 0 | 0 | 0 | 0 | 0 | 0 | 0 | 0 | 0 | 0 | 0 | 0 | 0 | 0 | 0 | 0 | 0 | 0 | 0 | 0 | 0 | 0 | 0 | 0 | 0 | 0 | 0 | 0 | 0 | 0 | 0 | 0 | 0 | 0 | 1 |
| virus_Sep8.50v_28128 | 1 | 0 | 0 | 0 | 0 | 0 | 0 | 0 | 0 | 0 | 0 | 0 | 0 | 0 | 0 | 0 | 0 | 0 | 0 | 0 | 0 | 0 | 0 | 0 | 0 | 0 | 0 | 0 | 0 | 0 | 0 | 0 | 0 | 0 | 0 | 0 | 0 | 0 | 0 | 0 | 0 | 0 | 0 | 0 | 0 | 0 | 0 | 0 | 0 | 0 | 0 | 0 | 0 | 0 | 1 |
| virus_Sep8.50v_28342 | 1 | 0 | 0 | 0 | 0 | 0 | 0 | 0 | 0 | 0 | 0 | 0 | 0 | 0 | 0 | 0 | 0 | 0 | 0 | 0 | 0 | 0 | 0 | 0 | 0 | 0 | 0 | 0 | 0 | 0 | 0 | 0 | 0 | 0 | 0 | 0 | 0 | 0 | 0 | 0 | 0 | 0 | 0 | 0 | 0 | 0 | 0 | 0 | 0 | 0 | 0 | 0 | 0 | 0 | 1 |
| virus_Sep8.50v_32606 | 1 | 0 | 0 | 0 | 0 | 0 | 0 | 0 | 0 | 0 | 0 | 0 | 0 | 0 | 0 | 0 | 0 | 0 | 0 | 0 | 0 | 0 | 0 | 0 | 0 | 0 | 0 | 0 | 0 | 0 | 0 | 0 | 0 | 0 | 0 | 0 | 0 | 0 | 0 | 0 | 0 | 0 | 0 | 0 | 0 | 0 | 0 | 0 | 0 | 0 | 0 | 0 | 0 | 0 | 1 |
| virus_Sep8.50v_35096 | 1 | 0 | 0 | 0 | 0 | 0 | 0 | 0 | 0 | 0 | 0 | 0 | 0 | 0 | 0 | 0 | 0 | 0 | 0 | 0 | 0 | 0 | 0 | 0 | 0 | 0 | 0 | 0 | 0 | 0 | 0 | 0 | 0 | 0 | 0 | 0 | 0 | 0 | 0 | 0 | 0 | 0 | 0 | 0 | 0 | 0 | 0 | 0 | 0 | 0 | 0 | 0 | 0 | 0 | 1 |
| virus_Sep8.50v_38342 | 1 | 0 | 0 | 0 | 0 | 0 | 0 | 0 | 0 | 0 | 0 | 0 | 0 | 0 | 0 | 0 | 0 | 0 | 0 | 0 | 0 | 0 | 0 | 0 | 0 | 0 | 0 | 0 | 0 | 0 | 0 | 0 | 0 | 0 | 0 | 0 | 0 | 0 | 0 | 0 | 0 | 0 | 0 | 0 | 0 | 0 | 0 | 0 | 0 | 0 | 0 | 0 | 0 | 0 | 1 |

**Supplementary Table S8:** Common RTPR 98% OTUs (>= 10 seqs in 27 samples with amplicons in both fractions)

| OTU_ID | fraction composition | clr(virioplankton)-clr(bacterioplankton) | clr(virioplankton)+clr(bacterioplankton) |
|---|---|---|---|
| virus_Aug5.75v_348 | only virioplankton | 5 | 1.111878605 |
| virus_Jul11.50v_15972 | only virioplankton | 5 | 0.603556111 |
| virus_Jul11.0v_7341 | only virioplankton | 5 | 0.113349775 |
| virus_Aug5.25v_30934 | only virioplankton | 5 | 0.050170873 |
| virus_May12.5v_21769 | only virioplankton | 5 | -0.140884364 |
| virus_Jun8.75v_1537 | only virioplankton | 5 | -0.377273142 |
| virus_Jun22.75v_24976 | only virioplankton | 5 | -0.600416693 |
| virus_Jun8.75v_10037 | only virioplankton | 5 | -0.600416693 |
| virus_Jul27.25v_13167 | only virioplankton | 5 | -0.68742807 |
| virus_Aug5.5v_13145 | only virioplankton | 5 | -0.78273825 |
| virus_Jul11.5v_10009 | only virioplankton | 5 | -0.834031544 |
| virus_Jul11.5v_17064 | only virioplankton | 5 | -0.834031544 |
| virus_Aug5.5v_32417 | only virioplankton | 5 | -0.888098766 |
| virus_Aug5.5v_13274 | only virioplankton | 5 | -1.070420322 |
| virus_Jun8.50v_27002 | only virioplankton | 5 | -1.139413194 |
| virus_Jul11.50v_16712 | only virioplankton | 5 | -1.213521166 |
| virus_Sep8.0v_3844 | only virioplankton | 5 | -1.293563874 |
| virus_Jul27.5v_1553 | only virioplankton | 5 | -1.47588543 |
| virus_Jun8.50v_40442 | only virioplankton | 5 | -1.47588543 |
| virus_Jun22.0v_12932 | only virioplankton | 5 | -1.47588543 |
| virus_Sep8.0v_10243 | only virioplankton | 5 | -1.47588543 |
| bacteria_Aug5d25b_1039 | both | 0.014897134 | 4.689742486 |
| bacteria_Jul11d0b_35947 | both | 0.027751265 | 4.317128178 |
| bacteria_Jul11d0b_349 | both | -0.692691665 | 4.18707505 |
| bacteria_Jul11d0b_5406 | both | 0.36440468 | 3.982422672 |
| bacteria_Aug5d25b_42861 | both | -0.774029677 | 3.876446524 |
| bacteria_Jul11d0b_35378 | both | 0.780412147 | 3.549966829 |
| bacteria_Jul11d0b_26490 | both | 0.155905927 | 3.535416308 |
| bacteria_Aug5d5b_22963 | both | 0.824043228 | 3.534749864 |
| bacteria_Jul11d0b_24544 | both | 0.267170825 | 3.449191975 |
| bacteria_Jul11d0b_624 | both | 0.852373735 | 3.045903147 |
| bacteria_Jul11d0b_26699 | both | 0.358228713 | 2.986568453 |
| bacteria_Aug5d25b_21123 | both | 0.786616089 | 2.970289024 |
| bacteria_May12d50b_25296 | both | 0.624551511 | 2.802168614 |
| virus_Jul11.25v_8557 | both | 0.518935214 | 2.620124411 |
| virus_Aug5.5v_32649 | both | 1.071576571 | 2.507527571 |
| virus_Jul11.25v_14520 | both | 0.356335812 | 2.39531558 |
| bacteria_Aug5d25b_14585 | both | 0.069560677 | 2.382736798 |
| bacteria_Jul27d5b_14425 | both | 0.692246198 | 2.259400396 |
| bacteria_Aug5d5b_43576 | both | 0.179418869 | 2.107633508 |
| virus_Aug5.25v_30214 | both | 0.878459404 | 2.104851865 |
| bacteria_Jul11d0b_19157 | both | -0.350388252 | 2.053411954 |
| bacteria_Aug5d25b_16734 | both | 0.53227421 | 2.020622131 |
| bacteria_Aug5d5b_15972 | both | 0.438202985 | 1.996081022 |
| bacteria_Aug5d5b_14294 | both | -0.148485046 | 1.925311951 |
| bacteria_Jul27d5b_24937 | both | 0.598799015 | 1.866976374 |
| bacteria_Jul11d25b_11656 | both | 0.109698918 | 1.778357538 |
| bacteria_Jul11d0b_27216 | both | 0.963689837 | 1.766706921 |
| bacteria_Jul11d0b_1759 | both | -0.056663865 | 1.697993029 |
| virus_Jul11.25v_9299 | both | 0.775468558 | 1.69380015 |
| bacteria_Jul27d75b_47191 | both | 0.082321377 | 1.663947187 |
| virus_Aug5.5v_9076 | both | 0.816053838 | 1.650875106 |
| bacteria_Jul11d0b_29643 | both | 0.912490356 | 1.559067556 |
| bacteria_Aug5d25b_11190 | both | -0.612278236 | 1.479024849 |
| bacteria_Jul11d0b_3527 | both | 0.673446944 | 1.420026508 |
| virus_Jul11.0v_6020 | both | 0.181066655 | 1.392013472 |
| virus_Aug5.5v_6951 | both | 4.755868861 | 1.386315451 |
| bacteria_Aug5d25b_52757 | both | 0.455475182 | 1.345493456 |
| bacteria_Jul11d0b_48732 | both | 0.608414474 | 1.32747495 |
| bacteria_Aug5d5b_65184 | both | 0.338750907 | 1.264954593 |
| bacteria_Jul11d0b_6084 | both | -4.295006736 | 1.232164771 |
| virus_Aug5.0v_4276 | both | 1.110107614 | 1.111878605 |
| bacteria_Jul11d0b_14765 | both | 1.319580159 | 1.104331399 |
| virus_Aug5.25v_38270 | both | 4.464348013 | 1.0967268 |
| virus_Aug5.0v_282 | both | 1.115279696 | 1.07355974 |
| virus_Jun22.50v_12875 | both | 0.314058294 | 1.049843214 |
| bacteria_Jul11d5b_2558 | both | -0.369284887 | 0.992214101 |
| bacteria_Aug5d0b_30617 | both | -0.518699325 | 0.865920376 |
| bacteria_Aug5d25b_41725 | both | -4.935785931 | 0.764824259 |
| bacteria_Aug5d5b_15792 | both | 0.695425851 | 0.743318054 |
| bacteria_Jun8d25b_45572 | both | 0.528371766 | 0.743318054 |
| bacteria_Jul11d0b_25777 | both | -0.357724064 | 0.698866291 |
| bacteria_Jul11d0b_9089 | both | 0.032131633 | 0.652346275 |
| virus_Aug5.0v_24 | both | 0.629828568 | 0.603556111 |
| virus_Aug5.0v_3150 | both | 0.890734603 | 0.590977329 |
| bacteria_Jul11d75b_8617 | both | 0.766884815 | 0.552262817 |
| virus_Aug5.5v_12684 | both | -0.269655046 | 0.53901759 |
| bacteria_Jun8d25b_21024 | both | 0.393144979 | 0.52559457 |
| bacteria_Jul11d0b_7291 | both | 0.188864626 | 0.511988918 |
| bacteria_Jul11d0b_47245 | both | -1.116952906 | 0.511988918 |
| bacteria_Jul11d0b_71717 | both | -0.009282152 | 0.498195596 |
| bacteria_Jul11d75b_24464 | both | -3.958534499 | 0.498195596 |
| bacteria_Jul11d0b_35117 | both | -3.899693999 | 0.441037182 |
| virus_Aug5.5v_4150 | both | 3.801506181 | 0.441037182 |
| bacteria_Jul11d0b_59045 | both | -0.962802226 | 0.411184219 |
| bacteria_Aug5d50b_2154 | both | 0.222519462 | 0.411184219 |
| bacteria_Jul11d75b_39799 | both | -0.061437144 | 0.395916746 |
| bacteria_May12d50b_8587 | both | -0.088105391 | 0.38041256 |
| virus_Jul11.50v_9120 | both | 0.082321378 | 0.364664203 |
| bacteria_Aug5d25b_46297 | both | -2.836799793 | 0.348663862 |
| bacteria_Aug5d50b_14505 | both | 0.570262708 | 0.348663862 |
| bacteria_Aug5d25b_11031 | both | -1.250484298 | 0.315874039 |
| bacteria_Jun8d25b_69463 | both | -0.851211161 | 0.29906692 |
| bacteria_Aug5d25b_32559 | both | -2.389101922 | 0.281972487 |
| virus_May12.75v_24636 | both | 0.395321258 | 0.281972487 |
| bacteria_Jul27d25b_10877 | both | 0.235893522 | 0.228862662 |
| bacteria_Aug5d0b_30481 | both | -0.32314373 | 0.133552482 |
| bacteria_Aug5d0b_47134 | both | 0.26010778 | 0.133552482 |
| bacteria_Jul11d25b_6636 | both | -0.103081846 | 0.071677078 |
| bacteria_Aug5d25b_33881 | both | 1.15495818 | 0.050170873 |
| virus_May12.75v_39998 | both | 1.288489573 | 0.028191966 |
| bacteria_Jun22d0b_13316 | both | 0.664654192 | -0.017270408 |
| bacteria_Jul11d25b_8174 | both | -0.990973103 | -0.040800905 |
| bacteria_Jul11d0b_58291 | both | -0.303102979 | -0.089591069 |
| bacteria_Jul27d5b_12006 | both | -2.915492062 | -0.089591069 |
| bacteria_Jul11d0b_58550 | both | 0.407743778 | -0.114908877 |
| virus_May12.50v_14428 | both | 0.066817191 | -0.114908877 |
| bacteria_Jul27d75b_43975 | both | 0.494755155 | -0.140884364 |
| bacteria_May12d50b_47769 | both | 0.626432979 | -0.140884364 |
| virus_Jul27.25v_12504 | both | 0.37000345 | -0.140884364 |
| bacteria_Jul11d75b_16607 | both | -0.020194186 | -0.167552611 |
| bacteria_Aug5d25b_53723 | both | -0.180042886 | -0.194951585 |
| bacteria_Aug5d25b_72218 | both | 0.048798685 | -0.194951585 |
| bacteria_Jul11d75b_30241 | both | 0.048798685 | -0.194951585 |
| bacteria_Jul11d0b_38787 | both | -3.929546963 | -0.223122462 |
| bacteria_Jun22d50b_26980 | both | 2.369402284 | -0.252109999 |
| bacteria_May12d25b_9336 | both | 0.775468558 | -0.252109999 |
| bacteria_Jun22d0b_13423 | both | 0.472282299 | -0.252109999 |
| bacteria_Jul11d75b_5405 | both | -3.868922341 | -0.281962962 |
| bacteria_Jul11d75b_6234 | both | -0.962802226 | -0.281962962 |
| bacteria_Jul11d75b_33761 | both | 0.15642935 | -0.281962962 |
| virus_Jun8.25v_17498 | both | 3.030800767 | -0.312734621 |
| bacteria_Sep8d5b_67580 | both | -1.592770505 | -0.377273142 |
| virus_Aug5.25v_27001 | both | 0.608414473 | -0.377273142 |
| bacteria_Jul27d75b_14154 | both | 1.794038139 | -0.377273142 |
| virus_Jul11.0v_1381 | both | 0.940548309 | -0.411174693 |
| virus_Jul11.0v_20945 | both | 1.429395026 | -0.411174693 |
| bacteria_Jul11d0b_2761 | both | 0.238667448 | -0.411174693 |
| bacteria_Aug5d25b_3402 | both | 0.184600227 | -0.446266013 |
| virus_Jul27.75v_9932 | both | 2.892650428 | -0.446266013 |

| OTU_ID | fraction composition | clr(virioplankton)-clr(bacterioplankton) | clr(virioplankton)+clr(bacterioplankton) |
|---|---|---|---|
| bacteria_Aug5d25b_35406 | both | -0.028492989 | -0.482633657 |
| bacteria_Aug5d50b_10806 | both | -0.329078466 | -0.482633657 |
| bacteria_Aug5d50b_38363 | both | 1.676255103 | -0.482633657 |
| bacteria_Jul27d25b_34258 | both | 0.127441813 | -0.482633657 |
| virus_Jul11.5v_18046 | both | 2.815689387 | -0.520373985 |
| virus_Jul11.0v_4093 | both | 0.172177707 | -0.559594699 |
| bacteria_Jul27d5b_12937 | both | 0.172177707 | -0.559594699 |
| virus_Jul11.50v_3426 | both | 0.983107924 | -0.559594699 |
| bacteria_Jul11d75b_22 | both | 0.931814629 | -0.600416693 |
| virus_Aug5.5v_14779 | both | -0.236132353 | -0.600416693 |
| bacteria_Aug5d0b_10997 | both | -2.754561695 | -0.642976307 |
| bacteria_Jul27d5b_7357 | both | 0.423492135 | -0.642976307 |
| virus_Jul11.0v_16718 | both | 2.687856015 | -0.642976307 |
| bacteria_Jul27d5b_8488 | both | 0.423492135 | -0.642976307 |
| virus_May12.50v_38831 | both | -0.490197815 | -0.642976307 |
| bacteria_Jun22d75b_73848 | both | 0.358953614 | -0.68742807 |
| virus_Jul11.5v_18386 | both | -0.403186438 | -0.68742807 |
| virus_Jun22.75v_32241 | both | 0.289960743 | -0.733948086 |
| virus_May12.5v_690 | both | 0.983107923 | -0.78273825 |
| bacteria_Jun22d0b_53542 | both | 0.00227867 | -0.78273825 |
| bacteria_Jul27d25b_21364 | both | -0.508546954 | -0.834031544 |
| bacteria_Jul11d0b_37708 | both | -1.358697883 | -0.888098766 |
| bacteria_Aug5d25b_41310 | both | 0.552325008 | -0.888098766 |
| bacteria_Jul11d5b_41219 | both | 1.137258603 | -0.945257179 |
| bacteria_Jul27d5b_12427 | both | 0.4722823 | -0.945257179 |
| bacteria_Sep8d5b_28522 | both | 0.775468559 | -0.945257179 |
| virus_Aug5.0v_3881 | both | -0.046511494 | -0.945257179 |
| virus_Jun8.50v_22514 | both | -0.285403403 | -0.945257179 |
| virus_May12.25v_28535 | both | 2.369402284 | -0.945257179 |
| bacteria_Aug5d25b_43739 | both | 0.775468559 | -0.945257179 |
| bacteria_Jun8d50b_165 | both | -0.15187201 | -1.005881801 |
| bacteria_Jun22d75b_72407 | both | 0.695425851 | -1.005881801 |
| bacteria_May12d50b_2360 | both | 0.695425851 | -1.005881801 |
| bacteria_Sep8d5b_207 | both | -0.403186438 | -1.005881801 |
| virus_Aug5.0v_4316 | both | -0.15187201 | -1.005881801 |
| bacteria_Jul27d75b_27465 | both | 1.468615739 | -1.070420322 |
| bacteria_Jun8d25b_22580 | both | -1.789480799 | -1.070420322 |
| bacteria_Jun8d25b_68299 | both | -2.274988615 | -1.070420322 |
| virus_Aug5.25v_14023 | both | 1.468615739 | -1.070420322 |
| virus_Jul11.50v_7164 | both | 0.983107923 | -1.070420322 |
| virus_Sep8.50v_12712 | both | -1.096333618 | -1.070420322 |
| bacteria_Jul11d25b_1285 | both | -1.789480799 | -1.070420322 |
| virus_May12.50v_5104 | both | 0.289960743 | -1.070420322 |
| bacteria_Jul27d25b_72494 | both | -0.690868511 | -1.139413194 |
| bacteria_Sep8d5b_60365 | both | 0.184600227 | -1.139413194 |
| virus_Aug5.0v_276 | both | 0.513104294 | -1.139413194 |
| virus_Jul27.5v_8523 | both | -2.194945907 | -1.139413194 |
| virus_Aug5.75v_7326 | both | 0.513104294 | -1.139413194 |
| virus_Jul11.50v_11743 | both | 0.184600227 | -1.139413194 |
| virus_Aug5.0v_1938 | both | -0.249035758 | -1.213521166 |
| bacteria_Jul27d75b_69505 | both | -0.249035758 | -1.213521166 |
| virus_Jul11.5v_8752 | both | -0.249035758 | -1.213521166 |
| bacteria_Aug5d0b_16335 | both | -0.557337118 | -1.213521166 |
| virus_Sep8.0v_17751 | both | 1.994708835 | -1.293563874 |
| bacteria_Jul11d0b_17993 | both | 0.69542585 | -1.293563874 |
| virus_Aug5.25v_16404 | both | 1.994708835 | -1.293563874 |
| virus_Aug5.25v_26024 | both | 0.289960743 | -1.293563874 |
| virus_Aug5.25v_40296 | both | 0.289960743 | -1.293563874 |
| virus_Jul27.25v_38635 | both | 0.69542585 | -1.293563874 |
| virus_Jul27.5v_8387 | both | 0.69542585 | -1.293563874 |
| virus_Jul27.5v_10015 | both | -1.096333619 | -1.293563874 |
| virus_Jul11.75v_30378 | both | 1.206251474 | -1.293563874 |
| bacteria_Jul11d5b_3980 | both | 0.15642935 | -1.380575251 |
| bacteria_Jun22d0b_31457 | both | 1.100890958 | -1.380575251 |
| virus_Aug5.0v_3331 | both | 1.100890958 | -1.380575251 |
| virus_Aug5.25v_22058 | both | 1.899398655 | -1.380575251 |
| virus_Sep8.0v_19674 | both | 1.100890958 | -1.380575251 |
| virus_Sep8.50v_1306 | both | -0.585507994 | -1.380575251 |
| bacteria_Jul11d75b_42061 | both | 1.100890958 | -1.380575251 |
| bacteria_Jul27d5b_7248 | both | 0.577642815 | -1.380575251 |
| virus_May12.75v_13478 | both | 0.577642815 | -1.380575251 |
| bacteria_Jul27d25b_32815 | both | 0.577642815 | -1.380575251 |
| bacteria_Aug5d25b_32181 | both | -0.808651546 | -1.47588543 |
| bacteria_Jul11d50b_41285 | both | -0.808651546 | -1.47588543 |
| bacteria_Jul27d75b_20546 | both | -1.789480799 | -1.47588543 |
| virus_Aug5.25v_20741 | both | 1.794038139 | -1.47588543 |
| virus_Aug5.25v_23422 | both | 0.983107923 | -1.47588543 |
| bacteria_Aug5d25b_10485 | both | 0.983107923 | -1.47588543 |
| bacteria_Aug5d25b_20975 | both | -1.250484298 | -1.47588543 |
| bacteria_Jul11d75b_15188 | both | -0.403186437 | -1.47588543 |
| bacteria_Jul11d75b_25852 | both | 0.983107923 | -1.47588543 |
| bacteria_Jul11d50b_13030 | both | 0.444111422 | -1.47588543 |
| virus_Jul11.25v_3423 | both | 0.983107923 | -1.47588543 |
| bacteria_Jul11d25b_4728 | both | -0.403186437 | -1.47588543 |
| bacteria_Aug5d50b_36848 | only bacterioplankton | -5 | 2.424064994 |
| bacteria_Aug5d50b_68297 | only bacterioplankton | -5 | 1.681114991 |
| bacteria_Jul11d25b_36968 | only bacterioplankton | -5 | 1.504733205 |
| bacteria_Jul11d5b_42515 | only bacterioplankton | -5 | 0.940028348 |
| bacteria_Jul11d0b_17411 | only bacterioplankton | -5 | 0.922009842 |
| bacteria_Sep8d5b_44832 | only bacterioplankton | -5 | 0.903660704 |
| bacteria_Aug5d25b_59498 | only bacterioplankton | -5 | 0.875489827 |
| bacteria_Jul27d5b_19668 | only bacterioplankton | -5 | 0.732388983 |
| bacteria_Jun8d25b_34384 | only bacterioplankton | -5 | 0.721339147 |
| bacteria_Aug5d0b_42381 | only bacterioplankton | -5 | 0.675876773 |
| bacteria_Jul11d50b_51257 | only bacterioplankton | -5 | 0.675876773 |
| bacteria_Jun8d25b_55773 | only bacterioplankton | -5 | 0.640370084 |
| bacteria_Jul11d50b_45782 | only bacterioplankton | -5 | 0.53901759 |
| bacteria_Jul11d5b_36272 | only bacterioplankton | -5 | 0.498195596 |
| bacteria_Jul11d0b_44865 | only bacterioplankton | -5 | -0.040800905 |
| bacteria_Jun8d25b_39854 | only bacterioplankton | -5 | -0.167552611 |
| bacteria_Jul11d25b_18912 | only bacterioplankton | -5 | -0.252109999 |
| bacteria_Aug5d5b_41735 | only bacterioplankton | -5 | -0.312734621 |
| bacteria_Aug5d50b_66004 | only bacterioplankton | -5 | -0.312734621 |
| bacteria_May12d25b_17650 | only bacterioplankton | -5 | -0.344483319 |
| bacteria_Aug5d5b_59932 | only bacterioplankton | -5 | -0.377273142 |
| bacteria_Aug5d25b_67931 | only bacterioplankton | -5 | -0.377273142 |
| bacteria_Aug5d0b_1353 | only bacterioplankton | -5 | -0.446266013 |
| bacteria_Jul11d0b_29913 | only bacterioplankton | -5 | -0.600416693 |
| bacteria_Jul11d75b_7370 | only bacterioplankton | -5 | -0.68742807 |
| bacteria_May12d25b_4727 | only bacterioplankton | -5 | -0.68742807 |
| bacteria_Jul11d75b_25529 | only bacterioplankton | -5 | -0.78273825 |
| bacteria_May12d25b_29611 | only bacterioplankton | -5 | -0.834031544 |
| bacteria_Jul11d0b_15322 | only bacterioplankton | -5 | -0.888098766 |
| bacteria_Sep8d25b_1335 | only bacterioplankton | -5 | -0.888098766 |
| bacteria_Jul27d5b_44900 | only bacterioplankton | -5 | -0.888098766 |
| bacteria_Jun8d50b_21017 | only bacterioplankton | -5 | -0.945257179 |
| bacteria_Jul11d0b_61222 | only bacterioplankton | -5 | -1.005881801 |
| bacteria_Jun8d75b_67192 | only bacterioplankton | -5 | -1.005881801 |
| bacteria_Jul27d75b_21145 | only bacterioplankton | -5 | -1.070420322 |
| bacteria_Jul11d5b_32250 | only bacterioplankton | -5 | -1.139413194 |
| bacteria_Aug5d50b_21866 | only bacterioplankton | -5 | -1.213521166 |
| bacteria_Jul27d5b_39021 | only bacterioplankton | -5 | -1.213521166 |
| bacteria_Jul11d50b_17953 | only bacterioplankton | -5 | -1.380575251 |
| bacteria_Sep8d5b_16061 | only bacterioplankton | -5 | -1.380575251 |

**Supplementary Table S9**. 29 August 5th 98% RTPR OTUs retained for assessing depth-resolved distribution patterns within both virio- and bacterioplankton fractions

98% RTPR OTUs from virio- and bacterioplankton occurring in Aug 5th samples, >50 total sequences and <=2 zero replacements

| Cluster repre | Aug 5th OTU | Aug5_0m | | | | Aug5_05m | | | | Aug5_25m | | | | Aug5_50m | | | | Aug5_75m | | | | total | number 0- |
|---|---|---|---|---|---|---|---|---|---|---|---|---|---|---|---|---|---|---|---|---|---|---|---|
| | | bacterioplankton | | virioplankton | | bacterioplankton | | virioplankton | | bacterioplankton | | virioplankton | | bacterioplankton | | virioplankton | | bacterioplankton | | virioplankton | | | |
| | | Sequences | CLR | Sequences | CLR | Sequences | CLR | Sequences | CLR | Sequences | CLR | Sequences | CLR | Sequences | CLR | Sequences | CLR | Sequences | CLR | Sequences | CLR | | |
| bacteria_Jul1 | 1 | 87 | 7.099 | 362 | 9.3 | 20 | 5.192 | 92 | 7.217 | 489 | 9.724 | 306 | 9.067 | 47 | 6.253 | 26 | 5.435 | 3 | 2.724 | 10 | 3.929 | 1442 | none |
| bacteria_Jul1 | 2 | 64 | 6.657 | 206 | 8.487 | 6 | 3.455 | 102 | 7.365 | 267 | 8.851 | 160 | 8.132 | 76 | 6.946 | 154 | 8.002 | 10 | 4.461 | 78 | 6.893 | 1123 | none |
| bacteria_Jul1 | 3 | 34 | 5.744 | 221 | 8.588 | 11 | 4.33 | 103 | 7.379 | 59 | 6.673 | 124 | 7.764 | 16 | 4.699 | 24 | 5.32 | 2 | 2.139 | 19 | 4.855 | 613 | none |
| bacteria_Jul1 | 4 | 41 | 6.014 | 126 | 7.778 | 14 | 4.678 | 67 | 6.759 | 50 | 6.434 | 67 | 6.876 | 4 | 2.699 | 11 | 4.194 | 0 | 2.006 | 12 | 4.192 | 392 | 1 |
| bacteria_Jul1 | 5 | 31 | 5.611 | 112 | 7.608 | 7 | 3.678 | 50 | 6.337 | 45 | 6.282 | 69 | 6.919 | 11 | 4.158 | 15 | 4.642 | 0 | 2.006 | 10 | 3.929 | 350 | 1 |
| bacteria_Aug | 6 | 4 | 2.657 | 23 | 5.324 | 3 | 2.455 | 12 | 4.278 | 44 | 6.25 | 22 | 5.269 | 64 | 6.699 | 22 | 5.194 | 75 | 7.368 | 55 | 6.389 | 324 | none |
| virus_Aug5.5 | 7 | 7 | 3.464 | 170 | 8.21 | 8 | 3.87 | 30 | 5.6 | 11 | 4.25 | 67 | 6.876 | 7 | 3.506 | 8 | 3.735 | 0 | 2.006 | 7 | 3.415 | 315 | 1 |
| bacteria_Jul1 | 8 | 16 | 4.657 | 62 | 6.755 | 3 | 2.455 | 33 | 5.737 | 38 | 6.038 | 21 | 5.202 | 4 | 2.699 | 15 | 4.642 | 1 | 1.139 | 22 | 5.067 | 215 | none |
| virus_Aug5.2 | 9 | 5 | 2.978 | 107 | 7.542 | 5 | 3.192 | 14 | 4.5 | 6 | 3.375 | 48 | 6.395 | 3 | 2.283 | 3 | 2.32 | 0 | 2.006 | 8 | 3.607 | 199 | 1 |
| bacteria_Aug | 10 | 1 | 0.657 | 12 | 4.385 | 0 | 1.525 | 11 | 4.152 | 16 | 4.791 | 19 | 5.058 | 62 | 6.653 | 29 | 5.593 | 23 | 5.662 | 24 | 5.192 | 197 | 1 |
| bacteria_Jul1 | 11 | 16 | 4.657 | 52 | 6.501 | 5 | 3.192 | 25 | 5.337 | 27 | 5.545 | 13 | 4.51 | 2 | 1.699 | 6 | 3.32 | 2 | 2.139 | 14 | 4.415 | 162 | none |
| bacteria_Jul1 | 12 | 10 | 3.978 | 34 | 5.888 | 4 | 2.87 | 16 | 4.693 | 48 | 6.375 | 23 | 5.334 | 2 | 1.699 | 3 | 2.32 | 0 | 0.92 | 1 | 0.607 | 141 | 1 |
| bacteria_Aug | 13 | 6 | 3.241 | 32 | 5.8 | 7 | 3.678 | 8 | 3.693 | 10 | 4.112 | 24 | 5.395 | 6 | 3.283 | 17 | 4.823 | 0 | 2.006 | 25 | 5.251 | 135 | 1 |
| bacteria_Jul2 | 14 | 4 | 2.657 | 37 | 6.01 | 2 | 1.87 | 20 | 5.015 | 17 | 4.878 | 21 | 5.202 | 4 | 2.699 | 11 | 4.194 | 0 | 2.006 | 17 | 4.695 | 133 | 1 |
| bacteria_Jul1 | 15 | 3 | 2.241 | 71 | 6.95 | 0 | 0.875 | 13 | 4.393 | 4 | 2.791 | 33 | 5.854 | 1 | 0.699 | 3 | 2.32 | 0 | 0.92 | 1 | 0.607 | 129 | 2 |
| virus_Jul11.2 | 16 | 2 | 1.657 | 66 | 6.845 | 3 | 2.455 | 12 | 4.278 | 3 | 2.375 | 29 | 5.668 | 0 | 0.292 | 6 | 3.32 | 0 | 1.01 | 3 | 2.192 | 124 | 2 |
| bacteria_Aug | 17 | 1 | 0.657 | 11 | 4.26 | 2 | 1.87 | 16 | 4.693 | 15 | 4.697 | 10 | 4.132 | 5 | 3.02 | 4 | 2.735 | 7 | 3.946 | 48 | 6.192 | 119 | none |
| bacteria_Aug | 18 | 5 | 2.978 | 35 | 5.93 | 2 | 1.87 | 18 | 4.863 | 21 | 5.183 | 12 | 4.395 | 1 | 0.699 | 7 | 3.542 | 0 | 1.23 | 15 | 4.514 | 116 | 1 |
| bacteria_Aug | 19 | 11 | 4.116 | 42 | 6.193 | 5 | 3.192 | 20 | 5.015 | 11 | 4.25 | 9 | 3.98 | 0 | 0.292 | 2 | 1.735 | 0 | 2.006 | 3 | 2.192 | 103 | 2 |
| virus_Aug5.5 | 20 | 3 | 2.241 | 36 | 5.97 | 4 | 2.87 | 19 | 4.941 | 8 | 3.791 | 15 | 4.717 | 2 | 1.699 | 5 | 3.057 | 0 | 2.006 | 9 | 3.777 | 101 | 1 |
| bacteria_Aug | 21 | 4 | 2.657 | 20 | 5.122 | 3 | 2.455 | 8 | 3.693 | 9 | 3.96 | 17 | 4.898 | 10 | 4.02 | 7 | 3.542 | 0 | 2.006 | 19 | 4.855 | 97 | 1 |
| bacteria_Jul1 | 22 | 5 | 2.978 | 18 | 4.97 | 2 | 1.87 | 6 | 3.278 | 17 | 4.878 | 25 | 5.454 | 3 | 2.283 | 13 | 4.435 | 0 | 2.006 | 7 | 3.415 | 96 | 1 |
| bacteria_May | 23 | 5 | 2.978 | 11 | 4.26 | 1 | 0.87 | 11 | 4.152 | 23 | 5.314 | 13 | 4.51 | 2 | 1.699 | 10 | 4.057 | 1 | 1.139 | 9 | 3.777 | 86 | none |
| bacteria_Jul1 | 24 | 10 | 3.978 | 17 | 4.888 | 2 | 1.87 | 19 | 4.941 | 12 | 4.375 | 5 | 3.132 | 2 | 1.699 | 13 | 4.435 | 0 | 2.006 | 4 | 2.607 | 84 | 1 |
| bacteria_Aug | 25 | 3 | 2.241 | 1 | 0.8 | 4 | 2.87 | 11 | 4.152 | 7 | 3.598 | 14 | 4.617 | 17 | 4.786 | 9 | 3.905 | 0 | -0.957 | 16 | 4.607 | 82 | 1 |
| virus_Aug5.0 | 26 | 1 | 0.657 | 46 | 6.324 | 1 | 0.87 | 7 | 3.5 | 1 | 0.791 | 15 | 4.717 | 2 | 1.699 | 2 | 1.735 | 0 | -0.575 | 1 | 0.607 | 76 | 1 |
| bacteria_Jul1 | 27 | 4 | 2.657 | 18 | 4.97 | 0 | 1.525 | 12 | 4.278 | 7 | 3.598 | 6 | 3.395 | 2 | 1.699 | 5 | 3.057 | 0 | 2.006 | 4 | 2.607 | 58 | 1 |
| virus_Jul11.2 | 28 | 0 | 0.273 | 4 | 2.8 | 0 | 1.525 | 9 | 3.863 | 14 | 4.598 | 6 | 3.395 | 5 | 3.02 | 6 | 3.32 | 3 | 2.724 | 7 | 3.415 | 54 | 2 |
| virus_Jul11.2 | 29 | 1 | 0.657 | 6 | 3.385 | 0 | 1.525 | 8 | 3.693 | 13 | 4.491 | 4 | 2.81 | 5 | 3.02 | 8 | 3.735 | 0 | 1.19 | 9 | 3.777 | 54 | 2 |
| total sequences each | | 384 | | 1958 | | 124 | | 772 | | 1292 | | 1197 | | 365 | | 444 | | 127 | | 457 | | | |
| total 0-replaced | | 1 | | none | | 4 | | none | | none | | none | | 2 | | none | | 19 | | none | | | |

Clustered 98% RTPR OTUs from virio- and bacterioplankton on August 5th (amplicons in both fractions at all depths)

| Cluster repre | Aug 5th OTU | Aug5_0m | | Aug5_05m | | Aug5_25m | | Aug5_50m | | Aug5_75m | | total sequences |
|---|---|---|---|---|---|---|---|---|---|---|---|---|
| | | bacterioplank | virioplankton | bacterioplank | virioplankton | bacterioplank | virioplankton | bacterioplank | virioplankton | bacterioplank | virioplankton | |
| bacteria_Jul1 | 1 | 87 | 362 | 20 | 92 | 489 | 306 | 47 | 26 | 3 | 10 | 1442 |
| bacteria_Jul1 | 11 | 16 | 52 | 5 | 25 | 27 | 13 | 2 | 6 | 2 | 14 | 162 |
| bacteria_Jul1 | 22 | 5 | 18 | 2 | 6 | 17 | 25 | 3 | 13 | 0 | 7 | 96 |
| bacteria_Jul11d0b_3527 | | 0 | 0 | 0 | 1 | 2 | 1 | 2 | 3 | 0 | 6 | 15 |
| bacteria_Jul1 | 3 | 34 | 221 | 11 | 103 | 59 | 124 | 16 | 24 | 2 | 19 | 613 |
| bacteria_Jul11d0b_6084 | | 2 | 0 | 1 | 0 | 6 | 0 | 6 | 0 | 0 | 0 | 15 |
| bacteria_Jul11d0b_7291 | | 0 | 4 | 0 | 2 | 3 | 2 | 0 | 0 | 1 | 2 | 14 |
| bacteria_Jul11d0b_9089 | | 2 | 2 | 0 | 1 | 0 | 0 | 1 | 0 | 0 | 1 | 7 |
| bacteria_Jul11d0b_11991 | | 0 | 0 | 0 | 0 | 0 | 0 | 0 | 0 | 0 | 0 | 0 |
| bacteria_Jul11d0b_14765 | | 0 | 0 | 0 | 2 | 1 | 4 | 2 | 2 | 0 | 2 | 13 |
| bacteria_Jul11d0b_15322 | | 0 | 0 | 0 | 0 | 2 | 0 | 2 | 0 | 0 | 0 | 4 |
| bacteria_Jul11d0b_17411 | | 4 | 0 | 2 | 0 | 7 | 0 | 3 | 0 | 1 | 0 | 17 |
| bacteria_Jul1 | 12 | 10 | 34 | 4 | 16 | 48 | 23 | 2 | 3 | 0 | 1 | 141 |
| bacteria_Jul1 | 5 | 31 | 112 | 7 | 50 | 45 | 69 | 11 | 15 | 0 | 10 | 350 |
| bacteria_Jul11d0b_25777 | | 1 | 3 | 0 | 5 | 11 | 0 | 0 | 1 | 1 | 2 | 24 |
| bacteria_Jul1 | 4 | 41 | 126 | 14 | 67 | 50 | 67 | 4 | 11 | 0 | 12 | 392 |
| bacteria_Jul11d0b_26699 | | 0 | 2 | 0 | 2 | 1 | 0 | 2 | 3 | 4 | 5 | 19 |
| bacteria_Jul11d0b_27216 | | 0 | 3 | 0 | 4 | 9 | 5 | 0 | 2 | 2 | 4 | 29 |
| bacteria_Jul1 | 15 | 3 | 71 | 0 | 13 | 4 | 33 | 1 | 3 | 0 | 1 | 129 |
| bacteria_Jul11d0b_29888 | | 0 | 0 | 0 | 0 | 0 | 0 | 0 | 0 | 0 | 0 | 0 |
| bacteria_Jul11d0b_29913 | | 0 | 0 | 1 | 0 | 4 | 0 | 1 | 0 | 0 | 0 | 6 |
| bacteria_Jul11d0b_35117 | | 2 | 0 | 2 | 0 | 5 | 0 | 6 | 0 | 0 | 0 | 15 |
| bacteria_Jul1 | 8 | 16 | 62 | 3 | 33 | 38 | 21 | 4 | 15 | 1 | 22 | 215 |
| bacteria_Jul1 | 2 | 64 | 206 | 6 | 102 | 267 | 160 | 76 | 154 | 10 | 78 | 1123 |
| bacteria_Jul11d0b_37708 | | 0 | 1 | 1 | 0 | 1 | 2 | 1 | 0 | 0 | 0 | 6 |
| bacteria_Jul11d0b_38787 | | 2 | 0 | 0 | 0 | 3 | 0 | 0 | 0 | 0 | 1 | 6 |
| bacteria_Jul11d0b_44865 | | 1 | 0 | 0 | 0 | 4 | 0 | 0 | 0 | 0 | 0 | 5 |
| bacteria_Jul11d0b_47245 | | 7 | 3 | 0 | 0 | 9 | 7 | 2 | 0 | 1 | 1 | 30 |
| bacteria_Jul1 | 27 | 4 | 18 | 0 | 12 | 7 | 6 | 2 | 5 | 0 | 4 | 58 |
| bacteria_Jul11d0b_58291 | | 1 | 1 | 0 | 3 | 4 | 1 | 1 | 0 | 0 | 0 | 11 |
| bacteria_Jul11d0b_58550 | | 1 | 2 | 0 | 3 | 0 | 1 | 0 | 1 | 0 | 0 | 8 |
| bacteria_Jul11d0b_59045 | | 3 | 3 | 0 | 0 | 0 | 2 | 0 | 3 | 1 | 2 | 14 |
| bacteria_Jul11d0b_61222 | | 1 | 0 | 0 | 0 | 1 | 0 | 0 | 0 | 0 | 0 | 2 |
| bacteria_Jul11d0b_71717 | | 3 | 2 | 0 | 2 | 3 | 1 | 0 | 0 | 0 | 1 | 12 |
| bacteria_Aug5d0b_10997 | | 8 | 1 | 0 | 0 | 8 | 0 | 0 | 0 | 0 | 0 | 17 |
| bacteria_Aug5d0b_30481 | | 5 | 7 | 2 | 1 | 5 | 4 | 0 | 0 | 0 | 1 | 25 |
| bacteria_Aug5d0b_30617 | | 6 | 7 | 0 | 3 | 7 | 7 | 6 | 4 | 0 | 1 | 41 |
| bacteria_Aug5d0b_32352 | | 1 | 0 | 0 | 0 | 0 | 0 | 0 | 0 | 0 | 0 | 1 |
| bacteria_Aug5d0b_42381 | | 3 | 0 | 0 | 0 | 3 | 0 | 2 | 0 | 0 | 0 | 8 |
| bacteria_Aug5d0b_47134 | | 1 | 3 | 2 | 2 | 4 | 3 | 0 | 0 | 0 | 2 | 17 |
| bacteria_Aug5d0b_57551 | | 1 | 0 | 0 | 0 | 0 | 0 | 1 | 0 | 0 | 0 | 2 |
| bacteria_Aug5d0b_63126 | | 1 | 0 | 0 | 0 | 0 | 0 | 0 | 0 | 0 | 0 | 1 |
| bacteria_Aug5d0b_73476 | | 1 | 0 | 0 | 0 | 0 | 0 | 0 | 0 | 0 | 0 | 1 |
| bacteria_Aug | 25 | 3 | 1 | 4 | 11 | 7 | 14 | 17 | 9 | 0 | 16 | 82 |
| bacteria_Aug5d5b_15792 | | 0 | 4 | 2 | 1 | 0 | 3 | 0 | 0 | 0 | 3 | 13 |
| bacteria_Aug | 21 | 4 | 20 | 3 | 8 | 9 | 17 | 10 | 7 | 0 | 19 | 97 |
| bacteria_Aug | 17 | 1 | 11 | 2 | 16 | 15 | 10 | 5 | 4 | 7 | 48 | 119 |
| bacteria_Aug5d5b_29960 | | 0 | 0 | 1 | 0 | 0 | 0 | 0 | 0 | 0 | 0 | 1 |
| bacteria_Aug5d5b_41001 | | 0 | 0 | 1 | 0 | 0 | 0 | 0 | 0 | 0 | 0 | 1 |
| bacteria_Aug5d5b_41735 | | 0 | 0 | 2 | 0 | 2 | 0 | 1 | 0 | 0 | 0 | 5 |
| bacteria_Aug | 19 | 11 | 42 | 5 | 20 | 11 | 9 | 0 | 2 | 0 | 3 | 103 |
| bacteria_Aug5d5b_58010 | | 0 | 0 | 1 | 0 | 0 | 0 | 0 | 0 | 0 | 0 | 1 |
| bacteria_Aug5d5b_59932 | | 0 | 0 | 1 | 0 | 0 | 0 | 1 | 0 | 0 | 0 | 2 |
| bacteria_Aug5d5b_65184 | | 1 | 17 | 1 | 0 | 3 | 4 | 3 | 1 | 1 | 2 | 33 |
| bacteria_Aug | 6 | 4 | 23 | 3 | 12 | 44 | 22 | 64 | 22 | 75 | 55 | 324 |
| bacteria_Aug5d25b_3402 | | 0 | 1 | 0 | 2 | 1 | 1 | 0 | 0 | 0 | 0 | 5 |
| bacteria_Aug5d25b_3440 | | 0 | 0 | 0 | 0 | 1 | 0 | 0 | 0 | 0 | 0 | 1 |
| bacteria_Aug5d25b_6187 | | 0 | 0 | 0 | 0 | 1 | 0 | 0 | 0 | 0 | 0 | 1 |
| bacteria_Aug5d25b_11031 | | 3 | 0 | 0 | 2 | 6 | 1 | 1 | 0 | 0 | 0 | 13 |
| bacteria_Aug5d25b_11190 | | 1 | 0 | 1 | 4 | 15 | 3 | 0 | 1 | 0 | 2 | 27 |
| bacteria_Aug5d25b_11412 | | 0 | 0 | 0 | 0 | 2 | 0 | 0 | 0 | 0 | 0 | 2 |
| bacteria_Aug5d25b_11494 | | 1 | 0 | 0 | 0 | 1 | 2 | 0 | 0 | 0 | 0 | 4 |
| bacteria_Aug | 13 | 6 | 32 | 7 | 8 | 10 | 24 | 6 | 17 | 0 | 25 | 135 |
| bacteria_Aug5d25b_16734 | | 0 | 8 | 0 | 10 | 9 | 4 | 1 | 2 | 2 | 8 | 44 |
| bacteria_Aug5d25b_16994 | | 0 | 0 | 0 | 0 | 2 | 0 | 0 | 0 | 0 | 0 | 2 |
| bacteria_Aug5d25b_19567 | | 0 | 0 | 0 | 0 | 1 | 0 | 0 | 0 | 0 | 0 | 1 |
| bacteria_Aug5d25b_20034 | | 0 | 0 | 0 | 0 | 1 | 0 | 0 | 0 | 0 | 0 | 1 |
| bacteria_Aug5d25b_20891 | | 0 | 0 | 0 | 0 | 1 | 0 | 0 | 0 | 0 | 0 | 1 |
| bacteria_Aug | 18 | 5 | 35 | 2 | 18 | 21 | 12 | 1 | 7 | 0 | 15 | 116 |
| bacteria_Aug5d25b_28995 | | 0 | 0 | 0 | 0 | 1 | 0 | 0 | 0 | 0 | 0 | 1 |
| bacteria_Aug5d25b_31207 | | 0 | 0 | 0 | 0 | 1 | 0 | 1 | 0 | 0 | 0 | 2 |
| bacteria_Aug5d25b_32181 | | 0 | 0 | 0 | 0 | 2 | 0 | 0 | 0 | 0 | 0 | 2 |
| bacteria_Aug5d25b_32559 | | 3 | 0 | 1 | 1 | 3 | 0 | 1 | 0 | 0 | 1 | 10 |
| bacteria_Aug5d25b_33881 | | 1 | 1 | 0 | 1 | 3 | 1 | 0 | 0 | 0 | 1 | 8 |
| bacteria_Aug5d25b_35406 | | 0 | 0 | 0 | 0 | 3 | 0 | 1 | 0 | 0 | 0 | 4 |
| bacteria_Aug5d25b_41725 | | 11 | 0 | 9 | 0 | 27 | 0 | 0 | 0 | 0 | 0 | 47 |
| bacteria_Aug | 10 | 1 | 12 | 0 | 11 | 16 | 19 | 62 | 29 | 23 | 24 | 197 |
| bacteria_Aug5d25b_46297 | | 1 | 0 | 0 | 0 | 3 | 0 | 1 | 0 | 0 | 0 | 5 |
| bacteria_Aug5d25b_47450 | | 0 | 0 | 0 | 0 | 1 | 0 | 0 | 0 | 0 | 0 | 1 |
| bacteria_Aug5d25b_52757 | | 0 | 9 | 0 | 4 | 4 | 11 | 2 | 7 | 0 | 14 | 51 |
| bacteria_Aug5d25b_53723 | | 0 | 0 | 0 | 2 | 3 | 0 | 1 | 0 | 0 | 0 | 6 |
| bacteria_Aug5d25b_56643 | | 0 | 0 | 0 | 0 | 1 | 0 | 0 | 0 | 0 | 0 | 1 |
| bacteria_Aug5d25b_59498 | | 1 | 0 | 0 | 0 | 6 | 0 | 4 | 0 | 0 | 0 | 11 |
| bacteria_Aug5d25b_64962 | | 0 | 0 | 0 | 0 | 1 | 0 | 0 | 0 | 0 | 0 | 1 |
| bacteria_Aug5d25b_67931 | | 0 | 0 | 0 | 0 | 4 | 0 | 0 | 0 | 0 | 0 | 4 |
| bacteria_Aug5d25b_68913 | | 0 | 0 | 0 | 0 | 3 | 0 | 0 | 0 | 0 | 0 | 3 |
| bacteria_Aug5d25b_72218 | | 0 | 0 | 0 | 0 | 4 | 1 | 1 | 0 | 0 | 0 | 6 |
| bacteria_Aug5d50b_2118 | | 0 | 0 | 0 | 1 | 0 | 0 | 1 | 0 | 0 | 0 | 2 |
| bacteria_Aug5d50b_2154 | | 0 | 0 | 0 | 0 | 2 | 4 | 4 | 2 | 0 | 1 | 13 |
| bacteria_Aug5d50b_8831 | | 0 | 0 | 0 | 0 | 0 | 0 | 1 | 0 | 0 | 0 | 1 |
| bacteria_Aug5d50b_10806 | | 1 | 0 | 0 | 1 | 3 | 0 | 1 | 0 | 0 | 0 | 6 |
| bacteria_Aug5d50b_13789 | | 0 | 0 | 0 | 0 | 1 | 0 | 1 | 0 | 0 | 0 | 2 |
| bacteria_Aug5d50b_14026 | | 1 | 0 | 0 | 0 | 0 | 0 | 1 | 0 | 0 | 0 | 2 |
| bacteria_Aug5d50b_14505 | | 0 | 0 | 0 | 0 | 1 | 0 | 1 | 0 | 0 | 2 | 4 |
| bacteria_Aug5d50b_17974 | | 0 | 0 | 0 | 0 | 0 | 0 | 1 | 0 | 0 | 0 | 1 |
| bacteria_Aug5d50b_21866 | | 0 | 0 | 0 | 0 | 0 | 0 | 1 | 0 | 0 | 0 | 1 |
| bacteria_Aug5d50b_28175 | | 0 | 0 | 0 | 0 | 0 | 0 | 1 | 0 | 0 | 0 | 1 |
| bacteria_Aug5d50b_36848 | | 0 | 0 | 0 | 0 | 4 | 0 | 2 | 0 | 0 | 0 | 6 |
| bacteria_Aug5d50b_38363 | | 0 | 1 | 0 | 2 | 1 | 0 | 2 | 0 | 0 | 0 | 6 |
| bacteria_Aug5d50b_64722 | | 0 | 0 | 0 | 0 | 0 | 0 | 1 | 0 | 0 | 0 | 1 |
| bacteria_Aug5d50b_66004 | | 0 | 0 | 0 | 0 | 1 | 0 | 1 | 0 | 0 | 0 | 2 |
| bacteria_Aug5d50b_68297 | | 3 | 0 | 5 | 0 | 15 | 0 | 4 | 0 | 2 | 0 | 29 |
| bacteria_Jul11d5b_2558 | | 1 | 3 | 0 | 2 | 9 | 1 | 1 | 1 | 0 | 3 | 21 |
| bacteria_Jul11d5b_3980 | | 0 | 0 | 0 | 0 | 0 | 1 | 0 | 0 | 0 | 0 | 1 |
| bacteria_Jul11d5b_20339 | | 0 | 0 | 0 | 0 | 0 | 0 | 0 | 0 | 0 | 0 | 0 |
| bacteria_Jul11d5b_25083 | | 0 | 0 | 0 | 0 | 0 | 0 | 0 | 0 | 0 | 0 | 0 |
| bacteria_Jul11d5b_32250 | | 0 | 0 | 1 | 0 | 0 | 0 | 1 | 0 | 0 | 0 | 2 |
| bacteria_Jul11d5b_36272 | | 0 | 0 | 0 | 0 | 5 | 0 | 3 | 0 | 0 | 0 | 8 |
| bacteria_Jul11d5b_41219 | | 0 | 0 | 1 | 0 | 0 | 0 | 0 | 0 | 0 | 1 | 2 |
| bacteria_Jul11d25b_195 | | 0 | 0 | 0 | 0 | 1 | 0 | 0 | 0 | 0 | 0 | 1 |
| bacteria_Jul11d25b_6636 | | 1 | 1 | 0 | 0 | 1 | 2 | 1 | 0 | 0 | 1 | 7 |
| bacteria_Jul11d25b_7263 | | 0 | 0 | 0 | 0 | 1 | 0 | 0 | 0 | 0 | 0 | 1 |
| bacteria_Jul11d25b_8174 | | 0 | 1 | 0 | 1 | 3 | 0 | 0 | 1 | 2 | 1 | 9 |
| bacteria_Jul1 | 24 | 10 | 17 | 2 | 19 | 12 | 5 | 2 | 13 | 0 | 4 | 84 |
| bacteria_Jul11d25b_18912 | | 0 | 0 | 0 | 0 | 0 | 0 | 0 | 0 | 0 | 0 | 0 |

| | | | | | | | | | | | |
|---|---|---|---|---|---|---|---|---|---|---|---|
| bacteria_Jul11d25b_20536 | 0 | 0 | 0 | 0 | 2 | 0 | 0 | 0 | 0 | 0 | 2 |
| bacteria_Jul11d25b_28609 | 0 | 0 | 0 | 0 | 0 | 0 | 0 | 0 | 0 | 0 | 0 |
| bacteria_Jul11d25b_30910 | 0 | 0 | 0 | 1 | 1 | 0 | 0 | 0 | 0 | 0 | 2 |
| bacteria_Jul11d25b_36968 | 2 | 0 | 1 | 0 | 9 | 0 | 2 | 0 | 2 | 0 | 16 |
| bacteria_Jul11d50b_3125 | 0 | 0 | 0 | 0 | 0 | 0 | 0 | 0 | 0 | 0 | 0 |
| bacteria_Jul11d50b_8402 | 0 | 0 | 0 | 0 | 0 | 0 | 0 | 0 | 0 | 1 | 1 |
| bacteria_Jul11d50b_9662 | 0 | 0 | 0 | 0 | 0 | 0 | 0 | 0 | 0 | 0 | 0 |
| bacteria_Jul11d50b_10402 | 0 | 0 | 0 | 0 | 0 | 0 | 0 | 0 | 0 | 0 | 0 |
| bacteria_Jul11d50b_15817 | 0 | 0 | 0 | 0 | 0 | 0 | 0 | 0 | 0 | 0 | 0 |
| bacteria_Jul11d50b_19565 | 0 | 0 | 0 | 0 | 0 | 1 | 0 | 0 | 0 | 0 | 1 |
| bacteria_Jul11d50b_23123 | 0 | 0 | 0 | 0 | 0 | 0 | 0 | 0 | 0 | 0 | 0 |
| bacteria_Jul11d50b_32886 | 0 | 0 | 0 | 0 | 0 | 0 | 0 | 0 | 0 | 0 | 0 |
| bacteria_Jul11d50b_41285 | 0 | 0 | 0 | 1 | 0 | 0 | 0 | 1 | 0 | 0 | 2 |
| bacteria_Jul11d50b_42829 | 0 | 0 | 0 | 0 | 0 | 1 | 0 | 0 | 0 | 0 | 1 |
| bacteria_Jul11d50b_45782 | 5 | 0 | 0 | 0 | 5 | 0 | 1 | 0 | 0 | 0 | 11 |
| bacteria_Jul11d50b_51257 | 2 | 0 | 3 | 0 | 7 | 0 | 3 | 0 | 0 | 0 | 15 |
| bacteria_Jul11d75b_22 | 0 | 3 | 0 | 0 | 0 | 3 | 0 | 0 | 0 | 0 | 6 |
| bacteria_Jul11d75b_2421 | 0 | 0 | 0 | 0 | 0 | 0 | 0 | 0 | 1 | 0 | 1 |
| bacteria_Jul11d75b_5405 | 0 | 0 | 0 | 0 | 3 | 0 | 0 | 0 | 0 | 0 | 3 |
| bacteria_Jul11d75b_6234 | 0 | 0 | 0 | 0 | 3 | 1 | 2 | 1 | 0 | 1 | 8 |
| bacteria_Jul11d75b_8044 | 0 | 0 | 0 | 0 | 0 | 0 | 0 | 0 | 0 | 0 | 0 |
| bacteria_Jul11d75b_9207 | 0 | 0 | 0 | 0 | 0 | 0 | 0 | 0 | 0 | 0 | 0 |
| bacteria_Jul11d75b_9365 | 0 | 0 | 0 | 0 | 0 | 0 | 0 | 0 | 0 | 0 | 0 |
| bacteria_Jul11d75b_10494 | 0 | 0 | 0 | 0 | 0 | 1 | 0 | 0 | 0 | 0 | 1 |
| bacteria_Jul11d75b_16607 | 0 | 2 | 0 | 1 | 1 | 1 | 0 | 4 | 0 | 0 | 9 |
| bacteria_Jul11d75b_17437 | 0 | 0 | 0 | 0 | 0 | 0 | 0 | 0 | 0 | 0 | 0 |
| bacteria_Jul11d75b_21130 | 0 | 0 | 0 | 0 | 0 | 0 | 0 | 0 | 0 | 0 | 0 |
| bacteria_Jul11d75b_24271 | 0 | 1 | 0 | 0 | 0 | 0 | 0 | 0 | 0 | 0 | 1 |
| bacteria_Jul11d75b_24464 | 0 | 0 | 1 | 0 | 3 | 0 | 0 | 0 | 0 | 0 | 4 |
| bacteria_Jul11d75b_24795 | 0 | 0 | 0 | 0 | 0 | 0 | 0 | 1 | 0 | 0 | 1 |
| bacteria_Jul11d75b_25529 | 0 | 0 | 0 | 0 | 0 | 0 | 1 | 0 | 0 | 0 | 1 |
| bacteria_Jul11d75b_26607 | 0 | 0 | 0 | 0 | 0 | 0 | 0 | 0 | 0 | 0 | 0 |
| bacteria_Jul11d75b_29776 | 0 | 0 | 0 | 0 | 0 | 0 | 0 | 0 | 0 | 0 | 0 |
| bacteria_Jul11d75b_30241 | 0 | 3 | 0 | 1 | 0 | 1 | 0 | 0 | 0 | 1 | 6 |
| bacteria_Jul11d75b_32574 | 0 | 0 | 0 | 0 | 0 | 0 | 1 | 0 | 0 | 0 | 1 |
| bacteria_Jul11d75b_33761 | 2 | 2 | 0 | 0 | 1 | 1 | 1 | 0 | 0 | 0 | 7 |
| bacteria_Jul11d75b_39799 | 0 | 2 | 0 | 1 | 0 | 1 | 0 | 0 | 0 | 0 | 4 |
| bacteria_Jul11d75b_52660 | 0 | 0 | 0 | 0 | 0 | 0 | 0 | 0 | 0 | 0 | 0 |
| bacteria_Jul27d5b_4608 | 0 | 0 | 0 | 0 | 0 | 0 | 0 | 0 | 0 | 0 | 0 |
| bacteria_Jul27d5b_7357 | 0 | 1 | 0 | 1 | 0 | 0 | 0 | 0 | 0 | 0 | 2 |
| bacteria_Jul27d5b_12006 | 1 | 0 | 0 | 1 | 1 | 0 | 0 | 0 | 2 | 0 | 5 |
| bacteria_Jul27d5b_12427 | 0 | 1 | 0 | 0 | 1 | 1 | 0 | 0 | 0 | 0 | 3 |
| bacteria_Jul27d5b_13272 | 0 | 0 | 0 | 0 | 0 | 0 | 0 | 0 | 0 | 0 | 0 |
| bacteria_Jul2 14 | 4 | 37 | 2 | 20 | 17 | 21 | 4 | 11 | 0 | 17 | 133 |
| bacteria_Jul27d5b_23117 | 0 | 0 | 0 | 0 | 0 | 1 | 0 | 0 | 0 | 0 | 1 |
| bacteria_Jul27d5b_24937 | 1 | 0 | 0 | 3 | 3 | 1 | 1 | 0 | 0 | 1 | 10 |
| bacteria_Jul27d5b_31534 | 0 | 0 | 0 | 0 | 0 | 0 | 0 | 0 | 0 | 0 | 0 |
| bacteria_Jul27d5b_32690 | 0 | 0 | 1 | 0 | 0 | 0 | 0 | 0 | 0 | 0 | 1 |
| bacteria_Jul27d5b_33739 | 0 | 0 | 0 | 0 | 0 | 0 | 0 | 0 | 0 | 0 | 0 |
| bacteria_Jul27d5b_35976 | 0 | 1 | 0 | 0 | 2 | 0 | 0 | 0 | 0 | 0 | 3 |
| bacteria_Jul27d5b_37077 | 0 | 0 | 0 | 0 | 0 | 0 | 0 | 0 | 0 | 0 | 0 |
| bacteria_Jul27d5b_39021 | 0 | 0 | 0 | 0 | 1 | 0 | 0 | 0 | 0 | 0 | 1 |
| bacteria_Jul27d5b_39765 | 0 | 0 | 0 | 0 | 0 | 0 | 0 | 0 | 0 | 0 | 0 |
| bacteria_Jul27d5b_39997 | 0 | 0 | 0 | 0 | 0 | 0 | 0 | 0 | 0 | 0 | 0 |
| bacteria_Jul27d5b_40094 | 0 | 1 | 0 | 0 | 0 | 0 | 0 | 0 | 0 | 0 | 1 |
| bacteria_Jul27d5b_43825 | 0 | 0 | 0 | 0 | 0 | 0 | 0 | 0 | 0 | 0 | 0 |
| bacteria_Jul27d5b_44206 | 0 | 0 | 0 | 0 | 0 | 0 | 0 | 0 | 0 | 0 | 0 |
| bacteria_Jul27d5b_45879 | 0 | 0 | 0 | 0 | 0 | 0 | 0 | 0 | 0 | 0 | 0 |
| bacteria_Jul27d5b_47803 | 0 | 0 | 0 | 0 | 0 | 0 | 0 | 0 | 0 | 0 | 0 |
| bacteria_Jul27d25b_1007 | 0 | 0 | 0 | 0 | 0 | 0 | 0 | 0 | 0 | 0 | 0 |
| bacteria_Jul27d25b_10877 | 0 | 0 | 0 | 0 | 2 | 2 | 0 | 0 | 0 | 0 | 4 |
| bacteria_Jul27d25b_11015 | 0 | 0 | 0 | 0 | 0 | 0 | 0 | 0 | 0 | 0 | 0 |
| bacteria_Jul27d25b_14037 | 0 | 0 | 0 | 0 | 0 | 0 | 0 | 0 | 0 | 0 | 0 |
| bacteria_Jul27d25b_17211 | 0 | 0 | 0 | 1 | 0 | 0 | 0 | 0 | 0 | 0 | 1 |
| bacteria_Jul27d25b_21364 | 0 | 2 | 0 | 1 | 0 | 0 | 0 | 1 | 0 | 1 | 5 |
| bacteria_Jul27d25b_34258 | 0 | 2 | 0 | 3 | 1 | 2 | 0 | 0 | 0 | 0 | 8 |
| bacteria_Jul27d25b_39850 | 0 | 0 | 0 | 0 | 0 | 0 | 0 | 0 | 0 | 0 | 0 |
| bacteria_Jul27d25b_40009 | 0 | 0 | 0 | 0 | 0 | 0 | 0 | 0 | 0 | 0 | 0 |
| bacteria_Jul27d25b_41610 | 0 | 0 | 0 | 0 | 0 | 0 | 0 | 0 | 0 | 0 | 0 |
| bacteria_Jul27d25b_42338 | 0 | 0 | 0 | 0 | 0 | 0 | 0 | 0 | 0 | 0 | 0 |
| bacteria_Jul27d25b_52979 | 0 | 0 | 0 | 0 | 0 | 0 | 1 | 0 | 0 | 0 | 1 |
| bacteria_Jul27d25b_57031 | 0 | 0 | 0 | 0 | 0 | 0 | 0 | 0 | 0 | 0 | 0 |
| bacteria_Jul27d25b_65153 | 0 | 0 | 0 | 0 | 0 | 0 | 0 | 0 | 0 | 0 | 0 |
| bacteria_Jul27d25b_72494 | 0 | 0 | 0 | 0 | 1 | 0 | 1 | 0 | 0 | 0 | 2 |
| bacteria_Jul27d75b_3332 | 0 | 0 | 0 | 0 | 0 | 0 | 0 | 0 | 0 | 0 | 0 |
| bacteria_Jul27d75b_3872 | 0 | 0 | 0 | 0 | 0 | 0 | 0 | 0 | 0 | 0 | 0 |
| bacteria_Jul27d75b_5063 | 0 | 0 | 0 | 0 | 1 | 0 | 0 | 1 | 0 | 0 | 2 |
| bacteria_Jul27d75b_8055 | 0 | 0 | 0 | 0 | 0 | 0 | 0 | 0 | 0 | 0 | 0 |
| bacteria_Jul27d75b_20546 | 0 | 0 | 0 | 1 | 0 | 0 | 1 | 0 | 0 | 0 | 2 |
| bacteria_Jul27d75b_21145 | 0 | 0 | 0 | 0 | 0 | 0 | 0 | 0 | 0 | 0 | 0 |
| bacteria_Jul27d75b_23045 | 0 | 0 | 0 | 0 | 0 | 0 | 0 | 0 | 0 | 0 | 0 |
| bacteria_Jul27d75b_27425 | 0 | 0 | 0 | 0 | 0 | 0 | 0 | 0 | 0 | 0 | 0 |
| bacteria_Jul27d75b_27465 | 0 | 0 | 0 | 1 | 0 | 0 | 0 | 0 | 0 | 1 | 2 |
| bacteria_Jul27d75b_27540 | 0 | 0 | 0 | 0 | 0 | 0 | 0 | 0 | 0 | 0 | 0 |
| bacteria_Jul27d75b_35331 | 0 | 1 | 0 | 0 | 0 | 0 | 0 | 0 | 0 | 0 | 1 |
| bacteria_Jul27d75b_42780 | 0 | 0 | 0 | 0 | 0 | 0 | 0 | 0 | 0 | 0 | 0 |
| bacteria_Jul27d75b_43975 | 0 | 0 | 0 | 0 | 0 | 1 | 0 | 0 | 0 | 3 | 4 |
| bacteria_Jul27d75b_45641 | 0 | 1 | 0 | 0 | 0 | 0 | 0 | 0 | 0 | 0 | 1 |
| bacteria_Jul27d75b_47191 | 4 | 32 | 1 | 10 | 34 | 20 | 0 | 0 | 0 | 0 | 101 |
| bacteria_Jul27d75b_53955 | 0 | 0 | 0 | 0 | 0 | 0 | 0 | 0 | 0 | 0 | 0 |
| bacteria_Jul27d75b_55377 | 0 | 0 | 0 | 0 | 0 | 0 | 0 | 0 | 0 | 0 | 0 |
| bacteria_Jul27d75b_56067 | 0 | 0 | 0 | 0 | 0 | 0 | 0 | 0 | 0 | 0 | 0 |
| bacteria_Jul27d75b_59703 | 0 | 0 | 0 | 0 | 0 | 0 | 0 | 0 | 0 | 0 | 0 |
| bacteria_Jul27d75b_69011 | 0 | 0 | 0 | 0 | 0 | 0 | 0 | 0 | 0 | 0 | 0 |
| bacteria_Jul27d75b_75175 | 0 | 0 | 0 | 0 | 0 | 0 | 0 | 0 | 0 | 0 | 0 |
| bacteria_Jun8d25b_1057 | 0 | 0 | 0 | 0 | 0 | 0 | 0 | 0 | 0 | 0 | 0 |
| bacteria_Jun8d25b_12904 | 0 | 0 | 0 | 0 | 0 | 0 | 0 | 0 | 0 | 0 | 0 |
| bacteria_Jun8d25b_21024 | 0 | 0 | 0 | 1 | 0 | 0 | 2 | 0 | 0 | 0 | 3 |
| bacteria_Jun8d25b_22580 | 0 | 0 | 0 | 0 | 0 | 1 | 0 | 0 | 0 | 0 | 1 |
| bacteria_Jun8d25b_25682 | 0 | 0 | 0 | 0 | 0 | 0 | 0 | 0 | 0 | 0 | 0 |
| bacteria_Jun8d25b_25810 | 0 | 0 | 0 | 0 | 0 | 0 | 0 | 0 | 0 | 0 | 0 |
| bacteria_Jun8d25b_26641 | 0 | 0 | 0 | 0 | 0 | 0 | 0 | 0 | 0 | 0 | 0 |
| bacteria_Jun8d25b_28680 | 0 | 0 | 0 | 0 | 0 | 0 | 0 | 0 | 0 | 0 | 0 |
| bacteria_Jun8d25b_29438 | 0 | 0 | 0 | 0 | 0 | 0 | 0 | 1 | 0 | 0 | 1 |
| bacteria_Jun8d25b_30221 | 0 | 0 | 0 | 0 | 0 | 0 | 0 | 0 | 0 | 0 | 0 |
| bacteria_Jun8d25b_34384 | 3 | 0 | 0 | 0 | 8 | 0 | 4 | 0 | 1 | 0 | 16 |
| bacteria_Jun8d25b_39854 | 0 | 0 | 0 | 0 | 0 | 0 | 0 | 0 | 0 | 0 | 0 |
| bacteria_Jun8d25b_43708 | 0 | 0 | 0 | 0 | 0 | 0 | 0 | 0 | 0 | 0 | 0 |
| bacteria_Jun8d25b_45572 | 0 | 1 | 0 | 0 | 0 | 0 | 1 | 3 | 0 | 1 | 6 |
| bacteria_Jun8d25b_55773 | 1 | 0 | 0 | 0 | 2 | 0 | 4 | 0 | 0 | 0 | 7 |
| bacteria_Jun8d25b_55947 | 0 | 0 | 0 | 0 | 0 | 0 | 0 | 0 | 0 | 0 | 0 |
| bacteria_Jun8d25b_59751 | 0 | 0 | 0 | 0 | 1 | 0 | 0 | 0 | 0 | 0 | 1 |
| bacteria_Jun8d25b_68299 | 0 | 0 | 0 | 0 | 3 | 0 | 0 | 0 | 0 | 0 | 3 |
| bacteria_Jun8d25b_68580 | 0 | 0 | 0 | 0 | 0 | 0 | 0 | 0 | 0 | 0 | 0 |
| bacteria_Jun8d25b_69463 | 0 | 0 | 0 | 0 | 2 | 0 | 0 | 2 | 0 | 1 | 5 |
| bacteria_Jun8d25b_71094 | 0 | 0 | 0 | 0 | 0 | 0 | 0 | 0 | 0 | 0 | 0 |
| bacteria_Jun8d25b_71830 | 0 | 0 | 0 | 0 | 0 | 0 | 0 | 0 | 0 | 0 | 0 |
| bacteria_Jun8d50b_165 | 0 | 0 | 0 | 0 | 0 | 0 | 0 | 0 | 0 | 0 | 0 |
| bacteria_Jun8d50b_1248 | 0 | 0 | 0 | 0 | 0 | 0 | 0 | 1 | 0 | 0 | 1 |
| bacteria_Jun8d50b_14625 | 0 | 0 | 0 | 0 | 0 | 0 | 0 | 0 | 0 | 0 | 0 |
| bacteria_Jun8d50b_18759 | 0 | 0 | 0 | 0 | 0 | 0 | 0 | 0 | 0 | 0 | 0 |
| bacteria_Jun8d50b_21017 | 1 | 0 | 0 | 0 | 1 | 0 | 0 | 0 | 0 | 0 | 2 |
| bacteria_Jun8d50b_30506 | 0 | 0 | 0 | 0 | 0 | 0 | 0 | 0 | 0 | 0 | 0 |
| bacteria_Jun8d50b_33815 | 0 | 0 | 0 | 0 | 0 | 0 | 0 | 0 | 0 | 0 | 0 |
| bacteria_Jun8d50b_34065 | 0 | 0 | 0 | 0 | 0 | 0 | 0 | 0 | 0 | 0 | 0 |
| bacteria_Jun8d75b_7768 | 0 | 0 | 0 | 0 | 0 | 0 | 0 | 0 | 0 | 0 | 0 |
| bacteria_Jun8d75b_67192 | 0 | 0 | 0 | 0 | 0 | 0 | 0 | 0 | 0 | 0 | 0 |
| bacteria_Jun22d0b_13316 | 1 | 2 | 0 | 0 | 0 | 0 | 0 | 0 | 0 | 2 | 5 |
| bacteria_Jun22d0b_31457 | 0 | 0 | 0 | 1 | 1 | 0 | 0 | 0 | 0 | 0 | 2 |
| bacteria_Jun22d0b_59869 | 0 | 0 | 1 | 0 | 0 | 0 | 0 | 0 | 0 | 0 | 1 |
| bacteria_Jun22d0b_69465 | 0 | 0 | 0 | 0 | 0 | 0 | 0 | 0 | 0 | 0 | 0 |
| bacteria_Jun22d0b_70419 | 0 | 0 | 0 | 0 | 0 | 0 | 0 | 0 | 0 | 0 | 0 |
| bacteria_Jun22d50b_2698 | 1 | 0 | 0 | 0 | 0 | 0 | 0 | 0 | 0 | 0 | 1 |
| bacteria_Jun22d50b_4570 | 0 | 0 | 0 | 0 | 0 | 0 | 0 | 0 | 0 | 0 | 0 |
| bacteria_Jun22d75b_2090 | 0 | 0 | 0 | 0 | 0 | 0 | 0 | 0 | 0 | 0 | 0 |
| bacteria_Jun22d75b_6264 | 0 | 0 | 0 | 0 | 1 | 0 | 0 | 0 | 0 | 0 | 1 |
| bacteria_Jun22d75b_1361 | 0 | 0 | 0 | 0 | 0 | 0 | 0 | 0 | 0 | 0 | 0 |
| bacteria_Jun22d75b_2486 | 0 | 0 | 0 | 0 | 0 | 0 | 0 | 0 | 0 | 0 | 0 |
| bacteria_Jun22d75b_2937 | 0 | 0 | 0 | 0 | 0 | 0 | 0 | 0 | 0 | 0 | 0 |
| bacteria_Jun22d75b_2965 | 0 | 0 | 0 | 0 | 0 | 0 | 0 | 0 | 0 | 0 | 0 |
| bacteria_Jun22d75b_3252 | 0 | 0 | 0 | 0 | 0 | 0 | 0 | 0 | 0 | 0 | 0 |
| bacteria_Jun22d75b_4450 | 0 | 0 | 0 | 0 | 0 | 0 | 0 | 0 | 0 | 0 | 0 |
| bacteria_Jun22d75b_4459 | 0 | 0 | 0 | 0 | 0 | 0 | 0 | 0 | 0 | 0 | 0 |
| bacteria_Jun22d75b_5443 | 1 | 1 | 0 | 0 | 0 | 2 | 0 | 0 | 0 | 0 | 4 |
| bacteria_Jun22d75b_5985 | 0 | 0 | 0 | 0 | 0 | 0 | 0 | 0 | 0 | 0 | 0 |
| bacteria_Jun22d75b_6571 | 0 | 0 | 0 | 0 | 0 | 0 | 0 | 0 | 0 | 0 | 0 |
| bacteria_Jun22d75b_7240 | 0 | 0 | 0 | 0 | 0 | 1 | 0 | 0 | 0 | 1 | 2 |
| bacteria_Jun22d75b_72651 | 0 | 0 | 0 | 0 | 0 | 0 | 0 | 0 | 0 | 0 | 0 |
| bacteria_Jun22d75b_7384 | 0 | 1 | 0 | 0 | 0 | 0 | 0 | 0 | 0 | 0 | 1 |
| bacteria_May12d25b_1528 | 0 | 0 | 0 | 0 | 0 | 0 | 0 | 0 | 0 | 0 | 0 |
| bacteria_May12d25b_4551 | 0 | 0 | 0 | 0 | 0 | 0 | 0 | 0 | 0 | 0 | 0 |
| bacteria_May12d25b_933 | 0 | 3 | 0 | 1 | 0 | 2 | 0 | 1 | 0 | 1 | 8 |
| bacteria_May12d25b_176 | 0 | 0 | 0 | 0 | 1 | 0 | 1 | 0 | 0 | 0 | 2 |
| bacteria_May12d25b_2961 | 0 | 0 | 0 | 0 | 0 | 0 | 0 | 0 | 0 | 0 | 0 |
| bacteria_May12d25b_3873 | 0 | 0 | 0 | 0 | 0 | 0 | 0 | 0 | 0 | 0 | 0 |
| bacteria_May12d25b_4768 | 0 | 0 | 0 | 0 | 0 | 0 | 0 | 0 | 0 | 0 | 0 |
| bacteria_May12d25b_479 | 0 | 0 | 0 | 0 | 0 | 0 | 0 | 0 | 0 | 0 | 0 |

| | | | | | | | | | | | | |
|---|---|---|---|---|---|---|---|---|---|---|---|---|
| bacteria_May12d25b_5404 | | 0 | 0 | 0 | 0 | 0 | 0 | 0 | 0 | 0 | 0 | 0 |
| bacteria_May12d50b_2360 | | 0 | 0 | 0 | 0 | 1 | 1 | 0 | 0 | 0 | 0 | 2 |
| bacteria_May12d50b_6049 | | 0 | 0 | 0 | 0 | 0 | 0 | 0 | 0 | 0 | 0 | 0 |
| bacteria_May12d50b_8587 | | 1 | 3 | 1 | 5 | 3 | 1 | 0 | 0 | 0 | 1 | 15 |
| bacteria_May12d50b_9610 | | 0 | 0 | 0 | 0 | 0 | 0 | 0 | 0 | 0 | 0 | 0 |
| bacteria_May12d50b_1244 | | 0 | 0 | 0 | 0 | 0 | 0 | 0 | 0 | 0 | 0 | 0 |
| bacteria_May12d50b_1817 | | 0 | 0 | 0 | 0 | 0 | 0 | 0 | 0 | 0 | 0 | 0 |
| bacteria_May12d50b_2099 | | 0 | 0 | 0 | 0 | 0 | 0 | 0 | 0 | 0 | 0 | 0 |
| bacteria_May12d50b_2225 | | 0 | 0 | 0 | 0 | 0 | 1 | 0 | 0 | 0 | 0 | 1 |
| bacteria_May12d50b_2364 | | 0 | 0 | 0 | 0 | 0 | 0 | 0 | 0 | 0 | 0 | 0 |
| bacteria_May | 23 | 5 | 11 | 1 | 11 | 23 | 13 | 2 | 10 | 1 | 9 | 86 |
| bacteria_May12d50b_2716 | | 0 | 0 | 0 | 0 | 0 | 0 | 0 | 0 | 0 | 0 | 0 |
| bacteria_May12d50b_3228 | | 0 | 0 | 0 | 0 | 0 | 0 | 0 | 0 | 0 | 0 | 0 |
| bacteria_May12d50b_4776 | | 0 | 1 | 0 | 1 | 2 | 1 | 0 | 0 | 0 | 0 | 5 |
| bacteria_May12d50b_4779 | | 0 | 0 | 0 | 0 | 0 | 0 | 0 | 0 | 0 | 0 | 0 |
| bacteria_May12d75b_2075 | | 0 | 0 | 0 | 0 | 0 | 0 | 0 | 0 | 0 | 0 | 0 |
| bacteria_May12d75b_2586 | | 0 | 0 | 0 | 0 | 0 | 0 | 0 | 0 | 0 | 0 | 0 |
| bacteria_May12d75b_3431 | | 0 | 0 | 0 | 0 | 0 | 0 | 0 | 0 | 0 | 0 | 0 |
| bacteria_Sep8d0b_31869 | | 0 | 0 | 0 | 0 | 0 | 0 | 0 | 0 | 0 | 0 | 0 |
| bacteria_Sep8d5b_207 | | 0 | 1 | 0 | 0 | 0 | 0 | 1 | 0 | 0 | 0 | 2 |
| bacteria_Sep8d5b_4491 | | 2 | 0 | 0 | 0 | 0 | 0 | 0 | 0 | 0 | 0 | 2 |
| bacteria_Sep8d5b_19262 | | 0 | 0 | 0 | 0 | 1 | 0 | 1 | 0 | 0 | 0 | 2 |
| bacteria_Sep8d5b_21974 | | 0 | 0 | 0 | 0 | 0 | 0 | 0 | 0 | 0 | 0 | 0 |
| bacteria_Sep8d5b_28522 | | 0 | 1 | 0 | 2 | 0 | 1 | 0 | 0 | 0 | 0 | 4 |
| bacteria_Sep8d5b_37775 | | 0 | 0 | 0 | 0 | 0 | 0 | 0 | 0 | 0 | 0 | 0 |
| bacteria_Sep8d5b_46368 | | 0 | 0 | 0 | 0 | 0 | 0 | 0 | 0 | 0 | 0 | 0 |
| bacteria_Sep8d5b_49913 | | 0 | 0 | 0 | 1 | 0 | 1 | 0 | 0 | 0 | 0 | 2 |
| bacteria_Sep8d5b_52820 | | 0 | 0 | 0 | 0 | 0 | 0 | 0 | 0 | 0 | 0 | 0 |
| bacteria_Sep8d5b_56440 | | 0 | 0 | 0 | 0 | 0 | 0 | 0 | 0 | 0 | 0 | 0 |
| bacteria_Sep8d5b_60365 | | 0 | 0 | 0 | 0 | 1 | 0 | 0 | 0 | 0 | 2 | 3 |
| bacteria_Sep8d5b_67580 | | 1 | 1 | 0 | 0 | 2 | 1 | 0 | 0 | 1 | 2 | 8 |
| bacteria_Sep8d5b_71861 | | 0 | 0 | 0 | 0 | 0 | 0 | 0 | 0 | 0 | 0 | 0 |
| bacteria_Sep8d25b_1335 | | 0 | 0 | 0 | 0 | 1 | 0 | 1 | 0 | 0 | 0 | 2 |
| bacteria_Sep8d25b_22520 | | 0 | 0 | 0 | 0 | 0 | 0 | 0 | 0 | 0 | 0 | 0 |
| virus_Aug5.0v_24 | | 1 | 3 | 0 | 5 | 4 | 0 | 0 | 1 | 0 | 0 | 14 |
| virus_Aug5.0v_276 | | 0 | 1 | 0 | 0 | 0 | 1 | 0 | 0 | 0 | 2 | 4 |
| virus_Aug5.0v_289 | | 0 | 1 | 0 | 0 | 0 | 0 | 0 | 0 | 0 | 0 | 1 |
| virus_Aug5.0v_581 | | 0 | 1 | 0 | 0 | 0 | 0 | 0 | 0 | 0 | 0 | 1 |
| virus_Aug5.0v_1733 | | 0 | 1 | 0 | 0 | 0 | 0 | 0 | 0 | 0 | 0 | 1 |
| virus_Aug5.0v_1938 | | 0 | 1 | 1 | 0 | 0 | 0 | 1 | 0 | 0 | 0 | 3 |
| virus_Aug5.0v_2681 | | 0 | 1 | 0 | 0 | 0 | 0 | 0 | 0 | 0 | 0 | 1 |
| virus_Aug5.0v_2811 | | 0 | 1 | 0 | 0 | 0 | 0 | 0 | 0 | 0 | 0 | 1 |
| virus_Aug5.0v_2845 | | 0 | 1 | 0 | 0 | 0 | 0 | 0 | 0 | 0 | 0 | 1 |
| virus_Aug5.0v_3150 | | 1 | 3 | 0 | 3 | 2 | 0 | 0 | 1 | 0 | 1 | 11 |
| virus_Aug5.0v_3331 | | 0 | 5 | 1 | 0 | 0 | 2 | 0 | 0 | 0 | 0 | 8 |
| virus_Aug5.0v_3417 | | 0 | 1 | 0 | 0 | 0 | 0 | 0 | 0 | 0 | 0 | 1 |
| virus_Aug5.0v_3516 | | 0 | 1 | 0 | 0 | 0 | 0 | 0 | 0 | 0 | 0 | 1 |
| virus_Aug5.0v_3629 | | 0 | 1 | 0 | 0 | 0 | 0 | 0 | 1 | 0 | 0 | 2 |
| virus_Aug5.0v_3862 | | 0 | 1 | 0 | 0 | 0 | 0 | 0 | 0 | 0 | 0 | 1 |
| virus_Aug5.0v_3863 | | 0 | 1 | 0 | 0 | 0 | 0 | 0 | 0 | 0 | 0 | 1 |
| virus_Aug5.0v_3881 | | 0 | 4 | 0 | 2 | 1 | 0 | 0 | 1 | 0 | 0 | 8 |
| virus_Aug5.0v_3883 | | 0 | 1 | 0 | 0 | 0 | 0 | 0 | 0 | 0 | 0 | 1 |
| virus_Aug5.0v_4047 | | 0 | 1 | 0 | 0 | 0 | 0 | 0 | 0 | 0 | 0 | 1 |
| virus_Aug5.0v_4121 | | 0 | 1 | 0 | 0 | 0 | 0 | 0 | 0 | 0 | 0 | 1 |
| virus_Aug5.0v_4275 | | 0 | 1 | 0 | 0 | 0 | 0 | 0 | 0 | 0 | 0 | 1 |
| virus_Aug5.0 | 26 | 1 | 46 | 1 | 7 | 1 | 15 | 2 | 2 | 0 | 1 | 76 |
| virus_Aug5.0v_4754 | | 0 | 1 | 0 | 0 | 0 | 0 | 0 | 0 | 0 | 0 | 1 |
| virus_Aug5.0v_5052 | | 0 | 1 | 0 | 0 | 0 | 0 | 0 | 0 | 0 | 0 | 1 |
| virus_Aug5.0v_5360 | | 0 | 1 | 0 | 0 | 0 | 0 | 0 | 0 | 0 | 0 | 1 |
| virus_Aug5.5v_3929 | | 0 | 0 | 0 | 1 | 0 | 0 | 0 | 0 | 0 | 0 | 1 |
| virus_Aug5.5v_4047 | | 0 | 0 | 0 | 1 | 0 | 0 | 0 | 0 | 0 | 0 | 1 |
| virus_Aug5.5v_4150 | | 0 | 0 | 0 | 3 | 0 | 1 | 0 | 1 | 0 | 0 | 5 |
| virus_Aug5.5v_6951 | | 0 | 0 | 0 | 12 | 0 | 10 | 0 | 16 | 0 | 7 | 45 |
| virus_Aug5.5v_12684 | | 5 | 1 | 2 | 3 | 3 | 2 | 0 | 0 | 0 | 0 | 16 |
| virus_Aug5.5v_13145 | | 0 | 4 | 0 | 5 | 0 | 1 | 0 | 1 | 0 | 0 | 11 |
| virus_Aug5.5v_13274 | | 0 | 0 | 0 | 3 | 0 | 0 | 0 | 0 | 0 | 0 | 3 |
| virus_Aug5.5v_14779 | | 1 | 2 | 0 | 1 | 1 | 0 | 0 | 0 | 0 | 0 | 5 |
| virus_Aug5.5v_15591 | | 0 | 0 | 0 | 1 | 0 | 0 | 0 | 0 | 0 | 0 | 1 |
| virus_Aug5.5v_15844 | | 0 | 0 | 0 | 1 | 0 | 0 | 0 | 1 | 0 | 1 | 3 |
| virus_Aug5.5v_19438 | | 0 | 1 | 0 | 1 | 0 | 1 | 0 | 0 | 0 | 0 | 3 |
| virus_Aug5.5v_26463 | | 0 | 0 | 0 | 1 | 0 | 0 | 0 | 0 | 0 | 0 | 1 |
| virus_Aug5.5v_27474 | | 0 | 0 | 0 | 1 | 0 | 0 | 0 | 0 | 0 | 0 | 1 |
| virus_Aug5.5v_32417 | | 0 | 0 | 0 | 1 | 0 | 1 | 0 | 1 | 0 | 0 | 3 |
| virus_Aug5.5 | 7 | 7 | 170 | 8 | 30 | 11 | 67 | 7 | 8 | 0 | 7 | 315 |
| virus_Aug5.5v_33459 | | 0 | 0 | 0 | 1 | 0 | 0 | 0 | 0 | 0 | 1 | 2 |
| virus_Aug5.5v_35202 | | 0 | 0 | 0 | 1 | 0 | 0 | 0 | 0 | 0 | 0 | 1 |
| virus_Aug5.25v_10994 | | 0 | 0 | 0 | 0 | 0 | 1 | 0 | 0 | 0 | 0 | 1 |
| virus_Aug5.25v_14023 | | 0 | 7 | 0 | 0 | 0 | 3 | 0 | 0 | 0 | 0 | 10 |
| virus_Aug5.25v_15437 | | 0 | 0 | 0 | 0 | 0 | 1 | 0 | 0 | 0 | 0 | 1 |
| virus_Aug5.25v_15975 | | 0 | 0 | 0 | 0 | 0 | 1 | 0 | 0 | 0 | 0 | 1 |
| virus_Aug5.25v_19753 | | 0 | 0 | 0 | 0 | 0 | 1 | 0 | 0 | 0 | 0 | 1 |
| virus_Aug5.25v_20741 | | 0 | 1 | 0 | 0 | 0 | 2 | 0 | 0 | 0 | 0 | 3 |
| virus_Aug5.25v_22058 | | 0 | 3 | 0 | 2 | 0 | 1 | 0 | 0 | 0 | 0 | 6 |
| virus_Aug5.25v_23422 | | 0 | 0 | 0 | 1 | 0 | 2 | 0 | 0 | 0 | 0 | 3 |
| virus_Aug5.25v_24147 | | 0 | 0 | 0 | 0 | 0 | 1 | 0 | 0 | 0 | 0 | 1 |
| virus_Aug5.25v_25126 | | 0 | 0 | 0 | 0 | 0 | 2 | 1 | 0 | 0 | 0 | 3 |
| virus_Aug5.25v_25628 | | 0 | 0 | 0 | 0 | 0 | 1 | 0 | 0 | 0 | 0 | 1 |
| virus_Aug5.25v_26534 | | 0 | 0 | 0 | 0 | 0 | 1 | 0 | 0 | 0 | 0 | 1 |
| virus_Aug5.25v_27001 | | 0 | 5 | 0 | 1 | 0 | 1 | 0 | 0 | 0 | 0 | 7 |
| virus_Aug5.25v_27584 | | 0 | 0 | 0 | 0 | 0 | 1 | 0 | 0 | 0 | 0 | 1 |
| virus_Aug5.25v_31270 | | 0 | 0 | 0 | 0 | 0 | 2 | 0 | 0 | 0 | 0 | 2 |
| virus_Aug5.25v_31556 | | 0 | 0 | 0 | 0 | 2 | 1 | 0 | 1 | 0 | 0 | 4 |
| virus_Aug5.25v_37532 | | 0 | 0 | 0 | 0 | 0 | 1 | 0 | 0 | 0 | 0 | 1 |
| virus_Aug5.25v_38270 | | 0 | 0 | 0 | 4 | 0 | 3 | 0 | 0 | 0 | 1 | 8 |
| virus_Aug5.50v_5434 | | 0 | 0 | 0 | 0 | 0 | 0 | 0 | 1 | 0 | 0 | 1 |
| virus_Aug5.50v_10921 | | 0 | 0 | 0 | 0 | 0 | 0 | 0 | 1 | 0 | 0 | 1 |
| virus_Aug5.50v_13857 | | 0 | 0 | 0 | 0 | 0 | 1 | 0 | 1 | 0 | 0 | 2 |
| virus_Aug5.50v_26485 | | 0 | 0 | 1 | 0 | 0 | 0 | 0 | 1 | 0 | 0 | 2 |
| virus_Aug5.50v_37878 | | 0 | 0 | 0 | 0 | 0 | 0 | 0 | 1 | 0 | 0 | 1 |
| virus_Aug5.75v_348 | | 0 | 0 | 0 | 0 | 0 | 0 | 0 | 0 | 0 | 4 | 4 |
| virus_Aug5.75v_10553 | | 0 | 0 | 0 | 0 | 0 | 0 | 0 | 0 | 0 | 1 | 1 |
| virus_Aug5.75v_14295 | | 0 | 0 | 0 | 0 | 0 | 0 | 0 | 0 | 0 | 1 | 1 |
| virus_Aug5.75v_18641 | | 0 | 0 | 0 | 0 | 0 | 0 | 0 | 0 | 0 | 1 | 1 |
| virus_Aug5.75v_25496 | | 0 | 0 | 0 | 0 | 0 | 0 | 0 | 0 | 0 | 1 | 1 |
| virus_Aug5.75v_34716 | | 0 | 0 | 0 | 0 | 0 | 0 | 0 | 0 | 0 | 1 | 1 |
| virus_Aug5.75v_35584 | | 0 | 0 | 0 | 0 | 0 | 0 | 0 | 0 | 0 | 2 | 2 |
| virus_Aug5.75v_36446 | | 0 | 0 | 0 | 0 | 0 | 0 | 0 | 0 | 0 | 1 | 1 |
| virus_Jul11.0v_1306 | | 0 | 0 | 0 | 0 | 0 | 0 | 0 | 0 | 0 | 0 | 0 |
| virus_Jul11.0v_1381 | | 0 | 0 | 0 | 0 | 2 | 0 | 0 | 0 | 0 | 1 | 3 |
| virus_Jul11.0v_4093 | | 0 | 1 | 0 | 0 | 2 | 1 | 0 | 2 | 0 | 0 | 6 |
| virus_Jul11.0v_6020 | | 3 | 3 | 1 | 5 | 8 | 3 | 0 | 0 | 0 | 1 | 24 |
| virus_Jul11.0v_7341 | | 0 | 0 | 0 | 1 | 0 | 1 | 0 | 1 | 0 | 0 | 3 |
| virus_Jul11.0v_9287 | | 0 | 0 | 0 | 0 | 0 | 0 | 0 | 0 | 0 | 0 | 0 |
| virus_Jul11.0v_11308 | | 0 | 0 | 0 | 0 | 0 | 0 | 0 | 0 | 0 | 0 | 0 |
| virus_Jul11.0v_11656 | | 0 | 0 | 0 | 0 | 0 | 0 | 0 | 0 | 0 | 0 | 0 |
| virus_Jul11.0v_16432 | | 0 | 0 | 0 | 0 | 0 | 0 | 0 | 0 | 0 | 0 | 0 |
| virus_Jul11.0v_16718 | | 0 | 1 | 0 | 1 | 0 | 1 | 0 | 0 | 0 | 0 | 3 |
| virus_Jul11.0v_19513 | | 0 | 1 | 0 | 0 | 0 | 0 | 0 | 0 | 0 | 0 | 1 |
| virus_Jul11.0v_19603 | | 0 | 0 | 0 | 0 | 0 | 0 | 0 | 0 | 0 | 0 | 0 |
| virus_Jul11.0v_20945 | | 0 | 1 | 0 | 2 | 0 | 0 | 0 | 0 | 0 | 0 | 3 |
| virus_Jul11.5v_2894 | | 0 | 0 | 0 | 0 | 0 | 0 | 0 | 0 | 0 | 0 | 0 |
| virus_Jul11.5v_3613 | | 0 | 0 | 0 | 1 | 0 | 0 | 0 | 0 | 0 | 0 | 1 |
| virus_Jul11.5v_4614 | | 0 | 0 | 0 | 0 | 0 | 0 | 0 | 0 | 0 | 0 | 0 |
| virus_Jul11.5v_6833 | | 0 | 0 | 0 | 1 | 0 | 1 | 0 | 0 | 0 | 0 | 2 |
| virus_Jul11.5v_9517 | | 0 | 0 | 0 | 0 | 0 | 0 | 0 | 0 | 0 | 0 | 0 |
| virus_Jul11.5v_10009 | | 0 | 0 | 0 | 3 | 0 | 0 | 0 | 0 | 0 | 1 | 4 |
| virus_Jul11.5v_11265 | | 0 | 0 | 0 | 0 | 0 | 0 | 0 | 0 | 0 | 0 | 0 |
| virus_Jul11.5v_12327 | | 0 | 0 | 0 | 0 | 0 | 0 | 0 | 0 | 0 | 0 | 0 |
| virus_Jul11.5v_12620 | | 0 | 0 | 0 | 0 | 0 | 0 | 0 | 0 | 0 | 0 | 0 |
| virus_Jul11.5v_13527 | | 0 | 0 | 0 | 0 | 0 | 0 | 0 | 0 | 0 | 0 | 0 |
| virus_Jul11.5v_13628 | | 0 | 0 | 0 | 0 | 0 | 0 | 0 | 0 | 0 | 0 | 0 |
| virus_Jul11.5v_15098 | | 0 | 0 | 0 | 0 | 0 | 0 | 0 | 0 | 0 | 0 | 0 |
| virus_Jul11.5v_15679 | | 0 | 0 | 0 | 0 | 0 | 0 | 0 | 0 | 0 | 0 | 0 |
| virus_Jul11.5v_17064 | | 0 | 0 | 0 | 1 | 0 | 0 | 0 | 0 | 0 | 1 | 2 |
| virus_Jul11.5v_18046 | | 0 | 0 | 0 | 2 | 0 | 6 | 0 | 0 | 0 | 0 | 8 |
| virus_Jul11.5v_18386 | | 0 | 0 | 1 | 0 | 1 | 0 | 0 | 1 | 0 | 1 | 4 |
| virus_Jul11.25v_5561 | | 0 | 0 | 0 | 0 | 0 | 0 | 0 | 0 | 0 | 0 | 0 |
| virus_Jul11.25v_6485 | | 0 | 0 | 0 | 0 | 0 | 0 | 0 | 0 | 0 | 0 | 0 |
| virus_Jul11.2 | 28 | 0 | 4 | 0 | 9 | 14 | 6 | 5 | 6 | 3 | 7 | 54 |
| virus_Jul11.25v_10897 | | 0 | 0 | 0 | 0 | 0 | 0 | 0 | 0 | 0 | 0 | 0 |
| virus_Jul11.25v_11955 | | 0 | 0 | 0 | 0 | 0 | 0 | 0 | 0 | 0 | 0 | 0 |
| virus_Jul11.25v_12861 | | 0 | 0 | 0 | 0 | 0 | 0 | 0 | 0 | 0 | 0 | 0 |
| virus_Jul11.25v_13450 | | 0 | 0 | 0 | 1 | 0 | 0 | 0 | 0 | 0 | 0 | 1 |
| virus_Jul11.2 | 29 | 1 | 6 | 0 | 8 | 13 | 4 | 5 | 8 | 0 | 9 | 54 |
| virus_Jul11.25v_15786 | | 0 | 0 | 0 | 0 | 0 | 0 | 0 | 0 | 0 | 0 | 0 |
| virus_Jul11.25v_16320 | | 0 | 0 | 0 | 0 | 0 | 0 | 0 | 0 | 0 | 0 | 0 |
| virus_Jul11.25v_16748 | | 0 | 0 | 0 | 0 | 0 | 0 | 0 | 0 | 0 | 0 | 0 |
| virus_Jul11.25v_17316 | | 0 | 0 | 0 | 0 | 0 | 0 | 0 | 0 | 0 | 0 | 0 |
| virus_Jul11.50v_1249 | | 0 | 0 | 0 | 0 | 0 | 0 | 0 | 0 | 0 | 0 | 0 |
| virus_Jul11.50v_5024 | | 0 | 0 | 0 | 0 | 0 | 0 | 0 | 0 | 0 | 0 | 0 |
| virus_Jul11.50v_5185 | | 0 | 0 | 0 | 0 | 0 | 0 | 0 | 0 | 0 | 0 | 0 |
| virus_Jul11.50v_7164 | | 0 | 2 | 0 | 1 | 0 | 0 | 0 | 0 | 0 | 0 | 3 |
| virus_Jul11.50v_8859 | | 0 | 0 | 0 | 0 | 0 | 0 | 0 | 0 | 0 | 0 | 0 |
| virus_Jul11.50v_9111 | | 0 | 0 | 0 | 0 | 0 | 0 | 0 | 0 | 0 | 0 | 0 |
| virus_Jul11.50v_9120 | | 5 | 0 | 0 | 0 | 1 | 0 | 0 | 1 | 0 | 0 | 7 |
| virus_Jul11.50v_12800 | | 0 | 0 | 0 | 0 | 0 | 0 | 0 | 0 | 0 | 0 | 0 |
| virus_Jul11.50v_13462 | | 0 | 0 | 0 | 0 | 0 | 0 | 0 | 0 | 0 | 0 | 0 |

| | | | | | | | | | | | |
|---|---|---|---|---|---|---|---|---|---|---|---|
| virus_Jul11.50v_13741 | 0 | 0 | 0 | 0 | 0 | 0 | 0 | 0 | 0 | 0 | 0 |
| virus_Jul11.50v_14684 | 0 | 0 | 0 | 0 | 0 | 0 | 0 | 0 | 0 | 0 | 0 |
| virus_Jul11.50v_15972 | 0 | 14 | 0 | 8 | 0 | 8 | 0 | 3 | 0 | 5 | 38 |
| virus_Jul11.50v_16904 | 0 | 0 | 0 | 0 | 0 | 0 | 0 | 0 | 0 | 0 | 0 |
| virus_Jul11.50v_17822 | 0 | 0 | 0 | 0 | 0 | 0 | 0 | 0 | 0 | 1 | 1 |
| virus_Jul11.50v_19933 | 0 | 0 | 0 | 0 | 0 | 0 | 0 | 0 | 0 | 0 | 0 |
| virus_Jul11.50v_20336 | 0 | 0 | 0 | 0 | 0 | 0 | 0 | 0 | 0 | 0 | 0 |
| virus_Jul11.75v_179 | 0 | 0 | 0 | 0 | 0 | 0 | 0 | 0 | 0 | 0 | 0 |
| virus_Jul11.75v_2729 | 0 | 0 | 0 | 0 | 1 | 0 | 0 | 0 | 0 | 0 | 1 |
| virus_Jul11.75v_5240 | 0 | 0 | 0 | 0 | 0 | 0 | 0 | 0 | 0 | 0 | 0 |
| virus_Jul11.75v_19905 | 0 | 0 | 0 | 0 | 0 | 0 | 0 | 0 | 0 | 0 | 0 |
| virus_Jul11.75v_23605 | 0 | 0 | 0 | 0 | 0 | 0 | 0 | 0 | 0 | 0 | 0 |
| virus_Jul11.75v_24726 | 0 | 0 | 0 | 0 | 0 | 0 | 0 | 0 | 0 | 0 | 0 |
| virus_Jul11.75v_35941 | 0 | 0 | 0 | 0 | 0 | 0 | 0 | 0 | 0 | 0 | 0 |
| virus_Jul11.75v_36621 | 0 | 0 | 0 | 0 | 0 | 0 | 0 | 0 | 0 | 0 | 0 |
| virus_Jul11.75v_36672 | 0 | 0 | 0 | 0 | 0 | 0 | 0 | 0 | 0 | 0 | 0 |
| virus_Jul11.75v_38959 | 0 | 0 | 0 | 0 | 0 | 0 | 0 | 0 | 0 | 0 | 0 |
| virus_Jul11.75v_40569 | 0 | 0 | 0 | 0 | 0 | 0 | 0 | 0 | 0 | 0 | 0 |
| virus_Jul27.5v_1426 | 0 | 0 | 0 | 0 | 0 | 0 | 0 | 0 | 0 | 0 | 0 |
| virus_Jul27.5v_1553 | 0 | 0 | 0 | 0 | 0 | 0 | 0 | 0 | 0 | 0 | 0 |
| virus_Jul27.5v_5032 | 0 | 0 | 0 | 0 | 0 | 0 | 0 | 0 | 0 | 0 | 0 |
| virus_Jul27.5v_5507 | 0 | 1 | 0 | 1 | 0 | 0 | 0 | 0 | 0 | 0 | 2 |
| virus_Jul27.5v_5739 | 0 | 0 | 0 | 0 | 0 | 0 | 0 | 0 | 0 | 0 | 0 |
| virus_Jul27.5v_5916 | 0 | 0 | 0 | 0 | 0 | 0 | 0 | 0 | 0 | 0 | 0 |
| virus_Jul27.5v_7114 | 0 | 0 | 0 | 0 | 0 | 0 | 0 | 0 | 0 | 0 | 0 |
| virus_Jul27.5v_7801 | 0 | 0 | 0 | 0 | 0 | 0 | 0 | 0 | 0 | 0 | 0 |
| virus_Jul27.5v_8304 | 0 | 0 | 0 | 0 | 0 | 0 | 0 | 0 | 0 | 0 | 0 |
| virus_Jul27.5v_8523 | 3 | 0 | 0 | 0 | 1 | 0 | 0 | 0 | 0 | 0 | 4 |
| virus_Jul27.5v_8690 | 0 | 0 | 0 | 0 | 0 | 1 | 0 | 0 | 0 | 0 | 1 |
| virus_Jul27.5v_9493 | 0 | 0 | 0 | 0 | 0 | 0 | 0 | 0 | 0 | 0 | 0 |
| virus_Jul27.5v_9941 | 0 | 0 | 0 | 0 | 0 | 0 | 0 | 0 | 0 | 0 | 0 |
| virus_Jul27.5v_10641 | 0 | 0 | 0 | 0 | 0 | 0 | 0 | 0 | 0 | 0 | 0 |
| virus_Jul27.5v_11478 | 0 | 0 | 0 | 0 | 0 | 0 | 0 | 0 | 0 | 0 | 0 |
| virus_Jul27.5v_12901 | 0 | 0 | 0 | 0 | 0 | 0 | 0 | 0 | 0 | 0 | 0 |
| virus_Jul27.5v_13266 | 0 | 0 | 0 | 0 | 0 | 0 | 0 | 0 | 0 | 0 | 0 |
| virus_Jul27.5v_13960 | 0 | 0 | 0 | 0 | 0 | 0 | 0 | 0 | 0 | 0 | 0 |
| virus_Jul27.5v_14499 | 0 | 0 | 0 | 0 | 0 | 0 | 0 | 0 | 0 | 0 | 0 |
| virus_Jul27.5v_15916 | 0 | 0 | 0 | 0 | 0 | 0 | 0 | 0 | 0 | 0 | 0 |
| virus_Jul27.5v_16280 | 0 | 0 | 0 | 0 | 1 | 0 | 0 | 0 | 0 | 0 | 1 |
| virus_Jul27.5v_16976 | 0 | 0 | 0 | 0 | 0 | 0 | 0 | 0 | 0 | 0 | 0 |
| virus_Jul27.5v_19523 | 0 | 0 | 0 | 0 | 0 | 0 | 0 | 0 | 0 | 0 | 0 |
| virus_Jul27.5v_20083 | 0 | 0 | 0 | 0 | 0 | 0 | 0 | 0 | 0 | 0 | 0 |
| virus_Jul27.5v_20148 | 0 | 0 | 0 | 0 | 0 | 0 | 0 | 0 | 0 | 0 | 0 |
| virus_Jul27.5v_20233 | 0 | 0 | 0 | 0 | 0 | 0 | 0 | 0 | 0 | 0 | 0 |
| virus_Jul27.25v_1517 | 0 | 0 | 0 | 0 | 0 | 0 | 0 | 0 | 0 | 0 | 0 |
| virus_Jul27.25v_6981 | 0 | 0 | 0 | 0 | 0 | 0 | 0 | 0 | 0 | 0 | 0 |
| virus_Jul27.25v_10497 | 0 | 0 | 0 | 0 | 0 | 0 | 0 | 0 | 0 | 0 | 0 |
| virus_Jul27.25v_13167 | 0 | 4 | 0 | 0 | 0 | 1 | 0 | 0 | 0 | 0 | 5 |
| virus_Jul27.25v_14448 | 0 | 0 | 0 | 0 | 0 | 0 | 0 | 0 | 0 | 0 | 0 |
| virus_Jul27.25v_15557 | 0 | 0 | 0 | 0 | 0 | 0 | 0 | 0 | 0 | 0 | 0 |
| virus_Jul27.25v_24394 | 0 | 0 | 0 | 0 | 0 | 0 | 0 | 0 | 0 | 0 | 0 |
| virus_Jul27.25v_27779 | 0 | 0 | 0 | 0 | 0 | 0 | 0 | 0 | 0 | 0 | 0 |
| virus_Jul27.25v_28260 | 0 | 0 | 0 | 0 | 1 | 0 | 0 | 1 | 0 | 0 | 2 |
| virus_Jul27.25v_28837 | 0 | 0 | 0 | 0 | 0 | 0 | 0 | 0 | 0 | 0 | 0 |
| virus_Jul27.25v_31066 | 0 | 0 | 0 | 0 | 0 | 0 | 0 | 0 | 0 | 0 | 0 |
| virus_Jul27.25v_32372 | 0 | 0 | 0 | 0 | 0 | 0 | 0 | 0 | 0 | 0 | 0 |
| virus_Jul27.25v_34410 | 0 | 0 | 0 | 0 | 0 | 0 | 0 | 0 | 0 | 0 | 0 |
| virus_Jul27.25v_35759 | 0 | 0 | 0 | 0 | 0 | 0 | 0 | 0 | 0 | 0 | 0 |
| virus_Jul27.25v_39770 | 0 | 0 | 0 | 0 | 0 | 0 | 0 | 0 | 0 | 0 | 0 |
| virus_Jul27.75v_1947 | 0 | 0 | 0 | 0 | 0 | 0 | 0 | 0 | 0 | 0 | 0 |
| virus_Jul27.75v_3620 | 0 | 0 | 0 | 0 | 0 | 0 | 0 | 0 | 0 | 0 | 0 |
| virus_Jul27.75v_12158 | 0 | 1 | 0 | 1 | 0 | 0 | 0 | 0 | 0 | 0 | 2 |
| virus_Jul27.75v_18162 | 0 | 0 | 0 | 0 | 0 | 0 | 0 | 0 | 0 | 0 | 0 |
| virus_Jul27.75v_18179 | 0 | 0 | 0 | 0 | 0 | 0 | 0 | 0 | 0 | 0 | 0 |
| virus_Jul27.75v_18645 | 0 | 0 | 0 | 0 | 0 | 0 | 0 | 0 | 0 | 0 | 0 |
| virus_Jun8.25v_1416 | 0 | 0 | 0 | 0 | 0 | 0 | 0 | 0 | 0 | 0 | 0 |
| virus_Jun8.25v_15884 | 0 | 0 | 0 | 0 | 0 | 0 | 0 | 0 | 0 | 0 | 0 |
| virus_Jun8.25v_17498 | 0 | 0 | 0 | 0 | 0 | 1 | 0 | 0 | 0 | 0 | 1 |
| virus_Jun8.25v_23058 | 0 | 0 | 0 | 0 | 0 | 0 | 0 | 0 | 0 | 0 | 0 |
| virus_Jun8.50v_219 | 0 | 0 | 0 | 0 | 0 | 0 | 0 | 0 | 0 | 0 | 0 |
| virus_Jun8.50v_3281 | 0 | 0 | 0 | 0 | 0 | 0 | 0 | 0 | 0 | 0 | 0 |
| virus_Jun8.50v_8532 | 0 | 0 | 0 | 0 | 0 | 0 | 0 | 0 | 0 | 0 | 0 |
| virus_Jun8.50v_22514 | 0 | 0 | 0 | 0 | 0 | 0 | 0 | 0 | 0 | 1 | 1 |
| virus_Jun8.50v_24075 | 0 | 0 | 0 | 0 | 0 | 0 | 0 | 0 | 0 | 0 | 0 |
| virus_Jun8.50v_24602 | 0 | 0 | 0 | 0 | 0 | 0 | 0 | 0 | 0 | 0 | 0 |
| virus_Jun8.50v_28378 | 0 | 0 | 0 | 0 | 0 | 0 | 0 | 0 | 0 | 0 | 0 |
| virus_Jun8.50v_36988 | 0 | 0 | 0 | 0 | 0 | 0 | 0 | 0 | 0 | 0 | 0 |
| virus_Jun8.50v_39951 | 0 | 0 | 0 | 0 | 0 | 0 | 0 | 0 | 0 | 0 | 0 |
| virus_Jun8.50v_40442 | 0 | 0 | 0 | 1 | 0 | 0 | 0 | 0 | 0 | 0 | 1 |
| virus_Jun8.75v_1576 | 0 | 0 | 0 | 0 | 0 | 0 | 0 | 0 | 0 | 0 | 0 |
| virus_Jun8.75v_3585 | 0 | 0 | 0 | 0 | 0 | 0 | 0 | 0 | 0 | 0 | 0 |
| virus_Jun22.0v_2504 | 0 | 0 | 0 | 0 | 0 | 0 | 0 | 0 | 0 | 0 | 0 |
| virus_Jun22.0v_3245 | 0 | 0 | 0 | 0 | 0 | 0 | 0 | 0 | 0 | 0 | 0 |
| virus_Jun22.0v_3466 | 0 | 0 | 0 | 0 | 0 | 0 | 0 | 0 | 0 | 0 | 0 |
| virus_Jun22.0v_6617 | 0 | 0 | 0 | 0 | 0 | 0 | 0 | 0 | 0 | 0 | 0 |
| virus_Jun22.0v_6931 | 0 | 0 | 0 | 0 | 0 | 0 | 0 | 0 | 0 | 0 | 0 |
| virus_Jun22.0v_11186 | 0 | 0 | 0 | 0 | 0 | 0 | 0 | 0 | 0 | 0 | 0 |
| virus_Jun22.0v_11829 | 0 | 0 | 0 | 0 | 0 | 0 | 0 | 0 | 0 | 0 | 0 |
| virus_Jun22.0v_12932 | 0 | 0 | 0 | 0 | 0 | 0 | 0 | 0 | 0 | 0 | 0 |
| virus_Jun22.0v_17823 | 0 | 0 | 0 | 0 | 0 | 0 | 0 | 0 | 0 | 0 | 0 |
| virus_Jun22.0v_17853 | 0 | 0 | 0 | 0 | 0 | 0 | 0 | 0 | 0 | 0 | 0 |
| virus_Jun22.50v_4162 | 0 | 0 | 0 | 0 | 0 | 0 | 0 | 0 | 0 | 0 | 0 |
| virus_Jun22.50v_12192 | 0 | 0 | 0 | 0 | 0 | 0 | 0 | 0 | 0 | 0 | 0 |
| virus_Jun22.50v_12875 | 3 | 4 | 0 | 2 | 6 | 4 | 0 | 1 | 0 | 1 | 21 |
| virus_Jun22.50v_13859 | 0 | 0 | 0 | 0 | 0 | 0 | 0 | 0 | 0 | 0 | 0 |
| virus_Jun22.50v_15780 | 0 | 0 | 0 | 0 | 0 | 0 | 0 | 0 | 0 | 0 | 0 |
| virus_Jun22.50v_15993 | 0 | 0 | 0 | 0 | 0 | 0 | 0 | 0 | 0 | 0 | 0 |
| virus_Jun22.50v_16105 | 0 | 0 | 0 | 0 | 0 | 0 | 0 | 0 | 0 | 0 | 0 |
| virus_Jun22.50v_17216 | 0 | 0 | 0 | 0 | 0 | 0 | 0 | 0 | 0 | 0 | 0 |
| virus_Jun22.50v_18262 | 0 | 0 | 0 | 0 | 0 | 0 | 0 | 0 | 0 | 0 | 0 |
| virus_Jun22.50v_27550 | 0 | 0 | 0 | 0 | 0 | 0 | 0 | 0 | 0 | 0 | 0 |
| virus_Jun22.50v_30159 | 0 | 0 | 0 | 0 | 0 | 0 | 0 | 0 | 0 | 0 | 0 |
| virus_Jun22.50v_31867 | 0 | 0 | 0 | 0 | 0 | 0 | 0 | 0 | 0 | 0 | 0 |
| virus_Jun22.50v_34570 | 0 | 0 | 0 | 0 | 0 | 0 | 0 | 0 | 0 | 0 | 0 |
| virus_Jun22.50v_35780 | 0 | 0 | 0 | 1 | 0 | 0 | 0 | 0 | 0 | 0 | 1 |
| virus_Jun22.50v_38582 | 0 | 0 | 0 | 0 | 0 | 0 | 0 | 0 | 0 | 0 | 0 |
| virus_Jun22.50v_38796 | 0 | 0 | 0 | 0 | 0 | 0 | 0 | 0 | 0 | 0 | 0 |
| virus_Jun22.50v_40223 | 0 | 0 | 0 | 0 | 0 | 0 | 0 | 0 | 0 | 0 | 0 |
| virus_Jun22.75v_2628 | 0 | 0 | 0 | 0 | 0 | 0 | 0 | 0 | 0 | 0 | 0 |
| virus_Jun22.75v_4010 | 0 | 0 | 0 | 0 | 0 | 0 | 0 | 0 | 0 | 0 | 0 |
| virus_Jun22.75v_5319 | 0 | 0 | 0 | 0 | 0 | 0 | 0 | 0 | 0 | 0 | 0 |
| virus_Jun22.75v_9284 | 0 | 0 | 0 | 0 | 0 | 0 | 0 | 0 | 0 | 0 | 0 |
| virus_Jun22.75v_13651 | 0 | 0 | 0 | 0 | 0 | 0 | 0 | 0 | 0 | 0 | 0 |
| virus_Jun22.75v_19461 | 0 | 0 | 0 | 0 | 0 | 0 | 0 | 0 | 0 | 0 | 0 |
| virus_Jun22.75v_19760 | 0 | 0 | 0 | 1 | 0 | 1 | 0 | 0 | 0 | 0 | 2 |
| virus_Jun22.75v_21291 | 0 | 0 | 0 | 0 | 0 | 0 | 0 | 0 | 0 | 0 | 0 |
| virus_Jun22.75v_23171 | 0 | 0 | 0 | 0 | 0 | 0 | 0 | 0 | 0 | 0 | 0 |
| virus_Jun22.75v_24976 | 0 | 3 | 0 | 1 | 0 | 1 | 0 | 2 | 0 | 1 | 8 |
| virus_Jun22.75v_32291 | 0 | 0 | 0 | 0 | 0 | 0 | 0 | 0 | 0 | 0 | 0 |
| virus_Jun22.75v_34153 | 0 | 0 | 0 | 0 | 0 | 0 | 0 | 0 | 0 | 0 | 0 |
| virus_Jun22.75v_37026 | 0 | 0 | 0 | 0 | 0 | 0 | 0 | 0 | 0 | 0 | 0 |
| virus_Jun22.75v_41066 | 0 | 0 | 0 | 0 | 0 | 0 | 0 | 0 | 0 | 0 | 0 |
| virus_May12.5v_690 | 0 | 1 | 0 | 0 | 0 | 2 | 0 | 0 | 0 | 4 | 7 |
| virus_May12.5v_11069 | 0 | 0 | 0 | 0 | 0 | 0 | 0 | 0 | 0 | 0 | 0 |
| virus_May12.5v_21702 | 0 | 0 | 0 | 0 | 0 | 0 | 0 | 0 | 0 | 0 | 0 |
| virus_May12.5v_21769 | 0 | 0 | 0 | 0 | 0 | 1 | 0 | 1 | 0 | 0 | 2 |
| virus_May12.5v_22134 | 0 | 0 | 0 | 0 | 0 | 0 | 0 | 0 | 0 | 0 | 0 |
| virus_May12.25v_935 | 0 | 0 | 0 | 0 | 0 | 0 | 0 | 0 | 0 | 0 | 0 |
| virus_May12.25v_2423 | 0 | 0 | 0 | 0 | 0 | 0 | 0 | 0 | 0 | 0 | 0 |
| virus_May12.25v_5937 | 0 | 0 | 0 | 0 | 0 | 0 | 0 | 0 | 0 | 0 | 0 |
| virus_May12.25v_13721 | 0 | 0 | 0 | 0 | 0 | 0 | 0 | 0 | 0 | 0 | 0 |
| virus_May12.25v_28535 | 0 | 0 | 0 | 0 | 0 | 0 | 0 | 0 | 0 | 0 | 0 |
| virus_May12.25v_29362 | 0 | 0 | 0 | 0 | 0 | 0 | 0 | 0 | 0 | 0 | 0 |
| virus_May12.25v_29365 | 0 | 1 | 0 | 1 | 1 | 0 | 0 | 0 | 0 | 0 | 3 |
| virus_May12.25v_31200 | 0 | 0 | 0 | 0 | 0 | 0 | 0 | 0 | 0 | 0 | 0 |
| virus_May12.25v_32088 | 0 | 0 | 0 | 0 | 0 | 0 | 0 | 0 | 0 | 0 | 0 |
| virus_May12.25v_33418 | 0 | 0 | 0 | 0 | 0 | 0 | 0 | 0 | 0 | 0 | 0 |
| virus_May12.25v_35142 | 0 | 0 | 0 | 0 | 0 | 0 | 0 | 0 | 0 | 0 | 0 |
| virus_May12.25v_35881 | 0 | 0 | 0 | 0 | 0 | 0 | 0 | 0 | 0 | 0 | 0 |
| virus_May12.25v_38963 | 0 | 0 | 0 | 0 | 0 | 0 | 0 | 0 | 0 | 0 | 0 |
| virus_May12.50v_8304 | 0 | 1 | 0 | 0 | 0 | 0 | 0 | 0 | 0 | 0 | 1 |
| virus_May12.50v_9895 | 0 | 0 | 0 | 0 | 0 | 0 | 0 | 0 | 0 | 0 | 0 |
| virus_May12.50v_9925 | 0 | 0 | 0 | 0 | 0 | 0 | 0 | 0 | 0 | 0 | 0 |
| virus_May12.50v_14428 | 0 | 0 | 0 | 0 | 0 | 0 | 0 | 1 | 0 | 1 | 2 |
| virus_May12.50v_26510 | 0 | 0 | 0 | 0 | 0 | 0 | 0 | 0 | 0 | 0 | 0 |
| virus_May12.50v_26875 | 0 | 0 | 0 | 0 | 0 | 0 | 0 | 0 | 0 | 0 | 0 |
| virus_May12.50v_29251 | 0 | 0 | 0 | 0 | 0 | 0 | 0 | 0 | 0 | 0 | 0 |
| virus_May12.50v_29669 | 0 | 0 | 0 | 0 | 0 | 0 | 0 | 0 | 0 | 0 | 0 |
| virus_May12.50v_35559 | 0 | 0 | 0 | 0 | 0 | 0 | 0 | 0 | 0 | 0 | 0 |
| virus_May12.50v_35624 | 0 | 0 | 0 | 0 | 0 | 0 | 0 | 0 | 0 | 0 | 0 |
| virus_May12.50v_36443 | 0 | 0 | 0 | 0 | 0 | 0 | 0 | 0 | 0 | 0 | 0 |
| virus_May12.50v_36683 | 0 | 0 | 0 | 0 | 0 | 0 | 0 | 0 | 0 | 0 | 0 |
| virus_May12.50v_36914 | 0 | 0 | 0 | 0 | 0 | 0 | 0 | 0 | 0 | 0 | 0 |
| virus_May12.50v_39522 | 0 | 0 | 0 | 0 | 0 | 0 | 0 | 0 | 0 | 0 | 0 |
| virus_May12.75v_1135 | 0 | 0 | 0 | 0 | 0 | 0 | 0 | 0 | 0 | 0 | 0 |
| virus_May12.75v_2430 | 0 | 0 | 0 | 0 | 0 | 0 | 0 | 0 | 0 | 0 | 0 |
| virus_May12.75v_4842 | 0 | 0 | 0 | 0 | 0 | 0 | 0 | 0 | 0 | 0 | 0 |

| | | | | | | | | | | | |
|---|---|---|---|---|---|---|---|---|---|---|---|
| virus_May12.75v_6881 | 0 | 0 | 0 | 0 | 0 | 0 | 0 | 0 | 0 | 0 | 0 |
| virus_May12.75v_11710 | 0 | 0 | 0 | 0 | 0 | 0 | 0 | 0 | 0 | 0 | 0 |
| virus_May12.75v_11814 | 0 | 0 | 0 | 0 | 0 | 0 | 0 | 0 | 0 | 0 | 0 |
| virus_May12.75v_14086 | 0 | 0 | 0 | 0 | 0 | 0 | 0 | 0 | 0 | 0 | 0 |
| virus_May12.75v_16053 | 0 | 0 | 0 | 0 | 0 | 0 | 0 | 0 | 0 | 0 | 0 |
| virus_May12.75v_16591 | 0 | 0 | 0 | 0 | 0 | 0 | 0 | 0 | 0 | 0 | 0 |
| virus_May12.75v_17774 | 0 | 0 | 0 | 0 | 0 | 0 | 0 | 0 | 0 | 0 | 0 |
| virus_May12.75v_18089 | 0 | 0 | 0 | 0 | 0 | 0 | 0 | 0 | 0 | 0 | 0 |
| virus_May12.75v_24636 | 0 | 4 | 2 | 1 | 0 | 6 | 1 | 1 | 0 | 4 | 19 |
| virus_May12.75v_24867 | 0 | 0 | 0 | 0 | 0 | 0 | 0 | 0 | 0 | 0 | 0 |
| virus_May12.75v_26070 | 0 | 0 | 0 | 0 | 0 | 0 | 0 | 0 | 0 | 0 | 0 |
| virus_May12.75v_26084 | 0 | 0 | 0 | 0 | 0 | 0 | 0 | 0 | 0 | 0 | 0 |
| virus_May12.75v_38498 | 0 | 0 | 0 | 0 | 0 | 0 | 0 | 0 | 0 | 0 | 0 |
| virus_May12.75v_39998 | 0 | 3 | 0 | 1 | 2 | 0 | 0 | 0 | 0 | 0 | 6 |
| virus_Sep8.0v_595 | 0 | 1 | 0 | 0 | 0 | 0 | 0 | 0 | 0 | 0 | 1 |
| virus_Sep8.0v_3393 | 0 | 0 | 0 | 1 | 0 | 0 | 0 | 0 | 0 | 0 | 1 |
| virus_Sep8.0v_4419 | 0 | 1 | 0 | 0 | 0 | 0 | 0 | 0 | 0 | 0 | 1 |
| virus_Sep8.0v_17751 | 0 | 1 | 0 | 1 | 0 | 0 | 0 | 0 | 0 | 1 | 3 |
| virus_Sep8.0v_19674 | 0 | 1 | 0 | 0 | 0 | 3 | 0 | 0 | 0 | 0 | 4 |
| virus_Sep8.0v_20500 | 0 | 0 | 0 | 0 | 0 | 0 | 0 | 0 | 0 | 0 | 0 |
| virus_Sep8.5v_498 | 0 | 0 | 0 | 0 | 0 | 0 | 0 | 0 | 0 | 0 | 0 |
| virus_Sep8.5v_2658 | 0 | 0 | 0 | 0 | 0 | 0 | 0 | 0 | 0 | 0 | 0 |
| virus_Sep8.5v_3819 | 0 | 0 | 0 | 0 | 0 | 0 | 0 | 0 | 0 | 0 | 0 |
| virus_Sep8.5v_5681 | 0 | 0 | 0 | 0 | 0 | 0 | 0 | 0 | 0 | 0 | 0 |
| virus_Sep8.5v_10931 | 0 | 0 | 0 | 0 | 0 | 0 | 0 | 0 | 0 | 0 | 0 |
| virus_Sep8.5v_11546 | 0 | 0 | 0 | 0 | 0 | 0 | 0 | 0 | 0 | 0 | 0 |
| virus_Sep8.5v_12583 | 0 | 0 | 0 | 0 | 0 | 0 | 0 | 0 | 0 | 0 | 0 |
| virus_Sep8.5v_17128 | 0 | 0 | 0 | 0 | 0 | 0 | 0 | 0 | 0 | 0 | 0 |
| virus_Sep8.5v_17395 | 0 | 0 | 0 | 0 | 0 | 0 | 0 | 0 | 0 | 0 | 0 |
| virus_Sep8.5v_17749 | 0 | 0 | 0 | 0 | 0 | 0 | 0 | 0 | 0 | 0 | 0 |
| virus_Sep8.25v_293 | 0 | 0 | 0 | 0 | 0 | 0 | 0 | 0 | 0 | 0 | 0 |
| virus_Sep8.25v_1130 | 0 | 0 | 0 | 0 | 0 | 0 | 0 | 0 | 0 | 0 | 0 |
| virus_Sep8.25v_6070 | 0 | 0 | 0 | 0 | 0 | 0 | 0 | 0 | 0 | 0 | 0 |
| virus_Sep8.25v_11003 | 0 | 0 | 0 | 0 | 0 | 0 | 0 | 0 | 0 | 0 | 0 |
| virus_Sep8.50v_906 | 0 | 0 | 0 | 0 | 0 | 0 | 0 | 0 | 0 | 0 | 0 |
| virus_Sep8.50v_1306 | 0 | 0 | 0 | 1 | 3 | 0 | 0 | 0 | 0 | 0 | 4 |
| virus_Sep8.50v_3182 | 0 | 1 | 0 | 0 | 0 | 0 | 0 | 0 | 0 | 0 | 1 |
| virus_Sep8.50v_5030 | 0 | 0 | 0 | 0 | 0 | 0 | 0 | 0 | 0 | 0 | 0 |
| virus_Sep8.50v_7061 | 0 | 0 | 0 | 0 | 0 | 0 | 0 | 0 | 0 | 0 | 0 |
| virus_Sep8.50v_8612 | 0 | 0 | 0 | 0 | 0 | 0 | 0 | 0 | 0 | 0 | 0 |
| virus_Sep8.50v_9119 | 0 | 0 | 0 | 0 | 0 | 0 | 0 | 0 | 0 | 0 | 0 |
| virus_Sep8.50v_9279 | 0 | 0 | 0 | 0 | 0 | 0 | 0 | 0 | 0 | 0 | 0 |
| virus_Sep8.50v_11061 | 0 | 0 | 0 | 0 | 0 | 0 | 0 | 0 | 0 | 0 | 0 |
| virus_Sep8.50v_11476 | 0 | 0 | 0 | 0 | 0 | 0 | 0 | 0 | 0 | 0 | 0 |
| virus_Sep8.50v_12712 | 0 | 1 | 0 | 1 | 2 | 0 | 1 | 1 | 0 | 0 | 6 |
| virus_Sep8.50v_14141 | 0 | 0 | 0 | 0 | 0 | 0 | 0 | 0 | 0 | 0 | 0 |
| virus_Sep8.50v_16464 | 0 | 0 | 0 | 0 | 0 | 0 | 0 | 0 | 0 | 0 | 0 |
| virus_Sep8.50v_19042 | 0 | 1 | 0 | 0 | 0 | 0 | 0 | 0 | 0 | 0 | 1 |
| virus_Sep8.50v_19346 | 0 | 0 | 0 | 0 | 0 | 0 | 0 | 0 | 0 | 0 | 0 |
| virus_Sep8.50v_20095 | 0 | 0 | 0 | 0 | 0 | 0 | 0 | 0 | 0 | 0 | 0 |
| virus_Sep8.50v_25302 | 0 | 0 | 0 | 0 | 0 | 0 | 0 | 0 | 0 | 0 | 0 |
| virus_Sep8.50v_29766 | 0 | 0 | 0 | 0 | 0 | 0 | 0 | 0 | 0 | 0 | 0 |
| virus_Sep8.50v_34884 | 0 | 0 | 0 | 0 | 0 | 0 | 0 | 0 | 0 | 0 | 0 |
| virus_Sep8.50v_38749 | 0 | 0 | 0 | 0 | 0 | 0 | 0 | 0 | 0 | 0 | 0 |
| virus_Sep8.50v_40247 | 0 | 0 | 0 | 0 | 0 | 0 | 0 | 0 | 0 | 0 | 0 |
| bacteria_Jul11d0b_1804 | 0 | 0 | 0 | 0 | 0 | 0 | 0 | 0 | 0 | 0 | 0 |
| bacteria_Jul11d0b_2761 | 0 | 1 | 0 | 0 | 0 | 0 | 0 | 0 | 0 | 0 | 1 |
| bacteria_Jul11d0b_14357 | 0 | 4 | 0 | 0 | 0 | 0 | 0 | 0 | 0 | 0 | 4 |
| bacteria_Jul11d0b_17993 | 0 | 2 | 0 | 0 | 0 | 1 | 0 | 0 | 0 | 1 | 4 |
| bacteria_Jul11d0b_38371 | 0 | 0 | 0 | 0 | 0 | 0 | 0 | 0 | 0 | 0 | 0 |
| bacteria_Jul11d0b_41269 | 0 | 0 | 0 | 0 | 0 | 0 | 0 | 0 | 0 | 0 | 0 |
| bacteria_Jul11d0b_57039 | 0 | 0 | 0 | 0 | 0 | 0 | 0 | 0 | 0 | 0 | 0 |
| bacteria_Jul11d0b_57590 | 0 | 0 | 0 | 0 | 0 | 0 | 0 | 0 | 0 | 0 | 0 |
| bacteria_Aug5d0b_1353 | 3 | 0 | 0 | 0 | 1 | 0 | 0 | 0 | 0 | 0 | 4 |
| bacteria_Aug5d0b_3944 | 1 | 0 | 0 | 0 | 0 | 0 | 0 | 0 | 0 | 0 | 1 |
| bacteria_Aug5d0b_10700 | 1 | 0 | 0 | 0 | 0 | 0 | 0 | 0 | 0 | 0 | 1 |
| bacteria_Aug5d0b_21195 | 1 | 0 | 0 | 0 | 0 | 0 | 0 | 0 | 0 | 0 | 1 |
| bacteria_Aug5d0b_27442 | 1 | 0 | 0 | 0 | 0 | 0 | 0 | 0 | 0 | 0 | 1 |
| bacteria_Aug5d0b_38140 | 1 | 0 | 0 | 0 | 0 | 0 | 0 | 0 | 0 | 0 | 1 |
| bacteria_Aug5d0b_45479 | 1 | 0 | 0 | 0 | 0 | 0 | 0 | 0 | 0 | 0 | 1 |
| bacteria_Aug5d0b_49411 | 1 | 0 | 0 | 0 | 0 | 0 | 0 | 0 | 0 | 0 | 1 |
| bacteria_Aug5d5b_4155 | 0 | 0 | 1 | 0 | 0 | 0 | 0 | 0 | 0 | 0 | 1 |
| bacteria_Aug5d5b_28863 | 0 | 0 | 1 | 0 | 0 | 0 | 0 | 0 | 0 | 0 | 1 |
| bacteria_Aug5d5b_68029 | 0 | 0 | 1 | 0 | 0 | 0 | 0 | 0 | 0 | 0 | 1 |
| bacteria_Aug5d5b_68781 | 0 | 0 | 1 | 0 | 0 | 0 | 0 | 0 | 0 | 0 | 1 |
| bacteria_Aug5d25b_543 | 0 | 0 | 0 | 0 | 1 | 0 | 0 | 0 | 0 | 0 | 1 |
| bacteria_Aug5d25b_5118 | 0 | 0 | 0 | 0 | 1 | 0 | 0 | 0 | 0 | 0 | 1 |
| bacteria_Aug5d25b_6004 | 0 | 0 | 0 | 0 | 1 | 0 | 0 | 0 | 0 | 0 | 1 |
| bacteria_Aug5d25b_9230 | 0 | 0 | 0 | 0 | 2 | 0 | 0 | 0 | 0 | 0 | 2 |
| bacteria_Aug5d25b_9853 | 0 | 0 | 0 | 0 | 1 | 0 | 0 | 0 | 0 | 0 | 1 |
| bacteria_Aug5d25b_9970 | 0 | 0 | 0 | 0 | 1 | 0 | 0 | 0 | 0 | 0 | 1 |
| bacteria_Aug5d25b_10011 | 0 | 0 | 0 | 0 | 2 | 0 | 0 | 0 | 0 | 0 | 2 |
| bacteria_Aug5d25b_10485 | 0 | 0 | 0 | 1 | 1 | 0 | 0 | 0 | 0 | 0 | 2 |
| bacteria_Aug5d25b_11145 | 0 | 0 | 0 | 0 | 1 | 0 | 0 | 0 | 0 | 0 | 1 |
| bacteria_Aug5d25b_11719 | 0 | 0 | 0 | 0 | 1 | 0 | 0 | 0 | 0 | 0 | 1 |
| bacteria_Aug5d25b_16162 | 0 | 0 | 0 | 0 | 1 | 0 | 0 | 1 | 0 | 0 | 2 |
| bacteria_Aug5d25b_17410 | 0 | 0 | 0 | 0 | 1 | 0 | 0 | 0 | 0 | 0 | 1 |
| bacteria_Aug5d25b_20975 | 0 | 0 | 0 | 0 | 2 | 0 | 1 | 0 | 0 | 0 | 3 |
| bacteria_Aug5d25b_22721 | 0 | 0 | 0 | 0 | 1 | 0 | 0 | 0 | 0 | 0 | 1 |
| bacteria_Aug5d25b_27681 | 0 | 0 | 0 | 0 | 1 | 0 | 0 | 0 | 0 | 0 | 1 |
| bacteria_Aug5d25b_28993 | 0 | 0 | 0 | 0 | 1 | 0 | 0 | 0 | 0 | 0 | 1 |
| bacteria_Aug5d25b_36633 | 0 | 0 | 0 | 0 | 1 | 0 | 0 | 0 | 0 | 0 | 1 |
| bacteria_Aug5d25b_41310 | 0 | 0 | 0 | 0 | 1 | 0 | 0 | 0 | 0 | 0 | 1 |
| bacteria_Aug5d25b_41414 | 0 | 0 | 0 | 0 | 1 | 0 | 0 | 0 | 0 | 0 | 1 |
| bacteria_Aug5d25b_42877 | 0 | 0 | 0 | 0 | 1 | 0 | 0 | 0 | 0 | 0 | 1 |
| bacteria_Aug5d25b_44898 | 0 | 0 | 0 | 0 | 1 | 0 | 0 | 0 | 0 | 0 | 1 |
| bacteria_Aug5d25b_46724 | 0 | 0 | 0 | 0 | 1 | 0 | 0 | 0 | 0 | 0 | 1 |
| bacteria_Aug5d25b_46892 | 0 | 0 | 0 | 0 | 1 | 0 | 0 | 0 | 0 | 0 | 1 |
| bacteria_Aug5d25b_51169 | 0 | 0 | 0 | 0 | 1 | 0 | 0 | 0 | 0 | 0 | 1 |
| bacteria_Aug5d25b_52110 | 0 | 0 | 0 | 0 | 1 | 0 | 0 | 0 | 0 | 0 | 1 |
| bacteria_Aug5d25b_52686 | 0 | 0 | 0 | 0 | 1 | 0 | 0 | 0 | 0 | 0 | 1 |
| bacteria_Aug5d25b_55450 | 0 | 0 | 0 | 0 | 1 | 0 | 0 | 0 | 0 | 0 | 1 |
| bacteria_Aug5d25b_58274 | 0 | 0 | 0 | 0 | 1 | 0 | 0 | 0 | 0 | 0 | 1 |
| bacteria_Aug5d25b_65829 | 0 | 0 | 0 | 0 | 1 | 0 | 0 | 0 | 0 | 0 | 1 |
| bacteria_Aug5d50b_10184 | 0 | 0 | 0 | 0 | 0 | 0 | 1 | 0 | 0 | 0 | 1 |
| bacteria_Aug5d50b_39196 | 0 | 0 | 0 | 0 | 0 | 0 | 1 | 0 | 0 | 0 | 1 |
| bacteria_Aug5d50b_67695 | 0 | 0 | 0 | 0 | 0 | 0 | 1 | 0 | 0 | 0 | 1 |
| bacteria_Aug5d75b_43040 | 0 | 0 | 0 | 0 | 0 | 0 | 0 | 0 | 1 | 0 | 1 |
| bacteria_Jul11d5b_15657 | 0 | 0 | 0 | 0 | 0 | 0 | 0 | 0 | 0 | 0 | 0 |
| bacteria_Jul11d5b_16587 | 0 | 0 | 0 | 0 | 0 | 0 | 0 | 0 | 0 | 0 | 0 |
| bacteria_Jul11d5b_42515 | 2 | 0 | 6 | 0 | 11 | 0 | 5 | 0 | 0 | 0 | 24 |
| bacteria_Jul11d25b_1285 | 0 | 1 | 0 | 0 | 2 | 0 | 1 | 0 | 0 | 1 | 5 |
| bacteria_Jul11d25b_23949 | 0 | 0 | 0 | 0 | 0 | 0 | 0 | 0 | 0 | 0 | 0 |
| bacteria_Jul11d25b_26453 | 0 | 0 | 0 | 0 | 0 | 0 | 0 | 0 | 0 | 0 | 0 |
| bacteria_Jul11d25b_27699 | 0 | 0 | 0 | 0 | 0 | 0 | 0 | 0 | 0 | 0 | 0 |
| bacteria_Jul11d25b_41160 | 0 | 0 | 0 | 0 | 0 | 0 | 0 | 0 | 0 | 0 | 0 |
| bacteria_Jul11d50b_4516 | 0 | 0 | 0 | 0 | 0 | 0 | 0 | 0 | 0 | 0 | 0 |
| bacteria_Jul11d50b_17953 | 0 | 0 | 0 | 0 | 0 | 0 | 1 | 0 | 1 | 0 | 2 |
| bacteria_Jul11d50b_18845 | 0 | 0 | 0 | 0 | 0 | 0 | 0 | 0 | 0 | 0 | 0 |
| bacteria_Jul11d50b_23065 | 0 | 0 | 0 | 0 | 0 | 0 | 0 | 0 | 0 | 0 | 0 |
| bacteria_Jul11d50b_43548 | 0 | 0 | 0 | 0 | 0 | 1 | 0 | 0 | 0 | 0 | 1 |
| bacteria_Jul11d50b_48235 | 0 | 0 | 0 | 0 | 0 | 0 | 0 | 0 | 0 | 0 | 0 |
| bacteria_Jul11d75b_3634 | 0 | 0 | 0 | 0 | 0 | 0 | 0 | 0 | 0 | 0 | 0 |
| bacteria_Jul11d75b_6691 | 0 | 0 | 0 | 0 | 0 | 0 | 0 | 0 | 0 | 0 | 0 |
| bacteria_Jul11d75b_7370 | 2 | 0 | 1 | 0 | 1 | 0 | 0 | 0 | 0 | 0 | 4 |
| bacteria_Jul11d75b_15188 | 0 | 1 | 0 | 1 | 3 | 0 | 0 | 0 | 0 | 0 | 5 |
| bacteria_Jul11d75b_25852 | 0 | 0 | 0 | 0 | 0 | 0 | 0 | 1 | 0 | 1 | 2 |
| bacteria_Jul11d75b_26807 | 0 | 0 | 0 | 0 | 0 | 0 | 0 | 0 | 0 | 0 | 0 |
| bacteria_Jul11d75b_27308 | 0 | 0 | 0 | 0 | 1 | 0 | 0 | 0 | 0 | 0 | 1 |
| bacteria_Jul11d75b_30582 | 0 | 0 | 0 | 0 | 0 | 0 | 0 | 0 | 0 | 0 | 0 |
| bacteria_Jul11d75b_31625 | 0 | 0 | 0 | 0 | 0 | 0 | 0 | 0 | 0 | 0 | 0 |
| bacteria_Jul11d75b_32512 | 0 | 0 | 0 | 0 | 0 | 0 | 0 | 0 | 0 | 0 | 0 |
| bacteria_Jul11d75b_36749 | 0 | 0 | 0 | 0 | 0 | 0 | 0 | 0 | 0 | 0 | 0 |
| bacteria_Jul11d75b_41365 | 0 | 0 | 0 | 0 | 0 | 0 | 0 | 0 | 0 | 0 | 0 |
| bacteria_Jul11d75b_41847 | 0 | 0 | 0 | 0 | 0 | 0 | 0 | 0 | 0 | 0 | 0 |
| bacteria_Jul11d75b_42061 | 0 | 0 | 0 | 1 | 0 | 0 | 0 | 0 | 0 | 0 | 1 |
| bacteria_Jul11d75b_48633 | 0 | 0 | 0 | 0 | 0 | 0 | 0 | 0 | 0 | 0 | 0 |
| bacteria_Jul11d75b_49870 | 0 | 0 | 0 | 0 | 0 | 0 | 0 | 0 | 0 | 0 | 0 |
| bacteria_Jul11d75b_50049 | 0 | 0 | 0 | 0 | 0 | 0 | 0 | 0 | 0 | 0 | 0 |
| bacteria_Jul27d5b_266 | 0 | 0 | 0 | 0 | 0 | 0 | 0 | 0 | 0 | 0 | 0 |
| bacteria_Jul27d5b_985 | 0 | 0 | 0 | 0 | 1 | 1 | 0 | 0 | 0 | 0 | 2 |
| bacteria_Jul27d5b_1964 | 0 | 0 | 0 | 0 | 0 | 0 | 0 | 0 | 0 | 0 | 0 |
| bacteria_Jul27d5b_7248 | 0 | 2 | 0 | 1 | 0 | 0 | 0 | 0 | 0 | 0 | 3 |
| bacteria_Jul27d5b_8488 | 0 | 1 | 0 | 0 | 0 | 1 | 0 | 1 | 0 | 1 | 4 |
| bacteria_Jul27d5b_12937 | 0 | 1 | 0 | 0 | 1 | 1 | 1 | 4 | 0 | 4 | 12 |
| bacteria_Jul27d5b_16291 | 0 | 0 | 0 | 0 | 0 | 0 | 0 | 0 | 0 | 0 | 0 |
| bacteria_Jul27d5b_19668 | 2 | 0 | 4 | 0 | 7 | 0 | 5 | 0 | 0 | 0 | 18 |
| bacteria_Jul27d5b_19715 | 0 | 1 | 0 | 0 | 0 | 0 | 0 | 0 | 0 | 0 | 1 |
| bacteria_Jul27d5b_20185 | 0 | 0 | 0 | 0 | 0 | 0 | 0 | 0 | 0 | 0 | 0 |
| bacteria_Jul27d5b_20306 | 0 | 0 | 0 | 0 | 0 | 0 | 0 | 0 | 0 | 0 | 0 |
| bacteria_Jul27d5b_23686 | 0 | 0 | 0 | 0 | 0 | 0 | 0 | 0 | 0 | 0 | 0 |
| bacteria_Jul27d5b_27118 | 0 | 0 | 0 | 0 | 0 | 0 | 0 | 0 | 0 | 0 | 0 |
| bacteria_Jul27d5b_29462 | 0 | 0 | 0 | 0 | 0 | 0 | 0 | 0 | 0 | 0 | 0 |
| bacteria_Jul27d5b_31136 | 0 | 0 | 0 | 0 | 0 | 0 | 0 | 0 | 0 | 0 | 0 |
| bacteria_Jul27d5b_35389 | 1 | 0 | 0 | 0 | 0 | 0 | 0 | 0 | 0 | 0 | 1 |
| bacteria_Jul27d5b_39640 | 0 | 0 | 0 | 0 | 0 | 0 | 0 | 0 | 0 | 0 | 0 |
| bacteria_Jul27d5b_39687 | 0 | 0 | 0 | 0 | 0 | 0 | 0 | 0 | 0 | 0 | 0 |

| | | | | | | | | | | | |
|---|---|---|---|---|---|---|---|---|---|---|---|
| bacteria_Jul27d5b_44472 | 0 | 0 | 0 | 0 | 0 | 0 | 0 | 0 | 0 | 0 | 0 |
| bacteria_Jul27d5b_44900 | 0 | 0 | 0 | 0 | 1 | 0 | 1 | 0 | 0 | 0 | 2 |
| bacteria_Jul27d25b_13646 | 0 | 1 | 0 | 1 | 1 | 1 | 0 | 0 | 0 | 0 | 4 |
| bacteria_Jul27d25b_19148 | 0 | 0 | 0 | 0 | 0 | 0 | 0 | 0 | 0 | 0 | 0 |
| bacteria_Jul27d25b_27671 | 0 | 0 | 0 | 0 | 0 | 0 | 0 | 0 | 0 | 0 | 0 |
| bacteria_Jul27d25b_31851 | 0 | 0 | 0 | 0 | 0 | 0 | 0 | 0 | 0 | 0 | 0 |
| bacteria_Jul27d25b_31859 | 0 | 0 | 0 | 0 | 0 | 0 | 0 | 0 | 0 | 0 | 0 |
| bacteria_Jul27d25b_31871 | 0 | 0 | 0 | 0 | 0 | 0 | 0 | 0 | 0 | 0 | 0 |
| bacteria_Jul27d25b_32000 | 0 | 0 | 0 | 0 | 0 | 0 | 0 | 0 | 0 | 0 | 0 |
| bacteria_Jul27d25b_36843 | 0 | 0 | 0 | 0 | 0 | 0 | 0 | 0 | 0 | 0 | 0 |
| bacteria_Jul27d25b_47179 | 0 | 0 | 0 | 0 | 0 | 0 | 0 | 0 | 0 | 0 | 0 |
| bacteria_Jul27d25b_48025 | 0 | 0 | 0 | 0 | 0 | 0 | 1 | 0 | 0 | 0 | 1 |
| bacteria_Jul27d25b_55213 | 0 | 0 | 0 | 0 | 0 | 0 | 0 | 0 | 0 | 0 | 0 |
| bacteria_Jul27d25b_68514 | 1 | 0 | 0 | 0 | 0 | 0 | 0 | 0 | 0 | 0 | 1 |
| bacteria_Jul27d25b_73665 | 0 | 0 | 0 | 0 | 0 | 0 | 0 | 0 | 0 | 0 | 0 |
| bacteria_Jul27d25b_75133 | 0 | 0 | 0 | 0 | 0 | 0 | 0 | 0 | 0 | 0 | 0 |
| bacteria_Jul27d25b_75238 | 0 | 0 | 0 | 0 | 0 | 0 | 1 | 0 | 0 | 0 | 1 |
| bacteria_Jul27d75b_1937 | 0 | 0 | 0 | 0 | 0 | 0 | 0 | 0 | 0 | 0 | 0 |
| bacteria_Jul27d75b_6107 | 0 | 0 | 0 | 0 | 1 | 0 | 0 | 0 | 0 | 0 | 1 |
| bacteria_Jul27d75b_12967 | 0 | 0 | 0 | 0 | 0 | 0 | 0 | 0 | 0 | 0 | 0 |
| bacteria_Jul27d75b_25053 | 0 | 0 | 1 | 0 | 0 | 0 | 0 | 0 | 0 | 0 | 1 |
| bacteria_Jul27d75b_35743 | 0 | 0 | 0 | 0 | 0 | 0 | 0 | 0 | 0 | 0 | 0 |
| bacteria_Jul27d75b_42798 | 0 | 0 | 0 | 0 | 0 | 0 | 0 | 0 | 0 | 0 | 0 |
| bacteria_Jul27d75b_51274 | 0 | 0 | 0 | 0 | 0 | 0 | 0 | 0 | 0 | 0 | 0 |
| bacteria_Jul27d75b_67015 | 0 | 0 | 0 | 0 | 0 | 0 | 0 | 0 | 0 | 0 | 0 |
| bacteria_Jul27d75b_69505 | 0 | 0 | 0 | 0 | 0 | 0 | 0 | 0 | 0 | 0 | 0 |
| bacteria_Jun8d25b_7303 | 0 | 0 | 0 | 0 | 0 | 0 | 0 | 0 | 0 | 0 | 0 |
| bacteria_Jun8d25b_7643 | 0 | 0 | 0 | 0 | 0 | 0 | 0 | 0 | 0 | 0 | 0 |
| bacteria_Jun8d25b_11474 | 0 | 0 | 0 | 0 | 0 | 0 | 0 | 0 | 0 | 0 | 0 |
| bacteria_Jun8d25b_15489 | 0 | 0 | 0 | 0 | 0 | 0 | 0 | 0 | 0 | 0 | 0 |
| bacteria_Jun8d25b_16198 | 0 | 0 | 0 | 0 | 0 | 0 | 0 | 0 | 0 | 0 | 0 |
| bacteria_Jun8d25b_18140 | 0 | 0 | 0 | 0 | 0 | 0 | 0 | 0 | 0 | 0 | 0 |
| bacteria_Jun8d25b_21457 | 0 | 0 | 0 | 0 | 1 | 0 | 0 | 1 | 0 | 0 | 2 |
| bacteria_Jun8d25b_37071 | 0 | 0 | 0 | 0 | 0 | 0 | 0 | 0 | 0 | 0 | 0 |
| bacteria_Jun8d25b_49086 | 0 | 0 | 0 | 0 | 0 | 0 | 0 | 1 | 0 | 0 | 1 |
| bacteria_Jun8d25b_57524 | 0 | 0 | 0 | 0 | 0 | 0 | 0 | 0 | 0 | 0 | 0 |
| bacteria_Jun8d25b_58090 | 0 | 0 | 0 | 0 | 0 | 0 | 0 | 0 | 0 | 0 | 0 |
| bacteria_Jun8d25b_59223 | 0 | 0 | 0 | 0 | 0 | 0 | 0 | 0 | 0 | 0 | 0 |
| bacteria_Jun8d25b_60249 | 0 | 0 | 0 | 0 | 0 | 0 | 0 | 0 | 0 | 0 | 0 |
| bacteria_Jun8d25b_70783 | 0 | 0 | 0 | 0 | 0 | 0 | 0 | 0 | 0 | 0 | 0 |
| bacteria_Jun8d25b_72033 | 0 | 0 | 0 | 0 | 0 | 0 | 0 | 0 | 0 | 0 | 0 |
| bacteria_Jun8d50b_14277 | 0 | 0 | 0 | 0 | 0 | 0 | 0 | 0 | 0 | 0 | 0 |
| bacteria_Jun8d50b_20532 | 0 | 0 | 0 | 0 | 1 | 0 | 0 | 0 | 0 | 0 | 1 |
| bacteria_Jun8d50b_26056 | 0 | 0 | 0 | 0 | 0 | 0 | 0 | 0 | 0 | 0 | 0 |
| bacteria_Jun8d50b_41294 | 0 | 0 | 0 | 0 | 0 | 0 | 0 | 0 | 0 | 0 | 0 |
| bacteria_Jun8d50b_47497 | 0 | 0 | 0 | 0 | 0 | 0 | 0 | 0 | 0 | 0 | 0 |
| bacteria_Jun8d75b_3886 | 0 | 0 | 0 | 0 | 0 | 0 | 0 | 0 | 0 | 0 | 0 |
| bacteria_Jun8d75b_42996 | 0 | 0 | 0 | 0 | 0 | 0 | 0 | 0 | 0 | 0 | 0 |
| bacteria_Jun8d75b_64247 | 0 | 0 | 0 | 0 | 0 | 0 | 0 | 0 | 0 | 0 | 0 |
| bacteria_Jun22d0b_11280 | 0 | 0 | 0 | 0 | 0 | 0 | 0 | 0 | 0 | 0 | 0 |
| bacteria_Jun22d0b_13423 | 0 | 4 | 0 | 2 | 1 | 0 | 0 | 0 | 0 | 0 | 7 |
| bacteria_Jun22d0b_15273 | 0 | 0 | 0 | 0 | 0 | 0 | 1 | 0 | 0 | 0 | 1 |
| bacteria_Jun22d0b_21445 | 0 | 0 | 0 | 0 | 0 | 0 | 0 | 0 | 0 | 0 | 0 |
| bacteria_Jun22d0b_21911 | 0 | 0 | 0 | 0 | 0 | 0 | 0 | 0 | 0 | 0 | 0 |
| bacteria_Jun22d0b_53542 | 0 | 0 | 0 | 0 | 0 | 1 | 0 | 0 | 0 | 0 | 1 |
| bacteria_Jun22d0b_57216 | 0 | 1 | 0 | 0 | 0 | 0 | 0 | 0 | 0 | 0 | 1 |
| bacteria_Jun22d0b_61197 | 0 | 0 | 0 | 0 | 0 | 0 | 0 | 0 | 0 | 0 | 0 |
| bacteria_Jun22d0b_62451 | 0 | 0 | 0 | 0 | 0 | 0 | 0 | 0 | 0 | 0 | 0 |
| bacteria_Jun22d50b_16203 | 0 | 0 | 0 | 0 | 0 | 0 | 0 | 0 | 0 | 0 | 0 |
| bacteria_Jun22d50b_22133 | 0 | 0 | 0 | 0 | 0 | 0 | 0 | 0 | 0 | 0 | 0 |
| bacteria_Jun22d50b_31671 | 0 | 0 | 0 | 0 | 0 | 0 | 0 | 0 | 0 | 0 | 0 |
| bacteria_Jun22d50b_42348 | 0 | 0 | 0 | 0 | 0 | 0 | 0 | 0 | 0 | 0 | 0 |
| bacteria_Jun22d75b_16420 | 0 | 0 | 0 | 0 | 0 | 0 | 0 | 0 | 0 | 0 | 0 |
| bacteria_Jun22d75b_29001 | 0 | 0 | 0 | 0 | 0 | 0 | 0 | 0 | 0 | 0 | 0 |
| bacteria_Jun22d75b_51022 | 0 | 0 | 0 | 0 | 0 | 0 | 0 | 0 | 0 | 0 | 0 |
| bacteria_Jun22d75b_58106 | 0 | 0 | 0 | 0 | 0 | 0 | 0 | 0 | 0 | 0 | 0 |
| bacteria_May12d25b_4727 | 0 | 0 | 0 | 0 | 2 | 0 | 1 | 0 | 0 | 0 | 3 |
| bacteria_May12d25b_6226 | 0 | 0 | 0 | 0 | 0 | 0 | 0 | 0 | 0 | 0 | 0 |
| bacteria_May12d25b_1011 | 0 | 0 | 0 | 0 | 0 | 0 | 0 | 0 | 0 | 0 | 0 |
| bacteria_May12d25b_1304 | 0 | 0 | 0 | 0 | 0 | 0 | 0 | 0 | 0 | 0 | 0 |
| bacteria_May12d25b_2195 | 0 | 0 | 0 | 0 | 0 | 0 | 0 | 0 | 0 | 0 | 0 |
| bacteria_May12d25b_5306 | 0 | 0 | 0 | 0 | 0 | 0 | 0 | 0 | 0 | 0 | 0 |
| bacteria_May12d25b_6229 | 0 | 0 | 0 | 0 | 0 | 0 | 0 | 0 | 0 | 0 | 0 |
| bacteria_May12d25b_6243 | 0 | 0 | 0 | 0 | 0 | 0 | 0 | 0 | 0 | 0 | 0 |
| bacteria_May12d25b_6370 | 0 | 0 | 0 | 0 | 0 | 0 | 0 | 0 | 0 | 0 | 0 |
| bacteria_May12d25b_6830 | 0 | 0 | 0 | 0 | 0 | 0 | 0 | 0 | 0 | 0 | 0 |
| bacteria_May12d50b_1715 | 0 | 0 | 0 | 0 | 0 | 0 | 0 | 0 | 0 | 1 | 1 |
| bacteria_May12d50b_1791 | 0 | 0 | 0 | 0 | 0 | 0 | 0 | 0 | 0 | 0 | 0 |
| bacteria_May12d50b_3993 | 0 | 0 | 0 | 0 | 0 | 0 | 0 | 0 | 0 | 0 | 0 |
| bacteria_May12d50b_1134 | 0 | 0 | 0 | 0 | 0 | 0 | 0 | 0 | 0 | 0 | 0 |
| bacteria_May12d50b_1266 | 0 | 0 | 0 | 0 | 1 | 0 | 1 | 0 | 0 | 0 | 2 |
| bacteria_May12d50b_1462 | 0 | 0 | 0 | 0 | 0 | 0 | 0 | 0 | 0 | 0 | 0 |
| bacteria_May12d50b_1526 | 0 | 0 | 0 | 0 | 0 | 0 | 0 | 0 | 0 | 0 | 0 |
| bacteria_May12d50b_1842 | 0 | 0 | 0 | 0 | 0 | 0 | 0 | 0 | 0 | 0 | 0 |
| bacteria_May12d50b_1859 | 0 | 0 | 0 | 0 | 0 | 0 | 0 | 0 | 0 | 0 | 0 |
| bacteria_May12d50b_1915 | 0 | 0 | 0 | 0 | 0 | 0 | 0 | 0 | 0 | 0 | 0 |
| bacteria_May12d50b_2285 | 0 | 0 | 0 | 0 | 0 | 0 | 0 | 0 | 0 | 0 | 0 |
| bacteria_May12d50b_4625 | 0 | 0 | 0 | 0 | 0 | 0 | 0 | 0 | 0 | 0 | 0 |
| bacteria_May12d75b_1590 | 0 | 0 | 0 | 0 | 0 | 0 | 0 | 0 | 0 | 0 | 0 |
| bacteria_Sep8d0b_37415 | 0 | 0 | 0 | 0 | 0 | 0 | 0 | 0 | 0 | 0 | 0 |
| bacteria_Sep8d0b_44016 | 0 | 0 | 0 | 0 | 0 | 0 | 0 | 0 | 0 | 0 | 0 |
| bacteria_Sep8d5b_15089 | 0 | 0 | 0 | 0 | 0 | 0 | 0 | 0 | 0 | 0 | 0 |
| bacteria_Sep8d5b_19474 | 0 | 0 | 0 | 0 | 0 | 0 | 0 | 0 | 0 | 0 | 0 |
| bacteria_Sep8d5b_25085 | 0 | 0 | 0 | 0 | 0 | 0 | 0 | 0 | 0 | 0 | 0 |
| bacteria_Sep8d5b_32964 | 0 | 0 | 0 | 0 | 1 | 0 | 0 | 0 | 0 | 0 | 1 |
| bacteria_Sep8d5b_34186 | 0 | 0 | 0 | 0 | 0 | 0 | 0 | 0 | 0 | 0 | 0 |
| bacteria_Sep8d5b_44398 | 0 | 0 | 0 | 0 | 0 | 0 | 0 | 0 | 0 | 0 | 0 |
| bacteria_Sep8d5b_44813 | 0 | 0 | 0 | 0 | 0 | 0 | 0 | 0 | 0 | 0 | 0 |
| bacteria_Sep8d5b_51298 | 0 | 0 | 0 | 0 | 0 | 0 | 0 | 0 | 0 | 0 | 0 |
| bacteria_Sep8d5b_53048 | 0 | 0 | 0 | 0 | 0 | 0 | 0 | 0 | 0 | 0 | 0 |
| bacteria_Sep8d5b_64021 | 0 | 0 | 0 | 0 | 0 | 0 | 0 | 0 | 0 | 0 | 0 |
| bacteria_Sep8d5b_69339 | 0 | 0 | 0 | 0 | 0 | 0 | 0 | 0 | 0 | 0 | 0 |
| bacteria_Sep8d25b_12041 | 0 | 0 | 0 | 0 | 0 | 0 | 0 | 0 | 0 | 0 | 0 |
| bacteria_Sep8d25b_24038 | 0 | 0 | 0 | 0 | 0 | 0 | 0 | 0 | 0 | 0 | 0 |
| bacteria_Sep8d25b_45808 | 0 | 0 | 0 | 0 | 0 | 0 | 0 | 0 | 0 | 0 | 0 |
| bacteria_Sep8d25b_51194 | 0 | 0 | 0 | 0 | 0 | 0 | 0 | 0 | 0 | 0 | 0 |
| bacteria_Sep8d50b_15224 | 0 | 0 | 0 | 0 | 0 | 0 | 0 | 0 | 0 | 0 | 0 |
| bacteria_Sep8d50b_37428 | 0 | 0 | 0 | 0 | 0 | 0 | 0 | 0 | 0 | 0 | 0 |
| bacteria_Sep8d50b_38337 | 0 | 0 | 0 | 0 | 0 | 0 | 0 | 0 | 0 | 0 | 0 |
| virus_Aug5.0v_202 | 0 | 1 | 0 | 0 | 0 | 0 | 0 | 0 | 0 | 0 | 1 |
| virus_Aug5.0v_282 | 1 | 11 | 0 | 8 | 7 | 5 | 1 | 2 | 1 | 3 | 39 |
| virus_Aug5.0v_793 | 0 | 1 | 0 | 0 | 0 | 0 | 0 | 0 | 0 | 0 | 1 |
| virus_Aug5.0v_1120 | 0 | 1 | 0 | 0 | 0 | 0 | 0 | 0 | 0 | 0 | 1 |
| virus_Aug5.0v_1193 | 0 | 1 | 0 | 0 | 0 | 0 | 0 | 0 | 0 | 0 | 1 |
| virus_Aug5.0v_1519 | 0 | 1 | 0 | 0 | 0 | 0 | 0 | 0 | 0 | 0 | 1 |
| virus_Aug5.0v_1537 | 0 | 1 | 0 | 0 | 0 | 0 | 0 | 0 | 0 | 0 | 1 |
| virus_Aug5.0v_1609 | 0 | 1 | 0 | 0 | 0 | 0 | 0 | 0 | 0 | 0 | 1 |
| virus_Aug5.0v_1760 | 0 | 3 | 0 | 0 | 0 | 0 | 0 | 0 | 0 | 0 | 3 |
| virus_Aug5.0v_2052 | 0 | 1 | 0 | 1 | 1 | 0 | 0 | 0 | 0 | 0 | 3 |
| virus_Aug5.0v_2246 | 0 | 1 | 0 | 0 | 0 | 0 | 0 | 0 | 0 | 0 | 1 |
| virus_Aug5.0v_2268 | 0 | 1 | 0 | 0 | 0 | 0 | 0 | 0 | 0 | 0 | 1 |
| virus_Aug5.0v_2799 | 0 | 1 | 0 | 0 | 0 | 0 | 0 | 0 | 0 | 0 | 1 |
| virus_Aug5.0v_3013 | 0 | 2 | 0 | 0 | 0 | 0 | 0 | 0 | 0 | 0 | 2 |
| virus_Aug5.0v_3099 | 0 | 2 | 0 | 0 | 0 | 2 | 0 | 0 | 0 | 0 | 4 |
| virus_Aug5.0v_3301 | 0 | 1 | 0 | 0 | 0 | 0 | 0 | 0 | 0 | 0 | 1 |
| virus_Aug5.0v_3389 | 0 | 1 | 0 | 0 | 0 | 0 | 0 | 0 | 0 | 0 | 1 |
| virus_Aug5.0v_3413 | 0 | 1 | 0 | 0 | 0 | 0 | 0 | 0 | 0 | 0 | 1 |
| virus_Aug5.0v_3520 | 0 | 1 | 0 | 0 | 0 | 0 | 0 | 0 | 0 | 0 | 1 |
| virus_Aug5.0v_4316 | 0 | 3 | 0 | 1 | 2 | 1 | 0 | 0 | 0 | 0 | 7 |
| virus_Aug5.0v_4562 | 0 | 1 | 0 | 0 | 0 | 0 | 0 | 0 | 0 | 0 | 1 |
| virus_Aug5.0v_4580 | 0 | 5 | 1 | 1 | 0 | 0 | 0 | 0 | 0 | 0 | 7 |
| virus_Aug5.0v_4614 | 0 | 1 | 0 | 0 | 0 | 0 | 0 | 0 | 0 | 0 | 1 |
| virus_Aug5.0v_5038 | 0 | 1 | 0 | 0 | 0 | 2 | 0 | 0 | 0 | 0 | 3 |
| virus_Aug5.0v_5057 | 0 | 1 | 0 | 0 | 0 | 0 | 0 | 0 | 0 | 0 | 1 |
| virus_Aug5.0v_5201 | 0 | 1 | 0 | 0 | 0 | 0 | 0 | 0 | 0 | 0 | 1 |
| virus_Aug5.0v_5281 | 0 | 1 | 0 | 0 | 0 | 0 | 0 | 0 | 0 | 0 | 1 |
| virus_Aug5.0v_5312 | 0 | 1 | 0 | 0 | 0 | 0 | 0 | 0 | 0 | 0 | 1 |
| virus_Aug5.0v_5323 | 0 | 1 | 0 | 0 | 0 | 0 | 0 | 0 | 0 | 0 | 1 |
| virus_Aug5.0v_5341 | 0 | 1 | 0 | 0 | 0 | 0 | 0 | 0 | 0 | 0 | 1 |
| virus_Aug5.5v_3959 | 0 | 0 | 0 | 1 | 0 | 0 | 0 | 0 | 0 | 0 | 1 |
| virus_Aug5.5v_4413 | 0 | 0 | 0 | 1 | 0 | 0 | 0 | 0 | 0 | 0 | 1 |
| virus_Aug5.5v_4974 | 0 | 0 | 0 | 1 | 0 | 0 | 0 | 0 | 0 | 0 | 1 |
| virus_Aug5.5v_11623 | 0 | 0 | 0 | 1 | 0 | 0 | 0 | 0 | 0 | 0 | 1 |
| virus_Aug5.5v_15186 | 0 | 0 | 0 | 1 | 0 | 0 | 0 | 0 | 0 | 0 | 1 |
| virus_Aug5.5v_17227 | 0 | 0 | 0 | 1 | 0 | 0 | 0 | 0 | 0 | 0 | 1 |
| virus_Aug5.5v_18550 | 0 | 0 | 0 | 1 | 0 | 0 | 0 | 0 | 0 | 0 | 1 |
| virus_Aug5.5v_33657 | 0 | 0 | 0 | 1 | 0 | 0 | 0 | 0 | 0 | 0 | 1 |
| virus_Aug5.5v_35015 | 0 | 0 | 0 | 1 | 0 | 0 | 0 | 0 | 0 | 0 | 1 |
| virus_Aug5.25v_86 | 0 | 0 | 0 | 0 | 0 | 1 | 0 | 0 | 0 | 0 | 1 |
| virus_Aug5.25v_552 | 0 | 0 | 0 | 0 | 0 | 1 | 0 | 0 | 0 | 0 | 1 |
| virus_Aug5.25v_2583 | 0 | 0 | 0 | 0 | 0 | 1 | 0 | 0 | 0 | 0 | 1 |
| virus_Aug5.25v_3738 | 0 | 0 | 0 | 0 | 0 | 1 | 0 | 0 | 0 | 0 | 1 |
| virus_Aug5.25v_4597 | 0 | 0 | 0 | 0 | 0 | 1 | 0 | 0 | 0 | 0 | 1 |
| virus_Aug5.25v_4730 | 0 | 0 | 0 | 0 | 0 | 1 | 0 | 0 | 0 | 0 | 1 |
| virus_Aug5.25v_8581 | 0 | 0 | 0 | 0 | 0 | 1 | 0 | 0 | 0 | 0 | 1 |
| virus_Aug5.25v_10776 | 0 | 0 | 0 | 0 | 0 | 1 | 0 | 0 | 0 | 0 | 1 |
| virus_Aug5.25v_16404 | 0 | 0 | 0 | 0 | 0 | 1 | 0 | 0 | 0 | 0 | 1 |

| | | | | | | | | | | | | |
|---|---|---|---|---|---|---|---|---|---|---|---|---|
| virus_Aug5.25v_20427 | | 0 | 0 | 0 | 0 | 0 | 1 | 0 | 0 | 0 | 0 | 1 |
| virus_Aug5.25v_20700 | | 0 | 0 | 0 | 0 | 0 | 1 | 0 | 0 | 0 | 0 | 1 |
| virus_Aug5.25v_21512 | | 0 | 0 | 0 | 0 | 0 | 1 | 0 | 0 | 0 | 0 | 1 |
| virus_Aug5.25v_24273 | | 0 | 0 | 0 | 0 | 0 | 1 | 0 | 0 | 0 | 0 | 1 |
| virus_Aug5.25v_25788 | | 0 | 0 | 0 | 0 | 0 | 1 | 0 | 0 | 0 | 0 | 1 |
| virus_Aug5.25v_25920 | | 0 | 0 | 0 | 0 | 0 | 1 | 0 | 0 | 0 | 0 | 1 |
| virus_Aug5.25v_26024 | | 0 | 2 | 0 | 1 | 0 | 1 | 0 | 0 | 0 | 0 | 4 |
| virus_Aug5.25v_28677 | | 0 | 0 | 0 | 0 | 0 | 1 | 0 | 0 | 0 | 0 | 1 |
| virus_Aug5.2 | 9 | 5 | 107 | 5 | 14 | 6 | 48 | 3 | 3 | 0 | 8 | 199 |
| virus_Aug5.25v_30934 | | 0 | 0 | 0 | 0 | 0 | 1 | 0 | 0 | 0 | 0 | 1 |
| virus_Aug5.25v_31488 | | 0 | 0 | 0 | 0 | 0 | 1 | 0 | 0 | 0 | 0 | 1 |
| virus_Aug5.25v_33587 | | 0 | 0 | 0 | 0 | 0 | 1 | 0 | 0 | 0 | 0 | 1 |
| virus_Aug5.25v_34030 | | 0 | 0 | 0 | 0 | 0 | 1 | 0 | 0 | 0 | 0 | 1 |
| virus_Aug5.25v_36661 | | 0 | 0 | 0 | 0 | 0 | 1 | 0 | 0 | 0 | 1 | 2 |
| virus_Aug5.25v_40296 | | 0 | 0 | 0 | 0 | 0 | 1 | 1 | 1 | 1 | 0 | 4 |
| virus_Aug5.25v_40869 | | 0 | 0 | 0 | 0 | 0 | 1 | 0 | 0 | 0 | 0 | 1 |
| virus_Aug5.50v_4473 | | 0 | 0 | 0 | 0 | 0 | 0 | 0 | 1 | 0 | 0 | 1 |
| virus_Aug5.50v_9849 | | 0 | 0 | 0 | 0 | 0 | 0 | 0 | 1 | 0 | 0 | 1 |
| virus_Aug5.50v_16031 | | 0 | 0 | 0 | 0 | 0 | 0 | 0 | 1 | 0 | 0 | 1 |
| virus_Aug5.50v_17646 | | 0 | 0 | 0 | 0 | 1 | 0 | 0 | 1 | 0 | 1 | 3 |
| virus_Aug5.50v_24085 | | 0 | 1 | 0 | 2 | 1 | 1 | 0 | 1 | 0 | 0 | 6 |
| virus_Aug5.50v_30098 | | 0 | 0 | 0 | 0 | 0 | 0 | 0 | 1 | 0 | 0 | 1 |
| virus_Aug5.50v_37809 | | 0 | 0 | 0 | 0 | 0 | 0 | 0 | 1 | 0 | 0 | 1 |
| virus_Aug5.75v_865 | | 0 | 0 | 0 | 0 | 0 | 0 | 0 | 0 | 0 | 1 | 1 |
| virus_Aug5.75v_2824 | | 0 | 0 | 0 | 0 | 0 | 0 | 0 | 0 | 0 | 1 | 1 |
| virus_Aug5.75v_7326 | | 0 | 1 | 0 | 2 | 0 | 1 | 0 | 0 | 0 | 1 | 5 |
| virus_Aug5.75v_14604 | | 0 | 0 | 0 | 0 | 0 | 1 | 0 | 0 | 0 | 1 | 2 |
| virus_Aug5.75v_17455 | | 0 | 0 | 0 | 0 | 0 | 0 | 0 | 0 | 0 | 1 | 1 |
| virus_Aug5.75v_17539 | | 0 | 0 | 0 | 0 | 0 | 0 | 0 | 0 | 0 | 1 | 1 |
| virus_Aug5.75v_28329 | | 0 | 0 | 0 | 0 | 0 | 0 | 0 | 0 | 0 | 1 | 1 |
| virus_Aug5.75v_34066 | | 0 | 0 | 0 | 0 | 0 | 0 | 0 | 0 | 0 | 1 | 1 |
| virus_Jul11.0v_7583 | | 0 | 0 | 0 | 0 | 0 | 0 | 0 | 0 | 0 | 0 | 0 |
| virus_Jul11.0v_13581 | | 0 | 1 | 0 | 1 | 0 | 1 | 0 | 0 | 0 | 0 | 3 |
| virus_Jul11.0v_16073 | | 0 | 0 | 0 | 0 | 0 | 0 | 0 | 0 | 0 | 0 | 0 |
| virus_Jul11.0v_17344 | | 0 | 1 | 0 | 0 | 0 | 0 | 0 | 0 | 0 | 0 | 1 |
| virus_Jul11.0v_17606 | | 0 | 0 | 0 | 0 | 0 | 0 | 0 | 0 | 0 | 0 | 0 |
| virus_Jul11.0v_20942 | | 0 | 0 | 0 | 0 | 0 | 0 | 0 | 0 | 0 | 0 | 0 |
| virus_Jul11.5v_1149 | | 0 | 1 | 0 | 0 | 0 | 0 | 0 | 0 | 0 | 0 | 1 |
| virus_Jul11.5v_2015 | | 0 | 0 | 0 | 0 | 0 | 0 | 0 | 0 | 0 | 0 | 0 |
| virus_Jul11.5v_3354 | | 0 | 0 | 0 | 0 | 0 | 0 | 0 | 0 | 0 | 0 | 0 |
| virus_Jul11.5v_6236 | | 0 | 0 | 0 | 0 | 0 | 0 | 0 | 0 | 0 | 0 | 0 |
| virus_Jul11.5v_8514 | | 0 | 0 | 0 | 0 | 0 | 0 | 0 | 0 | 0 | 0 | 0 |
| virus_Jul11.5v_8934 | | 0 | 0 | 0 | 0 | 0 | 0 | 0 | 0 | 0 | 0 | 0 |
| virus_Jul11.5v_9524 | | 0 | 0 | 0 | 0 | 0 | 0 | 0 | 0 | 0 | 0 | 0 |
| virus_Jul11.5v_11287 | | 0 | 0 | 0 | 0 | 0 | 0 | 0 | 0 | 0 | 0 | 0 |
| virus_Jul11.5v_11372 | | 0 | 0 | 0 | 0 | 0 | 0 | 0 | 0 | 0 | 0 | 0 |
| virus_Jul11.5v_13541 | | 0 | 0 | 0 | 0 | 0 | 0 | 0 | 0 | 0 | 0 | 0 |
| virus_Jul11.5v_13921 | | 0 | 0 | 0 | 0 | 0 | 0 | 0 | 0 | 0 | 0 | 0 |
| virus_Jul11.5v_15683 | | 0 | 0 | 0 | 0 | 0 | 0 | 0 | 0 | 0 | 0 | 0 |
| virus_Jul11.5v_16593 | | 0 | 0 | 0 | 0 | 0 | 0 | 0 | 0 | 0 | 0 | 0 |
| virus_Jul11.5v_18142 | | 0 | 0 | 0 | 0 | 0 | 0 | 0 | 0 | 0 | 0 | 0 |
| virus_Jul11.5v_18298 | | 0 | 0 | 0 | 0 | 0 | 0 | 0 | 0 | 0 | 0 | 0 |
| virus_Jul11.5v_18843 | | 0 | 0 | 0 | 0 | 0 | 0 | 0 | 0 | 0 | 0 | 0 |
| virus_Jul11.25v_400 | | 0 | 0 | 0 | 0 | 0 | 0 | 0 | 0 | 0 | 0 | 0 |
| virus_Jul11.25v_914 | | 0 | 0 | 0 | 0 | 0 | 0 | 0 | 0 | 0 | 0 | 0 |
| virus_Jul11.25v_2745 | | 0 | 0 | 0 | 0 | 0 | 0 | 0 | 0 | 0 | 0 | 0 |
| virus_Jul11.25v_2818 | | 0 | 0 | 0 | 0 | 0 | 0 | 0 | 0 | 0 | 0 | 0 |
| virus_Jul11.25v_3200 | | 0 | 0 | 0 | 0 | 0 | 0 | 0 | 0 | 0 | 0 | 0 |
| virus_Jul11.25v_5072 | | 0 | 0 | 0 | 0 | 0 | 0 | 0 | 0 | 0 | 0 | 0 |
| virus_Jul11.25v_5616 | | 0 | 0 | 0 | 0 | 0 | 0 | 0 | 0 | 0 | 0 | 0 |
| virus_Jul11.25v_6604 | | 0 | 0 | 0 | 0 | 0 | 0 | 0 | 0 | 0 | 0 | 0 |
| virus_Jul11.25v_7068 | | 0 | 0 | 0 | 0 | 0 | 0 | 0 | 0 | 0 | 0 | 0 |
| virus_Jul11.25v_7261 | | 0 | 0 | 0 | 0 | 0 | 0 | 0 | 0 | 0 | 0 | 0 |
| virus_Jul11.25v_8292 | | 0 | 0 | 0 | 0 | 0 | 1 | 0 | 0 | 0 | 0 | 1 |
| virus_Jul11.25v_9154 | | 0 | 0 | 0 | 0 | 0 | 0 | 0 | 0 | 0 | 0 | 0 |
| virus_Jul11.2 | 16 | 2 | 66 | 3 | 12 | 3 | 29 | 0 | 6 | 0 | 3 | 124 |
| virus_Jul11.25v_11137 | | 0 | 0 | 0 | 0 | 0 | 0 | 0 | 0 | 0 | 0 | 0 |
| virus_Jul11.25v_11902 | | 0 | 0 | 0 | 0 | 0 | 0 | 0 | 0 | 0 | 0 | 0 |
| virus_Jul11.25v_12432 | | 0 | 0 | 0 | 0 | 0 | 0 | 0 | 0 | 0 | 0 | 0 |
| virus_Jul11.25v_12881 | | 0 | 0 | 0 | 0 | 0 | 0 | 0 | 0 | 0 | 0 | 0 |
| virus_Jul11.25v_12958 | | 0 | 0 | 0 | 0 | 0 | 0 | 0 | 0 | 0 | 0 | 0 |
| virus_Jul11.25v_13332 | | 0 | 0 | 0 | 0 | 0 | 0 | 0 | 0 | 0 | 0 | 0 |
| virus_Jul11.25v_13997 | | 0 | 0 | 0 | 0 | 0 | 0 | 0 | 0 | 0 | 0 | 0 |
| virus_Jul11.25v_14963 | | 0 | 0 | 0 | 0 | 0 | 0 | 0 | 0 | 0 | 0 | 0 |
| virus_Jul11.25v_15112 | | 0 | 0 | 0 | 0 | 0 | 0 | 0 | 0 | 0 | 0 | 0 |
| virus_Jul11.25v_18292 | | 0 | 0 | 0 | 0 | 0 | 0 | 0 | 0 | 0 | 0 | 0 |
| virus_Jul11.25v_20780 | | 0 | 0 | 0 | 0 | 0 | 0 | 0 | 0 | 0 | 0 | 0 |
| virus_Jul11.50v_1243 | | 0 | 0 | 0 | 0 | 0 | 0 | 0 | 0 | 0 | 0 | 0 |
| virus_Jul11.50v_1408 | | 0 | 0 | 0 | 0 | 0 | 0 | 0 | 0 | 0 | 0 | 0 |
| virus_Jul11.50v_3426 | | 0 | 0 | 0 | 0 | 0 | 1 | 0 | 0 | 0 | 1 | 2 |
| virus_Jul11.50v_4166 | | 0 | 0 | 0 | 0 | 0 | 0 | 0 | 0 | 0 | 0 | 0 |
| virus_Jul11.50v_4788 | | 0 | 0 | 0 | 0 | 0 | 0 | 0 | 0 | 0 | 0 | 0 |
| virus_Jul11.50v_6014 | | 0 | 0 | 0 | 0 | 0 | 0 | 0 | 0 | 0 | 0 | 0 |
| virus_Jul11.50v_9303 | | 0 | 0 | 0 | 0 | 0 | 0 | 0 | 0 | 0 | 0 | 0 |
| virus_Jul11.50v_9390 | | 0 | 0 | 0 | 0 | 0 | 0 | 0 | 0 | 0 | 0 | 0 |
| virus_Jul11.50v_9468 | | 0 | 0 | 0 | 0 | 0 | 0 | 0 | 0 | 0 | 0 | 0 |
| virus_Jul11.50v_10110 | | 0 | 0 | 0 | 0 | 0 | 0 | 0 | 0 | 0 | 0 | 0 |
| virus_Jul11.50v_11281 | | 0 | 0 | 0 | 0 | 0 | 0 | 0 | 0 | 0 | 0 | 0 |
| virus_Jul11.50v_11743 | | 0 | 0 | 0 | 0 | 1 | 0 | 0 | 0 | 0 | 0 | 1 |
| virus_Jul11.50v_16973 | | 0 | 0 | 0 | 0 | 0 | 0 | 0 | 0 | 0 | 0 | 0 |
| virus_Jul11.75v_3084 | | 0 | 0 | 0 | 0 | 0 | 0 | 0 | 0 | 0 | 0 | 0 |
| virus_Jul11.75v_4190 | | 0 | 0 | 0 | 0 | 0 | 0 | 0 | 0 | 0 | 0 | 0 |
| virus_Jul11.75v_17025 | | 0 | 0 | 0 | 0 | 0 | 0 | 0 | 0 | 0 | 0 | 0 |
| virus_Jul11.75v_17405 | | 0 | 0 | 0 | 0 | 0 | 0 | 0 | 0 | 0 | 0 | 0 |
| virus_Jul11.75v_23737 | | 0 | 0 | 0 | 0 | 0 | 0 | 0 | 0 | 0 | 0 | 0 |
| virus_Jul11.75v_32525 | | 0 | 0 | 0 | 0 | 0 | 0 | 0 | 0 | 0 | 0 | 0 |
| virus_Jul11.75v_37018 | | 0 | 0 | 0 | 0 | 0 | 0 | 0 | 0 | 0 | 0 | 0 |
| virus_Jul11.75v_39594 | | 0 | 0 | 0 | 0 | 0 | 0 | 0 | 0 | 0 | 0 | 0 |
| virus_Jul11.75v_40073 | | 0 | 0 | 0 | 0 | 0 | 0 | 0 | 0 | 0 | 0 | 0 |
| virus_Jul27.5v_43 | | 0 | 0 | 0 | 0 | 0 | 0 | 0 | 0 | 0 | 0 | 0 |
| virus_Jul27.5v_380 | | 0 | 0 | 0 | 0 | 0 | 0 | 0 | 0 | 0 | 0 | 0 |
| virus_Jul27.5v_470 | | 0 | 0 | 0 | 0 | 0 | 0 | 0 | 0 | 0 | 0 | 0 |
| virus_Jul27.5v_1937 | | 0 | 0 | 0 | 0 | 0 | 0 | 0 | 0 | 0 | 0 | 0 |
| virus_Jul27.5v_2368 | | 0 | 0 | 0 | 0 | 0 | 0 | 0 | 0 | 0 | 0 | 0 |
| virus_Jul27.5v_3838 | | 0 | 0 | 0 | 0 | 0 | 0 | 0 | 0 | 0 | 0 | 0 |
| virus_Jul27.5v_3962 | | 0 | 2 | 0 | 0 | 0 | 0 | 0 | 0 | 0 | 0 | 2 |
| virus_Jul27.5v_5152 | | 0 | 0 | 0 | 0 | 0 | 0 | 0 | 0 | 0 | 0 | 0 |
| virus_Jul27.5v_5879 | | 0 | 1 | 0 | 0 | 1 | 1 | 0 | 0 | 0 | 0 | 3 |
| virus_Jul27.5v_7463 | | 0 | 0 | 0 | 0 | 0 | 0 | 0 | 0 | 0 | 0 | 0 |
| virus_Jul27.5v_7704 | | 0 | 0 | 0 | 0 | 0 | 0 | 0 | 0 | 0 | 0 | 0 |
| virus_Jul27.5v_8055 | | 0 | 0 | 0 | 0 | 0 | 0 | 0 | 0 | 0 | 0 | 0 |
| virus_Jul27.5v_8443 | | 0 | 0 | 0 | 0 | 0 | 0 | 0 | 0 | 0 | 0 | 0 |
| virus_Jul27.5v_12688 | | 0 | 0 | 0 | 0 | 0 | 0 | 0 | 0 | 0 | 0 | 0 |
| virus_Jul27.5v_13580 | | 0 | 0 | 0 | 0 | 0 | 0 | 0 | 0 | 0 | 0 | 0 |
| virus_Jul27.5v_13776 | | 0 | 0 | 0 | 0 | 0 | 0 | 0 | 0 | 0 | 0 | 0 |
| virus_Jul27.5v_13813 | | 0 | 0 | 0 | 1 | 0 | 1 | 0 | 0 | 1 | 0 | 3 |
| virus_Jul27.5v_14695 | | 0 | 0 | 0 | 0 | 0 | 0 | 0 | 0 | 0 | 0 | 0 |
| virus_Jul27.5v_16702 | | 0 | 0 | 0 | 0 | 0 | 0 | 0 | 0 | 0 | 0 | 0 |
| virus_Jul27.5v_16922 | | 0 | 0 | 0 | 0 | 0 | 0 | 0 | 0 | 0 | 0 | 0 |
| virus_Jul27.5v_18729 | | 0 | 0 | 0 | 0 | 0 | 0 | 0 | 0 | 0 | 0 | 0 |
| virus_Jul27.5v_19719 | | 0 | 0 | 0 | 0 | 0 | 0 | 0 | 0 | 0 | 0 | 0 |
| virus_Jul27.5v_19974 | | 0 | 0 | 0 | 0 | 0 | 0 | 0 | 0 | 0 | 0 | 0 |
| virus_Jul27.25v_3151 | | 0 | 1 | 0 | 0 | 0 | 1 | 0 | 0 | 0 | 0 | 2 |
| virus_Jul27.25v_5795 | | 0 | 0 | 0 | 0 | 0 | 0 | 0 | 0 | 0 | 0 | 0 |
| virus_Jul27.25v_7668 | | 0 | 0 | 0 | 0 | 0 | 0 | 0 | 0 | 0 | 0 | 0 |
| virus_Jul27.25v_8840 | | 0 | 0 | 0 | 0 | 0 | 0 | 0 | 0 | 0 | 0 | 0 |
| virus_Jul27.25v_9816 | | 0 | 0 | 0 | 0 | 0 | 0 | 0 | 0 | 0 | 0 | 0 |
| virus_Jul27.25v_10141 | | 0 | 0 | 0 | 0 | 0 | 0 | 0 | 0 | 0 | 0 | 0 |
| virus_Jul27.25v_13490 | | 0 | 0 | 0 | 0 | 0 | 0 | 0 | 0 | 0 | 0 | 0 |
| virus_Jul27.25v_15898 | | 0 | 0 | 0 | 0 | 0 | 0 | 0 | 0 | 0 | 0 | 0 |
| virus_Jul27.25v_17052 | | 0 | 0 | 0 | 0 | 0 | 0 | 0 | 0 | 0 | 0 | 0 |
| virus_Jul27.25v_20364 | | 0 | 0 | 0 | 0 | 0 | 0 | 0 | 0 | 0 | 0 | 0 |
| virus_Jul27.25v_24189 | | 0 | 0 | 0 | 0 | 0 | 0 | 0 | 0 | 0 | 0 | 0 |
| virus_Jul27.25v_25363 | | 0 | 0 | 0 | 0 | 0 | 0 | 0 | 0 | 0 | 0 | 0 |
| virus_Jul27.25v_27552 | | 0 | 0 | 0 | 0 | 0 | 0 | 0 | 0 | 0 | 0 | 0 |
| virus_Jul27.25v_34099 | | 0 | 0 | 0 | 0 | 0 | 0 | 0 | 0 | 0 | 0 | 0 |
| virus_Jul27.25v_34387 | | 0 | 0 | 0 | 0 | 0 | 0 | 0 | 0 | 0 | 0 | 0 |
| virus_Jul27.25v_38302 | | 0 | 0 | 0 | 0 | 0 | 0 | 0 | 0 | 0 | 0 | 0 |
| virus_Jul27.25v_40180 | | 0 | 0 | 0 | 0 | 0 | 0 | 0 | 0 | 0 | 0 | 0 |
| virus_Jul27.75v_1714 | | 0 | 0 | 0 | 0 | 0 | 0 | 0 | 0 | 0 | 0 | 0 |
| virus_Jul27.75v_2711 | | 0 | 0 | 0 | 0 | 0 | 0 | 0 | 0 | 0 | 0 | 0 |
| virus_Jul27.75v_10732 | | 0 | 0 | 0 | 0 | 0 | 0 | 0 | 0 | 0 | 0 | 0 |
| virus_Jun8.25v_9630 | | 0 | 0 | 0 | 0 | 0 | 0 | 0 | 0 | 0 | 0 | 0 |
| virus_Jun8.25v_9804 | | 0 | 0 | 0 | 0 | 0 | 0 | 0 | 0 | 0 | 0 | 0 |
| virus_Jun8.25v_16511 | | 0 | 0 | 0 | 0 | 0 | 0 | 0 | 0 | 0 | 0 | 0 |
| virus_Jun8.25v_20975 | | 0 | 0 | 0 | 0 | 0 | 0 | 0 | 0 | 0 | 0 | 0 |
| virus_Jun8.50v_7739 | | 0 | 0 | 0 | 0 | 0 | 0 | 0 | 0 | 0 | 0 | 0 |
| virus_Jun8.50v_16449 | | 0 | 0 | 0 | 0 | 0 | 0 | 0 | 0 | 0 | 0 | 0 |
| virus_Jun8.50v_19047 | | 0 | 0 | 0 | 0 | 0 | 0 | 0 | 0 | 0 | 0 | 0 |
| virus_Jun8.50v_20410 | | 0 | 0 | 0 | 0 | 0 | 0 | 0 | 0 | 0 | 0 | 0 |
| virus_Jun8.50v_28076 | | 0 | 0 | 0 | 0 | 0 | 0 | 0 | 0 | 0 | 0 | 0 |
| virus_Jun8.50v_30737 | | 0 | 0 | 0 | 0 | 0 | 0 | 0 | 0 | 0 | 0 | 0 |
| virus_Jun8.50v_32582 | | 0 | 0 | 0 | 0 | 0 | 0 | 0 | 0 | 0 | 0 | 0 |
| virus_Jun8.50v_40686 | | 0 | 0 | 0 | 0 | 0 | 0 | 0 | 0 | 0 | 0 | 0 |
| virus_Jun8.75v_561 | | 0 | 0 | 0 | 0 | 0 | 0 | 0 | 0 | 0 | 0 | 0 |
| virus_Jun8.75v_1537 | | 0 | 0 | 0 | 0 | 0 | 0 | 0 | 0 | 0 | 1 | 1 |
| virus_Jun8.75v_2728 | | 0 | 0 | 0 | 0 | 0 | 0 | 0 | 0 | 0 | 0 | 0 |

| | | | | | | | | | | | |
|---|---|---|---|---|---|---|---|---|---|---|---|
| virus_Jun8.75v_8412 | 0 | 0 | 0 | 0 | 0 | 0 | 0 | 0 | 0 | 0 | 0 |
| virus_Jun8.75v_10037 | 0 | 0 | 0 | 0 | 0 | 0 | 0 | 0 | 0 | 2 | 2 |
| virus_Jun8.75v_15985 | 0 | 0 | 0 | 0 | 0 | 0 | 0 | 0 | 0 | 0 | 0 |
| virus_Jun8.75v_18738 | 0 | 0 | 0 | 0 | 0 | 0 | 0 | 0 | 0 | 0 | 0 |
| virus_Jun8.75v_19828 | 0 | 0 | 0 | 0 | 0 | 0 | 0 | 0 | 0 | 0 | 0 |
| virus_Jun22.0v_1288 | 0 | 0 | 0 | 0 | 0 | 0 | 0 | 0 | 0 | 0 | 0 |
| virus_Jun22.0v_1661 | 0 | 0 | 0 | 0 | 0 | 0 | 0 | 0 | 0 | 0 | 0 |
| virus_Jun22.0v_2351 | 0 | 0 | 0 | 0 | 0 | 0 | 0 | 0 | 0 | 0 | 0 |
| virus_Jun22.0v_2549 | 0 | 0 | 0 | 0 | 0 | 0 | 0 | 0 | 0 | 0 | 0 |
| virus_Jun22.0v_8194 | 0 | 0 | 0 | 0 | 0 | 0 | 0 | 0 | 0 | 0 | 0 |
| virus_Jun22.0v_12992 | 0 | 0 | 0 | 0 | 0 | 0 | 0 | 0 | 0 | 0 | 0 |
| virus_Jun22.0v_17047 | 0 | 0 | 0 | 0 | 0 | 0 | 0 | 0 | 0 | 0 | 0 |
| virus_Jun22.0v_18187 | 0 | 0 | 0 | 0 | 0 | 0 | 0 | 0 | 0 | 0 | 0 |
| virus_Jun22.0v_18741 | 0 | 0 | 0 | 0 | 0 | 0 | 0 | 0 | 0 | 0 | 0 |
| virus_Jun22.0v_20897 | 0 | 0 | 0 | 0 | 0 | 0 | 0 | 0 | 0 | 0 | 0 |
| virus_Jun22.50v_2361 | 0 | 0 | 0 | 0 | 0 | 0 | 0 | 0 | 0 | 0 | 0 |
| virus_Jun22.50v_6516 | 0 | 0 | 0 | 0 | 0 | 0 | 0 | 0 | 0 | 0 | 0 |
| virus_Jun22.50v_8936 | 0 | 0 | 0 | 0 | 0 | 0 | 0 | 0 | 0 | 0 | 0 |
| virus_Jun22.50v_9120 | 0 | 0 | 0 | 0 | 0 | 0 | 0 | 0 | 0 | 0 | 0 |
| virus_Jun22.50v_9291 | 0 | 0 | 0 | 0 | 0 | 0 | 0 | 0 | 0 | 0 | 0 |
| virus_Jun22.50v_14807 | 0 | 0 | 0 | 0 | 0 | 0 | 0 | 0 | 0 | 0 | 0 |
| virus_Jun22.50v_17696 | 0 | 0 | 0 | 0 | 0 | 0 | 0 | 0 | 0 | 0 | 0 |
| virus_Jun22.50v_19008 | 0 | 0 | 0 | 0 | 0 | 0 | 0 | 0 | 0 | 0 | 0 |
| virus_Jun22.50v_20514 | 0 | 0 | 0 | 0 | 0 | 0 | 0 | 0 | 0 | 0 | 0 |
| virus_Jun22.50v_24843 | 0 | 0 | 0 | 0 | 0 | 0 | 0 | 0 | 0 | 0 | 0 |
| virus_Jun22.50v_26935 | 0 | 0 | 0 | 0 | 0 | 0 | 0 | 0 | 0 | 0 | 0 |
| virus_Jun22.50v_30078 | 0 | 0 | 0 | 0 | 0 | 0 | 0 | 0 | 0 | 0 | 0 |
| virus_Jun22.50v_31467 | 0 | 0 | 0 | 0 | 0 | 0 | 0 | 0 | 0 | 0 | 0 |
| virus_Jun22.50v_33089 | 0 | 0 | 0 | 0 | 0 | 0 | 0 | 0 | 0 | 0 | 0 |
| virus_Jun22.50v_33799 | 0 | 0 | 0 | 0 | 0 | 0 | 0 | 0 | 0 | 0 | 0 |
| virus_Jun22.50v_39024 | 0 | 0 | 0 | 0 | 0 | 0 | 0 | 0 | 0 | 0 | 0 |
| virus_Jun22.50v_39505 | 0 | 0 | 0 | 0 | 0 | 0 | 0 | 0 | 0 | 0 | 0 |
| virus_Jun22.50v_40677 | 0 | 0 | 0 | 0 | 0 | 0 | 0 | 0 | 0 | 0 | 0 |
| virus_Jun22.75v_3585 | 0 | 0 | 0 | 0 | 0 | 1 | 0 | 0 | 0 | 0 | 1 |
| virus_Jun22.75v_4346 | 0 | 0 | 0 | 0 | 0 | 0 | 0 | 0 | 0 | 0 | 0 |
| virus_Jun22.75v_5986 | 0 | 0 | 0 | 0 | 0 | 0 | 0 | 0 | 0 | 0 | 0 |
| virus_Jun22.75v_6745 | 0 | 0 | 0 | 0 | 0 | 1 | 1 | 1 | 0 | 0 | 3 |
| virus_Jun22.75v_12338 | 0 | 0 | 0 | 0 | 0 | 0 | 0 | 0 | 0 | 0 | 0 |
| virus_Jun22.75v_12846 | 0 | 0 | 0 | 0 | 0 | 0 | 0 | 0 | 0 | 0 | 0 |
| virus_Jun22.75v_18459 | 0 | 0 | 0 | 0 | 0 | 0 | 0 | 0 | 0 | 0 | 0 |
| virus_Jun22.75v_25850 | 0 | 0 | 0 | 0 | 0 | 0 | 0 | 0 | 0 | 0 | 0 |
| virus_Jun22.75v_29454 | 0 | 0 | 0 | 0 | 0 | 0 | 0 | 0 | 0 | 0 | 0 |
| virus_Jun22.75v_32241 | 0 | 1 | 0 | 0 | 0 | 0 | 0 | 0 | 0 | 2 | 3 |
| virus_Jun22.75v_36613 | 0 | 0 | 0 | 0 | 0 | 0 | 0 | 0 | 0 | 0 | 0 |
| virus_Jun22.75v_40825 | 0 | 0 | 0 | 0 | 0 | 0 | 0 | 0 | 0 | 0 | 0 |
| virus_May12.5v_2938 | 0 | 0 | 0 | 0 | 0 | 0 | 0 | 0 | 0 | 0 | 0 |
| virus_May12.5v_13330 | 0 | 0 | 0 | 0 | 0 | 0 | 0 | 0 | 0 | 0 | 0 |
| virus_May12.5v_21973 | 0 | 0 | 0 | 0 | 0 | 0 | 0 | 0 | 0 | 0 | 0 |
| virus_May12.5v_23168 | 0 | 0 | 0 | 0 | 0 | 0 | 0 | 0 | 0 | 0 | 0 |
| virus_May12.5v_27624 | 0 | 0 | 0 | 0 | 0 | 0 | 0 | 0 | 0 | 0 | 0 |
| virus_May12.5v_29202 | 0 | 0 | 0 | 0 | 0 | 0 | 0 | 0 | 0 | 0 | 0 |
| virus_May12.5v_29257 | 0 | 0 | 0 | 0 | 0 | 0 | 0 | 0 | 0 | 0 | 0 |
| virus_May12.5v_33029 | 0 | 0 | 0 | 0 | 0 | 0 | 0 | 0 | 0 | 0 | 0 |
| virus_May12.5v_33832 | 0 | 0 | 0 | 0 | 0 | 0 | 0 | 0 | 0 | 0 | 0 |
| virus_May12.5v_37184 | 0 | 0 | 0 | 0 | 0 | 0 | 0 | 0 | 0 | 0 | 0 |
| virus_May12.5v_41032 | 0 | 0 | 0 | 0 | 0 | 0 | 0 | 0 | 0 | 0 | 0 |
| virus_May12.25v_288 | 0 | 0 | 0 | 0 | 0 | 0 | 0 | 0 | 0 | 0 | 0 |
| virus_May12.25v_3047 | 0 | 0 | 0 | 0 | 0 | 0 | 0 | 0 | 0 | 0 | 0 |
| virus_May12.25v_8994 | 0 | 0 | 0 | 0 | 0 | 0 | 0 | 0 | 0 | 0 | 0 |
| virus_May12.25v_13023 | 0 | 0 | 0 | 0 | 0 | 0 | 0 | 0 | 0 | 0 | 0 |
| virus_May12.25v_16726 | 0 | 0 | 0 | 0 | 0 | 0 | 0 | 0 | 0 | 0 | 0 |
| virus_May12.25v_26714 | 0 | 0 | 0 | 0 | 0 | 0 | 0 | 0 | 0 | 0 | 0 |
| virus_May12.25v_28165 | 0 | 0 | 0 | 0 | 0 | 0 | 0 | 0 | 0 | 0 | 0 |
| virus_May12.25v_28620 | 0 | 0 | 0 | 0 | 0 | 0 | 0 | 0 | 0 | 0 | 0 |
| virus_May12.25v_35885 | 0 | 0 | 0 | 0 | 0 | 0 | 0 | 0 | 0 | 0 | 0 |
| virus_May12.50v_7190 | 0 | 0 | 0 | 0 | 0 | 0 | 0 | 0 | 0 | 0 | 0 |
| virus_May12.50v_8935 | 0 | 1 | 0 | 0 | 0 | 0 | 0 | 0 | 0 | 0 | 1 |
| virus_May12.50v_9400 | 0 | 0 | 0 | 0 | 0 | 0 | 0 | 0 | 0 | 0 | 0 |
| virus_May12.50v_9493 | 0 | 0 | 0 | 0 | 0 | 0 | 0 | 0 | 0 | 0 | 0 |
| virus_May12.50v_9880 | 0 | 0 | 0 | 0 | 0 | 0 | 0 | 0 | 0 | 0 | 0 |
| virus_May12.50v_11531 | 0 | 0 | 0 | 0 | 0 | 0 | 0 | 0 | 0 | 0 | 0 |
| virus_May12.50v_11953 | 0 | 0 | 0 | 0 | 0 | 0 | 0 | 0 | 0 | 0 | 0 |
| virus_May12.50v_19301 | 0 | 0 | 1 | 0 | 0 | 1 | 0 | 0 | 0 | 0 | 2 |
| virus_May12.50v_19340 | 0 | 0 | 0 | 0 | 0 | 0 | 0 | 0 | 0 | 0 | 0 |
| virus_May12.50v_22672 | 0 | 0 | 0 | 0 | 0 | 0 | 0 | 0 | 0 | 0 | 0 |
| virus_May12.50v_22804 | 0 | 0 | 0 | 0 | 0 | 0 | 0 | 0 | 0 | 0 | 0 |
| virus_May12.50v_22845 | 0 | 0 | 0 | 0 | 0 | 0 | 0 | 0 | 0 | 0 | 0 |
| virus_May12.50v_25185 | 0 | 0 | 0 | 0 | 0 | 0 | 0 | 0 | 0 | 0 | 0 |
| virus_May12.50v_26155 | 0 | 0 | 0 | 0 | 0 | 0 | 0 | 0 | 0 | 0 | 0 |
| virus_May12.50v_27974 | 0 | 0 | 0 | 0 | 0 | 0 | 0 | 0 | 0 | 0 | 0 |
| virus_May12.50v_32397 | 0 | 0 | 0 | 0 | 0 | 0 | 0 | 0 | 0 | 0 | 0 |
| virus_May12.50v_38464 | 0 | 0 | 0 | 0 | 0 | 0 | 0 | 0 | 0 | 0 | 0 |
| virus_May12.50v_38567 | 0 | 0 | 0 | 0 | 0 | 0 | 0 | 0 | 0 | 0 | 0 |
| virus_May12.50v_40245 | 0 | 0 | 0 | 0 | 0 | 0 | 0 | 0 | 0 | 0 | 0 |
| virus_May12.50v_40416 | 0 | 0 | 0 | 0 | 0 | 0 | 0 | 0 | 0 | 0 | 0 |
| virus_May12.75v_8355 | 0 | 0 | 0 | 0 | 0 | 0 | 0 | 0 | 0 | 0 | 0 |
| virus_May12.75v_10506 | 0 | 0 | 0 | 0 | 0 | 0 | 0 | 0 | 0 | 0 | 0 |
| virus_May12.75v_13478 | 0 | 1 | 0 | 0 | 0 | 0 | 0 | 1 | 0 | 0 | 2 |
| virus_May12.75v_13493 | 0 | 1 | 0 | 0 | 1 | 0 | 0 | 0 | 0 | 0 | 2 |
| virus_May12.75v_13516 | 0 | 0 | 0 | 0 | 0 | 0 | 0 | 0 | 0 | 0 | 0 |
| virus_May12.75v_13829 | 0 | 0 | 0 | 0 | 0 | 0 | 0 | 0 | 0 | 0 | 0 |
| virus_May12.75v_24463 | 0 | 0 | 0 | 0 | 0 | 0 | 0 | 0 | 0 | 0 | 0 |
| virus_May12.75v_27240 | 0 | 0 | 0 | 0 | 0 | 0 | 0 | 0 | 0 | 0 | 0 |
| virus_May12.75v_30830 | 0 | 0 | 0 | 0 | 0 | 0 | 0 | 0 | 0 | 0 | 0 |
| virus_May12.75v_31618 | 0 | 0 | 0 | 0 | 0 | 0 | 0 | 0 | 0 | 0 | 0 |
| virus_May12.75v_36787 | 0 | 0 | 0 | 0 | 0 | 0 | 0 | 0 | 0 | 0 | 0 |
| virus_May12.75v_39250 | 0 | 0 | 0 | 0 | 0 | 0 | 0 | 0 | 0 | 0 | 0 |
| virus_May12.75v_39704 | 0 | 0 | 0 | 0 | 0 | 0 | 0 | 0 | 0 | 0 | 0 |
| virus_Sep8.0v_1170 | 1 | 0 | 0 | 0 | 0 | 0 | 0 | 0 | 0 | 0 | 1 |
| virus_Sep8.0v_1807 | 0 | 1 | 0 | 0 | 0 | 0 | 0 | 0 | 0 | 0 | 1 |
| virus_Sep8.0v_2419 | 0 | 0 | 0 | 0 | 0 | 0 | 0 | 0 | 0 | 0 | 0 |
| virus_Sep8.0v_4952 | 0 | 0 | 0 | 0 | 0 | 0 | 0 | 0 | 0 | 1 | 1 |
| virus_Sep8.0v_5444 | 0 | 0 | 0 | 0 | 0 | 0 | 0 | 0 | 0 | 0 | 0 |
| virus_Sep8.0v_6319 | 0 | 0 | 0 | 0 | 0 | 0 | 0 | 0 | 0 | 0 | 0 |
| virus_Sep8.0v_7000 | 0 | 0 | 0 | 0 | 0 | 0 | 0 | 0 | 0 | 0 | 0 |
| virus_Sep8.0v_10367 | 0 | 0 | 0 | 0 | 0 | 0 | 0 | 0 | 0 | 0 | 0 |
| virus_Sep8.0v_10941 | 0 | 0 | 0 | 0 | 0 | 0 | 0 | 0 | 0 | 0 | 0 |
| virus_Sep8.0v_14363 | 0 | 0 | 0 | 0 | 0 | 0 | 0 | 0 | 0 | 0 | 0 |
| virus_Sep8.0v_15316 | 0 | 0 | 0 | 0 | 0 | 0 | 0 | 0 | 0 | 0 | 0 |
| virus_Sep8.0v_18231 | 0 | 0 | 0 | 0 | 0 | 0 | 0 | 0 | 0 | 0 | 0 |
| virus_Sep8.0v_18705 | 0 | 0 | 0 | 0 | 0 | 0 | 0 | 0 | 0 | 0 | 0 |
| virus_Sep8.0v_20528 | 0 | 0 | 0 | 0 | 0 | 0 | 0 | 0 | 0 | 0 | 0 |
| virus_Sep8.5v_143 | 0 | 0 | 0 | 0 | 0 | 0 | 0 | 0 | 0 | 0 | 0 |
| virus_Sep8.5v_5564 | 0 | 0 | 0 | 0 | 0 | 0 | 0 | 0 | 0 | 0 | 0 |
| virus_Sep8.5v_9257 | 0 | 0 | 0 | 0 | 0 | 0 | 0 | 0 | 0 | 0 | 0 |
| virus_Sep8.5v_10759 | 0 | 0 | 0 | 0 | 0 | 0 | 0 | 0 | 0 | 0 | 0 |
| virus_Sep8.5v_11860 | 0 | 0 | 0 | 0 | 0 | 0 | 0 | 0 | 0 | 0 | 0 |
| virus_Sep8.5v_13376 | 0 | 0 | 0 | 0 | 0 | 0 | 0 | 0 | 0 | 0 | 0 |
| virus_Sep8.5v_15067 | 0 | 0 | 0 | 0 | 0 | 0 | 0 | 0 | 0 | 0 | 0 |
| virus_Sep8.25v_7208 | 0 | 0 | 0 | 0 | 0 | 0 | 0 | 0 | 0 | 0 | 0 |
| virus_Sep8.25v_9123 | 0 | 0 | 0 | 0 | 0 | 0 | 0 | 0 | 0 | 0 | 0 |
| virus_Sep8.25v_12326 | 0 | 0 | 0 | 0 | 0 | 0 | 0 | 0 | 0 | 0 | 0 |
| virus_Sep8.25v_18163 | 0 | 0 | 0 | 0 | 0 | 0 | 0 | 0 | 0 | 0 | 0 |
| virus_Sep8.25v_20896 | 0 | 0 | 0 | 0 | 0 | 0 | 0 | 0 | 0 | 0 | 0 |
| virus_Sep8.25v_21004 | 0 | 0 | 0 | 0 | 0 | 0 | 0 | 0 | 0 | 0 | 0 |
| virus_Sep8.50v_3217 | 0 | 0 | 0 | 0 | 0 | 0 | 0 | 0 | 0 | 0 | 0 |
| virus_Sep8.50v_5179 | 0 | 0 | 0 | 0 | 0 | 0 | 0 | 0 | 0 | 0 | 0 |
| virus_Sep8.50v_11419 | 0 | 0 | 0 | 0 | 0 | 0 | 0 | 0 | 0 | 0 | 0 |
| virus_Sep8.50v_12252 | 0 | 0 | 0 | 0 | 0 | 0 | 0 | 0 | 0 | 0 | 0 |
| virus_Sep8.50v_12628 | 0 | 0 | 0 | 0 | 0 | 0 | 0 | 0 | 0 | 0 | 0 |
| virus_Sep8.50v_15779 | 0 | 0 | 0 | 0 | 0 | 0 | 0 | 0 | 0 | 0 | 0 |
| virus_Sep8.50v_18102 | 0 | 0 | 0 | 0 | 0 | 0 | 0 | 0 | 0 | 0 | 0 |
| virus_Sep8.50v_22893 | 0 | 0 | 0 | 0 | 0 | 0 | 0 | 0 | 0 | 0 | 0 |
| virus_Sep8.50v_27082 | 0 | 0 | 0 | 0 | 0 | 0 | 0 | 0 | 0 | 0 | 0 |
| virus_Sep8.50v_28279 | 0 | 0 | 0 | 0 | 0 | 0 | 0 | 0 | 0 | 0 | 0 |
| virus_Sep8.50v_30979 | 0 | 0 | 0 | 0 | 0 | 0 | 0 | 0 | 0 | 0 | 0 |
| virus_Sep8.50v_31633 | 0 | 0 | 0 | 0 | 0 | 0 | 0 | 0 | 0 | 0 | 0 |
| virus_Sep8.50v_32593 | 0 | 0 | 0 | 0 | 0 | 0 | 0 | 0 | 0 | 0 | 0 |
| virus_Sep8.50v_33778 | 0 | 1 | 0 | 0 | 0 | 0 | 0 | 0 | 0 | 0 | 1 |
| virus_Sep8.50v_34011 | 0 | 0 | 0 | 0 | 0 | 0 | 0 | 0 | 0 | 0 | 0 |
| virus_Sep8.50v_34791 | 0 | 0 | 0 | 0 | 0 | 0 | 0 | 0 | 0 | 0 | 0 |
| virus_Sep8.50v_35689 | 0 | 0 | 0 | 0 | 0 | 0 | 0 | 0 | 0 | 0 | 0 |
| virus_Sep8.50v_35750 | 0 | 0 | 0 | 0 | 0 | 0 | 0 | 0 | 0 | 0 | 0 |
| virus_Sep8.50v_37801 | 0 | 0 | 0 | 0 | 0 | 0 | 0 | 0 | 0 | 0 | 0 |
| virus_Sep8.50v_40086 | 0 | 0 | 0 | 0 | 0 | 0 | 0 | 0 | 0 | 0 | 0 |
| bacteria_Jul11d0b_5074 | 0 | 0 | 0 | 0 | 0 | 0 | 0 | 0 | 0 | 0 | 0 |
| bacteria_Jul11d0b_10479 | 0 | 0 | 0 | 0 | 0 | 0 | 0 | 0 | 0 | 0 | 0 |
| bacteria_Jul11d0b_11132 | 0 | 0 | 0 | 0 | 0 | 0 | 0 | 0 | 0 | 0 | 0 |
| bacteria_Jul11d0b_16778 | 0 | 0 | 0 | 0 | 0 | 0 | 0 | 0 | 0 | 0 | 0 |
| bacteria_Jul11d0b_29309 | 0 | 0 | 0 | 0 | 0 | 0 | 0 | 0 | 0 | 0 | 0 |
| bacteria_Jul11d0b_39221 | 0 | 0 | 0 | 0 | 0 | 0 | 0 | 0 | 0 | 0 | 0 |
| bacteria_Jul11d0b_50644 | 0 | 0 | 0 | 0 | 0 | 0 | 0 | 0 | 0 | 0 | 0 |
| bacteria_Aug5d0b_29542 | 1 | 0 | 0 | 0 | 0 | 0 | 0 | 0 | 0 | 0 | 1 |
| bacteria_Aug5d0b_36096 | 1 | 0 | 0 | 0 | 0 | 0 | 0 | 0 | 0 | 0 | 1 |
| bacteria_Aug5d0b_57201 | 1 | 0 | 0 | 0 | 0 | 0 | 0 | 0 | 0 | 0 | 1 |
| bacteria_Aug5d0b_57371 | 1 | 0 | 0 | 0 | 0 | 0 | 0 | 0 | 0 | 0 | 1 |
| bacteria_Aug5d0b_61259 | 1 | 0 | 0 | 0 | 0 | 0 | 0 | 0 | 0 | 0 | 1 |

| | | | | | | | | | | | |
|---|---|---|---|---|---|---|---|---|---|---|---|
| bacteria_Aug5d5b_2271 | 0 | 0 | 1 | 0 | 0 | 0 | 0 | 0 | 0 | 0 | 1 |
| bacteria_Aug5d5b_61768 | 0 | 0 | 1 | 0 | 0 | 0 | 0 | 0 | 0 | 0 | 1 |
| bacteria_Aug5d25b_3118 | 0 | 0 | 0 | 0 | 1 | 0 | 0 | 0 | 0 | 0 | 1 |
| bacteria_Aug5d25b_5836 | 0 | 0 | 0 | 0 | 1 | 0 | 1 | 0 | 0 | 0 | 2 |
| bacteria_Aug5d25b_16314 | 0 | 0 | 0 | 0 | 1 | 0 | 0 | 0 | 0 | 0 | 1 |
| bacteria_Aug5d25b_23204 | 0 | 0 | 0 | 0 | 1 | 0 | 0 | 0 | 0 | 0 | 1 |
| bacteria_Aug5d25b_25404 | 0 | 0 | 0 | 0 | 1 | 0 | 0 | 0 | 0 | 0 | 1 |
| bacteria_Aug5d25b_28476 | 0 | 0 | 0 | 0 | 1 | 0 | 0 | 0 | 0 | 0 | 1 |
| bacteria_Aug5d25b_29868 | 0 | 0 | 0 | 0 | 1 | 0 | 0 | 0 | 0 | 0 | 1 |
| bacteria_Aug5d25b_39761 | 0 | 0 | 0 | 0 | 1 | 0 | 0 | 0 | 0 | 0 | 1 |
| bacteria_Aug5d25b_40004 | 0 | 0 | 0 | 0 | 1 | 0 | 0 | 0 | 0 | 0 | 1 |
| bacteria_Aug5d25b_43739 | 0 | 4 | 0 | 1 | 2 | 1 | 1 | 0 | 0 | 3 | 12 |
| bacteria_Aug5d25b_45411 | 0 | 2 | 0 | 0 | 1 | 0 | 0 | 0 | 0 | 0 | 3 |
| bacteria_Aug5d25b_50585 | 0 | 0 | 0 | 0 | 1 | 0 | 0 | 0 | 0 | 0 | 1 |
| bacteria_Aug5d25b_57546 | 0 | 0 | 0 | 0 | 1 | 0 | 0 | 0 | 0 | 0 | 1 |
| bacteria_Aug5d25b_63069 | 0 | 0 | 0 | 0 | 1 | 0 | 1 | 0 | 0 | 0 | 2 |
| bacteria_Aug5d25b_64056 | 0 | 0 | 0 | 0 | 1 | 0 | 0 | 0 | 0 | 0 | 1 |
| bacteria_Aug5d25b_70143 | 0 | 0 | 0 | 0 | 1 | 0 | 0 | 0 | 0 | 0 | 1 |
| bacteria_Aug5d25b_70827 | 0 | 0 | 0 | 0 | 1 | 0 | 0 | 0 | 0 | 0 | 1 |
| bacteria_Aug5d25b_71726 | 0 | 0 | 0 | 0 | 1 | 0 | 0 | 1 | 0 | 0 | 2 |
| bacteria_Aug5d50b_2535 | 0 | 0 | 0 | 0 | 0 | 0 | 1 | 0 | 0 | 0 | 1 |
| bacteria_Aug5d50b_10508 | 0 | 0 | 0 | 0 | 0 | 0 | 1 | 0 | 0 | 0 | 1 |
| bacteria_Aug5d50b_23927 | 0 | 0 | 0 | 0 | 0 | 0 | 1 | 0 | 0 | 0 | 1 |
| bacteria_Aug5d50b_34659 | 0 | 0 | 0 | 0 | 0 | 0 | 1 | 0 | 0 | 0 | 1 |
| bacteria_Aug5d50b_41628 | 0 | 0 | 0 | 0 | 0 | 0 | 1 | 0 | 0 | 0 | 1 |
| bacteria_Aug5d50b_46055 | 0 | 0 | 0 | 0 | 0 | 0 | 1 | 0 | 0 | 0 | 1 |
| bacteria_Aug5d50b_47037 | 0 | 0 | 0 | 0 | 0 | 0 | 1 | 0 | 0 | 0 | 1 |
| bacteria_Aug5d50b_47622 | 0 | 0 | 0 | 0 | 0 | 0 | 1 | 0 | 0 | 0 | 1 |
| bacteria_Aug5d50b_62072 | 0 | 0 | 0 | 0 | 0 | 0 | 1 | 0 | 0 | 0 | 1 |
| bacteria_Aug5d50b_75188 | 0 | 0 | 0 | 0 | 0 | 0 | 1 | 0 | 0 | 0 | 1 |
| bacteria_Aug5d75b_2964 | 0 | 0 | 0 | 0 | 0 | 0 | 0 | 0 | 1 | 0 | 1 |
| bacteria_Jul11d5b_3608 | 0 | 0 | 0 | 0 | 0 | 0 | 0 | 0 | 0 | 0 | 0 |
| bacteria_Jul11d5b_8057 | 0 | 0 | 0 | 0 | 0 | 0 | 0 | 0 | 0 | 0 | 0 |
| bacteria_Jul11d5b_8474 | 0 | 0 | 0 | 0 | 0 | 0 | 0 | 0 | 0 | 0 | 0 |
| bacteria_Jul11d5b_10588 | 0 | 0 | 0 | 0 | 0 | 0 | 0 | 0 | 0 | 0 | 0 |
| bacteria_Jul11d5b_23901 | 0 | 0 | 0 | 0 | 0 | 0 | 0 | 0 | 0 | 0 | 0 |
| bacteria_Jul11d5b_41355 | 0 | 0 | 0 | 0 | 0 | 0 | 0 | 0 | 0 | 0 | 0 |
| bacteria_Jul11d25b_5650 | 0 | 0 | 0 | 0 | 0 | 0 | 0 | 0 | 0 | 0 | 0 |
| bacteria_Jul11d25b_11676 | 0 | 0 | 0 | 0 | 0 | 0 | 0 | 0 | 0 | 0 | 0 |
| bacteria_Jul11d25b_15790 | 0 | 0 | 0 | 0 | 0 | 0 | 0 | 0 | 0 | 0 | 0 |
| bacteria_Jul11d25b_18970 | 0 | 0 | 0 | 0 | 0 | 0 | 0 | 0 | 0 | 0 | 0 |
| bacteria_Jul11d25b_20521 | 0 | 0 | 0 | 0 | 0 | 0 | 0 | 0 | 0 | 0 | 0 |
| bacteria_Jul11d25b_25054 | 0 | 0 | 0 | 0 | 0 | 0 | 0 | 0 | 0 | 0 | 0 |
| bacteria_Jul11d25b_36298 | 0 | 0 | 0 | 0 | 0 | 0 | 0 | 0 | 0 | 0 | 0 |
| bacteria_Jul11d25b_38079 | 0 | 0 | 0 | 0 | 0 | 0 | 0 | 0 | 0 | 0 | 0 |
| bacteria_Jul11d25b_39933 | 0 | 0 | 0 | 0 | 0 | 0 | 0 | 0 | 0 | 0 | 0 |
| bacteria_Jul11d25b_40555 | 0 | 0 | 0 | 0 | 0 | 0 | 0 | 0 | 0 | 0 | 0 |
| bacteria_Jul11d25b_43790 | 0 | 0 | 0 | 0 | 0 | 0 | 0 | 0 | 0 | 0 | 0 |
| bacteria_Jul11d50b_13030 | 0 | 1 | 0 | 0 | 0 | 0 | 0 | 1 | 0 | 0 | 2 |
| bacteria_Jul11d50b_20957 | 0 | 0 | 0 | 0 | 0 | 0 | 0 | 0 | 0 | 0 | 0 |
| bacteria_Jul11d50b_22096 | 0 | 0 | 0 | 1 | 0 | 0 | 0 | 0 | 0 | 0 | 1 |
| bacteria_Jul11d50b_32360 | 0 | 0 | 0 | 0 | 0 | 0 | 0 | 0 | 0 | 0 | 0 |
| bacteria_Jul11d50b_43716 | 0 | 0 | 0 | 0 | 0 | 0 | 0 | 0 | 0 | 0 | 0 |
| bacteria_Jul11d75b_826 | 0 | 0 | 0 | 0 | 0 | 0 | 0 | 0 | 0 | 0 | 0 |
| bacteria_Jul11d75b_1469 | 0 | 0 | 0 | 0 | 0 | 0 | 0 | 0 | 0 | 0 | 0 |
| bacteria_Jul11d75b_2405 | 0 | 0 | 0 | 0 | 0 | 0 | 0 | 0 | 0 | 0 | 0 |
| bacteria_Jul11d75b_11767 | 0 | 0 | 0 | 0 | 1 | 0 | 0 | 0 | 0 | 0 | 1 |
| bacteria_Jul11d75b_14793 | 0 | 0 | 0 | 0 | 0 | 0 | 0 | 0 | 0 | 0 | 0 |
| bacteria_Jul11d75b_19147 | 0 | 0 | 0 | 0 | 0 | 0 | 0 | 0 | 0 | 0 | 0 |
| bacteria_Jul11d75b_21463 | 0 | 0 | 0 | 0 | 0 | 0 | 0 | 0 | 0 | 0 | 0 |
| bacteria_Jul11d75b_23919 | 0 | 0 | 0 | 0 | 0 | 0 | 0 | 0 | 0 | 0 | 0 |
| bacteria_Jul11d75b_31045 | 0 | 0 | 0 | 0 | 0 | 0 | 0 | 0 | 0 | 0 | 0 |
| bacteria_Jul11d75b_33654 | 0 | 0 | 0 | 0 | 0 | 0 | 0 | 0 | 0 | 0 | 0 |
| bacteria_Jul11d75b_38316 | 0 | 0 | 0 | 0 | 0 | 0 | 0 | 0 | 0 | 0 | 0 |
| bacteria_Jul11d75b_38912 | 0 | 0 | 0 | 0 | 0 | 0 | 0 | 0 | 0 | 0 | 0 |
| bacteria_Jul11d75b_38972 | 0 | 0 | 0 | 0 | 0 | 0 | 0 | 0 | 0 | 0 | 0 |
| bacteria_Jul11d75b_39322 | 0 | 0 | 0 | 0 | 0 | 0 | 0 | 0 | 0 | 0 | 0 |
| bacteria_Jul11d75b_43173 | 0 | 0 | 0 | 0 | 0 | 0 | 0 | 0 | 0 | 0 | 0 |
| bacteria_Jul11d75b_45433 | 0 | 0 | 0 | 0 | 0 | 0 | 0 | 0 | 0 | 0 | 0 |
| bacteria_Jul11d75b_48382 | 0 | 0 | 0 | 0 | 0 | 0 | 0 | 0 | 0 | 0 | 0 |
| bacteria_Jul27d5b_6962 | 0 | 0 | 0 | 0 | 0 | 0 | 0 | 0 | 0 | 0 | 0 |
| bacteria_Jul27d5b_8807 | 0 | 0 | 0 | 0 | 0 | 0 | 0 | 0 | 0 | 0 | 0 |
| bacteria_Jul27d5b_19865 | 0 | 0 | 0 | 0 | 0 | 0 | 0 | 0 | 0 | 0 | 0 |
| bacteria_Jul27d5b_20737 | 0 | 0 | 0 | 0 | 0 | 0 | 0 | 0 | 0 | 0 | 0 |
| bacteria_Jul27d5b_22999 | 0 | 0 | 0 | 0 | 0 | 0 | 0 | 0 | 0 | 0 | 0 |
| bacteria_Jul27d5b_25455 | 0 | 0 | 0 | 0 | 0 | 0 | 0 | 0 | 0 | 0 | 0 |
| bacteria_Jul27d5b_27004 | 0 | 0 | 0 | 0 | 0 | 0 | 0 | 0 | 0 | 0 | 0 |
| bacteria_Jul27d5b_33034 | 0 | 0 | 0 | 0 | 0 | 0 | 0 | 0 | 0 | 0 | 0 |
| bacteria_Jul27d5b_37905 | 0 | 0 | 0 | 0 | 0 | 0 | 0 | 0 | 0 | 0 | 0 |
| bacteria_Jul27d5b_40384 | 0 | 0 | 0 | 0 | 0 | 0 | 1 | 0 | 0 | 0 | 1 |
| bacteria_Jul27d5b_40933 | 0 | 0 | 0 | 0 | 0 | 0 | 0 | 0 | 0 | 0 | 0 |
| bacteria_Jul27d5b_43272 | 0 | 0 | 0 | 0 | 0 | 0 | 0 | 0 | 0 | 0 | 0 |
| bacteria_Jul27d5b_44170 | 0 | 0 | 0 | 0 | 0 | 0 | 0 | 0 | 0 | 0 | 0 |
| bacteria_Jul27d5b_49546 | 0 | 0 | 0 | 0 | 0 | 0 | 0 | 0 | 0 | 0 | 0 |
| bacteria_Jul27d5b_51156 | 0 | 0 | 0 | 0 | 0 | 0 | 0 | 0 | 0 | 0 | 0 |
| bacteria_Jul27d25b_6046 | 0 | 0 | 0 | 0 | 0 | 0 | 0 | 0 | 0 | 0 | 0 |
| bacteria_Jul27d25b_24951 | 0 | 0 | 0 | 0 | 0 | 0 | 0 | 0 | 0 | 0 | 0 |
| bacteria_Jul27d25b_25366 | 0 | 0 | 0 | 0 | 0 | 0 | 0 | 0 | 0 | 0 | 0 |
| bacteria_Jul27d25b_26214 | 0 | 0 | 0 | 0 | 0 | 0 | 0 | 0 | 0 | 0 | 0 |
| bacteria_Jul27d25b_27464 | 0 | 0 | 0 | 0 | 0 | 0 | 0 | 0 | 0 | 0 | 0 |
| bacteria_Jul27d25b_29312 | 0 | 0 | 0 | 0 | 0 | 0 | 0 | 0 | 0 | 0 | 0 |
| bacteria_Jul27d25b_31076 | 0 | 0 | 0 | 0 | 0 | 0 | 0 | 0 | 0 | 0 | 0 |
| bacteria_Jul27d25b_33502 | 0 | 0 | 0 | 0 | 0 | 0 | 0 | 0 | 0 | 0 | 0 |
| bacteria_Jul27d25b_47729 | 0 | 1 | 0 | 0 | 0 | 0 | 0 | 0 | 0 | 0 | 1 |
| bacteria_Jul27d25b_48531 | 0 | 0 | 0 | 0 | 0 | 0 | 0 | 0 | 0 | 0 | 0 |
| bacteria_Jul27d25b_54301 | 0 | 0 | 0 | 0 | 0 | 0 | 0 | 0 | 0 | 0 | 0 |
| bacteria_Jul27d25b_61671 | 0 | 0 | 0 | 0 | 0 | 0 | 0 | 0 | 0 | 0 | 0 |
| bacteria_Jul27d25b_65930 | 0 | 0 | 0 | 0 | 0 | 0 | 0 | 0 | 0 | 0 | 0 |
| bacteria_Jul27d25b_66926 | 0 | 0 | 0 | 0 | 0 | 0 | 0 | 0 | 0 | 0 | 0 |
| bacteria_Jul27d25b_69657 | 0 | 0 | 0 | 0 | 0 | 0 | 0 | 0 | 0 | 0 | 0 |
| bacteria_Jul27d75b_3855 | 0 | 0 | 0 | 0 | 0 | 0 | 1 | 0 | 0 | 0 | 1 |
| bacteria_Jul27d75b_5032 | 0 | 0 | 0 | 0 | 0 | 0 | 0 | 0 | 0 | 0 | 0 |
| bacteria_Jul27d75b_12770 | 0 | 0 | 0 | 0 | 0 | 0 | 0 | 0 | 0 | 0 | 0 |
| bacteria_Jul27d75b_13696 | 0 | 0 | 0 | 0 | 0 | 0 | 0 | 0 | 0 | 0 | 0 |
| bacteria_Jul27d75b_22239 | 0 | 0 | 0 | 0 | 0 | 0 | 0 | 0 | 0 | 0 | 0 |
| bacteria_Jul27d75b_30480 | 0 | 0 | 0 | 0 | 0 | 0 | 0 | 0 | 0 | 0 | 0 |
| bacteria_Jul27d75b_34331 | 0 | 0 | 0 | 0 | 0 | 0 | 0 | 0 | 0 | 0 | 0 |
| bacteria_Jul27d75b_51527 | 0 | 0 | 0 | 0 | 0 | 0 | 0 | 0 | 0 | 0 | 0 |
| bacteria_Jul27d75b_51955 | 0 | 0 | 0 | 0 | 0 | 0 | 0 | 0 | 0 | 0 | 0 |
| bacteria_Jul27d75b_57829 | 0 | 0 | 0 | 0 | 0 | 0 | 0 | 0 | 0 | 0 | 0 |
| bacteria_Jul27d75b_59189 | 0 | 0 | 0 | 0 | 0 | 0 | 0 | 0 | 0 | 0 | 0 |
| bacteria_Jul27d75b_61527 | 0 | 0 | 0 | 0 | 0 | 0 | 0 | 0 | 0 | 0 | 0 |
| bacteria_Jul27d75b_69037 | 0 | 0 | 0 | 0 | 0 | 0 | 0 | 0 | 0 | 0 | 0 |
| bacteria_Jul27d75b_71748 | 0 | 0 | 0 | 0 | 0 | 0 | 0 | 0 | 0 | 0 | 0 |
| bacteria_Jul27d75b_74013 | 0 | 0 | 0 | 0 | 0 | 0 | 0 | 0 | 0 | 0 | 0 |
| bacteria_Jun8d25b_2306 | 0 | 0 | 0 | 0 | 0 | 0 | 1 | 0 | 0 | 0 | 1 |
| bacteria_Jun8d25b_12666 | 0 | 0 | 0 | 0 | 0 | 0 | 0 | 0 | 0 | 0 | 0 |
| bacteria_Jun8d25b_31827 | 0 | 0 | 0 | 0 | 0 | 0 | 0 | 0 | 0 | 0 | 0 |
| bacteria_Jun8d25b_34108 | 0 | 0 | 0 | 0 | 0 | 0 | 0 | 0 | 0 | 0 | 0 |
| bacteria_Jun8d25b_39401 | 0 | 0 | 0 | 0 | 0 | 0 | 0 | 0 | 0 | 0 | 0 |
| bacteria_Jun8d25b_41365 | 0 | 0 | 0 | 0 | 0 | 0 | 0 | 0 | 0 | 0 | 0 |
| bacteria_Jun8d25b_46307 | 0 | 0 | 0 | 0 | 0 | 0 | 0 | 0 | 0 | 0 | 0 |
| bacteria_Jun8d25b_49754 | 0 | 0 | 0 | 0 | 0 | 0 | 0 | 0 | 0 | 0 | 0 |
| bacteria_Jun8d25b_54041 | 0 | 0 | 0 | 0 | 0 | 0 | 0 | 0 | 0 | 0 | 0 |
| bacteria_Jun8d25b_60553 | 0 | 0 | 0 | 0 | 0 | 0 | 0 | 0 | 0 | 0 | 0 |
| bacteria_Jun8d25b_64582 | 0 | 0 | 0 | 0 | 0 | 0 | 0 | 0 | 0 | 0 | 0 |
| bacteria_Jun8d25b_66773 | 0 | 0 | 0 | 0 | 0 | 0 | 0 | 0 | 0 | 0 | 0 |
| bacteria_Jun8d25b_68066 | 0 | 0 | 0 | 0 | 0 | 0 | 0 | 0 | 0 | 0 | 0 |
| bacteria_Jun8d25b_73990 | 0 | 0 | 0 | 0 | 0 | 0 | 0 | 0 | 0 | 0 | 0 |
| bacteria_Jun8d50b_4413 | 0 | 0 | 0 | 0 | 0 | 0 | 0 | 0 | 0 | 0 | 0 |
| bacteria_Jun8d50b_16298 | 0 | 0 | 0 | 0 | 0 | 0 | 0 | 0 | 0 | 0 | 0 |
| bacteria_Jun8d50b_25943 | 0 | 0 | 0 | 0 | 0 | 0 | 0 | 0 | 0 | 0 | 0 |
| bacteria_Jun8d50b_30780 | 0 | 0 | 0 | 0 | 0 | 0 | 0 | 0 | 0 | 0 | 0 |
| bacteria_Jun8d75b_53038 | 0 | 0 | 0 | 0 | 0 | 0 | 0 | 0 | 0 | 0 | 0 |
| bacteria_Jun22d0b_444 | 0 | 0 | 0 | 0 | 0 | 0 | 0 | 0 | 0 | 0 | 0 |
| bacteria_Jun22d0b_5193 | 0 | 0 | 0 | 0 | 0 | 0 | 0 | 0 | 0 | 0 | 0 |
| bacteria_Jun22d0b_14922 | 0 | 0 | 0 | 0 | 0 | 0 | 0 | 0 | 0 | 0 | 0 |
| bacteria_Jun22d0b_34569 | 0 | 0 | 0 | 0 | 0 | 0 | 0 | 0 | 0 | 0 | 0 |
| bacteria_Jun22d0b_73272 | 0 | 0 | 0 | 0 | 0 | 1 | 0 | 0 | 0 | 0 | 1 |
| bacteria_Jun22d50b_2567 | 0 | 0 | 0 | 0 | 0 | 0 | 0 | 0 | 0 | 0 | 0 |
| bacteria_Jun22d50b_13452 | 0 | 0 | 0 | 0 | 0 | 0 | 0 | 0 | 0 | 0 | 0 |
| bacteria_Jun22d50b_15377 | 0 | 0 | 0 | 0 | 0 | 0 | 0 | 0 | 0 | 0 | 0 |
| bacteria_Jun22d50b_26932 | 0 | 0 | 0 | 0 | 0 | 0 | 0 | 0 | 0 | 0 | 0 |
| bacteria_Jun22d50b_33190 | 0 | 0 | 0 | 0 | 0 | 0 | 0 | 0 | 0 | 0 | 0 |
| bacteria_Jun22d50b_36655 | 0 | 0 | 0 | 0 | 0 | 0 | 0 | 0 | 0 | 0 | 0 |
| bacteria_Jun22d75b_4712 | 0 | 0 | 0 | 0 | 0 | 0 | 0 | 0 | 0 | 0 | 0 |
| bacteria_Jun22d75b_40861 | 0 | 0 | 0 | 0 | 0 | 0 | 0 | 0 | 0 | 0 | 0 |
| bacteria_Jun22d75b_42560 | 0 | 0 | 0 | 0 | 0 | 0 | 0 | 0 | 0 | 0 | 0 |
| bacteria_Jun22d75b_45727 | 0 | 0 | 0 | 0 | 0 | 0 | 0 | 0 | 0 | 0 | 0 |
| bacteria_Jun22d75b_51343 | 0 | 0 | 0 | 0 | 0 | 0 | 0 | 0 | 0 | 0 | 0 |
| bacteria_Jun22d75b_62565 | 0 | 0 | 0 | 0 | 0 | 0 | 0 | 0 | 0 | 0 | 0 |
| bacteria_May12d5b_14865 | 0 | 0 | 0 | 0 | 0 | 0 | 0 | 0 | 0 | 0 | 0 |
| bacteria_May12d5b_47581 | 0 | 0 | 0 | 0 | 0 | 0 | 0 | 0 | 0 | 0 | 0 |
| bacteria_May12d25b_1475 | 0 | 0 | 0 | 0 | 0 | 0 | 0 | 0 | 0 | 0 | 0 |
| bacteria_May12d25b_3577 | 0 | 0 | 0 | 0 | 0 | 0 | 0 | 0 | 0 | 0 | 0 |
| bacteria_May12d25b_9512 | 0 | 0 | 0 | 0 | 0 | 0 | 0 | 0 | 0 | 0 | 0 |
| bacteria_May12d25b_1632 | 0 | 0 | 0 | 0 | 0 | 0 | 0 | 0 | 0 | 0 | 0 |

| | | | | | | | | | | | |
|---|---|---|---|---|---|---|---|---|---|---|---|
| bacteria_May12d25b_2397 | 0 | 0 | 0 | 0 | 0 | 0 | 0 | 0 | 0 | 0 | 0 |
| bacteria_May12d25b_5027 | 0 | 0 | 0 | 0 | 0 | 0 | 0 | 0 | 0 | 0 | 0 |
| bacteria_May12d25b_5646 | 0 | 0 | 0 | 0 | 0 | 0 | 0 | 0 | 0 | 0 | 0 |
| bacteria_May12d25b_5985 | 0 | 0 | 0 | 0 | 0 | 0 | 0 | 0 | 0 | 0 | 0 |
| bacteria_May12d25b_6845 | 0 | 0 | 0 | 0 | 0 | 0 | 0 | 0 | 0 | 0 | 0 |
| bacteria_May12d50b_3500 | 0 | 0 | 0 | 0 | 0 | 0 | 0 | 0 | 0 | 0 | 0 |
| bacteria_May12d50b_8013 | 0 | 0 | 0 | 0 | 0 | 0 | 0 | 0 | 0 | 0 | 0 |
| bacteria_May12d50b_1129 | 0 | 0 | 0 | 0 | 0 | 0 | 0 | 0 | 0 | 0 | 0 |
| bacteria_May12d50b_1685 | 0 | 0 | 0 | 0 | 0 | 0 | 0 | 0 | 0 | 0 | 0 |
| bacteria_May12d50b_2117 | 0 | 0 | 0 | 0 | 0 | 0 | 0 | 0 | 0 | 0 | 0 |
| bacteria_May12d50b_2171 | 0 | 0 | 0 | 0 | 0 | 0 | 0 | 0 | 0 | 0 | 0 |
| bacteria_May12d50b_2380 | 0 | 0 | 0 | 0 | 0 | 0 | 0 | 0 | 0 | 0 | 0 |
| bacteria_May12d50b_4723 | 0 | 0 | 0 | 0 | 0 | 0 | 0 | 0 | 0 | 0 | 0 |
| bacteria_May12d50b_5106 | 0 | 0 | 0 | 0 | 0 | 0 | 0 | 0 | 0 | 0 | 0 |
| bacteria_May12d75b_3880 | 0 | 0 | 0 | 0 | 0 | 0 | 0 | 0 | 0 | 0 | 0 |
| bacteria_Sep8d0b_25219 | 0 | 0 | 0 | 0 | 0 | 0 | 0 | 0 | 0 | 0 | 0 |
| bacteria_Sep8d0b_26669 | 0 | 0 | 0 | 0 | 0 | 0 | 0 | 0 | 0 | 0 | 0 |
| bacteria_Sep8d0b_36515 | 0 | 0 | 0 | 0 | 0 | 0 | 0 | 0 | 0 | 0 | 0 |
| bacteria_Sep8d5b_3542 | 0 | 0 | 0 | 0 | 0 | 0 | 0 | 0 | 0 | 0 | 0 |
| bacteria_Sep8d5b_10378 | 0 | 0 | 0 | 0 | 0 | 0 | 0 | 0 | 0 | 0 | 0 |
| bacteria_Sep8d5b_16031 | 0 | 0 | 0 | 0 | 0 | 0 | 0 | 0 | 0 | 0 | 0 |
| bacteria_Sep8d5b_16061 | 1 | 0 | 1 | 0 | 1 | 0 | 0 | 0 | 0 | 0 | 3 |
| bacteria_Sep8d5b_24502 | 0 | 0 | 0 | 0 | 0 | 0 | 0 | 0 | 0 | 0 | 0 |
| bacteria_Sep8d5b_39511 | 0 | 0 | 0 | 0 | 0 | 0 | 0 | 0 | 0 | 0 | 0 |
| bacteria_Sep8d5b_39919 | 0 | 0 | 0 | 0 | 0 | 0 | 1 | 0 | 0 | 0 | 1 |
| bacteria_Sep8d5b_42190 | 0 | 0 | 0 | 0 | 0 | 0 | 0 | 0 | 0 | 0 | 0 |
| bacteria_Sep8d5b_43629 | 0 | 0 | 0 | 0 | 0 | 0 | 0 | 0 | 0 | 0 | 0 |
| bacteria_Sep8d5b_43979 | 0 | 0 | 0 | 0 | 0 | 0 | 0 | 0 | 0 | 0 | 0 |
| bacteria_Sep8d5b_44832 | 5 | 0 | 0 | 0 | 9 | 0 | 1 | 0 | 2 | 0 | 17 |
| bacteria_Sep8d5b_45292 | 0 | 0 | 0 | 0 | 0 | 0 | 0 | 0 | 0 | 0 | 0 |
| bacteria_Sep8d5b_50208 | 0 | 0 | 0 | 0 | 0 | 0 | 0 | 0 | 0 | 0 | 0 |
| bacteria_Sep8d5b_70638 | 0 | 0 | 0 | 0 | 0 | 0 | 0 | 0 | 0 | 0 | 0 |
| bacteria_Sep8d5b_70779 | 0 | 0 | 0 | 0 | 0 | 0 | 0 | 0 | 0 | 0 | 0 |
| bacteria_Sep8d25b_10337 | 0 | 0 | 0 | 0 | 0 | 0 | 0 | 0 | 0 | 0 | 0 |
| bacteria_Sep8d25b_16917 | 0 | 0 | 0 | 0 | 0 | 0 | 0 | 0 | 0 | 0 | 0 |
| bacteria_Sep8d25b_24091 | 0 | 0 | 0 | 0 | 0 | 0 | 0 | 0 | 0 | 0 | 0 |
| bacteria_Sep8d25b_36037 | 0 | 0 | 0 | 0 | 0 | 0 | 0 | 0 | 0 | 0 | 0 |
| bacteria_Sep8d50b_3989 | 0 | 0 | 0 | 0 | 0 | 0 | 0 | 0 | 0 | 0 | 0 |
| bacteria_Sep8d50b_50416 | 0 | 0 | 0 | 0 | 0 | 0 | 0 | 0 | 0 | 0 | 0 |
| virus_Aug5.0v_789 | 0 | 1 | 0 | 0 | 0 | 0 | 0 | 0 | 0 | 0 | 1 |
| virus_Aug5.0v_1060 | 0 | 1 | 0 | 0 | 0 | 0 | 0 | 0 | 0 | 0 | 1 |
| virus_Aug5.0v_1297 | 0 | 1 | 0 | 0 | 0 | 0 | 0 | 0 | 0 | 0 | 1 |
| virus_Aug5.0v_1593 | 0 | 1 | 0 | 0 | 0 | 0 | 0 | 0 | 0 | 0 | 1 |
| virus_Aug5.0v_1658 | 0 | 1 | 0 | 0 | 0 | 0 | 0 | 0 | 0 | 0 | 1 |
| virus_Aug5.0v_2120 | 0 | 1 | 0 | 0 | 0 | 0 | 0 | 0 | 0 | 0 | 1 |
| virus_Aug5.0v_2155 | 0 | 1 | 0 | 0 | 0 | 0 | 0 | 0 | 0 | 0 | 1 |
| virus_Aug5.0v_2189 | 0 | 1 | 0 | 0 | 0 | 0 | 0 | 0 | 0 | 0 | 1 |
| virus_Aug5.0v_3190 | 0 | 1 | 0 | 0 | 0 | 0 | 0 | 0 | 0 | 0 | 1 |
| virus_Aug5.0v_3240 | 0 | 1 | 0 | 0 | 0 | 0 | 0 | 0 | 0 | 0 | 1 |
| virus_Aug5.0v_3433 | 0 | 2 | 0 | 1 | 1 | 0 | 0 | 0 | 0 | 0 | 4 |
| virus_Aug5.0v_3609 | 0 | 1 | 0 | 0 | 0 | 0 | 0 | 0 | 0 | 0 | 1 |
| virus_Aug5.0v_3795 | 0 | 1 | 0 | 0 | 0 | 0 | 0 | 0 | 0 | 0 | 1 |
| virus_Aug5.0v_4066 | 0 | 1 | 0 | 0 | 0 | 0 | 0 | 0 | 0 | 0 | 1 |
| virus_Aug5.0v_4136 | 0 | 1 | 0 | 0 | 0 | 0 | 0 | 0 | 0 | 0 | 1 |
| virus_Aug5.0v_4222 | 0 | 1 | 0 | 0 | 0 | 0 | 0 | 0 | 0 | 0 | 1 |
| virus_Aug5.0v_4306 | 0 | 1 | 0 | 0 | 0 | 0 | 0 | 0 | 0 | 0 | 1 |
| virus_Aug5.0v_4370 | 0 | 1 | 0 | 0 | 0 | 0 | 0 | 0 | 0 | 0 | 1 |
| virus_Aug5.0v_4499 | 0 | 1 | 0 | 0 | 0 | 0 | 0 | 0 | 0 | 0 | 1 |
| virus_Aug5.0v_4554 | 0 | 1 | 0 | 0 | 0 | 0 | 0 | 0 | 0 | 0 | 1 |
| virus_Aug5.0v_5152 | 0 | 1 | 0 | 0 | 0 | 0 | 0 | 0 | 0 | 0 | 1 |
| virus_Aug5.0v_5172 | 0 | 1 | 0 | 0 | 0 | 0 | 0 | 0 | 0 | 0 | 1 |
| virus_Aug5.0v_5204 | 0 | 1 | 0 | 0 | 0 | 0 | 0 | 0 | 0 | 0 | 1 |
| virus_Aug5.0v_5230 | 0 | 1 | 0 | 0 | 0 | 0 | 0 | 0 | 0 | 0 | 1 |
| virus_Aug5.0v_5247 | 0 | 1 | 0 | 0 | 0 | 0 | 0 | 0 | 0 | 0 | 1 |
| virus_Aug5.0v_5393 | 0 | 1 | 0 | 0 | 0 | 0 | 0 | 0 | 0 | 0 | 1 |
| virus_Aug5.5v_2571 | 0 | 0 | 0 | 1 | 0 | 0 | 0 | 0 | 0 | 0 | 1 |
| virus_Aug5.5v_3543 | 0 | 0 | 0 | 1 | 0 | 0 | 0 | 0 | 0 | 0 | 1 |
| virus_Aug5.5' 20 | 3 | 36 | 4 | 19 | 8 | 15 | 2 | 5 | 0 | 9 | 101 |
| virus_Aug5.5v_20294 | 0 | 1 | 0 | 1 | 0 | 0 | 0 | 0 | 0 | 0 | 2 |
| virus_Aug5.5v_20692 | 1 | 1 | 0 | 2 | 0 | 0 | 0 | 0 | 0 | 0 | 4 |
| virus_Aug5.5v_26014 | 0 | 0 | 0 | 1 | 0 | 0 | 0 | 0 | 0 | 0 | 1 |
| virus_Aug5.5v_34765 | 0 | 0 | 0 | 1 | 0 | 0 | 0 | 0 | 0 | 0 | 1 |
| virus_Aug5.5v_40500 | 0 | 0 | 0 | 1 | 0 | 0 | 0 | 0 | 0 | 0 | 1 |
| virus_Aug5.5v_40891 | 0 | 0 | 0 | 1 | 0 | 0 | 0 | 0 | 0 | 0 | 1 |
| virus_Aug5.25v_5930 | 0 | 0 | 0 | 0 | 0 | 1 | 0 | 0 | 0 | 0 | 1 |
| virus_Aug5.25v_6480 | 0 | 0 | 0 | 0 | 0 | 1 | 0 | 0 | 0 | 0 | 1 |
| virus_Aug5.25v_7470 | 0 | 0 | 0 | 0 | 0 | 1 | 0 | 0 | 0 | 0 | 1 |
| virus_Aug5.25v_11359 | 0 | 0 | 0 | 0 | 0 | 1 | 0 | 0 | 0 | 0 | 1 |
| virus_Aug5.25v_13128 | 0 | 0 | 0 | 0 | 0 | 1 | 0 | 0 | 0 | 0 | 1 |
| virus_Aug5.25v_13924 | 0 | 0 | 0 | 0 | 0 | 1 | 0 | 0 | 0 | 0 | 1 |
| virus_Aug5.25v_14876 | 0 | 0 | 0 | 0 | 0 | 1 | 0 | 0 | 0 | 0 | 1 |
| virus_Aug5.25v_18357 | 0 | 0 | 0 | 0 | 0 | 1 | 0 | 0 | 0 | 0 | 1 |
| virus_Aug5.25v_18580 | 0 | 0 | 0 | 0 | 0 | 1 | 0 | 0 | 0 | 0 | 1 |
| virus_Aug5.25v_18767 | 0 | 0 | 0 | 0 | 0 | 1 | 0 | 0 | 0 | 0 | 1 |
| virus_Aug5.25v_19083 | 0 | 0 | 0 | 0 | 0 | 1 | 0 | 0 | 0 | 0 | 1 |
| virus_Aug5.25v_22167 | 0 | 0 | 0 | 0 | 0 | 1 | 0 | 0 | 0 | 0 | 1 |
| virus_Aug5.25v_23454 | 0 | 0 | 0 | 0 | 0 | 1 | 0 | 0 | 0 | 0 | 1 |
| virus_Aug5.25v_27923 | 0 | 0 | 0 | 0 | 0 | 1 | 0 | 0 | 0 | 0 | 1 |
| virus_Aug5.25v_29513 | 0 | 0 | 0 | 0 | 0 | 1 | 0 | 0 | 0 | 0 | 1 |
| virus_Aug5.25v_35124 | 0 | 0 | 0 | 0 | 0 | 1 | 0 | 0 | 0 | 0 | 1 |
| virus_Aug5.25v_35953 | 0 | 0 | 0 | 0 | 0 | 1 | 0 | 0 | 0 | 0 | 1 |
| virus_Aug5.25v_37357 | 0 | 0 | 0 | 0 | 0 | 1 | 0 | 0 | 0 | 0 | 1 |
| virus_Aug5.25v_38436 | 0 | 0 | 0 | 0 | 0 | 1 | 0 | 0 | 0 | 0 | 1 |
| virus_Aug5.25v_38579 | 0 | 0 | 0 | 0 | 0 | 1 | 0 | 0 | 0 | 0 | 1 |
| virus_Aug5.50v_3350 | 0 | 0 | 0 | 0 | 0 | 1 | 0 | 1 | 0 | 0 | 2 |
| virus_Aug5.50v_6581 | 0 | 0 | 0 | 0 | 0 | 0 | 0 | 1 | 0 | 0 | 1 |
| virus_Aug5.50v_8883 | 0 | 0 | 0 | 0 | 0 | 0 | 0 | 1 | 0 | 0 | 1 |
| virus_Aug5.50v_12710 | 0 | 0 | 0 | 0 | 0 | 0 | 0 | 1 | 0 | 0 | 1 |
| virus_Aug5.50v_22260 | 0 | 0 | 0 | 0 | 1 | 0 | 0 | 2 | 0 | 0 | 3 |
| virus_Aug5.50v_24607 | 0 | 0 | 0 | 0 | 0 | 0 | 0 | 1 | 0 | 0 | 1 |
| virus_Aug5.50v_38847 | 0 | 0 | 0 | 0 | 0 | 0 | 0 | 1 | 0 | 0 | 1 |
| virus_Aug5.75v_739 | 0 | 0 | 0 | 0 | 0 | 0 | 0 | 0 | 0 | 1 | 1 |
| virus_Aug5.75v_13162 | 0 | 0 | 0 | 0 | 0 | 0 | 0 | 0 | 0 | 1 | 1 |
| virus_Aug5.75v_13845 | 0 | 0 | 0 | 0 | 0 | 0 | 0 | 0 | 0 | 1 | 1 |
| virus_Aug5.75v_21894 | 0 | 0 | 0 | 0 | 0 | 0 | 0 | 0 | 0 | 1 | 1 |
| virus_Aug5.75v_39888 | 0 | 0 | 0 | 0 | 0 | 0 | 0 | 0 | 0 | 1 | 1 |
| virus_Aug5.75v_41101 | 0 | 0 | 0 | 0 | 0 | 0 | 0 | 0 | 0 | 1 | 1 |
| virus_Jul11.0v_5281 | 0 | 0 | 0 | 0 | 0 | 0 | 0 | 0 | 0 | 0 | 0 |
| virus_Jul11.0v_14736 | 0 | 0 | 0 | 0 | 0 | 0 | 0 | 0 | 0 | 0 | 0 |
| virus_Jul11.0v_14923 | 0 | 0 | 0 | 0 | 0 | 0 | 0 | 0 | 0 | 0 | 0 |
| virus_Jul11.0v_17430 | 0 | 0 | 0 | 0 | 0 | 0 | 0 | 0 | 0 | 0 | 0 |
| virus_Jul11.0v_19171 | 0 | 0 | 0 | 0 | 0 | 0 | 0 | 0 | 0 | 0 | 0 |
| virus_Jul11.5v_1317 | 0 | 0 | 0 | 0 | 0 | 0 | 0 | 0 | 0 | 0 | 0 |
| virus_Jul11.5v_1898 | 0 | 0 | 0 | 0 | 0 | 0 | 0 | 0 | 0 | 0 | 0 |
| virus_Jul11.5v_2359 | 0 | 0 | 0 | 0 | 0 | 0 | 0 | 0 | 0 | 0 | 0 |
| virus_Jul11.5v_3203 | 0 | 0 | 0 | 0 | 0 | 0 | 0 | 0 | 0 | 0 | 0 |
| virus_Jul11.5v_4175 | 0 | 0 | 0 | 0 | 0 | 0 | 0 | 0 | 0 | 0 | 0 |
| virus_Jul11.5v_4210 | 0 | 0 | 0 | 0 | 0 | 0 | 0 | 0 | 0 | 0 | 0 |
| virus_Jul11.5v_5239 | 0 | 0 | 0 | 0 | 0 | 1 | 0 | 0 | 0 | 1 | 2 |
| virus_Jul11.5v_6190 | 0 | 1 | 0 | 0 | 0 | 0 | 0 | 0 | 0 | 0 | 1 |
| virus_Jul11.5v_8752 | 0 | 2 | 0 | 0 | 2 | 0 | 0 | 1 | 0 | 0 | 5 |
| virus_Jul11.5v_9168 | 0 | 0 | 0 | 0 | 0 | 0 | 0 | 0 | 0 | 0 | 0 |
| virus_Jul11.5v_9652 | 0 | 0 | 0 | 0 | 0 | 0 | 0 | 0 | 0 | 0 | 0 |
| virus_Jul11.5v_9989 | 0 | 0 | 0 | 0 | 0 | 0 | 0 | 0 | 0 | 0 | 0 |
| virus_Jul11.5v_11436 | 0 | 0 | 0 | 0 | 0 | 0 | 0 | 0 | 0 | 0 | 0 |
| virus_Jul11.5v_11629 | 0 | 0 | 0 | 0 | 0 | 0 | 0 | 0 | 0 | 0 | 0 |
| virus_Jul11.5v_12625 | 0 | 0 | 0 | 0 | 0 | 0 | 0 | 0 | 0 | 0 | 0 |
| virus_Jul11.5v_15222 | 0 | 0 | 0 | 0 | 0 | 0 | 0 | 0 | 0 | 0 | 0 |
| virus_Jul11.5v_17996 | 0 | 0 | 0 | 0 | 0 | 0 | 0 | 0 | 0 | 0 | 0 |
| virus_Jul11.5v_19242 | 0 | 0 | 0 | 0 | 0 | 0 | 0 | 0 | 0 | 0 | 0 |
| virus_Jul11.5v_20935 | 0 | 0 | 0 | 0 | 0 | 0 | 0 | 0 | 0 | 0 | 0 |
| virus_Jul11.25v_976 | 0 | 0 | 0 | 0 | 0 | 0 | 0 | 0 | 0 | 0 | 0 |
| virus_Jul11.25v_1782 | 0 | 0 | 0 | 0 | 0 | 0 | 0 | 0 | 0 | 0 | 0 |
| virus_Jul11.25v_3423 | 0 | 0 | 0 | 0 | 0 | 1 | 0 | 0 | 0 | 0 | 1 |
| virus_Jul11.25v_4514 | 0 | 0 | 0 | 0 | 1 | 0 | 0 | 0 | 0 | 0 | 1 |
| virus_Jul11.25v_4883 | 0 | 0 | 0 | 0 | 0 | 0 | 0 | 0 | 0 | 0 | 0 |
| virus_Jul11.25v_8380 | 0 | 0 | 0 | 0 | 0 | 0 | 0 | 0 | 0 | 0 | 0 |
| virus_Jul11.25v_10088 | 0 | 0 | 0 | 0 | 0 | 0 | 0 | 0 | 0 | 0 | 0 |
| virus_Jul11.25v_11920 | 0 | 0 | 0 | 0 | 0 | 0 | 0 | 0 | 0 | 0 | 0 |
| virus_Jul11.25v_14119 | 0 | 0 | 0 | 0 | 0 | 0 | 0 | 0 | 0 | 0 | 0 |
| virus_Jul11.25v_16470 | 0 | 0 | 0 | 0 | 0 | 0 | 0 | 0 | 0 | 0 | 0 |
| virus_Jul11.25v_19522 | 0 | 0 | 0 | 0 | 0 | 0 | 0 | 0 | 0 | 0 | 0 |
| virus_Jul11.25v_20568 | 0 | 0 | 0 | 0 | 0 | 0 | 0 | 0 | 0 | 0 | 0 |
| virus_Jul11.50v_1396 | 0 | 0 | 0 | 0 | 0 | 0 | 0 | 0 | 0 | 0 | 0 |
| virus_Jul11.50v_1579 | 0 | 0 | 0 | 0 | 0 | 0 | 0 | 0 | 0 | 0 | 0 |
| virus_Jul11.50v_2116 | 0 | 0 | 0 | 0 | 0 | 0 | 0 | 0 | 0 | 0 | 0 |
| virus_Jul11.50v_6507 | 0 | 0 | 0 | 0 | 0 | 0 | 0 | 0 | 0 | 0 | 0 |
| virus_Jul11.50v_9863 | 0 | 0 | 0 | 0 | 0 | 0 | 0 | 0 | 0 | 0 | 0 |
| virus_Jul11.50v_12178 | 1 | 0 | 0 | 0 | 0 | 0 | 0 | 0 | 0 | 0 | 1 |
| virus_Jul11.50v_12927 | 0 | 0 | 0 | 0 | 0 | 0 | 0 | 0 | 0 | 0 | 0 |
| virus_Jul11.50v_13993 | 0 | 0 | 0 | 0 | 0 | 0 | 0 | 0 | 0 | 0 | 0 |
| virus_Jul11.50v_14675 | 0 | 0 | 0 | 0 | 0 | 0 | 0 | 0 | 0 | 0 | 0 |
| virus_Jul11.50v_15423 | 0 | 0 | 0 | 0 | 0 | 0 | 0 | 0 | 0 | 0 | 0 |
| virus_Jul11.50v_17503 | 0 | 0 | 0 | 0 | 0 | 0 | 0 | 0 | 0 | 0 | 0 |
| virus_Jul11.50v_17530 | 0 | 0 | 0 | 0 | 0 | 0 | 0 | 0 | 0 | 0 | 0 |
| virus_Jul11.50v_17700 | 0 | 0 | 0 | 0 | 0 | 0 | 0 | 0 | 0 | 0 | 0 |
| virus_Jul11.50v_18986 | 0 | 0 | 0 | 0 | 0 | 0 | 0 | 0 | 0 | 0 | 0 |

| | | | | | | | | | | | |
|---|---|---|---|---|---|---|---|---|---|---|---|
| virus_Jul11.75v_6438 | 0 | 0 | 0 | 0 | 0 | 0 | 0 | 0 | 0 | 0 | 0 |
| virus_Jul11.75v_8152 | 0 | 0 | 0 | 0 | 0 | 0 | 0 | 0 | 0 | 0 | 0 |
| virus_Jul11.75v_10703 | 0 | 0 | 0 | 0 | 0 | 0 | 0 | 0 | 0 | 0 | 0 |
| virus_Jul11.75v_12927 | 0 | 0 | 0 | 0 | 0 | 0 | 0 | 0 | 0 | 0 | 0 |
| virus_Jul11.75v_23390 | 0 | 0 | 0 | 0 | 0 | 0 | 0 | 0 | 0 | 0 | 0 |
| virus_Jul11.75v_28208 | 0 | 0 | 0 | 0 | 0 | 0 | 0 | 0 | 0 | 0 | 0 |
| virus_Jul11.75v_36830 | 0 | 0 | 0 | 0 | 0 | 0 | 0 | 0 | 0 | 0 | 0 |
| virus_Jul11.75v_38785 | 0 | 0 | 0 | 0 | 0 | 0 | 0 | 0 | 0 | 0 | 0 |
| virus_Jul11.75v_38853 | 0 | 0 | 0 | 0 | 0 | 0 | 0 | 0 | 0 | 0 | 0 |
| virus_Jul27.5v_878 | 0 | 0 | 0 | 0 | 0 | 0 | 0 | 0 | 0 | 0 | 0 |
| virus_Jul27.5v_942 | 0 | 0 | 0 | 0 | 0 | 0 | 0 | 0 | 0 | 0 | 0 |
| virus_Jul27.5v_2522 | 0 | 0 | 0 | 0 | 0 | 0 | 0 | 0 | 0 | 0 | 0 |
| virus_Jul27.5v_7214 | 0 | 0 | 0 | 0 | 0 | 0 | 0 | 0 | 0 | 0 | 0 |
| virus_Jul27.5v_8478 | 0 | 0 | 0 | 0 | 1 | 0 | 0 | 0 | 0 | 0 | 1 |
| virus_Jul27.5v_8684 | 0 | 0 | 0 | 0 | 0 | 0 | 0 | 0 | 0 | 0 | 0 |
| virus_Jul27.5v_11959 | 0 | 0 | 0 | 0 | 0 | 0 | 0 | 0 | 0 | 0 | 0 |
| virus_Jul27.5v_12484 | 0 | 0 | 0 | 0 | 0 | 0 | 0 | 0 | 0 | 0 | 0 |
| virus_Jul27.5v_13978 | 0 | 0 | 0 | 0 | 0 | 0 | 0 | 1 | 0 | 0 | 1 |
| virus_Jul27.5v_14805 | 0 | 1 | 0 | 0 | 0 | 0 | 0 | 0 | 0 | 1 | 2 |
| virus_Jul27.5v_15103 | 0 | 0 | 0 | 0 | 0 | 0 | 0 | 0 | 0 | 0 | 0 |
| virus_Jul27.5v_15110 | 0 | 0 | 0 | 0 | 0 | 0 | 0 | 0 | 0 | 0 | 0 |
| virus_Jul27.5v_15470 | 0 | 0 | 0 | 0 | 0 | 0 | 0 | 0 | 0 | 0 | 0 |
| virus_Jul27.5v_17149 | 0 | 0 | 0 | 0 | 0 | 0 | 0 | 0 | 0 | 0 | 0 |
| virus_Jul27.5v_20206 | 0 | 0 | 0 | 0 | 0 | 0 | 0 | 0 | 0 | 0 | 0 |
| virus_Jul27.25v_1425 | 0 | 0 | 0 | 0 | 0 | 0 | 0 | 0 | 0 | 0 | 0 |
| virus_Jul27.25v_1689 | 0 | 0 | 0 | 0 | 0 | 0 | 0 | 0 | 0 | 0 | 0 |
| virus_Jul27.25v_2009 | 0 | 0 | 0 | 0 | 0 | 0 | 0 | 0 | 0 | 0 | 0 |
| virus_Jul27.25v_2238 | 0 | 1 | 0 | 0 | 0 | 0 | 0 | 0 | 0 | 0 | 1 |
| virus_Jul27.25v_5554 | 0 | 1 | 0 | 0 | 0 | 0 | 0 | 0 | 0 | 0 | 1 |
| virus_Jul27.25v_6038 | 0 | 0 | 0 | 0 | 0 | 0 | 0 | 0 | 0 | 0 | 0 |
| virus_Jul27.25v_11119 | 0 | 0 | 0 | 0 | 0 | 0 | 0 | 0 | 0 | 0 | 0 |
| virus_Jul27.25v_13759 | 0 | 0 | 0 | 0 | 0 | 0 | 0 | 0 | 0 | 0 | 0 |
| virus_Jul27.25v_20090 | 0 | 0 | 0 | 0 | 0 | 0 | 0 | 0 | 0 | 0 | 0 |
| virus_Jul27.25v_22399 | 0 | 0 | 0 | 0 | 0 | 0 | 0 | 0 | 0 | 0 | 0 |
| virus_Jul27.25v_29389 | 0 | 0 | 0 | 0 | 0 | 0 | 0 | 0 | 0 | 0 | 0 |
| virus_Jul27.25v_32053 | 0 | 0 | 0 | 0 | 0 | 0 | 0 | 0 | 0 | 0 | 0 |
| virus_Jul27.25v_38635 | 0 | 0 | 0 | 0 | 0 | 0 | 0 | 0 | 0 | 1 | 1 |
| virus_Jul27.25v_39795 | 0 | 0 | 0 | 0 | 0 | 0 | 0 | 0 | 0 | 0 | 0 |
| virus_Jul27.75v_56 | 0 | 0 | 0 | 0 | 0 | 0 | 0 | 0 | 0 | 0 | 0 |
| virus_Jul27.75v_735 | 0 | 0 | 0 | 0 | 0 | 0 | 0 | 0 | 0 | 0 | 0 |
| virus_Jul27.75v_975 | 0 | 0 | 0 | 0 | 0 | 0 | 0 | 0 | 0 | 0 | 0 |
| virus_Jul27.75v_3642 | 0 | 0 | 0 | 0 | 0 | 0 | 0 | 0 | 0 | 0 | 0 |
| virus_Jul27.75v_8888 | 0 | 0 | 0 | 0 | 0 | 0 | 0 | 0 | 0 | 0 | 0 |
| virus_Jul27.75v_9932 | 0 | 0 | 0 | 0 | 1 | 0 | 0 | 0 | 0 | 1 | 2 |
| virus_Jul27.75v_17542 | 0 | 0 | 0 | 0 | 0 | 0 | 0 | 0 | 0 | 0 | 0 |
| virus_Jun8.25v_10081 | 0 | 0 | 0 | 0 | 0 | 0 | 0 | 0 | 0 | 0 | 0 |
| virus_Jun8.25v_11433 | 0 | 0 | 0 | 0 | 0 | 0 | 0 | 0 | 0 | 0 | 0 |
| virus_Jun8.25v_35482 | 0 | 0 | 0 | 0 | 0 | 0 | 0 | 0 | 0 | 0 | 0 |
| virus_Jun8.50v_3611 | 0 | 0 | 0 | 0 | 0 | 0 | 0 | 0 | 0 | 0 | 0 |
| virus_Jun8.50v_3759 | 0 | 0 | 0 | 0 | 0 | 0 | 0 | 0 | 0 | 0 | 0 |
| virus_Jun8.50v_8834 | 0 | 0 | 0 | 0 | 0 | 0 | 0 | 0 | 0 | 0 | 0 |
| virus_Jun8.50v_12310 | 0 | 0 | 0 | 0 | 0 | 0 | 0 | 0 | 0 | 0 | 0 |
| virus_Jun8.50v_13238 | 0 | 0 | 0 | 0 | 0 | 0 | 0 | 0 | 0 | 0 | 0 |
| virus_Jun8.50v_17308 | 0 | 0 | 0 | 0 | 0 | 0 | 0 | 0 | 0 | 0 | 0 |
| virus_Jun8.50v_20040 | 0 | 0 | 0 | 0 | 0 | 0 | 0 | 0 | 0 | 0 | 0 |
| virus_Jun8.50v_22104 | 0 | 0 | 0 | 0 | 0 | 0 | 0 | 0 | 0 | 0 | 0 |
| virus_Jun8.50v_22877 | 0 | 0 | 0 | 0 | 0 | 0 | 0 | 0 | 0 | 0 | 0 |
| virus_Jun8.50v_25735 | 0 | 0 | 0 | 0 | 0 | 0 | 0 | 0 | 0 | 0 | 0 |
| virus_Jun8.50v_27002 | 0 | 0 | 0 | 0 | 0 | 1 | 0 | 0 | 0 | 2 | 3 |
| virus_Jun8.50v_34579 | 0 | 0 | 0 | 0 | 0 | 0 | 0 | 0 | 0 | 0 | 0 |
| virus_Jun8.50v_35343 | 0 | 0 | 0 | 0 | 0 | 0 | 0 | 0 | 0 | 0 | 0 |
| virus_Jun8.50v_40458 | 0 | 0 | 0 | 0 | 0 | 0 | 0 | 0 | 0 | 0 | 0 |
| virus_Jun8.75v_6822 | 0 | 1 | 0 | 0 | 0 | 0 | 0 | 0 | 0 | 0 | 1 |
| virus_Jun8.75v_12607 | 0 | 0 | 0 | 0 | 0 | 0 | 0 | 0 | 0 | 0 | 0 |
| virus_Jun8.75v_19896 | 0 | 0 | 0 | 0 | 0 | 0 | 0 | 1 | 0 | 0 | 1 |
| virus_Jun8.75v_20644 | 0 | 0 | 0 | 0 | 0 | 0 | 0 | 0 | 0 | 0 | 0 |
| virus_Jun22.0v_7656 | 0 | 0 | 0 | 0 | 0 | 0 | 0 | 0 | 0 | 0 | 0 |
| virus_Jun22.0v_9662 | 0 | 0 | 0 | 0 | 0 | 0 | 0 | 0 | 0 | 0 | 0 |
| virus_Jun22.0v_10421 | 0 | 0 | 0 | 0 | 0 | 0 | 0 | 0 | 0 | 0 | 0 |
| virus_Jun22.0v_11636 | 0 | 0 | 0 | 0 | 0 | 0 | 0 | 0 | 0 | 0 | 0 |
| virus_Jun22.0v_12525 | 0 | 0 | 0 | 0 | 0 | 0 | 0 | 0 | 0 | 0 | 0 |
| virus_Jun22.0v_12960 | 0 | 0 | 0 | 0 | 0 | 0 | 0 | 0 | 0 | 0 | 0 |
| virus_Jun22.0v_13049 | 0 | 0 | 0 | 0 | 0 | 0 | 0 | 0 | 0 | 0 | 0 |
| virus_Jun22.0v_13574 | 0 | 0 | 0 | 0 | 0 | 0 | 0 | 0 | 0 | 0 | 0 |
| virus_Jun22.0v_20743 | 0 | 0 | 0 | 0 | 0 | 0 | 0 | 0 | 0 | 0 | 0 |
| virus_Jun22.0v_20826 | 0 | 0 | 0 | 0 | 0 | 0 | 0 | 0 | 0 | 0 | 0 |
| virus_Jun22.50v_394 | 0 | 0 | 0 | 0 | 0 | 0 | 0 | 0 | 0 | 0 | 0 |
| virus_Jun22.50v_2786 | 0 | 0 | 0 | 0 | 0 | 0 | 0 | 0 | 0 | 0 | 0 |
| virus_Jun22.50v_3075 | 0 | 0 | 0 | 0 | 0 | 0 | 0 | 0 | 0 | 0 | 0 |
| virus_Jun22.50v_3567 | 0 | 0 | 0 | 0 | 0 | 0 | 0 | 0 | 0 | 0 | 0 |
| virus_Jun22.50v_3841 | 0 | 0 | 0 | 0 | 0 | 0 | 0 | 0 | 0 | 0 | 0 |
| virus_Jun22.50v_6918 | 0 | 0 | 0 | 0 | 0 | 0 | 0 | 0 | 0 | 0 | 0 |
| virus_Jun22.50v_9386 | 0 | 0 | 0 | 0 | 0 | 0 | 0 | 0 | 0 | 0 | 0 |
| virus_Jun22.50v_10904 | 0 | 0 | 0 | 0 | 0 | 0 | 0 | 0 | 0 | 0 | 0 |
| virus_Jun22.50v_11727 | 0 | 0 | 0 | 0 | 0 | 0 | 0 | 0 | 0 | 0 | 0 |
| virus_Jun22.50v_16876 | 0 | 0 | 0 | 0 | 0 | 0 | 0 | 0 | 0 | 0 | 0 |
| virus_Jun22.50v_17698 | 0 | 0 | 0 | 0 | 0 | 0 | 0 | 0 | 0 | 0 | 0 |
| virus_Jun22.50v_18819 | 0 | 0 | 0 | 0 | 0 | 0 | 0 | 0 | 0 | 0 | 0 |
| virus_Jun22.50v_23542 | 0 | 0 | 0 | 0 | 0 | 0 | 0 | 0 | 0 | 0 | 0 |
| virus_Jun22.50v_27048 | 0 | 0 | 0 | 0 | 0 | 0 | 0 | 0 | 0 | 0 | 0 |
| virus_Jun22.50v_31269 | 0 | 0 | 0 | 0 | 0 | 1 | 0 | 0 | 0 | 0 | 1 |
| virus_Jun22.50v_31296 | 0 | 0 | 0 | 0 | 0 | 0 | 0 | 0 | 0 | 0 | 0 |
| virus_Jun22.50v_33174 | 0 | 0 | 0 | 0 | 0 | 0 | 0 | 0 | 0 | 0 | 0 |
| virus_Jun22.50v_34549 | 0 | 0 | 0 | 0 | 0 | 0 | 0 | 0 | 0 | 0 | 0 |
| virus_Jun22.50v_39821 | 0 | 0 | 0 | 0 | 0 | 0 | 0 | 0 | 0 | 0 | 0 |
| virus_Jun22.75v_731 | 0 | 0 | 0 | 0 | 0 | 0 | 0 | 0 | 0 | 0 | 0 |
| virus_Jun22.75v_3480 | 0 | 0 | 0 | 0 | 0 | 0 | 0 | 0 | 0 | 0 | 0 |
| virus_Jun22.75v_5791 | 0 | 0 | 0 | 0 | 0 | 0 | 0 | 0 | 0 | 0 | 0 |
| virus_Jun22.75v_7618 | 0 | 0 | 0 | 0 | 0 | 0 | 0 | 0 | 0 | 0 | 0 |
| virus_Jun22.75v_14158 | 0 | 0 | 0 | 0 | 0 | 0 | 0 | 0 | 0 | 0 | 0 |
| virus_Jun22.75v_24388 | 0 | 0 | 0 | 0 | 0 | 0 | 0 | 0 | 0 | 0 | 0 |
| virus_Jun22.75v_24468 | 0 | 0 | 0 | 0 | 0 | 0 | 0 | 0 | 0 | 0 | 0 |
| virus_Jun22.75v_25721 | 0 | 0 | 0 | 0 | 0 | 0 | 0 | 0 | 0 | 0 | 0 |
| virus_Jun22.75v_27590 | 0 | 0 | 0 | 0 | 0 | 0 | 0 | 0 | 0 | 0 | 0 |
| virus_Jun22.75v_35587 | 0 | 0 | 0 | 0 | 0 | 0 | 0 | 0 | 0 | 0 | 0 |
| virus_Jun22.75v_39233 | 0 | 0 | 0 | 0 | 0 | 0 | 0 | 0 | 0 | 0 | 0 |
| virus_May12.5v_10392 | 0 | 0 | 0 | 0 | 0 | 0 | 0 | 0 | 0 | 0 | 0 |
| virus_May12.5v_16119 | 0 | 0 | 0 | 0 | 0 | 0 | 0 | 0 | 0 | 0 | 0 |
| virus_May12.5v_30925 | 0 | 0 | 0 | 0 | 0 | 0 | 0 | 0 | 0 | 0 | 0 |
| virus_May12.5v_38399 | 0 | 0 | 0 | 0 | 0 | 0 | 0 | 0 | 0 | 0 | 0 |
| virus_May12.5v_38602 | 0 | 0 | 0 | 0 | 0 | 0 | 0 | 0 | 0 | 0 | 0 |
| virus_May12.5v_39722 | 0 | 0 | 0 | 0 | 0 | 0 | 0 | 0 | 0 | 0 | 0 |
| virus_May12.25v_1198 | 0 | 0 | 0 | 0 | 0 | 0 | 0 | 0 | 0 | 0 | 0 |
| virus_May12.25v_1851 | 0 | 0 | 0 | 0 | 0 | 0 | 0 | 0 | 0 | 0 | 0 |
| virus_May12.25v_2853 | 0 | 0 | 0 | 0 | 0 | 0 | 0 | 0 | 0 | 0 | 0 |
| virus_May12.25v_6517 | 0 | 0 | 0 | 0 | 0 | 0 | 0 | 0 | 0 | 0 | 0 |
| virus_May12.25v_6591 | 0 | 0 | 0 | 0 | 0 | 0 | 0 | 0 | 0 | 0 | 0 |
| virus_May12.25v_13968 | 0 | 0 | 0 | 0 | 0 | 0 | 0 | 0 | 0 | 0 | 0 |
| virus_May12.25v_21905 | 0 | 0 | 0 | 0 | 0 | 0 | 0 | 0 | 0 | 0 | 0 |
| virus_May12.25v_24101 | 0 | 0 | 0 | 0 | 0 | 0 | 0 | 0 | 0 | 0 | 0 |
| virus_May12.25v_26384 | 0 | 0 | 0 | 0 | 0 | 0 | 0 | 0 | 0 | 0 | 0 |
| virus_May12.25v_33564 | 0 | 0 | 0 | 0 | 0 | 0 | 0 | 0 | 0 | 0 | 0 |
| virus_May12.25v_34745 | 0 | 0 | 0 | 0 | 0 | 0 | 0 | 0 | 0 | 0 | 0 |
| virus_May12.50v_254 | 0 | 0 | 0 | 0 | 0 | 0 | 0 | 0 | 0 | 0 | 0 |
| virus_May12.50v_2606 | 0 | 0 | 0 | 0 | 0 | 0 | 0 | 0 | 0 | 0 | 0 |
| virus_May12.50v_5052 | 0 | 0 | 0 | 0 | 0 | 0 | 0 | 0 | 0 | 0 | 0 |
| virus_May12.50v_5666 | 0 | 0 | 0 | 0 | 0 | 0 | 0 | 0 | 0 | 0 | 0 |
| virus_May12.50v_6376 | 0 | 0 | 0 | 0 | 0 | 0 | 0 | 0 | 0 | 0 | 0 |
| virus_May12.50v_13126 | 0 | 0 | 0 | 0 | 0 | 0 | 0 | 0 | 0 | 0 | 0 |
| virus_May12.50v_14452 | 0 | 0 | 0 | 0 | 0 | 0 | 0 | 0 | 0 | 0 | 0 |
| virus_May12.50v_21298 | 0 | 0 | 0 | 0 | 0 | 0 | 0 | 0 | 0 | 0 | 0 |
| virus_May12.50v_27028 | 0 | 0 | 0 | 0 | 0 | 0 | 0 | 0 | 0 | 0 | 0 |
| virus_May12.50v_29396 | 0 | 0 | 0 | 0 | 0 | 0 | 0 | 0 | 0 | 0 | 0 |
| virus_May12.50v_29767 | 0 | 0 | 0 | 0 | 0 | 0 | 0 | 0 | 0 | 0 | 0 |
| virus_May12.50v_31451 | 0 | 0 | 0 | 0 | 0 | 0 | 0 | 0 | 0 | 0 | 0 |
| virus_May12.50v_32115 | 0 | 0 | 0 | 0 | 0 | 0 | 0 | 0 | 0 | 0 | 0 |
| virus_May12.50v_33445 | 0 | 0 | 0 | 0 | 0 | 0 | 0 | 0 | 0 | 0 | 0 |
| virus_May12.50v_35098 | 0 | 0 | 0 | 0 | 0 | 0 | 0 | 0 | 0 | 0 | 0 |
| virus_May12.50v_35971 | 0 | 0 | 0 | 0 | 0 | 0 | 0 | 0 | 0 | 0 | 0 |
| virus_May12.50v_38831 | 0 | 1 | 0 | 0 | 1 | 0 | 1 | 1 | 0 | 0 | 4 |
| virus_May12.50v_39645 | 0 | 0 | 0 | 0 | 0 | 0 | 0 | 0 | 0 | 0 | 0 |
| virus_May12.50v_40170 | 0 | 0 | 0 | 0 | 0 | 0 | 0 | 0 | 0 | 0 | 0 |
| virus_May12.50v_40192 | 0 | 0 | 0 | 0 | 0 | 0 | 0 | 0 | 0 | 0 | 0 |
| virus_May12.75v_113 | 0 | 0 | 0 | 0 | 0 | 0 | 0 | 0 | 0 | 0 | 0 |
| virus_May12.75v_6261 | 0 | 0 | 0 | 0 | 0 | 0 | 0 | 0 | 0 | 0 | 0 |
| virus_May12.75v_9003 | 0 | 0 | 0 | 0 | 0 | 0 | 0 | 0 | 0 | 0 | 0 |
| virus_May12.75v_15769 | 0 | 0 | 0 | 0 | 0 | 0 | 0 | 0 | 0 | 0 | 0 |
| virus_May12.75v_16261 | 0 | 0 | 0 | 0 | 0 | 0 | 0 | 0 | 0 | 0 | 0 |
| virus_May12.75v_16954 | 0 | 0 | 0 | 0 | 0 | 0 | 0 | 0 | 0 | 0 | 0 |
| virus_May12.75v_20408 | 0 | 0 | 0 | 0 | 0 | 0 | 0 | 0 | 0 | 0 | 0 |
| virus_May12.75v_20647 | 0 | 0 | 0 | 0 | 0 | 0 | 0 | 0 | 0 | 0 | 0 |
| virus_May12.75v_20654 | 0 | 0 | 0 | 0 | 0 | 0 | 0 | 0 | 0 | 0 | 0 |
| virus_May12.75v_22749 | 0 | 0 | 0 | 0 | 0 | 0 | 0 | 0 | 0 | 0 | 0 |
| virus_May12.75v_29897 | 0 | 0 | 0 | 0 | 0 | 0 | 0 | 0 | 0 | 0 | 0 |
| virus_May12.75v_31871 | 0 | 0 | 0 | 0 | 0 | 0 | 0 | 0 | 0 | 0 | 0 |
| virus_May12.75v_37898 | 0 | 0 | 0 | 0 | 0 | 0 | 0 | 0 | 0 | 0 | 0 |
| virus_Sep8.0v_3844 | 0 | 0 | 0 | 0 | 0 | 1 | 0 | 2 | 0 | 0 | 3 |

| | | | | | | | | | | | |
|---|---|---|---|---|---|---|---|---|---|---|---|
| virus_Sep8.0v_3905 | 0 | 0 | 0 | 0 | 0 | 0 | 0 | 0 | 0 | 0 | 0 |
| virus_Sep8.0v_6064 | 0 | 0 | 0 | 0 | 0 | 0 | 0 | 0 | 0 | 0 | 0 |
| virus_Sep8.0v_6839 | 0 | 0 | 0 | 0 | 0 | 0 | 0 | 0 | 0 | 0 | 0 |
| virus_Sep8.0v_9317 | 0 | 0 | 0 | 0 | 0 | 0 | 0 | 0 | 0 | 0 | 0 |
| virus_Sep8.0v_10243 | 0 | 1 | 0 | 2 | 0 | 0 | 0 | 0 | 0 | 0 | 3 |
| virus_Sep8.0v_11663 | 0 | 0 | 0 | 0 | 0 | 0 | 0 | 0 | 0 | 0 | 0 |
| virus_Sep8.0v_13962 | 0 | 0 | 0 | 0 | 0 | 0 | 0 | 0 | 0 | 0 | 0 |
| virus_Sep8.0v_14253 | 0 | 0 | 0 | 0 | 0 | 0 | 0 | 0 | 0 | 0 | 0 |
| virus_Sep8.0v_15334 | 0 | 0 | 0 | 0 | 0 | 0 | 0 | 0 | 0 | 0 | 0 |
| virus_Sep8.0v_16261 | 0 | 0 | 0 | 0 | 0 | 0 | 0 | 0 | 0 | 0 | 0 |
| virus_Sep8.0v_17426 | 0 | 0 | 0 | 0 | 0 | 0 | 0 | 0 | 0 | 0 | 0 |
| virus_Sep8.0v_18398 | 1 | 1 | 0 | 0 | 0 | 1 | 0 | 0 | 0 | 1 | 4 |
| virus_Sep8.0v_18634 | 0 | 0 | 0 | 0 | 0 | 0 | 0 | 0 | 0 | 0 | 0 |
| virus_Sep8.0v_20158 | 0 | 0 | 0 | 0 | 0 | 0 | 0 | 0 | 0 | 0 | 0 |
| virus_Sep8.5v_7676 | 0 | 0 | 0 | 0 | 0 | 0 | 0 | 0 | 0 | 0 | 0 |
| virus_Sep8.5v_15643 | 0 | 0 | 0 | 0 | 0 | 0 | 0 | 0 | 0 | 0 | 0 |
| virus_Sep8.25v_4126 | 0 | 0 | 0 | 0 | 0 | 0 | 0 | 0 | 0 | 0 | 0 |
| virus_Sep8.25v_13893 | 0 | 0 | 0 | 0 | 0 | 0 | 0 | 0 | 0 | 0 | 0 |
| virus_Sep8.25v_16511 | 0 | 0 | 0 | 0 | 0 | 0 | 0 | 0 | 0 | 0 | 0 |
| virus_Sep8.25v_20096 | 0 | 0 | 0 | 0 | 0 | 0 | 0 | 0 | 0 | 0 | 0 |
| virus_Sep8.50v_1340 | 0 | 0 | 0 | 0 | 0 | 0 | 0 | 0 | 0 | 0 | 0 |
| virus_Sep8.50v_3956 | 0 | 0 | 0 | 0 | 0 | 0 | 0 | 0 | 0 | 0 | 0 |
| virus_Sep8.50v_7825 | 0 | 0 | 0 | 0 | 0 | 0 | 0 | 0 | 0 | 0 | 0 |
| virus_Sep8.50v_13124 | 0 | 0 | 0 | 0 | 0 | 0 | 0 | 0 | 0 | 0 | 0 |
| virus_Sep8.50v_15806 | 0 | 0 | 0 | 0 | 0 | 0 | 0 | 0 | 0 | 0 | 0 |
| virus_Sep8.50v_16868 | 0 | 0 | 0 | 0 | 0 | 0 | 0 | 0 | 0 | 0 | 0 |
| virus_Sep8.50v_20345 | 0 | 0 | 0 | 0 | 0 | 0 | 0 | 0 | 0 | 0 | 0 |
| virus_Sep8.50v_27827 | 0 | 0 | 0 | 0 | 0 | 0 | 0 | 0 | 0 | 0 | 0 |
| virus_Sep8.50v_30163 | 0 | 0 | 0 | 0 | 0 | 0 | 0 | 0 | 0 | 0 | 0 |
| virus_Sep8.50v_33206 | 0 | 0 | 0 | 0 | 0 | 0 | 0 | 0 | 0 | 0 | 0 |
| virus_Sep8.50v_36820 | 0 | 0 | 0 | 0 | 0 | 0 | 0 | 0 | 0 | 0 | 0 |
| virus_Sep8.50v_37938 | 0 | 0 | 0 | 0 | 0 | 0 | 0 | 0 | 0 | 0 | 0 |
| virus_Sep8.50v_39605 | 0 | 0 | 0 | 0 | 0 | 0 | 0 | 0 | 0 | 0 | 0 |
| bacteria_Jul11d0b_5361 | 0 | 0 | 0 | 0 | 0 | 0 | 0 | 0 | 0 | 0 | 0 |
| bacteria_Jul11d0b_30174 | 0 | 0 | 0 | 0 | 0 | 0 | 0 | 0 | 0 | 0 | 0 |
| bacteria_Jul11d0b_37224 | 0 | 0 | 0 | 0 | 0 | 0 | 0 | 0 | 0 | 0 | 0 |
| bacteria_Jul11d0b_38022 | 0 | 0 | 0 | 0 | 0 | 0 | 0 | 0 | 0 | 0 | 0 |
| bacteria_Jul11d0b_43588 | 0 | 0 | 0 | 0 | 0 | 0 | 0 | 0 | 0 | 0 | 0 |
| bacteria_Aug5d0b_16335 | 4 | 0 | 0 | 4 | 0 | 0 | 0 | 0 | 0 | 0 | 8 |
| bacteria_Aug5d0b_18242 | 1 | 0 | 0 | 0 | 0 | 0 | 0 | 0 | 0 | 0 | 1 |
| bacteria_Aug5d5b_43791 | 0 | 0 | 1 | 0 | 0 | 0 | 0 | 0 | 0 | 0 | 1 |
| bacteria_Aug5d5b_65398 | 0 | 0 | 1 | 0 | 0 | 0 | 0 | 0 | 0 | 0 | 1 |
| bacteria_Aug5d5b_67989 | 0 | 3 | 1 | 0 | 0 | 0 | 0 | 0 | 0 | 0 | 4 |
| bacteria_Aug5d25b_73 | 0 | 0 | 0 | 0 | 1 | 0 | 0 | 0 | 0 | 0 | 1 |
| bacteria_Aug5d25b_3701 | 0 | 0 | 0 | 0 | 1 | 0 | 0 | 0 | 0 | 0 | 1 |
| bacteria_Aug5d25b_7352 | 0 | 0 | 0 | 0 | 1 | 0 | 0 | 0 | 0 | 0 | 1 |
| bacteria_Aug5d25b_11334 | 0 | 0 | 0 | 0 | 1 | 0 | 0 | 0 | 0 | 0 | 1 |
| bacteria_Aug5d25b_15856 | 0 | 0 | 0 | 0 | 1 | 0 | 0 | 1 | 0 | 0 | 2 |
| bacteria_Aug5d25b_18616 | 0 | 0 | 0 | 0 | 1 | 0 | 0 | 0 | 0 | 0 | 1 |
| bacteria_Aug5d25b_21156 | 0 | 0 | 0 | 0 | 1 | 0 | 0 | 0 | 0 | 0 | 1 |
| bacteria_Aug5d25b_21262 | 0 | 0 | 0 | 0 | 1 | 0 | 0 | 0 | 0 | 0 | 1 |
| bacteria_Aug5d25b_23361 | 0 | 0 | 0 | 0 | 1 | 0 | 0 | 0 | 0 | 0 | 1 |
| bacteria_Aug5d25b_33607 | 0 | 0 | 0 | 0 | 1 | 0 | 0 | 0 | 0 | 0 | 1 |
| bacteria_Aug5d25b_33619 | 0 | 0 | 0 | 0 | 1 | 0 | 0 | 0 | 0 | 0 | 1 |
| bacteria_Aug5d25b_36648 | 0 | 0 | 0 | 0 | 1 | 0 | 0 | 0 | 0 | 0 | 1 |
| bacteria_Aug5d25b_37158 | 0 | 0 | 0 | 0 | 1 | 0 | 0 | 0 | 0 | 0 | 1 |
| bacteria_Aug5d25b_39375 | 0 | 0 | 0 | 0 | 1 | 0 | 0 | 0 | 0 | 0 | 1 |
| bacteria_Aug5d25b_44406 | 0 | 0 | 0 | 1 | 1 | 0 | 0 | 0 | 0 | 0 | 2 |
| bacteria_Aug5d25b_46040 | 0 | 0 | 0 | 0 | 1 | 0 | 0 | 0 | 0 | 0 | 1 |
| bacteria_Aug5d25b_50005 | 0 | 0 | 0 | 0 | 1 | 0 | 0 | 0 | 0 | 0 | 1 |
| bacteria_Aug5d25b_51012 | 0 | 0 | 0 | 0 | 1 | 0 | 0 | 0 | 0 | 0 | 1 |
| bacteria_Aug5d25b_67053 | 0 | 0 | 0 | 0 | 1 | 0 | 0 | 0 | 0 | 0 | 1 |
| bacteria_Aug5d25b_67151 | 0 | 0 | 0 | 0 | 1 | 0 | 0 | 0 | 0 | 0 | 1 |
| bacteria_Aug5d25b_68097 | 0 | 0 | 0 | 0 | 1 | 0 | 0 | 0 | 0 | 0 | 1 |
| bacteria_Aug5d50b_2721 | 0 | 0 | 0 | 0 | 0 | 0 | 1 | 0 | 0 | 0 | 1 |
| bacteria_Aug5d50b_3371 | 0 | 0 | 0 | 0 | 0 | 0 | 1 | 0 | 0 | 0 | 1 |
| bacteria_Aug5d50b_14373 | 0 | 0 | 0 | 0 | 0 | 0 | 1 | 0 | 0 | 0 | 1 |
| bacteria_Aug5d50b_19026 | 0 | 0 | 0 | 0 | 0 | 0 | 1 | 0 | 0 | 0 | 1 |
| bacteria_Aug5d50b_27643 | 0 | 0 | 0 | 0 | 0 | 0 | 1 | 0 | 0 | 0 | 1 |
| bacteria_Aug5d50b_52726 | 0 | 0 | 0 | 0 | 0 | 0 | 1 | 0 | 0 | 0 | 1 |
| bacteria_Aug5d50b_57824 | 0 | 0 | 0 | 0 | 0 | 0 | 1 | 0 | 0 | 0 | 1 |
| bacteria_Aug5d50b_70364 | 0 | 0 | 0 | 0 | 0 | 0 | 1 | 0 | 0 | 0 | 1 |
| bacteria_Jul11d5b_1349 | 0 | 0 | 0 | 0 | 0 | 0 | 0 | 0 | 0 | 0 | 0 |
| bacteria_Jul11d5b_23323 | 0 | 0 | 0 | 0 | 0 | 0 | 0 | 0 | 0 | 0 | 0 |
| bacteria_Jul11d5b_23680 | 0 | 0 | 0 | 0 | 0 | 0 | 0 | 0 | 0 | 0 | 0 |
| bacteria_Jul11d5b_27296 | 0 | 0 | 0 | 0 | 0 | 0 | 0 | 0 | 0 | 0 | 0 |
| bacteria_Jul11d5b_38272 | 0 | 0 | 0 | 0 | 0 | 0 | 0 | 0 | 0 | 0 | 0 |
| bacteria_Jul11d5b_45099 | 0 | 0 | 0 | 0 | 0 | 0 | 0 | 0 | 0 | 0 | 0 |
| bacteria_Jul11d25b_4728 | 0 | 0 | 0 | 0 | 0 | 1 | 0 | 0 | 0 | 0 | 1 |
| bacteria_Jul11d25b_10267 | 0 | 0 | 0 | 0 | 0 | 0 | 0 | 0 | 0 | 0 | 0 |
| bacteria_Jul11d25b_14302 | 0 | 0 | 0 | 0 | 0 | 0 | 0 | 0 | 0 | 0 | 0 |
| bacteria_Jul11d25b_19241 | 0 | 0 | 0 | 0 | 0 | 0 | 0 | 0 | 0 | 0 | 0 |
| bacteria_Jul11d25b_35754 | 0 | 0 | 0 | 0 | 0 | 0 | 0 | 0 | 0 | 0 | 0 |
| bacteria_Jul11d25b_41280 | 0 | 0 | 0 | 0 | 0 | 0 | 0 | 0 | 0 | 0 | 0 |
| bacteria_Jul11d25b_44295 | 0 | 1 | 0 | 0 | 0 | 0 | 0 | 0 | 0 | 0 | 1 |
| bacteria_Jul11d25b_50924 | 0 | 0 | 0 | 0 | 0 | 0 | 0 | 0 | 0 | 0 | 0 |
| bacteria_Jul11d25b_52368 | 0 | 1 | 0 | 1 | 0 | 0 | 0 | 0 | 0 | 0 | 2 |
| bacteria_Jul11d50b_10673 | 0 | 0 | 0 | 0 | 0 | 0 | 0 | 0 | 0 | 0 | 0 |
| bacteria_Jul11d50b_29403 | 0 | 0 | 0 | 0 | 0 | 0 | 0 | 0 | 0 | 0 | 0 |
| bacteria_Jul11d50b_44736 | 0 | 0 | 0 | 0 | 0 | 0 | 0 | 0 | 0 | 0 | 0 |
| bacteria_Jul11d50b_47362 | 0 | 0 | 0 | 0 | 0 | 0 | 0 | 0 | 0 | 0 | 0 |
| bacteria_Jul11d50b_52033 | 0 | 0 | 0 | 0 | 0 | 0 | 0 | 0 | 0 | 0 | 0 |
| bacteria_Jul11d75b_8184 | 0 | 0 | 0 | 0 | 0 | 0 | 0 | 0 | 0 | 0 | 0 |
| bacteria_Jul11d75b_8617 | 0 | 4 | 0 | 7 | 2 | 10 | 0 | 0 | 0 | 5 | 28 |
| bacteria_Jul11d75b_11391 | 0 | 0 | 0 | 0 | 0 | 0 | 0 | 0 | 0 | 0 | 0 |
| bacteria_Jul11d75b_16431 | 0 | 0 | 0 | 0 | 0 | 0 | 0 | 0 | 0 | 0 | 0 |
| bacteria_Jul11d75b_20357 | 0 | 0 | 0 | 0 | 0 | 0 | 0 | 0 | 0 | 0 | 0 |
| bacteria_Jul11d75b_23171 | 0 | 0 | 0 | 0 | 0 | 0 | 0 | 0 | 0 | 0 | 0 |
| bacteria_Jul11d75b_23633 | 0 | 0 | 0 | 0 | 0 | 0 | 0 | 0 | 0 | 0 | 0 |
| bacteria_Jul11d75b_25120 | 0 | 0 | 0 | 0 | 0 | 0 | 0 | 0 | 0 | 0 | 0 |
| bacteria_Jul11d75b_28025 | 0 | 0 | 0 | 0 | 0 | 0 | 0 | 0 | 0 | 0 | 0 |
| bacteria_Jul11d75b_29883 | 0 | 0 | 0 | 0 | 0 | 0 | 0 | 0 | 0 | 0 | 0 |
| bacteria_Jul11d75b_32862 | 0 | 0 | 0 | 0 | 0 | 0 | 0 | 0 | 0 | 0 | 0 |
| bacteria_Jul11d75b_33219 | 0 | 0 | 0 | 0 | 0 | 0 | 0 | 0 | 0 | 0 | 0 |
| bacteria_Jul11d75b_43477 | 0 | 0 | 0 | 0 | 0 | 0 | 0 | 0 | 0 | 0 | 0 |
| bacteria_Jul11d75b_46998 | 0 | 0 | 0 | 0 | 0 | 0 | 0 | 0 | 0 | 0 | 0 |
| bacteria_Jul11d75b_50166 | 0 | 0 | 0 | 0 | 0 | 0 | 0 | 0 | 0 | 0 | 0 |
| bacteria_Jul11d75b_51527 | 0 | 0 | 0 | 0 | 0 | 1 | 0 | 0 | 0 | 0 | 1 |
| bacteria_Jul27d5b_3120 | 0 | 0 | 0 | 0 | 0 | 0 | 0 | 0 | 0 | 0 | 0 |
| bacteria_Jul27d5b_5428 | 0 | 0 | 0 | 0 | 0 | 0 | 0 | 0 | 0 | 0 | 0 |
| bacteria_Jul27d5b_5943 | 0 | 0 | 0 | 0 | 0 | 0 | 0 | 0 | 0 | 0 | 0 |
| bacteria_Jul27d5b_10360 | 0 | 0 | 0 | 0 | 0 | 0 | 0 | 0 | 0 | 0 | 0 |
| bacteria_Jul27d5b_12140 | 0 | 0 | 0 | 0 | 0 | 0 | 0 | 0 | 0 | 0 | 0 |
| bacteria_Jul27d5b_15953 | 0 | 0 | 0 | 0 | 0 | 0 | 0 | 0 | 0 | 0 | 0 |
| bacteria_Jul27d5b_26185 | 0 | 0 | 0 | 0 | 0 | 0 | 0 | 0 | 0 | 0 | 0 |
| bacteria_Jul27d5b_26651 | 0 | 0 | 0 | 0 | 0 | 0 | 0 | 0 | 0 | 0 | 0 |
| bacteria_Jul27d5b_33096 | 0 | 0 | 0 | 0 | 0 | 0 | 0 | 0 | 0 | 0 | 0 |
| bacteria_Jul27d5b_33101 | 0 | 0 | 0 | 0 | 0 | 0 | 0 | 0 | 0 | 0 | 0 |
| bacteria_Jul27d5b_33613 | 0 | 1 | 0 | 0 | 0 | 0 | 0 | 0 | 0 | 0 | 1 |
| bacteria_Jul27d5b_35075 | 0 | 0 | 0 | 0 | 0 | 0 | 0 | 0 | 0 | 0 | 0 |
| bacteria_Jul27d5b_35447 | 0 | 0 | 0 | 0 | 1 | 0 | 0 | 0 | 0 | 0 | 1 |
| bacteria_Jul27d5b_35909 | 0 | 0 | 0 | 0 | 0 | 0 | 0 | 0 | 0 | 0 | 0 |
| bacteria_Jul27d5b_48077 | 0 | 0 | 0 | 0 | 0 | 0 | 0 | 0 | 0 | 0 | 0 |
| bacteria_Jul27d5b_48693 | 0 | 0 | 0 | 0 | 0 | 0 | 0 | 0 | 0 | 0 | 0 |
| bacteria_Jul27d5b_48854 | 0 | 0 | 0 | 0 | 0 | 0 | 0 | 0 | 0 | 0 | 0 |
| bacteria_Jul27d5b_49859 | 0 | 0 | 0 | 0 | 0 | 0 | 0 | 0 | 0 | 0 | 0 |
| bacteria_Jul27d25b_7972 | 0 | 0 | 0 | 0 | 0 | 0 | 0 | 0 | 0 | 0 | 0 |
| bacteria_Jul27d25b_9181 | 0 | 0 | 0 | 0 | 0 | 0 | 0 | 0 | 0 | 0 | 0 |
| bacteria_Jul27d25b_15951 | 0 | 0 | 0 | 0 | 0 | 0 | 0 | 0 | 0 | 0 | 0 |
| bacteria_Jul27d25b_20487 | 0 | 0 | 0 | 0 | 0 | 0 | 0 | 0 | 0 | 0 | 0 |
| bacteria_Jul27d25b_22563 | 0 | 0 | 0 | 0 | 0 | 0 | 0 | 0 | 0 | 0 | 0 |
| bacteria_Jul27d25b_25276 | 0 | 0 | 0 | 0 | 0 | 0 | 0 | 0 | 0 | 0 | 0 |
| bacteria_Jul27d25b_25284 | 0 | 0 | 0 | 0 | 0 | 0 | 0 | 0 | 0 | 0 | 0 |
| bacteria_Jul27d25b_26744 | 0 | 0 | 0 | 0 | 0 | 0 | 0 | 0 | 0 | 0 | 0 |
| bacteria_Jul27d25b_30235 | 0 | 0 | 0 | 0 | 0 | 0 | 0 | 0 | 0 | 0 | 0 |
| bacteria_Jul27d25b_30574 | 0 | 0 | 0 | 0 | 0 | 0 | 0 | 0 | 0 | 0 | 0 |
| bacteria_Jul27d25b_31099 | 0 | 0 | 0 | 0 | 0 | 0 | 0 | 0 | 0 | 0 | 0 |
| bacteria_Jul27d25b_32815 | 0 | 1 | 0 | 0 | 0 | 0 | 0 | 0 | 0 | 0 | 1 |
| bacteria_Jul27d25b_37694 | 0 | 0 | 0 | 0 | 0 | 0 | 0 | 0 | 0 | 0 | 0 |
| bacteria_Jul27d25b_51427 | 0 | 0 | 0 | 0 | 0 | 0 | 0 | 0 | 0 | 0 | 0 |
| bacteria_Jul27d25b_51956 | 0 | 0 | 0 | 0 | 0 | 0 | 0 | 0 | 0 | 0 | 0 |
| bacteria_Jul27d25b_60261 | 0 | 0 | 0 | 0 | 0 | 0 | 0 | 0 | 0 | 0 | 0 |
| bacteria_Jul27d25b_62967 | 0 | 0 | 0 | 0 | 0 | 0 | 0 | 0 | 0 | 0 | 0 |
| bacteria_Jul27d25b_64491 | 0 | 0 | 0 | 0 | 0 | 0 | 0 | 0 | 0 | 0 | 0 |
| bacteria_Jul27d25b_65588 | 0 | 2 | 0 | 0 | 0 | 0 | 0 | 0 | 0 | 0 | 2 |
| bacteria_Jul27d25b_73563 | 0 | 0 | 0 | 0 | 0 | 0 | 0 | 0 | 0 | 0 | 0 |
| bacteria_Jul27d75b_2253 | 0 | 0 | 0 | 0 | 0 | 0 | 0 | 0 | 0 | 0 | 0 |
| bacteria_Jul27d75b_14154 | 0 | 0 | 0 | 0 | 0 | 0 | 0 | 0 | 0 | 0 | 0 |
| bacteria_Jul27d75b_15243 | 0 | 0 | 0 | 0 | 0 | 0 | 0 | 0 | 0 | 0 | 0 |
| bacteria_Jul27d75b_16064 | 0 | 0 | 0 | 0 | 0 | 0 | 0 | 0 | 0 | 0 | 0 |
| bacteria_Jul27d75b_28594 | 0 | 0 | 0 | 0 | 0 | 0 | 0 | 0 | 0 | 0 | 0 |
| bacteria_Jul27d75b_52028 | 0 | 0 | 0 | 0 | 0 | 0 | 0 | 0 | 0 | 0 | 0 |
| bacteria_Jul27d75b_53903 | 0 | 0 | 0 | 0 | 0 | 0 | 0 | 0 | 0 | 0 | 0 |
| bacteria_Jul27d75b_55566 | 0 | 0 | 0 | 0 | 0 | 0 | 0 | 0 | 0 | 0 | 0 |
| bacteria_Jul27d75b_58121 | 0 | 0 | 0 | 0 | 0 | 0 | 0 | 0 | 0 | 0 | 0 |
| bacteria_Jun8d25b_7145 | 0 | 0 | 0 | 0 | 0 | 0 | 0 | 0 | 0 | 0 | 0 |
| bacteria_Jun8d25b_20564 | 0 | 0 | 0 | 0 | 0 | 0 | 0 | 0 | 0 | 0 | 0 |

| | | | | | | | | | | | |
|---|---|---|---|---|---|---|---|---|---|---|---|
| bacteria_Jun8d25b_21602 | 0 | 0 | 0 | 0 | 0 | 0 | 0 | 0 | 0 | 0 | 0 |
| bacteria_Jun8d25b_34360 | 0 | 0 | 0 | 0 | 0 | 0 | 0 | 0 | 0 | 0 | 0 |
| bacteria_Jun8d25b_34461 | 0 | 0 | 0 | 0 | 0 | 0 | 0 | 0 | 0 | 0 | 0 |
| bacteria_Jun8d25b_41983 | 0 | 0 | 0 | 0 | 0 | 0 | 0 | 0 | 0 | 0 | 0 |
| bacteria_Jun8d25b_45038 | 0 | 0 | 0 | 0 | 0 | 0 | 0 | 0 | 0 | 0 | 0 |
| bacteria_Jun8d25b_63896 | 0 | 0 | 0 | 0 | 0 | 0 | 0 | 0 | 0 | 0 | 0 |
| bacteria_Jun8d25b_72308 | 0 | 0 | 0 | 0 | 0 | 0 | 0 | 0 | 0 | 0 | 0 |
| bacteria_Jun8d25b_73418 | 0 | 0 | 0 | 0 | 0 | 0 | 0 | 0 | 0 | 0 | 0 |
| bacteria_Jun8d50b_825 | 0 | 0 | 0 | 0 | 0 | 0 | 0 | 0 | 0 | 0 | 0 |
| bacteria_Jun8d50b_20442 | 0 | 0 | 0 | 0 | 0 | 0 | 0 | 0 | 0 | 0 | 0 |
| bacteria_Jun8d50b_21639 | 0 | 0 | 0 | 0 | 0 | 0 | 0 | 0 | 0 | 0 | 0 |
| bacteria_Jun8d50b_34821 | 0 | 0 | 0 | 0 | 0 | 0 | 0 | 0 | 0 | 0 | 0 |
| bacteria_Jun8d50b_46464 | 0 | 0 | 0 | 0 | 0 | 0 | 0 | 0 | 0 | 0 | 0 |
| bacteria_Jun8d50b_50881 | 0 | 0 | 0 | 0 | 0 | 0 | 0 | 0 | 0 | 0 | 0 |
| bacteria_Jun8d75b_13815 | 0 | 0 | 0 | 0 | 0 | 0 | 0 | 0 | 0 | 0 | 0 |
| bacteria_Jun22d75b_545 | 0 | 0 | 0 | 0 | 0 | 0 | 0 | 0 | 0 | 0 | 0 |
| bacteria_Jun22d75b_27176 | 0 | 0 | 0 | 0 | 0 | 0 | 0 | 0 | 0 | 0 | 0 |
| bacteria_Jun22d75b_33698 | 0 | 0 | 0 | 0 | 0 | 0 | 0 | 0 | 0 | 0 | 0 |
| bacteria_Jun22d75b_69176 | 0 | 0 | 0 | 0 | 0 | 0 | 0 | 0 | 0 | 0 | 0 |
| bacteria_May12d25b_5131 | 0 | 0 | 0 | 0 | 0 | 0 | 0 | 0 | 0 | 0 | 0 |
| bacteria_May12d25b_1921 | 0 | 0 | 0 | 0 | 0 | 0 | 0 | 0 | 0 | 0 | 0 |
| bacteria_May12d25b_2388 | 0 | 0 | 0 | 0 | 0 | 0 | 0 | 0 | 0 | 0 | 0 |
| bacteria_May12d25b_2591 | 0 | 0 | 0 | 0 | 1 | 0 | 0 | 0 | 0 | 0 | 1 |
| bacteria_May12d25b_2703 | 0 | 0 | 0 | 0 | 0 | 0 | 0 | 0 | 0 | 0 | 0 |
| bacteria_May12d25b_2835 | 0 | 0 | 0 | 0 | 0 | 0 | 0 | 0 | 0 | 0 | 0 |
| bacteria_May12d25b_4039 | 0 | 0 | 0 | 0 | 0 | 0 | 0 | 0 | 0 | 0 | 0 |
| bacteria_May12d25b_4312 | 0 | 0 | 0 | 0 | 0 | 0 | 0 | 0 | 0 | 0 | 0 |
| bacteria_May12d25b_4780 | 0 | 0 | 0 | 0 | 0 | 0 | 0 | 0 | 0 | 0 | 0 |
| bacteria_May12d25b_4782 | 0 | 0 | 0 | 0 | 0 | 0 | 0 | 0 | 0 | 0 | 0 |
| bacteria_May12d25b_5111 | 0 | 0 | 0 | 0 | 0 | 0 | 0 | 0 | 0 | 0 | 0 |
| bacteria_May12d25b_5275 | 0 | 0 | 0 | 0 | 0 | 0 | 0 | 0 | 0 | 0 | 0 |
| bacteria_May12d50b_6923 | 0 | 0 | 0 | 0 | 0 | 0 | 0 | 0 | 0 | 0 | 0 |
| bacteria_May12d50b_1083 | 0 | 0 | 0 | 0 | 0 | 0 | 0 | 0 | 0 | 0 | 0 |
| bacteria_May12d50b_2322 | 0 | 0 | 0 | 0 | 0 | 0 | 0 | 0 | 0 | 0 | 0 |
| bacteria_May12d50b_2428 | 0 | 0 | 0 | 0 | 0 | 0 | 0 | 0 | 0 | 0 | 0 |
| bacteria_May12d50b_2771 | 0 | 0 | 0 | 0 | 0 | 0 | 0 | 0 | 0 | 0 | 0 |
| bacteria_May12d50b_4023 | 0 | 0 | 0 | 0 | 0 | 0 | 0 | 0 | 0 | 0 | 0 |
| bacteria_May12d50b_4198 | 0 | 0 | 0 | 0 | 0 | 0 | 0 | 0 | 0 | 0 | 0 |
| bacteria_May12d50b_4714 | 0 | 0 | 0 | 0 | 0 | 0 | 0 | 0 | 0 | 0 | 0 |
| bacteria_May12d50b_4792 | 0 | 0 | 0 | 0 | 0 | 0 | 0 | 0 | 0 | 0 | 0 |
| bacteria_Sep8d0b_352 | 0 | 0 | 0 | 0 | 0 | 0 | 0 | 0 | 0 | 0 | 0 |
| bacteria_Sep8d0b_46600 | 0 | 0 | 0 | 0 | 0 | 0 | 0 | 0 | 0 | 0 | 0 |
| bacteria_Sep8d0b_69954 | 0 | 0 | 0 | 0 | 0 | 0 | 0 | 0 | 0 | 0 | 0 |
| bacteria_Sep8d5b_3363 | 0 | 0 | 0 | 0 | 0 | 0 | 0 | 0 | 0 | 0 | 0 |
| bacteria_Sep8d5b_12958 | 0 | 0 | 0 | 0 | 0 | 0 | 0 | 0 | 0 | 0 | 0 |
| bacteria_Sep8d5b_26342 | 0 | 0 | 0 | 0 | 0 | 0 | 0 | 0 | 0 | 0 | 0 |
| bacteria_Sep8d5b_26406 | 0 | 0 | 0 | 0 | 0 | 0 | 0 | 0 | 0 | 0 | 0 |
| bacteria_Sep8d5b_26646 | 0 | 0 | 0 | 0 | 0 | 0 | 0 | 0 | 0 | 0 | 0 |
| bacteria_Sep8d5b_33456 | 0 | 0 | 0 | 0 | 0 | 0 | 0 | 0 | 0 | 0 | 0 |
| bacteria_Sep8d5b_35871 | 0 | 0 | 0 | 0 | 0 | 0 | 0 | 0 | 0 | 0 | 0 |
| bacteria_Sep8d5b_49612 | 0 | 0 | 0 | 0 | 0 | 0 | 0 | 0 | 0 | 0 | 0 |
| bacteria_Sep8d5b_53005 | 0 | 0 | 0 | 0 | 0 | 0 | 0 | 0 | 0 | 0 | 0 |
| bacteria_Sep8d5b_54964 | 0 | 0 | 0 | 0 | 0 | 0 | 0 | 0 | 0 | 0 | 0 |
| bacteria_Sep8d5b_58266 | 0 | 0 | 0 | 0 | 0 | 0 | 0 | 0 | 0 | 0 | 0 |
| bacteria_Sep8d5b_63780 | 0 | 0 | 0 | 0 | 0 | 0 | 0 | 0 | 0 | 0 | 0 |
| bacteria_Sep8d5b_66338 | 0 | 0 | 0 | 0 | 0 | 0 | 0 | 0 | 0 | 0 | 0 |
| bacteria_Sep8d5b_66689 | 0 | 0 | 0 | 0 | 0 | 0 | 0 | 0 | 0 | 0 | 0 |
| bacteria_Sep8d25b_899 | 0 | 0 | 0 | 0 | 0 | 0 | 0 | 0 | 0 | 0 | 0 |
| bacteria_Sep8d25b_4177 | 0 | 0 | 0 | 0 | 0 | 0 | 0 | 0 | 0 | 0 | 0 |
| bacteria_Sep8d25b_7083 | 0 | 0 | 0 | 0 | 0 | 0 | 0 | 0 | 0 | 0 | 0 |
| bacteria_Sep8d25b_17783 | 0 | 0 | 0 | 0 | 0 | 0 | 0 | 0 | 0 | 0 | 0 |
| bacteria_Sep8d25b_38783 | 0 | 0 | 0 | 0 | 0 | 0 | 0 | 0 | 0 | 0 | 0 |
| bacteria_Sep8d50b_12570 | 0 | 0 | 0 | 0 | 0 | 0 | 0 | 0 | 0 | 0 | 0 |
| virus_Aug5.0v_14 | 0 | 1 | 0 | 0 | 0 | 0 | 0 | 0 | 0 | 0 | 1 |
| virus_Aug5.0v_212 | 0 | 1 | 0 | 0 | 0 | 0 | 0 | 0 | 0 | 0 | 1 |
| virus_Aug5.0v_329 | 0 | 1 | 0 | 0 | 0 | 0 | 0 | 0 | 0 | 0 | 1 |
| virus_Aug5.0v_422 | 0 | 1 | 0 | 0 | 0 | 0 | 0 | 0 | 0 | 0 | 1 |
| virus_Aug5.0v_1810 | 0 | 1 | 0 | 0 | 0 | 0 | 0 | 0 | 0 | 0 | 1 |
| virus_Aug5.0v_1816 | 0 | 1 | 0 | 0 | 0 | 1 | 0 | 0 | 0 | 0 | 2 |
| virus_Aug5.0v_1821 | 0 | 1 | 0 | 0 | 0 | 0 | 0 | 0 | 0 | 0 | 1 |
| virus_Aug5.0v_2129 | 0 | 1 | 0 | 0 | 0 | 0 | 0 | 0 | 0 | 0 | 1 |
| virus_Aug5.0v_2186 | 0 | 1 | 0 | 0 | 0 | 0 | 0 | 0 | 0 | 0 | 1 |
| virus_Aug5.0v_2210 | 0 | 1 | 0 | 1 | 0 | 0 | 0 | 0 | 0 | 0 | 2 |
| virus_Aug5.0v_2365 | 0 | 1 | 0 | 0 | 0 | 0 | 0 | 0 | 0 | 0 | 1 |
| virus_Aug5.0v_2471 | 0 | 1 | 0 | 0 | 0 | 0 | 0 | 0 | 0 | 0 | 1 |
| virus_Aug5.0v_2521 | 0 | 1 | 0 | 0 | 0 | 0 | 0 | 0 | 0 | 0 | 1 |
| virus_Aug5.0v_2618 | 0 | 1 | 0 | 0 | 0 | 0 | 0 | 0 | 0 | 0 | 1 |
| virus_Aug5.0v_2756 | 0 | 1 | 0 | 0 | 0 | 0 | 0 | 0 | 0 | 0 | 1 |
| virus_Aug5.0v_2789 | 0 | 1 | 0 | 0 | 0 | 0 | 0 | 0 | 0 | 0 | 1 |
| virus_Aug5.0v_2905 | 0 | 1 | 0 | 0 | 0 | 0 | 0 | 0 | 0 | 0 | 1 |
| virus_Aug5.0v_2957 | 0 | 1 | 0 | 0 | 0 | 0 | 0 | 0 | 0 | 0 | 1 |
| virus_Aug5.0v_3560 | 0 | 1 | 0 | 0 | 0 | 0 | 0 | 0 | 0 | 0 | 1 |
| virus_Aug5.0v_3981 | 0 | 1 | 0 | 0 | 0 | 0 | 0 | 0 | 0 | 0 | 1 |
| virus_Aug5.0v_4043 | 0 | 1 | 0 | 0 | 0 | 0 | 0 | 0 | 0 | 0 | 1 |
| virus_Aug5.0v_4310 | 0 | 1 | 0 | 0 | 0 | 0 | 0 | 0 | 0 | 0 | 1 |
| virus_Aug5.0v_4450 | 0 | 1 | 0 | 0 | 0 | 0 | 0 | 0 | 0 | 0 | 1 |
| virus_Aug5.0v_4926 | 0 | 1 | 0 | 0 | 0 | 0 | 0 | 0 | 0 | 0 | 1 |
| virus_Aug5.0v_4936 | 0 | 1 | 0 | 0 | 0 | 0 | 0 | 0 | 0 | 0 | 1 |
| virus_Aug5.0v_4945 | 0 | 1 | 0 | 0 | 0 | 0 | 0 | 0 | 0 | 0 | 1 |
| virus_Aug5.0v_4969 | 0 | 1 | 0 | 0 | 0 | 0 | 0 | 0 | 0 | 0 | 1 |
| virus_Aug5.0v_5286 | 0 | 1 | 0 | 0 | 0 | 0 | 0 | 0 | 0 | 0 | 1 |
| virus_Aug5.0v_5428 | 0 | 1 | 0 | 0 | 0 | 0 | 0 | 0 | 0 | 0 | 1 |
| virus_Aug5.5v_357 | 0 | 0 | 0 | 1 | 0 | 0 | 0 | 0 | 0 | 0 | 1 |
| virus_Aug5.5v_7957 | 0 | 0 | 0 | 1 | 0 | 0 | 0 | 0 | 0 | 0 | 1 |
| virus_Aug5.5v_13568 | 0 | 0 | 0 | 1 | 0 | 0 | 0 | 0 | 0 | 0 | 1 |
| virus_Aug5.5v_14419 | 0 | 0 | 0 | 1 | 0 | 1 | 0 | 1 | 0 | 0 | 3 |
| virus_Aug5.5v_16378 | 0 | 0 | 0 | 1 | 0 | 0 | 0 | 0 | 0 | 0 | 1 |
| virus_Aug5.5v_18066 | 0 | 0 | 0 | 1 | 0 | 0 | 0 | 0 | 0 | 0 | 1 |
| virus_Aug5.5v_18078 | 0 | 0 | 0 | 1 | 0 | 0 | 0 | 0 | 0 | 0 | 1 |
| virus_Aug5.5v_20966 | 0 | 0 | 0 | 1 | 0 | 0 | 0 | 0 | 0 | 0 | 1 |
| virus_Aug5.5v_21274 | 0 | 0 | 0 | 1 | 0 | 0 | 0 | 0 | 0 | 0 | 1 |
| virus_Aug5.5v_23135 | 0 | 0 | 0 | 1 | 0 | 0 | 0 | 0 | 0 | 0 | 1 |
| virus_Aug5.5v_35539 | 0 | 0 | 0 | 1 | 0 | 0 | 0 | 0 | 0 | 0 | 1 |
| virus_Aug5.5v_36929 | 0 | 0 | 0 | 1 | 0 | 0 | 0 | 0 | 0 | 0 | 1 |
| virus_Aug5.5v_38127 | 0 | 0 | 0 | 1 | 0 | 0 | 0 | 0 | 0 | 0 | 1 |
| virus_Aug5.25v_2 | 0 | 0 | 0 | 0 | 0 | 1 | 0 | 0 | 0 | 0 | 1 |
| virus_Aug5.25v_1489 | 0 | 0 | 0 | 1 | 0 | 1 | 0 | 0 | 0 | 0 | 2 |
| virus_Aug5.25v_3142 | 0 | 0 | 0 | 0 | 0 | 1 | 0 | 0 | 0 | 0 | 1 |
| virus_Aug5.25v_3378 | 0 | 1 | 0 | 0 | 0 | 2 | 0 | 0 | 0 | 0 | 3 |
| virus_Aug5.25v_4313 | 0 | 0 | 0 | 0 | 0 | 1 | 0 | 0 | 0 | 0 | 1 |
| virus_Aug5.25v_7175 | 0 | 0 | 0 | 0 | 0 | 1 | 0 | 0 | 0 | 0 | 1 |
| virus_Aug5.25v_8171 | 0 | 0 | 0 | 0 | 0 | 1 | 0 | 0 | 0 | 0 | 1 |
| virus_Aug5.25v_9437 | 0 | 0 | 0 | 0 | 0 | 1 | 0 | 0 | 0 | 0 | 1 |
| virus_Aug5.25v_9868 | 0 | 0 | 0 | 0 | 0 | 1 | 0 | 0 | 0 | 0 | 1 |
| virus_Aug5.25v_11199 | 0 | 0 | 0 | 0 | 0 | 1 | 0 | 0 | 0 | 0 | 1 |
| virus_Aug5.25v_11227 | 0 | 0 | 0 | 0 | 0 | 1 | 0 | 0 | 0 | 0 | 1 |
| virus_Aug5.25v_17104 | 0 | 0 | 0 | 0 | 0 | 1 | 0 | 0 | 0 | 0 | 1 |
| virus_Aug5.25v_19259 | 0 | 0 | 0 | 0 | 0 | 1 | 0 | 0 | 0 | 0 | 1 |
| virus_Aug5.25v_20583 | 0 | 0 | 0 | 0 | 0 | 1 | 0 | 0 | 0 | 0 | 1 |
| virus_Aug5.25v_21003 | 0 | 0 | 0 | 0 | 0 | 1 | 0 | 0 | 0 | 0 | 1 |
| virus_Aug5.25v_21305 | 0 | 0 | 0 | 0 | 0 | 1 | 0 | 0 | 0 | 0 | 1 |
| virus_Aug5.25v_24050 | 0 | 0 | 0 | 0 | 0 | 1 | 0 | 0 | 0 | 0 | 1 |
| virus_Aug5.25v_25011 | 0 | 0 | 0 | 0 | 0 | 1 | 0 | 0 | 0 | 0 | 1 |
| virus_Aug5.25v_28937 | 0 | 0 | 0 | 0 | 0 | 1 | 0 | 0 | 0 | 0 | 1 |
| virus_Aug5.25v_36597 | 0 | 0 | 0 | 0 | 0 | 1 | 0 | 0 | 0 | 0 | 1 |
| virus_Aug5.25v_40304 | 0 | 0 | 0 | 0 | 0 | 1 | 0 | 0 | 0 | 0 | 1 |
| virus_Aug5.50v_4328 | 0 | 0 | 0 | 0 | 0 | 0 | 0 | 1 | 0 | 0 | 1 |
| virus_Aug5.50v_7119 | 0 | 0 | 0 | 0 | 0 | 0 | 0 | 1 | 0 | 0 | 1 |
| virus_Aug5.50v_11588 | 0 | 0 | 0 | 0 | 0 | 0 | 0 | 1 | 0 | 0 | 1 |
| virus_Aug5.50v_13084 | 0 | 0 | 0 | 0 | 0 | 0 | 0 | 1 | 0 | 0 | 1 |
| virus_Aug5.50v_13508 | 0 | 0 | 0 | 0 | 0 | 0 | 0 | 1 | 0 | 0 | 1 |
| virus_Aug5.50v_21345 | 0 | 0 | 0 | 0 | 0 | 0 | 0 | 1 | 0 | 0 | 1 |
| virus_Aug5.50v_22866 | 0 | 0 | 0 | 0 | 0 | 0 | 0 | 1 | 0 | 0 | 1 |
| virus_Aug5.50v_38232 | 0 | 0 | 0 | 0 | 0 | 0 | 0 | 1 | 0 | 0 | 1 |
| virus_Aug5.75v_1000 | 0 | 0 | 0 | 0 | 0 | 0 | 0 | 0 | 0 | 1 | 1 |
| virus_Aug5.75v_4421 | 0 | 0 | 0 | 0 | 0 | 0 | 0 | 0 | 0 | 1 | 1 |
| virus_Aug5.75v_6711 | 0 | 0 | 0 | 0 | 0 | 0 | 0 | 0 | 0 | 2 | 2 |
| virus_Aug5.75v_11084 | 0 | 0 | 0 | 0 | 0 | 0 | 0 | 0 | 0 | 1 | 1 |
| virus_Aug5.75v_18383 | 0 | 0 | 0 | 0 | 0 | 0 | 0 | 0 | 0 | 1 | 1 |
| virus_Aug5.75v_27104 | 0 | 0 | 0 | 0 | 0 | 0 | 0 | 0 | 0 | 1 | 1 |
| virus_Aug5.75v_28751 | 0 | 0 | 0 | 0 | 0 | 0 | 0 | 0 | 0 | 1 | 1 |
| virus_Aug5.75v_30233 | 0 | 0 | 0 | 0 | 0 | 0 | 0 | 0 | 0 | 1 | 1 |
| virus_Aug5.75v_30379 | 0 | 0 | 0 | 0 | 0 | 0 | 0 | 0 | 0 | 1 | 1 |
| virus_Aug5.75v_31313 | 0 | 0 | 0 | 0 | 0 | 0 | 0 | 0 | 0 | 1 | 1 |
| virus_Aug5.75v_32334 | 0 | 0 | 0 | 0 | 0 | 0 | 0 | 0 | 0 | 1 | 1 |
| virus_Aug5.75v_34269 | 0 | 0 | 0 | 0 | 0 | 0 | 0 | 0 | 0 | 1 | 1 |
| virus_Jul11.0v_1841 | 0 | 0 | 0 | 0 | 0 | 0 | 0 | 0 | 0 | 0 | 0 |
| virus_Jul11.0v_5433 | 0 | 0 | 0 | 0 | 0 | 0 | 0 | 0 | 0 | 0 | 0 |
| virus_Jul11.0v_5545 | 0 | 0 | 0 | 0 | 0 | 0 | 0 | 0 | 0 | 0 | 0 |
| virus_Jul11.0v_12161 | 0 | 0 | 0 | 0 | 0 | 0 | 0 | 0 | 0 | 0 | 0 |
| virus_Jul11.0v_16249 | 0 | 0 | 0 | 0 | 0 | 0 | 0 | 0 | 0 | 0 | 0 |
| virus_Jul11.0v_16850 | 0 | 0 | 0 | 0 | 0 | 0 | 0 | 0 | 0 | 0 | 0 |
| virus_Jul11.0v_17580 | 0 | 0 | 0 | 0 | 0 | 0 | 0 | 0 | 0 | 0 | 0 |
| virus_Jul11.0v_18060 | 0 | 0 | 0 | 0 | 0 | 0 | 0 | 0 | 0 | 0 | 0 |
| virus_Jul11.0v_18988 | 0 | 0 | 0 | 0 | 0 | 0 | 0 | 0 | 0 | 0 | 0 |
| virus_Jul11.5v_505 | 0 | 0 | 0 | 0 | 0 | 0 | 0 | 0 | 0 | 0 | 0 |
| virus_Jul11.5v_1015 | 0 | 0 | 0 | 0 | 0 | 0 | 0 | 0 | 0 | 0 | 0 |

| | | | | | | | | | | | |
|---|---|---|---|---|---|---|---|---|---|---|---|
| virus_Jul11.5v_2303 | 0 | 0 | 0 | 0 | 0 | 0 | 0 | 0 | 0 | 0 | 0 |
| virus_Jul11.5v_2601 | 0 | 0 | 0 | 0 | 0 | 0 | 0 | 0 | 0 | 0 | 0 |
| virus_Jul11.5v_3734 | 0 | 0 | 0 | 0 | 0 | 0 | 0 | 0 | 0 | 0 | 0 |
| virus_Jul11.5v_4525 | 0 | 0 | 0 | 0 | 0 | 0 | 0 | 0 | 0 | 0 | 0 |
| virus_Jul11.5v_4595 | 0 | 0 | 0 | 0 | 0 | 0 | 0 | 0 | 0 | 0 | 0 |
| virus_Jul11.5v_4718 | 0 | 0 | 0 | 0 | 0 | 0 | 0 | 0 | 0 | 0 | 0 |
| virus_Jul11.5v_8294 | 0 | 0 | 0 | 0 | 0 | 0 | 0 | 0 | 0 | 0 | 0 |
| virus_Jul11.5v_10738 | 0 | 0 | 0 | 0 | 0 | 0 | 0 | 0 | 0 | 0 | 0 |
| virus_Jul11.5v_13320 | 0 | 0 | 0 | 0 | 0 | 0 | 0 | 0 | 0 | 0 | 0 |
| virus_Jul11.5v_13543 | 0 | 0 | 0 | 0 | 0 | 0 | 0 | 0 | 0 | 0 | 0 |
| virus_Jul11.5v_14609 | 0 | 0 | 0 | 0 | 0 | 0 | 0 | 0 | 0 | 0 | 0 |
| virus_Jul11.5v_15561 | 0 | 0 | 0 | 0 | 0 | 0 | 0 | 0 | 0 | 0 | 0 |
| virus_Jul11.5v_16835 | 0 | 0 | 0 | 0 | 0 | 0 | 0 | 0 | 0 | 0 | 0 |
| virus_Jul11.5v_17515 | 0 | 0 | 0 | 0 | 0 | 0 | 0 | 0 | 0 | 0 | 0 |
| virus_Jul11.5v_20187 | 0 | 0 | 0 | 0 | 0 | 0 | 0 | 0 | 0 | 0 | 0 |
| virus_Jul11.25v_348 | 0 | 0 | 0 | 0 | 0 | 0 | 0 | 0 | 0 | 0 | 0 |
| virus_Jul11.25v_3684 | 0 | 0 | 0 | 0 | 0 | 0 | 0 | 0 | 0 | 0 | 0 |
| virus_Jul11.25v_3735 | 0 | 0 | 0 | 0 | 0 | 0 | 0 | 0 | 0 | 0 | 0 |
| virus_Jul11.25v_4792 | 0 | 0 | 0 | 0 | 0 | 0 | 0 | 0 | 0 | 0 | 0 |
| virus_Jul11.25v_6271 | 0 | 0 | 0 | 0 | 0 | 0 | 0 | 0 | 0 | 0 | 0 |
| virus_Jul11.25v_6769 | 0 | 0 | 0 | 0 | 0 | 0 | 0 | 0 | 0 | 0 | 0 |
| virus_Jul11.25v_9543 | 0 | 0 | 0 | 0 | 0 | 0 | 0 | 0 | 0 | 0 | 0 |
| virus_Jul11.25v_11244 | 0 | 0 | 0 | 0 | 0 | 0 | 0 | 0 | 0 | 0 | 0 |
| virus_Jul11.25v_14845 | 0 | 0 | 0 | 0 | 0 | 0 | 0 | 0 | 0 | 0 | 0 |
| virus_Jul11.25v_15608 | 0 | 0 | 0 | 0 | 0 | 0 | 0 | 0 | 0 | 0 | 0 |
| virus_Jul11.25v_18324 | 0 | 0 | 0 | 0 | 0 | 0 | 0 | 0 | 0 | 0 | 0 |
| virus_Jul11.25v_19146 | 0 | 0 | 0 | 0 | 0 | 0 | 0 | 0 | 0 | 0 | 0 |
| virus_Jul11.25v_19575 | 0 | 0 | 0 | 0 | 0 | 1 | 0 | 0 | 0 | 0 | 1 |
| virus_Jul11.50v_1478 | 0 | 0 | 0 | 0 | 0 | 0 | 0 | 0 | 0 | 0 | 0 |
| virus_Jul11.50v_1692 | 0 | 0 | 0 | 0 | 0 | 0 | 0 | 0 | 0 | 0 | 0 |
| virus_Jul11.50v_2377 | 0 | 0 | 0 | 0 | 0 | 0 | 0 | 0 | 0 | 0 | 0 |
| virus_Jul11.50v_8236 | 0 | 0 | 0 | 0 | 0 | 0 | 0 | 0 | 0 | 0 | 0 |
| virus_Jul11.50v_9034 | 0 | 0 | 0 | 0 | 0 | 0 | 0 | 0 | 0 | 0 | 0 |
| virus_Jul11.50v_9634 | 0 | 0 | 0 | 0 | 0 | 0 | 0 | 0 | 0 | 0 | 0 |
| virus_Jul11.50v_10452 | 0 | 0 | 0 | 0 | 0 | 0 | 0 | 0 | 0 | 0 | 0 |
| virus_Jul11.50v_10799 | 0 | 0 | 0 | 0 | 0 | 0 | 0 | 0 | 0 | 0 | 0 |
| virus_Jul11.50v_10873 | 0 | 0 | 0 | 0 | 0 | 0 | 0 | 0 | 0 | 0 | 0 |
| virus_Jul11.50v_11059 | 0 | 0 | 0 | 0 | 0 | 0 | 0 | 0 | 0 | 0 | 0 |
| virus_Jul11.50v_11565 | 0 | 0 | 0 | 0 | 0 | 0 | 0 | 0 | 0 | 0 | 0 |
| virus_Jul11.50v_13821 | 0 | 0 | 0 | 0 | 0 | 0 | 0 | 0 | 0 | 0 | 0 |
| virus_Jul11.50v_14199 | 0 | 0 | 0 | 0 | 0 | 0 | 0 | 0 | 0 | 0 | 0 |
| virus_Jul11.50v_16712 | 0 | 0 | 0 | 0 | 0 | 0 | 0 | 0 | 0 | 0 | 0 |
| virus_Jul11.50v_18621 | 0 | 0 | 0 | 0 | 0 | 0 | 0 | 0 | 0 | 0 | 0 |
| virus_Jul11.50v_19785 | 0 | 0 | 0 | 0 | 0 | 0 | 0 | 0 | 0 | 0 | 0 |
| virus_Jul11.50v_19926 | 0 | 0 | 0 | 0 | 0 | 0 | 0 | 0 | 0 | 0 | 0 |
| virus_Jul11.75v_3391 | 0 | 0 | 0 | 0 | 0 | 0 | 0 | 0 | 0 | 0 | 0 |
| virus_Jul11.75v_13608 | 0 | 0 | 0 | 0 | 0 | 0 | 0 | 0 | 0 | 0 | 0 |
| virus_Jul11.75v_24568 | 0 | 0 | 0 | 0 | 0 | 0 | 0 | 0 | 0 | 0 | 0 |
| virus_Jul11.75v_34355 | 0 | 0 | 0 | 0 | 0 | 0 | 0 | 0 | 0 | 0 | 0 |
| virus_Jul11.75v_37277 | 0 | 0 | 0 | 0 | 0 | 0 | 0 | 0 | 0 | 0 | 0 |
| virus_Jul27.5v_843 | 0 | 0 | 0 | 0 | 0 | 0 | 0 | 0 | 0 | 0 | 0 |
| virus_Jul27.5v_950 | 0 | 0 | 0 | 0 | 0 | 0 | 0 | 0 | 0 | 0 | 0 |
| virus_Jul27.5v_1171 | 0 | 0 | 0 | 0 | 0 | 0 | 0 | 0 | 0 | 0 | 0 |
| virus_Jul27.5v_1182 | 0 | 0 | 0 | 0 | 0 | 0 | 0 | 0 | 0 | 0 | 0 |
| virus_Jul27.5v_1870 | 0 | 0 | 0 | 0 | 0 | 0 | 0 | 0 | 0 | 0 | 0 |
| virus_Jul27.5v_2857 | 0 | 0 | 0 | 0 | 0 | 0 | 0 | 0 | 0 | 0 | 0 |
| virus_Jul27.5v_3275 | 0 | 0 | 0 | 0 | 0 | 0 | 0 | 0 | 0 | 0 | 0 |
| virus_Jul27.5v_3540 | 0 | 0 | 0 | 0 | 0 | 0 | 0 | 0 | 0 | 0 | 0 |
| virus_Jul27.5v_3810 | 0 | 0 | 0 | 0 | 0 | 0 | 0 | 0 | 0 | 0 | 0 |
| virus_Jul27.5v_4018 | 0 | 0 | 0 | 0 | 0 | 0 | 0 | 0 | 0 | 0 | 0 |
| virus_Jul27.5v_6689 | 0 | 0 | 0 | 0 | 0 | 0 | 0 | 0 | 0 | 0 | 0 |
| virus_Jul27.5v_8387 | 0 | 0 | 0 | 0 | 3 | 1 | 0 | 0 | 0 | 0 | 4 |
| virus_Jul27.5v_8906 | 0 | 0 | 0 | 0 | 0 | 0 | 0 | 0 | 0 | 0 | 0 |
| virus_Jul27.5v_9686 | 0 | 0 | 0 | 0 | 0 | 0 | 0 | 0 | 0 | 0 | 0 |
| virus_Jul27.5v_10015 | 0 | 1 | 1 | 0 | 0 | 0 | 0 | 0 | 0 | 0 | 2 |
| virus_Jul27.5v_10371 | 0 | 0 | 0 | 0 | 0 | 0 | 0 | 0 | 0 | 0 | 0 |
| virus_Jul27.5v_10375 | 0 | 0 | 0 | 0 | 0 | 0 | 0 | 0 | 0 | 0 | 0 |
| virus_Jul27.5v_10681 | 0 | 0 | 0 | 0 | 0 | 0 | 0 | 0 | 0 | 0 | 0 |
| virus_Jul27.5v_10725 | 0 | 0 | 0 | 0 | 0 | 0 | 0 | 0 | 0 | 0 | 0 |
| virus_Jul27.5v_11919 | 0 | 0 | 0 | 0 | 0 | 0 | 0 | 0 | 0 | 0 | 0 |
| virus_Jul27.5v_12166 | 0 | 0 | 0 | 0 | 0 | 0 | 0 | 0 | 0 | 0 | 0 |
| virus_Jul27.5v_13132 | 0 | 0 | 0 | 0 | 0 | 0 | 0 | 0 | 0 | 0 | 0 |
| virus_Jul27.5v_13408 | 0 | 0 | 0 | 0 | 0 | 0 | 0 | 0 | 0 | 0 | 0 |
| virus_Jul27.5v_13488 | 0 | 0 | 0 | 0 | 0 | 1 | 0 | 0 | 0 | 0 | 1 |
| virus_Jul27.5v_14080 | 0 | 0 | 0 | 0 | 0 | 0 | 0 | 0 | 0 | 0 | 0 |
| virus_Jul27.5v_14291 | 0 | 0 | 0 | 0 | 0 | 0 | 0 | 0 | 0 | 0 | 0 |
| virus_Jul27.5v_15997 | 0 | 0 | 0 | 0 | 0 | 0 | 0 | 0 | 0 | 0 | 0 |
| virus_Jul27.5v_16519 | 0 | 0 | 0 | 0 | 0 | 0 | 0 | 0 | 0 | 0 | 0 |
| virus_Jul27.5v_16814 | 0 | 0 | 0 | 0 | 0 | 0 | 0 | 0 | 0 | 0 | 0 |
| virus_Jul27.5v_18724 | 0 | 0 | 0 | 0 | 0 | 0 | 0 | 0 | 0 | 0 | 0 |
| virus_Jul27.5v_18780 | 0 | 0 | 0 | 0 | 0 | 0 | 0 | 0 | 0 | 0 | 0 |
| virus_Jul27.5v_20008 | 0 | 0 | 0 | 0 | 0 | 0 | 0 | 0 | 0 | 0 | 0 |
| virus_Jul27.5v_20962 | 0 | 0 | 0 | 0 | 0 | 0 | 0 | 0 | 0 | 0 | 0 |
| virus_Jul27.5v_21101 | 0 | 0 | 0 | 0 | 0 | 0 | 0 | 0 | 0 | 0 | 0 |
| virus_Jul27.25v_469 | 0 | 0 | 0 | 0 | 0 | 0 | 0 | 0 | 0 | 0 | 0 |
| virus_Jul27.25v_2097 | 0 | 0 | 0 | 0 | 0 | 0 | 0 | 0 | 0 | 0 | 0 |
| virus_Jul27.25v_3424 | 0 | 0 | 0 | 0 | 0 | 0 | 0 | 0 | 0 | 0 | 0 |
| virus_Jul27.25v_6615 | 0 | 0 | 0 | 0 | 0 | 0 | 0 | 0 | 0 | 0 | 0 |
| virus_Jul27.25v_6801 | 0 | 0 | 0 | 0 | 0 | 0 | 0 | 0 | 0 | 0 | 0 |
| virus_Jul27.25v_10565 | 0 | 0 | 0 | 0 | 0 | 0 | 0 | 0 | 0 | 0 | 0 |
| virus_Jul27.25v_12240 | 0 | 0 | 0 | 0 | 0 | 0 | 0 | 0 | 0 | 0 | 0 |
| virus_Jul27.25v_12504 | 1 | 1 | 0 | 3 | 4 | 0 | 0 | 0 | 0 | 0 | 9 |
| virus_Jul27.25v_13264 | 0 | 0 | 0 | 0 | 0 | 0 | 0 | 0 | 0 | 0 | 0 |
| virus_Jul27.25v_14754 | 0 | 0 | 0 | 0 | 0 | 0 | 0 | 0 | 0 | 0 | 0 |
| virus_Jul27.25v_22688 | 0 | 0 | 0 | 0 | 0 | 0 | 0 | 0 | 0 | 0 | 0 |
| virus_Jul27.25v_25885 | 0 | 0 | 0 | 0 | 0 | 0 | 0 | 0 | 0 | 0 | 0 |
| virus_Jul27.25v_32275 | 0 | 0 | 0 | 0 | 0 | 0 | 0 | 0 | 0 | 0 | 0 |
| virus_Jul27.25v_32554 | 0 | 0 | 0 | 0 | 0 | 0 | 0 | 0 | 0 | 0 | 0 |
| virus_Jul27.25v_35277 | 0 | 0 | 0 | 0 | 0 | 0 | 0 | 0 | 0 | 0 | 0 |
| virus_Jul27.75v_3174 | 0 | 0 | 0 | 0 | 0 | 0 | 0 | 0 | 0 | 0 | 0 |
| virus_Jul27.75v_11058 | 0 | 0 | 0 | 0 | 0 | 0 | 0 | 0 | 0 | 0 | 0 |
| virus_Jun8.25v_4189 | 0 | 0 | 0 | 0 | 0 | 0 | 0 | 0 | 0 | 0 | 0 |
| virus_Jun8.25v_4654 | 0 | 0 | 0 | 0 | 0 | 0 | 0 | 0 | 0 | 0 | 0 |
| virus_Jun8.25v_5811 | 0 | 0 | 0 | 0 | 0 | 0 | 0 | 0 | 0 | 0 | 0 |
| virus_Jun8.25v_37889 | 0 | 0 | 0 | 0 | 0 | 0 | 0 | 0 | 0 | 0 | 0 |
| virus_Jun8.50v_502 | 0 | 0 | 0 | 0 | 0 | 0 | 0 | 0 | 0 | 0 | 0 |
| virus_Jun8.50v_5068 | 0 | 0 | 0 | 0 | 0 | 0 | 0 | 0 | 0 | 0 | 0 |
| virus_Jun8.50v_7917 | 0 | 0 | 0 | 0 | 0 | 0 | 0 | 0 | 0 | 0 | 0 |
| virus_Jun8.50v_13174 | 0 | 0 | 0 | 0 | 0 | 0 | 0 | 0 | 0 | 0 | 0 |
| virus_Jun8.50v_17672 | 0 | 0 | 0 | 0 | 0 | 0 | 0 | 0 | 0 | 0 | 0 |
| virus_Jun8.50v_20182 | 0 | 0 | 0 | 0 | 0 | 0 | 0 | 0 | 0 | 0 | 0 |
| virus_Jun8.50v_24700 | 0 | 0 | 0 | 0 | 0 | 0 | 0 | 0 | 0 | 0 | 0 |
| virus_Jun8.50v_27475 | 0 | 0 | 0 | 0 | 0 | 0 | 0 | 0 | 0 | 0 | 0 |
| virus_Jun8.50v_39939 | 0 | 0 | 0 | 0 | 0 | 0 | 0 | 0 | 0 | 0 | 0 |
| virus_Jun8.75v_115 | 0 | 0 | 0 | 0 | 0 | 0 | 0 | 0 | 0 | 0 | 0 |
| virus_Jun8.75v_3911 | 0 | 0 | 0 | 0 | 0 | 0 | 0 | 0 | 0 | 0 | 0 |
| virus_Jun8.75v_10966 | 0 | 0 | 0 | 0 | 0 | 0 | 0 | 0 | 0 | 0 | 0 |
| virus_Jun8.75v_12670 | 0 | 0 | 0 | 0 | 0 | 0 | 0 | 0 | 0 | 0 | 0 |
| virus_Jun8.75v_14242 | 0 | 0 | 0 | 0 | 0 | 0 | 0 | 0 | 0 | 0 | 0 |
| virus_Jun8.75v_17613 | 0 | 0 | 0 | 0 | 0 | 0 | 0 | 0 | 0 | 0 | 0 |
| virus_Jun8.75v_18074 | 0 | 1 | 0 | 0 | 0 | 0 | 0 | 0 | 0 | 0 | 1 |
| virus_Jun22.0v_3590 | 0 | 0 | 0 | 0 | 0 | 0 | 0 | 0 | 0 | 0 | 0 |
| virus_Jun22.0v_5725 | 0 | 0 | 0 | 0 | 0 | 0 | 0 | 0 | 0 | 0 | 0 |
| virus_Jun22.0v_6413 | 0 | 0 | 0 | 0 | 0 | 0 | 0 | 0 | 0 | 0 | 0 |
| virus_Jun22.0v_8397 | 0 | 0 | 0 | 0 | 0 | 0 | 0 | 0 | 0 | 0 | 0 |
| virus_Jun22.0v_9242 | 0 | 0 | 0 | 0 | 0 | 0 | 0 | 0 | 0 | 0 | 0 |
| virus_Jun22.0v_9832 | 0 | 0 | 0 | 0 | 0 | 0 | 0 | 0 | 0 | 0 | 0 |
| virus_Jun22.0v_10533 | 0 | 0 | 0 | 0 | 0 | 0 | 0 | 0 | 0 | 0 | 0 |
| virus_Jun22.0v_10752 | 0 | 0 | 0 | 0 | 0 | 0 | 0 | 0 | 0 | 0 | 0 |
| virus_Jun22.0v_13380 | 0 | 0 | 0 | 0 | 0 | 0 | 0 | 0 | 0 | 0 | 0 |
| virus_Jun22.0v_15281 | 0 | 0 | 0 | 0 | 0 | 0 | 0 | 0 | 0 | 0 | 0 |
| virus_Jun22.0v_15382 | 0 | 0 | 0 | 0 | 0 | 1 | 0 | 0 | 0 | 0 | 1 |
| virus_Jun22.0v_15498 | 0 | 0 | 0 | 0 | 0 | 0 | 0 | 0 | 0 | 0 | 0 |
| virus_Jun22.0v_19381 | 0 | 0 | 0 | 0 | 0 | 0 | 0 | 0 | 0 | 0 | 0 |
| virus_Jun22.0v_20016 | 0 | 0 | 0 | 0 | 0 | 0 | 0 | 0 | 0 | 0 | 0 |
| virus_Jun22.50v_62 | 0 | 0 | 0 | 0 | 0 | 0 | 0 | 0 | 0 | 0 | 0 |
| virus_Jun22.50v_441 | 0 | 0 | 0 | 0 | 0 | 0 | 0 | 0 | 0 | 0 | 0 |
| virus_Jun22.50v_3420 | 0 | 0 | 0 | 0 | 0 | 0 | 0 | 0 | 0 | 0 | 0 |
| virus_Jun22.50v_4277 | 0 | 0 | 0 | 0 | 0 | 0 | 0 | 0 | 0 | 0 | 0 |
| virus_Jun22.50v_6586 | 0 | 0 | 0 | 0 | 0 | 0 | 0 | 0 | 0 | 0 | 0 |
| virus_Jun22.50v_9933 | 0 | 0 | 0 | 0 | 0 | 0 | 0 | 0 | 0 | 0 | 0 |
| virus_Jun22.50v_12043 | 0 | 0 | 0 | 0 | 0 | 0 | 0 | 0 | 0 | 0 | 0 |
| virus_Jun22.50v_14067 | 0 | 0 | 0 | 0 | 0 | 0 | 0 | 0 | 0 | 0 | 0 |
| virus_Jun22.50v_15762 | 0 | 0 | 0 | 0 | 0 | 0 | 0 | 0 | 0 | 0 | 0 |
| virus_Jun22.50v_16786 | 0 | 0 | 0 | 0 | 0 | 0 | 0 | 0 | 0 | 0 | 0 |
| virus_Jun22.50v_18709 | 0 | 0 | 0 | 0 | 0 | 0 | 0 | 0 | 0 | 0 | 0 |
| virus_Jun22.50v_19423 | 0 | 0 | 0 | 0 | 0 | 0 | 0 | 0 | 0 | 0 | 0 |
| virus_Jun22.50v_24832 | 0 | 0 | 0 | 1 | 0 | 0 | 0 | 0 | 0 | 0 | 1 |
| virus_Jun22.50v_26207 | 0 | 0 | 0 | 0 | 0 | 0 | 0 | 0 | 0 | 0 | 0 |
| virus_Jun22.50v_28387 | 0 | 0 | 0 | 0 | 0 | 0 | 0 | 0 | 0 | 0 | 0 |
| virus_Jun22.50v_31049 | 0 | 0 | 0 | 0 | 0 | 0 | 0 | 0 | 0 | 0 | 0 |
| virus_Jun22.50v_33443 | 0 | 0 | 0 | 0 | 0 | 0 | 0 | 0 | 0 | 0 | 0 |
| virus_Jun22.50v_34947 | 0 | 0 | 0 | 0 | 0 | 0 | 0 | 0 | 0 | 0 | 0 |
| virus_Jun22.50v_36431 | 0 | 0 | 0 | 0 | 0 | 0 | 0 | 0 | 0 | 0 | 0 |
| virus_Jun22.50v_38646 | 0 | 0 | 0 | 0 | 0 | 0 | 0 | 0 | 0 | 0 | 0 |
| virus_Jun22.50v_39947 | 0 | 0 | 0 | 0 | 0 | 0 | 0 | 0 | 0 | 0 | 0 |
| virus_Jun22.50v_40701 | 0 | 0 | 0 | 0 | 0 | 0 | 0 | 0 | 0 | 0 | 0 |

| | | | | | | | | | | | |
|---|---|---|---|---|---|---|---|---|---|---|---|
| virus_Jun22.75v_8444 | 0 | 0 | 0 | 0 | 0 | 0 | 0 | 0 | 0 | 0 | 0 |
| virus_Jun22.75v_21281 | 0 | 0 | 0 | 0 | 0 | 0 | 0 | 0 | 0 | 0 | 0 |
| virus_Jun22.75v_23329 | 0 | 0 | 0 | 0 | 0 | 0 | 0 | 0 | 0 | 0 | 0 |
| virus_Jun22.75v_30527 | 0 | 0 | 0 | 0 | 0 | 0 | 0 | 0 | 0 | 0 | 0 |
| virus_Jun22.75v_34611 | 0 | 0 | 0 | 0 | 0 | 0 | 0 | 0 | 0 | 0 | 0 |
| virus_Jun22.75v_36044 | 0 | 1 | 0 | 0 | 0 | 0 | 0 | 0 | 0 | 0 | 1 |
| virus_Jun22.75v_36530 | 0 | 0 | 0 | 0 | 0 | 0 | 0 | 0 | 0 | 0 | 0 |
| virus_May12.5v_10726 | 0 | 0 | 0 | 0 | 0 | 0 | 0 | 0 | 0 | 0 | 0 |
| virus_May12.5v_13906 | 0 | 0 | 0 | 0 | 0 | 0 | 0 | 0 | 0 | 0 | 0 |
| virus_May12.5v_20244 | 0 | 0 | 0 | 0 | 0 | 0 | 0 | 0 | 0 | 0 | 0 |
| virus_May12.5v_26333 | 0 | 0 | 0 | 0 | 0 | 0 | 0 | 0 | 0 | 0 | 0 |
| virus_May12.5v_29920 | 0 | 0 | 0 | 0 | 0 | 0 | 0 | 0 | 0 | 0 | 0 |
| virus_May12.5v_30341 | 0 | 0 | 0 | 0 | 0 | 0 | 0 | 0 | 0 | 0 | 0 |
| virus_May12.25v_2024 | 0 | 0 | 0 | 0 | 0 | 0 | 0 | 0 | 0 | 0 | 0 |
| virus_May12.25v_8993 | 0 | 0 | 0 | 0 | 0 | 0 | 0 | 0 | 0 | 0 | 0 |
| virus_May12.25v_10234 | 0 | 0 | 0 | 0 | 0 | 0 | 0 | 0 | 0 | 0 | 0 |
| virus_May12.25v_18546 | 0 | 0 | 0 | 0 | 0 | 0 | 0 | 0 | 0 | 0 | 0 |
| virus_May12.25v_19002 | 0 | 0 | 0 | 0 | 0 | 0 | 0 | 0 | 0 | 0 | 0 |
| virus_May12.25v_32287 | 0 | 0 | 0 | 0 | 0 | 0 | 0 | 0 | 0 | 0 | 0 |
| virus_May12.50v_2546 | 0 | 0 | 0 | 0 | 0 | 0 | 0 | 0 | 0 | 0 | 0 |
| virus_May12.50v_3821 | 0 | 0 | 0 | 0 | 0 | 0 | 0 | 0 | 0 | 0 | 0 |
| virus_May12.50v_5104 | 0 | 0 | 0 | 0 | 1 | 1 | 0 | 0 | 0 | 0 | 2 |
| virus_May12.50v_5278 | 0 | 0 | 0 | 0 | 0 | 0 | 0 | 0 | 0 | 0 | 0 |
| virus_May12.50v_7981 | 0 | 0 | 0 | 0 | 0 | 0 | 0 | 0 | 0 | 0 | 0 |
| virus_May12.50v_8873 | 0 | 0 | 0 | 0 | 0 | 0 | 0 | 0 | 0 | 0 | 0 |
| virus_May12.50v_10361 | 0 | 0 | 0 | 0 | 0 | 0 | 0 | 0 | 0 | 0 | 0 |
| virus_May12.50v_11921 | 0 | 0 | 0 | 0 | 0 | 0 | 0 | 0 | 0 | 0 | 0 |
| virus_May12.50v_12297 | 0 | 0 | 0 | 0 | 0 | 0 | 0 | 0 | 0 | 0 | 0 |
| virus_May12.50v_13239 | 0 | 0 | 0 | 0 | 0 | 0 | 0 | 0 | 0 | 0 | 0 |
| virus_May12.50v_18083 | 0 | 0 | 0 | 0 | 0 | 0 | 0 | 0 | 0 | 0 | 0 |
| virus_May12.50v_19435 | 0 | 0 | 0 | 0 | 0 | 0 | 0 | 0 | 0 | 0 | 0 |
| virus_May12.50v_26459 | 0 | 0 | 0 | 0 | 0 | 0 | 0 | 0 | 0 | 0 | 0 |
| virus_May12.50v_26716 | 0 | 0 | 0 | 0 | 0 | 0 | 0 | 0 | 0 | 0 | 0 |
| virus_May12.50v_26768 | 0 | 0 | 0 | 0 | 0 | 0 | 0 | 0 | 0 | 0 | 0 |
| virus_May12.50v_30347 | 0 | 0 | 0 | 0 | 0 | 0 | 0 | 0 | 0 | 0 | 0 |
| virus_May12.50v_30643 | 0 | 0 | 0 | 0 | 0 | 0 | 0 | 0 | 0 | 0 | 0 |
| virus_May12.50v_30988 | 0 | 0 | 0 | 0 | 0 | 0 | 0 | 0 | 0 | 0 | 0 |
| virus_May12.50v_32300 | 0 | 0 | 0 | 0 | 0 | 0 | 0 | 0 | 0 | 0 | 0 |
| virus_May12.50v_34468 | 0 | 0 | 0 | 0 | 0 | 0 | 0 | 0 | 0 | 0 | 0 |
| virus_May12.50v_37010 | 0 | 0 | 0 | 0 | 0 | 0 | 0 | 0 | 0 | 0 | 0 |
| virus_May12.50v_41150 | 0 | 0 | 0 | 0 | 0 | 0 | 0 | 0 | 0 | 0 | 0 |
| virus_May12.75v_1619 | 0 | 0 | 0 | 0 | 0 | 0 | 0 | 0 | 0 | 0 | 0 |
| virus_May12.75v_5902 | 0 | 0 | 0 | 0 | 0 | 0 | 0 | 0 | 0 | 0 | 0 |
| virus_May12.75v_7263 | 0 | 0 | 0 | 0 | 0 | 0 | 0 | 0 | 0 | 0 | 0 |
| virus_May12.75v_8282 | 0 | 0 | 0 | 0 | 0 | 0 | 0 | 0 | 0 | 0 | 0 |
| virus_May12.75v_12328 | 0 | 0 | 0 | 0 | 0 | 0 | 0 | 0 | 0 | 0 | 0 |
| virus_May12.75v_13736 | 0 | 0 | 0 | 0 | 0 | 0 | 0 | 0 | 0 | 0 | 0 |
| virus_May12.75v_15365 | 0 | 0 | 0 | 0 | 0 | 0 | 0 | 0 | 0 | 0 | 0 |
| virus_May12.75v_15510 | 0 | 0 | 0 | 0 | 0 | 0 | 0 | 0 | 0 | 0 | 0 |
| virus_May12.75v_16696 | 0 | 0 | 0 | 0 | 0 | 0 | 0 | 0 | 0 | 0 | 0 |
| virus_May12.75v_16871 | 0 | 0 | 0 | 0 | 0 | 0 | 0 | 0 | 0 | 0 | 0 |
| virus_May12.75v_23165 | 0 | 0 | 0 | 0 | 0 | 0 | 0 | 0 | 0 | 0 | 0 |
| virus_May12.75v_23945 | 0 | 0 | 0 | 0 | 0 | 0 | 0 | 0 | 0 | 0 | 0 |
| virus_May12.75v_26613 | 0 | 0 | 0 | 0 | 0 | 0 | 0 | 0 | 0 | 0 | 0 |
| virus_May12.75v_31817 | 0 | 0 | 0 | 0 | 0 | 0 | 0 | 0 | 0 | 0 | 0 |
| virus_May12.75v_35955 | 0 | 0 | 0 | 0 | 0 | 0 | 0 | 0 | 0 | 0 | 0 |
| virus_Sep8.0v_2763 | 0 | 0 | 0 | 0 | 0 | 0 | 0 | 0 | 0 | 0 | 0 |
| virus_Sep8.0v_2914 | 0 | 0 | 0 | 0 | 0 | 0 | 0 | 0 | 0 | 0 | 0 |
| virus_Sep8.0v_3366 | 0 | 0 | 0 | 0 | 0 | 0 | 0 | 0 | 0 | 0 | 0 |
| virus_Sep8.0v_6021 | 0 | 0 | 0 | 0 | 0 | 0 | 0 | 0 | 0 | 0 | 0 |
| virus_Sep8.0v_7698 | 0 | 0 | 0 | 0 | 0 | 0 | 0 | 0 | 0 | 0 | 0 |
| virus_Sep8.0v_12226 | 0 | 0 | 0 | 0 | 0 | 0 | 0 | 0 | 0 | 0 | 0 |
| virus_Sep8.0v_13681 | 0 | 0 | 0 | 0 | 0 | 0 | 0 | 0 | 0 | 0 | 0 |
| virus_Sep8.0v_14852 | 0 | 0 | 0 | 0 | 0 | 0 | 0 | 0 | 0 | 0 | 0 |
| virus_Sep8.0v_16804 | 0 | 0 | 0 | 0 | 0 | 0 | 0 | 0 | 0 | 0 | 0 |
| virus_Sep8.5v_4344 | 0 | 0 | 0 | 0 | 0 | 0 | 0 | 0 | 0 | 0 | 0 |
| virus_Sep8.5v_4658 | 0 | 0 | 0 | 0 | 0 | 0 | 0 | 0 | 0 | 0 | 0 |
| virus_Sep8.5v_6795 | 0 | 0 | 0 | 0 | 0 | 0 | 0 | 0 | 0 | 0 | 0 |
| virus_Sep8.5v_7952 | 0 | 0 | 0 | 0 | 0 | 0 | 0 | 0 | 0 | 0 | 0 |
| virus_Sep8.5v_9010 | 0 | 0 | 0 | 0 | 0 | 0 | 0 | 0 | 0 | 0 | 0 |
| virus_Sep8.5v_11686 | 0 | 0 | 0 | 0 | 0 | 0 | 0 | 0 | 0 | 0 | 0 |
| virus_Sep8.5v_14931 | 0 | 0 | 0 | 0 | 0 | 0 | 0 | 0 | 0 | 0 | 0 |
| virus_Sep8.5v_16321 | 0 | 0 | 0 | 0 | 0 | 0 | 0 | 0 | 0 | 0 | 0 |
| virus_Sep8.25v_10438 | 0 | 0 | 0 | 0 | 0 | 0 | 0 | 0 | 0 | 0 | 0 |
| virus_Sep8.25v_17802 | 0 | 0 | 0 | 0 | 0 | 0 | 0 | 0 | 0 | 0 | 0 |
| virus_Sep8.25v_20688 | 0 | 0 | 0 | 0 | 0 | 0 | 0 | 0 | 0 | 0 | 0 |
| virus_Sep8.50v_1914 | 0 | 0 | 0 | 0 | 0 | 0 | 0 | 0 | 0 | 0 | 0 |
| virus_Sep8.50v_1986 | 0 | 1 | 0 | 0 | 0 | 0 | 0 | 0 | 0 | 0 | 1 |
| virus_Sep8.50v_4677 | 0 | 0 | 0 | 0 | 0 | 0 | 0 | 0 | 0 | 0 | 0 |
| virus_Sep8.50v_5365 | 0 | 0 | 0 | 0 | 0 | 0 | 0 | 0 | 0 | 0 | 0 |
| virus_Sep8.50v_13663 | 0 | 0 | 0 | 0 | 0 | 0 | 0 | 0 | 0 | 0 | 0 |
| virus_Sep8.50v_17016 | 0 | 0 | 0 | 0 | 0 | 0 | 0 | 0 | 0 | 0 | 0 |
| virus_Sep8.50v_17084 | 0 | 0 | 0 | 0 | 0 | 0 | 0 | 0 | 0 | 0 | 0 |
| virus_Sep8.50v_17827 | 0 | 0 | 0 | 0 | 0 | 0 | 0 | 0 | 0 | 0 | 0 |
| virus_Sep8.50v_30838 | 0 | 0 | 0 | 0 | 0 | 0 | 0 | 0 | 0 | 0 | 0 |
| virus_Sep8.50v_32867 | 0 | 0 | 0 | 0 | 0 | 0 | 0 | 0 | 0 | 0 | 0 |
| virus_Sep8.50v_33302 | 0 | 0 | 0 | 0 | 0 | 0 | 0 | 0 | 0 | 0 | 0 |
| virus_Sep8.50v_36214 | 0 | 0 | 0 | 0 | 0 | 0 | 0 | 0 | 0 | 0 | 0 |
| virus_Sep8.50v_36971 | 0 | 0 | 0 | 0 | 0 | 0 | 0 | 0 | 0 | 0 | 0 |
| virus_Sep8.50v_38954 | 0 | 0 | 0 | 0 | 0 | 0 | 0 | 0 | 0 | 0 | 0 |
| virus_Sep8.50v_40927 | 0 | 0 | 0 | 0 | 0 | 0 | 0 | 0 | 0 | 0 | 0 |
| bacteria_Jul11d0b_3362 | 0 | 0 | 0 | 0 | 1 | 0 | 0 | 0 | 0 | 0 | 1 |
| bacteria_Jul11d0b_24044 | 0 | 0 | 0 | 0 | 0 | 0 | 0 | 0 | 0 | 0 | 0 |
| bacteria_Jul11d0b_30788 | 0 | 0 | 0 | 0 | 0 | 0 | 0 | 0 | 0 | 0 | 0 |
| bacteria_Jul11d0b_34563 | 0 | 0 | 0 | 0 | 0 | 0 | 0 | 0 | 0 | 0 | 0 |
| bacteria_Jul11d0b_43429 | 0 | 0 | 0 | 0 | 0 | 0 | 0 | 0 | 0 | 0 | 0 |
| bacteria_Jul11d0b_50464 | 0 | 0 | 0 | 0 | 0 | 0 | 0 | 0 | 0 | 0 | 0 |
| bacteria_Jul11d0b_53423 | 0 | 0 | 0 | 0 | 0 | 0 | 0 | 0 | 0 | 0 | 0 |
| bacteria_Jul11d0b_54836 | 0 | 0 | 0 | 0 | 0 | 0 | 0 | 0 | 0 | 0 | 0 |
| bacteria_Jul11d0b_63707 | 0 | 0 | 0 | 0 | 0 | 0 | 0 | 0 | 0 | 0 | 0 |
| bacteria_Jul11d0b_65078 | 0 | 0 | 0 | 0 | 0 | 0 | 0 | 0 | 0 | 0 | 0 |
| bacteria_Jul11d0b_71634 | 0 | 0 | 0 | 0 | 0 | 0 | 0 | 0 | 0 | 0 | 0 |
| bacteria_Aug5d0b_7396 | 1 | 0 | 0 | 0 | 0 | 0 | 0 | 0 | 0 | 0 | 1 |
| bacteria_Aug5d0b_11150 | 1 | 0 | 0 | 0 | 0 | 0 | 0 | 0 | 0 | 0 | 1 |
| bacteria_Aug5d0b_33754 | 1 | 0 | 0 | 0 | 0 | 0 | 0 | 0 | 0 | 0 | 1 |
| bacteria_Aug5d0b_37965 | 1 | 0 | 0 | 0 | 1 | 0 | 0 | 1 | 0 | 0 | 3 |
| bacteria_Aug5d0b_64213 | 1 | 0 | 0 | 0 | 0 | 0 | 0 | 0 | 0 | 0 | 1 |
| bacteria_Aug5d5b_2148 | 0 | 0 | 1 | 0 | 0 | 0 | 0 | 0 | 0 | 0 | 1 |
| bacteria_Aug5d5b_8186 | 0 | 0 | 1 | 0 | 0 | 0 | 0 | 0 | 0 | 0 | 1 |
| bacteria_Aug5d5b_26848 | 0 | 0 | 1 | 0 | 0 | 0 | 0 | 0 | 0 | 0 | 1 |
| bacteria_Aug5d5b_28101 | 0 | 0 | 1 | 0 | 0 | 0 | 0 | 0 | 0 | 0 | 1 |
| bacteria_Aug5d5b_55827 | 0 | 0 | 1 | 0 | 0 | 0 | 0 | 0 | 0 | 0 | 1 |
| bacteria_Aug5d5b_58626 | 0 | 0 | 1 | 0 | 0 | 0 | 0 | 0 | 0 | 0 | 1 |
| bacteria_Aug5d25b_6197 | 0 | 0 | 0 | 0 | 1 | 0 | 0 | 0 | 0 | 0 | 1 |
| bacteria_Aug5d25b_12313 | 0 | 0 | 0 | 0 | 1 | 0 | 0 | 0 | 0 | 0 | 1 |
| bacteria_Aug5d25b_13559 | 0 | 0 | 0 | 0 | 1 | 0 | 0 | 0 | 0 | 0 | 1 |
| bacteria_Aug5d25b_14783 | 0 | 0 | 0 | 0 | 1 | 0 | 0 | 0 | 0 | 0 | 1 |
| bacteria_Aug5d25b_16579 | 0 | 0 | 0 | 0 | 1 | 0 | 0 | 0 | 0 | 0 | 1 |
| bacteria_Aug5d25b_21783 | 0 | 0 | 0 | 0 | 1 | 0 | 0 | 0 | 0 | 0 | 1 |
| bacteria_Aug5d25b_25424 | 0 | 0 | 0 | 0 | 1 | 0 | 0 | 0 | 0 | 0 | 1 |
| bacteria_Aug5d25b_26381 | 0 | 0 | 0 | 0 | 1 | 0 | 0 | 0 | 0 | 0 | 1 |
| bacteria_Aug5d25b_30172 | 0 | 0 | 0 | 0 | 1 | 0 | 0 | 0 | 0 | 0 | 1 |
| bacteria_Aug5d25b_30549 | 0 | 0 | 0 | 0 | 1 | 0 | 0 | 0 | 0 | 0 | 1 |
| bacteria_Aug5d25b_44387 | 0 | 0 | 0 | 0 | 1 | 0 | 0 | 0 | 0 | 0 | 1 |
| bacteria_Aug5d25b_57302 | 0 | 0 | 0 | 0 | 1 | 0 | 0 | 0 | 0 | 0 | 1 |
| bacteria_Aug5d25b_72086 | 0 | 0 | 0 | 0 | 1 | 0 | 0 | 0 | 0 | 0 | 1 |
| bacteria_Aug5d25b_72845 | 0 | 0 | 0 | 0 | 1 | 0 | 0 | 0 | 0 | 0 | 1 |
| bacteria_Aug5d25b_73102 | 0 | 0 | 0 | 0 | 1 | 0 | 0 | 0 | 0 | 0 | 1 |
| bacteria_Aug5d25b_73860 | 0 | 0 | 0 | 0 | 1 | 0 | 0 | 0 | 0 | 0 | 1 |
| bacteria_Aug5d50b_37663 | 0 | 0 | 0 | 0 | 0 | 0 | 1 | 0 | 0 | 0 | 1 |
| bacteria_Aug5d50b_48721 | 0 | 0 | 0 | 0 | 0 | 0 | 1 | 0 | 0 | 0 | 1 |
| bacteria_Aug5d50b_60514 | 0 | 0 | 0 | 0 | 0 | 0 | 1 | 0 | 0 | 0 | 1 |
| bacteria_Aug5d50b_64610 | 0 | 0 | 0 | 0 | 0 | 0 | 1 | 0 | 0 | 0 | 1 |
| bacteria_Aug5d50b_65293 | 0 | 0 | 0 | 0 | 0 | 0 | 1 | 0 | 0 | 0 | 1 |
| bacteria_Jul11d5b_48719 | 0 | 0 | 0 | 0 | 0 | 0 | 0 | 0 | 0 | 0 | 0 |
| bacteria_Jul11d25b_1024 | 0 | 0 | 0 | 0 | 0 | 0 | 0 | 0 | 0 | 0 | 0 |
| bacteria_Jul11d25b_6161 | 0 | 0 | 0 | 0 | 0 | 0 | 0 | 0 | 0 | 0 | 0 |
| bacteria_Jul11d25b_10720 | 0 | 0 | 0 | 0 | 0 | 0 | 0 | 0 | 0 | 0 | 0 |
| bacteria_Jul11d25b_12879 | 0 | 0 | 0 | 0 | 0 | 0 | 0 | 0 | 0 | 0 | 0 |
| bacteria_Jul11d25b_13173 | 0 | 0 | 0 | 0 | 0 | 0 | 0 | 0 | 0 | 0 | 0 |
| bacteria_Jul11d25b_15054 | 0 | 0 | 0 | 0 | 0 | 0 | 0 | 0 | 0 | 0 | 0 |
| bacteria_Jul11d25b_15678 | 0 | 0 | 0 | 0 | 0 | 0 | 0 | 0 | 0 | 0 | 0 |
| bacteria_Jul11d25b_35128 | 0 | 0 | 0 | 0 | 0 | 0 | 0 | 0 | 0 | 0 | 0 |
| bacteria_Jul11d25b_49473 | 0 | 0 | 0 | 0 | 0 | 0 | 0 | 0 | 0 | 0 | 0 |
| bacteria_Jul11d25b_51857 | 0 | 0 | 0 | 0 | 0 | 0 | 0 | 0 | 0 | 0 | 0 |
| bacteria_Jul11d50b_3464 | 0 | 0 | 0 | 0 | 0 | 0 | 0 | 0 | 0 | 0 | 0 |
| bacteria_Jul11d50b_16219 | 0 | 0 | 0 | 0 | 0 | 0 | 0 | 0 | 0 | 0 | 0 |
| bacteria_Jul11d50b_32894 | 0 | 0 | 0 | 0 | 0 | 0 | 0 | 0 | 0 | 0 | 0 |
| bacteria_Jul11d50b_41228 | 0 | 0 | 0 | 0 | 0 | 0 | 0 | 0 | 0 | 0 | 0 |
| bacteria_Jul11d50b_42406 | 0 | 0 | 0 | 0 | 0 | 0 | 0 | 0 | 0 | 0 | 0 |
| bacteria_Jul11d50b_47170 | 0 | 0 | 0 | 0 | 0 | 0 | 0 | 0 | 0 | 0 | 0 |
| bacteria_Jul11d75b_2126 | 0 | 0 | 0 | 0 | 0 | 0 | 0 | 0 | 0 | 0 | 0 |
| bacteria_Jul11d75b_5378 | 0 | 0 | 0 | 0 | 0 | 0 | 0 | 0 | 0 | 0 | 0 |
| bacteria_Jul11d75b_7061 | 0 | 0 | 0 | 0 | 0 | 0 | 0 | 0 | 0 | 0 | 0 |
| bacteria_Jul11d75b_20424 | 0 | 0 | 0 | 0 | 0 | 0 | 0 | 0 | 0 | 0 | 0 |
| bacteria_Jul11d75b_22001 | 0 | 0 | 0 | 0 | 0 | 0 | 0 | 0 | 0 | 0 | 0 |
| bacteria_Jul11d75b_22382 | 0 | 0 | 0 | 0 | 0 | 0 | 0 | 0 | 0 | 0 | 0 |

| | | | | | | | | | | | |
|---|---|---|---|---|---|---|---|---|---|---|---|
| bacteria_Jul11d75b_40531 | 0 | 0 | 0 | 0 | 0 | 0 | 0 | 0 | 0 | 0 | 0 |
| bacteria_Jul11d75b_44131 | 0 | 0 | 0 | 0 | 0 | 0 | 0 | 0 | 0 | 0 | 0 |
| bacteria_Jul11d75b_44469 | 0 | 0 | 0 | 0 | 0 | 0 | 0 | 0 | 0 | 0 | 0 |
| bacteria_Jul11d75b_47026 | 0 | 0 | 0 | 0 | 0 | 0 | 0 | 0 | 0 | 0 | 0 |
| bacteria_Jul11d75b_48668 | 0 | 0 | 0 | 0 | 0 | 0 | 0 | 0 | 0 | 0 | 0 |
| bacteria_Jul11d75b_50693 | 0 | 0 | 0 | 0 | 0 | 0 | 0 | 0 | 0 | 0 | 0 |
| bacteria_Jul11d75b_51199 | 0 | 0 | 0 | 0 | 0 | 0 | 0 | 0 | 0 | 0 | 0 |
| bacteria_Jul27d5b_371 | 0 | 0 | 0 | 0 | 0 | 0 | 0 | 0 | 0 | 0 | 0 |
| bacteria_Jul27d5b_3836 | 0 | 0 | 0 | 0 | 0 | 0 | 0 | 0 | 0 | 0 | 0 |
| bacteria_Jul27d5b_4533 | 0 | 0 | 0 | 0 | 0 | 0 | 0 | 0 | 0 | 0 | 0 |
| bacteria_Jul27d5b_5095 | 0 | 0 | 0 | 0 | 0 | 0 | 0 | 0 | 0 | 0 | 0 |
| bacteria_Jul27d5b_8610 | 0 | 0 | 0 | 0 | 0 | 0 | 0 | 0 | 0 | 0 | 0 |
| bacteria_Jul27d5b_8809 | 0 | 0 | 0 | 0 | 0 | 0 | 0 | 0 | 0 | 0 | 0 |
| bacteria_Jul27d5b_11294 | 0 | 0 | 0 | 0 | 0 | 0 | 0 | 0 | 0 | 0 | 0 |
| bacteria_Jul27d5b_14307 | 0 | 0 | 0 | 0 | 0 | 0 | 0 | 0 | 0 | 0 | 0 |
| bacteria_Jul27d5b_16666 | 0 | 0 | 0 | 0 | 0 | 0 | 0 | 0 | 0 | 0 | 0 |
| bacteria_Jul27d5b_19792 | 0 | 0 | 0 | 0 | 0 | 0 | 0 | 0 | 0 | 0 | 0 |
| bacteria_Jul27d5b_20733 | 0 | 0 | 0 | 0 | 0 | 0 | 0 | 0 | 0 | 0 | 0 |
| bacteria_Jul27d5b_22807 | 0 | 0 | 0 | 0 | 0 | 0 | 0 | 0 | 0 | 0 | 0 |
| bacteria_Jul27d5b_27485 | 0 | 0 | 0 | 0 | 0 | 0 | 0 | 0 | 0 | 0 | 0 |
| bacteria_Jul27d5b_30402 | 0 | 0 | 0 | 0 | 0 | 0 | 0 | 0 | 0 | 0 | 0 |
| bacteria_Jul27d5b_31026 | 0 | 0 | 0 | 0 | 0 | 0 | 0 | 0 | 0 | 0 | 0 |
| bacteria_Jul27d5b_33079 | 0 | 0 | 0 | 0 | 0 | 0 | 0 | 0 | 0 | 0 | 0 |
| bacteria_Jul27d5b_34794 | 0 | 0 | 0 | 0 | 0 | 0 | 0 | 0 | 0 | 0 | 0 |
| bacteria_Jul27d5b_34881 | 0 | 0 | 0 | 0 | 0 | 0 | 0 | 0 | 0 | 0 | 0 |
| bacteria_Jul27d5b_35273 | 0 | 0 | 0 | 0 | 0 | 0 | 0 | 0 | 0 | 0 | 0 |
| bacteria_Jul27d5b_39022 | 0 | 0 | 0 | 0 | 0 | 0 | 0 | 0 | 0 | 0 | 0 |
| bacteria_Jul27d5b_39135 | 0 | 0 | 0 | 0 | 0 | 0 | 0 | 0 | 0 | 0 | 0 |
| bacteria_Jul27d5b_43532 | 0 | 0 | 0 | 0 | 0 | 0 | 0 | 0 | 0 | 0 | 0 |
| bacteria_Jul27d5b_49001 | 0 | 0 | 0 | 0 | 0 | 0 | 0 | 0 | 0 | 0 | 0 |
| bacteria_Jul27d5b_50003 | 0 | 0 | 0 | 0 | 0 | 0 | 0 | 0 | 0 | 0 | 0 |
| bacteria_Jul27d5b_51746 | 0 | 0 | 0 | 0 | 0 | 0 | 0 | 0 | 0 | 0 | 0 |
| bacteria_Jul27d5b_52090 | 0 | 0 | 0 | 0 | 0 | 0 | 0 | 0 | 0 | 0 | 0 |
| bacteria_Jul27d25b_1530 | 0 | 0 | 0 | 0 | 0 | 0 | 0 | 0 | 0 | 0 | 0 |
| bacteria_Jul27d25b_8485 | 0 | 0 | 0 | 0 | 0 | 0 | 0 | 0 | 0 | 0 | 0 |
| bacteria_Jul27d25b_11631 | 0 | 0 | 0 | 0 | 0 | 0 | 0 | 0 | 0 | 0 | 0 |
| bacteria_Jul27d25b_19745 | 0 | 0 | 0 | 0 | 0 | 0 | 0 | 0 | 0 | 0 | 0 |
| bacteria_Jul27d25b_25726 | 0 | 0 | 0 | 0 | 0 | 0 | 0 | 0 | 0 | 0 | 0 |
| bacteria_Jul27d25b_26121 | 0 | 0 | 0 | 0 | 0 | 0 | 0 | 0 | 0 | 0 | 0 |
| bacteria_Jul27d25b_28840 | 0 | 0 | 0 | 0 | 0 | 0 | 0 | 0 | 0 | 0 | 0 |
| bacteria_Jul27d25b_29152 | 0 | 0 | 0 | 0 | 0 | 0 | 0 | 0 | 0 | 0 | 0 |
| bacteria_Jul27d25b_30149 | 0 | 0 | 0 | 0 | 0 | 0 | 0 | 0 | 0 | 0 | 0 |
| bacteria_Jul27d25b_39334 | 0 | 0 | 0 | 0 | 0 | 0 | 0 | 0 | 0 | 0 | 0 |
| bacteria_Jul27d25b_40615 | 0 | 0 | 0 | 0 | 0 | 0 | 0 | 0 | 0 | 0 | 0 |
| bacteria_Jul27d25b_47385 | 0 | 0 | 0 | 0 | 0 | 0 | 0 | 0 | 0 | 0 | 0 |
| bacteria_Jul27d25b_55853 | 0 | 0 | 0 | 0 | 0 | 0 | 0 | 0 | 0 | 0 | 0 |
| bacteria_Jul27d25b_57018 | 0 | 0 | 0 | 0 | 0 | 0 | 0 | 0 | 0 | 0 | 0 |
| bacteria_Jul27d25b_58438 | 0 | 0 | 0 | 0 | 0 | 0 | 0 | 0 | 0 | 0 | 0 |
| bacteria_Jul27d25b_63781 | 0 | 0 | 0 | 0 | 0 | 0 | 0 | 0 | 0 | 0 | 0 |
| bacteria_Jul27d25b_71708 | 0 | 0 | 0 | 0 | 0 | 0 | 0 | 0 | 0 | 0 | 0 |
| bacteria_Jul27d25b_73132 | 0 | 0 | 0 | 0 | 0 | 0 | 0 | 0 | 0 | 0 | 0 |
| bacteria_Jul27d75b_18089 | 0 | 0 | 0 | 0 | 0 | 0 | 0 | 0 | 0 | 0 | 0 |
| bacteria_Jul27d75b_21158 | 0 | 0 | 0 | 0 | 0 | 0 | 0 | 0 | 0 | 0 | 0 |
| bacteria_Jul27d75b_22650 | 0 | 0 | 0 | 0 | 0 | 0 | 0 | 0 | 0 | 0 | 0 |
| bacteria_Jul27d75b_26096 | 0 | 0 | 0 | 0 | 0 | 0 | 0 | 0 | 0 | 0 | 0 |
| bacteria_Jul27d75b_31732 | 0 | 0 | 0 | 0 | 0 | 0 | 0 | 0 | 0 | 0 | 0 |
| bacteria_Jul27d75b_33579 | 0 | 0 | 0 | 0 | 0 | 0 | 0 | 0 | 0 | 0 | 0 |
| bacteria_Jul27d75b_41399 | 0 | 0 | 0 | 0 | 0 | 0 | 0 | 0 | 0 | 1 | 1 |
| bacteria_Jul27d75b_48580 | 0 | 0 | 0 | 0 | 0 | 0 | 0 | 0 | 0 | 0 | 0 |
| bacteria_Jul27d75b_53502 | 0 | 0 | 0 | 0 | 0 | 0 | 0 | 0 | 0 | 0 | 0 |
| bacteria_Jul27d75b_56542 | 0 | 0 | 0 | 0 | 0 | 0 | 0 | 0 | 0 | 0 | 0 |
| bacteria_Jul27d75b_63474 | 0 | 0 | 0 | 0 | 0 | 0 | 0 | 0 | 0 | 0 | 0 |
| bacteria_Jul27d75b_67767 | 0 | 0 | 0 | 0 | 0 | 0 | 0 | 0 | 0 | 0 | 0 |
| bacteria_Jul27d75b_68680 | 0 | 0 | 0 | 0 | 0 | 0 | 0 | 0 | 0 | 0 | 0 |
| bacteria_Jul27d75b_68855 | 0 | 0 | 0 | 0 | 0 | 0 | 0 | 0 | 0 | 0 | 0 |
| bacteria_Jul27d75b_72237 | 0 | 0 | 0 | 0 | 0 | 0 | 0 | 0 | 0 | 0 | 0 |
| bacteria_Jun8d25b_3103 | 0 | 0 | 0 | 0 | 0 | 0 | 0 | 0 | 0 | 0 | 0 |
| bacteria_Jun8d25b_34399 | 0 | 0 | 0 | 0 | 0 | 0 | 0 | 0 | 0 | 0 | 0 |
| bacteria_Jun8d25b_37545 | 0 | 0 | 0 | 0 | 0 | 0 | 0 | 0 | 0 | 0 | 0 |
| bacteria_Jun8d25b_39385 | 0 | 0 | 0 | 0 | 0 | 0 | 0 | 0 | 0 | 0 | 0 |
| bacteria_Jun8d25b_44318 | 0 | 0 | 0 | 0 | 0 | 0 | 0 | 0 | 0 | 0 | 0 |
| bacteria_Jun8d25b_52047 | 0 | 0 | 0 | 0 | 0 | 0 | 0 | 0 | 0 | 0 | 0 |
| bacteria_Jun8d25b_57777 | 0 | 0 | 0 | 0 | 0 | 0 | 0 | 0 | 0 | 0 | 0 |
| bacteria_Jun8d25b_62757 | 0 | 0 | 0 | 0 | 0 | 0 | 0 | 0 | 0 | 0 | 0 |
| bacteria_Jun8d25b_71380 | 0 | 0 | 0 | 0 | 0 | 0 | 0 | 0 | 0 | 0 | 0 |
| bacteria_Jun8d50b_341 | 0 | 0 | 0 | 0 | 0 | 0 | 0 | 0 | 0 | 0 | 0 |
| bacteria_Jun8d50b_4841 | 0 | 0 | 0 | 0 | 0 | 0 | 0 | 0 | 0 | 0 | 0 |
| bacteria_Jun8d50b_22493 | 0 | 0 | 0 | 0 | 0 | 0 | 0 | 0 | 0 | 0 | 0 |
| bacteria_Jun8d50b_24396 | 0 | 0 | 0 | 0 | 0 | 0 | 0 | 0 | 0 | 0 | 0 |
| bacteria_Jun8d50b_36238 | 0 | 0 | 0 | 0 | 0 | 0 | 0 | 0 | 0 | 0 | 0 |
| bacteria_Jun8d50b_45077 | 0 | 0 | 0 | 0 | 0 | 0 | 0 | 0 | 0 | 0 | 0 |
| bacteria_Jun8d50b_45883 | 0 | 0 | 0 | 0 | 0 | 0 | 0 | 0 | 0 | 0 | 0 |
| bacteria_Jun8d50b_46634 | 0 | 0 | 0 | 0 | 0 | 0 | 0 | 0 | 0 | 0 | 0 |
| bacteria_Jun8d75b_24040 | 0 | 0 | 0 | 0 | 0 | 0 | 0 | 0 | 0 | 0 | 0 |
| bacteria_Jun22d0b_1872 | 0 | 0 | 0 | 0 | 0 | 0 | 0 | 0 | 0 | 0 | 0 |
| bacteria_Jun22d0b_13561 | 0 | 0 | 0 | 0 | 0 | 0 | 0 | 0 | 0 | 0 | 0 |
| bacteria_Jun22d0b_72542 | 0 | 1 | 0 | 0 | 0 | 0 | 0 | 0 | 0 | 0 | 1 |
| bacteria_Jun22d50b_11173 | 0 | 0 | 0 | 0 | 0 | 0 | 0 | 0 | 0 | 0 | 0 |
| bacteria_Jun22d50b_37032 | 0 | 0 | 0 | 0 | 0 | 0 | 0 | 0 | 0 | 0 | 0 |
| bacteria_Jun22d50b_44843 | 0 | 0 | 0 | 0 | 0 | 0 | 0 | 0 | 0 | 0 | 0 |
| bacteria_Jun22d50b_49499 | 0 | 0 | 0 | 0 | 0 | 0 | 0 | 0 | 0 | 0 | 0 |
| bacteria_Jun22d75b_17053 | 0 | 0 | 0 | 0 | 0 | 0 | 0 | 0 | 0 | 0 | 0 |
| bacteria_Jun22d75b_41056 | 0 | 0 | 0 | 0 | 0 | 0 | 0 | 0 | 0 | 0 | 0 |
| bacteria_Jun22d75b_45229 | 0 | 0 | 0 | 0 | 0 | 0 | 0 | 0 | 0 | 1 | 1 |
| bacteria_Jun22d75b_61468 | 0 | 0 | 0 | 0 | 0 | 0 | 0 | 0 | 0 | 0 | 0 |
| bacteria_Jun22d75b_65430 | 0 | 0 | 0 | 0 | 0 | 0 | 0 | 0 | 0 | 0 | 0 |
| bacteria_Jun22d75b_72891 | 0 | 0 | 0 | 0 | 0 | 0 | 0 | 0 | 0 | 0 | 0 |
| bacteria_May12d25b_9976 | 0 | 0 | 0 | 0 | 0 | 0 | 0 | 0 | 0 | 0 | 0 |
| bacteria_May12d25b_1311 | 0 | 0 | 0 | 0 | 0 | 0 | 0 | 0 | 0 | 0 | 0 |
| bacteria_May12d25b_1315 | 0 | 0 | 0 | 0 | 0 | 0 | 0 | 0 | 0 | 0 | 0 |
| bacteria_May12d25b_1413 | 0 | 0 | 0 | 0 | 0 | 0 | 0 | 0 | 0 | 0 | 0 |
| bacteria_May12d25b_1852 | 0 | 0 | 0 | 0 | 1 | 1 | 0 | 0 | 0 | 0 | 2 |
| bacteria_May12d25b_4325 | 0 | 0 | 0 | 0 | 0 | 0 | 0 | 0 | 0 | 0 | 0 |
| bacteria_May12d25b_5412 | 0 | 0 | 0 | 0 | 0 | 0 | 0 | 0 | 0 | 0 | 0 |
| bacteria_May12d25b_6769 | 0 | 0 | 0 | 0 | 0 | 0 | 0 | 0 | 0 | 0 | 0 |
| bacteria_May12d25b_7124 | 0 | 0 | 0 | 0 | 0 | 0 | 0 | 0 | 0 | 0 | 0 |
| bacteria_May12d50b_505 | 0 | 0 | 0 | 0 | 0 | 0 | 0 | 0 | 0 | 0 | 0 |
| bacteria_May12d50b_8438 | 0 | 0 | 0 | 0 | 0 | 0 | 0 | 0 | 0 | 0 | 0 |
| bacteria_May12d50b_2063 | 0 | 0 | 0 | 0 | 0 | 0 | 0 | 0 | 0 | 0 | 0 |
| bacteria_May12d50b_2129 | 0 | 0 | 0 | 0 | 0 | 0 | 0 | 0 | 0 | 0 | 0 |
| bacteria_May12d50b_2729 | 0 | 0 | 0 | 0 | 0 | 0 | 0 | 0 | 0 | 0 | 0 |
| bacteria_May12d50b_3692 | 0 | 0 | 0 | 0 | 0 | 0 | 0 | 0 | 0 | 0 | 0 |
| bacteria_May12d50b_3851 | 0 | 0 | 0 | 0 | 0 | 0 | 0 | 0 | 0 | 0 | 0 |
| bacteria_May12d50b_4442 | 0 | 0 | 0 | 0 | 0 | 0 | 0 | 0 | 0 | 0 | 0 |
| bacteria_May12d50b_4471 | 0 | 0 | 0 | 0 | 0 | 0 | 0 | 0 | 0 | 0 | 0 |
| bacteria_May12d50b_4645 | 0 | 0 | 0 | 0 | 0 | 0 | 0 | 0 | 0 | 0 | 0 |
| bacteria_May12d50b_4892 | 0 | 0 | 0 | 0 | 0 | 0 | 0 | 0 | 0 | 0 | 0 |
| bacteria_May12d50b_5005 | 0 | 0 | 0 | 0 | 0 | 0 | 0 | 0 | 0 | 0 | 0 |
| bacteria_May12d50b_5105 | 0 | 0 | 0 | 0 | 0 | 0 | 0 | 0 | 0 | 0 | 0 |
| bacteria_Sep8d5b_2772 | 0 | 0 | 0 | 0 | 0 | 0 | 0 | 0 | 0 | 0 | 0 |
| bacteria_Sep8d5b_4362 | 0 | 0 | 0 | 0 | 0 | 0 | 0 | 0 | 0 | 0 | 0 |
| bacteria_Sep8d5b_4996 | 0 | 0 | 0 | 0 | 0 | 0 | 0 | 0 | 0 | 0 | 0 |
| bacteria_Sep8d5b_5046 | 0 | 0 | 0 | 0 | 0 | 0 | 0 | 0 | 0 | 0 | 0 |
| bacteria_Sep8d5b_5518 | 0 | 0 | 0 | 0 | 0 | 0 | 0 | 0 | 0 | 0 | 0 |
| bacteria_Sep8d5b_10019 | 0 | 0 | 0 | 0 | 0 | 0 | 0 | 0 | 0 | 0 | 0 |
| bacteria_Sep8d5b_10717 | 0 | 0 | 0 | 0 | 0 | 0 | 0 | 0 | 0 | 0 | 0 |
| bacteria_Sep8d5b_12119 | 0 | 0 | 0 | 0 | 0 | 0 | 0 | 0 | 0 | 0 | 0 |
| bacteria_Sep8d5b_18164 | 0 | 0 | 0 | 0 | 0 | 0 | 0 | 0 | 0 | 0 | 0 |
| bacteria_Sep8d5b_19593 | 0 | 0 | 0 | 0 | 0 | 0 | 0 | 0 | 0 | 0 | 0 |
| bacteria_Sep8d5b_24411 | 0 | 0 | 0 | 0 | 0 | 0 | 0 | 0 | 0 | 0 | 0 |
| bacteria_Sep8d5b_28614 | 0 | 0 | 0 | 0 | 0 | 0 | 0 | 0 | 0 | 0 | 0 |
| bacteria_Sep8d5b_30494 | 0 | 0 | 0 | 0 | 0 | 0 | 0 | 0 | 0 | 0 | 0 |
| bacteria_Sep8d5b_62465 | 0 | 0 | 0 | 0 | 0 | 0 | 0 | 0 | 0 | 0 | 0 |
| bacteria_Sep8d5b_64306 | 0 | 0 | 0 | 0 | 0 | 0 | 0 | 0 | 0 | 0 | 0 |
| bacteria_Sep8d5b_68076 | 0 | 0 | 0 | 1 | 0 | 0 | 0 | 0 | 0 | 0 | 1 |
| bacteria_Sep8d25b_18102 | 0 | 0 | 0 | 0 | 0 | 0 | 0 | 0 | 0 | 0 | 0 |
| bacteria_Sep8d25b_23922 | 0 | 0 | 0 | 0 | 0 | 0 | 0 | 0 | 0 | 0 | 0 |
| bacteria_Sep8d25b_35008 | 0 | 0 | 0 | 0 | 0 | 0 | 0 | 0 | 0 | 0 | 0 |
| bacteria_Sep8d25b_45121 | 0 | 0 | 0 | 0 | 0 | 0 | 0 | 0 | 0 | 0 | 0 |
| bacteria_Sep8d50b_5207 | 0 | 0 | 0 | 0 | 0 | 0 | 0 | 0 | 0 | 0 | 0 |
| bacteria_Sep8d50b_51343 | 0 | 0 | 0 | 0 | 0 | 0 | 0 | 0 | 0 | 0 | 0 |
| virus_Aug5.0v_393 | 0 | 1 | 0 | 0 | 0 | 0 | 0 | 0 | 0 | 0 | 1 |
| virus_Aug5.0v_397 | 0 | 1 | 0 | 0 | 0 | 0 | 0 | 0 | 0 | 0 | 1 |
| virus_Aug5.0v_705 | 0 | 1 | 0 | 0 | 0 | 0 | 0 | 0 | 0 | 0 | 1 |
| virus_Aug5.0v_722 | 0 | 1 | 0 | 0 | 0 | 0 | 0 | 0 | 0 | 0 | 1 |
| virus_Aug5.0v_1135 | 0 | 1 | 0 | 0 | 0 | 0 | 0 | 0 | 0 | 0 | 1 |
| virus_Aug5.0v_1409 | 0 | 1 | 0 | 0 | 0 | 0 | 0 | 0 | 0 | 0 | 1 |
| virus_Aug5.0v_1430 | 0 | 1 | 0 | 0 | 1 | 0 | 0 | 1 | 0 | 0 | 3 |
| virus_Aug5.0v_1774 | 0 | 3 | 0 | 0 | 0 | 0 | 0 | 0 | 0 | 0 | 3 |
| virus_Aug5.0v_1855 | 0 | 1 | 0 | 0 | 0 | 0 | 0 | 0 | 0 | 0 | 1 |
| virus_Aug5.0v_1978 | 0 | 1 | 0 | 0 | 0 | 0 | 0 | 0 | 0 | 0 | 1 |
| virus_Aug5.0v_2018 | 0 | 1 | 0 | 0 | 0 | 0 | 0 | 0 | 0 | 0 | 1 |
| virus_Aug5.0v_2075 | 0 | 1 | 0 | 0 | 0 | 0 | 0 | 0 | 0 | 0 | 1 |
| virus_Aug5.0v_2107 | 0 | 2 | 0 | 0 | 0 | 0 | 0 | 0 | 0 | 0 | 2 |
| virus_Aug5.0v_2760 | 0 | 1 | 0 | 0 | 0 | 0 | 0 | 0 | 0 | 0 | 1 |
| virus_Aug5.0v_3014 | 0 | 1 | 0 | 0 | 0 | 0 | 0 | 0 | 0 | 0 | 1 |
| virus_Aug5.0v_3711 | 0 | 1 | 0 | 0 | 0 | 0 | 0 | 0 | 0 | 0 | 1 |

| | | | | | | | | | | | |
|---|---|---|---|---|---|---|---|---|---|---|---|
| virus_Aug5.0v_3725 | 0 | 1 | 0 | 0 | 0 | 0 | 0 | 0 | 0 | 0 | 1 |
| virus_Aug5.0v_3810 | 0 | 1 | 0 | 0 | 0 | 0 | 0 | 0 | 0 | 0 | 1 |
| virus_Aug5.0v_3893 | 0 | 1 | 0 | 0 | 0 | 0 | 0 | 0 | 0 | 0 | 1 |
| virus_Aug5.0v_4361 | 0 | 1 | 0 | 0 | 0 | 0 | 0 | 0 | 0 | 0 | 1 |
| virus_Aug5.0v_4464 | 0 | 1 | 0 | 0 | 0 | 0 | 0 | 0 | 0 | 0 | 1 |
| virus_Aug5.0v_4490 | 0 | 1 | 0 | 0 | 0 | 0 | 0 | 0 | 0 | 0 | 1 |
| virus_Aug5.0v_4573 | 0 | 1 | 0 | 0 | 0 | 0 | 0 | 0 | 0 | 0 | 1 |
| virus_Aug5.0v_4613 | 0 | 1 | 0 | 0 | 0 | 0 | 0 | 0 | 0 | 0 | 1 |
| virus_Aug5.0v_4629 | 0 | 1 | 0 | 0 | 0 | 0 | 0 | 0 | 0 | 0 | 1 |
| virus_Aug5.0v_4761 | 0 | 1 | 0 | 0 | 0 | 0 | 0 | 0 | 0 | 0 | 1 |
| virus_Aug5.0v_4774 | 0 | 1 | 0 | 0 | 0 | 0 | 0 | 0 | 0 | 0 | 1 |
| virus_Aug5.0v_4925 | 0 | 1 | 0 | 0 | 0 | 0 | 0 | 0 | 0 | 1 | 2 |
| virus_Aug5.0v_5090 | 0 | 2 | 0 | 0 | 0 | 0 | 0 | 0 | 0 | 0 | 2 |
| virus_Aug5.0v_5373 | 0 | 1 | 0 | 0 | 0 | 0 | 0 | 0 | 0 | 0 | 1 |
| virus_Aug5.5v_7503 | 0 | 0 | 0 | 1 | 0 | 0 | 0 | 0 | 0 | 0 | 1 |
| virus_Aug5.5v_10513 | 0 | 0 | 0 | 1 | 0 | 0 | 0 | 0 | 0 | 0 | 1 |
| virus_Aug5.5v_15316 | 0 | 0 | 0 | 1 | 0 | 0 | 0 | 0 | 0 | 0 | 1 |
| virus_Aug5.5v_17722 | 0 | 0 | 0 | 1 | 0 | 0 | 0 | 0 | 0 | 0 | 1 |
| virus_Aug5.5v_22369 | 0 | 0 | 0 | 1 | 0 | 0 | 0 | 0 | 0 | 0 | 1 |
| virus_Aug5.5v_28614 | 0 | 0 | 0 | 1 | 0 | 0 | 0 | 0 | 0 | 0 | 1 |
| virus_Aug5.5v_28796 | 0 | 0 | 0 | 1 | 0 | 0 | 0 | 0 | 0 | 0 | 1 |
| virus_Aug5.5v_35563 | 0 | 0 | 0 | 1 | 0 | 0 | 0 | 0 | 0 | 0 | 1 |
| virus_Aug5.5v_39677 | 0 | 0 | 0 | 1 | 0 | 0 | 0 | 0 | 0 | 0 | 1 |
| virus_Aug5.5v_41029 | 0 | 0 | 0 | 1 | 0 | 0 | 0 | 0 | 0 | 0 | 1 |
| virus_Aug5.25v_1114 | 0 | 0 | 0 | 0 | 0 | 1 | 0 | 0 | 0 | 0 | 1 |
| virus_Aug5.25v_1472 | 0 | 0 | 0 | 0 | 0 | 1 | 0 | 0 | 0 | 0 | 1 |
| virus_Aug5.25v_2209 | 0 | 0 | 0 | 0 | 0 | 1 | 0 | 0 | 0 | 0 | 1 |
| virus_Aug5.25v_3366 | 0 | 0 | 0 | 0 | 0 | 1 | 0 | 0 | 0 | 0 | 1 |
| virus_Aug5.25v_6601 | 0 | 0 | 0 | 0 | 0 | 1 | 0 | 0 | 0 | 0 | 1 |
| virus_Aug5.25v_8939 | 0 | 0 | 0 | 0 | 0 | 1 | 0 | 0 | 0 | 0 | 1 |
| virus_Aug5.25v_9357 | 0 | 0 | 0 | 0 | 0 | 1 | 0 | 0 | 0 | 0 | 1 |
| virus_Aug5.25v_10907 | 0 | 0 | 0 | 0 | 0 | 1 | 0 | 0 | 0 | 0 | 1 |
| virus_Aug5.25v_12783 | 0 | 0 | 0 | 0 | 0 | 1 | 0 | 0 | 0 | 0 | 1 |
| virus_Aug5.25v_12787 | 0 | 0 | 0 | 0 | 0 | 1 | 0 | 0 | 0 | 0 | 1 |
| virus_Aug5.25v_14889 | 0 | 0 | 0 | 0 | 0 | 1 | 0 | 0 | 0 | 0 | 1 |
| virus_Aug5.25v_15452 | 0 | 0 | 0 | 0 | 0 | 1 | 0 | 0 | 0 | 0 | 1 |
| virus_Aug5.25v_19510 | 0 | 0 | 0 | 0 | 0 | 1 | 0 | 0 | 0 | 0 | 1 |
| virus_Aug5.25v_20101 | 0 | 0 | 0 | 0 | 0 | 1 | 0 | 0 | 0 | 0 | 1 |
| virus_Aug5.25v_20980 | 0 | 0 | 0 | 0 | 0 | 1 | 0 | 0 | 0 | 0 | 1 |
| virus_Aug5.25v_21421 | 0 | 0 | 0 | 0 | 0 | 1 | 0 | 0 | 0 | 0 | 1 |
| virus_Aug5.25v_26529 | 0 | 0 | 0 | 0 | 0 | 1 | 0 | 0 | 0 | 0 | 1 |
| virus_Aug5.25v_27987 | 0 | 0 | 0 | 0 | 0 | 1 | 0 | 0 | 0 | 0 | 1 |
| virus_Aug5.25v_33871 | 0 | 0 | 0 | 0 | 0 | 1 | 0 | 0 | 0 | 0 | 1 |
| virus_Aug5.25v_33981 | 0 | 0 | 0 | 0 | 0 | 1 | 0 | 0 | 0 | 0 | 1 |
| virus_Aug5.25v_34476 | 0 | 0 | 0 | 0 | 0 | 1 | 0 | 0 | 0 | 0 | 1 |
| virus_Aug5.25v_34563 | 0 | 0 | 0 | 0 | 0 | 1 | 0 | 0 | 0 | 0 | 1 |
| virus_Aug5.25v_40718 | 0 | 0 | 0 | 0 | 1 | 1 | 1 | 0 | 0 | 0 | 3 |
| virus_Aug5.50v_4994 | 0 | 0 | 0 | 0 | 0 | 0 | 0 | 1 | 0 | 0 | 1 |
| virus_Aug5.50v_15274 | 0 | 0 | 0 | 0 | 0 | 0 | 0 | 1 | 0 | 0 | 1 |
| virus_Aug5.50v_23067 | 0 | 0 | 0 | 0 | 0 | 0 | 0 | 1 | 0 | 0 | 1 |
| virus_Aug5.50v_25883 | 0 | 0 | 0 | 0 | 0 | 0 | 0 | 1 | 0 | 0 | 1 |
| virus_Aug5.50v_37660 | 0 | 0 | 0 | 0 | 0 | 0 | 0 | 1 | 0 | 0 | 1 |
| virus_Aug5.50v_40182 | 0 | 0 | 0 | 0 | 0 | 0 | 0 | 1 | 0 | 0 | 1 |
| virus_Aug5.75v_14261 | 0 | 0 | 0 | 0 | 0 | 0 | 0 | 0 | 0 | 1 | 1 |
| virus_Aug5.75v_14842 | 0 | 0 | 0 | 0 | 0 | 0 | 0 | 0 | 0 | 1 | 1 |
| virus_Aug5.75v_19722 | 0 | 0 | 0 | 0 | 0 | 0 | 0 | 0 | 0 | 1 | 1 |
| virus_Aug5.75v_26312 | 0 | 0 | 0 | 0 | 0 | 0 | 0 | 0 | 0 | 1 | 1 |
| virus_Aug5.75v_33388 | 0 | 0 | 0 | 0 | 0 | 0 | 0 | 0 | 0 | 1 | 1 |
| virus_Aug5.75v_36458 | 0 | 0 | 0 | 0 | 0 | 0 | 0 | 0 | 0 | 1 | 1 |
| virus_Aug5.75v_37349 | 0 | 0 | 0 | 0 | 0 | 0 | 0 | 0 | 0 | 1 | 1 |
| virus_Aug5.75v_38675 | 0 | 0 | 0 | 0 | 0 | 0 | 0 | 0 | 0 | 1 | 1 |
| virus_Jul11.0v_130 | 0 | 0 | 0 | 0 | 0 | 0 | 0 | 0 | 0 | 0 | 0 |
| virus_Jul11.0v_9445 | 0 | 0 | 0 | 0 | 0 | 0 | 0 | 0 | 0 | 0 | 0 |
| virus_Jul11.0v_11429 | 0 | 0 | 0 | 0 | 0 | 0 | 0 | 0 | 0 | 0 | 0 |
| virus_Jul11.0v_12253 | 0 | 0 | 0 | 0 | 0 | 0 | 0 | 0 | 0 | 0 | 0 |
| virus_Jul11.0v_12503 | 0 | 0 | 0 | 0 | 0 | 0 | 0 | 0 | 0 | 0 | 0 |
| virus_Jul11.0v_19984 | 0 | 0 | 0 | 0 | 0 | 0 | 0 | 0 | 0 | 0 | 0 |
| virus_Jul11.0v_20214 | 0 | 0 | 0 | 0 | 0 | 0 | 0 | 0 | 0 | 0 | 0 |
| virus_Jul11.5v_381 | 0 | 0 | 0 | 0 | 0 | 0 | 0 | 0 | 0 | 0 | 0 |
| virus_Jul11.5v_673 | 0 | 0 | 0 | 0 | 0 | 0 | 0 | 0 | 0 | 0 | 0 |
| virus_Jul11.5v_1191 | 0 | 0 | 0 | 0 | 0 | 0 | 0 | 0 | 0 | 0 | 0 |
| virus_Jul11.5v_2794 | 0 | 0 | 0 | 0 | 0 | 0 | 0 | 0 | 0 | 0 | 0 |
| virus_Jul11.5v_4074 | 0 | 0 | 0 | 0 | 0 | 0 | 0 | 0 | 0 | 0 | 0 |
| virus_Jul11.5v_5256 | 0 | 0 | 0 | 0 | 0 | 0 | 0 | 0 | 0 | 0 | 0 |
| virus_Jul11.5v_6463 | 0 | 0 | 0 | 0 | 0 | 0 | 0 | 0 | 0 | 0 | 0 |
| virus_Jul11.5v_7391 | 0 | 0 | 0 | 0 | 0 | 0 | 0 | 0 | 0 | 0 | 0 |
| virus_Jul11.5v_7560 | 0 | 0 | 0 | 0 | 0 | 0 | 0 | 0 | 0 | 0 | 0 |
| virus_Jul11.5v_8070 | 0 | 0 | 0 | 0 | 0 | 0 | 0 | 0 | 0 | 0 | 0 |
| virus_Jul11.5v_8177 | 0 | 0 | 0 | 0 | 0 | 0 | 0 | 0 | 0 | 0 | 0 |
| virus_Jul11.5v_11526 | 0 | 0 | 0 | 0 | 0 | 0 | 0 | 0 | 0 | 0 | 0 |
| virus_Jul11.5v_12569 | 0 | 0 | 0 | 0 | 0 | 0 | 0 | 0 | 0 | 0 | 0 |
| virus_Jul11.5v_13808 | 0 | 0 | 0 | 0 | 0 | 0 | 0 | 0 | 0 | 0 | 0 |
| virus_Jul11.5v_14896 | 0 | 0 | 0 | 0 | 0 | 0 | 0 | 0 | 0 | 0 | 0 |
| virus_Jul11.5v_15011 | 0 | 0 | 0 | 0 | 0 | 0 | 0 | 0 | 0 | 0 | 0 |
| virus_Jul11.5v_15540 | 0 | 0 | 0 | 0 | 0 | 0 | 0 | 0 | 0 | 0 | 0 |
| virus_Jul11.5v_17079 | 0 | 0 | 0 | 0 | 0 | 0 | 0 | 0 | 0 | 0 | 0 |
| virus_Jul11.5v_18487 | 0 | 0 | 0 | 0 | 0 | 0 | 0 | 0 | 0 | 0 | 0 |
| virus_Jul11.5v_19652 | 0 | 0 | 0 | 0 | 0 | 0 | 0 | 0 | 0 | 0 | 0 |
| virus_Jul11.5v_20584 | 0 | 0 | 0 | 0 | 0 | 0 | 0 | 0 | 0 | 0 | 0 |
| virus_Jul11.25v_543 | 0 | 0 | 0 | 0 | 0 | 0 | 0 | 0 | 0 | 0 | 0 |
| virus_Jul11.25v_3211 | 0 | 0 | 0 | 0 | 0 | 0 | 0 | 0 | 0 | 0 | 0 |
| virus_Jul11.25v_7446 | 0 | 0 | 0 | 0 | 0 | 0 | 0 | 0 | 0 | 0 | 0 |
| virus_Jul11.25v_8637 | 0 | 0 | 0 | 0 | 0 | 0 | 0 | 0 | 0 | 0 | 0 |
| virus_Jul11.25v_12490 | 0 | 0 | 0 | 0 | 0 | 0 | 0 | 0 | 0 | 0 | 0 |
| virus_Jul11.25v_16201 | 0 | 0 | 0 | 0 | 0 | 0 | 0 | 0 | 0 | 0 | 0 |
| virus_Jul11.25v_17500 | 0 | 0 | 0 | 0 | 0 | 0 | 0 | 0 | 0 | 0 | 0 |
| virus_Jul11.50v_5842 | 0 | 0 | 0 | 0 | 0 | 0 | 0 | 0 | 0 | 0 | 0 |
| virus_Jul11.50v_7997 | 0 | 0 | 0 | 0 | 0 | 0 | 0 | 0 | 0 | 0 | 0 |
| virus_Jul11.50v_9849 | 0 | 0 | 0 | 0 | 0 | 0 | 0 | 0 | 0 | 0 | 0 |
| virus_Jul11.50v_9853 | 0 | 0 | 0 | 0 | 0 | 0 | 0 | 0 | 0 | 0 | 0 |
| virus_Jul11.50v_10840 | 0 | 0 | 0 | 0 | 0 | 0 | 0 | 0 | 0 | 0 | 0 |
| virus_Jul11.50v_11789 | 0 | 0 | 0 | 0 | 0 | 0 | 0 | 0 | 0 | 0 | 0 |
| virus_Jul11.50v_11809 | 0 | 0 | 0 | 0 | 0 | 0 | 0 | 0 | 0 | 0 | 0 |
| virus_Jul11.50v_13214 | 0 | 0 | 0 | 0 | 0 | 0 | 0 | 0 | 0 | 0 | 0 |
| virus_Jul11.50v_14411 | 0 | 0 | 0 | 0 | 0 | 0 | 0 | 0 | 0 | 0 | 0 |
| virus_Jul11.50v_15139 | 0 | 0 | 0 | 0 | 0 | 1 | 0 | 0 | 0 | 0 | 1 |
| virus_Jul11.50v_16194 | 0 | 0 | 0 | 0 | 0 | 0 | 0 | 0 | 0 | 0 | 0 |
| virus_Jul11.50v_18630 | 0 | 0 | 0 | 0 | 0 | 0 | 0 | 0 | 0 | 0 | 0 |
| virus_Jul11.75v_2531 | 0 | 0 | 0 | 0 | 0 | 0 | 0 | 0 | 0 | 0 | 0 |
| virus_Jul11.75v_12470 | 0 | 0 | 0 | 0 | 0 | 0 | 0 | 0 | 0 | 0 | 0 |
| virus_Jul11.75v_23511 | 0 | 0 | 0 | 0 | 0 | 0 | 0 | 0 | 0 | 0 | 0 |
| virus_Jul11.75v_27674 | 0 | 0 | 0 | 0 | 0 | 0 | 0 | 0 | 0 | 0 | 0 |
| virus_Jul11.75v_30378 | 0 | 1 | 0 | 1 | 0 | 0 | 0 | 0 | 0 | 0 | 2 |
| virus_Jul11.75v_32918 | 0 | 0 | 0 | 0 | 0 | 0 | 0 | 0 | 0 | 0 | 0 |
| virus_Jul11.75v_34246 | 0 | 0 | 0 | 0 | 0 | 0 | 0 | 0 | 0 | 0 | 0 |
| virus_Jul11.75v_38552 | 0 | 0 | 0 | 0 | 0 | 0 | 0 | 0 | 0 | 0 | 0 |
| virus_Jul27.5v_370 | 0 | 0 | 0 | 0 | 0 | 0 | 0 | 0 | 0 | 0 | 0 |
| virus_Jul27.5v_1138 | 0 | 0 | 0 | 0 | 0 | 0 | 0 | 0 | 0 | 0 | 0 |
| virus_Jul27.5v_3305 | 0 | 1 | 0 | 0 | 0 | 0 | 0 | 0 | 0 | 0 | 1 |
| virus_Jul27.5v_3543 | 0 | 0 | 0 | 0 | 0 | 0 | 0 | 0 | 0 | 0 | 0 |
| virus_Jul27.5v_3646 | 0 | 0 | 0 | 0 | 0 | 0 | 0 | 0 | 0 | 0 | 0 |
| virus_Jul27.5v_6013 | 0 | 0 | 0 | 0 | 0 | 0 | 0 | 0 | 0 | 0 | 0 |
| virus_Jul27.5v_6125 | 0 | 0 | 0 | 0 | 0 | 0 | 0 | 0 | 0 | 0 | 0 |
| virus_Jul27.5v_8885 | 0 | 0 | 0 | 0 | 0 | 0 | 0 | 0 | 0 | 0 | 0 |
| virus_Jul27.5v_8922 | 0 | 0 | 0 | 0 | 0 | 0 | 0 | 0 | 0 | 0 | 0 |
| virus_Jul27.5v_9135 | 0 | 0 | 0 | 0 | 0 | 0 | 0 | 0 | 0 | 0 | 0 |
| virus_Jul27.5v_9777 | 0 | 0 | 0 | 0 | 0 | 0 | 0 | 0 | 0 | 0 | 0 |
| virus_Jul27.5v_9857 | 0 | 0 | 0 | 0 | 0 | 0 | 0 | 0 | 0 | 0 | 0 |
| virus_Jul27.5v_10389 | 0 | 0 | 0 | 0 | 0 | 0 | 0 | 0 | 0 | 0 | 0 |
| virus_Jul27.5v_11463 | 0 | 0 | 0 | 0 | 0 | 0 | 0 | 0 | 0 | 0 | 0 |
| virus_Jul27.5v_12791 | 0 | 0 | 0 | 0 | 0 | 0 | 0 | 0 | 0 | 0 | 0 |
| virus_Jul27.5v_14925 | 0 | 0 | 0 | 0 | 0 | 0 | 0 | 0 | 0 | 0 | 0 |
| virus_Jul27.5v_16264 | 0 | 0 | 0 | 0 | 0 | 0 | 0 | 0 | 0 | 0 | 0 |
| virus_Jul27.5v_16298 | 0 | 0 | 0 | 0 | 0 | 0 | 0 | 0 | 0 | 0 | 0 |
| virus_Jul27.5v_17000 | 0 | 0 | 0 | 0 | 0 | 0 | 0 | 0 | 0 | 0 | 0 |
| virus_Jul27.5v_17644 | 0 | 0 | 0 | 0 | 0 | 0 | 0 | 0 | 0 | 0 | 0 |
| virus_Jul27.5v_18255 | 0 | 0 | 0 | 0 | 0 | 0 | 0 | 0 | 0 | 0 | 0 |
| virus_Jul27.5v_18965 | 0 | 0 | 0 | 0 | 0 | 0 | 0 | 0 | 0 | 0 | 0 |
| virus_Jul27.5v_19824 | 0 | 0 | 0 | 0 | 0 | 0 | 0 | 0 | 0 | 0 | 0 |
| virus_Jul27.5v_20005 | 0 | 0 | 0 | 0 | 0 | 0 | 0 | 0 | 0 | 0 | 0 |
| virus_Jul27.5v_20218 | 0 | 0 | 0 | 0 | 0 | 0 | 0 | 0 | 0 | 0 | 0 |
| virus_Jul27.25v_193 | 0 | 0 | 0 | 0 | 0 | 0 | 0 | 0 | 0 | 0 | 0 |
| virus_Jul27.25v_412 | 0 | 0 | 0 | 0 | 0 | 0 | 0 | 0 | 0 | 0 | 0 |
| virus_Jul27.25v_861 | 0 | 0 | 0 | 0 | 0 | 0 | 0 | 0 | 0 | 0 | 0 |
| virus_Jul27.25v_9152 | 0 | 0 | 0 | 0 | 0 | 0 | 0 | 0 | 0 | 0 | 0 |
| virus_Jul27.25v_9384 | 0 | 0 | 0 | 0 | 0 | 0 | 0 | 0 | 0 | 0 | 0 |
| virus_Jul27.25v_10021 | 0 | 0 | 0 | 0 | 0 | 0 | 0 | 0 | 0 | 0 | 0 |
| virus_Jul27.25v_12717 | 0 | 0 | 0 | 0 | 0 | 0 | 0 | 0 | 0 | 0 | 0 |
| virus_Jul27.25v_16514 | 0 | 0 | 0 | 0 | 0 | 0 | 0 | 0 | 0 | 0 | 0 |
| virus_Jul27.25v_17182 | 0 | 0 | 0 | 0 | 0 | 0 | 0 | 0 | 0 | 0 | 0 |
| virus_Jul27.25v_19006 | 0 | 0 | 0 | 0 | 0 | 0 | 0 | 0 | 0 | 0 | 0 |
| virus_Jul27.25v_19311 | 0 | 0 | 0 | 0 | 0 | 0 | 0 | 0 | 0 | 0 | 0 |
| virus_Jul27.25v_25018 | 0 | 0 | 0 | 0 | 0 | 0 | 0 | 0 | 0 | 0 | 0 |
| virus_Jul27.25v_39273 | 0 | 0 | 0 | 0 | 0 | 0 | 0 | 0 | 0 | 0 | 0 |
| virus_Jul27.25v_39279 | 0 | 0 | 0 | 0 | 0 | 0 | 0 | 0 | 0 | 0 | 0 |
| virus_Jul27.75v_3872 | 0 | 0 | 0 | 0 | 0 | 0 | 0 | 0 | 0 | 0 | 0 |
| virus_Jun8.25v_468 | 0 | 0 | 0 | 0 | 0 | 0 | 0 | 0 | 0 | 0 | 0 |

| | | | | | | | | | | | |
|---|---|---|---|---|---|---|---|---|---|---|---|
| virus_Jun8.25v_2122 | 0 | 0 | 0 | 0 | 0 | 0 | 0 | 0 | 0 | 0 | 0 |
| virus_Jun8.25v_2418 | 0 | 0 | 0 | 0 | 0 | 0 | 0 | 0 | 0 | 0 | 0 |
| virus_Jun8.25v_36017 | 0 | 0 | 0 | 0 | 0 | 0 | 0 | 0 | 0 | 0 | 0 |
| virus_Jun8.50v_686 | 0 | 0 | 0 | 0 | 0 | 0 | 0 | 0 | 0 | 0 | 0 |
| virus_Jun8.50v_2793 | 0 | 0 | 0 | 0 | 0 | 0 | 0 | 0 | 0 | 0 | 0 |
| virus_Jun8.50v_4633 | 0 | 0 | 0 | 0 | 0 | 0 | 0 | 0 | 0 | 0 | 0 |
| virus_Jun8.50v_9271 | 0 | 0 | 0 | 0 | 0 | 0 | 0 | 0 | 0 | 0 | 0 |
| virus_Jun8.50v_11912 | 0 | 0 | 0 | 0 | 0 | 0 | 0 | 0 | 0 | 0 | 0 |
| virus_Jun8.50v_13861 | 0 | 0 | 0 | 0 | 0 | 0 | 0 | 0 | 0 | 0 | 0 |
| virus_Jun8.50v_18944 | 0 | 0 | 0 | 0 | 0 | 0 | 0 | 0 | 0 | 0 | 0 |
| virus_Jun8.50v_19908 | 0 | 0 | 0 | 0 | 0 | 0 | 0 | 0 | 0 | 0 | 0 |
| virus_Jun8.50v_23548 | 0 | 0 | 0 | 0 | 0 | 0 | 0 | 0 | 0 | 0 | 0 |
| virus_Jun8.50v_23587 | 0 | 0 | 0 | 0 | 0 | 0 | 0 | 0 | 0 | 0 | 0 |
| virus_Jun8.50v_32285 | 0 | 0 | 0 | 0 | 0 | 0 | 0 | 0 | 0 | 0 | 0 |
| virus_Jun8.50v_32983 | 0 | 0 | 0 | 0 | 0 | 0 | 0 | 0 | 0 | 0 | 0 |
| virus_Jun8.50v_37130 | 0 | 0 | 0 | 0 | 0 | 0 | 0 | 0 | 0 | 0 | 0 |
| virus_Jun8.50v_37568 | 0 | 0 | 0 | 0 | 1 | 0 | 0 | 0 | 0 | 0 | 1 |
| virus_Jun8.75v_6003 | 0 | 0 | 0 | 0 | 0 | 0 | 0 | 0 | 0 | 0 | 0 |
| virus_Jun8.75v_8467 | 0 | 0 | 0 | 0 | 0 | 0 | 0 | 0 | 0 | 0 | 0 |
| virus_Jun22.0v_839 | 0 | 0 | 0 | 0 | 0 | 0 | 0 | 0 | 0 | 0 | 0 |
| virus_Jun22.0v_2092 | 0 | 0 | 0 | 0 | 0 | 0 | 0 | 0 | 0 | 0 | 0 |
| virus_Jun22.0v_7724 | 0 | 0 | 0 | 0 | 0 | 0 | 0 | 0 | 0 | 0 | 0 |
| virus_Jun22.0v_13800 | 0 | 0 | 0 | 0 | 0 | 0 | 0 | 0 | 0 | 0 | 0 |
| virus_Jun22.0v_17340 | 0 | 0 | 0 | 0 | 0 | 0 | 0 | 0 | 0 | 0 | 0 |
| virus_Jun22.0v_19526 | 0 | 0 | 0 | 0 | 0 | 0 | 0 | 0 | 0 | 0 | 0 |
| virus_Jun22.50v_1067 | 0 | 0 | 0 | 0 | 0 | 0 | 0 | 0 | 0 | 0 | 0 |
| virus_Jun22.50v_1467 | 0 | 0 | 0 | 0 | 0 | 0 | 0 | 0 | 0 | 0 | 0 |
| virus_Jun22.50v_4776 | 0 | 0 | 0 | 0 | 0 | 0 | 0 | 0 | 0 | 0 | 0 |
| virus_Jun22.50v_7192 | 0 | 0 | 0 | 0 | 0 | 0 | 0 | 0 | 0 | 0 | 0 |
| virus_Jun22.50v_9359 | 0 | 0 | 0 | 0 | 0 | 0 | 0 | 0 | 0 | 0 | 0 |
| virus_Jun22.50v_9394 | 0 | 0 | 0 | 0 | 0 | 0 | 0 | 0 | 0 | 0 | 0 |
| virus_Jun22.50v_11217 | 0 | 0 | 0 | 0 | 0 | 0 | 0 | 0 | 0 | 0 | 0 |
| virus_Jun22.50v_13684 | 0 | 0 | 0 | 0 | 0 | 0 | 0 | 0 | 0 | 0 | 0 |
| virus_Jun22.50v_15620 | 0 | 0 | 0 | 0 | 0 | 0 | 0 | 0 | 0 | 0 | 0 |
| virus_Jun22.50v_15849 | 0 | 0 | 0 | 0 | 0 | 0 | 0 | 0 | 0 | 0 | 0 |
| virus_Jun22.50v_16076 | 0 | 0 | 0 | 0 | 0 | 0 | 0 | 0 | 0 | 0 | 0 |
| virus_Jun22.50v_16395 | 0 | 0 | 0 | 0 | 0 | 0 | 0 | 0 | 0 | 0 | 0 |
| virus_Jun22.50v_19149 | 0 | 0 | 0 | 0 | 0 | 0 | 0 | 0 | 0 | 0 | 0 |
| virus_Jun22.50v_23024 | 0 | 0 | 0 | 0 | 0 | 0 | 0 | 0 | 0 | 0 | 0 |
| virus_Jun22.50v_25119 | 0 | 0 | 0 | 0 | 0 | 0 | 0 | 0 | 0 | 0 | 0 |
| virus_Jun22.50v_25905 | 0 | 0 | 0 | 0 | 0 | 0 | 0 | 0 | 0 | 0 | 0 |
| virus_Jun22.50v_30485 | 0 | 0 | 0 | 0 | 0 | 0 | 0 | 0 | 0 | 0 | 0 |
| virus_Jun22.50v_31945 | 0 | 0 | 0 | 0 | 0 | 0 | 0 | 0 | 0 | 0 | 0 |
| virus_Jun22.50v_32309 | 0 | 0 | 0 | 0 | 0 | 0 | 0 | 0 | 0 | 0 | 0 |
| virus_Jun22.50v_33242 | 0 | 0 | 0 | 0 | 0 | 0 | 0 | 0 | 0 | 0 | 0 |
| virus_Jun22.50v_33321 | 0 | 0 | 0 | 0 | 0 | 0 | 0 | 0 | 0 | 0 | 0 |
| virus_Jun22.50v_35089 | 0 | 0 | 0 | 0 | 0 | 0 | 0 | 0 | 0 | 0 | 0 |
| virus_Jun22.50v_38892 | 0 | 0 | 0 | 0 | 0 | 0 | 0 | 0 | 0 | 0 | 0 |
| virus_Jun22.50v_39859 | 0 | 0 | 0 | 0 | 0 | 0 | 0 | 0 | 0 | 0 | 0 |
| virus_Jun22.50v_40139 | 0 | 0 | 0 | 0 | 0 | 0 | 0 | 0 | 0 | 0 | 0 |
| virus_Jun22.50v_40868 | 0 | 0 | 0 | 0 | 0 | 0 | 0 | 0 | 0 | 0 | 0 |
| virus_Jun22.75v_2594 | 0 | 0 | 0 | 0 | 0 | 0 | 0 | 0 | 0 | 0 | 0 |
| virus_Jun22.75v_7335 | 0 | 0 | 0 | 0 | 0 | 0 | 0 | 0 | 0 | 0 | 0 |
| virus_Jun22.75v_13569 | 0 | 0 | 0 | 0 | 0 | 0 | 0 | 0 | 0 | 0 | 0 |
| virus_Jun22.75v_14341 | 0 | 0 | 0 | 0 | 0 | 0 | 0 | 0 | 1 | 0 | 1 |
| virus_Jun22.75v_20936 | 0 | 0 | 0 | 0 | 0 | 0 | 0 | 0 | 0 | 0 | 0 |
| virus_Jun22.75v_28096 | 0 | 0 | 0 | 0 | 0 | 0 | 0 | 0 | 0 | 0 | 0 |
| virus_May12.5v_4693 | 0 | 0 | 0 | 0 | 0 | 0 | 0 | 0 | 0 | 0 | 0 |
| virus_May12.5v_7454 | 0 | 0 | 0 | 0 | 0 | 0 | 0 | 0 | 0 | 0 | 0 |
| virus_May12.5v_10900 | 0 | 0 | 0 | 0 | 0 | 0 | 0 | 0 | 0 | 0 | 0 |
| virus_May12.5v_11516 | 0 | 0 | 0 | 0 | 0 | 0 | 0 | 0 | 0 | 0 | 0 |
| virus_May12.5v_19292 | 0 | 0 | 0 | 0 | 0 | 0 | 0 | 0 | 0 | 0 | 0 |
| virus_May12.5v_29532 | 0 | 0 | 0 | 0 | 0 | 0 | 0 | 0 | 0 | 0 | 0 |
| virus_May12.5v_32445 | 0 | 0 | 0 | 0 | 0 | 0 | 0 | 0 | 0 | 0 | 0 |
| virus_May12.5v_38052 | 0 | 0 | 0 | 0 | 0 | 0 | 0 | 0 | 0 | 0 | 0 |
| virus_May12.25v_9039 | 0 | 0 | 0 | 0 | 0 | 0 | 0 | 0 | 0 | 0 | 0 |
| virus_May12.25v_9204 | 0 | 0 | 0 | 0 | 0 | 0 | 0 | 0 | 0 | 0 | 0 |
| virus_May12.25v_17390 | 0 | 0 | 0 | 0 | 0 | 0 | 0 | 0 | 0 | 0 | 0 |
| virus_May12.25v_22625 | 0 | 0 | 0 | 0 | 0 | 0 | 0 | 0 | 0 | 0 | 0 |
| virus_May12.25v_22808 | 0 | 0 | 0 | 0 | 0 | 0 | 0 | 0 | 0 | 0 | 0 |
| virus_May12.25v_31397 | 0 | 0 | 0 | 0 | 0 | 0 | 0 | 0 | 0 | 0 | 0 |
| virus_May12.25v_39988 | 0 | 0 | 0 | 0 | 0 | 0 | 0 | 0 | 0 | 0 | 0 |
| virus_May12.50v_1250 | 0 | 0 | 0 | 0 | 0 | 0 | 0 | 0 | 0 | 0 | 0 |
| virus_May12.50v_10400 | 0 | 0 | 0 | 0 | 0 | 0 | 0 | 0 | 0 | 0 | 0 |
| virus_May12.50v_12063 | 0 | 0 | 0 | 0 | 0 | 0 | 0 | 0 | 0 | 0 | 0 |
| virus_May12.50v_17053 | 0 | 0 | 0 | 0 | 0 | 0 | 0 | 0 | 0 | 0 | 0 |
| virus_May12.50v_20096 | 0 | 0 | 0 | 0 | 0 | 0 | 0 | 0 | 0 | 0 | 0 |
| virus_May12.50v_20803 | 0 | 0 | 0 | 0 | 0 | 0 | 0 | 0 | 0 | 0 | 0 |
| virus_May12.50v_23630 | 0 | 0 | 0 | 0 | 0 | 0 | 0 | 0 | 0 | 0 | 0 |
| virus_May12.50v_24948 | 0 | 0 | 0 | 0 | 0 | 0 | 0 | 0 | 0 | 0 | 0 |
| virus_May12.50v_27008 | 0 | 0 | 0 | 0 | 0 | 0 | 0 | 0 | 0 | 0 | 0 |
| virus_May12.50v_29488 | 0 | 0 | 0 | 0 | 0 | 0 | 0 | 0 | 0 | 0 | 0 |
| virus_May12.50v_30715 | 0 | 0 | 0 | 0 | 0 | 0 | 0 | 0 | 0 | 0 | 0 |
| virus_May12.50v_32540 | 0 | 0 | 0 | 0 | 0 | 0 | 0 | 0 | 0 | 0 | 0 |
| virus_May12.50v_33908 | 0 | 0 | 0 | 0 | 0 | 0 | 0 | 0 | 0 | 0 | 0 |
| virus_May12.50v_39712 | 0 | 0 | 0 | 0 | 0 | 0 | 0 | 0 | 0 | 0 | 0 |
| virus_May12.50v_39758 | 0 | 0 | 0 | 0 | 0 | 0 | 0 | 0 | 0 | 0 | 0 |
| virus_May12.75v_14789 | 0 | 0 | 0 | 0 | 0 | 0 | 0 | 0 | 0 | 0 | 0 |
| virus_May12.75v_20699 | 0 | 0 | 0 | 0 | 0 | 0 | 0 | 0 | 0 | 0 | 0 |
| virus_May12.75v_22093 | 0 | 0 | 0 | 0 | 0 | 0 | 0 | 0 | 0 | 0 | 0 |
| virus_May12.75v_26017 | 0 | 0 | 0 | 0 | 0 | 0 | 0 | 0 | 0 | 0 | 0 |
| virus_May12.75v_29072 | 0 | 0 | 0 | 0 | 0 | 0 | 0 | 0 | 0 | 0 | 0 |
| virus_May12.75v_32293 | 0 | 0 | 0 | 0 | 0 | 0 | 0 | 0 | 0 | 0 | 0 |
| virus_May12.75v_33047 | 0 | 0 | 0 | 0 | 0 | 0 | 0 | 0 | 0 | 0 | 0 |
| virus_May12.75v_39352 | 0 | 0 | 0 | 1 | 0 | 0 | 0 | 0 | 0 | 0 | 1 |
| virus_Sep8.0v_6322 | 0 | 0 | 0 | 0 | 0 | 0 | 0 | 0 | 0 | 0 | 0 |
| virus_Sep8.0v_6624 | 0 | 0 | 0 | 0 | 0 | 0 | 0 | 0 | 0 | 0 | 0 |
| virus_Sep8.0v_7899 | 0 | 0 | 0 | 0 | 0 | 0 | 0 | 0 | 0 | 0 | 0 |
| virus_Sep8.0v_8244 | 0 | 0 | 0 | 0 | 0 | 0 | 0 | 0 | 0 | 0 | 0 |
| virus_Sep8.0v_9273 | 0 | 0 | 0 | 0 | 0 | 0 | 0 | 0 | 0 | 0 | 0 |
| virus_Sep8.0v_10379 | 0 | 0 | 0 | 0 | 0 | 0 | 0 | 0 | 0 | 0 | 0 |
| virus_Sep8.0v_13182 | 0 | 0 | 0 | 0 | 0 | 0 | 0 | 0 | 0 | 0 | 0 |
| virus_Sep8.0v_13245 | 0 | 0 | 0 | 0 | 0 | 0 | 0 | 0 | 0 | 0 | 0 |
| virus_Sep8.0v_14474 | 0 | 0 | 0 | 0 | 0 | 0 | 0 | 0 | 0 | 0 | 0 |
| virus_Sep8.0v_15743 | 0 | 0 | 0 | 0 | 0 | 0 | 0 | 0 | 0 | 0 | 0 |
| virus_Sep8.0v_18564 | 0 | 0 | 0 | 0 | 0 | 0 | 0 | 0 | 0 | 0 | 0 |
| virus_Sep8.5v_616 | 0 | 0 | 0 | 0 | 0 | 0 | 0 | 0 | 0 | 0 | 0 |
| virus_Sep8.5v_5247 | 0 | 0 | 0 | 0 | 0 | 0 | 0 | 0 | 0 | 0 | 0 |
| virus_Sep8.5v_16613 | 0 | 0 | 0 | 0 | 0 | 0 | 0 | 0 | 0 | 0 | 0 |
| virus_Sep8.5v_17091 | 0 | 0 | 0 | 0 | 0 | 0 | 0 | 0 | 0 | 0 | 0 |
| virus_Sep8.25v_2227 | 0 | 0 | 0 | 0 | 0 | 0 | 0 | 0 | 0 | 0 | 0 |
| virus_Sep8.25v_9814 | 0 | 0 | 0 | 0 | 0 | 0 | 0 | 0 | 0 | 0 | 0 |
| virus_Sep8.25v_11905 | 0 | 0 | 0 | 0 | 0 | 0 | 0 | 0 | 0 | 0 | 0 |
| virus_Sep8.25v_16285 | 0 | 0 | 0 | 0 | 0 | 0 | 0 | 0 | 0 | 0 | 0 |
| virus_Sep8.25v_19115 | 0 | 0 | 0 | 0 | 0 | 0 | 0 | 0 | 0 | 0 | 0 |
| virus_Sep8.25v_19723 | 0 | 0 | 0 | 0 | 0 | 0 | 0 | 0 | 0 | 0 | 0 |
| virus_Sep8.25v_19816 | 0 | 0 | 0 | 0 | 0 | 0 | 0 | 0 | 0 | 0 | 0 |
| virus_Sep8.50v_128 | 0 | 0 | 0 | 0 | 0 | 0 | 0 | 0 | 0 | 0 | 0 |
| virus_Sep8.50v_1458 | 0 | 0 | 0 | 0 | 0 | 0 | 0 | 0 | 0 | 0 | 0 |
| virus_Sep8.50v_10422 | 0 | 0 | 0 | 0 | 0 | 0 | 0 | 0 | 0 | 0 | 0 |
| virus_Sep8.50v_13225 | 0 | 0 | 0 | 0 | 0 | 0 | 0 | 0 | 0 | 0 | 0 |
| virus_Sep8.50v_16320 | 0 | 0 | 0 | 0 | 0 | 0 | 0 | 0 | 0 | 0 | 0 |
| virus_Sep8.50v_21840 | 0 | 0 | 0 | 0 | 0 | 0 | 0 | 0 | 0 | 0 | 0 |
| virus_Sep8.50v_24019 | 0 | 0 | 0 | 0 | 0 | 0 | 0 | 0 | 0 | 0 | 0 |
| virus_Sep8.50v_24605 | 0 | 0 | 0 | 0 | 0 | 0 | 0 | 0 | 0 | 0 | 0 |
| virus_Sep8.50v_27465 | 0 | 0 | 0 | 0 | 0 | 0 | 0 | 0 | 0 | 0 | 0 |
| virus_Sep8.50v_30437 | 0 | 1 | 0 | 0 | 0 | 0 | 0 | 0 | 0 | 0 | 1 |
| virus_Sep8.50v_32091 | 0 | 0 | 0 | 0 | 0 | 0 | 0 | 0 | 0 | 0 | 0 |
| virus_Sep8.50v_32558 | 0 | 0 | 0 | 0 | 0 | 0 | 0 | 0 | 0 | 0 | 0 |
| virus_Sep8.50v_36039 | 0 | 0 | 0 | 0 | 0 | 0 | 0 | 0 | 0 | 0 | 0 |
| virus_Sep8.50v_38153 | 0 | 0 | 0 | 0 | 0 | 0 | 0 | 0 | 0 | 0 | 0 |
| virus_Sep8.50v_39493 | 0 | 0 | 0 | 0 | 0 | 0 | 0 | 0 | 0 | 0 | 0 |
| virus_Sep8.50v_40089 | 0 | 0 | 0 | 0 | 0 | 0 | 0 | 0 | 0 | 0 | 0 |
| bacteria_Jul11d0b_11488 | 0 | 0 | 0 | 0 | 0 | 0 | 0 | 0 | 0 | 0 | 0 |
| bacteria_Jul11d0b_13870 | 0 | 0 | 0 | 0 | 0 | 0 | 0 | 0 | 0 | 0 | 0 |
| bacteria_Jul11d0b_18094 | 0 | 0 | 0 | 0 | 0 | 0 | 0 | 0 | 0 | 0 | 0 |
| bacteria_Jul11d0b_24169 | 0 | 0 | 0 | 0 | 0 | 0 | 0 | 0 | 0 | 0 | 0 |
| bacteria_Jul11d0b_28117 | 0 | 0 | 0 | 0 | 0 | 0 | 0 | 0 | 0 | 0 | 0 |
| bacteria_Jul11d0b_38701 | 0 | 0 | 0 | 0 | 0 | 0 | 0 | 0 | 0 | 0 | 0 |
| bacteria_Jul11d0b_39470 | 0 | 0 | 0 | 0 | 0 | 0 | 0 | 0 | 0 | 0 | 0 |
| bacteria_Jul11d0b_39953 | 0 | 0 | 0 | 0 | 0 | 0 | 0 | 0 | 0 | 0 | 0 |
| bacteria_Jul11d0b_50725 | 0 | 0 | 0 | 0 | 0 | 0 | 0 | 0 | 0 | 0 | 0 |
| bacteria_Jul11d0b_50942 | 0 | 0 | 0 | 0 | 0 | 0 | 0 | 0 | 0 | 0 | 0 |
| bacteria_Jul11d0b_64156 | 0 | 0 | 0 | 0 | 0 | 0 | 0 | 0 | 0 | 0 | 0 |
| bacteria_Aug5d0b_32446 | 1 | 0 | 0 | 0 | 0 | 0 | 0 | 0 | 0 | 0 | 1 |
| bacteria_Aug5d0b_72428 | 1 | 0 | 0 | 0 | 0 | 0 | 0 | 0 | 0 | 0 | 1 |
| bacteria_Aug5d5b_10045 | 0 | 0 | 1 | 0 | 0 | 0 | 0 | 0 | 0 | 0 | 1 |
| bacteria_Aug5d5b_30134 | 0 | 0 | 1 | 0 | 0 | 0 | 0 | 0 | 0 | 0 | 1 |
| bacteria_Aug5d25b_9580 | 0 | 0 | 0 | 0 | 1 | 0 | 0 | 0 | 0 | 0 | 1 |
| bacteria_Aug5d25b_10628 | 0 | 0 | 0 | 0 | 1 | 0 | 0 | 0 | 0 | 0 | 1 |
| bacteria_Aug5d25b_12321 | 0 | 1 | 0 | 0 | 1 | 0 | 0 | 0 | 0 | 0 | 2 |
| bacteria_Aug5d25b_13592 | 0 | 0 | 0 | 0 | 1 | 0 | 0 | 0 | 0 | 0 | 1 |
| bacteria_Aug5d25b_16026 | 0 | 0 | 0 | 0 | 1 | 0 | 0 | 0 | 0 | 0 | 1 |
| bacteria_Aug5d25b_18342 | 0 | 0 | 0 | 0 | 1 | 0 | 0 | 0 | 0 | 0 | 1 |
| bacteria_Aug5d25b_18729 | 0 | 0 | 0 | 0 | 1 | 0 | 0 | 0 | 0 | 0 | 1 |
| bacteria_Aug5d25b_20128 | 0 | 0 | 0 | 0 | 1 | 0 | 0 | 0 | 0 | 0 | 1 |
| bacteria_Aug5d25b_22027 | 0 | 0 | 0 | 0 | 1 | 0 | 0 | 0 | 0 | 0 | 1 |

| | | | | | | | | | | | |
|---|---|---|---|---|---|---|---|---|---|---|---|
| bacteria_Aug5d25b_22709 | 0 | 0 | 0 | 0 | 1 | 0 | 0 | 0 | 0 | 0 | 1 |
| bacteria_Aug5d25b_23173 | 0 | 0 | 0 | 0 | 1 | 0 | 0 | 0 | 0 | 0 | 1 |
| bacteria_Aug5d25b_31402 | 0 | 0 | 0 | 0 | 1 | 0 | 0 | 0 | 0 | 0 | 1 |
| bacteria_Aug5d25b_46283 | 0 | 0 | 0 | 0 | 1 | 0 | 0 | 0 | 0 | 0 | 1 |
| bacteria_Aug5d25b_48257 | 0 | 0 | 0 | 0 | 1 | 0 | 0 | 0 | 0 | 0 | 1 |
| bacteria_Aug5d25b_50584 | 0 | 0 | 0 | 0 | 1 | 0 | 0 | 0 | 0 | 0 | 1 |
| bacteria_Aug5d50b_4093 | 0 | 0 | 0 | 0 | 0 | 0 | 1 | 0 | 0 | 0 | 1 |
| bacteria_Aug5d50b_11286 | 0 | 0 | 0 | 0 | 0 | 0 | 1 | 0 | 0 | 0 | 1 |
| bacteria_Aug5d50b_63025 | 0 | 0 | 0 | 0 | 0 | 0 | 1 | 0 | 0 | 0 | 1 |
| bacteria_Aug5d50b_70446 | 0 | 0 | 0 | 0 | 0 | 0 | 1 | 0 | 0 | 0 | 1 |
| bacteria_Jul11d5b_6987 | 0 | 0 | 0 | 0 | 0 | 0 | 0 | 0 | 0 | 0 | 0 |
| bacteria_Jul11d5b_13001 | 0 | 0 | 0 | 0 | 0 | 0 | 0 | 0 | 0 | 0 | 0 |
| bacteria_Jul11d5b_25213 | 0 | 0 | 0 | 0 | 0 | 0 | 0 | 0 | 0 | 0 | 0 |
| bacteria_Jul11d5b_37105 | 0 | 0 | 0 | 0 | 0 | 0 | 0 | 0 | 0 | 0 | 0 |
| bacteria_Jul11d5b_42893 | 0 | 0 | 0 | 0 | 0 | 0 | 0 | 0 | 0 | 0 | 0 |
| bacteria_Jul11d25b_3094 | 0 | 0 | 0 | 0 | 0 | 0 | 0 | 0 | 0 | 0 | 0 |
| bacteria_Jul11d25b_8817 | 0 | 0 | 0 | 0 | 0 | 0 | 0 | 0 | 0 | 0 | 0 |
| bacteria_Jul11d25b_11944 | 0 | 0 | 0 | 0 | 0 | 0 | 0 | 0 | 0 | 0 | 0 |
| bacteria_Jul11d25b_13107 | 0 | 0 | 0 | 0 | 0 | 0 | 0 | 0 | 0 | 0 | 0 |
| bacteria_Jul11d25b_14040 | 0 | 0 | 0 | 0 | 0 | 0 | 0 | 0 | 0 | 0 | 0 |
| bacteria_Jul11d25b_20003 | 0 | 0 | 0 | 0 | 0 | 0 | 0 | 0 | 0 | 0 | 0 |
| bacteria_Jul11d25b_26957 | 0 | 0 | 0 | 0 | 0 | 0 | 0 | 0 | 0 | 0 | 0 |
| bacteria_Jul11d25b_33937 | 0 | 1 | 0 | 0 | 0 | 0 | 0 | 0 | 0 | 0 | 1 |
| bacteria_Jul11d25b_34697 | 0 | 0 | 0 | 0 | 0 | 0 | 0 | 0 | 0 | 0 | 0 |
| bacteria_Jul11d25b_37794 | 0 | 0 | 0 | 0 | 0 | 0 | 0 | 0 | 0 | 0 | 0 |
| bacteria_Jul11d25b_38601 | 0 | 0 | 0 | 0 | 0 | 0 | 0 | 0 | 0 | 0 | 0 |
| bacteria_Jul11d25b_51429 | 0 | 0 | 0 | 0 | 0 | 0 | 0 | 0 | 0 | 0 | 0 |
| bacteria_Jul11d25b_51486 | 0 | 0 | 0 | 0 | 0 | 0 | 0 | 0 | 0 | 0 | 0 |
| bacteria_Jul11d25b_51509 | 0 | 0 | 0 | 0 | 0 | 0 | 0 | 0 | 0 | 0 | 0 |
| bacteria_Jul11d50b_1675 | 0 | 0 | 0 | 0 | 0 | 0 | 0 | 0 | 0 | 0 | 0 |
| bacteria_Jul11d50b_15432 | 0 | 0 | 0 | 0 | 0 | 0 | 0 | 0 | 0 | 0 | 0 |
| bacteria_Jul11d50b_33830 | 0 | 0 | 0 | 0 | 0 | 0 | 0 | 0 | 0 | 0 | 0 |
| bacteria_Jul11d50b_43519 | 0 | 0 | 0 | 0 | 0 | 0 | 0 | 0 | 0 | 0 | 0 |
| bacteria_Jul11d50b_46901 | 0 | 0 | 0 | 0 | 0 | 0 | 0 | 0 | 0 | 0 | 0 |
| bacteria_Jul11d50b_51108 | 0 | 0 | 0 | 0 | 0 | 0 | 0 | 0 | 0 | 0 | 0 |
| bacteria_Jul11d75b_3746 | 0 | 0 | 0 | 0 | 0 | 0 | 0 | 0 | 0 | 0 | 0 |
| bacteria_Jul11d75b_10346 | 0 | 0 | 0 | 0 | 0 | 0 | 0 | 0 | 0 | 0 | 0 |
| bacteria_Jul11d75b_13970 | 0 | 0 | 0 | 0 | 0 | 0 | 0 | 0 | 0 | 0 | 0 |
| bacteria_Jul11d75b_17694 | 0 | 0 | 0 | 0 | 0 | 0 | 0 | 0 | 0 | 0 | 0 |
| bacteria_Jul11d75b_17803 | 0 | 0 | 0 | 0 | 0 | 0 | 0 | 0 | 0 | 0 | 0 |
| bacteria_Jul11d75b_24366 | 0 | 0 | 0 | 0 | 0 | 0 | 0 | 0 | 0 | 0 | 0 |
| bacteria_Jul11d75b_29684 | 0 | 0 | 0 | 0 | 0 | 0 | 0 | 0 | 0 | 0 | 0 |
| bacteria_Jul11d75b_38527 | 0 | 0 | 0 | 0 | 0 | 0 | 0 | 0 | 0 | 0 | 0 |
| bacteria_Jul11d75b_39862 | 0 | 0 | 0 | 1 | 0 | 0 | 0 | 0 | 0 | 0 | 1 |
| bacteria_Jul11d75b_46826 | 0 | 0 | 0 | 0 | 0 | 0 | 0 | 0 | 0 | 0 | 0 |
| bacteria_Jul11d75b_49356 | 0 | 0 | 0 | 0 | 0 | 0 | 0 | 0 | 0 | 0 | 0 |
| bacteria_Jul11d75b_50938 | 0 | 0 | 0 | 0 | 0 | 0 | 0 | 0 | 0 | 0 | 0 |
| bacteria_Jul27d5b_2731 | 0 | 0 | 0 | 0 | 0 | 0 | 0 | 0 | 0 | 0 | 0 |
| bacteria_Jul27d5b_4398 | 0 | 0 | 0 | 0 | 0 | 0 | 0 | 0 | 0 | 0 | 0 |
| bacteria_Jul27d5b_12258 | 0 | 0 | 0 | 0 | 0 | 0 | 0 | 0 | 0 | 0 | 0 |
| bacteria_Jul27d5b_12885 | 0 | 0 | 0 | 0 | 0 | 0 | 0 | 0 | 0 | 0 | 0 |
| bacteria_Jul27d5b_15744 | 0 | 0 | 0 | 0 | 0 | 0 | 0 | 0 | 0 | 0 | 0 |
| bacteria_Jul27d5b_16149 | 0 | 0 | 0 | 0 | 0 | 0 | 0 | 0 | 0 | 0 | 0 |
| bacteria_Jul27d5b_18667 | 0 | 0 | 0 | 1 | 0 | 0 | 0 | 0 | 0 | 0 | 1 |
| bacteria_Jul27d5b_24390 | 0 | 0 | 0 | 0 | 0 | 0 | 0 | 0 | 0 | 0 | 0 |
| bacteria_Jul27d5b_24460 | 0 | 0 | 0 | 0 | 0 | 0 | 0 | 0 | 0 | 0 | 0 |
| bacteria_Jul27d5b_27471 | 0 | 0 | 0 | 0 | 0 | 0 | 0 | 0 | 0 | 0 | 0 |
| bacteria_Jul27d5b_28308 | 0 | 0 | 0 | 0 | 0 | 0 | 0 | 0 | 0 | 0 | 0 |
| bacteria_Jul27d5b_29681 | 0 | 0 | 0 | 0 | 0 | 0 | 0 | 0 | 0 | 0 | 0 |
| bacteria_Jul27d5b_29806 | 0 | 0 | 0 | 0 | 0 | 0 | 0 | 0 | 0 | 0 | 0 |
| bacteria_Jul27d5b_35891 | 0 | 0 | 0 | 0 | 0 | 0 | 0 | 0 | 0 | 0 | 0 |
| bacteria_Jul27d5b_37915 | 0 | 0 | 0 | 0 | 0 | 0 | 0 | 0 | 0 | 0 | 0 |
| bacteria_Jul27d5b_45730 | 0 | 0 | 0 | 0 | 0 | 0 | 0 | 0 | 0 | 0 | 0 |
| bacteria_Jul27d5b_45881 | 0 | 0 | 0 | 0 | 0 | 0 | 0 | 0 | 0 | 0 | 0 |
| bacteria_Jul27d5b_47242 | 0 | 0 | 0 | 0 | 0 | 0 | 0 | 0 | 0 | 0 | 0 |
| bacteria_Jul27d5b_50670 | 0 | 0 | 0 | 0 | 0 | 0 | 0 | 0 | 0 | 0 | 0 |
| bacteria_Jul27d5b_52174 | 0 | 0 | 0 | 0 | 0 | 0 | 0 | 0 | 0 | 0 | 0 |
| bacteria_Jul27d25b_11380 | 0 | 0 | 0 | 0 | 0 | 0 | 0 | 0 | 0 | 0 | 0 |
| bacteria_Jul27d25b_17921 | 0 | 0 | 0 | 0 | 0 | 0 | 0 | 0 | 0 | 0 | 0 |
| bacteria_Jul27d25b_20545 | 0 | 0 | 0 | 0 | 0 | 0 | 0 | 0 | 0 | 0 | 0 |
| bacteria_Jul27d25b_22389 | 0 | 0 | 0 | 0 | 0 | 0 | 0 | 0 | 0 | 0 | 0 |
| bacteria_Jul27d25b_26573 | 0 | 0 | 0 | 0 | 0 | 0 | 0 | 0 | 0 | 0 | 0 |
| bacteria_Jul27d25b_28741 | 0 | 0 | 0 | 0 | 0 | 0 | 0 | 0 | 0 | 0 | 0 |
| bacteria_Jul27d25b_30604 | 0 | 0 | 0 | 0 | 0 | 0 | 0 | 0 | 0 | 0 | 0 |
| bacteria_Jul27d25b_34265 | 0 | 0 | 0 | 0 | 0 | 0 | 0 | 0 | 0 | 0 | 0 |
| bacteria_Jul27d25b_34820 | 0 | 0 | 0 | 0 | 0 | 0 | 0 | 0 | 0 | 0 | 0 |
| bacteria_Jul27d25b_41223 | 0 | 0 | 0 | 0 | 0 | 0 | 0 | 0 | 0 | 0 | 0 |
| bacteria_Jul27d25b_45152 | 0 | 0 | 0 | 0 | 0 | 0 | 0 | 0 | 0 | 0 | 0 |
| bacteria_Jul27d25b_50037 | 0 | 0 | 0 | 0 | 0 | 0 | 0 | 0 | 0 | 0 | 0 |
| bacteria_Jul27d25b_50441 | 0 | 0 | 0 | 0 | 0 | 0 | 0 | 0 | 0 | 0 | 0 |
| bacteria_Jul27d25b_51397 | 0 | 0 | 0 | 0 | 0 | 0 | 0 | 0 | 0 | 0 | 0 |
| bacteria_Jul27d25b_67155 | 0 | 0 | 0 | 0 | 0 | 0 | 0 | 0 | 0 | 0 | 0 |
| bacteria_Jul27d25b_68270 | 0 | 0 | 0 | 0 | 0 | 0 | 0 | 0 | 0 | 0 | 0 |
| bacteria_Jul27d25b_69674 | 0 | 0 | 0 | 0 | 0 | 0 | 0 | 0 | 0 | 0 | 0 |
| bacteria_Jul27d25b_71275 | 0 | 0 | 0 | 0 | 0 | 0 | 0 | 0 | 0 | 0 | 0 |
| bacteria_Jul27d25b_72543 | 0 | 0 | 0 | 0 | 0 | 0 | 0 | 0 | 0 | 0 | 0 |
| bacteria_Jul27d25b_72811 | 0 | 0 | 0 | 0 | 0 | 0 | 0 | 0 | 0 | 0 | 0 |
| bacteria_Jul27d75b_1935 | 0 | 0 | 0 | 0 | 0 | 0 | 0 | 0 | 0 | 0 | 0 |
| bacteria_Jul27d75b_2931 | 0 | 0 | 0 | 0 | 0 | 0 | 0 | 0 | 0 | 0 | 0 |
| bacteria_Jul27d75b_4873 | 0 | 0 | 0 | 0 | 0 | 0 | 0 | 0 | 0 | 0 | 0 |
| bacteria_Jul27d75b_6449 | 0 | 0 | 0 | 0 | 0 | 0 | 0 | 0 | 0 | 0 | 0 |
| bacteria_Jul27d75b_6865 | 0 | 0 | 0 | 0 | 0 | 0 | 0 | 0 | 0 | 0 | 0 |
| bacteria_Jul27d75b_9384 | 0 | 0 | 0 | 0 | 0 | 0 | 0 | 0 | 0 | 0 | 0 |
| bacteria_Jul27d75b_9429 | 0 | 0 | 0 | 0 | 0 | 0 | 0 | 0 | 0 | 0 | 0 |
| bacteria_Jul27d75b_11468 | 0 | 0 | 0 | 0 | 0 | 0 | 0 | 0 | 0 | 0 | 0 |
| bacteria_Jul27d75b_12570 | 0 | 0 | 0 | 0 | 0 | 0 | 0 | 0 | 0 | 0 | 0 |
| bacteria_Jul27d75b_12648 | 0 | 0 | 0 | 0 | 0 | 0 | 0 | 0 | 0 | 0 | 0 |
| bacteria_Jul27d75b_12776 | 0 | 0 | 0 | 0 | 0 | 0 | 0 | 0 | 0 | 0 | 0 |
| bacteria_Jul27d75b_13527 | 0 | 0 | 0 | 0 | 0 | 0 | 0 | 0 | 0 | 0 | 0 |
| bacteria_Jul27d75b_15741 | 0 | 0 | 0 | 0 | 0 | 0 | 0 | 0 | 0 | 0 | 0 |
| bacteria_Jul27d75b_20602 | 0 | 0 | 0 | 0 | 0 | 0 | 0 | 0 | 0 | 0 | 0 |
| bacteria_Jul27d75b_22037 | 0 | 0 | 0 | 0 | 0 | 0 | 0 | 0 | 0 | 0 | 0 |
| bacteria_Jul27d75b_29716 | 0 | 0 | 0 | 0 | 0 | 0 | 0 | 0 | 0 | 0 | 0 |
| bacteria_Jul27d75b_34089 | 0 | 0 | 0 | 0 | 0 | 0 | 0 | 0 | 0 | 0 | 0 |
| bacteria_Jul27d75b_60772 | 0 | 0 | 0 | 0 | 0 | 0 | 0 | 0 | 0 | 0 | 0 |
| bacteria_Jul27d75b_66918 | 0 | 0 | 0 | 0 | 0 | 0 | 0 | 0 | 0 | 0 | 0 |
| bacteria_Jul27d75b_68002 | 0 | 0 | 0 | 0 | 0 | 0 | 0 | 0 | 0 | 0 | 0 |
| bacteria_Jun8d25b_5785 | 0 | 0 | 0 | 0 | 0 | 0 | 0 | 0 | 0 | 0 | 0 |
| bacteria_Jun8d25b_30135 | 0 | 0 | 0 | 0 | 0 | 0 | 0 | 0 | 0 | 0 | 0 |
| bacteria_Jun8d25b_38642 | 0 | 0 | 0 | 0 | 0 | 0 | 0 | 0 | 0 | 0 | 0 |
| bacteria_Jun8d25b_44065 | 0 | 0 | 0 | 0 | 0 | 0 | 0 | 0 | 0 | 0 | 0 |
| bacteria_Jun8d25b_48717 | 0 | 0 | 0 | 0 | 0 | 0 | 0 | 0 | 0 | 0 | 0 |
| bacteria_Jun8d25b_51731 | 0 | 0 | 0 | 0 | 0 | 0 | 0 | 0 | 0 | 0 | 0 |
| bacteria_Jun8d25b_57047 | 0 | 0 | 0 | 0 | 0 | 0 | 0 | 0 | 0 | 0 | 0 |
| bacteria_Jun8d25b_65983 | 0 | 0 | 0 | 0 | 0 | 0 | 0 | 0 | 0 | 0 | 0 |
| bacteria_Jun8d25b_73142 | 0 | 0 | 0 | 0 | 0 | 0 | 0 | 0 | 0 | 0 | 0 |
| bacteria_Jun8d50b_9073 | 0 | 0 | 0 | 0 | 0 | 0 | 0 | 0 | 0 | 0 | 0 |
| bacteria_Jun8d50b_23697 | 0 | 0 | 0 | 0 | 0 | 0 | 0 | 0 | 0 | 0 | 0 |
| bacteria_Jun8d50b_23918 | 0 | 0 | 0 | 0 | 0 | 0 | 0 | 0 | 0 | 0 | 0 |
| bacteria_Jun8d50b_37342 | 0 | 0 | 0 | 0 | 0 | 0 | 0 | 0 | 0 | 0 | 0 |
| bacteria_Jun8d50b_38853 | 0 | 0 | 0 | 0 | 0 | 0 | 0 | 0 | 0 | 0 | 0 |
| bacteria_Jun8d50b_40026 | 0 | 0 | 0 | 0 | 0 | 0 | 0 | 0 | 0 | 0 | 0 |
| bacteria_Jun8d50b_44799 | 0 | 0 | 0 | 0 | 0 | 0 | 0 | 0 | 0 | 0 | 0 |
| bacteria_Jun8d75b_16458 | 0 | 0 | 0 | 0 | 0 | 0 | 0 | 0 | 0 | 0 | 0 |
| bacteria_Jun8d75b_27066 | 0 | 0 | 0 | 0 | 0 | 0 | 0 | 0 | 0 | 0 | 0 |
| bacteria_Jun8d75b_72425 | 0 | 0 | 0 | 0 | 0 | 0 | 0 | 0 | 0 | 0 | 0 |
| bacteria_Jun22d0b_30541 | 0 | 0 | 0 | 0 | 0 | 0 | 0 | 0 | 0 | 0 | 0 |
| bacteria_Jun22d0b_55410 | 0 | 0 | 0 | 0 | 0 | 0 | 0 | 0 | 0 | 0 | 0 |
| bacteria_Jun22d0b_66924 | 0 | 0 | 0 | 0 | 0 | 0 | 0 | 0 | 0 | 0 | 0 |
| bacteria_Jun22d50b_4234 | 0 | 0 | 0 | 0 | 0 | 0 | 0 | 0 | 0 | 0 | 0 |
| bacteria_Jun22d75b_5705 | 0 | 0 | 0 | 0 | 0 | 0 | 0 | 0 | 0 | 0 | 0 |
| bacteria_Jun22d75b_3931 | 0 | 0 | 0 | 0 | 0 | 0 | 0 | 0 | 0 | 0 | 0 |
| bacteria_Jun22d75b_4748 | 0 | 0 | 0 | 0 | 0 | 0 | 0 | 0 | 0 | 0 | 0 |
| bacteria_Jun22d75b_5194 | 0 | 0 | 0 | 0 | 0 | 0 | 0 | 0 | 0 | 0 | 0 |
| bacteria_Jun22d75b_54631 | 0 | 0 | 0 | 0 | 0 | 0 | 0 | 0 | 0 | 0 | 0 |
| bacteria_Jun22d75b_5572 | 0 | 0 | 0 | 0 | 0 | 0 | 0 | 0 | 0 | 0 | 0 |
| bacteria_Jun22d75b_5755 | 0 | 0 | 0 | 0 | 0 | 0 | 0 | 0 | 0 | 0 | 0 |
| bacteria_Jun22d75b_6065 | 0 | 0 | 0 | 0 | 0 | 0 | 0 | 0 | 0 | 0 | 0 |
| bacteria_Jun22d75b_6400 | 0 | 0 | 0 | 0 | 0 | 0 | 0 | 0 | 0 | 0 | 0 |
| bacteria_Jun22d75b_6747 | 0 | 0 | 0 | 0 | 0 | 0 | 0 | 0 | 0 | 0 | 0 |
| bacteria_May12d5b_6614 | 0 | 0 | 0 | 0 | 0 | 0 | 0 | 0 | 0 | 0 | 0 |
| bacteria_May12d25b_4431 | 0 | 0 | 0 | 0 | 0 | 0 | 0 | 0 | 0 | 0 | 0 |
| bacteria_May12d25b_2134 | 0 | 0 | 0 | 0 | 0 | 0 | 0 | 0 | 0 | 0 | 0 |
| bacteria_May12d25b_3713 | 0 | 0 | 0 | 0 | 0 | 0 | 0 | 0 | 0 | 0 | 0 |
| bacteria_May12d25b_3954 | 0 | 0 | 0 | 0 | 0 | 0 | 0 | 0 | 0 | 0 | 0 |
| bacteria_May12d25b_6133 | 0 | 0 | 0 | 0 | 0 | 0 | 0 | 0 | 0 | 0 | 0 |
| bacteria_May12d25b_640 | 0 | 0 | 0 | 0 | 0 | 0 | 0 | 0 | 0 | 0 | 0 |
| bacteria_May12d25b_7221 | 0 | 0 | 0 | 0 | 0 | 0 | 0 | 0 | 0 | 0 | 0 |
| bacteria_May12d50b_133 | 0 | 0 | 0 | 0 | 0 | 0 | 0 | 0 | 0 | 0 | 0 |
| bacteria_May12d50b_1852 | 0 | 0 | 0 | 0 | 0 | 0 | 0 | 0 | 0 | 0 | 0 |
| bacteria_May12d50b_507 | 0 | 0 | 0 | 0 | 0 | 0 | 0 | 0 | 0 | 0 | 0 |
| bacteria_May12d50b_589 | 0 | 0 | 0 | 0 | 0 | 0 | 0 | 0 | 0 | 0 | 0 |
| bacteria_May12d50b_103 | 0 | 0 | 0 | 0 | 0 | 0 | 0 | 0 | 0 | 0 | 0 |
| bacteria_May12d50b_1125 | 0 | 0 | 0 | 0 | 0 | 0 | 0 | 0 | 0 | 0 | 0 |
| bacteria_May12d50b_118 | 0 | 0 | 0 | 0 | 0 | 0 | 0 | 0 | 0 | 0 | 0 |
| bacteria_May12d50b_1333 | 0 | 0 | 0 | 0 | 0 | 0 | 0 | 0 | 0 | 0 | 0 |
| bacteria_May12d50b_1493 | 0 | 0 | 0 | 0 | 0 | 0 | 0 | 0 | 0 | 0 | 0 |

| | | | | | | | | | | | |
|---|---|---|---|---|---|---|---|---|---|---|---|
| bacteria_May12d50b_3121 | 0 | 0 | 0 | 0 | 0 | 0 | 0 | 0 | 0 | 0 | 0 |
| bacteria_May12d50b_4003 | 0 | 0 | 0 | 0 | 0 | 0 | 0 | 0 | 0 | 0 | 0 |
| bacteria_May12d50b_4738 | 0 | 0 | 0 | 0 | 0 | 0 | 0 | 0 | 0 | 0 | 0 |
| bacteria_May12d75b_2274 | 0 | 0 | 0 | 0 | 0 | 0 | 0 | 0 | 0 | 0 | 0 |
| bacteria_Sep8d0b_54399 | 0 | 0 | 0 | 0 | 0 | 0 | 0 | 0 | 0 | 0 | 0 |
| bacteria_Sep8d5b_843 | 0 | 0 | 0 | 0 | 0 | 0 | 0 | 0 | 0 | 0 | 0 |
| bacteria_Sep8d5b_1310 | 0 | 0 | 0 | 0 | 0 | 0 | 0 | 0 | 0 | 0 | 0 |
| bacteria_Sep8d5b_12186 | 0 | 0 | 0 | 0 | 0 | 0 | 0 | 0 | 0 | 0 | 0 |
| bacteria_Sep8d5b_12323 | 0 | 0 | 0 | 0 | 0 | 0 | 0 | 0 | 0 | 0 | 0 |
| bacteria_Sep8d5b_16248 | 0 | 0 | 0 | 0 | 0 | 0 | 0 | 0 | 0 | 0 | 0 |
| bacteria_Sep8d5b_22301 | 0 | 0 | 0 | 0 | 0 | 0 | 0 | 0 | 0 | 0 | 0 |
| bacteria_Sep8d5b_25121 | 0 | 0 | 0 | 0 | 0 | 0 | 0 | 0 | 0 | 0 | 0 |
| bacteria_Sep8d5b_27115 | 0 | 0 | 0 | 0 | 0 | 0 | 0 | 0 | 0 | 0 | 0 |
| bacteria_Sep8d5b_27802 | 0 | 0 | 0 | 0 | 0 | 0 | 0 | 0 | 0 | 0 | 0 |
| bacteria_Sep8d5b_28161 | 0 | 0 | 0 | 0 | 0 | 0 | 0 | 0 | 0 | 0 | 0 |
| bacteria_Sep8d5b_29068 | 0 | 0 | 0 | 0 | 0 | 0 | 0 | 0 | 0 | 0 | 0 |
| bacteria_Sep8d5b_32864 | 0 | 0 | 0 | 0 | 0 | 0 | 0 | 0 | 0 | 0 | 0 |
| bacteria_Sep8d5b_39467 | 0 | 0 | 0 | 0 | 0 | 0 | 0 | 0 | 0 | 0 | 0 |
| bacteria_Sep8d5b_40166 | 0 | 0 | 0 | 0 | 0 | 0 | 0 | 0 | 0 | 0 | 0 |
| bacteria_Sep8d5b_42221 | 0 | 0 | 0 | 0 | 0 | 0 | 0 | 0 | 0 | 0 | 0 |
| bacteria_Sep8d5b_44029 | 0 | 0 | 0 | 0 | 0 | 0 | 0 | 0 | 0 | 0 | 0 |
| bacteria_Sep8d5b_44060 | 0 | 0 | 0 | 0 | 0 | 0 | 0 | 0 | 0 | 0 | 0 |
| bacteria_Sep8d5b_51717 | 0 | 0 | 0 | 0 | 0 | 0 | 0 | 0 | 0 | 0 | 0 |
| bacteria_Sep8d5b_57398 | 0 | 0 | 0 | 0 | 0 | 0 | 0 | 0 | 0 | 0 | 0 |
| bacteria_Sep8d5b_58396 | 0 | 0 | 0 | 0 | 0 | 0 | 0 | 0 | 0 | 0 | 0 |
| bacteria_Sep8d5b_62351 | 0 | 0 | 0 | 0 | 0 | 0 | 0 | 0 | 0 | 0 | 0 |
| bacteria_Sep8d5b_71080 | 0 | 0 | 0 | 0 | 0 | 0 | 0 | 0 | 0 | 0 | 0 |
| bacteria_Sep8d25b_2747 | 0 | 0 | 0 | 0 | 0 | 0 | 0 | 0 | 0 | 0 | 0 |
| bacteria_Sep8d25b_21835 | 0 | 0 | 0 | 0 | 0 | 0 | 0 | 0 | 0 | 0 | 0 |
| bacteria_Sep8d25b_40244 | 0 | 0 | 0 | 0 | 0 | 0 | 0 | 0 | 0 | 0 | 0 |
| bacteria_Sep8d25b_42928 | 0 | 0 | 0 | 0 | 0 | 0 | 0 | 0 | 0 | 0 | 0 |
| bacteria_Sep8d25b_45275 | 0 | 0 | 0 | 0 | 0 | 0 | 0 | 0 | 0 | 0 | 0 |
| bacteria_Sep8d50b_922 | 0 | 0 | 0 | 0 | 0 | 0 | 0 | 0 | 0 | 0 | 0 |
| bacteria_Sep8d50b_8892 | 0 | 0 | 0 | 0 | 0 | 0 | 0 | 0 | 0 | 0 | 0 |
| bacteria_Sep8d50b_20684 | 0 | 0 | 0 | 0 | 0 | 0 | 0 | 0 | 0 | 0 | 0 |
| bacteria_Sep8d50b_41897 | 0 | 0 | 0 | 0 | 0 | 0 | 0 | 0 | 0 | 0 | 0 |
| virus_Aug5.0v_144 | 0 | 1 | 0 | 0 | 0 | 0 | 0 | 0 | 0 | 0 | 1 |
| virus_Aug5.0v_357 | 0 | 1 | 0 | 0 | 0 | 0 | 0 | 0 | 0 | 0 | 1 |
| virus_Aug5.0v_508 | 0 | 1 | 0 | 0 | 0 | 0 | 0 | 0 | 0 | 0 | 1 |
| virus_Aug5.0v_648 | 0 | 1 | 0 | 0 | 0 | 0 | 0 | 0 | 0 | 0 | 1 |
| virus_Aug5.0v_809 | 0 | 1 | 0 | 0 | 0 | 0 | 0 | 0 | 0 | 0 | 1 |
| virus_Aug5.0v_1201 | 0 | 1 | 0 | 0 | 0 | 0 | 0 | 0 | 0 | 0 | 1 |
| virus_Aug5.0v_1242 | 0 | 1 | 0 | 0 | 0 | 0 | 0 | 0 | 0 | 0 | 1 |
| virus_Aug5.0v_1279 | 0 | 1 | 0 | 0 | 0 | 0 | 0 | 0 | 0 | 0 | 1 |
| virus_Aug5.0v_1502 | 0 | 1 | 0 | 0 | 0 | 0 | 0 | 0 | 0 | 0 | 1 |
| virus_Aug5.0v_1806 | 0 | 1 | 0 | 0 | 0 | 0 | 0 | 0 | 1 | 0 | 2 |
| virus_Aug5.0v_1839 | 0 | 1 | 0 | 0 | 0 | 0 | 0 | 0 | 0 | 0 | 1 |
| virus_Aug5.0v_1877 | 0 | 1 | 0 | 0 | 0 | 0 | 0 | 0 | 0 | 0 | 1 |
| virus_Aug5.0v_2151 | 0 | 1 | 0 | 0 | 0 | 0 | 0 | 0 | 0 | 0 | 1 |
| virus_Aug5.0v_2249 | 0 | 1 | 0 | 0 | 0 | 0 | 0 | 0 | 0 | 0 | 1 |
| virus_Aug5.0v_2548 | 0 | 1 | 0 | 0 | 0 | 0 | 0 | 0 | 0 | 0 | 1 |
| virus_Aug5.0v_2599 | 0 | 1 | 0 | 0 | 0 | 0 | 0 | 0 | 0 | 0 | 1 |
| virus_Aug5.0v_2941 | 0 | 1 | 0 | 0 | 0 | 0 | 0 | 0 | 0 | 0 | 1 |
| virus_Aug5.0v_3054 | 0 | 1 | 0 | 0 | 0 | 0 | 0 | 0 | 0 | 0 | 1 |
| virus_Aug5.0v_3215 | 0 | 1 | 0 | 0 | 0 | 0 | 0 | 0 | 0 | 0 | 1 |
| virus_Aug5.0v_3285 | 0 | 1 | 0 | 0 | 0 | 1 | 0 | 0 | 0 | 0 | 2 |
| virus_Aug5.0v_3596 | 0 | 1 | 0 | 0 | 0 | 0 | 0 | 0 | 0 | 0 | 1 |
| virus_Aug5.0v_3749 | 0 | 1 | 0 | 0 | 0 | 0 | 0 | 0 | 0 | 0 | 1 |
| virus_Aug5.0v_4149 | 0 | 1 | 0 | 0 | 0 | 0 | 0 | 0 | 0 | 0 | 1 |
| virus_Aug5.0v_4286 | 0 | 1 | 0 | 0 | 0 | 0 | 0 | 0 | 0 | 0 | 1 |
| virus_Aug5.0v_4436 | 0 | 1 | 0 | 0 | 0 | 0 | 0 | 0 | 0 | 0 | 1 |
| virus_Aug5.0v_5325 | 0 | 1 | 0 | 0 | 0 | 0 | 0 | 0 | 0 | 0 | 1 |
| virus_Aug5.0v_5390 | 0 | 1 | 0 | 0 | 0 | 0 | 0 | 0 | 0 | 0 | 1 |
| virus_Aug5.5v_1196 | 0 | 0 | 0 | 1 | 0 | 0 | 0 | 0 | 0 | 0 | 1 |
| virus_Aug5.5v_3201 | 0 | 0 | 0 | 1 | 0 | 1 | 0 | 0 | 0 | 0 | 2 |
| virus_Aug5.5v_14907 | 0 | 0 | 0 | 1 | 0 | 0 | 0 | 0 | 0 | 0 | 1 |
| virus_Aug5.5v_16686 | 0 | 0 | 0 | 1 | 0 | 0 | 0 | 0 | 0 | 0 | 1 |
| virus_Aug5.5v_17290 | 0 | 0 | 0 | 1 | 0 | 0 | 0 | 0 | 0 | 1 | 2 |
| virus_Aug5.5v_17871 | 0 | 0 | 0 | 1 | 0 | 0 | 0 | 0 | 0 | 0 | 1 |
| virus_Aug5.5v_20459 | 0 | 0 | 0 | 1 | 0 | 0 | 0 | 0 | 0 | 0 | 1 |
| virus_Aug5.5v_20826 | 0 | 0 | 0 | 1 | 0 | 0 | 1 | 0 | 0 | 0 | 2 |
| virus_Aug5.5v_27451 | 0 | 0 | 0 | 1 | 0 | 0 | 0 | 0 | 0 | 0 | 1 |
| virus_Aug5.5v_34422 | 0 | 0 | 0 | 1 | 0 | 0 | 0 | 0 | 0 | 0 | 1 |
| virus_Aug5.5v_36555 | 0 | 0 | 0 | 1 | 0 | 0 | 0 | 0 | 0 | 0 | 1 |
| virus_Aug5.5v_37292 | 0 | 0 | 0 | 1 | 0 | 0 | 0 | 0 | 0 | 0 | 1 |
| virus_Aug5.5v_37490 | 0 | 0 | 0 | 1 | 0 | 0 | 0 | 0 | 0 | 0 | 1 |
| virus_Aug5.5v_38688 | 0 | 0 | 0 | 1 | 0 | 0 | 0 | 0 | 0 | 0 | 1 |
| virus_Aug5.25v_4275 | 0 | 1 | 0 | 0 | 0 | 1 | 0 | 0 | 0 | 0 | 2 |
| virus_Aug5.25v_7723 | 0 | 0 | 0 | 0 | 0 | 1 | 0 | 0 | 0 | 0 | 1 |
| virus_Aug5.25v_10034 | 0 | 0 | 0 | 0 | 0 | 1 | 0 | 0 | 0 | 0 | 1 |
| virus_Aug5.25v_11212 | 0 | 0 | 0 | 0 | 0 | 1 | 0 | 0 | 0 | 0 | 1 |
| virus_Aug5.25v_14747 | 0 | 0 | 0 | 0 | 0 | 1 | 0 | 0 | 0 | 0 | 1 |
| virus_Aug5.25v_16298 | 0 | 0 | 0 | 0 | 0 | 1 | 0 | 0 | 0 | 0 | 1 |
| virus_Aug5.25v_17243 | 0 | 0 | 0 | 0 | 0 | 1 | 0 | 0 | 0 | 0 | 1 |
| virus_Aug5.25v_17930 | 0 | 0 | 0 | 0 | 0 | 1 | 0 | 0 | 0 | 0 | 1 |
| virus_Aug5.25v_19585 | 0 | 0 | 0 | 0 | 0 | 1 | 0 | 0 | 0 | 0 | 1 |
| virus_Aug5.25v_21808 | 0 | 0 | 0 | 0 | 0 | 1 | 0 | 0 | 0 | 0 | 1 |
| virus_Aug5.25v_21957 | 0 | 0 | 0 | 0 | 0 | 1 | 0 | 0 | 0 | 0 | 1 |
| virus_Aug5.25v_22285 | 0 | 0 | 0 | 0 | 0 | 1 | 0 | 0 | 0 | 0 | 1 |
| virus_Aug5.25v_23714 | 0 | 0 | 0 | 0 | 0 | 1 | 0 | 0 | 0 | 0 | 1 |
| virus_Aug5.25v_26919 | 0 | 0 | 0 | 0 | 0 | 1 | 0 | 0 | 0 | 0 | 1 |
| virus_Aug5.25v_30444 | 0 | 0 | 0 | 0 | 0 | 1 | 0 | 0 | 0 | 0 | 1 |
| virus_Aug5.25v_31255 | 0 | 0 | 0 | 0 | 0 | 1 | 0 | 0 | 0 | 0 | 1 |
| virus_Aug5.25v_32541 | 0 | 0 | 0 | 0 | 0 | 1 | 0 | 0 | 0 | 0 | 1 |
| virus_Aug5.25v_36318 | 0 | 0 | 0 | 0 | 0 | 1 | 0 | 0 | 0 | 0 | 1 |
| virus_Aug5.25v_36353 | 0 | 0 | 0 | 0 | 0 | 1 | 0 | 0 | 0 | 0 | 1 |
| virus_Aug5.25v_38368 | 0 | 0 | 0 | 0 | 0 | 1 | 0 | 0 | 0 | 0 | 1 |
| virus_Aug5.25v_38987 | 0 | 0 | 0 | 0 | 0 | 1 | 0 | 0 | 0 | 0 | 1 |
| virus_Aug5.25v_40327 | 0 | 0 | 0 | 0 | 0 | 1 | 0 | 0 | 0 | 0 | 1 |
| virus_Aug5.25v_40792 | 0 | 0 | 0 | 0 | 0 | 1 | 0 | 0 | 0 | 0 | 1 |
| virus_Aug5.25v_41255 | 0 | 0 | 0 | 0 | 0 | 1 | 0 | 0 | 0 | 0 | 1 |
| virus_Aug5.50v_5862 | 0 | 0 | 0 | 0 | 0 | 0 | 0 | 1 | 0 | 0 | 1 |
| virus_Aug5.50v_25014 | 0 | 0 | 0 | 0 | 0 | 0 | 0 | 1 | 0 | 0 | 1 |
| virus_Aug5.50v_25206 | 0 | 0 | 0 | 0 | 0 | 0 | 0 | 1 | 0 | 0 | 1 |
| virus_Aug5.50v_27756 | 0 | 0 | 0 | 0 | 0 | 0 | 0 | 1 | 0 | 0 | 1 |
| virus_Aug5.50v_40106 | 0 | 0 | 0 | 0 | 0 | 0 | 0 | 1 | 0 | 0 | 1 |
| virus_Aug5.75v_3968 | 0 | 0 | 0 | 0 | 0 | 0 | 0 | 0 | 0 | 1 | 1 |
| virus_Aug5.75v_5047 | 0 | 0 | 0 | 0 | 0 | 0 | 0 | 0 | 0 | 1 | 1 |
| virus_Jul11.0v_1613 | 0 | 0 | 0 | 0 | 0 | 0 | 0 | 0 | 0 | 0 | 0 |
| virus_Jul11.0v_3874 | 0 | 0 | 0 | 0 | 0 | 0 | 0 | 0 | 0 | 0 | 0 |
| virus_Jul11.0v_7815 | 0 | 0 | 0 | 0 | 0 | 0 | 0 | 0 | 0 | 0 | 0 |
| virus_Jul11.0v_14307 | 0 | 0 | 0 | 0 | 0 | 0 | 0 | 0 | 0 | 0 | 0 |
| virus_Jul11.0v_14910 | 0 | 0 | 0 | 0 | 0 | 0 | 0 | 0 | 0 | 0 | 0 |
| virus_Jul11.0v_18936 | 0 | 0 | 0 | 0 | 0 | 0 | 0 | 0 | 0 | 0 | 0 |
| virus_Jul11.0v_20475 | 0 | 0 | 0 | 0 | 0 | 0 | 0 | 0 | 0 | 0 | 0 |
| virus_Jul11.5v_948 | 0 | 0 | 0 | 0 | 0 | 0 | 0 | 0 | 0 | 0 | 0 |
| virus_Jul11.5v_1952 | 0 | 0 | 0 | 0 | 0 | 0 | 0 | 0 | 0 | 0 | 0 |
| virus_Jul11.5v_3073 | 0 | 0 | 0 | 0 | 0 | 0 | 0 | 0 | 0 | 0 | 0 |
| virus_Jul11.5v_3138 | 0 | 0 | 0 | 0 | 0 | 0 | 0 | 0 | 0 | 0 | 0 |
| virus_Jul11.5v_4412 | 0 | 0 | 0 | 0 | 0 | 0 | 0 | 0 | 0 | 0 | 0 |
| virus_Jul11.5v_6076 | 0 | 0 | 0 | 0 | 0 | 0 | 0 | 0 | 0 | 0 | 0 |
| virus_Jul11.5v_6230 | 0 | 0 | 0 | 0 | 0 | 0 | 0 | 0 | 0 | 0 | 0 |
| virus_Jul11.5v_6464 | 0 | 0 | 0 | 0 | 0 | 0 | 0 | 0 | 0 | 0 | 0 |
| virus_Jul11.5v_6977 | 0 | 0 | 0 | 0 | 0 | 0 | 0 | 0 | 0 | 0 | 0 |
| virus_Jul11.5v_9419 | 0 | 0 | 0 | 0 | 0 | 0 | 0 | 0 | 0 | 0 | 0 |
| virus_Jul11.5v_9901 | 0 | 0 | 0 | 0 | 0 | 0 | 0 | 0 | 0 | 0 | 0 |
| virus_Jul11.5v_13471 | 0 | 0 | 0 | 0 | 0 | 0 | 0 | 0 | 0 | 0 | 0 |
| virus_Jul11.5v_15182 | 0 | 0 | 0 | 0 | 0 | 0 | 0 | 0 | 0 | 0 | 0 |
| virus_Jul11.5v_17185 | 0 | 0 | 0 | 0 | 0 | 0 | 0 | 0 | 0 | 0 | 0 |
| virus_Jul11.5v_18128 | 0 | 0 | 0 | 0 | 0 | 0 | 0 | 0 | 0 | 0 | 0 |
| virus_Jul11.5v_18322 | 0 | 0 | 0 | 0 | 0 | 0 | 0 | 0 | 0 | 0 | 0 |
| virus_Jul11.25v_1151 | 0 | 0 | 0 | 0 | 0 | 0 | 0 | 0 | 0 | 0 | 0 |
| virus_Jul11.25v_1165 | 0 | 0 | 0 | 0 | 0 | 0 | 0 | 0 | 0 | 0 | 0 |
| virus_Jul11.25v_1628 | 0 | 0 | 0 | 0 | 0 | 0 | 0 | 0 | 0 | 0 | 0 |
| virus_Jul11.25v_3496 | 0 | 0 | 0 | 0 | 0 | 0 | 0 | 0 | 0 | 0 | 0 |
| virus_Jul11.25v_3823 | 0 | 0 | 0 | 0 | 0 | 0 | 0 | 0 | 0 | 0 | 0 |
| virus_Jul11.25v_4548 | 0 | 0 | 0 | 0 | 0 | 0 | 0 | 0 | 0 | 0 | 0 |
| virus_Jul11.25v_5123 | 0 | 0 | 0 | 0 | 0 | 0 | 0 | 0 | 0 | 0 | 0 |
| virus_Jul11.25v_5315 | 0 | 0 | 0 | 0 | 0 | 0 | 0 | 0 | 0 | 0 | 0 |
| virus_Jul11.25v_11011 | 0 | 0 | 0 | 0 | 0 | 0 | 0 | 0 | 0 | 0 | 0 |
| virus_Jul11.25v_11317 | 0 | 0 | 0 | 0 | 0 | 0 | 0 | 0 | 0 | 0 | 0 |
| virus_Jul11.25v_11801 | 0 | 0 | 0 | 0 | 0 | 0 | 0 | 0 | 0 | 0 | 0 |
| virus_Jul11.25v_12713 | 0 | 0 | 0 | 0 | 0 | 0 | 0 | 0 | 0 | 0 | 0 |
| virus_Jul11.25v_13785 | 0 | 0 | 0 | 0 | 0 | 0 | 0 | 0 | 0 | 0 | 0 |
| virus_Jul11.25v_16828 | 0 | 0 | 0 | 0 | 0 | 0 | 0 | 0 | 0 | 0 | 0 |
| virus_Jul11.25v_17692 | 0 | 0 | 0 | 0 | 0 | 0 | 0 | 0 | 0 | 0 | 0 |
| virus_Jul11.25v_20956 | 0 | 0 | 0 | 0 | 0 | 0 | 0 | 0 | 0 | 0 | 0 |
| virus_Jul11.50v_1658 | 0 | 0 | 0 | 0 | 0 | 0 | 0 | 0 | 0 | 0 | 0 |
| virus_Jul11.50v_3134 | 0 | 0 | 0 | 0 | 0 | 0 | 0 | 0 | 0 | 0 | 0 |
| virus_Jul11.50v_3570 | 0 | 0 | 0 | 0 | 0 | 0 | 0 | 0 | 0 | 0 | 0 |
| virus_Jul11.50v_5386 | 0 | 0 | 0 | 0 | 0 | 0 | 0 | 0 | 0 | 0 | 0 |
| virus_Jul11.50v_8132 | 0 | 0 | 0 | 0 | 0 | 0 | 0 | 0 | 0 | 0 | 0 |
| virus_Jul11.50v_9013 | 0 | 0 | 0 | 0 | 0 | 0 | 0 | 0 | 0 | 0 | 0 |
| virus_Jul11.50v_9145 | 0 | 0 | 0 | 0 | 0 | 0 | 0 | 0 | 0 | 0 | 0 |
| virus_Jul11.50v_10222 | 0 | 0 | 0 | 0 | 1 | 0 | 0 | 0 | 0 | 0 | 1 |
| virus_Jul11.50v_10292 | 0 | 0 | 0 | 0 | 0 | 0 | 0 | 0 | 0 | 0 | 0 |
| virus_Jul11.50v_14073 | 0 | 0 | 0 | 0 | 0 | 0 | 0 | 0 | 0 | 0 | 0 |

| | | | | | | | | | | | |
|---|---|---|---|---|---|---|---|---|---|---|---|
| virus_Jul11.50v_14817 | 0 | 0 | 0 | 0 | 0 | 0 | 0 | 0 | 0 | 0 | 0 |
| virus_Jul11.50v_15596 | 0 | 0 | 0 | 0 | 0 | 0 | 0 | 0 | 0 | 0 | 0 |
| virus_Jul11.50v_15921 | 0 | 0 | 0 | 0 | 0 | 0 | 0 | 0 | 0 | 0 | 0 |
| virus_Jul11.50v_16057 | 0 | 0 | 0 | 0 | 0 | 0 | 0 | 0 | 0 | 0 | 0 |
| virus_Jul11.50v_17405 | 0 | 0 | 0 | 0 | 0 | 0 | 0 | 0 | 0 | 0 | 0 |
| virus_Jul11.50v_18812 | 0 | 0 | 0 | 0 | 0 | 0 | 0 | 0 | 0 | 0 | 0 |
| virus_Jul11.50v_19569 | 0 | 0 | 0 | 0 | 0 | 0 | 0 | 0 | 0 | 0 | 0 |
| virus_Jul11.75v_20635 | 0 | 0 | 0 | 0 | 0 | 0 | 0 | 0 | 0 | 0 | 0 |
| virus_Jul11.75v_22505 | 0 | 0 | 0 | 0 | 0 | 0 | 0 | 0 | 0 | 0 | 0 |
| virus_Jul11.75v_23409 | 0 | 0 | 0 | 0 | 0 | 0 | 0 | 0 | 0 | 0 | 0 |
| virus_Jul11.75v_26829 | 0 | 0 | 0 | 0 | 0 | 0 | 0 | 0 | 0 | 0 | 0 |
| virus_Jul11.75v_27124 | 0 | 0 | 0 | 0 | 0 | 0 | 0 | 0 | 0 | 0 | 0 |
| virus_Jul11.75v_28798 | 0 | 0 | 0 | 0 | 0 | 0 | 0 | 0 | 0 | 0 | 0 |
| virus_Jul11.75v_31337 | 0 | 0 | 0 | 0 | 0 | 0 | 0 | 0 | 0 | 0 | 0 |
| virus_Jul11.75v_36835 | 0 | 0 | 0 | 0 | 0 | 0 | 0 | 0 | 0 | 0 | 0 |
| virus_Jul11.75v_37165 | 0 | 0 | 0 | 0 | 0 | 0 | 0 | 0 | 0 | 0 | 0 |
| virus_Jul27.5v_213 | 0 | 0 | 0 | 0 | 0 | 0 | 0 | 0 | 0 | 0 | 0 |
| virus_Jul27.5v_354 | 0 | 0 | 0 | 0 | 0 | 0 | 0 | 0 | 0 | 0 | 0 |
| virus_Jul27.5v_650 | 0 | 0 | 0 | 0 | 0 | 0 | 0 | 0 | 0 | 0 | 0 |
| virus_Jul27.5v_1734 | 0 | 0 | 0 | 0 | 0 | 0 | 0 | 0 | 0 | 0 | 0 |
| virus_Jul27.5v_2154 | 0 | 0 | 0 | 0 | 0 | 0 | 0 | 0 | 0 | 0 | 0 |
| virus_Jul27.5v_2593 | 0 | 0 | 0 | 0 | 0 | 0 | 0 | 0 | 0 | 0 | 0 |
| virus_Jul27.5v_3707 | 0 | 0 | 0 | 0 | 0 | 0 | 0 | 0 | 0 | 0 | 0 |
| virus_Jul27.5v_4775 | 0 | 0 | 0 | 0 | 0 | 0 | 0 | 0 | 0 | 0 | 0 |
| virus_Jul27.5v_5298 | 0 | 0 | 0 | 0 | 0 | 0 | 0 | 0 | 0 | 0 | 0 |
| virus_Jul27.5v_6476 | 0 | 0 | 0 | 0 | 0 | 0 | 0 | 0 | 0 | 0 | 0 |
| virus_Jul27.5v_6482 | 0 | 0 | 0 | 0 | 0 | 0 | 0 | 0 | 0 | 0 | 0 |
| virus_Jul27.5v_7913 | 0 | 0 | 0 | 0 | 0 | 0 | 0 | 0 | 0 | 0 | 0 |
| virus_Jul27.5v_8541 | 0 | 0 | 0 | 0 | 0 | 0 | 0 | 0 | 0 | 0 | 0 |
| virus_Jul27.5v_8588 | 0 | 0 | 0 | 0 | 0 | 0 | 0 | 0 | 0 | 0 | 0 |
| virus_Jul27.5v_9251 | 0 | 0 | 0 | 0 | 0 | 0 | 0 | 0 | 0 | 0 | 0 |
| virus_Jul27.5v_9916 | 0 | 0 | 0 | 0 | 0 | 0 | 0 | 0 | 0 | 0 | 0 |
| virus_Jul27.5v_10429 | 0 | 0 | 0 | 0 | 0 | 0 | 0 | 0 | 0 | 0 | 0 |
| virus_Jul27.5v_10453 | 0 | 0 | 0 | 0 | 0 | 0 | 0 | 0 | 0 | 0 | 0 |
| virus_Jul27.5v_11706 | 0 | 0 | 0 | 0 | 0 | 0 | 0 | 0 | 0 | 0 | 0 |
| virus_Jul27.5v_12269 | 0 | 0 | 0 | 0 | 0 | 0 | 0 | 0 | 0 | 0 | 0 |
| virus_Jul27.5v_12825 | 0 | 0 | 0 | 0 | 0 | 0 | 0 | 0 | 0 | 0 | 0 |
| virus_Jul27.5v_12858 | 0 | 0 | 0 | 0 | 0 | 0 | 0 | 0 | 0 | 0 | 0 |
| virus_Jul27.5v_13352 | 0 | 0 | 0 | 0 | 0 | 0 | 0 | 0 | 0 | 0 | 0 |
| virus_Jul27.5v_15851 | 0 | 0 | 0 | 0 | 0 | 0 | 0 | 0 | 0 | 0 | 0 |
| virus_Jul27.5v_16211 | 0 | 0 | 0 | 0 | 0 | 0 | 0 | 0 | 0 | 0 | 0 |
| virus_Jul27.5v_17045 | 0 | 0 | 0 | 0 | 0 | 0 | 0 | 0 | 0 | 0 | 0 |
| virus_Jul27.5v_17195 | 0 | 0 | 0 | 0 | 0 | 0 | 0 | 0 | 0 | 0 | 0 |
| virus_Jul27.5v_18023 | 0 | 0 | 0 | 0 | 0 | 0 | 0 | 0 | 0 | 0 | 0 |
| virus_Jul27.5v_18282 | 0 | 0 | 0 | 0 | 0 | 0 | 0 | 0 | 0 | 0 | 0 |
| virus_Jul27.5v_18332 | 0 | 0 | 0 | 0 | 0 | 0 | 0 | 0 | 0 | 0 | 0 |
| virus_Jul27.5v_18638 | 0 | 0 | 0 | 0 | 0 | 0 | 0 | 0 | 0 | 0 | 0 |
| virus_Jul27.5v_18651 | 0 | 0 | 0 | 0 | 0 | 0 | 0 | 0 | 0 | 0 | 0 |
| virus_Jul27.5v_19951 | 0 | 0 | 0 | 0 | 0 | 0 | 0 | 0 | 0 | 0 | 0 |
| virus_Jul27.5v_20309 | 0 | 0 | 0 | 1 | 0 | 0 | 0 | 0 | 0 | 0 | 1 |
| virus_Jul27.5v_20723 | 0 | 0 | 0 | 0 | 0 | 0 | 0 | 0 | 0 | 0 | 0 |
| virus_Jul27.5v_20949 | 0 | 0 | 0 | 0 | 0 | 0 | 0 | 0 | 0 | 0 | 0 |
| virus_Jul27.25v_4130 | 0 | 0 | 0 | 0 | 0 | 0 | 0 | 0 | 0 | 0 | 0 |
| virus_Jul27.25v_6746 | 0 | 0 | 0 | 0 | 0 | 0 | 0 | 0 | 0 | 0 | 0 |
| virus_Jul27.25v_9489 | 0 | 0 | 0 | 0 | 0 | 0 | 0 | 0 | 0 | 0 | 0 |
| virus_Jul27.25v_12888 | 0 | 0 | 0 | 0 | 0 | 0 | 0 | 0 | 0 | 0 | 0 |
| virus_Jul27.25v_13260 | 0 | 0 | 0 | 0 | 0 | 0 | 0 | 0 | 0 | 0 | 0 |
| virus_Jul27.25v_18132 | 0 | 0 | 0 | 0 | 0 | 0 | 0 | 0 | 0 | 0 | 0 |
| virus_Jul27.25v_19173 | 0 | 0 | 0 | 0 | 0 | 0 | 0 | 0 | 0 | 0 | 0 |
| virus_Jul27.25v_21996 | 0 | 0 | 0 | 0 | 0 | 0 | 0 | 0 | 0 | 0 | 0 |
| virus_Jul27.25v_22871 | 0 | 0 | 0 | 0 | 0 | 0 | 0 | 0 | 0 | 0 | 0 |
| virus_Jul27.25v_29083 | 0 | 0 | 0 | 0 | 0 | 0 | 0 | 0 | 0 | 0 | 0 |
| virus_Jul27.25v_29742 | 0 | 0 | 0 | 0 | 0 | 0 | 0 | 0 | 0 | 0 | 0 |
| virus_Jul27.25v_32099 | 0 | 0 | 0 | 0 | 0 | 0 | 0 | 0 | 0 | 0 | 0 |
| virus_Jul27.75v_15687 | 0 | 0 | 0 | 0 | 0 | 0 | 0 | 0 | 0 | 0 | 0 |
| virus_Jun8.25v_8775 | 0 | 0 | 0 | 0 | 0 | 0 | 0 | 0 | 0 | 0 | 0 |
| virus_Jun8.25v_13954 | 0 | 0 | 0 | 0 | 0 | 0 | 0 | 0 | 0 | 0 | 0 |
| virus_Jun8.25v_21043 | 0 | 0 | 0 | 0 | 0 | 0 | 0 | 0 | 0 | 0 | 0 |
| virus_Jun8.25v_24974 | 0 | 0 | 0 | 0 | 0 | 0 | 0 | 0 | 0 | 0 | 0 |
| virus_Jun8.25v_38200 | 0 | 0 | 0 | 0 | 0 | 0 | 0 | 0 | 0 | 0 | 0 |
| virus_Jun8.25v_39198 | 0 | 0 | 0 | 0 | 0 | 0 | 0 | 0 | 0 | 0 | 0 |
| virus_Jun8.50v_2049 | 0 | 0 | 0 | 0 | 0 | 0 | 0 | 0 | 0 | 0 | 0 |
| virus_Jun8.50v_4763 | 0 | 0 | 0 | 0 | 0 | 0 | 0 | 0 | 0 | 0 | 0 |
| virus_Jun8.50v_5089 | 0 | 0 | 0 | 0 | 0 | 0 | 0 | 0 | 0 | 0 | 0 |
| virus_Jun8.50v_6995 | 0 | 0 | 0 | 0 | 0 | 0 | 0 | 0 | 0 | 0 | 0 |
| virus_Jun8.50v_8716 | 0 | 0 | 0 | 0 | 0 | 0 | 0 | 0 | 0 | 0 | 0 |
| virus_Jun8.50v_13937 | 0 | 0 | 0 | 0 | 0 | 0 | 0 | 0 | 0 | 0 | 0 |
| virus_Jun8.50v_16334 | 0 | 0 | 0 | 0 | 0 | 0 | 0 | 0 | 0 | 0 | 0 |
| virus_Jun8.50v_17015 | 0 | 0 | 0 | 0 | 0 | 0 | 0 | 0 | 0 | 0 | 0 |
| virus_Jun8.50v_17419 | 0 | 0 | 0 | 0 | 0 | 0 | 0 | 0 | 0 | 0 | 0 |
| virus_Jun8.50v_17880 | 0 | 0 | 0 | 0 | 0 | 0 | 0 | 0 | 0 | 0 | 0 |
| virus_Jun8.50v_19106 | 0 | 0 | 0 | 0 | 0 | 0 | 0 | 0 | 0 | 0 | 0 |
| virus_Jun8.50v_19367 | 0 | 0 | 0 | 0 | 0 | 0 | 0 | 0 | 0 | 0 | 0 |
| virus_Jun8.50v_20991 | 0 | 0 | 0 | 0 | 0 | 0 | 0 | 0 | 0 | 0 | 0 |
| virus_Jun8.50v_26042 | 0 | 0 | 0 | 0 | 0 | 0 | 0 | 0 | 0 | 0 | 0 |
| virus_Jun8.50v_28327 | 0 | 0 | 0 | 0 | 0 | 0 | 0 | 0 | 0 | 0 | 0 |
| virus_Jun8.50v_28675 | 0 | 0 | 0 | 0 | 0 | 0 | 0 | 0 | 0 | 0 | 0 |
| virus_Jun8.50v_28682 | 0 | 0 | 0 | 0 | 0 | 0 | 0 | 0 | 0 | 0 | 0 |
| virus_Jun8.50v_30966 | 0 | 0 | 0 | 0 | 0 | 0 | 0 | 0 | 0 | 0 | 0 |
| virus_Jun8.50v_36205 | 0 | 0 | 0 | 0 | 0 | 0 | 0 | 0 | 0 | 0 | 0 |
| virus_Jun8.50v_40474 | 0 | 0 | 0 | 0 | 0 | 0 | 0 | 0 | 0 | 0 | 0 |
| virus_Jun8.75v_451 | 0 | 0 | 0 | 0 | 0 | 0 | 0 | 0 | 0 | 0 | 0 |
| virus_Jun8.75v_2736 | 0 | 0 | 0 | 0 | 0 | 0 | 0 | 0 | 0 | 0 | 0 |
| virus_Jun8.75v_6517 | 0 | 0 | 0 | 0 | 0 | 0 | 0 | 0 | 0 | 0 | 0 |
| virus_Jun8.75v_15662 | 0 | 0 | 0 | 0 | 0 | 0 | 0 | 0 | 0 | 0 | 0 |
| virus_Jun22.0v_1189 | 0 | 0 | 0 | 0 | 0 | 0 | 0 | 0 | 0 | 0 | 0 |
| virus_Jun22.0v_7061 | 0 | 0 | 0 | 0 | 0 | 0 | 0 | 0 | 0 | 0 | 0 |
| virus_Jun22.0v_7637 | 0 | 0 | 0 | 0 | 0 | 0 | 0 | 0 | 0 | 0 | 0 |
| virus_Jun22.0v_8025 | 0 | 0 | 0 | 0 | 0 | 0 | 0 | 0 | 0 | 0 | 0 |
| virus_Jun22.0v_9080 | 0 | 0 | 0 | 0 | 0 | 0 | 0 | 0 | 0 | 0 | 0 |
| virus_Jun22.0v_11812 | 0 | 0 | 0 | 0 | 0 | 0 | 0 | 0 | 0 | 0 | 0 |
| virus_Jun22.0v_12447 | 0 | 0 | 0 | 0 | 0 | 0 | 0 | 0 | 0 | 0 | 0 |
| virus_Jun22.0v_12683 | 0 | 0 | 0 | 0 | 0 | 0 | 0 | 0 | 0 | 0 | 0 |
| virus_Jun22.0v_13511 | 0 | 0 | 0 | 0 | 0 | 0 | 0 | 0 | 0 | 0 | 0 |
| virus_Jun22.0v_13900 | 0 | 0 | 0 | 0 | 0 | 0 | 0 | 0 | 0 | 0 | 0 |
| virus_Jun22.0v_15566 | 0 | 0 | 0 | 0 | 0 | 0 | 0 | 0 | 0 | 0 | 0 |
| virus_Jun22.0v_19488 | 0 | 0 | 0 | 0 | 0 | 0 | 0 | 0 | 0 | 0 | 0 |
| virus_Jun22.0v_19850 | 0 | 0 | 0 | 0 | 0 | 0 | 0 | 0 | 0 | 0 | 0 |
| virus_Jun22.50v_1367 | 0 | 0 | 0 | 0 | 0 | 0 | 0 | 0 | 0 | 0 | 0 |
| virus_Jun22.50v_1537 | 0 | 0 | 0 | 0 | 0 | 0 | 0 | 0 | 0 | 0 | 0 |
| virus_Jun22.50v_1966 | 0 | 0 | 0 | 0 | 0 | 0 | 0 | 0 | 0 | 0 | 0 |
| virus_Jun22.50v_2620 | 0 | 0 | 0 | 0 | 0 | 0 | 0 | 0 | 0 | 0 | 0 |
| virus_Jun22.50v_3703 | 0 | 0 | 0 | 0 | 0 | 0 | 0 | 0 | 0 | 0 | 0 |
| virus_Jun22.50v_4782 | 0 | 0 | 0 | 0 | 0 | 0 | 0 | 0 | 0 | 0 | 0 |
| virus_Jun22.50v_8144 | 0 | 0 | 0 | 0 | 0 | 0 | 0 | 0 | 0 | 0 | 0 |
| virus_Jun22.50v_9466 | 0 | 0 | 0 | 0 | 0 | 0 | 0 | 0 | 0 | 0 | 0 |
| virus_Jun22.50v_9692 | 0 | 0 | 0 | 0 | 0 | 0 | 0 | 0 | 0 | 0 | 0 |
| virus_Jun22.50v_13919 | 0 | 0 | 0 | 0 | 0 | 0 | 0 | 0 | 0 | 0 | 0 |
| virus_Jun22.50v_15738 | 0 | 0 | 0 | 0 | 0 | 0 | 0 | 0 | 0 | 0 | 0 |
| virus_Jun22.50v_16087 | 0 | 0 | 0 | 0 | 0 | 0 | 0 | 0 | 0 | 0 | 0 |
| virus_Jun22.50v_19633 | 0 | 0 | 0 | 0 | 0 | 0 | 0 | 0 | 0 | 0 | 0 |
| virus_Jun22.50v_20126 | 0 | 0 | 0 | 0 | 0 | 0 | 0 | 0 | 0 | 0 | 0 |
| virus_Jun22.50v_25683 | 0 | 0 | 0 | 0 | 0 | 0 | 0 | 0 | 0 | 0 | 0 |
| virus_Jun22.50v_27917 | 0 | 0 | 0 | 0 | 0 | 0 | 0 | 0 | 0 | 0 | 0 |
| virus_Jun22.50v_28817 | 0 | 0 | 0 | 0 | 0 | 0 | 0 | 0 | 0 | 0 | 0 |
| virus_Jun22.50v_32060 | 0 | 0 | 0 | 0 | 0 | 1 | 0 | 0 | 0 | 0 | 1 |
| virus_Jun22.50v_34629 | 0 | 0 | 0 | 0 | 0 | 0 | 0 | 0 | 0 | 0 | 0 |
| virus_Jun22.50v_35523 | 0 | 0 | 0 | 0 | 0 | 0 | 0 | 0 | 0 | 0 | 0 |
| virus_Jun22.50v_36714 | 0 | 0 | 0 | 0 | 0 | 0 | 0 | 0 | 0 | 0 | 0 |
| virus_Jun22.50v_38051 | 0 | 0 | 0 | 0 | 0 | 0 | 0 | 0 | 0 | 0 | 0 |
| virus_Jun22.75v_223 | 0 | 0 | 0 | 0 | 0 | 0 | 0 | 0 | 0 | 0 | 0 |
| virus_Jun22.75v_12929 | 0 | 0 | 0 | 0 | 0 | 0 | 0 | 0 | 0 | 0 | 0 |
| virus_Jun22.75v_20691 | 0 | 0 | 0 | 0 | 0 | 0 | 0 | 0 | 0 | 0 | 0 |
| virus_Jun22.75v_31125 | 0 | 0 | 0 | 0 | 0 | 0 | 0 | 0 | 0 | 0 | 0 |
| virus_Jun22.75v_31544 | 0 | 0 | 0 | 0 | 0 | 0 | 0 | 0 | 0 | 0 | 0 |
| virus_Jun22.75v_33788 | 0 | 0 | 0 | 0 | 0 | 0 | 0 | 0 | 0 | 0 | 0 |
| virus_Jun22.75v_35460 | 0 | 0 | 0 | 0 | 0 | 0 | 0 | 0 | 0 | 0 | 0 |
| virus_Jun22.75v_36894 | 0 | 0 | 0 | 0 | 0 | 0 | 0 | 0 | 0 | 0 | 0 |
| virus_Jun22.75v_39617 | 0 | 0 | 0 | 0 | 0 | 0 | 0 | 0 | 0 | 0 | 0 |
| virus_May12.5v_7184 | 0 | 0 | 0 | 0 | 0 | 0 | 0 | 0 | 0 | 0 | 0 |
| virus_May12.5v_8804 | 0 | 0 | 0 | 0 | 0 | 0 | 0 | 0 | 0 | 0 | 0 |
| virus_May12.5v_21886 | 0 | 0 | 0 | 0 | 0 | 0 | 0 | 0 | 0 | 0 | 0 |
| virus_May12.5v_34493 | 0 | 0 | 0 | 0 | 0 | 0 | 0 | 0 | 0 | 0 | 0 |
| virus_May12.5v_39318 | 0 | 0 | 0 | 0 | 0 | 0 | 0 | 0 | 0 | 0 | 0 |
| virus_May12.5v_40344 | 0 | 0 | 0 | 0 | 0 | 0 | 0 | 0 | 0 | 0 | 0 |
| virus_May12.25v_6074 | 0 | 0 | 0 | 0 | 0 | 0 | 0 | 0 | 0 | 0 | 0 |
| virus_May12.25v_13980 | 0 | 0 | 0 | 0 | 0 | 0 | 0 | 0 | 0 | 0 | 0 |
| virus_May12.25v_18305 | 0 | 0 | 0 | 0 | 0 | 0 | 0 | 0 | 0 | 0 | 0 |
| virus_May12.25v_20845 | 0 | 0 | 0 | 0 | 0 | 0 | 0 | 0 | 0 | 0 | 0 |
| virus_May12.25v_22797 | 0 | 0 | 0 | 0 | 0 | 0 | 0 | 0 | 0 | 0 | 0 |
| virus_May12.25v_33105 | 0 | 0 | 0 | 0 | 0 | 0 | 0 | 0 | 0 | 0 | 0 |
| virus_May12.25v_33383 | 0 | 0 | 0 | 0 | 0 | 0 | 0 | 0 | 0 | 0 | 0 |
| virus_May12.50v_11507 | 0 | 0 | 0 | 0 | 0 | 0 | 0 | 0 | 0 | 0 | 0 |
| virus_May12.50v_13686 | 0 | 0 | 0 | 0 | 0 | 0 | 0 | 0 | 0 | 0 | 0 |
| virus_May12.50v_14520 | 0 | 0 | 0 | 0 | 0 | 0 | 0 | 0 | 0 | 0 | 0 |
| virus_May12.50v_16193 | 0 | 0 | 0 | 0 | 0 | 0 | 0 | 0 | 0 | 0 | 0 |
| virus_May12.50v_18124 | 0 | 0 | 0 | 0 | 0 | 0 | 0 | 0 | 0 | 0 | 0 |

| | | | | | | | | | | | |
|---|---|---|---|---|---|---|---|---|---|---|---|
| virus_May12.50v_20309 | 0 | 0 | 0 | 0 | 0 | 0 | 0 | 0 | 0 | 0 | 0 |
| virus_May12.50v_21848 | 0 | 0 | 0 | 0 | 0 | 0 | 0 | 0 | 0 | 0 | 0 |
| virus_May12.50v_22108 | 0 | 0 | 0 | 0 | 0 | 0 | 0 | 0 | 0 | 0 | 0 |
| virus_May12.50v_22484 | 0 | 0 | 0 | 0 | 0 | 0 | 0 | 0 | 0 | 0 | 0 |
| virus_May12.50v_23050 | 0 | 0 | 0 | 0 | 0 | 0 | 0 | 0 | 0 | 0 | 0 |
| virus_May12.50v_24689 | 0 | 0 | 0 | 0 | 0 | 0 | 0 | 0 | 0 | 0 | 0 |
| virus_May12.50v_24998 | 0 | 0 | 0 | 0 | 0 | 0 | 0 | 0 | 0 | 0 | 0 |
| virus_May12.50v_28989 | 0 | 0 | 0 | 0 | 0 | 0 | 0 | 0 | 0 | 0 | 0 |
| virus_May12.50v_37033 | 0 | 0 | 0 | 0 | 0 | 0 | 0 | 0 | 0 | 0 | 0 |
| virus_May12.50v_38219 | 0 | 0 | 0 | 0 | 0 | 0 | 0 | 0 | 0 | 0 | 0 |
| virus_May12.50v_40959 | 0 | 0 | 0 | 0 | 0 | 0 | 0 | 0 | 0 | 0 | 0 |
| virus_May12.75v_5824 | 0 | 0 | 0 | 0 | 0 | 0 | 0 | 0 | 0 | 0 | 0 |
| virus_May12.75v_7621 | 0 | 0 | 0 | 0 | 0 | 0 | 0 | 0 | 0 | 0 | 0 |
| virus_May12.75v_8932 | 0 | 0 | 0 | 0 | 0 | 0 | 0 | 0 | 0 | 0 | 0 |
| virus_May12.75v_17207 | 0 | 0 | 0 | 0 | 0 | 0 | 0 | 0 | 0 | 0 | 0 |
| virus_May12.75v_19288 | 0 | 0 | 0 | 0 | 0 | 0 | 0 | 0 | 0 | 0 | 0 |
| virus_May12.75v_20779 | 0 | 0 | 0 | 0 | 0 | 0 | 0 | 0 | 0 | 0 | 0 |
| virus_May12.75v_26924 | 0 | 0 | 0 | 0 | 0 | 0 | 0 | 0 | 0 | 0 | 0 |
| virus_May12.75v_29765 | 0 | 0 | 0 | 0 | 0 | 0 | 0 | 0 | 0 | 0 | 0 |
| virus_Sep8.0v_344 | 0 | 0 | 0 | 0 | 0 | 0 | 0 | 0 | 0 | 0 | 0 |
| virus_Sep8.0v_4929 | 0 | 0 | 0 | 0 | 0 | 0 | 0 | 0 | 0 | 0 | 0 |
| virus_Sep8.0v_6668 | 0 | 0 | 0 | 0 | 0 | 0 | 0 | 0 | 0 | 0 | 0 |
| virus_Sep8.0v_8381 | 0 | 0 | 0 | 0 | 0 | 0 | 0 | 0 | 0 | 0 | 0 |
| virus_Sep8.0v_8949 | 0 | 0 | 0 | 1 | 0 | 0 | 0 | 0 | 0 | 1 | 2 |
| virus_Sep8.0v_14731 | 0 | 0 | 0 | 0 | 0 | 0 | 0 | 0 | 0 | 0 | 0 |
| virus_Sep8.0v_14734 | 0 | 0 | 0 | 0 | 0 | 0 | 0 | 0 | 0 | 0 | 0 |
| virus_Sep8.0v_15059 | 0 | 0 | 0 | 0 | 0 | 0 | 0 | 0 | 0 | 0 | 0 |
| virus_Sep8.0v_15617 | 0 | 0 | 0 | 0 | 0 | 0 | 0 | 0 | 0 | 0 | 0 |
| virus_Sep8.0v_16579 | 0 | 0 | 0 | 0 | 0 | 0 | 0 | 0 | 0 | 0 | 0 |
| virus_Sep8.0v_17533 | 0 | 0 | 0 | 0 | 0 | 0 | 0 | 0 | 0 | 0 | 0 |
| virus_Sep8.5v_784 | 0 | 0 | 0 | 0 | 0 | 0 | 0 | 0 | 0 | 0 | 0 |
| virus_Sep8.5v_4513 | 0 | 0 | 0 | 0 | 0 | 0 | 0 | 0 | 0 | 0 | 0 |
| virus_Sep8.5v_15177 | 0 | 0 | 0 | 0 | 0 | 0 | 0 | 0 | 0 | 0 | 0 |
| virus_Sep8.5v_19322 | 0 | 0 | 0 | 0 | 0 | 0 | 0 | 0 | 0 | 0 | 0 |
| virus_Sep8.5v_19493 | 0 | 0 | 0 | 0 | 0 | 0 | 0 | 0 | 0 | 0 | 0 |
| virus_Sep8.25v_2915 | 0 | 0 | 0 | 0 | 0 | 0 | 0 | 0 | 0 | 0 | 0 |
| virus_Sep8.25v_3482 | 0 | 0 | 0 | 0 | 0 | 0 | 0 | 0 | 0 | 0 | 0 |
| virus_Sep8.25v_8066 | 0 | 0 | 0 | 0 | 0 | 0 | 0 | 0 | 0 | 0 | 0 |
| virus_Sep8.25v_10270 | 0 | 0 | 0 | 0 | 0 | 0 | 0 | 0 | 0 | 0 | 0 |
| virus_Sep8.25v_13433 | 0 | 0 | 0 | 0 | 0 | 0 | 0 | 0 | 0 | 0 | 0 |
| virus_Sep8.25v_15394 | 0 | 0 | 0 | 0 | 0 | 0 | 0 | 0 | 0 | 0 | 0 |
| virus_Sep8.50v_1937 | 0 | 0 | 0 | 0 | 0 | 0 | 0 | 0 | 0 | 0 | 0 |
| virus_Sep8.50v_4736 | 0 | 0 | 0 | 0 | 0 | 0 | 0 | 0 | 0 | 0 | 0 |
| virus_Sep8.50v_8381 | 0 | 0 | 0 | 0 | 0 | 0 | 0 | 0 | 0 | 0 | 0 |
| virus_Sep8.50v_11308 | 0 | 0 | 0 | 0 | 0 | 0 | 0 | 0 | 0 | 0 | 0 |
| virus_Sep8.50v_12759 | 0 | 0 | 0 | 0 | 0 | 0 | 0 | 0 | 0 | 0 | 0 |
| virus_Sep8.50v_17838 | 0 | 0 | 0 | 0 | 0 | 0 | 0 | 0 | 0 | 0 | 0 |
| virus_Sep8.50v_18193 | 0 | 0 | 0 | 0 | 0 | 0 | 0 | 0 | 0 | 0 | 0 |
| virus_Sep8.50v_19332 | 0 | 1 | 0 | 0 | 0 | 0 | 0 | 0 | 0 | 0 | 1 |
| virus_Sep8.50v_19579 | 0 | 0 | 0 | 0 | 0 | 0 | 0 | 0 | 0 | 0 | 0 |
| virus_Sep8.50v_22015 | 0 | 0 | 0 | 0 | 0 | 0 | 0 | 0 | 0 | 0 | 0 |
| virus_Sep8.50v_22677 | 0 | 0 | 0 | 0 | 0 | 0 | 0 | 0 | 0 | 0 | 0 |
| virus_Sep8.50v_23280 | 0 | 0 | 0 | 0 | 0 | 0 | 0 | 0 | 0 | 0 | 0 |
| virus_Sep8.50v_24505 | 0 | 0 | 0 | 0 | 0 | 0 | 0 | 0 | 0 | 0 | 0 |
| virus_Sep8.50v_25933 | 0 | 0 | 0 | 0 | 0 | 0 | 0 | 0 | 0 | 0 | 0 |
| virus_Sep8.50v_27680 | 0 | 0 | 0 | 0 | 0 | 0 | 0 | 0 | 0 | 0 | 0 |
| virus_Sep8.50v_28215 | 0 | 0 | 0 | 0 | 0 | 0 | 0 | 0 | 0 | 0 | 0 |
| virus_Sep8.50v_36519 | 0 | 0 | 0 | 0 | 0 | 0 | 0 | 0 | 0 | 0 | 0 |
| virus_Sep8.50v_37232 | 0 | 0 | 0 | 0 | 0 | 0 | 0 | 0 | 0 | 0 | 0 |
| virus_Sep8.50v_39051 | 0 | 0 | 0 | 0 | 0 | 0 | 0 | 0 | 0 | 0 | 0 |
| virus_Sep8.50v_40957 | 0 | 0 | 0 | 1 | 0 | 0 | 0 | 0 | 0 | 0 | 1 |
| bacteria_Jul11d0b_8428 | 0 | 0 | 0 | 0 | 0 | 0 | 0 | 0 | 0 | 0 | 0 |
| bacteria_Jul11d0b_10918 | 0 | 0 | 0 | 0 | 0 | 0 | 0 | 0 | 0 | 0 | 0 |
| bacteria_Jul11d0b_12494 | 0 | 0 | 0 | 0 | 0 | 0 | 0 | 0 | 0 | 0 | 0 |
| bacteria_Jul11d0b_22352 | 0 | 0 | 0 | 0 | 0 | 0 | 0 | 0 | 0 | 0 | 0 |
| bacteria_Jul11d0b_24258 | 0 | 0 | 0 | 0 | 0 | 0 | 0 | 0 | 0 | 0 | 0 |
| bacteria_Jul11d0b_24428 | 0 | 0 | 0 | 0 | 0 | 0 | 0 | 0 | 0 | 0 | 0 |
| bacteria_Jul11d0b_38423 | 0 | 0 | 0 | 0 | 0 | 0 | 0 | 0 | 0 | 0 | 0 |
| bacteria_Jul11d0b_41022 | 0 | 0 | 0 | 0 | 0 | 0 | 0 | 0 | 0 | 0 | 0 |
| bacteria_Jul11d0b_41903 | 0 | 0 | 0 | 0 | 0 | 0 | 0 | 0 | 0 | 0 | 0 |
| bacteria_Jul11d0b_54474 | 0 | 0 | 0 | 0 | 0 | 0 | 0 | 0 | 0 | 0 | 0 |
| bacteria_Jul11d0b_58257 | 0 | 0 | 0 | 0 | 0 | 0 | 0 | 0 | 0 | 0 | 0 |
| bacteria_Jul11d0b_59524 | 0 | 0 | 0 | 0 | 0 | 0 | 0 | 0 | 0 | 0 | 0 |
| bacteria_Jul11d0b_61002 | 0 | 0 | 0 | 0 | 0 | 0 | 0 | 0 | 0 | 0 | 0 |
| bacteria_Jul11d0b_72150 | 0 | 0 | 0 | 0 | 0 | 0 | 0 | 0 | 0 | 0 | 0 |
| bacteria_Aug5d0b_35742 | 1 | 0 | 0 | 0 | 0 | 0 | 0 | 0 | 0 | 0 | 1 |
| bacteria_Aug5d0b_48375 | 1 | 0 | 0 | 0 | 0 | 0 | 0 | 0 | 0 | 0 | 1 |
| bacteria_Aug5d5b_5022 | 0 | 0 | 1 | 0 | 0 | 0 | 0 | 0 | 0 | 0 | 1 |
| bacteria_Aug5d5b_7350 | 0 | 0 | 1 | 0 | 0 | 0 | 0 | 0 | 0 | 0 | 1 |
| bacteria_Aug5d5b_36130 | 0 | 0 | 1 | 0 | 0 | 0 | 0 | 0 | 0 | 0 | 1 |
| bacteria_Aug5d25b_1978 | 0 | 0 | 0 | 0 | 1 | 0 | 0 | 0 | 0 | 0 | 1 |
| bacteria_Aug5d25b_3474 | 0 | 0 | 0 | 0 | 1 | 0 | 0 | 0 | 0 | 0 | 1 |
| bacteria_Aug5d25b_6289 | 0 | 0 | 0 | 0 | 1 | 0 | 0 | 0 | 0 | 0 | 1 |
| bacteria_Aug5d25b_20212 | 0 | 0 | 0 | 0 | 1 | 0 | 0 | 0 | 0 | 0 | 1 |
| bacteria_Aug5d25b_22280 | 0 | 0 | 0 | 0 | 1 | 0 | 0 | 0 | 0 | 0 | 1 |
| bacteria_Aug5d25b_25496 | 0 | 0 | 0 | 0 | 1 | 0 | 0 | 0 | 0 | 0 | 1 |
| bacteria_Aug5d25b_31735 | 0 | 0 | 0 | 0 | 1 | 0 | 0 | 0 | 0 | 0 | 1 |
| bacteria_Aug5d25b_33783 | 0 | 0 | 0 | 0 | 1 | 0 | 0 | 0 | 0 | 0 | 1 |
| bacteria_Aug5d25b_40949 | 0 | 0 | 0 | 0 | 1 | 0 | 0 | 0 | 0 | 0 | 1 |
| bacteria_Aug5d25b_42824 | 0 | 0 | 0 | 0 | 1 | 0 | 0 | 0 | 0 | 0 | 1 |
| bacteria_Aug5d25b_46191 | 0 | 0 | 0 | 0 | 1 | 0 | 0 | 0 | 0 | 0 | 1 |
| bacteria_Aug5d25b_47332 | 0 | 0 | 0 | 0 | 1 | 0 | 0 | 0 | 0 | 0 | 1 |
| bacteria_Aug5d25b_47879 | 0 | 0 | 0 | 0 | 1 | 0 | 0 | 0 | 0 | 0 | 1 |
| bacteria_Aug5d25b_48501 | 0 | 0 | 0 | 0 | 1 | 0 | 0 | 0 | 0 | 0 | 1 |
| bacteria_Aug5d25b_49248 | 0 | 0 | 0 | 0 | 1 | 0 | 0 | 0 | 0 | 0 | 1 |
| bacteria_Aug5d25b_56458 | 0 | 0 | 0 | 0 | 1 | 0 | 0 | 0 | 0 | 0 | 1 |
| bacteria_Aug5d25b_58151 | 0 | 0 | 0 | 0 | 1 | 0 | 0 | 0 | 0 | 0 | 1 |
| bacteria_Aug5d25b_59654 | 0 | 0 | 0 | 0 | 1 | 0 | 0 | 0 | 0 | 0 | 1 |
| bacteria_Aug5d25b_60503 | 0 | 0 | 0 | 0 | 1 | 0 | 0 | 0 | 0 | 0 | 1 |
| bacteria_Aug5d25b_62399 | 0 | 0 | 0 | 0 | 1 | 0 | 0 | 0 | 0 | 0 | 1 |
| bacteria_Aug5d25b_63085 | 0 | 0 | 0 | 0 | 1 | 0 | 0 | 0 | 0 | 0 | 1 |
| bacteria_Aug5d25b_66662 | 0 | 0 | 0 | 0 | 1 | 0 | 0 | 0 | 0 | 0 | 1 |
| bacteria_Aug5d25b_67028 | 0 | 0 | 0 | 0 | 1 | 0 | 0 | 0 | 0 | 0 | 1 |
| bacteria_Aug5d25b_72871 | 0 | 0 | 0 | 0 | 1 | 0 | 0 | 0 | 0 | 0 | 1 |
| bacteria_Aug5d50b_4420 | 0 | 0 | 0 | 0 | 0 | 0 | 1 | 0 | 0 | 0 | 1 |
| bacteria_Aug5d50b_6225 | 0 | 0 | 0 | 0 | 0 | 0 | 1 | 0 | 0 | 0 | 1 |
| bacteria_Aug5d50b_16661 | 0 | 0 | 0 | 0 | 0 | 0 | 1 | 0 | 0 | 0 | 1 |
| bacteria_Aug5d50b_33163 | 0 | 0 | 0 | 0 | 0 | 0 | 1 | 0 | 0 | 0 | 1 |
| bacteria_Aug5d50b_46306 | 0 | 0 | 0 | 0 | 0 | 0 | 1 | 0 | 0 | 0 | 1 |
| bacteria_Aug5d50b_65425 | 0 | 0 | 0 | 0 | 0 | 0 | 1 | 0 | 0 | 0 | 1 |
| bacteria_Aug5d50b_70461 | 0 | 0 | 0 | 0 | 0 | 0 | 1 | 0 | 0 | 0 | 1 |
| bacteria_Aug5d75b_33872 | 0 | 0 | 0 | 0 | 0 | 0 | 0 | 0 | 1 | 0 | 1 |
| bacteria_Aug5d75b_43197 | 0 | 0 | 0 | 0 | 0 | 0 | 0 | 0 | 1 | 0 | 1 |
| bacteria_Jul11d5b_5193 | 0 | 0 | 0 | 0 | 0 | 0 | 0 | 0 | 0 | 0 | 0 |
| bacteria_Jul11d5b_11014 | 0 | 0 | 0 | 0 | 0 | 0 | 0 | 0 | 0 | 0 | 0 |
| bacteria_Jul11d5b_11355 | 0 | 0 | 0 | 0 | 0 | 0 | 0 | 0 | 0 | 0 | 0 |
| bacteria_Jul11d5b_12692 | 0 | 0 | 0 | 0 | 0 | 0 | 0 | 0 | 0 | 0 | 0 |
| bacteria_Jul11d5b_22577 | 0 | 0 | 0 | 0 | 0 | 0 | 0 | 0 | 0 | 0 | 0 |
| bacteria_Jul11d5b_24110 | 0 | 0 | 0 | 0 | 0 | 0 | 0 | 0 | 0 | 0 | 0 |
| bacteria_Jul11d5b_30495 | 0 | 0 | 0 | 0 | 0 | 0 | 0 | 0 | 0 | 0 | 0 |
| bacteria_Jul11d5b_30935 | 0 | 0 | 0 | 0 | 0 | 0 | 0 | 0 | 0 | 0 | 0 |
| bacteria_Jul11d5b_43132 | 0 | 0 | 0 | 0 | 0 | 0 | 0 | 0 | 0 | 0 | 0 |
| bacteria_Jul11d5b_44630 | 0 | 0 | 0 | 0 | 0 | 0 | 0 | 0 | 0 | 0 | 0 |
| bacteria_Jul11d5b_46404 | 0 | 0 | 0 | 0 | 0 | 0 | 0 | 0 | 0 | 0 | 0 |
| bacteria_Jul11d5b_48870 | 0 | 0 | 0 | 0 | 0 | 0 | 0 | 0 | 0 | 0 | 0 |
| bacteria_Jul11d5b_50478 | 0 | 0 | 0 | 0 | 0 | 0 | 0 | 0 | 0 | 0 | 0 |
| bacteria_Jul11d25b_2700 | 0 | 0 | 0 | 0 | 0 | 0 | 0 | 0 | 0 | 0 | 0 |
| bacteria_Jul11d25b_15213 | 0 | 0 | 0 | 0 | 0 | 0 | 0 | 0 | 0 | 0 | 0 |
| bacteria_Jul11d25b_18916 | 0 | 0 | 0 | 0 | 0 | 0 | 0 | 0 | 0 | 0 | 0 |
| bacteria_Jul11d25b_22182 | 0 | 0 | 0 | 0 | 0 | 0 | 0 | 0 | 0 | 0 | 0 |
| bacteria_Jul11d25b_30071 | 0 | 0 | 0 | 0 | 0 | 0 | 0 | 0 | 0 | 0 | 0 |
| bacteria_Jul11d25b_34247 | 0 | 0 | 0 | 0 | 0 | 0 | 0 | 0 | 0 | 0 | 0 |
| bacteria_Jul11d25b_34379 | 0 | 0 | 0 | 0 | 0 | 0 | 0 | 0 | 0 | 0 | 0 |
| bacteria_Jul11d25b_38703 | 0 | 0 | 0 | 0 | 1 | 0 | 0 | 0 | 0 | 0 | 1 |
| bacteria_Jul11d50b_16078 | 0 | 0 | 0 | 0 | 0 | 0 | 0 | 0 | 0 | 0 | 0 |
| bacteria_Jul11d50b_23104 | 0 | 0 | 0 | 0 | 0 | 0 | 0 | 0 | 0 | 0 | 0 |
| bacteria_Jul11d50b_40497 | 0 | 0 | 0 | 0 | 0 | 0 | 0 | 0 | 0 | 0 | 0 |
| bacteria_Jul11d50b_47920 | 0 | 0 | 0 | 0 | 0 | 0 | 0 | 0 | 0 | 0 | 0 |
| bacteria_Jul11d50b_47967 | 0 | 0 | 0 | 0 | 0 | 0 | 0 | 0 | 0 | 0 | 0 |
| bacteria_Jul11d50b_48240 | 0 | 0 | 0 | 0 | 0 | 0 | 0 | 0 | 0 | 0 | 0 |
| bacteria_Jul11d75b_1312 | 0 | 0 | 0 | 0 | 0 | 0 | 0 | 0 | 0 | 0 | 0 |
| bacteria_Jul11d75b_10964 | 0 | 0 | 0 | 0 | 0 | 0 | 0 | 0 | 0 | 0 | 0 |
| bacteria_Jul11d75b_11531 | 0 | 0 | 0 | 0 | 0 | 0 | 0 | 0 | 0 | 0 | 0 |
| bacteria_Jul11d75b_16422 | 0 | 0 | 0 | 0 | 0 | 0 | 0 | 0 | 0 | 0 | 0 |
| bacteria_Jul11d75b_16739 | 0 | 0 | 0 | 0 | 0 | 0 | 0 | 0 | 0 | 0 | 0 |
| bacteria_Jul11d75b_17204 | 0 | 0 | 0 | 0 | 0 | 0 | 0 | 0 | 0 | 0 | 0 |
| bacteria_Jul11d75b_20620 | 0 | 0 | 0 | 0 | 0 | 0 | 0 | 0 | 0 | 0 | 0 |
| bacteria_Jul11d75b_20831 | 0 | 0 | 0 | 0 | 0 | 0 | 0 | 0 | 0 | 0 | 0 |
| bacteria_Jul11d75b_24101 | 0 | 0 | 0 | 0 | 0 | 0 | 0 | 0 | 0 | 0 | 0 |
| bacteria_Jul11d75b_36557 | 0 | 0 | 0 | 0 | 0 | 0 | 0 | 0 | 0 | 0 | 0 |
| bacteria_Jul11d75b_38056 | 0 | 0 | 0 | 0 | 0 | 0 | 0 | 0 | 0 | 0 | 0 |
| bacteria_Jul11d75b_38237 | 0 | 0 | 0 | 0 | 0 | 0 | 0 | 0 | 0 | 0 | 0 |
| bacteria_Jul11d75b_43467 | 0 | 0 | 0 | 0 | 0 | 0 | 0 | 0 | 0 | 0 | 0 |
| bacteria_Jul11d75b_47780 | 0 | 0 | 0 | 0 | 0 | 0 | 0 | 0 | 0 | 0 | 0 |
| bacteria_Jul11d75b_48402 | 0 | 0 | 0 | 0 | 0 | 0 | 0 | 0 | 0 | 0 | 0 |
| bacteria_Jul11d75b_51001 | 0 | 0 | 0 | 0 | 0 | 0 | 0 | 0 | 0 | 0 | 0 |
| bacteria_Jul27d5b_686 | 0 | 0 | 0 | 0 | 0 | 0 | 0 | 0 | 0 | 0 | 0 |

| | | | | | | | | | | | |
|---|---|---|---|---|---|---|---|---|---|---|---|
| bacteria_Jul27d5b_1259 | 0 | 0 | 0 | 0 | 0 | 0 | 0 | 0 | 0 | 0 | 0 |
| bacteria_Jul27d5b_12734 | 0 | 0 | 0 | 0 | 0 | 0 | 0 | 0 | 0 | 0 | 0 |
| bacteria_Jul27d5b_14326 | 0 | 0 | 0 | 0 | 0 | 0 | 0 | 0 | 0 | 0 | 0 |
| bacteria_Jul27d5b_16153 | 0 | 0 | 0 | 0 | 0 | 0 | 0 | 0 | 0 | 0 | 0 |
| bacteria_Jul27d5b_19267 | 0 | 0 | 0 | 0 | 0 | 0 | 0 | 0 | 0 | 0 | 0 |
| bacteria_Jul27d5b_19286 | 0 | 0 | 0 | 0 | 0 | 0 | 0 | 0 | 0 | 0 | 0 |
| bacteria_Jul27d5b_21977 | 0 | 0 | 0 | 0 | 0 | 0 | 0 | 0 | 0 | 0 | 0 |
| bacteria_Jul27d5b_27108 | 0 | 0 | 0 | 0 | 0 | 0 | 0 | 0 | 0 | 0 | 0 |
| bacteria_Jul27d5b_28753 | 0 | 0 | 0 | 0 | 0 | 0 | 0 | 0 | 0 | 0 | 0 |
| bacteria_Jul27d5b_29495 | 0 | 0 | 0 | 0 | 0 | 0 | 0 | 0 | 0 | 0 | 0 |
| bacteria_Jul27d5b_32759 | 0 | 0 | 0 | 0 | 0 | 0 | 0 | 0 | 0 | 0 | 0 |
| bacteria_Jul27d5b_35645 | 0 | 0 | 0 | 0 | 0 | 0 | 0 | 0 | 0 | 0 | 0 |
| bacteria_Jul27d5b_45172 | 0 | 0 | 0 | 0 | 0 | 0 | 0 | 0 | 0 | 0 | 0 |
| bacteria_Jul27d5b_50606 | 0 | 0 | 0 | 0 | 0 | 0 | 0 | 0 | 0 | 0 | 0 |
| bacteria_Jul27d5b_50718 | 0 | 0 | 0 | 0 | 0 | 0 | 0 | 0 | 0 | 0 | 0 |
| bacteria_Jul27d25b_502 | 0 | 0 | 0 | 0 | 0 | 0 | 0 | 0 | 0 | 0 | 0 |
| bacteria_Jul27d25b_5613 | 0 | 0 | 0 | 0 | 0 | 0 | 0 | 0 | 0 | 0 | 0 |
| bacteria_Jul27d25b_7793 | 0 | 0 | 0 | 0 | 0 | 0 | 0 | 0 | 0 | 0 | 0 |
| bacteria_Jul27d25b_10581 | 0 | 0 | 0 | 0 | 0 | 0 | 0 | 0 | 0 | 0 | 0 |
| bacteria_Jul27d25b_11446 | 0 | 0 | 0 | 0 | 0 | 0 | 0 | 0 | 0 | 0 | 0 |
| bacteria_Jul27d25b_18520 | 0 | 0 | 0 | 0 | 0 | 0 | 0 | 0 | 0 | 0 | 0 |
| bacteria_Jul27d25b_27633 | 0 | 0 | 0 | 0 | 0 | 0 | 0 | 0 | 0 | 1 | 1 |
| bacteria_Jul27d25b_28904 | 0 | 0 | 0 | 0 | 0 | 0 | 0 | 0 | 0 | 0 | 0 |
| bacteria_Jul27d25b_29322 | 0 | 0 | 0 | 0 | 0 | 0 | 0 | 0 | 0 | 0 | 0 |
| bacteria_Jul27d25b_30205 | 0 | 0 | 0 | 0 | 0 | 0 | 0 | 0 | 0 | 0 | 0 |
| bacteria_Jul27d25b_42959 | 0 | 0 | 0 | 0 | 0 | 0 | 0 | 0 | 0 | 0 | 0 |
| bacteria_Jul27d25b_47334 | 0 | 0 | 0 | 0 | 0 | 0 | 0 | 0 | 0 | 0 | 0 |
| bacteria_Jul27d25b_47345 | 0 | 0 | 0 | 0 | 0 | 0 | 0 | 0 | 0 | 0 | 0 |
| bacteria_Jul27d25b_51877 | 0 | 0 | 0 | 0 | 0 | 0 | 0 | 0 | 0 | 0 | 0 |
| bacteria_Jul27d25b_53264 | 0 | 0 | 0 | 0 | 0 | 0 | 0 | 0 | 0 | 0 | 0 |
| bacteria_Jul27d25b_58082 | 0 | 0 | 0 | 0 | 0 | 0 | 0 | 0 | 0 | 0 | 0 |
| bacteria_Jul27d25b_74608 | 0 | 0 | 0 | 0 | 0 | 0 | 0 | 0 | 0 | 0 | 0 |
| bacteria_Jul27d75b_8199 | 0 | 0 | 0 | 0 | 0 | 0 | 0 | 0 | 0 | 0 | 0 |
| bacteria_Jul27d75b_10648 | 0 | 0 | 0 | 0 | 0 | 0 | 0 | 1 | 0 | 0 | 1 |
| bacteria_Jul27d75b_15254 | 0 | 0 | 0 | 0 | 0 | 0 | 0 | 0 | 0 | 0 | 0 |
| bacteria_Jul27d75b_16713 | 0 | 0 | 0 | 0 | 0 | 0 | 0 | 0 | 0 | 0 | 0 |
| bacteria_Jul27d75b_25740 | 0 | 0 | 0 | 0 | 0 | 0 | 0 | 0 | 0 | 0 | 0 |
| bacteria_Jul27d75b_32002 | 0 | 0 | 0 | 0 | 0 | 0 | 0 | 0 | 0 | 0 | 0 |
| bacteria_Jul27d75b_35408 | 0 | 0 | 0 | 0 | 0 | 0 | 0 | 0 | 0 | 0 | 0 |
| bacteria_Jul27d75b_42054 | 0 | 0 | 0 | 0 | 0 | 0 | 0 | 0 | 0 | 0 | 0 |
| bacteria_Jul27d75b_48070 | 0 | 0 | 0 | 0 | 0 | 0 | 0 | 0 | 0 | 0 | 0 |
| bacteria_Jul27d75b_55336 | 0 | 0 | 0 | 0 | 0 | 0 | 0 | 0 | 0 | 0 | 0 |
| bacteria_Jul27d75b_73257 | 0 | 0 | 0 | 0 | 0 | 0 | 0 | 0 | 0 | 0 | 0 |
| bacteria_Jun8d25b_13733 | 0 | 0 | 0 | 0 | 0 | 0 | 0 | 0 | 0 | 0 | 0 |
| bacteria_Jun8d25b_19595 | 0 | 0 | 0 | 0 | 0 | 0 | 0 | 0 | 0 | 0 | 0 |
| bacteria_Jun8d25b_21178 | 0 | 0 | 2 | 0 | 0 | 0 | 0 | 0 | 0 | 0 | 2 |
| bacteria_Jun8d25b_34124 | 0 | 0 | 0 | 0 | 0 | 0 | 0 | 0 | 0 | 0 | 0 |
| bacteria_Jun8d25b_34838 | 0 | 0 | 0 | 0 | 0 | 0 | 0 | 0 | 0 | 0 | 0 |
| bacteria_Jun8d25b_36928 | 0 | 0 | 0 | 0 | 0 | 0 | 0 | 0 | 0 | 0 | 0 |
| bacteria_Jun8d25b_43589 | 0 | 0 | 0 | 0 | 0 | 0 | 0 | 0 | 0 | 0 | 0 |
| bacteria_Jun8d25b_46330 | 0 | 1 | 0 | 0 | 0 | 0 | 0 | 0 | 0 | 0 | 1 |
| bacteria_Jun8d25b_48941 | 0 | 0 | 0 | 0 | 0 | 0 | 0 | 0 | 0 | 0 | 0 |
| bacteria_Jun8d25b_58158 | 0 | 0 | 0 | 0 | 0 | 0 | 0 | 0 | 0 | 0 | 0 |
| bacteria_Jun8d25b_60913 | 0 | 0 | 0 | 0 | 0 | 0 | 0 | 0 | 0 | 0 | 0 |
| bacteria_Jun8d25b_64834 | 0 | 0 | 0 | 0 | 0 | 0 | 0 | 0 | 0 | 0 | 0 |
| bacteria_Jun8d25b_69873 | 0 | 0 | 0 | 0 | 0 | 0 | 0 | 0 | 0 | 0 | 0 |
| bacteria_Jun8d50b_775 | 0 | 0 | 0 | 0 | 0 | 0 | 0 | 0 | 0 | 0 | 0 |
| bacteria_Jun8d50b_1368 | 0 | 0 | 0 | 0 | 0 | 0 | 0 | 0 | 0 | 0 | 0 |
| bacteria_Jun8d50b_2490 | 0 | 0 | 0 | 0 | 0 | 0 | 0 | 0 | 0 | 0 | 0 |
| bacteria_Jun8d50b_4374 | 0 | 0 | 0 | 0 | 0 | 0 | 0 | 0 | 0 | 0 | 0 |
| bacteria_Jun8d50b_18591 | 0 | 0 | 0 | 0 | 0 | 0 | 0 | 0 | 0 | 0 | 0 |
| bacteria_Jun8d50b_48805 | 0 | 0 | 0 | 0 | 0 | 0 | 0 | 0 | 0 | 0 | 0 |
| bacteria_Jun8d50b_50538 | 0 | 0 | 0 | 0 | 0 | 0 | 0 | 0 | 0 | 0 | 0 |
| bacteria_Jun8d75b_57768 | 0 | 0 | 0 | 0 | 0 | 0 | 0 | 0 | 0 | 0 | 0 |
| bacteria_Jun8d75b_64161 | 0 | 0 | 0 | 0 | 0 | 0 | 0 | 0 | 0 | 0 | 0 |
| bacteria_Jun8d75b_73646 | 0 | 0 | 0 | 0 | 0 | 0 | 0 | 0 | 0 | 0 | 0 |
| bacteria_Jun22d0b_14396 | 0 | 0 | 0 | 0 | 0 | 0 | 0 | 0 | 0 | 0 | 0 |
| bacteria_Jun22d0b_29231 | 0 | 0 | 0 | 0 | 0 | 0 | 0 | 0 | 0 | 0 | 0 |
| bacteria_Jun22d50b_4723 | 0 | 0 | 0 | 0 | 0 | 0 | 0 | 0 | 0 | 0 | 0 |
| bacteria_Jun22d50b_14525 | 0 | 0 | 0 | 0 | 0 | 0 | 0 | 0 | 0 | 0 | 0 |
| bacteria_Jun22d50b_17170 | 0 | 0 | 0 | 0 | 0 | 0 | 0 | 0 | 0 | 0 | 0 |
| bacteria_Jun22d50b_17179 | 0 | 0 | 0 | 0 | 0 | 0 | 0 | 0 | 0 | 0 | 0 |
| bacteria_Jun22d50b_31234 | 0 | 0 | 0 | 0 | 0 | 0 | 0 | 0 | 0 | 0 | 0 |
| bacteria_Jun22d75b_9016 | 0 | 0 | 0 | 0 | 0 | 0 | 0 | 0 | 0 | 0 | 0 |
| bacteria_Jun22d75b_22034 | 0 | 0 | 0 | 0 | 0 | 0 | 0 | 0 | 0 | 0 | 0 |
| bacteria_Jun22d75b_22956 | 0 | 0 | 0 | 0 | 0 | 0 | 0 | 0 | 0 | 0 | 0 |
| bacteria_Jun22d75b_25067 | 0 | 0 | 0 | 0 | 0 | 0 | 0 | 0 | 0 | 0 | 0 |
| bacteria_Jun22d75b_30532 | 0 | 0 | 0 | 0 | 0 | 0 | 0 | 0 | 0 | 0 | 0 |
| bacteria_Jun22d75b_33826 | 0 | 0 | 0 | 0 | 0 | 0 | 0 | 0 | 0 | 0 | 0 |
| bacteria_Jun22d75b_35439 | 0 | 0 | 0 | 0 | 0 | 0 | 0 | 0 | 0 | 0 | 0 |
| bacteria_Jun22d75b_39195 | 0 | 0 | 0 | 0 | 0 | 0 | 0 | 0 | 0 | 0 | 0 |
| bacteria_Jun22d75b_47915 | 0 | 0 | 0 | 0 | 0 | 0 | 0 | 0 | 0 | 0 | 0 |
| bacteria_Jun22d75b_56874 | 0 | 0 | 0 | 0 | 0 | 0 | 0 | 0 | 0 | 0 | 0 |
| bacteria_Jun22d75b_61149 | 0 | 0 | 0 | 0 | 0 | 0 | 0 | 0 | 0 | 0 | 0 |
| bacteria_Jun22d75b_72028 | 0 | 0 | 0 | 0 | 0 | 0 | 0 | 0 | 0 | 0 | 0 |
| bacteria_May12d5b_3683 | 0 | 0 | 0 | 0 | 0 | 0 | 0 | 0 | 0 | 0 | 0 |
| bacteria_May12d25b_3351 | 0 | 0 | 0 | 0 | 0 | 0 | 0 | 0 | 0 | 0 | 0 |
| bacteria_May12d25b_3376 | 0 | 0 | 0 | 0 | 0 | 0 | 0 | 0 | 0 | 0 | 0 |
| bacteria_May12d25b_6292 | 0 | 0 | 0 | 0 | 0 | 0 | 0 | 0 | 0 | 0 | 0 |
| bacteria_May12d25b_1181 | 0 | 0 | 0 | 0 | 0 | 0 | 0 | 0 | 0 | 0 | 0 |
| bacteria_May12d25b_2149 | 0 | 0 | 0 | 0 | 0 | 0 | 0 | 0 | 0 | 0 | 0 |
| bacteria_May12d25b_2450 | 0 | 0 | 0 | 0 | 0 | 0 | 0 | 0 | 0 | 0 | 0 |
| bacteria_May12d25b_2547 | 0 | 0 | 0 | 0 | 0 | 0 | 0 | 0 | 0 | 0 | 0 |
| bacteria_May12d25b_4046 | 0 | 0 | 0 | 0 | 0 | 0 | 0 | 0 | 0 | 0 | 0 |
| bacteria_May12d25b_4839 | 0 | 0 | 0 | 0 | 0 | 0 | 0 | 0 | 0 | 0 | 0 |
| bacteria_May12d25b_6534 | 0 | 0 | 0 | 0 | 0 | 0 | 0 | 0 | 0 | 0 | 0 |
| bacteria_May12d25b_6889 | 0 | 0 | 0 | 0 | 0 | 0 | 0 | 0 | 0 | 0 | 0 |
| bacteria_May12d25b_7323 | 0 | 0 | 0 | 0 | 0 | 0 | 0 | 0 | 0 | 0 | 0 |
| bacteria_May12d25b_7520 | 0 | 0 | 0 | 0 | 0 | 0 | 0 | 0 | 0 | 0 | 0 |
| bacteria_May12d50b_4250 | 0 | 0 | 0 | 0 | 0 | 0 | 0 | 0 | 0 | 0 | 0 |
| bacteria_May12d50b_4868 | 0 | 0 | 0 | 0 | 0 | 0 | 0 | 0 | 0 | 0 | 0 |
| bacteria_May12d50b_5390 | 0 | 0 | 0 | 0 | 0 | 0 | 0 | 0 | 0 | 0 | 0 |
| bacteria_May12d50b_1071 | 0 | 0 | 0 | 0 | 0 | 0 | 0 | 0 | 0 | 0 | 0 |
| bacteria_May12d50b_1822 | 0 | 0 | 0 | 0 | 0 | 0 | 0 | 0 | 0 | 0 | 0 |
| bacteria_May12d50b_2804 | 0 | 0 | 0 | 0 | 0 | 0 | 0 | 0 | 0 | 0 | 0 |
| bacteria_May12d50b_2947 | 0 | 0 | 0 | 0 | 0 | 0 | 0 | 0 | 0 | 0 | 0 |
| bacteria_May12d50b_3046 | 0 | 0 | 0 | 0 | 0 | 0 | 0 | 0 | 0 | 0 | 0 |
| bacteria_May12d50b_3492 | 0 | 0 | 0 | 0 | 0 | 0 | 0 | 0 | 0 | 0 | 0 |
| bacteria_May12d50b_4365 | 0 | 0 | 0 | 0 | 0 | 0 | 0 | 0 | 0 | 0 | 0 |
| bacteria_May12d50b_4399 | 0 | 0 | 0 | 0 | 0 | 0 | 0 | 0 | 0 | 0 | 0 |
| bacteria_May12d50b_4483 | 0 | 0 | 0 | 0 | 0 | 0 | 0 | 0 | 0 | 0 | 0 |
| bacteria_May12d50b_4527 | 0 | 0 | 0 | 0 | 0 | 0 | 0 | 0 | 0 | 0 | 0 |
| bacteria_Sep8d0b_64274 | 0 | 0 | 0 | 0 | 0 | 0 | 0 | 0 | 0 | 0 | 0 |
| bacteria_Sep8d5b_2421 | 0 | 0 | 0 | 0 | 0 | 0 | 0 | 0 | 0 | 0 | 0 |
| bacteria_Sep8d5b_7498 | 0 | 0 | 0 | 0 | 0 | 0 | 0 | 0 | 0 | 0 | 0 |
| bacteria_Sep8d5b_9113 | 0 | 0 | 0 | 0 | 0 | 0 | 0 | 0 | 0 | 0 | 0 |
| bacteria_Sep8d5b_13510 | 0 | 0 | 0 | 0 | 0 | 0 | 0 | 0 | 0 | 0 | 0 |
| bacteria_Sep8d5b_18132 | 0 | 0 | 0 | 0 | 0 | 0 | 0 | 0 | 0 | 0 | 0 |
| bacteria_Sep8d5b_23650 | 0 | 0 | 0 | 0 | 0 | 0 | 0 | 0 | 0 | 0 | 0 |
| bacteria_Sep8d5b_30965 | 0 | 0 | 0 | 0 | 0 | 0 | 0 | 0 | 0 | 0 | 0 |
| bacteria_Sep8d5b_40204 | 0 | 0 | 0 | 0 | 0 | 0 | 0 | 0 | 0 | 0 | 0 |
| bacteria_Sep8d5b_49247 | 0 | 0 | 0 | 0 | 0 | 0 | 0 | 0 | 0 | 0 | 0 |
| bacteria_Sep8d5b_53742 | 0 | 0 | 0 | 0 | 0 | 0 | 0 | 0 | 0 | 0 | 0 |
| bacteria_Sep8d5b_54102 | 0 | 0 | 0 | 0 | 0 | 0 | 0 | 0 | 0 | 0 | 0 |
| bacteria_Sep8d5b_61998 | 0 | 0 | 0 | 0 | 0 | 0 | 0 | 0 | 0 | 0 | 0 |
| bacteria_Sep8d25b_5510 | 0 | 0 | 0 | 0 | 0 | 0 | 0 | 0 | 0 | 0 | 0 |
| bacteria_Sep8d25b_23440 | 0 | 0 | 0 | 0 | 0 | 0 | 0 | 0 | 0 | 0 | 0 |
| bacteria_Sep8d25b_27187 | 0 | 0 | 0 | 0 | 0 | 0 | 0 | 0 | 0 | 0 | 0 |
| bacteria_Sep8d25b_36989 | 0 | 0 | 0 | 0 | 0 | 0 | 0 | 0 | 0 | 0 | 0 |
| bacteria_Sep8d50b_37121 | 0 | 0 | 0 | 0 | 0 | 0 | 0 | 0 | 0 | 0 | 0 |
| virus_Aug5.0v_306 | 0 | 1 | 0 | 0 | 0 | 0 | 0 | 0 | 0 | 0 | 1 |
| virus_Aug5.0v_366 | 0 | 1 | 0 | 0 | 0 | 0 | 0 | 0 | 0 | 0 | 1 |
| virus_Aug5.0v_386 | 0 | 1 | 0 | 0 | 0 | 0 | 0 | 0 | 0 | 0 | 1 |
| virus_Aug5.0v_549 | 0 | 1 | 0 | 0 | 0 | 0 | 0 | 0 | 0 | 0 | 1 |
| virus_Aug5.0v_624 | 0 | 1 | 0 | 0 | 0 | 0 | 0 | 0 | 0 | 0 | 1 |
| virus_Aug5.0v_658 | 0 | 1 | 0 | 0 | 0 | 0 | 0 | 0 | 0 | 0 | 1 |
| virus_Aug5.0v_708 | 0 | 1 | 0 | 0 | 0 | 0 | 0 | 0 | 0 | 0 | 1 |
| virus_Aug5.0v_879 | 0 | 1 | 0 | 0 | 0 | 0 | 0 | 0 | 0 | 0 | 1 |
| virus_Aug5.0v_987 | 0 | 1 | 0 | 0 | 0 | 0 | 0 | 0 | 0 | 0 | 1 |
| virus_Aug5.0v_1092 | 0 | 1 | 0 | 0 | 0 | 0 | 0 | 0 | 0 | 0 | 1 |
| virus_Aug5.0v_1251 | 0 | 1 | 0 | 0 | 0 | 0 | 0 | 0 | 0 | 0 | 1 |
| virus_Aug5.0v_1426 | 0 | 1 | 0 | 0 | 0 | 0 | 0 | 0 | 0 | 0 | 1 |
| virus_Aug5.0v_1512 | 0 | 1 | 0 | 0 | 0 | 0 | 0 | 0 | 0 | 0 | 1 |
| virus_Aug5.0v_1608 | 0 | 1 | 0 | 0 | 0 | 0 | 0 | 0 | 0 | 0 | 1 |
| virus_Aug5.0v_1667 | 0 | 1 | 0 | 0 | 0 | 0 | 0 | 0 | 0 | 0 | 1 |
| virus_Aug5.0v_1708 | 0 | 1 | 0 | 0 | 0 | 0 | 0 | 0 | 0 | 0 | 1 |
| virus_Aug5.0v_1835 | 0 | 1 | 0 | 0 | 0 | 0 | 0 | 0 | 0 | 0 | 1 |
| virus_Aug5.0v_1954 | 0 | 2 | 0 | 0 | 0 | 0 | 0 | 0 | 0 | 0 | 2 |
| virus_Aug5.0v_2441 | 0 | 1 | 0 | 0 | 0 | 0 | 0 | 0 | 0 | 0 | 1 |
| virus_Aug5.0v_3382 | 0 | 1 | 0 | 0 | 0 | 0 | 0 | 0 | 0 | 1 | 2 |
| virus_Aug5.0v_3602 | 0 | 1 | 0 | 0 | 0 | 0 | 0 | 0 | 0 | 0 | 1 |
| virus_Aug5.0v_4028 | 0 | 1 | 0 | 0 | 0 | 0 | 0 | 0 | 0 | 0 | 1 |
| virus_Aug5.0v_4122 | 0 | 1 | 0 | 0 | 0 | 0 | 0 | 0 | 0 | 0 | 1 |
| virus_Aug5.0v_4191 | 0 | 1 | 0 | 0 | 0 | 0 | 0 | 0 | 0 | 0 | 1 |
| virus_Aug5.0v_4641 | 0 | 1 | 0 | 0 | 0 | 0 | 0 | 0 | 0 | 0 | 1 |
| virus_Aug5.0v_4883 | 0 | 1 | 0 | 0 | 0 | 0 | 0 | 0 | 0 | 0 | 1 |
| virus_Aug5.0v_4912 | 0 | 1 | 0 | 0 | 0 | 0 | 0 | 0 | 0 | 0 | 1 |

| | | | | | | | | | | | |
|---|---|---|---|---|---|---|---|---|---|---|---|
| virus_Aug5.0v_5136 | 0 | 1 | 0 | 0 | 0 | 0 | 0 | 0 | 0 | 0 | 1 |
| virus_Aug5.0v_5212 | 0 | 1 | 0 | 0 | 0 | 0 | 0 | 0 | 0 | 0 | 1 |
| virus_Aug5.0v_5430 | 0 | 1 | 0 | 0 | 0 | 0 | 0 | 0 | 0 | 0 | 1 |
| virus_Aug5.5v_427 | 0 | 0 | 0 | 1 | 0 | 0 | 0 | 0 | 0 | 0 | 1 |
| virus_Aug5.5v_8232 | 0 | 0 | 0 | 1 | 0 | 0 | 0 | 0 | 0 | 0 | 1 |
| virus_Aug5.5v_9607 | 0 | 0 | 0 | 1 | 0 | 0 | 0 | 0 | 0 | 0 | 1 |
| virus_Aug5.5v_11130 | 0 | 0 | 0 | 1 | 0 | 0 | 0 | 0 | 0 | 0 | 1 |
| virus_Aug5.5v_13719 | 0 | 0 | 0 | 1 | 0 | 0 | 0 | 0 | 0 | 0 | 1 |
| virus_Aug5.5v_15451 | 0 | 0 | 0 | 1 | 0 | 0 | 0 | 0 | 0 | 0 | 1 |
| virus_Aug5.5v_19338 | 0 | 0 | 0 | 1 | 0 | 0 | 0 | 0 | 0 | 0 | 1 |
| virus_Aug5.5v_24009 | 0 | 0 | 0 | 1 | 0 | 0 | 0 | 0 | 0 | 0 | 1 |
| virus_Aug5.5v_24892 | 0 | 0 | 0 | 1 | 0 | 0 | 0 | 0 | 0 | 0 | 1 |
| virus_Aug5.5v_24908 | 0 | 0 | 0 | 1 | 0 | 0 | 0 | 0 | 0 | 0 | 1 |
| virus_Aug5.5v_25676 | 0 | 0 | 0 | 1 | 0 | 0 | 0 | 0 | 0 | 0 | 1 |
| virus_Aug5.5v_27007 | 0 | 0 | 0 | 1 | 0 | 0 | 0 | 0 | 0 | 0 | 1 |
| virus_Aug5.5v_30894 | 0 | 0 | 0 | 1 | 0 | 0 | 0 | 0 | 0 | 0 | 1 |
| virus_Aug5.5v_31730 | 0 | 0 | 0 | 1 | 0 | 0 | 0 | 0 | 0 | 0 | 1 |
| virus_Aug5.5v_31869 | 0 | 0 | 0 | 1 | 0 | 0 | 0 | 0 | 0 | 0 | 1 |
| virus_Aug5.5v_33528 | 0 | 0 | 0 | 1 | 0 | 0 | 0 | 0 | 0 | 0 | 1 |
| virus_Aug5.25v_2561 | 0 | 1 | 0 | 0 | 0 | 1 | 0 | 0 | 0 | 0 | 2 |
| virus_Aug5.25v_7919 | 0 | 0 | 0 | 0 | 0 | 1 | 0 | 0 | 0 | 0 | 1 |
| virus_Aug5.25v_18348 | 0 | 0 | 0 | 0 | 0 | 1 | 0 | 0 | 0 | 0 | 1 |
| virus_Aug5.25v_19141 | 0 | 0 | 0 | 0 | 0 | 1 | 0 | 0 | 0 | 0 | 1 |
| virus_Aug5.25v_21303 | 0 | 0 | 0 | 0 | 0 | 1 | 0 | 0 | 0 | 0 | 1 |
| virus_Aug5.25v_22287 | 0 | 0 | 0 | 0 | 0 | 1 | 0 | 0 | 0 | 0 | 1 |
| virus_Aug5.25v_22964 | 0 | 0 | 0 | 0 | 0 | 1 | 0 | 0 | 0 | 0 | 1 |
| virus_Aug5.25v_24202 | 0 | 0 | 0 | 0 | 0 | 1 | 0 | 0 | 0 | 0 | 1 |
| virus_Aug5.25v_28461 | 0 | 0 | 0 | 0 | 0 | 1 | 0 | 0 | 0 | 0 | 1 |
| virus_Aug5.25v_30080 | 0 | 0 | 0 | 0 | 0 | 1 | 0 | 0 | 0 | 0 | 1 |
| virus_Aug5.25v_30997 | 0 | 0 | 0 | 0 | 0 | 1 | 0 | 0 | 0 | 0 | 1 |
| virus_Aug5.25v_35003 | 0 | 0 | 0 | 0 | 0 | 1 | 0 | 0 | 0 | 0 | 1 |
| virus_Aug5.25v_35301 | 0 | 0 | 0 | 0 | 0 | 1 | 0 | 0 | 0 | 0 | 1 |
| virus_Aug5.25v_38816 | 0 | 0 | 0 | 0 | 0 | 1 | 0 | 0 | 0 | 0 | 1 |
| virus_Aug5.25v_39418 | 0 | 0 | 0 | 0 | 0 | 1 | 0 | 0 | 0 | 0 | 1 |
| virus_Aug5.50v_7594 | 0 | 0 | 0 | 0 | 0 | 0 | 0 | 1 | 0 | 0 | 1 |
| virus_Aug5.50v_7694 | 0 | 0 | 0 | 0 | 0 | 0 | 0 | 1 | 0 | 0 | 1 |
| virus_Aug5.50v_18854 | 0 | 0 | 0 | 0 | 0 | 0 | 0 | 1 | 0 | 0 | 1 |
| virus_Aug5.50v_22967 | 0 | 0 | 0 | 0 | 0 | 0 | 0 | 1 | 0 | 0 | 1 |
| virus_Aug5.50v_26854 | 0 | 0 | 0 | 0 | 0 | 0 | 0 | 1 | 0 | 0 | 1 |
| virus_Aug5.50v_31053 | 0 | 0 | 0 | 0 | 0 | 0 | 0 | 1 | 0 | 0 | 1 |
| virus_Aug5.50v_31512 | 0 | 0 | 0 | 0 | 0 | 0 | 0 | 1 | 0 | 0 | 1 |
| virus_Aug5.50v_33561 | 0 | 0 | 0 | 0 | 0 | 0 | 0 | 1 | 0 | 0 | 1 |
| virus_Aug5.50v_36307 | 0 | 0 | 0 | 0 | 0 | 0 | 0 | 1 | 0 | 0 | 1 |
| virus_Aug5.75v_11363 | 0 | 0 | 0 | 0 | 0 | 0 | 0 | 0 | 0 | 1 | 1 |
| virus_Aug5.75v_24035 | 1 | 1 | 0 | 0 | 0 | 0 | 0 | 0 | 0 | 1 | 3 |
| virus_Aug5.75v_24045 | 0 | 0 | 0 | 0 | 0 | 0 | 0 | 0 | 0 | 1 | 1 |
| virus_Aug5.75v_27115 | 0 | 0 | 0 | 0 | 0 | 0 | 0 | 0 | 0 | 1 | 1 |
| virus_Aug5.75v_27904 | 0 | 0 | 0 | 0 | 0 | 0 | 0 | 0 | 0 | 1 | 1 |
| virus_Aug5.75v_28382 | 0 | 0 | 0 | 0 | 0 | 0 | 0 | 0 | 0 | 1 | 1 |
| virus_Aug5.75v_32948 | 0 | 0 | 0 | 0 | 0 | 0 | 0 | 0 | 0 | 1 | 1 |
| virus_Aug5.75v_37632 | 0 | 0 | 0 | 0 | 0 | 0 | 0 | 0 | 0 | 1 | 1 |
| virus_Aug5.75v_39058 | 0 | 0 | 0 | 0 | 0 | 0 | 0 | 0 | 0 | 1 | 1 |
| virus_Aug5.75v_39931 | 0 | 0 | 0 | 0 | 0 | 0 | 0 | 0 | 0 | 1 | 1 |
| virus_Jul11.0v_542 | 0 | 0 | 0 | 0 | 0 | 0 | 0 | 0 | 0 | 0 | 0 |
| virus_Jul11.0v_3981 | 0 | 0 | 0 | 0 | 0 | 0 | 0 | 0 | 0 | 0 | 0 |
| virus_Jul11.0v_5391 | 0 | 0 | 0 | 0 | 0 | 0 | 0 | 0 | 0 | 0 | 0 |
| virus_Jul11.0v_8193 | 0 | 0 | 0 | 0 | 0 | 0 | 0 | 0 | 0 | 0 | 0 |
| virus_Jul11.0v_9887 | 0 | 0 | 0 | 0 | 0 | 0 | 0 | 0 | 0 | 0 | 0 |
| virus_Jul11.0v_11131 | 0 | 0 | 0 | 0 | 0 | 0 | 0 | 0 | 0 | 0 | 0 |
| virus_Jul11.0v_13563 | 0 | 0 | 0 | 0 | 0 | 0 | 0 | 0 | 0 | 0 | 0 |
| virus_Jul11.0v_19224 | 0 | 0 | 0 | 0 | 0 | 0 | 0 | 0 | 0 | 0 | 0 |
| virus_Jul11.0v_19453 | 0 | 0 | 0 | 0 | 0 | 0 | 0 | 0 | 0 | 0 | 0 |
| virus_Jul11.5v_1837 | 0 | 0 | 0 | 0 | 0 | 0 | 0 | 0 | 0 | 0 | 0 |
| virus_Jul11.5v_2027 | 0 | 0 | 0 | 0 | 0 | 0 | 0 | 0 | 0 | 0 | 0 |
| virus_Jul11.5v_2104 | 0 | 0 | 0 | 0 | 0 | 0 | 0 | 0 | 0 | 0 | 0 |
| virus_Jul11.5v_2369 | 0 | 0 | 0 | 0 | 0 | 0 | 0 | 0 | 0 | 0 | 0 |
| virus_Jul11.5v_3342 | 0 | 0 | 0 | 0 | 0 | 0 | 0 | 0 | 0 | 0 | 0 |
| virus_Jul11.5v_3420 | 0 | 0 | 0 | 0 | 0 | 0 | 0 | 0 | 0 | 0 | 0 |
| virus_Jul11.5v_3575 | 0 | 0 | 0 | 0 | 0 | 0 | 0 | 0 | 0 | 0 | 0 |
| virus_Jul11.5v_5632 | 0 | 0 | 0 | 0 | 0 | 0 | 0 | 0 | 0 | 0 | 0 |
| virus_Jul11.5v_9729 | 0 | 0 | 0 | 0 | 0 | 0 | 0 | 0 | 0 | 0 | 0 |
| virus_Jul11.5v_12061 | 0 | 0 | 0 | 0 | 0 | 0 | 0 | 0 | 0 | 0 | 0 |
| virus_Jul11.5v_17463 | 0 | 0 | 0 | 0 | 0 | 0 | 0 | 0 | 0 | 0 | 0 |
| virus_Jul11.5v_18582 | 0 | 0 | 0 | 0 | 0 | 0 | 0 | 0 | 0 | 0 | 0 |
| virus_Jul11.5v_19033 | 0 | 0 | 0 | 0 | 0 | 0 | 0 | 0 | 0 | 0 | 0 |
| virus_Jul11.5v_19624 | 0 | 0 | 0 | 0 | 0 | 0 | 0 | 0 | 0 | 0 | 0 |
| virus_Jul11.5v_20599 | 0 | 0 | 0 | 0 | 0 | 0 | 0 | 0 | 0 | 0 | 0 |
| virus_Jul11.5v_21037 | 0 | 0 | 0 | 0 | 0 | 0 | 0 | 0 | 0 | 0 | 0 |
| virus_Jul11.25v_125 | 0 | 0 | 0 | 0 | 0 | 0 | 0 | 0 | 0 | 0 | 0 |
| virus_Jul11.25v_1142 | 0 | 0 | 0 | 0 | 0 | 0 | 0 | 0 | 0 | 0 | 0 |
| virus_Jul11.25v_1365 | 0 | 0 | 0 | 0 | 0 | 0 | 0 | 0 | 0 | 0 | 0 |
| virus_Jul11.25v_7826 | 0 | 0 | 0 | 0 | 0 | 0 | 0 | 0 | 0 | 0 | 0 |
| virus_Jul11.25v_9234 | 0 | 0 | 0 | 0 | 0 | 0 | 0 | 0 | 0 | 0 | 0 |
| virus_Jul11.25v_13802 | 0 | 0 | 0 | 0 | 0 | 0 | 0 | 0 | 0 | 0 | 0 |
| virus_Jul11.25v_16091 | 0 | 0 | 0 | 0 | 0 | 0 | 0 | 0 | 0 | 0 | 0 |
| virus_Jul11.25v_16658 | 0 | 0 | 0 | 0 | 0 | 0 | 0 | 0 | 0 | 0 | 0 |
| virus_Jul11.25v_16824 | 0 | 0 | 0 | 0 | 0 | 0 | 0 | 0 | 0 | 0 | 0 |
| virus_Jul11.25v_17732 | 0 | 0 | 0 | 0 | 0 | 0 | 0 | 0 | 0 | 0 | 0 |
| virus_Jul11.25v_19545 | 0 | 0 | 0 | 0 | 0 | 0 | 0 | 0 | 0 | 0 | 0 |
| virus_Jul11.50v_32 | 0 | 0 | 0 | 0 | 0 | 0 | 0 | 0 | 0 | 0 | 0 |
| virus_Jul11.50v_391 | 0 | 0 | 0 | 0 | 0 | 0 | 0 | 0 | 0 | 0 | 0 |
| virus_Jul11.50v_5094 | 0 | 0 | 0 | 0 | 0 | 0 | 0 | 0 | 0 | 0 | 0 |
| virus_Jul11.50v_6504 | 0 | 0 | 0 | 0 | 0 | 0 | 0 | 0 | 0 | 0 | 0 |
| virus_Jul11.50v_9430 | 0 | 0 | 0 | 0 | 0 | 0 | 0 | 0 | 0 | 0 | 0 |
| virus_Jul11.50v_9436 | 0 | 0 | 0 | 0 | 0 | 0 | 0 | 0 | 0 | 0 | 0 |
| virus_Jul11.50v_9556 | 0 | 0 | 0 | 0 | 0 | 0 | 0 | 0 | 0 | 0 | 0 |
| virus_Jul11.50v_14336 | 0 | 0 | 0 | 0 | 0 | 0 | 0 | 0 | 0 | 0 | 0 |
| virus_Jul11.50v_14475 | 0 | 0 | 0 | 0 | 0 | 0 | 0 | 0 | 0 | 0 | 0 |
| virus_Jul11.50v_14820 | 0 | 0 | 0 | 0 | 0 | 0 | 0 | 0 | 0 | 0 | 0 |
| virus_Jul11.50v_15682 | 0 | 0 | 0 | 0 | 0 | 0 | 0 | 0 | 0 | 0 | 0 |
| virus_Jul11.50v_15968 | 0 | 0 | 0 | 0 | 0 | 0 | 0 | 0 | 0 | 0 | 0 |
| virus_Jul11.50v_16447 | 0 | 0 | 0 | 0 | 0 | 0 | 0 | 0 | 0 | 0 | 0 |
| virus_Jul11.50v_17919 | 0 | 0 | 0 | 0 | 0 | 0 | 0 | 0 | 0 | 0 | 0 |
| virus_Jul11.75v_958 | 0 | 0 | 0 | 0 | 0 | 0 | 0 | 0 | 0 | 0 | 0 |
| virus_Jul11.75v_20354 | 0 | 0 | 0 | 0 | 0 | 0 | 0 | 0 | 0 | 0 | 0 |
| virus_Jul11.75v_23749 | 0 | 0 | 0 | 0 | 0 | 0 | 0 | 0 | 0 | 0 | 0 |
| virus_Jul11.75v_33809 | 0 | 0 | 0 | 0 | 0 | 0 | 0 | 0 | 0 | 0 | 0 |
| virus_Jul27.5v_170 | 0 | 0 | 0 | 0 | 0 | 0 | 0 | 0 | 0 | 0 | 0 |
| virus_Jul27.5v_179 | 0 | 0 | 0 | 0 | 0 | 0 | 0 | 0 | 0 | 0 | 0 |
| virus_Jul27.5v_364 | 0 | 0 | 0 | 0 | 0 | 0 | 0 | 0 | 0 | 0 | 0 |
| virus_Jul27.5v_519 | 0 | 0 | 0 | 0 | 0 | 0 | 0 | 0 | 0 | 0 | 0 |
| virus_Jul27.5v_3024 | 0 | 0 | 0 | 0 | 0 | 0 | 0 | 0 | 0 | 0 | 0 |
| virus_Jul27.5v_3767 | 0 | 0 | 0 | 0 | 0 | 0 | 0 | 0 | 0 | 0 | 0 |
| virus_Jul27.5v_5539 | 0 | 0 | 0 | 0 | 0 | 0 | 0 | 0 | 0 | 0 | 0 |
| virus_Jul27.5v_6121 | 0 | 0 | 0 | 0 | 0 | 0 | 0 | 0 | 0 | 0 | 0 |
| virus_Jul27.5v_7323 | 0 | 0 | 0 | 0 | 0 | 0 | 0 | 0 | 0 | 0 | 0 |
| virus_Jul27.5v_8652 | 0 | 0 | 0 | 0 | 0 | 0 | 0 | 0 | 0 | 0 | 0 |
| virus_Jul27.5v_8945 | 0 | 0 | 0 | 0 | 0 | 0 | 0 | 0 | 0 | 0 | 0 |
| virus_Jul27.5v_9148 | 0 | 0 | 0 | 0 | 0 | 0 | 0 | 0 | 0 | 0 | 0 |
| virus_Jul27.5v_11613 | 0 | 0 | 0 | 0 | 0 | 0 | 0 | 0 | 0 | 0 | 0 |
| virus_Jul27.5v_13084 | 0 | 0 | 0 | 0 | 0 | 0 | 0 | 0 | 0 | 0 | 0 |
| virus_Jul27.5v_13706 | 0 | 0 | 0 | 0 | 0 | 0 | 0 | 0 | 0 | 0 | 0 |
| virus_Jul27.5v_13910 | 0 | 0 | 0 | 0 | 0 | 0 | 0 | 0 | 0 | 0 | 0 |
| virus_Jul27.5v_15771 | 0 | 0 | 0 | 0 | 0 | 0 | 0 | 0 | 0 | 0 | 0 |
| virus_Jul27.5v_16350 | 0 | 0 | 0 | 0 | 0 | 0 | 0 | 0 | 0 | 0 | 0 |
| virus_Jul27.5v_16972 | 0 | 0 | 0 | 0 | 0 | 0 | 0 | 0 | 0 | 0 | 0 |
| virus_Jul27.5v_17153 | 0 | 0 | 0 | 0 | 0 | 0 | 0 | 0 | 0 | 0 | 0 |
| virus_Jul27.5v_17304 | 0 | 0 | 0 | 0 | 0 | 0 | 0 | 0 | 0 | 0 | 0 |
| virus_Jul27.5v_17778 | 0 | 0 | 0 | 0 | 0 | 0 | 0 | 0 | 0 | 0 | 0 |
| virus_Jul27.5v_17934 | 0 | 0 | 0 | 0 | 0 | 0 | 0 | 0 | 0 | 0 | 0 |
| virus_Jul27.5v_18031 | 0 | 0 | 0 | 0 | 0 | 0 | 0 | 0 | 0 | 0 | 0 |
| virus_Jul27.5v_18355 | 0 | 0 | 0 | 0 | 0 | 0 | 0 | 0 | 0 | 0 | 0 |
| virus_Jul27.5v_18500 | 0 | 0 | 0 | 0 | 0 | 0 | 0 | 0 | 0 | 0 | 0 |
| virus_Jul27.5v_18658 | 0 | 0 | 0 | 1 | 0 | 1 | 1 | 0 | 0 | 0 | 3 |
| virus_Jul27.5v_19079 | 0 | 0 | 0 | 0 | 0 | 0 | 0 | 0 | 0 | 0 | 0 |
| virus_Jul27.5v_19245 | 0 | 0 | 0 | 0 | 0 | 0 | 0 | 0 | 0 | 0 | 0 |
| virus_Jul27.25v_626 | 0 | 0 | 0 | 0 | 0 | 0 | 0 | 0 | 0 | 0 | 0 |
| virus_Jul27.25v_1516 | 0 | 0 | 0 | 0 | 0 | 0 | 0 | 0 | 0 | 0 | 0 |
| virus_Jul27.25v_3062 | 0 | 0 | 0 | 0 | 0 | 0 | 0 | 0 | 0 | 0 | 0 |
| virus_Jul27.25v_3573 | 0 | 0 | 0 | 0 | 0 | 0 | 0 | 0 | 0 | 0 | 0 |
| virus_Jul27.25v_3941 | 0 | 0 | 0 | 0 | 0 | 0 | 0 | 0 | 0 | 0 | 0 |
| virus_Jul27.25v_10421 | 0 | 0 | 0 | 0 | 0 | 0 | 0 | 0 | 0 | 0 | 0 |
| virus_Jul27.25v_17730 | 0 | 0 | 0 | 0 | 0 | 0 | 0 | 0 | 0 | 0 | 0 |
| virus_Jul27.25v_18805 | 0 | 0 | 0 | 0 | 0 | 0 | 0 | 0 | 0 | 0 | 0 |
| virus_Jul27.25v_29797 | 0 | 0 | 0 | 0 | 0 | 0 | 0 | 0 | 0 | 0 | 0 |
| virus_Jul27.25v_32502 | 0 | 0 | 0 | 0 | 0 | 0 | 0 | 0 | 0 | 0 | 0 |
| virus_Jul27.25v_35958 | 0 | 0 | 0 | 0 | 0 | 0 | 0 | 0 | 0 | 0 | 0 |
| virus_Jul27.25v_36115 | 0 | 0 | 0 | 0 | 0 | 0 | 0 | 0 | 0 | 0 | 0 |
| virus_Jul27.25v_37529 | 0 | 0 | 0 | 0 | 0 | 0 | 0 | 0 | 0 | 0 | 0 |
| virus_Jul27.25v_41111 | 0 | 0 | 0 | 0 | 0 | 0 | 0 | 0 | 0 | 0 | 0 |
| virus_Jul27.75v_5232 | 0 | 0 | 0 | 0 | 0 | 0 | 0 | 0 | 0 | 0 | 0 |
| virus_Jul27.75v_6139 | 0 | 0 | 0 | 0 | 0 | 0 | 0 | 0 | 0 | 0 | 0 |
| virus_Jul27.75v_18436 | 0 | 0 | 0 | 0 | 0 | 0 | 0 | 0 | 0 | 0 | 0 |
| virus_Jun8.25v_36706 | 0 | 0 | 0 | 0 | 0 | 0 | 0 | 0 | 0 | 0 | 0 |
| virus_Jun8.50v_8013 | 0 | 0 | 0 | 0 | 0 | 0 | 0 | 0 | 0 | 0 | 0 |
| virus_Jun8.50v_12566 | 0 | 0 | 0 | 0 | 0 | 0 | 0 | 0 | 0 | 0 | 0 |
| virus_Jun8.50v_15301 | 0 | 0 | 0 | 0 | 0 | 0 | 0 | 0 | 0 | 0 | 0 |

| | | | | | | | | | | | |
|---|---|---|---|---|---|---|---|---|---|---|---|
| virus_Jun8.50v_16840 | 0 | 0 | 0 | 0 | 0 | 0 | 0 | 0 | 0 | 0 | 0 |
| virus_Jun8.50v_25689 | 0 | 0 | 0 | 0 | 0 | 0 | 0 | 0 | 0 | 0 | 0 |
| virus_Jun8.50v_27280 | 0 | 0 | 0 | 0 | 0 | 0 | 0 | 0 | 0 | 0 | 0 |
| virus_Jun8.50v_29234 | 0 | 0 | 0 | 0 | 0 | 0 | 0 | 0 | 0 | 0 | 0 |
| virus_Jun8.50v_29243 | 0 | 0 | 0 | 0 | 0 | 0 | 0 | 0 | 0 | 0 | 0 |
| virus_Jun8.50v_30352 | 0 | 0 | 0 | 0 | 0 | 0 | 0 | 0 | 0 | 0 | 0 |
| virus_Jun8.50v_31585 | 0 | 0 | 0 | 0 | 0 | 0 | 0 | 0 | 0 | 0 | 0 |
| virus_Jun8.75v_6905 | 0 | 0 | 0 | 0 | 0 | 0 | 0 | 0 | 0 | 0 | 0 |
| virus_Jun8.75v_7324 | 0 | 0 | 0 | 0 | 0 | 0 | 0 | 0 | 0 | 0 | 0 |
| virus_Jun8.75v_17411 | 0 | 0 | 0 | 0 | 0 | 0 | 0 | 0 | 0 | 0 | 0 |
| virus_Jun22.0v_1940 | 0 | 0 | 0 | 1 | 0 | 1 | 0 | 0 | 0 | 0 | 2 |
| virus_Jun22.0v_2695 | 0 | 0 | 0 | 0 | 0 | 0 | 0 | 0 | 0 | 0 | 0 |
| virus_Jun22.0v_2799 | 0 | 0 | 0 | 0 | 0 | 0 | 0 | 0 | 0 | 0 | 0 |
| virus_Jun22.0v_5371 | 0 | 0 | 0 | 0 | 0 | 0 | 0 | 0 | 0 | 0 | 0 |
| virus_Jun22.0v_10868 | 0 | 0 | 0 | 0 | 0 | 0 | 0 | 0 | 0 | 0 | 0 |
| virus_Jun22.0v_12841 | 0 | 0 | 0 | 0 | 0 | 0 | 0 | 0 | 0 | 0 | 0 |
| virus_Jun22.0v_15014 | 0 | 0 | 0 | 0 | 0 | 0 | 0 | 0 | 0 | 0 | 0 |
| virus_Jun22.0v_16126 | 0 | 0 | 0 | 0 | 0 | 0 | 0 | 0 | 0 | 0 | 0 |
| virus_Jun22.0v_17581 | 0 | 0 | 0 | 0 | 0 | 0 | 0 | 0 | 0 | 0 | 0 |
| virus_Jun22.0v_18752 | 0 | 0 | 0 | 0 | 0 | 0 | 0 | 0 | 0 | 0 | 0 |
| virus_Jun22.0v_19176 | 0 | 0 | 0 | 0 | 0 | 0 | 0 | 0 | 0 | 0 | 0 |
| virus_Jun22.50v_1479 | 0 | 0 | 0 | 0 | 0 | 0 | 0 | 0 | 0 | 0 | 0 |
| virus_Jun22.50v_4224 | 0 | 0 | 0 | 0 | 0 | 0 | 0 | 0 | 0 | 0 | 0 |
| virus_Jun22.50v_4841 | 0 | 0 | 0 | 0 | 0 | 0 | 0 | 0 | 0 | 0 | 0 |
| virus_Jun22.50v_5419 | 0 | 0 | 0 | 0 | 0 | 0 | 0 | 0 | 0 | 0 | 0 |
| virus_Jun22.50v_7359 | 0 | 0 | 0 | 0 | 0 | 0 | 0 | 0 | 0 | 0 | 0 |
| virus_Jun22.50v_7769 | 0 | 0 | 0 | 0 | 0 | 0 | 0 | 0 | 0 | 0 | 0 |
| virus_Jun22.50v_8292 | 0 | 0 | 0 | 0 | 0 | 0 | 0 | 0 | 0 | 0 | 0 |
| virus_Jun22.50v_8773 | 0 | 0 | 0 | 0 | 0 | 0 | 0 | 0 | 0 | 0 | 0 |
| virus_Jun22.50v_9376 | 0 | 0 | 0 | 0 | 0 | 0 | 0 | 0 | 0 | 0 | 0 |
| virus_Jun22.50v_11549 | 0 | 0 | 0 | 0 | 0 | 0 | 0 | 0 | 0 | 0 | 0 |
| virus_Jun22.50v_12093 | 0 | 0 | 0 | 0 | 0 | 0 | 0 | 0 | 0 | 0 | 0 |
| virus_Jun22.50v_13394 | 0 | 0 | 0 | 0 | 0 | 0 | 0 | 0 | 0 | 0 | 0 |
| virus_Jun22.50v_15153 | 0 | 0 | 0 | 0 | 0 | 0 | 0 | 0 | 0 | 0 | 0 |
| virus_Jun22.50v_15625 | 0 | 0 | 0 | 0 | 0 | 0 | 0 | 0 | 0 | 0 | 0 |
| virus_Jun22.50v_18588 | 0 | 0 | 0 | 0 | 0 | 0 | 0 | 0 | 0 | 0 | 0 |
| virus_Jun22.50v_23531 | 0 | 0 | 0 | 0 | 0 | 0 | 0 | 0 | 0 | 0 | 0 |
| virus_Jun22.50v_28668 | 0 | 0 | 0 | 0 | 0 | 0 | 0 | 0 | 0 | 0 | 0 |
| virus_Jun22.50v_29227 | 0 | 0 | 0 | 0 | 0 | 0 | 0 | 0 | 0 | 0 | 0 |
| virus_Jun22.50v_31012 | 0 | 0 | 0 | 0 | 0 | 0 | 0 | 0 | 0 | 0 | 0 |
| virus_Jun22.50v_31086 | 0 | 0 | 0 | 0 | 0 | 0 | 0 | 0 | 0 | 0 | 0 |
| virus_Jun22.50v_31215 | 0 | 0 | 0 | 0 | 0 | 0 | 0 | 0 | 0 | 0 | 0 |
| virus_Jun22.50v_32786 | 0 | 0 | 0 | 0 | 0 | 0 | 0 | 0 | 0 | 0 | 0 |
| virus_Jun22.50v_33173 | 0 | 0 | 0 | 0 | 0 | 0 | 0 | 0 | 0 | 0 | 0 |
| virus_Jun22.50v_37659 | 0 | 0 | 0 | 0 | 0 | 0 | 0 | 0 | 0 | 0 | 0 |
| virus_Jun22.50v_40415 | 0 | 0 | 0 | 0 | 0 | 0 | 0 | 0 | 0 | 0 | 0 |
| virus_Jun22.75v_3347 | 0 | 0 | 0 | 0 | 0 | 0 | 0 | 0 | 0 | 0 | 0 |
| virus_Jun22.75v_9026 | 0 | 0 | 0 | 0 | 0 | 0 | 0 | 0 | 0 | 0 | 0 |
| virus_Jun22.75v_11856 | 0 | 0 | 0 | 0 | 0 | 0 | 0 | 0 | 0 | 0 | 0 |
| virus_Jun22.75v_28013 | 0 | 0 | 0 | 0 | 0 | 0 | 0 | 0 | 0 | 0 | 0 |
| virus_Jun22.75v_28044 | 0 | 0 | 0 | 0 | 0 | 0 | 0 | 0 | 0 | 0 | 0 |
| virus_Jun22.75v_35672 | 0 | 0 | 0 | 0 | 0 | 0 | 0 | 0 | 0 | 0 | 0 |
| virus_Jun22.75v_36577 | 0 | 0 | 0 | 0 | 0 | 0 | 0 | 0 | 0 | 0 | 0 |
| virus_Jun22.75v_37712 | 0 | 0 | 0 | 0 | 0 | 0 | 0 | 0 | 0 | 0 | 0 |
| virus_May12.5v_229 | 0 | 0 | 0 | 0 | 0 | 0 | 0 | 0 | 0 | 0 | 0 |
| virus_May12.5v_776 | 0 | 0 | 0 | 0 | 0 | 0 | 0 | 0 | 0 | 0 | 0 |
| virus_May12.5v_3677 | 0 | 0 | 0 | 0 | 0 | 0 | 0 | 0 | 0 | 0 | 0 |
| virus_May12.5v_5978 | 0 | 0 | 0 | 0 | 0 | 0 | 0 | 0 | 0 | 0 | 0 |
| virus_May12.5v_14498 | 0 | 0 | 0 | 0 | 0 | 0 | 0 | 0 | 0 | 0 | 0 |
| virus_May12.5v_20478 | 0 | 0 | 0 | 0 | 0 | 0 | 0 | 0 | 0 | 0 | 0 |
| virus_May12.5v_26165 | 0 | 0 | 0 | 0 | 0 | 0 | 0 | 0 | 0 | 0 | 0 |
| virus_May12.5v_30061 | 0 | 0 | 0 | 0 | 0 | 0 | 0 | 0 | 0 | 0 | 0 |
| virus_May12.5v_30272 | 0 | 0 | 0 | 0 | 0 | 0 | 0 | 0 | 0 | 0 | 0 |
| virus_May12.5v_34438 | 0 | 0 | 0 | 0 | 0 | 0 | 0 | 0 | 0 | 0 | 0 |
| virus_May12.5v_38517 | 0 | 0 | 0 | 0 | 0 | 0 | 0 | 0 | 0 | 0 | 0 |
| virus_May12.25v_6734 | 0 | 0 | 0 | 0 | 0 | 0 | 0 | 0 | 0 | 0 | 0 |
| virus_May12.25v_13063 | 0 | 0 | 0 | 0 | 0 | 0 | 0 | 0 | 0 | 0 | 0 |
| virus_May12.25v_18789 | 0 | 0 | 0 | 0 | 0 | 0 | 0 | 0 | 0 | 0 | 0 |
| virus_May12.25v_20561 | 0 | 0 | 0 | 0 | 0 | 0 | 0 | 0 | 0 | 0 | 0 |
| virus_May12.25v_22028 | 0 | 0 | 0 | 0 | 0 | 0 | 0 | 0 | 0 | 0 | 0 |
| virus_May12.25v_26376 | 0 | 0 | 0 | 0 | 0 | 0 | 0 | 0 | 0 | 0 | 0 |
| virus_May12.25v_37667 | 0 | 0 | 0 | 0 | 0 | 0 | 0 | 0 | 0 | 0 | 0 |
| virus_May12.25v_40561 | 0 | 0 | 0 | 0 | 0 | 0 | 0 | 0 | 0 | 0 | 0 |
| virus_May12.50v_517 | 0 | 0 | 0 | 0 | 0 | 0 | 0 | 0 | 0 | 0 | 0 |
| virus_May12.50v_1873 | 0 | 0 | 0 | 0 | 0 | 0 | 0 | 0 | 0 | 0 | 0 |
| virus_May12.50v_4359 | 0 | 0 | 0 | 0 | 0 | 0 | 0 | 0 | 0 | 0 | 0 |
| virus_May12.50v_6436 | 0 | 0 | 0 | 0 | 0 | 0 | 0 | 0 | 0 | 0 | 0 |
| virus_May12.50v_7012 | 0 | 0 | 0 | 0 | 0 | 0 | 0 | 0 | 0 | 0 | 0 |
| virus_May12.50v_7840 | 0 | 0 | 0 | 0 | 0 | 0 | 0 | 0 | 0 | 0 | 0 |
| virus_May12.50v_12855 | 0 | 0 | 0 | 0 | 0 | 0 | 0 | 0 | 0 | 0 | 0 |
| virus_May12.50v_14301 | 0 | 0 | 0 | 0 | 0 | 0 | 0 | 0 | 0 | 0 | 0 |
| virus_May12.50v_16267 | 0 | 0 | 0 | 0 | 0 | 0 | 0 | 0 | 0 | 0 | 0 |
| virus_May12.50v_19591 | 0 | 0 | 0 | 0 | 0 | 0 | 0 | 0 | 0 | 0 | 0 |
| virus_May12.50v_21730 | 0 | 0 | 0 | 0 | 0 | 0 | 0 | 0 | 0 | 0 | 0 |
| virus_May12.50v_22241 | 0 | 0 | 0 | 0 | 0 | 0 | 0 | 0 | 0 | 0 | 0 |
| virus_May12.50v_24086 | 0 | 0 | 0 | 0 | 0 | 0 | 0 | 0 | 0 | 0 | 0 |
| virus_May12.50v_24282 | 0 | 0 | 0 | 0 | 0 | 0 | 0 | 0 | 0 | 0 | 0 |
| virus_May12.50v_34656 | 0 | 0 | 0 | 0 | 0 | 0 | 0 | 0 | 0 | 0 | 0 |
| virus_May12.50v_37498 | 0 | 0 | 0 | 0 | 0 | 0 | 0 | 0 | 0 | 0 | 0 |
| virus_May12.50v_39241 | 0 | 0 | 0 | 0 | 0 | 0 | 0 | 0 | 0 | 0 | 0 |
| virus_May12.50v_40642 | 0 | 0 | 0 | 0 | 0 | 0 | 0 | 0 | 0 | 0 | 0 |
| virus_May12.75v_5955 | 0 | 0 | 0 | 0 | 0 | 0 | 0 | 0 | 0 | 0 | 0 |
| virus_May12.75v_6315 | 0 | 0 | 0 | 0 | 0 | 0 | 0 | 0 | 0 | 0 | 0 |
| virus_May12.75v_12526 | 0 | 0 | 0 | 0 | 0 | 0 | 0 | 0 | 0 | 0 | 0 |
| virus_May12.75v_13232 | 0 | 0 | 0 | 0 | 0 | 0 | 0 | 0 | 0 | 0 | 0 |
| virus_May12.75v_17992 | 0 | 0 | 0 | 0 | 0 | 0 | 0 | 0 | 0 | 0 | 0 |
| virus_May12.75v_19401 | 0 | 0 | 0 | 0 | 0 | 0 | 0 | 0 | 0 | 0 | 0 |
| virus_May12.75v_22279 | 0 | 0 | 0 | 0 | 0 | 0 | 0 | 0 | 0 | 0 | 0 |
| virus_May12.75v_23095 | 0 | 0 | 0 | 0 | 0 | 0 | 0 | 0 | 0 | 0 | 0 |
| virus_May12.75v_23746 | 0 | 0 | 0 | 0 | 0 | 0 | 0 | 0 | 0 | 0 | 0 |
| virus_May12.75v_26200 | 0 | 0 | 0 | 0 | 0 | 0 | 0 | 0 | 0 | 0 | 0 |
| virus_May12.75v_28437 | 0 | 0 | 0 | 0 | 0 | 0 | 0 | 0 | 0 | 0 | 0 |
| virus_May12.75v_33213 | 0 | 0 | 0 | 0 | 0 | 0 | 0 | 0 | 0 | 0 | 0 |
| virus_May12.75v_38720 | 0 | 0 | 0 | 0 | 0 | 0 | 0 | 0 | 0 | 0 | 0 |
| virus_May12.75v_38832 | 0 | 0 | 0 | 0 | 0 | 0 | 0 | 0 | 0 | 0 | 0 |
| virus_Sep8.0v_1101 | 0 | 0 | 0 | 0 | 0 | 0 | 0 | 0 | 0 | 0 | 0 |
| virus_Sep8.0v_1275 | 0 | 0 | 0 | 0 | 0 | 0 | 0 | 0 | 0 | 0 | 0 |
| virus_Sep8.0v_10736 | 0 | 0 | 0 | 0 | 0 | 0 | 0 | 0 | 0 | 0 | 0 |
| virus_Sep8.0v_13128 | 0 | 0 | 0 | 0 | 0 | 0 | 0 | 0 | 0 | 0 | 0 |
| virus_Sep8.0v_14349 | 0 | 0 | 0 | 0 | 0 | 0 | 0 | 0 | 0 | 0 | 0 |
| virus_Sep8.0v_15372 | 0 | 0 | 0 | 0 | 0 | 0 | 0 | 0 | 0 | 0 | 0 |
| virus_Sep8.0v_16220 | 0 | 0 | 0 | 0 | 0 | 0 | 0 | 0 | 0 | 0 | 0 |
| virus_Sep8.0v_17221 | 0 | 0 | 0 | 0 | 0 | 0 | 0 | 0 | 0 | 0 | 0 |
| virus_Sep8.0v_18019 | 0 | 0 | 0 | 0 | 0 | 0 | 0 | 0 | 0 | 0 | 0 |
| virus_Sep8.0v_19152 | 0 | 0 | 0 | 0 | 0 | 0 | 0 | 0 | 0 | 0 | 0 |
| virus_Sep8.0v_19482 | 0 | 0 | 0 | 0 | 0 | 0 | 0 | 0 | 0 | 0 | 0 |
| virus_Sep8.5v_1832 | 0 | 0 | 0 | 0 | 0 | 0 | 0 | 0 | 0 | 0 | 0 |
| virus_Sep8.5v_6144 | 0 | 0 | 0 | 0 | 0 | 0 | 0 | 0 | 0 | 0 | 0 |
| virus_Sep8.5v_8500 | 0 | 0 | 0 | 0 | 0 | 0 | 0 | 0 | 0 | 0 | 0 |
| virus_Sep8.5v_8644 | 0 | 0 | 0 | 0 | 0 | 0 | 0 | 0 | 0 | 0 | 0 |
| virus_Sep8.5v_17440 | 0 | 0 | 0 | 0 | 0 | 0 | 0 | 0 | 0 | 0 | 0 |
| virus_Sep8.5v_19205 | 0 | 0 | 0 | 0 | 0 | 0 | 0 | 0 | 0 | 0 | 0 |
| virus_Sep8.25v_1879 | 0 | 0 | 0 | 0 | 0 | 0 | 0 | 0 | 0 | 0 | 0 |
| virus_Sep8.25v_2068 | 0 | 0 | 0 | 0 | 0 | 0 | 0 | 0 | 0 | 0 | 0 |
| virus_Sep8.25v_5364 | 0 | 0 | 0 | 0 | 0 | 0 | 0 | 0 | 0 | 0 | 0 |
| virus_Sep8.25v_7572 | 0 | 0 | 0 | 0 | 0 | 0 | 0 | 0 | 0 | 0 | 0 |
| virus_Sep8.25v_8232 | 0 | 0 | 0 | 0 | 0 | 0 | 0 | 0 | 0 | 0 | 0 |
| virus_Sep8.25v_9026 | 0 | 0 | 0 | 0 | 0 | 0 | 0 | 0 | 0 | 0 | 0 |
| virus_Sep8.25v_14057 | 0 | 0 | 0 | 0 | 0 | 0 | 0 | 0 | 0 | 0 | 0 |
| virus_Sep8.25v_17217 | 0 | 0 | 0 | 0 | 0 | 0 | 0 | 0 | 0 | 0 | 0 |
| virus_Sep8.25v_17295 | 0 | 0 | 0 | 0 | 0 | 0 | 0 | 0 | 0 | 0 | 0 |
| virus_Sep8.25v_18280 | 0 | 0 | 0 | 0 | 0 | 0 | 0 | 0 | 0 | 0 | 0 |
| virus_Sep8.50v_1433 | 0 | 0 | 0 | 0 | 0 | 0 | 0 | 0 | 0 | 0 | 0 |
| virus_Sep8.50v_1984 | 0 | 0 | 0 | 0 | 0 | 0 | 0 | 0 | 0 | 0 | 0 |
| virus_Sep8.50v_7969 | 0 | 0 | 0 | 0 | 0 | 0 | 0 | 0 | 0 | 0 | 0 |
| virus_Sep8.50v_13391 | 0 | 0 | 0 | 0 | 0 | 0 | 0 | 0 | 0 | 0 | 0 |
| virus_Sep8.50v_20658 | 0 | 0 | 0 | 0 | 0 | 0 | 0 | 0 | 0 | 0 | 0 |
| virus_Sep8.50v_22833 | 0 | 0 | 0 | 0 | 0 | 0 | 0 | 0 | 0 | 0 | 0 |
| virus_Sep8.50v_25890 | 0 | 0 | 0 | 0 | 0 | 0 | 0 | 0 | 0 | 0 | 0 |
| virus_Sep8.50v_29542 | 0 | 0 | 0 | 0 | 0 | 0 | 0 | 0 | 0 | 0 | 0 |
| virus_Sep8.50v_32743 | 0 | 0 | 0 | 0 | 0 | 0 | 0 | 0 | 0 | 0 | 0 |
| virus_Sep8.50v_36946 | 0 | 0 | 0 | 0 | 0 | 0 | 0 | 0 | 0 | 0 | 0 |
| virus_Sep8.50v_39897 | 0 | 0 | 0 | 0 | 0 | 0 | 0 | 0 | 0 | 0 | 0 |
| bacteria_Jul11d0b_3604 | 0 | 0 | 0 | 0 | 0 | 0 | 0 | 0 | 0 | 0 | 0 |
| bacteria_Jul11d0b_5048 | 0 | 0 | 0 | 0 | 0 | 0 | 0 | 0 | 0 | 0 | 0 |
| bacteria_Jul11d0b_11384 | 0 | 0 | 0 | 0 | 0 | 0 | 0 | 0 | 0 | 0 | 0 |
| bacteria_Jul11d0b_15193 | 0 | 0 | 0 | 0 | 0 | 0 | 0 | 0 | 0 | 0 | 0 |
| bacteria_Jul11d0b_19957 | 0 | 0 | 0 | 0 | 0 | 0 | 0 | 0 | 0 | 0 | 0 |
| bacteria_Jul11d0b_23989 | 0 | 0 | 0 | 0 | 0 | 1 | 0 | 0 | 0 | 0 | 1 |
| bacteria_Jul11d0b_32605 | 0 | 0 | 0 | 0 | 0 | 0 | 0 | 0 | 0 | 0 | 0 |
| bacteria_Jul11d0b_34466 | 0 | 0 | 0 | 0 | 0 | 0 | 0 | 0 | 0 | 0 | 0 |
| bacteria_Jul11d0b_47693 | 0 | 0 | 0 | 0 | 0 | 0 | 0 | 0 | 0 | 0 | 0 |
| bacteria_Jul11d0b_59949 | 0 | 0 | 0 | 0 | 0 | 0 | 0 | 0 | 0 | 0 | 0 |
| bacteria_Jul11d0b_66100 | 0 | 0 | 0 | 0 | 0 | 0 | 0 | 0 | 0 | 0 | 0 |
| bacteria_Jul11d0b_74640 | 0 | 0 | 0 | 0 | 0 | 0 | 0 | 0 | 0 | 0 | 0 |
| bacteria_Aug5d0b_10414 | 1 | 0 | 0 | 0 | 0 | 0 | 0 | 0 | 0 | 0 | 1 |
| bacteria_Aug5d0b_15290 | 1 | 0 | 0 | 0 | 0 | 0 | 0 | 0 | 0 | 0 | 1 |

| | | | | | | | | | | | |
|---|---|---|---|---|---|---|---|---|---|---|---|
| bacteria_Aug5d0b_20121 | 1 | 0 | 0 | 0 | 0 | 0 | 0 | 0 | 0 | 0 | 1 |
| bacteria_Aug5d0b_25710 | 1 | 0 | 0 | 0 | 0 | 0 | 0 | 0 | 0 | 0 | 1 |
| bacteria_Aug5d0b_53114 | 1 | 0 | 0 | 0 | 0 | 0 | 0 | 0 | 0 | 0 | 1 |
| bacteria_Aug5d5b_14816 | 0 | 0 | 1 | 0 | 0 | 0 | 0 | 0 | 0 | 0 | 1 |
| bacteria_Aug5d5b_52619 | 0 | 0 | 1 | 0 | 0 | 0 | 0 | 0 | 0 | 0 | 1 |
| bacteria_Aug5d5b_60951 | 0 | 0 | 1 | 0 | 0 | 0 | 0 | 0 | 0 | 0 | 1 |
| bacteria_Aug5d25b_49 | 0 | 0 | 0 | 0 | 1 | 0 | 0 | 0 | 0 | 0 | 1 |
| bacteria_Aug5d25b_4558 | 0 | 0 | 0 | 0 | 1 | 0 | 0 | 0 | 0 | 0 | 1 |
| bacteria_Aug5d25b_5385 | 0 | 0 | 0 | 0 | 1 | 0 | 0 | 0 | 0 | 0 | 1 |
| bacteria_Aug5d25b_7514 | 0 | 0 | 0 | 0 | 1 | 0 | 0 | 0 | 0 | 0 | 1 |
| bacteria_Aug5d25b_7927 | 0 | 0 | 0 | 0 | 1 | 0 | 0 | 0 | 0 | 0 | 1 |
| bacteria_Aug5d25b_20832 | 0 | 0 | 0 | 0 | 1 | 0 | 0 | 0 | 0 | 0 | 1 |
| bacteria_Aug5d25b_21577 | 0 | 0 | 0 | 0 | 1 | 0 | 0 | 0 | 0 | 0 | 1 |
| bacteria_Aug5d25b_27355 | 0 | 0 | 0 | 0 | 1 | 0 | 0 | 0 | 0 | 0 | 1 |
| bacteria_Aug5d25b_28630 | 0 | 1 | 0 | 0 | 1 | 0 | 0 | 0 | 0 | 0 | 2 |
| bacteria_Aug5d25b_31703 | 0 | 0 | 0 | 0 | 1 | 0 | 0 | 0 | 0 | 0 | 1 |
| bacteria_Aug5d25b_33814 | 0 | 0 | 0 | 0 | 1 | 0 | 0 | 0 | 0 | 0 | 1 |
| bacteria_Aug5d25b_41695 | 0 | 0 | 0 | 0 | 1 | 0 | 0 | 0 | 0 | 0 | 1 |
| bacteria_Aug5d25b_46917 | 0 | 0 | 0 | 0 | 1 | 0 | 0 | 0 | 0 | 0 | 1 |
| bacteria_Aug5d25b_52810 | 0 | 0 | 0 | 0 | 1 | 0 | 0 | 0 | 0 | 0 | 1 |
| bacteria_Aug5d25b_56485 | 0 | 0 | 0 | 0 | 1 | 0 | 0 | 0 | 0 | 0 | 1 |
| bacteria_Aug5d25b_57550 | 0 | 0 | 0 | 0 | 1 | 0 | 0 | 0 | 0 | 0 | 1 |
| bacteria_Aug5d25b_59665 | 0 | 0 | 0 | 0 | 1 | 0 | 0 | 0 | 0 | 0 | 1 |
| bacteria_Aug5d25b_70648 | 0 | 0 | 0 | 0 | 1 | 0 | 0 | 0 | 0 | 0 | 1 |
| bacteria_Aug5d25b_75038 | 0 | 0 | 0 | 0 | 1 | 0 | 0 | 0 | 0 | 0 | 1 |
| bacteria_Aug5d50b_8022 | 0 | 0 | 0 | 0 | 0 | 0 | 1 | 0 | 0 | 0 | 1 |
| bacteria_Aug5d50b_25061 | 0 | 0 | 0 | 0 | 0 | 0 | 1 | 0 | 0 | 0 | 1 |
| bacteria_Aug5d50b_39269 | 0 | 0 | 0 | 0 | 0 | 0 | 1 | 0 | 0 | 0 | 1 |
| bacteria_Aug5d50b_49220 | 0 | 0 | 0 | 0 | 0 | 0 | 1 | 0 | 0 | 0 | 1 |
| bacteria_Aug5d50b_51216 | 0 | 0 | 0 | 0 | 0 | 0 | 1 | 0 | 0 | 0 | 1 |
| bacteria_Aug5d50b_51488 | 0 | 0 | 0 | 0 | 0 | 0 | 1 | 0 | 0 | 0 | 1 |
| bacteria_Aug5d50b_69459 | 0 | 0 | 0 | 0 | 0 | 0 | 1 | 0 | 0 | 0 | 1 |
| bacteria_Aug5d50b_70718 | 0 | 0 | 0 | 0 | 0 | 0 | 1 | 0 | 0 | 0 | 1 |
| bacteria_Aug5d75b_33598 | 0 | 0 | 0 | 0 | 0 | 0 | 0 | 0 | 1 | 0 | 1 |
| bacteria_Jul11d5b_18268 | 0 | 0 | 0 | 0 | 0 | 0 | 0 | 0 | 0 | 0 | 0 |
| bacteria_Jul11d5b_23101 | 0 | 0 | 0 | 0 | 0 | 0 | 0 | 0 | 0 | 0 | 0 |
| bacteria_Jul11d25b_4548 | 0 | 0 | 0 | 0 | 0 | 0 | 0 | 0 | 0 | 0 | 0 |
| bacteria_Jul11d25b_11310 | 0 | 0 | 0 | 0 | 0 | 0 | 0 | 0 | 0 | 0 | 0 |
| bacteria_Jul11d25b_18956 | 0 | 0 | 0 | 0 | 0 | 0 | 0 | 0 | 0 | 0 | 0 |
| bacteria_Jul11d25b_21450 | 0 | 0 | 0 | 0 | 0 | 0 | 0 | 0 | 0 | 0 | 0 |
| bacteria_Jul11d25b_25750 | 0 | 0 | 0 | 0 | 0 | 0 | 0 | 0 | 0 | 0 | 0 |
| bacteria_Jul11d25b_31720 | 0 | 0 | 0 | 0 | 0 | 0 | 0 | 0 | 0 | 0 | 0 |
| bacteria_Jul11d25b_32003 | 0 | 0 | 0 | 0 | 0 | 0 | 0 | 0 | 0 | 0 | 0 |
| bacteria_Jul11d25b_37071 | 0 | 0 | 0 | 0 | 0 | 0 | 0 | 0 | 0 | 0 | 0 |
| bacteria_Jul11d25b_42919 | 0 | 0 | 0 | 0 | 0 | 0 | 0 | 0 | 0 | 0 | 0 |
| bacteria_Jul11d25b_44777 | 0 | 0 | 0 | 0 | 0 | 0 | 0 | 0 | 0 | 0 | 0 |
| bacteria_Jul11d25b_47727 | 0 | 0 | 0 | 0 | 0 | 0 | 0 | 0 | 0 | 0 | 0 |
| bacteria_Jul11d50b_924 | 0 | 0 | 0 | 0 | 0 | 0 | 0 | 0 | 0 | 0 | 0 |
| bacteria_Jul11d50b_3387 | 0 | 0 | 0 | 0 | 0 | 0 | 0 | 0 | 0 | 0 | 0 |
| bacteria_Jul11d50b_7734 | 0 | 0 | 0 | 0 | 0 | 0 | 0 | 0 | 0 | 0 | 0 |
| bacteria_Jul11d50b_34795 | 0 | 0 | 0 | 0 | 0 | 0 | 0 | 0 | 0 | 0 | 0 |
| bacteria_Jul11d50b_38333 | 0 | 0 | 0 | 0 | 0 | 0 | 0 | 0 | 0 | 0 | 0 |
| bacteria_Jul11d50b_48480 | 0 | 0 | 0 | 0 | 0 | 0 | 0 | 0 | 0 | 0 | 0 |
| bacteria_Jul11d50b_52119 | 0 | 0 | 0 | 0 | 0 | 0 | 0 | 0 | 0 | 0 | 0 |
| bacteria_Jul11d75b_5601 | 0 | 0 | 0 | 0 | 0 | 0 | 0 | 0 | 0 | 0 | 0 |
| bacteria_Jul11d75b_10363 | 0 | 0 | 0 | 0 | 0 | 0 | 0 | 0 | 0 | 0 | 0 |
| bacteria_Jul11d75b_11150 | 0 | 0 | 0 | 0 | 0 | 0 | 0 | 0 | 0 | 0 | 0 |
| bacteria_Jul11d75b_13425 | 0 | 0 | 0 | 0 | 0 | 0 | 0 | 0 | 0 | 0 | 0 |
| bacteria_Jul11d75b_21186 | 0 | 0 | 0 | 0 | 0 | 0 | 0 | 0 | 0 | 0 | 0 |
| bacteria_Jul11d75b_26013 | 0 | 0 | 0 | 0 | 0 | 0 | 0 | 0 | 0 | 0 | 0 |
| bacteria_Jul11d75b_26160 | 0 | 0 | 0 | 0 | 0 | 0 | 0 | 0 | 0 | 0 | 0 |
| bacteria_Jul11d75b_28929 | 0 | 0 | 0 | 0 | 0 | 0 | 0 | 0 | 0 | 0 | 0 |
| bacteria_Jul11d75b_30008 | 0 | 0 | 0 | 0 | 0 | 0 | 0 | 0 | 0 | 0 | 0 |
| bacteria_Jul11d75b_30476 | 0 | 0 | 0 | 0 | 0 | 0 | 0 | 0 | 0 | 0 | 0 |
| bacteria_Jul11d75b_30545 | 0 | 0 | 0 | 0 | 0 | 0 | 0 | 0 | 0 | 0 | 0 |
| bacteria_Jul11d75b_37909 | 0 | 0 | 0 | 0 | 0 | 0 | 0 | 0 | 0 | 0 | 0 |
| bacteria_Jul11d75b_38354 | 0 | 0 | 0 | 0 | 0 | 0 | 0 | 0 | 0 | 0 | 0 |
| bacteria_Jul11d75b_44444 | 0 | 0 | 0 | 0 | 0 | 0 | 0 | 0 | 0 | 0 | 0 |
| bacteria_Jul11d75b_49794 | 0 | 0 | 0 | 0 | 0 | 0 | 0 | 0 | 0 | 0 | 0 |
| bacteria_Jul27d5b_39 | 0 | 0 | 0 | 0 | 0 | 0 | 0 | 0 | 0 | 0 | 0 |
| bacteria_Jul27d5b_4246 | 0 | 0 | 0 | 0 | 0 | 0 | 0 | 0 | 0 | 0 | 0 |
| bacteria_Jul27d5b_6030 | 0 | 0 | 0 | 0 | 0 | 1 | 0 | 0 | 0 | 0 | 1 |
| bacteria_Jul27d5b_6063 | 0 | 0 | 0 | 0 | 0 | 0 | 0 | 0 | 0 | 0 | 0 |
| bacteria_Jul27d5b_8869 | 0 | 0 | 0 | 0 | 0 | 0 | 0 | 0 | 0 | 0 | 0 |
| bacteria_Jul27d5b_8981 | 0 | 0 | 0 | 0 | 0 | 0 | 0 | 0 | 0 | 0 | 0 |
| bacteria_Jul27d5b_9683 | 0 | 0 | 0 | 0 | 0 | 0 | 0 | 0 | 0 | 0 | 0 |
| bacteria_Jul27d5b_11900 | 0 | 0 | 0 | 0 | 0 | 0 | 0 | 0 | 0 | 0 | 0 |
| bacteria_Jul27d5b_12737 | 0 | 0 | 0 | 0 | 0 | 0 | 0 | 0 | 0 | 0 | 0 |
| bacteria_Jul27d5b_17374 | 0 | 0 | 0 | 0 | 0 | 0 | 0 | 0 | 0 | 0 | 0 |
| bacteria_Jul27d5b_22266 | 0 | 0 | 0 | 0 | 0 | 0 | 0 | 0 | 0 | 0 | 0 |
| bacteria_Jul27d5b_25331 | 0 | 0 | 0 | 0 | 0 | 0 | 0 | 0 | 0 | 0 | 0 |
| bacteria_Jul27d5b_26540 | 0 | 0 | 0 | 0 | 0 | 0 | 0 | 0 | 0 | 0 | 0 |
| bacteria_Jul27d5b_29220 | 0 | 0 | 0 | 0 | 0 | 0 | 0 | 0 | 0 | 0 | 0 |
| bacteria_Jul27d5b_32879 | 0 | 0 | 0 | 0 | 0 | 0 | 0 | 0 | 0 | 0 | 0 |
| bacteria_Jul27d5b_34322 | 0 | 0 | 0 | 0 | 0 | 0 | 0 | 0 | 0 | 0 | 0 |
| bacteria_Jul27d5b_35460 | 0 | 0 | 0 | 0 | 0 | 0 | 0 | 0 | 0 | 0 | 0 |
| bacteria_Jul27d5b_40006 | 0 | 0 | 0 | 0 | 0 | 0 | 0 | 0 | 0 | 0 | 0 |
| bacteria_Jul27d5b_42270 | 0 | 0 | 0 | 0 | 0 | 0 | 0 | 0 | 0 | 0 | 0 |
| bacteria_Jul27d5b_42482 | 0 | 0 | 0 | 0 | 0 | 0 | 0 | 0 | 0 | 0 | 0 |
| bacteria_Jul27d5b_49113 | 0 | 0 | 0 | 0 | 0 | 0 | 0 | 0 | 0 | 0 | 0 |
| bacteria_Jul27d5b_49118 | 0 | 0 | 0 | 0 | 0 | 0 | 0 | 0 | 0 | 0 | 0 |
| bacteria_Jul27d25b_1325 | 0 | 0 | 0 | 0 | 0 | 0 | 0 | 0 | 0 | 0 | 0 |
| bacteria_Jul27d25b_5348 | 0 | 0 | 0 | 0 | 0 | 0 | 0 | 0 | 0 | 0 | 0 |
| bacteria_Jul27d25b_6232 | 0 | 0 | 0 | 0 | 0 | 0 | 0 | 0 | 0 | 0 | 0 |
| bacteria_Jul27d25b_10607 | 0 | 0 | 0 | 0 | 0 | 0 | 0 | 0 | 0 | 0 | 0 |
| bacteria_Jul27d25b_11196 | 0 | 0 | 0 | 0 | 0 | 0 | 0 | 0 | 0 | 0 | 0 |
| bacteria_Jul27d25b_15171 | 0 | 0 | 0 | 0 | 0 | 0 | 0 | 0 | 0 | 0 | 0 |
| bacteria_Jul27d25b_15238 | 0 | 0 | 0 | 0 | 0 | 0 | 0 | 0 | 0 | 0 | 0 |
| bacteria_Jul27d25b_19487 | 0 | 0 | 0 | 0 | 0 | 0 | 0 | 0 | 0 | 0 | 0 |
| bacteria_Jul27d25b_25567 | 0 | 0 | 0 | 0 | 0 | 0 | 0 | 0 | 0 | 0 | 0 |
| bacteria_Jul27d25b_27634 | 0 | 0 | 0 | 0 | 0 | 0 | 0 | 0 | 0 | 0 | 0 |
| bacteria_Jul27d25b_33142 | 0 | 0 | 0 | 0 | 0 | 0 | 0 | 0 | 0 | 0 | 0 |
| bacteria_Jul27d25b_34274 | 0 | 0 | 0 | 0 | 0 | 0 | 0 | 0 | 0 | 0 | 0 |
| bacteria_Jul27d25b_37741 | 0 | 0 | 0 | 0 | 0 | 0 | 0 | 0 | 0 | 0 | 0 |
| bacteria_Jul27d25b_38267 | 0 | 1 | 0 | 0 | 0 | 1 | 0 | 0 | 0 | 0 | 2 |
| bacteria_Jul27d25b_38444 | 0 | 0 | 0 | 0 | 0 | 0 | 0 | 0 | 0 | 0 | 0 |
| bacteria_Jul27d25b_43774 | 0 | 0 | 0 | 0 | 0 | 0 | 0 | 0 | 0 | 0 | 0 |
| bacteria_Jul27d25b_48027 | 0 | 0 | 0 | 0 | 0 | 0 | 0 | 0 | 0 | 0 | 0 |
| bacteria_Jul27d25b_52272 | 0 | 0 | 0 | 0 | 0 | 0 | 0 | 0 | 0 | 0 | 0 |
| bacteria_Jul27d25b_52879 | 0 | 0 | 0 | 0 | 0 | 0 | 0 | 0 | 0 | 0 | 0 |
| bacteria_Jul27d25b_57073 | 0 | 0 | 0 | 0 | 0 | 0 | 0 | 0 | 0 | 0 | 0 |
| bacteria_Jul27d25b_58761 | 0 | 0 | 0 | 0 | 0 | 0 | 0 | 0 | 0 | 0 | 0 |
| bacteria_Jul27d25b_66757 | 0 | 0 | 0 | 0 | 0 | 0 | 0 | 0 | 0 | 0 | 0 |
| bacteria_Jul27d25b_68366 | 0 | 0 | 0 | 0 | 0 | 0 | 0 | 0 | 0 | 0 | 0 |
| bacteria_Jul27d25b_69095 | 0 | 0 | 0 | 0 | 0 | 0 | 0 | 0 | 0 | 0 | 0 |
| bacteria_Jul27d25b_72755 | 0 | 0 | 0 | 0 | 0 | 0 | 0 | 0 | 0 | 0 | 0 |
| bacteria_Jul27d75b_533 | 0 | 0 | 0 | 0 | 0 | 0 | 0 | 0 | 0 | 0 | 0 |
| bacteria_Jul27d75b_1878 | 0 | 0 | 0 | 0 | 0 | 0 | 0 | 0 | 0 | 0 | 0 |
| bacteria_Jul27d75b_8694 | 0 | 0 | 0 | 0 | 0 | 0 | 0 | 0 | 0 | 0 | 0 |
| bacteria_Jul27d75b_14934 | 0 | 0 | 0 | 0 | 0 | 0 | 0 | 0 | 0 | 0 | 0 |
| bacteria_Jul27d75b_16893 | 0 | 0 | 0 | 0 | 0 | 0 | 0 | 0 | 0 | 0 | 0 |
| bacteria_Jul27d75b_17360 | 0 | 0 | 0 | 0 | 0 | 0 | 0 | 0 | 0 | 0 | 0 |
| bacteria_Jul27d75b_28236 | 0 | 0 | 0 | 0 | 0 | 0 | 0 | 0 | 0 | 0 | 0 |
| bacteria_Jul27d75b_29408 | 0 | 0 | 0 | 0 | 0 | 0 | 0 | 0 | 0 | 0 | 0 |
| bacteria_Jul27d75b_45905 | 0 | 0 | 0 | 0 | 0 | 0 | 0 | 0 | 0 | 0 | 0 |
| bacteria_Jul27d75b_47691 | 0 | 0 | 0 | 0 | 0 | 0 | 0 | 0 | 0 | 0 | 0 |
| bacteria_Jul27d75b_55391 | 0 | 0 | 0 | 0 | 0 | 0 | 0 | 0 | 0 | 0 | 0 |
| bacteria_Jul27d75b_57228 | 0 | 0 | 0 | 0 | 0 | 0 | 0 | 0 | 0 | 0 | 0 |
| bacteria_Jul27d75b_58869 | 0 | 0 | 0 | 0 | 0 | 0 | 0 | 0 | 0 | 0 | 0 |
| bacteria_Jul27d75b_64293 | 0 | 0 | 0 | 0 | 0 | 0 | 0 | 0 | 0 | 0 | 0 |
| bacteria_Jul27d75b_65719 | 0 | 0 | 0 | 0 | 0 | 0 | 0 | 0 | 0 | 0 | 0 |
| bacteria_Jul27d75b_68359 | 0 | 0 | 0 | 0 | 0 | 0 | 0 | 0 | 0 | 0 | 0 |
| bacteria_Jul27d75b_68843 | 0 | 0 | 0 | 0 | 0 | 0 | 0 | 0 | 0 | 0 | 0 |
| bacteria_Jun8d25b_1031 | 0 | 0 | 0 | 0 | 0 | 0 | 0 | 0 | 0 | 0 | 0 |
| bacteria_Jun8d25b_9502 | 0 | 0 | 0 | 0 | 0 | 0 | 0 | 0 | 0 | 0 | 0 |
| bacteria_Jun8d25b_21844 | 0 | 0 | 0 | 0 | 0 | 0 | 0 | 0 | 0 | 0 | 0 |
| bacteria_Jun8d25b_27181 | 0 | 0 | 0 | 0 | 0 | 0 | 0 | 0 | 0 | 0 | 0 |
| bacteria_Jun8d25b_31268 | 0 | 0 | 0 | 0 | 0 | 0 | 0 | 0 | 0 | 0 | 0 |
| bacteria_Jun8d25b_45582 | 0 | 0 | 0 | 0 | 0 | 0 | 0 | 0 | 0 | 0 | 0 |
| bacteria_Jun8d25b_53591 | 0 | 0 | 0 | 0 | 0 | 0 | 0 | 0 | 0 | 0 | 0 |
| bacteria_Jun8d25b_55128 | 0 | 0 | 0 | 0 | 0 | 0 | 0 | 0 | 0 | 0 | 0 |
| bacteria_Jun8d25b_59492 | 0 | 0 | 0 | 0 | 0 | 0 | 0 | 0 | 0 | 0 | 0 |
| bacteria_Jun8d25b_63591 | 0 | 0 | 0 | 0 | 0 | 0 | 0 | 0 | 0 | 0 | 0 |
| bacteria_Jun8d50b_9577 | 0 | 0 | 0 | 0 | 0 | 0 | 0 | 0 | 0 | 0 | 0 |
| bacteria_Jun8d50b_13763 | 0 | 0 | 0 | 0 | 0 | 0 | 0 | 0 | 0 | 0 | 0 |
| bacteria_Jun8d50b_40756 | 0 | 0 | 0 | 0 | 0 | 0 | 0 | 0 | 0 | 0 | 0 |
| bacteria_Jun8d75b_24465 | 0 | 0 | 0 | 0 | 0 | 0 | 0 | 0 | 0 | 0 | 0 |
| bacteria_Jun22d0b_29245 | 0 | 0 | 0 | 0 | 0 | 0 | 0 | 0 | 0 | 0 | 0 |
| bacteria_Jun22d0b_43496 | 0 | 0 | 0 | 0 | 0 | 0 | 0 | 0 | 0 | 0 | 0 |
| bacteria_Jun22d0b_66483 | 0 | 0 | 0 | 0 | 0 | 0 | 0 | 0 | 0 | 0 | 0 |
| bacteria_Jun22d0b_68415 | 0 | 0 | 0 | 0 | 0 | 0 | 0 | 0 | 0 | 0 | 0 |
| bacteria_Jun22d50b_4976 | 0 | 0 | 0 | 0 | 0 | 0 | 0 | 0 | 0 | 0 | 0 |
| bacteria_Jun22d50b_14818 | 0 | 0 | 0 | 0 | 0 | 0 | 0 | 0 | 0 | 0 | 0 |
| bacteria_Jun22d50b_24014 | 0 | 0 | 0 | 0 | 0 | 0 | 0 | 0 | 0 | 0 | 0 |
| bacteria_Jun22d50b_46532 | 0 | 0 | 0 | 0 | 0 | 0 | 0 | 0 | 0 | 0 | 0 |
| bacteria_Jun22d75b_1740 | 0 | 0 | 0 | 0 | 0 | 0 | 0 | 0 | 0 | 0 | 0 |
| bacteria_Jun22d75b_12653 | 0 | 0 | 0 | 0 | 0 | 0 | 0 | 0 | 0 | 0 | 0 |

| | | | | | | | | | | | |
|---|---|---|---|---|---|---|---|---|---|---|---|
| bacteria_Jun22d75b_13354 | 0 | 0 | 0 | 0 | 0 | 0 | 0 | 0 | 0 | 0 | 0 |
| bacteria_Jun22d75b_19843 | 0 | 0 | 0 | 0 | 0 | 0 | 0 | 0 | 0 | 0 | 0 |
| bacteria_Jun22d75b_28500 | 0 | 0 | 0 | 0 | 0 | 0 | 0 | 0 | 0 | 0 | 0 |
| bacteria_Jun22d75b_34927 | 0 | 0 | 0 | 0 | 0 | 0 | 0 | 0 | 0 | 0 | 0 |
| bacteria_Jun22d75b_71527 | 0 | 0 | 0 | 0 | 0 | 0 | 0 | 0 | 0 | 0 | 0 |
| bacteria_May12d25b_1893 | 0 | 0 | 0 | 0 | 0 | 0 | 0 | 0 | 0 | 0 | 0 |
| bacteria_May12d25b_2249 | 0 | 0 | 0 | 0 | 0 | 0 | 0 | 0 | 0 | 0 | 0 |
| bacteria_May12d25b_2738 | 0 | 0 | 0 | 0 | 0 | 0 | 0 | 0 | 0 | 0 | 0 |
| bacteria_May12d25b_3252 | 0 | 0 | 0 | 0 | 0 | 0 | 0 | 0 | 0 | 0 | 0 |
| bacteria_May12d25b_4353 | 0 | 0 | 0 | 0 | 0 | 0 | 0 | 0 | 0 | 0 | 0 |
| bacteria_May12d25b_4421 | 0 | 0 | 0 | 0 | 0 | 0 | 0 | 0 | 0 | 0 | 0 |
| bacteria_May12d25b_4471 | 0 | 0 | 0 | 0 | 0 | 0 | 0 | 0 | 0 | 0 | 0 |
| bacteria_May12d25b_4991 | 0 | 0 | 0 | 0 | 0 | 0 | 0 | 0 | 0 | 0 | 0 |
| bacteria_May12d25b_5814 | 0 | 0 | 0 | 0 | 0 | 0 | 0 | 0 | 0 | 0 | 0 |
| bacteria_May12d25b_6070 | 0 | 0 | 0 | 0 | 0 | 0 | 0 | 0 | 0 | 0 | 0 |
| bacteria_May12d25b_6835 | 0 | 0 | 0 | 0 | 0 | 0 | 0 | 0 | 0 | 0 | 0 |
| bacteria_May12d50b_1355 | 0 | 0 | 0 | 0 | 0 | 0 | 0 | 0 | 0 | 0 | 0 |
| bacteria_May12d50b_1469 | 0 | 0 | 0 | 0 | 0 | 0 | 0 | 0 | 0 | 0 | 0 |
| bacteria_May12d50b_2730 | 0 | 0 | 0 | 0 | 0 | 0 | 0 | 0 | 0 | 0 | 0 |
| bacteria_May12d50b_2745 | 0 | 0 | 0 | 0 | 0 | 0 | 0 | 0 | 0 | 0 | 0 |
| bacteria_May12d50b_3000 | 0 | 0 | 0 | 0 | 0 | 0 | 0 | 0 | 0 | 0 | 0 |
| bacteria_May12d50b_3081 | 0 | 0 | 0 | 0 | 0 | 0 | 0 | 0 | 0 | 0 | 0 |
| bacteria_May12d50b_3962 | 0 | 0 | 0 | 0 | 0 | 0 | 0 | 0 | 0 | 0 | 0 |
| bacteria_May12d50b_4838 | 0 | 0 | 0 | 0 | 0 | 0 | 0 | 0 | 0 | 0 | 0 |
| bacteria_May12d75b_5382 | 0 | 0 | 0 | 0 | 0 | 0 | 0 | 0 | 0 | 0 | 0 |
| bacteria_Sep8d0b_30961 | 0 | 0 | 0 | 0 | 0 | 0 | 0 | 0 | 0 | 0 | 0 |
| bacteria_Sep8d0b_35968 | 0 | 0 | 0 | 0 | 0 | 0 | 0 | 0 | 0 | 0 | 0 |
| bacteria_Sep8d5b_9448 | 0 | 0 | 0 | 0 | 0 | 0 | 0 | 0 | 0 | 0 | 0 |
| bacteria_Sep8d5b_12068 | 0 | 0 | 0 | 0 | 0 | 0 | 0 | 0 | 0 | 0 | 0 |
| bacteria_Sep8d5b_14374 | 0 | 0 | 0 | 0 | 0 | 0 | 0 | 0 | 0 | 0 | 0 |
| bacteria_Sep8d5b_19536 | 0 | 0 | 0 | 0 | 0 | 0 | 0 | 0 | 0 | 0 | 0 |
| bacteria_Sep8d5b_26480 | 0 | 0 | 0 | 0 | 0 | 0 | 0 | 0 | 0 | 0 | 0 |
| bacteria_Sep8d5b_29126 | 0 | 0 | 0 | 0 | 0 | 0 | 0 | 0 | 0 | 0 | 0 |
| bacteria_Sep8d5b_35637 | 0 | 0 | 0 | 0 | 0 | 0 | 0 | 0 | 0 | 0 | 0 |
| bacteria_Sep8d5b_36022 | 0 | 0 | 0 | 0 | 0 | 0 | 0 | 0 | 0 | 0 | 0 |
| bacteria_Sep8d5b_38265 | 0 | 0 | 0 | 0 | 0 | 0 | 0 | 0 | 0 | 0 | 0 |
| bacteria_Sep8d5b_41465 | 0 | 0 | 0 | 0 | 0 | 0 | 0 | 0 | 0 | 0 | 0 |
| bacteria_Sep8d5b_50190 | 0 | 0 | 0 | 0 | 0 | 0 | 0 | 0 | 0 | 0 | 0 |
| bacteria_Sep8d5b_52952 | 0 | 0 | 0 | 0 | 0 | 0 | 0 | 0 | 0 | 0 | 0 |
| bacteria_Sep8d5b_53435 | 0 | 0 | 0 | 0 | 0 | 0 | 0 | 0 | 0 | 0 | 0 |
| bacteria_Sep8d5b_55699 | 0 | 0 | 0 | 0 | 0 | 0 | 0 | 0 | 0 | 0 | 0 |
| bacteria_Sep8d5b_57191 | 0 | 0 | 0 | 0 | 0 | 0 | 0 | 0 | 0 | 0 | 0 |
| bacteria_Sep8d5b_59121 | 0 | 0 | 0 | 0 | 0 | 0 | 0 | 0 | 0 | 0 | 0 |
| bacteria_Sep8d5b_59920 | 0 | 0 | 0 | 0 | 0 | 0 | 0 | 0 | 0 | 0 | 0 |
| bacteria_Sep8d5b_60883 | 0 | 0 | 0 | 0 | 0 | 0 | 0 | 0 | 0 | 0 | 0 |
| bacteria_Sep8d5b_61021 | 0 | 0 | 0 | 0 | 0 | 0 | 0 | 0 | 0 | 0 | 0 |
| bacteria_Sep8d5b_63599 | 0 | 0 | 0 | 0 | 0 | 0 | 0 | 0 | 0 | 0 | 0 |
| bacteria_Sep8d5b_67463 | 0 | 0 | 0 | 0 | 0 | 0 | 0 | 0 | 0 | 0 | 0 |
| bacteria_Sep8d5b_68026 | 0 | 0 | 0 | 0 | 0 | 0 | 0 | 0 | 0 | 0 | 0 |
| bacteria_Sep8d5b_70353 | 0 | 0 | 0 | 0 | 0 | 0 | 0 | 0 | 0 | 0 | 0 |
| bacteria_Sep8d5b_74335 | 0 | 0 | 0 | 0 | 0 | 0 | 0 | 0 | 0 | 0 | 0 |
| bacteria_Sep8d25b_31062 | 0 | 0 | 0 | 0 | 0 | 0 | 0 | 0 | 0 | 0 | 0 |
| bacteria_Sep8d25b_43934 | 0 | 0 | 0 | 0 | 0 | 0 | 0 | 0 | 0 | 0 | 0 |
| bacteria_Sep8d50b_464 | 0 | 0 | 0 | 0 | 0 | 0 | 0 | 0 | 0 | 0 | 0 |
| virus_Aug5.0v_17 | 0 | 1 | 0 | 0 | 0 | 0 | 0 | 0 | 0 | 0 | 1 |
| virus_Aug5.0v_188 | 0 | 1 | 0 | 0 | 0 | 0 | 0 | 0 | 0 | 0 | 1 |
| virus_Aug5.0v_468 | 0 | 1 | 0 | 0 | 0 | 0 | 0 | 0 | 0 | 0 | 1 |
| virus_Aug5.0v_1188 | 0 | 1 | 0 | 0 | 0 | 0 | 0 | 0 | 0 | 0 | 1 |
| virus_Aug5.0v_1224 | 0 | 1 | 0 | 0 | 0 | 0 | 0 | 0 | 0 | 0 | 1 |
| virus_Aug5.0v_1237 | 0 | 1 | 0 | 0 | 0 | 0 | 0 | 0 | 0 | 0 | 1 |
| virus_Aug5.0v_1632 | 0 | 1 | 0 | 0 | 0 | 0 | 0 | 0 | 0 | 0 | 1 |
| virus_Aug5.0v_1656 | 0 | 1 | 0 | 0 | 0 | 0 | 0 | 0 | 0 | 0 | 1 |
| virus_Aug5.0v_2181 | 0 | 1 | 0 | 0 | 0 | 0 | 0 | 0 | 0 | 0 | 1 |
| virus_Aug5.0v_2555 | 0 | 1 | 0 | 0 | 0 | 0 | 0 | 0 | 0 | 0 | 1 |
| virus_Aug5.0v_2650 | 0 | 1 | 0 | 0 | 0 | 0 | 0 | 0 | 0 | 0 | 1 |
| virus_Aug5.0v_2672 | 0 | 1 | 0 | 0 | 0 | 0 | 0 | 0 | 0 | 0 | 1 |
| virus_Aug5.0v_2718 | 0 | 1 | 0 | 0 | 0 | 0 | 0 | 0 | 0 | 0 | 1 |
| virus_Aug5.0v_2990 | 0 | 1 | 0 | 0 | 0 | 0 | 0 | 0 | 0 | 0 | 1 |
| virus_Aug5.0v_3220 | 0 | 1 | 0 | 0 | 0 | 0 | 0 | 0 | 0 | 0 | 1 |
| virus_Aug5.0v_3323 | 0 | 1 | 0 | 0 | 0 | 0 | 0 | 0 | 0 | 0 | 1 |
| virus_Aug5.0v_3567 | 0 | 1 | 0 | 0 | 0 | 0 | 0 | 0 | 0 | 0 | 1 |
| virus_Aug5.0v_3574 | 0 | 1 | 0 | 0 | 0 | 0 | 0 | 0 | 0 | 0 | 1 |
| virus_Aug5.0v_4127 | 0 | 1 | 0 | 0 | 0 | 0 | 0 | 0 | 0 | 0 | 1 |
| virus_Aug5.0v_4322 | 0 | 1 | 0 | 0 | 0 | 0 | 0 | 0 | 0 | 0 | 1 |
| virus_Aug5.0v_4410 | 0 | 1 | 0 | 0 | 0 | 0 | 0 | 0 | 0 | 0 | 1 |
| virus_Aug5.0v_4542 | 0 | 1 | 0 | 0 | 0 | 0 | 0 | 0 | 0 | 0 | 1 |
| virus_Aug5.0v_4731 | 0 | 1 | 0 | 0 | 0 | 0 | 0 | 0 | 0 | 0 | 1 |
| virus_Aug5.0v_4824 | 0 | 1 | 0 | 0 | 0 | 0 | 0 | 0 | 0 | 0 | 1 |
| virus_Aug5.0v_4956 | 0 | 1 | 0 | 0 | 0 | 0 | 0 | 0 | 0 | 0 | 1 |
| virus_Aug5.0v_5074 | 0 | 1 | 0 | 0 | 0 | 0 | 0 | 0 | 0 | 0 | 1 |
| virus_Aug5.0v_5383 | 0 | 1 | 0 | 0 | 0 | 0 | 0 | 0 | 0 | 0 | 1 |
| virus_Aug5.5v_449 | 0 | 0 | 0 | 1 | 0 | 0 | 0 | 0 | 0 | 0 | 1 |
| virus_Aug5.5v_10746 | 0 | 0 | 0 | 1 | 0 | 0 | 0 | 0 | 0 | 0 | 1 |
| virus_Aug5.5v_12704 | 0 | 0 | 0 | 1 | 0 | 0 | 0 | 0 | 0 | 0 | 1 |
| virus_Aug5.5v_14009 | 0 | 0 | 0 | 1 | 0 | 0 | 0 | 0 | 0 | 0 | 1 |
| virus_Aug5.5v_23090 | 0 | 0 | 0 | 1 | 0 | 0 | 0 | 0 | 0 | 0 | 1 |
| virus_Aug5.5v_26095 | 0 | 0 | 0 | 1 | 0 | 0 | 0 | 0 | 0 | 0 | 1 |
| virus_Aug5.5v_26523 | 0 | 0 | 0 | 1 | 0 | 0 | 0 | 0 | 0 | 0 | 1 |
| virus_Aug5.5v_27701 | 0 | 0 | 0 | 1 | 0 | 0 | 0 | 0 | 0 | 0 | 1 |
| virus_Aug5.5v_28182 | 0 | 0 | 0 | 1 | 0 | 0 | 0 | 0 | 0 | 0 | 1 |
| virus_Aug5.5v_31422 | 0 | 0 | 0 | 1 | 0 | 0 | 0 | 0 | 0 | 0 | 1 |
| virus_Aug5.5v_35456 | 0 | 0 | 0 | 1 | 0 | 0 | 0 | 0 | 0 | 0 | 1 |
| virus_Aug5.5v_38006 | 0 | 0 | 0 | 1 | 0 | 0 | 0 | 0 | 0 | 0 | 1 |
| virus_Aug5.5v_40425 | 0 | 0 | 0 | 1 | 0 | 0 | 0 | 0 | 0 | 0 | 1 |
| virus_Aug5.25v_4744 | 0 | 0 | 0 | 0 | 0 | 1 | 0 | 0 | 0 | 0 | 1 |
| virus_Aug5.25v_6835 | 0 | 0 | 0 | 0 | 0 | 1 | 0 | 0 | 0 | 0 | 1 |
| virus_Aug5.25v_8038 | 0 | 0 | 0 | 0 | 0 | 1 | 0 | 0 | 0 | 0 | 1 |
| virus_Aug5.25v_8334 | 0 | 0 | 0 | 0 | 0 | 1 | 0 | 0 | 0 | 0 | 1 |
| virus_Aug5.25v_10634 | 0 | 0 | 0 | 0 | 0 | 1 | 0 | 0 | 0 | 0 | 1 |
| virus_Aug5.25v_12087 | 0 | 0 | 0 | 0 | 0 | 1 | 0 | 0 | 0 | 0 | 1 |
| virus_Aug5.25v_12350 | 0 | 0 | 0 | 0 | 0 | 1 | 0 | 0 | 0 | 0 | 1 |
| virus_Aug5.25v_14138 | 0 | 0 | 0 | 0 | 0 | 1 | 0 | 0 | 0 | 0 | 1 |
| virus_Aug5.25v_18338 | 0 | 0 | 0 | 0 | 0 | 1 | 0 | 0 | 0 | 0 | 1 |
| virus_Aug5.25v_20746 | 0 | 0 | 0 | 0 | 0 | 1 | 0 | 0 | 0 | 0 | 1 |
| virus_Aug5.25v_20934 | 0 | 0 | 0 | 0 | 0 | 1 | 0 | 0 | 0 | 0 | 1 |
| virus_Aug5.25v_22083 | 0 | 0 | 0 | 0 | 0 | 1 | 0 | 0 | 0 | 0 | 1 |
| virus_Aug5.25v_24775 | 0 | 0 | 0 | 0 | 0 | 1 | 0 | 0 | 0 | 0 | 1 |
| virus_Aug5.25v_27289 | 0 | 0 | 0 | 0 | 0 | 1 | 0 | 0 | 0 | 0 | 1 |
| virus_Aug5.25v_31155 | 0 | 0 | 0 | 0 | 0 | 1 | 0 | 0 | 0 | 0 | 1 |
| virus_Aug5.25v_32749 | 0 | 0 | 0 | 0 | 0 | 1 | 0 | 0 | 0 | 0 | 1 |
| virus_Aug5.25v_32847 | 0 | 0 | 0 | 0 | 0 | 1 | 0 | 0 | 0 | 0 | 1 |
| virus_Aug5.25v_32890 | 0 | 0 | 0 | 0 | 0 | 1 | 0 | 0 | 0 | 0 | 1 |
| virus_Aug5.25v_33426 | 0 | 0 | 0 | 0 | 0 | 1 | 0 | 0 | 0 | 0 | 1 |
| virus_Aug5.25v_35473 | 0 | 0 | 0 | 0 | 0 | 1 | 0 | 0 | 0 | 0 | 1 |
| virus_Aug5.25v_35775 | 0 | 0 | 0 | 0 | 0 | 1 | 0 | 0 | 0 | 0 | 1 |
| virus_Aug5.50v_78 | 0 | 0 | 0 | 0 | 0 | 0 | 0 | 1 | 0 | 0 | 1 |
| virus_Aug5.50v_7705 | 0 | 0 | 0 | 0 | 0 | 0 | 0 | 1 | 0 | 0 | 1 |
| virus_Aug5.50v_21180 | 0 | 0 | 0 | 0 | 0 | 0 | 0 | 1 | 0 | 0 | 1 |
| virus_Aug5.50v_22303 | 0 | 0 | 0 | 0 | 0 | 0 | 0 | 1 | 0 | 0 | 1 |
| virus_Aug5.50v_28901 | 0 | 0 | 0 | 0 | 0 | 0 | 0 | 1 | 0 | 0 | 1 |
| virus_Aug5.50v_31718 | 0 | 0 | 0 | 0 | 0 | 0 | 0 | 1 | 0 | 0 | 1 |
| virus_Aug5.50v_35175 | 0 | 0 | 0 | 0 | 0 | 0 | 0 | 1 | 0 | 0 | 1 |
| virus_Aug5.50v_38181 | 0 | 0 | 0 | 0 | 0 | 0 | 0 | 1 | 0 | 0 | 1 |
| virus_Aug5.75v_4962 | 0 | 0 | 0 | 0 | 0 | 0 | 0 | 0 | 0 | 1 | 1 |
| virus_Aug5.75v_7742 | 0 | 0 | 0 | 0 | 0 | 0 | 0 | 0 | 0 | 1 | 1 |
| virus_Aug5.75v_15194 | 0 | 0 | 0 | 0 | 0 | 0 | 0 | 0 | 0 | 1 | 1 |
| virus_Aug5.75v_16458 | 0 | 0 | 0 | 0 | 0 | 0 | 0 | 0 | 0 | 1 | 1 |
| virus_Aug5.75v_26667 | 0 | 0 | 0 | 0 | 0 | 0 | 0 | 0 | 0 | 1 | 1 |
| virus_Aug5.75v_31974 | 0 | 0 | 0 | 0 | 0 | 0 | 0 | 0 | 0 | 1 | 1 |
| virus_Aug5.75v_32037 | 0 | 0 | 0 | 0 | 0 | 0 | 0 | 0 | 0 | 1 | 1 |
| virus_Jul11.0v_1929 | 0 | 0 | 0 | 0 | 0 | 0 | 0 | 0 | 0 | 0 | 0 |
| virus_Jul11.0v_3574 | 0 | 0 | 0 | 0 | 0 | 0 | 0 | 0 | 0 | 0 | 0 |
| virus_Jul11.0v_3581 | 0 | 0 | 0 | 0 | 0 | 0 | 0 | 0 | 0 | 0 | 0 |
| virus_Jul11.0v_8858 | 0 | 0 | 0 | 1 | 0 | 0 | 0 | 0 | 0 | 0 | 1 |
| virus_Jul11.0v_10567 | 0 | 0 | 0 | 0 | 0 | 0 | 0 | 0 | 0 | 0 | 0 |
| virus_Jul11.0v_14863 | 0 | 0 | 0 | 0 | 0 | 0 | 0 | 0 | 0 | 0 | 0 |
| virus_Jul11.0v_19251 | 0 | 0 | 0 | 0 | 0 | 0 | 0 | 0 | 0 | 0 | 0 |
| virus_Jul11.0v_20973 | 0 | 0 | 0 | 0 | 0 | 0 | 0 | 0 | 0 | 0 | 0 |
| virus_Jul11.5v_3195 | 0 | 0 | 0 | 0 | 0 | 0 | 0 | 0 | 0 | 0 | 0 |
| virus_Jul11.5v_6366 | 0 | 0 | 0 | 0 | 0 | 0 | 0 | 0 | 0 | 0 | 0 |
| virus_Jul11.5v_8622 | 0 | 0 | 0 | 0 | 0 | 0 | 0 | 0 | 0 | 0 | 0 |
| virus_Jul11.5v_9693 | 0 | 0 | 0 | 0 | 0 | 0 | 0 | 0 | 0 | 0 | 0 |
| virus_Jul11.5v_10872 | 0 | 0 | 0 | 0 | 0 | 0 | 0 | 0 | 0 | 0 | 0 |
| virus_Jul11.5v_12056 | 0 | 0 | 0 | 0 | 0 | 0 | 0 | 0 | 0 | 0 | 0 |
| virus_Jul11.5v_12999 | 0 | 0 | 0 | 0 | 0 | 0 | 0 | 0 | 0 | 0 | 0 |
| virus_Jul11.5v_15655 | 0 | 0 | 0 | 0 | 0 | 0 | 0 | 0 | 0 | 0 | 0 |
| virus_Jul11.5v_18759 | 0 | 0 | 0 | 0 | 0 | 0 | 0 | 0 | 0 | 0 | 0 |
| virus_Jul11.5v_19132 | 0 | 0 | 0 | 0 | 0 | 0 | 0 | 0 | 0 | 0 | 0 |
| virus_Jul11.25v_858 | 0 | 0 | 0 | 0 | 0 | 0 | 0 | 0 | 0 | 0 | 0 |
| virus_Jul11.25v_4155 | 0 | 0 | 0 | 0 | 0 | 0 | 0 | 0 | 0 | 0 | 0 |
| virus_Jul11.25v_7357 | 0 | 0 | 0 | 0 | 0 | 0 | 0 | 0 | 0 | 0 | 0 |
| virus_Jul11.25v_9931 | 0 | 0 | 0 | 0 | 0 | 0 | 0 | 0 | 0 | 0 | 0 |
| virus_Jul11.25v_14451 | 0 | 0 | 0 | 0 | 0 | 0 | 0 | 0 | 0 | 0 | 0 |
| virus_Jul11.25v_15347 | 0 | 0 | 0 | 0 | 0 | 0 | 0 | 0 | 0 | 0 | 0 |
| virus_Jul11.25v_16147 | 0 | 0 | 0 | 0 | 0 | 0 | 0 | 0 | 0 | 0 | 0 |
| virus_Jul11.25v_17097 | 0 | 0 | 0 | 0 | 0 | 0 | 0 | 0 | 0 | 0 | 0 |
| virus_Jul11.25v_18940 | 0 | 0 | 0 | 0 | 0 | 0 | 0 | 0 | 0 | 0 | 0 |

| | | | | | | | | | | | |
|---|---|---|---|---|---|---|---|---|---|---|---|
| virus_Jul11.25v_19468 | 0 | 0 | 0 | 0 | 0 | 0 | 0 | 0 | 0 | 0 | 0 |
| virus_Jul11.50v_514 | 0 | 0 | 0 | 0 | 0 | 0 | 0 | 0 | 0 | 0 | 0 |
| virus_Jul11.50v_4248 | 0 | 0 | 0 | 0 | 0 | 0 | 0 | 0 | 0 | 0 | 0 |
| virus_Jul11.50v_4694 | 0 | 0 | 0 | 0 | 0 | 0 | 0 | 0 | 0 | 0 | 0 |
| virus_Jul11.50v_6169 | 0 | 0 | 0 | 0 | 0 | 0 | 0 | 0 | 0 | 0 | 0 |
| virus_Jul11.50v_6448 | 0 | 0 | 0 | 0 | 0 | 0 | 0 | 0 | 0 | 0 | 0 |
| virus_Jul11.50v_7910 | 0 | 0 | 0 | 0 | 0 | 0 | 0 | 0 | 0 | 0 | 0 |
| virus_Jul11.50v_8038 | 0 | 0 | 0 | 0 | 0 | 0 | 0 | 0 | 0 | 0 | 0 |
| virus_Jul11.50v_8950 | 0 | 0 | 0 | 0 | 0 | 0 | 0 | 0 | 0 | 0 | 0 |
| virus_Jul11.50v_10028 | 0 | 0 | 0 | 0 | 0 | 0 | 0 | 0 | 0 | 0 | 0 |
| virus_Jul11.50v_11322 | 0 | 0 | 0 | 0 | 0 | 0 | 0 | 0 | 0 | 0 | 0 |
| virus_Jul11.50v_12821 | 0 | 0 | 0 | 0 | 0 | 0 | 0 | 0 | 0 | 0 | 0 |
| virus_Jul11.50v_14197 | 0 | 0 | 0 | 0 | 0 | 0 | 0 | 0 | 0 | 0 | 0 |
| virus_Jul11.50v_14384 | 0 | 0 | 0 | 0 | 0 | 0 | 0 | 0 | 0 | 0 | 0 |
| virus_Jul11.50v_18584 | 0 | 0 | 0 | 0 | 0 | 0 | 0 | 0 | 0 | 0 | 0 |
| virus_Jul11.50v_18898 | 0 | 0 | 0 | 0 | 0 | 0 | 0 | 0 | 0 | 0 | 0 |
| virus_Jul11.75v_11827 | 0 | 0 | 0 | 0 | 0 | 0 | 0 | 0 | 0 | 0 | 0 |
| virus_Jul11.75v_18794 | 0 | 0 | 0 | 0 | 0 | 0 | 0 | 0 | 0 | 0 | 0 |
| virus_Jul11.75v_25365 | 0 | 0 | 0 | 0 | 0 | 0 | 0 | 0 | 0 | 0 | 0 |
| virus_Jul11.75v_26150 | 0 | 0 | 0 | 0 | 0 | 0 | 0 | 0 | 0 | 0 | 0 |
| virus_Jul11.75v_26801 | 0 | 0 | 0 | 0 | 0 | 0 | 0 | 0 | 0 | 0 | 0 |
| virus_Jul11.75v_36647 | 0 | 0 | 0 | 0 | 0 | 0 | 0 | 0 | 0 | 0 | 0 |
| virus_Jul27.5v_37 | 0 | 0 | 0 | 0 | 0 | 0 | 0 | 0 | 0 | 0 | 0 |
| virus_Jul27.5v_186 | 0 | 0 | 0 | 0 | 0 | 0 | 0 | 0 | 0 | 0 | 0 |
| virus_Jul27.5v_1242 | 0 | 0 | 0 | 0 | 0 | 0 | 0 | 0 | 0 | 0 | 0 |
| virus_Jul27.5v_2891 | 0 | 0 | 0 | 0 | 0 | 0 | 0 | 0 | 0 | 0 | 0 |
| virus_Jul27.5v_3224 | 0 | 0 | 0 | 0 | 0 | 0 | 0 | 0 | 0 | 0 | 0 |
| virus_Jul27.5v_3497 | 0 | 0 | 0 | 0 | 0 | 0 | 0 | 0 | 0 | 0 | 0 |
| virus_Jul27.5v_3834 | 0 | 0 | 0 | 0 | 0 | 0 | 0 | 0 | 0 | 0 | 0 |
| virus_Jul27.5v_3886 | 0 | 0 | 0 | 0 | 0 | 0 | 0 | 0 | 0 | 0 | 0 |
| virus_Jul27.5v_5273 | 0 | 0 | 0 | 0 | 0 | 0 | 0 | 0 | 0 | 0 | 0 |
| virus_Jul27.5v_6098 | 0 | 0 | 0 | 0 | 0 | 0 | 0 | 0 | 0 | 0 | 0 |
| virus_Jul27.5v_9291 | 0 | 0 | 0 | 0 | 0 | 0 | 0 | 0 | 0 | 0 | 0 |
| virus_Jul27.5v_10821 | 0 | 0 | 0 | 0 | 0 | 0 | 0 | 0 | 0 | 0 | 0 |
| virus_Jul27.5v_11134 | 0 | 0 | 0 | 0 | 0 | 0 | 0 | 0 | 0 | 0 | 0 |
| virus_Jul27.5v_11482 | 0 | 0 | 0 | 0 | 0 | 0 | 0 | 0 | 0 | 0 | 0 |
| virus_Jul27.5v_12822 | 0 | 0 | 0 | 0 | 0 | 0 | 0 | 0 | 0 | 0 | 0 |
| virus_Jul27.5v_12864 | 0 | 0 | 0 | 0 | 0 | 0 | 0 | 0 | 0 | 0 | 0 |
| virus_Jul27.5v_13223 | 0 | 0 | 0 | 0 | 0 | 0 | 0 | 0 | 0 | 0 | 0 |
| virus_Jul27.5v_13230 | 0 | 0 | 0 | 0 | 0 | 0 | 0 | 0 | 0 | 0 | 0 |
| virus_Jul27.5v_14174 | 0 | 0 | 0 | 0 | 0 | 0 | 0 | 0 | 0 | 0 | 0 |
| virus_Jul27.5v_17154 | 0 | 0 | 0 | 0 | 0 | 0 | 0 | 0 | 0 | 0 | 0 |
| virus_Jul27.5v_20216 | 0 | 0 | 0 | 0 | 0 | 0 | 0 | 0 | 0 | 0 | 0 |
| virus_Jul27.5v_21090 | 0 | 0 | 0 | 0 | 0 | 0 | 0 | 0 | 0 | 0 | 0 |
| virus_Jul27.25v_2860 | 0 | 0 | 0 | 0 | 0 | 0 | 0 | 0 | 0 | 0 | 0 |
| virus_Jul27.25v_5587 | 0 | 0 | 0 | 0 | 0 | 0 | 0 | 0 | 0 | 0 | 0 |
| virus_Jul27.25v_8794 | 0 | 0 | 0 | 1 | 0 | 0 | 0 | 0 | 0 | 0 | 1 |
| virus_Jul27.25v_13389 | 0 | 0 | 0 | 0 | 0 | 0 | 0 | 0 | 0 | 0 | 0 |
| virus_Jul27.25v_13523 | 0 | 0 | 0 | 0 | 0 | 0 | 0 | 0 | 0 | 0 | 0 |
| virus_Jul27.25v_14334 | 0 | 0 | 0 | 0 | 0 | 0 | 0 | 0 | 0 | 0 | 0 |
| virus_Jul27.25v_21311 | 0 | 0 | 0 | 0 | 0 | 0 | 0 | 0 | 0 | 0 | 0 |
| virus_Jul27.25v_21411 | 0 | 0 | 0 | 0 | 0 | 0 | 0 | 0 | 0 | 0 | 0 |
| virus_Jul27.25v_22225 | 0 | 0 | 0 | 0 | 0 | 0 | 0 | 0 | 0 | 0 | 0 |
| virus_Jul27.25v_26383 | 0 | 0 | 0 | 0 | 0 | 0 | 0 | 0 | 0 | 0 | 0 |
| virus_Jul27.25v_28250 | 0 | 0 | 0 | 0 | 0 | 0 | 0 | 0 | 0 | 0 | 0 |
| virus_Jul27.25v_29812 | 0 | 0 | 0 | 0 | 0 | 0 | 0 | 0 | 0 | 0 | 0 |
| virus_Jul27.25v_34443 | 0 | 0 | 0 | 0 | 0 | 0 | 0 | 0 | 0 | 0 | 0 |
| virus_Jul27.75v_898 | 0 | 0 | 0 | 0 | 0 | 0 | 0 | 0 | 0 | 0 | 0 |
| virus_Jul27.75v_8322 | 0 | 0 | 0 | 0 | 0 | 0 | 0 | 0 | 0 | 0 | 0 |
| virus_Jul27.75v_10122 | 0 | 0 | 0 | 0 | 0 | 0 | 0 | 0 | 0 | 0 | 0 |
| virus_Jul27.75v_10776 | 0 | 0 | 0 | 0 | 0 | 0 | 0 | 0 | 0 | 0 | 0 |
| virus_Jul27.75v_14092 | 0 | 0 | 0 | 0 | 0 | 0 | 0 | 0 | 0 | 0 | 0 |
| virus_Jul27.75v_18316 | 0 | 0 | 0 | 0 | 0 | 0 | 0 | 0 | 0 | 0 | 0 |
| virus_Jul27.75v_19148 | 0 | 0 | 0 | 0 | 0 | 0 | 0 | 0 | 0 | 0 | 0 |
| virus_Jun8.25v_16698 | 0 | 0 | 0 | 0 | 0 | 0 | 0 | 0 | 0 | 0 | 0 |
| virus_Jun8.25v_26408 | 0 | 0 | 0 | 0 | 0 | 0 | 0 | 0 | 0 | 0 | 0 |
| virus_Jun8.25v_27945 | 0 | 0 | 0 | 0 | 0 | 0 | 0 | 0 | 0 | 0 | 0 |
| virus_Jun8.25v_29954 | 0 | 0 | 0 | 0 | 0 | 0 | 0 | 0 | 0 | 0 | 0 |
| virus_Jun8.50v_8853 | 0 | 0 | 0 | 0 | 0 | 0 | 0 | 0 | 0 | 0 | 0 |
| virus_Jun8.50v_13689 | 0 | 0 | 0 | 0 | 0 | 0 | 0 | 0 | 0 | 0 | 0 |
| virus_Jun8.50v_17712 | 0 | 0 | 0 | 0 | 0 | 0 | 0 | 0 | 0 | 0 | 0 |
| virus_Jun8.50v_20481 | 0 | 0 | 0 | 0 | 0 | 0 | 0 | 0 | 0 | 0 | 0 |
| virus_Jun8.50v_27147 | 0 | 0 | 0 | 0 | 0 | 0 | 0 | 0 | 0 | 0 | 0 |
| virus_Jun8.50v_29052 | 0 | 0 | 0 | 0 | 0 | 0 | 0 | 0 | 0 | 0 | 0 |
| virus_Jun8.50v_29123 | 0 | 0 | 0 | 0 | 0 | 0 | 0 | 0 | 0 | 0 | 0 |
| virus_Jun8.50v_32615 | 0 | 0 | 0 | 0 | 0 | 0 | 0 | 0 | 0 | 0 | 0 |
| virus_Jun8.50v_35056 | 0 | 0 | 0 | 0 | 0 | 0 | 0 | 0 | 0 | 0 | 0 |
| virus_Jun8.50v_35836 | 0 | 0 | 0 | 0 | 0 | 0 | 0 | 0 | 0 | 0 | 0 |
| virus_Jun8.50v_36178 | 0 | 0 | 0 | 0 | 0 | 0 | 0 | 0 | 0 | 0 | 0 |
| virus_Jun8.75v_596 | 0 | 0 | 0 | 0 | 0 | 0 | 0 | 0 | 0 | 0 | 0 |
| virus_Jun8.75v_2130 | 0 | 0 | 0 | 0 | 0 | 0 | 0 | 0 | 0 | 0 | 0 |
| virus_Jun8.75v_2661 | 0 | 0 | 0 | 0 | 0 | 0 | 0 | 0 | 0 | 0 | 0 |
| virus_Jun8.75v_20642 | 0 | 0 | 0 | 0 | 0 | 0 | 0 | 0 | 0 | 0 | 0 |
| virus_Jun22.0v_397 | 0 | 0 | 0 | 0 | 0 | 0 | 0 | 0 | 0 | 0 | 0 |
| virus_Jun22.0v_2785 | 0 | 0 | 0 | 0 | 0 | 0 | 0 | 0 | 0 | 0 | 0 |
| virus_Jun22.0v_4319 | 0 | 0 | 0 | 0 | 0 | 0 | 0 | 0 | 0 | 0 | 0 |
| virus_Jun22.0v_5997 | 0 | 0 | 0 | 0 | 0 | 0 | 0 | 0 | 0 | 0 | 0 |
| virus_Jun22.0v_6860 | 0 | 0 | 0 | 0 | 0 | 0 | 0 | 0 | 0 | 0 | 0 |
| virus_Jun22.0v_7074 | 0 | 0 | 0 | 0 | 0 | 0 | 0 | 0 | 0 | 0 | 0 |
| virus_Jun22.0v_7652 | 0 | 0 | 0 | 0 | 0 | 0 | 0 | 0 | 0 | 0 | 0 |
| virus_Jun22.0v_12106 | 0 | 0 | 0 | 0 | 0 | 0 | 0 | 0 | 0 | 0 | 0 |
| virus_Jun22.0v_16886 | 0 | 0 | 0 | 0 | 0 | 0 | 0 | 0 | 0 | 0 | 0 |
| virus_Jun22.0v_16908 | 0 | 0 | 0 | 0 | 0 | 0 | 0 | 0 | 0 | 0 | 0 |
| virus_Jun22.0v_17757 | 0 | 0 | 0 | 0 | 0 | 0 | 0 | 0 | 0 | 0 | 0 |
| virus_Jun22.0v_17979 | 0 | 0 | 0 | 0 | 0 | 0 | 0 | 0 | 0 | 0 | 0 |
| virus_Jun22.0v_19010 | 0 | 0 | 0 | 0 | 0 | 0 | 0 | 0 | 0 | 0 | 0 |
| virus_Jun22.0v_19116 | 0 | 0 | 0 | 0 | 0 | 0 | 0 | 0 | 0 | 0 | 0 |
| virus_Jun22.0v_19212 | 0 | 0 | 0 | 0 | 0 | 0 | 0 | 0 | 0 | 0 | 0 |
| virus_Jun22.50v_170 | 0 | 0 | 0 | 0 | 0 | 0 | 0 | 0 | 0 | 0 | 0 |
| virus_Jun22.50v_754 | 0 | 0 | 0 | 0 | 0 | 0 | 0 | 0 | 0 | 0 | 0 |
| virus_Jun22.50v_9417 | 0 | 0 | 0 | 0 | 0 | 0 | 0 | 0 | 0 | 0 | 0 |
| virus_Jun22.50v_16142 | 0 | 0 | 0 | 0 | 0 | 0 | 0 | 0 | 0 | 0 | 0 |
| virus_Jun22.50v_16177 | 0 | 0 | 0 | 0 | 0 | 0 | 0 | 0 | 0 | 0 | 0 |
| virus_Jun22.50v_17142 | 0 | 0 | 0 | 0 | 0 | 0 | 0 | 0 | 0 | 0 | 0 |
| virus_Jun22.50v_18075 | 0 | 0 | 0 | 0 | 0 | 0 | 0 | 0 | 0 | 0 | 0 |
| virus_Jun22.50v_21417 | 0 | 0 | 0 | 0 | 0 | 0 | 0 | 0 | 0 | 0 | 0 |
| virus_Jun22.50v_22685 | 0 | 0 | 0 | 0 | 0 | 0 | 0 | 0 | 0 | 0 | 0 |
| virus_Jun22.50v_23272 | 0 | 0 | 0 | 0 | 0 | 0 | 0 | 0 | 0 | 0 | 0 |
| virus_Jun22.50v_27084 | 0 | 0 | 0 | 0 | 0 | 0 | 0 | 0 | 0 | 0 | 0 |
| virus_Jun22.50v_27535 | 0 | 0 | 0 | 0 | 0 | 0 | 0 | 0 | 0 | 0 | 0 |
| virus_Jun22.50v_29457 | 0 | 0 | 0 | 0 | 0 | 0 | 0 | 0 | 0 | 0 | 0 |
| virus_Jun22.50v_29617 | 0 | 0 | 0 | 0 | 0 | 0 | 0 | 0 | 0 | 0 | 0 |
| virus_Jun22.50v_32780 | 0 | 0 | 0 | 0 | 0 | 0 | 0 | 0 | 0 | 0 | 0 |
| virus_Jun22.50v_33897 | 0 | 0 | 0 | 0 | 0 | 1 | 0 | 0 | 0 | 0 | 1 |
| virus_Jun22.75v_2016 | 0 | 0 | 0 | 0 | 0 | 0 | 0 | 0 | 0 | 0 | 0 |
| virus_Jun22.75v_5295 | 0 | 0 | 0 | 0 | 0 | 0 | 0 | 0 | 0 | 0 | 0 |
| virus_Jun22.75v_5416 | 0 | 0 | 0 | 0 | 0 | 0 | 0 | 0 | 0 | 0 | 0 |
| virus_Jun22.75v_13437 | 0 | 0 | 0 | 0 | 0 | 0 | 0 | 0 | 0 | 0 | 0 |
| virus_Jun22.75v_21990 | 0 | 0 | 0 | 0 | 0 | 0 | 0 | 0 | 0 | 0 | 0 |
| virus_Jun22.75v_22208 | 0 | 0 | 0 | 0 | 0 | 0 | 0 | 0 | 0 | 0 | 0 |
| virus_Jun22.75v_23111 | 0 | 0 | 0 | 0 | 0 | 0 | 0 | 0 | 0 | 0 | 0 |
| virus_Jun22.75v_37444 | 0 | 0 | 0 | 0 | 0 | 0 | 0 | 0 | 0 | 0 | 0 |
| virus_Jun22.75v_37735 | 0 | 0 | 0 | 0 | 0 | 0 | 0 | 0 | 0 | 0 | 0 |
| virus_May12.5v_10352 | 0 | 0 | 0 | 0 | 0 | 0 | 0 | 0 | 0 | 0 | 0 |
| virus_May12.5v_12219 | 0 | 0 | 0 | 0 | 0 | 0 | 0 | 0 | 0 | 0 | 0 |
| virus_May12.5v_20341 | 0 | 0 | 0 | 0 | 0 | 0 | 0 | 0 | 0 | 0 | 0 |
| virus_May12.5v_24933 | 0 | 0 | 0 | 0 | 0 | 0 | 0 | 0 | 0 | 0 | 0 |
| virus_May12.5v_29781 | 0 | 0 | 0 | 0 | 0 | 0 | 0 | 0 | 0 | 0 | 0 |
| virus_May12.5v_33866 | 0 | 0 | 0 | 0 | 0 | 0 | 0 | 0 | 0 | 0 | 0 |
| virus_May12.5v_35782 | 0 | 0 | 0 | 0 | 0 | 0 | 0 | 0 | 0 | 0 | 0 |
| virus_May12.25v_3512 | 0 | 0 | 0 | 0 | 0 | 0 | 0 | 0 | 0 | 0 | 0 |
| virus_May12.25v_9922 | 0 | 0 | 0 | 0 | 0 | 0 | 0 | 0 | 0 | 0 | 0 |
| virus_May12.25v_13925 | 0 | 0 | 0 | 0 | 0 | 0 | 0 | 0 | 0 | 0 | 0 |
| virus_May12.25v_23099 | 0 | 0 | 0 | 0 | 0 | 0 | 0 | 0 | 0 | 0 | 0 |
| virus_May12.25v_23907 | 0 | 0 | 0 | 0 | 0 | 0 | 0 | 0 | 0 | 0 | 0 |
| virus_May12.25v_25260 | 0 | 0 | 0 | 0 | 0 | 0 | 0 | 0 | 0 | 0 | 0 |
| virus_May12.25v_26035 | 0 | 0 | 0 | 0 | 0 | 0 | 0 | 0 | 0 | 0 | 0 |
| virus_May12.50v_2829 | 0 | 0 | 0 | 0 | 0 | 0 | 0 | 0 | 0 | 0 | 0 |
| virus_May12.50v_3419 | 0 | 0 | 0 | 0 | 0 | 0 | 0 | 0 | 0 | 0 | 0 |
| virus_May12.50v_5753 | 0 | 0 | 0 | 0 | 0 | 0 | 0 | 0 | 0 | 0 | 0 |
| virus_May12.50v_6335 | 0 | 0 | 0 | 0 | 0 | 0 | 0 | 0 | 0 | 0 | 0 |
| virus_May12.50v_6736 | 0 | 0 | 0 | 0 | 0 | 0 | 0 | 0 | 0 | 0 | 0 |
| virus_May12.50v_10133 | 0 | 0 | 0 | 0 | 0 | 0 | 0 | 0 | 0 | 0 | 0 |
| virus_May12.50v_11498 | 0 | 0 | 0 | 0 | 0 | 0 | 0 | 0 | 0 | 0 | 0 |
| virus_May12.50v_16371 | 0 | 0 | 0 | 0 | 0 | 0 | 0 | 0 | 0 | 0 | 0 |
| virus_May12.50v_18048 | 0 | 0 | 0 | 0 | 0 | 0 | 0 | 0 | 0 | 0 | 0 |
| virus_May12.50v_19478 | 0 | 0 | 0 | 0 | 0 | 0 | 0 | 0 | 0 | 0 | 0 |
| virus_May12.50v_19686 | 0 | 0 | 0 | 0 | 0 | 0 | 0 | 0 | 0 | 0 | 0 |
| virus_May12.50v_22318 | 0 | 0 | 0 | 0 | 0 | 0 | 0 | 0 | 0 | 0 | 0 |
| virus_May12.50v_29146 | 0 | 0 | 0 | 0 | 0 | 0 | 0 | 0 | 0 | 0 | 0 |
| virus_May12.50v_32716 | 0 | 0 | 0 | 0 | 0 | 0 | 0 | 0 | 0 | 0 | 0 |
| virus_May12.50v_34294 | 0 | 0 | 0 | 0 | 0 | 0 | 0 | 0 | 0 | 0 | 0 |
| virus_May12.50v_38650 | 0 | 0 | 0 | 0 | 0 | 0 | 0 | 0 | 0 | 0 | 0 |
| virus_May12.75v_4038 | 0 | 0 | 0 | 0 | 0 | 0 | 0 | 0 | 0 | 0 | 0 |
| virus_May12.75v_5309 | 0 | 0 | 0 | 0 | 0 | 0 | 0 | 0 | 0 | 0 | 0 |
| virus_May12.75v_6936 | 0 | 0 | 0 | 0 | 0 | 0 | 0 | 0 | 0 | 0 | 0 |
| virus_May12.75v_21233 | 0 | 0 | 0 | 0 | 0 | 0 | 0 | 0 | 0 | 0 | 0 |

| | | | | | | | | | | | |
|---|---|---|---|---|---|---|---|---|---|---|---|
| virus_May12.75v_23655 | 0 | 0 | 0 | 0 | 0 | 0 | 0 | 0 | 0 | 0 | 0 |
| virus_May12.75v_26000 | 0 | 0 | 0 | 0 | 0 | 0 | 0 | 0 | 0 | 0 | 0 |
| virus_May12.75v_29927 | 0 | 0 | 0 | 0 | 0 | 0 | 0 | 0 | 0 | 0 | 0 |
| virus_May12.75v_31407 | 0 | 0 | 0 | 0 | 0 | 0 | 0 | 0 | 0 | 0 | 0 |
| virus_May12.75v_31866 | 0 | 0 | 0 | 0 | 0 | 0 | 0 | 0 | 0 | 0 | 0 |
| virus_May12.75v_34385 | 0 | 0 | 0 | 0 | 0 | 0 | 0 | 0 | 0 | 0 | 0 |
| virus_Sep8.0v_4725 | 0 | 0 | 0 | 0 | 0 | 0 | 0 | 0 | 0 | 0 | 0 |
| virus_Sep8.0v_11196 | 0 | 0 | 0 | 0 | 0 | 0 | 0 | 0 | 0 | 0 | 0 |
| virus_Sep8.0v_11708 | 0 | 0 | 0 | 0 | 0 | 0 | 0 | 0 | 0 | 0 | 0 |
| virus_Sep8.0v_13501 | 0 | 0 | 0 | 0 | 0 | 0 | 0 | 0 | 0 | 0 | 0 |
| virus_Sep8.0v_14400 | 0 | 0 | 0 | 0 | 0 | 0 | 0 | 0 | 0 | 0 | 0 |
| virus_Sep8.0v_15001 | 0 | 0 | 0 | 0 | 0 | 0 | 0 | 0 | 0 | 0 | 0 |
| virus_Sep8.0v_19519 | 0 | 0 | 0 | 0 | 0 | 0 | 0 | 0 | 0 | 0 | 0 |
| virus_Sep8.0v_20692 | 0 | 0 | 0 | 1 | 0 | 0 | 0 | 0 | 0 | 0 | 1 |
| virus_Sep8.5v_553 | 0 | 0 | 0 | 0 | 0 | 0 | 0 | 0 | 0 | 0 | 0 |
| virus_Sep8.5v_8152 | 0 | 0 | 0 | 0 | 0 | 0 | 0 | 0 | 0 | 0 | 0 |
| virus_Sep8.5v_10850 | 0 | 0 | 0 | 0 | 0 | 0 | 0 | 0 | 0 | 0 | 0 |
| virus_Sep8.5v_12898 | 0 | 0 | 0 | 0 | 0 | 0 | 0 | 0 | 0 | 0 | 0 |
| virus_Sep8.5v_14137 | 0 | 0 | 0 | 0 | 0 | 0 | 0 | 0 | 0 | 0 | 0 |
| virus_Sep8.5v_14369 | 0 | 0 | 0 | 0 | 0 | 0 | 0 | 0 | 0 | 0 | 0 |
| virus_Sep8.5v_18197 | 0 | 0 | 0 | 0 | 0 | 0 | 0 | 0 | 0 | 0 | 0 |
| virus_Sep8.5v_18285 | 0 | 0 | 0 | 0 | 0 | 0 | 0 | 0 | 0 | 0 | 0 |
| virus_Sep8.5v_19867 | 0 | 0 | 0 | 0 | 0 | 0 | 0 | 0 | 0 | 0 | 0 |
| virus_Sep8.5v_19912 | 0 | 0 | 0 | 0 | 0 | 0 | 0 | 0 | 0 | 0 | 0 |
| virus_Sep8.5v_20924 | 0 | 0 | 0 | 0 | 0 | 0 | 0 | 0 | 0 | 0 | 0 |
| virus_Sep8.25v_3414 | 0 | 0 | 0 | 0 | 0 | 0 | 0 | 0 | 0 | 0 | 0 |
| virus_Sep8.25v_4207 | 0 | 0 | 0 | 0 | 0 | 0 | 0 | 0 | 0 | 0 | 0 |
| virus_Sep8.25v_7191 | 0 | 0 | 0 | 0 | 0 | 0 | 0 | 0 | 0 | 0 | 0 |
| virus_Sep8.25v_17655 | 0 | 0 | 0 | 0 | 0 | 0 | 0 | 0 | 0 | 0 | 0 |
| virus_Sep8.25v_18733 | 0 | 0 | 0 | 0 | 0 | 0 | 0 | 0 | 0 | 0 | 0 |
| virus_Sep8.50v_376 | 0 | 0 | 0 | 0 | 0 | 0 | 0 | 0 | 0 | 0 | 0 |
| virus_Sep8.50v_3268 | 0 | 0 | 0 | 0 | 0 | 0 | 0 | 0 | 0 | 0 | 0 |
| virus_Sep8.50v_3315 | 0 | 0 | 0 | 0 | 0 | 0 | 0 | 0 | 0 | 0 | 0 |
| virus_Sep8.50v_4978 | 0 | 0 | 0 | 0 | 0 | 0 | 0 | 0 | 0 | 0 | 0 |
| virus_Sep8.50v_5872 | 0 | 0 | 0 | 0 | 0 | 0 | 0 | 0 | 0 | 0 | 0 |
| virus_Sep8.50v_6231 | 0 | 0 | 0 | 0 | 0 | 0 | 0 | 0 | 0 | 0 | 0 |
| virus_Sep8.50v_9900 | 0 | 0 | 0 | 0 | 0 | 0 | 0 | 0 | 0 | 0 | 0 |
| virus_Sep8.50v_16670 | 0 | 0 | 0 | 0 | 0 | 0 | 0 | 0 | 0 | 0 | 0 |
| virus_Sep8.50v_19482 | 0 | 0 | 0 | 0 | 0 | 0 | 0 | 0 | 0 | 0 | 0 |
| virus_Sep8.50v_22684 | 0 | 0 | 0 | 0 | 0 | 0 | 0 | 0 | 0 | 0 | 0 |
| virus_Sep8.50v_24010 | 0 | 0 | 0 | 0 | 0 | 0 | 0 | 0 | 0 | 0 | 0 |
| virus_Sep8.50v_24060 | 0 | 0 | 0 | 0 | 0 | 0 | 0 | 0 | 0 | 0 | 0 |
| virus_Sep8.50v_27169 | 0 | 0 | 0 | 0 | 0 | 0 | 0 | 0 | 0 | 0 | 0 |
| virus_Sep8.50v_30896 | 0 | 0 | 0 | 0 | 0 | 0 | 0 | 0 | 0 | 0 | 0 |
| virus_Sep8.50v_32747 | 0 | 0 | 0 | 0 | 0 | 0 | 0 | 0 | 0 | 0 | 0 |
| virus_Sep8.50v_40662 | 0 | 0 | 0 | 0 | 0 | 0 | 0 | 0 | 0 | 0 | 0 |
| bacteria_Jul11d0b_1559 | 0 | 0 | 0 | 0 | 0 | 0 | 0 | 0 | 0 | 0 | 0 |
| bacteria_Jul11d0b_17568 | 0 | 0 | 0 | 0 | 0 | 0 | 0 | 0 | 0 | 0 | 0 |
| bacteria_Jul11d0b_20622 | 0 | 0 | 0 | 0 | 0 | 0 | 0 | 0 | 0 | 0 | 0 |
| bacteria_Jul11d0b_37127 | 0 | 0 | 0 | 0 | 0 | 0 | 0 | 0 | 0 | 0 | 0 |
| bacteria_Jul11d0b_39804 | 0 | 0 | 0 | 0 | 0 | 0 | 0 | 0 | 0 | 0 | 0 |
| bacteria_Jul11d0b_43136 | 0 | 0 | 0 | 0 | 0 | 0 | 0 | 0 | 0 | 0 | 0 |
| bacteria_Jul11d0b_55109 | 0 | 0 | 0 | 0 | 0 | 0 | 0 | 0 | 0 | 0 | 0 |
| bacteria_Jul11d0b_59354 | 0 | 0 | 0 | 0 | 0 | 0 | 0 | 0 | 0 | 0 | 0 |
| bacteria_Aug5d0b_5736 | 1 | 0 | 0 | 0 | 0 | 0 | 0 | 0 | 0 | 0 | 1 |
| bacteria_Aug5d0b_22456 | 1 | 0 | 0 | 0 | 0 | 0 | 0 | 0 | 0 | 0 | 1 |
| bacteria_Aug5d0b_30877 | 1 | 0 | 0 | 0 | 0 | 0 | 0 | 0 | 0 | 0 | 1 |
| bacteria_Aug5d0b_59588 | 1 | 0 | 0 | 0 | 0 | 0 | 0 | 0 | 0 | 0 | 1 |
| bacteria_Aug5d5b_5262 | 0 | 0 | 1 | 0 | 0 | 0 | 0 | 0 | 0 | 0 | 1 |
| bacteria_Aug5d5b_31015 | 0 | 0 | 1 | 0 | 0 | 0 | 0 | 0 | 0 | 0 | 1 |
| bacteria_Aug5d25b_223 | 0 | 0 | 0 | 0 | 1 | 0 | 0 | 0 | 0 | 0 | 1 |
| bacteria_Aug5d25b_5676 | 0 | 0 | 0 | 0 | 1 | 0 | 0 | 0 | 0 | 0 | 1 |
| bacteria_Aug5d25b_6493 | 0 | 0 | 0 | 0 | 1 | 0 | 0 | 0 | 0 | 0 | 1 |
| bacteria_Aug5d25b_20095 | 0 | 0 | 0 | 0 | 1 | 0 | 0 | 0 | 0 | 0 | 1 |
| bacteria_Aug5d25b_20440 | 0 | 0 | 0 | 0 | 1 | 0 | 0 | 0 | 0 | 0 | 1 |
| bacteria_Aug5d25b_22051 | 0 | 0 | 0 | 0 | 1 | 0 | 0 | 0 | 0 | 0 | 1 |
| bacteria_Aug5d25b_24241 | 0 | 0 | 0 | 0 | 1 | 0 | 0 | 0 | 0 | 0 | 1 |
| bacteria_Aug5d25b_25889 | 0 | 0 | 0 | 0 | 1 | 0 | 0 | 0 | 0 | 0 | 1 |
| bacteria_Aug5d25b_31972 | 0 | 0 | 0 | 0 | 1 | 0 | 0 | 0 | 0 | 0 | 1 |
| bacteria_Aug5d25b_46756 | 0 | 0 | 0 | 0 | 2 | 0 | 0 | 0 | 0 | 0 | 2 |
| bacteria_Aug5d25b_47910 | 0 | 0 | 0 | 0 | 1 | 0 | 0 | 0 | 0 | 0 | 1 |
| bacteria_Aug5d25b_54161 | 0 | 0 | 0 | 0 | 1 | 0 | 0 | 0 | 0 | 0 | 1 |
| bacteria_Aug5d25b_55715 | 0 | 0 | 0 | 0 | 1 | 0 | 0 | 0 | 0 | 0 | 1 |
| bacteria_Aug5d25b_65492 | 0 | 0 | 0 | 0 | 1 | 0 | 0 | 0 | 0 | 0 | 1 |
| bacteria_Aug5d25b_65922 | 0 | 0 | 0 | 0 | 1 | 0 | 0 | 0 | 0 | 0 | 1 |
| bacteria_Aug5d25b_66302 | 0 | 0 | 0 | 0 | 1 | 0 | 0 | 0 | 0 | 0 | 1 |
| bacteria_Aug5d25b_67281 | 0 | 0 | 0 | 0 | 1 | 0 | 0 | 0 | 0 | 0 | 1 |
| bacteria_Aug5d25b_67714 | 0 | 0 | 0 | 0 | 1 | 0 | 0 | 0 | 0 | 0 | 1 |
| bacteria_Aug5d25b_68212 | 0 | 0 | 0 | 0 | 1 | 0 | 0 | 0 | 0 | 0 | 1 |
| bacteria_Aug5d25b_69711 | 0 | 0 | 0 | 0 | 1 | 0 | 0 | 0 | 0 | 0 | 1 |
| bacteria_Aug5d25b_74111 | 0 | 0 | 0 | 0 | 1 | 0 | 0 | 0 | 0 | 0 | 1 |
| bacteria_Aug5d50b_3303 | 0 | 0 | 0 | 0 | 0 | 0 | 1 | 0 | 0 | 0 | 1 |
| bacteria_Aug5d50b_30112 | 0 | 0 | 0 | 0 | 0 | 0 | 1 | 0 | 0 | 0 | 1 |
| bacteria_Aug5d50b_60497 | 0 | 0 | 0 | 0 | 0 | 0 | 1 | 0 | 0 | 0 | 1 |
| bacteria_Aug5d75b_30714 | 0 | 0 | 0 | 0 | 0 | 0 | 0 | 0 | 1 | 0 | 1 |
| bacteria_Jul11d5b_11533 | 0 | 0 | 0 | 0 | 0 | 0 | 0 | 0 | 0 | 0 | 0 |
| bacteria_Jul11d5b_23765 | 0 | 0 | 0 | 0 | 0 | 0 | 0 | 0 | 0 | 0 | 0 |
| bacteria_Jul11d5b_26487 | 0 | 0 | 0 | 0 | 0 | 0 | 0 | 0 | 0 | 0 | 0 |
| bacteria_Jul11d5b_34352 | 0 | 0 | 0 | 0 | 0 | 0 | 0 | 0 | 0 | 0 | 0 |
| bacteria_Jul11d5b_40809 | 0 | 0 | 0 | 0 | 0 | 0 | 0 | 0 | 0 | 0 | 0 |
| bacteria_Jul11d25b_337 | 0 | 0 | 0 | 0 | 0 | 0 | 0 | 0 | 0 | 0 | 0 |
| bacteria_Jul11d25b_14148 | 0 | 0 | 0 | 0 | 0 | 0 | 0 | 0 | 0 | 0 | 0 |
| bacteria_Jul11d25b_17876 | 0 | 0 | 0 | 0 | 1 | 0 | 0 | 0 | 0 | 0 | 1 |
| bacteria_Jul11d25b_40274 | 0 | 0 | 0 | 0 | 0 | 0 | 0 | 0 | 0 | 0 | 0 |
| bacteria_Jul11d25b_40724 | 0 | 0 | 0 | 0 | 0 | 0 | 0 | 0 | 0 | 0 | 0 |
| bacteria_Jul11d25b_43031 | 0 | 0 | 0 | 0 | 0 | 0 | 0 | 0 | 0 | 0 | 0 |
| bacteria_Jul11d50b_10166 | 0 | 0 | 0 | 0 | 0 | 0 | 0 | 0 | 0 | 0 | 0 |
| bacteria_Jul11d50b_30178 | 0 | 0 | 0 | 0 | 0 | 0 | 0 | 0 | 0 | 0 | 0 |
| bacteria_Jul11d50b_39541 | 0 | 0 | 0 | 0 | 0 | 0 | 0 | 0 | 0 | 0 | 0 |
| bacteria_Jul11d50b_41262 | 0 | 0 | 0 | 0 | 0 | 1 | 0 | 0 | 0 | 0 | 1 |
| bacteria_Jul11d50b_43330 | 0 | 0 | 0 | 0 | 0 | 0 | 0 | 0 | 0 | 0 | 0 |
| bacteria_Jul11d50b_48101 | 0 | 0 | 0 | 0 | 0 | 0 | 0 | 0 | 0 | 0 | 0 |
| bacteria_Jul11d50b_50386 | 0 | 0 | 0 | 0 | 0 | 0 | 0 | 0 | 0 | 0 | 0 |
| bacteria_Jul11d75b_964 | 0 | 0 | 0 | 0 | 0 | 0 | 0 | 0 | 0 | 0 | 0 |
| bacteria_Jul11d75b_3509 | 0 | 1 | 0 | 0 | 0 | 0 | 0 | 0 | 0 | 0 | 1 |
| bacteria_Jul11d75b_5615 | 0 | 0 | 0 | 0 | 0 | 0 | 0 | 0 | 0 | 0 | 0 |
| bacteria_Jul11d75b_8015 | 0 | 0 | 0 | 0 | 0 | 0 | 0 | 0 | 0 | 0 | 0 |
| bacteria_Jul11d75b_8648 | 0 | 0 | 0 | 0 | 0 | 0 | 0 | 0 | 0 | 0 | 0 |
| bacteria_Jul11d75b_14214 | 0 | 0 | 0 | 0 | 0 | 0 | 0 | 0 | 0 | 0 | 0 |
| bacteria_Jul11d75b_15766 | 0 | 0 | 0 | 0 | 0 | 0 | 0 | 0 | 0 | 0 | 0 |
| bacteria_Jul11d75b_20821 | 0 | 0 | 0 | 0 | 0 | 0 | 0 | 0 | 0 | 0 | 0 |
| bacteria_Jul11d75b_21556 | 0 | 0 | 0 | 0 | 0 | 0 | 0 | 0 | 0 | 0 | 0 |
| bacteria_Jul11d75b_31518 | 0 | 0 | 0 | 0 | 0 | 0 | 0 | 0 | 0 | 0 | 0 |
| bacteria_Jul11d75b_34393 | 0 | 0 | 0 | 0 | 0 | 0 | 0 | 0 | 0 | 0 | 0 |
| bacteria_Jul11d75b_34828 | 0 | 0 | 0 | 0 | 0 | 0 | 0 | 0 | 0 | 0 | 0 |
| bacteria_Jul11d75b_35503 | 0 | 0 | 0 | 0 | 0 | 0 | 0 | 0 | 0 | 0 | 0 |
| bacteria_Jul11d75b_37955 | 0 | 0 | 0 | 0 | 0 | 0 | 0 | 0 | 0 | 0 | 0 |
| bacteria_Jul11d75b_43478 | 0 | 0 | 0 | 0 | 0 | 0 | 0 | 0 | 0 | 0 | 0 |
| bacteria_Jul11d75b_44486 | 0 | 1 | 0 | 0 | 0 | 0 | 0 | 0 | 0 | 0 | 1 |
| bacteria_Jul11d75b_44982 | 0 | 0 | 0 | 0 | 0 | 0 | 0 | 0 | 0 | 0 | 0 |
| bacteria_Jul27d5b_2955 | 0 | 0 | 0 | 0 | 0 | 0 | 0 | 0 | 0 | 0 | 0 |
| bacteria_Jul27d5b_3738 | 0 | 0 | 0 | 0 | 0 | 0 | 0 | 0 | 0 | 0 | 0 |
| bacteria_Jul27d5b_4770 | 0 | 0 | 0 | 0 | 0 | 0 | 0 | 0 | 0 | 0 | 0 |
| bacteria_Jul27d5b_6501 | 0 | 0 | 0 | 0 | 0 | 0 | 0 | 0 | 0 | 1 | 1 |
| bacteria_Jul27d5b_6869 | 0 | 0 | 0 | 0 | 0 | 0 | 0 | 0 | 0 | 0 | 0 |
| bacteria_Jul27d5b_7754 | 0 | 0 | 0 | 0 | 0 | 0 | 0 | 0 | 0 | 0 | 0 |
| bacteria_Jul27d5b_9865 | 0 | 0 | 0 | 0 | 0 | 0 | 0 | 0 | 0 | 0 | 0 |
| bacteria_Jul27d5b_10093 | 0 | 0 | 0 | 0 | 0 | 0 | 0 | 0 | 0 | 0 | 0 |
| bacteria_Jul27d5b_13572 | 0 | 0 | 0 | 0 | 0 | 0 | 0 | 0 | 0 | 0 | 0 |
| bacteria_Jul27d5b_14333 | 0 | 0 | 0 | 0 | 0 | 0 | 0 | 0 | 0 | 0 | 0 |
| bacteria_Jul27d5b_14779 | 0 | 0 | 0 | 0 | 0 | 0 | 0 | 0 | 0 | 0 | 0 |
| bacteria_Jul27d5b_15894 | 0 | 0 | 0 | 0 | 0 | 0 | 0 | 0 | 0 | 0 | 0 |
| bacteria_Jul27d5b_19092 | 0 | 0 | 0 | 0 | 0 | 0 | 0 | 0 | 0 | 0 | 0 |
| bacteria_Jul27d5b_23521 | 0 | 0 | 0 | 0 | 0 | 0 | 0 | 0 | 0 | 0 | 0 |
| bacteria_Jul27d5b_23776 | 0 | 0 | 0 | 0 | 0 | 0 | 0 | 0 | 0 | 0 | 0 |
| bacteria_Jul27d5b_26535 | 0 | 0 | 0 | 0 | 0 | 0 | 0 | 0 | 0 | 0 | 0 |
| bacteria_Jul27d5b_31405 | 0 | 0 | 0 | 0 | 0 | 0 | 0 | 0 | 0 | 0 | 0 |
| bacteria_Jul27d5b_34275 | 0 | 0 | 0 | 0 | 0 | 0 | 0 | 0 | 0 | 0 | 0 |
| bacteria_Jul27d5b_44929 | 0 | 0 | 0 | 0 | 0 | 0 | 0 | 0 | 0 | 0 | 0 |
| bacteria_Jul27d5b_46842 | 0 | 0 | 0 | 0 | 0 | 0 | 0 | 0 | 0 | 0 | 0 |
| bacteria_Jul27d5b_49625 | 0 | 0 | 0 | 0 | 0 | 0 | 0 | 0 | 0 | 0 | 0 |
| bacteria_Jul27d5b_50810 | 0 | 0 | 0 | 0 | 0 | 0 | 0 | 0 | 0 | 0 | 0 |
| bacteria_Jul27d25b_3077 | 0 | 0 | 0 | 0 | 0 | 0 | 0 | 0 | 0 | 0 | 0 |
| bacteria_Jul27d25b_12212 | 0 | 0 | 0 | 0 | 0 | 0 | 0 | 0 | 0 | 0 | 0 |
| bacteria_Jul27d25b_17011 | 0 | 0 | 0 | 0 | 0 | 0 | 0 | 0 | 0 | 0 | 0 |
| bacteria_Jul27d25b_26053 | 0 | 0 | 0 | 0 | 0 | 0 | 0 | 0 | 0 | 0 | 0 |
| bacteria_Jul27d25b_31065 | 0 | 0 | 0 | 0 | 0 | 0 | 0 | 0 | 0 | 0 | 0 |
| bacteria_Jul27d25b_32437 | 0 | 0 | 0 | 0 | 0 | 0 | 0 | 0 | 0 | 0 | 0 |
| bacteria_Jul27d25b_32754 | 0 | 0 | 0 | 0 | 0 | 0 | 0 | 0 | 0 | 0 | 0 |
| bacteria_Jul27d25b_36728 | 0 | 0 | 0 | 0 | 0 | 0 | 0 | 0 | 0 | 0 | 0 |
| bacteria_Jul27d25b_46379 | 0 | 0 | 0 | 0 | 0 | 0 | 0 | 0 | 0 | 0 | 0 |
| bacteria_Jul27d25b_49428 | 0 | 0 | 0 | 0 | 0 | 0 | 0 | 0 | 0 | 0 | 0 |
| bacteria_Jul27d25b_49671 | 0 | 0 | 0 | 0 | 0 | 0 | 0 | 0 | 0 | 0 | 0 |
| bacteria_Jul27d25b_56762 | 0 | 0 | 0 | 0 | 0 | 0 | 0 | 0 | 0 | 0 | 0 |
| bacteria_Jul27d25b_59534 | 0 | 0 | 0 | 0 | 0 | 0 | 0 | 0 | 0 | 0 | 0 |
| bacteria_Jul27d25b_60387 | 0 | 0 | 0 | 0 | 0 | 0 | 0 | 0 | 0 | 0 | 0 |
| bacteria_Jul27d25b_61923 | 0 | 0 | 0 | 0 | 0 | 0 | 0 | 0 | 0 | 0 | 0 |

| | | | | | | | | | | | |
|---|---|---|---|---|---|---|---|---|---|---|---|
| bacteria_Jul27d25b_63149 | 0 | 0 | 0 | 0 | 0 | 0 | 0 | 0 | 0 | 0 | 0 |
| bacteria_Jul27d25b_63507 | 0 | 0 | 0 | 0 | 0 | 0 | 0 | 0 | 0 | 0 | 0 |
| bacteria_Jul27d25b_67608 | 0 | 0 | 0 | 0 | 0 | 0 | 0 | 0 | 0 | 0 | 0 |
| bacteria_Jul27d25b_69115 | 0 | 0 | 0 | 0 | 0 | 0 | 0 | 0 | 0 | 0 | 0 |
| bacteria_Jul27d25b_73855 | 0 | 0 | 0 | 0 | 0 | 0 | 0 | 0 | 0 | 0 | 0 |
| bacteria_Jul27d25b_74996 | 0 | 0 | 0 | 0 | 0 | 0 | 0 | 0 | 0 | 0 | 0 |
| bacteria_Jul27d75b_10084 | 0 | 1 | 0 | 0 | 0 | 0 | 0 | 0 | 0 | 0 | 1 |
| bacteria_Jul27d75b_16046 | 0 | 0 | 0 | 0 | 0 | 0 | 0 | 0 | 0 | 0 | 0 |
| bacteria_Jul27d75b_16153 | 0 | 0 | 0 | 0 | 0 | 0 | 0 | 0 | 0 | 0 | 0 |
| bacteria_Jul27d75b_20379 | 0 | 0 | 0 | 0 | 0 | 0 | 0 | 0 | 0 | 0 | 0 |
| bacteria_Jul27d75b_23081 | 0 | 0 | 0 | 0 | 0 | 0 | 0 | 0 | 0 | 0 | 0 |
| bacteria_Jul27d75b_39181 | 0 | 0 | 0 | 0 | 0 | 0 | 0 | 0 | 0 | 0 | 0 |
| bacteria_Jul27d75b_52832 | 0 | 0 | 0 | 0 | 0 | 0 | 0 | 0 | 0 | 0 | 0 |
| bacteria_Jul27d75b_64220 | 0 | 0 | 0 | 0 | 0 | 0 | 0 | 0 | 0 | 0 | 0 |
| bacteria_Jul27d75b_72128 | 0 | 0 | 0 | 0 | 0 | 0 | 0 | 0 | 0 | 0 | 0 |
| bacteria_Jun8d25b_26857 | 0 | 0 | 0 | 0 | 0 | 0 | 0 | 0 | 0 | 0 | 0 |
| bacteria_Jun8d25b_32993 | 0 | 0 | 0 | 0 | 0 | 0 | 0 | 0 | 0 | 0 | 0 |
| bacteria_Jun8d25b_33746 | 0 | 0 | 0 | 0 | 0 | 0 | 0 | 0 | 0 | 0 | 0 |
| bacteria_Jun8d25b_36164 | 0 | 0 | 0 | 0 | 0 | 0 | 0 | 0 | 0 | 0 | 0 |
| bacteria_Jun8d25b_37526 | 0 | 0 | 0 | 0 | 0 | 0 | 0 | 0 | 0 | 0 | 0 |
| bacteria_Jun8d25b_45101 | 0 | 0 | 0 | 0 | 0 | 0 | 0 | 0 | 0 | 0 | 0 |
| bacteria_Jun8d25b_46474 | 0 | 0 | 0 | 0 | 0 | 0 | 0 | 0 | 0 | 0 | 0 |
| bacteria_Jun8d25b_50962 | 0 | 0 | 0 | 0 | 0 | 0 | 0 | 0 | 0 | 0 | 0 |
| bacteria_Jun8d25b_55397 | 0 | 0 | 0 | 0 | 0 | 0 | 0 | 0 | 0 | 0 | 0 |
| bacteria_Jun8d25b_61830 | 0 | 0 | 0 | 0 | 0 | 0 | 0 | 0 | 0 | 0 | 0 |
| bacteria_Jun8d25b_64064 | 0 | 0 | 0 | 0 | 0 | 0 | 0 | 0 | 0 | 0 | 0 |
| bacteria_Jun8d25b_71606 | 0 | 0 | 0 | 0 | 0 | 0 | 0 | 0 | 0 | 0 | 0 |
| bacteria_Jun8d25b_74871 | 0 | 0 | 0 | 0 | 0 | 0 | 0 | 0 | 0 | 0 | 0 |
| bacteria_Jun8d50b_5228 | 0 | 0 | 0 | 0 | 0 | 0 | 0 | 0 | 0 | 0 | 0 |
| bacteria_Jun8d50b_18155 | 0 | 0 | 0 | 0 | 0 | 0 | 0 | 0 | 0 | 0 | 0 |
| bacteria_Jun8d50b_35654 | 0 | 0 | 0 | 0 | 0 | 0 | 0 | 0 | 0 | 0 | 0 |
| bacteria_Jun8d75b_5458 | 0 | 0 | 0 | 0 | 0 | 0 | 0 | 0 | 0 | 0 | 0 |
| bacteria_Jun8d75b_65005 | 0 | 0 | 0 | 0 | 0 | 0 | 0 | 0 | 0 | 0 | 0 |
| bacteria_Jun22d50b_6099 | 0 | 0 | 0 | 0 | 0 | 0 | 0 | 0 | 0 | 0 | 0 |
| bacteria_Jun22d75b_7275 | 0 | 0 | 0 | 0 | 0 | 0 | 0 | 0 | 0 | 0 | 0 |
| bacteria_Jun22d75b_8202 | 0 | 0 | 0 | 0 | 0 | 0 | 0 | 0 | 0 | 0 | 0 |
| bacteria_Jun22d75b_9524 | 0 | 0 | 0 | 0 | 0 | 0 | 0 | 0 | 0 | 0 | 0 |
| bacteria_Jun22d75b_9806 | 0 | 0 | 0 | 0 | 0 | 0 | 0 | 0 | 0 | 0 | 0 |
| bacteria_Jun22d75b_11079 | 0 | 0 | 0 | 0 | 0 | 0 | 0 | 0 | 0 | 0 | 0 |
| bacteria_Jun22d75b_19778 | 0 | 0 | 0 | 0 | 0 | 0 | 0 | 0 | 0 | 0 | 0 |
| bacteria_Jun22d75b_30180 | 0 | 0 | 0 | 0 | 0 | 0 | 0 | 0 | 0 | 0 | 0 |
| bacteria_Jun22d75b_42184 | 0 | 0 | 0 | 0 | 0 | 0 | 0 | 0 | 0 | 0 | 0 |
| bacteria_Jun22d75b_57662 | 0 | 0 | 0 | 0 | 0 | 0 | 0 | 0 | 0 | 0 | 0 |
| bacteria_Jun22d75b_59900 | 0 | 0 | 0 | 0 | 0 | 0 | 0 | 0 | 0 | 0 | 0 |
| bacteria_May12d25b_723 | 0 | 0 | 0 | 0 | 0 | 0 | 0 | 0 | 0 | 0 | 0 |
| bacteria_May12d25b_2113 | 0 | 0 | 0 | 0 | 0 | 0 | 0 | 0 | 0 | 0 | 0 |
| bacteria_May12d25b_2300 | 0 | 0 | 0 | 0 | 0 | 0 | 0 | 0 | 0 | 0 | 0 |
| bacteria_May12d25b_2926 | 0 | 0 | 0 | 0 | 0 | 0 | 0 | 0 | 0 | 0 | 0 |
| bacteria_May12d25b_4842 | 0 | 0 | 0 | 0 | 0 | 0 | 0 | 0 | 0 | 0 | 0 |
| bacteria_May12d25b_5661 | 0 | 0 | 0 | 0 | 0 | 0 | 0 | 0 | 0 | 0 | 0 |
| bacteria_May12d25b_5973 | 0 | 0 | 0 | 0 | 0 | 0 | 0 | 0 | 0 | 0 | 0 |
| bacteria_May12d25b_6037 | 0 | 0 | 0 | 0 | 0 | 0 | 0 | 0 | 0 | 0 | 0 |
| bacteria_May12d25b_6141 | 0 | 0 | 0 | 0 | 0 | 0 | 0 | 0 | 0 | 0 | 0 |
| bacteria_May12d25b_6784 | 0 | 0 | 0 | 0 | 0 | 0 | 0 | 0 | 0 | 0 | 0 |
| bacteria_May12d25b_7448 | 0 | 0 | 0 | 0 | 0 | 0 | 0 | 0 | 0 | 0 | 0 |
| bacteria_May12d50b_2570 | 0 | 0 | 0 | 0 | 0 | 0 | 0 | 0 | 0 | 0 | 0 |
| bacteria_May12d50b_9866 | 0 | 0 | 0 | 0 | 0 | 0 | 0 | 0 | 0 | 0 | 0 |
| bacteria_May12d50b_1296 | 0 | 0 | 0 | 0 | 0 | 0 | 0 | 0 | 0 | 0 | 0 |
| bacteria_May12d50b_1887 | 0 | 0 | 0 | 0 | 0 | 0 | 0 | 0 | 0 | 0 | 0 |
| bacteria_May12d50b_3563 | 0 | 0 | 0 | 0 | 0 | 0 | 0 | 0 | 0 | 0 | 0 |
| bacteria_May12d50b_3632 | 0 | 0 | 0 | 0 | 0 | 0 | 0 | 0 | 0 | 0 | 0 |
| bacteria_May12d50b_4768 | 0 | 0 | 0 | 0 | 0 | 0 | 0 | 0 | 0 | 0 | 0 |
| bacteria_Sep8d5b_10520 | 0 | 0 | 0 | 0 | 0 | 0 | 0 | 0 | 0 | 0 | 0 |
| bacteria_Sep8d5b_15212 | 0 | 0 | 0 | 0 | 0 | 0 | 0 | 0 | 0 | 0 | 0 |
| bacteria_Sep8d5b_16078 | 0 | 0 | 0 | 0 | 0 | 0 | 0 | 0 | 0 | 0 | 0 |
| bacteria_Sep8d5b_19656 | 0 | 0 | 0 | 0 | 0 | 0 | 0 | 0 | 0 | 0 | 0 |
| bacteria_Sep8d5b_20813 | 0 | 0 | 0 | 0 | 0 | 0 | 0 | 0 | 0 | 0 | 0 |
| bacteria_Sep8d5b_23945 | 0 | 0 | 0 | 0 | 0 | 0 | 0 | 0 | 0 | 0 | 0 |
| bacteria_Sep8d5b_27058 | 0 | 0 | 0 | 0 | 0 | 0 | 0 | 0 | 0 | 0 | 0 |
| bacteria_Sep8d5b_37485 | 0 | 0 | 0 | 0 | 0 | 0 | 0 | 0 | 0 | 0 | 0 |
| bacteria_Sep8d5b_43068 | 0 | 0 | 0 | 0 | 0 | 0 | 0 | 0 | 0 | 0 | 0 |
| bacteria_Sep8d5b_48305 | 0 | 0 | 0 | 0 | 0 | 0 | 0 | 0 | 0 | 0 | 0 |
| bacteria_Sep8d5b_69488 | 0 | 0 | 0 | 0 | 0 | 0 | 0 | 0 | 0 | 0 | 0 |
| bacteria_Sep8d5b_72192 | 0 | 0 | 0 | 0 | 0 | 0 | 0 | 0 | 0 | 0 | 0 |
| bacteria_Sep8d25b_1641 | 0 | 0 | 0 | 0 | 0 | 0 | 0 | 0 | 0 | 0 | 0 |
| bacteria_Sep8d25b_22164 | 0 | 0 | 0 | 0 | 0 | 0 | 0 | 0 | 0 | 0 | 0 |
| bacteria_Sep8d25b_27591 | 0 | 0 | 0 | 0 | 0 | 0 | 0 | 0 | 0 | 0 | 0 |
| bacteria_Sep8d25b_42892 | 0 | 0 | 0 | 0 | 0 | 0 | 0 | 0 | 0 | 0 | 0 |
| bacteria_Sep8d50b_9349 | 0 | 0 | 0 | 0 | 0 | 0 | 0 | 0 | 0 | 0 | 0 |
| virus_Aug5.0v_146 | 0 | 1 | 0 | 0 | 0 | 0 | 0 | 0 | 0 | 0 | 1 |
| virus_Aug5.0v_475 | 0 | 1 | 0 | 0 | 0 | 0 | 0 | 0 | 0 | 0 | 1 |
| virus_Aug5.0v_482 | 0 | 1 | 0 | 0 | 0 | 0 | 0 | 0 | 0 | 0 | 1 |
| virus_Aug5.0v_519 | 0 | 1 | 0 | 0 | 0 | 0 | 0 | 0 | 0 | 0 | 1 |
| virus_Aug5.0v_829 | 0 | 1 | 0 | 0 | 0 | 0 | 0 | 0 | 0 | 0 | 1 |
| virus_Aug5.0v_910 | 0 | 1 | 0 | 0 | 0 | 0 | 0 | 0 | 0 | 0 | 1 |
| virus_Aug5.0v_2014 | 0 | 1 | 0 | 0 | 0 | 0 | 0 | 0 | 0 | 0 | 1 |
| virus_Aug5.0v_2035 | 0 | 1 | 0 | 0 | 0 | 0 | 0 | 0 | 0 | 0 | 1 |
| virus_Aug5.0v_2209 | 0 | 1 | 0 | 0 | 0 | 0 | 0 | 0 | 0 | 0 | 1 |
| virus_Aug5.0v_2290 | 0 | 1 | 0 | 0 | 0 | 0 | 0 | 0 | 0 | 0 | 1 |
| virus_Aug5.0v_2334 | 0 | 1 | 0 | 0 | 0 | 0 | 0 | 0 | 0 | 0 | 1 |
| virus_Aug5.0v_2629 | 0 | 1 | 0 | 0 | 0 | 0 | 0 | 0 | 0 | 0 | 1 |
| virus_Aug5.0v_2801 | 0 | 1 | 0 | 0 | 0 | 0 | 0 | 0 | 0 | 0 | 1 |
| virus_Aug5.0v_2804 | 0 | 1 | 0 | 0 | 0 | 0 | 0 | 0 | 0 | 0 | 1 |
| virus_Aug5.0v_3270 | 0 | 1 | 0 | 0 | 0 | 0 | 0 | 0 | 0 | 0 | 1 |
| virus_Aug5.0v_3405 | 0 | 1 | 0 | 0 | 0 | 0 | 0 | 0 | 0 | 0 | 1 |
| virus_Aug5.0v_3473 | 0 | 1 | 0 | 0 | 0 | 0 | 0 | 0 | 0 | 0 | 1 |
| virus_Aug5.0v_3709 | 0 | 1 | 0 | 0 | 0 | 0 | 0 | 0 | 0 | 0 | 1 |
| virus_Aug5.0v_4007 | 0 | 1 | 0 | 0 | 0 | 0 | 0 | 0 | 0 | 0 | 1 |
| virus_Aug5.0v_4045 | 0 | 1 | 0 | 0 | 0 | 0 | 0 | 0 | 0 | 0 | 1 |
| virus_Aug5.0v_4098 | 0 | 1 | 0 | 0 | 0 | 0 | 0 | 0 | 0 | 0 | 1 |
| virus_Aug5.0v_4228 | 0 | 1 | 0 | 0 | 0 | 0 | 0 | 0 | 0 | 0 | 1 |
| virus_Aug5.0v_4777 | 0 | 1 | 0 | 0 | 0 | 0 | 0 | 0 | 0 | 0 | 1 |
| virus_Aug5.0v_4813 | 0 | 1 | 0 | 0 | 0 | 0 | 0 | 0 | 0 | 0 | 1 |
| virus_Aug5.0v_5214 | 0 | 1 | 0 | 0 | 0 | 0 | 0 | 0 | 0 | 0 | 1 |
| virus_Aug5.0v_5277 | 0 | 1 | 0 | 0 | 0 | 0 | 0 | 0 | 0 | 0 | 1 |
| virus_Aug5.5v_1027 | 0 | 0 | 0 | 1 | 0 | 0 | 0 | 0 | 0 | 0 | 1 |
| virus_Aug5.5v_11011 | 0 | 0 | 0 | 1 | 0 | 0 | 0 | 0 | 0 | 0 | 1 |
| virus_Aug5.5v_16881 | 0 | 0 | 0 | 1 | 0 | 0 | 0 | 0 | 0 | 0 | 1 |
| virus_Aug5.5v_19223 | 0 | 0 | 0 | 1 | 0 | 0 | 0 | 0 | 0 | 0 | 1 |
| virus_Aug5.5v_19590 | 0 | 0 | 0 | 1 | 0 | 0 | 0 | 0 | 0 | 0 | 1 |
| virus_Aug5.5v_21555 | 0 | 0 | 0 | 1 | 0 | 0 | 0 | 0 | 0 | 0 | 1 |
| virus_Aug5.5v_27059 | 0 | 0 | 0 | 1 | 0 | 0 | 0 | 0 | 0 | 0 | 1 |
| virus_Aug5.5v_34126 | 0 | 0 | 0 | 1 | 0 | 0 | 0 | 0 | 0 | 0 | 1 |
| virus_Aug5.5v_36038 | 0 | 0 | 0 | 1 | 0 | 0 | 0 | 0 | 0 | 0 | 1 |
| virus_Aug5.5v_36607 | 0 | 0 | 0 | 1 | 0 | 0 | 0 | 0 | 0 | 0 | 1 |
| virus_Aug5.5v_39651 | 0 | 0 | 0 | 1 | 0 | 0 | 0 | 0 | 0 | 0 | 1 |
| virus_Aug5.25v_5018 | 0 | 0 | 0 | 0 | 0 | 1 | 0 | 0 | 0 | 0 | 1 |
| virus_Aug5.25v_5611 | 0 | 0 | 0 | 0 | 0 | 1 | 0 | 0 | 0 | 0 | 1 |
| virus_Aug5.25v_6389 | 0 | 0 | 0 | 0 | 0 | 1 | 0 | 0 | 0 | 0 | 1 |
| virus_Aug5.25v_8358 | 0 | 0 | 0 | 0 | 0 | 1 | 0 | 0 | 0 | 0 | 1 |
| virus_Aug5.25v_9617 | 0 | 0 | 0 | 0 | 0 | 1 | 0 | 0 | 0 | 0 | 1 |
| virus_Aug5.25v_13579 | 0 | 0 | 0 | 0 | 0 | 1 | 0 | 0 | 0 | 0 | 1 |
| virus_Aug5.25v_15035 | 0 | 0 | 0 | 0 | 0 | 1 | 0 | 0 | 0 | 0 | 1 |
| virus_Aug5.25v_15566 | 0 | 0 | 0 | 0 | 0 | 1 | 0 | 0 | 0 | 0 | 1 |
| virus_Aug5.25v_16100 | 0 | 0 | 0 | 0 | 0 | 1 | 0 | 0 | 0 | 0 | 1 |
| virus_Aug5.25v_19477 | 0 | 0 | 0 | 0 | 0 | 1 | 0 | 0 | 0 | 0 | 1 |
| virus_Aug5.25v_22020 | 0 | 0 | 0 | 0 | 0 | 1 | 0 | 0 | 0 | 0 | 1 |
| virus_Aug5.25v_22567 | 0 | 0 | 0 | 0 | 0 | 1 | 0 | 0 | 0 | 0 | 1 |
| virus_Aug5.25v_23323 | 0 | 0 | 0 | 0 | 0 | 1 | 0 | 0 | 0 | 0 | 1 |
| virus_Aug5.25v_23484 | 0 | 0 | 0 | 0 | 0 | 1 | 0 | 0 | 0 | 0 | 1 |
| virus_Aug5.25v_23672 | 0 | 0 | 0 | 0 | 0 | 1 | 0 | 0 | 0 | 0 | 1 |
| virus_Aug5.25v_25630 | 0 | 0 | 0 | 0 | 0 | 1 | 0 | 0 | 0 | 0 | 1 |
| virus_Aug5.25v_26039 | 0 | 0 | 0 | 0 | 0 | 1 | 0 | 0 | 0 | 0 | 1 |
| virus_Aug5.25v_27480 | 0 | 0 | 0 | 0 | 0 | 1 | 0 | 0 | 0 | 0 | 1 |
| virus_Aug5.25v_27726 | 0 | 0 | 0 | 0 | 0 | 1 | 0 | 0 | 0 | 0 | 1 |
| virus_Aug5.25v_28987 | 0 | 0 | 0 | 0 | 0 | 1 | 0 | 0 | 0 | 0 | 1 |
| virus_Aug5.25v_29117 | 0 | 0 | 0 | 0 | 0 | 1 | 0 | 0 | 0 | 0 | 1 |
| virus_Aug5.25v_33610 | 0 | 0 | 0 | 0 | 0 | 1 | 0 | 0 | 0 | 0 | 1 |
| virus_Aug5.25v_34955 | 0 | 0 | 0 | 0 | 0 | 1 | 0 | 0 | 0 | 0 | 1 |
| virus_Aug5.25v_39753 | 0 | 0 | 0 | 0 | 0 | 1 | 0 | 0 | 0 | 0 | 1 |
| virus_Aug5.25v_40751 | 0 | 0 | 0 | 0 | 0 | 1 | 0 | 0 | 0 | 0 | 1 |
| virus_Aug5.50v_720 | 0 | 0 | 0 | 0 | 0 | 0 | 0 | 1 | 0 | 0 | 1 |
| virus_Aug5.50v_14187 | 0 | 0 | 0 | 0 | 0 | 0 | 0 | 1 | 0 | 0 | 1 |
| virus_Aug5.50v_15857 | 0 | 0 | 0 | 0 | 0 | 0 | 0 | 1 | 0 | 0 | 1 |
| virus_Aug5.50v_18806 | 0 | 0 | 0 | 0 | 0 | 0 | 0 | 1 | 0 | 0 | 1 |
| virus_Aug5.50v_25476 | 0 | 0 | 0 | 0 | 0 | 0 | 0 | 1 | 0 | 0 | 1 |
| virus_Aug5.50v_26588 | 0 | 0 | 0 | 0 | 0 | 0 | 0 | 1 | 0 | 0 | 1 |
| virus_Aug5.50v_32577 | 0 | 0 | 0 | 0 | 0 | 0 | 0 | 1 | 0 | 0 | 1 |
| virus_Aug5.50v_32970 | 0 | 0 | 0 | 0 | 0 | 0 | 0 | 1 | 0 | 0 | 1 |
| virus_Aug5.50v_37664 | 0 | 0 | 0 | 0 | 0 | 0 | 0 | 1 | 0 | 0 | 1 |
| virus_Aug5.50v_41091 | 0 | 0 | 0 | 0 | 0 | 0 | 0 | 1 | 0 | 0 | 1 |
| virus_Aug5.75v_9587 | 0 | 0 | 0 | 0 | 0 | 0 | 0 | 0 | 0 | 1 | 1 |
| virus_Aug5.75v_16573 | 0 | 0 | 0 | 0 | 0 | 0 | 0 | 0 | 0 | 1 | 1 |
| virus_Aug5.75v_16676 | 0 | 0 | 0 | 0 | 0 | 0 | 0 | 0 | 0 | 1 | 1 |
| virus_Aug5.75v_17768 | 0 | 0 | 0 | 0 | 0 | 0 | 0 | 0 | 0 | 1 | 1 |
| virus_Aug5.75v_19491 | 0 | 0 | 0 | 0 | 0 | 0 | 0 | 0 | 0 | 1 | 1 |
| virus_Jul11.0v_442 | 0 | 0 | 0 | 0 | 0 | 0 | 0 | 0 | 0 | 0 | 0 |

| | | | | | | | | | | | |
|---|---|---|---|---|---|---|---|---|---|---|---|
| virus_Jul11.0v_4604 | 0 | 0 | 0 | 0 | 0 | 0 | 0 | 0 | 0 | 0 | 0 |
| virus_Jul11.0v_5551 | 0 | 0 | 0 | 0 | 0 | 0 | 0 | 0 | 0 | 0 | 0 |
| virus_Jul11.0v_7048 | 0 | 0 | 0 | 0 | 0 | 0 | 0 | 0 | 0 | 0 | 0 |
| virus_Jul11.0v_8126 | 0 | 0 | 0 | 0 | 0 | 0 | 0 | 0 | 0 | 0 | 0 |
| virus_Jul11.0v_9962 | 0 | 0 | 0 | 0 | 0 | 0 | 0 | 0 | 0 | 0 | 0 |
| virus_Jul11.0v_10133 | 0 | 0 | 0 | 0 | 0 | 0 | 0 | 0 | 0 | 0 | 0 |
| virus_Jul11.0v_14097 | 0 | 0 | 0 | 0 | 0 | 0 | 0 | 0 | 0 | 0 | 0 |
| virus_Jul11.5v_96 | 0 | 0 | 0 | 0 | 0 | 0 | 0 | 0 | 0 | 0 | 0 |
| virus_Jul11.5v_2249 | 0 | 0 | 0 | 0 | 0 | 0 | 0 | 0 | 0 | 0 | 0 |
| virus_Jul11.5v_2297 | 0 | 0 | 0 | 0 | 0 | 0 | 0 | 0 | 0 | 0 | 0 |
| virus_Jul11.5v_3433 | 0 | 0 | 0 | 0 | 0 | 0 | 0 | 0 | 0 | 0 | 0 |
| virus_Jul11.5v_4199 | 0 | 0 | 0 | 0 | 0 | 0 | 0 | 0 | 0 | 0 | 0 |
| virus_Jul11.5v_5665 | 0 | 0 | 0 | 0 | 0 | 0 | 0 | 0 | 0 | 0 | 0 |
| virus_Jul11.5v_8581 | 0 | 0 | 0 | 0 | 0 | 0 | 0 | 0 | 0 | 0 | 0 |
| virus_Jul11.5v_9486 | 0 | 0 | 0 | 0 | 0 | 0 | 0 | 0 | 0 | 0 | 0 |
| virus_Jul11.5v_12100 | 0 | 0 | 0 | 0 | 0 | 0 | 0 | 0 | 0 | 0 | 0 |
| virus_Jul11.5v_12724 | 0 | 0 | 0 | 0 | 0 | 0 | 0 | 0 | 0 | 0 | 0 |
| virus_Jul11.5v_13492 | 0 | 0 | 0 | 0 | 0 | 0 | 0 | 0 | 0 | 0 | 0 |
| virus_Jul11.5v_14178 | 0 | 0 | 0 | 0 | 0 | 0 | 0 | 0 | 0 | 0 | 0 |
| virus_Jul11.5v_14940 | 0 | 0 | 0 | 0 | 0 | 0 | 0 | 0 | 0 | 0 | 0 |
| virus_Jul11.5v_15366 | 0 | 0 | 0 | 0 | 0 | 0 | 0 | 0 | 0 | 0 | 0 |
| virus_Jul11.5v_15420 | 0 | 0 | 0 | 0 | 0 | 0 | 0 | 0 | 0 | 0 | 0 |
| virus_Jul11.5v_17790 | 0 | 0 | 0 | 0 | 0 | 0 | 0 | 0 | 0 | 0 | 0 |
| virus_Jul11.5v_18748 | 0 | 0 | 0 | 0 | 0 | 0 | 0 | 0 | 0 | 0 | 0 |
| virus_Jul11.5v_20090 | 0 | 0 | 0 | 0 | 0 | 0 | 0 | 0 | 0 | 0 | 0 |
| virus_Jul11.5v_20509 | 0 | 0 | 0 | 0 | 0 | 0 | 0 | 0 | 0 | 0 | 0 |
| virus_Jul11.25v_1083 | 0 | 0 | 0 | 0 | 0 | 0 | 0 | 0 | 0 | 0 | 0 |
| virus_Jul11.25v_3761 | 0 | 0 | 0 | 0 | 0 | 0 | 0 | 0 | 0 | 0 | 0 |
| virus_Jul11.25v_4477 | 0 | 0 | 0 | 0 | 0 | 0 | 0 | 0 | 0 | 0 | 0 |
| virus_Jul11.25v_6383 | 0 | 0 | 0 | 0 | 0 | 0 | 0 | 0 | 0 | 0 | 0 |
| virus_Jul11.25v_8611 | 0 | 0 | 0 | 0 | 0 | 0 | 0 | 0 | 0 | 0 | 0 |
| virus_Jul11.25v_13188 | 0 | 0 | 0 | 0 | 0 | 0 | 0 | 0 | 0 | 0 | 0 |
| virus_Jul11.25v_15692 | 0 | 0 | 0 | 0 | 0 | 0 | 0 | 0 | 0 | 0 | 0 |
| virus_Jul11.25v_17401 | 0 | 0 | 0 | 0 | 0 | 0 | 0 | 0 | 0 | 0 | 0 |
| virus_Jul11.25v_18199 | 0 | 0 | 0 | 0 | 0 | 0 | 0 | 0 | 0 | 0 | 0 |
| virus_Jul11.25v_19237 | 0 | 1 | 0 | 0 | 0 | 0 | 0 | 0 | 0 | 0 | 1 |
| virus_Jul11.25v_19730 | 0 | 0 | 0 | 0 | 0 | 0 | 0 | 0 | 0 | 0 | 0 |
| virus_Jul11.50v_760 | 0 | 0 | 0 | 0 | 0 | 0 | 0 | 0 | 0 | 0 | 0 |
| virus_Jul11.50v_1622 | 0 | 0 | 0 | 0 | 0 | 0 | 0 | 0 | 0 | 0 | 0 |
| virus_Jul11.50v_2621 | 0 | 0 | 0 | 0 | 0 | 0 | 0 | 0 | 0 | 0 | 0 |
| virus_Jul11.50v_3228 | 0 | 0 | 0 | 0 | 0 | 0 | 0 | 0 | 0 | 0 | 0 |
| virus_Jul11.50v_9019 | 0 | 0 | 0 | 0 | 0 | 0 | 0 | 0 | 0 | 0 | 0 |
| virus_Jul11.50v_9942 | 0 | 0 | 0 | 0 | 0 | 0 | 0 | 0 | 0 | 0 | 0 |
| virus_Jul11.50v_10932 | 0 | 0 | 0 | 0 | 0 | 0 | 0 | 0 | 0 | 0 | 0 |
| virus_Jul11.50v_12102 | 0 | 0 | 0 | 0 | 0 | 0 | 0 | 0 | 0 | 0 | 0 |
| virus_Jul11.50v_12580 | 0 | 0 | 0 | 0 | 0 | 0 | 0 | 0 | 0 | 0 | 0 |
| virus_Jul11.50v_12998 | 0 | 0 | 0 | 0 | 0 | 0 | 0 | 0 | 0 | 0 | 0 |
| virus_Jul11.50v_14281 | 0 | 0 | 0 | 0 | 0 | 0 | 0 | 0 | 0 | 0 | 0 |
| virus_Jul11.50v_19238 | 0 | 0 | 0 | 0 | 0 | 0 | 0 | 0 | 0 | 0 | 0 |
| virus_Jul11.75v_310 | 0 | 0 | 0 | 0 | 0 | 0 | 0 | 0 | 0 | 0 | 0 |
| virus_Jul11.75v_5099 | 0 | 0 | 0 | 0 | 0 | 0 | 0 | 0 | 0 | 0 | 0 |
| virus_Jul11.75v_20708 | 0 | 0 | 0 | 0 | 0 | 0 | 0 | 0 | 0 | 0 | 0 |
| virus_Jul11.75v_22172 | 0 | 0 | 0 | 0 | 0 | 0 | 0 | 0 | 0 | 0 | 0 |
| virus_Jul11.75v_28690 | 0 | 0 | 0 | 0 | 0 | 0 | 0 | 0 | 0 | 0 | 0 |
| virus_Jul11.75v_30859 | 0 | 0 | 0 | 0 | 0 | 0 | 0 | 0 | 0 | 0 | 0 |
| virus_Jul11.75v_32281 | 0 | 0 | 0 | 0 | 0 | 0 | 0 | 0 | 0 | 0 | 0 |
| virus_Jul11.75v_32550 | 0 | 0 | 0 | 0 | 0 | 0 | 0 | 0 | 0 | 0 | 0 |
| virus_Jul11.75v_35753 | 0 | 0 | 0 | 0 | 0 | 0 | 0 | 0 | 0 | 0 | 0 |
| virus_Jul11.75v_38733 | 0 | 0 | 0 | 0 | 0 | 0 | 0 | 0 | 0 | 0 | 0 |
| virus_Jul11.75v_40626 | 0 | 0 | 0 | 0 | 0 | 0 | 0 | 0 | 0 | 0 | 0 |
| virus_Jul27.5v_612 | 0 | 0 | 0 | 0 | 0 | 0 | 0 | 0 | 0 | 0 | 0 |
| virus_Jul27.5v_1625 | 0 | 0 | 0 | 0 | 0 | 0 | 0 | 0 | 0 | 0 | 0 |
| virus_Jul27.5v_1963 | 0 | 0 | 0 | 0 | 0 | 0 | 0 | 0 | 0 | 0 | 0 |
| virus_Jul27.5v_2044 | 0 | 0 | 0 | 0 | 0 | 0 | 0 | 0 | 0 | 0 | 0 |
| virus_Jul27.5v_2291 | 0 | 0 | 0 | 0 | 0 | 0 | 0 | 0 | 0 | 0 | 0 |
| virus_Jul27.5v_2985 | 0 | 0 | 0 | 0 | 0 | 0 | 0 | 0 | 0 | 0 | 0 |
| virus_Jul27.5v_4301 | 0 | 0 | 0 | 0 | 0 | 0 | 0 | 0 | 0 | 0 | 0 |
| virus_Jul27.5v_5003 | 0 | 0 | 0 | 0 | 0 | 0 | 0 | 0 | 0 | 0 | 0 |
| virus_Jul27.5v_5430 | 0 | 0 | 0 | 0 | 0 | 0 | 0 | 0 | 0 | 0 | 0 |
| virus_Jul27.5v_5955 | 0 | 0 | 0 | 0 | 0 | 0 | 0 | 0 | 0 | 0 | 0 |
| virus_Jul27.5v_7634 | 0 | 0 | 0 | 0 | 0 | 0 | 0 | 0 | 0 | 0 | 0 |
| virus_Jul27.5v_8937 | 0 | 0 | 0 | 0 | 0 | 0 | 0 | 0 | 0 | 0 | 0 |
| virus_Jul27.5v_10211 | 0 | 0 | 0 | 0 | 0 | 0 | 0 | 0 | 0 | 0 | 0 |
| virus_Jul27.5v_11104 | 0 | 0 | 0 | 0 | 0 | 0 | 0 | 0 | 0 | 0 | 0 |
| virus_Jul27.5v_11617 | 0 | 0 | 0 | 0 | 0 | 0 | 0 | 0 | 0 | 0 | 0 |
| virus_Jul27.5v_11739 | 0 | 0 | 0 | 0 | 0 | 0 | 0 | 0 | 0 | 0 | 0 |
| virus_Jul27.5v_12604 | 0 | 0 | 0 | 0 | 0 | 0 | 0 | 0 | 0 | 0 | 0 |
| virus_Jul27.5v_13178 | 0 | 0 | 0 | 0 | 0 | 0 | 0 | 0 | 0 | 0 | 0 |
| virus_Jul27.5v_13640 | 0 | 0 | 0 | 0 | 0 | 0 | 0 | 0 | 0 | 0 | 0 |
| virus_Jul27.5v_14443 | 0 | 0 | 0 | 0 | 0 | 0 | 0 | 0 | 0 | 0 | 0 |
| virus_Jul27.5v_15185 | 0 | 0 | 0 | 0 | 0 | 0 | 0 | 0 | 0 | 0 | 0 |
| virus_Jul27.5v_15390 | 0 | 0 | 0 | 0 | 0 | 0 | 0 | 0 | 0 | 0 | 0 |
| virus_Jul27.5v_15609 | 0 | 0 | 0 | 0 | 0 | 0 | 0 | 0 | 0 | 0 | 0 |
| virus_Jul27.5v_15809 | 0 | 0 | 0 | 0 | 0 | 0 | 0 | 0 | 0 | 0 | 0 |
| virus_Jul27.5v_16884 | 0 | 0 | 0 | 0 | 0 | 0 | 0 | 0 | 0 | 0 | 0 |
| virus_Jul27.5v_16984 | 0 | 0 | 0 | 0 | 0 | 0 | 0 | 0 | 0 | 0 | 0 |
| virus_Jul27.5v_19180 | 0 | 0 | 0 | 0 | 0 | 0 | 0 | 0 | 0 | 0 | 0 |
| virus_Jul27.5v_19187 | 0 | 0 | 0 | 0 | 0 | 0 | 0 | 0 | 0 | 0 | 0 |
| virus_Jul27.5v_20806 | 0 | 0 | 0 | 0 | 0 | 0 | 0 | 0 | 0 | 0 | 0 |
| virus_Jul27.5v_21031 | 0 | 0 | 0 | 0 | 0 | 0 | 0 | 0 | 0 | 0 | 0 |
| virus_Jul27.25v_578 | 0 | 0 | 0 | 0 | 0 | 0 | 0 | 0 | 0 | 0 | 0 |
| virus_Jul27.25v_765 | 0 | 0 | 0 | 0 | 0 | 0 | 0 | 0 | 0 | 0 | 0 |
| virus_Jul27.25v_3153 | 0 | 0 | 0 | 0 | 0 | 0 | 0 | 0 | 0 | 0 | 0 |
| virus_Jul27.25v_4579 | 0 | 0 | 0 | 0 | 0 | 0 | 0 | 0 | 0 | 0 | 0 |
| virus_Jul27.25v_17282 | 0 | 0 | 0 | 0 | 0 | 0 | 0 | 0 | 0 | 0 | 0 |
| virus_Jul27.25v_18466 | 0 | 0 | 0 | 0 | 0 | 0 | 0 | 0 | 0 | 0 | 0 |
| virus_Jul27.25v_21341 | 0 | 0 | 0 | 0 | 0 | 0 | 0 | 0 | 0 | 0 | 0 |
| virus_Jul27.25v_28362 | 0 | 0 | 0 | 0 | 0 | 0 | 0 | 0 | 0 | 0 | 0 |
| virus_Jul27.25v_31892 | 0 | 0 | 0 | 0 | 0 | 0 | 0 | 0 | 0 | 0 | 0 |
| virus_Jul27.25v_32817 | 0 | 0 | 0 | 0 | 0 | 0 | 0 | 0 | 0 | 0 | 0 |
| virus_Jul27.25v_33044 | 0 | 0 | 0 | 0 | 0 | 0 | 0 | 0 | 0 | 0 | 0 |
| virus_Jul27.25v_33364 | 0 | 0 | 0 | 0 | 0 | 0 | 0 | 0 | 0 | 0 | 0 |
| virus_Jul27.25v_34509 | 0 | 0 | 0 | 0 | 0 | 0 | 0 | 0 | 0 | 0 | 0 |
| virus_Jul27.25v_36312 | 0 | 0 | 0 | 0 | 0 | 0 | 0 | 0 | 0 | 0 | 0 |
| virus_Jul27.25v_37678 | 0 | 0 | 0 | 0 | 0 | 0 | 0 | 0 | 0 | 0 | 0 |
| virus_Jul27.25v_39511 | 0 | 0 | 0 | 0 | 0 | 0 | 0 | 0 | 0 | 0 | 0 |
| virus_Jul27.75v_1676 | 0 | 0 | 0 | 0 | 0 | 0 | 0 | 0 | 0 | 0 | 0 |
| virus_Jul27.75v_12255 | 0 | 0 | 0 | 0 | 0 | 0 | 0 | 0 | 0 | 0 | 0 |
| virus_Jun8.25v_4802 | 0 | 0 | 0 | 0 | 0 | 0 | 0 | 0 | 0 | 0 | 0 |
| virus_Jun8.25v_5246 | 0 | 0 | 0 | 0 | 0 | 0 | 0 | 0 | 0 | 0 | 0 |
| virus_Jun8.25v_23093 | 0 | 0 | 0 | 0 | 0 | 0 | 0 | 0 | 0 | 0 | 0 |
| virus_Jun8.25v_27165 | 0 | 0 | 0 | 0 | 0 | 0 | 0 | 0 | 0 | 0 | 0 |
| virus_Jun8.50v_4437 | 0 | 0 | 0 | 0 | 0 | 0 | 0 | 0 | 0 | 0 | 0 |
| virus_Jun8.50v_5126 | 0 | 0 | 0 | 0 | 0 | 0 | 0 | 0 | 0 | 0 | 0 |
| virus_Jun8.50v_13375 | 0 | 0 | 0 | 0 | 0 | 0 | 0 | 0 | 0 | 0 | 0 |
| virus_Jun8.50v_16842 | 0 | 0 | 0 | 0 | 0 | 0 | 0 | 0 | 0 | 0 | 0 |
| virus_Jun8.50v_17840 | 0 | 0 | 0 | 0 | 0 | 0 | 0 | 0 | 0 | 0 | 0 |
| virus_Jun8.50v_20729 | 0 | 0 | 0 | 0 | 0 | 0 | 0 | 0 | 0 | 0 | 0 |
| virus_Jun8.50v_30353 | 0 | 0 | 0 | 0 | 0 | 0 | 0 | 0 | 0 | 0 | 0 |
| virus_Jun8.50v_37590 | 0 | 0 | 0 | 0 | 0 | 0 | 0 | 0 | 0 | 0 | 0 |
| virus_Jun8.50v_37882 | 0 | 0 | 0 | 0 | 0 | 0 | 0 | 0 | 0 | 0 | 0 |
| virus_Jun8.75v_3226 | 0 | 0 | 0 | 0 | 0 | 0 | 0 | 0 | 0 | 0 | 0 |
| virus_Jun8.75v_18911 | 0 | 0 | 0 | 0 | 0 | 0 | 0 | 0 | 0 | 0 | 0 |
| virus_Jun22.0v_556 | 0 | 0 | 0 | 0 | 0 | 0 | 0 | 0 | 0 | 0 | 0 |
| virus_Jun22.0v_861 | 0 | 0 | 0 | 0 | 0 | 0 | 0 | 0 | 0 | 0 | 0 |
| virus_Jun22.0v_1274 | 0 | 0 | 0 | 0 | 0 | 0 | 0 | 0 | 0 | 0 | 0 |
| virus_Jun22.0v_3429 | 0 | 0 | 0 | 0 | 0 | 0 | 0 | 0 | 0 | 0 | 0 |
| virus_Jun22.0v_4953 | 0 | 0 | 0 | 0 | 0 | 0 | 0 | 0 | 0 | 0 | 0 |
| virus_Jun22.0v_8658 | 0 | 0 | 0 | 0 | 0 | 0 | 0 | 0 | 0 | 0 | 0 |
| virus_Jun22.0v_11126 | 0 | 0 | 0 | 0 | 0 | 0 | 0 | 0 | 0 | 0 | 0 |
| virus_Jun22.0v_11947 | 0 | 0 | 0 | 0 | 0 | 0 | 0 | 0 | 0 | 0 | 0 |
| virus_Jun22.0v_15447 | 0 | 0 | 0 | 0 | 0 | 0 | 0 | 0 | 0 | 0 | 0 |
| virus_Jun22.0v_17818 | 0 | 0 | 0 | 0 | 0 | 0 | 0 | 0 | 0 | 0 | 0 |
| virus_Jun22.0v_19577 | 0 | 0 | 0 | 0 | 0 | 0 | 0 | 0 | 0 | 0 | 0 |
| virus_Jun22.50v_1210 | 0 | 0 | 0 | 0 | 0 | 0 | 0 | 0 | 0 | 0 | 0 |
| virus_Jun22.50v_4968 | 0 | 0 | 0 | 0 | 0 | 0 | 0 | 0 | 0 | 0 | 0 |
| virus_Jun22.50v_8234 | 0 | 0 | 0 | 0 | 0 | 0 | 0 | 0 | 0 | 0 | 0 |
| virus_Jun22.50v_8302 | 0 | 0 | 0 | 0 | 0 | 0 | 0 | 0 | 0 | 0 | 0 |
| virus_Jun22.50v_11598 | 0 | 0 | 0 | 0 | 0 | 0 | 0 | 0 | 0 | 0 | 0 |
| virus_Jun22.50v_15338 | 0 | 0 | 0 | 0 | 0 | 0 | 0 | 0 | 0 | 0 | 0 |
| virus_Jun22.50v_15599 | 0 | 0 | 0 | 0 | 0 | 0 | 0 | 0 | 0 | 0 | 0 |
| virus_Jun22.50v_19630 | 0 | 0 | 0 | 0 | 0 | 0 | 0 | 0 | 0 | 0 | 0 |
| virus_Jun22.50v_19641 | 0 | 0 | 0 | 0 | 0 | 0 | 0 | 0 | 0 | 0 | 0 |
| virus_Jun22.50v_20864 | 0 | 0 | 0 | 0 | 0 | 0 | 0 | 0 | 0 | 0 | 0 |
| virus_Jun22.50v_23741 | 0 | 0 | 0 | 0 | 0 | 0 | 0 | 0 | 0 | 0 | 0 |
| virus_Jun22.50v_25125 | 0 | 0 | 0 | 0 | 0 | 0 | 0 | 0 | 0 | 0 | 0 |
| virus_Jun22.50v_27762 | 0 | 0 | 0 | 0 | 0 | 0 | 0 | 0 | 0 | 0 | 0 |
| virus_Jun22.50v_28667 | 0 | 0 | 0 | 0 | 0 | 0 | 0 | 0 | 0 | 0 | 0 |
| virus_Jun22.50v_29776 | 0 | 0 | 0 | 0 | 0 | 0 | 0 | 0 | 0 | 0 | 0 |
| virus_Jun22.50v_31917 | 0 | 0 | 0 | 0 | 0 | 0 | 0 | 0 | 0 | 0 | 0 |
| virus_Jun22.50v_32459 | 0 | 0 | 0 | 0 | 0 | 0 | 0 | 0 | 0 | 0 | 0 |
| virus_Jun22.50v_32722 | 0 | 0 | 0 | 0 | 0 | 0 | 0 | 0 | 0 | 0 | 0 |
| virus_Jun22.50v_38736 | 0 | 0 | 0 | 0 | 0 | 0 | 0 | 0 | 0 | 0 | 0 |
| virus_Jun22.75v_2967 | 0 | 0 | 0 | 0 | 0 | 0 | 0 | 0 | 0 | 0 | 0 |
| virus_Jun22.75v_3511 | 0 | 0 | 0 | 0 | 0 | 0 | 0 | 0 | 0 | 0 | 0 |
| virus_Jun22.75v_4608 | 0 | 0 | 0 | 0 | 0 | 0 | 0 | 0 | 0 | 0 | 0 |
| virus_Jun22.75v_8296 | 0 | 0 | 0 | 0 | 0 | 0 | 0 | 0 | 0 | 0 | 0 |

| | | | | | | | | | | | |
|---|---|---|---|---|---|---|---|---|---|---|---|
| virus_Jun22.75v_9197 | 0 | 0 | 0 | 0 | 0 | 0 | 0 | 0 | 0 | 0 | 0 |
| virus_Jun22.75v_11180 | 0 | 0 | 0 | 0 | 0 | 0 | 0 | 0 | 0 | 0 | 0 |
| virus_Jun22.75v_15887 | 0 | 0 | 0 | 0 | 0 | 0 | 0 | 0 | 0 | 0 | 0 |
| virus_Jun22.75v_17235 | 0 | 0 | 0 | 0 | 0 | 0 | 0 | 0 | 0 | 0 | 0 |
| virus_Jun22.75v_17750 | 0 | 0 | 0 | 0 | 0 | 0 | 0 | 0 | 0 | 0 | 0 |
| virus_Jun22.75v_19113 | 0 | 0 | 0 | 0 | 0 | 0 | 0 | 0 | 0 | 0 | 0 |
| virus_Jun22.75v_21972 | 0 | 0 | 0 | 0 | 0 | 0 | 0 | 0 | 0 | 0 | 0 |
| virus_Jun22.75v_25000 | 0 | 0 | 0 | 0 | 0 | 0 | 0 | 0 | 0 | 0 | 0 |
| virus_Jun22.75v_27225 | 0 | 0 | 0 | 0 | 0 | 0 | 0 | 0 | 0 | 0 | 0 |
| virus_Jun22.75v_30843 | 0 | 0 | 0 | 0 | 0 | 0 | 0 | 0 | 0 | 0 | 0 |
| virus_Jun22.75v_30907 | 0 | 0 | 0 | 0 | 0 | 0 | 0 | 0 | 0 | 0 | 0 |
| virus_Jun22.75v_32988 | 0 | 0 | 0 | 0 | 0 | 0 | 0 | 0 | 0 | 0 | 0 |
| virus_Jun22.75v_39688 | 0 | 0 | 0 | 0 | 0 | 0 | 0 | 0 | 0 | 0 | 0 |
| virus_Jun22.75v_40555 | 0 | 0 | 0 | 0 | 0 | 0 | 0 | 0 | 0 | 0 | 0 |
| virus_May12.5v_19120 | 0 | 0 | 0 | 0 | 0 | 0 | 0 | 0 | 0 | 0 | 0 |
| virus_May12.5v_19860 | 0 | 0 | 0 | 0 | 0 | 0 | 0 | 0 | 0 | 0 | 0 |
| virus_May12.5v_19934 | 0 | 0 | 0 | 0 | 0 | 0 | 0 | 0 | 0 | 0 | 0 |
| virus_May12.5v_25221 | 0 | 0 | 0 | 0 | 0 | 0 | 0 | 0 | 0 | 0 | 0 |
| virus_May12.5v_26224 | 0 | 0 | 0 | 0 | 0 | 0 | 0 | 0 | 0 | 0 | 0 |
| virus_May12.5v_31990 | 0 | 0 | 0 | 0 | 0 | 0 | 0 | 0 | 0 | 0 | 0 |
| virus_May12.5v_39748 | 0 | 0 | 0 | 0 | 0 | 0 | 0 | 0 | 0 | 0 | 0 |
| virus_May12.5v_40204 | 0 | 0 | 0 | 0 | 0 | 0 | 0 | 0 | 0 | 0 | 0 |
| virus_May12.5v_40693 | 0 | 0 | 0 | 0 | 0 | 0 | 0 | 0 | 0 | 0 | 0 |
| virus_May12.25v_3798 | 0 | 0 | 0 | 0 | 0 | 0 | 0 | 0 | 0 | 0 | 0 |
| virus_May12.25v_8321 | 0 | 0 | 0 | 0 | 0 | 0 | 0 | 0 | 0 | 0 | 0 |
| virus_May12.25v_9339 | 0 | 0 | 0 | 0 | 0 | 0 | 0 | 0 | 0 | 0 | 0 |
| virus_May12.25v_17687 | 0 | 0 | 0 | 0 | 0 | 0 | 0 | 0 | 0 | 0 | 0 |
| virus_May12.25v_18938 | 0 | 0 | 0 | 0 | 0 | 0 | 0 | 0 | 0 | 0 | 0 |
| virus_May12.25v_24167 | 0 | 0 | 0 | 0 | 0 | 0 | 0 | 0 | 0 | 0 | 0 |
| virus_May12.25v_31660 | 0 | 0 | 0 | 0 | 0 | 0 | 0 | 0 | 0 | 0 | 0 |
| virus_May12.25v_31949 | 0 | 0 | 0 | 0 | 0 | 0 | 0 | 0 | 0 | 0 | 0 |
| virus_May12.25v_33909 | 0 | 0 | 0 | 0 | 0 | 0 | 0 | 0 | 0 | 0 | 0 |
| virus_May12.50v_5143 | 0 | 0 | 0 | 0 | 0 | 0 | 0 | 0 | 0 | 0 | 0 |
| virus_May12.50v_11557 | 0 | 0 | 0 | 0 | 0 | 0 | 0 | 0 | 0 | 0 | 0 |
| virus_May12.50v_11845 | 0 | 0 | 0 | 0 | 0 | 0 | 0 | 0 | 0 | 0 | 0 |
| virus_May12.50v_12016 | 0 | 0 | 0 | 0 | 0 | 0 | 0 | 0 | 0 | 0 | 0 |
| virus_May12.50v_12239 | 0 | 0 | 0 | 0 | 0 | 0 | 0 | 0 | 0 | 0 | 0 |
| virus_May12.50v_12696 | 0 | 0 | 0 | 0 | 0 | 0 | 0 | 0 | 0 | 0 | 0 |
| virus_May12.50v_19565 | 0 | 0 | 0 | 0 | 0 | 0 | 0 | 0 | 0 | 0 | 0 |
| virus_May12.50v_19765 | 0 | 0 | 0 | 0 | 0 | 0 | 0 | 0 | 0 | 0 | 0 |
| virus_May12.50v_33817 | 0 | 0 | 0 | 0 | 0 | 0 | 0 | 0 | 0 | 0 | 0 |
| virus_May12.50v_36826 | 0 | 0 | 0 | 0 | 0 | 0 | 0 | 0 | 0 | 0 | 0 |
| virus_May12.75v_19610 | 0 | 0 | 0 | 0 | 0 | 0 | 0 | 0 | 0 | 0 | 0 |
| virus_May12.75v_21318 | 0 | 0 | 0 | 0 | 0 | 0 | 0 | 0 | 0 | 0 | 0 |
| virus_May12.75v_22671 | 0 | 0 | 0 | 0 | 0 | 0 | 0 | 0 | 0 | 0 | 0 |
| virus_May12.75v_23551 | 0 | 0 | 0 | 0 | 0 | 0 | 0 | 0 | 0 | 0 | 0 |
| virus_May12.75v_33699 | 0 | 0 | 0 | 0 | 0 | 0 | 0 | 0 | 0 | 0 | 0 |
| virus_May12.75v_35375 | 0 | 0 | 0 | 0 | 0 | 0 | 0 | 0 | 0 | 0 | 0 |
| virus_Sep8.0v_1296 | 0 | 0 | 0 | 0 | 0 | 0 | 0 | 0 | 0 | 0 | 0 |
| virus_Sep8.0v_5587 | 0 | 0 | 0 | 0 | 0 | 0 | 0 | 0 | 0 | 0 | 0 |
| virus_Sep8.0v_6401 | 0 | 0 | 0 | 0 | 0 | 0 | 0 | 0 | 0 | 0 | 0 |
| virus_Sep8.0v_7485 | 0 | 0 | 0 | 0 | 0 | 0 | 0 | 0 | 0 | 0 | 0 |
| virus_Sep8.0v_8165 | 0 | 0 | 0 | 0 | 0 | 0 | 0 | 0 | 0 | 0 | 0 |
| virus_Sep8.0v_9852 | 0 | 0 | 0 | 0 | 0 | 0 | 0 | 0 | 0 | 0 | 0 |
| virus_Sep8.0v_12375 | 0 | 0 | 0 | 0 | 0 | 0 | 0 | 0 | 0 | 0 | 0 |
| virus_Sep8.0v_12732 | 0 | 0 | 0 | 0 | 0 | 0 | 0 | 0 | 0 | 0 | 0 |
| virus_Sep8.0v_14290 | 0 | 0 | 0 | 0 | 0 | 0 | 0 | 0 | 0 | 0 | 0 |
| virus_Sep8.0v_15345 | 0 | 0 | 0 | 0 | 0 | 0 | 0 | 0 | 0 | 0 | 0 |
| virus_Sep8.0v_15453 | 0 | 0 | 0 | 0 | 0 | 0 | 0 | 0 | 0 | 0 | 0 |
| virus_Sep8.0v_16923 | 0 | 0 | 0 | 0 | 0 | 0 | 0 | 0 | 0 | 0 | 0 |
| virus_Sep8.5v_1908 | 0 | 0 | 0 | 0 | 0 | 0 | 0 | 0 | 0 | 0 | 0 |
| virus_Sep8.5v_3741 | 0 | 0 | 0 | 0 | 0 | 0 | 0 | 0 | 0 | 0 | 0 |
| virus_Sep8.5v_8003 | 0 | 0 | 0 | 0 | 0 | 0 | 0 | 0 | 0 | 0 | 0 |
| virus_Sep8.5v_8909 | 0 | 0 | 0 | 0 | 0 | 0 | 0 | 0 | 0 | 0 | 0 |
| virus_Sep8.5v_12833 | 0 | 0 | 0 | 0 | 0 | 0 | 0 | 0 | 0 | 0 | 0 |
| virus_Sep8.5v_13874 | 0 | 0 | 0 | 0 | 0 | 0 | 0 | 0 | 0 | 0 | 0 |
| virus_Sep8.5v_13987 | 0 | 0 | 0 | 0 | 0 | 0 | 0 | 0 | 0 | 0 | 0 |
| virus_Sep8.5v_15731 | 0 | 0 | 0 | 0 | 0 | 0 | 0 | 0 | 0 | 0 | 0 |
| virus_Sep8.25v_3817 | 0 | 0 | 0 | 0 | 0 | 0 | 0 | 0 | 0 | 0 | 0 |
| virus_Sep8.25v_4686 | 0 | 0 | 0 | 0 | 0 | 0 | 0 | 0 | 0 | 0 | 0 |
| virus_Sep8.25v_9252 | 0 | 0 | 0 | 0 | 0 | 0 | 0 | 0 | 0 | 0 | 0 |
| virus_Sep8.25v_11089 | 0 | 0 | 0 | 0 | 0 | 0 | 0 | 0 | 0 | 0 | 0 |
| virus_Sep8.25v_11794 | 0 | 0 | 0 | 0 | 0 | 0 | 0 | 0 | 0 | 0 | 0 |
| virus_Sep8.25v_13890 | 0 | 0 | 0 | 0 | 0 | 0 | 0 | 0 | 0 | 0 | 0 |
| virus_Sep8.25v_15359 | 0 | 0 | 0 | 0 | 0 | 0 | 0 | 0 | 0 | 0 | 0 |
| virus_Sep8.25v_17635 | 0 | 0 | 0 | 0 | 0 | 0 | 0 | 0 | 0 | 0 | 0 |
| virus_Sep8.50v_1245 | 0 | 0 | 0 | 0 | 0 | 0 | 0 | 0 | 0 | 0 | 0 |
| virus_Sep8.50v_2515 | 0 | 0 | 0 | 0 | 0 | 0 | 0 | 0 | 0 | 0 | 0 |
| virus_Sep8.50v_6130 | 0 | 0 | 0 | 0 | 0 | 0 | 0 | 0 | 0 | 0 | 0 |
| virus_Sep8.50v_6763 | 0 | 0 | 0 | 0 | 0 | 0 | 0 | 0 | 0 | 0 | 0 |
| virus_Sep8.50v_8608 | 0 | 0 | 0 | 0 | 0 | 0 | 0 | 0 | 0 | 0 | 0 |
| virus_Sep8.50v_9662 | 0 | 0 | 0 | 0 | 0 | 0 | 0 | 0 | 0 | 0 | 0 |
| virus_Sep8.50v_10971 | 0 | 0 | 0 | 0 | 0 | 0 | 0 | 0 | 0 | 0 | 0 |
| virus_Sep8.50v_11571 | 0 | 0 | 0 | 0 | 0 | 0 | 0 | 0 | 0 | 0 | 0 |
| virus_Sep8.50v_14560 | 0 | 0 | 0 | 0 | 0 | 0 | 0 | 0 | 0 | 0 | 0 |
| virus_Sep8.50v_14612 | 0 | 0 | 0 | 0 | 0 | 0 | 0 | 0 | 0 | 0 | 0 |
| virus_Sep8.50v_18336 | 0 | 0 | 0 | 0 | 0 | 0 | 0 | 0 | 0 | 0 | 0 |
| virus_Sep8.50v_19636 | 0 | 0 | 0 | 0 | 0 | 0 | 0 | 0 | 0 | 0 | 0 |
| virus_Sep8.50v_21101 | 0 | 0 | 0 | 0 | 0 | 0 | 0 | 0 | 0 | 0 | 0 |
| virus_Sep8.50v_21263 | 0 | 0 | 0 | 0 | 0 | 0 | 0 | 0 | 0 | 0 | 0 |
| virus_Sep8.50v_28086 | 0 | 0 | 0 | 0 | 0 | 0 | 0 | 0 | 0 | 0 | 0 |
| virus_Sep8.50v_37486 | 0 | 0 | 0 | 0 | 0 | 0 | 0 | 0 | 0 | 0 | 0 |
| virus_Sep8.50v_37785 | 0 | 0 | 0 | 0 | 0 | 0 | 0 | 0 | 0 | 0 | 0 |
| bacteria_Jul11d0b_4064 | 0 | 0 | 0 | 0 | 0 | 0 | 0 | 0 | 0 | 0 | 0 |
| bacteria_Jul11d0b_7450 | 0 | 0 | 0 | 0 | 0 | 0 | 0 | 0 | 0 | 0 | 0 |
| bacteria_Jul11d0b_10933 | 0 | 0 | 0 | 0 | 0 | 0 | 0 | 0 | 0 | 0 | 0 |
| bacteria_Jul11d0b_17925 | 1 | 0 | 0 | 0 | 0 | 0 | 0 | 0 | 0 | 0 | 1 |
| bacteria_Jul11d0b_25686 | 0 | 0 | 0 | 0 | 0 | 0 | 0 | 0 | 0 | 0 | 0 |
| bacteria_Jul11d0b_31332 | 0 | 0 | 0 | 0 | 0 | 0 | 0 | 0 | 0 | 0 | 0 |
| bacteria_Jul11d0b_33463 | 0 | 0 | 0 | 0 | 0 | 0 | 0 | 0 | 0 | 0 | 0 |
| bacteria_Jul11d0b_33558 | 0 | 0 | 0 | 0 | 0 | 0 | 0 | 0 | 0 | 0 | 0 |
| bacteria_Jul11d0b_54020 | 0 | 0 | 0 | 0 | 0 | 0 | 0 | 0 | 0 | 0 | 0 |
| bacteria_Aug5d0b_33601 | 1 | 0 | 0 | 0 | 0 | 0 | 0 | 0 | 0 | 0 | 1 |
| bacteria_Aug5d0b_54816 | 1 | 0 | 0 | 0 | 0 | 0 | 0 | 0 | 0 | 0 | 1 |
| bacteria_Aug5d0b_56553 | 1 | 0 | 0 | 0 | 0 | 0 | 0 | 0 | 0 | 0 | 1 |
| bacteria_Aug5d0b_60065 | 1 | 0 | 0 | 0 | 0 | 0 | 0 | 0 | 0 | 0 | 1 |
| bacteria_Aug5d0b_60239 | 1 | 0 | 0 | 0 | 0 | 0 | 0 | 0 | 0 | 0 | 1 |
| bacteria_Aug5d0b_60568 | 1 | 0 | 0 | 0 | 0 | 0 | 0 | 0 | 0 | 0 | 1 |
| bacteria_Aug5d5b_5724 | 0 | 0 | 1 | 0 | 0 | 0 | 0 | 0 | 0 | 0 | 1 |
| bacteria_Aug5d5b_19122 | 0 | 0 | 1 | 0 | 0 | 0 | 0 | 0 | 0 | 0 | 1 |
| bacteria_Aug5d5b_61659 | 0 | 0 | 1 | 0 | 0 | 0 | 0 | 0 | 0 | 0 | 1 |
| bacteria_Aug5d25b_791 | 0 | 0 | 0 | 0 | 1 | 0 | 0 | 0 | 0 | 0 | 1 |
| bacteria_Aug5d25b_2222 | 0 | 0 | 0 | 0 | 1 | 0 | 0 | 0 | 0 | 0 | 1 |
| bacteria_Aug5d25b_2981 | 0 | 0 | 0 | 0 | 1 | 0 | 0 | 0 | 0 | 0 | 1 |
| bacteria_Aug5d25b_7759 | 0 | 0 | 0 | 0 | 1 | 0 | 0 | 0 | 0 | 0 | 1 |
| bacteria_Aug5d25b_12071 | 0 | 0 | 0 | 0 | 1 | 0 | 0 | 0 | 0 | 0 | 1 |
| bacteria_Aug5d25b_23554 | 0 | 0 | 0 | 0 | 1 | 0 | 0 | 0 | 0 | 0 | 1 |
| bacteria_Aug5d25b_25050 | 0 | 0 | 0 | 0 | 1 | 0 | 0 | 0 | 0 | 0 | 1 |
| bacteria_Aug5d25b_25665 | 0 | 0 | 0 | 0 | 1 | 0 | 0 | 0 | 0 | 0 | 1 |
| bacteria_Aug5d25b_26824 | 0 | 0 | 0 | 0 | 1 | 0 | 0 | 0 | 0 | 0 | 1 |
| bacteria_Aug5d25b_28046 | 0 | 0 | 0 | 0 | 1 | 0 | 0 | 0 | 0 | 0 | 1 |
| bacteria_Aug5d25b_31025 | 0 | 0 | 0 | 0 | 1 | 0 | 0 | 0 | 0 | 0 | 1 |
| bacteria_Aug5d25b_35349 | 0 | 0 | 0 | 0 | 1 | 0 | 0 | 0 | 0 | 0 | 1 |
| bacteria_Aug5d25b_35959 | 0 | 0 | 0 | 0 | 1 | 0 | 0 | 0 | 0 | 0 | 1 |
| bacteria_Aug5d25b_39343 | 0 | 0 | 0 | 0 | 1 | 0 | 0 | 0 | 0 | 0 | 1 |
| bacteria_Aug5d25b_46921 | 0 | 0 | 0 | 0 | 1 | 0 | 0 | 0 | 0 | 0 | 1 |
| bacteria_Aug5d25b_48644 | 0 | 0 | 0 | 0 | 1 | 0 | 0 | 0 | 0 | 0 | 1 |
| bacteria_Aug5d25b_60264 | 0 | 0 | 0 | 0 | 1 | 0 | 0 | 0 | 0 | 0 | 1 |
| bacteria_Aug5d25b_61264 | 0 | 0 | 0 | 0 | 1 | 0 | 0 | 0 | 0 | 0 | 1 |
| bacteria_Aug5d25b_61912 | 0 | 0 | 0 | 0 | 1 | 0 | 0 | 0 | 0 | 0 | 1 |
| bacteria_Aug5d25b_64549 | 0 | 0 | 0 | 0 | 1 | 0 | 0 | 0 | 0 | 0 | 1 |
| bacteria_Aug5d25b_70162 | 0 | 0 | 0 | 0 | 1 | 0 | 0 | 0 | 0 | 0 | 1 |
| bacteria_Aug5d50b_8785 | 0 | 0 | 0 | 0 | 0 | 0 | 1 | 0 | 0 | 0 | 1 |
| bacteria_Aug5d50b_26982 | 0 | 0 | 0 | 0 | 0 | 0 | 1 | 0 | 0 | 0 | 1 |
| bacteria_Aug5d50b_41700 | 0 | 0 | 0 | 0 | 0 | 0 | 1 | 0 | 0 | 0 | 1 |
| bacteria_Aug5d50b_62509 | 0 | 0 | 0 | 0 | 0 | 0 | 1 | 0 | 0 | 0 | 1 |
| bacteria_Aug5d75b_33727 | 0 | 0 | 0 | 0 | 0 | 0 | 0 | 0 | 1 | 0 | 1 |
| bacteria_Jul11d5b_31800 | 0 | 0 | 0 | 0 | 0 | 0 | 0 | 0 | 0 | 0 | 0 |
| bacteria_Jul11d5b_36453 | 0 | 0 | 0 | 0 | 0 | 0 | 0 | 0 | 0 | 0 | 0 |
| bacteria_Jul11d5b_38728 | 0 | 0 | 0 | 0 | 0 | 0 | 0 | 0 | 0 | 0 | 0 |
| bacteria_Jul11d5b_42764 | 0 | 0 | 0 | 0 | 0 | 0 | 0 | 0 | 0 | 0 | 0 |
| bacteria_Jul11d5b_47970 | 0 | 0 | 0 | 0 | 0 | 0 | 0 | 0 | 0 | 0 | 0 |
| bacteria_Jul11d5b_51347 | 0 | 0 | 0 | 0 | 0 | 0 | 0 | 0 | 0 | 0 | 0 |
| bacteria_Jul11d25b_5809 | 0 | 0 | 0 | 0 | 0 | 0 | 0 | 0 | 0 | 0 | 0 |
| bacteria_Jul11d25b_12907 | 0 | 0 | 0 | 0 | 0 | 0 | 0 | 0 | 0 | 0 | 0 |
| bacteria_Jul11d25b_13423 | 0 | 0 | 0 | 0 | 0 | 0 | 0 | 0 | 0 | 0 | 0 |
| bacteria_Jul11d25b_14185 | 0 | 0 | 0 | 0 | 0 | 0 | 0 | 0 | 0 | 0 | 0 |
| bacteria_Jul11d50b_10287 | 0 | 0 | 0 | 0 | 0 | 0 | 0 | 0 | 0 | 0 | 0 |
| bacteria_Jul11d50b_15301 | 0 | 0 | 0 | 0 | 0 | 0 | 0 | 0 | 0 | 0 | 0 |
| bacteria_Jul11d50b_16716 | 0 | 0 | 0 | 0 | 0 | 0 | 0 | 0 | 0 | 0 | 0 |
| bacteria_Jul11d50b_20983 | 0 | 0 | 0 | 0 | 0 | 0 | 0 | 0 | 0 | 0 | 0 |
| bacteria_Jul11d50b_40527 | 0 | 0 | 0 | 0 | 0 | 0 | 0 | 0 | 0 | 0 | 0 |
| bacteria_Jul11d50b_41113 | 0 | 0 | 0 | 0 | 0 | 0 | 0 | 0 | 0 | 0 | 0 |
| bacteria_Jul11d50b_51144 | 0 | 0 | 0 | 0 | 0 | 0 | 0 | 0 | 0 | 0 | 0 |
| bacteria_Jul11d75b_6043 | 0 | 0 | 0 | 0 | 0 | 0 | 0 | 0 | 0 | 0 | 0 |
| bacteria_Jul11d75b_15586 | 0 | 0 | 0 | 0 | 0 | 0 | 0 | 0 | 0 | 0 | 0 |
| bacteria_Jul11d75b_19273 | 0 | 0 | 0 | 0 | 0 | 0 | 0 | 0 | 0 | 0 | 0 |

| | | | | | | | | | | | |
|---|---|---|---|---|---|---|---|---|---|---|---|
| bacteria_Jul11d75b_22805 | 0 | 0 | 0 | 0 | 0 | 0 | 0 | 0 | 0 | 0 | 0 |
| bacteria_Jul11d75b_22856 | 0 | 0 | 0 | 0 | 0 | 0 | 0 | 0 | 0 | 0 | 0 |
| bacteria_Jul11d75b_23815 | 0 | 0 | 0 | 0 | 0 | 0 | 0 | 0 | 0 | 0 | 0 |
| bacteria_Jul11d75b_25694 | 0 | 0 | 0 | 0 | 0 | 0 | 0 | 0 | 0 | 0 | 0 |
| bacteria_Jul11d75b_35410 | 0 | 0 | 0 | 0 | 0 | 0 | 0 | 0 | 0 | 0 | 0 |
| bacteria_Jul11d75b_43579 | 0 | 0 | 0 | 0 | 0 | 0 | 0 | 0 | 0 | 0 | 0 |
| bacteria_Jul11d75b_47361 | 0 | 0 | 0 | 0 | 0 | 0 | 0 | 0 | 0 | 0 | 0 |
| bacteria_Jul11d75b_50279 | 0 | 0 | 0 | 0 | 0 | 0 | 0 | 0 | 0 | 0 | 0 |
| bacteria_Jul11d75b_50788 | 0 | 0 | 0 | 0 | 0 | 0 | 0 | 0 | 0 | 0 | 0 |
| bacteria_Jul27d5b_1828 | 0 | 0 | 0 | 0 | 0 | 0 | 0 | 0 | 0 | 0 | 0 |
| bacteria_Jul27d5b_2410 | 0 | 0 | 0 | 0 | 0 | 0 | 0 | 0 | 0 | 0 | 0 |
| bacteria_Jul27d5b_5023 | 0 | 0 | 0 | 0 | 0 | 0 | 0 | 0 | 0 | 0 | 0 |
| bacteria_Jul27d5b_5198 | 0 | 0 | 0 | 0 | 0 | 0 | 0 | 0 | 0 | 0 | 0 |
| bacteria_Jul27d5b_9568 | 0 | 0 | 0 | 0 | 0 | 0 | 0 | 0 | 0 | 0 | 0 |
| bacteria_Jul27d5b_10242 | 0 | 0 | 0 | 0 | 0 | 0 | 0 | 0 | 0 | 0 | 0 |
| bacteria_Jul27d5b_16318 | 0 | 0 | 0 | 0 | 0 | 0 | 0 | 0 | 0 | 0 | 0 |
| bacteria_Jul27d5b_16574 | 0 | 0 | 0 | 0 | 0 | 0 | 0 | 0 | 0 | 0 | 0 |
| bacteria_Jul27d5b_22465 | 0 | 0 | 0 | 0 | 0 | 0 | 0 | 0 | 0 | 0 | 0 |
| bacteria_Jul27d5b_27923 | 0 | 0 | 0 | 0 | 0 | 0 | 0 | 0 | 0 | 0 | 0 |
| bacteria_Jul27d5b_29555 | 0 | 0 | 0 | 0 | 0 | 0 | 0 | 0 | 0 | 0 | 0 |
| bacteria_Jul27d5b_33466 | 0 | 0 | 0 | 0 | 0 | 0 | 0 | 0 | 0 | 0 | 0 |
| bacteria_Jul27d5b_34460 | 0 | 0 | 0 | 0 | 0 | 0 | 0 | 0 | 0 | 0 | 0 |
| bacteria_Jul27d5b_35314 | 0 | 0 | 0 | 0 | 0 | 0 | 0 | 0 | 0 | 0 | 0 |
| bacteria_Jul27d5b_36799 | 0 | 0 | 0 | 0 | 0 | 0 | 0 | 0 | 0 | 0 | 0 |
| bacteria_Jul27d5b_36998 | 0 | 0 | 0 | 0 | 0 | 0 | 0 | 0 | 0 | 0 | 0 |
| bacteria_Jul27d5b_39938 | 0 | 0 | 0 | 0 | 0 | 0 | 0 | 0 | 0 | 0 | 0 |
| bacteria_Jul27d5b_39975 | 0 | 0 | 0 | 0 | 0 | 0 | 0 | 0 | 0 | 0 | 0 |
| bacteria_Jul27d5b_40256 | 0 | 0 | 0 | 0 | 0 | 0 | 0 | 0 | 0 | 0 | 0 |
| bacteria_Jul27d5b_41588 | 0 | 0 | 0 | 0 | 0 | 0 | 0 | 0 | 0 | 0 | 0 |
| bacteria_Jul27d5b_42493 | 0 | 0 | 0 | 0 | 0 | 0 | 0 | 0 | 0 | 0 | 0 |
| bacteria_Jul27d5b_43443 | 0 | 0 | 0 | 0 | 0 | 0 | 0 | 0 | 0 | 0 | 0 |
| bacteria_Jul27d5b_44981 | 0 | 0 | 0 | 0 | 0 | 0 | 0 | 0 | 0 | 0 | 0 |
| bacteria_Jul27d5b_50313 | 0 | 0 | 0 | 0 | 0 | 0 | 0 | 0 | 0 | 0 | 0 |
| bacteria_Jul27d25b_2841 | 0 | 0 | 0 | 0 | 0 | 0 | 0 | 0 | 0 | 0 | 0 |
| bacteria_Jul27d25b_8443 | 0 | 0 | 0 | 0 | 0 | 0 | 0 | 0 | 0 | 0 | 0 |
| bacteria_Jul27d25b_8988 | 0 | 0 | 0 | 0 | 0 | 0 | 0 | 0 | 0 | 0 | 0 |
| bacteria_Jul27d25b_9583 | 0 | 0 | 0 | 0 | 0 | 0 | 0 | 0 | 0 | 0 | 0 |
| bacteria_Jul27d25b_9926 | 0 | 0 | 0 | 0 | 0 | 0 | 0 | 0 | 0 | 0 | 0 |
| bacteria_Jul27d25b_16877 | 0 | 0 | 0 | 0 | 0 | 0 | 0 | 0 | 0 | 0 | 0 |
| bacteria_Jul27d25b_21227 | 0 | 0 | 0 | 0 | 0 | 0 | 0 | 0 | 0 | 0 | 0 |
| bacteria_Jul27d25b_28632 | 0 | 0 | 0 | 0 | 0 | 0 | 0 | 0 | 0 | 0 | 0 |
| bacteria_Jul27d25b_31910 | 0 | 0 | 0 | 0 | 0 | 0 | 0 | 0 | 0 | 0 | 0 |
| bacteria_Jul27d25b_31926 | 0 | 0 | 0 | 0 | 0 | 0 | 0 | 0 | 0 | 0 | 0 |
| bacteria_Jul27d25b_37673 | 0 | 0 | 0 | 0 | 0 | 0 | 0 | 0 | 0 | 0 | 0 |
| bacteria_Jul27d25b_40645 | 0 | 0 | 0 | 0 | 0 | 0 | 0 | 0 | 0 | 0 | 0 |
| bacteria_Jul27d25b_42445 | 0 | 0 | 0 | 0 | 0 | 0 | 0 | 0 | 0 | 0 | 0 |
| bacteria_Jul27d25b_53675 | 0 | 0 | 0 | 0 | 0 | 0 | 0 | 0 | 0 | 0 | 0 |
| bacteria_Jul27d25b_59509 | 0 | 0 | 0 | 0 | 0 | 0 | 0 | 0 | 0 | 0 | 0 |
| bacteria_Jul27d25b_61017 | 0 | 0 | 0 | 0 | 0 | 0 | 0 | 0 | 0 | 0 | 0 |
| bacteria_Jul27d25b_64458 | 0 | 0 | 0 | 0 | 0 | 0 | 0 | 0 | 0 | 0 | 0 |
| bacteria_Jul27d25b_64687 | 0 | 0 | 0 | 0 | 0 | 0 | 0 | 0 | 0 | 0 | 0 |
| bacteria_Jul27d25b_67245 | 0 | 0 | 0 | 0 | 0 | 0 | 0 | 0 | 0 | 0 | 0 |
| bacteria_Jul27d25b_70877 | 0 | 0 | 0 | 0 | 0 | 0 | 0 | 0 | 0 | 0 | 0 |
| bacteria_Jul27d75b_5075 | 0 | 0 | 0 | 0 | 0 | 0 | 0 | 0 | 0 | 0 | 0 |
| bacteria_Jul27d75b_9102 | 0 | 0 | 0 | 0 | 0 | 0 | 0 | 0 | 0 | 0 | 0 |
| bacteria_Jul27d75b_11137 | 0 | 0 | 0 | 0 | 0 | 0 | 0 | 0 | 0 | 0 | 0 |
| bacteria_Jul27d75b_20773 | 0 | 0 | 0 | 0 | 0 | 0 | 0 | 0 | 0 | 0 | 0 |
| bacteria_Jul27d75b_21908 | 0 | 0 | 0 | 0 | 0 | 0 | 0 | 0 | 0 | 0 | 0 |
| bacteria_Jul27d75b_26709 | 0 | 0 | 0 | 0 | 0 | 0 | 0 | 0 | 0 | 0 | 0 |
| bacteria_Jul27d75b_27684 | 0 | 0 | 0 | 0 | 0 | 0 | 0 | 0 | 0 | 0 | 0 |
| bacteria_Jul27d75b_30548 | 0 | 0 | 0 | 0 | 0 | 0 | 0 | 0 | 0 | 0 | 0 |
| bacteria_Jul27d75b_35681 | 0 | 0 | 0 | 0 | 0 | 0 | 0 | 0 | 0 | 0 | 0 |
| bacteria_Jul27d75b_42367 | 0 | 0 | 0 | 0 | 0 | 0 | 0 | 0 | 0 | 0 | 0 |
| bacteria_Jul27d75b_67952 | 0 | 0 | 0 | 0 | 0 | 0 | 0 | 0 | 0 | 0 | 0 |
| bacteria_Jul27d75b_70493 | 0 | 0 | 0 | 0 | 0 | 0 | 0 | 0 | 0 | 0 | 0 |
| bacteria_Jun8d25b_2004 | 0 | 0 | 0 | 0 | 0 | 0 | 0 | 0 | 0 | 0 | 0 |
| bacteria_Jun8d25b_3182 | 0 | 0 | 0 | 0 | 0 | 0 | 0 | 0 | 0 | 0 | 0 |
| bacteria_Jun8d25b_3988 | 0 | 0 | 0 | 0 | 0 | 0 | 0 | 0 | 0 | 0 | 0 |
| bacteria_Jun8d25b_4019 | 0 | 0 | 0 | 0 | 0 | 0 | 0 | 0 | 0 | 0 | 0 |
| bacteria_Jun8d25b_5230 | 0 | 0 | 0 | 0 | 0 | 0 | 0 | 0 | 0 | 0 | 0 |
| bacteria_Jun8d25b_10147 | 0 | 0 | 0 | 0 | 0 | 0 | 0 | 0 | 0 | 0 | 0 |
| bacteria_Jun8d25b_12595 | 0 | 0 | 0 | 0 | 0 | 0 | 0 | 0 | 0 | 0 | 0 |
| bacteria_Jun8d25b_30233 | 0 | 0 | 0 | 0 | 0 | 0 | 0 | 0 | 0 | 0 | 0 |
| bacteria_Jun8d25b_41464 | 0 | 0 | 0 | 0 | 0 | 0 | 0 | 0 | 0 | 0 | 0 |
| bacteria_Jun8d25b_45795 | 0 | 0 | 0 | 0 | 0 | 0 | 0 | 0 | 0 | 0 | 0 |
| bacteria_Jun8d25b_46747 | 0 | 0 | 0 | 0 | 0 | 0 | 0 | 0 | 0 | 0 | 0 |
| bacteria_Jun8d25b_70968 | 0 | 0 | 0 | 0 | 0 | 0 | 0 | 0 | 0 | 0 | 0 |
| bacteria_Jun8d50b_6659 | 0 | 0 | 0 | 0 | 0 | 0 | 0 | 0 | 0 | 0 | 0 |
| bacteria_Jun8d75b_66505 | 0 | 0 | 0 | 0 | 0 | 0 | 0 | 0 | 0 | 0 | 0 |
| bacteria_Jun22d0b_13028 | 0 | 0 | 0 | 0 | 0 | 0 | 0 | 0 | 0 | 0 | 0 |
| bacteria_Jun22d0b_21865 | 0 | 0 | 0 | 0 | 0 | 0 | 0 | 0 | 0 | 0 | 0 |
| bacteria_Jun22d0b_23038 | 0 | 0 | 0 | 0 | 0 | 0 | 0 | 0 | 0 | 0 | 0 |
| bacteria_Jun22d0b_42214 | 0 | 0 | 0 | 0 | 0 | 0 | 0 | 0 | 0 | 0 | 0 |
| bacteria_Jun22d50b_3291 | 0 | 0 | 0 | 0 | 0 | 0 | 0 | 0 | 0 | 0 | 0 |
| bacteria_Jun22d50b_1156 | 0 | 0 | 0 | 0 | 0 | 0 | 0 | 0 | 0 | 0 | 0 |
| bacteria_Jun22d50b_2111 | 0 | 0 | 0 | 0 | 0 | 0 | 0 | 0 | 0 | 0 | 0 |
| bacteria_Jun22d50b_3249 | 0 | 0 | 0 | 0 | 0 | 0 | 0 | 0 | 0 | 0 | 0 |
| bacteria_Jun22d50b_3415 | 0 | 0 | 0 | 0 | 0 | 0 | 0 | 0 | 0 | 0 | 0 |
| bacteria_Jun22d75b_2954 | 0 | 0 | 0 | 0 | 0 | 0 | 0 | 0 | 0 | 0 | 0 |
| bacteria_Jun22d75b_3312 | 0 | 0 | 0 | 0 | 0 | 0 | 0 | 0 | 0 | 0 | 0 |
| bacteria_Jun22d75b_8601 | 0 | 0 | 0 | 0 | 0 | 0 | 0 | 0 | 0 | 0 | 0 |
| bacteria_Jun22d75b_1278 | 0 | 0 | 0 | 0 | 0 | 0 | 0 | 0 | 0 | 0 | 0 |
| bacteria_Jun22d75b_1526 | 0 | 0 | 0 | 0 | 0 | 0 | 0 | 0 | 0 | 0 | 0 |
| bacteria_Jun22d75b_2272 | 0 | 0 | 0 | 0 | 0 | 0 | 0 | 0 | 0 | 0 | 0 |
| bacteria_Jun22d75b_2979 | 0 | 0 | 0 | 0 | 0 | 0 | 0 | 0 | 0 | 0 | 0 |
| bacteria_Jun22d75b_3525 | 0 | 0 | 0 | 0 | 0 | 0 | 0 | 0 | 0 | 0 | 0 |
| bacteria_Jun22d75b_4262 | 0 | 0 | 0 | 0 | 0 | 0 | 0 | 0 | 0 | 0 | 0 |
| bacteria_Jun22d75b_4726 | 0 | 0 | 0 | 0 | 0 | 0 | 0 | 0 | 0 | 0 | 0 |
| bacteria_Jun22d75b_5584 | 0 | 0 | 0 | 0 | 0 | 0 | 0 | 0 | 0 | 0 | 0 |
| bacteria_Jun22d75b_5889 | 0 | 0 | 0 | 0 | 0 | 0 | 0 | 0 | 0 | 0 | 0 |
| bacteria_May12d5b_6907 | 0 | 0 | 0 | 0 | 0 | 0 | 0 | 0 | 0 | 0 | 0 |
| bacteria_May12d5b_4683 | 0 | 0 | 0 | 0 | 0 | 0 | 0 | 0 | 0 | 0 | 0 |
| bacteria_May12d25b_207 | 0 | 0 | 0 | 0 | 0 | 0 | 0 | 0 | 0 | 0 | 0 |
| bacteria_May12d25b_608 | 0 | 0 | 0 | 0 | 0 | 0 | 0 | 0 | 0 | 0 | 0 |
| bacteria_May12d25b_712 | 0 | 0 | 0 | 0 | 0 | 0 | 0 | 0 | 0 | 0 | 0 |
| bacteria_May12d25b_1214 | 0 | 0 | 0 | 0 | 0 | 0 | 0 | 0 | 0 | 0 | 0 |
| bacteria_May12d25b_1242 | 0 | 0 | 0 | 0 | 0 | 0 | 0 | 0 | 0 | 0 | 0 |
| bacteria_May12d25b_317 | 0 | 0 | 0 | 0 | 0 | 0 | 0 | 0 | 0 | 0 | 0 |
| bacteria_May12d25b_322 | 0 | 0 | 0 | 0 | 0 | 0 | 0 | 0 | 0 | 0 | 0 |
| bacteria_May12d25b_3821 | 0 | 0 | 0 | 0 | 0 | 0 | 0 | 0 | 0 | 0 | 0 |
| bacteria_May12d25b_408 | 0 | 0 | 0 | 0 | 0 | 0 | 0 | 0 | 0 | 0 | 0 |
| bacteria_May12d25b_4602 | 0 | 0 | 0 | 0 | 0 | 0 | 0 | 0 | 0 | 0 | 0 |
| bacteria_May12d50b_158 | 0 | 0 | 0 | 0 | 0 | 0 | 0 | 0 | 0 | 0 | 0 |
| bacteria_May12d50b_239 | 0 | 0 | 0 | 0 | 0 | 0 | 0 | 0 | 0 | 0 | 0 |
| bacteria_May12d50b_256 | 0 | 0 | 0 | 0 | 0 | 0 | 0 | 0 | 0 | 0 | 0 |
| bacteria_May12d50b_459 | 0 | 0 | 0 | 0 | 0 | 0 | 0 | 0 | 0 | 0 | 0 |
| bacteria_May12d50b_5114 | 0 | 0 | 0 | 0 | 0 | 0 | 0 | 0 | 0 | 0 | 0 |
| bacteria_May12d75b_601 | 0 | 0 | 0 | 0 | 0 | 0 | 0 | 0 | 0 | 0 | 0 |
| bacteria_Sep8d5b_2294 | 0 | 0 | 0 | 0 | 0 | 0 | 0 | 0 | 0 | 0 | 0 |
| bacteria_Sep8d5b_12590 | 0 | 0 | 0 | 0 | 0 | 0 | 0 | 0 | 0 | 0 | 0 |
| bacteria_Sep8d5b_14868 | 0 | 0 | 0 | 0 | 0 | 0 | 0 | 0 | 0 | 0 | 0 |
| bacteria_Sep8d5b_17067 | 0 | 0 | 0 | 0 | 0 | 0 | 0 | 0 | 0 | 0 | 0 |
| bacteria_Sep8d5b_18883 | 0 | 0 | 0 | 0 | 0 | 0 | 0 | 0 | 0 | 0 | 0 |
| bacteria_Sep8d5b_23224 | 0 | 0 | 0 | 0 | 0 | 0 | 0 | 0 | 0 | 0 | 0 |
| bacteria_Sep8d5b_27106 | 0 | 0 | 0 | 0 | 0 | 0 | 0 | 0 | 0 | 0 | 0 |
| bacteria_Sep8d5b_30260 | 0 | 0 | 0 | 0 | 0 | 0 | 0 | 0 | 0 | 0 | 0 |
| bacteria_Sep8d5b_32539 | 0 | 0 | 0 | 0 | 0 | 0 | 0 | 0 | 0 | 0 | 0 |
| bacteria_Sep8d5b_33226 | 0 | 0 | 0 | 0 | 0 | 0 | 0 | 0 | 0 | 0 | 0 |
| bacteria_Sep8d5b_39706 | 0 | 0 | 0 | 0 | 0 | 0 | 0 | 0 | 0 | 0 | 0 |
| bacteria_Sep8d5b_43802 | 0 | 0 | 0 | 0 | 0 | 0 | 0 | 0 | 0 | 0 | 0 |
| bacteria_Sep8d5b_45766 | 0 | 0 | 0 | 0 | 0 | 0 | 0 | 0 | 0 | 0 | 0 |
| bacteria_Sep8d5b_46599 | 0 | 0 | 0 | 0 | 0 | 0 | 0 | 0 | 0 | 0 | 0 |
| bacteria_Sep8d5b_53057 | 0 | 0 | 0 | 0 | 0 | 0 | 0 | 0 | 0 | 0 | 0 |
| bacteria_Sep8d5b_57581 | 0 | 0 | 0 | 0 | 0 | 0 | 0 | 0 | 0 | 0 | 0 |
| bacteria_Sep8d5b_62136 | 0 | 0 | 0 | 0 | 0 | 0 | 0 | 0 | 0 | 0 | 0 |
| bacteria_Sep8d5b_62790 | 0 | 0 | 0 | 0 | 0 | 0 | 0 | 0 | 0 | 0 | 0 |
| bacteria_Sep8d5b_65244 | 0 | 0 | 0 | 0 | 0 | 0 | 0 | 0 | 0 | 0 | 0 |
| bacteria_Sep8d5b_67161 | 0 | 0 | 0 | 0 | 0 | 0 | 0 | 0 | 0 | 0 | 0 |
| bacteria_Sep8d5b_68717 | 0 | 0 | 0 | 0 | 0 | 0 | 0 | 0 | 0 | 0 | 0 |
| bacteria_Sep8d5b_69240 | 0 | 0 | 0 | 0 | 0 | 0 | 0 | 0 | 0 | 0 | 0 |
| bacteria_Sep8d25b_13259 | 0 | 0 | 0 | 0 | 0 | 0 | 0 | 0 | 0 | 0 | 0 |
| bacteria_Sep8d25b_16890 | 0 | 0 | 0 | 0 | 0 | 0 | 0 | 0 | 0 | 0 | 0 |
| bacteria_Sep8d25b_47178 | 0 | 0 | 0 | 0 | 0 | 0 | 0 | 0 | 0 | 0 | 0 |
| bacteria_Sep8d50b_29588 | 0 | 0 | 0 | 0 | 0 | 0 | 0 | 0 | 0 | 0 | 0 |
| virus_Aug5.0v_290 | 0 | 1 | 0 | 0 | 0 | 0 | 0 | 0 | 0 | 0 | 1 |
| virus_Aug5.0v_348 | 0 | 1 | 0 | 0 | 0 | 0 | 0 | 0 | 0 | 0 | 1 |
| virus_Aug5.0v_390 | 0 | 1 | 0 | 0 | 0 | 0 | 0 | 0 | 0 | 0 | 1 |
| virus_Aug5.0v_626 | 0 | 1 | 0 | 0 | 0 | 0 | 0 | 0 | 0 | 0 | 1 |
| virus_Aug5.0v_695 | 0 | 1 | 0 | 0 | 0 | 0 | 0 | 0 | 0 | 0 | 1 |
| virus_Aug5.0v_845 | 0 | 1 | 0 | 0 | 0 | 0 | 0 | 0 | 0 | 0 | 1 |
| virus_Aug5.0v_1048 | 0 | 1 | 0 | 0 | 0 | 0 | 0 | 0 | 0 | 0 | 1 |
| virus_Aug5.0v_1451 | 0 | 1 | 0 | 0 | 0 | 0 | 0 | 0 | 0 | 0 | 1 |
| virus_Aug5.0v_1478 | 0 | 1 | 0 | 0 | 0 | 0 | 0 | 0 | 0 | 0 | 1 |
| virus_Aug5.0v_1688 | 0 | 1 | 0 | 0 | 0 | 0 | 0 | 0 | 0 | 0 | 1 |
| virus_Aug5.0v_2536 | 0 | 1 | 0 | 0 | 0 | 0 | 0 | 0 | 0 | 0 | 1 |
| virus_Aug5.0v_3006 | 0 | 1 | 0 | 0 | 0 | 0 | 0 | 0 | 0 | 0 | 1 |
| virus_Aug5.0v_3052 | 0 | 1 | 0 | 0 | 0 | 0 | 0 | 0 | 0 | 0 | 1 |

| | | | | | | | | | | | |
|---|---|---|---|---|---|---|---|---|---|---|---|
| virus_Aug5.0v_3248 | 0 | 1 | 0 | 0 | 0 | 0 | 0 | 0 | 0 | 0 | 1 |
| virus_Aug5.0v_3442 | 0 | 1 | 0 | 0 | 0 | 0 | 0 | 0 | 0 | 0 | 1 |
| virus_Aug5.0v_3538 | 0 | 1 | 0 | 0 | 0 | 0 | 0 | 0 | 0 | 0 | 1 |
| virus_Aug5.0v_3549 | 0 | 1 | 0 | 0 | 0 | 0 | 0 | 0 | 0 | 0 | 1 |
| virus_Aug5.0v_3564 | 0 | 1 | 0 | 0 | 0 | 0 | 0 | 0 | 0 | 0 | 1 |
| virus_Aug5.0v_3625 | 0 | 1 | 0 | 0 | 0 | 0 | 0 | 0 | 0 | 0 | 1 |
| virus_Aug5.0v_3985 | 0 | 1 | 0 | 0 | 0 | 0 | 0 | 0 | 0 | 0 | 1 |
| virus_Aug5.0v_4021 | 0 | 1 | 0 | 0 | 0 | 0 | 0 | 0 | 0 | 0 | 1 |
| virus_Aug5.0v_4206 | 0 | 1 | 0 | 0 | 0 | 0 | 0 | 0 | 0 | 0 | 1 |
| virus_Aug5.0v_4337 | 0 | 1 | 0 | 0 | 0 | 0 | 0 | 0 | 0 | 0 | 1 |
| virus_Aug5.0v_4514 | 0 | 1 | 0 | 0 | 0 | 0 | 0 | 0 | 0 | 0 | 1 |
| virus_Aug5.0v_4642 | 0 | 1 | 0 | 0 | 0 | 0 | 0 | 0 | 0 | 0 | 1 |
| virus_Aug5.0v_4674 | 0 | 1 | 0 | 0 | 0 | 0 | 0 | 0 | 0 | 0 | 1 |
| virus_Aug5.0v_4814 | 0 | 1 | 0 | 0 | 0 | 0 | 0 | 0 | 0 | 0 | 1 |
| virus_Aug5.0v_5066 | 0 | 1 | 0 | 0 | 0 | 0 | 0 | 0 | 0 | 0 | 1 |
| virus_Aug5.0v_5222 | 0 | 1 | 0 | 0 | 0 | 0 | 0 | 0 | 0 | 0 | 1 |
| virus_Aug5.0v_5270 | 0 | 1 | 0 | 0 | 0 | 0 | 0 | 0 | 0 | 0 | 1 |
| virus_Aug5.5v_4964 | 0 | 0 | 0 | 1 | 0 | 0 | 0 | 0 | 0 | 0 | 1 |
| virus_Aug5.5v_4987 | 0 | 0 | 0 | 1 | 0 | 0 | 0 | 0 | 0 | 0 | 1 |
| virus_Aug5.5v_5667 | 0 | 0 | 0 | 1 | 0 | 0 | 0 | 0 | 0 | 0 | 1 |
| virus_Aug5.5v_6323 | 0 | 0 | 0 | 1 | 0 | 0 | 0 | 0 | 0 | 0 | 1 |
| virus_Aug5.5v_6613 | 0 | 0 | 0 | 1 | 0 | 0 | 0 | 0 | 0 | 0 | 1 |
| virus_Aug5.5v_7763 | 0 | 0 | 0 | 1 | 0 | 0 | 0 | 0 | 0 | 0 | 1 |
| virus_Aug5.5v_21254 | 0 | 0 | 0 | 1 | 0 | 0 | 0 | 0 | 0 | 0 | 1 |
| virus_Aug5.5v_23129 | 0 | 0 | 0 | 1 | 0 | 0 | 0 | 0 | 0 | 0 | 1 |
| virus_Aug5.5v_28046 | 0 | 0 | 0 | 1 | 0 | 0 | 0 | 0 | 0 | 0 | 1 |
| virus_Aug5.5v_33291 | 0 | 0 | 0 | 1 | 0 | 0 | 0 | 0 | 0 | 0 | 1 |
| virus_Aug5.5v_36082 | 0 | 0 | 0 | 1 | 0 | 0 | 0 | 0 | 0 | 0 | 1 |
| virus_Aug5.5v_36926 | 0 | 0 | 0 | 1 | 0 | 0 | 0 | 0 | 0 | 0 | 1 |
| virus_Aug5.5v_36992 | 0 | 0 | 0 | 1 | 0 | 0 | 0 | 0 | 0 | 0 | 1 |
| virus_Aug5.25v_652 | 0 | 0 | 0 | 0 | 0 | 1 | 0 | 0 | 0 | 0 | 1 |
| virus_Aug5.25v_1084 | 0 | 0 | 0 | 0 | 0 | 1 | 0 | 0 | 0 | 0 | 1 |
| virus_Aug5.25v_1465 | 0 | 0 | 0 | 0 | 0 | 1 | 0 | 0 | 0 | 0 | 1 |
| virus_Aug5.25v_5891 | 0 | 0 | 0 | 0 | 0 | 1 | 0 | 0 | 0 | 0 | 1 |
| virus_Aug5.25v_6260 | 0 | 0 | 0 | 0 | 0 | 1 | 0 | 0 | 0 | 0 | 1 |
| virus_Aug5.25v_12315 | 0 | 0 | 0 | 0 | 0 | 1 | 0 | 0 | 0 | 0 | 1 |
| virus_Aug5.25v_13513 | 0 | 0 | 0 | 0 | 0 | 1 | 0 | 0 | 0 | 0 | 1 |
| virus_Aug5.25v_17057 | 0 | 0 | 0 | 0 | 0 | 1 | 0 | 0 | 0 | 0 | 1 |
| virus_Aug5.25v_18271 | 0 | 0 | 0 | 0 | 0 | 1 | 0 | 0 | 0 | 0 | 1 |
| virus_Aug5.25v_18561 | 0 | 0 | 0 | 0 | 0 | 1 | 0 | 0 | 0 | 0 | 1 |
| virus_Aug5.25v_18955 | 0 | 0 | 0 | 0 | 0 | 1 | 0 | 0 | 0 | 0 | 1 |
| virus_Aug5.25v_21604 | 0 | 0 | 0 | 0 | 0 | 1 | 0 | 0 | 0 | 0 | 1 |
| virus_Aug5.25v_22660 | 0 | 0 | 0 | 0 | 0 | 1 | 0 | 0 | 0 | 0 | 1 |
| virus_Aug5.25v_22819 | 0 | 0 | 0 | 0 | 0 | 1 | 0 | 0 | 0 | 0 | 1 |
| virus_Aug5.25v_24413 | 0 | 0 | 0 | 0 | 0 | 1 | 0 | 0 | 0 | 0 | 1 |
| virus_Aug5.25v_24767 | 0 | 0 | 0 | 0 | 0 | 1 | 0 | 0 | 0 | 0 | 1 |
| virus_Aug5.25v_27579 | 0 | 0 | 0 | 0 | 0 | 1 | 0 | 0 | 0 | 0 | 1 |
| virus_Aug5.25v_29883 | 0 | 0 | 0 | 0 | 0 | 1 | 0 | 0 | 0 | 0 | 1 |
| virus_Aug5.25v_32838 | 0 | 0 | 0 | 0 | 0 | 1 | 0 | 0 | 0 | 0 | 1 |
| virus_Aug5.25v_33658 | 0 | 0 | 0 | 0 | 0 | 1 | 0 | 0 | 0 | 0 | 1 |
| virus_Aug5.25v_33983 | 0 | 0 | 0 | 0 | 0 | 1 | 0 | 0 | 0 | 0 | 1 |
| virus_Aug5.25v_35907 | 0 | 0 | 0 | 0 | 0 | 1 | 0 | 0 | 0 | 0 | 1 |
| virus_Aug5.25v_37846 | 0 | 0 | 0 | 0 | 0 | 1 | 0 | 0 | 0 | 0 | 1 |
| virus_Aug5.25v_37860 | 0 | 0 | 0 | 0 | 0 | 1 | 0 | 0 | 0 | 0 | 1 |
| virus_Aug5.25v_38021 | 0 | 0 | 0 | 0 | 0 | 1 | 0 | 0 | 0 | 0 | 1 |
| virus_Aug5.50v_8035 | 0 | 0 | 0 | 0 | 0 | 0 | 0 | 1 | 0 | 0 | 1 |
| virus_Aug5.50v_28836 | 0 | 0 | 0 | 0 | 0 | 0 | 0 | 1 | 0 | 0 | 1 |
| virus_Aug5.50v_31030 | 0 | 0 | 0 | 0 | 0 | 0 | 0 | 1 | 0 | 0 | 1 |
| virus_Aug5.50v_31317 | 0 | 0 | 0 | 0 | 0 | 0 | 0 | 1 | 0 | 0 | 1 |
| virus_Aug5.50v_32846 | 0 | 0 | 0 | 0 | 0 | 0 | 0 | 1 | 0 | 0 | 1 |
| virus_Aug5.50v_39142 | 0 | 0 | 0 | 0 | 0 | 0 | 0 | 1 | 0 | 0 | 1 |
| virus_Aug5.75v_499 | 0 | 0 | 0 | 0 | 0 | 0 | 0 | 0 | 0 | 1 | 1 |
| virus_Aug5.75v_2997 | 0 | 0 | 0 | 0 | 0 | 0 | 0 | 0 | 0 | 1 | 1 |
| virus_Aug5.75v_11712 | 0 | 0 | 0 | 0 | 0 | 0 | 0 | 0 | 0 | 1 | 1 |
| virus_Aug5.75v_16093 | 0 | 0 | 0 | 0 | 0 | 0 | 0 | 0 | 0 | 1 | 1 |
| virus_Aug5.75v_19284 | 0 | 0 | 0 | 0 | 0 | 0 | 0 | 0 | 0 | 1 | 1 |
| virus_Aug5.75v_26254 | 0 | 0 | 0 | 0 | 0 | 0 | 0 | 0 | 0 | 1 | 1 |
| virus_Aug5.75v_26509 | 0 | 0 | 0 | 0 | 0 | 0 | 0 | 0 | 0 | 1 | 1 |
| virus_Aug5.75v_26937 | 0 | 0 | 0 | 0 | 0 | 0 | 0 | 0 | 0 | 1 | 1 |
| virus_Aug5.75v_40920 | 0 | 0 | 0 | 0 | 0 | 0 | 0 | 0 | 0 | 1 | 1 |
| virus_Jul11.0v_583 | 0 | 0 | 0 | 0 | 0 | 0 | 0 | 0 | 0 | 0 | 0 |
| virus_Jul11.0v_2963 | 0 | 0 | 0 | 0 | 0 | 0 | 0 | 0 | 0 | 0 | 0 |
| virus_Jul11.0v_5945 | 0 | 0 | 0 | 0 | 0 | 0 | 0 | 0 | 0 | 0 | 0 |
| virus_Jul11.0v_7008 | 0 | 0 | 0 | 0 | 0 | 0 | 0 | 0 | 0 | 0 | 0 |
| virus_Jul11.0v_8246 | 0 | 0 | 0 | 0 | 0 | 0 | 0 | 0 | 0 | 0 | 0 |
| virus_Jul11.0v_9478 | 0 | 0 | 0 | 0 | 0 | 0 | 0 | 0 | 0 | 0 | 0 |
| virus_Jul11.0v_9971 | 0 | 0 | 0 | 0 | 0 | 0 | 0 | 0 | 0 | 0 | 0 |
| virus_Jul11.0v_10217 | 0 | 0 | 0 | 0 | 0 | 0 | 0 | 0 | 0 | 0 | 0 |
| virus_Jul11.0v_13147 | 0 | 0 | 0 | 0 | 0 | 0 | 0 | 0 | 0 | 0 | 0 |
| virus_Jul11.0v_17969 | 0 | 0 | 0 | 0 | 0 | 0 | 0 | 0 | 0 | 0 | 0 |
| virus_Jul11.5v_412 | 0 | 0 | 0 | 0 | 0 | 0 | 0 | 0 | 0 | 0 | 0 |
| virus_Jul11.5v_3527 | 0 | 0 | 0 | 0 | 0 | 0 | 0 | 0 | 0 | 0 | 0 |
| virus_Jul11.5v_4015 | 0 | 0 | 0 | 0 | 0 | 0 | 0 | 0 | 0 | 0 | 0 |
| virus_Jul11.5v_4145 | 0 | 0 | 0 | 0 | 0 | 0 | 0 | 0 | 0 | 0 | 0 |
| virus_Jul11.5v_9420 | 0 | 0 | 0 | 0 | 0 | 0 | 0 | 0 | 0 | 0 | 0 |
| virus_Jul11.5v_11138 | 0 | 0 | 0 | 0 | 0 | 0 | 0 | 0 | 0 | 0 | 0 |
| virus_Jul11.5v_14679 | 0 | 0 | 0 | 0 | 0 | 0 | 0 | 0 | 0 | 0 | 0 |
| virus_Jul11.5v_20285 | 0 | 0 | 0 | 0 | 0 | 0 | 0 | 0 | 0 | 0 | 0 |
| virus_Jul11.25v_89 | 0 | 0 | 0 | 0 | 0 | 0 | 0 | 0 | 0 | 0 | 0 |
| virus_Jul11.25v_670 | 0 | 0 | 0 | 0 | 0 | 0 | 0 | 0 | 0 | 0 | 0 |
| virus_Jul11.25v_1359 | 0 | 0 | 0 | 0 | 0 | 0 | 0 | 0 | 0 | 0 | 0 |
| virus_Jul11.25v_2817 | 0 | 0 | 0 | 0 | 0 | 0 | 0 | 0 | 0 | 0 | 0 |
| virus_Jul11.25v_8754 | 0 | 0 | 0 | 0 | 0 | 0 | 0 | 0 | 0 | 0 | 0 |
| virus_Jul11.25v_10130 | 0 | 0 | 0 | 0 | 0 | 0 | 0 | 0 | 0 | 0 | 0 |
| virus_Jul11.25v_12389 | 0 | 0 | 0 | 0 | 0 | 0 | 0 | 0 | 0 | 0 | 0 |
| virus_Jul11.25v_14332 | 0 | 0 | 0 | 0 | 0 | 0 | 0 | 0 | 0 | 0 | 0 |
| virus_Jul11.25v_14388 | 0 | 0 | 0 | 0 | 0 | 0 | 0 | 0 | 0 | 0 | 0 |
| virus_Jul11.25v_17868 | 0 | 0 | 0 | 0 | 0 | 0 | 0 | 0 | 0 | 0 | 0 |
| virus_Jul11.50v_1076 | 0 | 0 | 0 | 0 | 0 | 0 | 0 | 0 | 0 | 0 | 0 |
| virus_Jul11.50v_1938 | 0 | 0 | 0 | 0 | 0 | 0 | 0 | 0 | 0 | 0 | 0 |
| virus_Jul11.50v_3146 | 0 | 0 | 0 | 0 | 0 | 0 | 0 | 0 | 0 | 0 | 0 |
| virus_Jul11.50v_7126 | 0 | 0 | 0 | 0 | 0 | 0 | 0 | 0 | 0 | 0 | 0 |
| virus_Jul11.50v_8009 | 0 | 0 | 0 | 0 | 0 | 0 | 0 | 0 | 0 | 0 | 0 |
| virus_Jul11.50v_8700 | 0 | 0 | 0 | 0 | 0 | 0 | 0 | 0 | 0 | 0 | 0 |
| virus_Jul11.50v_11036 | 0 | 0 | 0 | 0 | 0 | 0 | 0 | 0 | 0 | 0 | 0 |
| virus_Jul11.50v_11517 | 0 | 0 | 0 | 0 | 0 | 0 | 0 | 0 | 0 | 0 | 0 |
| virus_Jul11.50v_18833 | 0 | 0 | 0 | 0 | 0 | 0 | 0 | 0 | 0 | 0 | 0 |
| virus_Jul11.50v_20177 | 0 | 0 | 0 | 0 | 0 | 0 | 0 | 0 | 0 | 0 | 0 |
| virus_Jul11.50v_20537 | 0 | 0 | 0 | 0 | 0 | 0 | 0 | 0 | 0 | 0 | 0 |
| virus_Jul11.75v_3327 | 0 | 0 | 0 | 0 | 0 | 0 | 0 | 0 | 0 | 0 | 0 |
| virus_Jul11.75v_6584 | 0 | 0 | 0 | 0 | 0 | 0 | 0 | 0 | 0 | 0 | 0 |
| virus_Jul11.75v_11755 | 0 | 0 | 0 | 0 | 0 | 0 | 0 | 0 | 0 | 0 | 0 |
| virus_Jul11.75v_20801 | 0 | 0 | 0 | 0 | 0 | 0 | 0 | 0 | 0 | 0 | 0 |
| virus_Jul11.75v_24996 | 0 | 0 | 0 | 0 | 0 | 0 | 0 | 0 | 0 | 0 | 0 |
| virus_Jul11.75v_29943 | 0 | 0 | 0 | 0 | 0 | 0 | 0 | 0 | 0 | 0 | 0 |
| virus_Jul11.75v_35859 | 0 | 0 | 0 | 0 | 0 | 0 | 0 | 0 | 0 | 0 | 0 |
| virus_Jul27.5v_1048 | 0 | 0 | 0 | 0 | 0 | 0 | 0 | 0 | 0 | 0 | 0 |
| virus_Jul27.5v_1642 | 0 | 0 | 0 | 0 | 0 | 0 | 0 | 0 | 0 | 0 | 0 |
| virus_Jul27.5v_2167 | 0 | 0 | 0 | 0 | 0 | 0 | 0 | 0 | 0 | 0 | 0 |
| virus_Jul27.5v_2735 | 0 | 0 | 0 | 0 | 0 | 0 | 0 | 0 | 0 | 0 | 0 |
| virus_Jul27.5v_4455 | 0 | 0 | 0 | 0 | 0 | 0 | 0 | 0 | 0 | 0 | 0 |
| virus_Jul27.5v_7648 | 0 | 0 | 0 | 0 | 0 | 0 | 0 | 0 | 0 | 0 | 0 |
| virus_Jul27.5v_8002 | 0 | 0 | 0 | 0 | 0 | 0 | 0 | 0 | 0 | 0 | 0 |
| virus_Jul27.5v_9780 | 0 | 0 | 0 | 0 | 0 | 0 | 0 | 0 | 0 | 0 | 0 |
| virus_Jul27.5v_12543 | 0 | 0 | 0 | 0 | 0 | 0 | 0 | 0 | 0 | 0 | 0 |
| virus_Jul27.5v_13117 | 0 | 0 | 0 | 0 | 0 | 0 | 0 | 0 | 0 | 0 | 0 |
| virus_Jul27.5v_13619 | 0 | 0 | 0 | 0 | 0 | 0 | 0 | 0 | 0 | 0 | 0 |
| virus_Jul27.5v_14056 | 0 | 0 | 0 | 0 | 0 | 0 | 0 | 0 | 0 | 0 | 0 |
| virus_Jul27.5v_14572 | 0 | 0 | 0 | 0 | 0 | 0 | 0 | 0 | 0 | 0 | 0 |
| virus_Jul27.5v_14825 | 0 | 0 | 0 | 0 | 0 | 0 | 0 | 0 | 0 | 0 | 0 |
| virus_Jul27.5v_17228 | 0 | 0 | 0 | 0 | 0 | 0 | 0 | 0 | 0 | 0 | 0 |
| virus_Jul27.5v_18212 | 0 | 0 | 0 | 0 | 0 | 0 | 0 | 0 | 0 | 0 | 0 |
| virus_Jul27.5v_19511 | 0 | 0 | 0 | 0 | 0 | 0 | 0 | 0 | 0 | 0 | 0 |
| virus_Jul27.5v_20574 | 0 | 0 | 0 | 0 | 0 | 0 | 0 | 0 | 0 | 0 | 0 |
| virus_Jul27.25v_5603 | 0 | 0 | 0 | 0 | 0 | 0 | 0 | 0 | 0 | 0 | 0 |
| virus_Jul27.25v_11841 | 0 | 0 | 0 | 0 | 0 | 0 | 0 | 0 | 0 | 0 | 0 |
| virus_Jul27.25v_13112 | 0 | 0 | 0 | 0 | 0 | 0 | 0 | 0 | 0 | 0 | 0 |
| virus_Jul27.25v_16854 | 0 | 0 | 0 | 0 | 0 | 0 | 0 | 0 | 0 | 0 | 0 |
| virus_Jul27.25v_19127 | 0 | 0 | 0 | 0 | 0 | 0 | 0 | 0 | 0 | 0 | 0 |
| virus_Jul27.25v_27739 | 0 | 0 | 0 | 0 | 0 | 0 | 0 | 0 | 0 | 0 | 0 |
| virus_Jul27.25v_28745 | 0 | 0 | 0 | 0 | 0 | 0 | 0 | 0 | 0 | 0 | 0 |
| virus_Jul27.25v_32390 | 0 | 0 | 0 | 0 | 0 | 0 | 0 | 0 | 0 | 0 | 0 |
| virus_Jul27.25v_32724 | 0 | 0 | 0 | 0 | 0 | 0 | 0 | 0 | 0 | 0 | 0 |
| virus_Jul27.25v_33018 | 0 | 0 | 0 | 0 | 0 | 0 | 0 | 0 | 0 | 0 | 0 |
| virus_Jul27.25v_35410 | 0 | 0 | 0 | 0 | 0 | 0 | 0 | 0 | 0 | 0 | 0 |
| virus_Jul27.25v_37043 | 0 | 0 | 0 | 0 | 0 | 0 | 0 | 0 | 0 | 0 | 0 |
| virus_Jul27.25v_39824 | 0 | 0 | 0 | 0 | 0 | 0 | 0 | 0 | 0 | 0 | 0 |
| virus_Jul27.75v_2284 | 0 | 0 | 0 | 0 | 0 | 0 | 0 | 0 | 0 | 0 | 0 |
| virus_Jul27.75v_2820 | 0 | 0 | 0 | 0 | 0 | 0 | 0 | 0 | 0 | 0 | 0 |
| virus_Jul27.75v_4233 | 0 | 0 | 0 | 0 | 0 | 0 | 0 | 0 | 0 | 0 | 0 |
| virus_Jul27.75v_6963 | 0 | 0 | 0 | 0 | 0 | 0 | 0 | 0 | 0 | 0 | 0 |
| virus_Jul27.75v_18485 | 0 | 0 | 0 | 0 | 0 | 0 | 0 | 0 | 0 | 0 | 0 |
| virus_Jun8.25v_21223 | 0 | 0 | 0 | 0 | 0 | 0 | 0 | 0 | 0 | 0 | 0 |
| virus_Jun8.25v_27543 | 0 | 0 | 0 | 0 | 0 | 0 | 0 | 0 | 0 | 0 | 0 |
| virus_Jun8.50v_610 | 0 | 0 | 0 | 0 | 0 | 0 | 0 | 0 | 0 | 0 | 0 |
| virus_Jun8.50v_663 | 0 | 0 | 0 | 0 | 0 | 0 | 0 | 0 | 0 | 0 | 0 |
| virus_Jun8.50v_5606 | 0 | 0 | 0 | 0 | 0 | 0 | 0 | 0 | 0 | 0 | 0 |

| | | | | | | | | | | | |
|---|---|---|---|---|---|---|---|---|---|---|---|
| virus_Jun8.50v_9744 | 0 | 0 | 0 | 0 | 0 | 0 | 0 | 0 | 0 | 0 | 0 |
| virus_Jun8.50v_12412 | 0 | 0 | 0 | 0 | 0 | 0 | 0 | 0 | 0 | 0 | 0 |
| virus_Jun8.50v_16697 | 0 | 0 | 0 | 0 | 0 | 0 | 0 | 0 | 0 | 0 | 0 |
| virus_Jun8.50v_17108 | 0 | 0 | 0 | 0 | 0 | 0 | 0 | 0 | 0 | 0 | 0 |
| virus_Jun8.50v_19258 | 0 | 0 | 0 | 0 | 0 | 0 | 0 | 0 | 0 | 0 | 0 |
| virus_Jun8.50v_21260 | 0 | 0 | 0 | 0 | 0 | 0 | 0 | 0 | 0 | 0 | 0 |
| virus_Jun8.50v_23094 | 0 | 0 | 0 | 0 | 0 | 0 | 0 | 0 | 0 | 0 | 0 |
| virus_Jun8.50v_32175 | 0 | 0 | 0 | 0 | 0 | 0 | 0 | 0 | 0 | 0 | 0 |
| virus_Jun8.50v_37894 | 0 | 0 | 0 | 0 | 0 | 0 | 0 | 0 | 0 | 0 | 0 |
| virus_Jun8.75v_14531 | 0 | 0 | 0 | 0 | 0 | 0 | 0 | 0 | 0 | 0 | 0 |
| virus_Jun8.75v_16443 | 0 | 0 | 0 | 0 | 0 | 0 | 0 | 0 | 0 | 0 | 0 |
| virus_Jun8.75v_17650 | 0 | 0 | 0 | 0 | 0 | 0 | 0 | 0 | 0 | 0 | 0 |
| virus_Jun22.0v_628 | 0 | 0 | 0 | 0 | 0 | 0 | 0 | 0 | 0 | 0 | 0 |
| virus_Jun22.0v_6661 | 0 | 0 | 0 | 0 | 0 | 0 | 0 | 0 | 0 | 0 | 0 |
| virus_Jun22.0v_8077 | 0 | 0 | 0 | 0 | 0 | 0 | 0 | 0 | 0 | 0 | 0 |
| virus_Jun22.0v_10341 | 0 | 0 | 0 | 0 | 0 | 0 | 0 | 0 | 0 | 0 | 0 |
| virus_Jun22.0v_11443 | 0 | 0 | 0 | 0 | 0 | 0 | 0 | 0 | 0 | 0 | 0 |
| virus_Jun22.0v_13622 | 0 | 0 | 0 | 0 | 0 | 0 | 0 | 0 | 0 | 0 | 0 |
| virus_Jun22.0v_14147 | 0 | 0 | 0 | 0 | 0 | 0 | 0 | 0 | 0 | 0 | 0 |
| virus_Jun22.0v_15267 | 0 | 0 | 0 | 0 | 0 | 0 | 0 | 0 | 0 | 0 | 0 |
| virus_Jun22.0v_15322 | 0 | 0 | 0 | 0 | 0 | 0 | 0 | 0 | 0 | 0 | 0 |
| virus_Jun22.0v_17184 | 0 | 0 | 0 | 0 | 0 | 0 | 0 | 0 | 0 | 0 | 0 |
| virus_Jun22.0v_18743 | 0 | 0 | 0 | 0 | 0 | 0 | 0 | 0 | 0 | 0 | 0 |
| virus_Jun22.0v_19420 | 0 | 0 | 0 | 0 | 0 | 0 | 0 | 0 | 0 | 0 | 0 |
| virus_Jun22.0v_19636 | 0 | 0 | 0 | 0 | 0 | 0 | 0 | 0 | 0 | 0 | 0 |
| virus_Jun22.50v_1003 | 0 | 0 | 0 | 0 | 0 | 0 | 0 | 0 | 0 | 0 | 0 |
| virus_Jun22.50v_3776 | 0 | 0 | 0 | 0 | 0 | 0 | 0 | 0 | 0 | 0 | 0 |
| virus_Jun22.50v_5212 | 0 | 0 | 0 | 0 | 0 | 0 | 0 | 0 | 0 | 0 | 0 |
| virus_Jun22.50v_5520 | 0 | 0 | 0 | 0 | 0 | 0 | 0 | 0 | 0 | 0 | 0 |
| virus_Jun22.50v_10823 | 0 | 0 | 0 | 0 | 0 | 0 | 0 | 0 | 0 | 0 | 0 |
| virus_Jun22.50v_16755 | 0 | 0 | 0 | 0 | 0 | 0 | 0 | 0 | 0 | 0 | 0 |
| virus_Jun22.50v_20221 | 0 | 0 | 0 | 0 | 0 | 0 | 0 | 0 | 0 | 0 | 0 |
| virus_Jun22.50v_24243 | 0 | 0 | 0 | 0 | 0 | 0 | 0 | 0 | 0 | 0 | 0 |
| virus_Jun22.50v_29480 | 0 | 0 | 0 | 0 | 0 | 0 | 0 | 0 | 0 | 0 | 0 |
| virus_Jun22.50v_36324 | 0 | 0 | 0 | 0 | 0 | 0 | 0 | 0 | 0 | 0 | 0 |
| virus_Jun22.50v_36973 | 0 | 0 | 0 | 0 | 0 | 0 | 0 | 0 | 0 | 0 | 0 |
| virus_Jun22.50v_39809 | 0 | 0 | 0 | 0 | 0 | 0 | 0 | 0 | 0 | 0 | 0 |
| virus_Jun22.50v_41054 | 0 | 0 | 0 | 0 | 0 | 0 | 0 | 0 | 0 | 0 | 0 |
| virus_Jun22.75v_705 | 0 | 0 | 0 | 0 | 0 | 0 | 0 | 0 | 0 | 0 | 0 |
| virus_Jun22.75v_1417 | 0 | 0 | 0 | 0 | 0 | 0 | 0 | 0 | 0 | 0 | 0 |
| virus_Jun22.75v_3936 | 0 | 0 | 0 | 0 | 0 | 0 | 0 | 0 | 0 | 0 | 0 |
| virus_Jun22.75v_4550 | 0 | 0 | 0 | 0 | 0 | 0 | 0 | 0 | 0 | 0 | 0 |
| virus_Jun22.75v_20711 | 0 | 0 | 0 | 0 | 0 | 0 | 0 | 0 | 0 | 0 | 0 |
| virus_Jun22.75v_26391 | 0 | 0 | 0 | 0 | 0 | 0 | 0 | 0 | 0 | 0 | 0 |
| virus_Jun22.75v_28430 | 0 | 0 | 0 | 0 | 0 | 0 | 0 | 0 | 0 | 0 | 0 |
| virus_Jun22.75v_36860 | 0 | 0 | 0 | 0 | 0 | 0 | 0 | 0 | 0 | 0 | 0 |
| virus_Jun22.75v_38535 | 0 | 0 | 0 | 0 | 0 | 0 | 0 | 0 | 0 | 0 | 0 |
| virus_May12.5v_5981 | 0 | 0 | 0 | 0 | 0 | 0 | 0 | 0 | 0 | 0 | 0 |
| virus_May12.5v_9250 | 0 | 0 | 0 | 0 | 0 | 0 | 0 | 0 | 0 | 0 | 0 |
| virus_May12.5v_14281 | 0 | 0 | 0 | 0 | 0 | 0 | 0 | 0 | 0 | 0 | 0 |
| virus_May12.5v_16513 | 0 | 0 | 0 | 0 | 0 | 0 | 0 | 0 | 0 | 0 | 0 |
| virus_May12.5v_20899 | 0 | 0 | 0 | 0 | 0 | 0 | 0 | 0 | 0 | 0 | 0 |
| virus_May12.5v_35100 | 0 | 0 | 0 | 0 | 0 | 0 | 0 | 0 | 0 | 0 | 0 |
| virus_May12.5v_35952 | 0 | 0 | 0 | 0 | 0 | 0 | 0 | 0 | 0 | 0 | 0 |
| virus_May12.25v_3321 | 0 | 0 | 0 | 0 | 0 | 0 | 0 | 0 | 0 | 0 | 0 |
| virus_May12.25v_6494 | 0 | 0 | 0 | 0 | 0 | 0 | 0 | 0 | 0 | 0 | 0 |
| virus_May12.25v_12960 | 0 | 0 | 0 | 0 | 0 | 0 | 0 | 0 | 0 | 0 | 0 |
| virus_May12.25v_13959 | 0 | 0 | 0 | 0 | 0 | 0 | 0 | 0 | 0 | 0 | 0 |
| virus_May12.25v_18041 | 0 | 0 | 0 | 0 | 0 | 0 | 0 | 0 | 0 | 0 | 0 |
| virus_May12.25v_24338 | 0 | 0 | 0 | 0 | 0 | 0 | 0 | 0 | 0 | 0 | 0 |
| virus_May12.25v_32267 | 0 | 0 | 0 | 0 | 0 | 0 | 0 | 0 | 0 | 0 | 0 |
| virus_May12.25v_32412 | 0 | 0 | 0 | 0 | 0 | 0 | 0 | 0 | 0 | 0 | 0 |
| virus_May12.50v_2088 | 0 | 0 | 0 | 0 | 0 | 0 | 0 | 0 | 0 | 0 | 0 |
| virus_May12.50v_2141 | 0 | 0 | 0 | 0 | 0 | 0 | 0 | 0 | 0 | 0 | 0 |
| virus_May12.50v_3367 | 0 | 0 | 0 | 0 | 0 | 0 | 0 | 0 | 0 | 0 | 0 |
| virus_May12.50v_4477 | 0 | 0 | 0 | 0 | 0 | 0 | 0 | 0 | 0 | 0 | 0 |
| virus_May12.50v_5150 | 0 | 0 | 0 | 0 | 0 | 0 | 0 | 0 | 0 | 0 | 0 |
| virus_May12.50v_5341 | 0 | 0 | 0 | 0 | 0 | 0 | 0 | 0 | 0 | 0 | 0 |
| virus_May12.50v_9190 | 0 | 0 | 0 | 0 | 0 | 0 | 0 | 0 | 0 | 0 | 0 |
| virus_May12.50v_10911 | 0 | 0 | 0 | 0 | 0 | 0 | 0 | 0 | 0 | 0 | 0 |
| virus_May12.50v_14924 | 0 | 0 | 0 | 0 | 0 | 0 | 0 | 0 | 0 | 0 | 0 |
| virus_May12.50v_16415 | 0 | 0 | 0 | 0 | 0 | 0 | 0 | 0 | 0 | 0 | 0 |
| virus_May12.50v_16743 | 0 | 0 | 0 | 0 | 0 | 0 | 0 | 0 | 0 | 0 | 0 |
| virus_May12.50v_23648 | 0 | 0 | 0 | 0 | 0 | 0 | 0 | 0 | 0 | 0 | 0 |
| virus_May12.50v_35285 | 0 | 0 | 0 | 0 | 0 | 0 | 0 | 0 | 0 | 0 | 0 |
| virus_May12.75v_5202 | 0 | 0 | 0 | 0 | 0 | 0 | 0 | 0 | 0 | 0 | 0 |
| virus_May12.75v_7025 | 0 | 0 | 0 | 0 | 0 | 0 | 0 | 0 | 0 | 0 | 0 |
| virus_May12.75v_7338 | 0 | 0 | 0 | 0 | 0 | 0 | 0 | 0 | 0 | 0 | 0 |
| virus_May12.75v_7417 | 0 | 0 | 0 | 0 | 0 | 0 | 0 | 0 | 0 | 0 | 0 |
| virus_May12.75v_13775 | 0 | 0 | 0 | 0 | 0 | 0 | 0 | 0 | 0 | 0 | 0 |
| virus_May12.75v_13877 | 0 | 0 | 0 | 0 | 0 | 0 | 0 | 0 | 0 | 0 | 0 |
| virus_May12.75v_14700 | 0 | 0 | 0 | 0 | 0 | 0 | 0 | 0 | 0 | 0 | 0 |
| virus_May12.75v_21511 | 0 | 0 | 0 | 0 | 0 | 0 | 0 | 0 | 0 | 0 | 0 |
| virus_May12.75v_24328 | 0 | 0 | 0 | 0 | 0 | 0 | 0 | 0 | 0 | 0 | 0 |
| virus_May12.75v_26799 | 0 | 0 | 0 | 0 | 0 | 0 | 0 | 0 | 0 | 0 | 0 |
| virus_May12.75v_26962 | 0 | 0 | 0 | 0 | 0 | 0 | 0 | 0 | 0 | 0 | 0 |
| virus_May12.75v_38312 | 0 | 0 | 0 | 0 | 0 | 0 | 0 | 0 | 0 | 0 | 0 |
| virus_Sep8.0v_205 | 0 | 0 | 0 | 0 | 0 | 0 | 0 | 0 | 0 | 0 | 0 |
| virus_Sep8.0v_2148 | 0 | 0 | 0 | 0 | 0 | 0 | 0 | 0 | 0 | 0 | 0 |
| virus_Sep8.0v_3627 | 0 | 0 | 0 | 0 | 0 | 0 | 0 | 0 | 0 | 0 | 0 |
| virus_Sep8.0v_3910 | 0 | 0 | 0 | 0 | 0 | 0 | 0 | 0 | 0 | 0 | 0 |
| virus_Sep8.0v_4218 | 0 | 0 | 0 | 0 | 0 | 0 | 0 | 0 | 0 | 0 | 0 |
| virus_Sep8.0v_4467 | 0 | 0 | 0 | 0 | 0 | 0 | 0 | 0 | 0 | 0 | 0 |
| virus_Sep8.0v_6813 | 0 | 0 | 0 | 0 | 0 | 0 | 0 | 0 | 0 | 0 | 0 |
| virus_Sep8.0v_11375 | 0 | 0 | 0 | 0 | 0 | 0 | 0 | 0 | 0 | 0 | 0 |
| virus_Sep8.0v_11931 | 0 | 0 | 0 | 0 | 0 | 0 | 0 | 0 | 0 | 0 | 0 |
| virus_Sep8.0v_15950 | 0 | 0 | 0 | 0 | 0 | 0 | 0 | 0 | 0 | 0 | 0 |
| virus_Sep8.0v_17805 | 0 | 0 | 0 | 0 | 0 | 0 | 0 | 0 | 0 | 0 | 0 |
| virus_Sep8.5v_460 | 0 | 0 | 0 | 0 | 0 | 0 | 0 | 0 | 0 | 0 | 0 |
| virus_Sep8.5v_4351 | 0 | 0 | 0 | 0 | 0 | 0 | 0 | 0 | 0 | 0 | 0 |
| virus_Sep8.5v_15891 | 0 | 0 | 0 | 0 | 0 | 0 | 0 | 0 | 0 | 0 | 0 |
| virus_Sep8.5v_20490 | 0 | 0 | 0 | 0 | 0 | 0 | 0 | 0 | 0 | 0 | 0 |
| virus_Sep8.25v_19340 | 0 | 0 | 0 | 0 | 0 | 0 | 0 | 0 | 0 | 0 | 0 |
| virus_Sep8.50v_1272 | 0 | 0 | 0 | 0 | 0 | 0 | 0 | 0 | 0 | 0 | 0 |
| virus_Sep8.50v_5442 | 0 | 0 | 0 | 0 | 0 | 0 | 0 | 0 | 0 | 0 | 0 |
| virus_Sep8.50v_7321 | 0 | 0 | 0 | 0 | 0 | 0 | 0 | 0 | 0 | 0 | 0 |
| virus_Sep8.50v_9303 | 0 | 0 | 0 | 0 | 0 | 0 | 0 | 0 | 0 | 0 | 0 |
| virus_Sep8.50v_11296 | 0 | 0 | 0 | 0 | 0 | 0 | 0 | 0 | 0 | 0 | 0 |
| virus_Sep8.50v_17089 | 0 | 0 | 0 | 0 | 0 | 0 | 0 | 0 | 0 | 0 | 0 |
| virus_Sep8.50v_21946 | 0 | 0 | 0 | 0 | 0 | 0 | 0 | 0 | 0 | 0 | 0 |
| virus_Sep8.50v_28128 | 0 | 0 | 0 | 0 | 0 | 0 | 0 | 0 | 0 | 0 | 0 |
| virus_Sep8.50v_28342 | 0 | 0 | 0 | 0 | 0 | 0 | 0 | 0 | 0 | 0 | 0 |
| virus_Sep8.50v_32606 | 0 | 0 | 0 | 0 | 0 | 0 | 0 | 0 | 0 | 0 | 0 |
| virus_Sep8.50v_35096 | 0 | 0 | 0 | 0 | 0 | 0 | 0 | 0 | 0 | 0 | 0 |
| virus_Sep8.50v_38342 | 0 | 0 | 0 | 0 | 0 | 0 | 0 | 0 | 0 | 0 | 0 |

**Supplementary Table S10**. Amplicon sequencing overview

| Sample (Date.Depth-fraction) | Date | Depth (m) | Amplicon type | Fraction | Number of QC sequences |
|---|---|---|---|---|---|
| May12.0v | 2016-05-12 | 0 | RTPR | 0.02 - 0.22 µm | NA |
| May12.5v | 2016-05-12 | 5 | RTPR | 0.02 - 0.22 µm | 973 |
| May12.25v | 2016-05-12 | 25 | RTPR | 0.02 - 0.22 µm | 1034 |
| May12.50v | 2016-05-12 | 50 | RTPR | 0.02 - 0.22 µm | 1559 |
| May12.75v | 2016-05-12 | 75 | RTPR | 0.02 - 0.22 µm | 1514 |
| Jun8.0v | 2016-06-08 | 0 | RTPR | 0.02 - 0.22 µm | NA |
| Jun8.5v | 2016-06-08 | 5 | RTPR | 0.02 - 0.22 µm | NA |
| Jun8.25v | 2016-06-08 | 25 | RTPR | 0.02 - 0.22 µm | 492 |
| Jun8.50v | 2016-06-08 | 50 | RTPR | 0.02 - 0.22 µm | 1571 |
| Jun8.75v | 2016-06-08 | 75 | RTPR | 0.02 - 0.22 µm | 367 |
| Jun22.0v | 2016-06-22 | 0 | RTPR | 0.02 - 0.22 µm | 714 |
| Jun22.5v | 2016-06-22 | 5 | RTPR | 0.02 - 0.22 µm | 618 |
| Jun22.25v | 2016-06-22 | 25 | RTPR | 0.02 - 0.22 µm | NA |
| Jun22.50v | 2016-06-22 | 50 | RTPR | 0.02 - 0.22 µm | 1852 |
| Jun22.75v | 2016-06-22 | 75 | RTPR | 0.02 - 0.22 µm | 1326 |
| Jul11.0v | 2016-07-11 | 0 | RTPR | 0.02 - 0.22 µm | 641 |
| Jul11.5v | 2016-07-11 | 5 | RTPR | 0.02 - 0.22 µm | 1094 |
| Jul11.25v | 2016-07-11 | 25 | RTPR | 0.02 - 0.22 µm | 960 |
| Jul11.50v | 2016-07-11 | 50 | RTPR | 0.02 - 0.22 µm | 854 |
| Jul11.75v | 2016-07-11 | 75 | RTPR | 0.02 - 0.22 µm | 838 |
| Jul27.0v | 2016-07-27 | 5 | RTPR | 0.02 - 0.22 µm | NA |
| Jul27.5v | 2016-07-27 | 5 | RTPR | 0.02 - 0.22 µm | 1523 |
| Jul27.25v | 2016-07-27 | 25 | RTPR | 0.02 - 0.22 µm | 1045 |
| Jul27.50v | 2016-07-27 | 50 | RTPR | 0.02 - 0.22 µm | 1112 |
| Jul27.75v | 2016-07-27 | 75 | RTPR | 0.02 - 0.22 µm | 249 |
| Aug5.0v | 2016-08-05 | 0 | RTPR | 0.02 - 0.22 µm | 2604 |
| Aug5.5v | 2016-08-05 | 5 | RTPR | 0.02 - 0.22 µm | 1143 |
| Aug5.25v | 2016-08-05 | 25 | RTPR | 0.02 - 0.22 µm | 1684 |
| Aug5.50v | 2016-08-05 | 50 | RTPR | 0.02 - 0.22 µm | 635 |
| Aug5.75v | 2016-08-05 | 75 | RTPR | 0.02 - 0.22 µm | 709 |
| Sep8.0v | 2016-09-08 | 0 | RTPR | 0.02 - 0.22 µm | 666 |
| Sep8.5v | 2016-09-08 | 5 | RTPR | 0.02 - 0.22 µm | 640 |
| Sep8.25v | 2016-09-08 | 25 | RTPR | 0.02 - 0.22 µm | 425 |
| Sep8.50v | 2016-09-08 | 50 | RTPR | 0.02 - 0.22 µm | 1631 |
| Sep8.75v | 2016-09-08 | 50 | RTPR | 0.02 - 0.22 µm | NA |
| May12d0b | 2016-05-12 | 0 | RTPR | > 0.22 µm | 3703 |
| May12d5b | 2016-05-12 | 5 | RTPR | > 0.22 µm | 57 |
| May12d25b | 2016-05-12 | 25 | RTPR | > 0.22 µm | 999 |
| May12d50b | 2016-05-12 | 50 | RTPR | > 0.22 µm | 623 |
| May12d75b | 2016-05-12 | 75 | RTPR | > 0.22 µm | 85 |
| Jun8d25b | 2016-06-08 | 25 | RTPR | > 0.22 µm | 1846 |
| Jun8d50b | 2016-06-08 | 50 | RTPR | > 0.22 µm | 676 |
| Jun8d75b | 2016-06-08 | 75 | RTPR | > 0.22 µm | 188 |
| Jun22d0b | 2016-06-22 | 0 | RTPR | > 0.22 µm | 436 |
| Jun22d25b | 2016-06-22 | 25 | RTPR | > 0.22 µm | 1289 |
| Jun22d50b | 2016-06-22 | 50 | RTPR | > 0.22 µm | 453 |
| Jun22d75b | 2016-06-22 | 75 | RTPR | > 0.22 µm | 695 |
| Jul11d0b | 2016-07-11 | 0 | RTPR | > 0.22 µm | 756 |
| Jul11d5b | 2016-07-11 | 5 | RTPR | > 0.22 µm | 366 |
| Jul11d25b | 2016-07-11 | 25 | RTPR | > 0.22 µm | 570 |
| Jul11d50b | 2016-07-11 | 50 | RTPR | > 0.22 µm | 351 |
| Jul11d75b | 2016-07-11 | 75 | RTPR | > 0.22 µm | 764 |
| Jul27d0b | 2016-07-27 | 0 | RTPR | > 0.22 µm | 4091 |
| Jul27d5b | 2016-07-27 | 5 | RTPR | > 0.22 µm | 2013 |
| Jul27d25b | 2016-07-27 | 25 | RTPR | > 0.22 µm | 1411 |
| Jul27d75b | 2016-07-27 | 75 | RTPR | > 0.22 µm | 1332 |
| Aug5d0b | 2016-08-05 | 0 | RTPR | > 0.22 µm | 588 |
| Aug5d5b | 2016-08-05 | 5 | RTPR | > 0.22 µm | 226 |
| Aug5d25b | 2016-08-05 | 25 | RTPR | > 0.22 µm | 1999 |
| Aug5d50b | 2016-08-05 | 50 | RTPR | > 0.22 µm | 572 |
| Aug5d75b | 2016-08-05 | 75 | RTPR | > 0.22 µm | 167 |
| Sep8d0b | 2016-09-08 | 0 | RTPR | > 0.22 µm | 164 |
| Sep8d5b | 2016-09-08 | 5 | RTPR | > 0.22 µm | 1607 |
| Sep8d25b | 2016-09-08 | 25 | RTPR | > 0.22 µm | 189 |
| Sep8d50b | 2016-09-08 | 50 | RTPR | > 0.22 µm | 121 |
| Sep8d75b | 2016-09-08 | 75 | RTPR | > 0.22 µm | 1508 |
| May12d0 | 2016-05-12 | 0 | 16S rDNA | > 0.22 µm | 21121 |
| May12d5 | 2016-05-12 | 5 | 16S rDNA | > 0.22 µm | 15579 |
| May12d25 | 2016-05-12 | 25 | 16S rDNA | > 0.22 µm | 56214 |
| May12d50 | 2016-05-12 | 50 | 16S rDNA | > 0.22 µm | 31693 |
| May12d75 | 2016-05-12 | 75 | 16S rDNA | > 0.22 µm | 44420 |
| Jun8d0 | 2016-06-08 | 0 | 16S rDNA | > 0.22 µm | 43527 |
| Jun8d5 | 2016-06-08 | 5 | 16S rDNA | > 0.22 µm | 17396 |
| Jun8d25 | 2016-06-08 | 25 | 16S rDNA | > 0.22 µm | 25563 |
| Jun8d50 | 2016-06-08 | 50 | 16S rDNA | > 0.22 µm | 17422 |
| Jun8d75 | 2016-06-08 | 75 | 16S rDNA | > 0.22 µm | 19713 |
| Jun22d0 | 2016-06-22 | 0 | 16S rDNA | > 0.22 µm | 8994 |
| Jun22d5 | 2016-06-22 | 5 | 16S rDNA | > 0.22 µm | 31765 |
| Jun22d25 | 2016-06-22 | 25 | 16S rDNA | > 0.22 µm | 17107 |
| Jun22d50 | 2016-06-22 | 50 | 16S rDNA | > 0.22 µm | 22436 |
| Jun22d75 | 2016-06-22 | 75 | 16S rDNA | > 0.22 µm | 35561 |
| Jul11d0 | 2016-07-11 | 0 | 16S rDNA | > 0.22 µm | 41201 |
| Jul11d5 | 2016-07-11 | 5 | 16S rDNA | > 0.22 µm | 62296 |
| Jul11d25 | 2016-07-11 | 25 | 16S rDNA | > 0.22 µm | 19278 |
| Jul11d50 | 2016-07-11 | 50 | 16S rDNA | > 0.22 µm | 17046 |
| Jul11d75 | 2016-07-11 | 75 | 16S rDNA | > 0.22 µm | 9061 |
| Jul27d0 | 2016-07-27 | 0 | 16S rDNA | > 0.22 µm | 22915 |
| Jul27d5 | 2016-07-27 | 5 | 16S rDNA | > 0.22 µm | 15373 |
| Jul27d25 | 2016-07-27 | 25 | 16S rDNA | > 0.22 µm | 12357 |
| Jul27d50 | 2016-07-27 | 50 | 16S rDNA | > 0.22 µm | 19990 |
| Jul27d75 | 2016-07-27 | 75 | 16S rDNA | > 0.22 µm | 21001 |
| Aug5d0 | 2016-08-05 | 0 | 16S rDNA | > 0.22 µm | 26995 |
| Aug5d5 | 2016-08-05 | 5 | 16S rDNA | > 0.22 µm | 16138 |
| Aug5d25 | 2016-08-05 | 25 | 16S rDNA | > 0.22 µm | 47639 |
| Aug5d50 | 2016-08-05 | 50 | 16S rDNA | > 0.22 µm | 47639 |
| Aug5d75 | 2016-08-05 | 75 | 16S rDNA | > 0.22 µm | 21963 |
| Sep8d0 | 2016-09-08 | 0 | 16S rDNA | > 0.22 µm | 42591 |
| Sep8d5 | 2016-09-08 | 5 | 16S rDNA | > 0.22 µm | 20200 |
| Sep8d25 | 2016-09-08 | 25 | 16S rDNA | > 0.22 µm | 15533 |
| Sep8d50 | 2016-09-08 | 50 | 16S rDNA | > 0.22 µm | 22105 |
| Sep8d75 | 2016-09-08 | 75 | 16S rDNA | > 0.22 µm | 17313 |

**Supplementary Table S11**. Determination of environmental factors correlated with community composition

(A) Mantel test between virioplankton composition similarity matrix based on RTPR phylogenetic clades and metadata similarity matrices

| Tested metadata | Spearman | | Pearson | |
|---|---|---|---|---|
| | r | p-value | r | p-value |
| Day | 0.42 | 0.001* | 0.42 | 0.001* |
| Depth | 0.19 | 0.008* | 0.22 | 0.006* |
| Temperature | 0.48 | 0.001* | 0.51 | 0.001* |
| Density | 0.44 | 0.001* | 0.48 | 0.001* |
| Current Velocity | 0.07 | 0.211 | -0.01 | 0.496 |
| Salinity | 0.16 | 0.029* | 0.12 | 0.091 |
| Nanoeukaryote Abundance | 0.06 | 0.226 | 0.02 | 0.376 |
| Picoeukaryote Abundance | 0.10 | 0.085 | 0.05 | 0.256 |
| Prochlorococcus Abundance | -0.08 | 0.861 | -0.09 | 0.877 |
| Synechococcus Abundance | 0.12 | 0.096 | 0.02 | 0.398 |
| Total Autotroph Abundance | 0.05 | 0.276 | 0.01 | 0.413 |
| Viral abundance | 0.03 | 0.315 | 0.04 | 0.297 |
| Bacterial abundance | -0.08 | 0.911 | -0.10 | 0.951 |

(B) Mantel test between virioplankton composition similarity matrix based on RTPR 98% OTUs and metadata similarity matrices

| Tested metadata | Spearman | | Pearson | |
|---|---|---|---|---|
| | r | p-value | r | p-value |
| Day | 0.30 | 0.001* | 0.27 | 0.001* |
| Depth | 0.15 | 0.014* | 0.17 | 0.005* |
| Temperature | 0.38 | 0.001* | 0.44 | 0.001* |
| Density | 0.35 | 0.001* | 0.43 | 0.001* |
| Current Velocity | 0.11 | 0.143 | 0.06 | 0.289 |
| Salinity | 0.14 | 0.081 | 0.09 | 0.197 |
| Nanoeukaryote | 0.14 | 0.065 | 0.09 | 0.197 |
| Picoeukaryote Abundance | 0.14 | 0.079 | 0.07 | 0.216 |
| Prochlorococcus | -0.13 | 0.938 | -0.16 | 0.967 |
| Synechococcus Abundance | 0.09 | 0.159 | 0.02 | 0.387 |
| Total Autotroph | 0.09 | 0.154 | 0.02 | 0.387 |
| Viral abundance | 0.08 | 0.152 | 0.08 | 0.194 |
| Bacterial abundance | -0.03 | 0.627 | -0.04 | 0.694 |

(C) Mantel test between bacterioplankton composition similarity matrix and metadata similarity matrices

| Tested metadata | Spearman | | Pearson | |
|---|---|---|---|---|
| | r | p-value | r | p-value |
| Day | 0.35 | 0.001* | 0.37 | 0.001* |
| Depth | 0.45 | 0.001* | 0.47 | 0.001* |
| Temperature | 0.42 | 0.001* | 0.46 | 0.001* |
| Density | 0.46 | 0.001* | 0.50 | 0.001* |
| Current Velocity | -0.04 | 0.690 | -0.02 | 0.878 |
| Salinity | 0.15 | 0.103 | 0.15 | 0.145 |
| Nanoeukaryote Abundance | 0.10 | 0.326 | 0.08 | 0.493 |
| Picoeukaryote Abundance | 0.06 | 0.549 | 0.04 | 0.733 |
| Prochlorococcus Abundance | -0.01 | 0.945 | -0.08 | 0.493 |
| Synechococcus Abundance | 0.11 | 0.282 | 0.03 | 0.745 |
| Total Autotroph Abundance | 0.08 | 0.441 | 0.05 | 0.642 |
| Viral abundance | 0.11 | 0.122 | 0.10 | 0.248 |
| Bacterial abundance | 0.06 | 0.432 | 0.04 | 0.643 |

**p*-value <.05 permutation = 999

**Supplementary Table S12.** Beta group diversity significance based on day and depth

(A) PERMANOVA test of virioplankton community beta group diversity significance based on RTPR phylogenetic clades across sampling day

| Group 1 | Group2 | Sample | pseudo-F | p-value | q-value |
|---|---|---|---|---|---|
| 12-May | 8-Jun | 7 | 1.272 | 0.334 | 0.438 |
| 12-May | 22-Jun | 8 | 4.340 | 0.024* | 0.091 |
| 12-May | Jul11 | 9 | 5.096 | 0.019* | 0.091 |
| 12-May | Jul27 | 8 | 9.294 | 0.026* | 0.091 |
| 12-May | Aug5 | 9 | 15.921 | 0.012* | 0.091 |
| 12-May | Sep8 | 8 | 3.191 | 0.025* | 0.091 |
| 8-Jun | Jun22 | 7 | 1.950 | 0.097 | 0.185 |
| 8-Jun | Jul11 | 8 | 2.453 | 0.052 | 0.109 |
| 8-Jun | Jul27 | 7 | 4.299 | 0.032* | 0.096 |
| 8-Jun | Aug5 | 8 | 8.345 | 0.022* | 0.091 |
| 8-Jun | Sep8 | 7 | 1.586 | 0.261 | 0.365 |
| 22-Jun | Jul11 | 9 | 0.574 | 0.750 | 0.788 |
| 22-Jun | 27-Jul | 8 | 1.398 | 0.246 | 0.365 |
| 22-Jun | Aug5 | 9 | 3.985 | 0.037* | 0.097 |
| 22-Jun | Sep8 | 8 | 0.558 | 0.823 | 0.823 |
| 11-Jul | Jul27 | 9 | 1.030 | 0.428 | 0.529 |
| 11-Jul | Aug5 | 10 | 3.476 | 0.042* | 0.098 |
| 11-Jul | Sep8 | 9 | 0.638 | 0.728 | 0.788 |
| 27-Jul | Aug5 | 9 | 1.777 | 0.141 | 0.228 |
| 27-Jul | Sep8 | 8 | 0.830 | 0.555 | 0.648 |
| 5-Aug | Sep8 | 9 | 1.610 | 0.133 | 0.228 |

(B) PERMANOVA test of virioplankton community beta group diversity significance based on RTPR 98% OTUs across sampling day

| Group 1 | Group2 | Sample | pseudo-F | p-value | q-value |
|---|---|---|---|---|---|
| 12-May | 8-Jun | 7 | 1.827 | 0.04* | 0.095 |
| 12-May | 22-Jun | 8 | 1.263 | 0.153 | 0.262 |
| 12-May | Jul11 | 9 | 2.101 | 0.016* | 0.090 |
| 12-May | Jul27 | 8 | 2.437 | 0.025* | 0.090 |
| 12-May | Aug5 | 9 | 3.749 | 0.008* | 0.090 |
| 12-May | Sep8 | 8 | 1.983 | 0.03* | 0.090 |
| 8-Jun | Jun22 | 7 | 1.365 | 0.140 | 0.262 |
| 8-Jun | Jul11 | 8 | 1.611 | 0.045* | 0.105 |
| 8-Jun | Jul27 | 7 | 1.989 | 0.056 | 0.118 |
| 8-Jun | Aug5 | 8 | 3.882 | 0.019* | 0.090 |
| 8-Jun | Sep8 | 7 | 1.458 | 0.176 | 0.264 |
| 22-Jun | Jul11 | 9 | 0.890 | 0.640 | 0.704 |
| 22-Jun | 27-Jul | 8 | 1.103 | 0.256 | 0.358 |
| 22-Jun | Aug5 | 9 | 2.019 | 0.026* | 0.090 |
| 22-Jun | Sep8 | 8 | 0.915 | 0.505 | 0.589 |
| 11-Jul | Jul27 | 9 | 1.017 | 0.369 | 0.456 |
| 11-Jul | Aug5 | 10 | 2.164 | 0.029* | 0.090 |
| 11-Jul | Sep8 | 9 | 0.869 | 0.670 | 0.704 |
| 27-Jul | Aug5 | 9 | 1.066 | 0.281 | 0.369 |
| 27-Jul | Sep8 | 8 | 0.761 | 0.770 | 0.770 |
| 5-Aug | Sep8 | 9 | 1.480 | 0.162 | 0.262 |

(C) PERMANOVA test of bacterioplankton community beta group diversity significance across sampling day

| Group 1 | Group2 | Sample | pseudo-F | p-value | q-value |
|---|---|---|---|---|---|
| 12-May | 8-Jun | 10 | 2.066 | 0.005* | 0.042* |
| 12-May | 22-Jun | 10 | 2.229 | 0.023* | 0.069 |
| 12-May | Jul11 | 10 | 3.922 | 0.007* | 0.042* |
| 12-May | Jul27 | 10 | 3.548 | 0.009* | 0.042* |
| 12-May | Aug5 | 10 | 3.141 | 0.012* | 0.042* |
| 12-May | Sep8 | 10 | 4.365 | 0.008* | 0.042* |
| 8-Jun | Jun22 | 10 | 1.147 | 0.263 | 0.29 |
| 8-Jun | Jul11 | 10 | 2.371 | 0.036* | 0.084 |
| 8-Jun | Jul27 | 10 | 1.891 | 0.042* | 0.088 |
| 8-Jun | Aug5 | 10 | 2.046 | 0.031* | 0.081 |
| 8-Jun | Sep8 | 10 | 2.988 | 0.012* | 0.042* |
| 22-Jun | Jul11 | 10 | 1.586 | 0.136 | 0.196 |
| 22-Jun | 27-Jul | 10 | 1.328 | 0.169 | 0.209 |
| 22-Jun | Aug5 | 10 | 1.449 | 0.147 | 0.196 |
| 22-Jun | Sep8 | 10 | 2.291 | 0.058 | 0.111 |
| 11-Jul | Jul27 | 10 | 1.13 | 0.277 | 0.291 |
| 11-Jul | Aug5 | 10 | 1.191 | 0.236 | 0.275 |
| 11-Jul | Sep8 | 10 | 1.548 | 0.113 | 0.196 |
| 27-Jul | Aug5 | 10 | 0.929 | 0.464 | 0.464 |
| 27-Jul | Sep8 | 10 | 1.604 | 0.131 | 0.196 |
| 5-Aug | Sep8 | 10 | 1.486 | 0.149 | 0.196 |

(D) PERMANOVA test of virioplankton community beta group diversity significance based on RTPR phylogenetic clades across sampling depths

| Group 1 | Group2 | Sample | pseudo-F | p-value | q-value |
|---|---|---|---|---|---|
| 0m | 5m | 10 | 2.268 | 0.101 | 0.165 |
| 0m | 25m | 10 | 2.026 | 0.115 | 0.165 |
| 0m | 50m | 11 | 3.105 | 0.039* | 0.125 |
| 0m | 75m | 10 | 6.688 | 0.01* | 0.100 |
| 5m | 25m | 12 | 1.318 | 0.237 | 0.263 |
| 5m | 50m | 13 | 2.503 | 0.050 | 0.125 |
| 5m | 75m | 12 | 3.492 | 0.025* | 0.125 |
| 25m | 50m | 13 | 0.557 | 0.685 | 0.685 |
| 25m | 75m | 12 | 1.760 | 0.121 | 0.165 |
| 50m | 75m | 13 | 1.769 | 0.132 | 0.165 |

(D) PERMANOVA test of virioplankton community beta group diversity significance based on RTPR 98% OTUs across sampling depths

| Group 1 | Group2 | Sample | pseudo-F | p-value | q-value |
|---|---|---|---|---|---|
| 0m | 5m | 10 | 1.104 | 0.294 | 0.368 |
| 0m | 25m | 10 | 1.246 | 0.217 | 0.362 |
| 0m | 50m | 11 | 1.608 | 0.067 | 0.223 |
| 0m | 75m | 10 | 2.370 | 0.013* | 0.130 |
| 5m | 25m | 12 | 0.944 | 0.458 | 0.509 |
| 5m | 50m | 13 | 1.125 | 0.279 | 0.368 |
| 5m | 75m | 12 | 0.869 | 0.571 | 0.571 |
| 25m | 50m | 13 | 1.255 | 0.146 | 0.362 |
| 25m | 75m | 12 | 1.738 | 0.055 | 0.223 |
| 50m | 75m | 13 | 1.242 | 0.183 | 0.362 |

(F) PERMANOVA test of bacterioplankton community beta group diversity significance across sampling depths

| Group 1 | Group2 | Sample | pseudo-F | p-value | q-value |
|---|---|---|---|---|---|
| 0m | 5m | 14 | 0.412 | 0.993 | 0.993 |
| 0m | 25m | 14 | 0.862 | 0.561 | 0.701 |
| 0m | 50m | 14 | 2.318 | 0.013* | 0.026* |
| 0m | 75m | 14 | 4.469 | 0.001* | 0.001* |
| 5m | 25m | 14 | 0.847 | 0.634 | 0.704 |
| 5m | 50m | 14 | 2.388 | 0.007* | 0.018* |
| 5m | 75m | 14 | 4.596 | 0.002* | 0.007* |
| 25m | 50m | 14 | 1.22 | 0.171 | 0.244 |
| 25m | 75m | 14 | 3.183 | 0.002* | 0.007* |
| 50m | 75m | 14 | 1.519 | 0.040* | 0.067 |

Permutation time: 999; three decimal places were kept.
*p-value or q-value < 0.05

**Supplementary Table S13.** Proportionality tests within and between 221 16S ASVs (in samples with RTPR viral amplicons) and 17 RTPR phylogenetic clades

| Feature_ID | Feature_number | ASV1 | ASV2 | ASV3 | ASV4 | ASV5 | ASV6 | ASV7 | ASV8 | ASV9 | [illegible] |
|---|---|---|---|---|---|---|---|---|---|---|---|
| e5cad0f83c5da3a8e4f138aac9423460 | ASV1 | 1 | 0.213557 | -0.273029 | 0.853361 | 0.636629 | 0.568959 | 0.105416 | 0.326286 | 0.722327 | [illegible] |
| 36b651dfb1ecf5381fe35ea3c4a1061f | ASV2 | 0.213557 | 1 | 0.122933 | 0.327105 | 0.376278 | 0.340636 | 0.49118 | 0.52808 | 0.486706 | [illegible] |
| 1d72f3f992bcd714b2a0c1b37d614dd2 | ASV3 | -0.273029 | 0.122933 | 1 | -0.293201 | -0.401793 | -0.523347 | -0.210983 | -0.25139 | 0.147816 | [illegible] |
| 936be975fb97fd3b6b5829e1c83be6bd | ASV4 | 0.853361 | 0.327105 | -0.293201 | 1 | 0.829239 | 0.740229 | 0.283642 | 0.520759 | 0.604348 | [illegible] |
| a6a79b8a486f3cdaf899162d34b8035a | ASV5 | 0.636629 | 0.376278 | -0.401793 | 0.829239 | 1 | 0.824233 | 0.600304 | 0.804157 | 0.381932 | [illegible] |
| 1122e6b7ea8ece945016e945302ee1ad | ASV6 | 0.568959 | 0.340636 | -0.523347 | 0.740229 | 0.824233 | 1 | 0.576123 | 0.66936 | 0.223503 | [illegible] |
| a3d4f90910b762e33dd4d344f63357f8 | ASV7 | 0.105416 | 0.49118 | -0.210983 | 0.283642 | 0.600304 | 0.576123 | 1 | 0.905548 | 0.000332 | [illegible] |
| a756a146af3f113055fcd3b5ebebe18f | ASV8 | 0.326286 | 0.52808 | -0.25139 | 0.520759 | 0.804157 | 0.66936 | 0.905548 | 1 | 0.187709 | [illegible] |
| b77bf87ad1c17314baa9d7c9acc10a3b | ASV9 | 0.722327 | 0.486706 | 0.147816 | 0.604348 | 0.381932 | 0.223503 | 0.000332 | 0.187709 | 1 | [illegible] |
| 2d4bbf1e16d0492f979b8c5510decc4c | ASV10 | 0.600098 | 0.299019 | -0.515818 | 0.764539 | 0.805707 | 0.973417 | 0.551696 | 0.653143 | 0.195258 | [illegible] |
| 0194fcb71fb1f171d26db7ddaeb20811 | ASV11 | 0.768423 | 0.211869 | -0.405603 | 0.937266 | 0.849397 | 0.784731 | 0.275415 | 0.500493 | 0.394688 | [illegible] |
| 2b7c6eeca2efe4e826759346836efc05 | ASV12 | 0.649281 | 0.06765 | -0.058932 | 0.484115 | 0.256515 | 0.202906 | -0.016416 | 0.093348 | 0.383525 | [illegible] |
| c1b1851dde5aaca57cc59dc350c37be7 | ASV13 | 0.664495 | 0.396404 | -0.17621 | 0.69601 | 0.735626 | 0.473136 | 0.419215 | 0.664858 | 0.63893 | [illegible] |
| 8b0192de06c3b10f85bc482236c92366 | ASV14 | 0.507592 | -0.003027 | -0.082089 | 0.442888 | 0.239126 | 0.178499 | -0.080895 | 0.046096 | 0.261687 | [illegible] |
| c7d8e6d1fd5ae641fff2b96fa1112d7b | ASV15 | -0.644334 | -0.021605 | 0.663593 | -0.728386 | -0.571013 | -0.641557 | -0.069683 | -0.249827 | -0.188031 | [illegible] |
| 81092899f1c0c2db02ec2ab6e65d86f2 | ASV16 | 0.277282 | 0.496506 | -0.294134 | 0.494574 | 0.787355 | 0.72852 | 0.937318 | 0.966597 | 0.099418 | [illegible] |
| 1c54ed87c1705610cad2896671526efe | ASV17 | -0.418853 | -0.129931 | 0.039872 | -0.505828 | -0.302301 | -0.238257 | -0.014952 | -0.125197 | -0.343423 | [illegible] |
| a5e6de786770e07687674cc32d097319 | ASV18 | 0.878261 | 0.401507 | -0.316679 | 0.855085 | 0.73701 | 0.716493 | 0.275546 | 0.486415 | 0.748891 | [illegible] |
| b4aaa30f8f64f72ba6cfd139f2023a8d | ASV19 | -0.177234 | 0.56352 | 0.208248 | -0.139604 | -0.058598 | -0.028721 | 0.411554 | 0.219019 | 0.092856 | [illegible] |
| e43703ac625ad9fda149e97e827b0553 | ASV20 | 0.199744 | -0.157187 | -0.442085 | 0.221039 | 0.542969 | 0.508777 | 0.593307 | 0.600008 | -0.150145 | [illegible] |
| a35da92fe92a502bc206941a91a633bc | ASV21 | -0.121531 | 0.063048 | 0.009377 | -0.015449 | 0.071859 | 0.214478 | 0.488439 | 0.348831 | -0.317323 | [illegible] |
| 91f687fdb62026d040baa83be4ad4556 | ASV22 | -0.19823 | -0.090344 | -0.12054 | -0.330432 | -0.173074 | -0.084636 | 0.045059 | -0.05286 | -0.20476 | [illegible] |
| 5234f65d442af2c647f49e161bb35ffa | ASV23 | 0.319207 | -0.194873 | -0.082958 | 0.390175 | 0.21664 | 0.091778 | -0.201461 | -0.000481 | 0.027352 | [illegible] |
| 8941b912fff03d55d958a89a8510b9c2 | ASV24 | 0.798045 | 0.13182 | -0.408601 | 0.9192 | 0.788867 | 0.689447 | 0.16769 | 0.418977 | 0.424259 | [illegible] |
| 3ab38fc647166c931ad18a473daa0560 | ASV25 | 0.548103 | 0.100289 | -0.042558 | 0.56019 | 0.262827 | 0.229925 | -0.135838 | 0.029173 | 0.452974 | [illegible] |
| 46eb9145a13d564c00f01e905866be62 | ASV26 | -0.485266 | 0.013922 | 0.644327 | -0.606075 | -0.536242 | -0.582499 | -0.128425 | -0.280356 | -0.003422 | [illegible] |
| 1b980624ac540cbc5036300a8919d6da | ASV27 | 0.533886 | 0.653298 | 0.104977 | 0.571565 | 0.384172 | 0.35224 | 0.060963 | 0.221708 | 0.775419 | [illegible] |
| 05932a0c91ddcdbca8bcd2c6ad0849b7 | ASV28 | 0.114917 | 0.741415 | 0.251145 | 0.131904 | 0.137221 | 0.037177 | 0.221353 | 0.217603 | 0.512635 | [illegible] |
| e057f66ccacf6c694b1b69b11f19533f | ASV29 | 0.737182 | 0.289253 | -0.242786 | 0.680786 | 0.699513 | 0.629448 | 0.261229 | 0.473692 | 0.596272 | [illegible] |
| 2c6e619dbaab11dccebfaa0c0bc5d1d3 | ASV30 | 0.28091 | 0.243652 | 0.071053 | 0.278029 | 0.129098 | -0.010579 | -0.010136 | 0.13438 | 0.441748 | [illegible] |
| [illegible] | [illegible] | [illegible] | [illegible] | [illegible] | [illegible] | [illegible] | [illegible] | [illegible] | [illegible] | [illegible] | [illegible] |

**Supplementary Table S13.** Proportionality tests within and

| Feature_ID | Feature_number | ASV86 | ASV87 | ASV88 | ASV89 | ASV90 | ASV91 | ASV92 | ASV93 | ASV94 | ASV95 | ASV96 | ASV97 | ASV98 | ASV99 |
|---|---|---|---|---|---|---|---|---|---|---|---|---|---|---|---|
| e5cad0f83c5da3a8e4f138aac9423460 | ASV1 | 0.380389 | -0.11986 | 0.23092 | 0.037125 | 0.724351 | 0.501586 | -0.501782 | -0.278032 | 0.363876 | 0.454073 | -0.598193 | -0.440946 | -0.536355 | -0.495318 |
| 36b651dfb1ecf5381fe35ea3c4a1061f | ASV2 | -0.048871 | -0.157949 | 0.177552 | 0.405559 | 0.068042 | 0.024535 | 0.072302 | 0.315827 | 0.122532 | 0.015422 | -0.160406 | 0.041707 | -0.037458 | -0.254939 |
| 1d72f3f992bcd714b2a0c1b37d614dd2 | ASV3 | -0.225337 | -0.546222 | 0.343355 | 0.228405 | -0.239731 | -0.376918 | 0.619854 | 0.219128 | 0.071997 | -0.05027 | 0.058692 | 0.203318 | 0.310873 | 0.255342 |
| 936be975fb97fd3b6b5829e1c83be6bd | ASV4 | 0.393766 | 0.002533 | 0.249483 | -0.009694 | 0.600751 | 0.442055 | -0.602777 | -0.275078 | 0.390167 | 0.470514 | -0.596975 | -0.30871 | -0.544615 | -0.531089 |
| a6a79b8a486f3cdaf899162d34b8035a | ASV5 | 0.269141 | 0.162196 | 0.139119 | -0.099952 | 0.368411 | 0.347942 | -0.608517 | -0.216118 | 0.197093 | 0.190589 | -0.366451 | -0.213494 | -0.404463 | -0.450864 |
| 1122e6b7ea8ece945016e945302ee1ad | ASV6 | 0.158702 | 0.350676 | -0.035498 | -0.231063 | 0.273073 | 0.349498 | -0.536739 | -0.28629 | 0.006799 | 0.139329 | -0.298664 | -0.21411 | -0.323921 | -0.499276 |
| a3d4f90910b762e33dd4d344f63357f8 | ASV7 | -0.089614 | 0.190608 | -0.087117 | 0.011426 | -0.076723 | 0.048271 | -0.204286 | 0.099398 | -0.138448 | -0.133165 | -0.003939 | -0.011575 | -0.034955 | -0.209094 |
| a756a146af3f113055fcd3b5ebebe18f | ASV8 | 0.051091 | 0.132501 | 0.014126 | -0.003778 | 0.091465 | 0.125198 | -0.319314 | -0.022042 | 0.016085 | 0.004023 | -0.137189 | -0.088363 | -0.144433 | -0.260444 |
| b77bf87ad1c17314baa9d7c9acc10a3b | ASV9 | 0.211765 | -0.417013 | 0.376073 | 0.42717 | 0.577489 | 0.292301 | -0.12258 | 0.146194 | 0.34856 | 0.32749 | -0.475103 | -0.16918 | -0.303932 | -0.367245 |
| 2d4bbf1e16d0492f979b8c5510decc4c | ASV10 | 0.155738 | 0.329596 | -0.087049 | -0.248867 | 0.255372 | 0.356216 | -0.541722 | -0.312531 | 0.025111 | 0.142256 | -0.307817 | -0.24013 | -0.355542 | -0.480117 |
| 0194fcb71fb1f171d26db7ddaeb20811 | ASV11 | 0.433802 | 0.163633 | 0.178924 | -0.169963 | 0.534004 | 0.480062 | -0.703556 | -0.356617 | 0.353076 | 0.370369 | -0.5287 | -0.314315 | -0.571101 | -0.538713 |
| 2b7c6eeca2efe4e826759346836efc05 | ASV12 | 0.212452 | -0.16027 | 0.376944 | 0.036293 | 0.653071 | 0.404084 | -0.253425 | -0.19167 | 0.466839 | 0.588227 | -0.636121 | -0.42868 | -0.483965 | -0.414575 |
| c1b1851dde5aaca57cc59dc350c37be7 | ASV13 | 0.236867 | -0.090819 | 0.20234 | 0.201676 | 0.381548 | 0.219808 | -0.373898 | -0.046091 | 0.228701 | 0.257133 | -0.310238 | -0.198937 | -0.269762 | -0.300159 |
| 8b0192de06c3b10f85bc482236c92366 | ASV14 | 0.525935 | -0.143584 | 0.154333 | -0.089918 | 0.705276 | 0.577515 | -0.399682 | -0.277075 | 0.423594 | 0.433926 | -0.718619 | -0.619612 | -0.507038 | -0.206215 |
| c7d8e6d1fd5ae641fff2b96fa1112d7b | ASV15 | -0.463476 | -0.282742 | -0.007031 | 0.160905 | -0.558429 | -0.516536 | 0.639011 | 0.339924 | -0.336873 | -0.462841 | 0.496122 | 0.360863 | 0.546446 | 0.439227 |
| 81092899f1c0c2db02ec2ab6e65d86f2 | ASV16 | 0.012337 | 0.220752 | -0.027645 | -0.061483 | 0.040991 | 0.120364 | -0.330762 | -0.048731 | -0.04399 | -0.010459 | -0.102342 | -0.047381 | -0.145435 | -0.281107 |
| 1c54ed87c1705610cad2896671526efe | ASV17 | -0.450166 | 0.184409 | -0.516782 | -0.038429 | -0.623781 | -0.546002 | 0.347836 | 0.119423 | -0.571196 | -0.551979 | 0.891371 | 0.289555 | 0.591054 | 0.433551 |
| a5e6de786770e07687674cc32d097319 | ASV18 | 0.246029 | -0.024407 | 0.195194 | 0.114071 | 0.562124 | 0.398492 | -0.483905 | -0.152378 | 0.262056 | 0.379701 | -0.463407 | -0.248725 | -0.43063 | -0.513517 |
| b4aaa30f8f64f72ba6cfd139f2023a8d | ASV19 | -0.158961 | -0.141699 | -0.029996 | 0.271029 | -0.203108 | -0.087275 | 0.182022 | 0.479118 | -0.138738 | -0.2489 | 0.09529 | 0.099798 | 0.117677 | -0.03366 |
| e43703ac625ad9fda149e97e827b0553 | ASV20 | 0.0293 | 0.273694 | -0.21841 | -0.368297 | 0.039146 | 0.189724 | -0.416592 | -0.213817 | -0.178084 | -0.200128 | 0.018645 | -0.112958 | -0.168515 | -0.128879 |
| a35da92fe92a502bc206941a91a633bc | ASV21 | -0.121017 | 0.084254 | -0.222755 | -0.155484 | -0.22581 | -0.077193 | 0.009289 | -0.01338 | -0.20605 | -0.183401 | 0.118162 | -0.025009 | 0.082286 | 0.03106 |
| 91f687fdb62026d040baa83be4ad4556 | ASV22 | -0.416007 | 0.201966 | -0.494855 | -0.008781 | -0.422966 | -0.192964 | 0.136851 | 0.10279 | -0.548838 | -0.496323 | 0.647708 | 0.138836 | 0.327452 | 0.139374 |
| 5234f65d442af2c647f49e161bb35ffa | ASV23 | 0.576284 | -0.136806 | 0.157243 | -0.300572 | 0.434875 | 0.253455 | -0.291027 | -0.471377 | 0.357059 | 0.420165 | -0.400115 | -0.448101 | -0.30842 | 0.14984 |
| 8941b912fff03d55d958a89a8510b9c2 | ASV24 | 0.536503 | 0.062214 | 0.254936 | -0.215044 | 0.664762 | 0.496221 | -0.711766 | -0.459848 | 0.443345 | 0.521266 | -0.604392 | -0.435551 | -0.551148 | -0.443485 |
| 3ab38fc647166c931ad18a473daa0560 | ASV25 | 0.641347 | -0.311696 | 0.518111 | -0.009523 | 0.808962 | 0.438965 | -0.273157 | -0.287858 | 0.624575 | 0.789541 | -0.841918 | -0.498583 | -0.444938 | -0.23004 |
| 46eb9145a13d564c00f01e905866be62 | ASV26 | -0.436626 | -0.389098 | 0.050275 | 0.484846 | -0.427546 | -0.383473 | 0.615288 | 0.566145 | -0.185353 | -0.452215 | 0.398441 | 0.252176 | 0.446462 | 0.24838 |
| 1b980624ac540cbc5036300a8919d6da | ASV27 | 0.089725 | -0.263894 | 0.232389 | 0.419263 | 0.340897 | 0.208607 | -0.082083 | 0.16686 | 0.286345 | 0.231229 | -0.353475 | -0.089245 | -0.229756 | -0.333766 |

**Supplementary Table S13.** Proportionality tests within and

| Feature_ID | Feature_number | ASV171 | ASV172 | ASV173 | ASV174 | ASV175 | ASV176 | ASV177 | ASV178 | ASV179 | ASV180 | ASV181 | ASV182 | ASV183 | ASV184 | ASV185 | ASV186 | ASV187 | ASV188 | ASV189 | ASV190 | ASV191 | ASV192 | ASV193 | ASV194 | ASV195 | ASV196 | ASV197 | ASV198 | ASV199 | ASV200 | ASV201 | ASV202 | ASV203 | ASV204 | ASV205 | ASV206 | ASV207 | ASV208 | ASV209 | ASV210 | ASV211 | ASV212 | ASV213 | ASV214 | ASV215 | ASV216 | ASV217 | ASV218 | ASV219 | ASV220 | ASV221 | clade1 | clade2 | clade3 | clade4 | clade5 | clade6 | clade7 | clade8 | clade9 | clade10 | clade11 | clade12 | clade13 | clade14 |
|---|---|---|---|---|---|---|---|---|---|---|---|---|---|---|---|---|---|---|---|---|---|---|---|---|---|---|---|---|---|---|---|---|---|---|---|---|---|---|---|---|---|---|---|---|---|---|---|---|---|---|---|---|---|---|---|---|---|---|---|---|---|---|---|---|---|---|
| e5cad0f83c5da3a8e4f138aac9423460 | ASV1 | -0.618674 | 0.193581 | 0.442808 | -0.736729 | 0.289409 | 0.510954 | 0.222376 | 0.266638 | -0.275722 | 0.645812 | 0.381175 | -0.73455 | 0.632208 | -0.634714 | -0.472245 | -0.519396 | -0.457954 | 0.412065 | -0.324867 | 0.585122 | 0.126778 | -0.707429 | -0.538329 | -0.280514 | -0.335736 | -0.757131 | 0.447956 | -0.478239 | -0.672928 | 0.496502 | -0.606222 | 0.499937 | -0.380067 | 0.421764 | -0.513169 | 0.481519 | -0.441376 | -0.08298 | -0.162881 | 0.567855 | -0.426187 | 0.30259 | -0.572406 | -0.486253 | 0.036299 | 0.385448 | 0.022455 | -0.522139 | -0.558604 | 0.623204 | -0.113706 | 0.521232 | 0.43575 | 0.507991 | 0.3722 | -0.558165 | -0.449467 | 0.317638 | -0.030818 | -0.480961 | -0.174465 | -0.290177 | -0.557606 | 0.36218 | -0.140765 |
| 36b651dfb1ecf5381fe35ea3c4a1061f | ASV2 | -0.031924 | -0.125786 | 0.060971 | -0.182004 | 0.154757 | 0.064313 | -0.070252 | 0.286023 | 0.047806 | 0.082791 | 0.170113 | -0.084976 | 0.036807 | -0.032235 | -0.051578 | -0.142725 | -0.247778 | 0.132943 | -0.085948 | 0.087412 | -0.09164 | -0.19708 | -0.084678 | 0.009988 | -0.267942 | -0.16779 | 0.056263 | -0.071432 | -0.15309 | 0.167053 | -0.00819 | -0.01322 | -0.135942 | 0.177882 | -0.024229 | 0.091369 | -0.089669 | 0.115792 | 0.110405 | 0.055872 | -0.204433 | 0.232726 | -0.053712 | -0.08984 | 0.106976 | 0.09188 | -0.099019 | -0.082209 | -0.096991 | 0.102577 | -0.144934 | -0.07098 | -0.01956 | 0.133244 | 0.250322 | -0.108634 | 0.148677 | 0.04406 | -0.213465 | -0.520885 | -0.139011 | 0.228428 | -0.209357 | -0.171003 | 0.265399 |
| 1d72f3f992bcd714b2a0c1b37d614dd2 | ASV3 | 0.329467 | -0.390205 | -0.264751 | 0.113754 | 0.190565 | -0.424338 | 0.136471 | 0.385118 | 0.519973 | -0.460961 | -0.067291 | 0.282169 | -0.504633 | 0.298987 | 0.390286 | 0.288856 | 0.30099 | -0.18528 | -0.129268 | -0.424181 | 0.260871 | 0.206161 | 0.33825 | -0.133333 | -0.139474 | 0.324773 | -0.331342 | 0.39142 | 0.338393 | -0.144988 | 0.405256 | -0.456984 | -0.332465 | 0.088059 | 0.233065 | -0.139031 | -0.128745 | 0.386448 | 0.217332 | -0.429977 | -0.174928 | 0.221314 | 0.159976 | 0.058962 | 0.375412 | -0.284964 | -0.396962 | 0.317637 | 0.035739 | -0.416957 | -0.443398 | -0.08875 | -0.127691 | -0.129687 | 0.071295 | 0.042068 | -0.05689 | 0.035177 | 0.037238 | 0.198933 | -0.099622 | 0.314452 | 0.132215 | -0.134288 | 0.000814 |
| 936be975fb97fd3b6b5829e1c83be6bd | ASV4 | -0.51044 | 0.078389 | 0.433972 | -0.754868 | 0.393287 | 0.538193 | 0.202272 | 0.291123 | -0.199726 | 0.624385 | 0.498726 | -0.623677 | 0.627447 | -0.527674 | -0.564057 | -0.548979 | -0.66078 | 0.480759 | -0.376141 | 0.588049 | 0.181596 | -0.737313 | -0.54839 | -0.192028 | -0.283735 | -0.710176 | 0.560399 | -0.423659 | -0.61546 | 0.645658 | -0.468425 | 0.419185 | -0.300335 | 0.292654 | -0.483334 | 0.343451 | -0.451519 | 0.115005 | -0.314421 | 0.606338 | -0.479715 | 0.467979 | -0.430081 | -0.513038 | 0.195561 | 0.460423 | 0.041726 | -0.512373 | -0.491795 | 0.550419 | -0.02566 | 0.451674 | 0.146235 | 0.283714 | 0.35042 | -0.419641 | -0.258615 | 0.229051 | -0.193839 | -0.573154 | -0.081046 | -0.12427 | -0.531778 | 0.257101 | 0.011114 |
| a6a79b8a486f3cdaf899162d34b8035a | ASV5 | -0.34128 | 0.093652 | 0.387883 | -0.456623 | 0.23429 | 0.41406 | 0.106895 | 0.076596 | -0.318348 | 0.512986 | 0.406695 | -0.388205 | 0.477254 | -0.322726 | -0.444608 | -0.410268 | -0.621884 | 0.261658 | -0.18723 | 0.512914 | 0.030743 | -0.466724 | -0.359332 | -0.022396 | -0.182249 | -0.481479 | 0.461506 | -0.367847 | -0.38986 | 0.435166 | -0.356256 | 0.304936 | -0.170928 | 0.201148 | -0.277906 | 0.318309 | -0.208122 | 0.049403 | -0.245138 | 0.481544 | -0.299379 | 0.265338 | -0.222212 | -0.334242 | 0.059536 | 0.375982 | 0.092056 | -0.464868 | -0.301689 | 0.488675 | 0.065085 | 0.449425 | 0.044185 | 0.186267 | 0.324612 | -0.290783 | -0.145986 | 0.325003 | -0.353892 | -0.628672 | -0.02392 | -0.056016 | -0.551271 | 0.382883 | 0.023258 |
| 1122e6b7ea8ece945016e945302ee1ad | ASV6 | -0.275441 | 0.218534 | 0.279644 | -0.367614 | 0.038353 | 0.445878 | -0.155117 | -0.090083 | -0.319794 | 0.491883 | 0.294212 | -0.329919 | 0.451374 | -0.26461 | -0.39729 | -0.363968 | -0.597602 | 0.406583 | -0.039178 | 0.481616 | -0.217908 | -0.486098 | -0.404827 | 0.096398 | -0.228038 | -0.456407 | 0.334809 | -0.393851 | -0.435323 | 0.337014 | -0.315471 | 0.315969 | 0.125921 | 0.050455 | -0.250088 | 0.231825 | -0.0699 | -0.174105 | -0.179049 | 0.483031 | -0.211467 | 0.157042 | -0.221911 | -0.169117 | -0.064575 | 0.40631 | 0.25812 | -0.346176 | -0.220899 | 0.512601 | 0.137327 | 0.211671 | 0.218734 | 0.188918 | 0.071858 | -0.145646 | -0.069291 | 0.017953 | -0.25935 | -0.465025 | -0.136846 | 0.068314 | -0.36931 | 0.119435 | 0.070573 |
| a3d4f90910b762e33dd4d344f63357f8 | ASV7 | -0.013361 | -0.136981 | 0.095628 | -0.024606 | 0.007243 | 0.10064 | -0.169717 | -0.099865 | -0.240287 | 0.125501 | 0.129712 | 0.019642 | 0.075566 | 0.068585 | -0.098016 | -0.08222 | -0.276988 | 0.104033 | 0.138709 | 0.1546 | -0.20266 | -0.035679 | -0.097297 | 0.157056 | -0.006969 | -0.074008 | 0.065637 | -0.077914 | 0.005022 | 0.04777 | -0.079565 | 0.044492 | 0.150558 | -0.080994 | 0.025395 | 0.020197 | 0.1574 | 0.01967 | 0.013621 | 0.12308 | 0.023235 | 0.06815 | 0.090002 | 0.012017 | -0.04285 | 0.183098 | 0.135945 | -0.129581 | 0.097051 | 0.147145 | 0.153291 | -0.157137 | -0.109864 | -0.019153 | 0.013454 | 0.119884 | 0.248786 | -0.051578 | -0.394627 | -0.360902 | 0.043972 | 0.200568 | -0.106557 | -0.07139 | 0.29895 |
| a756a146af3f113055fcd3b5ebebe18f | ASV8 | -0.117027 | -0.06147 | 0.174992 | -0.195889 | 0.089634 | 0.179803 | -0.03907 | 0.008172 | -0.218886 | 0.233734 | 0.222217 | -0.118918 | 0.204326 | -0.059466 | -0.202082 | -0.192007 | -0.369022 | 0.147526 | -0.008953 | 0.247502 | -0.093216 | -0.175153 | -0.153896 | 0.061427 | -0.042472 | -0.210938 | 0.200251 | -0.141514 | -0.125312 | 0.2063 | -0.158603 | 0.126463 | 0.01967 | 0.023502 | -0.067905 | 0.095077 | -0.000205 | 0.083525 | -0.063315 | 0.224258 | -0.100625 | 0.150192 | -0.025551 | -0.130747 | 0.038619 | 0.229452 | 0.07222 | -0.227628 | -0.046858 | 0.246654 | 0.121117 | 0.131889 | -0.107165 | 0.065129 | 0.176644 | -0.013704 | 0.135734 | 0.089403 | -0.463199 | -0.582757 | 0.058698 | 0.09009 | -0.247653 | 0.089651 | 0.222042 |
| b77bf87ad1c17314baa9d7c9acc10a3b | ASV9 | -0.449798 | -0.048767 | 0.323793 | -0.568478 | 0.404127 | 0.219578 | 0.31801 | 0.535965 | -0.05503 | 0.35484 | 0.369301 | -0.543911 | 0.333724 | -0.435776 | -0.359611 | -0.454886 | -0.297524 | 0.244587 | -0.333702 | 0.304848 | 0.106399 | -0.538112 | -0.393471 | -0.208386 | -0.510508 | -0.54786 | 0.254835 | -0.339697 | -0.466695 | 0.326588 | -0.341591 | 0.314864 | -0.558041 | 0.547847 | -0.386238 | 0.443389 | -0.407527 | 0.048708 | -0.050538 | 0.268083 | -0.363913 | 0.427568 | -0.464201 | -0.389575 | 0.139351 | 0.172723 | -0.247519 | -0.284847 | -0.473843 | 0.314753 | -0.295884 | 0.349356 | 0.405484 | 0.556759 | 0.4986 | -0.668626 | -0.439911 | 0.463007 | 0.155467 | -0.352941 | -0.293112 | -0.116829 | -0.535791 | 0.227103 | -0.226241 |
| 2d4bbf1e16d0492f979b8c5510decc4c | ASV10 | -0.252054 | 0.178802 | 0.258208 | -0.381423 | 0.015749 | 0.456096 | -0.180847 | -0.111764 | -0.309439 | 0.492092 | 0.267148 | -0.338089 | 0.450374 | -0.247686 | -0.377388 | -0.327409 | -0.579581 | 0.426116 | -0.033673 | 0.48514 | -0.199269 | -0.499746 | -0.397383 | 0.088824 | -0.184956 | -0.453109 | 0.314074 | -0.330899 | -0.424379 | 0.341572 | -0.343988 | 0.313417 | 0.143732 | -0.019002 | -0.258399 | 0.194996 | -0.061004 | -0.14121 | -0.139344 | 0.49428 | -0.18901 | 0.13305 | -0.187728 | -0.141684 | -0.06097 | 0.447326 | 0.275189 | -0.330681 | -0.20631 | 0.52986 | 0.172533 | 0.201892 | 0.166429 | 0.223405 | 0.074551 | -0.087633 | -0.005835 | -0.030301 | -0.284351 | -0.473668 | -0.139777 | 0.005896 | -0.329704 | 0.081711 | 0.105739 |
| 0194fcb71fb1f171d26db7ddaeb20811 | ASV11 | -0.462309 | 0.20225 | 0.458139 | -0.650614 | 0.303682 | 0.58345 | 0.139385 | 0.108545 | -0.257763 | 0.652762 | 0.463372 | -0.570807 | 0.634499 | -0.522441 | -0.589849 | -0.519654 | -0.683802 | 0.433153 | -0.318058 | 0.633126 | 0.160605 | -0.695514 | -0.488931 | -0.13002 | -0.248508 | -0.644992 | 0.575846 | -0.48976 | -0.587719 | 0.601777 | -0.476177 | 0.418439 | -0.225191 | 0.203411 | -0.423397 | 0.380966 | -0.375577 | 0.022359 | -0.406187 | 0.647853 | -0.441645 | 0.326848 | -0.36371 | -0.419259 | 0.094339 | 0.464418 | 0.142171 | -0.565458 | -0.464981 | 0.585392 | 0.051845 | 0.535737 | 0.129233 | 0.179287 | 0.280016 | -0.378824 | -0.288839 | 0.251054 | -0.235396 | -0.534133 | -0.024111 | -0.152033 | -0.556296 | 0.379975 | 0.012781 |
| 2b7c6eeca2efe4e826759346836efc05 | ASV12 | -0.486035 | 0.050973 | 0.402358 | -0.691681 | 0.230873 | 0.455646 | 0.136496 | 0.33135 | -0.109423 | 0.562913 | 0.318727 | -0.67304 | 0.582025 | -0.628339 | -0.136575 | -0.26113 | -0.275855 | 0.328876 | -0.496435 | 0.499501 | 0.292813 | -0.558374 | -0.437126 | -0.517256 | -0.130664 | -0.678855 | 0.452504 | -0.245987 | -0.584713 | 0.541277 | -0.454679 | 0.303216 | -0.383854 | 0.478243 | -0.492428 | 0.314011 | -0.588432 | 0.114331 | 0.04707 | 0.47116 | -0.500486 | 0.273288 | -0.508412 | -0.613015 | 0.191831 | 0.244198 | -0.001179 | -0.466076 | -0.487862 | 0.492178 | -0.078787 | 0.220377 | 0.257356 | 0.257627 | 0.213566 | -0.286868 | -0.255083 | 0.100124 | -0.02843 | -0.257516 | -0.085327 | -0.194572 | -0.22161 | 0.144409 | -0.012558 |
| c1b1851dde5aaca57cc59dc350c37be7 | ASV13 | -0.395587 | -0.076373 | 0.299842 | -0.436287 | 0.300077 | 0.259839 | 0.281488 | 0.233055 | -0.195339 | 0.347296 | 0.366574 | -0.390316 | 0.353746 | -0.325276 | -0.32915 | -0.37318 | -0.353544 | 0.193822 | -0.176701 | 0.319694 | 0.145206 | -0.350106 | -0.327359 | -0.121846 | -0.118532 | -0.440983 | 0.327409 | -0.289342 | -0.358553 | 0.328568 | -0.354058 | 0.249544 | -0.260458 | 0.232503 | -0.284135 | 0.224238 | -0.207635 | 0.06214 | -0.160386 | 0.307308 | -0.194945 | 0.290239 | -0.266019 | -0.360524 | 0.075804 | 0.196204 | -0.033662 | -0.418455 | -0.276291 | 0.297814 | -2.05E-05 | 0.470638 | 0.052014 | 0.317556 | 0.548406 | -0.391225 | -0.207354 | 0.431118 | -0.172422 | -0.59212 | 0.079872 | -0.127236 | -0.53042 | 0.301262 | -0.066856 |
| 8b0192de06c3b10f85bc482236c92366 | ASV14 | -0.447212 | 0.383077 | 0.309288 | -0.612017 | 0.251476 | 0.554105 | 0.29612 | 0.2374 | -0.032925 | 0.546926 | 0.140785 | -0.492503 | 0.539612 | -0.551165 | -0.33394 | -0.434371 | -0.204875 | 0.378129 | -0.586822 | 0.578813 | 0.360312 | -0.493003 | -0.36305 | -0.375938 | -0.2157 | -0.429651 | 0.552203 | -0.250975 | -0.363031 | 0.543224 | -0.29199 | 0.331241 | -0.371045 | 0.3763 | -0.296326 | 0.447145 | -0.614356 | 0.147051 | -0.388575 | 0.569811 | -0.520897 | 0.253863 | -0.490662 | -0.325219 | 0.261342 | 0.307156 | -0.227365 | -0.232932 | -0.586227 | 0.515521 | -0.185809 | 0.33104 | 0.193772 | 0.159622 | 0.096269 | -0.300398 | -0.324362 | 0.100344 | -0.029732 | -0.190748 | -0.054879 | -0.194941 | -0.199752 | 0.231735 | -0.067885 |
| c7d8e6d1fd5ae641fff2b96fa1112d7b | ASV15 | 0.433689 | -0.36812 | -0.348138 | 0.578646 | -0.148638 | -0.602006 | -0.053302 | 0.035047 | 0.250131 | -0.632794 | -0.272904 | 0.541961 | -0.667767 | 0.540058 | 0.530928 | 0.446893 | 0.517812 | -0.446707 | 0.303016 | -0.587353 | -0.121346 | 0.658127 | 0.503578 | 0.147099 | 0.130272 | 0.584365 | -0.536577 | 0.439256 | 0.589195 | -0.545713 | 0.414971 | -0.507284 | 0.079962 | -0.127198 | 0.470123 | -0.291782 | 0.36135 | 0.126147 | 0.418185 | -0.616618 | 0.3625 | -0.137474 | 0.39963 | 0.327522 | 0.050839 | -0.427054 | -0.279791 | 0.419516 | 0.441756 | -0.545085 | -0.138608 | -0.373397 | -0.164162 | -0.173711 | -0.121974 | 0.320721 | 0.194264 | -0.056531 | 0.058129 | 0.39226 | -0.037087 | 0.328099 | 0.356778 | -0.227056 | -0.046895 |
| 81092899f1c0c2db02ec2ab6e65d86f2 | ASV16 | -0.096377 | -0.073844 | 0.168055 | -0.15716 | 0.059155 | 0.191932 | -0.096688 | -0.05399 | -0.248747 | 0.232076 | 0.208637 | -0.090726 | 0.199219 | -0.036047 | -0.204642 | -0.16371 | -0.407933 | 0.176692 | 0.043655 | 0.249935 | -0.129097 | -0.177833 | -0.180027 | 0.111841 | -0.045765 | -0.196582 | 0.181846 | -0.150793 | -0.123458 | 0.180599 | -0.161378 | 0.127117 | 0.084308 | -0.02356 | -0.058476 | 0.07364 | 0.05178 | 0.03413 | -0.074512 | 0.222172 | -0.078275 | 0.134909 | 0.002254 | -0.087589 | -0.009281 | 0.236457 | 0.143824 | -0.221398 | -0.010168 | 0.243546 | 0.140128 | 0.025413 | -0.094491 | 0.011031 | 0.111288 | 0.036709 | 0.169822 | 0.035084 | -0.448758 | -0.50769 | 0.051474 | 0.15134 | -0.228904 | 0.02936 | 0.271361 |
| 1c54ed87c1705610cad2896671526efe | ASV17 | 0.475689 | -0.107945 | -0.484655 | 0.750184 | -0.556415 | -0.609126 | -0.372363 | -0.474124 | -0.057334 | -0.609711 | -0.58805 | 0.552483 | -0.612945 | 0.563277 | 0.407515 | 0.558576 | 0.495478 | -0.489383 | 0.73902 | -0.593729 | -0.35082 | 0.681461 | 0.495087 | 0.329927 | 0.348633 | 0.577831 | -0.646621 | 0.285669 | 0.516751 | -0.711078 | 0.194379 | -0.314276 | 0.450402 | -0.401767 | 0.542454 | -0.354001 | 0.705331 | -0.259263 | 0.245389 | -0.625618 | 0.603403 | -0.554442 | 0.437239 | 0.55663 | -0.34071 | -0.339363 | 0.192757 | 0.319343 | 0.529026 | -0.441406 | 0.215422 | -0.193175 | -0.074718 | -0.109578 | -0.16033 | 0.282103 | 0.161084 | -0.086404 | 0.065189 | 0.223844 | 0.049259 | 0.049343 | 0.246034 | -0.102708 | 0.023809 |

| Supplementary Table S14. Virioplankton RTPR barcodes | | |
|---|---|---|
| SampleID | one strand of barcode sequences | sequencing library |
| May12_05m | ACACGCATGACACACTT | 1st |
| May12_25m | ATGATGTGCTACATCTT | 1st |
| May12_50m | CTGCTAGAGTCTACAGT | 1st |
| May12_75m | CTGCGTGCTCTACGACT | 1st |
| Jun8_25m | TATACATACACGCACTT | 1st |
| Jun8_50m | GCTGTCATGTACTAGCT | 1st |
| Jun8_75m | ACACGCATGACACACTT | 2nd |
| Jun22_0m | CTGCTAGAGTCTACAGT | 2nd |
| Jun22_05m | CGCATCTGTGCATGCAT | 2nd |
| Jun22_50m | TAGATGCGAGAGTAGAT | 1st |
| Jun22_75m | CATCACTACGCTAGATT | 1st |
| Jul11_0m | GTGTCTAGACAGCTGTT | 2nd |
| Jul11_05m | CTGCGTGCTCTACGACT | 2nd |
| Jul11_25m | TAGATGCGAGAGTAGAT | 2nd |
| Jul11_50m | CACTAGCTCTGACTACT | 2nd |
| Jul11_75m | GCGCGATACGATGACTT | 1st |
| Jul27_05m | GCTGTCATGTACTAGCT | 2nd |
| Jul27_25m | GACTGAGATCATGATCT | 1st |
| Jul27_50m | CGATCATCTATAGACAT | 1st |
| Jul27_75m | CGAGCTATCTCATACTT | 2nd |
| Aug5_0m | ATGATGTGCTACATCTT | 2nd |
| Aug5_05m | GTGTCTAGACAGCTGTT | 1st |
| Aug5_25m | CACTAGCTCTGACTACT | 1st |
| Aug5_50m | CGAGCTATCTCATACTT | 1st |
| Aug5_75m | ATAGAGACTCAGAGCTT | 1st |
| Sep8_0m | CGATCATCTATAGACAT | 2nd |
| Sep8_05m | CATCACTACGCTAGATT | 2nd |
| Sep8_25m | GCGCGATACGATGACTT | 2nd |
| Sep8_50m | CGCATCTGTGCATGCAT | 1st |

| **Supplementary Table S15.** Bacterioplankton 16S barcodes | | | | |
|---|---|---|---|---|
| | 1st sequencing library | | 2nd sequencing library | |
| SampleID | i7 Index | i5 Index | i7 Index | i5 Index |
| May12_0m | ACTCGCTA | TCGACTAG | CTCTCTAC | AAGGAGTA |
| May12_05m | GGAGCTAC | TCGACTAG | CGAGGCTG | AAGGAGTA |
| May12_25m | GCGTAGTA | TCGACTAG | AAGAGGCA | AAGGAGTA |
| May12_50m | CGGAGCCT | TCGACTAG | GTAGAGGA | AAGGAGTA |
| May12_75m | TACGCTGC | TCGACTAG | GCTCATGA | AAGGAGTA |
| Jun8_0m | ACTCGCTA | TTCTAGCT | ATCTCAGG | AAGGAGTA |
| Jun8_05m | GGAGCTAC | TTCTAGCT | TAAGGCGA | CTAAGCCT |
| Jun8_25m | GCGTAGTA | TTCTAGCT | CGTACTAG | CTAAGCCT |
| Jun8_50m | CGGAGCCT | TTCTAGCT | AGGCAGAA | CTAAGCCT |
| Jun8_75m | TACGCTGC | TTCTAGCT | TCCTGAGC | CTAAGCCT |
| Jun22_0m | ACTCGCTA | CCTAGAGT | GGACTCCT | CTAAGCCT |
| Jun22_05m | GGAGCTAC | CCTAGAGT | TAGGCATG | CTAAGCCT |
| Jun22_25m | GCGTAGTA | CCTAGAGT | CTCTCTAC | CTAAGCCT |
| Jun22_50m | CGGAGCCT | CCTAGAGT | CGAGGCTG | CTAAGCCT |
| Jun22_75m | TACGCTGC | CCTAGAGT | AAGAGGCA | CTAAGCCT |
| Jul11_0m | ACTCGCTA | GCGTAAGA | GTAGAGGA | CTAAGCCT |
| Jul11_05m | GGAGCTAC | GCGTAAGA | GCTCATGA | CTAAGCCT |
| Jul11_25m | GCGTAGTA | GCGTAAGA | ATCTCAGG | CTAAGCCT |
| Jul11_50m | CGGAGCCT | GCGTAAGA | TAAGGCGA | CGTCTAAT |
| Jul11_75m | TACGCTGC | GCGTAAGA | CGTACTAG | CGTCTAAT |
| Jul27_0m | ACTCGCTA | CTATTAAG | AGGCAGAA | CGTCTAAT |
| Jul27_05m | GGAGCTAC | CTATTAAG | TCCTGAGC | CGTCTAAT |
| Jul27_25m | GCGTAGTA | CTATTAAG | GGACTCCT | CGTCTAAT |
| Jul27_50m | CGGAGCCT | CTATTAAG | TAGGCATG | CGTCTAAT |
| Jul27_75m | TACGCTGC | CTATTAAG | CTCTCTAC | CGTCTAAT |
| Aug5_0m | ACTCGCTA | AAGGCTAT | CGAGGCTG | CGTCTAAT |
| Aug5_05m | GGAGCTAC | AAGGCTAT | AAGAGGCA | CGTCTAAT |
| Aug5_25m | GCGTAGTA | AAGGCTAT | GTAGAGGA | CGTCTAAT |
| Aug5_50m | CGGAGCCT | AAGGCTAT | GCTCATGA | CGTCTAAT |
| Aug5_75m | TACGCTGC | AAGGCTAT | ATCTCAGG | CGTCTAAT |
| Sep8_0m | ACTCGCTA | GAGCCTTA | TAAGGCGA | TCTCTCCG |
| Sep8_05m | GGAGCTAC | GAGCCTTA | CGTACTAG | TCTCTCCG |
| Sep8_25m | GCGTAGTA | GAGCCTTA | AGGCAGAA | TCTCTCCG |
| Sep8_50m | CGGAGCCT | GAGCCTTA | TCCTGAGC | TCTCTCCG |
| Sep8_75m | TACGCTGC | GAGCCTTA | GGACTCCT | TCTCTCCG |

| **Supplementary Table S16.** Bacterioplankton RTPR barcodes | | |
|---|---|---|
| SampleID | One strand of barcode sequences | sequencing library |
| May12_0m | GCTGTCATGTACTAGCT | 1st |
| May12_05m | ACACGCATGACACACTT | 2nd |
| May12_25m | CTGCTAGAGTCTACAGT | 1st |
| May12_50m | CTGCTAGAGTCTACAGT | 2nd |
| May12_75m | CTGCGTGCTCTACGACT | 1st |
| Jun8_0m | ATAGAGACTCAGAGCTT | 1st |
| Jun8_05m | TATACATACACGCACTT | 2nd |
| Jun8_25m | TATACATACACGCACTT | 1st |
| Jun8_50m | GTGTCTAGACAGCTGTT | 2nd |
| Jun8_75m | TGTGTATCAGTACATGT | 1st |
| Jun22_0m | ACACGCATGACACACTT | 1st |
| Jun22_05m | CATCACTACGCTAGATT | 2nd |
| Jun22_25m | GCTGTCATGTACTAGCT | 2nd |
| Jun22_50m | TAGATGCGAGAGTAGAT | 2nd |
| Jun22_75m | CATCACTACGCTAGATT | 1st |
| Jul11_0m | GATCTCTACTATATGCT | 1st |
| Jul11_05m | CACTAGCTCTGACTACT | 2nd |
| Jul11_25m | TGTGTATCAGTACATGT | 2nd |
| Jul11_50m | GATCTCTACTATATGCT | 2nd |
| Jul11_75m | GCGCGATACGATGACTT | 2nd |
| Jul27_0m | ATGATGTGCTACATCTT | 1st |
| Jul27_05m | CTGCGTGCTCTACGACT | 2nd |
| Jul27_25m | CACTAGCTCTGACTACT | 1st |
| Jul27_50m | CGATCATCTATAGACAT | 2nd |
| Jul27_75m | TACAGATAGTGTAGCGT | 1st |
| Aug5_0m | GCGCGATACGATGACTT | 1st |
| Aug5_05m | GTGTCTAGACAGCTGTT | 1st |
| Aug5_25m | CGATCATCTATAGACAT | 1st |
| Aug5_50m | CGAGCTATCTCATACTT | 1st |
| Aug5_75m | ATAGAGACTCAGAGCTT | 2nd |
| Sep8_0m | TAGATGCGAGAGTAGAT | 1st |
| Sep8_05m | CGCATCTGTGCATGCAT | 1st |
| Sep8_25m | TACAGATAGTGTAGCGT | 2nd |
| Sep8_50m | CGCATCTGTGCATGCAT | 2nd |
| Sep8_75m | CGAGCTATCTCATACTT | 2nd |
