## supplementary figures for "Ubiquitous, B_12_-dependent virioplankton utilizing ribonucleotide triphosphate reductase demonstrate interseasonal dynamics and associate with a diverse range of bacterial hosts in the pelagic ocean"

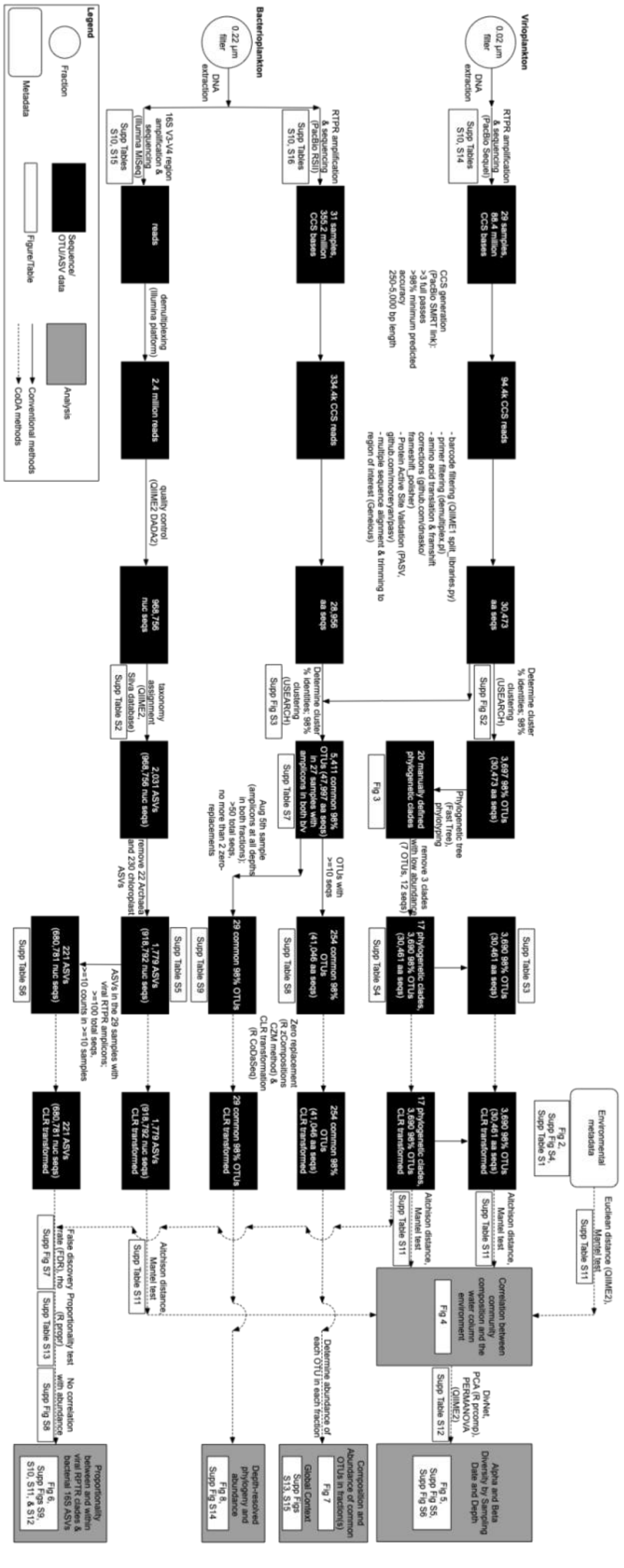


**Supplementary Figure S1.** Provenance diagram of pipeline and analyses.

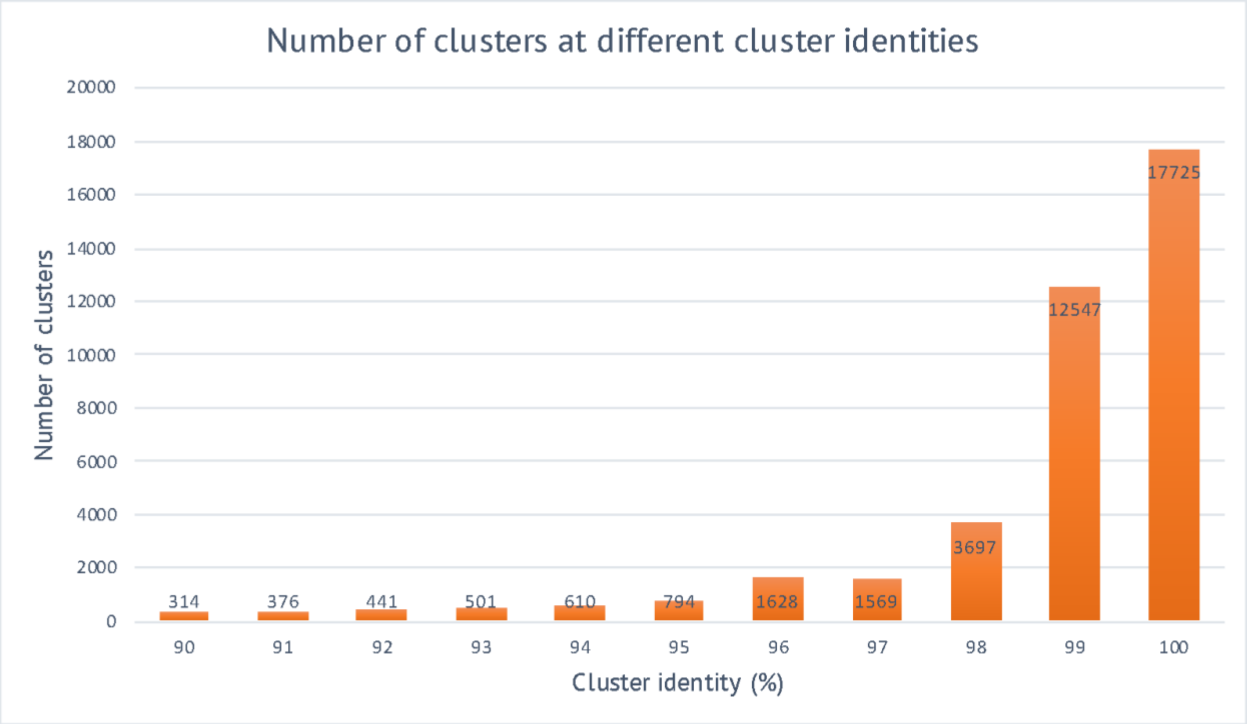


**Supplementary Figure S2.** Number of virioplankton RTPR sequence OTU clusters changed with percent cluster identity. Clustering percent identity selection was guided by the number of clusters generated at each identity.  Clustering at 98% identity was selected for subsequent analyses as this level showed a dramatic drop in the overall number of clusters, thus balancing data reduction and data loss (i.e., lumping and splitting of clusters).

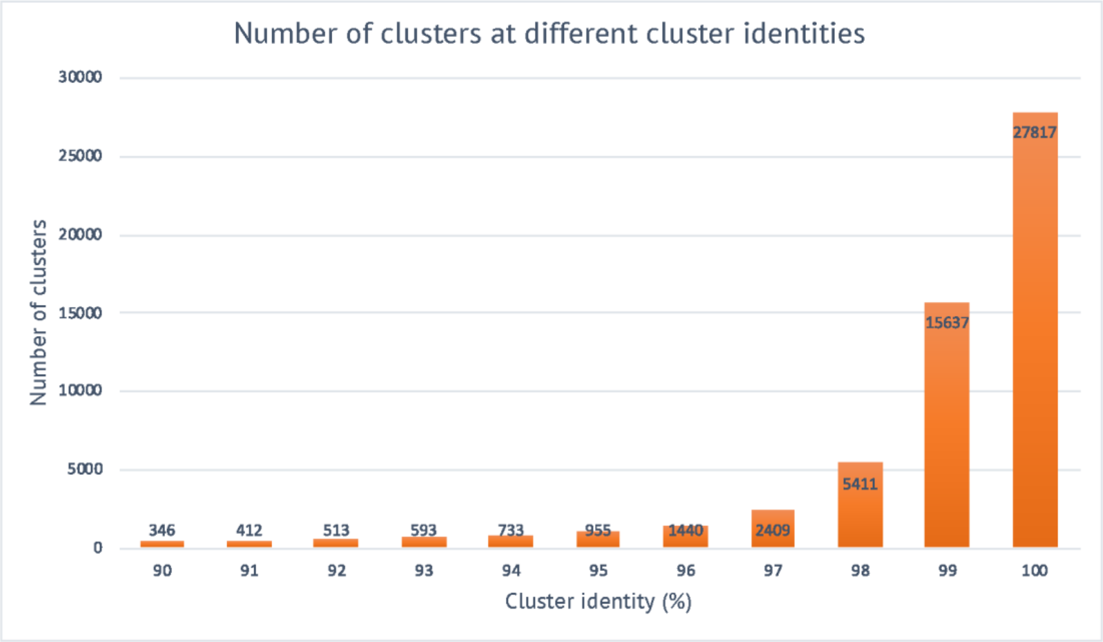


**Supplementary Figure S3.** Number of virio- and bacterioplankton common RTPR sequence OTU clusters changed with percent cluster identity. Clustering percent identity selection was guided by the number of clusters generated at each identity. Clustering at 98% identity was selected for subsequent analyses as this level showed a dramatic drop in the overall number of clusters, thus balancing data reduction and data loss (i.e., lumping and splitting of clusters).

A

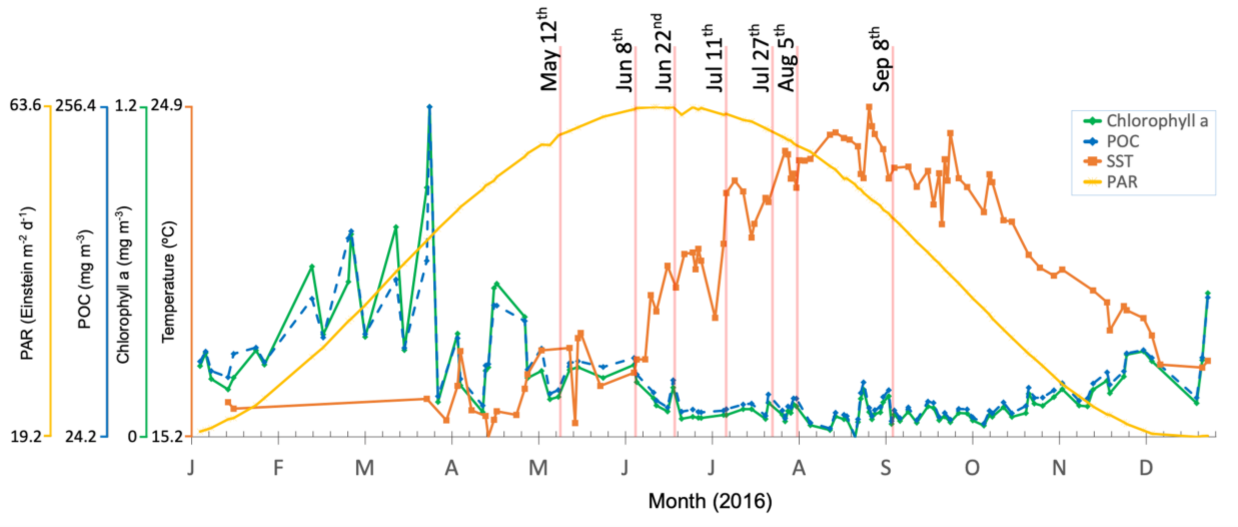


B

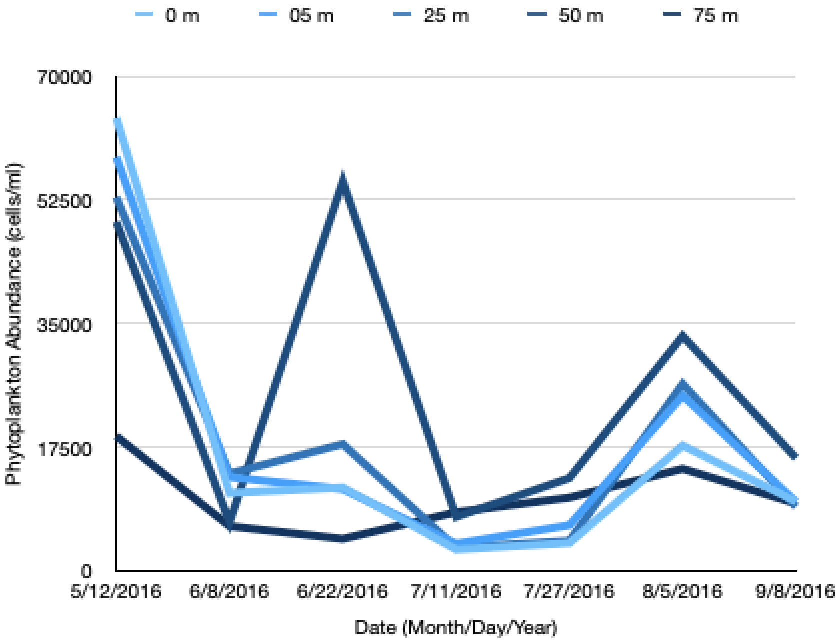


**Supplementary Figure S4.** Satellite remote sensing data and phytoplankton abundance demonstrated dramatic seasonality. **(A)** Line plots of photosynthetically available radiance (PAR) at the ocean surface, particulate organic carbon (POC), chlorophyll a, and sea surface temperature (SST) taken from satellite imagery data of the 9 $km^2$ area surrounding the sampling site. **(B)** Line plots of overall phytoplankton abundance at all sampling depths across sampling dates.

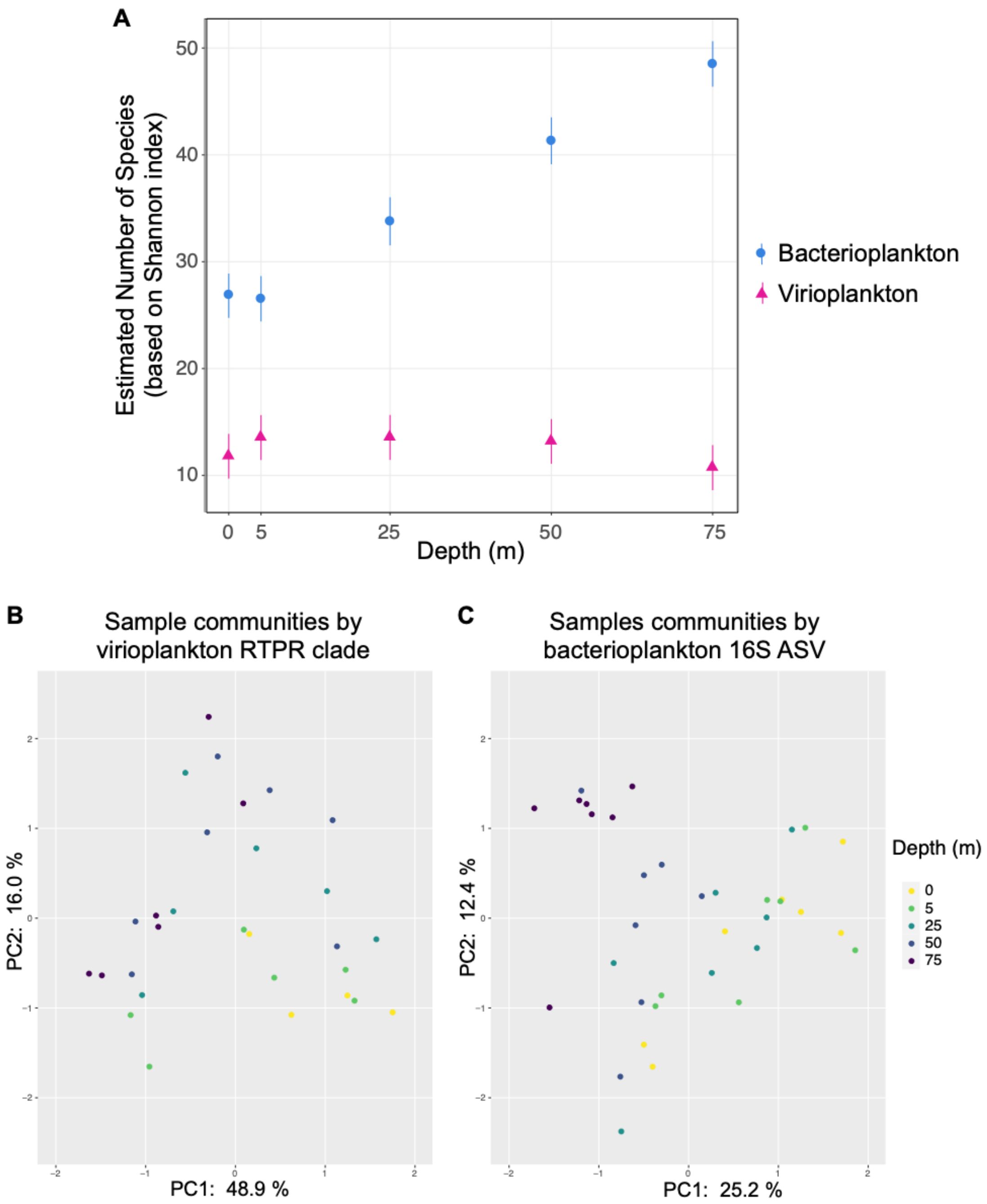


**Supplementary Figure S5.** Sampling depth influenced virioplankton (RTPR phylogenetic clade) and bacterioplankton (16S ASV) community alpha and beta diversity. **(A)** Effective number of species (ENS, based on Shannon indices) of virioplankton communities by sampling depth. Error bars represent two standard deviations of the estimates. Principal component analysis (PCA) plots of beta diversity based on centered-log ratio (clr) transformed abundance of **(B)** virioplankton RPTR phylogenetic clade communities or **(C)** bacterioplankton 16S ASV communities. Samples (circles) are colored according to sampling depth.

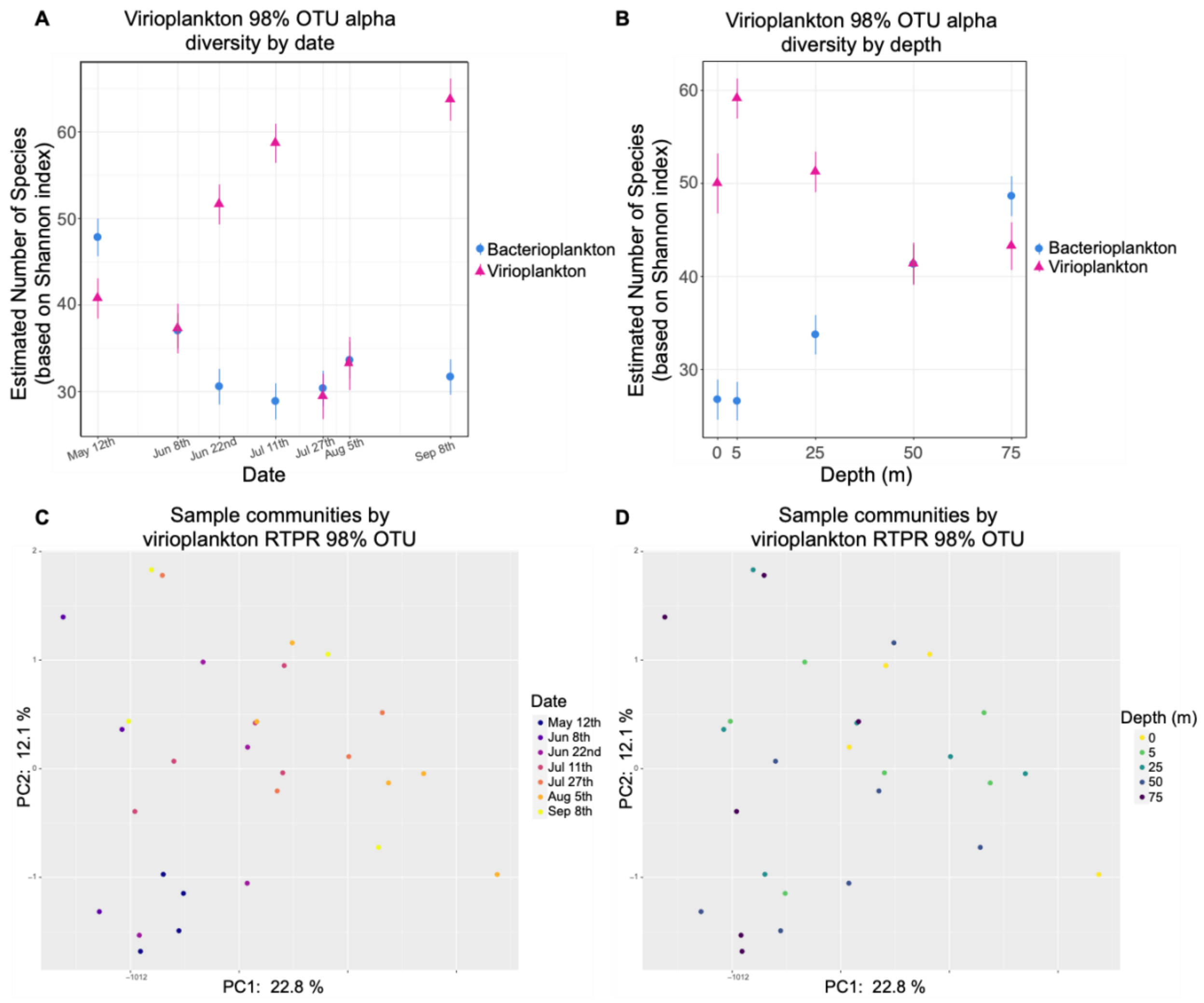


**Supplementary Figure S6.** Sampling date and depth influenced virioplankton (RTPR 98% OTUs) community alpha and beta diversity. Effective number of species (ENS, based on Shannon indices) of virioplankton communities by **(A)** sampling date or **(B)** depth. Error bars represent two standard deviations of the estimates. Principal component analysis (PCA) plots of virioplankton RPTR 98% OTU beta diversity based on centered-log ratio transformed (clr) abundance, with samples (circles) colored according to **(C)** sampling date or **(D)** depth.

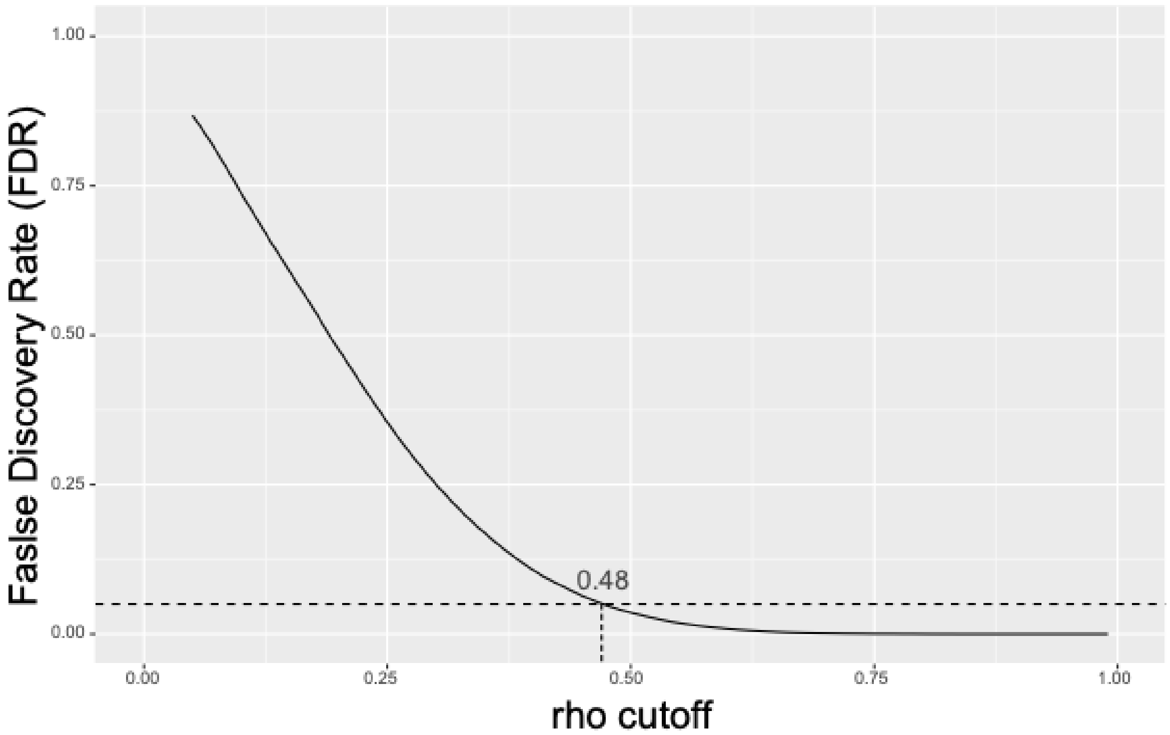


**Supplementary Figure S7.** False discovery rate (FDR) of proportionality test. A rho (ρ) cutoff of 0.48 was determined for positive associations between virioplankton or/and bacterioplankton populations in the proportionality test based on a false discovery rate less than 0.05.

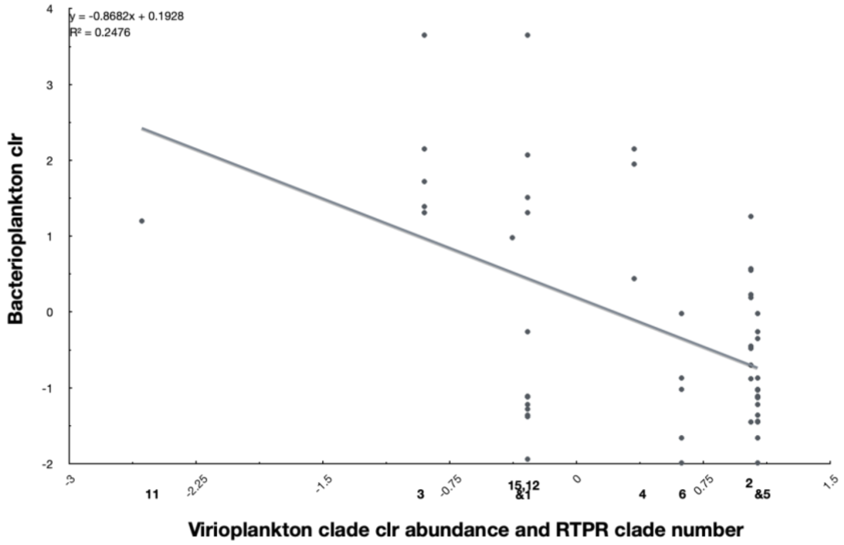


**Supplementary Figure S8**. Bacterioplankton–virioplankton associations are not driven by observed 16S ASV or RTPR clade clr abundance. RTPR clade numbers shown in bold along x-axis corresponding with their clr value.

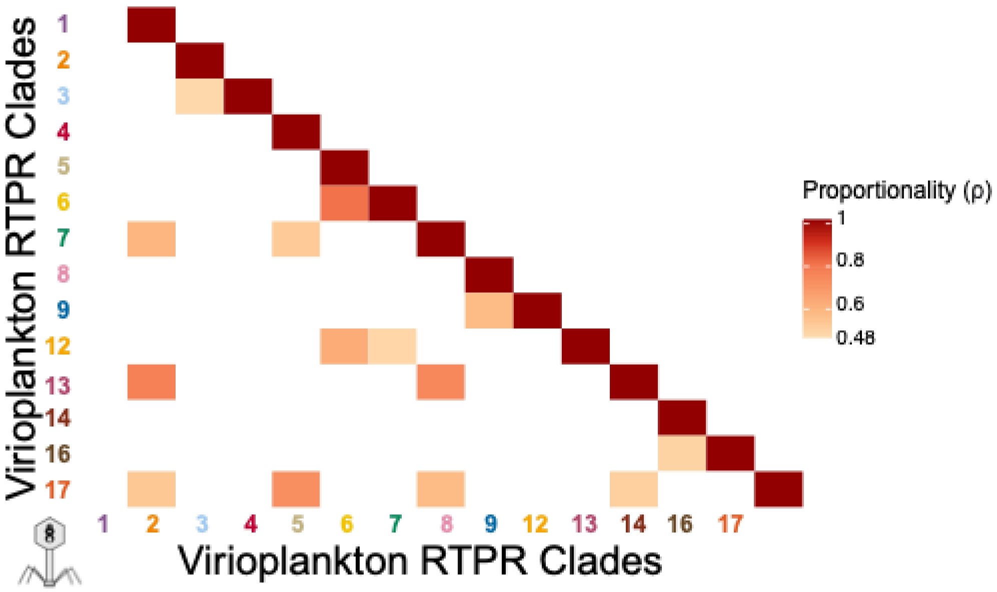


**Supplementary Figure S9**. Association matrix (ρ proportionality) within the virioplankton RTPR phylogenetic clades identified on the 98% OTU phylogenetic tree (**Fig. 3**). Only those clades with a positive non-self association (ρ ≥.48) are shown.

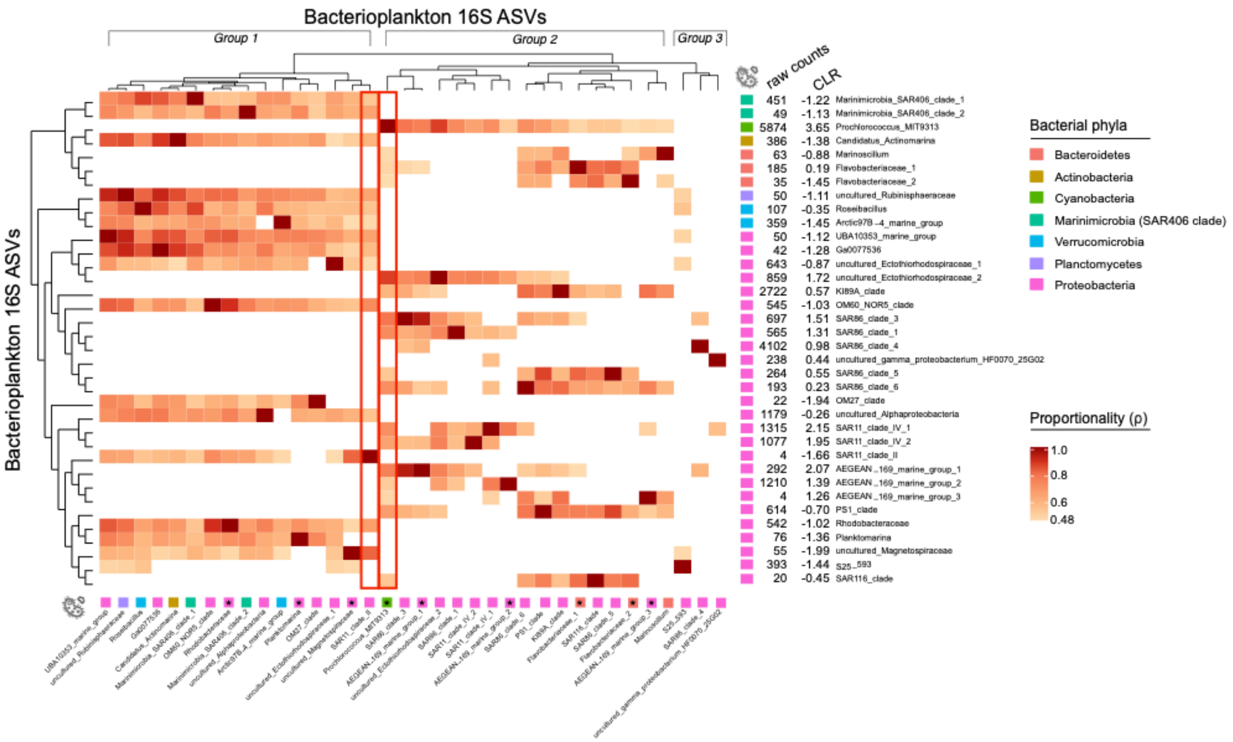


**Supplementary Figure S10**. Association matrix (ρ proportionality) within the bacterioplankton 16S ASVs. The y-axis dendrogram shows phylogenetic relationships between ASVs. The x-axis dendrogram is based upon the association patterns. Three major groups according to association patterns are shown above the x-axis dendrogram. Vertical red boxes indicate the positive associations with SAR11_clade_II and Prochlorococcus_MIT9313. Asterisk indicates those ASVs within orders, families, or genera containing species capable of $B_{12}$ synthesis according to a pangenomic survey by Heal et al. 2017 [90].

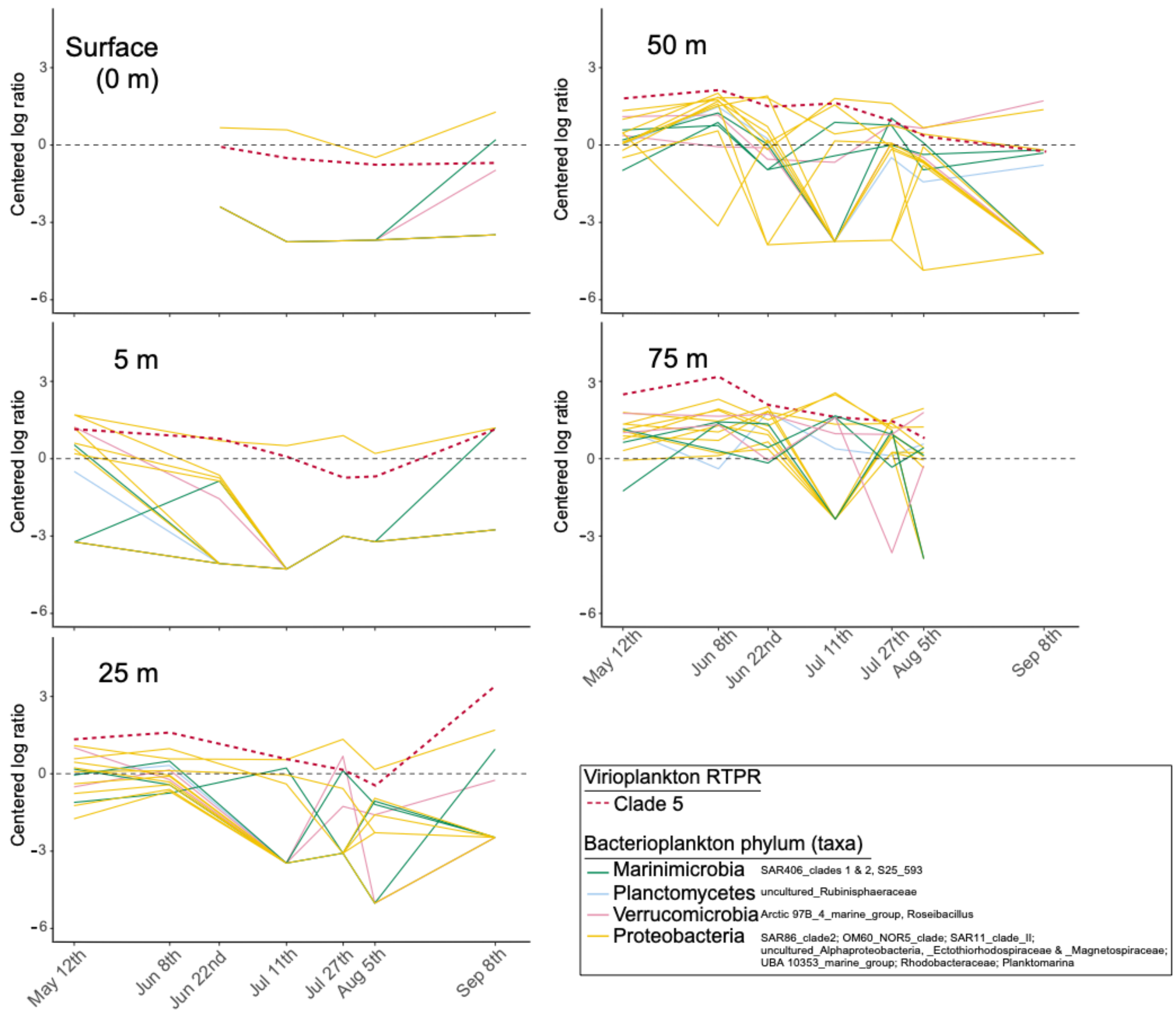


**Supplementary Figure S11**. Changes in centered log ratio abundance of virioplankton RTPR phylogenetic clade 5 and associating bacterioplankton 16S ASVs along sampling dates at each sampling depth.

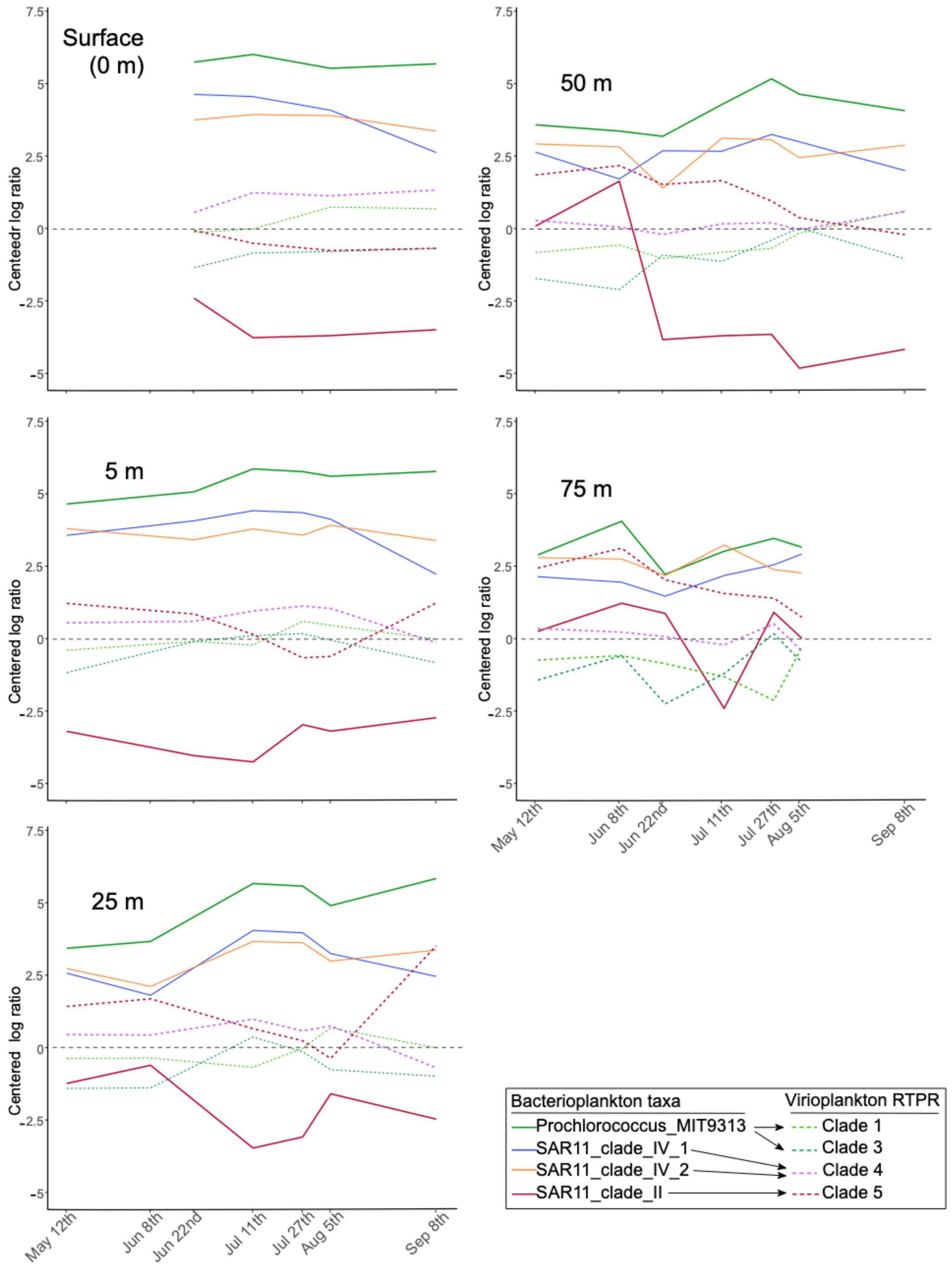


**Supplementary Figure S12**. Changes in centered log ratio abundance of Prochlorococcus_MIT9313, SAR11 clades, and associating virioplankton RTPR phylogenetic clades (1, 3, 4, and 5) along sampling dates at each sampling depth. Arrows in legend indicate positive associations between bacterioplankton taxa and virioplankton RTPR clades.

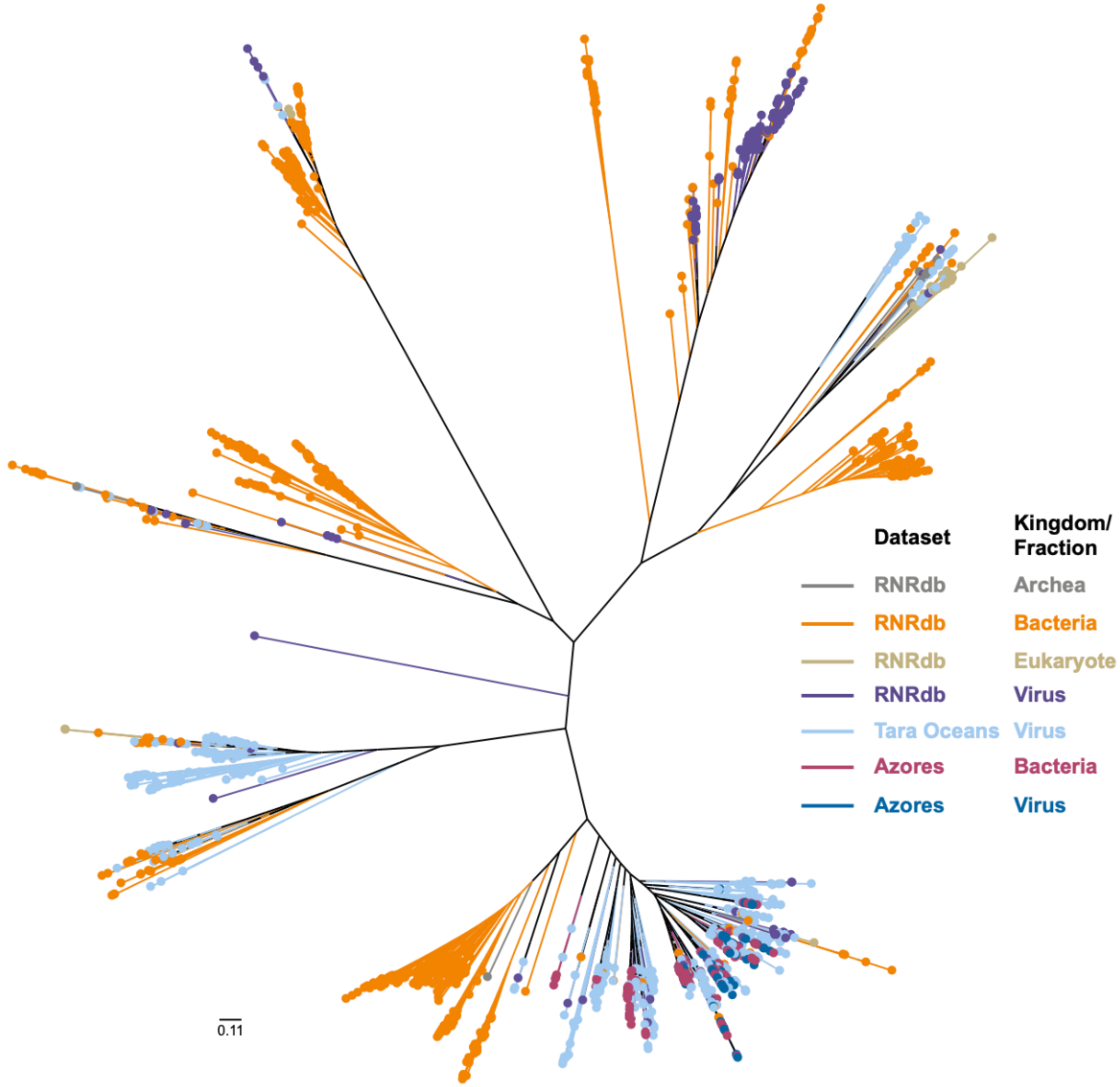


**Supplementary Figure S13**. A phylogenetic tree of RTPR 98% OTUs from the virioplankton and bacterioplankton fraction in the context of RTPR sequences from the *Tara* Ocean virome and RNRdb databases. Scale bar indicates amino acid substitutions per site.

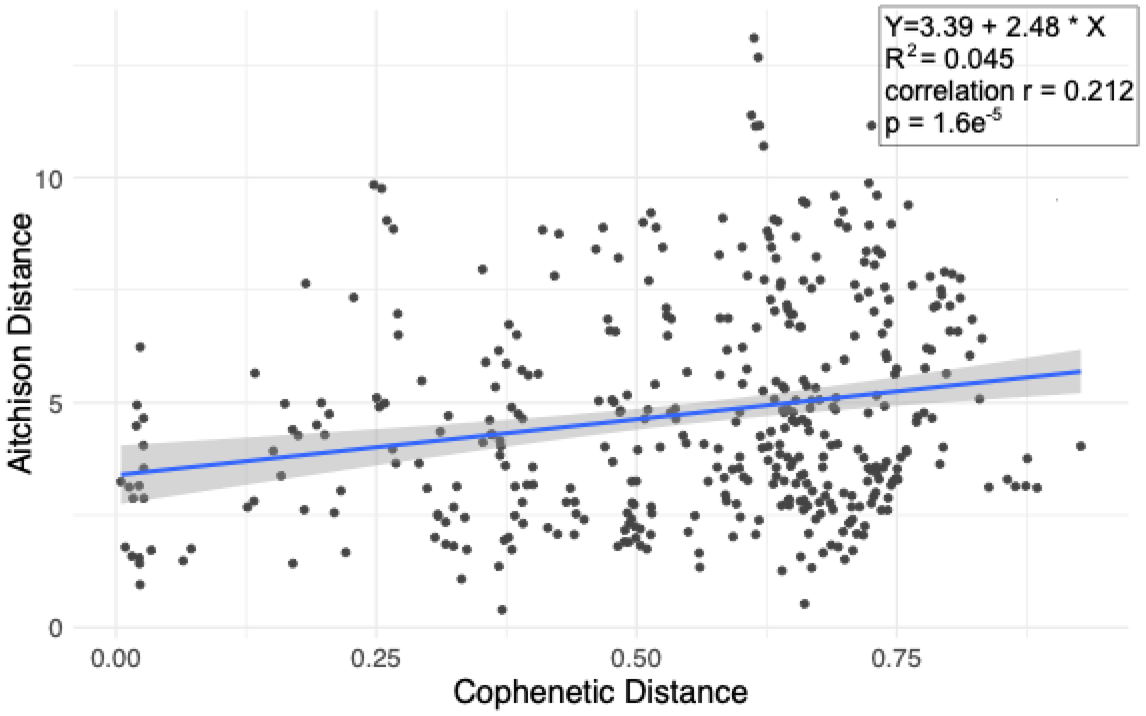


**Supplementary Figure S14**. Scatterplot of pairwise comparisons between 29 RTPR 98% OTUs occurring in both the bacterioplankton and virioplankton fractions on the August 5th sampling date. x-axis: cophenetic distance between OTU reference sequences. y-axis: Aitchison distance of OTUs in each sample. Pairwise self-comparisons were removed.

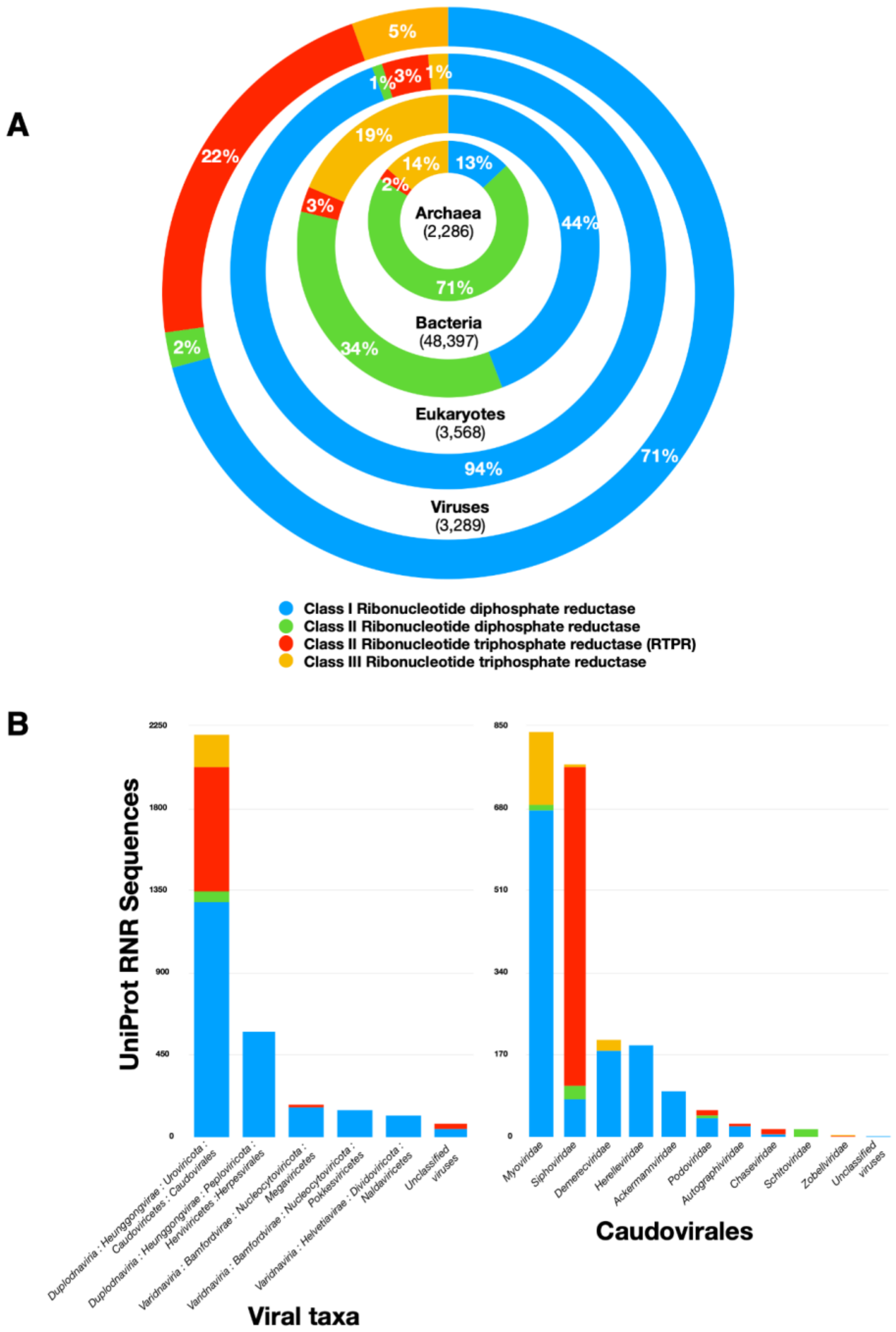


**Supplementary Figure S15**. Frequency of ribonucleotide reductase proteins in UniProt Knowledge Base (Jan. 5, 2022). **(A)** RNRs according to taxonomic kingdom. **(B)** RNRs according to viral taxa (left) and Caudovirales family (right). Numbers in parentheses indicate the total number of RNR proteins in each kingdom.
