## supplementary methods for "Ubiquitous, B_12_-dependent virioplankton utilizing ribonucleotide triphosphate reductase demonstrate interseasonal dynamics and associate with a diverse range of bacterial hosts in the pelagic ocean"

### Molecular genetic methods and sequencing

#### DNA Extraction from the Virioplankton Fraction and Tag-Encoded RTPR Amplicon Sequencing

DNA was extracted from the 0.02 µm Anotop filters (virioplankton fraction) using the MasterPure complete DNA/RNA purification kit (Epicentre, Madison, WI). The Qubit™ dsDNA HS Assay kit (Invitrogen, Carlsbad, CA) measured resulting DNA concentration. RTPR genes within the virioplankton fraction were PCR-amplified using a set of degenerate primers developed from RTPR sequence alignments derived from uncultivated virus populations observed within marine viromes. Forward primers (F1: 5'-CCAAGWCKGGYSARTGGTGGG-3'; F2: 5'-CTAAATCAGGTGAKTGGTGGG-3'; F3: 5'-CTAAGWCAGGWSANTGGTGGR-3' and F4: 5'-GTAARTCWGGYAAYTGGTGGG-3') were mixed together equimolar to a final concentration of 10 µM each. Reverse primers (R1: 5'-GTAACMGAKGGCTTGTGYTC-3'; R2: 5'-GTCAYAGAWGGCTTRTGYTC-3'; R3: 5'-GTGRCAGAARSWTTGTGWKY-3' and R4 5'-GTTAYASWAGGTTTRTGTTC-3') were mixed together equimolar to a final concentration of 10 µM each. PCR assays were performed in a 50 µL reaction volume using 3 ng of extracted DNA, 25 µL of Dream Taq 2x concentrated master mix (Thermo Fisher Scientific, Waltham, MA), 1 µL of BSA (bovine serum albumin, Thermo Fisher Scientific), 0.5 µL 10 µM forward primer mixture, 0.5 µL 10 µM reverse primer mixture, and Ambion Nuclease-Free Water (Ambion, Inc, Austin, TX) under the following cycling conditions: initial denaturation step of 95 °C for 5 min; 35 cycles of 95 ºC for 45 s, 52 ºC for 45 s, and 72 ºC for 90 s; and a final elongation step of 72 ºC for 4 min. Reactions were held at 4 ºC until further processing. PCR amplification products (amplicons) were purified using AMPure XP Magnetic Beads (Beckman Coulter, Brea, CA). A unique barcode sequence was ligated to purified amplicons from each sample (**Supplementary Table S15**). Ligation reactions were performed in 10 µL reaction volume using 10 ng of purified amplicons, 1 µL 10X concentrated ligase buffer (Thermo Fisher Scientific), 2 µL dNTPs (Thermo Fisher Scientific), 1 µL Promega™ T4 DNA Ligase (Thermo Fisher Scientific), 1 µL of 10 mM barcode oligonucleotides and Ambion Nuclease-Free Water at 15 °C for 12 h, and then moved to -20 °C for 2 h to fully stop the reaction. Ligated products were purified using Zymo DNA Clean & Concentrator kit (Zymo Research, Irvine, CA) and enriched by a limited cycle PCR using barcodes as primers (**Supplementary Table S15**). Reactions were performed in 50 µL reaction volume using 2 ng of barcoded amplicons, 5 µL 10X concentrated Taq DNA polymerase buffer (Takara Bio, Japan), 5 µL dNTPs (Thermo Fisher Scientific), 0.5 µL TaKaRa Ex Taq DNA Polymerase Hot-Start Version (Takara Bio), 1 µL single strand barcode oligonucleotides and Ambion Nuclease-Free Water under the following cycling conditions: initial denaturation step of 95 °C for five min; 22 cycles of 95 °C for 30 s, 52 °C for 60 s, and 72 °C for 90 s; and a final elongation step of 72 ºC for 7 min. Reactions were held at 4 ºC until further processing. Enriched ligated products were purified using Zymo DNA Clean & Concentrator kit. All PCR and ligation reactions were performed on a Bio-Rad MJ Research PTC 200 Peltier Thermal Cycler (Bio-Rad Laboratories, Hercules, CA). Final DNA concentration of each barcoded sample was determined using the Qubit™ dsDNA HS Assay kit. Barcoded samples were pooled together in equal proportions based on their molecular weight and DNA concentrations, and sequenced on a PacBio RSII sequencer (Pacific Biosciences, Menlo Park, CA) at the University of Delaware Sequencing and Genotyping Center (UDSGC), Newark, Delaware. Two RTPR sequencing libraries were prepared in total for the virioplankton fraction and one of the libraries (1st sequencing library) was sequenced twice to improve sequence recovery (**Supplementary Table S15**).

**RTPR primer design**

Primers were developed to amplify an approximately 750 bp fragment from RTPR RNR alpha-subunit sequences identified in marine viromes [77]. Briefly, RTPR RNR alpha-subunit protein sequences from reference viruses and Chesapeake Bay, Gulf of Maine, and Dry Tortugas marine viral metagenomes were aligned with MAFFT (version 7.450) using the FFT-NS-i x1000 algorithm [111]. Alignments were visualized in Geneious (version 5.6.2) [112] to identify regions with high conservation, which were used to design potential primers. Putative primer sequences were screened for melting temperature and dimerization using an online primer calculator (www.thermofisher.com), and primer specificity was assessed with the NCBI Primer-BLAST tool against viruses in the nr database. Four primer pairs were chosen to minimize degeneracy while maximizing targeted viral diversity: Forward primers F1 (5’ – CCAAGWCKGGYSARTGGTGGG – 3’), F2 (5’ – CTAAATCAGGTGAKTGGTGGG – 3’), F3 (5’ – CTAAGWCAGGWSANTGGTGGR – 3’), and F4 (5’ – GTAARTCWGGYAAYTGGTGGG – 3’); reverse primers R1 (5’ – GTAACMGAKGGCTTGTGYTC – 3’), R2 (5’ – GTCAYAGAWGGCTTRTGYTC – 3’), R3 (5’ – GTGRCAGAARSWTTGTGWKY – 3’), and R4 (5’ – GTTAYASWAGGTTTRTGTTC – 3’).

**DNA Extraction from the Bacterioplankton Fraction, and 16S rRNA Amplicon and Tag-Encoded RTPR Sequencing**

DNA was extracted from the 0.22 µm Sterivex (Merck Millipore, Darmstadt, Germany) filters (bacterioplankton fraction) using the phenol/chloroform method [53] and quantified with a Qubit™ dsDNA HS Assay kit. The Sterivex filters were thawed to room temperature, after which the housing was cut open with a razor blade. The filter was removed and cut into small pieces (~4x4 mm) using sterile technique. Cells captured on the filter were lysed, first by incubating the filter pieces at room temperature with agitation in 1000 µL of lysis buffer (40 mM glucose, 20 mM Tris-HCl, 60 mM EDTA, 1 mg/mL lysozyme) for 2 h and then for an additional 10 min after adding 200 µL of 10 % sodium dodecyl sulfate (1.6% final concentration). DNA was purified by two sequential organic solvent extractions. The first extraction was performed with buffer-saturated phenol pH >7.4 (Thermo Fisher Scientific) in a solvent:sample ratio of 1:1.4 (840 µL). The extraction phases were separated by a 10 min centrifugation at 21,000 × g at 4 °C. The aqueous phase (1 mL) was transferred to a new 2 mL centrifuge tube and an equal volume (1 mL) of 24:1 chloroform/isoamyl alcohol was added to the sample. After mixing by inversion 15 times, the aqueous phase was separated by a 10 min centrifugation at 21,000 × g at 4 °C. The aqueous phase (0.8–1mL) was kept and a half volume of 7.4 M ammonium acetate was added, followed by a 30 min incubation at room temperature and pelleting of the precipitate (21,000 × g for 10 min at 4 °C). The pellet was discarded and the supernatant split into three new 2 mL tubes. The DNA was precipitated by adding 2.5X of 100% ice cold ethanol and incubating overnight at 4 °C. The precipitated DNA was pelleted into a single tube by 3 subsequent centrifugations at 21,000 × g for 30 min at 4 °C (the supernatant was discarded). The resulting DNA pellet was washed by adding 300 µL of 70% ethanol, followed by centrifugation at 21,000 × g for 5 min at 4 °C and discarding of the supernatant. The DNA pellet was dried at room temperature for 1–2 h and resuspended in 50 µL of 55 °C elution buffer (10 mM Tris-HCl, pH 8.5). Purified DNA was quantified with a QubitTM dsDNA HS Assay kit and stored at -80 °C until further analysis. All centrifugation steps were performed in an Eppendorf centrifuge 5417 R (Eppendorf AG, Hamburg, Germany) equipped with a 30 place aerosol tight rotor with a diameter of 9.5 cm (FA-45-30-11).

The V3–V4 hypervariable region of the 16S rRNA gene was PCR-amplified and sequenced from the purified bacterioplankton fraction DNA on the Illumina MiSeq (Illumina, San Diego, CA) using a Nextera XT DNA Library Preparation Kit (Illumina), which exploits a dual-indexing strategy for multiplexed sequencing [54]. Each sample was associated with a unique pair of indices (**Supplementary Table S16**) from Nextera XT DNA Library Preparation Kit. Two bacterial 16S rRNA libraries consisting of all 35 samples were prepared equilmolar and sequenced independently using the same protocol to obtain greater read depth.

RTPR genes within the bacterioplankton fraction were PCR-amplified, barcoded, and enriched using the same protocol as described for virioplankton. Purified RTPR amplicons were ligated with a unique barcode per sample (**Supplementary Table S17**) and then pooled into one of two RTPR sequencing libraries in equal proportions based on their molecular weight and DNA concentrations. Each library was ligated with a distinct PacBio index enabling subsequent sequence deconvolution. Finally, the two sequence libraries were pooled together in equal proportions and sequenced on a PacBio Sequel (Pacific Biosciences) at the UDSGC.

### Bioinformatic methods

#### RTPR amplicon quality control

General and detailed analysis pipelines are summarized in **Fig. 1** and **Supplementary Fig. S1**, respectively. Briefly, RTPR gene amplicon sequences from the virioplankton fraction (PacBio RSII sequencer) and the bacterioplankton fraction (PacBio Sequel sequencer) were initially screened for low quality bases and read length using the PacBio Data Repository of the Delaware Biotechnology Institute (http://pacifica.dbi.udel.edu) and Delaware Biotechnology Institute's SMRT Link 6.0.0.47841 (https://sequel.dbi.udel.edu), respectively. Circular consensus sequence (CCS) reads were generated. Reads with less than three full passes, less than 98% minimum predicted accuracy, and a length shorter than 250 bp or longer than 5,000 bp were excluded. Generated CCS reads were demultiplexed using the split_libraries.py script from the QIIME 1 package version 1.7.0 [113].

Demultiplexed CCS nucleotide sequence reads were screened for forward and reverse degenerate primer sequences using a custom demultiplex.pl script (https://github.com/dnasko/biobin/tree/master/pcr products/adapter searching). Reads containing both forward and reverse primer sequences with at least 85% similarity in their first 60 base pairs at either end were retained. Barcode and primer sequences were trimmed from the reads.

Demultiplexed and trimmed nucleotide sequence reads were translated into predicted amino acid sequences using a custom frameshift polishing pipeline (https://github.com/dnasko/frameshift_polisher). Briefly, the frameshift polishing pipeline performs a six-frame translated BLASTx [114] search of each CCS read against a curated database of reference peptides from the *Tara* Oceans [75] and RNRdb [76] databases (https://doi.org/10.5281/zenodo.3756785). Alignments between each of the translated CCS reads and reference peptides were then parsed to identify instances in which multiple alignments were reported between a CCS read and a reference peptide in multiple frames. In the event a frameshift was detected the corrected peptide was produced in FASTA format. Additional frameshift polishing pipeline details are provided in the next section**.**

Key catalytic residues in translated RTPR amino acid sequences [10] were confirmed by multiple sequence alignments using Protein Active Site Validation (PASV) (https://github.com/mooreryan/pasv) (Moore et al. 2021). RTPR amino acid sequences without the key residues (C408, E410, and C418 in *Lactobacillus leichmannii* monomeric class 2 adenosylcobalamin-dependent ribonucleoside-triphosphate reductase (UniProt acc. Q59490 (RTPR_LACLE)) were removed. All RTPR amino acid sequences with key residues were aligned using MAFFT (version 7.450) (FFT-NS-1) [115] in Geneious (version 10.0.9) and subsequently trimmed to the region of interest (H346 to S643 in *Lactobacillus leichmannii* monomeric class 2 adenosylcobalamin-dependent ribonucleoside-triphosphate reductase).

**Frameshift polishing pipeline**

Despite improvements in PacBio DNA sequencing accuracy and read correction algorithms, frameshifts can still occur in 3X circular consensus sequences. These errors must be identified and corrected prior to subsequent bioinformatic analyses of peptide sequences. A custom frameshift polishing pipeline (https://github.com/dnasko/frameshift_polisher) is a BLAST-based pipeline capable of detecting and correcting potential frameshifts in coding genes. Users provide a FASTA file of nucleotide sequences to be corrected and the pipeline searches these sequences against a user-defined protein database.

The protein database should include a set of proteins related to those in the survey. For example, if a user sequenced amplicons of the ribonucleotide triphosphate reductase (RTPR) gene, the database should include a diverse set of RTPR peptide sequences. Users can generate a custom database of proteins by searching for them on UniProt (https://www.uniprot.org) and downloading the set of relevant sequences as a FASTA file. A protein BLAST database of these sequences must then be created using the command line BLAST software (https://blast.ncbi.nlm.nih.gov/). Importantly, the sequences in the reference database must all be in-frame (i.e., contain no frameshifts).

The frame shift polishing pipeline will search the input nucleotide sequences against the protein database in all six frames and produce alignments with an e value ≤1 and a percent similarity >30%. Results of this search are then parsed, searching for instances where a query sequence (a translated nucleotide amplicon sequence in the input FASTA file) aligns with a subject sequence (a sequence in the protein database):

1. multiple times, and
2. in multiple frames, and
3. in non-overlapping regions of the protein

The multiple non-overlapping alignments are then concatenated together to form one frameshift-corrected protein sequence. Ambiguous amino acids (denoted by an ‘X’) are inserted in regions where the multi-frame alignments do not overlap.
